# HIV-1 protein coding sequences are present in relevant bacteria

**DOI:** 10.1101/2024.12.04.626764

**Authors:** Hector F. Pelaez-Prestel, Juan Mozas-Gutierrez, Esther M. Lafuente, Pedro A. Reche

## Abstract

Human Immunodeficiency Virus (HIV) is a retrovirus that attacks the immune system, causing acquired immunodeficiency syndrome (AIDS). Early diagnosis and treatment of HIV infected individuals are considered key to reduce HIV transmission and developing AIDS. Therefore, HIV diagnostics play an important role in the battle against AIDS. HIV tests are regarded as reliable and very specific. However, false-positive results are known to occur, usually caused by infections with unrelated pathogens leading to cross-reactive antibodies. In this work, we found through TBLASTN searches that the genome of several bacterial species, most frequently *Klebsiella pneumoniae and Escherichia coli*, contain segments matching HIV-1 proteins, including p24 and other proteins relevant for HIV testing, with a high degree of similarity (sequence identity > 95 %). The presence of HIV-1 in these bacteria of the human microbiota does not appear to be an artifact, since HIV-1 proteins were detected in different isolates of the same species. The proteome of other common viruses, particularly Influenza A virus and Hepatitis B virus, was also detected in bacterial genomes, but to a much lesser extent. Overall, our findings support that some bacteria can acquire HIV-1 genetic material and could interfere with HIV testing, causing false-positives.

## 1. INTRODUCTION

Human Immunodeficiency Virus (HIV) is an enveloped retrovirus that attacks CD4^+^ T cells, leading to their gradual depletion and the development of acquired immunodeficiency syndrome (AIDS) (1,2). HIV transmission occurs mainly through sexual contact, often between men who have sex with men but in Africa, where heterosexual sex transmission is dominant (3). The virus can be categorized into 2 distinct subtypes: HIV type 1 (HIV-1) and HIV type 2 (HIV-2). Both viruses are transmitted similarly, show the same tropism and cause AIDS (4). However, infections by HIV-1 are much more common and progress quicker to AIDS than HIV-2 infections (4).

AIDS is the last stage of HIV infection, where the clinical symptoms of a weakened immune system manifest (5,6). The first stage is the acute phase, which can pass unnoticed or manifest with flu-like symptoms. During this phase, lasting 2-4 weeks after infection, the virus replicates quickly, spreading through the body, and HIV genome and antigens, notably HIV-1 p24, can be detected in blood. The second stage is the chronic or asymptomatic phase, in which the host starts producing antibodies (Ab) and becomes HIV seropositive. In this stage, HIV antigens drop and may be undetectable, yet viral replication and destruction of CD4^+^ T cells continue and patients eventually enter the AIDS stage unless they receive antiretroviral therapy (ART) (5,6).

There is evidence that starting ART early in the course of HIV infection has long-term health benefits for patients and decreases the risk of transmission (7,8). Therefore, ART is currently recommended for all HIV positive people, regardless of CD4^+^ T cell count (9). Moreover, in order to decrease the risk of infection, antiretroviral prophylaxis is also recommended for perinatally-exposed newborns and people who has been exposed to contagious HIV-positive material (9). Despite the noted benefits, ARTs do not eliminate the virus and have a considerably degree of toxicity and long-term adverse effects (10). Thereby, patients in ART are continuously monitored, and may require adjustments of the ART regime and additional supportive treatments, all of which represent a considerable burden to both, patients and health care systems (10).

HIV diagnostics play a relevant role in the battle against HIV/AIDS, facilitating early initiation of HIV treatments. Moreover, they are also useful to stage and monitor patients, and for outbreak surveillance. HIV tests have historically relied on commercial enzyme immunoassays (EIAs) that detect antibodies to HIV antigens in serum (5,11). Current generation of commercial HIV EIA tests (4-th and 5-th generation), detects Abs against HIV-1 and HIV-2 as well as HIV-1 p24 antigen, which allows for earlier diagnosis of HIV-1 infection (5). A list of FDA approved laboratory Antibody/Antigen (Ab/Ag) HIV tests are available at https://www.cdc.gov/hiv/partners/testing/laboratorytests.html (accessed on October/10/2024). In these tests, HIV-1 p24 is detected by using Abs generated against this protein or fragments derived from it, while HIV antibodies are detected by using HIV antigens and/or antigen fragment cocktails. The following antigens are generally considered for HIV-1 testing (coding gene in italic): gp160 (envelope protein precursor, *env*), gp120 (extracellular region of envelope protein), gp41 (transmembrane region of envelope protein), p31 (integrase, *pol*), p51 (reverse transcriptase, *pol*), p24 (core protein, *gag*) and p17 (matrix protein, *gag*). In the laboratory, detection of Ab is sufficient to confirm HIV infection but differentiation IEA tests may be necessary to distinguish between HIV-1 and HIV-2 infection. If the Ag test for HIV-1 p24 is positive but the Ab is negative, supplemental confirmatory IEA tests (e.g. western-blotting), or RNA tests will be necessary to confirm or negate HIV-1 infection (12). HIV RNA tests involve reverse transcription of HIV genetic material into DNA and are aimed to detect conserved regions in *gag* or *pol* genes (5).

HIV tests are regarded as very reliable and specific. However, false-positives (FPs) can occur due to technical reasons (e.g. misinterpretation and clerical errors) and more importantly under certain biological or medical conditions (13–15). Relevant conditions that have been reported to interfere with HIV IEA tests and produce FP results include: pregnancy (16), recent vaccination (e.g. flu, Tdap or Hepatitis B vaccine) (17–19), cancer (19), and autoimmune diseases like lupus (20) or rheumatoid arthritis (20). Likewise, various infectious diseases (e.g. schistosomiasis, malaria, COVID-19 and Epstein-Barr virus mononucleosis) have been shown to produce FPs (21–24), likely due to cross-reactivity.

In this work, we investigated potential cross-reactivity between bacteria and HIV-1, using TBLASTN searches against bacterial genomes with the HIV-1 proteome. Interestingly, we detected the presence of HIV-1 protein coding sequences in many bacterial species but more frequently in *Klebsiella pneumoniae* and *Escherichia coli,* which are part of the human gut microbiota. Sequence identity between HIV-1 and bacterial genomes was not only high (up to 100%) but reached the entire HIV-1 proteome, including p24 and other relevant proteins in HIV-1 testing. Other common viruses were also detected in bacterial genomes, but to a much lesser degree. These results support that some bacteria can acquire HIV-1 sequences, which could potentially induce immune cross-reactivity to HIV-1, causing FP results in HIV tests. Ruling out this possibility will reduce the chance of putting healthy individuals on ART and alleviate the economic burden of HIV treatments on health care systems.

## 2. METHODS

### Genomes and proteins

The HIV-1 reference genome (HIV-1 group M isotype B, isolate HXB2) was obtained from NCBI (Accession: NC_001802), and amino acid coding sequences (CDS) and mature proteins were collected upon it in a FASTA file. The entire proteomes of nine common human viruses, consisting of the CDS in the relevant NCBI genomic sequences, were also collected in FASTA files (Table 1).

**Table 1.** Proteomes of common human viruses considered in this study.

| Accession | Virus | CDS | Amino Acids |
| --- | --- | --- | --- |
| NC_038311 | Human rhinovirus A | 1 | 2157 |
| NC_009996 | Human rhinovirus C | 1 | 2144 |
| NC_038235 | Respiratory Syncytial virus A | 11 | 4536 |
| NC_001781 | Respiratory Syncytial virus B | 11 | 4540 |
| NC_026422_29* | Influenza A virus | 12 | 4784 |
| NC_002204_11* | Influenza B virus | 10 | 4609 |
| NC_003977 | Hepatitis B virus | 8 | 2299 |
| M62321 | Hepatitis C virus | 1 | 3011 |
| NC_027779 | Papilloma virus | 6 | 2309 |

\* Proteome is contained in various independent accessions varying the last two digits and the range is indicated. The corresponding author will provide the FASTA files upon written request.

### Microbial genome BLAST searches

Sequence similarity searches of viral proteins in microbial genomes were carried online at NCBI BLAST site (https://blast.ncbi.nlm.nih.gov/Blast.cgi), selecting TBLASTN. The amino acid sequences of viral proteomes/proteins were entered as query and the RefSeq Genome Database was selected as the target database, restricting the search to Bacteria (taxid:2). TBLASTN searches were carried out with default settings but increasing the maximum target to 1000. The raw output from searches was downloaded as text and BLAST summary results with taxonomic information were saved in Excel (Microsoft). The taxonomic TBLASTN information tabulated the bacterial species with TBLASTN hits, indicating the number of hits and top bit-score of all hits *per* bacteria as well.

Similar TBLASTN searches were carried out to determine bit-scores of viral proteomes/proteins with themselves but restricting the search in RefSeq Genome Database to Viruses (taxid:10239). As result, for each protein a self bit-score resulting from a hit alignment with itself was obtained and recorded. Bit-scores only depend on the hit sequence alignment, not the size of the database. Hence, the self bit- score (sBS) of a viral sequence query corresponds to the maximum bit-score (mBS) that the query sequence can reach.

### Relative bit-scores

Relative bit-scores (RBSs) of TBLASTN hits between viral protein queries and subject bacterial genomic sequences were obtained by dividing bit-scores (BS) with the corresponding mBS, as indicated in Equation 1.

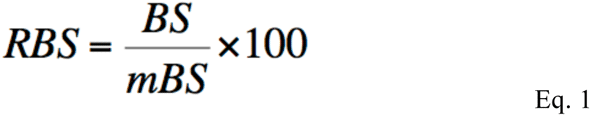

Where mBS is the maximum bit-score of the relevant viral protein sequence obtained from a self-hit alignment, as defined previously. Since a given viral protein query sequence can produce numerous TBLASTN hits to bacterial sequences from the same species, the top RBS for each viral protein per bacterial species was computed after the best hit, that with the top bit-score.

### Viral proteome coverage

The viral proteome coverage (VPC) was introduced as an estimation of the overall presence and similarity of an entire viral proteome (*p*) in a given bacterial species (*b*). For any given viral proteome used in TBLASTN searches, VPC (*p,b*) was obtained using Equation 2, which tallies the top RBS (tRBS) from the best hit of each protein in the viral proteome (*p*:*1 .. N*) *per* bacterial species (b), divided by the total number of proteins, N, in the viral proteome.

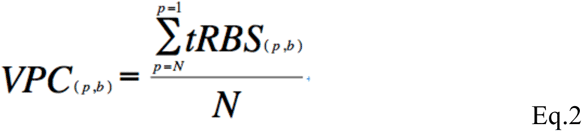

### Other procedures

TBLASTN results were parsed using PERL scripts to compute RBSs of hit alignments and data was processed in Excel (Microsoft, Corp.) Multiple sequence alignments between HIV-1 proteins and bacterial TBLASTN hits were generated using T-COFFEE (25). All graphics were generated using GraphPad Prism (GraphPad Software, Inc.).

## 3. RESULTS

### Detection of HIV-1 proteins in bacterial genomes

We investigated bacteria as potential sources of immune cross-reactivity with HIV-1 by queering bacterial genomes with HIV-1 protein sequences, using TBLASTN (details in Methods). The results revealed that all HIV-1 proteins produced highly significant TBLASTN hits in the genome of various bacterial species (TBLASTN output is provided in Supplementary File 1). The single most significant hits with bacteria were produced by p24, Vpr, Integrase, Protease and Reverse Transcriptase (RT) p51, which reached a relative bit-score (RBS) of > 97 % (Fig 1). The RBS of the hits between HIV-1 proteins and bacterial genomes were computed by dividing the bit-score of the hit alignments with the maximum possible bit-score of the relevant HIV-1 proteins, obtained from a self-hit (details in Methods). An RBS of 100% for a given HIV-1 protein indicates that the protein produces a hit alignment over their entire length with 100% sequence identity. As shown in Figure 1A, all HIV-1 proteins produced a considerable number of hits with bacteria genomes with RBS ≥ 50 % and the RBS of the best scoring hit was ≥ 58 % for all HIV-1 proteins. Interestingly, only a handful of bacterial species, including *Escherichia coli* and *Klebsiella pneumoniae*, were the subject of the best scoring hits with HIV-1 proteins (Fig.1). These best scoring hits between HIV-1 proteins and bacteria had a sequence identity ≥ 63 % (value from Tat) and covered at least 65% of the length of query (value from Vpu) (Table 2). Multiple sequence alignments of HIV-1 proteins with bacterial genomes are provided in Supplementary File 2.

**Figure 1.**
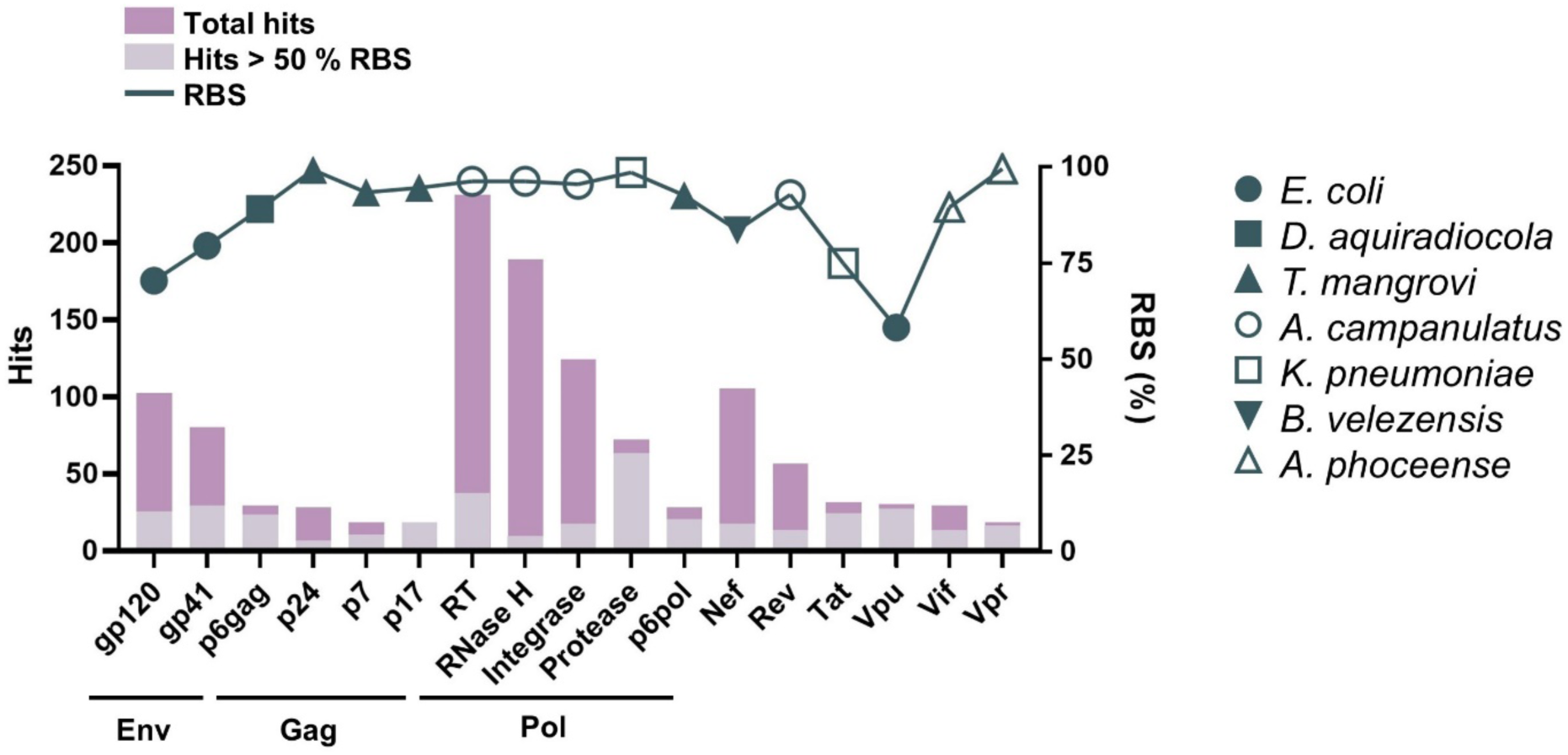
**Presence of HIV-1 proteins in bacterial genomes**. Chart showing for each HIV-1 protein (x-axis) the number of **TBLASTN** hits with bacterial genomes (left y-axis). Stacked columns in light grey show the number of hits with RBS ≥ 50%. The RBS (right y-axis) of the best hit for each HIV-1 protein is also plotted in the chart, using different symbols that represent the bacterial species, indicated on the right, subject of the hit. HIV-1 proteins that derive from Env, Gag and Pol are underlined.

**Table 2.**
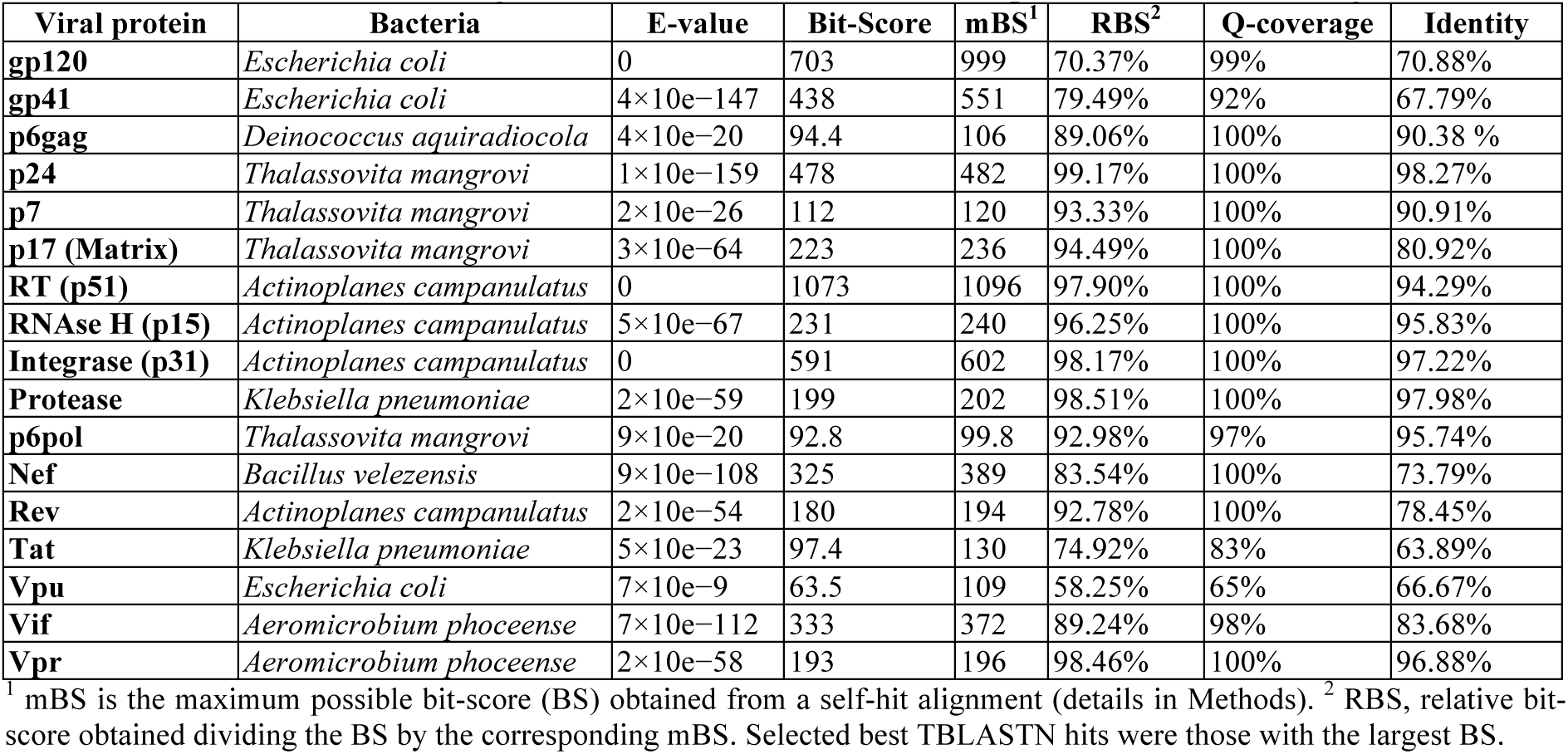
Statistics of best scoring TBLASTN hits between HIV-1 proteins and bacterial genomes.

| Viral protein | Bacteria | E-value | Bit-Score | mBS<sup>1</sup> | RBS<sup>2</sup> | Q-coverage | Identity |
| --- | --- | --- | --- | --- | --- | --- | --- |
| gp120 | *Escherichia coli* | 0 | 703 | 999 | 70.37% | 99% | 70.88% |
| gp41 | *Escherichia coli* | $4 \times 10^{-147}$ | 438 | 551 | 79.49% | 92% | 67.79% |
| p6gag | *Deinococcus aquiradiocola* | $4 \times 10^{-20}$ | 94.4 | 106 | 89.06% | 100% | 90.38 % |
| p24 | *Thalassovita mangrovi* | $1 \times 10^{-159}$ | 478 | 482 | 99.17% | 100% | 98.27% |
| p7 | *Thalassovita mangrovi* | $2 \times 10^{-26}$ | 112 | 120 | 93.33% | 100% | 90.91% |
| p17 (Matrix) | *Thalassovita mangrovi* | $3 \times 10^{-64}$ | 223 | 236 | 94.49% | 100% | 80.92% |
| RT (p51) | *Actinoplanes campanulatus* | 0 | 1073 | 1096 | 97.90% | 100% | 94.29% |
| RNase H (p15) | *Actinoplanes campanulatus* | $5 \times 10^{-67}$ | 231 | 240 | 96.25% | 100% | 95.83% |
| Integrase (p31) | *Actinoplanes campanulatus* | 0 | 591 | 602 | 98.17% | 100% | 97.22% |
| Protease | *Klebsiella pneumoniae* | $2 \times 10^{-59}$ | 199 | 202 | 98.51% | 100% | 97.98% |
| p6pol | *Thalassovita mangrovi* | $9 \times 10^{-20}$ | 92.8 | 99.8 | 92.98% | 97% | 95.74% |
| Nef | *Bacillus velezensis* | $9 \times 10^{-108}$ | 325 | 389 | 83.54% | 100% | 73.79% |
| Rev | *Actinoplanes campanulatus* | $2 \times 10^{-54}$ | 180 | 194 | 92.78% | 100% | 78.45% |
| Tat | *Klebsiella pneumoniae* | $5 \times 10^{-23}$ | 97.4 | 130 | 74.92% | 83% | 63.89% |
| Vpu | *Escherichia coli* | $7 \times 10^{-9}$ | 63.5 | 109 | 58.25% | 65% | 66.67% |
| Vif | *Aeromicrobium phoceense* | $7 \times 10^{-112}$ | 333 | 372 | 89.24% | 98% | 83.68% |
| Vpr | *Aeromicrobium phoceense* | $2 \times 10^{-58}$ | 193 | 196 | 98.46% | 100% | 96.88% |

<sup>1</sup> mBS is the maximum possible bit-score (BS) obtained from a self-hit alignment (details in Methods). <sup>2</sup> RBS, relative bit-score obtained dividing the BS by the corresponding mBS. Selected best TBLASTN hits were those with the largest BS.

The genome database used in TBLASTN searches included not one but many genomes from the same bacterial species, usually different strains/isolates. Therefore, it is worth noting that HIV-1 proteins produced very similar hits in different representative genomes of a particular bacterial species (see Supplementary file 1). The number of hits per bacterial species with RBS ≥ 50% corresponding to HIV-1 proteins relevant for HIV-1 testing is shown in Figure 2 along with the RBS of the best hit.

**Figure 2.**
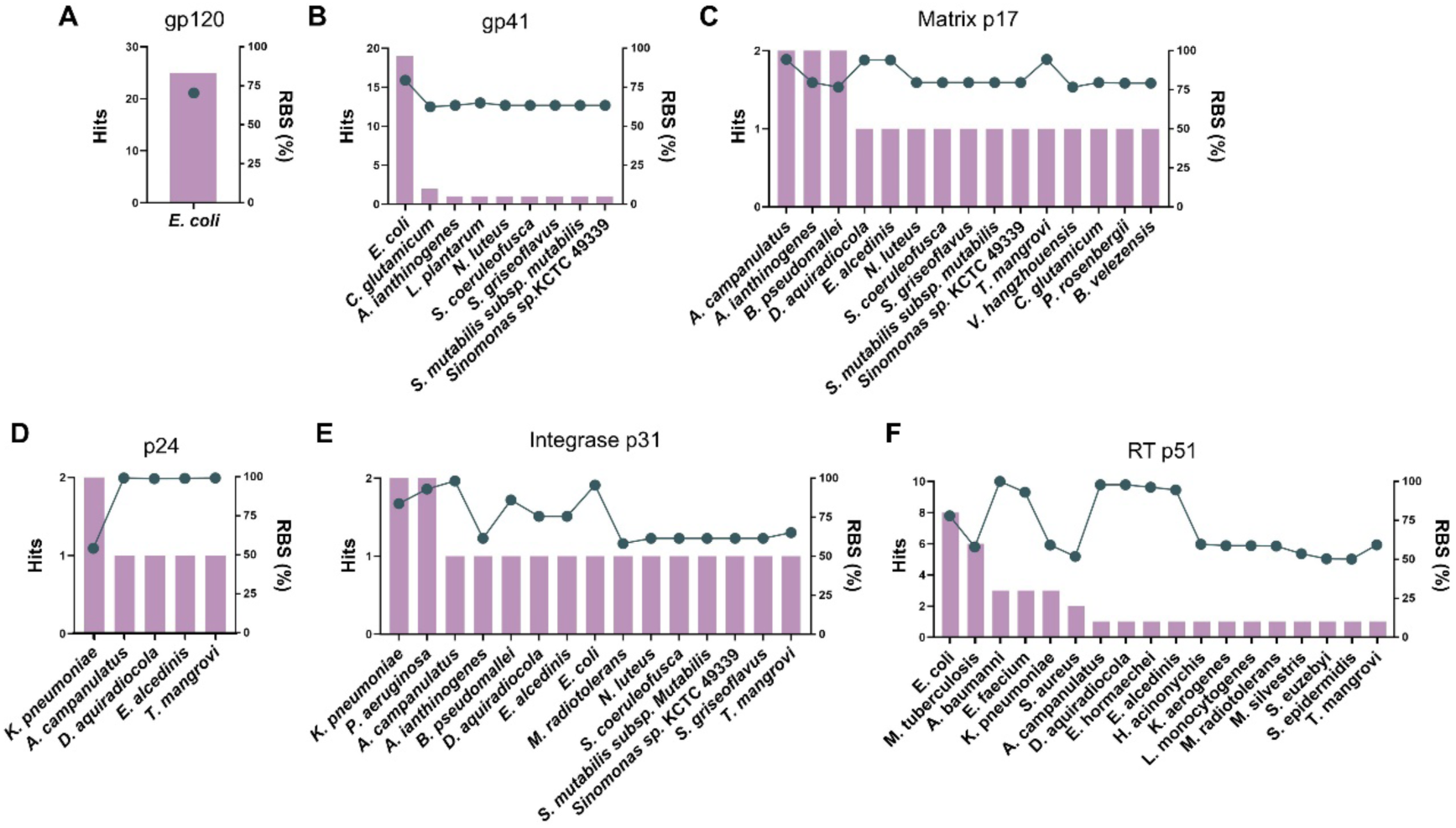
HIV-1 protein hits per bacterial species. The graphs show the number of hits on bacterial genomes (RBS ≥ 50%) of gp120 (**A**), gp41 (**B**), p17 (**C**), p24 (**D**), p51 (**E**), and p31 (**F**), stratified by bacterial species. The graphs also depict the RBS (line and dots) of the best scoring hit per bacterial species.

The selected proteins include gp120 and gp41, p24 and p17, p31 and p51. The first two are part of the HIV-1 envelope glycoprotein (gp160 or Env) and derive by cleavage, while p24 and p17 (matrix) derive from Gag protein, and p31 (integrase) and p51 (reverse transcriptase, RT) from Pol. The bacterial species whose genomes resulted in significant hits varied with the distinct HIV-1 proteins (Fig.2). However, *E. coli* and *K. pneumoniae* were the subject of the most significant hits with the majority of HIV-1 proteins. Only p17 (matrix) did not yield any significant hit with these two bacterial species.

### HIV-1 proteome coverage in bacterial genomes

Since many HIV-1 proteins produced significant hits with the same bacterial species, we introduced the viral proteome coverage (VPC) to estimate how much of the HIV-1 proteome, not just a single protein, is present in the genome of a specific bacterial species. The VPC was computed by adding up the RBS value from the best scoring hit of each viral protein to the genome of a specific bacterial species and dividing by the total number of viral proteins, 17 for HIV-1, used in TBLASTN searches (details in Methods). A VPC of 100% will indicate that all proteins included in the viral proteome produce 100% identity hit alignments spanning their entire length. As shown in Figure 3, HIV-1 reached the largest VPC in *K*. *pneumoniae* (67.74%), followed by *E. coli* (63.72%), *Actinoplanes campanulatus* (60.12 %), *Enterococcus alcedinis* (59.91%) and *Deinococcus aquiradiocola* (59.87%) However, while *A. campanulatus*, *E. alcedinis* and *D. aquiradiocola* were the subject of only a few, yet highly significant hits*, K*. *pneumoniae and E. coli* were the subject of the majority of significant hits, indicating that HIV-1 proteome appears repeatedly in these two species and this presence is unlikely the result of contamination or experimental artifacts. The data used to compute VPC values is provided in Supplementary Table S1.

**Figure 3.**
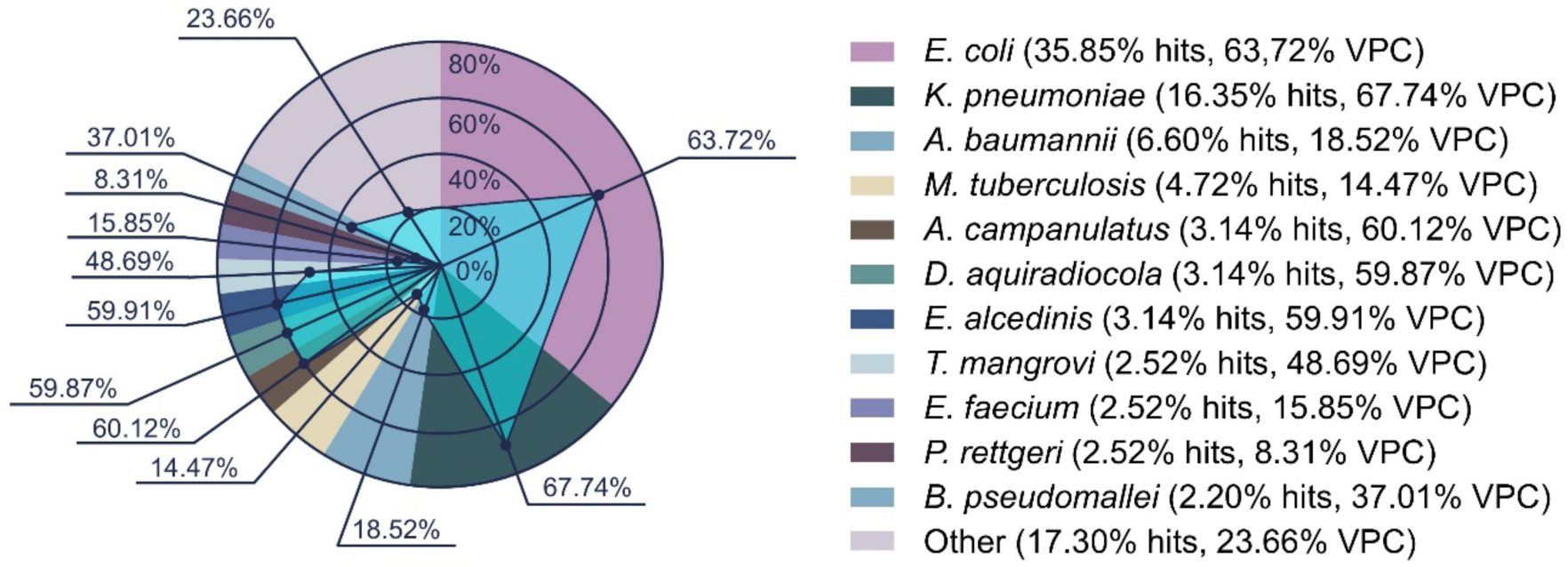
HIV-1 proteome in bacterial species. Parts-to-whole chart show the percentage of significant TBLASTN hits (RBS ≥ 50%) to each bacterial species considering all HIV-1 proteins. Total number of hits with ≥ 50% was 318. The spider charts represent the VPC per bacterial species computed after the best hits. The VPC shown in *Other* corresponds to the VPC determined on *Pseudomonas aeruginosa*, which reaches the largest VPC of all bacterial species included under *Other*.

### Proteome coverage of common viruses in bacterial genomes

Given the extensive presence of HIV-1 in bacterial genomes, we examined whether the same was true for other viruses. To that end, we carried out TBLASTN searches against bacterial genomes with the proteome of common viruses and analyzed hits computing their RBS. The VPC of the viruses per bacterial species was also computed after the RBS of the best scoring hits. The virus proteomes included Human Rhinovirus A (HRVA) and C (HRVC), Respiratory Syncitial virus A (RSVA) and B (RSVB), Influenza A virus (IAV), Influenza B virus (IBV), Hepatitis B virus (HBV), Hepatitis C virus (HCV) and Human Papiloma virus (HPV). The results revealed that, all these viruses, with the exception of IAV, produced much fewer significant hits (RBS ≥ 50%) in bacterial genomes than HIV- 1 (Fig.4A). IAV produced a considerable number of highly significant hits and reached a VPC of ≥ 75% in *Listeria monocytogenes* (Fig.4A). HBV produced few but yet highly significant hits and actually reached a VPC of 100% in *Streptomyces sp. 2BB-J2* (Fig.4). Supplementary Table S2 and S3 collect the statistics of the best TBLASTN hits between IAV and HBV proteins with bacterial genomes, respectively. Interestingly, we also noted that IAV produced a significant number of hits with *E. coli* reaching a VPC ∼60% (Fig.3B). Likewise, HBV reached a VPC of ∼60% in *K. pneumoniae* (Fig.4C). In supplementary Table S4 and S5, we provide the VPC data of IAV with *E. coli* and of HBV with *K. pneumoniae*, respectively.

**Fig. 4.**
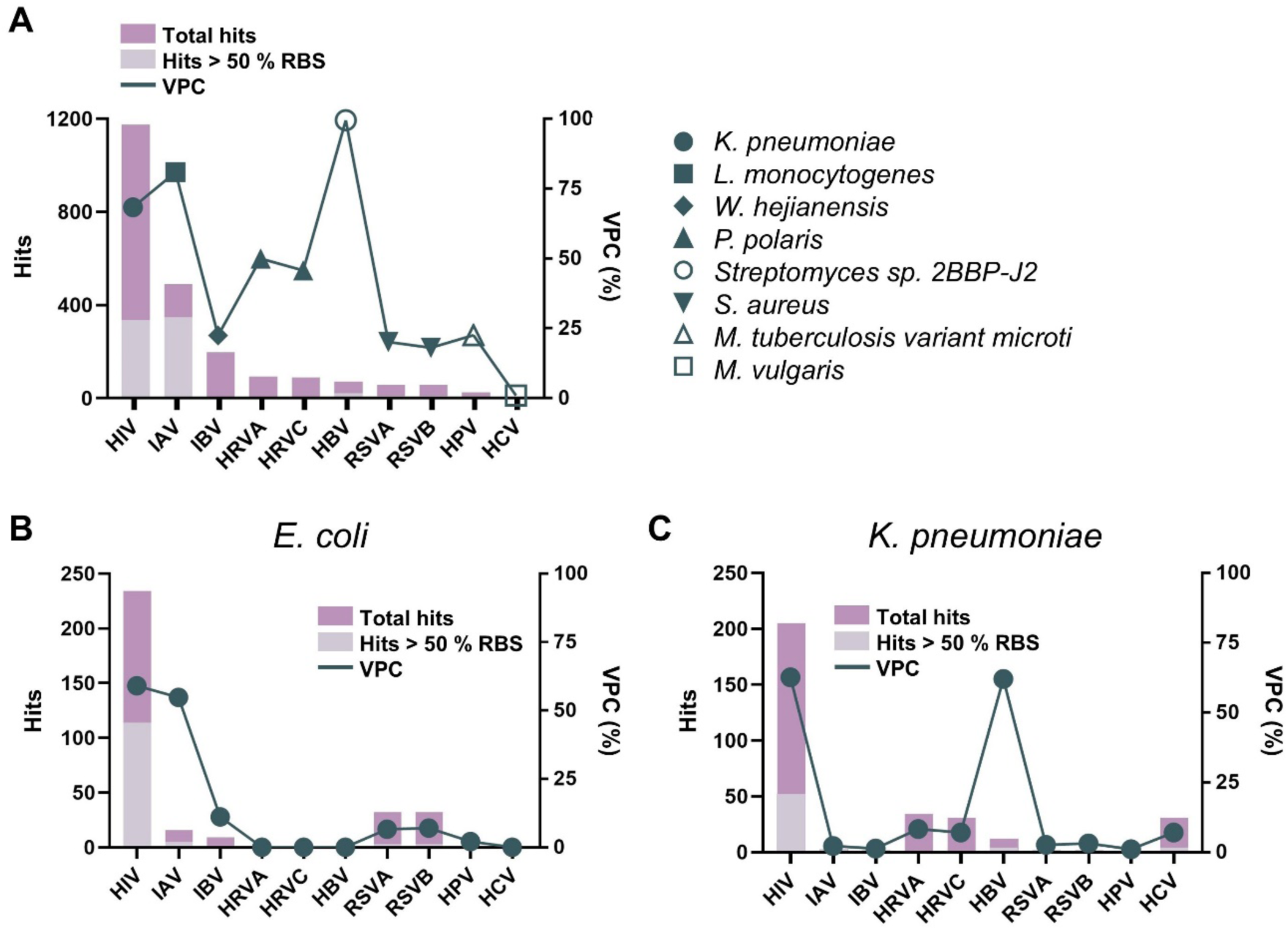
Presence of common virus proteomes in bacterial species. **A)** Chart showing for each virus (HIV-1, IAV IBV, HRVA, HRVC, RSVA, RSVB, HPV and HCV) (x-axis) the total number of TBLASTN hits with bacterial genomes (left y-axis). Stacked columns in light grey show the number of hits with RBS ≥ 50%. The largest VPC value (right y-axis) reached by the viruses is also plotted in the chart (right y-axis), using different symbols that represent the bacterial species (shown on the right) where that particular value was attained. **B**) and **C**) show the same data as in panel A but limited to TBLASTN hits with *E. coli* and *K. pneumoniae*, respectively.

## 4. DISCUSSION

HIV testing is an important part in the fight against AIDS since early diagnosis and treatment maximize the benefits of ART therapy (7,8). Thereby, major attention is paid to minimize false-negatives in HIV tests results that could delay ART therapy. However, false-positive (FP) results in HIV testing are not without consequences, including those derived from exposing healthy individuals to the adverse effects of lifelong treatments, not to mention the significant social and emotional impact on a person. Fortunately, FPs are considered rare, yet they are known to occur. Therefore, it is important to investigate and know the circumstances under which they may arise.

FP HIV results are more commonly reported in relation to infections that induce antibodies cross-reacting with HIV antigens (15). These cross-reactive antibodies likely result from antigens showing molecular mimicry with HIV-1 proteins, which can be readily detected by sequence similarity (26–28). However, the infectious agents that can potentially induce cross-reactive antibodies with HIV remain to be investigated and are unknown. In this work, we investigated bacteria as potential sources of immune cross-reactivity to HIV-1 due to molecular mimicry, using TBLASTN searches with HIV- 1 proteins against bacterial genomes. The results were surprising and unexpected.

We found that HIV-1 proteins appear to be present in quite a few bacterial species (Fig. 1, Fig. 2 and Fig. 3). Proteins like p24, Integrase, Protease, R and Vpr and were found to produce hit alignments with ≥ 96% sequence identity spanning their entire length (relative bit-score, RBS, > 97%). Moreover, hit alignments produced by other HIV-1 proteins were also highly significant (RBS ≥ 58 %) (Fig. 1 and Table 1). *K. pneumoniae* and *E. coli* were the subject of the best hits with HIV-1 proteins that are relevant for HIV testing like gp120, gp41, p24, p31 and p51 (Fig. 2). *K. pneumoniae* and *E. coli* also produced highly significant hits with the remaining HIV-1 proteins, reaching a VPC of 67.74% and 63.72%, respectively. Only three other bacterial species, *A. campanulatus, E. alcedinis and D. aquiradiocola*, reached a comparable VPC of 60.12 %, 59.91% and 59.87%, respectively.

Arguably, the detection of bacterial genomic sequences that are highly similar (if not identical) to HIV-1 proteins or derived fragments could be attributed to contamination or some other experimental artifact. However, the fact that the same/similar highly significant HIV-1 hits were detected in different strains/isolates of the same bacterial species, such as in the case of *K. pneumoniae* and *E. coli,* makes this possibility unlikely. Furthermore, the presence of HIV-1 related sequences in bacterial genomes might be greater if sequences that are not aligned with the relevant taxa are excluded during annotation and curation processes. In comparison with HIV-1, the proteome of other common viruses was seldom detected in bacterial genomes with the exception of Influenza virus A (IAV) and Hepatitis B virus (HBV) (Fig. 4). Interestingly, *K. pneumoniae* and *E. coli* also capitalized a relevant number of highly significant hits with IAV and HBV but with differences. Thus, IAV reached a VPC of ∼ 60% in *E. coli* and of 20% in *K. pneumoniae.* In contrast, HPV reached a VPC of ∼60% in *K. pneumoniae*, while in *E. coli* was ≤ 20%. Overall, these data support that some bacteria (*e.g. K. pneumoniae and E. coli*) can acquire viral genomic sequences, with perhaps certain preferences. The presence of HIV-1 like sequences have been indeed confirmed in bacteria isolated from the gut and respiratory tract of HIV/AIDS patients (29,30). Of note, *E. coli* and *K. pneumoniae* are both part of the human gut microbiota, where they are usually harmless (31,32). However, there are strains of *E. coli* that are pathogenic (33) and *K. pneumoniae* can colonize other tissues and produce an ample range of diseases. Moreover, bacteria in the gut microbiota can cause harm if they enter the organism, in which case the immune system responds and eliminates them. Indeed, antibodies against the gut microbiota are very common in human sera (34), which surely result after occasional breaches in the gut mucosa leading to the entrance of bacteria and subsequent immune recognition.

Viruses, particularly retroviruses, are well known vectors of horizontal gene transfer, posing an important evolutionary force in most life forms, including bacteria (35,36). However, how bacteria can acquire sequences from viruses that infect humans is unclear –more so in the case of RNA viruses– and deserves to be investigated. Nonetheless, human commensal bacteria, including *E. coli,* are known to interact with enteric viruses (37) and likely with other viruses. Hence, we can speculate that such close interactions at the gut, or sites of infection, may also facilitate gene transfer from viruses to bacteria. Alternatively, viral gene transfer may occur between bacteria and virus-infected cells (35). Integration of genomic material from RNA viruses into bacterial genome will require prior reverse transcription to DNA. Incidentally, *E. coli* has been found to express a functional reverse transcriptase (38) and likewise *K. pneumoniae* (39), which is part of a viral defense mechanism (40).

The presence of HIV-1 sequences in *K. pneumoniae* and *E. coli* can have wide implications, since those bacteria are part the gut microbiota and can be transmitted through person-to-person contact or contact with contaminated sources. Bacteria carrying HIV-1 sequences are very unlikely to shed infectious viral particles and contribute to HIV-1 transmission. However, *K. pneumoniae* and *E. coli* specimens bearing HIV-1 sequences could serve as transmission vectors of HIV seropositivity, provided that the bacteria can express HIV-1 proteins. Interestingly, this appears to be the case. By using HIV-1 antibodies against various Gag derived proteins, Hainova *et al.* (41) detected a protein in bacteria isolated from HIV-1 patients with a size comparable to the Gag precursor. It could be argued that bacteria expressing HIV-1 proteins will unlikely secrete them, which may be anticipated as necessary to induce HIV-1 antibodies and transmit HIV seropositivity. However, once bacteria enter the body, there are processes that destroy them and result in the release of hidden antigens and epitopes, which can then induce specific antibodies. Indeed, a recent work has revealed that the majority of antibodies elicited during infection are directed against hidden antigens and epitopes (42). Relevant immune processes that would lead to particulate/degrade bacteria and antigens include complement activation (43) and phagocytosis (44).

## 5. CONCLUSION

Our work supports that highly common and clinically relevant bacteria such as *E. coli* and *K. pneumoniae* can acquire HIV-1 sequences and could transmit HIV seropositivity to people. Since these bacteria are present in the gut microbiota, transmission of HIV positivity without actual HIV infection might not be uncommon: it could occur through person-to-person contacts, from contaminated sources and procedures such as microbiota transplantation, which demands testing the presence of HIV sequences in microbiota samples. The possibility that microbiota bacteria could transmit HIV seropositivity deserves serious consideration in order to prevent treating healthy individuals with life-long medications that carry significant toxicity and take a heavy toll on health care systems. Therefore, it is crucial to count with accurate HIV diagnostics guaranteeing that life-saving ART is only received by HIV infected patients.

## ACKNOWLEDGMENTS

We wish to thank to PAR lab members for comments and critical reading

## AUTHOR CONTRIBUTIONS

Conceptualization: P.A.R; Methodology: H.F.P.-P. & J.M.-G.; Investigation: P.A.R & E.M.L; Data Analysis: H.F.P.-P. & J.M.-G; Writing-Original Draft: P.A.R; Final Writing & Editing: H.F.P.-P., J.M.-G, E.M.L & P.A.R. All authors have read and approved the final manuscript.

## STATEMENTS

Institutional Review Board Statement: Not applicable. Informed Consent Statement: Not applicable.

## Author Approvals

All the individuals named as authors of this manuscript have approved its submission.

## SUPPLEMENTARY MATERIALS

**Supplementary File 1:** Output of TBLASTN searches with HIV-1 proteins against bacterial genomes

**Supplementary File 2**: Supplementary File 2: Multiple Sequence Alignments of HIV-1 proteins and top TBLASTN hits to bacterial genomes

**Supplementary Table S1:** HIV-1 proteome coverage statistics in *E. coli* and *K. pneumoniae*

**Supplementary Table S2**: IAV proteome coverage in *L. monocytogenes*.

**Supplementary Table S3:** HBV proteome coverage in *Streptomyces sp. 2BBP J2*.

**Supplementary Table S4**: IAV proteome coverage in *E. coli*.

**Supplementary Table S5:** HBV proteome coverage in *K. pneumoniae*

