## Supplementary File 1 for "HIV-1 protein coding sequences are present in relevant bacteria"

### Supplementary File 1: Output of TBLASTN searches with HIV-1 proteins against bacterial genomes

RID: 5YZRAEE2016
Job Title:HIV Mature Proteins tblastn Bacteria
Program: TBLASTN
Database: refseq_genomes (99 databases)
Query #1: sp|P04578|ENV_HV1H2 Envelope glycoprotein gp160 OS=Human immunodeficiency virus type 1 group M subtype B (isolate HXB2) OX=11706 GN=env PE=1 SV=2 Query ID: lcl|Query_2387706 Length: 856

Sequences producing significant alignments:

| Description | Scientific Name | Common Name | Taxid | Max Score | Total Score | Query cover | E Value | Per. Ident | Acc. Len | Accession |
|---|---|---|---|---|---|---|---|---|---|---|
| Escherichia coli strain EN5177 contig00087, whole genome shotg... | Escherichia ... | NA | 562 | 1082 | 1082 | 100% | 0.0 | 61.15 | 3403 | NZ_JAPMNK0100... |
| Escherichia coli strain NBE0004 contig00090, whole genome... | Escherichia ... | NA | 562 | 936 | 936 | 84% | 0.0 | 61.99 | 2818 | NZ_JARBDH0100... |
| Escherichia coli strain NBE0006 contig00106, whole genome... | Escherichia ... | NA | 562 | 857 | 857 | 79% | 0.0 | 61.42 | 2090 | NZ_JARBDI0100... |
| Escherichia coli strain NBE0009 contig00112, whole genome... | Escherichia ... | NA | 562 | 743 | 743 | 63% | 0.0 | 66.79 | 2003 | NZ_JARBDL0100... |
| Escherichia coli strain NBE0009 contig00111, whole genome... | Escherichia ... | NA | 562 | 740 | 740 | 63% | 0.0 | 65.95 | 2027 | NZ_JARBDL0100... |
| Escherichia coli strain EN5076 contig00066, whole genome shotg... | Escherichia ... | NA | 562 | 659 | 911 | 83% | 0.0 | 67.36 | 2732 | NZ_JAODTC0100... |
| Escherichia coli strain EN5374 contig00094, whole genome shotg... | Escherichia ... | NA | 562 | 656 | 908 | 83% | 0.0 | 66.74 | 2720 | NZ_JAPTGI0100... |
| Escherichia coli strain EN5090 contig00081, whole genome shotg... | Escherichia ... | NA | 562 | 655 | 954 | 83% | 0.0 | 67.43 | 2798 | NZ_JAODTD0100... |
| Escherichia coli strain NBE0010 contig00080, whole genome... | Escherichia ... | NA | 562 | 654 | 907 | 83% | 0.0 | 66.53 | 2743 | NZ_JARBDM0100... |
| Escherichia coli strain EN5177 contig00105, whole genome shotg... | Escherichia ... | NA | 562 | 652 | 652 | 59% | 0.0 | 64.93 | 1783 | NZ_JAPMNK0100... |
| Escherichia coli strain EN5379 contig00092, whole genome shotg... | Escherichia ... | NA | 562 | 646 | 939 | 83% | 0.0 | 67.50 | 2592 | NZ_JAPTGJ0100... |
| Escherichia coli strain EN5317 contig00057, whole genome shotg... | Escherichia ... | NA | 562 | 642 | 946 | 84% | 0.0 | 66.25 | 2867 | NZ_JARBDO0100... |
| Escherichia coli strain EN5169 contig00082, whole genome shotg... | Escherichia ... | NA | 562 | 634 | 934 | 83% | 0.0 | 65.98 | 2837 | NZ_JAPMNJ0100... |
| Escherichia coli strain NBE0003 contig00096, whole genome... | Escherichia ... | NA | 562 | 634 | 934 | 83% | 0.0 | 65.98 | 2837 | NZ_JARBDG0100... |
| Escherichia coli strain EN5143 contig00069, whole genome shotg... | Escherichia ... | NA | 562 | 629 | 885 | 76% | 0.0 | 66.53 | 2688 | NZ_JAPMNI0100... |
| Escherichia coli strain NBE0002 contig00097, whole genome... | Escherichia ... | NA | 562 | 629 | 884 | 76% | 0.0 | 66.53 | 2717 | NZ_JARBDF0100... |
| Escherichia coli strain EN5308 contig00099, whole genome shotg... | Escherichia ... | NA | 562 | 626 | 916 | 83% | 0.0 | 64.95 | 2827 | NZ_JARBDE0100... |
| Escherichia coli strain NBE0009 contig00116, whole genome... | Escherichia ... | NA | 562 | 612 | 612 | 54% | 0.0 | 65.61 | 1713 | NZ_JARBDL0100... |
| Escherichia coli strain EN5286 contig00120, whole genome shotg... | Escherichia ... | NA | 562 | 603 | 937 | 85% | 0.0 | 60.50 | 3033 | NZ_JAPMNM0100... |
| Escherichia coli strain NBE0007 contig00093, whole genome... | Escherichia ... | NA | 562 | 599 | 937 | 85% | 0.0 | 59.73 | 3013 | NZ_JARBDJ0100... |
| Escherichia coli strain EN5192 contig00108, whole genome shotg... | Escherichia ... | NA | 562 | 599 | 925 | 86% | 0.0 | 58.69 | 3032 | NZ_JAPMNL0100... |
| Escherichia coli strain VRES0669, whole genome shotgun sequence | Escherichia ... | NA | 562 | 593 | 593 | 54% | 0.0 | 63.71 | 5544 | NZ_UINV010000... |
| Escherichia coli strain EN5308 contig00096, whole genome shotg... | Escherichia ... | NA | 562 | 591 | 920 | 85% | 0.0 | 60.08 | 3024 | NZ_JARBDE0100... |
| Escherichia coli strain EN5143 contig00071, whole genome shotg... | Escherichia ... | NA | 562 | 590 | 914 | 84% | 0.0 | 59.03 | 2774 | NZ_JAPMNI0100... |
| Escherichia coli strain EN5383 contig00096, whole genome shotg... | Escherichia ... | NA | 562 | 564 | 864 | 75% | 0.0 | 67.16 | 2210 | NZ_JAPTZN0100... |
| Escherichia coli strain EN5192 contig00149, whole genome shotg... | Escherichia ... | NA | 562 | 538 | 538 | 45% | 4e-180 | 66.83 | 1397 | NZ_JAPMNL0100... |
| Escherichia coli strain NBE0006 contig00124, whole genome... | Escherichia ... | NA | 562 | 529 | 529 | 44% | 4e-177 | 68.30 | 1306 | NZ_JARBDI0100... |
| Klebsiella pneumoniae strain EuSCAPE_IL004, whole genome shotg... | Klebsiella p... | NA | 573 | 495 | 495 | 43% | 4e-165 | 65.69 | 1081 | NZ_UKJP020000... |
| Escherichia coli strain NBE0010 contig00098, whole genome... | Escherichia ... | NA | 562 | 504 | 850 | 71% | 3e-164 | 68.87 | 1912 | NZ_JARBDM0100... |
| Pseudomonas aeruginosa strain VET-121... | Pseudomonas ... | NA | 287 | 449 | 449 | 37% | 2e-147 | 66.87 | 1017 | NZ_QZXB010001... |
| Escherichia coli strain NBE0002 contig00149, whole genome... | Escherichia ... | NA | 562 | 431 | 431 | 36% | 2e-140 | 65.47 | 996 | NZ_JARBDF0100... |
| Escherichia coli strain EN5308 contig00145, whole genome shotg... | Escherichia ... | NA | 562 | 430 | 430 | 38% | 3e-140 | 64.41 | 1005 | NZ_JARBDE0100... |
| Escherichia coli strain NBE0002 contig00132, whole genome... | Escherichia ... | NA | 562 | 431 | 431 | 37% | 9e-139 | 65.14 | 1381 | NZ_JARBDF0100... |
| Escherichia coli strain EN5383 contig00089, whole genome shotg... | Escherichia ... | NA | 562 | 438 | 531 | 48% | 1e-138 | 64.91 | 1970 | NZ_JAPTZN0100... |
| Escherichia coli strain EN5095 contig00078, whole genome shotg... | Escherichia ... | NA | 562 | 428 | 721 | 68% | 9e-135 | 64.83 | 1972 | NZ_JAPYLY0100... |
| Escherichia coli strain EN5132 contig00077, whole genome shotg... | Escherichia ... | NA | 562 | 428 | 720 | 65% | 1e-132 | 66.46 | 2498 | NZ_JAPYLZ0100... |
| Nocardioides luteus strain JCM 3358 sequence39, whole genome... | Nocardioides... | NA | 1844 | 446 | 446 | 33% | 2e-131 | 79.37 | 5754 | NZ_BMRK010000... |
| Streptomyces griseoflavus strain JCM 4479 sequence37, whole... | Streptomyces... | NA | 35619 | 446 | 446 | 33% | 2e-131 | 79.37 | 5775 | NZ_BMUC010000... |
| Saccharothrix coeruleofusca strain JCM 3313 sequence45, whole... | Saccharothri... | NA | 33919 | 446 | 446 | 33% | 2e-131 | 79.37 | 5775 | NZ_BMRG010000... |
| Actinoplanes ianthinogenes strain JCM 3249 sequence037, whole... | Actinoplanes... | NA | 122358 | 446 | 446 | 33% | 2e-131 | 79.37 | 5815 | NZ_BMQZ010000... |
| Sinomonas sp. KCTC 49339 sequence28, whole genome shotgun... | Sinomonas sp... | NA | 2769422 | 446 | 446 | 33% | 2e-131 | 79.37 | 5849 | NZ_BNCM010000... |
| Saccharothrix mutabilis subsp. mutabilis strain JCM 3380... | Saccharothri... | NA | 66855 | 446 | 446 | 33% | 2e-131 | 79.37 | 5899 | NZ_BAAABU0100... |
| Corynebacterium glutamicum strain 39 Scaffolds_32_9166, whole... | Corynebacter... | NA | 1718 | 441 | 441 | 33% | 5e-129 | 79.37 | 8862 | NZ_JACEQW0100... |
| Corynebacterium glutamicum strain 37 Scaffolds_33_8928, whole... | Corynebacter... | NA | 1718 | 441 | 441 | 33% | 5e-129 | 79.37 | 8928 | NZ_JACENO0100... |
| Methyloversatilis universalis FAM5 contig00006, whole genome... | Methyloversa... | NA | 1000565 | 380 | 380 | 25% | 1e-122 | 82.65 | 649 | NZ_AFHG010000... |
| Klebsiella pneumoniae strain P7701... | Klebsiella p... | NA | 573 | 378 | 378 | 22% | 3e-122 | 96.30 | 570 | NZ_JACAWM0100... |
| Escherichia coli strain NBE0004 contig00110, whole genome... | Escherichia ... | NA | 562 | 388 | 388 | 41% | 8e-122 | 59.45 | 1390 | NZ_JARBDH0100... |
| Escherichia coli strain NBE0004 contig00131, whole genome... | Escherichia ... | NA | 562 | 375 | 375 | 32% | 2e-119 | 66.90 | 847 | NZ_JARBDH0100... |
| Pseudomonas aeruginosa strain VET-42... | Pseudomonas ... | NA | 287 | 357 | 357 | 28% | 4e-112 | 66.93 | 963 | NZ_RACX010002... |
| Escherichia coli strain EN5143 contig00088, whole genome shotg... | Escherichia ... | NA | 562 | 319 | 319 | 32% | 4e-98 | 60.50 | 833 | NZ_JAPMNI0100... |
| Escherichia coli strain EN5192 contig00135, whole genome shotg... | Escherichia ... | NA | 562 | 269 | 431 | 44% | 6e-77 | 55.56 | 1341 | NZ_JAPMNL0100... |
| Klebsiella pneumoniae strain EuSCAPE_IL001, whole genome shotg... | Klebsiella p... | NA | 573 | 243 | 243 | 21% | 2e-70 | 64.55 | 599 | NZ_UKJN020000... |
| Neisseria gonorrhoeae strain GC-206... | Neisseria go... | NA | 485 | 240 | 240 | 19% | 6e-70 | 71.08 | 488 | NZ_SAWN010001... |
| Klebsiella pneumoniae strain OK8 jcf7180000003506, whole genom... | Klebsiella p... | NA | 573 | 232 | 232 | 20% | 2e-66 | 64.48 | 577 | NZ_MZXR010000... |
| Klebsiella pneumoniae strain EuSCAPE_IL006, whole genome shotg... | Klebsiella p... | NA | 573 | 230 | 230 | 20% | 5e-66 | 63.59 | 560 | NZ_UKJR020001... |
| Aggregatibacter actinomycetemcomitans serotype d str. SA269... | Aggregatibac... | NA | 1434259 | 227 | 227 | 18% | 4e-65 | 74.69 | 504 | NZ_AZTX010004... |
| Burkholderia pseudomallei strain 2604STDY5519032... | Burkholderia... | NA | 28450 | 225 | 225 | 19% | 2e-64 | 66.86 | 499 | NZ_UVID020000... |
| Escherichia coli strain EN5192 contig00156, whole genome shotg... | Escherichia ... | NA | 562 | 226 | 226 | 22% | 3e-64 | 61.93 | 562 | NZ_JAPMNL0100... |
| Klebsiella pneumoniae strain PN085E1IA... | Klebsiella p... | NA | 573 | 224 | 224 | 17% | 5e-64 | 72.26 | 474 | NZ_PDVD010000... |
| Pseudomonas aeruginosa strain VET-42... | Pseudomonas ... | NA | 287 | 215 | 215 | 21% | 2e-60 | 58.03 | 548 | NZ_RACX010001... |
| Klebsiella pneumoniae strain ST-234:K080... | Klebsiella p... | NA | 573 | 206 | 206 | 18% | 2e-57 | 68.15 | 474 | NZ_NXJY010001... |
| Escherichia coli strain EN5177 contig00148, whole genome shotg... | Escherichia ... | NA | 562 | 208 | 208 | 21% | 1e-56 | 58.95 | 850 | NZ_JAPMNK0100... |
| Enterobacter hormaechei strain ST-234:K063... | Enterobacter... | NA | 158836 | 202 | 202 | 18% | 3e-56 | 66.46 | 475 | NZ_NXJN010001... |
| Klebsiella pneumoniae strain ST-1552:K089... | Klebsiella p... | NA | 573 | 202 | 202 | 17% | 5e-56 | 68.39 | 474 | NZ_NXIW010001... |
| Escherichia coli strain NBE0009 contig00153, whole genome... | Escherichia ... | NA | 562 | 204 | 204 | 18% | 1e-55 | 62.13 | 753 | NZ_JARBDL0100... |
| Escherichia coli strain E003 NODE_241_length_474_cov_17.072,... | Escherichia ... | NA | 562 | 200 | 200 | 17% | 3e-55 | 67.10 | 474 | NZ_NXIZ010002... |
| Cronobacter sakazakii strain cro4114A1 contig126, whole genome... | Cronobacter ... | NA | 28141 | 192 | 192 | 16% | 8e-53 | 67.13 | 413 | NZ_NRKJ010001... |
| Cronobacter malonaticus strain cro3263A1 contig69, whole genom... | Cronobacter ... | NA | 413503 | 192 | 192 | 16% | 1e-52 | 67.13 | 413 | NZ_NRMM010000... |
| Citrobacter freundii strain CF003... | Citrobacter ... | NA | 546 | 192 | 192 | 17% | 2e-52 | 66.23 | 474 | NZ_NXKE010001... |
| Escherichia coli strain NBE0009 contig00151, whole genome... | Escherichia ... | NA | 562 | 194 | 194 | 18% | 1e-51 | 59.76 | 802 | NZ_JARBDL0100... |
| Klebsiella pneumoniae strain EuSCAPE_IL005, whole genome shotg... | Klebsiella p... | NA | 573 | 188 | 188 | 16% | 6e-51 | 63.27 | 455 | NZ_UKJL020000... |
| Klebsiella pneumoniae strain EuSCAPE_IL004, whole genome shotg... | Klebsiella p... | NA | 573 | 186 | 186 | 12% | 1e-50 | 80.00 | 368 | NZ_UKJP020001... |
| Klebsiella pneumoniae strain EuSCAPE_MT003, whole genome shotg... | Klebsiella p... | NA | 573 | 193 | 193 | 18% | 2e-50 | 60.87 | 964 | NZ_UKIK020000... |
| Escherichia coli strain NBE0006 contig00145, whole genome... | Escherichia ... | NA | 562 | 186 | 186 | 19% | 7e-49 | 56.98 | 738 | NZ_JARBDI0100... |
| Cronobacter malonaticus strain cro3263A1 contig68, whole genom... | Cronobacter ... | NA | 413503 | 179 | 179 | 16% | 7e-48 | 64.79 | 413 | NZ_NRMM010000... |
| Escherichia coli strain NBE0006 contig00147, whole genome... | Escherichia ... | NA | 562 | 182 | 182 | 17% | 1e-47 | 60.76 | 694 | NZ_JARBDI0100... |
| Actinoplanes campanulatus strain JCM 3059 sequence095, whole... | Actinoplanes... | NA | 113559 | 179 | 179 | 13% | 3e-47 | 81.03 | 582 | NZ_BMPW010000... |
| Cronobacter sakazakii strain cro4215W contig103, whole genome... | Cronobacter ... | NA | 28141 | 176 | 176 | 16% | 7e-47 | 64.08 | 413 | NZ_NRKF010001... |
| Deinococcus aquiradiocola strain JCM 14371 sequence41, whole... | Deinococcus ... | NA | 393059 | 179 | 179 | 13% | 8e-47 | 81.03 | 690 | NZ_BMOE010000... |
| Klebsiella pneumoniae strain EuSCAPE_IL004, whole genome shotg... | Klebsiella p... | NA | 573 | 174 | 174 | 10% | 1e-46 | 84.78 | 277 | NZ_UKJP020001... |
| Enterococcus alcedinis strain CCM 8433 sequence26, whole genom... | Enterococcus... | NA | 1274384 | 179 | 179 | 13% | 4e-46 | 81.03 | 853 | NZ_BMDT010000... |
| Cronobacter dublinensis strain cro981C2 contig304, whole genom... | Cronobacter ... | NA | 413497 | 174 | 174 | 16% | 4e-46 | 63.38 | 413 | NZ_NROB010003... |
| Cronobacter sakazakii strain cro4114B1 contig109, whole genome... | Cronobacter ... | NA | 28141 | 174 | 174 | 16% | 4e-46 | 63.38 | 413 | NZ_NRKI010001... |
| Cronobacter sakazakii strain cro4114A1 contig127, whole genome... | Cronobacter ... | NA | 28141 | 174 | 174 | 16% | 4e-46 | 63.38 | 413 | NZ_NRKJ010001... |
| Cronobacter sakazakii strain cro611A3 contig61, whole genome... | Cronobacter ... | NA | 28141 | 172 | 172 | 16% | 1e-45 | 62.94 | 407 | NZ_NRMC010000... |
| Enterobacter hormaechei strain ST-234:K063... | Enterobacter... | NA | 158836 | 173 | 173 | 15% | 2e-45 | 65.15 | 480 | NZ_NXJN010001... |
| Cronobacter sakazakii strain cro3040W contig81, whole genome... | Cronobacter ... | NA | 28141 | 169 | 169 | 16% | 2e-44 | 62.68 | 404 | NZ_NRJO010000... |
| Cronobacter sakazakii strain cro4114A1 contig132, whole genome... | Cronobacter ... | NA | 28141 | 167 | 167 | 16% | 6e-44 | 61.97 | 392 | NZ_NRKJ010001... |
| Klebsiella pneumoniae strain EuSCAPE_IL022, whole genome shotg... | Klebsiella p... | NA | 573 | 167 | 167 | 12% | 1e-43 | 73.87 | 411 | NZ_UKKD020000... |
| Cronobacter sakazakii strain cro4215W contig104, whole genome... | Cronobacter ... | NA | 28141 | 166 | 166 | 16% | 2e-43 | 61.97 | 405 | NZ_NRKF010001... |
| Escherichia coli strain NBE0009 contig00162, whole genome... | Escherichia ... | NA | 562 | 165 | 165 | 13% | 3e-43 | 65.52 | 350 | NZ_JARBDL0100... |
| Cronobacter sakazakii strain cro4114A1 contig129, whole genome... | Cronobacter ... | NA | 28141 | 165 | 165 | 16% | 6e-43 | 61.27 | 404 | NZ_NRKJ010001... |
| Cronobacter malonaticus strain cro3263A1 contig74, whole genom... | Cronobacter ... | NA | 413503 | 165 | 165 | 16% | 8e-43 | 58.45 | 404 | NZ_NRMM010000... |
| Cronobacter sakazakii strain cro4114B1 contig117, whole genome... | Cronobacter ... | NA | 28141 | 164 | 164 | 16% | 1e-42 | 61.27 | 404 | NZ_NRKI010001... |
| Cronobacter sakazakii strain cro4114B1 contig115, whole genome... | Cronobacter ... | NA | 28141 | 163 | 163 | 16% | 2e-42 | 57.75 | 404 | NZ_NRKI010001... |
| Corynebacterium glutamicum strain 38 Scaffolds_31_4109, whole... | Corynebacter... | NA | 1718 | 181 | 181 | 13% | 3e-42 | 80.51 | 3118 | NZ_JACENN0100... |
| Cronobacter sakazakii strain cro4114A1 contig128, whole genome... | Cronobacter ... | NA | 28141 | 163 | 163 | 16% | 3e-42 | 57.75 | 404 | NZ_NRKJ010001... |
| Lactiplantibacillus plantarum strain MH-301 Scaffolds_50_4236,... | Lactiplantib... | NA | 1590 | 181 | 181 | 13% | 4e-42 | 80.51 | 4236 | NZ_JAOYUG0100... |
| Corynebacterium glutamicum strain 51 Scaffolds_32_4236, whole... | Corynebacter... | NA | 1718 | 181 | 181 | 13% | 4e-42 | 80.51 | 4236 | NZ_JACENK0100... |
| Cronobacter sakazakii strain cro4215W contig105, whole genome... | Cronobacter ... | NA | 28141 | 162 | 162 | 16% | 5e-42 | 57.75 | 404 | NZ_NRKF010001... |
| Cronobacter malonaticus strain cro3263A1 contig76, whole genom... | Cronobacter ... | NA | 413503 | 162 | 162 | 16% | 5e-42 | 59.86 | 392 | NZ_NRMM010000... |
| Cronobacter dublinensis strain cro981C2 contig308, whole genom... | Cronobacter ... | NA | 413497 | 161 | 161 | 16% | 1e-41 | 60.99 | 405 | NZ_NROB010003... |
| Deinococcus aquiradiocola strain JCM 14371 sequence36, whole... | Deinococcus ... | NA | 393059 | 166 | 166 | 12% | 2e-41 | 79.28 | 869 | NZ_BMOE010000... |
| Cronobacter sakazakii strain cro4215W contig109, whole genome... | Cronobacter ... | NA | 28141 | 160 | 160 | 16% | 2e-41 | 57.75 | 389 | NZ_NRKF010001... |
| Cronobacter sakazakii strain cro4114B1 contig105, whole genome... | Cronobacter ... | NA | 28141 | 159 | 159 | 16% | 9e-41 | 59.03 | 416 | NZ_NRKI010001... |
| Thalassovita mangrovi strain GS-10 Scaffold21_1, whole genome... | Thalassovita... | NA | 2692236 | 169 | 169 | 12% | 2e-40 | 79.28 | 1440 | NZ_WWEN010000... |
| Klebsiella pneumoniae strain EuSCAPE_MT015, whole genome shotg... | Klebsiella p... | NA | 573 | 156 | 156 | 13% | 3e-40 | 64.29 | 338 | NZ_UKIW020001... |
| Escherichia coli strain EN5308 contig00153, whole genome shotg... | Escherichia ... | NA | 562 | 162 | 162 | 19% | 4e-40 | 54.29 | 786 | NZ_JARBDE0100... |
| Cronobacter sakazakii strain cro4114A1 contig133, whole genome... | Cronobacter ... | NA | 28141 | 157 | 157 | 16% | 4e-40 | 58.45 | 389 | NZ_NRKJ010001... |
| Cronobacter sakazakii strain cro4191B1-2 contig85, whole genom... | Cronobacter ... | NA | 28141 | 156 | 156 | 16% | 6e-40 | 58.45 | 392 | NZ_NRKG010000... |
| Cronobacter sakazakii strain cro3040W contig84, whole genome... | Cronobacter ... | NA | 28141 | 156 | 156 | 16% | 6e-40 | 58.45 | 392 | NZ_NRJO010000... |

| | | | | | | | | | | |
|---|---|---|---|---|---|---|---|---|---|---|
| Planctopirus hydrillae strain JC280 Contig113, whole genome... | Planctopirus... | NA | 1841610 | 157 | 157 | 17% | 7e-40 | 59.86 | 437 | NZ_LYDR010000... |
| Cronobacter sakazakii strain cro4114B1 contig122, whole genome... | Cronobacter ... | NA | 28141 | 155 | 155 | 16% | 9e-40 | 57.04 | 383 | NZ_NRKI010001... |
| Cronobacter malonaticus strain cro3263A1 contig75, whole genom... | Cronobacter ... | NA | 413503 | 154 | 154 | 16% | 5e-39 | 57.75 | 392 | NZ_NRMM010000... |
| Cronobacter sakazakii strain cro4114B1 contig124, whole genome... | Cronobacter ... | NA | 28141 | 154 | 154 | 16% | 5e-39 | 57.75 | 392 | NZ_NRKI010001... |
| Cronobacter sakazakii strain cro4191B1-2 contig83, whole genom... | Cronobacter ... | NA | 28141 | 154 | 154 | 16% | 6e-39 | 56.64 | 410 | NZ_NRKG010000... |
| Cronobacter sakazakii strain cro4114A1 contig131, whole genome... | Cronobacter ... | NA | 28141 | 153 | 153 | 16% | 7e-39 | 57.75 | 392 | NZ_NRKJ010001... |
| Cronobacter sakazakii strain cro3040W contig83, whole genome... | Cronobacter ... | NA | 28141 | 152 | 152 | 16% | 2e-38 | 54.93 | 392 | NZ_NRJO010000... |
| Klebsiella pneumoniae strain EuSCAPE_MT017, whole genome shotg... | Klebsiella p... | NA | 573 | 153 | 153 | 12% | 3e-38 | 67.83 | 513 | NZ_UKJA020000... |
| Cronobacter dublinensis strain cro3540W contig49, whole genome... | Cronobacter ... | NA | 413497 | 151 | 151 | 16% | 6e-38 | 54.23 | 392 | NZ_NRNN010000... |
| Actinoplanes campanulatus strain JCM 3059 sequence058, whole... | Actinoplanes... | NA | 113559 | 167 | 167 | 12% | 2e-37 | 79.28 | 4754 | NZ_BMPW010000... |
| Photobacterium rosenbergii strain DP6N14-7 DP6N14-7_ctg0065,... | Photobacteri... | NA | 294936 | 147 | 147 | 12% | 1e-35 | 63.64 | 564 | NZ_JAHVIB0100... |
| Cronobacter sakazakii strain cro4191B1-2 contig84, whole genom... | Cronobacter ... | NA | 28141 | 127 | 185 | 15% | 1e-35 | 60.58 | 402 | NZ_NRKG010000... |
| Klebsiella pneumoniae strain EuSCAPE_NO017, whole genome shotg... | Klebsiella p... | NA | 573 | 146 | 146 | 12% | 2e-35 | 66.10 | 607 | NZ_UKIO020000... |
| Cronobacter malonaticus strain cro4164A3 contig145, whole geno... | Cronobacter ... | NA | 413503 | 120 | 181 | 15% | 2e-34 | 61.62 | 405 | NZ_NRNF010001... |
| Klebsiella pneumoniae strain EuSCAPE_IL018, whole genome shotg... | Klebsiella p... | NA | 573 | 139 | 139 | 14% | 1e-33 | 56.25 | 388 | NZ_UKJU020001... |
| Klebsiella pneumoniae strain EuSCAPE_IL015, whole genome shotg... | Klebsiella p... | NA | 573 | 140 | 140 | 11% | 1e-33 | 63.21 | 532 | NZ_UKJV020000... |
| Klebsiella pneumoniae strain EuSCAPE_DE091, whole genome shotg... | Klebsiella p... | NA | 573 | 140 | 140 | 12% | 4e-33 | 63.41 | 618 | NZ_UKMA020000... |
| Neisseria gonorrhoeae strain GC-206... | Neisseria go... | NA | 485 | 135 | 135 | 9% | 5e-33 | 74.70 | 243 | NZ_SAWN010001... |
| Bacillus velezensis strain TN14-11 TN14-11_ctg0032, whole geno... | Bacillus vel... | NA | 492670 | 144 | 144 | 12% | 1e-32 | 63.64 | 1125 | NZ_JAHVIU0100... |
| Burkholderia pseudomallei strain 2604STDY5518971... | Burkholderia... | NA | 28450 | 134 | 134 | 12% | 2e-32 | 60.55 | 313 | NZ_UVFZ020001... |
| Cronobacter malonaticus strain cro3263A1 contig73, whole genom... | Cronobacter ... | NA | 413503 | 116 | 173 | 16% | 6e-32 | 52.89 | 406 | NZ_NRMM010000... |
| Klebsiella pneumoniae strain EuSCAPE_MT015, whole genome shotg... | Klebsiella p... | NA | 573 | 130 | 130 | 12% | 6e-31 | 60.55 | 328 | NZ_UKIW020001... |
| Klebsiella pneumoniae strain EuSCAPE_MT009, whole genome shotg... | Klebsiella p... | NA | 573 | 129 | 129 | 11% | 2e-30 | 66.00 | 324 | NZ_UKIV020000... |
| Acinetobacter baumannii strain P4 contig_P4_92, whole genome... | Acinetobacte... | NA | 470 | 134 | 134 | 10% | 5e-29 | 76.40 | 1226 | NZ_JBBXLU0100... |
| Staphylococcus aureus strain SA3... | Staphylococc... | NA | 1280 | 122 | 122 | 7% | 1e-28 | 80.95 | 205 | NZ_RQSW010007... |
| Klebsiella pneumoniae strain EuSCAPE_IL009, whole genome shotg... | Klebsiella p... | NA | 573 | 119 | 119 | 10% | 3e-27 | 63.74 | 274 | NZ_UKJO020000... |
| Klebsiella pneumoniae strain EuSCAPE_MT009, whole genome shotg... | Klebsiella p... | NA | 573 | 119 | 119 | 12% | 1e-26 | 56.52 | 398 | NZ_UKIV020000... |
| Klebsiella pneumoniae strain EuSCAPE_MT006, whole genome shotg... | Klebsiella p... | NA | 573 | 115 | 115 | 9% | 4e-26 | 70.59 | 250 | NZ_UKIU020000... |
| Klebsiella pneumoniae strain EuSCAPE_NO001, whole genome shotg... | Klebsiella p... | NA | 573 | 113 | 113 | 10% | 3e-25 | 69.15 | 289 | NZ_UKMF020000... |
| Klebsiella pneumoniae strain EuSCAPE_IL022, whole genome shotg... | Klebsiella p... | NA | 573 | 112 | 112 | 10% | 1e-24 | 58.89 | 290 | NZ_UKKD020000... |
| Acinetobacter baumannii strain AB49... | Acinetobacte... | NA | 470 | 111 | 111 | 11% | 2e-24 | 54.90 | 263 | NZ_JAAXKW0100... |
| Klebsiella pneumoniae strain EuSCAPE_IL004, whole genome shotg... | Klebsiella p... | NA | 573 | 110 | 110 | 10% | 4e-24 | 58.62 | 327 | NZ_UKJP020001... |
| Klebsiella pneumoniae strain EuSCAPE_NO001, whole genome shotg... | Klebsiella p... | NA | 573 | 109 | 109 | 8% | 6e-24 | 68.83 | 245 | NZ_UKMF020000... |
| Klebsiella pneumoniae strain EuSCAPE_MT002, whole genome shotg... | Klebsiella p... | NA | 573 | 106 | 106 | 10% | 1e-22 | 63.64 | 265 | NZ_UKIM020000... |
| Klebsiella pneumoniae strain EuSCAPE_IL016, whole genome shotg... | Klebsiella p... | NA | 573 | 106 | 106 | 11% | 2e-22 | 55.67 | 345 | NZ_UKKA020000... |
| Neisseria gonorrhoeae strain GC-206... | Neisseria go... | NA | 485 | 104 | 104 | 9% | 3e-22 | 59.30 | 258 | NZ_SAWN010001... |
| Pseudomonas aeruginosa strain VET-121... | Pseudomonas ... | NA | 287 | 99.0 | 99.0 | 9% | 3e-20 | 66.67 | 275 | NZ_QZXB010002... |
| Klebsiella pneumoniae strain EuSCAPE_IL019, whole genome shotg... | Klebsiella p... | NA | 573 | 96.7 | 96.7 | 9% | 2e-19 | 62.20 | 244 | NZ_UKJZ020000... |
| Photobacterium rosenbergii strain DP6N14-7 DP6N14-7_ctg0054,... | Photobacteri... | NA | 294936 | 96.7 | 96.7 | 7% | 5e-17 | 70.15 | 960 | NZ_JAHVIB0100... |
| Klebsiella pneumoniae strain EuSCAPE_LT003, whole genome shotg... | Klebsiella p... | NA | 573 | 83.2 | 83.2 | 7% | 2e-14 | 65.08 | 317 | NZ_UKJH020000... |
| Bacillus velezensis strain TN14-11 TN14-11_ctg0036, whole geno... | Bacillus vel... | NA | 492670 | 70.9 | 70.9 | 5% | 3e-08 | 70.59 | 1101 | NZ_JAHVIU0100... |
| Klebsiella pneumoniae strain EuSCAPE_IL005, whole genome shotg... | Klebsiella p... | NA | 573 | 61.2 | 61.2 | 4% | 3e-06 | 73.17 | 442 | NZ_UKJL020000... |

Alignments:

>Escherichia coli strain EN5177 contig00087, whole genome shotgun sequence
Sequence ID: NZ_JAPMNK010000087.1 Length: 3403
Range 1: 352 to 3099

Score:1082 bits(2797), Expect:0.0,
Method:Compositional matrix adjust.,
Identities:606/924(66%), Positives:711/924(76%), Gaps:76/924(8%)

```
Query  1     MRVK---EKYQHLWRWGWRWGTMLLG--MLMICSATEKLWVTVYYGVPVWKEATTTLFCA  55
             MRV+     YQH W WG      +LG  MLM C+    LWVTVYYGVPVW++A  TLFCA
Sbjct  352   MRVRGILRSYQHWWIWG------ILGFWMLMTCNVAGNLWVTVYYGVPVWRDANATLFCA  513

Query  56    SDAKAYDTEVHNVWATHACVPTDPNPQEVVLVNVTENFNMWKNDMVEQMHEDIISLWDQS  115
             SDAK Y TEVHNVWATHACVPTDPNPQE+VL NVTENFNMW+NDMVEQMH+D+IS+WDQ
Sbjct  514   SDAKVYKTEVHNVWATHACVPTDPNPQELVLENVTENFNMWENDMVEQMHQDVISIWDQG  693

Query  116   LKPCVKLTPLCVSLKCTDLK------NDTNTNSSSGRMIMEK--------GEIKNCSFNI  161
             LKPCVKLTPLCV+L CT  +      ++ NT + S    ++          EI NCSFN
Sbjct  694   LKPCVKLTPLCVTLNCTTYEPKKVTADNNNTCAPSQNTTVKNLTIDPEAVKEISNCSFNT  873

Query  162   STSIRGKVQKEYAFFYKLDIIPIDNDTTS-----YKLTSCNTSVITQACPKVSFEPIPIH  216
             +T +R K +K +A FYK+D++ +  ++T+      Y+L +CNTS ITQACPKVSFEPIPIH
Sbjct  874   TTELRDKTRKVHALFYKIDMVELGENSTAPNFTLYRLINCNTSAITQACPKVSFEPIPIH  1053

Query  217   YCAPAGFAILKCNNKTFNGTGPCTNVSTVQCTHGIRPVVSTQLLLNGSLAEEEVVIRSVN  276
             YCAPAG+A+LKCNN TFNGTGPC N+STVQCTHGIRPVVSTQLLLNGSLAE  V IRS N
Sbjct  1054  YCAPAGYALLKCNNNTFNGTGPCRNISTVQCTHGIRPVVSTQLLLNGSLAEGNVTIRSEN  1233

Query  277   FTDNAKTIIVQLNTSVEINCTRPNNNTRKRIRIQRGPGRAFVTIGK-IGNMRQAHCNISR  335
              TDNAKTII  LN SVEI CTRPNNNTR+ +RI  GPG+ F   G  IG++R+AHCN+S
Sbjct  1234  LTDNAKTIIAHLNQSVEILCTRPNNNTRQSVRI--GPGQTFFATGDIIGDIRRAHCNVSA  1407

Query  336   AKWNNTLKQIASKLREQFGNNKTIIFKQSSGGDPEIVTHSFNCGGEFFYCNSTQLFNS--  393
               W+  L+ ++ +L + F NNKTI F   SGGD EI THSFNC GEFFYCN++QLFNS
Sbjct  1408  KHWHTALRNVSIELAKLFPNNKTINFTSPSGGDLEITTHSFNCRGEFFYCNTSQLFNSLY  1587

Query  394   -------TWFNSTWSTEGSNNTE----GSD-------------------TITLPCRIKQ  422
                    W+N    +    N T+    GS                     ITLPCRIKQ
Sbjct  1588  MPNNDTYIWYNENHTYVWHNGTQYVINGSTQYVTLKNSAQNVTINNITDFNITLPCRIKQ  1767

Query  423   IINMWQKVGKAMYAPPISGQIRCSSNITGLLLTRDGGNSN---NESEIFRPGGGDMRDNW  479
             IINMWQ+VGKAMYAPPI+G I C SNITGLLLTRDGG  +   N++EIFRPGGG+M+DNW
Sbjct  1768  IINMWQEVGKAMYAPPIAGNITCLSNITGLLLTRDGGKGDEIRNDTEIFRPGGGNMKDNW  1947

Query  480   RSELYKYKVVKIEPLGVAPTKAKRRVVQREKRAVGIgalflgflgaagstmgaasmtLTV  539
             RSELYKYKVV+I+PLGVAPT A+RRVV RE+RAVG+GA+F GFLGAAGSTMGAAS+ LTV
Sbjct  1948  RSELYKYKVVEIKPLGVAPTPARRRVVGRERRAVGLGAVFFGFLGAAGSTMGAASVALTV  2127

Query  540   QARQLLSGIVQQQNNLLRAIEAQQHLLQLTVWGIKQLQARILAVERYLKDQQLLGIWGCS  599
             QARQLLSGIVQQQNNLLRAIEAQQ++L+LTVWGIKQLQ R+LA+ERYL++Q+LLG+WGCS
Sbjct  2128  QARQLLSGIVQQQNNLLRAIEAQQNMLRLTVWGIKQLQTRVLAIERYLREQRLLGLWGCS  2307

Query  600   GKLICTTAVPWNASWSNKSLEQIWNHTTWMEWDREINNYTSLIHSLIeesqnqqekneqe  659
              K+ICTT VPWN+SWSNKS   IWN+ TWMEWD+EI  +T+ I+ L+EESQ QQE+NE++
Sbjct  2308  EKIICTTNVPWNSSWSNKSYSDIWNNLTWMEWDKEIEQHTNTIYQLLEESQGQQEQNEKD  2487

Query  660   llelDKWASLWNWFNITNWLWYIKLFIMIVGGLVGLRIVFAVLSIVNRVRQGYSPLSFQT  719
             LL LDKW  LW+WF+ITNWLWYIK+FIMIVGGL+GLRIVFAVLSIVNRVRQGYSPLSFQT
Sbjct  2488  LLALDKWNDLWSWFDITNWLWYIKIFIMIVGGLIGLRIVFAVLSIVNRVRQGYSPLSFQT  2667

Query  720   HLPTprgpdrpegieeeggerdrdrSIRLVNGSLALIWDDLRSLCLFSYHRLRDLLLIVT  779
               P+P G DRP  I+EEGGE+ R+ SIRLV+G LAL WDDLRSL F YHRL+DL+ +
Sbjct  2668  LAPSPGGLDRPGRIDEEGGEQGRNSSIRLVSGFLALAWDDLRSLFLFCYHRLKDLISVTA  2847

Query  780   RIVELLGR-------RGWEALKYWWNLLQYWSQELKNSAVSLLNATAIAVAEGTDRVIEV  832
             R VELLGR       RGWEALKY  +L+QYW  ELK SA+SLL+  AIAV EGTDR+IE
Sbjct  2848  RAVELLGRSSLKGLQRGWEALKYLGSLVQYWGLELKKSAISLLDTIAIAVGEGTDRIIEG  3027

Query  833   VQGACRAIRHIPRRIRQGLERILL  856
             +QG CRAIR+IPRR+RQG E  LL
Sbjct  3028  IQGLCRAIRNIPRRVRQGFEAALL  3099
```

>Escherichia coli strain NBE0004 contig00090, whole genome shotgun sequence
Sequence ID: NZ_JARBDH010000090.1 Length: 2818
Range 1: 140 to 2467

```
Score:936 bits(2418), Expect:0.0,
Method:Compositional matrix adjust.,
Identities:515/784(66%), Positives:605/784(77%), Gaps:69/784(8%)

Query  1     MRVK---EKYQHLWRWGWRWGTMLLG--MLMICSATEKLWVTVYYGVPVWKEATTTLFCA  55
             MRV+     YQH W WG      +LG  MLM C+    LWVTVYYGVPVW++A  TLFCA
Sbjct  2467  MRVRGILRSYQHWWIWG------ILGFWMLMTCNVAGNLWVTVYYGVPVWRDANATLFCA  2306

Query  56    SDAKAYDTEVHNVWATHACVPTDPNPQEVVLVNVTENFNMWKNDMVEQMHEDIISLWDQS  115
             SDAK Y TEVHNVWATHACVPTDPNPQE+VL NVTENFNMW+NDMVEQMH+D+IS+WDQ
Sbjct  2305  SDAKVYKTEVHNVWATHACVPTDPNPQELVLENVTENFNMWENDMVEQMHQDVISIWDQG  2126

Query  116   LKPCVKLTPLCVSLKCTDLK------NDTNTNSSSGRMIMEK--------GEIKNCSFNI  161
             LKPCVKLTPLCV+L CT  +      ++ NT + S    ++          EI NCSFN
Sbjct  2125  LKPCVKLTPLCVTLNCTTYEPKKVTADNNNTCAPSQNTTVKNLTIDPEAVKEISNCSFNT  1946

Query  162   STSIRGKVQKEYAFFYKLDIIPIDNDTTS-----YKLTSCNTSVITQACPKVSFEPIPIH  216
             +T +R K +K +A FYK+D++ +  ++T+     Y+L +CNTS ITQACPKVSFEPIPIH
Sbjct  1945  TTELRDKTRKVHALFYKIDMVELGENSTAPNFTLYRLINCNTSAITQACPKVSFEPIPIH  1766

Query  217   YCAPAGFAILKCNNKTFNGTGPCTNVSTVQCTHGIRPVVSTQLLLNGSLAEEEVVIRSVN  276
             YCAPAG+A+LKCNN TFNGTGPC N+STVQCTHGIRPVVSTQLLLNGSLAE  V IRS N
Sbjct  1765  YCAPAGYALLKCNNNTFNGTGPCRNISTVQCTHGIRPVVSTQLLLNGSLAEGNVTIRSEN  1586

Query  277   FTDNAKTIIVQLNTSVEINCTRPNNNTRKRIRIQRGPGRAFVTIGK-IGNMRQAHCNISR  335
              TDNAKTII  LN SVEI CTRPNNNTR+ +RI  GPG+ F   G  IG++R+AHCN+S
Sbjct  1585  LTDNAKTIIAHLNQSVEILCTRPNNNTRQSVRI--GPGQTFFATGDIIGDIRRAHCNVSA  1412

Query  336   AKWNNTLKQIASKLREQFGNNKTIIFKQSSGGDPEIVTHSFNCGGEFFYCNSTQLFNS--  393
               W+  L+ ++ +L + F NNKTI F   SGGD EI THSFNC GEFFYCN++QLFNS
Sbjct  1411  KHWHTALRNVSIELAKLFPNNKTINFTSPSGGDLEITTHSFNCRGEFFYCNTSQLFNSLY  1232

Query  394   -------TWFNSTWSTEGSNNTE----GSD--------------------TITLPCRIKQ  422
                     W+N   +    N T+    GS                      ITLPCRIKQ
Sbjct  1231  MPNNDTYIWYNENHTYVWHNGTQYVINGSTQYVTLKNSAQNVTINNITDFNITLPCRIKQ  1052

Query  423   IINMWQKVGKAMYAPPISGQIRCSSNITGLLLTRDGGNSN---NESEIFRPGGGDMRDNW  479
             IINMWQ+VGKAMYAPPI+G I C SNITGLLLTRDGG  +   N++EIFRPGGG+M+DNW
Sbjct  1051  IINMWQEVGKAMYAPPIAGNITCLSNITGLLLTRDGGKGDEIRNDTEIFRPGGGNMKDNW  872

Query  480   RSELYKYKVVKIEPLGVAPTKAKRRVVQREKRAVGIgalflgflgaagstmgaasmtLTV  539
             RSELYKYKVV+I+PLGVAPT A+RRVV RE+RAVG+GA+F GFLGAAGSTMGAAS+ LTV
Sbjct  871   RSELYKYKVVEIKPLGVAPTPARRRVVGRERRAVGLGAVFFGFLGAAGSTMGAASVALTV  692

Query  540   QARQLLSGIVQQQNNLLRAIEAQQHLLQLTVWGIKQLQARILAVERYLKDQQLLGIWGCS  599
             QARQLLSGIVQQQNNLLRAIEAQQ++L+LTVWGIKQLQ R+LA+ERYL++Q+LLG+WGCS
Sbjct  691   QARQLLSGIVQQQNNLLRAIEAQQNMLRLTVWGIKQLQTRVLAIERYLREQRLLGLWGCS  512

Query  600   GKLICTTAVPWNASWSNKSLEQIWNHTTWMEWDREINNYTSLIHSLIeesqnqqekneqe  659
              K+ICTT VPWN+SWSNKS   IWN+ TWMEWD+EI  +T+ I+ L+EESQ QQE+NE++
Sbjct  511   EKIICTTNVPWNSSWSNKSYSDIWNNLTWMEWDKEIEQHTNTIYQLLEESQGQQEQNEKD  332

Query  660   llelDKWASLWNWFNITNWLWYIKLFIMIVGGLVGLRIVFAVLSIVNRVRQGYSPLSFQT  719
             LL LDKW  LW+WF+ITNWLWYIK+FIMIVGGL+GLRIVFAVLSIVNRVRQGYSPLSFQT
Sbjct  331   LLALDKWNDLWSWFDITNWLWYIKIFIMIVGGLIGLRIVFAVLSIVNRVRQGYSPLSFQT  152

Query  720   HLPT  723
               P+
Sbjct  151   LAPS  140



>Escherichia coli strain NBE0006 contig00106, whole genome shotgun sequence
Sequence ID: NZ_JARBDI010000106.1 Length: 2090
Range 1: 1 to 2088

Score:857 bits(2214), Expect:0.0,
Method:Compositional matrix adjust.,
Identities:476/705(68%), Positives:563/705(79%), Gaps:36/705(5%)

Query  107   DIISLWDQSLKPCVKLTPLCVSLKCT--DLKNDTNTNSSSGR--MIMEKG--EIKNCSFN  160
             D+IS+WDQ LKPCVKLTPLCV+L CT  + +N+T + +S+ +   I E+   EI NCSF
Sbjct  2088  DVISIWDQGLKPCVKLTPLCVTLNCTKYEPRNETVSGNSTAKSPTIDEEALKEISNCSFE  1909

Query  161   ISTSIRGKVQKEYAFFYKLDIIPIDNDTTS------------YKLTSCNTSVITQACPKV  208
              +T +R K +K +A FYK+D+I +  ++T             Y+L +CNTS ITQACPKV
Sbjct  1908  TTTELRDKTRKVHALFYKIDMIELGQNSTCQKPENSTSECTMYRLINCNTSAITQACPKV  1729

Query  209   SFEPIPIHYCAPAGFAILKCNNKTFNGTGPCTNVSTVQCTHGIRPVVSTQLLLNGSLAEE  268
             SFEPIPIHYCAPAG+A+LKCNN TFNGTGPC N+STVQCTHGIRPVVSTQLLLNGSLAE
Sbjct  1728  SFEPIPIHYCAPAGYALLKCNNNTFNGTGPCRNISTVQCTHGIRPVVSTQLLLNGSLAEG  1549

Query  269   EVVIRSVNFTDNAKTIIVQLNTSVEINCTRPNNNTRKRIRIQRGPGRAFVTIGK-IGNMR  327
              + IRS N TDNAKTII  LN SVEINCTRPNNNTRK IRI  GPG+ F   G+ IGN+R
Sbjct  1548  NITIRSENLTDNAKTIIAHLNESVEINCTRPNNNTRKSIRI--GPGQTFYATGEIIGNIR  1375

Query  328   QAHCNISRAKWNNTLKQIASKLREQFGNNKTIIFKQSSGGDPEIVTHSFNCGGEFFYCNS  387
             QAHCNIS   WN TL+++  KL E F  NKTI F  SSGGD EI THSFNC GEFFYC++
Sbjct  1374  QAHCNISENMWNKTLQRVGKKLAEHFP-NKTIKFASSSGGDLEITTHSFNCRGEFFYCST  1198

Query  388   TQLFNSTWFNSTWSTEGSNNTEGSDTITLPCRIKQIINMWQKVGKAMYAPPISGQIRCSS  447
             ++LFNST+            ++ + TIT+PC IKQI+NMWQ+VG+AMYAPPI G I C S
Sbjct  1197  SRLFNSTYM------PNDTKSKSNKTITIPCSIKQIVNMWQEVGRAMYAPPIEGSITCES  1036

Query  448   NITGLLLTRDGG---NSN-----NESEIFRPGGGDMRDNWRSELYKYKVVKIEPLGVAPT  499
             NITGLLL RDGG    SN     N +E FRP GGDMR+NWRSELYKYKVV+I+PLG+APT
Sbjct  1035  NITGLLLVRDGGIESESNCTETLNCTETFRPEGGDMRNNWRSELYKYKVVEIKPLGIAPT  856

Query  500   KAKRRVVQREKRAVGIgalflgflgaagstmgaasmtLTVQARQLLSGIVQQQNNLLRAI  559
              AKRRVV+REKRAVG+GA+FLGFLGAAGSTMGAASM LTVQARQLLSGIVQQQNNLLRAI
Sbjct  855   AAKRRVVEREKRAVGLGAVFLGFLGAAGSTMGAASMALTVQARQLLSGIVQQQNNLLRAI  676

Query  560   EAQQHLLQLTVWGIKQLQARILAVERYLKDQQLLGIWGCSGKLICTTAVPWNASWSNKSL  619
             EAQQ++L+LTVWGIKQLQ R+LA+ERYL++Q+LLG+WGCS + ICTT VPWN+SWSN+S
Sbjct  675   EAQQNMLRLTVWGIKQLQTRVLAIERYLREQRLLGLWGCSERTICTTNVPWNSSWSNRSY  496

Query  620   EQIWNHTTWMEWDREINNYTSLIHSLIeesqnqqekneqellelDKWASLWNWFNITNWL  679
               IWN+ TWMEWD+EI  +T+ I+ L+EESQ QQE+NE++LL LDKW  LW+WF+ITNWL
Sbjct  495   SDIWNNLTWMEWDKEIEQHTNTIYQLLEESQGQQEQNEKDLLALDKWNDLWSWFDITNWL  316

Query  680   WYIKLFIMIVGGLVGLRIVFAVLSIVNRVRQGYSPLSFQTHLPTprgpdrpegieeegge  739
             WYI++FIMIVGGL+GLRIVFAVLSIVNRVRQGYSPLSFQT  P+P G DRP  I+EEGGE
Sbjct  315   WYIRIFIMIVGGLIGLRIVFAVLSIVNRVRQGYSPLSFQTLAPSPGGLDRPGRIDEEGGE  136

Query  740   rdrdrSIRLVNGSLALIWDDLRSLCLFSYHRLRDLLLIVTRIVEL  784
             + R+ S RLV+G LAL W+DLRSL LF YHRL+DL+ +  R VEL
Sbjct  135   QGRNNSTRLVSGFLALAWEDLRSLFLFCYHRLKDLISVTARAVEL  1



>Escherichia coli strain NBE0009 contig00112, whole genome shotgun sequence
```

```
Sequence ID: NZ_JARBDL010000112.1 Length: 2003
Range 1: 3 to 1646

Score:743 bits(1919), Expect:0.0,
Method:Compositional matrix adjust.,
Identities:393/560(70%), Positives:455/560(81%), Gaps:25/560(4%)

Query  1     MRVKEKYQHLWRWGWRWGTMLLGMLMICSATEKLWVTVYYGVPVWKEATTTLFCASDAKA  60
             MRV+   ++  +W W WG +   MLM C     LWVTVYYGVPVW EA TTLFCASDAKA
Sbjct  1646  MRVRGILRNYQQW-WIWGVLGFWMLMNCKVGGNLWVTVYYGVPVWTEAKTTLFCASDAKA  1470

Query  61    YDTEVHNVWATHACVPTDPNPQEVVLVNVTENFNMWKNDMVEQMHEDIISLWDQSLKPCV  120
             Y+ EVHNVWATHACVPTDPNPQE+VL NVTENFNMWKNDMV QMHED+ISLWDQSLKPCV
Sbjct  1469  YEKEVHNVWATHACVPTDPNPQEMVLENVTENFNMWKNDMVNQMHEDVISLWDQSLKPCV  1290

Query  121   KLTPLCVSLKCTDLKNDTNTNSSSGRMIMEKGEIKNCSFNISTSIRGKVQKEYAFFYKLD  180
             KLTPLCV+L+C    N +  N SS R+     E+KNCSFN +T +R K  K  A F++LD
Sbjct  1289  KLTPLCVTLEC----NSSTYNQSSVRV----HEVKNCSFNATTELRDKKHKVQALFHRLD  1134

Query  181   IIPIDN------DTTSYKLTSCNTSVITQACPKVSFEPIPIHYCAPAGFAILKCNNKTFN  234
             ++P+DN       T  Y+L +CNTSVITQACPKV+F+PIPIHYC PAG+AIL+CN+KTFN
Sbjct  1133  VVPLDNRSSNENSTGYYRLINCNTSVITQACPKVTFDPIPIHYCTPAGYAILQCNDKTFN  954

Query  235   GTGPCTNVSTVQCTHGIRPVVSTQLLLNGSLAEEEVVIRSVNFTDNAKTIIVQLNTSVEI  294
             GTGPC NVSTVQCTHGI+PVVSTQLLLNGSLAEEE+VIRS N T+NAKTIIV LN SV+I
Sbjct  953   GTGPCNNVSTVQCTHGIKPVVSTQLLLNGSLAEEEIVIRSENLTNNAKTIIVHLNKSVDI  774

Query  295   NCTRPNNNTRKRIRIQRGPGRAFVTIGK-IGNMRQAHCNISRAKWNNTLKQIASKLREQF  353
              CTRPNNNTRK IRI  GPG+ F   G  IG++R+AHCNIS  +WN+T+ ++  KL E F
Sbjct  773   VCTRPNNNTRKSIRI--GPGQTFYATGDIIGDIRKAHCNISLNRWNDTILKVREKLAEHF  600

Query  354   GNNKTIIFKQSSGGDPEIVTHSFNCGGEFFYCNSTQLFNSTWF--NSTWSTEGS--NNTE  409
               N+TI F+ SSGGD EI THSFNCGGEFFYCN++ LFN T++  N T+   G+  N+T+
Sbjct  599   P-NRTIEFQPSSGGDLEITTHSFNCGGEFFYCNTSNLFNFTYYYDNGTYHYNGTYYNSTQ  423

Query  410   GSDT--ITLPCRIKQIINMWQKVGKAMYAPPISGQIRCSSNITGLLLTRDGGNSNNESEI  467
              + T  IT+PCRIKQIINMWQ+VG+AMYAPPI+G I C SNITGLLL RDGG S+NE+E
Sbjct  422   YNSTANITIPCRIKQIINMWQEVGRAMYAPPIAGNITCRSNITGLLLVRDGGPSSNETET  243

Query  468   FRPGGGDMRDNWRSELYKYKVVKIEPLGVAPTKAKRRVVQREKRAVGIgalflgflgaag  527
             FRPGGGDMRDNWRSELYKYKVV+I+PLG+APT AKRRVV REKRAVGIGA+FLGFLGAAG
Sbjct  242   FRPGGGDMRDNWRSELYKYKVVEIKPLGIAPTGAKRRVVGREKRAVGIGAVFLGFLGAAG  63

Query  528   stmgaasmtLTVQARQLLSG  547
             STMGAAS+TLTVQARQLLSG
Sbjct  62    STMGAASITLTVQARQLLSG  3



>Escherichia coli strain NBE0009 contig00111, whole genome shotgun sequence
Sequence ID: NZ_JARBDL010000111.1 Length: 2027
Range 1: 358 to 2025

Score:740 bits(1911), Expect:0.0,
Method:Compositional matrix adjust.,
Identities:388/561(69%), Positives:450/561(80%), Gaps:19/561(3%)

Query  1     MRVKEKYQHLWRWGWRWGTMLLGMLMICSATEKLWVTVYYGVPVWKEATTTLFCASDAKA  60
             MRV+   +   +W W WG +   MLMIC+  + LWVTVYYGVPVWKEA TTLFCASDAK
Sbjct  358   MRVRGTPRSYPQW-WIWGVLGFWMLMICNMKKDLWVTVYYGVPVWKEAKTTLFCASDAKG  534

Query  61    YDTEVHNVWATHACVPTDPNPQEVVLVNVTENFNMWKNDMVEQMHEDIISLWDQSLKPCV  120
             Y+TEVHNVWATHACVPTDP+PQE+ L NVTENFNMW+N+MV+QMHEDIISLWDQSLKPC+
Sbjct  535   YETEVHNVWATHACVPTDPSPQEMRLDNVTENFNMWENNMVDQMHEDIISLWDQSLKPCI  714

Query  121   KLTPLCVSLKCTDLKNDTNTNSSSGRMIMEKGEIKNCSFNISTSIRGKVQKEYAFFYKLD  180
             KLTPLCV+L+C +  N T TN  +      K EI+NCSFN +T I  K +KEYA FY+LD
Sbjct  715   KLTPLCVTLEC-EKANRTVTNILN-ETYDSKNEIQNCSFNATTEIEDKKKKEYALFYRLD  888

Query  181   IIPIDN-------DTTSYKLTSCNTSVITQACPKVSFEPIPIHYCAPAGFAILKCNNKTF  233
             I+ +DN       D+  Y+L +CNTS ITQACPKVSF+PIPIHYCAPAG+AILKCNNKTF
Sbjct  889   IVSLDNENNSSGNDSRPYRLINCNTSAITQACPKVSFDPIPIHYCAPAGYAILKCNNKTF  1068

Query  234   NGTGPCTNVSTVQCTHGIRPVVSTQLLLNGSLAEEEVVIRSVNFTDNAKTIIVQLNTSVE  293
             NGTGPC NVSTVQCTHGI+PVVSTQLL+NGSLAEEE++IRS N TDNAKTIIVQLN SV
Sbjct  1069  NGTGPCNNVSTVQCTHGIKPVVSTQLLVNGSLAEEEIIIRSENLTDNAKTIIVQLNKSVP  1248

Query  294   INCTRPNNNTRKRIRIQRGPGRA-FVTIGKIGNMRQAHCNISRAKWNNTLKQIASKLREQ  352
             I C RPNNNTRK IRI  GPG+   + T   IG++RQAHCN+SR  W  TL  +  KL+E
Sbjct  1249  ITCIRPNNNTRKSIRI--GPGQTVYATGDSIGDIRQAHCNVSRKDWEGTLYNVNKKLKEH  1422

Query  353   FGNNKTIIFKQSSGGDPEIVTHSFNCGGEFFYCNSTQLFNSTW---FNSTWSTEGSNNTE  409
             F N  TI FK SSGGD EI THSFNC GEFFYCN++ LFN T+   +  T++  G N++
Sbjct  1423  FPNITTIKFKPSSGGDLEITTHSFNCRGEFFYCNTSDLFNGTYNSTYKGTYNNTGGNSSA  1602

Query  410   GSDTITLPCRIKQIINMWQKVGKAMYAPPISGQIRCSSNITGLLLTRDGGN--SNNESEI  467
                 ITL CRIKQIINMWQ+VGKAMYAPPI+G I C SNITGLLL RDGG   ++N++E
Sbjct  1603  ADPNITLQCRIKQIINMWQQVGKAMYAPPIAGNITCISNITGLLLVRDGGTNRTDNDTET  1782

Query  468   FRPGGGDMRDNWRSELYKYKVVKIEPLGVAPTKAKRRVVQREKRAV-GIgalflgflgaa  526
             FRPGGG+M+DNWRSELYKYKVV+I+PLGVAPT AKRRVV+REKRA+ G+GA+FLG LGAA
Sbjct  1783  FRPGGGNMKDNWRSELYKYKVVEIQPLGVAPTGAKRRVVEREKRAITGLGAVFLGILGAA  1962

Query  527   gstmgaasmtLTVQARQLLSG  547
             GSTMGAAS+TLTVQARQLLSG
Sbjct  1963  GSTMGAASITLTVQARQLLSG  2025



>Escherichia coli strain EN5076 contig00066, whole genome shotgun sequence
Sequence ID: NZ_JAODTC010000066.1 Length: 2732
Range 1: 957 to 2375

Score:659 bits(1700), Expect:0.0,
Method:Compositional matrix adjust.,
Identities:322/478(67%), Positives:378/478(79%), Gaps:13/478(2%)

Query  1     MRVKEKYQHLWRWGWRWGTMLLGMLMICSATEKLWVTVYYGVPVWKEATTTLFCASDAKA  60
             MRV+   ++  +W W WG +   MLM C     LWVTVYYGVPVWKEA TTLFCASDAKA
Sbjct  2375  MRVRGILRNYQQW-WIWGILGFWMLMNCKGKGNLWVTVYYGVPVWKEAKTTLFCASDAKA  2199

Query  61    YDTEVHNVWATHACVPTDPNPQEVVLVNVTENFNMWKNDMVEQMHEDIISLWDQSLKPCV  120
             Y TEVHNVWATHACVPTDPNPQE+VL NVTENFNMWKNDMV QMHED+ISLWDQSLKPCV
Sbjct  2198  YKTEVHNVWATHACVPTDPNPQEMVLENVTENFNMWKNDMVNQMHEDVISLWDQSLKPCV  2019

Query  121   KLTPLCVSLKCTDLKNDTNTNSSSGRMIMEKGEIKNCSFNISTSIRGKVQKEYAFFYKLD  180
             KLTPLCV+LKC    ++  ++++         E+KNCSFNI+T +R + QK YA FYKLD
Sbjct  2018  KLTPLCVTLKCNATTHNGTSDAADNDTSSGGEEMKNCSFNITTELRDRKQKVYALFYKLD  1839

Query  181   IIPID-----NDTTSYKLTSCNTSVITQACPKVSFEPIPIHYCAPAGFAILKCNNKTFNG  235
             ++P+D     N + SY+L +CNTS ITQACPKVSF+PIPIHYCAPAG+AIL+CN+KTFNG
Sbjct  1838  VVPLDNTNGSNSSRSYRLINCNTSAITQACPKVSFDPIPIHYCAPAGYAILRCNDKTFNG  1659
```

```
Query  236   TGPCTNVSTVQCTHGIRPVVSTQLLLNGSLAEEEVVIRSVNFTDNAKTIIVQLNTSVEIN  295
             TGPC NVSTVQCTHGI+PVVSTQLLLNGSLAEEE++IRS N +DNAKTIIV LN SVEI
Sbjct  1658  TGPCHNVSTVQCTHGIKPVVSTQLLLNGSLAEEEIIIRSQNLSDNAKTIIVHLNQSVEIV  1479

Query  296   CTRPNNNTRKRIRIQRGPGRAFVTIGK-IGNMRQAHCNISRAKWNNTLKQIASKLREQFG  354
             CTRP+NNTRK IRI  GPG+ F   G  IGN+RQA+C+I+ +KW+ TL  ++ KL E+F
Sbjct  1478  CTRPHNNTRKSIRI--GPGQTFYATGAIIGNIRQAYCSINISKWSETLHNVSKKLAERFP  1305

Query  355   NNKTIIFKQSSGGDPEIVTHSFNCGGEFFYCNSTQLFNST--WFNSTWSTEGSNNTEGSD  412
              NKTI F  SSGGD E+ THSFNC GEFFYCN+++LFNST  + N T+   G+N T  ++
Sbjct  1304  -NKTINFAASSGGDLEVTTHSFNCRGEFFYCNTSELFNSTYIYLNRTYMFNGTNGT-STN  1131

Query  413   TITLPCRIKQIINMWQKVGKAMYAPPISGQIRCSSNITGLLLTRDGGNSNNESEIFRP  470
               T+PCRIKQIINMWQ+VG+AMYAPPI G I C SNITGLLL RDGG  N  +EIFRP
Sbjct  1130  NFTIPCRIKQIINMWQEVGRAMYAPPIEGNITCKSNITGLLLERDGGEENRTTEIFRP  957



Range 2: 282 to 950

Score:251 bits(642), Expect:2e-66,
Method:Compositional matrix adjust.,
Identities:160/243(66%), Positives:186/243(76%), Gaps:20/243(8%)

Query  613  SWSNKSLEQIWNHTTWMEWDREINNYTSLIHSLIeesqnqqekneqellelDKWASLWNW  672
            SWSN+S   IW++ TWM+WDREI+NYT  I+ L+EESQNQQEKNE+ELL LD W SLWNW
Sbjct  950  SWSNRSQTDIWDNMTWMQWDREISNYTDTIYKLLEESQNQQEKNEKELLALDSWESLWNW  771

Query  673  FNITNWLWYIKLFIMIVGGLVGLRIVFAVLSIVNRVRQGYSPLSFQTHLPTprgpdrpeg  732
            F+IT WLWYIK+FIMIVGG                    YSPLSFQT  P PRGPDR
Sbjct  770  FSITKWLWYIKIFIMIVGG--------------------YSPLSFQTLTPNPRGPDRLGR  651

Query  733  ieeeggerdrdrSIRLVNGSLALIWDDLRSLCLFSYHRLRDLLLIVTRIVELLGRRGWEA  792
            IEEEGGE+D+DRSIRLV+G LAL WDDLRSLCLFSYH+LR+ +L+  R+VELLGRRGWEA
Sbjct  650  IEEEGGEQDKDRSIRLVSGFLALAWDDLRSLCLFSYHQLRNFILVTARVVELLGRRGWEA  471

Query  793  LKYWWNLLQYWSQELKNSAVSLLNATAIAVAEGTDRVIEVVQGACRAIRHIPRRIRQGLE  852
            LKY  NL+ YW  ELK SA+ LL+  AIAVAEGTDR+I++Q  CRAI +IPRRIRQG E
Sbjct  470  LKYLGNLVLYWGLELKKSAIGLLDVIAIAVAEGTDRLIDLIQRICRAICNIPRRIRQGFE  291

Query  853  RIL  855
              L
Sbjct  290  AAL  282




>Escherichia coli strain EN5374 contig00094, whole genome shotgun sequence
Sequence ID: NZ_JAPTGI010000094.1 Length: 2720
Range 1: 352 to 1770

Score:656 bits(1693), Expect:0.0,
Method:Compositional matrix adjust.,
Identities:319/478(67%), Positives:379/478(79%), Gaps:13/478(2%)

Query  1     MRVKEKYQHLWRWGWRWGTMLLGMLMICSATEKLWVTVYYGVPVWKEATTTLFCASDAKA  60
             MRV+   ++  +W W WG +   MLM C+    LWVTVYYGVPVW++A TTLFCASDAKA
Sbjct  352   MRVRGMLRNYQQW-WIWGVLGFWMLMNCNGGGNLWVTVYYGVPVWRDAKTTLFCASDAKA  528

Query  61    YDTEVHNVWATHACVPTDPNPQEVVLVNVTENFNMWKNDMVEQMHEDIISLWDQSLKPCV  120
             Y+ EVHNVWATHACVPTDPNPQE+VL NVTENFNMWKNDMV QMHED+ISLWDQSLKPCV
Sbjct  529   YEKEVHNVWATHACVPTDPNPQEMVLENVTENFNMWKNDMVNQMHEDVISLWDQSLKPCV  708

Query  121   KLTPLCVSLKCTDLKNDTNTNSSSGRMIMEKGEIKNCSFNISTSIRGKVQKEYAFFYKLD  180
             KLTPLCV+LKC    ++  ++++          E+KNCSFNI+T +R + QK YA FYKLD
Sbjct  709   KLTPLCVTLKCNATTHNGTSDAADNDTSSGGEEMKNCSFNITTELRDRKQKVYALFYKLD  888

Query  181   IIPID-----NDTTSYKLTSCNTSVITQACPKVSFEPIPIHYCAPAGFAILKCNNKTFNG  235
             ++P+D     N + SY+L +CNTS ITQACPKVSF+PIPIHYCAPAG+AIL+CN+KTFNG
Sbjct  889   VVPLDNTNGSNSSRSYRLINCNTSAITQACPKVSFDPIPIHYCAPAGYAILRCNDKTFNG  1068

Query  236   TGPCTNVSTVQCTHGIRPVVSTQLLLNGSLAEEEVVIRSVNFTDNAKTIIVQLNTSVEIN  295
             TGPC NVSTVQCTHGI+PVVSTQLLLNGSLAEEE++IRS N +DNAKTIIV LN SVEI
Sbjct  1069  TGPCHNVSTVQCTHGIKPVVSTQLLLNGSLAEEEIIIRSQNLSDNAKTIIVHLNQSVEIV  1248

Query  296   CTRPNNNTRKRIRIQRGPGRAFVTIGK-IGNMRQAHCNISRAKWNNTLKQIASKLREQFG  354
             CTRP+NNTRK IRI  GPG+ F   G  IGN+RQA+C+I+ +KW+ TL  ++ KL E+F
Sbjct  1249  CTRPHNNTRKSIRI--GPGQTFYATGAIIGNIRQAYCSINISKWSETLHNVSKKLAERFP  1422

Query  355   NNKTIIFKQSSGGDPEIVTHSFNCGGEFFYCNSTQLFNST--WFNSTWSTEGSNNTEGSD  412
              NKTI F  SSGGD E+ THSFNC GEFFYCN+++LFNST  + N T+   G+N T  ++
Sbjct  1423  -NKTINFAASSGGDLEVTTHSFNCRGEFFYCNTSELFNSTYIYLNRTYMFNGTNGT-STN  1596

Query  413   TITLPCRIKQIINMWQKVGKAMYAPPISGQIRCSSNITGLLLTRDGGNSNNESEIFRP  470
               T+PCRIKQIINMWQ+VG+AMYAPPI G I C SNITGLLL RDGG  N  +EIFRP
Sbjct  1597  NFTIPCRIKQIINMWQEVGRAMYAPPIEGNITCKSNITGLLLERDGGEENRTTEIFRP  1770



Range 2: 1777 to 2445

Score:251 bits(642), Expect:2e-66,
Method:Compositional matrix adjust.,
Identities:160/243(66%), Positives:186/243(76%), Gaps:20/243(8%)

Query  613   SWSNKSLEQIWNHTTWMEWDREINNYTSLIHSLIeesqnqqekneqellelDKWASLWNW  672
             SWSN+S   IW++ TWM+WDREI+NYT  I+ L+EESQNQQEKNE+ELL LD W SLWNW
Sbjct  1777  SWSNRSQTDIWDNMTWMQWDREISNYTDTIYKLLEESQNQQEKNEKELLALDSWESLWNW  1956

Query  673   FNITNWLWYIKLFIMIVGGLVGLRIVFAVLSIVNRVRQGYSPLSFQTHLPTprgpdrpeg  732
             F+IT WLWYIK+FIMIVGG                    YSPLSFQT  P PRGPDR
Sbjct  1957  FSITKWLWYIKIFIMIVGG--------------------YSPLSFQTLTPNPRGPDRLGR  2076

Query  733   ieeeggerdrdrSIRLVNGSLALIWDDLRSLCLFSYHRLRDLLLIVTRIVELLGRRGWEA  792
             IEEEGGE+D+DRSIRLV+G LAL WDDLRSLCLFSYH+LR+ +L+  R+VELLGRRGWEA
Sbjct  2077  IEEEGGEQDKDRSIRLVSGFLALAWDDLRSLCLFSYHQLRNFILVTARVVELLGRRGWEA  2256

Query  793   LKYWWNLLQYWSQELKNSAVSLLNATAIAVAEGTDRVIEVVQGACRAIRHIPRRIRQGLE  852
             LKY  NL+ YW  ELK SA+ LL+  AIAVAEGTDR+I++Q  CRAI +IPRRIRQG E
Sbjct  2257  LKYLGNLVLYWGLELKKSAIGLLDVIAIAVAEGTDRLIDLIQRICRAICNIPRRIRQGFE  2436

Query  853   RIL  855
               L
Sbjct  2437  AAL  2445



>Escherichia coli strain EN5090 contig00081, whole genome shotgun sequence
Sequence ID: NZ_JAODTD010000081.1 Length: 2798
Range 1: 1020 to 2441
```

```
Score:655 bits(1691), Expect:0.0,
Method:Compositional matrix adjust.,
Identities:323/479(67%), Positives:377/479(78%), Gaps:14/479(2%)

Query  1     MRVKEKYQHLWRWGWRWGTMLLGMLMICSATEKLWVTVYYGVPVWKEATTTLFCASDAKA  60
             MRV+   +   +W W WG +   MLM C+    LWVTVYYGVPVWKEA TTLFCASDAKA
Sbjct  2441  MRVRGILRSYQQW-WIWGILGFWMLMNCNGKGNLWVTVYYGVPVWKEAKTTLFCASDAKA  2265

Query  61    YDTEVHNVWATHACVPTDPNPQEVVLVNVTENFNMWKNDMVEQMHEDIISLWDQSLKPCV  120
             Y TEVHNVWATHACVPTDPNPQE+VL NVTENFNMWKNDMV QMHED+ISLWDQSLKPCV
Sbjct  2264  YKTEVHNVWATHACVPTDPNPQEMVLENVTENFNMWKNDMVNQMHEDVISLWDQSLKPCV  2085

Query  121   KLTPLCVSLKCTDLKNDTNTNSSSGRMIMEKGEIKNCSFNISTSIRGKVQKEYAFFYKLD  180
             KLTPLCV+LKCT   ++   +++         E+KNCSFNI+T +R + QK YA FYKLD
Sbjct  2084  KLTPLCVTLKCTATTHNGTRDAADNDTSSGGEEMKNCSFNITTELRDRKQKVYALFYKLD  1905

Query  181   IIPIDNDTTS-----YKLTSCNTSVITQACPKVSFEPIPIHYCAPAGFAILKCNNKTFNG  235
             ++P+DN   S     Y+L +CNTS ITQACPKVSF+PIPIHYCAPAG+AIL+CN+KTFNG
Sbjct  1904  VVPLDNTNGSNSSRPYRLINCNTSAITQACPKVSFDPIPIHYCAPAGYAILRCNDKTFNG  1725

Query  236   TGPCTNVSTVQCTHGIRPVVSTQLLLNGSLAEEEVVIRSVNFTDNAKTIIVQLNTSVEIN  295
             TGPC NVSTVQCTHGI+PVVSTQLLLNGSLAEEE++IRS N +DNAKTIIV LN SVEI
Sbjct  1724  TGPCHNVSTVQCTHGIKPVVSTQLLLNGSLAEEEIIIRSQNLSDNAKTIIVHLNQSVEIV  1545

Query  296   CTRPNNNTRKRIRIQRGPGRAFVTIGK-IGNMRQAHCNISRAKWNNTLKQIASKLREQFG  354
             CTRP+NNTRK IRI  GPG+ F   G  IGN+RQA+C+I+ +KW+ TL  ++ KL E+F
Sbjct  1544  CTRPHNNTRKSIRI--GPGQTFYATGAIIGNIRQAYCSINISKWSETLHNVSKKLAERFP  1371

Query  355   NNKTIIFKQSSGGDPEIVTHSFNCGGEFFYCNSTQLFNSTWF--NSTWSTEGSNNTEGSD  412
              NKTI F  SSGGD E+ THSFNC GEFFYCN+++LFNST+   + T+   G+N T  ++
Sbjct  1370  -NKTINFAASSGGDLEVTTHSFNCRGEFFYCNTSELFNSTYIYPDRTYMFNGTNGT-STN  1197

Query  413   TITLPCRIKQIINMWQKVGKAMYAPPISGQIRCSSNITGLLLTRDGGNSNNE-SEIFRP  470
               T+PCRIKQIINMWQKVG+AMYAPPI G I C SNITGLLL RDGG   N  +EIFRP
Sbjct  1196  NFTIPCRIKQIINMWQKVGRAMYAPPIEGNITCKSNITGLLLERDGGEEENRTTEIFRP  1020



Range 2: 285 to 1013

Score:298 bits(764), Expect:4e-83,
Method:Compositional matrix adjust.,
Identities:177/243(73%), Positives:205/243(84%), Gaps:0/243(0%)

Query  613   SWSNKSLEQIWNHTTWMEWDREINNYTSLIHSLIeesqnqqekneqellelDKWASLWNW  672
             SWSN+S   IW++ TWM+WDREI+NYT  I+ L+EESQNQQEKNE+ELL L  W SLWNW
Sbjct  1013  SWSNRSQTDIWDNMTWMQWDREISNYTDTIYKLLEESQNQQEKNEKELLALGSWESLWNW  834

Query  673   FNITNWLWYIKLFIMIVGGLVGLRIVFAVLSIVNRVRQGYSPLSFQTHLPTprgpdrpeg  732
             F+IT WLWYIK+FIMIVGGL+GLRI+FAVLSIVNRVRQGYSPLSFQT  P PRG DR
Sbjct  833   FSITKWLWYIKIFIMIVGGLIGLRIIFAVLSIVNRVRQGYSPLSFQTLTPNPRGLDRLRR  654

Query  733   ieeeggerdrdrSIRLVNGSLALIWDDLRSLCLFSYHRLRDLLLIVTRIVELLGRRGWEA  792
             IEEEGGE+D+DRSIRLV+G LAL WDDLRSLCLFSYH+LR+ +L+  R+VELLGRRGWEA
Sbjct  653   IEEEGGEQDKDRSIRLVSGFLALAWDDLRSLCLFSYHQLRNFILVTARVVELLGRRGWEA  474

Query  793   LKYWWNLLQYWSQELKNSAVSLLNATAIAVAEGTDRVIEVVQGACRAIRHIPRRIRQGLE  852
             LKY  NL+ YW  ELK SA+SLL+  AIAVAEGTDR+I+++Q  CRAI +IPRRIRQG E
Sbjct  473   LKYLGNLVLYWGLELKKSAISLLDVIAIAVAEGTDRLIDLIQRICRAICNIPRRIRQGFE  294

Query  853   RIL  855
               L
Sbjct  293   AAL  285



>Escherichia coli strain NBE0010 contig00080, whole genome shotgun sequence
Sequence ID: NZ_JARBDM010000080.1 Length: 2743
Range 1: 53 to 1471

Score:654 bits(1687), Expect:0.0,
Method:Compositional matrix adjust.,
Identities:318/478(67%), Positives:378/478(79%), Gaps:13/478(2%)

Query  1     MRVKEKYQHLWRWGWRWGTMLLGMLMICSATEKLWVTVYYGVPVWKEATTTLFCASDAKA  60
             MRV+   ++  +W   WG +   ML+ C   E LWVTVYYGVPVW++A TTLFCASDAKA
Sbjct  53    MRVRGILRNYQQW*-IWGVLSF*MLITCKVEENLWVTVYYGVPVWRDAKTTLFCASDAKA  229

Query  61    YDTEVHNVWATHACVPTDPNPQEVVLVNVTENFNMWKNDMVEQMHEDIISLWDQSLKPCV  120
             Y+ EVHNVWATHACVPTDPNPQE+VL NVTENFNMWKNDMV QMHED+ISLWDQSLKPCV
Sbjct  230   YEKEVHNVWATHACVPTDPNPQEMVLENVTENFNMWKNDMVNQMHEDVISLWDQSLKPCV  409

Query  121   KLTPLCVSLKCTDLKNDTNTNSSSGRMIMEKGEIKNCSFNISTSIRGKVQKEYAFFYKLD  180
             KLTPLCV+LKC    ++  ++++         E+KNCSFNI+T +R + QK YA FYKLD
Sbjct  410   KLTPLCVTLKCNATTHNGTSDAADNDTSSGGEEMKNCSFNITTELRDRKQKVYALFYKLD  589

Query  181   IIPID-----NDTTSYKLTSCNTSVITQACPKVSFEPIPIHYCAPAGFAILKCNNKTFNG  235
             ++P+D     N + SY+L +CNTS ITQACPKVSF+PIPIHYCAPAG+AIL+CN+KTFNG
Sbjct  590   VVPLDNTNGSNSSRSYRLINCNTSAITQACPKVSFDPIPIHYCAPAGYAILRCNDKTFNG  769

Query  236   TGPCTNVSTVQCTHGIRPVVSTQLLLNGSLAEEEVVIRSVNFTDNAKTIIVQLNTSVEIN  295
             TGPC NVSTVQCTHGI+PVVSTQLLLNGSLAEEE++IRS N +DNAKTIIV LN SVEI
Sbjct  770   TGPCHNVSTVQCTHGIKPVVSTQLLLNGSLAEEEIIIRSQNLSDNAKTIIVHLNQSVEIV  949

Query  296   CTRPNNNTRKRIRIQRGPGRAFVTIGK-IGNMRQAHCNISRAKWNNTLKQIASKLREQFG  354
             CTRP+NNTRK IRI  GPG+ F   G  IGN+RQA+C+I+ +KW+ TL  ++ KL E+F
Sbjct  950   CTRPHNNTRKSIRI--GPGQTFYATGAIIGNIRQAYCSINISKWSETLHNVSKKLAERFP  1123

Query  355   NNKTIIFKQSSGGDPEIVTHSFNCGGEFFYCNSTQLFNST--WFNSTWSTEGSNNTEGSD  412
              NKTI F  SSGGD E+ THSFNC GEFFYCN+++LFNST  + N T+   G+N T  ++
Sbjct  1124  -NKTINFAASSGGDLEVTTHSFNCRGEFFYCNTSELFNSTYIYLNRTYMFNGTNGT-STN  1297

Query  413   TITLPCRIKQIINMWQKVGKAMYAPPISGQIRCSSNITGLLLTRDGGNSNNESEIFRP  470
               T+PCRIKQIINMWQ+VG+AMYAPPI G I C SNITGLLL RDGG  N  +EIFRP
Sbjct  1298  NFTIPCRIKQIINMWQEVGRAMYAPPIEGNITCKSNITGLLLERDGGEENRTTEIFRP  1471



Range 2: 1478 to 2146

Score:253 bits(646), Expect:8e-67,
Method:Compositional matrix adjust.,
Identities:160/243(66%), Positives:186/243(76%), Gaps:20/243(8%)

Query  613   SWSNKSLEQIWNHTTWMEWDREINNYTSLIHSLIeesqnqqekneqellelDKWASLWNW  672
             SWSN+S   IW++ TWM+WDREI+NYT  I+ L+EESQNQQEKNE+ELL LD W SLWNW
Sbjct  1478  SWSNRSQTDIWDNMTWMQWDREISNYTDTIYKLLEESQNQQEKNEKELLALDSWESLWNW  1657

Query  673   FNITNWLWYIKLFIMIVGGLVGLRIVFAVLSIVNRVRQGYSPLSFQTHLPTprgpdrpeg  732
             F+IT WLWYIK+FIMIVGG                    YSPLSFQT  P PRGPDR
```

```
Sbjct  1658  FSITKWLWYIKIFIMIVGG--------------------YSPLSFQTLTPNPRGPDRLGR  1777

Query  733   ieeeggerdrdrSIRLVNGSLALIWDDLRSLCLFSYHRLRDLLLIVTRIVELLGRRGWEA  792
             IEEEGGE+D+DRSIRLV+G LAL WDDLRSLCLFSYH+LR+ +L+  R+VELLGRRGWEA
Sbjct  1778  IEEEGGEQDKDRSIRLVSGFLALAWDDLRSLCLFSYHQLRNFILVTARVVELLGRRGWEA  1957

Query  793   LKYWWNLLQYWSQELKNSAVSLLNATAIAVAEGTDRVIEVVQGACRAIRHIPRRIRQGLE  852
             LKY  NL+ YW  ELK SA+ LL+  AIAVAEGTDR+I+++Q  CRAI +IPRRIRQG E
Sbjct  1958  LKYLGNLVLYWGLELKKSAIGLLDVIAIAVAEGTDRLIDLIQRICRAICNIPRRIRQGFE  2137

Query  853   RIL  855
               L
Sbjct  2138  AAL  2146
```

>Escherichia coli strain EN5177 contig00105, whole genome shotgun sequence
Sequence ID: NZ_JAPMNK010000105.1 Length: 1783
Range 1: 176 to 1681

Score:652 bits(1681), Expect:0.0,
Method:Compositional matrix adjust.,
Identities:337/519(65%), Positives:389/519(74%), Gaps:30/519(5%)

```
Query  1     MRVKEKYQHLWRWGWRWGTMLLGMLMICSATEKLWVTVYYGVPVWKEATTTLFCASDAKA  60
             MRVK   ++  +W W  G +   MLMIC+      VTVYY VPVWKEA  TLFCASDAKA
Sbjct  176   MRVKGTLRNYQQW-WI*GVLGF*MLMICNKGGDS*VTVYYEVPVWKEAKATLFCASDAKA  352

Query  61    YDTEVHNVWATHACVPTDPNPQEVVLVNVTENFNMWKNDMVEQMHEDIISLWDQSLKPCV  120
             ++ EVHNVWATHACVPTDPNPQE+VL NVTENFNMWK+DMV+QMHED+ISL DQSLKPCV
Sbjct  353   HEREVHNVWATHACVPTDPNPQEIVLENVTENFNMWKSDMVDQMHEDVISL*DQSLKPCV  532

Query  121   KLTPLCVSLKCTDLKNDTNTNSSSGRMIMEKGEIKNCSFNISTSIRGKVQKEYAFFYKLD  180
             KLTPLCV+L+C +     N          E  E+K C FN ST +  K QK  A FY LD
Sbjct  533   KLTPLCVTLECDNETVSVNHTD-------ESKELKRCFFNASTEVEDKKQKVSALFYSLD  691

Query  181   IIPIDNDTT---------SYKLTSCNTSVITQACPKVSFEPIPIHYCAPAGFAILKCNNK  231
             I P++  ++         +Y L SCNTSVI QACPKV+F+PIPIHYCAPAG+AILKCNN+
Sbjct  692   IRPLNKSSSRNSSINSNETYVLISCNTSVIKQACPKVTFDPIPIHYCAPAGYAILKCNNE  871

Query  232   TFNGTGPCTNVSTVQCTHGIRPVVSTQLLLNGSLAEEEVVIRSVNFTDNAKTIIVQLNTS  291
             TFN TGPC NVSTVQCTH I+PVVSTQLLLN SLA+ E++IRS N TDNAKTIIV LN S
Sbjct  872   TFNRTGPCNNVSTVQCTHRIKPVVSTQLLLNSSLAKGEIIIRSENLTDNAKTIIVHLNES  1051

Query  292   VEINCTRPNNNTRKRIRIQRGPGRAFVTIGKI-GNMRQAHCNISRAKWNNTLKQIASKLR  350
             VEINCTRPNNNTRK IRI  GPG+ F   G+I  N+RQAHCNIS  + N TL+++  KL
Sbjct  1052  VEINCTRPNNNTRKSIRI--GPGQTFYATGEIVRNIRQAHCNISEKE*NKTLQKVGIKLA  1225

Query  351   EQFGNNKTIIFKQSSGGDPEIVTHSFNCGGEFFYCNSTQLFNSTWFNSTWSTEGSNNTEG  410
             E F  NKTI F   SGGD EI+THSFNC GEFFYCN++ LFNST +NST+    S
Sbjct  1226  EHFP-NKTINFTSHSGGDLEIMTHSFNCRGEFFYCNTSSLFNST-YNSTYIRNNS-----  1384

Query  411   SDTITLPCRIKQIINMWQKVGKAMYAPPISGQIRCSSNITGLLLTRDGGNSN---NESEI  467
             S  IT+ CRIKQIINMWQ+VG+AMYAPPI+G I C SNITGLLL RD G+ N   N +E
Sbjct  1385  SSIITIQCRIKQIINMWQEVGRAMYAPPIAGNITCQSNITGLLLVRDRGHRNETSNSTET  1564

Query  468   FRPGGGDMRDNWRSELYKYKVVKIEPLGVAPTKAKRRVV  506
             FRP GGDM++N RSELYKYKVV+I+PLG+APT AKRRVV
Sbjct  1565  FRPKGGDMKNN*RSELYKYKVVEIKPLGIAPTAAKRRVV  1681
```

>Escherichia coli strain EN5379 contig00092, whole genome shotgun sequence
Sequence ID: NZ_JAPTGJ010000092.1 Length: 2592
Range 1: 1019 to 2434

Score:646 bits(1667), Expect:0.0,
Method:Compositional matrix adjust.,
Identities:324/480(68%), Positives:377/480(78%), Gaps:18/480(3%)

```
Query  1     MRVKEKYQHLWRWGWRWGTMLLGMLMICSATEKLWVTVYYGVPVWKEATTTLFCASDAKA  60
             MRV+   +   +W W WG +   MLM C+    LWVTVYYGVPVWKEA TTLFCASDAKA
Sbjct  2434  MRVRGILRSYQQW-WIWGILGFWMLMNCNGKGNLWVTVYYGVPVWKEAKTTLFCASDAKA  2258

Query  61    YDTEVHNVWATHACVPTDPNPQEVVLVNVTENFNMWKNDMVEQMHEDIISLWDQSLKPCV  120
             Y TEVHNVWATHACVPTDPNPQE+VL NVTENFNMWKNDMV QMHED+ISLWD+SLKPCV
Sbjct  2257  YKTEVHNVWATHACVPTDPNPQEMVLENVTENFNMWKNDMVNQMHEDVISLWDESLKPCV  2078

Query  121   KLTPLCVSLKC-TDLKNDTNTNSSSGRMIMEKGEIKNCSFNISTSIRGKVQKEYAFFYKL  179
             KLTPLCV+LKC     N T   + +G +     E+KNCSFNI+T +R + QK YA FYKL
Sbjct  2077  KLTPLCVTLKCNATASNGTAAGNGTGSV---GEEMKNCSFNITTELRDRKQKVYALFYKL  1907

Query  180   DIIPIDNDTTS-----YKLTSCNTSVITQACPKVSFEPIPIHYCAPAGFAILKCNNKTFN  234
             D++P+DN   S     Y+L +CNTS ITQACPKVSF+PIPIHYCAPAG+AIL+CN+KTFN
Sbjct  1906  DVVPLDNTNGSNSSRPYRLINCNTSAITQACPKVSFDPIPIHYCAPAGYAILRCNDKTFN  1727

Query  235   GTGPCTNVSTVQCTHGIRPVVSTQLLLNGSLAEEEVVIRSVNFTDNAKTIIVQLNTSVEI  294
             GTGPC NVSTVQCTHGI+PVVSTQLLLNGSLAEEE++IRS N +DNAKTIIV LN SVEI
Sbjct  1726  GTGPCHNVSTVQCTHGIKPVVSTQLLLNGSLAEEEIIIRSQNLSDNAKTIIVHLNQSVEI  1547

Query  295   NCTRPNNNTRKRIRIQRGPGRAFVTIGK-IGNMRQAHCNISRAKWNNTLKQIASKLREQF  353
              CTRP+NNTRK IRI  GPG+ F   G  IGN+RQA+C+I+ +KW+ TL  ++ KL E+F
Sbjct  1546  VCTRPHNNTRKSIRI--GPGQTFYATGAIIGNIRQAYCSINISKWSETLHNVSKKLAERF  1373

Query  354   GNNKTIIFKQSSGGDPEIVTHSFNCGGEFFYCNSTQLFNSTWF--NSTWSTEGSNNTEGS  411
               NKTI F  SSGGD E+ THSFNC GEFFYCN+++LFNST+   + T+   G+N T  +
Sbjct  1372  P-NKTINFAASSGGDLEVTTHSFNCRGEFFYCNTSELFNSTYIYPDRTYMFNGTNGT-ST  1199

Query  412   DTITLPCRIKQIINMWQKVGKAMYAPPISGQIRCSSNITGLLLTRDGGNSNNE-SEIFRP  470
             +  T+PCRIKQIINMWQKVG+AMYAPPI G I C SNITGLLL RDGG   N  +EIFRP
Sbjct  1198  NNFTIPCRIKQIINMWQKVGRAMYAPPIEGNITCKSNITGLLLERDGGEEENRTTEIFRP  1019
```

Range 2: 263 to 1012

Score:292 bits(748), Expect:3e-81,
Method:Compositional matrix adjust.,
Identities:180/250(72%), Positives:204/250(81%), Gaps:7/250(2%)

```
Query  613   SWSNKSLEQIWNHTTWMEWDREINNYTSLIHSLIeesqnqqekneqellelDKWASLWNW  672
             SWSN+S   IW++ TWM+WDREI+NYT  I+ L+EESQNQQEKNE+ELL L  W SLWNW
Sbjct  1012  SWSNRSQTDIWDNMTWMQWDREISNYTDTIYKLLEESQNQQEKNEKELLALGSWESLWNW  833

Query  673   FNITNWLWYIKLFIMIVGGLVGLRIVFAVLSIVNRVRQGYSPLSFQTHLPTprgpdrpeg  732
             F+IT WLWYIK+FIMIVGGL+GLRI+FAVLSIVNRVRQGYSPLSFQT  P PRG DR
Sbjct  832   FSITKWLWYIKIFIMIVGGLIGLRIIFAVLSIVNRVRQGYSPLSFQTLTPNPRGLDRLRR  653

Query  733   ieeeggerdrdrSIRLVNGSLALIWDDLRSLCLFSYHRLRDLLLIVTRIVELLGR-----  787
             IEEEGGE+DRDRSIRLVNG L+L  DDLR+LCLFSYHRLRD +L+  R+VELLGR
Sbjct  652   IEEEGGEQDRDRSIRLVNGFLSLA*DDLRNLCLFSYHRLRDFILLTARVVELLGRSSLRG  473
```

```
Query  788   --RGWEALKYWWNLLQYWSQELKNSAVSLLNATAIAVAEGTDRVIEVVQGACRAIRHIPR  845
               RGWEALKY  +L+QYW  ELK SA+SLL+  AIAVAEGTDR+I VVQ  CRAI +IPR
Sbjct  472   LQRGWEALKYLGSLVQYWGLELKKSAISLLDTIAIAVAEGTDRIISVVQRICRAIYNIPR  293

Query  846   RIRQGLERIL  855
             RIRQG E  L
Sbjct  292   RIRQGFEAAL  263
```

```
>Escherichia coli strain EN5317 contig00057, whole genome shotgun sequence
Sequence ID: NZ_JARBDO010000057.1 Length: 2867
Range 1: 1077 to 2510

Score:642 bits(1657), Expect:0.0,
Method:Compositional matrix adjust.,
Identities:320/483(66%), Positives:373/483(77%), Gaps:18/483(3%)

Query  1     MRVKEKYQHLWRWGWRWGTMLLGMLMICSATEKLWVTVYYGVPVWKEATTTLFCASDAKA  60
             MRV+   +   +W W WG +   MLM C+  + LWVTVYYGVPVWKEA TTLFCASDAK
Sbjct  2510  MRVRGTPRSYQQW-WIWGVLGFWMLMNCNMKKDLWVTVYYGVPVWKEAKTTLFCASDAKG  2334

Query  61    YDTEVHNVWATHACVPTDPNPQEVVLVNVTENFNMWKNDMVEQMHEDIISLWDQSLKPCV  120
             Y+TEVHNVWATHACVPTDP+PQE+ L NVTENFNMW+N+MV+QMHEDIISLWDQSLKPC+
Sbjct  2333  YETEVHNVWATHACVPTDPSPQEMRLDNVTENFNMWENNMVDQMHEDIISLWDQSLKPCI  2154

Query  121   KLTPLCVSLKCTDLKNDTNTNSSSGRMIMEKGEIKNCSFNISTSIRGKVQKEYAFFYKLD  180
             KLTPLCV+L+C +  N T TN  +      K EI+NCSFN +T I  K +KEYA FY+LD
Sbjct  2153  KLTPLCVTLEC-EKANRTVTNILN-ETYDSKNEIQNCSFNATTEIEDKKKKEYALFYRLD  1980

Query  181   IIPIDN-------DTTSYKLTSCNTSVITQACPKVSFEPIPIHYCAPAGFAILKCNNKTF  233
             I+ +DN       D+  Y+L +CNTS ITQACPKVSF+PIPIHYCAPAG+AILKCNNKTF
Sbjct  1979  IVSLDNENNSSGNDSRPYRLINCNTSAITQACPKVSFDPIPIHYCAPAGYAILKCNNKTF  1800

Query  234   NGTGPCTNVSTVQCTHGIRPVVSTQLLLNGSLAEEEVVIRSVNFTDNAKTIIVQLNTSVE  293
             NGTGPC NVSTVQCTHGI+PVVSTQLL+NGSLAEEE++IRS N TDNAKTIIVQLN SV
Sbjct  1799  NGTGPCNNVSTVQCTHGIKPVVSTQLLVNGSLAEEEIIIRSENLTDNAKTIIVQLNKSVP  1620

Query  294   INCTRPNNNTRKRIRIQRGPGRA-FVTIGKIGNMRQAHCNISRAKWNNTLKQIASKLREQ  352
             I C RPNNNTRK IRI  GPG+  + T   IG++RQAHCN+SR  W  TL  +  KL+E
Sbjct  1619  ITCIRPNNNTRKSIRI--GPGQTVYATGDSIGDIRQAHCNVSRKDWEGTLYNVNKKLKEH  1446

Query  353   FGNNKTIIFKQSSGGDPEIVTHSFNCGGEFFYCNSTQLFNSTW---FNSTWSTEGSNNTE  409
             F N  TI FK SSGGD EI THSFNC GEFFYCN++ LFN T+   +  T++  G N++
Sbjct  1445  FPNITTIKFKPSSGGDLEITTHSFNCRGEFFYCNTSDLFNGTYNSTYKGTYNNTGGNSSA  1266

Query  410   GSDTITLPCRIKQIINMWQKVGKAMYAPPISGQIRCSSNITGLLLTRDGGN--SNNESEI  467
                 ITL CRIKQIINMWQ+VGKAMYAPPI+G I C SNITGLLL RDGG   ++N++E
Sbjct  1265  ADPNITLQCRIKQIINMWQQVGKAMYAPPIAGNITCISNITGLLLVRDGGTNRTDNDTET  1086

Query  468   FRP  470
             F P
Sbjct  1085  FLP  1077
```

```
Range 2: 305 to 1084

Score:303 bits(775), Expect:2e-84,
Method:Compositional matrix adjust.,
Identities:181/260(70%), Positives:213/260(81%), Gaps:7/260(2%)

Query  603   ICTTAVPWNASWSNKSLEQIWNHTTWMEWDREINNYTSLIHSLIeesqnqqekneqelle  662
              C TAVPWN+SWSNK+ +QIWN+ TWM+WD+EI+NYT  I+ L+E+SQ QQEKNE++LL
Sbjct  1084  FCPTAVPWNSSWSNKTQDQIWNNMTWMQWDKEISNYTYTIYRLLEDSQTQQEKNEKDLLA  905

Query  663   lDKWASLWNWFNITNWLWYIKLFIMIVGGLVGLRIVFAVLSIVNRVRQGYSPLSFQTHLP  722
             LD WASLWNWF+IT WLWYIK+FIMIVGGL+GLRI+FAVLSIVNRVRQGYSPLSFQT  P
Sbjct  904   LDSWASLWNWFDITKWLWYIKIFIMIVGGLIGLRIIFAVLSIVNRVRQGYSPLSFQTLTP  725

Query  723   TprgpdrpegieeeggerdrdrSIRLVNGSLALIWDDLRSLCLFSYHRLRDLLLIVTRIV  782
             TP  PDR  GIEEEGGE+DR RS+RLV+G LA+ WDDLR+LCLFSYHRLRDLLL++ R V
Sbjct  724   TPGEPDRLRGIEEEGGEQDRVRSVRLVSGFLAIAWDDLRNLCLFSYHRLRDLLLVIARAV  545

Query  783   ELLG-------RRGWEALKYWWNLLQYWSQELKNSAVSLLNATAIAVAEGTDRVIEVVQG  835
             ELLG       +RG EALK+   ++QYW  ELK SA+SL +  AIAVAEGTDR+I  +Q
Sbjct  544   ELLGHSCLVGLQRGREALKHLGGIVQYWGLELKKSAISLFDTIAIAVAEGTDRIIIGIQR  365

Query  836   ACRAIRHIPRRIRQGLERIL  855
              CRAI +IPRRIRQG E  L
Sbjct  364   LCRAICNIPRRIRQGFEAAL  305
```

```
>Escherichia coli strain EN5169 contig00082, whole genome shotgun sequence
Sequence ID: NZ_JAPMNJ010000082.1 Length: 2837
Range 1: 1062 to 2480

Score:634 bits(1634), Expect:0.0,
Method:Compositional matrix adjust.,
Identities:320/485(66%), Positives:373/485(76%), Gaps:28/485(5%)

Query  1     MRVKEKYQHLWRWGWRWGTMLLGMLMICSATEKLWVTVYYGVPVWKEATTTLFCASDAKA  60
             MRVK   ++  +W W WG +   MLMIC+    LWVTVYYGVPVWKEA TTLFCAS+AKA
Sbjct  2480  MRVKGILRNYRQW-WIWGILGFWMLMICNG--NLWVTVYYGVPVWKEAKTTLFCASNAKA  2310

Query  61    YDTEVHNVWATHACVPTDPNPQEVVLVNVTENFNMWKNDMVEQMHEDIISLWDQSLKPCV  120
             Y+ EVHNVWATHACVPTDPNPQE+VL NVTENFNMWKNDMV QMHED+ISLWDQSLKPCV
Sbjct  2309  YEKEVHNVWATHACVPTDPNPQEMVLENVTENFNMWKNDMVNQMHEDVISLWDQSLKPCV  2130

Query  121   KLTPLCVSLKCTDLK---NDTNTNSSSGRMIMEKGEIKNCSFNISTSIRGKVQKEYAFFY  177
             KLTPLCV+L+C  +    N TN   ++  +     EI+NCSFN +T IR + Q+ YA FY
Sbjct  2129  KLTPLCVTLECRKVNATHNATNNGDATHNVTNNGQEIQNCSFNATTEIRDRKQRVYALFY  1950

Query  178   KLDIIPIDNDTTS-------YKLTSCNTSVITQACPKVSFEPIPIHYCAPAGFAILKCNN  230
             +LDI+P+D + +S       Y+L +CNTS ITQACPKVSF+PIPIHYCAPAG+AILKCNN
Sbjct  1949  RLDIVPLDKNNSSKNNSSEYYRLINCNTSAITQACPKVSFDPIPIHYCAPAGYAILKCNN  1770

Query  231   KTFNGTGPCTNVSTVQCTHGIRPVVSTQLLLNGSLAEEEVVIRSVNFTDNAKTIIVQLNT  290
             KTFNGTGPC NVSTVQCTHGI+PVVSTQLLLNGSLAE E++IRS N TDN KTIIV L+
Sbjct  1769  KTFNGTGPCNNVSTVQCTHGIKPVVSTQLLLNGSLAEGEIIIRSENLTDNVKTIIVHLDQ  1590

Query  291   SVEINCTRPNNNTRKRIRIQRGPGRAF-VTIGKIGNMRQAHCNISRAKWNNTLKQIASKL  349
             SVEI CTRPNNNTRK IRI  GPG+ F  T G IGN+RQAHCNIS  KWN TL+++  KL
Sbjct  1589  SVEIVCTRPNNNTRKSIRI--GPGQTFYATGGIIGNIRQAHCNISEDKWNETLQRVGKKL  1416

Query  350   REQFGNNKTIIFKQSSGGDPEIVTHSFNCGGEFFYCNSTQLFNSTWFNSTWSTEGSNNTE  409
              E F  NKTI F  SSGGD EI THSFNC GEFFYC++++LFNST+            ++
Sbjct  1415  VEHFP-NKTIKFAPSSGGDLEITTHSFNCRGEFFYCSTSRLFNSTYM------PNDTKSK  1257
```

```
Query  410   GSDTITLPCRIKQIINMWQKVGKAMYAPPISGQIRCSSNITGLLLTRDGG-----NSNNE  464
              + TIT+PC IKQI+NMWQ+VG+AMYAPPI G I C SNITG+LL RDGG       NN+
Sbjct  1256  SNKTITIPCSIKQIVNMWQEVGRAMYAPPIEGNITCRSNITGILLVRDGGVDSEDPENNK  1077

Query  465   SEIFR  469
             +E F+
Sbjct  1076  TETFQ  1062



Range 2: 296 to 1060

Score:300 bits(767), Expect:2e-83,
Method:Compositional matrix adjust.,
Identities:180/255(71%), Positives:208/255(81%), Gaps:7/255(2%)

Query  608   VPWNASWSNKSLEQIWNHTTWMEWDREINNYTSLIHSLIeesqnqqekneqellelDKWA  667
             VPWN+SWSNKS   IW + TWM+WD+E++NYT  I+ L+E+SQ QQE+NE++LL LD W
Sbjct  1060  VPWNSSWSNKSQTDIWENMTWMQWDKEVSNYTDTIYRLLEDSQTQQERNEKDLLALDNWK  881

Query  668   SLWNWFNITNWLWYIKLFIMIVGGLVGLRIVFAVLSIVNRVRQGYSPLSFQTHLPTprgp  727
             +LWNWF+ITNWLWYIK+FIMIVGGL+GLRI+FAVLSIVNRVRQGYSPLSFQT  P PRGP
Sbjct  880   NLWNWFSITNWLWYIKIFIMIVGGLIGLRIIFAVLSIVNRVRQGYSPLSFQTLTPNPRGP  701

Query  728   drpegieeeggerdrdrSIRLVNGSLALIWDDLRSLCLFSYHRLRDLLLIVTRIVELLG-  786
             DR   IEEEGG +DRDRSIRLVNG LAL WDDL SLCLFSYHRLRDL+L+  R VELLG
Sbjct  700   DRLGRIEEEGGGQDRDRSIRLVNGFLALAWDDLWSLCLFSYHRLRDLILVTARAVELLGH  521

Query  787   ------RRGWEALKYWWNLLQYWSQELKNSAVSLLNATAIAVAEGTDRVIEVVQGACRAI  840
                   +RGWEALKY   ++QYW  ELK  A+SLL+  AIAVAEGTDR+IE +Q  CRAI
Sbjct  520   SSLRGLQRGWEALKYLGGIVQYWGLELKKRAISLLDTVAIAVAEGTDRIIEFLQRICRAI  341

Query  841   RHIPRRIRQGLERIL  855
             R+IPRRIRQG E  L
Sbjct  340   RNIPRRIRQGFEAAL  296



>Escherichia coli strain NBE0003 contig00096, whole genome shotgun sequence
Sequence ID: NZ_JARBDG010000096.1 Length: 2837
Range 1: 1062 to 2480

Score:634 bits(1634), Expect:0.0,
Method:Compositional matrix adjust.,
Identities:320/485(66%), Positives:373/485(76%), Gaps:28/485(5%)

Query  1     MRVKEKYQHLWRWGWRWGTMLLGMLMICSATEKLWVTVYYGVPVWKEATTTLFCASDAKA  60
             MRVK   ++  +W W WG +   MLMIC+    LWVTVYYGVPVWKEA TTLFCAS+AKA
Sbjct  2480  MRVKGILRNYRQW-WIWGILGFWMLMICNG--NLWVTVYYGVPVWKEAKTTLFCASNAKA  2310

Query  61    YDTEVHNVWATHACVPTDPNPQEVVLVNVTENFNMWKNDMVEQMHEDIISLWDQSLKPCV  120
             Y+ EVHNVWATHACVPTDPNPQE+VL NVTENFNMWKNDMV QMHED+ISLWDQSLKPCV
Sbjct  2309  YEKEVHNVWATHACVPTDPNPQEMVLENVTENFNMWKNDMVNQMHEDVISLWDQSLKPCV  2130

Query  121   KLTPLCVSLKCTDLK---NDTNTNSSSGRMIMEKGEIKNCSFNISTSIRGKVQKEYAFFY  177
             KLTPLCV+L+C  +    N TN   ++  +     EI+NCSFN +T IR + Q+ YA FY
Sbjct  2129  KLTPLCVTLECRKVNATHNATNNGDATHNVTNNGQEIQNCSFNATTEIRDRKQRVYALFY  1950

Query  178   KLDIIPIDNDTTS-------YKLTSCNTSVITQACPKVSFEPIPIHYCAPAGFAILKCNN  230
             +LDI+P+D + +S       Y+L +CNTS ITQACPKVSF+PIPIHYCAPAG+AILKCNN
Sbjct  1949  RLDIVPLDKNNSSKNNSSEYRLINCNTSAITQACPKVSFDPIPIHYCAPAGYAILKCNN  1770

Query  231   KTFNGTGPCTNVSTVQCTHGIRPVVSTQLLLNGSLAEEEVVIRSVNFTDNAKTIIVQLNT  290
             KTFNGTGPC NVSTVQCTHGI+PVVSTQLLLNGSLAE E++IRS N TDN KTIIV L+
Sbjct  1769  KTFNGTGPCNNVSTVQCTHGIKPVVSTQLLLNGSLAEGEIIIRSENLTDNVKTIIVHLDQ  1590

Query  291   SVEINCTRPNNNTRKRIRIQRGPGRAF-VTIGKIGNMRQAHCNISRAKWNNTLKQIASKL  349
             SVEI CTRPNNNTRK IRI  GPG+ F  T G IGN+RQAHCNIS  KWN TL+++  KL
Sbjct  1589  SVEIVCTRPNNNTRKSIRI--GPGQTFYATGGIIGNIRQAHCNISEDKWNETLQRVGKKL  1416

Query  350   REQFGNNKTIIFKQSSGGDPEIVTHSFNCGGEFFYCNSTQLFNSTWFNSTWSTEGSNNTE  409
              E F  NKTI F  SSGGD EI THSFNC GEFFYC++++LFNST+            ++
Sbjct  1415  VEHFP-NKTIKFAPSSGGDLEITTHSFNCRGEFFYCSTSRLFNSTYM------PNDTKSK  1257

Query  410   GSDTITLPCRIKQIINMWQKVGKAMYAPPISGQIRCSSNITGLLLTRDGG-----NSNNE  464
              + TIT+PC IKQI+NMWQ+VG+AMYAPPI G I C SNITG+LL RDGG       NN+
Sbjct  1256  SNKTITIPCSIKQIVNMWQEVGRAMYAPPIEGNITCRSNITGILLVRDGGVDSEDPENNK  1077

Query  465   SEIFR  469
             +E F+
Sbjct  1076  TETFQ  1062



Range 2: 296 to 1060

Score:300 bits(767), Expect:2e-83,
Method:Compositional matrix adjust.,
Identities:180/255(71%), Positives:208/255(81%), Gaps:7/255(2%)

Query  608   VPWNASWSNKSLEQIWNHTTWMEWDREINNYTSLIHSLIeesqnqqekneqellelDKWA  667
             VPWN+SWSNKS   IW + TWM+WD+E++NYT  I+ L+E+SQ QQE+NE++LL LD W
Sbjct  1060  VPWNSSWSNKSQTDIWENMTWMQWDKEVSNYTDTIYRLLEDSQTQQERNEKDLLALDNWK  881

Query  668   SLWNWFNITNWLWYIKLFIMIVGGLVGLRIVFAVLSIVNRVRQGYSPLSFQTHLPTprgp  727
             +LWNWF+ITNWLWYIK+FIMIVGGL+GLRI+FAVLSIVNRVRQGYSPLSFQT  P PRGP
Sbjct  880   NLWNWFSITNWLWYIKIFIMIVGGLIGLRIIFAVLSIVNRVRQGYSPLSFQTLTPNPRGP  701

Query  728   drpegieeeggerdrdrSIRLVNGSLALIWDDLRSLCLFSYHRLRDLLLIVTRIVELLG-  786
             DR   IEEEGG +DRDRSIRLVNG LAL WDDL SLCLFSYHRLRDL+L+  R VELLG
Sbjct  700   DRLGRIEEEGGGQDRDRSIRLVNGFLALAWDDLWSLCLFSYHRLRDLILVTARAVELLGH  521

Query  787   ------RRGWEALKYWWNLLQYWSQELKNSAVSLLNATAIAVAEGTDRVIEVVQGACRAI  840
                   +RGWEALKY   ++QYW  ELK  A+SLL+  AIAVAEGTDR+IE +Q  CRAI
Sbjct  520   SSLRGLQRGWEALKYLGGIVQYWGLELKKRAISLLDTVAIAVAEGTDRIIEFLQRICRAI  341

Query  841   RHIPRRIRQGLERIL  855
             R+IPRRIRQG E  L
Sbjct  340   RNIPRRIRQGFEAAL  296



>Escherichia coli strain EN5143 contig00069, whole genome shotgun sequence
Sequence ID: NZ_JAPMNI010000069.1 Length: 2688
Range 1: 358 to 1731

Score:629 bits(1622), Expect:0.0,
Method:Compositional matrix adjust.,
Identities:316/475(67%), Positives:366/475(77%), Gaps:27/475(5%)
```

```
Query  1     MRVKEKYQHLWRWGWRWGTMLLGMLMICSATEKLWVTVYYGVPVWKEATTTLFCASDAKA  60
             MRV+   ++  +W W WG +   MLMIC+     WVTVYYGVPVWKEA TTLFCASDAKA
Sbjct  358   MRVRGTLRNYQQW-WIWGVLGFWMLMICNGGGDSWVTVYYGVPVWKEAKTTLFCASDAKA  534

Query  61    YDTEVHNVWATHACVPTDPNPQEVVLVNVTENFNMWKNDMVEQMHEDIISLWDQSLKPCV  120
             ++ EVHNVWATHACVPTDPNPQE+VL NVTENFNMW+NDMV+QMHED+ISLWDQSLKPCV
Sbjct  535   HEREVHNVWATHACVPTDPNPQEMVLENVTENFNMWRNDMVDQMHEDVISLWDQSLKPCV  714

Query  121   KLTPLCVSLKCTDLKNDTNTNSSSGRMIMEKGEIKNCSFNISTSIRGKVQKEYAFFYKLD  180
             KLTPLCV+L+C       N N++      E  E+K C FN ST +  K QK  A FY LD
Sbjct  715   KLTPLCVTLEC-------NNNTTFKNYTDESKELKRCFFNASTEVEDKKQKVSALFYSLD  873

Query  181   IIPIDNDTT---------SYKLTSCNTSVITQACPKVSFEPIPIHYCAPAGFAILKCNNK  231
             I P++  ++         +Y L SCNTSVI QACPKV+F+PIPIHYCAPAG+AILKCNN+
Sbjct  874   IRPLNKSSSGNSSINSNETYVLISCNTSVIKQACPKVTFDPIPIHYCAPAGYAILKCNNE  1053

Query  232   TFNGTGPCTNVSTVQCTHGIRPVVSTQLLLNGSLAEEEVVIRSVNFTDNAKTIIVQLNTS  291
             TFNGTGPC NVSTVQCTHGI+PVVSTQLLLNGSLA+ E+VIRS N TDNAKTIIV LN S
Sbjct  1054  TFNGTGPCNNVSTVQCTHGIKPVVSTQLLLNGSLAKGEIVIRSENLTDNAKTIIVHLNES  1233

Query  292   VEINCTRPNNNTRKRIRIQRGPGRAFVTIGK-IGNMRQAHCNISRAKWNNTLKQIASKLR  350
             VEINCTRPNNNTRK IRI  GPG+ F   G+ IGN+RQAHCNIS +KWN TL+++  KL
Sbjct  1234  VEINCTRPNNNTRKSIRI--GPGQTFYATGEIIGNIRQAHCNISESKWNETLQRVGIKLA  1407

Query  351   EQFGNNKTIIFKQSSGGDPEIVTHSFNCGGEFFYCNSTQLFNSTWFNSTWSTEGSNNTEG  410
             E F  NKTI F   SGGD EI+THSFNCGGEFFYCN++ LFN T +N T+    S
Sbjct  1408  EHFP-NKTINFTSHSGGDLEIMTHSFNCGGEFFYCNTSSLFNGT-YNGTYIRNNS-----  1566

Query  411   SDTITLPCRIKQIINMWQKVGKAMYAPPISGQIRCSSNITGLLLTRDGGNSNNES  465
             S  IT+ CRIKQIINMWQ+VG+AMYAPPI+G+I C+S+ITG+LLTRDGG  N  S
Sbjct  1567  SSIITIQCRIKQIINMWQEVGRAMYAPPIAGKITCNSSITGILLTRDGGTGNETS  1731



Range 2: 1819 to 2418

Score:256 bits(654), Expect:4e-68,
Method:Compositional matrix adjust.,
Identities:142/200(71%), Positives:163/200(81%), Gaps:8/200(4%)

Query  664   DKWASLWNWFNITNWLWYIKLFIMIVGGLVGLRIVFAVLSIVNRVRQGYSPLSFQTHLP-  722
             D W +LWNWF IT WLWYIK+FIMIVGGL+GLRI+FAV+SIVNR RQGYSPLSFQT  P
Sbjct  1819  DSWNNLWNWFGITKWLWYIKIFIMIVGGLIGLRILFAVISIVNRFRQGYSPLSFQTLTPN  1998

Query  723   TprgpdrpegieeeggerdrdrSIRLVNGSLALIWDDLRSLCLFSYHRLRDLLLIVTRIV  782
             +   PDR   IEEEGGE+DR+RSIRLV+G LAL WDDLR+LCLFSYHRLRD +L+  R++
Sbjct  1999  SRGPPDRLGRIEEEGGEQDRNRSIRLVSGFLALAWDDLRNLCLFSYHRLRDFILVTARVL  2178

Query  783   ELLGR-------RGWEALKYWWNLLQYWSQELKNSAVSLLNATAIAVAEGTDRVIEVVQG  835
             ELLGR       RGWEALKY  +L+QYW  ELKNSA+SLL+  AIAVAEGTDR+I +VQG
Sbjct  2179  ELLGRSSLKGLQRGWEALKYLGSLVQYWGLELKNSAISLLDTIAIAVAEGTDRIINLVQG  2358

Query  836   ACRAIRHIPRRIRQGLERIL  855
              CRAIR+IPRRIRQG E  L
Sbjct  2359  ICRAIRNIPRRIRQGFEAAL  2418



>Escherichia coli strain NBE0002 contig00097, whole genome shotgun sequence
Sequence ID: NZ_JARBDF010000097.1 Length: 2717
Range 1: 987 to 2360

Score:629 bits(1622), Expect:0.0,
Method:Compositional matrix adjust.,
Identities:316/475(67%), Positives:366/475(77%), Gaps:27/475(5%)

Query  1     MRVKEKYQHLWRWGWRWGTMLLGMLMICSATEKLWVTVYYGVPVWKEATTTLFCASDAKA  60
             MRV+   ++  +W W WG +   MLMIC+     WVTVYYGVPVWKEA TTLFCASDAKA
Sbjct  2360  MRVRGTLRNYQQW-WIWGVLGFWMLMICNGGGDSWVTVYYGVPVWKEAKTTLFCASDAKA  2184

Query  61    YDTEVHNVWATHACVPTDPNPQEVVLVNVTENFNMWKNDMVEQMHEDIISLWDQSLKPCV  120
             ++ EVHNVWATHACVPTDPNPQE+VL NVTENFNMW+NDMV+QMHED+ISLWDQSLKPCV
Sbjct  2183  HEREVHNVWATHACVPTDPNPQEMVLENVTENFNMWRNDMVDQMHEDVISLWDQSLKPCV  2004

Query  121   KLTPLCVSLKCTDLKNDTNTNSSSGRMIMEKGEIKNCSFNISTSIRGKVQKEYAFFYKLD  180
             KLTPLCV+L+C       N N++      E  E+K C FN ST +  K QK  A FY LD
Sbjct  2003  KLTPLCVTLEC-------NNNTTFKNYTDESKELKRCFFNASTEVEDKKQKVSALFYSLD  1845

Query  181   IIPIDNDTT---------SYKLTSCNTSVITQACPKVSFEPIPIHYCAPAGFAILKCNNK  231
             I P++  ++         +Y L SCNTSVI QACPKV+F+PIPIHYCAPAG+AILKCNN+
Sbjct  1844  IRPLNKSSSGNSSINSNETYVLISCNTSVIKQACPKVTFDPIPIHYCAPAGYAILKCNNE  1665

Query  232   TFNGTGPCTNVSTVQCTHGIRPVVSTQLLLNGSLAEEEVVIRSVNFTDNAKTIIVQLNTS  291
             TFNGTGPC NVSTVQCTHGI+PVVSTQLLLNGSLA+ E+VIRS N TDNAKTIIV LN S
Sbjct  1664  TFNGTGPCNNVSTVQCTHGIKPVVSTQLLLNGSLAKGEIVIRSENLTDNAKTIIVHLNES  1485

Query  292   VEINCTRPNNNTRKRIRIQRGPGRAFVTIGK-IGNMRQAHCNISRAKWNNTLKQIASKLR  350
             VEINCTRPNNNTRK IRI  GPG+ F   G+ IGN+RQAHCNIS +KWN TL+++  KL
Sbjct  1484  VEINCTRPNNNTRKSIRI--GPGQTFYATGEIIGNIRQAHCNISESKWNETLQRVGIKLA  1311

Query  351   EQFGNNKTIIFKQSSGGDPEIVTHSFNCGGEFFYCNSTQLFNSTWFNSTWSTEGSNNTEG  410
             E F  NKTI F   SGGD EI+THSFNCGGEFFYCN++ LFN T +N T+    S
Sbjct  1310  EHFP-NKTINFTSHSGGDLEIMTHSFNCGGEFFYCNTSSLFNGT-YNGTYIRNNS-----  1152

Query  411   SDTITLPCRIKQIINMWQKVGKAMYAPPISGQIRCSSNITGLLLTRDGGNSNNES  465
             S  IT+ CRIKQIINMWQ+VG+AMYAPPI+G+I C+S+ITG+LLTRDGG  N  S
Sbjct  1151  SSIITIQCRIKQIINMWQEVGRAMYAPPIAGKITCNSSITGILLTRDGGTGNETS  987



Range 2: 300 to 899

Score:255 bits(651), Expect:1e-67,
Method:Compositional matrix adjust.,
Identities:141/200(71%), Positives:160/200(80%), Gaps:7/200(3%)

Query  664  DKWASLWNWFNITNWLWYIKLFIMIVGGLVGLRIVFAVLSIVNRVRQGYSPLSFQTHLPT  723
            D W +LWNWF IT WLWYIK+FIMIVGGL+GLRIVFAVLSIVNRVRQGYSPLSFQT  P+
Sbjct  899  DSWNNLWNWFGITKWLWYIKIFIMIVGGLIGLRIVFAVLSIVNRVRQGYSPLSFQTLAPS  720

Query  724  prgpdrpegieeeggerdrdrSIRLVNGSLALIWDDLRSLCLFSYHRLRDLLLIVTRIVE  783
            P G DRP  I+EEGGE+ R+ S RLV+G LAL WDDLRSL LF YHRL+DL+ +  R+VE
Sbjct  719  PGGLDRPGRIDEEGGEQGRNNSTRLVSGFLALAWDDLRSLFLFCYHRLKDLISVTARVVE  540

Query  784  LLGR-------RGWEALKYWWNLLQYWSQELKNSAVSLLNATAIAVAEGTDRVIEVVQGA  836
            LLGR       RGWEALKY  +L+QYW  ELK SA+SLL+  AIAV EGTDR+IE VQG
Sbjct  539  LLGRSSLKGLQRGWEALKYLGSLVQYWGLELKKSAISLLDTIAIAVGEGTDRIIEGVQGL  360

Query  837  CRAIRHIPRRIRQGLERILL  856
            CRAIR+IPRR+RQG E  LL
Sbjct  359  CRAIRNIPRRVRQGFEAALL  300
```

>Escherichia coli strain EN5308 contig00099, whole genome shotgun sequence
Sequence ID: NZ_JARBDE010000099.1 Length: 2827
Range 1: 1127 to 2524

Score:626 bits(1614), Expect:0.0,
Method:Compositional matrix adjust.,
Identities:315/485(65%), Positives:375/485(77%), Gaps:32/485(6%)

```
Query  1     MRVKEKYQHLWRWGWRWGTMLLGMLMICSATEKLWVTVYYGVPVWKEATTTLFCASDAKA  60
             MRV+   ++   W W W ++   MLMIC+  E LWVTVYYGVPVWKEA   LFCASDA+A
Sbjct  2524  MRVRGILRNYPPW-WIWASLGFWMLMICNGGENLWVTVYYGVPVWKEAKAPLFCASDARA  2348

Query  61    YDTEVHNVWATHACVPTDPNPQEVVLVNVTENFNMWKNDMVEQMHEDIISLWDQSLKPCV  120
             YD EVHNVWATHACVPTDP+PQE+VL NVTENFNMW N MV+QMHED+ISLWDQSLKPCV
Sbjct  2347  YDREVHNVWATHACVPTDPSPQEMVLENVTENFNMWNNYMVDQMHEDVISLWDQSLKPCV  2168

Query  121   KLTPLCVSLKCTDLKNDTNTNSSSGRMIMEKGEIKNCSFNISTSIRGKVQKEYAFFYKLD  180
             KLTPLCV+L C ++     T +++ R      E+KNCSFN +T +  K QK YA FY++D
Sbjct  2167  KLTPLCVTLNCVNVTKTNVTVNNTNRQ-----EMKNCSFNATTELVDKKQKVYALFYRVD  2003

Query  181   IIPI----------DNDTTSYKLTSCNTSVITQACPKVSFEPIPIHYCAPAGFAILKCNN  230
             I+P+          +N+++ Y+L +CNTS ITQACPKV+F+PIPIHYCAPAGFAILKCNN
Sbjct  2002  IVPLNKENNSKSNEENNSSEYRLINCNTSAITQACPKVTFDPIPIHYCAPAGFAILKCNN  1823

Query  231   KTFNGTGPCTNVSTVQCTHGIRPVVSTQLLLNGSLAEEEVVIRSVNFTDNAKTIIVQLNT  290
             KTFN T PC NVSTVQCTHGI+PVVSTQLLLNGSLAEEE++IRS N T+NAK IIVQLN
Sbjct  1822  KTFNKTRPCHNVSTVQCTHGIKPVVSTQLLLNGSLAEEEIIIRSENLTNNAKIIIVQLNQ  1643

Query  291   SVEINCTRPNNNTRKRIRIQRGPGRAFVTIGK-IGNMRQAHCNISRAKWNNTLKQIASKL  349
             SVEI C RP NNTRK IRI  GPG+ F   G  IGN+R+AHCNIS   W NTL++++ KL
Sbjct  1642  SVEIECIRPGNNTRKSIRI--GPGQTFYATGDIIGNIREAHCNIS--NWTNTLERVSGKL  1475

Query  350   REQFGNNKTIIFKQSSGGDPEIVTHSFNCGGEFFYCNSTQLFNSTWFNSTWSTEGSNNTE  409
             RE F  NKTIIFK +SGGDPE+ THSFNC GEFFYCN+++LFN T+        G++N E
Sbjct  1474  REHFP-NKTIIFKPASGGDPEVTTHSFNCRGEFFYCNTSKLFNGTY-------NGTSN-E  1322

Query  410   GSDTITLPCRIKQIINMWQKVGKAMYAPPISGQIRCSSNITGLLLTRDGGNSNNES--EI  467
              S  IT+PC+I+Q +N+WQ+VG+AMYAPPI+G I C SNITGLLL RDGG+ NN S  EI
Sbjct  1321  SSSNITIPCKIRQFVNLWQEVGRAMYAPPIAGNITCKSNITGLLLVRDGGSGNNSSNDEI  1142

Query  468   FRPGG  472
             FRP G
Sbjct  1141  FRPIG  1127
```

Range 2: 373 to 1125

Score:290 bits(741), Expect:8e-80,
Method:Compositional matrix adjust.,
Identities:176/251(70%), Positives:204/251(81%), Gaps:7/251(2%)

```
Query  612   ASWSNKSLEQIWNHTTWMEWDREINNYTSLIHSLIeesqnqqekneqellelDKWASLWN  671
             +SWSNKS   IW + TWM+WD+E++NYT  I+ L+E+SQ QQE+NE++LL LD W +LWN
Sbjct  1125  SSWSNKSQTDIWENMTWMQWDKEVSNYTDTIYRLLEDSQTQQERNEKDLLALDNWKNLWN  946

Query  672   WFNITNWLWYIKLFIMIVGGLVGLRIVFAVLSIVNRVRQGYSPLSFQTHLPTprgpdrpe  731
             WF+ITNWLWYIK+FIMIVGGL+GLRI+FAVLSIVNRVRQGYSPLSFQT  P PRGPDR
Sbjct  945   WFSITNWLWYIKIFIMIVGGLIGLRIIFAVLSIVNRVRQGYSPLSFQTLTPNPRGPDRLG  766

Query  732   gieeeggerdrdrSIRLVNGSLALIWDDLRSLCLFSYHRLRDLLLIVTRIVELLG-----  786
              IEEEGG +DRDRSIRLVNG LAL WDDL SLCLFSYHRLRDL+L+  R VELLG
Sbjct  765   RIEEEGGGQDRDRSIRLVNGFLALAWDDLWSLCLFSYHRLRDLILVTARAVELLGHSSLR  586

Query  787   --RRGWEALKYWWNLLQYWSQELKNSAVSLLNATAIAVAEGTDRVIEVVQGACRAIRHIP  844
               +RGWEALKY   ++QYW  ELK  A+SLL+  AIAVAEGTDR+IE +Q  CRAIR+IP
Sbjct  585   GLQRGWEALKYLGGIVQYWGLELKKRAISLLDTVAIAVAEGTDRIIEFLQRICRAIRNIP  406

Query  845   RRIRQGLERIL  855
             RRIRQG E  L
Sbjct  405   RRIRQGFEAAL  373
```

>Escherichia coli strain NBE0009 contig00116, whole genome shotgun sequence
Sequence ID: NZ_JARBDL010000116.1 Length: 1713
Range 1: 308 to 1705

Score:612 bits(1578), Expect:0.0,
Method:Compositional matrix adjust.,
Identities:357/474(75%), Positives:406/474(85%), Gaps:17/474(3%)

```
Query  391   FNSTWFNSTWSTEGSNNTEGSDTITLPCRIKQIINMWQKVGKAMYAPPISGQIRCSSNIT  450
             +N T++N T     +N       IT+PCRIKQIINMWQKVG+AMYAPPI+G I C+S IT
Sbjct  1705  YNGTYYNRTQYNSTAN-------ITIPCRIKQIINMWQKVGRAMYAPPIAGNITCNSTIT  1547

Query  451   GLLLTRDGGNSN--NESEIFRPGGGDMRDNWRSELYKYKVVKIEPLGVAPTKAKRRVVQR  508
             GLLL RDGGN+N  N +EIFRPGGGDMRDNWRSELYKYKVV+I+PLG+APT+AKRRVV R
Sbjct  1546  GLLLLRDGGNTNDTNYTEIFRPGGGDMRDNWRSELYKYKVVEIKPLGIAPTRAKRRVVGR  1367

Query  509   EKRAVGIgalflgflgaagstmgaasmtLTVQARQLLSGIVQQQNNLLRAIEAQQHLLQL  568
             EKRA GIGA+FLGFLGAAGSTMGAAS+TLTVQARQLLSGIVQQQ+NLLRAIEAQQHLLQL
Sbjct  1366  EKRA-GIGAVFLGFLGAAGSTMGAASITLTVQARQLLSGIVQQQSNLLRAIEAQQHLLQL  1190

Query  569   TVWGIKQLQARILAVERYLKDQQLLGIWGCSGKLICTTAVPWNASWSNKSLEQIWNHTTW  628
             TVWGIKQLQ R+LA+ERYLKDQQLLGIWGCSGKLIC TAVPWN+SWSNK+  +IW++ TW
Sbjct  1189  TVWGIKQLQTRVLALERYLKDQQLLGIWGCSGKLICPTAVPWNSSWSNKNQSEIWDNMTW  1010

Query  629   MEWDREINNYTSLIHSLIeesqnqqekneqellelDKWASLWNWFNITNWLWYIKLFIMI  688
             M+WDREI+NYT  I+ L+EESQNQQEKNE++LL LD WASLWNWFNIT WLWYIK+FIMI
Sbjct  1009  MQWDREISNYTYTIYRLLEESQNQQEKNEEDLLALDSWASLWNWFNITQWLWYIKIFIMI  830

Query  689   VGGLVGLRIVFAVLSIVNRVRQGYSPLSFQTHLPTprgpdrpegieeeggerdrdrSIRL  748
             VGGL+GLRI+FAVLSIVNRVRQGYSPLSFQT +P+P   D+  GIEEEGGE+DR+RSIRL
Sbjct  829   VGGLIGLRIIFAVLSIVNRVRQGYSPLSFQTPIPSPGELDKLRGIEEEGGEQDRNRSIRL  650

Query  749   VNGSLALIWDDLRSLCLFSYHRLRDLLLIVTRIVELLGR-------RGWEALKYWWNLLQ  801
             VNG LAL WDDLR+LCLFSYHRLRD +L+  R+VELLGR       RGWEALKY  +L+Q
Sbjct  649   VNGFLALAWDDLRNLCLFSYHRLRDFILLTARVVELLGRSSLRGLQRGWEALKYLGSLVQ  470

Query  802   YWSQELKNSAVSLLNATAIAVAEGTDRVIEVVQGACRAIRHIPRRIRQGLERIL  855
             YW  ELK SA+SL +  AI VAEGTDR+I VVQ  CRAI +IPRR+RQG E  L
Sbjct  469   YWGLELKKSAISLFDTIAITVAEGTDRIISVVQRICRAIYNIPRRLRQGFEAAL  308
```

>Escherichia coli strain EN5286 contig00120, whole genome shotgun sequence

```
Sequence ID: NZ_JAPMNM010000120.1 Length: 3033
Range 1: 352 to 1884

Score:603 bits(1555), Expect:0.0,
Method:Compositional matrix adjust.,
Identities:314/519(61%), Positives:364/519(70%), Gaps:65/519(12%)

Query  1      MRVK---EKYQHLWRWGWRWGTMLLG--MLMICSATEKLWVTVYYGVPVWKEATTTLFCA  55
              MRV+     YQH W WG      +LG  MLMIC+    LWVTVYYGVPVWK+A  TLFCA
Sbjct  352    MRVRGILRSYQHWWIWG------ILGFWMLMICNVVGNLWVTVYYGVPVWKDANATLFCA  513

Query  56     SDAKAYDTEVHNVWATHACVPTDPNPQEVVLVNVTENFNMWKNDMVEQMHEDIISLWDQS  115
              SDAK Y TEVHNVWATHACVPTDPNPQE+VL NVTENFNMWKN+MVEQMHED+IS+WDQ
Sbjct  514    SDAKVYKTEVHNVWATHACVPTDPNPQELVLGNVTENFNMWKNNMVEQMHEDVISIWDQG  693

Query  116    LKPCVKLTPLCVSLKCTDLK------NDTNTNSSSGRMIMEKG-------EIKNCSFNIS  162
              LKPCVKLTPLCV+L CT  +      +  NT ++S  M +          EI NCSF  +
Sbjct  694    LKPCVKLTPLCVTLNCTKYEPRKVTVDKNNTGATSQNMTVNLTIDKEALEEISNCSFETT  873

Query  163    TSIRGKVQKEYAFFYKLDIIPI-----DNDTTSYKLTSCNTSVITQACPKVSFEPIPIHY  217
              T +R K +K +A FYK+D++ I     D+  T Y+L +CNTS +TQACPKVSFEPIPIHY
Sbjct  874    TELRDKTRKVHALFYKIDMVEISKIGNDSTITRYRLINCNTSALTQACPKVSFEPIPIHY  1053

Query  218    CAPAGFAILKCNNKTFNGTGPCTNVSTVQCTHGIRPVVSTQLLLNGSLAEEEVVIRSVNF  277
              CAPAG+A+LKCNN TFNGTGPC N+STVQCTHGIRPVVSTQLLLNGSLAE  + IRS N
Sbjct  1054   CAPAGYALLKCNNNTFNGTGPCRNISTVQCTHGIRPVVSTQLLLNGSLAEGNITIRSENL  1233

Query  278    TDNAKTIIVQLNTSVEINCTRPNNNTRKRIRIQRGPGRAFVTIGKI-GNMRQAHCNISRA  336
              T+NAKTIIV LN SVEI C RPNNNTR+ +RI  GPG+ F   G I G++R+AHCNIS
Sbjct  1234   TNNAKTIIVHLNQSVEILCVRPNNNTRQSVRI--GPGQTFFATGDIIGDIRRAHCNISAK  1407

Query  337    KWNNTLKQIASKLREQFGNNKTIIFKQSSGGDPEIVTHSFNCGGEFFYCNSTQLFNS---  393
               W+  L+ ++ +L + F NNKTI F   SGGD EI THSFNC GEFFYCN++QLFNS
Sbjct  1408   DWHTALRNVSIELAKLFPNNKTINFTSPSGGDLEITTHSFNCRGEFFYCNTSQLFNSLYM  1587

Query  394    ------TWFNSTWSTEGSNNTE---GSDT---------------------ITLPCRIKQI  423
                     W+N   +    N T+    S T                     ITLPC+IKQI
Sbjct  1588   PDNDTYIWYNGNHTYVWHNGTQYVINSSTQYVTINNSAQNVAINNMTDFNITLPCKIKQI  1767

Query  424    INMWQKVGKAMYAPPISGQIRCSSNITGLLLTRDGGNSN  462
              INMWQ+VGKAMYAPPI+G I C SNITGLLLTRDGG  N
Sbjct  1768   INMWQEVGKAMYAPPIAGNITCLSNITGLLLTRDGGKGN  1884



Range 2: 1893 to 2729

Score:334 bits(856), Expect:2e-95,
Method:Compositional matrix adjust.,
Identities:194/280(69%), Positives:227/280(81%), Gaps:8/280(2%)

Query  584    ERYLKDQQLLGIWGCSGKLICTTAVPWNASWSNKSLEQIWNHTTWMEWDREINNYTSLIH  643
              ER+ ++Q+LLG+WGCS + ICTT VPWN+SWSNKS   IWN+ TWMEWD+EI  +T+ I+
Sbjct  1893   ERH-REQRLLGLWGCSERTICTTNVPWNSSWSNKSYSDIWNNLTWMEWDKEIEKHTNTIY  2069

Query  644    SLIeesqnqqekneqellelDKWASLWNWFNITNWLWYIKLFIMIVGGLVGLRIVFAVLS  703
               L+EESQ QQE+NE++LL LDKW  LW+WF+ITNWLWYIK+FIMIVGGL+GLRIVFAVLS
Sbjct  2070   QLLEESQGQQEQNEKDLLALDKWNDLWSWFDITNWLWYIKIFIMIVGGLIGLRIVFAVLS  2249

Query  704    IVNRVRQGYSPLSFQTHLPTprgpdrpegieeeggerdrdrSIRLVNGSLALIWDDLRSL  763
              IVNRVRQGYSPLSFQT  P+P G DRP  IEE GGE+ R  SIRLV+G LAL WDDLRSL
Sbjct  2250   IVNRVRQGYSPLSFQTLAPSPGGLDRPGRIEEGGGEQGRSSSIRLVSGFLALAWDDLRSL  2429

Query  764    CLFSYHRLRDLLLIVTRIVELLGR-------RGWEALKYWWNLLQYWSQELKNSAVSLLN  816
              CLF YHRL+DL+ +  R VELLGR       RGWEALKY  +L+QYW  ELK SA+SLL+
Sbjct  2430   CLFCYHRLKDLISVTARAVELLGRSSLKGLQRGWEALKYLGSLVQYWGLELKKSAISLLD  2609

Query  817    ATAIAVAEGTDRVIEVVQGACRAIRHIPRRIRQGLERILL  856
                AIAV EGTDR+IE VQG CRAIR+IPRR+RQG E  LL
Sbjct  2610   TIAIAVGEGTDRIIEGVQGLCRAIRNIPRRVRQGFEAALL  2729



>Escherichia coli strain NBE0007 contig00093, whole genome shotgun sequence
Sequence ID: NZ_JARBDJ010000093.1 Length: 3013
Range 1: 1130 to 2662

Score:599 bits(1545), Expect:0.0,
Method:Compositional matrix adjust.,
Identities:310/519(60%), Positives:363/519(69%), Gaps:65/519(12%)

Query  1      MRVK---EKYQHLWRWGWRWGTMLLG--MLMICSATEKLWVTVYYGVPVWKEATTTLFCA  55
              MRV+     YQH W WG      +LG  MLMIC+    LWVTVYYGVPVW++A  TLFCA
Sbjct  2662   MRVRGILRSYQHWWIWG------ILGFWMLMICNVVGNLWVTVYYGVPVWRDANATLFCA  2501

Query  56     SDAKAYDTEVHNVWATHACVPTDPNPQEVVLVNVTENFNMWKNDMVEQMHEDIISLWDQS  115
              SDAK Y TEVHNVWATHACVPTDPNPQE+VL NVTENFNMW+N+MVEQMHED+IS+WDQ
Sbjct  2500   SDAKVYKTEVHNVWATHACVPTDPNPQELVLENVTENFNMWENNMVEQMHEDVISIWDQG  2321

Query  116    LKPCVKLTPLCVSLKCTDLK------NDTNTNSSSGRMIMEKG-------EIKNCSFNIS  162
              LKPCVKLTPLCV+L CT  +      +  NT ++S  M +          EI NCSF  +
Sbjct  2320   LKPCVKLTPLCVTLNCTKYEPRKVTVDKNNTGATSQNMTVNPTIEKEALEEISNCSFETT  2141

Query  163    TSIRGKVQKEYAFFYKLDIIPI-----DNDTTSYKLTSCNTSVITQACPKVSFEPIPIHY  217
              T +R K +K +A FYK+D++ +     D+  T Y+L +CNTS +TQACPKVSFEPIPIHY
Sbjct  2140   TELRDKTRKVHALFYKIDMVELGKNGNDSTITRYRLINCNTSALTQACPKVSFEPIPIHY  1961

Query  218    CAPAGFAILKCNNKTFNGTGPCTNVSTVQCTHGIRPVVSTQLLLNGSLAEEEVVIRSVNF  277
              CAPAG+A+LKCNN TFNGTGPC N+STVQCTHGIRPVVSTQLLLNGSLAE  + IRS N
Sbjct  1960   CAPAGYALLKCNNNTFNGTGPCRNISTVQCTHGIRPVVSTQLLLNGSLAEGNITIRSENL  1781

Query  278    TDNAKTIIVQLNTSVEINCTRPNNNTRKRIRIQRGPGRAFVTIGKI-GNMRQAHCNISRA  336
              T+NAKTII  LN SVEI C RPNNNTR+ +RI  GPG+ F   G I G++R+AHCNIS
Sbjct  1780   TNNAKTIIAHLNKSVEILCARPNNNTRQSVRI--GPGQTFFATGDIIGDIRRAHCNISAK  1607

Query  337    KWNNTLKQIASKLREQFGNNKTIIFKQSSGGDPEIVTHSFNCGGEFFYCNSTQLFNST--  394
               W+  L+ ++ +L + F NNKTI F   SGGD EI THSFNC GEFFYCN++QLFNS
Sbjct  1606   DWHTALRNVSIELAKLFPNNKTINFTSPSGGDLEITTHSFNCRGEFFYCNTSQLFNSLYM  1427

Query  395    -------WFNSTWSTEGSNNTE---GSDT---------------------ITLPCRIKQI  423
                     W+N   +    N T+    S T                     ITLPC+IKQI
Sbjct  1426   PDNDTYLWYNGNHTYVWHNGTQYVINSSTQYVTINNSAQNVAINNMTDFNITLPCKIKQI  1247

Query  424    INMWQKVGKAMYAPPISGQIRCSSNITGLLLTRDGGNSN  462
              INMWQ+VGKAMYAPPI+G I C SNITGLLLTRDGG  N
Sbjct  1246   INMWQEVGKAMYAPPIAGNITCLSNITGLLLTRDGGKGN  1130



Range 2: 289 to 1110
```

```
Score:337 bits(864), Expect:1e-96,
Method:Compositional matrix adjust.,
Identities:196/274(72%), Positives:230/274(83%), Gaps:7/274(2%)

Query  590   QQLLGIWGCSGKLICTTAVPWNASWSNKSLEQIWNHTTWMEWDREINNYTSLIHSLIees  649
             Q+LLG+WGCS K+ICTT VPWN+SWSNKS   IWN+ TWMEWD+EI+ +T+ I+ L+EES
Sbjct  1110  QRLLGLWGCSEKIICTTNVPWNSSWSNKSYSDIWNNLTWMEWDKEIDKHTNTIYQLLEES  931

Query  650   qnqqekneqellelDKWASLWNWFNITNWLWYIKLFIMIVGGLVGLRIVFAVLSIVNRVR  709
             Q QQE+NE++LL LDKW  LW+WF+ITNWLWYIK+FIMIVGGL+GLRIVFAVLSIVNRVR
Sbjct  930   QGQQEQNEKDLLALDKWNDLWSWFDITNWLWYIKIFIMIVGGLIGLRIVFAVLSIVNRVR  751

Query  710   QGYSPLSFQTHLPTprgpdrpegieeeggerdrdrSIRLVNGSLALIWDDLRSLCLFSYH  769
             QGYSPLSFQT  P+P G DRP  IEEEGGE+D+DRSIRLV+G LAL WDDLR+LCLFSYH
Sbjct  750   QGYSPLSFQTLAPSPGGLDRPGRIEEEGGEQDKDRSIRLVSGFLALAWDDLRNLCLFSYH  571

Query  770   RLRDLLLIVTRIVELLGR-------RGWEALKYWWNLLQYWSQELKNSAVSLLNATAIAV  822
             RLRD +L+  R+VELLGR       +GWEALKY  +L+QYW  ELK SA+SLL+  AIAV
Sbjct  570   RLRDCILVTARVVELLGRNSLRGLQKGWEALKYLGSLVQYWGLELKKSAISLLDTIAIAV  391

Query  823   AEGTDRVIEVVQGACRAIRHIPRRIRQGLERILL  856
              EGTDR+IE VQG CRAIR+IPRR+RQG E  LL
Sbjct  390   GEGTDRIIEGVQGLCRAIRNIPRRVRQGFEAALL  289



>Escherichia coli strain EN5192 contig00108, whole genome shotgun sequence
Sequence ID: NZ_JAPMNL010000108.1 Length: 3032
Range 1: 353 to 1942

Score:599 bits(1544), Expect:0.0,
Method:Compositional matrix adjust.,
Identities:314/535(59%), Positives:369/535(68%), Gaps:66/535(12%)

Query  1     MRVKEKYQHLWRWGWRWGTMLLGMLMICSATEKLWVTVYYGVPVWKEATTTLFCASDAKA  60
             M+V+   ++   W W WG +   M MIC+    LWVTVYYGVPVWK+A  TLFCASDAK
Sbjct  353   MKVRGILRNYQHW-WIWGILGFWMFMICNVVGNLWVTVYYGVPVWKDANATLFCASDAKV  529

Query  61    YDTEVHNVWATHACVPTDPNPQEVVLVNVTENFNMWKNDMVEQMHEDIISLWDQSLKPCV  120
             Y TEVHNVWATHACVPTDPNPQE+VL NVTENFNMWKN+MVEQMHED+IS+WDQ LKPCV
Sbjct  530   YKTEVHNVWATHACVPTDPNPQELVLENVTENFNMWKNNMVEQMHEDVISIWDQGLKPCV  709

Query  121   KLTPLCVSLKCTDLK--NDTNTNSSSGRMIMEKG-------------EIKNCSFNISTSI  165
             KLTPLCV+L CT  K  N+T   +S+G                    EI NCSF+ +T +
Sbjct  710   KLTPLCVTLNCTKYKPRNETVDGTSTGNTGQNNTGKLNLTIDPEALEEISNCSFDTTTEL  889

Query  166   RGKVQKEYAFFYKLDIIPIDNDTTS---------YKLTSCNTSVITQACPKVSFEPIPIH  216
             R K +K +A FYK+D++ +D ++T          Y+L +CNTS ITQACPKVSFEPIPIH
Sbjct  890   RDKTRKVHALFYKIDMVELDKNSTCQTPTPNFTLYRLINCNTSAITQACPKVSFEPIPIH  1069

Query  217   YCAPAGFAILKCNNKTFNGTGPCTNVSTVQCTHGIRPVVSTQLLLNGSLAEEEVVIRSVN  276
             YCAPAG+A+LKCNN TFNGTGPC N+STVQCTHGIRPVVSTQLLLNGSLAE  + IRS N
Sbjct  1070  YCAPAGYALLKCNNNTFNGTGPCRNISTVQCTHGIRPVVSTQLLLNGSLAEGNITIRSEN  1249

Query  277   FTDNAKTIIVQLNTSVEINCTRPNNNTRKRIRIQRGPGRAFVTIGK-IGNMRQAHCNISR  335
              TDNAKTII  LN SVEI C RPNNNTR+ +RI  GPG+ F   G  IG++R+AHCNIS
Sbjct  1250  LTDNAKTIIAHLNQSVEILCVRPNNNTRQSVRI--GPGQTFFATGDIIGDIRRAHCNISA  1423

Query  336   AKWNNTLKQIASKLREQFGNNKTIIFKQSSGGDPEIVTHSFNCGGEFFYCNSTQLFNS--  393
               W+  L+ ++ +L + +  NKTI F   SGGD EI THSFNC GEFFYCN++QLFNS
Sbjct  1424  KDWHTALRNVSIELAKLY--NKTINFTSHSGGDLEITTHSFNCRGEFFYCNTSQLFNSLY  1597

Query  394   ----------------TWFNST-WSTEGS------NNTEGSDT--------ITLPCRIKQ  422
                              W N T +   GS      NN+  + T        ITLPC+IKQ
Sbjct  1598  MPNNDTYIWYNGNHTHVWHNGTQYVINGSTQYVTLNNSAQNVTINNITDFNITLPCKIKQ  1777

Query  423   IINMWQKVGKAMYAPPISGQIRCSSNITGLLLTRDGGNSN---NESEIFRPGGGD  474
             IINMWQ+VGKAMYAPPI+G I C SNITGLLLTRD G      N +EIFRP G +
Sbjct  1778  IINMWQEVGKAMYAPPIAGNITCLSNITGLLLTRDRGKGKEVRNNTEIFRPEGNN  1942



Range 2: 1939 to 2760

Score:326 bits(836), Expect:1e-92,
Method:Compositional matrix adjust.,
Identities:191/274(70%), Positives:223/274(81%), Gaps:7/274(2%)

Query  590   QQLLGIWGCSGKLICTTAVPWNASWSNKSLEQIWNHTTWMEWDREINNYTSLIHSLIees  649
             Q+LLG+WGCS + ICTT VPWN+SWSN+S   IWN+ TWMEWD+EI  +T+ I+ L+EES
Sbjct  1939  QRLLGLWGCSERTICTTNVPWNSSWSNRSYSDIWNNLTWMEWDKEIEQHTNTIYQLLEES  2118

Query  650   qnqqekneqellelDKWASLWNWFNITNWLWYIKLFIMIVGGLVGLRIVFAVLSIVNRVR  709
             Q QQE+NE++LL LDKW  LW+WF+ITNWLWYIK+FIMIVGGL+GLRIVFAVLSIVNRVR
Sbjct  2119  QGQQEQNEKDLLALDKWNDLWSWFDITNWLWYIKIFIMIVGGLIGLRIVFAVLSIVNRVR  2298

Query  710   QGYSPLSFQTHLPTprgpdrpegieeeggerdrdrSIRLVNGSLALIWDDLRSLCLFSYH  769
             QGYSPLSFQT  P+P G DRP  IEE GGE+ R+ SIRLV+G LAL W+DLRSL LF YH
Sbjct  2299  QGYSPLSFQTLAPSPGGLDRPGRIEEGGGEQGRNSSIRLVSGFLALAWEDLRSLFLFCYH  2478

Query  770   RLRDLLLIVTRIVELLGR-------RGWEALKYWWNLLQYWSQELKNSAVSLLNATAIAV  822
             RL+DL+ + TR VELLGR       RGWEALKY  +L+QYW  ELK SA+SLL+  AIAV
Sbjct  2479  RLKDLISVTTRAVELLGRSSLKGLQRGWEALKYLGSLVQYWVLELKKSAISLLDTIAIAV  2658

Query  823   AEGTDRVIEVVQGACRAIRHIPRRIRQGLERILL  856
              EGTDR+IE VQG CRAIR+IPRRIRQG E  LL
Sbjct  2659  GEGTDRIIEGVQGFCRAIRNIPRRIRQGFEAALL  2760



>Escherichia coli strain VRES0669, whole genome shotgun sequence
Sequence ID: NZ_UINV01000086.1 Length: 5544
Range 1: 6 to 1391

Score:593 bits(1529), Expect:0.0,
Method:Compositional matrix adjust.,
Identities:302/474(64%), Positives:365/474(77%), Gaps:21/474(4%)

Query  1     MRVKEKYQHLWRWGWRWGTMLLGMLMICSATEKLWVTVYYGVPVWKEATTTLFCASDAKA  60
             MRVK   Q  W+  W WG  +L   +IC+     WVTVYYGVPVWKEA  TLFCASDAK
Sbjct  1391  MRVK-GIQRNWQQWWIWG--ILSFWIICNVAGNRWVTVYYGVPVWKEAKATLFCASDAKT  1221

Query  61    YDTEVHNVWATHACVPTDPNPQEVVLVNVTENFNMWKNDMVEQMHEDIISLWDQSLKPCV  120
                EVHN+WATHACVPT+PNPQE+VL NVTENFNMWKNDMV+QMHEDIISLWDQSLKPCV
Sbjct  1220  QAEEVHNIWATHACVPTNPNPQEIVLENVTENFNMWKNDMVDQMHEDIISLWDQSLKPCV  1041

Query  121   KLTPLCVSLKCTDLK-NDTNTNSSSGRMIMEKGEIKNCSFNISTSIRGKVQKEYAFFYKL  179
             KLT LCV+L C+++K N T  N +  R      ++KNCSFN++T +R K QK +A FY+L
Sbjct  1040  KLTSLCVTLNCSNVKINVTAINGTYLR--NSTSDMKNCSFNVTTELRDKRQKAHALFYRL  867

Query  180   DIIPIDND----TTSYKLTSCNTSVITQACPKVSFEPIPIHYCAPAGFAILKCNNKTFNG  235
```

```
             D++P++ +    ++SY+L +CNTS I+QACPK SF+P+PIHYCAPAG+AILKCNNKTFNG
Sbjct  866   DVVPLEEEKNSSSSSYRLINCNTSTISQACPKTSFDPLPIHYCAPAGYAILKCNNKTFNG  687

Query  236   TGPCTNVSTVQCTHGIRPVVSTQLLLNGSLAEEEVVIRSVNFTDNAKTIIVQLNTSVEIN  295
             TGPC NVSTVQCTHGI+PVVSTQLLLNGSLA E+++IRS N T N +TIIV L   VEI
Sbjct  686   TGPCQNVSTVQCTHGIKPVVSTQLLLNGSLA-EDIIIRSENLTSNTRTIIVHLKDPVEIR  510

Query  296   CTRPNNNTRKRIRIQRGPGRAFVTIGKI-GNMRQAHCNISRAKWNNTLKQIASKLREQFG  354
             C RPNNNTR  IRI  G G+ F   G+I G++RQAHCNI+  KWN TL+++  KL+E F
Sbjct  509   CIRPNNNTRHSIRIGIGRGQTFYATGRIVGDIRQAHCNITERKWNETLQKVVEKLKEHFP  330

Query  355   NNKTIIFKQS-SGGDPEIVTHSFNCGGEFFYCNSTQLFNSTWFNST--WSTEGSNNTEGS  411
              NKTI FK +  GGD E+  H+FNC GEFFYCN++QLF +  FN T  +++ G+N T
Sbjct  329   -NKTIQFKPAIPGGDLEVTMHTFNCRGEFFYCNTSQLF-TYMFNDTDMFNSTGNNATN--  162

Query  412   DTITLPCRIKQIINMWQKVGKAMYAPPISGQIRCSSNITGLLLTRDGGNSNNES  465
               ITLPCRIKQI+NMWQ+VG+A+YAPPI+G I C+SNITGLLL RDGGNS N S
Sbjct  161   --ITLPCRIKQIVNMWQEVGRAIYAPPITGIINCTSNITGLLLIRDGGNSGNGS  6
```

>Escherichia coli strain EN5308 contig00096, whole genome shotgun sequence
Sequence ID: NZ_JARBDE010000096.1 Length: 3024
Range 1: 1156 to 2673

Score:591 bits(1524), Expect:0.0,
Method:Compositional matrix adjust.,
Identities:310/516(60%), Positives:366/516(70%), Gaps:67/516(12%)

```
Query  1     MRVK---EKYQHLWRWGWRWGTMLLG--MLMICSATEKLWVTVYYGVPVWKEATTTLFCA  55
             MRV+     YQH W WG      +LG  MLMIC+    LWVTVYYGVPVWK+A  TLFCA
Sbjct  2673  MRVRGILRSYQHWWIWG------ILGFWMLMICNVVGNLWVTVYYGVPVWKDANATLFCA  2512

Query  56    SDAKAYDTEVHNVWATHACVPTDPNPQEVVLVNVTENFNMWKNDMVEQMHEDIISLWDQS  115
             SDAK Y TEVHNVWATHACVPTDPNPQE+VL NVTENFNMW+N+MVEQMHED+IS+WDQ
Sbjct  2511  SDAKVYKTEVHNVWATHACVPTDPNPQELVLGNVTENFNMWENNMVEQMHEDVISIWDQG  2332

Query  116   LKPCVKLTPLCVSLKCT--DLKNDT-NTNSSSGRMIMEK---GEIKNCSFNISTSIRGKV  169
             LKPCVKLTPLCV+L CT  + +N T N NS++    +++    EI NCSF  +T +R K
Sbjct  2331  LKPCVKLTPLCVTLNCTKYEPRNKTVNGNSTAKSPTIDEEALKEISNCSFETTTELRDKT  2152

Query  170   QKEYAFFYKLDIIPIDNDTTS------------YKLTSCNTSVITQACPKVSFEPIPIHY  217
             +K +A FYK+D+I +  ++T              Y+L +CNTS ITQACPKVSFEPIPIHY
Sbjct  2151  RKVHALFYKIDMIELGQNSTCQNPENSTSECTMYRLINCNTSAITQACPKVSFEPIPIHY  1972

Query  218   CAPAGFAILKCNNKTFNGTGPCTNVSTVQCTHGIRPVVSTQLLLNGSLAEEEVVIRSVNF  277
             CAPAG+A+LKCNN TFNGTGPC N+STVQCTHGIRPVVSTQLLLNGSLAE  + IRS N
Sbjct  1971  CAPAGYALLKCNNNTFNGTGPCRNISTVQCTHGIRPVVSTQLLLNGSLAEGNITIRSENL  1792

Query  278   TDNAKTIIVQLNTSVEINCTRPNNNTRKRIRIQRGPGRAFVTIGK-IGNMRQAHCNISRA  336
             T+NAKTII  LN SVEI C RPNNNTR+ +RI  GPG+ F   G  IG++R+AHCNIS
Sbjct  1791  TNNAKTIIAHLNKSVEILCVRPNNNTRQSVRI--GPGQTFFATGDIIGDIRRAHCNISAK  1618

Query  337   KWNNTLKQIASKLREQFGNNKTIIFKQSSGGDPEIVTHSFNCGGEFFYCNSTQLFNS---  393
              W+  L+ ++ +L + +  NKTI F   SGGD EI THSFNC GEFFYCN++QLFNS
Sbjct  1617  HWHTALRNVSIELAKLY--NKTINFTSHSGGDLEITTHSFNCRGEFFYCNTSQLFNSLYM  1444

Query  394   ------TWFNSTWSTEGSNNTE----GSD--------------------TITLPCRIKQI  423
                     W+N + +    N+T+    GS                         ITLPC+IKQI
Sbjct  1443  PDNDTYIWYNESHTHVWHNDTQYVINGSTQYVTLNKSAQNVTINNITDFNITLPCKIKQI  1264

Query  424   INMWQKVGKAMYAPPISGQIRCSSNITGLLLTRDGG  459
             INMWQ+VGKAMYAPPI+G I C SNITGLLLTRDGG
Sbjct  1263  INMWQEVGKAMYAPPIAGNITCLSNITGLLLTRDGG  1156
```

Range 2: 302 to 1138

Score:328 bits(841), Expect:3e-93,
Method:Compositional matrix adjust.,
Identities:190/280(68%), Positives:227/280(81%), Gaps:8/280(2%)

```
Query  584   ERYLKDQQLLGIWGCSGKLICTTAVPWNASWSNKSLEQIWNHTTWMEWDREINNYTSLIH  643
             ER+ ++Q+LLG+WGCS + ICTT VPWN+SWSNKS   IWN+ TWMEWD+EI  +T+ I+
Sbjct  1138  ERH-REQRLLGLWGCSERTICTTNVPWNSSWSNKSYSDIWNNLTWMEWDKEIEQHTNTIY  962

Query  644   SLIeesqnqqekneqellelDKWASLWNWFNITNWLWYIKLFIMIVGGLVGLRIVFAVLS  703
              L+EESQ QQE+NE++LL LDKW  LW+WF+ITNWLWYI++FIMIVGGL+GLRIVFAVLS
Sbjct  961   QLLEESQGQQEQNEKDLLALDKWNDLWSWFDITNWLWYIRIFIMIVGGLIGLRIVFAVLS  782

Query  704   IVNRVRQGYSPLSFQTHLPTprgpdrpegieeeggerdrdrSIRLVNGSLALIWDDLRSL  763
             IVNRVRQGYSPLSFQT  P+P G DRP  I+ EGGE+ R+ S RLV+G LAL WDDLRSL
Sbjct  781   IVNRVRQGYSPLSFQTLAPSPGGLDRPGRIDGEGGEQGRNNSTRLVSGFLALAWDDLRSL  602

Query  764   CLFSYHRLRDLLLIVTRIVELLGR-------RGWEALKYWWNLLQYWSQELKNSAVSLLN  816
              LF YHRL+DL+ +  R+VELLGR       RGWEALKY  +L+QYW  ELK SA+SLL+
Sbjct  601   FLFCYHRLKDLISVTARVVELLGRSSLKGLQRGWEALKYLGSLVQYWGLELKKSAISLLD  422

Query  817   ATAIAVAEGTDRVIEVVQGACRAIRHIPRRIRQGLERILL  856
               AIAV EGTDR+IE VQG CRAIR+IPRR+RQG E  LL
Sbjct  421   TIAIAVGEGTDRIIEGVQGLCRAIRNIPRRVRQGFEAALL  302
```

>Escherichia coli strain EN5143 contig00071, whole genome shotgun sequence
Sequence ID: NZ_JAPMNI010000071.1 Length: 2774
Range 1: 109 to 1623

Score:590 bits(1520), Expect:0.0,
Method:Compositional matrix adjust.,
Identities:304/515(59%), Positives:357/515(69%), Gaps:66/515(12%)

```
Query  1     MRVK---EKYQHLWRWGWRWGTMLLGMLMICSATEKLWVTVYYGVPVWKEATTTLFCASD  57
             MRV+     YQH W WG      +LG  MIC+    LWVTVYYGVPVW++A  TLFCASD
Sbjct  109   MRVRGILRSYQHWWIWG------ILGFWMICNVVGNLWVTVYYGVPVWRDANATLFCASD  270

Query  58    AKAYDTEVHNVWATHACVPTDPNPQEVVLVNVTENFNMWKNDMVEQMHEDIISLWDQSLK  117
             AK Y TEVHNVWATHACVPTDPNPQE+VL NVTENFNMW+NDMVEQMH+D+IS+WDQ LK
Sbjct  271   AKVYKTEVHNVWATHACVPTDPNPQELVLGNVTENFNMWENDMVEQMHQDVISIWDQGLK  450

Query  118   PCVKLTPLCVSLKCTDLKNDTNTNSSSGRMIME-------KGEIKNCSFNISTSIRGKVQ  170
             PCVKLTPLCV+L CT  +    T   + ++ +           EI NCSF  +T +R K +
Sbjct  451   PCVKLTPLCVTLNCTKYEPRNETKKDNIKISLNLTIDQEAVDEINNCSFETTTELRDKTR  630

Query  171   KEYAFFYKLDIIPIDNDTTS------------YKLTSCNTSVITQACPKVSFEPIPIHYC  218
             K +A FYK+D+I +  ++T              Y+L +CNTS ITQACPKVSFEPIPIHYC
Sbjct  631   KVHALFYKIDMIELGQNSTCQKPENSTSKCTMYRLINCNTSAITQACPKVSFEPIPIHYC  810

Query  219   APAGFAILKCNNKTFNGTGPCTNVSTVQCTHGIRPVVSTQLLLNGSLAEEEVVIRSVNFT  278
             APAG+A+LKCNN TFNGTGPC N+STVQCTHGIRPVVSTQLLLNGSLAE  + IRS N T
```

```
Sbjct  811   APAGYALLKCNNNTFNGTGPCRNISTVQCTHGIRPVVSTQLLLNGSLAEGNITIRSENLT  990

Query  279   DNAKTIIVQLNTSVEINCTRPNNNTRKRIRIQRGPGRAFVTIGK-IGNMRQAHCNISRAK  337
             DNAKTII  LN SVEI C RPNNNTR+ +RI  GPG+ F   G  IG++R+AHCNIS
Sbjct  991   DNAKTIIAHLNQSVEILCVRPNNNTRQSVRI--GPGQTFFATGDIIGDIRRAHCNISAKH  1164

Query  338   WNNTLKQIASKLREQFGNNKTIIFKQSSGGDPEIVTHSFNCGGEFFYCNSTQLFNS----  393
             W+  L+ ++ +L + +  NKTI F   SGGD EI THSFNC GEFFYCN++QLFNS
Sbjct  1165  WHTALRNVSIELAKLY--NKTINFTSHSGGDLEITTHSFNCRGEFFYCNTSQLFNSLYMP  1338

Query  394   -----TWFNSTWSTEGSNNTE----GSD--------------------TITLPCRIKQII  424
                   W+N   +    N+T+    GS                      ITLPC+IKQII
Sbjct  1339  DNDTYIWYNGNHTHVWHNDTQYVINGSTQYVTLKNSAQNVTINNITDFNITLPCKIKQII  1518

Query  425   NMWQKVGKAMYAPPISGQIRCSSNITGLLLTRDGG  459
             NMWQ+VGKAMYAPPI+G I C SNITGLLLTRDGG
Sbjct  1519  NMWQEVGKAMYAPPIAGNITCLSNITGLLLTRDGG  1623
```

Range 2: 1652 to 2473

Score:323 bits(829), Expect:2e-92,
Method:Compositional matrix adjust.,
Identities:188/274(69%), Positives:222/274(81%), Gaps:7/274(2%)

```
Query  590   QQLLGIWGCSGKLICTTAVPWNASWSNKSLEQIWNHTTWMEWDREINNYTSLIHSLIees  649
             Q+LLG+WGCS + ICTT VPWN+SWSN+S   IWN+ TWMEWD+EI  +T+ I+ L+EES
Sbjct  1652  QRLLGLWGCSERTICTTNVPWNSSWSNRSYSDIWNNLTWMEWDKEIEQHTNTIYQLLEES  1831

Query  650   qnqqekneqellelDKWASLWNWFNITNWLWYIKLFIMIVGGLVGLRIVFAVLSIVNRVR  709
             Q QQE+NE++LL LDKW  LW+WF+ITNWLWYIK+FIMIVGGL+GLRIVFAVLSIVNRVR
Sbjct  1832  QGQQEQNEKDLLALDKWNDLWSWFDITNWLWYIKIFIMIVGGLIGLRIVFAVLSIVNRVR  2011

Query  710   QGYSPLSFQTHLPTprgpdrpegieeeggerdrdrSIRLVNGSLALIWDDLRSLCLFSYH  769
             QGYSPLSFQT  P+P G DRP  I+EEGGE+ R+ S RLV+G LAL WDDLRSL F YH
Sbjct  2012  QGYSPLSFQTLAPSPGGLDRPGRIDEEGGEQGRNNSTRLVSGFLALAWDDLRSLFLFCYH  2191

Query  770   RLRDLLLIVTRIVELLGR-------RGWEALKYWWNLLQYWSQELKNSAVSLLNATAIAV  822
             RL+DL+ +  R VELLGR       RGWEALKY  +L+QYW  ELK SA+SLL+  AIAV
Sbjct  2192  RLKDLISVTARAVELLGRSSLKGLQRGWEALKYLGSLVQYWVLELKKSAISLLDTIAIAV  2371

Query  823   AEGTDRVIEVVQGACRAIRHIPRRIRQGLERILL  856
              EGTDR+IE +QG CRAIR+IPRR+RQG E  LL
Sbjct  2372  GEGTDRIIEGIQGLCRAIRNIPRRVRQGFEAALL  2473
```

>Escherichia coli strain EN5383 contig00096, whole genome shotgun sequence
Sequence ID: NZ_JAPTZN010000096.1 Length: 2210
Range 1: 1 to 1206

Score:564 bits(1453), Expect:0.0,
Method:Compositional matrix adjust.,
Identities:274/408(67%), Positives:328/408(80%), Gaps:14/408(3%)

```
Query  71    THACVPTDPNPQEVVLVNVTENFNMWKNDMVEQMHEDIISLWDQSLKPCVKLTPLCVSLK  130
             THACVPTDP+PQE+VLVNVTENFNMWKNDM+ QMHED+ISLWDQSLKPCVKLTPLCV+LK
Sbjct  1     THACVPTDPSPQEMVLVNVTENFNMWKNDMMNQMHEDVISLWDQSLKPCVKLTPLCVTLK  180

Query  131   CTDLKNDTNTNSSSGRMIMEKGEIKNCSFNISTSIRGKVQKEYAFFYKLDIIPIDN----  186
             C        T +S+G   + + E+KNCSFNI+T +R + QK YA FYKLD++P+DN
Sbjct  181   CNAAIARNGTAASNGTSSVRE-EMKNCSFNITTELRDRKQKVYALFYKLDVVPLDNTNGS  357

Query  187   -DTTSYKLTSCNTSVITQACPKVSFEPIPIHYCAPAGFAILKCNNKTFNGTGPCTNVSTV  245
               + SY+L +CNTS ITQACPKVSF+PIPIHYCAPAG+AIL+CN+KTFNGTGPC NVSTV
Sbjct  358   NSSRSYRLINCNTSAITQACPKVSFDPIPIHYCAPAGYAILRCNDKTFNGTGPCHNVSTV  537

Query  246   QCTHGIRPVVSTQLLLNGSLAEEEVVIRSVNFTDNAKTIIVQLNTSVEINCTRPNNNTRK  305
             QCTHGI+PVVSTQLLLNGSLAEEE++IRS N +D+ K IIV LN SVEI CTRPN N RK
Sbjct  538   QCTHGIKPVVSTQLLLNGSLAEEEIMIRSQNLSDDTKIIIVHLNQSVEIVCTRPNVNKRK  717

Query  306   RIRIQRGPGRAFVTIGK-IGNMRQAHCNISRAKWNNTLKQIASKLREQFGNNKTIIFKQS  364
              IRI  GPG+ F   G+ IG++RQA+C+I+ +KW+ TL  ++ KL E+F  NKTI F  S
Sbjct  718   SIRI--GPGQTFYATGEIIGDIRQAYCSINISKWSETLYNVSKKLAERF--NKTINFASS  885

Query  365   SGGDPEIVTHSFNCGGEFFYCNSTQLFNSTWF--NSTWSTEGSNNTEGSDTITLPCRIKQ  422
             SGGD E++THSFNCGGEFFYCN+++LFNST+   N T+   G+N T  ++  T+PCRIKQ
Sbjct  886   SGGDLEVITHSFNCGGEFFYCNTSELFNSTYIYPNRTYMFNGTNGT-STNNFTIPCRIKQ  1062

Query  423   IINMWQKVGKAMYAPPISGQIRCSSNITGLLLTRDGGNSNNESEIFRP  470
             IINMWQ+VG+AMYAPPI G I C SNITGLLL RDGG  N  +EIFRP
Sbjct  1063  IINMWQEVGRAMYAPPIEGNITCKSNITGLLLERDGGEENKTTEIFRP  1206
```

Range 2: 1213 to 1941

Score:300 bits(767), Expect:4e-85,
Method:Compositional matrix adjust.,
Identities:177/243(73%), Positives:205/243(84%), Gaps:0/243(0%)

```
Query  613   SWSNKSLEQIWNHTTWMEWDREINNYTSLIHSLIeesqnqqekneqellelDKWASLWNW  672
             SWSN+S   IW++ TWM+WDREI NYT  I+ L+EESQNQQEKNE+ELL L  W SLWNW
Sbjct  1213  SWSNRSQTDIWDNMTWMQWDREISNYTDTIYKLLEESQNQQEKNEKELLALGSWESLWNW  1392

Query  673   FNITNWLWYIKLFIMIVGGLVGLRIVFAVLSIVNRVRQGYSPLSFQTHLPTprgpdrpeg  732
             F+IT WLWYIK+FIMIVGGL+GLRI+FAVLSIVNRVRQGYSPLSFQT  P PRGPDR
Sbjct  1393  FSITKWLWYIKIFIMIVGGLIGLRIIFAVLSIVNRVRQGYSPLSFQTLTPNPRGPDRLGR  1572

Query  733   ieeeggerdrdrSIRLVNGSLALIWDDLRSLCLFSYHRLRDLLLIVTRIVELLGRRGWEA  792
             IEEEGGE+D+DRSIRLV+G LAL WDDLRSLCLFSYH+LR+ +L+  R+VELLGRRGWEA
Sbjct  1573  IEEEGGEQDKDRSIRLVSGFLALAWDDLRSLCLFSYHQLRNFILVTARVVELLGRRGWEA  1752

Query  793   LKYWWNLLQYWSQELKNSAVSLLNATAIAVAEGTDRVIEVVQGACRAIRHIPRRIRQGLE  852
             LKY  NL+ YW  ELK SA+ LL+  AIAVAEGTDR+I+++Q  CRAI +IPRRIRQG E
Sbjct  1753  LKYLGNLVLYWGLELKKSAIGLLDVIAIAVAEGTDRLIDLIQRICRAIYNIPRRIRQGFE  1932

Query  853   RIL  855
               L
Sbjct  1933  AAL  1941
```

>Escherichia coli strain EN5192 contig00149, whole genome shotgun sequence
Sequence ID: NZ_JAPMNL010000149.1 Length: 1397
Range 1: 223 to 1392

Score:538 bits(1386), Expect:4e-180,
Method:Compositional matrix adjust.,

```
Identities:268/401(67%), Positives:315/401(78%), Gaps:20/401(4%)

Query  1     MRVKEKYQHLWRWGWRWGTMLLGMLMICSATEKLWVTVYYGVPVWKEATTTLFCASDAKA  60
             MRV+   ++  +W W WG +   MLMIC+     WVTVYYGVPVWKEA TTLFCASDAKA
Sbjct  223   MRVRGTLRNYQQW-WIWGILGFWMLMICNGGRDSWVTVYYGVPVWKEAKTTLFCASDAKA  399

Query  61    YDTEVHNVWATHACVPTDPNPQEVVLVNVTENFNMWKNDMVEQMHEDIISLWDQSLKPCV  120
             ++ EVHNVWATHACVPTDPNPQE+VL NVTENFNMWK+DMV+QMHED+ISLWDQSLKPCV
Sbjct  400   HEREVHNVWATHACVPTDPNPQEMVLENVTENFNMWKSDMVDQMHEDVISLWDQSLKPCV  579

Query  121   KLTPLCVSLKCTDLKNDTNTNSSSGRMIMEKGEIKNCSFNISTSIRGKVQKEYAFFYKLD  180
             KLTPLCV+L CT+ K    TN S  + + +K E+KNCSF  +T ++ K Q+  A FY LD
Sbjct  580   KLTPLCVTLNCTNAK----TNVS--KQVTDK-EVKNCSFKANTEVQDKRQRVSALFYSLD  738

Query  181   IIPIDNDTT--------SYKLTSCNTSVITQACPKVSFEPIPIHYCAPAGFAILKCNNKT  232
             I P++  ++        +Y L SCNTSVI QACPKV+F+PIPIHYCAPAGFAILKCNN+T
Sbjct  739   IRPLNKSSSGNSSENYENYVLISCNTSVIKQACPKVTFDPIPIHYCAPAGFAILKCNNET  918

Query  233   FNGTGPCTNVSTVQCTHGIRPVVSTQLLLNGSLAEEEVVIRSVNFTDNAKTIIVQLNTSV  292
             FNGTGPC NVSTVQCTHGI+PVVSTQLLLNGSLA+ E++IRS N T+N KTIIV LN SV
Sbjct  919   FNGTGPCNNVSTVQCTHGIKPVVSTQLLLNGSLAKGEIIIRSENLTNNVKTIIVHLNESV  1098

Query  293   EINCTRPNNNTRKRIRIQRGPGRAFVTIGK-IGNMRQAHCNISRAKWNNTLKQIASKLRE  351
             +I C RP+NNTRK IRI  GPG+ F   G  IG++RQAHCNIS   WN TL+++  KL E
Sbjct  1099  KIECIRPHNNTRKSIRI--GPGQTFYATGDIIGDIRQAHCNISENMWNKTLQRVGKKLAE  1272

Query  352   QFGNNKTIIFKQSSGGDPEIVTHSFNCGGEFFYCNSTQLFN  392
              F  NKTI F+ SSGGD EI+THSFNCGGEFFYCN++ LFN
Sbjct  1273  HFP-NKTIKFESSSGGDLEIMTHSFNCGGEFFYCNTSGLFN  1392



>Escherichia coli strain NBE0006 contig00124, whole genome shotgun sequence
Sequence ID: NZ_JARBDI010000124.1 Length: 1306
Range 1: 1 to 1152

Score:529 bits(1363), Expect:4e-177,
Method:Compositional matrix adjust.,
Identities:265/388(68%), Positives:305/388(78%), Gaps:15/388(3%)

Query  1     MRVKEKYQHLWRWGWRWGTMLLGMLMICSATEKLWVTVYYGVPVWKEATTTLFCASDAKA  60
             MRV+   ++  +W W WG +   MLMIC+     WVTVYYGVPVWKEA TTLFCASDAKA
Sbjct  1152  MRVRGTLRNYQQW-WIWGVLGFWMLMICNGGGDSWVTVYYGVPVWKEAKTTLFCASDAKA  976

Query  61    YDTEVHNVWATHACVPTDPNPQEVVLVNVTENFNMWKNDMVEQMHEDIISLWDQSLKPCV  120
             ++ EVHNVWATHACVPTDPNPQE+VL NVTENFNMW+NDMV+QMHED+ISLWDQSLKPCV
Sbjct  975   HEREVHNVWATHACVPTDPNPQEMVLENVTENFNMWRNDMVDQMHEDVISLWDQSLKPCV  796

Query  121   KLTPLCVSLKCTDL---KNDTNTNSSSGRMIMEKGEIKNCSFNISTSIRGKVQKEYAFFY  177
             KLTPLCV+L+C  +    N TN   ++  +     EI+NCSFN +T IR + Q+ YA FY
Sbjct  795   KLTPLCVTLECRKVNATHNATNNGDATHNVTNNGQEIQNCSFNATTEIRDRKQRVYALFY  616

Query  178   KLDIIPIDNDTTS-------YKLTSCNTSVITQACPKVSFEPIPIHYCAPAGFAILKCNN  230
             +LDI+P+D + +S       Y+L +CNTS ITQACPKVSF+PIPIHYCAPAG+AILKCNN
Sbjct  615   RLDIVPLDKNNSSKNNSSEYYRLINCNTSAITQACPKVSFDPIPIHYCAPAGYAILKCNN  436

Query  231   KTFNGTGPCTNVSTVQCTHGIRPVVSTQLLLNGSLAEEEVVIRSVNFTDNAKTIIVQLNT  290
             KTFNGTGPC NVSTVQCTHGI+PVVSTQLLLNGSLAE E++IRS N TDN KTIIV L+
Sbjct  435   KTFNGTGPCNNVSTVQCTHGIKPVVSTQLLLNGSLAEGEIIIRSENLTDNVKTIIVHLDQ  256

Query  291   SVEINCTRPNNNTRKRIRIQRGPGRAF-VTIGKIGNMRQAHCNISRAKWNNTLKQIASKL  349
             SVEI CTRPNNNTRK IRI  GPG+ F  T G IGN+RQAHCNIS  KWN TL+++  KL
Sbjct  255   SVEIVCTRPNNNTRKSIRI--GPGQTFYATGGIIGNIRQAHCNISEDKWNETLQRVGKKL  82

Query  350   REQFGNNKTIIFKQSSGGDPEIVTHSFN  377
              E F  NKTI F  SSGGD EI THSFN
Sbjct  81    VEHFP-NKTIKFAPSSGGDLEITTHSFN  1



>Klebsiella pneumoniae strain EuSCAPE_IL004, whole genome shotgun sequence
Sequence ID: NZ_UKJP02000086.1 Length: 1081
Range 1: 1 to 1080

Score:495 bits(1275), Expect:4e-165,
Method:Compositional matrix adjust.,
Identities:266/376(71%), Positives:314/376(83%), Gaps:20/376(5%)

Query  176   FYKLDIIPI---DNDTTSYKLTSCNTSVITQACPKVSFEPIPIHYCAPAGFAILKCNNKT  232
             FY+LDI+PI   +++ + Y+L +CNTS ++QACPKVSF+PIPIHYCAPAG+AILKCN+KT
Sbjct  1080  FYRLDIVPIPGENSNISKYRLINCNTSAMSQACPKVSFDPIPIHYCAPAGYAILKCNDKT  901

Query  233   FNGTGPCTNVSTVQCTHGIRPVVSTQLLLNGSLAEEEVVIRSVNFTDNAKTIIVQLNTSV  292
             FNGTGPC NVSTVQCTHGI+PVVSTQLLLNGSL+EE ++IRS N T+N KTIIV L+  V
Sbjct  900   FNGTGPCLNVSTVQCTHGIKPVVSTQLLLNGSLSEEGIIIRSENLTNNVKTIIVHLDKPV  721

Query  293   EINCTRPNNNTRKRIRIQRGPGRAFVTIGKIGNMRQAHCNISRAKWNNTLKQIASKLREQ  352
             +I CTRPNNNTRK IRI  GPG+ F     IG++RQA+CNIS  KW +TL+ + +KLR+
Sbjct  720   DIMCTRPNNNTRKSIRI--GPGQTFYATDIIGDIRQAYCNISGKKWYDTLQGVKNKLRKH  547

Query  353   FGNNKTIIFKQSSGGDPEIVTHSFNCGGEFFYCNSTQLFNSTWFNSTWSTEGSNNTEGSD  412
             F  +KTI F+ SSGGD EI THSFNC GEFFYC++++LFNST  +++
Sbjct  546   F-PSKTIQFEPSSGGDLEITTHSFNCMGEFFYCDTSKLFNSTKRDNS-------------  409

Query  413   TITLPCRIKQIINMWQKVGKAMYAPPISGQIRCSSNITGLLLTRD-GGNSNNESEIFRPG  471
             TITLPCRIKQIINMWQ VG+AMYAPPI+G I C SNITGLLLTRD GGNS+  +E FRPG
Sbjct  408   TITLPCRIKQIINMWQGVGRAMYAPPIAGNITCRSNITGLLLTRDGGGNSSATTETFRPG  229

Query  472   GGDMRDNWRSELYKYKVVKIEPLGVAPTKAKRRVVQREKRAVGIgalflgflgaagstmg  531
             GGDMRDNWRSELYKYKVV+I+PLGVAPT+AKRRVV+R KRAVGIGA+FLGFLGAAGSTMG
Sbjct  228   GGDMRDNWRSELYKYKVVEIKPLGVAPTEAKRRVVERGKRAVGIGAVFLGFLGAAGSTMG  49

Query  532   aasmtLTVQARQLLSG  547
             AAS+TLTVQARQLLSG
Sbjct  48    AASITLTVQARQLLSG  1



>Escherichia coli strain NBE0010 contig00098, whole genome shotgun sequence
Sequence ID: NZ_JARBDM010000098.1 Length: 1912
Range 1: 825 to 1910

Score:504 bits(1297), Expect:3e-164,
Method:Compositional matrix adjust.,
Identities:297/363(82%), Positives:330/363(90%), Gaps:1/363(0%)

Query  424   INMWQKVGKAMYAPPISGQIRCSSNITGLLLTRDGGNSNNESEIFRPGGGDMRDNWRSEL  483
             INMWQ+VG+AMYAPPI+G I+C SNITGLLL RDGG  +N +E FRPGGGDMRDNWRSEL
Sbjct  1910  INMWQEVGRAMYAPPIAGNIKCGSNITGLLLVRDGG-PDNGTETFRPGGGDMRDNWRSEL  1734

Query  484   YKYKVVKIEPLGVAPTKAKRRVVQREKRAVGIgalflgflgaagstmgaasmtLTVQARQ  543
```

```
             YKYKVV+I+PLGVAPT+AKRRVV R KRAVGIGA+FLGFLGAAGSTMGAAS+TLTVQARQ
Sbjct  1733  YKYKVVEIKPLGVAPTEAKRRVVGRGKRAVGIGAVFLGFLGAAGSTMGAASITLTVQARQ  1554

Query  544   LLSGIVQQQNNLLRAIEAQQHLLQLTVWGIKQLQARILAVERYLKDQQLLGIWGCSGKLI  603
             LLSGIVQQQ+NLLRAIEAQQHLLQLTVWGIKQLQ R+LA+ERYLKDQQLLGIWGCSGKLI
Sbjct  1553  LLSGIVQQQSNLLRAIEAQQHLLQLTVWGIKQLQTRVLAIERYLKDQQLLGIWGCSGKLI  1374

Query  604   CTTAVPWNASWSNKSLEQIWNHTTWMEWDREINNYTSLIHSLIeesqnqqekneqellel  663
             C TAVPWN+SWSNK+  +IWN+ TWM+WDREI+NYT  I+ L+EESQNQQEKNE++LL L
Sbjct  1373  CPTAVPWNSSWSNKNQSEIWNNMTWMQWDREISNYTYTIYRLLEESQNQQEKNEEDLLAL  1194

Query  664   DKWASLWNWFNITNWLWYIKLFIMIVGGLVGLRIVFAVLSIVNRVRQGYSPLSFQTHLPT  723
             D WASLWNWFNIT WLWYIK+FIMIVGGL+GLRI+FAVLSIVNRVRQGYSPLSFQT  P
Sbjct  1193  DSWASLWNWFNITQWLWYIKIFIMIVGGLIGLRIIFAVLSIVNRVRQGYSPLSFQTPTPN  1014

Query  724   prgpdrpegieeeggerdrdrSIRLVNGSLALIWDDLRSLCLFSYHRLRDLLLIVTRIVE  783
             P   DR  GIEEEGGE+DR+RSIRLVNG LAL WDDLR+LCLFSYHRLRD +L+  R+VE
Sbjct  1013  PGELDRLRGIEEEGGEQDRNRSIRLVNGFLALAWDDLRNLCLFSYHRLRDFILLTARVVE  834

Query  784   LLG  786
             LLG
Sbjct  833   LLG  825



Range 2: 47 to 793

Score:346 bits(888), Expect:1e-103,
Method:Compositional matrix adjust.,
Identities:170/252(67%), Positives:204/252(80%), Gaps:7/252(2%)

Query  153   EIKNCSFNISTSIRGKVQKEYAFFYKLDIIPIDNDTT-SYKLTSCNTSVITQACPKVSFE  211
             E+KNCSFN +T +R K QK  A F++LDI+P+ N +  +Y+L +CNTS I QACPKVSF+
Sbjct  47    EVKNCSFNATTELRDKKQKVQALFHRLDIMPLTNGSDGNYRLINCNTSTIAQACPKVSFD  226

Query  212   PIPIHYCAPAGFAILKCNNKTFNGTGPCTNVSTVQCTHGIRPVVSTQLLLNGSLAEEEVV  271
             PIPIHYC PAG+AIL+CNNKTFNGTGPC NVSTVQCTHGI+PVVSTQLLLNGSLA+EE+V
Sbjct  227   PIPIHYCTPAGYAILQCNNKTFNGTGPCNNVSTVQCTHGIKPVVSTQLLLNGSLAKEEIV  406

Query  272   IRSVNFTDNAKTIIVQLNTSVEINCTRPNNNTRKRIRIQRGPGRAFVTIGK-IGNMRQAH  330
             IRS N T+NAK IIV LN SV+I CTRPNNNTRK IRI  GPG+ F   G  IG++R+AH
Sbjct  407   IRSENLTNNAKIIIVHLNKSVDIVCTRPNNNTRKSIRI--GPGQTFYATGDIIGDIRKAH  580

Query  331   CNISRAKWNNTLKQIASKLREQFGNNKTIIFKQSSGGDPEIVTHSFNCGGEFFYCNSTQL  390
             CNI+  +WN TL +++ KL E+F  NKTI FK SSGGD EI THSFNCGGEFFYCN++ L
Sbjct  581   CNININRWNETLLKVSKKLAERFP-NKTIQFKPSSGGDLEITTHSFNCGGEFFYCNTSNL  757

Query  391   FNSTWF--NSTW  400
             FN T++  N T+
Sbjct  758   FNFTYYYVNGTY  793



>Pseudomonas aeruginosa strain VET-121 NODE_76_length_1017_cov_1.90426_ID_151, whole genome shotgun sequence
Sequence ID: NZ_QZXB01000124.1 Length: 1017
Range 1: 61 to 981

Score:449 bits(1156), Expect:2e-147,
Method:Compositional matrix adjust.,
Identities:216/323(67%), Positives:259/323(80%), Gaps:21/323(6%)

Query  189   TSYKLTSCNTSVITQACPKVSFEPIPIHYCAPAGFAILKCNNKTFNGTGPCTNVSTVQCT  248
             +SY L +CN + I QACPKVSFEPIPIHYCAPAGFAILKC +K FNGTGPC NVSTVQCT
Sbjct  61    SSYMLINCNATTIKQACPKVSFEPIPIHYCAPAGFAILKCRDKNFNGTGPCKNVSTVQCT  240

Query  249   HGIRPVVSTQLLLNGSLAEEEVVIRSVNFTDNAKTIIVQLNTSVEINCTRPNNNTRKRIR  308
             HGI+PVVSTQLLLNGSLAEE+++IRS N ++NAKTIIVQ N SVEINCTRPNNNTRK I
Sbjct  241   HGIKPVVSTQLLLNGSLAEEDIIIRSENISNNAKTIIVQFNQSVEINCTRPNNNTRKSIH  420

Query  309   IQRGPGRAFVTIGK-IGNMRQAHCNISRAKWNNTLKQIASKLREQFGNNKTIIFKQSSGG  367
             +  GPG+AF   G  IG++RQA+CNIS  KW++ L ++ +KL+E F  N  I FK S+GG
Sbjct  421   L--GPGQAFFATGDIIGSIRQAYCNISGDKWDDMLNKVKAKLKEFFPGN--ISFKSSAGG  588

Query  368   DPEIVTHSFNCGGEFFYCNSTQLFNSTWFNSTWSTEGSNNTEGSDTITLPCRIKQIINMW  427
             D EI+TH+FNCGGEFFYCN++ LFNS+  N+               I LPC+I+QI+ MW
Sbjct  589   DLEIITHTFNCGGEFFYCNTSDLFNSSSKNN------------GTIIKLPCKIRQIVRMW  732

Query  428   QKVGKAMYAPPISGQIRCSSNITGLLLTRD----GGNSNNESEIFRPGGGDMRDNWRSEL  483
             Q+V +AMYAPPI+G I C SNITGLLLTRD    GGN+ N++EIFRPGGGDM+DNWRSEL
Sbjct  733   QRVAQAMYAPPIAGIITCRSNITGLLLTRDGSSKGGNNTNDTEIFRPGGGDMKDNWRSEL  912

Query  484   YKYKVVKIEPLGVAPTKAKRRVV  506
             YKYK+VKI+PLG+AP+KA+RR V
Sbjct  913   YKYKIVKIKPLGIAPSKARRRAV  981



>Escherichia coli strain NBE0002 contig00149, whole genome shotgun sequence
Sequence ID: NZ_JARBDF010000149.1 Length: 996
Range 1: 11 to 991

Score:431 bits(1107), Expect:2e-140,
Method:Compositional matrix adjust.,
Identities:218/333(65%), Positives:250/333(75%), Gaps:30/333(9%)

Query  1     MRVK---EKYQHLWRWGWRWGTMLLG--MLMICSATEKLWVTVYYGVPVWKEATTTLFCA  55
             MRV+     YQH W WG      +LG  M MIC+    LWVTVYYGVPVWK+A  TLFCA
Sbjct  11    MRVRGVLRSYQHWWIWG------ILGFWMFMICNVVGNLWVTVYYGVPVWKDANATLFCA  172

Query  56    SDAKAYDTEVHNVWATHACVPTDPNPQEVVLVNVTENFNMWKNDMVEQMHEDIISLWDQS  115
             SDAK Y TEVHNVWATHACVPTDPNPQE+VL NVTENFNMW+NDMVEQMH+D+IS+WDQ
Sbjct  173   SDAKVYKTEVHNVWATHACVPTDPNPQELVLGNVTENFNMWENDMVEQMHQDVISIWDQG  352

Query  116   LKPCVKLTPLCVSLKCT--DLKNDT---NTNSSSGRMIMEKG--EIKNCSFNISTSIRGK  168
             LKPCVKLTPLCV+L CT  + +N+T   N   S    I ++   EI NCSF  +T +R K
Sbjct  353   LKPCVKLTPLCVTLNCTKYEPRNETKKDNIKISLNLTIDQEAVEEINNCSFETTTELRDK  532

Query  169   VQKEYAFFYKLDIIPIDNDTTS------------YKLTSCNTSVITQACPKVSFEPIPIH  216
              +K +A FYK+D+I +  ++T             Y+L +CNTS ITQACPKVSFEPIPIH
Sbjct  533   TRKVHALFYKIDMIELGQNSTCQKPENSTSECTMYRLINCNTSAITQACPKVSFEPIPIH  712

Query  217   YCAPAGFAILKCNNKTFNGTGPCTNVSTVQCTHGIRPVVSTQLLLNGSLAEEEVVIRSVN  276
             YCAPAG+A+LKCNN TFNGTGPC N+STVQCTHGIRPVVSTQLLLNGSLAE  + IRS N
Sbjct  713   YCAPAGYALLKCNNNTFNGTGPCRNISTVQCTHGIRPVVSTQLLLNGSLAEGNITIRSEN  892

Query  277   FTDNAKTIIVQLNTSVEINCTRPNNNTRKRIRI  309
              TDNAKTII  LN SVEI C RPNNNTR+ +RI
Sbjct  893   LTDNAKTIIAHLNQSVEILCVRPNNNTRQSVRI  991
```

```
>Escherichia coli strain EN5308 contig00145, whole genome shotgun sequence
Sequence ID: NZ_JARBDE010000145.1 Length: 1005
Range 1: 5 to 994

Score:430 bits(1106), Expect:3e-140,
Method:Compositional matrix adjust.,
Identities:219/340(64%), Positives:258/340(75%), Gaps:19/340(5%)

Query  136  NDTNTNSSSGRMIMEKGEIKNCSFNISTSIRGKVQKEYAFFYKLDIIPIDNDTTS-----  190
            N TN   ++  +     EI+NCSFN +T IR + Q+ YA FY+LDI+P+D + +S
Sbjct  5    NATNNGDATHNVTNNGQEIQNCSFNATTEIRDRKQRVYALFYRLDIVPLDKNNSSKNNSS  184

Query  191  --YKLTSCNTSVITQACPKVSFEPIPIHYCAPAGFAILKCNNKTFNGTGPCTNVSTVQCT  248
              Y+L +CNTS ITQACPKVSF+PIPIHYCAPAG+AILKCNNKTFNGTGPC NVSTVQCT
Sbjct  185  EYYRLINCNTSAITQACPKVSFDPIPIHYCAPAGYAILKCNNKTFNGTGPCNNVSTVQCT  364

Query  249  HGIRPVVSTQLLLNGSLAEEEVVIRSVNFTDNAKTIIVQLNTSVEINCTRPNNNTRKRIR  308
            HGI+PVVSTQLLLNGSLAE E++IRS N TDN KTIIV L+ SVEI CTRPNNNTRK IR
Sbjct  365  HGIKPVVSTQLLLNGSLAEGEIIIRSENLTDNVKTIIVHLDQSVEIVCTRPNNNTRKSIR  544

Query  309  IQRGPGRAF-VTIGKIGNMRQAHCNISRAKWNNTLKQIASKLREQFGNNKTIIFKQSSGG  367
            I  GPG+ F  T G IGN+RQAHCNIS  KWN TL+++  KL E F  NKTI F  SSGG
Sbjct  545  I--GPGQTFYATGGIIGNIRQAHCNISEDKWNETLQRVGKKLVEHFP-NKTIKFAPSSGG  715

Query  368  DPEIVTHSFNCGGEFFYCNSTQLFNSTWF-NSTWSTEGSNNTEGSDTITLPCRIKQIINM  426
            D EI THSFNC GEFFYC++++LFNST+  N T S       + + TIT+PC IKQI+NM
Sbjct  716  DLEITTHSFNCRGEFFYCSTSRLFNSTYMPNDTKS-------KSNKTITIPCSIKQIVNM  874

Query  427  WQKVGKAMYAPPISGQIRCSSNITGLLLTRDGGNSNNESE  466
            WQ+VG+AMYAPPI G I C SNITG+LL RDGG  + + E
Sbjct  875  WQEVGRAMYAPPIEGNITCRSNITGILLVRDGGVDSEDPE  994



>Escherichia coli strain NBE0002 contig00132, whole genome shotgun sequence
Sequence ID: NZ_JARBDF010000132.1 Length: 1381
Range 1: 2 to 976

Score:431 bits(1109), Expect:9e-139,
Method:Compositional matrix adjust.,
Identities:213/327(65%), Positives:251/327(76%), Gaps:11/327(3%)

Query  18   GTMLLGMLMICSATEKLWVTVYYGVPVWKEATTTLFCASDAKAYDTEVHNVWATHACVPT  77
            G +   MLM C     L VTVYY VPVW EA TTLFCASDAKAY+ EVHN+WATHACVPT
Sbjct  976  GVLGF*MLMNCKVGRNL*VTVYYEVPVWTEAKTTLFCASDAKAYEKEVHNIWATHACVPT  797

Query  78   DPNPQEVVLVNVTENFNMWKNDMVEQMHEDIISLWDQSLKPCVKLTPLCVSLKCTDLK-N  136
            DPNPQE+VL NVTE FNMWKND+V+QMHED+ISL DQSLKPCVKLTPLCV+L+C     N
Sbjct  796  DPNPQEMVLKNVTEYFNMWKNDIVDQMHEDVISL*DQSLKPCVKLTPLCVTLECNSTTYN  617

Query  137  DTNTNSSSGRMIMEKGEIKNCSFNISTSIRGKVQKEYAFFYKLDIIPIDND-------TT  189
            +T T  +   ++ +  E+KNCSFN +T +R K  K  A F++LDI+PIDN        T
Sbjct  616  NTYTYKNKSSVLEQVHEVKNCSFNATTELRDKKNKV*ALFHRLDIVPIDNKENFNENIT*  437

Query  190  SYKLTSCNTSVITQACPKVSFEPIPIHYCAPAGFAILKCNNKTFNGTGPCTNVSTVQCTH  249
            SY+L +CNTSVITQACPKV+F+PIPIHYC PA FAIL+CNNKTF+  GPC NVSTVQCTH
Sbjct  436  SYRLINCNTSVITQACPKVNFDPIPIHYCTPASFAILQCNNKTFSKKGPCNNVSTVQCTH  257

Query  250  GIRPVVSTQLLLNGSLAEEEVVIRSVNFTDNAKTIIVQLNTSVEINCTRPNNNTRKRIRI  309
             I+PVVSTQLLLN SLAEEE++IRS N TDN KTIIV LN SV I C RP NNTRK ++I
Sbjct  256  RIKPVVSTQLLLNSSLAEEEIIIRSENLTDNVKTIIVNLNESVVIECMRPGNNTRKSVKI  77

Query  310  QRGPGRA-FVTIGKIGNMRQAHCNISR  335
            +  PG+  F T   + ++RQAHCN+S+
Sbjct  76   R--PGQTFFATEDIVEDIRQAHCNVSK  2



>Escherichia coli strain EN5383 contig00089, whole genome shotgun sequence
Sequence ID: NZ_JAPTZN010000089.1 Length: 1970
Range 1: 448 to 1443

Score:438 bits(1127), Expect:1e-138,
Method:Compositional matrix adjust.,
Identities:222/342(65%), Positives:265/342(77%), Gaps:18/342(5%)

Query  128   SLKCTDLK-NDTNTNSSSGRMIMEKGEIKNCSFNISTSIRGKVQKEYAFFYKLDIIPIDN  186
             +L CTD   N T TN+++   +  + EIKNCSFN +T I+ K +K  A FYKLDI+PID
Sbjct  448   TLDCTDANTNRTMTNANTTGTVTNREEIKNCSFNATTEIKDKKRKVQALFYKLDIVPIDK  627

Query  187   DTT-SYKLTSCNTSVITQACPKVSFEPIPIHYCAPAGFAILKCNNKTFNGTGPCTNVSTV  245
             D   SY+L +CNTS I QACPKV+F+PIPIHYCAPAG+AILKCNNKTFNGTGPC NVSTV
Sbjct  628   DNNNSYRLINCNTSTIAQACPKVNFDPIPIHYCAPAGYAILKCNNKTFNGTGPCNNVSTV  807

Query  246   QCTHGIRPVVSTQLLLNGSLAEEEVVIRSVNFTDNAKTIIVQLNTSVEINCTRPNNNTRK  305
             QCTHGI+PVVSTQLLLNGSLAEE+++IRS N TDNAKTIIV LN SVEI CTRPNNNTRK
Sbjct  808   QCTHGIKPVVSTQLLLNGSLAEEDIIIRSKNLTDNAKTIIVHLNESVEIECTRPNNNTRK  987

Query  306   RIRIQRGPGRAFVTIGK-IGNMRQAHCNISRAKWNNTLKQIASKLREQFGNNKTIIFKQS  364
              IRI  GPG+ F   G  IG++RQAHCNIS  +WN TL+++  KL+E F  NKTI F++S
Sbjct  988   SIRI--GPGQTFYATGDIIGDIRQAHCNISAERWNRTLQRVRKKLKEHFP-NKTINFEKS  1158

Query  365   SGGDPEIVTHSFNCGGEFFYCNSTQLFNSTWFNSTWSTEGS---NNTEGSDT--ITLPCR  419
             SGGD EI  HSFNC GEFFYCN++ LF       T++  G+   N TEG+ +  IT+PC+
Sbjct  1159  SGGDLEITMHSFNCRGEFFYCNTSGLF-------TYNPNGTYNPNGTEGNSSSFITIPCK  1317

Query  420   IKQIINMWQKVGKAMYAPPISGQIRCSSNITGLLLTRDGGNS  461
             +KQ+IN+WQ+VG+AMYAPPI G I C SNITGLLL RDGGN
Sbjct  1318  LKQVINLWQEVGRAMYAPPIEGNITCKSNITGLLLVRDGGND  1443



Range 2: 1454 to 1708

Score:92.4 bits(228), Expect:1e-14,
Method:Compositional matrix adjust.,
Identities:49/85(58%), Positives:60/85(70%), Gaps:7/85(8%)

Query  775   LLIVTRIVELLG-------RRGWEALKYWWNLLQYWSQELKNSAVSLLNATAIAVAEGTD  827
             +L+  R VELLG       +RGW  LKY   L+QYW  ELK S +SL+++ AIAVAEGTD
Sbjct  1454  ILVTVRGVELLGHSILKGLQRGWRVLKYLGGLVQYWGLELKKSTISLVDSIAIAVAEGTD  1633

Query  828   RVIEVVQGACRAIRHIPRRIRQGLE  852
             R+IEV+Q   RAI +IPRRIRQG E
Sbjct  1634  RIIEVLQRIGRAICNIPRRIRQGFE  1708



>Escherichia coli strain EN5095 contig00078, whole genome shotgun sequence
```

Sequence ID: NZ_JAPYLY010000078.1 Length: 1972
Range 1: 1002 to 1970

Score:428 bits(1101), Expect:9e-135,
Method:Compositional matrix adjust.,
Identities:223/344(65%), Positives:265/344(77%), Gaps:27/344(7%)

```
Query  125   LCVSLKCTDLKNDTNTNSSSGRMIMEKGEIKNCSFNISTSIRGKVQKEYAFFYKLDIIPI  184
             LCV+L C++    T  NSSS        EIKNC+FN +T IR K QK  A FYKLD++P+
Sbjct  1970  LCVTLHCSNAT-STFGNSSS--------EIKNCTFNATTEIRDKKQKVNALFYKLDVVPL  1818

Query  185   -DNDTTSYKLTSCNTSVITQACPKVSFEPIPIHYCAPAGFAILKCNNKTFNGTGPCTNVS  243
              DN+ + Y+L +CNTS +TQACPK+SF+PIPIHYCAPAG+AILKCN+KTFNGTGPC NVS
Sbjct  1817  NDNNNSYYRLINCNTSTVTQACPKISFDPIPIHYCAPAGYAILKCNDKTFNGTGPCKNVS  1638

Query  244   TVQCTHGIRPVVSTQLLLNGSLAE-EEVVIRSVNFTDNAKTIIVQLNTSVEINCTRPNNN  302
             TVQCTHGI+PVVSTQLLLNGSLAE  E++IRS N T+NAKTIIV LN SV+I C RPNNN
Sbjct  1637  TVQCTHGIKPVVSTQLLLNGSLAEGGEIIIRSENITNNAKTIIVHLNESVQIRCIRPNNN  1458

Query  303   TRKRIRIQRGPGRAF-VTIGKIGNMRQAHCNISRAKWNNTLKQIASKLREQFGNNKTIIF  361
             TRK IRI  GPG+ F  T G IGN+R+A C+I+ ++WN TL+ +  KLRE F  N+TI F
Sbjct  1457  TRKSIRI--GPGQTFYATGGIIGNIREAQCSINGSEWNKTLQGVREKLREHFP-NRTITF  1287

Query  362   KQSSGGDPEIVTHSFNCGGEFFYCNSTQLFNS---TWFNSTWSTEGSNNTEGSDTITLPC  418
               SSGGD EI THSFNC GEFFYCN++ LFNS   T  NSTW          + T+ LPC
Sbjct  1286  NSSSGGDLEITTHSFNCRGEFFYCNTSSLFNSTNGTESNSTWV---------NGTLILPC  1134

Query  419   RIKQIINMWQKVGKAMYAPPISGQIRCSSNITGLLLTRDGGNSN  462
             RIKQIINMWQ+VG+AMYAPPI+G I C+S+ITGLLLTRDGGN+N
Sbjct  1133  RIKQIINMWQEVGRAMYAPPIAGNITCNSSITGLLLTRDGGNTN  1002
```

Range 2: 239 to 994

Score:292 bits(748), Expect:3e-83,
Method:Compositional matrix adjust.,
Identities:176/252(70%), Positives:206/252(81%), Gaps:7/252(2%)

```
Query  611   NASWSNKSLEQIWNHTTWMEWDREINNYTSLIHSLIeesqnqqekneqellelDKWASLW  670
             +ASWSN++ + IW++ TWM+WDREI+NYT  I+ L+E+SQNQQ+KNE++LL LD W +LW
Sbjct  994   SASWSNRTKKDIWDNMTWMQWDREISNYTETIYRLLEDSQNQQDKNEKDLLALDSWKNLW  815

Query  671   NWFNITNWLWYIKLFIMIVGGLVGLRIVFAVLSIVNRVRQGYSPLSFQTHLPTprgpdrp  730
             NWF+I NWLWYIK+FIMIVGGL+GLRI+FAVLSIVNRVRQGYSPLSFQT  P P G DR
Sbjct  814   NWFDIANWLWYIKIFIMIVGGLIGLRIIFAVLSIVNRVRQGYSPLSFQTLTPNPGGLDRL  635

Query  731   egieeeggerdrdrSIRLVNGSLALIWDDLRSLCLFSYHRLRDLLLIVTRIVELLGR---  787
               IEEEGGE+DR RSIRLV G LAL WDDLRSLCLFSYHRLRDLLL+ TR VE+LGR
Sbjct  634   GRIEEEGGEQDRGRSIRLVQGFLALAWDDLRSLCLFSYHRLRDLLLVTTRAVEILGRSSL  455

Query  788   ----RGWEALKYWWNLLQYWSQELKNSAVSLLNATAIAVAEGTDRVIEVVQGACRAIRHI  843
                 RGWEALKY  +++QYW  ELK SA+SLL+ TAI VAEGTDR IE+VQ   RAI ++
Sbjct  454   RGLQRGWEALKYLGSIVQYWVLELKKSAISLLDTTAIVVAEGTDRTIELVQRTYRAICNV  275

Query  844   PRRIRQGLERIL  855
             PRRIRQG E  L
Sbjct  274   PRRIRQGFEAAL  239
```

>Escherichia coli strain EN5132 contig00077, whole genome shotgun sequence
Sequence ID: NZ_JAPYLZ010000077.1 Length: 2498
Range 1: 500 to 1456

Score:428 bits(1101), Expect:1e-132,
Method:Compositional matrix adjust.,
Identities:218/328(66%), Positives:251/328(76%), Gaps:19/328(5%)

```
Query  150   EKGEIKNCSFNISTSIRGKVQKEYAFFYKLDIIPIDND---------TTSYKLTSCNTSV  200
             E  E+K C FN ST +  K QK  A FY LDI P++           T +Y LTSCNTSV
Sbjct  500   ETKELKRCFFNASTEVEDKKQKVSALFYSLDIRPLNKSSSGNSSINSTETYVLTSCNTSV  679

Query  201   ITQACPKVSFEPIPIHYCAPAGFAILKCNNKTFNGTGPCTNVSTVQCTHGIRPVVSTQLL  260
             I QACPKV+F+PIPIHYCAPAG+AILKCNN+TFNGTGPC NVSTVQCTHGI+PVVSTQLL
Sbjct  680   IKQACPKVTFDPIPIHYCAPAGYAILKCNNETFNGTGPCNNVSTVQCTHGIKPVVSTQLL  859

Query  261   LNGSLAEEEVVIRSVNFTDNAKTIIVQLNTSVEINCTRPNNNTRKRIRIQRGPGRAFVTI  320
             LNGSLA+ ++VIRS N TDNAKTIIV LN SVEINCTRPNNNTRK IRI  GPG+ F
Sbjct  860   LNGSLAKGDIVIRSENLTDNAKTIIVHLNESVEINCTRPNNNTRKSIRI--GPGQTFYAT  1033

Query  321   GK-IGNMRQAHCNISRAKWNNTLKQIASKLREQFGNNKTIIFKQSSGGDPEIVTHSFNCG  379
             G+ IG++RQAHCNIS   WN TL+++  KL E F  NKTI F+ SSGGD EI+THSFNCG
Sbjct  1034  GEIIGDIRQAHCNISENMWNKTLQRVGKKLAEHFP-NKTIKFEPSSGGDLEIMTHSFNCG  1210

Query  380   GEFFYCNSTQLFNSTWFNSTWSTEGSNNTEGSDTITLPCRIKQIINMWQKVGKAMYAPPI  439
             GEFFYCN++ LFN T+ N T+  + S     S  IT+ CRIKQIINMWQ+VG+AMYAPPI
Sbjct  1211  GEFFYCNTSSLFNGTY-NGTYIEKNS-----SSIITIQCRIKQIINMWQEVGRAMYAPPI  1372

Query  440   SGQIRCSSNITGLLLTRDGGNSNNESEI  467
             +G I C SNITGLLL RDGG  +N +EI
Sbjct  1373  AGNITCRSNITGLLLVRDGGTGSNSTEI  1456
```

Range 2: 1473 to 2228

Score:291 bits(745), Expect:3e-81,
Method:Compositional matrix adjust.,
Identities:172/252(68%), Positives:206/252(81%), Gaps:7/252(2%)

```
Query  611   NASWSNKSLEQIWNHTTWMEWDREINNYTSLIHSLIeesqnqqekneqellelDKWASLW  670
             ++SWSNKS ++IW++ TWM+WD+EI+NYT  I+ L+E+SQ QQEKNE +LL LD W +LW
Sbjct  1473  HSSWSNKSRDEIWDNMTWMQWDKEISNYTDTIYRLLEDSQIQQEKNETDLLALDSWNNLW  1652

Query  671   NWFNITNWLWYIKLFIMIVGGLVGLRIVFAVLSIVNRVRQGYSPLSFQTHLPTprgpdrp  730
             NWF IT WLWYIK+FIMIVGGL+GLRI+FAV+SIVNR RQGYSPLSFQT  P P GPDR
Sbjct  1653  NWFGITKWLWYIKIFIMIVGGLIGLRIIFAVISIVNRFRQGYSPLSFQTLTPNPGGPDRL  1832

Query  731   egieeeggerdrdrSIRLVNGSLALIWDDLRSLCLFSYHRLRDLLLIVTRIVELLGR---  787
               IEEEGGE+D++RSIRLVNG LAL W+DLR+LCLFSYHRLRD +L+  R++ELLGR
Sbjct  1833  GRIEEEGGEQDKNRSIRLVNGFLALAWEDLRNLCLFSYHRLRDFILVTARVLELLGRSSL  2012

Query  788   ----RGWEALKYWWNLLQYWSQELKNSAVSLLNATAIAVAEGTDRVIEVVQGACRAIRHI  843
                 R WEALKY  +L+QYW  ELK SA+SLL+  AIAV EGTDR+I +VQG CRAIR+I
Sbjct  2013  KGLQRVWEALKYLGSLVQYWGLELKESAISLLDTIAIAVGEGTDRIISLVQGICRAIRNI  2192

Query  844   PRRIRQGLERIL  855
             PRRIRQG E  L
Sbjct  2193  PRRIRQGFEAAL  2228
```

```
>Nocardioides luteus strain JCM 3358 sequence39, whole genome shotgun sequence
Sequence ID: NZ_BMRK01000039.1 Length: 5754
Range 1: 566 to 1423

Score:446 bits(1147), Expect:2e-131,
Method:Compositional matrix adjust.,
Identities:286/286(100%), Positives:286/286(100%), Gaps:0/286(0%)

Query  467   IFRPGGGDMRDNWRSELYKYKVVKIEPLGVAPTKAKRRVVQREKRAVGIgalflgflgaa  526
             IFRPGGGDMRDNWRSELYKYKVVKIEPLGVAPTKAKRRVVQREKRAVGIGALFLGFLGAA
Sbjct  566   IFRPGGGDMRDNWRSELYKYKVVKIEPLGVAPTKAKRRVVQREKRAVGIGALFLGFLGAA  745

Query  527   gstmgaasmtLTVQARQLLSGIVQQQNNLLRAIEAQQHLLQLTVWGIKQLQARILAVERY  586
             GSTMGAASMTLTVQARQLLSGIVQQQNNLLRAIEAQQHLLQLTVWGIKQLQARILAVERY
Sbjct  746   GSTMGAASMTLTVQARQLLSGIVQQQNNLLRAIEAQQHLLQLTVWGIKQLQARILAVERY  925

Query  587   LKDQQLLGIWGCSGKLICTTAVPWNASWSNKSLEQIWNHTTWMEWDREINNYTSLIHSLI  646
             LKDQQLLGIWGCSGKLICTTAVPWNASWSNKSLEQIWNHTTWMEWDREINNYTSLIHSLI
Sbjct  926   LKDQQLLGIWGCSGKLICTTAVPWNASWSNKSLEQIWNHTTWMEWDREINNYTSLIHSLI  1105

Query  647   eesqnqqekneqellelDKWASLWNWFNITNWLWYIKLFIMIVGGLVGLRIVFAVLSIVN  706
             EESQNQQEKNEQELLELDKWASLWNWFNITNWLWYIKLFIMIVGGLVGLRIVFAVLSIVN
Sbjct  1106  EESQNQQEKNEQELLELDKWASLWNWFNITNWLWYIKLFIMIVGGLVGLRIVFAVLSIVN  1285

Query  707   RVRQGYSPLSFQTHLPTprgpdrpegieeeggerdrdrSIRLVNGS  752
             RVRQGYSPLSFQTHLPTPRGPDRPEGIEEEGGERDRDRSIRLVNGS
Sbjct  1286  RVRQGYSPLSFQTHLPTPRGPDRPEGIEEEGGERDRDRSIRLVNGS  1423
```

```
>Streptomyces griseoflavus strain JCM 4479 sequence37, whole genome shotgun sequence
Sequence ID: NZ_BMUC01000037.1 Length: 5775
Range 1: 4325 to 5182

Score:446 bits(1147), Expect:2e-131,
Method:Compositional matrix adjust.,
Identities:286/286(100%), Positives:286/286(100%), Gaps:0/286(0%)

Query  467   IFRPGGGDMRDNWRSELYKYKVVKIEPLGVAPTKAKRRVVQREKRAVGIgalflgflgaa  526
             IFRPGGGDMRDNWRSELYKYKVVKIEPLGVAPTKAKRRVVQREKRAVGIGALFLGFLGAA
Sbjct  5182  IFRPGGGDMRDNWRSELYKYKVVKIEPLGVAPTKAKRRVVQREKRAVGIGALFLGFLGAA  5003

Query  527   gstmgaasmtLTVQARQLLSGIVQQQNNLLRAIEAQQHLLQLTVWGIKQLQARILAVERY  586
             GSTMGAASMTLTVQARQLLSGIVQQQNNLLRAIEAQQHLLQLTVWGIKQLQARILAVERY
Sbjct  5002  GSTMGAASMTLTVQARQLLSGIVQQQNNLLRAIEAQQHLLQLTVWGIKQLQARILAVERY  4823

Query  587   LKDQQLLGIWGCSGKLICTTAVPWNASWSNKSLEQIWNHTTWMEWDREINNYTSLIHSLI  646
             LKDQQLLGIWGCSGKLICTTAVPWNASWSNKSLEQIWNHTTWMEWDREINNYTSLIHSLI
Sbjct  4822  LKDQQLLGIWGCSGKLICTTAVPWNASWSNKSLEQIWNHTTWMEWDREINNYTSLIHSLI  4643

Query  647   eesqnqqekneqellelDKWASLWNWFNITNWLWYIKLFIMIVGGLVGLRIVFAVLSIVN  706
             EESQNQQEKNEQELLELDKWASLWNWFNITNWLWYIKLFIMIVGGLVGLRIVFAVLSIVN
Sbjct  4642  EESQNQQEKNEQELLELDKWASLWNWFNITNWLWYIKLFIMIVGGLVGLRIVFAVLSIVN  4463

Query  707   RVRQGYSPLSFQTHLPTprgpdrpegieeeggerdrdrSIRLVNGS  752
             RVRQGYSPLSFQTHLPTPRGPDRPEGIEEEGGERDRDRSIRLVNGS
Sbjct  4462  RVRQGYSPLSFQTHLPTPRGPDRPEGIEEEGGERDRDRSIRLVNGS  4325
```

```
>Saccharothrix coeruleofusca strain JCM 3313 sequence45, whole genome shotgun sequence
Sequence ID: NZ_BMRG01000045.1 Length: 5775
Range 1: 4325 to 5182

Score:446 bits(1147), Expect:2e-131,
Method:Compositional matrix adjust.,
Identities:286/286(100%), Positives:286/286(100%), Gaps:0/286(0%)

Query  467   IFRPGGGDMRDNWRSELYKYKVVKIEPLGVAPTKAKRRVVQREKRAVGIgalflgflgaa  526
             IFRPGGGDMRDNWRSELYKYKVVKIEPLGVAPTKAKRRVVQREKRAVGIGALFLGFLGAA
Sbjct  5182  IFRPGGGDMRDNWRSELYKYKVVKIEPLGVAPTKAKRRVVQREKRAVGIGALFLGFLGAA  5003

Query  527   gstmgaasmtLTVQARQLLSGIVQQQNNLLRAIEAQQHLLQLTVWGIKQLQARILAVERY  586
             GSTMGAASMTLTVQARQLLSGIVQQQNNLLRAIEAQQHLLQLTVWGIKQLQARILAVERY
Sbjct  5002  GSTMGAASMTLTVQARQLLSGIVQQQNNLLRAIEAQQHLLQLTVWGIKQLQARILAVERY  4823

Query  587   LKDQQLLGIWGCSGKLICTTAVPWNASWSNKSLEQIWNHTTWMEWDREINNYTSLIHSLI  646
             LKDQQLLGIWGCSGKLICTTAVPWNASWSNKSLEQIWNHTTWMEWDREINNYTSLIHSLI
Sbjct  4822  LKDQQLLGIWGCSGKLICTTAVPWNASWSNKSLEQIWNHTTWMEWDREINNYTSLIHSLI  4643

Query  647   eesqnqqekneqellelDKWASLWNWFNITNWLWYIKLFIMIVGGLVGLRIVFAVLSIVN  706
             EESQNQQEKNEQELLELDKWASLWNWFNITNWLWYIKLFIMIVGGLVGLRIVFAVLSIVN
Sbjct  4642  EESQNQQEKNEQELLELDKWASLWNWFNITNWLWYIKLFIMIVGGLVGLRIVFAVLSIVN  4463

Query  707   RVRQGYSPLSFQTHLPTprgpdrpegieeeggerdrdrSIRLVNGS  752
             RVRQGYSPLSFQTHLPTPRGPDRPEGIEEEGGERDRDRSIRLVNGS
Sbjct  4462  RVRQGYSPLSFQTHLPTPRGPDRPEGIEEEGGERDRDRSIRLVNGS  4325
```

```
>Actinoplanes ianthinogenes strain JCM 3249 sequence037, whole genome shotgun sequence
Sequence ID: NZ_BMQZ01000037.1 Length: 5815
Range 1: 4345 to 5202

Score:446 bits(1147), Expect:2e-131,
Method:Compositional matrix adjust.,
Identities:286/286(100%), Positives:286/286(100%), Gaps:0/286(0%)

Query  467   IFRPGGGDMRDNWRSELYKYKVVKIEPLGVAPTKAKRRVVQREKRAVGIgalflgflgaa  526
             IFRPGGGDMRDNWRSELYKYKVVKIEPLGVAPTKAKRRVVQREKRAVGIGALFLGFLGAA
Sbjct  5202  IFRPGGGDMRDNWRSELYKYKVVKIEPLGVAPTKAKRRVVQREKRAVGIGALFLGFLGAA  5023

Query  527   gstmgaasmtLTVQARQLLSGIVQQQNNLLRAIEAQQHLLQLTVWGIKQLQARILAVERY  586
             GSTMGAASMTLTVQARQLLSGIVQQQNNLLRAIEAQQHLLQLTVWGIKQLQARILAVERY
Sbjct  5022  GSTMGAASMTLTVQARQLLSGIVQQQNNLLRAIEAQQHLLQLTVWGIKQLQARILAVERY  4843

Query  587   LKDQQLLGIWGCSGKLICTTAVPWNASWSNKSLEQIWNHTTWMEWDREINNYTSLIHSLI  646
             LKDQQLLGIWGCSGKLICTTAVPWNASWSNKSLEQIWNHTTWMEWDREINNYTSLIHSLI
Sbjct  4842  LKDQQLLGIWGCSGKLICTTAVPWNASWSNKSLEQIWNHTTWMEWDREINNYTSLIHSLI  4663

Query  647   eesqnqqekneqellelDKWASLWNWFNITNWLWYIKLFIMIVGGLVGLRIVFAVLSIVN  706
             EESQNQQEKNEQELLELDKWASLWNWFNITNWLWYIKLFIMIVGGLVGLRIVFAVLSIVN
Sbjct  4662  EESQNQQEKNEQELLELDKWASLWNWFNITNWLWYIKLFIMIVGGLVGLRIVFAVLSIVN  4483

Query  707   RVRQGYSPLSFQTHLPTprgpdrpegieeeggerdrdrSIRLVNGS  752
             RVRQGYSPLSFQTHLPTPRGPDRPEGIEEEGGERDRDRSIRLVNGS
Sbjct  4482  RVRQGYSPLSFQTHLPTPRGPDRPEGIEEEGGERDRDRSIRLVNGS  4345
```

```
>Sinomonas sp. KCTC 49339 sequence28, whole genome shotgun sequence
Sequence ID: NZ_BNCM01000028.1 Length: 5849
Range 1: 4316 to 5173

Score:446 bits(1147), Expect:2e-131,
Method:Compositional matrix adjust.,
Identities:286/286(100%), Positives:286/286(100%), Gaps:0/286(0%)

Query  467   IFRPGGGDMRDNWRSELYKYKVVKIEPLGVAPTKAKRRVVQREKRAVGIgalflgflgaa  526
             IFRPGGGDMRDNWRSELYKYKVVKIEPLGVAPTKAKRRVVQREKRAVGIGALFLGFLGAA
Sbjct  5173  IFRPGGGDMRDNWRSELYKYKVVKIEPLGVAPTKAKRRVVQREKRAVGIGALFLGFLGAA  4994

Query  527   gstmgaasmtLTVQARQLLSGIVQQQNNLLRAIEAQQHLLQLTVWGIKQLQARILAVERY  586
             GSTMGAASMTLTVQARQLLSGIVQQQNNLLRAIEAQQHLLQLTVWGIKQLQARILAVERY
Sbjct  4993  GSTMGAASMTLTVQARQLLSGIVQQQNNLLRAIEAQQHLLQLTVWGIKQLQARILAVERY  4814

Query  587   LKDQQLLGIWGCSGKLICTTAVPWNASWSNKSLEQIWNHTTWMEWDREINNYTSLIHSLI  646
             LKDQQLLGIWGCSGKLICTTAVPWNASWSNKSLEQIWNHTTWMEWDREINNYTSLIHSLI
Sbjct  4813  LKDQQLLGIWGCSGKLICTTAVPWNASWSNKSLEQIWNHTTWMEWDREINNYTSLIHSLI  4634

Query  647   eesqnqqekneqellelDKWASLWNWFNITNWLWYIKLFIMIVGGLVGLRIVFAVLSIVN  706
             EESQNQQEKNEQELLELDKWASLWNWFNITNWLWYIKLFIMIVGGLVGLRIVFAVLSIVN
Sbjct  4633  EESQNQQEKNEQELLELDKWASLWNWFNITNWLWYIKLFIMIVGGLVGLRIVFAVLSIVN  4454

Query  707   RVRQGYSPLSFQTHLPTprgpdrpegieeeggerdrdrSIRLVNGS  752
             RVRQGYSPLSFQTHLPTPRGPDRPEGIEEEGGERDRDRSIRLVNGS
Sbjct  4453  RVRQGYSPLSFQTHLPTPRGPDRPEGIEEEGGERDRDRSIRLVNGS  4316



>Saccharothrix mutabilis subsp. mutabilis strain JCM 3380 sequence46, whole genome shotgun sequence
Sequence ID: NZ_BAAABU010000046.1 Length: 5899
Range 1: 614 to 1471

Score:446 bits(1147), Expect:2e-131,
Method:Compositional matrix adjust.,
Identities:286/286(100%), Positives:286/286(100%), Gaps:0/286(0%)

Query  467   IFRPGGGDMRDNWRSELYKYKVVKIEPLGVAPTKAKRRVVQREKRAVGIgalflgflgaa  526
             IFRPGGGDMRDNWRSELYKYKVVKIEPLGVAPTKAKRRVVQREKRAVGIGALFLGFLGAA
Sbjct  614   IFRPGGGDMRDNWRSELYKYKVVKIEPLGVAPTKAKRRVVQREKRAVGIGALFLGFLGAA  793

Query  527   gstmgaasmtLTVQARQLLSGIVQQQNNLLRAIEAQQHLLQLTVWGIKQLQARILAVERY  586
             GSTMGAASMTLTVQARQLLSGIVQQQNNLLRAIEAQQHLLQLTVWGIKQLQARILAVERY
Sbjct  794   GSTMGAASMTLTVQARQLLSGIVQQQNNLLRAIEAQQHLLQLTVWGIKQLQARILAVERY  973

Query  587   LKDQQLLGIWGCSGKLICTTAVPWNASWSNKSLEQIWNHTTWMEWDREINNYTSLIHSLI  646
             LKDQQLLGIWGCSGKLICTTAVPWNASWSNKSLEQIWNHTTWMEWDREINNYTSLIHSLI
Sbjct  974   LKDQQLLGIWGCSGKLICTTAVPWNASWSNKSLEQIWNHTTWMEWDREINNYTSLIHSLI  1153

Query  647   eesqnqqekneqellelDKWASLWNWFNITNWLWYIKLFIMIVGGLVGLRIVFAVLSIVN  706
             EESQNQQEKNEQELLELDKWASLWNWFNITNWLWYIKLFIMIVGGLVGLRIVFAVLSIVN
Sbjct  1154  EESQNQQEKNEQELLELDKWASLWNWFNITNWLWYIKLFIMIVGGLVGLRIVFAVLSIVN  1333

Query  707   RVRQGYSPLSFQTHLPTprgpdrpegieeeggerdrdrSIRLVNGS  752
             RVRQGYSPLSFQTHLPTPRGPDRPEGIEEEGGERDRDRSIRLVNGS
Sbjct  1334  RVRQGYSPLSFQTHLPTPRGPDRPEGIEEEGGERDRDRSIRLVNGS  1471



>Corynebacterium glutamicum strain 39 Scaffolds_32_9166, whole genome shotgun sequence
Sequence ID: NZ_JACEQW010000032.1 Length: 8862
Range 1: 4105 to 4953

Score:441 bits(1134), Expect:5e-129,
Method:Compositional matrix adjust.,
Identities:273/286(95%), Positives:276/286(96%), Gaps:3/286(1%)

Query  467   IFRPGGGDMRDNWRSELYKYKVVKIEPLGVAPTKAKRRVVQREKRAVGIgalflgflgaa  526
             IFRPGGGDMRDNWRSELYKYKVVKIEPLGVAPTKAKRRVVQREKRAVGIGALFLGFLGAA
Sbjct  4953  IFRPGGGDMRDNWRSELYKYKVVKIEPLGVAPTKAKRRVVQREKRAVGIGALFLGFLGAA  4774

Query  527   gstmgaasmtLTVQARQLLSGIVQQQNNLLRAIEAQQHLLQLTVWGIKQLQARILAVERY  586
             GSTMGAASMTLTVQARQLLSGIVQQQNNLLRAIEAQQHLLQLTVWGIKQLQARILAVERY
Sbjct  4773  GSTMGAASMTLTVQARQLLSGIVQQQNNLLRAIEAQQHLLQLTVWGIKQLQARILAVERY  4594

Query  587   LKDQQLLGIWGCSGKLICTTAVPWNASWSNKSLEQIWNHTTWMEWDREINNYTSLIHSLI  646
             LKDQQLLGIWGCSGKLICTTAVPWNASWSNKSLEQIWNHTTWMEWDREINNYTSLIHSLI
Sbjct  4593  LKDQQLLGIWGCSGKLICTTAVPWNASWSNKSLEQIWNHTTWMEWDREINNYTSLIHSLI  4414

Query  647   eesqnqqekneqellelDKWASLWNWFNITNWLWYIKLFIMIVGGLVGLRIVFAVLSIVN  706
                +  Q + E+   ELDKWASLWNWFNITNWLWYIKLFIMIVGGLVGLRIVFAVLSIVN
Sbjct  4413  ---EESQNQQEKNEQELDKWASLWNWFNITNWLWYIKLFIMIVGGLVGLRIVFAVLSIVN  4243

Query  707   RVRQGYSPLSFQTHLPTprgpdrpegieeeggerdrdrSIRLVNGS  752
             RVRQGYSPLSFQTHLPTPRGPDRPEGIEEEGGERDRDRSIRLVNGS
Sbjct  4242  RVRQGYSPLSFQTHLPTPRGPDRPEGIEEEGGERDRDRSIRLVNGS  4105



>Corynebacterium glutamicum strain 37 Scaffolds_33_8928, whole genome shotgun sequence
Sequence ID: NZ_JACENO010000033.1 Length: 8928
Range 1: 4079 to 4927

Score:441 bits(1134), Expect:5e-129,
Method:Compositional matrix adjust.,
Identities:273/286(95%), Positives:276/286(96%), Gaps:3/286(1%)

Query  467   IFRPGGGDMRDNWRSELYKYKVVKIEPLGVAPTKAKRRVVQREKRAVGIgalflgflgaa  526
             IFRPGGGDMRDNWRSELYKYKVVKIEPLGVAPTKAKRRVVQREKRAVGIGALFLGFLGAA
Sbjct  4927  IFRPGGGDMRDNWRSELYKYKVVKIEPLGVAPTKAKRRVVQREKRAVGIGALFLGFLGAA  4748

Query  527   gstmgaasmtLTVQARQLLSGIVQQQNNLLRAIEAQQHLLQLTVWGIKQLQARILAVERY  586
             GSTMGAASMTLTVQARQLLSGIVQQQNNLLRAIEAQQHLLQLTVWGIKQLQARILAVERY
Sbjct  4747  GSTMGAASMTLTVQARQLLSGIVQQQNNLLRAIEAQQHLLQLTVWGIKQLQARILAVERY  4568

Query  587   LKDQQLLGIWGCSGKLICTTAVPWNASWSNKSLEQIWNHTTWMEWDREINNYTSLIHSLI  646
             LKDQQLLGIWGCSGKLICTTAVPWNASWSNKSLEQIWNHTTWMEWDREINNYTSLIHSLI
Sbjct  4567  LKDQQLLGIWGCSGKLICTTAVPWNASWSNKSLEQIWNHTTWMEWDREINNYTSLIHSLI  4388

Query  647   eesqnqqekneqellelDKWASLWNWFNITNWLWYIKLFIMIVGGLVGLRIVFAVLSIVN  706
                +  Q + E+   ELDKWASLWNWFNITNWLWYIKLFIMIVGGLVGLRIVFAVLSIVN
Sbjct  4387  ---EESQNQQEKNEQELDKWASLWNWFNITNWLWYIKLFIMIVGGLVGLRIVFAVLSIVN  4217

Query  707   RVRQGYSPLSFQTHLPTprgpdrpegieeeggerdrdrSIRLVNGS  752
             RVRQGYSPLSFQTHLPTPRGPDRPEGIEEEGGERDRDRSIRLVNGS
Sbjct  4216  RVRQGYSPLSFQTHLPTPRGPDRPEGIEEEGGERDRDRSIRLVNGS  4079
```

```
>Methyloversatilis universalis FAM5 contig00006, whole genome shotgun sequence
Sequence ID: NZ_AFHG01000004.1 Length: 649
Range 1: 2 to 649

Score:380 bits(976), Expect:1e-122,
Method:Compositional matrix adjust.,
Identities:181/219(83%), Positives:193/219(88%), Gaps:8/219(3%)

Query  23   GMLMICSATEKLWVTVYYGVPVWKEATTTLFCASDAKAYDTEVHNVWATHACVPTDPNPQ  82
            GMLMIC+ATEKLWVTVYYGVPVWKEATTTLFCASD KAYDTEVHNVWATHACVPTDPNPQ
Sbjct  2    GMLMICNATEKLWVTVYYGVPVWKEATTTLFCASDTKAYDTEVHNVWATHACVPTDPNPQ  181

Query  83   EVVLVNVTENFNMWKNDMVEQMHEDIISLWDQSLKPCVKLTPLCVSLKCTDLKNDTNT--  140
            EVVL NVTENFNMW+NDMVEQMHEDIISLWDQSLKPCVKLTPLCV+L C D
Sbjct  182  EVVLGNVTENFNMWRNDMVEQMHEDIISLWDQSLKPCVKLTPLCVTLNCNDYNGTNTNGT  361

Query  141  ---NSSSGRMIMEKGEIKNCSFNISTSIRGKVQKEYAFFYKLDIIPIDNDTTSYKLTSCN  197
               NSS+  MI   GEIKNCSFNI+T+IR K++KEYA FYKLD++PIDND TSY+L SCN
Sbjct  362  NTKNSSNVEMI---GEIKNCSFNITTNIRDKIKKEYALFYKLDVVPIDNDNTSYRLISCN  532

Query  198  TSVITQACPKVSFEPIPIHYCAPAGFAILKCNNKTFNGT  236
            TSVITQACPK+SFEPIPIHYCAPAGFAILKCNNK FNGT
Sbjct  533  TSVITQACPKISFEPIPIHYCAPAGFAILKCNNKKFNGT  649



>Klebsiella pneumoniae strain P7701 NODE_62_length_570_cov_92.6095, whole genome shotgun sequence
Sequence ID: NZ_JACAWM010000060.1 Length: 570
Range 1: 3 to 569

Score:378 bits(971), Expect:3e-122,
Method:Compositional matrix adjust.,
Identities:182/189(96%), Positives:184/189(97%), Gaps:0/189(0%)

Query  248  THGIRPVVSTQLLLNGSLAEEEVVIRSVNFTDNAKTIIVQLNTSVEINCTRPNNNTRKRI  307
            THGIRPV STQLLLNGSLAEE+VVIRS NFTDNAKTIIVQLNTSVEINCTRPNNNTRK I
Sbjct  3    THGIRPVASTQLLLNGSLAEEDVVIRSANFTDNAKTIIVQLNTSVEINCTRPNNNTRKSI  182

Query  308  RIQRGPGRAFVTIGKIGNMRQAHCNISRAKWNNTLKQIASKLREQFGNNKTIIFKQSSGG  367
            RIQRGPGRAFVTIGKIGNMRQAHCNISRAKWN TLKQIASKLREQFGNNKTIIFKQSSGG
Sbjct  183  RIQRGPGRAFVTIGKIGNMRQAHCNISRAKWNATLKQIASKLREQFGNNKTIIFKQSSGG  362

Query  368  DPEIVTHSFNCGGEFFYCNSTQLFNSTWFNSTWSTEGSNNTEGSDTITLPCRIKQIINMW  427
            DPEIVTHSFNCGGEFFYCNSTQLFNSTWFNSTWSTEGSNNTEGSDTITLPCRIKQ INMW
Sbjct  363  DPEIVTHSFNCGGEFFYCNSTQLFNSTWFNSTWSTEGSNNTEGSDTITLPCRIKQFINMW  542

Query  428  QKVGKAMYA  436
            Q+VGKAMYA
Sbjct  543  QEVGKAMYA  569



>Escherichia coli strain NBE0004 contig00110, whole genome shotgun sequence
Sequence ID: NZ_JARBDH010000110.1 Length: 1390
Range 1: 294 to 1388

Score:388 bits(996), Expect:8e-122,
Method:Compositional matrix adjust.,
Identities:258/365(71%), Positives:300/365(82%), Gaps:7/365(1%)

Query  498   PTKAKRRVVQREKRAVGIgalflgflgaagstmgaasmtLTVQARQLLSGIVQQQNNLLR  557
             PT AKRRVV+REKRAVGIGA+FL FLG AGST+GAAS+ LTVQ RQLLSGIVQQQ+NLL
Sbjct  1388  PTAAKRRVVEREKRAVGIGAVFLRFLGTAGSTIGAASIALTVQTRQLLSGIVQQQSNLLM  1209

Query  558   AIEAQQHLLQLTVWGIKQLQARILAVERYLKDQQLLGIWGCSGKLICTTAVPWNASWSNK  617
             AIEAQQHLLQLTV GIKQLQ R+LA++RYLKDQQLLG+WGCSGKLICTTAVP N+S SNK
Sbjct  1208  AIEAQQHLLQLTV*GIKQLQTRVLALKRYLKDQQLLGMWGCSGKLICTTAVP*NSS*SNK  1029

Query  618   SLEQIWNHTTWMEWDREINNYTSLIHSLIeesqnqqekneqellelDKWASLWNWFNITN  677
             S ++I ++ T M+ DREI+NYT  I+ L+E+SQ QQE+NE++LL LD   +LWN F IT
Sbjct  1028  SRDEI*DNMT*MQ*DREISNYTDTIYRLLEDSQIQQERNEKDLLALDS*NNLWN*FGITK  849

Query  678   WLWYIKLFIMIVGGLVGLRIVFAVLSIVNRVRQGYSPLSFQTHLPTprgpdrpegieeeg  737
             WLWYIK+FIMIVGGL+ LRI+FAV+SIVNR RQ YSPLSFQT  P PRGPDR   IEEEG
Sbjct  848   WLWYIKIFIMIVGGLISLRIIFAVISIVNRFRQRYSPLSFQTLTPNPRGPDRLGRIEEEG  669

Query  738   gerdrdrSIRLVNGSLALIWDDLRSLCLFSYHRLRDLLLIVTRIVELLGR-------RGW  790
              E+DR+RSIRLVNG LAL  DDL++LCLFSYHRLRD +L+  R+VELLGR       R
Sbjct  668   REQDRNRSIRLVNGFLALA*DDLQNLCLFSYHRLRDFILVAARVVELLGRSSLKGLQRR*  489

Query  791   EALKYWWNLLQYWSQELKNSAVSLLNATAIAVAEGTDRVIEVVQGACRAIRHIPRRIRQG  850
             EALKY  +L+QY   ELK SA+SLL+  AIAV EGTDR+I +VQG CRA+R+IPRRIRQG
Sbjct  488   EALKYLGSLVQY*GLELKGSAISLLDTIAIAVGEGTDRIISLVQGICRAVRNIPRRIRQG  309

Query  851   LERIL  855
              E  L
Sbjct  308   FEAAL  294



>Escherichia coli strain NBE0004 contig00131, whole genome shotgun sequence
Sequence ID: NZ_JARBDH010000131.1 Length: 847
Range 1: 2 to 841

Score:375 bits(962), Expect:2e-119,
Method:Compositional matrix adjust.,
Identities:194/290(67%), Positives:219/290(75%), Gaps:22/290(7%)

Query  15   WRWGTMLLGMLMICSATEKLWVTVYYGVPVWKEATTTLFCASDAKAYDTEVHNVWATHAC  74
            W  G +   MLMIC+      VTVYY VPVWKEA  TLFCASDAKA++ EVHNV ATHAC
Sbjct  841  WI*GVLGF*MLMICNKGGDS*VTVYYEVPVWKEAKATLFCASDAKAHEREVHNV*ATHAC  662

Query  75   VPTDPNPQEVVLVNVTENFNMWKNDMVEQMHEDIISLWDQSLKPCVKLTPLCVSLKC---  131
            VPTDPNPQE+VL NVTENFNMWKNDMV+QMHED+ISL DQSLKPCVKLTPLCV+L+C
Sbjct  661  VPTDPNPQEMVLENVTENFNMWKNDMVDQMHEDVISL*DQSLKPCVKLTPLCVTLECNNY  482

Query  132  TDLKNDTNTNSSSGRMIMEKGEIKNCSFNISTSIRGKVQKEYAFFYKLDIIPIDNDTT--  189
            T  KN T+          E  E+K C FN ST +  K QK  A FY LDI P++  ++
Sbjct  481  TTYKNYTD----------ETKELKRCFFNASTEVEDKKQKVSALFYSLDIRPLNKSSSRN  332

Query  190  -------SYKLTSCNTSVITQACPKVSFEPIPIHYCAPAGFAILKCNNKTFNGTGPCTNV  242
                   +Y LTSCNTSVI QACPKV+F+PIPIHYCAPAG+AILKCNN+TFN TGPC NV
Sbjct  331  SSINSNGTYVLTSCNTSVIKQACPKVTFDPIPIHYCAPAGYAILKCNNETFNRTGPCNNV  152

Query  243  STVQCTHGIRPVVSTQLLLNGSLAEEEVVIRSVNFTDNAKTIIVQLNTSV  292
            STVQCTH I+PVVSTQLLLNGSLA+ ++VIRS N  DNAKTIIV LN SV
Sbjct  151  STVQCTHRIKPVVSTQLLLNGSLAKGDIVIRSENLADNAKTIIVHLNESV  2
```

>Pseudomonas aeruginosa strain VET-42 NODE_89_length_1008_cov_1.38346_ID_177, whole genome shotgun sequence
Sequence ID: NZ_RACX01000220.1 Length: 963
Range 1: 207 to 956

Score:357 bits(916), Expect:4e-112,
Method:Compositional matrix adjust.,
Identities:168/251(67%), Positives:201/251(80%), Gaps:6/251(2%)

```
Query  1    MRVKEKYQHLWRWGWRWGTMLLGMLMICSATEKLWVTVYYGVPVWKEATTTLFCASDAKA  60
            MRV+   Q  W   W+WGT++LG+++IC+A++ LWVTVYYGVP W++A T LFCASDAKA
Sbjct  207  MRVRGT-QMNWLNLWKWGTLMLGLVIICNASKDLWVTVYYGVPAWEDADTILFCASDAKA  383

Query  61   YDTEVHNVWATHACVPTDPNPQEVVLVNVTENFNMWKNDMVEQMHEDIISLWDQSLKPCV  120
            + TE HN+WATHACVPTDPNPQE+ L NVTENFNMWKN+MVEQMHEDIISLWD+SLK CV
Sbjct  384  HSTESHNIWATHACVPTDPNPQEIPLENVTENFNMWKNNMVEQMHEDIISLWDESLKRCV  563

Query  121  KLTPLCVSLKCTDL-KNDTNTNSSSGRMIMEKGEIKNCSFNISTSIRGKVQKEYAFFYKL  179
            KLTPLCV+L C ++ K D +TN      ++ KGEI NCSFNI+T IR + + EYA FYKL
Sbjct  564  KLTPLCVTLYCENVNKTDNSTNGEDNSTVINKGEITNCSFNITTEIRDRKKTEYALFYKL  743

Query  180  DIIPID----NDTTSYKLTSCNTSVITQACPKVSFEPIPIHYCAPAGFAILKCNNKTFNG  235
            D+IPI+    +  + Y L +CN + I QACPKVSFEPIPIHYCAPAGFAILKC +K FNG
Sbjct  744  DVIPIEGRDRSSNSKYMLINCNATTIKQACPKVSFEPIPIHYCAPAGFAILKCRDKNFNG  923

Query  236  TGPCTNVSTVQ  246
            TGPC NVSTVQ
Sbjct  924  TGPCKNVSTVQ  956
```

>Escherichia coli strain EN5143 contig00088, whole genome shotgun sequence
Sequence ID: NZ_JAPMNI010000088.1 Length: 833
Range 1: 1 to 831

Score:319 bits(817), Expect:4e-98,
Method:Compositional matrix adjust.,
Identities:197/281(70%), Positives:230/281(81%), Gaps:4/281(1%)

```
Query  391  FNSTWFNSTWSTEGSNNTEGSDTITLPCRIKQIINMWQKVGKAMYAPPISGQIRCSSNIT  450
            FNST +N T      N T     IT+PCRIKQIINMWQKV +AMYAPPI+G I C SNIT
Sbjct  831  FNSTLYNETID----NGTHLISNITIPCRIKQIINMWQKVRRAMYAPPIAGDITCRSNIT  664

Query  451  GLLLTRDGGNSNNESEIFRPGGGDMRDNWRSELYKYKVVKIEPLGVAPTKAKRRVVQREK  510
            GLLL RD G  +NE+EIFRP   DM+DN RS+LYKYKVV+I+PL +APT+AKRRVV+REK
Sbjct  663  GLLLLRDRGVGDNETEIFRPRERDMKDN*RSKLYKYKVVEIKPLRIAPTRAKRRVVKREK  484

Query  511  RAVGIgalflgflgaagstmgaasmtLTVQARQLLSGIVQQQNNLLRAIEAQQHLLQLTV  570
            RAV I A+FL FLGAA ST+GAAS+TLTVQARQLLSGIVQQQ+NLL+AIEAQQHLLQLTV
Sbjct  483  RAVRIRAVFLKFLGAARSTIGAASITLTVQARQLLSGIVQQQSNLLKAIEAQQHLLQLTV  304

Query  571  WGIKQLQARILAVERYLKDQQLLGIWGCSGKLICTTAVPWNASWSNKSLEQIWNHTTWME  630
             GIKQLQ R+LA+ERYLKDQQLL I GCS KLIC TA+PWN+S SNK+ +QI N+ T M+
Sbjct  303  *GIKQLQTRVLAIERYLKDQQLLRI*GCSRKLICPTAIPWNSS*SNKTYDQI*NNMT*MQ  124

Query  631  WDREINNYTSLIHSLIeesqnqqekneqellelDKWASLWN  671
             D+EI+NYT  I+ L+E+SQ QQEKNE++LL LD  ASL N
Sbjct  123  *DKEISNYTYTIYRLLEDSQTQQEKNEKDLLALDS*ASL*N  1
```

>Escherichia coli strain EN5192 contig00135, whole genome shotgun sequence
Sequence ID: NZ_JAPMNL010000135.1 Length: 1341
Range 1: 243 to 965

Score:269 bits(688), Expect:6e-77,
Method:Compositional matrix adjust.,
Identities:166/252(66%), Positives:196/252(77%), Gaps:18/252(7%)

```
Query  611  NASWSNKSLEQIWNHTTWMEWDREINNYTSLIHSLIeesqnqqekneqellelDKWASLW  670
            ++SWSNKS  +IW++ TWM+WDREI+NYT  I+ L+E+SQ QQEKNE++LL LD W +LW
Sbjct  965  HSSWSNKSRSEIWDNMTWMQWDREISNYTDTIYRLLEDSQIQQEKNEKDLLALDSWNNLW  786

Query  671  NWFNITNWLWYIKLFIMIVGGLVGLRIVFAVLSIVNRVRQGYSPLSFQTHLPTprgpdrp  730
            NWF IT WLWYIK+FIMIVGGL+GLRI+FAV+SIVNR RQGYSPLSFQT  P P GPDR
Sbjct  785  NWFGITKWLWYIKIFIMIVGGLIGLRIIFAVISIVNRFRQGYSPLSFQTLTPNPGGPDRL  606

Query  731  egieeeggerdrdrSIRLVNGSLALIWDDLRSLCLFSYHRLRDLLLIVTRIVELLGR---  787
              IEEEGGE+ R+RSIRLVNG LAL W+DLR+LCLFSYHRLRD +L+  R+VELLGR
Sbjct  605  GRIEEEGGEQGRNRSIRLVNGFLALAWEDLRNLCLFSYHRLRDFILVTARVVELLGRNSL  426

Query  788  ----RGWEALKYWWNLLQYWSQELKNSAVSLLNATAIAVAEGTDRVIEVVQGACRAIRHI  843
                RGWEALKY  +L+QYW            +  AIAVAEGTDR+I + QG CRAI +I
Sbjct  425  KGLQRGWEALKYLGSLVQYW-----------FDTIAIAVAEGTDRIINLAQGICRAICNI  279

Query  844  PRRIRQGLERIL  855
            PRRIRQG E  L
Sbjct  278  PRRIRQGFEAAL  243
```

Range 2: 958 to 1335

Score:162 bits(409), Expect:3e-38,
Method:Compositional matrix adjust.,
Identities:83/135(61%), Positives:95/135(70%), Gaps:11/135(8%)

```
Query  337   KWNNTLKQIASKLREQFGNNKTIIFKQSSGGDPEIVTHSFNCGGEFFYCNSTQLFNSTWF  396
             KWN TL++++ KL E F N KTI F  SSGGD EI THSFNC GEFFYCN++ LFN T+
Sbjct  1335  KWNETLQRVSKKLAEHFPN-KTIKFASSSGGDLEITTHSFNCRGEFFYCNTSGLFNGTYM  1159

Query  397   NSTWSTEGSNNTEG--SDTITLPCRIKQIINMWQKVGKAMYAPPISGQIRCSSNITGLLL  454
                     SN TE   S  IT+PCRIKQIINMWQ+VG+AMYAPPI G I C SNITGLLL
Sbjct  1158  --------SNGTESNSSSIITIPCRIKQIINMWQEVGRAMYAPPIEGNITCKSNITGLLL  1003

Query  455   TRDGGNSNNESEIFR  469
              RDGG   N +E  +
Sbjct  1002  VRDGGTEANTTETLQ  958
```

>Klebsiella pneumoniae strain EuSCAPE_IL001, whole genome shotgun sequence
Sequence ID: NZ_UKJN02000084.1 Length: 599
Range 1: 32 to 598

Score:243 bits(620), Expect:2e-70,
Method:Compositional matrix adjust.,
Identities:139/189(74%), Positives:155/189(82%), Gaps:7/189(3%)

```
Query  664  DKWASLWNWFNITNWLWYIKLFIMIVGGLVGLRIVFAVLSIVNRVRQGYSPLSFQTHLPT  723
            DKW  LW+WF+IT WLWYI++FIMIVGGL+GLRI+FAV SIV RVRQGYSPLSFQT +P
Sbjct  32   DKWNDLWSWFSITQWLWYIRIFIMIVGGLIGLRIIFAVFSIVKRVRQGYSPLSFQTLIPN  211

Query  724  prgpdrpegieeeggerdrdrSIRLVNGSLALIWDDLRSLCLFSYHRLRDLLLIVTRIVE  783
            PR PDR  GIEEEGGE+DRDRSIRLVNG LAL WDDLRSLCLF YHRLRD +L+  R  E
Sbjct  212  PRDPDRHGGIEEEGGEQDRDRSIRLVNGFLALFWDDLRSLCLFCYHRLRDFILVTARAAE  391

Query  784  LLGR-------RGWEALKYWWNLLQYWSQELKNSAVSLLNATAIAVAEGTDRVIEVVQGA  836
            LLGR       RGWE LKY  NL+QYW  ELK SA+SLL+ TAIAVAEGTDR+IE+VQ
Sbjct  392  LLGRSSLRGLQRGWEILKYLGNLVQYWGLELKKSAISLLDTTAIAVAEGTDRIIEIVQAT  571

Query  837  CRAIRHIPR  845
            CRAIR+IPR
Sbjct  572  CRAIRNIPR  598
```

>Neisseria gonorrhoeae strain GC-206 NODE_120_length_488_cov_1.24711_ID_20412, whole genome shotgun sequence
Sequence ID: NZ_SAWN01000120.1 Length: 488
Range 1: 2 to 487

Score:240 bits(612), Expect:6e-70,
Method:Compositional matrix adjust.,
Identities:118/166(71%), Positives:134/166(80%), Gaps:5/166(3%)

```
Query  197  NTSVITQACPKVSFEPIPIHYCAPAGFAILKCNNKTFNGTGPCTNVSTVQCTHGIRPVVS  256
            NTS I QACPK++F+PIPIHYCAPAG+AILKCNNKTFNGTGPC NVSTVQCTHGI+PVVS
Sbjct  487  NTSAIAQACPKITFDPIPIHYCAPAGYAILKCNNKTFNGTGPCQNVSTVQCTHGIKPVVS  308

Query  257  TQLLLNGSLAEEEVVIRSVNFTDNAKTIIVQLNTSVEINCTRPNNNTRKRIRIQRGPGRA  316
            TQLLLNGSLAE +++IRS N TD+ KTIIV LN SV I CTRP NNTRK +RI  GPG+A
Sbjct  307  TQLLLNGSLAEGDIIIRSENITDSIKTIIVHLNESVYIACTRPGNNTRKSVRI--GPGQA  134

Query  317  FVTIGK-IGNMRQAHCNISRAKWNNTLKQIASKLREQFGNNKTIIF  361
            F   G  IG++RQAHCNISR  W+ TL ++  KL   F  NKTI F
Sbjct  133  FYATGDVIGDIRQAHCNISRGNWSRTLGKVRDKLAAHF--NKTIEF  2
```

>Klebsiella pneumoniae strain OK8 jcf7180000003506, whole genome shotgun sequence
Sequence ID: NZ_MZXR01000036.1 Length: 577
Range 1: 1 to 549

Score:232 bits(591), Expect:2e-66,
Method:Composition-based stats.,
Identities:130/183(71%), Positives:148/183(80%), Gaps:8/183(4%)

```
Query  681  YIKLFIMIVGGLVGLRIVFAVLSIVNRVRQGYSPLSFQTHLPTprgp-drpegieeegge  739
            YIK+FIMIVGGL+GLRI+FAVLSIV RVRQGYSPLSFQT  PT  G  DR   IEEEGGE
Sbjct  1    YIKIFIMIVGGLIGLRIIFAVLSIVKRVRQGYSPLSFQTLTPTSGGRPDRLGRIEEEGGE  180

Query  740  rdrdrSIRLVNGSLALIWDDLRSLCLFSYHRLRDLLLIVTRIVELLGRR-------GWEA  792
            +D+ RSIRLV+G LAL WDDLR+LCLFSYHRLRD +L+  R+VELLGR       GWEA
Sbjct  181  QDKGRSIRLVSGFLALAWDDLRNLCLFSYHRLRDFILVAARVVELLGRSSLRGLQTGWEA  360

Query  793  LKYWWNLLQYWSQELKNSAVSLLNATAIAVAEGTDRVIEVVQGACRAIRHIPRRIRQGLE  852
            LKY  +L+QYW QELK SA++LL+  AIAVAEGTDR+IE+ QG CRAIR+IPRRIRQG E
Sbjct  361  LKYLGSLVQYWGQELKKSAINLLDTVAIAVAEGTDRIIELAQGICRAIRNIPRRIRQGFE  540

Query  853  RIL  855
              L
Sbjct  541  AAL  549
```

>Klebsiella pneumoniae strain EuSCAPE_IL006, whole genome shotgun sequence
Sequence ID: NZ_UKJR02000114.1 Length: 560
Range 1: 5 to 556

Score:230 bits(587), Expect:5e-66,
Method:Compositional matrix adjust.,
Identities:136/184(74%), Positives:156/184(84%), Gaps:7/184(3%)

```
Query  664  DKWASLWNWFNITNWLWYIKLFIMIVGGLVGLRIVFAVLSIVNRVRQGYSPLSFQTHLPT  723
            D W SLW+WF+I+NWLWYI++FIMIVGGL+GLRI+FAV+S+VNRVRQGYSPLS QT +P
Sbjct  556  DSWNSLWSWFSISNWLWYIRIFIMIVGGLIGLRIIFAVISVVNRVRQGYSPLSLQTLIPN  377

Query  724  prgpdrpegieeeggerdrdrSIRLVNGSLALIWDDLRSLCLFSYHRLRDLLLIVTRIVE  783
            PRGPDRP GIEEEGGE+DRDRSIRLVNG LA+ WDDLRSLCLFSYHRLRDL L+  R VE
Sbjct  376  PRGPDRPGGIEEEGGEQDRDRSIRLVNGFLAIAWDDLRSLCLFSYHRLRDLTLVTARAVE  197

Query  784  LLGR-------RGWEALKYWWNLLQYWSQELKNSAVSLLNATAIAVAEGTDRVIEVVQGA  836
            LLGR       RGWEALKY  +++QYW  ELK SA+SLL+  AIAVAEGTDR+IEV+Q
Sbjct  196  LLGRSCLKGLQRGWEALKYLGSIVQYWGLELKKSAISLLDTIAIAVAEGTDRIIEVIQRI  17

Query  837  CRAI  840
            CRAI
Sbjct  16   CRAI  5
```

>Aggregatibacter actinomycetemcomitans serotype d str. SA269 contig0497, whole genome shotgun sequence
Sequence ID: NZ_AZTX01000497.1 Length: 504
Range 1: 1 to 486

Score:227 bits(579), Expect:4e-65,
Method:Compositional matrix adjust.,
Identities:141/162(87%), Positives:146/162(90%), Gaps:1/162(0%)

```
Query  679  LWYIKLFIMIVGGLVGLRIVFAVLSIVNRVRQGYSPLSFQTHLPTprgpdrpegieeegg  738
            LWYIK+FIMIVGGL+GLRI+FAVLSIVNRVRQGYSPLSFQTHLP PRG DRPEGIEEEGG
Sbjct  1    LWYIKIFIMIVGGLIGLRIIFAVLSIVNRVRQGYSPLSFQTHLPLPRGADRPEGIEEEGG  180

Query  739  erdrdrSIRLVNGSLALIWDDLRSLCLFSYHRLRDLLLIVTRIVELLGRRGWEALKYWWN  798
            ERDRDRSIRLV GSLALIWDDLRSLCLFSYHRLRDLLLIVTR VELLGRRGWEALKYWWN
Sbjct  181  ERDRDRSIRLVTGSLALIWDDLRSLCLFSYHRLRDLLLIVTRTVELLGRRGWEALKYWWN  360

Query  799  LLQYWSQELKNSAVSLLNATAIAVAE-GTDRVIEVVQGACRA  839
            LL YWSQELKNSAVSLLNATAIAV + G     E VQ   R+
Sbjct  361  LLLYWSQELKNSAVSLLNATAIAVRQYGWSYFHEAVQAVWRS  486
```

>Burkholderia pseudomallei strain 2604STDY5519032 ERS240165SCcontig000080, whole genome shotgun sequence
Sequence ID: NZ_UVID02000080.1 Length: 499
Range 1: 8 to 490

```
Score:225 bits(574), Expect:2e-64,
Method:Compositional matrix adjust.,
Identities:115/172(67%), Positives:132/172(76%), Gaps:12/172(6%)

Query  186  NDTTSYKLTSCNTSVITQACPKVSFEPIPIHYCAPAGFAILKCNNKTFNGTGPCTNVSTV  245
            N++  Y+L +CNTS I QACPKVSF+PIPIHYCAPAG+AILKC +KTFNGTGPC NVSTV
Sbjct  490  NESKEYRLINCNTSAIKQACPKVSFDPIPIHYCAPAGYAILKCKDKTFNGTGPCHNVSTV  311

Query  246  QCTHGIRPVVSTQLLLNGSLAEEEVVIRSVNFTDNAKTIIVQLNTSVEINCTRPNNNTRK  305
            QCTHGI+PVVSTQLLLNGS        RS N TDNAKTIIV LN SV+I CTRP NNTR
Sbjct  310  QCTHGIKPVVSTQLLLNGSF-------RSENLTDNAKTIIVHLNESVDIVCTRPGNNTRT  152

Query  306  RIRIQRGPGRAFVTIGK-IGNMRQAHCNISRAKWNNTLKQIASKLREQFGNN  356
             IRI  GPG+ F   G  IG++RQAHCNIS   W  TL+++ +KL + F NN
Sbjct  151  SIRI--GPGQTFYARGAVIGDIRQAHCNIS--NWYKTLQRVGNKLAKHFPNN  8



>Escherichia coli strain EN5192 contig00156, whole genome shotgun sequence
Sequence ID: NZ_JAPMNL010000156.1 Length: 562
Range 1: 2 to 562

Score:226 bits(575), Expect:3e-64,
Method:Compositional matrix adjust.,
Identities:122/197(62%), Positives:142/197(72%), Gaps:17/197(8%)

Query  279  DNAKTIIVQLNTSVEINCTRPNNNTRKRIRIQRGPGRAF-VTIGKIGNMRQAHCNISRAK  337
            DN KTIIV L+ SVEI CT PNNNTRK IRI  GPG+ F  T G IGN+RQAHCNIS  K
Sbjct  562  DNVKTIIVHLDQSVEIVCTGPNNNTRKSIRI--GPGQTFYATGGIIGNIRQAHCNISEDK  389

Query  338  WNNTLKQIASKLREQFGNNKTIIFKQSSGGDPEIVTHSFNCGGEFFYCNSTQLFNSTWF-  396
            WN TL+++  KL E F N KTI F  SSGGD EI THSFNC GEFFYC++++LFNST+
Sbjct  388  WNETLQRVGKKLVEHFPN-KTIKFAPSSGGDLEITTHSFNCRGEFFYCSTSRLFNSTYMP  212

Query  397  NSTWSTEGSNNTEGSDTITLPCRIKQIINMWQKVGKAMYAPPISGQIRCSSNITGLLLTR  456
            N T S       + + TIT+PC IKQI+NMWQ+VG+AMYAPPI G I C SNITG+LL R
Sbjct  211  NDTKS-------KSNKTITIPCSIKQIVNMWQEVGRAMYAPPIEGNITCRSNITGILLVR  53

Query  457  DGG-----NSNNESEIF  468
            DGG       NN++E F
Sbjct  52   DGGVDSEDPENNKTETF  2



>Klebsiella pneumoniae strain PN085E1IA NODE_127_length_474_cov_225.069_ID_253, whole genome shotgun sequence
Sequence ID: NZ_PDVD01000016.1 Length: 474
Range 1: 15 to 467

Score:224 bits(571), Expect:5e-64,
Method:Compositional matrix adjust.,
Identities:112/155(72%), Positives:125/155(80%), Gaps:5/155(3%)

Query  221  AGFAILKCNNKTFNGTGPCTNVSTVQCTHGIRPVVSTQLLLNGSLAEEEVVIRSVNFTDN  280
            AG+AILKCNNKTFNGTGPC NVSTVQCTHGI+PVVSTQLLLNGSLAEEE++IRS N T+N
Sbjct  467  AGYAILKCNNKTFNGTGPCNNVSTVQCTHGIKPVVSTQLLLNGSLAEEEIIIRSENLTNN  288

Query  281  AKTIIVQLNTSVEINCTRPNNNTRKRIRIQRGPGRAFVTIGK-IGNMRQAHCNISRAKWN  339
            AKTIIV  N SVEI C RP NNTRK +RI  GPG+ F   G  IG++RQAHCNIS A WN
Sbjct  287  AKTIIVHFNESVEITCIRPGNNTRKSVRI--GPGQTFYATGDIIGDIRQAHCNISEADWN  114

Query  340  NTLKQIASKLREQFGNNKTIIFKQSSGGDPEIVTH  374
             TL ++  KL+E F  NKTIIF+ SSGGD EI TH
Sbjct  113  KTLARVVKKLKEHF--NKTIIFQPSSGGDLEITTH  15



>Pseudomonas aeruginosa strain VET-42 NODE_97_length_548_cov_2.70913_ID_193, whole genome shotgun sequence
Sequence ID: NZ_RACX01000130.1 Length: 548
Range 1: 3 to 533

Score:215 bits(548), Expect:2e-60,
Method:Compositional matrix adjust.,
Identities:112/193(58%), Positives:139/193(72%), Gaps:21/193(10%)

Query  278  TDNAKTIIVQLNTSVEINCTRPNNNTRKRIRIQRGPGRAFVTIGKI-GNMRQAHCNISRA  336
            TDN KTIIVQ N SVEINCTRPNNNTRK I +  GPG+AF   G I G++RQA+CNIS
Sbjct  533  TDNTKTIIVQFNQSVEINCTRPNNNTRKSIHL--GPGQAFFATGDIIGSIRQAYCNISGD  360

Query  337  KWNNTLKQIASKLREQFGNNKTIIFKQSSGGDPEIVTHSFNCGGEFFYCNSTQLFNSTWF  396
            KW++ L ++ ++ +KL+E F  N  I FK S+GGD EI+TH+FNCGGEFFYCN++ LFNS+
Sbjct  359  KWDDMLNKVKAKLKEFFPGN--ISFKSSAGGDLEIITHTFNCGGEFFYCNTSDLFNSSSK  186

Query  397  NSTWSTEGSNNTEGSDTITLPCRIKQIINMWQKVGKAMYAPPISGQIRCSSNITGLLLTR  456
            N+               I LPC+I+QI+ MWQ+V +AMYAPPI+G I C SNITGLLLTR
Sbjct  185  NN------------GTIIKLPCKIRQIVRMWQRVAQAMYAPPIAGIITCRSNITGLLLTR  42

Query  457  D----GGNSNNES  465
            D    GGN+ N++
Sbjct  41   DGSSKGGNNTNDT  3



>Klebsiella pneumoniae strain ST-234:K080 NODE_195_length_474_cov_25.8761, whole genome shotgun sequence
Sequence ID: NZ_NXJY01000191.1 Length: 474
Range 1: 8 to 466

Score:206 bits(525), Expect:2e-57,
Method:Compositional matrix adjust.,
Identities:107/157(68%), Positives:123/157(78%), Gaps:5/157(3%)

Query  221  AGFAILKCNNKTFNGTGPCTNVSTVQCTHGIRPVVSTQLLLNGSLAEEEVVIRSVNFTDN  280
            AG+AILKCNN+TFNG GPC NVSTVQCTHGI+PVVSTQLLLNGSLA +EVVIRS N T+N
Sbjct  8    AGYAILKCNNETFNGKGPCNNVSTVQCTHGIKPVVSTQLLLNGSLA-KEVVIRSENLTNN  184

Query  281  AKTIIVQLNTSVEINCTRPNNNTRKRIRIQRGPGRAFVTIGK-IGNMRQAHCNISRAKWN  339
            AK IIV LN SV I C RP+NNTRK +RI  GPG+ F   G  IG++R+AHCNIS+ +WN
Sbjct  185  AKIIIVHLNKSVTIECIRPSNNTRKSVRI--GPGQTFYATGDIIGDIREAHCNISKQEWN  358

Query  340  NTLKQIASKLREQFGNNKTIIFKQSSGGDPEIVTHSF  376
             TL+ +  KL + F  NKTI F QSSGGD EI THS
Sbjct  359  KTLQLVGEKLAKHFP-NKTITFNQSSGGDLEITTHSL  466



>Escherichia coli strain EN5177 contig00148, whole genome shotgun sequence
Sequence ID: NZ_JAPMNK010000148.1 Length: 850
Range 1: 279 to 848

Score:208 bits(530), Expect:1e-56,
```

```
Method:Composition-based stats.,
Identities:127/190(67%), Positives:146/190(76%), Gaps:7/190(3%)

Query  673  FNITNWLWYIKLFIMIVGGLVGLRIVFAVLSIVNRVRQGYSPLSFQTHLPTprgpdrpeg  732
            F IT WL YIK+FIMIVGGL+ LRI+FAV+SIVNR RQ YSPLSFQT  P PRGPDR
Sbjct  848  FGITKWL*YIKIFIMIVGGLISLRIIFAVISIVNRFRQRYSPLSFQTLTPNPRGPDRLGR  669

Query  733  ieeeggerdrdrSIRLVNGSLALIWDDLRSLCLFSYHRLRDLLLIVTRIVELLGR-----  787
            IEEEG E+DR+RSIRLVNG LAL  DDL++LCLFSYHRLRD +L+  R+VELLGR
Sbjct  668  IEEEGREQDRNRSIRLVNGFLALA*DDLQNLCLFSYHRLRDFILVAARVVELLGRSSLKG  489

Query  788  --RGWEALKYWWNLLQYWSQELKNSAVSLLNATAIAVAEGTDRVIEVVQGACRAIRHIPR  845
              R  EALKY  +L+QY   ELK SA+SLL+  AIAVAE TD++I +VQ  CRAIR+IPR
Sbjct  488  LQRR*EALKYLGSLVQY*GLELKGSAISLLDTIAIAVAERTDKIINLVQRICRAIRNIPR  309

Query  846  RIRQGLERIL  855
            RIRQ  E  L
Sbjct  308  RIRQSFEAAL  279



>Enterobacter hormaechei strain ST-234:K063 NODE_159_length_475_cov_27.7644, whole genome shotgun sequence
Sequence ID: NZ_NXJN01000152.1 Length: 475
Range 1: 6 to 467

Score:202 bits(515), Expect:3e-56,
Method:Compositional matrix adjust.,
Identities:105/158(66%), Positives:123/158(77%), Gaps:5/158(3%)

Query  220  PAGFAILKCNNKTFNGTGPCTNVSTVQCTHGIRPVVSTQLLLNGSLAEEEVVIRSVNFTD  279
            PAG+AILKCNN+TFNG GPC NVSTVQCTHGI+PVVSTQLLLNGSLA +EVVIRS N T+
Sbjct  6    PAGYAILKCNNETFNGKGPCNNVSTVQCTHGIKPVVSTQLLLNGSLA-KEVVIRSENLTN  182

Query  280  NAKTIIVQLNTSVEINCTRPNNNTRKRIRIQRGPGRAFVTIGK-IGNMRQAHCNISRAKW  338
            NAK IIVQLN SV I C RP+NNTRK +RI  GPG+ F   GK  GN R+A+CNI++ +W
Sbjct  183  NAKIIIVQLNESVTIECIRPSNNTRKSVRI--GPGQTFYATGKRRGNRREAYCNITKQEW  356

Query  339  NNTLKQIASKLREQFGNNKTIIFKQSSGGDPEIVTHSF  376
            + TL+ +  KL + F  NK I F QSSGGD E+ THS
Sbjct  357  SRTLQLVGEKLAKHFP-NKAITFNQSSGGDLEVTTHSI  467



>Klebsiella pneumoniae strain ST-1552:K089 NODE_185_length_474_cov_21.4899, whole genome shotgun sequence
Sequence ID: NZ_NXIW01000184.1 Length: 474
Range 1: 15 to 467

Score:202 bits(514), Expect:5e-56,
Method:Compositional matrix adjust.,
Identities:106/155(68%), Positives:121/155(78%), Gaps:5/155(3%)

Query  221  AGFAILKCNNKTFNGTGPCTNVSTVQCTHGIRPVVSTQLLLNGSLAEEEVVIRSVNFTDN  280
            AG+AILKCNN+TFNG GPC NVSTVQCTHGI+PVVSTQLLLNGSLA +EVVIRS N T+N
Sbjct  467  AGYAILKCNNETFNGKGPCNNVSTVQCTHGIKPVVSTQLLLNGSLA-KEVVIRSENLTNN  291

Query  281  AKTIIVQLNTSVEINCTRPNNNTRKRIRIQRGPGRAFVTIGK-IGNMRQAHCNISRAKWN  339
            AK IIV LN SV I C RP+NNTRK +RI  GPG+ F   G  IGN+R+A+CNISR +WN
Sbjct  290  AKIIIVHLNKSVTIECIRPSNNTRKSVRI--GPGQTFYATGDIIGNIREAYCNISRQEWN  117

Query  340  NTLKQIASKLREQFGNNKTIIFKQSSGGDPEIVTH  374
             TL+ +  KL + F  NKTI F QSSGGD EI T
Sbjct  116  KTLQLVGEKLAKHFP-NKTITFNQSSGGDLEITTQ  15



>Escherichia coli strain NBE0009 contig00153, whole genome shotgun sequence
Sequence ID: NZ_JARBDL010000153.1 Length: 753
Range 1: 1 to 507

Score:204 bits(520), Expect:1e-55,
Method:Composition-based stats.,
Identities:120/169(71%), Positives:136/169(80%), Gaps:7/169(4%)

Query  694  GLRIVFAVLSIVNRVRQGYSPLSFQTHLPTprgpdrpegieeeggerdrdrSIRLVNGSL  753
            GLRI+FAVLSIVNRVRQGYSPLSFQT +P P G DR   IEEEGGE+DRDRSIRLVNG L
Sbjct  1    GLRIIFAVLSIVNRVRQGYSPLSFQTPIPNPGGLDRLGRIEEEGGEQDRDRSIRLVNGFL  180

Query  754  ALIWDDLRSLCLFSYHRLRDLLLIVTRIVELLG-------RRGWEALKYWWNLLQYWSQE  806
            +L WDDLR+LCLFSYHRLRD +L+  R+VELLG       +RGW+ALKY  +L+QYW  E
Sbjct  181  SLAWDDLRNLCLFSYHRLRDFILVTARVVELLGHSSLRGLQRGWKALKYLRSLVQYWGLE  360

Query  807  LKNSAVSLLNATAIAVAEGTDRVIEVVQGACRAIRHIPRRIRQGLERIL  855
            LK SA+SLL+  AIAVAEGTDR+I VVQ  CRAI +IPRRIRQG E  L
Sbjct  361  LKRSAISLLDTIAIAVAEGTDRIISVVQRICRAIYNIPRRIRQGFEAAL  507



>Escherichia coli strain E003 NODE_241_length_474_cov_17.072, whole genome shotgun sequence
Sequence ID: NZ_NXIZ01000238.1 Length: 474
Range 1: 15 to 467

Score:200 bits(508), Expect:3e-55,
Method:Compositional matrix adjust.,
Identities:104/155(67%), Positives:121/155(78%), Gaps:5/155(3%)

Query  221  AGFAILKCNNKTFNGTGPCTNVSTVQCTHGIRPVVSTQLLLNGSLAEEEVVIRSVNFTDN  280
            AG+AILKCNN+TFNG GPC NVSTVQCTHGI+PVVSTQLLLNGSLA +EVVIRS N T+N
Sbjct  467  AGYAILKCNNETFNGKGPCNNVSTVQCTHGIKPVVSTQLLLNGSLA-KEVVIRSENLTNN  291

Query  281  AKTIIVQLNTSVEINCTRPNNNTRKRIRIQRGPGRAFVTIGK-IGNMRQAHCNISRAKWN  339
            AK IIV LN SV I C RP+NNTRK +RI  GPG+ F   G  IGN+R+A+CNIS+ +WN
Sbjct  290  AKIIIVHLNKSVTIECIRPSNNTRKSVRI--GPGQTFYATGDIIGNIREAYCNISKQEWN  117

Query  340  NTLKQIASKLREQFGNNKTIIFKQSSGGDPEIVTH  374
             TL+ +  KL + F  NKTI F QSSGGD E+ T
Sbjct  116  KTLQLVGEKLAKHFP-NKTITFNQSSGGDLEVTTQ  15



>Cronobacter sakazakii strain cro4114A1 contig126, whole genome shotgun sequence
Sequence ID: NZ_NRKJ01000126.1 Length: 413
Range 1: 3 to 413

Score:192 bits(489), Expect:8e-53,
Method:Compositional matrix adjust.,
Identities:96/143(67%), Positives:111/143(77%), Gaps:8/143(5%)

Query  298  RPNNNTRKRIRIQRGPGRAFVTIGKI-GNMRQAHCNISRAKWNNTLKQIASKLREQFGNN  356
            RP+NNTRK I    GPGR+    G I G++RQAHCNIS  KWNNTLK++A KLREQFGNN
```

```
Sbjct  413  RPDNNTRKSIHF--GPGRSLYATGNIIGDIRQAHCNISGEKWNNTLKRVAIKLREQFGNN  240

Query  357  KTIIFKQSSGGDPEIVTHSFNCGGEFFYCNSTQLFNSTWF-NSTWSTEGSNNTEGSDTIT  415
            KTI+F+Q SGGDPE+V H  NCGGEFFYCN+T LFNST   N TW++ G    +GS  IT
Sbjct  239  KTIVFQQPSGGDPEVVMHILNCGGEFFYCNTTNLFNSTLLANDTWNSTG----QGSGNIT  72

Query  416  LPCRIKQIINMWQKVGKAMYAPP  438
            + CRIKQIIN WQ+VGKAMYAPP
Sbjct  71   IQCRIKQIINRWQEVGKAMYAPP  3
```

>Cronobacter malonaticus strain cro3263A1 contig69, whole genome shotgun sequence
Sequence ID: NZ_NRMM01000069.1 Length: 413
Range 1: 1 to 411

Score:192 bits(488), Expect:1e-52,
Method:Compositional matrix adjust.,
Identities:96/143(67%), Positives:111/143(77%), Gaps:8/143(5%)

```
Query  298  RPNNNTRKRIRIQRGPGRAFVTIGKI-GNMRQAHCNISRAKWNNTLKQIASKLREQFGNN  356
            RP+NNTRK I    GPGR+    G I G++RQAHCNIS  KWNNTLK++A KLREQFGNN
Sbjct  1    RPSNNTRKSIHF--GPGRSLYATGNIIGDIRQAHCNISGEKWNNTLKRVAIKLREQFGNN  174

Query  357  KTIIFKQSSGGDPEIVTHSFNCGGEFFYCNSTQLFNSTWF-NSTWSTEGSNNTEGSDTIT  415
            KTI+F+Q SGGDPE+V H  NCGGEFFYCN+T LFNST   N TW++ G    +GS  IT
Sbjct  175  KTIVFQQPSGGDPEVVMHILNCGGEFFYCNTTNLFNSTLLANDTWNSTG----QGSGNIT  342

Query  416  LPCRIKQIINMWQKVGKAMYAPP  438
            + CRIKQIIN WQ+VGKAMYAPP
Sbjct  343  IQCRIKQIINRWQEVGKAMYAPP  411
```

>Citrobacter freundii strain CF003 NODE_156_length_474_cov_23.0173, whole genome shotgun sequence
Sequence ID: NZ_NXKE01000152.1 Length: 474
Range 1: 8 to 457

Score:192 bits(489), Expect:2e-52,
Method:Compositional matrix adjust.,
Identities:102/154(66%), Positives:118/154(76%), Gaps:5/154(3%)

```
Query  221  AGFAILKCNNKTFNGTGPCTNVSTVQCTHGIRPVVSTQLLLNGSLAEEEVVIRSVNFTDN  280
            AG+AILKCNN+TFNG GPC NVSTVQCTHGI+PVVSTQLLLNGSLA +EVVIRS N T+N
Sbjct  8    AGYAILKCNNETFNGKGPCNNVSTVQCTHGIKPVVSTQLLLNGSLA-KEVVIRSENLTNN  184

Query  281  AKTIIVQLNTSVEINCTRPNNNTRKRIRIQRGPGRAFVTIGK-IGNMRQAHCNISRAKWN  339
            AK IIV LN SV I C RP+NNTRK +RI  GPG+ F   G  IG +R+A+CNISR + N
Sbjct  185  AKIIIVHLNKSVTIECIRPSNNTRKSVRI--GPGQTFYATGYIIGEIREAYCNISRQELN  358

Query  340  NTLKQIASKLREQFGNNKTIIFKQSSGGDPEIVT  373
             TL+ +  KL +    NKTI F QSSGGD E+ T
Sbjct  359  KTLQLVGEKLAKHVP-NKTITFNQSSGGDLEVTT  457
```

>Escherichia coli strain NBE0009 contig00151, whole genome shotgun sequence
Sequence ID: NZ_JARBDL010000151.1 Length: 802
Range 1: 296 to 802

Score:194 bits(493), Expect:1e-51,
Method:Composition-based stats.,
Identities:116/169(69%), Positives:133/169(78%), Gaps:7/169(4%)

```
Query  694  GLRIVFAVLSIVNRVRQGYSPLSFQTHLPTprgpdrpegieeeggerdrdrSIRLVNGSL  753
            GLRI+FAVLSIVNRVRQGYSPLSFQT  PTP  PDR  GIEEEGGE+DR RS+RLV+G L
Sbjct  802  GLRIIFAVLSIVNRVRQGYSPLSFQTLTPTPGEPDRLRGIEEEGGEQDRVRSVRLVSGFL  623

Query  754  ALIWDDLRSLCLFSYHRLRDLLLIVTRIVELLG-------RRGWEALKYWWNLLQYWSQE  806
            A+ WDDLR+LCLFSYHRLRDLLL++ R VELLG       +RG EALK+   ++QYW  E
Sbjct  622  AIAWDDLRNLCLFSYHRLRDLLLVIARAVELLGHSCLVGLQRGREALKHLGGIVQYWGLE  443

Query  807  LKNSAVSLLNATAIAVAEGTDRVIEVVQGACRAIRHIPRRIRQGLERIL  855
            LK SA+SL +  AIAVAEGTDR+I  +Q  CRAI +IPRRIRQG E  L
Sbjct  442  LKKSAISLFDTIAIAVAEGTDRIIIGIQRLCRAICNIPRRIRQGFEAAL  296
```

>Klebsiella pneumoniae strain EuSCAPE_IL005, whole genome shotgun sequence
Sequence ID: NZ_UKJL02000055.1 Length: 455
Range 1: 15 to 455

Score:188 bits(477), Expect:6e-51,
Method:Composition-based stats.,
Identities:110/147(75%), Positives:125/147(85%), Gaps:7/147(4%)

```
Query  664  DKWASLWNWFNITNWLWYIKLFIMIVGGLVGLRIVFAVLSIVNRVRQGYSPLSFQTHLPT  723
            DKW +LW+WF+ITNWLWYI++FIMIVGGL+GLRI+FAVLS+VNRVRQGYSPLSFQT +P+
Sbjct  15   DKWGNLWSWFDITNWLWYIRIFIMIVGGLIGLRIIFAVLSMVNRVRQGYSPLSFQTLIPS  194

Query  724  prgpdrpegieeeggerdrdrSIRLVNGSLALIWDDLRSLCLFSYHRLRDLLLIVTRIVE  783
            PRGPDR E IEEEGGE+DR+RSIRLV+G LAL WDDLRSLCLF YHRLRD +L+  R VE
Sbjct  195  PRGPDRLERIEEEGGEQDRNRSIRLVSGFLALAWDDLRSLCLFCYHRLRDFILVTVRAVE  374

Query  784  LLGR-------RGWEALKYWWNLLQYW  803
            LLGR       RGWE LKY  NL QYW
Sbjct  375  LLGRSSLKGLQRGWEILKYLGNLAQYW  455
```

>Klebsiella pneumoniae strain EuSCAPE_IL004, whole genome shotgun sequence
Sequence ID: NZ_UKJP02000107.1 Length: 368
Range 1: 14 to 343

Score:186 bits(472), Expect:1e-50,
Method:Compositional matrix adjust.,
Identities:88/110(80%), Positives:99/110(90%), Gaps:0/110(0%)

```
Query  537  LTVQARQLLSGIVQQQNNLLRAIEAQQHLLQLTVWGIKQLQARILAVERYLKDQQLLGIW  596
            LTVQARQLLSGIVQQQ NLLRAIEAQQH+LQLTVWGIKQLQ R+LA+ERYLKDQQLLGIW
Sbjct  343  LTVQARQLLSGIVQQQGNLLRAIEAQQHMLQLTVWGIKQLQTRVLAIERYLKDQQLLGIW  164

Query  597  GCSGKLICTTAVPWNASWSNKSLEQIWNHTTWMEWDREINNYTSLIHSLI  646
            GCSGKLICTTAVPWN++WS KS   IW + TWM+WDREI+N+T  I+ L+
Sbjct  163  GCSGKLICTTAVPWNSTWSKKSQNDIWKNMTWMQWDREISNHTDTIYRLL  14
```

```
>Klebsiella pneumoniae strain EuSCAPE_MT003, whole genome shotgun sequence
Sequence ID: NZ_UKIK02000085.1 Length: 964
Range 1: 480 to 962

Score:193 bits(490), Expect:2e-50,
Method:Compositional matrix adjust.,
Identities:113/161(70%), Positives:129/161(80%), Gaps:0/161(0%)

Query  696  RIVFAVLSIVNRVRQGYSPLSFQTHLPTprgpdrpegieeeggerdrdrSIRLVNGSLAL  755
            RI+FAVLSIVNRVRQGYSPLS QT    P G DRP  IEEEGGE+D+DRSIRLVNG  +L
Sbjct  962  RIIFAVLSIVNRVRQGYSPLSLQTLTQNPGGLDRPGRIEEEGGEQDKDRSIRLVNGFFSL  783

Query  756  IWDDLRSLCLFSYHRLRDLLLIVTRIVELLGRRGWEALKYWWNLLQYWSQELKNSAVSLL  815
             W+DLRSLCLFSY +LRD  LI  R VE+LGRRGWE LKY  +L+QYW  ELK SA+SLL
Sbjct  782  AWEDLRSLCLFSYRQLRDFTLIAARAVEILGRRGWETLKYLGSLIQYWGLELKKSAISLL  603

Query  816  NATAIAVAEGTDRVIEVVQGACRAIRHIPRRIRQGLERILL  856
            +  AI VAEGTDR++E +QG CRAIR++PRRIRQG E  LL
Sbjct  602  DTIAITVAEGTDRILEFIQGICRAIRNVPRRIRQGFEAALL  480



>Escherichia coli strain NBE0006 contig00145, whole genome shotgun sequence
Sequence ID: NZ_JARBDI010000145.1 Length: 738
Range 1: 1 to 483

Score:186 bits(472), Expect:7e-49,
Method:Composition-based stats.,
Identities:113/172(66%), Positives:129/172(75%), Gaps:18/172(10%)

Query  691  GLVGLRIVFAVLSIVNRVRQGYSPLSFQTHLPTprgpdrpegieeeggerdrdrSIRLVN  750
            GL+GLRI+FAV+SIVNR RQGYSPLSFQT  P P GPDR   IEEEGGE+DR+RSIRLVN
Sbjct  1    GLIGLRIIFAVISIVNRFRQGYSPLSFQTLTPNPGGPDRLGRIEEEGGEQDRNRSIRLVN  180

Query  751  GSLALIWDDLRSLCLFSYHRLRDLLLIVTRIVELLGR-------RGWEALKYWWNLLQYW  803
            G LAL WDDLR+LCLFSYH+LRD +L+  R+VELLGR       RGWEALKY  +L+QYW
Sbjct  181  GFLALAWDDLRNLCLFSYHQLRDFILVTARVVELLGRNSLKGLQRGWEALKYLGSLVQYW  360

Query  804  SQELKNSAVSLLNATAIAVAEGTDRVIEVVQGACRAIRHIPRRIRQGLERIL  855
                        +  AIAVAEGTDR+I + QG CRAI +IPRRIRQG E  L
Sbjct  361  -----------FDTIAIAVAEGTDRIINLAQGICRAICNIPRRIRQGFEAAL  483



>Cronobacter malonaticus strain cro3263A1 contig68, whole genome shotgun sequence
Sequence ID: NZ_NRMM01000068.1 Length: 413
Range 1: 1 to 411

Score:179 bits(454), Expect:7e-48,
Method:Compositional matrix adjust.,
Identities:92/142(65%), Positives:107/142(75%), Gaps:6/142(4%)

Query  298  RPNNNTRKRIRIQRGPGRAFVTIGKI-GNMRQAHCNISRAKWNNTLKQIASKLREQFGNN  356
            RP+NNTRK IRI  GPG+ F   G I G++RQAHCNISR  W  TL  ++ KL E F N
Sbjct  1    RPDNNTRKSIRI--GPGQTFYATGDIIGDIRQAHCNISRTDWERTLHNVSKKLAEHFPN-  171

Query  357  KTIIFKQSSGGDPEIVTHSFNCGGEFFYCNSTQLFNSTWFNSTWSTEGSNNTEGSDTITL  416
            KTI F  SSGGD EI THSFNC GEFFYCN++ LFNST+  +T  T  SN T+G+ TIT+
Sbjct  172  KTIRFTNSSGGDLEITTHSFNCRGEFFYCNTSGLFNSTY--TTNGTFESNETKGNSTITI  345

Query  417  PCRIKQIINMWQKVGKAMYAPP  438
            PCRIKQIINMWQ+VG+AMYAPP
Sbjct  346  PCRIKQIINMWQEVGRAMYAPP  411



>Escherichia coli strain NBE0006 contig00147, whole genome shotgun sequence
Sequence ID: NZ_JARBDI010000147.1 Length: 694
Range 1: 219 to 692

Score:182 bits(461), Expect:1e-47,
Method:Composition-based stats.,
Identities:112/158(71%), Positives:123/158(77%), Gaps:7/158(4%)

Query  705  VNRVRQGYSPLSFQTHLPTprgpdrpegieeeggerdrdrSIRLVNGSLALIWDDLRSLC  764
            VNRVRQGYSPLSFQT  P PRGPDR   IEEEGG +DRDRSIRLVNG LAL WDDL SLC
Sbjct  692  VNRVRQGYSPLSFQTLTPNPRGPDRLGRIEEEGGGQDRDRSIRLVNGFLALAWDDLWSLC  513

Query  765  LFSYHRLRDLLLIVTRIVELLG-------RRGWEALKYWWNLLQYWSQELKNSAVSLLNA  817
            LFSYHRLRDL+L+  R VELLG       +RGWEALKY   ++QYW  ELK  A+SLL+
Sbjct  512  LFSYHRLRDLILVTARAVELLGHSSLRGLQRGWEALKYLGGIVQYWGLELKKRAISLLDT  333

Query  818  TAIAVAEGTDRVIEVVQGACRAIRHIPRRIRQGLERIL  855
             AIAVAEGTDR+IE +Q  CRAIR+IPRRIRQG E  L
Sbjct  332  VAIAVAEGTDRIIEFLQRICRAIRNIPRRIRQGFEAAL  219



>Actinoplanes campanulatus strain JCM 3059 sequence095, whole genome shotgun sequence
Sequence ID: NZ_BMPW01000095.1 Length: 582
Range 1: 111 to 458

Score:179 bits(455), Expect:3e-47,
Method:Compositional matrix adjust.,
Identities:115/116(99%), Positives:115/116(99%), Gaps:0/116(0%)

Query  719  THLPTprgpdrpegieeeggerdrdrSIRLVNGSLALIWDDLRSLCLFSYHRLRDLLLIV  778
            THLP PRGPDRPEGIEEEGGERDRDRSIRLVNGSLALIWDDLRSLCLFSYHRLRDLLLIV
Sbjct  111  THLPIPRGPDRPEGIEEEGGERDRDRSIRLVNGSLALIWDDLRSLCLFSYHRLRDLLLIV  290

Query  779  TRIVELLGRRGWEALKYWWNLLQYWSQELKNSAVSLLNATAIAVAEGTDRVIEVVQ  834
            TRIVELLGRRGWEALKYWWNLLQYWSQELKNSAVSLLNATAIAVAEGTDRVIEVVQ
Sbjct  291  TRIVELLGRRGWEALKYWWNLLQYWSQELKNSAVSLLNATAIAVAEGTDRVIEVVQ  458



>Cronobacter sakazakii strain cro4215W contig103, whole genome shotgun sequence
Sequence ID: NZ_NRKF01000103.1 Length: 413
Range 1: 1 to 411

Score:176 bits(446), Expect:7e-47,
Method:Compositional matrix adjust.,
Identities:91/142(64%), Positives:106/142(74%), Gaps:6/142(4%)

Query  298  RPNNNTRKRIRIQRGPGRAFVTIGKI-GNMRQAHCNISRAKWNNTLKQIASKLREQFGNN  356
            RP+NNTRK IRI  GPG+ F   G I G++RQAHCNISR  W  TL  ++ KL E F N
Sbjct  1    RPSNNTRKSIRI--GPGQTFYATGDIIGDIRQAHCNISRTDWERTLHNVSKKLAEHFPN-  171

Query  357  KTIIFKQSSGGDPEIVTHSFNCGGEFFYCNSTQLFNSTWFNSTWSTEGSNNTEGSDTITL  416
```

```
            KTI F  SSGGD EI THSFNC GEFFYCN++ LFNST+  +T  T  SN T+ + TIT+
Sbjct  172  KTIRFTNSSGGDLEITTHSFNCRGEFFYCNTSGLFNSTY--TTNGTFESNETKSNSTITI  345

Query  417  PCRIKQIINMWQKVGKAMYAPP  438
            PCRIKQIINMWQ+VG+AMYAPP
Sbjct  346  PCRIKQIINMWQEVGRAMYAPP  411
```

>Deinococcus aquiradiocola strain JCM 14371 sequence41, whole genome shotgun sequence
Sequence ID: NZ_BMOE01000041.1 Length: 690
Range 1: 107 to 454

Score:179 bits(455), Expect:8e-47,
Method:Compositional matrix adjust.,
Identities:115/116(99%), Positives:115/116(99%), Gaps:0/116(0%)

```
Query  719  THLPTprgpdrpegieeeggerdrdrSIRLVNGSLALIWDDLRSLCLFSYHRLRDLLLIV  778
            THLP PRGPDRPEGIEEEGGERDRDRSIRLVNGSLALIWDDLRSLCLFSYHRLRDLLLIV
Sbjct  107  THLPIPRGPDRPEGIEEEGGERDRDRSIRLVNGSLALIWDDLRSLCLFSYHRLRDLLLIV  286

Query  779  TRIVELLGRRGWEALKYWWNLLQYWSQELKNSAVSLLNATAIAVAEGTDRVIEVVQ  834
            TRIVELLGRRGWEALKYWWNLLQYWSQELKNSAVSLLNATAIAVAEGTDRVIEVVQ
Sbjct  287  TRIVELLGRRGWEALKYWWNLLQYWSQELKNSAVSLLNATAIAVAEGTDRVIEVVQ  454
```

>Klebsiella pneumoniae strain EuSCAPE_IL004, whole genome shotgun sequence
Sequence ID: NZ_UKJP02000113.1 Length: 277
Range 1: 1 to 276

Score:174 bits(440), Expect:1e-46,
Method:Composition-based stats.,
Identities:78/92(85%), Positives:83/92(90%), Gaps:0/92(0%)

```
Query  48   ATTTLFCASDAKAYDTEVHNVWATHACVPTDPNPQEVVLVNVTENFNMWKNDMVEQMHED  107
            A   LFCASDAKAYD EVHNVWATHACVPTDPNPQE+ L NVTENFNMWKNDMVEQMH+D
Sbjct  1    AKAPLFCASDAKAYDKEVHNVWATHACVPTDPNPQEMFLENVTENFNMWKNDMVEQMHKD  180

Query  108  IISLWDQSLKPCVKLTPLCVSLKCTDLKNDTN  139
            IISLWD+SLKPCVKLTPLCV+L CTD+ N TN
Sbjct  181  IISLWDESLKPCVKLTPLCVTLNCTDVDNSTN  276
```

>Enterococcus alcedinis strain CCM 8433 sequence26, whole genome shotgun sequence
Sequence ID: NZ_BMDT01000026.1 Length: 853
Range 1: 129 to 476

Score:179 bits(455), Expect:4e-46,
Method:Compositional matrix adjust.,
Identities:115/116(99%), Positives:115/116(99%), Gaps:0/116(0%)

```
Query  719  THLPTprgpdrpegieeeggerdrdrSIRLVNGSLALIWDDLRSLCLFSYHRLRDLLLIV  778
            THLP PRGPDRPEGIEEEGGERDRDRSIRLVNGSLALIWDDLRSLCLFSYHRLRDLLLIV
Sbjct  129  THLPIPRGPDRPEGIEEEGGERDRDRSIRLVNGSLALIWDDLRSLCLFSYHRLRDLLLIV  308

Query  779  TRIVELLGRRGWEALKYWWNLLQYWSQELKNSAVSLLNATAIAVAEGTDRVIEVVQ  834
            TRIVELLGRRGWEALKYWWNLLQYWSQELKNSAVSLLNATAIAVAEGTDRVIEVVQ
Sbjct  309  TRIVELLGRRGWEALKYWWNLLQYWSQELKNSAVSLLNATAIAVAEGTDRVIEVVQ  476
```

>Cronobacter dublinensis strain cro981C2 contig304, whole genome shotgun sequence
Sequence ID: NZ_NROB01000304.1 Length: 413
Range 1: 1 to 411

Score:174 bits(441), Expect:4e-46,
Method:Compositional matrix adjust.,
Identities:90/142(63%), Positives:105/142(73%), Gaps:6/142(4%)

```
Query  298  RPNNNTRKRIRIQRGPGRAFVTIGKI-GNMRQAHCNISRAKWNNTLKQIASKLREQFGNN  356
            RP+NNTRK IRI  GPG+ F   G I G++RQAHCNISR  W  TL  ++ KL E F
Sbjct  1    RPSNNTRKSIRI--GPGQTFYATGDIIGDIRQAHCNISRTDWERTLHNVSKKLAEHFPK-  171

Query  357  KTIIFKQSSGGDPEIVTHSFNCGGEFFYCNSTQLFNSTWFNSTWSTEGSNNTEGSDTITL  416
            KTI F  SSGGD EI THSFNC GEFFYCN++ LFNST+  +T  T  SN T+ + TIT+
Sbjct  172  KTIRFTNSSGGDLEITTHSFNCRGEFFYCNTSGLFNSTY--TTNGTFESNETKSNSTITI  345

Query  417  PCRIKQIINMWQKVGKAMYAPP  438
            PCRIKQIINMWQ+VG+AMYAPP
Sbjct  346  PCRIKQIINMWQEVGRAMYAPP  411
```

>Cronobacter sakazakii strain cro4114B1 contig109, whole genome shotgun sequence
Sequence ID: NZ_NRKI01000109.1 Length: 413
Range 1: 1 to 411

Score:174 bits(441), Expect:4e-46,
Method:Compositional matrix adjust.,
Identities:90/142(63%), Positives:105/142(73%), Gaps:6/142(4%)

```
Query  298  RPNNNTRKRIRIQRGPGRAFVTIGKI-GNMRQAHCNISRAKWNNTLKQIASKLREQFGNN  356
            RP+NNTRK IRI  GPG+ F   G I G++RQAHCNISR  W  TL  ++ KL E F
Sbjct  1    RPSNNTRKSIRI--GPGQTFYATGDIIGDIRQAHCNISRTDWERTLHNVSKKLAEHFPK-  171

Query  357  KTIIFKQSSGGDPEIVTHSFNCGGEFFYCNSTQLFNSTWFNSTWSTEGSNNTEGSDTITL  416
            KTI F  SSGGD EI THSFNC GEFFYCN++ LFNST+  +T  T  SN T+ + TIT+
Sbjct  172  KTIRFTNSSGGDLEITTHSFNCRGEFFYCNTSGLFNSTY--TTNGTFESNETKSNSTITI  345

Query  417  PCRIKQIINMWQKVGKAMYAPP  438
            PCRIKQIINMWQ+VG+AMYAPP
Sbjct  346  PCRIKQIINMWQEVGRAMYAPP  411
```

>Cronobacter sakazakii strain cro4114A1 contig127, whole genome shotgun sequence
Sequence ID: NZ_NRKJ01000127.1 Length: 413
Range 1: 1 to 411

Score:174 bits(441), Expect:4e-46,
Method:Compositional matrix adjust.,
Identities:90/142(63%), Positives:105/142(73%), Gaps:6/142(4%)

```
Query  298  RPNNNTRKRIRIQRGPGRAFVTIGKI-GNMRQAHCNISRAKWNNTLKQIASKLREQFGNN  356
            RP+NNTRK IRI  GPG+ F   G I G++RQAHCNISR  W  TL  ++ KL E F
Sbjct  1    RPSNNTRKSIRI--GPGQTFYATGDIIGDIRQAHCNISRTDWERTLHNVSKKLAEHFPK-  171
```

```
Query  357  KTIIFKQSSGGDPEIVTHSFNCGGEFFYCNSTQLFNSTWFNSTWSTEGSNNTEGSDTITL  416
            KTI F  SSGGD EI THSFNC GEFFYCN++ LFNST+  +T  T  SN T+ + TIT+
Sbjct  172  KTIRFTNSSGGDLEITTHSFNCRGEFFYCNTSGLFNSTY--TTNGTFESNETKSNSTITI  345

Query  417  PCRIKQIINMWQKVGKAMYAPP  438
            PCRIKQIINMWQ+VG+AMYAPP
Sbjct  346  PCRIKQIINMWQEVGRAMYAPP  411
```

```
>Cronobacter sakazakii strain cro611A3 contig61, whole genome shotgun sequence
Sequence ID: NZ_NRMC01000061.1 Length: 407
Range 1: 3 to 407

Score:172 bits(437), Expect:1e-45,
Method:Compositional matrix adjust.,
Identities:90/143(63%), Positives:106/143(74%), Gaps:10/143(6%)

Query  298  RPNNNTRKRIRIQRGPGRAFVTIGKI-GNMRQAHCNISRAKWNNTLKQIASKLREQFGNN  356
            RP+NNTRK IRI  GPG+ F   G I G++RQAHCNIS+  W  TL +++ KL E F  N
Sbjct  407  RPSNNTRKSIRI--GPGQTFYATGDIIGDIRQAHCNISKRDWEETLYKVSKKLAEHF-QN  237

Query  357  KTIIFKQSSGGDPEIVTHSFNCGGEFFYCNSTQLFNSTWF-NSTWSTEGSNNTEGSDTIT  415
            KTI F+ SSGGD EI THSFNC GEFFYCN++ LFN T+  N T ++ GSN      TIT
Sbjct  236  KTIQFEPSSGGDLEITTHSFNCRGEFFYCNTSGLFNKTYMHNGTLNSTGSNG-----TIT  72

Query  416  LPCRIKQIINMWQKVGKAMYAPP  438
            +PCRIKQIINMWQKVG+AMYAPP
Sbjct  71   IPCRIKQIINMWQKVGQAMYAPP  3
```

```
>Enterobacter hormaechei strain ST-234:K063 NODE_153_length_480_cov_7.37677, whole genome shotgun sequence
Sequence ID: NZ_NXJN01000148.1 Length: 480
Range 1: 84 to 473

Score:173 bits(438), Expect:2e-45,
Method:Compositional matrix adjust.,
Identities:86/132(65%), Positives:103/132(78%), Gaps:3/132(2%)

Query  221  AGFAILKCNNKTFNGTGPCTNVSTVQCTHGIRPVVSTQLLLNGSLAEEEVVIRSVNFTDN  280
            AG+AILKCNNKTFNGTGPC NVSTVQCTHGI+PVVSTQLLLNGSLAE E++IRS +  +N
Sbjct  473  AGYAILKCNNKTFNGTGPCNNVSTVQCTHGIKPVVSTQLLLNGSLAEGEIIIRSESLVEN  294

Query  281  AKTIIVQLNTSVEINCTRPNNNTRKRIRIQRGPGRAFVTIGK-IGNMRQAHCNISRAKWN  339
              TII  LN SV+I C RPNNNTRK I I  GP +AF   G  IGN+RQA+CN++   WN
Sbjct  293  TNTIIAHLNDSVDIMCIRPNNNTRKSIGI--GPVQAFYATGDVIGNIRQAYCNVTTKPWN  120

Query  340  NTLKQIASKLRE  351
             TL+++  ++ E
Sbjct  119  ETLERVKIEIEE  84
```

```
>Cronobacter sakazakii strain cro3040W contig81, whole genome shotgun sequence
Sequence ID: NZ_NRJO01000081.1 Length: 404
Range 1: 1 to 402

Score:169 bits(429), Expect:2e-44,
Method:Compositional matrix adjust.,
Identities:89/142(63%), Positives:103/142(72%), Gaps:9/142(6%)

Query  298  RPNNNTRKRIRIQRGPGRAFVTIGKI-GNMRQAHCNISRAKWNNTLKQIASKLREQFGNN  356
            RP+NNTRK IRI  GPG+ F   G I G++RQAHCNISR  W  TL  ++ KL E F  N
Sbjct  1    RPSNNTRKSIRI--GPGQTFYATGDIIGDIRQAHCNISRKDWEETLHNVSKKLAEHF-QN  171

Query  357  KTIIFKQSSGGDPEIVTHSFNCGGEFFYCNSTQLFNSTWFNSTWSTEGSNNTEGSDTITL  416
            K I F  SSGGD EI THSFNC GEFFYCN++ LFN T+  +   TE SN+T    TIT+
Sbjct  172  KKIQFASSSGGDLEITTHSFNCRGEFFYCNTSGLFNGTYDGTLNGTE-SNST----TITI  336

Query  417  PCRIKQIINMWQKVGKAMYAPP  438
            PCRIKQIINMWQ+VG+AMYAPP
Sbjct  337  PCRIKQIINMWQEVGRAMYAPP  402
```

```
>Cronobacter sakazakii strain cro4114A1 contig132, whole genome shotgun sequence
Sequence ID: NZ_NRKJ01000132.1 Length: 392
Range 1: 1 to 390

Score:167 bits(424), Expect:6e-44,
Method:Compositional matrix adjust.,
Identities:88/142(62%), Positives:99/142(69%), Gaps:13/142(9%)

Query  298  RPNNNTRKRIRIQRGPGRAFVTIGKI-GNMRQAHCNISRAKWNNTLKQIASKLREQFGNN  356
            RP NNTR  I I  GPGRAF  IG I G++RQAHC I+  KWN TLKQ+  KL+E F  N
Sbjct  1    RPTNNTRTSIPI--GPGRAFYRIGDIIGDIRQAHCKINGPKWNETLKQVVRKLKEHF--N  168

Query  357  KTIIFKQSSGGDPEIVTHSFNCGGEFFYCNSTQLFNSTWFNSTWSTEGSNNTEGSDTITL  416
            KTI F+  SGGD EI  H FNC GEFFYCN+ +LFNSTW          N T  +DTI L
Sbjct  169  KTIKFQPPSGGDLEITMHHFNCRGEFFYCNTAKLFNSTWI--------GNATMENDTIIL  324

Query  417  PCRIKQIINMWQKVGKAMYAPP  438
            PCRIKQIIN+WQ VG+AMYAPP
Sbjct  325  PCRIKQIINLWQGVGQAMYAPP  390
```

```
>Klebsiella pneumoniae strain EuSCAPE_IL022, whole genome shotgun sequence
Sequence ID: NZ_UKKD02000023.1 Length: 411
Range 1: 46 to 378

Score:167 bits(422), Expect:1e-43,
Method:Composition-based stats.,
Identities:82/111(74%), Positives:92/111(82%), Gaps:0/111(0%)

Query  745  SIRLVNGSLALIWDDLRSLCLFSYHRLRDLLLIVTRIVELLGRRGWEALKYWWNLLQYWS  804
            SIRLV+G LAL W+DLRSLCLFSYHRLRDLLLIVTR VELLG+RGWE LKY  +L+QYW
Sbjct  378  SIRLVSGFLALAWEDLRSLCLFSYHRLRDLLLIVTRAVELLGQRGWEILKYLGSLIQYWG  199

Query  805  QELKNSAVSLLNATAIAVAEGTDRVIEVVQGACRAIRHIPRRIRQGLERIL  855
             ELK SA+SLL+  AIA AEGTDR+IE +Q  CRAI +IPRRIRQG E  L
Sbjct  198  LELKKSAISLLDTIAIATAEGTDRIIEGIQRICRAICNIPRRIRQGFEAAL  46
```

```
>Cronobacter sakazakii strain cro4215W contig104, whole genome shotgun sequence
```

```
Sequence ID: NZ_NRKF01000104.1 Length: 405
Range 1: 1 to 402

Score:166 bits(421), Expect:2e-43,
Method:Compositional matrix adjust.,
Identities:88/142(62%), Positives:102/142(71%), Gaps:9/142(6%)

Query  298  RPNNNTRKRIRIQRGPGRAFVTIGKI-GNMRQAHCNISRAKWNNTLKQIASKLREQFGNN  356
            RPNNNTR  IRI  GPG+ F   G I G++RQA+C I+  KWN TL+Q+A KL+EQF  N
Sbjct  1    RPNNNTRTSIRI--GPGQTFYRTGDIIGDIRQAYCEINGTKWNETLRQVAKKLKEQF--N  168

Query  357  KTIIFKQSSGGDPEIVTHSFNCGGEFFYCNSTQLFNSTWFNSTWSTEGSNNTEGSDTITL  416
            KTI F+  SGGD EI    FNC GEFFYCN+T+LFNSTW     + +G N   GSDT  L
Sbjct  169  KTIKFQPPSGGDLEITMLHFNCRGEFFYCNTTKLFNSTW-EGNETIKGGN---GSDTTIL  336

Query  417  PCRIKQIINMWQKVGKAMYAPP  438
            PCRIKQIINMWQ  G+AMYAPP
Sbjct  337  PCRIKQIINMWQGAGQAMYAPP  402



>Escherichia coli strain NBE0009 contig00162, whole genome shotgun sequence
Sequence ID: NZ_JARBDL010000162.1 Length: 350
Range 1: 2 to 349

Score:165 bits(418), Expect:3e-43,
Method:Compositional matrix adjust.,
Identities:88/116(76%), Positives:105/116(90%), Gaps:0/116(0%)

Query  604  CTTAVPWNASWSNKSLEQIWNHTTWMEWDREINNYTSLIHSLIeesqnqqekneqellel  663
            C TAVPWN+SWSNK+ E+IW++ TWM+WDREINNYT  I+ L+EESQ QQE+NE++LL L
Sbjct  349  CPTAVPWNSSWSNKTQEEIWDNMTWMQWDREINNYTKTIYDLLEESQYQQERNEKDLLAL  170

Query  664  DKWASLWNWFNITNWLWYIKLFIMIVGGLVGLRIVFAVLSIVNRVRQGYSPLSFQT  719
            D W +LWNWF+I+ WLWYIK+FIMIVGGL+GLRI+FAVLSIVNRVRQGYSPLSFQT
Sbjct  169  DSWNNLWNWFDISKWLWYIKIFIMIVGGLIGLRIIFAVLSIVNRVRQGYSPLSFQT  2



>Cronobacter sakazakii strain cro4114A1 contig129, whole genome shotgun sequence
Sequence ID: NZ_NRKJ01000129.1 Length: 404
Range 1: 1 to 402

Score:165 bits(417), Expect:6e-43,
Method:Compositional matrix adjust.,
Identities:87/142(61%), Positives:102/142(71%), Gaps:9/142(6%)

Query  298  RPNNNTRKRIRIQRGPGRAFVTIGKI-GNMRQAHCNISRAKWNNTLKQIASKLREQFGNN  356
            RP+NNTR  IRI  GPG+ F   G I G++RQA+C I+  KWN TL+Q+A KL+EQF  N
Sbjct  1    RPDNNTRTSIRI--GPGQTFYRTGDIIGDIRQAYCEINGTKWNETLRQVAKKLKEQF--N  168

Query  357  KTIIFKQSSGGDPEIVTHSFNCGGEFFYCNSTQLFNSTWFNSTWSTEGSNNTEGSDTITL  416
            KTI F+  SGGD EI    FNC GEFFYCN+T+LFNSTW     + +G N   GSDT  L
Sbjct  169  KTIKFQPPSGGDLEITMLHFNCRGEFFYCNTTKLFNSTW-EGNETIKGGN---GSDTTIL  336

Query  417  PCRIKQIINMWQKVGKAMYAPP  438
            PCRIKQIINMWQ  G+AMYAPP
Sbjct  337  PCRIKQIINMWQGAGQAMYAPP  402



>Cronobacter malonaticus strain cro3263A1 contig74, whole genome shotgun sequence
Sequence ID: NZ_NRMM01000074.1 Length: 404
Range 1: 3 to 404

Score:165 bits(417), Expect:8e-43,
Method:Compositional matrix adjust.,
Identities:83/142(58%), Positives:99/142(69%), Gaps:9/142(6%)

Query  298  RPNNNTRKRIRIQRGPGRAFVTIGKI-GNMRQAHCNISRAKWNNTLKQIASKLREQFGNN  356
            RPNNNTRK I +    GRA    G I G++RQAHCN+S+  W  TL Q+A+KLREQF  N
Sbjct  404  RPNNNTRKSISL--AGGRALYATGDIIGDIRQAHCNVSKGNWTETLGQVATKLREQF-KN  234

Query  357  KTIIFKQSSGGDPEIVTHSFNCGGEFFYCNSTQLFNSTWFNSTWSTEGSNNTEGSDTITL  416
            KTI+F  SSGGDPE+    FNCGGEFFYCN+++LFN T+        G+     +  ITL
Sbjct  233  KTIVFNSSSGGDPEVEMFIFNCGGEFFYCNTSKLFNYTFL-----ANGTGQNYSTGNITL  69

Query  417  PCRIKQIINMWQKVGKAMYAPP  438
             CRIKQIIN WQ+VGKAMYAPP
Sbjct  68   QCRIKQIINRWQEVGKAMYAPP  3



>Cronobacter sakazakii strain cro4114B1 contig117, whole genome shotgun sequence
Sequence ID: NZ_NRKI01000117.1 Length: 404
Range 1: 1 to 402

Score:164 bits(416), Expect:1e-42,
Method:Compositional matrix adjust.,
Identities:87/142(61%), Positives:102/142(71%), Gaps:9/142(6%)

Query  298  RPNNNTRKRIRIQRGPGRAFVTIGKI-GNMRQAHCNISRAKWNNTLKQIASKLREQFGNN  356
            RP+NNTR  IRI  GPG+ F   G I G++RQA+C I+  KWN TL+Q+A KL+EQF  N
Sbjct  1    RPSNNTRTSIRI--GPGQTFYRTGDIIGDIRQAYCEINGTKWNETLRQVAKKLKEQF--N  168

Query  357  KTIIFKQSSGGDPEIVTHSFNCGGEFFYCNSTQLFNSTWFNSTWSTEGSNNTEGSDTITL  416
            KTI F+  SGGD EI    FNC GEFFYCN+T+LFNSTW     + +G N   GSDT  L
Sbjct  169  KTIKFQPPSGGDLEITMLHFNCRGEFFYCNTTKLFNSTW-EGNETIKGGN---GSDTTIL  336

Query  417  PCRIKQIINMWQKVGKAMYAPP  438
            PCRIKQIINMWQ  G+AMYAPP
Sbjct  337  PCRIKQIINMWQGAGQAMYAPP  402



>Cronobacter sakazakii strain cro4114B1 contig115, whole genome shotgun sequence
Sequence ID: NZ_NRKI01000115.1 Length: 404
Range 1: 1 to 402

Score:163 bits(413), Expect:2e-42,
Method:Compositional matrix adjust.,
Identities:82/142(58%), Positives:99/142(69%), Gaps:9/142(6%)

Query  298  RPNNNTRKRIRIQRGPGRAFVTIGKI-GNMRQAHCNISRAKWNNTLKQIASKLREQFGNN  356
            RPNNNTRK I +    GRA    G I G++RQAHCN+S+  W  TL Q+A+KLREQF  N
Sbjct  1    RPNNNTRKSISL--AGGRALYATGNIIGDIRQAHCNVSKGNWTETLGQVATKLREQF-KN  171

Query  357  KTIIFKQSSGGDPEIVTHSFNCGGEFFYCNSTQLFNSTWFNSTWSTEGSNNTEGSDTITL  416
            KTI+F  SSGGDPE+    FNCGGEFFYCN+++LFN T+        G+     +  IT+
```

```
Sbjct  172  KTIVFNSSSGGDPEVEMFIFNCGGEFFYCNTSKLFNYTFL-----ANGTGQNYSTGNITI  336

Query  417  PCRIKQIINMWQKVGKAMYAPP  438
             CRIKQIIN WQ+VGKAMYAPP
Sbjct  337  QCRIKQIINRWQEVGKAMYAPP  402
```

```
>Corynebacterium glutamicum strain 38 Scaffolds_31_4109, whole genome shotgun sequence
Sequence ID: NZ_JACENN010000031.1 Length: 3118
Range 1: 2381 to 2734

Score:181 bits(459), Expect:3e-42,
Method:Compositional matrix adjust.,
Identities:116/118(98%), Positives:116/118(98%), Gaps:0/118(0%)

Query  719   THLPTprgpdrpegieeeggerdrdrSIRLVNGSLALIWDDLRSLCLFSYHRLRDLLLIV  778
             THLP PRGPDRPEGIEEEGGERDRDRSIRLVNGSLALIWDDLRSLCLFSYHRLRDLLLIV
Sbjct  2381  THLPIPRGPDRPEGIEEEGGERDRDRSIRLVNGSLALIWDDLRSLCLFSYHRLRDLLLIV  2560

Query  779   TRIVELLGRRGWEALKYWWNLLQYWSQELKNSAVSLLNATAIAVAEGTDRVIEVVQGA  836
             TRIVELLGRRGWEALKYWWNLLQYWSQELKNSAVSLLNATAIAVAEGTDRVIEVVQ A
Sbjct  2561  TRIVELLGRRGWEALKYWWNLLQYWSQELKNSAVSLLNATAIAVAEGTDRVIEVVQEA  2734
```

```
>Cronobacter sakazakii strain cro4114A1 contig128, whole genome shotgun sequence
Sequence ID: NZ_NRKJ01000128.1 Length: 404
Range 1: 1 to 402

Score:163 bits(412), Expect:3e-42,
Method:Compositional matrix adjust.,
Identities:82/142(58%), Positives:99/142(69%), Gaps:9/142(6%)

Query  298  RPNNNTRKRIRIQRGPGRAFVTIGKI-GNMRQAHCNISRAKWNNTLKQIASKLREQFGNN  356
            RP+NNTRK I +    GRA    G I G++RQAHCN+S+  W  TL Q+A+KLREQF  N
Sbjct  1    RPSNNTRKSISL--AGGRALYATGDIIGDIRQAHCNVSKGNWTETLGQVATKLREQF-KN  171

Query  357  KTIIFKQSSGGDPEIVTHSFNCGGEFFYCNSTQLFNSTWFNSTWSTEGSNNTEGSDTITL  416
            KTI+F  SSGGDPE+    FNCGGEFFYCN+++LFN T+        G+     +  ITL
Sbjct  172  KTIVFNSSSGGDPEVEMFIFNCGGEFFYCNTSKLFNYTFL-----ANGTGQNYSTGNITL  336

Query  417  PCRIKQIINMWQKVGKAMYAPP  438
             CRIKQIIN WQ+VGKAMYAPP
Sbjct  337  QCRIKQIINRWQEVGKAMYAPP  402
```

```
>Lactiplantibacillus plantarum strain MH-301 Scaffolds_50_4236, whole genome shotgun sequence
Sequence ID: NZ_JAOYUG010000050.1 Length: 4236
Range 1: 196 to 549

Score:181 bits(459), Expect:4e-42,
Method:Compositional matrix adjust.,
Identities:116/118(98%), Positives:116/118(98%), Gaps:0/118(0%)

Query  719  THLPTprgpdrpegieeeggerdrdrSIRLVNGSLALIWDDLRSLCLFSYHRLRDLLLIV  778
            THLP PRGPDRPEGIEEEGGERDRDRSIRLVNGSLALIWDDLRSLCLFSYHRLRDLLLIV
Sbjct  196  THLPIPRGPDRPEGIEEEGGERDRDRSIRLVNGSLALIWDDLRSLCLFSYHRLRDLLLIV  375

Query  779  TRIVELLGRRGWEALKYWWNLLQYWSQELKNSAVSLLNATAIAVAEGTDRVIEVVQGA  836
            TRIVELLGRRGWEALKYWWNLLQYWSQELKNSAVSLLNATAIAVAEGTDRVIEVVQ A
Sbjct  376  TRIVELLGRRGWEALKYWWNLLQYWSQELKNSAVSLLNATAIAVAEGTDRVIEVVQEA  549
```

```
>Corynebacterium glutamicum strain 51 Scaffolds_32_4236, whole genome shotgun sequence
Sequence ID: NZ_JACENK010000032.1 Length: 4236
Range 1: 2963 to 3316

Score:181 bits(459), Expect:4e-42,
Method:Compositional matrix adjust.,
Identities:116/118(98%), Positives:116/118(98%), Gaps:0/118(0%)

Query  719   THLPTprgpdrpegieeeggerdrdrSIRLVNGSLALIWDDLRSLCLFSYHRLRDLLLIV  778
             THLP PRGPDRPEGIEEEGGERDRDRSIRLVNGSLALIWDDLRSLCLFSYHRLRDLLLIV
Sbjct  2963  THLPIPRGPDRPEGIEEEGGERDRDRSIRLVNGSLALIWDDLRSLCLFSYHRLRDLLLIV  3142

Query  779   TRIVELLGRRGWEALKYWWNLLQYWSQELKNSAVSLLNATAIAVAEGTDRVIEVVQGA  836
             TRIVELLGRRGWEALKYWWNLLQYWSQELKNSAVSLLNATAIAVAEGTDRVIEVVQ A
Sbjct  3143  TRIVELLGRRGWEALKYWWNLLQYWSQELKNSAVSLLNATAIAVAEGTDRVIEVVQEA  3316
```

```
>Cronobacter sakazakii strain cro4215W contig105, whole genome shotgun sequence
Sequence ID: NZ_NRKF01000105.1 Length: 404
Range 1: 1 to 402

Score:162 bits(411), Expect:5e-42,
Method:Compositional matrix adjust.,
Identities:82/142(58%), Positives:98/142(69%), Gaps:9/142(6%)

Query  298  RPNNNTRKRIRIQRGPGRAFVTIGKI-GNMRQAHCNISRAKWNNTLKQIASKLREQFGNN  356
            RP NNTRK I +    GRA    G I G++RQAHCN+S+  W  TL Q+A+KLREQF  N
Sbjct  1    RPTNNTRKSISL--AGGRALYATGDIIGDIRQAHCNVSKGNWTETLGQVATKLREQF-KN  171

Query  357  KTIIFKQSSGGDPEIVTHSFNCGGEFFYCNSTQLFNSTWFNSTWSTEGSNNTEGSDTITL  416
            KTI+F  SSGGDPE+    FNCGGEFFYCN+++LFN T+        G+     +  ITL
Sbjct  172  KTIVFNSSSGGDPEVEMFIFNCGGEFFYCNTSKLFNYTFL-----ANGTGQNYSTGNITL  336

Query  417  PCRIKQIINMWQKVGKAMYAPP  438
             CRIKQIIN WQ+VGKAMYAPP
Sbjct  337  QCRIKQIINRWQEVGKAMYAPP  402
```

```
>Cronobacter malonaticus strain cro3263A1 contig76, whole genome shotgun sequence
Sequence ID: NZ_NRMM01000076.1 Length: 392
Range 1: 1 to 390

Score:162 bits(411), Expect:5e-42,
Method:Compositional matrix adjust.,
Identities:85/142(60%), Positives:100/142(70%), Gaps:13/142(9%)

Query  298  RPNNNTRKRIRIQRGPGRAFVTIGKI-GNMRQAHCNISRAKWNNTLKQIASKLREQFGNN  356
            RP+NNTR  IRI  GPG+ F   G I G++RQA+C I+  KWN TL+Q+A KL+EQF  N
Sbjct  1    RPDNNTRTSIRI--GPGQTFYRTGDIIGDIRQAYCEINGTKWNETLRQVAKKLKEQF--N  168
```

```
Query  357  KTIIFKQSSGGDPEIVTHSFNCGGEFFYCNSTQLFNSTWFNSTWSTEGSNNTEGSDTITL  416
            KTI F+  SGGD EI    FNC GEFFYCN+ +LFNSTW          N T  +DTI L
Sbjct  169  KTIKFQPPSGGDLEITMLHFNCRGEFFYCNTAKLFNSTWI--------GNATMENDTIIL  324

Query  417  PCRIKQIINMWQKVGKAMYAPP  438
            PCRIKQIIN+WQ VG+AMYAPP
Sbjct  325  PCRIKQIINLWQGVGQAMYAPP  390
```

```
>Cronobacter dublinensis strain cro981C2 contig308, whole genome shotgun sequence
Sequence ID: NZ_NROB01000308.1 Length: 405
Range 1: 1 to 399

Score:161 bits(408), Expect:1e-41,
Method:Compositional matrix adjust.,
Identities:86/141(61%), Positives:101/141(71%), Gaps:9/141(6%)

Query  298  RPNNNTRKRIRIQRGPGRAFVTIGKI-GNMRQAHCNISRAKWNNTLKQIASKLREQFGNN  356
            RP+NNTR  IRI  GPG+ F   G I G++RQA+C I+  KWN TL+Q+A KL+EQF  N
Sbjct  1    RPSNNTRTSIRI--GPGQTFYRTGDIIGDIRQAYCEINGTKWNETLRQVAKKLKEQF--N  168

Query  357  KTIIFKQSSGGDPEIVTHSFNCGGEFFYCNSTQLFNSTWFNSTWSTEGSNNTEGSDTITL  416
            KTI F+  SGGD EI    FNC GEFFYCN+T+LFNSTW     + +G N   GSDT  L
Sbjct  169  KTIKFQPPSGGDLEITMLHFNCRGEFFYCNTTKLFNSTW-EGNETIKGGN---GSDTTIL  336

Query  417  PCRIKQIINMWQKVGKAMYAP  437
            PCRIKQIINMWQ  G+AMYAP
Sbjct  337  PCRIKQIINMWQGAGQAMYAP  399
```

```
>Deinococcus aquiradiocola strain JCM 14371 sequence36, whole genome shotgun sequence
Sequence ID: NZ_BMOE01000036.1 Length: 869
Range 1: 321 to 653

Score:166 bits(421), Expect:2e-41,
Method:Compositional matrix adjust.,
Identities:109/111(98%), Positives:109/111(98%), Gaps:0/111(0%)

Query  499  TKAKRRVVQREKRAVGIgalflgflgaagstmgaasmtLTVQARQLLSGIVQQQNNLLRA  558
             KAKRRVVQREKRAVGIGALFLGFLGAAGSTMGAASMTLTVQARQLLS IVQQQNNLLRA
Sbjct  321  AKAKRRVVQREKRAVGIGALFLGFLGAAGSTMGAASMTLTVQARQLLSDIVQQQNNLLRA  500

Query  559  IEAQQHLLQLTVWGIKQLQARILAVERYLKDQQLLGIWGCSGKLICTTAVP  609
            IEAQQHLLQLTVWGIKQLQARILAVERYLKDQQLLGIWGCSGKLICTTAVP
Sbjct  501  IEAQQHLLQLTVWGIKQLQARILAVERYLKDQQLLGIWGCSGKLICTTAVP  653
```

```
>Cronobacter sakazakii strain cro4215W contig109, whole genome shotgun sequence
Sequence ID: NZ_NRKF01000109.1 Length: 389
Range 1: 3 to 389

Score:160 bits(406), Expect:2e-41,
Method:Composition-based stats.,
Identities:82/142(58%), Positives:98/142(69%), Gaps:14/142(9%)

Query  298  RPNNNTRKRIRIQRGPGRAFVTIGKI-GNMRQAHCNISRAKWNNTLKQIASKLREQFGNN  356
            RP+NNTR  I I  GPGRAF   G I G++RQA+C ++R +WN  L Q+  KL E F  N
Sbjct  389  RPDNNTRTSINI--GPGRAFYRTGDIIGDIRQAYCEVNRTEWNEILGQVIKKLEEHF-KN  219

Query  357  KTIIFKQSSGGDPEIVTHSFNCGGEFFYCNSTQLFNSTWFNSTWSTEGSNNTEGSDTITL  416
            KTI F+  +GGD EI  H FNC GEFFYCN+T+L     FNS+ + +GSN     DTI L
Sbjct  218  KTIEFQPPAGGDLEITMHHFNCRGEFFYCNTTKL-----FNSSETGKGSN-----DTIIL  69

Query  417  PCRIKQIINMWQKVGKAMYAPP  438
            PCRIKQIINMWQ  G+AMYAPP
Sbjct  68   PCRIKQIINMWQGAGQAMYAPP  3
```

```
>Cronobacter sakazakii strain cro4114B1 contig105, whole genome shotgun sequence
Sequence ID: NZ_NRKI01000105.1 Length: 416
Range 1: 3 to 416

Score:159 bits(402), Expect:9e-41,
Method:Compositional matrix adjust.,
Identities:85/144(59%), Positives:105/144(72%), Gaps:9/144(6%)

Query  298  RPNNNTRKRIRIQRGPGRAFVTIGKI-GNMRQAHCNISRAKWNNTLKQIASKLREQFGNN  356
            RP NNTRK IRI  GPG+ F   G+I G++RQA+CNIS  +W  TL  ++ KL E F  N
Sbjct  416  RPGNNTRKSIRI--GPGQTFYATGEIIGDIRQAYCNISGKEWKETLHNVSKKLAEYF-PN  246

Query  357  KTIIFKQSSGGDPEIVTHSFNCGGEFFYCNSTQLFNSTW-FNSTWSTEGSNNTEGSDT-I  414
            KTI +  SSGGD EI THSFNC GEFFYCN+++LFN T+  N T++  G   TE   T I
Sbjct  245  KTINWTSSSGGDLEITTHSFNCRGEFFYCNTSELFNGTYMLNGTFTFNG---TESEPTNI  75

Query  415  TLPCRIKQIINMWQKVGKAMYAPP  438
            T+PC+IKQ+INMWQ+VG+AMYAPP
Sbjct  74   TIPCKIKQVINMWQEVGRAMYAPP  3
```

```
>Thalassovita mangrovi strain GS-10 Scaffold21_1, whole genome shotgun sequence
Sequence ID: NZ_WWEN01000021.1 Length: 1440
Range 1: 752 to 1084

Score:169 bits(427), Expect:2e-40,
Method:Compositional matrix adjust.,
Identities:109/111(98%), Positives:109/111(98%), Gaps:0/111(0%)

Query  499   TKAKRRVVQREKRAVGIgalflgflgaagstmgaasmtLTVQARQLLSGIVQQQNNLLRA  558
              KAKRRVVQREKRAVGIGALFLGFLGAAGSTMGAASMTLTVQARQLLS IVQQQNNLLRA
Sbjct  1084  AKAKRRVVQREKRAVGIGALFLGFLGAAGSTMGAASMTLTVQARQLLSDIVQQQNNLLRA  905

Query  559   IEAQQHLLQLTVWGIKQLQARILAVERYLKDQQLLGIWGCSGKLICTTAVP  609
             IEAQQHLLQLTVWGIKQLQARILAVERYLKDQQLLGIWGCSGKLICTTAVP
Sbjct  904   IEAQQHLLQLTVWGIKQLQARILAVERYLKDQQLLGIWGCSGKLICTTAVP  752
```

```
>Klebsiella pneumoniae strain EuSCAPE_MT015, whole genome shotgun sequence
Sequence ID: NZ_UKIW02000108.1 Length: 338
Range 1: 2 to 337

Score:156 bits(395), Expect:3e-40,
Method:Compositional matrix adjust.,
```

```
Identities:85/112(76%), Positives:104/112(92%), Gaps:0/112(0%)

Query  594  GIWGCSGKLICTTAVPWNASWSNKSLEQIWNHTTWMEWDREINNYTSLIHSLIeesqnqq  653
            GIWGCSGKLICTT VPWN+SWSNKSL  IW++ TWM+WDRE++NYT+ I+ L+E+SQNQQ
Sbjct  337  GIWGCSGKLICTTNVPWNSSWSNKSLTDIWDNMTWMQWDREVSNYTNTIYKLLEDSQNQQ  158

Query  654  ekneqellelDKWASLWNWFNITNWLWYIKLFIMIVGGLVGLRIVFAVLSIV  705
            +KNEQ+LL LDKW +LWNWFNIT+WLWYI++FIMIVGGL+GLRI+FAV+SIV
Sbjct  157  DKNEQDLLALDKWDNLWNWFNITSWLWYIRIFIMIVGGLIGLRIIFAVISIV  2




>Escherichia coli strain EN5308 contig00153, whole genome shotgun sequence
Sequence ID: NZ_JARBDE010000153.1 Length: 786
Range 1: 262 to 786

Score:162 bits(409), Expect:4e-40,
Method:Composition-based stats.,
Identities:109/175(62%), Positives:128/175(73%), Gaps:7/175(4%)

Query  688  IVGGLVGLRIVFAVLSIVNRVRQGYSPLSFQTHLPTprgpdrpegieeeggerdrdrSIR  747
            IVGGL+ LRI+FAVLSIVN+VRQ YSPLSFQT  P P GPDR   IEEE GE+D+DRSI+
Sbjct  786  IVGGLIKLRIIFAVLSIVNKVRQRYSPLSFQTLTPNPGGPDRLRRIEEESGEQDKDRSIQ  607

Query  748  LVNGSLALIWDDLRSLCLFSYHRLRDLLLIVTRIVELL---GRRGW----EALKYWWNLL  800
            LV+G LAL  DDL++LCLFSYHRLRD + +  R VELL     RG     EALKY  +L+
Sbjct  606  LVSGFLALA*DDLQNLCLFSYHRLRDFISVTARTVELLRHHSLRGLQKR*EALKYLRSLV  427

Query  801  QYWSQELKNSAVSLLNATAIAVAEGTDRVIEVVQGACRAIRHIPRRIRQGLERIL  855
            QY   ELK SA+SLL+  AIAVAE TD++IE+ Q   RA  +IPRRIRQ  E  L
Sbjct  426  QY*GLELKKSAISLLDTIAIAVAERTDKIIELGQRIYRAFCNIPRRIRQSFEAAL  262




>Cronobacter sakazakii strain cro4114A1 contig133, whole genome shotgun sequence
Sequence ID: NZ_NRKJ01000133.1 Length: 389
Range 1: 3 to 389

Score:157 bits(396), Expect:4e-40,
Method:Compositional matrix adjust.,
Identities:83/142(58%), Positives:98/142(69%), Gaps:14/142(9%)

Query  298  RPNNNTRKRIRIQRGPGRAFVTIGKI-GNMRQAHCNISRAKWNNTLKQIASKLREQFGNN  356
            RPNNNTR  I I  GPGRAF   G I G++RQA+C ++R +WN  L Q+  KL E F  N
Sbjct  389  RPNNNTRTSINI--GPGRAFYRTGDIIGDIRQAYCEVNRTEWNEILGQVIKKLEEHF-KN  219

Query  357  KTIIFKQSSGGDPEIVTHSFNCGGEFFYCNSTQLFNSTWFNSTWSTEGSNNTEGSDTITL  416
            KTI F+  +GGD EI  H FNC GEFFYCN+T+L     FNS+ + +GSN     DTI L
Sbjct  218  KTIEFQPPAGGDLEITMHHFNCRGEFFYCNTTKL-----FNSSETGKGSN-----DTIIL  69

Query  417  PCRIKQIINMWQKVGKAMYAPP  438
            PCRIKQIINMWQ  G+AMYAPP
Sbjct  68   PCRIKQIINMWQGAGQAMYAPP  3




>Cronobacter sakazakii strain cro4191B1-2 contig85, whole genome shotgun sequence
Sequence ID: NZ_NRKG01000085.1 Length: 392
Range 1: 1 to 390

Score:156 bits(395), Expect:6e-40,
Method:Compositional matrix adjust.,
Identities:83/142(58%), Positives:97/142(68%), Gaps:13/142(9%)

Query  298  RPNNNTRKRIRIQRGPGRAFVTIGKI-GNMRQAHCNISRAKWNNTLKQIASKLREQFGNN  356
            RP NNTRK IRI  GPG+ F   G I G++RQAHC ++   +WN TLKQ+  KL+E F  N
Sbjct  1    RPTNNTRKSIRI--GPGQMFYRTGDIIGDIRQAHCEVNGTEWNETLKQVTKKLKEYF--N  168

Query  357  KTIIFKQSSGGDPEIVTHSFNCGGEFFYCNSTQLFNSTWFNSTWSTEGSNNTEGSDTITL  416
            KTI+F+  SGGD E     FNC GEFFYCN+T LFNST      + EG N      TI L
Sbjct  169  KTIVFRPPSGGDLETTMLHFNCRGEFFYCNTTNLFNST---ENGTMEGLNT-----TIIL  324

Query  417  PCRIKQIINMWQKVGKAMYAPP  438
            PCRIKQI+NMWQ VG+AMYAPP
Sbjct  325  PCRIKQIVNMWQGVGQAMYAPP  390




>Cronobacter sakazakii strain cro3040W contig84, whole genome shotgun sequence
Sequence ID: NZ_NRJO01000084.1 Length: 392
Range 1: 1 to 390

Score:156 bits(395), Expect:6e-40,
Method:Compositional matrix adjust.,
Identities:83/142(58%), Positives:97/142(68%), Gaps:13/142(9%)

Query  298  RPNNNTRKRIRIQRGPGRAFVTIGKI-GNMRQAHCNISRAKWNNTLKQIASKLREQFGNN  356
            RP NNTRK IRI  GPG+ F   G I G++RQAHC ++   +WN TLKQ+  KL+E F  N
Sbjct  1    RPTNNTRKSIRI--GPGQMFYRTGDIIGDIRQAHCEVNGTEWNETLKQVTKKLKEYF--N  168

Query  357  KTIIFKQSSGGDPEIVTHSFNCGGEFFYCNSTQLFNSTWFNSTWSTEGSNNTEGSDTITL  416
            KTI+F+  SGGD E     FNC GEFFYCN+T LFNST      + EG N      TI L
Sbjct  169  KTIVFRPPSGGDLETTMLHFNCRGEFFYCNTTNLFNST---ENGTMEGLNT-----TIIL  324

Query  417  PCRIKQIINMWQKVGKAMYAPP  438
            PCRIKQI+NMWQ VG+AMYAPP
Sbjct  325  PCRIKQIVNMWQGVGQAMYAPP  390




>Planctopirus hydrillae strain JC280 Contig113, whole genome shotgun sequence
Sequence ID: NZ_LYDR01000013.1 Length: 437
Range 1: 11 to 436

Score:157 bits(396), Expect:7e-40,
Method:Compositional matrix adjust.,
Identities:88/147(60%), Positives:107/147(72%), Gaps:6/147(4%)

Query  292  VEINCTRPNNNTRKRIRIQRGPGRAFVTIGKI-GNMRQAHCNISRAKWNNTLKQIASKLR  350
            VEI C RPNNNTRK +RI  GPG+ F   G+I G++R A+CNI+  KWN TL+ ++ KL
Sbjct  11   VEIVCIRPNNNTRKSMRI--GPGQTFYATGEIIGDIRYAYCNINEQKWNATLQNVSRKLL  184

Query  351  EQFGNNKTIIFKQSSGGDPEIVTHSFNCGGEFFYCNSTQLFNSTWFNSTWSTEGSNNTEG  410
            E F N KTI F++SSGGD EI THSFNC GEFFYCN++ LFN T+  S      +N+T
Sbjct  185  EHFPN-KTISFQRSSGGDLEITTHSFNCRGEFFYCNTSNLFNRTF--SELVNNSTNSTNS  355

Query  411  SDTITLPCRIKQIINMWQKVGKAMYAP  437
            +  ITL CRIKQIINMWQ VG+AMYAP
Sbjct  356  TSIITLQCRIKQIINMWQGVGRAMYAP  436
```

```
>Cronobacter sakazakii strain cro4114B1 contig122, whole genome shotgun sequence
Sequence ID: NZ_NRKI01000122.1 Length: 383
Range 1: 1 to 381

Score:155 bits(393), Expect:9e-40,
Method:Compositional matrix adjust.,
Identities:81/142(57%), Positives:95/142(66%), Gaps:16/142(11%)

Query  298  RPNNNTRKRIRIQRGPGRAFVTIGKI-GNMRQAHCNISRAKWNNTLKQIASKLREQFGNN  356
            RPNNNTR  I I  GPGRAF   G I G++RQA+C ++R +WN  L Q+  KL E F  N
Sbjct  1    RPNNNTRTSINI--GPGRAFYRTGDIIGDIRQAYCEVNRTEWNEILGQVIKKLEEHF-KN  171

Query  357  KTIIFKQSSGGDPEIVTHSFNCGGEFFYCNSTQLFNSTWFNSTWSTEGSNNTEGSDTITL  416
            KTI F+  +GGD EI  H FNC GEFFYCN+T+LFNS+             T  +DTI L
Sbjct  172  KTIEFQPPAGGDLEITMHHFNCRGEFFYCNTTKLFNSS------------ETRENDTIIL  315

Query  417  PCRIKQIINMWQKVGKAMYAPP  438
            PCRIKQIINMWQ  G+AMYAPP
Sbjct  316  PCRIKQIINMWQGAGQAMYAPP  381



>Cronobacter malonaticus strain cro3263A1 contig75, whole genome shotgun sequence
Sequence ID: NZ_NRMM01000075.1 Length: 392
Range 1: 1 to 390

Score:154 bits(389), Expect:5e-39,
Method:Compositional matrix adjust.,
Identities:82/142(58%), Positives:96/142(67%), Gaps:13/142(9%)

Query  298  RPNNNTRKRIRIQRGPGRAFVTIGKI-GNMRQAHCNISRAKWNNTLKQIASKLREQFGNN  356
            RP NNTRK IRI  GPG+ F   G I G++R AHC ++  +WN TLKQ+  KL+E F  N
Sbjct  1    RPTNNTRKSIRI--GPGQMFYRTGDIIGDIRNAHCEVNGTEWNETLKQVTKKLKEYF--N  168

Query  357  KTIIFKQSSGGDPEIVTHSFNCGGEFFYCNSTQLFNSTWFNSTWSTEGSNNTEGSDTITL  416
            KTI+F+  SGGD E     FNC GEFFYCN+T LFNST      + EG N      TI L
Sbjct  169  KTIVFRPPSGGDLETTMLHFNCRGEFFYCNTTNLFNST---ENGTMEGLNT-----TIIL  324

Query  417  PCRIKQIINMWQKVGKAMYAPP  438
            PCRIKQI+NMWQ VG+AMYAPP
Sbjct  325  PCRIKQIVNMWQGVGQAMYAPP  390



>Cronobacter sakazakii strain cro4114B1 contig124, whole genome shotgun sequence
Sequence ID: NZ_NRKI01000124.1 Length: 392
Range 1: 1 to 390

Score:154 bits(389), Expect:5e-39,
Method:Compositional matrix adjust.,
Identities:82/142(58%), Positives:96/142(67%), Gaps:13/142(9%)

Query  298  RPNNNTRKRIRIQRGPGRAFVTIGKI-GNMRQAHCNISRAKWNNTLKQIASKLREQFGNN  356
            RP NNTRK IRI  GPG+ F   G I G++R AHC ++  +WN TLKQ+  KL+E F  N
Sbjct  1    RPTNNTRKSIRI--GPGQMFYRTGDIIGDIRNAHCEVNGTEWNETLKQVTKKLKEYF--N  168

Query  357  KTIIFKQSSGGDPEIVTHSFNCGGEFFYCNSTQLFNSTWFNSTWSTEGSNNTEGSDTITL  416
            KTI+F+  SGGD E     FNC GEFFYCN+T LFNST      + EG N      TI L
Sbjct  169  KTIVFRPPSGGDLETTMLHFNCRGEFFYCNTTNLFNST---ENGTMEGLNT-----TIIL  324

Query  417  PCRIKQIINMWQKVGKAMYAPP  438
            PCRIKQI+NMWQ VG+AMYAPP
Sbjct  325  PCRIKQIVNMWQGVGQAMYAPP  390



>Cronobacter sakazakii strain cro4191B1-2 contig83, whole genome shotgun sequence
Sequence ID: NZ_NRKG01000083.1 Length: 410
Range 1: 1 to 408

Score:154 bits(388), Expect:6e-39,
Method:Compositional matrix adjust.,
Identities:81/143(57%), Positives:97/143(67%), Gaps:9/143(6%)

Query  298  RPNNNTRKRIRIQRGPGRAFVTIGKI-GNMRQAHCNISRAKWNNTLKQIASKLREQFGNN  356
            RP NNTR  I I  GPG+ F   G I GN+R+A C I++ KW   LKQ+A+KL+  F N
Sbjct  1    RPYNNTRTGITI--GPGQVFYRTGDIIGNIRKAFCEINKTKWKEVLKQVATKLKGHF-NT  171

Query  357  KTIIFKQSSGGDPEIVTHSFNCGGEFFYCNSTQLFNSTWF-NSTWSTEGSNNTEGSDTIT  415
              II +  SGGD EI TH F+C GEFFYCN+T+LFNST   N TW T      EG+ TIT
Sbjct  172  TKIILQPPSGGDLEITTHHFSCRGEFFYCNTTELFNSTLMENGTWET----TKEGNGTIT  339

Query  416  LPCRIKQIINMWQKVGKAMYAPP  438
            +PCRIKQI+ MWQ  G+AMYAPP
Sbjct  340  IPCRIKQIVRMWQGTGQAMYAPP  408



>Cronobacter sakazakii strain cro4114A1 contig131, whole genome shotgun sequence
Sequence ID: NZ_NRKJ01000131.1 Length: 392
Range 1: 1 to 390

Score:153 bits(387), Expect:7e-39,
Method:Compositional matrix adjust.,
Identities:82/142(58%), Positives:96/142(67%), Gaps:13/142(9%)

Query  298  RPNNNTRKRIRIQRGPGRAFVTIGKI-GNMRQAHCNISRAKWNNTLKQIASKLREQFGNN  356
            RP NNTRK IRI  GPG+ F   G I G++R AHC ++  +WN TLKQ+  KL+E F  N
Sbjct  1    RPYNNTRKSIRI--GPGQMFYRTGDIIGDIRNAHCEVNGTEWNETLKQVTKKLKEYF--N  168

Query  357  KTIIFKQSSGGDPEIVTHSFNCGGEFFYCNSTQLFNSTWFNSTWSTEGSNNTEGSDTITL  416
            KTI+F+  SGGD E     FNC GEFFYCN+T LFNST      + EG N      TI L
Sbjct  169  KTIVFRPPSGGDLETTMLHFNCRGEFFYCNTTNLFNST---ENGTMEGLNT-----TIIL  324

Query  417  PCRIKQIINMWQKVGKAMYAPP  438
            PCRIKQI+NMWQ VG+AMYAPP
Sbjct  325  PCRIKQIVNMWQGVGQAMYAPP  390



>Cronobacter sakazakii strain cro3040W contig83, whole genome shotgun sequence
Sequence ID: NZ_NRJO01000083.1 Length: 392
Range 1: 1 to 390

Score:152 bits(383), Expect:2e-38,
```

```
Method:Compositional matrix adjust.,
Identities:78/142(55%), Positives:99/142(69%), Gaps:13/142(9%)

Query  298  RPNNNTRKRIRIQRGPGRAFVTIGKI-GNMRQAHCNISRAKWNNTLKQIASKLREQFGNN  356
            RPNNNTR  +RI  GPG+ F     I G++R+A+C I+  +W  TL+Q+ +KL+E F NN
Sbjct  1    RPNNNTRTSVRI--GPGQMFYRTEDITGDIRKAYCEINGTEWTETLEQVTAKLKEHF-NN  171

Query  357  KTIIFKQSSGGDPEIVTHSFNCGGEFFYCNSTQLFNSTWFNSTWSTEGSNNTEGSDTITL  416
            KTI+F+  SGGD EI  H FNC GEFFYCN+T+LFNST      + EG N      TI L
Sbjct  172  KTILFRPPSGGDLEITMHHFNCKGEFFYCNTTKLFNST----NGTMEGLNG-----TIIL  324

Query  417  PCRIKQIINMWQKVGKAMYAPP  438
            PCRI+QI+ MWQ+ G+AMYAPP
Sbjct  325  PCRIRQIVRMWQETGQAMYAPP  390
```

```
>Klebsiella pneumoniae strain EuSCAPE_MT017, whole genome shotgun sequence
Sequence ID: NZ_UKJA02000086.1 Length: 513
Range 1: 155 to 499

Score:153 bits(387), Expect:3e-38,
Method:Composition-based stats.,
Identities:78/115(68%), Positives:85/115(73%), Gaps:7/115(6%)

Query  745  SIRLVNGSLALIWDDLRSLCLFSYHRLRDLLLIVTRIVELLGR-------RGWEALKYWW  797
            SIRLV G LAL WDDLRSLCLFSYHRLRDL+L+  R  ELLGR       RGWE LKY
Sbjct  499  SIRLVQGFLALAWDDLRSLCLFSYHRLRDLILVTARAGELLGRSSLKGLQRGWEILKYLG  320

Query  798  NLLQYWSQELKNSAVSLLNATAIAVAEGTDRVIEVVQGACRAIRHIPRRIRQGLE  852
            +L+QYW  ELK SA SLL+  AIAVA GTDR+IEV+Q   RAI HIP RIRQG E
Sbjct  319  SLIQYWGLELKKSATSLLDTIAIAVATGTDRIIEVLQAIGRAIYHIPTRIRQGFE  155
```

```
>Cronobacter dublinensis strain cro3540W contig49, whole genome shotgun sequence
Sequence ID: NZ_NRNN01000049.1 Length: 392
Range 1: 1 to 390

Score:151 bits(381), Expect:6e-38,
Method:Compositional matrix adjust.,
Identities:77/142(54%), Positives:99/142(69%), Gaps:13/142(9%)

Query  298  RPNNNTRKRIRIQRGPGRAFVTIGKI-GNMRQAHCNISRAKWNNTLKQIASKLREQFGNN  356
            RP+NNTR  +RI  GPG+ F     I G++R+A+C I+  +W  TL+Q+ +KL+E F NN
Sbjct  1    RPDNNTRTSVRI--GPGQMFYRTEDITGDIRKAYCEINGTEWTETLEQVTAKLKEHF-NN  171

Query  357  KTIIFKQSSGGDPEIVTHSFNCGGEFFYCNSTQLFNSTWFNSTWSTEGSNNTEGSDTITL  416
            KTI+F+  SGGD EI  H FNC GEFFYCN+T+LFNST      + EG N      TI L
Sbjct  172  KTILFRPPSGGDLEITMHHFNCKGEFFYCNTTKLFNST----NGTMEGLNG-----TIIL  324

Query  417  PCRIKQIINMWQKVGKAMYAPP  438
            PCRI+QI+ MWQ+ G+AMYAPP
Sbjct  325  PCRIRQIVRMWQETGQAMYAPP  390
```

```
>Actinoplanes campanulatus strain JCM 3059 sequence058, whole genome shotgun sequence
Sequence ID: NZ_BMPW01000058.1 Length: 4754
Range 1: 4417 to 4749

Score:167 bits(422), Expect:2e-37,
Method:Compositional matrix adjust.,
Identities:109/111(98%), Positives:109/111(98%), Gaps:0/111(0%)

Query  499   TKAKRRVVQREKRAVGIgalflgflgaagstmgaasmtLTVQARQLLSGIVQQQNNLLRA  558
              KAKRRVVQREKRAVGIGALFLGFLGAAGSTMGAASMTLTVQARQLLS IVQQQNNLLRA
Sbjct  4417  AKAKRRVVQREKRAVGIGALFLGFLGAAGSTMGAASMTLTVQARQLLSDIVQQQNNLLRA  4596

Query  559   IEAQQHLLQLTVWGIKQLQARILAVERYLKDQQLLGIWGCSGKLICTTAVP  609
             IEAQQHLLQLTVWGIKQLQARILAVERYLKDQQLLGIWGCSGKLICTTAVP
Sbjct  4597  IEAQQHLLQLTVWGIKQLQARILAVERYLKDQQLLGIWGCSGKLICTTAVP  4749
```

```
>Photobacterium rosenbergii strain DP6N14-7 DP6N14-7_ctg0065, whole genome shotgun sequence
Sequence ID: NZ_JAHVIB010000057.1 Length: 564
Range 1: 116 to 445

Score:147 bits(370), Expect:1e-35,
Method:Composition-based stats.,
Identities:84/110(76%), Positives:97/110(88%), Gaps:0/110(0%)

Query  664  DKWASLWNWFNITNWLWYIKLFIMIVGGLVGLRIVFAVLSIVNRVRQGYSPLSFQTHLPT  723
            D W +LWNWF+I+NWLWYI++FIMIVGGL+GLRIVFAVLSIVNRVRQGYSPLS QT +P
Sbjct  116  DSWKNLWNWFDISNWLWYIRIFIMIVGGLIGLRIVFAVLSIVNRVRQGYSPLSLQTLIPN  295

Query  724  prgpdrpegieeeggerdrdrSIRLVNGSLALIWDDLRSLCLFSYHRLRD  773
            P GPDR   I+EEGGE+ RDRS+RLV+G LAL W+DLR+LCLFSYH LRD
Sbjct  296  PVGPDRLGRIDEEGGEQGRDRSVRLVSGFLALAWEDLRNLCLFSYHHLRD  445
```

```
>Cronobacter sakazakii strain cro4191B1-2 contig84, whole genome shotgun sequence
Sequence ID: NZ_NRKG01000084.1 Length: 402
Range 1: 1 to 303

Score:127 bits(318), Expect:1e-35,
Method:Composition-based stats.,
Identities:63/104(61%), Positives:72/104(69%), Gaps:4/104(3%)

Query  298  RPNNNTRKRIRIQRGPGRAFVTIGKI-GNMRQAHCNISRAKWNNTLKQIASKLREQFGNN  356
            RPNNNTRK IRI  GPG+ F   G I G++RQAHCNISR  W  TL  ++ KL E F  N
Sbjct  1    RPNNNTRKSIRI--GPGQTFYATGDIIGDIRQAHCNISRKDWEETLHNVSKKLAEHF-QN  171

Query  357  KTIIFKQSSGGDPEIVTHSFNCGGEFFYCNSTQLFNSTWFNSTW  400
            K I F  SSGGD EI THSFNC GEFFYCN++ LFN T+  + W
Sbjct  172  KIIQFASSSGGDLEITTHSFNCRGEFFYCNTSGLFNGTYRYT*W  303
```

```
Range 2: 299 to 400

Score:58.5 bits(140), Expect:1e-35,
Method:Compositional matrix adjust.,
Identities:27/34(79%), Positives:30/34(88%), Gaps:1/34(2%)

Query  406  NNTEG-SDTITLPCRIKQIINMWQKVGKAMYAPP  438
```

```
            N TE  S TIT+PCRIKQIINMWQ+VG+AMYAPP
Sbjct  299  NGTESNSTTITIPCRIKQIINMWQEVGRAMYAPP  400
```

```
>Klebsiella pneumoniae strain EuSCAPE_NO017, whole genome shotgun sequence
Sequence ID: NZ_UKIO02000061.1 Length: 607
Range 1: 4 to 357

Score:146 bits(369), Expect:2e-35,
Method:Composition-based stats.,
Identities:78/118(66%), Positives:88/118(74%), Gaps:7/118(5%)

Query  745  SIRLVNGSLALIWDDLRSLCLFSYHRLRDLLLIVTRIVELLG-------RRGWEALKYWW  797
            SIRLV+G LAL+WDDLRSLCLFSYHRLRD +L+  R  ELLG       +RGWE LKY
Sbjct  4    SIRLVSGFLALVWDDLRSLCLFSYHRLRDFILVTARAAELLGRNLLRGLQRGWEILKYLG  183

Query  798  NLLQYWSQELKNSAVSLLNATAIAVAEGTDRVIEVVQGACRAIRHIPRRIRQGLERIL  855
            +L QYW  ELK SA SLL+A AIA AEGTDR+IE +Q   RAI +IPRRIRQGLE  L
Sbjct  184  SLAQYWGTELKKSATSLLDAIAIATAEGTDRIIEAIQRIGRAIYNIPRRIRQGLELAL  357
```

```
>Cronobacter malonaticus strain cro4164A3 contig145, whole genome shotgun sequence
Sequence ID: NZ_NRNF01000145.1 Length: 405
Range 1: 118 to 405

Score:120 bits(301), Expect:2e-34,
Method:Compositional matrix adjust.,
Identities:61/99(62%), Positives:70/99(70%), Gaps:4/99(4%)

Query  298  RPNNNTRKRIRIQRGPGRAFVTIGKI-GNMRQAHCNISRAKWNNTLKQIASKLREQFGNN  356
            RP+NNTRK IRI  GPG+ F   G I G++RQAHCNISR  W  TL  ++ KL E F  N
Sbjct  405  RPSNNTRKSIRI--GPGQTFYATGDIIGDIRQAHCNISRKDWEETLHNVSKKLAEHF-QN  235

Query  357  KTIIFKQSSGGDPEIVTHSFNCGGEFFYCNSTQLFNSTW  395
            K I F  SSGGD EI THSFNC GEFFYCN++ LFN T+
Sbjct  234  KKIQFASSSGGDLEITTHSFNCRGEFFYCNTSGLFNKTY  118
```

```
Range 2: 3 to 101

Score:60.8 bits(146), Expect:2e-34,
Method:Compositional matrix adjust.,
Identities:26/33(79%), Positives:30/33(90%), Gaps:0/33(0%)

Query  406  NNTEGSDTITLPCRIKQIINMWQKVGKAMYAPP  438
            N TE + TIT+PCRIKQIINMWQ+VG+AMYAPP
Sbjct  101  NGTESNGTITIPCRIKQIINMWQEVGQAMYAPP  3
```

```
>Klebsiella pneumoniae strain EuSCAPE_IL018, whole genome shotgun sequence
Sequence ID: NZ_UKJU02000101.1 Length: 388
Range 1: 3 to 386

Score:139 bits(349), Expect:1e-33,
Method:Composition-based stats.,
Identities:89/128(70%), Positives:101/128(78%), Gaps:7/128(5%)

Query  710  QGYSPLSFQTHLPTprgpdrpegieeeggerdrdrSIRLVNGSLALIWDDLRSLCLFSYH  769
            QGYSP+SFQT  P PRGPDR   IEEEGGE+DRDRS+RLV+G LAL WDDLRSLCLF YH
Sbjct  386  QGYSPVSFQTLTPNPRGPDRLGRIEEEGGEQDRDRSVRLVSGFLALAWDDLRSLCLFCYH  207

Query  770  RLRDLLLIVTRIVELLG-------RRGWEALKYWWNLLQYWSQELKNSAVSLLNATAIAV  822
            RLRD +L+  R VELLG       +RGWE LKY  +L+QYW  ELK SA+SLL+ TAIAV
Sbjct  206  RLRDFILVTARAVELLGHSSLKGLQRGWEILKYLGSLVQYWGLELKRSAISLLDTTAIAV  27

Query  823  AEGTDRVI  830
            AEGTDR+I
Sbjct  26   AEGTDRII  3
```

```
>Klebsiella pneumoniae strain EuSCAPE_IL015, whole genome shotgun sequence
Sequence ID: NZ_UKJV02000040.1 Length: 532
Range 1: 4 to 321

Score:140 bits(353), Expect:1e-33,
Method:Composition-based stats.,
Identities:67/106(63%), Positives:80/106(75%), Gaps:7/106(6%)

Query  757  WDDLRSLCLFSYHRLRDLLLIVTRIVELLGR-------RGWEALKYWWNLLQYWSQELKN  809
            WDDLR+LCLFSYHRLRDL L+  R+VELLGR       +GW+ALKY  +L+QYW  ELK
Sbjct  4    WDDLRNLCLFSYHRLRDLTLVTARVVELLGRSSLRGLQKGWDALKYLGSLVQYWGLELKK  183

Query  810  SAVSLLNATAIAVAEGTDRVIEVVQGACRAIRHIPRRIRQGLERIL  855
            SA+SL +  AI VAEGTDR+IEV+Q  CRAI ++P RIRQG E  L
Sbjct  184  SAISLFDTIAIVVAEGTDRIIEVIQRFCRAIYNVPTRIRQGFEAAL  321
```

```
>Klebsiella pneumoniae strain EuSCAPE_DE091, whole genome shotgun sequence
Sequence ID: NZ_UKMA02000049.1 Length: 618
Range 1: 202 to 570

Score:140 bits(352), Expect:4e-33,
Method:Compositional matrix adjust.,
Identities:78/123(63%), Positives:88/123(71%), Gaps:16/123(13%)

Query  745  SIRLVNGSLALIWDDLRSLCLFSYHRLRDLLLIVTRIVELLGR-------RGWEALKYWW  797
            S+RLVNG L L+W+DLRSLCLFSYHRLRDLLLI  R VELLGR       R WEALKY
Sbjct  570  SVRLVNGFLTLVWNDLRSLCLFSYHRLRDLLLITARAVELLGRCSLRGLQRVWEALKYLG  391

Query  798  NLLQYWSQELKNSAVSLLNATAIAVAEGTDRVIEVVQGACRAIRH---------IPRRIR  848
            +L+QYW  ELK SA+SLL+  AIAVAEGTDR+IEV+Q  CRA R+           P RIR
Sbjct  390  SLVQYWGLELKKSAISLLDTIAIAVAEGTDRIIEVIQIICRANRNNIDNL*GYPQPTRIR  211

Query  849  QGL  851
            Q L
Sbjct  210  QKL  202
```

```
>Neisseria gonorrhoeae strain GC-206 NODE_162_length_243_cov_2.87234_ID_20496, whole genome shotgun sequence
Sequence ID: NZ_SAWN01000162.1 Length: 243
```

```
Range 1: 1 to 243

Score:135 bits(339), Expect:5e-33,
Method:Compositional matrix adjust.,
Identities:62/83(75%), Positives:74/83(89%), Gaps:2/83(2%)

Query  424  INMWQKVGKAMYAPPISGQIRCSSNITGLLLTRDGGNSNNESEIFRPGGGDMRDNWRSEL  483
            INMW +VG+A+YAPPI+G I C+S+ITG+LLTRDG    N++E FRPGGGDMRDNWRSEL
Sbjct  243  INMWHEVGQAIYAPPIAGNITCNSSITGILLTRDG--DKNDTETFRPGGGDMRDNWRSEL  70

Query  484  YKYKVVKIEPLGVAPTKAKRRVV  506
            YKYKVV+I+PLGVAPT+AKR VV
Sbjct  69   YKYKVVEIKPLGVAPTEAKRNVV  1




>Bacillus velezensis strain TN14-11 TN14-11_ctg0032, whole genome shotgun sequence
Sequence ID: NZ_JAHVIU010000017.1 Length: 1125
Range 1: 116 to 445

Score:144 bits(363), Expect:1e-32,
Method:Compositional matrix adjust.,
Identities:84/110(76%), Positives:97/110(88%), Gaps:0/110(0%)

Query  664  DKWASLWNWFNITNWLWYIKLFIMIVGGLVGLRIVFAVLSIVNRVRQGYSPLSFQTHLPT  723
            D W +LWNWF+I+NWLWYI++FIMIVGGL+GLRIVFAVLSIVNRVRQGYSPLS QT +P
Sbjct  116  DSWKNLWNWFDISNWLWYIRIFIMIVGGLIGLRIVFAVLSIVNRVRQGYSPLSLQTLIPN  295

Query  724  prgpdrpegieeeggerdrdrSIRLVNGSLALIWDDLRSLCLFSYHRLRD  773
            P GPDR   I+EEGGE+ RDRS+RLV+G LAL W+DLR+LCLFSYH LRD
Sbjct  296  PVGPDRLGRIDEEGGEQGRDRSVRLVSGFLALAWEDLRNLCLFSYHHLRD  445




>Burkholderia pseudomallei strain 2604STDY5518971 ERS240105SCcontig000123, whole genome shotgun sequence
Sequence ID: NZ_UVFZ02000123.1 Length: 313
Range 1: 2 to 310

Score:134 bits(338), Expect:2e-32,
Method:Compositional matrix adjust.,
Identities:66/109(61%), Positives:82/109(75%), Gaps:6/109(5%)

Query  358  TIIFKQSSGGDPEIVTHSFNCGGEFFYCNSTQLFNSTWFNSTWSTEGSNNTEGSDTITLP  417
            TI F+  SGGD EI TH FNC GEFFYCN+T+LF +   ++  ST G N+T    TI +P
Sbjct  2    TISFEPHSGGDLEITTHHFNCRGEFFYCNTTKLFTNNCTDN--STGGCNDT----TIIIP  163

Query  418  CRIKQIINMWQKVGKAMYAPPISGQIRCSSNITGLLLTRDGGNSNNESE  466
            C+I+QI+ MWQ VG+AMYAPPISG+I+C SNITG+LLTRDGG   + SE
Sbjct  164  CKIRQIVRMWQGVGQAMYAPPISGEIKCESNITGILLTRDGGQDKSTSE  310




>Cronobacter malonaticus strain cro3263A1 contig73, whole genome shotgun sequence
Sequence ID: NZ_NRMM01000073.1 Length: 406
Range 1: 1 to 333

Score:116 bits(290), Expect:6e-32,
Method:Compositional matrix adjust.,
Identities:64/121(53%), Positives:78/121(64%), Gaps:11/121(9%)

Query  298  RPNNNTRKRIRIQRGPGRAFVTIGKI-GNMRQAHCNISRAKWNNTLKQIASKLREQFGNN  356
            RP+NNTR  I I  GPGRAF   G I G++RQA+C ++R +WN  L Q+  KL E F  N
Sbjct  1    RPDNNTRTSINI--GPGRAFYRTGDIIGDIRQAYCEVNRTEWNEILGQVIKKLEEHF-KN  171

Query  357  KTIIFKQSSGGDPEIVTHSFNCGGEFFYCNSTQLFNSTWFNSTWSTEGSNNTEGSDTITL  416
            KTI F+  SGGD EI    FNC GEFFYCN+T+LFNSTW       EG+   +G   +TL
Sbjct  172  KTIKFQPPSGGDLEITMLHFNCRGEFFYCNTTKLFNSTW-------EGNEP*KGVMAVTL  330

Query  417  P  417
            P
Sbjct  331  P  333




Range 2: 318 to 404

Score:57.4 bits(137), Expect:6e-32,
Method:Composition-based stats.,
Identities:24/29(83%), Positives:25/29(86%), Gaps:0/29(0%)

Query  410  GSDTITLPCRIKQIINMWQKVGKAMYAPP  438
            GSDT  LPCRIKQIINMWQ  G+AMYAPP
Sbjct  318  GSDTTILPCRIKQIINMWQGAGQAMYAPP  404




>Klebsiella pneumoniae strain EuSCAPE_MT015, whole genome shotgun sequence
Sequence ID: NZ_UKIW02000110.1 Length: 328
Range 1: 1 to 327

Score:130 bits(327), Expect:6e-31,
Method:Composition-based stats.,
Identities:79/109(72%), Positives:89/109(81%), Gaps:0/109(0%)

Query  708  VRQGYSPLSFQTHLPTprgpdrpegieeeggerdrdrSIRLVNGSLALIWDDLRSLCLFS  767
            VRQGYSPLSFQT  P+PR  DR   IEE GGE+D+DRSIRLVNG LAL W+DLRSLCLFS
Sbjct  1    VRQGYSPLSFQTLTPSPRELDRLGRIEEGGGEQDKDRSIRLVNGFLALAWEDLRSLCLFS  180

Query  768  YHRLRDLLLIVTRIVELLGRRGWEALKYWWNLLQYWSQELKNSAVSLLN  816
            YH+LRD +LIV R VELLGRRGW+ LKY  +L QYW  ELK SA+SLL+
Sbjct  181  YHQLRDFILIVVRAVELLGRRGWDTLKYLGSLGQYWGLELKKSAISLLD  327




>Klebsiella pneumoniae strain EuSCAPE_MT009, whole genome shotgun sequence
Sequence ID: NZ_UKIV02000095.1 Length: 324
Range 1: 3 to 302

Score:129 bits(323), Expect:2e-30,
Method:Compositional matrix adjust.,
Identities:66/100(66%), Positives:82/100(82%), Gaps:0/100(0%)

Query  547  GIVQQQNNLLRAIEAQQHLLQLTVWGIKQLQARILAVERYLKDQQLLGIWGCSGKLICTT  606
            GIVQQQ+NLLRAIEAQ H+LQLTV GIKQLQAR+LA+E+YLKDQQLL I  CS KLICTT
Sbjct  3    GIVQQQSNLLRAIEAQHHMLQLTV*GIKQLQARVLALEKYLKDQQLLKI*SCSRKLICTT  182

Query  607  AVPWNASWSNKSLEQIWNHTTWMEWDREINNYTSLIHSLI  646
             VP N+S SN++ + I ++ T M+ D+EI+NYT+ I  L+
```

Sbjct  183  NVP*NSS*SNRTYDDI*DNMT*MQ*DKEIDNYTNTIFQLL  302

>Acinetobacter baumannii strain P4 contig_P4_92, whole genome shotgun sequence
Sequence ID: NZ_JBBXLU010000071.1 Length: 1226
Range 1: 53 to 319

Score:134 bits(337), Expect:5e-29,
Method:Compositional matrix adjust.,
Identities:89/89(100%), Positives:89/89(100%), Gaps:0/89(0%)

Query  664  DKWASLWNWFNITNWLWYIKLFIMIVGGLVGLRIVFAVLSIVNRVRQGYSPLSFQTHLPT  723
            DKWASLWNWFNITNWLWYIKLFIMIVGGLVGLRIVFAVLSIVNRVRQGYSPLSFQTHLPT
Sbjct  53   DKWASLWNWFNITNWLWYIKLFIMIVGGLVGLRIVFAVLSIVNRVRQGYSPLSFQTHLPT  232

Query  724  prgpdrpegieeeggerdrdrSIRLVNGS  752
            PRGPDRPEGIEEEGGERDRDRSIRLVNGS
Sbjct  233  PRGPDRPEGIEEEGGERDRDRSIRLVNGS  319

>Staphylococcus aureus strain SA3 SA3_S21_L001_R1_001__paired__sampled_738, whole genome shotgun sequence
Sequence ID: NZ_RQSW01000723.1 Length: 205
Range 1: 5 to 193

Score:122 bits(306), Expect:1e-28,
Method:Composition-based stats.,
Identities:51/63(81%), Positives:57/63(90%), Gaps:0/63(0%)

Query  198  TSVITQACPKVSFEPIPIHYCAPAGFAILKCNNKTFNGTGPCTNVSTVQCTHGIRPVVST  257
            TS ITQACPKVSF+PIPIHYCAPAG+AILKCN+K FNGTGPC NVSTVQCTHGI+P  +T
Sbjct  5    TSAITQACPKVSFDPIPIHYCAPAGYAILKCNDKMFNGTGPCKNVSTVQCTHGIKPRGNT  184

Query  258  QLL  260
             +L
Sbjct  185  HVL  193

>Klebsiella pneumoniae strain EuSCAPE_IL009, whole genome shotgun sequence
Sequence ID: NZ_UKJO02000079.1 Length: 274
Range 1: 2 to 274

Score:119 bits(297), Expect:3e-27,
Method:Composition-based stats.,
Identities:74/91(81%), Positives:83/91(91%), Gaps:0/91(0%)

Query  687  MIVGGLVGLRIVFAVLSIVNRVRQGYSPLSFQTHLPTprgpdrpegieeeggerdrdrSI  746
            MIVGGL+GLRI+FAVLS+VNRVRQGYSPLSFQT +PTPR  DRP GIEEEGG +DRDRSI
Sbjct  274  MIVGGLIGLRIIFAVLSVVNRVRQGYSPLSFQTLIPTPRELDRPGGIEEEGGQDRDRSI  95

Query  747  RLVNGSLALIWDDLRSLCLFSYHRLRDLLLI  777
            RLV+G LAL WDDLR+LCLFSYH+LRD +LI
Sbjct  94   RLVSGFLALAWDDLRNLCLFSYHQLRDFILI  2

>Klebsiella pneumoniae strain EuSCAPE_MT009, whole genome shotgun sequence
Sequence ID: NZ_UKIV02000092.1 Length: 398
Range 1: 1 to 345

Score:119 bits(297), Expect:1e-26,
Method:Composition-based stats.,
Identities:65/115(57%), Positives:79/115(68%), Gaps:7/115(6%)

Query  748  LVNGSLALIWDDLRSLCLFSYHRLRDLLLIVTRIVELLG-------RRGWEALKYWWNLL  800
            LV+  LAL  DDLRSLCLFSYHRLRD +LI  R VELL        ++  E LKY  +L
Sbjct  1    LVSRFLALA*DDLRSLCLFSYHRLRDFILIAARAVELLRHSILRRLQKK*EILKYLRSLA  180

Query  801  QYWSQELKNSAVSLLNATAIAVAEGTDRVIEVVQGACRAIRHIPRRIRQGLERIL  855
            QY S ELK SA+SL +  AIAVAE TD++IE+ Q  C++I +IP+RIRQG E  L
Sbjct  181  QY*SVELKKSAISLFDTIAIAVAEKTDKIIELAQRICKSILNIPKRIRQGFEAAL  345

>Klebsiella pneumoniae strain EuSCAPE_MT006, whole genome shotgun sequence
Sequence ID: NZ_UKIU02000080.1 Length: 250
Range 1: 1 to 249

Score:115 bits(288), Expect:4e-26,
Method:Compositional matrix adjust.,
Identities:60/85(71%), Positives:68/85(80%), Gaps:3/85(3%)

Query  244  TVQCTHGIRPVVSTQLLLNGSLAEEEVVIRSVNFTDNAKTIIVQLNTSVEINCTRPNNNT  303
            TV CTHGI+PVVSTQLLLNGSLAE+E+VIRS N TDNAKTIIV LN SVEI C RP NNT
Sbjct  249  TVLCTHGIKPVVSTQLLLNGSLAEKEIVIRSENLTDNAKTIIVHLNESVEIVCMRPGNNT  70

Query  304  RKRIRIQRGPGRAFVTIGK-IGNMR  327
            R+ +RI  GPG+ F   G  IG++R
Sbjct  69   RRSMRI--GPGQTFYATGDIIGDIR  1

>Klebsiella pneumoniae strain EuSCAPE_NO001, whole genome shotgun sequence
Sequence ID: NZ_UKMF02000044.1 Length: 289
Range 1: 2 to 283

Score:113 bits(283), Expect:3e-25,
Method:Composition-based stats.,
Identities:84/94(89%), Positives:93/94(98%), Gaps:0/94(0%)

Query  465  SEIFRPGGGDMRDNWRSELYKYKVVKIEPLGVAPTKAKRRVVQREKRAVGIgalflgflg  524
            +E FRPGGGDMRDNWRSELYKYKVV+I+PLG+APTKA+RRVV+REKRAVGIGA+FLGFLG
Sbjct  283  NETFRPGGGDMRDNWRSELYKYKVVEIKPLGIAPTKARRRVVEREKRAVGIGAVFLGFLG  104

Query  525  aagstmgaasmtLTVQARQLLSGIVQQQNNLLRA  558
            AAGSTMGAAS+TLTVQARQLLSGIVQQQ+NLLRA
Sbjct  103  AAGSTMGAASITLTVQARQLLSGIVQQQSNLLRA  2

>Klebsiella pneumoniae strain EuSCAPE_IL022, whole genome shotgun sequence
Sequence ID: NZ_UKKD02000026.1 Length: 290
Range 1: 20 to 289

Score:112 bits(279), Expect:1e-24,

```
Method:Compositional matrix adjust.,
Identities:66/90(73%), Positives:81/90(90%), Gaps:0/90(0%)

Query  633  REINNYTSLIHSLIeesqnqqekneqellelDKWASLWNWFNITNWLWYIKLFIMIVGGL  692
            REI+NYT +I++L+E SQNQQEKNE++LL L+ W SLW+WF+IT WLWYI++FI+IVGGL
Sbjct  289  REISNYTEIIYNLLEVSQNQQEKNEKDLLALNSWDSLWSWFDITKWLWYIRIFIIIVGGL  110

Query  693  VGLRIVFAVLSIVNRVRQGYSPLSFQTHLP  722
            +GLRI+FAVLSIVNRVRQGYSPLS QT  P
Sbjct  109  IGLRIIFAVLSIVNRVRQGYSPLSLQTLNP  20



>Acinetobacter baumannii strain AB49 NODE_71_length_263_cov_0.536585_pilon, whole genome shotgun sequence
Sequence ID: NZ_JAAXKW010000068.1 Length: 263
Range 1: 2 to 262

Score:111 bits(277), Expect:2e-24,
Method:Compositional matrix adjust.,
Identities:56/102(55%), Positives:66/102(64%), Gaps:15/102(14%)

Query  347  SKLREQFGNNKTIIFKQSSGGDPEIVTHSFNCGGEFFYCNSTQLFNSTWFNSTWSTEGSN  406
            +KL+  F N  TI F  SSGGDPEI  HSFNC GEFFYCN++ LF +   N
Sbjct  262  TKLQSYFPN-ATINFTSSSGGDPEITMHSFNCRGEFFYCNTSNLFTNNITN---------  113

Query  407  NTEGSDTITLPCRIKQIINMWQKVGKAMYAPPISGQIRCSSN  448
                 + ITLPCRIKQIINMWQ+VG+AMY  P++G I C SN
Sbjct  112  -----ENITLPCRIKQIINMWQEVGRAMYTAPVAGSITCRSN  2



>Klebsiella pneumoniae strain EuSCAPE_IL004, whole genome shotgun sequence
Sequence ID: NZ_UKJP02000111.1 Length: 327
Range 1: 65 to 325

Score:110 bits(276), Expect:4e-24,
Method:Compositional matrix adjust.,
Identities:64/87(74%), Positives:78/87(89%), Gaps:0/87(0%)

Query  636  NNYTSLIHSLIeesqnqqekneqellelDKWASLWNWFNITNWLWYIKLFIMIVGGLVGL  695
            +N+T  I+ L+E SQNQQEKNE++LL L+ W +LWNWF+IT WLWYIK+FIMIVGGL+GL
Sbjct  325  SNHTDTIYRLLEISQNQQEKNEKDLLALNSWQNLWNWFDITQWLWYIKIFIMIVGGLIGL  146

Query  696  RIVFAVLSIVNRVRQGYSPLSFQTHLP  722
            RI+FAV+SIVNRVRQGYSPLSFQT +P
Sbjct  145  RIIFAVISIVNRVRQGYSPLSFQTLIP  65



>Klebsiella pneumoniae strain EuSCAPE_NO001, whole genome shotgun sequence
Sequence ID: NZ_UKMF02000047.1 Length: 245
Range 1: 3 to 230

Score:109 bits(272), Expect:6e-24,
Method:Compositional matrix adjust.,
Identities:53/77(69%), Positives:60/77(77%), Gaps:1/77(1%)

Query  1    MRVKEKYQHLWRWGWRWGTMLLGMLMICSATEKLWVTVYYGVPVWKEATTTLFCASDAKA  60
            M+VK   ++  +W W WG +   MLMIC+  EK WVTVYYGVPVWKEA TTLFCASDAKA
Sbjct  230  MKVKGMLRNYPQW-WIWGILGFWMLMICNGKEKSWVTVYYGVPVWKEAKTTLFCASDAKA  54

Query  61   YDTEVHNVWATHACVPT  77
            Y  EVHN+WATHACVPT
Sbjct  53   YSKEVHNIWATHACVPT  3



>Klebsiella pneumoniae strain EuSCAPE_MT002, whole genome shotgun sequence
Sequence ID: NZ_UKIM02000087.1 Length: 265
Range 1: 2 to 265

Score:106 bits(264), Expect:1e-22,
Method:Composition-based stats.,
Identities:71/88(81%), Positives:78/88(88%), Gaps:1/88(1%)

Query  676  TNWLWYIKLFIMIVGGLVGLRIVFAVLSIVNRVRQGYSPLSFQTHLP-Tprgpdrpegie  734
            TNWLWYI++FIMIVGGL+GLRI+FAVLSIVNRVRQGYSPLSFQT  P +   PDR E IE
Sbjct  265  TNWLWYIRIFIMIVGGLIGLRIIFAVLSIVNRVRQGYSPLSFQTLAPNSRGRPDRLERIE  86

Query  735  eeggerdrdrSIRLVNGSLALIWDDLRS  762
            EEGGE+DRDRSIRLV+G LAL WDDLRS
Sbjct  85   EEGGEQDRDRSIRLVSGFLALAWDDLRS  2



>Klebsiella pneumoniae strain EuSCAPE_IL016, whole genome shotgun sequence
Sequence ID: NZ_UKKA02000070.1 Length: 345
Range 1: 53 to 343

Score:106 bits(264), Expect:2e-22,
Method:Compositional matrix adjust.,
Identities:67/97(69%), Positives:83/97(85%), Gaps:0/97(0%)

Query  627  TWMEWDREINNYTSLIHSLIeesqnqqekneqellelDKWASLWNWFNITNWLWYIKLFI  686
            T M+ DREINNYT+ I+ L+EESQNQQEKN+++LL LD W +L + F+IT WL YIK+FI
Sbjct  343  T*MQ*DREINNYTNTIYRLLEESQNQQEKNKKDLLALDSWQNL*S*FSITKWL*YIKIFI  164

Query  687  MIVGGLVGLRIVFAVLSIVNRVRQGYSPLSFQTHLPT  723
            +IV GL+ LRI+FAVLSIVNRVRQGYSPLSFQT +P+
Sbjct  163  IIVRGLISLRIIFAVLSIVNRVRQGYSPLSFQTLIPS  53



>Neisseria gonorrhoeae strain GC-206 NODE_155_length_258_cov_1.77833_ID_20482, whole genome shotgun sequence
Sequence ID: NZ_SAWN01000155.1 Length: 258
Range 1: 1 to 258

Score:104 bits(260), Expect:3e-22,
Method:Compositional matrix adjust.,
Identities:51/86(59%), Positives:65/86(75%), Gaps:1/86(1%)

Query  81   PQEVVLVNVTENFNMWKNDMVEQMHEDIISLWDQSLKPCVKLTPLCVSLKCTDLKNDTNT  140
            PQE+VL NVTENFNMWKN MV+QMHED++SLWDQSLKPCVKLT +CV+L CT +K    T
Sbjct  258  PQEIVLGNVTENFNMWKNYMVDQMHEDVVSLWDQSLKPCVKLTTVCVTLNCTSVKVTNVT  79

Query  141  NSSSGRMIME-KGEIKNCSFNISTSI  165
             S+S +       +++NC+F  +T I
Sbjct  78   ASASVKGTDNITNDMQNCTFTATTEI  1
```

```
>Pseudomonas aeruginosa strain VET-121 NODE_144_length_275_cov_1.53535_ID_287, whole genome shotgun sequence
Sequence ID: NZ_QZXB01000257.1 Length: 275
Range 1: 40 to 273

Score:99.0 bits(245), Expect:3e-20,
Method:Composition-based stats.,
Identities:70/78(90%), Positives:75/78(96%), Gaps:0/78(0%)

Query  503  RRVVQREKRAVGIgalflgflgaagstmgaasmtLTVQARQLLSGIVQQQNNLLRAIEAQ  562
            RRVV REKRAVG+GA+FLGFLG AGSTMGAAS+TLTVQ RQLLSGIVQQQ+NLLRAIEAQ
Sbjct  273  RRVVGREKRAVGLGAVFLGFLGTAGSTMGAASITLTVQVRQLLSGIVQQQSNLLRAIEAQ  94

Query  563  QHLLQLTVWGIKQLQARI  580
            QHLLQLTVWGIKQLQAR+
Sbjct  93   QHLLQLTVWGIKQLQARV  40
```

```
>Klebsiella pneumoniae strain EuSCAPE_IL019, whole genome shotgun sequence
Sequence ID: NZ_UKJZ02000064.1 Length: 244
Range 1: 3 to 242

Score:96.7 bits(239), Expect:2e-19,
Method:Compositional matrix adjust.,
Identities:51/82(62%), Positives:58/82(70%), Gaps:2/82(2%)

Query  272  IRSVNFTDNAKTIIVQLNTSVEINCTRPNNNTRKRIRIQRGPGRAFVTIGKIGNMRQAHC  331
            I S N TDNAKTIIV LN SV I CTRP NNTRK IRI  GPG+ F     IG++R+A+C
Sbjct  242  ISSKNLTDNAKTIIVHLNESVTIICTRPGNNTRKSIRI--GPGQTFYANKIIGDIRRAYC  69

Query  332  NISRAKWNNTLKQIASKLREQF  353
            NIS   WN TLKQ+  KL+E F
Sbjct  68   NISGEAWNKTLKQVEEKLKEYF  3
```

```
>Photobacterium rosenbergii strain DP6N14-7 DP6N14-7_ctg0054, whole genome shotgun sequence
Sequence ID: NZ_JAHVIB010000049.1 Length: 960
Range 1: 14 to 214

Score:96.7 bits(239), Expect:5e-17,
Method:Composition-based stats.,
Identities:47/67(70%), Positives:53/67(79%), Gaps:0/67(0%)

Query  789  GWEALKYWWNLLQYWSQELKNSAVSLLNATAIAVAEGTDRVIEVVQGACRAIRHIPRRIR  848
            GWEALKY  +L+QYW QELK SA+SL +A AI VAEGTDR+IEV Q  CRAI +IPRRIR
Sbjct  14   GWEALKYLGSLVQYWGQELKKSAISLFDALAITVAEGTDRIIEVGQRLCRAICNIPRRIR  193

Query  849  QGLERIL  855
            QG E  L
Sbjct  194  QGFEAAL  214
```

```
>Klebsiella pneumoniae strain EuSCAPE_LT003, whole genome shotgun sequence
Sequence ID: NZ_UKJH02000068.1 Length: 317
Range 1: 126 to 314

Score:83.2 bits(204), Expect:2e-14,
Method:Composition-based stats.,
Identities:41/63(65%), Positives:48/63(76%), Gaps:0/63(0%)

Query  793  LKYWWNLLQYWSQELKNSAVSLLNATAIAVAEGTDRVIEVVQGACRAIRHIPRRIRQGLE  852
            LKY  + +QYW  ELK SAVSL +  AIAVAEGTDR+IEVVQG  +AI +IPRRIRQG +
Sbjct  314  LKYLGSRVQYWGLELKKSAVSLFDTLAIAVAEGTDRIIEVVQGISKAILNIPRRIRQGFK  135

Query  853  RIL  855
              L
Sbjct  134  AAL  126
```

```
>Bacillus velezensis strain TN14-11 TN14-11_ctg0036, whole genome shotgun sequence
Sequence ID: NZ_JAHVIU010000019.1 Length: 1101
Range 1: 203 to 355

Score:70.9 bits(172), Expect:3e-08,
Method:Compositional matrix adjust.,
Identities:36/51(71%), Positives:40/51(78%), Gaps:0/51(0%)

Query  805  QELKNSAVSLLNATAIAVAEGTDRVIEVVQGACRAIRHIPRRIRQGLERIL  855
            QELK SA+SL +A AI VAEGTDR+IEV Q  CRAI +IPRRIRQG E  L
Sbjct  203  QELKKSAISLFDALAITVAEGTDRIIEVGQRLCRAICNIPRRIRQGFEAAL  355
```

```
>Klebsiella pneumoniae strain EuSCAPE_IL005, whole genome shotgun sequence
Sequence ID: NZ_UKJL02000056.1 Length: 442
Range 1: 318 to 440

Score:61.2 bits(147), Expect:3e-06,
Method:Composition-based stats.,
Identities:30/41(73%), Positives:34/41(82%), Gaps:0/41(0%)

Query  816  NATAIAVAEGTDRVIEVVQGACRAIRHIPRRIRQGLERILL  856
            +ATAIAVAEGTDR+IE+VQ   RAI +IPRRIRQG E  LL
Sbjct  440  DATAIAVAEGTDRIIELVQRIGRAICNIPRRIRQGFETALL  318
```

Query #2: SP|P04578|GP120 Query ID: lcl|Query_4184771 Length: 479

Sequences producing significant alignments:

| Description | Scientific Name | Common Name | Taxid | Max Score | Total Score | Query cover | E Value | Per. Ident | Acc. Len | Accession |
|---|---|---|---|---|---|---|---|---|---|---|
| Escherichia coli strain NBE0009 contig00112, whole genome... | Escherichia ... | NA | 562 | 703 | 703 | 99% | 0.0 | 70.88 | 2003 | NZ_JARBDL0100... |
| Escherichia coli strain NBE0009 contig00111, whole genome... | Escherichia ... | NA | 562 | 701 | 701 | 99% | 0.0 | 70.26 | 2027 | NZ_JARBDL0100... |
| Escherichia coli strain EN5177 contig00087, whole genome shotg... | Escherichia ... | NA | 562 | 648 | 648 | 97% | 0.0 | 62.64 | 3403 | NZ_JAPMNK0100... |
| Escherichia coli strain NBE0004 contig00090, whole genome... | Escherichia ... | NA | 562 | 648 | 648 | 97% | 0.0 | 62.64 | 2818 | NZ_JARBDH0100... |
| Escherichia coli strain EN5076 contig00066, whole genome shotg... | Escherichia ... | NA | 562 | 639 | 639 | 91% | 0.0 | 69.89 | 2732 | NZ_JAODTC0100... |
| Escherichia coli strain NBE0010 contig00080, whole genome... | Escherichia ... | NA | 562 | 637 | 637 | 91% | 0.0 | 69.21 | 2743 | NZ_JARBDM0100... |
| Escherichia coli strain EN5374 contig00094, whole genome shotg... | Escherichia ... | NA | 562 | 636 | 636 | 91% | 0.0 | 69.21 | 2720 | NZ_JAPTGI0100... |
| Escherichia coli strain EN5177 contig00105, whole genome shotg... | Escherichia ... | NA | 562 | 636 | 636 | 98% | 0.0 | 67.15 | 1783 | NZ_JAPMNK0100... |

| | | | | | | | | | | |
|---|---|---|---|---|---|---|---|---|---|---|
| Escherichia coli strain EN5090 contig00081, whole genome shotg... | Escherichia ... | NA | 562 | 634 | 634 | 91% | 0.0 | 69.96 | 2798 | NZ_JAODTD0100... |
| Escherichia coli strain EN5379 contig00092, whole genome shotg... | Escherichia ... | NA | 562 | 629 | 629 | 91% | 0.0 | 70.02 | 2592 | NZ_JAPTGJ0100... |
| Escherichia coli strain EN5317 contig00057, whole genome shotg... | Escherichia ... | NA | 562 | 620 | 620 | 91% | 0.0 | 68.67 | 2867 | NZ_JARBDO0100... |
| Escherichia coli strain NBE0003 contig00096, whole genome... | Escherichia ... | NA | 562 | 613 | 613 | 91% | 0.0 | 67.92 | 2837 | NZ_JARBDG0100... |
| Escherichia coli strain EN5169 contig00082, whole genome shotg... | Escherichia ... | NA | 562 | 613 | 613 | 91% | 0.0 | 67.92 | 2837 | NZ_JAPMNJ0100... |
| Escherichia coli strain NBE0002 contig00097, whole genome... | Escherichia ... | NA | 562 | 605 | 605 | 89% | 0.0 | 68.93 | 2717 | NZ_JARBDF0100... |
| Escherichia coli strain EN5143 contig00069, whole genome shotg... | Escherichia ... | NA | 562 | 605 | 605 | 89% | 0.0 | 68.93 | 2688 | NZ_JAPMNI0100... |
| Escherichia coli strain EN5308 contig00099, whole genome shotg... | Escherichia ... | NA | 562 | 603 | 603 | 91% | 0.0 | 67.04 | 2827 | NZ_JARBDE0100... |
| Escherichia coli strain EN5286 contig00120, whole genome shotg... | Escherichia ... | NA | 562 | 582 | 582 | 89% | 0.0 | 61.95 | 3033 | NZ_JAPMNM0100... |
| Escherichia coli strain EN5192 contig00108, whole genome shotg... | Escherichia ... | NA | 562 | 579 | 579 | 92% | 0.0 | 60.16 | 3032 | NZ_JAPMNL0100... |
| Escherichia coli strain VRES0669, whole genome shotgun sequence | Escherichia ... | NA | 562 | 578 | 578 | 89% | 0.0 | 65.91 | 5544 | NZ_UINV010000... |
| Escherichia coli strain NBE0007 contig00093, whole genome... | Escherichia ... | NA | 562 | 578 | 578 | 89% | 0.0 | 61.54 | 3013 | NZ_JARBDJ0100... |
| Escherichia coli strain EN5308 contig00096, whole genome shotg... | Escherichia ... | NA | 562 | 570 | 570 | 88% | 0.0 | 61.51 | 3024 | NZ_JARBDE0100... |
| Escherichia coli strain EN5143 contig00071, whole genome shotg... | Escherichia ... | NA | 562 | 570 | 570 | 88% | 0.0 | 61.38 | 2774 | NZ_JAPMNI0100... |
| Escherichia coli strain NBE0006 contig00106, whole genome... | Escherichia ... | NA | 562 | 533 | 533 | 84% | 1e-180 | 63.43 | 2090 | NZ_JARBDI0100... |
| Escherichia coli strain EN5192 contig00149, whole genome shotg... | Escherichia ... | NA | 562 | 514 | 514 | 74% | 2e-176 | 69.75 | 1397 | NZ_JAPMNL0100... |
| Escherichia coli strain NBE0006 contig00124, whole genome... | Escherichia ... | NA | 562 | 508 | 508 | 71% | 4e-174 | 71.47 | 1306 | NZ_JARBDI0100... |
| Klebsiella pneumoniae strain EuSCAPE_IL004, whole genome shotg... | Klebsiella p... | NA | 573 | 471 | 471 | 70% | 1e-160 | 68.24 | 1081 | NZ_UKJP020000... |
| Pseudomonas aeruginosa strain VET-121... | Pseudomonas ... | NA | 287 | 445 | 445 | 66% | 5e-151 | 66.87 | 1017 | NZ_QZXB010001... |
| Escherichia coli strain EN5308 contig00145, whole genome shotg... | Escherichia ... | NA | 562 | 427 | 427 | 69% | 7e-144 | 64.41 | 1005 | NZ_JARBDE0100... |
| Escherichia coli strain NBE0002 contig00132, whole genome... | Escherichia ... | NA | 562 | 421 | 421 | 63% | 1e-139 | 66.88 | 1381 | NZ_JARBDF0100... |
| Escherichia coli strain EN5095 contig00078, whole genome shotg... | Escherichia ... | NA | 562 | 425 | 425 | 70% | 1e-138 | 65.69 | 1972 | NZ_JAPYLY0100... |
| Escherichia coli strain NBE0002 contig00149, whole genome... | Escherichia ... | NA | 562 | 412 | 412 | 57% | 6e-138 | 68.81 | 996 | NZ_JARBDF0100... |
| Escherichia coli strain EN5132 contig00077, whole genome shotg... | Escherichia ... | NA | 562 | 424 | 424 | 66% | 3e-136 | 66.46 | 2498 | NZ_JAPYLZ0100... |
| Klebsiella pneumoniae strain P7701... | Klebsiella p... | NA | 573 | 378 | 378 | 39% | 8e-127 | 96.30 | 570 | NZ_JACAWM0100... |
| Escherichia coli strain NBE0004 contig00131, whole genome... | Escherichia ... | NA | 562 | 365 | 365 | 53% | 2e-120 | 69.52 | 847 | NZ_JARBDH0100... |
| Methyloversatilis universalis FAM5 contig00006, whole genome... | Methyloversa... | NA | 1000565 | 359 | 359 | 42% | 4e-119 | 82.30 | 649 | NZ_AFHG010000... |
| Escherichia coli strain NBE0010 contig00098, whole genome... | Escherichia ... | NA | 562 | 342 | 494 | 69% | 9e-107 | 68.29 | 1912 | NZ_JARBDM0100... |
| Pseudomonas aeruginosa strain VET-42... | Pseudomonas ... | NA | 287 | 328 | 328 | 44% | 3e-105 | 70.45 | 963 | NZ_RACX010002... |
| Neisseria gonorrhoeae strain GC-206... | Neisseria go... | NA | 485 | 238 | 238 | 34% | 2e-72 | 71.08 | 488 | NZ_SAWN010001... |
| Burkholderia pseudomallei strain 2604STDY5519032... | Burkholderia... | NA | 28450 | 223 | 223 | 35% | 1e-66 | 66.86 | 499 | NZ_UVID020000... |
| Escherichia coli strain EN5192 contig00156, whole genome shotg... | Escherichia ... | NA | 562 | 223 | 223 | 39% | 2e-66 | 61.93 | 562 | NZ_JAPMNL0100... |
| Klebsiella pneumoniae strain PN085E1IA... | Klebsiella p... | NA | 573 | 222 | 222 | 32% | 2e-66 | 72.26 | 474 | NZ_PDVD010000... |
| Pseudomonas aeruginosa strain VET-42... | Pseudomonas ... | NA | 287 | 214 | 214 | 39% | 7e-63 | 58.03 | 548 | NZ_RACX010001... |
| Klebsiella pneumoniae strain ST-234:K080... | Klebsiella p... | NA | 573 | 205 | 205 | 32% | 8e-60 | 68.15 | 474 | NZ_NXJY010001... |
| Enterobacter hormaechei strain ST-234:K063... | Enterobacter... | NA | 158836 | 201 | 201 | 32% | 2e-58 | 66.46 | 475 | NZ_NXJN010001... |
| Klebsiella pneumoniae strain ST-1552:K089... | Klebsiella p... | NA | 573 | 201 | 201 | 32% | 4e-58 | 68.39 | 474 | NZ_NXIW010001... |
| Escherichia coli strain E003 NODE_241_length_474_cov_17.072,... | Escherichia ... | NA | 562 | 198 | 198 | 32% | 3e-57 | 67.10 | 474 | NZ_NXIZ010002... |
| Cronobacter sakazakii strain cro4114A1 contig126, whole genome... | Cronobacter ... | NA | 28141 | 192 | 192 | 29% | 4e-55 | 67.13 | 413 | NZ_NRKJ010001... |
| Cronobacter malonaticus strain cro3263A1 contig69, whole genom... | Cronobacter ... | NA | 413503 | 192 | 192 | 29% | 6e-55 | 67.13 | 413 | NZ_NRMM010000... |
| Citrobacter freundii strain CF003... | Citrobacter ... | NA | 546 | 191 | 191 | 31% | 2e-54 | 66.23 | 474 | NZ_NXKE010001... |
| Cronobacter malonaticus strain cro3263A1 contig68, whole genom... | Cronobacter ... | NA | 413503 | 177 | 177 | 29% | 2e-49 | 64.79 | 413 | NZ_NRMM010000... |
| Cronobacter sakazakii strain cro4215W contig103, whole genome... | Cronobacter ... | NA | 28141 | 174 | 174 | 29% | 2e-48 | 64.08 | 413 | NZ_NRKF010001... |
| Cronobacter dublinensis strain cro981C2 contig304, whole genom... | Cronobacter ... | NA | 413497 | 172 | 172 | 29% | 2e-47 | 63.38 | 413 | NZ_NROB010003... |
| Cronobacter sakazakii strain cro4114B1 contig109, whole genome... | Cronobacter ... | NA | 28141 | 172 | 172 | 29% | 2e-47 | 63.38 | 413 | NZ_NRKI010001... |
| Cronobacter sakazakii strain cro4114A1 contig127, whole genome... | Cronobacter ... | NA | 28141 | 172 | 172 | 29% | 2e-47 | 63.38 | 413 | NZ_NRKJ010001... |
| Cronobacter sakazakii strain cro611A3 contig61, whole genome... | Cronobacter ... | NA | 28141 | 171 | 171 | 29% | 3e-47 | 62.94 | 407 | NZ_NRMC010000... |
| Enterobacter hormaechei strain ST-234:K063... | Enterobacter... | NA | 158836 | 171 | 171 | 27% | 7e-47 | 65.15 | 480 | NZ_NXJN010001... |
| Klebsiella pneumoniae strain EuSCAPE_IL004, whole genome shotg... | Klebsiella p... | NA | 573 | 169 | 169 | 19% | 1e-46 | 84.78 | 277 | NZ_UKJP020001... |
| Cronobacter sakazakii strain cro3040W contig81, whole genome... | Cronobacter ... | NA | 28141 | 168 | 168 | 29% | 8e-46 | 62.68 | 404 | NZ_NRJO010000... |
| Escherichia coli strain NBE0009 contig00116, whole genome... | Escherichia ... | NA | 562 | 180 | 180 | 25% | 9e-46 | 70.73 | 1713 | NZ_JARBDL0100... |
| Cronobacter sakazakii strain cro4114A1 contig132, whole genome... | Cronobacter ... | NA | 28141 | 167 | 167 | 29% | 1e-45 | 61.97 | 392 | NZ_NRKJ010001... |
| Cronobacter sakazakii strain cro4215W contig104, whole genome... | Cronobacter ... | NA | 28141 | 165 | 165 | 29% | 8e-45 | 61.97 | 405 | NZ_NRKF010001... |
| Cronobacter malonaticus strain cro3263A1 contig74, whole genom... | Cronobacter ... | NA | 413503 | 164 | 164 | 29% | 2e-44 | 58.45 | 404 | NZ_NRMM010000... |
| Cronobacter sakazakii strain cro4114A1 contig129, whole genome... | Cronobacter ... | NA | 28141 | 164 | 164 | 29% | 2e-44 | 61.27 | 404 | NZ_NRKJ010001... |
| Cronobacter sakazakii strain cro4114B1 contig117, whole genome... | Cronobacter ... | NA | 28141 | 163 | 163 | 29% | 4e-44 | 61.27 | 404 | NZ_NRKI010001... |
| Cronobacter sakazakii strain cro4114B1 contig115, whole genome... | Cronobacter ... | NA | 28141 | 163 | 163 | 29% | 7e-44 | 57.75 | 404 | NZ_NRKI010001... |
| Cronobacter sakazakii strain cro4114A1 contig128, whole genome... | Cronobacter ... | NA | 28141 | 162 | 162 | 29% | 9e-44 | 57.75 | 404 | NZ_NRKJ010001... |
| Cronobacter malonaticus strain cro3263A1 contig76, whole genom... | Cronobacter ... | NA | 413503 | 162 | 162 | 29% | 1e-43 | 59.86 | 392 | NZ_NRMM010000... |
| Cronobacter sakazakii strain cro4215W contig105, whole genome... | Cronobacter ... | NA | 28141 | 162 | 162 | 29% | 1e-43 | 57.75 | 404 | NZ_NRKF010001... |
| Cronobacter dublinensis strain cro981C2 contig308, whole genom... | Cronobacter ... | NA | 413497 | 160 | 160 | 29% | 4e-43 | 60.99 | 405 | NZ_NROB010003... |
| Escherichia coli strain EN5143 contig00088, whole genome shotg... | Escherichia ... | NA | 562 | 166 | 166 | 25% | 4e-43 | 66.94 | 833 | NZ_JAPMNI0100... |
| Cronobacter sakazakii strain cro4114B1 contig105, whole genome... | Cronobacter ... | NA | 28141 | 157 | 157 | 29% | 6e-42 | 59.03 | 416 | NZ_NRKI010001... |
| Planctopirus hydrillae strain JC280 Contig113, whole genome... | Planctopirus... | NA | 1841610 | 157 | 157 | 30% | 2e-41 | 59.86 | 437 | NZ_LYDR010000... |
| Cronobacter sakazakii strain cro4114A1 contig133, whole genome... | Cronobacter ... | NA | 28141 | 155 | 155 | 29% | 3e-41 | 58.45 | 389 | NZ_NRKJ010001... |
| Cronobacter sakazakii strain cro4191B1-2 contig85, whole genom... | Cronobacter ... | NA | 28141 | 155 | 155 | 29% | 3e-41 | 58.45 | 392 | NZ_NRKG010000... |
| Cronobacter sakazakii strain cro3040W contig84, whole genome... | Cronobacter ... | NA | 28141 | 155 | 155 | 29% | 3e-41 | 58.45 | 392 | NZ_NRJO010000... |
| Cronobacter sakazakii strain cro4114B1 contig122, whole genome... | Cronobacter ... | NA | 28141 | 155 | 155 | 29% | 5e-41 | 57.04 | 383 | NZ_NRKI010001... |
| Cronobacter sakazakii strain cro4215W contig109, whole genome... | Cronobacter ... | NA | 28141 | 154 | 154 | 29% | 9e-41 | 57.75 | 389 | NZ_NRKF010001... |
| Cronobacter sakazakii strain cro4191B1-2 contig83, whole genom... | Cronobacter ... | NA | 28141 | 153 | 153 | 29% | 3e-40 | 56.64 | 410 | NZ_NRKG010000... |
| Cronobacter malonaticus strain cro3263A1 contig75, whole genom... | Cronobacter ... | NA | 413503 | 153 | 153 | 29% | 3e-40 | 57.75 | 392 | NZ_NRMM010000... |
| Cronobacter sakazakii strain cro4114B1 contig124, whole genome... | Cronobacter ... | NA | 28141 | 153 | 153 | 29% | 3e-40 | 57.75 | 392 | NZ_NRKI010001... |
| Cronobacter sakazakii strain cro4114A1 contig131, whole genome... | Cronobacter ... | NA | 28141 | 152 | 152 | 29% | 4e-40 | 57.75 | 392 | NZ_NRKJ010001... |
| Escherichia coli strain EN5192 contig00135, whole genome shotg... | Escherichia ... | NA | 562 | 161 | 161 | 27% | 8e-40 | 61.48 | 1341 | NZ_JAPMNL0100... |
| Cronobacter sakazakii strain cro3040W contig83, whole genome... | Cronobacter ... | NA | 28141 | 151 | 151 | 29% | 1e-39 | 54.93 | 392 | NZ_NRJO010000... |
| Cronobacter dublinensis strain cro3540W contig49, whole genome... | Cronobacter ... | NA | 413497 | 150 | 150 | 29% | 4e-39 | 55.63 | 392 | NZ_NRNN010000... |
| Cronobacter sakazakii strain cro4191B1-2 contig84, whole genom... | Cronobacter ... | NA | 28141 | 122 | 180 | 28% | 2e-34 | 60.58 | 402 | NZ_NRKG010000... |
| Cronobacter malonaticus strain cro4164A3 contig145, whole geno... | Cronobacter ... | NA | 413503 | 119 | 180 | 27% | 2e-34 | 61.62 | 405 | NZ_NRNF010001... |
| Neisseria gonorrhoeae strain GC-206... | Neisseria go... | NA | 485 | 134 | 134 | 17% | 6e-34 | 74.70 | 243 | NZ_SAWN010001... |
| Burkholderia pseudomallei strain 2604STDY5518971... | Burkholderia... | NA | 28450 | 133 | 133 | 22% | 5e-33 | 60.55 | 313 | NZ_UVFZ020001... |
| Cronobacter malonaticus strain cro3263A1 contig73, whole genom... | Cronobacter ... | NA | 413503 | 115 | 173 | 29% | 2e-32 | 52.89 | 406 | NZ_NRMM010000... |
| Klebsiella pneumoniae strain EuSCAPE_MT006, whole genome shotg... | Klebsiella p... | NA | 573 | 114 | 114 | 17% | 2e-26 | 70.59 | 250 | NZ_UKIU020000... |
| Neisseria gonorrhoeae strain GC-206... | Neisseria go... | NA | 485 | 104 | 104 | 17% | 1e-22 | 59.30 | 258 | NZ_SAWN010001... |
| Klebsiella pneumoniae strain EuSCAPE_IL019, whole genome shotg... | Klebsiella p... | NA | 573 | 96.7 | 96.7 | 17% | 8e-20 | 62.20 | 244 | NZ_UKJZ020000... |
| Nocardioides luteus strain JCM 3358 sequence39, whole genome... | Nocardioides... | NA | 1844 | 101 | 101 | 9% | 8e-18 | 100.00 | 5754 | NZ_BMRK010000... |
| Streptomyces griseoflavus strain JCM 4479 sequence37, whole... | Streptomyces... | NA | 35619 | 101 | 101 | 9% | 8e-18 | 100.00 | 5775 | NZ_BMUC010000... |
| Saccharothrix coeruleofusca strain JCM 3313 sequence45, whole... | Saccharothri... | NA | 33919 | 101 | 101 | 9% | 8e-18 | 100.00 | 5775 | NZ_BMRG010000... |
| Streptomyces griseoloalbus strain JCM 4480 sequence054, whole... | Streptomyces... | NA | 67303 | 101 | 101 | 9% | 8e-18 | 100.00 | 5815 | NZ_BMUD010000... |
| Actinoplanes ianthinogenes strain JCM 3249 sequence037, whole... | Actinoplanes... | NA | 122358 | 101 | 101 | 9% | 8e-18 | 100.00 | 5815 | NZ_BMQZ010000... |
| Sinomonas sp. KCTC 49339 sequence28, whole genome shotgun... | Sinomonas sp... | NA | 2769422 | 101 | 101 | 9% | 8e-18 | 100.00 | 5849 | NZ_BNCM010000... |
| Saccharothrix mutabilis subsp. mutabilis strain JCM 3380... | Saccharothri... | NA | 66855 | 101 | 101 | 9% | 8e-18 | 100.00 | 5899 | NZ_BAAABU0100... |
| Corynebacterium glutamicum strain 39 Scaffolds_32_9166, whole... | Corynebacter... | NA | 1718 | 101 | 101 | 9% | 1e-17 | 100.00 | 8862 | NZ_JACEQW0100... |
| Corynebacterium glutamicum strain 37 Scaffolds_33_8928, whole... | Corynebacter... | NA | 1718 | 101 | 101 | 9% | 1e-17 | 100.00 | 8928 | NZ_JACENO0100... |
| Klebsiella pneumoniae strain EuSCAPE_NO001, whole genome shotg... | Klebsiella p... | NA | 573 | 89.7 | 89.7 | 9% | 2e-17 | 88.89 | 245 | NZ_UKMF020000... |
| Klebsiella pneumoniae strain EuSCAPE_NO001, whole genome shotg... | Klebsiella p... | NA | 573 | 78.6 | 78.6 | 7% | 3e-13 | 86.49 | 289 | NZ_UKMF020000... |

Alignments:

```
>Escherichia coli strain NBE0009 contig00112, whole genome shotgun sequence
Sequence ID: NZ_JARBDL010000112.1 Length: 2003
Range 1: 111 to 1550

Score:703 bits(1815), Expect:0.0,
Method:Compositional matrix adjust.,
Identities:348/491(71%), Positives:403/491(82%), Gaps:24/491(4%)

Query  2     LWVTVYYGVPVWKEATTTLFCASDAKAYDTEVHNVWATHACVPTDPNPQEVVLVNVTENF  61
             LWVTVYYGVPVW EA TTLFCASDAKAY+ EVHNVWATHACVPTDPNPQE+VL NVTENF
Sbjct  1550  LWVTVYYGVPVWTEAKTTLFCASDAKAYEKEVHNVWATHACVPTDPNPQEMVLENVTENF  1371

Query  62    NMWKNDMVEQMHEDIISLWDQSLKPCVKLTPLCVSLKCTDLKNDTNTNSSSGRMIMEKGE  121
             NMWKNDMV QMHED+ISLWDQSLKPCVKLTPLCV+L+C    N +  N SS R+     E
Sbjct  1370  NMWKNDMVNQMHEDVISLWDQSLKPCVKLTPLCVTLEC----NSSTYNQSSVRV----HE  1215

Query  122   IKNCSFNISTSIRGKVQKEYAFFYKLDIIPIDN------DTTSYKLTSCNTSVITQACPK  175
             +KNCSFN +T +R K  K  A F++LD++P+DN       T  Y+L +CNTSVITQACPK
Sbjct  1214  VKNCSFNATTELRDKKHKVQALFHRLDVVPLDNRSSNENSTGYYRLINCNTSVITQACPK  1035

Query  176   VSFEPIPIHYCAPAGFAILKCNNKTFNGTGPCTNVSTVQCTHGIRPVVSTQLLLNGSLAE  235
             V+F+PIPIHYC PAG+AIL+CN+KTFNGTGPC NVSTVQCTHGI+PVVSTQLLLNGSLAE
Sbjct  1034  VTFDPIPIHYCTPAGYAILQCNDKTFNGTGPCNNVSTVQCTHGIKPVVSTQLLLNGSLAE  855

Query  236   EEVVIRSVNFTDNAKTIIVQLNTSVEINCTRPNNNTRKRIRIQRGPGRAFVTIGK-IGNM  294
```

```
             EE+VIRS N T+NAKTIIV LN SV+I CTRPNNNTRK IRI  GPG+ F   G  IG++
Sbjct  854   EEIVIRSENLTNNAKTIIVHLNKSVDIVCTRPNNNTRKSIRI--GPGQTFYATGDIIGDI  681

Query  295   RQAHCNISRAKWNNTLKQIASKLREQFGNNKTIIFKQSSGGDPEIVTHSFNCGGEFFYCN  354
             R+AHCNIS  +WN+T+ ++  KL E F  N+TI F+ SSGGD EI THSFNCGGEFFYCN
Sbjct  680   RKAHCNISLNRWNDTILKVREKLAEHFP-NRTIEFQPSSGGDLEITTHSFNCGGEFFYCN  504

Query  355   STQLFNSTWF--NSTWSTEGS--NNTEGSDT--ITLPCRIKQIINMWQKVGKAMYAPPIS  408
             ++ LFN T++  N T+   G+  N+T+ + T  IT+PCRIKQIINMWQ+VG+AMYAPPI+
Sbjct  503   TSNLFNFTYYYDNGTYHYNGTYYNSTQYNSTANITIPCRIKQIINMWQEVGRAMYAPPIA  324

Query  409   GQIRCSSNITGLLLTRDGGNSNNESEIFRPGGGDMRDNWRSELYKYKVVKIEPLGVAPTK  468
             G I C SNITGLLL RDGG S+NE+E FRPGGGDMRDNWRSELYKYKVV+I+PLG+APT
Sbjct  323   GNITCRSNITGLLLVRDGGPSSNETETFRPGGGDMRDNWRSELYKYKVVEIKPLGIAPTG  144

Query  469   AKRRVVQREKR  479
             AKRRVV REKR
Sbjct  143   AKRRVVGREKR  111
```

>Escherichia coli strain NBE0009 contig00111, whole genome shotgun sequence
Sequence ID: NZ_JARBDL010000111.1 Length: 2027
Range 1: 454 to 1914

Score:701 bits(1810), Expect:0.0,
Method:Compositional matrix adjust.,
Identities:345/491(70%), Positives:397/491(80%), Gaps:17/491(3%)

```
Query  2     LWVTVYYGVPVWKEATTTLFCASDAKAYDTEVHNVWATHACVPTDPNPQEVVLVNVTENF  61
             LWVTVYYGVPVWKEA TTLFCASDAK Y+TEVHNVWATHACVPTDP+PQE+ L NVTENF
Sbjct  454   LWVTVYYGVPVWKEAKTTLFCASDAKGYETEVHNVWATHACVPTDPSPQEMRLDNVTENF  633

Query  62    NMWKNDMVEQMHEDIISLWDQSLKPCVKLTPLCVSLKCTDLKNDTNTNSSSGRMIMEKGE  121
             NMW+N+MV+QMHEDIISLWDQSLKPC+KLTPLCV+L+C +  N T TN  +      K E
Sbjct  634   NMWENNMVDQMHEDIISLWDQSLKPCIKLTPLCVTLEC-EKANRTVTNILN-ETYDSKNE  807

Query  122   IKNCSFNISTSIRGKVQKEYAFFYKLDIIPIDN-------DTTSYKLTSCNTSVITQACP  174
             I+NCSFN +T I  K +KEYA FY+LDI+ +DN       D+  Y+L +CNTS ITQACP
Sbjct  808   IQNCSFNATTEIEDKKKKEYALFYRLDIVSLDNENNSSGNDSRPYRLINCNTSAITQACP  987

Query  175   KVSFEPIPIHYCAPAGFAILKCNNKTFNGTGPCTNVSTVQCTHGIRPVVSTQLLLNGSLA  234
             KVSF+PIPIHYCAPAG+AILKCNNKTFNGTGPC NVSTVQCTHGI+PVVSTQLL+NGSLA
Sbjct  988   KVSFDPIPIHYCAPAGYAILKCNNKTFNGTGPCNNVSTVQCTHGIKPVVSTQLLVNGSLA  1167

Query  235   EEEVVIRSVNFTDNAKTIIVQLNTSVEINCTRPNNNTRKRIRIQRGPGR-AFVTIGKIGN  293
             EEE++IRS N TDNAKTIIVQLN SV I C RPNNNTRK IRI  GPG+  + T   IG+
Sbjct  1168  EEEIIIRSENLTDNAKTIIVQLNKSVPITCIRPNNNTRKSIRI--GPGQTVYATGDSIGD  1341

Query  294   MRQAHCNISRAKWNNTLKQIASKLREQFGNNKTIIFKQSSGGDPEIVTHSFNCGGEFFYC  353
             +RQAHCN+SR  W  TL  +  KL+E F N  TI FK SSGGD EI THSFNC GEFFYC
Sbjct  1342  IRQAHCNVSRKDWEGTLYNVNKKLKEHFPNITTIKFKPSSGGDLEITTHSFNCRGEFFYC  1521

Query  354   NSTQLFNSTW---FNSTWSTEGSNNTEGSDTITLPCRIKQIINMWQKVGKAMYAPPISGQ  410
             N++ LFN T+   +  T++  G N++      ITL CRIKQIINMWQ+VGKAMYAPPI+G
Sbjct  1522  NTSDLFNGTYNSTYKGTYNNTGGNSSAADPNITLQCRIKQIINMWQQVGKAMYAPPIAGN  1701

Query  411   IRCSSNITGLLLTRDGGN--SNNESEIFRPGGGDMRDNWRSELYKYKVVKIEPLGVAPTK  468
             I C SNITGLLL RDGG   ++N++E FRPGGG+M+DNWRSELYKYKVV+I+PLGVAPT
Sbjct  1702  ITCISNITGLLLVRDGGTNRTDNDTETFRPGGGNMKDNWRSELYKYKVVEIQPLGVAPTG  1881

Query  469   AKRRVVQREKR  479
             AKRRVV+REKR
Sbjct  1882  AKRRVVEREKR  1914
```

>Escherichia coli strain EN5177 contig00087, whole genome shotgun sequence
Sequence ID: NZ_JAPMNK010000087.1 Length: 3403
Range 1: 448 to 2007

Score:648 bits(1671), Expect:0.0,
Method:Compositional matrix adjust.,
Identities:327/522(63%), Positives:384/522(73%), Gaps:58/522(11%)

```
Query  2     LWVTVYYGVPVWKEATTTLFCASDAKAYDTEVHNVWATHACVPTDPNPQEVVLVNVTENF  61
             LWVTVYYGVPVW++A  TLFCASDAK Y TEVHNVWATHACVPTDPNPQE+VL NVTENF
Sbjct  448   LWVTVYYGVPVWRDANATLFCASDAKVYKTEVHNVWATHACVPTDPNPQELVLENVTENF  627

Query  62    NMWKNDMVEQMHEDIISLWDQSLKPCVKLTPLCVSLKCTDLK------NDTNTNSSSGRM  115
             NMW+NDMVEQMH+D+IS+WDQ LKPCVKLTPLCV+L CT  +      ++ NT + S
Sbjct  628   NMWENDMVEQMHQDVISIWDQGLKPCVKLTPLCVTLNCTTYEPKKVTADNNNTCAPSQNT  807

Query  116   IMEK--------GEIKNCSFNISTSIRGKVQKEYAFFYKLDIIPIDNDTTS-----YKLT  162
              ++          EI NCSFN +T +R K +K +A FYK+D++ +  ++T+     Y+L
Sbjct  808   TVKNLTIDPEAVKEISNCSFNTTTELRDKTRKVHALFYKIDMVELGENSTAPNFTLYRLI  987

Query  163   SCNTSVITQACPKVSFEPIPIHYCAPAGFAILKCNNKTFNGTGPCTNVSTVQCTHGIRPV  222
             +CNTS ITQACPKVSFEPIPIHYCAPAG+A+LKCNN TFNGTGPC N+STVQCTHGIRPV
Sbjct  988   NCNTSAITQACPKVSFEPIPIHYCAPAGYALLKCNNNTFNGTGPCRNISTVQCTHGIRPV  1167

Query  223   VSTQLLLNGSLAEEEVVIRSVNFTDNAKTIIVQLNTSVEINCTRPNNNTRKRIRIQRGPG  282
             VSTQLLLNGSLAE  V IRS N TDNAKTII  LN SVEI CTRPNNNTR+ +RI  GPG
Sbjct  1168  VSTQLLLNGSLAEGNVTIRSENLTDNAKTIIAHLNQSVEILCTRPNNNTRQSVRI--GPG  1341

Query  283   RAFVTIGK-IGNMRQAHCNISRAKWNNTLKQIASKLREQFGNNKTIIFKQSSGGDPEIVT  341
             + F   G  IG++R+AHCN+S   W+  L+ ++ +L + F NNKTI F   SGGD EI T
Sbjct  1342  QTFFATGDIIGDIRRAHCNVSAKHWHTALRNVSIELAKLFPNNKTINFTSPSGGDLEITT  1521

Query  342   HSFNCGGEFFYCNSTQLFNS---------TWFNSTWSTEGSNNTE----GSD--------  380
             HSFNC GEFFYCN++QLFNS          W+N   +    N T+    GS
Sbjct  1522  HSFNCRGEFFYCNTSQLFNSLYMPNNDTYIWYNENHTYVWHNGTQYVINGSTQYVTLKNS  1701

Query  381   ------------TITLPCRIKQIINMWQKVGKAMYAPPISGQIRCSSNITGLLLTRDGGN  428
                          ITLPCRIKQIINMWQ+VGKAMYAPPI+G I C SNITGLLLTRDGG
Sbjct  1702  AQNVTINNITDFNITLPCRIKQIINMWQEVGKAMYAPPIAGNITCLSNITGLLLTRDGGK  1881

Query  429   SN---NESEIFRPGGGDMRDNWRSELYKYKVVKIEPLGVAPT  467
              +   N++EIFRPGGG+M+DNWRSELYKYKVV+I+PLGVAPT
Sbjct  1882  GDEIRNDTEIFRPGGGNMKDNWRSELYKYKVVEIKPLGVAPT  2007
```

>Escherichia coli strain NBE0004 contig00090, whole genome shotgun sequence
Sequence ID: NZ_JARBDH010000090.1 Length: 2818
Range 1: 812 to 2371

Score:648 bits(1671), Expect:0.0,
Method:Compositional matrix adjust.,
Identities:327/522(63%), Positives:384/522(73%), Gaps:58/522(11%)

```
Query  2      LWVTVYYGVPVWKEATTTLFCASDAKAYDTEVHNVWATHACVPTDPNPQEVVLVNVTENF  61
              LWVTVYYGVPVW++A  TLFCASDAK Y TEVHNVWATHACVPTDPNPQE+VL NVTENF
Sbjct  2371   LWVTVYYGVPVWRDANATLFCASDAKVYKTEVHNVWATHACVPTDPNPQELVLENVTENF  2192

Query  62     NMWKNDMVEQMHEDIISLWDQSLKPCVKLTPLCVSLKCTDLK------NDTNTNSSSGRM  115
              NMW+NDMVEQMH+D+IS+WDQ LKPCVKLTPLCV+L CT  +      ++ NT + S
Sbjct  2191   NMWENDMVEQMHQDVISIWDQGLKPCVKLTPLCVTLNCTTYEPKKVTADNNNTCAPSQNT  2012

Query  116    IMEK--------GEIKNCSFNISTSIRGKVQKEYAFFYKLDIIPIDNDTTS-----YKLT  162
               ++          EI NCSFN +T +R K +K +A FYK+D++ +  ++T+     Y+L
Sbjct  2011   TVKNLTIDPEAVKEISNCSFNTTTELRDKTRKVHALFYKIDMVELGENSTAPNFTLYRLI  1832

Query  163    SCNTSVITQACPKVSFEPIPIHYCAPAGFAILKCNNKTFNGTGPCTNVSTVQCTHGIRPV  222
              +CNTS ITQACPKVSFEPIPIHYCAPAG+A+LKCNN TFNGTGPC N+STVQCTHGIRPV
Sbjct  1831   NCNTSAITQACPKVSFEPIPIHYCAPAGYALLKCNNNTFNGTGPCRNISTVQCTHGIRPV  1652

Query  223    VSTQLLLNGSLAEEEVVIRSVNFTDNAKTIIVQLNTSVEINCTRPNNNTRKRIRIQRGPG  282
              VSTQLLLNGSLAE  V IRS N TDNAKTII  LN SVEI CTRPNNNTR+ +RI  GPG
Sbjct  1651   VSTQLLLNGSLAEGNVTIRSENLTDNAKTIIAHLNQSVEILCTRPNNNTRQSVRI--GPG  1478

Query  283    RAFVTIGK-IGNMRQAHCNISRAKWNNTLKQIASKLREQFGNNKTIIFKQSSGGDPEIVT  341
              + F   G  IG++R+AHCN+S   W+  L+ ++ +L + F NNKTI F   SGGD EI T
Sbjct  1477   QTFFATGDIIGDIRRAHCNVSAKHWHTALRNVSIELAKLFPNNKTINFTSPSGGDLEITT  1298

Query  342    HSFNCGGEFFYCNSTQLFNS---------TWFNSTWSTEGSNNTE----GSD--------  380
              HSFNC GEFFYCN++QLFNS          W+N   +    N T+    GS
Sbjct  1297   HSFNCRGEFFYCNTSQLFNSLYMPNNDTYIWYNENHTYVWHNGTQYVINGSTQYVTLKNS  1118

Query  381    ------------TITLPCRIKQIINMWQKVGKAMYAPPISGQIRCSSNITGLLLTRDGGN  428
                           ITLPCRIKQIINMWQ+VGKAMYAPPI+G I C SNITGLLLTRDGG
Sbjct  1117   AQNVTINNITDFNITLPCRIKQIINMWQEVGKAMYAPPIAGNITCLSNITGLLLTRDGGK  938

Query  429    SN---NESEIFRPGGGDMRDNWRSELYKYKVVKIEPLGVAPT  467
               +   N++EIFRPGGG+M+DNWRSELYKYKVV+I+PLGVAPT
Sbjct  937    GDEIRNDTEIFRPGGGNMKDNWRSELYKYKVVEIKPLGVAPT  812
```

```
>Escherichia coli strain EN5076 contig00066, whole genome shotgun sequence
Sequence ID: NZ_JAODTC010000066.1 Length: 2732
Range 1: 957 to 2279

Score:639 bits(1648), Expect:0.0,
Method:Compositional matrix adjust.,
Identities:311/445(70%), Positives:362/445(81%), Gaps:12/445(2%)

Query  2      LWVTVYYGVPVWKEATTTLFCASDAKAYDTEVHNVWATHACVPTDPNPQEVVLVNVTENF  61
              LWVTVYYGVPVWKEA TTLFCASDAKAY TEVHNVWATHACVPTDPNPQE+VL NVTENF
Sbjct  2279   LWVTVYYGVPVWKEAKTTLFCASDAKAYKTEVHNVWATHACVPTDPNPQEMVLENVTENF  2100

Query  62     NMWKNDMVEQMHEDIISLWDQSLKPCVKLTPLCVSLKCTDLKNDTNTNSSSGRMIMEKGE  121
              NMWKNDMV QMHED+ISLWDQSLKPCVKLTPLCV+LKC    ++  ++++         E
Sbjct  2099   NMWKNDMVNQMHEDVISLWDQSLKPCVKLTPLCVTLKCNATTHNGTSDAADNDTSSGGEE  1920

Query  122    IKNCSFNISTSIRGKVQKEYAFFYKLDIIPID-----NDTTSYKLTSCNTSVITQACPKV  176
              +KNCSFNI+T +R + QK YA FYKLD++P+D     N + SY+L +CNTS ITQACPKV
Sbjct  1919   MKNCSFNITTELRDRKQKVYALFYKLDVVPLDNTNGSNSSRSYRLINCNTSAITQACPKV  1740

Query  177    SFEPIPIHYCAPAGFAILKCNNKTFNGTGPCTNVSTVQCTHGIRPVVSTQLLLNGSLAEE  236
              SF+PIPIHYCAPAG+AIL+CN+KTFNGTGPC NVSTVQCTHGI+PVVSTQLLLNGSLAEE
Sbjct  1739   SFDPIPIHYCAPAGYAILRCNDKTFNGTGPCHNVSTVQCTHGIKPVVSTQLLLNGSLAEE  1560

Query  237    EVVIRSVNFTDNAKTIIVQLNTSVEINCTRPNNNTRKRIRIQRGPGRAFVTIGK-IGNMR  295
              E++IRS N +DNAKTIIV LN SVEI CTRP+NNTRK IRI  GPG+ F   G  IGN+R
Sbjct  1559   EIIIRSQNLSDNAKTIIVHLNQSVEIVCTRPHNNTRKSIRI--GPGQTFYATGAIIGNIR  1386

Query  296    QAHCNISRAKWNNTLKQIASKLREQFGNNKTIIFKQSSGGDPEIVTHSFNCGGEFFYCNS  355
              QA+C+I+ +KW+ TL  ++ KL E+F  NKTI F  SSGGD E+ THSFNC GEFFYCN+
Sbjct  1385   QAYCSINISKWSETLHNVSKKLAERFP-NKTINFAASSGGDLEVTTHSFNCRGEFFYCNT  1209

Query  356    TQLFNST--WFNSTWSTEGSNNTEGSDTITLPCRIKQIINMWQKVGKAMYAPPISGQIRC  413
              ++LFNST  + N T+   G+N T  ++  T+PCRIKQIINMWQ+VG+AMYAPPI G I C
Sbjct  1208   SELFNSTYIYLNRTYMFNGTNGT-STNNFTIPCRIKQIINMWQEVGRAMYAPPIEGNITC  1032

Query  414    SSNITGLLLTRDGGNSNNESEIFRP  438
               SNITGLLL RDGG  N  +EIFRP
Sbjct  1031   KSNITGLLLERDGGEENRTTEIFRP  957
```

```
>Escherichia coli strain NBE0010 contig00080, whole genome shotgun sequence
Sequence ID: NZ_JARBDM010000080.1 Length: 2743
Range 1: 149 to 1471

Score:637 bits(1644), Expect:0.0,
Method:Compositional matrix adjust.,
Identities:308/445(69%), Positives:362/445(81%), Gaps:12/445(2%)

Query  2      LWVTVYYGVPVWKEATTTLFCASDAKAYDTEVHNVWATHACVPTDPNPQEVVLVNVTENF  61
              LWVTVYYGVPVW++A TTLFCASDAKAY+ EVHNVWATHACVPTDPNPQE+VL NVTENF
Sbjct  149    LWVTVYYGVPVWRDAKTTLFCASDAKAYEKEVHNVWATHACVPTDPNPQEMVLENVTENF  328

Query  62     NMWKNDMVEQMHEDIISLWDQSLKPCVKLTPLCVSLKCTDLKNDTNTNSSSGRMIMEKGE  121
              NMWKNDMV QMHED+ISLWDQSLKPCVKLTPLCV+LKC    ++  ++++         E
Sbjct  329    NMWKNDMVNQMHEDVISLWDQSLKPCVKLTPLCVTLKCNATTHNGTSDAADNDTSSGGEE  508

Query  122    IKNCSFNISTSIRGKVQKEYAFFYKLDIIPID-----NDTTSYKLTSCNTSVITQACPKV  176
              +KNCSFNI+T +R + QK YA FYKLD++P+D     N + SY+L +CNTS ITQACPKV
Sbjct  509    MKNCSFNITTELRDRKQKVYALFYKLDVVPLDNTNGSNSSRSYRLINCNTSAITQACPKV  688

Query  177    SFEPIPIHYCAPAGFAILKCNNKTFNGTGPCTNVSTVQCTHGIRPVVSTQLLLNGSLAEE  236
              SF+PIPIHYCAPAG+AIL+CN+KTFNGTGPC NVSTVQCTHGI+PVVSTQLLLNGSLAEE
Sbjct  689    SFDPIPIHYCAPAGYAILRCNDKTFNGTGPCHNVSTVQCTHGIKPVVSTQLLLNGSLAEE  868

Query  237    EVVIRSVNFTDNAKTIIVQLNTSVEINCTRPNNNTRKRIRIQRGPGRAFVTIGK-IGNMR  295
              E++IRS N +DNAKTIIV LN SVEI CTRP+NNTRK IRI  GPG+ F   G  IGN+R
Sbjct  869    EIIIRSQNLSDNAKTIIVHLNQSVEIVCTRPHNNTRKSIRI--GPGQTFYATGAIIGNIR  1042

Query  296    QAHCNISRAKWNNTLKQIASKLREQFGNNKTIIFKQSSGGDPEIVTHSFNCGGEFFYCNS  355
              QA+C+I+ +KW+ TL  ++ KL E+F  NKTI F  SSGGD E+ THSFNC GEFFYCN+
Sbjct  1043   QAYCSINISKWSETLHNVSKKLAERFP-NKTINFAASSGGDLEVTTHSFNCRGEFFYCNT  1219

Query  356    TQLFNST--WFNSTWSTEGSNNTEGSDTITLPCRIKQIINMWQKVGKAMYAPPISGQIRC  413
              ++LFNST  + N T+   G+N T  ++  T+PCRIKQIINMWQ+VG+AMYAPPI G I C
Sbjct  1220   SELFNSTYIYLNRTYMFNGTNGT-STNNFTIPCRIKQIINMWQEVGRAMYAPPIEGNITC  1396

Query  414    SSNITGLLLTRDGGNSNNESEIFRP  438
               SNITGLLL RDGG  N  +EIFRP
Sbjct  1397   KSNITGLLLERDGGEENRTTEIFRP  1471
```

```
>Escherichia coli strain EN5374 contig00094, whole genome shotgun sequence
Sequence ID: NZ_JAPTGI010000094.1 Length: 2720
Range 1: 448 to 1770

Score:636 bits(1641), Expect:0.0,
Method:Compositional matrix adjust.,
Identities:308/445(69%), Positives:362/445(81%), Gaps:12/445(2%)

Query  2     LWVTVYYGVPVWKEATTTLFCASDAKAYDTEVHNVWATHACVPTDPNPQEVVLVNVTENF  61
             LWVTVYYGVPVW++A TTLFCASDAKAY+ EVHNVWATHACVPTDPNPQE+VL NVTENF
Sbjct  448   LWVTVYYGVPVWRDAKTTLFCASDAKAYEKEVHNVWATHACVPTDPNPQEMVLENVTENF  627

Query  62    NMWKNDMVEQMHEDIISLWDQSLKPCVKLTPLCVSLKCTDLKNDTNTNSSSGRMIMEKGE  121
             NMWKNDMV QMHED+ISLWDQSLKPCVKLTPLCV+LKC    ++  ++++         E
Sbjct  628   NMWKNDMVNQMHEDVISLWDQSLKPCVKLTPLCVTLKCNATTHNGTSDAADNDTSSGGEE  807

Query  122   IKNCSFNISTSIRGKVQKEYAFFYKLDIIPID-----NDTTSYKLTSCNTSVITQACPKV  176
             +KNCSFNI+T +R + QK YA FYKLD++P+D     N + SY+L +CNTS ITQACPKV
Sbjct  808   MKNCSFNITTELRDRKQKVYALFYKLDVVPLDNTNGSNSSRSYRLINCNTSAITQACPKV  987

Query  177   SFEPIPIHYCAPAGFAILKCNNKTFNGTGPCTNVSTVQCTHGIRPVVSTQLLLNGSLAEE  236
             SF+PIPIHYCAPAG+AIL+CN+KTFNGTGPC NVSTVQCTHGI+PVVSTQLLLNGSLAEE
Sbjct  988   SFDPIPIHYCAPAGYAILRCNDKTFNGTGPCHNVSTVQCTHGIKPVVSTQLLLNGSLAEE  1167

Query  237   EVVIRSVNFTDNAKTIIVQLNTSVEINCTRPNNNTRKRIRIQRGPGRAFVTIGK-IGNMR  295
             E++IRS N +DNAKTIIV LN SVEI CTRP+NNTRK IRI  GPG+ F   G  IGN+R
Sbjct  1168  EIIIRSQNLSDNAKTIIVHLNQSVEIVCTRPHNNTRKSIRI--GPGQTFYATGAIIGNIR  1341

Query  296   QAHCNISRAKWNNTLKQIASKLREQFGNNKTIIFKQSSGGDPEIVTHSFNCGGEFFYCNS  355
             QA+C+I+ +KW+ TL  ++ KL E+F  NKTI F  SSGGD E+ THSFNC GEFFYCN+
Sbjct  1342  QAYCSINISKWSETLHNVSKKLAERFP-NKTINFAASSGGDLEVTTHSFNCRGEFFYCNT  1518

Query  356   TQLFNST--WFNSTWSTEGSNNTEGSDTITLPCRIKQIINMWQKVGKAMYAPPISGQIRC  413
             ++LFNST  + N T+   G+N T  ++  T+PCRIKQIINMWQ+VG+AMYAPPI G I C
Sbjct  1519  SELFNSTYIYLNRTYMFNGTNGT-STNNFTIPCRIKQIINMWQEVGRAMYAPPIEGNITC  1695

Query  414   SSNITGLLLTRDGGNSNNESEIFRP  438
              SNITGLLL RDGG  N  +EIFRP
Sbjct  1696  KSNITGLLLERDGGEENRTTEIFRP  1770



>Escherichia coli strain EN5177 contig00105, whole genome shotgun sequence
Sequence ID: NZ_JAPMNK010000105.1 Length: 1783
Range 1: 278 to 1681

Score:636 bits(1640), Expect:0.0,
Method:Compositional matrix adjust.,
Identities:325/484(67%), Positives:372/484(76%), Gaps:29/484(5%)

Query  4     VTVYYGVPVWKEATTTLFCASDAKAYDTEVHNVWATHACVPTDPNPQEVVLVNVTENFNM  63
             VTVYY VPVWKEA  TLFCASDAKA++ EVHNVWATHACVPTDPNPQE+VL NVTENFNM
Sbjct  278   VTVYYEVPVWKEAKATLFCASDAKAHEREVHNVWATHACVPTDPNPQEIVLENVTENFNM  457

Query  64    WKNDMVEQMHEDIISLWDQSLKPCVKLTPLCVSLKCTDLKNDTNTNSSSGRMIMEKGEIK  123
             WK+DMV+QMHED+ISL DQSLKPCVKLTPLCV+L+C +     N          E  E+K
Sbjct  458   WKSDMVDQMHEDVISL*DQSLKPCVKLTPLCVTLECDNETVSVNHTD-------ESKELK  616

Query  124   NCSFNISTSIRGKVQKEYAFFYKLDIIPIDNDTT---------SYKLTSCNTSVITQACP  174
              C FN ST +  K QK  A FY LDI P++  ++         +Y L SCNTSVI QACP
Sbjct  617   RCFFNASTEVEDKKQKVSALFYSLDIRPLNKSSSRNSSINSNETYVLISCNTSVIKQACP  796

Query  175   KVSFEPIPIHYCAPAGFAILKCNNKTFNGTGPCTNVSTVQCTHGIRPVVSTQLLLNGSLA  234
             KV+F+PIPIHYCAPAG+AILKCNN+TFN TGPC NVSTVQCTH I+PVVSTQLLLN SLA
Sbjct  797   KVTFDPIPIHYCAPAGYAILKCNNETFNRTGPCNNVSTVQCTHRIKPVVSTQLLLNSSLA  976

Query  235   EEEVVIRSVNFTDNAKTIIVQLNTSVEINCTRPNNNTRKRIRIQRGPGRAFVTIGKI-GN  293
             + E++IRS N TDNAKTIIV LN SVEINCTRPNNNTRK IRI  GPG+ F   G+I  N
Sbjct  977   KGEIIIRSENLTDNAKTIIVHLNESVEINCTRPNNNTRKSIRI--GPGQTFYATGEIVRN  1150

Query  294   MRQAHCNISRAKWNNTLKQIASKLREQFGNNKTIIFKQSSGGDPEIVTHSFNCGGEFFYC  353
             +RQAHCNIS  + N TL+++  KL E F  NKTI F   SGGD EI+THSFNC GEFFYC
Sbjct  1151  IRQAHCNISEKE*NKTLQKVGIKLAEHFP-NKTINFTSHSGGDLEIMTHSFNCRGEFFYC  1327

Query  354   NSTQLFNSTWFNSTWSTEGSNNTEGSDTITLPCRIKQIINMWQKVGKAMYAPPISGQIRC  413
             N++ LFNST +NST+    S     S  IT+ CRIKQIINMWQ+VG+AMYAPPI+G I C
Sbjct  1328  NTSSLFNST-YNSTYIRNNS-----SSIITIQCRIKQIINMWQEVGRAMYAPPIAGNITC  1489

Query  414   SSNITGLLLTRDGGNSN---NESEIFRPGGGDMRDNWRSELYKYKVVKIEPLGVAPTKAK  470
              SNITGLLL RD G+ N   N +E FRP GGDM++N RSELYKYKVV+I+PLG+APT AK
Sbjct  1490  QSNITGLLLVRDRGHRNETSNSTETFRPKGGDMKNN*RSELYKYKVVEIKPLGIAPTAAK  1669

Query  471   RRVV  474
             RRVV
Sbjct  1670  RRVV  1681



>Escherichia coli strain EN5090 contig00081, whole genome shotgun sequence
Sequence ID: NZ_JAODTD010000081.1 Length: 2798
Range 1: 1020 to 2345

Score:634 bits(1635), Expect:0.0,
Method:Compositional matrix adjust.,
Identities:312/446(70%), Positives:361/446(80%), Gaps:13/446(2%)

Query  2     LWVTVYYGVPVWKEATTTLFCASDAKAYDTEVHNVWATHACVPTDPNPQEVVLVNVTENF  61
             LWVTVYYGVPVWKEA TTLFCASDAKAY TEVHNVWATHACVPTDPNPQE+VL NVTENF
Sbjct  2345  LWVTVYYGVPVWKEAKTTLFCASDAKAYKTEVHNVWATHACVPTDPNPQEMVLENVTENF  2166

Query  62    NMWKNDMVEQMHEDIISLWDQSLKPCVKLTPLCVSLKCTDLKNDTNTNSSSGRMIMEKGE  121
             NMWKNDMV QMHED+ISLWDQSLKPCVKLTPLCV+LKCT   ++   +++         E
Sbjct  2165  NMWKNDMVNQMHEDVISLWDQSLKPCVKLTPLCVTLKCTATTHNGTRDAADNDTSSGGEE  1986

Query  122   IKNCSFNISTSIRGKVQKEYAFFYKLDIIPIDNDTTS-----YKLTSCNTSVITQACPKV  176
             +KNCSFNI+T +R + QK YA FYKLD++P+DN   S     Y+L +CNTS ITQACPKV
Sbjct  1985  MKNCSFNITTELRDRKQKVYALFYKLDVVPLDNTNGSNSSRPYRLINCNTSAITQACPKV  1806

Query  177   SFEPIPIHYCAPAGFAILKCNNKTFNGTGPCTNVSTVQCTHGIRPVVSTQLLLNGSLAEE  236
             SF+PIPIHYCAPAG+AIL+CN+KTFNGTGPC NVSTVQCTHGI+PVVSTQLLLNGSLAEE
Sbjct  1805  SFDPIPIHYCAPAGYAILRCNDKTFNGTGPCHNVSTVQCTHGIKPVVSTQLLLNGSLAEE  1626

Query  237   EVVIRSVNFTDNAKTIIVQLNTSVEINCTRPNNNTRKRIRIQRGPGRAFVTIGK-IGNMR  295
             E++IRS N +DNAKTIIV LN SVEI CTRP+NNTRK IRI  GPG+ F   G  IGN+R
Sbjct  1625  EIIIRSQNLSDNAKTIIVHLNQSVEIVCTRPHNNTRKSIRI--GPGQTFYATGAIIGNIR  1452

Query  296   QAHCNISRAKWNNTLKQIASKLREQFGNNKTIIFKQSSGGDPEIVTHSFNCGGEFFYCNS  355
             QA+C+I+ +KW+ TL  ++ KL E+F  NKTI F  SSGGD E+ THSFNC GEFFYCN+
Sbjct  1451  QAYCSINISKWSETLHNVSKKLAERFP-NKTINFAASSGGDLEVTTHSFNCRGEFFYCNT  1275

Query  356   TQLFNSTWF--NSTWSTEGSNNTEGSDTITLPCRIKQIINMWQKVGKAMYAPPISGQIRC  413
```

```
             ++LFNST+   + T+   G+N T  ++  T+PCRIKQIINMWQKVG+AMYAPPI G I C
Sbjct  1274  SELFNSTYIYPDRTYMFNGTNGT-STNNFTIPCRIKQIINMWQKVGRAMYAPPIEGNITC  1098

Query  414   SSNITGLLLTRDGGNSNNE-SEIFRP  438
              SNITGLLL RDGG   N  +EIFRP
Sbjct  1097  KSNITGLLLERDGGEEENRTTEIFRP  1020
```

>Escherichia coli strain EN5379 contig00092, whole genome shotgun sequence
Sequence ID: NZ_JAPTGJ010000092.1 Length: 2592
Range 1: 1019 to 2338

Score:629 bits(1621), Expect:0.0,
Method:Compositional matrix adjust.,
Identities:313/447(70%), Positives:361/447(80%), Gaps:17/447(3%)

```
Query  2     LWVTVYYGVPVWKEATTTLFCASDAKAYDTEVHNVWATHACVPTDPNPQEVVLVNVTENF  61
             LWVTVYYGVPVWKEA TTLFCASDAKAY TEVHNVWATHACVPTDPNPQE+VL NVTENF
Sbjct  2338  LWVTVYYGVPVWKEAKTTLFCASDAKAYKTEVHNVWATHACVPTDPNPQEMVLENVTENF  2159

Query  62    NMWKNDMVEQMHEDIISLWDQSLKPCVKLTPLCVSLKC-TDLKNDTNTNSSSGRMIMEKG  120
             NMWKNDMV QMHED+ISLWD+SLKPCVKLTPLCV+LKC     N T   + +G +
Sbjct  2158  NMWKNDMVNQMHEDVISLWDESLKPCVKLTPLCVTLKCNATASNGTAAGNGTGSV---GE  1988

Query  121   EIKNCSFNISTSIRGKVQKEYAFFYKLDIIPIDNDTTS-----YKLTSCNTSVITQACPK  175
             E+KNCSFNI+T +R + QK YA FYKLD++P+DN   S      Y+L +CNTS ITQACPK
Sbjct  1987  EMKNCSFNITTELRDRKQKVYALFYKLDVVPLDNTNGSNSSRPYRLINCNTSAITQACPK  1808

Query  176   VSFEPIPIHYCAPAGFAILKCNNKTFNGTGPCTNVSTVQCTHGIRPVVSTQLLLNGSLAE  235
             VSF+PIPIHYCAPAG+AIL+CN+KTFNGTGPC NVSTVQCTHGI+PVVSTQLLLNGSLAE
Sbjct  1807  VSFDPIPIHYCAPAGYAILRCNDKTFNGTGPCHNVSTVQCTHGIKPVVSTQLLLNGSLAE  1628

Query  236   EEVVIRSVNFTDNAKTIIVQLNTSVEINCTRPNNNTRKRIRIQRGPGRAFVTIGK-IGNM  294
             EE++IRS N +DNAKTIIV LN SVEI CTRP+NNTRK IRI  GPG+ F   G  IGN+
Sbjct  1627  EEIIIRSQNLSDNAKTIIVHLNQSVEIVCTRPHNNTRKSIRI--GPGQTFYATGAIIGNI  1454

Query  295   RQAHCNISRAKWNNTLKQIASKLREQFGNNKTIIFKQSSGGDPEIVTHSFNCGGEFFYCN  354
             RQA+C+I+ +KW+ TL  ++ KL E+F  NKTI F  SSGGD E+ THSFNC GEFFYCN
Sbjct  1453  RQAYCSINISKWSETLHNVSKKLAERFP-NKTINFAASSGGDLEVTTHSFNCRGEFFYCN  1277

Query  355   STQLFNSTWF--NSTWSTEGSNNTEGSDTITLPCRIKQIINMWQKVGKAMYAPPISGQIR  412
             +++LFNST+   + T+   G+N T  ++  T+PCRIKQIINMWQKVG+AMYAPPI G I
Sbjct  1276  TSELFNSTYIYPDRTYMFNGTNGT-STNNFTIPCRIKQIINMWQKVGRAMYAPPIEGNIT  1100

Query  413   CSSNITGLLLTRDGGNSNNE-SEIFRP  438
             C SNITGLLL RDGG   N  +EIFRP
Sbjct  1099  CKSNITGLLLERDGGEEENRTTEIFRP  1019
```

>Escherichia coli strain EN5317 contig00057, whole genome shotgun sequence
Sequence ID: NZ_JARBDO010000057.1 Length: 2867
Range 1: 1077 to 2414

Score:620 bits(1600), Expect:0.0,
Method:Compositional matrix adjust.,
Identities:309/450(69%), Positives:356/450(79%), Gaps:17/450(3%)

```
Query  2     LWVTVYYGVPVWKEATTTLFCASDAKAYDTEVHNVWATHACVPTDPNPQEVVLVNVTENF  61
             LWVTVYYGVPVWKEA TTLFCASDAK Y+TEVHNVWATHACVPTDP+PQE+ L NVTENF
Sbjct  2414  LWVTVYYGVPVWKEAKTTLFCASDAKGYETEVHNVWATHACVPTDPSPQEMRLDNVTENF  2235

Query  62    NMWKNDMVEQMHEDIISLWDQSLKPCVKLTPLCVSLKCTDLKNDTNTNSSSGRMIMEKGE  121
             NMW+N+MV+QMHEDIISLWDQSLKPC+KLTPLCV+L+C +  N T TN  +      K E
Sbjct  2234  NMWENNMVDQMHEDIISLWDQSLKPCIKLTPLCVTLEC-EKANRTVTNILN-ETYDSKNE  2061

Query  122   IKNCSFNISTSIRGKVQKEYAFFYKLDIIPIDN-------DTTSYKLTSCNTSVITQACP  174
             I+NCSFN +T I  K +KEYA FY+LDI+ +DN       D+  Y+L +CNTS ITQACP
Sbjct  2060  IQNCSFNATTEIEDKKKKEYALFYRLDIVSLDNENNSSGNDSRPYRLINCNTSAITQACP  1881

Query  175   KVSFEPIPIHYCAPAGFAILKCNNKTFNGTGPCTNVSTVQCTHGIRPVVSTQLLLNGSLA  234
             KVSF+PIPIHYCAPAG+AILKCNNKTFNGTGPC NVSTVQCTHGI+PVVSTQLL+NGSLA
Sbjct  1880  KVSFDPIPIHYCAPAGYAILKCNNKTFNGTGPCNNVSTVQCTHGIKPVVSTQLLVNGSLA  1701

Query  235   EEEVVIRSVNFTDNAKTIIVQLNTSVEINCTRPNNNTRKRIRIQRGPGR-AFVTIGKIGN  293
             EEE++IRS N TDNAKTIIVQLN SV I C RPNNNTRK IRI  GPG+  + T   IG+
Sbjct  1700  EEEIIIRSENLTDNAKTIIVQLNKSVPITCIRPNNNTRKSIRI--GPGQTVYATGDSIGD  1527

Query  294   MRQAHCNISRAKWNNTLKQIASKLREQFGNNKTIIFKQSSGGDPEIVTHSFNCGGEFFYC  353
             +RQAHCN+SR  W  TL  +  KL+E F N  TI FK SSGGD EI THSFNC GEFFYC
Sbjct  1526  IRQAHCNVSRKDWEGTLYNVNKKLKEHFPNITTIKFKPSSGGDLEITTHSFNCRGEFFYC  1347

Query  354   NSTQLFNSTW---FNSTWSTEGSNNTEGSDTITLPCRIKQIINMWQKVGKAMYAPPISGQ  410
             N++ LFN T+   +  T++  G N++     ITL CRIKQIINMWQ+VGKAMYAPPI+G
Sbjct  1346  NTSDLFNGTYNSTYKGTYNNTGGNSSAADPNITLQCRIKQIINMWQQVGKAMYAPPIAGN  1167

Query  411   IRCSSNITGLLLTRDGGN--SNNESEIFRP  438
             I C SNITGLLL RDGG   ++N++E F P
Sbjct  1166  ITCISNITGLLLVRDGGTNRTDNDTETFLP  1077
```

>Escherichia coli strain NBE0003 contig00096, whole genome shotgun sequence
Sequence ID: NZ_JARBDG010000096.1 Length: 2837
Range 1: 1062 to 2390

Score:613 bits(1580), Expect:0.0,
Method:Compositional matrix adjust.,
Identities:307/452(68%), Positives:355/452(78%), Gaps:25/452(5%)

```
Query  2     LWVTVYYGVPVWKEATTTLFCASDAKAYDTEVHNVWATHACVPTDPNPQEVVLVNVTENF  61
             LWVTVYYGVPVWKEA TTLFCAS+AKAY+ EVHNVWATHACVPTDPNPQE+VL NVTENF
Sbjct  2390  LWVTVYYGVPVWKEAKTTLFCASNAKAYEKEVHNVWATHACVPTDPNPQEMVLENVTENF  2211

Query  62    NMWKNDMVEQMHEDIISLWDQSLKPCVKLTPLCVSLKCTDL---KNDTNTNSSSGRMIME  118
             NMWKNDMV QMHED+ISLWDQSLKPCVKLTPLCV+L+C  +    N TN   ++  +
Sbjct  2210  NMWKNDMVNQMHEDVISLWDQSLKPCVKLTPLCVTLECRKVNATHNATNNGDATHNVTNN  2031

Query  119   KGEIKNCSFNISTSIRGKVQKEYAFFYKLDIIPIDNDTTS-------YKLTSCNTSVITQ  171
               EI+NCSFN +T IR + Q+ YA FY+LDI+P+D + +S       Y+L +CNTS ITQ
Sbjct  2030  GQEIQNCSFNATTEIRDRKQRVYALFYRLDIVPLDKNNSSKNNSSEYYRLINCNTSAITQ  1851

Query  172   ACPKVSFEPIPIHYCAPAGFAILKCNNKTFNGTGPCTNVSTVQCTHGIRPVVSTQLLLNG  231
             ACPKVSF+PIPIHYCAPAG+AILKCNNKTFNGTGPC NVSTVQCTHGI+PVVSTQLLLNG
Sbjct  1850  ACPKVSFDPIPIHYCAPAGYAILKCNNKTFNGTGPCNNVSTVQCTHGIKPVVSTQLLLNG  1671

Query  232   SLAEEEVVIRSVNFTDNAKTIIVQLNTSVEINCTRPNNNTRKRIRIQRGPGRAF-VTIGK  290
             SLAE E++IRS N TDN KTIIV L+ SVEI CTRPNNNTRK IRI  GPG+ F  T G
Sbjct  1670  SLAEGEIIIRSENLTDNVKTIIVHLDQSVEIVCTRPNNNTRKSIRI--GPGQTFYATGGI  1497
```

```
Query  291   IGNMRQAHCNISRAKWNNTLKQIASKLREQFGNNKTIIFKQSSGGDPEIVTHSFNCGGEF  350
             IGN+RQAHCNIS  KWN TL+++  KL E F  NKTI F  SSGGD EI THSFNC GEF
Sbjct  1496  IGNIRQAHCNISEDKWNETLQRVGKKLVEHFP-NKTIKFAPSSGGDLEITTHSFNCRGEF  1320

Query  351   FYCNSTQLFNSTWFNSTWSTEGSNNTEGSDTITLPCRIKQIINMWQKVGKAMYAPPISGQ  410
             FYC++++LFNST+            ++ + TIT+PC IKQI+NMWQ+VG+AMYAPPI G
Sbjct  1319  FYCSTSRLFNSTYM------PNDTKSKSNKTITIPCSIKQIVNMWQEVGRAMYAPPIEGN  1158

Query  411   IRCSSNITGLLLTRDGG-----NSNNESEIFR  437
             I C SNITG+LL RDGG       NN++E F+
Sbjct  1157  ITCRSNITGILLVRDGGVDSEDPENNKTETFQ  1062
```

>Escherichia coli strain EN5169 contig00082, whole genome shotgun sequence
Sequence ID: NZ_JAPMNJ010000082.1 Length: 2837
Range 1: 1062 to 2390

Score:613 bits(1580), Expect:0.0,
Method:Compositional matrix adjust.,
Identities:307/452(68%), Positives:355/452(78%), Gaps:25/452(5%)

```
Query  2     LWVTVYYGVPVWKEATTTLFCASDAKAYDTEVHNVWATHACVPTDPNPQEVVLVNVTENF  61
             LWVTVYYGVPVWKEA TTLFCAS+AKAY+ EVHNVWATHACVPTDPNPQE+VL NVTENF
Sbjct  2390  LWVTVYYGVPVWKEAKTTLFCASNAKAYEKEVHNVWATHACVPTDPNPQEMVLENVTENF  2211

Query  62    NMWKNDMVEQMHEDIISLWDQSLKPCVKLTPLCVSLKCTDL---KNDTNTNSSSGRMIME  118
             NMWKNDMV QMHED+ISLWDQSLKPCVKLTPLCV+L+C  +    N TN   ++  +
Sbjct  2210  NMWKNDMVNQMHEDVISLWDQSLKPCVKLTPLCVTLECRKVNATHNATNNGDATHNVTNN  2031

Query  119   KGEIKNCSFNISTSIRGKVQKEYAFFYKLDIIPIDNDTTS-------YKLTSCNTSVITQ  171
               EI+NCSFN +T IR + Q+ YA FY+LDI+P+D + +S       Y+L +CNTS ITQ
Sbjct  2030  GQEIQNCSFNATTEIRDRKQRVYALFYRLDIVPLDKNNSSKNNSSEYYRLINCNTSAITQ  1851

Query  172   ACPKVSFEPIPIHYCAPAGFAILKCNNKTFNGTGPCTNVSTVQCTHGIRPVVSTQLLLNG  231
             ACPKVSF+PIPIHYCAPAG+AILKCNNKTFNGTGPC NVSTVQCTHGI+PVVSTQLLLNG
Sbjct  1850  ACPKVSFDPIPIHYCAPAGYAILKCNNKTFNGTGPCNNVSTVQCTHGIKPVVSTQLLLNG  1671

Query  232   SLAEEEVVIRSVNFTDNAKTIIVQLNTSVEINCTRPNNNTRKRIRIQRGPGRAF-VTIGK  290
             SLAE E++IRS N TDN KTIIV L+ SVEI CTRPNNNTRK IRI  GPG+ F  T G
Sbjct  1670  SLAEGEIIIRSENLTDNVKTIIVHLDQSVEIVCTRPNNNTRKSIRI--GPGQTFYATGGI  1497

Query  291   IGNMRQAHCNISRAKWNNTLKQIASKLREQFGNNKTIIFKQSSGGDPEIVTHSFNCGGEF  350
             IGN+RQAHCNIS  KWN TL+++  KL E F  NKTI F  SSGGD EI THSFNC GEF
Sbjct  1496  IGNIRQAHCNISEDKWNETLQRVGKKLVEHFP-NKTIKFAPSSGGDLEITTHSFNCRGEF  1320

Query  351   FYCNSTQLFNSTWFNSTWSTEGSNNTEGSDTITLPCRIKQIINMWQKVGKAMYAPPISGQ  410
             FYC++++LFNST+            ++ + TIT+PC IKQI+NMWQ+VG+AMYAPPI G
Sbjct  1319  FYCSTSRLFNSTYM------PNDTKSKSNKTITIPCSIKQIVNMWQEVGRAMYAPPIEGN  1158

Query  411   IRCSSNITGLLLTRDGG-----NSNNESEIFR  437
             I C SNITG+LL RDGG       NN++E F+
Sbjct  1157  ITCRSNITGILLVRDGGVDSEDPENNKTETFQ  1062
```

>Escherichia coli strain NBE0002 contig00097, whole genome shotgun sequence
Sequence ID: NZ_JARBDF010000097.1 Length: 2717
Range 1: 987 to 2261

Score:605 bits(1561), Expect:0.0,
Method:Compositional matrix adjust.,
Identities:304/441(69%), Positives:348/441(78%), Gaps:26/441(5%)

```
Query  3     WVTVYYGVPVWKEATTTLFCASDAKAYDTEVHNVWATHACVPTDPNPQEVVLVNVTENFN  62
             WVTVYYGVPVWKEA TTLFCASDAKA++ EVHNVWATHACVPTDPNPQE+VL NVTENFN
Sbjct  2261  WVTVYYGVPVWKEAKTTLFCASDAKAHEREVHNVWATHACVPTDPNPQEMVLENVTENFN  2082

Query  63    MWKNDMVEQMHEDIISLWDQSLKPCVKLTPLCVSLKCTDLKNDTNTNSSSGRMIMEKGEI  122
             MW+NDMV+QMHED+ISLWDQSLKPCVKLTPLCV+L+C       N N++      E  E+
Sbjct  2081  MWRNDMVDQMHEDVISLWDQSLKPCVKLTPLCVTLEC-------NNNTTFKNYTDESKEL  1923

Query  123   KNCSFNISTSIRGKVQKEYAFFYKLDIIPIDNDTT---------SYKLTSCNTSVITQAC  173
             K C FN ST +  K QK  A FY LDI P++  ++         +Y L SCNTSVI QAC
Sbjct  1922  KRCFFNASTEVEDKKQKVSALFYSLDIRPLNKSSSGNSSINSNETYVLISCNTSVIKQAC  1743

Query  174   PKVSFEPIPIHYCAPAGFAILKCNNKTFNGTGPCTNVSTVQCTHGIRPVVSTQLLLNGSL  233
             PKV+F+PIPIHYCAPAG+AILKCNN+TFNGTGPC NVSTVQCTHGI+PVVSTQLLLNGSL
Sbjct  1742  PKVTFDPIPIHYCAPAGYAILKCNNETFNGTGPCNNVSTVQCTHGIKPVVSTQLLLNGSL  1563

Query  234   AEEEVVIRSVNFTDNAKTIIVQLNTSVEINCTRPNNNTRKRIRIQRGPGRAFVTIGK-IG  292
             A+ E+VIRS N TDNAKTIIV LN SVEINCTRPNNNTRK IRI  GPG+ F   G+ IG
Sbjct  1562  AKGEIVIRSENLTDNAKTIIVHLNESVEINCTRPNNNTRKSIRI--GPGQTFYATGEIIG  1389

Query  293   NMRQAHCNISRAKWNNTLKQIASKLREQFGNNKTIIFKQSSGGDPEIVTHSFNCGGEFFY  352
             N+RQAHCNIS +KWN TL+++  KL E F  NKTI F   SGGD EI+THSFNCGGEFFY
Sbjct  1388  NIRQAHCNISESKWNETLQRVGIKLAEHFP-NKTINFTSHSGGDLEIMTHSFNCGGEFFY  1212

Query  353   CNSTQLFNSTWFNSTWSTEGSNNTEGSDTITLPCRIKQIINMWQKVGKAMYAPPISGQIR  412
             CN++ LFN T +N T+    S     S  IT+ CRIKQIINMWQ+VG+AMYAPPI+G+I
Sbjct  1211  CNTSSLFNGT-YNGTYIRNNS-----SSIITIQCRIKQIINMWQEVGRAMYAPPIAGKIT  1050

Query  413   CSSNITGLLLTRDGGNSNNES  433
             C+S+ITG+LLTRDGG  N  S
Sbjct  1049  CNSSITGILLTRDGGTGNETS  987
```

>Escherichia coli strain EN5143 contig00069, whole genome shotgun sequence
Sequence ID: NZ_JAPMNI010000069.1 Length: 2688
Range 1: 457 to 1731

Score:605 bits(1561), Expect:0.0,
Method:Compositional matrix adjust.,
Identities:304/441(69%), Positives:348/441(78%), Gaps:26/441(5%)

```
Query  3     WVTVYYGVPVWKEATTTLFCASDAKAYDTEVHNVWATHACVPTDPNPQEVVLVNVTENFN  62
             WVTVYYGVPVWKEA TTLFCASDAKA++ EVHNVWATHACVPTDPNPQE+VL NVTENFN
Sbjct  457   WVTVYYGVPVWKEAKTTLFCASDAKAHEREVHNVWATHACVPTDPNPQEMVLENVTENFN  636

Query  63    MWKNDMVEQMHEDIISLWDQSLKPCVKLTPLCVSLKCTDLKNDTNTNSSSGRMIMEKGEI  122
             MW+NDMV+QMHED+ISLWDQSLKPCVKLTPLCV+L+C       N N++      E  E+
Sbjct  637   MWRNDMVDQMHEDVISLWDQSLKPCVKLTPLCVTLEC-------NNNTTFKNYTDESKEL  795

Query  123   KNCSFNISTSIRGKVQKEYAFFYKLDIIPIDNDTT---------SYKLTSCNTSVITQAC  173
             K C FN ST +  K QK  A FY LDI P++  ++         +Y L SCNTSVI QAC
Sbjct  796   KRCFFNASTEVEDKKQKVSALFYSLDIRPLNKSSSGNSSINSNETYVLISCNTSVIKQAC  975

Query  174   PKVSFEPIPIHYCAPAGFAILKCNNKTFNGTGPCTNVSTVQCTHGIRPVVSTQLLLNGSL  233
             PKV+F+PIPIHYCAPAG+AILKCNN+TFNGTGPC NVSTVQCTHGI+PVVSTQLLLNGSL
Sbjct  976   PKVTFDPIPIHYCAPAGYAILKCNNETFNGTGPCNNVSTVQCTHGIKPVVSTQLLLNGSL  1155
```

```
Query  234   AEEEVVIRSVNFTDNAKTIIVQLNTSVEINCTRPNNNTRKRIRIQRGPGRAFVTIGK-IG  292
             A+ E+VIRS N TDNAKTIIV LN SVEINCTRPNNNTRK IRI  GPG+ F   G+ IG
Sbjct  1156  AKGEIVIRSENLTDNAKTIIVHLNESVEINCTRPNNNTRKSIRI--GPGQTFYATGEIIG  1329

Query  293   NMRQAHCNISRAKWNNTLKQIASKLREQFGNNKTIIFKQSSGGDPEIVTHSFNCGGEFFY  352
             N+RQAHCNIS +KWN TL+++  KL E F  NKTI F   SGGD EI+THSFNCGGEFFY
Sbjct  1330  NIRQAHCNISESKWNETLQRVGIKLAEHFP-NKTINFTSHSGGDLEIMTHSFNCGGEFFY  1506

Query  353   CNSTQLFNSTWFNSTWSTEGSNNTEGSDTITLPCRIKQIINMWQKVGKAMYAPPISGQIR  412
             CN++ LFN T +N T+    S     S  IT+ CRIKQIINMWQ+VG+AMYAPPI+G+I
Sbjct  1507  CNTSSLFNGT-YNGTYIRNNS-----SSIITIQCRIKQIINMWQEVGRAMYAPPIAGKIT  1668

Query  413   CSSNITGLLLTRDGGNSNNES  433
             C+S+ITG+LLTRDGG  N  S
Sbjct  1669  CNSSITGILLTRDGGTGNETS  1731
```

```
>Escherichia coli strain EN5308 contig00099, whole genome shotgun sequence
Sequence ID: NZ_JARBDE010000099.1 Length: 2827
Range 1: 1127 to 2428

Score:603 bits(1555), Expect:0.0,
Method:Compositional matrix adjust.,
Identities:303/452(67%), Positives:355/452(78%), Gaps:31/452(6%)

Query  2     LWVTVYYGVPVWKEATTTLFCASDAKAYDTEVHNVWATHACVPTDPNPQEVVLVNVTENF  61
             LWVTVYYGVPVWKEA   LFCASDA+AYD EVHNVWATHACVPTDP+PQE+VL NVTENF
Sbjct  2428  LWVTVYYGVPVWKEAKAPLFCASDARAYDREVHNVWATHACVPTDPSPQEMVLENVTENF  2249

Query  62    NMWKNDMVEQMHEDIISLWDQSLKPCVKLTPLCVSLKCTDLKNDTNTNSSSGRMIMEKGE  121
             NMW N MV+QMHED+ISLWDQSLKPCVKLTPLCV+L C ++     T +++ R      E
Sbjct  2248  NMWNNYMVDQMHEDVISLWDQSLKPCVKLTPLCVTLNCVNVTKTNVTVNNTNRQ-----E  2084

Query  122   IKNCSFNISTSIRGKVQKEYAFFYKLDIIPIDNDTTS----------YKLTSCNTSVITQ  171
             +KNCSFN +T +  K QK YA FY++DI+P++ +  S          Y+L +CNTS ITQ
Sbjct  2083  MKNCSFNATTELVDKKQKVYALFYRVDIVPLNKENNSKSNEENNSSEYRLINCNTSAITQ  1904

Query  172   ACPKVSFEPIPIHYCAPAGFAILKCNNKTFNGTGPCTNVSTVQCTHGIRPVVSTQLLLNG  231
             ACPKV+F+PIPIHYCAPAGFAILKCNNKTFN T PC NVSTVQCTHGI+PVVSTQLLLNG
Sbjct  1903  ACPKVTFDPIPIHYCAPAGFAILKCNNKTFNKTRPCHNVSTVQCTHGIKPVVSTQLLLNG  1724

Query  232   SLAEEEVVIRSVNFTDNAKTIIVQLNTSVEINCTRPNNNTRKRIRIQRGPGRAFVTIGK-  290
             SLAEEE++IRS N T+NAK IIVQLN SVEI C RP NNTRK IRI  GPG+ F   G
Sbjct  1723  SLAEEEIIIRSENLTNNAKIIIVQLNQSVEIECIRPGNNTRKSIRI--GPGQTFYATGDI  1550

Query  291   IGNMRQAHCNISRAKWNNTLKQIASKLREQFGNNKTIIFKQSSGGDPEIVTHSFNCGGEF  350
             IGN+R+AHCNIS   W NTL++++ KLRE F  NKTIIFK +SGGDPE+ THSFNC GEF
Sbjct  1549  IGNIREAHCNIS--NWTNTLERVSGKLREHFP-NKTIIFKPASGGDPEVTTHSFNCRGEF  1379

Query  351   FYCNSTQLFNSTWFNSTWSTEGSNNTEGSDTITLPCRIKQIINMWQKVGKAMYAPPISGQ  410
             FYCN+++LFN T+       G++N E S  IT+PC+I+Q +N+WQ+VG+AMYAPPI+G
Sbjct  1378  FYCNTSKLFNGTY-------NGTSN-ESSSNITIPCKIRQFVNLWQEVGRAMYAPPIAGN  1223

Query  411   IRCSSNITGLLLTRDGGNSNNES--EIFRPGG  440
             I C SNITGLLL RDGG+ NN S  EIFRP G
Sbjct  1222  ITCKSNITGLLLVRDGGSGNNSSNDEIFRPIG  1127
```

```
>Escherichia coli strain EN5286 contig00120, whole genome shotgun sequence
Sequence ID: NZ_JAPMNM010000120.1 Length: 3033
Range 1: 448 to 1884

Score:582 bits(1501), Expect:0.0,
Method:Compositional matrix adjust.,
Identities:298/481(62%), Positives:345/481(71%), Gaps:54/481(11%)

Query  2     LWVTVYYGVPVWKEATTTLFCASDAKAYDTEVHNVWATHACVPTDPNPQEVVLVNVTENF  61
             LWVTVYYGVPVWK+A  TLFCASDAK Y TEVHNVWATHACVPTDPNPQE+VL NVTENF
Sbjct  448   LWVTVYYGVPVWKDANATLFCASDAKVYKTEVHNVWATHACVPTDPNPQELVLGNVTENF  627

Query  62    NMWKNDMVEQMHEDIISLWDQSLKPCVKLTPLCVSLKCTDLK------NDTNTNSSSGRM  115
             NMWKN+MVEQMHED+IS+WDQ LKPCVKLTPLCV+L CT  +      +  NT ++S  M
Sbjct  628   NMWKNNMVEQMHEDVISIWDQGLKPCVKLTPLCVTLNCTKYEPRKVTVDKNNTGATSQNM  807

Query  116   IMEKG-------EIKNCSFNISTSIRGKVQKEYAFFYKLDIIPI-----DNDTTSYKLTS  163
              +           EI NCSF  +T +R K +K +A FYK+D++ I     D+  T Y+L +
Sbjct  808   TVNLTIDKEALEEISNCSFETTTELRDKTRKVHALFYKIDMVEISKIGNDSTITRYRLIN  987

Query  164   CNTSVITQACPKVSFEPIPIHYCAPAGFAILKCNNKTFNGTGPCTNVSTVQCTHGIRPVV  223
             CNTS +TQACPKVSFEPIPIHYCAPAG+A+LKCNN TFNGTGPC N+STVQCTHGIRPVV
Sbjct  988   CNTSALTQACPKVSFEPIPIHYCAPAGYALLKCNNNTFNGTGPCRNISTVQCTHGIRPVV  1167

Query  224   STQLLLNGSLAEEEVVIRSVNFTDNAKTIIVQLNTSVEINCTRPNNNTRKRIRIQRGPGR  283
             STQLLLNGSLAE  + IRS N T+NAKTIIV LN SVEI C RPNNNTR+ +RI  GPG+
Sbjct  1168  STQLLLNGSLAEGNITIRSENLTNNAKTIIVHLNQSVEILCVRPNNNTRQSVRI--GPGQ  1341

Query  284   AFVTIGKI-GNMRQAHCNISRAKWNNTLKQIASKLREQFGNNKTIIFKQSSGGDPEIVTH  342
              F   G I G++R+AHCNIS   W+  L+ ++ +L + F NNKTI F   SGGD EI TH
Sbjct  1342  TFFATGDIIGDIRRAHCNISAKDWHTALRNVSIELAKLFPNNKTINFTSPSGGDLEITTH  1521

Query  343   SFNCGGEFFYCNSTQLFNS---------TWFNSTWSTEGSNNTE---GSDT---------  381
             SFNC GEFFYCN++QLFNS          W+N   +    N T+    S T
Sbjct  1522  SFNCRGEFFYCNTSQLFNSLYMPDNDTYIWYNGNHTYVWHNGTQYVINSSTQYVTINNSA  1701

Query  382   ------------ITLPCRIKQIINMWQKVGKAMYAPPISGQIRCSSNITGLLLTRDGGNS  429
                         ITLPC+IKQIINMWQ+VGKAMYAPPI+G I C SNITGLLLTRDGG
Sbjct  1702  QNVAINNMTDFNITLPCKIKQIINMWQEVGKAMYAPPIAGNITCLSNITGLLLTRDGGKG  1881

Query  430   N  430
             N
Sbjct  1882  N  1884
```

```
>Escherichia coli strain EN5192 contig00108, whole genome shotgun sequence
Sequence ID: NZ_JAPMNL010000108.1 Length: 3032
Range 1: 449 to 1942

Score:579 bits(1493), Expect:0.0,
Method:Compositional matrix adjust.,
Identities:302/502(60%), Positives:351/502(69%), Gaps:65/502(12%)

Query  2     LWVTVYYGVPVWKEATTTLFCASDAKAYDTEVHNVWATHACVPTDPNPQEVVLVNVTENF  61
             LWVTVYYGVPVWK+A  TLFCASDAK Y TEVHNVWATHACVPTDPNPQE+VL NVTENF
Sbjct  449   LWVTVYYGVPVWKDANATLFCASDAKVYKTEVHNVWATHACVPTDPNPQELVLENVTENF  628

Query  62    NMWKNDMVEQMHEDIISLWDQSLKPCVKLTPLCVSLKCTDLK--NDTNTNSSSGRMIMEK  119
             NMWKN+MVEQMHED+IS+WDQ LKPCVKLTPLCV+L CT  K  N+T   +S+G
```

```
Sbjct  629   NMWKNNMVEQMHEDVISIWDQGLKPCVKLTPLCVTLNCTKYKPRNETVDGTSTGNTGQNN  808

Query  120   G-------------EIKNCSFNISTSIRGKVQKEYAFFYKLDIIPIDNDTTS--------  158
                           EI NCSF+ +T +R K +K +A FYK+D++ +D ++T
Sbjct  809   TGKLNLTIDPEALEEISNCSFDTTTELRDKTRKVHALFYKIDMVELDKNSTCQTPTPNFT  988

Query  159   -YKLTSCNTSVITQACPKVSFEPIPIHYCAPAGFAILKCNNKTFNGTGPCTNVSTVQCTH  217
              Y+L +CNTS ITQACPKVSFEPIPIHYCAPAG+A+LKCNN TFNGTGPC N+STVQCTH
Sbjct  989   LYRLINCNTSAITQACPKVSFEPIPIHYCAPAGYALLKCNNNTFNGTGPCRNISTVQCTH  1168

Query  218   GIRPVVSTQLLLNGSLAEEEVVIRSVNFTDNAKTIIVQLNTSVEINCTRPNNNTRKRIRI  277
             GIRPVVSTQLLLNGSLAE  + IRS N TDNAKTII  LN SVEI C RPNNNTR+ +RI
Sbjct  1169  GIRPVVSTQLLLNGSLAEGNITIRSENLTDNAKTIIAHLNQSVEILCVRPNNNTRQSVRI  1348

Query  278   QRGPGRAFVTIGK-IGNMRQAHCNISRAKWNNTLKQIASKLREQFGNNKTIIFKQSSGGD  336
               GPG+ F   G  IG++R+AHCNIS   W+  L+ ++ +L + +  NKTI F   SGGD
Sbjct  1349  --GPGQTFFATGDIIGDIRRAHCNISAKDWHTALRNVSIELAKLY--NKTINFTSHSGGD  1516

Query  337   PEIVTHSFNCGGEFFYCNSTQLFNS---------TWFNSTWSTEGSNNTE----GSD---  380
              EI THSFNC GEFFYCN++QLFNS          W+N   +    N T+    GS
Sbjct  1517  LEITTHSFNCRGEFFYCNTSQLFNSLYMPNNDTYIWYNGNHTHVWHNGTQYVINGSTQYV  1696

Query  381   -----------------TITLPCRIKQIINMWQKVGKAMYAPPISGQIRCSSNITGLLLT  423
                               ITLPC+IKQIINMWQ+VGKAMYAPPI+G I C SNITGLLLT
Sbjct  1697  TLNNSAQNVTINNITDFNITLPCKIKQIINMWQEVGKAMYAPPIAGNITCLSNITGLLLT  1876

Query  424   RDGGNSN---NESEIFRPGGGD  442
             RD G      N +EIFRP G +
Sbjct  1877  RDRGKGKEVRNNTEIFRPEGNN  1942
```

>Escherichia coli strain VRES0669, whole genome shotgun sequence
Sequence ID: NZ_UINV01000086.1 Length: 5544
Range 1: 6 to 1298

Score:578 bits(1491), Expect:0.0,
Method:Compositional matrix adjust.,
Identities:290/440(66%), Positives:349/440(79%), Gaps:18/440(4%)

```
Query  3     WVTVYYGVPVWKEATTTLFCASDAKAYDTEVHNVWATHACVPTDPNPQEVVLVNVTENFN  62
             WVTVYYGVPVWKEA  TLFCASDAK    EVHN+WATHACVPT+PNPQE+VL NVTENFN
Sbjct  1298  WVTVYYGVPVWKEAKATLFCASDAKTQAEEVHNIWATHACVPTNPNPQEIVLENVTENFN  1119

Query  63    MWKNDMVEQMHEDIISLWDQSLKPCVKLTPLCVSLKCTDLK-NDTNTNSSSGRMIMEKGE  121
             MWKNDMV+QMHEDIISLWDQSLKPCVKLT LCV+L C+++K N T  N +  R      +
Sbjct  1118  MWKNDMVDQMHEDIISLWDQSLKPCVKLTSLCVTLNCSNVKINVTAINGTYLR--NSTSD  945

Query  122   IKNCSFNISTSIRGKVQKEYAFFYKLDIIPIDND----TTSYKLTSCNTSVITQACPKVS  177
             +KNCSFN++T +R K QK +A FY+LD++P++ +    ++SY+L +CNTS I+QACPK S
Sbjct  944   MKNCSFNVTTELRDKRQKAHALFYRLDVVPLEEEKNSSSSSYRLINCNTSTISQACPKTS  765

Query  178   FEPIPIHYCAPAGFAILKCNNKTFNGTGPCTNVSTVQCTHGIRPVVSTQLLLNGSLAEEE  237
             F+P+PIHYCAPAG+AILKCNNKTFNGTGPC NVSTVQCTHGI+PVVSTQLLLNGSLA E+
Sbjct  764   FDPLPIHYCAPAGYAILKCNNKTFNGTGPCQNVSTVQCTHGIKPVVSTQLLLNGSLA-ED  588

Query  238   VVIRSVNFTDNAKTIIVQLNTSVEINCTRPNNNTRKRIRIQRGPGRAFVTIGKI-GNMRQ  296
             ++IRS N T N +TIIV L   VEI C RPNNNTR  IRI  G G+ F   G+I G++RQ
Sbjct  587   IIIRSENLTSNTRTIIVHLKDPVEIRCIRPNNNTRHSIRIGIGRGQTFYATGRIVGDIRQ  408

Query  297   AHCNISRAKWNNTLKQIASKLREQFGNNKTIIFKQS-SGGDPEIVTHSFNCGGEFFYCNS  355
             AHCNI+  KWN TL+++  KL+E F  NKTI FK +  GGD E+  H+FNC GEFFYCN+
Sbjct  407   AHCNITERKWNETLQKVVEKLKEHFP-NKTIQFKPAIPGGDLEVTMHTFNCRGEFFYCNT  231

Query  356   TQLFNSTWFNST--WSTEGSNNTEGSDTITLPCRIKQIINMWQKVGKAMYAPPISGQIRC  413
             +QLF +  FN T  +++ G+N T     ITLPCRIKQI+NMWQ+VG+A+YAPPI+G I C
Sbjct  230   SQLF-TYMFNDTDMFNSTGNNATN----ITLPCRIKQIVNMWQEVGRAIYAPPITGIINC  66

Query  414   SSNITGLLLTRDGGNSNNES  433
             +SNITGLLL RDGGNS N S
Sbjct  65    TSNITGLLLIRDGGNSGNGS  6
```

>Escherichia coli strain NBE0007 contig00093, whole genome shotgun sequence
Sequence ID: NZ_JARBDJ010000093.1 Length: 3013
Range 1: 1130 to 2566

Score:578 bits(1491), Expect:0.0,
Method:Compositional matrix adjust.,
Identities:296/481(62%), Positives:346/481(71%), Gaps:54/481(11%)

```
Query  2     LWVTVYYGVPVWKEATTTLFCASDAKAYDTEVHNVWATHACVPTDPNPQEVVLVNVTENF  61
             LWVTVYYGVPVW++A  TLFCASDAK Y TEVHNVWATHACVPTDPNPQE+VL NVTENF
Sbjct  2566  LWVTVYYGVPVWRDANATLFCASDAKVYKTEVHNVWATHACVPTDPNPQELVLENVTENF  2387

Query  62    NMWKNDMVEQMHEDIISLWDQSLKPCVKLTPLCVSLKCTDLK------NDTNTNSSSGRM  115
             NMW+N+MVEQMHED+IS+WDQ LKPCVKLTPLCV+L CT  +      +  NT ++S  M
Sbjct  2386  NMWENNMVEQMHEDVISIWDQGLKPCVKLTPLCVTLNCTKYEPRKVTVDKNNTGATSQNM  2207

Query  116   I----MEKG---EIKNCSFNISTSIRGKVQKEYAFFYKLDIIPI-----DNDTTSYKLTS  163
                  +EK    EI NCSF  +T +R K +K +A FYK+D++ +     D+  T Y+L +
Sbjct  2206  TVNPTIEKEALEEISNCSFETTTELRDKTRKVHALFYKIDMVELGKNGNDSTITRYRLIN  2027

Query  164   CNTSVITQACPKVSFEPIPIHYCAPAGFAILKCNNKTFNGTGPCTNVSTVQCTHGIRPVV  223
             CNTS +TQACPKVSFEPIPIHYCAPAG+A+LKCNN TFNGTGPC N+STVQCTHGIRPVV
Sbjct  2026  CNTSALTQACPKVSFEPIPIHYCAPAGYALLKCNNNTFNGTGPCRNISTVQCTHGIRPVV  1847

Query  224   STQLLLNGSLAEEEVVIRSVNFTDNAKTIIVQLNTSVEINCTRPNNNTRKRIRIQRGPGR  283
             STQLLLNGSLAE  + IRS N T+NAKTII  LN SVEI C RPNNNTR+ +RI  GPG+
Sbjct  1846  STQLLLNGSLAEGNITIRSENLTNNAKTIIAHLNKSVEILCARPNNNTRQSVRI--GPGQ  1673

Query  284   AFVTIGKI-GNMRQAHCNISRAKWNNTLKQIASKLREQFGNNKTIIFKQSSGGDPEIVTH  342
              F   G I G++R+AHCNIS   W+  L+ ++ +L + F NNKTI F   SGGD EI TH
Sbjct  1672  TFFATGDIIGDIRRAHCNISAKDWHTALRNVSIELAKLFPNNKTINFTSPSGGDLEITTH  1493

Query  343   SFNCGGEFFYCNSTQLFNST---------WFNSTWSTEGSNNTE---GSDT---------  381
             SFNC GEFFYCN++QLFNS          W+N   +    N T+    S T
Sbjct  1492  SFNCRGEFFYCNTSQLFNSLYMPDNDTYLWYNGNHTYVWHNGTQYVINSSTQYVTINNSA  1313

Query  382   ------------ITLPCRIKQIINMWQKVGKAMYAPPISGQIRCSSNITGLLLTRDGGNS  429
                         ITLPC+IKQIINMWQ+VGKAMYAPPI+G I C SNITGLLLTRDGG
Sbjct  1312  QNVAINNMTDFNITLPCKIKQIINMWQEVGKAMYAPPIAGNITCLSNITGLLLTRDGGKG  1133

Query  430   N  430
             N
Sbjct  1132  N  1130
```

>Escherichia coli strain EN5308 contig00096, whole genome shotgun sequence

```
Sequence ID: NZ_JARBDE010000096.1 Length: 3024
Range 1: 1156 to 2577

Score:570 bits(1470), Expect:0.0,
Method:Compositional matrix adjust.,
Identities:294/478(62%), Positives:347/478(72%), Gaps:56/478(11%)

Query  2     LWVTVYYGVPVWKEATTTLFCASDAKAYDTEVHNVWATHACVPTDPNPQEVVLVNVTENF  61
             LWVTVYYGVPVWK+A  TLFCASDAK Y TEVHNVWATHACVPTDPNPQE+VL NVTENF
Sbjct  2577  LWVTVYYGVPVWKDANATLFCASDAKVYKTEVHNVWATHACVPTDPNPQELVLGNVTENF  2398

Query  62    NMWKNDMVEQMHEDIISLWDQSLKPCVKLTPLCVSLKCT--DLKNDT-NTNSSSGRMIME  118
             NMW+N+MVEQMHED+IS+WDQ LKPCVKLTPLCV+L CT  + +N T N NS++    ++
Sbjct  2397  NMWENNMVEQMHEDVISIWDQGLKPCVKLTPLCVTLNCTKYEPRNKTVNGNSTAKSPTID  2218

Query  119   K---GEIKNCSFNISTSIRGKVQKEYAFFYKLDIIPIDNDTTS------------YKLTS  163
             +    EI NCSF  +T +R K +K +A FYK+D+I +  ++T             Y+L +
Sbjct  2217  EEALKEISNCSFETTTELRDKTRKVHALFYKIDMIELGQNSTCQNPENSTSECTMYRLIN  2038

Query  164   CNTSVITQACPKVSFEPIPIHYCAPAGFAILKCNNKTFNGTGPCTNVSTVQCTHGIRPVV  223
             CNTS ITQACPKVSFEPIPIHYCAPAG+A+LKCNN TFNGTGPC N+STVQCTHGIRPVV
Sbjct  2037  CNTSAITQACPKVSFEPIPIHYCAPAGYALLKCNNNTFNGTGPCRNISTVQCTHGIRPVV  1858

Query  224   STQLLLNGSLAEEEVVIRSVNFTDNAKTIIVQLNTSVEINCTRPNNNTRKRIRIQRGPGR  283
             STQLLLNGSLAE  + IRS N T+NAKTII  LN SVEI C RPNNNTR+ +RI  GPG+
Sbjct  1857  STQLLLNGSLAEGNITIRSENLTNNAKTIIAHLNKSVEILCVRPNNNTRQSVRI--GPGQ  1684

Query  284   AFVTIGK-IGNMRQAHCNISRAKWNNTLKQIASKLREQFGNNKTIIFKQSSGGDPEIVTH  342
              F   G  IG++R+AHCNIS   W+  L+ ++ +L + +  NKTI F   SGGD EI TH
Sbjct  1683  TFFATGDIIGDIRRAHCNISAKHWHTALRNVSIELAKLY--NKTINFTSHSGGDLEITTH  1510

Query  343   SFNCGGEFFYCNSTQLFNS---------TWFNSTWSTEGSNNTE----GSD---------  380
             SFNC GEFFYCN++QLFNS          W+N + +    N+T+    GS
Sbjct  1509  SFNCRGEFFYCNTSQLFNSLYMPDNDTYIWYNESHTHVWHNDTQYVINGSTQYVTLNKSA  1330

Query  381   -----------TITLPCRIKQIINMWQKVGKAMYAPPISGQIRCSSNITGLLLTRDGG  427
                         ITLPC+IKQIINMWQ+VGKAMYAPPI+G I C SNITGLLLTRDGG
Sbjct  1329  QNVTINNITDFNITLPCKIKQIINMWQEVGKAMYAPPIAGNITCLSNITGLLLTRDGG  1156



>Escherichia coli strain EN5143 contig00071, whole genome shotgun sequence
Sequence ID: NZ_JAPMNI010000071.1 Length: 2774
Range 1: 199 to 1623

Score:570 bits(1468), Expect:0.0,
Method:Compositional matrix adjust.,
Identities:294/479(61%), Positives:343/479(71%), Gaps:57/479(11%)

Query  2     LWVTVYYGVPVWKEATTTLFCASDAKAYDTEVHNVWATHACVPTDPNPQEVVLVNVTENF  61
             LWVTVYYGVPVW++A  TLFCASDAK Y TEVHNVWATHACVPTDPNPQE+VL NVTENF
Sbjct  199   LWVTVYYGVPVWRDANATLFCASDAKVYKTEVHNVWATHACVPTDPNPQELVLGNVTENF  378

Query  62    NMWKNDMVEQMHEDIISLWDQSLKPCVKLTPLCVSLKCTDL--KNDT---NTNSSSGRMI  116
             NMW+NDMVEQMH+D+IS+WDQ LKPCVKLTPLCV+L CT    +N+T   N   S    I
Sbjct  379   NMWENDMVEQMHQDVISIWDQGLKPCVKLTPLCVTLNCTKYEPRNETKKDNIKISLNLTI  558

Query  117   MEKG--EIKNCSFNISTSIRGKVQKEYAFFYKLDIIPIDNDTTS------------YKLT  162
              ++   EI NCSF  +T +R K +K +A FYK+D+I +  ++T             Y+L
Sbjct  559   DQEAVDEINNCSFETTTELRDKTRKVHALFYKIDMIELGQNSTCQKPENSTSKCTMYRLI  738

Query  163   SCNTSVITQACPKVSFEPIPIHYCAPAGFAILKCNNKTFNGTGPCTNVSTVQCTHGIRPV  222
             +CNTS ITQACPKVSFEPIPIHYCAPAG+A+LKCNN TFNGTGPC N+STVQCTHGIRPV
Sbjct  739   NCNTSAITQACPKVSFEPIPIHYCAPAGYALLKCNNNTFNGTGPCRNISTVQCTHGIRPV  918

Query  223   VSTQLLLNGSLAEEEVVIRSVNFTDNAKTIIVQLNTSVEINCTRPNNNTRKRIRIQRGPG  282
             VSTQLLLNGSLAE  + IRS N TDNAKTII  LN SVEI C RPNNNTR+ +RI  GPG
Sbjct  919   VSTQLLLNGSLAEGNITIRSENLTDNAKTIIAHLNQSVEILCVRPNNNTRQSVRI--GPG  1092

Query  283   RAFVTIGK-IGNMRQAHCNISRAKWNNTLKQIASKLREQFGNNKTIIFKQSSGGDPEIVT  341
             + F   G  IG++R+AHCNIS   W+  L+ ++ +L + +  NKTI F   SGGD EI T
Sbjct  1093  QTFFATGDIIGDIRRAHCNISAKHWHTALRNVSIELAKLY--NKTINFTSHSGGDLEITT  1266

Query  342   HSFNCGGEFFYCNSTQLFNS---------TWFNSTWSTEGSNNTE----GSD--------  380
             HSFNC GEFFYCN++QLFNS          W+N   +    N+T+    GS
Sbjct  1267  HSFNCRGEFFYCNTSQLFNSLYMPDNDTYIWYNGNHTHVWHNDTQYVINGSTQYVTLKNS  1446

Query  381   ------------TITLPCRIKQIINMWQKVGKAMYAPPISGQIRCSSNITGLLLTRDGG  427
                          ITLPC+IKQIINMWQ+VGKAMYAPPI+G I C SNITGLLLTRDGG
Sbjct  1447  AQNVTINNITDFNITLPCKIKQIINMWQEVGKAMYAPPIAGNITCLSNITGLLLTRDGG  1623



>Escherichia coli strain NBE0006 contig00106, whole genome shotgun sequence
Sequence ID: NZ_JARBDI010000106.1 Length: 2090
Range 1: 820 to 2088

Score:533 bits(1374), Expect:1e-180,
Method:Compositional matrix adjust.,
Identities:274/432(63%), Positives:323/432(74%), Gaps:36/432(8%)

Query  75    DIISLWDQSLKPCVKLTPLCVSLKCTDL--KNDTNTNSSSGR--MIMEKG--EIKNCSFN  128
             D+IS+WDQ LKPCVKLTPLCV+L CT    +N+T + +S+ +   I E+   EI NCSF
Sbjct  2088  DVISIWDQGLKPCVKLTPLCVTLNCTKYEPRNETVSGNSTAKSPTIDEEALKEISNCSFE  1909

Query  129   ISTSIRGKVQKEYAFFYKLDIIPIDNDTT------------SYKLTSCNTSVITQACPKV  176
              +T +R K +K +A FYK+D+I +  ++T             Y+L +CNTS ITQACPKV
Sbjct  1908  TTTELRDKTRKVHALFYKIDMIELGQNSTCQKPENSTSECTMYRLINCNTSAITQACPKV  1729

Query  177   SFEPIPIHYCAPAGFAILKCNNKTFNGTGPCTNVSTVQCTHGIRPVVSTQLLLNGSLAEE  236
             SFEPIPIHYCAPAG+A+LKCNN TFNGTGPC N+STVQCTHGIRPVVSTQLLLNGSLAE
Sbjct  1728  SFEPIPIHYCAPAGYALLKCNNNTFNGTGPCRNISTVQCTHGIRPVVSTQLLLNGSLAEG  1549

Query  237   EVVIRSVNFTDNAKTIIVQLNTSVEINCTRPNNNTRKRIRIQRGPGRAFVTIGK-IGNMR  295
              + IRS N TDNAKTII  LN SVEINCTRPNNNTRK IRI  GPG+ F   G+ IGN+R
Sbjct  1548  NITIRSENLTDNAKTIIAHLNESVEINCTRPNNNTRKSIRI--GPGQTFYATGEIIGNIR  1375

Query  296   QAHCNISRAKWNNTLKQIASKLREQFGNNKTIIFKQSSGGDPEIVTHSFNCGGEFFYCNS  355
             QAHCNIS   WN TL+++  KL E F  NKTI F  SSGGD EI THSFNC GEFFYC++
Sbjct  1374  QAHCNISENMWNKTLQRVGKKLAEHFP-NKTIKFASSSGGDLEITTHSFNCRGEFFYCST  1198

Query  356   TQLFNSTWFNSTWSTEGSNNTEGSDTITLPCRIKQIINMWQKVGKAMYAPPISGQIRCSS  415
             ++LFNST+            ++ + TIT+PC IKQI+NMWQ+VG+AMYAPPI G I C S
Sbjct  1197  SRLFNSTYM------PNDTKSKSNKTITIPCSIKQIVNMWQEVGRAMYAPPIEGSITCES  1036

Query  416   NITGLLLTRDGG---NSN-----NESEIFRPGGGDMRDNWRSELYKYKVVKIEPLGVAPT  467
             NITGLLL RDGG    SN     N +E FRP GGDMR+NWRSELYKYKVV+I+PLG+APT
Sbjct  1035  NITGLLLVRDGGIESESNCTETLNCTETFRPEGGDMRNNWRSELYKYKVVEIKPLGIAPT  856

Query  468   KAKRRVVQREKR  479
              AKRRVV+REKR
Sbjct  855   AAKRRVVEREKR  820
```

>Escherichia coli strain EN5192 contig00149, whole genome shotgun sequence
Sequence ID: NZ_JAPMNL010000149.1 Length: 1397
Range 1: 322 to 1392

Score:514 bits(1325), Expect:2e-176,
Method:Compositional matrix adjust.,
Identities:256/367(70%), Positives:297/367(80%), Gaps:19/367(5%)

```
Query  3     WVTVYYGVPVWKEATTTLFCASDAKAYDTEVHNVWATHACVPTDPNPQEVVLVNVTENFN  62
             WVTVYYGVPVWKEA TTLFCASDAKA++ EVHNVWATHACVPTDPNPQE+VL NVTENFN
Sbjct  322   WVTVYYGVPVWKEAKTTLFCASDAKAHEREVHNVWATHACVPTDPNPQEMVLENVTENFN  501

Query  63    MWKNDMVEQMHEDIISLWDQSLKPCVKLTPLCVSLKCTDLKNDTNTNSSSGRMIMEKGEI  122
             MWK+DMV+QMHED+ISLWDQSLKPCVKLTPLCV+L CT+ K    TN S  + + +K E+
Sbjct  502   MWKSDMVDQMHEDVISLWDQSLKPCVKLTPLCVTLNCTNAK----TNVS--KQVTDK-EV  660

Query  123   KNCSFNISTSIRGKVQKEYAFFYKLDIIPIDNDTT--------SYKLTSCNTSVITQACP  174
             KNCSF  +T ++ K Q+  A FY LDI P++  ++        +Y L SCNTSVI QACP
Sbjct  661   KNCSFKANTEVQDKRQRVSALFYSLDIRPLNKSSSGNSSENYENYVLISCNTSVIKQACP  840

Query  175   KVSFEPIPIHYCAPAGFAILKCNNKTFNGTGPCTNVSTVQCTHGIRPVVSTQLLLNGSLA  234
             KV+F+PIPIHYCAPAGFAILKCNN+TFNGTGPC NVSTVQCTHGI+PVVSTQLLLNGSLA
Sbjct  841   KVTFDPIPIHYCAPAGFAILKCNNETFNGTGPCNNVSTVQCTHGIKPVVSTQLLLNGSLA  1020

Query  235   EEEVVIRSVNFTDNAKTIIVQLNTSVEINCTRPNNNTRKRIRIQRGPGRAFVTIGK-IGN  293
             + E++IRS N T+N KTIIV LN SV+I C RP+NNTRK IRI  GPG+ F   G  IG+
Sbjct  1021  KGEIIIRSENLTNNVKTIIVHLNESVKIECIRPHNNTRKSIRI--GPGQTFYATGDIIGD  1194

Query  294   MRQAHCNISRAKWNNTLKQIASKLREQFGNNKTIIFKQSSGGDPEIVTHSFNCGGEFFYC  353
             +RQAHCNIS   WN TL+++  KL E F  NKTI F+ SSGGD EI+THSFNCGGEFFYC
Sbjct  1195  IRQAHCNISENMWNKTLQRVGKKLAEHFP-NKTIKFESSSGGDLEIMTHSFNCGGEFFYC  1371

Query  354   NSTQLFN  360
             N++ LFN
Sbjct  1372  NTSGLFN  1392
```

>Escherichia coli strain NBE0006 contig00124, whole genome shotgun sequence
Sequence ID: NZ_JARBDI010000124.1 Length: 1306
Range 1: 1 to 1053

Score:508 bits(1307), Expect:4e-174,
Method:Compositional matrix adjust.,
Identities:253/354(71%), Positives:287/354(81%), Gaps:14/354(3%)

```
Query  3     WVTVYYGVPVWKEATTTLFCASDAKAYDTEVHNVWATHACVPTDPNPQEVVLVNVTENFN  62
             WVTVYYGVPVWKEA TTLFCASDAKA++ EVHNVWATHACVPTDPNPQE+VL NVTENFN
Sbjct  1053  WVTVYYGVPVWKEAKTTLFCASDAKAHEREVHNVWATHACVPTDPNPQEMVLENVTENFN  874

Query  63    MWKNDMVEQMHEDIISLWDQSLKPCVKLTPLCVSLKCTDL---KNDTNTNSSSGRMIMEK  119
             MW+NDMV+QMHED+ISLWDQSLKPCVKLTPLCV+L+C  +    N TN   ++  +
Sbjct  873   MWRNDMVDQMHEDVISLWDQSLKPCVKLTPLCVTLECRKVNATHNATNNGDATHNVTNNG  694

Query  120   GEIKNCSFNISTSIRGKVQKEYAFFYKLDIIPIDNDTTS-------YKLTSCNTSVITQA  172
              EI+NCSFN +T IR + Q+ YA FY+LDI+P+D + +S       Y+L +CNTS ITQA
Sbjct  693   QEIQNCSFNATTEIRDRKQRVYALFYRLDIVPLDKNNSSKNNSSEYYRLINCNTSAITQA  514

Query  173   CPKVSFEPIPIHYCAPAGFAILKCNNKTFNGTGPCTNVSTVQCTHGIRPVVSTQLLLNGS  232
             CPKVSF+PIPIHYCAPAG+AILKCNNKTFNGTGPC NVSTVQCTHGI+PVVSTQLLLNGS
Sbjct  513   CPKVSFDPIPIHYCAPAGYAILKCNNKTFNGTGPCNNVSTVQCTHGIKPVVSTQLLLNGS  334

Query  233   LAEEEVVIRSVNFTDNAKTIIVQLNTSVEINCTRPNNNTRKRIRIQRGPGRAF-VTIGKI  291
             LAE E++IRS N TDN KTIIV L+ SVEI CTRPNNNTRK IRI  GPG+ F  T G I
Sbjct  333   LAEGEIIIRSENLTDNVKTIIVHLDQSVEIVCTRPNNNTRKSIRI--GPGQTFYATGGII  160

Query  292   GNMRQAHCNISRAKWNNTLKQIASKLREQFGNNKTIIFKQSSGGDPEIVTHSFN  345
             GN+RQAHCNIS  KWN TL+++  KL E F  NKTI F  SSGGD EI THSFN
Sbjct  159   GNIRQAHCNISEDKWNETLQRVGKKLVEHFP-NKTIKFAPSSGGDLEITTHSFN  1
```

>Klebsiella pneumoniae strain EuSCAPE_IL004, whole genome shotgun sequence
Sequence ID: NZ_UKJP02000086.1 Length: 1081
Range 1: 109 to 1080

Score:471 bits(1211), Expect:1e-160,
Method:Compositional matrix adjust.,
Identities:232/340(68%), Positives:278/340(81%), Gaps:20/340(5%)

```
Query  144   FYKLDIIPI---DNDTTSYKLTSCNTSVITQACPKVSFEPIPIHYCAPAGFAILKCNNKT  200
             FY+LDI+PI   +++ + Y+L +CNTS ++QACPKVSF+PIPIHYCAPAG+AILKCN+KT
Sbjct  1080  FYRLDIVPIPGENSNISKYRLINCNTSAMSQACPKVSFDPIPIHYCAPAGYAILKCNDKT  901

Query  201   FNGTGPCTNVSTVQCTHGIRPVVSTQLLLNGSLAEEEVVIRSVNFTDNAKTIIVQLNTSV  260
             FNGTGPC NVSTVQCTHGI+PVVSTQLLLNGSL+EE ++IRS N T+N KTIIV L+  V
Sbjct  900   FNGTGPCLNVSTVQCTHGIKPVVSTQLLLNGSLSEEGIIIRSENLTNNVKTIIVHLDKPV  721

Query  261   EINCTRPNNNTRKRIRIQRGPGRAFVTIGKIGNMRQAHCNISRAKWNNTLKQIASKLREQ  320
             +I CTRPNNNTRK IRI  GPG+ F     IG++RQA+CNIS  KW +TL+ + +KLR+
Sbjct  720   DIMCTRPNNNTRKSIRI--GPGQTFYATDIIGDIRQAYCNISGKKWYDTLQGVKNKLRKH  547

Query  321   FGNNKTIIFKQSSGGDPEIVTHSFNCGGEFFYCNSTQLFNSTWFNSTWSTEGSNNTEGSD  380
             F  +KTI F+ SSGGD EI THSFNC GEFFYC++++LFNST  +++
Sbjct  546   F-PSKTIQFEPSSGGDLEITTHSFNCMGEFFYCDTSKLFNSTKRDNS-------------  409

Query  381   TITLPCRIKQIINMWQKVGKAMYAPPISGQIRCSSNITGLLLTRD-GGNSNNESEIFRPG  439
             TITLPCRIKQIINMWQ VG+AMYAPPI+G I C SNITGLLLTRD GGNS+  +E FRPG
Sbjct  408   TITLPCRIKQIINMWQGVGRAMYAPPIAGNITCRSNITGLLLTRDGGGNSSATTETFRPG  229

Query  440   GGDMRDNWRSELYKYKVVKIEPLGVAPTKAKRRVVQREKR  479
             GGDMRDNWRSELYKYKVV+I+PLGVAPT+AKRRVV+R KR
Sbjct  228   GGDMRDNWRSELYKYKVVEIKPLGVAPTEAKRRVVERGKR  109
```

>Pseudomonas aeruginosa strain VET-121 NODE_76_length_1017_cov_1.90426_ID_151, whole genome shotgun sequence
Sequence ID: NZ_QZXB01000124.1 Length: 1017
Range 1: 61 to 981

Score:445 bits(1145), Expect:5e-151,
Method:Compositional matrix adjust.,
Identities:216/323(67%), Positives:259/323(80%), Gaps:21/323(6%)

```
Query  157  TSYKLTSCNTSVITQACPKVSFEPIPIHYCAPAGFAILKCNNKTFNGTGPCTNVSTVQCT  216
            +SY L +CN + I QACPKVSFEPIPIHYCAPAGFAILKC +K FNGTGPC NVSTVQCT
Sbjct  61   SSYMLINCNATTIKQACPKVSFEPIPIHYCAPAGFAILKCRDKNFNGTGPCKNVSTVQCT  240
```

```
Query  217  HGIRPVVSTQLLLNGSLAEEEVVIRSVNFTDNAKTIIVQLNTSVEINCTRPNNNTRKRIR  276
            HGI+PVVSTQLLLNGSLAEE+++IRS N ++NAKTIIVQ N SVEINCTRPNNNTRK I
Sbjct  241  HGIKPVVSTQLLLNGSLAEEDIIIRSENISNNAKTIIVQFNQSVEINCTRPNNNTRKSIH  420

Query  277  IQRGPGRAFVTIGK-IGNMRQAHCNISRAKWNNTLKQIASKLREQFGNNKTIIFKQSSGG  335
            +  GPG+AF   G  IG++RQA+CNIS  KW++ L ++ +KL+E F  N  I FK S+GG
Sbjct  421  L--GPGQAFFATGDIIGSIRQAYCNISGDKWDDMLNKVKAKLKEFFPGN--ISFKSSAGG  588

Query  336  DPEIVTHSFNCGGEFFYCNSTQLFNSTWFNSTWSTEGSNNTEGSDTITLPCRIKQIINMW  395
            D EI+TH+FNCGGEFFYCN++ LFNS+  N+               I LPC+I+QI+ MW
Sbjct  589  DLEIITHTFNCGGEFFYCNTSDLFNSSSKNN------------GTIIKLPCKIRQIVRMW  732

Query  396  QKVGKAMYAPPISGQIRCSSNITGLLLTRD----GGNSNNESEIFRPGGGDMRDNWRSEL  451
            Q+V +AMYAPPI+G I C SNITGLLLTRD    GGN+ N++EIFRPGGGDM+DNWRSEL
Sbjct  733  QRVAQAMYAPPIAGIITCRSNITGLLLTRDGSSKGGNNTNDTEIFRPGGGDMKDNWRSEL  912

Query  452  YKYKVVKIEPLGVAPTKAKRRVV  474
            YKYK+VKI+PLG+AP+KA+RR V
Sbjct  913  YKYKIVKIKPLGIAPSKARRRAV  981



>Escherichia coli strain EN5308 contig00145, whole genome shotgun sequence
Sequence ID: NZ_JARBDE010000145.1 Length: 1005
Range 1: 5 to 994

Score:427 bits(1097), Expect:7e-144,
Method:Compositional matrix adjust.,
Identities:219/340(64%), Positives:258/340(75%), Gaps:19/340(5%)

Query  104  NDTNTNSSSGRMIMEKGEIKNCSFNISTSIRGKVQKEYAFFYKLDIIPIDNDTTS-----  158
            N TN   ++   +    EI+NCSFN +T IR + Q+ YA FY+LDI+P+D + +S
Sbjct  5    NATNNGDATHNVTNNGQEIQNCSFNATTEIRDRKQRVYALFYRLDIVPLDKNNSSKNNSS  184

Query  159  --YKLTSCNTSVITQACPKVSFEPIPIHYCAPAGFAILKCNNKTFNGTGPCTNVSTVQCT  216
              Y+L +CNTS ITQACPKVSF+PIPIHYCAPAG+AILKCNNKTFNGTGPC NVSTVQCT
Sbjct  185  EYYRLINCNTSAITQACPKVSFDPIPIHYCAPAGYAILKCNNKTFNGTGPCNNVSTVQCT  364

Query  217  HGIRPVVSTQLLLNGSLAEEEVVIRSVNFTDNAKTIIVQLNTSVEINCTRPNNNTRKRIR  276
            HGI+PVVSTQLLLNGSLAE E++IRS N TDN KTIIV L+ SVEI CTRPNNNTRK IR
Sbjct  365  HGIKPVVSTQLLLNGSLAEGEIIIRSENLTDNVKTIIVHLDQSVEIVCTRPNNNTRKSIR  544

Query  277  IQRGPGRAF-VTIGKIGNMRQAHCNISRAKWNNTLKQIASKLREQFGNNKTIIFKQSSGG  335
            I  GPG+ F  T G IGN+RQAHCNIS  KWN TL+++  KL E F  NKTI F  SSGG
Sbjct  545  I--GPGQTFYATGGIIGNIRQAHCNISEDKWNETLQRVGKKLVEHFP-NKTIKFAPSSGG  715

Query  336  DPEIVTHSFNCGGEFFYCNSTQLFNSTWF-NSTWSTEGSNNTEGSDTITLPCRIKQIINM  394
            D EI THSFNC GEFFYC++++LFNST+  N T S       + + TIT+PC IKQI+NM
Sbjct  716  DLEITTHSFNCRGEFFYCSTSRLFNSTYMPNDTKS-------KSNKTITIPCSIKQIVNM  874

Query  395  WQKVGKAMYAPPISGQIRCSSNITGLLLTRDGGNSNNESE  434
            WQ+VG+AMYAPPI G I C SNITG+LL RDGG  + + E
Sbjct  875  WQEVGRAMYAPPIEGNITCRSNITGILLVRDGGVDSEDPE  994



>Escherichia coli strain NBE0002 contig00132, whole genome shotgun sequence
Sequence ID: NZ_JARBDF010000132.1 Length: 1381
Range 1: 2 to 928

Score:421 bits(1082), Expect:1e-139,
Method:Compositional matrix adjust.,
Identities:208/311(67%), Positives:246/311(79%), Gaps:11/311(3%)

Query  2    LWVTVYYGVPVWKEATTTLFCASDAKAYDTEVHNVWATHACVPTDPNPQEVVLVNVTENF  61
            L VTVYY VPVW EA TTLFCASDAKAY+ EVHN+WATHACVPTDPNPQE+VL NVTE F
Sbjct  928  L*VTVYYEVPVWTEAKTTLFCASDAKAYEKEVHNIWATHACVPTDPNPQEMVLKNVTEYF  749

Query  62   NMWKNDMVEQMHEDIISLWDQSLKPCVKLTPLCVSLKC-TDLKNDTNTNSSSGRMIMEKG  120
            NMWKND+V+QMHED+ISL DQSLKPCVKLTPLCV+L+C +   N+T T  +   ++ +
Sbjct  748  NMWKNDIVDQMHEDVISL*DQSLKPCVKLTPLCVTLECNSTTYNNTYTYKNKSSVLEQVH  569

Query  121  EIKNCSFNISTSIRGKVQKEYAFFYKLDIIPIDND-------TTSYKLTSCNTSVITQAC  173
            E+KNCSFN +T +R K  K  A F++LDI+PIDN        T SY+L +CNTSVITQAC
Sbjct  568  EVKNCSFNATTELRDKKNKV*ALFHRLDIVPIDNKENFNENIT*SYRLINCNTSVITQAC  389

Query  174  PKVSFEPIPIHYCAPAGFAILKCNNKTFNGTGPCTNVSTVQCTHGIRPVVSTQLLLNGSL  233
            PKV+F+PIPIHYC PA FAIL+CNNKTF+  GPC NVSTVQCTH I+PVVSTQLLLN SL
Sbjct  388  PKVNFDPIPIHYCTPASFAILQCNNKTFSKKGPCNNVSTVQCTHRIKPVVSTQLLLNSSL  209

Query  234  AEEEVVIRSVNFTDNAKTIIVQLNTSVEINCTRPNNNTRKRIRIQRGPGRA-FVTIGKIG  292
            AEEE++IRS N TDN KTIIV LN SV I C RP NNTRK ++I+  PG+  F T   +
Sbjct  208  AEEEIIIRSENLTDNVKTIIVNLNESVVIECMRPGNNTRKSVKIR--PGQTFFATEDIVE  35

Query  293  NMRQAHCNISR  303
            ++RQAHCN+S+
Sbjct  34   DIRQAHCNVSK  2



>Escherichia coli strain EN5095 contig00078, whole genome shotgun sequence
Sequence ID: NZ_JAPYLY010000078.1 Length: 1972
Range 1: 1002 to 1970

Score:425 bits(1092), Expect:1e-138,
Method:Compositional matrix adjust.,
Identities:224/341(66%), Positives:267/341(78%), Gaps:21/341(6%)

Query  93    LCVSLKCTDLKNDTNTNSSSGRMIMEKGEIKNCSFNISTSIRGKVQKEYAFFYKLDIIPI  152
             LCV+L C++    T  NSSS        EIKNC+FN +T IR K QK  A FYKLD++P+
Sbjct  1970  LCVTLHCSN-ATSTFGNSSS--------EIKNCTFNATTEIRDKKQKVNALFYKLDVVPL  1818

Query  153   -DNDTTSYKLTSCNTSVITQACPKVSFEPIPIHYCAPAGFAILKCNNKTFNGTGPCTNVS  211
              DN+ + Y+L +CNTS +TQACPK+SF+PIPIHYCAPAG+AILKCN+KTFNGTGPC NVS
Sbjct  1817  NDNNNSYYRLINCNTSTVTQACPKISFDPIPIHYCAPAGYAILKCNDKTFNGTGPCKNVS  1638

Query  212   TVQCTHGIRPVVSTQLLLNGSLAE-EEVVIRSVNFTDNAKTIIVQLNTSVEINCTRPNNN  270
             TVQCTHGI+PVVSTQLLLNGSLAE  E++IRS N T+NAKTIIV LN SV+I C RPNNN
Sbjct  1637  TVQCTHGIKPVVSTQLLLNGSLAEGGEIIIRSENITNNAKTIIVHLNESVQIRCIRPNNN  1458

Query  271   TRKRIRIQRGPGRAF-VTIGKIGNMRQAHCNISRAKWNNTLKQIASKLREQFGNNKTIIF  329
             TRK IRI  GPG+ F  T G IGN+R+A C+I+ ++WN TL+ +  KLRE F  N+TI F
Sbjct  1457  TRKSIRI--GPGQTFYATGGIIGNIREAQCSINGSEWNKTLQGVREKLREHFP-NRTITF  1287

Query  330   KQSSGGDPEIVTHSFNCGGEFFYCNSTQLFNSTWFNSTWSTEGSNNTEGSDTITLPCRIK  389
               SSGGD EI THSFNC GEFFYCN++ LFNST  N T     SN+T  + T+ LPCRIK
Sbjct  1286  NSSSGGDLEITTHSFNCRGEFFYCNTSSLFNST--NGTE----SNSTWVNGTLILPCRIK  1125

Query  390   QIINMWQKVGKAMYAPPISGQIRCSSNITGLLLTRDGGNSN  430
             QIINMWQ+VG+AMYAPPI+G I C+S+ITGLLLTRDGGN+N
```

```
Sbjct  1124  QIINMWQEVGRAMYAPPIAGNITCNSSITGLLLTRDGGNTN  1002



>Escherichia coli strain NBE0002 contig00149, whole genome shotgun sequence
Sequence ID: NZ_JARBDF010000149.1 Length: 996
Range 1: 107 to 991

Score:412 bits(1058), Expect:6e-138,
Method:Compositional matrix adjust.,
Identities:203/295(69%), Positives:232/295(78%), Gaps:19/295(6%)

Query  2    LWVTVYYGVPVWKEATTTLFCASDAKAYDTEVHNVWATHACVPTDPNPQEVVLVNVTENF  61
            LWVTVYYGVPVWK+A  TLFCASDAK Y TEVHNVWATHACVPTDPNPQE+VL NVTENF
Sbjct  107  LWVTVYYGVPVWKDANATLFCASDAKVYKTEVHNVWATHACVPTDPNPQELVLGNVTENF  286

Query  62   NMWKNDMVEQMHEDIISLWDQSLKPCVKLTPLCVSLKCT--DLKNDT---NTNSSSGRMI  116
            NMW+NDMVEQMH+D+IS+WDQ LKPCVKLTPLCV+L CT  + +N+T   N   S    I
Sbjct  287  NMWENDMVEQMHQDVISIWDQGLKPCVKLTPLCVTLNCTKYEPRNETKKDNIKISLNLTI  466

Query  117  MEKG--EIKNCSFNISTSIRGKVQKEYAFFYKLDIIPIDNDTTS------------YKLT  162
             ++   EI NCSF  +T +R K +K +A FYK+D+I +  ++T             Y+L
Sbjct  467  DQEAVEEINNCSFETTTELRDKTRKVHALFYKIDMIELGQNSTCQKPENSTSECTMYRLI  646

Query  163  SCNTSVITQACPKVSFEPIPIHYCAPAGFAILKCNNKTFNGTGPCTNVSTVQCTHGIRPV  222
            +CNTS ITQACPKVSFEPIPIHYCAPAG+A+LKCNN TFNGTGPC N+STVQCTHGIRPV
Sbjct  647  NCNTSAITQACPKVSFEPIPIHYCAPAGYALLKCNNNTFNGTGPCRNISTVQCTHGIRPV  826

Query  223  VSTQLLLNGSLAEEEVVIRSVNFTDNAKTIIVQLNTSVEINCTRPNNNTRKRIRI  277
            VSTQLLLNGSLAE  + IRS N TDNAKTII  LN SVEI C RPNNNTR+ +RI
Sbjct  827  VSTQLLLNGSLAEGNITIRSENLTDNAKTIIAHLNQSVEILCVRPNNNTRQSVRI  991



>Escherichia coli strain EN5132 contig00077, whole genome shotgun sequence
Sequence ID: NZ_JAPYLZ010000077.1 Length: 2498
Range 1: 500 to 1456

Score:424 bits(1091), Expect:3e-136,
Method:Compositional matrix adjust.,
Identities:218/328(66%), Positives:251/328(76%), Gaps:19/328(5%)

Query  118   EKGEIKNCSFNISTSIRGKVQKEYAFFYKLDIIPIDND---------TTSYKLTSCNTSV  168
             E  E+K C FN ST +  K QK  A FY LDI P++           T +Y LTSCNTSV
Sbjct  500   ETKELKRCFFNASTEVEDKKQKVSALFYSLDIRPLNKSSSGNSSINSTETYVLTSCNTSV  679

Query  169   ITQACPKVSFEPIPIHYCAPAGFAILKCNNKTFNGTGPCTNVSTVQCTHGIRPVVSTQLL  228
             I QACPKV+F+PIPIHYCAPAG+AILKCNN+TFNGTGPC NVSTVQCTHGI+PVVSTQLL
Sbjct  680   IKQACPKVTFDPIPIHYCAPAGYAILKCNNETFNGTGPCNNVSTVQCTHGIKPVVSTQLL  859

Query  229   LNGSLAEEEVVIRSVNFTDNAKTIIVQLNTSVEINCTRPNNNTRKRIRIQRGPGRAFVTI  288
             LNGSLA+ ++VIRS N TDNAKTIIV LN SVEINCTRPNNNTRK IRI  GPG+ F
Sbjct  860   LNGSLAKGDIVIRSENLTDNAKTIIVHLNESVEINCTRPNNNTRKSIRI--GPGQTFYAT  1033

Query  289   GK-IGNMRQAHCNISRAKWNNTLKQIASKLREQFGNNKTIIFKQSSGGDPEIVTHSFNCG  347
             G+ IG++RQAHCNIS   WN TL+++  KL E F  NKTI F+ SSGGD EI+THSFNCG
Sbjct  1034  GEIIGDIRQAHCNISENMWNKTLQRVGKKLAEHFP-NKTIKFEPSSGGDLEIMTHSFNCG  1210

Query  348   GEFFYCNSTQLFNSTWFNSTWSTEGSNNTEGSDTITLPCRIKQIINMWQKVGKAMYAPPI  407
             GEFFYCN++ LFN T+ N T+  + S     S  IT+ CRIKQIINMWQ+VG+AMYAPPI
Sbjct  1211  GEFFYCNTSSLFNGTY-NGTYIEKNS-----SSIITIQCRIKQIINMWQEVGRAMYAPPI  1372

Query  408   SGQIRCSSNITGLLLTRDGGNSNNESEI  435
             +G I C SNITGLLL RDGG  +N +EI
Sbjct  1373  AGNITCRSNITGLLLVRDGGTGSNSTEI  1456



>Klebsiella pneumoniae strain P7701 NODE_62_length_570_cov_92.6095, whole genome shotgun sequence
Sequence ID: NZ_JACAWM010000060.1 Length: 570
Range 1: 3 to 569

Score:378 bits(970), Expect:8e-127,
Method:Compositional matrix adjust.,
Identities:182/189(96%), Positives:184/189(97%), Gaps:0/189(0%)

Query  216  THGIRPVVSTQLLLNGSLAEEEVVIRSVNFTDNAKTIIVQLNTSVEINCTRPNNNTRKRI  275
            THGIRPV STQLLLNGSLAEE+VVIRS NFTDNAKTIIVQLNTSVEINCTRPNNNTRK I
Sbjct  3    THGIRPVASTQLLLNGSLAEEDVVIRSANFTDNAKTIIVQLNTSVEINCTRPNNNTRKSI  182

Query  276  RIQRGPGRAFVTIGKIGNMRQAHCNISRAKWNNTLKQIASKLREQFGNNKTIIFKQSSGG  335
            RIQRGPGRAFVTIGKIGNMRQAHCNISRAKWN TLKQIASKLREQFGNNKTIIFKQSSGG
Sbjct  183  RIQRGPGRAFVTIGKIGNMRQAHCNISRAKWNATLKQIASKLREQFGNNKTIIFKQSSGG  362

Query  336  DPEIVTHSFNCGGEFFYCNSTQLFNSTWFNSTWSTEGSNNTEGSDTITLPCRIKQIINMW  395
            DPEIVTHSFNCGGEFFYCNSTQLFNSTWFNSTWSTEGSNNTEGSDTITLPCRIKQ INMW
Sbjct  363  DPEIVTHSFNCGGEFFYCNSTQLFNSTWFNSTWSTEGSNNTEGSDTITLPCRIKQFINMW  542

Query  396  QKVGKAMYA  404
            Q+VGKAMYA
Sbjct  543  QEVGKAMYA  569



>Escherichia coli strain NBE0004 contig00131, whole genome shotgun sequence
Sequence ID: NZ_JARBDH010000131.1 Length: 847
Range 1: 2 to 778

Score:365 bits(937), Expect:2e-120,
Method:Compositional matrix adjust.,
Identities:187/269(70%), Positives:210/269(78%), Gaps:22/269(8%)

Query  4    VTVYYGVPVWKEATTTLFCASDAKAYDTEVHNVWATHACVPTDPNPQEVVLVNVTENFNM  63
            VTVYY VPVWKEA  TLFCASDAKA++ EVHNV ATHACVPTDPNPQE+VL NVTENFNM
Sbjct  778  VTVYYEVPVWKEAKATLFCASDAKAHEREVHNV*ATHACVPTDPNPQEMVLENVTENFNM  599

Query  64   WKNDMVEQMHEDIISLWDQSLKPCVKLTPLCVSLKC---TDLKNDTNTNSSSGRMIMEKG  120
            WKNDMV+QMHED+ISL DQSLKPCVKLTPLCV+L+C   T  KN T+          E
Sbjct  598  WKNDMVDQMHEDVISL*DQSLKPCVKLTPLCVTLECNNYTTYKNYTD----------ETK  449

Query  121  EIKNCSFNISTSIRGKVQKEYAFFYKLDIIPIDNDTT---------SYKLTSCNTSVITQ  171
            E+K C FN ST +  K QK  A FY LDI P++  ++         +Y LTSCNTSVI Q
Sbjct  448  ELKRCFFNASTEVEDKKQKVSALFYSLDIRPLNKSSSRNSSINSNGTYVLTSCNTSVIKQ  269

Query  172  ACPKVSFEPIPIHYCAPAGFAILKCNNKTFNGTGPCTNVSTVQCTHGIRPVVSTQLLLNG  231
            ACPKV+F+PIPIHYCAPAG+AILKCNN+TFN TGPC NVSTVQCTH I+PVVSTQLLLNG
Sbjct  268  ACPKVTFDPIPIHYCAPAGYAILKCNNETFNRTGPCNNVSTVQCTHRIKPVVSTQLLLNG  89

Query  232  SLAEEEVVIRSVNFTDNAKTIIVQLNTSV  260
            SLA+ ++VIRS N  DNAKTIIV LN SV
```

```
Sbjct  88   SLAKGDIVIRSENLADNAKTIIVHLNESV  2




>Methyloversatilis universalis FAM5 contig00006, whole genome shotgun sequence
Sequence ID: NZ_AFHG01000004.1 Length: 649
Range 1: 32 to 649

Score:359 bits(922), Expect:4e-119,
Method:Compositional matrix adjust.,
Identities:172/209(82%), Positives:183/209(87%), Gaps:8/209(3%)

Query  1    KLWVTVYYGVPVWKEATTTLFCASDAKAYDTEVHNVWATHACVPTDPNPQEVVLVNVTEN  60
            KLWVTVYYGVPVWKEATTTLFCASD KAYDTEVHNVWATHACVPTDPNPQEVVL NVTEN
Sbjct  32   KLWVTVYYGVPVWKEATTTLFCASDTKAYDTEVHNVWATHACVPTDPNPQEVVLGNVTEN  211

Query  61   FNMWKNDMVEQMHEDIISLWDQSLKPCVKLTPLCVSLKCTDLKNDTNT-----NSSSGRM  115
            FNMW+NDMVEQMHEDIISLWDQSLKPCVKLTPLCV+L C D            NSS+  M
Sbjct  212  FNMWRNDMVEQMHEDIISLWDQSLKPCVKLTPLCVTLNCNDYNGTNTNGTNTKNSSNVEM  391

Query  116  IMEKGEIKNCSFNISTSIRGKVQKEYAFFYKLDIIPIDNDTTSYKLTSCNTSVITQACPK  175
            I   GEIKNCSFNI+T+IR K++KEYA FYKLD++PIDND TSY+L SCNTSVITQACPK
Sbjct  392  I---GEIKNCSFNITTNIRDKIKKEYALFYKLDVVPIDNDNTSYRLISCNTSVITQACPK  562

Query  176  VSFEPIPIHYCAPAGFAILKCNNKTFNGT  204
            +SFEPIPIHYCAPAGFAILKCNNK FNGT
Sbjct  563  ISFEPIPIHYCAPAGFAILKCNNKKFNGT  649




>Escherichia coli strain NBE0010 contig00098, whole genome shotgun sequence
Sequence ID: NZ_JARBDM010000098.1 Length: 1912
Range 1: 47 to 775

Score:342 bits(878), Expect:9e-107,
Method:Compositional matrix adjust.,
Identities:168/246(68%), Positives:201/246(81%), Gaps:5/246(2%)

Query  121  EIKNCSFNISTSIRGKVQKEYAFFYKLDIIPIDNDTT-SYKLTSCNTSVITQACPKVSFE  179
            E+KNCSFN +T +R K QK  A F++LDI+P+ N +  +Y+L +CNTS I QACPKVSF+
Sbjct  47   EVKNCSFNATTELRDKKQKVQALFHRLDIMPLTNGSDGNYRLINCNTSTIAQACPKVSFD  226

Query  180  PIPIHYCAPAGFAILKCNNKTFNGTGPCTNVSTVQCTHGIRPVVSTQLLLNGSLAEEEVV  239
            PIPIHYC PAG+AIL+CNNKTFNGTGPC NVSTVQCTHGI+PVVSTQLLLNGSLA+EE+V
Sbjct  227  PIPIHYCTPAGYAILQCNNKTFNGTGPCNNVSTVQCTHGIKPVVSTQLLLNGSLAKEEIV  406

Query  240  IRSVNFTDNAKTIIVQLNTSVEINCTRPNNNTRKRIRIQRGPGRAFVTIGK-IGNMRQAH  298
            IRS N T+NAK IIV LN SV+I CTRPNNNTRK IRI  GPG+ F   G  IG++R+AH
Sbjct  407  IRSENLTNNAKIIIVHLNKSVDIVCTRPNNNTRKSIRI--GPGQTFYATGDIIGDIRKAH  580

Query  299  CNISRAKWNNTLKQIASKLREQFGNNKTIIFKQSSGGDPEIVTHSFNCGGEFFYCNSTQL  358
            CNI+  +WN TL +++ KL E+F  NKTI FK SSGGD EI THSFNCGGEFFYCN++ L
Sbjct  581  CNININRWNETLLKVSKKLAERFP-NKTIQFKPSSGGDLEITTHSFNCGGEFFYCNTSNL  757

Query  359  FNSTWF  364
            FN T++
Sbjct  758  FNFTYY  775




Range 2: 1650 to 1910

Score:152 bits(383), Expect:3e-35,
Method:Compositional matrix adjust.,
Identities:70/88(80%), Positives:79/88(89%), Gaps:1/88(1%)

Query  392   INMWQKVGKAMYAPPISGQIRCSSNITGLLLTRDGGNSNNESEIFRPGGGDMRDNWRSEL  451
             INMWQ+VG+AMYAPPI+G I+C SNITGLLL RDGG  +N +E FRPGGGDMRDNWRSEL
Sbjct  1910  INMWQEVGRAMYAPPIAGNIKCGSNITGLLLVRDGG-PDNGTETFRPGGGDMRDNWRSEL  1734

Query  452   YKYKVVKIEPLGVAPTKAKRRVVQREKR  479
             YKYKVV+I+PLGVAPT+AKRRVV R KR
Sbjct  1733  YKYKVVEIKPLGVAPTEAKRRVVGRGKR  1650




>Pseudomonas aeruginosa strain VET-42 NODE_89_length_1008_cov_1.38346_ID_177, whole genome shotgun sequence
Sequence ID: NZ_RACX01000220.1 Length: 963
Range 1: 303 to 962

Score:328 bits(840), Expect:3e-105,
Method:Compositional matrix adjust.,
Identities:155/220(70%), Positives:178/220(80%), Gaps:5/220(2%)

Query  2    LWVTVYYGVPVWKEATTTLFCASDAKAYDTEVHNVWATHACVPTDPNPQEVVLVNVTENF  61
            LWVTVYYGVP W++A T LFCASDAKA+ TE HN+WATHACVPTDPNPQE+ L NVTENF
Sbjct  303  LWVTVYYGVPAWEDADTILFCASDAKAHSTESHNIWATHACVPTDPNPQEIPLENVTENF  482

Query  62   NMWKNDMVEQMHEDIISLWDQSLKPCVKLTPLCVSLKCTDL-KNDTNTNSSSGRMIMEKG  120
            NMWKN+MVEQMHEDIISLWD+SLK CVKLTPLCV+L C ++ K D +TN      ++ KG
Sbjct  483  NMWKNNMVEQMHEDIISLWDESLKRCVKLTPLCVTLYCENVNKTDNSTNGEDNSTVINKG  662

Query  121  EIKNCSFNISTSIRGKVQKEYAFFYKLDIIPID----NDTTSYKLTSCNTSVITQACPKV  176
            EI NCSFNI+T IR + + EYA FYKLD+IPI+    +  + Y L +CN + I QACPKV
Sbjct  663  EITNCSFNITTEIRDRKKTEYALFYKLDVIPIEGRDRSSNSKYMLINCNATTIKQACPKV  842

Query  177  SFEPIPIHYCAPAGFAILKCNNKTFNGTGPCTNVSTVQCT  216
            SFEPIPIHYCAPAGFAILKC +K FNGTGPC NVSTVQ T
Sbjct  843  SFEPIPIHYCAPAGFAILKCRDKNFNGTGPCKNVSTVQWT  962




>Neisseria gonorrhoeae strain GC-206 NODE_120_length_488_cov_1.24711_ID_20412, whole genome shotgun sequence
Sequence ID: NZ_SAWN01000120.1 Length: 488
Range 1: 2 to 487

Score:238 bits(606), Expect:2e-72,
Method:Compositional matrix adjust.,
Identities:118/166(71%), Positives:134/166(80%), Gaps:5/166(3%)

Query  165  NTSVITQACPKVSFEPIPIHYCAPAGFAILKCNNKTFNGTGPCTNVSTVQCTHGIRPVVS  224
            NTS I QACPK++F+PIPIHYCAPAG+AILKCNNKTFNGTGPC NVSTVQCTHGI+PVVS
Sbjct  487  NTSAIAQACPKITFDPIPIHYCAPAGYAILKCNNKTFNGTGPCQNVSTVQCTHGIKPVVS  308

Query  225  TQLLLNGSLAEEEVVIRSVNFTDNAKTIIVQLNTSVEINCTRPNNNTRKRIRIQRGPGRA  284
            TQLLLNGSLAE +++IRS N TD+ KTIIV LN SV I CTRP NNTRK +RI  GPG+A
Sbjct  307  TQLLLNGSLAEGDIIIRSENITDSIKTIIVHLNESVYIACTRPGNNTRKSVRI--GPGQA  134

Query  285  FVTIGK-IGNMRQAHCNISRAKWNNTLKQIASKLREQFGNNKTIIF  329
```

```
            F   G  IG++RQAHCNISR  W+ TL ++  KL   F  NKTI F
Sbjct  133  FYATGDVIGDIRQAHCNISRGNWSRTLGKVRDKLAAHF--NKTIEF  2
```

>Burkholderia pseudomallei strain 2604STDY5519032 ERS240165SCcontig000080, whole genome shotgun sequence
Sequence ID: NZ_UVID02000080.1 Length: 499
Range 1: 8 to 490

Score:223 bits(568), Expect:1e-66,
Method:Compositional matrix adjust.,
Identities:115/172(67%), Positives:132/172(76%), Gaps:12/172(6%)

```
Query  154  NDTTSYKLTSCNTSVITQACPKVSFEPIPIHYCAPAGFAILKCNNKTFNGTGPCTNVSTV  213
            N++  Y+L +CNTS I QACPKVSF+PIPIHYCAPAG+AILKC +KTFNGTGPC NVSTV
Sbjct  490  NESKEYRLINCNTSAIKQACPKVSFDPIPIHYCAPAGYAILKCKDKTFNGTGPCHNVSTV  311

Query  214  QCTHGIRPVVSTQLLLNGSLAEEEVVIRSVNFTDNAKTIIVQLNTSVEINCTRPNNNTRK  273
            QCTHGI+PVVSTQLLLNGS        RS N TDNAKTIIV LN SV+I CTRP NNTR
Sbjct  310  QCTHGIKPVVSTQLLLNGSF-------RSENLTDNAKTIIVHLNESVDIVCTRPGNNTRT  152

Query  274  RIRIQRGPGRAFVTIGK-IGNMRQAHCNISRAKWNNTLKQIASKLREQFGNN  324
             IRI  GPG+ F   G  IG++RQAHCNIS   W  TL+++ +KL + F NN
Sbjct  151  SIRI--GPGQTFYARGAVIGDIRQAHCNIS--NWYKTLQRVGNKLAKHFPNN  8
```

>Escherichia coli strain EN5192 contig00156, whole genome shotgun sequence
Sequence ID: NZ_JAPMNL010000156.1 Length: 562
Range 1: 2 to 562

Score:223 bits(569), Expect:2e-66,
Method:Compositional matrix adjust.,
Identities:122/197(62%), Positives:142/197(72%), Gaps:17/197(8%)

```
Query  247  DNAKTIIVQLNTSVEINCTRPNNNTRKRIRIQRGPGRAF-VTIGKIGNMRQAHCNISRAK  305
            DN KTIIV L+ SVEI CT PNNNTRK IRI  GPG+ F  T G IGN+RQAHCNIS  K
Sbjct  562  DNVKTIIVHLDQSVEIVCTGPNNNTRKSIRI--GPGQTFYATGGIIGNIRQAHCNISEDK  389

Query  306  WNNTLKQIASKLREQFGNNKTIIFKQSSGGDPEIVTHSFNCGGEFFYCNSTQLFNSTWF-  364
            WN TL+++  KL E F N KTI F  SSGGD EI THSFNC GEFFYC++++LFNST+
Sbjct  388  WNETLQRVGKKLVEHFPN-KTIKFAPSSGGDLEITTHSFNCRGEFFYCSTSRLFNSTYMP  212

Query  365  NSTWSTEGSNNTEGSDTITLPCRIKQIINMWQKVGKAMYAPPISGQIRCSSNITGLLLTR  424
            N T S       + + TIT+PC IKQI+NMWQ+VG+AMYAPPI G I C SNITG+LL R
Sbjct  211  NDTKS-------KSNKTITIPCSIKQIVNMWQEVGRAMYAPPIEGNITCRSNITGILLVR  53

Query  425  DGG-----NSNNESEIF  436
            DGG       NN++E F
Sbjct  52   DGGVDSEDPENNKTETF  2
```

>Klebsiella pneumoniae strain PN085E1IA NODE_127_length_474_cov_225.069_ID_253, whole genome shotgun sequence
Sequence ID: NZ_PDVD01000016.1 Length: 474
Range 1: 15 to 467

Score:222 bits(566), Expect:2e-66,
Method:Compositional matrix adjust.,
Identities:112/155(72%), Positives:125/155(80%), Gaps:5/155(3%)

```
Query  189  AGFAILKCNNKTFNGTGPCTNVSTVQCTHGIRPVVSTQLLLNGSLAEEEVVIRSVNFTDN  248
            AG+AILKCNNKTFNGTGPC NVSTVQCTHGI+PVVSTQLLLNGSLAEEE++IRS N T+N
Sbjct  467  AGYAILKCNNKTFNGTGPCNNVSTVQCTHGIKPVVSTQLLLNGSLAEEEIIIRSENLTNN  288

Query  249  AKTIIVQLNTSVEINCTRPNNNTRKRIRIQRGPGRAFVTIGK-IGNMRQAHCNISRAKWN  307
            AKTIIV  N SVEI C RP NNTRK +RI  GPG+ F   G  IG++RQAHCNIS A WN
Sbjct  287  AKTIIVHFNESVEITCIRPGNNTRKSVRI--GPGQTFYATGDIIGDIRQAHCNISEADWN  114

Query  308  NTLKQIASKLREQFGNNKTIIFKQSSGGDPEIVTH  342
             TL ++  KL+E F  NKTIIF+ SSGGD EI TH
Sbjct  113  KTLARVVKKLKEHF--NKTIIFQPSSGGDLEITTH  15
```

>Pseudomonas aeruginosa strain VET-42 NODE_97_length_548_cov_2.70913_ID_193, whole genome shotgun sequence
Sequence ID: NZ_RACX01000130.1 Length: 548
Range 1: 3 to 533

Score:214 bits(545), Expect:7e-63,
Method:Compositional matrix adjust.,
Identities:112/193(58%), Positives:139/193(72%), Gaps:21/193(10%)

```
Query  246  TDNAKTIIVQLNTSVEINCTRPNNNTRKRIRIQRGPGRAFVTIGKI-GNMRQAHCNISRA  304
            TDN KTIIVQ N SVEINCTRPNNNTRK I +  GPG+AF   G I G++RQA+CNIS
Sbjct  533  TDNTKTIIVQFNQSVEINCTRPNNNTRKSIHL--GPGQAFFATGDIIGSIRQAYCNISGD  360

Query  305  KWNNTLKQIASKLREQFGNNKTIIFKQSSGGDPEIVTHSFNCGGEFFYCNSTQLFNSTWF  364
            KW++ L ++ +KL+E F  N   I FK S+GGD EI+TH+FNCGGEFFYCN++ LFNS+
Sbjct  359  KWDDMLNKVKAKLKEFFPGN--ISFKSSAGGDLEIITHTFNCGGEFFYCNTSDLFNSSSK  186

Query  365  NSTWSTEGSNNTEGSDTITLPCRIKQIINMWQKVGKAMYAPPISGQIRCSSNITGLLLTR  424
            N+               I LPC+I+QI+ MWQ+V +AMYAPPI+G I C SNITGLLLTR
Sbjct  185  NN------------GTIIKLPCKIRQIVRMWQRVAQAMYAPPIAGIITCRSNITGLLLTR  42

Query  425  D----GGNSNNES  433
            D    GGN+ N++
Sbjct  41   DGSSKGGNNTNDT  3
```

>Klebsiella pneumoniae strain ST-234:K080 NODE_195_length_474_cov_25.8761, whole genome shotgun sequence
Sequence ID: NZ_NXJY01000191.1 Length: 474
Range 1: 8 to 466

Score:205 bits(522), Expect:8e-60,
Method:Compositional matrix adjust.,
Identities:107/157(68%), Positives:123/157(78%), Gaps:5/157(3%)

```
Query  189  AGFAILKCNNKTFNGTGPCTNVSTVQCTHGIRPVVSTQLLLNGSLAEEEVVIRSVNFTDN  248
            AG+AILKCNN+TFNG GPC NVSTVQCTHGI+PVVSTQLLLNGSLA +EVVIRS N T+N
Sbjct  8    AGYAILKCNNETFNGKGPCNNVSTVQCTHGIKPVVSTQLLLNGSLA-KEVVIRSENLTNN  184

Query  249  AKTIIVQLNTSVEINCTRPNNNTRKRIRIQRGPGRAFVTIGK-IGNMRQAHCNISRAKWN  307
            AK IIV LN SV I C RP+NNTRK +RI  GPG+ F   G  IG++R+AHCNIS+ +WN
Sbjct  185  AKIIIVHLNKSVTIECIRPSNNTRKSVRI--GPGQTFYATGDIIGDIREAHCNISKQEWN  358

Query  308  NTLKQIASKLREQFGNNKTIIFKQSSGGDPEIVTHSF  344
             TL+ +  KL + F  NKTI F QSSGGD EI THS
```

```
Sbjct  359  KTLQLVGEKLAKHFP-NKTITFNQSSGGDLEITTHSL  466
```

>Enterobacter hormaechei strain ST-234:K063 NODE_159_length_475_cov_27.7644, whole genome shotgun sequence
Sequence ID: NZ_NXJN01000152.1 Length: 475
Range 1: 6 to 467

Score:201 bits(512), Expect:2e-58,
Method:Compositional matrix adjust.,
Identities:105/158(66%), Positives:123/158(77%), Gaps:5/158(3%)

```
Query  188  PAGFAILKCNNKTFNGTGPCTNVSTVQCTHGIRPVVSTQLLLNGSLAEEEVVIRSVNFTD  247
            PAG+AILKCNN+TFNG GPC NVSTVQCTHGI+PVVSTQLLLNGSLA +EVVIRS N T+
Sbjct  6    PAGYAILKCNNETFNGKGPCNNVSTVQCTHGIKPVVSTQLLLNGSLA-KEVVIRSENLTN  182

Query  248  NAKTIIVQLNTSVEINCTRPNNNTRKRIRIQRGPGRAFVTIGK-IGNMRQAHCNISRAKW  306
            NAK IIVQLN SV I C RP+NNTRK +RI  GPG+ F   GK  GN R+A+CNI++ +W
Sbjct  183  NAKIIIVQLNESVTIECIRPSNNTRKSVRI--GPGQTFYATGKRRGNRREAYCNITKQEW  356

Query  307  NNTLKQIASKLREQFGNNKTIIFKQSSGGDPEIVTHSF  344
            + TL+ +  KL + F  NK I F QSSGGD E+ THS
Sbjct  357  SRTLQLVGEKLAKHFP-NKAITFNQSSGGDLEVTTHSI  467
```

>Klebsiella pneumoniae strain ST-1552:K089 NODE_185_length_474_cov_21.4899, whole genome shotgun sequence
Sequence ID: NZ_NXIW01000184.1 Length: 474
Range 1: 15 to 467

Score:201 bits(510), Expect:4e-58,
Method:Compositional matrix adjust.,
Identities:106/155(68%), Positives:121/155(78%), Gaps:5/155(3%)

```
Query  189  AGFAILKCNNKTFNGTGPCTNVSTVQCTHGIRPVVSTQLLLNGSLAEEEVVIRSVNFTDN  248
            AG+AILKCNN+TFNG GPC NVSTVQCTHGI+PVVSTQLLLNGSLA +EVVIRS N T+N
Sbjct  467  AGYAILKCNNETFNGKGPCNNVSTVQCTHGIKPVVSTQLLLNGSLA-KEVVIRSENLTNN  291

Query  249  AKTIIVQLNTSVEINCTRPNNNTRKRIRIQRGPGRAFVTIGK-IGNMRQAHCNISRAKWN  307
            AK IIV LN SV I C RP+NNTRK +RI  GPG+ F   G  IGN+R+A+CNISR +WN
Sbjct  290  AKIIIVHLNKSVTIECIRPSNNTRKSVRI--GPGQTFYATGDIIGNIREAYCNISRQEWN  117

Query  308  NTLKQIASKLREQFGNNKTIIFKQSSGGDPEIVTH  342
             TL+ +  KL + F  NKTI F QSSGGD EI T
Sbjct  116  KTLQLVGEKLAKHFP-NKTITFNQSSGGDLEITTQ  15
```

>Escherichia coli strain E003 NODE_241_length_474_cov_17.072, whole genome shotgun sequence
Sequence ID: NZ_NXIZ01000238.1 Length: 474
Range 1: 15 to 467

Score:198 bits(504), Expect:3e-57,
Method:Compositional matrix adjust.,
Identities:104/155(67%), Positives:121/155(78%), Gaps:5/155(3%)

```
Query  189  AGFAILKCNNKTFNGTGPCTNVSTVQCTHGIRPVVSTQLLLNGSLAEEEVVIRSVNFTDN  248
            AG+AILKCNN+TFNG GPC NVSTVQCTHGI+PVVSTQLLLNGSLA +EVVIRS N T+N
Sbjct  467  AGYAILKCNNETFNGKGPCNNVSTVQCTHGIKPVVSTQLLLNGSLA-KEVVIRSENLTNN  291

Query  249  AKTIIVQLNTSVEINCTRPNNNTRKRIRIQRGPGRAFVTIGK-IGNMRQAHCNISRAKWN  307
            AK IIV LN SV I C RP+NNTRK +RI  GPG+ F   G  IGN+R+A+CNIS+ +WN
Sbjct  290  AKIIIVHLNKSVTIECIRPSNNTRKSVRI--GPGQTFYATGDIIGNIREAYCNISKQEWN  117

Query  308  NTLKQIASKLREQFGNNKTIIFKQSSGGDPEIVTH  342
             TL+ +  KL + F  NKTI F QSSGGD E+ T
Sbjct  116  KTLQLVGEKLAKHFP-NKTITFNQSSGGDLEVTTQ  15
```

>Cronobacter sakazakii strain cro4114A1 contig126, whole genome shotgun sequence
Sequence ID: NZ_NRKJ01000126.1 Length: 413
Range 1: 3 to 413

Score:192 bits(488), Expect:4e-55,
Method:Compositional matrix adjust.,
Identities:96/143(67%), Positives:111/143(77%), Gaps:8/143(5%)

```
Query  266  RPNNNTRKRIRIQRGPGRAFVTIGKI-GNMRQAHCNISRAKWNNTLKQIASKLREQFGNN  324
            RP+NNTRK I    GPGR+    G I G++RQAHCNIS  KWNNTLK++A KLREQFGNN
Sbjct  413  RPDNNTRKSIHF--GPGRSLYATGNIIGDIRQAHCNISGEKWNNTLKRVAIKLREQFGNN  240

Query  325  KTIIFKQSSGGDPEIVTHSFNCGGEFFYCNSTQLFNSTWF-NSTWSTEGSNNTEGSDTIT  383
            KTI+F+Q SGGDPE+V H  NCGGEFFYCN+T LFNST   N TW++ G    +GS  IT
Sbjct  239  KTIVFQQPSGGDPEVVMHILNCGGEFFYCNTTNLFNSTLLANDTWNSTG----QGSGNIT  72

Query  384  LPCRIKQIINMWQKVGKAMYAPP  406
            + CRIKQIIN WQ+VGKAMYAPP
Sbjct  71   IQCRIKQIINRWQEVGKAMYAPP  3
```

>Cronobacter malonaticus strain cro3263A1 contig69, whole genome shotgun sequence
Sequence ID: NZ_NRMM01000069.1 Length: 413
Range 1: 1 to 411

Score:192 bits(487), Expect:6e-55,
Method:Compositional matrix adjust.,
Identities:96/143(67%), Positives:111/143(77%), Gaps:8/143(5%)

```
Query  266  RPNNNTRKRIRIQRGPGRAFVTIGKI-GNMRQAHCNISRAKWNNTLKQIASKLREQFGNN  324
            RP+NNTRK I    GPGR+    G I G++RQAHCNIS  KWNNTLK++A KLREQFGNN
Sbjct  1    RPSNNTRKSIHF--GPGRSLYATGNIIGDIRQAHCNISGEKWNNTLKRVAIKLREQFGNN  174

Query  325  KTIIFKQSSGGDPEIVTHSFNCGGEFFYCNSTQLFNSTWF-NSTWSTEGSNNTEGSDTIT  383
            KTI+F+Q SGGDPE+V H  NCGGEFFYCN+T LFNST   N TW++ G    +GS  IT
Sbjct  175  KTIVFQQPSGGDPEVVMHILNCGGEFFYCNTTNLFNSTLLANDTWNSTG----QGSGNIT  342

Query  384  LPCRIKQIINMWQKVGKAMYAPP  406
            + CRIKQIIN WQ+VGKAMYAPP
Sbjct  343  IQCRIKQIINRWQEVGKAMYAPP  411
```

>Citrobacter freundii strain CF003 NODE_156_length_474_cov_23.0173, whole genome shotgun sequence
Sequence ID: NZ_NXKE01000152.1 Length: 474
Range 1: 8 to 457

```
Score:191 bits(486), Expect:2e-54,
Method:Compositional matrix adjust.,
Identities:102/154(66%), Positives:118/154(76%), Gaps:5/154(3%)

Query  189  AGFAILKCNNKTFNGTGPCTNVSTVQCTHGIRPVVSTQLLLNGSLAEEEVVIRSVNFTDN  248
            AG+AILKCNN+TFNG GPC NVSTVQCTHGI+PVVSTQLLLNGSLA +EVVIRS N T+N
Sbjct  8    AGYAILKCNNETFNGKGPCNNVSTVQCTHGIKPVVSTQLLLNGSLA-KEVVIRSENLTNN  184

Query  249  AKTIIVQLNTSVEINCTRPNNNTRKRIRIQRGPGRAFVTIGK-IGNMRQAHCNISRAKWN  307
            AK IIV LN SV I C RP+NNTRK +RI  GPG+ F   G  IG +R+A+CNISR + N
Sbjct  185  AKIIIVHLNKSVTIECIRPSNNTRKSVRI--GPGQTFYATGYIIGEIREAYCNISRQELN  358

Query  308  NTLKQIASKLREQFGNNKTIIFKQSSGGDPEIVT  341
             TL+ +  KL +    NKTI F QSSGGD E+ T
Sbjct  359  KTLQLVGEKLAKHVP-NKTITFNQSSGGDLEVTT  457



>Cronobacter malonaticus strain cro3263A1 contig68, whole genome shotgun sequence
Sequence ID: NZ_NRMM01000068.1 Length: 413
Range 1: 1 to 411

Score:177 bits(449), Expect:2e-49,
Method:Compositional matrix adjust.,
Identities:92/142(65%), Positives:107/142(75%), Gaps:6/142(4%)

Query  266  RPNNNTRKRIRIQRGPGRAFVTIGKI-GNMRQAHCNISRAKWNNTLKQIASKLREQFGNN  324
            RP+NNTRK IRI  GPG+ F   G I G++RQAHCNISR  W  TL  ++ KL E F N
Sbjct  1    RPDNNTRKSIRI--GPGQTFYATGDIIGDIRQAHCNISRTDWERTLHNVSKKLAEHFPN-  171

Query  325  KTIIFKQSSGGDPEIVTHSFNCGGEFFYCNSTQLFNSTWFNSTWSTEGSNNTEGSDTITL  384
            KTI F  SSGGD EI THSFNC GEFFYCN++ LFNST+  +T  T  SN T+G+ TIT+
Sbjct  172  KTIRFTNSSGGDLEITTHSFNCRGEFFYCNTSGLFNSTY--TTNGTFESNETKGNSTITI  345

Query  385  PCRIKQIINMWQKVGKAMYAPP  406
            PCRIKQIINMWQ+VG+AMYAPP
Sbjct  346  PCRIKQIINMWQEVGRAMYAPP  411



>Cronobacter sakazakii strain cro4215W contig103, whole genome shotgun sequence
Sequence ID: NZ_NRKF01000103.1 Length: 413
Range 1: 1 to 411

Score:174 bits(442), Expect:2e-48,
Method:Compositional matrix adjust.,
Identities:91/142(64%), Positives:106/142(74%), Gaps:6/142(4%)

Query  266  RPNNNTRKRIRIQRGPGRAFVTIGKI-GNMRQAHCNISRAKWNNTLKQIASKLREQFGNN  324
            RP+NNTRK IRI  GPG+ F   G I G++RQAHCNISR  W  TL  ++ KL E F N
Sbjct  1    RPSNNTRKSIRI--GPGQTFYATGDIIGDIRQAHCNISRTDWERTLHNVSKKLAEHFPN-  171

Query  325  KTIIFKQSSGGDPEIVTHSFNCGGEFFYCNSTQLFNSTWFNSTWSTEGSNNTEGSDTITL  384
            KTI F  SSGGD EI THSFNC GEFFYCN++ LFNST+  +T  T  SN T+ + TIT+
Sbjct  172  KTIRFTNSSGGDLEITTHSFNCRGEFFYCNTSGLFNSTY--TTNGTFESNETKSNSTITI  345

Query  385  PCRIKQIINMWQKVGKAMYAPP  406
            PCRIKQIINMWQ+VG+AMYAPP
Sbjct  346  PCRIKQIINMWQEVGRAMYAPP  411



>Cronobacter dublinensis strain cro981C2 contig304, whole genome shotgun sequence
Sequence ID: NZ_NROB01000304.1 Length: 413
Range 1: 1 to 411

Score:172 bits(437), Expect:2e-47,
Method:Compositional matrix adjust.,
Identities:90/142(63%), Positives:105/142(73%), Gaps:6/142(4%)

Query  266  RPNNNTRKRIRIQRGPGRAFVTIGKI-GNMRQAHCNISRAKWNNTLKQIASKLREQFGNN  324
            RP+NNTRK IRI  GPG+ F   G I G++RQAHCNISR  W  TL  ++ KL E F
Sbjct  1    RPSNNTRKSIRI--GPGQTFYATGDIIGDIRQAHCNISRTDWERTLHNVSKKLAEHFPK-  171

Query  325  KTIIFKQSSGGDPEIVTHSFNCGGEFFYCNSTQLFNSTWFNSTWSTEGSNNTEGSDTITL  384
            KTI F  SSGGD EI THSFNC GEFFYCN++ LFNST+  +T  T  SN T+ + TIT+
Sbjct  172  KTIRFTNSSGGDLEITTHSFNCRGEFFYCNTSGLFNSTY--TTNGTFESNETKSNSTITI  345

Query  385  PCRIKQIINMWQKVGKAMYAPP  406
            PCRIKQIINMWQ+VG+AMYAPP
Sbjct  346  PCRIKQIINMWQEVGRAMYAPP  411



>Cronobacter sakazakii strain cro4114B1 contig109, whole genome shotgun sequence
Sequence ID: NZ_NRKI01000109.1 Length: 413
Range 1: 1 to 411

Score:172 bits(437), Expect:2e-47,
Method:Compositional matrix adjust.,
Identities:90/142(63%), Positives:105/142(73%), Gaps:6/142(4%)

Query  266  RPNNNTRKRIRIQRGPGRAFVTIGKI-GNMRQAHCNISRAKWNNTLKQIASKLREQFGNN  324
            RP+NNTRK IRI  GPG+ F   G I G++RQAHCNISR  W  TL  ++ KL E F
Sbjct  1    RPSNNTRKSIRI--GPGQTFYATGDIIGDIRQAHCNISRTDWERTLHNVSKKLAEHFPK-  171

Query  325  KTIIFKQSSGGDPEIVTHSFNCGGEFFYCNSTQLFNSTWFNSTWSTEGSNNTEGSDTITL  384
            KTI F  SSGGD EI THSFNC GEFFYCN++ LFNST+  +T  T  SN T+ + TIT+
Sbjct  172  KTIRFTNSSGGDLEITTHSFNCRGEFFYCNTSGLFNSTY--TTNGTFESNETKSNSTITI  345

Query  385  PCRIKQIINMWQKVGKAMYAPP  406
            PCRIKQIINMWQ+VG+AMYAPP
Sbjct  346  PCRIKQIINMWQEVGRAMYAPP  411



>Cronobacter sakazakii strain cro4114A1 contig127, whole genome shotgun sequence
Sequence ID: NZ_NRKJ01000127.1 Length: 413
Range 1: 1 to 411

Score:172 bits(437), Expect:2e-47,
Method:Compositional matrix adjust.,
Identities:90/142(63%), Positives:105/142(73%), Gaps:6/142(4%)

Query  266  RPNNNTRKRIRIQRGPGRAFVTIGKI-GNMRQAHCNISRAKWNNTLKQIASKLREQFGNN  324
            RP+NNTRK IRI  GPG+ F   G I G++RQAHCNISR  W  TL  ++ KL E F
Sbjct  1    RPSNNTRKSIRI--GPGQTFYATGDIIGDIRQAHCNISRTDWERTLHNVSKKLAEHFPK-  171

Query  325  KTIIFKQSSGGDPEIVTHSFNCGGEFFYCNSTQLFNSTWFNSTWSTEGSNNTEGSDTITL  384
```

```
            KTI F  SSGGD EI THSFNC GEFFYCN++ LFNST+  +T  T  SN T+ + TIT+
Sbjct  172  KTIRFTNSSGGDLEITTHSFNCRGEFFYCNTSGLFNSTY--TTNGTFESNETKSNSTITI  345

Query  385  PCRIKQIINMWQKVGKAMYAPP  406
            PCRIKQIINMWQ+VG+AMYAPP
Sbjct  346  PCRIKQIINMWQEVGRAMYAPP  411
```

```
>Cronobacter sakazakii strain cro611A3 contig61, whole genome shotgun sequence
Sequence ID: NZ_NRMC01000061.1 Length: 407
Range 1: 3 to 407

Score:171 bits(434), Expect:3e-47,
Method:Compositional matrix adjust.,
Identities:90/143(63%), Positives:106/143(74%), Gaps:10/143(6%)

Query  266  RPNNNTRKRIRIQRGPGRAFVTIGKI-GNMRQAHCNISRAKWNNTLKQIASKLREQFGNN  324
            RP+NNTRK IRI  GPG+ F   G I G++RQAHCNIS+  W  TL +++ KL E F  N
Sbjct  407  RPSNNTRKSIRI--GPGQTFYATGDIIGDIRQAHCNISKRDWEETLYKVSKKLAEHF-QN  237

Query  325  KTIIFKQSSGGDPEIVTHSFNCGGEFFYCNSTQLFNSTWF-NSTWSTEGSNNTEGSDTIT  383
            KTI F+ SSGGD EI THSFNC GEFFYCN++ LFN T+  N T ++ GSN      TIT
Sbjct  236  KTIQFEPSSGGDLEITTHSFNCRGEFFYCNTSGLFNKTYMHNGTLNSTGSNG-----TIT  72

Query  384  LPCRIKQIINMWQKVGKAMYAPP  406
            +PCRIKQIINMWQKVG+AMYAPP
Sbjct  71   IPCRIKQIINMWQKVGQAMYAPP  3
```

```
>Enterobacter hormaechei strain ST-234:K063 NODE_153_length_480_cov_7.37677, whole genome shotgun sequence
Sequence ID: NZ_NXJN01000148.1 Length: 480
Range 1: 84 to 473

Score:171 bits(434), Expect:7e-47,
Method:Compositional matrix adjust.,
Identities:86/132(65%), Positives:103/132(78%), Gaps:3/132(2%)

Query  189  AGFAILKCNNKTFNGTGPCTNVSTVQCTHGIRPVVSTQLLLNGSLAEEEVVIRSVNFTDN  248
            AG+AILKCNNKTFNGTGPC NVSTVQCTHGI+PVVSTQLLLNGSLAE E++IRS +  +N
Sbjct  473  AGYAILKCNNKTFNGTGPCNNVSTVQCTHGIKPVVSTQLLLNGSLAEGEIIIRSESLVEN  294

Query  249  AKTIIVQLNTSVEINCTRPNNNTRKRIRIQRGPGRAFVTIGK-IGNMRQAHCNISRAKWN  307
              TII  LN SV+I C RPNNNTRK I I  GP +AF   G  IGN+RQA+CN++   WN
Sbjct  293  TNTIIAHLNDSVDIMCIRPNNNTRKSIGI--GPVQAFYATGDVIGNIRQAYCNVTTKPWN  120

Query  308  NTLKQIASKLRE  319
             TL+++  ++ E
Sbjct  119  ETLERVKIEIEE  84
```

```
>Klebsiella pneumoniae strain EuSCAPE_IL004, whole genome shotgun sequence
Sequence ID: NZ_UKJP02000113.1 Length: 277
Range 1: 1 to 276

Score:169 bits(427), Expect:1e-46,
Method:Composition-based stats.,
Identities:78/92(85%), Positives:83/92(90%), Gaps:0/92(0%)

Query  16   ATTTLFCASDAKAYDTEVHNVWATHACVPTDPNPQEVVLVNVTENFNMWKNDMVEQMHED  75
            A   LFCASDAKAYD EVHNVWATHACVPTDPNPQE+ L NVTENFNMWKNDMVEQMH+D
Sbjct  1    AKAPLFCASDAKAYDKEVHNVWATHACVPTDPNPQEMFLENVTENFNMWKNDMVEQMHKD  180

Query  76   IISLWDQSLKPCVKLTPLCVSLKCTDLKNDTN  107
            IISLWD+SLKPCVKLTPLCV+L CTD+ N TN
Sbjct  181  IISLWDESLKPCVKLTPLCVTLNCTDVDNSTN  276
```

```
>Cronobacter sakazakii strain cro3040W contig81, whole genome shotgun sequence
Sequence ID: NZ_NRJO01000081.1 Length: 404
Range 1: 1 to 402

Score:168 bits(425), Expect:8e-46,
Method:Compositional matrix adjust.,
Identities:89/142(63%), Positives:103/142(72%), Gaps:9/142(6%)

Query  266  RPNNNTRKRIRIQRGPGRAFVTIGKI-GNMRQAHCNISRAKWNNTLKQIASKLREQFGNN  324
            RP+NNTRK IRI  GPG+ F   G I G++RQAHCNISR  W  TL  ++ KL E F  N
Sbjct  1    RPSNNTRKSIRI--GPGQTFYATGDIIGDIRQAHCNISRKDWEETLHNVSKKLAEHF-QN  171

Query  325  KTIIFKQSSGGDPEIVTHSFNCGGEFFYCNSTQLFNSTWFNSTWSTEGSNNTEGSDTITL  384
            K I F  SSGGD EI THSFNC GEFFYCN++ LFN T+  +   TE SN+T    TIT+
Sbjct  172  KKIQFASSSGGDLEITTHSFNCRGEFFYCNTSGLFNGTYDGTLNGTE-SNST----TITI  336

Query  385  PCRIKQIINMWQKVGKAMYAPP  406
            PCRIKQIINMWQ+VG+AMYAPP
Sbjct  337  PCRIKQIINMWQEVGRAMYAPP  402
```

```
>Escherichia coli strain NBE0009 contig00116, whole genome shotgun sequence
Sequence ID: NZ_JARBDL010000116.1 Length: 1713
Range 1: 1358 to 1705

Score:180 bits(457), Expect:9e-46,
Method:Compositional matrix adjust.,
Identities:87/123(71%), Positives:101/123(82%), Gaps:9/123(7%)

Query  359   FNSTWFNSTWSTEGSNNTEGSDTITLPCRIKQIINMWQKVGKAMYAPPISGQIRCSSNIT  418
             +N T++N T     +N       IT+PCRIKQIINMWQKVG+AMYAPPI+G I C+S IT
Sbjct  1705  YNGTYYNRTQYNSTAN-------ITIPCRIKQIINMWQKVGRAMYAPPIAGNITCNSTIT  1547

Query  419   GLLLTRDGGNSN--NESEIFRPGGGDMRDNWRSELYKYKVVKIEPLGVAPTKAKRRVVQR  476
             GLLL RDGGN+N  N +EIFRPGGGDMRDNWRSELYKYKVV+I+PLG+APT+AKRRVV R
Sbjct  1546  GLLLLRDGGNTNDTNYTEIFRPGGGDMRDNWRSELYKYKVVEIKPLGIAPTRAKRRVVGR  1367

Query  477   EKR  479
             EKR
Sbjct  1366  EKR  1358
```

```
>Cronobacter sakazakii strain cro4114A1 contig132, whole genome shotgun sequence
Sequence ID: NZ_NRKJ01000132.1 Length: 392
Range 1: 1 to 390
```

Score:167 bits(423), Expect:1e-45,
Method:Compositional matrix adjust.,
Identities:88/142(62%), Positives:99/142(69%), Gaps:13/142(9%)

```
Query  266  RPNNNTRKRIRIQRGPGRAFVTIGKI-GNMRQAHCNISRAKWNNTLKQIASKLREQFGNN  324
            RP NNTR  I I  GPGRAF  IG I G++RQAHC I+  KWN TLKQ+  KL+E F  N
Sbjct  1    RPTNNTRTSIPI--GPGRAFYRIGDIIGDIRQAHCKINGPKWNETLKQVVRKLKEHF--N  168

Query  325  KTIIFKQSSGGDPEIVTHSFNCGGEFFYCNSTQLFNSTWFNSTWSTEGSNNTEGSDTITL  384
            KTI F+  SGGD EI  H FNC GEFFYCN+ +LFNSTW          N T  +DTI L
Sbjct  169  KTIKFQPPSGGDLEITMHHFNCRGEFFYCNTAKLFNSTWI--------GNATMENDTIIL  324

Query  385  PCRIKQIINMWQKVGKAMYAPP  406
            PCRIKQIIN+WQ VG+AMYAPP
Sbjct  325  PCRIKQIINLWQGVGQAMYAPP  390
```

>Cronobacter sakazakii strain cro4215W contig104, whole genome shotgun sequence
Sequence ID: NZ_NRKF01000104.1 Length: 405
Range 1: 1 to 402

Score:165 bits(418), Expect:8e-45,
Method:Compositional matrix adjust.,
Identities:88/142(62%), Positives:102/142(71%), Gaps:9/142(6%)

```
Query  266  RPNNNTRKRIRIQRGPGRAFVTIGKI-GNMRQAHCNISRAKWNNTLKQIASKLREQFGNN  324
            RPNNNTR  IRI  GPG+ F   G I G++RQA+C I+  KWN TL+Q+A KL+EQF  N
Sbjct  1    RPNNNTRTSIRI--GPGQTFYRTGDIIGDIRQAYCEINGTKWNETLRQVAKKLKEQF--N  168

Query  325  KTIIFKQSSGGDPEIVTHSFNCGGEFFYCNSTQLFNSTWFNSTWSTEGSNNTEGSDTITL  384
            KTI F+  SGGD EI    FNC GEFFYCN+T+LFNSTW     + +G N   GSDT  L
Sbjct  169  KTIKFQPPSGGDLEITMLHFNCRGEFFYCNTTKLFNSTW-EGNETIKGGN---GSDTTIL  336

Query  385  PCRIKQIINMWQKVGKAMYAPP  406
            PCRIKQIINMWQ  G+AMYAPP
Sbjct  337  PCRIKQIINMWQGAGQAMYAPP  402
```

>Cronobacter malonaticus strain cro3263A1 contig74, whole genome shotgun sequence
Sequence ID: NZ_NRMM01000074.1 Length: 404
Range 1: 3 to 404

Score:164 bits(415), Expect:2e-44,
Method:Compositional matrix adjust.,
Identities:83/142(58%), Positives:99/142(69%), Gaps:9/142(6%)

```
Query  266  RPNNNTRKRIRIQRGPGRAFVTIGKI-GNMRQAHCNISRAKWNNTLKQIASKLREQFGNN  324
            RPNNNTRK I +    GRA    G I G++RQAHCN+S+  W  TL Q+A+KLREQF  N
Sbjct  404  RPNNNTRKSISL--AGGRALYATGDIIGDIRQAHCNVSKGNWTETLGQVATKLREQF-KN  234

Query  325  KTIIFKQSSGGDPEIVTHSFNCGGEFFYCNSTQLFNSTWFNSTWSTEGSNNTEGSDTITL  384
            KTI+F  SSGGDPE+    FNCGGEFFYCN+++LFN T+        G+     +  ITL
Sbjct  233  KTIVFNSSSGGDPEVEMFIFNCGGEFFYCNTSKLFNYTFL-----ANGTGQNYSTGNITL  69

Query  385  PCRIKQIINMWQKVGKAMYAPP  406
             CRIKQIIN WQ+VGKAMYAPP
Sbjct  68   QCRIKQIINRWQEVGKAMYAPP  3
```

>Cronobacter sakazakii strain cro4114A1 contig129, whole genome shotgun sequence
Sequence ID: NZ_NRKJ01000129.1 Length: 404
Range 1: 1 to 402

Score:164 bits(415), Expect:2e-44,
Method:Compositional matrix adjust.,
Identities:87/142(61%), Positives:102/142(71%), Gaps:9/142(6%)

```
Query  266  RPNNNTRKRIRIQRGPGRAFVTIGKI-GNMRQAHCNISRAKWNNTLKQIASKLREQFGNN  324
            RP+NNTR  IRI  GPG+ F   G I G++RQA+C I+  KWN TL+Q+A KL+EQF  N
Sbjct  1    RPDNNTRTSIRI--GPGQTFYRTGDIIGDIRQAYCEINGTKWNETLRQVAKKLKEQF--N  168

Query  325  KTIIFKQSSGGDPEIVTHSFNCGGEFFYCNSTQLFNSTWFNSTWSTEGSNNTEGSDTITL  384
            KTI F+  SGGD EI    FNC GEFFYCN+T+LFNSTW     + +G N   GSDT  L
Sbjct  169  KTIKFQPPSGGDLEITMLHFNCRGEFFYCNTTKLFNSTW-EGNETIKGGN---GSDTTIL  336

Query  385  PCRIKQIINMWQKVGKAMYAPP  406
            PCRIKQIINMWQ  G+AMYAPP
Sbjct  337  PCRIKQIINMWQGAGQAMYAPP  402
```

>Cronobacter sakazakii strain cro4114B1 contig117, whole genome shotgun sequence
Sequence ID: NZ_NRKI01000117.1 Length: 404
Range 1: 1 to 402

Score:163 bits(413), Expect:4e-44,
Method:Compositional matrix adjust.,
Identities:87/142(61%), Positives:102/142(71%), Gaps:9/142(6%)

```
Query  266  RPNNNTRKRIRIQRGPGRAFVTIGKI-GNMRQAHCNISRAKWNNTLKQIASKLREQFGNN  324
            RP+NNTR  IRI  GPG+ F   G I G++RQA+C I+  KWN TL+Q+A KL+EQF  N
Sbjct  1    RPSNNTRTSIRI--GPGQTFYRTGDIIGDIRQAYCEINGTKWNETLRQVAKKLKEQF--N  168

Query  325  KTIIFKQSSGGDPEIVTHSFNCGGEFFYCNSTQLFNSTWFNSTWSTEGSNNTEGSDTITL  384
            KTI F+  SGGD EI    FNC GEFFYCN+T+LFNSTW     + +G N   GSDT  L
Sbjct  169  KTIKFQPPSGGDLEITMLHFNCRGEFFYCNTTKLFNSTW-EGNETIKGGN---GSDTTIL  336

Query  385  PCRIKQIINMWQKVGKAMYAPP  406
            PCRIKQIINMWQ  G+AMYAPP
Sbjct  337  PCRIKQIINMWQGAGQAMYAPP  402
```

>Cronobacter sakazakii strain cro4114B1 contig115, whole genome shotgun sequence
Sequence ID: NZ_NRKI01000115.1 Length: 404
Range 1: 1 to 402

Score:163 bits(412), Expect:7e-44,
Method:Compositional matrix adjust.,
Identities:82/142(58%), Positives:99/142(69%), Gaps:9/142(6%)

```
Query  266  RPNNNTRKRIRIQRGPGRAFVTIGKI-GNMRQAHCNISRAKWNNTLKQIASKLREQFGNN  324
            RPNNNTRK I +    GRA    G I G++RQAHCN+S+  W  TL Q+A+KLREQF  N
Sbjct  1    RPNNNTRKSISL--AGGRALYATGNIIGDIRQAHCNVSKGNWTETLGQVATKLREQF-KN  171
```

```
Query  325  KTIIFKQSSGGDPEIVTHSFNCGGEFFYCNSTQLFNSTWFNSTWSTEGSNNTEGSDTITL  384
            KTI+F  SSGGDPE+    FNCGGEFFYCN+++LFN T+        G+     +  IT+
Sbjct  172  KTIVFNSSSGGDPEVEMFIFNCGGEFFYCNTSKLFNYTFL-----ANGTGQNYSTGNITI  336

Query  385  PCRIKQIINMWQKVGKAMYAPP  406
             CRIKQIIN WQ+VGKAMYAPP
Sbjct  337  QCRIKQIINRWQEVGKAMYAPP  402
```

```
>Cronobacter sakazakii strain cro4114A1 contig128, whole genome shotgun sequence
Sequence ID: NZ_NRKJ01000128.1 Length: 404
Range 1: 1 to 402

Score:162 bits(411), Expect:9e-44,
Method:Compositional matrix adjust.,
Identities:82/142(58%), Positives:99/142(69%), Gaps:9/142(6%)

Query  266  RPNNNTRKRIRIQRGPGRAFVTIGKI-GNMRQAHCNISRAKWNNTLKQIASKLREQFGNN  324
            RP+NNTRK I +    GRA    G I G++RQAHCN+S+  W  TL Q+A+KLREQF  N
Sbjct  1    RPSNNTRKSISL--AGGRALYATGDIIGDIRQAHCNVSKGNWTETLGQVATKLREQF-KN  171

Query  325  KTIIFKQSSGGDPEIVTHSFNCGGEFFYCNSTQLFNSTWFNSTWSTEGSNNTEGSDTITL  384
            KTI+F  SSGGDPE+    FNCGGEFFYCN+++LFN T+        G+     +  ITL
Sbjct  172  KTIVFNSSSGGDPEVEMFIFNCGGEFFYCNTSKLFNYTFL-----ANGTGQNYSTGNITL  336

Query  385  PCRIKQIINMWQKVGKAMYAPP  406
             CRIKQIIN WQ+VGKAMYAPP
Sbjct  337  QCRIKQIINRWQEVGKAMYAPP  402
```

```
>Cronobacter malonaticus strain cro3263A1 contig76, whole genome shotgun sequence
Sequence ID: NZ_NRMM01000076.1 Length: 392
Range 1: 1 to 390

Score:162 bits(410), Expect:1e-43,
Method:Compositional matrix adjust.,
Identities:85/142(60%), Positives:100/142(70%), Gaps:13/142(9%)

Query  266  RPNNNTRKRIRIQRGPGRAFVTIGKI-GNMRQAHCNISRAKWNNTLKQIASKLREQFGNN  324
            RP+NNTR  IRI  GPG+ F   G I G++RQA+C I+  KWN TL+Q+A KL+EQF  N
Sbjct  1    RPDNNTRTSIRI--GPGQTFYRTGDIIGDIRQAYCEINGTKWNETLRQVAKKLKEQF--N  168

Query  325  KTIIFKQSSGGDPEIVTHSFNCGGEFFYCNSTQLFNSTWFNSTWSTEGSNNTEGSDTITL  384
            KTI F+  SGGD EI    FNC GEFFYCN+ +LFNSTW          N T  +DTI L
Sbjct  169  KTIKFQPPSGGDLEITMLHFNCRGEFFYCNTAKLFNSTWI--------GNATMENDTIIL  324

Query  385  PCRIKQIINMWQKVGKAMYAPP  406
            PCRIKQIIN+WQ VG+AMYAPP
Sbjct  325  PCRIKQIINLWQGVGQAMYAPP  390
```

```
>Cronobacter sakazakii strain cro4215W contig105, whole genome shotgun sequence
Sequence ID: NZ_NRKF01000105.1 Length: 404
Range 1: 1 to 402

Score:162 bits(410), Expect:1e-43,
Method:Compositional matrix adjust.,
Identities:82/142(58%), Positives:98/142(69%), Gaps:9/142(6%)

Query  266  RPNNNTRKRIRIQRGPGRAFVTIGKI-GNMRQAHCNISRAKWNNTLKQIASKLREQFGNN  324
            RP NNTRK I +    GRA    G I G++RQAHCN+S+  W  TL Q+A+KLREQF  N
Sbjct  1    RPTNNTRKSISL--AGGRALYATGDIIGDIRQAHCNVSKGNWTETLGQVATKLREQF-KN  171

Query  325  KTIIFKQSSGGDPEIVTHSFNCGGEFFYCNSTQLFNSTWFNSTWSTEGSNNTEGSDTITL  384
            KTI+F  SSGGDPE+    FNCGGEFFYCN+++LFN T+        G+     +  ITL
Sbjct  172  KTIVFNSSSGGDPEVEMFIFNCGGEFFYCNTSKLFNYTFL-----ANGTGQNYSTGNITL  336

Query  385  PCRIKQIINMWQKVGKAMYAPP  406
             CRIKQIIN WQ+VGKAMYAPP
Sbjct  337  QCRIKQIINRWQEVGKAMYAPP  402
```

```
>Cronobacter dublinensis strain cro981C2 contig308, whole genome shotgun sequence
Sequence ID: NZ_NROB01000308.1 Length: 405
Range 1: 1 to 399

Score:160 bits(406), Expect:4e-43,
Method:Compositional matrix adjust.,
Identities:86/141(61%), Positives:101/141(71%), Gaps:9/141(6%)

Query  266  RPNNNTRKRIRIQRGPGRAFVTIGKI-GNMRQAHCNISRAKWNNTLKQIASKLREQFGNN  324
            RP+NNTR  IRI  GPG+ F   G I G++RQA+C I+  KWN TL+Q+A KL+EQF  N
Sbjct  1    RPSNNTRTSIRI--GPGQTFYRTGDIIGDIRQAYCEINGTKWNETLRQVAKKLKEQF--N  168

Query  325  KTIIFKQSSGGDPEIVTHSFNCGGEFFYCNSTQLFNSTWFNSTWSTEGSNNTEGSDTITL  384
            KTI F+  SGGD EI    FNC GEFFYCN+T+LFNSTW     + +G N   GSDT  L
Sbjct  169  KTIKFQPPSGGDLEITMLHFNCRGEFFYCNTTKLFNSTW-EGNETIKGGN---GSDTTIL  336

Query  385  PCRIKQIINMWQKVGKAMYAP  405
            PCRIKQIINMWQ  G+AMYAP
Sbjct  337  PCRIKQIINMWQGAGQAMYAP  399
```

```
>Escherichia coli strain EN5143 contig00088, whole genome shotgun sequence
Sequence ID: NZ_JAPMNI010000088.1 Length: 833
Range 1: 481 to 831

Score:166 bits(419), Expect:4e-43,
Method:Compositional matrix adjust.,
Identities:81/121(67%), Positives:94/121(77%), Gaps:4/121(3%)

Query  359  FNSTWFNSTWSTEGSNNTEGSDTITLPCRIKQIINMWQKVGKAMYAPPISGQIRCSSNIT  418
            FNST +N T      N T     IT+PCRIKQIINMWQKV +AMYAPPI+G I C SNIT
Sbjct  831  FNSTLYNETID----NGTHLISNITIPCRIKQIINMWQKVRRAMYAPPIAGDITCRSNIT  664

Query  419  GLLLTRDGGNSNNESEIFRPGGGDMRDNWRSELYKYKVVKIEPLGVAPTKAKRRVVQREK  478
            GLLL RD G  +NE+EIFRP   DM+DN RS+LYKYKVV+I+PL +APT+AKRRVV+REK
Sbjct  663  GLLLLRDRGVGDNETEIFRPRERDMKDN*RSKLYKYKVVEIKPLRIAPTRAKRRVVKREK  484

Query  479  R  479
            R
Sbjct  483  R  481
```

```
>Cronobacter sakazakii strain cro4114B1 contig105, whole genome shotgun sequence
Sequence ID: NZ_NRKI01000105.1 Length: 416
Range 1: 3 to 416

Score:157 bits(398), Expect:6e-42,
Method:Compositional matrix adjust.,
Identities:85/144(59%), Positives:105/144(72%), Gaps:9/144(6%)

Query  266  RPNNNTRKRIRIQRGPGRAFVTIGKI-GNMRQAHCNISRAKWNNTLKQIASKLREQFGNN  324
            RP NNTRK IRI  GPG+ F   G+I G++RQA+CNIS  +W  TL  ++ KL E F  N
Sbjct  416  RPGNNTRKSIRI--GPGQTFYATGEIIGDIRQAYCNISGKEWKETLHNVSKKLAEYF-PN  246

Query  325  KTIIFKQSSGGDPEIVTHSFNCGGEFFYCNSTQLFNSTW-FNSTWSTEGSNNTEGSDT-I  382
            KTI +  SSGGD EI THSFNC GEFFYCN+++LFN T+  N T++  G   TE   T I
Sbjct  245  KTINWTSSSGGDLEITTHSFNCRGEFFYCNTSELFNGTYMLNGTFTFNG---TESEPTNI  75

Query  383  TLPCRIKQIINMWQKVGKAMYAPP  406
            T+PC+IKQ+INMWQ+VG+AMYAPP
Sbjct  74   TIPCKIKQVINMWQEVGRAMYAPP  3




>Planctopirus hydrillae strain JC280 Contig113, whole genome shotgun sequence
Sequence ID: NZ_LYDR01000013.1 Length: 437
Range 1: 11 to 436

Score:157 bits(396), Expect:2e-41,
Method:Compositional matrix adjust.,
Identities:88/147(60%), Positives:107/147(72%), Gaps:6/147(4%)

Query  260  VEINCTRPNNNTRKRIRIQRGPGRAFVTIGKI-GNMRQAHCNISRAKWNNTLKQIASKLR  318
            VEI C RPNNNTRK +RI  GPG+ F   G+I G++R A+CNI+  KWN TL+ ++ KL
Sbjct  11   VEIVCIRPNNNTRKSMRI--GPGQTFYATGEIIGDIRYAYCNINEQKWNATLQNVSRKLL  184

Query  319  EQFGNNKTIIFKQSSGGDPEIVTHSFNCGGEFFYCNSTQLFNSTWFNSTWSTEGSNNTEG  378
            E F N KTI F++SSGGD EI THSFNC GEFFYCN++ LFN T+  S      +N+T
Sbjct  185  EHFPN-KTISFQRSSGGDLEITTHSFNCRGEFFYCNTSNLFNRTF--SELVNNSTNSTNS  355

Query  379  SDTITLPCRIKQIINMWQKVGKAMYAP  405
            +  ITL CRIKQIINMWQ VG+AMYAP
Sbjct  356  TSIITLQCRIKQIINMWQGVGRAMYAP  436




>Cronobacter sakazakii strain cro4114A1 contig133, whole genome shotgun sequence
Sequence ID: NZ_NRKJ01000133.1 Length: 389
Range 1: 3 to 389

Score:155 bits(393), Expect:3e-41,
Method:Compositional matrix adjust.,
Identities:83/142(58%), Positives:98/142(69%), Gaps:14/142(9%)

Query  266  RPNNNTRKRIRIQRGPGRAFVTIGKI-GNMRQAHCNISRAKWNNTLKQIASKLREQFGNN  324
            RPNNNTR  I I  GPGRAF   G I G++RQA+C ++R +WN  L Q+  KL E F  N
Sbjct  389  RPNNNTRTSINI--GPGRAFYRTGDIIGDIRQAYCEVNRTEWNEILGQVIKKLEEHF-KN  219

Query  325  KTIIFKQSSGGDPEIVTHSFNCGGEFFYCNSTQLFNSTWFNSTWSTEGSNNTEGSDTITL  384
            KTI F+  +GGD EI  H FNC GEFFYCN+T+L     FNS+ + +GSN     DTI L
Sbjct  218  KTIEFQPPAGGDLEITMHHFNCRGEFFYCNTTKL-----FNSSETGKGSN-----DTIIL  69

Query  385  PCRIKQIINMWQKVGKAMYAPP  406
            PCRIKQIINMWQ  G+AMYAPP
Sbjct  68   PCRIKQIINMWQGAGQAMYAPP  3




>Cronobacter sakazakii strain cro4191B1-2 contig85, whole genome shotgun sequence
Sequence ID: NZ_NRKG01000085.1 Length: 392
Range 1: 1 to 390

Score:155 bits(393), Expect:3e-41,
Method:Compositional matrix adjust.,
Identities:83/142(58%), Positives:97/142(68%), Gaps:13/142(9%)

Query  266  RPNNNTRKRIRIQRGPGRAFVTIGKI-GNMRQAHCNISRAKWNNTLKQIASKLREQFGNN  324
            RP NNTRK IRI  GPG+ F   G I G++RQAHC ++  +WN TLKQ+  KL+E F  N
Sbjct  1    RPTNNTRKSIRI--GPGQMFYRTGDIIGDIRQAHCEVNGTEWNETLKQVTKKLKEYF--N  168

Query  325  KTIIFKQSSGGDPEIVTHSFNCGGEFFYCNSTQLFNSTWFNSTWSTEGSNNTEGSDTITL  384
            KTI+F+  SGGD E     FNC GEFFYCN+T LFNST      + EG N      TI L
Sbjct  169  KTIVFRPPSGGDLETTMLHFNCRGEFFYCNTTNLFNST---ENGTMEGLNT-----TIIL  324

Query  385  PCRIKQIINMWQKVGKAMYAPP  406
            PCRIKQI+NMWQ VG+AMYAPP
Sbjct  325  PCRIKQIVNMWQGVGQAMYAPP  390




>Cronobacter sakazakii strain cro3040W contig84, whole genome shotgun sequence
Sequence ID: NZ_NRJO01000084.1 Length: 392
Range 1: 1 to 390

Score:155 bits(393), Expect:3e-41,
Method:Compositional matrix adjust.,
Identities:83/142(58%), Positives:97/142(68%), Gaps:13/142(9%)

Query  266  RPNNNTRKRIRIQRGPGRAFVTIGKI-GNMRQAHCNISRAKWNNTLKQIASKLREQFGNN  324
            RP NNTRK IRI  GPG+ F   G I G++RQAHC ++  +WN TLKQ+  KL+E F  N
Sbjct  1    RPTNNTRKSIRI--GPGQMFYRTGDIIGDIRQAHCEVNGTEWNETLKQVTKKLKEYF--N  168

Query  325  KTIIFKQSSGGDPEIVTHSFNCGGEFFYCNSTQLFNSTWFNSTWSTEGSNNTEGSDTITL  384
            KTI+F+  SGGD E     FNC GEFFYCN+T LFNST      + EG N      TI L
Sbjct  169  KTIVFRPPSGGDLETTMLHFNCRGEFFYCNTTNLFNST---ENGTMEGLNT-----TIIL  324

Query  385  PCRIKQIINMWQKVGKAMYAPP  406
            PCRIKQI+NMWQ VG+AMYAPP
Sbjct  325  PCRIKQIVNMWQGVGQAMYAPP  390




>Cronobacter sakazakii strain cro4114B1 contig122, whole genome shotgun sequence
Sequence ID: NZ_NRKI01000122.1 Length: 383
Range 1: 1 to 381

Score:155 bits(391), Expect:5e-41,
Method:Compositional matrix adjust.,
Identities:81/142(57%), Positives:95/142(66%), Gaps:16/142(11%)
```

```
Query  266  RPNNNTRKRIRIQRGPGRAFVTIGKI-GNMRQAHCNISRAKWNNTLKQIASKLREQFGNN  324
            RPNNNTR  I I  GPGRAF   G I G++RQA+C ++R +WN  L Q+  KL E F  N
Sbjct  1    RPNNNTRTSINI--GPGRAFYRTGDIIGDIRQAYCEVNRTEWNEILGQVIKKLEEHF-KN  171

Query  325  KTIIFKQSSGGDPEIVTHSFNCGGEFFYCNSTQLFNSTWFNSTWSTEGSNNTEGSDTITL  384
            KTI F+  +GGD EI  H FNC GEFFYCN+T+LFNS+             T  +DTI L
Sbjct  172  KTIEFQPPAGGDLEITMHHFNCRGEFFYCNTTKLFNSS------------ETRENDTIIL  315

Query  385  PCRIKQIINMWQKVGKAMYAPP  406
            PCRIKQIINMWQ  G+AMYAPP
Sbjct  316  PCRIKQIINMWQGAGQAMYAPP  381
```

>Cronobacter sakazakii strain cro4215W contig109, whole genome shotgun sequence
Sequence ID: NZ_NRKF01000109.1 Length: 389
Range 1: 3 to 389

Score:154 bits(390), Expect:9e-41,
Method:Compositional matrix adjust.,
Identities:82/142(58%), Positives:98/142(69%), Gaps:14/142(9%)

```
Query  266  RPNNNTRKRIRIQRGPGRAFVTIGKI-GNMRQAHCNISRAKWNNTLKQIASKLREQFGNN  324
            RP+NNTR  I I  GPGRAF   G I G++RQA+C ++R +WN  L Q+  KL E F  N
Sbjct  389  RPDNNTRTSINI--GPGRAFYRTGDIIGDIRQAYCEVNRTEWNEILGQVIKKLEEHF-KN  219

Query  325  KTIIFKQSSGGDPEIVTHSFNCGGEFFYCNSTQLFNSTWFNSTWSTEGSNNTEGSDTITL  384
            KTI F+  +GGD EI  H FNC GEFFYCN+T+L     FNS+ + +GSN     DTI L
Sbjct  218  KTIEFQPPAGGDLEITMHHFNCRGEFFYCNTTKL-----FNSSETGKGSN-----DTIIL  69

Query  385  PCRIKQIINMWQKVGKAMYAPP  406
            PCRIKQIINMWQ  G+AMYAPP
Sbjct  68   PCRIKQIINMWQGAGQAMYAPP  3
```

>Cronobacter sakazakii strain cro4191B1-2 contig83, whole genome shotgun sequence
Sequence ID: NZ_NRKG01000083.1 Length: 410
Range 1: 1 to 408

Score:153 bits(387), Expect:3e-40,
Method:Compositional matrix adjust.,
Identities:81/143(57%), Positives:97/143(67%), Gaps:9/143(6%)

```
Query  266  RPNNNTRKRIRIQRGPGRAFVTIGKI-GNMRQAHCNISRAKWNNTLKQIASKLREQFGNN  324
            RP NNTR  I I  GPG+ F   G I GN+R+A C I++ KW   LKQ+A+KL+  F N
Sbjct  1    RPYNNTRTGITI--GPGQVFYRTGDIIGNIRKAFCEINKTKWKEVLKQVATKLKGHF-NT  171

Query  325  KTIIFKQSSGGDPEIVTHSFNCGGEFFYCNSTQLFNSTWF-NSTWSTEGSNNTEGSDTIT  383
              II +  SGGD EI TH F+C GEFFYCN+T+LFNST   N TW T      EG+ TIT
Sbjct  172  TKIILQPPSGGDLEITTHHFSCRGEFFYCNTTELFNSTLMENGTWET----TKEGNGTIT  339

Query  384  LPCRIKQIINMWQKVGKAMYAPP  406
            +PCRIKQI+ MWQ  G+AMYAPP
Sbjct  340  IPCRIKQIVRMWQGTGQAMYAPP  408
```

>Cronobacter malonaticus strain cro3263A1 contig75, whole genome shotgun sequence
Sequence ID: NZ_NRMM01000075.1 Length: 392
Range 1: 1 to 390

Score:153 bits(386), Expect:3e-40,
Method:Compositional matrix adjust.,
Identities:82/142(58%), Positives:96/142(67%), Gaps:13/142(9%)

```
Query  266  RPNNNTRKRIRIQRGPGRAFVTIGKI-GNMRQAHCNISRAKWNNTLKQIASKLREQFGNN  324
            RP NNTRK IRI  GPG+ F   G I G++R AHC ++  +WN TLKQ+  KL+E F  N
Sbjct  1    RPTNNTRKSIRI--GPGQMFYRTGDIIGDIRNAHCEVNGTEWNETLKQVTKKLKEYF--N  168

Query  325  KTIIFKQSSGGDPEIVTHSFNCGGEFFYCNSTQLFNSTWFNSTWSTEGSNNTEGSDTITL  384
            KTI+F+  SGGD E     FNC GEFFYCN+T LFNST      + EG N      TI L
Sbjct  169  KTIVFRPPSGGDLETTMLHFNCRGEFFYCNTTNLFNST---ENGTMEGLNT-----TIIL  324

Query  385  PCRIKQIINMWQKVGKAMYAPP  406
            PCRIKQI+NMWQ VG+AMYAPP
Sbjct  325  PCRIKQIVNMWQGVGQAMYAPP  390
```

>Cronobacter sakazakii strain cro4114B1 contig124, whole genome shotgun sequence
Sequence ID: NZ_NRKI01000124.1 Length: 392
Range 1: 1 to 390

Score:153 bits(386), Expect:3e-40,
Method:Compositional matrix adjust.,
Identities:82/142(58%), Positives:96/142(67%), Gaps:13/142(9%)

```
Query  266  RPNNNTRKRIRIQRGPGRAFVTIGKI-GNMRQAHCNISRAKWNNTLKQIASKLREQFGNN  324
            RP NNTRK IRI  GPG+ F   G I G++R AHC ++  +WN TLKQ+  KL+E F  N
Sbjct  1    RPTNNTRKSIRI--GPGQMFYRTGDIIGDIRNAHCEVNGTEWNETLKQVTKKLKEYF--N  168

Query  325  KTIIFKQSSGGDPEIVTHSFNCGGEFFYCNSTQLFNSTWFNSTWSTEGSNNTEGSDTITL  384
            KTI+F+  SGGD E     FNC GEFFYCN+T LFNST      + EG N      TI L
Sbjct  169  KTIVFRPPSGGDLETTMLHFNCRGEFFYCNTTNLFNST---ENGTMEGLNT-----TIIL  324

Query  385  PCRIKQIINMWQKVGKAMYAPP  406
            PCRIKQI+NMWQ VG+AMYAPP
Sbjct  325  PCRIKQIVNMWQGVGQAMYAPP  390
```

>Cronobacter sakazakii strain cro4114A1 contig131, whole genome shotgun sequence
Sequence ID: NZ_NRKJ01000131.1 Length: 392
Range 1: 1 to 390

Score:152 bits(385), Expect:4e-40,
Method:Compositional matrix adjust.,
Identities:82/142(58%), Positives:96/142(67%), Gaps:13/142(9%)

```
Query  266  RPNNNTRKRIRIQRGPGRAFVTIGKI-GNMRQAHCNISRAKWNNTLKQIASKLREQFGNN  324
            RP NNTRK IRI  GPG+ F   G I G++R AHC ++  +WN TLKQ+  KL+E F  N
Sbjct  1    RPYNNTRKSIRI--GPGQMFYRTGDIIGDIRNAHCEVNGTEWNETLKQVTKKLKEYF--N  168

Query  325  KTIIFKQSSGGDPEIVTHSFNCGGEFFYCNSTQLFNSTWFNSTWSTEGSNNTEGSDTITL  384
            KTI+F+  SGGD E     FNC GEFFYCN+T LFNST      + EG N      TI L
Sbjct  169  KTIVFRPPSGGDLETTMLHFNCRGEFFYCNTTNLFNST---ENGTMEGLNT-----TIIL  324
```

```
Query  385  PCRIKQIINMWQKVGKAMYAPP  406
            PCRIKQI+NMWQ VG+AMYAPP
Sbjct  325  PCRIKQIVNMWQGVGQAMYAPP  390
```

>Escherichia coli strain EN5192 contig00135, whole genome shotgun sequence

Sequence ID: NZ_JAPMNL010000135.1 Length: 1341

Range 1: 958 to 1335

Score:161 bits(408), Expect:8e-40,

Method:Compositional matrix adjust.,

Identities:83/135(61%), Positives:95/135(70%), Gaps:11/135(8%)

```
Query  305   KWNNTLKQIASKLREQFGNNKTIIFKQSSGGDPEIVTHSFNCGGEFFYCNSTQLFNSTWF  364
             KWN TL++++ KL E F  NKTI F  SSGGD EI THSFNC GEFFYCN++ LFN T+
Sbjct  1335  KWNETLQRVSKKLAEHF-PNKTIKFASSSGGDLEITTHSFNCRGEFFYCNTSGLFNGTYM  1159

Query  365   NSTWSTEGSNNTEG--SDTITLPCRIKQIINMWQKVGKAMYAPPISGQIRCSSNITGLLL  422
                     SN TE   S  IT+PCRIKQIINMWQ+VG+AMYAPPI G I C SNITGLLL
Sbjct  1158  --------SNGTESNSSSIITIPCRIKQIINMWQEVGRAMYAPPIEGNITCKSNITGLLL  1003

Query  423   TRDGGNSNNESEIFR  437
              RDGG   N +E  +
Sbjct  1002  VRDGGTEANTTETLQ  958
```

>Cronobacter sakazakii strain cro3040W contig83, whole genome shotgun sequence

Sequence ID: NZ_NRJO01000083.1 Length: 392

Range 1: 1 to 390

Score:151 bits(382), Expect:1e-39,

Method:Compositional matrix adjust.,

Identities:78/142(55%), Positives:99/142(69%), Gaps:13/142(9%)

```
Query  266  RPNNNTRKRIRIQRGPGRAFVTIGKI-GNMRQAHCNISRAKWNNTLKQIASKLREQFGNN  324
            RPNNNTR  +RI  GPG+ F     I G++R+A+C I+  +W  TL+Q+ +KL+E F NN
Sbjct  1    RPNNNTRTSVRI--GPGQMFYRTEDITGDIRKAYCEINGTEWTETLEQVTAKLKEHF-NN  171

Query  325  KTIIFKQSSGGDPEIVTHSFNCGGEFFYCNSTQLFNSTWFNSTWSTEGSNNTEGSDTITL  384
            KTI+F+  SGGD EI  H FNC GEFFYCN+T+LFNST      + EG N      TI L
Sbjct  172  KTILFRPPSGGDLEITMHHFNCKGEFFYCNTTKLFNST----NGTMEGLNG-----TIIL  324

Query  385  PCRIKQIINMWQKVGKAMYAPP  406
            PCRI+QI+ MWQ+ G+AMYAPP
Sbjct  325  PCRIRQIVRMWQETGQAMYAPP  390
```

>Cronobacter dublinensis strain cro3540W contig49, whole genome shotgun sequence

Sequence ID: NZ_NRNN01000049.1 Length: 392

Range 1: 1 to 390

Score:150 bits(379), Expect:4e-39,

Method:Compositional matrix adjust.,

Identities:79/142(56%), Positives:100/142(70%), Gaps:13/142(9%)

```
Query  266  RPNNNTRKRIRIQRGPGRAFVTIGKI-GNMRQAHCNISRAKWNNTLKQIASKLREQFGNN  324
            RP+NNTR  +RI  GPG+ F     I G++R+A+C I+  +W  TL+Q+ +KL+E F NN
Sbjct  1    RPDNNTRTSVRI--GPGQMFYRTEDITGDIRKAYCEINGTEWTETLEQVTAKLKEHF-NN  171

Query  325  KTIIFKQSSGGDPEIVTHSFNCGGEFFYCNSTQLFNSTWFNSTWSTEGSNNTEGSDTITL  384
            KTI+F+  SGGD EI  H FNC GEFFYCN+T+LFNST  N T   EG N      TI L
Sbjct  172  KTILFRPPSGGDLEITMHHFNCKGEFFYCNTTKLFNST--NGT--MEGLNG-----TIIL  324

Query  385  PCRIKQIINMWQKVGKAMYAPP  406
            PCRI+QI+ MWQ+ G+AMYAPP
Sbjct  325  PCRIRQIVRMWQETGQAMYAPP  390
```

>Cronobacter sakazakii strain cro4191B1-2 contig84, whole genome shotgun sequence

Sequence ID: NZ_NRKG01000084.1 Length: 402

Range 1: 1 to 303

Score:122 bits(306), Expect:2e-34,

Method:Compositional matrix adjust.,

Identities:63/104(61%), Positives:72/104(69%), Gaps:4/104(3%)

```
Query  266  RPNNNTRKRIRIQRGPGRAFVTIGKI-GNMRQAHCNISRAKWNNTLKQIASKLREQFGNN  324
            RPNNNTRK IRI  GPG+ F   G I G++RQAHCNISR  W  TL  ++ KL E F  N
Sbjct  1    RPNNNTRKSIRI--GPGQTFYATGDIIGDIRQAHCNISRKDWEETLHNVSKKLAEHF-QN  171

Query  325  KTIIFKQSSGGDPEIVTHSFNCGGEFFYCNSTQLFNSTWFNSTW  368
            K I F  SSGGD EI THSFNC GEFFYCN++ LFN T+  + W
Sbjct  172  KIIQFASSSGGDLEITTHSFNCRGEFFYCNTSGLFNGTYRYT*W  303
```

Range 2: 299 to 400

Score:58.2 bits(139), Expect:2e-34,

Method:Compositional matrix adjust.,

Identities:27/34(79%), Positives:30/34(88%), Gaps:1/34(2%)

```
Query  374  NNTEG-SDTITLPCRIKQIINMWQKVGKAMYAPP  406
            N TE  S TIT+PCRIKQIINMWQ+VG+AMYAPP
Sbjct  299  NGTESNSTTITIPCRIKQIINMWQEVGRAMYAPP  400
```

>Cronobacter malonaticus strain cro4164A3 contig145, whole genome shotgun sequence

Sequence ID: NZ_NRNF01000145.1 Length: 405

Range 1: 118 to 405

Score:119 bits(299), Expect:2e-34,

Method:Compositional matrix adjust.,

Identities:61/99(62%), Positives:70/99(70%), Gaps:4/99(4%)

```
Query  266  RPNNNTRKRIRIQRGPGRAFVTIGKI-GNMRQAHCNISRAKWNNTLKQIASKLREQFGNN  324
            RP+NNTRK IRI  GPG+ F   G I G++RQAHCNISR  W  TL  ++ KL E F  N
Sbjct  405  RPSNNTRKSIRI--GPGQTFYATGDIIGDIRQAHCNISRKDWEETLHNVSKKLAEHF-QN  235

Query  325  KTIIFKQSSGGDPEIVTHSFNCGGEFFYCNSTQLFNSTW  363
            K I F  SSGGD EI THSFNC GEFFYCN++ LFN T+
Sbjct  234  KKIQFASSSGGDLEITTHSFNCRGEFFYCNTSGLFNKTY  118
```

Range 2: 3 to 101

Score:60.8 bits(146), Expect:2e-34,
Method:Compositional matrix adjust.,
Identities:26/33(79%), Positives:30/33(90%), Gaps:0/33(0%)

```
Query  374  NNTEGSDTITLPCRIKQIINMWQKVGKAMYAPP  406
            N TE + TIT+PCRIKQIINMWQ+VG+AMYAPP
Sbjct  101  NGTESNGTITIPCRIKQIINMWQEVGQAMYAPP  3
```

>Neisseria gonorrhoeae strain GC-206 NODE_162_length_243_cov_2.87234_ID_20496, whole genome shotgun sequence
Sequence ID: NZ_SAWN01000162.1 Length: 243
Range 1: 1 to 243

Score:134 bits(338), Expect:6e-34,
Method:Compositional matrix adjust.,
Identities:62/83(75%), Positives:74/83(89%), Gaps:2/83(2%)

```
Query  392  INMWQKVGKAMYAPPISGQIRCSSNITGLLLTRDGGNSNNESEIFRPGGGDMRDNWRSEL  451
            INMW +VG+A+YAPPI+G I C+S+ITG+LLTRDG    N++E FRPGGGDMRDNWRSEL
Sbjct  243  INMWHEVGQAIYAPPIAGNITCNSSITGILLTRDG--DKNDTETFRPGGGDMRDNWRSEL  70

Query  452  YKYKVVKIEPLGVAPTKAKRRVV  474
            YKYKVV+I+PLGVAPT+AKR VV
Sbjct  69   YKYKVVEIKPLGVAPTEAKRNVV  1
```

>Burkholderia pseudomallei strain 2604STDY5518971 ERS240105SCcontig000123, whole genome shotgun sequence
Sequence ID: NZ_UVFZ02000123.1 Length: 313
Range 1: 2 to 310

Score:133 bits(334), Expect:5e-33,
Method:Compositional matrix adjust.,
Identities:66/109(61%), Positives:82/109(75%), Gaps:6/109(5%)

```
Query  326  TIIFKQSSGGDPEIVTHSFNCGGEFFYCNSTQLFNSTWFNSTWSTEGSNNTEGSDTITLP  385
            TI F+  SGGD EI TH FNC GEFFYCN+T+LF +   ++  ST G N+T    TI +P
Sbjct  2    TISFEPHSGGDLEITTHHFNCRGEFFYCNTTKLFTNNCTDN--STGGCNDT----TIIIP  163

Query  386  CRIKQIINMWQKVGKAMYAPPISGQIRCSSNITGLLLTRDGGNSNNESE  434
            C+I+QI+ MWQ VG+AMYAPPISG+I+C SNITG+LLTRDGG   + SE
Sbjct  164  CKIRQIVRMWQGVGQAMYAPPISGEIKCESNITGILLTRDGGQDKSTSE  310
```

>Cronobacter malonaticus strain cro3263A1 contig73, whole genome shotgun sequence
Sequence ID: NZ_NRMM01000073.1 Length: 406
Range 1: 1 to 333

Score:115 bits(289), Expect:2e-32,
Method:Compositional matrix adjust.,
Identities:64/121(53%), Positives:78/121(64%), Gaps:11/121(9%)

```
Query  266  RPNNNTRKRIRIQRGPGRAFVTIGKI-GNMRQAHCNISRAKWNNTLKQIASKLREQFGNN  324
            RP+NNTR  I I  GPGRAF   G I G++RQA+C ++R +WN  L Q+  KL E F  N
Sbjct  1    RPDNNTRTSINI--GPGRAFYRTGDIIGDIRQAYCEVNRTEWNEILGQVIKKLEEHF-KN  171

Query  325  KTIIFKQSSGGDPEIVTHSFNCGGEFFYCNSTQLFNSTWFNSTWSTEGSNNTEGSDTITL  384
            KTI F+  SGGD EI    FNC GEFFYCN+T+LFNSTW       EG+   +G   +TL
Sbjct  172  KTIKFQPPSGGDLEITMLHFNCRGEFFYCNTTKLFNSTW-------EGNEP*KGVMAVTL  330

Query  385  P  385
            P
Sbjct  331  P  333
```

Range 2: 318 to 404

Score:57.8 bits(138), Expect:2e-32,
Method:Composition-based stats.,
Identities:24/29(83%), Positives:25/29(86%), Gaps:0/29(0%)

```
Query  378  GSDTITLPCRIKQIINMWQKVGKAMYAPP  406
            GSDT  LPCRIKQIINMWQ  G+AMYAPP
Sbjct  318  GSDTTILPCRIKQIINMWQGAGQAMYAPP  404
```

>Klebsiella pneumoniae strain EuSCAPE_MT006, whole genome shotgun sequence
Sequence ID: NZ_UKIU02000080.1 Length: 250
Range 1: 1 to 249

Score:114 bits(286), Expect:2e-26,
Method:Compositional matrix adjust.,
Identities:60/85(71%), Positives:68/85(80%), Gaps:3/85(3%)

```
Query  212  TVQCTHGIRPVVSTQLLLNGSLAEEEVVIRSVNFTDNAKTIIVQLNTSVEINCTRPNNNT  271
            TV CTHGI+PVVSTQLLLNGSLAE+E+VIRS N TDNAKTIIV LN SVEI C RP NNT
Sbjct  249  TVLCTHGIKPVVSTQLLLNGSLAEKEIVIRSENLTDNAKTIIVHLNESVEIVCMRPGNNT  70

Query  272  RKRIRIQRGPGRAFVTIGK-IGNMR  295
            R+ +RI  GPG+ F   G  IG++R
Sbjct  69   RRSMRI--GPGQTFYATGDIIGDIR  1
```

>Neisseria gonorrhoeae strain GC-206 NODE_155_length_258_cov_1.77833_ID_20482, whole genome shotgun sequence
Sequence ID: NZ_SAWN01000155.1 Length: 258
Range 1: 1 to 258

Score:104 bits(259), Expect:1e-22,
Method:Compositional matrix adjust.,
Identities:51/86(59%), Positives:65/86(75%), Gaps:1/86(1%)

```
Query  49   PQEVVLVNVTENFNMWKNDMVEQMHEDIISLWDQSLKPCVKLTPLCVSLKCTDLKNDTNT  108
            PQE+VL NVTENFNMWKN MV+QMHED++SLWDQSLKPCVKLT +CV+L CT +K    T
Sbjct  258  PQEIVLGNVTENFNMWKNYMVDQMHEDVVSLWDQSLKPCVKLTTVCVTLNCTSVKVTNVT  79

Query  109  NSSSGRMIME-KGEIKNCSFNISTSI  133
             S+S +       +++NC+F  +T I
Sbjct  78   ASASVKGTDNITNDMQNCTFTATTEI  1
```

```
>Klebsiella pneumoniae strain EuSCAPE_IL019, whole genome shotgun sequence
Sequence ID: NZ_UKJZ02000064.1 Length: 244
Range 1: 3 to 242

Score:96.7 bits(239), Expect:8e-20,
Method:Compositional matrix adjust.,
Identities:51/82(62%), Positives:58/82(70%), Gaps:2/82(2%)

Query  240  IRSVNFTDNAKTIIVQLNTSVEINCTRPNNNTRKRIRIQRGPGRAFVTIGKIGNMRQAHC  299
            I S N TDNAKTIIV LN SV I CTRP NNTRK IRI  GPG+ F     IG++R+A+C
Sbjct  242  ISSKNLTDNAKTIIVHLNESVTIICTRPGNNTRKSIRI--GPGQTFYANKIIGDIRRAYC  69

Query  300  NISRAKWNNTLKQIASKLREQF  321
            NIS   WN TLKQ+  KL+E F
Sbjct  68   NISGEAWNKTLKQVEEKLKEYF  3




>Nocardioides luteus strain JCM 3358 sequence39, whole genome shotgun sequence
Sequence ID: NZ_BMRK01000039.1 Length: 5754
Range 1: 566 to 700

Score:101 bits(251), Expect:8e-18,
Method:Compositional matrix adjust.,
Identities:45/45(100%), Positives:45/45(100%), Gaps:0/45(0%)

Query  435  IFRPGGGDMRDNWRSELYKYKVVKIEPLGVAPTKAKRRVVQREKR  479
            IFRPGGGDMRDNWRSELYKYKVVKIEPLGVAPTKAKRRVVQREKR
Sbjct  566  IFRPGGGDMRDNWRSELYKYKVVKIEPLGVAPTKAKRRVVQREKR  700




>Streptomyces griseoflavus strain JCM 4479 sequence37, whole genome shotgun sequence
Sequence ID: NZ_BMUC01000037.1 Length: 5775
Range 1: 5048 to 5182

Score:101 bits(251), Expect:8e-18,
Method:Compositional matrix adjust.,
Identities:45/45(100%), Positives:45/45(100%), Gaps:0/45(0%)

Query  435   IFRPGGGDMRDNWRSELYKYKVVKIEPLGVAPTKAKRRVVQREKR  479
             IFRPGGGDMRDNWRSELYKYKVVKIEPLGVAPTKAKRRVVQREKR
Sbjct  5182  IFRPGGGDMRDNWRSELYKYKVVKIEPLGVAPTKAKRRVVQREKR  5048




>Saccharothrix coeruleofusca strain JCM 3313 sequence45, whole genome shotgun sequence
Sequence ID: NZ_BMRG01000045.1 Length: 5775
Range 1: 5048 to 5182

Score:101 bits(251), Expect:8e-18,
Method:Compositional matrix adjust.,
Identities:45/45(100%), Positives:45/45(100%), Gaps:0/45(0%)

Query  435   IFRPGGGDMRDNWRSELYKYKVVKIEPLGVAPTKAKRRVVQREKR  479
             IFRPGGGDMRDNWRSELYKYKVVKIEPLGVAPTKAKRRVVQREKR
Sbjct  5182  IFRPGGGDMRDNWRSELYKYKVVKIEPLGVAPTKAKRRVVQREKR  5048




>Streptomyces griseoloalbus strain JCM 4480 sequence054, whole genome shotgun sequence
Sequence ID: NZ_BMUD01000054.1 Length: 5815
Range 1: 5068 to 5202

Score:101 bits(251), Expect:8e-18,
Method:Compositional matrix adjust.,
Identities:45/45(100%), Positives:45/45(100%), Gaps:0/45(0%)

Query  435   IFRPGGGDMRDNWRSELYKYKVVKIEPLGVAPTKAKRRVVQREKR  479
             IFRPGGGDMRDNWRSELYKYKVVKIEPLGVAPTKAKRRVVQREKR
Sbjct  5202  IFRPGGGDMRDNWRSELYKYKVVKIEPLGVAPTKAKRRVVQREKR  5068




>Actinoplanes ianthinogenes strain JCM 3249 sequence037, whole genome shotgun sequence
Sequence ID: NZ_BMQZ01000037.1 Length: 5815
Range 1: 5068 to 5202

Score:101 bits(251), Expect:8e-18,
Method:Compositional matrix adjust.,
Identities:45/45(100%), Positives:45/45(100%), Gaps:0/45(0%)

Query  435   IFRPGGGDMRDNWRSELYKYKVVKIEPLGVAPTKAKRRVVQREKR  479
             IFRPGGGDMRDNWRSELYKYKVVKIEPLGVAPTKAKRRVVQREKR
Sbjct  5202  IFRPGGGDMRDNWRSELYKYKVVKIEPLGVAPTKAKRRVVQREKR  5068




>Sinomonas sp. KCTC 49339 sequence28, whole genome shotgun sequence
Sequence ID: NZ_BNCM01000028.1 Length: 5849
Range 1: 5039 to 5173

Score:101 bits(251), Expect:8e-18,
Method:Compositional matrix adjust.,
Identities:45/45(100%), Positives:45/45(100%), Gaps:0/45(0%)

Query  435   IFRPGGGDMRDNWRSELYKYKVVKIEPLGVAPTKAKRRVVQREKR  479
             IFRPGGGDMRDNWRSELYKYKVVKIEPLGVAPTKAKRRVVQREKR
Sbjct  5173  IFRPGGGDMRDNWRSELYKYKVVKIEPLGVAPTKAKRRVVQREKR  5039




>Saccharothrix mutabilis subsp. mutabilis strain JCM 3380 sequence46, whole genome shotgun sequence
Sequence ID: NZ_BAAABU010000046.1 Length: 5899
Range 1: 614 to 748

Score:101 bits(251), Expect:8e-18,
Method:Compositional matrix adjust.,
Identities:45/45(100%), Positives:45/45(100%), Gaps:0/45(0%)

Query  435  IFRPGGGDMRDNWRSELYKYKVVKIEPLGVAPTKAKRRVVQREKR  479
            IFRPGGGDMRDNWRSELYKYKVVKIEPLGVAPTKAKRRVVQREKR
Sbjct  614  IFRPGGGDMRDNWRSELYKYKVVKIEPLGVAPTKAKRRVVQREKR  748
```

```
>Corynebacterium glutamicum strain 39 Scaffolds_32_9166, whole genome shotgun sequence
Sequence ID: NZ_JACEQW010000032.1 Length: 8862
Range 1: 4819 to 4953

Score:101 bits(251), Expect:1e-17,
Method:Compositional matrix adjust.,
Identities:45/45(100%), Positives:45/45(100%), Gaps:0/45(0%)

Query  435   IFRPGGGDMRDNWRSELYKYKVVKIEPLGVAPTKAKRRVVQREKR  479
             IFRPGGGDMRDNWRSELYKYKVVKIEPLGVAPTKAKRRVVQREKR
Sbjct  4953  IFRPGGGDMRDNWRSELYKYKVVKIEPLGVAPTKAKRRVVQREKR  4819
```

```
>Corynebacterium glutamicum strain 37 Scaffolds_33_8928, whole genome shotgun sequence
Sequence ID: NZ_JACENO010000033.1 Length: 8928
Range 1: 4793 to 4927

Score:101 bits(251), Expect:1e-17,
Method:Compositional matrix adjust.,
Identities:45/45(100%), Positives:45/45(100%), Gaps:0/45(0%)

Query  435   IFRPGGGDMRDNWRSELYKYKVVKIEPLGVAPTKAKRRVVQREKR  479
             IFRPGGGDMRDNWRSELYKYKVVKIEPLGVAPTKAKRRVVQREKR
Sbjct  4927  IFRPGGGDMRDNWRSELYKYKVVKIEPLGVAPTKAKRRVVQREKR  4793
```

```
>Klebsiella pneumoniae strain EuSCAPE_NO001, whole genome shotgun sequence
Sequence ID: NZ_UKMF02000047.1 Length: 245
Range 1: 3 to 137

Score:89.7 bits(221), Expect:2e-17,
Method:Compositional matrix adjust.,
Identities:40/45(89%), Positives:41/45(91%), Gaps:0/45(0%)

Query  1    KLWVTVYYGVPVWKEATTTLFCASDAKAYDTEVHNVWATHACVPT  45
            K WVTVYYGVPVWKEA TTLFCASDAKAY  EVHN+WATHACVPT
Sbjct  137  KSWVTVYYGVPVWKEAKTTLFCASDAKAYSKEVHNIWATHACVPT  3
```

```
>Klebsiella pneumoniae strain EuSCAPE_NO001, whole genome shotgun sequence
Sequence ID: NZ_UKMF02000044.1 Length: 289
Range 1: 173 to 283

Score:78.6 bits(192), Expect:3e-13,
Method:Composition-based stats.,
Identities:32/37(86%), Positives:36/37(97%), Gaps:0/37(0%)

Query  433  SEIFRPGGGDMRDNWRSELYKYKVVKIEPLGVAPTKA  469
            +E FRPGGGDMRDNWRSELYKYKVV+I+PLG+APTKA
Sbjct  283  NETFRPGGGDMRDNWRSELYKYKVVEIKPLGIAPTKA  173
```

Query #3: SP|P04578|GP41 Query ID: lcl|Query_4184773 Length: 345

Sequences producing significant alignments:

| Description | Scientific Name | Common Name | Taxid | Max Score | Total Score | Query cover | E Value | Per. Ident | Acc. Len | Accession |
|---|---|---|---|---|---|---|---|---|---|---|
| Escherichia coli strain NBE0009 contig00116, whole genome... | Escherichia ... | NA | 562 | 438 | 438 | 92% | 4e-147 | 67.79 | 1713 | NZ_JARBDL0100... |
| Escherichia coli strain EN5177 contig00087, whole genome shotg... | Escherichia ... | NA | 562 | 419 | 419 | 92% | 2e-133 | 64.83 | 3403 | NZ_JAPMNK0100... |
| Escherichia coli strain NBE0004 contig00110, whole genome... | Escherichia ... | NA | 562 | 364 | 364 | 92% | 2e-119 | 61.66 | 1390 | NZ_JARBDH0100... |
| Escherichia coli strain NBE0010 contig00098, whole genome... | Escherichia ... | NA | 562 | 355 | 355 | 79% | 1e-113 | 65.45 | 1912 | NZ_JARBDM0100... |
| Nocardioides luteus strain JCM 3358 sequence39, whole genome... | Nocardioides... | NA | 1844 | 349 | 349 | 62% | 2e-104 | 82.41 | 5754 | NZ_BMRK010000... |
| Streptomyces griseoflavus strain JCM 4479 sequence37, whole... | Streptomyces... | NA | 35619 | 349 | 349 | 62% | 2e-104 | 82.41 | 5775 | NZ_BMUC010000... |
| Saccharothrix coeruleofusca strain JCM 3313 sequence45, whole... | Saccharothri... | NA | 33919 | 349 | 349 | 62% | 2e-104 | 82.41 | 5775 | NZ_BMRG010000... |
| Streptomyces griseoloalbus strain JCM 4480 sequence054, whole... | Streptomyces... | NA | 67303 | 349 | 349 | 62% | 2e-104 | 82.41 | 5815 | NZ_BMUD010000... |
| Actinoplanes ianthinogenes strain JCM 3249 sequence037, whole... | Actinoplanes... | NA | 122358 | 349 | 349 | 62% | 2e-104 | 82.41 | 5815 | NZ_BMQZ010000... |
| Sinomonas sp. KCTC 49339 sequence28, whole genome shotgun... | Sinomonas sp... | NA | 2769422 | 349 | 349 | 62% | 2e-104 | 82.41 | 5849 | NZ_BNCM010000... |
| Saccharothrix mutabilis subsp. mutabilis strain JCM 3380... | Saccharothri... | NA | 66855 | 349 | 349 | 62% | 2e-104 | 82.41 | 5899 | NZ_BAAABU0100... |
| Escherichia coli strain NBE0007 contig00093, whole genome... | Escherichia ... | NA | 562 | 337 | 337 | 77% | 4e-103 | 61.68 | 3013 | NZ_JARBDJ0100... |
| Corynebacterium glutamicum strain 39 Scaffolds_32_9166, whole... | Corynebacter... | NA | 1718 | 344 | 344 | 62% | 1e-102 | 82.41 | 8862 | NZ_JACEQW0100... |
| Corynebacterium glutamicum strain 37 Scaffolds_33_8928, whole... | Corynebacter... | NA | 1718 | 344 | 344 | 62% | 1e-102 | 82.41 | 8928 | NZ_JACENO0100... |
| Escherichia coli strain EN5286 contig00120, whole genome shotg... | Escherichia ... | NA | 562 | 334 | 334 | 79% | 5e-102 | 60.71 | 3033 | NZ_JAPMNM0100... |
| Escherichia coli strain EN5308 contig00096, whole genome shotg... | Escherichia ... | NA | 562 | 327 | 327 | 79% | 2e-99 | 59.64 | 3024 | NZ_JARBDE0100... |
| Escherichia coli strain EN5192 contig00108, whole genome shotg... | Escherichia ... | NA | 562 | 325 | 325 | 77% | 1e-98 | 60.95 | 3032 | NZ_JAPMNL0100... |
| Escherichia coli strain EN5143 contig00071, whole genome shotg... | Escherichia ... | NA | 562 | 323 | 323 | 77% | 2e-98 | 59.85 | 2774 | NZ_JAPMNI0100... |
| Escherichia coli strain NBE0006 contig00106, whole genome... | Escherichia ... | NA | 562 | 317 | 317 | 71% | 3e-98 | 62.90 | 2090 | NZ_JARBDI0100... |
| Escherichia coli strain EN5317 contig00057, whole genome shotg... | Escherichia ... | NA | 562 | 303 | 303 | 73% | 1e-90 | 59.23 | 2867 | NZ_JARBDO0100... |
| Escherichia coli strain NBE0003 contig00096, whole genome... | Escherichia ... | NA | 562 | 298 | 298 | 72% | 1e-88 | 59.53 | 2837 | NZ_JARBDG0100... |
| Escherichia coli strain EN5169 contig00082, whole genome shotg... | Escherichia ... | NA | 562 | 298 | 298 | 72% | 1e-88 | 59.53 | 2837 | NZ_JAPMNJ0100... |
| Escherichia coli strain EN5095 contig00078, whole genome shotg... | Escherichia ... | NA | 562 | 291 | 291 | 71% | 1e-88 | 59.52 | 1972 | NZ_JAPYLY0100... |
| Escherichia coli strain EN5090 contig00081, whole genome shotg... | Escherichia ... | NA | 562 | 293 | 293 | 70% | 4e-87 | 60.49 | 2798 | NZ_JAODTD0100... |
| Escherichia coli strain EN5132 contig00077, whole genome shotg... | Escherichia ... | NA | 562 | 289 | 289 | 71% | 2e-86 | 57.94 | 2498 | NZ_JAPYLZ0100... |
| Escherichia coli strain EN5379 contig00092, whole genome shotg... | Escherichia ... | NA | 562 | 288 | 288 | 70% | 7e-86 | 59.60 | 2592 | NZ_JAPTGJ0100... |
| Escherichia coli strain EN5308 contig00099, whole genome shotg... | Escherichia ... | NA | 562 | 288 | 288 | 70% | 4e-85 | 59.36 | 2827 | NZ_JARBDE0100... |
| Escherichia coli strain EN5192 contig00135, whole genome shotg... | Escherichia ... | NA | 562 | 268 | 268 | 71% | 3e-82 | 55.56 | 1341 | NZ_JAPMNL0100... |
| Escherichia coli strain NBE0004 contig00090, whole genome... | Escherichia ... | NA | 562 | 276 | 276 | 54% | 9e-81 | 71.12 | 2818 | NZ_JARBDH0100... |
| Klebsiella pneumoniae strain EuSCAPE_IL001, whole genome shotg... | Klebsiella p... | NA | 573 | 242 | 242 | 52% | 2e-75 | 64.55 | 599 | NZ_UKJN020000... |
| Escherichia coli strain EN5143 contig00069, whole genome shotg... | Escherichia ... | NA | 562 | 253 | 253 | 55% | 2e-72 | 64.50 | 2688 | NZ_JAPMNI0100... |
| Escherichia coli strain NBE0002 contig00097, whole genome... | Escherichia ... | NA | 562 | 253 | 253 | 55% | 3e-72 | 64.50 | 2717 | NZ_JARBDF0100... |
| Klebsiella pneumoniae strain EuSCAPE_IL006, whole genome shotg... | Klebsiella p... | NA | 573 | 229 | 229 | 51% | 1e-70 | 63.59 | 560 | NZ_UKJR020001... |
| Escherichia coli strain EN5374 contig00094, whole genome shotg... | Escherichia ... | NA | 562 | 248 | 248 | 70% | 2e-70 | 53.09 | 2720 | NZ_JAPTGI0100... |
| Escherichia coli strain EN5076 contig00066, whole genome shotg... | Escherichia ... | NA | 562 | 248 | 248 | 70% | 2e-70 | 53.09 | 2732 | NZ_JAODTC0100... |
| Escherichia coli strain NBE0010 contig00080, whole genome... | Escherichia ... | NA | 562 | 248 | 248 | 70% | 3e-70 | 53.09 | 2743 | NZ_JARBDM0100... |
| Klebsiella pneumoniae strain OK8 jcf7180000003506, whole genom... | Klebsiella p... | NA | 573 | 227 | 227 | 50% | 1e-69 | 64.48 | 577 | NZ_MZXR010000... |
| Aggregatibacter actinomycetemcomitans serotype d str. SA269... | Aggregatibac... | NA | 1434259 | 223 | 223 | 46% | 2e-68 | 74.69 | 504 | NZ_AZTX010004... |
| Escherichia coli strain EN5177 contig00148, whole genome shotg... | Escherichia ... | NA | 562 | 204 | 204 | 52% | 4e-59 | 59.36 | 850 | NZ_JAPMNK0100... |
| Escherichia coli strain NBE0009 contig00153, whole genome... | Escherichia ... | NA | 562 | 199 | 199 | 46% | 8e-58 | 62.65 | 753 | NZ_JARBDL0100... |
| Klebsiella pneumoniae strain EuSCAPE_MT003, whole genome shotg... | Klebsiella p... | NA | 573 | 193 | 193 | 46% | 1e-54 | 60.87 | 964 | NZ_UKIK020000... |
| Klebsiella pneumoniae strain EuSCAPE_IL004, whole genome shotg... | Klebsiella p... | NA | 573 | 184 | 184 | 31% | 9e-54 | 80.00 | 368 | NZ_UKJP020001... |
| Escherichia coli strain NBE0009 contig00151, whole genome... | Escherichia ... | NA | 562 | 189 | 189 | 46% | 1e-53 | 60.24 | 802 | NZ_JARBDL0100... |
| Klebsiella pneumoniae strain EuSCAPE_IL005, whole genome shotg... | Klebsiella p... | NA | 573 | 181 | 181 | 40% | 3e-52 | 63.27 | 455 | NZ_UKJL020000... |
| Escherichia coli strain NBE0006 contig00145, whole genome... | Escherichia ... | NA | 562 | 182 | 182 | 47% | 4e-51 | 56.98 | 738 | NZ_JARBDI0100... |
| Actinoplanes campanulatus strain JCM 3059 sequence095, whole... | Actinoplanes... | NA | 113559 | 177 | 177 | 33% | 3e-50 | 81.03 | 582 | NZ_BMPW010000... |
| Deinococcus aquiradiocola strain JCM 14371 sequence41, whole... | Deinococcus ... | NA | 393059 | 177 | 177 | 33% | 1e-49 | 81.03 | 690 | NZ_BMOE010000... |
| Escherichia coli strain NBE0006 contig00147, whole genome... | Escherichia ... | NA | 562 | 177 | 177 | 43% | 2e-49 | 60.76 | 694 | NZ_JARBDI0100... |
| Enterococcus alcedinis strain CCM 8433 sequence26, whole genom... | Enterococcus... | NA | 1274384 | 177 | 177 | 33% | 5e-49 | 81.03 | 853 | NZ_BMDT010000... |
| Escherichia coli strain NBE0009 contig00162, whole genome... | Escherichia ... | NA | 562 | 165 | 165 | 33% | 2e-46 | 65.52 | 350 | NZ_JARBDL0100... |
| Corynebacterium glutamicum strain 38 Scaffolds_31_4109, whole... | Corynebacter... | NA | 1718 | 179 | 179 | 34% | 2e-45 | 80.51 | 3118 | NZ_JACENN0100... |
| Corynebacterium glutamicum strain 51 Scaffolds_32_4236, whole... | Corynebacter... | NA | 1718 | 179 | 179 | 34% | 2e-45 | 80.51 | 4236 | NZ_JACENK0100... |
| Lactiplantibacillus plantarum strain MH-301 Scaffolds_50_4236,... | Lactiplantib... | NA | 1590 | 179 | 358 | 34% | 2e-45 | 80.51 | 4236 | NZ_JAOYUG0100... |
| Klebsiella pneumoniae strain EuSCAPE_IL022, whole genome shotg... | Klebsiella p... | NA | 573 | 161 | 161 | 32% | 1e-44 | 73.87 | 411 | NZ_UKKD020000... |

| | | | | | | | | | | |
|---|---|---|---|---|---|---|---|---|---|---|
| Klebsiella pneumoniae strain EuSCAPE_MT015, whole genome shotg... | Klebsiella p... | NA | 573 | 155 | 155 | 32% | 8e-43 | 64.29 | 338 | NZ_UKIW020001... |
| Escherichia coli strain EN5308 contig00153, whole genome shotg... | Escherichia ... | NA | 562 | 160 | 160 | 47% | 1e-42 | 54.65 | 786 | NZ_JARBDE0100... |
| Escherichia coli strain EN5143 contig00088, whole genome shotg... | Escherichia ... | NA | 562 | 160 | 160 | 39% | 1e-42 | 63.70 | 833 | NZ_JAPMNI0100... |
| Klebsiella pneumoniae strain EuSCAPE_MT017, whole genome shotg... | Klebsiella p... | NA | 573 | 149 | 149 | 31% | 2e-39 | 67.83 | 513 | NZ_UKJA020000... |
| Klebsiella pneumoniae strain EuSCAPE_NO017, whole genome shotg... | Klebsiella p... | NA | 573 | 143 | 143 | 31% | 8e-37 | 66.96 | 607 | NZ_UKIO020000... |
| Photobacterium rosenbergii strain DP6N14-7 DP6N14-7_ctg0065,... | Photobacteri... | NA | 294936 | 143 | 143 | 31% | 8e-37 | 63.64 | 564 | NZ_JAHVIB0100... |
| Deinococcus aquiradiocola strain JCM 14371 sequence36, whole... | Deinococcus ... | NA | 393059 | 144 | 144 | 21% | 6e-36 | 98.63 | 869 | NZ_BMOE010000... |
| Thalassovita mangrovi strain GS-10 Scaffold21_1, whole genome... | Thalassovita... | NA | 2692236 | 147 | 147 | 21% | 2e-35 | 98.63 | 1440 | NZ_WWEN010000... |
| Klebsiella pneumoniae strain EuSCAPE_DE091, whole genome shotg... | Klebsiella p... | NA | 573 | 139 | 139 | 31% | 3e-35 | 63.41 | 618 | NZ_UKMA020000... |
| Klebsiella pneumoniae strain EuSCAPE_IL018, whole genome shotg... | Klebsiella p... | NA | 573 | 136 | 136 | 35% | 5e-35 | 56.25 | 388 | NZ_UKJU020001... |
| Klebsiella pneumoniae strain EuSCAPE_IL015, whole genome shotg... | Klebsiella p... | NA | 573 | 137 | 137 | 28% | 7e-35 | 63.21 | 532 | NZ_UKJV020000... |
| Bacillus velezensis strain TN14-11 TN14-11_ctg0032, whole geno... | Bacillus vel... | NA | 492670 | 143 | 143 | 31% | 9e-35 | 63.64 | 1125 | NZ_JAHVIU0100... |
| Actinoplanes campanulatus strain JCM 3059 sequence058, whole... | Actinoplanes... | NA | 113559 | 143 | 143 | 21% | 4e-33 | 98.63 | 4754 | NZ_BMPW010000... |
| Klebsiella pneumoniae strain EuSCAPE_MT009, whole genome shotg... | Klebsiella p... | NA | 573 | 130 | 130 | 28% | 8e-33 | 66.00 | 324 | NZ_UKIV020000... |
| Klebsiella pneumoniae strain EuSCAPE_MT015, whole genome shotg... | Klebsiella p... | NA | 573 | 127 | 127 | 31% | 1e-31 | 60.55 | 328 | NZ_UKIW020001... |
| Acinetobacter baumannii strain P4 contig_P4_92, whole genome... | Acinetobacte... | NA | 470 | 133 | 133 | 25% | 6e-31 | 76.40 | 1226 | NZ_JBBXLU0100... |
| Klebsiella pneumoniae strain EuSCAPE_MT009, whole genome shotg... | Klebsiella p... | NA | 573 | 119 | 119 | 30% | 2e-28 | 57.14 | 398 | NZ_UKIV020000... |
| Klebsiella pneumoniae strain EuSCAPE_IL009, whole genome shotg... | Klebsiella p... | NA | 573 | 113 | 113 | 26% | 9e-27 | 63.74 | 274 | NZ_UKJO020000... |
| Klebsiella pneumoniae strain EuSCAPE_IL022, whole genome shotg... | Klebsiella p... | NA | 573 | 110 | 110 | 26% | 1e-25 | 58.24 | 290 | NZ_UKKD020000... |
| Klebsiella pneumoniae strain EuSCAPE_IL004, whole genome shotg... | Klebsiella p... | NA | 573 | 109 | 109 | 25% | 6e-25 | 57.95 | 327 | NZ_UKJP020001... |
| Klebsiella pneumoniae strain EuSCAPE_MT002, whole genome shotg... | Klebsiella p... | NA | 573 | 104 | 104 | 25% | 2e-23 | 63.64 | 265 | NZ_UKIM020000... |
| Klebsiella pneumoniae strain EuSCAPE_IL016, whole genome shotg... | Klebsiella p... | NA | 573 | 105 | 105 | 28% | 3e-23 | 55.67 | 345 | NZ_UKKA020000... |
| Photobacterium rosenbergii strain DP6N14-7 DP6N14-7_ctg0054,... | Photobacteri... | NA | 294936 | 95.9 | 95.9 | 19% | 8e-18 | 70.15 | 960 | NZ_JAHVIB0100... |
| Klebsiella pneumoniae strain EuSCAPE_LT003, whole genome shotg... | Klebsiella p... | NA | 573 | 83.2 | 83.2 | 18% | 3e-15 | 65.08 | 317 | NZ_UKJH020000... |
| Pseudomonas aeruginosa strain VET-121... | Pseudomonas ... | NA | 287 | 82.0 | 82.0 | 12% | 7e-15 | 93.18 | 275 | NZ_QZXB010002... |
| Bacillus velezensis strain TN14-11 TN14-11_ctg0036, whole geno... | Bacillus vel... | NA | 492670 | 71.6 | 71.6 | 14% | 3e-09 | 70.59 | 1101 | NZ_JAHVIU0100... |
| Klebsiella pneumoniae strain EuSCAPE_IL005, whole genome shotg... | Klebsiella p... | NA | 573 | 57.8 | 57.8 | 11% | 1e-05 | 73.17 | 442 | NZ_UKJL020000... |

Alignments:

```
>Escherichia coli strain NBE0009 contig00116, whole genome shotgun sequence
Sequence ID: NZ_JARBDL010000116.1 Length: 1713
Range 1: 308 to 1285

Score:438 bits(1127), Expect:4e-147,
Method:Compositional matrix adjust.,
Identities:248/326(76%), Positives:281/326(86%), Gaps:7/326(2%)

Query  26    LTVQARQLLSGIVQQQNNLLRAIEAQQHLLQLTVWGIKQLQARILAVERYLKDQQLLGIW  85
             LTVQARQLLSGIVQQQ+NLLRAIEAQQHLLQLTVWGIKQLQ R+LA+ERYLKDQQLLGIW
Sbjct  1285  LTVQARQLLSGIVQQQSNLLRAIEAQQHLLQLTVWGIKQLQTRVLALERYLKDQQLLGIW  1106

Query  86    GCSGKLICTTAVPWNASWSNKSLEQIWNHTTWMEWDREINNYTSLIHSLIeesqnqqekn  145
             GCSGKLIC TAVPWN+SWSNK+  +IW++ TWM+WDREI+NYT  I+ L+EESQNQQEKN
Sbjct  1105  GCSGKLICPTAVPWNSSWSNKNQSEIWDNMTWMQWDREISNYTYTIYRLLEESQNQQEKN  926

Query  146   eqellelDKWASLWNWFNITNWLWYIKLFIMIVGGLVGLRIVFAVLSIVNRVRQGYSPLS  205
             E++LL LD WASLWNWFNIT WLWYIK+FIMIVGGL+GLRI+FAVLSIVNRVRQGYSPLS
Sbjct  925   EEDLLALDSWASLWNWFNITQWLWYIKIFIMIVGGLIGLRIIFAVLSIVNRVRQGYSPLS  746

Query  206   FQTHLPTprgpdrpegieeeggerdrdrSIRLVNGSLALIWDDLRSLCLFSYHRLRDLLL  265
             FQT +P+P   D+  GIEEEGGE+DR+RSIRLVNG LAL WDDLR+LCLFSYHRLRD +L
Sbjct  745   FQTPIPSPGELDKLRGIEEEGGEQDRNRSIRLVNGFLALAWDDLRNLCLFSYHRLRDFIL  566

Query  266   IVTRIVELLGR-------RGWEALKYWWNLLQYWSQELKNSAVSLLNATAIAVAEGTDRV  318
             +  R+VELLGR       RGWEALKY  +L+QYW  ELK SA+SL +  AI VAEGTDR+
Sbjct  565   LTARVVELLGRSSLRGLQRGWEALKYLGSLVQYWGLELKKSAISLFDTIAITVAEGTDRI  386

Query  319   IEVVQGACRAIRHIPRRIRQGLERIL  344
             I VVQ  CRAI +IPRR+RQG E  L
Sbjct  385   ISVVQRICRAIYNIPRRLRQGFEAAL  308



>Escherichia coli strain EN5177 contig00087, whole genome shotgun sequence
Sequence ID: NZ_JAPMNK010000087.1 Length: 3403
Range 1: 2119 to 3099

Score:419 bits(1078), Expect:2e-133,
Method:Compositional matrix adjust.,
Identities:236/327(72%), Positives:276/327(84%), Gaps:7/327(2%)

Query  26    LTVQARQLLSGIVQQQNNLLRAIEAQQHLLQLTVWGIKQLQARILAVERYLKDQQLLGIW  85
             LTVQARQLLSGIVQQQNNLLRAIEAQQ++L+LTVWGIKQLQ R+LA+ERYL++Q+LLG+W
Sbjct  2119  LTVQARQLLSGIVQQQNNLLRAIEAQQNMLRLTVWGIKQLQTRVLAIERYLREQRLLGLW  2298

Query  86    GCSGKLICTTAVPWNASWSNKSLEQIWNHTTWMEWDREINNYTSLIHSLIeesqnqqekn  145
             GCS K+ICTT VPWN+SWSNKS   IWN+ TWMEWD+EI  +T+ I+ L+EESQ QQE+N
Sbjct  2299  GCSEKIICTTNVPWNSSWSNKSYSDIWNNLTWMEWDKEIEQHTNTIYQLLEESQGQQEQN  2478

Query  146   eqellelDKWASLWNWFNITNWLWYIKLFIMIVGGLVGLRIVFAVLSIVNRVRQGYSPLS  205
             E++LL LDKW  LW+WF+ITNWLWYIK+FIMIVGGL+GLRIVFAVLSIVNRVRQGYSPLS
Sbjct  2479  EKDLLALDKWNDLWSWFDITNWLWYIKIFIMIVGGLIGLRIVFAVLSIVNRVRQGYSPLS  2658

Query  206   FQTHLPTprgpdrpegieeeggerdrdrSIRLVNGSLALIWDDLRSLCLFSYHRLRDLLL  265
             FQT  P+P G DRP  I+EEGGE+ R+ SIRLV+G LAL WDDLRSL LF YHRL+DL+
Sbjct  2659  FQTLAPSPGGLDRPGRIDEEGGEQGRNSSIRLVSGFLALAWDDLRSLFLFCYHRLKDLIS  2838

Query  266   IVTRIVELLGR-------RGWEALKYWWNLLQYWSQELKNSAVSLLNATAIAVAEGTDRV  318
             +  R VELLGR       RGWEALKY  +L+QYW  ELK SA+SLL+  AIAV EGTDR+
Sbjct  2839  VTARAVELLGRSSLKGLQRGWEALKYLGSLVQYWGLELKKSAISLLDTIAIAVGEGTDRI  3018

Query  319   IEVVQGACRAIRHIPRRIRQGLERILL  345
             IE +QG CRAIR+IPRR+RQG E  LL
Sbjct  3019  IEGIQGLCRAIRNIPRRVRQGFEAALL  3099



>Escherichia coli strain NBE0004 contig00110, whole genome shotgun sequence
Sequence ID: NZ_JARBDH010000110.1 Length: 1390
Range 1: 294 to 1271

Score:364 bits(934), Expect:2e-119,
Method:Compositional matrix adjust.,
Identities:227/326(70%), Positives:265/326(81%), Gaps:7/326(2%)

Query  26    LTVQARQLLSGIVQQQNNLLRAIEAQQHLLQLTVWGIKQLQARILAVERYLKDQQLLGIW  85
             LTVQ RQLLSGIVQQQ+NLL AIEAQQHLLQLTV GIKQLQ R+LA++RYLKDQQLLG+W
Sbjct  1271  LTVQTRQLLSGIVQQQSNLLMAIEAQQHLLQLTV*GIKQLQTRVLALKRYLKDQQLLGMW  1092

Query  86    GCSGKLICTTAVPWNASWSNKSLEQIWNHTTWMEWDREINNYTSLIHSLIeesqnqqekn  145
             GCSGKLICTTAVP N+S SNKS ++I ++ T M+ DREI+NYT  I+ L+E+SQ QQE+N
Sbjct  1091  GCSGKLICTTAVP*NSS*SNKSRDEI*DNMT*MQ*DREISNYTDTIYRLLEDSQIQQERN  912

Query  146   eqellelDKWASLWNWFNITNWLWYIKLFIMIVGGLVGLRIVFAVLSIVNRVRQGYSPLS  205
             E++LL LD   +LWN F IT WLWYIK+FIMIVGGL+ LRI+FAV+SIVNR RQ YSPLS
Sbjct  911   EKDLLALDS*NNLWN*FGITKWLWYIKIFIMIVGGLISLRIIFAVISIVNRFRQRYSPLS  732

Query  206   FQTHLPTprgpdrpegieeeggerdrdrSIRLVNGSLALIWDDLRSLCLFSYHRLRDLLL  265
             FQT  P PRGPDR   IEEEG E+DR+RSIRLVNG LAL  DDL++LCLFSYHRLRD +L
Sbjct  731   FQTLTPNPRGPDRLGRIEEEGREQDRNRSIRLVNGFLALA*DDLQNLCLFSYHRLRDFIL  552
```

```
Query  266   IVTRIVELLGR-------RGWEALKYWWNLLQYWSQELKNSAVSLLNATAIAVAEGTDRV  318
             +  R+VELLGR       R  EALKY  +L+QY   ELK SA+SLL+  AIAV EGTDR+
Sbjct  551   VAARVVELLGRSSLKGLQRR*EALKYLGSLVQY*GLELKGSAISLLDTIAIAVGEGTDRI  372

Query  319   IEVVQGACRAIRHIPRRIRQGLERIL  344
             I +VQG CRA+R+IPRRIRQG E  L
Sbjct  371   ISLVQGICRAVRNIPRRIRQGFEAAL  294
```

>Escherichia coli strain NBE0010 contig00098, whole genome shotgun sequence
Sequence ID: NZ_JARBDM010000098.1 Length: 1912
Range 1: 825 to 1649

Score:355 bits(911), Expect:1e-113,
Method:Compositional matrix adjust.,
Identities:227/275(83%), Positives:251/275(91%), Gaps:0/275(0%)

```
Query  1     AVGIgalflgflgaagstmgaasmtLTVQARQLLSGIVQQQNNLLRAIEAQQHLLQLTVW  60
             AVGIGA+FLGFLGAAGSTMGAAS+TLTVQARQLLSGIVQQQ+NLLRAIEAQQHLLQLTVW
Sbjct  1649  AVGIGAVFLGFLGAAGSTMGAASITLTVQARQLLSGIVQQQSNLLRAIEAQQHLLQLTVW  1470

Query  61    GIKQLQARILAVERYLKDQQLLGIWGCSGKLICTTAVPWNASWSNKSLEQIWNHTTWMEW  120
             GIKQLQ R+LA+ERYLKDQQLLGIWGCSGKLIC TAVPWN+SWSNK+  +IWN+ TWM+W
Sbjct  1469  GIKQLQTRVLAIERYLKDQQLLGIWGCSGKLICPTAVPWNSSWSNKNQSEIWNNMTWMQW  1290

Query  121   DREINNYTSLIHSLIeesqnqqekneqellelDKWASLWNWFNITNWLWYIKLFIMIVGG  180
             DREI+NYT  I+ L+EESQNQQEKNE++LL LD WASLWNWFNIT WLWYIK+FIMIVGG
Sbjct  1289  DREISNYTYTIYRLLEESQNQQEKNEEDLLALDSWASLWNWFNITQWLWYIKIFIMIVGG  1110

Query  181   LVGLRIVFAVLSIVNRVRQGYSPLSFQTHLPTprgpdrpegieeeggerdrdrSIRLVNG  240
             L+GLRI+FAVLSIVNRVRQGYSPLSFQT  P P   DR  GIEEEGGE+DR+RSIRLVNG
Sbjct  1109  LIGLRIIFAVLSIVNRVRQGYSPLSFQTPTPNPGELDRLRGIEEEGGEQDRNRSIRLVNG  930

Query  241   SLALIWDDLRSLCLFSYHRLRDLLLIVTRIVELLG  275
              LAL WDDLR+LCLFSYHRLRD +L+  R+VELLG
Sbjct  929   FLALAWDDLRNLCLFSYHRLRDFILLTARVVELLG  825
```

>Nocardioides luteus strain JCM 3358 sequence39, whole genome shotgun sequence
Sequence ID: NZ_BMRK01000039.1 Length: 5754
Range 1: 776 to 1423

Score:349 bits(896), Expect:2e-104,
Method:Compositional matrix adjust.,
Identities:216/216(100%), Positives:216/216(100%), Gaps:0/216(0%)

```
Query  26    LTVQARQLLSGIVQQQNNLLRAIEAQQHLLQLTVWGIKQLQARILAVERYLKDQQLLGIW  85
             LTVQARQLLSGIVQQQNNLLRAIEAQQHLLQLTVWGIKQLQARILAVERYLKDQQLLGIW
Sbjct  776   LTVQARQLLSGIVQQQNNLLRAIEAQQHLLQLTVWGIKQLQARILAVERYLKDQQLLGIW  955

Query  86    GCSGKLICTTAVPWNASWSNKSLEQIWNHTTWMEWDREINNYTSLIHSLIeesqnqqekn  145
             GCSGKLICTTAVPWNASWSNKSLEQIWNHTTWMEWDREINNYTSLIHSLIEESQNQQEKN
Sbjct  956   GCSGKLICTTAVPWNASWSNKSLEQIWNHTTWMEWDREINNYTSLIHSLIEESQNQQEKN  1135

Query  146   eqellelDKWASLWNWFNITNWLWYIKLFIMIVGGLVGLRIVFAVLSIVNRVRQGYSPLS  205
             EQELLELDKWASLWNWFNITNWLWYIKLFIMIVGGLVGLRIVFAVLSIVNRVRQGYSPLS
Sbjct  1136  EQELLELDKWASLWNWFNITNWLWYIKLFIMIVGGLVGLRIVFAVLSIVNRVRQGYSPLS  1315

Query  206   FQTHLPTprgpdrpegieeeggerdrdrSIRLVNGS  241
             FQTHLPTPRGPDRPEGIEEEGGERDRDRSIRLVNGS
Sbjct  1316  FQTHLPTPRGPDRPEGIEEEGGERDRDRSIRLVNGS  1423
```

>Streptomyces griseoflavus strain JCM 4479 sequence37, whole genome shotgun sequence
Sequence ID: NZ_BMUC01000037.1 Length: 5775
Range 1: 4325 to 4972

Score:349 bits(896), Expect:2e-104,
Method:Compositional matrix adjust.,
Identities:216/216(100%), Positives:216/216(100%), Gaps:0/216(0%)

```
Query  26    LTVQARQLLSGIVQQQNNLLRAIEAQQHLLQLTVWGIKQLQARILAVERYLKDQQLLGIW  85
             LTVQARQLLSGIVQQQNNLLRAIEAQQHLLQLTVWGIKQLQARILAVERYLKDQQLLGIW
Sbjct  4972  LTVQARQLLSGIVQQQNNLLRAIEAQQHLLQLTVWGIKQLQARILAVERYLKDQQLLGIW  4793

Query  86    GCSGKLICTTAVPWNASWSNKSLEQIWNHTTWMEWDREINNYTSLIHSLIeesqnqqekn  145
             GCSGKLICTTAVPWNASWSNKSLEQIWNHTTWMEWDREINNYTSLIHSLIEESQNQQEKN
Sbjct  4792  GCSGKLICTTAVPWNASWSNKSLEQIWNHTTWMEWDREINNYTSLIHSLIEESQNQQEKN  4613

Query  146   eqellelDKWASLWNWFNITNWLWYIKLFIMIVGGLVGLRIVFAVLSIVNRVRQGYSPLS  205
             EQELLELDKWASLWNWFNITNWLWYIKLFIMIVGGLVGLRIVFAVLSIVNRVRQGYSPLS
Sbjct  4612  EQELLELDKWASLWNWFNITNWLWYIKLFIMIVGGLVGLRIVFAVLSIVNRVRQGYSPLS  4433

Query  206   FQTHLPTprgpdrpegieeeggerdrdrSIRLVNGS  241
             FQTHLPTPRGPDRPEGIEEEGGERDRDRSIRLVNGS
Sbjct  4432  FQTHLPTPRGPDRPEGIEEEGGERDRDRSIRLVNGS  4325
```

>Saccharothrix coeruleofusca strain JCM 3313 sequence45, whole genome shotgun sequence
Sequence ID: NZ_BMRG01000045.1 Length: 5775
Range 1: 4325 to 4972

Score:349 bits(896), Expect:2e-104,
Method:Compositional matrix adjust.,
Identities:216/216(100%), Positives:216/216(100%), Gaps:0/216(0%)

```
Query  26    LTVQARQLLSGIVQQQNNLLRAIEAQQHLLQLTVWGIKQLQARILAVERYLKDQQLLGIW  85
             LTVQARQLLSGIVQQQNNLLRAIEAQQHLLQLTVWGIKQLQARILAVERYLKDQQLLGIW
Sbjct  4972  LTVQARQLLSGIVQQQNNLLRAIEAQQHLLQLTVWGIKQLQARILAVERYLKDQQLLGIW  4793

Query  86    GCSGKLICTTAVPWNASWSNKSLEQIWNHTTWMEWDREINNYTSLIHSLIeesqnqqekn  145
             GCSGKLICTTAVPWNASWSNKSLEQIWNHTTWMEWDREINNYTSLIHSLIEESQNQQEKN
Sbjct  4792  GCSGKLICTTAVPWNASWSNKSLEQIWNHTTWMEWDREINNYTSLIHSLIEESQNQQEKN  4613

Query  146   eqellelDKWASLWNWFNITNWLWYIKLFIMIVGGLVGLRIVFAVLSIVNRVRQGYSPLS  205
             EQELLELDKWASLWNWFNITNWLWYIKLFIMIVGGLVGLRIVFAVLSIVNRVRQGYSPLS
Sbjct  4612  EQELLELDKWASLWNWFNITNWLWYIKLFIMIVGGLVGLRIVFAVLSIVNRVRQGYSPLS  4433

Query  206   FQTHLPTprgpdrpegieeeggerdrdrSIRLVNGS  241
             FQTHLPTPRGPDRPEGIEEEGGERDRDRSIRLVNGS
Sbjct  4432  FQTHLPTPRGPDRPEGIEEEGGERDRDRSIRLVNGS  4325
```

```
>Streptomyces griseoloalbus strain JCM 4480 sequence054, whole genome shotgun sequence
Sequence ID: NZ_BMUD01000054.1 Length: 5815
Range 1: 4345 to 4992

Score:349 bits(896), Expect:2e-104,
Method:Compositional matrix adjust.,
Identities:216/216(100%), Positives:216/216(100%), Gaps:0/216(0%)

Query  26    LTVQARQLLSGIVQQQNNLLRAIEAQQHLLQLTVWGIKQLQARILAVERYLKDQQLLGIW  85
             LTVQARQLLSGIVQQQNNLLRAIEAQQHLLQLTVWGIKQLQARILAVERYLKDQQLLGIW
Sbjct  4992  LTVQARQLLSGIVQQQNNLLRAIEAQQHLLQLTVWGIKQLQARILAVERYLKDQQLLGIW  4813

Query  86    GCSGKLICTTAVPWNASWSNKSLEQIWNHTTWMEWDREINNYTSLIHSLIeesqnqqekn  145
             GCSGKLICTTAVPWNASWSNKSLEQIWNHTTWMEWDREINNYTSLIHSLIEESQNQQEKN
Sbjct  4812  GCSGKLICTTAVPWNASWSNKSLEQIWNHTTWMEWDREINNYTSLIHSLIEESQNQQEKN  4633

Query  146   eqellelDKWASLWNWFNITNWLWYIKLFIMIVGGLVGLRIVFAVLSIVNRVRQGYSPLS  205
             EQELLELDKWASLWNWFNITNWLWYIKLFIMIVGGLVGLRIVFAVLSIVNRVRQGYSPLS
Sbjct  4632  EQELLELDKWASLWNWFNITNWLWYIKLFIMIVGGLVGLRIVFAVLSIVNRVRQGYSPLS  4453

Query  206   FQTHLPTprgpdrpegieeeggerdrdrSIRLVNGS  241
             FQTHLPTPRGPDRPEGIEEEGGERDRDRSIRLVNGS
Sbjct  4452  FQTHLPTPRGPDRPEGIEEEGGERDRDRSIRLVNGS  4345
```

```
>Actinoplanes ianthinogenes strain JCM 3249 sequence037, whole genome shotgun sequence
Sequence ID: NZ_BMQZ01000037.1 Length: 5815
Range 1: 4345 to 4992

Score:349 bits(896), Expect:2e-104,
Method:Compositional matrix adjust.,
Identities:216/216(100%), Positives:216/216(100%), Gaps:0/216(0%)

Query  26    LTVQARQLLSGIVQQQNNLLRAIEAQQHLLQLTVWGIKQLQARILAVERYLKDQQLLGIW  85
             LTVQARQLLSGIVQQQNNLLRAIEAQQHLLQLTVWGIKQLQARILAVERYLKDQQLLGIW
Sbjct  4992  LTVQARQLLSGIVQQQNNLLRAIEAQQHLLQLTVWGIKQLQARILAVERYLKDQQLLGIW  4813

Query  86    GCSGKLICTTAVPWNASWSNKSLEQIWNHTTWMEWDREINNYTSLIHSLIeesqnqqekn  145
             GCSGKLICTTAVPWNASWSNKSLEQIWNHTTWMEWDREINNYTSLIHSLIEESQNQQEKN
Sbjct  4812  GCSGKLICTTAVPWNASWSNKSLEQIWNHTTWMEWDREINNYTSLIHSLIEESQNQQEKN  4633

Query  146   eqellelDKWASLWNWFNITNWLWYIKLFIMIVGGLVGLRIVFAVLSIVNRVRQGYSPLS  205
             EQELLELDKWASLWNWFNITNWLWYIKLFIMIVGGLVGLRIVFAVLSIVNRVRQGYSPLS
Sbjct  4632  EQELLELDKWASLWNWFNITNWLWYIKLFIMIVGGLVGLRIVFAVLSIVNRVRQGYSPLS  4453

Query  206   FQTHLPTprgpdrpegieeeggerdrdrSIRLVNGS  241
             FQTHLPTPRGPDRPEGIEEEGGERDRDRSIRLVNGS
Sbjct  4452  FQTHLPTPRGPDRPEGIEEEGGERDRDRSIRLVNGS  4345
```

```
>Sinomonas sp. KCTC 49339 sequence28, whole genome shotgun sequence
Sequence ID: NZ_BNCM01000028.1 Length: 5849
Range 1: 4316 to 4963

Score:349 bits(896), Expect:2e-104,
Method:Compositional matrix adjust.,
Identities:216/216(100%), Positives:216/216(100%), Gaps:0/216(0%)

Query  26    LTVQARQLLSGIVQQQNNLLRAIEAQQHLLQLTVWGIKQLQARILAVERYLKDQQLLGIW  85
             LTVQARQLLSGIVQQQNNLLRAIEAQQHLLQLTVWGIKQLQARILAVERYLKDQQLLGIW
Sbjct  4963  LTVQARQLLSGIVQQQNNLLRAIEAQQHLLQLTVWGIKQLQARILAVERYLKDQQLLGIW  4784

Query  86    GCSGKLICTTAVPWNASWSNKSLEQIWNHTTWMEWDREINNYTSLIHSLIeesqnqqekn  145
             GCSGKLICTTAVPWNASWSNKSLEQIWNHTTWMEWDREINNYTSLIHSLIEESQNQQEKN
Sbjct  4783  GCSGKLICTTAVPWNASWSNKSLEQIWNHTTWMEWDREINNYTSLIHSLIEESQNQQEKN  4604

Query  146   eqellelDKWASLWNWFNITNWLWYIKLFIMIVGGLVGLRIVFAVLSIVNRVRQGYSPLS  205
             EQELLELDKWASLWNWFNITNWLWYIKLFIMIVGGLVGLRIVFAVLSIVNRVRQGYSPLS
Sbjct  4603  EQELLELDKWASLWNWFNITNWLWYIKLFIMIVGGLVGLRIVFAVLSIVNRVRQGYSPLS  4424

Query  206   FQTHLPTprgpdrpegieeeggerdrdrSIRLVNGS  241
             FQTHLPTPRGPDRPEGIEEEGGERDRDRSIRLVNGS
Sbjct  4423  FQTHLPTPRGPDRPEGIEEEGGERDRDRSIRLVNGS  4316
```

```
>Saccharothrix mutabilis subsp. mutabilis strain JCM 3380 sequence46, whole genome shotgun sequence
Sequence ID: NZ_BAAABU010000046.1 Length: 5899
Range 1: 824 to 1471

Score:349 bits(896), Expect:2e-104,
Method:Compositional matrix adjust.,
Identities:216/216(100%), Positives:216/216(100%), Gaps:0/216(0%)

Query  26    LTVQARQLLSGIVQQQNNLLRAIEAQQHLLQLTVWGIKQLQARILAVERYLKDQQLLGIW  85
             LTVQARQLLSGIVQQQNNLLRAIEAQQHLLQLTVWGIKQLQARILAVERYLKDQQLLGIW
Sbjct  824   LTVQARQLLSGIVQQQNNLLRAIEAQQHLLQLTVWGIKQLQARILAVERYLKDQQLLGIW  1003

Query  86    GCSGKLICTTAVPWNASWSNKSLEQIWNHTTWMEWDREINNYTSLIHSLIeesqnqqekn  145
             GCSGKLICTTAVPWNASWSNKSLEQIWNHTTWMEWDREINNYTSLIHSLIEESQNQQEKN
Sbjct  1004  GCSGKLICTTAVPWNASWSNKSLEQIWNHTTWMEWDREINNYTSLIHSLIEESQNQQEKN  1183

Query  146   eqellelDKWASLWNWFNITNWLWYIKLFIMIVGGLVGLRIVFAVLSIVNRVRQGYSPLS  205
             EQELLELDKWASLWNWFNITNWLWYIKLFIMIVGGLVGLRIVFAVLSIVNRVRQGYSPLS
Sbjct  1184  EQELLELDKWASLWNWFNITNWLWYIKLFIMIVGGLVGLRIVFAVLSIVNRVRQGYSPLS  1363

Query  206   FQTHLPTprgpdrpegieeeggerdrdrSIRLVNGS  241
             FQTHLPTPRGPDRPEGIEEEGGERDRDRSIRLVNGS
Sbjct  1364  FQTHLPTPRGPDRPEGIEEEGGERDRDRSIRLVNGS  1471
```

```
>Escherichia coli strain NBE0007 contig00093, whole genome shotgun sequence
Sequence ID: NZ_JARBDJ010000093.1 Length: 3013
Range 1: 289 to 1110

Score:337 bits(864), Expect:4e-103,
Method:Compositional matrix adjust.,
Identities:196/274(72%), Positives:230/274(83%), Gaps:7/274(2%)

Query  79    QQLLGIWGCSGKLICTTAVPWNASWSNKSLEQIWNHTTWMEWDREINNYTSLIHSLIees  138
             Q+LLG+WGCS K+ICTT VPWN+SWSNKS   IWN+ TWMEWD+EI+ +T+ I+ L+EES
Sbjct  1110  QRLLGLWGCSEKIICTTNVPWNSSWSNKSYSDIWNNLTWMEWDKEIDKHTNTIYQLLEES  931

Query  139   qnqqekneqellelDKWASLWNWFNITNWLWYIKLFIMIVGGLVGLRIVFAVLSIVNRVR  198
             Q QQE+NE++LL LDKW  LW+WF+ITNWLWYIK+FIMIVGGL+GLRIVFAVLSIVNRVR
Sbjct  930   QGQQEQNEKDLLALDKWNDLWSWFDITNWLWYIKIFIMIVGGLIGLRIVFAVLSIVNRVR  751
```

```
Query  199   QGYSPLSFQTHLPTprgpdrpegieeeggerdrdrSIRLVNGSLALIWDDLRSLCLFSYH  258
             QGYSPLSFQT  P+P G DRP  IEEEGGE+D+DRSIRLV+G LAL WDDLR+LCLFSYH
Sbjct  750   QGYSPLSFQTLAPSPGGLDRPGRIEEEGGEQDKDRSIRLVSGFLALAWDDLRNLCLFSYH  571

Query  259   RLRDLLLIVTRIVELLGR-------RGWEALKYWWNLLQYWSQELKNSAVSLLNATAIAV  311
             RLRD +L+  R+VELLGR       +GWEALKY  +L+QYW  ELK SA+SLL+  AIAV
Sbjct  570   RLRDCILVTARVVELLGRNSLRGLQKGWEALKYLGSLVQYWGLELKKSAISLLDTIAIAV  391

Query  312   AEGTDRVIEVVQGACRAIRHIPRRIRQGLERILL  345
              EGTDR+IE VQG CRAIR+IPRR+RQG E  LL
Sbjct  390   GEGTDRIIEGVQGLCRAIRNIPRRVRQGFEAALL  289
```

>Corynebacterium glutamicum strain 39 Scaffolds_32_9166, whole genome shotgun sequence
Sequence ID: NZ_JACEQW010000032.1 Length: 8862
Range 1: 4105 to 4743

Score:344 bits(883), Expect:1e-102,
Method:Compositional matrix adjust.,
Identities:203/216(94%), Positives:206/216(95%), Gaps:3/216(1%)

```
Query  26    LTVQARQLLSGIVQQQNNLLRAIEAQQHLLQLTVWGIKQLQARILAVERYLKDQQLLGIW  85
             LTVQARQLLSGIVQQQNNLLRAIEAQQHLLQLTVWGIKQLQARILAVERYLKDQQLLGIW
Sbjct  4743  LTVQARQLLSGIVQQQNNLLRAIEAQQHLLQLTVWGIKQLQARILAVERYLKDQQLLGIW  4564

Query  86    GCSGKLICTTAVPWNASWSNKSLEQIWNHTTWMEWDREINNYTSLIHSLIeesqnqqekn  145
             GCSGKLICTTAVPWNASWSNKSLEQIWNHTTWMEWDREINNYTSLIHSLI   +  Q +
Sbjct  4563  GCSGKLICTTAVPWNASWSNKSLEQIWNHTTWMEWDREINNYTSLIHSLI---EESQNQQ  4393

Query  146   eqellelDKWASLWNWFNITNWLWYIKLFIMIVGGLVGLRIVFAVLSIVNRVRQGYSPLS  205
             E+   ELDKWASLWNWFNITNWLWYIKLFIMIVGGLVGLRIVFAVLSIVNRVRQGYSPLS
Sbjct  4392  EKNEQELDKWASLWNWFNITNWLWYIKLFIMIVGGLVGLRIVFAVLSIVNRVRQGYSPLS  4213

Query  206   FQTHLPTprgpdrpegieeeggerdrdrSIRLVNGS  241
             FQTHLPTPRGPDRPEGIEEEGGERDRDRSIRLVNGS
Sbjct  4212  FQTHLPTPRGPDRPEGIEEEGGERDRDRSIRLVNGS  4105
```

>Corynebacterium glutamicum strain 37 Scaffolds_33_8928, whole genome shotgun sequence
Sequence ID: NZ_JACENO010000033.1 Length: 8928
Range 1: 4079 to 4717

Score:344 bits(883), Expect:1e-102,
Method:Compositional matrix adjust.,
Identities:203/216(94%), Positives:206/216(95%), Gaps:3/216(1%)

```
Query  26    LTVQARQLLSGIVQQQNNLLRAIEAQQHLLQLTVWGIKQLQARILAVERYLKDQQLLGIW  85
             LTVQARQLLSGIVQQQNNLLRAIEAQQHLLQLTVWGIKQLQARILAVERYLKDQQLLGIW
Sbjct  4717  LTVQARQLLSGIVQQQNNLLRAIEAQQHLLQLTVWGIKQLQARILAVERYLKDQQLLGIW  4538

Query  86    GCSGKLICTTAVPWNASWSNKSLEQIWNHTTWMEWDREINNYTSLIHSLIeesqnqqekn  145
             GCSGKLICTTAVPWNASWSNKSLEQIWNHTTWMEWDREINNYTSLIHSLI   +  Q +
Sbjct  4537  GCSGKLICTTAVPWNASWSNKSLEQIWNHTTWMEWDREINNYTSLIHSLI---EESQNQQ  4367

Query  146   eqellelDKWASLWNWFNITNWLWYIKLFIMIVGGLVGLRIVFAVLSIVNRVRQGYSPLS  205
             E+   ELDKWASLWNWFNITNWLWYIKLFIMIVGGLVGLRIVFAVLSIVNRVRQGYSPLS
Sbjct  4366  EKNEQELDKWASLWNWFNITNWLWYIKLFIMIVGGLVGLRIVFAVLSIVNRVRQGYSPLS  4187

Query  206   FQTHLPTprgpdrpegieeeggerdrdrSIRLVNGS  241
             FQTHLPTPRGPDRPEGIEEEGGERDRDRSIRLVNGS
Sbjct  4186  FQTHLPTPRGPDRPEGIEEEGGERDRDRSIRLVNGS  4079
```

>Escherichia coli strain EN5286 contig00120, whole genome shotgun sequence
Sequence ID: NZ_JAPMNM010000120.1 Length: 3033
Range 1: 1893 to 2729

Score:334 bits(856), Expect:5e-102,
Method:Compositional matrix adjust.,
Identities:194/280(69%), Positives:227/280(81%), Gaps:8/280(2%)

```
Query  73    ERYLKDQQLLGIWGCSGKLICTTAVPWNASWSNKSLEQIWNHTTWMEWDREINNYTSLIH  132
             ER+ ++Q+LLG+WGCS + ICTT VPWN+SWSNKS   IWN+ TWMEWD+EI  +T+ I+
Sbjct  1893  ERH-REQRLLGLWGCSERTICTTNVPWNSSWSNKSYSDIWNNLTWMEWDKEIEKHTNTIY  2069

Query  133   SLIeesqnqqekneqellelDKWASLWNWFNITNWLWYIKLFIMIVGGLVGLRIVFAVLS  192
              L+EESQ QQE+NE++LL LDKW  LW+WF+ITNWLWYIK+FIMIVGGL+GLRIVFAVLS
Sbjct  2070  QLLEESQGQQEQNEKDLLALDKWNDLWSWFDITNWLWYIKIFIMIVGGLIGLRIVFAVLS  2249

Query  193   IVNRVRQGYSPLSFQTHLPTprgpdrpegieeeggerdrdrSIRLVNGSLALIWDDLRSL  252
             IVNRVRQGYSPLSFQT  P+P G DRP  IEE GGE+ R  SIRLV+G LAL WDDLRSL
Sbjct  2250  IVNRVRQGYSPLSFQTLAPSPGGLDRPGRIEEGGGEQGRSSSIRLVSGFLALAWDDLRSL  2429

Query  253   CLFSYHRLRDLLLIVTRIVELLGR-------RGWEALKYWWNLLQYWSQELKNSAVSLLN  305
             CLF YHRL+DL+ +  R VELLGR       RGWEALKY  +L+QYW  ELK SA+SLL+
Sbjct  2430  CLFCYHRLKDLISVTARAVELLGRSSLKGLQRGWEALKYLGSLVQYWGLELKKSAISLLD  2609

Query  306   ATAIAVAEGTDRVIEVVQGACRAIRHIPRRIRQGLERILL  345
               AIAV EGTDR+IE VQG CRAIR+IPRR+RQG E  LL
Sbjct  2610  TIAIAVGEGTDRIIEGVQGLCRAIRNIPRRVRQGFEAALL  2729
```

>Escherichia coli strain EN5308 contig00096, whole genome shotgun sequence
Sequence ID: NZ_JARBDE010000096.1 Length: 3024
Range 1: 302 to 1138

Score:327 bits(838), Expect:2e-99,
Method:Compositional matrix adjust.,
Identities:190/280(68%), Positives:227/280(81%), Gaps:8/280(2%)

```
Query  73    ERYLKDQQLLGIWGCSGKLICTTAVPWNASWSNKSLEQIWNHTTWMEWDREINNYTSLIH  132
             ER+ ++Q+LLG+WGCS + ICTT VPWN+SWSNKS   IWN+ TWMEWD+EI  +T+ I+
Sbjct  1138  ERH-REQRLLGLWGCSERTICTTNVPWNSSWSNKSYSDIWNNLTWMEWDKEIEQHTNTIY  962

Query  133   SLIeesqnqqekneqellelDKWASLWNWFNITNWLWYIKLFIMIVGGLVGLRIVFAVLS  192
              L+EESQ QQE+NE++LL LDKW  LW+WF+ITNWLWYI++FIMIVGGL+GLRIVFAVLS
Sbjct  961   QLLEESQGQQEQNEKDLLALDKWNDLWSWFDITNWLWYIRIFIMIVGGLIGLRIVFAVLS  782

Query  193   IVNRVRQGYSPLSFQTHLPTprgpdrpegieeeggerdrdrSIRLVNGSLALIWDDLRSL  252
             IVNRVRQGYSPLSFQT  P+P G DRP  I+ EGGE+ R+ S RLV+G LAL WDDLRSL
Sbjct  781   IVNRVRQGYSPLSFQTLAPSPGGLDRPGRIDEGGEQGRNNSTRLVSGFLALAWDDLRSL  602

Query  253   CLFSYHRLRDLLLIVTRIVELLGR-------RGWEALKYWWNLLQYWSQELKNSAVSLLN  305
              LF YHRL+DL+ +  R+VELLGR       RGWEALKY  +L+QYW  ELK SA+SLL+
Sbjct  601   FLFCYHRLKDLISVTARVVELLGRSSLKGLQRGWEALKYLGSLVQYWGLELKKSAISLLD  422
```

```
Query  306   ATAIAVAEGTDRVIEVVQGACRAIRHIPRRIRQGLERILL  345
               AIAV EGTDR+IE VQG CRAIR+IPRR+RQG E  LL
Sbjct  421   TIAIAVGEGTDRIIEGVQGLCRAIRNIPRRVRQGFEAALL  302
```

>Escherichia coli strain EN5192 contig00108, whole genome shotgun sequence
Sequence ID: NZ_JAPMNL010000108.1 Length: 3032
Range 1: 1939 to 2760

Score:325 bits(834), Expect:1e-98,
Method:Compositional matrix adjust.,
Identities:191/274(70%), Positives:223/274(81%), Gaps:7/274(2%)

```
Query  79    QQLLGIWGCSGKLICTTAVPWNASWSNKSLEQIWNHTTWMEWDREINNYTSLIHSLIees  138
             Q+LLG+WGCS + ICTT VPWN+SWSN+S   IWN+ TWMEWD+EI  +T+ I+ L+EES
Sbjct  1939  QRLLGLWGCSERTICTTNVPWNSSWSNRSYSDIWNNLTWMEWDKEIEQHTNTIYQLLEES  2118

Query  139   qnqqekneqellelDKWASLWNWFNITNWLWYIKLFIMIVGGLVGLRIVFAVLSIVNRVR  198
             Q QQE+NE++LL LDKW  LW+WF+ITNWLWYIK+FIMIVGGL+GLRIVFAVLSIVNRVR
Sbjct  2119  QGQQEQNEKDLLALDKWNDLWSWFDITNWLWYIKIFIMIVGGLIGLRIVFAVLSIVNRVR  2298

Query  199   QGYSPLSFQTHLPTprgpdrpegieeeggerdrdrSIRLVNGSLALIWDDLRSLCLFSYH  258
             QGYSPLSFQT  P+P G DRP  IEE GGE+ R+ SIRLV+G LAL W+DLRSL LF YH
Sbjct  2299  QGYSPLSFQTLAPSPGGLDRPGRIEEGGGEQGRNSSIRLVSGFLALAWEDLRSLFLFCYH  2478

Query  259   RLRDLLLIVTRIVELLGR-------RGWEALKYWWNLLQYWSQELKNSAVSLLNATAIAV  311
             RL+DL+ + TR VELLGR       RGWEALKY  +L+QYW  ELK SA+SLL+  AIAV
Sbjct  2479  RLKDLISVTTRAVELLGRSSLKGLQRGWEALKYLGSLVQYWVLELKKSAISLLDTIAIAV  2658

Query  312   AEGTDRVIEVVQGACRAIRHIPRRIRQGLERILL  345
              EGTDR+IE VQG CRAIR+IPRRIRQG E  LL
Sbjct  2659  GEGTDRIIEGVQGFCRAIRNIPRRIRQGFEAALL  2760
```

>Escherichia coli strain EN5143 contig00071, whole genome shotgun sequence
Sequence ID: NZ_JAPMNI010000071.1 Length: 2774
Range 1: 1652 to 2473

Score:323 bits(827), Expect:2e-98,
Method:Compositional matrix adjust.,
Identities:188/274(69%), Positives:222/274(81%), Gaps:7/274(2%)

```
Query  79    QQLLGIWGCSGKLICTTAVPWNASWSNKSLEQIWNHTTWMEWDREINNYTSLIHSLIees  138
             Q+LLG+WGCS + ICTT VPWN+SWSN+S   IWN+ TWMEWD+EI  +T+ I+ L+EES
Sbjct  1652  QRLLGLWGCSERTICTTNVPWNSSWSNRSYSDIWNNLTWMEWDKEIEQHTNTIYQLLEES  1831

Query  139   qnqqekneqellelDKWASLWNWFNITNWLWYIKLFIMIVGGLVGLRIVFAVLSIVNRVR  198
             Q QQE+NE++LL LDKW  LW+WF+ITNWLWYIK+FIMIVGGL+GLRIVFAVLSIVNRVR
Sbjct  1832  QGQQEQNEKDLLALDKWNDLWSWFDITNWLWYIKIFIMIVGGLIGLRIVFAVLSIVNRVR  2011

Query  199   QGYSPLSFQTHLPTprgpdrpegieeeggerdrdrSIRLVNGSLALIWDDLRSLCLFSYH  258
             QGYSPLSFQT  P+P G DRP  I+EEGGE+ R+ S RLV+G LAL WDDLRSL LF YH
Sbjct  2012  QGYSPLSFQTLAPSPGGLDRPGRIDEEGGEQGRNNSTRLVSGFLALAWDDLRSLFLFCYH  2191

Query  259   RLRDLLLIVTRIVELLGR-------RGWEALKYWWNLLQYWSQELKNSAVSLLNATAIAV  311
             RL+DL+ +  R VELLGR       RGWEALKY  +L+QYW  ELK SA+SLL+  AIAV
Sbjct  2192  RLKDLISVTARAVELLGRSSLKGLQRGWEALKYLGSLVQYWVLELKKSAISLLDTIAIAV  2371

Query  312   AEGTDRVIEVVQGACRAIRHIPRRIRQGLERILL  345
              EGTDR+IE +QG CRAIR+IPRR+RQG E  LL
Sbjct  2372  GEGTDRIIEGIQGLCRAIRNIPRRVRQGFEAALL  2473
```

>Escherichia coli strain NBE0006 contig00106, whole genome shotgun sequence
Sequence ID: NZ_JARBDI010000106.1 Length: 2090
Range 1: 1 to 744

Score:317 bits(812), Expect:3e-98,
Method:Compositional matrix adjust.,
Identities:180/248(73%), Positives:215/248(86%), Gaps:0/248(0%)

```
Query  26   LTVQARQLLSGIVQQQNNLLRAIEAQQHLLQLTVWGIKQLQARILAVERYLKDQQLLGIW  85
            LTVQARQLLSGIVQQQNNLLRAIEAQQ++L+LTVWGIKQLQ R+LA+ERYL++Q+LLG+W
Sbjct  744  LTVQARQLLSGIVQQQNNLLRAIEAQQNMLRLTVWGIKQLQTRVLAIERYLREQRLLGLW  565

Query  86   GCSGKLICTTAVPWNASWSNKSLEQIWNHTTWMEWDREINNYTSLIHSLIeesqnqqekn  145
            GCS + ICTT VPWN+SWSN+S   IWN+ TWMEWD+EI  +T+ I+ L+EESQ QQE+N
Sbjct  564  GCSERTICTTNVPWNSSWSNRSYSDIWNNLTWMEWDKEIEQHTNTIYQLLEESQGQQEQN  385

Query  146  eqellelDKWASLWNWFNITNWLWYIKLFIMIVGGLVGLRIVFAVLSIVNRVRQGYSPLS  205
            E++LL LDKW  LW+WF+ITNWLWYI++FIMIVGGL+GLRIVFAVLSIVNRVRQGYSPLS
Sbjct  384  EKDLLALDKWNDLWSWFDITNWLWYIRIFIMIVGGLIGLRIVFAVLSIVNRVRQGYSPLS  205

Query  206  FQTHLPTprgpdrpegieeeggerdrdrSIRLVNGSLALIWDDLRSLCLFSYHRLRDLLL  265
            FQT  P+P G DRP  I+EEGGE+ R+ S RLV+G LAL W+DLRSL LF YHRL+DL+
Sbjct  204  FQTLAPSPGGLDRPGRIDEEGGEQGRNNSTRLVSGFLALAWEDLRSLFLFCYHRLKDLIS  25

Query  266  IVTRIVEL  273
            +  R VEL
Sbjct  24   VTARAVEL  1
```

>Escherichia coli strain EN5317 contig00057, whole genome shotgun sequence
Sequence ID: NZ_JARBDO010000057.1 Length: 2867
Range 1: 305 to 1084

Score:303 bits(775), Expect:1e-90,
Method:Compositional matrix adjust.,
Identities:181/260(70%), Positives:213/260(81%), Gaps:7/260(2%)

```
Query  92    ICTTAVPWNASWSNKSLEQIWNHTTWMEWDREINNYTSLIHSLIeesqnqqekneqelle  151
              C TAVPWN+SWSNK+ +QIWN+ TWM+WD+EI+NYT  I+ L+E+SQ QQEKNE++LL
Sbjct  1084  FCPTAVPWNSSWSNKTQDQIWNNMTWMQWDKEISNYTYTIYRLLEDSQTQQEKNEKDLLA  905

Query  152   lDKWASLWNWFNITNWLWYIKLFIMIVGGLVGLRIVFAVLSIVNRVRQGYSPLSFQTHLP  211
             LD WASLWNWF+IT WLWYIK+FIMIVGGL+GLRI+FAVLSIVNRVRQGYSPLSFQT  P
Sbjct  904   LDSWASLWNWFDITKWLWYIKIFIMIVGGLIGLRIIFAVLSIVNRVRQGYSPLSFQTLTP  725

Query  212   TprgpdrpegieeeggerdrdrSIRLVNGSLALIWDDLRSLCLFSYHRLRDLLLIVTRIV  271
             TP  PDR  GIEEEGGE+DR RS+RLV+G LA+ WDDLR+LCLFSYHRLRDLLL++ R V
Sbjct  724   TPGEPDRLRGIEEEGGEQDRVRSVRLVSGFLAIAWDDLRNLCLFSYHRLRDLLLVIARAV  545

Query  272   ELLG-------RRGWEALKYWWNLLQYWSQELKNSAVSLLNATAIAVAEGTDRVIEVVQG  324
             ELLG       +RG EALK+    ++QYW  ELK SA+SL +  AIAVAEGTDR+I  +Q
Sbjct  544   ELLGHSCLVGLQRGREALKHLGGIVQYWGLELKKSAISLFDTIAIAVAEGTDRIIGIQR  365
```

```
Query  325   ACRAIRHIPRRIRQGLERIL  344
              CRAI +IPRRIRQG E  L
Sbjct  364   LCRAICNIPRRIRQGFEAAL  305
```

>Escherichia coli strain NBE0003 contig00096, whole genome shotgun sequence
Sequence ID: NZ_JARBDG010000096.1 Length: 2837
Range 1: 296 to 1066

Score:298 bits(762), Expect:1e-88,
Method:Compositional matrix adjust.,
Identities:180/257(70%), Positives:209/257(81%), Gaps:7/257(2%)

```
Query  95    TAVPWNASWSNKSLEQIWNHTTWMEWDREINNYTSLIHSLIeesqnqqekneqellelDK  154
             + VPWN+SWSNKS   IW + TWM+WD+E++NYT  I+ L+E+SQ QQE+NE++LL LD
Sbjct  1066  SNVPWNSSWSNKSQTDIWENMTWMQWDKEVSNYTDTIYRLLEDSQTQQERNEKDLLALDN  887

Query  155   WASLWNWFNITNWLWYIKLFIMIVGGLVGLRIVFAVLSIVNRVRQGYSPLSFQTHLPTpr  214
             W +LWNWF+ITNWLWYIK+FIMIVGGL+GLRI+FAVLSIVNRVRQGYSPLSFQT  P PR
Sbjct  886   WKNLWNWFSITNWLWYIKIFIMIVGGLIGLRIIFAVLSIVNRVRQGYSPLSFQTLTPNPR  707

Query  215   gpdrpegieeeggerdrdrSIRLVNGSLALIWDDLRSLCLFSYHRLRDLLLIVTRIVELL  274
             GPDR   IEEEGG +DRDRSIRLVNG LAL WDDL SLCLFSYHRLRDL+L+  R VELL
Sbjct  706   GPDRLGRIEEEGGGQDRDRSIRLVNGFLALAWDDLWSLCLFSYHRLRDLILVTARAVELL  527

Query  275   G-------RRGWEALKYWWNLLQYWSQELKNSAVSLLNATAIAVAEGTDRVIEVVQGACR  327
             G       +RGWEALKY   ++QYW  ELK  A+SLL+  AIAVAEGTDR+IE +Q  CR
Sbjct  526   GHSSLRGLQRGWEALKYLGGIVQYWGLELKKRAISLLDTVAIAVAEGTDRIIEFLQRICR  347

Query  328   AIRHIPRRIRQGLERIL  344
             AIR+IPRRIRQG E  L
Sbjct  346   AIRNIPRRIRQGFEAAL  296
```

>Escherichia coli strain EN5169 contig00082, whole genome shotgun sequence
Sequence ID: NZ_JAPMNJ010000082.1 Length: 2837
Range 1: 296 to 1066

Score:298 bits(762), Expect:1e-88,
Method:Compositional matrix adjust.,
Identities:180/257(70%), Positives:209/257(81%), Gaps:7/257(2%)

```
Query  95    TAVPWNASWSNKSLEQIWNHTTWMEWDREINNYTSLIHSLIeesqnqqekneqellelDK  154
             + VPWN+SWSNKS   IW + TWM+WD+E++NYT  I+ L+E+SQ QQE+NE++LL LD
Sbjct  1066  SNVPWNSSWSNKSQTDIWENMTWMQWDKEVSNYTDTIYRLLEDSQTQQERNEKDLLALDN  887

Query  155   WASLWNWFNITNWLWYIKLFIMIVGGLVGLRIVFAVLSIVNRVRQGYSPLSFQTHLPTpr  214
             W +LWNWF+ITNWLWYIK+FIMIVGGL+GLRI+FAVLSIVNRVRQGYSPLSFQT  P PR
Sbjct  886   WKNLWNWFSITNWLWYIKIFIMIVGGLIGLRIIFAVLSIVNRVRQGYSPLSFQTLTPNPR  707

Query  215   gpdrpegieeeggerdrdrSIRLVNGSLALIWDDLRSLCLFSYHRLRDLLLIVTRIVELL  274
             GPDR   IEEEGG +DRDRSIRLVNG LAL WDDL SLCLFSYHRLRDL+L+  R VELL
Sbjct  706   GPDRLGRIEEEGGGQDRDRSIRLVNGFLALAWDDLWSLCLFSYHRLRDLILVTARAVELL  527

Query  275   G-------RRGWEALKYWWNLLQYWSQELKNSAVSLLNATAIAVAEGTDRVIEVVQGACR  327
             G       +RGWEALKY   ++QYW  ELK  A+SLL+  AIAVAEGTDR+IE +Q  CR
Sbjct  526   GHSSLRGLQRGWEALKYLGGIVQYWGLELKKRAISLLDTVAIAVAEGTDRIIEFLQRICR  347

Query  328   AIRHIPRRIRQGLERIL  344
             AIR+IPRRIRQG E  L
Sbjct  346   AIRNIPRRIRQGFEAAL  296
```

>Escherichia coli strain EN5095 contig00078, whole genome shotgun sequence
Sequence ID: NZ_JAPYLY010000078.1 Length: 1972
Range 1: 239 to 994

Score:291 bits(745), Expect:1e-88,
Method:Compositional matrix adjust.,
Identities:176/252(70%), Positives:206/252(81%), Gaps:7/252(2%)

```
Query  100  NASWSNKSLEQIWNHTTWMEWDREINNYTSLIHSLIeesqnqqekneqellelDKWASLW  159
            +ASWSN++ + IW++ TWM+WDREI+NYT  I+ L+E+SQNQQ+KNE++LL LD W +LW
Sbjct  994  SASWSNRTKKDIWDNMTWMQWDREISNYTETIYRLLEDSQNQQDKNEKDLLALDSWKNLW  815

Query  160  NWFNITNWLWYIKLFIMIVGGLVGLRIVFAVLSIVNRVRQGYSPLSFQTHLPTprgpdrp  219
            NWF+I NWLWYIK+FIMIVGGL+GLRI+FAVLSIVNRVRQGYSPLSFQT  P P G DR
Sbjct  814  NWFDIANWLWYIKIFIMIVGGLIGLRIIFAVLSIVNRVRQGYSPLSFQTLTPNPGGLDRL  635

Query  220  egieeeggerdrdrSIRLVNGSLALIWDDLRSLCLFSYHRLRDLLLIVTRIVELLGR---  276
              IEEEGGE+DR RSIRLV G LAL WDDLRSLCLFSYHRLRDLLL+ TR VE+LGR
Sbjct  634  GRIEEEGGEQDRGRSIRLVQGFLALAWDDLRSLCLFSYHRLRDLLLVTTRAVEILGRSSL  455

Query  277  ----RGWEALKYWWNLLQYWSQELKNSAVSLLNATAIAVAEGTDRVIEVVQGACRAIRHI  332
                RGWEALKY  +++QYW  ELK SA+SLL+ TAI VAEGTDR IE+VQ   RAI ++
Sbjct  454  RGLQRGWEALKYLGSIVQYWVLELKKSAISLLDTTAIVVAEGTDRTIELVQRTYRAICNV  275

Query  333  PRRIRQGLERIL  344
            PRRIRQG E  L
Sbjct  274  PRRIRQGFEAAL  239
```

>Escherichia coli strain EN5090 contig00081, whole genome shotgun sequence
Sequence ID: NZ_JAODTD010000081.1 Length: 2798
Range 1: 285 to 1013

Score:293 bits(750), Expect:4e-87,
Method:Compositional matrix adjust.,
Identities:177/243(73%), Positives:205/243(84%), Gaps:0/243(0%)

```
Query  102   SWSNKSLEQIWNHTTWMEWDREINNYTSLIHSLIeesqnqqekneqellelDKWASLWNW  161
             SWSN+S   IW++ TWM+WDREI+NYT  I+ L+EESQNQQEKNE+ELL L  W SLWNW
Sbjct  1013  SWSNRSQTDIWDNMTWMQWDREISNYTDTIYKLLEESQNQQEKNEKELLALGSWESLWNW  834

Query  162   FNITNWLWYIKLFIMIVGGLVGLRIVFAVLSIVNRVRQGYSPLSFQTHLPTprgpdrpeg  221
             F+IT WLWYIK+FIMIVGGL+GLRI+FAVLSIVNRVRQGYSPLSFQT  P PRG DR
Sbjct  833   FSITKWLWYIKIFIMIVGGLIGLRIIFAVLSIVNRVRQGYSPLSFQTLTPNPRGLDRLRR  654

Query  222   ieeeggerdrdrSIRLVNGSLALIWDDLRSLCLFSYHRLRDLLLIVTRIVELLGRRGWEA  281
             IEEEGGE+D+DRSIRLV+G LAL WDDLRSLCLFSYH+LR+ +L+  R+VELLGRRGWEA
Sbjct  653   IEEEGGEQDKDRSIRLVSGFLALAWDDLRSLCLFSYHQLRNFILVTARVVELLGRRGWEA  474

Query  282   LKYWWNLLQYWSQELKNSAVSLLNATAIAVAEGTDRVIEVVQGACRAIRHIPRRIRQGLE  341
             LKY  NL+ YW  ELK SA+SLL+  AIAVAEGTDR+I+++Q  CRAI +IPRRIRQG E
Sbjct  473   LKYLGNLVLYWGLELKKSAISLLDVIAIAVAEGTDRLIDLIQRICRAICNIPRRIRQGFE  294
```

```
Query  342   RIL  344
               L
Sbjct  293   AAL  285
```

>Escherichia coli strain EN5132 contig00077, whole genome shotgun sequence
Sequence ID: NZ_JAPYLZ010000077.1 Length: 2498
Range 1: 1473 to 2228

Score:289 bits(740), Expect:2e-86,
Method:Compositional matrix adjust.,
Identities:172/252(68%), Positives:206/252(81%), Gaps:7/252(2%)

```
Query  100   NASWSNKSLEQIWNHTTWMEWDREINNYTSLIHSLIeesqnqqekneqellelDKWASLW  159
             ++SWSNKS ++IW++ TWM+WD+EI+NYT  I+ L+E+SQ QQEKNE +LL LD W +LW
Sbjct  1473  HSSWSNKSRDEIWDNMTWMQWDKEISNYTDTIYRLLEDSQIQQEKNETDLLALDSWNNLW  1652

Query  160   NWFNITNWLWYIKLFIMIVGGLVGLRIVFAVLSIVNRVRQGYSPLSFQTHLPTprgpdrp  219
             NWF IT WLWYIK+FIMIVGGL+GLRI+FAV+SIVNR RQGYSPLSFQT  P P GPDR
Sbjct  1653  NWFGITKWLWYIKIFIMIVGGLIGLRIIFAVISIVNRFRQGYSPLSFQTLTPNPGGPDRL  1832

Query  220   egieeeggerdrdrSIRLVNGSLALIWDDLRSLCLFSYHRLRDLLLIVTRIVELLGR---  276
               IEEEGGE+D++RSIRLVNG LAL W+DLR+LCLFSYHRLRD +L+  R++ELLGR
Sbjct  1833  GRIEEEGGEQDKNRSIRLVNGFLALAWEDLRNLCLFSYHRLRDFILVTARVLELLGRSSL  2012

Query  277   ----RGWEALKYWWNLLQYWSQELKNSAVSLLNATAIAVAEGTDRVIEVVQGACRAIRHI  332
                 R WEALKY  +L+QYW  ELK SA+SLL+  AIAV EGTDR+I +VQG CRAIR+I
Sbjct  2013  KGLQRVWEALKYLGSLVQYWGLELKESAISLLDTIAIAVGEGTDRIISLVQGICRAIRNI  2192

Query  333   PRRIRQGLERIL  344
             PRRIRQG E  L
Sbjct  2193  PRRIRQGFEAAL  2228
```

>Escherichia coli strain EN5379 contig00092, whole genome shotgun sequence
Sequence ID: NZ_JAPTGJ010000092.1 Length: 2592
Range 1: 263 to 1012

Score:288 bits(738), Expect:7e-86,
Method:Compositional matrix adjust.,
Identities:180/250(72%), Positives:204/250(81%), Gaps:7/250(2%)

```
Query  102   SWSNKSLEQIWNHTTWMEWDREINNYTSLIHSLIeesqnqqekneqellelDKWASLWNW  161
             SWSN+S   IW++ TWM+WDREI+NYT  I+ L+EESQNQQEKNE+ELL L  W SLWNW
Sbjct  1012  SWSNRSQTDIWDNMTWMQWDREISNYTDTIYKLLEESQNQQEKNEKELLALGSWESLWNW  833

Query  162   FNITNWLWYIKLFIMIVGGLVGLRIVFAVLSIVNRVRQGYSPLSFQTHLPTprgpdrpeg  221
             F+IT WLWYIK+FIMIVGGL+GLRI+FAVLSIVNRVRQGYSPLSFQT  P PRG DR
Sbjct  832   FSITKWLWYIKIFIMIVGGLIGLRIIFAVLSIVNRVRQGYSPLSFQTLTPNPRGLDRLRR  653

Query  222   ieeeggerdrdrSIRLVNGSLALIWDDLRSLCLFSYHRLRDLLLIVTRIVELLGR-----  276
             IEEEGGE+DRDRSIRLVNG L+L  DDLR+LCLFSYHRLRD +L+  R+VELLGR
Sbjct  652   IEEEGGEQDRDRSIRLVNGFLSLA*DDLRNLCLFSYHRLRDFILLTARVVELLGRSSLRG  473

Query  277   --RGWEALKYWWNLLQYWSQELKNSAVSLLNATAIAVAEGTDRVIEVVQGACRAIRHIPR  334
               RGWEALKY  +L+QYW  ELK SA+SLL+  AIAVAEGTDR+I VVQ  CRAI +IPR
Sbjct  472   LQRGWEALKYLGSLVQYWGLELKKSAISLLDTIAIAVAEGTDRIISVVQRICRAIYNIPR  293

Query  335   RIRQGLERIL  344
             RIRQG E  L
Sbjct  292   RIRQGFEAAL  263
```

>Escherichia coli strain EN5308 contig00099, whole genome shotgun sequence
Sequence ID: NZ_JARBDE010000099.1 Length: 2827
Range 1: 373 to 1125

Score:288 bits(737), Expect:4e-85,
Method:Compositional matrix adjust.,
Identities:176/251(70%), Positives:204/251(81%), Gaps:7/251(2%)

```
Query  101   ASWSNKSLEQIWNHTTWMEWDREINNYTSLIHSLIeesqnqqekneqellelDKWASLWN  160
             +SWSNKS   IW + TWM+WD+E++NYT  I+ L+E+SQ QQE+NE++LL LD W +LWN
Sbjct  1125  SSWSNKSQTDIWENMTWMQWDKEVSNYTDTIYRLLEDSQTQQERNEKDLLALDNWKNLWN  946

Query  161   WFNITNWLWYIKLFIMIVGGLVGLRIVFAVLSIVNRVRQGYSPLSFQTHLPTprgpdrpe  220
             WF+ITNWLWYIK+FIMIVGGL+GLRI+FAVLSIVNRVRQGYSPLSFQT  P PRGPDR
Sbjct  945   WFSITNWLWYIKIFIMIVGGLIGLRIIFAVLSIVNRVRQGYSPLSFQTLTPNPRGPDRLG  766

Query  221   gieeeggerdrdrSIRLVNGSLALIWDDLRSLCLFSYHRLRDLLLIVTRIVELLG-----  275
              IEEGG +DRDRSIRLVNG LAL WDDL SLCLFSYHRLRDL+L+  R VELLG
Sbjct  765   RIEEEGGGQDRDRSIRLVNGFLALAWDDLWSLCLFSYHRLRDLILVTARAVELLGHSSLR  586

Query  276   --RRGWEALKYWWNLLQYWSQELKNSAVSLLNATAIAVAEGTDRVIEVVQGACRAIRHIP  333
               +RGWEALKY   ++QYW  ELK  A+SLL+  AIAVAEGTDR+IE +Q  CRAIR+IP
Sbjct  585   GLQRGWEALKYLGGIVQYWGLELKKRAISLLDTVAIAVAEGTDRIIEFLQRICRAIRNIP  406

Query  334   RRIRQGLERIL  344
             RRIRQG E  L
Sbjct  405   RRIRQGFEAAL  373
```

>Escherichia coli strain EN5192 contig00135, whole genome shotgun sequence
Sequence ID: NZ_JAPMNL010000135.1 Length: 1341
Range 1: 243 to 965

Score:268 bits(686), Expect:3e-82,
Method:Compositional matrix adjust.,
Identities:166/252(66%), Positives:196/252(77%), Gaps:18/252(7%)

```
Query  100  NASWSNKSLEQIWNHTTWMEWDREINNYTSLIHSLIeesqnqqekneqellelDKWASLW  159
            ++SWSNKS  +IW++ TWM+WDREI+NYT  I+ L+E+SQ QQEKNE++LL LD W +LW
Sbjct  965  HSSWSNKSRSEIWDNMTWMQWDREISNYTDTIYRLLEDSQIQQEKNEKDLLALDSWNNLW  786

Query  160  NWFNITNWLWYIKLFIMIVGGLVGLRIVFAVLSIVNRVRQGYSPLSFQTHLPTprgpdrp  219
            NWF IT WLWYIK+FIMIVGGL+GLRI+FAV+SIVNR RQGYSPLSFQT  P P GPDR
Sbjct  785  NWFGITKWLWYIKIFIMIVGGLIGLRIIFAVISIVNRFRQGYSPLSFQTLTPNPGGPDRL  606

Query  220  egieeeggerdrdrSIRLVNGSLALIWDDLRSLCLFSYHRLRDLLLIVTRIVELLGR---  276
              IEEEGGE+ R+RSIRLVNG LAL W+DLR+LCLFSYHRLRD +L+  R+VELLGR
Sbjct  605  GRIEEEGGEQGRNRSIRLVNGFLALAWEDLRNLCLFSYHRLRDFILVTARVVELLGRNSL  426

Query  277  ----RGWEALKYWWNLLQYWSQELKNSAVSLLNATAIAVAEGTDRVIEVVQGACRAIRHI  332
                RGWEALKY  +L+QYW            +  AIAVAEGTDR+I + QG CRAI +I
Sbjct  425  KGLQRGWEALKYLGSLVQYW-----------FDTIAIAVAEGTDRIINLAQGICRAICNI  279
```

```
Query  333  PRRIRQGLERIL  344
            PRRIRQG E  L
Sbjct  278  PRRIRQGFEAAL  243
```

>Escherichia coli strain NBE0004 contig00090, whole genome shotgun sequence
Sequence ID: NZ_JARBDH010000090.1 Length: 2818
Range 1: 140 to 700

Score:276 bits(706), Expect:9e-81,
Method:Compositional matrix adjust.,
Identities:145/187(78%), Positives:170/187(90%), Gaps:0/187(0%)

```
Query  26   LTVQARQLLSGIVQQQNNLLRAIEAQQHLLQLTVWGIKQLQARILAVERYLKDQQLLGIW  85
            LTVQARQLLSGIVQQQNNLLRAIEAQQ++L+LTVWGIKQLQ R+LA+ERYL++Q+LLG+W
Sbjct  700  LTVQARQLLSGIVQQQNNLLRAIEAQQNMLRLTVWGIKQLQTRVLAIERYLREQRLLGLW  521

Query  86   GCSGKLICTTAVPWNASWSNKSLEQIWNHTTWMEWDREINNYTSLIHSLIeesqnqqekn  145
            GCS K+ICTT VPWN+SWSNKS   IWN+ TWMEWD+EI  +T+ I+ L+EESQ QQE+N
Sbjct  520  GCSEKIICTTNVPWNSSWSNKSYSDIWNNLTWMEWDKEIEQHTNTIYQLLEESQGQQEQN  341

Query  146  eqellelDKWASLWNWFNITNWLWYIKLFIMIVGGLVGLRIVFAVLSIVNRVRQGYSPLS  205
            E++LL LDKW  LW+WF+ITNWLWYIK+FIMIVGGL+GLRIVFAVLSIVNRVRQGYSPLS
Sbjct  340  EKDLLALDKWNDLWSWFDITNWLWYIKIFIMIVGGLIGLRIVFAVLSIVNRVRQGYSPLS  161

Query  206  FQTHLPT  212
            FQT  P+
Sbjct  160  FQTLAPS  140
```

>Klebsiella pneumoniae strain EuSCAPE_IL001, whole genome shotgun sequence
Sequence ID: NZ_UKJN02000084.1 Length: 599
Range 1: 32 to 598

Score:242 bits(618), Expect:2e-75,
Method:Compositional matrix adjust.,
Identities:139/189(74%), Positives:155/189(82%), Gaps:7/189(3%)

```
Query  153  DKWASLWNWFNITNWLWYIKLFIMIVGGLVGLRIVFAVLSIVNRVRQGYSPLSFQTHLPT  212
            DKW  LW+WF+IT WLWYI++FIMIVGGL+GLRI+FAV SIV RVRQGYSPLSFQT +P
Sbjct  32   DKWNDLWSWFSITQWLWYIRIFIMIVGGLIGLRIIFAVFSIVKRVRQGYSPLSFQTLIPN  211

Query  213  prgpdrpegieeeggerdrdrSIRLVNGSLALIWDDLRSLCLFSYHRLRDLLLIVTRIVE  272
            PR PDR  GIEEEGGE+DRDRSIRLVNG LAL WDDLRSLCLF YHRLRD +L+  R  E
Sbjct  212  PRDPDRHGGIEEEGGEQDRDRSIRLVNGFLALFWDDLRSLCLFCYHRLRDFILVTARAAE  391

Query  273  LLGR-------RGWEALKYWWNLLQYWSQELKNSAVSLLNATAIAVAEGTDRVIEVVQGA  325
            LLGR       RGWE LKY  NL+QYW  ELK SA+SLL+ TAIAVAEGTDR+IE+VQ
Sbjct  392  LLGRSSLRGLQRGWEILKYLGNLVQYWGLELKKSAISLLDTTAIAVAEGTDRIIEIVQAT  571

Query  326  CRAIRHIPR  334
            CRAIR+IPR
Sbjct  572  CRAIRNIPR  598
```

>Escherichia coli strain EN5143 contig00069, whole genome shotgun sequence
Sequence ID: NZ_JAPMNI010000069.1 Length: 2688
Range 1: 1819 to 2418

Score:253 bits(647), Expect:2e-72,
Method:Compositional matrix adjust.,
Identities:142/200(71%), Positives:163/200(81%), Gaps:8/200(4%)

```
Query  153   DKWASLWNWFNITNWLWYIKLFIMIVGGLVGLRIVFAVLSIVNRVRQGYSPLSFQTHLP-  211
             D W +LWNWF IT WLWYIK+FIMIVGGL+GLRI+FAV+SIVNR RQGYSPLSFQT  P
Sbjct  1819  DSWNNLWNWFGITKWLWYIKIFIMIVGGLIGLRILFAVISIVNRFRQGYSPLSFQTLTPN  1998

Query  212   TprgpdrpegieeeggerdrdrSIRLVNGSLALIWDDLRSLCLFSYHRLRDLLLIVTRIV  271
             +   PDR   IEEEGGE+DR+RSIRLV+G LAL WDDLR+LCLFSYHRLRD +L+  R++
Sbjct  1999  SRGPPDRLGRIEEEGGEQDRNRSIRLVSGFLALAWDDLRNLCLFSYHRLRDFILVTARVL  2178

Query  272   ELLGR-------RGWEALKYWWNLLQYWSQELKNSAVSLLNATAIAVAEGTDRVIEVVQG  324
             ELLGR       RGWEALKY  +L+QYW  ELKNSA+SLL+  AIAVAEGTDR+I +VQG
Sbjct  2179  ELLGRSSLKGLQRGWEALKYLGSLVQYWGLELKNSAISLLDTIAIAVAEGTDRIINLVQG  2358

Query  325   ACRAIRHIPRRIRQGLERIL  344
              CRAIR+IPRRIRQG E  L
Sbjct  2359  ICRAIRNIPRRIRQGFEAAL  2418
```

>Escherichia coli strain NBE0002 contig00097, whole genome shotgun sequence
Sequence ID: NZ_JARBDF010000097.1 Length: 2717
Range 1: 300 to 899

Score:253 bits(645), Expect:3e-72,
Method:Compositional matrix adjust.,
Identities:141/200(71%), Positives:160/200(80%), Gaps:7/200(3%)

```
Query  153  DKWASLWNWFNITNWLWYIKLFIMIVGGLVGLRIVFAVLSIVNRVRQGYSPLSFQTHLPT  212
            D W +LWNWF IT WLWYIK+FIMIVGGL+GLRIVFAVLSIVNRVRQGYSPLSFQT  P+
Sbjct  899  DSWNNLWNWFGITKWLWYIKIFIMIVGGLIGLRIVFAVLSIVNRVRQGYSPLSFQTLAPS  720

Query  213  prgpdrpegieeeggerdrdrSIRLVNGSLALIWDDLRSLCLFSYHRLRDLLLIVTRIVE  272
            P G DRP  I+EEGGE+ R+ S RLV+G LAL WDDLRSL LF YHRL+DL+ +  R+VE
Sbjct  719  PGGLDRPGRIDEEGGEQGRNNSTRLVSGFLALAWDDLRSLFLFCYHRLKDLISVTARVVE  540

Query  273  LLGR-------RGWEALKYWWNLLQYWSQELKNSAVSLLNATAIAVAEGTDRVIEVVQGA  325
            LLGR       RGWEALKY  +L+QYW  ELK SA+SLL+  AIAV EGTDR+IE VQG
Sbjct  539  LLGRSSLKGLQRGWEALKYLGSLVQYWGLELKKSAISLLDTIAIAVGEGTDRIIEGVQGL  360

Query  326  CRAIRHIPRRIRQGLERILL  345
            CRAIR+IPRR+RQG E  LL
Sbjct  359  CRAIRNIPRRVRQGFEAALL  300
```

>Klebsiella pneumoniae strain EuSCAPE_IL006, whole genome shotgun sequence
Sequence ID: NZ_UKJR02000114.1 Length: 560
Range 1: 5 to 556

Score:229 bits(585), Expect:1e-70,
Method:Compositional matrix adjust.,
Identities:136/184(74%), Positives:156/184(84%), Gaps:7/184(3%)

```
Query  153  DKWASLWNWFNITNWLWYIKLFIMIVGGLVGLRIVFAVLSIVNRVRQGYSPLSFQTHLPT  212
```

```
            D W SLW+WF+I+NWLWYI++FIMIVGGL+GLRI+FAV+S+VNRVRQGYSPLS QT +P
Sbjct  556  DSWNSLWSWFSISNWLWYIRIFIMIVGGLIGLRIIFAVISVVNRVRQGYSPLSLQTLIPN  377

Query  213  prgpdrpegieeeggerdrdrSIRLVNGSLALIWDDLRSLCLFSYHRLRDLLLIVTRIVE  272
            PRGPDRP GIEEEGGE+DRDRSIRLVNG LA+ WDDLRSLCLFSYHRLRDL L+  R VE
Sbjct  376  PRGPDRPGGIEEEGGEQDRDRSIRLVNGFLAIAWDDLRSLCLFSYHRLRDLTLVTARAVE  197

Query  273  LLGR-------RGWEALKYWWNLLQYWSQELKNSAVSLLNATAIAVAEGTDRVIEVVQGA  325
            LLGR       RGWEALKY  +++QYW  ELK SA+SLL+  AIAVAEGTDR+IEV+Q
Sbjct  196  LLGRSCLKGLQRGWEALKYLGSIVQYWGLELKKSAISLLDTIAIAVAEGTDRIIEVIQRI  17

Query  326  CRAI  329
            CRAI
Sbjct  16   CRAI  5
```

>Escherichia coli strain EN5374 contig00094, whole genome shotgun sequence
Sequence ID: NZ_JAPTGI010000094.1 Length: 2720
Range 1: 1777 to 2445

Score:248 bits(632), Expect:2e-70,
Method:Compositional matrix adjust.,
Identities:160/243(66%), Positives:186/243(76%), Gaps:20/243(8%)

```
Query  102   SWSNKSLEQIWNHTTWMEWDREINNYTSLIHSLIeesqnqqekneqellelDKWASLWNW  161
             SWSN+S   IW++ TWM+WDREI+NYT  I+ L+EESQNQQEKNE+ELL LD W SLWNW
Sbjct  1777  SWSNRSQTDIWDNMTWMQWDREISNYTDTIYKLLEESQNQQEKNEKELLALDSWESLWNW  1956

Query  162   FNITNWLWYIKLFIMIVGGLVGLRIVFAVLSIVNRVRQGYSPLSFQTHLPTprgpdrpeg  221
             F+IT WLWYIK+FIMIVGG                    YSPLSFQT  P PRGPDR
Sbjct  1957  FSITKWLWYIKIFIMIVGG--------------------YSPLSFQTLTPNPRGPDRLGR  2076

Query  222   ieeeggerdrdrSIRLVNGSLALIWDDLRSLCLFSYHRLRDLLLIVTRIVELLGRRGWEA  281
             IEEEGGE+D+DRSIRLV+G LAL WDDLRSLCLFSYH+LR+ +L+  R+VELLGRRGWEA
Sbjct  2077  IEEEGGEQDKDRSIRLVSGFLALAWDDLRSLCLFSYHQLRNFILVTARVVELLGRRGWEA  2256

Query  282   LKYWWNLLQYWSQELKNSAVSLLNATAIAVAEGTDRVIEVVQGACRAIRHIPRRIRQGLE  341
             LKY  NL+ YW  ELK SA+ LL+  AIAVAEGTDR+I++Q  CRAI +IPRRIRQG E
Sbjct  2257  LKYLGNLVLYWGLELKKSAIGLLDVIAIAVAEGTDRLIDLIQRICRAICNIPRRIRQGFE  2436

Query  342   RIL  344
               L
Sbjct  2437  AAL  2445
```

>Escherichia coli strain EN5076 contig00066, whole genome shotgun sequence
Sequence ID: NZ_JAODTC010000066.1 Length: 2732
Range 1: 282 to 950

Score:248 bits(632), Expect:2e-70,
Method:Compositional matrix adjust.,
Identities:160/243(66%), Positives:186/243(76%), Gaps:20/243(8%)

```
Query  102  SWSNKSLEQIWNHTTWMEWDREINNYTSLIHSLIeesqnqqekneqellelDKWASLWNW  161
            SWSN+S   IW++ TWM+WDREI+NYT  I+ L+EESQNQQEKNE+ELL LD W SLWNW
Sbjct  950  SWSNRSQTDIWDNMTWMQWDREISNYTDTIYKLLEESQNQQEKNEKELLALDSWESLWNW  771

Query  162  FNITNWLWYIKLFIMIVGGLVGLRIVFAVLSIVNRVRQGYSPLSFQTHLPTprgpdrpeg  221
            F+IT WLWYIK+FIMIVGG                    YSPLSFQT  P PRGPDR
Sbjct  770  FSITKWLWYIKIFIMIVGG--------------------YSPLSFQTLTPNPRGPDRLGR  651

Query  222  ieeeggerdrdrSIRLVNGSLALIWDDLRSLCLFSYHRLRDLLLIVTRIVELLGRRGWEA  281
            IEEEGGE+D+DRSIRLV+G LAL WDDLRSLCLFSYH+LR+ +L+  R+VELLGRRGWEA
Sbjct  650  IEEEGGEQDKDRSIRLVSGFLALAWDDLRSLCLFSYHQLRNFILVTARVVELLGRRGWEA  471

Query  282  LKYWWNLLQYWSQELKNSAVSLLNATAIAVAEGTDRVIEVVQGACRAIRHIPRRIRQGLE  341
            LKY  NL+ YW  ELK SA+ LL+  AIAVAEGTDR+I++Q  CRAI +IPRRIRQG E
Sbjct  470  LKYLGNLVLYWGLELKKSAIGLLDVIAIAVAEGTDRLIDLIQRICRAICNIPRRIRQGFE  291

Query  342  RIL  344
              L
Sbjct  290  AAL  282
```

>Escherichia coli strain NBE0010 contig00080, whole genome shotgun sequence
Sequence ID: NZ_JARBDM010000080.1 Length: 2743
Range 1: 1478 to 2146

Score:248 bits(632), Expect:3e-70,
Method:Compositional matrix adjust.,
Identities:160/243(66%), Positives:186/243(76%), Gaps:20/243(8%)

```
Query  102   SWSNKSLEQIWNHTTWMEWDREINNYTSLIHSLIeesqnqqekneqellelDKWASLWNW  161
             SWSN+S   IW++ TWM+WDREI+NYT  I+ L+EESQNQQEKNE+ELL LD W SLWNW
Sbjct  1478  SWSNRSQTDIWDNMTWMQWDREISNYTDTIYKLLEESQNQQEKNEKELLALDSWESLWNW  1657

Query  162   FNITNWLWYIKLFIMIVGGLVGLRIVFAVLSIVNRVRQGYSPLSFQTHLPTprgpdrpeg  221
             F+IT WLWYIK+FIMIVGG                    YSPLSFQT  P PRGPDR
Sbjct  1658  FSITKWLWYIKIFIMIVGG--------------------YSPLSFQTLTPNPRGPDRLGR  1777

Query  222   ieeeggerdrdrSIRLVNGSLALIWDDLRSLCLFSYHRLRDLLLIVTRIVELLGRRGWEA  281
             IEEEGGE+D+DRSIRLV+G LAL WDDLRSLCLFSYH+LR+ +L+  R+VELLGRRGWEA
Sbjct  1778  IEEEGGEQDKDRSIRLVSGFLALAWDDLRSLCLFSYHQLRNFILVTARVVELLGRRGWEA  1957

Query  282   LKYWWNLLQYWSQELKNSAVSLLNATAIAVAEGTDRVIEVVQGACRAIRHIPRRIRQGLE  341
             LKY  NL+ YW  ELK SA+ LL+  AIAVAEGTDR+I++Q  CRAI +IPRRIRQG E
Sbjct  1958  LKYLGNLVLYWGLELKKSAIGLLDVIAIAVAEGTDRLIDLIQRICRAICNIPRRIRQGFE  2137

Query  342   RIL  344
               L
Sbjct  2138  AAL  2146
```

>Klebsiella pneumoniae strain OK8 jcf7180000003506, whole genome shotgun sequence
Sequence ID: NZ_MZXR01000036.1 Length: 577
Range 1: 1 to 549

Score:227 bits(579), Expect:1e-69,
Method:Compositional matrix adjust.,
Identities:128/183(70%), Positives:146/183(79%), Gaps:8/183(4%)

```
Query  170  YIKLFIMIVGGLVGLRIVFAVLSIVNRVRQGYSPLSFQTHLPTprgpdrpegi-eeegge  228
            YIK+FIMIVGGL+GLRI+FAVLSIV RVRQGYSPLSFQT  PT G     G  EEEGGE
Sbjct  1    YIKIFIMIVGGLIGLRIIFAVLSIVKRVRQGYSPLSFQTLTPTSGGRPDRLGRIEEEGGE  180

Query  229  rdrdrSIRLVNGSLALIWDDLRSLCLFSYHRLRDLLLIVTRIVELLGRR-------GWEA  281
```

```
            +D+ RSIRLV+G LAL WDDLR+LCLFSYHRLRD +L+  R+VELLGR        GWEA
Sbjct  181  QDKGRSIRLVSGFLALAWDDLRNLCLFSYHRLRDFILVAARVVELLGRSSLRGLQTGWEA  360

Query  282  LKYWWNLLQYWSQELKNSAVSLLNATAIAVAEGTDRVIEVVQGACRAIRHIPRRIRQGLE  341
            LKY  +L+QYW QELK SA++LL+  AIAVAEGTDR+IE+ QG CRAIR+IPRRIRQG E
Sbjct  361  LKYLGSLVQYWGQELKKSAINLLDTVAIAVAEGTDRIIELAQGICRAIRNIPRRIRQGFE  540

Query  342  RIL  344
              L
Sbjct  541  AAL  549
```

>Aggregatibacter actinomycetemcomitans serotype d str. SA269 contig0497, whole genome shotgun sequence
Sequence ID: NZ_AZTX01000497.1 Length: 504
Range 1: 1 to 486

Score:223 bits(569), Expect:2e-68,
Method:Compositional matrix adjust.,
Identities:141/162(87%), Positives:146/162(90%), Gaps:1/162(0%)

```
Query  168  LWYIKLFIMIVGGLVGLRIVFAVLSIVNRVRQGYSPLSFQTHLPTprgpdrpegieeegg  227
            LWYIK+FIMIVGGL+GLRI+FAVLSIVNRVRQGYSPLSFQTHLP PRG DRPEGIEEEGG
Sbjct  1    LWYIKIFIMIVGGLIGLRIIFAVLSIVNRVRQGYSPLSFQTHLPLPRGADRPEGIEEEGG  180

Query  228  erdrdrSIRLVNGSLALIWDDLRSLCLFSYHRLRDLLLIVTRIVELLGRRGWEALKYWWN  287
            ERDRDRSIRLV GSLALIWDDLRSLCLFSYHRLRDLLLIVTR VELLGRRGWEALKYWWN
Sbjct  181  ERDRDRSIRLVTGSLALIWDDLRSLCLFSYHRLRDLLLIVTRTVELLGRRGWEALKYWWN  360

Query  288  LLQYWSQELKNSAVSLLNATAIAVAE-GTDRVIEVVQGACRA  328
            LL YWSQELKNSAVSLLNATAIAV + G     E VQ   R+
Sbjct  361  LLLYWSQELKNSAVSLLNATAIAVRQYGWSYFHEAVQAVWRS  486
```

>Escherichia coli strain EN5177 contig00148, whole genome shotgun sequence
Sequence ID: NZ_JAPMNK010000148.1 Length: 850
Range 1: 288 to 848

Score:204 bits(518), Expect:4e-59,
Method:Compositional matrix adjust.,
Identities:126/187(67%), Positives:145/187(77%), Gaps:7/187(3%)

```
Query  162  FNITNWLWYIKLFIMIVGGLVGLRIVFAVLSIVNRVRQGYSPLSFQTHLPTprgpdrpeg  221
            F IT WL YIK+FIMIVGGL+ LRI+FAV+SIVNR RQ YSPLSFQT  P PRGPDR
Sbjct  848  FGITKWL*YIKIFIMIVGGLISLRIIFAVISIVNRFRQRYSPLSFQTLTPNPRGPDRLGR  669

Query  222  ieeeggerdrdrSIRLVNGSLALIWDDLRSLCLFSYHRLRDLLLIVTRIVELLGR-----  276
            IEEEG E+DR+RSIRLVNG LAL  DDL++LCLFSYHRLRD +L+  R+VELLGR
Sbjct  668  IEEEGREQDRNRSIRLVNGFLALA*DDLQNLCLFSYHRLRDFILVAARVVELLGRSSLKG  489

Query  277  --RGWEALKYWWNLLQYWSQELKNSAVSLLNATAIAVAEGTDRVIEVVQGACRAIRHIPR  334
              R  EALKY  +L+QY   ELK SA+SLL+  AIAVAE TD++I +VQ  CRAIR+IPR
Sbjct  488  LQRR*EALKYLGSLVQY*GLELKGSAISLLDTIAIAVAERTDKIINLVQRICRAIRNIPR  309

Query  335  RIRQGLE  341
            RIRQ  E
Sbjct  308  RIRQSFE  288
```

>Escherichia coli strain NBE0009 contig00153, whole genome shotgun sequence
Sequence ID: NZ_JARBDL010000153.1 Length: 753
Range 1: 1 to 498

Score:199 bits(506), Expect:8e-58,
Method:Compositional matrix adjust.,
Identities:119/166(72%), Positives:135/166(81%), Gaps:7/166(4%)

```
Query  183  GLRIVFAVLSIVNRVRQGYSPLSFQTHLPTprgpdrpegieeeggerdrdrSIRLVNGSL  242
            GLRI+FAVLSIVNRVRQGYSPLSFQT +P P G DR   IEEEGGE+DRDRSIRLVNG L
Sbjct  1    GLRIIFAVLSIVNRVRQGYSPLSFQTPIPNPGGLDRLGRIEEEGGEQDRDRSIRLVNGFL  180

Query  243  ALIWDDLRSLCLFSYHRLRDLLLIVTRIVELLG-------RRGWEALKYWWNLLQYWSQE  295
            +L WDDLR+LCLFSYHRLRD +L+  R+VELLG       +RGW+ALKY  +L+QYW  E
Sbjct  181  SLAWDDLRNLCLFSYHRLRDFILVTARVVELLGHSSLRGLQRGWKALKYLRSLVQYWGLE  360

Query  296  LKNSAVSLLNATAIAVAEGTDRVIEVVQGACRAIRHIPRRIRQGLE  341
            LK SA+SLL+  AIAVAEGTDR+I VVQ  CRAI +IPRRIRQG E
Sbjct  361  LKRSAISLLDTIAIAVAEGTDRIISVVQRICRAIYNIPRRIRQGFE  498
```

>Klebsiella pneumoniae strain EuSCAPE_MT003, whole genome shotgun sequence
Sequence ID: NZ_UKIK02000085.1 Length: 964
Range 1: 480 to 962

Score:193 bits(491), Expect:1e-54,
Method:Compositional matrix adjust.,
Identities:113/161(70%), Positives:129/161(80%), Gaps:0/161(0%)

```
Query  185  RIVFAVLSIVNRVRQGYSPLSFQTHLPTprgpdrpegieeeggerdrdrSIRLVNGSLAL  244
            RI+FAVLSIVNRVRQGYSPLS QT    P G DRP  IEEEGGE+D+DRSIRLVNG  +L
Sbjct  962  RIIFAVLSIVNRVRQGYSPLSLQTLTQNPGGLDRPGRIEEEGGEQDKDRSIRLVNGFFSL  783

Query  245  IWDDLRSLCLFSYHRLRDLLLIVTRIVELLGRRGWEALKYWWNLLQYWSQELKNSAVSLL  304
             W+DLRSLCLFSY +LRD  LI  R VE+LGRRGWE LKY  +L+QYW  ELK SA+SLL
Sbjct  782  AWEDLRSLCLFSYRQLRDFTLIAARAVEILGRRGWETLKYLGSLIQYWGLELKKSAISLL  603

Query  305  NATAIAVAEGTDRVIEVVQGACRAIRHIPRRIRQGLERILL  345
            +  AI VAEGTDR++E +QG CRAIR++PRRIRQG E  LL
Sbjct  602  DTIAITVAEGTDRILEFIQGICRAIRNVPRRIRQGFEAALL  480
```

>Klebsiella pneumoniae strain EuSCAPE_IL004, whole genome shotgun sequence
Sequence ID: NZ_UKJP02000107.1 Length: 368
Range 1: 14 to 343

Score:184 bits(467), Expect:9e-54,
Method:Compositional matrix adjust.,
Identities:88/110(80%), Positives:99/110(90%), Gaps:0/110(0%)

```
Query  26   LTVQARQLLSGIVQQQNNLLRAIEAQQHLLQLTVWGIKQLQARILAVERYLKDQQLLGIW  85
            LTVQARQLLSGIVQQQ NLLRAIEAQQH+LQLTVWGIKQLQ R+LA+ERYLKDQQLLGIW
Sbjct  343  LTVQARQLLSGIVQQQGNLLRAIEAQQHMLQLTVWGIKQLQTRVLAIERYLKDQQLLGIW  164

Query  86   GCSGKLICTTAVPWNASWSNKSLEQIWNHTTWMEWDREINNYTSLIHSLI  135
            GCSGKLICTTAVPWN++WS KS   IW + TWM+WDREI+N+T  I+ L+
```

```
Sbjct  163  GCSGKLICTTAVPWNSTWSKKSQNDIWKNMTWMQWDREISNHTDTIYRLL  14



>Escherichia coli strain NBE0009 contig00151, whole genome shotgun sequence
Sequence ID: NZ_JARBDL010000151.1 Length: 802
Range 1: 305 to 802

Score:189 bits(479), Expect:1e-53,
Method:Compositional matrix adjust.,
Identities:115/166(69%), Positives:132/166(79%), Gaps:7/166(4%)

Query  183  GLRIVFAVLSIVNRVRQGYSPLSFQTHLPTprgpdrpegieeeggerdrdrSIRLVNGSL  242
            GLRI+FAVLSIVNRVRQGYSPLSFQT  PTP  PDR  GIEEEGGE+DR RS+RLV+G L
Sbjct  802  GLRIIFAVLSIVNRVRQGYSPLSFQTLTPTPGEPDRLRGIEEEGGEQDRVRSVRLVSGFL  623

Query  243  ALIWDDLRSLCLFSYHRLRDLLLIVTRIVELLG-------RRGWEALKYWWNLLQYWSQE  295
            A+ WDDLR+LCLFSYHRLRDLLL++ R VELLG       +RG EALK+   ++QYW  E
Sbjct  622  AIAWDDLRNLCLFSYHRLRDLLLVIARAVELLGHSCLVGLQRGREALKHLGGIVQYWGLE  443

Query  296  LKNSAVSLLNATAIAVAEGTDRVIEVVQGACRAIRHIPRRIRQGLE  341
            LK SA+SL +  AIAVAEGTDR+I  +Q  CRAI +IPRRIRQG E
Sbjct  442  LKKSAISLFDTIAIAVAEGTDRIIIGIQRLCRAICNIPRRIRQGFE  305



>Klebsiella pneumoniae strain EuSCAPE_IL005, whole genome shotgun sequence
Sequence ID: NZ_UKJL02000055.1 Length: 455
Range 1: 15 to 455

Score:181 bits(460), Expect:3e-52,
Method:Compositional matrix adjust.,
Identities:110/147(75%), Positives:125/147(85%), Gaps:7/147(4%)

Query  153  DKWASLWNWFNITNWLWYIKLFIMIVGGLVGLRIVFAVLSIVNRVRQGYSPLSFQTHLPT  212
            DKW +LW+WF+ITNWLWYI++FIMIVGGL+GLRI+FAVLS+VNRVRQGYSPLSFQT +P+
Sbjct  15   DKWGNLWSWFDITNWLWYIRIFIMIVGGLIGLRIIFAVLSMVNRVRQGYSPLSFQTLIPS  194

Query  213  prgpdrpegieeeggerdrdrSIRLVNGSLALIWDDLRSLCLFSYHRLRDLLLIVTRIVE  272
            PRGPDR E IEEEGGE+DR+RSIRLV+G LAL WDDLRSLCLF YHRLRD +L+  R VE
Sbjct  195  PRGPDRLERIEEEGGEQDRNRSIRLVSGFLALAWDDLRSLCLFCYHRLRDFILVTVRAVE  374

Query  273  LLGR-------RGWEALKYWWNLLQYW  292
            LLGR       RGWE LKY  NL QYW
Sbjct  375  LLGRSSLKGLQRGWEILKYLGNLAQYW  455



>Escherichia coli strain NBE0006 contig00145, whole genome shotgun sequence
Sequence ID: NZ_JARBDI010000145.1 Length: 738
Range 1: 1 to 483

Score:182 bits(461), Expect:4e-51,
Method:Compositional matrix adjust.,
Identities:113/172(66%), Positives:129/172(75%), Gaps:18/172(10%)

Query  180  GLVGLRIVFAVLSIVNRVRQGYSPLSFQTHLPTprgpdrpegieeeggerdrdrSIRLVN  239
            GL+GLRI+FAV+SIVNR RQGYSPLSFQT  P P GPDR   IEEEGGE+DR+RSIRLVN
Sbjct  1    GLIGLRIIFAVISIVNRFRQGYSPLSFQTLTPNPGGPDRLGRIEEEGGEQDRNRSIRLVN  180

Query  240  GSLALIWDDLRSLCLFSYHRLRDLLLIVTRIVELLGR-------RGWEALKYWWNLLQYW  292
            G LAL WDDLR+LCLFSYH+LRD +L+  R+VELLGR       RGWEALKY  +L+QYW
Sbjct  181  GFLALAWDDLRNLCLFSYHQLRDFILVTARVVELLGRNSLKGLQRGWEALKYLGSLVQYW  360

Query  293  SQELKNSAVSLLNATAIAVAEGTDRVIEVVQGACRAIRHIPRRIRQGLERIL  344
                       +  AIAVAEGTDR+I + QG CRAI +IPRRIRQG E  L
Sbjct  361  -----------FDTIAIAVAEGTDRIINLAQGICRAICNIPRRIRQGFEAAL  483



>Actinoplanes campanulatus strain JCM 3059 sequence095, whole genome shotgun sequence
Sequence ID: NZ_BMPW01000095.1 Length: 582
Range 1: 111 to 458

Score:177 bits(450), Expect:3e-50,
Method:Compositional matrix adjust.,
Identities:115/116(99%), Positives:115/116(99%), Gaps:0/116(0%)

Query  208  THLPTprgpdrpegieeeggerdrdrSIRLVNGSLALIWDDLRSLCLFSYHRLRDLLLIV  267
            THLP PRGPDRPEGIEEEGGERDRDRSIRLVNGSLALIWDDLRSLCLFSYHRLRDLLLIV
Sbjct  111  THLPIPRGPDRPEGIEEEGGERDRDRSIRLVNGSLALIWDDLRSLCLFSYHRLRDLLLIV  290

Query  268  TRIVELLGRRGWEALKYWWNLLQYWSQELKNSAVSLLNATAIAVAEGTDRVIEVVQ  323
            TRIVELLGRRGWEALKYWWNLLQYWSQELKNSAVSLLNATAIAVAEGTDRVIEVVQ
Sbjct  291  TRIVELLGRRGWEALKYWWNLLQYWSQELKNSAVSLLNATAIAVAEGTDRVIEVVQ  458



>Deinococcus aquiradiocola strain JCM 14371 sequence41, whole genome shotgun sequence
Sequence ID: NZ_BMOE01000041.1 Length: 690
Range 1: 107 to 454

Score:177 bits(450), Expect:1e-49,
Method:Compositional matrix adjust.,
Identities:115/116(99%), Positives:115/116(99%), Gaps:0/116(0%)

Query  208  THLPTprgpdrpegieeeggerdrdrSIRLVNGSLALIWDDLRSLCLFSYHRLRDLLLIV  267
            THLP PRGPDRPEGIEEEGGERDRDRSIRLVNGSLALIWDDLRSLCLFSYHRLRDLLLIV
Sbjct  107  THLPIPRGPDRPEGIEEEGGERDRDRSIRLVNGSLALIWDDLRSLCLFSYHRLRDLLLIV  286

Query  268  TRIVELLGRRGWEALKYWWNLLQYWSQELKNSAVSLLNATAIAVAEGTDRVIEVVQ  323
            TRIVELLGRRGWEALKYWWNLLQYWSQELKNSAVSLLNATAIAVAEGTDRVIEVVQ
Sbjct  287  TRIVELLGRRGWEALKYWWNLLQYWSQELKNSAVSLLNATAIAVAEGTDRVIEVVQ  454



>Escherichia coli strain NBE0006 contig00147, whole genome shotgun sequence
Sequence ID: NZ_JARBDI010000147.1 Length: 694
Range 1: 219 to 692

Score:177 bits(449), Expect:2e-49,
Method:Compositional matrix adjust.,
Identities:112/158(71%), Positives:123/158(77%), Gaps:7/158(4%)

Query  194  VNRVRQGYSPLSFQTHLPTprgpdrpegieeeggerdrdrSIRLVNGSLALIWDDLRSLC  253
            VNRVRQGYSPLSFQT  P PRGPDR   IEEEGG +DRDRSIRLVNG LAL WDDL SLC
Sbjct  692  VNRVRQGYSPLSFQTLTPNPRGPDRLGRIEEEGGGQDRDRSIRLVNGFLALAWDDLWSLC  513
```

```
Query  254  LFSYHRLRDLLLIVTRIVELLG-------RRGWEALKYWWNLLQYWSQELKNSAVSLLNA  306
            LFSYHRLRDL+L+  R VELLG       +RGWEALKY   ++QYW  ELK  A+SLL+
Sbjct  512  LFSYHRLRDLILVTARAVELLGHSSLRGLQRGWEALKYLGGIVQYWGLELKKRAISLLDT  333

Query  307  TAIAVAEGTDRVIEVVQGACRAIRHIPRRIRQGLERIL  344
             AIAVAEGTDR+IE +Q  CRAIR+IPRRIRQG E  L
Sbjct  332  VAIAVAEGTDRIIEFLQRICRAIRNIPRRIRQGFEAAL  219
```

>Enterococcus alcedinis strain CCM 8433 sequence26, whole genome shotgun sequence
Sequence ID: NZ_BMDT01000026.1 Length: 853
Range 1: 129 to 476

Score:177 bits(450), Expect:5e-49,
Method:Compositional matrix adjust.,
Identities:115/116(99%), Positives:115/116(99%), Gaps:0/116(0%)

```
Query  208  THLPTprgpdrpegieeeggerdrdrSIRLVNGSLALIWDDLRSLCLFSYHRLRDLLLIV  267
            THLP PRGPDRPEGIEEEGGERDRDRSIRLVNGSLALIWDDLRSLCLFSYHRLRDLLLIV
Sbjct  129  THLPIPRGPDRPEGIEEEGGERDRDRSIRLVNGSLALIWDDLRSLCLFSYHRLRDLLLIV  308

Query  268  TRIVELLGRRGWEALKYWWNLLQYWSQELKNSAVSLLNATAIAVAEGTDRVIEVVQ  323
            TRIVELLGRRGWEALKYWWNLLQYWSQELKNSAVSLLNATAIAVAEGTDRVIEVVQ
Sbjct  309  TRIVELLGRRGWEALKYWWNLLQYWSQELKNSAVSLLNATAIAVAEGTDRVIEVVQ  476
```

>Escherichia coli strain NBE0009 contig00162, whole genome shotgun sequence
Sequence ID: NZ_JARBDL010000162.1 Length: 350
Range 1: 2 to 349

Score:165 bits(417), Expect:2e-46,
Method:Compositional matrix adjust.,
Identities:88/116(76%), Positives:105/116(90%), Gaps:0/116(0%)

```
Query  93   CTTAVPWNASWSNKSLEQIWNHTTWMEWDREINNYTSLIHSLIeesqnqqekneqellel  152
            C TAVPWN+SWSNK+ E+IW++ TWM+WDREINNYT  I+ L+EESQ QQE+NE++LL L
Sbjct  349  CPTAVPWNSSWSNKTQEEIWDNMTWMQWDREINNYTKTIYDLLEESQYQQERNEKDLLAL  170

Query  153  DKWASLWNWFNITNWLWYIKLFIMIVGGLVGLRIVFAVLSIVNRVRQGYSPLSFQT  208
            D W +LWNWF+I+ WLWYIK+FIMIVGGL+GLRI+FAVLSIVNRVRQGYSPLSFQT
Sbjct  169  DSWNNLWNWFDISKWLWYIKIFIMIVGGLIGLRIIFAVLSIVNRVRQGYSPLSFQT  2
```

>Corynebacterium glutamicum strain 38 Scaffolds_31_4109, whole genome shotgun sequence
Sequence ID: NZ_JACENN010000031.1 Length: 3118
Range 1: 2381 to 2734

Score:179 bits(454), Expect:2e-45,
Method:Compositional matrix adjust.,
Identities:116/118(98%), Positives:116/118(98%), Gaps:0/118(0%)

```
Query  208   THLPTprgpdrpegieeeggerdrdrSIRLVNGSLALIWDDLRSLCLFSYHRLRDLLLIV  267
             THLP PRGPDRPEGIEEEGGERDRDRSIRLVNGSLALIWDDLRSLCLFSYHRLRDLLLIV
Sbjct  2381  THLPIPRGPDRPEGIEEEGGERDRDRSIRLVNGSLALIWDDLRSLCLFSYHRLRDLLLIV  2560

Query  268   TRIVELLGRRGWEALKYWWNLLQYWSQELKNSAVSLLNATAIAVAEGTDRVIEVVQGA  325
             TRIVELLGRRGWEALKYWWNLLQYWSQELKNSAVSLLNATAIAVAEGTDRVIEVVQ A
Sbjct  2561  TRIVELLGRRGWEALKYWWNLLQYWSQELKNSAVSLLNATAIAVAEGTDRVIEVVQEA  2734
```

>Corynebacterium glutamicum strain 51 Scaffolds_32_4236, whole genome shotgun sequence
Sequence ID: NZ_JACENK010000032.1 Length: 4236
Range 1: 2963 to 3316

Score:179 bits(454), Expect:2e-45,
Method:Compositional matrix adjust.,
Identities:116/118(98%), Positives:116/118(98%), Gaps:0/118(0%)

```
Query  208   THLPTprgpdrpegieeeggerdrdrSIRLVNGSLALIWDDLRSLCLFSYHRLRDLLLIV  267
             THLP PRGPDRPEGIEEEGGERDRDRSIRLVNGSLALIWDDLRSLCLFSYHRLRDLLLIV
Sbjct  2963  THLPIPRGPDRPEGIEEEGGERDRDRSIRLVNGSLALIWDDLRSLCLFSYHRLRDLLLIV  3142

Query  268   TRIVELLGRRGWEALKYWWNLLQYWSQELKNSAVSLLNATAIAVAEGTDRVIEVVQGA  325
             TRIVELLGRRGWEALKYWWNLLQYWSQELKNSAVSLLNATAIAVAEGTDRVIEVVQ A
Sbjct  3143  TRIVELLGRRGWEALKYWWNLLQYWSQELKNSAVSLLNATAIAVAEGTDRVIEVVQEA  3316
```

>Lactiplantibacillus plantarum strain MH-301 Scaffolds_50_4236, whole genome shotgun sequence
Sequence ID: NZ_JAOYUG010000050.1 Length: 4236
Range 1: 196 to 549

Score:179 bits(454), Expect:2e-45,
Method:Compositional matrix adjust.,
Identities:116/118(98%), Positives:116/118(98%), Gaps:0/118(0%)

```
Query  208  THLPTprgpdrpegieeeggerdrdrSIRLVNGSLALIWDDLRSLCLFSYHRLRDLLLIV  267
            THLP PRGPDRPEGIEEEGGERDRDRSIRLVNGSLALIWDDLRSLCLFSYHRLRDLLLIV
Sbjct  196  THLPIPRGPDRPEGIEEEGGERDRDRSIRLVNGSLALIWDDLRSLCLFSYHRLRDLLLIV  375

Query  268  TRIVELLGRRGWEALKYWWNLLQYWSQELKNSAVSLLNATAIAVAEGTDRVIEVVQGA  325
            TRIVELLGRRGWEALKYWWNLLQYWSQELKNSAVSLLNATAIAVAEGTDRVIEVVQ A
Sbjct  376  TRIVELLGRRGWEALKYWWNLLQYWSQELKNSAVSLLNATAIAVAEGTDRVIEVVQEA  549
```

Range 2: 196 to 549

Score:179 bits(454), Expect:2e-45,
Method:Compositional matrix adjust.,
Identities:116/118(98%), Positives:116/118(98%), Gaps:0/118(0%)

```
Query  208  THLPTprgpdrpegieeeggerdrdrSIRLVNGSLALIWDDLRSLCLFSYHRLRDLLLIV  267
            THLP PRGPDRPEGIEEEGGERDRDRSIRLVNGSLALIWDDLRSLCLFSYHRLRDLLLIV
Sbjct  196  THLPIPRGPDRPEGIEEEGGERDRDRSIRLVNGSLALIWDDLRSLCLFSYHRLRDLLLIV  375

Query  268  TRIVELLGRRGWEALKYWWNLLQYWSQELKNSAVSLLNATAIAVAEGTDRVIEVVQGA  325
            TRIVELLGRRGWEALKYWWNLLQYWSQELKNSAVSLLNATAIAVAEGTDRVIEVVQ A
Sbjct  376  TRIVELLGRRGWEALKYWWNLLQYWSQELKNSAVSLLNATAIAVAEGTDRVIEVVQEA  549
```

>Klebsiella pneumoniae strain EuSCAPE_IL022, whole genome shotgun sequence

Sequence ID: NZ_UKKD02000023.1 Length: 411
Range 1: 46 to 378

Score:161 bits(408), Expect:1e-44,
Method:Compositional matrix adjust.,
Identities:82/111(74%), Positives:92/111(82%), Gaps:0/111(0%)

```
Query  234  SIRLVNGSLALIWDDLRSLCLFSYHRLRDLLLIVTRIVELLGRRGWEALKYWWNLLQYWS  293
            SIRLV+G LAL W+DLRSLCLFSYHRLRDLLLIVTR VELLG+RGWE LKY  +L+QYW
Sbjct  378  SIRLVSGFLALAWEDLRSLCLFSYHRLRDLLLIVTRAVELLGQRGWEILKYLGSLIQYWG  199

Query  294  QELKNSAVSLLNATAIAVAEGTDRVIEVVQGACRAIRHIPRRIRQGLERIL  344
             ELK SA+SLL+  AIA AEGTDR+IE +Q  CRAI +IPRRIRQG E  L
Sbjct  198  LELKKSAISLLDTIAIATAEGTDRIIEGIQRICRAICNIPRRIRQGFEAAL  46
```

>Klebsiella pneumoniae strain EuSCAPE_MT015, whole genome shotgun sequence
Sequence ID: NZ_UKIW02000108.1 Length: 338
Range 1: 2 to 337

Score:155 bits(393), Expect:8e-43,
Method:Compositional matrix adjust.,
Identities:85/112(76%), Positives:104/112(92%), Gaps:0/112(0%)

```
Query  83   GIWGCSGKLICTTAVPWNASWSNKSLEQIWNHTTWMEWDREINNYTSLIHSLIeesqnqq  142
            GIWGCSGKLICTT VPWN+SWSNKSL  IW++ TWM+WDRE++NYT+ I+ L+E+SQNQQ
Sbjct  337  GIWGCSGKLICTTNVPWNSSWSNKSLTDIWDNMTWMQWDREVSNYTNTIYKLLEDSQNQQ  158

Query  143  ekneqellelDKWASLWNWFNITNWLWYIKLFIMIVGGLVGLRIVFAVLSIV  194
            +KNEQ+LL LDKW +LWNWFNIT+WLWYI++FIMIVGGL+GLRI+FAV+SIV
Sbjct  157  DKNEQDLLALDKWDNLWNWFNITSWLWYIRIFIMIVGGLIGLRIIFAVISIV  2
```

>Escherichia coli strain EN5308 contig00153, whole genome shotgun sequence
Sequence ID: NZ_JARBDE010000153.1 Length: 786
Range 1: 271 to 786

Score:160 bits(405), Expect:1e-42,
Method:Compositional matrix adjust.,
Identities:108/172(63%), Positives:127/172(73%), Gaps:7/172(4%)

```
Query  177  IVGGLVGLRIVFAVLSIVNRVRQGYSPLSFQTHLPTprgpdrpegieeeggerdrdrSIR  236
            IVGGL+ LRI+FAVLSIVN+VRQ YSPLSFQT  P P GPDR   IEEE GE+D+DRSI+
Sbjct  786  IVGGLIKLRIIFAVLSIVNKVRQRYSPLSFQTLTPNPGGPDRLRRIEEESGEQDKDRSIQ  607

Query  237  LVNGSLALIWDDLRSLCLFSYHRLRDLLLIVTRIVELLGR---RGW----EALKYWWNLL  289
            LV+G LAL  DDL++LCLFSYHRLRD + +  R VELL     RG     EALKY  +L+
Sbjct  606  LVSGFLALA*DDLQNLCLFSYHRLRDFISVTARTVELLRHHSLRGLQKR*EALKYLRSLV  427

Query  290  QYWSQELKNSAVSLLNATAIAVAEGTDRVIEVVQGACRAIRHIPRRIRQGLE  341
            QY   ELK SA+SLL+  AIAVAE TD++IE+ Q   RA  +IPRRIRQ  E
Sbjct  426  QY*GLELKKSAISLLDTIAIAVAERTDKIIELGQRIYRAFCNIPRRIRQSFE  271
```

>Escherichia coli strain EN5143 contig00088, whole genome shotgun sequence
Sequence ID: NZ_JAPMNI010000088.1 Length: 833
Range 1: 1 to 405

Score:160 bits(406), Expect:1e-42,
Method:Compositional matrix adjust.,
Identities:98/135(73%), Positives:115/135(85%), Gaps:0/135(0%)

```
Query  26   LTVQARQLLSGIVQQQNNLLRAIEAQQHLLQLTVWGIKQLQARILAVERYLKDQQLLGIW  85
            LTVQARQLLSGIVQQQ+NLL+AIEAQQHLLQLTV GIKQLQ R+LA+ERYLKDQQLL I
Sbjct  405  LTVQARQLLSGIVQQQSNLLKAIEAQQHLLQLTV*GIKQLQTRVLAIERYLKDQQLLRI*  226

Query  86   GCSGKLICTTAVPWNASWSNKSLEQIWNHTTWMEWDREINNYTSLIHSLIeesqnqqekn  145
            GCS KLIC TA+PWN+S SNK+ +QI N+ T M+ D+EI+NYT  I+ L+E+SQ QQEKN
Sbjct  225  GCSRKLICPTAIPWNSS*SNKTYDQI*NNMT*MQ*DKEISNYTYTIYRLLEDSQTQQEKN  46

Query  146  eqellelDKWASLWN  160
            E++LL LD  ASL N
Sbjct  45   EKDLLALDS*ASL*N  1
```

>Klebsiella pneumoniae strain EuSCAPE_MT017, whole genome shotgun sequence
Sequence ID: NZ_UKJA02000086.1 Length: 513
Range 1: 155 to 499

Score:149 bits(376), Expect:2e-39,
Method:Compositional matrix adjust.,
Identities:78/115(68%), Positives:85/115(73%), Gaps:7/115(6%)

```
Query  234  SIRLVNGSLALIWDDLRSLCLFSYHRLRDLLLIVTRIVELLGR-------RGWEALKYWW  286
            SIRLV G LAL WDDLRSLCLFSYHRLRDL+L+  R  ELLGR       RGWE LKY
Sbjct  499  SIRLVQGFLALAWDDLRSLCLFSYHRLRDLILVTARAGELLGRSSLKGLQRGWEILKYLG  320

Query  287  NLLQYWSQELKNSAVSLLNATAIAVAEGTDRVIEVVQGACRAIRHIPRRIRQGLE  341
            +L+QYW  ELK SA SLL+  AIAVA GTDR+IEV+Q   RAI HIP RIRQG E
Sbjct  319  SLIQYWGLELKKSATSLLDTIAIAVATGTDRIIEVLQAIGRAIYHIPTRIRQGFE  155
```

>Klebsiella pneumoniae strain EuSCAPE_NO017, whole genome shotgun sequence
Sequence ID: NZ_UKIO02000061.1 Length: 607
Range 1: 4 to 348

Score:143 bits(361), Expect:8e-37,
Method:Compositional matrix adjust.,
Identities:77/115(67%), Positives:87/115(75%), Gaps:7/115(6%)

```
Query  234  SIRLVNGSLALIWDDLRSLCLFSYHRLRDLLLIVTRIVELLG-------RRGWEALKYWW  286
            SIRLV+G LAL+WDDLRSLCLFSYHRLRD +L+  R  ELLG       +RGWE LKY
Sbjct  4    SIRLVSGFLALVWDDLRSLCLFSYHRLRDFILVTARAAELLGRNLLRGLQRGWEILKYLG  183

Query  287  NLLQYWSQELKNSAVSLLNATAIAVAEGTDRVIEVVQGACRAIRHIPRRIRQGLE  341
            +L QYW  ELK SA SLL+A AIA AEGTDR+IE +Q   RAI +IPRRIRQGLE
Sbjct  184  SLAQYWGTELKKSATSLLDAIAIATAEGTDRIIEAIQRIGRAIYNIPRRIRQGLE  348
```

>Photobacterium rosenbergii strain DP6N14-7 DP6N14-7_ctg0065, whole genome shotgun sequence
Sequence ID: NZ_JAHVIB010000057.1 Length: 564
Range 1: 116 to 445

```
Score:143 bits(360), Expect:8e-37,
Method:Compositional matrix adjust.,
Identities:84/110(76%), Positives:97/110(88%), Gaps:0/110(0%)

Query  153  DKWASLWNWFNITNWLWYIKLFIMIVGGLVGLRIVFAVLSIVNRVRQGYSPLSFQTHLPT  212
            D W +LWNWF+I+NWLWYI++FIMIVGGL+GLRIVFAVLSIVNRVRQGYSPLS QT +P
Sbjct  116  DSWKNLWNWFDISNWLWYIRIFIMIVGGLIGLRIVFAVLSIVNRVRQGYSPLSLQTLIPN  295

Query  213  prgpdrpegieeeggerdrdrSIRLVNGSLALIWDDLRSLCLFSYHRLRD  262
            P GPDR   I+EEGGE+ RDRS+RLV+G LAL W+DLR+LCLFSYH LRD
Sbjct  296  PVGPDRLGRIDEEGGEQGRDRSVRLVSGFLALAWEDLRNLCLFSYHHLRD  445
```

>Deinococcus aquiradiocola strain JCM 14371 sequence36, whole genome shotgun sequence
Sequence ID: NZ_BMOE01000036.1 Length: 869
Range 1: 435 to 653

```
Score:144 bits(362), Expect:6e-36,
Method:Compositional matrix adjust.,
Identities:72/73(99%), Positives:72/73(98%), Gaps:0/73(0%)

Query  26   LTVQARQLLSGIVQQQNNLLRAIEAQQHLLQLTVWGIKQLQARILAVERYLKDQQLLGIW  85
            LTVQARQLLS IVQQQNNLLRAIEAQQHLLQLTVWGIKQLQARILAVERYLKDQQLLGIW
Sbjct  435  LTVQARQLLSDIVQQQNNLLRAIEAQQHLLQLTVWGIKQLQARILAVERYLKDQQLLGIW  614

Query  86   GCSGKLICTTAVP  98
            GCSGKLICTTAVP
Sbjct  615  GCSGKLICTTAVP  653
```

>Thalassovita mangrovi strain GS-10 Scaffold21_1, whole genome shotgun sequence
Sequence ID: NZ_WWEN01000021.1 Length: 1440
Range 1: 752 to 970

```
Score:147 bits(370), Expect:2e-35,
Method:Compositional matrix adjust.,
Identities:72/73(99%), Positives:72/73(98%), Gaps:0/73(0%)

Query  26   LTVQARQLLSGIVQQQNNLLRAIEAQQHLLQLTVWGIKQLQARILAVERYLKDQQLLGIW  85
            LTVQARQLLS IVQQQNNLLRAIEAQQHLLQLTVWGIKQLQARILAVERYLKDQQLLGIW
Sbjct  970  LTVQARQLLSDIVQQQNNLLRAIEAQQHLLQLTVWGIKQLQARILAVERYLKDQQLLGIW  791

Query  86   GCSGKLICTTAVP  98
            GCSGKLICTTAVP
Sbjct  790  GCSGKLICTTAVP  752
```

>Klebsiella pneumoniae strain EuSCAPE_DE091, whole genome shotgun sequence
Sequence ID: NZ_UKMA02000049.1 Length: 618
Range 1: 202 to 570

```
Score:139 bits(351), Expect:3e-35,
Method:Compositional matrix adjust.,
Identities:78/123(63%), Positives:88/123(71%), Gaps:16/123(13%)

Query  234  SIRLVNGSLALIWDDLRSLCLFSYHRLRDLLLIVTRIVELLGR-------RGWEALKYWW  286
            S+RLVNG L L+W+DLRSLCLFSYHRLRDLLLI  R VELLGR       R WEALKY
Sbjct  570  SVRLVNGFLTLVWNDLRSLCLFSYHRLRDLLLITARAVELLGRCSLRGLQRVWEALKYLG  391

Query  287  NLLQYWSQELKNSAVSLLNATAIAVAEGTDRVIEVVQGACRAIRH---------IPRRIR  337
            +L+QYW  ELK SA+SLL+  AIAVAEGTDR+IEV+Q  CRA R+           P RIR
Sbjct  390  SLVQYWGLELKKSAISLLDTIAIAVAEGTDRIIEVIQIICRANRNNIDNL*GYPQPTRIR  211

Query  338  QGL  340
            Q L
Sbjct  210  QKL  202
```

>Klebsiella pneumoniae strain EuSCAPE_IL018, whole genome shotgun sequence
Sequence ID: NZ_UKJU02000101.1 Length: 388
Range 1: 3 to 386

```
Score:136 bits(342), Expect:5e-35,
Method:Compositional matrix adjust.,
Identities:89/128(70%), Positives:101/128(78%), Gaps:7/128(5%)

Query  199  QGYSPLSFQTHLPTprgpdrpegieeeggerdrdrSIRLVNGSLALIWDDLRSLCLFSYH  258
            QGYSP+SFQT  P PRGPDR   IEEGGE+DRDRS+RLV+G LAL WDDLRSLCLF YH
Sbjct  386  QGYSPVSFQTLTPNPRGPDRLGRIEEEGGEQDRDRSVRLVSGFLALAWDDLRSLCLFCYH  207

Query  259  RLRDLLLIVTRIVELLG-------RRGWEALKYWWNLLQYWSQELKNSAVSLLNATAIAV  311
            RLRD +L+  R VELLG       +RGWE LKY  +L+QYW  ELK SA+SLL+ TAIAV
Sbjct  206  RLRDFILVTARAVELLGHSSLKGLQRGWEILKYLGSLVQYWGLELKRSAISLLDTTAIAV  27

Query  312  AEGTDRVI  319
            AEGTDR+I
Sbjct  26   AEGTDRII  3
```

>Klebsiella pneumoniae strain EuSCAPE_IL015, whole genome shotgun sequence
Sequence ID: NZ_UKJV02000040.1 Length: 532
Range 1: 4 to 321

```
Score:137 bits(346), Expect:7e-35,
Method:Compositional matrix adjust.,
Identities:67/106(63%), Positives:80/106(75%), Gaps:7/106(6%)

Query  246  WDDLRSLCLFSYHRLRDLLLIVTRIVELLGR-------RGWEALKYWWNLLQYWSQELKN  298
            WDDLR+LCLFSYHRLRDL L+  R+VELLGR       +GW+ALKY  +L+QYW  ELK
Sbjct  4    WDDLRNLCLFSYHRLRDLTLVTARVVELLGRSSLRGLQKGWDALKYLGSLVQYWGLELKK  183

Query  299  SAVSLLNATAIAVAEGTDRVIEVVQGACRAIRHIPRRIRQGLERIL  344
            SA+SL +  AI VAEGTDR+IEV+Q  CRAI ++P RIRQG E  L
Sbjct  184  SAISLFDTIAIVVAEGTDRIIEVIQRFCRAIYNVPTRIRQGFEAAL  321
```

>Bacillus velezensis strain TN14-11 TN14-11_ctg0032, whole genome shotgun sequence
Sequence ID: NZ_JAHVIU010000017.1 Length: 1125
Range 1: 116 to 445

Score:143 bits(360), Expect:9e-35,

```
Method:Compositional matrix adjust.,
Identities:84/110(76%), Positives:97/110(88%), Gaps:0/110(0%)

Query  153  DKWASLWNWFNITNWLWYIKLFIMIVGGLVGLRIVFAVLSIVNRVRQGYSPLSFQTHLPT  212
            D W +LWNWF+I+NWLWYI++FIMIVGGL+GLRIVFAVLSIVNRVRQGYSPLS QT +P
Sbjct  116  DSWKNLWNWFDISNWLWYIRIFIMIVGGLIGLRIVFAVLSIVNRVRQGYSPLSLQTLIPN  295

Query  213  prgpdrpegieeeggerdrdrSIRLVNGSLALIWDDLRSLCLFSYHRLRD  262
            P GPDR   I+EEGGE+ RDRS+RLV+G LAL W+DLR+LCLFSYH LRD
Sbjct  296  PVGPDRLGRIDEEGGEQGRDRSVRLVSGFLALAWEDLRNLCLFSYHHLRD  445




>Actinoplanes campanulatus strain JCM 3059 sequence058, whole genome shotgun sequence
Sequence ID: NZ_BMPW01000058.1 Length: 4754
Range 1: 4531 to 4749

Score:143 bits(361), Expect:4e-33,
Method:Compositional matrix adjust.,
Identities:72/73(99%), Positives:72/73(98%), Gaps:0/73(0%)

Query  26    LTVQARQLLSGIVQQQNNLLRAIEAQQHLLQLTVWGIKQLQARILAVERYLKDQQLLGIW  85
             LTVQARQLLS IVQQQNNLLRAIEAQQHLLQLTVWGIKQLQARILAVERYLKDQQLLGIW
Sbjct  4531  LTVQARQLLSDIVQQQNNLLRAIEAQQHLLQLTVWGIKQLQARILAVERYLKDQQLLGIW  4710

Query  86    GCSGKLICTTAVP  98
             GCSGKLICTTAVP
Sbjct  4711  GCSGKLICTTAVP  4749




>Klebsiella pneumoniae strain EuSCAPE_MT009, whole genome shotgun sequence
Sequence ID: NZ_UKIV02000095.1 Length: 324
Range 1: 3 to 302

Score:130 bits(326), Expect:8e-33,
Method:Compositional matrix adjust.,
Identities:66/100(66%), Positives:82/100(82%), Gaps:0/100(0%)

Query  36   GIVQQQNNLLRAIEAQQHLLQLTVWGIKQLQARILAVERYLKDQQLLGIWGCSGKLICTT  95
            GIVQQQ+NLLRAIEAQ H+LQLTV GIKQLQAR+LA+E+YLKDQQLL I  CS KLICTT
Sbjct  3    GIVQQQSNLLRAIEAQHHMLQLTV*GIKQLQARVLALEKYLKDQQLLKI*SCSRKLICTT  182

Query  96   AVPWNASWSNKSLEQIWNHTTWMEWDREINNYTSLIHSLI  135
             VP N+S SN++ + I ++ T M+ D+EI+NYT+ I  L+
Sbjct  183  NVP*NSS*SNRTYDDI*DNMT*MQ*DKEIDNYTNTIFQLL  302




>Klebsiella pneumoniae strain EuSCAPE_MT015, whole genome shotgun sequence
Sequence ID: NZ_UKIW02000110.1 Length: 328
Range 1: 1 to 327

Score:127 bits(318), Expect:1e-31,
Method:Compositional matrix adjust.,
Identities:79/109(72%), Positives:89/109(81%), Gaps:0/109(0%)

Query  197  VRQGYSPLSFQTHLPTprgpdrpegieeeggerdrdrSIRLVNGSLALIWDDLRSLCLFS  256
            VRQGYSPLSFQT  P+PR  DR   IEE GGE+D+DRSIRLVNG LAL W+DLRSLCLFS
Sbjct  1    VRQGYSPLSFQTLTPSPRELDRLGRIEEGGGEQDKDRSIRLVNGFLALAWEDLRSLCLFS  180

Query  257  YHRLRDLLLIVTRIVELLGRRGWEALKYWWNLLQYWSQELKNSAVSLLN  305
            YH+LRD +LIV R VELLGRRGW+ LKY  +L QYW  ELK SA+SLL+
Sbjct  181  YHQLRDFILIVVRAVELLGRRGWDTLKYLGSLGQYWGLELKKSAISLLD  327




>Acinetobacter baumannii strain P4 contig_P4_92, whole genome shotgun sequence
Sequence ID: NZ_JBBXLU010000071.1 Length: 1226
Range 1: 53 to 319

Score:133 bits(335), Expect:6e-31,
Method:Compositional matrix adjust.,
Identities:89/89(100%), Positives:89/89(100%), Gaps:0/89(0%)

Query  153  DKWASLWNWFNITNWLWYIKLFIMIVGGLVGLRIVFAVLSIVNRVRQGYSPLSFQTHLPT  212
            DKWASLWNWFNITNWLWYIKLFIMIVGGLVGLRIVFAVLSIVNRVRQGYSPLSFQTHLPT
Sbjct  53   DKWASLWNWFNITNWLWYIKLFIMIVGGLVGLRIVFAVLSIVNRVRQGYSPLSFQTHLPT  232

Query  213  prgpdrpegieeeggerdrdrSIRLVNGS  241
            PRGPDRPEGIEEEGGERDRDRSIRLVNGS
Sbjct  233  PRGPDRPEGIEEEGGERDRDRSIRLVNGS  319




>Klebsiella pneumoniae strain EuSCAPE_MT009, whole genome shotgun sequence
Sequence ID: NZ_UKIV02000092.1 Length: 398
Range 1: 1 to 336

Score:119 bits(298), Expect:2e-28,
Method:Compositional matrix adjust.,
Identities:64/112(57%), Positives:78/112(69%), Gaps:7/112(6%)

Query  237  LVNGSLALIWDDLRSLCLFSYHRLRDLLLIVTRIVELLG-------RRGWEALKYWWNLL  289
            LV+  LAL  DDLRSLCLFSYHRLRD +LI  R VELL        ++  E LKY  +L
Sbjct  1    LVSRFLALA*DDLRSLCLFSYHRLRDFILIAARAVELLRHSILRRLQKK*EILKYLRSLA  180

Query  290  QYWSQELKNSAVSLLNATAIAVAEGTDRVIEVVQGACRAIRHIPRRIRQGLE  341
            QY S ELK SA+SL +  AIAVAE TD++IE+ Q  C++I +IP+RIRQG E
Sbjct  181  QY*SVELKKSAISLFDTIAIAVAEKTDKIIELAQRICKSILNIPKRIRQGFE  336




>Klebsiella pneumoniae strain EuSCAPE_IL009, whole genome shotgun sequence
Sequence ID: NZ_UKJO02000079.1 Length: 274
Range 1: 2 to 274

Score:113 bits(283), Expect:9e-27,
Method:Compositional matrix adjust.,
Identities:74/91(81%), Positives:83/91(91%), Gaps:0/91(0%)

Query  176  MIVGGLVGLRIVFAVLSIVNRVRQGYSPLSFQTHLPTprgpdrpegieeeggerdrdrSI  235
            MIVGGL+GLRI+FAVLS+VNRVRQGYSPLSFQT +PTPR  DRP GIEEEGG +DRDRSI
Sbjct  274  MIVGGLIGLRIIFAVLSVVNRVRQGYSPLSFQTLIPTPRELDRPGGIEEEGGGQDRDRSI  95

Query  236  RLVNGSLALIWDDLRSLCLFSYHRLRDLLLI  266
            RLV+G LAL WDDLR+LCLFSYH+LRD +LI
Sbjct  94   RLVSGFLALAWDDLRNLCLFSYHQLRDFILI  2
```

>Klebsiella pneumoniae strain EuSCAPE_IL022, whole genome shotgun sequence
Sequence ID: NZ_UKKD02000026.1 Length: 290
Range 1: 17 to 289

Score:110 bits(275), Expect:1e-25,
Method:Compositional matrix adjust.,
Identities:66/91(73%), Positives:81/91(89%), Gaps:0/91(0%)

```
Query  122  REINNYTSLIHSLIeesqnqqekneqellelDKWASLWNWFNITNWLWYIKLFIMIVGGL  181
            REI+NYT +I++L+E SQNQQEKNE++LL L+ W SLW+WF+IT WLWYI++FI+IVGGL
Sbjct  289  REISNYTEIIYNLLEVSQNQQEKNEKDLLALNSWDSLWSWFDITKWLWYIRIFIIIVGGL  110

Query  182  VGLRIVFAVLSIVNRVRQGYSPLSFQTHLPT  212
            +GLRI+FAVLSIVNRVRQGYSPLS QT  P
Sbjct  109  IGLRIIFAVLSIVNRVRQGYSPLSLQTLNPN  17
```

>Klebsiella pneumoniae strain EuSCAPE_IL004, whole genome shotgun sequence
Sequence ID: NZ_UKJP02000111.1 Length: 327
Range 1: 62 to 325

Score:109 bits(273), Expect:6e-25,
Method:Compositional matrix adjust.,
Identities:64/88(73%), Positives:78/88(88%), Gaps:0/88(0%)

```
Query  125  NNYTSLIHSLIeesqnqqekneqellelDKWASLWNWFNITNWLWYIKLFIMIVGGLVGL  184
            +N+T  I+ L+E SQNQQEKNE++LL L+ W +LWNWF+IT WLWYIK+FIMIVGGL+GL
Sbjct  325  SNHTDTIYRLLEISQNQQEKNEKDLLALNSWQNLWNWFDITQWLWYIKIFIMIVGGLIGL  146

Query  185  RIVFAVLSIVNRVRQGYSPLSFQTHLPT  212
            RI+FAV+SIVNRVRQGYSPLSFQT +P
Sbjct  145  RIIFAVISIVNRVRQGYSPLSFQTLIPN  62
```

>Klebsiella pneumoniae strain EuSCAPE_MT002, whole genome shotgun sequence
Sequence ID: NZ_UKIM02000087.1 Length: 265
Range 1: 2 to 265

Score:104 bits(260), Expect:2e-23,
Method:Compositional matrix adjust.,
Identities:71/88(81%), Positives:78/88(88%), Gaps:1/88(1%)

```
Query  165  TNWLWYIKLFIMIVGGLVGLRIVFAVLSIVNRVRQGYSPLSFQTHLP-Tprgpdrpegie  223
            TNWLWYI++FIMIVGGL+GLRI+FAVLSIVNRVRQGYSPLSFQT  P +   PDR E IE
Sbjct  265  TNWLWYIRIFIMIVGGLIGLRIIFAVLSIVNRVRQGYSPLSFQTLAPNSRGRPDRLERIE  86

Query  224  eeggerdrdrSIRLVNGSLALIWDDLRS  251
            EEGGE+DRDRSIRLV+G LAL WDDLRS
Sbjct  85   EEGGEQDRDRSIRLVSGFLALAWDDLRS  2
```

>Klebsiella pneumoniae strain EuSCAPE_IL016, whole genome shotgun sequence
Sequence ID: NZ_UKKA02000070.1 Length: 345
Range 1: 53 to 343

Score:105 bits(262), Expect:3e-23,
Method:Compositional matrix adjust.,
Identities:67/97(69%), Positives:83/97(85%), Gaps:0/97(0%)

```
Query  116  TWMEWDREINNYTSLIHSLIeesqnqqekneqellelDKWASLWNWFNITNWLWYIKLFI  175
            T M+ DREINNYT+ I+ L+EESQNQQEKN+++LL LD W +L + F+IT WL YIK+FI
Sbjct  343  T*MQ*DREINNYTNTIYRLLEESQNQQEKNKKDLLALDSWQNL*S*FSITKWL*YIKIFI  164

Query  176  MIVGGLVGLRIVFAVLSIVNRVRQGYSPLSFQTHLPT  212
            +IV GL+ LRI+FAVLSIVNRVRQGYSPLSFQT +P+
Sbjct  163  IIVRGLISLRIIFAVLSIVNRVRQGYSPLSFQTLIPS  53
```

>Photobacterium rosenbergii strain DP6N14-7 DP6N14-7_ctg0054, whole genome shotgun sequence
Sequence ID: NZ_JAHVIB010000049.1 Length: 960
Range 1: 14 to 214

Score:95.9 bits(237), Expect:8e-18,
Method:Compositional matrix adjust.,
Identities:47/67(70%), Positives:53/67(79%), Gaps:0/67(0%)

```
Query  278  GWEALKYWWNLLQYWSQELKNSAVSLLNATAIAVAEGTDRVIEVVQGACRAIRHIPRRIR  337
            GWEALKY  +L+QYW QELK SA+SL +A AI VAEGTDR+IEV Q  CRAI +IPRRIR
Sbjct  14   GWEALKYLGSLVQYWGQELKKSAISLFDALAITVAEGTDRIIEVGQRLCRAICNIPRRIR  193

Query  338  QGLERIL  344
            QG E  L
Sbjct  194  QGFEAAL  214
```

>Klebsiella pneumoniae strain EuSCAPE_LT003, whole genome shotgun sequence
Sequence ID: NZ_UKJH02000068.1 Length: 317
Range 1: 126 to 314

Score:83.2 bits(204), Expect:3e-15,
Method:Compositional matrix adjust.,
Identities:41/63(65%), Positives:48/63(76%), Gaps:0/63(0%)

```
Query  282  LKYWWNLLQYWSQELKNSAVSLLNATAIAVAEGTDRVIEVVQGACRAIRHIPRRIRQGLE  341
            LKY  + +QYW  ELK SAVSL +  AIAVAEGTDR+IEVVQG  +AI +IPRRIRQG +
Sbjct  314  LKYLGSRVQYWGLELKKSAVSLFDTLAIAVAEGTDRIIEVVQGISKAILNIPRRIRQGFK  135

Query  342  RIL  344
              L
Sbjct  134  AAL  126
```

>Pseudomonas aeruginosa strain VET-121 NODE_144_length_275_cov_1.53535_ID_287, whole genome shotgun sequence
Sequence ID: NZ_QZXB01000257.1 Length: 275
Range 1: 40 to 171

Score:82.0 bits(201), Expect:7e-15,
Method:Compositional matrix adjust.,

Identities:41/44(93%), Positives:43/44(97%), Gaps:0/44(0%)

```
Query  26   LTVQARQLLSGIVQQQNNLLRAIEAQQHLLQLTVWGIKQLQARI  69
            LTVQ RQLLSGIVQQQ+NLLRAIEAQQHLLQLTVWGIKQLQAR+
Sbjct  171  LTVQVRQLLSGIVQQQSNLLRAIEAQQHLLQLTVWGIKQLQARV  40
```

>Bacillus velezensis strain TN14-11 TN14-11_ctg0036, whole genome shotgun sequence
Sequence ID: NZ_JAHVIU010000019.1 Length: 1101
Range 1: 203 to 355

Score:71.6 bits(174), Expect:3e-09,
Method:Compositional matrix adjust.,
Identities:36/51(71%), Positives:40/51(78%), Gaps:0/51(0%)

```
Query  294  QELKNSAVSLLNATAIAVAEGTDRVIEVVQGACRAIRHIPRRIRQGLERIL  344
            QELK SA+SL +A AI VAEGTDR+IEV Q  CRAI +IPRRIRQG E  L
Sbjct  203  QELKKSAISLFDALAITVAEGTDRIIEVGQRLCRAICNIPRRIRQGFEAAL  355
```

>Klebsiella pneumoniae strain EuSCAPE_IL005, whole genome shotgun sequence
Sequence ID: NZ_UKJL02000056.1 Length: 442
Range 1: 318 to 440

Score:57.8 bits(138), Expect:1e-05,
Method:Compositional matrix adjust.,
Identities:30/41(73%), Positives:34/41(82%), Gaps:0/41(0%)

```
Query  305  NATAIAVAEGTDRVIEVVQGACRAIRHIPRRIRQGLERILL  345
            +ATAIAVAEGTDR+IE+VQ   RAI +IPRRIRQG E  LL
Sbjct  440  DATAIAVAEGTDRIIELVQRIGRAICNIPRRIRQGFETALL  318
```

Query #4: sp|P04591|GAG_HV1H2 Gag polyprotein OS=Human immunodeficiency virus type 1 group M subtype B (isolate HXB2) OX=11706 GN=gag PE=1 SV=3 Query ID: lcl|Query_837153
Length: 500

Sequences producing significant alignments:

| Description | Scientific Name | Common Name | Taxid | Max Score | Total Score | Query cover | E Value | Per. Ident | Acc. Len | Accession |
|---|---|---|---|---|---|---|---|---|---|---|
| Thalassovita mangrovi strain GS-10 Scaffold19_1, whole genome... | Thalassovita... | NA | 2692236 | 969 | 969 | 100% | 0.0 | 91.80 | 2900 | NZ_WWEN010000... |
| Actinoplanes campanulatus strain JCM 3059 sequence058, whole... | Actinoplanes... | NA | 113559 | 967 | 967 | 100% | 0.0 | 91.80 | 4754 | NZ_BMPW010000... |
| Enterococcus alcedinis strain CCM 8433 sequence23, whole genom... | Enterococcus... | NA | 1274384 | 966 | 966 | 100% | 0.0 | 91.80 | 6075 | NZ_BMDT010000... |
| Deinococcus aquiradiocola strain JCM 14371 sequence30, whole... | Deinococcus ... | NA | 393059 | 966 | 966 | 100% | 0.0 | 91.80 | 6128 | NZ_BMOE010000... |
| Burkholderia pseudomallei strain 2604STDY5519032... | Burkholderia... | NA | 28450 | 381 | 381 | 44% | 2e-127 | 81.17 | 656 | NZ_UVID020000... |
| Klebsiella pneumoniae strain EuSCAPE_IL020, whole genome shotg... | Klebsiella p... | NA | 573 | 278 | 278 | 31% | 8e-88 | 82.80 | 469 | NZ_UKKB020000... |
| Bacillus velezensis strain TN14-11 TN14-11_ctg0032, whole geno... | Bacillus vel... | NA | 492670 | 278 | 278 | 36% | 1e-84 | 74.03 | 1125 | NZ_JAHVIU0100... |
| Klebsiella pneumoniae strain EuSCAPE_IL017, whole genome shotg... | Klebsiella p... | NA | 573 | 266 | 266 | 26% | 2e-83 | 91.79 | 403 | NZ_UKKG020000... |
| Klebsiella pneumoniae strain EuSCAPE_LT003, whole genome shotg... | Klebsiella p... | NA | 573 | 266 | 266 | 28% | 2e-83 | 85.71 | 419 | NZ_UKJH020000... |
| Photobacterium rosenbergii strain DP6N14-7 DP6N14-7_ctg0069,... | Photobacteri... | NA | 294936 | 230 | 230 | 31% | 2e-68 | 69.81 | 667 | NZ_JAHVIB0100... |
| Vibrio hangzhouensis strain DP3N7-4 DP3N7-4_ctg0051, whole... | Vibrio hangz... | NA | 462991 | 230 | 230 | 31% | 5e-68 | 69.81 | 742 | NZ_JAHVKH0100... |
| Bacillus velezensis strain TN14-11 TN14-11_ctg0135, whole geno... | Bacillus vel... | NA | 492670 | 225 | 225 | 30% | 1e-66 | 70.86 | 633 | NZ_JAHVIU0100... |
| Photobacterium rosenbergii strain DP6N14-7 DP6N14-7_ctg0073,... | Photobacteri... | NA | 294936 | 225 | 225 | 30% | 1e-66 | 70.86 | 641 | NZ_JAHVIB0100... |
| Burkholderia pseudomallei strain 2604STDY5518971... | Burkholderia... | NA | 28450 | 217 | 217 | 21% | 6e-65 | 92.66 | 336 | NZ_UVFZ020001... |
| Klebsiella pneumoniae strain EuSCAPE_MT003, whole genome shotg... | Klebsiella p... | NA | 573 | 209 | 209 | 21% | 9e-62 | 90.48 | 318 | NZ_UKIK020001... |
| Klebsiella pneumoniae strain EuSCAPE_NO017, whole genome shotg... | Klebsiella p... | NA | 573 | 209 | 209 | 22% | 2e-61 | 84.35 | 348 | NZ_UKIO020000... |
| Klebsiella pneumoniae strain EuSCAPE_IL019, whole genome shotg... | Klebsiella p... | NA | 573 | 208 | 208 | 20% | 2e-61 | 93.20 | 312 | NZ_UKJZ020000... |
| Neisseria gonorrhoeae strain GC-206... | Neisseria go... | NA | 485 | 206 | 206 | 22% | 9e-61 | 85.71 | 334 | NZ_SAWN010001... |
| Neisseria gonorrhoeae strain GC-191... | Neisseria go... | NA | 485 | 186 | 186 | 18% | 3e-53 | 89.36 | 284 | NZ_SAWA010001... |
| Klebsiella pneumoniae strain EuSCAPE_IL005, whole genome shotg... | Klebsiella p... | NA | 573 | 183 | 183 | 20% | 5e-52 | 84.47 | 308 | NZ_UKJL020000... |
| Burkholderia pseudomallei strain 2604STDY5519032... | Burkholderia... | NA | 28450 | 182 | 182 | 20% | 3e-51 | 81.55 | 362 | NZ_UVID020000... |
| Neisseria gonorrhoeae strain gc-168... | Neisseria go... | NA | 485 | 180 | 180 | 18% | 8e-51 | 89.01 | 274 | NZ_SAVF010001... |
| Klebsiella pneumoniae strain EuSCAPE_MT002, whole genome shotg... | Klebsiella p... | NA | 573 | 172 | 172 | 17% | 6e-48 | 91.76 | 257 | NZ_UKIM020000... |
| Nocardioides luteus strain JCM 3358 sequence39, whole genome... | Nocardioides... | NA | 1844 | 189 | 189 | 20% | 3e-47 | 92.00 | 5754 | NZ_BMRK010000... |
| Streptomyces griseoflavus strain JCM 4479 sequence37, whole... | Streptomyces... | NA | 35619 | 189 | 189 | 20% | 3e-47 | 92.00 | 5775 | NZ_BMUC010000... |
| Saccharothrix coeruleofusca strain JCM 3313 sequence45, whole... | Saccharothri... | NA | 33919 | 189 | 189 | 20% | 3e-47 | 92.00 | 5775 | NZ_BMRG010000... |
| Streptomyces griseoloalbus strain JCM 4480 sequence054, whole... | Streptomyces... | NA | 67303 | 189 | 189 | 20% | 3e-47 | 92.00 | 5815 | NZ_BMUD010000... |
| Actinoplanes ianthinogenes strain JCM 3249 sequence037, whole... | Actinoplanes... | NA | 122358 | 189 | 189 | 20% | 3e-47 | 92.00 | 5815 | NZ_BMQZ010000... |
| Sinomonas sp. KCTC 49339 sequence28, whole genome shotgun... | Sinomonas sp... | NA | 2769422 | 189 | 189 | 20% | 3e-47 | 92.00 | 5849 | NZ_BNCM010000... |
| Saccharothrix mutabilis subsp. mutabilis strain JCM 3380... | Saccharothri... | NA | 66855 | 189 | 189 | 20% | 3e-47 | 92.00 | 5899 | NZ_BAAABU0100... |
| Corynebacterium glutamicum strain 39 Scaffolds_32_9166, whole... | Corynebacter... | NA | 1718 | 189 | 189 | 20% | 4e-47 | 92.00 | 8862 | NZ_JACEQW0100... |
| Corynebacterium glutamicum strain 37 Scaffolds_33_8928, whole... | Corynebacter... | NA | 1718 | 189 | 189 | 20% | 4e-47 | 92.00 | 8928 | NZ_JACENO0100... |
| Klebsiella pneumoniae strain EuSCAPE_LT007, whole genome shotg... | Klebsiella p... | NA | 573 | 166 | 166 | 16% | 1e-45 | 90.36 | 252 | NZ_UKJI020000... |
| Klebsiella pneumoniae strain EuSCAPE_IL005, whole genome shotg... | Klebsiella p... | NA | 573 | 166 | 166 | 16% | 2e-45 | 91.57 | 250 | NZ_UKJL020000... |
| Klebsiella pneumoniae strain EuSCAPE_IL019, whole genome shotg... | Klebsiella p... | NA | 573 | 157 | 157 | 17% | 4e-42 | 83.53 | 254 | NZ_UKJZ020000... |
| Klebsiella pneumoniae strain B2 isolate B2, whole genome shotg... | Klebsiella p... | NA | 573 | 154 | 154 | 82% | 2e-35 | 27.55 | 7306 | NZ_CABWNT0100... |
| Klebsiella quasipneumoniae strain EuSCAPE_IL014, whole genome... | Klebsiella q... | NA | 1463165 | 137 | 137 | 16% | 9e-35 | 79.52 | 269 | NZ_UJYK020000... |
| Klebsiella pneumoniae strain EuSCAPE_MT015, whole genome shotg... | Klebsiella p... | NA | 573 | 118 | 118 | 18% | 2e-27 | 73.20 | 281 | NZ_UKIW020001... |
| Mycobacterium tuberculosis variant bovis strain CSURQ1132, who... | Mycobacteriu... | NA | 1765 | 125 | 125 | 16% | 2e-26 | 78.82 | 1524 | NZ_CAJEUK0100... |
| Mycobacterium tuberculosis variant bovis strain CSURQ1134, who... | Mycobacteriu... | NA | 1765 | 125 | 125 | 17% | 5e-26 | 77.01 | 1697 | NZ_CAJEUT0100... |
| Klebsiella pneumoniae strain EuSCAPE_IL017, whole genome shotg... | Klebsiella p... | NA | 573 | 114 | 114 | 16% | 6e-26 | 73.49 | 325 | NZ_UKKG020000... |
| Enterococcus faecium strain A209R1B0C1... | Enterococcus... | NA | 1352 | 123 | 123 | 16% | 3e-25 | 77.78 | 1819 | NZ_PGCR010000... |
| Klebsiella pneumoniae strain EuSCAPE_LT007, whole genome shotg... | Klebsiella p... | NA | 573 | 106 | 106 | 14% | 9e-23 | 77.46 | 392 | NZ_UKJI020000... |
| Escherichia coli strain VRES0669, whole genome shotgun sequence | Escherichia ... | NA | 562 | 113 | 113 | 14% | 1e-21 | 78.57 | 5544 | NZ_UINV010000... |
| Cellulophaga baltica strain 14 NODE_63_length_422_cov_10.20678... | Cellulophaga... | NA | 76594 | 99.8 | 99.8 | 14% | 4e-20 | 75.00 | 422 | NZ_VIQP010000... |
| Enterococcus faecium strain A206R2B0... | Enterococcus... | NA | 1352 | 103 | 103 | 15% | 8e-19 | 73.08 | 1761 | NZ_PGCU010000... |
| Listeria monocytogenes strain YA00082404... | Listeria mon... | NA | 1639 | 103 | 103 | 15% | 1e-18 | 72.29 | 1816 | NZ_QEXE010000... |
| Rhizobium bangladeshense strain 1017 scf43_contig_2, whole... | Rhizobium ba... | NA | 1138189 | 93.6 | 168 | 9% | 1e-18 | 91.49 | 288 | NZ_LYPK010000... |
| Burkholderia pseudomallei strain A-175-05-2-02, whole genome... | Burkholderia... | NA | 28450 | 81.3 | 81.3 | 11% | 2e-13 | 71.19 | 455 | NZ_UVFW010001... |
| Pseudomonas sp. MPR-LB3 NODE_257_length_230_cov_0.882353, whol... | Pseudomonas ... | NA | 2070625 | 78.6 | 78.6 | 7% | 2e-13 | 97.37 | 230 | NZ_POFY010002... |
| Escherichia coli strain YA00028237 contig_187, whole genome... | Escherichia ... | NA | 562 | 86.3 | 86.3 | 15% | 3e-13 | 68.24 | 1828 | NZ_NGBP020001... |
| Photobacterium rosenbergii strain DP6N14-7 DP6N14-7_ctg0065,... | Photobacteri... | NA | 294936 | 79.7 | 79.7 | 7% | 1e-12 | 94.87 | 564 | NZ_JAHVIB0100... |
| Klebsiella pneumoniae strain P7701... | Klebsiella p... | NA | 573 | 81.6 | 81.6 | 9% | 6e-12 | 88.89 | 1305 | NZ_JACAWM0100... |
| Acinetobacter baumannii strain ACM-35... | Acinetobacte... | NA | 470 | 75.5 | 75.5 | 8% | 7e-10 | 84.09 | 1409 | NZ_QEHN010000... |
| Chlamydia trachomatis strain R4175extract, whole genome shotgu... | Chlamydia tr... | NA | 813 | 74.3 | 147 | 31% | 3e-09 | 33.33 | 2947 | NZ_CVNX010000... |
| Acinetobacter baumannii strain ACM-36... | Acinetobacte... | NA | 470 | 68.6 | 68.6 | 7% | 1e-08 | 86.84 | 567 | NZ_QAGO010000... |
| Chlamydia abortus strain 11_01, whole genome shotgun sequence | Chlamydia ab... | NA | 83555 | 72.4 | 72.4 | 29% | 1e-08 | 33.55 | 5240 | NZ_FQLK010000... |
| Chlamydia abortus strain 09_02, whole genome shotgun sequence | Chlamydia ab... | NA | 83555 | 72.0 | 72.0 | 29% | 2e-08 | 33.55 | 4507 | NZ_FQLN010000... |
| Klebsiella pneumoniae strain NUBRI-K 73, whole genome shotgun... | Klebsiella p... | NA | 573 | 69.3 | 69.3 | 12% | 8e-08 | 64.18 | 1717 | NZ_SOYT010000... |
| Salirhabdus euzebyi strain Marseille-Q1438, whole genome shotg... | Salirhabdus ... | NA | 394506 | 68.9 | 68.9 | 8% | 9e-08 | 80.49 | 1279 | NZ_CADDWK0100... |
| Pseudomonas aromaticivorans strain MAP12 42, whole genome... | Pseudomonas ... | NA | 2849492 | 69.3 | 69.3 | 29% | 1e-07 | 30.46 | 2454 | NZ_JAHRGL0100... |
| Enterobacter hormaechei strain AUH-ENM6... | Enterobacter... | NA | 158836 | 68.6 | 68.6 | 8% | 1e-07 | 82.50 | 1394 | NZ_PYHB010000... |
| Pseudomonas sp. MPR-R2A3 NODE_254_length_244_cov_0.832335, who... | Pseudomonas ... | NA | 2070623 | 62.0 | 62.0 | 6% | 1e-07 | 84.38 | 244 | NZ_POFW010002... |
| Acinetobacter baumannii strain ACM-14... | Acinetobacte... | NA | 470 | 65.9 | 65.9 | 8% | 2e-07 | 77.50 | 787 | NZ_PYDA010000... |
| Helicobacter acinonychis isolate 212_10, whole genome shotgun... | Helicobacter... | NA | 212 | 63.5 | 63.5 | 8% | 5e-06 | 78.05 | 1526 | NZ_FZLW010000... |
| Listeria monocytogenes strain CrieF364... | Listeria mon... | NA | 1639 | 61.6 | 61.6 | 6% | 6e-06 | 90.00 | 769 | NZ_JASJTP0100... |
| Escherichia coli strain 8374wH11... | Escherichia ... | NA | 562 | 62.4 | 62.4 | 8% | 1e-05 | 75.00 | 1397 | NZ_WSJI010000... |
| Methylocella silvestris strain TVC... | Methylocella... | NA | 199596 | 60.5 | 60.5 | 7% | 4e-05 | 82.86 | 1453 | NZ_PDZR010000... |
| Enterococcus faecalis strain 44.1... | Enterococcus... | NA | 1351 | 58.5 | 58.5 | 6% | 8e-05 | 84.85 | 806 | NZ_NSJR010000... |
| Mycobacterium tuberculosis variant bovis strain CSURQ1140, who... | Mycobacteriu... | NA | 1765 | 56.6 | 56.6 | 9% | 8e-04 | 70.83 | 1286 | NZ_CAJEUP0100... |
| Escherichia coli strain HU8 NODE_685_length_270_cov_0.756477,... | Escherichia ... | NA | 562 | 51.6 | 51.6 | 7% | 8e-04 | 48.65 | 270 | NZ_JAHCTM0100... |
| Salmonella enterica subsp. enterica serovar Typhimurium strain... | Salmonella e... | NA | 90371 | 50.4 | 50.4 | 7% | 0.001 | 44.74 | 211 | NZ_JALDNE0100... |
| Klebsiella pneumoniae strain EuSCAPE_IL004, whole genome shotg... | Klebsiella p... | NA | 573 | 51.2 | 51.2 | 9% | 0.002 | 65.96 | 356 | NZ_UKJP020001... |
| Escherichia coli strain 8374wG10... | Escherichia ... | NA | 562 | 54.3 | 54.3 | 6% | 0.004 | 78.79 | 1373 | NZ_WSJQ010000... |
| Klebsiella pneumoniae strain EuSCAPE_IL005, whole genome shotg... | Klebsiella p... | NA | 573 | 48.9 | 48.9 | 10% | 0.011 | 62.75 | 323 | NZ_UKJL020000... |
| Rickettsia endosymbiont of Ceutorhynchus assimilis isolate... | Rickettsia e... | NA | 2884482 | 52.4 | 52.4 | 29% | 0.027 | 23.98 | 4063491 | NZ_OU906081.1 |
| Geminicoccus harenae strain CPCC 101083 Scaffold77_1, whole... | Geminicoccus... | NA | 2498453 | 52.0 | 52.0 | 7% | 0.035 | 48.65 | 7497 | NZ_JABGCL0100... |
| Modestobacter marinus strain CPCC 205245 Scaffold62_1, whole... | Modestobacte... | NA | 477641 | 52.0 | 52.0 | 7% | 0.036 | 48.65 | 7666 | NZ_JABGCH0100... |

```
Modestobacter italicus strain CPCC 205242 Scaffold14_1, whole...  Modestobacte... NA             2732864    52.0    52.0  7%    0.037  48.65  15567      NZ_JABGCJ0100...
```

Alignments:

```
>Thalassovita mangrovi strain GS-10 Scaffold19_1, whole genome shotgun sequence
Sequence ID: NZ_WWEN01000019.1 Length: 2900
Range 1: 1040 to 2539

Score:969 bits(2505), Expect:0.0,
Method:Compositional matrix adjust.,
Identities:475/500(95%), Positives:491/500(98%), Gaps:0/500(0%)

Query  1     MGARASVLSGGELDRWEKIRLRPGGKKKYKLKHIVWASRELERFAVNPGLLETSEGCRQI  60
             MGARASVLSGGELD+WEKIRLRPGGKK+YKLKHIVWASRELERFAVNPGLLETSEGCRQI
Sbjct  1040  MGARASVLSGGELDKWEKIRLRPGGKKQYKLKHIVWASRELERFAVNPGLLETSEGCRQI  1219

Query  61    LGQLQPSLQTGSEELRSLYNTVATLYCVHQRIEIKDTKEALDKIeeeqnkskkkaqqaaa  120
             LGQLQPSLQTGSEELRSLYNT+A LYCVHQRI++KDTKEALDKIEEEQNKSKKKAQQAAA
Sbjct  1220  LGQLQPSLQTGSEELRSLYNTIAVLYCVHQRIDVKDTKEALDKIEEEQNKSKKKAQQAAA  1399

Query  121   DTGHSNQVSQNYPIVQNIQGQMVHQAISPRTLNAWVKVVEEKAFSPEVIPMFSALSEGAT  180
             DTG+++QVSQNYPIVQN+QGQMVHQAISPRTLNAWVKVVEEKAFSPEVIPMFSALSEGAT
Sbjct  1400  DTGNNSQVSQNYPIVQNLQGQMVHQAISPRTLNAWVKVVEEKAFSPEVIPMFSALSEGAT  1579

Query  181   PQDLNTMLNTVGGHQAAMQMLKETINEEAAEWDRVHPVHAGPIAPGQMREPRGSDIAGTT  240
             PQDLNTMLNTVGGHQAAMQMLKETINEEAAEWDR+HPVHAGPIAPGQMREPRGSDIAGTT
Sbjct  1580  PQDLNTMLNTVGGHQAAMQMLKETINEEAAEWDRLHPVHAGPIAPGQMREPRGSDIAGTT  1759

Query  241   STLQEQIGWMTNNPPIPVGEIYKRWIILGLNKIVRMYSPTSILDIRQGPKEPFRDYVDRF  300
             STLQEQIGWMT+NPPIPVGEIYKRWIILGLNKIVRMYSPTSILDIRQGPKEPFRDYVDRF
Sbjct  1760  STLQEQIGWMTHNPPIPVGEIYKRWIILGLNKIVRMYSPTSILDIRQGPKEPFRDYVDRF  1939

Query  301   YKTLRAEQASQEVKNWMTETLLVQNANPDCKTILKALGPAATLEEMMTACQGVGGPGHKA  360
             YKTLRAEQASQEVKNWMTETLLVQNANPDCKTILKALGP ATLEEMMTACQGVGGPGHKA
Sbjct  1940  YKTLRAEQASQEVKNWMTETLLVQNANPDCKTILKALGPGATLEEMMTACQGVGGPGHKA  2119

Query  361   RVLAEAMSQVTNSATIMMQRGNFRNQRKIVKCFNCGKEGHTARNCRAPRKKGCWKCGKEG  420
             RVLAEAMSQVTN ATIM+Q+GNFRNQRK VKCFNCGKEGH A+NCRAPRKKGCWKCGKEG
Sbjct  2120  RVLAEAMSQVTNPATIMIQKGNFRNQRKTVKCFNCGKEGHIAKNCRAPRKKGCWKCGKEG  2299

Query  421   HQMKDCTERQANFLGKIWPSYKGRPGNFLQSRPEPTAPPEESFRSGVETTTPPQKQEPID  480
             HQMKDCTERQANFLGKIWPS+KGRPGNFLQSRPEPTAPPEESFR G ETTTP QKQEPID
Sbjct  2300  HQMKDCTERQANFLGKIWPSHKGRPGNFLQSRPEPTAPPEESFRFGEETTTPSQKQEPID  2479

Query  481   KELYPLTSLRSLFGNDPSSQ  500
             KELYPL SLRSLFG+DPSSQ
Sbjct  2480  KELYPLASLRSLFGSDPSSQ  2539


>Actinoplanes campanulatus strain JCM 3059 sequence058, whole genome shotgun sequence
Sequence ID: NZ_BMPW01000058.1 Length: 4754
Range 1: 97 to 1596

Score:967 bits(2501), Expect:0.0,
Method:Compositional matrix adjust.,
Identities:475/500(95%), Positives:491/500(98%), Gaps:0/500(0%)

Query  1     MGARASVLSGGELDRWEKIRLRPGGKKKYKLKHIVWASRELERFAVNPGLLETSEGCRQI  60
             MGARASVLSGGELD+WEKIRLRPGGKK+YKLKHIVWASRELERFAVNPGLLETSEGCRQI
Sbjct  97    MGARASVLSGGELDKWEKIRLRPGGKKQYKLKHIVWASRELERFAVNPGLLETSEGCRQI  276

Query  61    LGQLQPSLQTGSEELRSLYNTVATLYCVHQRIEIKDTKEALDKIeeeqnkskkkaqqaaa  120
             LGQLQPSLQTGSEELRSLYNT+A LYCVHQRI++KDTKEALDKIEEEQNKSKKKAQQAAA
Sbjct  277   LGQLQPSLQTGSEELRSLYNTIAVLYCVHQRIDVKDTKEALDKIEEEQNKSKKKAQQAAA  456

Query  121   DTGHSNQVSQNYPIVQNIQGQMVHQAISPRTLNAWVKVVEEKAFSPEVIPMFSALSEGAT  180
             DTG+++QVSQNYPIVQN+QGQMVHQAISPRTLNAWVKVVEEKAFSPEVIPMFSALSEGAT
Sbjct  457   DTGNNSQVSQNYPIVQNLQGQMVHQAISPRTLNAWVKVVEEKAFSPEVIPMFSALSEGAT  636

Query  181   PQDLNTMLNTVGGHQAAMQMLKETINEEAAEWDRVHPVHAGPIAPGQMREPRGSDIAGTT  240
             PQDLNTMLNTVGGHQAAMQMLKETINEEAAEWDR+HPVHAGPIAPGQMREPRGSDIAGTT
Sbjct  637   PQDLNTMLNTVGGHQAAMQMLKETINEEAAEWDRLHPVHAGPIAPGQMREPRGSDIAGTT  816

Query  241   STLQEQIGWMTNNPPIPVGEIYKRWIILGLNKIVRMYSPTSILDIRQGPKEPFRDYVDRF  300
             STLQEQIGWMT+NPPIPVGEIYKRWIILGLNKIVRMYSPTSILDIRQGPKEPFRDYVDRF
Sbjct  817   STLQEQIGWMTHNPPIPVGEIYKRWIILGLNKIVRMYSPTSILDIRQGPKEPFRDYVDRF  996

Query  301   YKTLRAEQASQEVKNWMTETLLVQNANPDCKTILKALGPAATLEEMMTACQGVGGPGHKA  360
             YKTLRAEQASQEVKNWMTETLLVQNANPDCKTILKALGP ATLEEMMTACQGVGGPGHKA
Sbjct  997   YKTLRAEQASQEVKNWMTETLLVQNANPDCKTILKALGPGATLEEMMTACQGVGGPGHKA  1176

Query  361   RVLAEAMSQVTNSATIMMQRGNFRNQRKIVKCFNCGKEGHTARNCRAPRKKGCWKCGKEG  420
             RVLAEAMSQVTN ATIM+Q+GNFRNQRK VKCFNCGKEGH A+NCRAPRKKGCWKCGKEG
Sbjct  1177  RVLAEAMSQVTNPATIMIQKGNFRNQRKTVKCFNCGKEGHIAKNCRAPRKKGCWKCGKEG  1356

Query  421   HQMKDCTERQANFLGKIWPSYKGRPGNFLQSRPEPTAPPEESFRSGVETTTPPQKQEPID  480
             HQMKDCTERQANFLGKIWPS+KGRPGNFLQSRPEPTAPPEESFR G ETTTP QKQEPID
Sbjct  1357  HQMKDCTERQANFLGKIWPSHKGRPGNFLQSRPEPTAPPEESFRFGEETTTPSQKQEPID  1536

Query  481   KELYPLTSLRSLFGNDPSSQ  500
             KELYPL SLRSLFG+DPSSQ
Sbjct  1537  KELYPLASLRSLFGSDPSSQ  1596


>Enterococcus alcedinis strain CCM 8433 sequence23, whole genome shotgun sequence
Sequence ID: NZ_BMDT01000023.1 Length: 6075
Range 1: 55 to 1554

Score:966 bits(2498), Expect:0.0,
Method:Compositional matrix adjust.,
Identities:475/500(95%), Positives:491/500(98%), Gaps:0/500(0%)

Query  1     MGARASVLSGGELDRWEKIRLRPGGKKKYKLKHIVWASRELERFAVNPGLLETSEGCRQI  60
             MGARASVLSGGELD+WEKIRLRPGGKK+YKLKHIVWASRELERFAVNPGLLETSEGCRQI
Sbjct  55    MGARASVLSGGELDKWEKIRLRPGGKKQYKLKHIVWASRELERFAVNPGLLETSEGCRQI  234

Query  61    LGQLQPSLQTGSEELRSLYNTVATLYCVHQRIEIKDTKEALDKIeeeqnkskkkaqqaaa  120
             LGQLQPSLQTGSEELRSLYNT+A LYCVHQRI++KDTKEALDKIEEEQNKSKKKAQQAAA
Sbjct  235   LGQLQPSLQTGSEELRSLYNTIAVLYCVHQRIDVKDTKEALDKIEEEQNKSKKKAQQAAA  414

Query  121   DTGHSNQVSQNYPIVQNIQGQMVHQAISPRTLNAWVKVVEEKAFSPEVIPMFSALSEGAT  180
             DTG+++QVSQNYPIVQN+QGQMVHQAISPRTLNAWVKVVEEKAFSPEVIPMFSALSEGAT
Sbjct  415   DTGNNSQVSQNYPIVQNLQGQMVHQAISPRTLNAWVKVVEEKAFSPEVIPMFSALSEGAT  594

Query  181   PQDLNTMLNTVGGHQAAMQMLKETINEEAAEWDRVHPVHAGPIAPGQMREPRGSDIAGTT  240
             PQDLNTMLNTVGGHQAAMQMLKETINEEAAEWDR+HPVHAGPIAPGQMREPRGSDIAGTT
Sbjct  595   PQDLNTMLNTVGGHQAAMQMLKETINEEAAEWDRLHPVHAGPIAPGQMREPRGSDIAGTT  774

Query  241   STLQEQIGWMTNNPPIPVGEIYKRWIILGLNKIVRMYSPTSILDIRQGPKEPFRDYVDRF  300
```

```
              STLQEQIGWMT+NPPIPVGEIYKRWIILGLNKIVRMYSPTSILDIRQGPKEPFRDYVDRF
Sbjct  775   STLQEQIGWMTHNPPIPVGEIYKRWIILGLNKIVRMYSPTSILDIRQGPKEPFRDYVDRF  954

Query  301   YKTLRAEQASQEVKNWMTETLLVQNANPDCKTILKALGPAATLEEMMTACQGVGGPGHKA  360
              YKTLRAEQASQEVKNWMTETLLVQNANPDCKTILKALGP ATLEEMMTACQGVGGPGHKA
Sbjct  955   YKTLRAEQASQEVKNWMTETLLVQNANPDCKTILKALGPGATLEEMMTACQGVGGPGHKA  1134

Query  361   RVLAEAMSQVTNSATIMMQRGNFRNQRKIVKCFNCGKEGHTARNCRAPRKKGCWKCGKEG  420
              RVLAEAMSQVTN ATIM+Q+GNFRNQRK VKCFNCGKEGH A+NCRAPRKKGCWKCGKEG
Sbjct  1135  RVLAEAMSQVTNPATIMIQKGNFRNQRKTVKCFNCGKEGHIAKNCRAPRKKGCWKCGKEG  1314

Query  421   HQMKDCTERQANFLGKIWPSYKGRPGNFLQSRPEPTAPPEESFRSGVETTTPPQKQEPID  480
              HQMKDCTERQANFLGKIWPS+KGRPGNFLQSRPEPTAPPEESFR G ETTTP QKQEPID
Sbjct  1315  HQMKDCTERQANFLGKIWPSHKGRPGNFLQSRPEPTAPPEESFRFGEETTTPSQKQEPID  1494

Query  481   KELYPLTSLRSLFGNDPSSQ  500
              KELYPL SLRSLFG+DPSSQ
Sbjct  1495  KELYPLASLRSLFGSDPSSQ  1554
```

>Deinococcus aquiradiocola strain JCM 14371 sequence30, whole genome shotgun sequence
Sequence ID: NZ_BMOE01000030.1 Length: 6128
Range 1: 108 to 1607

Score:966 bits(2497), Expect:0.0,
Method:Compositional matrix adjust.,
Identities:475/500(95%), Positives:491/500(98%), Gaps:0/500(0%)

```
Query  1     MGARASVLSGGELDRWEKIRLRPGGKKKYKLKHIVWASRELERFAVNPGLLETSEGCRQI  60
              MGARASVLSGGELD+WEKIRLRPGGKK+YKLKHIVWASRELERFAVNPGLLETSEGCRQI
Sbjct  108   MGARASVLSGGELDKWEKIRLRPGGKKQYKLKHIVWASRELERFAVNPGLLETSEGCRQI  287

Query  61    LGQLQPSLQTGSEELRSLYNTVATLYCVHQRIEIKDTKEALDKIeeeqnkskkkaqqaaa  120
              LGQLQPSLQTGSEELRSLYNT+A LYCVHQRI++KDTKEALDKIEEEQNKSKKKAQQAAA
Sbjct  288   LGQLQPSLQTGSEELRSLYNTIAVLYCVHQRIDVKDTKEALDKIEEEQNKSKKKAQQAAA  467

Query  121   DTGHSNQVSQNYPIVQNIQGQMVHQAISPRTLNAWVKVVEEKAFSPEVIPMFSALSEGAT  180
              DTG+++QVSQNYPIVQN+QGQMVHQAISPRTLNAWVKVVEEKAFSPEVIPMFSALSEGAT
Sbjct  468   DTGNNSQVSQNYPIVQNLQGQMVHQAISPRTLNAWVKVVEEKAFSPEVIPMFSALSEGAT  647

Query  181   PQDLNTMLNTVGGHQAAMQMLKETINEEAAEWDRVHPVHAGPIAPGQMREPRGSDIAGTT  240
              PQDLNTMLNTVGGHQAAMQMLKETINEEAAEWDR+HPVHAGPIAPGQMREPRGSDIAGTT
Sbjct  648   PQDLNTMLNTVGGHQAAMQMLKETINEEAAEWDRLHPVHAGPIAPGQMREPRGSDIAGTT  827

Query  241   STLQEQIGWMTNNPPIPVGEIYKRWIILGLNKIVRMYSPTSILDIRQGPKEPFRDYVDRF  300
              STLQEQIGWMT+NPPIPVGEIYKRWIILGLNKIVRMYSPTSILDIRQGPKEPFRDYVDRF
Sbjct  828   STLQEQIGWMTHNPPIPVGEIYKRWIILGLNKIVRMYSPTSILDIRQGPKEPFRDYVDRF  1007

Query  301   YKTLRAEQASQEVKNWMTETLLVQNANPDCKTILKALGPAATLEEMMTACQGVGGPGHKA  360
              YKTLRAEQASQEVKNWMTETLLVQNANPDCKTILKALGP ATLEEMMTACQGVGGPGHKA
Sbjct  1008  YKTLRAEQASQEVKNWMTETLLVQNANPDCKTILKALGPGATLEEMMTACQGVGGPGHKA  1187

Query  361   RVLAEAMSQVTNSATIMMQRGNFRNQRKIVKCFNCGKEGHTARNCRAPRKKGCWKCGKEG  420
              RVLAEAMSQVTN ATIM+Q+GNFRNQRK VKCFNCGKEGH A+NCRAPRKKGCWKCGKEG
Sbjct  1188  RVLAEAMSQVTNPATIMIQKGNFRNQRKTVKCFNCGKEGHIAKNCRAPRKKGCWKCGKEG  1367

Query  421   HQMKDCTERQANFLGKIWPSYKGRPGNFLQSRPEPTAPPEESFRSGVETTTPPQKQEPID  480
              HQMKDCTERQANFLGKIWPS+KGRPGNFLQSRPEPTAPPEESFR G ETTTP QKQEPID
Sbjct  1368  HQMKDCTERQANFLGKIWPSHKGRPGNFLQSRPEPTAPPEESFRFGEETTTPSQKQEPID  1547

Query  481   KELYPLTSLRSLFGNDPSSQ  500
              KELYPL SLRSLFG+DPSSQ
Sbjct  1548  KELYPLASLRSLFGSDPSSQ  1607
```

>Burkholderia pseudomallei strain 2604STDY5519032 ERS240165SCcontig000074, whole genome shotgun sequence
Sequence ID: NZ_UVID02000074.1 Length: 656
Range 1: 1 to 654

Score:381 bits(979), Expect:2e-127,
Method:Compositional matrix adjust.,
Identities:181/223(81%), Positives:197/223(88%), Gaps:5/223(2%)

```
Query  274  VRMYSPTSILDIRQGPKEPFRDYVDRFYKTLRAEQASQEVKNWMTETLLVQNANPDCKTI  333
            VRMYSP SILDI+QGPKEPFRDYVDRF+KTLRAEQA+QEVKNWMT+TLLVQNANPDCKTI
Sbjct  654  VRMYSPVSILDIKQGPKEPFRDYVDRFFKTLRAEQATQEVKNWMTDTLLVQNANPDCKTI  475

Query  334  LKALGPAATLEEMMTACQGVGGPGHKARVLAEAMSQVTNSATIMMQRGNFRNQRKIVKCF  393
            L+ LGP ATLEEMMTACQGVGGP HKARVLAEAMSQ  N+ T+MMQ+ NF+  R+ VKCF
Sbjct  474  LRGLGPGATLEEMMTACQGVGGPSHKARVLAEAMSQANNT-TVMMQKNNFKGPRRTVKCF  298

Query  394  NCGKEGHTARNCRAPRKKGCWKCGKEGHQMKDCTERQANFLGKIWPSYKGRPGNFLQSRP  453
            NCGKEGH A+NCRAPRKKGCWKCGKEGHQMKDCTERQANFLGKIWPS+KGRPGNF+QSR
Sbjct  297  NCGKEGHIAKNCRAPRKKGCWKCGKEGHQMKDCTERQANFLGKIWPSHKGRPGNFIQSRL  118

Query  454  EPTAPPEESFRSGVETTTPPQKQEPIDKELYPLTSLRSLFGND  496
            EPTAPP ESF+   E  TP  +QE   KE  PLTSL+SLFGND
Sbjct  117  EPTAPPAESFK--FEEATPAPRQE--SKEREPLTSLKSLFGND  1
```

>Klebsiella pneumoniae strain EuSCAPE_IL020, whole genome shotgun sequence
Sequence ID: NZ_UKKB02000090.1 Length: 469
Range 1: 1 to 468

Score:278 bits(710), Expect:8e-88,
Method:Compositional matrix adjust.,
Identities:130/157(83%), Positives:142/157(90%), Gaps:1/157(0%)

```
Query  233  GSDIAGTTSTLQEQIGWMTNNPPIPVGEIYKRWIILGLNKIVRMYSPTSILDIRQGPKEP  292
            GSDIAGTTSTLQEQ+ W+T NP IPVGEIYKRWIILGLNKIVRMYSP SILDI+QGPKEP
Sbjct  468  GSDIAGTTSTLQEQVAWITGNPAIPVGEIYKRWIILGLNKIVRMYSPVSILDIKQGPKEP  289

Query  293  FRDYVDRFYKTLRAEQASQEVKNWMTETLLVQNANPDCKTILKALGPAATLEEMMTACQG  352
            FRDYVDRF+K LRAEQA+QEVKNWMT+TLLVQNANPDCK IL+ALGP A LEEMMTACQG
Sbjct  288  FRDYVDRFFKCLRAEQATQEVKNWMTDTLLVQNANPDCKAILRALGPGANLEEMMTACQG  109

Query  353  VGGPGHKARVLAEAMSQVTNSATIMMQRGNFRNQRKI  389
            VGGP HKARVLAEAMSQ  N+A IMMQR NF+  +++
Sbjct  108  VGGPSHKARVLAEAMSQ-ANNANIMMQRNNFKGIKRV  1
```

>Bacillus velezensis strain TN14-11 TN14-11_ctg0032, whole genome shotgun sequence
Sequence ID: NZ_JAHVIU010000017.1 Length: 1125
Range 1: 446 to 979

Score:278 bits(710), Expect:1e-84,

```
Method:Compositional matrix adjust.,
Identities:145/181(80%), Positives:164/181(90%), Gaps:3/181(1%)

Query  1    MGARASVLSGGELDRWEKIRLRPGGKKKYKLKHIVWASRELERFAVNPGLLETSEGCRQI  60
            +GARAS+L GG+L +WEKI+LRP GKK Y LKH+VWASRELERFAVNPGLLET+EGC+QI
Sbjct  979  IGARASILRGGKLGKWEKIKLRPVGKKHYMLKHLVWASRELERFAVNPGLLETAEGCKQI  800

Query  61   LGQLQPSLQTGSEELRSLYNTVATLYCVHQRIEIKDTKEALDKIeeeqnkskkkaqqaaa  120
            + QLQP+LQTG+EELRSLYNTVA LYCVH+ IE+KDTKEALDKIEEEQNKS++K QQA
Sbjct  799  INQLQPALQTGTEELRSLYNTVAVLYCVHKGIEVKDTKEALDKIEEEQNKSQQKTQQAKE  620

Query  121  DTGHSNQVSQNYPIVQNIQGQMVHQAISPRTLNAWVKVVEEKAFSPEVIPMFSALSEGAT  180
                  +VSQNYPIVQN+QGQMVHQ++SPRTLNAWVKVVEEKAFSPEVIPMFSALSEGAT
Sbjct  619  A---DMKVSQNYPIVQNLQGQMVHQSLSPRTLNAWVKVVEEKAFSPEVIPMFSALSEGAT  449

Query  181  P  181
            P
Sbjct  448  P  446
```

>Klebsiella pneumoniae strain EuSCAPE_IL017, whole genome shotgun sequence
Sequence ID: NZ_UKKG02000043.1 Length: 403
Range 1: 1 to 402

```
Score:266 bits(679), Expect:2e-83,
Method:Compositional matrix adjust.,
Identities:123/134(92%), Positives:131/134(97%), Gaps:0/134(0%)

Query  233  GSDIAGTTSTLQEQIGWMTNNPPIPVGEIYKRWIILGLNKIVRMYSPTSILDIRQGPKEP  292
            GSDIAGTTSTLQEQI WMT+NPP+PVG+IYKRWIILGLNKIVRMYSP SILDIRQGPKEP
Sbjct  402  GSDIAGTTSTLQEQITWMTSNPPVPVGDIYKRWIILGLNKIVRMYSPVSILDIRQGPKEP  223

Query  293  FRDYVDRFYKTLRAEQASQEVKNWMTETLLVQNANPDCKTILKALGPAATLEEMMTACQG  352
            FRDYVDRF+KTLRAEQA+Q+VKNWMT+TLLVQNANPDCKTIL+ALGP ATLEEMMTACQG
Sbjct  222  FRDYVDRFFKTLRAEQATQDVKNWMTDTLLVQNANPDCKTILRALGPGATLEEMMTACQG  43

Query  353  VGGPGHKARVLAEA  366
            VGGPGHKARVLAEA
Sbjct  42   VGGPGHKARVLAEA  1
```

>Klebsiella pneumoniae strain EuSCAPE_LT003, whole genome shotgun sequence
Sequence ID: NZ_UKJH02000061.1 Length: 419
Range 1: 2 to 418

```
Score:266 bits(679), Expect:2e-83,
Method:Compositional matrix adjust.,
Identities:120/140(86%), Positives:133/140(95%), Gaps:1/140(0%)

Query  284  DIRQGPKEPFRDYVDRFYKTLRAEQASQEVKNWMTETLLVQNANPDCKTILKALGPAATL  343
            DI+QGPKEPFRDYVDRF+K LRAEQA+Q+VKNWMT+TLLVQNANPDCKTIL+ALGPAA+L
Sbjct  418  DIKQGPKEPFRDYVDRFFKVLRAEQATQDVKNWMTDTLLVQNANPDCKTILRALGPAASL  239

Query  344  EEMMTACQGVGGPGHKARVLAEAMSQVTNSATIMMQRGNFRNQRKIVKCFNCGKEGHTAR  403
            EEMMTACQGVGGP HKARVLAEAMSQV ++A +MMQ+ NF+N RKIVKCFNCGKEGH AR
Sbjct  238  EEMMTACQGVGGPSHKARVLAEAMSQVNSNAQVMMQKSNFKN-RKIVKCFNCGKEGHIAR  62

Query  404  NCRAPRKKGCWKCGKEGHQM  423
            NCRAPRKKGCWKCG+EGHQM
Sbjct  61   NCRAPRKKGCWKCGREGHQM  2
```

>Photobacterium rosenbergii strain DP6N14-7 DP6N14-7_ctg0069, whole genome shotgun sequence
Sequence ID: NZ_JAHVIB010000058.1 Length: 667
Range 1: 169 to 636

```
Score:230 bits(586), Expect:2e-68,
Method:Compositional matrix adjust.,
Identities:122/159(77%), Positives:141/159(88%), Gaps:3/159(1%)

Query  1    MGARASVLSGGELDRWEKIRLRPGGKKKYKLKHIVWASRELERFAVNPGLLETSEGCRQI  60
            +GARAS+L GG+L +WEKI+LRP GKK Y LKH+VWASRELERFAVNPGLLET+EGCRQI
Sbjct  169  IGARASILRGGKLGKWEKIKLRPVGKKHYMLKHLVWASRELERFAVNPGLLETAEGCRQI  348

Query  61   LGQLQPSLQTGSEELRSLYNTVATLYCVHQRIEIKDTKEALDKIeeeqnkskkkaqqaaa  120
            + QLQP+LQTG+EELRSLYNTVA LYCVH+ IE++DTKEALDKIEEEQNKS++K QQA
Sbjct  349  INQLQPALQTGTEELRSLYNTVAVLYCVHKEIEVRDTKEALDKIEEEQNKSQQKTQQAKE  528

Query  121  DTGHSNQVSQNYPIVQNIQGQMVHQAISPRTLNAWVKVV  159
                  +VSQNYPIVQN+QGQMVHQ+ISPRTLNA +KVV
Sbjct  529  A---DMKVSQNYPIVQNLQGQMVHQSISPRTLNA*IKVV  636
```

>Vibrio hangzhouensis strain DP3N7-4 DP3N7-4_ctg0051, whole genome shotgun sequence
Sequence ID: NZ_JAHVKH010000037.1 Length: 742
Range 1: 77 to 544

```
Score:230 bits(586), Expect:5e-68,
Method:Compositional matrix adjust.,
Identities:122/159(77%), Positives:141/159(88%), Gaps:3/159(1%)

Query  1    MGARASVLSGGELDRWEKIRLRPGGKKKYKLKHIVWASRELERFAVNPGLLETSEGCRQI  60
            +GARAS+L GG+L +WEKI+LRP GKK Y LKH+VWASRELERFAVNPGLLET+EGCRQI
Sbjct  544  IGARASILRGGKLGKWEKIKLRPVGKKHYMLKHLVWASRELERFAVNPGLLETAEGCRQI  365

Query  61   LGQLQPSLQTGSEELRSLYNTVATLYCVHQRIEIKDTKEALDKIeeeqnkskkkaqqaaa  120
            + QLQP+LQTG+EELRSLYNTVA LYCVH+ IE++DTKEALDKIEEEQNKS++K QQA
Sbjct  364  INQLQPALQTGTEELRSLYNTVAVLYCVHKEIEVRDTKEALDKIEEEQNKSQQKTQQAKE  185

Query  121  DTGHSNQVSQNYPIVQNIQGQMVHQAISPRTLNAWVKVV  159
                  +VSQNYPIVQN+QGQMVHQ+ISPRTLNA +KVV
Sbjct  184  A---DMKVSQNYPIVQNLQGQMVHQSISPRTLNA*IKVV  77
```

>Bacillus velezensis strain TN14-11 TN14-11_ctg0135, whole genome shotgun sequence
Sequence ID: NZ_JAHVIU010000020.1 Length: 633
Range 1: 188 to 631

```
Score:225 bits(574), Expect:1e-66,
Method:Compositional matrix adjust.,
Identities:118/151(78%), Positives:136/151(90%), Gaps:3/151(1%)

Query  1    MGARASVLSGGELDRWEKIRLRPGGKKKYKLKHIVWASRELERFAVNPGLLETSEGCRQI  60
            MGARAS+L GG+LD+WEKIRLRPGGKK Y LKH+VWASRELERFAVNPGLLET+EGC+QI
```

```
Sbjct  188  MGARASILRGGKLDKWEKIRLRPGGKKHYMLKHLVWASRELERFAVNPGLLETAEGCKQI  367

Query  61   LGQLQPSLQTGSEELRSLYNTVATLYCVHQRIEIKDTKEALDKIeeeqnkskkkaqqaaa  120
            + QLQP+LQTG+EELRSLYNTVA LYCVH+ IE++DTKEALDKIEEEQNKS++K QQA
Sbjct  368  INQLQPALQTGTEELRSLYNTVAVLYCVHKGIEVRDTKEALDKIEEEQNKSQQKTQQAKE  547

Query  121  DTGHSNQVSQNYPIVQNIQGQMVHQAISPRT  151
                    +VSQNYPIVQN+QGQMVHQ++SPRT
Sbjct  548  A---DMKVSQNYPIVQNLQGQMVHQSLSPRT  631
```

```
>Photobacterium rosenbergii strain DP6N14-7 DP6N14-7_ctg0073, whole genome shotgun sequence
Sequence ID: NZ_JAHVIB010000060.1 Length: 641
Range 1: 196 to 639

Score:225 bits(574), Expect:1e-66,
Method:Compositional matrix adjust.,
Identities:118/151(78%), Positives:136/151(90%), Gaps:3/151(1%)

Query  1    MGARASVLSGGELDRWEKIRLRPGGKKKYKLKHIVWASRELERFAVNPGLLETSEGCRQI  60
            MGARAS+L GG+LD+WEKIRLRPGGKK Y LKH+VWASRELERFAVNPGLLET+EGC+QI
Sbjct  196  MGARASILRGGKLDKWEKIRLRPGGKKHYMLKHLVWASRELERFAVNPGLLETAEGCKQI  375

Query  61   LGQLQPSLQTGSEELRSLYNTVATLYCVHQRIEIKDTKEALDKIeeeqnkskkkaqqaaa  120
            + QLQP+LQTG+EELRSLYNTVA LYCVH+ IE++DTKEALDKIEEEQNKS++K QQA
Sbjct  376  INQLQPALQTGTEELRSLYNTVAVLYCVHKGIEVRDTKEALDKIEEEQNKSQQKTQQAKE  555

Query  121  DTGHSNQVSQNYPIVQNIQGQMVHQAISPRT  151
                    +VSQNYPIVQN+QGQMVHQ++SPRT
Sbjct  556  A---DMKVSQNYPIVQNLQGQMVHQSLSPRT  639
```

```
>Burkholderia pseudomallei strain 2604STDY5518971 ERS240105SCcontig000120, whole genome shotgun sequence
Sequence ID: NZ_UVFZ02000120.1 Length: 336
Range 1: 9 to 335

Score:217 bits(553), Expect:6e-65,
Method:Compositional matrix adjust.,
Identities:101/109(93%), Positives:105/109(96%), Gaps:0/109(0%)

Query  261  IYKRWIILGLNKIVRMYSPTSILDIRQGPKEPFRDYVDRFYKTLRAEQASQEVKNWMTET  320
            IYKRWIILGLNKIVRMYSP SILDI+QGPKEPFRDYVDRFY+TLRAEQA+QEVKNWMTET
Sbjct  335  IYKRWIILGLNKIVRMYSPVSILDIKQGPKEPFRDYVDRFYRTLRAEQATQEVKNWMTET  156

Query  321  LLVQNANPDCKTILKALGPAATLEEMMTACQGVGGPGHKARVLAEAMSQ  369
            LLVQNANPDCK+ILKALG  ATLEEMMTACQGVGGP HKARVLAEAMSQ
Sbjct  155  LLVQNANPDCKSILKALGTGATLEEMMTACQGVGGPSHKARVLAEAMSQ  9
```

```
>Klebsiella pneumoniae strain EuSCAPE_MT003, whole genome shotgun sequence
Sequence ID: NZ_UKIK02000113.1 Length: 318
Range 1: 3 to 317

Score:209 bits(531), Expect:9e-62,
Method:Compositional matrix adjust.,
Identities:95/105(90%), Positives:104/105(99%), Gaps:0/105(0%)

Query  233  GSDIAGTTSTLQEQIGWMTNNPPIPVGEIYKRWIILGLNKIVRMYSPTSILDIRQGPKEP  292
            GSDIAGTTS+LQEQIGWMT+NPPIPVG+IYKRWIILGLNKIVRMYSP SILDI+QGPKEP
Sbjct  317  GSDIAGTTSSLQEQIGWMTSNPPIPVGDIYKRWIILGLNKIVRMYSPVSILDIKQGPKEP  138

Query  293  FRDYVDRFYKTLRAEQASQEVKNWMTETLLVQNANPDCKTILKAL  337
            FRDYVDRF+KTLRAEQA+Q+VKNWMT+TLLVQNANPDCKTIL+AL
Sbjct  137  FRDYVDRFFKTLRAEQATQDVKNWMTDTLLVQNANPDCKTILRAL  3
```

```
>Klebsiella pneumoniae strain EuSCAPE_NO017, whole genome shotgun sequence
Sequence ID: NZ_UKIO02000074.1 Length: 348
Range 1: 2 to 346

Score:209 bits(531), Expect:2e-61,
Method:Compositional matrix adjust.,
Identities:97/115(84%), Positives:105/115(91%), Gaps:1/115(0%)

Query  312  EVKNWMTETLLVQNANPDCKTILKALGPAATLEEMMTACQGVGGPGHKARVLAEAMSQVT  371
            EVKNWMT+TLLVQNANPDCKTIL+ALGP A+LEEMMTACQGVGGPGHKARVLAEAMSQV
Sbjct  346  EVKNWMTDTLLVQNANPDCKTILRALGPGASLEEMMTACQGVGGPGHKARVLAEAMSQVN  167

Query  372  NSA-TIMMQRGNFRNQRKIVKCFNCGKEGHTARNCRAPRKKGCWKCGKEGHQMKD  425
            N+   IMMQ+ NF+  R+ VKCFNCGKEGH ARNCRAPRKKGCWKCG+EGHQMKD
Sbjct  166  NNNPAIMMQKSNFKGPRRAVKCFNCGKEGHIARNCRAPRKKGCWKCGREGHQMKD  2
```

```
>Klebsiella pneumoniae strain EuSCAPE_IL019, whole genome shotgun sequence
Sequence ID: NZ_UKJZ02000061.1 Length: 312
Range 1: 2 to 310

Score:208 bits(529), Expect:2e-61,
Method:Compositional matrix adjust.,
Identities:96/103(93%), Positives:101/103(98%), Gaps:0/103(0%)

Query  233  GSDIAGTTSTLQEQIGWMTNNPPIPVGEIYKRWIILGLNKIVRMYSPTSILDIRQGPKEP  292
            GSDIAGTTSTLQEQI WMT+NPPIPVG+IYKRWIILGLNKIVRMYSP SILDIRQGPKEP
Sbjct  2    GSDIAGTTSTLQEQIAWMTSNPPIPVGDIYKRWIILGLNKIVRMYSPVSILDIRQGPKEP  181

Query  293  FRDYVDRFYKTLRAEQASQEVKNWMTETLLVQNANPDCKTILK  335
            FRDYVDRF+KTLRAEQA+QEVKNWMTETLLVQNANPDCKTIL+
Sbjct  182  FRDYVDRFFKTLRAEQATQEVKNWMTETLLVQNANPDCKTILR  310
```

```
>Neisseria gonorrhoeae strain GC-206 NODE_136_length_334_cov_1.12903_ID_20444, whole genome shotgun sequence
Sequence ID: NZ_SAWN01000136.1 Length: 334
Range 1: 2 to 334

Score:206 bits(525), Expect:9e-61,
Method:Compositional matrix adjust.,
Identities:96/112(86%), Positives:104/112(92%), Gaps:1/112(0%)

Query  311  QEVKNWMTETLLVQNANPDCKTILKALGPAATLEEMMTACQGVGGPGHKARVLAEAMSQV  370
            Q+VKNWMTETLLVQNANPDCK IL+ALGP ATLEEMMTACQGVGGPGHKARVLAEAMSQV
Sbjct  2    QDVKNWMTETLLVQNANPDCKNILRALGPGATLEEMMTACQGVGGPGHKARVLAEAMSQV  181
```

```
Query  371  TNSATIMMQRGNFRNQRKIVKCFNCGKEGHTARNCRAPRKKGCWKCGKEGHQ  422
             N+  IMMQRGNF+  ++I+KCFNCGKEGH A+NCRAPRKKGCWKCGKEGHQ
Sbjct  182  -NNPNIMMQRGNFKGPKRIIKCFNCGKEGHIAKNCRAPRKKGCWKCGKEGHQ  334




>Neisseria gonorrhoeae strain GC-191 NODE_147_length_284_cov_0.751092_ID_22618, whole genome shotgun sequence
Sequence ID: NZ_SAWA01000147.1 Length: 284
Range 1: 1 to 282

Score:186 bits(473), Expect:3e-53,
Method:Compositional matrix adjust.,
Identities:84/94(89%), Positives:92/94(97%), Gaps:0/94(0%)

Query  259  GEIYKRWIILGLNKIVRMYSPTSILDIRQGPKEPFRDYVDRFYKTLRAEQASQEVKNWMT  318
            G+IYKRWIILGLNKIVRMYSP SILDI+QGPKEPFRDYVDRF+KTLRAEQA+Q+VKNWMT
Sbjct  282  GDIYKRWIILGLNKIVRMYSPVSILDIKQGPKEPFRDYVDRFFKTLRAEQATQDVKNWMT  103

Query  319  ETLLVQNANPDCKTILKALGPAATLEEMMTACQG  352
            +TLLVQNANPDCKTIL+ALGP A+LEEMMTACQG
Sbjct  102  DTLLVQNANPDCKTILRALGPGASLEEMMTACQG  1




>Klebsiella pneumoniae strain EuSCAPE_IL005, whole genome shotgun sequence
Sequence ID: NZ_UKJL02000060.1 Length: 308
Range 1: 1 to 306

Score:183 bits(465), Expect:5e-52,
Method:Compositional matrix adjust.,
Identities:87/103(84%), Positives:93/103(90%), Gaps:1/103(0%)

Query  307  EQASQEVKNWMTETLLVQNANPDCKTILKALGPAATLEEMMTACQGVGGPGHKARVLAEA  366
            EQ+SQEVKNWMT+TLLVQNANPDCKTILKALG  ATLEEMMTACQGVGGP HKARVLAEA
Sbjct  306  EQSSQEVKNWMTDTLLVQNANPDCKTILKALGTGATLEEMMTACQGVGGPSHKARVLAEA  127

Query  367  MSQVTNSATIMMQRGNFRNQRKIVKCFNCGKEGHTARNCRAPR  409
            MSQ TN+  +MMQ+ NF+  RKIVKCFNCGKEGH ARNCRAPR
Sbjct  126  MSQ-TNNINVMMQKSNFKGPRKIVKCFNCGKEGHIARNCRAPR  1




>Burkholderia pseudomallei strain 2604STDY5519032 ERS240165SCcontig000086, whole genome shotgun sequence
Sequence ID: NZ_UVID02000086.1 Length: 362
Range 1: 3 to 311

Score:182 bits(461), Expect:3e-51,
Method:Compositional matrix adjust.,
Identities:84/103(82%), Positives:96/103(93%), Gaps:0/103(0%)

Query  2    GARASVLSGGELDRWEKIRLRPGGKKKYKLKHIVWASRELERFAVNPGLLETSEGCRQIL  61
            GARAS+L GG+LD WEKIRLRPGGKK Y LKH+VWASRELERFA+NPGLLETSEGC+QI+
Sbjct  3    GARASILRGGKLDTWEKIRLRPGGKKHYMLKHLVWASRELERFALNPGLLETSEGCKQII  182

Query  62   GQLQPSLQTGSEELRSLYNTVATLYCVHQRIEIKDTKEALDKI  104
             QLQP+LQTG+EELRSL+NTVATLYCVH+ I++ DTKEALDKI
Sbjct  183  QQLQPALQTGTEELRSLFNTVATLYCVHKDIKVADTKEALDKI  311




>Neisseria gonorrhoeae strain gc-168 NODE_130_length_274_cov_0.69863_ID_20302, whole genome shotgun sequence
Sequence ID: NZ_SAVF01000130.1 Length: 274
Range 1: 1 to 273

Score:180 bits(456), Expect:8e-51,
Method:Compositional matrix adjust.,
Identities:81/91(89%), Positives:89/91(97%), Gaps:0/91(0%)

Query  233  GSDIAGTTSTLQEQIGWMTNNPPIPVGEIYKRWIILGLNKIVRMYSPTSILDIRQGPKEP  292
            GSDIAGTTSTLQEQI WMT+NPP+PVG+IYKRWIILGLNKIVRMYSP SILDI+QGPKEP
Sbjct  273  GSDIAGTTSTLQEQIAWMTSNPPVPVGDIYKRWIILGLNKIVRMYSPVSILDIKQGPKEP  94

Query  293  FRDYVDRFYKTLRAEQASQEVKNWMTETLLV  323
            FRDYVDRF+KTLRAEQ++QEVKNWMT+TLLV
Sbjct  93   FRDYVDRFFKTLRAEQSTQEVKNWMTDTLLV  1




>Klebsiella pneumoniae strain EuSCAPE_MT002, whole genome shotgun sequence
Sequence ID: NZ_UKIM02000089.1 Length: 257
Range 1: 2 to 256

Score:172 bits(436), Expect:6e-48,
Method:Compositional matrix adjust.,
Identities:78/85(92%), Positives:83/85(97%), Gaps:0/85(0%)

Query  233  GSDIAGTTSTLQEQIGWMTNNPPIPVGEIYKRWIILGLNKIVRMYSPTSILDIRQGPKEP  292
            GSDIAGTTSTLQEQI WMT+NPP+PVGEIYKRWIILGLNKIVRMYSP SILDIRQGPKEP
Sbjct  2    GSDIAGTTSTLQEQIAWMTSNPPVPVGEIYKRWIILGLNKIVRMYSPVSILDIRQGPKEP  181

Query  293  FRDYVDRFYKTLRAEQASQEVKNWM  317
            FRDYVDRF+KTLRAEQA+Q+VKNWM
Sbjct  182  FRDYVDRFFKTLRAEQATQDVKNWM  256




>Nocardioides luteus strain JCM 3358 sequence39, whole genome shotgun sequence
Sequence ID: NZ_BMRK01000039.1 Length: 5754
Range 1: 217 to 516

Score:189 bits(481), Expect:3e-47,
Method:Compositional matrix adjust.,
Identities:92/100(92%), Positives:93/100(93%), Gaps:0/100(0%)

Query  5    ASVLSGGELDRWEKIRLRPGGKKKYKLKHIVWASRELERFAVNPGLLETSEGCRQILGQL  64
             S+  G    RWEKIRLRPGGKKKYKLKHIVWASRELERFAVNPGLLETSEGCRQILGQL
Sbjct  217  VSIKRGRIRSRWEKIRLRPGGKKKYKLKHIVWASRELERFAVNPGLLETSEGCRQILGQL  396

Query  65   QPSLQTGSEELRSLYNTVATLYCVHQRIEIKDTKEALDKI  104
            QPSLQTGSEELRSLYNTVATLYCVHQRIEIKDTKEALDKI
Sbjct  397  QPSLQTGSEELRSLYNTVATLYCVHQRIEIKDTKEALDKI  516




>Streptomyces griseoflavus strain JCM 4479 sequence37, whole genome shotgun sequence
Sequence ID: NZ_BMUC01000037.1 Length: 5775
Range 1: 5232 to 5531
```

```
Score:189 bits(481), Expect:3e-47,
Method:Compositional matrix adjust.,
Identities:92/100(92%), Positives:93/100(93%), Gaps:0/100(0%)

Query  5     ASVLSGGELDRWEKIRLRPGGKKKYKLKHIVWASRELERFAVNPGLLETSEGCRQILGQL  64
              S+  G    RWEKIRLRPGGKKKYKLKHIVWASRELERFAVNPGLLETSEGCRQILGQL
Sbjct  5531  VSIKRGRIRSRWEKIRLRPGGKKKYKLKHIVWASRELERFAVNPGLLETSEGCRQILGQL  5352

Query  65    QPSLQTGSEELRSLYNTVATLYCVHQRIEIKDTKEALDKI  104
             QPSLQTGSEELRSLYNTVATLYCVHQRIEIKDTKEALDKI
Sbjct  5351  QPSLQTGSEELRSLYNTVATLYCVHQRIEIKDTKEALDKI  5232



>Saccharothrix coeruleofusca strain JCM 3313 sequence45, whole genome shotgun sequence
Sequence ID: NZ_BMRG01000045.1 Length: 5775
Range 1: 5232 to 5531

Score:189 bits(481), Expect:3e-47,
Method:Compositional matrix adjust.,
Identities:92/100(92%), Positives:93/100(93%), Gaps:0/100(0%)

Query  5     ASVLSGGELDRWEKIRLRPGGKKKYKLKHIVWASRELERFAVNPGLLETSEGCRQILGQL  64
              S+  G    RWEKIRLRPGGKKKYKLKHIVWASRELERFAVNPGLLETSEGCRQILGQL
Sbjct  5531  VSIKRGRIRSRWEKIRLRPGGKKKYKLKHIVWASRELERFAVNPGLLETSEGCRQILGQL  5352

Query  65    QPSLQTGSEELRSLYNTVATLYCVHQRIEIKDTKEALDKI  104
             QPSLQTGSEELRSLYNTVATLYCVHQRIEIKDTKEALDKI
Sbjct  5351  QPSLQTGSEELRSLYNTVATLYCVHQRIEIKDTKEALDKI  5232



>Streptomyces griseoloalbus strain JCM 4480 sequence054, whole genome shotgun sequence
Sequence ID: NZ_BMUD01000054.1 Length: 5815
Range 1: 5252 to 5551

Score:189 bits(481), Expect:3e-47,
Method:Compositional matrix adjust.,
Identities:92/100(92%), Positives:93/100(93%), Gaps:0/100(0%)

Query  5     ASVLSGGELDRWEKIRLRPGGKKKYKLKHIVWASRELERFAVNPGLLETSEGCRQILGQL  64
              S+  G    RWEKIRLRPGGKKKYKLKHIVWASRELERFAVNPGLLETSEGCRQILGQL
Sbjct  5551  VSIKRGRIRSRWEKIRLRPGGKKKYKLKHIVWASRELERFAVNPGLLETSEGCRQILGQL  5372

Query  65    QPSLQTGSEELRSLYNTVATLYCVHQRIEIKDTKEALDKI  104
             QPSLQTGSEELRSLYNTVATLYCVHQRIEIKDTKEALDKI
Sbjct  5371  QPSLQTGSEELRSLYNTVATLYCVHQRIEIKDTKEALDKI  5252



>Actinoplanes ianthinogenes strain JCM 3249 sequence037, whole genome shotgun sequence
Sequence ID: NZ_BMQZ01000037.1 Length: 5815
Range 1: 5252 to 5551

Score:189 bits(481), Expect:3e-47,
Method:Compositional matrix adjust.,
Identities:92/100(92%), Positives:93/100(93%), Gaps:0/100(0%)

Query  5     ASVLSGGELDRWEKIRLRPGGKKKYKLKHIVWASRELERFAVNPGLLETSEGCRQILGQL  64
              S+  G    RWEKIRLRPGGKKKYKLKHIVWASRELERFAVNPGLLETSEGCRQILGQL
Sbjct  5551  VSIKRGRIRSRWEKIRLRPGGKKKYKLKHIVWASRELERFAVNPGLLETSEGCRQILGQL  5372

Query  65    QPSLQTGSEELRSLYNTVATLYCVHQRIEIKDTKEALDKI  104
             QPSLQTGSEELRSLYNTVATLYCVHQRIEIKDTKEALDKI
Sbjct  5371  QPSLQTGSEELRSLYNTVATLYCVHQRIEIKDTKEALDKI  5252



>Sinomonas sp. KCTC 49339 sequence28, whole genome shotgun sequence
Sequence ID: NZ_BNCM01000028.1 Length: 5849
Range 1: 5223 to 5522

Score:189 bits(481), Expect:3e-47,
Method:Compositional matrix adjust.,
Identities:92/100(92%), Positives:93/100(93%), Gaps:0/100(0%)

Query  5     ASVLSGGELDRWEKIRLRPGGKKKYKLKHIVWASRELERFAVNPGLLETSEGCRQILGQL  64
              S+  G    RWEKIRLRPGGKKKYKLKHIVWASRELERFAVNPGLLETSEGCRQILGQL
Sbjct  5522  VSIKRGRIRSRWEKIRLRPGGKKKYKLKHIVWASRELERFAVNPGLLETSEGCRQILGQL  5343

Query  65    QPSLQTGSEELRSLYNTVATLYCVHQRIEIKDTKEALDKI  104
             QPSLQTGSEELRSLYNTVATLYCVHQRIEIKDTKEALDKI
Sbjct  5342  QPSLQTGSEELRSLYNTVATLYCVHQRIEIKDTKEALDKI  5223



>Saccharothrix mutabilis subsp. mutabilis strain JCM 3380 sequence46, whole genome shotgun sequence
Sequence ID: NZ_BAAABU010000046.1 Length: 5899
Range 1: 265 to 564

Score:189 bits(481), Expect:3e-47,
Method:Compositional matrix adjust.,
Identities:92/100(92%), Positives:93/100(93%), Gaps:0/100(0%)

Query  5    ASVLSGGELDRWEKIRLRPGGKKKYKLKHIVWASRELERFAVNPGLLETSEGCRQILGQL  64
             S+  G    RWEKIRLRPGGKKKYKLKHIVWASRELERFAVNPGLLETSEGCRQILGQL
Sbjct  265  VSIKRGRIRSRWEKIRLRPGGKKKYKLKHIVWASRELERFAVNPGLLETSEGCRQILGQL  444

Query  65   QPSLQTGSEELRSLYNTVATLYCVHQRIEIKDTKEALDKI  104
            QPSLQTGSEELRSLYNTVATLYCVHQRIEIKDTKEALDKI
Sbjct  445  QPSLQTGSEELRSLYNTVATLYCVHQRIEIKDTKEALDKI  564



>Corynebacterium glutamicum strain 39 Scaffolds_32_9166, whole genome shotgun sequence
Sequence ID: NZ_JACEQW010000032.1 Length: 8862
Range 1: 5003 to 5302

Score:189 bits(481), Expect:4e-47,
Method:Compositional matrix adjust.,
Identities:92/100(92%), Positives:93/100(93%), Gaps:0/100(0%)

Query  5     ASVLSGGELDRWEKIRLRPGGKKKYKLKHIVWASRELERFAVNPGLLETSEGCRQILGQL  64
              S+  G    RWEKIRLRPGGKKKYKLKHIVWASRELERFAVNPGLLETSEGCRQILGQL
Sbjct  5302  VSIKRGRIRSRWEKIRLRPGGKKKYKLKHIVWASRELERFAVNPGLLETSEGCRQILGQL  5123

Query  65    QPSLQTGSEELRSLYNTVATLYCVHQRIEIKDTKEALDKI  104
```

```
             QPSLQTGSEELRSLYNTVATLYCVHQRIEIKDTKEALDKI
Sbjct  5122  QPSLQTGSEELRSLYNTVATLYCVHQRIEIKDTKEALDKI  5003
```

>Corynebacterium glutamicum strain 37 Scaffolds_33_8928, whole genome shotgun sequence
Sequence ID: NZ_JACENO010000033.1 Length: 8928
Range 1: 4977 to 5276

Score:189 bits(481), Expect:4e-47,
Method:Compositional matrix adjust.,
Identities:92/100(92%), Positives:93/100(93%), Gaps:0/100(0%)

```
Query  5     ASVLSGGELDRWEKIRLRPGGKKKYKLKHIVWASRELERFAVNPGLLETSEGCRQILGQL  64
              S+  G    RWEKIRLRPGGKKKYKLKHIVWASRELERFAVNPGLLETSEGCRQILGQL
Sbjct  5276  VSIKRGRIRSRWEKIRLRPGGKKKYKLKHIVWASRELERFAVNPGLLETSEGCRQILGQL  5097

Query  65    QPSLQTGSEELRSLYNTVATLYCVHQRIEIKDTKEALDKI  104
             QPSLQTGSEELRSLYNTVATLYCVHQRIEIKDTKEALDKI
Sbjct  5096  QPSLQTGSEELRSLYNTVATLYCVHQRIEIKDTKEALDKI  4977
```

>Klebsiella pneumoniae strain EuSCAPE_LT007, whole genome shotgun sequence
Sequence ID: NZ_UKJI02000076.1 Length: 252
Range 1: 2 to 250

Score:166 bits(420), Expect:1e-45,
Method:Compositional matrix adjust.,
Identities:75/83(90%), Positives:82/83(98%), Gaps:0/83(0%)

```
Query  233  GSDIAGTTSTLQEQIGWMTNNPPIPVGEIYKRWIILGLNKIVRMYSPTSILDIRQGPKEP  292
            GSDIAGTTSTLQEQ+GWMT+NPPIPVG+IYKRWIILGLNKIVRMYSP SILDI+QGPKEP
Sbjct  2    GSDIAGTTSTLQEQVGWMTSNPPIPVGDIYKRWIILGLNKIVRMYSPVSILDIKQGPKEP  181

Query  293  FRDYVDRFYKTLRAEQASQEVKN  315
            FRDYVDRF+KTLRAEQA+Q+VKN
Sbjct  182  FRDYVDRFFKTLRAEQATQDVKN  250
```

>Klebsiella pneumoniae strain EuSCAPE_IL005, whole genome shotgun sequence
Sequence ID: NZ_UKJL02000066.1 Length: 250
Range 1: 1 to 249

Score:166 bits(419), Expect:2e-45,
Method:Compositional matrix adjust.,
Identities:76/83(92%), Positives:81/83(97%), Gaps:0/83(0%)

```
Query  233  GSDIAGTTSTLQEQIGWMTNNPPIPVGEIYKRWIILGLNKIVRMYSPTSILDIRQGPKEP  292
            GSDIAGTTSTLQEQI WMT+NPPIPVG+IYKRWIILGLNKIVRMYSP SILDI+QGPKEP
Sbjct  249  GSDIAGTTSTLQEQIAWMTSNPPIPVGDIYKRWIILGLNKIVRMYSPVSILDIKQGPKEP  70

Query  293  FRDYVDRFYKTLRAEQASQEVKN  315
            FRDYVDRF+KTLRAEQ+SQEVKN
Sbjct  69   FRDYVDRFFKTLRAEQSSQEVKN  1
```

>Klebsiella pneumoniae strain EuSCAPE_IL019, whole genome shotgun sequence
Sequence ID: NZ_UKJZ02000062.1 Length: 254
Range 1: 1 to 252

Score:157 bits(396), Expect:4e-42,
Method:Compositional matrix adjust.,
Identities:71/85(84%), Positives:78/85(91%), Gaps:1/85(1%)

```
Query  366  AMSQVTNSATIMMQRGNFRNQRKIVKCFNCGKEGHTARNCRAPRKKGCWKCGKEGHQMKD  425
            AMSQ  N+  IMMQ+GNF+  ++I+KCFNCGKEGH ARNCRAPRKKGCWKCGKEGHQMKD
Sbjct  1    AMSQA-NNINIMMQKGNFKGPKRIIKCFNCGKEGHIARNCRAPRKKGCWKCGKEGHQMKD  177

Query  426  CTERQANFLGKIWPSYKGRPGNFLQ  450
            CTERQANFLGKIWPS+KGRPGNFLQ
Sbjct  178  CTERQANFLGKIWPSHKGRPGNFLQ  252
```

>Klebsiella pneumoniae strain B2 isolate B2, whole genome shotgun sequence
Sequence ID: NZ_CABWNT010000058.1 Length: 7306
Range 1: 2421 to 3641

Score:154 bits(390), Expect:2e-35,
Method:Compositional matrix adjust.,
Identities:118/421(28%), Positives:201/421(47%), Gaps:21/421(4%)

```
Query  13    LDRWEKIRLRPGGKKKYKLKHIVWASRELERFAVNPGLLETSEGCRQILGQLQPSLQTGS  72
             L + EK+ ++  G +K    +  WA   ++ F       E     R ++  L+   QT S
Sbjct  2421  LKKLEKVTVQ--GSQKLTSGNCNWALNLVDLFHDTNFCKEKDWQLRDVIPLLEDVSQTLS  2594

Query  73    -EELRSLYNTVATLYCVHQRIEIKDTKEALDKIeeeqnkskkkaqqaaaDTGHSNQVSQN  131
              +E  +   T   +  V   ++I    +A       + K ++K       T   ++  +
Sbjct  2595  GQEREAFEKTWWAIAAVKMGLQINTVNDAKSTFSILKAKFERKTANT---TKKQSEPEEE  2765

Query  132   YPIVQNIQGQMVHQAISPRTLNAWVKVVEEKAFSPEV-IPMFSALSEGATPQDLNTMLNT  190
             YPI+ +  G    + ++PR    WV  +++     E  + +F  LS   T +++N  L+
Sbjct  2766  YPIMIDGAGNRNFRPLTPRGYTTWVNTIQQNNLLNEASVNLFGILSVDCTSEEMNAFLDV  2945

Query  191   VGGHQAAMQMLKETINEEAAEWDRVHPVHAGP-IAPGQMREPRGSD-IAGTTSTLQEQIG  248
             V G     Q+L + +++ A EWDR HP+   P +AP Q   P  +  I G     + Q+
Sbjct  2946  VPGQAGQKQVLLDNLDKIAEEWDRRHPMPNPPLVAPPQGPIPMTARFIRGLGVPRERQME  3125

Query  249   WMTNNPPIPVGEIYKRWIILGLNKIVR-MYSPTSILDIRQGPKEPFRDYVDRFYKTLRAE  307
                +       + Y++WII  + + ++ M       +IRQGPKEP+ ++VDR    +++E
Sbjct  3126  PAFDQ----FRQTYRQWIIEAMTEGIKIMIGKPKAQNIRQGPKEPYPEFVDRLLSQIKSE  3293

Query  308   QASQEVKNWMTETLLVQNANPDCKTILKALGPAATLEEMMTACQGVGGPGHKARVLAEAM  367
                 ++  ++T+TL +QNAN +CK  ++ L P  TLEE M AC+ +G    K  +LA+A+
Sbjct  3294  GHPADITKFLTDTLTIQNANDECKNAMRHLRPEDTLEEKMYACRDIGTMKQKMALLAKAL  3473

Query  368   SQVTNSATIMMQRGNFRNQRKIVK--CFNCGKEGHTARNCRAPRKKGCWKCGKEGHQMKD  425
                       M+ G F+      K  C+NCGK GH +  C+AP  K C+KC + GH  K
Sbjct  3474  Q---TGLAGPMKGGIFKGGPLGAKQTCYNCGKPGHLSSQCKAP--KVCFKCKQPGHFSKQ  3638

Query  426   C  426
             C
Sbjct  3639  C  3641
```

```
>Klebsiella quasipneumoniae strain EuSCAPE_IL014, whole genome shotgun sequence
Sequence ID: NZ_UJYK02000052.1 Length: 269
Range 1: 3 to 245

Score:137 bits(346), Expect:9e-35,
Method:Compositional matrix adjust.,
Identities:66/83(80%), Positives:74/83(89%), Gaps:2/83(2%)

Query  292  PFRDYVDRFYKTLRAEQASQEVKNWMTETLLVQNANPDCKTILKALGPAATLEEMMTACQ  351
            PFRDYVDRF+KTLRAEQA+QEVKNWMTETLLVQNANPDCKTIL+ALGP ATLEEMMTACQ
Sbjct  3    PFRDYVDRFFKTLRAEQATQEVKNWMTETLLVQNANPDCKTILRALGPGATLEEMMTACQ  182

Query  352  GVGGPGHKARVLAEAMSQVTNSA  374
            GVG  GH+A  +  A+S  T+S+
Sbjct  183  GVG--GHQAASIKLALSGGTSSS  245




>Klebsiella pneumoniae strain EuSCAPE_MT015, whole genome shotgun sequence
Sequence ID: NZ_UKIW02000113.1 Length: 281
Range 1: 3 to 281

Score:118 bits(295), Expect:2e-27,
Method:Compositional matrix adjust.,
Identities:71/97(73%), Positives:75/97(77%), Gaps:11/97(11%)

Query  411  KGCWKCGKEGHQMKDCTERQANFLGKIWPSYKGRPGNFLQSR-------PEPTAPPEESF  463
            KGCWKCGKEGHQMKDCTERQANFLGK+WPS KGRPGNFLQSR        EPTAPP ESF
Sbjct  3    KGCWKCGKEGHQMKDCTERQANFLGKVWPSNKGRPGNFLQSRPEPTAPPAEPTAPPAESF  182

Query  464  RSGVETTTPPQKQEPIDKELYPLTSLRSLFGNDPSSQ  500
            R   E TTP  KQE  D+E  PL SL+SLFG+DP SQ
Sbjct  183  R--FEETTPALKQEVKDRE--PLISLKSLFGSDPLSQ  281




>Mycobacterium tuberculosis variant bovis strain CSURQ1132, whole genome shotgun sequence
Sequence ID: NZ_CAJEUK010000155.1 Length: 1524
Range 1: 1271 to 1522

Score:125 bits(314), Expect:2e-26,
Method:Compositional matrix adjust.,
Identities:67/85(79%), Positives:69/85(81%), Gaps:2/85(2%)

Query  417   GKEGHQMKDCTERQANFLGKIWPSYKGRPGNFLQSRPEPTAPPEESFRSGVETTTPPQKQ  476
             GKEGHQMKDCTERQANFLGKIWPS KGRPGNF Q R EPTAPP ESF  G E  +  QKQ
Sbjct  1522  GKEGHQMKDCTERQANFLGKIWPSSKGRPGNFPQGRLEPTAPPAESFGMGEEIPS-SQKQ  1346

Query  477   EPIDKELY-PLTSLRSLFGNDPSSQ  500
             EP DK LY PLTSL+SLFGNDP SQ
Sbjct  1345  EPGDKGLYPPLTSLKSLFGNDP*SQ  1271




>Mycobacterium tuberculosis variant bovis strain CSURQ1134, whole genome shotgun sequence
Sequence ID: NZ_CAJEUT010000109.1 Length: 1697
Range 1: 1233 to 1490

Score:125 bits(313), Expect:5e-26,
Method:Compositional matrix adjust.,
Identities:67/87(77%), Positives:69/87(79%), Gaps:2/87(2%)

Query  415   KCGKEGHQMKDCTERQANFLGKIWPSYKGRPGNFLQSRPEPTAPPEESFRSGVETTTPPQ  474
             K  KEGHQMKDCTERQANFLGKIWPS KGRPGNF Q R EPTAPP ESF  G E  +  Q
Sbjct  1490  KQAKEGHQMKDCTERQANFLGKIWPSSKGRPGNFPQGRLEPTAPPAESFGMGEEIPS-SQ  1314

Query  475   KQEPIDKELY-PLTSLRSLFGNDPSSQ  500
             KQEP DK LY PLTSL+SLFGNDP SQ
Sbjct  1313  KQEPGDKGLYPPLTSLKSLFGNDP*SQ  1233




>Klebsiella pneumoniae strain EuSCAPE_IL017, whole genome shotgun sequence
Sequence ID: NZ_UKKG02000044.1 Length: 325
Range 1: 2 to 238

Score:114 bits(286), Expect:6e-26,
Method:Compositional matrix adjust.,
Identities:61/83(73%), Positives:66/83(79%), Gaps:6/83(7%)

Query  420  GHQMKDCT--ERQANFLGKIWPSYKGRPGNFLQSRPEPTAPPEESFRSGVETTTPPQKQE  477
            GHQMKDC   ERQANFLGKIWPS KGRPGNFLQ+RPEP+APP ESFR   E TTP  KQE
Sbjct  2    GHQMKDCNNNERQANFLGKIWPSSKGRPGNFLQNRPEPSAPPAESFR--FEETTPAPKQE  175

Query  478  PIDKELYPLTSLRSLFGNDPSSQ  500
              D+E   LTSL+SLFG+DP SQ
Sbjct  176  SKDREA--LTSLKSLFGSDPLSQ  238




>Enterococcus faecium strain A209R1B0C1 NODE_70_length_1819_cov_3.01832_ID_139, whole genome shotgun sequence
Sequence ID: NZ_PGCR01000088.1 Length: 1819
Range 1: 1 to 225

Score:123 bits(308), Expect:3e-25,
Method:Compositional matrix adjust.,
Identities:63/81(78%), Positives:68/81(83%), Gaps:6/81(7%)

Query  420  GHQMKDCTERQANFLGKIWPSYKGRPGNFLQSRPEPTAPPEESFRSGVETTTPPQKQEPI  479
            GHQMKDCTERQANFLGKIWPS+KGRPGNFLQ+RPEPTAPP ESFR   E TTP QKQE
Sbjct  1    GHQMKDCTERQANFLGKIWPSHKGRPGNFLQNRPEPTAPPAESFR--FEETTPAQKQESR  174

Query  480  DKELYPLTSLRSLFGNDPSSQ  500
            +    PLTSL+SLFG+DP SQ
Sbjct  175  E----PLTSLKSLFGSDPLSQ  225




>Klebsiella pneumoniae strain EuSCAPE_LT007, whole genome shotgun sequence
Sequence ID: NZ_UKJI02000072.1 Length: 392
Range 1: 192 to 392

Score:106 bits(265), Expect:9e-23,
Method:Compositional matrix adjust.,
Identities:55/71(77%), Positives:59/71(83%), Gaps:4/71(5%)

Query  426  CTERQANFLGKIWPSYKGRPGNFLQSRPEPTAPPEESFRSGVETTTPPQKQEPIDKELYP  485
```

```
            CTERQANFLGKIWPS+KGRPGNFLQSRPEP+APP ESFR   E TTP  KQE  DKE
Sbjct  392  CTERQANFLGKIWPSHKGRPGNFLQSRPEPSAPPAESFR--FEETTPAPKQESKDKEA--  225

Query  486  LTSLRSLFGND  496
            LTSL+SLFG+D
Sbjct  224  LTSLKSLFGSD  192
```

```
>Escherichia coli strain VRES0669, whole genome shotgun sequence
Sequence ID: NZ_UINV01000086.1 Length: 5544
Range 1: 5338 to 5544

Score:113 bits(283), Expect:1e-21,
Method:Compositional matrix adjust.,
Identities:55/70(79%), Positives:60/70(85%), Gaps:1/70(1%)

Query  321   LLVQNANPDCKTILKALGPAATLEEMMTACQGVGGPGHKARVLAEAMSQVTNSATIMMQR  380
             LLVQNANPDCKTIL+ALGP ATLEEMMTACQGVGGP HKARVLAEAMSQ  N A IMMQR
Sbjct  5544  LLVQNANPDCKTILRALGPGATLEEMMTACQGVGGPSHKARVLAEAMSQ-ANPANIMMQR  5368

Query  381   GNFRNQRKIV  390
              NF+  ++ +
Sbjct  5367  NNFKGPKRTI  5338
```

```
>Cellulophaga baltica strain 14 NODE_63_length_422_cov_10.206780, whole genome shotgun sequence
Sequence ID: NZ_VIQP01000062.1 Length: 422
Range 1: 210 to 422

Score:99.8 bits(247), Expect:4e-20,
Method:Compositional matrix adjust.,
Identities:54/72(75%), Positives:58/72(80%), Gaps:2/72(2%)

Query  430  QANFLGKIWPSYKGRPGNFLQSRPEPTAPPEESFRSGVETTTPPQKQEPIDKELY-PLTS  488
            QANFLG++WPS KGRPGNF QSRPEPTAPP ES   G E T+ PQ QEP DK LY PLTS
Sbjct  422  QANFLGRLWPSNKGRPGNFPQSRPEPTAPPAESMGMGEEITSSPQ-QEPRDKGLYPPLTS  246

Query  489  LRSLFGNDPSSQ  500
            L+SLFGNDP SQ
Sbjct  245  LKSLFGNDP*SQ  210
```

```
>Enterococcus faecium strain A206R2B0 NODE_116_length_1761_cov_4.01102_ID_231, whole genome shotgun sequence
Sequence ID: NZ_PGCU01000052.1 Length: 1761
Range 1: 3 to 209

Score:103 bits(258), Expect:8e-19,
Method:Compositional matrix adjust.,
Identities:57/78(73%), Positives:60/78(76%), Gaps:9/78(11%)

Query  423  MKDCTERQANFLGKIWPSYKGRPGNFLQSRPEPTAPPEESFRSGVETTTPPQKQEPIDKE  482
            MKDCTERQANFLGKIWPS KGRPGNF QSR EPTAPP ESFR         +KQE  D+E
Sbjct  3    MKDCTERQANFLGKIWPSQKGRPGNFHQSRLEPTAPPAESFRF--------EKQEMEDRE  158

Query  483  LYPLTSLRSLFGNDPSSQ  500
              PLTSL+SLFGNDP SQ
Sbjct  159  -PPLTSLKSLFGNDPLSQ  209
```

```
>Listeria monocytogenes strain YA00082404 NODE_73_length_1816_cov_5.664890, whole genome shotgun sequence
Sequence ID: NZ_QEXE01000073.1 Length: 1816
Range 1: 2 to 238

Score:103 bits(257), Expect:1e-18,
Method:Compositional matrix adjust.,
Identities:60/83(72%), Positives:64/83(77%), Gaps:12/83(14%)

Query  426  CT-ERQANFLGKIWPSYKGRPGNFLQSRP-------EPTAPPEESFRSGVETTTPPQKQE  477
            CT ERQANFLGKIWPS+KGRPGNFLQ+RP       EPTAPP ESFR G   TTP  KQE
Sbjct  2    CTNERQANFLGKIWPSHKGRPGNFLQNRPEPTAPPVEPTAPPAESFRFG--ETTPAPKQE  175

Query  478  PIDKELYPLTSLRSLFGNDPSSQ  500
            P D+E  PLTSL+SLFGNDP SQ
Sbjct  176  PKDRE--PLTSLKSLFGNDPWSQ  238
```

```
>Rhizobium bangladeshense strain 1017 scf43_contig_2, whole genome shotgun sequence
Sequence ID: NZ_LYPK01000043.1 Length: 288
Range 1: 13 to 153

Score:93.6 bits(231), Expect:1e-18,
Method:Compositional matrix adjust.,
Identities:43/47(91%), Positives:47/47(100%), Gaps:0/47(0%)

Query  121  DTGHSNQVSQNYPIVQNIQGQMVHQAISPRTLNAWVKVVEEKAFSPE  167
            DTG+++QVSQNYPIVQN+QGQMVHQAISPRTLNAWVKVVEEKAFSPE
Sbjct  13   DTGNNSQVSQNYPIVQNLQGQMVHQAISPRTLNAWVKVVEEKAFSPE  153
```

```
Range 2: 174 to 287

Score:75.1 bits(183), Expect:5e-12,
Method:Compositional matrix adjust.,
Identities:34/38(89%), Positives:38/38(100%), Gaps:0/38(0%)

Query  121  DTGHSNQVSQNYPIVQNIQGQMVHQAISPRTLNAWVKV  158
            DTG+++QVSQNYPIVQN+QGQMVHQAISPRTLNAWVKV
Sbjct  174  DTGNNSQVSQNYPIVQNLQGQMVHQAISPRTLNAWVKV  287
```

```
>Burkholderia pseudomallei strain A-175-05-2-02, whole genome shotgun sequence
Sequence ID: NZ_UVFW01000138.1 Length: 455
Range 1: 281 to 454

Score:81.3 bits(199), Expect:2e-13,
Method:Compositional matrix adjust.,
Identities:42/59(71%), Positives:47/59(79%), Gaps:1/59(1%)

Query  424  KDCTERQANFLGKIWPSYKGRPGNFLQSRPEPTAPPEESFRSGVETTTPPQKQEPIDKE  482
            KDCTERQANFLG+IWPS +GRPGNF QSR EPTAPP E++  G ETT+   KQE  DKE
```

```
Sbjct  454  KDCTERQANFLGRIWPSNRGRPGNFPQSRTEPTAPPAENWGMGEETTS-LLKQEQKDKE  281
```

```
>Pseudomonas sp. MPR-LB3 NODE_257_length_230_cov_0.882353, whole genome shotgun sequence
Sequence ID: NZ_POFY01000255.1 Length: 230
Range 1: 52 to 165

Score:78.6 bits(192), Expect:2e-13,
Method:Compositional matrix adjust.,
Identities:37/38(97%), Positives:37/38(97%), Gaps:0/38(0%)

Query  147  ISPRTLNAWVKVVEEKAFSPEVIPMFSALSEGATPQDL  184
            ISPRTLNAWVKVVEEKAFSPEVIPMFSALSEGATPQD
Sbjct  165  ISPRTLNAWVKVVEEKAFSPEVIPMFSALSEGATPQDF  52
```

```
>Escherichia coli strain YA00028237 contig_187, whole genome shotgun sequence
Sequence ID: NZ_NGBP02000187.1 Length: 1828
Range 1: 1586 to 1828

Score:86.3 bits(212), Expect:3e-13,
Method:Compositional matrix adjust.,
Identities:58/85(68%), Positives:61/85(71%), Gaps:11/85(12%)

Query  423   MKDCTERQANFLGKIWPSYKGRPGNFLQSR-------PEPTAPPEESFRSGVETTTPPQK  475
              KDC ERQ NFLGKIWPS+KGRPGNFLQSR        EPTAPP ESFR   E TTP  K
Sbjct  1828  YKDCAERQVNFLGKIWPSHKGRPGNFLQSRPEPTAPPAEPTAPPAESFR--FEGTTPTPK  1655

Query  476   QEPIDKELYPLTSLRSLFGNDPSSQ  500
             QE  D+E  PL SL+SLFGNDP SQ
Sbjct  1654  QEKKDRE--PLISLKSLFGNDPLSQ  1586
```

```
>Photobacterium rosenbergii strain DP6N14-7 DP6N14-7_ctg0065, whole genome shotgun sequence
Sequence ID: NZ_JAHVIB010000057.1 Length: 564
Range 1: 446 to 562

Score:79.7 bits(195), Expect:1e-12,
Method:Compositional matrix adjust.,
Identities:37/39(95%), Positives:39/39(100%), Gaps:0/39(0%)

Query  143  VHQAISPRTLNAWVKVVEEKAFSPEVIPMFSALSEGATP  181
            VHQ++SPRTLNAWVKVVEEKAFSPEVIPMFSALSEGATP
Sbjct  562  VHQSLSPRTLNAWVKVVEEKAFSPEVIPMFSALSEGATP  446
```

```
>Klebsiella pneumoniae strain P7701 NODE_50_length_1305_cov_481.863, whole genome shotgun sequence
Sequence ID: NZ_JACAWM010000050.1 Length: 1305
Range 1: 1169 to 1303

Score:81.6 bits(200), Expect:6e-12,
Method:Compositional matrix adjust.,
Identities:40/45(89%), Positives:41/45(91%), Gaps:0/45(0%)

Query  456   TAPPEESFRSGVETTTPPQKQEPIDKELYPLTSLRSLFGNDPSSQ  500
             TAPPEESFR G ETTTP QKQEPIDKELYPL SLRSLFG+DPSSQ
Sbjct  1303  TAPPEESFRFGEETTTPSQKQEPIDKELYPLASLRSLFGSDPSSQ  1169
```

```
>Acinetobacter baumannii strain ACM-35 NODE_65_length_1409_cov_2.30031, whole genome shotgun sequence
Sequence ID: NZ_QEHN01000065.1 Length: 1409
Range 1: 1 to 132

Score:75.5 bits(184), Expect:7e-10,
Method:Compositional matrix adjust.,
Identities:37/44(84%), Positives:37/44(84%), Gaps:0/44(0%)

Query  453  PEPTAPPEESFRSGVETTTPPQKQEPIDKELYPLTSLRSLFGND  496
            PEPTAPPEES R G ET TP QKQEP DKELYPL SLRSLFGND
Sbjct  1    PEPTAPPEESLRLGGETATPSQKQEPTDKELYPLASLRSLFGND  132
```

```
>Chlamydia trachomatis strain R4175extract, whole genome shotgun sequence
Sequence ID: NZ_CVNX01000009.1 Length: 2947
Range 1: 706 to 1158

Score:74.3 bits(181), Expect:3e-09,
Method:Compositional matrix adjust.,
Identities:55/165(33%), Positives:79/165(47%), Gaps:21/165(12%)

Query  282   ILDIRQGPKEPFRDYVDRFYKTLRAEQASQEVKNWMTETLLVQNANPDCKTILKALGPAA  341
             +  I QGP+E F D+V R  +         E    + E L+ + A  +C+    A+ P
Sbjct  706   LTKIIQGPQESFSDFVARMTEAAERIFGESEQAAPLIEQLIYEQATKECRA---AIAPRK  876

Query  342   T--LEEMMTACQGVGGPGHKARVLAEAMSQVTNSATIMMQRGNFRNQRKIVKCFNCGKEG  399
                L++ +  C+ +GGP   A  LA A+ Q  N +   M R N R       CFNCGK G
Sbjct  877   NKGLQDWLRVCRELGGPLSNA-GLAAAILQSQNRS---MGRNNQRT------CFNCGKPG  1026

Query  400   HTARNCRAPRKKG-----CWKCGKEGHQMKDCTERQANFLGKIWP  439
             H  ++CRAP K+G     C KCGK  H+   C    + +  G+I P
Sbjct  1027  HFKKDCRAPDKQGGTLTLCSKCGKGYHRADQCRSVR-DIKGRILP  1158
```

```
Range 2: 1680 to 2132

Score:73.6 bits(179), Expect:5e-09,
Method:Compositional matrix adjust.,
Identities:53/165(32%), Positives:79/165(47%), Gaps:21/165(12%)

Query  282   ILDIRQGPKEPFRDYVDRFYKTLRAEQASQEVKNWMTETLLVQNANPDCKTILKALGPAA  341
             +  I QGP+E F D+V R  +         E    + E L+ + A  +C+    A+ P
Sbjct  1680  LTKIIQGPQESFSDFVARMTEAAERIFGESEQAAPLIEQLIYEQATKECRA---AIAPRK  1850

Query  342   T--LEEMMTACQGVGGPGHKARVLAEAMSQVTNSATIMMQRGNFRNQRKIVKCFNCGKEG  399
                L++ +  C+ +GGP   A  LA A+ Q  N +   M R + R       CFNCGK G
Sbjct  1851  NKGLQDWLRVCRELGGPLTNAG-LAAAILQSQNRS---MSRNDQRT------CFNCGKPG  2000

Query  400   HTARNCRAPRKKG-----CWKCGKEGHQMKDCTERQANFLGKIWP  439
             H  ++CRAP K+G     C KCGK  H+   C    + +  G++ P
Sbjct  2001  HFKKDCRAPDKQGGTLTLCSKCGKGYHRADQCRSVR-DIKGRVLP  2132
```

```
>Acinetobacter baumannii strain ACM-36 NODE_67_length_567_cov_1.13409, whole genome shotgun sequence
Sequence ID: NZ_QAGO01000067.1 Length: 567
Range 1: 1 to 114

Score:68.6 bits(166), Expect:1e-08,
Method:Compositional matrix adjust.,
Identities:33/38(87%), Positives:33/38(86%), Gaps:0/38(0%)

Query  459  PEESFRSGVETTTPPQKQEPIDKELYPLTSLRSLFGND  496
            PEESFR G ETTTP QKQEP DKELYPL SLRSLFGND
Sbjct  1    PEESFRLGGETTTPSQKQEPTDKELYPLASLRSLFGND  114
```

```
>Chlamydia abortus strain 11_01, whole genome shotgun sequence
Sequence ID: NZ_FQLK01000002.1 Length: 5240
Range 1: 4532 to 4948

Score:72.4 bits(176), Expect:1e-08,
Method:Compositional matrix adjust.,
Identities:51/152(34%), Positives:73/152(48%), Gaps:20/152(13%)

Query  282   ILDIRQGPKEPFRDYVDRFYKTLRAEQASQEVKNWMTETLLVQNANPDCKTILKALGPAA  341
             +  I QGP+E F D+V R  +          E    + E L+ + A  +C+    A+ P
Sbjct  4948  LTKIIQGPQESFSDFVARMTEAAERIFGESEQAAPLIEQLIYEQATKECRA---AIAPRK  4778

Query  342   T--LEEMMTACQGVGGPGHKARVLAEAMSQVTNSATIMMQRGNFRNQRKIVKCFNCGKEG  399
                L++ +  C+ +GGP   A  LA A+ Q  N +   M R + R       CFNCGK G
Sbjct  4777  NKGLQDWLRVCRELGGPLTNA-GLAAAILQSQNRS---MSRNDQRT------CFNCGKPG  4628

Query  400   HTARNCRAPRKKG-----CWKCGKEGHQMKDC  426
             H  ++CRAP K+G     C KCGK  H+   C
Sbjct  4627  HFKKDCRAPDKQGGTLTLCSKCGKGYHRADQC  4532
```

```
>Chlamydia abortus strain 09_02, whole genome shotgun sequence
Sequence ID: NZ_FQLN01000004.1 Length: 4507
Range 1: 934 to 1350

Score:72.0 bits(175), Expect:2e-08,
Method:Compositional matrix adjust.,
Identities:51/152(34%), Positives:73/152(48%), Gaps:20/152(13%)

Query  282   ILDIRQGPKEPFRDYVDRFYKTLRAEQASQEVKNWMTETLLVQNANPDCKTILKALGPAA  341
             +  I QGP+E F D+V R  +          E    + E L+ + A  +C+    A+ P
Sbjct  934   LTKIIQGPQESFSDFVARMTEAAERIFGESEQAAPLIEQLIYEQATKECRA---AIAPRK  1104

Query  342   T--LEEMMTACQGVGGPGHKARVLAEAMSQVTNSATIMMQRGNFRNQRKIVKCFNCGKEG  399
                L++ +  C+ +GGP   A  LA A+ Q  N +   M R + R       CFNCGK G
Sbjct  1105  NKGLQDWLRVCRELGGPLTNA-GLAAAILQSQNRS---MSRNDQRT------CFNCGKPG  1254

Query  400   HTARNCRAPRKKG-----CWKCGKEGHQMKDC  426
             H  ++CRAP K+G     C KCGK  H+   C
Sbjct  1255  HFKKDCRAPDKQGGTLTLCSKCGKGYHRADQC  1350
```

```
>Klebsiella pneumoniae strain NUBRI-K 73, whole genome shotgun sequence
Sequence ID: NZ_SOYT01000073.1 Length: 1717
Range 1: 3 to 191

Score:69.3 bits(168), Expect:8e-08,
Method:Compositional matrix adjust.,
Identities:43/67(64%), Positives:48/67(71%), Gaps:11/67(16%)

Query  441  YKGRPGNFLQSRP-------EPTAPPEESFRSGVETTTPPQKQEPIDKELYPLTSLRSLF  493
            +KGRPGNFLQ+RP       EPTAPP ESFR   E TT   KQEP D+E  PLTSL+SLF
Sbjct  3    HKGRPGNFLQNRPEPTAPPVEPTAPPAESFR--FEETTLAPKQEPKDRE--PLTSLKSLF  170

Query  494  GNDPSSQ  500
            G+DP SQ
Sbjct  171  GSDPLSQ  191
```

```
>Salirhabdus euzebyi strain Marseille-Q1438, whole genome shotgun sequence
Sequence ID: NZ_CADDWK010000035.1 Length: 1279
Range 1: 1157 to 1279

Score:68.9 bits(167), Expect:9e-08,
Method:Compositional matrix adjust.,
Identities:33/41(80%), Positives:35/41(85%), Gaps:0/41(0%)

Query  460   EESFRSGVETTTPPQKQEPIDKELYPLTSLRSLFGNDPSSQ  500
             EES R G ETTTP QKQE IDK++YPL SLRSLFGNDPSSQ
Sbjct  1279  EESLRLGEETTTPSQKQEQIDKDMYPLASLRSLFGNDPSSQ  1157
```

```
>Pseudomonas aromaticivorans strain MAP12 42, whole genome shotgun sequence
Sequence ID: NZ_JAHRGL010000043.1 Length: 2454
Range 1: 2005 to 2406

Score:69.3 bits(168), Expect:1e-07,
Method:Compositional matrix adjust.,
Identities:46/151(30%), Positives:72/151(47%), Gaps:23/151(15%)

Query  282   ILDIRQGPKEPFRDYVDRFYKTLRAEQASQEVKNWMTETLLVQNANPDCKTILKALGPAA  341
             +  I QGP+EPF D+V R  +          E    + E L+ + A  +C+    A+ P
Sbjct  2005  LTKIVQGPQEPFSDFVARMTEAASRIFGDAEQAMPLIEQLVFEQATQECRA---AIAPRK  2175

Query  342   T--LEEMMTACQGVGGPGHKARVLAEAMSQVTNSATIMMQRGNFRNQRKIVKCFNCGKEG  399
             +  L++ +  C+ +GGP   A + A          A +  QR     +R    CFNCGK G
Sbjct  2176  SKGLQDWLKICRELGGPLTNAGLAA---------AILQTQR-----RRNTSACFNCGKTG  2313

Query  400   HTARNCRAPRK----KGCWKCGKEGHQMKDC  426
             H  ++CRAP +    + C +CGK  H+  +C
Sbjct  2314  HLKKDCRAPERMREVELCRRCGKGYHRASEC  2406
```

```
>Enterobacter hormaechei strain AUH-ENM6 NODE_27_length_1394_cov_0.980268, whole genome shotgun sequence
Sequence ID: NZ_PYHB01000027.1 Length: 1394
```

Range 1: 2 to 121

Score:68.6 bits(166), Expect:1e-07,
Method:Compositional matrix adjust.,
Identities:33/40(83%), Positives:33/40(82%), Gaps:0/40(0%)

```
Query  457  APPEESFRSGVETTTPPQKQEPIDKELYPLTSLRSLFGND  496
            APPEES R G ET TP QKQEP DKELYPL SLRSLFGND
Sbjct  2    APPEESLRLGGETATPSQKQEPTDKELYPLASLRSLFGND  121
```

>Pseudomonas sp. MPR-R2A3 NODE_254_length_244_cov_0.832335, whole genome shotgun sequence
Sequence ID: NZ_POFW01000253.1 Length: 244
Range 1: 23 to 118

Score:62.0 bits(149), Expect:1e-07,
Method:Compositional matrix adjust.,
Identities:27/32(84%), Positives:30/32(93%), Gaps:0/32(0%)

```
Query  430  QANFLGKIWPSYKGRPGNFLQSRPEPTAPPEE  461
            Q+  LGKIWPS+KGRPGNFLQSRPEPTAPPE+
Sbjct  23   QSQHLGKIWPSHKGRPGNFLQSRPEPTAPPED  118
```

>Acinetobacter baumannii strain ACM-14 NODE_74_length_787_cov_0.637879, whole genome shotgun sequence
Sequence ID: NZ_PYDA01000074.1 Length: 787
Range 1: 665 to 784

Score:65.9 bits(159), Expect:2e-07,
Method:Compositional matrix adjust.,
Identities:31/40(78%), Positives:34/40(85%), Gaps:0/40(0%)

```
Query  461  ESFRSGVETTTPPQKQEPIDKELYPLTSLRSLFGNDPSSQ  500
            ESF+ G ETTTPPQKQE  DKE+YPL SL+SLFGNDP SQ
Sbjct  784  ESFKFGEETTTPPQKQEEKDKEMYPLASLKSLFGNDPLSQ  665
```

>Helicobacter acinonychis isolate 212_10, whole genome shotgun sequence
Sequence ID: NZ_FZLW01000041.1 Length: 1526
Range 1: 1365 to 1487

Score:63.5 bits(153), Expect:5e-06,
Method:Compositional matrix adjust.,
Identities:32/41(78%), Positives:32/41(78%), Gaps:0/41(0%)

```
Query  460   EESFRSGVETTTPPQKQEPIDKELYPLTSLRSLFGNDPSSQ  500
              ESFR G ET TP QKQE IDKE YPL SLRSLFGNDP SQ
Sbjct  1487  VESFRFGEETATPHQKQEQIDKETYPLASLRSLFGNDPLSQ  1365
```

>Listeria monocytogenes strain CrieF364 NODE_22_length_769_cov_13.562305, whole genome shotgun sequence
Sequence ID: NZ_JASJTP010000022.1 Length: 769
Range 1: 82 to 171

Score:61.6 bits(148), Expect:6e-06,
Method:Compositional matrix adjust.,
Identities:27/30(90%), Positives:28/30(93%), Gaps:0/30(0%)

```
Query  188  LNTVGGHQAAMQMLKETINEEAAEWDRVHP  217
            +  VGGHQAAMQMLKETINEEAAEWDRVHP
Sbjct  171  IRLVGGHQAAMQMLKETINEEAAEWDRVHP  82
```

>Escherichia coli strain 8374wH11 NODE_36_length_1397_cov_3.54567, whole genome shotgun sequence
Sequence ID: NZ_WSJI01000036.1 Length: 1397
Range 1: 5 to 124

Score:62.4 bits(150), Expect:1e-05,
Method:Compositional matrix adjust.,
Identities:30/40(75%), Positives:33/40(82%), Gaps:0/40(0%)

```
Query  461  ESFRSGVETTTPPQKQEPIDKELYPLTSLRSLFGNDPSSQ  500
            ESF+ G ETTT PQKQE  DKE+YPL SL+SLFGNDP SQ
Sbjct  5    ESFKFGEETTTLPQKQEEKDKEMYPLASLKSLFGNDPLSQ  124
```

>Methylocella silvestris strain TVC NODE_58_length_1453_cov_3.56561, whole genome shotgun sequence
Sequence ID: NZ_PDZR01000058.1 Length: 1453
Range 1: 2 to 106

Score:60.5 bits(145), Expect:4e-05,
Method:Compositional matrix adjust.,
Identities:29/35(83%), Positives:29/35(82%), Gaps:0/35(0%)

```
Query  462  SFRSGVETTTPPQKQEPIDKELYPLTSLRSLFGND  496
            S R G ETTTP QKQEP DKELYPL SLRSLFGND
Sbjct  2    SLRLGGETTTPSQKQEPTDKELYPLASLRSLFGND  106
```

>Enterococcus faecalis strain 44.1 NODE_33_length_806_cov_1.20619, whole genome shotgun sequence
Sequence ID: NZ_NSJR01000033.1 Length: 806
Range 1: 43 to 141

Score:58.5 bits(140), Expect:8e-05,
Method:Compositional matrix adjust.,
Identities:28/33(85%), Positives:28/33(84%), Gaps:0/33(0%)

```
Query  464  RSGVETTTPPQKQEPIDKELYPLTSLRSLFGND  496
            R G ETTTP QKQEP DKELYPL SLRSLFGND
Sbjct  43   RLGGETTTPSQKQEPTDKELYPLASLRSLFGND  141
```

>Mycobacterium tuberculosis variant bovis strain CSURQ1140, whole genome shotgun sequence
Sequence ID: NZ_CAJEUP010000157.1 Length: 1286
Range 1: 1141 to 1281

Score:56.6 bits(135), Expect:8e-04,
Method:Compositional matrix adjust.,

```
Identities:34/48(71%), Positives:36/48(75%), Gaps:2/48(4%)

Query  454   EPTAPPEESFRSGVETTTPPQKQEPIDKELYP-LTSLRSLFGNDPSSQ  500
             EPTAPP ESF  G E  +  QKQEP DK LYP LTSL+SLFGNDP SQ
Sbjct  1281  EPTAPPAESFGMGEEIPSS-QKQEPGDKGLYPPLTSLKSLFGNDP*SQ  1141



>Escherichia coli strain HU8 NODE_685_length_270_cov_0.756477, whole genome shotgun sequence
Sequence ID: NZ_JAHCTM010000685.1 Length: 270
Range 1: 21 to 131

Score:51.6 bits(122), Expect:8e-04,
Method:Composition-based stats.,
Identities:18/37(49%), Positives:25/37(67%), Gaps:0/37(0%)

Query  390  VKCFNCGKEGHTARNCRAPRKKGCWKCGKEGHQMKDC  426
            V C+NCG++GH + +C  PR+  C+ CG  GH  KDC
Sbjct  21   VTCYNCGEKGHISTSCPKPRRLTCYNCGGVGHISKDC  131



>Salmonella enterica subsp. enterica serovar Typhimurium strain SH6609 NODE_771_length_211_cov_1.211538, whole genome shotgun sequence
Sequence ID: NZ_JALDNE010000502.1 Length: 211
Range 1: 86 to 199

Score:50.4 bits(119), Expect:0.001,
Method:Composition-based stats.,
Identities:17/38(45%), Positives:25/38(65%), Gaps:0/38(0%)

Query  390  VKCFNCGKEGHTARNCRAPRKKGCWKCGKEGHQMKDCT  427
            VKC+ CG+ GH A NC    +  C++CG+ GH  ++CT
Sbjct  86   VKCYRCGETGHVAINCSKTSEDICYRCGESGHLARECT  199



>Klebsiella pneumoniae strain EuSCAPE_IL004, whole genome shotgun sequence
Sequence ID: NZ_UKJP02000108.1 Length: 356
Range 1: 204 to 332

Score:51.2 bits(121), Expect:0.002,
Method:Compositional matrix adjust.,
Identities:31/47(66%), Positives:35/47(74%), Gaps:4/47(8%)

Query  454  EPTAPPEESFRSGVETTTPPQKQEPIDKELYPLTSLRSLFGNDPSSQ  500
            EPTAPP ESF+   E TTP  +QE  D+E  PLTSL+SLFGNDP SQ
Sbjct  332  EPTAPPAESFK--FEETTPAPRQEQKDRE--PLTSLKSLFGNDPLSQ  204



>Escherichia coli strain 8374wG10 NODE_34_length_1373_cov_2.57865, whole genome shotgun sequence
Sequence ID: NZ_WSJQ01000034.1 Length: 1373
Range 1: 1274 to 1372

Score:54.3 bits(129), Expect:0.004,
Method:Compositional matrix adjust.,
Identities:26/33(79%), Positives:28/33(84%), Gaps:0/33(0%)

Query  468   ETTTPPQKQEPIDKELYPLTSLRSLFGNDPSSQ  500
             ETTT PQKQE  DKE+YPL SL+SLFGNDP SQ
Sbjct  1372  ETTTLPQKQEEKDKEMYPLASLKSLFGNDPLSQ  1274



>Klebsiella pneumoniae strain EuSCAPE_IL005, whole genome shotgun sequence
Sequence ID: NZ_UKJL02000059.1 Length: 323
Range 1: 5 to 139

Score:48.9 bits(115), Expect:0.011,
Method:Compositional matrix adjust.,
Identities:32/51(63%), Positives:36/51(70%), Gaps:6/51(11%)

Query  450  QSRPEPTAPPEESFRSGVETTTPPQKQEPIDKELYPLTSLRSLFGNDPSSQ  500
            QSR EPTAPP +SF    E TTP  KQE  D+E  PL SL+SLFG+DP SQ
Sbjct  5    QSRLEPTAPPADSF----EETTPAPKQEMKDRE--PLISLKSLFGSDPLSQ  139



>Rickettsia endosymbiont of Ceutorhynchus assimilis isolate Rickettsia_endosymbiont_Ceutorhynchus_assimilis chromosome 1
Sequence ID: NZ_OU906081.1 Length: 4063491
Range 1: 350606 to 351085

Score:52.4 bits(124), Expect:0.027,
Method:Compositional matrix adjust.,
Identities:41/171(24%), Positives:64/171(37%), Gaps:37/171(21%)

Query  282     ILDIRQGPKEPFRDYVDRFYKTLR-----AEQASQEVKNWMTETLLVQNANPDCKTILKA  336
               ILD  Q P E   DY+       R      E+A  E+        +++N  P   T L
Sbjct  350606  ILDRTQDPSEGIVDYLTTMNVLFRRYLNVPEEAQLEI--------IMRNLAPFYTTQLPV  350761

Query  337     LGPAATLEEMMTACQGVGGPGHKAR-----------------VLAEAMSQVTNSATIMMQ  379
               +    TLEE+   C  +    ++A                   E    V + + + +
Sbjct  350762  VN---TLEELEVECLKLEAKKYRAEHYVPPPRRRVNLVEPDFTFVETSDTVPDVSALRLN  350932

Query  380     RG----NFRNQRKIVKCFNCGKEGHTARNCRAPRKKGCWKCGKEGHQMKDC  426
                G    N    R    C+NC K GH  R C+ P++  C++CG  G   ++C
Sbjct  350933  EGSGAENSNRSRSSALCWNCQKPGHLNRECKQPKRLHCYRCGHSGSTSRNC  351085



>Geminicoccus harenae strain CPCC 101083 Scaffold77_1, whole genome shotgun sequence
Sequence ID: NZ_JABGCL010000077.1 Length: 7497
Range 1: 1986 to 2096

Score:52.0 bits(123), Expect:0.035,
Method:Compositional matrix adjust.,
Identities:18/37(49%), Positives:25/37(67%), Gaps:0/37(0%)

Query  390   VKCFNCGKEGHTARNCRAPRKKGCWKCGKEGHQMKDC  426
             V C+NCG++GH + +C  PR+  C+ CG  GH  KDC
Sbjct  1986  VTCYNCGEKGHISTSCPKPRRLTCYNCGGVGHISKDC  2096



>Modestobacter marinus strain CPCC 205245 Scaffold62_1, whole genome shotgun sequence
Sequence ID: NZ_JABGCH010000063.1 Length: 7666
```

```
Range 1: 1990 to 2100

Score:52.0 bits(123), Expect:0.036,
Method:Compositional matrix adjust.,
Identities:18/37(49%), Positives:25/37(67%), Gaps:0/37(0%)

Query  390   VKCFNCGKEGHTARNCRAPRKKGCWKCGKEGHQMKDC  426
             V C+NCG++GH + +C  PR+  C+ CG  GH  KDC
Sbjct  1990  VTCYNCGEKGHISTSCPKPRRLTCYNCGGVGHISKDC  2100
```

```
>Modestobacter italicus strain CPCC 205242 Scaffold14_1, whole genome shotgun sequence
Sequence ID: NZ_JABGCJ010000014.1 Length: 15567
Range 1: 10921 to 11031

Score:52.0 bits(123), Expect:0.037,
Method:Compositional matrix adjust.,
Identities:18/37(49%), Positives:25/37(67%), Gaps:0/37(0%)

Query  390    VKCFNCGKEGHTARNCRAPRKKGCWKCGKEGHQMKDC  426
              V C+NCG++GH + +C  PR+  C+ CG  GH  KDC
Sbjct  11031  VTCYNCGEKGHISTSCPKPRRLTCYNCGGVGHISKDC  10921
```

Query #5: sp|P04585|p6_gag_HV1H2 Gag-Pol polyprotein OS=Human immunodeficiency virus type 1 group M subtype B (isolate HXB2) OX=11706 GN=gag-pol PE=1 SV=4 Query ID: lcl|Query_2387715 Length: 52

Sequences producing significant alignments:

| Description | Scientific Name | Common Name | Taxid | Max Score | Total Score | Query cover | E Value | Per. Ident | Acc. Len | Accession |
|---|---|---|---|---|---|---|---|---|---|---|
| Deinococcus aquiradiocola strain JCM 14371 sequence30, whole... | Deinococcus ... | NA | 393059 | 94.4 | 94.4 | 100% | 4e-20 | 90.38 | 6128 | NZ_BMOE010000... |
| Enterococcus alcedinis strain CCM 8433 sequence23, whole genom... | Enterococcus... | NA | 1274384 | 94.4 | 94.4 | 100% | 4e-20 | 90.38 | 6075 | NZ_BMDT010000... |
| Actinoplanes campanulatus strain JCM 3059 sequence058, whole... | Actinoplanes... | NA | 113559 | 94.4 | 94.4 | 100% | 4e-20 | 90.38 | 4754 | NZ_BMPW010000... |
| Thalassovita mangrovi strain GS-10 Scaffold19_1, whole genome... | Thalassovita... | NA | 2692236 | 94.4 | 94.4 | 100% | 4e-20 | 90.38 | 2900 | NZ_WWEN010000... |
| Klebsiella pneumoniae strain P7701... | Klebsiella p... | NA | 573 | 79.0 | 79.0 | 86% | 7e-15 | 88.89 | 1305 | NZ_JACAWM0100... |
| Acinetobacter baumannii strain ACM-35... | Acinetobacte... | NA | 470 | 73.9 | 73.9 | 84% | 5e-13 | 84.09 | 1409 | NZ_QEHN010000... |
| Acinetobacter baumannii strain ACM-36... | Acinetobacte... | NA | 470 | 67.0 | 67.0 | 73% | 3e-11 | 86.84 | 567 | NZ_QAGO010000... |
| Salirhabdus euzebyi strain Marseille-Q1438, whole genome shotg... | Salirhabdus ... | NA | 394506 | 68.2 | 68.2 | 78% | 5e-11 | 80.49 | 1279 | NZ_CADDWK0100... |
| Enterobacter hormaechei strain AUH-ENM6... | Enterobacter... | NA | 158836 | 67.0 | 67.0 | 76% | 1e-10 | 82.50 | 1394 | NZ_PYHB010000... |
| Cellulophaga baltica strain 14 NODE_63_length_422_cov_10.20678... | Cellulophaga... | NA | 76594 | 62.4 | 62.4 | 98% | 6e-10 | 73.08 | 422 | NZ_VIQP010000... |
| Acinetobacter baumannii strain ACM-14... | Acinetobacte... | NA | 470 | 64.3 | 64.3 | 76% | 6e-10 | 77.50 | 787 | NZ_PYDA010000... |
| Helicobacter acinonychis isolate 212_10, whole genome shotgun... | Helicobacter... | NA | 212 | 63.2 | 63.2 | 78% | 3e-09 | 78.05 | 1526 | NZ_FZLW010000... |
| Escherichia coli strain 8374wH11... | Escherichia ... | NA | 562 | 61.6 | 61.6 | 76% | 9e-09 | 75.00 | 1397 | NZ_WSJI010000... |
| Methylocella silvestris strain TVC... | Methylocella... | NA | 199596 | 59.7 | 59.7 | 67% | 4e-08 | 82.86 | 1453 | NZ_PDZR010000... |
| Klebsiella pneumoniae strain EuSCAPE_IL017, whole genome shotg... | Klebsiella p... | NA | 573 | 56.2 | 56.2 | 100% | 8e-08 | 65.38 | 325 | NZ_UKKG020000... |
| Mycobacterium tuberculosis variant bovis strain CSURQ1134, who... | Mycobacteriu... | NA | 1765 | 58.9 | 117 | 98% | 1e-07 | 69.23 | 1697 | NZ_CAJEUT0100... |
| Mycobacterium tuberculosis variant bovis strain CSURQ1132, who... | Mycobacteriu... | NA | 1765 | 58.5 | 117 | 98% | 1e-07 | 69.23 | 1524 | NZ_CAJEUK0100... |
| Enterococcus faecalis strain 44.1... | Enterococcus... | NA | 1351 | 57.8 | 57.8 | 63% | 2e-07 | 84.85 | 806 | NZ_NSJR010000... |
| Enterococcus faecium strain A209R1B0C1... | Enterococcus... | NA | 1352 | 57.4 | 57.4 | 100% | 3e-07 | 67.31 | 1819 | NZ_PGCR010000... |
| Listeria monocytogenes strain YA00082404... | Listeria mon... | NA | 1639 | 57.0 | 57.0 | 100% | 4e-07 | 64.41 | 1816 | NZ_QEXE010000... |
| Klebsiella pneumoniae strain EuSCAPE_LT007, whole genome shotg... | Klebsiella p... | NA | 573 | 54.3 | 54.3 | 92% | 8e-07 | 68.75 | 392 | NZ_UKJI020000... |
| Mycobacterium tuberculosis strain 32845_S29 214, whole genome... | Mycobacteriu... | NA | 1773 | 50.8 | 50.8 | 96% | 4e-06 | 62.96 | 223 | NZ_QTBV010000... |
| Mycobacterium tuberculosis strain 32847_S31 462, whole genome... | Mycobacteriu... | NA | 1773 | 53.9 | 53.9 | 100% | 5e-06 | 62.50 | 893 | NZ_QTBX010001... |
| Escherichia coli strain 8374wG10... | Escherichia ... | NA | 562 | 53.9 | 53.9 | 63% | 5e-06 | 78.79 | 1373 | NZ_WSJQ010000... |
| Mycobacterium tuberculosis strain 32846_S30 476, whole genome... | Mycobacteriu... | NA | 1773 | 51.6 | 51.6 | 100% | 3e-05 | 61.54 | 861 | NZ_QTBW010001... |
| Klebsiella pneumoniae strain EuSCAPE_IL004, whole genome shotg... | Klebsiella p... | NA | 573 | 49.7 | 49.7 | 90% | 4e-05 | 65.96 | 356 | NZ_UKJP020001... |
| Burkholderia pseudomallei strain 2604STDY5519032... | Burkholderia... | NA | 28450 | 50.4 | 50.4 | 92% | 7e-05 | 60.42 | 656 | NZ_UVID020000... |
| Klebsiella pneumoniae strain NUBRI-K 73, whole genome shotgun... | Klebsiella p... | NA | 573 | 50.4 | 50.4 | 100% | 8e-05 | 61.02 | 1717 | NZ_SOYT010000... |
| Klebsiella pneumoniae strain EuSCAPE_IL005, whole genome shotg... | Klebsiella p... | NA | 573 | 48.1 | 48.1 | 98% | 1e-04 | 62.75 | 323 | NZ_UKJL020000... |
| Enterococcus faecium strain A206R2B0... | Enterococcus... | NA | 1352 | 48.1 | 48.1 | 98% | 7e-04 | 62.75 | 1761 | NZ_PGCU010000... |

Alignments:

```
>Deinococcus aquiradiocola strain JCM 14371 sequence30, whole genome shotgun sequence
Sequence ID: NZ_BMOE01000030.1 Length: 6128
Range 1: 1452 to 1607

Score:94.4 bits(233), Expect:4e-20,
Method:Compositional matrix adjust.,
Identities:47/52(90%), Positives:48/52(92%), Gaps:0/52(0%)

Query  1     LQSRPEPTAPPEESFRSGVETTTPPQKQEPIDKELYPLTSLRSLFGNDPSSQ  52
             LQSRPEPTAPPEESFR G ETTTP QKQEPIDKELYPL SLRSLFG+DPSSQ
Sbjct  1452  LQSRPEPTAPPEESFRFGEETTTPSQKQEPIDKELYPLASLRSLFGSDPSSQ  1607
```

```
>Enterococcus alcedinis strain CCM 8433 sequence23, whole genome shotgun sequence
Sequence ID: NZ_BMDT01000023.1 Length: 6075
Range 1: 1399 to 1554

Score:94.4 bits(233), Expect:4e-20,
Method:Compositional matrix adjust.,
Identities:47/52(90%), Positives:48/52(92%), Gaps:0/52(0%)

Query  1     LQSRPEPTAPPEESFRSGVETTTPPQKQEPIDKELYPLTSLRSLFGNDPSSQ  52
             LQSRPEPTAPPEESFR G ETTTP QKQEPIDKELYPL SLRSLFG+DPSSQ
Sbjct  1399  LQSRPEPTAPPEESFRFGEETTTPSQKQEPIDKELYPLASLRSLFGSDPSSQ  1554
```

```
>Actinoplanes campanulatus strain JCM 3059 sequence058, whole genome shotgun sequence
Sequence ID: NZ_BMPW01000058.1 Length: 4754
Range 1: 1441 to 1596

Score:94.4 bits(233), Expect:4e-20,
Method:Compositional matrix adjust.,
Identities:47/52(90%), Positives:48/52(92%), Gaps:0/52(0%)

Query  1     LQSRPEPTAPPEESFRSGVETTTPPQKQEPIDKELYPLTSLRSLFGNDPSSQ  52
             LQSRPEPTAPPEESFR G ETTTP QKQEPIDKELYPL SLRSLFG+DPSSQ
Sbjct  1441  LQSRPEPTAPPEESFRFGEETTTPSQKQEPIDKELYPLASLRSLFGSDPSSQ  1596
```

```
>Thalassovita mangrovi strain GS-10 Scaffold19_1, whole genome shotgun sequence
Sequence ID: NZ_WWEN01000019.1 Length: 2900
Range 1: 2384 to 2539

Score:94.4 bits(233), Expect:4e-20,
Method:Compositional matrix adjust.,
Identities:47/52(90%), Positives:48/52(92%), Gaps:0/52(0%)

Query  1     LQSRPEPTAPPEESFRSGVETTTPPQKQEPIDKELYPLTSLRSLFGNDPSSQ  52
             LQSRPEPTAPPEESFR G ETTTP QKQEPIDKELYPL SLRSLFG+DPSSQ
```

```
Sbjct  2384  LQSRPEPTAPPEESFRFGEETTTPSQKQEPIDKELYPLASLRSLFGSDPSSQ  2539
```

>Klebsiella pneumoniae strain P7701 NODE_50_length_1305_cov_481.863, whole genome shotgun sequence
Sequence ID: NZ_JACAWM010000050.1 Length: 1305
Range 1: 1169 to 1303

Score:79.0 bits(193), Expect:7e-15,
Method:Compositional matrix adjust.,
Identities:40/45(89%), Positives:41/45(91%), Gaps:0/45(0%)

```
Query  8     TAPPEESFRSGVETTTPPQKQEPIDKELYPLTSLRSLFGNDPSSQ  52
             TAPPEESFR G ETTTP QKQEPIDKELYPL SLRSLFG+DPSSQ
Sbjct  1303  TAPPEESFRFGEETTTPSQKQEPIDKELYPLASLRSLFGSDPSSQ  1169
```

>Acinetobacter baumannii strain ACM-35 NODE_65_length_1409_cov_2.30031, whole genome shotgun sequence
Sequence ID: NZ_QEHN01000065.1 Length: 1409
Range 1: 1 to 132

Score:73.9 bits(180), Expect:5e-13,
Method:Compositional matrix adjust.,
Identities:37/44(84%), Positives:37/44(84%), Gaps:0/44(0%)

```
Query  5    PEPTAPPEESFRSGVETTTPPQKQEPIDKELYPLTSLRSLFGND  48
            PEPTAPPEES R G ET TP QKQEP DKELYPL SLRSLFGND
Sbjct  1    PEPTAPPEESLRLGGETATPSQKQEPTDKELYPLASLRSLFGND  132
```

>Acinetobacter baumannii strain ACM-36 NODE_67_length_567_cov_1.13409, whole genome shotgun sequence
Sequence ID: NZ_QAGO01000067.1 Length: 567
Range 1: 1 to 114

Score:67.0 bits(162), Expect:3e-11,
Method:Compositional matrix adjust.,
Identities:33/38(87%), Positives:33/38(86%), Gaps:0/38(0%)

```
Query  11   PEESFRSGVETTTPPQKQEPIDKELYPLTSLRSLFGND  48
            PEESFR G ETTTP QKQEP DKELYPL SLRSLFGND
Sbjct  1    PEESFRLGGETTTPSQKQEPTDKELYPLASLRSLFGND  114
```

>Salirhabdus euzebyi strain Marseille-Q1438, whole genome shotgun sequence
Sequence ID: NZ_CADDWK010000035.1 Length: 1279
Range 1: 1157 to 1279

Score:68.2 bits(165), Expect:5e-11,
Method:Compositional matrix adjust.,
Identities:33/41(80%), Positives:35/41(85%), Gaps:0/41(0%)

```
Query  12    EESFRSGVETTTPPQKQEPIDKELYPLTSLRSLFGNDPSSQ  52
             EES R G ETTTP QKQE IDK++YPL SLRSLFGNDPSSQ
Sbjct  1279  EESLRLGEETTTPSQKQEQIDKDMYPLASLRSLFGNDPSSQ  1157
```

>Enterobacter hormaechei strain AUH-ENM6 NODE_27_length_1394_cov_0.980268, whole genome shotgun sequence
Sequence ID: NZ_PYHB01000027.1 Length: 1394
Range 1: 2 to 121

Score:67.0 bits(162), Expect:1e-10,
Method:Compositional matrix adjust.,
Identities:33/40(83%), Positives:33/40(82%), Gaps:0/40(0%)

```
Query  9    APPEESFRSGVETTTPPQKQEPIDKELYPLTSLRSLFGND  48
            APPEES R G ET TP QKQEP DKELYPL SLRSLFGND
Sbjct  2    APPEESLRLGGETATPSQKQEPTDKELYPLASLRSLFGND  121
```

>Cellulophaga baltica strain 14 NODE_63_length_422_cov_10.206780, whole genome shotgun sequence
Sequence ID: NZ_VIQP01000062.1 Length: 422
Range 1: 210 to 362

Score:62.4 bits(150), Expect:6e-10,
Method:Compositional matrix adjust.,
Identities:38/52(73%), Positives:40/52(76%), Gaps:2/52(3%)

```
Query  2    QSRPEPTAPPEESFRSGVETTTPPQKQEPIDKELY-PLTSLRSLFGNDPSSQ  52
            QSRPEPTAPP ES   G E T+ PQ QEP DK LY PLTSL+SLFGNDP SQ
Sbjct  362  QSRPEPTAPPAESMGMGEEITSSPQ-QEPRDKGLYPPLTSLKSLFGNDP*SQ  210
```

>Acinetobacter baumannii strain ACM-14 NODE_74_length_787_cov_0.637879, whole genome shotgun sequence
Sequence ID: NZ_PYDA01000074.1 Length: 787
Range 1: 665 to 784

Score:64.3 bits(155), Expect:6e-10,
Method:Compositional matrix adjust.,
Identities:31/40(78%), Positives:34/40(85%), Gaps:0/40(0%)

```
Query  13   ESFRSGVETTTPPQKQEPIDKELYPLTSLRSLFGNDPSSQ  52
            ESF+ G ETTTPPQKQE  DKE+YPL SL+SLFGNDP SQ
Sbjct  784  ESFKFGEETTTPPQKQEEKDKEMYPLASLKSLFGNDPLSQ  665
```

>Helicobacter acinonychis isolate 212_10, whole genome shotgun sequence
Sequence ID: NZ_FZLW01000041.1 Length: 1526
Range 1: 1365 to 1487

Score:63.2 bits(152), Expect:3e-09,
Method:Compositional matrix adjust.,
Identities:32/41(78%), Positives:32/41(78%), Gaps:0/41(0%)

```
Query  12    EESFRSGVETTTPPQKQEPIDKELYPLTSLRSLFGNDPSSQ  52
              ESFR G ET TP QKQE IDKE YPL SLRSLFGNDP SQ
Sbjct  1487  VESFRFGEETATPHQKQEQIDKETYPLASLRSLFGNDPLSQ  1365
```

>Escherichia coli strain 8374wH11 NODE_36_length_1397_cov_3.54567, whole genome shotgun sequence
Sequence ID: NZ_WSJI01000036.1 Length: 1397
Range 1: 5 to 124

Score:61.6 bits(148), Expect:9e-09,
Method:Compositional matrix adjust.,
Identities:30/40(75%), Positives:33/40(82%), Gaps:0/40(0%)

```
Query  13   ESFRSGVETTTPPQKQEPIDKELYPLTSLRSLFGNDPSSQ  52
            ESF+ G ETTT PQKQE  DKE+YPL SL+SLFGNDP SQ
Sbjct  5    ESFKFGEETTTLPQKQEEKDKEMYPLASLKSLFGNDPLSQ  124
```

>Methylocella silvestris strain TVC NODE_58_length_1453_cov_3.56561, whole genome shotgun sequence
Sequence ID: NZ_PDZR01000058.1 Length: 1453
Range 1: 2 to 106

Score:59.7 bits(143), Expect:4e-08,
Method:Compositional matrix adjust.,
Identities:29/35(83%), Positives:29/35(82%), Gaps:0/35(0%)

```
Query  14   SFRSGVETTTPPQKQEPIDKELYPLTSLRSLFGND  48
            S R G ETTTP QKQEP DKELYPL SLRSLFGND
Sbjct  2    SLRLGGETTTPSQKQEPTDKELYPLASLRSLFGND  106
```

>Klebsiella pneumoniae strain EuSCAPE_IL017, whole genome shotgun sequence
Sequence ID: NZ_UKKG02000044.1 Length: 325
Range 1: 95 to 238

Score:56.2 bits(134), Expect:8e-08,
Method:Compositional matrix adjust.,
Identities:34/52(65%), Positives:39/52(75%), Gaps:4/52(7%)

```
Query  1    LQSRPEPTAPPEESFRSGVETTTPPQKQEPIDKELYPLTSLRSLFGNDPSSQ  52
            LQ+RPEP+APP ESFR   E TTP  KQE  D+E   LTSL+SLFG+DP SQ
Sbjct  95   LQNRPEPSAPPAESFR--FEETTPAPKQESKDREA--LTSLKSLFGSDPLSQ  238
```

>Mycobacterium tuberculosis variant bovis strain CSURQ1134, whole genome shotgun sequence
Sequence ID: NZ_CAJEUT010000109.1 Length: 1697
Range 1: 1233 to 1385

Score:58.9 bits(141), Expect:1e-07,
Method:Compositional matrix adjust.,
Identities:36/52(69%), Positives:38/52(73%), Gaps:2/52(3%)

```
Query  2     QSRPEPTAPPEESFRSGVETTTPPQKQEPIDKELYP-LTSLRSLFGNDPSSQ  52
             Q R EPTAPP ESF  G E  +  QKQEP DK LYP LTSL+SLFGNDP SQ
Sbjct  1385  QGRLEPTAPPAESFGMGEEIPSS-QKQEPGDKGLYPPLTSLKSLFGNDP*SQ  1233
```

Range 2: 1233 to 1385

Score:58.9 bits(141), Expect:1e-07,
Method:Compositional matrix adjust.,
Identities:36/52(69%), Positives:38/52(73%), Gaps:2/52(3%)

```
Query  2     QSRPEPTAPPEESFRSGVETTTPPQKQEPIDKELYP-LTSLRSLFGNDPSSQ  52
             Q R EPTAPP ESF  G E  +  QKQEP DK LYP LTSL+SLFGNDP SQ
Sbjct  1385  QGRLEPTAPPAESFGMGEEIPSS-QKQEPGDKGLYPPLTSLKSLFGNDP*SQ  1233
```

>Mycobacterium tuberculosis variant bovis strain CSURQ1132, whole genome shotgun sequence
Sequence ID: NZ_CAJEUK010000155.1 Length: 1524
Range 1: 1271 to 1423

Score:58.5 bits(140), Expect:1e-07,
Method:Compositional matrix adjust.,
Identities:36/52(69%), Positives:38/52(73%), Gaps:2/52(3%)

```
Query  2     QSRPEPTAPPEESFRSGVETTTPPQKQEPIDKELYP-LTSLRSLFGNDPSSQ  52
             Q R EPTAPP ESF  G E  +  QKQEP DK LYP LTSL+SLFGNDP SQ
Sbjct  1423  QGRLEPTAPPAESFGMGEEIPSS-QKQEPGDKGLYPPLTSLKSLFGNDP*SQ  1271
```

Range 2: 1271 to 1423

Score:58.5 bits(140), Expect:1e-07,
Method:Compositional matrix adjust.,
Identities:36/52(69%), Positives:38/52(73%), Gaps:2/52(3%)

```
Query  2     QSRPEPTAPPEESFRSGVETTTPPQKQEPIDKELYP-LTSLRSLFGNDPSSQ  52
             Q R EPTAPP ESF  G E  +  QKQEP DK LYP LTSL+SLFGNDP SQ
Sbjct  1423  QGRLEPTAPPAESFGMGEEIPSS-QKQEPGDKGLYPPLTSLKSLFGNDP*SQ  1271
```

>Enterococcus faecalis strain 44.1 NODE_33_length_806_cov_1.20619, whole genome shotgun sequence
Sequence ID: NZ_NSJR01000033.1 Length: 806
Range 1: 43 to 141

Score:57.8 bits(138), Expect:2e-07,
Method:Compositional matrix adjust.,
Identities:28/33(85%), Positives:28/33(84%), Gaps:0/33(0%)

```
Query  16   RSGVETTTPPQKQEPIDKELYPLTSLRSLFGND  48
            R G ETTTP QKQEP DKELYPL SLRSLFGND
Sbjct  43   RLGGETTTPSQKQEPTDKELYPLASLRSLFGND  141
```

>Enterococcus faecium strain A209R1B0C1 NODE_70_length_1819_cov_3.01832_ID_139, whole genome shotgun sequence
Sequence ID: NZ_PGCR01000088.1 Length: 1819
Range 1: 88 to 225

Score:57.4 bits(137), Expect:3e-07,
Method:Compositional matrix adjust.,
Identities:35/52(67%), Positives:39/52(75%), Gaps:6/52(11%)

```
Query  1    LQSRPEPTAPPEESFRSGVETTTPPQKQEPIDKELYPLTSLRSLFGNDPSSQ  52
            LQ+RPEPTAPP ESFR   E TTP QKQE  +    PLTSL+SLFG+DP SQ
Sbjct  88   LQNRPEPTAPPAESFR--FEETTPAQKQESRE----PLTSLKSLFGSDPLSQ  225
```

>Listeria monocytogenes strain YA00082404 NODE_73_length_1816_cov_5.664890, whole genome shotgun sequence
Sequence ID: NZ_QEXE01000073.1 Length: 1816
Range 1: 74 to 238

Score:57.0 bits(136), Expect:4e-07,
Method:Compositional matrix adjust.,
Identities:38/59(64%), Positives:41/59(69%), Gaps:11/59(18%)

```
Query  1    LQSRPEPTAPP-------EESFRSGVETTTPPQKQEPIDKELYPLTSLRSLFGNDPSSQ  52
            LQ+RPEPTAPP        ESFR G   TTP  KQEP D+E  PLTSL+SLFGNDP SQ
Sbjct  74   LQNRPEPTAPPVEPTAPPAESFRFG--ETTPAPKQEPKDRE--PLTSLKSLFGNDPWSQ  238
```

>Klebsiella pneumoniae strain EuSCAPE_LT007, whole genome shotgun sequence
Sequence ID: NZ_UKJI02000072.1 Length: 392
Range 1: 192 to 323

Score:54.3 bits(129), Expect:8e-07,
Method:Compositional matrix adjust.,
Identities:33/48(69%), Positives:36/48(75%), Gaps:4/48(8%)

```
Query  1    LQSRPEPTAPPEESFRSGVETTTPPQKQEPIDKELYPLTSLRSLFGND  48
            LQSRPEP+APP ESFR   E TTP  KQE  DKE   LTSL+SLFG+D
Sbjct  323  LQSRPEPSAPPAESFR--FEETTPAPKQESKDKEA--LTSLKSLFGSD  192
```

>Mycobacterium tuberculosis strain 32845_S29 214, whole genome shotgun sequence
Sequence ID: NZ_QTBV01000045.1 Length: 223
Range 1: 2 to 157

Score:50.8 bits(120), Expect:4e-06,
Method:Compositional matrix adjust.,
Identities:34/54(63%), Positives:40/54(74%), Gaps:6/54(11%)

```
Query  3    SRPEPTAPPEESF-RSGVETTTPPQ---KQEPIDKELYPLTSLRSLFGNDPSSQ  52
            SRPEPTAPP ESF ++  E T PP    +QEP D+E  PL SL+SLFG+DPSSQ
Sbjct  2    SRPEPTAPPAESFLQNRPEPTAPPAESFRQEPKDRE--PLISLKSLFGSDPSSQ  157
```

>Mycobacterium tuberculosis strain 32847_S31 462, whole genome shotgun sequence
Sequence ID: NZ_QTBX01000137.1 Length: 893
Range 1: 69 to 230

Score:53.9 bits(128), Expect:5e-06,
Method:Compositional matrix adjust.,
Identities:35/56(63%), Positives:42/56(75%), Gaps:6/56(10%)

```
Query  1    LQSRPEPTAPPEESF-RSGVETTTPPQ---KQEPIDKELYPLTSLRSLFGNDPSSQ  52
            +QSRPEPTAPP ESF ++  E T PP    +QEP D+E  PL SL+SLFG+DPSSQ
Sbjct  69   IQSRPEPTAPPAESFLQNRPEPTAPPAESFRQEPKDRE--PLISLKSLFGSDPSSQ  230
```

>Escherichia coli strain 8374wG10 NODE_34_length_1373_cov_2.57865, whole genome shotgun sequence
Sequence ID: NZ_WSJQ01000034.1 Length: 1373
Range 1: 1274 to 1372

Score:53.9 bits(128), Expect:5e-06,
Method:Compositional matrix adjust.,
Identities:26/33(79%), Positives:28/33(84%), Gaps:0/33(0%)

```
Query  20    ETTTPPQKQEPIDKELYPLTSLRSLFGNDPSSQ  52
             ETTT PQKQE  DKE+YPL SL+SLFGNDP SQ
Sbjct  1372  ETTTLPQKQEEKDKEMYPLASLKSLFGNDPLSQ  1274
```

>Mycobacterium tuberculosis strain 32846_S30 476, whole genome shotgun sequence
Sequence ID: NZ_QTBW01000113.1 Length: 861
Range 1: 744 to 860

Score:51.6 bits(122), Expect:3e-05,
Method:Compositional matrix adjust.,
Identities:32/52(62%), Positives:36/52(69%), Gaps:13/52(25%)

```
Query  1    LQSRPEPTAPPEESFRSGVETTTPPQKQEPIDKELYPLTSLRSLFGNDPSSQ  52
            LQ+RPEPTAPP ESFR           QEP D+E  PL SL+SLFG+DPSSQ
Sbjct  860  LQNRPEPTAPPAESFR-----------QEPKDRE--PLISLKSLFGSDPSSQ  744
```

>Klebsiella pneumoniae strain EuSCAPE_IL004, whole genome shotgun sequence
Sequence ID: NZ_UKJP02000108.1 Length: 356
Range 1: 204 to 332

Score:49.7 bits(117), Expect:4e-05,
Method:Compositional matrix adjust.,
Identities:31/47(66%), Positives:35/47(74%), Gaps:4/47(8%)

```
Query  6    EPTAPPEESFRSGVETTTPPQKQEPIDKELYPLTSLRSLFGNDPSSQ  52
            EPTAPP ESF+   E TTP  +QE  D+E  PLTSL+SLFGNDP SQ
Sbjct  332  EPTAPPAESFK--FEETTPAPRQEQKDRE--PLTSLKSLFGNDPLSQ  204
```

>Burkholderia pseudomallei strain 2604STDY5519032 ERS240165SCcontig000074, whole genome shotgun sequence
Sequence ID: NZ_UVID02000074.1 Length: 656
Range 1: 1 to 132

Score:50.4 bits(119), Expect:7e-05,
Method:Compositional matrix adjust.,
Identities:29/48(60%), Positives:35/48(72%), Gaps:4/48(8%)

```
Query  1    LQSRPEPTAPPEESFRSGVETTTPPQKQEPIDKELYPLTSLRSLFGND  48
            +QSR EPTAPP ESF+   E  TP  +QE  ++E  PLTSL+SLFGND
Sbjct  132  IQSRLEPTAPPAESFK--FEEATPAPRQESKERE--PLTSLKSLFGND  1
```

```
>Klebsiella pneumoniae strain NUBRI-K 73, whole genome shotgun sequence
Sequence ID: NZ_SOYT01000073.1 Length: 1717
Range 1: 27 to 191

Score:50.4 bits(119), Expect:8e-05,
Method:Compositional matrix adjust.,
Identities:36/59(61%), Positives:40/59(67%), Gaps:11/59(18%)

Query  1    LQSRPEPTAPP-------EESFRSGVETTTPPQKQEPIDKELYPLTSLRSLFGNDPSSQ  52
            LQ+RPEPTAPP        ESFR   E TT   KQEP D+E  PLTSL+SLFG+DP SQ
Sbjct  27   LQNRPEPTAPPVEPTAPPAESFR--FEETTLAPKQEPKDRE--PLTSLKSLFGSDPLSQ  191
```

```
>Klebsiella pneumoniae strain EuSCAPE_IL005, whole genome shotgun sequence
Sequence ID: NZ_UKJL02000059.1 Length: 323
Range 1: 5 to 139

Score:48.1 bits(113), Expect:1e-04,
Method:Compositional matrix adjust.,
Identities:32/51(63%), Positives:36/51(70%), Gaps:6/51(11%)

Query  2    QSRPEPTAPPEESFRSGVETTTPPQKQEPIDKELYPLTSLRSLFGNDPSSQ  52
            QSR EPTAPP +SF    E TTP  KQE  D+E  PL SL+SLFG+DP SQ
Sbjct  5    QSRLEPTAPPADSF----EETTPAPKQEMKDRE--PLISLKSLFGSDPLSQ  139
```

```
>Enterococcus faecium strain A206R2B0 NODE_116_length_1761_cov_4.01102_ID_231, whole genome shotgun sequence
Sequence ID: NZ_PGCU01000052.1 Length: 1761
Range 1: 84 to 209

Score:48.1 bits(113), Expect:7e-04,
Method:Compositional matrix adjust.,
Identities:32/51(63%), Positives:35/51(68%), Gaps:9/51(17%)

Query  2    QSRPEPTAPPEESFRSGVETTTPPQKQEPIDKELYPLTSLRSLFGNDPSSQ  52
            QSR EPTAPP ESFR         +KQE  D+E  PLTSL+SLFGNDP SQ
Sbjct  84   QSRLEPTAPPAESFRF--------EKQEMEDRE-PPLTSLKSLFGNDPLSQ  209
```

Query #6: sp|P04585|Capsid_Protein_p24_HV1H2 Gag-Pol polyprotein OS=Human immunodeficiency virus type 1 group M subtype B (isolate HXB2) OX=11706 GN=gag-pol PE=1 SV=4 Query ID: lcl|Query_2387713 Length: 231

Sequences producing significant alignments:

| Description | Scientific Name | Common Name | Taxid | Max Score | Total Score | Query cover | E Value | Per. Ident | Acc. Len | Accession |
|---|---|---|---|---|---|---|---|---|---|---|
| Thalassovita mangrovi strain GS-10 Scaffold19_1, whole genome... | Thalassovita... | NA | 2692236 | 478 | 478 | 100% | 1e-159 | 98.27 | 2900 | NZ_WWEN010000... |
| Actinoplanes campanulatus strain JCM 3059 sequence058, whole... | Actinoplanes... | NA | 113559 | 478 | 478 | 100% | 4e-154 | 98.27 | 4754 | NZ_BMPW010000... |
| Enterococcus alcedinis strain CCM 8433 sequence23, whole genom... | Enterococcus... | NA | 1274384 | 477 | 477 | 100% | 1e-151 | 98.27 | 6075 | NZ_BMDT010000... |
| Deinococcus aquiradiocola strain JCM 14371 sequence30, whole... | Deinococcus ... | NA | 393059 | 476 | 476 | 100% | 2e-151 | 98.27 | 6128 | NZ_BMOE010000... |
| Klebsiella pneumoniae strain EuSCAPE_IL017, whole genome shotg... | Klebsiella p... | NA | 573 | 261 | 261 | 56% | 3e-85 | 91.60 | 403 | NZ_UKKG020000... |
| Klebsiella pneumoniae strain EuSCAPE_IL020, whole genome shotg... | Klebsiella p... | NA | 573 | 251 | 251 | 56% | 3e-81 | 87.79 | 469 | NZ_UKKB020000... |
| Klebsiella pneumoniae strain EuSCAPE_MT003, whole genome shotg... | Klebsiella p... | NA | 573 | 208 | 208 | 45% | 4e-65 | 90.48 | 318 | NZ_UKIK020001... |
| Klebsiella pneumoniae strain EuSCAPE_IL019, whole genome shotg... | Klebsiella p... | NA | 573 | 208 | 208 | 44% | 6e-65 | 93.20 | 312 | NZ_UKJZ020000... |
| Burkholderia pseudomallei strain 2604STDY5518971... | Burkholderia... | NA | 28450 | 207 | 207 | 44% | 1e-64 | 92.23 | 336 | NZ_UVFZ020001... |
| Neisseria gonorrhoeae strain GC-191... | Neisseria go... | NA | 485 | 187 | 187 | 40% | 7e-57 | 89.36 | 284 | NZ_SAWA010001... |
| Neisseria gonorrhoeae strain gc-168... | Neisseria go... | NA | 485 | 179 | 179 | 39% | 1e-53 | 89.01 | 274 | NZ_SAVF010001... |
| Burkholderia pseudomallei strain 2604STDY5519032... | Burkholderia... | NA | 28450 | 178 | 178 | 38% | 1e-51 | 90.00 | 656 | NZ_UVID020000... |
| Klebsiella pneumoniae strain EuSCAPE_MT002, whole genome shotg... | Klebsiella p... | NA | 573 | 171 | 171 | 36% | 7e-51 | 91.76 | 257 | NZ_UKIM020000... |
| Klebsiella pneumoniae strain EuSCAPE_IL005, whole genome shotg... | Klebsiella p... | NA | 573 | 166 | 166 | 35% | 2e-48 | 91.57 | 250 | NZ_UKJL020000... |
| Klebsiella pneumoniae strain EuSCAPE_LT007, whole genome shotg... | Klebsiella p... | NA | 573 | 166 | 166 | 35% | 2e-48 | 90.36 | 252 | NZ_UKJI020000... |
| Klebsiella pneumoniae strain EuSCAPE_LT003, whole genome shotg... | Klebsiella p... | NA | 573 | 160 | 160 | 34% | 1e-45 | 88.75 | 419 | NZ_UKJH020000... |
| Klebsiella quasipneumoniae strain EuSCAPE_IL014, whole genome... | Klebsiella q... | NA | 1463165 | 135 | 135 | 29% | 2e-36 | 89.86 | 269 | NZ_UJYK020000... |
| Klebsiella pneumoniae strain EuSCAPE_IL005, whole genome shotg... | Klebsiella p... | NA | 573 | 112 | 112 | 24% | 3e-27 | 91.23 | 308 | NZ_UKJL020000... |
| Neisseria gonorrhoeae strain GC-206... | Neisseria go... | NA | 485 | 106 | 106 | 22% | 5e-25 | 92.45 | 334 | NZ_SAWN010001... |
| Klebsiella pneumoniae strain B2 isolate B2, whole genome shotg... | Klebsiella p... | NA | 573 | 115 | 115 | 100% | 1e-24 | 30.21 | 7306 | NZ_CABWNT0100... |
| Klebsiella pneumoniae strain EuSCAPE_NO017, whole genome shotg... | Klebsiella p... | NA | 573 | 105 | 105 | 22% | 1e-24 | 92.31 | 348 | NZ_UKIO020000... |
| Bacillus velezensis strain TN14-11 TN14-11_ctg0032, whole geno... | Bacillus vel... | NA | 492670 | 100 | 100 | 21% | 3e-20 | 93.88 | 1125 | NZ_JAHVIU0100... |
| Escherichia coli strain VRES0669, whole genome shotgun sequence | Escherichia ... | NA | 562 | 87.8 | 87.8 | 18% | 3e-15 | 93.02 | 5544 | NZ_UINV010000... |
| Pseudomonas sp. MPR-LB3 NODE_257_length_230_cov_0.882353, whol... | Pseudomonas ... | NA | 2070625 | 78.2 | 78.2 | 16% | 2e-14 | 97.37 | 230 | NZ_POFY010002... |
| Photobacterium rosenbergii strain DP6N14-7 DP6N14-7_ctg0065,... | Photobacteri... | NA | 294936 | 79.3 | 79.3 | 16% | 1e-13 | 94.87 | 564 | NZ_JAHVIB0100... |
| Rhizobium bangladeshense strain 1017 scf43_contig_2, whole... | Rhizobium ba... | NA | 1138189 | 73.9 | 129 | 15% | 1e-12 | 97.14 | 288 | NZ_LYPK010000... |
| Listeria monocytogenes strain CrieF364... | Listeria mon... | NA | 1639 | 61.6 | 61.6 | 13% | 9e-07 | 87.10 | 769 | NZ_JASJTP0100... |
| Photobacterium rosenbergii strain DP6N14-7 DP6N14-7_ctg0069,... | Photobacteri... | NA | 294936 | 51.2 | 51.2 | 11% | 0.003 | 85.19 | 667 | NZ_JAHVIB0100... |
| Vibrio hangzhouensis strain DP3N7-4 DP3N7-4_ctg0051, whole... | Vibrio hangz... | NA | 462991 | 51.2 | 51.2 | 11% | 0.004 | 85.19 | 742 | NZ_JAHVKH0100... |

Alignments:

```
>Thalassovita mangrovi strain GS-10 Scaffold19_1, whole genome shotgun sequence
Sequence ID: NZ_WWEN01000019.1 Length: 2900
Range 1: 1436 to 2128

Score:478 bits(1230), Expect:1e-159,
Method:Compositional matrix adjust.,
Identities:227/231(98%), Positives:230/231(99%), Gaps:0/231(0%)

Query  1     PIVQNIQGQMVHQAISPRTLNAWVKVVEEKAFSPEVIPMFSALSEGATPQDLNTMLNTVG  60
             PIVQN+QGQMVHQAISPRTLNAWVKVVEEKAFSPEVIPMFSALSEGATPQDLNTMLNTVG
Sbjct  1436  PIVQNLQGQMVHQAISPRTLNAWVKVVEEKAFSPEVIPMFSALSEGATPQDLNTMLNTVG  1615

Query  61    GHQAAMQMLKETINEEAAEWDRVHPVHAGPIAPGQMREPRGSDIAGTTSTLQEQIGWMTN  120
             GHQAAMQMLKETINEEAAEWDR+HPVHAGPIAPGQMREPRGSDIAGTTSTLQEQIGWMT+
Sbjct  1616  GHQAAMQMLKETINEEAAEWDRLHPVHAGPIAPGQMREPRGSDIAGTTSTLQEQIGWMTH  1795

Query  121   NPPIPVGEIYKRWIILGLNKIVRMYSPTSILDIRQGPKEPFRDYVDRFYKTLRAEQASQE  180
             NPPIPVGEIYKRWIILGLNKIVRMYSPTSILDIRQGPKEPFRDYVDRFYKTLRAEQASQE
Sbjct  1796  NPPIPVGEIYKRWIILGLNKIVRMYSPTSILDIRQGPKEPFRDYVDRFYKTLRAEQASQE  1975

Query  181   VKNWMTETLLVQNANPDCKTILKALGPAATLEEMMTACQGVGGPGHKARVL  231
             VKNWMTETLLVQNANPDCKTILKALGP ATLEEMMTACQGVGGPGHKARVL
Sbjct  1976  VKNWMTETLLVQNANPDCKTILKALGPGATLEEMMTACQGVGGPGHKARVL  2128
```

```
>Actinoplanes campanulatus strain JCM 3059 sequence058, whole genome shotgun sequence
Sequence ID: NZ_BMPW01000058.1 Length: 4754
Range 1: 493 to 1185

Score:478 bits(1230), Expect:4e-154,
Method:Compositional matrix adjust.,
Identities:227/231(98%), Positives:230/231(99%), Gaps:0/231(0%)
```

```
Query  1     PIVQNIQGQMVHQAISPRTLNAWVKVVEEKAFSPEVIPMFSALSEGATPQDLNTMLNTVG  60
             PIVQN+QGQMVHQAISPRTLNAWVKVVEEKAFSPEVIPMFSALSEGATPQDLNTMLNTVG
Sbjct  493   PIVQNLQGQMVHQAISPRTLNAWVKVVEEKAFSPEVIPMFSALSEGATPQDLNTMLNTVG  672

Query  61    GHQAAMQMLKETINEEAAEWDRVHPVHAGPIAPGQMREPRGSDIAGTTSTLQEQIGWMTN  120
             GHQAAMQMLKETINEEAAEWDR+HPVHAGPIAPGQMREPRGSDIAGTTSTLQEQIGWMT+
Sbjct  673   GHQAAMQMLKETINEEAAEWDRLHPVHAGPIAPGQMREPRGSDIAGTTSTLQEQIGWMTH  852

Query  121   NPPIPVGEIYKRWIILGLNKIVRMYSPTSILDIRQGPKEPFRDYVDRFYKTLRAEQASQE  180
             NPPIPVGEIYKRWIILGLNKIVRMYSPTSILDIRQGPKEPFRDYVDRFYKTLRAEQASQE
Sbjct  853   NPPIPVGEIYKRWIILGLNKIVRMYSPTSILDIRQGPKEPFRDYVDRFYKTLRAEQASQE  1032

Query  181   VKNWMTETLLVQNANPDCKTILKALGPAATLEEMMTACQGVGGPGHKARVL  231
             VKNWMTETLLVQNANPDCKTILKALGP ATLEEMMTACQGVGGPGHKARVL
Sbjct  1033  VKNWMTETLLVQNANPDCKTILKALGPGATLEEMMTACQGVGGPGHKARVL  1185
```

>Enterococcus alcedinis strain CCM 8433 sequence23, whole genome shotgun sequence
Sequence ID: NZ_BMDT01000023.1 Length: 6075
Range 1: 451 to 1143

Score:477 bits(1227), Expect:1e-151,
Method:Compositional matrix adjust.,
Identities:227/231(98%), Positives:230/231(99%), Gaps:0/231(0%)

```
Query  1     PIVQNIQGQMVHQAISPRTLNAWVKVVEEKAFSPEVIPMFSALSEGATPQDLNTMLNTVG  60
             PIVQN+QGQMVHQAISPRTLNAWVKVVEEKAFSPEVIPMFSALSEGATPQDLNTMLNTVG
Sbjct  451   PIVQNLQGQMVHQAISPRTLNAWVKVVEEKAFSPEVIPMFSALSEGATPQDLNTMLNTVG  630

Query  61    GHQAAMQMLKETINEEAAEWDRVHPVHAGPIAPGQMREPRGSDIAGTTSTLQEQIGWMTN  120
             GHQAAMQMLKETINEEAAEWDR+HPVHAGPIAPGQMREPRGSDIAGTTSTLQEQIGWMT+
Sbjct  631   GHQAAMQMLKETINEEAAEWDRLHPVHAGPIAPGQMREPRGSDIAGTTSTLQEQIGWMTH  810

Query  121   NPPIPVGEIYKRWIILGLNKIVRMYSPTSILDIRQGPKEPFRDYVDRFYKTLRAEQASQE  180
             NPPIPVGEIYKRWIILGLNKIVRMYSPTSILDIRQGPKEPFRDYVDRFYKTLRAEQASQE
Sbjct  811   NPPIPVGEIYKRWIILGLNKIVRMYSPTSILDIRQGPKEPFRDYVDRFYKTLRAEQASQE  990

Query  181   VKNWMTETLLVQNANPDCKTILKALGPAATLEEMMTACQGVGGPGHKARVL  231
             VKNWMTETLLVQNANPDCKTILKALGP ATLEEMMTACQGVGGPGHKARVL
Sbjct  991   VKNWMTETLLVQNANPDCKTILKALGPGATLEEMMTACQGVGGPGHKARVL  1143
```

>Deinococcus aquiradiocola strain JCM 14371 sequence30, whole genome shotgun sequence
Sequence ID: NZ_BMOE01000030.1 Length: 6128
Range 1: 504 to 1196

Score:476 bits(1226), Expect:2e-151,
Method:Compositional matrix adjust.,
Identities:227/231(98%), Positives:230/231(99%), Gaps:0/231(0%)

```
Query  1     PIVQNIQGQMVHQAISPRTLNAWVKVVEEKAFSPEVIPMFSALSEGATPQDLNTMLNTVG  60
             PIVQN+QGQMVHQAISPRTLNAWVKVVEEKAFSPEVIPMFSALSEGATPQDLNTMLNTVG
Sbjct  504   PIVQNLQGQMVHQAISPRTLNAWVKVVEEKAFSPEVIPMFSALSEGATPQDLNTMLNTVG  683

Query  61    GHQAAMQMLKETINEEAAEWDRVHPVHAGPIAPGQMREPRGSDIAGTTSTLQEQIGWMTN  120
             GHQAAMQMLKETINEEAAEWDR+HPVHAGPIAPGQMREPRGSDIAGTTSTLQEQIGWMT+
Sbjct  684   GHQAAMQMLKETINEEAAEWDRLHPVHAGPIAPGQMREPRGSDIAGTTSTLQEQIGWMTH  863

Query  121   NPPIPVGEIYKRWIILGLNKIVRMYSPTSILDIRQGPKEPFRDYVDRFYKTLRAEQASQE  180
             NPPIPVGEIYKRWIILGLNKIVRMYSPTSILDIRQGPKEPFRDYVDRFYKTLRAEQASQE
Sbjct  864   NPPIPVGEIYKRWIILGLNKIVRMYSPTSILDIRQGPKEPFRDYVDRFYKTLRAEQASQE  1043

Query  181   VKNWMTETLLVQNANPDCKTILKALGPAATLEEMMTACQGVGGPGHKARVL  231
             VKNWMTETLLVQNANPDCKTILKALGP ATLEEMMTACQGVGGPGHKARVL
Sbjct  1044  VKNWMTETLLVQNANPDCKTILKALGPGATLEEMMTACQGVGGPGHKARVL  1196
```

>Klebsiella pneumoniae strain EuSCAPE_IL017, whole genome shotgun sequence
Sequence ID: NZ_UKKG02000043.1 Length: 403
Range 1: 10 to 402

Score:261 bits(666), Expect:3e-85,
Method:Compositional matrix adjust.,
Identities:120/131(92%), Positives:128/131(97%), Gaps:0/131(0%)

```
Query  101  GSDIAGTTSTLQEQIGWMTNNPPIPVGEIYKRWIILGLNKIVRMYSPTSILDIRQGPKEP  160
            GSDIAGTTSTLQEQI WMT+NPP+PVG+IYKRWIILGLNKIVRMYSP SILDIRQGPKEP
Sbjct  402  GSDIAGTTSTLQEQITWMTSNPPVPVGDIYKRWIILGLNKIVRMYSPVSILDIRQGPKEP  223

Query  161  FRDYVDRFYKTLRAEQASQEVKNWMTETLLVQNANPDCKTILKALGPAATLEEMMTACQG  220
            FRDYVDRF+KTLRAEQA+Q+VKNWMT+TLLVQNANPDCKTIL+ALGP ATLEEMMTACQG
Sbjct  222  FRDYVDRFFKTLRAEQATQDVKNWMTDTLLVQNANPDCKTILRALGPGATLEEMMTACQG  43

Query  221  VGGPGHKARVL  231
            VGGPGHKARVL
Sbjct  42   VGGPGHKARVL  10
```

>Klebsiella pneumoniae strain EuSCAPE_IL020, whole genome shotgun sequence
Sequence ID: NZ_UKKB02000090.1 Length: 469
Range 1: 76 to 468

Score:251 bits(641), Expect:3e-81,
Method:Compositional matrix adjust.,
Identities:115/131(88%), Positives:122/131(93%), Gaps:0/131(0%)

```
Query  101  GSDIAGTTSTLQEQIGWMTNNPPIPVGEIYKRWIILGLNKIVRMYSPTSILDIRQGPKEP  160
            GSDIAGTTSTLQEQ+ W+T NP IPVGEIYKRWIILGLNKIVRMYSP SILDI+QGPKEP
Sbjct  468  GSDIAGTTSTLQEQVAWITGNPAIPVGEIYKRWIILGLNKIVRMYSPVSILDIKQGPKEP  289

Query  161  FRDYVDRFYKTLRAEQASQEVKNWMTETLLVQNANPDCKTILKALGPAATLEEMMTACQG  220
            FRDYVDRF+K LRAEQA+QEVKNWMT+TLLVQNANPDCK IL+ALGP A LEEMMTACQG
Sbjct  288  FRDYVDRFFKCLRAEQATQEVKNWMTDTLLVQNANPDCKAILRALGPGANLEEMMTACQG  109

Query  221  VGGPGHKARVL  231
            VGGP HKARVL
Sbjct  108  VGGPSHKARVL  76
```

>Klebsiella pneumoniae strain EuSCAPE_MT003, whole genome shotgun sequence
Sequence ID: NZ_UKIK02000113.1 Length: 318
Range 1: 3 to 317

Score:208 bits(530), Expect:4e-65,

```
Method:Compositional matrix adjust.,
Identities:95/105(90%), Positives:104/105(99%), Gaps:0/105(0%)

Query  101  GSDIAGTTSTLQEQIGWMTNNPPIPVGEIYKRWIILGLNKIVRMYSPTSILDIRQGPKEP  160
             GSDIAGTTS+LQEQIGWMT+NPPIPVG+IYKRWIILGLNKIVRMYSP SILDI+QGPKEP
Sbjct  317  GSDIAGTTSSLQEQIGWMTSNPPIPVGDIYKRWIILGLNKIVRMYSPVSILDIKQGPKEP  138

Query  161  FRDYVDRFYKTLRAEQASQEVKNWMTETLLVQNANPDCKTILKAL  205
             FRDYVDRF+KTLRAEQA+Q+VKNWMT+TLLVQNANPDCKTIL+AL
Sbjct  137  FRDYVDRFFKTLRAEQATQDVKNWMTDTLLVQNANPDCKTILRAL  3
```

```
>Klebsiella pneumoniae strain EuSCAPE_IL019, whole genome shotgun sequence
Sequence ID: NZ_UKJZ02000061.1 Length: 312
Range 1: 2 to 310

Score:208 bits(529), Expect:6e-65,
Method:Compositional matrix adjust.,
Identities:96/103(93%), Positives:101/103(98%), Gaps:0/103(0%)

Query  101  GSDIAGTTSTLQEQIGWMTNNPPIPVGEIYKRWIILGLNKIVRMYSPTSILDIRQGPKEP  160
             GSDIAGTTSTLQEQI WMT+NPPIPVG+IYKRWIILGLNKIVRMYSP SILDIRQGPKEP
Sbjct  2    GSDIAGTTSTLQEQIAWMTSNPPIPVGDIYKRWIILGLNKIVRMYSPVSILDIRQGPKEP  181

Query  161  FRDYVDRFYKTLRAEQASQEVKNWMTETLLVQNANPDCKTILK  203
             FRDYVDRF+KTLRAEQA+QEVKNWMTETLLVQNANPDCKTIL+
Sbjct  182  FRDYVDRFFKTLRAEQATQEVKNWMTETLLVQNANPDCKTILR  310
```

```
>Burkholderia pseudomallei strain 2604STDY5518971 ERS240105SCcontig000120, whole genome shotgun sequence
Sequence ID: NZ_UVFZ02000120.1 Length: 336
Range 1: 27 to 335

Score:207 bits(528), Expect:1e-64,
Method:Compositional matrix adjust.,
Identities:95/103(92%), Positives:99/103(96%), Gaps:0/103(0%)

Query  129  IYKRWIILGLNKIVRMYSPTSILDIRQGPKEPFRDYVDRFYKTLRAEQASQEVKNWMTET  188
             IYKRWIILGLNKIVRMYSP SILDI+QGPKEPFRDYVDRFY+TLRAEQA+QEVKNWMTET
Sbjct  335  IYKRWIILGLNKIVRMYSPVSILDIKQGPKEPFRDYVDRFYRTLRAEQATQEVKNWMTET  156

Query  189  LLVQNANPDCKTILKALGPAATLEEMMTACQGVGGPGHKARVL  231
             LLVQNANPDCK+ILKALG  ATLEEMMTACQGVGGP HKARVL
Sbjct  155  LLVQNANPDCKSILKALGTGATLEEMMTACQGVGGPSHKARVL  27
```

```
>Neisseria gonorrhoeae strain GC-191 NODE_147_length_284_cov_0.751092_ID_22618, whole genome shotgun sequence
Sequence ID: NZ_SAWA01000147.1 Length: 284
Range 1: 1 to 282

Score:187 bits(475), Expect:7e-57,
Method:Compositional matrix adjust.,
Identities:84/94(89%), Positives:92/94(97%), Gaps:0/94(0%)

Query  127  GEIYKRWIILGLNKIVRMYSPTSILDIRQGPKEPFRDYVDRFYKTLRAEQASQEVKNWMT  186
             G+IYKRWIILGLNKIVRMYSP SILDI+QGPKEPFRDYVDRF+KTLRAEQA+Q+VKNWMT
Sbjct  282  GDIYKRWIILGLNKIVRMYSPVSILDIKQGPKEPFRDYVDRFFKTLRAEQATQDVKNWMT  103

Query  187  ETLLVQNANPDCKTILKALGPAATLEEMMTACQG  220
             +TLLVQNANPDCKTIL+ALGP A+LEEMMTACQG
Sbjct  102  DTLLVQNANPDCKTILRALGPGASLEEMMTACQG  1
```

```
>Neisseria gonorrhoeae strain gc-168 NODE_130_length_274_cov_0.69863_ID_20302, whole genome shotgun sequence
Sequence ID: NZ_SAVF01000130.1 Length: 274
Range 1: 1 to 273

Score:179 bits(454), Expect:1e-53,
Method:Compositional matrix adjust.,
Identities:81/91(89%), Positives:89/91(97%), Gaps:0/91(0%)

Query  101  GSDIAGTTSTLQEQIGWMTNNPPIPVGEIYKRWIILGLNKIVRMYSPTSILDIRQGPKEP  160
             GSDIAGTTSTLQEQI WMT+NPP+PVG+IYKRWIILGLNKIVRMYSP SILDI+QGPKEP
Sbjct  273  GSDIAGTTSTLQEQIAWMTSNPPVPVGDIYKRWIILGLNKIVRMYSPVSILDIKQGPKEP  94

Query  161  FRDYVDRFYKTLRAEQASQEVKNWMTETLLV  191
             FRDYVDRF+KTLRAEQ++QEVKNWMT+TLLV
Sbjct  93   FRDYVDRFFKTLRAEQSTQEVKNWMTDTLLV  1
```

```
>Burkholderia pseudomallei strain 2604STDY5519032 ERS240165SCcontig000074, whole genome shotgun sequence
Sequence ID: NZ_UVID02000074.1 Length: 656
Range 1: 385 to 654

Score:178 bits(452), Expect:1e-51,
Method:Compositional matrix adjust.,
Identities:81/90(90%), Positives:86/90(95%), Gaps:0/90(0%)

Query  142  VRMYSPTSILDIRQGPKEPFRDYVDRFYKTLRAEQASQEVKNWMTETLLVQNANPDCKTI  201
             VRMYSP SILDI+QGPKEPFRDYVDRF+KTLRAEQA+QEVKNWMT+TLLVQNANPDCKTI
Sbjct  654  VRMYSPVSILDIKQGPKEPFRDYVDRFFKTLRAEQATQEVKNWMTDTLLVQNANPDCKTI  475

Query  202  LKALGPAATLEEMMTACQGVGGPGHKARVL  231
             L+ LGP ATLEEMMTACQGVGGP HKARVL
Sbjct  474  LRGLGPGATLEEMMTACQGVGGPSHKARVL  385
```

```
>Klebsiella pneumoniae strain EuSCAPE_MT002, whole genome shotgun sequence
Sequence ID: NZ_UKIM02000089.1 Length: 257
Range 1: 2 to 256

Score:171 bits(434), Expect:7e-51,
Method:Compositional matrix adjust.,
Identities:78/85(92%), Positives:83/85(97%), Gaps:0/85(0%)

Query  101  GSDIAGTTSTLQEQIGWMTNNPPIPVGEIYKRWIILGLNKIVRMYSPTSILDIRQGPKEP  160
             GSDIAGTTSTLQEQI WMT+NPP+PVGEIYKRWIILGLNKIVRMYSP SILDIRQGPKEP
Sbjct  2    GSDIAGTTSTLQEQIAWMTSNPPVPVGEIYKRWIILGLNKIVRMYSPVSILDIRQGPKEP  181

Query  161  FRDYVDRFYKTLRAEQASQEVKNWM  185
             FRDYVDRF+KTLRAEQA+Q+VKNWM
Sbjct  182  FRDYVDRFFKTLRAEQATQDVKNWM  256
```

>Klebsiella pneumoniae strain EuSCAPE_IL005, whole genome shotgun sequence
Sequence ID: NZ_UKJL02000066.1 Length: 250
Range 1: 1 to 249

Score:166 bits(419), Expect:2e-48,
Method:Compositional matrix adjust.,
Identities:76/83(92%), Positives:81/83(97%), Gaps:0/83(0%)

```
Query  101  GSDIAGTTSTLQEQIGWMTNNPPIPVGEIYKRWIILGLNKIVRMYSPTSILDIRQGPKEP  160
            GSDIAGTTSTLQEQI WMT+NPPIPVG+IYKRWIILGLNKIVRMYSP SILDI+QGPKEP
Sbjct  249  GSDIAGTTSTLQEQIAWMTSNPPIPVGDIYKRWIILGLNKIVRMYSPVSILDIKQGPKEP  70

Query  161  FRDYVDRFYKTLRAEQASQEVKN  183
            FRDYVDRF+KTLRAEQ+SQEVKN
Sbjct  69   FRDYVDRFFKTLRAEQSSQEVKN  1
```

>Klebsiella pneumoniae strain EuSCAPE_LT007, whole genome shotgun sequence
Sequence ID: NZ_UKJI02000076.1 Length: 252
Range 1: 2 to 250

Score:166 bits(419), Expect:2e-48,
Method:Compositional matrix adjust.,
Identities:75/83(90%), Positives:82/83(98%), Gaps:0/83(0%)

```
Query  101  GSDIAGTTSTLQEQIGWMTNNPPIPVGEIYKRWIILGLNKIVRMYSPTSILDIRQGPKEP  160
            GSDIAGTTSTLQEQ+GWMT+NPPIPVG+IYKRWIILGLNKIVRMYSP SILDI+QGPKEP
Sbjct  2    GSDIAGTTSTLQEQVGWMTSNPPIPVGDIYKRWIILGLNKIVRMYSPVSILDIKQGPKEP  181

Query  161  FRDYVDRFYKTLRAEQASQEVKN  183
            FRDYVDRF+KTLRAEQA+Q+VKN
Sbjct  182  FRDYVDRFFKTLRAEQATQDVKN  250
```

>Klebsiella pneumoniae strain EuSCAPE_LT003, whole genome shotgun sequence
Sequence ID: NZ_UKJH02000061.1 Length: 419
Range 1: 179 to 418

Score:160 bits(404), Expect:1e-45,
Method:Compositional matrix adjust.,
Identities:71/80(89%), Positives:78/80(97%), Gaps:0/80(0%)

```
Query  152  DIRQGPKEPFRDYVDRFYKTLRAEQASQEVKNWMTETLLVQNANPDCKTILKALGPAATL  211
            DI+QGPKEPFRDYVDRF+K LRAEQA+Q+VKNWMT+TLLVQNANPDCKTIL+ALGPAA+L
Sbjct  418  DIKQGPKEPFRDYVDRFFKVLRAEQATQDVKNWMTDTLLVQNANPDCKTILRALGPAASL  239

Query  212  EEMMTACQGVGGPGHKARVL  231
            EEMMTACQGVGGP HKARVL
Sbjct  238  EEMMTACQGVGGPSHKARVL  179
```

>Klebsiella quasipneumoniae strain EuSCAPE_IL014, whole genome shotgun sequence
Sequence ID: NZ_UJYK02000052.1 Length: 269
Range 1: 3 to 203

Score:135 bits(339), Expect:2e-36,
Method:Compositional matrix adjust.,
Identities:62/69(90%), Positives:66/69(95%), Gaps:2/69(2%)

```
Query  160  PFRDYVDRFYKTLRAEQASQEVKNWMTETLLVQNANPDCKTILKALGPAATLEEMMTACQ  219
            PFRDYVDRF+KTLRAEQA+QEVKNWMTETLLVQNANPDCKTIL+ALGP ATLEEMMTACQ
Sbjct  3    PFRDYVDRFFKTLRAEQATQEVKNWMTETLLVQNANPDCKTILRALGPGATLEEMMTACQ  182

Query  220  GVGGPGHKA  228
            GVG  GH+A
Sbjct  183  GVG--GHQA  203
```

>Klebsiella pneumoniae strain EuSCAPE_IL005, whole genome shotgun sequence
Sequence ID: NZ_UKJL02000060.1 Length: 308
Range 1: 136 to 306

Score:112 bits(279), Expect:3e-27,
Method:Compositional matrix adjust.,
Identities:52/57(91%), Positives:54/57(94%), Gaps:0/57(0%)

```
Query  175  EQASQEVKNWMTETLLVQNANPDCKTILKALGPAATLEEMMTACQGVGGPGHKARVL  231
            EQ+SQEVKNWMT+TLLVQNANPDCKTILKALG  ATLEEMMTACQGVGGP HKARVL
Sbjct  306  EQSSQEVKNWMTDTLLVQNANPDCKTILKALGTGATLEEMMTACQGVGGPSHKARVL  136
```

>Neisseria gonorrhoeae strain GC-206 NODE_136_length_334_cov_1.12903_ID_20444, whole genome shotgun sequence
Sequence ID: NZ_SAWN01000136.1 Length: 334
Range 1: 2 to 160

Score:106 bits(265), Expect:5e-25,
Method:Compositional matrix adjust.,
Identities:49/53(92%), Positives:51/53(96%), Gaps:0/53(0%)

```
Query  179  QEVKNWMTETLLVQNANPDCKTILKALGPAATLEEMMTACQGVGGPGHKARVL  231
            Q+VKNWMTETLLVQNANPDCK IL+ALGP ATLEEMMTACQGVGGPGHKARVL
Sbjct  2    QDVKNWMTETLLVQNANPDCKNILRALGPGATLEEMMTACQGVGGPGHKARVL  160
```

>Klebsiella pneumoniae strain B2 isolate B2, whole genome shotgun sequence
Sequence ID: NZ_CABWNT010000058.1 Length: 7306
Range 1: 2769 to 3461

Score:115 bits(288), Expect:1e-24,
Method:Compositional matrix adjust.,
Identities:71/235(30%), Positives:122/235(51%), Gaps:8/235(3%)

```
Query  1     PIVQNIQGQMVHQAISPRTLNAWVKVVEEKAFSPEV-IPMFSALSEGATPQDLNTMLNTV  59
             PI+ +  G    + ++PR    WV  +++     E  + +F  LS   T +++N  L+ V
Sbjct  2769  PIMIDGAGNRNFRPLTPRGYTTWVNTIQQNNLLNEASVNLFGILSVDCTSEEMNAFLDVV  2948

Query  60    GGHQAAMQMLKETINEEAAEWDRVHPVHAGP-IAPGQMREPRGSD-IAGTTSTLQEQIGW  117
              G     Q+L + +++ A EWDR HP+   P +AP Q   P  +  I G     + Q+
```

```
Sbjct  2949  PGQAGQKQVLLDNLDKIAEEWDRRHPMPNPPLVAPPQGPIPMTARFIRGLGVPRERQMEP  3128

Query  118   MTNNPPIPVGEIYKRWIILGLNKIVR-MYSPTSILDIRQGPKEPFRDYVDRFYKTLRAEQ  176
               +       + Y++WII  + + ++ M       +IRQGPKEP+ ++VDR    +++E
Sbjct  3129  AFDQ----FRQTYRQWIIEAMTEGIKIMIGKPKAQNIRQGPKEPYPEFVDRLLSQIKSEG  3296

Query  177   ASQEVKNWMTETLLVQNANPDCKTILKALGPAATLEEMMTACQGVGGPGHKARVL  231
                ++   ++T+TL +QNAN +CK  ++ L P  TLEE M AC+ +G    K  +L
Sbjct  3297  HPADITKFLTDTLTIQNANDECKNAMRHLRPEDTLEEKMYACRDIGTMKQKMALL  3461
```

>Klebsiella pneumoniae strain EuSCAPE_NO017, whole genome shotgun sequence
Sequence ID: NZ_UKIO02000074.1 Length: 348
Range 1: 191 to 346

Score:105 bits(263), Expect:1e-24,
Method:Compositional matrix adjust.,
Identities:48/52(92%), Positives:51/52(98%), Gaps:0/52(0%)

```
Query  180  EVKNWMTETLLVQNANPDCKTILKALGPAATLEEMMTACQGVGGPGHKARVL  231
            EVKNWMT+TLLVQNANPDCKTIL+ALGP A+LEEMMTACQGVGGPGHKARVL
Sbjct  346  EVKNWMTDTLLVQNANPDCKTILRALGPGASLEEMMTACQGVGGPGHKARVL  191
```

>Bacillus velezensis strain TN14-11 TN14-11_ctg0032, whole genome shotgun sequence
Sequence ID: NZ_JAHVIU010000017.1 Length: 1125
Range 1: 446 to 592

Score:100 bits(249), Expect:3e-20,
Method:Compositional matrix adjust.,
Identities:46/49(94%), Positives:49/49(100%), Gaps:0/49(0%)

```
Query  1    PIVQNIQGQMVHQAISPRTLNAWVKVVEEKAFSPEVIPMFSALSEGATP  49
            PIVQN+QGQMVHQ++SPRTLNAWVKVVEEKAFSPEVIPMFSALSEGATP
Sbjct  592  PIVQNLQGQMVHQSLSPRTLNAWVKVVEEKAFSPEVIPMFSALSEGATP  446
```

>Escherichia coli strain VRES0669, whole genome shotgun sequence
Sequence ID: NZ_UINV01000086.1 Length: 5544
Range 1: 5416 to 5544

Score:87.8 bits(216), Expect:3e-15,
Method:Compositional matrix adjust.,
Identities:40/43(93%), Positives:41/43(95%), Gaps:0/43(0%)

```
Query  189   LLVQNANPDCKTILKALGPAATLEEMMTACQGVGGPGHKARVL  231
             LLVQNANPDCKTIL+ALGP ATLEEMMTACQGVGGP HKARVL
Sbjct  5544  LLVQNANPDCKTILRALGPGATLEEMMTACQGVGGPSHKARVL  5416
```

>Pseudomonas sp. MPR-LB3 NODE_257_length_230_cov_0.882353, whole genome shotgun sequence
Sequence ID: NZ_POFY01000255.1 Length: 230
Range 1: 52 to 165

Score:78.2 bits(191), Expect:2e-14,
Method:Compositional matrix adjust.,
Identities:37/38(97%), Positives:37/38(97%), Gaps:0/38(0%)

```
Query  15   ISPRTLNAWVKVVEEKAFSPEVIPMFSALSEGATPQDL  52
            ISPRTLNAWVKVVEEKAFSPEVIPMFSALSEGATPQD
Sbjct  165  ISPRTLNAWVKVVEEKAFSPEVIPMFSALSEGATPQDF  52
```

>Photobacterium rosenbergii strain DP6N14-7 DP6N14-7_ctg0065, whole genome shotgun sequence
Sequence ID: NZ_JAHVIB010000057.1 Length: 564
Range 1: 446 to 562

Score:79.3 bits(194), Expect:1e-13,
Method:Compositional matrix adjust.,
Identities:37/39(95%), Positives:39/39(100%), Gaps:0/39(0%)

```
Query  11   VHQAISPRTLNAWVKVVEEKAFSPEVIPMFSALSEGATP  49
            VHQ++SPRTLNAWVKVVEEKAFSPEVIPMFSALSEGATP
Sbjct  562  VHQSLSPRTLNAWVKVVEEKAFSPEVIPMFSALSEGATP  446
```

>Rhizobium bangladeshense strain 1017 scf43_contig_2, whole genome shotgun sequence
Sequence ID: NZ_LYPK01000043.1 Length: 288
Range 1: 49 to 153

Score:73.9 bits(180), Expect:1e-12,
Method:Compositional matrix adjust.,
Identities:34/35(97%), Positives:35/35(100%), Gaps:0/35(0%)

```
Query  1    PIVQNIQGQMVHQAISPRTLNAWVKVVEEKAFSPE  35
            PIVQN+QGQMVHQAISPRTLNAWVKVVEEKAFSPE
Sbjct  49   PIVQNLQGQMVHQAISPRTLNAWVKVVEEKAFSPE  153
```

Range 2: 210 to 287

Score:55.8 bits(133), Expect:9e-06,
Method:Compositional matrix adjust.,
Identities:25/26(96%), Positives:26/26(100%), Gaps:0/26(0%)

```
Query  1    PIVQNIQGQMVHQAISPRTLNAWVKV  26
            PIVQN+QGQMVHQAISPRTLNAWVKV
Sbjct  210  PIVQNLQGQMVHQAISPRTLNAWVKV  287
```

>Listeria monocytogenes strain CrieF364 NODE_22_length_769_cov_13.562305, whole genome shotgun sequence
Sequence ID: NZ_JASJTP010000022.1 Length: 769
Range 1: 79 to 171

Score:61.6 bits(148), Expect:9e-07,
Method:Compositional matrix adjust.,
Identities:27/31(87%), Positives:28/31(90%), Gaps:0/31(0%)

```
Query  56   LNTVGGHQAAMQMLKETINEEAAEWDRVHPV  86
            +  VGGHQAAMQMLKETINEEAAEWDRVHP
Sbjct  171  IRLVGGHQAAMQMLKETINEEAAEWDRVHPC  79
```

>Photobacterium rosenbergii strain DP6N14-7 DP6N14-7_ctg0069, whole genome shotgun sequence
Sequence ID: NZ_JAHVIB010000058.1 Length: 667
Range 1: 556 to 636

Score:51.2 bits(121), Expect:0.003,
Method:Compositional matrix adjust.,
Identities:23/27(85%), Positives:26/27(96%), Gaps:0/27(0%)

```
Query  1    PIVQNIQGQMVHQAISPRTLNAWVKVV  27
            PIVQN+QGQMVHQ+ISPRTLNA +KVV
Sbjct  556  PIVQNLQGQMVHQSISPRTLNA*IKVV  636
```

>Vibrio hangzhouensis strain DP3N7-4 DP3N7-4_ctg0051, whole genome shotgun sequence
Sequence ID: NZ_JAHVKH010000037.1 Length: 742
Range 1: 77 to 157

Score:51.2 bits(121), Expect:0.004,
Method:Compositional matrix adjust.,
Identities:23/27(85%), Positives:26/27(96%), Gaps:0/27(0%)

```
Query  1    PIVQNIQGQMVHQAISPRTLNAWVKVV  27
            PIVQN+QGQMVHQ+ISPRTLNA +KVV
Sbjct  157  PIVQNLQGQMVHQSISPRTLNA*IKVV  77
```

Query #7: sp|P04585|Nucleocapsid_Protein__p7_HV1H2 Gag-Pol polyprotein OS=Human immunodeficiency virus type 1 group M subtype B (isolate HXB2) OX=11706 GN=gag-pol PE=1 SV=4
Query ID: lcl|Query_2387714 Length: 55

Sequences producing significant alignments:

| Description | Scientific Name | Common Name | Taxid | Max Score | Total Score | Query cover | E Value | Per. Ident | Acc. Len | Accession |
|---|---|---|---|---|---|---|---|---|---|---|
| Klebsiella pneumoniae strain EuSCAPE_IL019, whole genome shotg... | Klebsiella p... | NA | 573 | 103 | 103 | 100% | 1e-26 | 85.45 | 254 | NZ_UKJZ020000... |
| Thalassovita mangrovi strain GS-10 Scaffold19_1, whole genome... | Thalassovita... | NA | 2692236 | 112 | 112 | 100% | 2e-26 | 90.91 | 2900 | NZ_WWEN010000... |
| Actinoplanes campanulatus strain JCM 3059 sequence058, whole... | Actinoplanes... | NA | 113559 | 112 | 112 | 100% | 2e-26 | 90.91 | 4754 | NZ_BMPW010000... |
| Enterococcus alcedinis strain CCM 8433 sequence23, whole genom... | Enterococcus... | NA | 1274384 | 112 | 112 | 100% | 2e-26 | 90.91 | 6075 | NZ_BMDT010000... |
| Deinococcus aquiradiocola strain JCM 14371 sequence30, whole... | Deinococcus ... | NA | 393059 | 112 | 112 | 100% | 2e-26 | 90.91 | 6128 | NZ_BMOE010000... |
| Burkholderia pseudomallei strain 2604STDY5519032... | Burkholderia... | NA | 28450 | 103 | 103 | 100% | 3e-25 | 83.64 | 656 | NZ_UVID020000... |
| Klebsiella pneumoniae strain EuSCAPE_NO017, whole genome shotg... | Klebsiella p... | NA | 573 | 88.6 | 88.6 | 87% | 2e-20 | 81.25 | 348 | NZ_UKIO020000... |
| Klebsiella pneumoniae strain EuSCAPE_LT003, whole genome shotg... | Klebsiella p... | NA | 573 | 87.0 | 87.0 | 83% | 1e-19 | 86.96 | 419 | NZ_UKJH020000... |
| Neisseria gonorrhoeae strain GC-206... | Neisseria go... | NA | 485 | 85.5 | 85.5 | 81% | 2e-19 | 82.22 | 334 | NZ_SAWN010001... |
| Klebsiella pneumoniae strain EuSCAPE_IL005, whole genome shotg... | Klebsiella p... | NA | 573 | 60.5 | 60.5 | 58% | 2e-09 | 81.25 | 308 | NZ_UKJL020000... |
| Klebsiella pneumoniae strain EuSCAPE_MT015, whole genome shotg... | Klebsiella p... | NA | 573 | 51.6 | 51.6 | 40% | 5e-06 | 100.00 | 281 | NZ_UKIW020001... |
| Modestobacter marinus strain CPCC 205245 Scaffold62_1, whole... | Modestobacte... | NA | 477641 | 51.2 | 51.2 | 67% | 6e-05 | 48.65 | 7666 | NZ_JABGCH0100... |
| Geminicoccus harenae strain CPCC 101083 Scaffold77_1, whole... | Geminicoccus... | NA | 2498453 | 50.8 | 50.8 | 67% | 7e-05 | 48.65 | 7497 | NZ_JABGCL0100... |
| Modestobacter italicus strain CPCC 205242 Scaffold14_1, whole... | Modestobacte... | NA | 2732864 | 50.8 | 50.8 | 67% | 7e-05 | 48.65 | 15567 | NZ_JABGCJ0100... |
| Escherichia coli strain HU8 NODE_685_length_270_cov_0.756477,... | Escherichia ... | NA | 562 | 47.4 | 47.4 | 67% | 2e-04 | 48.65 | 270 | NZ_JAHCTM0100... |
| Salmonella enterica subsp. enterica serovar Typhimurium strain... | Salmonella e... | NA | 90371 | 43.1 | 43.1 | 85% | 0.005 | 38.30 | 211 | NZ_JALDNE0100... |
| Chlamydia trachomatis strain R4175extract, whole genome shotgu... | Chlamydia tr... | NA | 813 | 45.1 | 45.1 | 85% | 0.007 | 46.15 | 2947 | NZ_CVNX010000... |
| Insolitispirillum peregrinum strain DSM 11589, whole genome... | Insolitispir... | NA | 80876 | 44.7 | 44.7 | 87% | 0.011 | 45.83 | 474983 | NZ_FTOA010000... |
| Rickettsia endosymbiont of Ceutorhynchus assimilis isolate... | Rickettsia e... | NA | 2884482 | 43.9 | 43.9 | 81% | 0.020 | 35.56 | 4063491 | NZ_OU906081.1 |

Alignments:

>Klebsiella pneumoniae strain EuSCAPE_IL019, whole genome shotgun sequence
Sequence ID: NZ_UKJZ02000062.1 Length: 254
Range 1: 34 to 198

Score:103 bits(257), Expect:1e-26,
Method:Compositional matrix adjust.,
Identities:47/55(85%), Positives:52/55(94%), Gaps:0/55(0%)

```
Query  1    MQRGNFRNQRKIVKCFNCGKEGHTARNCRAPRKKGCWKCGKEGHQMKDCTERQAN  55
            MQ+GNF+  ++I+KCFNCGKEGH ARNCRAPRKKGCWKCGKEGHQMKDCTERQAN
Sbjct  34   MQKGNFKGPKRIIKCFNCGKEGHIARNCRAPRKKGCWKCGKEGHQMKDCTERQAN  198
```

>Thalassovita mangrovi strain GS-10 Scaffold19_1, whole genome shotgun sequence
Sequence ID: NZ_WWEN01000019.1 Length: 2900
Range 1: 2171 to 2335

Score:112 bits(280), Expect:2e-26,
Method:Compositional matrix adjust.,
Identities:50/55(91%), Positives:53/55(96%), Gaps:0/55(0%)

```
Query  1     MQRGNFRNQRKIVKCFNCGKEGHTARNCRAPRKKGCWKCGKEGHQMKDCTERQAN  55
             +Q+GNFRNQRK VKCFNCGKEGH A+NCRAPRKKGCWKCGKEGHQMKDCTERQAN
Sbjct  2171  IQKGNFRNQRKTVKCFNCGKEGHIAKNCRAPRKKGCWKCGKEGHQMKDCTERQAN  2335
```

>Actinoplanes campanulatus strain JCM 3059 sequence058, whole genome shotgun sequence
Sequence ID: NZ_BMPW01000058.1 Length: 4754
Range 1: 1228 to 1392

Score:112 bits(280), Expect:2e-26,
Method:Compositional matrix adjust.,
Identities:50/55(91%), Positives:53/55(96%), Gaps:0/55(0%)

```
Query  1     MQRGNFRNQRKIVKCFNCGKEGHTARNCRAPRKKGCWKCGKEGHQMKDCTERQAN  55
             +Q+GNFRNQRK VKCFNCGKEGH A+NCRAPRKKGCWKCGKEGHQMKDCTERQAN
Sbjct  1228  IQKGNFRNQRKTVKCFNCGKEGHIAKNCRAPRKKGCWKCGKEGHQMKDCTERQAN  1392
```

>Enterococcus alcedinis strain CCM 8433 sequence23, whole genome shotgun sequence
Sequence ID: NZ_BMDT01000023.1 Length: 6075
Range 1: 1186 to 1350

Score:112 bits(280), Expect:2e-26,
Method:Compositional matrix adjust.,
Identities:50/55(91%), Positives:53/55(96%), Gaps:0/55(0%)

```
Query  1     MQRGNFRNQRKIVKCFNCGKEGHTARNCRAPRKKGCWKCGKEGHQMKDCTERQAN  55
             +Q+GNFRNQRK VKCFNCGKEGH A+NCRAPRKKGCWKCGKEGHQMKDCTERQAN
Sbjct  1186  IQKGNFRNQRKTVKCFNCGKEGHIAKNCRAPRKKGCWKCGKEGHQMKDCTERQAN  1350
```

```
>Deinococcus aquiradiocola strain JCM 14371 sequence30, whole genome shotgun sequence
Sequence ID: NZ_BMOE01000030.1 Length: 6128
Range 1: 1239 to 1403

Score:112 bits(280), Expect:2e-26,
Method:Compositional matrix adjust.,
Identities:50/55(91%), Positives:53/55(96%), Gaps:0/55(0%)

Query  1     MQRGNFRNQRKIVKCFNCGKEGHTARNCRAPRKKGCWKCGKEGHQMKDCTERQAN  55
             +Q+GNFRNQRK VKCFNCGKEGH A+NCRAPRKKGCWKCGKEGHQMKDCTERQAN
Sbjct  1239  IQKGNFRNQRKTVKCFNCGKEGHIAKNCRAPRKKGCWKCGKEGHQMKDCTERQAN  1403
```

```
>Burkholderia pseudomallei strain 2604STDY5519032 ERS240165SCcontig000074, whole genome shotgun sequence
Sequence ID: NZ_UVID02000074.1 Length: 656
Range 1: 181 to 345

Score:103 bits(258), Expect:3e-25,
Method:Compositional matrix adjust.,
Identities:46/55(84%), Positives:50/55(90%), Gaps:0/55(0%)

Query  1    MQRGNFRNQRKIVKCFNCGKEGHTARNCRAPRKKGCWKCGKEGHQMKDCTERQAN  55
            MQ+ NF+  R+ VKCFNCGKEGH A+NCRAPRKKGCWKCGKEGHQMKDCTERQAN
Sbjct  345  MQKNNFKGPRRTVKCFNCGKEGHIAKNCRAPRKKGCWKCGKEGHQMKDCTERQAN  181
```

```
>Klebsiella pneumoniae strain EuSCAPE_NO017, whole genome shotgun sequence
Sequence ID: NZ_UKIO02000074.1 Length: 348
Range 1: 2 to 145

Score:88.6 bits(218), Expect:2e-20,
Method:Compositional matrix adjust.,
Identities:39/48(81%), Positives:43/48(89%), Gaps:0/48(0%)

Query  1    MQRGNFRNQRKIVKCFNCGKEGHTARNCRAPRKKGCWKCGKEGHQMKD  48
            MQ+ NF+  R+ VKCFNCGKEGH ARNCRAPRKKGCWKCG+EGHQMKD
Sbjct  145  MQKSNFKGPRRAVKCFNCGKEGHIARNCRAPRKKGCWKCGREGHQMKD  2
```

```
>Klebsiella pneumoniae strain EuSCAPE_LT003, whole genome shotgun sequence
Sequence ID: NZ_UKJH02000061.1 Length: 419
Range 1: 2 to 136

Score:87.0 bits(214), Expect:1e-19,
Method:Compositional matrix adjust.,
Identities:40/46(87%), Positives:43/46(93%), Gaps:1/46(2%)

Query  1    MQRGNFRNQRKIVKCFNCGKEGHTARNCRAPRKKGCWKCGKEGHQM  46
            MQ+ NF+N RKIVKCFNCGKEGH ARNCRAPRKKGCWKCG+EGHQM
Sbjct  136  MQKSNFKN-RKIVKCFNCGKEGHIARNCRAPRKKGCWKCGREGHQM  2
```

```
>Neisseria gonorrhoeae strain GC-206 NODE_136_length_334_cov_1.12903_ID_20444, whole genome shotgun sequence
Sequence ID: NZ_SAWN01000136.1 Length: 334
Range 1: 200 to 334

Score:85.5 bits(210), Expect:2e-19,
Method:Compositional matrix adjust.,
Identities:37/45(82%), Positives:42/45(93%), Gaps:0/45(0%)

Query  1    MQRGNFRNQRKIVKCFNCGKEGHTARNCRAPRKKGCWKCGKEGHQ  45
            MQRGNF+  ++I+KCFNCGKEGH A+NCRAPRKKGCWKCGKEGHQ
Sbjct  200  MQRGNFKGPKRIIKCFNCGKEGHIAKNCRAPRKKGCWKCGKEGHQ  334
```

```
>Klebsiella pneumoniae strain EuSCAPE_IL005, whole genome shotgun sequence
Sequence ID: NZ_UKJL02000060.1 Length: 308
Range 1: 1 to 96

Score:60.5 bits(145), Expect:2e-09,
Method:Compositional matrix adjust.,
Identities:26/32(81%), Positives:28/32(87%), Gaps:0/32(0%)

Query  1   MQRGNFRNQRKIVKCFNCGKEGHTARNCRAPR  32
           MQ+ NF+  RKIVKCFNCGKEGH ARNCRAPR
Sbjct  96  MQKSNFKGPRKIVKCFNCGKEGHIARNCRAPR  1
```

```
>Klebsiella pneumoniae strain EuSCAPE_MT015, whole genome shotgun sequence
Sequence ID: NZ_UKIW02000113.1 Length: 281
Range 1: 3 to 68

Score:51.6 bits(122), Expect:5e-06,
Method:Compositional matrix adjust.,
Identities:22/22(100%), Positives:22/22(100%), Gaps:0/22(0%)

Query  34  KGCWKCGKEGHQMKDCTERQAN  55
           KGCWKCGKEGHQMKDCTERQAN
Sbjct  3   KGCWKCGKEGHQMKDCTERQAN  68
```

```
>Modestobacter marinus strain CPCC 205245 Scaffold62_1, whole genome shotgun sequence
Sequence ID: NZ_JABGCH010000063.1 Length: 7666
Range 1: 1990 to 2100

Score:51.2 bits(121), Expect:6e-05,
Method:Composition-based stats.,
Identities:18/37(49%), Positives:25/37(67%), Gaps:0/37(0%)

Query  13    VKCFNCGKEGHTARNCRAPRKKGCWKCGKEGHQMKDC  49
             V C+NCG++GH + +C  PR+  C+ CG  GH  KDC
Sbjct  1990  VTCYNCGEKGHISTSCPKPRRLTCYNCGGVGHISKDC  2100
```

```
>Geminicoccus harenae strain CPCC 101083 Scaffold77_1, whole genome shotgun sequence
```

Sequence ID: NZ_JABGCL010000077.1 Length: 7497
Range 1: 1986 to 2096

Score:50.8 bits(120), Expect:7e-05,
Method:Composition-based stats.,
Identities:18/37(49%), Positives:25/37(67%), Gaps:0/37(0%)

```
Query  13    VKCFNCGKEGHTARNCRAPRKKGCWKCGKEGHQMKDC  49
             V C+NCG++GH + +C  PR+  C+ CG  GH  KDC
Sbjct  1986  VTCYNCGEKGHISTSCPKPRRLTCYNCGGVGHISKDC  2096
```

>Modestobacter italicus strain CPCC 205242 Scaffold14_1, whole genome shotgun sequence
Sequence ID: NZ_JABGCJ010000014.1 Length: 15567
Range 1: 10921 to 11031

Score:50.8 bits(120), Expect:7e-05,
Method:Composition-based stats.,
Identities:18/37(49%), Positives:25/37(67%), Gaps:0/37(0%)

```
Query  13     VKCFNCGKEGHTARNCRAPRKKGCWKCGKEGHQMKDC  49
              V C+NCG++GH + +C  PR+  C+ CG  GH  KDC
Sbjct  11031  VTCYNCGEKGHISTSCPKPRRLTCYNCGGVGHISKDC  10921
```

>Escherichia coli strain HU8 NODE_685_length_270_cov_0.756477, whole genome shotgun sequence
Sequence ID: NZ_JAHCTM010000685.1 Length: 270
Range 1: 21 to 131

Score:47.4 bits(111), Expect:2e-04,
Method:Compositional matrix adjust.,
Identities:18/37(49%), Positives:25/37(67%), Gaps:0/37(0%)

```
Query  13   VKCFNCGKEGHTARNCRAPRKKGCWKCGKEGHQMKDC  49
            V C+NCG++GH + +C  PR+  C+ CG  GH  KDC
Sbjct  21   VTCYNCGEKGHISTSCPKPRRLTCYNCGGVGHISKDC  131
```

>Salmonella enterica subsp. enterica serovar Typhimurium strain SH6609 NODE_771_length_211_cov_1.211538, whole genome shotgun sequence
Sequence ID: NZ_JALDNE010000502.1 Length: 211
Range 1: 59 to 199

Score:43.1 bits(100), Expect:0.005,
Method:Compositional matrix adjust.,
Identities:18/47(38%), Positives:28/47(59%), Gaps:0/47(0%)

```
Query  4    GNFRNQRKIVKCFNCGKEGHTARNCRAPRKKGCWKCGKEGHQMKDCT  50
            G+ +     VKC+ CG+ GH A NC    +  C++CG+ GH  ++CT
Sbjct  59   GHIQKDCTKVKCYRCGETGHVAINCSKTSEDICYRCGESGHLARECT  199
```

>Chlamydia trachomatis strain R4175extract, whole genome shotgun sequence
Sequence ID: NZ_CVNX01000009.1 Length: 2947
Range 1: 1947 to 2096

Score:45.1 bits(105), Expect:0.007,
Method:Composition-based stats.,
Identities:24/52(46%), Positives:30/52(57%), Gaps:7/52(13%)

```
Query  3     RGNFRNQRKIVKCFNCGKEGHTARNCRAPRKKG-----CWKCGKEGHQMKDC  49
             R   RN ++   CFNCGK GH  ++CRAP K+G     C KCGK  H+   C
Sbjct  1947  RSMSRNDQR--TCFNCGKPGHFKKDCRAPDKQGGTLTLCSKCGKGYHRADQC  2096
```

>Insolitispirillum peregrinum strain DSM 11589, whole genome shotgun sequence
Sequence ID: NZ_FTOA01000005.1 Length: 474983
Range 1: 265843 to 265980

Score:44.7 bits(104), Expect:0.011,
Method:Compositional matrix adjust.,
Identities:22/48(46%), Positives:29/48(60%), Gaps:2/48(4%)

```
Query  5       NFRNQRKIVKCFNCGKEGHTARNCRAPRKKGCWKCGKEGHQMKDCTER  52
               + RN   + +C  CGK GH  RNC  P  + C KCGK GH +++C ER
Sbjct  265843  DIRNCPNVRRCSGCGKPGHDIRNC--PNVRRCSKCGKPGHDLRNCPER  265980
```

>Rickettsia endosymbiont of Ceutorhynchus assimilis isolate Rickettsia_endosymbiont_Ceutorhynchus_assimilis chromosome 1
Sequence ID: NZ_OU906081.1 Length: 4063491
Range 1: 350951 to 351085

Score:43.9 bits(102), Expect:0.020,
Method:Composition-based stats.,
Identities:16/45(36%), Positives:24/45(53%), Gaps:0/45(0%)

```
Query  5       NFRNQRKIVKCFNCGKEGHTARNCRAPRKKGCWKCGKEGHQMKDC  49
               N    R    C+NC K GH  R C+ P++  C++CG  G   ++C
Sbjct  350951  NSNRSRSSALCWNCQKPGHLNRECKQPKRLHCYRCGHSGSTSRNC  351085
```

Query #8: sp|P04585|Matrix_Protein_p17_HV1H2 Gag-Pol polyprotein OS=Human immunodeficiency virus type 1 group M subtype B (isolate HXB2) OX=11706 GN=gag-pol PE=1 SV=4 Query ID: lcl|Query_2387712 Length: 131

Sequences producing significant alignments:

| Description | Scientific Name | Common Name | Taxid | Max Score | Total Score | Query cover | E Value | Per. Ident | Acc. Len | Accession |
|---|---|---|---|---|---|---|---|---|---|---|
| Thalassovita mangrovi strain GS-10 Scaffold19_1, whole genome... | Thalassovita... | NA | 2692236 | 223 | 223 | 100% | 3e-64 | 80.92 | 2900 | NZ_WWEN010000... |
| Actinoplanes campanulatus strain JCM 3059 sequence058, whole... | Actinoplanes... | NA | 113559 | 223 | 223 | 100% | 9e-64 | 80.92 | 4754 | NZ_BMPW010000... |
| Enterococcus alcedinis strain CCM 8433 sequence23, whole genom... | Enterococcus... | NA | 1274384 | 222 | 222 | 100% | 9e-64 | 80.92 | 6075 | NZ_BMDT010000... |
| Deinococcus aquiradiocola strain JCM 14371 sequence30, whole... | Deinococcus ... | NA | 393059 | 222 | 222 | 100% | 1e-63 | 80.92 | 6128 | NZ_BMOE010000... |
| Bacillus velezensis strain TN14-11 TN14-11_ctg0135, whole geno... | Bacillus vel... | NA | 492670 | 187 | 187 | 78% | 8e-57 | 82.52 | 633 | NZ_JAHVIU0100... |
| Photobacterium rosenbergii strain DP6N14-7 DP6N14-7_ctg0073,... | Photobacteri... | NA | 294936 | 187 | 187 | 78% | 8e-57 | 82.52 | 641 | NZ_JAHVIB0100... |
| Burkholderia pseudomallei strain 2604STDY5519032... | Burkholderia... | NA | 28450 | 181 | 181 | 78% | 7e-56 | 81.55 | 362 | NZ_UVID020000... |
| Photobacterium rosenbergii strain DP6N14-7 DP6N14-7_ctg0069,... | Photobacteri... | NA | 294936 | 181 | 181 | 78% | 2e-54 | 80.58 | 667 | NZ_JAHVIB0100... |
| Vibrio hangzhouensis strain DP3N7-4 DP3N7-4_ctg0051, whole... | Vibrio hangz... | NA | 462991 | 181 | 181 | 78% | 4e-54 | 80.58 | 742 | NZ_JAHVKH0100... |
| Bacillus velezensis strain TN14-11 TN14-11_ctg0032, whole geno... | Bacillus vel... | NA | 492670 | 181 | 181 | 78% | 1e-52 | 80.58 | 1125 | NZ_JAHVIU0100... |
| Nocardioides luteus strain JCM 3358 sequence39, whole genome... | Nocardioides... | NA | 1844 | 188 | 188 | 76% | 8e-52 | 92.00 | 5754 | NZ_BMRK010000... |
| Streptomyces griseoflavus strain JCM 4479 sequence37, whole... | Streptomyces... | NA | 35619 | 188 | 188 | 76% | 8e-52 | 92.00 | 5775 | NZ_BMUC010000... |
| Saccharothrix coeruleofusca strain JCM 3313 sequence45, whole... | Saccharothri... | NA | 33919 | 188 | 188 | 76% | 8e-52 | 92.00 | 5775 | NZ_BMRG010000... |

```
Actinoplanes ianthinogenes strain JCM 3249 sequence037, whole...  Actinoplanes... NA              122358     188    188    76%    8e-52 92.00  5815       NZ_BMQZ010000...
Sinomonas sp. KCTC 49339 sequence28, whole genome shotgun...      Sinomonas sp... NA              2769422    188    188    76%    8e-52 92.00  5849       NZ_BNCM010000...
Saccharothrix mutabilis subsp. mutabilis strain JCM 3380...       Saccharothri... NA              66855      188    188    76%    8e-52 92.00  5899       NZ_BAAABU0100...
Corynebacterium glutamicum strain 39 Scaffolds_32_9166, whole...  Corynebacter... NA              1718       188    188    76%    8e-52 92.00  8862       NZ_JACEQW0100...
Corynebacterium glutamicum strain 37 Scaffolds_33_8928, whole...  Corynebacter... NA              1718       188    188    76%    8e-52 92.00  8928       NZ_JACENO0100...
```

Alignments:

```
>Thalassovita mangrovi strain GS-10 Scaffold19_1, whole genome shotgun sequence
Sequence ID: NZ_WWEN01000019.1 Length: 2900
Range 1: 1043 to 1435

Score:223 bits(567), Expect:3e-64,
Method:Compositional matrix adjust.,
Identities:122/131(93%), Positives:130/131(99%), Gaps:0/131(0%)

Query  1     GARASVLSGGELDRWEKIRLRPGGKKKYKLKHIVWASRELERFAVNPGLLETSEGCRQIL  60
             GARASVLSGGELD+WEKIRLRPGGKK+YKLKHIVWASRELERFAVNPGLLETSEGCRQIL
Sbjct  1043  GARASVLSGGELDKWEKIRLRPGGKKQYKLKHIVWASRELERFAVNPGLLETSEGCRQIL  1222

Query  61    GQLQPSLQTGSEELRSLYNTVATLYCVHQRIEIKDTKEALDKIeeeqnkskkkaqqaaaD  120
             GQLQPSLQTGSEELRSLYNT+A LYCVHQRI++KDTKEALDKIEEEQNKSKKKAQQAAAD
Sbjct  1223  GQLQPSLQTGSEELRSLYNTIAVLYCVHQRIDVKDTKEALDKIEEEQNKSKKKAQQAAAD  1402

Query  121   TGHSNQVSQNY  131
             TG+++QVSQNY
Sbjct  1403  TGNNSQVSQNY  1435



>Actinoplanes campanulatus strain JCM 3059 sequence058, whole genome shotgun sequence
Sequence ID: NZ_BMPW01000058.1 Length: 4754
Range 1: 100 to 492

Score:223 bits(567), Expect:9e-64,
Method:Compositional matrix adjust.,
Identities:122/131(93%), Positives:130/131(99%), Gaps:0/131(0%)

Query  1    GARASVLSGGELDRWEKIRLRPGGKKKYKLKHIVWASRELERFAVNPGLLETSEGCRQIL  60
            GARASVLSGGELD+WEKIRLRPGGKK+YKLKHIVWASRELERFAVNPGLLETSEGCRQIL
Sbjct  100  GARASVLSGGELDKWEKIRLRPGGKKQYKLKHIVWASRELERFAVNPGLLETSEGCRQIL  279

Query  61   GQLQPSLQTGSEELRSLYNTVATLYCVHQRIEIKDTKEALDKIeeeqnkskkkaqqaaaD  120
            GQLQPSLQTGSEELRSLYNT+A LYCVHQRI++KDTKEALDKIEEEQNKSKKKAQQAAAD
Sbjct  280  GQLQPSLQTGSEELRSLYNTIAVLYCVHQRIDVKDTKEALDKIEEEQNKSKKKAQQAAAD  459

Query  121  TGHSNQVSQNY  131
            TG+++QVSQNY
Sbjct  460  TGNNSQVSQNY  492



>Enterococcus alcedinis strain CCM 8433 sequence23, whole genome shotgun sequence
Sequence ID: NZ_BMDT01000023.1 Length: 6075
Range 1: 58 to 450

Score:222 bits(566), Expect:9e-64,
Method:Compositional matrix adjust.,
Identities:122/131(93%), Positives:130/131(99%), Gaps:0/131(0%)

Query  1    GARASVLSGGELDRWEKIRLRPGGKKKYKLKHIVWASRELERFAVNPGLLETSEGCRQIL  60
            GARASVLSGGELD+WEKIRLRPGGKK+YKLKHIVWASRELERFAVNPGLLETSEGCRQIL
Sbjct  58   GARASVLSGGELDKWEKIRLRPGGKKQYKLKHIVWASRELERFAVNPGLLETSEGCRQIL  237

Query  61   GQLQPSLQTGSEELRSLYNTVATLYCVHQRIEIKDTKEALDKIeeeqnkskkkaqqaaaD  120
            GQLQPSLQTGSEELRSLYNT+A LYCVHQRI++KDTKEALDKIEEEQNKSKKKAQQAAAD
Sbjct  238  GQLQPSLQTGSEELRSLYNTIAVLYCVHQRIDVKDTKEALDKIEEEQNKSKKKAQQAAAD  417

Query  121  TGHSNQVSQNY  131
            TG+++QVSQNY
Sbjct  418  TGNNSQVSQNY  450



>Deinococcus aquiradiocola strain JCM 14371 sequence30, whole genome shotgun sequence
Sequence ID: NZ_BMOE01000030.1 Length: 6128
Range 1: 111 to 503

Score:222 bits(566), Expect:1e-63,
Method:Compositional matrix adjust.,
Identities:122/131(93%), Positives:130/131(99%), Gaps:0/131(0%)

Query  1    GARASVLSGGELDRWEKIRLRPGGKKKYKLKHIVWASRELERFAVNPGLLETSEGCRQIL  60
            GARASVLSGGELD+WEKIRLRPGGKK+YKLKHIVWASRELERFAVNPGLLETSEGCRQIL
Sbjct  111  GARASVLSGGELDKWEKIRLRPGGKKQYKLKHIVWASRELERFAVNPGLLETSEGCRQIL  290

Query  61   GQLQPSLQTGSEELRSLYNTVATLYCVHQRIEIKDTKEALDKIeeeqnkskkkaqqaaaD  120
            GQLQPSLQTGSEELRSLYNT+A LYCVHQRI++KDTKEALDKIEEEQNKSKKKAQQAAAD
Sbjct  291  GQLQPSLQTGSEELRSLYNTIAVLYCVHQRIDVKDTKEALDKIEEEQNKSKKKAQQAAAD  470

Query  121  TGHSNQVSQNY  131
            TG+++QVSQNY
Sbjct  471  TGNNSQVSQNY  503



>Bacillus velezensis strain TN14-11 TN14-11_ctg0135, whole genome shotgun sequence
Sequence ID: NZ_JAHVIU010000020.1 Length: 633
Range 1: 191 to 499

Score:187 bits(475), Expect:8e-57,
Method:Compositional matrix adjust.,
Identities:85/103(83%), Positives:97/103(94%), Gaps:0/103(0%)

Query  1    GARASVLSGGELDRWEKIRLRPGGKKKYKLKHIVWASRELERFAVNPGLLETSEGCRQIL  60
            GARAS+L GG+LD+WEKIRLRPGGKK Y LKH+VWASRELERFAVNPGLLET+EGC+QI+
Sbjct  191  GARASILRGGKLDKWEKIRLRPGGKKHYMLKHLVWASRELERFAVNPGLLETAEGCKQII  370

Query  61   GQLQPSLQTGSEELRSLYNTVATLYCVHQRIEIKDTKEALDKI  103
             QLQP+LQTG+EELRSLYNTVA LYCVH+ IE++DTKEALDKI
Sbjct  371  NQLQPALQTGTEELRSLYNTVAVLYCVHKGIEVRDTKEALDKI  499



>Photobacterium rosenbergii strain DP6N14-7 DP6N14-7_ctg0073, whole genome shotgun sequence
Sequence ID: NZ_JAHVIB010000060.1 Length: 641
Range 1: 199 to 507
```

```
Score:187 bits(475), Expect:8e-57,
Method:Compositional matrix adjust.,
Identities:85/103(83%), Positives:97/103(94%), Gaps:0/103(0%)

Query  1    GARASVLSGGELDRWEKIRLRPGGKKKYKLKHIVWASRELERFAVNPGLLETSEGCRQIL  60
            GARAS+L GG+LD+WEKIRLRPGGKK Y LKH+VWASRELERFAVNPGLLET+EGC+QI+
Sbjct  199  GARASILRGGKLDKWEKIRLRPGGKKHYMLKHLVWASRELERFAVNPGLLETAEGCKQII  378

Query  61   GQLQPSLQTGSEELRSLYNTVATLYCVHQRIEIKDTKEALDKI  103
             QLQP+LQTG+EELRSLYNTVA LYCVH+ IE++DTKEALDKI
Sbjct  379  NQLQPALQTGTEELRSLYNTVAVLYCVHKGIEVRDTKEALDKI  507
```

```
>Burkholderia pseudomallei strain 2604STDY5519032 ERS240165SCcontig000086, whole genome shotgun sequence
Sequence ID: NZ_UVID02000086.1 Length: 362
Range 1: 3 to 311

Score:181 bits(460), Expect:7e-56,
Method:Compositional matrix adjust.,
Identities:84/103(82%), Positives:96/103(93%), Gaps:0/103(0%)

Query  1    GARASVLSGGELDRWEKIRLRPGGKKKYKLKHIVWASRELERFAVNPGLLETSEGCRQIL  60
            GARAS+L GG+LD WEKIRLRPGGKK Y LKH+VWASRELERFA+NPGLLETSEGC+QI+
Sbjct  3    GARASILRGGKLDTWEKIRLRPGGKKHYMLKHLVWASRELERFALNPGLLETSEGCKQII  182

Query  61   GQLQPSLQTGSEELRSLYNTVATLYCVHQRIEIKDTKEALDKI  103
             QLQP+LQTG+EELRSL+NTVATLYCVH+ I++ DTKEALDKI
Sbjct  183  QQLQPALQTGTEELRSLFNTVATLYCVHKDIKVADTKEALDKI  311
```

```
>Photobacterium rosenbergii strain DP6N14-7 DP6N14-7_ctg0069, whole genome shotgun sequence
Sequence ID: NZ_JAHVIB010000058.1 Length: 667
Range 1: 172 to 480

Score:181 bits(460), Expect:2e-54,
Method:Compositional matrix adjust.,
Identities:83/103(81%), Positives:95/103(92%), Gaps:0/103(0%)

Query  1    GARASVLSGGELDRWEKIRLRPGGKKKYKLKHIVWASRELERFAVNPGLLETSEGCRQIL  60
            GARAS+L GG+L +WEKI+LRP GKK Y LKH+VWASRELERFAVNPGLLET+EGCRQI+
Sbjct  172  GARASILRGGKLGKWEKIKLRPVGKKHYMLKHLVWASRELERFAVNPGLLETAEGCRQII  351

Query  61   GQLQPSLQTGSEELRSLYNTVATLYCVHQRIEIKDTKEALDKI  103
             QLQP+LQTG+EELRSLYNTVA LYCVH+ IE++DTKEALDKI
Sbjct  352  NQLQPALQTGTEELRSLYNTVAVLYCVHKEIEVRDTKEALDKI  480
```

```
>Vibrio hangzhouensis strain DP3N7-4 DP3N7-4_ctg0051, whole genome shotgun sequence
Sequence ID: NZ_JAHVKH010000037.1 Length: 742
Range 1: 233 to 541

Score:181 bits(460), Expect:4e-54,
Method:Compositional matrix adjust.,
Identities:83/103(81%), Positives:95/103(92%), Gaps:0/103(0%)

Query  1    GARASVLSGGELDRWEKIRLRPGGKKKYKLKHIVWASRELERFAVNPGLLETSEGCRQIL  60
            GARAS+L GG+L +WEKI+LRP GKK Y LKH+VWASRELERFAVNPGLLET+EGCRQI+
Sbjct  541  GARASILRGGKLGKWEKIKLRPVGKKHYMLKHLVWASRELERFAVNPGLLETAEGCRQII  362

Query  61   GQLQPSLQTGSEELRSLYNTVATLYCVHQRIEIKDTKEALDKI  103
             QLQP+LQTG+EELRSLYNTVA LYCVH+ IE++DTKEALDKI
Sbjct  361  NQLQPALQTGTEELRSLYNTVAVLYCVHKEIEVRDTKEALDKI  233
```

```
>Bacillus velezensis strain TN14-11 TN14-11_ctg0032, whole genome shotgun sequence
Sequence ID: NZ_JAHVIU010000017.1 Length: 1125
Range 1: 668 to 976

Score:181 bits(459), Expect:1e-52,
Method:Compositional matrix adjust.,
Identities:83/103(81%), Positives:95/103(92%), Gaps:0/103(0%)

Query  1    GARASVLSGGELDRWEKIRLRPGGKKKYKLKHIVWASRELERFAVNPGLLETSEGCRQIL  60
            GARAS+L GG+L +WEKI+LRP GKK Y LKH+VWASRELERFAVNPGLLET+EGC+QI+
Sbjct  976  GARASILRGGKLGKWEKIKLRPVGKKHYMLKHLVWASRELERFAVNPGLLETAEGCKQII  797

Query  61   GQLQPSLQTGSEELRSLYNTVATLYCVHQRIEIKDTKEALDKI  103
             QLQP+LQTG+EELRSLYNTVA LYCVH+ IE+KDTKEALDKI
Sbjct  796  NQLQPALQTGTEELRSLYNTVAVLYCVHKGIEVKDTKEALDKI  668
```

```
>Nocardioides luteus strain JCM 3358 sequence39, whole genome shotgun sequence
Sequence ID: NZ_BMRK01000039.1 Length: 5754
Range 1: 217 to 516

Score:188 bits(478), Expect:8e-52,
Method:Compositional matrix adjust.,
Identities:92/100(92%), Positives:93/100(93%), Gaps:0/100(0%)

Query  4    ASVLSGGELDRWEKIRLRPGGKKKYKLKHIVWASRELERFAVNPGLLETSEGCRQILGQL  63
             S+  G    RWEKIRLRPGGKKKYKLKHIVWASRELERFAVNPGLLETSEGCRQILGQL
Sbjct  217  VSIKRGRIRSRWEKIRLRPGGKKKYKLKHIVWASRELERFAVNPGLLETSEGCRQILGQL  396

Query  64   QPSLQTGSEELRSLYNTVATLYCVHQRIEIKDTKEALDKI  103
            QPSLQTGSEELRSLYNTVATLYCVHQRIEIKDTKEALDKI
Sbjct  397  QPSLQTGSEELRSLYNTVATLYCVHQRIEIKDTKEALDKI  516
```

```
>Streptomyces griseoflavus strain JCM 4479 sequence37, whole genome shotgun sequence
Sequence ID: NZ_BMUC01000037.1 Length: 5775
Range 1: 5232 to 5531

Score:188 bits(478), Expect:8e-52,
Method:Compositional matrix adjust.,
Identities:92/100(92%), Positives:93/100(93%), Gaps:0/100(0%)

Query  4     ASVLSGGELDRWEKIRLRPGGKKKYKLKHIVWASRELERFAVNPGLLETSEGCRQILGQL  63
              S+  G    RWEKIRLRPGGKKKYKLKHIVWASRELERFAVNPGLLETSEGCRQILGQL
Sbjct  5531  VSIKRGRIRSRWEKIRLRPGGKKKYKLKHIVWASRELERFAVNPGLLETSEGCRQILGQL  5352

Query  64    QPSLQTGSEELRSLYNTVATLYCVHQRIEIKDTKEALDKI  103
             QPSLQTGSEELRSLYNTVATLYCVHQRIEIKDTKEALDKI
```

```
Sbjct  5351  QPSLQTGSEELRSLYNTVATLYCVHQRIEIKDTKEALDKI  5232
```

>Saccharothrix coeruleofusca strain JCM 3313 sequence45, whole genome shotgun sequence
Sequence ID: NZ_BMRG01000045.1 Length: 5775
Range 1: 5232 to 5531

Score:188 bits(478), Expect:8e-52,
Method:Compositional matrix adjust.,
Identities:92/100(92%), Positives:93/100(93%), Gaps:0/100(0%)

```
Query  4     ASVLSGGELDRWEKIRLRPGGKKKYKLKHIVWASRELERFAVNPGLLETSEGCRQILGQL  63
              S+  G    RWEKIRLRPGGKKKYKLKHIVWASRELERFAVNPGLLETSEGCRQILGQL
Sbjct  5531  VSIKRGRIRSRWEKIRLRPGGKKKYKLKHIVWASRELERFAVNPGLLETSEGCRQILGQL  5352

Query  64    QPSLQTGSEELRSLYNTVATLYCVHQRIEIKDTKEALDKI  103
             QPSLQTGSEELRSLYNTVATLYCVHQRIEIKDTKEALDKI
Sbjct  5351  QPSLQTGSEELRSLYNTVATLYCVHQRIEIKDTKEALDKI  5232
```

>Actinoplanes ianthinogenes strain JCM 3249 sequence037, whole genome shotgun sequence
Sequence ID: NZ_BMQZ01000037.1 Length: 5815
Range 1: 5252 to 5551

Score:188 bits(478), Expect:8e-52,
Method:Compositional matrix adjust.,
Identities:92/100(92%), Positives:93/100(93%), Gaps:0/100(0%)

```
Query  4     ASVLSGGELDRWEKIRLRPGGKKKYKLKHIVWASRELERFAVNPGLLETSEGCRQILGQL  63
              S+  G    RWEKIRLRPGGKKKYKLKHIVWASRELERFAVNPGLLETSEGCRQILGQL
Sbjct  5551  VSIKRGRIRSRWEKIRLRPGGKKKYKLKHIVWASRELERFAVNPGLLETSEGCRQILGQL  5372

Query  64    QPSLQTGSEELRSLYNTVATLYCVHQRIEIKDTKEALDKI  103
             QPSLQTGSEELRSLYNTVATLYCVHQRIEIKDTKEALDKI
Sbjct  5371  QPSLQTGSEELRSLYNTVATLYCVHQRIEIKDTKEALDKI  5252
```

>Sinomonas sp. KCTC 49339 sequence28, whole genome shotgun sequence
Sequence ID: NZ_BNCM01000028.1 Length: 5849
Range 1: 5223 to 5522

Score:188 bits(478), Expect:8e-52,
Method:Compositional matrix adjust.,
Identities:92/100(92%), Positives:93/100(93%), Gaps:0/100(0%)

```
Query  4     ASVLSGGELDRWEKIRLRPGGKKKYKLKHIVWASRELERFAVNPGLLETSEGCRQILGQL  63
              S+  G    RWEKIRLRPGGKKKYKLKHIVWASRELERFAVNPGLLETSEGCRQILGQL
Sbjct  5522  VSIKRGRIRSRWEKIRLRPGGKKKYKLKHIVWASRELERFAVNPGLLETSEGCRQILGQL  5343

Query  64    QPSLQTGSEELRSLYNTVATLYCVHQRIEIKDTKEALDKI  103
             QPSLQTGSEELRSLYNTVATLYCVHQRIEIKDTKEALDKI
Sbjct  5342  QPSLQTGSEELRSLYNTVATLYCVHQRIEIKDTKEALDKI  5223
```

>Saccharothrix mutabilis subsp. mutabilis strain JCM 3380 sequence46, whole genome shotgun sequence
Sequence ID: NZ_BAAABU010000046.1 Length: 5899
Range 1: 265 to 564

Score:188 bits(478), Expect:8e-52,
Method:Compositional matrix adjust.,
Identities:92/100(92%), Positives:93/100(93%), Gaps:0/100(0%)

```
Query  4    ASVLSGGELDRWEKIRLRPGGKKKYKLKHIVWASRELERFAVNPGLLETSEGCRQILGQL  63
             S+  G    RWEKIRLRPGGKKKYKLKHIVWASRELERFAVNPGLLETSEGCRQILGQL
Sbjct  265  VSIKRGRIRSRWEKIRLRPGGKKKYKLKHIVWASRELERFAVNPGLLETSEGCRQILGQL  444

Query  64   QPSLQTGSEELRSLYNTVATLYCVHQRIEIKDTKEALDKI  103
            QPSLQTGSEELRSLYNTVATLYCVHQRIEIKDTKEALDKI
Sbjct  445  QPSLQTGSEELRSLYNTVATLYCVHQRIEIKDTKEALDKI  564
```

>Corynebacterium glutamicum strain 39 Scaffolds_32_9166, whole genome shotgun sequence
Sequence ID: NZ_JACEQW010000032.1 Length: 8862
Range 1: 5003 to 5302

Score:188 bits(478), Expect:8e-52,
Method:Compositional matrix adjust.,
Identities:92/100(92%), Positives:93/100(93%), Gaps:0/100(0%)

```
Query  4     ASVLSGGELDRWEKIRLRPGGKKKYKLKHIVWASRELERFAVNPGLLETSEGCRQILGQL  63
              S+  G    RWEKIRLRPGGKKKYKLKHIVWASRELERFAVNPGLLETSEGCRQILGQL
Sbjct  5302  VSIKRGRIRSRWEKIRLRPGGKKKYKLKHIVWASRELERFAVNPGLLETSEGCRQILGQL  5123

Query  64    QPSLQTGSEELRSLYNTVATLYCVHQRIEIKDTKEALDKI  103
             QPSLQTGSEELRSLYNTVATLYCVHQRIEIKDTKEALDKI
Sbjct  5122  QPSLQTGSEELRSLYNTVATLYCVHQRIEIKDTKEALDKI  5003
```

>Corynebacterium glutamicum strain 37 Scaffolds_33_8928, whole genome shotgun sequence
Sequence ID: NZ_JACENO010000033.1 Length: 8928
Range 1: 4977 to 5276

Score:188 bits(478), Expect:8e-52,
Method:Compositional matrix adjust.,
Identities:92/100(92%), Positives:93/100(93%), Gaps:0/100(0%)

```
Query  4     ASVLSGGELDRWEKIRLRPGGKKKYKLKHIVWASRELERFAVNPGLLETSEGCRQILGQL  63
              S+  G    RWEKIRLRPGGKKKYKLKHIVWASRELERFAVNPGLLETSEGCRQILGQL
Sbjct  5276  VSIKRGRIRSRWEKIRLRPGGKKKYKLKHIVWASRELERFAVNPGLLETSEGCRQILGQL  5097

Query  64    QPSLQTGSEELRSLYNTVATLYCVHQRIEIKDTKEALDKI  103
             QPSLQTGSEELRSLYNTVATLYCVHQRIEIKDTKEALDKI
Sbjct  5096  QPSLQTGSEELRSLYNTVATLYCVHQRIEIKDTKEALDKI  4977
```

Query #9: SP|P04585|POL POLYPROTEIN OS=Human immunodeficiency virus type 1 group M subtype B (isolate HXB2) OX=11706 GN=GAG-POL PE=1 SV=4 Query ID: lcl|Query_1446707 Length: 1115

Sequences producing significant alignments:

| Description | Scientific Name | Common Name | Taxid | Max Score | Total Score | Query cover | E Value | Per. Ident | Acc. Len | Accession |
|---|---|---|---|---|---|---|---|---|---|---|
| Deinococcus aquiradiocola strain JCM 14371 sequence30, whole... | Deinococcus ... | NA | 393059 | 1381 | 2057 | 94% | 0.0 | 94.65 | 6128 | NZ_BMOE010000... |
| Enterococcus alcedinis strain CCM 8433 sequence23, whole genom... | Enterococcus... | NA | 1274384 | 1381 | 2057 | 94% | 0.0 | 94.65 | 6075 | NZ_BMDT010000... |
| Actinoplanes campanulatus strain JCM 3059 sequence058, whole... | Actinoplanes... | NA | 113559 | 1380 | 2192 | 99% | 0.0 | 94.65 | 4754 | NZ_BMPW010000... |
| Klebsiella pneumoniae strain ARGID_31501, whole genome shotgun... | Klebsiella p... | NA | 573 | 1208 | 1934 | 90% | 0.0 | 92.01 | 2655 | NZ_CAKLPD0100... |
| Enterococcus faecium strain A209R1B0C1... | Enterococcus... | NA | 1352 | 1050 | 1050 | 52% | 0.0 | 86.44 | 1819 | NZ_PGCR010000... |
| Listeria monocytogenes strain YA00082404... | Listeria mon... | NA | 1639 | 1038 | 1038 | 52% | 0.0 | 85.14 | 1816 | NZ_QEXE010000... |
| Escherichia coli strain YA00028237 contig_187, whole genome... | Escherichia ... | NA | 562 | 1026 | 1026 | 52% | 0.0 | 82.49 | 1828 | NZ_NGBP020001... |
| Enterococcus faecium strain A206R2B0... | Enterococcus... | NA | 1352 | 1020 | 1020 | 51% | 0.0 | 86.11 | 1761 | NZ_PGCU010000... |
| Klebsiella pneumoniae strain NUBRI-K 73, whole genome shotgun... | Klebsiella p... | NA | 573 | 1001 | 1001 | 50% | 0.0 | 84.52 | 1717 | NZ_SOYT010000... |
| Enterococcus faecium strain A201R2B0... | Enterococcus... | NA | 1352 | 975 | 975 | 47% | 0.0 | 89.02 | 1589 | NZ_PGCT010000... |
| Thalassovita mangrovi strain GS-10 Scaffold20_1, whole genome... | Thalassovita... | NA | 2692236 | 950 | 1145 | 53% | 0.0 | 93.54 | 1488 | NZ_WWEN010000... |
| Helicobacter acinonychis isolate 212_10, whole genome shotgun... | Helicobacter... | NA | 212 | 901 | 964 | 42% | 0.0 | 90.70 | 1526 | NZ_FZLW010000... |
| Escherichia coli strain 8374wH11... | Escherichia ... | NA | 562 | 891 | 891 | 41% | 0.0 | 92.24 | 1397 | NZ_WSJI010000... |
| Methylocella silvestris strain TVC... | Methylocella... | NA | 199596 | 887 | 887 | 42% | 0.0 | 90.02 | 1453 | NZ_PDZR010000... |
| Acinetobacter baumannii strain ACM-35... | Acinetobacte... | NA | 470 | 882 | 882 | 42% | 0.0 | 90.62 | 1409 | NZ_QEHN010000... |
| Enterobacter hormaechei strain AUH-ENM6... | Enterobacter... | NA | 158836 | 878 | 878 | 41% | 0.0 | 90.73 | 1394 | NZ_PYHB010000... |
| Escherichia coli strain 8374wG10... | Escherichia ... | NA | 562 | 875 | 875 | 40% | 0.0 | 92.32 | 1373 | NZ_WSJQ010000... |
| Klebsiella pneumoniae strain P7701... | Klebsiella p... | NA | 573 | 869 | 869 | 38% | 0.0 | 97.66 | 1305 | NZ_JACAWM0100... |
| Streptococcus agalactiae strain JH02623 contig_36, whole genom... | Streptococcu... | NA | 1311 | 857 | 857 | 42% | 0.0 | 86.01 | 1479 | NZ_RPAW010000... |
| Escherichia coli strain 8374wB4 NODE_60_length_1385_cov_3.2655... | Escherichia ... | NA | 562 | 855 | 855 | 41% | 0.0 | 90.63 | 1385 | NZ_WSPX010000... |
| Klebsiella aerogenes strain AUH-KAM5... | Klebsiella a... | NA | 548 | 848 | 848 | 40% | 0.0 | 90.47 | 1427 | NZ_PYHJ010000... |
| Streptococcus agalactiae strain JH02623 contig_37, whole genom... | Streptococcu... | NA | 1311 | 848 | 848 | 42% | 0.0 | 85.42 | 1467 | NZ_RPAW010000... |
| Salirhabdus euzebyi strain Marseille-Q1438, whole genome shotg... | Salirhabdus ... | NA | 394506 | 842 | 842 | 37% | 0.0 | 95.51 | 1279 | NZ_CADDWK0100... |
| Escherichia coli strain 8374wB11... | Escherichia ... | NA | 562 | 825 | 825 | 39% | 0.0 | 90.83 | 1310 | NZ_WSJO010000... |
| Mycobacterium tuberculosis variant bovis strain CSURQ1134, who... | Mycobacteriu... | NA | 1765 | 812 | 1625 | 39% | 0.0 | 88.49 | 1697 | NZ_CAJEUT0100... |
| Mycobacterium tuberculosis variant bovis strain CSURQ1132, who... | Mycobacteriu... | NA | 1765 | 811 | 1623 | 39% | 0.0 | 88.49 | 1524 | NZ_CAJEUK0100... |
| Acinetobacter baumannii strain ACM-5... | Acinetobacte... | NA | 470 | 811 | 811 | 38% | 0.0 | 91.51 | 1274 | NZ_PYCX010000... |
| Mycobacterium tuberculosis variant bovis strain CSURQ1140, who... | Mycobacteriu... | NA | 1765 | 788 | 1577 | 38% | 0.0 | 88.58 | 1286 | NZ_CAJEUP0100... |
| Escherichia coli strain VRES0669, whole genome shotgun sequence | Escherichia ... | NA | 562 | 777 | 1831 | 88% | 0.0 | 90.51 | 5544 | NZ_UINV010000... |
| Burkholderia pseudomallei strain 2604STDY5518971... | Burkholderia... | NA | 28450 | 733 | 1070 | 51% | 0.0 | 91.82 | 1345 | NZ_UVFZ020000... |
| Providencia rettgeri strain AFE652_S1 38_contig_28, whole geno... | Providencia ... | NA | 587 | 724 | 724 | 33% | 0.0 | 92.95 | 1127 | NZ_SHDK010000... |
| Staphylococcus aureus strain G703N1B1... | Staphylococ... | NA | 1280 | 722 | 722 | 34% | 0.0 | 88.66 | 1173 | NZ_PGXA010000... |
| Levilactobacillus brevis strain CRL 2013 scaffold12, whole... | Levilactobac... | NA | 1580 | 697 | 697 | 31% | 0.0 | 93.82 | 1126 | NZ_MZMW010000... |
| Pseudomonas aeruginosa strain VET-121... | Pseudomonas ... | NA | 287 | 686 | 686 | 31% | 0.0 | 91.81 | 1158 | NZ_QZXB010001... |
| Pseudomonas aeruginosa strain VET-42... | Pseudomonas ... | NA | 287 | 683 | 683 | 31% | 0.0 | 91.27 | 1166 | NZ_RACX010001... |
| Streptococcus agalactiae strain JH02623 contig_39, whole genom... | Streptococcu... | NA | 1311 | 676 | 791 | 36% | 0.0 | 92.75 | 1204 | NZ_RPAW010000... |
| Levilactobacillus brevis strain CRL 2013 scaffold13, whole... | Levilactobac... | NA | 1580 | 668 | 668 | 30% | 0.0 | 93.29 | 1030 | NZ_MZMW010000... |
| Pseudomonas aeruginosa strain VET-42... | Pseudomonas ... | NA | 287 | 664 | 774 | 36% | 0.0 | 90.38 | 1154 | NZ_RACX010001... |
| Providencia rettgeri strain AFE96_S1 41_contig_60, whole genom... | Providencia ... | NA | 587 | 654 | 654 | 30% | 0.0 | 92.54 | 1101 | NZ_SHDP010000... |
| Staphylococcus epidermidis strain PR246B0... | Staphylococc... | NA | 1282 | 651 | 651 | 32% | 0.0 | 85.01 | 1113 | NZ_PCFD010000... |
| Pseudomonas aeruginosa strain ENV-297... | Pseudomonas ... | NA | 287 | 635 | 716 | 32% | 0.0 | 91.08 | 1100 | NZ_RADD010000... |
| Staphylococcus aureus strain PN235B0... | Staphylococc... | NA | 1280 | 621 | 621 | 31% | 0.0 | 83.76 | 1083 | NZ_PDUY010001... |
| Enterococcus durans strain E115_10... | Enterococcus... | NA | 53345 | 612 | 612 | 28% | 0.0 | 90.22 | 955 | NZ_JAANZI0100... |
| Escherichia coli strain 8374wA11 NODE_37_length_940_cov_2.7589... | Escherichia ... | NA | 562 | 609 | 609 | 28% | 0.0 | 91.05 | 940 | NZ_WSPR010000... |
| Escherichia coli strain TzEc080 NODE_37_length_940_cov_2.75892... | Escherichia ... | NA | 562 | 609 | 609 | 28% | 0.0 | 91.05 | 940 | NZ_WSFZ010000... |
| Methylobacterium radiotolerans strain MAMP 4754... | Methylobacte... | NA | 31998 | 599 | 1157 | 58% | 0.0 | 84.84 | 1636 | NZ_NKQS010006... |
| Providencia rettgeri strain APW139_S1 24_contig_65, whole geno... | Providencia ... | NA | 587 | 588 | 588 | 27% | 0.0 | 89.39 | 941 | NZ_SHDG010000... |
| Providencia rettgeri strain APW139_S1 24_contig_64, whole geno... | Providencia ... | NA | 587 | 578 | 578 | 26% | 0.0 | 90.33 | 902 | NZ_SHDG010000... |
| Acinetobacter baumannii strain ACM-29... | Acinetobacte... | NA | 470 | 568 | 568 | 26% | 0.0 | 91.53 | 888 | NZ_PYDO010001... |
| Mycobacterium tuberculosis variant bovis strain CSURQ1139, who... | Mycobacteriu... | NA | 1765 | 563 | 1126 | 25% | 0.0 | 93.62 | 848 | NZ_CAJEUO0100... |
| Mycobacterium tuberculosis strain 32846_S30 184, whole genome... | Mycobacteriu... | NA | 1773 | 558 | 558 | 27% | 0.0 | 85.76 | 932 | NZ_QTBW010000... |
| Acinetobacter baumannii strain ACM-36... | Acinetobacte... | NA | 470 | 554 | 554 | 26% | 0.0 | 89.15 | 953 | NZ_QAGO010000... |
| Providencia rettgeri strain AFE97_S1 21_contig_71, whole genom... | Providencia ... | NA | 587 | 482 | 724 | 34% | 0.0 | 86.52 | 1157 | NZ_SHDO010000... |
| Providencia rettgeri strain AFE422_S1 40_contig_34, whole geno... | Providencia ... | NA | 587 | 468 | 689 | 31% | 0.0 | 89.88 | 1058 | NZ_SHDL010000... |
| Mycobacterium tuberculosis strain 32846_S30 476, whole genome... | Mycobacteriu... | NA | 1773 | 536 | 536 | 26% | 5e-179 | 87.58 | 861 | NZ_QTBW010001... |
| Mycobacterium tuberculosis variant bovis strain CSURQ1134, who... | Mycobacteriu... | NA | 1765 | 530 | 1060 | 25% | 2e-176 | 89.44 | 910 | NZ_CAJEUT0100... |
| Providencia rettgeri strain AFE97_S1 21_contig_72, whole genom... | Providencia ... | NA | 587 | 528 | 528 | 24% | 5e-176 | 89.44 | 854 | NZ_SHDO010000... |
| Enterobacter hormaechei strain AUH-ENM4... | Enterobacter... | NA | 158836 | 525 | 525 | 24% | 6e-175 | 92.19 | 809 | NZ_PYGZ010000... |
| Candidatus Tokpelaia sp. strain RhiAcro1 isolate RhiAcro1_RAo1... | Candidatus T... | NA | 2233777 | 514 | 514 | 24% | 1e-170 | 89.45 | 826 | NZ_QLZD010000... |
| Acinetobacter baumannii strain ACM-27... | Acinetobacte... | NA | 470 | 507 | 507 | 26% | 1e-167 | 82.53 | 895 | NZ_PYDM010000... |
| Acinetobacter baumannii strain ACM-14... | Acinetobacte... | NA | 470 | 497 | 497 | 23% | 3e-164 | 91.57 | 787 | NZ_PYDA010000... |
| Klebsiella pneumoniae strain EuSCAPE_DE086, whole genome shotg... | Klebsiella p... | NA | 573 | 499 | 580 | 34% | 6e-164 | 76.44 | 995 | NZ_UKLW020000... |
| Mycobacterium tuberculosis strain 32847_S31 462, whole genome... | Mycobacteriu... | NA | 1773 | 496 | 496 | 25% | 3e-163 | 84.86 | 893 | NZ_QTBX010001... |
| Enterococcus faecalis strain 44.1... | Enterococcus... | NA | 1351 | 490 | 490 | 22% | 1e-161 | 91.76 | 806 | NZ_NSJR010000... |
| Mycobacterium tuberculosis strain 32845_S29 549, whole genome... | Mycobacteriu... | NA | 1773 | 486 | 486 | 24% | 7e-160 | 84.31 | 825 | NZ_QTBV010001... |
| Klebsiella pneumoniae strain P7701... | Klebsiella p... | NA | 573 | 484 | 484 | 21% | 8e-160 | 97.05 | 711 | NZ_JACAWM0100... |
| Thalassovita mangrovi strain GS-10 Scaffold30_1, whole genome... | Thalassovita... | NA | 2692236 | 464 | 545 | 25% | 5e-152 | 91.91 | 722 | NZ_WWEN010000... |
| Escherichia coli strain 8374wB4 NODE_74_length_738_cov_3.41735... | Escherichia ... | NA | 562 | 464 | 464 | 21% | 6e-152 | 87.76 | 738 | NZ_WSPX010000... |
| Acinetobacter baumannii strain ACM-17... | Acinetobacte... | NA | 470 | 458 | 458 | 22% | 2e-149 | 86.80 | 757 | NZ_PYDD010000... |
| Acinetobacter baumannii strain ACM-41... | Acinetobacte... | NA | 470 | 454 | 454 | 21% | 4e-148 | 89.96 | 718 | NZ_QAGT010000... |
| Acinetobacter baumannii strain ACM-22... | Acinetobacte... | NA | 470 | 452 | 452 | 21% | 4e-147 | 88.43 | 726 | NZ_PYDI010000... |
| Acinetobacter baumannii strain ACM-13... | Acinetobacte... | NA | 470 | 438 | 438 | 20% | 4e-142 | 89.27 | 701 | NZ_PYCZ010000... |
| Escherichia coli strain ECM-2 NODE_127_length_636_cov_2.0668,... | Escherichia ... | NA | 562 | 421 | 421 | 18% | 1e-135 | 95.26 | 636 | NZ_PXIA010001... |
| Acinetobacter baumannii strain ACM-14... | Acinetobacte... | NA | 470 | 416 | 416 | 19% | 8e-134 | 90.41 | 665 | NZ_PYDA010000... |
| Acinetobacter baumannii strain ACM-11... | Acinetobacte... | NA | 470 | 413 | 413 | 21% | 3e-132 | 83.83 | 706 | NZ_PYCW010000... |
| Mycobacterium tuberculosis strain 32847_S31 312, whole genome... | Mycobacteriu... | NA | 1773 | 409 | 409 | 20% | 5e-131 | 83.33 | 704 | NZ_QTBX010000... |
| Providencia rettgeri strain AFE85_S2 19_contig_46, whole genom... | Providencia ... | NA | 587 | 409 | 491 | 18% | 5e-130 | 93.33 | 846 | NZ_SHDQ010000... |
| Mycobacterium tuberculosis variant bovis strain CSURQ1132, who... | Mycobacteriu... | NA | 1765 | 406 | 812 | 20% | 9e-129 | 87.01 | 862 | NZ_CAJEUK0100... |
| Acinetobacter baumannii strain ACM-25... | Acinetobacte... | NA | 470 | 399 | 399 | 19% | 2e-127 | 86.70 | 655 | NZ_PXYP010000... |
| Levilactobacillus brevis strain CRL 2013 scaffold15, whole... | Levilactobac... | NA | 1580 | 396 | 396 | 16% | 2e-126 | 98.94 | 566 | NZ_MZMW010000... |
| Pseudomonas aeruginosa strain ENV-297... | Pseudomonas ... | NA | 287 | 394 | 394 | 17% | 3e-125 | 93.37 | 638 | NZ_RADD010001... |
| Mycobacterium tuberculosis strain 32845_S29 230, whole genome... | Mycobacteriu... | NA | 1773 | 391 | 391 | 17% | 2e-124 | 94.00 | 601 | NZ_QTBV010000... |
| Acinetobacter baumannii strain ACM-38... | Acinetobacte... | NA | 470 | 391 | 391 | 19% | 2e-124 | 86.64 | 652 | NZ_QAGQ010000... |
| Acinetobacter baumannii strain ACM-32... | Acinetobacte... | NA | 470 | 389 | 389 | 19% | 2e-123 | 86.05 | 654 | NZ_PYDR010000... |
| Providencia rettgeri strain AMW697_S1 18_contig_57, whole geno... | Providencia ... | NA | 587 | 384 | 384 | 17% | 7e-122 | 93.30 | 585 | NZ_SHDI010000... |
| Levilactobacillus brevis strain CRL 2013 scaffold16, whole... | Levilactobac... | NA | 1580 | 376 | 376 | 16% | 5e-119 | 97.25 | 547 | NZ_MZMW010000... |
| Enterococcus faecalis strain 44.1... | Enterococcus... | NA | 1351 | 367 | 367 | 18% | 4e-115 | 85.22 | 611 | NZ_NSJR010000... |
| Cuspidothrix issatschenkoi CHARLIE-1 contig_175, whole genome... | Cuspidothrix... | NA | 2052836 | 385 | 456 | 60% | 1e-114 | 41.62 | 2237 | NZ_PGEM010001... |
| Acinetobacter baumannii strain ACM-36... | Acinetobacte... | NA | 470 | 363 | 363 | 16% | 4e-114 | 91.49 | 567 | NZ_QAGO010000... |
| Enterobacter hormaechei strain 2011WA-SCV PROKKA_contig000046,... | Enterobacter... | NA | 158836 | 352 | 352 | 16% | 4e-110 | 92.22 | 544 | NZ_SJEC010000... |
| Levilactobacillus brevis strain CRL 2013 scaffold14, whole... | Levilactobac... | NA | 1580 | 352 | 405 | 15% | 1e-109 | 94.29 | 588 | NZ_MZMW010000... |
| Acinetobacter baumannii strain ACM-38... | Acinetobacte... | NA | 470 | 351 | 351 | 16% | 1e-109 | 91.11 | 542 | NZ_QAGQ010000... |
| Cellulophaga baltica strain 14 NODE_54_length_581_cov_63.51321... | Cellulophaga... | NA | 76594 | 344 | 344 | 17% | 7e-107 | 84.97 | 581 | NZ_VIQP010000... |
| Shigella sonnei strain 07401ss_1, whole genome shotgun sequence | Shigella sonnei | NA | 624 | 342 | 342 | 15% | 3e-106 | 93.49 | 509 | NZ_CWST010003... |
| Mycobacterium tuberculosis variant bovis strain CSURQ1131, who... | Mycobacteriu... | NA | 1765 | 343 | 865 | 19% | 7e-105 | 80.98 | 888 | NZ_CAJEUJ0100... |
| Enterococcus faecium strain INF91_9... | Enterococcus... | NA | 1352 | 338 | 338 | 15% | 1e-104 | 92.40 | 515 | NZ_JAANZG0100... |
| Klebsiella aerogenes strain AUH-KAM3... | Klebsiella a... | NA | 548 | 338 | 338 | 16% | 1e-104 | 85.48 | 558 | NZ_PYHH010000... |
| Thalassovita mangrovi strain GS-10 Scaffold19_1, whole genome... | Thalassovita... | NA | 2692236 | 360 | 360 | 16% | 4e-103 | 97.21 | 2900 | NZ_WWEN010000... |
| Nocardioides luteus strain JCM 3358 sequence39, whole genome... | Nocardioides... | NA | 1844 | 371 | 371 | 16% | 7e-103 | 95.08 | 5754 | NZ_BMRK010000... |
| Streptomyces griseoflavus strain JCM 4479 sequence37, whole... | Streptomyces... | NA | 35619 | 371 | 371 | 16% | 7e-103 | 95.08 | 5775 | NZ_BMUC010000... |
| Saccharothrix coeruleofusca strain JCM 3313 sequence45, whole... | Saccharothri... | NA | 33919 | 371 | 371 | 16% | 7e-103 | 95.08 | 5775 | NZ_BMRG010000... |
| Streptomyces griseoloalbus strain JCM 4480 sequence054, whole... | Streptomyces... | NA | 67303 | 371 | 371 | 16% | 7e-103 | 95.08 | 5815 | NZ_BMUD010000... |
| Actinoplanes ianthinogenes strain JCM 3249 sequence037, whole... | Actinoplanes... | NA | 122358 | 371 | 371 | 16% | 7e-103 | 95.08 | 5815 | NZ_BMQZ010000... |
| Sinomonas cellulolyticus strain KCTC 49339 sequence28, whole... | Sinomonas ce... | NA | 2801916 | 371 | 371 | 16% | 7e-103 | 95.08 | 5849 | NZ_BNCM010000... |
| Saccharothrix mutabilis subsp. mutabilis strain JCM 3380... | Saccharothri... | NA | 66855 | 371 | 371 | 16% | 7e-103 | 95.08 | 5899 | NZ_BAAABU0100... |
| Neisseria gonorrhoeae strain GC-206... | Neisseria go... | NA | 485 | 332 | 332 | 14% | 2e-102 | 91.62 | 504 | NZ_SAWN010001... |
| Providencia rettgeri strain AFE96_S1 41_contig_58, whole genom... | Providencia ... | NA | 587 | 330 | 330 | 15% | 8e-102 | 89.71 | 546 | NZ_SHDP010000... |
| Providencia rettgeri strain AFE422_S1 40_contig_33, whole geno... | Providencia ... | NA | 587 | 329 | 329 | 14% | 1e-101 | 91.62 | 501 | NZ_SHDL010000... |
| Providencia rettgeri strain AFE220_S1 26_contig_66, whole geno... | Providencia ... | NA | 587 | 326 | 326 | 14% | 2e-100 | 91.57 | 502 | NZ_SHDM010000... |
| Acinetobacter baumannii strain ACM-23... | Acinetobacte... | NA | 470 | 326 | 326 | 15% | 2e-100 | 93.45 | 505 | NZ_PYDJ010000... |
| Providencia rettgeri strain AFE220_S1 26_contig_67, whole geno... | Providencia ... | NA | 587 | 325 | 325 | 15% | 6e-100 | 90.59 | 512 | NZ_SHDM010000... |
| Acinetobacter baumannii strain ACM-24... | Acinetobacte... | NA | 470 | 324 | 324 | 14% | 1e-99 | 93.41 | 503 | NZ_PYDK010000... |
| Escherichia coli strain 8374wG10 NODE_51_length_497_cov_1.1680... | Escherichia ... | NA | 562 | 323 | 323 | 14% | 2e-99 | 92.12 | 497 | NZ_WSJQ010000... |
| Escherichia coli strain 8374wH11 NODE_48_length_500_cov_2.0026... | Escherichia ... | NA | 562 | 320 | 320 | 14% | 3e-98 | 91.52 | 500 | NZ_WSJI010000... |
| Acinetobacter baumannii strain ACM-35... | Acinetobacte... | NA | 470 | 317 | 317 | 13% | 5e-97 | 97.42 | 466 | NZ_QEHN010000... |
| Klebsiella pneumoniae strain B2 isolate B2, whole genome shotg... | Klebsiella p... | NA | 573 | 350 | 564 | 94% | 2e-95 | 38.27 | 7306 | NZ_CABWNT0100... |
| Escherichia coli strain 8374wH11 NODE_51_length_464_cov_1.9881... | Escherichia ... | NA | 562 | 310 | 310 | 13% | 2e-94 | 92.86 | 464 | NZ_WSJI010000... |
| Escherichia coli strain 8374wB4 NODE_86_length_485_cov_1.49162... | Escherichia ... | NA | 562 | 309 | 309 | 14% | 3e-94 | 93.04 | 485 | NZ_WSPX010000... |
| Escherichia coli strain 8374wA11 NODE_44_length_523_cov_1.7197... | Escherichia ... | NA | 562 | 308 | 308 | 15% | 9e-94 | 82.66 | 523 | NZ_WSPR010000... |
| Escherichia coli strain TzEc080 NODE_44_length_523_cov_1.7197,... | Escherichia ... | NA | 562 | 308 | 308 | 15% | 9e-94 | 82.66 | 523 | NZ_WSFZ010000... |
| Enterococcus faecium strain E21_6... | Enterococcus... | NA | 1352 | 305 | 305 | 15% | 1e-92 | 85.38 | 518 | NZ_JAANZJ0100... |
| Escherichia coli strain P010... | Escherichia ... | NA | 562 | 301 | 301 | 16% | 9e-91 | 79.46 | 556 | NZ_VKOQ010001... |

| | | | | | | | | | | |
|---|---|---|---|---|---|---|---|---|---|---|
| Methylobacterium radiotolerans strain MAMP 4754... | Methylobacte... | NA | 31998 | 298 | 298 | 14% | 6e-90 | 88.12 | 483 | NZ_NKQS010002... |
| Klebsiella pneumoniae strain EuSCAPE_MT006, whole genome shotg... | Klebsiella p... | NA | 573 | 296 | 296 | 13% | 1e-89 | 94.04 | 455 | NZ_UKIU020000... |
| Cronobacter sakazakii strain cro4215W contig98, whole genome... | Cronobacter ... | NA | 28141 | 294 | 294 | 12% | 7e-89 | 100.00 | 429 | NZ_NRKF010000... |
| Klebsiella pneumoniae strain EuSCAPE_IL016, whole genome shotg... | Klebsiella p... | NA | 573 | 295 | 295 | 15% | 1e-88 | 83.91 | 523 | NZ_UKKA020000... |
| Cronobacter sakazakii strain cro4114A1 contig119, whole genome... | Cronobacter ... | NA | 28141 | 293 | 293 | 12% | 3e-88 | 99.30 | 429 | NZ_NRKJ010001... |
| Escherichia coli strain 8374wB11 NODE_49_length_463_cov_1.7708... | Escherichia ... | NA | 562 | 293 | 293 | 13% | 4e-88 | 88.89 | 463 | NZ_WSJO010000... |
| Klebsiella pneumoniae strain EuSCAPE_IL021, whole genome shotg... | Klebsiella p... | NA | 573 | 291 | 291 | 12% | 7e-88 | 97.89 | 426 | NZ_UKJX020001... |
| Cronobacter sakazakii strain cro4114B1 contig94, whole genome... | Cronobacter ... | NA | 28141 | 291 | 291 | 12% | 8e-88 | 97.89 | 429 | NZ_NRKI010000... |
| Escherichia coli strain 8374wA11 NODE_51_length_444_cov_2.5520... | Escherichia ... | NA | 562 | 290 | 290 | 13% | 2e-87 | 91.22 | 444 | NZ_WSPR010000... |
| Escherichia coli strain TzEc080 NODE_51_length_444_cov_2.55205... | Escherichia ... | NA | 562 | 290 | 290 | 13% | 2e-87 | 91.22 | 444 | NZ_WSFZ010000... |
| Cronobacter malonaticus strain cro3263A1 contig63, whole genom... | Cronobacter ... | NA | 413503 | 286 | 286 | 12% | 7e-86 | 95.77 | 429 | NZ_NRMM010000... |
| Cronobacter dublinensis strain cro981C2 contig293, whole genom... | Cronobacter ... | NA | 413497 | 285 | 285 | 12% | 1e-85 | 96.48 | 429 | NZ_NROB010002... |
| Klebsiella pneumoniae strain EuSCAPE_MT015, whole genome shotg... | Klebsiella p... | NA | 573 | 285 | 285 | 12% | 1e-85 | 95.14 | 435 | NZ_UKIW020000... |
| Burkholderia pseudomallei strain 2604STDY5518971... | Burkholderia... | NA | 28450 | 285 | 285 | 12% | 2e-85 | 95.07 | 427 | NZ_UVFZ020001... |
| Cronobacter sakazakii strain cro3040W contig78, whole genome... | Cronobacter ... | NA | 28141 | 285 | 285 | 12% | 2e-85 | 96.48 | 429 | NZ_NRJO010000... |
| Cronobacter sakazakii strain cro4215W contig97, whole genome... | Cronobacter ... | NA | 28141 | 284 | 284 | 12% | 5e-85 | 95.07 | 429 | NZ_NRKF010000... |
| Cronobacter sakazakii strain cro4114B1 contig95, whole genome... | Cronobacter ... | NA | 28141 | 284 | 284 | 12% | 5e-85 | 95.07 | 429 | NZ_NRKI010000... |
| Cronobacter sakazakii strain cro4114A1 contig120, whole genome... | Cronobacter ... | NA | 28141 | 284 | 284 | 12% | 5e-85 | 95.07 | 429 | NZ_NRKJ010001... |
| Cronobacter sakazakii strain cro4191B1-2 contig80, whole genom... | Cronobacter ... | NA | 28141 | 284 | 284 | 12% | 5e-85 | 95.77 | 429 | NZ_NRKG010000... |
| Escherichia coli strain 8374wB11 NODE_47_length_476_cov_2.5988... | Escherichia ... | NA | 562 | 283 | 283 | 14% | 1e-84 | 83.54 | 476 | NZ_WSJO010000... |
| Burkholderia pseudomallei strain 2604STDY5518971... | Burkholderia... | NA | 28450 | 281 | 281 | 13% | 6e-84 | 87.50 | 457 | NZ_UVFZ020001... |
| Cronobacter malonaticus strain cro4164A3 contig124, whole geno... | Cronobacter ... | NA | 413503 | 280 | 280 | 12% | 9e-84 | 94.37 | 429 | NZ_NRNF010001... |
| Escherichia coli strain 8374wB11 NODE_56_length_419_cov_1.4349... | Escherichia ... | NA | 562 | 280 | 280 | 12% | 1e-83 | 94.24 | 419 | NZ_WSJO010000... |
| Cronobacter sakazakii strain cro3040W contig77, whole genome... | Cronobacter ... | NA | 28141 | 280 | 280 | 12% | 2e-83 | 94.37 | 429 | NZ_NRJO010000... |
| Levilactobacillus brevis strain CRL 2013 scaffold20, whole... | Levilactobac... | NA | 1580 | 271 | 271 | 11% | 9e-81 | 96.97 | 398 | NZ_MZMW010000... |
| Escherichia coli strain 8374wB11... | Escherichia ... | NA | 562 | 271 | 271 | 12% | 2e-80 | 87.14 | 422 | NZ_WSJO010000... |
| Klebsiella pneumoniae strain EuSCAPE_IL018, whole genome shotg... | Klebsiella p... | NA | 573 | 267 | 267 | 11% | 5e-79 | 94.62 | 392 | NZ_UKJU020001... |
| Klebsiella pneumoniae strain EuSCAPE_NO017, whole genome shotg... | Klebsiella p... | NA | 573 | 266 | 266 | 13% | 3e-78 | 84.21 | 458 | NZ_UKIO020000... |
| Acinetobacter baumannii strain ACM-28... | Acinetobacte... | NA | 470 | 263 | 263 | 11% | 2e-77 | 93.13 | 395 | NZ_PYDN010000... |
| Klebsiella pneumoniae strain EuSCAPE_LT003, whole genome shotg... | Klebsiella p... | NA | 573 | 262 | 262 | 12% | 3e-77 | 91.91 | 422 | NZ_UKJH020000... |
| Klebsiella pneumoniae strain EuSCAPE_MT012, whole genome shotg... | Klebsiella p... | NA | 573 | 261 | 440 | 22% | 8e-77 | 85.33 | 452 | NZ_UKIX020000... |
| Klebsiella pneumoniae strain EuSCAPE_IL004, whole genome shotg... | Klebsiella p... | NA | 573 | 255 | 255 | 11% | 7e-75 | 92.91 | 382 | NZ_UKJP020001... |
| Acinetobacter baumannii strain ACM-15... | Acinetobacte... | NA | 470 | 255 | 255 | 12% | 2e-74 | 87.41 | 432 | NZ_PYDB010000... |
| Acinetobacter baumannii strain ACM-29... | Acinetobacte... | NA | 470 | 254 | 254 | 12% | 5e-74 | 86.23 | 447 | NZ_PYDO010001... |
| Escherichia coli strain 8374wH11 NODE_62_length_437_cov_1.9548... | Escherichia ... | NA | 562 | 252 | 252 | 13% | 1e-73 | 82.07 | 437 | NZ_WSJI010000... |
| Escherichia coli strain 8374wA11 NODE_52_length_437_cov_2.2451... | Escherichia ... | NA | 562 | 247 | 247 | 13% | 2e-71 | 80.00 | 437 | NZ_WSPR010000... |
| Escherichia coli strain TzEc080 NODE_52_length_437_cov_2.24516... | Escherichia ... | NA | 562 | 247 | 247 | 13% | 2e-71 | 80.00 | 437 | NZ_WSFZ010000... |
| Escherichia coli strain 8374wB4 NODE_91_length_444_cov_2.17035... | Escherichia ... | NA | 562 | 246 | 305 | 13% | 2e-71 | 92.62 | 444 | NZ_WSPX010000... |
| Klebsiella aerogenes strain AUH-KAM4... | Klebsiella a... | NA | 548 | 244 | 244 | 11% | 5e-71 | 91.27 | 381 | NZ_PYHI010000... |
| Pseudomonas aeruginosa strain P60_London_6_VIM_2_11_13, whole... | Pseudomonas ... | NA | 287 | 243 | 243 | 11% | 2e-70 | 87.69 | 395 | NZ_CVXD010000... |
| Klebsiella pneumoniae strain EuSCAPE_MT020, whole genome shotg... | Klebsiella p... | NA | 573 | 243 | 243 | 10% | 3e-70 | 95.90 | 368 | NZ_UKJB020000... |
| Escherichia coli strain 8374wH11... | Escherichia ... | NA | 562 | 242 | 242 | 12% | 6e-70 | 81.75 | 417 | NZ_WSJI010000... |
| Escherichia coli strain 8374wA11 NODE_70_length_381_cov_1.2598... | Escherichia ... | NA | 562 | 238 | 238 | 11% | 2e-68 | 90.48 | 381 | NZ_WSPR010000... |
| Escherichia coli strain TzEc080 NODE_70_length_381_cov_1.25984... | Escherichia ... | NA | 562 | 238 | 238 | 11% | 2e-68 | 90.48 | 381 | NZ_WSFZ010000... |
| Acinetobacter baumannii strain ACM-37... | Acinetobacte... | NA | 470 | 238 | 238 | 12% | 2e-68 | 83.45 | 418 | NZ_QAGP010000... |
| Mycobacterium tuberculosis strain 32845_S29 147, whole genome... | Mycobacteriu... | NA | 1773 | 237 | 237 | 10% | 3e-68 | 92.37 | 357 | NZ_QTBV010000... |
| Mycobacterium tuberculosis strain 32847_S31 147, whole genome... | Mycobacteriu... | NA | 1773 | 237 | 237 | 10% | 3e-68 | 92.37 | 357 | NZ_QTBX010000... |
| Escherichia coli strain 8374wG10 NODE_65_length_433_cov_1.7679... | Escherichia ... | NA | 562 | 237 | 237 | 12% | 5e-68 | 77.08 | 433 | NZ_WSJQ010000... |
| Neisseria gonorrhoeae strain GC-206... | Neisseria go... | NA | 485 | 234 | 234 | 10% | 3e-67 | 92.98 | 344 | NZ_SAWN010001... |
| Klebsiella pneumoniae strain EuSCAPE_MT010, whole genome shotg... | Klebsiella p... | NA | 573 | 233 | 233 | 10% | 8e-67 | 92.62 | 367 | NZ_UKIT020000... |
| Klebsiella aerogenes strain AUH-KAM2... | Klebsiella a... | NA | 548 | 233 | 233 | 12% | 1e-66 | 81.34 | 404 | NZ_PYHG010000... |
| Methylobacterium radiotolerans strain MAMP 4754... | Methylobacte... | NA | 31998 | 232 | 232 | 10% | 1e-65 | 90.68 | 539 | NZ_NKQS010010... |
| Escherichia coli strain ECM-2 NODE_147_length_382_cov_1.52941,... | Escherichia ... | NA | 562 | 223 | 223 | 11% | 3e-63 | 84.80 | 382 | NZ_PXIA010001... |
| Klebsiella pneumoniae strain EuSCAPE_IL007, whole genome shotg... | Klebsiella p... | NA | 573 | 221 | 221 | 10% | 8e-63 | 92.92 | 343 | NZ_UKJT020000... |
| Escherichia coli strain 8374wB11 NODE_46_length_516_cov_2.1516... | Escherichia ... | NA | 562 | 218 | 348 | 15% | 8e-61 | 91.96 | 516 | NZ_WSJO010000... |
| Thalassovita mangrovi strain GS-10 Scaffold21_1, whole genome... | Thalassovita... | NA | 2692236 | 228 | 228 | 10% | 2e-60 | 97.35 | 1440 | NZ_WWEN010000... |
| Klebsiella pneumoniae strain EuSCAPE_LT004, whole genome shotg... | Klebsiella p... | NA | 573 | 202 | 202 | 9% | 5e-56 | 86.49 | 335 | NZ_UKJE020000... |
| Klebsiella pneumoniae strain EuSCAPE_IL005, whole genome shotg... | Klebsiella p... | NA | 573 | 199 | 199 | 8% | 4e-55 | 96.94 | 298 | NZ_UKJL020000... |
| Klebsiella pneumoniae strain EuSCAPE_LT003, whole genome shotg... | Klebsiella p... | NA | 573 | 198 | 198 | 8% | 1e-54 | 92.86 | 297 | NZ_UKJH020000... |
| Klebsiella pneumoniae strain EuSCAPE_LT004, whole genome shotg... | Klebsiella p... | NA | 573 | 197 | 197 | 10% | 3e-54 | 77.05 | 367 | NZ_UKJE020000... |
| Deinococcus aquiradiocola strain JCM 14371 sequence36, whole... | Deinococcus ... | NA | 393059 | 204 | 204 | 9% | 4e-54 | 98.02 | 869 | NZ_BMOE010000... |
| Klebsiella pneumoniae strain EuSCAPE_LT003, whole genome shotg... | Klebsiella p... | NA | 573 | 194 | 194 | 8% | 2e-53 | 89.90 | 305 | NZ_UKJH020000... |
| Klebsiella pneumoniae strain EuSCAPE_IL005, whole genome shotg... | Klebsiella p... | NA | 573 | 197 | 197 | 9% | 8e-53 | 92.23 | 636 | NZ_UKJL020000... |
| Cellulophaga baltica strain 14 NODE_63_length_422_cov_10.20678... | Cellulophaga... | NA | 76594 | 192 | 192 | 11% | 3e-52 | 76.15 | 422 | NZ_VIQP010000... |
| Klebsiella pneumoniae strain EuSCAPE_IL004, whole genome shotg... | Klebsiella p... | NA | 573 | 191 | 191 | 10% | 3e-52 | 80.00 | 356 | NZ_UKJP020001... |
| Klebsiella pneumoniae strain EuSCAPE_IL005, whole genome shotg... | Klebsiella p... | NA | 573 | 187 | 187 | 8% | 6e-51 | 92.47 | 283 | NZ_UKJL020000... |
| Klebsiella pneumoniae strain EuSCAPE_LT007, whole genome shotg... | Klebsiella p... | NA | 573 | 186 | 186 | 10% | 3e-50 | 78.99 | 392 | NZ_UKJI020000... |
| Klebsiella pneumoniae strain EuSCAPE_DE082, whole genome shotg... | Klebsiella p... | NA | 573 | 178 | 178 | 8% | 9e-48 | 88.66 | 295 | NZ_UKLV020000... |
| Klebsiella pneumoniae strain EuSCAPE_IL005, whole genome shotg... | Klebsiella p... | NA | 573 | 167 | 167 | 10% | 1e-43 | 75.00 | 323 | NZ_UKJL020000... |
| Salmonella enterica subsp. enterica serovar Anatum strain... | Salmonella e... | NA | 58712 | 164 | 164 | 7% | 5e-43 | 89.66 | 264 | NZ_CALPAM0100... |
| Klebsiella pneumoniae strain EuSCAPE_IL004, whole genome shotg... | Klebsiella p... | NA | 573 | 163 | 163 | 7% | 1e-42 | 95.06 | 244 | NZ_UKJP020001... |
| Burkholderia pseudomallei strain A-175-05-2-02, whole genome... | Burkholderia... | NA | 28450 | 160 | 214 | 11% | 7e-41 | 91.67 | 455 | NZ_UVFW010001... |
| Mycobacterium tuberculosis strain 32847_S31 146, whole genome... | Mycobacteriu... | NA | 1773 | 154 | 154 | 6% | 1e-39 | 89.74 | 235 | NZ_QTBX010000... |
| Methylobacterium radiotolerans strain MAMP 4754... | Methylobacte... | NA | 31998 | 151 | 151 | 7% | 2e-38 | 88.89 | 289 | NZ_NKQS010005... |
| Chlamydia abortus strain 09_01, whole genome shotgun sequence | Chlamydia ab... | NA | 83555 | 155 | 155 | 24% | 2e-37 | 37.55 | 847 | NZ_FQLJ010000... |
| Chlamydia trachomatis strain R33512extract, whole genome shotg... | Chlamydia tr... | NA | 813 | 159 | 159 | 24% | 4e-36 | 37.14 | 1732 | NZ_CVNW010000... |
| Neisseria gonorrhoeae strain GC-206... | Neisseria go... | NA | 485 | 144 | 144 | 6% | 5e-36 | 91.89 | 223 | NZ_SAWN010001... |
| Chlamydia abortus strain 09_03, whole genome shotgun sequence | Chlamydia ab... | NA | 83555 | 163 | 344 | 59% | 7e-36 | 36.56 | 4440 | NZ_FQLL010000... |
| Chlamydia trachomatis strain R4175extract, whole genome shotgu... | Chlamydia tr... | NA | 813 | 160 | 160 | 25% | 1e-35 | 36.43 | 2328 | NZ_CVNX010000... |
| Chlamydia abortus strain 11_01, whole genome shotgun sequence | Chlamydia ab... | NA | 83555 | 162 | 261 | 60% | 1e-35 | 36.43 | 5240 | NZ_FQLK010000... |
| Chlamydia abortus strain 15a, whole genome shotgun sequence | Chlamydia ab... | NA | 83555 | 160 | 160 | 25% | 3e-35 | 36.43 | 3222 | NZ_FQLM010000... |
| Chlamydia abortus strain 09_02, whole genome shotgun sequence | Chlamydia ab... | NA | 83555 | 160 | 160 | 25% | 6e-35 | 36.43 | 4507 | NZ_FQLN010000... |
| Chlamydia abortus strain 09_05, whole genome shotgun sequence | Chlamydia ab... | NA | 83555 | 158 | 259 | 59% | 3e-34 | 37.55 | 4831 | NZ_FQLG010000... |
| Methylobacterium radiotolerans strain MAMP 4754... | Methylobacte... | NA | 31998 | 136 | 136 | 7% | 3e-33 | 80.95 | 279 | NZ_NKQS010012... |
| Pseudomonas aromaticivorans strain MAP12 34, whole genome... | Pseudomonas ... | NA | 2849492 | 143 | 330 | 69% | 1e-29 | 33.58 | 7031 | NZ_JAHRGL0100... |
| Klebsiella pneumoniae strain EuSCAPE_IL017, whole genome shotg... | Klebsiella p... | NA | 573 | 125 | 125 | 7% | 5e-29 | 72.73 | 325 | NZ_UKKG020000... |
| Klebsiella pneumoniae strain EuSCAPE_MT012, whole genome shotg... | Klebsiella p... | NA | 573 | 105 | 105 | 4% | 9e-22 | 96.23 | 385 | NZ_UKIX020000... |
| Mycobacterium tuberculosis strain 32845_S29 214, whole genome... | Mycobacteriu... | NA | 1773 | 94.4 | 94.4 | 6% | 1e-18 | 61.54 | 223 | NZ_QTBV010000... |
| Acinetobacter baumannii strain P4 contig_P4_92, whole genome... | Acinetobacte... | NA | 470 | 103 | 103 | 4% | 2e-18 | 92.31 | 1226 | NZ_JBBXLU0100... |
| Chlamydia trachomatis strain R33512extract, whole genome shotg... | Chlamydia tr... | NA | 813 | 95.9 | 95.9 | 33% | 7e-16 | 25.56 | 1310 | NZ_CVNW010000... |
| Acinetobacter baumannii strain E12 contig_E12_75, whole genome... | Acinetobacte... | NA | 470 | 82.8 | 82.8 | 3% | 2e-13 | 90.48 | 514 | NZ_JBBXKS0100... |
| Tumebacillus flagellatus strain GST4 Contig077, whole genome... | Tumebacillus... | NA | 1157490 | 89.0 | 89.0 | 15% | 3e-13 | 29.67 | 2091 | NZ_JMIR010000... |
| Listeria monocytogenes strain 1827/2,1... | Listeria mon... | NA | 1639 | 89.0 | 89.0 | 26% | 5e-13 | 25.71 | 4618 | NZ_JAENKW0100... |
| Chlamydia abortus strain 09_01, whole genome shotgun sequence | Chlamydia ab... | NA | 83555 | 85.5 | 85.5 | 26% | 2e-12 | 27.59 | 1528 | NZ_FQLJ010000... |
| Uncultured Roseobacter sp. isolate 53.fasta NODE_38, whole... | uncultured R... | NA | 114847 | 75.9 | 75.9 | 3% | 1e-11 | 100.00 | 315 | NZ_CANMKI0100... |
| Cellulophaga baltica isolate 925.fasta NODE_42, whole genome... | Cellulophaga... | NA | 76594 | 74.7 | 74.7 | 3% | 3e-11 | 100.00 | 316 | NZ_CANLPS0100... |
| Rickettsia endosymbiont of Ceutorhynchus assimilis isolate... | Rickettsia e... | NA | 2884482 | 83.6 | 165 | 35% | 3e-11 | 23.98 | 4063491 | NZ_OU906081.1 |
| Klebsiella pneumoniae strain EuSCAPE_MT015, whole genome shotg... | Klebsiella p... | NA | 573 | 73.9 | 73.9 | 5% | 3e-11 | 65.15 | 281 | NZ_UKIW020001... |
| Uncultured Dokdonia sp. isolate 746.fasta NODE_102, whole geno... | uncultured D... | NA | 575653 | 75.9 | 75.9 | 3% | 3e-11 | 100.00 | 440 | NZ_CANLOB0100... |
| Klebsiella pneumoniae strain BL2026... | Klebsiella p... | NA | 573 | 81.6 | 81.6 | 16% | 5e-11 | 31.77 | 1934 | NZ_VICY010000... |
| Streptococcus pneumoniae strain NN2021-1 12_contig_107, whole... | Streptococcu... | NA | 1313 | 80.1 | 80.1 | 17% | 2e-10 | 30.77 | 1657 | NZ_JAGYHD0100... |
| Klebsiella pneumoniae strain BL1427... | Klebsiella p... | NA | 573 | 78.2 | 78.2 | 16% | 2e-10 | 30.73 | 1114 | NZ_VICX010000... |
| Salmonella enterica subsp. enterica serovar Typhi strain 7-sc-... | Salmonella e... | NA | 90370 | 71.2 | 71.2 | 2% | 3e-10 | 100.00 | 289 | NZ_JAEPKN0100... |
| Acinetobacter baumannii strain E1 contig_E1_51, whole genome... | Acinetobacte... | NA | 470 | 68.9 | 68.9 | 3% | 6e-09 | 88.57 | 412 | NZ_JBBXKX0100... |
| Burkholderia pseudomallei strain 2604STDY5519032... | Burkholderia... | NA | 28450 | 70.5 | 70.5 | 4% | 1e-08 | 67.27 | 656 | NZ_UVID020000... |
| Wolbachia endosymbiont of Psylliodes chrysocephala isolate... | Wolbachia en... | NA | 2883236 | 73.9 | 73.9 | 20% | 3e-08 | 26.89 | 1438854 | NZ_OU909470.1 |
| Gemmatimonas sp. isolate Sample-104-S78-bin-40... | Gemmatimonas... | NA | 1962908 | 73.2 | 73.2 | 15% | 4e-08 | 28.42 | 8099 | NZ_JBEILP0100... |
| Corynebacterium glutamicum strain 39 Scaffolds_32_9166, whole... | Corynebacter... | NA | 1718 | 72.8 | 72.8 | 5% | 6e-08 | 66.07 | 8862 | NZ_JACEQW0100... |
| Corynebacterium glutamicum strain 37 Scaffolds_33_8928, whole... | Corynebacter... | NA | 1718 | 72.8 | 72.8 | 5% | 6e-08 | 66.07 | 8928 | NZ_JACENO0100... |
| Klebsiella pneumoniae strain BL2026... | Klebsiella p... | NA | 573 | 66.2 | 66.2 | 13% | 3e-07 | 31.88 | 659 | NZ_VICY010000... |
| Sphingobacterium bovisgrunnientis strain KCTC 52685... | Sphingobacte... | NA | 1874697 | 64.3 | 64.3 | 14% | 5e-07 | 26.38 | 503 | NZ_WUQY010001... |
| Acinetobacter baumannii strain BM7085... | Acinetobacte... | NA | 470 | 63.5 | 63.5 | 12% | 5e-07 | 28.47 | 446 | NZ_JBBONG0100... |
| Uncultured Roseobacter sp. isolate 930.fasta NODE_31, whole... | uncultured R... | NA | 114847 | 60.1 | 60.1 | 2% | 2e-06 | 100.00 | 257 | NZ_CANLQB0100... |
| Brachybacterium paraconglomeratum isolate 859.fasta NODE_71,... | Brachybacter... | NA | 173362 | 60.1 | 60.1 | 2% | 2e-06 | 100.00 | 258 | NZ_CANLNX0100... |
| Streptococcus anginosus strain San49... | Streptococcu... | NA | 1328 | 59.3 | 59.3 | 7% | 5e-06 | 35.80 | 269 | NZ_JAXIIG0100... |
| Klebsiella pneumoniae strain c10 contig85, whole genome shotgu... | Klebsiella p... | NA | 573 | 66.2 | 66.2 | 21% | 5e-06 | 26.94 | 6264 | NZ_JAIBKQ0100... |
| Enterococcus faecalis strain 15224... | Enterococcus... | NA | 1351 | 62.4 | 62.4 | 11% | 5e-06 | 29.66 | 632 | NZ_QNGP010000... |
| Enterococcus faecalis strain AKSZ-144... | Enterococcus... | NA | 1351 | 58.5 | 58.5 | 2% | 6e-06 | 96.15 | 234 | NZ_JAOXSP0100... |
| Roseobacter litoralis isolate 289-G.fasta NODE_32, whole genom... | Roseobacter ... | NA | 42443 | 58.2 | 58.2 | 2% | 8e-06 | 96.15 | 249 | NZ_CANMVH0100... |
| Uncultured Pseudoteredinibacter sp. isolate 208.fasta NODE_42,... | uncultured P... | NA | 1641701 | 58.2 | 58.2 | 2% | 8e-06 | 96.15 | 253 | NZ_CANMSS0100... |
| Uncultured Tateyamaria sp. isolate 203.fasta NODE_54, whole... | uncultured T... | NA | 455651 | 58.2 | 58.2 | 2% | 9e-06 | 96.15 | 263 | NZ_CANLVZ0100... |
| Uncultured Tateyamaria sp. isolate 159-G.fasta NODE_34, whole... | uncultured T... | NA | 455651 | 58.2 | 58.2 | 2% | 1e-05 | 96.15 | 252 | NZ_CANNDY0100... |
| Uncultured Tenacibaculum sp. isolate 838.fasta NODE_42, whole... | uncultured T... | NA | 174713 | 58.2 | 58.2 | 2% | 1e-05 | 96.15 | 262 | NZ_CANMMT0100... |
| Pseudomonas sp. QS1027... | Pseudomonas ... | NA | 2056244 | 57.8 | 57.8 | 2% | 1e-05 | 100.00 | 240 | NZ_PHSU010000... |

| | | | | | | | | | | |
|---|---|---|---|---|---|---|---|---|---|---|
| Uncultured Roseobacter sp. isolate 908.fasta NODE_28, whole... | uncultured R... | NA | 114847 | 57.8 | 57.8 | 2% | 1e-05 | 100.00 | 244 | NZ_CANMNK0100... |
| Uncultured Roseibium sp. isolate 190.fasta NODE_26, whole geno... | uncultured R... | NA | 1936171 | 57.8 | 57.8 | 2% | 1e-05 | 100.00 | 248 | NZ_CANNEN0100... |
| Uncultured Roseobacter sp. isolate 407-3Z.fasta NODE_45, whole... | uncultured R... | NA | 114847 | 57.8 | 57.8 | 2% | 1e-05 | 100.00 | 252 | NZ_CANMKD0100... |
| Uncultured Winogradskyella sp. isolate 408-3Z.fasta NODE_32,... | uncultured W... | NA | 395353 | 57.8 | 57.8 | 2% | 1e-05 | 100.00 | 256 | NZ_CANMJY0100... |
| Uncultured Ruegeria sp. isolate 409.fasta NODE_44, whole genom... | uncultured R... | NA | 259304 | 57.8 | 57.8 | 2% | 1e-05 | 100.00 | 256 | NZ_CANLNV0100... |
| Epibacterium ulvae isolate 20.fasta NODE_62, whole genome... | Epibacterium... | NA | 1156985 | 57.8 | 57.8 | 2% | 1e-05 | 100.00 | 257 | NZ_CANLDO0100... |
| Roseibium album isolate 321-3Z.fasta NODE_38, whole genome... | Roseibium album | NA | 311410 | 57.8 | 57.8 | 2% | 1e-05 | 100.00 | 255 | NZ_CANMGD0100... |
| Proteus mirabilis strain PM2-4 PM2-4_scaffold37, whole genome... | Proteus mira... | NA | 584 | 64.7 | 64.7 | 20% | 1e-05 | 25.40 | 5277 | NZ_JAMKNT0100... |
| Polyangium sp. y55x31 NODE_26_length_169233_cov_17.164843, who... | Polyangium s... | NA | 3042688 | 63.5 | 125 | 21% | 4e-05 | 33.33 | 169233 | NZ_JARZHG0100... |
| Stenotrophomonas sp. isolate AP15.006 scaffold_7194, whole... | Stenotrophom... | NA | 69392 | 62.0 | 62.0 | 12% | 4e-05 | 31.06 | 1073 | NZ_JACXUC0100... |
| Thalassolituus oleivorans isolate... | Thalassolitu... | NA | 187493 | 63.2 | 63.2 | 24% | 4e-05 | 26.76 | 10375 | NZ_CAWUIR0100... |
| Klebsiella pneumoniae strain BL198... | Klebsiella p... | NA | 573 | 59.3 | 59.3 | 11% | 5e-05 | 29.50 | 611 | NZ_VICV010000... |
| Escherichia coli strain w183 NODE_188_length_269_cov_26.989,... | Escherichia ... | NA | 562 | 56.2 | 56.2 | 2% | 5e-05 | 100.00 | 269 | NZ_JAJNDO0100... |
| Pseudomonas nitroreducens strain FY43... | Pseudomonas ... | NA | 46680 | 56.2 | 56.2 | 2% | 6e-05 | 96.15 | 306 | NZ_JAJISQ0100... |
| Polyangium spumosum strain DSM 14734 Scaffold16, whole genome... | Polyangium s... | NA | 889282 | 62.4 | 123 | 21% | 9e-05 | 33.33 | 180314 | NZ_WJIE010000... |
| Salmonella sp. gx-f5 contig70, whole genome shotgun sequence | Salmonella s... | NA | 2582605 | 56.2 | 56.2 | 8% | 1e-04 | 34.78 | 380 | NZ_VCKP010000... |
| Klebsiella pneumoniae strain AS012315 AS012315_632, whole geno... | Klebsiella p... | NA | 573 | 54.3 | 54.3 | 2% | 1e-04 | 100.00 | 234 | NZ_VKXG010006... |
| Klebsiella pneumoniae isolate ena-yuan-GCF_010597935.1... | Klebsiella p... | NA | 573 | 54.3 | 54.3 | 2% | 1e-04 | 100.00 | 234 | NZ_CAWPJY0100... |
| Proteus mirabilis strain PM2-4 PM2-4_scaffold43, whole genome... | Proteus mira... | NA | 584 | 61.2 | 61.2 | 13% | 2e-04 | 26.90 | 3726 | NZ_JAMKNT0100... |
| Escherichia coli strain w189 NODE_129_length_265_cov_25.6333,... | Escherichia ... | NA | 562 | 54.3 | 54.3 | 2% | 2e-04 | 100.00 | 265 | NZ_JAJNDQ0100... |
| Escherichia coli strain w190 NODE_198_length_265_cov_25.5778,... | Escherichia ... | NA | 562 | 54.3 | 54.3 | 2% | 2e-04 | 100.00 | 265 | NZ_JAJNDR0100... |
| Escherichia coli strain w176 NODE_127_length_264_cov_65.95,... | Escherichia ... | NA | 562 | 54.3 | 54.3 | 2% | 2e-04 | 100.00 | 264 | NZ_JAJNDN0100... |
| Acinetobacter baumannii strain BM4942... | Acinetobacte... | NA | 470 | 55.1 | 55.1 | 8% | 2e-04 | 28.26 | 338 | NZ_JBBONW0100... |
| Escherichia coli strain w202 NODE_136_length_268_cov_26.95,... | Escherichia ... | NA | 562 | 54.3 | 54.3 | 2% | 2e-04 | 100.00 | 268 | NZ_JAJNDS0100... |
| Escherichia coli strain w184 NODE_200_length_268_cov_29.5537,... | Escherichia ... | NA | 562 | 54.3 | 54.3 | 2% | 2e-04 | 100.00 | 268 | NZ_JAJNDP0100... |
| Escherichia coli strain w208 NODE_126_length_276_cov_44.5, who... | Escherichia ... | NA | 562 | 54.3 | 54.3 | 2% | 2e-04 | 100.00 | 276 | NZ_JAJNDT0100... |
| Streptomyces albidochromogenes strain DSM 41800... | Streptomyces... | NA | 329524 | 56.6 | 56.6 | 16% | 2e-04 | 27.96 | 521 | NZ_VBVX010007... |
| Polyangium sp. 15x6 NODE_112_length_36528_cov_15.618939, whole... | Polyangium s... | NA | 3042687 | 60.5 | 119 | 21% | 3e-04 | 32.54 | 36528 | NZ_JASBQE0100... |
| Acinetobacter baumannii strain BM4825... | Acinetobacte... | NA | 470 | 54.7 | 54.7 | 10% | 4e-04 | 29.84 | 342 | NZ_JBBOMF0100... |
| Thiolapillus sp. isolate Ifr-SOX-238... | Thiolapillus... | NA | 2017437 | 58.9 | 58.9 | 17% | 6e-04 | 28.92 | 2294 | NZ_JANEJA0100... |
| Klebsiella pneumoniae strain BL3427... | Klebsiella p... | NA | 573 | 57.8 | 57.8 | 11% | 6e-04 | 30.08 | 966 | NZ_VIDH010000... |
| Polyangium sp. 6x1 NODE_22_length_166486_cov_22.101912, whole... | Polyangium s... | NA | 3042689 | 59.3 | 117 | 21% | 8e-04 | 33.33 | 166486 | NZ_JARZHH0100... |
| Penaeicola halotolerans strain LMIT005... | Penaeicola h... | NA | 2793196 | 53.1 | 53.1 | 5% | 0.001 | 45.90 | 336 | NZ_JAEAGO0100... |
| Candidatus Erwinia dacicola strain IL contig_608, whole genome... | Candidatus E... | NA | 252393 | 57.8 | 57.8 | 9% | 0.001 | 30.84 | 1576 | NZ_MAYS010005... |
| Escherichia coli strain EC53 NODE_385_length_1072_cov_0.533668... | Escherichia ... | NA | 562 | 56.6 | 56.6 | 11% | 0.002 | 29.25 | 1072 | NZ_RDNE010003... |
| Enterococcus faecalis strain 17838... | Enterococcus... | NA | 1351 | 52.0 | 52.0 | 8% | 0.002 | 33.70 | 288 | NZ_QNGS010001... |
| Piscirickettsia salmonis strain FIUCHILE-89L01, whole genome... | Piscirickett... | NA | 1238 | 57.4 | 57.4 | 12% | 0.002 | 29.08 | 2765 | NZ_LELB010002... |
| Escherichia coli strain 8551F 237, whole genome shotgun sequence | Escherichia ... | NA | 562 | 54.7 | 54.7 | 18% | 0.003 | 27.78 | 689 | NZ_JAFCNN0100... |
| Pseudomonas aeruginosa isolate PSA-13, whole genome shotgun... | Pseudomonas ... | NA | 287 | 53.5 | 53.5 | 12% | 0.003 | 30.66 | 505 | NZ_FNME010000... |
| Polyangium jinanense strain SDU14 Scaffold9, whole genome... | Polyangium j... | NA | 2829994 | 57.0 | 112 | 20% | 0.004 | 32.23 | 231060 | NZ_JAGTJJ0100... |
| Polyangium jinanense strain SDU13 Scaffold15, whole genome... | Polyangium j... | NA | 2829994 | 57.0 | 112 | 20% | 0.004 | 32.23 | 231263 | NZ_JAGTJK0100... |
| Bradyrhizobium sp. 33ap4 NODE_290_length_534_cov_0.849015, who... | Bradyrhizobi... | NA | 3061630 | 53.1 | 53.1 | 12% | 0.004 | 29.37 | 534 | NZ_JAUMTJ0100... |
| Eikenella corrodens strain 491_NMEN, whole genome shotgun... | Eikenella co... | NA | 539 | 53.9 | 53.9 | 10% | 0.005 | 28.46 | 659 | NZ_JVFA010000... |
| Enterobacter cloacae complex sp. GF14B... | Enterobacter... | NA | 2511982 | 55.8 | 55.8 | 21% | 0.006 | 27.00 | 4351 | NZ_QFRV010001... |
| Methylobacterium radiotolerans strain MAMP 4754... | Methylobacte... | NA | 31998 | 50.1 | 50.1 | 4% | 0.007 | 57.45 | 253 | NZ_NKQS010008... |
| Proteus terrae subsp. cibarius strain pt-OS9-1... | Proteus terr... | NA | 626774 | 50.1 | 50.1 | 8% | 0.007 | 35.11 | 267 | NZ_JBBEZU0100... |
| Eikenella corrodens strain EI_05 NODE_86_length_856_cov_136.29... | Eikenella co... | NA | 539 | 53.9 | 53.9 | 10% | 0.007 | 28.46 | 856 | NZ_VYYO010000... |
| Devosia insulae DS-56 contig234, whole genome shotgun sequence | Devosia insu... | NA | 1116389 | 55.8 | 55.8 | 9% | 0.008 | 30.53 | 24799 | NZ_LAJE020001... |
| Escherichia coli strain RF108-1... | Escherichia ... | NA | 562 | 53.1 | 53.1 | 6% | 0.008 | 34.94 | 640 | NZ_JAAQGD0100... |
| Streptococcus mutans strain PKUSS-9 C209, whole genome shotgun... | Streptococcu... | NA | 1309 | 50.1 | 50.1 | 6% | 0.009 | 38.67 | 307 | NZ_NBFI010000... |
| Klebsiella pneumoniae strain A13... | Klebsiella p... | NA | 573 | 50.1 | 50.1 | 9% | 0.014 | 30.36 | 333 | NZ_JARWSQ0100... |
| Escherichia coli strain EC86 NODE_205_length_902_cov_0.835152,... | Escherichia ... | NA | 562 | 53.1 | 53.1 | 8% | 0.019 | 32.98 | 902 | NZ_RDNB010002... |
| Klebsiella pneumoniae strain kp110... | Klebsiella p... | NA | 573 | 52.4 | 52.4 | 13% | 0.019 | 25.64 | 728 | NZ_JAPMWX0100... |
| Enterococcus faecalis strain 15407... | Enterococcus... | NA | 1351 | 48.9 | 48.9 | 7% | 0.022 | 32.58 | 285 | NZ_QNGQ010002... |
| Marmoricola endophyticus strain CGMCC 1.16067 sequence01, whol... | Marmoricola ... | NA | 2040280 | 54.3 | 54.3 | 12% | 0.023 | 27.92 | 3374816 | NZ_BMKQ010000... |
| Escherichia coli strain 90 NODE_134_length_1268_cov_0.816121,... | Escherichia ... | NA | 562 | 53.1 | 53.1 | 6% | 0.025 | 33.77 | 1268 | NZ_JACAMA0100... |
| Breoghania sp. isolate MO_035.B22 MO_035.c_6474, whole genome... | Breoghania sp. | NA | 2065378 | 53.9 | 53.9 | 20% | 0.025 | 25.51 | 5920 | NZ_JASJAV0100... |
| Devosia sp. Root413D1 contig_4, whole genome shotgun sequence | Devosia sp. ... | NA | 1736531 | 54.3 | 54.3 | 9% | 0.025 | 29.63 | 575790 | NZ_LMEA010000... |
| Breoghania sp. isolate MO_035.B22 MO_035.c_2803, whole genome... | Breoghania sp. | NA | 2065378 | 53.9 | 53.9 | 11% | 0.027 | 28.37 | 6353 | NZ_JASJAV0100... |
| Clostridium paraputrificum strain 28-X28 Scaffold60_1, whole... | Clostridium ... | NA | 29363 | 51.6 | 51.6 | 12% | 0.030 | 30.37 | 680 | NZ_RAYJ010000... |
| Agreia sp. PsM10 NODE_68_length_253_cov_0.818182, whole genome... | Agreia sp. P... | NA | 3030533 | 48.1 | 48.1 | 5% | 0.031 | 43.86 | 253 | NZ_JARUXC0100... |
| Campylobacter sp. JMF_08 NE1 NODE_13_length_3482_cov_0.037836,... | Campylobacte... | NA | 2983821 | 53.5 | 53.5 | 15% | 0.031 | 29.28 | 3482 | NZ_JAOVUC0100... |
| Campylobacter sp. JMF_10 EL2 NODE_13_length_1688_cov_0.027690,... | Campylobacte... | NA | 2983828 | 53.1 | 53.1 | 14% | 0.033 | 30.30 | 1688 | NZ_JAOVTV0100... |
| Campylobacter sp. CS_ED1 strain Cs ED1... | Campylobacte... | NA | 2984140 | 53.5 | 53.5 | 15% | 0.035 | 29.28 | 5400 | NZ_JAOVUL0100... |
| Klebsiella pneumoniae strain BL198... | Klebsiella p... | NA | 573 | 50.1 | 50.1 | 7% | 0.035 | 34.94 | 484 | NZ_VICV010000... |
| Campylobacter sp. CN_NE4 strain Cn NE4... | Campylobacte... | NA | 2984148 | 53.5 | 53.5 | 15% | 0.037 | 29.28 | 5576 | NZ_JAOVUF0100... |
| Pseudomonas aromaticivorans strain MAP12 47, whole genome... | Pseudomonas ... | NA | 2849492 | 53.1 | 53.1 | 10% | 0.037 | 27.78 | 2177 | NZ_JAHRGL0100... |
| Devosia sp. Root105 contig_7, whole genome shotgun sequence | Devosia sp. ... | NA | 1736423 | 53.5 | 53.5 | 8% | 0.038 | 31.86 | 620452 | NZ_LMCR010000... |
| Eikenella corrodens strain KCOM 1378 B230_1_2, whole genome... | Eikenella co... | NA | 539 | 53.5 | 53.5 | 10% | 0.040 | 28.46 | 572658 | NZ_NHRS010000... |
| Wolbachia endosymbiont of Corcyra cephalonica isolate SYLI2103... | Wolbachia en... | NA | 218111 | 53.5 | 53.5 | 13% | 0.040 | 29.11 | 5332 | NZ_CP087956.1 |
| Eikenella corrodens ATCC 23834 Scfld2, whole genome shotgun... | Eikenella co... | NA | 546274 | 53.5 | 53.5 | 10% | 0.040 | 28.46 | 131618 | NZ_EQ973318.1 |
| Eikenella corrodens strain FDAARGOS_1474 chromosome, complete... | Eikenella co... | NA | 539 | 53.5 | 53.5 | 10% | 0.041 | 28.46 | 2192571 | NZ_CP082861.1 |
| Eikenella corrodens strain NCTC10596 chromosome 1, complete... | Eikenella co... | NA | 539 | 53.5 | 53.5 | 10% | 0.041 | 28.46 | 2192577 | NZ_LT906482.1 |
| Lachnoclostridium phocaeense strain Marseille-P3177T chromosom... | Lachnoclostr... | NA | 1871021 | 53.5 | 53.5 | 14% | 0.042 | 27.17 | 3500754 | NZ_LT635479.1 |
| Roseiconus lacunae strain JC742 soapdenovo_K_105_82, whole... | Roseiconus l... | NA | 2605694 | 53.5 | 53.5 | 10% | 0.044 | 28.77 | 35968 | NZ_JAJMQV0100... |
| Klebsiella pneumoniae strain KP231... | Klebsiella p... | NA | 573 | 47.4 | 47.4 | 4% | 0.045 | 46.94 | 233 | NZ_JAUUHQ0100... |
| Eikenella corrodens strain NML01-0040 Eikcor_contig000008, who... | Eikenella co... | NA | 539 | 53.1 | 53.1 | 10% | 0.046 | 28.46 | 65735 | NZ_LXSE010000... |
| Eikenella corrodens strain EI_05... | Eikenella co... | NA | 539 | 53.1 | 53.1 | 10% | 0.050 | 28.46 | 165032 | NZ_VYYO010000... |

Alignments:

```
>Deinococcus aquiradiocola strain JCM 14371 sequence30, whole genome shotgun sequence
Sequence ID: NZ_BMOE01000030.1 Length: 6128
Range 1: 1430 to 3559

Score:1381 bits(3574), Expect:0.0,
Method:Compositional matrix adjust.,
Identities:689/710(97%), Positives:703/710(99%), Gaps:0/710(0%)

Query  2     QGKAREFSSEQTRANSPTRRELQVWGRDNNSPSEAGADRQGTVSFNFPQVTLWQRPLVTI   61
             QGKAREFSSEQTRANSPTRRELQVWGRDNNS SEAGADRQGTVSF+FPQ+TLWQRPLVTI
Sbjct  1430  QGKAREFSSEQTRANSPTRRELQVWGRDNNSLSEAGADRQGTVSFSFPQITLWQRPLVTI   1609

Query  62    KIGGQLKEALLDTGADDTVLEEMSLPGRWKPKMIGGIGGFIKVRQYDQILIEICGHKAIG   121
             KIGGQLKEALLDTGADDTVLEEM+LPGRWKPKMIGGIGGFIKVRQYDQILIEICGHKAIG
Sbjct  1610  KIGGQLKEALLDTGADDTVLEEMNLPGRWKPKMIGGIGGFIKVRQYDQILIEICGHKAIG   1789

Query  122   TVLVGPTPVNIIGRNLLTQIGCTLNFPISPIETVPVKLKPGMDGPKVKQWPLTEEKIKAL   181
             TVLVGPTPVNIIGRNLLTQIGCTLNFPISPIETVPVKLKPGMDGPKVKQWPLTEEKIKAL
Sbjct  1790  TVLVGPTPVNIIGRNLLTQIGCTLNFPISPIETVPVKLKPGMDGPKVKQWPLTEEKIKAL   1969

Query  182   VEICTEMEKEGKISKIGPENPYNTPVFAIKKKDSTKWRKLVDFRELNKRTQDFWEVQLGI   241
             VEICTEMEKEGKISKIGPENPYNTPVFAIKKKDSTKWRKLVDFRELNKRTQDFWEVQLGI
Sbjct  1970  VEICTEMEKEGKISKIGPENPYNTPVFAIKKKDSTKWRKLVDFRELNKRTQDFWEVQLGI   2149

Query  242   PHPAGLKKKKSVTVLDVGDAYFSVPLDEDFRKYTAFTIPSINNETPGIRYQYNVLPQGWK   301
             PHPAGLK+KKSVTVLDVGDAYFSVPLD+DFRKYTAFTIPSINNETPGIRYQYNVLPQGWK
Sbjct  2150  PHPAGLKQKKSVTVLDVGDAYFSVPLDKDFRKYTAFTIPSINNETPGIRYQYNVLPQGWK   2329

Query  302   GSPAIFQSSMTKILEPFRKQNPDIVIYQYMDDLYVGSDLEIGQHRTKIEELRQHLLRWGL   361
             GSPAIFQ SMTKILEPFRKQNPDIVIYQYMDDLYVGSDLEIGQHRTKIEELRQHLLRWG
Sbjct  2330  GSPAIFQCSMTKILEPFRKQNPDIVIYQYMDDLYVGSDLEIGQHRTKIEELRQHLLRWGF   2509

Query  362   TTPDKKHQKEPPFLWMGYELHPDKWTVQPIVLPEKDSWTVNDIQKLVGKLNWASQIYPGI   421
             TTPDKKHQKEPPFLWMGYELHPDKWTVQPIVLPEKDSWTVNDIQKLVGKLNWASQIY GI
Sbjct  2510  TTPDKKHQKEPPFLWMGYELHPDKWTVQPIVLPEKDSWTVNDIQKLVGKLNWASQIYAGI   2689

Query  422   KVRQLCKLLRGTKALTEVIPLTeeaelelaeNREILKEPVHGVYYDPSKDLIAEIQKQGQ   481
             KVRQLCKLLRGTKALTEV+PLTEEAELELAENREILKEPVHGVYYDPSKDLIAEIQKQGQ
Sbjct  2690  KVRQLCKLLRGTKALTEVVPLTEEAELELAENREILKEPVHGVYYDPSKDLIAEIQKQGQ   2869

Query  482   GQWTYQIYQEPFKNLKTGKYARMRGAHTNDVKQLTEAVQKITTESIVIWGKTPKFKLPIQ   541
             GQWTYQIYQEPFKNLKTGKYARM+GAHTNDVKQLTEAVQKI TESIVIWGKTPKFKLPIQ
```

```
Sbjct  2870  GQWTYQIYQEPFKNLKTGKYARMKGAHTNDVKQLTEAVQKIATESIVIWGKTPKFKLPIQ  3049

Query  542   KetwetwwteYWQATWIPEWEFVNTPPLVKLWYQLEKEPIVGAETFYVDGAANRETKLGK  601
             KETWE WWTEYWQATWIPEWEFVNTPPLVKLWYQLEKEPI+GAETFYVDGAANRETKLGK
Sbjct  3050  KETWEAWWTEYWQATWIPEWEFVNTPPLVKLWYQLEKEPIIGAETFYVDGAANRETKLGK  3229

Query  602   AGYVTNRGRQKVVTLTDTTNQKTELQAIYLALQDSGLEVNIVTDSQYALGIIQAQPDQSE  661
             AGYVT+RGRQKVV LTDTTNQKTELQAI+LALQDSGLEVNIVTDSQYALGIIQAQPD+SE
Sbjct  3230  AGYVTDRGRQKVVPLTDTTNQKTELQAIHLALQDSGLEVNIVTDSQYALGIIQAQPDKSE  3409

Query  662   SELVNQIIEQLIKKEKVYLAWVPAHKGIGGNEQVDKLVSAGIRKVLYVDG  711
             SELV+QIIEQLIKKEKVYLAWVPAHKGIGGNEQVDKLVSAGIRKVL++DG
Sbjct  3410  SELVSQIIEQLIKKEKVYLAWVPAHKGIGGNEQVDKLVSAGIRKVLFLDG  3559
```

Range 2: 3179 to 4216

Score:676 bits(1744), Expect:0.0,
Method:Compositional matrix adjust.,
Identities:331/346(96%), Positives:340/346(98%), Gaps:0/346(0%)

```
Query  705   KVLYVDGAANRETKLGKAGYVTNRGRQKVVTLTDTTNQKTELQAIYLALQDSGLEVNIVT  764
             +  YVDGAANRETKLGKAGYVT+RGRQKVV LTDTTNQKTELQAI+LALQDSGLEVNIVT
Sbjct  3179  ETFYVDGAANRETKLGKAGYVTDRGRQKVVPLTDTTNQKTELQAIHLALQDSGLEVNIVT  3358

Query  765   DSQYALGIIQAQPDQSESELVNQIIEQLIKKEKVYLAWVPAHKGIGGNEQVDKLVSAGIR  824
             DSQYALGIIQAQPD+SESELV+QIIEQLIKKEKVYLAWVPAHKGIGGNEQVDKLVSAGIR
Sbjct  3359  DSQYALGIIQAQPDKSESELVSQIIEQLIKKEKVYLAWVPAHKGIGGNEQVDKLVSAGIR  3538

Query  825   KVLFLDGIDKAQDEHEKYHSNWRAMASDFNLPPVVAKEIVASCDKCQLKGEAMHGQVDCS  884
             KVLFLDGIDKAQ+EHEKYHSNWRAMASDFNLPPVVAKEIVASCDKCQLKGEAMHGQVDCS
Sbjct  3539  KVLFLDGIDKAQEEHEKYHSNWRAMASDFNLPPVVAKEIVASCDKCQLKGEAMHGQVDCS  3718

Query  885   PGIWQLDCTHLEGKVILVAVHVASGYIEAEVIPAETGQETAYFLLKLAGRWPVKTIHTDN  944
             PGIWQLDCTHLEGKVILVAVHVASGYIEAEVIPAETGQETAYFLLKLAGRWPVKT+HTDN
Sbjct  3719  PGIWQLDCTHLEGKVILVAVHVASGYIEAEVIPAETGQETAYFLLKLAGRWPVKTVHTDN  3898

Query  945   GSNFTGATVRAACWWAGIKQEFGIPYNPQSQGVVESMNKELKKIIGQVRDQAEHLKTAVQ  1004
             GSNFT  TV+AACWWAGIKQEFGIPYNPQSQGV+ESMNKELKKIIGQVRDQAEHLKTAVQ
Sbjct  3899  GSNFTSTTVKAACWWAGIKQEFGIPYNPQSQGVIESMNKELKKIIGQVRDQAEHLKTAVQ  4078

Query  1005  MAVFIHNFKRKGGIGGYSAGERIVDIIATDIQTKELQKQITKIQNF  1050
             MAVFIHNFKRKGGIGGYSAGERIVDIIATDIQTKELQKQITKIQ F
Sbjct  4079  MAVFIHNFKRKGGIGGYSAGERIVDIIATDIQTKELQKQITKIQIF  4216
```

>Enterococcus alcedinis strain CCM 8433 sequence23, whole genome shotgun sequence
Sequence ID: NZ_BMDT01000023.1 Length: 6075
Range 1: 1377 to 3506

Score:1381 bits(3574), Expect:0.0,
Method:Compositional matrix adjust.,
Identities:689/710(97%), Positives:703/710(99%), Gaps:0/710(0%)

```
Query  2     QGKAREFSSEQTRANSPTRRELQVWGRDNNSPSEAGADRQGTVSFNFPQVTLWQRPLVTI  61
             QGKAREFSSEQTRANSPTRRELQVWGRDNNS SEAGADRQGTVSF+FPQ+TLWQRPLVTI
Sbjct  1377  QGKAREFSSEQTRANSPTRRELQVWGRDNNSLSEAGADRQGTVSFSFPQITLWQRPLVTI  1556

Query  62    KIGGQLKEALLDTGADDTVLEEMSLPGRWKPKMIGGIGGFIKVRQYDQILIEICGHKAIG  121
             KIGGQLKEALLDTGADDTVLEEM+LPGRWKPKMIGGIGGFIKVRQYDQILIEICGHKAIG
Sbjct  1557  KIGGQLKEALLDTGADDTVLEEMNLPGRWKPKMIGGIGGFIKVRQYDQILIEICGHKAIG  1736

Query  122   TVLVGPTPVNIIGRNLLTQIGCTLNFPISPIETVPVKLKPGMDGPKVKQWPLTEEKIKAL  181
             TVLVGPTPVNIIGRNLLTQIGCTLNFPISPIETVPVKLKPGMDGPKVKQWPLTEEKIKAL
Sbjct  1737  TVLVGPTPVNIIGRNLLTQIGCTLNFPISPIETVPVKLKPGMDGPKVKQWPLTEEKIKAL  1916

Query  182   VEICTEMEKEGKISKIGPENPYNTPVFAIKKKDSTKWRKLVDFRELNKRTQDFWEVQLGI  241
             VEICTEMEKEGKISKIGPENPYNTPVFAIKKKDSTKWRKLVDFRELNKRTQDFWEVQLGI
Sbjct  1917  VEICTEMEKEGKISKIGPENPYNTPVFAIKKKDSTKWRKLVDFRELNKRTQDFWEVQLGI  2096

Query  242   PHPAGLKKKKSVTVLDVGDAYFSVPLDEDFRKYTAFTIPSINNETPGIRYQYNVLPQGWK  301
             PHPAGLK+KKSVTVLDVGDAYFSVPLD+DFRKYTAFTIPSINNETPGIRYQYNVLPQGWK
Sbjct  2097  PHPAGLKQKKSVTVLDVGDAYFSVPLDKDFRKYTAFTIPSINNETPGIRYQYNVLPQGWK  2276

Query  302   GSPAIFQSSMTKILEPFRKQNPDIVIYQYMDDLYVGSDLEIGQHRTKIEELRQHLLRWGL  361
             GSPAIFQ SMTKILEPFRKQNPDIVIYQYMDDLYVGSDLEIGQHRTKIEELRQHLLRWG
Sbjct  2277  GSPAIFQCSMTKILEPFRKQNPDIVIYQYMDDLYVGSDLEIGQHRTKIEELRQHLLRWGF  2456

Query  362   TTPDKKHQKEPPFLWMGYELHPDKWTVQPIVLPEKDSWTVNDIQKLVGKLNWASQIYPGI  421
             TTPDKKHQKEPPFLWMGYELHPDKWTVQPIVLPEKDSWTVNDIQKLVGKLNWASQIY GI
Sbjct  2457  TTPDKKHQKEPPFLWMGYELHPDKWTVQPIVLPEKDSWTVNDIQKLVGKLNWASQIYAGI  2636

Query  422   KVRQLCKLLRGTKALTEVIPLTeeaelelaeNREILKEPVHGVYYDPSKDLIAEIQKQGQ  481
             KVRQLCKLLRGTKALTEV+PLTEEAELELAENREILKEPVHGVYYDPSKDLIAEIQKQGQ
Sbjct  2637  KVRQLCKLLRGTKALTEVVPLTEEAELELAENREILKEPVHGVYYDPSKDLIAEIQKQGQ  2816

Query  482   GQWTYQIYQEPFKNLKTGKYARMRGAHTNDVKQLTEAVQKITTESIVIWGKTPKFKLPIQ  541
             GQWTYQIYQEPFKNLKTGKYARM+GAHTNDVKQLTEAVQKI TESIVIWGKTPKFKLPIQ
Sbjct  2817  GQWTYQIYQEPFKNLKTGKYARMKGAHTNDVKQLTEAVQKIATESIVIWGKTPKFKLPIQ  2996

Query  542   KetwetwwteYWQATWIPEWEFVNTPPLVKLWYQLEKEPIVGAETFYVDGAANRETKLGK  601
             KETWE WWTEYWQATWIPEWEFVNTPPLVKLWYQLEKEPI+GAETFYVDGAANRETKLGK
Sbjct  2997  KETWEAWWTEYWQATWIPEWEFVNTPPLVKLWYQLEKEPIIGAETFYVDGAANRETKLGK  3176

Query  602   AGYVTNRGRQKVVTLTDTTNQKTELQAIYLALQDSGLEVNIVTDSQYALGIIQAQPDQSE  661
             AGYVT+RGRQKVV LTDTTNQKTELQAI+LALQDSGLEVNIVTDSQYALGIIQAQPD+SE
Sbjct  3177  AGYVTDRGRQKVVPLTDTTNQKTELQAIHLALQDSGLEVNIVTDSQYALGIIQAQPDKSE  3356

Query  662   SELVNQIIEQLIKKEKVYLAWVPAHKGIGGNEQVDKLVSAGIRKVLYVDG  711
             SELV+QIIEQLIKKEKVYLAWVPAHKGIGGNEQVDKLVSAGIRKVL++DG
Sbjct  3357  SELVSQIIEQLIKKEKVYLAWVPAHKGIGGNEQVDKLVSAGIRKVLFLDG  3506
```

Range 2: 3126 to 4163

Score:676 bits(1744), Expect:0.0,
Method:Compositional matrix adjust.,
Identities:331/346(96%), Positives:340/346(98%), Gaps:0/346(0%)

```
Query  705   KVLYVDGAANRETKLGKAGYVTNRGRQKVVTLTDTTNQKTELQAIYLALQDSGLEVNIVT  764
             +  YVDGAANRETKLGKAGYVT+RGRQKVV LTDTTNQKTELQAI+LALQDSGLEVNIVT
Sbjct  3126  ETFYVDGAANRETKLGKAGYVTDRGRQKVVPLTDTTNQKTELQAIHLALQDSGLEVNIVT  3305

Query  765   DSQYALGIIQAQPDQSESELVNQIIEQLIKKEKVYLAWVPAHKGIGGNEQVDKLVSAGIR  824
             DSQYALGIIQAQPD+SESELV+QIIEQLIKKEKVYLAWVPAHKGIGGNEQVDKLVSAGIR
Sbjct  3306  DSQYALGIIQAQPDKSESELVSQIIEQLIKKEKVYLAWVPAHKGIGGNEQVDKLVSAGIR  3485

Query  825   KVLFLDGIDKAQDEHEKYHSNWRAMASDFNLPPVVAKEIVASCDKCQLKGEAMHGQVDCS  884
```

```
             KVLFLDGIDKAQ+EHEKYHSNWRAMASDFNLPPVVAKEIVASCDKCQLKGEAMHGQVDCS
Sbjct  3486  KVLFLDGIDKAQEEHEKYHSNWRAMASDFNLPPVVAKEIVASCDKCQLKGEAMHGQVDCS  3665

Query  885   PGIWQLDCTHLEGKVILVAVHVASGYIEAEVIPAETGQETAYFLLKLAGRWPVKTIHTDN  944
             PGIWQLDCTHLEGKVILVAVHVASGYIEAEVIPAETGQETAYFLLKLAGRWPVKT+HTDN
Sbjct  3666  PGIWQLDCTHLEGKVILVAVHVASGYIEAEVIPAETGQETAYFLLKLAGRWPVKTVHTDN  3845

Query  945   GSNFTGATVRAACWWAGIKQEFGIPYNPQSQGVVESMNKELKKIIGQVRDQAEHLKTAVQ  1004
             GSNFT  TV+AACWWAGIKQEFGIPYNPQSQGV+ESMNKELKKIIGQVRDQAEHLKTAVQ
Sbjct  3846  GSNFTSTTVKAACWWAGIKQEFGIPYNPQSQGVIESMNKELKKIIGQVRDQAEHLKTAVQ  4025

Query  1005  MAVFIHNFKRKGGIGGYSAGERIVDIIATDIQTKELQKQITKIQNF  1050
             MAVFIHNFKRKGGIGGYSAGERIVDIIATDIQTKELQKQITKIQ F
Sbjct  4026  MAVFIHNFKRKGGIGGYSAGERIVDIIATDIQTKELQKQITKIQIF  4163
```

>Actinoplanes campanulatus strain JCM 3059 sequence058, whole genome shotgun sequence
Sequence ID: NZ_BMPW01000058.1 Length: 4754
Range 1: 1419 to 3548

Score:1380 bits(3571), Expect:0.0,
Method:Compositional matrix adjust.,
Identities:689/710(97%), Positives:703/710(99%), Gaps:0/710(0%)

```
Query  2     QGKAREFSSEQTRANSPTRRELQVWGRDNNSPSEAGADRQGTVSFNFPQVTLWQRPLVTI  61
             QGKAREFSSEQTRANSPTRRELQVWGRDNNS SEAGADRQGTVSF+FPQ+TLWQRPLVTI
Sbjct  1419  QGKAREFSSEQTRANSPTRRELQVWGRDNNSLSEAGADRQGTVSFSFPQITLWQRPLVTI  1598

Query  62    KIGGQLKEALLDTGADDTVLEEMSLPGRWKPKMIGGIGGFIKVRQYDQILIEICGHKAIG  121
             KIGGQLKEALLDTGADDTVLEEM+LPGRWKPKMIGGIGGFIKVRQYDQILIEICGHKAIG
Sbjct  1599  KIGGQLKEALLDTGADDTVLEEMNLPGRWKPKMIGGIGGFIKVRQYDQILIEICGHKAIG  1778

Query  122   TVLVGPTPVNIIGRNLLTQIGCTLNFPISPIETVPVKLKPGMDGPKVKQWPLTEEKIKAL  181
             TVLVGPTPVNIIGRNLLTQIGCTLNFPISPIETVPVKLKPGMDGPKVKQWPLTEEKIKAL
Sbjct  1779  TVLVGPTPVNIIGRNLLTQIGCTLNFPISPIETVPVKLKPGMDGPKVKQWPLTEEKIKAL  1958

Query  182   VEICTEMEKEGKISKIGPENPYNTPVFAIKKKDSTKWRKLVDFRELNKRTQDFWEVQLGI  241
             VEICTEMEKEGKISKIGPENPYNTPVFAIKKKDSTKWRKLVDFRELNKRTQDFWEVQLGI
Sbjct  1959  VEICTEMEKEGKISKIGPENPYNTPVFAIKKKDSTKWRKLVDFRELNKRTQDFWEVQLGI  2138

Query  242   PHPAGLKKKKSVTVLDVGDAYFSVPLDEDFRKYTAFTIPSINNETPGIRYQYNVLPQGWK  301
             PHPAGLK+KKSVTVLDVGDAYFSVPLD+DFRKYTAFTIPSINNETPGIRYQYNVLPQGWK
Sbjct  2139  PHPAGLKQKKSVTVLDVGDAYFSVPLDKDFRKYTAFTIPSINNETPGIRYQYNVLPQGWK  2318

Query  302   GSPAIFQSSMTKILEPFRKQNPDIVIYQYMDDLYVGSDLEIGQHRTKIEELRQHLLRWGL  361
             GSPAIFQ SMTKILEPFRKQNPDIVIYQYMDDLYVGSDLEIGQHRTKIEELRQHLLRWG
Sbjct  2319  GSPAIFQCSMTKILEPFRKQNPDIVIYQYMDDLYVGSDLEIGQHRTKIEELRQHLLRWGF  2498

Query  362   TTPDKKHQKEPPFLWMGYELHPDKWTVQPIVLPEKDSWTVNDIQKLVGKLNWASQIYPGI  421
             TTPDKKHQKEPPFLWMGYELHPDKWTVQPIVLPEKDSWTVNDIQKLVGKLNWASQIY GI
Sbjct  2499  TTPDKKHQKEPPFLWMGYELHPDKWTVQPIVLPEKDSWTVNDIQKLVGKLNWASQIYAGI  2678

Query  422   KVRQLCKLLRGTKALTEVIPLTeeaelelaeNREILKEPVHGVYYDPSKDLIAEIQKQGQ  481
             KVRQLCKLLRGTKALTEV+PLTEEAELELAENREILKEPVHGVYYDPSKDLIAEIQKQGQ
Sbjct  2679  KVRQLCKLLRGTKALTEVVPLTEEAELELAENREILKEPVHGVYYDPSKDLIAEIQKQGQ  2858

Query  482   GQWTYQIYQEPFKNLKTGKYARMRGAHTNDVKQLTEAVQKITTESIVIWGKTPKFKLPIQ  541
             GQWTYQIYQEPFKNLKTGKYARM+GAHTNDVKQLTEAVQKI TESIVIWGKTPKFKLPIQ
Sbjct  2859  GQWTYQIYQEPFKNLKTGKYARMKGAHTNDVKQLTEAVQKIATESIVIWGKTPKFKLPIQ  3038

Query  542   KetwetwwteYWQATWIPEWEFVNTPPLVKLWYQLEKEPIVGAETFYVDGAANRETKLGK  601
             KETWE WWTEYWQATWIPEWEFVNTPPLVKLWYQLEKEPI+GAETFYVDGAANRETKLGK
Sbjct  3039  KETWEAWWTEYWQATWIPEWEFVNTPPLVKLWYQLEKEPIIGAETFYVDGAANRETKLGK  3218

Query  602   AGYVTNRGRQKVVTLTDTTNQKTELQAIYLALQDSGLEVNIVTDSQYALGIIQAQPDQSE  661
             AGYVT+RGRQKVV LTDTTNQKTELQAI+LALQDSGLEVNIVTDSQYALGIIQAQPD+SE
Sbjct  3219  AGYVTDRGRQKVVPLTDTTNQKTELQAIHLALQDSGLEVNIVTDSQYALGIIQAQPDKSE  3398

Query  662   SELVNQIIEQLIKKEKVYLAWVPAHKGIGGNEQVDKLVSAGIRKVLYVDG  711
             SELV+QIIEQLIKKEKVYLAWVPAHKGIGGNEQVDKLVSAGIRKVL++DG
Sbjct  3399  SELVSQIIEQLIKKEKVYLAWVPAHKGIGGNEQVDKLVSAGIRKVLFLDG  3548
```

Range 2: 3168 to 4400

Score:811 bits(2096), Expect:0.0,
Method:Compositional matrix adjust.,
Identities:395/411(96%), Positives:406/411(98%), Gaps:0/411(0%)

```
Query  705   KVLYVDGAANRETKLGKAGYVTNRGRQKVVTLTDTTNQKTELQAIYLALQDSGLEVNIVT  764
             +  YVDGAANRETKLGKAGYVT+RGRQKVV LTDTTNQKTELQAI+LALQDSGLEVNIVT
Sbjct  3168  ETFYVDGAANRETKLGKAGYVTDRGRQKVVPLTDTTNQKTELQAIHLALQDSGLEVNIVT  3347

Query  765   DSQYALGIIQAQPDQSESELVNQIIEQLIKKEKVYLAWVPAHKGIGGNEQVDKLVSAGIR  824
             DSQYALGIIQAQPD+SESELV+QIIEQLIKKEKVYLAWVPAHKGIGGNEQVDKLVSAGIR
Sbjct  3348  DSQYALGIIQAQPDKSESELVSQIIEQLIKKEKVYLAWVPAHKGIGGNEQVDKLVSAGIR  3527

Query  825   KVLFLDGIDKAQDEHEKYHSNWRAMASDFNLPPVVAKEIVASCDKCQLKGEAMHGQVDCS  884
             KVLFLDGIDKAQ+EHEKYHSNWRAMASDFNLPPVVAKEIVASCDKCQLKGEAMHGQVDCS
Sbjct  3528  KVLFLDGIDKAQEEHEKYHSNWRAMASDFNLPPVVAKEIVASCDKCQLKGEAMHGQVDCS  3707

Query  885   PGIWQLDCTHLEGKVILVAVHVASGYIEAEVIPAETGQETAYFLLKLAGRWPVKTIHTDN  944
             PGIWQLDCTHLEGKVILVAVHVASGYIEAEVIPAETGQETAYFLLKLAGRWPVKT+HTDN
Sbjct  3708  PGIWQLDCTHLEGKVILVAVHVASGYIEAEVIPAETGQETAYFLLKLAGRWPVKTVHTDN  3887

Query  945   GSNFTGATVRAACWWAGIKQEFGIPYNPQSQGVVESMNKELKKIIGQVRDQAEHLKTAVQ  1004
             GSNFT  TV+AACWWAGIKQEFGIPYNPQSQGV+ESMNKELKKIIGQVRDQAEHLKTAVQ
Sbjct  3888  GSNFTSTTVKAACWWAGIKQEFGIPYNPQSQGVIESMNKELKKIIGQVRDQAEHLKTAVQ  4067

Query  1005  MAVFIHNFKRKGGIGGYSAGERIVDIIATDIQTKELQKQITKIQNFRVYYRDSRNPLWKG  1064
             MAVFIHNFKRKGGIGGYSAGERIVDIIATDIQTKELQKQITKIQNFRVYYRDSR+P+WKG
Sbjct  4068  MAVFIHNFKRKGGIGGYSAGERIVDIIATDIQTKELQKQITKIQNFRVYYRDSRDPVWKG  4247

Query  1065  PAKLLWKGEGAVVIQDNSDIKVVPRRKAKIIRDYGKQMAGDDCVASRQDED  1115
             PAKLLWKGEGAVVIQDNSDIKVVPRRKAKIIRDYGKQMAGDDCVASRQDED
Sbjct  4248  PAKLLWKGEGAVVIQDNSDIKVVPRRKAKIIRDYGKQMAGDDCVASRQDED  4400
```

>Klebsiella pneumoniae strain ARGID_31501, whole genome shotgun sequence
Sequence ID: NZ_CAKLPD010000006.1 Length: 2655
Range 1: 2 to 1915

Score:1208 bits(3126), Expect:0.0,
Method:Compositional matrix adjust.,
Identities:603/638(95%), Positives:621/638(97%), Gaps:0/638(0%)

```
Query  74    TGADDTVLEEMSLPGRWKPKMIGGIGGFIKVRQYDQILIEICGHKAIGTVLVGPTPVNII  133
             TGAD TVLE+M L G+WKPKMIGGIGGFIKVRQYDQI IEICGHKAIGTVLVGPTPVNII
```

```
Sbjct  2     TGADGTVLEDMQLSGKWKPKMIGGIGGFIKVRQYDQIPIEICGHKAIGTVLVGPTPVNII  181

Query  134   GRNLLTQIGCTLNFPISPIETVPVKLKPGMDGPKVKQWPLTEEKIKALVEICTEMEKEGK  193
             GRNLLTQ+GCTLNFPISPIETVPVKLKPGMDGPKVKQWPLTEEKIKALVEICTEMEKEGK
Sbjct  182   GRNLLTQLGCTLNFPISPIETVPVKLKPGMDGPKVKQWPLTEEKIKALVEICTEMEKEGK  361

Query  194   ISKIGPENPYNTPVFAIKKKDSTKWRKLVDFRELNKRTQDFWEVQLGIPHPAGLKKKKSV  253
             ISKIGPENPYNTP+FAIKKKDSTKWRKLVDFRELNKRTQDFWEVQLGIPHPAGLKKKKSV
Sbjct  362   ISKIGPENPYNTPIFAIKKKDSTKWRKLVDFRELNKRTQDFWEVQLGIPHPAGLKKKKSV  541

Query  254   TVLDVGDAYFSVPLDEDFRKYTAFTIPSINNETPGIRYQYNVLPQGWKGSPAIFQSSMTK  313
             TVLDVGDAYFSVPLD+DFRKYTAFTIPSINNETPG+RYQYNVLPQGWKGSPAIFQSSMTK
Sbjct  542   TVLDVGDAYFSVPLDQDFRKYTAFTIPSINNETPGVRYQYNVLPQGWKGSPAIFQSSMTK  721

Query  314   ILEPFRKQNPDIVIYQYMDDLYVGSDLEIGQHRTKIEELRQHLLRWGLTTPDKKHQKEPP  373
             ILEPFRKQNPDIVIYQYMDDLYVGSDLEIGQHR KIEELRQHLL+WG TTPDKKHQKEPP
Sbjct  722   ILEPFRKQNPDIVIYQYMDDLYVGSDLEIGQHRIKIEELRQHLLKWGFTTPDKKHQKEPP  901

Query  374   FLWMGYELHPDKWTVQPIVLPEKDSWTVNDIQKLVGKLNWASQIYPGIKVRQLCKLLRGT  433
             FLWMGYELHPDKWTVQPI LPEKDSWTVNDIQKLVGKLNWASQIY GIKVRQLCKLLRG
Sbjct  902   FLWMGYELHPDKWTVQPIKLPEKDSWTVNDIQKLVGKLNWASQIYAGIKVRQLCKLLRGA  1081

Query  434   KALTEVIPLTeeaelelaeNREILKEPVHGVYYDPSKDLIAEIQKQGQGQWTYQIYQEPF  493
             KALTEV+PLT+EAELELAENREILKEPVHGVYYDPSK+LIAEIQKQG+GQWTYQIYQE F
Sbjct  1082  KALTEVVPLTKEAELELAENREILKEPVHGVYYDPSKELIAEIQKQGEGQWTYQIYQEQF  1261

Query  494   KNLKTGKYARMRGAHTNDVKQLTEAVQKITTESIVIWGKTPKFKLPIQKetwetwwteYW  553
             KNLKTGKYARMRG HTNDVKQLTEAVQKI TESIVIWGKTPKF+LPI KETWETWWT+YW
Sbjct  1262  KNLKTGKYARMRGTHTNDVKQLTEAVQKIATESIVIWGKTPKFRLPIHKETWETWWTDYW  1441

Query  554   QATWIPEWEFVNTPPLVKLWYQLEKEPIVGAETFYVDGAANRETKLGKAGYVTNRGRQKV  613
             QATWIPEWEFVNTPPLVKLWYQLEKEPI GAETFYVDGAANRETK+GKAGYVTNRGRQKV
Sbjct  1442  QATWIPEWEFVNTPPLVKLWYQLEKEPIEGAETFYVDGAANRETKIGKAGYVTNRGRQKV  1621

Query  614   VTLTDTTNQKTELQAIYLALQDSGLEVNIVTDSQYALGIIQAQPDQSESELVNQIIEQLI  673
             V LTDTTNQKTELQAIYLALQDS LEVNIVTDSQYALGIIQAQPD+SESELVNQIIEQLI
Sbjct  1622  VPLTDTTNQKTELQAIYLALQDSELEVNIVTDSQYALGIIQAQPDKSESELVNQIIEQLI  1801

Query  674   KKEKVYLAWVPAHKGIGGNEQVDKLVSAGIRKVLYVDG  711
             KKEK+YLAWVPAHKGIGGNEQVDKLVS GIRKVL++DG
Sbjct  1802  KKEKIYLAWVPAHKGIGGNEQVDKLVSTGIRKVLFLDG  1915
```

Range 2: 1535 to 2653

Score:726 bits(1873), Expect:0.0,
Method:Compositional matrix adjust.,
Identities:350/373(94%), Positives:364/373(97%), Gaps:0/373(0%)

```
Query  705   KVLYVDGAANRETKLGKAGYVTNRGRQKVVTLTDTTNQKTELQAIYLALQDSGLEVNIVT  764
             +  YVDGAANRETK+GKAGYVTNRGRQKVV LTDTTNQKTELQAIYLALQDS LEVNIVT
Sbjct  1535  ETFYVDGAANRETKIGKAGYVTNRGRQKVVPLTDTTNQKTELQAIYLALQDSELEVNIVT  1714

Query  765   DSQYALGIIQAQPDQSESELVNQIIEQLIKKEKVYLAWVPAHKGIGGNEQVDKLVSAGIR  824
             DSQYALGIIQAQPD+SESELVNQIIEQLIKKEK+YLAWVPAHKGIGGNEQVDKLVS GIR
Sbjct  1715  DSQYALGIIQAQPDKSESELVNQIIEQLIKKEKIYLAWVPAHKGIGGNEQVDKLVSTGIR  1894

Query  825   KVLFLDGIDKAQDEHEKYHSNWRAMASDFNLPPVVAKEIVASCDKCQLKGEAMHGQVDCS  884
             KVLFLDGIDKAQ++H+KYHSNWRAMASDFNLPP+VAKEIVASCDKCQLKGEAMHGQVDCS
Sbjct  1895  KVLFLDGIDKAQEDHDKYHSNWRAMASDFNLPPIVAKEIVASCDKCQLKGEAMHGQVDCS  2074

Query  885   PGIWQLDCTHLEGKVILVAVHVASGYIEAEVIPAETGQETAYFLLKLAGRWPVKTIHTDN  944
             PGIWQLDCTHLEGK+ILVAVHVASGYIEAEVIPAETGQETAYF+LKLAGRWPV TIHTDN
Sbjct  2075  PGIWQLDCTHLEGKIILVAVHVASGYIEAEVIPAETGQETAYFILKLAGRWPVTTIHTDN  2254

Query  945   GSNFTGATVRAACWWAGIKQEFGIPYNPQSQGVVESMNKELKKIIGQVRDQAEHLKTAVQ  1004
             G NF  +TV+AACWWAGIKQEFGIPYNPQSQGVVESMNKELKKIIGQVRDQAEHL+TAVQ
Sbjct  2255  GPNFISSTVKAACWWAGIKQEFGIPYNPQSQGVVESMNKELKKIIGQVRDQAEHLRTAVQ  2434

Query  1005  MAVFIHNFKRKGGIGGYSAGERIVDIIATDIQTKELQKQITKIQNFRVYYRDSRNPLWKG  1064
             MAVFIHNFKRKGGIGGYSAGERIVDIIATDIQTKELQKQITKIQNFRVYYRDSR+PLWKG
Sbjct  2435  MAVFIHNFKRKGGIGGYSAGERIVDIIATDIQTKELQKQITKIQNFRVYYRDSRDPLWKG  2614

Query  1065  PAKLLWKGEGAVV  1077
             PAKLLWKGEGAVV
Sbjct  2615  PAKLLWKGEGAVV  2653
```

>Enterococcus faecium strain A209R1B0C1 NODE_70_length_1819_cov_3.01832_ID_139, whole genome shotgun sequence
Sequence ID: NZ_PGCR01000088.1 Length: 1819
Range 1: 66 to 1817

Score:1050 bits(2716), Expect:0.0,
Method:Compositional matrix adjust.,
Identities:527/590(89%), Positives:559/590(94%), Gaps:6/590(1%)

```
Query  2     QGKAREFSSEQTRANSPTRRELQVWGRDNNSPSEAGADRQGTVSFNFPQVTLWQRPLVTI  61
             QG+AREF SEQTRANSPT  ELQV  R +N  SE G   +GT+  NFPQ+TLWQRPLVTI
Sbjct  66    QGEAREFPSEQTRANSPTSGELQV--RGDNPCSETGV--EGTL--NFPQITLWQRPLVTI  227

Query  62    KIGGQLKEALLDTGADDTVLEEMSLPGRWKPKMIGGIGGFIKVRQYDQILIEICGHKAIG  121
             ++GGQ+KEALLDTGADDTVLE+++LPG+WKPKMIGGIGGFIKVRQYDQI IEICG KAIG
Sbjct  228   RVGGQIKEALLDTGADDTVLEDINLPGKWKPKMIGGIGGFIKVRQYDQIPIEICGKKAIG  407

Query  122   TVLVGPTPVNIIGRNLLTQIGCTLNFPISPIETVPVKLKPGMDGPKVKQWPLTEEKIKAL  181
             TVLVGPTPVNIIGRNLLTQ+GCTLNFPISPIETVPVKLKPGMDGPKVKQWPLTEEKIKAL
Sbjct  408   TVLVGPTPVNIIGRNLLTQLGCTLNFPISPIETVPVKLKPGMDGPKVKQWPLTEEKIKAL  587

Query  182   VEICTEMEKEGKISKIGPENPYNTPVFAIKKKDSTKWRKLVDFRELNKRTQDFWEVQLGI  241
              EIC EMEKEGKI+KIGPENPYNTP+FAIKKKDSTKWRKLVDFRELNKRTQDFWEVQLGI
Sbjct  588   TEICNEMEKEGKITKIGPENPYNTPIFAIKKKDSTKWRKLVDFRELNKRTQDFWEVQLGI  767

Query  242   PHPAGLKKKKSVTVLDVGDAYFSVPLDEDFRKYTAFTIPSINNETPGIRYQYNVLPQGWK  301
             PHPAGLKKKKSVTVLDVGDAYFSVPLD++FRKYTAFTIPSINNETPGIRYQYNVLPQGWK
Sbjct  768   PHPAGLKKKKSVTVLDVGDAYFSVPLDKEFRKYTAFTIPSINNETPGIRYQYNVLPQGWK  947

Query  302   GSPAIFQSSMTKILEPFRKQNPDIVIYQYMDDLYVGSDLEIGQHRTKIEELRQHLLRWGL  361
             GSP+IFQSSM KILEPFR +NP++VIYQYMDDLYVGSDLEIGQHR KIEELR+HLLRWG
Sbjct  948   GSPSIFQSSMIKILEPFRAKNPEMVIYQYMDDLYVGSDLEIGQHRAKIEELREHLLRWGF  1127

Query  362   TTPDKKHQKEPPFLWMGYELHPDKWTVQPIVLPEKDSWTVNDIQKLVGKLNWASQIYPGI  421
             TTPDKKHQKEPPFLWMGYELHPDKWTVQPI LP+KDSWTVNDIQKLVGKLNWASQIYPGI
Sbjct  1128  TTPDKKHQKEPPFLWMGYELHPDKWTVQPIQLPDKDSWTVNDIQKLVGKLNWASQIYPGI  1307

Query  422   KVRQLCKLLRGTKALTEVIPLTeeaelelaeNREILKEPVHGVYYDPSKDLIAEIQKQGQ  481
             KVRQLCKLLRG KALT+++PLTEEAELELAENREILKEPVHGVYYDPSKDLIAEIQKQG
Sbjct  1308  KVRQLCKLLRGAKALTDIVPLTEEAELELAENREILKEPVHGVYYDPSKDLIAEIQKQGH  1487

Query  482   GQWTYQIYQEPFKNLKTGKYARMRGAHTNDVKQLTEAVQKITTESIVIWGKTPKFKLPIQ  541
              QWTYQIYQ+PFKNLKTGKYA+MR AHTNDVKQL EAVQKI  ESIVIWGKTPKF+LPIQ
```

```
Sbjct  1488   DQWTYQIYQDPFKNLKTGKYAKMRTAHTNDVKQLAEAVQKIAIESIVIWGKTPKFRLPIQ  1667

Query  542    KetwetwwteYWQATWIPEWEFVNTPPLVKLWYQLEKEPIVGAETFYVDG  591
              KETWETWWT+YWQATWIP+WEFVNTPPLVKLWYQL KEPI+GAETFYVDG
Sbjct  1668   KETWETWWTDYWQATWIPDWEFVNTPPLVKLWYQLVKEPIIGAETFYVDG  1817



>Listeria monocytogenes strain YA00082404 NODE_73_length_1816_cov_5.664890, whole genome shotgun sequence
Sequence ID: NZ_QEXE01000073.1 Length: 1816
Range 1: 52 to 1815

Score:1038 bits(2685), Expect:0.0,
Method:Compositional matrix adjust.,
Identities:520/592(88%), Positives:553/592(93%), Gaps:11/592(1%)

Query  2      QGKAREFSSEQ-------TRANSPTRRELQVWGRDNNSPSEAGADRQGTVSFNFPQVTLW  54
              QG+AREF SEQ       +RANSPT RELQV  R +N  SE GA+ QGT  FNFPQ+TLW
Sbjct  52     QGEAREFPSEQARTNSPTSRANSPTGRELQV--RGDNPRSETGAEGQGT--FNFPQITLW  219

Query  55     QRPLVTIKIGGQLKEALLDTGADDTVLEEMSLPGRWKPKMIGGIGGFIKVRQYDQILIEI  114
              QRP+VTIK+GGQ+KEALLDTGADDTVLE+M LPGRWKPKMIGGIGGFIKVRQYDQI+IEI
Sbjct  220    QRPMVTIKVGGQVKEALLDTGADDTVLEDMELPGRWKPKMIGGIGGFIKVRQYDQIVIEI  399

Query  115    CGHKAIGTVLVGPTPVNIIGRNLLTQIGCTLNFPISPIETVPVKLKPGMDGPKVKQWPLT  174
              CG KAIGTVLVGPTPVNIIGRN+LTQ+GCTLNFPISPIET+PVKLKPGMDGP+VKQWPLT
Sbjct  400    CGKKAIGTVLVGPTPVNIIGRNMLTQLGCTLNFPISPIETIPVKLKPGMDGPRVKQWPLT  579

Query  175    EEKIKALVEICTEMEKEGKISKIGPENPYNTPVFAIKKKDSTKWRKLVDFRELNKRTQDF  234
              EEKIKAL EIC +MEKEGKI++IGPENPYNTP+FAIK+KD TKWRKLVDFRELNKRTQDF
Sbjct  580    EEKIKALTEICEDMEKEGKITRIGPENPYNTPIFAIKRKDGTKWRKLVDFRELNKRTQDF  759

Query  235    WEVQLGIPHPAGLKKKKSVTVLDVGDAYFSVPLDEDFRKYTAFTIPSINNETPGIRYQYN  294
              WEVQLGIPHPAGLKKKKS+TVLDVGDAYFSVPLDE FRKYTAFTIPSINN TPGIRYQYN
Sbjct  760    WEVQLGIPHPAGLKKKKSMTVLDVGDAYFSVPLDESFRKYTAFTIPSINNSTPGIRYQYN  939

Query  295    VLPQGWKGSPAIFQSSMTKILEPFRKQNPDIVIYQYMDDLYVGSDLEIGQHRTKIEELRQ  354
              VLPQGWKGSPAIFQSSMTKIL+PFR +NPDIVIYQY+DDL VGSDLEIGQHR KIE+LR+
Sbjct  940    VLPQGWKGSPAIFQSSMTKILQPFRDENPDIVIYQYVDDLLVGSDLEIGQHRAKIEQLRE  1119

Query  355    HLLRWGLTTPDKKHQKEPPFLWMGYELHPDKWTVQPIVLPEKDSWTVNDIQKLVGKLNWA  414
              HLLRWGLTTPDKKHQKEPPFLWMGYELHPDKWTVQPI LPEKDSWTVNDIQKLVGKLNWA
Sbjct  1120   HLLRWGLTTPDKKHQKEPPFLWMGYELHPDKWTVQPIQLPEKDSWTVNDIQKLVGKLNWA  1299

Query  415    SQIYPGIKVRQLCKLLRGTKALTEVIPLTeeaelelaeNREILKEPVHGVYYDPSKDLIA  474
              SQIYPGI  +QLC+LLRGTKALT+++PLTEEAELELAENREILKEPVHGVYYDPSKDLIA
Sbjct  1300   SQIYPGICTKQLCRLLRGTKALTDIVPLTEEAELELAENREILKEPVHGVYYDPSKDLIA  1479

Query  475    EIQKQGQGQWTYQIYQEPFKNLKTGKYARMRGAHTNDVKQLTEAVQKITTESIVIWGKTP  534
              EIQKQG  QWTYQIYQEPFKNLKTGKYA+MR AHTNDVKQLTEAVQKI  ESIVIWGKTP
Sbjct  1480   EIQKQGDDQWTYQIYQEPFKNLKTGKYAKMRTAHTNDVKQLTEAVQKIALESIVIWGKTP  1659

Query  535    KFKLPIQKetwetwwteYWQATWIPEWEFVNTPPLVKLWYQLEKEPIVGAET  586
              KF+LPIQKETWE WWT+YWQATWIPEWEFVNTPPLVKLWYQLEKEPI  AET
Sbjct  1660   KFRLPIQKETWEIWWTDYWQATWIPEWEFVNTPPLVKLWYQLEKEPIAEAET  1815



>Escherichia coli strain YA00028237 contig_187, whole genome shotgun sequence
Sequence ID: NZ_NGBP02000187.1 Length: 1828
Range 1: 3 to 1772

Score:1026 bits(2652), Expect:0.0,
Method:Compositional matrix adjust.,
Identities:506/594(85%), Positives:554/594(93%), Gaps:11/594(1%)

Query  2      QGKAREFSSEQTRANSPTRR-------ELQVWGRDNNSPSEAGADRQGTVSFNFPQVTLW  54
              QG+AREF SEQTRANSPT R       ELQV  R +NS SEAG +RQGT  FNFPQ+TLW
Sbjct  1772   QGEAREFPSEQTRANSPTSRANSPTGRELQV--RGDNSHSEAGEERQGT--FNFPQITLW  1605

Query  55     QRPLVTIKIGGQLKEALLDTGADDTVLEEMSLPGRWKPKMIGGIGGFIKVRQYDQILIEI  114
              QRP VT+K+GGQ++EAL+DTGADDTVLE+++LPG+W PK+IGGIGGF+KV+QYD I+IE+
Sbjct  1604   QRPFVTVKVGGQIREALIDTGADDTVLEDINLPGKWTPKIIGGIGGFVKVKQYDHIVIEM  1425

Query  115    CGHKAIGTVLVGPTPVNIIGRNLLTQIGCTLNFPISPIETVPVKLKPGMDGPKVKQWPLT  174
              CG KAIG+VLVGPTP NIIGRN+L+Q+GCTLNFPISPIETVPVKLKPGMDGP+VKQWPLT
Sbjct  1424   CGKKAIGSVLVGPTPANIIGRNMLSQLGCTLNFPISPIETVPVKLKPGMDGPRVKQWPLT  1245

Query  175    EEKIKALVEICTEMEKEGKISKIGPENPYNTPVFAIKKKDSTKWRKLVDFRELNKRTQDF  234
              EEKIKAL EIC EM+KEGKI+KIGPENPYNTP+FAIKKK+STKWRKLVDFRELNKRTQDF
Sbjct  1244   EEKIKALTEICDEMKKEGKITKIGPENPYNTPIFAIKKKNSTKWRKLVDFRELNKRTQDF  1065

Query  235    WEVQLGIPHPAGLKKKKSVTVLDVGDAYFSVPLDEDFRKYTAFTIPSINNETPGIRYQYN  294
              WEVQLGIPHPAGLKK KSVTVLDVGDAYFSVPLDE+FRKYTAFTIPS+NNETPGIRYQYN
Sbjct  1064   WEVQLGIPHPAGLKKNKSVTVLDVGDAYFSVPLDENFRKYTAFTIPSVNNETPGIRYQYN  885

Query  295    VLPQGWKGSPAIFQSSMTKILEPFRKQNPDIVIYQYMDDLYVGSDLEIGQHRTKIEELRQ  354
              VLPQGWKGSPAIFQSSMTKILEPFR QNP+IVIYQY+DDLYVGSDLEIGQHR K+E+LR
Sbjct  884    VLPQGWKGSPAIFQSSMTKILEPFRAQNPEIVIYQYVDDLYVGSDLEIGQHRAKVEKLRD  705

Query  355    HLLRWGLTTPDKKHQKEPPFLWMGYELHPDKWTVQPIVLPEKDSWTVNDIQKLVGKLNWA  414
              HLL+WG TTPDKKHQKEPPFLWMGYELHPDKWTVQ I LPEK+SWTVNDIQKLVGKLNWA
Sbjct  704    HLLKWGFTTPDKKHQKEPPFLWMGYELHPDKWTVQQIQLPEKESWTVNDIQKLVGKLNWA  525

Query  415    SQIYPGIKVRQLCKLLRGTKALTEVIPLTeeaelelaeNREILKEPVHGVYYDPSKDLIA  474
              SQIY GIKV+QLC+LLRG KALT+++PLTEEAELELAENREILKEPVHGVYYDPSKDL+A
Sbjct  524    SQIYAGIKVKQLCRLLRGAKALTDIVPLTEEAELELAENREILKEPVHGVYYDPSKDLVA  345

Query  475    EIQKQGQGQWTYQIYQEPFKNLKTGKYARMRGAHTNDVKQLTEAVQKITTESIVIWGKTP  534
              EIQKQG  QWTYQIYQEPFKNLKTGKYA+ R AHTNDV+QLTE VQKI+ ESIVIWGKTP
Sbjct  344    EIQKQGNDQWTYQIYQEPFKNLKTGKYAKRRTAHTNDVQQLTEVVQKISRESIVIWGKTP  165

Query  535    KFKLPIQKetwetwwteYWQATWIPEWEFVNTPPLVKLWYQLEKEPIVGAETFY  588
              KF+LPIQKETWETWW +YWQATWIP+WEFVNTPPLVKLWYQLEK+PI GAETFY
Sbjct  164    KFRLPIQKETWETWWADYWQATWIPDWEFVNTPPLVKLWYQLEKDPIAGAETFY  3



>Enterococcus faecium strain A206R2B0 NODE_116_length_1761_cov_4.01102_ID_231, whole genome shotgun sequence
Sequence ID: NZ_PGCU01000052.1 Length: 1761
Range 1: 59 to 1759

Score:1020 bits(2637), Expect:0.0,
Method:Compositional matrix adjust.,
Identities:512/576(89%), Positives:540/576(93%), Gaps:9/576(1%)

Query  2      QGKAREFSSEQTRANSPTRRELQVWGRDNNSPSEAGADRQGTVSFNFPQVTLWQRPLVTI  61
              +G+A EF SEQTRANSPT RELQV        EAG   QG  + NFPQ+TLWQRPLV+I
Sbjct  59     EGEAGEFPSEQTRANSPTSRELQV--------REAGNGGQG-ATLNFPQITLWQRPLVSI  211

Query  62     KIGGQLKEALLDTGADDTVLEEMSLPGRWKPKMIGGIGGFIKVRQYDQILIEICGHKAIG  121
```

```
             KI GQ+KEALLDTGADDTVLEE++LPG+WKPKMIGGIGGFIKVRQY++ILIEICG KA G
Sbjct  212   KIEGQVKEALLDTGADDTVLEEINLPGKWKPKMIGGIGGFIKVRQYEEILIEICGKKARG  391

Query  122   TVLVGPTPVNIIGRNLLTQIGCTLNFPISPIETVPVKLKPGMDGPKVKQWPLTEEKIKAL  181
             TVLVGPTPVNIIGRN+LTQ+GCTLNFPISPI TVPVKLKPGMDGPKVKQWPLTEEKIKAL
Sbjct  392   TVLVGPTPVNIIGRNMLTQLGCTLNFPISPIGTVPVKLKPGMDGPKVKQWPLTEEKIKAL  571

Query  182   VEICTEMEKEGKISKIGPENPYNTPVFAIKKKDSTKWRKLVDFRELNKRTQDFWEVQLGI  241
               IC EMEKEGKISKIGPENPYNTPVFAIKKKDSTKWRKLVDFRELNKRTQDFWEVQLGI
Sbjct  572   TAICEEMEKEGKISKIGPENPYNTPVFAIKKKDSTKWRKLVDFRELNKRTQDFWEVQLGI  751

Query  242   PHPAGLKKKKSVTVLDVGDAYFSVPLDEDFRKYTAFTIPSINNETPGIRYQYNVLPQGWK  301
             PHPAGLKKKKSVTVLDVGDAYFSVPLDE FR+YTAFTIPSINNETPGIRYQYNVLPQGWK
Sbjct  752   PHPAGLKKKKSVTVLDVGDAYFSVPLDEGFRRYTAFTIPSINNETPGIRYQYNVLPQGWK  931

Query  302   GSPAIFQSSMTKILEPFRKQNPDIVIYQYMDDLYVGSDLEIGQHRTKIEELRQHLLRWGL  361
             GSPAIFQSSMTKILEPFR +NPDI+IYQYMDDLYVGSDLEIGQHR KIEELR+HLLRWGL
Sbjct  932   GSPAIFQSSMTKILEPFRTRNPDIIIYQYMDDLYVGSDLEIGQHRAKIEELREHLLRWGL  1111

Query  362   TTPDKKHQKEPPFLWMGYELHPDKWTVQPIVLPEKDSWTVNDIQKLVGKLNWASQIYPGI  421
             TTPDKKHQKEPPFLWMGYELHPDKWTVQPIVLPEKDSWTVNDIQKLVGKLNWASQIY GI
Sbjct  1112  TTPDKKHQKEPPFLWMGYELHPDKWTVQPIVLPEKDSWTVNDIQKLVGKLNWASQIYQGI  1291

Query  422   KVRQLCKLLRGTKALTEVIPLTeeaelelaeNREILKEPVHGVYYDPSKDLIAEIQKQGQ  481
             KV+QLCKLLRG KALT+++PLTEEAELELAENREILKEPVHGVYYDPSKDLIAEIQKQG
Sbjct  1292  KVKQLCKLLRGAKALTDIVPLTEEAELELAENREILKEPVHGVYYDPSKDLIAEIQKQGH  1471

Query  482   GQWTYQIYQEPFKNLKTGKYARMRGAHTNDVKQLTEAVQKITTESIVIWGKTPKFKLPIQ  541
              QWTYQIYQEPFKNLKTGKYA+MR AHTND+KQLTEAVQKI  ESIVIWGKTPKF+LPIQ
Sbjct  1472  DQWTYQIYQEPFKNLKTGKYAKMRTAHTNDIKQLTEAVQKIAMESIVIWGKTPKFRLPIQ  1651

Query  542   KetwetwwteYWQATWIPEWEFVNTPPLVKLWYQLE  577
             K TWETWWT+YWQATWIP+WEFVNTPP++KLWYQLE
Sbjct  1652  KGTWETWWTDYWQATWIPDWEFVNTPPIIKLWYQLE  1759
```

>Klebsiella pneumoniae strain NUBRI-K 73, whole genome shotgun sequence
Sequence ID: NZ_SOYT01000073.1 Length: 1717
Range 1: 5 to 1717

Score:1001 bits(2589), Expect:0.0,
Method:Compositional matrix adjust.,
Identities:503/575(87%), Positives:539/575(93%), Gaps:11/575(1%)

```
Query  2     QGKAREFSSEQ-------TRANSPTRRELQVWGRDNNSPSEAGADRQGTVSFNFPQVTLW  54
             QGKAREF  EQ       +RANSPT RELQV  R +NS SEAGA RQGT+  NFPQ+TLW
Sbjct  5     QGKAREFPPEQARTNSPTSRANSPTSRELQV--RGDNSRSEAGAKRQGTL--NFPQITLW  172

Query  55    QRPLVTIKIGGQLKEALLDTGADDTVLEEMSLPGRWKPKMIGGIGGFIKVRQYDQILIEI  114
             QRPLV+IK+GGQ++EALLDTGADDTVLE+++LPG+WKPKMIGGIGGFIKVRQYD+ILIEI
Sbjct  173   QRPLVSIKVGGQIREALLDTGADDTVLEDINLPGKWKPKMIGGIGGFIKVRQYDEILIEI  352

Query  115   CGHKAIGTVLVGPTPVNIIGRNLLTQIGCTLNFPISPIETVPVKLKPGMDGPKVKQWPLT  174
             CG KAIG+VLVGPTPVNIIGRN+LTQ+GCTLNFPISPIETVPV+LKPGMDGPKVKQWPLT
Sbjct  353   CGKKAIGSVLVGPTPVNIIGRNMLTQLGCTLNFPISPIETVPVQLKPGMDGPKVKQWPLT  532

Query  175   EEKIKALVEICTEMEKEGKISKIGPENPYNTPVFAIKKKDSTKWRKLVDFRELNKRTQDF  234
             EEKIKAL  IC EMEKEGKISKIGPENPYNTPVFAIKKKDSTKWRK+VDFRELNKRTQDF
Sbjct  533   EEKIKALTAICEEMEKEGKISKIGPENPYNTPVFAIKKKDSTKWRKVVDFRELNKRTQDF  712

Query  235   WEVQLGIPHPAGLKKKKSVTVLDVGDAYFSVPLDEDFRKYTAFTIPSINNETPGIRYQYN  294
             WEVQLGIPHPAGLKKKKSVTVLDVGDA+FSVPL E FRKYTAFTIPSINNETPGIRYQYN
Sbjct  713   WEVQLGIPHPAGLKKKKSVTVLDVGDAFFSVPLYEGFRKYTAFTIPSINNETPGIRYQYN  892

Query  295   VLPQGWKGSPAIFQSSMTKILEPFRKQNPDIVIYQYMDDLYVGSDLEIGQHRTKIEELRQ  354
             VLPQGWKGSPAIFQ SM KILEPFR +NP+IVIYQY+DDLYV SDLEIGQHR ++EELR+
Sbjct  893   VLPQGWKGSPAIFQCSMIKILEPFRARNPEIVIYQYVDDLYVASDLEIGQHRARLEELRE  1072

Query  355   HLLRWGLTTPDKKHQKEPPFLWMGYELHPDKWTVQPIVLPEKDSWTVNDIQKLVGKLNWA  414
             HLL+WG TTPDKKHQKEPPFLWMGYELHPDKWTVQPI LPEK+SWTVNDIQKLVGKLNWA
Sbjct  1073  HLLKWGFTTPDKKHQKEPPFLWMGYELHPDKWTVQPIQLPEKESWTVNDIQKLVGKLNWA  1252

Query  415   SQIYPGIKVRQLCKLLRGTKALTEVIPLTeeaelelaeNREILKEPVHGVYYDPSKDLIA  474
             SQIYPGIKV+QLCKLLRGTKALT+++PLTEEAELELAENREILKEPVHGVYYDPSKDLIA
Sbjct  1253  SQIYPGIKVKQLCKLLRGTKALTDIVPLTEEAELELAENREILKEPVHGVYYDPSKDLIA  1432

Query  475   EIQKQGQGQWTYQIYQEPFKNLKTGKYARMRGAHTNDVKQLTEAVQKITTESIVIWGKTP  534
             EIQKQG  QWTYQIYQEPFKNLKTGKYA+ + AHTNDVKQLTEAVQKI+ ESIVIWGK P
Sbjct  1433  EIQKQGNDQWTYQIYQEPFKNLKTGKYAKRKSAHTNDVKQLTEAVQKISMESIVIWGKIP  1612

Query  535   KFKLPIQKetwetwwteYWQATWIPEWEFVNTPPL  569
             KF+LPIQKETWETWWT+YWQATWIPEWEFVNTPPL
Sbjct  1613  KFRLPIQKETWETWWTDYWQATWIPEWEFVNTPPL  1717
```

>Enterococcus faecium strain A201R2B0 NODE_60_length_1589_cov_4.20178_ID_119, whole genome shotgun sequence
Sequence ID: NZ_PGCT01000032.1 Length: 1589
Range 1: 2 to 1585

Score:975 bits(2520), Expect:0.0,
Method:Compositional matrix adjust.,
Identities:486/528(92%), Positives:508/528(96%), Gaps:0/528(0%)

```
Query  63    IGGQLKEALLDTGADDTVLEEMSLPGRWKPKMIGGIGGFIKVRQYDQILIEICGHKAIGT  122
             I GQ+KEALL+TG DDTVLEE++LPG+WKPKMIGGIGGFIKVRQY++ILIEICG KA GT
Sbjct  1585  IEGQVKEALLETGEDDTVLEEINLPGKWKPKMIGGIGGFIKVRQYEEILIEICGKKARGT  1406

Query  123   VLVGPTPVNIIGRNLLTQIGCTLNFPISPIETVPVKLKPGMDGPKVKQWPLTEEKIKALV  182
             VLVGPTPVNIIGRN+LTQ+GCTLNFPISPI TVPVKLKPGMDGPKVKQWPLTEEKIKAL
Sbjct  1405  VLVGPTPVNIIGRNMLTQLGCTLNFPISPIGTVPVKLKPGMDGPKVKQWPLTEEKIKALT  1226

Query  183   EICTEMEKEGKISKIGPENPYNTPVFAIKKKDSTKWRKLVDFRELNKRTQDFWEVQLGIP  242
              IC EMEKEGKISKIGPENPYNTPVFAIKKKDSTKWRKLVDFRELNKRTQDFWEVQLGIP
Sbjct  1225  AICEEMEKEGKISKIGPENPYNTPVFAIKKKDSTKWRKLVDFRELNKRTQDFWEVQLGIP  1046

Query  243   HPAGLKKKKSVTVLDVGDAYFSVPLDEDFRKYTAFTIPSINNETPGIRYQYNVLPQGWKG  302
             HPAGLKKKKSVTVLDVGDAYFSVPLDE FR+YTAFTIPSINNETPGIRYQYNVLPQGWKG
Sbjct  1045  HPAGLKKKKSVTVLDVGDAYFSVPLDEGFRRYTAFTIPSINNETPGIRYQYNVLPQGWKG  866

Query  303   SPAIFQSSMTKILEPFRKQNPDIVIYQYMDDLYVGSDLEIGQHRTKIEELRQHLLRWGLT  362
             SPAIFQSSMTKILEPFR +NPDI+IYQYMDDLYVGSDLEIGQHR KIEELR+HLLRWGLT
Sbjct  865   SPAIFQSSMTKILEPFRTRNPDIIIYQYMDDLYVGSDLEIGQHRAKIEELREHLLRWGLT  686

Query  363   TPDKKHQKEPPFLWMGYELHPDKWTVQPIVLPEKDSWTVNDIQKLVGKLNWASQIYPGIK  422
             TPDKKHQKEPPFLWMGYELHPDKWTVQPIVLPEKDSWTVNDIQKLVGKLNWASQIY GIK
Sbjct  685   TPDKKHQKEPPFLWMGYELHPDKWTVQPIVLPEKDSWTVNDIQKLVGKLNWASQIYQGIK  506

Query  423   VRQLCKLLRGTKALTEVIPLTeeaelelaeNREILKEPVHGVYYDPSKDLIAEIQKQGQG  482
             V+QLCKLLRG KALT+++PLTEEAELELAENREILKEPVHGVYYDPSKDLIAEIQKQG
Sbjct  505   VKQLCKLLRGAKALTDIVPLTEEAELELAENREILKEPVHGVYYDPSKDLIAEIQKQGHD  326
```

```
Query  483   QWTYQIYQEPFKNLKTGKYARMRGAHTNDVKQLTEAVQKITTESIVIWGKTPKFKLPIQK  542
             QWTYQIYQEPFKNLKTGKYA+MR AHTND+KQLTEAVQKI  ESIVIWGKTPKF+LPIQK
Sbjct  325   QWTYQIYQEPFKNLKTGKYAKMRTAHTNDIKQLTEAVQKIAMESIVIWGKTPKFRLPIQK  146

Query  543   etwetwwteYWQATWIPEWEFVNTPPLVKLWYQLEKEPIVGAETFYVD  590
              TWETWWT+YWQATWIP+WEFVNTPPLVKLWYQLEKEPI GAETFYVD
Sbjct  145   GTWETWWTDYWQATWIPDWEFVNTPPLVKLWYQLEKEPIAGAETFYVD  2



>Thalassovita mangrovi strain GS-10 Scaffold20_1, whole genome shotgun sequence
Sequence ID: NZ_WWEN01000020.1 Length: 1488
Range 1: 2 to 1486

Score:950 bits(2456), Expect:0.0,
Method:Compositional matrix adjust.,
Identities:480/495(97%), Positives:489/495(98%), Gaps:0/495(0%)

Query  196   KIGPENPYNTPVFAIKKKDSTKWRKLVDFRELNKRTQDFWEVQLGIPHPAGLKKKKSVTV  255
             KIGPENPYNTPVFAIKKKDSTKWRKLVDFRELNKRTQDFWEVQLGIPHPAGLK+KKSVTV
Sbjct  1486  KIGPENPYNTPVFAIKKKDSTKWRKLVDFRELNKRTQDFWEVQLGIPHPAGLKQKKSVTV  1307

Query  256   LDVGDAYFSVPLDEDFRKYTAFTIPSINNETPGIRYQYNVLPQGWKGSPAIFQSSMTKIL  315
             LDVGDAYFSVPLD+DFRKYTAFTIPSINNETPGIRYQYNVLPQGWKGSPAIFQ SMTKIL
Sbjct  1306  LDVGDAYFSVPLDKDFRKYTAFTIPSINNETPGIRYQYNVLPQGWKGSPAIFQCSMTKIL  1127

Query  316   EPFRKQNPDIVIYQYMDDLYVGSDLEIGQHRTKIEELRQHLLRWGLTTPDKKHQKEPPFL  375
             EPFRKQNPDIVIYQYMDDLYVGSDLEIGQHRTKIEELRQHLLRWG TTPDKKHQKEPPFL
Sbjct  1126  EPFRKQNPDIVIYQYMDDLYVGSDLEIGQHRTKIEELRQHLLRWGFTTPDKKHQKEPPFL  947

Query  376   WMGYELHPDKWTVQPIVLPEKDSWTVNDIQKLVGKLNWASQIYPGIKVRQLCKLLRGTKA  435
             WMGYELHPDKWTVQPIVLPEKDSWTVNDIQKLVGKLNWASQIY GIKVRQLCKLLRGTKA
Sbjct  946   WMGYELHPDKWTVQPIVLPEKDSWTVNDIQKLVGKLNWASQIYAGIKVRQLCKLLRGTKA  767

Query  436   LTEVIPLTeeaelelaeNREILKEPVHGVYYDPSKDLIAEIQKQGQGQWTYQIYQEPFKN  495
             LTEV+PLTEEAELELAENREILKEPVHGVYYDPSKDLIAEIQKQGQGQWTYQIYQEPFKN
Sbjct  766   LTEVVPLTEEAELELAENREILKEPVHGVYYDPSKDLIAEIQKQGQGQWTYQIYQEPFKN  587

Query  496   LKTGKYARMRGAHTNDVKQLTEAVQKITTESIVIWGKTPKFKLPIQKetwetwwteYWQA  555
             LKTGKYARM+GAHTNDVKQLTEAVQKI TESIVIWGKTPKFKLPIQKETWE WWTEYWQA
Sbjct  586   LKTGKYARMKGAHTNDVKQLTEAVQKIATESIVIWGKTPKFKLPIQKETWEAWWTEYWQA  407

Query  556   TWIPEWEFVNTPPLVKLWYQLEKEPIVGAETFYVDGAANRETKLGKAGYVTNRGRQKVVT  615
             TWIPEWEFVNTPPLVKLWYQLEKEPI+GAETFYVDGAANRETKLGKAGYVT+RGRQKVV
Sbjct  406   TWIPEWEFVNTPPLVKLWYQLEKEPIIGAETFYVDGAANRETKLGKAGYVTDRGRQKVVP  227

Query  616   LTDTTNQKTELQAIYLALQDSGLEVNIVTDSQYALGIIQAQPDQSESELVNQIIEQLIKK  675
             LTDTTNQKTELQAI+LALQDSGLEVNIVTDSQYALGIIQAQPD+SESELV+QIIEQLIKK
Sbjct  226   LTDTTNQKTELQAIHLALQDSGLEVNIVTDSQYALGIIQAQPDKSESELVSQIIEQLIKK  47

Query  676   EKVYLAWVPAHKGIG  690
             EKVYLAWVPAHKGIG
Sbjct  46    EKVYLAWVPAHKGIG  2



Range 2: 2 to 319

Score:194 bits(493), Expect:1e-48,
Method:Compositional matrix adjust.,
Identities:98/106(92%), Positives:103/106(97%), Gaps:0/106(0%)

Query  705  KVLYVDGAANRETKLGKAGYVTNRGRQKVVTLTDTTNQKTELQAIYLALQDSGLEVNIVT  764
            +  YVDGAANRETKLGKAGYVT+RGRQKVV LTDTTNQKTELQAI+LALQDSGLEVNIVT
Sbjct  319  ETFYVDGAANRETKLGKAGYVTDRGRQKVVPLTDTTNQKTELQAIHLALQDSGLEVNIVT  140

Query  765  DSQYALGIIQAQPDQSESELVNQIIEQLIKKEKVYLAWVPAHKGIG  810
            DSQYALGIIQAQPD+SESELV+QIIEQLIKKEKVYLAWVPAHKGIG
Sbjct  139  DSQYALGIIQAQPDKSESELVSQIIEQLIKKEKVYLAWVPAHKGIG  2



>Helicobacter acinonychis isolate 212_10, whole genome shotgun sequence
Sequence ID: NZ_FZLW01000041.1 Length: 1526
Range 1: 103 to 1521

Score:901 bits(2329), Expect:0.0,
Method:Compositional matrix adjust.,
Identities:437/473(92%), Positives:460/473(97%), Gaps:0/473(0%)

Query  9     SSEQTRANSPTRRELQVWGRDNNSPSEAGADRQGTVSFNFPQVTLWQRPLVTIKIGGQLK  68
             +S  +RANSPT RELQVWG D+NSPSEAGADRQG +SF+FPQ+TLWQRP+VTIKIGGQLK
Sbjct  1521  NSPTSRANSPTSRELQVWGGDSNSPSEAGADRQGDISFSFPQITLWQRPIVTIKIGGQLK  1342

Query  69    EALLDTGADDTVLEEMSLPGRWKPKMIGGIGGFIKVRQYDQILIEICGHKAIGTVLVGPT  128
             EALLDTGADDTVLEEMSLPG+WKPKMIGGIGGFIKVRQY+++ IEICGHKA GTVL+GPT
Sbjct  1341  EALLDTGADDTVLEEMSLPGKWKPKMIGGIGGFIKVRQYEEVPIEICGHKATGTVLIGPT  1162

Query  129   PVNIIGRNLLTQIGCTLNFPISPIETVPVKLKPGMDGPKVKQWPLTEEKIKALVEICTEM  188
             PVNIIGRNLLTQ+GCTLNFPISPIETVPVKLKPGMDGPKVKQWPLTEEKIKAL+EICTEM
Sbjct  1161  PVNIIGRNLLTQLGCTLNFPISPIETVPVKLKPGMDGPKVKQWPLTEEKIKALIEICTEM  982

Query  189   EKEGKISKIGPENPYNTPVFAIKKKDSTKWRKLVDFRELNKRTQDFWEVQLGIPHPAGLK  248
             EKEGKISKIGPENPYNTPVFAIKKKDSTKWRKLVDFRELNK+TQDFWEVQLGIPHP+GLK
Sbjct  981   EKEGKISKIGPENPYNTPVFAIKKKDSTKWRKLVDFRELNKKTQDFWEVQLGIPHPSGLK  802

Query  249   KKKSVTVLDVGDAYFSVPLDEDFRKYTAFTIPSINNETPGIRYQYNVLPQGWKGSPAIFQ  308
             KKKSVTVLDVGDAYFSVPLD+DFRKYTAFTIPS+NNETPGIRYQYNVLPQGWKGSPAIFQ
Sbjct  801   KKKSVTVLDVGDAYFSVPLDKDFRKYTAFTIPSMNNETPGIRYQYNVLPQGWKGSPAIFQ  622

Query  309   SSMTKILEPFRKQNPDIVIYQYMDDLYVGSDLEIGQHRTKIEELRQHLLRWGLTTPDKKH  368
              SMTKILEPFRKQNP+IVIYQYMDDLYVGSDLEI QHRTKIEELRQHLL WG +TPDKKH
Sbjct  621   HSMTKILEPFRKQNPNIVIYQYMDDLYVGSDLEIEQHRTKIEELRQHLLGWGFSTPDKKH  442

Query  369   QKEPPFLWMGYELHPDKWTVQPIVLPEKDSWTVNDIQKLVGKLNWASQIYPGIKVRQLCK  428
             QKEPPFLWMGYELHPDKWTVQPI+LPEKDSWTVNDIQKLVGKLNWASQIYPGIKVRQLCK
Sbjct  441   QKEPPFLWMGYELHPDKWTVQPIMLPEKDSWTVNDIQKLVGKLNWASQIYPGIKVRQLCK  262

Query  429   LLRGTKALTEVIPLTeeaelelaeNREILKEPVHGVYYDPSKDLIAEIQKQGQ  481
             LLRGTKALTEV+PLT EAELELAENREILKEPVHGVYYDPSKDLIAE+QKQ Q
Sbjct  261   LLRGTKALTEVVPLTAEAELELAENREILKEPVHGVYYDPSKDLIAELQKQEQ  103



Range 2: 1 to 102

Score:63.2 bits(152), Expect:3e-05,
Method:Compositional matrix adjust.,
Identities:28/34(82%), Positives:31/34(91%), Gaps:0/34(0%)
```

```
Query  40   RQGTVSFNFPQVTLWQRPLVTIKIGGQLKEALLD  73
            R   +SFNFPQ+TLWQRP+VTIKIGGQLKEALLD
Sbjct  102  RPMDISFNFPQITLWQRPIVTIKIGGQLKEALLD  1
```

>Escherichia coli strain 8374wH11 NODE_36_length_1397_cov_3.54567, whole genome shotgun sequence
Sequence ID: NZ_WSJI01000036.1 Length: 1397
Range 1: 4 to 1395

Score:891 bits(2302), Expect:0.0,
Method:Compositional matrix adjust.,
Identities:436/464(94%), Positives:451/464(97%), Gaps:0/464(0%)

```
Query  21    RELQVWGRDNNSPSEAGADRQGTVSFNFPQVTLWQRPLVTIKIGGQLKEALLDTGADDTV  80
             RELQVWG DNNSPSEAG +RQG VSF+ PQ+TLWQRPLVTIKIGGQLKEALLDTGADDTV
Sbjct  4     RELQVWGGDNNSPSEAGGERQGDVSFSLPQITLWQRPLVTIKIGGQLKEALLDTGADDTV  183

Query  81    LEEMSLPGRWKPKMIGGIGGFIKVRQYDQILIEICGHKAIGTVLVGPTPVNIIGRNLLTQ  140
             LEEM+LPGRWKPKMIGGIGGFIKVRQYDQI IEICGHKAIGTVLVGPTPVNIIGRN+LTQ
Sbjct  184   LEEMNLPGRWKPKMIGGIGGFIKVRQYDQIPIEICGHKAIGTVLVGPTPVNIIGRNMLTQ  363

Query  141   IGCTLNFPISPIETVPVKLKPGMDGPKVKQWPLTEEKIKALVEICTEMEKEGKISKIGPE  200
             +GCTLNFPISPI+TVPVKLKPGMDGPKVKQWPLTEEKIKALVEICTEMEKEGKISKIGPE
Sbjct  364   LGCTLNFPISPIDTVPVKLKPGMDGPKVKQWPLTEEKIKALVEICTEMEKEGKISKIGPE  543

Query  201   NPYNTPVFAIKKKDSTKWRKLVDFRELNKRTQDFWEVQLGIPHPAGLKKKKSVTVLDVGD  260
             NPYNTPVFAIKKKDSTKWRKLVDFRELNKRTQDFWEVQLGIPHPAGLKKKKSVTVLDVGD
Sbjct  544   NPYNTPVFAIKKKDSTKWRKLVDFRELNKRTQDFWEVQLGIPHPAGLKKKKSVTVLDVGD  723

Query  261   AYFSVPLDEDFRKYTAFTIPSINNETPGIRYQYNVLPQGWKGSPAIFQSSMTKILEPFRK  320
             AYFSVPLD++FRKYTAFTIPS NNETPGIRYQYNVLPQGWKGSPAIFQ SMTKILEPFRK
Sbjct  724   AYFSVPLDKEFRKYTAFTIPSTNNETPGIRYQYNVLPQGWKGSPAIFQCSMTKILEPFRK  903

Query  321   QNPDIVIYQYMDDLYVGSDLEIGQHRTKIEELRQHLLRWGLTTPDKKHQKEPPFLWMGYE  380
             QNPDIVIYQY+DDLYVGSDLEI QHRTK+EELRQHLL WG TTPDKKHQKEPPFLWMGYE
Sbjct  904   QNPDIVIYQYVDDLYVGSDLEIEQHRTKVEELRQHLLTWGFTTPDKKHQKEPPFLWMGYE  1083

Query  381   LHPDKWTVQPIVLPEKDSWTVNDIQKLVGKLNWASQIYPGIKVRQLCKLLRGTKALTEVI  440
             LHPDKWTVQPIVLPEKDSWTVNDIQKLVGKLNWASQIYPGIKVRQLCKLLRG KALTEV+
Sbjct  1084  LHPDKWTVQPIVLPEKDSWTVNDIQKLVGKLNWASQIYPGIKVRQLCKLLRGAKALTEVV  1263

Query  441   PLTeeaelelaeNREILKEPVHGVYYDPSKDLIAEIQKQGQGQW  484
              LT+EAELELAENREI+KEPVHGVYYDP+KDLIAEIQKQGQ QW
Sbjct  1264  TLTKEAELELAENREIIKEPVHGVYYDPTKDLIAEIQKQGQDQW  1395
```

>Methylocella silvestris strain TVC NODE_58_length_1453_cov_3.56561, whole genome shotgun sequence
Sequence ID: NZ_PDZR01000058.1 Length: 1453
Range 1: 1 to 1413

Score:887 bits(2291), Expect:0.0,
Method:Compositional matrix adjust.,
Identities:433/471(92%), Positives:455/471(96%), Gaps:0/471(0%)

```
Query  22    ELQVWGRDNNSPSEAGADRQGTVSFNFPQVTLWQRPLVTIKIGGQLKEALLDTGADDTVL  81
             E Q WG DNN SEAGADRQGTVSF+FPQ+TLWQRP+V I+IGGQLKEALLDTGADDTVL
Sbjct  1     EPQAWGGDNNSLSEAGADRQGTVSFSFPQITLWQRPVVEIRIGGQLKEALLDTGADDTVL  180

Query  82    EEMSLPGRWKPKMIGGIGGFIKVRQYDQILIEICGHKAIGTVLVGPTPVNIIGRNLLTQI  141
             E+M LPGRWKPKMIGGIGGFIKVRQYD++ IEICGHKAIGTVLVGPTPVNIIGRNLLTQ+
Sbjct  181   EDMQLPGRWKPKMIGGIGGFIKVRQYDEVPIEICGHKAIGTVLVGPTPVNIIGRNLLTQL  360

Query  142   GCTLNFPISPIETVPVKLKPGMDGPKVKQWPLTEEKIKALVEICTEMEKEGKISKIGPEN  201
             GCTLNFPISPIETVPVKLKPGMDGPKVKQWPLTEEKIKALVEICTEMEKEGKI+KIGPEN
Sbjct  361   GCTLNFPISPIETVPVKLKPGMDGPKVKQWPLTEEKIKALVEICTEMEKEGKITKIGPEN  540

Query  202   PYNTPVFAIKKKDSTKWRKLVDFRELNKRTQDFWEVQLGIPHPAGLKKKKSVTVLDVGDA  261
             PYNTP+FAIKKK+S KWRKLVDFRELNKRTQ+FWEVQLGIPHPAGLKKKKSVTVLDVGDA
Sbjct  541   PYNTPIFAIKKKNSNKWRKLVDFRELNKRTQEFWEVQLGIPHPAGLKKKKSVTVLDVGDA  720

Query  262   YFSVPLDEDFRKYTAFTIPSINNETPGIRYQYNVLPQGWKGSPAIFQSSMTKILEPFRKQ  321
             YFSVPLD DFRKYTAFTIPS NNETPGIRYQYNVLPQGWKGSPAIFQSSMTKILEPFRKQ
Sbjct  721   YFSVPLDPDFRKYTAFTIPSRNNETPGIRYQYNVLPQGWKGSPAIFQSSMTKILEPFRKQ  900

Query  322   NPDIVIYQYMDDLYVGSDLEIGQHRTKIEELRQHLLRWGLTTPDKKHQKEPPFLWMGYEL  381
             NPD+VIYQYMDDLYVGSDLE+ QHRTK+EELR HLL+ G +TPD+KHQKEPPFLWMGYEL
Sbjct  901   NPDMVIYQYMDDLYVGSDLELEQHRTKVEELRAHLLK*GFSTPDQKHQKEPPFLWMGYEL  1080

Query  382   HPDKWTVQPIVLPEKDSWTVNDIQKLVGKLNWASQIYPGIKVRQLCKLLRGTKALTEVIP  441
             HPDKWTVQPIVLPEKDSWTVNDIQKLVGKLNWASQIY GIKV+QLCKLLRGTKALTEV+P
Sbjct  1081  HPDKWTVQPIVLPEKDSWTVNDIQKLVGKLNWASQIYAGIKVKQLCKLLRGTKALTEVVP  1260

Query  442   LTeeaelelaeNREILKEPVHGVYYDPSKDLIAEIQKQGQGQWTYQIYQEP  492
             L+EEAELELAENREILKEPVHGVYYDPSKDL+AEIQKQ QGQWTYQIYQEP
Sbjct  1261  LSEEAELELAENREILKEPVHGVYYDPSKDLVAEIQKQEQGQWTYQIYQEP  1413
```

>Acinetobacter baumannii strain ACM-35 NODE_65_length_1409_cov_2.30031, whole genome shotgun sequence
Sequence ID: NZ_QEHN01000065.1 Length: 1409
Range 1: 3 to 1409

Score:882 bits(2278), Expect:0.0,
Method:Compositional matrix adjust.,
Identities:434/469(93%), Positives:453/469(96%), Gaps:0/469(0%)

```
Query  14    RANSPTRRELQVWGRDNNSPSEAGADRQGTVSFNFPQVTLWQRPLVTIKIGGQLKEALLD  73
             RANSPTRRE Q WG D+NS SEAGADRQGTVSF+FPQ+TLWQRP+VTIK+GGQLKEALLD
Sbjct  3     RANSPTRREPQAWGGDSNSLSEAGADRQGTVSFSFPQITLWQRPVVTIKVGGQLKEALLD  182

Query  74    TGADDTVLEEMSLPGRWKPKMIGGIGGFIKVRQYDQILIEICGHKAIGTVLVGPTPVNII  133
             TGADDTVLEEM+LPGRWKPKMIGGIGGFIKVRQYDQI IEICGHK IGTVL+GPTPVNII
Sbjct  183   TGADDTVLEEMNLPGRWKPKMIGGIGGFIKVRQYDQIPIEICGHKVIGTVLIGPTPVNII  362

Query  134   GRNLLTQIGCTLNFPISPIETVPVKLKPGMDGPKVKQWPLTEEKIKALVEICTEMEKEGK  193
             GRNLLTQ+GCTLNFPISPIETVPVKLKPGMDGPKVKQWPLTEEKIKALVEICTEMEKEGK
Sbjct  363   GRNLLTQLGCTLNFPISPIETVPVKLKPGMDGPKVKQWPLTEEKIKALVEICTEMEKEGK  542

Query  194   ISKIGPENPYNTPVFAIKKKDSTKWRKLVDFRELNKRTQDFWEVQLGIPHPAGLKKKKSV  253
             ISKIGPENPYNTP+FAIKKK+S KWRKLVDFRELNKRTQ+FWEVQLGIPHPAGLKKKKSV
Sbjct  543   ISKIGPENPYNTPIFAIKKKNSDKWRKLVDFRELNKRTQEFWEVQLGIPHPAGLKKKKSV  722

Query  254   TVLDVGDAYFSVPLDEDFRKYTAFTIPSINNETPGIRYQYNVLPQGWKGSPAIFQSSMTK  313
             TVLDVGDAYFSVPLD DFRKYTAFTIPS NNETPGIRYQYNVLPQGWKGSPAIFQSSMTK
Sbjct  723   TVLDVGDAYFSVPLDPDFRKYTAFTIPSRNNETPGIRYQYNVLPQGWKGSPAIFQSSMTK  902

Query  314   ILEPFRKQNPDIVIYQYMDDLYVGSDLEIGQHRTKIEELRQHLLRWGLTTPDKKHQKEPP  373
             ILEPFRKQNPD+VIYQYMDDLYVGSDLEI QHRTK+EELR HLL+WG +TPD+KHQKEPP
```

```
Sbjct  903   ILEPFRKQNPDMVIYQYMDDLYVGSDLEIEQHRTKVEELRAHLLKWGFSTPDQKHQKEPP  1082

Query  374   FLWMGYELHPDKWTVQPIVLPEKDSWTVNDIQKLVGKLNWASQIYPGIKVRQLCKLLRGT  433
             FLWMGYELHPDKWTVQPIVLPEKDSWTVNDIQKLVGKLNWASQIY GIKV+QLCKLLRG
Sbjct  1083  FLWMGYELHPDKWTVQPIVLPEKDSWTVNDIQKLVGKLNWASQIYAGIKVKQLCKLLRGA  1262

Query  434   KALTEVIPLTeeaelelaeNREILKEPVHGVYYDPSKDLIAEIQKQGQG  482
             KALTEV+ LTEEAELELAENREILKEPVHGVYYDPSKDL+AEIQKQG G
Sbjct  1263  KALTEVVTLTEEAELELAENREILKEPVHGVYYDPSKDLVAEIQKQGLG  1409
```

```
>Enterobacter hormaechei strain AUH-ENM6 NODE_27_length_1394_cov_0.980268, whole genome shotgun sequence
Sequence ID: NZ_PYHB01000027.1 Length: 1394
Range 1: 1 to 1392

Score:878 bits(2269), Expect:0.0,
Method:Compositional matrix adjust.,
Identities:430/464(93%), Positives:449/464(96%), Gaps:0/464(0%)

Query  17    SPTRRELQVWGRDNNSPSEAGADRQGTVSFNFPQVTLWQRPLVTIKIGGQLKEALLDTGA  76
             SPTRRE Q WG D NS SEAGADRQGTVSF+FPQ+TLWQRP+VTIK+GGQLKEALLDTGA
Sbjct  1     SPTRREPQAWGGDRNSLSEAGADRQGTVSFSFPQITLWQRPVVTIKVGGQLKEALLDTGA  180

Query  77    DDTVLEEMSLPGRWKPKMIGGIGGFIKVRQYDQILIEICGHKAIGTVLVGPTPVNIIGRN  136
             DDTVLE+M LPGRWKPKMIGGIGGFIKVRQYD++ IEICGHKAIGTVLVGPTPVNIIGRN
Sbjct  181   DDTVLEDMHLPGRWKPKMIGGIGGFIKVRQYDEVPIEICGHKAIGTVLVGPTPVNIIGRN  360

Query  137   LLTQIGCTLNFPISPIETVPVKLKPGMDGPKVKQWPLTEEKIKALVEICTEMEKEGKISK  196
             LLTQ+GCTLNFPISPIETVPVKLKPGMDGPKVKQWPLTEEKIKALVEICTEMEKEGKISK
Sbjct  361   LLTQLGCTLNFPISPIETVPVKLKPGMDGPKVKQWPLTEEKIKALVEICTEMEKEGKISK  540

Query  197   IGPENPYNTPVFAIKKKDSTKWRKLVDFRELNKRTQDFWEVQLGIPHPAGLKKKKSVTVL  256
             IGPENPYNTP+FAIKKK+S KWRKLVDFRELNKRTQ+FWEVQLGIPHPAGLKKKKSVTVL
Sbjct  541   IGPENPYNTPIFAIKKKNSDKWRKLVDFRELNKRTQEFWEVQLGIPHPAGLKKKKSVTVL  720

Query  257   DVGDAYFSVPLDEDFRKYTAFTIPSINNETPGIRYQYNVLPQGWKGSPAIFQSSMTKILE  316
             DVGDAYFSVPLD DFRKYTAFTIPS NNETPGIRYQYNVLPQGWKGSPAIFQSSMTKILE
Sbjct  721   DVGDAYFSVPLDPDFRKYTAFTIPSRNNETPGIRYQYNVLPQGWKGSPAIFQSSMTKILE  900

Query  317   PFRKQNPDIVIYQYMDDLYVGSDLEIGQHRTKIEELRQHLLRWGLTTPDKKHQKEPPFLW  376
             PFRKQNPDIVIYQYMDDLYVGSDLE+ QHRTK+EELR HLL+WG +TPD+KHQKEPPFLW
Sbjct  901   PFRKQNPDIVIYQYMDDLYVGSDLELEQHRTKVEELRAHLLKWGFSTPDQKHQKEPPFLW  1080

Query  377   MGYELHPDKWTVQPIVLPEKDSWTVNDIQKLVGKLNWASQIYPGIKVRQLCKLLRGTKAL  436
             MGYELHPDKWTVQPIVLPEKDSWTVNDIQKLVGKLNWASQIY  IKV+QLCKLLRGTKAL
Sbjct  1081  MGYELHPDKWTVQPIVLPEKDSWTVNDIQKLVGKLNWASQIYAEIKVKQLCKLLRGTKAL  1260

Query  437   TEVIPLTeeaelelaeNREILKEPVHGVYYDPSKDLIAEIQKQG  480
             TEV+PL+EEAELELAENREILKEPVHGVYYDPSKDLIAEIQKQG
Sbjct  1261  TEVVPLSEEAELELAENREILKEPVHGVYYDPSKDLIAEIQKQG  1392
```

```
>Escherichia coli strain 8374wG10 NODE_34_length_1373_cov_2.57865, whole genome shotgun sequence
Sequence ID: NZ_WSJQ01000034.1 Length: 1373
Range 1: 3 to 1370

Score:875 bits(2261), Expect:0.0,
Method:Compositional matrix adjust.,
Identities:430/456(94%), Positives:444/456(97%), Gaps:0/456(0%)

Query  29    DNNSPSEAGADRQGTVSFNFPQVTLWQRPLVTIKIGGQLKEALLDTGADDTVLEEMSLPG  88
             DNNSPSEAG +RQG VSF+ PQ+TLWQRPLVTIKIGGQLKEALLDTGADDTVLEEM+LPG
Sbjct  1370  DNNSPSEAGGERQGDVSFSLPQITLWQRPLVTIKIGGQLKEALLDTGADDTVLEEMNLPG  1191

Query  89    RWKPKMIGGIGGFIKVRQYDQILIEICGHKAIGTVLVGPTPVNIIGRNLLTQIGCTLNFP  148
             RWKPKMIGGIGGFIKVRQYDQI IEICGHK IGTVL+G TPVNIIGRNLLTQ+GCTLNFP
Sbjct  1190  RWKPKMIGGIGGFIKVRQYDQIPIEICGHKVIGTVLIGSTPVNIIGRNLLTQLGCTLNFP  1011

Query  149   ISPIETVPVKLKPGMDGPKVKQWPLTEEKIKALVEICTEMEKEGKISKIGPENPYNTPVF  208
             ISPIETVPVKLKPGMDGPKVKQWPLTEEKIKALVEICTEMEKEGKISKIGPENPYNTPVF
Sbjct  1010  ISPIETVPVKLKPGMDGPKVKQWPLTEEKIKALVEICTEMEKEGKISKIGPENPYNTPVF  831

Query  209   AIKKKDSTKWRKLVDFRELNKRTQDFWEVQLGIPHPAGLKKKKSVTVLDVGDAYFSVPLD  268
             AIKKKDSTKWRKLVDFRELNKRTQDFWEVQLGIPHPAGLKKKKSVTVLDVGDAYFSVPLD
Sbjct  830   AIKKKDSTKWRKLVDFRELNKRTQDFWEVQLGIPHPAGLKKKKSVTVLDVGDAYFSVPLD  651

Query  269   EDFRKYTAFTIPSINNETPGIRYQYNVLPQGWKGSPAIFQSSMTKILEPFRKQNPDIVIY  328
             ++FRKYTAFTIPS NNETPGIRYQYNVLPQGWKGSPAIFQ SMTKILEPFRKQNPDIVIY
Sbjct  650   KEFRKYTAFTIPSTNNETPGIRYQYNVLPQGWKGSPAIFQCSMTKILEPFRKQNPDIVIY  471

Query  329   QYMDDLYVGSDLEIGQHRTKIEELRQHLLRWGLTTPDKKHQKEPPFLWMGYELHPDKWTV  388
             QY+DDLYVGSDLEIGQHR K+EELRQHLL WGLTTPDKKHQKEPPFLWMGYELHPDKWTV
Sbjct  470   QYVDDLYVGSDLEIGQHRAKVEELRQHLLTWGLTTPDKKHQKEPPFLWMGYELHPDKWTV  291

Query  389   QPIVLPEKDSWTVNDIQKLVGKLNWASQIYPGIKVRQLCKLLRGTKALTEVIPLTeeael  448
             QPI LPEKDSWTVNDIQKLVGKLNWASQIYPGIKV+QLCKLLRG KALT+VIPLTEEAEL
Sbjct  290   QPIQLPEKDSWTVNDIQKLVGKLNWASQIYPGIKVKQLCKLLRGAKALTDVIPLTEEAEL  111

Query  449   elaeNREILKEPVHGVYYDPSKDLIAEIQKQGQGQW  484
             ELAENREI+KEPVHGVYYDP+KDLIAEIQKQGQGQW
Sbjct  110   ELAENREIIKEPVHGVYYDPTKDLIAEIQKQGQGQW  3
```

```
>Klebsiella pneumoniae strain P7701 NODE_50_length_1305_cov_481.863, whole genome shotgun sequence
Sequence ID: NZ_JACAWM010000050.1 Length: 1305
Range 1: 21 to 1304

Score:869 bits(2246), Expect:0.0,
Method:Compositional matrix adjust.,
Identities:418/428(98%), Positives:424/428(99%), Gaps:0/428(0%)

Query  16    NSPTRRELQVWGRDNNSPSEAGADRQGTVSFNFPQVTLWQRPLVTIKIGGQLKEALLDTG  75
             NSPTRRELQVWGRDNNS SEAGADRQGTVSF+FPQ+TLWQRPLVTIKIGGQLKEALLDTG
Sbjct  1304  NSPTRRELQVWGRDNNSLSEAGADRQGTVSFSFPQITLWQRPLVTIKIGGQLKEALLDTG  1125

Query  76    ADDTVLEEMSLPGRWKPKMIGGIGGFIKVRQYDQILIEICGHKAIGTVLVGPTPVNIIGR  135
             ADDTVLEEM+LPGRWKPKMIGGIGGFIKVRQYDQILIEICGHKAIGTVLVGPTPVNIIGR
Sbjct  1124  ADDTVLEEMNLPGRWKPKMIGGIGGFIKVRQYDQILIEICGHKAIGTVLVGPTPVNIIGR  945

Query  136   NLLTQIGCTLNFPISPIETVPVKLKPGMDGPKVKQWPLTEEKIKALVEICTEMEKEGKIS  195
             NLLTQIGCTLNFPISPIETVPVKLKPGMDGPKVKQWPLTEEKIKALVEICTEMEKEGKIS
Sbjct  944   NLLTQIGCTLNFPISPIETVPVKLKPGMDGPKVKQWPLTEEKIKALVEICTEMEKEGKIS  765

Query  196   KIGPENPYNTPVFAIKKKDSTKWRKLVDFRELNKRTQDFWEVQLGIPHPAGLKKKKSVTV  255
             KIGPENPYNTPVFAIKKKDSTKWRKLVDFRELNKRTQDFWEVQLGIPHPAGLK+KKSVTV
Sbjct  764   KIGPENPYNTPVFAIKKKDSTKWRKLVDFRELNKRTQDFWEVQLGIPHPAGLKQKKSVTV  585

Query  256   LDVGDAYFSVPLDEDFRKYTAFTIPSINNETPGIRYQYNVLPQGWKGSPAIFQSSMTKIL  315
```

```
             LDVGDAYFSVPLD+DFRKYTAFTIPSINNETPGIRYQYNVLPQGWKGSPAIFQ SMTKIL
Sbjct  584   LDVGDAYFSVPLDKDFRKYTAFTIPSINNETPGIRYQYNVLPQGWKGSPAIFQCSMTKIL  405

Query  316   EPFRKQNPDIVIYQYMDDLYVGSDLEIGQHRTKIEELRQHLLRWGLTTPDKKHQKEPPFL  375
             EPFRKQNPDIVIYQYMDDLYVGSDLEIGQHRTKIEELRQHLLRWG TTPDKKHQKEPPFL
Sbjct  404   EPFRKQNPDIVIYQYMDDLYVGSDLEIGQHRTKIEELRQHLLRWGFTTPDKKHQKEPPFL  225

Query  376   WMGYELHPDKWTVQPIVLPEKDSWTVNDIQKLVGKLNWASQIYPGIKVRQLCKLLRGTKA  435
             WMGYELHPDKWTVQPIVLPEKDSWTVNDIQKLVGKLNWASQIY GIKVRQLCKLLRGTKA
Sbjct  224   WMGYELHPDKWTVQPIVLPEKDSWTVNDIQKLVGKLNWASQIYAGIKVRQLCKLLRGTKA  45

Query  436   LTEVIPLT  443
             LTEV+PLT
Sbjct  44    LTEVVPLT  21
```

>Streptococcus agalactiae strain JH02623 contig_36, whole genome shotgun sequence
Sequence ID: NZ_RPAW01000036.1 Length: 1479
Range 1: 3 to 1448

Score:857 bits(2213), Expect:0.0,
Method:Compositional matrix adjust.,
Identities:427/486(88%), Positives:449/486(92%), Gaps:17/486(3%)

```
Query  1     LQGKAREFSSEQTRANSPTR-------------RELQVWGRDNNSPSEAGADRQGTVSFN  47
             LQG+AR+  SEQTRANSPT              RELQV  R ++  SEAGA+RQGT+  N
Sbjct  1448  LQGEARKLPSEQTRANSPTSSPPEQTRANSPTSRELQV--RGDSPLSEAGAERQGTL--N  1281

Query  48    FPQVTLWQRPLVTIKIGGQLKEALLDTGADDTVLEEMSLPGRWKPKMIGGIGGFIKVRQY  107
             FPQ+TLWQRPLV I++GGQLKEALLDTGADDTVLEE+SLPG+WKPKMIGGIGGFIKVRQY
Sbjct  1280  FPQITLWQRPLVAIRVGGQLKEALLDTGADDTVLEEISLPGKWKPKMIGGIGGFIKVRQY  1101

Query  108   DQILIEICGHKAIGTVLVGPTPVNIIGRNLLTQIGCTLNFPISPIETVPVKLKPGMDGPK  167
             DQI IEICG KAIGTVLVGPTPVNIIGRN+LTQ+GCTLNFPISPIETVPVKLKPGMDGPK
Sbjct  1100  DQIPIEICGKKAIGTVLVGPTPVNIIGRNMLTQLGCTLNFPISPIETVPVKLKPGMDGPK  921

Query  168   VKQWPLTEEKIKALVEICTEMEKEGKISKIGPENPYNTPVFAIKKKDSTKWRKLVDFREL  227
             VKQWPLTEEKIKAL EIC EMEKEGKI+KIGPENPYNTPVFAIKKKDSTKWRKLVDFREL
Sbjct  920   VKQWPLTEEKIKALTEICEEMEKEGKITKIGPENPYNTPVFAIKKKDSTKWRKLVDFREL  741

Query  228   NKRTQDFWEVQLGIPHPAGLKKKKSVTVLDVGDAYFSVPLDEDFRKYTAFTIPSINNETP  287
             NKRTQDFWEVQLGIPHPAGLKK KSVTVLDVGDAYFSVPL E+FRKYTAFTIPS NNETP
Sbjct  740   NKRTQDFWEVQLGIPHPAGLKKNKSVTVLDVGDAYFSVPLHEEFRKYTAFTIPSTNNETP  561

Query  288   GIRYQYNVLPQGWKGSPAIFQSSMTKILEPFRKQNPDIVIYQYMDDLYVGSDLEIGQHRT  347
             GIRYQYNVLPQGWKGSPAIFQ SMTKILEPFR QNP++VIYQYMDDLYVGSDLEIGQHR
Sbjct  560   GIRYQYNVLPQGWKGSPAIFQCSMTKILEPFRTQNPELVIYQYMDDLYVGSDLEIGQHRA  381

Query  348   KIEELRQHLLRWGLTTPDKKHQKEPPFLWMGYELHPDKWTVQPIVLPEKDSWTVNDIQKL  407
             KIEELR+HLL+WG TTPDKKHQKEPPFLWMGYELHPDKWTVQPI LPEKDSWTVNDIQKL
Sbjct  380   KIEELREHLLKWGFTTPDKKHQKEPPFLWMGYELHPDKWTVQPIQLPEKDSWTVNDIQKL  201

Query  408   VGKLNWASQIYPGIKVRQLCKLLRGTKALTEVIPLTeeaelelaeNREILKEPVHGVYYD  467
             VGKLNWASQIYPGIKVRQLCKLLRGTKALT+++PLTEEAELELAENREILKEPVHG YYD
Sbjct  200   VGKLNWASQIYPGIKVRQLCKLLRGTKALTDIVPLTEEAELELAENREILKEPVHGAYYD  21

Query  468   PSKDLI  473
             PSKDLI
Sbjct  20    PSKDLI  3
```

>Escherichia coli strain 8374wB4 NODE_60_length_1385_cov_3.2655, whole genome shotgun sequence
Sequence ID: NZ_WSPX01000060.1 Length: 1385
Range 1: 9 to 1385

Score:855 bits(2210), Expect:0.0,
Method:Compositional matrix adjust.,
Identities:424/459(92%), Positives:441/459(96%), Gaps:1/459(0%)

```
Query  31    NSPSEAGADRQGTV-SFNFPQVTLWQRPLVTIKIGGQLKEALLDTGADDTVLEEMSLPGR  89
             N   +AG    GT+ SFNFPQ+TLWQRPLVTIKIGGQLKEALLDTGADDTVLEE++LPG+
Sbjct  9     NLLPKAGDKGHGTLPSFNFPQITLWQRPLVTIKIGGQLKEALLDTGADDTVLEEINLPGK  188

Query  90    WKPKMIGGIGGFIKVRQYDQILIEICGHKAIGTVLVGPTPVNIIGRNLLTQIGCTLNFPI  149
             WKPKMIGGIGGFIKVRQYDQI IEICG KAIGTVLVGPTPVNIIGRNLLTQ+GCTLNFPI
Sbjct  189   WKPKMIGGIGGFIKVRQYDQIPIEICGKKAIGTVLVGPTPVNIIGRNLLTQLGCTLNFPI  368

Query  150   SPIETVPVKLKPGMDGPKVKQWPLTEEKIKALVEICTEMEKEGKISKIGPENPYNTPVFA  209
             SPIETVPVKLKP MDGPKVKQWPLTEEKIKALVEICTEMEKEGKISKIGPENPYNTPVFA
Sbjct  369   SPIETVPVKLKPRMDGPKVKQWPLTEEKIKALVEICTEMEKEGKISKIGPENPYNTPVFA  548

Query  210   IKKKDSTKWRKLVDFRELNKRTQDFWEVQLGIPHPAGLKKKKSVTVLDVGDAYFSVPLDE  269
             IKKKDSTKWRKLVDFRELNKRTQDFWEVQLGIPHPAGLKKKKSVTVLDVGDAYFSVPLD+
Sbjct  549   IKKKDSTKWRKLVDFRELNKRTQDFWEVQLGIPHPAGLKKKKSVTVLDVGDAYFSVPLDK  728

Query  270   DFRKYTAFTIPSINNETPGIRYQYNVLPQGWKGSPAIFQSSMTKILEPFRKQNPDIVIYQ  329
             +FRKYTAFTIPS NN+TPGIRYQYNVLPQGWKGSPAIFQ SMTKILEPFRKQNPDIVIYQ
Sbjct  729   EFRKYTAFTIPSTNNKTPGIRYQYNVLPQGWKGSPAIFQCSMTKILEPFRKQNPDIVIYQ  908

Query  330   YMDDLYVGSDLEIGQHRTKIEELRQHLLRWGLTTPDKKHQKEPPFLWMGYELHPDKWTVQ  389
             Y+DDLYVGSDLEI QHRTK+EELRQHLL WG TTPDKK+QKEPPFLWMGYELHPDKWTVQ
Sbjct  909   YVDDLYVGSDLEIEQHRTKVEELRQHLLTWGFTTPDKKYQKEPPFLWMGYELHPDKWTVQ  1088

Query  390   PIVLPEKDSWTVNDIQKLVGKLNWASQIYPGIKVRQLCKLLRGTKALTEVIPLTeeaele  449
             PIVLPEKDSWTVNDIQKLVGKLNWASQIYPGIKVRQLCKLLRG KALTEV+ LT+EAELE
Sbjct  1089  PIVLPEKDSWTVNDIQKLVGKLNWASQIYPGIKVRQLCKLLRGAKALTEVVTLTKEAELE  1268

Query  450   laeNREILKEPVHGVYYDPSKDLIAEIQKQGQGQWTYQI  488
             LAENREI+KEPVHGVYYDP+KDLIAEIQKQGQGQWTYQI
Sbjct  1269  LAENREIIKEPVHGVYYDPTKDLIAEIQKQGQGQWTYQI  1385
```

>Klebsiella aerogenes strain AUH-KAM5 NODE_25_length_1427_cov_1.66077, whole genome shotgun sequence
Sequence ID: NZ_PYHJ01000025.1 Length: 1427
Range 1: 45 to 1397

Score:848 bits(2191), Expect:0.0,
Method:Compositional matrix adjust.,
Identities:417/451(92%), Positives:435/451(96%), Gaps:0/451(0%)

```
Query  36    AGADRQGTVSFNFPQVTLWQRPLVTIKIGGQLKEALLDTGADDTVLEEMSLPGRWKPKMI  95
             AG +RQG VSF+ PQ+TLWQRPLVTIKIGGQLKEALLDTGADDTVLEEM+LPGRWKPKMI
Sbjct  45    AGGERQGDVSFSLPQITLWQRPLVTIKIGGQLKEALLDTGADDTVLEEMNLPGRWKPKMI  224

Query  96    GGIGGFIKVRQYDQILIEICGHKAIGTVLVGPTPVNIIGRNLLTQIGCTLNFPISPIETV  155
             GGIGGFIKVRQYDQI IEICGHK IGTVL+G TPVNIIGRNLLTQ+GCTLNFPISPIETV
Sbjct  225   GGIGGFIKVRQYDQIPIEICGHKVIGTVLIGSTPVNIIGRNLLTQLGCTLNFPISPIETV  404
```

```
Query  156   PVKLKPGMDGPKVKQWPLTEEKIKALVEICTEMEKEGKISKIGPENPYNTPVFAIKKKDS  215
             PVKLKPGMDGPKVKQWPLTEEKIKALVEICTEMEKEGKISKIGPENPYNTPVFAIKKKDS
Sbjct  405   PVKLKPGMDGPKVKQWPLTEEKIKALVEICTEMEKEGKISKIGPENPYNTPVFAIKKKDS  584

Query  216   TKWRKLVDFRELNKRTQDFWEVQLGIPHPAGLKKKKSVTVLDVGDAYFSVPLDEDFRKYT  275
             TKWRKLVDFRELNKRTQDFWEVQLGIPHPAGLKKKKSVTVLDVGDAYFSVPLD++FRKYT
Sbjct  585   TKWRKLVDFRELNKRTQDFWEVQLGIPHPAGLKKKKSVTVLDVGDAYFSVPLDKEFRKYT  764

Query  276   AFTIPSINNETPGIRYQYNVLPQGWKGSPAIFQSSMTKILEPFRKQNPDIVIYQYMDDLY  335
             AFTIPS NNETPGIRYQYNVLPQGWKGSPAIFQSSMTKILEPFRKQNPDIVIYQY+DDLY
Sbjct  765   AFTIPSRNNETPGIRYQYNVLPQGWKGSPAIFQSSMTKILEPFRKQNPDIVIYQYIDDLY  944

Query  336   VGSDLEIGQHRTKIEELRQHLLRWGLTTPDKKHQKEPPFLWMGYELHPDKWTVQPIVLPE  395
             VGSDLE+ QHRTK+EELRQHLL WG +TPD+KHQKEPPFLWMGYELHPDKWTVQPIVLPE
Sbjct  945   VGSDLELEQHRTKVEELRQHLLTWGFSTPDQKHQKEPPFLWMGYELHPDKWTVQPIVLPE  1124

Query  396   KDSWTVNDIQKLVGKLNWASQIYPGIKVRQLCKLLRGTKALTEVIPLTeeaelelaeNRE  455
             KDSWTVNDIQKLVGKLNWASQIY GIKV+QLCKLL+GTKALTEV+PL+EEAELELAENRE
Sbjct  1125  KDSWTVNDIQKLVGKLNWASQIYAGIKVKQLCKLLKGTKALTEVVPLSEEAELELAENRE  1304

Query  456   ILKEPVHGVYYDPSKDLIAEIQKQGQGQWTY  486
             ILKEPVHGVYYDPSKDL+AEIQKQ  G   Y
Sbjct  1305  ILKEPVHGVYYDPSKDLVAEIQKQEHGPVPY  1397



>Streptococcus agalactiae strain JH02623 contig_37, whole genome shotgun sequence
Sequence ID: NZ_RPAW01000037.1 Length: 1467
Range 1: 3 to 1439

Score:848 bits(2190), Expect:0.0,
Method:Compositional matrix adjust.,
Identities:419/480(87%), Positives:446/480(92%), Gaps:8/480(1%)

Query  1     LQGKAREFSSEQTRA-------NSPTRRELQVWGRDNNSPSEAGADRQGTVSFNFPQVTL  53
              QGKAREFSSEQ RA       NSPT REL+V   DN  P EA  +  G +S NFPQ+TL
Sbjct  1439  FQGKAREFSSEQARASSPQARANSPTCRELRVRRGDNPLP-EARTEEGGAISSNFPQITL  1263

Query  54    WQRPLVTIKIGGQLKEALLDTGADDTVLEEMSLPGRWKPKMIGGIGGFIKVRQYDQILIE  113
             WQRPLVT++IGGQL EALLDTGADDTVLEE++LPG+WKPKMIGGIGGFIKVRQYDQILIE
Sbjct  1262  WQRPLVTVRIGGQLIEALLDTGADDTVLEEINLPGKWKPKMIGGIGGFIKVRQYDQILIE  1083

Query  114   ICGHKAIGTVLVGPTPVNIIGRNLLTQIGCTLNFPISPIETVPVKLKPGMDGPKVKQWPL  173
             I G KAIGTVLVGPTP+NIIGRN+LTQIGCTLNFPISPIETVPVKLKPGMDGP+VKQWPL
Sbjct  1082  IEGKKAIGTVLVGPTPINIIGRNMLTQIGCTLNFPISPIETVPVKLKPGMDGPRVKQWPL  903

Query  174   TEEKIKALVEICTEMEKEGKISKIGPENPYNTPVFAIKKKDSTKWRKLVDFRELNKRTQD  233
             TEEKIKAL EIC EMEKEGKISKIGPENPYNTP+FAIKKKDST WRKLVDFRELNKRTQD
Sbjct  902   TEEKIKALTEICIEMEKEGKISKIGPENPYNTPIFAIKKKDSTTWRKLVDFRELNKRTQD  723

Query  234   FWEVQLGIPHPAGLKKKKSVTVLDVGDAYFSVPLDEDFRKYTAFTIPSINNETPGIRYQY  293
             FWE+QLGIPHPAGLKKKKS+TVLDVGDAYFS+PLDE+FRKYTAFTIPSINNETPGIRYQY
Sbjct  722   FWEIQLGIPHPAGLKKKKSITVLDVGDAYFSIPLDENFRKYTAFTIPSINNETPGIRYQY  543

Query  294   NVLPQGWKGSPAIFQSSMTKILEPFRKQNPDIVIYQYMDDLYVGSDLEIGQHRTKIEELR  353
             NVLPQGWKGSPAIFQSSMTKILEPFR +NP++VIYQYMDDLYVGSDLEI QHR KIEELR
Sbjct  542   NVLPQGWKGSPAIFQSSMTKILEPFRAKNPEMVIYQYMDDLYVGSDLEIKQHRAKIEELR  363

Query  354   QHLLRWGLTTPDKKHQKEPPFLWMGYELHPDKWTVQPIVLPEKDSWTVNDIQKLVGKLNW  413
             +HLL+WG TTPDKKHQKEPPFLWMGYELHPDKWTVQPI LP+KDSWTVNDIQKLVGKLNW
Sbjct  362   EHLLKWGFTTPDKKHQKEPPFLWMGYELHPDKWTVQPIQLPDKDSWTVNDIQKLVGKLNW  183

Query  414   ASQIYPGIKVRQLCKLLRGTKALTEVIPLTeeaelelaeNREILKEPVHGVYYDPSKDLI  473
             ASQIYPGIKV+QLCKLLRG KALT++IPLTEEAELELAENREILK+PVHGVYYDPSKDLI
Sbjct  182   ASQIYPGIKVKQLCKLLRGAKALTDIIPLTEEAELELAENREILKQPVHGVYYDPSKDLI  3



>Salirhabdus euzebyi strain Marseille-Q1438, whole genome shotgun sequence
Sequence ID: NZ_CADDWK010000035.1 Length: 1279
Range 1: 9 to 1277

Score:842 bits(2175), Expect:0.0,
Method:Compositional matrix adjust.,
Identities:404/423(96%), Positives:416/423(98%), Gaps:0/423(0%)

Query  21    RELQVWGRDNNSPSEAGADRQGTVSFNFPQVTLWQRPLVTIKIGGQLKEALLDTGADDTV  80
             R+ QV GRDNNS SEAGADRQG VSF+FPQ+TLWQRPLV IKIGGQLKEALLDTGADDTV
Sbjct  1277  RKPQVGGRDNNSLSEAGADRQGHVSFSFPQITLWQRPLVAIKIGGQLKEALLDTGADDTV  1098

Query  81    LEEMSLPGRWKPKMIGGIGGFIKVRQYDQILIEICGHKAIGTVLVGPTPVNIIGRNLLTQ  140
             LEEM+LPGRWKPKMIGGIGGFIKVRQYDQI IEICGHKAIGTVL+GPTPVNIIGRNLLTQ
Sbjct  1097  LEEMNLPGRWKPKMIGGIGGFIKVRQYDQIPIEICGHKAIGTVLIGPTPVNIIGRNLLTQ  918

Query  141   IGCTLNFPISPIETVPVKLKPGMDGPKVKQWPLTEEKIKALVEICTEMEKEGKISKIGPE  200
             IGCTLNFPISPIETVPVKLKPGMDGPKVKQWPLTEEKIKALVEICTEMEKEGKIS++GPE
Sbjct  917   IGCTLNFPISPIETVPVKLKPGMDGPKVKQWPLTEEKIKALVEICTEMEKEGKISRVGPE  738

Query  201   NPYNTPVFAIKKKDSTKWRKLVDFRELNKRTQDFWEVQLGIPHPAGLKKKKSVTVLDVGD  260
             NPYNTP+FAIKKKDSTKWRKLVDFRELNK+TQDFWEVQLGIPHPAGLKKKKSVTVLDVGD
Sbjct  737   NPYNTPIFAIKKKDSTKWRKLVDFRELNKKTQDFWEVQLGIPHPAGLKKKKSVTVLDVGD  558

Query  261   AYFSVPLDEDFRKYTAFTIPSINNETPGIRYQYNVLPQGWKGSPAIFQSSMTKILEPFRK  320
             AYFSVPLD+DFRKYTAFTIPS+NNETPGIRYQYNVLPQGWKGSPAIFQSSMTKILEPFRK
Sbjct  557   AYFSVPLDKDFRKYTAFTIPSVNNETPGIRYQYNVLPQGWKGSPAIFQSSMTKILEPFRK  378

Query  321   QNPDIVIYQYMDDLYVGSDLEIGQHRTKIEELRQHLLRWGLTTPDKKHQKEPPFLWMGYE  380
             QNPDIVIYQYMDDLYVGSDLEIGQHRTKIEELRQHLLRWG TTPDKKHQKEPPFLWMGYE
Sbjct  377   QNPDIVIYQYMDDLYVGSDLEIGQHRTKIEELRQHLLRWGFTTPDKKHQKEPPFLWMGYE  198

Query  381   LHPDKWTVQPIVLPEKDSWTVNDIQKLVGKLNWASQIYPGIKVRQLCKLLRGTKALTEVI  440
             LHPDKWTVQPIVLPEKDSWTVNDIQKLVGKLNWASQIYPGIKV+QLCKLLRGTKALTEVI
Sbjct  197   LHPDKWTVQPIVLPEKDSWTVNDIQKLVGKLNWASQIYPGIKVKQLCKLLRGTKALTEVI  18

Query  441   PLT  443
             PLT
Sbjct  17    PLT  9



>Escherichia coli strain 8374wB11 NODE_34_length_1310_cov_3.21048, whole genome shotgun sequence
Sequence ID: NZ_WSJO01000034.1 Length: 1310
Range 1: 1 to 1308

Score:825 bits(2132), Expect:0.0,
Method:Compositional matrix adjust.,
Identities:404/436(93%), Positives:420/436(96%), Gaps:0/436(0%)

Query  53    LWQRPLVTIKIGGQLKEALLDTGADDTVLEEMSLPGRWKPKMIGGIGGFIKVRQYDQILI  112
             LW RP V ++IGGQ +EALLD GADD VLEE++LPG+WKPKMIGGIGGFIKVRQYDQI I
Sbjct  1308  LWHRPFVPVRIGGQPREALLDNGADDPVLEEINLPGKWKPKMIGGIGGFIKVRQYDQIPI  1129
```

```
Query  113   EICGHKAIGTVLVGPTPVNIIGRNLLTQIGCTLNFPISPIETVPVKLKPGMDGPKVKQWP  172
             EICGHK IGTVL+G TPVNIIGRNLLTQ+GCTLNFPISPIETVPVKLKPGMDGPKVKQWP
Sbjct  1128  EICGHKVIGTVLIGSTPVNIIGRNLLTQLGCTLNFPISPIETVPVKLKPGMDGPKVKQWP  949

Query  173   LTEEKIKALVEICTEMEKEGKISKIGPENPYNTPVFAIKKKDSTKWRKLVDFRELNKRTQ  232
             LTEEKIKALVEICTEMEKEGKISKIGPENPYNTPVFAIKKKDSTKWRKLVDFRELNKRTQ
Sbjct  948   LTEEKIKALVEICTEMEKEGKISKIGPENPYNTPVFAIKKKDSTKWRKLVDFRELNKRTQ  769

Query  233   DFWEVQLGIPHPAGLKKKKSVTVLDVGDAYFSVPLDEDFRKYTAFTIPSINNETPGIRYQ  292
             DFWEVQLGIPHPAGLKKKKSVTVLDVGDAYFSVPLD++FRKYTAFTIPS NNETPGIRYQ
Sbjct  768   DFWEVQLGIPHPAGLKKKKSVTVLDVGDAYFSVPLDKEFRKYTAFTIPSTNNETPGIRYQ  589

Query  293   YNVLPQGWKGSPAIFQSSMTKILEPFRKQNPDIVIYQYMDDLYVGSDLEIGQHRTKIEEL  352
             YNVLPQGWKGSPAIFQ SMTKILEPFRKQNPDIVIYQY+DDLYVGSDLEI QHRTK+EEL
Sbjct  588   YNVLPQGWKGSPAIFQCSMTKILEPFRKQNPDIVIYQYVDDLYVGSDLEIEQHRTKVEEL  409

Query  353   RQHLLRWGLTTPDKKHQKEPPFLWMGYELHPDKWTVQPIVLPEKDSWTVNDIQKLVGKLN  412
             RQHLL WG TTPDKKHQKEPPFLWMGYELHPDKWTVQPIVLPEKDSWTVNDIQKLVGKLN
Sbjct  408   RQHLLTWGFTTPDKKHQKEPPFLWMGYELHPDKWTVQPIVLPEKDSWTVNDIQKLVGKLN  229

Query  413   WASQIYPGIKVRQLCKLLRGTKALTEVIPLTeeaelelaeNREILKEPVHGVYYDPSKDL  472
             WASQIYPGIKVRQLCKLLRG KALTEV+ LT+EAELELAENREI+KEPVHGVYYDP+KDL
Sbjct  228   WASQIYPGIKVRQLCKLLRGAKALTEVVTLTKEAELELAENREIIKEPVHGVYYDPTKDL  49

Query  473   IAEIQKQGQGQWTYQI  488
             IAEIQKQGQGQWTYQI
Sbjct  48    IAEIQKQGQGQWTYQI  1
```

>Mycobacterium tuberculosis variant bovis strain CSURQ1134, whole genome shotgun sequence
Sequence ID: NZ_CAJEUT010000109.1 Length: 1697
Range 1: 85 to 1410

Score:812 bits(2098), Expect:0.0,
Method:Compositional matrix adjust.,
Identities:392/443(88%), Positives:416/443(93%), Gaps:2/443(0%)

```
Query  2     QGKAREFSSEQTRANSPTRRELQVWGRDNNSPSEAGADRQGTVS-FNFPQVTLWQRPLVT  60
             QG+AR+FSS QT  NSPT REL   GRD   P EAG   QG +S FNFPQ+TLWQRPLVT
Sbjct  1410  QGEARKFSSGQTGTNSPTSRELWNGGRDTILP-EAGTGGQGPISSFNFPQITLWQRPLVT  1234

Query  61    IKIGGQLKEALLDTGADDTVLEEMSLPGRWKPKMIGGIGGFIKVRQYDQILIEICGHKAI  120
             ++I GQL EALLDTGADDTVLEE++LPGRWKPKMIGGIGGFIKVRQYDQI IEICG KAI
Sbjct  1233  VRIEGQLIEALLDTGADDTVLEEINLPGRWKPKMIGGIGGFIKVRQYDQISIEICGKKAI  1054

Query  121   GTVLVGPTPVNIIGRNLLTQIGCTLNFPISPIETVPVKLKPGMDGPKVKQWPLTEEKIKA  180
             GTVLVGPTPVNIIGRNLLTQIGCTLNFPISPIETVPVKLKPGMDGPKVKQWPLTEEKIKA
Sbjct  1053  GTVLVGPTPVNIIGRNLLTQIGCTLNFPISPIETVPVKLKPGMDGPKVKQWPLTEEKIKA  874

Query  181   LVEICTEMEKEGKISKIGPENPYNTPVFAIKKKDSTKWRKLVDFRELNKRTQDFWEVQLG  240
             L EICTEMEKEGKISKIGPENPYNTPVFAI+KKDSTKWRKLVDFRELNKRTQDFWE+QLG
Sbjct  873   LTEICTEMEKEGKISKIGPENPYNTPVFAIRKKDSTKWRKLVDFRELNKRTQDFWEIQLG  694

Query  241   IPHPAGLKKKKSVTVLDVGDAYFSVPLDEDFRKYTAFTIPSINNETPGIRYQYNVLPQGW  300
             IPHPAGLKKKKSVTVLDVGDAYFSVPLD+DFRKYTAFTIPSINNETPGIRYQYNVLPQGW
Sbjct  693   IPHPAGLKKKKSVTVLDVGDAYFSVPLDKDFRKYTAFTIPSINNETPGIRYQYNVLPQGW  514

Query  301   KGSPAIFQSSMTKILEPFRKQNPDIVIYQYMDDLYVGSDLEIGQHRTKIEELRQHLLRWG  360
             KGSPAIFQ+SMTKILEPFR +NP++VIYQYMDDLY+GSDLEIGQHRTKIEELR HLLRWG
Sbjct  513   KGSPAIFQASMTKILEPFRTKNPELVIYQYMDDLYIGSDLEIGQHRTKIEELRAHLLRWG  334

Query  361   LTTPDKKHQKEPPFLWMGYELHPDKWTVQPIVLPEKDSWTVNDIQKLVGKLNWASQIYPG  420
              TTPDKKHQKEPPFLWMGYELHPDKWTVQP+ LP+K++WTVNDIQKLVGKLNWASQIY G
Sbjct  333   FTTPDKKHQKEPPFLWMGYELHPDKWTVQPVQLPDKENWTVNDIQKLVGKLNWASQIYAG  154

Query  421   IKVRQLCKLLRGTKALTEVIPLT  443
             IKV+QLCKLLRG KALT+++PLT
Sbjct  153   IKVKQLCKLLRGAKALTDIVPLT  85
```

Range 2: 85 to 1410

Score:812 bits(2098), Expect:0.0,
Method:Compositional matrix adjust.,
Identities:392/443(88%), Positives:416/443(93%), Gaps:2/443(0%)

```
Query  2     QGKAREFSSEQTRANSPTRRELQVWGRDNNSPSEAGADRQGTVS-FNFPQVTLWQRPLVT  60
             QG+AR+FSS QT  NSPT REL   GRD   P EAG   QG +S FNFPQ+TLWQRPLVT
Sbjct  1410  QGEARKFSSGQTGTNSPTSRELWNGGRDTILP-EAGTGGQGPISSFNFPQITLWQRPLVT  1234

Query  61    IKIGGQLKEALLDTGADDTVLEEMSLPGRWKPKMIGGIGGFIKVRQYDQILIEICGHKAI  120
             ++I GQL EALLDTGADDTVLEE++LPGRWKPKMIGGIGGFIKVRQYDQI IEICG KAI
Sbjct  1233  VRIEGQLIEALLDTGADDTVLEEINLPGRWKPKMIGGIGGFIKVRQYDQISIEICGKKAI  1054

Query  121   GTVLVGPTPVNIIGRNLLTQIGCTLNFPISPIETVPVKLKPGMDGPKVKQWPLTEEKIKA  180
             GTVLVGPTPVNIIGRNLLTQIGCTLNFPISPIETVPVKLKPGMDGPKVKQWPLTEEKIKA
Sbjct  1053  GTVLVGPTPVNIIGRNLLTQIGCTLNFPISPIETVPVKLKPGMDGPKVKQWPLTEEKIKA  874

Query  181   LVEICTEMEKEGKISKIGPENPYNTPVFAIKKKDSTKWRKLVDFRELNKRTQDFWEVQLG  240
             L EICTEMEKEGKISKIGPENPYNTPVFAI+KKDSTKWRKLVDFRELNKRTQDFWE+QLG
Sbjct  873   LTEICTEMEKEGKISKIGPENPYNTPVFAIRKKDSTKWRKLVDFRELNKRTQDFWEIQLG  694

Query  241   IPHPAGLKKKKSVTVLDVGDAYFSVPLDEDFRKYTAFTIPSINNETPGIRYQYNVLPQGW  300
             IPHPAGLKKKKSVTVLDVGDAYFSVPLD+DFRKYTAFTIPSINNETPGIRYQYNVLPQGW
Sbjct  693   IPHPAGLKKKKSVTVLDVGDAYFSVPLDKDFRKYTAFTIPSINNETPGIRYQYNVLPQGW  514

Query  301   KGSPAIFQSSMTKILEPFRKQNPDIVIYQYMDDLYVGSDLEIGQHRTKIEELRQHLLRWG  360
             KGSPAIFQ+SMTKILEPFR +NP++VIYQYMDDLY+GSDLEIGQHRTKIEELR HLLRWG
Sbjct  513   KGSPAIFQASMTKILEPFRTKNPELVIYQYMDDLYIGSDLEIGQHRTKIEELRAHLLRWG  334

Query  361   LTTPDKKHQKEPPFLWMGYELHPDKWTVQPIVLPEKDSWTVNDIQKLVGKLNWASQIYPG  420
              TTPDKKHQKEPPFLWMGYELHPDKWTVQP+ LP+K++WTVNDIQKLVGKLNWASQIY G
Sbjct  333   FTTPDKKHQKEPPFLWMGYELHPDKWTVQPVQLPDKENWTVNDIQKLVGKLNWASQIYAG  154

Query  421   IKVRQLCKLLRGTKALTEVIPLT  443
             IKV+QLCKLLRG KALT+++PLT
Sbjct  153   IKVKQLCKLLRGAKALTDIVPLT  85
```

>Mycobacterium tuberculosis variant bovis strain CSURQ1132, whole genome shotgun sequence
Sequence ID: NZ_CAJEUK010000155.1 Length: 1524
Range 1: 123 to 1448

Score:811 bits(2096), Expect:0.0,
Method:Compositional matrix adjust.,
Identities:392/443(88%), Positives:416/443(93%), Gaps:2/443(0%)

```
Query  2     QGKAREFSSEQTRANSPTRRELQVWGRDNNSPSEAGADRQGTVS-FNFPQVTLWQRPLVT  60
             QG+AR+FSS QT  NSPT REL   GRD   P EAG   QG +S FNFPQ+TLWQRPLVT
Sbjct  1448  QGEARKFSSGQTGTNSPTSRELWNGGRDTILP-EAGTGGQGPISSFNFPQITLWQRPLVT  1272

Query  61    IKIGGQLKEALLDTGADDTVLEEMSLPGRWKPKMIGGIGGFIKVRQYDQILIEICGHKAI  120
             ++I GQL EALLDTGADDTVLEE++LPGRWKPKMIGGIGGFIKVRQYDQI IEICG KAI
Sbjct  1271  VRIEGQLIEALLDTGADDTVLEEINLPGRWKPKMIGGIGGFIKVRQYDQISIEICGKKAI  1092

Query  121   GTVLVGPTPVNIIGRNLLTQIGCTLNFPISPIETVPVKLKPGMDGPKVKQWPLTEEKIKA  180
             GTVLVGPTPVNIIGRNLLTQIGCTLNFPISPIETVPVKLKPGMDGPKVKQWPLTEEKIKA
Sbjct  1091  GTVLVGPTPVNIIGRNLLTQIGCTLNFPISPIETVPVKLKPGMDGPKVKQWPLTEEKIKA  912

Query  181   LVEICTEMEKEGKISKIGPENPYNTPVFAIKKKDSTKWRKLVDFRELNKRTQDFWEVQLG  240
             L EICTEMEKEGKISKIGPENPYNTPVFAI+KKDSTKWRKLVDFRELNKRTQDFWE+QLG
Sbjct  911   LTEICTEMEKEGKISKIGPENPYNTPVFAIRKKDSTKWRKLVDFRELNKRTQDFWEIQLG  732

Query  241   IPHPAGLKKKKSVTVLDVGDAYFSVPLDEDFRKYTAFTIPSINNETPGIRYQYNVLPQGW  300
             IPHPAGLKKKKSVTVLDVGDAYFSVPLD+DFRKYTAFTIPSINNETPGIRYQYNVLPQGW
Sbjct  731   IPHPAGLKKKKSVTVLDVGDAYFSVPLDKDFRKYTAFTIPSINNETPGIRYQYNVLPQGW  552

Query  301   KGSPAIFQSSMTKILEPFRKQNPDIVIYQYMDDLYVGSDLEIGQHRTKIEELRQHLLRWG  360
             KGSPAIFQ+SMTKILEPFR +NP++VIYQYMDDLY+GSDLEIGQHRTKIEELR HLLRWG
Sbjct  551   KGSPAIFQASMTKILEPFRTKNPELVIYQYMDDLYIGSDLEIGQHRTKIEELRAHLLRWG  372

Query  361   LTTPDKKHQKEPPFLWMGYELHPDKWTVQPIVLPEKDSWTVNDIQKLVGKLNWASQIYPG  420
              TTPDKKHQKEPPFLWMGYELHPDKWTVQP+ LP+K++WTVNDIQKLVGKLNWASQIY G
Sbjct  371   FTTPDKKHQKEPPFLWMGYELHPDKWTVQPVQLPDKENWTVNDIQKLVGKLNWASQIYAG  192

Query  421   IKVRQLCKLLRGTKALTEVIPLT  443
             IKV+QLCKLLRG KALT+++PLT
Sbjct  191   IKVKQLCKLLRGAKALTDIVPLT  123
```

Range 2: 123 to 1448

Score:811 bits(2096), Expect:0.0,
Method:Compositional matrix adjust.,
Identities:392/443(88%), Positives:416/443(93%), Gaps:2/443(0%)

```
Query  2     QGKAREFSSEQTRANSPTRRELQVWGRDNNSPSEAGADRQGTVS-FNFPQVTLWQRPLVT  60
             QG+AR+FSS QT  NSPT REL   GRD   P EAG   QG +S FNFPQ+TLWQRPLVT
Sbjct  1448  QGEARKFSSGQTGTNSPTSRELWNGGRDTILP-EAGTGGQGPISSFNFPQITLWQRPLVT  1272

Query  61    IKIGGQLKEALLDTGADDTVLEEMSLPGRWKPKMIGGIGGFIKVRQYDQILIEICGHKAI  120
             ++I GQL EALLDTGADDTVLEE++LPGRWKPKMIGGIGGFIKVRQYDQI IEICG KAI
Sbjct  1271  VRIEGQLIEALLDTGADDTVLEEINLPGRWKPKMIGGIGGFIKVRQYDQISIEICGKKAI  1092

Query  121   GTVLVGPTPVNIIGRNLLTQIGCTLNFPISPIETVPVKLKPGMDGPKVKQWPLTEEKIKA  180
             GTVLVGPTPVNIIGRNLLTQIGCTLNFPISPIETVPVKLKPGMDGPKVKQWPLTEEKIKA
Sbjct  1091  GTVLVGPTPVNIIGRNLLTQIGCTLNFPISPIETVPVKLKPGMDGPKVKQWPLTEEKIKA  912

Query  181   LVEICTEMEKEGKISKIGPENPYNTPVFAIKKKDSTKWRKLVDFRELNKRTQDFWEVQLG  240
             L EICTEMEKEGKISKIGPENPYNTPVFAI+KKDSTKWRKLVDFRELNKRTQDFWE+QLG
Sbjct  911   LTEICTEMEKEGKISKIGPENPYNTPVFAIRKKDSTKWRKLVDFRELNKRTQDFWEIQLG  732

Query  241   IPHPAGLKKKKSVTVLDVGDAYFSVPLDEDFRKYTAFTIPSINNETPGIRYQYNVLPQGW  300
             IPHPAGLKKKKSVTVLDVGDAYFSVPLD+DFRKYTAFTIPSINNETPGIRYQYNVLPQGW
Sbjct  731   IPHPAGLKKKKSVTVLDVGDAYFSVPLDKDFRKYTAFTIPSINNETPGIRYQYNVLPQGW  552

Query  301   KGSPAIFQSSMTKILEPFRKQNPDIVIYQYMDDLYVGSDLEIGQHRTKIEELRQHLLRWG  360
             KGSPAIFQ+SMTKILEPFR +NP++VIYQYMDDLY+GSDLEIGQHRTKIEELR HLLRWG
Sbjct  551   KGSPAIFQASMTKILEPFRTKNPELVIYQYMDDLYIGSDLEIGQHRTKIEELRAHLLRWG  372

Query  361   LTTPDKKHQKEPPFLWMGYELHPDKWTVQPIVLPEKDSWTVNDIQKLVGKLNWASQIYPG  420
              TTPDKKHQKEPPFLWMGYELHPDKWTVQP+ LP+K++WTVNDIQKLVGKLNWASQIY G
Sbjct  371   FTTPDKKHQKEPPFLWMGYELHPDKWTVQPVQLPDKENWTVNDIQKLVGKLNWASQIYAG  192

Query  421   IKVRQLCKLLRGTKALTEVIPLT  443
             IKV+QLCKLLRG KALT+++PLT
Sbjct  191   IKVKQLCKLLRGAKALTDIVPLT  123
```

>Acinetobacter baumannii strain ACM-5 NODE_55_length_1274_cov_1.90584, whole genome shotgun sequence
Sequence ID: NZ_PYCX01000055.1 Length: 1274
Range 1: 2 to 1273

Score:811 bits(2094), Expect:0.0,
Method:Compositional matrix adjust.,
Identities:396/424(93%), Positives:411/424(96%), Gaps:0/424(0%)

```
Query  63    IGGQLKEALLDTGADDTVLEEMSLPGRWKPKMIGGIGGFIKVRQYDQILIEICGHKAIGT  122
             + GQLKEALLDTGA+DTVLE+M LPGRWKPKMIGGI GFIKVRQYD++ IEICGHKAIGT
Sbjct  1273  VEGQLKEALLDTGAEDTVLEDMQLPGRWKPKMIGGIRGFIKVRQYDEVPIEICGHKAIGT  1094

Query  123   VLVGPTPVNIIGRNLLTQIGCTLNFPISPIETVPVKLKPGMDGPKVKQWPLTEEKIKALV  182
             VL+GPTPVNIIGRNLLTQ+GCTLNFPISPIETVPVKLKPGMDGPKVKQWPLTEEKIKALV
Sbjct  1093  VLIGPTPVNIIGRNLLTQLGCTLNFPISPIETVPVKLKPGMDGPKVKQWPLTEEKIKALV  914

Query  183   EICTEMEKEGKISKIGPENPYNTPVFAIKKKDSTKWRKLVDFRELNKRTQDFWEVQLGIP  242
             EICTEMEKEGKISKIGPENPYNTP+FAIKKK+S KWRKLVDFRELNKRTQ+FWEVQLGIP
Sbjct  913   EICTEMEKEGKISKIGPENPYNTPIFAIKKKNSDKWRKLVDFRELNKRTQEFWEVQLGIP  734

Query  243   HPAGLKKKKSVTVLDVGDAYFSVPLDEDFRKYTAFTIPSINNETPGIRYQYNVLPQGWKG  302
             HPAGLKKKKSVTVLDVGDAYFSVPLD DFRKYTAFTIPS NNETPGIRYQYNVLPQGWKG
Sbjct  733   HPAGLKKKKSVTVLDVGDAYFSVPLDPDFRKYTAFTIPSTNNETPGIRYQYNVLPQGWKG  554

Query  303   SPAIFQSSMTKILEPFRKQNPDIVIYQYMDDLYVGSDLEIGQHRTKIEELRQHLLRWGLT  362
             SPAIFQ SMTKILEPFRKQNPDIVIYQYMDDLYVGSDLEI QHRTK+EELRQHLL WG T
Sbjct  553   SPAIFQCSMTKILEPFRKQNPDIVIYQYMDDLYVGSDLEIEQHRTKVEELRQHLLTWGFT  374

Query  363   TPDKKHQKEPPFLWMGYELHPDKWTVQPIVLPEKDSWTVNDIQKLVGKLNWASQIYPGIK  422
             TPDKKHQKEPPFLWMGYELHPDKWTVQPIVLPEKDSWTVNDIQKLVGKLNWASQIYPGIK
Sbjct  373   TPDKKHQKEPPFLWMGYELHPDKWTVQPIVLPEKDSWTVNDIQKLVGKLNWASQIYPGIK  194

Query  423   VRQLCKLLRGTKALTEVIPLTeeaelelaeNREILKEPVHGVYYDPSKDLIAEIQKQGQG  482
             VRQLCKLLRG KALTEV+ LT+EAELELAENREI+KEPVHGVYYDP+KDLIAEIQKQGQG
Sbjct  193   VRQLCKLLRGAKALTEVVTLTKEAELELAENREIIKEPVHGVYYDPTKDLIAEIQKQGQG  14

Query  483   QWTY  486
             QWTY
Sbjct  13    QWTY  2
```

>Mycobacterium tuberculosis variant bovis strain CSURQ1140, whole genome shotgun sequence
Sequence ID: NZ_CAJEUP010000157.1 Length: 1286
Range 1: 2 to 1285

Score:788 bits(2036), Expect:0.0,

```
Method:Compositional matrix adjust.,
Identities:380/429(89%), Positives:403/429(93%), Gaps:2/429(0%)

Query  13    TRANSPTRRELQVWGRDNNSPSEAGADRQGTVS-FNFPQVTLWQRPLVTIKIGGQLKEAL  71
             T  NSPT REL   GRD   P EAG   QG +S FNFPQ+TLWQRPLVT++I GQL EAL
Sbjct  1285  TGTNSPTSRELWNGGRDTILP-EAGTGGQGPISSFNFPQITLWQRPLVTVRIEGQLIEAL  1109

Query  72    LDTGADDTVLEEMSLPGRWKPKMIGGIGGFIKVRQYDQILIEICGHKAIGTVLVGPTPVN  131
             LDTGADDTVLEE++LPGRWKPKMIGGIGGFIKVRQYDQI IEICG KAIGTVLVGPTPVN
Sbjct  1108  LDTGADDTVLEEINLPGRWKPKMIGGIGGFIKVRQYDQISIEICGKKAIGTVLVGPTPVN  929

Query  132   IIGRNLLTQIGCTLNFPISPIETVPVKLKPGMDGPKVKQWPLTEEKIKALVEICTEMEKE  191
             IIGRNLLTQIGCTLNFPISPIETVPVKLKPGMDGPK+KQWPLTEEKIKAL EICTEMEKE
Sbjct  928   IIGRNLLTQIGCTLNFPISPIETVPVKLKPGMDGPKIKQWPLTEEKIKALTEICTEMEKE  749

Query  192   GKISKIGPENPYNTPVFAIKKKDSTKWRKLVDFRELNKRTQDFWEVQLGIPHPAGLKKKK  251
             GKISKIGPENPYNTPVFAI+KKDSTKWRKLVDFRELNKRTQDFWE+QLGIPHPAGLKKKK
Sbjct  748   GKISKIGPENPYNTPVFAIRKKDSTKWRKLVDFRELNKRTQDFWEIQLGIPHPAGLKKKK  569

Query  252   SVTVLDVGDAYFSVPLDEDFRKYTAFTIPSINNETPGIRYQYNVLPQGWKGSPAIFQSSM  311
             SVTVLDVGDAYFSVPLD+DFRKYTAFTIPSINNETPGIRYQYNVLPQGWKGSPAIFQ+SM
Sbjct  568   SVTVLDVGDAYFSVPLDKDFRKYTAFTIPSINNETPGIRYQYNVLPQGWKGSPAIFQASM  389

Query  312   TKILEPFRKQNPDIVIYQYMDDLYVGSDLEIGQHRTKIEELRQHLLRWGLTTPDKKHQKE  371
             TKILEPFR +NP++VIYQYMDDLY+GSDLEIGQHRTKIEELR HLLRWG TTPDKKHQKE
Sbjct  388   TKILEPFRTKNPELVIYQYMDDLYIGSDLEIGQHRTKIEELRAHLLRWGFTTPDKKHQKE  209

Query  372   PPFLWMGYELHPDKWTVQPIVLPEKDSWTVNDIQKLVGKLNWASQIYPGIKVRQLCKLLR  431
             PPFLWMGYELHPDKWTVQP+ LP+K++WTVNDIQKLVGKLNWASQIY GIKV+QLCKLLR
Sbjct  208   PPFLWMGYELHPDKWTVQPVQLPDKENWTVNDIQKLVGKLNWASQIYAGIKVKQLCKLLR  29

Query  432   GTKALTEVI  440
             G KALT+++
Sbjct  28    GAKALTDIV  2



Range 2: 2 to 1285

Score:788 bits(2036), Expect:0.0,
Method:Compositional matrix adjust.,
Identities:380/429(89%), Positives:403/429(93%), Gaps:2/429(0%)

Query  13    TRANSPTRRELQVWGRDNNSPSEAGADRQGTVS-FNFPQVTLWQRPLVTIKIGGQLKEAL  71
             T  NSPT REL   GRD   P EAG   QG +S FNFPQ+TLWQRPLVT++I GQL EAL
Sbjct  1285  TGTNSPTSRELWNGGRDTILP-EAGTGGQGPISSFNFPQITLWQRPLVTVRIEGQLIEAL  1109

Query  72    LDTGADDTVLEEMSLPGRWKPKMIGGIGGFIKVRQYDQILIEICGHKAIGTVLVGPTPVN  131
             LDTGADDTVLEE++LPGRWKPKMIGGIGGFIKVRQYDQI IEICG KAIGTVLVGPTPVN
Sbjct  1108  LDTGADDTVLEEINLPGRWKPKMIGGIGGFIKVRQYDQISIEICGKKAIGTVLVGPTPVN  929

Query  132   IIGRNLLTQIGCTLNFPISPIETVPVKLKPGMDGPKVKQWPLTEEKIKALVEICTEMEKE  191
             IIGRNLLTQIGCTLNFPISPIETVPVKLKPGMDGPK+KQWPLTEEKIKAL EICTEMEKE
Sbjct  928   IIGRNLLTQIGCTLNFPISPIETVPVKLKPGMDGPKIKQWPLTEEKIKALTEICTEMEKE  749

Query  192   GKISKIGPENPYNTPVFAIKKKDSTKWRKLVDFRELNKRTQDFWEVQLGIPHPAGLKKKK  251
             GKISKIGPENPYNTPVFAI+KKDSTKWRKLVDFRELNKRTQDFWE+QLGIPHPAGLKKKK
Sbjct  748   GKISKIGPENPYNTPVFAIRKKDSTKWRKLVDFRELNKRTQDFWEIQLGIPHPAGLKKKK  569

Query  252   SVTVLDVGDAYFSVPLDEDFRKYTAFTIPSINNETPGIRYQYNVLPQGWKGSPAIFQSSM  311
             SVTVLDVGDAYFSVPLD+DFRKYTAFTIPSINNETPGIRYQYNVLPQGWKGSPAIFQ+SM
Sbjct  568   SVTVLDVGDAYFSVPLDKDFRKYTAFTIPSINNETPGIRYQYNVLPQGWKGSPAIFQASM  389

Query  312   TKILEPFRKQNPDIVIYQYMDDLYVGSDLEIGQHRTKIEELRQHLLRWGLTTPDKKHQKE  371
             TKILEPFR +NP++VIYQYMDDLY+GSDLEIGQHRTKIEELR HLLRWG TTPDKKHQKE
Sbjct  388   TKILEPFRTKNPELVIYQYMDDLYIGSDLEIGQHRTKIEELRAHLLRWGFTTPDKKHQKE  209

Query  372   PPFLWMGYELHPDKWTVQPIVLPEKDSWTVNDIQKLVGKLNWASQIYPGIKVRQLCKLLR  431
             PPFLWMGYELHPDKWTVQP+ LP+K++WTVNDIQKLVGKLNWASQIY GIKV+QLCKLLR
Sbjct  208   PPFLWMGYELHPDKWTVQPVQLPDKENWTVNDIQKLVGKLNWASQIYAGIKVKQLCKLLR  29

Query  432   GTKALTEVI  440
             G KALT+++
Sbjct  28    GAKALTDIV  2



>Escherichia coli strain VRES0669, whole genome shotgun sequence
Sequence ID: NZ_UINV01000086.1 Length: 5544
Range 1: 2544 to 3776

Score:777 bits(2006), Expect:0.0,
Method:Compositional matrix adjust.,
Identities:372/411(91%), Positives:394/411(95%), Gaps:0/411(0%)

Query  705   KVLYVDGAANRETKLGKAGYVTNRGRQKVVTLTDTTNQKTELQAIYLALQDSGLEVNIVT  764
             +  YVDGAANRETK GKAGYVT+RGRQKV++ T+TTNQK ELQAI LALQDSG EVNIVT
Sbjct  3776  ETFYVDGAANRETKAGKAGYVTDRGRQKVISFTETTNQKAELQAIQLALQDSGSEVNIVT  3597

Query  765   DSQYALGIIQAQPDQSESELVNQIIEQLIKKEKVYLAWVPAHKGIGGNEQVDKLVSAGIR  824
             DSQYALGII A PD+SESELVN+IIE LI+KE+VYL+WVPAHKGIGGNEQVDKLVS GIR
Sbjct  3596  DSQYALGIIHALPDKSESELVNKIIELLIQKERVYLSWVPAHKGIGGNEQVDKLVSTGIR  3417

Query  825   KVLFLDGIDKAQDEHEKYHSNWRAMASDFNLPPVVAKEIVASCDKCQLKGEAMHGQVDCS  884
             KVLFLDGIDKAQ+EHEKYHSNWRAMAS+FNLPP+VAKEIVASCDKCQLKGEAMHGQVDCS
Sbjct  3416  KVLFLDGIDKAQEEHEKYHSNWRAMASEFNLPPIVAKEIVASCDKCQLKGEAMHGQVDCS  3237

Query  885   PGIWQLDCTHLEGKVILVAVHVASGYIEAEVIPAETGQETAYFLLKLAGRWPVKTIHTDN  944
             PGIWQLDCTHLEGKVILVAVHVASGY+EAEVIPAETGQETAYF+LKLAGRWPVKTIHTDN
Sbjct  3236  PGIWQLDCTHLEGKVILVAVHVASGYMEAEVIPAETGQETAYFILKLAGRWPVKTIHTDN  3057

Query  945   GSNFTGATVRAACWWAGIKQEFGIPYNPQSQGVVESMNKELKKIIGQVRDQAEHLKTAVQ  1004
             GSNFT   V+AACWWAGI+QEFGIPYNPQSQGVVESMNKELKKII Q+RDQAEHLKTAVQ
Sbjct  3056  GSNFTSTVVKAACWWAGIQQEFGIPYNPQSQGVVESMNKELKKIIEQLRDQAEHLKTAVQ  2877

Query  1005  MAVFIHNFKRKGGIGGYSAGERIVDIIATDIQTKELQKQITKIQNFRVYYRDSRNPLWKG  1064
             MAVF+HNFKRKGGIGGYSAGERI+DIIATDIQTKELQKQITKIQNFRVYYRDSR+P+WKG
Sbjct  2876  MAVFVHNFKRKGGIGGYSAGERIIDIIATDIQTKELQKQITKIQNFRVYYRDSRDPIWKG  2697

Query  1065  PAKLLWKGEGAVVIQDNSDIKVVPRRKAKIIRDYGKQMAGDDCVASRQDED  1115
             PAKLLWKGEGAVVIQDNSDIKVVPRRKAKIIRDYGKQMAG DCVA RQDED
Sbjct  2696  PAKLLWKGEGAVVIQDNSDIKVVPRRKAKIIRDYGKQMAGTDCVAGRQDED  2544



Range 2: 4301 to 5194

Score:571 bits(1472), Expect:2e-174,
Method:Compositional matrix adjust.,
Identities:271/298(91%), Positives:288/298(96%), Gaps:0/298(0%)
```

```
Query  42    GTVSFNFPQVTLWQRPLVTIKIGGQLKEALLDTGADDTVLEEMSLPGRWKPKMIGGIGGF  101
             G  + N PQ+TLWQRPLV+IK+GGQ+KEALLDTGADDTVLEE++LPG+WKPKMIGGIGGF
Sbjct  5194  GQGTLNLPQITLWQRPLVSIKVGGQIKEALLDTGADDTVLEEINLPGKWKPKMIGGIGGF  5015

Query  102   IKVRQYDQILIEICGHKAIGTVLVGPTPVNIIGRNLLTQIGCTLNFPISPIETVPVKLKP  161
             IKVRQYDQI IEICG KAIGTVLVGPTPVNIIGRN+LTQ+GCTLNFPISPIETVPVKLKP
Sbjct  5014  IKVRQYDQIPIEICGKKAIGTVLVGPTPVNIIGRNMLTQLGCTLNFPISPIETVPVKLKP  4835

Query  162   GMDGPKVKQWPLTEEKIKALVEICTEMEKEGKISKIGPENPYNTPVFAIKKKDSTKWRKL  221
             GMDGPKVKQWPLTEEKIKAL EIC EMEKEGKI+KIGP+NPYNTPVFAIKKKDSTKWRKL
Sbjct  4834  GMDGPKVKQWPLTEEKIKALTEICEEMEKEGKITKIGPDNPYNTPVFAIKKKDSTKWRKL  4655

Query  222   VDFRELNKRTQDFWEVQLGIPHPAGLKKKKSVTVLDVGDAYFSVPLDEDFRKYTAFTIPS  281
             +DFRELNKRTQDFWEVQLGIPHPAGLKKKKSVTVLDVGDAYFSVPLDE+FRKYTAFTIPS
Sbjct  4654  IDFRELNKRTQDFWEVQLGIPHPAGLKKKKSVTVLDVGDAYFSVPLDENFRKYTAFTIPS  4475

Query  282   INNETPGIRYQYNVLPQGWKGSPAIFQSSMTKILEPFRKQNPDIVIYQYMDDLYVGSD  339
              NNETPGIRYQYNVLPQGWKGSPAIFQSSMT+ILEPFR QNP+IVIYQY+DDLYVGSD
Sbjct  4474  TNNETPGIRYQYNVLPQGWKGSPAIFQSSMTRILEPFRAQNPEIVIYQYVDDLYVGSD  4301



Range 3: 3396 to 4259

Score:482 bits(1241), Expect:7e-142,
Method:Compositional matrix adjust.,
Identities:253/288(88%), Positives:271/288(94%), Gaps:0/288(0%)

Query  424   RQLCKLLRGTKALTEVIPLTeeaelelaeNREILKEPVHGVYYDPSKDLIAEIQKQGQGQ  483
             RQLC+LLRGTKALT+++PLTEEAELELAENREILKEPVHGVYYDPSKDLIAEIQKQG  Q
Sbjct  4259  RQLCRLLRGTKALTDIVPLTEEAELELAENREILKEPVHGVYYDPSKDLIAEIQKQGNDQ  4080

Query  484   WTYQIYQEPFKNLKTGKYARMRGAHTNDVKQLTEAVQKITTESIVIWGKTPKFKLPIQKe  543
             WTYQIYQEPFKNLKTGKYARMR AHTNDVKQLTEAVQKI  ESIVIWGK PKF+LPIQKE
Sbjct  4079  WTYQIYQEPFKNLKTGKYARMRTAHTNDVKQLTEAVQKIVMESIVIWGKPPKFRLPIQKE  3900

Query  544   twetwwteYWQATWIPEWEFVNTPPLVKLWYQLEKEPIVGAETFYVDGAANRETKLGKAG  603
             TWETWW++YWQATWIPEWEFVNTPPLVKLWYQLEKEPI G ETFYVDGAANRETK GKAG
Sbjct  3899  TWETWWSDYWQATWIPEWEFVNTPPLVKLWYQLEKEPIEGVETFYVDGAANRETKAGKAG  3720

Query  604   YVTNRGRQKVVTLTDTTNQKTELQAIYLALQDSGLEVNIVTDSQYALGIIQAQPDQSESE  663
             YVT+RGRQKV++ T+TTNQK ELQAI LALQDSG EVNIVTDSQYALGII A PD+SESE
Sbjct  3719  YVTDRGRQKVISFTETTNQKAELQAIQLALQDSGSEVNIVTDSQYALGIIHALPDKSESE  3540

Query  664   LVNQIIEQLIKKEKVYLAWVPAHKGIGGNEQVDKLVSAGIRKVLYVDG  711
             LVN+IIE LI+KE+VYL+WVPAHKGIGGNEQVDKLVS GIRKVL++DG
Sbjct  3539  LVNKIIELLIQKERVYLSWVPAHKGIGGNEQVDKLVSTGIRKVLFLDG  3396



>Burkholderia pseudomallei strain 2604STDY5518971 ERS240105SCcontig000097, whole genome shotgun sequence
Sequence ID: NZ_UVFZ02000097.1 Length: 1345
Range 1: 208 to 1344

Score:733 bits(1892), Expect:0.0,
Method:Compositional matrix adjust.,
Identities:348/379(92%), Positives:370/379(97%), Gaps:0/379(0%)

Query  708   YVDGAANRETKLGKAGYVTNRGRQKVVTLTDTTNQKTELQAIYLALQDSGLEVNIVTDSQ  767
             YVDGAA+RETKLGKAGYVT+RGRQKVV+LT+TTNQKTEL AI+LALQDSG EVNIVTDSQ
Sbjct  208   YVDGAASRETKLGKAGYVTDRGRQKVVSLTETTNQKTELHAIHLALQDSGSEVNIVTDSQ  387

Query  768   YALGIIQAQPDQSESELVNQIIEQLIKKEKVYLAWVPAHKGIGGNEQVDKLVSAGIRKVL  827
             YALGIIQAQPD SESE+VNQIIE+LIKKE+VYL+WVPAHKGIGGNEQVDKLVS+GIRKVL
Sbjct  388   YALGIIQAQPDMSESEVVNQIIEELIKKERVYLSWVPAHKGIGGNEQVDKLVSSGIRKVL  567

Query  828   FLDGIDKAQDEHEKYHSNWRAMASDFNLPPVVAKEIVASCDKCQLKGEAMHGQVDCSPGI  887
             FLDGIDKAQ++HE+YHSNWR MASDFNLPP+VAKEIVA+CDKCQLKGEAMHGQVDCSPGI
Sbjct  568   FLDGIDKAQEDHERYHSNWRTMASDFNLPPIVAKEIVANCDKCQLKGEAMHGQVDCSPGI  747

Query  888   WQLDCTHLEGKVILVAVHVASGYIEAEVIPAETGQETAYFLLKLAGRWPVKTIHTDNGSN  947
             WQLDCTHLEGKVILVAVHVASGYIEAEVIPAETGQETAYFLLKLAGRWPVK IHTDNGSN
Sbjct  748   WQLDCTHLEGKVILVAVHVASGYIEAEVIPAETGQETAYFLLKLAGRWPVKVIHTDNGSN  927

Query  948   FTGATVRAACWWAGIKQEFGIPYNPQSQGVVESMNKELKKIIGQVRDQAEHLKTAVQMAV  1007
             FT   V+AACWWA ++QEFGIPYNPQSQGVVESMNKELKKIIGQVR+QAEHLKTAVQMAV
Sbjct  928   FTSTAVKAACWWANVQQEFGIPYNPQSQGVVESMNKELKKIIGQVREQAEHLKTAVQMAV  1107

Query  1008  FIHNFKRKGGIGGYSAGERIVDIIATDIQTKELQKQITKIQNFRVYYRDSRNPLWKGPAK  1067
             FIHNFKRKGGIGGYSAGERI+DIIATDIQTKELQKQITKIQNFRVYYRDSR+P+WKGPAK
Sbjct  1108  FIHNFKRKGGIGGYSAGERIIDIIATDIQTKELQKQITKIQNFRVYYRDSRDPIWKGPAK  1287

Query  1068  LLWKGEGAVVIQDNSDIKV  1086
             LLWKGEGAVVIQDNSDIKV
Sbjct  1288  LLWKGEGAVVIQDNSDIKV  1344



Range 2: 205 to 579

Score:223 bits(569), Expect:7e-81,
Method:Compositional matrix adjust.,
Identities:109/125(87%), Positives:122/125(97%), Gaps:0/125(0%)

Query  587  FYVDGAANRETKLGKAGYVTNRGRQKVVTLTDTTNQKTELQAIYLALQDSGLEVNIVTDS  646
            +YVDGAA+RETKLGKAGYVT+RGRQKVV+LT+TTNQKTEL AI+LALQDSG EVNIVTDS
Sbjct  205  YYVDGAASRETKLGKAGYVTDRGRQKVVSLTETTNQKTELHAIHLALQDSGSEVNIVTDS  384

Query  647  QYALGIIQAQPDQSESELVNQIIEQLIKKEKVYLAWVPAHKGIGGNEQVDKLVSAGIRKV  706
            QYALGIIQAQPD SESE+VNQIIE+LIKKE+VYL+WVPAHKGIGGNEQVDKLVS+GIRKV
Sbjct  385  QYALGIIQAQPDMSESEVVNQIIEELIKKERVYLSWVPAHKGIGGNEQVDKLVSSGIRKV  564

Query  707  LYVDG  711
            L++DG
Sbjct  565  LFLDG  579



Range 3: 2 to 208

Score:113 bits(283), Expect:7e-81,
Method:Compositional matrix adjust.,
Identities:61/69(88%), Positives:67/69(97%), Gaps:0/69(0%)

Query  498  TGKYARMRGAHTNDVKQLTEAVQKITTESIVIWGKTPKFKLPIQKetwetwwteYWQATW  557
            TGKYA++RGAHTNDVK LTEAVQKI TESIVIWGKTP+F+LPIQ+ETW+TWWTEYWQATW
Sbjct  2    TGKYAKIRGAHTNDVKLLTEAVQKIATESIVIWGKTPRFRLPIQRETWDTWWTEYWQATW  181

Query  558  IPEWEFVNT  566
            IPEWEFVNT
```

```
Sbjct  182  IPEWEFVNT  208
```

>Providencia rettgeri strain AFE652_S1 38_contig_28, whole genome shotgun sequence
Sequence ID: NZ_SHDK01000028.1 Length: 1127
Range 1: 1 to 1107

Score:724 bits(1868), Expect:0.0,
Method:Compositional matrix adjust.,
Identities:343/369(93%), Positives:360/369(97%), Gaps:0/369(0%)

```
Query  49    PQVTLWQRPLVTIKIGGQLKEALLDTGADDTVLEEMSLPGRWKPKMIGGIGGFIKVRQYD  108
             PQ+TLWQRPLV+IK+GGQ+KEALLDTGADDTVLEE++LPG+WKPKMIGGIGGFIKVRQYD
Sbjct  1107  PQITLWQRPLVSIKVGGQIKEALLDTGADDTVLEEINLPGKWKPKMIGGIGGFIKVRQYD  928

Query  109   QILIEICGHKAIGTVLVGPTPVNIIGRNLLTQIGCTLNFPISPIETVPVKLKPGMDGPKV  168
              ILIEICG KAIGTVLVGPTPVNIIGRN+LTQ+GCTLNFPISPIETVPVKLKPGMDGP+V
Sbjct  927   HILIEICGKKAIGTVLVGPTPVNIIGRNMLTQLGCTLNFPISPIETVPVKLKPGMDGPRV  748

Query  169   KQWPLTEEKIKALVEICTEMEKEGKISKIGPENPYNTPVFAIKKKDSTKWRKLVDFRELN  228
             KQWPLTEEKIKAL EIC EMEKEGKI +IGPENPYNTPVFAIKKKDSTKWRKLVDFRELN
Sbjct  747   KQWPLTEEKIKALTEICEEMEKEGKIERIGPENPYNTPVFAIKKKDSTKWRKLVDFRELN  568

Query  229   KRTQDFWEVQLGIPHPAGLKKKKSVTVLDVGDAYFSVPLDEDFRKYTAFTIPSINNETPG  288
             KRTQDFWEVQLGIPHPAGLKKKKSVTVLDVGDAYFSVPLDEDFRKYTAFTIPSINNETPG
Sbjct  567   KRTQDFWEVQLGIPHPAGLKKKKSVTVLDVGDAYFSVPLDEDFRKYTAFTIPSINNETPG  388

Query  289   IRYQYNVLPQGWKGSPAIFQSSMTKILEPFRKQNPDIVIYQYMDDLYVGSDLEIGQHRTK  348
             IRYQYNVLPQGWKGSPAIFQSSM +ILEPFR +NP+IVIYQYMDDLYVGSDLEIGQHR K
Sbjct  387   IRYQYNVLPQGWKGSPAIFQSSMIRILEPFRAKNPEIVIYQYMDDLYVGSDLEIGQHRAK  208

Query  349   IEELRQHLLRWGLTTPDKKHQKEPPFLWMGYELHPDKWTVQPIVLPEKDSWTVNDIQKLV  408
             IE+LR+HLL+WGLTTPDKKHQKEPPFLWMGYELHPDKWTVQPI LPEKDSWTVNDIQKLV
Sbjct  207   IEKLREHLLKWGLTTPDKKHQKEPPFLWMGYELHPDKWTVQPIQLPEKDSWTVNDIQKLV  28

Query  409   GKLNWASQI  417
             GKLNWASQI
Sbjct  27    GKLNWASQI  1
```

>Staphylococcus aureus strain G703N1B1 NODE_79_length_1173_cov_3.01147_ID_157, whole genome shotgun sequence
Sequence ID: NZ_PGXA01000017.1 Length: 1173
Range 1: 3 to 1166

Score:722 bits(1864), Expect:0.0,
Method:Compositional matrix adjust.,
Identities:361/388(93%), Positives:375/388(96%), Gaps:0/388(0%)

```
Query  202   PYNTPVFAIKKKDSTKWRKLVDFRELNKRTQDFWEVQLGIPHPAGLKKKKSVTVLDVGDA  261
             PYNTP+FAIKKKDSTKWRKLVDFRELNKRTQDFWEVQLGIPHPAGLKKKKSVTVLDVGDA
Sbjct  1166  PYNTPIFAIKKKDSTKWRKLVDFRELNKRTQDFWEVQLGIPHPAGLKKKKSVTVLDVGDA  987

Query  262   YFSVPLDEDFRKYTAFTIPSINNETPGIRYQYNVLPQGWKGSPAIFQSSMTKILEPFRKQ  321
             YFSVPLD+DFRKYTAFTIPSINNETPGIRYQYNVLPQGWKGSP+IFQSSMT+ILEPFR Q
Sbjct  986   YFSVPLDKDFRKYTAFTIPSINNETPGIRYQYNVLPQGWKGSPSIFQSSMTRILEPFRAQ  807

Query  322   NPDIVIYQYMDDLYVGSDLEIGQHRTKIEELRQHLLRWGLTTPDKKHQKEPPFLWMGYEL  381
             NP+IVIYQYMDDLYVGSDLEIGQHR KIEELR+HLL+WG TTPDKKHQKEPPFLWMGYEL
Sbjct  806   NPEIVIYQYMDDLYVGSDLEIGQHRAKIEELREHLLKWGFTTPDKKHQKEPPFLWMGYEL  627

Query  382   HPDKWTVQPIVLPEKDSWTVNDIQKLVGKLNWASQIYPGIKVRQLCKLLRGTKALTEVIP  441
             HPDKWTVQPI LPEKDSWTVNDIQKLVGKLNWASQIY GIKV+QLC+LLRG KALT+V+P
Sbjct  626   HPDKWTVQPIQLPEKDSWTVNDIQKLVGKLNWASQIYAGIKVKQLCRLLRGAKALTDVVP  447

Query  442   LTeeaelelaeNREILKEPVHGVYYDPSKDLIAEIQKQGQGQWTYQIYQEPFKNLKTGKY  501
             LTEEAELELAENREILKEPVHGVYYDPSKDLIAEIQKQG  QWTYQIYQEPFKNLKTGKY
Sbjct  446   LTEEAELELAENREILKEPVHGVYYDPSKDLIAEIQKQGHDQWTYQIYQEPFKNLKTGKY  267

Query  502   ARMRGAHTNDVKQLTEAVQKITTESIVIWGKTPKFKLPIQKetwetwwteYWQATWIPEW  561
             A+MR AHTNDVKQLTEAVQKI  E IVIWGKTPKF+LPIQKETWETWWT+YWQATWIPEW
Sbjct  266   AKMRSAHTNDVKQLTEAVQKIAMECIVIWGKTPKFRLPIQKETWETWWTDYWQATWIPEW  87

Query  562   EFVNTPPLVKLWYQLEKEPIVGAETFYV  589
             EFVNTPPLVKLWYQLEKEPI GAETFYV
Sbjct  86    EFVNTPPLVKLWYQLEKEPIAGAETFYV  3
```

>Levilactobacillus brevis strain CRL 2013 scaffold12, whole genome shotgun sequence
Sequence ID: NZ_MZMW01000016.1 Length: 1126
Range 1: 59 to 1126

Score:697 bits(1800), Expect:0.0,
Method:Compositional matrix adjust.,
Identities:334/356(94%), Positives:344/356(96%), Gaps:0/356(0%)

```
Query  48    FPQVTLWQRPLVTIKIGGQLKEALLDTGADDTVLEEMSLPGRWKPKMIGGIGGFIKVRQY  107
             +PQ+TLWQRPLVT K+GGQL EALLDTGADDTVLE+++L G+WKPKMIGGIGGFIKV+QY
Sbjct  59    WPQITLWQRPLVTXKVGGQLXEALLDTGADDTVLEDINLXGKWKPKMIGGIGGFIKVKQY  238

Query  108   DQILIEICGHKAIGTVLVGPTPVNIIGRNLLTQIGCTLNFPISPIETVPVKLKPGMDGPK  167
             D ILIEICGHKAIGTVLVGPTP NIIGRNLLTQ+GCTLNFPISPIETVPVKLKPGMDGPK
Sbjct  239   DNILIEICGHKAIGTVLVGPTPXNIIGRNLLTQLGCTLNFPISPIETVPVKLKPGMDGPK  418

Query  168   VKQWPLTEEKIKALVEICTEMEKEGKISKIGPENPYNTPVFAIKKKDSTKWRKLVDFREL  227
             VKQWPLTEEKIKAL EICTEMEKEGKISKIGPENPYNTP FAIKKKDSTKWRKLVDFREL
Sbjct  419   VKQWPLTEEKIKALXEICTEMEKEGKISKIGPENPYNTPXFAIKKKDSTKWRKLVDFREL  598

Query  228   NKRTQDFWEVQLGIPHPAGLKKKKSVTVLDVGDAYFSVPLDEDFRKYTAFTIPSINNETP  287
             NKRTQDFWEVQLGIPHPAGLK+ KSVTVLDVGDAYFSVPLDE FRKYTAFTIPSINNETP
Sbjct  599   NKRTQDFWEVQLGIPHPAGLKQNKSVTVLDVGDAYFSVPLDEAFRKYTAFTIPSINNETP  778

Query  288   GIRYQYNVLPQGWKGSPAIFQSSMTKILEPFRKQNPDIVIYQYMDDLYVGSDLEIGQHRT  347
             GIRYQYNVLPQGWKGSPAIFQSSMTKILEPFRKQNPDIVIYQYMDDLYVGSDLEIGQHR
Sbjct  779   GIRYQYNVLPQGWKGSPAIFQSSMTKILEPFRKQNPDIVIYQYMDDLYVGSDLEIGQHRA  958

Query  348   KIEELRQHLLRWGLTTPDKKHQKEPPFLWMGYELHPDKWTVQPIVLPEKDSWTVND  403
             KIEELRQHLLRWG TTPDKKHQKE PFLWMGYELHPDKWTVQPIVLPEKDSWTVND
Sbjct  959   KIEELRQHLLRWGFTTPDKKHQKEHPFLWMGYELHPDKWTVQPIVLPEKDSWTVND  1126
```

>Pseudomonas aeruginosa strain VET-121 NODE_73_length_1192_cov_8.02063_ID_145, whole genome shotgun sequence
Sequence ID: NZ_QZXB01000185.1 Length: 1158
Range 1: 57 to 1118

Score:686 bits(1770), Expect:0.0,
Method:Compositional matrix adjust.,

```
Identities:325/354(92%), Positives:342/354(96%), Gaps:0/354(0%)

Query  50    QVTLWQRPLVTIKIGGQLKEALLDTGADDTVLEEMSLPGRWKPKMIGGIGGFIKVRQYDQ  109
             Q+TLWQRPLVT++I  QL EALLDTGADDTVLE++ LPG+WKPKMIGGIGGFIKVRQYDQ
Sbjct  1118  QITLWQRPLVTVRIEEQLIEALLDTGADDTVLEDIDLPGKWKPKMIGGIGGFIKVRQYDQ  939

Query  110   ILIEICGHKAIGTVLVGPTPVNIIGRNLLTQIGCTLNFPISPIETVPVKLKPGMDGPKVK  169
             I IEICG KAIGTVLVGPTPVNIIGRN+LT+IGCTLNFPISPIETVPVKLKPGMDGPKVK
Sbjct  938   IHIEICGKKAIGTVLVGPTPVNIIGRNILTKIGCTLNFPISPIETVPVKLKPGMDGPKVK  759

Query  170   QWPLTEEKIKALVEICTEMEKEGKISKIGPENPYNTPVFAIKKKDSTKWRKLVDFRELNK  229
             QWPLTEEKIKAL EICTEMEKEGKISKIGPENPYNTP+FAIKKKDSTKWRKLVDFRELNK
Sbjct  758   QWPLTEEKIKALTEICTEMEKEGKISKIGPENPYNTPIFAIKKKDSTKWRKLVDFRELNK  579

Query  230   RTQDFWEVQLGIPHPAGLKKKKSVTVLDVGDAYFSVPLDEDFRKYTAFTIPSINNETPGI  289
             RTQDFWEVQLGIPHPAGLKKKKSVT+LDVGDAYFSVPLD DFRKYTAFTIPSINNETPGI
Sbjct  578   RTQDFWEVQLGIPHPAGLKKKKSVTILDVGDAYFSVPLDPDFRKYTAFTIPSINNETPGI  399

Query  290   RYQYNVLPQGWKGSPAIFQSSMTKILEPFRKQNPDIVIYQYMDDLYVGSDLEIGQHRTKI  349
             RYQYNVLPQGWKGSPAIFQ SMTKILEPFRKQNP++VIYQYMDDLY+GSDLEIGQHR KI
Sbjct  398   RYQYNVLPQGWKGSPAIFQCSMTKILEPFRKQNPELVIYQYMDDLYIGSDLEIGQHRAKI  219

Query  350   EELRQHLLRWGLTTPDKKHQKEPPFLWMGYELHPDKWTVQPIVLPEKDSWTVND  403
             EELR+HLL+WG TTPDKKHQKEPPFLWMGYELHPDKWTVQPI LP+K+SWTVND
Sbjct  218   EELREHLLKWGFTTPDKKHQKEPPFLWMGYELHPDKWTVQPIELPDKESWTVND  57
```

>Pseudomonas aeruginosa strain VET-42 NODE_85_length_1204_cov_6.50932_ID_169, whole genome shotgun sequence
Sequence ID: NZ_RACX01000100.1 Length: 1166
Range 1: 31 to 1095

Score:683 bits(1763), Expect:0.0,
Method:Compositional matrix adjust.,
Identities:324/355(91%), Positives:341/355(96%), Gaps:0/355(0%)

```
Query  50    QVTLWQRPLVTIKIGGQLKEALLDTGADDTVLEEMSLPGRWKPKMIGGIGGFIKVRQYDQ  109
             +  LWQRPLVT++I  QL EALLDTGADDTVLE++ LPG+WKPKMIGGIGGFIKVRQYDQ
Sbjct  31    ETALWQRPLVTVRIEEQLIEALLDTGADDTVLEDIDLPGKWKPKMIGGIGGFIKVRQYDQ  210

Query  110   ILIEICGHKAIGTVLVGPTPVNIIGRNLLTQIGCTLNFPISPIETVPVKLKPGMDGPKVK  169
             I IEICG KAIGTVLVGPTPVNIIGRN+LT+IGCTLNFPISPIETVPVKLKPGMDGPKVK
Sbjct  211   IHIEICGKKAIGTVLVGPTPVNIIGRNILTKIGCTLNFPISPIETVPVKLKPGMDGPKVK  390

Query  170   QWPLTEEKIKALVEICTEMEKEGKISKIGPENPYNTPVFAIKKKDSTKWRKLVDFRELNK  229
             QWPLTEEKIKAL EICTEMEKEGKISKIGPENPYNTP+FAIKKKDSTKWRKLVDFRELNK
Sbjct  391   QWPLTEEKIKALTEICTEMEKEGKISKIGPENPYNTPIFAIKKKDSTKWRKLVDFRELNK  570

Query  230   RTQDFWEVQLGIPHPAGLKKKKSVTVLDVGDAYFSVPLDEDFRKYTAFTIPSINNETPGI  289
             RTQDFWEVQLGIPHPAGLKKKKSVT+LDVGDAYFSVPLD DFRKYTAFTIPSINNETPGI
Sbjct  571   RTQDFWEVQLGIPHPAGLKKKKSVTILDVGDAYFSVPLDPDFRKYTAFTIPSINNETPGI  750

Query  290   RYQYNVLPQGWKGSPAIFQSSMTKILEPFRKQNPDIVIYQYMDDLYVGSDLEIGQHRTKI  349
             RYQYNVLPQGWKGSPAIFQ SMTKILEPFRKQNP++VIYQYMDDLY+GSDLEIGQHR KI
Sbjct  751   RYQYNVLPQGWKGSPAIFQCSMTKILEPFRKQNPELVIYQYMDDLYIGSDLEIGQHRAKI  930

Query  350   EELRQHLLRWGLTTPDKKHQKEPPFLWMGYELHPDKWTVQPIVLPEKDSWTVNDI  404
             EELR+HLL+WG TTPDKKHQKEPPFLWMGYELHPDKWTVQPI LP+K+SWTVNDI
Sbjct  931   EELREHLLKWGFTTPDKKHQKEPPFLWMGYELHPDKWTVQPIELPDKESWTVNDI  1095
```

>Streptococcus agalactiae strain JH02623 contig_39, whole genome shotgun sequence
Sequence ID: NZ_RPAW01000039.1 Length: 1204
Range 1: 170 to 1204

Score:676 bits(1744), Expect:0.0,
Method:Compositional matrix adjust.,
Identities:320/345(93%), Positives:337/345(97%), Gaps:0/345(0%)

```
Query  771   GIIQAQPDQSESELVNQIIEQLIKKEKVYLAWVPAHKGIGGNEQVDKLVSAGIRKVLFLD  830
             GIIQAQPD+SESELVNQIIEQLIKKE+VYLAWVPAHKGIGGNEQVDKLVS GIRKVLFLD
Sbjct  1204  GIIQAQPDKSESELVNQIIEQLIKKERVYLAWVPAHKGIGGNEQVDKLVSNGIRKVLFLD  1025

Query  831   GIDKAQDEHEKYHSNWRAMASDFNLPPVVAKEIVASCDKCQLKGEAMHGQVDCSPGIWQL  890
             GIDKAQ+EHEKYH+NW+AMASDFNLPP+VAKEIVASCDKCQLKGEAMHGQVDCSPGIWQL
Sbjct  1024  GIDKAQEEHEKYHNNWKAMASDFNLPPIVAKEIVASCDKCQLKGEAMHGQVDCSPGIWQL  845

Query  891   DCTHLEGKVILVAVHVASGYIEAEVIPAETGQETAYFLLKLAGRWPVKTIHTDNGSNFTG  950
             DCTHLEGK+ILVAVHVASGYIEAEVIPAETGQETAYF+LKLAGRWPVK IHTDNGSNFT
Sbjct  844   DCTHLEGKIILVAVHVASGYIEAEVIPAETGQETAYFILKLAGRWPVKVIHTDNGSNFTS  665

Query  951   ATVRAACWWAGIKQEFGIPYNPQSQGVVESMNKELKKIIGQVRDQAEHLKTAVQMAVFIH  1010
             A V+AACWWA I QEFGIPYNPQSQGVVESMNKELK+II QVRDQAEHLKTAVQMAVFIH
Sbjct  664   AAVKAACWWANITQEFGIPYNPQSQGVVESMNKELKRIIQQVRDQAEHLKTAVQMAVFIH  485

Query  1011  NFKRKGGIGGYSAGERIVDIIATDIQTKELQKQITKIQNFRVYYRDSRNPLWKGPAKLLW  1070
             NFK+KGGIGGYSAGERI+DIIA+DIQTKELQKQITKIQNFRVYYRDSR+P+WKGPAKLLW
Sbjct  484   NFKKKGGIGGYSAGERIIDIIASDIQTKELQKQITKIQNFRVYYRDSRDPIWKGPAKLLW  305

Query  1071  KGEGAVVIQDNSDIKVVPRRKAKIIRDYGKQMAGDDCVASRQDED  1115
             KGEGAVVIQDN++IKVVPRRKAKIIRDYGKQMAGDDCVA RQDED
Sbjct  304   KGEGAVVIQDNNEIKVVPRRKAKIIRDYGKQMAGDDCVAGRQDED  170
```

Range 2: 1022 to 1204

Score:115 bits(288), Expect:1e-22,
Method:Compositional matrix adjust.,
Identities:56/61(92%), Positives:60/61(98%), Gaps:0/61(0%)

```
Query  651   GIIQAQPDQSESELVNQIIEQLIKKEKVYLAWVPAHKGIGGNEQVDKLVSAGIRKVLYVD  710
             GIIQAQPD+SESELVNQIIEQLIKKE+VYLAWVPAHKGIGGNEQVDKLVS GIRKVL++D
Sbjct  1204  GIIQAQPDKSESELVNQIIEQLIKKERVYLAWVPAHKGIGGNEQVDKLVSNGIRKVLFLD  1025

Query  711   G  711
             G
Sbjct  1024  G  1022
```

>Levilactobacillus brevis strain CRL 2013 scaffold13, whole genome shotgun sequence
Sequence ID: NZ_MZMW01000017.1 Length: 1030
Range 1: 1 to 1029

Score:668 bits(1724), Expect:0.0,
Method:Compositional matrix adjust.,
Identities:320/343(93%), Positives:329/343(95%), Gaps:0/343(0%)

```
Query  49    PQVTLWQRPLVTIKIGGQLKEALLDTGADDTVLEEMSLPGRWKPKMIGGIGGFIKVRQYD  108
             PQ+TLWQRPLVTIK+G QLKEALLDTGADDTVLE ++LPG WKPKMIGGIGGFIKV QYD
Sbjct  1     PQITLWQRPLVTIKVGXQLKEALLDTGADDTVLEXINLPGXWKPKMIGGIGGFIKVXQYD  180

Query  109   QILIEICGHKAIGTVLVGPTPVNIIGRNLLTQIGCTLNFPISPIETVPVKLKPGMDGPKV  168
              I IEICGH A GTVLVGPTPVNIIGRNLLTQ+GCTLNFPISPIETVPVKLKPGMDGPKV
Sbjct  181   XIPIEICGHXAXGTVLVGPTPVNIIGRNLLTQLGCTLNFPISPIETVPVKLKPGMDGPKV  360

Query  169   KQWPLTEEKIKALVEICTEMEKEGKISKIGPENPYNTPVFAIKKKDSTKWRKLVDFRELN  228
             KQWPLTEEKI AL EIC EMEKEGKISKIGPENPYNTPVFAIKKKDSTKWRKLVDFRELN
Sbjct  361   KQWPLTEEKIXALTEICXEMEKEGKISKIGPENPYNTPVFAIKKKDSTKWRKLVDFRELN  540

Query  229   KRTQDFWEVQLGIPHPAGLKKKKSVTVLDVGDAYFSVPLDEDFRKYTAFTIPSINNETPG  288
             KRTQDFWEVQLGIPHPAGLK+KKSVTVLDVGDAYFSVPLDE+FRKYTAFTIPS NNETPG
Sbjct  541   KRTQDFWEVQLGIPHPAGLKRKKSVTVLDVGDAYFSVPLDENFRKYTAFTIPSTNNETPG  720

Query  289   IRYQYNVLPQGWKGSPAIFQSSMTKILEPFRKQNPDIVIYQYMDDLYVGSDLEIGQHRTK  348
             IRYQYNVLPQGWKGSPAIFQSSMTKILEPFRKQNPDI IYQYMDDLYVGSDLEIGQHR K
Sbjct  721   IRYQYNVLPQGWKGSPAIFQSSMTKILEPFRKQNPDIDIYQYMDDLYVGSDLEIGQHRAK  900

Query  349   IEELRQHLLRWGLTTPDKKHQKEPPFLWMGYELHPDKWTVQPI  391
             IEELR+HLLRWGLTTPDKKHQK+PPFLWMGYELHPDKWTVQPI
Sbjct  901   IEELREHLLRWGLTTPDKKHQKKPPFLWMGYELHPDKWTVQPI  1029



>Pseudomonas aeruginosa strain VET-42 NODE_86_length_1154_cov_8.30919_ID_171, whole genome shotgun sequence
Sequence ID: NZ_RACX01000118.1 Length: 1154
Range 1: 75 to 1103

Score:664 bits(1712), Expect:0.0,
Method:Compositional matrix adjust.,
Identities:310/343(90%), Positives:333/343(97%), Gaps:0/343(0%)

Query  773   IQAQPDQSESELVNQIIEQLIKKEKVYLAWVPAHKGIGGNEQVDKLVSAGIRKVLFLDGI  832
             +QAQPD+SESE+VNQIIEQLIKKEKVYL+WVPAHKGIGGNEQVDKLVS+GIRKVLFLDGI
Sbjct  1103  VQAQPDKSESEVVNQIIEQLIKKEKVYLSWVPAHKGIGGNEQVDKLVSSGIRKVLFLDGI  924

Query  833   DKAQDEHEKYHSNWRAMASDFNLPPVVAKEIVASCDKCQLKGEAMHGQVDCSPGIWQLDC  892
             DKAQ+EHE+YH+NWRAMASDFNLPP+VAKEIVASCDKCQ KGEAMHGQVDCSPGIWQ+DC
Sbjct  923   DKAQEEHERYHNNWRAMASDFNLPPIVAKEIVASCDKCQQKGEAMHGQVDCSPGIWQIDC  744

Query  893   THLEGKVILVAVHVASGYIEAEVIPAETGQETAYFLLKLAGRWPVKTIHTDNGSNFTGAT  952
             THLEGK+I+VAVHVASGYIEAEVIPAETGQETAYF+LKLA RWPVK IHTDNGSNFT A
Sbjct  743   THLEGKIIIVAVHVASGYIEAEVIPAETGQETAYFILKLAARWPVKVIHTDNGSNFTSAA  564

Query  953   VRAACWWAGIKQEFGIPYNPQSQGVVESMNKELKKIIGQVRDQAEHLKTAVQMAVFIHNF  1012
             V+AACWWA + QEFGIPYNPQSQGVVESMNKELKKIIGQ+RDQAEHLKTAVQMAVFIHNF
Sbjct  563   VKAACWWANVTQEFGIPYNPQSQGVVESMNKELKKIIGQIRDQAEHLKTAVQMAVFIHNF  384

Query  1013  KRKGGIGGYSAGERIVDIIATDIQTKELQKQITKIQNFRVYYRDSRNPLWKGPAKLLWKG  1072
             KRKGGIG YSAGERI+DIIA+DIQTKELQKQI KIQNFRVYYRDSR+P+WKGPAKLLWKG
Sbjct  383   KRKGGIGRYSAGERIIDIIASDIQTKELQKQIIKIQNFRVYYRDSRDPIWKGPAKLLWKG  204

Query  1073  EGAVVIQDNSDIKVVPRRKAKIIRDYGKQMAGDDCVASRQDED  1115
             EGAVVIQDNS+IKV+PRRKAKIIRDYGKQMAGDDCVA RQDE+
Sbjct  203   EGAVVIQDNSEIKVIPRRKAKIIRDYGKQMAGDDCVAGRQDEN  75



Range 2: 927 to 1103

Score:110 bits(275), Expect:5e-21,
Method:Compositional matrix adjust.,
Identities:52/59(88%), Positives:59/59(100%), Gaps:0/59(0%)

Query  653   IQAQPDQSESELVNQIIEQLIKKEKVYLAWVPAHKGIGGNEQVDKLVSAGIRKVLYVDG  711
             +QAQPD+SESE+VNQIIEQLIKKEKVYL+WVPAHKGIGGNEQVDKLVS+GIRKVL++DG
Sbjct  1103  VQAQPDKSESEVVNQIIEQLIKKEKVYLSWVPAHKGIGGNEQVDKLVSSGIRKVLFLDG  927



>Providencia rettgeri strain AFE96_S1 41_contig_60, whole genome shotgun sequence
Sequence ID: NZ_SHDP01000060.1 Length: 1101
Range 1: 96 to 1100

Score:654 bits(1688), Expect:0.0,
Method:Compositional matrix adjust.,
Identities:310/335(93%), Positives:325/335(97%), Gaps:0/335(0%)

Query  62    KIGGQLKEALLDTGADDTVLEEMSLPGRWKPKMIGGIGGFIKVRQYDQILIEICGHKAIG  121
             K+GGQ+KEALLDTGADDTVLE+++LPG+WKP+MIGGIGGFIKVRQYDQILIEICG KAIG
Sbjct  96    KVGGQVKEALLDTGADDTVLEDINLPGKWKPRMIGGIGGFIKVRQYDQILIEICGKKAIG  275

Query  122   TVLVGPTPVNIIGRNLLTQIGCTLNFPISPIETVPVKLKPGMDGPKVKQWPLTEEKIKAL  181
             TVLVGPTPVNIIGRN+LTQ+GCTLNFPISPIETVPVKLKPGMDGPKVKQWPLTEEKIKAL
Sbjct  276   TVLVGPTPVNIIGRNMLTQLGCTLNFPISPIETVPVKLKPGMDGPKVKQWPLTEEKIKAL  455

Query  182   VEICTEMEKEGKISKIGPENPYNTPVFAIKKKDSTKWRKLVDFRELNKRTQDFWEVQLGI  241
               IC EMEKEGKI+KIGPENPYNTP+FAIKKKDSTKWRKLVDFRELNKRTQDFWEVQLGI
Sbjct  456   TAICKEMEKEGKITKIGPENPYNTPIFAIKKKDSTKWRKLVDFRELNKRTQDFWEVQLGI  635

Query  242   PHPAGLKKKKSVTVLDVGDAYFSVPLDEDFRKYTAFTIPSINNETPGIRYQYNVLPQGWK  301
             PHPAGLKKKKSVTVLDVGDAYFSVPLDE FRKYTAFTIPSINNETPGIRYQYNVLPQGWK
Sbjct  636   PHPAGLKKKKSVTVLDVGDAYFSVPLDEGFRKYTAFTIPSINNETPGIRYQYNVLPQGWK  815

Query  302   GSPAIFQSSMTKILEPFRKQNPDIVIYQYMDDLYVGSDLEIGQHRTKIEELRQHLLRWGL  361
             GSPAIFQSSMTKILEPFR QNP+IVIYQY+DDLYVGSDLEIGQHR KIEELR+HLL+WG
Sbjct  816   GSPAIFQSSMTKILEPFRAQNPEIVIYQYVDDLYVGSDLEIGQHRAKIEELREHLLKWGF  995

Query  362   TTPDKKHQKEPPFLWMGYELHPDKWTVQPIVLPEK  396
             TTPDKKHQKEPPFLWMGYELHPDKWTVQPI LP K
Sbjct  996   TTPDKKHQKEPPFLWMGYELHPDKWTVQPIQLPTK  1100



>Staphylococcus epidermidis strain PR246B0 NODE_70_length_1113_cov_1.18763_ID_139, whole genome shotgun sequence
Sequence ID: NZ_PCFD01000070.1 Length: 1113
Range 1: 3 to 1103

Score:651 bits(1679), Expect:0.0,
Method:Compositional matrix adjust.,
Identities:329/367(90%), Positives:349/367(95%), Gaps:0/367(0%)

Query  219   RKLVDFRELNKRTQDFWEVQLGIPHPAGLKKKKSVTVLDVGDAYFSVPLDEDFRKYTAFT  278
             +K+VDFRELNKRTQDFWEVQLGIPHPAGLKKKKSVTVLDVGDA+FSVPLDE FRKYTAFT
Sbjct  1103  KKVVDFRELNKRTQDFWEVQLGIPHPAGLKKKKSVTVLDVGDAFFSVPLDESFRKYTAFT  924

Query  279   IPSINNETPGIRYQYNVLPQGWKGSPAIFQSSMTKILEPFRKQNPDIVIYQYMDDLYVGS  338
             IPSINN TPGIRYQYNVLPQGWKGSPAIFQSSMTKILEPFR +NP++VIYQY+DDLYV S
```

```
Sbjct  923   IPSINNATPGIRYQYNVLPQGWKGSPAIFQSSMTKILEPFRTKNPEMVIYQYVDDLYVES  744

Query  339   DLEIGQHRTKIEELRQHLLRWGLTTPDKKHQKEPPFLWMGYELHPDKWTVQPIVLPEKDS  398
             DLEI QHR K+EELR+HLL+WG TTPD+KHQKEPPFLWMGYELHPDKWTVQPI LPEKD
Sbjct  743   DLEIMQHRAKVEELREHLLKWGFTTPDEKHQKEPPFLWMGYELHPDKWTVQPIQLPEKDI  564

Query  399   WTVNDIQKLVGKLNWASQIYPGIKVRQLCKLLRGTKALTEVIPLTeeaelelaeNREILK  458
             WTVNDIQKLVGKLNWASQIYP IKV+QLCKLLRGTKALT+++ LTEEAELELAENREILK
Sbjct  563   WTVNDIQKLVGKLNWASQIYPEIKVKQLCKLLRGTKALTDIVTLTEEAELELAENREILK  384

Query  459   EPVHGVYYDPSKDLIAEIQKQGQGQWTYQIYQEPFKNLKTGKYARMRGAHTNDVKQLTEA  518
             EPVHGVYYDPSKDLIAEIQKQG  QWTYQIYQEPFKNLKTGKYA+MR AHTNDV+QLTEA
Sbjct  383   EPVHGVYYDPSKDLIAEIQKQGNDQWTYQIYQEPFKNLKTGKYAKMRTAHTNDVRQLTEA  204

Query  519   VQKITTESIVIWGKTPKFKLPIQKetwetwwteYWQATWIPEWEFVNTPPLVKLWYQLEK  578
             VQKI  ESIVIWGK PKF+LPIQKETWETWWT+YWQATWIP+WEFVNTPPLVKLWYQLEK
Sbjct  203   VQKIARESIVIWGKIPKFRLPIQKETWETWWTDYWQATWIPDWEFVNTPPLVKLWYQLEK  24

Query  579   EPIVGAE  585
             EPI GAE
Sbjct  23    EPIAGAE  3



>Pseudomonas aeruginosa strain ENV-297 NODE_79_length_1100_cov_9.24438_ID_157, whole genome shotgun sequence
Sequence ID: NZ_RADD01000036.1 Length: 1100
Range 1: 81 to 1055

Score:635 bits(1637), Expect:0.0,
Method:Compositional matrix adjust.,
Identities:296/325(91%), Positives:315/325(96%), Gaps:0/325(0%)

Query  791   QLIKKEKVYLAWVPAHKGIGGNEQVDKLVSAGIRKVLFLDGIDKAQDEHEKYHSNWRAMA  850
             QLIKKEKVYL+WVPAHKGIGGNEQVDKLVS+GIRKVLFLDGIDKAQ+EHE+YH+NWRAMA
Sbjct  1055  QLIKKEKVYLSWVPAHKGIGGNEQVDKLVSSGIRKVLFLDGIDKAQEEHERYHNNWRAMA  876

Query  851   SDFNLPPVVAKEIVASCDKCQLKGEAMHGQVDCSPGIWQLDCTHLEGKVILVAVHVASGY  910
             SDFNLPP+VAKEIVASCDKCQ KGEAMHGQVDCSPGIWQ+DCTHLEGK+I+VAVHVASGY
Sbjct  875   SDFNLPPIVAKEIVASCDKCQQKGEAMHGQVDCSPGIWQIDCTHLEGKIIIVAVHVASGY  696

Query  911   IEAEVIPAETGQETAYFLLKLAGRWPVKTIHTDNGSNFTGATVRAACWWAGIKQEFGIPY  970
             IEAEVIPAETGQETAYF+LKLA RWPVK IHTDNGSNFT A V+AACWWA + QEFGIPY
Sbjct  695   IEAEVIPAETGQETAYFILKLAARWPVKVIHTDNGSNFTSAAVKAACWWANVTQEFGIPY  516

Query  971   NPQSQGVVESMNKELKKIIGQVRDQAEHLKTAVQMAVFIHNFKRKGGIGGYSAGERIVDI  1030
             NPQSQGVVESMNKELKKIIGQ+RDQAEHLKTAVQMAVFIHNFKRKGGIG YSAGERI+DI
Sbjct  515   NPQSQGVVESMNKELKKIIGQIRDQAEHLKTAVQMAVFIHNFKRKGGIGRYSAGERIIDI  336

Query  1031  IATDIQTKELQKQITKIQNFRVYYRDSRNPLWKGPAKLLWKGEGAVVIQDNSDIKVVPRR  1090
             IA+DIQTKELQKQI KIQNFRVYYRDSR+P+WKGPAKLLWKGEGAVVIQDNS+IKV+PRR
Sbjct  335   IASDIQTKELQKQIIKIQNFRVYYRDSRDPIWKGPAKLLWKGEGAVVIQDNSEIKVIPRR  156

Query  1091  KAKIIRDYGKQMAGDDCVASRQDED  1115
             KAKIIRDYGKQMAGDDCVA RQDED
Sbjct  155   KAKIIRDYGKQMAGDDCVAGRQDED  81




Range 2: 933 to 1055

Score:80.9 bits(198), Expect:3e-11,
Method:Compositional matrix adjust.,
Identities:37/41(90%), Positives:41/41(100%), Gaps:0/41(0%)

Query  671   QLIKKEKVYLAWVPAHKGIGGNEQVDKLVSAGIRKVLYVDG  711
             QLIKKEKVYL+WVPAHKGIGGNEQVDKLVS+GIRKVL++DG
Sbjct  1055  QLIKKEKVYLSWVPAHKGIGGNEQVDKLVSSGIRKVLFLDG  933



>Staphylococcus aureus strain PN235B0 NODE_123_length_1117_cov_0.929293_ID_245, whole genome shotgun sequence
Sequence ID: NZ_PDUY01000159.1 Length: 1083
Range 1: 30 to 1082

Score:621 bits(1601), Expect:0.0,
Method:Compositional matrix adjust.,
Identities:311/351(89%), Positives:331/351(94%), Gaps:0/351(0%)

Query  238   QLGIPHPAGLKKKKSVTVLDVGDAYFSVPLDEDFRKYTAFTIPSINNETPGIRYQYNVLP  297
             Q+GIPHPAG+KK KSVTVLDVGDAYFSVPL EDFRKYTAFTIPS+NNE PGIRYQYNVLP
Sbjct  30    QIGIPHPAGIKKNKSVTVLDVGDAYFSVPLHEDFRKYTAFTIPSVNNEAPGIRYQYNVLP  209

Query  298   QGWKGSPAIFQSSMTKILEPFRKQNPDIVIYQYMDDLYVGSDLEIGQHRTKIEELRQHLL  357
             QGWKGSPAIFQSSMTKILEPFR +NP++VIYQY+DDLY+ SDLEI QHR K+EELR+HLL
Sbjct  210   QGWKGSPAIFQSSMTKILEPFRTKNPEVVIYQYVDDLYIESDLEIMQHRAKVEELREHLL  389

Query  358   RWGLTTPDKKHQKEPPFLWMGYELHPDKWTVQPIVLPEKDSWTVNDIQKLVGKLNWASQI  417
             +WG TTPD+KHQKEPPFLWMGYELHPDKWTVQPI LPEKD WTVNDIQKLVGKLNWASQI
Sbjct  390   KWGFTTPDEKHQKEPPFLWMGYELHPDKWTVQPIQLPEKDIWTVNDIQKLVGKLNWASQI  569

Query  418   YPGIKVRQLCKLLRGTKALTEVIPLTeeaelelaeNREILKEPVHGVYYDPSKDLIAEIQ  477
             YP IKV+QLCKLLRGTKALT+++PLTEEAELELAENREILKEPVHG YYDPSKDLIAEIQ
Sbjct  570   YPEIKVKQLCKLLRGTKALTDIVPLTEEAELELAENREILKEPVHGAYYDPSKDLIAEIQ  749

Query  478   KQGQGQWTYQIYQEPFKNLKTGKYARMRGAHTNDVKQLTEAVQKITTESIVIWGKTPKFK  537
             KQG  QWTYQIYQEPFKNLKTGKYA+MR AHTNDVKQLTEAVQKI  E IVIWGK PKF+
Sbjct  750   KQGNDQWTYQIYQEPFKNLKTGKYAKMRTAHTNDVKQLTEAVQKIARECIVIWGKIPKFR  929

Query  538   LPIQKetwetwwteYWQATWIPEWEFVNTPPLVKLWYQLEKEPIVGAETFY  588
             LPIQKETWETWWT+YWQATWIP+WEFVNTPPLVKLWYQLEKEPI GAETFY
Sbjct  930   LPIQKETWETWWTDYWQATWIPDWEFVNTPPLVKLWYQLEKEPIAGAETFY  1082



>Enterococcus durans strain E115_10 NODE_63_length_955_cov_1.30556, whole genome shotgun sequence
Sequence ID: NZ_JAANZI010000063.1 Length: 955
Range 1: 3 to 953

Score:612 bits(1578), Expect:0.0,
Method:Compositional matrix adjust.,
Identities:286/317(90%), Positives:305/317(96%), Gaps:0/317(0%)

Query  75   GADDTVLEEMSLPGRWKPKMIGGIGGFIKVRQYDQILIEICGHKAIGTVLVGPTPVNIIG  134
            GADDT+LE+++LPG+WKPKM GGIGGFIKV+QYDQ+L+EICG K IGTVLVGPTP+NIIG
Sbjct  953  GADDTILEDINLPGKWKPKMAGGIGGFIKVKQYDQVLVEICGQKVIGTVLVGPTPINIIG  774

Query  135  RNLLTQIGCTLNFPISPIETVPVKLKPGMDGPKVKQWPLTEEKIKALVEICTEMEKEGKI  194
            RNL+TQ+GCTLNFPISPIETVPVKLKPGMDGPKVKQWPLTEEKIKAL EIC EMEKEGKI
Sbjct  773  RNLMTQLGCTLNFPISPIETVPVKLKPGMDGPKVKQWPLTEEKIKALTEICEEMEKEGKI  594
```

```
Query  195  SKIGPENPYNTPVFAIKKKDSTKWRKLVDFRELNKRTQDFWEVQLGIPHPAGLKKKKSVT  254
            +KIGPENPYNTPVFAIKKKDSTKWRKLVDFRELNKRTQDFWEVQLGIPHPAGLK+ KSVT
Sbjct  593  TKIGPENPYNTPVFAIKKKDSTKWRKLVDFRELNKRTQDFWEVQLGIPHPAGLKQNKSVT  414

Query  255  VLDVGDAYFSVPLDEDFRKYTAFTIPSINNETPGIRYQYNVLPQGWKGSPAIFQSSMTKI  314
            VLDVGDAYFSVPLDE FRKYTAFTIPSINNETPGIRYQYNVLPQGWKGSPAIFQ SMTKI
Sbjct  413  VLDVGDAYFSVPLDEGFRKYTAFTIPSINNETPGIRYQYNVLPQGWKGSPAIFQCSMTKI  234

Query  315  LEPFRKQNPDIVIYQYMDDLYVGSDLEIGQHRTKIEELRQHLLRWGLTTPDKKHQKEPPF  374
            LEPFR +NP++VIYQY+DDLYVGSDLEIGQHR KIEELR+HLL+WG TTPDKKHQKE PF
Sbjct  233  LEPFRAKNPELVIYQYVDDLYVGSDLEIGQHRAKIEELREHLLKWGFTTPDKKHQKEHPF  54

Query  375  LWMGYELHPDKWTVQPI  391
            LWMGYELHPDKWTVQPI
Sbjct  53   LWMGYELHPDKWTVQPI  3
```

>Escherichia coli strain 8374wA11 NODE_37_length_940_cov_2.75892, whole genome shotgun sequence
Sequence ID: NZ_WSPR01000036.1 Length: 940
Range 1: 2 to 940

Score:609 bits(1570), Expect:0.0,
Method:Compositional matrix adjust.,
Identities:285/313(91%), Positives:304/313(97%), Gaps:0/313(0%)

```
Query  129  PVNIIGRNLLTQIGCTLNFPISPIETVPVKLKPGMDGPKVKQWPLTEEKIKALVEICTEM  188
            P+NIIGRN+ TQ+GCTLNFPISPIETVPVKLKPGMDGP+VKQWPLTEEKIKAL  IC EM
Sbjct  940  PINIIGRNMSTQLGCTLNFPISPIETVPVKLKPGMDGPRVKQWPLTEEKIKALQAICEEM  761

Query  189  EKEGKISKIGPENPYNTPVFAIKKKDSTKWRKLVDFRELNKRTQDFWEVQLGIPHPAGLK  248
            EKEGKI+KIGPENPYNTPVFAIKKKDSTKWRKLVDFRELNKRTQDFWEVQLGIPHPAGLK
Sbjct  760  EKEGKITKIGPENPYNTPVFAIKKKDSTKWRKLVDFRELNKRTQDFWEVQLGIPHPAGLK  581

Query  249  KKKSVTVLDVGDAYFSVPLDEDFRKYTAFTIPSINNETPGIRYQYNVLPQGWKGSPAIFQ  308
            KKKSVTVLDVGDAYFSVPLDE+FRKYTAFTIPSINN TPGIRYQYNVLPQGWKGSPAIFQ
Sbjct  580  KKKSVTVLDVGDAYFSVPLDENFRKYTAFTIPSINNATPGIRYQYNVLPQGWKGSPAIFQ  401

Query  309  SSMTKILEPFRKQNPDIVIYQYMDDLYVGSDLEIGQHRTKIEELRQHLLRWGLTTPDKKH  368
            S+MTKIL+PFR +NPDIVIYQYMDDLYVGSDLEIGQHR KIEELR+HLL+WGLTTPDKKH
Sbjct  400  STMTKILQPFRTRNPDIVIYQYMDDLYVGSDLEIGQHRAKIEELREHLLKWGLTTPDKKH  221

Query  369  QKEPPFLWMGYELHPDKWTVQPIVLPEKDSWTVNDIQKLVGKLNWASQIYPGIKVRQLCK  428
            QKEPPFLWMGYELHPDKWTVQPI LP+K+SWTVNDIQKLVGKLNWASQIYPGI+V++LC+
Sbjct  220  QKEPPFLWMGYELHPDKWTVQPIQLPDKESWTVNDIQKLVGKLNWASQIYPGIRVKRLCR  41

Query  429  LLRGTKALTEVIP  441
            LLRG KALT++IP
Sbjct  40   LLRGAKALTDIIP  2
```

>Escherichia coli strain TzEc080 NODE_37_length_940_cov_2.75892, whole genome shotgun sequence
Sequence ID: NZ_WSFZ01000036.1 Length: 940
Range 1: 2 to 940

Score:609 bits(1570), Expect:0.0,
Method:Compositional matrix adjust.,
Identities:285/313(91%), Positives:304/313(97%), Gaps:0/313(0%)

```
Query  129  PVNIIGRNLLTQIGCTLNFPISPIETVPVKLKPGMDGPKVKQWPLTEEKIKALVEICTEM  188
            P+NIIGRN+ TQ+GCTLNFPISPIETVPVKLKPGMDGP+VKQWPLTEEKIKAL  IC EM
Sbjct  940  PINIIGRNMSTQLGCTLNFPISPIETVPVKLKPGMDGPRVKQWPLTEEKIKALQAICEEM  761

Query  189  EKEGKISKIGPENPYNTPVFAIKKKDSTKWRKLVDFRELNKRTQDFWEVQLGIPHPAGLK  248
            EKEGKI+KIGPENPYNTPVFAIKKKDSTKWRKLVDFRELNKRTQDFWEVQLGIPHPAGLK
Sbjct  760  EKEGKITKIGPENPYNTPVFAIKKKDSTKWRKLVDFRELNKRTQDFWEVQLGIPHPAGLK  581

Query  249  KKKSVTVLDVGDAYFSVPLDEDFRKYTAFTIPSINNETPGIRYQYNVLPQGWKGSPAIFQ  308
            KKKSVTVLDVGDAYFSVPLDE+FRKYTAFTIPSINN TPGIRYQYNVLPQGWKGSPAIFQ
Sbjct  580  KKKSVTVLDVGDAYFSVPLDENFRKYTAFTIPSINNATPGIRYQYNVLPQGWKGSPAIFQ  401

Query  309  SSMTKILEPFRKQNPDIVIYQYMDDLYVGSDLEIGQHRTKIEELRQHLLRWGLTTPDKKH  368
            S+MTKIL+PFR +NPDIVIYQYMDDLYVGSDLEIGQHR KIEELR+HLL+WGLTTPDKKH
Sbjct  400  STMTKILQPFRTRNPDIVIYQYMDDLYVGSDLEIGQHRAKIEELREHLLKWGLTTPDKKH  221

Query  369  QKEPPFLWMGYELHPDKWTVQPIVLPEKDSWTVNDIQKLVGKLNWASQIYPGIKVRQLCK  428
            QKEPPFLWMGYELHPDKWTVQPI LP+K+SWTVNDIQKLVGKLNWASQIYPGI+V++LC+
Sbjct  220  QKEPPFLWMGYELHPDKWTVQPIQLPDKESWTVNDIQKLVGKLNWASQIYPGIRVKRLCR  41

Query  429  LLRGTKALTEVIP  441
            LLRG KALT++IP
Sbjct  40   LLRGAKALTDIIP  2
```

>Methylobacterium radiotolerans strain MAMP 4754 MP-M-radiotolerans_contig_704, whole genome shotgun sequence
Sequence ID: NZ_NKQS01000691.1 Length: 1636
Range 1: 606 to 1634

Score:599 bits(1544), Expect:0.0,
Method:Compositional matrix adjust.,
Identities:307/343(90%), Positives:325/343(94%), Gaps:0/343(0%)

```
Query  369   QKEPPFLWMGYELHPDKWTVQPIVLPEKDSWTVNDIQKLVGKLNWASQIYPGIKVRQLCK  428
             QKEPPFLWMGYELHPDKWTVQPIVLPEK+SWTVNDIQKLVGKLNWASQIYPGIKVRQ+CK
Sbjct  1634  QKEPPFLWMGYELHPDKWTVQPIVLPEKESWTVNDIQKLVGKLNWASQIYPGIKVRQMCK  1455

Query  429   LLRGTKALTEVIPLTeeaelelaeNREILKEPVHGVYYDPSKDLIAEIQKQGQGQWTYQI  488
             LLRGTKALT+++PLT EAELELAENREILKEPVHG YYDPSKDLIAEIQKQG  QWTYQI
Sbjct  1454  LLRGTKALTDIVPLTTEAELELAENREILKEPVHGAYYDPSKDLIAEIQKQGHDQWTYQI  1275

Query  489   YQEPFKNLKTGKYARMRGAHTNDVKQLTEAVQKITTESIVIWGKTPKFKLPIQKetwetw  548
             YQEPFKNLKTGKYA M+ A+ NDVKQLTEAVQKI  ESIVIWGKTPKF+LPIQKETWETW
Sbjct  1274  YQEPFKNLKTGKYA*MKSANINDVKQLTEAVQKIALESIVIWGKTPKFRLPIQKETWETW  1095

Query  549   wteYWQATWIPEWEFVNTPPLVKLWYQLEKEPIVGAETFYVDGAANRETKLGKAGYVTNR  608
             WT+YWQATWIPEWEFVNTPPLVKLWYQLEKEPI GAETFYVDGAANRETK GKAGYVT+R
Sbjct  1094  WTDYWQATWIPEWEFVNTPPLVKLWYQLEKEPIAGAETFYVDGAANRETKRGKAGYVTDR  915

Query  609   GRQKVVTLTDTTNQKTELQAIYLALQDSGLEVNIVTDSQYALGIIQAQPDQSESELVNQI  668
             GRQK+V L +TTNQK ELQAI LALQDSG EVNIVTDSQYALGIIQAQPD+SESELVNQI
Sbjct  914   GRQKIVPLNETTNQKAELQAIQLALQDSGSEVNIVTDSQYALGIIQAQPDKSESELVNQI  735

Query  669   IEQLIKKEKVYLAWVPAHKGIGGNEQVDKLVSAGIRKVLYVDG  711
             IEQLI KE++YL+WVPAHKGIGGNEQVDKLVS GIRKVL++DG
Sbjct  734   IEQLINKERIYLSWVPAHKGIGGNEQVDKLVSTGIRKVLFLDG  606
```

Range 2: 42 to 1016

Score:558 bits(1438), Expect:0.0,
Method:Compositional matrix adjust.,
Identities:273/326(84%), Positives:297/326(91%), Gaps:3/326(0%)

```
Query  694   QVDKLVSAGIRKVLYVDGAANRETKLGKAGYVTNRGRQKVVTLTDTTNQKTELQAIYLAL  753
             Q++K   AG  +  YVDGAANRETK GKAGYVT+RGRQK+V L +TTNQK ELQAI LAL
Sbjct  1016  QLEKEPIAG-AETFYVDGAANRETKRGKAGYVTDRGRQKIVPLNETTNQKAELQAIQLAL  840

Query  754   QDSGLEVNIVTDSQYALGIIQAQPDQSESELVNQIIEQLIKKEKVYLAWVPAHKGIGGNE  813
             QDSG EVNIVTDSQYALGIIQAQPD+SESELVNQIIEQLI KE++YL+WVPAHKGIGGNE
Sbjct  839   QDSGSEVNIVTDSQYALGIIQAQPDKSESELVNQIIEQLINKERIYLSWVPAHKGIGGNE  660

Query  814   QVDKLVSAGIRKVLFLDGIDKAQDEHEKYHSNWRAMASDFNLPPVVAKEIVASCDKCQLK  873
             QVDKLVS GIRKVLFLDGIDKAQ+EHEKYHSNWRAMAS+FNLPP+VAKEIVASCDKCQLK
Sbjct  659   QVDKLVSTGIRKVLFLDGIDKAQEEHEKYHSNWRAMASEFNLPPIVAKEIVASCDKCQLK  480

Query  874   GEAMHGQVDCSPGIWQLDCTHLEGKVILVAVHVASGYIEAEVIPAETGQETAYFLLKLAG  933
             GEAMHGQVDCSPGIWQLDCTHLEGK+ILVAVHVASGYIEAEVIPAETGQETAYF+LKLAG
Sbjct  479   GEAMHGQVDCSPGIWQLDCTHLEGKIILVAVHVASGYIEAEVIPAETGQETAYFILKLAG  300

Query  934   RWPVKTIHTDNGSNFTGATVRAACWWAGIKQEFGIPYNPQSQGVVESMNKELKKIIGQVR  993
             RWPVK IHTDNGSNFT A V+AACWWAGI+QEFGI YNPQ +GVV+SM+ EL KIIGQ R
Sbjct  299   RWPVKVIHTDNGSNFTSAAVKAACWWAGIQQEFGIHYNPQRKGVVDSMHTELTKIIGQER  120

Query  994   DQAEHLKTAVQMAVFIHNF--KRKGG  1017
             D+ EH+KTAV MAV+IHN   KR+ G
Sbjct  119   DKDEHIKTAVLMAVYIHNCSSKREAG  42
```

>Providencia rettgeri strain APW139_S1 24_contig_65, whole genome shotgun sequence
Sequence ID: NZ_SHDG01000065.1 Length: 941
Range 1: 9 to 941

Score:588 bits(1516), Expect:0.0,
Method:Compositional matrix adjust.,
Identities:278/311(89%), Positives:298/311(95%), Gaps:0/311(0%)

```
Query  50   QVTLWQRPLVTIKIGGQLKEALLDTGADDTVLEEMSLPGRWKPKMIGGIGGFIKVRQYDQ  109
            ++TLW+RPL TIK+G Q+KEALLDTGADDTVLEE++LPG+WKPKMIGGIGGFIKVRQYD
Sbjct  9    RITLWRRPLDTIKVGSQIKEALLDTGADDTVLEEINLPGKWKPKMIGGIGGFIKVRQYDH  188

Query  110  ILIEICGHKAIGTVLVGPTPVNIIGRNLLTQIGCTLNFPISPIETVPVKLKPGMDGPKVK  169
            I IEICG KAIGTVLVGPTPVNIIGRNLLTQ+GCTLNFPIS IETVPVKLKPGMDGP+VK
Sbjct  189  IPIEICGKKAIGTVLVGPTPVNIIGRNLLTQLGCTLNFPISSIETVPVKLKPGMDGPRVK  368

Query  170  QWPLTEEKIKALVEICTEMEKEGKISKIGPENPYNTPVFAIKKKDSTKWRKLVDFRELNK  229
            QWPLTEEKIKAL EIC EMEKEGKI+KIGPENPYNTPVFAIKKKDSTKWRK+VDFRELNK
Sbjct  369  QWPLTEEKIKALTEICEEMEKEGKITKIGPENPYNTPVFAIKKKDSTKWRKVVDFRELNK  548

Query  230  RTQDFWEVQLGIPHPAGLKKKKSVTVLDVGDAYFSVPLDEDFRKYTAFTIPSINNETPGI  289
            RTQDFWEVQLGIPHPAG+KK KSVTVLDVGDA+FSVPLDE FRKYTAFTIPSINNE PGI
Sbjct  549  RTQDFWEVQLGIPHPAGIKKNKSVTVLDVGDAFFSVPLDESFRKYTAFTIPSINNEAPGI  728

Query  290  RYQYNVLPQGWKGSPAIFQSSMTKILEPFRKQNPDIVIYQYMDDLYVGSDLEIGQHRTKI  349
            RYQYNVLPQGWKGSPAIFQSSMT+ILEPFR +NP+IVIYQY+DDLYVGSDLEIGQHR KI
Sbjct  729  RYQYNVLPQGWKGSPAIFQSSMTRILEPFRAKNPEIVIYQYVDDLYVGSDLEIGQHRAKI  908

Query  350  EELRQHLLRWG  360
            EELR+HLL+WG
Sbjct  909  EELREHLLKWG  941
```

>Providencia rettgeri strain APW139_S1 24_contig_64, whole genome shotgun sequence
Sequence ID: NZ_SHDG01000064.1 Length: 902
Range 1: 3 to 902

Score:578 bits(1490), Expect:0.0,
Method:Compositional matrix adjust.,
Identities:271/300(90%), Positives:287/300(95%), Gaps:0/300(0%)

```
Query  104  VRQYDQILIEICGHKAIGTVLVGPTPVNIIGRNLLTQIGCTLNFPISPIETVPVKLKPGM  163
            VRQYDQI IEICG KAIGTVLVGPTPVNIIGRNLLTQ+GCTLNFPISPI+TVPVKLKPGM
Sbjct  3    VRQYDQIPIEICGKKAIGTVLVGPTPVNIIGRNLLTQLGCTLNFPISPIKTVPVKLKPGM  182

Query  164  DGPKVKQWPLTEEKIKALVEICTEMEKEGKISKIGPENPYNTPVFAIKKKDSTKWRKLVD  223
            DGPKVKQWPLTEEKIKAL  IC EMEKEGKI+KIGPENPYNTP+FAIKKK STKWRK+VD
Sbjct  183  DGPKVKQWPLTEEKIKALTAICEEMEKEGKITKIGPENPYNTPIFAIKKKGSTKWRKVVD  362

Query  224  FRELNKRTQDFWEVQLGIPHPAGLKKKKSVTVLDVGDAYFSVPLDEDFRKYTAFTIPSIN  283
            FRELNKRTQDFWEVQLGIPHPAGLK+ KS+TVLDVGDA+FSVPLD+DFRKYTAFTIPS+N
Sbjct  363  FRELNKRTQDFWEVQLGIPHPAGLKRNKSMTVLDVGDAFFSVPLDKDFRKYTAFTIPSVN  542

Query  284  NETPGIRYQYNVLPQGWKGSPAIFQSSMTKILEPFRKQNPDIVIYQYMDDLYVGSDLEIG  343
            N TPGIRYQYNVLPQGWKGSPAIFQ SMTKILEPFR QNPDIVIYQY+DDLYVGSDLEIG
Sbjct  543  NATPGIRYQYNVLPQGWKGSPAIFQCSMTKILEPFRTQNPDIVIYQYVDDLYVGSDLEIG  722

Query  344  QHRTKIEELRQHLLRWGLTTPDKKHQKEPPFLWMGYELHPDKWTVQPIVLPEKDSWTVND  403
            QHR K+EELR+HLL+WG TTPDKKHQKEPPFLWMGYELHPDKWTVQPI LP+K+SWTVND
Sbjct  723  QHRAKVEELRKHLLKWGFTTPDKKHQKEPPFLWMGYELHPDKWTVQPIQLPDKESWTVND  902
```

>Acinetobacter baumannii strain ACM-29 NODE_109_length_888_cov_0.827858, whole genome shotgun sequence
Sequence ID: NZ_PYDO01000109.1 Length: 888
Range 1: 2 to 886

Score:568 bits(1465), Expect:0.0,
Method:Compositional matrix adjust.,
Identities:270/295(92%), Positives:286/295(96%), Gaps:0/295(0%)

```
Query  61   IKIGGQLKEALLDTGADDTVLEEMSLPGRWKPKMIGGIGGFIKVRQYDQILIEICGHKAI  120
            IK+GGQLKEALLDTGA+DTVLE+M LPGRWKPKMIGGI GFIKVRQYD++ IEICGHKAI
Sbjct  886  IKVGGQLKEALLDTGAEDTVLEDMQLPGRWKPKMIGGIRGFIKVRQYDEVPIEICGHKAI  707

Query  121  GTVLVGPTPVNIIGRNLLTQIGCTLNFPISPIETVPVKLKPGMDGPKVKQWPLTEEKIKA  180
            GTVL+GPTPVNIIGRNLLTQ+GCTLNFPISPIETVPVKLKPGMDGPKVKQWPLTEEKIKA
Sbjct  706  GTVLIGPTPVNIIGRNLLTQLGCTLNFPISPIETVPVKLKPGMDGPKVKQWPLTEEKIKA  527

Query  181  LVEICTEMEKEGKISKIGPENPYNTPVFAIKKKDSTKWRKLVDFRELNKRTQDFWEVQLG  240
            LVEICTEMEKEGKI+KIGPENPYNTP+FAIKKK+S KWRKLVDFRELNKRTQ+FWEVQLG
Sbjct  526  LVEICTEMEKEGKITKIGPENPYNTPIFAIKKKNSDKWRKLVDFRELNKRTQEFWEVQLG  347

Query  241  IPHPAGLKKKKSVTVLDVGDAYFSVPLDEDFRKYTAFTIPSINNETPGIRYQYNVLPQGW  300
            IPHPAGLKKKKSVTVLDVGDAYFSVPLD DFRKYTAFT PS NNETPGIRYQYNVLPQGW
Sbjct  346  IPHPAGLKKKKSVTVLDVGDAYFSVPLDPDFRKYTAFTTPSRNNETPGIRYQYNVLPQGW  167

Query  301  KGSPAIFQSSMTKILEPFRKQNPDIVIYQYMDDLYVGSDLEIGQHRTKIEELRQH  355
```

```
           KGSPAIFQSSMTKILEPFRKQNPD+VIYQY+DDLYVGSDLE+ QHRTK+EELR +
Sbjct  166  KGSPAIFQSSMTKILEPFRKQNPDMVIYQYIDDLYVGSDLELEQHRTKVEELRAY  2
```

```
>Mycobacterium tuberculosis variant bovis strain CSURQ1139, whole genome shotgun sequence
Sequence ID: NZ_CAJEUO010000157.1 Length: 848
Range 1: 1 to 846

Score:563 bits(1450), Expect:0.0,
Method:Compositional matrix adjust.,
Identities:264/282(94%), Positives:277/282(98%), Gaps:0/282(0%)

Query  125  VGPTPVNIIGRNLLTQIGCTLNFPISPIETVPVKLKPGMDGPKVKQWPLTEEKIKALVEI  184
            VGPTPVNIIGRNLLTQIGCTLNFPISPIETVPVKLKPGMDGPKVKQWPLTEEKIKAL EI
Sbjct  846  VGPTPVNIIGRNLLTQIGCTLNFPISPIETVPVKLKPGMDGPKVKQWPLTEEKIKALTEI  667

Query  185  CTEMEKEGKISKIGPENPYNTPVFAIKKKDSTKWRKLVDFRELNKRTQDFWEVQLGIPHP  244
            CTEMEKEGKISKIGPENPYNTPVFAI+KKDSTKWRKLVDFRELNKRTQDFWE+QLGIPHP
Sbjct  666  CTEMEKEGKISKIGPENPYNTPVFAIRKKDSTKWRKLVDFRELNKRTQDFWEIQLGIPHP  487

Query  245  AGLKKKKSVTVLDVGDAYFSVPLDEDFRKYTAFTIPSINNETPGIRYQYNVLPQGWKGSP  304
            AGLKKKKSVTVLDVGDAYFSVPLD+DFRKYTAFTIPSINNETPGIRYQYNVLPQGWKGSP
Sbjct  486  AGLKKKKSVTVLDVGDAYFSVPLDKDFRKYTAFTIPSINNETPGIRYQYNVLPQGWKGSP  307

Query  305  AIFQSSMTKILEPFRKQNPDIVIYQYMDDLYVGSDLEIGQHRTKIEELRQHLLRWGLTTP  364
            AIFQ+SMTKILEPFR +NP++VIYQYMDDLY+GSDLEIGQHRTKIEELR HLLRWG TTP
Sbjct  306  AIFQASMTKILEPFRTKNPELVIYQYMDDLYIGSDLEIGQHRTKIEELRAHLLRWGFTTP  127

Query  365  DKKHQKEPPFLWMGYELHPDKWTVQPIVLPEKDSWTVNDIQK  406
            DKKHQKEPPFLWMGYELHPDKWTVQP+ LP+K++WTVNDI+K
Sbjct  126  DKKHQKEPPFLWMGYELHPDKWTVQPVQLPDKENWTVNDIRK  1
```

```
Range 2: 1 to 846

Score:563 bits(1450), Expect:0.0,
Method:Compositional matrix adjust.,
Identities:264/282(94%), Positives:277/282(98%), Gaps:0/282(0%)

Query  125  VGPTPVNIIGRNLLTQIGCTLNFPISPIETVPVKLKPGMDGPKVKQWPLTEEKIKALVEI  184
            VGPTPVNIIGRNLLTQIGCTLNFPISPIETVPVKLKPGMDGPKVKQWPLTEEKIKAL EI
Sbjct  846  VGPTPVNIIGRNLLTQIGCTLNFPISPIETVPVKLKPGMDGPKVKQWPLTEEKIKALTEI  667

Query  185  CTEMEKEGKISKIGPENPYNTPVFAIKKKDSTKWRKLVDFRELNKRTQDFWEVQLGIPHP  244
            CTEMEKEGKISKIGPENPYNTPVFAI+KKDSTKWRKLVDFRELNKRTQDFWE+QLGIPHP
Sbjct  666  CTEMEKEGKISKIGPENPYNTPVFAIRKKDSTKWRKLVDFRELNKRTQDFWEIQLGIPHP  487

Query  245  AGLKKKKSVTVLDVGDAYFSVPLDEDFRKYTAFTIPSINNETPGIRYQYNVLPQGWKGSP  304
            AGLKKKKSVTVLDVGDAYFSVPLD+DFRKYTAFTIPSINNETPGIRYQYNVLPQGWKGSP
Sbjct  486  AGLKKKKSVTVLDVGDAYFSVPLDKDFRKYTAFTIPSINNETPGIRYQYNVLPQGWKGSP  307

Query  305  AIFQSSMTKILEPFRKQNPDIVIYQYMDDLYVGSDLEIGQHRTKIEELRQHLLRWGLTTP  364
            AIFQ+SMTKILEPFR +NP++VIYQYMDDLY+GSDLEIGQHRTKIEELR HLLRWG TTP
Sbjct  306  AIFQASMTKILEPFRTKNPELVIYQYMDDLYIGSDLEIGQHRTKIEELRAHLLRWGFTTP  127

Query  365  DKKHQKEPPFLWMGYELHPDKWTVQPIVLPEKDSWTVNDIQK  406
            DKKHQKEPPFLWMGYELHPDKWTVQP+ LP+K++WTVNDI+K
Sbjct  126  DKKHQKEPPFLWMGYELHPDKWTVQPVQLPDKENWTVNDIRK  1
```

```
>Mycobacterium tuberculosis strain 32846_S30 184, whole genome shotgun sequence
Sequence ID: NZ_QTBW01000041.1 Length: 932
Range 1: 4 to 930

Score:558 bits(1438), Expect:0.0,
Method:Compositional matrix adjust.,
Identities:283/309(92%), Positives:297/309(96%), Gaps:0/309(0%)

Query  273  KYTAFTIPSINNETPGIRYQYNVLPQGWKGSPAIFQSSMTKILEPFRKQNPDIVIYQYMD  332
            KYTAFTIPSINNETPGIRYQYNVLPQGWKGSPAIFQ SMTKILEPFR +NP+IVIYQYMD
Sbjct  930  KYTAFTIPSINNETPGIRYQYNVLPQGWKGSPAIFQYSMTKILEPFRAKNPEIVIYQYMD  751

Query  333  DLYVGSDLEIGQHRTKIEELRQHLLRWGLTTPDKKHQKEPPFLWMGYELHPDKWTVQPIV  392
            DLYVGSDLEIGQHR KIEELR HLL+WG TTPDKKHQKEPPFLWMGYELHPDKWTVQPI
Sbjct  750  DLYVGSDLEIGQHRVKIEELRGHLLKWGFTTPDKKHQKEPPFLWMGYELHPDKWTVQPIQ  571

Query  393  LPEKDSWTVNDIQKLVGKLNWASQIYPGIKVRQLCKLLRGTKALTEVIPLTeeaelelae  452
            LPEKDSWTVNDIQKLVGKLNWASQIYPGIKV+QLC+LLRG KALT+++PLTEEAELELAE
Sbjct  570  LPEKDSWTVNDIQKLVGKLNWASQIYPGIKVKQLCRLLRGAKALTDIVPLTEEAELELAE  391

Query  453  NREILKEPVHGVYYDPSKDLIAEIQKQGQGQWTYQIYQEPFKNLKTGKYARMRGAHTNDV  512
            NREILKEPVHGVYYDP+KDLIAEIQKQ  GQWTYQIYQEPFKNLKTGKYA+MR AHTNDV
Sbjct  390  NREILKEPVHGVYYDPAKDLIAEIQKQDHGQWTYQIYQEPFKNLKTGKYAKMRTAHTNDV  211

Query  513  KQLTEAVQKITTESIVIWGKTPKFKLPIQKetwetwwteYWQATWIPEWEFVNTPPLVKL  572
            KQLTEAVQKI  ESIV+WGKTPKF+LPIQKETWETWWTEYWQATWIPEWEFVNTPPLVKL
Sbjct  210  KQLTEAVQKIAMESIVVWGKTPKFRLPIQKETWETWWTEYWQATWIPEWEFVNTPPLVKL  31

Query  573  WYQLEKEPI  581
            WYQLEK+P+
Sbjct  30   WYQLEKDPL  4
```

```
>Acinetobacter baumannii strain ACM-36 NODE_62_length_953_cov_1.8184, whole genome shotgun sequence
Sequence ID: NZ_QAGO01000062.1 Length: 953
Range 1: 69 to 953

Score:554 bits(1427), Expect:0.0,
Method:Compositional matrix adjust.,
Identities:272/295(92%), Positives:285/295(96%), Gaps:0/295(0%)

Query  189  EKEGKISKIGPENPYNTPVFAIKKKDSTKWRKLVDFRELNKRTQDFWEVQLGIPHPAGLK  248
            EKEGKISKIG E PYNTP+FAIKKK+S KWRKLVDFRELNKRTQ+FWEVQLGIPHPAGLK
Sbjct  953  EKEGKISKIGSETPYNTPIFAIKKKNSDKWRKLVDFRELNKRTQEFWEVQLGIPHPAGLK  774

Query  249  KKKSVTVLDVGDAYFSVPLDEDFRKYTAFTIPSINNETPGIRYQYNVLPQGWKGSPAIFQ  308
            KKKSVTVLDVGDAYFSVPLD DFRKYTAFTIPS NNETPGIRYQYNVLPQGWKGSPAIFQ
Sbjct  773  KKKSVTVLDVGDAYFSVPLDPDFRKYTAFTIPSRNNETPGIRYQYNVLPQGWKGSPAIFQ  594

Query  309  SSMTKILEPFRKQNPDIVIYQYMDDLYVGSDLEIGQHRTKIEELRQHLLRWGLTTPDKKH  368
            SSMTKILEPFRKQNPD+VIYQYMDDLYVGSDLE+ QHRTK+EELR HLL+WG +TPD+KH
Sbjct  593  SSMTKILEPFRKQNPDMVIYQYMDDLYVGSDLELEQHRTKVEELRAHLLKWGFSTPDQKH  414

Query  369  QKEPPFLWMGYELHPDKWTVQPIVLPEKDSWTVNDIQKLVGKLNWASQIYPGIKVRQLCK  428
            QKEPPFLWMGYELHPDKWTVQPIVLPEKDSWTVNDIQKLVGKLNWASQIY GIKV+QLCK
Sbjct  413  QKEPPFLWMGYELHPDKWTVQPIVLPEKDSWTVNDIQKLVGKLNWASQIYAGIKVKQLCK  234
```

```
Query  429  LLRGTKALTEVIPLTeeaelelaeNREILKEPVHGVYYDPSKDLIAEIQKQGQGQ  483
            LLRGTKALTEV+PL+EEAELELAENREILKEPVHGVYYDPSKDL+AEIQKQG GQ
Sbjct  233  LLRGTKALTEVVPLSEEAELELAENREILKEPVHGVYYDPSKDLVAEIQKQGLGQ  69
```

```
>Providencia rettgeri strain AFE97_S1 21_contig_71, whole genome shotgun sequence
Sequence ID: NZ_SHDO01000071.1 Length: 1157
Range 1: 10 to 804

Score:482 bits(1240), Expect:0.0,
Method:Compositional matrix adjust.,
Identities:231/267(87%), Positives:249/267(93%), Gaps:2/267(0%)

Query  28   RDNNSPSEAGADRQGTVSFNFPQVTLWQRPLVTIKIGGQLKEALLDTGADDTVLEEMSLP  87
            R +   SEAG + QGT+  NFPQ+TLWQRPLV+IK+GGQ+KEALLDTGADDTVLEE++LP
Sbjct  10   RGDKPRSEAGVEGQGTI--NFPQITLWQRPLVSIKVGGQVKEALLDTGADDTVLEEINLP  183

Query  88   GRWKPKMIGGIGGFIKVRQYDQILIEICGHKAIGTVLVGPTPVNIIGRNLLTQIGCTLNF  147
            G+WKPKMIGGIGGFIKVRQYDQI IEICG KAIGTVLVGPTPVNIIGRN+LTQ+GCTLNF
Sbjct  184  GKWKPKMIGGIGGFIKVRQYDQIPIEICGKKAIGTVLVGPTPVNIIGRNMLTQLGCTLNF  363

Query  148  PISPIETVPVKLKPGMDGPKVKQWPLTEEKIKALVEICTEMEKEGKISKIGPENPYNTPV  207
            PISPIETVPVKLKPGMDGPKVKQWPLTEEKIKAL  IC EMEKEGKI+KIGPENPYNTP+
Sbjct  364  PISPIETVPVKLKPGMDGPKVKQWPLTEEKIKALTAICEEMEKEGKITKIGPENPYNTPI  543

Query  208  FAIKKKDSTKWRKLVDFRELNKRTQDFWEVQLGIPHPAGLKKKKSVTVLDVGDAYFSVPL  267
            FAIKKKDSTKWRKLVDFRELNKRTQDFWEVQLGIPHPAGLK+ KS+TVLDVGDAYFSVPL
Sbjct  544  FAIKKKDSTKWRKLVDFRELNKRTQDFWEVQLGIPHPAGLKRNKSMTVLDVGDAYFSVPL  723

Query  268  DEDFRKYTAFTIPSINNETPGIRYQYN  294
            DE FRKYTAFTIPSINNETPGI +  +
Sbjct  724  DEGFRKYTAFTIPSINNETPGISFHRD  804



Range 2: 792 to 1157

Score:242 bits(618), Expect:0.0,
Method:Compositional matrix adjust.,
Identities:113/122(93%), Positives:116/122(95%), Gaps:0/122(0%)

Query  296   LPQGWKGSPAIFQSSMTKILEPFRKQNPDIVIYQYMDDLYVGSDLEIGQHRTKIEELRQH  355
             LPQGWKGSPAIFQSSMTKILEPFR QNP++VIYQYMDDLYVGSDLEIGQHR KIEELR H
Sbjct  792   LPQGWKGSPAIFQSSMTKILEPFRTQNPEMVIYQYMDDLYVGSDLEIGQHRAKIEELRAH  971

Query  356   LLRWGLTTPDKKHQKEPPFLWMGYELHPDKWTVQPIVLPEKDSWTVNDIQKLVGKLNWAS  415
             LL+WG TTPDKKHQKEPPFLWMGYELHPDKWTVQPI LPEKDSWTVNDIQKL GKLNWAS
Sbjct  972   LLKWGFTTPDKKHQKEPPFLWMGYELHPDKWTVQPIKLPEKDSWTVNDIQKLGGKLNWAS  1151

Query  416   QI  417
             QI
Sbjct  1152  QI  1157
```

```
>Providencia rettgeri strain AFE422_S1 40_contig_34, whole genome shotgun sequence
Sequence ID: NZ_SHDL01000034.1 Length: 1058
Range 1: 3 to 743

Score:468 bits(1205), Expect:0.0,
Method:Compositional matrix adjust.,
Identities:222/247(90%), Positives:237/247(95%), Gaps:0/247(0%)

Query  48   FPQVTLWQRPLVTIKIGGQLKEALLDTGADDTVLEEMSLPGRWKPKMIGGIGGFIKVRQY  107
            FPQ+TLWQRPLV+IK+GGQ+KEALLDTGADDTVLEE++LPG+WKPKMIGGIGGFIKVRQY
Sbjct  3    FPQITLWQRPLVSIKVGGQVKEALLDTGADDTVLEEINLPGKWKPKMIGGIGGFIKVRQY  182

Query  108  DQILIEICGHKAIGTVLVGPTPVNIIGRNLLTQIGCTLNFPISPIETVPVKLKPGMDGPK  167
            DQI IEICG KAIGTVLVGPTPVNIIGRN+LTQ+GCTLNFPISPIETVPVKLKPGMDGPK
Sbjct  183  DQIPIEICGKKAIGTVLVGPTPVNIIGRNMLTQLGCTLNFPISPIETVPVKLKPGMDGPK  362

Query  168  VKQWPLTEEKIKALVEICTEMEKEGKISKIGPENPYNTPVFAIKKKDSTKWRKLVDFREL  227
            VKQWPLTEEKIKAL  IC EMEKEGKI+KIGPENPYNTP+FAIKKKDSTKWRKLVDFREL
Sbjct  363  VKQWPLTEEKIKALTAICEEMEKEGKITKIGPENPYNTPIFAIKKKDSTKWRKLVDFREL  542

Query  228  NKRTQDFWEVQLGIPHPAGLKKKKSVTVLDVGDAYFSVPLDEDFRKYTAFTIPSINNETP  287
            NKRTQDFWEVQLGIPHPAGLK+ KS+TVLDVGDAYFSVPLDE FRKYTAFTIPSINNETP
Sbjct  543  NKRTQDFWEVQLGIPHPAGLKRNKSMTVLDVGDAYFSVPLDEGFRKYTAFTIPSINNETP  722

Query  288  GIRYQYN  294
            GI +  +
Sbjct  723  GISFHRD  743



Range 2: 731 to 1057

Score:220 bits(560), Expect:0.0,
Method:Compositional matrix adjust.,
Identities:101/109(93%), Positives:104/109(95%), Gaps:0/109(0%)

Query  296   LPQGWKGSPAIFQSSMTKILEPFRKQNPDIVIYQYMDDLYVGSDLEIGQHRTKIEELRQH  355
             LPQGWKGSPAIFQSSMTKILEPFR QNP++VIYQYMDDLYVGSDLEIGQHR KIEELR H
Sbjct  731   LPQGWKGSPAIFQSSMTKILEPFRTQNPEMVIYQYMDDLYVGSDLEIGQHRAKIEELRAH  910

Query  356   LLRWGLTTPDKKHQKEPPFLWMGYELHPDKWTVQPIVLPEKDSWTVNDI  404
             LL+WG TTPDKKHQKEPPFLWMGYELHPDKWTVQPI LPEKDSWTVNDI
Sbjct  911   LLKWGFTTPDKKHQKEPPFLWMGYELHPDKWTVQPIKLPEKDSWTVNDI  1057
```

```
>Mycobacterium tuberculosis strain 32846_S30 476, whole genome shotgun sequence
Sequence ID: NZ_QTBW01000113.1 Length: 861
Range 1: 1 to 855

Score:536 bits(1381), Expect:5e-179,
Method:Compositional matrix adjust.,
Identities:261/298(88%), Positives:275/298(92%), Gaps:13/298(4%)

Query  11   EQTRANSPTRRELQVWGRDNNSPSEAGADRQGTVSFNFPQVTLWQRPLVTIKIGGQLKEA  70
            EQ RA+SPT RELQ           AGA+RQG  + N PQ+TLWQRPLV+IK+GGQ++EA
Sbjct  855  EQARADSPTSRELQ-----------AGAERQG--ALNLPQITLWQRPLVSIKVGGQIREA  715

Query  71   LLDTGADDTVLEEMSLPGRWKPKMIGGIGGFIKVRQYDQILIEICGHKAIGTVLVGPTPV  130
            LLDTGADDTVLE+++LPG+WKPKMIGGIGGFIKVRQYDQILIEICG KAIGTVLVGPTPV
Sbjct  714  LLDTGADDTVLEDINLPGKWKPKMIGGIGGFIKVRQYDQILIEICGKKAIGTVLVGPTPV  535
```

```
Query  131  NIIGRNLLTQIGCTLNFPISPIETVPVKLKPGMDGPKVKQWPLTEEKIKALVEICTEMEK  190
            NIIGRN+LTQIGCTLNFPISPIETVPVKLKPGMDGPKVKQWPLTEEKIKAL  IC EMEK
Sbjct  534  NIIGRNMLTQIGCTLNFPISPIETVPVKLKPGMDGPKVKQWPLTEEKIKALTAICEEMEK  355

Query  191  EGKISKIGPENPYNTPVFAIKKKDSTKWRKLVDFRELNKRTQDFWEVQLGIPHPAGLKKK  250
            EGKI+KIGPENPYNTPVFAIKKKDSTKWRKLVDFRELNKRTQDFWEVQLGIPHPAGLKK
Sbjct  354  EGKITKIGPENPYNTPVFAIKKKDSTKWRKLVDFRELNKRTQDFWEVQLGIPHPAGLKKN  175

Query  251  KSVTVLDVGDAYFSVPLDEDFRKYTAFTIPSINNETPGIRYQYNVLPQGWKGSPAIFQ  308
            KSVTVLDVGDAYFSVPLDE FRKYTAFTIPSINNETPGIRYQYNVLPQGWKGSPAIFQ
Sbjct  174  KSVTVLDVGDAYFSVPLDESFRKYTAFTIPSINNETPGIRYQYNVLPQGWKGSPAIFQ  1
```

```
>Mycobacterium tuberculosis variant bovis strain CSURQ1134, whole genome shotgun sequence
Sequence ID: NZ_CAJEUT010000134.1 Length: 910
Range 1: 54 to 905

Score:530 bits(1365), Expect:2e-176,
Method:Compositional matrix adjust.,
Identities:254/284(89%), Positives:264/284(92%), Gaps:0/284(0%)

Query  143  CTLNFPISPIETVPVKLKPGMDGPKVKQWPLTEEKIKALVEICTEMEKEGKISKIGPENP  202
             +LNFP  P ET+ VKLK  M GP+ KQWPLT   I +L EICTEMEKEGKISKIGPENP
Sbjct  905  MSLNFPSRPTETLRVKLKR*MVGPQAKQWPLTA*TINSLTEICTEMEKEGKISKIGPENP  726

Query  203  YNTPVFAIKKKDSTKWRKLVDFRELNKRTQDFWEVQLGIPHPAGLKKKKSVTVLDVGDAY  262
            YNTPVFAIKKK+STKWRKLVDFRELNK+TQDFWEVQLGIPHP+GLKKKKSVTVLDVGDAY
Sbjct  725  YNTPVFAIKKKNSTKWRKLVDFRELNKKTQDFWEVQLGIPHPSGLKKKKSVTVLDVGDAY  546

Query  263  FSVPLDEDFRKYTAFTIPSINNETPGIRYQYNVLPQGWKGSPAIFQSSMTKILEPFRKQN  322
            FSVPLDEDFRKYTAFTIPS+NN TPGIRYQYNVLPQGWKGSPAIFQSSMTKILEPFRKQN
Sbjct  545  FSVPLDEDFRKYTAFTIPSVNNATPGIRYQYNVLPQGWKGSPAIFQSSMTKILEPFRKQN  366

Query  323  PDIVIYQYMDDLYVGSDLEIGQHRTKIEELRQHLLRWGLTTPDKKHQKEPPFLWMGYELH  382
            PDIVIYQY+DDLYVGSDLEI QHRTK+EELRQHL RWG  TPDKKHQKEPPFLWMGYELH
Sbjct  365  PDIVIYQYVDDLYVGSDLEIEQHRTKVEELRQHLWRWGFDTPDKKHQKEPPFLWMGYELH  186

Query  383  PDKWTVQPIVLPEKDSWTVNDIQKLVGKLNWASQIYPGIKVRQL  426
            PDKWTVQPIVLPEKDSWTVNDIQKLVGKLNWASQIYPGIKV QL
Sbjct  185  PDKWTVQPIVLPEKDSWTVNDIQKLVGKLNWASQIYPGIKVNQL  54
```

```
Range 2: 54 to 905

Score:530 bits(1365), Expect:2e-176,
Method:Compositional matrix adjust.,
Identities:254/284(89%), Positives:264/284(92%), Gaps:0/284(0%)

Query  143  CTLNFPISPIETVPVKLKPGMDGPKVKQWPLTEEKIKALVEICTEMEKEGKISKIGPENP  202
             +LNFP  P ET+ VKLK  M GP+ KQWPLT   I +L EICTEMEKEGKISKIGPENP
Sbjct  905  MSLNFPSRPTETLRVKLKR*MVGPQAKQWPLTA*TINSLTEICTEMEKEGKISKIGPENP  726

Query  203  YNTPVFAIKKKDSTKWRKLVDFRELNKRTQDFWEVQLGIPHPAGLKKKKSVTVLDVGDAY  262
            YNTPVFAIKKK+STKWRKLVDFRELNK+TQDFWEVQLGIPHP+GLKKKKSVTVLDVGDAY
Sbjct  725  YNTPVFAIKKKNSTKWRKLVDFRELNKKTQDFWEVQLGIPHPSGLKKKKSVTVLDVGDAY  546

Query  263  FSVPLDEDFRKYTAFTIPSINNETPGIRYQYNVLPQGWKGSPAIFQSSMTKILEPFRKQN  322
            FSVPLDEDFRKYTAFTIPS+NN TPGIRYQYNVLPQGWKGSPAIFQSSMTKILEPFRKQN
Sbjct  545  FSVPLDEDFRKYTAFTIPSVNNATPGIRYQYNVLPQGWKGSPAIFQSSMTKILEPFRKQN  366

Query  323  PDIVIYQYMDDLYVGSDLEIGQHRTKIEELRQHLLRWGLTTPDKKHQKEPPFLWMGYELH  382
            PDIVIYQY+DDLYVGSDLEI QHRTK+EELRQHL RWG  TPDKKHQKEPPFLWMGYELH
Sbjct  365  PDIVIYQYVDDLYVGSDLEIEQHRTKVEELRQHLWRWGFDTPDKKHQKEPPFLWMGYELH  186

Query  383  PDKWTVQPIVLPEKDSWTVNDIQKLVGKLNWASQIYPGIKVRQL  426
            PDKWTVQPIVLPEKDSWTVNDIQKLVGKLNWASQIYPGIKV QL
Sbjct  185  PDKWTVQPIVLPEKDSWTVNDIQKLVGKLNWASQIYPGIKVNQL  54
```

```
>Providencia rettgeri strain AFE97_S1 21_contig_72, whole genome shotgun sequence
Sequence ID: NZ_SHDO01000072.1 Length: 854
Range 1: 3 to 854

Score:528 bits(1361), Expect:5e-176,
Method:Compositional matrix adjust.,
Identities:254/284(89%), Positives:271/284(95%), Gaps:7/284(2%)

Query  61   IKIGGQLKEALLDTGADDTVLEEMSLPGRWKPKMIGGIGGFIKVRQYDQILIEICGHKAI  120
            IKIGGQ+KEALLDTGADDTVLEE++LPG+WKPKMIGGIGGFIKVRQY++I IEICG KAI
Sbjct  3    IKIGGQIKEALLDTGADDTVLEEINLPGKWKPKMIGGIGGFIKVRQYEEIPIEICGKKAI  182

Query  121  GTVLVGPTPVNIIGRNLLTQIGCTLNFPISP-------IETVPVKLKPGMDGPKVKQWPL  173
            G+VLVGPTPVNIIGRN+LTQ+GCTLNFPISP       I+TVPVKLKPGMDGPKVKQWPL
Sbjct  183  GSVLVGPTPVNIIGRNMLTQLGCTLNFPISPPFSFKSPIKTVPVKLKPGMDGPKVKQWPL  362

Query  174  TEEKIKALVEICTEMEKEGKISKIGPENPYNTPVFAIKKKDSTKWRKLVDFRELNKRTQD  233
            TEEKIKAL EIC EMEKEGKI+KIGPENPYNTP+FAIK+KDSTKWRKLVDFRELNKRTQD
Sbjct  363  TEEKIKALTEICDEMEKEGKITKIGPENPYNTPIFAIKRKDSTKWRKLVDFRELNKRTQD  542

Query  234  FWEVQLGIPHPAGLKKKKSVTVLDVGDAYFSVPLDEDFRKYTAFTIPSINNETPGIRYQY  293
            FWEVQLGIPHPAGLKKKKS+TVLDVGDAYFSVPLDE FRKYTAFTIPSINNETPGIRYQY
Sbjct  543  FWEVQLGIPHPAGLKKKKSMTVLDVGDAYFSVPLDESFRKYTAFTIPSINNETPGIRYQY  722

Query  294  NVLPQGWKGSPAIFQSSMTKILEPFRKQNPDIVIYQYMDDLYVG  337
            NVLPQGWKGSPAIFQSSMTKILEPFR +NP+IVIYQY+DDLYVG
Sbjct  723  NVLPQGWKGSPAIFQSSMTKILEPFRAKNPEIVIYQYVDDLYVG  854
```

```
>Enterobacter hormaechei strain AUH-ENM4 NODE_27_length_809_cov_0.64956, whole genome shotgun sequence
Sequence ID: NZ_PYGZ01000027.1 Length: 809
Range 1: 3 to 809

Score:525 bits(1352), Expect:6e-175,
Method:Compositional matrix adjust.,
Identities:248/269(92%), Positives:261/269(97%), Gaps:0/269(0%)

Query  123  VLVGPTPVNIIGRNLLTQIGCTLNFPISPIETVPVKLKPGMDGPKVKQWPLTEEKIKALV  182
            VLVGPTPVNIIGRNLLTQ+GCTLNFPISPIETVPVKLKPGMDGPKVKQWPLTEEKIKALV
Sbjct  809  VLVGPTPVNIIGRNLLTQLGCTLNFPISPIETVPVKLKPGMDGPKVKQWPLTEEKIKALV  630

Query  183  EICTEMEKEGKISKIGPENPYNTPVFAIKKKDSTKWRKLVDFRELNKRTQDFWEVQLGIP  242
            EICTEMEKEGKISKIGPENPYNTP+FAIKKK+S KWRKLV FRELNKRTQ+FWE+QLGIP
Sbjct  629  EICTEMEKEGKISKIGPENPYNTPIFAIKKKNSDKWRKLVGFRELNKRTQEFWEIQLGIP  450

Query  243  HPAGLKKKKSVTVLDVGDAYFSVPLDEDFRKYTAFTIPSINNETPGIRYQYNVLPQGWKG  302
            HPAGLKKKKSVTVLDVGDAYFSVPLD++FRKYTAFTIPS NN+TPGIRYQYNVLPQGWKG
```

```
Sbjct  449  HPAGLKKKKSVTVLDVGDAYFSVPLDKEFRKYTAFTIPSTNNKTPGIRYQYNVLPQGWKG  270

Query  303  SPAIFQSSMTKILEPFRKQNPDIVIYQYMDDLYVGSDLEIGQHRTKIEELRQHLLRWGLT  362
            SPAIFQSSMTKILEPFRKQNPD+VI QYMDDLYVGSDLEI Q+R K+EELR HLL+WG T
Sbjct  269  SPAIFQSSMTKILEPFRKQNPDMVICQYMDDLYVGSDLEIEQNRIKVEELRAHLLKWGFT  90

Query  363  TPDKKHQKEPPFLWMGYELHPDKWTVQPI  391
            TPD+KHQKEPPFLWMGYELHPDKWTVQPI
Sbjct  89   TPDQKHQKEPPFLWMGYELHPDKWTVQPI  3
```

>Candidatus Tokpelaia sp. strain RhiAcro1 isolate RhiAcro1_RAo1 NODE_51_length_826_cov_3.34846_ID_101, whole genome shotgun sequence
Sequence ID: NZ_QLZD01000026.1 Length: 826
Range 1: 2 to 826

Score:514 bits(1324), Expect:1e-170,
Method:Compositional matrix adjust.,
Identities:255/275(93%), Positives:267/275(97%), Gaps:0/275(0%)

```
Query  209  AIKKKDSTKWRKLVDFRELNKRTQDFWEVQLGIPHPAGLKKKKSVTVLDVGDAYFSVPLD  268
            AIKKKDSTKWRKLVDFRELNKRTQDFWEVQLGIPHPAGLKKKKSVTVLDVGDAYFSVPLD
Sbjct  826  AIKKKDSTKWRKLVDFRELNKRTQDFWEVQLGIPHPAGLKKKKSVTVLDVGDAYFSVPLD  647

Query  269  EDFRKYTAFTIPSINNETPGIRYQYNVLPQGWKGSPAIFQSSMTKILEPFRKQNPDIVIY  328
            ++FRKYT FTIPSINNETPGIRYQYNVLPQGWKGSPAIFQ SMTKILEPFRKQNP++VIY
Sbjct  646  KEFRKYTTFTIPSINNETPGIRYQYNVLPQGWKGSPAIFQCSMTKILEPFRKQNPEMVIY  467

Query  329  QYMDDLYVGSDLEIGQHRTKIEELRQHLLRWGLTTPDKKHQKEPPFLWMGYELHPDKWTV  388
            QYMDDLYVGSDLE+GQHRTKIEELR+HLL WG TTPDKKHQKEPPFLWMGYELHPDKWTV
Sbjct  466  QYMDDLYVGSDLELGQHRTKIEELRKHLLAWGFTTPDKKHQKEPPFLWMGYELHPDKWTV  287

Query  389  QPIVLPEKDSWTVNDIQKLVGKLNWASQIYPGIKVRQLCKLLRGTKALTEVIPLTeeael  448
            QPI+LPEKDSWTVNDIQKLVGKLNWASQIY GIKV+QLCKLLRG K+LTEVI LTEEAEL
Sbjct  286  QPIMLPEKDSWTVNDIQKLVGKLNWASQIYAGIKVKQLCKLLRGAKSLTEVITLTEEAEL  107

Query  449  elaeNREILKEPVHGVYYDPSKDLIAEIQKQGQGQ  483
            ELAENREILKEPVHG YYDP+KDL+AE+QKQGQGQ
Sbjct  106  ELAENREILKEPVHGAYYDPTKDLVAELQKQGQGQ  2
```

>Acinetobacter baumannii strain ACM-27 NODE_68_length_895_cov_1.29167, whole genome shotgun sequence
Sequence ID: NZ_PYDM01000068.1 Length: 895
Range 1: 15 to 890

Score:507 bits(1306), Expect:1e-167,
Method:Compositional matrix adjust.,
Identities:241/292(83%), Positives:267/292(91%), Gaps:0/292(0%)

```
Query  152  IETVPVKLKPGMDGPKVKQWPLTEEKIKALVEICTEMEKEGKISKIGPENPYNTPVFAIK  211
            IET  VKL  GM GPK+KQW      I AL EI TE++++GKISKIGPENP NTP+FAIK
Sbjct  890  IETNKVKL*SGMYGPKIKQWS*ITHIIVALTEIFTEVKQDGKISKIGPENPCNTPIFAIK  711

Query  212  KKDSTKWRKLVDFRELNKRTQDFWEVQLGIPHPAGLKKKKSVTVLDVGDAYFSVPLDEDF  271
            KK+ST WRKL+DFRELNKRTQDFWEVQLGIPHPAGL+KKKSVTVLDVGDAYFS+PLDEDF
Sbjct  710  KKNSTGWRKLMDFRELNKRTQDFWEVQLGIPHPAGLEKKKSVTVLDVGDAYFSIPLDEDF  531

Query  272  RKYTAFTIPSINNETPGIRYQYNVLPQGWKGSPAIFQSSMTKILEPFRKQNPDIVIYQYM  331
            RKYTAFTIPS NNETPG+RYQYNVLPQGWKGSPAIFQSSMT+ILEPFR +NP+IVIYQ +
Sbjct  530  RKYTAFTIPSTNNETPGVRYQYNVLPQGWKGSPAIFQSSMTRILEPFRAKNPEIVIYQSV  351

Query  332  DDLYVGSDLEIGQHRTKIEELRQHLLRWGLTTPDKKHQKEPPFLWMGYELHPDKWTVQPI  391
            DDLYVGSDLEIGQHR KI+ELR++L +WG  TPDKKHQ+EPPFLWMGY+LHPDKWTVQPI
Sbjct  350  DDLYVGSDLEIGQHRAKIKELREYLWKWGFYTPDKKHQEEPPFLWMGYKLHPDKWTVQPI  171

Query  392  VLPEKDSWTVNDIQKLVGKLNWASQIYPGIKVRQLCKLLRGTKALTEVIPLT  443
             LP K+SWTVNDIQKLVGKLNWASQIYPGIKV+QLCKLLRGTKALT+++PLT
Sbjct  170  QLPNKESWTVNDIQKLVGKLNWASQIYPGIKVKQLCKLLRGTKALTDIVPLT  15
```

>Acinetobacter baumannii strain ACM-14 NODE_74_length_787_cov_0.637879, whole genome shotgun sequence
Sequence ID: NZ_PYDA01000074.1 Length: 787
Range 1: 3 to 785

Score:497 bits(1280), Expect:3e-164,
Method:Compositional matrix adjust.,
Identities:239/261(92%), Positives:251/261(96%), Gaps:0/261(0%)

```
Query  21   RELQVWGRDNNSPSEAGADRQGTVSFNFPQVTLWQRPLVTIKIGGQLKEALLDTGADDTV  80
            RELQVWG DNNSPSEAG +RQG VSF+ PQ+TLWQRPLVTIKIGGQLKEALLDTGADDTV
Sbjct  785  RELQVWGGDNNSPSEAGGERQGDVSFSLPQITLWQRPLVTIKIGGQLKEALLDTGADDTV  606

Query  81   LEEMSLPGRWKPKMIGGIGGFIKVRQYDQILIEICGHKAIGTVLVGPTPVNIIGRNLLTQ  140
            LE+M LP RWKPKMIGGI GFIKVRQYD++ IEICGHKAIGTVL+GPTPVNIIGRNLLTQ
Sbjct  605  LEDMQLPERWKPKMIGGIRGFIKVRQYDEVPIEICGHKAIGTVLIGPTPVNIIGRNLLTQ  426

Query  141  IGCTLNFPISPIETVPVKLKPGMDGPKVKQWPLTEEKIKALVEICTEMEKEGKISKIGPE  200
            +GCTLNFPISPIETVPVKLKPGMDGPKVKQWPLTEEKIKALVEICTEMEKEGKISKIGPE
Sbjct  425  LGCTLNFPISPIETVPVKLKPGMDGPKVKQWPLTEEKIKALVEICTEMEKEGKISKIGPE  246

Query  201  NPYNTPVFAIKKKDSTKWRKLVDFRELNKRTQDFWEVQLGIPHPAGLKKKKSVTVLDVGD  260
            NPYNTP+FAIKKK+S KWRKLVDFRELNKRTQDFWEVQLGIPHPAGLKKKKSVTVLDVGD
Sbjct  245  NPYNTPIFAIKKKNSNKWRKLVDFRELNKRTQDFWEVQLGIPHPAGLKKKKSVTVLDVGD  66

Query  261  AYFSVPLDEDFRKYTAFTIPS  281
            AYFSVPLD++FRKYTA TIPS
Sbjct  65   AYFSVPLDKEFRKYTASTIPS  3
```

>Klebsiella pneumoniae strain EuSCAPE_DE086, whole genome shotgun sequence
Sequence ID: NZ_UKLW02000059.1 Length: 995
Range 1: 2 to 994

Score:499 bits(1285), Expect:6e-164,
Method:Compositional matrix adjust.,
Identities:267/331(81%), Positives:289/331(87%), Gaps:0/331(0%)

```
Query  310  SMTKILEPFRKQNPDIVIYQYMDDLYVGSDLEIGQHRTKIEELRQHLLRWGLTTPDKKHQ  369
            SMTKILEPF+ QNP+IVIYQY+DDLYV SDL+I QHR KIEELR+HLLR   TTPDKKHQ
Sbjct  2    SMTKILEPFKAQNPEIVIYQYIDDLYVRSDLKIRQHRAKIEELREHLLR*RFTTPDKKHQ  181

Query  370  KEPPFLWMGYELHPDKWTVQPIVLPEKDSWTVNDIQKLVGKLNWASQIYPGIKVRQLCKL  429
            KEPPFL + YELHPDK TVQPI LPEK+S TVNDIQKLV KLN ASQIY  IKV+QLCKL
Sbjct  182  KEPPFL*IKYELHPDK*TVQPIKLPEKES*TVNDIQKLVRKLN*ASQIYSKIKVKQLCKL  361

Query  430  LRGTKALTEVIPLTeeaelelaeNREILKEPVHGVYYDPSKDLIAEIQKQGQGQWTYQIY  489
            LR  KALT+++PLTEEAELELAENREILKEPVH  YYDPSKDLIAE+QKQG  QWTYQIY
```

```
Sbjct  362  LREAKALTDIVPLTEEAELELAENREILKEPVHEAYYDPSKDLIAELQKQGHDQWTYQIY  541

Query  490  QEPFKNLKTGKYARMRGAHTNDVKQLTEAVQKITTESIVIWGKTPKFKLPIQKetwetww  549
            Q+PFKNLKTGKYA+MR AHTNDVKQLTEAVQKI  E IVI  KTPKFKLPIQKET E  W
Sbjct  542  QKPFKNLKTGKYAKMRTAHTNDVKQLTEAVQKIAIECIVI*RKTPKFKLPIQKET*EL*W  721

Query  550  teYWQATWIPEWEFVNTPPLVKLWYQLEKEPIVGAETFYVDGAANRETKLGKAGYVTNRG  609
            T+YWQAT IPE EFVNTPPLVKL YQLEKEPIV AETFYVD AAN+ETK+ KA YVT++
Sbjct  722  TDYWQAT*IPE*EFVNTPPLVKL*YQLEKEPIVRAETFYVDRAANKETKIRKARYVTDKR  901

Query  610  RQKVVTLTDTTNQKTELQAIYLALQDSGLEV  640
            RQK+V+LT+TTNQKTELQ I LALQDSG EV
Sbjct  902  RQKIVSLTETTNQKTELQTIQLALQDSGSEV  994




Range 2: 830 to 994

Score:80.9 bits(198), Expect:2e-11,
Method:Compositional matrix adjust.,
Identities:39/55(71%), Positives:46/55(83%), Gaps:0/55(0%)

Query  706  VLYVDGAANRETKLGKAGYVTNRGRQKVVTLTDTTNQKTELQAIYLALQDSGLEV  760
              YVD AAN+ETK+ KA YVT++ RQK+V+LT+TTNQKTELQ I LALQDSG EV
Sbjct  830  TFYVDRAANKETKIRKARYVTDKRRQKIVSLTETTNQKTELQTIQLALQDSGSEV  994




>Mycobacterium tuberculosis strain 32847_S31 462, whole genome shotgun sequence
Sequence ID: NZ_QTBX01000137.1 Length: 893
Range 1: 47 to 892

Score:496 bits(1276), Expect:3e-163,
Method:Compositional matrix adjust.,
Identities:241/284(85%), Positives:260/284(91%), Gaps:6/284(2%)

Query  2    QGKAREFSSEQTRANSPTRRELQVWGRDNNSPS----EAGADRQGTVSFNFPQVTLWQRP  57
            QG+AREF  EQ RANSPT R+L       +SP+    +AGA+RQG  + N PQ+TLWQRP
Sbjct  47   QGEAREFYPEQARANSPTSRKLPPEQARADSPTSRELQAGAERQG--ALNLPQITLWQRP  220

Query  58   LVTIKIGGQLKEALLDTGADDTVLEEMSLPGRWKPKMIGGIGGFIKVRQYDQILIEICGH  117
            LV+IK+GGQ++EALLDTGADDTVLE+++LPG+WKPKMIGGIGGFIKVRQYDQILIEICG
Sbjct  221  LVSIKVGGQIREALLDTGADDTVLEDINLPGKWKPKMIGGIGGFIKVRQYDQILIEICGK  400

Query  118  KAIGTVLVGPTPVNIIGRNLLTQIGCTLNFPISPIETVPVKLKPGMDGPKVKQWPLTEEK  177
            KAIGTVLVGPTPVNIIGRN+LTQIGCTLNFPISPIETVPVKLKPGMDGPKVKQWPLTEEK
Sbjct  401  KAIGTVLVGPTPVNIIGRNMLTQIGCTLNFPISPIETVPVKLKPGMDGPKVKQWPLTEEK  580

Query  178  IKALVEICTEMEKEGKISKIGPENPYNTPVFAIKKKDSTKWRKLVDFRELNKRTQDFWEV  237
            IKAL  IC EMEKEGKI+KIGPENPYNTPVFAIKKKDSTKWRKLVDFRELNKRTQDFWE+
Sbjct  581  IKALTAICEEMEKEGKITKIGPENPYNTPVFAIKKKDSTKWRKLVDFRELNKRTQDFWEI  760

Query  238  QLGIPHPAGLKKKKSVTVLDVGDAYFSVPLDEDFRKYTAFTIPS  281
            QLGIPHPAGLKKKKSVTVLDVGDAYFSVPLDE FRKYTAFTIPS
Sbjct  761  QLGIPHPAGLKKKKSVTVLDVGDAYFSVPLDESFRKYTAFTIPS  892




>Enterococcus faecalis strain 44.1 NODE_33_length_806_cov_1.20619, whole genome shotgun sequence
Sequence ID: NZ_NSJR01000033.1 Length: 806
Range 1: 42 to 806

Score:490 bits(1262), Expect:1e-161,
Method:Compositional matrix adjust.,
Identities:234/255(92%), Positives:245/255(96%), Gaps:0/255(0%)

Query  24   QVWGRDNNSPSEAGADRQGTVSFNFPQVTLWQRPLVTIKIGGQLKEALLDTGADDTVLEE  83
            Q WG DNNS SEAGADRQGTVSF+FPQ+TLWQRP+VTIK+GGQLKEALLDTGADDTVLEE
Sbjct  42   QAWGGDNNSLSEAGADRQGTVSFSFPQITLWQRPVVTIKVGGQLKEALLDTGADDTVLEE  221

Query  84   MSLPGRWKPKMIGGIGGFIKVRQYDQILIEICGHKAIGTVLVGPTPVNIIGRNLLTQIGC  143
            M+LPGRWKPKMIGGIGGFIKVRQYD++ IEICGHK IGTVL+G TPVNIIGRNLLTQ+GC
Sbjct  222  MNLPGRWKPKMIGGIGGFIKVRQYDEVPIEICGHKVIGTVLIGSTPVNIIGRNLLTQLGC  401

Query  144  TLNFPISPIETVPVKLKPGMDGPKVKQWPLTEEKIKALVEICTEMEKEGKISKIGPENPY  203
            TLNFPISPIETVPVKLKPGMDGPKVKQWPLTEEKIKALVEICTEMEKEGKISKIGPENPY
Sbjct  402  TLNFPISPIETVPVKLKPGMDGPKVKQWPLTEEKIKALVEICTEMEKEGKISKIGPENPY  581

Query  204  NTPVFAIKKKDSTKWRKLVDFRELNKRTQDFWEVQLGIPHPAGLKKKKSVTVLDVGDAYF  263
            NTP+FAIKKK+S KWRKLVDFR LNKRTQ FWEVQLGIPHPAGLKKKKSVTVLDVGDAYF
Sbjct  582  NTPIFAIKKKNSNKWRKLVDFRGLNKRTQGFWEVQLGIPHPAGLKKKKSVTVLDVGDAYF  761

Query  264  SVPLDEDFRKYTAFT  278
            SVPLD DFRKYTAFT
Sbjct  762  SVPLDPDFRKYTAFT  806




>Mycobacterium tuberculosis strain 32845_S29 549, whole genome shotgun sequence
Sequence ID: NZ_QTBV01000198.1 Length: 825
Range 1: 2 to 823

Score:486 bits(1251), Expect:7e-160,
Method:Compositional matrix adjust.,
Identities:249/274(91%), Positives:262/274(95%), Gaps:0/274(0%)

Query  310  SMTKILEPFRKQNPDIVIYQYMDDLYVGSDLEIGQHRTKIEELRQHLLRWGLTTPDKKHQ  369
            SMTKILEPFR +NP+IVIYQYMDDLYVGSDLEIGQHR KIEELR HLL+WG TTPDKKHQ
Sbjct  823  SMTKILEPFRTENPEIVIYQYMDDLYVGSDLEIGQHRVKIEELRGHLLKWGFTTPDKKHQ  644

Query  370  KEPPFLWMGYELHPDKWTVQPIVLPEKDSWTVNDIQKLVGKLNWASQIYPGIKVRQLCKL  429
            KEPPFLWMGYELHPDKWTVQPI LPEKDSWTVNDIQKLVGKLNWASQIYPGIKV+QLC+L
Sbjct  643  KEPPFLWMGYELHPDKWTVQPIQLPEKDSWTVNDIQKLVGKLNWASQIYPGIKVKQLCRL  464

Query  430  LRGTKALTEVIPLTeeaelelaeNREILKEPVHGVYYDPSKDLIAEIQKQGQGQWTYQIY  489
            LRG KALT+++PLTEEAELELAENREILKEPVHGVYYDP+KDLIAEIQKQ  GQWTYQIY
Sbjct  463  LRGAKALTDIVPLTEEAELELAENREILKEPVHGVYYDPAKDLIAEIQKQDHGQWTYQIY  284

Query  490  QEPFKNLKTGKYARMRGAHTNDVKQLTEAVQKITTESIVIWGKTPKFKLPIQKetwetww  549
            QEPFKNLKTGKYA+MR AHTNDVKQLTEAVQKI  ESIV+WGKTPKF+LPIQKETWETWW
Sbjct  283  QEPFKNLKTGKYAKMRTAHTNDVKQLTEAVQKIAMESIVVWGKTPKFRLPIQKETWETWW  104

Query  550  teYWQATWIPEWEFVNTPPLVKLWYQLEKEPIVG  583
            TEYWQATWIPEWEFVNTPPLVKLWYQLEK+PI G
Sbjct  103  TEYWQATWIPEWEFVNTPPLVKLWYQLEKDPIAG  2




>Klebsiella pneumoniae strain P7701 NODE_60_length_711_cov_294.76, whole genome shotgun sequence
```

```
Sequence ID: NZ_JACAWM010000058.1 Length: 711
Range 1: 1 to 711

Score:484 bits(1247), Expect:8e-160,
Method:Compositional matrix adjust.,
Identities:230/237(97%), Positives:235/237(99%), Gaps:0/237(0%)

Query  877   MHGQVDCSPGIWQLDCTHLEGKVILVAVHVASGYIEAEVIPAETGQETAYFLLKLAGRWP  936
             MHGQVDCSPGIWQLDCTHLEGKVILVAVHVASGYIEAEVIPAETGQETAYFLLKLAGRWP
Sbjct  1     MHGQVDCSPGIWQLDCTHLEGKVILVAVHVASGYIEAEVIPAETGQETAYFLLKLAGRWP  180

Query  937   VKTIHTDNGSNFTGATVRAACWWAGIKQEFGIPYNPQSQGVVESMNKELKKIIGQVRDQA  996
             VKT+HTDNGSNFT  TV+AACWWAGIKQEFGIPYNPQSQGV+ESMNKELKKIIGQVRDQA
Sbjct  181   VKTVHTDNGSNFTSTTVKAACWWAGIKQEFGIPYNPQSQGVIESMNKELKKIIGQVRDQA  360

Query  997   EHLKTAVQMAVFIHNFKRKGGIGGYSAGERIVDIIATDIQTKELQKQITKIQNFRVYYRD  1056
             EHLKTAVQMAVFIHNFKRKGGIGGYSAGERIVDIIATDIQTKELQKQITKIQNFRVYYRD
Sbjct  361   EHLKTAVQMAVFIHNFKRKGGIGGYSAGERIVDIIATDIQTKELQKQITKIQNFRVYYRD  540

Query  1057  SRNPLWKGPAKLLWKGEGAVVIQDNSDIKVVPRRKAKIIRDYGKQMAGDDCVASRQD  1113
             SR+P+WKGPAKLLWKGEGAVVIQDNSDIKVVPRRKAKIIRDYGKQMAGDDCVASRQD
Sbjct  541   SRDPVWKGPAKLLWKGEGAVVIQDNSDIKVVPRRKAKIIRDYGKQMAGDDCVASRQD  711



>Thalassovita mangrovi strain GS-10 Scaffold30_1, whole genome shotgun sequence
Sequence ID: NZ_WWEN01000028.1 Length: 722
Range 1: 2 to 706

Score:464 bits(1195), Expect:5e-152,
Method:Compositional matrix adjust.,
Identities:216/235(92%), Positives:228/235(97%), Gaps:0/235(0%)

Query  782   SELVNQIIEQLIKKEKVYLAWVPAHKGIGGNEQVDKLVSAGIRKVLFLDGIDKAQDEHEK  841
             ++  +++IEQ+ KK + Y+AWVPAHKGIGGNEQVDKLVSAGIRKVLFLDGIDKAQ+EHEK
Sbjct  706   AKAASKVIEQITKKAEAYMAWVPAHKGIGGNEQVDKLVSAGIRKVLFLDGIDKAQEEHEK  527

Query  842   YHSNWRAMASDFNLPPVVAKEIVASCDKCQLKGEAMHGQVDCSPGIWQLDCTHLEGKVIL  901
             YHSNWRAMASDFNLPPVVAKEIVASCDKCQLKGEAMHGQVDCSPGIWQLDCTHLEGKVIL
Sbjct  526   YHSNWRAMASDFNLPPVVAKEIVASCDKCQLKGEAMHGQVDCSPGIWQLDCTHLEGKVIL  347

Query  902   VAVHVASGYIEAEVIPAETGQETAYFLLKLAGRWPVKTIHTDNGSNFTGATVRAACWWAG  961
             VAVHVASGYIEAEVIPAETGQETAYFLLKLAGRWPVKT+HTDNGSNFT  TV+AACWWAG
Sbjct  346   VAVHVASGYIEAEVIPAETGQETAYFLLKLAGRWPVKTVHTDNGSNFTSTTVKAACWWAG  167

Query  962   IKQEFGIPYNPQSQGVVESMNKELKKIIGQVRDQAEHLKTAVQMAVFIHNFKRKG  1016
             IKQEFGIPYNPQSQGV+ESMNKELKKIIGQVRDQAEHLKTAVQMAVFIHNFKRKG
Sbjct  166   IKQEFGIPYNPQSQGVIESMNKELKKIIGQVRDQAEHLKTAVQMAVFIHNFKRKG  2




Range 2: 557 to 706

Score:80.1 bits(196), Expect:8e-12,
Method:Compositional matrix adjust.,
Identities:35/50(70%), Positives:45/50(90%), Gaps:0/50(0%)

Query  662  SELVNQIIEQLIKKEKVYLAWVPAHKGIGGNEQVDKLVSAGIRKVLYVDG  711
            ++  +++IEQ+ KK + Y+AWVPAHKGIGGNEQVDKLVSAGIRKVL++DG
Sbjct  706  AKAASKVIEQITKKAEAYMAWVPAHKGIGGNEQVDKLVSAGIRKVLFLDG  557




>Escherichia coli strain 8374wB4 NODE_74_length_738_cov_3.41735, whole genome shotgun sequence
Sequence ID: NZ_WSPX01000074.1 Length: 738
Range 1: 3 to 737

Score:464 bits(1195), Expect:6e-152,
Method:Compositional matrix adjust.,
Identities:215/245(88%), Positives:236/245(96%), Gaps:0/245(0%)

Query  147  FPISPIETVPVKLKPGMDGPKVKQWPLTEEKIKALVEICTEMEKEGKISKIGPENPYNTP  206
            FPISPIETVPVKLKPGMDGPK+KQWPLT+EKI+AL EICTE+E++GKISKIGPENPYNTP
Sbjct  3    FPISPIETVPVKLKPGMDGPKIKQWPLTKEKIEALTEICTELEQDGKISKIGPENPYNTP  182

Query  207  VFAIKKKDSTKWRKLVDFRELNKRTQDFWEVQLGIPHPAGLKKKKSVTVLDVGDAYFSVP  266
            +FAIKKK+ST WRKL+DFRELNKRTQDFWEVQLGIPHPAGL+KKKSVTVLDVGDAYFS+P
Sbjct  183  IFAIKKKNSTGWRKLMDFRELNKRTQDFWEVQLGIPHPAGLEKKKSVTVLDVGDAYFSIP  362

Query  267  LDEDFRKYTAFTIPSINNETPGIRYQYNVLPQGWKGSPAIFQSSMTKILEPFRKQNPDIV  326
            LDEDFRKYTAFTIPS NNETPG+RYQYNVLPQGWKGSPAIFQSSMT+ILEPFR +NP+IV
Sbjct  363  LDEDFRKYTAFTIPSTNNETPGVRYQYNVLPQGWKGSPAIFQSSMTRILEPFRAKNPEIV  542

Query  327  IYQYMDDLYVGSDLEIGQHRTKIEELRQHLLRWGLTTPDKKHQKEPPFLWMGYELHPDKW  386
            IYQY+DDLYVGSDLEIGQHR KI+ELR++L +WG  TPDKKHQ+ PPFLWMGYELHPDKW
Sbjct  543  IYQYVDDLYVGSDLEIGQHRAKIKELREYLWKWGFYTPDKKHQEGPPFLWMGYELHPDKW  722

Query  387  TVQPI  391
            TVQPI
Sbjct  723  TVQPI  737




>Acinetobacter baumannii strain ACM-17 NODE_68_length_757_cov_0.560317, whole genome shotgun sequence
Sequence ID: NZ_PYDD01000068.1 Length: 757
Range 1: 8 to 757

Score:458 bits(1179), Expect:2e-149,
Method:Compositional matrix adjust.,
Identities:226/250(90%), Positives:237/250(94%), Gaps:0/250(0%)

Query  253  VTVLDVGDAYFSVPLDEDFRKYTAFTIPSINNETPGIRYQYNVLPQGWKGSPAIFQSSMT  312
            VTV+DVGDAYFSVPLD DFRKYTAFTIPS NNETPGIRYQYNVLPQGWKGSPAIFQSSMT
Sbjct  757  VTVMDVGDAYFSVPLDPDFRKYTAFTIPSKNNETPGIRYQYNVLPQGWKGSPAIFQSSMT  578

Query  313  KILEPFRKQNPDIVIYQYMDDLYVGSDLEIGQHRTKIEELRQHLLRWGLTTPDKKHQKEP  372
            KILEPFRKQNPDIVI QYMDDLYVGSDLEI QHRTK+EELR +LL+WG +TPDKKHQKEP
Sbjct  577  KILEPFRKQNPDIVICQYMDDLYVGSDLEIEQHRTKVEELRAYLLKWGFSTPDKKHQKEP  398

Query  373  PFLWMGYELHPDKWTVQPIVLPEKDSWTVNDIQKLVGKLNWASQIYPGIKVRQLCKLLRG  432
            PFLWMGYELHPDKWTVQPIVLPEKDSWTVNDIQKLVGKLNWASQIY GIKV+QLCKLLRG
Sbjct  397  PFLWMGYELHPDKWTVQPIVLPEKDSWTVNDIQKLVGKLNWASQIYAGIKVKQLCKLLRG  218

Query  433  TKALTEVIPLTeeaelelaeNREILKEPVHGVYYDPSKDLIAEIQKQGQGQWTYQIYQEP  492
            TKALTEV+PL+EEAELELAENREILKEPVHGVYYDPSKDL+AEIQKQGQGQWTYQIYQEP
Sbjct  217  TKALTEVVPLSEEAELELAENREILKEPVHGVYYDPSKDLVAEIQKQGQGQWTYQIYQEP  38

Query  493  FKNLKTGKYA  502
                ++  YA
Sbjct  37   AHETRSSSYA  8
```

```
>Acinetobacter baumannii strain ACM-41 NODE_63_length_718_cov_0.852792, whole genome shotgun sequence
Sequence ID: NZ_QAGT01000063.1 Length: 718
Range 1: 2 to 718

Score:454 bits(1168), Expect:4e-148,
Method:Compositional matrix adjust.,
Identities:215/239(90%), Positives:225/239(94%), Gaps:0/239(0%)

Query  136  NLLTQIGCTLNFPISPIETVPVKLKPGMDGPKVKQWPLTEEKIKALVEICTEMEKEGKIS  195
            NLLTQ+GCTLNFPISPIETVPVKLKPGMDGPKVKQWPL+EEKIKAL EIC EMEKEGKIS
Sbjct  718  NLLTQLGCTLNFPISPIETVPVKLKPGMDGPKVKQWPLSEEKIKALTEICEEMEKEGKIS  539

Query  196  KIGPENPYNTPVFAIKKKDSTKWRKLVDFRELNKRTQDFWEVQLGIPHPAGLKKKKSVTV  255
            KIGPENPYNTPVFAIK++DSTKWRKLVDFRELNKRTQDFWEVQLGIPHPAGLKKKKS+TV
Sbjct  538  KIGPENPYNTPVFAIKRRDSTKWRKLVDFRELNKRTQDFWEVQLGIPHPAGLKKKKSMTV  359

Query  256  LDVGDAYFSVPLDEDFRKYTAFTIPSINNETPGIRYQYNVLPQGWKGSPAIFQSSMTKIL  315
            LDVGDAYFSVPLDE FRKYTAFTIPS NNETPGIR QYNVLPQGWKGSPAIF+ SMTK L
Sbjct  358  LDVGDAYFSVPLDEGFRKYTAFTIPSRNNETPGIRKQYNVLPQGWKGSPAIFKYSMTKTL  179

Query  316  EPFRKQNPDIVIYQYMDDLYVGSDLEIGQHRTKIEELRQHLLRWGLTTPDKKHQKEPPF  374
            EP R +NP+I+I QYMDDLYVGSDLEIGQHR KIEELR HLL+WG TTPDKKHQKEPP
Sbjct  178  EPIRAKNPEIIICQYMDDLYVGSDLEIGQHRAKIEELRDHLLKWGFTTPDKKHQKEPPL  2
```

```
>Acinetobacter baumannii strain ACM-22 NODE_78_length_726_cov_1.48748, whole genome shotgun sequence
Sequence ID: NZ_PYDI01000078.1 Length: 726
Range 1: 1 to 726

Score:452 bits(1162), Expect:4e-147,
Method:Compositional matrix adjust.,
Identities:223/242(92%), Positives:232/242(95%), Gaps:0/242(0%)

Query  226  ELNKRTQDFWEVQLGIPHPAGLKKKKSVTVLDVGDAYFSVPLDEDFRKYTAFTIPSINNE  285
            ELNKRTQDFWEVQLGIPHPAGLKKKKS+TVLDVGDAYFSVPLDE FRKYTAFTIPS NNE
Sbjct  1    ELNKRTQDFWEVQLGIPHPAGLKKKKSMTVLDVGDAYFSVPLDEGFRKYTAFTIPSRNNE  180

Query  286  TPGIRYQYNVLPQGWKGSPAIFQSSMTKILEPFRKQNPDIVIYQYMDDLYVGSDLEIGQH  345
            TPGIRYQYNVLPQGWKGSPAIFQ SMTKILEPFR +NP+I+I QYMDDLYVGSDLEIGQH
Sbjct  181  TPGIRYQYNVLPQGWKGSPAIFQCSMTKILEPFRAKNPEIIICQYMDDLYVGSDLEIGQH  360

Query  346  RTKIEELRQHLLRWGLTTPDKKHQKEPPFLWMGYELHPDKWTVQPIVLPEKDSWTVNDIQ  405
            R KIEELR HLL+WG TTPDKKHQKEPPFLWMGYELHPDKWTVQPI LPEKDSWTVNDIQ
Sbjct  361  RAKIEELRDHLLKWGFTTPDKKHQKEPPFLWMGYELHPDKWTVQPIQLPEKDSWTVNDIQ  540

Query  406  KLVGKLNWASQIYPGIKVRQLCKLLRGTKALTEVIPLTeeaelelaeNREILKEPVHGVY  465
            KLVGKLNWASQIYPGIKV+QLCKLLRG KALT+++PLTEEAELELAENREILKEPVHGVY
Sbjct  541  KLVGKLNWASQIYPGIKVKQLCKLLRGAKALTDIVPLTEEAELELAENREILKEPVHGVY  720

Query  466  YD  467
            YD
Sbjct  721  YD  726
```

```
>Acinetobacter baumannii strain ACM-13 NODE_69_length_701_cov_0.510453, whole genome shotgun sequence
Sequence ID: NZ_PYCZ01000069.1 Length: 701
Range 1: 3 to 701

Score:438 bits(1127), Expect:4e-142,
Method:Compositional matrix adjust.,
Identities:208/233(89%), Positives:219/233(93%), Gaps:0/233(0%)

Query  190  KEGKISKIGPENPYNTPVFAIKKKDSTKWRKLVDFRELNKRTQDFWEVQLGIPHPAGLKK  249
            KE K SK GPENPYNTP+FAIKKK+S KWRKLVDFRELNKRTQ+FWEVQLGIPHPAGLKK
Sbjct  3    KEWKTSKSGPENPYNTPIFAIKKKNSDKWRKLVDFRELNKRTQEFWEVQLGIPHPAGLKK  182

Query  250  KKSVTVLDVGDAYFSVPLDEDFRKYTAFTIPSINNETPGIRYQYNVLPQGWKGSPAIFQS  309
            K SVTVL VGDAYFSVP D DFRKYTAF+IPS NN+TPGIRYQYNVLPQGWKGSPAIFQS
Sbjct  183  KISVTVLCVGDAYFSVP*DPDFRKYTAFSIPSRNNKTPGIRYQYNVLPQGWKGSPAIFQS  362

Query  310  SMTKILEPFRKQNPDIVIYQYMDDLYVGSDLEIGQHRTKIEELRQHLLRWGLTTPDKKHQ  369
            SMTKILEPFRKQNPD+VI QYMDDLYVGSDLEI QHR K+EELR HLL+WG +TPD+KHQ
Sbjct  363  SMTKILEPFRKQNPDMVICQYMDDLYVGSDLEIEQHRIKVEELRAHLLKWGFSTPDQKHQ  542

Query  370  KEPPFLWMGYELHPDKWTVQPIVLPEKDSWTVNDIQKLVGKLNWASQIYPGIK  422
            KEPPFLWMGYELHPDKWTVQPIVLPEKDSWTVNDIQKLVGKLNWASQIYPG+K
Sbjct  543  KEPPFLWMGYELHPDKWTVQPIVLPEKDSWTVNDIQKLVGKLNWASQIYPGVK  701
```

```
>Escherichia coli strain ECM-2 NODE_127_length_636_cov_2.0668, whole genome shotgun sequence
Sequence ID: NZ_PXIA01000127.1 Length: 636
Range 1: 3 to 635

Score:421 bits(1082), Expect:1e-135,
Method:Compositional matrix adjust.,
Identities:201/211(95%), Positives:206/211(97%), Gaps:0/211(0%)

Query  151  PIETVPVKLKPGMDGPKVKQWPLTEEKIKALVEICTEMEKEGKISKIGPENPYNTPVFAI  210
            PIETVPVKLKPGMDGPKVKQWPLTEEKIKALVEICTEMEKEGKISKIGPENPYNTP+FAI
Sbjct  3    PIETVPVKLKPGMDGPKVKQWPLTEEKIKALVEICTEMEKEGKISKIGPENPYNTPIFAI  182

Query  211  KKKDSTKWRKLVDFRELNKRTQDFWEVQLGIPHPAGLKKKKSVTVLDVGDAYFSVPLDED  270
            KKK+S KWRKLVDFRELNKRTQ+FWEVQLGIPHPAGLKKKKSVTVLDVGDAYFSVPLD D
Sbjct  183  KKKNSDKWRKLVDFRELNKRTQEFWEVQLGIPHPAGLKKKKSVTVLDVGDAYFSVPLDPD  362

Query  271  FRKYTAFTIPSINNETPGIRYQYNVLPQGWKGSPAIFQSSMTKILEPFRKQNPDIVIYQY  330
            FRKYTAFTIPS NNETPGIRYQYNVLPQGWKGSPAIFQSSMTKILEPFRKQNPDIVIYQY
Sbjct  363  FRKYTAFTIPSRNNETPGIRYQYNVLPQGWKGSPAIFQSSMTKILEPFRKQNPDIVIYQY  542

Query  331  MDDLYVGSDLEIGQHRTKIEELRQHLLRWGL  361
            MDDLYVGSDLEI QHRTK+EELRQHLL+WG
Sbjct  543  MDDLYVGSDLEIEQHRTKVEELRQHLLKWGF  635
```

```
>Acinetobacter baumannii strain ACM-14 NODE_75_length_665_cov_0.98513, whole genome shotgun sequence
Sequence ID: NZ_PYDA01000075.1 Length: 665
Range 1: 1 to 657

Score:416 bits(1070), Expect:8e-134,
Method:Compositional matrix adjust.,
Identities:207/219(95%), Positives:213/219(97%), Gaps:0/219(0%)
```

```
Query  261  AYFSVPLDEDFRKYTAFTIPSINNETPGIRYQYNVLPQGWKGSPAIFQSSMTKILEPFRK  320
            AYFSVPLD DFRKYTAFTIPS NNETPGIRYQYNVLPQGWKGSPAIFQSSMTKILEPFRK
Sbjct  1    AYFSVPLDPDFRKYTAFTIPSRNNETPGIRYQYNVLPQGWKGSPAIFQSSMTKILEPFRK  180

Query  321  QNPDIVIYQYMDDLYVGSDLEIGQHRTKIEELRQHLLRWGLTTPDKKHQKEPPFLWMGYE  380
            QNPDIVIYQY+DDLYVGSDLEI QHRTK+EELRQHLL WG TTPDKKHQKEPPFLWMGYE
Sbjct  181  QNPDIVIYQYVDDLYVGSDLEIEQHRTKVEELRQHLLTWGFTTPDKKHQKEPPFLWMGYE  360

Query  381  LHPDKWTVQPIVLPEKDSWTVNDIQKLVGKLNWASQIYPGIKVRQLCKLLRGTKALTEVI  440
            LHPDKWTVQPIVLPEKDSWTVNDIQKLVGKLNWASQIY GIKV+QLCKLLRGTKALTEV+
Sbjct  361  LHPDKWTVQPIVLPEKDSWTVNDIQKLVGKLNWASQIYAGIKVKQLCKLLRGTKALTEVV  540

Query  441  PLTeeaelelaeNREILKEPVHGVYYDPSKDLIAEIQKQ  479
            PL+EEAELELAENREILKEPVHGVYYDPSKDL+AEIQKQ
Sbjct  541  PLSEEAELELAENREILKEPVHGVYYDPSKDLVAEIQKQ  657
```

```
>Acinetobacter baumannii strain ACM-11 NODE_65_length_706_cov_0.799655, whole genome shotgun sequence
Sequence ID: NZ_PYCW01000065.1 Length: 706
Range 1: 2 to 706

Score:413 bits(1061), Expect:3e-132,
Method:Compositional matrix adjust.,
Identities:206/235(88%), Positives:221/235(94%), Gaps:0/235(0%)

Query  237  VQLGIPHPAGLKKKKSVTVLDVGDAYFSVPLDEDFRKYTAFTIPSINNETPGIRYQYNVL  296
            V+LG      LK K S+++LDVGDAYFSVPLDE+FRKYTAFTIPS+NNETPGIRYQYNVL
Sbjct  2    VKLGKLQQPRLKHK*SLSILDVGDAYFSVPLDENFRKYTAFTIPSVNNETPGIRYQYNVL  181

Query  297  PQGWKGSPAIFQSSMTKILEPFRKQNPDIVIYQYMDDLYVGSDLEIGQHRTKIEELRQHL  356
            PQGWKGSPAIFQ+SMTKILEPFR +NPDIVIYQYMDDLYVGSDLEIGQHR KIEELR+HL
Sbjct  182  PQGWKGSPAIFQASMTKILEPFRLKNPDIVIYQYMDDLYVGSDLEIGQHRAKIEELREHL  361

Query  357  LRWGLTTPDKKHQKEPPFLWMGYELHPDKWTVQPIVLPEKDSWTVNDIQKLVGKLNWASQ  416
            L+WGLTTPDKKHQKEPPFLWMGYELHPDKWTVQPI LPEKDSWTVNDIQKLVGKL WASQ
Sbjct  362  LKWGLTTPDKKHQKEPPFLWMGYELHPDKWTVQPIQLPEKDSWTVNDIQKLVGKLIWASQ  541

Query  417  IYPGIKVRQLCKLLRGTKALTEVIPLTeeaelelaeNREILKEPVHGVYYDPSKD  471
            IY GIKV+QLCKLLRG KALT+VIPLTEEAELELAENREIL+EPVHGVYYDP++D
Sbjct  542  IYAGIKVKQLCKLLRGAKALTDVIPLTEEAELELAENREILREPVHGVYYDPTRD  706
```

```
>Mycobacterium tuberculosis strain 32847_S31 312, whole genome shotgun sequence
Sequence ID: NZ_QTBX01000084.1 Length: 704
Range 1: 3 to 704

Score:409 bits(1052), Expect:5e-131,
Method:Compositional matrix adjust.,
Identities:213/234(91%), Positives:224/234(95%), Gaps:0/234(0%)

Query  349  IEELRQHLLRWGLTTPDKKHQKEPPFLWMGYELHPDKWTVQPIVLPEKDSWTVNDIQKLV  408
            IEELR HLL+WG TTPDKKHQKEPPFLWMGYELHPDKWTVQPI LPEKDSWTVNDIQKLV
Sbjct  704  IEELRGHLLKWGFTTPDKKHQKEPPFLWMGYELHPDKWTVQPIQLPEKDSWTVNDIQKLV  525

Query  409  GKLNWASQIYPGIKVRQLCKLLRGTKALTEVIPLTeeaelelaeNREILKEPVHGVYYDP  468
            GKLNWASQIYPGIKV+QLC+LLRG KALT+++PLTEEAELELAENREILKEPVHGVYYDP
Sbjct  524  GKLNWASQIYPGIKVKQLCRLLRGAKALTDIVPLTEEAELELAENREILKEPVHGVYYDP  345

Query  469  SKDLIAEIQKQGQGQWTYQIYQEPFKNLKTGKYARMRGAHTNDVKQLTEAVQKITTESIV  528
            +KDLIAEIQKQ  GQWTYQIYQEPFKNLKTGKYA+MR AHTNDVKQLTEAVQKI  ESIV
Sbjct  344  AKDLIAEIQKQDHGQWTYQIYQEPFKNLKTGKYAKMRTAHTNDVKQLTEAVQKIAMESIV  165

Query  529  IWGKTPKFKLPIQKetwetwwteYWQATWIPEWEFVNTPPLVKLWYQLEKEPIV  582
            +WGKTPKF+LPIQKETWETWWTEYWQATWIPEWEFVNTPPLVKLWYQLEK+PI
Sbjct  164  VWGKTPKFRLPIQKETWETWWTEYWQATWIPEWEFVNTPPLVKLWYQLEKDPIA  3
```

```
>Providencia rettgeri strain AFE85_S2 19_contig_46, whole genome shotgun sequence
Sequence ID: NZ_SHDQ01000046.1 Length: 846
Range 1: 3 to 632

Score:409 bits(1051), Expect:5e-130,
Method:Compositional matrix adjust.,
Identities:196/210(93%), Positives:205/210(97%), Gaps:0/210(0%)

Query  62   KIGGQLKEALLDTGADDTVLEEMSLPGRWKPKMIGGIGGFIKVRQYDQILIEICGHKAIG  121
            ++GGQ+KEALLDTGADDTVLEE++LPG+WKPKMIGGIGGFIKVRQYD ILIEICG KAIG
Sbjct  632  EVGGQIKEALLDTGADDTVLEEINLPGKWKPKMIGGIGGFIKVRQYDHILIEICGKKAIG  453

Query  122  TVLVGPTPVNIIGRNLLTQIGCTLNFPISPIETVPVKLKPGMDGPKVKQWPLTEEKIKAL  181
            TVLVGPTPVNIIGRN+LTQ+GCTLNFPISPIETVPVKLKPGMDGP+VKQWPLTEEKIKAL
Sbjct  452  TVLVGPTPVNIIGRNMLTQLGCTLNFPISPIETVPVKLKPGMDGPRVKQWPLTEEKIKAL  273

Query  182  VEICTEMEKEGKISKIGPENPYNTPVFAIKKKDSTKWRKLVDFRELNKRTQDFWEVQLGI  241
             EIC EMEKEGKI KIGPENPYNTPVFAIKKKDSTKWRKLVDFRELNKRTQDFWEVQLGI
Sbjct  272  TEICEEMEKEGKIEKIGPENPYNTPVFAIKKKDSTKWRKLVDFRELNKRTQDFWEVQLGI  93

Query  242  PHPAGLKKKKSVTVLDVGDAYFSVPLDEDF  271
            PHPAGLKKKKSVTVLDVGDAYFSVPLDEDF
Sbjct  92   PHPAGLKKKKSVTVLDVGDAYFSVPLDEDF  3
```

```
Range 2: 680 to 844

Score:82.0 bits(201), Expect:4e-12,
Method:Composition-based stats.,
Identities:39/55(71%), Positives:42/55(76%), Gaps:0/55(0%)

Query  160  KPGMDGPKVKQWPLTEEKIKALVEICTEMEKEGKISKIGPENPYNTPVFAIKKKD  214
            KP     K +  PLTEEKIKA+ EIC EMEKEGKI+KIGPEN YNTP FAIKK D
Sbjct  680  KPAFA*KKKRH*PLTEEKIKAITEICKEMEKEGKITKIGPENQYNTPGFAIKKND  844
```

```
>Mycobacterium tuberculosis variant bovis strain CSURQ1132, whole genome shotgun sequence
Sequence ID: NZ_CAJEUK010000179.1 Length: 862
Range 1: 47 to 739

Score:406 bits(1043), Expect:9e-129,
Method:Compositional matrix adjust.,
Identities:201/231(87%), Positives:214/231(92%), Gaps:0/231(0%)

Query  38   ADRQGTVSFNFPQVTLWQRPLVTIKIGGQLKEALLDTGADDTVLEEMSLPGRWKPKMIGG  97
            AD QG   F + Q T+ +  +V IK GGQ+KEALLDTGADDTVLEE+ LPGRWKPKMIGG
Sbjct  47   ADGQGRDCF*YTQDTVEEGEIV*IKRGGQVKEALLDTGADDTVLEEIDLPGRWKPKMIGG  226
```

```
Query  98   IGGFIKVRQYDQILIEICGHKAIGTVLVGPTPVNIIGRNLLTQIGCTLNFPISPIETVPV  157
            IGGFIKVRQYDQ+ +EICGHKAIGTVLVGPTPVNIIGRNLLTQIGCTLNFPISPIETVPV
Sbjct  227  IGGFIKVRQYDQVQVEICGHKAIGTVLVGPTPVNIIGRNLLTQIGCTLNFPISPIETVPV  406

Query  158  KLKPGMDGPKVKQWPLTEEKIKALVEICTEMEKEGKISKIGPENPYNTPVFAIKKKDSTK  217
            KLKPGMDGPKVKQWPLTEEKIKAL+EICTEMEKEGKISKIGPENPYNTPVFAIKKK+STK
Sbjct  407  KLKPGMDGPKVKQWPLTEEKIKALIEICTEMEKEGKISKIGPENPYNTPVFAIKKKNSTK  586

Query  218  WRKLVDFRELNKRTQDFWEVQLGIPHPAGLKKKKSVTVLDVGDAYFSVPLD  268
            WRKLVDFRELNK+TQDFWEVQLGIPHP+GLKKKKSVTVLDVGDA  SV +D
Sbjct  587  WRKLVDFRELNKKTQDFWEVQLGIPHPSGLKKKKSVTVLDVGDA*SSVHVD  739
```

Range 2: 47 to 739

Score:406 bits(1043), Expect:9e-129,
Method:Compositional matrix adjust.,
Identities:201/231(87%), Positives:214/231(92%), Gaps:0/231(0%)

```
Query  38   ADRQGTVSFNFPQVTLWQRPLVTIKIGGQLKEALLDTGADDTVLEEMSLPGRWKPKMIGG  97
            AD QG   F + Q T+ +  +V IK GGQ+KEALLDTGADDTVLEE+ LPGRWKPKMIGG
Sbjct  47   ADGQGRDCF*YTQDTVEEGEIV*IKRGGQVKEALLDTGADDTVLEEIDLPGRWKPKMIGG  226

Query  98   IGGFIKVRQYDQILIEICGHKAIGTVLVGPTPVNIIGRNLLTQIGCTLNFPISPIETVPV  157
            IGGFIKVRQYDQ+ +EICGHKAIGTVLVGPTPVNIIGRNLLTQIGCTLNFPISPIETVPV
Sbjct  227  IGGFIKVRQYDQVQVEICGHKAIGTVLVGPTPVNIIGRNLLTQIGCTLNFPISPIETVPV  406

Query  158  KLKPGMDGPKVKQWPLTEEKIKALVEICTEMEKEGKISKIGPENPYNTPVFAIKKKDSTK  217
            KLKPGMDGPKVKQWPLTEEKIKAL+EICTEMEKEGKISKIGPENPYNTPVFAIKKK+STK
Sbjct  407  KLKPGMDGPKVKQWPLTEEKIKALIEICTEMEKEGKISKIGPENPYNTPVFAIKKKNSTK  586

Query  218  WRKLVDFRELNKRTQDFWEVQLGIPHPAGLKKKKSVTVLDVGDAYFSVPLD  268
            WRKLVDFRELNK+TQDFWEVQLGIPHP+GLKKKKSVTVLDVGDA  SV +D
Sbjct  587  WRKLVDFRELNKKTQDFWEVQLGIPHPSGLKKKKSVTVLDVGDA*SSVHVD  739
```

>Acinetobacter baumannii strain ACM-25 NODE_71_length_740_cov_1.2447, whole genome shotgun sequence
Sequence ID: NZ_PXYP01000071.1 Length: 655
Range 1: 1 to 654

Score:399 bits(1026), Expect:2e-127,
Method:Compositional matrix adjust.,
Identities:198/218(91%), Positives:206/218(94%), Gaps:0/218(0%)

```
Query  273  KYTAFTIPSINNETPGIRYQYNVLPQGWKGSPAIFQSSMTKILEPFRKQNPDIVIYQYMD  332
            KYTAFTIPS NNETPGIRYQYNVLPQGWKGSPAIFQ SMTKILEPFR +NP+I+I QYMD
Sbjct  654  KYTAFTIPSRNNETPGIRYQYNVLPQGWKGSPAIFQCSMTKILEPFRAKNPEIIICQYMD  475

Query  333  DLYVGSDLEIGQHRTKIEELRQHLLRWGLTTPDKKHQKEPPFLWMGYELHPDKWTVQPIV  392
            DLYVGSDLEIGQHR KIEELR HLL+WG TTPDKKHQKEPPFLWMGYELHPDKWTVQPI
Sbjct  474  DLYVGSDLEIGQHRAKIEELRDHLLKWGFTTPDKKHQKEPPFLWMGYELHPDKWTVQPIQ  295

Query  393  LPEKDSWTVNDIQKLVGKLNWASQIYPGIKVRQLCKLLRGTKALTEVIPLTeeaelelae  452
            LPEKDSWTVNDIQKLVGKLNWASQIYPGIKV+QLCKLLRG KALT+++PLTEEAELELAE
Sbjct  294  LPEKDSWTVNDIQKLVGKLNWASQIYPGIKVKQLCKLLRGAKALTDIVPLTEEAELELAE  115

Query  453  NREILKEPVHGVYYDPSKDLIAEIQKQGQGQWTYQIYQ  490
            NREILKEPVHG YYDPSKDLIAEIQKQG  QWTYQIYQ
Sbjct  114  NREILKEPVHGAYYDPSKDLIAEIQKQGNDQWTYQIYQ  1
```

>Levilactobacillus brevis strain CRL 2013 scaffold15, whole genome shotgun sequence
Sequence ID: NZ_MZMW01000019.1 Length: 566
Range 1: 3 to 566

Score:396 bits(1017), Expect:2e-126,
Method:Compositional matrix adjust.,
Identities:186/188(99%), Positives:187/188(99%), Gaps:0/188(0%)

```
Query  201  NPYNTPVFAIKKKDSTKWRKLVDFRELNKRTQDFWEVQLGIPHPAGLKKKKSVTVLDVGD  260
            NPYNTPVFAIKKKDSTKWRKLVDFRELNKRTQDFWEVQLGIPHPAGLKKKKSVTVLDVGD
Sbjct  566  NPYNTPVFAIKKKDSTKWRKLVDFRELNKRTQDFWEVQLGIPHPAGLKKKKSVTVLDVGD  387

Query  261  AYFSVPLDEDFRKYTAFTIPSINNETPGIRYQYNVLPQGWKGSPAIFQSSMTKILEPFRK  320
            AYFSVPLD+DFRKYTAFTIPSINNETPGIRYQYNVLPQGWKGSPAIFQSSMTKILEPFRK
Sbjct  386  AYFSVPLDKDFRKYTAFTIPSINNETPGIRYQYNVLPQGWKGSPAIFQSSMTKILEPFRK  207

Query  321  QNPDIVIYQYMDDLYVGSDLEIGQHRTKIEELRQHLLRWGLTTPDKKHQKEPPFLWMGYE  380
            QNPDIVIYQYMDDLYVGSDLEIGQHRTKIEELRQHLLRWG TTPDKKHQKEPPFLWMGYE
Sbjct  206  QNPDIVIYQYMDDLYVGSDLEIGQHRTKIEELRQHLLRWGFTTPDKKHQKEPPFLWMGYE  27

Query  381  LHPDKWTV  388
            LHPDKWTV
Sbjct  26   LHPDKWTV  3
```

>Pseudomonas aeruginosa strain ENV-297 NODE_95_length_638_cov_6.84848_ID_189, whole genome shotgun sequence
Sequence ID: NZ_RADD01000130.1 Length: 638
Range 1: 2 to 589

Score:394 bits(1011), Expect:3e-125,
Method:Compositional matrix adjust.,
Identities:183/196(93%), Positives:191/196(97%), Gaps:0/196(0%)

```
Query  208  FAIKKKDSTKWRKLVDFRELNKRTQDFWEVQLGIPHPAGLKKKKSVTVLDVGDAYFSVPL  267
            FAIKKKDSTKWRKLVDFRELNKRTQDFWEVQLGIPHPAGLKKKKSVT+LDVGDAYFSVPL
Sbjct  2    FAIKKKDSTKWRKLVDFRELNKRTQDFWEVQLGIPHPAGLKKKKSVTILDVGDAYFSVPL  181

Query  268  DEDFRKYTAFTIPSINNETPGIRYQYNVLPQGWKGSPAIFQSSMTKILEPFRKQNPDIVI  327
            D DFRKYTAFTIPSINNETPGIRYQYNVLPQGWKGSPAIFQ SMTKILEPFRKQNP++VI
Sbjct  182  DPDFRKYTAFTIPSINNETPGIRYQYNVLPQGWKGSPAIFQCSMTKILEPFRKQNPELVI  361

Query  328  YQYMDDLYVGSDLEIGQHRTKIEELRQHLLRWGLTTPDKKHQKEPPFLWMGYELHPDKWT  387
            YQYMDDLY+GSDLEIGQHR KIEELR+HLL+WG TTPDKKHQKEPPFLWMGYELHPDKWT
Sbjct  362  YQYMDDLYIGSDLEIGQHRAKIEELREHLLKWGFTTPDKKHQKEPPFLWMGYELHPDKWT  541

Query  388  VQPIVLPEKDSWTVND  403
            VQPI LP+K+SWTVND
Sbjct  542  VQPIELPDKESWTVND  589
```

>Mycobacterium tuberculosis strain 32845_S29 230, whole genome shotgun sequence
Sequence ID: NZ_QTBV01000049.1 Length: 601

Range 1: 1 to 600

Score:391 bits(1004), Expect:2e-124,
Method:Compositional matrix adjust.,
Identities:188/200(94%), Positives:195/200(97%), Gaps:0/200(0%)

```
Query  70   ALLDTGADDTVLEEMSLPGRWKPKMIGGIGGFIKVRQYDQILIEICGHKAIGTVLVGPTP  129
            ALLDTGADDTVLE+++LPG+WKPKMIGGIGGFIKVRQYDQILIEICG KAIGTVLVGPTP
Sbjct  600  ALLDTGADDTVLEDINLPGKWKPKMIGGIGGFIKVRQYDQILIEICGKKAIGTVLVGPTP  421

Query  130  VNIIGRNLLTQIGCTLNFPISPIETVPVKLKPGMDGPKVKQWPLTEEKIKALVEICTEME  189
            VNIIGRN+LTQIGCTLNFPISPIETVPVKLKPGMDGPKVKQWPLTEEKIKAL  IC EME
Sbjct  420  VNIIGRNMLTQIGCTLNFPISPIETVPVKLKPGMDGPKVKQWPLTEEKIKALTAICEEME  241

Query  190  KEGKISKIGPENPYNTPVFAIKKKDSTKWRKLVDFRELNKRTQDFWEVQLGIPHPAGLKK  249
            KEGKI+KIGPENPYNTPVFAIKKKDSTKWRKLVDFRELNKRTQDFWE+QLGIPHPAGLKK
Sbjct  240  KEGKITKIGPENPYNTPVFAIKKKDSTKWRKLVDFRELNKRTQDFWEIQLGIPHPAGLKK  61

Query  250  KKSVTVLDVGDAYFSVPLDE  269
             KSVTVLDVGDAYFSVPLDE
Sbjct  60   NKSVTVLDVGDAYFSVPLDE  1
```

>Acinetobacter baumannii strain ACM-38 NODE_58_length_652_cov_1.55048, whole genome shotgun sequence
Sequence ID: NZ_QAGQ01000058.1 Length: 652
Range 1: 1 to 651

Score:391 bits(1005), Expect:2e-124,
Method:Compositional matrix adjust.,
Identities:188/217(87%), Positives:205/217(94%), Gaps:0/217(0%)

```
Query  32   SPSEAGADRQGTVSFNFPQVTLWQRPLVTIKIGGQLKEALLDTGADDTVLEEMSLPGRWK  91
            S S+AG++RQGTV+ +FPQ+T+W  P+VTI++G QLK ALLDTG  DTVLE+M LPGRWK
Sbjct  651  SRSKAGSNRQGTVTCHFPQITIWLLPVVTIQVGRQLKVALLDTGEYDTVLEDMQLPGRWK  472

Query  92   PKMIGGIGGFIKVRQYDQILIEICGHKAIGTVLVGPTPVNIIGRNLLTQIGCTLNFPISP  151
            PKMIGGI GFIKVRQYD++ IEICGHKAIGTVLVGPTPVNIIGRNLLTQ+GCTLNFPISP
Sbjct  471  PKMIGGIRGFIKVRQYDEVPIEICGHKAIGTVLVGPTPVNIIGRNLLTQLGCTLNFPISP  292

Query  152  IETVPVKLKPGMDGPKVKQWPLTEEKIKALVEICTEMEKEGKISKIGPENPYNTPVFAIK  211
            IETVPVKLKPGMDGPKVKQWPLTEEKIKALVEICTEMEKEGKISKIGPENPYNTP+FAIK
Sbjct  291  IETVPVKLKPGMDGPKVKQWPLTEEKIKALVEICTEMEKEGKISKIGPENPYNTPIFAIK  112

Query  212  KKDSTKWRKLVDFRELNKRTQDFWEVQLGIPHPAGLK  248
            KK+S KWRKLVDFRELNKRTQ+FWEVQLGIPHPAGLK
Sbjct  111  KKNSDKWRKLVDFRELNKRTQEFWEVQLGIPHPAGLK  1
```

>Acinetobacter baumannii strain ACM-32 NODE_81_length_654_cov_1.01708, whole genome shotgun sequence
Sequence ID: NZ_PYDR01000081.1 Length: 654
Range 1: 9 to 653

Score:389 bits(999), Expect:2e-123,
Method:Compositional matrix adjust.,
Identities:185/215(86%), Positives:206/215(95%), Gaps:0/215(0%)

```
Query  45   SFNFPQVTLWQRPLVTIKIGGQLKEALLDTGADDTVLEEMSLPGRWKPKMIGGIGGFIKV  104
            +F+FPQ+T WQRPLV IK+GGQ+KEALLDTGADDTVLE+++LPG+WKP+MIGGIGGFIKV
Sbjct  9    AFDFPQITPWQRPLVPIKVGGQIKEALLDTGADDTVLEDINLPGKWKPRMIGGIGGFIKV  188

Query  105  RQYDQILIEICGHKAIGTVLVGPTPVNIIGRNLLTQIGCTLNFPISPIETVPVKLKPGMD  164
            RQY+QI IEICG KAIGT+LVGPTP+NIIGRN+LTQ+GCTLNFPISPIETVPVKLKPGMD
Sbjct  189  RQYEQIPIEICGKKAIGTILVGPTPINIIGRNMLTQLGCTLNFPISPIETVPVKLKPGMD  368

Query  165  GPKVKQWPLTEEKIKALVEICTEMEKEGKISKIGPENPYNTPVFAIKKKDSTKWRKLVDF  224
            GP+VKQWPLTEEKIKAL  IC EMEKEGKI+KIGPENPYNTPVFAIKKK+STK RKL+DF
Sbjct  369  GPRVKQWPLTEEKIKALQAICEEMEKEGKITKIGPENPYNTPVFAIKKKNSTK*RKLIDF  548

Query  225  RELNKRTQDFWEVQLGIPHPAGLKKKKSVTVLDVG  259
            R+LNK +QDFWEVQLGIPHPAGLKKKKSVTVLDVG
Sbjct  549  RKLNKISQDFWEVQLGIPHPAGLKKKKSVTVLDVG  653
```

>Providencia rettgeri strain AMW697_S1 18_contig_57, whole genome shotgun sequence
Sequence ID: NZ_SHDI01000057.1 Length: 585
Range 1: 2 to 583

Score:384 bits(986), Expect:7e-122,
Method:Compositional matrix adjust.,
Identities:181/194(93%), Positives:186/194(95%), Gaps:0/194(0%)

```
Query  170  QWPLTEEKIKALVEICTEMEKEGKISKIGPENPYNTPVFAIKKKDSTKWRKLVDFRELNK  229
            QWPLTEEKIKAL EIC EMEKEGKI+KIGPENPYNTP+FAIKKKDSTKWRKLVDFRELNK
Sbjct  2    QWPLTEEKIKALTEICEEMEKEGKITKIGPENPYNTPIFAIKKKDSTKWRKLVDFRELNK  181

Query  230  RTQDFWEVQLGIPHPAGLKKKKSVTVLDVGDAYFSVPLDEDFRKYTAFTIPSINNETPGI  289
            RTQDFWEVQLGIPHPAGLKK KSVTVLDVGDAYFSVPLDE FRKYTAFTIPSINNETPGI
Sbjct  182  RTQDFWEVQLGIPHPAGLKKNKSVTVLDVGDAYFSVPLDEGFRKYTAFTIPSINNETPGI  361

Query  290  RYQYNVLPQGWKGSPAIFQSSMTKILEPFRKQNPDIVIYQYMDDLYVGSDLEIGQHRTKI  349
            RYQYNVLPQGWKGSPAIFQSSMTKILEPFR QNP++VIYQYMDDLYVGSDLEIGQHR KI
Sbjct  362  RYQYNVLPQGWKGSPAIFQSSMTKILEPFRAQNPELVIYQYMDDLYVGSDLEIGQHRAKI  541

Query  350  EELRQHLLRWGLTT  363
            EELR HLL+WG TT
Sbjct  542  EELRNHLLKWGFTT  583
```

>Levilactobacillus brevis strain CRL 2013 scaffold16, whole genome shotgun sequence
Sequence ID: NZ_MZMW01000020.1 Length: 547
Range 1: 2 to 547

Score:376 bits(965), Expect:5e-119,
Method:Compositional matrix adjust.,
Identities:177/182(97%), Positives:179/182(98%), Gaps:0/182(0%)

```
Query  210  IKKKDSTKWRKLVDFRELNKRTQDFWEVQLGIPHPAGLKKKKSVTVLDVGDAYFSVPLDE  269
            IKKKDSTKWRKLVDFRELNKRTQDFWEVQLGIPHPAGLKKKK+VTVLDVGDAYFSVPLD+
Sbjct  547  IKKKDSTKWRKLVDFRELNKRTQDFWEVQLGIPHPAGLKKKKAVTVLDVGDAYFSVPLDK  368

Query  270  DFRKYTAFTIPSINNETPGIRYQYNVLPQGWKGSPAIFQSSMTKILEPFRKQNPDIVIYQ  329
            DFRKYTAFTIPSINNETPGIRYQYNVLPQGWKGSPAIFQSSMTKILEPFRKQNPDIVIYQ
Sbjct  367  DFRKYTAFTIPSINNETPGIRYQYNVLPQGWKGSPAIFQSSMTKILEPFRKQNPDIVIYQ  188

Query  330  YMDDLYVGSDLEIGQHRTKIEELRQHLLRWGLTTPDKKHQKEPPFLWMGYELHPDKWTVQ  389
            YMDDLYVGSDLEIGQHR KIEELRQHLL WG TTPDKKHQKEPPFLWMGYELHPDKWTVQ
```

Sbjct  187  YMDDLYVGSDLEIGQHRAKIEELRQHLLEWGFTTPDKKHQKEPPFLWMGYELHPDKWTVQ  8

Query  390  PI  391
            PI
Sbjct  7    PI  2

>Enterococcus faecalis strain 44.1 NODE_36_length_611_cov_1.15083, whole genome shotgun sequence
Sequence ID: NZ_NSJR01000036.1 Length: 611
Range 1: 2 to 610

Score:367 bits(941), Expect:4e-115,
Method:Compositional matrix adjust.,
Identities:182/203(90%), Positives:194/203(95%), Gaps:0/203(0%)

Query  274  YTAFTIPSINNETPGIRYQYNVLPQGWKGSPAIFQSSMTKILEPFRKQNPDIVIYQYMDD  333
            YTAFTI S  NET GI+Y YN+LPQGWKGSPAIFQSSMTKILEPFRKQNPDIVIYQY+DD
Sbjct  610  YTAFTILSRKNETSGIKY*YNILPQGWKGSPAIFQSSMTKILEPFRKQNPDIVIYQYVDD  431

Query  334  LYVGSDLEIGQHRTKIEELRQHLLRWGLTTPDKKHQKEPPFLWMGYELHPDKWTVQPIVL  393
            LYVGSDLE+ QHRTK+EELR HLL+WG +TPD+KHQKEPPFLWMGYELHPDKWTVQPIVL
Sbjct  430  LYVGSDLELEQHRTKVEELRAHLLKWGFSTPDQKHQKEPPFLWMGYELHPDKWTVQPIVL  251

Query  394  PEKDSWTVNDIQKLVGKLNWASQIYPGIKVRQLCKLLRGTKALTEVIPLTeeaelelaeN  453
            PEKDSWTVNDIQKLVGKLNWASQIY GIKV+QLCKLLRGTKALTEV+PL+EEAELELAEN
Sbjct  250  PEKDSWTVNDIQKLVGKLNWASQIYAGIKVKQLCKLLRGTKALTEVVPLSEEAELELAEN  71

Query  454  REILKEPVHGVYYDPSKDLIAEI  476
            REILKEPVHGVYYDPSKDL+AEI
Sbjct  70   REILKEPVHGVYYDPSKDLVAEI  2

>Cuspidothrix issatschenkoi CHARLIE-1 contig_175, whole genome shotgun sequence
Sequence ID: NZ_PGEM01000173.1 Length: 2237
Range 1: 286 to 1899

Score:385 bits(989), Expect:1e-114,
Method:Compositional matrix adjust.,
Identities:233/543(43%), Positives:321/543(59%), Gaps:7/543(1%)

Query  157   VKLKPGMDGPKVKQWPLTEEKIKALVEICTEMEKEGKISKIGPENPYNTPVFAIKKKDST  216
             ++LK G  GPK+ QWPLT+EK++   EI   +  EGKIS+    NPYN+P+F IKK+ S
Sbjct  286   IELKEGTMGPKIPQWPLTKEKLEGAKEIVQRLLSEGKISEASDNNPYNSPIFVIKKR-SG  462

Query  217   KWRKLVDFRELNKRTQDFWEVQLGIPHPAGLKKKKSVTVLDVGDAYFSVPLDEDFRKYTA  276
             KWR L D RELNK  Q   E+  G+PHP GL K K +TVLD+GDAYF++PLD +FR YTA
Sbjct  463   KWRLLQDLRELNKTVQVGTEISRGLPHPGGLIKCKHMTVLDIGDAYFTIPLDPEFRPYTA  642

Query  277   FTIPSINNETPGIRYQYNVLPQGWKGSPAIFQSSMTKILEPFRKQNPDIVIYQYMDDLYV  336
             FTIPSIN++ P  RY +N LPQG+  SP I+Q ++ +IL+PFR++ P++ +YQYMDDL+V
Sbjct  643   FTIPSINHQEPDKRYVWNCLPQGFVLSPYIYQKTLQEILQPFRERYPEVQLYQYMDDLFV  822

Query  337   GSDLEIGQHRTKIEELRQHLLRWGLTTPDKKHQKEPPFLWMGYELHPDKWTVQPIVLPEK  396
             GS+    QH+  I ELR  LL  G  TPD K Q+ PP+ W+GY+L P+ W VQ + L
Sbjct  823   GSNGSKKQHKELIIELRAILLEKGFETPDDKLQEVPPYSWLGYQLCPENWKVQKMQLDMV  1002

Query  397   DSWTVNDIQKLVGKLNWASQIYPGIKVRQLCKLLRGTKALTEVIPLTeeaelelaeNREI  456
              + T+ND+QKL+G + W S   PG+ V+ +    +G   L + +  TEEA+ EL EN E
Sbjct  1003  KNPTLNDVQKLMGNITWMSSGVPGLTVKHIAATTKGCLELNQKVIWTEEAQKELEENNEK  1182

Query  457   LKEPVHGVYYDPSKDLIAEIQKQGQGQWTYQIYQEPFKNLKTGKYARMRGAHTNDVKQLT  516
             +K      YY+P ++++ E++     + TY I Q     L  GK          + VK L
Sbjct  1183  IKNAQGLQYYNPEEEMLCEVEITKNYEATYVIKQSQ-GILWAGKKIMKANKGWSTVKNLM  1359

Query  517   EAVQKITTESIVIWGKTPKFKLPIQKetwetwwteYWQATWIPEWEFVNTPPLVKLWYQL  576
               +Q + TESI   GK P FK+P  KE       + W  +W+PE  + +         +L
Sbjct  1360  LLLQHVATESITRVGKCPTFKVPFTKEQVMWEMQKGWYYSWLPEIVYTHQVVHDDWRMKL  1539

Query  577   EKEPIVGAETFYVDGAANRETKLGKAGYVTNRGRQKVVTLTDTTNQKTELQAIYLALQDS  636
              +EP  G  T Y DG   ++   G A YVT+ GR K   L   T+Q  E  AI +AL+D+
Sbjct  1540  VEEPTSGI-TIYTDGG--KQNGEGIAAYVTSNGRTKQKRLGPVTHQVAERMAIQMALEDT  1710

Query  637   -GLEVNIVTDSQYAL-GIIQAQPDQSESELVNQIIEQLIKKEKVYLAWVPAHKGIGGNEQ  694
                +VNIVTDS Y    I +    +        II+ + +KE VY AWVP HKGI GN+
Sbjct  1711  RDKQVNIVTDSYYCWKNITEGLGLEGPQSPWWPIIQNIREKEIVYFAWVPGHKGICGNQL  1890

Query  695   VDK  697
              D+
Sbjct  1891  ADE  1899

Range 2: 1471 to 1929

Score:70.5 bits(171), Expect:2e-07,
Method:Compositional matrix adjust.,
Identities:58/160(36%), Positives:87/160(54%), Gaps:16/160(10%)

Query  679   YLAWVP----AHKGIGGNEQVDKLV---SAGIRKVLYVDGAANRETKLGKAGYVTNRGRQ  731
             Y +W+P     H+ +  + ++ KLV   ++GI   +Y DG   ++   G A YVT+ GR
Sbjct  1471  YYSWLPEIVYTHQVVHDDWRM-KLVEEPTSGI--TIYTDGG--KQNGEGIAAYVTSNGRT  1635

Query  732   KVVTLTDTTNQKTELQAIYLALQDS-GLEVNIVTDSQYAL-GIIQAQPDQSESELVNQII  789
             K   L   T+Q  E  AI +AL+D+   +VNIVTDS Y    I +    +        II
Sbjct  1636  KQKRLGPVTHQVAERMAIQMALEDTRDKQVNIVTDSYYCWKNITEGLGLEGPQSPWWPII  1815

Query  790   EQLIKKEKVYLAWVPAHKGIGGNEQVDKLVSAGIRKVLFL  829
             + + +KE VY AWVP HKGI GN+  D+  +A I++ + L
Sbjct  1816  QNIREKEIVYFAWVPGHKGICGNQLADE--AAKIKEEIVL  1929

>Acinetobacter baumannii strain ACM-36 NODE_67_length_567_cov_1.13409, whole genome shotgun sequence
Sequence ID: NZ_QAGO01000067.1 Length: 567
Range 1: 3 to 566

Score:363 bits(932), Expect:4e-114,
Method:Compositional matrix adjust.,
Identities:172/188(91%), Positives:183/188(97%), Gaps:0/188(0%)

Query  20   RRELQVWGRDNNSPSEAGADRQGTVSFNFPQVTLWQRPLVTIKIGGQLKEALLDTGADDT  79
            RRELQ WG DNNS SEAGADRQGTVSF+FPQ+TLWQRP+VTIK+GGQLKEALLDTGADDT
Sbjct  3    RRELQAWGGDNNSLSEAGADRQGTVSFSFPQITLWQRPVVTIKVGGQLKEALLDTGADDT  182

Query  80   VLEEMSLPGRWKPKMIGGIGGFIKVRQYDQILIEICGHKAIGTVLVGPTPVNIIGRNLLT  139
            VLE+M LPGRWKPKMIGGIGGFIKVRQYD++ IEICGHKAIGTVL+GPTPVNIIGRNLLT
Sbjct  183  VLEDMQLPGRWKPKMIGGIGGFIKVRQYDEVPIEICGHKAIGTVLIGPTPVNIIGRNLLT  362

```
Query  140  QIGCTLNFPISPIETVPVKLKPGMDGPKVKQWPLTEEKIKALVEICTEMEKEGKISKIGP  199
            Q+GCTLNFPISPIETVPVKLKPGMDGPKVKQWPLTEEKIKALVEICTEMEKEGKI+KIGP
Sbjct  363  QLGCTLNFPISPIETVPVKLKPGMDGPKVKQWPLTEEKIKALVEICTEMEKEGKITKIGP  542

Query  200  ENPYNTPV  207
            ENPYNTP+
Sbjct  543  ENPYNTPI  566
```

>Enterobacter hormaechei strain 2011WA-SCV PROKKA_contig000046, whole genome shotgun sequence
Sequence ID: NZ_SJEC01000046.1 Length: 544
Range 1: 4 to 543

Score:352 bits(904), Expect:4e-110,
Method:Compositional matrix adjust.,
Identities:166/180(92%), Positives:172/180(95%), Gaps:0/180(0%)

```
Query  213  KDSTKWRKLVDFRELNKRTQDFWEVQLGIPHPAGLKKKKSVTVLDVGDAYFSVPLDEDFR  272
            KDST+W KL DFRELNKR QDFWEVQLGIPHPAGLKKKKSVTVLDVGDAYFSVPLD+DFR
Sbjct  4    KDSTQWGKLGDFRELNKRIQDFWEVQLGIPHPAGLKKKKSVTVLDVGDAYFSVPLDKDFR  183

Query  273  KYTAFTIPSINNETPGIRYQYNVLPQGWKGSPAIFQSSMTKILEPFRKQNPDIVIYQYMD  332
            KYTAFTIPS NNETPG+RYQYNVLPQGWKGSPAIFQSSM KILEPFRKQNP+I I QYMD
Sbjct  184  KYTAFTIPSRNNETPGMRYQYNVLPQGWKGSPAIFQSSMIKILEPFRKQNPNIEINQYMD  363

Query  333  DLYVGSDLEIGQHRTKIEELRQHLLRWGLTTPDKKHQKEPPFLWMGYELHPDKWTVQPIV  392
            DLYVGSDLEIGQHRTK+EELRQHLL+WGLTTPDKKHQKEPPFLWMGYELHPDKWTVQPI
Sbjct  364  DLYVGSDLEIGQHRTKVEELRQHLLKWGLTTPDKKHQKEPPFLWMGYELHPDKWTVQPIT  543
```

>Levilactobacillus brevis strain CRL 2013 scaffold14, whole genome shotgun sequence
Sequence ID: NZ_MZMW01000018.1 Length: 588
Range 1: 62 to 586

Score:352 bits(903), Expect:1e-109,
Method:Compositional matrix adjust.,
Identities:165/175(94%), Positives:167/175(95%), Gaps:0/175(0%)

```
Query  229  KRTQDFWEVQLGIPHPAGLKKKKSVTVLDVGDAYFSVPLDEDFRKYTAFTIPSINNETPG  288
            KRTQDFWEVQLGIPHPAGLKKKKSVTVLDVGDAYFSVPLD DFRKYTAFTIPS+NNETPG
Sbjct  586  KRTQDFWEVQLGIPHPAGLKKKKSVTVLDVGDAYFSVPLDXDFRKYTAFTIPSVNNETPG  407

Query  289  IRYQYNVLPQGWKGSPAIFQSSMTKILEPFRKQNPDIVIYQYMDDLYVGSDLEIGQHRTK  348
            +RYQYNVLPQGWKGSPAIFQSSMTKILEPFRK  PDIVIYQYMDDLYVGSDLEI QHR K
Sbjct  406  VRYQYNVLPQGWKGSPAIFQSSMTKILEPFRKXXPDIVIYQYMDDLYVGSDLEIEQHRAK  227

Query  349  IEELRQHLLRWGLTTPDKKHQKEPPFLWMGYELHPDKWTVQPIVLPEKDSWTVND  403
            IEELR HLL WG TTPDKKHQKEPPFLWMGYELHPDKWTVQPIVLPEKDSWTVND
Sbjct  226  IEELRXHLLXWGFTTPDKKHQKEPPFLWMGYELHPDKWTVQPIVLPEKDSWTVND  62
```

Range 2: 2 to 70

Score:52.8 bits(125), Expect:0.008,
Method:Composition-based stats.,
Identities:22/23(96%), Positives:22/23(95%), Gaps:0/23(0%)

```
Query  381  LHPDKWTVQPIVLPEKDSWTVND  403
            LHPDKWTVQPIVLPEKDSWTV D
Sbjct  2    LHPDKWTVQPIVLPEKDSWTVID  70
```

>Acinetobacter baumannii strain ACM-38 NODE_62_length_542_cov_0.942169, whole genome shotgun sequence
Sequence ID: NZ_QAGQ01000062.1 Length: 542
Range 1: 2 to 541

Score:351 bits(900), Expect:1e-109,
Method:Compositional matrix adjust.,
Identities:164/180(91%), Positives:172/180(95%), Gaps:0/180(0%)

```
Query  245  AGLKKKKSVTVLDVGDAYFSVPLDEDFRKYTAFTIPSINNETPGIRYQYNVLPQGWKGSP  304
            A LKKK S+TVLDVGDAYFSVPLD DFRKYTAFTIPS NNETPGIRYQYNVLPQGWKGSP
Sbjct  541  AELKKKNSLTVLDVGDAYFSVPLDPDFRKYTAFTIPSRNNETPGIRYQYNVLPQGWKGSP  362

Query  305  AIFQSSMTKILEPFRKQNPDIVIYQYMDDLYVGSDLEIGQHRTKIEELRQHLLRWGLTTP  364
            AIFQSSMTKILEPFRKQNPD+VIYQYMDDLYVGSDLE+ QHRTK+EELR HLL+WG +TP
Sbjct  361  AIFQSSMTKILEPFRKQNPDMVIYQYMDDLYVGSDLELEQHRTKVEELRAHLLKWGFSTP  182

Query  365  DKKHQKEPPFLWMGYELHPDKWTVQPIVLPEKDSWTVNDIQKLVGKLNWASQIYPGIKVR  424
            D+KHQKEPPFLWMGYELHPDKWTVQPIVLPEKDSWTVNDIQKLVGKLNWASQIY GIKV+
Sbjct  181  DQKHQKEPPFLWMGYELHPDKWTVQPIVLPEKDSWTVNDIQKLVGKLNWASQIYAGIKVK  2
```

>Cellulophaga baltica strain 14 NODE_54_length_581_cov_63.513216, whole genome shotgun sequence
Sequence ID: NZ_VIQP01000054.1 Length: 581
Range 1: 1 to 579

Score:344 bits(883), Expect:7e-107,
Method:Compositional matrix adjust.,
Identities:173/193(90%), Positives:184/193(95%), Gaps:0/193(0%)

```
Query  298  QGWKGSPAIFQSSMTKILEPFRKQNPDIVIYQYMDDLYVGSDLEIGQHRTKIEELRQHLL  357
            QGWKGSPAIFQ+SMTKIL+PFR++NP+IVIYQYMDDLYVGSDLEIGQHR KIEELR HLL
Sbjct  1    QGWKGSPAIFQASMTKILDPFRRKNPEIVIYQYMDDLYVGSDLEIGQHRAKIEELRAHLL  180

Query  358  RWGLTTPDKKHQKEPPFLWMGYELHPDKWTVQPIVLPEKDSWTVNDIQKLVGKLNWASQI  417
             WG TTPDKKHQKEPPFLWMGYELHPDKWTVQPI LP+KDSWTVNDIQKLVGKLNWASQI
Sbjct  181  NWGFTTPDKKHQKEPPFLWMGYELHPDKWTVQPIQLPDKDSWTVNDIQKLVGKLNWASQI  360

Query  418  YPGIKVRQLCKLLRGTKALTEVIPLTeeaelelaeNREILKEPVHGVYYDPSKDLIAEIQ  477
            YPGIKV +LCKLLRG KALT+++ LTEEAELELAENREILKEPVHGVYYDPSKDL+AEIQ
Sbjct  361  YPGIKVTRLCKLLRGAKALTDIVTLTEEAELELAENREILKEPVHGVYYDPSKDLVAEIQ  540

Query  478  KQGQGQWTYQIYQ  490
            KQGQ QWTYQIYQ
Sbjct  541  KQGQDQWTYQIYQ  579
```

>Shigella sonnei strain 07401ss_1, whole genome shotgun sequence
Sequence ID: NZ_CWST01000362.1 Length: 509
Range 1: 3 to 509

```
Score:342 bits(876), Expect:3e-106,
Method:Compositional matrix adjust.,
Identities:158/169(93%), Positives:165/169(97%), Gaps:0/169(0%)

Query  227  LNKRTQDFWEVQLGIPHPAGLKKKKSVTVLDVGDAYFSVPLDEDFRKYTAFTIPSINNET  286
            LNKRTQDFWEVQLGIPHPAGLKKKKSVTVLDVGDAYFSVPLD+DFRKYTAFTIPS+NNET
Sbjct  509  LNKRTQDFWEVQLGIPHPAGLKKKKSVTVLDVGDAYFSVPLDKDFRKYTAFTIPSVNNET  330

Query  287  PGIRYQYNVLPQGWKGSPAIFQSSMTKILEPFRKQNPDIVIYQYMDDLYVGSDLEIGQHR  346
            PGIRYQYNVLPQGWKGSPAIFQ+SMTKILEPFR +NP+IVIYQYMDDLYVGSDLEIGQHR
Sbjct  329  PGIRYQYNVLPQGWKGSPAIFQASMTKILEPFRIKNPEIVIYQYMDDLYVGSDLEIGQHR  150

Query  347  TKIEELRQHLLRWGLTTPDKKHQKEPPFLWMGYELHPDKWTVQPIVLPE  395
             KIEELR+HLL+WG TTPDKKHQKEPPFLWMGYELHPDKWTVQPI LPE
Sbjct  149  AKIEELREHLLKWGFTTPDKKHQKEPPFLWMGYELHPDKWTVQPIALPE  3
```

>Mycobacterium tuberculosis variant bovis strain CSURQ1131, whole genome shotgun sequence
Sequence ID: NZ_CAJEUJ010000145.1 Length: 888
Range 1: 114 to 701

```
Score:343 bits(880), Expect:7e-105,
Method:Compositional matrix adjust.,
Identities:166/205(81%), Positives:177/205(86%), Gaps:9/205(4%)

Query  188  MEKEGKISKIGPENPYNTPVFAIKKKDSTKWRKLVDFRELNKRTQDFWEVQLGIPHPAGL  247
            +E EG +S IGPENPYNTPVFAIKKK+STKWRKLVDFRELNK+TQDFWEVQLGIPHP+GL
Sbjct  114  VEHEGSLSNIGPENPYNTPVFAIKKKNSTKWRKLVDFRELNKKTQDFWEVQLGIPHPSGL  293

Query  248  KKKKSVTVLDVGDAYFSVPLDEDFRKYTAFTIPSINNETPGIRYQYNVLPQGWKGSPAIF  307
            KKKKSVTVLDVGDAYFSVPLDEDFRKYTAFTIPS+NN TPGIRYQYNVLPQGWKGSPAIF
Sbjct  294  KKKKSVTVLDVGDAYFSVPLDEDFRKYTAFTIPSVNNTTPGIRYQYNVLPQGWKGSPAIF  473

Query  308  QSSMTKILEPFRKQNPDIVIYQYMDDLYVGSDLEIGQHRTKIEELRQHLLRWGLTTPDKK  367
            QSSMTKILEPFRKQNPDIVIYQY+DDLYVGSDLEI QHR K+EELRQHL RWGLT   K
Sbjct  474  QSSMTKILEPFRKQNPDIVIYQYVDDLYVGSDLEIEQHRAKVEELRQHLWRWGLTHQTKN  653

Query  368  HQKEPPFLWMGYELHPDKWTVQPIV  392
             +K          LH   W +  I+
Sbjct  654  IRK---------NLHSFGWVMNSIL  701
```

Range 2: 629 to 769

```
Score:89.4 bits(220), Expect:2e-14,
Method:Compositional matrix adjust.,
Identities:39/47(83%), Positives:43/47(91%), Gaps:0/47(0%)

Query  360  GLTTPDKKHQKEPPFLWMGYELHPDKWTVQPIVLPEKDSWTVNDIQK  406
            G+ TPDKKHQKEPPFLWMGYELHPDKWTVQPIVLPEKDSW V+ + +
Sbjct  629  GIDTPDKKHQKEPPFLWMGYELHPDKWTVQPIVLPEKDSWNVHRLMQ  769
```

Range 3: 114 to 701

```
Score:343 bits(880), Expect:7e-105,
Method:Compositional matrix adjust.,
Identities:166/205(81%), Positives:177/205(86%), Gaps:9/205(4%)

Query  188  MEKEGKISKIGPENPYNTPVFAIKKKDSTKWRKLVDFRELNKRTQDFWEVQLGIPHPAGL  247
            +E EG +S IGPENPYNTPVFAIKKK+STKWRKLVDFRELNK+TQDFWEVQLGIPHP+GL
Sbjct  114  VEHEGSLSNIGPENPYNTPVFAIKKKNSTKWRKLVDFRELNKKTQDFWEVQLGIPHPSGL  293

Query  248  KKKKSVTVLDVGDAYFSVPLDEDFRKYTAFTIPSINNETPGIRYQYNVLPQGWKGSPAIF  307
            KKKKSVTVLDVGDAYFSVPLDEDFRKYTAFTIPS+NN TPGIRYQYNVLPQGWKGSPAIF
Sbjct  294  KKKKSVTVLDVGDAYFSVPLDEDFRKYTAFTIPSVNNTTPGIRYQYNVLPQGWKGSPAIF  473

Query  308  QSSMTKILEPFRKQNPDIVIYQYMDDLYVGSDLEIGQHRTKIEELRQHLLRWGLTTPDKK  367
            QSSMTKILEPFRKQNPDIVIYQY+DDLYVGSDLEI QHR K+EELRQHL RWGLT   K
Sbjct  474  QSSMTKILEPFRKQNPDIVIYQYVDDLYVGSDLEIEQHRAKVEELRQHLWRWGLTHQTKN  653

Query  368  HQKEPPFLWMGYELHPDKWTVQPIV  392
             +K          LH   W +  I+
Sbjct  654  IRK---------NLHSFGWVMNSIL  701
```

Range 4: 629 to 769

```
Score:89.4 bits(220), Expect:2e-14,
Method:Compositional matrix adjust.,
Identities:39/47(83%), Positives:43/47(91%), Gaps:0/47(0%)

Query  360  GLTTPDKKHQKEPPFLWMGYELHPDKWTVQPIVLPEKDSWTVNDIQK  406
            G+ TPDKKHQKEPPFLWMGYELHPDKWTVQPIVLPEKDSW V+ + +
Sbjct  629  GIDTPDKKHQKEPPFLWMGYELHPDKWTVQPIVLPEKDSWNVHRLMQ  769
```

>Enterococcus faecium strain INF91_9 NODE_617_length_515_cov_1.76289, whole genome shotgun sequence
Sequence ID: NZ_JAANZG010000617.1 Length: 515
Range 1: 2 to 514

```
Score:338 bits(866), Expect:1e-104,
Method:Compositional matrix adjust.,
Identities:158/171(92%), Positives:162/171(94%), Gaps:0/171(0%)

Query  234  FWEVQLGIPHPAGLKKKKSVTVLDVGDAYFSVPLDEDFRKYTAFTIPSINNETPGIRYQY  293
            FWEVQLGIPHP GLKKKKSVTVLDVGDAYFSVPLDE FRKYTAFTIPS NNETPGIRYQY
Sbjct  2    FWEVQLGIPHPGGLKKKKSVTVLDVGDAYFSVPLDEGFRKYTAFTIPSTNNETPGIRYQY  181

Query  294  NVLPQGWKGSPAIFQSSMTKILEPFRKQNPDIVIYQYMDDLYVGSDLEIGQHRTKIEELR  353
            NVLPQGWKGSPAIFQSSMTKILEPFR +NPDIVIYQY+DDL VGSDLEIGQHR KIEELR
Sbjct  182  NVLPQGWKGSPAIFQSSMTKILEPFRAKNPDIVIYQYVDDLLVGSDLEIGQHRAKIEELR  361

Query  354  QHLLRWGLTTPDKKHQKEPPFLWMGYELHPDKWTVQPIVLPEKDSWTVNDI  404
            +HLLRWG  TPDKKHQKEPPFLWMGYELHPDKWT+QPI LPEKDSWTVNDI
Sbjct  362  EHLLRWGFYTPDKKHQKEPPFLWMGYELHPDKWTIQPIQLPEKDSWTVNDI  514
```

>Klebsiella aerogenes strain AUH-KAM3 NODE_59_length_558_cov_0.865429, whole genome shotgun sequence
Sequence ID: NZ_PYHH01000059.1 Length: 558
Range 1: 1 to 558

```
Score:338 bits(866), Expect:1e-104,
Method:Compositional matrix adjust.,
Identities:168/186(90%), Positives:179/186(96%), Gaps:0/186(0%)

Query  282  INNETPGIRYQYNVLPQGWKGSPAIFQSSMTKILEPFRKQNPDIVIYQYMDDLYVGSDLE  341
            +NNETPGIRYQYNVLPQGWKGSPAIFQ+SMTKILEPFR +NPDIVIYQYMDDLYVGSDLE
Sbjct  558  VNNETPGIRYQYNVLPQGWKGSPAIFQASMTKILEPFRLKNPDIVIYQYMDDLYVGSDLE  379

Query  342  IGQHRTKIEELRQHLLRWGLTTPDKKHQKEPPFLWMGYELHPDKWTVQPIVLPEKDSWTV  401
            IGQHRTK+EELR HLL+ G +TPD+KHQKEPPFLW+GYELHPDKWTVQPI LPEKDSWTV
Sbjct  378  IGQHRTKVEELRAHLLK*GFSTPDQKHQKEPPFLWIGYELHPDKWTVQPIQLPEKDSWTV  199

Query  402  NDIQKLVGKLNWASQIYPGIKVRQLCKLLRGTKALTEVIPLTeeaelelaeNREILKEPV  461
            NDIQKLVGKLNWASQIY GIKV+QLCKLLRG KALT+VIPLTEEAELELAENREIL+EPV
Sbjct  198  NDIQKLVGKLNWASQIYAGIKVKQLCKLLRGAKALTDVIPLTEEAELELAENREILREPV  19

Query  462  HGVYYD  467
            HGVYYD
Sbjct  18   HGVYYD  1



>Thalassovita mangrovi strain GS-10 Scaffold19_1, whole genome shotgun sequence
Sequence ID: NZ_WWEN01000019.1 Length: 2900
Range 1: 2362 to 2898

Score:360 bits(924), Expect:4e-103,
Method:Compositional matrix adjust.,
Identities:174/179(97%), Positives:178/179(99%), Gaps:0/179(0%)

Query  2     QGKAREFSSEQTRANSPTRRELQVWGRDNNSPSEAGADRQGTVSFNFPQVTLWQRPLVTI  61
             QGKAREFSSEQTRANSPTRRELQVWGRDNNS SEAGADRQGTVSF+FPQ+TLWQRPLVTI
Sbjct  2362  QGKAREFSSEQTRANSPTRRELQVWGRDNNSLSEAGADRQGTVSFSFPQITLWQRPLVTI  2541

Query  62    KIGGQLKEALLDTGADDTVLEEMSLPGRWKPKMIGGIGGFIKVRQYDQILIEICGHKAIG  121
             KIGGQLKEALLDTGADDTVLEEM+LPGRWKPKMIGGIGGFIKVRQYDQILIEICGHKAIG
Sbjct  2542  KIGGQLKEALLDTGADDTVLEEMNLPGRWKPKMIGGIGGFIKVRQYDQILIEICGHKAIG  2721

Query  122   TVLVGPTPVNIIGRNLLTQIGCTLNFPISPIETVPVKLKPGMDGPKVKQWPLTEEKIKA  180
             TVLVGPTPVNIIGRNLLTQIGCTLNFPISPIETVPVKLKPGMDGP+VKQWPLTEEKIKA
Sbjct  2722  TVLVGPTPVNIIGRNLLTQIGCTLNFPISPIETVPVKLKPGMDGPQVKQWPLTEEKIKA  2898



>Nocardioides luteus strain JCM 3358 sequence39, whole genome shotgun sequence
Sequence ID: NZ_BMRK01000039.1 Length: 5754
Range 1: 1433 to 1981

Score:371 bits(953), Expect:7e-103,
Method:Compositional matrix adjust.,
Identities:174/183(95%), Positives:181/183(98%), Gaps:0/183(0%)

Query  880   QVDCSPGIWQLDCTHLEGKVILVAVHVASGYIEAEVIPAETGQETAYFLLKLAGRWPVKT  939
             ++DCSPGIWQLDCTHLEGKVILVAVHVASGYIEAEVIPAETGQETAYFLLKLAGRWPVKT
Sbjct  1433  RLDCSPGIWQLDCTHLEGKVILVAVHVASGYIEAEVIPAETGQETAYFLLKLAGRWPVKT  1612

Query  940   IHTDNGSNFTGATVRAACWWAGIKQEFGIPYNPQSQGVVESMNKELKKIIGQVRDQAEHL  999
             +HTDNGSNFT  TV+AACWWAGIKQEFGIPYNPQSQGV+ESMNKELKKIIGQVRDQAEHL
Sbjct  1613  VHTDNGSNFTSTTVKAACWWAGIKQEFGIPYNPQSQGVIESMNKELKKIIGQVRDQAEHL  1792

Query  1000  KTAVQMAVFIHNFKRKGGIGGYSAGERIVDIIATDIQTKELQKQITKIQNFRVYYRDSRN  1059
             KTAVQMAVFIHNFKRKGGIGGYSAGERIVDIIATDIQTKELQKQITKIQNFRVYYRDSR+
Sbjct  1793  KTAVQMAVFIHNFKRKGGIGGYSAGERIVDIIATDIQTKELQKQITKIQNFRVYYRDSRD  1972

Query  1060  PLW  1062
             P+W
Sbjct  1973  PVW  1981



>Streptomyces griseoflavus strain JCM 4479 sequence37, whole genome shotgun sequence
Sequence ID: NZ_BMUC01000037.1 Length: 5775
Range 1: 3767 to 4315

Score:371 bits(953), Expect:7e-103,
Method:Compositional matrix adjust.,
Identities:174/183(95%), Positives:181/183(98%), Gaps:0/183(0%)

Query  880   QVDCSPGIWQLDCTHLEGKVILVAVHVASGYIEAEVIPAETGQETAYFLLKLAGRWPVKT  939
             ++DCSPGIWQLDCTHLEGKVILVAVHVASGYIEAEVIPAETGQETAYFLLKLAGRWPVKT
Sbjct  4315  RLDCSPGIWQLDCTHLEGKVILVAVHVASGYIEAEVIPAETGQETAYFLLKLAGRWPVKT  4136

Query  940   IHTDNGSNFTGATVRAACWWAGIKQEFGIPYNPQSQGVVESMNKELKKIIGQVRDQAEHL  999
             +HTDNGSNFT  TV+AACWWAGIKQEFGIPYNPQSQGV+ESMNKELKKIIGQVRDQAEHL
Sbjct  4135  VHTDNGSNFTSTTVKAACWWAGIKQEFGIPYNPQSQGVIESMNKELKKIIGQVRDQAEHL  3956

Query  1000  KTAVQMAVFIHNFKRKGGIGGYSAGERIVDIIATDIQTKELQKQITKIQNFRVYYRDSRN  1059
             KTAVQMAVFIHNFKRKGGIGGYSAGERIVDIIATDIQTKELQKQITKIQNFRVYYRDSR+
Sbjct  3955  KTAVQMAVFIHNFKRKGGIGGYSAGERIVDIIATDIQTKELQKQITKIQNFRVYYRDSRD  3776

Query  1060  PLW  1062
             P+W
Sbjct  3775  PVW  3767



>Saccharothrix coeruleofusca strain JCM 3313 sequence45, whole genome shotgun sequence
Sequence ID: NZ_BMRG01000045.1 Length: 5775
Range 1: 3767 to 4315

Score:371 bits(953), Expect:7e-103,
Method:Compositional matrix adjust.,
Identities:174/183(95%), Positives:181/183(98%), Gaps:0/183(0%)

Query  880   QVDCSPGIWQLDCTHLEGKVILVAVHVASGYIEAEVIPAETGQETAYFLLKLAGRWPVKT  939
             ++DCSPGIWQLDCTHLEGKVILVAVHVASGYIEAEVIPAETGQETAYFLLKLAGRWPVKT
Sbjct  4315  RLDCSPGIWQLDCTHLEGKVILVAVHVASGYIEAEVIPAETGQETAYFLLKLAGRWPVKT  4136

Query  940   IHTDNGSNFTGATVRAACWWAGIKQEFGIPYNPQSQGVVESMNKELKKIIGQVRDQAEHL  999
             +HTDNGSNFT  TV+AACWWAGIKQEFGIPYNPQSQGV+ESMNKELKKIIGQVRDQAEHL
Sbjct  4135  VHTDNGSNFTSTTVKAACWWAGIKQEFGIPYNPQSQGVIESMNKELKKIIGQVRDQAEHL  3956

Query  1000  KTAVQMAVFIHNFKRKGGIGGYSAGERIVDIIATDIQTKELQKQITKIQNFRVYYRDSRN  1059
             KTAVQMAVFIHNFKRKGGIGGYSAGERIVDIIATDIQTKELQKQITKIQNFRVYYRDSR+
Sbjct  3955  KTAVQMAVFIHNFKRKGGIGGYSAGERIVDIIATDIQTKELQKQITKIQNFRVYYRDSRD  3776

Query  1060  PLW  1062
             P+W
Sbjct  3775  PVW  3767
```

```
>Streptomyces griseoloalbus strain JCM 4480 sequence054, whole genome shotgun sequence
Sequence ID: NZ_BMUD01000054.1 Length: 5815
Range 1: 3787 to 4335

Score:371 bits(953), Expect:7e-103,
Method:Compositional matrix adjust.,
Identities:174/183(95%), Positives:181/183(98%), Gaps:0/183(0%)

Query  880   QVDCSPGIWQLDCTHLEGKVILVAVHVASGYIEAEVIPAETGQETAYFLLKLAGRWPVKT  939
             ++DCSPGIWQLDCTHLEGKVILVAVHVASGYIEAEVIPAETGQETAYFLLKLAGRWPVKT
Sbjct  4335  RLDCSPGIWQLDCTHLEGKVILVAVHVASGYIEAEVIPAETGQETAYFLLKLAGRWPVKT  4156

Query  940   IHTDNGSNFTGATVRAACWWAGIKQEFGIPYNPQSQGVVESMNKELKKIIGQVRDQAEHL  999
             +HTDNGSNFT  TV+AACWWAGIKQEFGIPYNPQSQGV+ESMNKELKKIIGQVRDQAEHL
Sbjct  4155  VHTDNGSNFTSTTVKAACWWAGIKQEFGIPYNPQSQGVIESMNKELKKIIGQVRDQAEHL  3976

Query  1000  KTAVQMAVFIHNFKRKGGIGGYSAGERIVDIIATDIQTKELQKQITKIQNFRVYYRDSRN  1059
             KTAVQMAVFIHNFKRKGGIGGYSAGERIVDIIATDIQTKELQKQITKIQNFRVYYRDSR+
Sbjct  3975  KTAVQMAVFIHNFKRKGGIGGYSAGERIVDIIATDIQTKELQKQITKIQNFRVYYRDSRD  3796

Query  1060  PLW  1062
             P+W
Sbjct  3795  PVW  3787
```

```
>Actinoplanes ianthinogenes strain JCM 3249 sequence037, whole genome shotgun sequence
Sequence ID: NZ_BMQZ01000037.1 Length: 5815
Range 1: 3787 to 4335

Score:371 bits(953), Expect:7e-103,
Method:Compositional matrix adjust.,
Identities:174/183(95%), Positives:181/183(98%), Gaps:0/183(0%)

Query  880   QVDCSPGIWQLDCTHLEGKVILVAVHVASGYIEAEVIPAETGQETAYFLLKLAGRWPVKT  939
             ++DCSPGIWQLDCTHLEGKVILVAVHVASGYIEAEVIPAETGQETAYFLLKLAGRWPVKT
Sbjct  4335  RLDCSPGIWQLDCTHLEGKVILVAVHVASGYIEAEVIPAETGQETAYFLLKLAGRWPVKT  4156

Query  940   IHTDNGSNFTGATVRAACWWAGIKQEFGIPYNPQSQGVVESMNKELKKIIGQVRDQAEHL  999
             +HTDNGSNFT  TV+AACWWAGIKQEFGIPYNPQSQGV+ESMNKELKKIIGQVRDQAEHL
Sbjct  4155  VHTDNGSNFTSTTVKAACWWAGIKQEFGIPYNPQSQGVIESMNKELKKIIGQVRDQAEHL  3976

Query  1000  KTAVQMAVFIHNFKRKGGIGGYSAGERIVDIIATDIQTKELQKQITKIQNFRVYYRDSRN  1059
             KTAVQMAVFIHNFKRKGGIGGYSAGERIVDIIATDIQTKELQKQITKIQNFRVYYRDSR+
Sbjct  3975  KTAVQMAVFIHNFKRKGGIGGYSAGERIVDIIATDIQTKELQKQITKIQNFRVYYRDSRD  3796

Query  1060  PLW  1062
             P+W
Sbjct  3795  PVW  3787
```

```
>Sinomonas cellulolyticus strain KCTC 49339 sequence28, whole genome shotgun sequence
Sequence ID: NZ_BNCM01000028.1 Length: 5849
Range 1: 3758 to 4306

Score:371 bits(953), Expect:7e-103,
Method:Compositional matrix adjust.,
Identities:174/183(95%), Positives:181/183(98%), Gaps:0/183(0%)

Query  880   QVDCSPGIWQLDCTHLEGKVILVAVHVASGYIEAEVIPAETGQETAYFLLKLAGRWPVKT  939
             ++DCSPGIWQLDCTHLEGKVILVAVHVASGYIEAEVIPAETGQETAYFLLKLAGRWPVKT
Sbjct  4306  RLDCSPGIWQLDCTHLEGKVILVAVHVASGYIEAEVIPAETGQETAYFLLKLAGRWPVKT  4127

Query  940   IHTDNGSNFTGATVRAACWWAGIKQEFGIPYNPQSQGVVESMNKELKKIIGQVRDQAEHL  999
             +HTDNGSNFT  TV+AACWWAGIKQEFGIPYNPQSQGV+ESMNKELKKIIGQVRDQAEHL
Sbjct  4126  VHTDNGSNFTSTTVKAACWWAGIKQEFGIPYNPQSQGVIESMNKELKKIIGQVRDQAEHL  3947

Query  1000  KTAVQMAVFIHNFKRKGGIGGYSAGERIVDIIATDIQTKELQKQITKIQNFRVYYRDSRN  1059
             KTAVQMAVFIHNFKRKGGIGGYSAGERIVDIIATDIQTKELQKQITKIQNFRVYYRDSR+
Sbjct  3946  KTAVQMAVFIHNFKRKGGIGGYSAGERIVDIIATDIQTKELQKQITKIQNFRVYYRDSRD  3767

Query  1060  PLW  1062
             P+W
Sbjct  3766  PVW  3758
```

```
>Saccharothrix mutabilis subsp. mutabilis strain JCM 3380 sequence46, whole genome shotgun sequence
Sequence ID: NZ_BAAABU010000046.1 Length: 5899
Range 1: 1481 to 2029

Score:371 bits(953), Expect:7e-103,
Method:Compositional matrix adjust.,
Identities:174/183(95%), Positives:181/183(98%), Gaps:0/183(0%)

Query  880   QVDCSPGIWQLDCTHLEGKVILVAVHVASGYIEAEVIPAETGQETAYFLLKLAGRWPVKT  939
             ++DCSPGIWQLDCTHLEGKVILVAVHVASGYIEAEVIPAETGQETAYFLLKLAGRWPVKT
Sbjct  1481  RLDCSPGIWQLDCTHLEGKVILVAVHVASGYIEAEVIPAETGQETAYFLLKLAGRWPVKT  1660

Query  940   IHTDNGSNFTGATVRAACWWAGIKQEFGIPYNPQSQGVVESMNKELKKIIGQVRDQAEHL  999
             +HTDNGSNFT  TV+AACWWAGIKQEFGIPYNPQSQGV+ESMNKELKKIIGQVRDQAEHL
Sbjct  1661  VHTDNGSNFTSTTVKAACWWAGIKQEFGIPYNPQSQGVIESMNKELKKIIGQVRDQAEHL  1840

Query  1000  KTAVQMAVFIHNFKRKGGIGGYSAGERIVDIIATDIQTKELQKQITKIQNFRVYYRDSRN  1059
             KTAVQMAVFIHNFKRKGGIGGYSAGERIVDIIATDIQTKELQKQITKIQNFRVYYRDSR+
Sbjct  1841  KTAVQMAVFIHNFKRKGGIGGYSAGERIVDIIATDIQTKELQKQITKIQNFRVYYRDSRD  2020

Query  1060  PLW  1062
             P+W
Sbjct  2021  PVW  2029
```

```
>Neisseria gonorrhoeae strain GC-206 NODE_117_length_504_cov_1.30067_ID_20406, whole genome shotgun sequence
Sequence ID: NZ_SAWN01000117.1 Length: 504
Range 1: 2 to 502

Score:332 bits(850), Expect:2e-102,
Method:Compositional matrix adjust.,
Identities:153/167(92%), Positives:160/167(95%), Gaps:0/167(0%)

Query  261  AYFSVPLDEDFRKYTAFTIPSINNETPGIRYQYNVLPQGWKGSPAIFQSSMTKILEPFRK  320
            AYFSVPLDE+FRKYTAFTIPSINNETPGIRYQYNVLPQGWKGSP+IFQSSM KILEPFR
Sbjct  502  AYFSVPLDENFRKYTAFTIPSINNETPGIRYQYNVLPQGWKGSPSIFQSSMIKILEPFRA  323

Query  321  QNPDIVIYQYMDDLYVGSDLEIGQHRTKIEELRQHLLRWGLTTPDKKHQKEPPFLWMGYE  380
```

```
            +NP++VIYQY+DDLYVGSDLEIGQHR KIEELR HLL+WG TTPDKKHQKEPPFLWMGYE
Sbjct  322  ENPELVIYQYIDDLYVGSDLEIGQHRAKIEELRDHLLKWGFTTPDKKHQKEPPFLWMGYE  143

Query  381  LHPDKWTVQPIVLPEKDSWTVNDIQKLVGKLNWASQIYPGIKVRQLC  427
            LHPDKWTVQPI LP KDSWTVNDIQKLVGKLNWASQIYPGIKVRQLC
Sbjct  142  LHPDKWTVQPIQLPTKDSWTVNDIQKLVGKLNWASQIYPGIKVRQLC  2
```

>Providencia rettgeri strain AFE96_S1 41_contig_58, whole genome shotgun sequence
Sequence ID: NZ_SHDP01000058.1 Length: 546
Range 1: 20 to 544

Score:330 bits(847), Expect:8e-102,
Method:Compositional matrix adjust.,
Identities:157/175(90%), Positives:165/175(94%), Gaps:0/175(0%)

```
Query  187  EMEKEGKISKIGPENPYNTPVFAIKKKDSTKWRKLVDFRELNKRTQDFWEVQLGIPHPAG  246
            EMEKE KI+K G ENPYNTPVFAIKKKDSTKWRKLVD RELNKRT+D WEVQLGIPHPAG
Sbjct  20   EMEKEEKITKNGTENPYNTPVFAIKKKDSTKWRKLVDVRELNKRTKDMWEVQLGIPHPAG  199

Query  247  LKKKKSVTVLDVGDAYFSVPLDEDFRKYTAFTIPSINNETPGIRYQYNVLPQGWKGSPAI  306
            LKK KSVT+LDVGDAYFSVPLDE+FRKYTAFTIPSINN TPGIRYQYNVLPQGWKGSPAI
Sbjct  200  LKKNKSVTILDVGDAYFSVPLDENFRKYTAFTIPSINNATPGIRYQYNVLPQGWKGSPAI  379

Query  307  FQSSMTKILEPFRKQNPDIVIYQYMDDLYVGSDLEIGQHRTKIEELRQHLLRWGL  361
            FQSSMTKILEPF  +NPDIVIYQY+DDLYVGSDLEIGQHR KIEELR+HLL+WGL
Sbjct  380  FQSSMTKILEPFIAKNPDIVIYQYVDDLYVGSDLEIGQHRRKIEELREHLLKWGL  544
```

>Providencia rettgeri strain AFE422_S1 40_contig_33, whole genome shotgun sequence
Sequence ID: NZ_SHDL01000033.1 Length: 501
Range 1: 1 to 501

Score:329 bits(844), Expect:1e-101,
Method:Compositional matrix adjust.,
Identities:153/167(92%), Positives:162/167(97%), Gaps:0/167(0%)

```
Query  206  PVFAIKKKDSTKWRKLVDFRELNKRTQDFWEVQLGIPHPAGLKKKKSVTVLDVGDAYFSV  265
            P+FAIK+KDSTKWRKLVDFRELNKRTQDFWEVQLGIPHPAGLKKKKS+TVLDVGDAYFSV
Sbjct  501  PIFAIKRKDSTKWRKLVDFRELNKRTQDFWEVQLGIPHPAGLKKKKSMTVLDVGDAYFSV  322

Query  266  PLDEDFRKYTAFTIPSINNETPGIRYQYNVLPQGWKGSPAIFQSSMTKILEPFRKQNPDI  325
            PLDE FRKYTAFTIPSINNETPGIRYQYNVLPQGWKGSPAIFQSSMT+ILEPFR +NP+I
Sbjct  321  PLDEGFRKYTAFTIPSINNETPGIRYQYNVLPQGWKGSPAIFQSSMTRILEPFRAKNPEI  142

Query  326  VIYQYMDDLYVGSDLEIGQHRTKIEELRQHLLRWGLTTPDKKHQKEP  372
            VIYQY+DDLYVGSDLEIGQHR K+EELR+HLL WG TTPDKKHQKEP
Sbjct  141  VIYQYVDDLYVGSDLEIGQHRAKVEELRKHLLMWGFTTPDKKHQKEP  1
```

>Providencia rettgeri strain AFE220_S1 26_contig_66, whole genome shotgun sequence
Sequence ID: NZ_SHDM01000066.1 Length: 502
Range 1: 5 to 502

Score:326 bits(836), Expect:2e-100,
Method:Compositional matrix adjust.,
Identities:152/166(92%), Positives:160/166(96%), Gaps:0/166(0%)

```
Query  237  VQLGIPHPAGLKKKKSVTVLDVGDAYFSVPLDEDFRKYTAFTIPSINNETPGIRYQYNVL  296
            VQLGIPHPAGLKKKKSVTVLDVGDAYFSVPLDE+FRKYTAFTIPSINNETPGIRYQYNVL
Sbjct  502  VQLGIPHPAGLKKKKSVTVLDVGDAYFSVPLDENFRKYTAFTIPSINNETPGIRYQYNVL  323

Query  297  PQGWKGSPAIFQSSMTKILEPFRKQNPDIVIYQYMDDLYVGSDLEIGQHRTKIEELRQHL  356
            PQGWKGSPAIFQ SMTKILEPFR +NP+IVIYQYMDDLYVGSDLEIGQHR KIEELR+HL
Sbjct  322  PQGWKGSPAIFQCSMTKILEPFRAKNPEIVIYQYMDDLYVGSDLEIGQHRAKIEELREHL  143

Query  357  LRWGLTTPDKKHQKEPPFLWMGYELHPDKWTVQPIVLPEKDSWTVN  402
            L+WGLTTPDKKHQKEPPFLWMGYELHPDKWTVQPI LPEK+ + +N
Sbjct  142  LKWGLTTPDKKHQKEPPFLWMGYELHPDKWTVQPIQLPEKELYLIN  5
```

>Acinetobacter baumannii strain ACM-23 NODE_77_length_505_cov_0.960317, whole genome shotgun sequence
Sequence ID: NZ_PYDJ01000077.1 Length: 505
Range 1: 2 to 505

Score:326 bits(836), Expect:2e-100,
Method:Compositional matrix adjust.,
Identities:157/168(93%), Positives:165/168(98%), Gaps:0/168(0%)

```
Query  58   LVTIKIGGQLKEALLDTGADDTVLEEMSLPGRWKPKMIGGIGGFIKVRQYDQILIEICGH  117
            +V I+IGGQLKEALLDTGADDTVLE+M LPGRWKPKMIGGIGGFIKVRQYD++ IEICGH
Sbjct  505  VVEIRIGGQLKEALLDTGADDTVLEDMQLPGRWKPKMIGGIGGFIKVRQYDEVPIEICGH  326

Query  118  KAIGTVLVGPTPVNIIGRNLLTQIGCTLNFPISPIETVPVKLKPGMDGPKVKQWPLTEEK  177
            KAIGTVL+GPTPVNIIGRNLLTQ+GCTLNFPISPIETVPVKLKPGMDGPKVKQWPLTEEK
Sbjct  325  KAIGTVLIGPTPVNIIGRNLLTQLGCTLNFPISPIETVPVKLKPGMDGPKVKQWPLTEEK  146

Query  178  IKALVEICTEMEKEGKISKIGPENPYNTPVFAIKKKDSTKWRKLVDFR  225
            IKALVEICTEMEK+GKISKIGPENPYNTPVFAIKKKDSTKWRKLVDFR
Sbjct  145  IKALVEICTEMEKKGKISKIGPENPYNTPVFAIKKKDSTKWRKLVDFR  2
```

>Providencia rettgeri strain AFE220_S1 26_contig_67, whole genome shotgun sequence
Sequence ID: NZ_SHDM01000067.1 Length: 512
Range 1: 1 to 510

Score:325 bits(833), Expect:6e-100,
Method:Compositional matrix adjust.,
Identities:154/170(91%), Positives:167/170(98%), Gaps:0/170(0%)

```
Query  51   VTLWQRPLVTIKIGGQLKEALLDTGADDTVLEEMSLPGRWKPKMIGGIGGFIKVRQYDQI  110
            +TLWQRPLV+IK+GGQ+KEALLDTGADDTVLE+++LPG+WKPKMIGGIGGFIKVRQYDQI
Sbjct  510  ITLWQRPLVSIKVGGQIKEALLDTGADDTVLEDINLPGKWKPKMIGGIGGFIKVRQYDQI  331

Query  111  LIEICGHKAIGTVLVGPTPVNIIGRNLLTQIGCTLNFPISPIETVPVKLKPGMDGPKVKQ  170
            LIEICG KAIGTVLVGPTPVNIIGRN+LTQ+GCTLNFPISPIETVPVKLKPGMDGPKVKQ
Sbjct  330  LIEICGKKAIGTVLVGPTPVNIIGRNMLTQLGCTLNFPISPIETVPVKLKPGMDGPKVKQ  151

Query  171  WPLTEEKIKALVEICTEMEKEGKISKIGPENPYNTPVFAIKKKDSTKWRK  220
            WPLTEEKIKAL EIC EME+EGKI+KIGPENPYNTP+FAIKKKDSTKWRK
Sbjct  150  WPLTEEKIKALTEICDEMEREGKITKIGPENPYNTPIFAIKKKDSTKWRK  1
```

```
>Acinetobacter baumannii strain ACM-24 NODE_72_length_503_cov_0.531915, whole genome shotgun sequence
Sequence ID: NZ_PYDK01000072.1 Length: 503
Range 1: 1 to 501

Score:324 bits(831), Expect:1e-99,
Method:Compositional matrix adjust.,
Identities:156/167(93%), Positives:162/167(97%), Gaps:0/167(0%)

Query  104  VRQYDQILIEICGHKAIGTVLVGPTPVNIIGRNLLTQIGCTLNFPISPIETVPVKLKPGM  163
            VRQY+QI IEICG K IGTVLVGPTPVNIIGRN+LTQIGCTLNFPISPIETVPV LKPGM
Sbjct  501  VRQYNQIPIEICGKKTIGTVLVGPTPVNIIGRNILTQIGCTLNFPISPIETVPVTLKPGM  322

Query  164  DGPKVKQWPLTEEKIKALVEICTEMEKEGKISKIGPENPYNTPVFAIKKKDSTKWRKLVD  223
            DGPKVKQWPLTEEKIKAL +ICTEME+EGKISKIGPENPYNTPVFAIKKKDSTKWRKLVD
Sbjct  321  DGPKVKQWPLTEEKIKALTDICTEMEQEGKISKIGPENPYNTPVFAIKKKDSTKWRKLVD  142

Query  224  FRELNKRTQDFWEVQLGIPHPAGLKKKKSVTVLDVGDAYFSVPLDED  270
            FRELNKRTQDFWEVQLGIPHPAGLK+KKSVTVLDVGDAYFSVPLDE+
Sbjct  141  FRELNKRTQDFWEVQLGIPHPAGLKQKKSVTVLDVGDAYFSVPLDEN  1
```

```
>Escherichia coli strain 8374wG10 NODE_51_length_497_cov_1.16802, whole genome shotgun sequence
Sequence ID: NZ_WSJQ01000051.1 Length: 497
Range 1: 3 to 497

Score:323 bits(828), Expect:2e-99,
Method:Compositional matrix adjust.,
Identities:152/165(92%), Positives:160/165(96%), Gaps:0/165(0%)

Query  167  KVKQWPLTEEKIKALVEICTEMEKEGKISKIGPENPYNTPVFAIKKKDSTKWRKLVDFRE  226
            ++KQWPLTEEKIKAL  IC EMEKEGKI+KIGPENPYNTPVFAIKKKDSTKWRKLVDFRE
Sbjct  497  RIKQWPLTEEKIKALQAICEEMEKEGKITKIGPENPYNTPVFAIKKKDSTKWRKLVDFRE  318

Query  227  LNKRTQDFWEVQLGIPHPAGLKKKKSVTVLDVGDAYFSVPLDEDFRKYTAFTIPSINNET  286
            LNKRTQDFWEVQLGIPHPAGLKKKKSVTVLDVGDAYFSVPLDE+FRKYTAFTIPSINN T
Sbjct  317  LNKRTQDFWEVQLGIPHPAGLKKKKSVTVLDVGDAYFSVPLDENFRKYTAFTIPSINNAT  138

Query  287  PGIRYQYNVLPQGWKGSPAIFQSSMTKILEPFRKQNPDIVIYQYM  331
            PGIRYQYNVLPQGWKGSPAIFQS+MTKIL+PFR +NPD+VIYQYM
Sbjct  137  PGIRYQYNVLPQGWKGSPAIFQSTMTKILQPFRTRNPDLVIYQYM  3
```

```
>Escherichia coli strain 8374wH11 NODE_48_length_500_cov_2.00268, whole genome shotgun sequence
Sequence ID: NZ_WSJI01000048.1 Length: 500
Range 1: 1 to 495

Score:320 bits(821), Expect:3e-98,
Method:Compositional matrix adjust.,
Identities:151/165(92%), Positives:160/165(96%), Gaps:0/165(0%)

Query  173  LTEEKIKALVEICTEMEKEGKISKIGPENPYNTPVFAIKKKDSTKWRKLVDFRELNKRTQ  232
            L+EEKIKAL  IC EMEKEGKISKIGPENPYNTPVFAIK+KDSTKWRKLVDFRELNKRTQ
Sbjct  495  LSEEKIKALTGICEEMEKEGKISKIGPENPYNTPVFAIKRKDSTKWRKLVDFRELNKRTQ  316

Query  233  DFWEVQLGIPHPAGLKKKKSVTVLDVGDAYFSVPLDEDFRKYTAFTIPSINNETPGIRYQ  292
            DFWEVQLGIPHPAGLKKKKS+TVLDVGDAYFSVPLDE+FRKYTAFTIPSINN TPGIRYQ
Sbjct  315  DFWEVQLGIPHPAGLKKKKSMTVLDVGDAYFSVPLDENFRKYTAFTIPSINNATPGIRYQ  136

Query  293  YNVLPQGWKGSPAIFQSSMTKILEPFRKQNPDIVIYQYMDDLYVG  337
            YNVLPQGWKGSPAIFQS+MTKIL+PFR +NPDIVIYQY++DLYVG
Sbjct  135  YNVLPQGWKGSPAIFQSTMTKILQPFRTRNPDIVIYQYVEDLYVG  1
```

```
>Acinetobacter baumannii strain ACM-35 NODE_89_length_466_cov_2.51917, whole genome shotgun sequence
Sequence ID: NZ_QEHN01000089.1 Length: 466
Range 1: 2 to 466

Score:317 bits(811), Expect:5e-97,
Method:Compositional matrix adjust.,
Identities:151/155(97%), Positives:154/155(99%), Gaps:0/155(0%)

Query  139  TQIGCTLNFPISPIETVPVKLKPGMDGPKVKQWPLTEEKIKALVEICTEMEKEGKISKIG  198
            TQ+GCTLNFPISPIETVPVKLKPGMDGPKVKQWPLTEEKIKALVEICTEMEKEGKISKIG
Sbjct  2    TQLGCTLNFPISPIETVPVKLKPGMDGPKVKQWPLTEEKIKALVEICTEMEKEGKISKIG  181

Query  199  PENPYNTPVFAIKKKDSTKWRKLVDFRELNKRTQDFWEVQLGIPHPAGLKKKKSVTVLDV  258
            PENPYNTPVFAIKKKDSTKWRKLVDFRELNKRTQDFWEVQLGIPHPAGLKKKKSVTVLDV
Sbjct  182  PENPYNTPVFAIKKKDSTKWRKLVDFRELNKRTQDFWEVQLGIPHPAGLKKKKSVTVLDV  361

Query  259  GDAYFSVPLDEDFRKYTAFTIPSINNETPGIRYQY  293
            GDAYFSVPLD++FRKYTAFTIPS NNETPGIRYQY
Sbjct  362  GDAYFSVPLDKEFRKYTAFTIPSTNNETPGIRYQY  466
```

```
>Klebsiella pneumoniae strain B2 isolate B2, whole genome shotgun sequence
Sequence ID: NZ_CABWNT010000058.1 Length: 7306
Range 1: 3707 to 5398

Score:350 bits(897), Expect:2e-95,
Method:Compositional matrix adjust.,
Identities:224/567(40%), Positives:320/567(56%), Gaps:14/567(2%)

Query  5     AREFSSEQTRANSPTRRELQVWGRDNNSPSEAGADRQG-TVSFNFPQVT--LWQRPLVTI  61
             A   + + T      R  +  +  D     + G  + G + S  F  VT  L +RP   +
Sbjct  3707  AEGVNGQNTNGGETARNLISRFESDETGVQDQGRGKSGGSQSEQFVGVTYNLEKRPTTIV  3886

Query  62    KIGGQLKEALLDTGADDTVL-----EEMSLPGR-WKPKMIGGIGGFIKVRQYDQILIEIC  115
              I       LLDTGAD +VL       +   GR ++   I G+GG ++      + ++
Sbjct  3887  LINDTPLNVLLDTGADTSVLTIAHYNRLKYRGRKYQGTGIVGVGGNVETFS-TPVTVKKK  4063

Query  116   GHKAIGTVLVGPTPVNIIGRNLLTQIGCTLNFPISPIETVP--VKLKPGMDGPKVKQWPL  173
             G +    +LV   PV I+GR++L ++G  L       E  P  +KLK G  GPKV QWPL
Sbjct  4064  GKQIKTRMLVADIPVTILGRDILQELGAQLIMAQLSKEITPREIKLKTGTVGPKVPQWPL  4243

Query  174   TEEKIKALVEICTEMEKEGKISKIGPENPYNTPVFAIKKKDSTKWRKLVDFRELNKRTQD  233
             T+EK+    EI  ++  EGKIS+   +NPYN+P+F IKKK S KWR L D RELNK  Q
Sbjct  4244  TKEKLLGAKEIVKKLLDEGKISEASDDNPYNSPIFVIKKK-SGKWRLLQDLRELNKVVQV  4420

Query  234   FWEVQLGIPHPAGLKKKKSVTVLDVGDAYFSVPLDEDFRKYTAFTIPSINNETPGIRYQY  293
               E+  G+PHP GL K   +TVLD+GDAYF++PLD  FR+YTAFT+PSIN++ P  RY +
Sbjct  4421  GTEISRGLPHPGGLIKCNHMTVLDIGDAYFTIPLDPKFRQYTAFTVPSINHQEPDKRYVW  4600

Query  294   NVLPQGWKGSPAIFQSSMTKILEPFRKQNPDIVIYQYMDDLYVGSDLEIGQHRTKIEELR  353
             N LPQG+  SP I+Q ++  IL+ FR+++PD+ +YQYMDDL++GS+    QH+  +EELR
Sbjct  4601  NCLPQGFVLSPYIYQKTLQDILQAFRERHPDVQLYQYMDDLFIGSNESKRQHKELVEELR  4780
```

```
Query  354   QHLLRWGLTTPDKKHQKEPPFLWMGYELHPDKWTVQPIVLPEKDSWTVNDIQKLVGKLNW  413
               LL  G  TP+ K Q+E P+ W+GY+L P  W VQ + L      T+ND+QKL+G + W
Sbjct  4781  AILLEKGFETPEDKLQEEAPYNWLGYQLSPGNWKVQKMQLELVKEPTLNDVQKLMGNITW  4960

Query  414   ASQIYPGIKVRQLCKLLRGTKALTEVIPLTeeaelelaeNREILKEPVHGVYYDPSKDLI  473
              S   PG+ V+Q+    +G   L + +  TEEA+ EL EN + ++E     YY+P +++I
Sbjct  4961  MSSGVPGLTVKQIAATTKGCLDLNQKVVWTEEAQKELEENNKKIQEAQGLQYYNPEEEVI  5140

Query  474   AEIQKQGQGQWTYQIYQEPFKNLKTGKYARMRGAHTNDVKQLTEAVQKITTESIVIWGKT  533
              EI+     + TY I Q     L  GK         +  K L   +Q + TESIV  G
Sbjct  5141  CEIEITKNYEATYIIKQSQ-GILWAGKKIMRANKGWSAAKNLMLLLQHVATESIVRIGTC  5317

Query  534   PKFKLPIQKetwetwwteYWQATWIPE  560
             PKFK+P  KE  +    + W  +W+P+
Sbjct  5318  PKFKVPFTKEQVKWEMEKGWYYSWLPD  5398
```

Range 2: 5654 to 7180

Score:214 bits(545), Expect:1e-51,
Method:Compositional matrix adjust.,
Identities:163/516(32%), Positives:251/516(48%), Gaps:27/516(5%)

```
Query  600   GKAGYVTNRGRQKVVTLTDTTNQKTELQAIYLALQDSG-LEVNIVTDSQYAL-GIIQAQP  657
             G A YVT+ G+ K   L   T+Q  E  AI +AL+D+    VNIVTDS Y    I +
Sbjct  5654  GVAAYVTSNGKTKQKRLGPVTHQTAERIAIQMALEDTEETLVNIVTDSYYCWKNITEGLG  5833

Query  658   DQSESELVNQIIEQLIKKEKVYLAWVPAHKGIGGNEQVDKLVSAGIRKVLYVDGAANRET  717
              +        II+ +  KE VY AWVP HKGI GN+  D+        +L   G   +E
Sbjct  5834  LEGPDSPWWPIIQNIRAKEMVYFAWVPGHKGIYGNQLADEATKITEEIMLAYQGTQIKEK  6013

Query  718   KLGKAGYVTNRGRQKVVTLTDT----TNQKTELQAIYL-------ALQDSGLEVN-IVTD  765
             +   AG+        ++ +++T    T+ K ++            ++   GL +N  + D
Sbjct  6014  RDEDAGFDLCIPYDMIIPVSETKVIPTDVKIQVPHKCFGWVTGKSSMAKQGLLINGGIID  6193

Query  766   SQYA--LGIIQAQPDQSESELV-NQIIEQLIKKEKVYLAWVPAHKGIGGNEQVDKLVSAG  822
               Y   + +I     +S  +L   Q   QLI      L      K I    +  +    G
Sbjct  6194  EGYTGEIQVICTNIGKSNMKLREGQKFAQLI-----ILQHRSNDKQIWDENKTSQRGDKG  6358

Query  823   IRK--VLFLDGIDKAQDEHEKYHSNWRAMASDFNLPPVVAKEIVASCDKCQLKGEAMHGQ  880
                  + +++ I +AQDEHE +H++ + +A  + LP  VAK+I   C  C  +G    G
Sbjct  6359  FGSTGIFWVENIQEAQDEHENWHTSPKILAKRYGLPLTVAKQITQECPHCTKQGSGPAGC  6538

Query  881   VDCSPGIWQLDCTHLEGKVILVAVHVASGYIEAEVIPAETGQETAYFLLKLAGRWPVKTI  940
             V  SP  WQ DCTHLE +VI+  V   SGYI A ++  E    T+  +L+    +  K++
Sbjct  6539  VMRSPNHWQADCTHLENRVIMTFVESNSGYIHATLLSKENALCTSLAILEWVRLFSPKSL  6718

Query  941   HTDNGSNFTGATVRAACWWAGIKQEFGIPYNPQSQGVVESMNKELKKIIGQVRDQAEHLK  1000
             HTDNG+NF   +V     +  +    GIPY+P+SQG+VE  N+ LK+ I   R+  + L+
Sbjct  6719  HTDNGTNFVAESVANLLKFLKVTHTTGIPYHPESQGIVERANRTLKEKIKSHRENTQTLE  6898

Query  1001  TAVQMAVFIHNFKRKGGIGGYSAGERIVDIIATDIQTKELQKQITKIQNFRVYYRDSRNP  1060
              A+Q+A+   N K +  +GG +  E  +   A  I  + L +Q    + F  +Y+
Sbjct  6899  AALQLALITCN-KGRESMGGQTPWEVFITNQAQTIHEELLLQQAQSSKKF-CFYKIPGEH  7072

Query  1061  LWKGPAKLLWKGEGAVVIQDNSD-IKVVPRRKAKII  1095
              WKGP ++LWKG+GAVV+ D    I  VP  + K++
Sbjct  7073  NWKGPTRVLWKGDGAVVVNDEKKGIIAVPLTRTKLL  7180
```

>Escherichia coli strain 8374wH11 NODE_51_length_464_cov_1.98813, whole genome shotgun sequence
Sequence ID: NZ_WSJI01000051.1 Length: 464
Range 1: 1 to 462

Score:310 bits(793), Expect:2e-94,
Method:Compositional matrix adjust.,
Identities:143/154(93%), Positives:150/154(97%), Gaps:0/154(0%)

```
Query  250  KKSVTVLDVGDAYFSVPLDEDFRKYTAFTIPSINNETPGIRYQYNVLPQGWKGSPAIFQS  309
            KK+VTVLDVGDAYFSVPLDE FRKYTAFTIPS+NNETPGIRYQYNVLPQGWKGSPAIFQ+
Sbjct  462  KKAVTVLDVGDAYFSVPLDESFRKYTAFTIPSVNNETPGIRYQYNVLPQGWKGSPAIFQA  283

Query  310  SMTKILEPFRKQNPDIVIYQYMDDLYVGSDLEIGQHRTKIEELRQHLLRWGLTTPDKKHQ  369
            SMTKILEPFR +NPDIVIYQYM+DLYVGSDLEIGQHR KIEELR+HLL+WGLTTPDKKHQ
Sbjct  282  SMTKILEPFRLKNPDIVIYQYMEDLYVGSDLEIGQHRAKIEELREHLLKWGLTTPDKKHQ  103

Query  370  KEPPFLWMGYELHPDKWTVQPIVLPEKDSWTVND  403
            KEPPFLWMGYELHPDKWTVQPI LPEKDSWTVND
Sbjct  102  KEPPFLWMGYELHPDKWTVQPIQLPEKDSWTVND  1
```

>Escherichia coli strain 8374wB4 NODE_86_length_485_cov_1.49162, whole genome shotgun sequence
Sequence ID: NZ_WSPX01000086.1 Length: 485
Range 1: 12 to 485

Score:309 bits(792), Expect:3e-94,
Method:Compositional matrix adjust.,
Identities:147/158(93%), Positives:153/158(96%), Gaps:0/158(0%)

```
Query  126  GPTPVNIIGRNLLTQIGCTLNFPISPIETVPVKLKPGMDGPKVKQWPLTEEKIKALVEIC  185
            GP P+ IIGRN+LTQ+GCTLNFPISPIETVPVKLKPGMDGP+VKQWPLTEEKIKAL  IC
Sbjct  12   GPKPIEIIGRNMLTQLGCTLNFPISPIETVPVKLKPGMDGPRVKQWPLTEEKIKALQAIC  191

Query  186  TEMEKEGKISKIGPENPYNTPVFAIKKKDSTKWRKLVDFRELNKRTQDFWEVQLGIPHPA  245
             EMEKEGKI+KIGPENPYNTPVFAIKKKDSTKWRKLVDFRELNKRTQDFWEVQLGIPHPA
Sbjct  192  EEMEKEGKITKIGPENPYNTPVFAIKKKDSTKWRKLVDFRELNKRTQDFWEVQLGIPHPA  371

Query  246  GLKKKKSVTVLDVGDAYFSVPLDEDFRKYTAFTIPSIN  283
            GLKKKKSVTVLDVGDAYFSVPLDE+FRKYTAFTIPSIN
Sbjct  372  GLKKKKSVTVLDVGDAYFSVPLDENFRKYTAFTIPSIN  485
```

>Escherichia coli strain 8374wA11 NODE_44_length_523_cov_1.7197, whole genome shotgun sequence
Sequence ID: NZ_WSPR01000043.1 Length: 523
Range 1: 4 to 522

Score:308 bits(790), Expect:9e-94,
Method:Compositional matrix adjust.,
Identities:143/173(83%), Positives:164/173(94%), Gaps:0/173(0%)

```
Query  84   MSLPGRWKPKMIGGIGGFIKVRQYDQILIEICGHKAIGTVLVGPTPVNIIGRNLLTQIGC  143
            ++LPG+W PK+IGG+GG ++V+QYD ILIE  G KAIG+VLVGPTP NIIGRN++TQIGC
Sbjct  4    INLPGKWTPKLIGGVGGLVRVKQYDNILIEFEGRKAIGSVLVGPTPSNIIGRNMMTQIGC  183

Query  144  TLNFPISPIETVPVKLKPGMDGPKVKQWPLTEEKIKALVEICTEMEKEGKISKIGPENPY  203
```

```
           TLNFPISPIETVPVKLKPGMDGPK+KQWPLT+EKI+AL EICTE+E++GKISKIGPENPY
Sbjct  184  TLNFPISPIETVPVKLKPGMDGPKIKQWPLTKEKIEALTEICTELEQDGKISKIGPENPY  363

Query  204  NTPVFAIKKKDSTKWRKLVDFRELNKRTQDFWEVQLGIPHPAGLKKKKSVTVL  256
            NTP+FAIKKK+ST WRKL+DFRELNKRTQDFWEVQLGIPHPAGL+KKKSVTVL
Sbjct  364  NTPIFAIKKKNSTGWRKLMDFRELNKRTQDFWEVQLGIPHPAGLEKKKSVTVL  522



>Escherichia coli strain TzEc080 NODE_44_length_523_cov_1.7197, whole genome shotgun sequence
Sequence ID: NZ_WSFZ01000043.1 Length: 523
Range 1: 4 to 522

Score:308 bits(790), Expect:9e-94,
Method:Compositional matrix adjust.,
Identities:143/173(83%), Positives:164/173(94%), Gaps:0/173(0%)

Query  84   MSLPGRWKPKMIGGIGGFIKVRQYDQILIEICGHKAIGTVLVGPTPVNIIGRNLLTQIGC  143
            ++LPG+W PK+IGG+GG ++V+QYD ILIE  G KAIG+VLVGPTP NIIGRN++TQIGC
Sbjct  4    INLPGKWTPKLIGGVGGLVRVKQYDNILIEFEGRKAIGSVLVGPTPSNIIGRNMMTQIGC  183

Query  144  TLNFPISPIETVPVKLKPGMDGPKVKQWPLTEEKIKALVEICTEMEKEGKISKIGPENPY  203
            TLNFPISPIETVPVKLKPGMDGPK+KQWPLT+EKI+AL EICTE+E++GKISKIGPENPY
Sbjct  184  TLNFPISPIETVPVKLKPGMDGPKIKQWPLTKEKIEALTEICTELEQDGKISKIGPENPY  363

Query  204  NTPVFAIKKKDSTKWRKLVDFRELNKRTQDFWEVQLGIPHPAGLKKKKSVTVL  256
            NTP+FAIKKK+ST WRKL+DFRELNKRTQDFWEVQLGIPHPAGL+KKKSVTVL
Sbjct  364  NTPIFAIKKKNSTGWRKLMDFRELNKRTQDFWEVQLGIPHPAGLEKKKSVTVL  522



>Enterococcus faecium strain E21_6 NODE_88_length_518_cov_1.29923, whole genome shotgun sequence
Sequence ID: NZ_JAANZJ010000088.1 Length: 518
Range 1: 2 to 514

Score:305 bits(782), Expect:1e-92,
Method:Compositional matrix adjust.,
Identities:146/171(85%), Positives:163/171(95%), Gaps:2/171(1%)

Query  49   PQVTLWQRPLVTIKIGGQLKEALLDTGADDTVLEEMSL--PGRWKPKMIGGIGGFIKVRQ  106
            PQ+TLWQRPLV+IKIGGQ+KEALLDTGADDTVLEE+ L  PG+WKPKMIGGIGGFIKV+Q
Sbjct  2    PQITLWQRPLVSIKIGGQIKEALLDTGADDTVLEELGLELPGKWKPKMIGGIGGFIKVKQ  181

Query  107  YDQILIEICGHKAIGTVLVGPTPVNIIGRNLLTQIGCTLNFPISPIETVPVKLKPGMDGP  166
            Y++I IEICG +AIGTVLVGPTPVNIIGRN+LTQ+GCTLNFPIS IETVPVKLKPGMDGP
Sbjct  182  YEEIPIEICGKRAIGTVLVGPTPVNIIGRNMLTQLGCTLNFPISTIETVPVKLKPGMDGP  361

Query  167  KVKQWPLTEEKIKALVEICTEMEKEGKISKIGPENPYNTPVFAIKKKDSTK  217
            +VKQWPLTEEKIKAL EIC EME+EGKI+KIGPENPYNTP+FAIKKK+ST+
Sbjct  362  RVKQWPLTEEKIKALTEICAEMEREGKITKIGPENPYNTPIFAIKKKNSTE  514



>Escherichia coli strain P010 NODE_181_length_556_cov_0.773893_ID_361, whole genome shotgun sequence
Sequence ID: NZ_VKOQ01000154.1 Length: 556
Range 1: 2 to 556

Score:301 bits(770), Expect:9e-91,
Method:Compositional matrix adjust.,
Identities:163/185(88%), Positives:173/185(93%), Gaps:0/185(0%)

Query  388  VQPIVLPEKDSWTVNDIQKLVGKLNWASQIYPGIKVRQLCKLLRGTKALTEVIPLTeeae  447
            VQPI LPEKD WTVNDIQKLVGKLNWASQIYP IKV+QLCKLLRGTKALT+++PLTEEAE
Sbjct  556  VQPIQLPEKDIWTVNDIQKLVGKLNWASQIYPEIKVKQLCKLLRGTKALTDIVPLTEEAE  377

Query  448  lelaeNREILKEPVHGVYYDPSKDLIAEIQKQGQGQWTYQIYQEPFKNLKTGKYARMRGA  507
            L LAE REILKEPVHG YYDPS+DLIAEIQKQG  +WTYQIYQEPFKNLKTGKYA+MR A
Sbjct  376  LALAEIREILKEPVHGAYYDPSEDLIAEIQKQGNDKWTYQIYQEPFKNLKTGKYAKMRTA  197

Query  508  HTNDVKQLTEAVQKITTESIVIWGKTPKFKLPIQKetwetwwteYWQATWIPEWEFVNTP  567
            HTNDVKQLTEAVQKI  ESIVIWGK PKF+LPIQKETWETWWT+YWQATWIP+WEFVNTP
Sbjct  196  HTNDVKQLTEAVQKIARESIVIWGKIPKFRLPIQKETWETWWTDYWQATWIPDWEFVNTP  17

Query  568  PLVKL  572
            PLVKL
Sbjct  16   PLVKL  2



>Methylobacterium radiotolerans strain MAMP 4754 MP-M-radiotolerans_contig_206, whole genome shotgun sequence
Sequence ID: NZ_NKQS01000204.1 Length: 483
Range 1: 3 to 482

Score:298 bits(762), Expect:6e-90,
Method:Compositional matrix adjust.,
Identities:141/160(88%), Positives:155/160(96%), Gaps:0/160(0%)

Query  51   VTLWQRPLVTIKIGGQLKEALLDTGADDTVLEEMSLPGRWKPKMIGGIGGFIKVRQYDQI  110
            +TLWQRPLV+IK+ GQ+KEALLDTGADDTVLE+M+LPG+WKP+MIGGIGGFIKVRQY+QI
Sbjct  482  ITLWQRPLVSIKVEGQIKEALLDTGADDTVLEDMNLPGKWKPRMIGGIGGFIKVRQYEQI  303

Query  111  LIEICGHKAIGTVLVGPTPVNIIGRNLLTQIGCTLNFPISPIETVPVKLKPGMDGPKVKQ  170
             IEICG KAIGTVLVGPTP+NIIGRN+LTQ+GCTLNFPISPI+TVPVKLKPGMDGPKVKQ
Sbjct  302  NIEICGKKAIGTVLVGPTPINIIGRNMLTQLGCTLNFPISPIDTVPVKLKPGMDGPKVKQ  123

Query  171  WPLTEEKIKALVEICTEMEKEGKISKIGPENPYNTPVFAI  210
            WPLTEEKIKAL EIC +MEKEGKISKIGPENPYNTPVFAI
Sbjct  122  WPLTEEKIKALTEICEDMEKEGKISKIGPENPYNTPVFAI  3



>Klebsiella pneumoniae strain EuSCAPE_MT006, whole genome shotgun sequence
Sequence ID: NZ_UKIU02000071.1 Length: 455
Range 1: 3 to 455

Score:296 bits(758), Expect:1e-89,
Method:Compositional matrix adjust.,
Identities:142/151(94%), Positives:146/151(96%), Gaps:0/151(0%)

Query  98   IGGFIKVRQYDQILIEICGHKAIGTVLVGPTPVNIIGRNLLTQIGCTLNFPISPIETVPV  157
            IGGFIKVRQYDQI IEICG KAIGTVL+GPTPVNIIGRN+LTQ+GCTLNFPISPIETVPV
Sbjct  455  IGGFIKVRQYDQIPIEICGKKAIGTVLIGPTPVNIIGRNMLTQLGCTLNFPISPIETVPV  276

Query  158  KLKPGMDGPKVKQWPLTEEKIKALVEICTEMEKEGKISKIGPENPYNTPVFAIKKKDSTK  217
            KLKPGMDGPKVKQWPLTEEKIKAL  IC EMEKEGKI+KIGPENPYNTPVFAIKKKDSTK
Sbjct  275  KLKPGMDGPKVKQWPLTEEKIKALTAICEEMEKEGKITKIGPENPYNTPVFAIKKKDSTK  96

Query  218  WRKLVDFRELNKRTQDFWEVQLGIPHPAGLK  248
            WRKLVDFRELNKRTQDFWEVQLGIPHPAGLK
```

```
Sbjct  95   WRKLVDFRELNKRTQDFWEVQLGIPHPAGLK  3



>Cronobacter sakazakii strain cro4215W contig98, whole genome shotgun sequence
Sequence ID: NZ_NRKF01000098.1 Length: 429
Range 1: 3 to 428

Score:294 bits(753), Expect:7e-89,
Method:Compositional matrix adjust.,
Identities:142/142(100%), Positives:142/142(100%), Gaps:0/142(0%)

Query  139  TQIGCTLNFPISPIETVPVKLKPGMDGPKVKQWPLTEEKIKALVEICTEMEKEGKISKIG  198
            TQIGCTLNFPISPIETVPVKLKPGMDGPKVKQWPLTEEKIKALVEICTEMEKEGKISKIG
Sbjct  428  TQIGCTLNFPISPIETVPVKLKPGMDGPKVKQWPLTEEKIKALVEICTEMEKEGKISKIG  249

Query  199  PENPYNTPVFAIKKKDSTKWRKLVDFRELNKRTQDFWEVQLGIPHPAGLKKKKSVTVLDV  258
            PENPYNTPVFAIKKKDSTKWRKLVDFRELNKRTQDFWEVQLGIPHPAGLKKKKSVTVLDV
Sbjct  248  PENPYNTPVFAIKKKDSTKWRKLVDFRELNKRTQDFWEVQLGIPHPAGLKKKKSVTVLDV  69

Query  259  GDAYFSVPLDEDFRKYTAFTIP  280
            GDAYFSVPLDEDFRKYTAFTIP
Sbjct  68   GDAYFSVPLDEDFRKYTAFTIP  3



>Klebsiella pneumoniae strain EuSCAPE_IL016, whole genome shotgun sequence
Sequence ID: NZ_UKKA02000066.1 Length: 523
Range 1: 2 to 523

Score:295 bits(754), Expect:1e-88,
Method:Compositional matrix adjust.,
Identities:146/174(84%), Positives:159/174(91%), Gaps:0/174(0%)

Query  836   QDEHEKYHSNWRAMASDFNLPPVVAKEIVASCDKCQLKGEAMHGQVDCSPGIWQLDCTHL  895
             Q+EHEKYHSN RAMAS+FNLPP+VAKEIVAS DKCQLK EA+ GQVDCSP IWQLDCTHL
Sbjct  523   QEEHEKYHSN*RAMASEFNLPPIVAKEIVASYDKCQLKKEAIQGQVDCSPRIWQLDCTHL  344

Query  896   EGKVILVAVHVASGYIEAEVIPAETGQETAYFLLKLAGRWPVKTIHTDNGSNFTGATVRA  955
             E K+ILVAVHVASGY+EAEVIPA+TGQETAY++LKLAGRWPVK IHTDN SNFT A V+A
Sbjct  343   ERKIILVAVHVASGYLEAEVIPAKTGQETAYYILKLAGRWPVKVIHTDNSSNFTSAAVKA  164

Query  956   ACWWAGIKQEFGIPYNPQSQGVVESMNKELKKIIGQVRDQAEHLKTAVQMAVFI  1009
             AC  AGI+QEF IPYNPQSQ VVESMNKELKKII QV++QAEHLKTAVQMAVFI
Sbjct  163   AC**AGIQQEFRIPYNPQSQRVVESMNKELKKIIRQVKEQAEHLKTAVQMAVFI  2



>Cronobacter sakazakii strain cro4114A1 contig119, whole genome shotgun sequence
Sequence ID: NZ_NRKJ01000119.1 Length: 429
Range 1: 3 to 428

Score:293 bits(749), Expect:3e-88,
Method:Compositional matrix adjust.,
Identities:141/142(99%), Positives:142/142(100%), Gaps:0/142(0%)

Query  139  TQIGCTLNFPISPIETVPVKLKPGMDGPKVKQWPLTEEKIKALVEICTEMEKEGKISKIG  198
            TQIGCTLNFPISPIETVPVKLKPGMDGPKVKQWPLTEEKIKALVEICTEMEKEGKISKIG
Sbjct  428  TQIGCTLNFPISPIETVPVKLKPGMDGPKVKQWPLTEEKIKALVEICTEMEKEGKISKIG  249

Query  199  PENPYNTPVFAIKKKDSTKWRKLVDFRELNKRTQDFWEVQLGIPHPAGLKKKKSVTVLDV  258
            PENPYNTPVFAIKKKDSTKWRKLVDFRELNKRTQDFWEVQLGIPHPAGLKKKKSVTVLDV
Sbjct  248  PENPYNTPVFAIKKKDSTKWRKLVDFRELNKRTQDFWEVQLGIPHPAGLKKKKSVTVLDV  69

Query  259  GDAYFSVPLDEDFRKYTAFTIP  280
            GDAYFSVPLDE+FRKYTAFTIP
Sbjct  68   GDAYFSVPLDEEFRKYTAFTIP  3



>Escherichia coli strain 8374wB11 NODE_49_length_463_cov_1.77083, whole genome shotgun sequence
Sequence ID: NZ_WSJO01000049.1 Length: 463
Range 1: 1 to 459

Score:293 bits(749), Expect:4e-88,
Method:Compositional matrix adjust.,
Identities:136/153(89%), Positives:149/153(97%), Gaps:0/153(0%)

Query  129  PVNIIGRNLLTQIGCTLNFPISPIETVPVKLKPGMDGPKVKQWPLTEEKIKALVEICTEM  188
            P NIIGRN++TQIGCTLNFPISPIETVPVKLKPGMDGPK+KQWPLT+EKI+AL EICTE+
Sbjct  1    PCNIIGRNMMTQIGCTLNFPISPIETVPVKLKPGMDGPKIKQWPLTKEKIEALTEICTEL  180

Query  189  EKEGKISKIGPENPYNTPVFAIKKKDSTKWRKLVDFRELNKRTQDFWEVQLGIPHPAGLK  248
            E++GKISKIGPENPYNTP+FAIKKK+ST WRKL+DFRELNKRTQDFWEVQLGIPHPAGL+
Sbjct  181  EQDGKISKIGPENPYNTPIFAIKKKNSTGWRKLMDFRELNKRTQDFWEVQLGIPHPAGLE  360

Query  249  KKKSVTVLDVGDAYFSVPLDEDFRKYTAFTIPS  281
            KKKSVTVLDVGDAYFS+PLDEDFRK TAFTIPS
Sbjct  361  KKKSVTVLDVGDAYFSIPLDEDFRKDTAFTIPS  459



>Klebsiella pneumoniae strain EuSCAPE_IL021, whole genome shotgun sequence
Sequence ID: NZ_UKJX02000112.1 Length: 426
Range 1: 1 to 426

Score:291 bits(746), Expect:7e-88,
Method:Compositional matrix adjust.,
Identities:139/142(98%), Positives:139/142(97%), Gaps:0/142(0%)

Query  211  KKKDSTKWRKLVDFRELNKRTQDFWEVQLGIPHPAGLKKKKSVTVLDVGDAYFSVPLDED  270
            KKKDSTKWRKLVDFRELNKRTQDFWEVQLGIPHPAGLKKKKSVTVLDVGDAYFSVPLDED
Sbjct  1    KKKDSTKWRKLVDFRELNKRTQDFWEVQLGIPHPAGLKKKKSVTVLDVGDAYFSVPLDED  180

Query  271  FRKYTAFTIPSINNETPGIRYQYNVLPQGWKGSPAIFQSSMTKILEPFRKQNPDIVIYQY  330
            FRKYTAFTIPSINNETPGIRYQYNVLPQGWKGSPAIFQSSMTKILEPFR QNP IVIYQY
Sbjct  181  FRKYTAFTIPSINNETPGIRYQYNVLPQGWKGSPAIFQSSMTKILEPFRAQNPQIVIYQY  360

Query  331  MDDLYVGSDLEIGQHRTKIEEL  352
            MDDLYVGSDLEIGQHR KIEEL
Sbjct  361  MDDLYVGSDLEIGQHRAKIEEL  426



>Cronobacter sakazakii strain cro4114B1 contig94, whole genome shotgun sequence
Sequence ID: NZ_NRKI01000094.1 Length: 429
Range 1: 2 to 427
```

```
Score:291 bits(745), Expect:8e-88,
Method:Compositional matrix adjust.,
Identities:139/142(98%), Positives:142/142(100%), Gaps:0/142(0%)

Query  139  TQIGCTLNFPISPIETVPVKLKPGMDGPKVKQWPLTEEKIKALVEICTEMEKEGKISKIG  198
            TQIGCTLNFPISPIETVPVKLKPGMDGPKVKQWPLTEEKIKALVEICTEMEKEGKISKIG
Sbjct  2    TQIGCTLNFPISPIETVPVKLKPGMDGPKVKQWPLTEEKIKALVEICTEMEKEGKISKIG  181

Query  199  PENPYNTPVFAIKKKDSTKWRKLVDFRELNKRTQDFWEVQLGIPHPAGLKKKKSVTVLDV  258
            PENPYNTPVFAIKKKDSTKWRKLVDFRELNKRTQDFWEVQLGIPHPAGLK+KKSVTVLDV
Sbjct  182  PENPYNTPVFAIKKKDSTKWRKLVDFRELNKRTQDFWEVQLGIPHPAGLKRKKSVTVLDV  361

Query  259  GDAYFSVPLDEDFRKYTAFTIP  280
            GDAYFS+PLDE+FRKYTAFTIP
Sbjct  362  GDAYFSIPLDEEFRKYTAFTIP  427
```

```
>Escherichia coli strain 8374wA11 NODE_51_length_444_cov_2.55205, whole genome shotgun sequence
Sequence ID: NZ_WSPR01000049.1 Length: 444
Range 1: 1 to 444

Score:290 bits(742), Expect:2e-87,
Method:Compositional matrix adjust.,
Identities:135/148(91%), Positives:139/148(93%), Gaps:0/148(0%)

Query  259  GDAYFSVPLDEDFRKYTAFTIPSINNETPGIRYQYNVLPQGWKGSPAIFQSSMTKILEPF  318
            GDAYFSVPLDE FRKYTAFTIPS NNETPGIRYQYNVLPQGWKGSPAIFQ SMTKILEPF
Sbjct  444  GDAYFSVPLDEGFRKYTAFTIPSRNNETPGIRYQYNVLPQGWKGSPAIFQCSMTKILEPF  265

Query  319  RKQNPDIVIYQYMDDLYVGSDLEIGQHRTKIEELRQHLLRWGLTTPDKKHQKEPPFLWMG  378
            R +NP+I+I QYMDDLYVGSDLEIGQHR KIEELR HLL+WG TTPDKKHQKEPPFLWMG
Sbjct  264  RAKNPEIIICQYMDDLYVGSDLEIGQHRAKIEELRDHLLKWGFTTPDKKHQKEPPFLWMG  85

Query  379  YELHPDKWTVQPIVLPEKDSWTVNDIQK  406
            YELHPDKWTVQPI LPEKDSWTVNDIQK
Sbjct  84   YELHPDKWTVQPIQLPEKDSWTVNDIQK  1
```

```
>Escherichia coli strain TzEc080 NODE_51_length_444_cov_2.55205, whole genome shotgun sequence
Sequence ID: NZ_WSFZ01000049.1 Length: 444
Range 1: 1 to 444

Score:290 bits(742), Expect:2e-87,
Method:Compositional matrix adjust.,
Identities:135/148(91%), Positives:139/148(93%), Gaps:0/148(0%)

Query  259  GDAYFSVPLDEDFRKYTAFTIPSINNETPGIRYQYNVLPQGWKGSPAIFQSSMTKILEPF  318
            GDAYFSVPLDE FRKYTAFTIPS NNETPGIRYQYNVLPQGWKGSPAIFQ SMTKILEPF
Sbjct  444  GDAYFSVPLDEGFRKYTAFTIPSRNNETPGIRYQYNVLPQGWKGSPAIFQCSMTKILEPF  265

Query  319  RKQNPDIVIYQYMDDLYVGSDLEIGQHRTKIEELRQHLLRWGLTTPDKKHQKEPPFLWMG  378
            R +NP+I+I QYMDDLYVGSDLEIGQHR KIEELR HLL+WG TTPDKKHQKEPPFLWMG
Sbjct  264  RAKNPEIIICQYMDDLYVGSDLEIGQHRAKIEELRDHLLKWGFTTPDKKHQKEPPFLWMG  85

Query  379  YELHPDKWTVQPIVLPEKDSWTVNDIQK  406
            YELHPDKWTVQPI LPEKDSWTVNDIQK
Sbjct  84   YELHPDKWTVQPIQLPEKDSWTVNDIQK  1
```

```
>Cronobacter malonaticus strain cro3263A1 contig63, whole genome shotgun sequence
Sequence ID: NZ_NRMM01000063.1 Length: 429
Range 1: 2 to 427

Score:286 bits(732), Expect:7e-86,
Method:Compositional matrix adjust.,
Identities:136/142(96%), Positives:141/142(99%), Gaps:0/142(0%)

Query  139  TQIGCTLNFPISPIETVPVKLKPGMDGPKVKQWPLTEEKIKALVEICTEMEKEGKISKIG  198
            TQIGCTLNFPISPI+TVPVKLKPGMDGPKVKQWPLTEEKIKAL+EICTEMEKEGKISKIG
Sbjct  2    TQIGCTLNFPISPIDTVPVKLKPGMDGPKVKQWPLTEEKIKALIEICTEMEKEGKISKIG  181

Query  199  PENPYNTPVFAIKKKDSTKWRKLVDFRELNKRTQDFWEVQLGIPHPAGLKKKKSVTVLDV  258
            PENPYNTPVFAIKKKDSTKWRKLVDFRELNKRTQDF EVQLGIPHPAGLK+KKSVTVLDV
Sbjct  182  PENPYNTPVFAIKKKDSTKWRKLVDFRELNKRTQDFSEVQLGIPHPAGLKRKKSVTVLDV  361

Query  259  GDAYFSVPLDEDFRKYTAFTIP  280
            GDAYFS+PLDE+FRKYTAFTIP
Sbjct  362  GDAYFSIPLDEEFRKYTAFTIP  427
```

```
>Cronobacter dublinensis strain cro981C2 contig293, whole genome shotgun sequence
Sequence ID: NZ_NROB01000293.1 Length: 429
Range 1: 2 to 427

Score:285 bits(730), Expect:1e-85,
Method:Compositional matrix adjust.,
Identities:137/142(96%), Positives:139/142(97%), Gaps:0/142(0%)

Query  139  TQIGCTLNFPISPIETVPVKLKPGMDGPKVKQWPLTEEKIKALVEICTEMEKEGKISKIG  198
            TQIGCTLNFPISPI+TVPVKLKPGMDGPKVKQWPLTEEKIKAL EIC EMEKEGKISKIG
Sbjct  2    TQIGCTLNFPISPIDTVPVKLKPGMDGPKVKQWPLTEEKIKALTEICKEMEKEGKISKIG  181

Query  199  PENPYNTPVFAIKKKDSTKWRKLVDFRELNKRTQDFWEVQLGIPHPAGLKKKKSVTVLDV  258
            PENPYNTPVFAIKKKDS KWRKLVDFRELNKRTQDFWEVQLGIPHPAGLKKKKSVTVLDV
Sbjct  182  PENPYNTPVFAIKKKDSNKWRKLVDFRELNKRTQDFWEVQLGIPHPAGLKKKKSVTVLDV  361

Query  259  GDAYFSVPLDEDFRKYTAFTIP  280
            GDAYFSVPLDE+FRKYTAFTIP
Sbjct  362  GDAYFSVPLDENFRKYTAFTIP  427
```

```
>Klebsiella pneumoniae strain EuSCAPE_MT015, whole genome shotgun sequence
Sequence ID: NZ_UKIW02000099.1 Length: 435
Range 1: 3 to 434

Score:285 bits(730), Expect:1e-85,
Method:Compositional matrix adjust.,
Identities:137/144(95%), Positives:139/144(96%), Gaps:0/144(0%)

Query  103  KVRQYDQILIEICGHKAIGTVLVGPTPVNIIGRNLLTQIGCTLNFPISPIETVPVKLKPG  162
            KVRQYDQI IEICG KAIGTVLVGPTPVNIIGRNLLTQ+GCTLNFPISPIETVPVKLKPG
Sbjct  434  KVRQYDQITIEICGKKAIGTVLVGPTPVNIIGRNLLTQLGCTLNFPISPIETVPVKLKPG  255

Query  163  MDGPKVKQWPLTEEKIKALVEICTEMEKEGKISKIGPENPYNTPVFAIKKKDSTKWRKLV  222
```

```
             MDGPKVKQWPLTEEKIKAL  IC EMEKEGKI+KIGPENPYNTPVFAIKKKDSTKWRKLV
Sbjct  254  MDGPKVKQWPLTEEKIKALTAICEEMEKEGKITKIGPENPYNTPVFAIKKKDSTKWRKLV  75

Query  223  DFRELNKRTQDFWEVQLGIPHPAG  246
            DFRELNKRTQDFWEVQLGIPHPAG
Sbjct  74   DFRELNKRTQDFWEVQLGIPHPAG  3
```

>Burkholderia pseudomallei strain 2604STDY5518971 ERS240105SCcontig000116, whole genome shotgun sequence
Sequence ID: NZ_UVFZ02000116.1 Length: 427
Range 1: 1 to 426

Score:285 bits(729), Expect:2e-85,
Method:Compositional matrix adjust.,
Identities:135/142(95%), Positives:139/142(97%), Gaps:0/142(0%)

```
Query  173  LTEEKIKALVEICTEMEKEGKISKIGPENPYNTPVFAIKKKDSTKWRKLVDFRELNKRTQ  232
            LTEEKIKAL EIC +MEKEGKISKIGPENPYNTP+FAIKKKDSTKWRKLVDFRELNKRTQ
Sbjct  426  LTEEKIKALTEICNDMEKEGKISKIGPENPYNTPIFAIKKKDSTKWRKLVDFRELNKRTQ  247

Query  233  DFWEVQLGIPHPAGLKKKKSVTVLDVGDAYFSVPLDEDFRKYTAFTIPSINNETPGIRYQ  292
            DFWEVQLGIPHPAGLKKKKSVTVLDVGDAYFSVPLDE+FRKYTAFTIPS NNETPG+RYQ
Sbjct  246  DFWEVQLGIPHPAGLKKKKSVTVLDVGDAYFSVPLDENFRKYTAFTIPSRNNETPGVRYQ  67

Query  293  YNVLPQGWKGSPAIFQSSMTKI  314
            YNVLPQGWKGSPAIFQSSMTKI
Sbjct  66   YNVLPQGWKGSPAIFQSSMTKI  1
```

>Cronobacter sakazakii strain cro3040W contig78, whole genome shotgun sequence
Sequence ID: NZ_NRJO01000078.1 Length: 429
Range 1: 2 to 427

Score:285 bits(729), Expect:2e-85,
Method:Compositional matrix adjust.,
Identities:137/142(96%), Positives:138/142(97%), Gaps:0/142(0%)

```
Query  139  TQIGCTLNFPISPIETVPVKLKPGMDGPKVKQWPLTEEKIKALVEICTEMEKEGKISKIG  198
            TQIGCTLNFPISPI+TVPV LKPGMDGPKVKQWPLTEEKIKAL EIC EMEKEGKISKIG
Sbjct  2    TQIGCTLNFPISPIDTVPVTLKPGMDGPKVKQWPLTEEKIKALTEICREMEKEGKISKIG  181

Query  199  PENPYNTPVFAIKKKDSTKWRKLVDFRELNKRTQDFWEVQLGIPHPAGLKKKKSVTVLDV  258
            PENPYNTPVFAIKKKDSTKWRKLVDFRELNKRTQDFWEVQLGIPHPAGLKKKKSVTVLDV
Sbjct  182  PENPYNTPVFAIKKKDSTKWRKLVDFRELNKRTQDFWEVQLGIPHPAGLKKKKSVTVLDV  361

Query  259  GDAYFSVPLDEDFRKYTAFTIP  280
            GDAYFSVPLDE FRKYTAFTIP
Sbjct  362  GDAYFSVPLDESFRKYTAFTIP  427
```

>Cronobacter sakazakii strain cro4215W contig97, whole genome shotgun sequence
Sequence ID: NZ_NRKF01000097.1 Length: 429
Range 1: 2 to 427

Score:284 bits(726), Expect:5e-85,
Method:Compositional matrix adjust.,
Identities:135/142(95%), Positives:138/142(97%), Gaps:0/142(0%)

```
Query  139  TQIGCTLNFPISPIETVPVKLKPGMDGPKVKQWPLTEEKIKALVEICTEMEKEGKISKIG  198
            TQIGCTLNFPISPI+T+PV LKPGMDGPKVKQWPLTEEKIKAL EIC EMEKEGKISKIG
Sbjct  2    TQIGCTLNFPISPIDTIPVTLKPGMDGPKVKQWPLTEEKIKALTEICKEMEKEGKISKIG  181

Query  199  PENPYNTPVFAIKKKDSTKWRKLVDFRELNKRTQDFWEVQLGIPHPAGLKKKKSVTVLDV  258
            PENPYNTPVFAIKKKDSTKWRKLVDFRELNKRTQDFWEVQLGIPHPAGLKKKKSVTVLD+
Sbjct  182  PENPYNTPVFAIKKKDSTKWRKLVDFRELNKRTQDFWEVQLGIPHPAGLKKKKSVTVLDI  361

Query  259  GDAYFSVPLDEDFRKYTAFTIP  280
            GDAYFSVPLDE FRKYTAFTIP
Sbjct  362  GDAYFSVPLDESFRKYTAFTIP  427
```

>Cronobacter sakazakii strain cro4114B1 contig95, whole genome shotgun sequence
Sequence ID: NZ_NRKI01000095.1 Length: 429
Range 1: 2 to 427

Score:284 bits(726), Expect:5e-85,
Method:Compositional matrix adjust.,
Identities:135/142(95%), Positives:138/142(97%), Gaps:0/142(0%)

```
Query  139  TQIGCTLNFPISPIETVPVKLKPGMDGPKVKQWPLTEEKIKALVEICTEMEKEGKISKIG  198
            TQIGCTLNFPISPI+T+PV LKPGMDGPKVKQWPLTEEKIKAL EIC EMEKEGKISKIG
Sbjct  2    TQIGCTLNFPISPIDTIPVTLKPGMDGPKVKQWPLTEEKIKALTEICKEMEKEGKISKIG  181

Query  199  PENPYNTPVFAIKKKDSTKWRKLVDFRELNKRTQDFWEVQLGIPHPAGLKKKKSVTVLDV  258
            PENPYNTPVFAIKKKDSTKWRKLVDFRELNKRTQDFWEVQLGIPHPAGLKKKKSVTVLD+
Sbjct  182  PENPYNTPVFAIKKKDSTKWRKLVDFRELNKRTQDFWEVQLGIPHPAGLKKKKSVTVLDI  361

Query  259  GDAYFSVPLDEDFRKYTAFTIP  280
            GDAYFSVPLDE FRKYTAFTIP
Sbjct  362  GDAYFSVPLDESFRKYTAFTIP  427
```

>Cronobacter sakazakii strain cro4114A1 contig120, whole genome shotgun sequence
Sequence ID: NZ_NRKJ01000120.1 Length: 429
Range 1: 2 to 427

Score:284 bits(726), Expect:5e-85,
Method:Compositional matrix adjust.,
Identities:135/142(95%), Positives:138/142(97%), Gaps:0/142(0%)

```
Query  139  TQIGCTLNFPISPIETVPVKLKPGMDGPKVKQWPLTEEKIKALVEICTEMEKEGKISKIG  198
            TQIGCTLNFPISPI+T+PV LKPGMDGPKVKQWPLTEEKIKAL EIC EMEKEGKISKIG
Sbjct  2    TQIGCTLNFPISPIDTIPVTLKPGMDGPKVKQWPLTEEKIKALTEICKEMEKEGKISKIG  181

Query  199  PENPYNTPVFAIKKKDSTKWRKLVDFRELNKRTQDFWEVQLGIPHPAGLKKKKSVTVLDV  258
            PENPYNTPVFAIKKKDSTKWRKLVDFRELNKRTQDFWEVQLGIPHPAGLKKKKSVTVLD+
Sbjct  182  PENPYNTPVFAIKKKDSTKWRKLVDFRELNKRTQDFWEVQLGIPHPAGLKKKKSVTVLDI  361

Query  259  GDAYFSVPLDEDFRKYTAFTIP  280
            GDAYFSVPLDE FRKYTAFTIP
Sbjct  362  GDAYFSVPLDESFRKYTAFTIP  427
```

```
>Cronobacter sakazakii strain cro4191B1-2 contig80, whole genome shotgun sequence
Sequence ID: NZ_NRKG01000080.1 Length: 429
Range 1: 2 to 427

Score:284 bits(726), Expect:5e-85,
Method:Compositional matrix adjust.,
Identities:136/142(96%), Positives:138/142(97%), Gaps:0/142(0%)

Query  139  TQIGCTLNFPISPIETVPVKLKPGMDGPKVKQWPLTEEKIKALVEICTEMEKEGKISKIG  198
            TQIGCTLNFPISPI+TVPV LKPGMDGPKVKQWPLTEEKIKAL EIC EMEKEGKI+KIG
Sbjct  2    TQIGCTLNFPISPIDTVPVTLKPGMDGPKVKQWPLTEEKIKALTEICREMEKEGKITKIG  181

Query  199  PENPYNTPVFAIKKKDSTKWRKLVDFRELNKRTQDFWEVQLGIPHPAGLKKKKSVTVLDV  258
            PENPYNTPVFAIKKKDSTKWRKLVDFRELNKRTQDFWEVQLGIPHPAGLKKKKSVTVLDV
Sbjct  182  PENPYNTPVFAIKKKDSTKWRKLVDFRELNKRTQDFWEVQLGIPHPAGLKKKKSVTVLDV  361

Query  259  GDAYFSVPLDEDFRKYTAFTIP  280
            GDAYFSVPLDE FRKYTAFTIP
Sbjct  362  GDAYFSVPLDESFRKYTAFTIP  427



>Escherichia coli strain 8374wB11 NODE_47_length_476_cov_2.59885, whole genome shotgun sequence
Sequence ID: NZ_WSJO01000047.1 Length: 476
Range 1: 1 to 474

Score:283 bits(725), Expect:1e-84,
Method:Compositional matrix adjust.,
Identities:141/158(89%), Positives:153/158(96%), Gaps:0/158(0%)

Query  314  ILEPFRKQNPDIVIYQYMDDLYVGSDLEIGQHRTKIEELRQHLLRWGLTTPDKKHQKEPP  373
            IL+PFR +NPDIVIYQY+DDLYVGSDLEIGQHR KIEELR+HLL+WGLTTPDKKHQKEPP
Sbjct  474  ILQPFRTRNPDIVIYQYVDDLYVGSDLEIGQHRAKIEELREHLLKWGLTTPDKKHQKEPP  295

Query  374  FLWMGYELHPDKWTVQPIVLPEKDSWTVNDIQKLVGKLNWASQIYPGIKVRQLCKLLRGT  433
            FLWMGYELHPDKWTVQPI LP+K+SWTVNDIQKLVGKLNWASQIYPGI+V+ LCKLLRG
Sbjct  294  FLWMGYELHPDKWTVQPIQLPDKESWTVNDIQKLVGKLNWASQIYPGIRVKHLCKLLRGA  115

Query  434  KALTEVIPLTeeaelelaeNREILKEPVHGVYYDPSKD  471
            KALT++IPLTEEAELELAENREILKEPVHGVYYDPS+D
Sbjct  114  KALTDIIPLTEEAELELAENREILKEPVHGVYYDPSRD  1



>Burkholderia pseudomallei strain 2604STDY5518971 ERS240105SCcontig000113, whole genome shotgun sequence
Sequence ID: NZ_UVFZ02000113.1 Length: 457
Range 1: 2 to 457

Score:281 bits(719), Expect:6e-84,
Method:Compositional matrix adjust.,
Identities:141/152(93%), Positives:146/152(96%), Gaps:0/152(0%)

Query  335  YVGSDLEIGQHRTKIEELRQHLLRWGLTTPDKKHQKEPPFLWMGYELHPDKWTVQPIVLP  394
            YVGSDLEI QHRTKIEELR HLL+WG TTPDKKHQKEPPFLWMGYELHPDKWTVQPI LP
Sbjct  2    YVGSDLEIEQHRTKIEELRHHLLKWGFTTPDKKHQKEPPFLWMGYELHPDKWTVQPIKLP  181

Query  395  EKDSWTVNDIQKLVGKLNWASQIYPGIKVRQLCKLLRGTKALTEVIPLTeeaelelaeNR  454
            EKDSWTVNDIQKLVGKLNWASQIYPGIKV+QLCKL+RGTKALTEV+PLT+EAELELAENR
Sbjct  182  EKDSWTVNDIQKLVGKLNWASQIYPGIKVKQLCKLIRGTKALTEVVPLTKEAELELAENR  361

Query  455  EILKEPVHGVYYDPSKDLIAEIQKQGQGQWTY  486
            EILKEPVHG YYDPSKDLIAEIQKQG GQWTY
Sbjct  362  EILKEPVHGTYYDPSKDLIAEIQKQGNGQWTY  457



>Cronobacter malonaticus strain cro4164A3 contig124, whole genome shotgun sequence
Sequence ID: NZ_NRNF01000124.1 Length: 429
Range 1: 2 to 427

Score:280 bits(717), Expect:9e-84,
Method:Compositional matrix adjust.,
Identities:134/142(94%), Positives:138/142(97%), Gaps:0/142(0%)

Query  139  TQIGCTLNFPISPIETVPVKLKPGMDGPKVKQWPLTEEKIKALVEICTEMEKEGKISKIG  198
            TQIGCTLNFPISPI+TVPVKLKPGMDGPKVKQWPLT+EKI+AL  IC EMEKEGKI+KIG
Sbjct  2    TQIGCTLNFPISPIDTVPVKLKPGMDGPKVKQWPLTKEKIEALTAICDEMEKEGKITKIG  181

Query  199  PENPYNTPVFAIKKKDSTKWRKLVDFRELNKRTQDFWEVQLGIPHPAGLKKKKSVTVLDV  258
            PENPYNTPVFAIKKKDSTKWRKLVDFRELNKRTQDFWEVQLGIPHPAGLKKKKSVTVLDV
Sbjct  182  PENPYNTPVFAIKKKDSTKWRKLVDFRELNKRTQDFWEVQLGIPHPAGLKKKKSVTVLDV  361

Query  259  GDAYFSVPLDEDFRKYTAFTIP  280
            GDAYFSVPLDE FRKYTAFTIP
Sbjct  362  GDAYFSVPLDESFRKYTAFTIP  427



>Escherichia coli strain 8374wB11 NODE_56_length_419_cov_1.43493, whole genome shotgun sequence
Sequence ID: NZ_WSJO01000056.1 Length: 419
Range 1: 2 to 418

Score:280 bits(716), Expect:1e-83,
Method:Compositional matrix adjust.,
Identities:131/139(94%), Positives:136/139(97%), Gaps:0/139(0%)

Query  212  KKDSTKWRKLVDFRELNKRTQDFWEVQLGIPHPAGLKKKKSVTVLDVGDAYFSVPLDEDF  271
            K+DSTKWRKLVDFRELNKRTQDFWEVQLGIPHPAGLK+KKSVTVLDVGDAYFSVPLDE F
Sbjct  418  KRDSTKWRKLVDFRELNKRTQDFWEVQLGIPHPAGLKQKKSVTVLDVGDAYFSVPLDESF  239

Query  272  RKYTAFTIPSINNETPGIRYQYNVLPQGWKGSPAIFQSSMTKILEPFRKQNPDIVIYQYM  331
            RKYTAFTIPS+NNETPGIRYQYNVLPQGWKGSPAIFQ+SMTKILEPFR +NPDIVIYQYM
Sbjct  238  RKYTAFTIPSVNNETPGIRYQYNVLPQGWKGSPAIFQASMTKILEPFRLKNPDIVIYQYM  59

Query  332  DDLYVGSDLEIGQHRTKIE  350
            DDLYVGSDLEIGQHR KIE
Sbjct  58   DDLYVGSDLEIGQHRAKIE  2



>Cronobacter sakazakii strain cro3040W contig77, whole genome shotgun sequence
Sequence ID: NZ_NRJO01000077.1 Length: 429
Range 1: 3 to 428

Score:280 bits(715), Expect:2e-83,
Method:Compositional matrix adjust.,
Identities:134/142(94%), Positives:138/142(97%), Gaps:0/142(0%)
```

```
Query  139  TQIGCTLNFPISPIETVPVKLKPGMDGPKVKQWPLTEEKIKALVEICTEMEKEGKISKIG  198
            TQIGCTLNFPISPIETVPVKLKPGMDGPKVKQWPLT+EKI+AL  IC EMEKEGKI+KIG
Sbjct  428  TQIGCTLNFPISPIETVPVKLKPGMDGPKVKQWPLTKEKIEALTAICDEMEKEGKITKIG  249

Query  199  PENPYNTPVFAIKKKDSTKWRKLVDFRELNKRTQDFWEVQLGIPHPAGLKKKKSVTVLDV  258
            PENPYNTP+FAIKKKDSTKWRKLVDFRELNKRTQDFWEVQLGIPHPAGLKKKKSVTVLDV
Sbjct  248  PENPYNTPIFAIKKKDSTKWRKLVDFRELNKRTQDFWEVQLGIPHPAGLKKKKSVTVLDV  69

Query  259  GDAYFSVPLDEDFRKYTAFTIP  280
            GDAYFSVPL EDFRKYTAFTIP
Sbjct  68   GDAYFSVPLYEDFRKYTAFTIP  3
```

```
>Levilactobacillus brevis strain CRL 2013 scaffold20, whole genome shotgun sequence
Sequence ID: NZ_MZMW01000021.1 Length: 398
Range 1: 2 to 397

Score:271 bits(694), Expect:9e-81,
Method:Compositional matrix adjust.,
Identities:128/132(97%), Positives:129/132(97%), Gaps:0/132(0%)

Query  232  QDFWEVQLGIPHPAGLKKKKSVTVLDVGDAYFSVPLDEDFRKYTAFTIPSINNETPGIRY  291
            QDFWEVQLGIPHPAGLKKKKSVTVLDVGDAYFSVPLD+DFRKYTAFTIPSINNETPGIRY
Sbjct  2    QDFWEVQLGIPHPAGLKKKKSVTVLDVGDAYFSVPLDKDFRKYTAFTIPSINNETPGIRY  181

Query  292  QYNVLPQGWKGSPAIFQSSMTKILEPFRKQNPDIVIYQYMDDLYVGSDLEIGQHRTKIEE  351
            QYNVLPQGWKGSPAIFQSSMTKILEPFRKQNPDIVIYQYMDDLYVGSDLEIGQHR KIEE
Sbjct  182  QYNVLPQGWKGSPAIFQSSMTKILEPFRKQNPDIVIYQYMDDLYVGSDLEIGQHRAKIEE  361

Query  352  LRQHLLRWGLTT  363
            LRQHLL WG TT
Sbjct  362  LRQHLLTWGFTT  397
```

```
>Escherichia coli strain 8374wB11 NODE_55_length_422_cov_0.938983, whole genome shotgun sequence
Sequence ID: NZ_WSJO01000055.1 Length: 422
Range 1: 2 to 421

Score:271 bits(693), Expect:2e-80,
Method:Compositional matrix adjust.,
Identities:122/140(87%), Positives:134/140(95%), Gaps:0/140(0%)

Query  252  SVTVLDVGDAYFSVPLDEDFRKYTAFTIPSINNETPGIRYQYNVLPQGWKGSPAIFQSSM  311
            SVTV+DVGDAYFS+PLDEDFRKYTAFTIPS NNETPG+RYQYNVLPQGWKGSPAIFQSSM
Sbjct  421  SVTVMDVGDAYFSIPLDEDFRKYTAFTIPSTNNETPGVRYQYNVLPQGWKGSPAIFQSSM  242

Query  312  TKILEPFRKQNPDIVIYQYMDDLYVGSDLEIGQHRTKIEELRQHLLRWGLTTPDKKHQKE  371
            T+ILEPFR +NP+IVIYQY+DDLYVGSDLEIGQHR KI+ELR++L +WG  TPDKKHQ+E
Sbjct  241  TRILEPFRAKNPEIVIYQYVDDLYVGSDLEIGQHRAKIKELREYLWKWGFYTPDKKHQEE  62

Query  372  PPFLWMGYELHPDKWTVQPI  391
            PPFLWMGYELHPDKWTVQPI
Sbjct  61   PPFLWMGYELHPDKWTVQPI  2
```

```
>Klebsiella pneumoniae strain EuSCAPE_IL018, whole genome shotgun sequence
Sequence ID: NZ_UKJU02000100.1 Length: 392
Range 1: 3 to 392

Score:267 bits(682), Expect:5e-79,
Method:Compositional matrix adjust.,
Identities:123/130(95%), Positives:126/130(96%), Gaps:0/130(0%)

Query  262  YFSVPLDEDFRKYTAFTIPSINNETPGIRYQYNVLPQGWKGSPAIFQSSMTKILEPFRKQ  321
            YFSVPL EDFRKYTAFTIPSINNETPGIRYQYNVLPQGWKGSPAIFQSSMTKILEPFR Q
Sbjct  392  YFSVPLYEDFRKYTAFTIPSINNETPGIRYQYNVLPQGWKGSPAIFQSSMTKILEPFRTQ  213

Query  322  NPDIVIYQYMDDLYVGSDLEIGQHRTKIEELRQHLLRWGLTTPDKKHQKEPPFLWMGYEL  381
            NP+IVIYQYMDDLYVGSDLEIGQHR KIEELR+HLL+WG TTPDKKHQKEPPFLWMGYEL
Sbjct  212  NPEIVIYQYMDDLYVGSDLEIGQHRAKIEELREHLLKWGFTTPDKKHQKEPPFLWMGYEL  33

Query  382  HPDKWTVQPI  391
            HPDKWTVQPI
Sbjct  32   HPDKWTVQPI  3
```

```
>Klebsiella pneumoniae strain EuSCAPE_NO017, whole genome shotgun sequence
Sequence ID: NZ_UKIO02000069.1 Length: 458
Range 1: 6 to 455

Score:266 bits(679), Expect:3e-78,
Method:Compositional matrix adjust.,
Identities:128/152(84%), Positives:141/152(92%), Gaps:2/152(1%)

Query  30   NNSPSEAGADRQGTVSFNFPQVTLWQRPLVTIKIGGQLKEALLDTGADDTVLEEMSLPGR  89
            +N  SEAG  RQG    N PQ+TLWQRPLV+IK+GGQ++EALLDTGADDTVLE+++LPG+
Sbjct  6    DNPHSEAGVKRQG--GLNLPQITLWQRPLVSIKVGGQIREALLDTGADDTVLEDINLPGK  179

Query  90   WKPKMIGGIGGFIKVRQYDQILIEICGHKAIGTVLVGPTPVNIIGRNLLTQIGCTLNFPI  149
            WKPKMIGGIGGFIKVRQY+ ILIEICG KAIGTVLVGPTPVNIIGRN+LTQ+GCTLNFPI
Sbjct  180  WKPKMIGGIGGFIKVRQYEDILIEICGKKAIGTVLVGPTPVNIIGRNMLTQLGCTLNFPI  359

Query  150  SPIETVPVKLKPGMDGPKVKQWPLTEEKIKAL  181
            SPIETVPVKLKPGMDGPKVKQWPLTEEKIKAL
Sbjct  360  SPIETVPVKLKPGMDGPKVKQWPLTEEKIKAL  455
```

```
>Acinetobacter baumannii strain ACM-28 NODE_88_length_395_cov_2.15672, whole genome shotgun sequence
Sequence ID: NZ_PYDN01000087.1 Length: 395
Range 1: 3 to 395

Score:263 bits(671), Expect:2e-77,
Method:Compositional matrix adjust.,
Identities:122/131(93%), Positives:127/131(96%), Gaps:0/131(0%)

Query  282  INNETPGIRYQYNVLPQGWKGSPAIFQSSMTKILEPFRKQNPDIVIYQYMDDLYVGSDLE  341
            +NNETPGIRYQYNVLPQGWKGSPAIFQ+SMTKILEP R +NPDIVIYQYMDDLYVGSDLE
Sbjct  3    VNNETPGIRYQYNVLPQGWKGSPAIFQASMTKILEPLRLKNPDIVIYQYMDDLYVGSDLE  182

Query  342  IGQHRTKIEELRQHLLRWGLTTPDKKHQKEPPFLWMGYELHPDKWTVQPIVLPEKDSWTV  401
            IGQHR KIEELR+HLL+WGLTTPDKKHQKEPPFLWMGYELHPDKWTVQPI LPEKDSWTV
Sbjct  183  IGQHRAKIEELREHLLKWGLTTPDKKHQKEPPFLWMGYELHPDKWTVQPIQLPEKDSWTV  362

Query  402  NDIQKLVGKLN  412
```

```
            NDIQKLVGKLN
Sbjct  363  NDIQKLVGKLN  395




>Klebsiella pneumoniae strain EuSCAPE_LT003, whole genome shotgun sequence
Sequence ID: NZ_UKJH02000060.1 Length: 422
Range 1: 15 to 422

Score:262 bits(670), Expect:3e-77,
Method:Compositional matrix adjust.,
Identities:125/136(92%), Positives:133/136(97%), Gaps:0/136(0%)

Query  980   SMNKELKKIIGQVRDQAEHLKTAVQMAVFIHNFKRKGGIGGYSAGERIVDIIATDIQTKE  1039
             SMNKELKKIIGQVRDQAE LKTAVQMAVFIHNFKRKGGIGGYSAGERI+DIIA+DIQTKE
Sbjct  422   SMNKELKKIIGQVRDQAEQLKTAVQMAVFIHNFKRKGGIGGYSAGERIIDIIASDIQTKE  243

Query  1040  LQKQITKIQNFRVYYRDSRNPLWKGPAKLLWKGEGAVVIQDNSDIKVVPRRKAKIIRDYG  1099
             LQKQITK+QNFRVYYRDSR+P+WKGPAKLLWKGEGAVV+QDNSDIKVVPRRKAKII+DYG
Sbjct  242   LQKQITKLQNFRVYYRDSRDPIWKGPAKLLWKGEGAVVLQDNSDIKVVPRRKAKIIKDYG  63

Query  1100  KQMAGDDCVASRQDED  1115
             KQMAG DCVA RQ+ED
Sbjct  62    KQMAGADCVAGRQNED  15




>Klebsiella pneumoniae strain EuSCAPE_MT012, whole genome shotgun sequence
Sequence ID: NZ_UKIX02000064.1 Length: 452
Range 1: 3 to 452

Score:261 bits(668), Expect:8e-77,
Method:Compositional matrix adjust.,
Identities:136/150(91%), Positives:147/150(98%), Gaps:0/150(0%)

Query  538  LPIQKetwetwwteYWQATWIPEWEFVNTPPLVKLWYQLEKEPIVGAETFYVDGAANRET  597
            LPIQKETWETWWT+YWQATWIPEWEFVNTPPLVKLWYQLEKEPI+GAETFYVDGAANR+T
Sbjct  452  LPIQKETWETWWTDYWQATWIPEWEFVNTPPLVKLWYQLEKEPIIGAETFYVDGAANRDT  273

Query  598  KLGKAGYVTNRGRQKVVTLTDTTNQKTELQAIYLALQDSGLEVNIVTDSQYALGIIQAQP  657
            K+GKAGYVT++GRQKVV+LT+TTNQKTEL AI LALQDSG EVNIVTDSQYALGIIQAQP
Sbjct  272  KMGKAGYVTDKGRQKVVSLTETTNQKTELHAIQLALQDSGPEVNIVTDSQYALGIIQAQP  93

Query  658  DQSESELVNQIIEQLIKKEKVYLAWVPAHK  687
            D+SESELVNQIIEQLIKKE+VYL+WVPAHK
Sbjct  92   DKSESELVNQIIEQLIKKERVYLSWVPAHK  3




Range 2: 3 to 311

Score:178 bits(452), Expect:3e-47,
Method:Compositional matrix adjust.,
Identities:88/103(85%), Positives:98/103(95%), Gaps:0/103(0%)

Query  705  KVLYVDGAANRETKLGKAGYVTNRGRQKVVTLTDTTNQKTELQAIYLALQDSGLEVNIVT  764
            +  YVDGAANR+TK+GKAGYVT++GRQKVV+LT+TTNQKTEL AI LALQDSG EVNIVT
Sbjct  311  ETFYVDGAANRDTKMGKAGYVTDKGRQKVVSLTETTNQKTELHAIQLALQDSGPEVNIVT  132

Query  765  DSQYALGIIQAQPDQSESELVNQIIEQLIKKEKVYLAWVPAHK  807
            DSQYALGIIQAQPD+SESELVNQIIEQLIKKE+VYL+WVPAHK
Sbjct  131  DSQYALGIIQAQPDKSESELVNQIIEQLIKKERVYLSWVPAHK  3




>Klebsiella pneumoniae strain EuSCAPE_IL004, whole genome shotgun sequence
Sequence ID: NZ_UKJP02000105.1 Length: 382
Range 1: 2 to 382

Score:255 bits(652), Expect:7e-75,
Method:Compositional matrix adjust.,
Identities:118/127(93%), Positives:123/127(96%), Gaps:0/127(0%)

Query  855  LPPVVAKEIVASCDKCQLKGEAMHGQVDCSPGIWQLDCTHLEGKVILVAVHVASGYIEAE  914
            LPPVVAKEIVASCDKCQLKG AMHGQVDCSPGIWQLDCTHLEGK+ILVAVHVASGYIEAE
Sbjct  382  LPPVVAKEIVASCDKCQLKGGAMHGQVDCSPGIWQLDCTHLEGKIILVAVHVASGYIEAE  203

Query  915  VIPAETGQETAYFLLKLAGRWPVKTIHTDNGSNFTGATVRAACWWAGIKQEFGIPYNPQS  974
            VIPAETGQETAY++LKLAGRWPVK IHTDNGSNFT A V+AACWWAGI+QEFGIPYNPQS
Sbjct  202  VIPAETGQETAYYILKLAGRWPVKVIHTDNGSNFTSAAVKAACWWAGIQQEFGIPYNPQS  23

Query  975  QGVVESM  981
            QGVVESM
Sbjct  22   QGVVESM  2




>Acinetobacter baumannii strain ACM-15 NODE_85_length_432_cov_0.596721, whole genome shotgun sequence
Sequence ID: NZ_PYDB01000084.1 Length: 432
Range 1: 3 to 431

Score:255 bits(651), Expect:2e-74,
Method:Compositional matrix adjust.,
Identities:125/143(87%), Positives:134/143(93%), Gaps:0/143(0%)

Query  46   FNFPQVTLWQRPLVTIKIGGQLKEALLDTGADDTVLEEMSLPGRWKPKMIGGIGGFIKVR  105
            FNFPQ+TLWQRPLV+IK+GGQ+KEALLDTGADDTVLEE+SLPG+WKPKMIGGIGGFIKVR
Sbjct  3    FNFPQITLWQRPLVSIKVGGQIKEALLDTGADDTVLEEISLPGKWKPKMIGGIGGFIKVR  182

Query  106  QYDQILIEICGHKAIGTVLVGPTPVNIIGRNLLTQIGCTLNFPISPIETVPVKLKPGMDG  165
            QY QI IEICG KAIGTVLVGPTPVNIIGRNLLTQ+GCTLNFPISPIETVPVKLK GMDG
Sbjct  183  QYGQIPIEICGKKAIGTVLVGPTPVNIIGRNLLTQLGCTLNFPISPIETVPVKLKQGMDG  362

Query  166  PKVKQWPLTEEKIKALVEICTEM  188
            PKVKQW LTE+K+KA VE   EM
Sbjct  363  PKVKQWLLTEDKLKA*VET*KEM  431




>Acinetobacter baumannii strain ACM-29 NODE_131_length_447_cov_1.05937, whole genome shotgun sequence
Sequence ID: NZ_PYDO01000131.1 Length: 447
Range 1: 29 to 442

Score:254 bits(648), Expect:5e-74,
Method:Compositional matrix adjust.,
Identities:128/138(93%), Positives:135/138(97%), Gaps:0/138(0%)

Query  355  HLLRWGLTTPDKKHQKEPPFLWMGYELHPDKWTVQPIVLPEKDSWTVNDIQKLVGKLNWA  414
```

```
          HLL+WG +TPD+KHQKEPPFLWMGYELHPDKWTVQPIVLPEKDSWTVNDIQKLVGKLNWA
Sbjct  442  HLLKWGFSTPDQKHQKEPPFLWMGYELHPDKWTVQPIVLPEKDSWTVNDIQKLVGKLNWA  263

Query  415  SQIYPGIKVRQLCKLLRGTKALTEVIPLTeeaelelaeNREILKEPVHGVYYDPSKDLIA  474
            SQIY GIKV+QLCKLLRGTKALTEV+PL+EEAELELAENREILKEPVHGVYYDPSKDL+A
Sbjct  262  SQIYAGIKVKQLCKLLRGTKALTEVVPLSEEAELELAENREILKEPVHGVYYDPSKDLVA  83

Query  475  EIQKQGQGQWTYQIYQEP  492
            EIQKQG GQWTYQIYQEP
Sbjct  82   EIQKQGLGQWTYQIYQEP  29
```

>Escherichia coli strain 8374wH11 NODE_62_length_437_cov_1.95484, whole genome shotgun sequence
Sequence ID: NZ_WSJI01000059.1 Length: 437
Range 1: 2 to 436

Score:252 bits(644), Expect:1e-73,
Method:Compositional matrix adjust.,
Identities:128/145(88%), Positives:139/145(95%), Gaps:0/145(0%)

```
Query  336  VGSDLEIGQHRTKIEELRQHLLRWGLTTPDKKHQKEPPFLWMGYELHPDKWTVQPIVLPE  395
            VGSDLEIGQHR KI+ELR++L +WG  TPDKKHQ+EPPFLWMGYELHPDKWTVQPI LP
Sbjct  436  VGSDLEIGQHRAKIKELREYLWKWGFYTPDKKHQEEPPFLWMGYELHPDKWTVQPIQLPN  257

Query  396  KDSWTVNDIQKLVGKLNWASQIYPGIKVRQLCKLLRGTKALTEVIPLTeeaelelaeNRE  455
            K+SWTVNDIQKLVGKLNWASQIYPGIKV+QLCKLLRGTKALT+++PLTEEAELELAENRE
Sbjct  256  KESWTVNDIQKLVGKLNWASQIYPGIKVKQLCKLLRGTKALTDIVPLTEEAELELAENRE  77

Query  456  ILKEPVHGVYYDPSKDLIAEIQKQG  480
            ILKEPVHGVYYDPSKDLIAE+QKQG
Sbjct  76   ILKEPVHGVYYDPSKDLIAEVQKQG  2
```

>Escherichia coli strain 8374wA11 NODE_52_length_437_cov_2.24516, whole genome shotgun sequence
Sequence ID: NZ_WSPR01000050.1 Length: 437
Range 1: 1 to 435

Score:247 bits(630), Expect:2e-71,
Method:Compositional matrix adjust.,
Identities:125/145(86%), Positives:137/145(94%), Gaps:0/145(0%)

```
Query  340  LEIGQHRTKIEELRQHLLRWGLTTPDKKHQKEPPFLWMGYELHPDKWTVQPIVLPEKDSW  399
            LEIGQHR KI+ELR++L +WG  TPDKKHQ+EPPFLWMGYELHPDKWTVQPI LP K+SW
Sbjct  1    LEIGQHRAKIKELREYLWKWGFYTPDKKHQEEPPFLWMGYELHPDKWTVQPIQLPNKESW  180

Query  400  TVNDIQKLVGKLNWASQIYPGIKVRQLCKLLRGTKALTEVIPLTeeaelelaeNREILKE  459
            TVNDIQKLVGKLNWASQIYPGIKV+QLCKLLRGTKALT+++PLTEEAELELAENREILKE
Sbjct  181  TVNDIQKLVGKLNWASQIYPGIKVKQLCKLLRGTKALTDIVPLTEEAELELAENREILKE  360

Query  460  PVHGVYYDPSKDLIAEIQKQGQGQW  484
            PVHGVYYDPSK+LIAE+QKQG  QW
Sbjct  361  PVHGVYYDPSKELIAEVQKQGLDQW  435
```

>Escherichia coli strain TzEc080 NODE_52_length_437_cov_2.24516, whole genome shotgun sequence
Sequence ID: NZ_WSFZ01000050.1 Length: 437
Range 1: 1 to 435

Score:247 bits(630), Expect:2e-71,
Method:Compositional matrix adjust.,
Identities:125/145(86%), Positives:137/145(94%), Gaps:0/145(0%)

```
Query  340  LEIGQHRTKIEELRQHLLRWGLTTPDKKHQKEPPFLWMGYELHPDKWTVQPIVLPEKDSW  399
            LEIGQHR KI+ELR++L +WG  TPDKKHQ+EPPFLWMGYELHPDKWTVQPI LP K+SW
Sbjct  1    LEIGQHRAKIKELREYLWKWGFYTPDKKHQEEPPFLWMGYELHPDKWTVQPIQLPNKESW  180

Query  400  TVNDIQKLVGKLNWASQIYPGIKVRQLCKLLRGTKALTEVIPLTeeaelelaeNREILKE  459
            TVNDIQKLVGKLNWASQIYPGIKV+QLCKLLRGTKALT+++PLTEEAELELAENREILKE
Sbjct  181  TVNDIQKLVGKLNWASQIYPGIKVKQLCKLLRGTKALTDIVPLTEEAELELAENREILKE  360

Query  460  PVHGVYYDPSKDLIAEIQKQGQGQW  484
            PVHGVYYDPSK+LIAE+QKQG  QW
Sbjct  361  PVHGVYYDPSKELIAEVQKQGLDQW  435
```

>Escherichia coli strain 8374wB4 NODE_91_length_444_cov_2.17035, whole genome shotgun sequence
Sequence ID: NZ_WSPX01000090.1 Length: 444
Range 1: 78 to 443

Score:246 bits(628), Expect:2e-71,
Method:Compositional matrix adjust.,
Identities:113/122(93%), Positives:118/122(96%), Gaps:0/122(0%)

```
Query  259  GDAYFSVPLDEDFRKYTAFTIPSINNETPGIRYQYNVLPQGWKGSPAIFQSSMTKILEPF  318
            GDAYFSVPLDE FRKYTAFTIPS+NNETPGIRYQYNVLPQGWKGSPAIFQ+SMTKILEPF
Sbjct  78   GDAYFSVPLDESFRKYTAFTIPSVNNETPGIRYQYNVLPQGWKGSPAIFQASMTKILEPF  257

Query  319  RKQNPDIVIYQYMDDLYVGSDLEIGQHRTKIEELRQHLLRWGLTTPDKKHQKEPPFLWMG  378
            R +NPDIVIYQYMDDLYVGSDLEIGQHR KIEELR+HLL+WGLTTPDKKHQKEPPFLWMG
Sbjct  258  RLKNPDIVIYQYMDDLYVGSDLEIGQHRAKIEELREHLLKWGLTTPDKKHQKEPPFLWMG  437

Query  379  YE  380
             E
Sbjct  438  CE  443
```

Range 2: 2 to 103

Score:59.3 bits(142), Expect:2e-71,
Method:Composition-based stats.,
Identities:28/34(82%), Positives:30/34(88%), Gaps:1/34(2%)

```
Query  234  FWEVQLGIPHPAGLKKKKSVTVLDVGDA-YFSVP  266
            FWEVQLGIPHPAGLK+KKSVTVLDVG   +FS P
Sbjct  2    FWEVQLGIPHPAGLKQKKSVTVLDVGGCIFFSSP  103
```

>Klebsiella aerogenes strain AUH-KAM4 NODE_62_length_381_cov_0.925197, whole genome shotgun sequence
Sequence ID: NZ_PYHI01000061.1 Length: 381
Range 1: 2 to 379

Score:244 bits(624), Expect:5e-71,

```
Method:Compositional matrix adjust.,
Identities:115/126(91%), Positives:119/126(94%), Gaps:0/126(0%)

Query  238   QLGIPHPAGLKKKKSVTVLDVGDAYFSVPLDEDFRKYTAFTIPSINNETPGIRYQYNVLP  297
             QLGIPHPAGLKKKKSVTVLDVGDAYFSVPLDE+FRKYTAFTIPSINN TPGIRYQYNVLP
Sbjct  379   QLGIPHPAGLKKKKSVTVLDVGDAYFSVPLDENFRKYTAFTIPSINNATPGIRYQYNVLP  200

Query  298   QGWKGSPAIFQSSMTKILEPFRKQNPDIVIYQYMDDLYVGSDLEIGQHRTKIEELRQHLL  357
             QGWKGSPAIFQSSMTKILEPFRKQNPDIVIYQY DDLYVGS LE+ QHR K+EELR HLL
Sbjct  199   QGWKGSPAIFQSSMTKILEPFRKQNPDIVIYQYRDDLYVGSGLELEQHRIKVEELRAHLL  20

Query  358   RWGLTT  363
             +WG TT
Sbjct  19    KWGFTT  2
```

>Pseudomonas aeruginosa strain P60_London_6_VIM_2_11_13, whole genome shotgun sequence
Sequence ID: NZ_CVXD01000084.1 Length: 395
Range 1: 3 to 392

```
Score:243 bits(621), Expect:2e-70,
Method:Compositional matrix adjust.,
Identities:114/130(88%), Positives:127/130(97%), Gaps:0/130(0%)

Query  51    VTLWQRPLVTIKIGGQLKEALLDTGADDTVLEEMSLPGRWKPKMIGGIGGFIKVRQYDQI  110
             +TLWQRP+V +K+GGQL+EALLDTGADDTVLE+++LPG+WKPKMIGGIGGFIKV+QYDQI
Sbjct  392   ITLWQRPVVKVKVGGQLREALLDTGADDTVLEDINLPGKWKPKMIGGIGGFIKVKQYDQI  213

Query  111   LIEICGHKAIGTVLVGPTPVNIIGRNLLTQIGCTLNFPISPIETVPVKLKPGMDGPKVKQ  170
              I+ICGH+A GTVLVGPTPVNIIGRN+LTQIGCTLNFPISPIETVPVKLKPGMDGPKVKQ
Sbjct  212   TIDICGHRATGTVLVGPTPVNIIGRNMLTQIGCTLNFPISPIETVPVKLKPGMDGPKVKQ  33

Query  171   WPLTEEKIKA  180
             WPLTEEKIKA
Sbjct  32    WPLTEEKIKA  3
```

>Klebsiella pneumoniae strain EuSCAPE_MT020, whole genome shotgun sequence
Sequence ID: NZ_UKJB02000092.1 Length: 368
Range 1: 1 to 366

```
Score:243 bits(619), Expect:3e-70,
Method:Compositional matrix adjust.,
Identities:117/122(96%), Positives:119/122(97%), Gaps:0/122(0%)

Query  151   PIETVPVKLKPGMDGPKVKQWPLTEEKIKALVEICTEMEKEGKISKIGPENPYNTPVFAI  210
             PIETVPVKLKPGMDGPKVKQWPLTEEKIKAL  IC EMEKEGKI+KIGPENPYNTPVFAI
Sbjct  366   PIETVPVKLKPGMDGPKVKQWPLTEEKIKALTAICEEMEKEGKITKIGPENPYNTPVFAI  187

Query  211   KKKDSTKWRKLVDFRELNKRTQDFWEVQLGIPHPAGLKKKKSVTVLDVGDAYFSVPLDED  270
             KKKDSTKWRKLVDFRELNKRTQDFWEVQLGIPHPAGLKKKKSVTVLDVGDAYFSVPLDED
Sbjct  186   KKKDSTKWRKLVDFRELNKRTQDFWEVQLGIPHPAGLKKKKSVTVLDVGDAYFSVPLDED  7

Query  271   FR  272
             F+
Sbjct  6     FK  1
```

>Escherichia coli strain 8374wH11 NODE_67_length_417_cov_0.882759, whole genome shotgun sequence
Sequence ID: NZ_WSJI01000064.1 Length: 417
Range 1: 3 to 413

```
Score:242 bits(618), Expect:6e-70,
Method:Compositional matrix adjust.,
Identities:121/137(88%), Positives:133/137(97%), Gaps:0/137(0%)

Query  336   VGSDLEIGQHRTKIEELRQHLLRWGLTTPDKKHQKEPPFLWMGYELHPDKWTVQPIVLPE  395
             VGSDLEIGQHR KIEE+R+HLL+WGLTTPD+KHQKEPPFLWMGYELHPDKWTVQPI LP+
Sbjct  413   VGSDLEIGQHRAKIEEIREHLLKWGLTTPDRKHQKEPPFLWMGYELHPDKWTVQPIQLPD  234

Query  396   KDSWTVNDIQKLVGKLNWASQIYPGIKVRQLCKLLRGTKALTEVIPLTeeaelelaeNRE  455
             K+SWTVNDIQKLVGKLNWASQIYPGI+V+ LCKLLRG KALT+++PLTEEAELELAENRE
Sbjct  233   KESWTVNDIQKLVGKLNWASQIYPGIRVKHLCKLLRGAKALTDIMPLTEEAELELAENRE  54

Query  456   ILKEPVHGVYYDPSKDL  472
             ILKEPVHGVYYDPSK+L
Sbjct  53    ILKEPVHGVYYDPSKEL  3
```

>Escherichia coli strain 8374wA11 NODE_70_length_381_cov_1.25984, whole genome shotgun sequence
Sequence ID: NZ_WSPR01000067.1 Length: 381
Range 1: 1 to 378

```
Score:238 bits(606), Expect:2e-68,
Method:Compositional matrix adjust.,
Identities:114/126(90%), Positives:118/126(93%), Gaps:0/126(0%)

Query  161   PGMDGPKVKQWPLTEEKIKALVEICTEMEKEGKISKIGPENPYNTPVFAIKKKDSTKWRK  220
             PGMDGP VKQWP +EE IKAL E C +MEKEGKISKIGPENPYNTPVFAIK+KDSTKWRK
Sbjct  1     PGMDGP*VKQWPWSEETIKALTESCEDMEKEGKISKIGPENPYNTPVFAIKRKDSTKWRK  180

Query  221   LVDFRELNKRTQDFWEVQLGIPHPAGLKKKKSVTVLDVGDAYFSVPLDEDFRKYTAFTIP  280
             LVDFRELNKRTQDFWEVQLGIPHPAGLKKKKS+TVLDVGDAYFSVPLDE FRKYTAFTIP
Sbjct  181   LVDFRELNKRTQDFWEVQLGIPHPAGLKKKKSMTVLDVGDAYFSVPLDEGFRKYTAFTIP  360

Query  281   SINNET  286
             S NNET
Sbjct  361   SRNNET  378
```

>Escherichia coli strain TzEc080 NODE_70_length_381_cov_1.25984, whole genome shotgun sequence
Sequence ID: NZ_WSFZ01000067.1 Length: 381
Range 1: 1 to 378

```
Score:238 bits(606), Expect:2e-68,
Method:Compositional matrix adjust.,
Identities:114/126(90%), Positives:118/126(93%), Gaps:0/126(0%)

Query  161   PGMDGPKVKQWPLTEEKIKALVEICTEMEKEGKISKIGPENPYNTPVFAIKKKDSTKWRK  220
             PGMDGP VKQWP +EE IKAL E C +MEKEGKISKIGPENPYNTPVFAIK+KDSTKWRK
Sbjct  1     PGMDGP*VKQWPWSEETIKALTESCEDMEKEGKISKIGPENPYNTPVFAIKRKDSTKWRK  180

Query  221   LVDFRELNKRTQDFWEVQLGIPHPAGLKKKKSVTVLDVGDAYFSVPLDEDFRKYTAFTIP  280
             LVDFRELNKRTQDFWEVQLGIPHPAGLKKKKS+TVLDVGDAYFSVPLDE FRKYTAFTIP
```

```
Sbjct  181  LVDFRELNKRTQDFWEVQLGIPHPAGLKKKKSMTVLDVGDAYFSVPLDEGFRKYTAFTIP  360

Query  281  SINNET  286
            S NNET
Sbjct  361  SRNNET  378
```

>Acinetobacter baumannii strain ACM-37 NODE_88_length_418_cov_0.570447, whole genome shotgun sequence
Sequence ID: NZ_QAGP01000088.1 Length: 418
Range 1: 1 to 417

Score:238 bits(607), Expect:2e-68,
Method:Compositional matrix adjust.,
Identities:116/139(83%), Positives:124/139(89%), Gaps:0/139(0%)

```
Query  152  IETVPVKLKPGMDGPKVKQWPLTEEKIKALVEICTEMEKEGKISKIGPENPYNTPVFAIK  211
            IET  ++LK G DGP++KQW LTE KIK+L  IC EMEK GKI+KIGPENPYNTPVFAIK
Sbjct  417  IETETIRLKLGRDGPRIKQWALTEGKIKSLQAICEEMEKVGKITKIGPENPYNTPVFAIK  238

Query  212  KKDSTKWRKLVDFRELNKRTQDFWEVQLGIPHPAGLKKKKSVTVLDVGDAYFSVPLDEDF  271
            KK STKWRKLVDFRELNKRTQDFWEVQLGIPHPAGLKKKKSVTVLDV DAYFSV LDE+F
Sbjct  237  KKASTKWRKLVDFRELNKRTQDFWEVQLGIPHPAGLKKKKSVTVLDVVDAYFSVLLDENF  58

Query  272  RKYTAFTIPSINNETPGIR  290
             KYTAFTIPSINN TPG+R
Sbjct  57   MKYTAFTIPSINNATPGVR  1
```

>Mycobacterium tuberculosis strain 32845_S29 147, whole genome shotgun sequence
Sequence ID: NZ_QTBV01000025.1 Length: 357
Range 1: 3 to 356

Score:237 bits(604), Expect:3e-68,
Method:Compositional matrix adjust.,
Identities:109/118(92%), Positives:112/118(94%), Gaps:0/118(0%)

```
Query  262  YFSVPLDEDFRKYTAFTIPSINNETPGIRYQYNVLPQGWKGSPAIFQSSMTKILEPFRKQ  321
            YFSVPLDE FRKYTAFTIPSINNETPGIRYQYNVLPQGWKGSPAIFQ SMTKILEPFR +
Sbjct  3    YFSVPLDESFRKYTAFTIPSINNETPGIRYQYNVLPQGWKGSPAIFQYSMTKILEPFRAK  182

Query  322  NPDIVIYQYMDDLYVGSDLEIGQHRTKIEELRQHLLRWGLTTPDKKHQKEPPFLWMGY  379
            NP+IVIYQYMDDLYVGSDLEIGQHR KIEELR HLL+WG TTPDKKHQKEPPFLWMGY
Sbjct  183  NPEIVIYQYMDDLYVGSDLEIGQHRVKIEELRGHLLKWGFTTPDKKHQKEPPFLWMGY  356
```

>Mycobacterium tuberculosis strain 32847_S31 147, whole genome shotgun sequence
Sequence ID: NZ_QTBX01000037.1 Length: 357
Range 1: 2 to 355

Score:237 bits(604), Expect:3e-68,
Method:Compositional matrix adjust.,
Identities:109/118(92%), Positives:112/118(94%), Gaps:0/118(0%)

```
Query  262  YFSVPLDEDFRKYTAFTIPSINNETPGIRYQYNVLPQGWKGSPAIFQSSMTKILEPFRKQ  321
            YFSVPLDE FRKYTAFTIPSINNETPGIRYQYNVLPQGWKGSPAIFQ SMTKILEPFR +
Sbjct  355  YFSVPLDESFRKYTAFTIPSINNETPGIRYQYNVLPQGWKGSPAIFQYSMTKILEPFRAK  176

Query  322  NPDIVIYQYMDDLYVGSDLEIGQHRTKIEELRQHLLRWGLTTPDKKHQKEPPFLWMGY  379
            NP+IVIYQYMDDLYVGSDLEIGQHR KIEELR HLL+WG TTPDKKHQKEPPFLWMGY
Sbjct  175  NPEIVIYQYMDDLYVGSDLEIGQHRVKIEELRGHLLKWGFTTPDKKHQKEPPFLWMGY  2
```

>Escherichia coli strain 8374wG10 NODE_65_length_433_cov_1.76797, whole genome shotgun sequence
Sequence ID: NZ_WSJQ01000065.1 Length: 433
Range 1: 1 to 432

Score:237 bits(604), Expect:5e-68,
Method:Compositional matrix adjust.,
Identities:111/144(77%), Positives:127/144(88%), Gaps:0/144(0%)

```
Query  71   LLDTGADDTVLEEMSLPGRWKPKMIGGIGGFIKVRQYDQILIEICGHKAIGTVLVGPTPV  130
            +LDTGADDTV E ++LPG+W PK+IGG+GG ++V+QYD ILIE  G KAIG+VLVGPTP
Sbjct  1    ILDTGADDTVFENINLPGKWTPKLIGGVGGLVRVKQYDNILIEFEGRKAIGSVLVGPTPS  180

Query  131  NIIGRNLLTQIGCTLNFPISPIETVPVKLKPGMDGPKVKQWPLTEEKIKALVEICTEMEK  190
            NIIGRN++TQIGCTLNFPISPIETVPVKLKPGMDGPK+KQWPLT+EKI+AL E C E EK
Sbjct  181  NIIGRNMMTQIGCTLNFPISPIETVPVKLKPGMDGPKIKQWPLTKEKIEALTESCEEREK  360

Query  191  EGKISKIGPENPYNTPVFAIKKKD  214
            EGKISKIGPENP N PVF  K+KD
Sbjct  361  EGKISKIGPENPSNPPVFDRKRKD  432
```

>Neisseria gonorrhoeae strain GC-206 NODE_134_length_344_cov_1.3045_ID_20440, whole genome shotgun sequence
Sequence ID: NZ_SAWN01000134.1 Length: 344
Range 1: 1 to 342

Score:234 bits(596), Expect:3e-67,
Method:Compositional matrix adjust.,
Identities:106/114(93%), Positives:114/114(100%), Gaps:0/114(0%)

```
Query  825  KVLFLDGIDKAQDEHEKYHSNWRAMASDFNLPPVVAKEIVASCDKCQLKGEAMHGQVDCS  884
            KVLFLDGIDKAQ+EHE+YHSNWRAMAS+FNLPP+VAKEIVASCDKCQLKGEA+HGQVDCS
Sbjct  342  KVLFLDGIDKAQEEHERYHSNWRAMASEFNLPPIVAKEIVASCDKCQLKGEAIHGQVDCS  163

Query  885  PGIWQLDCTHLEGKVILVAVHVASGYIEAEVIPAETGQETAYFLLKLAGRWPVK  938
            PGIWQLDCTHLEGK+ILVAVHVASGY+EAEVIPAETGQETAYF+LKLAGRWPVK
Sbjct  162  PGIWQLDCTHLEGKIILVAVHVASGYMEAEVIPAETGQETAYFILKLAGRWPVK  1
```

>Klebsiella pneumoniae strain EuSCAPE_MT010, whole genome shotgun sequence
Sequence ID: NZ_UKIT02000046.1 Length: 367
Range 1: 1 to 366

Score:233 bits(594), Expect:8e-67,
Method:Compositional matrix adjust.,
Identities:113/122(93%), Positives:119/122(97%), Gaps:0/122(0%)

```
Query  982   NKELKKIIGQVRDQAEHLKTAVQMAVFIHNFKRKGGIGGYSAGERIVDIIATDIQTKELQ  1041
             NKELKKII QV+DQAEHLKTAVQMAVFIHNFKRKGGI GYSAGERI+DIIATD+QTKELQ
Sbjct  1     NKELKKIIRQVKDQAEHLKTAVQMAVFIHNFKRKGGIRGYSAGERIIDIIATDLQTKELQ  180
```

```
Query  1042  KQITKIQNFRVVYYRDSRNPLWKGPAKLLWKGEGAVVIQDNSDIKVVPRRKAKIIRDYGKQ  1101
             KQI KIQNFRVVYYRDSR+P+WKGPAKLLWKGEGAVVIQDNSDIKVVPRRKAKII+DYGKQ
Sbjct  181   KQIIKIQNFRVVYYRDSRDPIWKGPAKLLWKGEGAVVIQDNSDIKVVPRRKAKIIKDYGKQ  360

Query  1102  MA  1103
             MA
Sbjct  361   MA  366
```

>Klebsiella aerogenes strain AUH-KAM2 NODE_49_length_404_cov_0.407942, whole genome shotgun sequence
Sequence ID: NZ_PYHG01000048.1 Length: 404
Range 1: 3 to 404

Score:233 bits(593), Expect:1e-66,
Method:Compositional matrix adjust.,
Identities:118/134(88%), Positives:126/134(94%), Gaps:0/134(0%)

```
Query  339   DLEIGQHRTKIEELRQHLLRWGLTTPDKKHQKEPPFLWMGYELHPDKWTVQPIVLPEKDS  398
             DLEIGQHR KIEELR HLL+WG TTPDKKHQKEPPFLWMGYELHPDKWTV+PI LPEKDS
Sbjct  404   DLEIGQHRAKIEELRDHLLKWGFTTPDKKHQKEPPFLWMGYELHPDKWTVKPIQLPEKDS  225

Query  399   WTVNDIQKLVGKLNWASQIYPGIKVRQLCKLLRGTKALTEVIPLTeeaelelaeNREILK  458
             WTVN IQKLVGKLNWASQIYPGIK++QLCKLLRG KALT+++ LTEEAELELAENREILK
Sbjct  224   WTVNGIQKLVGKLNWASQIYPGIKIKQLCKLLRGAKALTDIVQLTEEAELELAENREILK  45

Query  459   EPVHGVYYDPSKDL  472
             EP HGVYYDPSK+L
Sbjct  44    EPAHGVYYDPSKEL  3
```

>Methylobacterium radiotolerans strain MAMP 4754 MP-M-radiotolerans_contig_1065, whole genome shotgun sequence
Sequence ID: NZ_NKQS01001052.1 Length: 539
Range 1: 166 to 519

Score:232 bits(591), Expect:1e-65,
Method:Compositional matrix adjust.,
Identities:107/118(91%), Positives:113/118(95%), Gaps:0/118(0%)

```
Query  326   VIYQYMDDLYVGSDLEIGQHRTKIEELRQHLLRWGLTTPDKKHQKEPPFLWMGYELHPDK  385
              +YQYMDDLYVGSDLEIGQHR KIEELR+HLLRWGLTTPDKKHQKEPPFLWMGYELHPDK
Sbjct  519   CVYQYMDDLYVGSDLEIGQHRAKIEELRKHLLRWGLTTPDKKHQKEPPFLWMGYELHPDK  340

Query  386   WTVQPIVLPEKDSWTVNDIQKLVGKLNWASQIYPGIKVRQLCKLLRGTKALTEVIPLT  443
             WTVQPIVLPEK+SWTVNDIQKLVGKLNWASQIY GIKV Q+CKLLR TKALT++IPLT
Sbjct  339   WTVQPIVLPEKESWTVNDIQKLVGKLNWASQIYTGIKVTQMCKLLR*TKALTDIIPLT  166
```

>Escherichia coli strain ECM-2 NODE_147_length_382_cov_1.52941, whole genome shotgun sequence
Sequence ID: NZ_PXIA01000147.1 Length: 382
Range 1: 1 to 375

Score:223 bits(568), Expect:3e-63,
Method:Compositional matrix adjust.,
Identities:115/125(92%), Positives:121/125(96%), Gaps:0/125(0%)

```
Query  360   GLTTPDKKHQKEPPFLWMGYELHPDKWTVQPIVLPEKDSWTVNDIQKLVGKLNWASQIYP  419
             GL+TPD+KHQKEPPFLWMGYEL PDKWTVQPIVLPEKDSWTVNDIQKLVGKLNWASQIY
Sbjct  375   GLSTPDQKHQKEPPFLWMGYELRPDKWTVQPIVLPEKDSWTVNDIQKLVGKLNWASQIYA  196

Query  420   GIKVRQLCKLLRGTKALTEVIPLTeeaelelaeNREILKEPVHGVYYDPSKDLIAEIQKQ  479
             GIKV+QLCKLLRG KALTEV+PL+EEAELELAENREILKEPVHGVYYDPSKDL+AEIQKQ
Sbjct  195   GIKVKQLCKLLRGAKALTEVVPLSEEAELELAENREILKEPVHGVYYDPSKDLVAEIQKQ  16

Query  480   GQGQW  484
             G GQW
Sbjct  15    GLGQW  1
```

>Klebsiella pneumoniae strain EuSCAPE_IL007, whole genome shotgun sequence
Sequence ID: NZ_UKJT02000028.1 Length: 343
Range 1: 3 to 341

Score:221 bits(564), Expect:8e-63,
Method:Compositional matrix adjust.,
Identities:105/113(93%), Positives:110/113(97%), Gaps:0/113(0%)

```
Query  996   AEHLKTAVQMAVFIHNFKRKGGIGGYSAGERIVDIIATDIQTKELQKQITKIQNFRVVYYR  1055
             AEHL+TAVQMAVFIHNFKRKGGIGGYSAGERI+DIIATDIQTKELQKQI KIQNFRVVYYR
Sbjct  3     AEHLRTAVQMAVFIHNFKRKGGIGGYSAGERIIDIIATDIQTKELQKQIIKIQNFRVVYYR  182

Query  1056  DSRNPLWKGPAKLLWKGEGAVVIQDNSDIKVVPRRKAKIIRDYGKQMAGDDCV  1108
             DSR+P+WKGPAKLLWKGEGAVVIQDNSDIKVVPRRK KII+DYGKQMAG DCV
Sbjct  183   DSRDPIWKGPAKLLWKGEGAVVIQDNSDIKVVPRRKVKIIKDYGKQMAGADCV  341
```

>Escherichia coli strain 8374wB11 NODE_46_length_516_cov_2.15167, whole genome shotgun sequence
Sequence ID: NZ_WSJO01000046.1 Length: 516
Range 1: 181 to 516

Score:218 bits(555), Expect:8e-61,
Method:Compositional matrix adjust.,
Identities:103/112(92%), Positives:109/112(97%), Gaps:0/112(0%)

```
Query  123   VLVGPTPVNIIGRNLLTQIGCTLNFPISPIETVPVKLKPGMDGPKVKQWPLTEEKIKALV  182
             VLVGPTP+NIIGRN+LTQ+GCTLNFPISPIETVPVKLKPGMDGP+VKQWPLTEEKIKAL
Sbjct  516   VLVGPTPINIIGRNMLTQLGCTLNFPISPIETVPVKLKPGMDGPRVKQWPLTEEKIKALQ  337

Query  183   EICTEMEKEGKISKIGPENPYNTPVFAIKKKDSTKWRKLVDFRELNKRTQDF  234
              IC EMEKEGKI+KIGPENPYNTPVFAIKKK+STKWRKLVDFRELNKRTQDF
Sbjct  336   AICEEMEKEGKITKIGPENPYNTPVFAIKKKNSTKWRKLVDFRELNKRTQDF  181
```

Range 2: 2 to 253

Score:130 bits(326), Expect:6e-30,
Method:Compositional matrix adjust.,
Identities:63/84(75%), Positives:69/84(82%), Gaps:0/84(0%)

```
Query  211   KKKDSTKWRKLVDFRELNKRTQDFWEVQLGIPHPAGLKKKKSVTVLDVGDAYFSVPLDED  270
             KK++     K+  F+    +   FWEVQLGIPHPAGLKKKKSVTVLDVGDAYFSVPLDE+
Sbjct  253   KKEEQY*VEKISRFQGTQ*KNSRFWEVQLGIPHPAGLKKKKSVTVLDVGDAYFSVPLDEN  74
```

```
Query  271  FRKYTAFTIPSINNETPGIRYQYN  294
            FRKYTAFTIPSINN TPGIRYQYN
Sbjct  73   FRKYTAFTIPSINNATPGIRYQYN  2



>Thalassovita mangrovi strain GS-10 Scaffold21_1, whole genome shotgun sequence
Sequence ID: NZ_WWEN01000021.1 Length: 1440
Range 1: 1101 to 1439

Score:228 bits(581), Expect:2e-60,
Method:Compositional matrix adjust.,
Identities:110/113(97%), Positives:112/113(99%), Gaps:0/113(0%)

Query  1003  VQMAVFIHNFKRKGGIGGYSAGERIVDIIATDIQTKELQKQITKIQNFRVYYRDSRNPLW  1062
             VQMAV IHNFKRKGGIGGYSAGERIVDIIATDIQTKELQKQITKIQNFRVYYRDSR+P+W
Sbjct  1439  VQMAVLIHNFKRKGGIGGYSAGERIVDIIATDIQTKELQKQITKIQNFRVYYRDSRDPVW  1260

Query  1063  KGPAKLLWKGEGAVVIQDNSDIKVVPRRKAKIIRDYGKQMAGDDCVASRQDED  1115
             KGPAKLLWKGEGAVVIQDNSDIKVVPRRKAKIIRDYGKQMAGDDCVASRQDED
Sbjct  1259  KGPAKLLWKGEGAVVIQDNSDIKVVPRRKAKIIRDYGKQMAGDDCVASRQDED  1101



>Klebsiella pneumoniae strain EuSCAPE_LT004, whole genome shotgun sequence
Sequence ID: NZ_UKJE02000070.1 Length: 335
Range 1: 1 to 333

Score:202 bits(514), Expect:5e-56,
Method:Compositional matrix adjust.,
Identities:96/111(86%), Positives:103/111(92%), Gaps:0/111(0%)

Query  221  LVDFRELNKRTQDFWEVQLGIPHPAGLKKKKSVTVLDVGDAYFSVPLDEDFRKYTAFTIP  280
            LVDF+ELNKRTQDFWEVQLGIPHPA LKKKKSVTVLDV DAYFSVPLD+ F+KYTAFTIP
Sbjct  333  LVDFKELNKRTQDFWEVQLGIPHPAELKKKKSVTVLDVKDAYFSVPLDKGFKKYTAFTIP  154

Query  281  SINNETPGIRYQYNVLPQGWKGSPAIFQSSMTKILEPFRKQNPDIVIYQYM  331
            SINNETP IRYQYNVLPQ  K SPAIFQSSMT+ILEPFR +NP+IVIYQY+
Sbjct  153  SINNETPEIRYQYNVLPQR*KRSPAIFQSSMTRILEPFRAKNPEIVIYQYI  1



>Klebsiella pneumoniae strain EuSCAPE_IL005, whole genome shotgun sequence
Sequence ID: NZ_UKJL02000063.1 Length: 298
Range 1: 3 to 296

Score:199 bits(506), Expect:4e-55,
Method:Compositional matrix adjust.,
Identities:95/98(97%), Positives:96/98(97%), Gaps:0/98(0%)

Query  184  ICTEMEKEGKISKIGPENPYNTPVFAIKKKDSTKWRKLVDFRELNKRTQDFWEVQLGIPH  243
            IC EMEKEGKI+KIGPENPYNTPVFAIKKKDSTKWRKLVDFRELNKRTQDFWEVQLGIPH
Sbjct  3    ICEEMEKEGKITKIGPENPYNTPVFAIKKKDSTKWRKLVDFRELNKRTQDFWEVQLGIPH  182

Query  244  PAGLKKKKSVTVLDVGDAYFSVPLDEDFRKYTAFTIPS  281
            PAGLKKKKSVTVLDVGDAYFSVPLDE FRKYTAFTIPS
Sbjct  183  PAGLKKKKSVTVLDVGDAYFSVPLDEGFRKYTAFTIPS  296



>Klebsiella pneumoniae strain EuSCAPE_LT003, whole genome shotgun sequence
Sequence ID: NZ_UKJH02000070.1 Length: 297
Range 1: 2 to 295

Score:198 bits(503), Expect:1e-54,
Method:Compositional matrix adjust.,
Identities:91/98(93%), Positives:94/98(95%), Gaps:0/98(0%)

Query  880  QVDCSPGIWQLDCTHLEGKVILVAVHVASGYIEAEVIPAETGQETAYFLLKLAGRWPVKT  939
            QVDCSPGIWQLDCTHLEGKVILVAVHVASGYIEAEVIPAETGQETAYF+LKLAGRWPVK
Sbjct  295  QVDCSPGIWQLDCTHLEGKVILVAVHVASGYIEAEVIPAETGQETAYFILKLAGRWPVKV  116

Query  940  IHTDNGSNFTGATVRAACWWAGIKQEFGIPYNPQSQGV  977
            IHTDNGSNFT   V+AACWWAGI+QEFGIPYNPQSQGV
Sbjct  115  IHTDNGSNFTSNAVKAACWWAGIQQEFGIPYNPQSQGV  2



>Klebsiella pneumoniae strain EuSCAPE_LT004, whole genome shotgun sequence
Sequence ID: NZ_UKJE02000066.1 Length: 367
Range 1: 1 to 366

Score:197 bits(502), Expect:3e-54,
Method:Compositional matrix adjust.,
Identities:102/122(84%), Positives:113/122(92%), Gaps:0/122(0%)

Query  407  LVGKLNWASQIYPGIKVRQLCKLLRGTKALTEVIPLTeeaelelaeNREILKEPVHGVYY  466
            LVGKLNWASQIYPGIKVRQLC+LLRG KALT+V+PLTEEAEL+LAEN+EILKEPVH VYY
Sbjct  1    LVGKLNWASQIYPGIKVRQLCRLLRGAKALTDVVPLTEEAELKLAENKEILKEPVHKVYY  180

Query  467  DPSKDLIAEIQKQGQGQWTYQIYQEPFKNLKTGKYARMRGAHTNDVKQLTEAVQKITTES  526
            DPSK+LIAEIQKQ   Q TYQ++QEPFKNLKTGKYA+M+ AHTNDVKQLTEAVQKI  E+
Sbjct  181  DPSKELIAEIQKQKNDQ*TYQVFQEPFKNLKTGKYAKMKSAHTNDVKQLTEAVQKIALEA  360

Query  527  IV  528
            IV
Sbjct  361  IV  366



>Deinococcus aquiradiocola strain JCM 14371 sequence36, whole genome shotgun sequence
Sequence ID: NZ_BMOE01000036.1 Length: 869
Range 1: 2 to 304

Score:204 bits(518), Expect:4e-54,
Method:Compositional matrix adjust.,
Identities:99/101(98%), Positives:101/101(100%), Gaps:0/101(0%)

Query  1015  KGGIGGYSAGERIVDIIATDIQTKELQKQITKIQNFRVYYRDSRNPLWKGPAKLLWKGEG  1074
             KGGIGGYSAGERIVDIIATDIQTKELQKQITKIQNFRVYYRDSR+P+WKGPAKLLWKGEG
Sbjct  2     KGGIGGYSAGERIVDIIATDIQTKELQKQITKIQNFRVYYRDSRDPVWKGPAKLLWKGEG  181

Query  1075  AVVIQDNSDIKVVPRRKAKIIRDYGKQMAGDDCVASRQDED  1115
             AVVIQDNSDIKVVPRRKAKIIRDYGKQMAGDDCVASRQDED
Sbjct  182   AVVIQDNSDIKVVPRRKAKIIRDYGKQMAGDDCVASRQDED  304
```

```
>Klebsiella pneumoniae strain EuSCAPE_LT003, whole genome shotgun sequence
Sequence ID: NZ_UKJH02000069.1 Length: 305
Range 1: 9 to 305

Score:194 bits(493), Expect:2e-53,
Method:Composition-based stats.,
Identities:89/99(90%), Positives:95/99(95%), Gaps:0/99(0%)

Query  803  VPAHKGIGGNEQVDKLVSAGIRKVLFLDGIDKAQDEHEKYHSNWRAMASDFNLPPVVAKE  862
            VPAHK I GNEQV+KLVS+ IRKVLFLD IDKAQ+EHEKYH+NWRA+ASDFNLPP+VAKE
Sbjct  9    VPAHKRIRGNEQVNKLVSSRIRKVLFLDRIDKAQEEHEKYHNNWRAIASDFNLPPIVAKE  188

Query  863  IVASCDKCQLKGEAMHGQVDCSPGIWQLDCTHLEGKVIL  901
            IVASCDKCQLKGEAMHGQVDCSPGIWQLDCTHLEGKVIL
Sbjct  189  IVASCDKCQLKGEAMHGQVDCSPGIWQLDCTHLEGKVIL  305



>Klebsiella pneumoniae strain EuSCAPE_IL005, whole genome shotgun sequence
Sequence ID: NZ_UKJL02000050.1 Length: 636
Range 1: 326 to 634

Score:197 bits(501), Expect:8e-53,
Method:Compositional matrix adjust.,
Identities:95/103(92%), Positives:99/103(96%), Gaps:0/103(0%)

Query  1013  KRKGGIGGYSAGERIVDIIATDIQTKELQKQITKIQNFRVYYRDSRNPLWKGPAKLLWKG  1072
             KRKGGIGGYSAGERI+DIIATDIQTKELQKQI KIQNFRVYYRDSR+P+WKGPAKLLWKG
Sbjct  634   KRKGGIGGYSAGERIIDIIATDIQTKELQKQIIKIQNFRVYYRDSRDPIWKGPAKLLWKG  455

Query  1073  EGAVVIQDNSDIKVVPRRKAKIIRDYGKQMAGDDCVASRQDED  1115
             EGAVV+QDNSDIKVVPRRK KIIRDYGKQMAG DCVA RQDED
Sbjct  454   EGAVVLQDNSDIKVVPRRKVKIIRDYGKQMAGADCVAGRQDED  326



>Cellulophaga baltica strain 14 NODE_63_length_422_cov_10.206780, whole genome shotgun sequence
Sequence ID: NZ_VIQP01000062.1 Length: 422
Range 1: 1 to 387

Score:192 bits(489), Expect:3e-52,
Method:Compositional matrix adjust.,
Identities:99/130(76%), Positives:111/130(85%), Gaps:2/130(1%)

Query  2    QGKAREFSSEQTRANSPTRRELQVWGRDNNSPSEAGADRQGTVS-FNFPQVTLWQRPLVT  60
            Q +AR+FSSEQTR N PT RE    GRDN   S AGA+ QG +S FNFPQ+TLWQRPLVT
Sbjct  387  QREARKFSSEQTRTNCPTSREHGDGGRDN-LLSAAGAEGQGAISSFNFPQITLWQRPLVT  211

Query  61   IKIGGQLKEALLDTGADDTVLEEMSLPGRWKPKMIGGIGGFIKVRQYDQILIEICGHKAI  120
            ++I GQL EALLDTGADDTVLE+++LPG+WKPKMIGGIGGFIKVRQYDQIL+EI G KAI
Sbjct  210  VRIEGQLIEALLDTGADDTVLEDINLPGKWKPKMIGGIGGFIKVRQYDQILMEIHGKKAI  31

Query  121  GTVLVGPTPV  130
            GTVLVGPTPV
Sbjct  30   GTVLVGPTPV  1



>Klebsiella pneumoniae strain EuSCAPE_IL004, whole genome shotgun sequence
Sequence ID: NZ_UKJP02000108.1 Length: 356
Range 1: 1 to 348

Score:191 bits(486), Expect:3e-52,
Method:Compositional matrix adjust.,
Identities:96/120(80%), Positives:106/120(88%), Gaps:4/120(3%)

Query  9    SSEQTRANSPTRRELQVWGRDNNSPSEAGADRQGTVSFNFPQVTLWQRPLVTIKIGGQLK  68
            +S  +RANSPT RELQV  R +N  SEAGA+ QGT   NFPQ+TLWQRPLV+IK+GGQ K
Sbjct  348  NSPTSRANSPTSRELQV--RGDNPCSEAGAEGQGT--LNFPQITLWQRPLVSIKVGGQTK  181

Query  69   EALLDTGADDTVLEEMSLPGRWKPKMIGGIGGFIKVRQYDQILIEICGHKAIGTVLVGPT  128
            EALLDTGADDTVLEE++LPG+WKPKMIGGIGGFIKVRQYDQILIEICG KAIGTVLVGPT
Sbjct  180  EALLDTGADDTVLEEINLPGKWKPKMIGGIGGFIKVRQYDQILIEICGKKAIGTVLVGPT  1



>Klebsiella pneumoniae strain EuSCAPE_IL005, whole genome shotgun sequence
Sequence ID: NZ_UKJL02000064.1 Length: 283
Range 1: 5 to 283

Score:187 bits(474), Expect:6e-51,
Method:Compositional matrix adjust.,
Identities:86/93(92%), Positives:88/93(94%), Gaps:0/93(0%)

Query  271  FRKYTAFTIPSINNETPGIRYQYNVLPQGWKGSPAIFQSSMTKILEPFRKQNPDIVIYQY  330
            F KYTAFTIPS NNETPGIRYQYNVLPQGWKGSPAIFQSSMTKILEPFR QNPDIVIYQY
Sbjct  5    FMKYTAFTIPSTNNETPGIRYQYNVLPQGWKGSPAIFQSSMTKILEPFRAQNPDIVIYQY  184

Query  331  MDDLYVGSDLEIGQHRTKIEELRQHLLRWGLTT  363
            MDDLYVGSDLEIGQHR KIEELR+HLL+WG TT
Sbjct  185  MDDLYVGSDLEIGQHRAKIEELREHLLKWGFTT  283



>Klebsiella pneumoniae strain EuSCAPE_LT007, whole genome shotgun sequence
Sequence ID: NZ_UKJI02000072.1 Length: 392
Range 1: 1 to 345

Score:186 bits(472), Expect:3e-50,
Method:Compositional matrix adjust.,
Identities:94/119(79%), Positives:104/119(87%), Gaps:4/119(3%)

Query  2    QGKAREFSSEQTRANSPTRRELQVWGRDNNSPSEAGADRQGTVSFNFPQVTLWQRPLVTI  61
            QG+AREF SEQ RA SPT RELQV  R +N  SEAG + QG  SFNFPQ+TLWQRP+VTI
Sbjct  345  QGEAREFPSEQARAISPTSRELQV--RGDNPCSEAGVEGQG--SFNFPQITLWQRPVVTI  178

Query  62   KIGGQLKEALLDTGADDTVLEEMSLPGRWKPKMIGGIGGFIKVRQYDQILIEICGHKAI  120
            K+GGQ+KEALLDTGADDTVLEE++LPG+WKPKMIGGIGGFIKVRQYDQI IEICG KAI
Sbjct  177  KVGGQVKEALLDTGADDTVLEEINLPGKWKPKMIGGIGGFIKVRQYDQIPIEICGKKAI  1



>Klebsiella pneumoniae strain EuSCAPE_DE082, whole genome shotgun sequence
Sequence ID: NZ_UKLV02000054.1 Length: 295
Range 1: 3 to 293

Score:178 bits(451), Expect:9e-48,
```

```
Method:Compositional matrix adjust.,
Identities:86/97(89%), Positives:89/97(91%), Gaps:0/97(0%)

Query  180  ALVEICTEMEKEGKISKIGPENPYNTPVFAIKKKDSTKWRKLVDFRELNKRTQDFWEVQL  239
            AL  IC EME EGKI+KI PENPYNTPVFAIKKKDSTKWRKLVDFRELNKRTQDF EVQL
Sbjct  3    ALTAICEEMENEGKITKIRPENPYNTPVFAIKKKDSTKWRKLVDFRELNKRTQDF*EVQL  182

Query  240  GIPHPAGLKKKKSVTVLDVGDAYFSVPLDEDFRKYTA  276
             IPHPAGLKKKKSVTVLDV DAYFSVPLD++FRKYTA
Sbjct  183  RIPHPAGLKKKKSVTVLDVEDAYFSVPLDKNFRKYTA  293



>Klebsiella pneumoniae strain EuSCAPE_IL005, whole genome shotgun sequence
Sequence ID: NZ_UKJL02000059.1 Length: 323
Range 1: 4 to 321

Score:167 bits(422), Expect:1e-43,
Method:Compositional matrix adjust.,
Identities:84/112(75%), Positives:95/112(84%), Gaps:6/112(5%)

Query  10   SEQTRANSPTRRELQVWGRDNNSPSEAGADRQGTVSFNFPQVTLWQRPLVTIKIGGQLKE  69
            SEQT A+S T R+L    R +N  SEAG +RQGT   NFPQ+TLWQRPLV+IK+GGQ KE
Sbjct  4    SEQTGADSSTSRQL----RGDNPCSEAGDERQGT--LNFPQITLWQRPLVSIKVGGQTKE  165

Query  70   ALLDTGADDTVLEEMSLPGRWKPKMIGGIGGFIKVRQYDQILIEICGHKAIG  121
            ALLDTGADDTVLEE++LPG+WKPKMIGGIGGFIKVRQYD+I IEICG KAIG
Sbjct  166  ALLDTGADDTVLEEINLPGKWKPKMIGGIGGFIKVRQYDEITIEICGKKAIG  321



>Salmonella enterica subsp. enterica serovar Anatum strain 2014LSAL01667 2014LSAL01667_120, whole genome shotgun sequence
Sequence ID: NZ_CALPAM010000120.1 Length: 264
Range 1: 2 to 262

Score:164 bits(415), Expect:5e-43,
Method:Compositional matrix adjust.,
Identities:78/87(90%), Positives:85/87(97%), Gaps:0/87(0%)

Query  1026  RIVDIIATDIQTKELQKQITKIQNFRVYYRDSRNPLWKGPAKLLWKGEGAVVIQDNSDIK  1085
             RI+DIIA+DIQTKELQKQITKIQ FRVYYRDSR+P+WKGPAKLLWKGEGAVVI DNSDIK
Sbjct  262   RIIDIIASDIQTKELQKQITKIQKFRVYYRDSRDPIWKGPAKLLWKGEGAVVIPDNSDIK  83

Query  1086  VVPRRKAKIIRDYGKQMAGDDCVASRQ  1112
             VVPRRKAKI+RDYG+QMAG+DCVASRQ
Sbjct  82    VVPRRKAKILRDYGQQMAGNDCVASRQ  2



>Klebsiella pneumoniae strain EuSCAPE_IL004, whole genome shotgun sequence
Sequence ID: NZ_UKJP02000116.1 Length: 244
Range 1: 2 to 244

Score:163 bits(412), Expect:1e-42,
Method:Compositional matrix adjust.,
Identities:77/81(95%), Positives:81/81(100%), Gaps:0/81(0%)

Query  984   ELKKIIGQVRDQAEHLKTAVQMAVFIHNFKRKGGIGGYSAGERIVDIIATDIQTKELQKQ  1043
             ELKKIIGQVRDQAEHLKTAVQMAVFIHNFKR+GGIGGYSAGERI+DIIATDIQTKELQKQ
Sbjct  2     ELKKIIGQVRDQAEHLKTAVQMAVFIHNFKRRGGIGGYSAGERIIDIIATDIQTKELQKQ  181

Query  1044  ITKIQNFRVYYRDSRNPLWKG  1064
             ITKIQNFRVYYRDSR+P+WKG
Sbjct  182   ITKIQNFRVYYRDSRDPIWKG  244



>Burkholderia pseudomallei strain A-175-05-2-02, whole genome shotgun sequence
Sequence ID: NZ_UVFW01000138.1 Length: 455
Range 1: 1 to 252

Score:160 bits(405), Expect:7e-41,
Method:Compositional matrix adjust.,
Identities:77/84(92%), Positives:80/84(95%), Gaps:0/84(0%)

Query  110  ILIEICGHKAIGTVLVGPTPVNIIGRNLLTQIGCTLNFPISPIETVPVKLKPGMDGPKVK  169
            + IEICG KAIGTVLVGPTPVNIIGRN+LTQIGCTLNFPISPIETVPVKLKPGMDGPKVK
Sbjct  252  LCIEICGKKAIGTVLVGPTPVNIIGRNMLTQIGCTLNFPISPIETVPVKLKPGMDGPKVK  73

Query  170  QWPLTEEKIKALVEICTEMEKEGK  193
            QWPLTEEKIKAL EIC +MEKEGK
Sbjct  72   QWPLTEEKIKALTEICNDMEKEGK  1



Range 2: 273 to 401

Score:54.3 bits(129), Expect:0.001,
Method:Composition-based stats.,
Identities:31/44(70%), Positives:33/44(75%), Gaps:1/44(2%)

Query  2    QGKAREFSSEQTRANSPTRRELQVWGRDNNSPSEAGADRQGTVS  45
            Q KAREFSSEQ RANSPT REL   GRD N  +EAGA+ QGT S
Sbjct  401  QRKAREFSSEQNRANSPTSRELGDGGRD-NLLTEAGAEGQGTSS  273



>Mycobacterium tuberculosis strain 32847_S31 146, whole genome shotgun sequence
Sequence ID: NZ_QTBX01000036.1 Length: 235
Range 1: 2 to 235

Score:154 bits(389), Expect:1e-39,
Method:Compositional matrix adjust.,
Identities:70/78(90%), Positives:73/78(93%), Gaps:0/78(0%)

Query  302  GSPAIFQSSMTKILEPFRKQNPDIVIYQYMDDLYVGSDLEIGQHRTKIEELRQHLLRWGL  361
            GSPAIFQ SMTKILEPFR +NP+IVIYQYMDDLYVGSDLEIGQHR KIEELR HLL+WG
Sbjct  235  GSPAIFQHSMTKILEPFRTENPEIVIYQYMDDLYVGSDLEIGQHRVKIEELRGHLLKWGF  56

Query  362  TTPDKKHQKEPPFLWMGY  379
            TTPDKKHQKEPPFLWMGY
Sbjct  55   TTPDKKHQKEPPFLWMGY  2



>Methylobacterium radiotolerans strain MAMP 4754 MP-M-radiotolerans_contig_571, whole genome shotgun sequence
Sequence ID: NZ_NKQS01000559.1 Length: 289
```

```
Range 1: 2 to 244

Score:151 bits(382), Expect:2e-38,
Method:Compositional matrix adjust.,
Identities:72/81(89%), Positives:76/81(93%), Gaps:0/81(0%)

Query  1035  IQTKELQKQITKIQNFRVYYRDSRNPLWKGPAKLLWKGEGAVVIQDNSDIKVVPRRKAKI  1094
             IQTK +QKQI KIQNFRVYYRDSR+P+WKGPAKLLWKGEGAVVIQDNSDIKVVPRRK KI
Sbjct  2     IQTKAVQKQIIKIQNFRVYYRDSRDPIWKGPAKLLWKGEGAVVIQDNSDIKVVPRRKVKI  181

Query  1095  IRDYGKQMAGDDCVASRQDED  1115
             I+DYGKQMAG DCVA RQDED
Sbjct  182   IKDYGKQMAGADCVAGRQDED  244



>Chlamydia abortus strain 09_01, whole genome shotgun sequence
Sequence ID: NZ_FQLJ01000007.1 Length: 847
Range 1: 31 to 810

Score:155 bits(392), Expect:2e-37,
Method:Compositional matrix adjust.,
Identities:101/269(38%), Positives:149/269(55%), Gaps:10/269(3%)

Query  156  PVKLKPGMDGPKVKQWPLTEEKIKALVEICTEMEKEGKISKIGPENPYNTPVFAIKKKDS  215
            P+  K G D   V QW L+ EK++A++++  E  K G I      +P+NTP+F IKKK S
Sbjct  810  PIPWKTG-DPVWVPQWHLSSEKLEAVIQLVEEQLKLGHIEP--STSPWNTPIFVIKKK-S  643

Query  216  TKWRKLVDFRELNKRTQDFWEVQLGIPHPAGLKKKKSVTVLDVGDAYFSVPLDEDFRKYT  275
             KWR L D R +N++   F  VQ G+P  + L +  ++ ++D+ D +FS+PL    R
Sbjct  642  GKWRLLHDLRAINEQMNLFGPVQRGLPVLSALPRGWNLIIIDIKDCFFSIPLCPRDRPRF  463

Query  276  AFTIPSINNETPGIRYQYNVLPQGWKGSPAIFQSSMTKILEPFRKQNPDIVIYQYMDD-L  334
            AFTIPSIN+  P  RYQ+ VLPQG   SP + Q  + + L P R+Q P +++  YMDD L
Sbjct  462  AFTIPSINHMEPDKRYQWKVLPQGMSNSPTMCQLYVQEALLPVREQFPSLILLLYMDDIL  283

Query  335  YVGSDLEIGQHRTKIEELRQHLLRWGLTTPDKKHQKEPPFLWMGYELHPDKWTVQPIVLP  394
                DL + Q       L + L +WGL    +K Q      ++G  + PDK   Q + +
Sbjct  282  LCHKDLTMLQKAYPF--LLKTLSQWGLQIATEKVQISDTGQFLGSVVSPDKIVPQKVEIR  109

Query  395  EKDSWTVNDIQKLVGKLNWASQIYPGIKV  423
                 T+ND QKL+G +NW   + P +K+
Sbjct  108  RDHLHTLNDFQKLLGDINW---LRPFLKI  31



>Chlamydia trachomatis strain R33512extract, whole genome shotgun sequence
Sequence ID: NZ_CVNW01000006.1 Length: 1732
Range 1: 563 to 1384

Score:159 bits(402), Expect:4e-36,
Method:Compositional matrix adjust.,
Identities:104/280(37%), Positives:152/280(54%), Gaps:9/280(3%)

Query  156   PVKLKPGMDGPKVKQWPLTEEKIKALVEICTEMEKEGKISKIGPENPYNTPVFAIKKKDS  215
             P+  K G D   V QW L+ EK++A++++  E  K G I      +P+NTP+F IKKK S
Sbjct  563   PIPWKTG-DPVWVPQWHLSSEKLEAVIQLVEEQLKLGHIEP--STSPWNTPIFVIKKK-S  730

Query  216   TKWRKLVDFRELNKRTQDFWEVQLGIPHPAGLKKKKSVTVLDVGDAYFSVPLDEDFRKYT  275
              KWR L D R +N++   F  VQ G+P  + L +  ++ ++D+ D +FS+PL    R
Sbjct  731   GKWRLLHDLRAINEQMNLFGPVQRGLPVLSALPRGWNLIIIDIKDCFFSIPLCPRDRPRF  910

Query  276   AFTIPSINNETPGIRYQYNVLPQGWKGSPAIFQSSMTKILEPFRKQNPDIVIYQYMDD-L  334
             AFTIPSIN+  P  RYQ+ VLPQG   SP + Q  + + L P R+Q P +++  YMDD L
Sbjct  911   AFTIPSINHMEPDKRYQWKVLPQGMSNSPTMCQLYVQEALLPVREQFPSLILLLYMDDIL  1090

Query  335   YVGSDLEIGQHRTKIEELRQHLLRWGLTTPDKKHQKEPPFLWMGYELHPDKWTVQPIVLP  394
                 DL + Q       L + L +WGL    +K Q      ++G  + PDK   Q + +
Sbjct  1091  LCHKDLTMLQKAYPF--LLKTLSQWGLQIATEKVQISDTGQFLGSVVSPDKIVPQKVEIR  1264

Query  395   EKDSWTVNDIQKLVGKLNWASQIY--PGIKVRQLCKLLRG  432
                  T+ND QKL+G +NW       P  ++R L  +L G
Sbjct  1265  RDHLHTLNDFQKLLGDINWLRPFLKIPSAELRPLFGILEG  1384



>Neisseria gonorrhoeae strain GC-206 NODE_177_length_223_cov_0.803571_ID_20526, whole genome shotgun sequence
Sequence ID: NZ_SAWN01000176.1 Length: 223
Range 1: 2 to 223

Score:144 bits(362), Expect:5e-36,
Method:Compositional matrix adjust.,
Identities:68/74(92%), Positives:71/74(95%), Gaps:0/74(0%)

Query  140  QIGCTLNFPISPIETVPVKLKPGMDGPKVKQWPLTEEKIKALVEICTEMEKEGKISKIGP  199
            Q+GCTLNFPISPIETVPVKLKPGMDGPKVKQWPLTEEKIKAL  IC EMEKEGKI+KIGP
Sbjct  2    QLGCTLNFPISPIETVPVKLKPGMDGPKVKQWPLTEEKIKALTAICEEMEKEGKITKIGP  181

Query  200  ENPYNTPVFAIKKK  213
            ENPYNTPVFAI+KK
Sbjct  182  ENPYNTPVFAIRKK  223



>Chlamydia abortus strain 09_03, whole genome shotgun sequence
Sequence ID: NZ_FQLL01000003.1 Length: 4440
Range 1: 80 to 901

Score:163 bits(412), Expect:7e-36,
Method:Compositional matrix adjust.,
Identities:102/279(37%), Positives:151/279(54%), Gaps:8/279(2%)

Query  168  VKQWPLTEEKIKALVEICTEMEKEGKISKIGPENPYNTPVFAIKKKDSTKWRKLVDFREL  227
            V QW L+ EK++A++++  E  K G I      +P+NTP+F IKKK S KWR L D R +
Sbjct  80   VPQWHLSSEKLEAVIQLVEEQLKLGHIEP--STSPWNTPIFVIKKK-SGKWRLLHDLRAI  250

Query  228  NKRTQDFWEVQLGIPHPAGLKKKKSVTVLDVGDAYFSVPLDEDFRKYTAFTIPSINNETP  287
            N++   F  VQ G+P  + L +  ++ ++D+ D +FS+PL    R   AFTIPSIN+  P
Sbjct  251  NEQMNLFGPVQRGLPVLSALPRGWNLIIIDIKDCFFSIPLCPRDRPRFAFTIPSINHMEP  430

Query  288  GIRYQYNVLPQGWKGSPAIFQSSMTKILEPFRKQNPDIVIYQYMDD-LYVGSDLEIGQHR  346
              RYQ+ VLPQG   SP + Q  + + L P R+Q P +++  YMDD L    DL + Q
Sbjct  431  DKRYQWKVLPQGMSNSPTMCQLYVQEALLPVREQFPSLILLLYMDDILLCHKDLTMLQKA  610

Query  347  TKIEELRQHLLRWGLTTPDKKHQKEPPFLWMGYELHPDKWTVQPIVLPEKDSWTVNDIQK  406
                 L + L +WGL    +K Q      ++G  + PDK   Q + +      T+ND QK
Sbjct  611  YPF--LLKTLSQWGLQIATEKVQISDTGQFLGSVVSPDKIVPQKVEIRRDHLHTLNDFQK  784

Query  407  LVGKLNWASQIY--PGIKVRQLCKLLRGTKALTEVIPLT  443
            L+G +NW       P  ++R L  +L G   ++    LT
```

```
Sbjct  785   LLGDINWLRPFLKIPSAELRPLFSILEGDPHISSPRTLT  901



Range 2: 1364 to 2416

Score:92.4 bits(228), Expect:4e-14,
Method:Compositional matrix adjust.,
Identities:94/371(25%), Positives:153/371(41%), Gaps:45/371(12%)

Query  706   VLYVDGAANRETKLGKAGYVTNRGRQKVVTLTDTTNQKTELQAIYLALQDSGLEVNIVTD  765
             V+Y DG     +K G   YV N G+       + + Q  E   +   L+     +NIV+D
Sbjct  1364  VVYTDG-----SKTGIGAYVAN-GKVVSKQYNENSPQVVECLVVLEVLKTFLEPLNIVSD  1525

Query  766   SQYALGIIQA-------QPDQSESELVNQI-IEQLIKKEKVYLAWVPAHKGIGG-----N  812
             S Y +  +         +P    + +  QI +  L ++  VY+  V AH G+ G     N
Sbjct  1526  SCYVVNAVNLLEVAGVIKPSSRVANIFQQIQLVLLSRRSPVYITHVRAHSGLPGPMALGN  1705

Query  813   EQVDKLVSAGIRKVLFLDGIDKAQDEHEKYHSNWRAMASDFNLPPVVAKEIVASCDKCQL  872
             +  DK  +  +        ++ A++ H  +H     + S F+L    A++IV  C  C
Sbjct  1706  DLADK--ATKVVAAALSSPVEAARNFHNNFHVTAETLRSRFSLTRKEARDIVTQCQSCCE  1879

Query  873   KGEAMHGQVDCSP------GIWQLDCTHLE--GKV--ILVAVHVASGYIEAEVIPAETGQ  922
                  H  V  +P       +WQ+D TH+   GK+  + V++   SG + A  +  E
Sbjct  1880  FLPVPH--VGINPRGIRPLQVWQMDVTHVSSFGKLQYLHVSIDTCSGIMFASPLTGEKAS  2053

Query  923   ETAYFLLKLAGRW-PVKTIHTDNGSNFTGATVRAACWWAGIKQEFGIPYNPQSQGVVESM  981
                   L+    W   K + TDNG  +T    +  C    +    G+PYNPQ QG+VE
Sbjct  2054  HVIQHCLEAWSAWGKPKLLKTDNGPAYTSQKFQQFCRQMDVTHLTGLPYNPQGQGIVERA  2233

Query  982   NKELKKIIGQVRDQAEHLKTAVQMAVFIHNFKRKGGIGGYSA-GERIVDIIATDIQTKEL  1040
             ++ LK  +         LK      V+I + +   G+ G  A G  + D  AT +    L
Sbjct  2234  HRTLKSYL---------LKQKRGXPVYITHVRAHSGLPGPMALGNDLAD-KATKVVAAAL  2383

Query  1041  QKQITKIQNFR  1051
                +   +NF
Sbjct  2384  SSPVEAARNFH  2416



Range 3: 2255 to 3193

Score:88.6 bits(218), Expect:7e-13,
Method:Compositional matrix adjust.,
Identities:85/332(26%), Positives:143/332(43%), Gaps:46/332(13%)

Query  792   LIKKEK---VYLAWVPAHKGIGG-----NEQVDKLVSAGIRKVLFLDGIDKAQDEHEKYH  843
             L+K+++   VY+  V AH G+ G     N+  DK  +  +        ++ A++ H  +H
Sbjct  2255  LLKQKRGXPVYITHVRAHSGLPGPMALGNDLADK--ATKVVAAALSSPVEAARNFHNNFH  2428

Query  844   SNWRAMASDFNLPPVVAKEIVASCDKCQLKGEAMHGQVDCSP------GIWQLDCTHLE-  896
                  + S F+L    A++IV  C  C       H  V  +P       +WQ+D TH+
Sbjct  2429  VTAETLRSRFSLTRKEARDIVTQCQSCCEFLPVPH--VGINPRGIRPLQVWQMDVTHVSS  2602

Query  897   -GKV--ILVAVHVASGYIEAEVIPAETGQETAYFLLKLAGRW-PVKTIHTDNGSNFTGAT  952
              GK+  + V++   SG + A  +  E         L+    W   + + TDNG  +T
Sbjct  2603  FGKLQYLHVSIDTCSGIMFASPLTGEKASHVIQHCLEAWSAWGKPRLLKTDNGPAYTSQK  2782

Query  953   VRAACWWAGIKQEFGIPYNPQSQGVVESMNKELKK-IIGQVRDQAEHL-----KTAVQMA  1006
              +  C    +    G+PYNPQ QG+VE  ++ LK  +I Q R   E       + +V +A
Sbjct  2783  FQQFCRQMDVTHLTGLPYNPQGQGIVERAHRTLKAYLIKQKRGTFEETVPRAPRVSVSLA  2962

Query  1007  VFIHNFKRKGGIGGYSAGERIVDIIATDIQTKELQKQITKIQNFRVYYRDSRNPLWKGPA  1066
             +F  NF       G++A ER              E  +      N  V +++  +  W GP
Sbjct  2963  LFTLNFLNIDA-HGHTAAER---------HCSEPDR-----PNEMVKWKNVLDNKWYGPD  3097

Query  1067  KLLWKGEGAVVI--QDNSDIKVVPRRKAKIIR  1096
              +L +  GA+ +  Q+  +   VP R  + I+
Sbjct  3098  PILIRSRGAICVFPQNEDNPFWVPERLTRKIQ  3193



>Chlamydia trachomatis strain R4175extract, whole genome shotgun sequence
Sequence ID: NZ_CVNX01000006.1 Length: 2328
Range 1: 1202 to 2056

Score:160 bits(404), Expect:1e-35,
Method:Compositional matrix adjust.,
Identities:106/291(36%), Positives:156/291(53%), Gaps:9/291(3%)

Query  156   PVKLKPGMDGPKVKQWPLTEEKIKALVEICTEMEKEGKISKIGPENPYNTPVFAIKKKDS  215
             P+  K G D   V QW L+ EK++A++++  E  K G I      +P+NTP+F IKKK S
Sbjct  2056  PIPWKTG-DPVWVPQWHLSSEKLEAVIQLVEEQLKLGHIEP--STSPWNTPIFVIKKK-S  1889

Query  216   TKWRKLVDFRELNKRTQDFWEVQLGIPHPAGLKKKKSVTVLDVGDAYFSVPLDEDFRKYT  275
              KWR L D R +N++   F  VQ G+P  + L +  ++ ++D+ D +FS+PL    R
Sbjct  1888  GKWRLLHDLRAINEQMNLFGPVQRGLPVLSALPRGWNLIIIDIKDCFFSIPLCPRDRPRF  1709

Query  276   AFTIPSINNETPGIRYQYNVLPQGWKGSPAIFQSSMTKILEPFRKQNPDIVIYQYMDD-L  334
             AFTIPSIN+  P  RYQ+ VLPQG   SP + Q  + + L P R+Q P +++  YMDD L
Sbjct  1708  AFTIPSINHMEPDKRYQWKVLPQGMSNSPTMCQLYVQEALLPVREQFPSLILLLYMDDIL  1529

Query  335   YVGSDLEIGQHRTKIEELRQHLLRWGLTTPDKKHQKEPPFLWMGYELHPDKWTVQPIVLP  394
                 DL + Q       L + L +WGL    +K Q      ++G  + PDK   Q + +
Sbjct  1528  LCHKDLTMLQKAYPF--LLKTLSQWGLQIATEKVQISDTGQFLGSVVSPDKIVPQKVEIR  1355

Query  395   EKDSWTVNDIQKLVGKLNWASQIY--PGIKVRQLCKLLRGTKALTEVIPLT  443
                 T+ND QKL+G +NW       P  ++R L  +L G   ++    LT
Sbjct  1354  RDHLHTLNDFQKLLGDINWLRPFLKIPSAELRPLFGILEGDPHISSPRTLT  1202



>Chlamydia abortus strain 11_01, whole genome shotgun sequence
Sequence ID: NZ_FQLK01000002.1 Length: 5240
Range 1: 2674 to 3528

Score:162 bits(410), Expect:1e-35,
Method:Compositional matrix adjust.,
Identities:106/291(36%), Positives:156/291(53%), Gaps:9/291(3%)

Query  156   PVKLKPGMDGPKVKQWPLTEEKIKALVEICTEMEKEGKISKIGPENPYNTPVFAIKKKDS  215
             P+  K G D   V QW L+ EK++A++++  E  K G I      +P+NTP+F IKKK S
Sbjct  3528  PIPWKTG-DPVWVPQWHLSSEKLEAVIQLVEEQLKLGHIEP--STSPWNTPIFVIKKK-S  3361

Query  216   TKWRKLVDFRELNKRTQDFWEVQLGIPHPAGLKKKKSVTVLDVGDAYFSVPLDEDFRKYT  275
              KWR L D R +N++   F  VQ G+P  + L +  ++ ++D+ D +FS+PL    R
Sbjct  3360  GKWRLLHDLRAINEQMNLFGPVQRGLPVLSALPRGWNLIIIDIKDCFFSIPLCPRDRPRF  3181

Query  276   AFTIPSINNETPGIRYQYNVLPQGWKGSPAIFQSSMTKILEPFRKQNPDIVIYQYMDD-L  334
             AFTIPSIN+  P  RYQ+ VLPQG   SP + Q  + + L P R+Q P +++  YMDD L
```

```
Sbjct  3180  AFTIPSINHMEPDKRYQWKVLPQGMSNSPTMCQLYVQEALLPVREQFPSLILLLYMDDIL  3001

Query  335   YVGSDLEIGQHRTKIEELRQHLLRWGLTTPDKKHQKEPPFLWMGYELHPDKWTVQPIVLP  394
                 DL + Q       L + L +WGL    +K Q      ++G  + PDK   Q + +
Sbjct  3000  LCHKDLTMLQKAYPF--LLKTLSQWGLQIATEKVQISDTGQFLGSVVSPDKIVPQKVEIR  2827

Query  395   EKDSWTVNDIQKLVGKLNWASQIY--PGIKVRQLCKLLRGTKALTEVIPLT  443
                  T+ND QKL+G +NW       P  ++R L  +L G   ++    LT
Sbjct  2826  RDHLHTLNDFQKLLGDINWLRPFLKIPSAELRPLFSILEGDPHISSPRTLT  2674
```

Range 2: 1018 to 2211

Score:99.0 bits(245), Expect:4e-16,
Method:Compositional matrix adjust.,
Identities:105/423(25%), Positives:177/423(41%), Gaps:57/423(13%)

```
Query  706   VLYVDGAANRETKLGKAGYVTNRGRQKVVTLTDTTNQKTELQAIYLALQDSGLEVNIVTD  765
             V+Y DG     +K G   YV N G+       + + Q  E   +   L+     +NIV+D
Sbjct  2211  VVYTDG-----SKTGIGAYVAN-GKVVSKQYNENSPQVVECLVVLEVLKTFLEPLNIVSD  2050

Query  766   SQYALGIIQA-------QPDQSESELVNQI-IEQLIKKEKVYLAWVPAHKGIGG-----N  812
             S Y +  +         +P    + +  QI +  L ++  VY+  V AH G+ G     N
Sbjct  2049  SCYVVNAVNLLEVAGVIKPSSRVANIFQQIQLVLLSRRSPVYITHVRAHSGLPGPMALGN  1870

Query  813   EQVDKLVSAGIRKVLFLDGIDKAQDEHEKYHSNWRAMASDFNLPPVVAKEIVASCDKCQL  872
             +  DK  +  +        ++ A++ H  +H     + S F+L    A++IV  C  C
Sbjct  1869  DLADK--ATKVVAAALSSPVEAARNFHNNFHVTAETLRSRFSLTRKEARDIVTQCQSCCE  1696

Query  873   KGEAMHGQVDCSP------GIWQLDCTHLE--GKV--ILVAVHVASGYIEAEVIPAETGQ  922
                  H  V  +P       +WQ+D TH+   GK+  + V++   SG + A  +  E
Sbjct  1695  FLPVPH--VGINPRGIRPLQVWQMDVTHVSSFGKLQYLHVSIDTCSGIMFASPLTGEKAS  1522

Query  923   ETAYFLLKLAGRW-PVKTIHTDNGSNFTGATVRAACWWAGIKQEFGIPYNPQSQGVVESM  981
                   L+    W   + + TDNG  +T    +  C    +    G+PYNPQ QG+VE
Sbjct  1521  HVIQHCLEAWSAWGKPRLLKTDNGPAYTSQKFQQFCRQMDVTHLTGLPYNPQGQGIVERA  1342

Query  982   NKELKK-IIGQVRDQAEHL-----KTAVQMAVFIHNFKRKGGIGGYSAGERIVDIIATDI  1035
             ++ LK  +I Q R   E       + +V +A+F  NF       G++A ER
Sbjct  1341  HRTLKAYLIKQKRGTFEETVPRAPRVSVSLALFTLNFLNIDA-HGHTAAER---------  1192

Query  1036  QTKELQKQITKIQNFRVYYRDSRNPLWKGPAKLLWKGEGAVVI--QDNSDIKVVPRRKAK  1093
                E  +      N  V +++  +  W GP  +L +  GA+ +  Q+  +   +P R  +
Sbjct  1191  HCSEPDR-----PNEMVKWKNVLDNKWYGPDPILIRSRGAICVFPQNEDNPFWIPERLTR  1027

Query  1094  IIR  1096
              I+
Sbjct  1026  KIQ  1018
```

>Chlamydia abortus strain 15a, whole genome shotgun sequence
Sequence ID: NZ_FQLM01000004.1 Length: 3222
Range 1: 1236 to 2090

Score:160 bits(406), Expect:3e-35,
Method:Compositional matrix adjust.,
Identities:106/291(36%), Positives:156/291(53%), Gaps:9/291(3%)

```
Query  156   PVKLKPGMDGPKVKQWPLTEEKIKALVEICTEMEKEGKISKIGPENPYNTPVFAIKKKDS  215
             P+  K G D   V QW L+ EK++A++++  E  K G I      +P+NTP+F IKKK S
Sbjct  2090  PIPWKTG-DPVWVPQWHLSSEKLEAVIQLVEEQLKLGHIEP--STSPWNTPIFVIKKK-S  1923

Query  216   TKWRKLVDFRELNKRTQDFWEVQLGIPHPAGLKKKKSVTVLDVGDAYFSVPLDEDFRKYT  275
              KWR L D R +N++   F  VQ G+P  + L +  ++ ++D+ D +FS+PL    R
Sbjct  1922  GKWRLLHDLRAINEQMNLFGPVQRGLPVLSALPRGWNLIIIDIKDCFFSIPLCPRDRPRF  1743

Query  276   AFTIPSINNETPGIRYQYNVLPQGWKGSPAIFQSSMTKILEPFRKQNPDIVIYQYMDD-L  334
             AFTIPSIN+  P  RYQ+ VLPQG   SP + Q  + + L P R+Q P +++  YMDD L
Sbjct  1742  AFTIPSINHMEPDKRYQWKVLPQGMSNSPTMCQLYVQEALLPVREQFPSLILLLYMDDIL  1563

Query  335   YVGSDLEIGQHRTKIEELRQHLLRWGLTTPDKKHQKEPPFLWMGYELHPDKWTVQPIVLP  394
                 DL + Q       L + L +WGL    +K Q      ++G  + PDK   Q + +
Sbjct  1562  LCHKDLTMLQKAYPF--LLKTLSQWGLQIATEKVQISDTGQFLGSVVSPDKIVPQKVEIR  1389

Query  395   EKDSWTVNDIQKLVGKLNWASQIY--PGIKVRQLCKLLRGTKALTEVIPLT  443
                  T+ND QKL+G +NW       P  ++R L  +L G   ++    LT
Sbjct  1388  RDHLHTLNDFQKLLGDINWLRPFLKIPSAELRPLFSILEGDPHISSPRTLT  1236
```

>Chlamydia abortus strain 09_02, whole genome shotgun sequence
Sequence ID: NZ_FQLN01000004.1 Length: 4507
Range 1: 2379 to 3233

Score:160 bits(404), Expect:6e-35,
Method:Compositional matrix adjust.,
Identities:106/291(36%), Positives:156/291(53%), Gaps:9/291(3%)

```
Query  156   PVKLKPGMDGPKVKQWPLTEEKIKALVEICTEMEKEGKISKIGPENPYNTPVFAIKKKDS  215
             P+  K G D   V QW L+ EK++A++++  E  K G I      +P+NTP+F IKKK S
Sbjct  2379  PIPWKTG-DPVWVPQWHLSSEKLEAVIQLVEEQLKLGHIEP--STSPWNTPIFVIKKK-S  2546

Query  216   TKWRKLVDFRELNKRTQDFWEVQLGIPHPAGLKKKKSVTVLDVGDAYFSVPLDEDFRKYT  275
              KWR L D R +N++   F  VQ G+P  + L +  ++ ++D+ D +FS+PL    R
Sbjct  2547  GKWRLLHDLRAINEQMNLFGPVQRGLPVLSALPRGWNLIIIDIKDCFFSIPLCPRDRPRF  2726

Query  276   AFTIPSINNETPGIRYQYNVLPQGWKGSPAIFQSSMTKILEPFRKQNPDIVIYQYMDD-L  334
             AFTIPSIN+  P  RYQ+ VLPQG   SP + Q  + + L P R+Q P +++  YMDD L
Sbjct  2727  AFTIPSINHMEPDKRYQWKVLPQGMSNSPTMCQLYVQEALLPVREQFPSLILLLYMDDIL  2906

Query  335   YVGSDLEIGQHRTKIEELRQHLLRWGLTTPDKKHQKEPPFLWMGYELHPDKWTVQPIVLP  394
                 DL + Q       L + L +WGL    +K Q      ++G  + PDK   Q + +
Sbjct  2907  LCHKDLTMLQKAYPF--LLKTLSQWGLQIATEKVQISDTGQFLGSVVSPDKIVPQKVEIR  3080

Query  395   EKDSWTVNDIQKLVGKLNWASQIY--PGIKVRQLCKLLRGTKALTEVIPLT  443
                  T+ND QKL+G +NW       P  ++R L  +L G   ++    LT
Sbjct  3081  RDHLHTLNDFQKLLGDINWLRPFLKIPSAELRPLFGILEGDPHISSPRTLT  3233
```

>Chlamydia abortus strain 09_05, whole genome shotgun sequence
Sequence ID: NZ_FQLG01000005.1 Length: 4831
Range 1: 3055 to 3834

Score:158 bits(399), Expect:3e-34,
Method:Compositional matrix adjust.,
Identities:101/269(38%), Positives:149/269(55%), Gaps:10/269(3%)

```
Query  156   PVKLKPGMDGPKVKQWPLTEEKIKALVEICTEMEKEGKISKIGPENPYNTPVFAIKKKDS  215
             P+  K G D   V QW L+ EK++A++++  E  K G I      +P+NTP+F IKKK S
Sbjct  3834  PIPWKTG-DPVWVPQWHLSSEKLEAVIQLVEEQLKLGHIEP--STSPWNTPIFVIKKK-S  3667

Query  216   TKWRKLVDFRELNKRTQDFWEVQLGIPHPAGLKKKKSVTVLDVGDAYFSVPLDEDFRKYT  275
              KWR L D R +N++   F  VQ G+P  + L +  ++ ++D+ D +FS+PL    R
Sbjct  3666  GKWRLLHDLRAINEQMNLFGPVQRGLPVLSALPRGWNLIIIDIKDCFFSIPLCPRDRPRF  3487

Query  276   AFTIPSINNETPGIRYQYNVLPQGWKGSPAIFQSSMTKILEPFRKQNPDIVIYQYMDD-L  334
             AFTIPSIN+  P  RYQ+ VLPQG   SP + Q  + + L P R+Q P +++  YMDD L
Sbjct  3486  AFTIPSINHMEPDKRYQWKVLPQGMSNSPTMCQLYVQEALLPVREQFPSLILLLYMDDIL  3307

Query  335   YVGSDLEIGQHRTKIEELRQHLLRWGLTTPDKKHQKEPPFLWMGYELHPDKWTVQPIVLP  394
                 DL + Q       L + L +WGL    +K Q      ++G  + PDK   Q + +
Sbjct  3306  LCHKDLTMLQKAYPF--LLKTLSQWGLQIATEKVQISDTGQFLGSVVSPDKIVPQKVEIR  3133

Query  395   EKDSWTVNDIQKLVGKLNWASQIYPGIKV  423
                  T+ND QKL+G +NW   + P +K+
Sbjct  3132  RDHLHTLNDFQKLLGDINW---LRPFLKI  3055
```

Range 2: 1318 to 2511

Score:101 bits(251), Expect:1e-16,
Method:Compositional matrix adjust.,
Identities:107/423(25%), Positives:177/423(41%), Gaps:57/423(13%)

```
Query  706   VLYVDGAANRETKLGKAGYVTNRGRQKVVTLTDTTNQKTELQAIYLALQDSGLEVNIVTD  765
             V+Y DG     +K G   YV N G+       + + Q  E   +   L+     +NIV+D
Sbjct  2511  VVYTDG-----SKTGIGAYVAN-GKVVSKQYNENSPQVVECLVVLEVLKTFLEPLNIVSD  2350

Query  766   SQYALGIIQA-------QPDQSESELVNQI-IEQLIKKEKVYLAWVPAHKGIGG-----N  812
             S Y +  +         +P    + +  QI +  L ++  VY+  V AH G+ G     N
Sbjct  2349  SCYVVNAVNLLEVAGVIKPSSRVANIFQQIQLVLLSRRSPVYITHVRAHSGLPGPMALGN  2170

Query  813   EQVDKLVSAGIRKVLFLDGIDKAQDEHEKYHSNWRAMASDFNLPPVVAKEIVASCDKCQL  872
             +  DK  +  +        ++ A++ H  +H     + S F+L    A++IV  C  C
Sbjct  2169  DLADK--ATKVVAAALSSPVEAARNFHNNFHVTAETLRSRFSLTRKEARDIVTQCQSCCE  1996

Query  873   KGEAMHGQVDCSP------GIWQLDCTHLE--GKV--ILVAVHVASGYIEAEVIPAETGQ  922
                  H  V  +P       +WQ+D TH+   GK+  + V++   SG + A  +  E
Sbjct  1995  FLPVPH--VGINPRGIRPLQVWQMDVTHVSSFGKLQYLHVSIDTCSGIMFASPLTGEKAS  1822

Query  923   ETAYFLLKLAGRW-PVKTIHTDNGSNFTGATVRAACWWAGIKQEFGIPYNPQSQGVVESM  981
                   L+    W   + + TDNG  +T    +  C    +    G+PYNPQ QG+VE
Sbjct  1821  HVIQHCLEAWSAWGKPRLLKTDNGPAYTSQKFQQFCRQMDVTHLTGLPYNPQGQGIVERA  1642

Query  982   NKELKK-IIGQVRDQAEHL-----KTAVQMAVFIHNFKRKGGIGGYSAGERIVDIIATDI  1035
             ++ LK  +I Q R   E       + +V MA+F  NF       G++A ER
Sbjct  1641  HRTLKAYLIKQKRGTFEETVPRAPRVSVSMALFTLNFLNIDA-HGHTAAER---------  1492

Query  1036  QTKELQKQITKIQNFRVYYRDSRNPLWKGPAKLLWKGEGAVVI--QDNSDIKVVPRRKAK  1093
                E  +      N  V +++  +  W GP  +L +  GA+ +  Q+  +   VP  R  +
Sbjct  1491  HCSEPDR-----PNEMVKWKNVLDNKWYGPDPILIRSRGAICVFPQNEDNPFWVPERLTR  1327

Query  1094  IIR  1096
              I+
Sbjct  1326  KIQ  1318
```

>Methylobacterium radiotolerans strain MAMP 4754 MP-M-radiotolerans_contig_1227, whole genome shotgun sequence
Sequence ID: NZ_NKQS01001213.1 Length: 279
Range 1: 3 to 254

Score:136 bits(343), Expect:3e-33,
Method:Compositional matrix adjust.,
Identities:68/84(81%), Positives:74/84(88%), Gaps:0/84(0%)

```
Query  995   QAEHLKTAVQMAVFIHNFKRKGGIGGYSAGERIVDIIATDIQTKELQKQITKIQNFRVYY  1054
             QAEHLKTAV MAVFIHNFKRKGGIGGYSAGERI+DIIATDIQTKELQKQ  KIQN++V Y
Sbjct  3     QAEHLKTAVLMAVFIHNFKRKGGIGGYSAGERIIDIIATDIQTKELQKQNIKIQNYQVDY  182

Query  1055  RDSRNPLWKGPAKLLWKGEGAVVI  1078
             RDS +   KG AKL+WKG+GAV I
Sbjct  183   RDSIDHSRKGSAKLIWKGKGAVGI  254
```

>Pseudomonas aromaticivorans strain MAP12 34, whole genome shotgun sequence
Sequence ID: NZ_JAHRGL010000035.1 Length: 7031
Range 1: 3 to 803

Score:143 bits(361), Expect:1e-29,
Method:Compositional matrix adjust.,
Identities:91/271(34%), Positives:147/271(54%), Gaps:6/271(2%)

```
Query  168  VKQWPLTEEKIKALVEICTEMEKEGKISKIGPENPYNTPVFAIKKKDSTKWRKLVDFREL  227
            V QWP+  E++ +L ++         + +     P+N+ VF IKKK S KWR L D R +
Sbjct  3    VGQWPMNSEELVSLEKLVH*QLDA*HVEETT--FPWNSTVFVIKKK-SGKWRMLTDLRAI  173

Query  228  NKRTQDFWEVQLGIPHPAGLKKKKSVTVLDVGDAYFSVPLDEDFRKYTAFTIPSINNETP  287
            NK  Q    +Q G+P P+ + K   + V+D+ D +F++PL E+ R+  AFT+P++NN  P
Sbjct  174  NKVIQPMGSLQPGMPLPSLIPKGWPIIVIDLKDCFFTIPLQENDREKFAFTVPTLNNSQP  353

Query  288  GIRYQYNVLPQGWKGSPAIFQSSMTKILEPFRKQNPDIVIYQYMDDLYVGSDLEIGQHRT  347
              RYQ+ VLPQG   SP + Q  + + LE  RK+ P  ++Y YMDD+ + SD
Sbjct  354  VRRYQWRVLPQGMLNSPTLCQHFVQQPLEIIRKKLPQSLVYHYMDDILL-SDSNKETLER  530

Query  348  KIEELRQHLLRWGLTTPDKKHQKEPPFLWMGYELHPDKWTVQPIVLPEKDSWTVNDIQKL  407
              E +++ L RWGL    +K Q+      ++GY++   +    Q + +       T+N +QKL
Sbjct  531  MFEMVKEVLPRWGLQIAPEKIQRGDSINYLGYKIDLQRIRPQKVQIRRDRYQTLNSLQKL  710

Query  408  VGKLNWASQI--YPGIKVRQLCKLLRGTKAL  436
            +G+++    I    G  ++ L   L+G K L
Sbjct  711  LGEISQLQTIIGVEGHDLKHLKMALKGDKDL  803
```

Range 2: 1278 to 2393

Score:114 bits(285), Expect:1e-20,
Method:Compositional matrix adjust.,
Identities:115/400(29%), Positives:176/400(44%), Gaps:53/400(13%)

```
Query  703   IRKVLYVDGAANRETKLG-KAGYVTNRGRQKVVTLTDTTNQKTELQAIYLALQDSGLEVN  761
             I  VL     AN+  K G KAG V+     KVV    T+ QK EL AI + L D    +N
Sbjct  1278  ISGVLTFYTDANKSGKAGYKAGEVS-----KVVQSPYTSVQKAELYAILMVLMDFTEPLN  1442
```

```
Query  762   IVTDSQYA----LGIIQAQ--PDQSE-SELVNQIIEQLIKKEK-VYLAWVPAHKGI----  809
             IVTDSQYA    L I  A+  PD +E + L  Q+ E +  +   +Y+  + +H G+
Sbjct  1443  IVTDSQYAERVVLHIETAEFIPDNTELTSLFLQLQETIRNRSNPIYITHIRSHTGLPGPL  1622

Query  810   -GGNEQVDKLVSAGIRKVLFLDGIDKAQDEHEKYHSNWRAMASDFNLPPVVAKEIVASCD  868
               GN+++D+L+         +  + +A + H+K+H N + +  DF++    AKEIV +C
Sbjct  1623  AQGNDEIDRLL---------IGSVLEASEFHKKHHVNSKGLKKDFSITWQQAKEIVRNCP  1775

Query  869   KCQLKGEAMHGQVDCSPG------IWQLDCTHLEGKVILVAVH----VASGYIEAEVIPA  918
              C    +       C+P       +WQ+D  H      L  VH      SG+  A  + +
Sbjct  1776  TCSFYNQT-PLPAGCNPKGIRRNEVWQMDVFHFAEFGNLKYVHHTIDTFSGFQWATALNS  1952

Query  919   ETGQETAYFLLKLAGRWPVKT-IHTDNGSNFTGATVRAACWWAGIKQEFGIPYNPQSQGV  977
             E         LL++     +   I TDN   +    +     +  IK   GIP+NP  Q V
Sbjct  1953  EKADSVITHLLEVMAVMGIPAQIKTDNAPAYVSTKLEQFFKYYNIKHVTGIPHNPTGQAV  2132

Query  978   VESMNKELKKIIGQVRDQAEHLKTAVQMAVFIHNFKRKGGIGGYSAGERIVDIIATDIQT  1037
             VE  N+ LK+++ +   + +  K  +  A+   NF      G  +A         T  +T
Sbjct  2133  VERSNRTLKEMLNKQAWKTKPPKHRLHNALLTLNFLNANEKGQTAAKRH-----WTREKT  2297

Query  1038  KELQKQITKIQNFRVYYRDSRNPLWKGPAKLLWKGEGAVV  1077
              EL        N  VY++D    +WK    L W    A+V
Sbjct  2298  AEL--------NQPVYFKDVLTSVWKSGHVL*WCRGFALV  2393
```

Range 3: 6287 to 7015

Score:72.0 bits(175), Expect:1e-07,
Method:Compositional matrix adjust.,
Identities:70/248(28%), Positives:106/248(42%), Gaps:58/248(23%)

```
Query  38    ADRQGTVSFNFPQVTLWQ------RPLVTIKIGGQLKEALLDTGADDTVLEEMSLPGRWK  91
             +D + +++    +   WQ       RP + +KI  ++   L+DTGAD T++ + S P +W
Sbjct  6287  SDEESSMNGQNEKRIFWQASINDKRPQLKVKINNKVISGLVDTGADVTIITQKSWPQKWP  6466

Query  92    PKMIG----GIGGFIKVRQYDQILIEIC--GHKAIGTVLVGPTPVNIIGRNLLTQIGCTL  145
              +       GIG   +VRQ    ++ I   G K I     V    +N+ GR+LL Q    +
Sbjct  6467  LREANVQFLGIGTLSRVRQSVNSVVCIGPEGQKGILKPYVADIAINLWGRDLLQQWNTQI  6646

Query  146   NFPISPIE-----------------------TVPVKLKPGM-DGPK--------------  167
             N P  P+                        T+    K G  DGP
Sbjct  6647  NIP--PVSDTNYVQSLDSRKDLVRRYGKRLPTIHAVQKLGTNDGPSEEPKALPLKWLTDE  6820

Query  168   ---VKQWPLTEEKIKALVEICTEMEKEGKISKIGPENPYNTPVFAIKKKDSTKWRKLVDF  224
                V QWP+T EK++AL ++  E    G I +    + +++PVF  KKK S K R L D
Sbjct  6821  PVWVGQWPMTSEKLEALEKLVQEQLDAGHIEE--STSSWSSPVFVTKKK-SDK*RMLTDL  6991

Query  225   RELNKRTQ  232
             R +NK  Q
Sbjct  6992  RVINKAIQ  7015
```

>Klebsiella pneumoniae strain EuSCAPE_IL017, whole genome shotgun sequence
Sequence ID: NZ_UKKG02000044.1 Length: 325
Range 1: 73 to 324

Score:125 bits(313), Expect:5e-29,
Method:Compositional matrix adjust.,
Identities:64/88(73%), Positives:73/88(82%), Gaps:4/88(4%)

```
Query  2    QGKAREFSSEQTRANSPTRRELQVWGRDNNSPSEAGADRQGTVSFNFPQVTLWQRPLVTI  61
            QG+AREF  EQ RA SPT RELQV G  +N  SEAG +RQG  S NFPQ+TLWQRPLV+I
Sbjct  73   QGEAREFPPEQARAISPTSRELQVRG--DNPRSEAGVERQG--SLNFPQITLWQRPLVSI  240

Query  62   KIGGQLKEALLDTGADDTVLEEMSLPGR  89
            K+GGQ+KEALLDTGADDTVLE+M LPG+
Sbjct  241  KVGGQIKEALLDTGADDTVLEDMHLPGK  324
```

>Klebsiella pneumoniae strain EuSCAPE_MT012, whole genome shotgun sequence
Sequence ID: NZ_UKIX02000067.1 Length: 385
Range 1: 226 to 384

Score:105 bits(262), Expect:9e-22,
Method:Compositional matrix adjust.,
Identities:51/53(96%), Positives:51/53(96%), Gaps:0/53(0%)

```
Query  1063  KGPAKLLWKGEGAVVIQDNSDIKVVPRRKAKIIRDYGKQMAGDDCVASRQDED  1115
             KGPAKLLWKGEGAVVIQDNSDIKVVPRRKAKIIRDYGKQMAG DCVA RQDED
Sbjct  384   KGPAKLLWKGEGAVVIQDNSDIKVVPRRKAKIIRDYGKQMAGADCVAGRQDED  226
```

>Mycobacterium tuberculosis strain 32845_S29 214, whole genome shotgun sequence
Sequence ID: NZ_QTBV01000045.1 Length: 223
Range 1: 28 to 222

Score:94.4 bits(233), Expect:1e-18,
Method:Compositional matrix adjust.,
Identities:48/78(62%), Positives:58/78(74%), Gaps:13/78(16%)

```
Query  5    AREFSSEQTRANSPTRRELQVWGRDNNSPSEAGADRQGTVSFNFPQVTLWQRPLVTIKIG  64
            +R+   EQ RA+SPT RELQ           AGA+RQG  + N PQ+TLWQRPLV+IK+G
Sbjct  28   SRKLPPEQARADSPTSRELQ-----------AGAERQG--ALNLPQITLWQRPLVSIKVG  168

Query  65   GQLKEALLDTGADDTVLE  82
            GQ++EALLDTGADDTVLE
Sbjct  169  GQIREALLDTGADDTVLE  222
```

>Acinetobacter baumannii strain P4 contig_P4_92, whole genome shotgun sequence
Sequence ID: NZ_JBBXLU010000071.1 Length: 1226
Range 1: 333 to 488

Score:103 bits(257), Expect:2e-18,
Method:Compositional matrix adjust.,
Identities:48/52(92%), Positives:52/52(100%), Gaps:0/52(0%)

```
Query  1012  FKRKGGIGGYSAGERIVDIIATDIQTKELQKQITKIQNFRVYYRDSRNPLWK  1063
             FKRKGGIGGYSAGERIVDIIATDIQTKELQKQITKIQNFRVYYRDSR+P+++
Sbjct  333   FKRKGGIGGYSAGERIVDIIATDIQTKELQKQITKIQNFRVYYRDSRDPVYR  488
```

```
>Chlamydia trachomatis strain R33512extract, whole genome shotgun sequence
Sequence ID: NZ_CVNW01000009.1 Length: 1310
Range 1: 31 to 1164

Score:95.9 bits(237), Expect:7e-16,
Method:Compositional matrix adjust.,
Identities:103/403(26%), Positives:169/403(41%), Gaps:55/403(13%)

Query  706   VLYVDGAANRETKLGKAGYVTNRGRQKVVTLTDTTNQKTELQAIYLALQDSGLEVNIVTD  765
             V+Y DG     +K G   YV N G+       + + Q  E   +   L+     +NIV+D
Sbjct  1164  VVYTDG-----SKTGIGAYVAN-GKVVSKQYNENSPQVVECLVVLEVLKTFLEPLNIVSD  1003

Query  766   SQY---ALGIIQAQPDQSESELVNQIIEQ-----LIKKEKVYLAWVPAHKGIGG-----N  812
             S Y   A+ +++       S  V  I +Q     L ++  VY+  V AH G+ G     N
Sbjct  1002  SCYVVNAVNLLEVAGVIKPSSRVANIFQQIQLVLLSRRFPVYITHVRAHSGLPGPMALGN  823

Query  813   EQVDKLVSAGIRKVLFLDGIDKAQDEHEKYHSNWRAMASDFNLPPVVAKEIVASCDKCQL  872
             +  DK  +  +        ++ A++ H  +H     + S F+L    A++IV  C  C
Sbjct  822   DLADK--ATKVVAAALSSPVEAARNFHNNFHVTAETLRSRFSLTRKEARDIVTQCQSCCE  649

Query  873   KGEAMHGQVDCSPG------IWQLDCTHLE--GKV--ILVAVHVASGYIEAEVIPAETGQ  922
                  H  V  +P       +WQ+D TH+   GK+  + V++   SG + A  +  E
Sbjct  648   FLPVPH--VGINPRGIRPLQVWQMDVTHVSSFGKLQYLHVSIDTCSGIMFASPLTGEKAS  475

Query  923   ETAYFLLKLAGRW-PVKTIHTDNGSNFTGATVRAACWWAGIKQEFGIPYNPQSQGVVESM  981
                   L+    W   + + TDNG  +T    +  C    +    G+PYNPQ QG+VE
Sbjct  474   HVIQHCLEAWSAWGKPRLLKTDNGPAYTSQKFQQFCRQMDVTHLTGLPYNPQGQGIVERA  295

Query  982   NKELKK-IIGQVRDQAEHL-----KTAVQMAVFIHNFKRKGGIGGYSAGERIVDIIATDI  1035
             ++ LK  +I Q R   E       + +V +A+F  NF       G++A ER
Sbjct  294   HRTLKAYLIKQKRGTFEETVPRAPRVSVSLALFTLNFLNIDA-HGHTAAER---------  145

Query  1036  QTKELQKQITKIQNFRVYYRDSRNPLWKGPAKLLWKGEGAVVI  1078
                E  +      N  V +++  +  W GP  +L +  GA+ +
Sbjct  144   HCSEPDR-----PNEMVKWKNVLDNKWYGPDPILIRSRGAICV  31



>Acinetobacter baumannii strain E12 contig_E12_75, whole genome shotgun sequence
Sequence ID: NZ_JBBXKS010000057.1 Length: 514
Range 1: 2 to 127

Score:82.8 bits(203), Expect:2e-13,
Method:Compositional matrix adjust.,
Identities:38/42(90%), Positives:42/42(100%), Gaps:0/42(0%)

Query  1022  SAGERIVDIIATDIQTKELQKQITKIQNFRVYYRDSRNPLWK  1063
             SAGERIVDIIATDIQTKELQKQITKIQNFRVYYRDSR+P+++
Sbjct  2     SAGERIVDIIATDIQTKELQKQITKIQNFRVYYRDSRDPVYR  127



>Tumebacillus flagellatus strain GST4 Contig077, whole genome shotgun sequence
Sequence ID: NZ_JMIR01000077.1 Length: 2091
Range 1: 1295 to 1816

Score:89.0 bits(219), Expect:3e-13,
Method:Compositional matrix adjust.,
Identities:54/182(30%), Positives:97/182(53%), Gaps:15/182(8%)

Query  168   VKQWPLTEEKIKALVEICTEMEKEGKISKIGPENPYNTPVFAIKKKDSTKWRKLVDFREL  227
             +KQ+P++++    +      +  +G +     ++P+NTP+  +KK  +  +R + D RE+
Sbjct  1816  IKQYPMSQKARLGIKPHIQRLLDQGILVPC--QSPWNTPLLPVKKPGTNDYRPVQDLREV  1643

Query  228   NKRTQDFWEVQLGIPHPAGL-----KKKKSVTVLDVGDAYFSVPLDEDFRKYTAFTIPSI  282
             NKR +D   +   +P+P  L        +  TVLD+ DA+F + L    +   AF
Sbjct  1642  NKRVED---IHPTVPNPYNLLSGLPPSHQWYTVLDLKDAFFCLRLHPTSQPLFAF---EW  1481

Query  283   NNETPGIRYQ--YNVLPQGWKGSPAIFQSSMTKILEPFRKQNPDIVIYQYMDDLYVGSDL  340
              +   GI  Q  +  LPQG+K SP +F  ++ + L  FR Q+PD+++ QY+DDL + +
Sbjct  1480  RDPEMGISGQLTWTRLPQGFKNSPTLFDEALHRDLADFRIQHPDLILLQYVDDLLLAATS  1301

Query  341   EI  342
             E+
Sbjct  1300  EL  1295



>Listeria monocytogenes strain 1827/2,1 NODE_48_length_4618_cov_6.857588, whole genome shotgun sequence
Sequence ID: NZ_JAENKW010000028.1 Length: 4618
Range 1: 1715 to 2731

Score:89.0 bits(219), Expect:5e-13,
Method:Compositional matrix adjust.,
Identities:91/354(26%), Positives:153/354(43%), Gaps:73/354(20%)

Query  57    PLVTIKIGGQLKEALLDTGADDTVLEEMSLPGRWKPKMIGGIGGFIKVRQYDQILI----  112
             P V + + G+ +  L+DTGA  +VL   S     +   I G  G    +++ + L+
Sbjct  1715  PRVQLDVAGRSENFLVDTGATYSVLTSYSRAFSSQTCTILGATGKTTTKRFTRALLCCWD  1894

Query  113   -EICGHKAIGTVLVGPTPVNIIGRNLLTQIGCTLNF-----------------PISP---  151
              +I  H+ +   +V   P  ++GR++LT++G TL                   PI+P
Sbjct  1895  GQIFSHQFL---VVPECPTPLLGRDILTKLGTTLVMGSFSAPRALQLLVTTEEPITPSIE  2065

Query  152   -------------------------IETVPVKLKPGMDGPKVKQWPLTEEKIKALVEICT  186
                                       E V + L+     P  KQ+PL  E  + L  +
Sbjct  2066  RDQKLWEDKINPQVWDQGIPGRAHQAEPVIIVLRDPTRFPNRKQYPLKREAREGLQPLIN  2245

Query  187   EMEKEGKISKIGPENPYNTPVFAIKKKDSTKWRKLVDFRELNKRTQDFWEVQLGIPHPAG  246
             +    G +  +   +P NTP+ ++KKKD T W  + D R +N+           +P+P
Sbjct  2246  KFLACGLL--VPTSSPCNTPILSVKKKDGT-W*MVQDLRIINEAVVPLHPT---VPNPYV  2407

Query  247   L-----KKKKSVTVLDVGDAYFSVPLDEDFRKYTAFTIPSINNETPGIRYQ---YNVLPQ  298
             +        K  TVLD+ DA+F +PL ++ +   AF        E PG ++Q   + VLPQ
Sbjct  2408  ILGEIPPSAKWFTVLDLKDAFFCIPLAKESQYLFAF-----EWEAPGEKHQQMTWTVLPQ  2572

Query  299   GWKGSPAIFQSSMTKILEPFRKQNPDIVIYQYMDDLYVGSDLEIGQHRTKIEEL  352
             G++ SP +F  ++++ L       P+  I QY+DDL + S  E    +  I+ L
Sbjct  2573  GFRDSPHLFGQALSQDLL-DLDLGPNGKILQYVDDLLICSPDEKSAQQHAIQVL  2731



>Chlamydia abortus strain 09_01, whole genome shotgun sequence
Sequence ID: NZ_FQLJ01000005.1 Length: 1528
Range 1: 357 to 1277

Score:85.5 bits(210), Expect:2e-12,
Method:Compositional matrix adjust.,
Identities:88/319(28%), Positives:139/319(43%), Gaps:39/319(12%)
```

```
Query  706   VLYVDGAANRETKLGKAGYVTNRGRQKVVTLTDTTNQ-KTELQAIYLALQDSGLE-VNIV  763
             V+Y DG     +K G   YV N    KVV+     N  +     + L +  + LE +NIV
Sbjct  357   VVYTDG-----SKTGIGAYVAN---GKVVSKQYNENSPQVVECLVVLEVLKTFLEPLNIV  512

Query  764   TDSQY---ALGIIQAQPDQSESELVNQIIEQ-----LIKKEKVYLAWVPAHKGIGG----  811
             +DS Y   A+ +++       S  V  I +Q     L ++  VY+  V AH G+ G
Sbjct  513   SDSCYVVNAVNLLEVAGVIKPSSRVANIFQQIQLVLLSRRSPVYITHVRAHSGLPGPMAL  692

Query  812   -NEQVDKLVSAGIRKVLFLDGIDKAQDEHEKYHSNWRAMASDFNLPPVVAKEIVASCDKC  870
              N+  DK  +  +        ++ A++ H  +H     + S F+L    A++IV  C  C
Sbjct  693   GNDLADK--ATKVVAAALSSPVEAARNFHNNFHVTAETLRSRFSLTRKEARDIVTQCQSC  866

Query  871   QLKGEAMHGQVDCSPG------IWQLDCTHLE--GKV--ILVAVHVASGYIEAEVIPAET  920
                    H  V  +P       +WQ+D TH+   GK+  + V++   SG + A  +  E
Sbjct  867   CEFLPVPH--VGINPRGIRPLQVWQMDVTHVSSFGKLQYLHVSIDTCSGIMFASPLTGEK  1040

Query  921   GQETAYFLLKLAGRW-PVKTIHTDNGSNFTGATVRAACWWAGIKQEFGIPYNPQSQGVVE  979
                     L+    W   + + TDNG  +T    +  C    +    G+PYNPQ QG+VE
Sbjct  1041  ASHVIQHCLEAWSAWGKPRLLKTDNGPAYTSQKFQQFCRQMDVTHLTGLPYNPQGQGIVE  1220

Query  980   SMNKELKK-IIGQVRDQAE  997
               ++ LK  +I Q R   E
Sbjct  1221  RAHRTLKAYLIKQKRGTFE  1277
```

>Uncultured Roseobacter sp. isolate 53.fasta NODE_38, whole genome shotgun sequence
Sequence ID: NZ_CANMKI010000038.1 Length: 315
Range 1: 210 to 314

Score:75.9 bits(185), Expect:1e-11,
Method:Composition-based stats.,
Identities:35/35(100%), Positives:35/35(100%), Gaps:0/35(0%)

```
Query  1004  QMAVFIHNFKRKGGIGGYSAGERIVDIIATDIQTK  1038
             QMAVFIHNFKRKGGIGGYSAGERIVDIIATDIQTK
Sbjct  210   QMAVFIHNFKRKGGIGGYSAGERIVDIIATDIQTK  314
```

>Cellulophaga baltica isolate 925.fasta NODE_42, whole genome shotgun sequence
Sequence ID: NZ_CANLPS010000042.1 Length: 316
Range 1: 35 to 136

Score:74.7 bits(182), Expect:3e-11,
Method:Composition-based stats.,
Identities:34/34(100%), Positives:34/34(100%), Gaps:0/34(0%)

```
Query  1004  QMAVFIHNFKRKGGIGGYSAGERIVDIIATDIQT  1037
             QMAVFIHNFKRKGGIGGYSAGERIVDIIATDIQT
Sbjct  136   QMAVFIHNFKRKGGIGGYSAGERIVDIIATDIQT  35
```

>Rickettsia endosymbiont of Ceutorhynchus assimilis isolate Rickettsia_endosymbiont_Ceutorhynchus_assimilis chromosome 1
Sequence ID: NZ_OU906081.1 Length: 4063491
Range 1: 42499 to 43827

Score:83.6 bits(205), Expect:3e-11,
Method:Compositional matrix adjust.,
Identities:112/467(24%), Positives:189/467(40%), Gaps:99/467(21%)

```
Query  30     NNSPSEAGADRQGTVSFNFPQVTLWQRPLVTIKIGGQLKEALLDTGAD-DTVLEEMSLPG  88
              +++P   G     + S N        RP++TI++ G+   AL+DTGAD + + + +   G
Sbjct  42499  HSTPGHVGLCTNSSDSHN------TNRPIITIRLNGKPIPALIDTGADVNLISDSLITRG  42660

Query  89     RWKP----KMIGGIGGFIKVRQYDQILIEICGHKAIGTVLVGP--TPVNIIGRNLLTQIG  142
               +KP      +   G   KV     + I+I G        LV P   P  I+G   L
Sbjct  42661  SYKPVNTRLQLACEGESAKVIGNADLTIDIEGIVISAKFLVVPGIEPRIILGMPFLQDNQ  42840

Query  143    CTLNFP----------------ISPIETVPVKLK-PGMDG--------------------  165
                L+F                S + TVP +++ P +D
Sbjct  42841  TILDFTRGCAYLGTKARKTIFWSSKLRTVPGEIQLPPLDNHLPQEFQDLVEEYSDLFDTG  43020

Query  166    -----------------PKV---KQWPLTEEKIKALVEICTEMEKEGKISKIGPENPYNT  205
                               PK+   + +P++ EK K L     +M + G I      + Y +
Sbjct  43021  LRQPTTRTTLHKIDLKEPKIVNKRCYPMSPEKRKILYGQLNQMLQAGVIEPTT--SAYAS  43194

Query  206    PVFAIKKKDSTKWRKLVDFRELNKRTQD----FWEVQLGIPHPAGLKKKKSVTVLDVGDA  261
              P   + + D  + R  VD+R +N  T D       +Q  +     L + K  T+LD+
Sbjct  43195  PPVLVTRPDK-EPRFCVDYRAVNDITLDESTTLPRIQDCV---KPLAQAKIFTLLDLKSG  43362

Query  262    YFSVPLDEDFRKYTAFTIPSINNETPGIRYQYNVLPQGWKGSPAIFQSSMTKILEPFRKQ  321
              Y+ +P+DE  +KYTAF+ P       G  YQ+ V+P G K +P+ FQ  MT+ +     Q
Sbjct  43363  YWQIPMDEQSKKYTAFSTPD------GASYQFTVMPFGLKNAPSTFQKLMTQEVLVGHLQ  43524

Query  322    NPDIVIYQYMDDLYVGSDL---EIGQHRTKIEELRQHLLRWGLTTPDKKHQKEPPFLWMG  378
              +    ++ Y+DD+ + S+      +   R   E LR H L+           Q+     ++G
Sbjct  43525  H---FVHVYLDDIIIYSETMEEHVNHVRKVFERLRIHKLQVSAAKCQIATQE---LDYLG  43686

Query  379    YELHPDKWTVQPIVLPEKDSWT--VN--DIQKLVGKLNWASQIYPGI  421
              Y +  D    Q   L + ++++  VN   +Q  +G  NW  +  P +
Sbjct  43687  YHIKGDITVPQNKHLKQIETFSRPVNKKQLQSFIGTCNWLREHVPHM  43827
```

Range 2: 14359 to 15687

Score:82.4 bits(202), Expect:6e-11,
Method:Compositional matrix adjust.,
Identities:111/467(24%), Positives:188/467(40%), Gaps:99/467(21%)

```
Query  30     NNSPSEAGADRQGTVSFNFPQVTLWQRPLVTIKIGGQLKEALLDTGAD-DTVLEEMSLPG  88
              +++P   G     + S N        RP++TI++ G+   A +DTGAD + + + +   G
Sbjct  15687  HSTPGHVGLCTNSSDSHN------TNRPIITIRLNGKPIPAFIDTGADVNLISDSLITRG  15526

Query  89     RWKP----KMIGGIGGFIKVRQYDQILIEICGHKAIGTVLVGP--TPVNIIGRNLLTQIG  142
               +KP      +   G   KV     + I+I G        LV P   P  I+G   L
Sbjct  15525  SYKPVNTRLQLACEGESAKVIGNADLTIDIEGIVISAKFLVVPGIEPRIILGMPFLQDNQ  15346

Query  143    CTLNFP----------------ISPIETVPVKLK-PGMDG--------------------  165
                L+F                S + TVP +++ P +D
Sbjct  15345  TILDFTRGCAYLGTKTRKTIFWSSKLRTVPGEIQLPPLDNHLPQEFQDLVEEYSDLFDTG  15166

Query  166    -----------------PKV---KQWPLTEEKIKALVEICTEMEKEGKISKIGPENPYNT  205
                               PK+   + +P++ EK K L     +M + G I      + Y +
Sbjct  15165  LRQPTTRTTLHKIDLKEPKIVNKRCYPMSPEKRKILYGQLNQMLQAGVIEPTT--SAYAS  14992

Query  206    PVFAIKKKDSTKWRKLVDFRELNKRTQD----FWEVQLGIPHPAGLKKKKSVTVLDVGDA  261
              P   + + D  + R  VD+R +N  T D       +Q  +     L + K  T+LD+
```

```
Sbjct  14991  PPVLVTRPDK-EPRFCVDYRAVNDITLDESTTLPRIQDCV---KPLAQAKIFTLLDLKSG  14824

Query  262    YFSVPLDEDFRKYTAFTIPSINNETPGIRYQYNVLPQGWKGSPAIFQSSMTKILEPFRKQ  321
              Y+ +P+DE  +KYTAF+ P       G  YQ+ V+P G K +P+ FQ  MT+ +     Q
Sbjct  14823  YWQIPMDEQSKKYTAFSTPD------GASYQFTVMPFGLKNAPSTFQKLMTQEVLVGHLQ  14662

Query  322    NPDIVIYQYMDDLYVGSDL---EIGQHRTKIEELRQHLLRWGLTTPDKKHQKEPPFLWMG  378
              +    ++ Y+DD+ + S+     +   R   E LR H L+           Q+     ++G
Sbjct  14661  H---FVHVYLDDIIIYSETMEEHVNHVRKVFERLRIHKLQVSAAKCQIATQE---LDYLG  14500

Query  379    YELHPDKWTVQPIVLPEKDSWT--VN--DIQKLVGKLNWASQIYPGI  421
              Y +  D    Q   L + ++++  VN   +Q  +G  NW  +  P +
Sbjct  14499  YHIKGDITVPQTKHLKQIETFSRPVNKKQLQSFIGTCNWLREHVPHM  14359
```

>Klebsiella pneumoniae strain EuSCAPE_MT015, whole genome shotgun sequence
Sequence ID: NZ_UKIW02000113.1 Length: 281
Range 1: 95 to 280

Score:73.9 bits(180), Expect:3e-11,
Method:Composition-based stats.,
Identities:43/66(65%), Positives:48/66(72%), Gaps:11/66(16%)

```
Query  2    QGKAREFSSEQTRAN-------SPTRRELQVWGRDNNSPSEAGADRQGTVSFNFPQVTLW  54
            QG+AREF  EQTRAN       SPT RELQV  R NN  SEAG +RQGT+  NFPQ+TLW
Sbjct  95   QGEAREFPPEQTRANSPTSRANSPTSRELQV--RGNNPGSEAGGERQGTL--NFPQITLW  262

Query  55   QRPLVT  60
            QRPLV+
Sbjct  263  QRPLVS  280
```

>Uncultured Dokdonia sp. isolate 746.fasta NODE_102, whole genome shotgun sequence
Sequence ID: NZ_CANLOB010000102.1 Length: 440
Range 1: 2 to 106

Score:75.9 bits(185), Expect:3e-11,
Method:Composition-based stats.,
Identities:35/35(100%), Positives:35/35(100%), Gaps:0/35(0%)

```
Query  1004  QMAVFIHNFKRKGGIGGYSAGERIVDIIATDIQTK  1038
             QMAVFIHNFKRKGGIGGYSAGERIVDIIATDIQTK
Sbjct  106   QMAVFIHNFKRKGGIGGYSAGERIVDIIATDIQTK  2
```

>Klebsiella pneumoniae strain BL2026 NODE_25_length_1934_cov_0.482568, whole genome shotgun sequence
Sequence ID: NZ_VICY01000025.1 Length: 1934
Range 1: 1208 to 1768

Score:81.6 bits(200), Expect:5e-11,
Method:Compositional matrix adjust.,
Identities:61/192(32%), Positives:102/192(53%), Gaps:10/192(5%)

```
Query  153   ETVPVKLKPGMDGPKVKQWPLTEEKIKALVEICTEMEKEGKISKIGPENPYNTPVFAIKK  212
             E V + LK      + KQ+PL  E  + L ++     K+ ++  +  E+ +NTP+  +KK
Sbjct  1208  EPVKIFLKESTKLVRQKQYPLKLEARRGLEKMIENFLKKYRLL-VECESEFNTPILPVKK  1384

Query  213   KDSTKWRKLVDFRELNKRTQDFWEVQLGIPHP--AGLKKK-KSVTVLDVGDAYFSVPLDE  269
              +S ++R + DF   NK     +    G P      LK+  K  TVL++ DA+F + LD+
Sbjct  1385  PESEEYRLVQDFNS-NKFNSSRYSPLSGKPVSLLTALKENHKWFTVLNL*DAFFCILLDK  1561

Query  270   DFRKYTAFTIPSINNETPGIRYQYN--VLPQGWKGSPAIFQSSMTKILEPFRKQNPDIVI  327
             + + Y AF    + N T G + Q    VLPQG+K SP IF + + K LE  +   P+ ++
Sbjct  1562  NSQAYFAF---ELQNPTTGQKTQLTWAVLPQGFKNSPMIFGNQLAKELETCKIIYPERLV  1732

Query  328   YQYMDDLYVGSD  339
              QY+DD+ + ++
Sbjct  1733  LQYVDDILIATE  1768
```

>Streptococcus pneumoniae strain NN2021-1 12_contig_107, whole genome shotgun sequence
Sequence ID: NZ_JAGYHD010000101.1 Length: 1657
Range 1: 54 to 641

Score:80.1 bits(196), Expect:2e-10,
Method:Compositional matrix adjust.,
Identities:64/208(31%), Positives:103/208(49%), Gaps:20/208(9%)

```
Query  153  ETVPVKLKPGMDGPKVKQWPLTEEKIKALVEICTEMEKEGKISKIGPENPYNTPVFAIKK  212
            E V + L+     P  KQ+PL  E  + L  +  +    G +  +   +P NTP+ ++KK
Sbjct  54   ELVIIVLQDPTRFPNQKQYPLKREAREGLQPLINKFLACGLL--VPTSSPCNTPILSVKK  227

Query  213  KDSTKWRKLVDFRELNKRTQDFWEVQLGIPHPAG-----LKKKKSVTVLDVGDAYFSVPL  267
            KD T WR + D R +N+       +   +P+P       L   K  TVLD+ DA+F +PL
Sbjct  228  KDRT-WRMVQDLRIINEAVVP---LHPTVPNPYVILGEILPSAKWFTVLDLKDAFFCIPL  395

Query  268  DEDFRKYTAFTIPSINNETPGIRYQ---YNVLPQGWKGSPAIFQSSMTKILEPFRKQNPD  324
             ++ +   AF       E PG ++Q   + VLPQG++ SP +F  ++++ L       P+
Sbjct  396  AKESQYLFAF-----EWEAPGEKHQQMTWTVLPQGFRDSPHLFGQALSRDLL-DLDLGPN  557

Query  325  IVIYQYMDDLYVGSDLEIGQHRTKIEEL  352
              I QY+DDL + S  E    +  I+ L
Sbjct  558  GKILQYVDDLLICSPDEKSAQQHAIQVL  641
```

>Klebsiella pneumoniae strain BL1427 NODE_43_length_1114_cov_0.430598, whole genome shotgun sequence
Sequence ID: NZ_VICX01000043.1 Length: 1114
Range 1: 49 to 609

Score:78.2 bits(191), Expect:2e-10,
Method:Composition-based stats.,
Identities:59/192(31%), Positives:101/192(52%), Gaps:10/192(5%)

```
Query  153  ETVPVKLKPGMDGPKVKQWPLTEEKIKALVEICTEMEKEGKISKIGPENPYNTPVFAIKK  212
            E V + LK      +  Q+P   E  + L ++     K+ ++  +  E+ +NTP+  +KK
Sbjct  609  EPVKIFLKESTKLVRQNQYPSKLEARRGLEKMIENFLKKYRLL-VECESEFNTPILPVKK  433

Query  213  KDSTKWRKLVDFRELNKRTQDFWEVQLGIPHP--AGLKKK-KSVTVLDVGDAYFSVPLDE  269
             +S ++R + DF   NK     +    G P      LK+  K  TVL++ DA+F + LD+
Sbjct  432  PESEEYRLVQDFNS-NKFNSSRYSPLSGKPVSLLTALKENHKWFTVLNL*DAFFCILLDK  256

Query  270  DFRKYTAFTIPSINNETPGIRYQ--YNVLPQGWKGSPAIFQSSMTKILEPFRKQNPDIVI  327
            + + Y AF    + N T G + Q  + VLPQG+K SP IF + + K LE  +   P+ ++
Sbjct  255  NSQAYFAF---ELQNPTTGQKTQLTWAVLPQGFKNSPMIFGNQLAKELETCKIIYPERLV  85
```

```
Query  328  YQYMDDLYVGSD  339
            QY+DD+ + ++
Sbjct  84   LQYVDDILIATE  49
```

>Salmonella enterica subsp. enterica serovar Typhi strain 7-sc-cl NODE_243_length_289_cov_0.314815, whole genome shotgun sequence
Sequence ID: NZ_JAEPKN010000213.1 Length: 289
Range 1: 1 to 96

Score:71.2 bits(173), Expect:3e-10,
Method:Composition-based stats.,
Identities:32/32(100%), Positives:32/32(100%), Gaps:0/32(0%)

```
Query  1004  QMAVFIHNFKRKGGIGGYSAGERIVDIIATDI  1035
              QMAVFIHNFKRKGGIGGYSAGERIVDIIATDI
Sbjct  96    QMAVFIHNFKRKGGIGGYSAGERIVDIIATDI  1
```

>Acinetobacter baumannii strain E1 contig_E1_51, whole genome shotgun sequence
Sequence ID: NZ_JBBXKX010000035.1 Length: 412
Range 1: 2 to 106

Score:68.9 bits(167), Expect:6e-09,
Method:Compositional matrix adjust.,
Identities:31/35(89%), Positives:35/35(100%), Gaps:0/35(0%)

```
Query  1029  DIIATDIQTKELQKQITKIQNFRVYYRDSRNPLWK  1063
              DIIATDIQTKELQKQITKIQNFRVYYRDSR+P+++
Sbjct  2     DIIATDIQTKELQKQITKIQNFRVYYRDSRDPVYR  106
```

>Burkholderia pseudomallei strain 2604STDY5519032 ERS240165SCcontig000074, whole genome shotgun sequence
Sequence ID: NZ_UVID02000074.1 Length: 656
Range 1: 2 to 154

Score:70.5 bits(171), Expect:1e-08,
Method:Composition-based stats.,
Identities:37/55(67%), Positives:42/55(76%), Gaps:4/55(7%)

```
Query  2    QGKAREFSSEQTRANSPTRRELQVWGRDNNSPSEAGADRQGTVSFNFPQVTLWQR  56
            QG+AREF  EQTRANSPT RELQV  R  N  +EAG + +GT  FNFPQ+TLWQR
Sbjct  154  QGEAREFHPEQTRANSPTSRELQV--RGGNPCAEAGVEGEGT--FNFPQITLWQR  2
```

>Wolbachia endosymbiont of Psylliodes chrysocephala isolate Wolbachia_endosymbiont_of_Psylliodes chrysocephala_genome chromosome 1, complete sequence
Sequence ID: NZ_OU909470.1 Length: 1438854
Range 1: 1002729 to 1003400

Score:73.9 bits(180), Expect:3e-08,
Method:Compositional matrix adjust.,
Identities:64/238(27%), Positives:114/238(47%), Gaps:22/238(9%)

```
Query  202      PYNTPVFAIKKKDSTKWRKLVDFRELNKRTQDFWEVQLGIPHPAG-LKKKKSVTVLDVGD  260
                PY  P+  +KKKD    R  +D RELNKR    +E  L I    G +   K  T LD+
Sbjct  1002729  PYINPIVIVKKKDGD-IRICLDARELNKRCIPEYEAPLTIEAILGRISHAKIFTKLDLRH  1002905

Query  261      AYFSVPLDEDFRKYTAFTIPSINNETPGIRYQYNVLPQGWKGSPAIFQSSMTKILEPFRK  320
                +++ +PLD+D R YT F I        G+ Y++ V+P G + + +    ++ K+L  F +
Sbjct  1002906  SFWLIPLDKDSRDYTGFMI-------DGVVYRFRVVPFGTQSATSALLKALHKVLNKFEQ  1003064

Query  321      QNPDIVIYQYMDDLYVGSDLEIGQHRTKIEELRQHLLRWGL-TTPDKKHQKEPPFLWMGY  379
                         Y+DD+ + S+ E  +H   IE +   L   GL    +K    +   +++GY
Sbjct  1003065  YT-----LHYIDDILIFSENE-EEHWKHIEIILHALDEAGLKLNINKCEFFKKEVIYLGY  1003226

Query  380      ELHPDKWTVQP----IVLPEKDSWTVNDIQKLVGKLNWASQIYPGIKVRQ--LCKLLR  431
                ++  D  ++      I+   K    +  ++  +G L++  ++ P +  ++  L +LLR
Sbjct  1003227  KIDTDGVSMDANRIKIIKDYKRPTNLKTLRGFLGILHYFKRMVPDLTKKEIPLIELLR  1003400
```

>MAG: Gemmatimonas sp. isolate Sample-104-S78-bin-40 Sample_104_S78-bin_40-k141_92450, whole genome shotgun sequence
Sequence ID: NZ_JBEILP010000206.1 Length: 8099
Range 1: 4123 to 4653

Score:73.2 bits(178), Expect:4e-08,
Method:Compositional matrix adjust.,
Identities:52/183(28%), Positives:87/183(47%), Gaps:19/183(10%)

```
Query  153   ETVPVKLKPGMDGPKVKQWPLTEEKIKALVEICTEMEKEGKISKIGPENPYNTPVFAIKK  212
             E   ++L+PG +    KQ+ + +     L +   E+   G I ++   +PYN+P+    K
Sbjct  4123  EYARLRLQPGAEPRSRKQYRIPDALRPQLQKTIEELAAHGLI-EVQSGSPYNSPILFAPK  4299

Query  213   KDSTKWRKLVDFRELNK----------RTQDFWEVQLGIPHP---AGLKKKKSVTVLDVG  259
              +ST+ R   D RELNK           T++ ++    + H    AG++     +  D
Sbjct  4300  PNSTEMRFCFDSRELNKVLLDHPYPAPTTEELFDRVARLQHDAKLAGVEAPLWFSKTDAR  4479

Query  260   DAYFSVPLDEDFRKYTAFTIPSINNETPGIRYQYNVLPQGWKGSPAIFQSSMTKILEPFR  319
               Y+ + +  D R   AFT+P ++       YQ+ V+P G K S AIFQ +M ++L PF
Sbjct  4480  HGYWQIEVHPDDRPLLAFTVPVLD-----CSYQWCVMPMGTKSSAAIFQRAMDQVLAPFS  4644

Query  320   KQN  322
               N
Sbjct  4645  NTN  4653
```

>Corynebacterium glutamicum strain 39 Scaffolds_32_9166, whole genome shotgun sequence
Sequence ID: NZ_JACEQW010000032.1 Length: 8862
Range 1: 3976 to 4143

Score:72.8 bits(177), Expect:6e-08,
Method:Compositional matrix adjust.,
Identities:37/56(66%), Positives:42/56(75%), Gaps:0/56(0%)

```
Query  993   RDQAEHLKTAVQMAVFIHNFKRKGGIGGYSAGERIVDIIATDIQTKELQKQITKIQ  1048
             RD+   ++       +   FKRKGGIGGYSAGERIVDIIATDIQTKELQKQITK +
Sbjct  4143  RDRDRSIRLVNGSRRYRLTFKRKGGIGGYSAGERIVDIIATDIQTKELQKQITKFK  3976
```

>Corynebacterium glutamicum strain 37 Scaffolds_33_8928, whole genome shotgun sequence
Sequence ID: NZ_JACENO010000033.1 Length: 8928

```
Range 1: 3950 to 4117

Score:72.8 bits(177), Expect:6e-08,
Method:Compositional matrix adjust.,
Identities:37/56(66%), Positives:42/56(75%), Gaps:0/56(0%)

Query  993   RDQAEHLKTAVQMAVFIHNFKRKGGIGGYSAGERIVDIIATDIQTKELQKQITKIQ  1048
             RD+   ++       +   FKRKGGIGGYSAGERIVDIIATDIQTKELQKQITK +
Sbjct  4117  RDRDRSIRLVNGSRRYRLTFKRKGGIGGYSAGERIVDIIATDIQTKELQKQITKFK  3950




>Klebsiella pneumoniae strain BL2026 NODE_43_length_659_cov_0.466165, whole genome shotgun sequence
Sequence ID: NZ_VICY01000043.1 Length: 659
Range 1: 198 to 650

Score:66.2 bits(160), Expect:3e-07,
Method:Compositional matrix adjust.,
Identities:51/160(32%), Positives:84/160(52%), Gaps:16/160(10%)

Query  155  VPVKLKPGMDGPKVKQWPLTEEKIKALVEICTEMEKEGKISKIGPENPYNTPVFAIKKKD  214
            V VK+K G    ++KQ+PL +E  + +  +  +  + G + +   E  +NTP+  ++K D
Sbjct  198  VEVKVKEGRRPVRIKQYPLKKEDREGIRPVIEKFLQLGLLKE*VSE--FNTPILPVRKPD  371

Query  215  STKWRKLVDFRELNKRTQDFWEVQLGIPHPAGLKKKKS-----VTVLDVGDAYFSVPLDE  269
             + +R + D R +NK T+D + V   + +P  L    +      TVLD+ DA+F +PL E
Sbjct  372  GS-YRVVQDLRAVNKITEDLYPV---VANPYTLLTVLTPELIWFTVLDLKDAFFCLPLHE  539

Query  270  DFRKYTAFTIPSINNETPGIRYQ--YNVLPQGWKGSPAIF  307
              +K  AF      N   G + Q  +  LPQG+K SP +F
Sbjct  540  ASQKLFAF---EWENPKSGRKTQLTWTALPQGFKNSPTLF  650




>Sphingobacterium bovisgrunnientis strain KCTC 52685 scaffold_128, whole genome shotgun sequence
Sequence ID: NZ_WUQY01000124.1 Length: 503
Range 1: 14 to 448

Score:64.3 bits(155), Expect:5e-07,
Method:Composition-based stats.,
Identities:43/163(26%), Positives:81/163(49%), Gaps:22/163(13%)

Query  201  NPYNTPVFAIKKKDSTK---WRKLVDFRELNKRT-QDFWEVQLGIPHPAGLKKKKSVTVL  256
            +PYN+P F ++  +  K    R ++D+RELN +T  + +++ +       ++  K  +
Sbjct  448  SPYNSPAFMVRNHNEIKRGKARMVIDYRELNSKTIPNSYKLPIKDLLIKQIQGSKIYSKF  269

Query  257  DVGDAYFSVPLDEDFRKYTAFTIPSINNETPGIRYQYNVLPQGWKGSPAIFQSSMTKILE  316
            D    Y+ + LDE  +  T+FT P+     P  ++++NV+  G   +PAIFQ  M KI+
Sbjct  268  DCKSGYWQIKLDEISKPLTSFTTPN-----PLGQFEWNVMSFGLMNAPAIFQQRMDKIIR  104

Query  317  PFRKQNPDIVIYQYMDDLYVGSDLEIGQHRTKIEELRQHLLRW  359
            P +          Y+DD+ +        H   ++E  +HL ++
Sbjct  103  PIQT-----FCIAYVDDILI--------HSKNLKEHIEHLKKF  14




>Acinetobacter baumannii strain BM7085 NODE_482_length_446_cov_1.108401, whole genome shotgun sequence
Sequence ID: NZ_JBBONG010000155.1 Length: 446
Range 1: 55 to 435

Score:63.5 bits(153), Expect:5e-07,
Method:Composition-based stats.,
Identities:41/144(28%), Positives:72/144(50%), Gaps:23/144(15%)

Query  201  NPYNTPVFAIKKKDSTKWRKLVDFRELNKRTQDFWEVQLGIPHP------AGLKKKKSVT  254
            +PY+     ++K +  + R  +DFR LN  TQ   E     P P      A L++ K  +
Sbjct  55   SPYSAGTVLLRKPNG-EHRLCIDFRLLNAVTQFQPE-----PLPDQTDLFAKLQRAKYFS  216

Query  255  VLDVGDAYFSVPLDEDFRKYTAFTIPSINNETPGIRYQYNVLPQGWKGSPAIFQSSMTKI  314
             +D+   Y+ +P+    RKY AF+       T    +++NVLP G   +P+IF + M K+
Sbjct  217  RIDLSKGYYQIPVQPQVRKYLAFS-------TMEGHFEFNVLPFGLHNAPSIFSAMMRKL  375

Query  315  LEPFRKQNPDIVIYQYMDDLYVGS  338
            L P   ++    ++ +MDD+ + +
Sbjct  376  LAPLSSEH----LHSFMDDILIAT  435




>Uncultured Roseobacter sp. isolate 930.fasta NODE_31, whole genome shotgun sequence
Sequence ID: NZ_CANLQB010000031.1 Length: 257
Range 1: 180 to 257

Score:60.1 bits(144), Expect:2e-06,
Method:Composition-based stats.,
Identities:26/26(100%), Positives:26/26(100%), Gaps:0/26(0%)

Query  1004  QMAVFIHNFKRKGGIGGYSAGERIVD  1029
             QMAVFIHNFKRKGGIGGYSAGERIVD
Sbjct  180   QMAVFIHNFKRKGGIGGYSAGERIVD  257




>Brachybacterium paraconglomeratum isolate 859.fasta NODE_71, whole genome shotgun sequence
Sequence ID: NZ_CANLNX010000071.1 Length: 258
Range 1: 181 to 258

Score:60.1 bits(144), Expect:2e-06,
Method:Composition-based stats.,
Identities:26/26(100%), Positives:26/26(100%), Gaps:0/26(0%)

Query  1004  QMAVFIHNFKRKGGIGGYSAGERIVD  1029
             QMAVFIHNFKRKGGIGGYSAGERIVD
Sbjct  181   QMAVFIHNFKRKGGIGGYSAGERIVD  258




>Streptococcus anginosus strain San49 NODE_781_length_269_cov_0.734375, whole genome shotgun sequence
Sequence ID: NZ_JAXIIG010000228.1 Length: 269
Range 1: 6 to 245

Score:59.3 bits(142), Expect:5e-06,
Method:Composition-based stats.,
Identities:29/81(36%), Positives:49/81(60%), Gaps:1/81(1%)

Query  201  NPYNTPVFAIKKKDSTKWRKLVDFRELNKRTQDFWEVQLGIPHPAGLKKKKSVTVLDVGD  260
            +P+NTP+F IKK+ S KWR L D + +N + Q    +Q G+P  + +  +  + ++++ D
Sbjct  245  SPWNTPIFVIKKQ-SGKWRILHDLKAINSQMQIMGPIQRGLPLLSSIPTEWLIIIINIKD  69
```

```
Query  261  AYFSVPLDEDFRKYTAFTIPS  281
             +FS+ L    R   AFT+PS
Sbjct  68   CFFSISLAPQDRVRFAFTLPS  6
```

```
>Klebsiella pneumoniae strain c10 contig85, whole genome shotgun sequence
Sequence ID: NZ_JAIBKQ010000085.1 Length: 6264
Range 1: 3409 to 4095

Score:66.2 bits(160), Expect:5e-06,
Method:Compositional matrix adjust.,
Identities:66/245(27%), Positives:113/245(46%), Gaps:23/245(9%)

Query  184   ICTEMEKEGKISKIGPENPYNTPVFAIKKKDSTKWRKLVDFRELNKRT-QDFWEVQLGIP  242
             I  EM KEG I   G  +P+ +PV  +KKKD +  R  VD+R+LN  T +D + +
Sbjct  4095  IVQEMAKEGVIEPSG--SPWASPVVLVKKKDGST-RFCVDYRQLNNVTKKDSYPLPRIDD  3925

Query  243   HPAGLKKKKSVTVLDVGDAYFSVPLDEDFRKYTAFTIPSINNETPGIRYQYNVLPQGWKG  302
                 L   +  + LD+   Y+ V L  + ++ TAFT+ +      G+ +Q+ V+P G
Sbjct  3924  TLDTLAGSRIFSTLDLKSGYWQVELAPEDKEKTAFTVGT------GL-WQFTVMPFGLCN  3766

Query  303   SPAIFQSSMTKILEPFRKQNPDIVIYQYMDD-LYVGSDLEIGQHRTKIEELRQHLLRWGL  361
             +PA F+  M  +L+    +   +    Y+DD + VG   E   H   +EE+ Q L   GL
Sbjct  3765  APATFERLMDTVLKGLSWKTCLV----YLDDIIVVGKTFE--DHLRNLEEVFQRLRNSGL  3604

Query  362   TTPDKK-HQKEPPFLWMGYELHPDKWTVQPIVLPEKDSWTV----NDIQKLVGKLNWASQ  416
                 KK H  +    ++G+ +  +   V    +     W V    ++++  +G   +  +
Sbjct  3603  RLSSKKCHLFQREVQYLGHVVSSEGVAVDKKKVESVKEWPVPTDKHELRSFLGLCTYYRR  3424

Query  417   IYPGI  421
               PG
Sbjct  3423  FVPGF  3409
```

```
>Enterococcus faecalis strain 15224 NODE_126_length_632_cov_0.918812, whole genome shotgun sequence
Sequence ID: NZ_QNGP01000071.1 Length: 632
Range 1: 177 to 611

Score:62.4 bits(150), Expect:5e-06,
Method:Compositional matrix adjust.,
Identities:43/145(30%), Positives:68/145(46%), Gaps:15/145(10%)

Query  860  AKEIVASCDKCQLKGEAMHGQVDCSPGIWQLDCTHLEG-------------KVILVAVHV  906
            ++E V +CD CQ  G   HG      GI +++   + G               ILVAV
Sbjct  177  SREFVRNCDSCQRAGNLPHGYAMPQQGILEIELFDVWGIDFMGPFPPSYSNTYILVAVDY  356

Query  907  ASGYIEAEVIPAETGQETAYFLLK-LAGRWPV-KTIHTDNGSNFTGATVRAACWWAGIKQ  964
             S ++EA   P    +    FL K +  R+ V + + +D G++F    + +A    GI
Sbjct  357  VSKWVEAIATPTNDTKTVLKFLQKHIFSRFGVPRALISDGGTHFCNKQLYSAMVRYGISH  536

Query  965  EFGIPYNPQSQGVVESMNKELKKII  989
            +   PY+PQ+ G  E  N+ELK+I+
Sbjct  537  KVATPYHPQTNGQAEVSNRELKRIL  611
```

```
>Enterococcus faecalis strain AKSZ-144 NODE_42_length_234_cov_1622.847134, whole genome shotgun sequence
Sequence ID: NZ_JAOXSP010000040.1 Length: 234
Range 1: 153 to 230

Score:58.5 bits(140), Expect:6e-06,
Method:Composition-based stats.,
Identities:25/26(96%), Positives:26/26(100%), Gaps:0/26(0%)

Query  1004  QMAVFIHNFKRKGGIGGYSAGERIVD  1029
             QMAVFIHNFKRKGGIGGYSAGERIV+
Sbjct  153   QMAVFIHNFKRKGGIGGYSAGERIVE  230
```

```
>Roseobacter litoralis isolate 289-G.fasta NODE_32, whole genome shotgun sequence
Sequence ID: NZ_CANMVH010000032.1 Length: 249
Range 1: 2 to 79

Score:58.2 bits(139), Expect:8e-06,
Method:Composition-based stats.,
Identities:25/26(96%), Positives:26/26(100%), Gaps:0/26(0%)

Query  1004  QMAVFIHNFKRKGGIGGYSAGERIVD  1029
             QMAVFIHNFKRKGGIGGYSAGERIV+
Sbjct  79    QMAVFIHNFKRKGGIGGYSAGERIVE  2
```

```
>Uncultured Pseudoteredinibacter sp. isolate 208.fasta NODE_42, whole genome shotgun sequence
Sequence ID: NZ_CANMSS010000042.1 Length: 253
Range 1: 174 to 251

Score:58.2 bits(139), Expect:8e-06,
Method:Composition-based stats.,
Identities:25/26(96%), Positives:26/26(100%), Gaps:0/26(0%)

Query  1004  QMAVFIHNFKRKGGIGGYSAGERIVD  1029
             QMAVFIHNFKRKGGIGGYSAGERIV+
Sbjct  174   QMAVFIHNFKRKGGIGGYSAGERIVE  251
```

```
>Uncultured Tateyamaria sp. isolate 203.fasta NODE_54, whole genome shotgun sequence
Sequence ID: NZ_CANLVZ010000054.1 Length: 263
Range 1: 185 to 262

Score:58.2 bits(139), Expect:9e-06,
Method:Composition-based stats.,
Identities:25/26(96%), Positives:26/26(100%), Gaps:0/26(0%)

Query  1004  QMAVFIHNFKRKGGIGGYSAGERIVD  1029
             QMAVFIHNFKRKGGIGGYSAGERIV+
Sbjct  185   QMAVFIHNFKRKGGIGGYSAGERIVE  262
```

```
>Uncultured Tateyamaria sp. isolate 159-G.fasta NODE_34, whole genome shotgun sequence
Sequence ID: NZ_CANNDY010000034.1 Length: 252
Range 1: 168 to 245
```

```
Score:58.2 bits(139), Expect:1e-05,
Method:Composition-based stats.,
Identities:25/26(96%), Positives:26/26(100%), Gaps:0/26(0%)

Query  1004  QMAVFIHNFKRKGGIGGYSAGERIVD  1029
             QMAVFIHNFKRKGGIGGYSAGERIV+
Sbjct  168   QMAVFIHNFKRKGGIGGYSAGERIVE  245



>Uncultured Tenacibaculum sp. isolate 838.fasta NODE_42, whole genome shotgun sequence
Sequence ID: NZ_CANMMT010000042.1 Length: 262
Range 1: 180 to 257

Score:58.2 bits(139), Expect:1e-05,
Method:Composition-based stats.,
Identities:25/26(96%), Positives:26/26(100%), Gaps:0/26(0%)

Query  1004  QMAVFIHNFKRKGGIGGYSAGERIVD  1029
             QMAVFIHNFKRKGGIGGYSAGERIV+
Sbjct  180   QMAVFIHNFKRKGGIGGYSAGERIVE  257



>Pseudomonas sp. QS1027 NG-9811_QS116_lib127709_4633_4663_4674_contig_78, whole genome shotgun sequence
Sequence ID: NZ_PHSU01000078.1 Length: 240
Range 1: 9 to 83

Score:57.8 bits(138), Expect:1e-05,
Method:Composition-based stats.,
Identities:25/25(100%), Positives:25/25(100%), Gaps:0/25(0%)

Query  1004  QMAVFIHNFKRKGGIGGYSAGERIV  1028
             QMAVFIHNFKRKGGIGGYSAGERIV
Sbjct  83    QMAVFIHNFKRKGGIGGYSAGERIV  9



>Uncultured Roseobacter sp. isolate 908.fasta NODE_28, whole genome shotgun sequence
Sequence ID: NZ_CANMNK010000028.1 Length: 244
Range 1: 168 to 242

Score:57.8 bits(138), Expect:1e-05,
Method:Composition-based stats.,
Identities:25/25(100%), Positives:25/25(100%), Gaps:0/25(0%)

Query  1004  QMAVFIHNFKRKGGIGGYSAGERIV  1028
             QMAVFIHNFKRKGGIGGYSAGERIV
Sbjct  168   QMAVFIHNFKRKGGIGGYSAGERIV  242



>Uncultured Roseibium sp. isolate 190.fasta NODE_26, whole genome shotgun sequence
Sequence ID: NZ_CANNEN010000026.1 Length: 248
Range 1: 172 to 246

Score:57.8 bits(138), Expect:1e-05,
Method:Composition-based stats.,
Identities:25/25(100%), Positives:25/25(100%), Gaps:0/25(0%)

Query  1004  QMAVFIHNFKRKGGIGGYSAGERIV  1028
             QMAVFIHNFKRKGGIGGYSAGERIV
Sbjct  172   QMAVFIHNFKRKGGIGGYSAGERIV  246



>Uncultured Roseobacter sp. isolate 407-3Z.fasta NODE_45, whole genome shotgun sequence
Sequence ID: NZ_CANMKD010000045.1 Length: 252
Range 1: 3 to 77

Score:57.8 bits(138), Expect:1e-05,
Method:Composition-based stats.,
Identities:25/25(100%), Positives:25/25(100%), Gaps:0/25(0%)

Query  1004  QMAVFIHNFKRKGGIGGYSAGERIV  1028
             QMAVFIHNFKRKGGIGGYSAGERIV
Sbjct  77    QMAVFIHNFKRKGGIGGYSAGERIV  3



>Uncultured Winogradskyella sp. isolate 408-3Z.fasta NODE_32, whole genome shotgun sequence
Sequence ID: NZ_CANMJY010000032.1 Length: 256
Range 1: 180 to 254

Score:57.8 bits(138), Expect:1e-05,
Method:Composition-based stats.,
Identities:25/25(100%), Positives:25/25(100%), Gaps:0/25(0%)

Query  1004  QMAVFIHNFKRKGGIGGYSAGERIV  1028
             QMAVFIHNFKRKGGIGGYSAGERIV
Sbjct  180   QMAVFIHNFKRKGGIGGYSAGERIV  254



>Uncultured Ruegeria sp. isolate 409.fasta NODE_44, whole genome shotgun sequence
Sequence ID: NZ_CANLNV010000044.1 Length: 256
Range 1: 3 to 77

Score:57.8 bits(138), Expect:1e-05,
Method:Composition-based stats.,
Identities:25/25(100%), Positives:25/25(100%), Gaps:0/25(0%)

Query  1004  QMAVFIHNFKRKGGIGGYSAGERIV  1028
             QMAVFIHNFKRKGGIGGYSAGERIV
Sbjct  77    QMAVFIHNFKRKGGIGGYSAGERIV  3



>Epibacterium ulvae isolate 20.fasta NODE_62, whole genome shotgun sequence
Sequence ID: NZ_CANLDO010000062.1 Length: 257
Range 1: 181 to 255

Score:57.8 bits(138), Expect:1e-05,
Method:Composition-based stats.,
Identities:25/25(100%), Positives:25/25(100%), Gaps:0/25(0%)
```

```
Query  1004  QMAVFIHNFKRKGGIGGYSAGERIV  1028
             QMAVFIHNFKRKGGIGGYSAGERIV
Sbjct  181   QMAVFIHNFKRKGGIGGYSAGERIV  255
```

>Roseibium album isolate 321-3Z.fasta NODE_38, whole genome shotgun sequence
Sequence ID: NZ_CANMGD010000038.1 Length: 255
Range 1: 3 to 77

Score:57.8 bits(138), Expect:1e-05,
Method:Composition-based stats.,
Identities:25/25(100%), Positives:25/25(100%), Gaps:0/25(0%)

```
Query  1004  QMAVFIHNFKRKGGIGGYSAGERIV  1028
             QMAVFIHNFKRKGGIGGYSAGERIV
Sbjct  77    QMAVFIHNFKRKGGIGGYSAGERIV  3
```

>Proteus mirabilis strain PM2-4 PM2-4_scaffold37, whole genome shotgun sequence
Sequence ID: NZ_JAMKNT010000037.1 Length: 5277
Range 1: 2285 to 2986

Score:64.7 bits(156), Expect:1e-05,
Method:Compositional matrix adjust.,
Identities:64/252(25%), Positives:104/252(41%), Gaps:37/252(14%)

```
Query  191   EGKISKIGPENPYNTPVFAIKKKD-----STKWRKLVDFRELNKRTQDFWEVQLGIPHPA  245
             +G I    P   +N+P+  ++KKD     S   R  +D R +N         +  +P  A
Sbjct  2986  DGVIEIAPPNTAFNSPLLTVRKKDANGNYSDDVRVCIDPRHINAILIKDRTDRFQLPLIA  2807

Query  246   GLKKKKS----VTVLDVGDAYFSVPLDEDFRKYTAFTIPSINNETPGIRYQYNVLPQGWK  301
              L +K S     TVLD+  A+  +PL E+ R  TAFT         G +YQ+   P G
Sbjct  2806  ELHQKMSAATLFTVLDLKQAFHRLPLAEEHRALTAFTF-------NGQQYQFCRAPFGLT  2648

Query  302   GSPAIFQSSMTKILE--PFRKQNPDIVIYQYMDDLYVGSDLEIGQH----RTKIEELRQH  355
               P  FQ  +T +L   P+        +  ++DDL +G+  +I +H     T IE L
Sbjct  2647  PIPNHFQRVLTTLLHDLPY--------VTCFIDDLTIGTGPDIEEHIKCVTTVIERLTSA  2492

Query  356   LLRWGLTTPDKKHQKEPPFLWMGYELHPDKWTVQPIVLPEKDSWTV----NDIQKLVGKL  411
                  +   DK H  +     +G+ +  +  ++    +    +W V     DIQ+ +G
Sbjct  2491  KFILNV---DKCHFMQTSVNILGFTISHNSLSLDSRKVANALAWPVPKTGKDIQRFLGLA  2321

Query  412   NWASQIYPGIKV  423
             N+     P
Sbjct  2320  NYFRSHLPNFST  2285
```

>Polyangium sp. y55x31 NODE_26_length_169233_cov_17.164843, whole genome shotgun sequence
Sequence ID: NZ_JARZHG010000026.1 Length: 169233
Range 1: 66608 to 66982

Score:63.5 bits(153), Expect:4e-05,
Method:Compositional matrix adjust.,
Identities:42/126(33%), Positives:67/126(53%), Gaps:8/126(6%)

```
Query  706    VLYVDGAANRETKLGKAGYVTNRGRQKVVT---LTDTTNQKTELQAIYLAL---QDSGLE  759
              V Y DGA          G V   G +++     L D TN   EL AI  A+   +D+G
Sbjct  66982  VAYADGACTGNPGPAGLGVVVVDGDRRIEISEYLGDGTNNIAELTAILRAVGAVEDTGKP  66803

Query  760    VNIVTDSQYALGIIQAQ-PDQSESELVNQIIEQLIKKEKVYLAWVPAHKGIGGNEQVDKL  818
              + + TDSQYA+G++Q     ++ ++LV Q ++ ++K+ K  L +VP H G+  NE+ D+L
Sbjct  66802  LLVRTDSQYAIGVLQKGWKAKANTDLVAQ-VKDVLKRNKTRLEYVPGHSGVALNERADEL  66626

Query  819    VSAGIR  824
                  +R
Sbjct  66625  AREAVR  66608
```

Range 2: 66608 to 66976

Score:62.4 bits(150), Expect:9e-05,
Method:Compositional matrix adjust.,
Identities:41/124(33%), Positives:66/124(53%), Gaps:8/124(6%)

```
Query  588    YVDGAANRETKLGKAGYVTNRGRQKVVT---LTDTTNQKTELQAIYLAL---QDSGLEVN  641
              Y DGA          G V   G +++     L D TN   EL AI  A+   +D+G  +
Sbjct  66976  YADGACTGNPGPAGLGVVVVDGDRRIEISEYLGDGTNNIAELTAILRAVGAVEDTGKPLL  66797

Query  642    IVTDSQYALGIIQAQ-PDQSESELVNQIIEQLIKKEKVYLAWVPAHKGIGGNEQVDKLVS  700
              + TDSQYA+G++Q     ++ ++LV Q ++ ++K+ K  L +VP H G+  NE+ D+L
Sbjct  66796  VRTDSQYAIGVLQKGWKAKANTDLVAQ-VKDVLKRNKTRLEYVPGHSGVALNERADELAR  66620

Query  701    AGIR  704
                +R
Sbjct  66619  EAVR  66608
```

>MAG: Stenotrophomonas sp. isolate AP15.006 scaffold_7194, whole genome shotgun sequence
Sequence ID: NZ_JACXUC010000133.1 Length: 1073
Range 1: 556 to 1038

Score:62.0 bits(149), Expect:4e-05,
Method:Compositional matrix adjust.,
Identities:50/161(31%), Positives:70/161(43%), Gaps:21/161(13%)

```
Query  860   AKEIVASCDKCQLKGE-AMHGQVDCSPGI-------WQLDCT------HLEGKVILVAVH  905
             AK  V  CD CQ  G   +  ++   P +       W LD          + K ILV
Sbjct  1038  AKHYVKRCDSCQRIGRPTLSSEMPLQPQVLIEPFEKWALDFIGPINPPSKQKKYILVCTD  859

Query  906   VASGYIEAEVIPAETGQETAYFLLK-LAGRWPV-KTIHTDNGSNFTGATVRAACWWAGIK  963
               + ++EA+ + + T      FL + +  R+ V + I TD GS FT   +        IK
Sbjct  858   YVTKWVEAKALSSATENSVVQFLFEDIFTRFGVPREIVTDQGSQFTSNLMEKLMEEYKIK  679

Query  964   QEFGIPYNPQSQGVVESMNKELKKIIGQV-----RDQAEHL  999
                  PY+PQ+ G VES NK L+ II +      RD AE L
Sbjct  678   HRTSTPYHPQANGQVESTNKVLENIITKTVHLHRRDWAERL  556
```

>MAG: Thalassolituus oleivorans isolate SRR17458625_maxbin2_1_1705322953 ERZ21854034.981, whole genome shotgun sequence
Sequence ID: NZ_CAWUIR010000113.1 Length: 10375
Range 1: 4275 to 5099

Score:63.2 bits(152), Expect:4e-05,
Method:Compositional matrix adjust.,
Identities:80/299(27%), Positives:124/299(41%), Gaps:45/299(15%)

```
Query  149   ISPIETVPVKLKPGMDGPK-------VKQWPLTEEKIKALVEICTEMEKEGKISKIGPEN  201
             +S ++ + V+LKPG + P         +Q     +KIK L++I    E    I
Sbjct  5099  MSKLDPIEVELKPGHEPPNSYGRTLGYEQRMFLNDKIKDLIKIGILKESANPI-------  4941

Query  202   PYNTPVFAIKKKDSTKWRKLVDFRELNKRT-QDFWEVQLGIPHPAGLKKKKSVTVLDVGD  260
              Y +P+F + KK   KWR +VD R LN  T +   E+ L       +K  K     DV
Sbjct  4940  -YGSPIFIVPKKGPKKWRMVVDMRALNTITIKTAMEMPLLDEQIDWIKGAKVFGTFDVLS  4764

Query  261   AYFSVPLDEDFRKYTAFTIPSINNETPGIRYQYNVLPQGWKGSPAIFQSSMTK-ILEPFR  319
              +  +P  E+ +KY  FT+  I  E     Y     P GW  +PA+F   M + IL+P
Sbjct  4763  GFDFLPTAENSQKY--FTL--ITKEGA---YTMCGAPMGWVNTPALFMDRMVRNILKP--  4611

Query  320   KQNPDIVIY--------QYMDDLYVGSDLEIGQHRTKIEELRQHLLRWGLTTPDKKHQKE  371
                  I IY        Q++DD  + +D      +   + L Q + R    + DK H
Sbjct  4610  -----IGIYGIATNGAIQWLDDTCLYADSFDRYAKLLDKFLAQMIKRAVRLSIDKCHLYG  4446

Query  372   PPFLWMGYELHPDKWTVQP----IVLPEKDSWTVNDIQKLVGKLNWASQIYPGIKVRQL  426
                  W G  ++   W  Q      +L  +    V ++ +++   NW S   P  K+ QL
Sbjct  4445  EEVEWCGRRINAKGWNFQSKFYDKILTVQKPQRVWEMAQVIYLCNWLSPAIP--KLSQL  4275
```

>Klebsiella pneumoniae strain BL198 NODE_37_length_611_cov_0.421488, whole genome shotgun sequence
Sequence ID: NZ_VICV01000037.1 Length: 611
Range 1: 59 to 448

Score:59.3 bits(142), Expect:5e-05,
Method:Compositional matrix adjust.,
Identities:41/139(29%), Positives:75/139(53%), Gaps:17/139(12%)

```
Query  251  KSVTVLDVGDAYFSVPLDEDFRKYTAFTI--PSINNETPGIRYQYNVLPQGWKGSPAIFQ  308
            K  TVL++ DA+F + LD++ + Y AF +   P+   +T   +  + VLPQG+K SP IF
Sbjct  59   KWFTVLNL*DAFFCILLDKNSQAYFAFELQNPTTRQKT---QLTWAVLPQGFKNSPMIFG  229

Query  309  SSMTKILEPFRKQNPDIVIYQYMDDLYVGSDLEIGQHRTKIEELRQHLLRWGLTTPDKKH  368
            + + K LE  +   P+ ++ QY+DD+ + ++      R +   L  +LL + L   D +
Sbjct  230  NQLAKELETCKIIYPERLVLQYVDDILIATE-----TREEYFNLTTNLLNF-LGQSDNRF  391

Query  369  QKEPP------FLWMGYEL  381
             K+         +++G+E+
Sbjct  392  SKKKAQIGKETVMYLGFEI  448
```

>Escherichia coli strain w183 NODE_188_length_269_cov_26.989, whole genome shotgun sequence
Sequence ID: NZ_JAJNDO010000188.1 Length: 269
Range 1: 198 to 269

Score:56.2 bits(134), Expect:5e-05,
Method:Composition-based stats.,
Identities:24/24(100%), Positives:24/24(100%), Gaps:0/24(0%)

```
Query  1004  QMAVFIHNFKRKGGIGGYSAGERI  1027
             QMAVFIHNFKRKGGIGGYSAGERI
Sbjct  198   QMAVFIHNFKRKGGIGGYSAGERI  269
```

>Pseudomonas nitroreducens strain FY43 NODE_35_length_306_cov_242.462882, whole genome shotgun sequence
Sequence ID: NZ_JAJISQ010000035.1 Length: 306
Range 1: 161 to 238

Score:56.2 bits(134), Expect:6e-05,
Method:Composition-based stats.,
Identities:25/26(96%), Positives:26/26(100%), Gaps:0/26(0%)

```
Query  1004  QMAVFIHNFKRKGGIGGYSAGERIVD  1029
             QMAVFIHNFKRKGGIGGYSAGERIV+
Sbjct  161   QMAVFIHNFKRKGGIGGYSAGERIVE  238
```

>Polyangium spumosum strain DSM 14734 Scaffold16, whole genome shotgun sequence
Sequence ID: NZ_WJIE01000016.1 Length: 180314
Range 1: 106383 to 106757

Score:62.4 bits(150), Expect:9e-05,
Method:Compositional matrix adjust.,
Identities:42/126(33%), Positives:67/126(53%), Gaps:8/126(6%)

```
Query  706     VLYVDGAANRETKLGKAGYVTNRGRQKVVT---LTDTTNQKTELQAIYLAL---QDSGLE  759
               V Y DGA           G V   G +++     L D TN   EL AI  A+   Q++G
Sbjct  106383  VAYADGACTGNPGPAGLGVVVVDGDRRIEVSEYLGDGTNNIAELTAILRAVEAVQETGKP  106562

Query  760     VNIVTDSQYALGIIQAQ-PDQSESELVNQIIEQLIKKEKVYLAWVPAHKGIGGNEQVDKL  818
               + + TDSQYA+G++Q     ++ ++LV Q+ + ++K+ K  L +VP H G+  NE+ D+L
Sbjct  106563  LLVRTDSQYAIGVLQKGWKAKANTDLVAQV-KDVLKRNKTRLEYVPGHSGVALNERADEL  106739

Query  819     VSAGIR  824
                   +R
Sbjct  106740  AREAVR  106757
```

Range 2: 106389 to 106757

Score:61.2 bits(147), Expect:2e-04,
Method:Compositional matrix adjust.,
Identities:41/124(33%), Positives:66/124(53%), Gaps:8/124(6%)

```
Query  588     YVDGAANRETKLGKAGYVTNRGRQKVVT---LTDTTNQKTELQAIYLAL---QDSGLEVN  641
               Y DGA           G V   G +++     L D TN   EL AI  A+   Q++G  +
Sbjct  106389  YADGACTGNPGPAGLGVVVVDGDRRIEVSEYLGDGTNNIAELTAILRAVEAVQETGKPLL  106568

Query  642     IVTDSQYALGIIQAQ-PDQSESELVNQIIEQLIKKEKVYLAWVPAHKGIGGNEQVDKLVS  700
               + TDSQYA+G++Q     ++ ++LV Q+ + ++K+ K  L +VP H G+  NE+ D+L
Sbjct  106569  VRTDSQYAIGVLQKGWKAKANTDLVAQV-KDVLKRNKTRLEYVPGHSGVALNERADELAR  106745

Query  701     AGIR  704
                 +R
Sbjct  106746  EAVR  106757
```

>Salmonella sp. gx-f5 contig70, whole genome shotgun sequence
Sequence ID: NZ_VCKP01000064.1 Length: 380
Range 1: 127 to 369

Score:56.2 bits(134), Expect:1e-04,
Method:Compositional matrix adjust.,
Identities:32/92(35%), Positives:49/92(53%), Gaps:11/92(11%)

```
Query  248  KKKKSVTVLDVGDAYFSVPLDEDFRKYTAFTIPSINNETPGIRYQYNVLPQGWKGSPAIF  307
            +  K    +D+ +A+FS+PL  + R   AFT         G++Y +N LPQGWK SP I
Sbjct  369  RAAKWYATIDIANAFFSIPLAAECRPQFAFT-------WRGVQYTWNRLPQGWKHSPTIC  211

Query  308  QSSMTKILEPFRKQNPDIVIYQYMDDLYVGSD  339
               +   LE  + + P+ +  QY+DD+ V  D
Sbjct  210  HGLIQSALE--QGEAPEHL--QYIDDIIVWGD  127
```

>Klebsiella pneumoniae strain AS012315 AS012315_632, whole genome shotgun sequence
Sequence ID: NZ_VKXG01000632.1 Length: 234
Range 1: 166 to 234

Score:54.3 bits(129), Expect:1e-04,
Method:Composition-based stats.,
Identities:23/23(100%), Positives:23/23(100%), Gaps:0/23(0%)

```
Query  1004  QMAVFIHNFKRKGGIGGYSAGER  1026
             QMAVFIHNFKRKGGIGGYSAGER
Sbjct  166   QMAVFIHNFKRKGGIGGYSAGER  234
```

>Klebsiella pneumoniae isolate ena-yuan-GCF_010597935.1 NZ_VKXG01000632.1, whole genome shotgun sequence
Sequence ID: NZ_CAWPJY010000593.1 Length: 234
Range 1: 166 to 234

Score:54.3 bits(129), Expect:1e-04,
Method:Composition-based stats.,
Identities:23/23(100%), Positives:23/23(100%), Gaps:0/23(0%)

```
Query  1004  QMAVFIHNFKRKGGIGGYSAGER  1026
             QMAVFIHNFKRKGGIGGYSAGER
Sbjct  166   QMAVFIHNFKRKGGIGGYSAGER  234
```

>Proteus mirabilis strain PM2-4 PM2-4_scaffold43, whole genome shotgun sequence
Sequence ID: NZ_JAMKNT010000043.1 Length: 3726
Range 1: 2922 to 3395

Score:61.2 bits(147), Expect:2e-04,
Method:Compositional matrix adjust.,
Identities:46/171(27%), Positives:71/171(41%), Gaps:32/171(18%)

```
Query  846   WRAMASDFNLPPVVAKEIVASCDKCQLKGEAMHGQVDCSPGIW------------QLDCT  893
             W  M  D        +E    C +CQ     MHG       ++             LD
Sbjct  3395  WENMTHDI-------QEYTRKCSRCQ-----MHGSKSLREELYPIPVSTKPFDRIALDVK  3252

Query  894   HLEG-----KVILVAVHVASGYIEAEVIPAETGQETAYFLL-KLAGRWPVKTI-HTDNGS  946
             H++      + I+  +   + Y+EA  +  +T  E A FL  ++  R    TI  +DNG
Sbjct  3251  HVQASRAGYRYIIAGIDYLTKYVEARPMRFQTSSEIALFLYDEIICRHGCPTIIVSDNGK  3072

Query  947   NFTGATVRAACWWAGIKQEFGIPYNPQSQGVVESMNKELKKIIGQVRDQAE  997
              F    +R  C    I  +   PYNPQS G++E  N+ L +I+ Q R+ +E
Sbjct  3071  PFLSDLIRQVCRNYSIIHKTTTPYNPQSNGLIERFNRTLGQIL-QRREDSE  2922
```

>Escherichia coli strain w189 NODE_129_length_265_cov_25.6333, whole genome shotgun sequence
Sequence ID: NZ_JAJNDQ010000122.1 Length: 265
Range 1: 195 to 263

Score:54.3 bits(129), Expect:2e-04,
Method:Composition-based stats.,
Identities:23/23(100%), Positives:23/23(100%), Gaps:0/23(0%)

```
Query  1004  QMAVFIHNFKRKGGIGGYSAGER  1026
             QMAVFIHNFKRKGGIGGYSAGER
Sbjct  195   QMAVFIHNFKRKGGIGGYSAGER  263
```

>Escherichia coli strain w190 NODE_198_length_265_cov_25.5778, whole genome shotgun sequence
Sequence ID: NZ_JAJNDR010000198.1 Length: 265
Range 1: 3 to 71

Score:54.3 bits(129), Expect:2e-04,
Method:Composition-based stats.,
Identities:23/23(100%), Positives:23/23(100%), Gaps:0/23(0%)

```
Query  1004  QMAVFIHNFKRKGGIGGYSAGER  1026
             QMAVFIHNFKRKGGIGGYSAGER
Sbjct  71    QMAVFIHNFKRKGGIGGYSAGER  3
```

>Escherichia coli strain w176 NODE_127_length_264_cov_65.95, whole genome shotgun sequence
Sequence ID: NZ_JAJNDN010000127.1 Length: 264
Range 1: 3 to 71

Score:54.3 bits(129), Expect:2e-04,
Method:Composition-based stats.,
Identities:23/23(100%), Positives:23/23(100%), Gaps:0/23(0%)

```
Query  1004  QMAVFIHNFKRKGGIGGYSAGER  1026
             QMAVFIHNFKRKGGIGGYSAGER
Sbjct  71    QMAVFIHNFKRKGGIGGYSAGER  3
```

>Acinetobacter baumannii strain BM4942 NODE_1051_length_338_cov_0.601533, whole genome shotgun sequence
Sequence ID: NZ_JBBONW010000251.1 Length: 338
Range 1: 5 to 247

Score:55.1 bits(131), Expect:2e-04,
Method:Composition-based stats.,

```
Identities:26/92(28%), Positives:50/92(54%), Gaps:11/92(11%)

Query  247  LKKKKSVTVLDVGDAYFSVPLDEDFRKYTAFTIPSINNETPGIRYQYNVLPQGWKGSPAI  306
            L++ K  + +D+   Y+ +P+    RKY AF+       T    +++NVLP G   +P+I
Sbjct  5    LQRAKYFSRIDLSKGYYQIPVQPQVRKYLAFS-------TMEGHFEFNVLPFGLHNAPSI  163

Query  307  FQSSMTKILEPFRKQNPDIVIYQYMDDLYVGS  338
            F + M K+L P   ++    ++ +MDD+ + +
Sbjct  164  FSAMMGKLLAPLSSEH----LHSFMDDILIAT  247




>Escherichia coli strain w202 NODE_136_length_268_cov_26.95, whole genome shotgun sequence
Sequence ID: NZ_JAJNDS010000125.1 Length: 268
Range 1: 198 to 266

Score:54.3 bits(129), Expect:2e-04,
Method:Composition-based stats.,
Identities:23/23(100%), Positives:23/23(100%), Gaps:0/23(0%)

Query  1004  QMAVFIHNFKRKGGIGGYSAGER  1026
             QMAVFIHNFKRKGGIGGYSAGER
Sbjct  198   QMAVFIHNFKRKGGIGGYSAGER  266




>Escherichia coli strain w184 NODE_200_length_268_cov_29.5537, whole genome shotgun sequence
Sequence ID: NZ_JAJNDP010000200.1 Length: 268
Range 1: 3 to 71

Score:54.3 bits(129), Expect:2e-04,
Method:Composition-based stats.,
Identities:23/23(100%), Positives:23/23(100%), Gaps:0/23(0%)

Query  1004  QMAVFIHNFKRKGGIGGYSAGER  1026
             QMAVFIHNFKRKGGIGGYSAGER
Sbjct  71    QMAVFIHNFKRKGGIGGYSAGER  3




>Escherichia coli strain w208 NODE_126_length_276_cov_44.5, whole genome shotgun sequence
Sequence ID: NZ_JAJNDT010000124.1 Length: 276
Range 1: 3 to 71

Score:54.3 bits(129), Expect:2e-04,
Method:Composition-based stats.,
Identities:23/23(100%), Positives:23/23(100%), Gaps:0/23(0%)

Query  1004  QMAVFIHNFKRKGGIGGYSAGER  1026
             QMAVFIHNFKRKGGIGGYSAGER
Sbjct  71    QMAVFIHNFKRKGGIGGYSAGER  3




>Streptomyces albidochromogenes strain DSM 41800 NODE_804_length_521_cov_1.81218, whole genome shotgun sequence
Sequence ID: NZ_VBVX01000736.1 Length: 521
Range 1: 4 to 498

Score:56.6 bits(135), Expect:2e-04,
Method:Compositional matrix adjust.,
Identities:52/186(28%), Positives:82/186(44%), Gaps:27/186(14%)

Query  157  VKLKPGMDGPKVKQWPLTEEKIKALVEICTEMEKEGKISKIGPENPYNTPVFAIKKKDST  216
            + L PG        + +++ + + L +   E+ ++G I K    +PY  PV  +KKKD
Sbjct  498  IHLIPGAKPTARPPYRISQFETEELKKQIEELLQQGFIQK--SNSPYAAPVLFVKKKDKA  325

Query  217  KWRKLVDFRELNKRTQDFWEVQLGIPHP------AGLKKKKSVTVLDVGDAYFSVPLDED  270
              R  VDFR LN  T     ++   P P        +   K  + LD+   YF + +D
Sbjct  324  -LRLCVDFRLLNINT-----IKNAFPIPLIEDLFMKIGTAKVFSKLDLMSGYFQIRMDPR  163

Query  271  FRKYTAFTIPSINNETPGIRYQYNVLPQGWKGSPAIFQSSMTKILEPFRKQNPDIVIYQY  330
              + T F        TP   Y + V+P G    PA FQS M  IL       P++++  Y
Sbjct  162  DEEKTGFV-------TPFGHYHWRVMPFGVVNGPATFQSFMNDIL----GDTPNVLV--Y  22

Query  331  MDDLYV  336
            +DD+ +
Sbjct  21   LDDILI  4




>Polyangium sp. 15x6 NODE_112_length_36528_cov_15.618939, whole genome shotgun sequence
Sequence ID: NZ_JASBQE010000112.1 Length: 36528
Range 1: 5134 to 5508

Score:60.5 bits(145), Expect:3e-04,
Method:Compositional matrix adjust.,
Identities:41/126(33%), Positives:67/126(53%), Gaps:8/126(6%)

Query  706   VLYVDGAANRETKLGKAGYVTNRGRQKVVT---LTDTTNQKTELQAIYLAL---QDSGLE  759
             V Y DGA          G V   G +++     L + TN   EL AI  A+   +D+G
Sbjct  5508  VAYADGACTGNPGPAGLGVVVVDGDRRIEISEYLGNGTNNIAELTAILRAVGAVEDTGKP  5329

Query  760   VNIVTDSQYALGIIQAQ-PDQSESELVNQIIEQLIKKEKVYLAWVPAHKGIGGNEQVDKL  818
             + + TDSQYA+G++Q     ++ +ELV + ++ ++K+ K  L +VP H G+  NE+ D+L
Sbjct  5328  LLVRTDSQYAIGVLQKGWKAKANTELVAE-VKDVLKRNKTRLEYVPGHSGVALNERADEL  5152

Query  819   VSAGIR  824
                 +R
Sbjct  5151  AREAVR  5134




Range 2: 5134 to 5502

Score:58.9 bits(141), Expect:8e-04,
Method:Compositional matrix adjust.,
Identities:40/124(32%), Positives:66/124(53%), Gaps:8/124(6%)

Query  588   YVDGAANRETKLGKAGYVTNRGRQKVVT---LTDTTNQKTELQAIYLAL---QDSGLEVN  641
             Y DGA          G V   G +++     L + TN   EL AI  A+   +D+G  +
Sbjct  5502  YADGACTGNPGPAGLGVVVVDGDRRIEISEYLGNGTNNIAELTAILRAVGAVEDTGKPLL  5323

Query  642   IVTDSQYALGIIQAQ-PDQSESELVNQIIEQLIKKEKVYLAWVPAHKGIGGNEQVDKLVS  700
             + TDSQYA+G++Q     ++ +ELV + ++ ++K+ K  L +VP H G+  NE+ D+L
Sbjct  5322  VRTDSQYAIGVLQKGWKAKANTELVAE-VKDVLKRNKTRLEYVPGHSGVALNERADELAR  5146

Query  701   AGIR  704
               +R
```

```
Sbjct  5145  EAVR  5134
```

>Acinetobacter baumannii strain BM4825 NODE_352_length_342_cov_0.830189, whole genome shotgun sequence
Sequence ID: NZ_JBBOMF010000201.1 Length: 342
Range 1: 1 to 333

Score:54.7 bits(130), Expect:4e-04,
Method:Composition-based stats.,
Identities:37/124(30%), Positives:59/124(47%), Gaps:19/124(15%)

```
Query  201  NPYNTPVFAIKKKDSTKWRKLVDFRELNKRTQDFWEVQLGIPHP------AGLKKKKSVT  254
            +PY+     ++K +       +DFR LN  TQ   E     P P      A L++ K  +
Sbjct  333  SPYSAGTVLLRKPNGEHLL-CIDFRLLNAVTQFQPE-----PLPDQTDLFAKLQRAKYFS  172

Query  255  VLDVGDAYFSVPLDEDFRKYTAFTIPSINNETPGIRYQYNVLPQGWKGSPAIFQSSMTKI  314
             +D+   Y+ +P+    RKY AF+       T    +++NVLP G   +P+IF + M K+
Sbjct  171  RIDLSKGYYQIPVQPQVRKYLAFS-------TMEGHFEFNVLPFGLHNAPSIFSAMMRKL  13

Query  315  LEPF  318
            L P
Sbjct  12   LAPL  1
```

>MAG: Thiolapillus sp. isolate Ifr-SOX-238 NODE_5703_length_2294_cov_6.676024, whole genome shotgun sequence
Sequence ID: NZ_JANEJA010000497.1 Length: 2294
Range 1: 1495 to 2034

Score:58.9 bits(141), Expect:6e-04,
Method:Compositional matrix adjust.,
Identities:59/204(29%), Positives:96/204(47%), Gaps:32/204(15%)

```
Query  167   KVKQWPLTEEKIKALVEICTEMEKEGKISKIGPE-NPYNTPVFAIKKKDSTKWRKLVDFR  225
             ++K +P+   K +   EI  E++    +  I P  + Y++P+  +KKKD T  R  +DFR
Sbjct  1495  RLKSYPIPFAKRE---EIRKEVKSMLDLGVIEPSTSSYSSPIVLVKKKDGTN-RFCIDFR  1662

Query  226   ELNKRTQDFWEVQLGIPHP------AGLKKKKSVTVLDVGDAYFSVPLDEDFRKYTAFTI  279
              LNK T     V    P P      A L K +  T +D+   Y+ +P+ ++ R  TAF
Sbjct  1663  ALNKVT-----VFDAEPIPDTEELFARLTKARFFTKIDLAKGYWQIPVKQEDRHKTAF--  1821

Query  280   PSINNETPGIRYQYNVLPQGWKGSPAIFQSSMTKI-LEPFRKQNPDIVIYQYMDDLYVGS  338
                  +TP   YQ+  +P G   +PA F   M K+ LE     N       + DD+ + +
Sbjct  1822  -----QTPLGLYQWTKMPFGLVSAPASFARMMRKLKLEENSALN-------FFDDILIAT  1965

Query  339   DLEIGQHRTKIEELRQHLLRWGLT  362
             +     H   ++ + Q L ++GLT
Sbjct  1966  E-RWDDHLESVKGVLQKLQKFGLT  2034
```

>Klebsiella pneumoniae strain BL3427 NODE_36_length_966_cov_0.293206, whole genome shotgun sequence
Sequence ID: NZ_VIDH01000036.1 Length: 966
Range 1: 228 to 599

Score:57.8 bits(138), Expect:6e-04,
Method:Compositional matrix adjust.,
Identities:40/133(30%), Positives:70/133(52%), Gaps:17/133(12%)

```
Query  257  DVGDAYFSVPLDEDFRKYTAFTIPSINNETPGIRYQYN--VLPQGWKGSPAIFQSSMTKI  314
             V DA+F + LD++ + Y AF    + N T G + Q +  VLPQG+K SP IF + + K
Sbjct  228  SV*DAFFCILLDKNSQAYFAF---ELQNPTTGQKTQLSWAVLPQGFKNSPMIFGNQLAKE  398

Query  315  LEPFRKQNPDIVIYQYMDDLYVGSDLEIGQHRTKIEELRQHLLRWGLTTPDKKHQKEPP-  373
            LE  +   P+ ++ QY+DD+ + ++      R +   L  +LL + L   D +  K+
Sbjct  399  LETCKIIYPERLVLQYVDDILIATE-----TREEYFNLTTNLLNF-LGQSDNRFSKKKAQ  560

Query  374  -----FLWMGYEL  381
                 +++G+E+
Sbjct  561  IGKETVMYLGFEI  599
```

>Polyangium sp. 6x1 NODE_22_length_166486_cov_22.101912, whole genome shotgun sequence
Sequence ID: NZ_JARZHH010000022.1 Length: 166486
Range 1: 110098 to 110472

Score:59.3 bits(142), Expect:8e-04,
Method:Compositional matrix adjust.,
Identities:42/126(33%), Positives:63/126(50%), Gaps:8/126(6%)

```
Query  706     VLYVDGAANRETKLGKAGYVTNRGRQKVVT---LTDTTNQKTELQAIYLALQ---DSGLE  759
               V Y DGA           G +   G +++     L   TN   EL AI  A++    +G
Sbjct  110098  VAYADGACTGNPGPAGLGVLVEDGDRRIEISEYLGQGTNNIAELTAILRAVEAVEGTGKS  110277

Query  760     VNIVTDSQYALGIIQAQ-PDQSESELVNQIIEQLIKKEKVYLAWVPAHKGIGGNEQVDKL  818
               + + TDSQYA+G++Q     ++ +ELV  + + L KK K  L +VP H G+ GNE+ D L
Sbjct  110278  LLVRTDSQYAIGVLQKGWKAKANTELVAAVKDAL-KKSKARLEYVPGHSGVRGNERADAL  110454

Query  819     VSAGIR  824
                   +R
Sbjct  110455  AREAVR  110472
```

Range 2: 110104 to 110472

Score:57.8 bits(138), Expect:0.002,
Method:Compositional matrix adjust.,
Identities:41/124(33%), Positives:62/124(50%), Gaps:8/124(6%)

```
Query  588     YVDGAANRETKLGKAGYVTNRGRQKVVT---LTDTTNQKTELQAIYLALQ---DSGLEVN  641
               Y DGA           G +   G +++     L   TN   EL AI  A++    +G  +
Sbjct  110104  YADGACTGNPGPAGLGVLVEDGDRRIEISEYLGQGTNNIAELTAILRAVEAVEGTGKSLL  110283

Query  642     IVTDSQYALGIIQAQ-PDQSESELVNQIIEQLIKKEKVYLAWVPAHKGIGGNEQVDKLVS  700
               + TDSQYA+G++Q     ++ +ELV  + + L KK K  L +VP H G+ GNE+ D L
Sbjct  110284  VRTDSQYAIGVLQKGWKAKANTELVAAVKDAL-KKSKARLEYVPGHSGVRGNERADALAR  110460

Query  701     AGIR  704
                 +R
Sbjct  110461  EAVR  110472
```

>Penaeicola halotolerans strain LMIT005 NODE_33_length_336_cov_0.594595, whole genome shotgun sequence
Sequence ID: NZ_JAEAGO010000029.1 Length: 336

```
Range 1: 3 to 176

Score:53.1 bits(126), Expect:0.001,
Method:Compositional matrix adjust.,
Identities:28/61(46%), Positives:39/61(63%), Gaps:3/61(4%)

Query  169  KQWPLTEEKIKALVEICTEMEKEGKISKIGPENPYNTPVFAIKKKDSTKWRKLVDFRELN  228
            KQWPL EEK++AL ++  E    G I +    +P+N+PVF +KKK   KWR + D R +N
Sbjct  176  KQWPLAEEKLQALEQLVQEQLDAGHIEE--STSPWNSPVFVVKKKFG-KWRMVTDLRAIN  6

Query  229  K  229
            K
Sbjct  5    K  3




>Candidatus Erwinia dacicola strain IL contig_608, whole genome shotgun sequence
Sequence ID: NZ_MAYS01000566.1 Length: 1576
Range 1: 725 to 1012

Score:57.8 bits(138), Expect:0.001,
Method:Compositional matrix adjust.,
Identities:33/107(31%), Positives:56/107(52%), Gaps:17/107(15%)

Query  217   KWRKLVDFRELNKRTQDFWEVQLGIPHPA------GLKKKKSVTVLDVGDAYFSVPLDED  270
             K RK  D+R+LN R+     V    P P        L++ + ++ LD+ + Y+ +PL++
Sbjct  725   KRRKCTDYRQLNARS-----VPDAYPLPRIQHILDKLRQARYISSLDLKNGYWQIPLEQG  889

Query  271   FRKYTAFTIPSINNETPGIRYQYNVLPQGWKGSPAIFQSSMTKILEP  317
              R YTAFT+P           +Q+ V+P G   +PA FQ  + +++ P
Sbjct  890   SRPYTAFTVPGRG------LFQWRVMPFGLHSAPATFQRVLDQVIGP  1012




>Escherichia coli strain EC53 NODE_385_length_1072_cov_0.533668, whole genome shotgun sequence
Sequence ID: NZ_RDNE01000376.1 Length: 1072
Range 1: 139 to 579

Score:56.6 bits(135), Expect:0.002,
Method:Compositional matrix adjust.,
Identities:43/147(29%), Positives:68/147(46%), Gaps:15/147(10%)

Query  860  AKEIVASCDKCQLKGEAMHGQ-VDCSP-------GIWQLD-----CTHLEGKVILVAVHV  906
            A +   SCD CQ +G     Q +  +P        +W +D      +    K ILVAV
Sbjct  139  AHDFAKSCDCCQREGGISKRQELPMNPILVIELFDVWGIDFMGPFVSSHGMKYILVAVDY  318

Query  907  ASGYIEAEVIPAETGQETAYFLLK-LAGRWPV-KTIHTDNGSNFTGATVRAACWWAGIKQ  964
             S ++EA  +    G+    FL K +  R+   + I +D GS+F     +A     G+
Sbjct  319  VSKWVEAVALSNNEGKSVTAFLKKNIFSRFGTPRAIISDGGSHFCNKLFKALLEKYGVCH  498

Query  965  EFGIPYNPQSQGVVESMNKELKKIIGQ  991
                PY+PQ+ G VE  N+E+K+I+ +
Sbjct  499  NVATPYHPQTSGQVEVSNREIKQILAK  579




>Enterococcus faecalis strain 17838 NODE_651_length_288_cov_1.37267, whole genome shotgun sequence
Sequence ID: NZ_QNGS01000142.1 Length: 288
Range 1: 6 to 281

Score:52.0 bits(123), Expect:0.002,
Method:Compositional matrix adjust.,
Identities:31/92(34%), Positives:50/92(54%), Gaps:2/92(2%)

Query  900  ILVAVHVASGYIEAEVIPAETGQETAYFLLK-LAGRWPV-KTIHTDNGSNFTGATVRAAC  957
            ILVAV   S ++EA   P    +    FL K +  R+ V + + +D G++F    + +A
Sbjct  6    ILVAVDYVSKWVEAIATPNNDTKTVLKFLQKHIFSRFGVPRVLISDGGTHFCNKQLYSAM  185

Query  958  WWAGIKQEFGIPYNPQSQGVVESMNKELKKII  989
               GI+ +   PY+PQ+ G  E  N+ELK+I+
Sbjct  186  VRYGIRHKVATPYHPQTNGQAEVSNRELKRIL  281




>Piscirickettsia salmonis strain FIUCHILE-89L01, whole genome shotgun sequence
Sequence ID: NZ_LELB01000266.1 Length: 2765
Range 1: 476 to 865

Score:57.4 bits(137), Expect:0.002,
Method:Compositional matrix adjust.,
Identities:41/141(29%), Positives:67/141(47%), Gaps:12/141(8%)

Query  200  ENPYNTPVFAIKKKDSTKWRKLVDFRELNKRT-QDFWEVQLGIPHPAGLKKKKSVTVLDV  258
            E+ Y +PV  +KK+D +  R  VDFR LN+ T +D + + L   H   L   K  + LD+
Sbjct  865  ESEYASPVLLVKKRDGSD-RLCVDFRALNRITVKDRYPLPLIDDHIDRLGSSKYFSCLDM  689

Query  259  GDAYFSVPLDEDFRKYTAFTIPSINNETPGIRYQYNVLPQGWKGSPAIFQSSMTKILEPF  318
               +  +P+ ED    T F        TP   ++Y  +P G   SP ++Q  +   L
Sbjct  688  ASGFHQIPIHEDSIHKTGFV-------TPEGHFEYLKMPFGLCNSPTVYQRIINGTLRRL  530

Query  319  RKQNPDIVIYQYMDDLYVGSD  339
             +    +V   Y+DD+ + S+
Sbjct  529  IETGNVLV---YVDDVLLVSE  476




>Escherichia coli strain 8551F 237, whole genome shotgun sequence
Sequence ID: NZ_JAFCNN010000170.1 Length: 689
Range 1: 93 to 665

Score:54.7 bits(130), Expect:0.003,
Method:Compositional matrix adjust.,
Identities:60/216(28%), Positives:96/216(44%), Gaps:32/216(14%)

Query  176  EKIKALVEICTEMEKEGKISKIGPENPYNTPVFAIKKKDSTKWRKLVDFRELNKRTQDFW  235
            E++K  +E   E+ ++G I K    +P   PV  +KKKD    R  VDFR LN  T
Sbjct  665  EELKKQIE---ELLQQGFIQK--SNSPXAAPVLFVKKKDKA-LRLCVDFRLLNINT----  516

Query  236  EVQLGIPHP------AGLKKKKSVTVLDVGDAYFSVPLDEDFRKYTAFTIPSINNETPGI  289
             ++   P P        +   K  + LD+   YF + +D    + T F        TP
Sbjct  515  -IKNAFPIPLIEDLFMKIGTAKVFSKLDLMSGYFQIRMDPRDEEKTGFV-------TPFG  360

Query  290  RYQYNVLPQGWKGSPAIFQSSMTKILEPFRKQNPDIVIYQYMDDLYVGSDLEIGQHRTKI  349
             Y + V+P G    PA FQS M  IL       P++++  Y+DD+ + S      +H   +
Sbjct  359  HYHWRVMPFGVVNGPATFQSFMNDIL----GDTPNVLV--YLDDILIYSH-NRKEHERDL  201

Query  350  EELRQHLLRWGLTTPDKKHQKEPPFL-WMGYELHPD  384
            + + + L    L    +KK +     L ++G+ + PD
Sbjct  200  KRVLRVLRENKLIAKEKKCEFFKDTLDFLGHTITPD  93
```

```
>Pseudomonas aeruginosa isolate PSA-13, whole genome shotgun sequence
Sequence ID: NZ_FNME01000065.1 Length: 505
Range 1: 74 to 448

Score:53.5 bits(127), Expect:0.003,
Method:Composition-based stats.,
Identities:42/137(31%), Positives:68/137(49%), Gaps:13/137(9%)

Query  201  NPYNTPVFAIKKKDSTKWRKLVDFRELNKRT-QDFWEVQLGIPHPAGLKKKKSVTVLDVG  259
            +P     F +KKKD T  R  VD+R LN  T ++ + + L       L+  +  T LD+
Sbjct  448  SPAGAGFFFVKKKDGT-LRPCVDYRGLNDITVKNRYPLLLMSSAFEILQGARFFTKLDLR  272

Query  260  DAYFSVPLDEDFRKYTAFTIPSINNETPGIRYQYNVLPQGWKGSPAIFQSSMTKILEPFR  319
            +AY  V + E     TAF        TP   ++Y VLP G   +PA+FQ+ +   +L
Sbjct  271  NAYHLVRIREGDEWKTAFN-------TPLGHFEYRVLPFGLANAPAVFQALVNDVL----  125

Query  320  KQNPDIVIYQYMDDLYV  336
            +   +I ++ Y+DD+ +
Sbjct  124  RDMLNIFVFVYLDDILI  74




>Polyangium jinanense strain SDU14 Scaffold9, whole genome shotgun sequence
Sequence ID: NZ_JAGTJJ010000009.1 Length: 231060
Range 1: 11606 to 11965

Score:57.0 bits(136), Expect:0.004,
Method:Compositional matrix adjust.,
Identities:39/121(32%), Positives:65/121(53%), Gaps:8/121(6%)

Query  706    VLYVDGAANRETKLGKAGYVTNRGRQKVVT---LTDTTNQKTELQAIYLAL---QDSGLE  759
              V Y DGA          G V   G +++     L + TN   EL AI  A+   +D+G
Sbjct  11606  VAYADGACTGNPGPAGLGVVVVDGDRRIEISEYLGNGTNNIAELTAILRAVGAVEDTGKP  11785

Query  760    VNIVTDSQYALGIIQAQ-PDQSESELVNQIIEQLIKKEKVYLAWVPAHKGIGGNEQVDKL  818
              + + TDSQYA+G++Q     ++ ++LV + ++ ++K+ K  L +VP H G+  NE+ D+L
Sbjct  11786  LLVRTDSQYAIGVLQKGWKAKANTDLVAE-VKDVLKRNKTRLEYVPGHSGVALNERADEL  11962

Query  819    V  819

Sbjct  11963  A  11965




Range 2: 11612 to 11965

Score:55.8 bits(133), Expect:0.009,
Method:Compositional matrix adjust.,
Identities:38/119(32%), Positives:64/119(53%), Gaps:8/119(6%)

Query  588    YVDGAANRETKLGKAGYVTNRGRQKVVT---LTDTTNQKTELQAIYLAL---QDSGLEVN  641
              Y DGA          G V   G +++     L + TN   EL AI  A+   +D+G  +
Sbjct  11612  YADGACTGNPGPAGLGVVVVDGDRRIEISEYLGNGTNNIAELTAILRAVGAVEDTGKPLL  11791

Query  642    IVTDSQYALGIIQAQ-PDQSESELVNQIIEQLIKKEKVYLAWVPAHKGIGGNEQVDKLV  699
              + TDSQYA+G++Q     ++ ++LV + ++ ++K+ K  L +VP H G+  NE+ D+L
Sbjct  11792  VRTDSQYAIGVLQKGWKAKANTDLVAE-VKDVLKRNKTRLEYVPGHSGVALNERADELA  11965




>Polyangium jinanense strain SDU13 Scaffold15, whole genome shotgun sequence
Sequence ID: NZ_JAGTJK010000015.1 Length: 231263
Range 1: 11759 to 12118

Score:57.0 bits(136), Expect:0.004,
Method:Compositional matrix adjust.,
Identities:39/121(32%), Positives:65/121(53%), Gaps:8/121(6%)

Query  706    VLYVDGAANRETKLGKAGYVTNRGRQKVVT---LTDTTNQKTELQAIYLAL---QDSGLE  759
              V Y DGA          G V   G +++     L + TN   EL AI  A+   +D+G
Sbjct  11759  VAYADGACTGNPGPAGLGVVVVDGDRRIEISEYLGNGTNNIAELTAILRAVGAVEDTGKP  11938

Query  760    VNIVTDSQYALGIIQAQ-PDQSESELVNQIIEQLIKKEKVYLAWVPAHKGIGGNEQVDKL  818
              + + TDSQYA+G++Q     ++ ++LV + ++ ++K+ K  L +VP H G+  NE+ D+L
Sbjct  11939  LLVRTDSQYAIGVLQKGWKAKANTDLVAE-VKDVLKRNKTRLEYVPGHSGVALNERADEL  12115

Query  819    V  819

Sbjct  12116  A  12118




Range 2: 11765 to 12118

Score:55.8 bits(133), Expect:0.009,
Method:Compositional matrix adjust.,
Identities:38/119(32%), Positives:64/119(53%), Gaps:8/119(6%)

Query  588    YVDGAANRETKLGKAGYVTNRGRQKVVT---LTDTTNQKTELQAIYLAL---QDSGLEVN  641
              Y DGA          G V   G +++     L + TN   EL AI  A+   +D+G  +
Sbjct  11765  YADGACTGNPGPAGLGVVVVDGDRRIEISEYLGNGTNNIAELTAILRAVGAVEDTGKPLL  11944

Query  642    IVTDSQYALGIIQAQ-PDQSESELVNQIIEQLIKKEKVYLAWVPAHKGIGGNEQVDKLV  699
              + TDSQYA+G++Q     ++ ++LV + ++ ++K+ K  L +VP H G+  NE+ D+L
Sbjct  11945  VRTDSQYAIGVLQKGWKAKANTDLVAE-VKDVLKRNKTRLEYVPGHSGVALNERADELA  12118




>Bradyrhizobium sp. 33ap4 NODE_290_length_534_cov_0.849015, whole genome shotgun sequence
Sequence ID: NZ_JAUMTJ010000287.1 Length: 534
Range 1: 107 to 481

Score:53.1 bits(126), Expect:0.004,
Method:Composition-based stats.,
Identities:42/143(29%), Positives:72/143(50%), Gaps:25/143(17%)

Query  201  NPYNTPVFAIKKKDSTKWRKLVDFRELNKRTQDFWEVQLGI-PHPA------GLKKKKSV  253
            +P+++PV  +KKKD T  R  VD+R+LN  T+       G+ P P        L++ + +
Sbjct  107  SPWSSPVVLVKKKDGT-LRFCVDYRKLNSVTKK------GVCPLPRIDDTLDRLRRSRYL  265

Query  254  TVLDVGDAYFSVPLDEDFRKYTAFTIPSINNETPGIRYQYNVLPQGWKGSPAIFQSSMTK  313
            + +D+   ++ + +DE  R+ TAF        TP   Y++ V+P G    PA FQ  +
Sbjct  266  SSMDLKSGHWQIEVDERDREKTAFI-------TPYGLYEFKVMPSGLCTPPATFQRVIDT  424

Query  314  ILEPFRKQNPDIVIYQYMDDLYV  336
```

```
            +L   + Q   +    Y+DD+ V
Sbjct  425  VLAGPKWQTCLV----YLDDVVV  481
```

>Eikenella corrodens strain 491_NMEN, whole genome shotgun sequence
Sequence ID: NZ_JVFA01000031.1 Length: 659
Range 1: 60 to 449

Score:53.9 bits(128), Expect:0.005,
Method:Compositional matrix adjust.,
Identities:37/130(28%), Positives:60/130(46%), Gaps:9/130(6%)

```
Query  900   ILVAVHVASGYIEAEVIPAETGQETAYFLLKLA---GRWPVKTIHTDNGSNFTGATVRA-  955
             + VA+   S  + A ++P +T    A FL +       + ++ +++DNGS + G    A
Sbjct  449   LFVAIDDFSRELYAAILPDKTADSAAKFLAEQVIEPCPYLIECVYSDNGSEYKGGANHAF  270

Query  956   --ACWWAGIKQEFGIPYNPQSQGVVESMNKELKKI--IGQVRDQAEHL-KTAVQMAVFIH  1010
                 CW  GI Q+F  P  PQ+ G  E + + L ++    Q  D  EH  K   +   F +
Sbjct  269   GVTCWENGINQKFTRPARPQTNGKAERVIRTLMEMWHEKQSFDSPEHRQKELCRFVNFYN  90

Query  1011  NFKRKGGIGG  1020
               K  G +GG
Sbjct  89    TVKPHGSLGG  60
```

>Enterobacter cloacae complex sp. GF14B NODE_151_length_4351_cov_0.721117, whole genome shotgun sequence
Sequence ID: NZ_QFRV01000155.1 Length: 4351
Range 1: 3283 to 3981

Score:55.8 bits(133), Expect:0.006,
Method:Compositional matrix adjust.,
Identities:71/263(27%), Positives:109/263(41%), Gaps:52/263(19%)

```
Query  153   ETVPVKLKPGMDGPKVKQWPLTEEKIKALVEICTEMEKEGKISKIGPENPYNTPVFAIKK  212
             E + ++LK G    + +   + +E+++AL E   ++ K G I  +     + +PV    K
Sbjct  3283  EEMHIELKEGAKPVRQRLRRMGQEQMEALREEVDKLLKAGFIVPV-ETTEWVSPVVVSPK  3459

Query  213   KDSTKWRKLVDFRELNKRTQDFWEVQLGIPHP-----------AGLKKKKSVTVLDVGDA  261
             KD  KWR  +DF+ LN  T+          P+P            AG ++    +V D
Sbjct  3460  KDG-KWRICIDFKPLNAATKKN-------PYPLPFIDQILDSVAGYER---YSVCDGFSG  3606

Query  262   YFSVPLDEDFRKYTAFTIPSINNETPGIRYQYNVLPQGWKGSPAIFQSSMTKILEPFRKQ  321
             YF + +    R  T F        TP   Y Y VLP G    P+ +Q     +L PF
Sbjct  3607  YFQLKIALKDRTKTTFI-------TPWGCYCYTVLPFGLSNGPSFYQERANWVLAPFIGS  3765

Query  322   NPDIVIYQYMDDLYVGSDLEIGQHRTKIEELRQHLLRW----GLTTPDKKHQKEPPFLWM  377
                  +  +MDD  V S     +H    E+L+  L R+    G   P K H  +P    +
Sbjct  3766  ----CVKAFMDDFCVYSSR--AEHH---EKLQMVLARYDECGGQLNPKKCHLAQPRVKLL  3918

Query  378   GY-------ELHPDKWTVQPIVL  393
             G+       E  PDK  V+ I+L
Sbjct  3919  GHMVSENGIEADPDK--VKSIIL  3981
```

>Methylobacterium radiotolerans strain MAMP 4754 MP-M-radiotolerans_contig_818, whole genome shotgun sequence
Sequence ID: NZ_NKQS01000805.1 Length: 253
Range 1: 15 to 155

Score:50.1 bits(118), Expect:0.007,
Method:Compositional matrix adjust.,
Identities:27/47(57%), Positives:33/47(70%), Gaps:0/47(0%)

```
Query  245  AGLKKKKSVTVLDVGDAYFSVPLDEDFRKYTAFTIPSINNETPGIRY  291
            AGLKKKK+ TVLDVGD   S  + E+FR+ T FT P+ +NET  I Y
Sbjct  15   AGLKKKKTETVLDVGDT*MSDTVTENFRQDTEFTRPNRHNETTWIIY  155
```

>Proteus terrae subsp. cibarius strain pt-OS9-1 NODE_353_length_267_cov_0.768421, whole genome shotgun sequence
Sequence ID: NZ_JBBEZU010000351.1 Length: 267
Range 1: 13 to 267

Score:50.1 bits(118), Expect:0.007,
Method:Composition-based stats.,
Identities:33/94(35%), Positives:50/94(53%), Gaps:12/94(12%)

```
Query  225  RELNKRTQDFWE---VQLGIPHPAGLKKKKSVTVLDVGDAYFSVPLDEDFRKYTAFTIPS  281
            RE+NK      E   +QL + +    +  K ++++D+ +AYF +P+ +  +KYT F   S
Sbjct  267  REINKHMISETEKPPLQLDVLN--SFQGAKFISIVDLNNAYFQIPISDSSKKYTGF---S  103

Query  282  INNETPGIRYQYNVLPQGWKGSPAIFQSSMTKIL  315
             N    G  Y YNVLPQG K S   F  +M  I+
Sbjct  102  FN----GKSYVYNVLPQGLKTSVGSFSRAMDFII  13
```

>Eikenella corrodens strain EI_05 NODE_86_length_856_cov_136.295, whole genome shotgun sequence
Sequence ID: NZ_VYYO01000085.1 Length: 856
Range 1: 408 to 797

Score:53.9 bits(128), Expect:0.007,
Method:Compositional matrix adjust.,
Identities:37/130(28%), Positives:61/130(46%), Gaps:9/130(6%)

```
Query  900   ILVAVHVASGYIEAEVIPAETGQETAYFLLKLA---GRWPVKTIHTDNGSNFTGATVRA-  955
             + VA+   S  + A ++P +T    A FL +       + ++ +++DNGS + G    A
Sbjct  408   LFVAIDDFSRELYAAILPDKTADSAAKFLAEQVIEPCPYLIECVYSDNGSEYKGGANHAF  587

Query  956   --ACWWAGIKQEFGIPYNPQSQGVVESMNKELKKI--IGQVRDQAEHL-KTAVQMAVFIH  1010
                 CW  GI Q+F  P  PQ+ G  E + + L ++    Q  D +EH  K   +   F +
Sbjct  588   GVTCWENGINQKFTRPARPQTNGKAERVIRTLMEMWHEKQSFDSSEHRQKELCRFVNFYN  767

Query  1011  NFKRKGGIGG  1020
               K  G +GG
Sbjct  768   TVKPHGSLGG  797
```

>Devosia insulae DS-56 contig234, whole genome shotgun sequence
Sequence ID: NZ_LAJE02000151.1 Length: 24799
Range 1: 24261 to 24653

Score:55.8 bits(133), Expect:0.008,
Method:Compositional matrix adjust.,
Identities:40/131(31%), Positives:59/131(45%), Gaps:24/131(18%)

```
Query  879    GQVDCSP-GIWQLDCTHL---EG--KVILVAVHVASGYIEAEVIPAETGQETAYFLLKL-  931
              G+ + +P G   +D  HL    EG    + VA+  A+ ++  E+I         A  L++L
Sbjct  24653  GRFEPAPVGFIHIDLKHLTRLEGHPSYVFVAIDRATRFVHIEIIERRDASTIAACLVRLL  24474

Query  932    -AGRWPVKTIHTDNGSNFT---GAT-------------VRAACWWAGIKQEFGIPYNPQS  974
               A  +PV TI TDNG+ FT   GA                   C W GI       P++PQ+
Sbjct  24473  TAFPYPVHTILTDNGAEFTDRFGAARWRNPSRPTGKHAFDLICRWYGIDHRLTRPFHPQT  24294

Query  975    QGVVESMNKEL  985
               G+ E  N+ L
Sbjct  24293  NGMAERFNRRL  24261
```

>Escherichia coli strain RF108-1 NODE_139_length_640_cov_0.833037, whole genome shotgun sequence
Sequence ID: NZ_JAAQGD010000139.1 Length: 640
Range 1: 47 to 286

Score:53.1 bits(126), Expect:0.008,
Method:Compositional matrix adjust.,
Identities:29/83(35%), Positives:42/83(50%), Gaps:8/83(9%)

```
Query  937   VKTIHTDNGSNFTGATVRAACWWAGIKQEFGIPYNPQSQGVVESMNKELKKIIGQVRDQA  996
             VK + +DNG  +T    +  C   GIK+EF  PYNPQ  GV E MN+ +++   +VR
Sbjct  286   VKCLRSDNGGEYTSKAFQDFCDMKGIKREFTAPYNPQQNGVAERMNRTIQE---KVRSML  116

Query  997   EHLKT-----AVQMAVFIHNFKR  1014
              +        A  +A+ IH   R
Sbjct  115   SYASLPNGF*AEALAIAIHLINR  47
```

>Streptococcus mutans strain PKUSS-9 C209, whole genome shotgun sequence
Sequence ID: NZ_NBFI01000031.1 Length: 307
Range 1: 12 to 230

Score:50.1 bits(118), Expect:0.009,
Method:Composition-based stats.,
Identities:29/75(39%), Positives:43/75(57%), Gaps:3/75(4%)

```
Query  296  LPQGWKGSPAIFQSSMTKILEPFRKQNPDIVIYQYMDDLYVGSDLEIG-QHRTKIEELRQ  354
            LPQG+K SP +F  ++ + L  FR Q+PD+++ QY+DDL + +  E+  Q  T+   L Q
Sbjct  12   LPQGFKNSPTLFDEALHRDLADFRIQHPDLILLQYVDDLLLAATSELDCQQGTR--ALLQ  185

Query  355  HLLRWGLTTPDKKHQ  369
             L   G     KK Q
Sbjct  186  TLGNLGYRASAKKAQ  230
```

>Klebsiella pneumoniae strain A13 NODE_160_length_467_cov_0.764103, whole genome shotgun sequence
Sequence ID: NZ_JARWSQ010000155.1 Length: 333
Range 1: 5 to 316

Score:50.1 bits(118), Expect:0.014,
Method:Composition-based stats.,
Identities:34/112(30%), Positives:56/112(50%), Gaps:9/112(8%)

```
Query  201  NPYNTPVFAIKKKDSTKWRKLVDFRELNKRT-QDFWEVQLGIPHPAGLKKKKSVTVLDVG  259
            +PY++PV  ++KKD T WR  +D++ LN  T +D + + +        LK  +  T LD+
Sbjct  316  SPYSSPVLLVRKKDGT-WRMCIDYQALNSMTVKDKYPIPVVDELLDELKGARVFTKLDLR  140

Query  260  DAYFSVPLDEDFRKYTAFTIPSINNETPGIRYQYNVLPQGWKGSPAIFQSSM  311
              Y  + + +D    TAF        T    Y++ V+  G   +P+ FQS M
Sbjct  139  SGYHQIRVCDDGIPKTAF-------RTHDGHYEFLVMSFGLTNAPSTFQSLM  5
```

>Escherichia coli strain EC86 NODE_205_length_902_cov_0.835152, whole genome shotgun sequence
Sequence ID: NZ_RDNB01000202.1 Length: 902
Range 1: 619 to 900

Score:53.1 bits(126), Expect:0.019,
Method:Compositional matrix adjust.,
Identities:31/94(33%), Positives:50/94(53%), Gaps:2/94(2%)

```
Query  900  ILVAVHVASGYIEAEVIPAETGQETAYFLLK-LAGRWPV-KTIHTDNGSNFTGATVRAAC  957
            ILVAV   S ++EA  +    G+    FL K +  R+   + I +D GS+F     +A
Sbjct  900  ILVAVDYVSKWVEAVALSNNEGKSVTAFLKKNIFSRFGTPRAIISDGGSHFCNKLFKALL  721

Query  958  WWAGIKQEFGIPYNPQSQGVVESMNKELKKIIGQ  991
              G++     PY+PQ+ G VE  N+E+K+I+ +
Sbjct  720  EKYGVRHNVATPYHPQTSGQVEVSNREIKQILAK  619
```

>Klebsiella pneumoniae strain kp110 NODE_70_length_728_cov_1.213518, whole genome shotgun sequence
Sequence ID: NZ_JAPMWX010000070.1 Length: 728
Range 1: 10 to 441

Score:52.4 bits(124), Expect:0.019,
Method:Compositional matrix adjust.,
Identities:40/156(26%), Positives:74/156(47%), Gaps:16/156(10%)

```
Query  157  VKLKPGMDGPKVKQWPLTEEKIKALVEICTEMEKEGKISKIGPENPYNTPVFAIKKKDST  216
            ++LKPG +   +KQWPL  E++ A+ +   +    G + +    +P ++P F ++K  +
Sbjct  441  IELKPGSNYCVMKQWPLPREQVLAIDKFFADRLAAGHVRE--STSPRSSPTFCVRKA-TG  271

Query  217  KWRKLVDFRELNKRTQDFWEVQLGIPHP----AGLKKKKSVTVLDVGDAYFSVPLDEDFR  272
             W  +  F +LN  T      Q  IP       G+ K    + +D+ D ++ + + E
Sbjct  270  GWWIVHAFNKLNAATV---PAQTPIPRKDVIIDGMSKITIFSSMDLMDGFYQILMRERDV  100

Query  273  KYTAFTIPSINNETPGIRYQYNVLPQGWKGSPAIFQ  308
             +TA + PS      G+ ++  V+P G   +PA F
Sbjct  99   PFTAVSTPS------GMLWERLVMP*GLSNAPATFN  10
```

>Enterococcus faecalis strain 15407 NODE_710_length_285_cov_0.436709, whole genome shotgun sequence
Sequence ID: NZ_QNGQ01000207.1 Length: 285
Range 1: 17 to 283

Score:48.9 bits(115), Expect:0.022,
Method:Compositional matrix adjust.,
Identities:29/89(33%), Positives:46/89(51%), Gaps:2/89(2%)

```
Query  903  AVHVASGYIEAEVIPAETGQETAYFLLK-LAGRWPV-KTIHTDNGSNFTGATVRAACWWA  960
```

```
           AV   S ++EA   P    +    FL K +  R+ V + + +D GS+F    + A
Sbjct  283  AVDYVSKWVEAIATPTNDNKVVMNFLRKNIFSRFGVPRALISDGGSHFCNRPLEALLRY  104

Query  961  GIKQEFGIPYNPQSQGVVESMNKELKKII  989
            G+K +   PY+PQ+ G  E  N+ELK+I+
Sbjct  103  GVKHKVATPYHPQTSGQTEISNRELKRIL  17
```

```
>Marmoricola endophyticus strain CGMCC 1.16067 sequence01, whole genome shotgun sequence
Sequence ID: NZ_BMKQ01000001.1 Length: 3374816
Range 1: 339118 to 339573

Score:54.3 bits(129), Expect:0.023,
Method:Compositional matrix adjust.,
Identities:43/154(28%), Positives:65/154(42%), Gaps:17/154(11%)

Query  683     VPAHKGIGGNEQVDKLVSAGIRKVLYVDGAANRETKLGKAGYVTNRGRQKVVTLTDTTNQ  742
               VPA  G  G  +V   VSA I   +Y DGA       G   +    G Q       TTNQ
Sbjct  339118  VPARLGAAGGSRVPHRVSAVIE--VYTDGACKGNPGPGGWAWAVRDGPQGSGGDPHTTNQ  339291

Query  743     KTELQAIYLALQDSGLEVNIVTDSQYALGIIQ---------------AQPDQSESELVNQ  787
               + E++A Y A+Q     + +V+DS Y +   +                A+   +  +L
Sbjct  339292  RMEIRAAYEAVQTLEGPLRVVSDSTYVVNCFRQGWWKGWHKRGWRNAAKQPVANRDLWEP  339471

Query  788     IIEQLIKKEKVYLAWVPAHKGIGGNEQVDKLVSA  821
                ++ + ++  V  AWV  H G   N+ VD+L  A
Sbjct  339472  FVDLVNERGDVEFAWVKGHSGHPMNDLVDRLAVA  339573
```

```
>Escherichia coli strain 90 NODE_134_length_1268_cov_0.816121, whole genome shotgun sequence
Sequence ID: NZ_JACAMA010000134.1 Length: 1268
Range 1: 537 to 767

Score:53.1 bits(126), Expect:0.025,
Method:Compositional matrix adjust.,
Identities:26/77(34%), Positives:40/77(51%), Gaps:8/77(10%)

Query  937   VKTIHTDNGSNFTGATVRAACWWAGIKQEFGIPYNPQSQGVVESMNKELKKIIGQVRDQ-  995
             VK + +DNG  +T    +  C   GIK+EF  PYNPQ  G+ E MN+ +++ +  +
Sbjct  767   VKCLRSDNGGEYTSKAFQDFCDMKGIKREFTAPYNPQQNGIAERMNRTIQEKVRSMLSNA  588

Query  996   -------AEHLKTAVQM  1005
                     AE L TA+ +
Sbjct  587   SLPNGFWAEALATAIHL  537
```

```
>MAG: Breoghania sp. isolate MO_035.B22 MO_035.c_6474, whole genome shotgun sequence
Sequence ID: NZ_JASJAV010000052.1 Length: 5920
Range 1: 2426 to 3088

Score:53.9 bits(128), Expect:0.025,
Method:Compositional matrix adjust.,
Identities:63/247(26%), Positives:104/247(42%), Gaps:50/247(20%)

Query  162   GMDGPKVKQWPLTEEKIKA-----LVEICTEMEKEG--KISKIGPEN-PYNTPVFAIKKK  213
             G   P  ++W   E          + EI T + K+   K+S+  PE+  + + +F + KK
Sbjct  3088  GTRPPLTRRWSAHESATSGNRAVLIQEIQTLLRKQAVEKVSR--PESLGFYSHIFLVPKK  2915

Query  214   DSTKWRKLVDFRELNKRTQDFWEVQLGIPH---------PAGLKKKKSVTVLDVGDAYFS  264
             +  + R +++ R LN+          L  PH           A ++     T +D+ DAYF
Sbjct  2914  NG-QLRPVINLRPLNR--------VLLCPHFQMETVGTIAAAIQPGDWATSVDLTDAYFH  2762

Query  265   VPLDEDFRKYTAFTIPSINNETPGIRYQYNVLPQGWKGSPAIFQSSMTKILEPF--RKQN  322
             +P+   FRKY  F +        G  +Q+  LP G   +P +     T++L P
Sbjct  2761  IPIAPWFRKYLRFVV-------AGEVFQFRALPFGLAPAPLV----CTRLLAPLAIHLHT  2615

Query  323   PDIVIYQYMDDLYV---GSDLEIGQHRTKIEELRQHLLRWGLTTPDKKHQKEPP--FLWM  377
               ++ ++Y+DDL +    SDL      T I      L R GL     K + EP   F+++
Sbjct  2614  QGVLFHRYLDDLLIRAPSSDLCRRWTDTTISL----LYRMGLGVNLDKSELEPSRDFVYV  2447

Query  378   GYELHPD  384
             G     D
Sbjct  2446  GVRFRTD  2426
```

```
>Devosia sp. Root413D1 contig_4, whole genome shotgun sequence
Sequence ID: NZ_LMEA01000009.1 Length: 575790
Range 1: 574945 to 575349

Score:54.3 bits(129), Expect:0.025,
Method:Compositional matrix adjust.,
Identities:40/135(30%), Positives:60/135(44%), Gaps:24/135(17%)

Query  879     GQVDCSP-GIWQLDCTHL---EG--KVILVAVHVASGYIEAEVIPAETGQETAYFLLKL-  931
               G+ + +P G   +D  HL   EG    + VA+  A+ ++  E+I        A  L++L
Sbjct  575349  GRFEPAPVGFIHIDLKHLTRLEGHPSYVFVAIDRATRFVHIEIIERRDASTIAACLVRLL  575170

Query  932      -AGRWPVKTIHTDNGSNFT---GAT-------------VRAACWWAGIKQEFGIPYNPQS  974
                A  +PV TI TDNG+ FT   GA                   C W GI      P++PQ+
Sbjct  575169  TAFPYPVHTILTDNGAEFTDRFGAARWRNPSKPTGKHAFDLVCRWYGIDHRLTRPFHPQT  574990

Query  975     QGVVESMNKELKKII  989
                G+ E  N+ L   +
Sbjct  574989  NGMAERFNRRLADAL  574945
```

```
>MAG: Breoghania sp. isolate MO_035.B22 MO_035.c_2803, whole genome shotgun sequence
Sequence ID: NZ_JASJAV010000047.1 Length: 6353
Range 1: 3126 to 3488

Score:53.9 bits(128), Expect:0.027,
Method:Compositional matrix adjust.,
Identities:40/141(28%), Positives:64/141(45%), Gaps:31/141(21%)

Query  207   VFAIKKKDSTKWRKLVDFRELNKRTQDFWEVQLGIPH---------PAGLKKKKSVTVLD  257
             VF + KK+  K R +++ R LN+         L  PH           A ++     T LD
Sbjct  3488  VFLVPKKNG-KLRPVINLRCLNR--------VLHCPHFQMETVGSISAAIQPGDWATSLD  3336

Query  258   VGDAYFSVPLDEDFRKYTAFTIPSINNETPGIRYQYNVLPQGWKGSPAIFQSSMTKILEP  317
             + DAYF +P+   FRKY  F +        G  +Q+  LP G   +P +     T++L P
Sbjct  3335  LTDAYFHIPIAPWFRKYLRFVV-------AGQVFQFRALPFGLAPAPLV----CTRLLAP  3189

Query  318   F--RKQNPDIVIYQYMDDLYV  336
                   +  I+ ++Y+DDL +
```

```
Sbjct  3188  LAIHLHSQGILFHRYLDDLLI  3126



>Clostridium paraputrificum strain 28-X28 Scaffold60_1, whole genome shotgun sequence
Sequence ID: NZ_RAYJ01000063.1 Length: 680
Range 1: 1 to 363

Score:51.6 bits(122), Expect:0.030,
Method:Compositional matrix adjust.,
Identities:41/135(30%), Positives:61/135(45%), Gaps:15/135(11%)

Query  206  PVFAIKKKDSTKWRKLVDFRELNKRTQ-DFWEVQLGIPHPAGLKKKKSVTVLDVGDAYFS  264
            P   + K D T +R   DFR++N  T+ D + +         +   K V+  D+   Y+
Sbjct  1    PCILVPKPDGT-YRLCTDFRKVNSVTKTDSYPIPRIDDCIDKIGSAKFVSKFDLLKGYWQ  177

Query  265  VPLDEDFRKYTAFTIPSINNETPGIRYQYNVLPQGWKGSPAIFQSSMTKILEPFRKQNPD  324
            VPL E  R+ +AF        TP   YQY V+P G K +PA FQ  +  +L
Sbjct  178  VPLTERAREISAFA-------TPDGLYQYKVMPFGMKNAPATFQRMIHSLLNHLEGCE--  330

Query  325  IVIYQYMDDLYVGSD  339
                 Y+DD+ + SD
Sbjct  331  ----AYIDDVIIYSD  363



>Agreia sp. PsM10 NODE_68_length_253_cov_0.818182, whole genome shotgun sequence
Sequence ID: NZ_JARUXC010000068.1 Length: 253
Range 1: 78 to 245

Score:48.1 bits(113), Expect:0.031,
Method:Composition-based stats.,
Identities:25/57(44%), Positives:33/57(57%), Gaps:1/57(1%)

Query  936  PVKTIHTDNGSNFTGATVRAACWWAGIKQEFGIPYNPQSQGVVESMNKELKKIIGQV  992
            P K I TDNGSNF    +R  C    I+     PY P+  GVVE+ NK +KKI+ ++
Sbjct  245  PSKII-TDNGSNFNNKMMRELCEEFKIEHHNSSPYRPKMNGVVEAANKNIKKIVQKM  78



>Campylobacter sp. JMF_08 NE1 NODE_13_length_3482_cov_0.037836, whole genome shotgun sequence
Sequence ID: NZ_JAOVUC010000013.1 Length: 3482
Range 1: 873 to 1355

Score:53.5 bits(127), Expect:0.031,
Method:Compositional matrix adjust.,
Identities:53/181(29%), Positives:86/181(47%), Gaps:25/181(13%)

Query  153   ETVPVKLKPGMDGPKVKQWPLTEEKIKALVEICTEMEKEGKISKIGP-ENPYNTPVFAIK  211
             +T P+K  P    P VKQ  L +E +K L+E             I P ++ Y++P   +
Sbjct  1355  DTQPIKQHPYRMNP-VKQQCLRDE-VKYLLEN----------DFIEPSQSNYSSPCILVP  1212

Query  212   KKDSTKWRKLVDFRELNKRTQ-DFWEVQLGIPHPAGLKKKKSVTVLDVGDAYFSVPLDED  270
             K + T +R   D+R++N  T+ D + +         + K K VT  D+   ++ VPL +
Sbjct  1211  KSNGT-YRMCTDYRKINAVTKTDSFPIPRIDDCIDNVGKAKYVTKFDLLKGFWQVPLTDR  1035

Query  271   FRKYTAFTIPSINNETPGIRYQYNVLPQGWKGSPAIFQSSMTKI---LEPFRKQNPDIVI  327
              ++ +AF        TP   +QY V+P G K SPA FQ  +  I   L+  +    D++I
Sbjct  1034  AKEISAFV-------TPDGLFQYKVMPFGMKNSPATFQRLINSILAGLDNCKAYIDDVII  876

Query  328   Y  328
             Y
Sbjct  875   Y  873



>Campylobacter sp. JMF_10 EL2 NODE_13_length_1688_cov_0.027690, whole genome shotgun sequence
Sequence ID: NZ_JAOVTV010000013.1 Length: 1688
Range 1: 397 to 831

Score:53.1 bits(126), Expect:0.033,
Method:Compositional matrix adjust.,
Identities:50/165(30%), Positives:79/165(47%), Gaps:22/165(13%)

Query  153  ETVPVKLKPGMDGPKVKQWPLTEEKIKALVEICTEMEKEGKISKIGP-ENPYNTPVFAIK  211
            +T P+K  P    P VKQ  L +E +K L+E             I P ++ Y++P   +
Sbjct  831  DTQPIKQHPYRMNP-VKQQCLKDE-VKYLLEN----------DFIEPSQSNYSSPCILVP  688

Query  212  KKDSTKWRKLVDFRELNKRTQ-DFWEVQLGIPHPAGLKKKKSVTVLDVGDAYFSVPLDED  270
            K + T +R   D+R++N  T+ D + +         + K K VT  D+   ++ VPL +
Sbjct  687  KSNGT-YRMCTDYRKINAVTKTDSFPIPRIDDCIDNVGKAKYVTKFDLLKGFWQVPLTDR  511

Query  271  FRKYTAFTIPSINNETPGIRYQYNVLPQGWKGSPAIFQSSMTKIL  315
             ++ +AF        TP   +QY V+P G K SPA FQ  +  IL
Sbjct  510  AKEISAFV-------TPDGLFQYKVMPFGMKNSPATFQRLINSIL  397



>Campylobacter sp. CS_ED1 strain Cs ED1 NODE_9_length_5400_cov_0.175250, whole genome shotgun sequence
Sequence ID: NZ_JAOVUL010000009.1 Length: 5400
Range 1: 3715 to 4197

Score:53.5 bits(127), Expect:0.035,
Method:Compositional matrix adjust.,
Identities:53/181(29%), Positives:86/181(47%), Gaps:25/181(13%)

Query  153   ETVPVKLKPGMDGPKVKQWPLTEEKIKALVEICTEMEKEGKISKIGP-ENPYNTPVFAIK  211
             +T P+K  P    P VKQ  L +E +K L+E             I P ++ Y++P   +
Sbjct  3715  DTQPIKQHPYRMNP-VKQQCLKDE-VKYLLEN----------DFIEPSQSNYSSPCILVP  3858

Query  212   KKDSTKWRKLVDFRELNKRTQ-DFWEVQLGIPHPAGLKKKKSVTVLDVGDAYFSVPLDED  270
             K + T +R   D+R++N  T+ D + +         + K K VT  D+   ++ VPL +
Sbjct  3859  KSNGT-YRMCTDYRKINAVTKTDSFPIPRIDDCIDNVGKAKYVTKFDLLKGFWQVPLTDR  4035

Query  271   FRKYTAFTIPSINNETPGIRYQYNVLPQGWKGSPAIFQSSMTKI---LEPFRKQNPDIVI  327
              ++ +AF        TP   +QY V+P G K SPA FQ  +  I   L+  +    D++I
Sbjct  4036  AKEISAFV-------TPDGLFQYKVMPFGMKNSPATFQRLINSILAGLDNCKAYIDDVII  4194

Query  328   Y  328
             Y
Sbjct  4195  Y  4197



>Klebsiella pneumoniae strain BL198 NODE_55_length_484_cov_0.434174, whole genome shotgun sequence
Sequence ID: NZ_VICV01000055.1 Length: 484
Range 1: 243 to 482
```

```
Score:50.1 bits(118), Expect:0.035,
Method:Compositional matrix adjust.,
Identities:29/83(35%), Positives:46/83(55%), Gaps:5/83(6%)

Query  261  AYFSVPLDEDFRKYTAFTIPSINNETPGIRYQ--YNVLPQGWKGSPAIFQSSMTKILEPF  318
            A+F +PL E  +K  AF     +N   G + Q  + VLPQG+K SP IF + + K LE +
Sbjct  482  AFFCLPLHEASQKLFAF---EWDNPKSGRKTQLTWTVLPQGFKNSPTIFGNQLAKDLESW  312

Query  319  RKQNPDIVIYQYMDDLYVGSDLE  341
               + +  + QY+DD+ + +  E
Sbjct  311  ETPSGEGKLLQYVDDILIATKTE  243




>Campylobacter sp. CN_NE4 strain Cn NE4 NODE_12_length_5576_cov_0.042351, whole genome shotgun sequence
Sequence ID: NZ_JAOVUF010000012.1 Length: 5576
Range 1: 3483 to 3965

Score:53.5 bits(127), Expect:0.037,
Method:Compositional matrix adjust.,
Identities:53/181(29%), Positives:86/181(47%), Gaps:25/181(13%)

Query  153   ETVPVKLKPGMDGPKVKQWPLTEEKIKALVEICTEMEKEGKISKIGP-ENPYNTPVFAIK  211
             +T P+K  P    P VKQ  L +E +K L+E             I P ++ Y++P   +
Sbjct  3483  DTQPIKQHPYRMNP-VKQQCLKDE-VKYLLEN----------DFIEPSQSNYSSPCILVP  3626

Query  212   KKDSTKWRKLVDFRELNKRTQ-DFWEVQLGIPHPAGLKKKKSVTVLDVGDAYFSVPLDED  270
             K + T +R   D+R++N  T+ D + +          + K K VT  D+   ++ VPL +
Sbjct  3627  KSNGT-YRMCTDYRKINAVTKTDSFPIPRIDDCIDNVGKAKYVTKFDLLKGFWQVPLTDR  3803

Query  271   FRKYTAFTIPSINNETPGIRYQYNVLPQGWKGSPAIFQSSMTKI---LEPFRKQNPDIVI  327
              ++ +AF        TP   +QY V+P G K SPA FQ  +  I   L+  +    D++I
Sbjct  3804  AKEISAFV-------TPDGLFQYKVMPFGMKNSPATFQRLINSILAGLDNCKAYIDDVII  3962

Query  328   Y  328
             Y
Sbjct  3963  Y  3965




>Pseudomonas aromaticivorans strain MAP12 47, whole genome shotgun sequence
Sequence ID: NZ_JAHRGL010000047.1 Length: 2177
Range 1: 5 to 367

Score:53.1 bits(126), Expect:0.037,
Method:Compositional matrix adjust.,
Identities:35/126(28%), Positives:66/126(52%), Gaps:10/126(7%)

Query  157  VKLKPGMDGPKVKQWPLTEEKIKALVEICTEMEKEGKISKIGPENPYNTPVFAIKKKDST  216
            V LK  +    V+Q+P+ +E  + +      + ++G +     ++P+NTP+  ++K  +
Sbjct  367  VTLKAAILPASVRQYPMPKEAREGIRPHIKRLLEQGILVPC--KSPWNTPLLPVRKPGTN  194

Query  217  KWRKLVDFRELNKRTQDFWEVQLGIPHP----AGLKKKKS-VTVLDVGDAYFSVPLDEDF  271
             +R + D RE+NKR +D   +   +P+P    +GL    +  TVLD+ DA+F + L
Sbjct  193  DYRPVQDLREVNKRIED---MHPTVPNPYNLLSGLPPNYTWYTVLDLKDAFFCLRLHPTN  23

Query  272  RKYTAF  277
            +   AF
Sbjct  22   QPIFAF  5




>Devosia sp. Root105 contig_7, whole genome shotgun sequence
Sequence ID: NZ_LMCR01000019.1 Length: 620452
Range 1: 183 to 521

Score:53.5 bits(127), Expect:0.038,
Method:Compositional matrix adjust.,
Identities:36/113(32%), Positives:51/113(45%), Gaps:20/113(17%)

Query  893  THLEG--KVILVAVHVASGYIEAEVIPAETGQETAYFLLKL--AGRWPVKTIHTDNGSNF  948
            T LEG    + VA+  A+ ++  E+I        A  L++L  A  +PV TI TDNG+ F
Sbjct  521  TRLEGHPSYVFVAIDRATRFVHIEIIERRDASTIAACLVRLLTAFPYPVHTILTDNGAEF  342

Query  949  T---GAT-------------VRAACWWAGIKQEFGIPYNPQSQGVVESMNKEL  985
            T   GA                   C W GI      P++PQ+ G+ E  N+ L
Sbjct  341  TDRFGAARWRNPSKPTGKHAFDLVCRWYGIDHRLTRPFHPQTNGMAERFNRRL  183




>Eikenella corrodens strain KCOM 1378 B230_1_2, whole genome shotgun sequence
Sequence ID: NZ_NHRS01000002.1 Length: 572658
Range 1: 511602 to 511991

Score:53.5 bits(127), Expect:0.040,
Method:Compositional matrix adjust.,
Identities:37/130(28%), Positives:60/130(46%), Gaps:9/130(6%)

Query  900     ILVAVHVASGYIEAEVIPAETGQETAYFLLKLA---GRWPVKTIHTDNGSNFTGATVRA-  955
               + VA+   S  + A ++P +T    A FL +       + ++ +++DNGS + G    A
Sbjct  511602  LFVAIDDFSRELYAAILPDKTADSAAKFLAEQVIEPCPYLIECVYSDNGSEYKGGANHAF  511781

Query  956     --ACWWAGIKQEFGIPYNPQSQGVVESMNKELKKI--IGQVRDQAEHL-KTAVQMAVFIH  1010
                 CW  GI Q+F  P  PQ+ G  E + + L ++    Q  D  EH  K   +   F +
Sbjct  511782  GVTCWENGINQKFTRPARPQTNGKAERVIRTLMEMWHEKQSFDSPEHRQKELCRFVNFYN  511961

Query  1011    NFKRKGGIGG  1020
                 K  G +GG
Sbjct  511962  TVKPHGSLGG  511991




>Wolbachia endosymbiont of Corcyra cephalonica isolate SYLI2103 plasmid unnamed3, complete sequence
Sequence ID: NZ_CP087956.1 Length: 5332
Range 1: 1719 to 2156

Score:53.5 bits(127), Expect:0.040,
Method:Compositional matrix adjust.,
Identities:46/158(29%), Positives:81/158(51%), Gaps:14/158(8%)

Query  201   NPYNTPVFAIKKKDSTKWRKLVDFRELNKRTQ-DFWEVQLGIPHPAGLKKKKSVTVLDVG  259
             +P+++PV   +KKD T  R  VD+R LN  T+ D + +          L   K  + +D+
Sbjct  1719  SPWSSPVVLVKKKDGTT-RFCVDYRMLNNVTKKDSYPLPRIDDTLDMLSGMKWFSTIDLK  1895

Query  260   DAYFSVPLDEDFRKYTAFTIPSINNETPGIRYQYNVLPQGWKGSPAIFQSSMTKILEPFR  319
               Y+ V +  + ++ TAF+   I N   G+ +Q+NV+P G   +PA F+  M +++
Sbjct  1896  SGYWQVEISPEDKEKTAFST-GIGN---GL-WQFNVMPFGLCNAPATFERLMERVMVGLL  2060

Query  320   KQNPDIVIYQYMDDLYV-GSDLEIGQHRTKIEELRQHL  356
```

```
                  V   Y+DD+ + G D E  +H   +E++ Q +
Sbjct  2061  GD----VCLVYLDDVIIMGRDFE--EHLKNLEKVFQRI  2156
```

>Eikenella corrodens ATCC 23834 Scfld2, whole genome shotgun sequence
Sequence ID: NZ_EQ973318.1 Length: 131618
Range 1: 59282 to 59671

Score:53.5 bits(127), Expect:0.040,
Method:Compositional matrix adjust.,
Identities:37/130(28%), Positives:60/130(46%), Gaps:9/130(6%)

```
Query  900    ILVAVHVASGYIEAEVIPAETGQETAYFLLKLA---GRWPVKTIHTDNGSNFTGATVRA-  955
              + VA+   S  + A ++P +T    A FL +       + ++ +++DNGS + G    A
Sbjct  59282  LFVAIDDFSRELYAAILPDKTADSAAKFLAEQVIEPCPYLIECVYSDNGSEYKGGANHAF  59461

Query  956    --ACWWAGIKQEFGIPYNPQSQGVVESMNKELKKI--IGQVRDQAEHL-KTAVQMAVFIH  1010
                 CW  GI Q+F  P  PQ+ G  E + + L ++    Q  D  EH  K   +    F +
Sbjct  59462  GVTCWENGINQKFTRPARPQTNGKAERVIRTLMEMWHEKQSFDSPEHRQKELCRFVNFYN  59641

Query  1011   NFKRKGGIGG  1020
                K  G +GG
Sbjct  59642  TVKPHGSLGG  59671
```

>Eikenella corrodens strain FDAARGOS_1474 chromosome, complete genome
Sequence ID: NZ_CP082861.1 Length: 2192571
Range 1: 344094 to 344483

Score:53.5 bits(127), Expect:0.041,
Method:Compositional matrix adjust.,
Identities:37/130(28%), Positives:60/130(46%), Gaps:9/130(6%)

```
Query  900     ILVAVHVASGYIEAEVIPAETGQETAYFLLKLA---GRWPVKTIHTDNGSNFTGATVRA-  955
               + VA+   S  + A ++P +T    A FL +       + ++ +++DNGS + G    A
Sbjct  344483  LFVAIDDFSRELYAAILPDKTADSAAKFLAEQVIEPCPYLIECVYSDNGSEYKGGANHAF  344304

Query  956     --ACWWAGIKQEFGIPYNPQSQGVVESMNKELKKI--IGQVRDQAEHL-KTAVQMAVFIH  1010
                  CW  GI Q+F  P  PQ+ G  E + + L ++    Q  D  EH  K   +    F +
Sbjct  344303  GVTCWENGINQKFTRPARPQTNGKAERVIRTLMEMWHEKQSFDSPEHRQKELCRFVNFYN  344124

Query  1011    NFKRKGGIGG  1020
                 K  G +GG
Sbjct  344123  TVKPHGSLGG  344094
```

>Eikenella corrodens strain NCTC10596 chromosome 1, complete sequence
Sequence ID: NZ_LT906482.1 Length: 2192577
Range 1: 716530 to 716919

Score:53.5 bits(127), Expect:0.041,
Method:Compositional matrix adjust.,
Identities:37/130(28%), Positives:60/130(46%), Gaps:9/130(6%)

```
Query  900     ILVAVHVASGYIEAEVIPAETGQETAYFLLKLA---GRWPVKTIHTDNGSNFTGATVRA-  955
               + VA+   S  + A ++P +T    A FL +       + ++ +++DNGS + G    A
Sbjct  716530  LFVAIDDFSRELYAAILPDKTADSAAKFLAEQVIEPCPYLIECVYSDNGSEYKGGANHAF  716709

Query  956     --ACWWAGIKQEFGIPYNPQSQGVVESMNKELKKI--IGQVRDQAEHL-KTAVQMAVFIH  1010
                  CW  GI Q+F  P  PQ+ G  E + + L ++    Q  D  EH  K   +    F +
Sbjct  716710  GVTCWENGINQKFTRPARPQTNGKAERVIRTLMEMWHEKQSFDSPEHRQKELCRFVNFYN  716889

Query  1011    NFKRKGGIGG  1020
                 K  G +GG
Sbjct  716890  TVKPHGSLGG  716919
```

>Lachnoclostridium phocaeense strain Marseille-P3177T chromosome contig00001
Sequence ID: NZ_LT635479.1 Length: 3500754
Range 1: 1457999 to 1458487

Score:53.5 bits(127), Expect:0.042,
Method:Compositional matrix adjust.,
Identities:47/173(27%), Positives:80/173(46%), Gaps:18/173(10%)

```
Query  569      LVKLWYQLEKEPIVGAETFYVDGAAN-RETKLGKAGYVTNRGRQKVVT-----LTDTTNQ  622
                +++L+ +L KE + G  T Y DG         G  G V      KV+         +TN
Sbjct  1458487  ILELFPELAKEKLPGKYTVYTDGGCGFNPGGPGGIGAVILDVEGKVIQEISKGFHASTNN  1458308

Query  623      KTELQAIYLALQD--SGLEVNIVTDSQYALGIIQAQPDQSESELVNQIIEQLIKKEKVYL  680
                + EL A   AL+   +G EV + +DSQY +  ++    ++++  + + +++ ++ +KV
Sbjct  1458307  RMELLAAIEALKTIPAGAEVKVKSDSQYLVQTMEEGWSRNKNHDLWKQLDEALRGKKVSF  1458128

Query  681      AWVPAHKGIGGNEQVDKLVSAGIRKVLYVDGAANRETKLGKAGYVTNRGRQKV  733
                 WV  H G   NE+ D+L +AGI          N    K    GY+  R   +V
Sbjct  1458127  KWVRGHNGNQWNERCDELATAGI----------NSSDKEEDTGYIPTRNASQV  1457999
```

>Roseiconus lacunae strain JC742 soapdenovo_K_105_82, whole genome shotgun sequence
Sequence ID: NZ_JAJMQV010000082.1 Length: 35968
Range 1: 13242 to 13676

Score:53.5 bits(127), Expect:0.044,
Method:Compositional matrix adjust.,
Identities:42/146(29%), Positives:70/146(47%), Gaps:26/146(17%)

```
Query  707    LYVDGAANRETKLGKAGYVT-------NRGRQKVVTLTDTTNQKTELQAIYLALQDSGLE  759
              LY DG+++ ++ +G   Y+        +   Q+  +    +N + EL+A+   L+  G E
Sbjct  13242  LYTDGSSDPDSGVGGWAYLIINEDSPLDLSDQQAGSHPRASNNRMELEAVLQGLRAIGEE  13421

Query  760    ---VNIVTDSQYA-------LGI-IQAQPDQ-------SESELVNQIIEQLIKKEKVYLA  801
                 + I TDS+Y        L I + ++PD        S   L  ++IE LI + +VY
Sbjct  13422  PACITIKTDSEYVERGLNEYLPIWLSSRPDSRFKDPIPSHPRLWRRVIE-LINRHEVYCI  13598

Query  802    WVPAHKGIGGNEQVDKLVSAGIRKVL  827
              W+P H G   NE VD+L  + +R  L
Sbjct  13599  WIPGHSGHTENEIVDRLAGSAVRDHL  13676
```

>Klebsiella pneumoniae strain KP231 NODE_2205_length_233_cov_2.807692, whole genome shotgun sequence
Sequence ID: NZ_JAUUHQ010001574.1 Length: 233
Range 1: 22 to 168

```
Score:47.4 bits(111), Expect:0.045,
Method:Compositional matrix adjust.,
Identities:23/49(47%), Positives:26/49(53%), Gaps:0/49(0%)

Query  937   VKTIHTDNGSNFTGATVRAACWWAGIKQEFGIPYNPQSQGVVESMNKEL  985
             VK I +DNGS F    V       GIK +F  PY PQ  GVVE  N+ L
Sbjct  22    VKKIRSDNGSEFKNLQVEEYLEEEGIKHKFSAPYTPQQNGVVERKNRTL  168
```

>Eikenella corrodens strain NML01-0040 Eikcor_contig000008, whole genome shotgun sequence
Sequence ID: NZ_LXSE01000008.1 Length: 65735
Range 1: 45607 to 45996

```
Score:53.1 bits(126), Expect:0.046,
Method:Compositional matrix adjust.,
Identities:37/130(28%), Positives:60/130(46%), Gaps:9/130(6%)

Query  900    ILVAVHVASGYIEAEVIPAETGQETAYFLLKLA---GRWPVKTIHTDNGSNFTGATVRA-  955
              + VA+    S   + A ++P +T    A FL +       + ++ +++DNGS + G    A
Sbjct  45607  LFVAIDDFSRELYAAILPDKTADSAAKFLAEQVIEPCPYLIECVYSDNGSEYKGGANHAF  45786

Query  956    --ACWWAGIKQEFGIPYNPQSQGVVESMNKELKKI--IGQVRDQAEHL-KTAVQMAVFIH  1010
                  CW  GI Q+F  P  PQ+ G  E + + L ++    Q  D  EH  K   +   F +
Sbjct  45787  GVTCWENGINQKFTRPARPQTNGKAERVIRTLMEMWHEKQSFDSPEHRQKELCRFVNFYN  45966

Query  1011   NFKRKGGIGG  1020
                 K  G +GG
Sbjct  45967  TVKPHGSLGG  45996
```

>Eikenella corrodens strain EI_05 NODE_1_length_165032_cov_49.9348, whole genome shotgun sequence
Sequence ID: NZ_VYYO01000001.1 Length: 165032
Range 1: 50396 to 50785

```
Score:53.1 bits(126), Expect:0.050,
Method:Compositional matrix adjust.,
Identities:37/130(28%), Positives:60/130(46%), Gaps:9/130(6%)

Query  900    ILVAVHVASGYIEAEVIPAETGQETAYFLLKLA---GRWPVKTIHTDNGSNFTGATVRA-  955
              + VA+    S   + A ++P +T    A FL +       + ++ +++DNGS + G    A
Sbjct  50396  LFVAIDDFSRELYAAILPDKTADSAAKFLAEQVIEPCPYLIECVYSDNGSEYKGGANHAF  50575

Query  956    --ACWWAGIKQEFGIPYNPQSQGVVESMNKELKKI--IGQVRDQAEHL-KTAVQMAVFIH  1010
                  CW  GI Q+F  P  PQ+ G  E + + L ++    Q  D  EH  K   +   F +
Sbjct  50576  GVTCWENGINQKFTRPARPQTNGKAERVIRTLMEMWHEKQSFDSPEHRQKELCRFVNFYN  50755

Query  1011   NFKRKGGIGG  1020
                 K  G +GG
Sbjct  50756  TVKPHGSLGG  50785
```

Query #10: sp|P04585|p51_RT_HV1H2 Gag-Pol polyprotein OS=Human immunodeficiency virus type 1 group M subtype B (isolate HXB2) OX=11706 GN=gag-pol PE=1 SV=4 Query ID: lcl|Query_2387718 Length: 560

Sequences producing significant alignments:

| Description | Scientific Name | Common Name | Taxid | Max Score | Total Score | Query cover | E Value | Per. Ident | Acc. Len | Accession |
|---|---|---|---|---|---|---|---|---|---|---|
| Enterococcus alcedinis strain CCM 8433 sequence23, whole genom... | Enterococcus... | NA | 1274384 | 1073 | 1073 | 100% | 0.0 | 94.29 | 6075 | NZ_BMDT010000... |
| Deinococcus aquiradiocola strain JCM 14371 sequence30, whole... | Deinococcus ... | NA | 393059 | 1073 | 1073 | 100% | 0.0 | 94.29 | 6128 | NZ_BMOE010000... |
| Actinoplanes campanulatus strain JCM 3059 sequence058, whole... | Actinoplanes... | NA | 113559 | 1073 | 1073 | 100% | 0.0 | 94.29 | 4754 | NZ_BMPW010000... |
| Klebsiella pneumoniae strain ARGID_31501, whole genome shotgun... | Klebsiella p... | NA | 573 | 1055 | 1055 | 100% | 0.0 | 92.50 | 2655 | NZ_CAKLPD0100... |
| Thalassovita mangrovi strain GS-10 Scaffold20_1, whole genome... | Thalassovita... | NA | 2692236 | 941 | 941 | 88% | 0.0 | 93.54 | 1488 | NZ_WWEN010000... |
| Enterococcus faecium strain A201R2B0... | Enterococcus... | NA | 1352 | 818 | 818 | 79% | 0.0 | 89.84 | 1589 | NZ_PGCT010000... |
| Enterococcus faecium strain A209R1B0C1... | Enterococcus... | NA | 1352 | 814 | 814 | 79% | 0.0 | 88.51 | 1819 | NZ_PGCR010000... |
| Listeria monocytogenes strain YA00082404... | Listeria mon... | NA | 1639 | 795 | 795 | 78% | 0.0 | 87.24 | 1816 | NZ_QEXE010000... |
| Escherichia coli strain YA00028237 contig_187, whole genome... | Escherichia ... | NA | 562 | 793 | 793 | 78% | 0.0 | 86.17 | 1828 | NZ_NGBP020001... |
| Enterococcus faecium strain A206R2B0... | Enterococcus... | NA | 1352 | 793 | 793 | 76% | 0.0 | 89.30 | 1761 | NZ_PGCU010000... |
| Klebsiella pneumoniae strain NUBRI-K 73, whole genome shotgun... | Klebsiella p... | NA | 573 | 759 | 759 | 75% | 0.0 | 86.97 | 1717 | NZ_SOYT010000... |
| Staphylococcus aureus strain G703N1B1... | Staphylococc... | NA | 1280 | 715 | 715 | 69% | 0.0 | 88.66 | 1173 | NZ_PGXA010000... |
| Escherichia coli strain 8374wB11... | Escherichia ... | NA | 562 | 654 | 654 | 60% | 0.0 | 93.26 | 1310 | NZ_WSJO010000... |
| Acinetobacter baumannii strain ACM-5... | Acinetobacte... | NA | 470 | 650 | 650 | 60% | 0.0 | 92.63 | 1274 | NZ_PYCX010000... |
| Escherichia coli strain 8374wB4 NODE_60_length_1385_cov_3.2655... | Escherichia ... | NA | 562 | 649 | 649 | 60% | 0.0 | 92.38 | 1385 | NZ_WSPX010000... |
| Escherichia coli strain 8374wG10... | Escherichia ... | NA | 562 | 648 | 648 | 60% | 0.0 | 93.18 | 1373 | NZ_WSJQ010000... |
| Methylocella silvestris strain TVC... | Methylocella... | NA | 199596 | 647 | 647 | 61% | 0.0 | 90.72 | 1453 | NZ_PDZR010000... |
| Escherichia coli strain 8374wH11... | Escherichia ... | NA | 562 | 644 | 644 | 60% | 0.0 | 92.58 | 1397 | NZ_WSJI010000... |
| Staphylococcus epidermidis strain PR246B0... | Staphylococc... | NA | 1282 | 644 | 644 | 65% | 0.0 | 85.01 | 1113 | NZ_PCFD010000... |
| Helicobacter acinonychis isolate 212_10, whole genome shotgun... | Helicobacter... | NA | 212 | 642 | 642 | 59% | 0.0 | 92.81 | 1526 | NZ_FZLW010000... |
| Klebsiella aerogenes strain AUH-KAM5... | Klebsiella a... | NA | 548 | 635 | 635 | 60% | 0.0 | 90.86 | 1427 | NZ_PYHJ010000... |
| Enterobacter hormaechei strain AUH-ENM6... | Enterobacter... | NA | 158836 | 629 | 629 | 59% | 0.0 | 91.59 | 1394 | NZ_PYHB010000... |
| Acinetobacter baumannii strain ACM-35... | Acinetobacte... | NA | 470 | 626 | 626 | 59% | 0.0 | 91.04 | 1409 | NZ_QEHN010000... |
| Staphylococcus aureus strain PN235B0... | Staphylococc... | NA | 1280 | 614 | 614 | 62% | 0.0 | 83.76 | 1083 | NZ_PDUY010001... |
| Klebsiella pneumoniae strain P7701... | Klebsiella p... | NA | 573 | 601 | 601 | 52% | 0.0 | 97.97 | 1305 | NZ_JACAWM0100... |
| Salirhabdus euzebyi strain Marseille-Q1438, whole genome shotg... | Salirhabdus ... | NA | 394506 | 600 | 600 | 52% | 0.0 | 97.30 | 1279 | NZ_CADDWK0100... |
| Methylobacterium radiotolerans strain MAMP 4754... | Methylobacte... | NA | 31998 | 588 | 588 | 60% | 0.0 | 85.25 | 1636 | NZ_NKQS010006... |
| Mycobacterium tuberculosis variant bovis strain CSURQ1132, who... | Mycobacteriu... | NA | 1765 | 578 | 1157 | 52% | 0.0 | 92.23 | 1524 | NZ_CAJEUK0100... |
| Mycobacterium tuberculosis variant bovis strain CSURQ1134, who... | Mycobacteriu... | NA | 1765 | 578 | 1157 | 52% | 0.0 | 92.23 | 1697 | NZ_CAJEUT0100... |
| Escherichia coli strain 8374wA11 NODE_37_length_940_cov_2.7589... | Escherichia ... | NA | 562 | 569 | 569 | 52% | 0.0 | 91.84 | 940 | NZ_WSPR010000... |
| Escherichia coli strain TzEc080 NODE_37_length_940_cov_2.75892... | Escherichia ... | NA | 562 | 569 | 569 | 52% | 0.0 | 91.84 | 940 | NZ_WSFZ010000... |
| Mycobacterium tuberculosis strain 32846_S30 184, whole genome... | Mycobacteriu... | NA | 1773 | 552 | 552 | 55% | 0.0 | 85.76 | 932 | NZ_QTBW010000... |
| Acinetobacter baumannii strain ACM-36... | Acinetobacte... | NA | 470 | 550 | 550 | 52% | 0.0 | 89.15 | 953 | NZ_QAGO010000... |
| Providencia rettgeri strain AFE652_S1 38_contig_28, whole geno... | Providencia ... | NA | 587 | 533 | 533 | 48% | 0.0 | 94.44 | 1127 | NZ_SHDK010000... |
| Mycobacterium tuberculosis variant bovis strain CSURQ1134, who... | Mycobacteriu... | NA | 1765 | 518 | 1036 | 49% | 5e-179 | 89.96 | 910 | NZ_CAJEUT0100... |
| Candidatus Tokpelaia sp. strain RhiAcro1 isolate RhiAcro1_RAo1... | Candidatus T... | NA | 2233777 | 511 | 511 | 49% | 8e-177 | 89.45 | 826 | NZ_QLZD010000... |
| Mycobacterium tuberculosis variant bovis strain CSURQ1139, who... | Mycobacteriu... | NA | 1765 | 510 | 1020 | 46% | 4e-176 | 93.05 | 848 | NZ_CAJEUO0100... |
| Levilactobacillus brevis strain CRL 2013 scaffold12, whole... | Levilactobac... | NA | 1580 | 513 | 513 | 45% | 4e-176 | 96.88 | 1126 | NZ_MZMW010000... |
| Pseudomonas aeruginosa strain VET-42... | Pseudomonas ... | NA | 287 | 510 | 510 | 46% | 1e-174 | 93.46 | 1166 | NZ_RACX010001... |
| Pseudomonas aeruginosa strain VET-121... | Pseudomonas ... | NA | 287 | 508 | 508 | 45% | 1e-173 | 94.14 | 1158 | NZ_QZXB010001... |
| Acinetobacter baumannii strain ACM-27... | Acinetobacte... | NA | 470 | 504 | 504 | 52% | 2e-173 | 82.53 | 895 | NZ_PYDM010000... |
| Klebsiella pneumoniae strain EuSCAPE_DE086, whole genome shotg... | Klebsiella p... | NA | 573 | 495 | 495 | 59% | 1e-169 | 76.44 | 995 | NZ_UKLW020000... |
| Providencia rettgeri strain APW139_S1 24_contig_64, whole geno... | Providencia ... | NA | 587 | 490 | 490 | 45% | 4e-168 | 89.84 | 902 | NZ_SHDG010000... |
| Levilactobacillus brevis strain CRL 2013 scaffold13, whole... | Levilactobac... | NA | 1580 | 488 | 488 | 43% | 2e-166 | 95.90 | 1030 | NZ_MZMW010000... |
| Providencia rettgeri strain AFE96_S1 41_contig_60, whole genom... | Providencia ... | NA | 587 | 488 | 488 | 44% | 2e-166 | 93.98 | 1101 | NZ_SHDP010000... |
| Mycobacterium tuberculosis strain 32845_S29 549, whole genome... | Mycobacteriu... | NA | 1773 | 481 | 481 | 48% | 7e-165 | 84.31 | 825 | NZ_QTBV010001... |
| Enterococcus durans strain E115_10... | Enterococcus... | NA | 53345 | 476 | 476 | 43% | 2e-162 | 93.03 | 955 | NZ_JAANZI0100... |
| Enterobacter hormaechei strain AUH-ENM4... | Enterobacter... | NA | 158836 | 471 | 471 | 43% | 5e-161 | 91.80 | 809 | NZ_PYGZ010000... |
| Klebsiella oxytoca strain WCHKO085006... | Klebsiella o... | NA | 571 | 462 | 462 | 40% | 4e-158 | 98.22 | 682 | NZ_NWES010001... |
| Escherichia coli strain 8374wB4 NODE_74_length_738_cov_3.41735... | Escherichia ... | NA | 562 | 458 | 458 | 43% | 3e-156 | 87.70 | 738 | NZ_WSPX010000... |
| Acinetobacter baumannii strain ACM-17... | Acinetobacte... | NA | 470 | 455 | 455 | 44% | 3e-155 | 86.80 | 757 | NZ_PYDD010000... |
| Acinetobacter baumannii strain ACM-22... | Acinetobacte... | NA | 470 | 449 | 449 | 43% | 5e-153 | 88.43 | 726 | NZ_PYDI010000... |
| Acinetobacter baumannii strain ACM-13... | Acinetobacte... | NA | 470 | 435 | 435 | 41% | 2e-147 | 89.27 | 701 | NZ_PYCZ010000... |
| Escherichia coli strain VRES0669, whole genome shotgun sequence | Escherichia ... | NA | 562 | 474 | 854 | 85% | 4e-146 | 82.75 | 5544 | NZ_UINV010000... |
| Acinetobacter baumannii strain ACM-41... | Acinetobacte... | NA | 470 | 427 | 427 | 40% | 4e-144 | 89.87 | 718 | NZ_QAGT010000... |
| Escherichia coli strain ECM-2 NODE_127_length_636_cov_2.0668,... | Escherichia ... | NA | 562 | 417 | 417 | 37% | 8e-141 | 95.26 | 636 | NZ_PXIA010001... |
| Acinetobacter baumannii strain ACM-14... | Acinetobacte... | NA | 470 | 412 | 412 | 39% | 1e-138 | 90.41 | 665 | NZ_PYDA010000... |
| Acinetobacter baumannii strain ACM-11... | Acinetobacte... | NA | 470 | 410 | 410 | 41% | 1e-137 | 83.83 | 706 | NZ_PYCW010000... |
| Providencia rettgeri strain APW139_S1 24_contig_65, whole geno... | Providencia ... | NA | 587 | 410 | 410 | 38% | 2e-136 | 91.08 | 941 | NZ_SHDG010000... |

| | | | | | | | | | | |
|---|---|---|---|---|---|---|---|---|---|---|
| Providencia rettgeri strain AFE97_S1 21_contig_71, whole genom... | Providencia ... | NA | 587 | 279 | 519 | 48% | 5e-136 | 90.48 | 1157 | NZ_SHDO010000... |
| Mycobacterium tuberculosis strain 32847_S31 312, whole genome... | Mycobacteriu... | NA | 1773 | 404 | 404 | 41% | 2e-135 | 83.33 | 704 | NZ_QTBX010000... |
| Acinetobacter baumannii strain ACM-29... | Acinetobacte... | NA | 470 | 402 | 402 | 37% | 1e-133 | 92.79 | 888 | NZ_PYDO010001... |
| Acinetobacter baumannii strain ACM-25... | Acinetobacte... | NA | 470 | 397 | 397 | 38% | 6e-133 | 86.70 | 655 | NZ_PXYP010000... |
| Levilactobacillus brevis strain CRL 2013 scaffold15, whole... | Levilactobac... | NA | 1580 | 391 | 391 | 33% | 5e-131 | 98.94 | 566 | NZ_MZMW010000... |
| Pseudomonas aeruginosa strain ENV-297... | Pseudomonas ... | NA | 287 | 390 | 390 | 35% | 5e-130 | 93.37 | 638 | NZ_RADD010001... |
| Providencia rettgeri strain AFE422_S1 40_contig_34, whole geno... | Providencia ... | NA | 587 | 278 | 496 | 45% | 6e-129 | 90.48 | 1058 | NZ_SHDL010000... |
| Providencia rettgeri strain AMW697_S1 18_contig_57, whole geno... | Providencia ... | NA | 587 | 382 | 382 | 34% | 4e-127 | 93.30 | 585 | NZ_SHDI010000... |
| Levilactobacillus brevis strain CRL 2013 scaffold16, whole... | Levilactobac... | NA | 1580 | 372 | 372 | 32% | 1e-123 | 97.25 | 547 | NZ_MZMW010000... |
| Providencia rettgeri strain AFE97_S1 21_contig_72, whole genom... | Providencia ... | NA | 587 | 369 | 369 | 33% | 9e-121 | 93.62 | 854 | NZ_SHDO010000... |
| Cuspidothrix issatschenkoi CHARLIE-1 contig_175, whole genome... | Cuspidothrix... | NA | 2052836 | 385 | 385 | 96% | 1e-120 | 41.62 | 2237 | NZ_PGEM010001... |
| Enterococcus faecalis strain 44.1... | Enterococcus... | NA | 1351 | 364 | 364 | 36% | 5e-120 | 85.22 | 611 | NZ_NSJR010000... |
| Enterobacter hormaechei strain 2011WA-SCV PROKKA_contig000046,... | Enterobacter... | NA | 158836 | 349 | 349 | 32% | 1e-114 | 92.22 | 544 | NZ_SJEC010000... |
| Levilactobacillus brevis strain CRL 2013 scaffold14, whole... | Levilactobac... | NA | 1580 | 348 | 398 | 31% | 6e-114 | 94.29 | 588 | NZ_MZMW010000... |
| Acinetobacter baumannii strain ACM-38... | Acinetobacte... | NA | 470 | 347 | 347 | 32% | 7e-114 | 91.11 | 542 | NZ_QAGQ010000... |
| Cellulophaga baltica strain 14 NODE_54_length_581_cov_63.51321... | Cellulophaga... | NA | 76594 | 342 | 342 | 34% | 1e-111 | 84.97 | 581 | NZ_VIQP010000... |
| Shigella sonnei strain 07401ss_1, whole genome shotgun sequence | Shigella sonnei | NA | 624 | 338 | 338 | 30% | 2e-110 | 93.49 | 509 | NZ_CWST010003... |
| Mycobacterium tuberculosis variant bovis strain CSURQ1131, who... | Mycobacteriu... | NA | 1765 | 341 | 859 | 39% | 1e-109 | 80.98 | 888 | NZ_CAJEUJ0100... |
| Klebsiella aerogenes strain AUH-KAM3... | Klebsiella a... | NA | 548 | 336 | 336 | 33% | 3e-109 | 85.48 | 558 | NZ_PYHH010000... |
| Enterococcus faecium strain INF91_9... | Enterococcus... | NA | 1352 | 334 | 334 | 30% | 6e-109 | 92.40 | 515 | NZ_JAANZG0100... |
| Neisseria gonorrhoeae strain GC-206... | Neisseria go... | NA | 485 | 329 | 329 | 29% | 7e-107 | 91.62 | 504 | NZ_SAWN010001... |
| Providencia rettgeri strain AFE96_S1 41_contig_58, whole genom... | Providencia ... | NA | 587 | 328 | 328 | 31% | 2e-106 | 89.71 | 546 | NZ_SHDP010000... |
| Providencia rettgeri strain AFE422_S1 40_contig_33, whole geno... | Providencia ... | NA | 587 | 327 | 327 | 29% | 4e-106 | 91.62 | 501 | NZ_SHDL010000... |
| Providencia rettgeri strain AFE220_S1 26_contig_66, whole geno... | Providencia ... | NA | 587 | 323 | 323 | 29% | 9e-105 | 91.57 | 502 | NZ_SHDM010000... |
| Escherichia coli strain 8374wG10 NODE_51_length_497_cov_1.1680... | Escherichia ... | NA | 562 | 321 | 321 | 29% | 7e-104 | 92.12 | 497 | NZ_WSJQ010000... |
| Klebsiella oxytoca strain WCHKO085010... | Klebsiella o... | NA | 571 | 320 | 320 | 27% | 1e-103 | 96.79 | 472 | NZ_NWER010001... |
| Escherichia coli strain 8374wH11 NODE_48_length_500_cov_2.0026... | Escherichia ... | NA | 562 | 319 | 319 | 29% | 5e-103 | 91.52 | 500 | NZ_WSJI010000... |
| Mycobacterium tuberculosis strain 32846_S30 476, whole genome... | Mycobacteriu... | NA | 1773 | 322 | 322 | 28% | 3e-102 | 96.27 | 861 | NZ_QTBW010001... |
| Escherichia coli strain 8374wH11 NODE_51_length_464_cov_1.9881... | Escherichia ... | NA | 562 | 306 | 306 | 27% | 3e-98 | 92.86 | 464 | NZ_WSJI010000... |
| Escherichia coli strain P010... | Escherichia ... | NA | 562 | 298 | 298 | 33% | 2e-94 | 79.46 | 556 | NZ_VKOQ010001... |
| Acinetobacter baumannii strain ACM-35... | Acinetobacte... | NA | 470 | 295 | 295 | 26% | 1e-93 | 97.95 | 466 | NZ_QEHN010000... |
| Klebsiella pneumoniae strain EuSCAPE_IL021, whole genome shotg... | Klebsiella p... | NA | 573 | 289 | 289 | 25% | 8e-92 | 97.89 | 426 | NZ_UKJX020001... |
| Escherichia coli strain 8374wA11 NODE_51_length_444_cov_2.5520... | Escherichia ... | NA | 562 | 288 | 288 | 26% | 2e-91 | 91.22 | 444 | NZ_WSPR010000... |
| Escherichia coli strain TzEc080 NODE_51_length_444_cov_2.55205... | Escherichia ... | NA | 562 | 288 | 288 | 26% | 2e-91 | 91.22 | 444 | NZ_WSFZ010000... |
| Klebsiella pneumoniae strain B2 isolate B2, whole genome shotg... | Klebsiella p... | NA | 573 | 315 | 375 | 91% | 2e-89 | 43.20 | 7306 | NZ_CABWNT0100... |
| Burkholderia pseudomallei strain 2604STDY5518971... | Burkholderia... | NA | 28450 | 283 | 283 | 25% | 2e-89 | 95.07 | 427 | NZ_UVFZ020001... |
| Escherichia coli strain 8374wB11 NODE_47_length_476_cov_2.5988... | Escherichia ... | NA | 562 | 281 | 281 | 28% | 2e-88 | 83.54 | 476 | NZ_WSJO010000... |
| Escherichia coli strain 8374wB11 NODE_56_length_419_cov_1.4349... | Escherichia ... | NA | 562 | 278 | 278 | 24% | 1e-87 | 94.24 | 419 | NZ_WSJO010000... |
| Burkholderia pseudomallei strain 2604STDY5518971... | Burkholderia... | NA | 28450 | 278 | 278 | 27% | 2e-87 | 87.50 | 457 | NZ_UVFZ020001... |
| Cronobacter sakazakii strain cro4215W contig98, whole genome... | Cronobacter ... | NA | 28141 | 272 | 272 | 23% | 4e-85 | 100.00 | 429 | NZ_NRKF010000... |
| Levilactobacillus brevis strain CRL 2013 scaffold20, whole... | Levilactobac... | NA | 1580 | 270 | 270 | 23% | 3e-84 | 96.97 | 398 | NZ_MZMW010000... |
| Cronobacter sakazakii strain cro4114A1 contig119, whole genome... | Cronobacter ... | NA | 28141 | 270 | 270 | 23% | 3e-84 | 99.25 | 429 | NZ_NRKJ010001... |
| Cronobacter sakazakii strain cro4114B1 contig94, whole genome... | Cronobacter ... | NA | 28141 | 269 | 269 | 23% | 8e-84 | 97.74 | 429 | NZ_NRKI010000... |
| Escherichia coli strain 8374wB11... | Escherichia ... | NA | 562 | 268 | 268 | 25% | 1e-83 | 87.14 | 422 | NZ_WSJO010000... |
| Escherichia coli strain 8374wB4 NODE_86_length_485_cov_1.49162... | Escherichia ... | NA | 562 | 267 | 267 | 24% | 5e-83 | 95.59 | 485 | NZ_WSPX010000... |
| Klebsiella pneumoniae strain EuSCAPE_IL018, whole genome shotg... | Klebsiella p... | NA | 573 | 264 | 264 | 23% | 4e-82 | 94.62 | 392 | NZ_UKJU020001... |
| Cronobacter malonaticus strain cro3263A1 contig63, whole genom... | Cronobacter ... | NA | 413503 | 264 | 264 | 23% | 6e-82 | 95.49 | 429 | NZ_NRMM010000... |
| Cronobacter dublinensis strain cro981C2 contig293, whole genom... | Cronobacter ... | NA | 413497 | 263 | 263 | 23% | 1e-81 | 96.24 | 429 | NZ_NROB010002... |
| Cronobacter sakazakii strain cro3040W contig78, whole genome... | Cronobacter ... | NA | 28141 | 263 | 263 | 23% | 2e-81 | 96.24 | 429 | NZ_NRJO010000... |
| Cronobacter sakazakii strain cro4215W contig97, whole genome... | Cronobacter ... | NA | 28141 | 262 | 262 | 23% | 4e-81 | 94.74 | 429 | NZ_NRKF010000... |
| Cronobacter sakazakii strain cro4114B1 contig95, whole genome... | Cronobacter ... | NA | 28141 | 262 | 262 | 23% | 4e-81 | 94.74 | 429 | NZ_NRKI010000... |
| Cronobacter sakazakii strain cro4114A1 contig120, whole genome... | Cronobacter ... | NA | 28141 | 262 | 262 | 23% | 4e-81 | 94.74 | 429 | NZ_NRKJ010001... |
| Cronobacter sakazakii strain cro4191B1-2 contig80, whole genom... | Cronobacter ... | NA | 28141 | 262 | 262 | 23% | 4e-81 | 95.49 | 429 | NZ_NRKG010000... |
| Acinetobacter baumannii strain ACM-28... | Acinetobacte... | NA | 470 | 260 | 260 | 23% | 1e-80 | 93.13 | 395 | NZ_PYDN010000... |
| Mycobacterium tuberculosis strain 32847_S31 462, whole genome... | Mycobacteriu... | NA | 1773 | 266 | 266 | 23% | 2e-80 | 95.52 | 893 | NZ_QTBX010001... |
| Acinetobacter baumannii strain ACM-14... | Acinetobacte... | NA | 470 | 263 | 263 | 23% | 6e-80 | 95.52 | 787 | NZ_PYDA010000... |
| Klebsiella pneumoniae strain EuSCAPE_MT012, whole genome shotg... | Klebsiella p... | NA | 573 | 259 | 259 | 26% | 9e-80 | 85.33 | 452 | NZ_UKIX020000... |
| Cronobacter malonaticus strain cro4164A3 contig124, whole geno... | Cronobacter ... | NA | 413503 | 258 | 258 | 23% | 9e-80 | 93.98 | 429 | NZ_NRNF010001... |
| Burkholderia pseudomallei strain 2604STDY5518971... | Burkholderia... | NA | 28450 | 219 | 332 | 33% | 1e-79 | 88.43 | 1345 | NZ_UVFZ020000... |
| Cronobacter sakazakii strain cro3040W contig77, whole genome... | Cronobacter ... | NA | 28141 | 258 | 258 | 23% | 2e-79 | 93.98 | 429 | NZ_NRJO010000... |
| Escherichia coli strain 8374wB11 NODE_49_length_463_cov_1.7708... | Escherichia ... | NA | 562 | 254 | 254 | 23% | 6e-78 | 89.55 | 463 | NZ_WSJO010000... |
| Enterococcus faecalis strain 44.1... | Enterococcus... | NA | 1351 | 258 | 258 | 23% | 7e-78 | 95.42 | 806 | NZ_NSJR010000... |
| Acinetobacter baumannii strain ACM-29... | Acinetobacte... | NA | 470 | 251 | 251 | 24% | 5e-77 | 86.23 | 447 | NZ_PYDO010001... |
| Escherichia coli strain 8374wH11 NODE_62_length_437_cov_1.9548... | Escherichia ... | NA | 562 | 250 | 250 | 25% | 1e-76 | 82.07 | 437 | NZ_WSJI010000... |
| Escherichia coli strain 8374wB4 NODE_91_length_444_cov_2.17035... | Escherichia ... | NA | 562 | 245 | 303 | 26% | 2e-74 | 92.62 | 444 | NZ_WSPX010000... |
| Klebsiella aerogenes strain AUH-KAM4... | Klebsiella a... | NA | 548 | 243 | 243 | 22% | 4e-74 | 91.27 | 381 | NZ_PYHI010000... |
| Escherichia coli strain 8374wA11 NODE_52_length_437_cov_2.2451... | Escherichia ... | NA | 562 | 244 | 244 | 25% | 4e-74 | 80.00 | 437 | NZ_WSPR010000... |
| Escherichia coli strain TzEc080 NODE_52_length_437_cov_2.24516... | Escherichia ... | NA | 562 | 244 | 244 | 25% | 4e-74 | 80.00 | 437 | NZ_WSFZ010000... |
| Acinetobacter baumannii strain ACM-24... | Acinetobacte... | NA | 470 | 242 | 242 | 21% | 3e-73 | 95.12 | 503 | NZ_PYDK010000... |
| Klebsiella pneumoniae strain EuSCAPE_MT020, whole genome shotg... | Klebsiella p... | NA | 573 | 240 | 240 | 21% | 4e-73 | 95.90 | 368 | NZ_UKJB020000... |
| Providencia rettgeri strain AFE85_S2 19_contig_46, whole genom... | Providencia ... | NA | 587 | 246 | 324 | 22% | 4e-73 | 96.77 | 846 | NZ_SHDQ010000... |
| Escherichia coli strain 8374wH11... | Escherichia ... | NA | 562 | 241 | 241 | 24% | 6e-73 | 81.75 | 417 | NZ_WSJI010000... |
| Acinetobacter baumannii strain ACM-37... | Acinetobacte... | NA | 470 | 238 | 238 | 24% | 9e-72 | 83.45 | 418 | NZ_QAGP010000... |
| Mycobacterium tuberculosis strain 32845_S29 230, whole genome... | Mycobacteriu... | NA | 1773 | 239 | 239 | 21% | 1e-71 | 95.08 | 601 | NZ_QTBV010000... |
| Escherichia coli strain 8374wA11 NODE_70_length_381_cov_1.2598... | Escherichia ... | NA | 562 | 236 | 236 | 22% | 2e-71 | 90.48 | 381 | NZ_WSPR010000... |
| Escherichia coli strain TzEc080 NODE_70_length_381_cov_1.25984... | Escherichia ... | NA | 562 | 236 | 236 | 22% | 2e-71 | 90.48 | 381 | NZ_WSFZ010000... |
| Mycobacterium tuberculosis strain 32845_S29 147, whole genome... | Mycobacteriu... | NA | 1773 | 234 | 234 | 21% | 7e-71 | 92.37 | 357 | NZ_QTBV010000... |
| Mycobacterium tuberculosis strain 32847_S31 147, whole genome... | Mycobacteriu... | NA | 1773 | 234 | 234 | 21% | 7e-71 | 92.37 | 357 | NZ_QTBX010000... |
| Klebsiella aerogenes strain AUH-KAM2... | Klebsiella a... | NA | 548 | 231 | 231 | 23% | 3e-69 | 81.34 | 404 | NZ_PYHG010000... |
| Methylobacterium radiotolerans strain MAMP 4754... | Methylobacte... | NA | 31998 | 231 | 231 | 21% | 1e-68 | 90.68 | 539 | NZ_NKQS010010... |
| Mycobacterium tuberculosis variant bovis strain CSURQ1132, who... | Mycobacteriu... | NA | 1765 | 229 | 459 | 21% | 2e-66 | 93.39 | 862 | NZ_CAJEUK0100... |
| Escherichia coli strain ECM-2 NODE_147_length_382_cov_1.52941,... | Escherichia ... | NA | 562 | 222 | 222 | 22% | 8e-66 | 84.80 | 382 | NZ_PXIA010001... |
| Escherichia coli strain 8374wA11 NODE_44_length_523_cov_1.7197... | Escherichia ... | NA | 562 | 205 | 205 | 19% | 1e-58 | 88.99 | 523 | NZ_WSPR010000... |
| Escherichia coli strain TzEc080 NODE_44_length_523_cov_1.7197,... | Escherichia ... | NA | 562 | 205 | 205 | 19% | 1e-58 | 88.99 | 523 | NZ_WSFZ010000... |
| Acinetobacter baumannii strain ACM-32... | Acinetobacte... | NA | 470 | 204 | 204 | 20% | 7e-58 | 90.18 | 654 | NZ_PYDR010000... |
| Klebsiella pneumoniae strain EuSCAPE_LT004, whole genome shotg... | Klebsiella p... | NA | 573 | 200 | 200 | 19% | 1e-57 | 86.49 | 335 | NZ_UKJE020000... |
| Klebsiella pneumoniae strain EuSCAPE_MT006, whole genome shotg... | Klebsiella p... | NA | 573 | 201 | 201 | 18% | 2e-57 | 96.04 | 455 | NZ_UKIU020000... |
| Klebsiella pneumoniae strain EuSCAPE_IL005, whole genome shotg... | Klebsiella p... | NA | 573 | 198 | 198 | 17% | 4e-57 | 96.94 | 298 | NZ_UKJL020000... |
| Acinetobacter baumannii strain ACM-38... | Acinetobacte... | NA | 470 | 202 | 202 | 18% | 6e-57 | 96.04 | 652 | NZ_QAGQ010000... |
| Klebsiella pneumoniae strain EuSCAPE_LT004, whole genome shotg... | Klebsiella p... | NA | 573 | 197 | 197 | 21% | 2e-56 | 77.05 | 367 | NZ_UKJE020000... |
| Klebsiella pneumoniae strain EuSCAPE_MT015, whole genome shotg... | Klebsiella p... | NA | 573 | 198 | 198 | 17% | 2e-56 | 95.96 | 435 | NZ_UKIW020000... |
| Klebsiella pneumoniae strain EuSCAPE_IL005, whole genome shotg... | Klebsiella p... | NA | 573 | 186 | 186 | 16% | 2e-52 | 92.47 | 283 | NZ_UKJL020000... |
| Klebsiella oxytoca strain WCHKO085010... | Klebsiella o... | NA | 571 | 183 | 183 | 16% | 1e-51 | 97.80 | 275 | NZ_NWER010001... |
| Klebsiella pneumoniae strain EuSCAPE_DE082, whole genome shotg... | Klebsiella p... | NA | 573 | 177 | 177 | 17% | 4e-49 | 88.66 | 295 | NZ_UKLV020000... |
| Klebsiella oxytoca strain WCHKO085010... | Klebsiella o... | NA | 571 | 174 | 174 | 15% | 6e-48 | 97.65 | 262 | NZ_NWER010001... |
| Escherichia coli strain 8374wB11 NODE_46_length_516_cov_2.1516... | Escherichia ... | NA | 562 | 168 | 298 | 26% | 1e-44 | 93.10 | 516 | NZ_WSJO010000... |
| Acinetobacter baumannii strain ACM-23... | Acinetobacte... | NA | 470 | 158 | 158 | 13% | 5e-41 | 98.72 | 505 | NZ_PYDJ010000... |
| Mycobacterium tuberculosis strain 32847_S31 146, whole genome... | Mycobacteriu... | NA | 1773 | 153 | 153 | 13% | 3e-40 | 89.74 | 235 | NZ_QTBX010000... |
| Chlamydia abortus strain 09_01, whole genome shotgun sequence | Chlamydia ab... | NA | 83555 | 156 | 156 | 43% | 7e-39 | 38.46 | 847 | NZ_FQLJ010000... |
| Chlamydia trachomatis strain R33512extract, whole genome shotg... | Chlamydia tr... | NA | 813 | 159 | 159 | 47% | 1e-37 | 37.31 | 1732 | NZ_CVNW010000... |
| Chlamydia trachomatis strain R4175extract, whole genome shotgu... | Chlamydia tr... | NA | 813 | 160 | 160 | 49% | 4e-37 | 36.56 | 2328 | NZ_CVNX010000... |
| Chlamydia abortus strain 09_03, whole genome shotgun sequence | Chlamydia ab... | NA | 83555 | 161 | 161 | 49% | 4e-37 | 36.56 | 4440 | NZ_FQLL010000... |
| Chlamydia abortus strain 15a, whole genome shotgun sequence | Chlamydia ab... | NA | 83555 | 160 | 160 | 49% | 4e-37 | 36.56 | 3222 | NZ_FQLM010000... |
| Chlamydia abortus strain 11_01, whole genome shotgun sequence | Chlamydia ab... | NA | 83555 | 160 | 160 | 49% | 5e-37 | 36.56 | 5240 | NZ_FQLK010000... |
| Chlamydia abortus strain 09_02, whole genome shotgun sequence | Chlamydia ab... | NA | 83555 | 160 | 160 | 49% | 8e-37 | 36.56 | 4507 | NZ_FQLN010000... |
| Chlamydia abortus strain 09_05, whole genome shotgun sequence | Chlamydia ab... | NA | 83555 | 157 | 157 | 43% | 1e-35 | 38.46 | 4831 | NZ_FQLG010000... |
| Providencia rettgeri strain AFE220_S1 26_contig_67, whole geno... | Providencia ... | NA | 587 | 142 | 142 | 13% | 4e-35 | 93.15 | 512 | NZ_SHDM010000... |
| Pseudomonas aromaticivorans strain MAP12 34, whole genome... | Pseudomonas ... | NA | 2849492 | 143 | 143 | 48% | 5e-31 | 33.46 | 7031 | NZ_JAHRGL0100... |
| Neisseria gonorrhoeae strain GC-206... | Neisseria go... | NA | 485 | 125 | 125 | 11% | 4e-30 | 92.42 | 223 | NZ_SAWN010001... |
| Enterococcus faecium strain E21_6... | Enterococcus... | NA | 1352 | 128 | 128 | 12% | 4e-30 | 87.14 | 518 | NZ_JAANZJ0100... |
| Methylobacterium radiotolerans strain MAMP 4754... | Methylobacte... | NA | 31998 | 122 | 122 | 11% | 4e-28 | 93.65 | 483 | NZ_NKQS010002... |
| Acinetobacter baumannii strain ACM-36... | Acinetobacte... | NA | 470 | 122 | 122 | 10% | 2e-27 | 96.67 | 567 | NZ_QAGO010000... |
| Escherichia coli strain 8374wG10 NODE_65_length_433_cov_1.7679... | Escherichia ... | NA | 562 | 112 | 112 | 11% | 2e-24 | 82.09 | 433 | NZ_WSJQ010000... |
| Pseudomonas aeruginosa strain VET-42... | Pseudomonas ... | NA | 287 | 104 | 104 | 9% | 1e-19 | 90.91 | 1154 | NZ_RACX010001... |
| Burkholderia pseudomallei strain A-175-05-2-02, whole genome... | Burkholderia... | NA | 28450 | 89.7 | 89.7 | 8% | 2e-16 | 93.48 | 455 | NZ_UVFW010001... |
| Tumebacillus flagellatus strain GST4 Contig077, whole genome... | Tumebacillus... | NA | 1157490 | 87.8 | 87.8 | 45% | 2e-13 | 26.59 | 2091 | NZ_JMIR010000... |
| Klebsiella pneumoniae strain BL2026... | Klebsiella p... | NA | 573 | 81.6 | 81.6 | 40% | 2e-11 | 29.17 | 1934 | NZ_VICY010000... |
| Streptococcus pneumoniae strain NN2021-1 12_contig_107, whole... | Streptococcu... | NA | 1313 | 79.3 | 79.3 | 35% | 6e-11 | 30.77 | 1657 | NZ_JAGYHD0100... |
| Klebsiella pneumoniae strain BL1427... | Klebsiella p... | NA | 573 | 76.6 | 76.6 | 33% | 2e-10 | 30.73 | 1114 | NZ_VICX010000... |
| Escherichia coli B7A gcontig_1112495760658, whole genome shotg... | Escherichia ... | NA | 340184 | 76.3 | 76.3 | 43% | 3e-10 | 26.17 | 1060 | NZ_AAJT020002... |
| Thalassovita mangrovi strain GS-10 Scaffold30_1, whole genome... | Thalassovita... | NA | 2692236 | 73.9 | 73.9 | 8% | 4e-10 | 71.74 | 722 | NZ_WWEN010000... |
| Rickettsia endosymbiont of Ceutorhynchus assimilis isolate... | Rickettsia e... | NA | 2884482 | 77.4 | 207 | 45% | 6e-10 | 26.52 | 4063491 | NZ_OU906081.1 |
| Klebsiella pneumoniae strain EuSCAPE_NO017, whole genome shotg... | Klebsiella p... | NA | 573 | 70.9 | 70.9 | 6% | 9e-10 | 100.00 | 458 | NZ_UKIO020000... |
| Pseudomonas aeruginosa strain ENV-297... | Pseudomonas ... | NA | 287 | 74.7 | 74.7 | 6% | 1e-09 | 94.59 | 1100 | NZ_RADD010000... |

| | | | | | | | | | | |
|---|---|---|---|---|---|---|---|---|---|---|
| Pseudomonas aeruginosa strain P60_London_6_VIM_2_11_13, whole... | Pseudomonas ... | NA | 287 | 68.6 | 68.6 | 5% | 3e-09 | 100.00 | 395 | NZ_CVXD010000... |
| Listeria monocytogenes strain 1827/2,1... | Listeria mon... | NA | 1639 | 73.9 | 73.9 | 35% | 6e-09 | 29.81 | 4618 | NZ_JAENKW0100... |
| Klebsiella pneumoniae strain BL2026... | Klebsiella p... | NA | 573 | 66.2 | 66.2 | 27% | 1e-07 | 31.88 | 659 | NZ_VICY010000... |
| Thalassovita mangrovi strain GS-10 Scaffold19_1, whole genome... | Thalassovita... | NA | 2692236 | 68.2 | 68.2 | 5% | 3e-07 | 96.97 | 2900 | NZ_WWEN010000... |
| Sphingobacterium bovisgrunnientis strain KCTC 52685... | Sphingobacte... | NA | 1874697 | 63.5 | 63.5 | 28% | 4e-07 | 26.22 | 503 | NZ_WUQY010001... |
| Acinetobacter baumannii strain ACM-15... | Acinetobacte... | NA | 470 | 62.8 | 62.8 | 7% | 5e-07 | 80.49 | 432 | NZ_PYDB010000... |
| Proteus mirabilis strain PM2-4 PM2-4_scaffold37, whole genome... | Proteus mira... | NA | 584 | 66.6 | 66.6 | 41% | 1e-06 | 24.60 | 5277 | NZ_JAMKNT0100... |
| Klebsiella pneumoniae strain c10 contig85, whole genome shotgu... | Klebsiella p... | NA | 573 | 66.2 | 66.2 | 42% | 2e-06 | 26.94 | 6264 | NZ_JAIBKQ0100... |
| Streptococcus anginosus strain San49... | Streptococcu... | NA | 1328 | 57.4 | 57.4 | 14% | 1e-05 | 35.80 | 269 | NZ_JAXIIG0100... |
| Klebsiella pneumoniae strain BL198... | Klebsiella p... | NA | 573 | 60.1 | 60.1 | 23% | 1e-05 | 29.20 | 611 | NZ_VICV010000... |
| Thalassolituus oleivorans isolate... | Thalassolitu... | NA | 187493 | 62.8 | 62.8 | 49% | 2e-05 | 26.76 | 10375 | NZ_CAWUIR0100... |
| Polyangium sp. y55x31 NODE_26_length_169233_cov_17.164843, who... | Polyangium s... | NA | 3042688 | 62.4 | 62.4 | 20% | 3e-05 | 33.06 | 169233 | NZ_JARZHG0100... |
| Polyangium spumosum strain DSM 14734 Scaffold16, whole genome... | Polyangium s... | NA | 889282 | 61.2 | 61.2 | 20% | 6e-05 | 33.06 | 180314 | NZ_WJIE010000... |
| Streptomyces albidochromogenes strain DSM 41800... | Streptomyces... | NA | 329524 | 57.4 | 57.4 | 32% | 7e-05 | 27.96 | 521 | NZ_VBVX010007... |
| Klebsiella pneumoniae strain BL3427... | Klebsiella p... | NA | 573 | 58.2 | 58.2 | 22% | 2e-04 | 30.08 | 966 | NZ_VIDH010000... |
| Thiolapillus sp. isolate Ifr-SOX-238... | Thiolapillus... | NA | 2017437 | 59.3 | 59.3 | 40% | 2e-04 | 28.33 | 2294 | NZ_JANEJA0100... |
| Polyangium sp. 15x6 NODE_112_length_36528_cov_15.618939, whole... | Polyangium s... | NA | 3042687 | 58.5 | 58.5 | 20% | 4e-04 | 32.26 | 36528 | NZ_JASBQE0100... |
| Penaeicola halotolerans strain LMIT005... | Penaeicola h... | NA | 2793196 | 53.1 | 53.1 | 10% | 6e-04 | 45.90 | 336 | NZ_JAEAGO0100... |
| Piscirickettsia salmonis strain FIUCHILE-89L01, whole genome... | Piscirickett... | NA | 1238 | 57.8 | 57.8 | 25% | 6e-04 | 29.08 | 2765 | NZ_LELB010002... |
| Escherichia coli strain 8551F 237, whole genome shotgun sequence | Escherichia ... | NA | 562 | 55.5 | 55.5 | 37% | 7e-04 | 27.78 | 689 | NZ_JAFCNN0100... |
| Candidatus Erwinia dacicola strain IL contig_608, whole genome... | Candidatus E... | NA | 252393 | 57.0 | 57.0 | 18% | 7e-04 | 31.78 | 1576 | NZ_MAYS010005... |
| Polyangium sp. 6x1 NODE_22_length_166486_cov_22.101912, whole... | Polyangium s... | NA | 3042689 | 57.8 | 57.8 | 20% | 7e-04 | 33.06 | 166486 | NZ_JARZHH0100... |
| Pseudomonas aeruginosa isolate PSA-13, whole genome shotgun... | Pseudomonas ... | NA | 287 | 53.5 | 53.5 | 24% | 0.001 | 30.66 | 505 | NZ_FNME010000... |
| Enterobacter cloacae complex sp. GF14B... | Enterobacter... | NA | 2511982 | 56.2 | 56.2 | 43% | 0.002 | 27.00 | 4351 | NZ_QFRV010001... |
| Methylobacterium radiotolerans strain MAMP 4754... | Methylobacte... | NA | 31998 | 50.1 | 50.1 | 8% | 0.003 | 56.25 | 253 | NZ_NKQS010008... |
| Polyangium jinanense strain SDU14 Scaffold9, whole genome... | Polyangium j... | NA | 2829994 | 55.1 | 55.1 | 20% | 0.004 | 31.93 | 231060 | NZ_JAGTJJ0100... |
| Polyangium jinanense strain SDU13 Scaffold15, whole genome... | Polyangium j... | NA | 2829994 | 55.1 | 55.1 | 20% | 0.004 | 31.93 | 231263 | NZ_JAGTJK0100... |
| Proteus terrae subsp. cibarius strain pt-OS9-1... | Proteus terr... | NA | 626774 | 49.3 | 49.3 | 16% | 0.006 | 32.98 | 267 | NZ_JBBEZU0100... |
| Streptococcus mutans strain PKUSS-9 C209, whole genome shotgun... | Streptococcu... | NA | 1309 | 49.7 | 49.7 | 13% | 0.006 | 38.67 | 307 | NZ_NBFI010000... |
| Bradyrhizobium sp. 33ap4 NODE_290_length_534_cov_0.849015, who... | Bradyrhizobi... | NA | 3061630 | 51.2 | 51.2 | 24% | 0.008 | 29.37 | 534 | NZ_JAUMTJ0100... |
| Breoghania sp. isolate MO_035.B22 MO_035.c_2803, whole genome... | Breoghania sp. | NA | 2065378 | 53.9 | 53.9 | 23% | 0.010 | 27.59 | 6353 | NZ_JASJAV0100... |
| Klebsiella pneumoniae strain BL198... | Klebsiella p... | NA | 573 | 50.4 | 50.4 | 27% | 0.011 | 25.00 | 484 | NZ_VICV010000... |
| Breoghania sp. isolate MO_035.B22 MO_035.c_6474, whole genome... | Breoghania sp. | NA | 2065378 | 53.9 | 53.9 | 39% | 0.011 | 25.51 | 5920 | NZ_JASJAV0100... |
| Wolbachia endosymbiont of Corcyra cephalonica isolate SYLI2103... | Wolbachia en... | NA | 218111 | 53.5 | 53.5 | 29% | 0.012 | 28.99 | 5332 | NZ_CP087956.1 |
| Klebsiella pneumoniae strain A13... | Klebsiella p... | NA | 573 | 49.3 | 49.3 | 19% | 0.012 | 30.36 | 333 | NZ_JARWSQ0100... |
| Clostridium paraputrificum strain 28-X28 Scaffold60_1, whole... | Clostridium ... | NA | 29363 | 51.6 | 51.6 | 23% | 0.013 | 30.37 | 680 | NZ_RAYJ010000... |
| Campylobacter sp. JMF_08 NE1 NODE_13_length_3482_cov_0.037836,... | Campylobacte... | NA | 2983821 | 53.5 | 53.5 | 31% | 0.014 | 29.28 | 3482 | NZ_JAOVUC0100... |
| Campylobacter sp. JMF_10 EL2 NODE_13_length_1688_cov_0.027690,... | Campylobacte... | NA | 2983828 | 52.8 | 52.8 | 31% | 0.015 | 29.28 | 1688 | NZ_JAOVTV0100... |
| Campylobacter sp. CS_ED1 strain Cs ED1... | Campylobacte... | NA | 2984140 | 53.1 | 53.1 | 31% | 0.016 | 29.28 | 5400 | NZ_JAOVUL0100... |
| Campylobacter sp. CN_NE4 strain Cn NE4... | Campylobacte... | NA | 2984148 | 53.1 | 53.1 | 31% | 0.017 | 29.28 | 5576 | NZ_JAOVUF0100... |
| Klebsiella pneumoniae strain BL1427... | Klebsiella p... | NA | 573 | 48.1 | 48.1 | 18% | 0.026 | 25.96 | 321 | NZ_VICX010001... |
| Pseudomonas aromaticivorans strain MAP12 47, whole genome... | Pseudomonas ... | NA | 2849492 | 52.4 | 52.4 | 21% | 0.028 | 27.78 | 2177 | NZ_JAHRGL0100... |
| Lachnoclostridium phocaeense strain Marseille-P3177T chromosom... | Lachnoclostr... | NA | 1871021 | 52.4 | 52.4 | 24% | 0.030 | 28.67 | 3500754 | NZ_LT635479.1 |
| Kribbella italica strain DSM 28967 Ga0415174_01, whole genome... | Kribbella it... | NA | 1540520 | 52.4 | 52.4 | 22% | 0.032 | 28.28 | 8885108 | NZ_JACHMY0100... |
| Campylobacter sp. JMF_03 NE3 NODE_16_length_1924_cov_0.035616,... | Campylobacte... | NA | 2983831 | 52.0 | 52.0 | 23% | 0.035 | 28.78 | 1924 | NZ_JAOVTS0100... |
| Campylobacter jejuni strain CJJD2010009 CJJD2010009_contig_657... | Campylobacte... | NA | 197 | 47.0 | 47.0 | 7% | 0.035 | 52.27 | 251 | NZ_JAJUND0100... |
| Saccharospirillum mangrovi strain HK-33 Scaffold2, whole genom... | Saccharospir... | NA | 2161747 | 52.4 | 52.4 | 25% | 0.035 | 26.54 | 517067 | NZ_QAPE010000... |
| Saccharospirillum mangrovi strain HK-33 chromosome, complete... | Saccharospir... | NA | 2161747 | 52.4 | 52.4 | 25% | 0.035 | 26.54 | 3686911 | NZ_CP031415.1 |
| Pseudomonas morbosilactucae strain MAFF 302046 302046_contig_4... | Pseudomonas ... | NA | 2938197 | 52.4 | 52.4 | 20% | 0.035 | 30.99 | 73784 | NZ_JALQCX0100... |

Alignments:

>Enterococcus alcedinis strain CCM 8433 sequence23, whole genome shotgun sequence
Sequence ID: NZ_BMDT01000023.1 Length: 6075
Range 1: 1815 to 3494

Score:1073 bits(2775), Expect:0.0,
Method:Compositional matrix adjust.,
Identities:545/560(97%), Positives:554/560(98%), Gaps:0/560(0%)

```
Query  1     PISPIETVPVKLKPGMDGPKVKQWPLTEEKIKALVEICTEMEKEGKISKIGPENPYNTPV  60
             PISPIETVPVKLKPGMDGPKVKQWPLTEEKIKALVEICTEMEKEGKISKIGPENPYNTPV
Sbjct  1815  PISPIETVPVKLKPGMDGPKVKQWPLTEEKIKALVEICTEMEKEGKISKIGPENPYNTPV  1994

Query  61    FAIKKKDSTKWRKLVDFRELNKRTQDFWEVQLGIPHPAGLKKKKSVTVLDVGDAYFSVPL  120
             FAIKKKDSTKWRKLVDFRELNKRTQDFWEVQLGIPHPAGLK+KKSVTVLDVGDAYFSVPL
Sbjct  1995  FAIKKKDSTKWRKLVDFRELNKRTQDFWEVQLGIPHPAGLKQKKSVTVLDVGDAYFSVPL  2174

Query  121   DEDFRKYTAFTIPSINNETPGIRYQYNVLPQGWKGSPAIFQSSMTKILEPFRKQNPDIVI  180
             D+DFRKYTAFTIPSINNETPGIRYQYNVLPQGWKGSPAIFQ SMTKILEPFRKQNPDIVI
Sbjct  2175  DKDFRKYTAFTIPSINNETPGIRYQYNVLPQGWKGSPAIFQCSMTKILEPFRKQNPDIVI  2354

Query  181   YQYMDDLYVGSDLEIGQHRTKIEELRQHLLRWGLTTPDKKHQKEPPFLWMGYELHPDKWT  240
             YQYMDDLYVGSDLEIGQHRTKIEELRQHLLRWG TTPDKKHQKEPPFLWMGYELHPDKWT
Sbjct  2355  YQYMDDLYVGSDLEIGQHRTKIEELRQHLLRWGFTTPDKKHQKEPPFLWMGYELHPDKWT  2534

Query  241   VQPIVLPEKDSWTVNDIQKLVGKLNWASQIYPGIKVRQLCKLLRGTKALTEVIPLTeeae  300
             VQPIVLPEKDSWTVNDIQKLVGKLNWASQIY GIKVRQLCKLLRGTKALTEV+PLTEEAE
Sbjct  2535  VQPIVLPEKDSWTVNDIQKLVGKLNWASQIYAGIKVRQLCKLLRGTKALTEVVPLTEEAE  2714

Query  301   lelaeNREILKEPVHGVYYDPSKDLIAEIQKQGQGQWTYQIYQEPFKNLKTGKYARMRGA  360
             LELAENREILKEPVHGVYYDPSKDLIAEIQKQGQGQWTYQIYQEPFKNLKTGKYARM+GA
Sbjct  2715  LELAENREILKEPVHGVYYDPSKDLIAEIQKQGQGQWTYQIYQEPFKNLKTGKYARMKGA  2894

Query  361   HTNDVKQLTEAVQKITTESIVIWGKTPKFKLPIQKetwetwwteYWQATWIPEWEFVNTP  420
             HTNDVKQLTEAVQKI TESIVIWGKTPKFKLPIQKETWE WWTEYWQATWIPEWEFVNTP
Sbjct  2895  HTNDVKQLTEAVQKIATESIVIWGKTPKFKLPIQKETWEAWWTEYWQATWIPEWEFVNTP  3074

Query  421   PLVKLWYQLEKEPIVGAETFYVDGAANRETKLGKAGYVTNRGRQKVVTLTDTTNQKTELQ  480
             PLVKLWYQLEKEPI+GAETFYVDGAANRETKLGKAGYVT+RGRQKVV LTDTTNQKTELQ
Sbjct  3075  PLVKLWYQLEKEPIIGAETFYVDGAANRETKLGKAGYVTDRGRQKVVPLTDTTNQKTELQ  3254

Query  481   AIYLALQDSGLEVNIVTDSQYALGIIQAQPDQSESELVNQIIEQLIKKEKVYLAWVPAHK  540
             AI+LALQDSGLEVNIVTDSQYALGIIQAQPD+SESELV+QIIEQLIKKEKVYLAWVPAHK
Sbjct  3255  AIHLALQDSGLEVNIVTDSQYALGIIQAQPDKSESELVSQIIEQLIKKEKVYLAWVPAHK  3434

Query  541   GIGGNEQVDKLVSAGIRKVL  560
             GIGGNEQVDKLVSAGIRKVL
Sbjct  3435  GIGGNEQVDKLVSAGIRKVL  3494
```

>Deinococcus aquiradiocola strain JCM 14371 sequence30, whole genome shotgun sequence
Sequence ID: NZ_BMOE01000030.1 Length: 6128
Range 1: 1868 to 3547

Score:1073 bits(2775), Expect:0.0,
Method:Compositional matrix adjust.,
Identities:545/560(97%), Positives:554/560(98%), Gaps:0/560(0%)

```
Query  1     PISPIETVPVKLKPGMDGPKVKQWPLTEEKIKALVEICTEMEKEGKISKIGPENPYNTPV  60
             PISPIETVPVKLKPGMDGPKVKQWPLTEEKIKALVEICTEMEKEGKISKIGPENPYNTPV
Sbjct  1868  PISPIETVPVKLKPGMDGPKVKQWPLTEEKIKALVEICTEMEKEGKISKIGPENPYNTPV  2047

Query  61    FAIKKKDSTKWRKLVDFRELNKRTQDFWEVQLGIPHPAGLKKKKSVTVLDVGDAYFSVPL  120
             FAIKKKDSTKWRKLVDFRELNKRTQDFWEVQLGIPHPAGLK+KKSVTVLDVGDAYFSVPL
Sbjct  2048  FAIKKKDSTKWRKLVDFRELNKRTQDFWEVQLGIPHPAGLKQKKSVTVLDVGDAYFSVPL  2227

Query  121   DEDFRKYTAFTIPSINNETPGIRYQYNVLPQGWKGSPAIFQSSMTKILEPFRKQNPDIVI  180
             D+DFRKYTAFTIPSINNETPGIRYQYNVLPQGWKGSPAIFQ SMTKILEPFRKQNPDIVI
Sbjct  2228  DKDFRKYTAFTIPSINNETPGIRYQYNVLPQGWKGSPAIFQCSMTKILEPFRKQNPDIVI  2407
```

```
Query  181   YQYMDDLYVGSDLEIGQHRTKIEELRQHLLRWGLTTPDKKHQKEPPFLWMGYELHPDKWT  240
             YQYMDDLYVGSDLEIGQHRTKIEELRQHLLRWG TTPDKKHQKEPPFLWMGYELHPDKWT
Sbjct  2408  YQYMDDLYVGSDLEIGQHRTKIEELRQHLLRWGFTTPDKKHQKEPPFLWMGYELHPDKWT  2587

Query  241   VQPIVLPEKDSWTVNDIQKLVGKLNWASQIYPGIKVRQLCKLLRGTKALTEVIPLTeeae  300
             VQPIVLPEKDSWTVNDIQKLVGKLNWASQIY GIKVRQLCKLLRGTKALTEV+PLTEEAE
Sbjct  2588  VQPIVLPEKDSWTVNDIQKLVGKLNWASQIYAGIKVRQLCKLLRGTKALTEVVPLTEEAE  2767

Query  301   lelaeNREILKEPVHGVYYDPSKDLIAEIQKQGQGQWTYQIYQEPFKNLKTGKYARMRGA  360
             LELAENREILKEPVHGVYYDPSKDLIAEIQKQGQGQWTYQIYQEPFKNLKTGKYARM+GA
Sbjct  2768  LELAENREILKEPVHGVYYDPSKDLIAEIQKQGQGQWTYQIYQEPFKNLKTGKYARMKGA  2947

Query  361   HTNDVKQLTEAVQKITTESIVIWGKTPKFKLPIQKetwetwwteYWQATWIPEWEFVNTP  420
             HTNDVKQLTEAVQKI TESIVIWGKTPKFKLPIQKETWE WWTEYWQATWIPEWEFVNTP
Sbjct  2948  HTNDVKQLTEAVQKIATESIVIWGKTPKFKLPIQKETWEAWWTEYWQATWIPEWEFVNTP  3127

Query  421   PLVKLWYQLEKEPIVGAETFYVDGAANRETKLGKAGYVTNRGRQKVVTLTDTTNQKTELQ  480
             PLVKLWYQLEKEPI+GAETFYVDGAANRETKLGKAGYVT+RGRQKVV LTDTTNQKTELQ
Sbjct  3128  PLVKLWYQLEKEPIIGAETFYVDGAANRETKLGKAGYVTDRGRQKVVPLTDTTNQKTELQ  3307

Query  481   AIYLALQDSGLEVNIVTDSQYALGIIQAQPDQSESELVNQIIEQLIKKEKVYLAWVPAHK  540
             AI+LALQDSGLEVNIVTDSQYALGIIQAQPD+SESELV+QIIEQLIKKEKVYLAWVPAHK
Sbjct  3308  AIHLALQDSGLEVNIVTDSQYALGIIQAQPDKSESELVSQIIEQLIKKEKVYLAWVPAHK  3487

Query  541   GIGGNEQVDKLVSAGIRKVL  560
             GIGGNEQVDKLVSAGIRKVL
Sbjct  3488  GIGGNEQVDKLVSAGIRKVL  3547
```

>Actinoplanes campanulatus strain JCM 3059 sequence058, whole genome shotgun sequence
Sequence ID: NZ_BMPW01000058.1 Length: 4754
Range 1: 1857 to 3536

Score:1073 bits(2775), Expect:0.0,
Method:Compositional matrix adjust.,
Identities:545/560(97%), Positives:554/560(98%), Gaps:0/560(0%)

```
Query  1     PISPIETVPVKLKPGMDGPKVKQWPLTEEKIKALVEICTEMEKEGKISKIGPENPYNTPV  60
             PISPIETVPVKLKPGMDGPKVKQWPLTEEKIKALVEICTEMEKEGKISKIGPENPYNTPV
Sbjct  1857  PISPIETVPVKLKPGMDGPKVKQWPLTEEKIKALVEICTEMEKEGKISKIGPENPYNTPV  2036

Query  61    FAIKKKDSTKWRKLVDFRELNKRTQDFWEVQLGIPHPAGLKKKKSVTVLDVGDAYFSVPL  120
             FAIKKKDSTKWRKLVDFRELNKRTQDFWEVQLGIPHPAGLK+KKSVTVLDVGDAYFSVPL
Sbjct  2037  FAIKKKDSTKWRKLVDFRELNKRTQDFWEVQLGIPHPAGLKQKKSVTVLDVGDAYFSVPL  2216

Query  121   DEDFRKYTAFTIPSINNETPGIRYQYNVLPQGWKGSPAIFQSSMTKILEPFRKQNPDIVI  180
             D+DFRKYTAFTIPSINNETPGIRYQYNVLPQGWKGSPAIFQ SMTKILEPFRKQNPDIVI
Sbjct  2217  DKDFRKYTAFTIPSINNETPGIRYQYNVLPQGWKGSPAIFQCSMTKILEPFRKQNPDIVI  2396

Query  181   YQYMDDLYVGSDLEIGQHRTKIEELRQHLLRWGLTTPDKKHQKEPPFLWMGYELHPDKWT  240
             YQYMDDLYVGSDLEIGQHRTKIEELRQHLLRWG TTPDKKHQKEPPFLWMGYELHPDKWT
Sbjct  2397  YQYMDDLYVGSDLEIGQHRTKIEELRQHLLRWGFTTPDKKHQKEPPFLWMGYELHPDKWT  2576

Query  241   VQPIVLPEKDSWTVNDIQKLVGKLNWASQIYPGIKVRQLCKLLRGTKALTEVIPLTeeae  300
             VQPIVLPEKDSWTVNDIQKLVGKLNWASQIY GIKVRQLCKLLRGTKALTEV+PLTEEAE
Sbjct  2577  VQPIVLPEKDSWTVNDIQKLVGKLNWASQIYAGIKVRQLCKLLRGTKALTEVVPLTEEAE  2756

Query  301   lelaeNREILKEPVHGVYYDPSKDLIAEIQKQGQGQWTYQIYQEPFKNLKTGKYARMRGA  360
             LELAENREILKEPVHGVYYDPSKDLIAEIQKQGQGQWTYQIYQEPFKNLKTGKYARM+GA
Sbjct  2757  LELAENREILKEPVHGVYYDPSKDLIAEIQKQGQGQWTYQIYQEPFKNLKTGKYARMKGA  2936

Query  361   HTNDVKQLTEAVQKITTESIVIWGKTPKFKLPIQKetwetwwteYWQATWIPEWEFVNTP  420
             HTNDVKQLTEAVQKI TESIVIWGKTPKFKLPIQKETWE WWTEYWQATWIPEWEFVNTP
Sbjct  2937  HTNDVKQLTEAVQKIATESIVIWGKTPKFKLPIQKETWEAWWTEYWQATWIPEWEFVNTP  3116

Query  421   PLVKLWYQLEKEPIVGAETFYVDGAANRETKLGKAGYVTNRGRQKVVTLTDTTNQKTELQ  480
             PLVKLWYQLEKEPI+GAETFYVDGAANRETKLGKAGYVT+RGRQKVV LTDTTNQKTELQ
Sbjct  3117  PLVKLWYQLEKEPIIGAETFYVDGAANRETKLGKAGYVTDRGRQKVVPLTDTTNQKTELQ  3296

Query  481   AIYLALQDSGLEVNIVTDSQYALGIIQAQPDQSESELVNQIIEQLIKKEKVYLAWVPAHK  540
             AI+LALQDSGLEVNIVTDSQYALGIIQAQPD+SESELV+QIIEQLIKKEKVYLAWVPAHK
Sbjct  3297  AIHLALQDSGLEVNIVTDSQYALGIIQAQPDKSESELVSQIIEQLIKKEKVYLAWVPAHK  3476

Query  541   GIGGNEQVDKLVSAGIRKVL  560
             GIGGNEQVDKLVSAGIRKVL
Sbjct  3477  GIGGNEQVDKLVSAGIRKVL  3536
```

>Klebsiella pneumoniae strain ARGID_31501, whole genome shotgun sequence
Sequence ID: NZ_CAKLPD010000006.1 Length: 2655
Range 1: 224 to 1903

Score:1055 bits(2729), Expect:0.0,
Method:Compositional matrix adjust.,
Identities:534/560(95%), Positives:547/560(97%), Gaps:0/560(0%)

```
Query  1     PISPIETVPVKLKPGMDGPKVKQWPLTEEKIKALVEICTEMEKEGKISKIGPENPYNTPV  60
             PISPIETVPVKLKPGMDGPKVKQWPLTEEKIKALVEICTEMEKEGKISKIGPENPYNTP+
Sbjct  224   PISPIETVPVKLKPGMDGPKVKQWPLTEEKIKALVEICTEMEKEGKISKIGPENPYNTPI  403

Query  61    FAIKKKDSTKWRKLVDFRELNKRTQDFWEVQLGIPHPAGLKKKKSVTVLDVGDAYFSVPL  120
             FAIKKKDSTKWRKLVDFRELNKRTQDFWEVQLGIPHPAGLKKKKSVTVLDVGDAYFSVPL
Sbjct  404   FAIKKKDSTKWRKLVDFRELNKRTQDFWEVQLGIPHPAGLKKKKSVTVLDVGDAYFSVPL  583

Query  121   DEDFRKYTAFTIPSINNETPGIRYQYNVLPQGWKGSPAIFQSSMTKILEPFRKQNPDIVI  180
             D+DFRKYTAFTIPSINNETPG+RYQYNVLPQGWKGSPAIFQSSMTKILEPFRKQNPDIVI
Sbjct  584   DQDFRKYTAFTIPSINNETPGVRYQYNVLPQGWKGSPAIFQSSMTKILEPFRKQNPDIVI  763

Query  181   YQYMDDLYVGSDLEIGQHRTKIEELRQHLLRWGLTTPDKKHQKEPPFLWMGYELHPDKWT  240
             YQYMDDLYVGSDLEIGQHR KIEELRQHLL+WG TTPDKKHQKEPPFLWMGYELHPDKWT
Sbjct  764   YQYMDDLYVGSDLEIGQHRIKIEELRQHLLKWGFTTPDKKHQKEPPFLWMGYELHPDKWT  943

Query  241   VQPIVLPEKDSWTVNDIQKLVGKLNWASQIYPGIKVRQLCKLLRGTKALTEVIPLTeeae  300
             VQPI LPEKDSWTVNDIQKLVGKLNWASQIY GIKVRQLCKLLRG KALTEV+PLT+EAE
Sbjct  944   VQPIKLPEKDSWTVNDIQKLVGKLNWASQIYAGIKVRQLCKLLRGAKALTEVVPLTKEAE  1123

Query  301   lelaeNREILKEPVHGVYYDPSKDLIAEIQKQGQGQWTYQIYQEPFKNLKTGKYARMRGA  360
             LELAENREILKEPVHGVYYDPSK+LIAEIQKQG+GQWTYQIYQE FKNLKTGKYARMRG
Sbjct  1124  LELAENREILKEPVHGVYYDPSKELIAEIQKQGEGQWTYQIYQEQFKNLKTGKYARMRGT  1303

Query  361   HTNDVKQLTEAVQKITTESIVIWGKTPKFKLPIQKetwetwwteYWQATWIPEWEFVNTP  420
             HTNDVKQLTEAVQKI TESIVIWGKTPKF+LPI KETWETWWT+YWQATWIPEWEFVNTP
Sbjct  1304  HTNDVKQLTEAVQKIATESIVIWGKTPKFRLPIHKETWETWWTDYWQATWIPEWEFVNTP  1483

Query  421   PLVKLWYQLEKEPIVGAETFYVDGAANRETKLGKAGYVTNRGRQKVVTLTDTTNQKTELQ  480
             PLVKLWYQLEKEPI GAETFYVDGAANRETK+GKAGYVTNRGRQKVV LTDTTNQKTELQ
Sbjct  1484  PLVKLWYQLEKEPIEGAETFYVDGAANRETKIGKAGYVTNRGRQKVVPLTDTTNQKTELQ  1663

Query  481   AIYLALQDSGLEVNIVTDSQYALGIIQAQPDQSESELVNQIIEQLIKKEKVYLAWVPAHK  540
             AIYLALQDS LEVNIVTDSQYALGIIQAQPD+SESELVNQIIEQLIKKEK+YLAWVPAHK
```

```
Sbjct  1664  AIYLALQDSELEVNIVTDSQYALGIIQAQPDKSESELVNQIIEQLIKKEKIYLAWVPAHK  1843

Query  541   GIGGNEQVDKLVSAGIRKVL  560
             GIGGNEQVDKLVS GIRKVL
Sbjct  1844  GIGGNEQVDKLVSTGIRKVL  1903
```

```
>Thalassovita mangrovi strain GS-10 Scaffold20_1, whole genome shotgun sequence
Sequence ID: NZ_WWEN01000020.1 Length: 1488
Range 1: 2 to 1486

Score:941 bits(2433), Expect:0.0,
Method:Compositional matrix adjust.,
Identities:480/495(97%), Positives:489/495(98%), Gaps:0/495(0%)

Query  49    KIGPENPYNTPVFAIKKKDSTKWRKLVDFRELNKRTQDFWEVQLGIPHPAGLKKKKSVTV  108
             KIGPENPYNTPVFAIKKKDSTKWRKLVDFRELNKRTQDFWEVQLGIPHPAGLK+KKSVTV
Sbjct  1486  KIGPENPYNTPVFAIKKKDSTKWRKLVDFRELNKRTQDFWEVQLGIPHPAGLKQKKSVTV  1307

Query  109   LDVGDAYFSVPLDEDFRKYTAFTIPSINNETPGIRYQYNVLPQGWKGSPAIFQSSMTKIL  168
             LDVGDAYFSVPLD+DFRKYTAFTIPSINNETPGIRYQYNVLPQGWKGSPAIFQ SMTKIL
Sbjct  1306  LDVGDAYFSVPLDKDFRKYTAFTIPSINNETPGIRYQYNVLPQGWKGSPAIFQCSMTKIL  1127

Query  169   EPFRKQNPDIVIYQYMDDLYVGSDLEIGQHRTKIEELRQHLLRWGLTTPDKKHQKEPPFL  228
             EPFRKQNPDIVIYQYMDDLYVGSDLEIGQHRTKIEELRQHLLRWG TTPDKKHQKEPPFL
Sbjct  1126  EPFRKQNPDIVIYQYMDDLYVGSDLEIGQHRTKIEELRQHLLRWGFTTPDKKHQKEPPFL  947

Query  229   WMGYELHPDKWTVQPIVLPEKDSWTVNDIQKLVGKLNWASQIYPGIKVRQLCKLLRGTKA  288
             WMGYELHPDKWTVQPIVLPEKDSWTVNDIQKLVGKLNWASQIY GIKVRQLCKLLRGTKA
Sbjct  946   WMGYELHPDKWTVQPIVLPEKDSWTVNDIQKLVGKLNWASQIYAGIKVRQLCKLLRGTKA  767

Query  289   LTEVIPLTeeaelelaeNREILKEPVHGVYYDPSKDLIAEIQKQGQGQWTYQIYQEPFKN  348
             LTEV+PLTEEAELELAENREILKEPVHGVYYDPSKDLIAEIQKQGQGQWTYQIYQEPFKN
Sbjct  766   LTEVVPLTEEAELELAENREILKEPVHGVYYDPSKDLIAEIQKQGQGQWTYQIYQEPFKN  587

Query  349   LKTGKYARMRGAHTNDVKQLTEAVQKITTESIVIWGKTPKFKLPIQKetwetwwteYWQA  408
             LKTGKYARM+GAHTNDVKQLTEAVQKI TESIVIWGKTPKFKLPIQKETWE WWTEYWQA
Sbjct  586   LKTGKYARMKGAHTNDVKQLTEAVQKIATESIVIWGKTPKFKLPIQKETWEAWWTEYWQA  407

Query  409   TWIPEWEFVNTPPLVKLWYQLEKEPIVGAETFYVDGAANRETKLGKAGYVTNRGRQKVVT  468
             TWIPEWEFVNTPPLVKLWYQLEKEPI+GAETFYVDGAANRETKLGKAGYVT+RGRQKVV
Sbjct  406   TWIPEWEFVNTPPLVKLWYQLEKEPIIGAETFYVDGAANRETKLGKAGYVTDRGRQKVVP  227

Query  469   LTDTTNQKTELQAIYLALQDSGLEVNIVTDSQYALGIIQAQPDQSESELVNQIIEQLIKK  528
             LTDTTNQKTELQAI+LALQDSGLEVNIVTDSQYALGIIQAQPD+SESELV+QIIEQLIKK
Sbjct  226   LTDTTNQKTELQAIHLALQDSGLEVNIVTDSQYALGIIQAQPDKSESELVSQIIEQLIKK  47

Query  529   EKVYLAWVPAHKGIG  543
             EKVYLAWVPAHKGIG
Sbjct  46    EKVYLAWVPAHKGIG  2
```

```
>Enterococcus faecium strain A201R2B0 NODE_60_length_1589_cov_4.20178_ID_119, whole genome shotgun sequence
Sequence ID: NZ_PGCT01000032.1 Length: 1589
Range 1: 2 to 1330

Score:818 bits(2113), Expect:0.0,
Method:Compositional matrix adjust.,
Identities:414/443(93%), Positives:427/443(96%), Gaps:0/443(0%)

Query  1     PISPIETVPVKLKPGMDGPKVKQWPLTEEKIKALVEICTEMEKEGKISKIGPENPYNTPV  60
             PISPI TVPVKLKPGMDGPKVKQWPLTEEKIKAL  IC EMEKEGKISKIGPENPYNTPV
Sbjct  1330  PISPIGTVPVKLKPGMDGPKVKQWPLTEEKIKALTAICEEMEKEGKISKIGPENPYNTPV  1151

Query  61    FAIKKKDSTKWRKLVDFRELNKRTQDFWEVQLGIPHPAGLKKKKSVTVLDVGDAYFSVPL  120
             FAIKKKDSTKWRKLVDFRELNKRTQDFWEVQLGIPHPAGLKKKKSVTVLDVGDAYFSVPL
Sbjct  1150  FAIKKKDSTKWRKLVDFRELNKRTQDFWEVQLGIPHPAGLKKKKSVTVLDVGDAYFSVPL  971

Query  121   DEDFRKYTAFTIPSINNETPGIRYQYNVLPQGWKGSPAIFQSSMTKILEPFRKQNPDIVI  180
             DE FR+YTAFTIPSINNETPGIRYQYNVLPQGWKGSPAIFQSSMTKILEPFR +NPDI+I
Sbjct  970   DEGFRRYTAFTIPSINNETPGIRYQYNVLPQGWKGSPAIFQSSMTKILEPFRTRNPDIII  791

Query  181   YQYMDDLYVGSDLEIGQHRTKIEELRQHLLRWGLTTPDKKHQKEPPFLWMGYELHPDKWT  240
             YQYMDDLYVGSDLEIGQHR KIEELR+HLLRWGLTTPDKKHQKEPPFLWMGYELHPDKWT
Sbjct  790   YQYMDDLYVGSDLEIGQHRAKIEELREHLLRWGLTTPDKKHQKEPPFLWMGYELHPDKWT  611

Query  241   VQPIVLPEKDSWTVNDIQKLVGKLNWASQIYPGIKVRQLCKLLRGTKALTEVIPLTeeae  300
             VQPIVLPEKDSWTVNDIQKLVGKLNWASQIY GIKV+QLCKLLRG KALT+++PLTEEAE
Sbjct  610   VQPIVLPEKDSWTVNDIQKLVGKLNWASQIYQGIKVKQLCKLLRGAKALTDIVPLTEEAE  431

Query  301   lelaeNREILKEPVHGVYYDPSKDLIAEIQKQGQGQWTYQIYQEPFKNLKTGKYARMRGA  360
             LELAENREILKEPVHGVYYDPSKDLIAEIQKQG  QWTYQIYQEPFKNLKTGKYA+MR A
Sbjct  430   LELAENREILKEPVHGVYYDPSKDLIAEIQKQGHDQWTYQIYQEPFKNLKTGKYAKMRTA  251

Query  361   HTNDVKQLTEAVQKITTESIVIWGKTPKFKLPIQKetwetwwteYWQATWIPEWEFVNTP  420
             HTND+KQLTEAVQKI  ESIVIWGKTPKF+LPIQK TWETWWT+YWQATWIP+WEFVNTP
Sbjct  250   HTNDIKQLTEAVQKIAMESIVIWGKTPKFRLPIQKGTWETWWTDYWQATWIPDWEFVNTP  71

Query  421   PLVKLWYQLEKEPIVGAETFYVD  443
             PLVKLWYQLEKEPI GAETFYVD
Sbjct  70    PLVKLWYQLEKEPIAGAETFYVD  2
```

```
>Enterococcus faecium strain A209R1B0C1 NODE_70_length_1819_cov_3.01832_ID_139, whole genome shotgun sequence
Sequence ID: NZ_PGCR01000088.1 Length: 1819
Range 1: 486 to 1817

Score:814 bits(2102), Expect:0.0,
Method:Compositional matrix adjust.,
Identities:410/444(92%), Positives:429/444(96%), Gaps:0/444(0%)

Query  1     PISPIETVPVKLKPGMDGPKVKQWPLTEEKIKALVEICTEMEKEGKISKIGPENPYNTPV  60
             PISPIETVPVKLKPGMDGPKVKQWPLTEEKIKAL EIC EMEKEGKI+KIGPENPYNTP+
Sbjct  486   PISPIETVPVKLKPGMDGPKVKQWPLTEEKIKALTEICNEMEKEGKITKIGPENPYNTPI  665

Query  61    FAIKKKDSTKWRKLVDFRELNKRTQDFWEVQLGIPHPAGLKKKKSVTVLDVGDAYFSVPL  120
             FAIKKKDSTKWRKLVDFRELNKRTQDFWEVQLGIPHPAGLKKKKSVTVLDVGDAYFSVPL
Sbjct  666   FAIKKKDSTKWRKLVDFRELNKRTQDFWEVQLGIPHPAGLKKKKSVTVLDVGDAYFSVPL  845

Query  121   DEDFRKYTAFTIPSINNETPGIRYQYNVLPQGWKGSPAIFQSSMTKILEPFRKQNPDIVI  180
             D++FRKYTAFTIPSINNETPGIRYQYNVLPQGWKGSP+IFQSSM KILEPFR +NP++VI
Sbjct  846   DKEFRKYTAFTIPSINNETPGIRYQYNVLPQGWKGSPSIFQSSMIKILEPFRAKNPEMVI  1025

Query  181   YQYMDDLYVGSDLEIGQHRTKIEELRQHLLRWGLTTPDKKHQKEPPFLWMGYELHPDKWT  240
             YQYMDDLYVGSDLEIGQHR KIEELR+HLLRWG TTPDKKHQKEPPFLWMGYELHPDKWT
Sbjct  1026  YQYMDDLYVGSDLEIGQHRAKIEELREHLLRWGFTTPDKKHQKEPPFLWMGYELHPDKWT  1205

Query  241   VQPIVLPEKDSWTVNDIQKLVGKLNWASQIYPGIKVRQLCKLLRGTKALTEVIPLTeeae  300
```

```
                VQPI LP+KDSWTVNDIQKLVGKLNWASQIYPGIKVRQLCKLLRG KALT+++PLTEEAE
Sbjct  1206   VQPIQLPDKDSWTVNDIQKLVGKLNWASQIYPGIKVRQLCKLLRGAKALTDIVPLTEEAE  1385

Query  301    lelaeNREILKEPVHGVYYDPSKDLIAEIQKQGQGQWTYQIYQEPFKNLKTGKYARMRGA  360
              LELAENREILKEPVHGVYYDPSKDLIAEIQKQG  QWTYQIYQ+PFKNLKTGKYA+MR A
Sbjct  1386   LELAENREILKEPVHGVYYDPSKDLIAEIQKQGHDQWTYQIYQDPFKNLKTGKYAKMRTA  1565

Query  361    HTNDVKQLTEAVQKITTESIVIWGKTPKFKLPIQKetwetwwteYWQATWIPEWEFVNTP  420
              HTNDVKQL EAVQKI  ESIVIWGKTPKF+LPIQKETWETWWT+YWQATWIP+WEFVNTP
Sbjct  1566   HTNDVKQLAEAVQKIAIESIVIWGKTPKFRLPIQKETWETWWTDYWQATWIPDWEFVNTP  1745

Query  421    PLVKLWYQLEKEPIVGAETFYVDG  444
              PLVKLWYQL KEPI+GAETFYVDG
Sbjct  1746   PLVKLWYQLVKEPIIGAETFYVDG  1817
```

>Listeria monocytogenes strain YA00082404 NODE_73_length_1816_cov_5.664890, whole genome shotgun sequence
Sequence ID: NZ_QEXE01000073.1 Length: 1816
Range 1: 499 to 1815

Score:795 bits(2053), Expect:0.0,
Method:Compositional matrix adjust.,
Identities:399/439(91%), Positives:420/439(95%), Gaps:0/439(0%)

```
Query  1      PISPIETVPVKLKPGMDGPKVKQWPLTEEKIKALVEICTEMEKEGKISKIGPENPYNTPV  60
              PISPIET+PVKLKPGMDGP+VKQWPLTEEKIKAL EIC +MEKEGKI++IGPENPYNTP+
Sbjct  499    PISPIETIPVKLKPGMDGPRVKQWPLTEEKIKALTEICEDMEKEGKITRIGPENPYNTPI  678

Query  61     FAIKKKDSTKWRKLVDFRELNKRTQDFWEVQLGIPHPAGLKKKKSVTVLDVGDAYFSVPL  120
              FAIK+KD TKWRKLVDFRELNKRTQDFWEVQLGIPHPAGLKKKKS+TVLDVGDAYFSVPL
Sbjct  679    FAIKRKDGTKWRKLVDFRELNKRTQDFWEVQLGIPHPAGLKKKKSMTVLDVGDAYFSVPL  858

Query  121    DEDFRKYTAFTIPSINNETPGIRYQYNVLPQGWKGSPAIFQSSMTKILEPFRKQNPDIVI  180
              DE FRKYTAFTIPSINN TPGIRYQYNVLPQGWKGSPAIFQSSMTKIL+PFR +NPDIVI
Sbjct  859    DESFRKYTAFTIPSINNSTPGIRYQYNVLPQGWKGSPAIFQSSMTKILQPFRDENPDIVI  1038

Query  181    YQYMDDLYVGSDLEIGQHRTKIEELRQHLLRWGLTTPDKKHQKEPPFLWMGYELHPDKWT  240
              YQY+DDL VGSDLEIGQHR KIE+LR+HLLRWGLTTPDKKHQKEPPFLWMGYELHPDKWT
Sbjct  1039   YQYVDDLLVGSDLEIGQHRAKIEQLREHLLRWGLTTPDKKHQKEPPFLWMGYELHPDKWT  1218

Query  241    VQPIVLPEKDSWTVNDIQKLVGKLNWASQIYPGIKVRQLCKLLRGTKALTEVIPLTeeae  300
              VQPI LPEKDSWTVNDIQKLVGKLNWASQIYPGI  +QLC+LLRGTKALT+++PLTEEAE
Sbjct  1219   VQPIQLPEKDSWTVNDIQKLVGKLNWASQIYPGICTKQLCRLLRGTKALTDIVPLTEEAE  1398

Query  301    lelaeNREILKEPVHGVYYDPSKDLIAEIQKQGQGQWTYQIYQEPFKNLKTGKYARMRGA  360
              LELAENREILKEPVHGVYYDPSKDLIAEIQKQG  QWTYQIYQEPFKNLKTGKYA+MR A
Sbjct  1399   LELAENREILKEPVHGVYYDPSKDLIAEIQKQGDDQWTYQIYQEPFKNLKTGKYAKMRTA  1578

Query  361    HTNDVKQLTEAVQKITTESIVIWGKTPKFKLPIQKetwetwwteYWQATWIPEWEFVNTP  420
              HTNDVKQLTEAVQKI  ESIVIWGKTPKF+LPIQKETWE WT+YWQATWIPEWEFVNTP
Sbjct  1579   HTNDVKQLTEAVQKIALESIVIWGKTPKFRLPIQKETWEIWWTDYWQATWIPEWEFVNTP  1758

Query  421    PLVKLWYQLEKEPIVGAET  439
              PLVKLWYQLEKEPI  AET
Sbjct  1759   PLVKLWYQLEKEPIAEAET  1815
```

>Escherichia coli strain YA00028237 contig_187, whole genome shotgun sequence
Sequence ID: NZ_NGBP02000187.1 Length: 1828
Range 1: 3 to 1325

Score:793 bits(2047), Expect:0.0,
Method:Compositional matrix adjust.,
Identities:396/441(90%), Positives:422/441(95%), Gaps:0/441(0%)

```
Query  1      PISPIETVPVKLKPGMDGPKVKQWPLTEEKIKALVEICTEMEKEGKISKIGPENPYNTPV  60
              PISPIETVPVKLKPGMDGP+VKQWPLTEEKIKAL EIC EM+KEGKI+KIGPENPYNTP+
Sbjct  1325   PISPIETVPVKLKPGMDGPRVKQWPLTEEKIKALTEICDEMKKEGKITKIGPENPYNTPI  1146

Query  61     FAIKKKDSTKWRKLVDFRELNKRTQDFWEVQLGIPHPAGLKKKKSVTVLDVGDAYFSVPL  120
              FAIKKK+STKWRKLVDFRELNKRTQDFWEVQLGIPHPAGLKK KSVTVLDVGDAYFSVPL
Sbjct  1145   FAIKKKNSTKWRKLVDFRELNKRTQDFWEVQLGIPHPAGLKKNKSVTVLDVGDAYFSVPL  966

Query  121    DEDFRKYTAFTIPSINNETPGIRYQYNVLPQGWKGSPAIFQSSMTKILEPFRKQNPDIVI  180
              DE+FRKYTAFTIPS+NNETPGIRYQYNVLPQGWKGSPAIFQSSMTKILEPFR QNP+IVI
Sbjct  965    DENFRKYTAFTIPSVNNETPGIRYQYNVLPQGWKGSPAIFQSSMTKILEPFRAQNPEIVI  786

Query  181    YQYMDDLYVGSDLEIGQHRTKIEELRQHLLRWGLTTPDKKHQKEPPFLWMGYELHPDKWT  240
              YQY+DDLYVGSDLEIGQHR K+E+LR HLL+WG TTPDKKHQKEPPFLWMGYELHPDKWT
Sbjct  785    YQYVDDLYVGSDLEIGQHRAKVEKLRDHLLKWGFTTPDKKHQKEPPFLWMGYELHPDKWT  606

Query  241    VQPIVLPEKDSWTVNDIQKLVGKLNWASQIYPGIKVRQLCKLLRGTKALTEVIPLTeeae  300
              VQ I LPEK+SWTVNDIQKLVGKLNWASQIY GIKV+QLC+LLRG KALT+++PLTEEAE
Sbjct  605    VQQIQLPEKESWTVNDIQKLVGKLNWASQIYAGIKVKQLCRLLRGAKALTDIVPLTEEAE  426

Query  301    lelaeNREILKEPVHGVYYDPSKDLIAEIQKQGQGQWTYQIYQEPFKNLKTGKYARMRGA  360
              LELAENREILKEPVHGVYYDPSKDL+AEIQKQG  QWTYQIYQEPFKNLKTGKYA+ R A
Sbjct  425    LELAENREILKEPVHGVYYDPSKDLVAEIQKQGNDQWTYQIYQEPFKNLKTGKYAKRRTA  246

Query  361    HTNDVKQLTEAVQKITTESIVIWGKTPKFKLPIQKetwetwwteYWQATWIPEWEFVNTP  420
              HTNDV+QLTE VQKI+ ESIVIWGKTPKF+LPIQKETWETWW +YWQATWIP+WEFVNTP
Sbjct  245    HTNDVQQLTEVVQKISRESIVIWGKTPKFRLPIQKETWETWWADYWQATWIPDWEFVNTP  66

Query  421    PLVKLWYQLEKEPIVGAETFY  441
              PLVKLWYQLEK+PI GAETFY
Sbjct  65     PLVKLWYQLEKDPIAGAETFY  3
```

>Enterococcus faecium strain A206R2B0 NODE_116_length_1761_cov_4.01102_ID_231, whole genome shotgun sequence
Sequence ID: NZ_PGCU01000052.1 Length: 1761
Range 1: 470 to 1759

Score:793 bits(2047), Expect:0.0,
Method:Compositional matrix adjust.,
Identities:400/430(93%), Positives:415/430(96%), Gaps:0/430(0%)

```
Query  1      PISPIETVPVKLKPGMDGPKVKQWPLTEEKIKALVEICTEMEKEGKISKIGPENPYNTPV  60
              PISPI TVPVKLKPGMDGPKVKQWPLTEEKIKAL  IC EMEKEGKISKIGPENPYNTPV
Sbjct  470    PISPIGTVPVKLKPGMDGPKVKQWPLTEEKIKALTAICEEMEKEGKISKIGPENPYNTPV  649

Query  61     FAIKKKDSTKWRKLVDFRELNKRTQDFWEVQLGIPHPAGLKKKKSVTVLDVGDAYFSVPL  120
              FAIKKKDSTKWRKLVDFRELNKRTQDFWEVQLGIPHPAGLKKKKSVTVLDVGDAYFSVPL
Sbjct  650    FAIKKKDSTKWRKLVDFRELNKRTQDFWEVQLGIPHPAGLKKKKSVTVLDVGDAYFSVPL  829

Query  121    DEDFRKYTAFTIPSINNETPGIRYQYNVLPQGWKGSPAIFQSSMTKILEPFRKQNPDIVI  180
              DE FR+YTAFTIPSINNETPGIRYQYNVLPQGWKGSPAIFQSSMTKILEPFR +NPDI+I
Sbjct  830    DEGFRRYTAFTIPSINNETPGIRYQYNVLPQGWKGSPAIFQSSMTKILEPFRTRNPDIII  1009
```

```
Query  181   YQYMDDLYVGSDLEIGQHRTKIEELRQHLLRWGLTTPDKKHQKEPPFLWMGYELHPDKWT  240
             YQYMDDLYVGSDLEIGQHR KIEELR+HLLRWGLTTPDKKHQKEPPFLWMGYELHPDKWT
Sbjct  1010  YQYMDDLYVGSDLEIGQHRAKIEELREHLLRWGLTTPDKKHQKEPPFLWMGYELHPDKWT  1189

Query  241   VQPIVLPEKDSWTVNDIQKLVGKLNWASQIYPGIKVRQLCKLLRGTKALTEVIPLTeeae  300
             VQPIVLPEKDSWTVNDIQKLVGKLNWASQIY GIKV+QLCKLLRG KALT+++PLTEEAE
Sbjct  1190  VQPIVLPEKDSWTVNDIQKLVGKLNWASQIYQGIKVKQLCKLLRGAKALTDIVPLTEEAE  1369

Query  301   lelaeNREILKEPVHGVYYDPSKDLIAEIQKQGQGQWTYQIYQEPFKNLKTGKYARMRGA  360
             LELAENREILKEPVHGVYYDPSKDLIAEIQKQG  QWTYQIYQEPFKNLKTGKYA+MR A
Sbjct  1370  LELAENREILKEPVHGVYYDPSKDLIAEIQKQGHDQWTYQIYQEPFKNLKTGKYAKMRTA  1549

Query  361   HTNDVKQLTEAVQKITTESIVIWGKTPKFKLPIQKetwetwwteYWQATWIPEWEFVNTP  420
             HTND+KQLTEAVQKI  ESIVIWGKTPKF+LPIQK TWETWWT+YWQATWIP+WEFVNTP
Sbjct  1550  HTNDIKQLTEAVQKIAMESIVIWGKTPKFRLPIQKGTWETWWTDYWQATWIPDWEFVNTP  1729

Query  421   PLVKLWYQLE  430
             P++KLWYQLE
Sbjct  1730  PIIKLWYQLE  1759
```

```
>Klebsiella pneumoniae strain NUBRI-K 73, whole genome shotgun sequence
Sequence ID: NZ_SOYT01000073.1 Length: 1717
Range 1: 452 to 1717

Score:759 bits(1961), Expect:0.0,
Method:Compositional matrix adjust.,
Identities:384/422(91%), Positives:404/422(95%), Gaps:0/422(0%)

Query  1     PISPIETVPVKLKPGMDGPKVKQWPLTEEKIKALVEICTEMEKEGKISKIGPENPYNTPV  60
             PISPIETVPV+LKPGMDGPKVKQWPLTEEKIKAL  IC EMEKEGKISKIGPENPYNTPV
Sbjct  452   PISPIETVPVQLKPGMDGPKVKQWPLTEEKIKALTAICEEMEKEGKISKIGPENPYNTPV  631

Query  61    FAIKKKDSTKWRKLVDFRELNKRTQDFWEVQLGIPHPAGLKKKKSVTVLDVGDAYFSVPL  120
             FAIKKKDSTKWRK+VDFRELNKRTQDFWEVQLGIPHPAGLKKKKSVTVLDVGDA+FSVPL
Sbjct  632   FAIKKKDSTKWRKVVDFRELNKRTQDFWEVQLGIPHPAGLKKKKSVTVLDVGDAFFSVPL  811

Query  121   DEDFRKYTAFTIPSINNETPGIRYQYNVLPQGWKGSPAIFQSSMTKILEPFRKQNPDIVI  180
              E FRKYTAFTIPSINNETPGIRYQYNVLPQGWKGSPAIFQ SM KILEPFR +NP+IVI
Sbjct  812   YEGFRKYTAFTIPSINNETPGIRYQYNVLPQGWKGSPAIFQCSMIKILEPFRARNPEIVI  991

Query  181   YQYMDDLYVGSDLEIGQHRTKIEELRQHLLRWGLTTPDKKHQKEPPFLWMGYELHPDKWT  240
             YQY+DDLYV SDLEIGQHR ++EELR+HLL+WG TTPDKKHQKEPPFLWMGYELHPDKWT
Sbjct  992   YQYVDDLYVASDLEIGQHRARLEELREHLLKWGFTTPDKKHQKEPPFLWMGYELHPDKWT  1171

Query  241   VQPIVLPEKDSWTVNDIQKLVGKLNWASQIYPGIKVRQLCKLLRGTKALTEVIPLTeeae  300
             VQPI LPEK+SWTVNDIQKLVGKLNWASQIYPGIKV+QLCKLLRGTKALT+++PLTEEAE
Sbjct  1172  VQPIQLPEKESWTVNDIQKLVGKLNWASQIYPGIKVKQLCKLLRGTKALTDIVPLTEEAE  1351

Query  301   lelaeNREILKEPVHGVYYDPSKDLIAEIQKQGQGQWTYQIYQEPFKNLKTGKYARMRGA  360
             LELAENREILKEPVHGVYYDPSKDLIAEIQKQG  QWTYQIYQEPFKNLKTGKYA+ + A
Sbjct  1352  LELAENREILKEPVHGVYYDPSKDLIAEIQKQGNDQWTYQIYQEPFKNLKTGKYAKRKSA  1531

Query  361   HTNDVKQLTEAVQKITTESIVIWGKTPKFKLPIQKetwetwwteYWQATWIPEWEFVNTP  420
             HTNDVKQLTEAVQKI+ ESIVIWGK PKF+LPIQKETWETWWT+YWQATWIPEWEFVNTP
Sbjct  1532  HTNDVKQLTEAVQKISMESIVIWGKIPKFRLPIQKETWETWWTDYWQATWIPEWEFVNTP  1711

Query  421   PL  422
             PL
Sbjct  1712  PL  1717
```

```
>Staphylococcus aureus strain G703N1B1 NODE_79_length_1173_cov_3.01147_ID_157, whole genome shotgun sequence
Sequence ID: NZ_PGXA01000017.1 Length: 1173
Range 1: 3 to 1166

Score:715 bits(1846), Expect:0.0,
Method:Compositional matrix adjust.,
Identities:361/388(93%), Positives:375/388(96%), Gaps:0/388(0%)

Query  55    PYNTPVFAIKKKDSTKWRKLVDFRELNKRTQDFWEVQLGIPHPAGLKKKKSVTVLDVGDA  114
             PYNTP+FAIKKKDSTKWRKLVDFRELNKRTQDFWEVQLGIPHPAGLKKKKSVTVLDVGDA
Sbjct  1166  PYNTPIFAIKKKDSTKWRKLVDFRELNKRTQDFWEVQLGIPHPAGLKKKKSVTVLDVGDA  987

Query  115   YFSVPLDEDFRKYTAFTIPSINNETPGIRYQYNVLPQGWKGSPAIFQSSMTKILEPFRKQ  174
             YFSVPLD+DFRKYTAFTIPSINNETPGIRYQYNVLPQGWKGSP+IFQSSMT+ILEPFR Q
Sbjct  986   YFSVPLDKDFRKYTAFTIPSINNETPGIRYQYNVLPQGWKGSPSIFQSSMTRILEPFRAQ  807

Query  175   NPDIVIYQYMDDLYVGSDLEIGQHRTKIEELRQHLLRWGLTTPDKKHQKEPPFLWMGYEL  234
             NP+IVIYQYMDDLYVGSDLEIGQHR KIEELR+HLL+WG TTPDKKHQKEPPFLWMGYEL
Sbjct  806   NPEIVIYQYMDDLYVGSDLEIGQHRAKIEELREHLLKWGFTTPDKKHQKEPPFLWMGYEL  627

Query  235   HPDKWTVQPIVLPEKDSWTVNDIQKLVGKLNWASQIYPGIKVRQLCKLLRGTKALTEVIP  294
             HPDKWTVQPI LPEKDSWTVNDIQKLVGKLNWASQIY GIKV+QLC+LLRG KALT+V+P
Sbjct  626   HPDKWTVQPIQLPEKDSWTVNDIQKLVGKLNWASQIYAGIKVKQLCRLLRGAKALTDVVP  447

Query  295   LTeeaelelaeNREILKEPVHGVYYDPSKDLIAEIQKQGQGQWTYQIYQEPFKNLKTGKY  354
             LTEEAELELAENREILKEPVHGVYYDPSKDLIAEIQKQG  QWTYQIYQEPFKNLKTGKY
Sbjct  446   LTEEAELELAENREILKEPVHGVYYDPSKDLIAEIQKQGHDQWTYQIYQEPFKNLKTGKY  267

Query  355   ARMRGAHTNDVKQLTEAVQKITTESIVIWGKTPKFKLPIQKetwetwwteYWQATWIPEW  414
             A+MR AHTNDVKQLTEAVQKI  E IVIWGKTPKF+LPIQKETWETWWT+YWQATWIPEW
Sbjct  266   AKMRSAHTNDVKQLTEAVQKIAMECIVIWGKTPKFRLPIQKETWETWWTDYWQATWIPEW  87

Query  415   EFVNTPPLVKLWYQLEKEPIVGAETFYV  442
             EFVNTPPLVKLWYQLEKEPI GAETFYV
Sbjct  86    EFVNTPPLVKLWYQLEKEPIAGAETFYV  3
```

```
>Escherichia coli strain 8374wB11 NODE_34_length_1310_cov_3.21048, whole genome shotgun sequence
Sequence ID: NZ_WSJO01000034.1 Length: 1310
Range 1: 1 to 1023

Score:654 bits(1688), Expect:0.0,
Method:Compositional matrix adjust.,
Identities:326/341(96%), Positives:334/341(97%), Gaps:0/341(0%)

Query  1     PISPIETVPVKLKPGMDGPKVKQWPLTEEKIKALVEICTEMEKEGKISKIGPENPYNTPV  60
             PISPIETVPVKLKPGMDGPKVKQWPLTEEKIKALVEICTEMEKEGKISKIGPENPYNTPV
Sbjct  1023  PISPIETVPVKLKPGMDGPKVKQWPLTEEKIKALVEICTEMEKEGKISKIGPENPYNTPV  844

Query  61    FAIKKKDSTKWRKLVDFRELNKRTQDFWEVQLGIPHPAGLKKKKSVTVLDVGDAYFSVPL  120
             FAIKKKDSTKWRKLVDFRELNKRTQDFWEVQLGIPHPAGLKKKKSVTVLDVGDAYFSVPL
Sbjct  843   FAIKKKDSTKWRKLVDFRELNKRTQDFWEVQLGIPHPAGLKKKKSVTVLDVGDAYFSVPL  664

Query  121   DEDFRKYTAFTIPSINNETPGIRYQYNVLPQGWKGSPAIFQSSMTKILEPFRKQNPDIVI  180
             D++FRKYTAFTIPS NNETPGIRYQYNVLPQGWKGSPAIFQ SMTKILEPFRKQNPDIVI
Sbjct  663   DKEFRKYTAFTIPSTNNETPGIRYQYNVLPQGWKGSPAIFQCSMTKILEPFRKQNPDIVI  484
```

```
Query  181   YQYMDDLYVGSDLEIGQHRTKIEELRQHLLRWGLTTPDKKHQKEPPFLWMGYELHPDKWT  240
             YQY+DDLYVGSDLEI QHRTK+EELRQHLL WG TTPDKKHQKEPPFLWMGYELHPDKWT
Sbjct  483   YQYVDDLYVGSDLEIEQHRTKVEELRQHLLTWGFTTPDKKHQKEPPFLWMGYELHPDKWT  304

Query  241   VQPIVLPEKDSWTVNDIQKLVGKLNWASQIYPGIKVRQLCKLLRGTKALTEVIPLTeeae  300
             VQPIVLPEKDSWTVNDIQKLVGKLNWASQIYPGIKVRQLCKLLRG KALTEV+ LT+EAE
Sbjct  303   VQPIVLPEKDSWTVNDIQKLVGKLNWASQIYPGIKVRQLCKLLRGAKALTEVVTLTKEAE  124

Query  301   lelaeNREILKEPVHGVYYDPSKDLIAEIQKQGQGQWTYQI  341
             LELAENREI+KEPVHGVYYDP+KDLIAEIQKQGQGQWTYQI
Sbjct  123   LELAENREIIKEPVHGVYYDPTKDLIAEIQKQGQGQWTYQI  1
```

>Acinetobacter baumannii strain ACM-5 NODE_55_length_1274_cov_1.90584, whole genome shotgun sequence
Sequence ID: NZ_PYCX01000055.1 Length: 1274
Range 1: 2 to 1018

Score:650 bits(1676), Expect:0.0,
Method:Compositional matrix adjust.,
Identities:322/339(95%), Positives:330/339(97%), Gaps:0/339(0%)

```
Query  1     PISPIETVPVKLKPGMDGPKVKQWPLTEEKIKALVEICTEMEKEGKISKIGPENPYNTPV  60
             PISPIETVPVKLKPGMDGPKVKQWPLTEEKIKALVEICTEMEKEGKISKIGPENPYNTP+
Sbjct  1018  PISPIETVPVKLKPGMDGPKVKQWPLTEEKIKALVEICTEMEKEGKISKIGPENPYNTPI  839

Query  61    FAIKKKDSTKWRKLVDFRELNKRTQDFWEVQLGIPHPAGLKKKKSVTVLDVGDAYFSVPL  120
             FAIKKK+S KWRKLVDFRELNKRTQ+FWEVQLGIPHPAGLKKKKSVTVLDVGDAYFSVPL
Sbjct  838   FAIKKKNSDKWRKLVDFRELNKRTQEFWEVQLGIPHPAGLKKKKSVTVLDVGDAYFSVPL  659

Query  121   DEDFRKYTAFTIPSINNETPGIRYQYNVLPQGWKGSPAIFQSSMTKILEPFRKQNPDIVI  180
             D DFRKYTAFTIPS NNETPGIRYQYNVLPQGWKGSPAIFQ SMTKILEPFRKQNPDIVI
Sbjct  658   DPDFRKYTAFTIPSTNNETPGIRYQYNVLPQGWKGSPAIFQCSMTKILEPFRKQNPDIVI  479

Query  181   YQYMDDLYVGSDLEIGQHRTKIEELRQHLLRWGLTTPDKKHQKEPPFLWMGYELHPDKWT  240
             YQYMDDLYVGSDLEI QHRTK+EELRQHLL WG TTPDKKHQKEPPFLWMGYELHPDKWT
Sbjct  478   YQYMDDLYVGSDLEIEQHRTKVEELRQHLLTWGFTTPDKKHQKEPPFLWMGYELHPDKWT  299

Query  241   VQPIVLPEKDSWTVNDIQKLVGKLNWASQIYPGIKVRQLCKLLRGTKALTEVIPLTeeae  300
             VQPIVLPEKDSWTVNDIQKLVGKLNWASQIYPGIKVRQLCKLLRG KALTEV+ LT+EAE
Sbjct  298   VQPIVLPEKDSWTVNDIQKLVGKLNWASQIYPGIKVRQLCKLLRGAKALTEVVTLTKEAE  119

Query  301   lelaeNREILKEPVHGVYYDPSKDLIAEIQKQGQGQWTY  339
             LELAENREI+KEPVHGVYYDP+KDLIAEIQKQGQGQWTY
Sbjct  118   LELAENREIIKEPVHGVYYDPTKDLIAEIQKQGQGQWTY  2
```

>Escherichia coli strain 8374wB4 NODE_60_length_1385_cov_3.2655, whole genome shotgun sequence
Sequence ID: NZ_WSPX01000060.1 Length: 1385
Range 1: 363 to 1385

Score:649 bits(1673), Expect:0.0,
Method:Compositional matrix adjust.,
Identities:323/341(95%), Positives:333/341(97%), Gaps:0/341(0%)

```
Query  1     PISPIETVPVKLKPGMDGPKVKQWPLTEEKIKALVEICTEMEKEGKISKIGPENPYNTPV  60
             PISPIETVPVKLKP MDGPKVKQWPLTEEKIKALVEICTEMEKEGKISKIGPENPYNTPV
Sbjct  363   PISPIETVPVKLKPRMDGPKVKQWPLTEEKIKALVEICTEMEKEGKISKIGPENPYNTPV  542

Query  61    FAIKKKDSTKWRKLVDFRELNKRTQDFWEVQLGIPHPAGLKKKKSVTVLDVGDAYFSVPL  120
             FAIKKKDSTKWRKLVDFRELNKRTQDFWEVQLGIPHPAGLKKKKSVTVLDVGDAYFSVPL
Sbjct  543   FAIKKKDSTKWRKLVDFRELNKRTQDFWEVQLGIPHPAGLKKKKSVTVLDVGDAYFSVPL  722

Query  121   DEDFRKYTAFTIPSINNETPGIRYQYNVLPQGWKGSPAIFQSSMTKILEPFRKQNPDIVI  180
             D++FRKYTAFTIPS NN+TPGIRYQYNVLPQGWKGSPAIFQ SMTKILEPFRKQNPDIVI
Sbjct  723   DKEFRKYTAFTIPSTNNKTPGIRYQYNVLPQGWKGSPAIFQCSMTKILEPFRKQNPDIVI  902

Query  181   YQYMDDLYVGSDLEIGQHRTKIEELRQHLLRWGLTTPDKKHQKEPPFLWMGYELHPDKWT  240
             YQY+DDLYVGSDLEI QHRTK+EELRQHLL WG TTPDKK+QKEPPFLWMGYELHPDKWT
Sbjct  903   YQYVDDLYVGSDLEIEQHRTKVEELRQHLLTWGFTTPDKKYQKEPPFLWMGYELHPDKWT  1082

Query  241   VQPIVLPEKDSWTVNDIQKLVGKLNWASQIYPGIKVRQLCKLLRGTKALTEVIPLTeeae  300
             VQPIVLPEKDSWTVNDIQKLVGKLNWASQIYPGIKVRQLCKLLRG KALTEV+ LT+EAE
Sbjct  1083  VQPIVLPEKDSWTVNDIQKLVGKLNWASQIYPGIKVRQLCKLLRGAKALTEVVTLTKEAE  1262

Query  301   lelaeNREILKEPVHGVYYDPSKDLIAEIQKQGQGQWTYQI  341
             LELAENREI+KEPVHGVYYDP+KDLIAEIQKQGQGQWTYQI
Sbjct  1263  LELAENREIIKEPVHGVYYDPTKDLIAEIQKQGQGQWTYQI  1385
```

>Escherichia coli strain 8374wG10 NODE_34_length_1373_cov_2.57865, whole genome shotgun sequence
Sequence ID: NZ_WSJQ01000034.1 Length: 1373
Range 1: 3 to 1013

Score:648 bits(1671), Expect:0.0,
Method:Compositional matrix adjust.,
Identities:323/337(96%), Positives:331/337(98%), Gaps:0/337(0%)

```
Query  1     PISPIETVPVKLKPGMDGPKVKQWPLTEEKIKALVEICTEMEKEGKISKIGPENPYNTPV  60
             PISPIETVPVKLKPGMDGPKVKQWPLTEEKIKALVEICTEMEKEGKISKIGPENPYNTPV
Sbjct  1013  PISPIETVPVKLKPGMDGPKVKQWPLTEEKIKALVEICTEMEKEGKISKIGPENPYNTPV  834

Query  61    FAIKKKDSTKWRKLVDFRELNKRTQDFWEVQLGIPHPAGLKKKKSVTVLDVGDAYFSVPL  120
             FAIKKKDSTKWRKLVDFRELNKRTQDFWEVQLGIPHPAGLKKKKSVTVLDVGDAYFSVPL
Sbjct  833   FAIKKKDSTKWRKLVDFRELNKRTQDFWEVQLGIPHPAGLKKKKSVTVLDVGDAYFSVPL  654

Query  121   DEDFRKYTAFTIPSINNETPGIRYQYNVLPQGWKGSPAIFQSSMTKILEPFRKQNPDIVI  180
             D++FRKYTAFTIPS NNETPGIRYQYNVLPQGWKGSPAIFQ SMTKILEPFRKQNPDIVI
Sbjct  653   DKEFRKYTAFTIPSTNNETPGIRYQYNVLPQGWKGSPAIFQCSMTKILEPFRKQNPDIVI  474

Query  181   YQYMDDLYVGSDLEIGQHRTKIEELRQHLLRWGLTTPDKKHQKEPPFLWMGYELHPDKWT  240
             YQY+DDLYVGSDLEIGQHR K+EELRQHLL WGLTTPDKKHQKEPPFLWMGYELHPDKWT
Sbjct  473   YQYVDDLYVGSDLEIGQHRAKVEELRQHLLTWGLTTPDKKHQKEPPFLWMGYELHPDKWT  294

Query  241   VQPIVLPEKDSWTVNDIQKLVGKLNWASQIYPGIKVRQLCKLLRGTKALTEVIPLTeeae  300
             VQPI LPEKDSWTVNDIQKLVGKLNWASQIYPGIKV+QLCKLLRG KALT+VIPLTEEAE
Sbjct  293   VQPIQLPEKDSWTVNDIQKLVGKLNWASQIYPGIKVKQLCKLLRGAKALTDVIPLTEEAE  114

Query  301   lelaeNREILKEPVHGVYYDPSKDLIAEIQKQGQGQW  337
             LELAENREI+KEPVHGVYYDP+KDLIAEIQKQGQGQW
Sbjct  113   LELAENREIIKEPVHGVYYDPTKDLIAEIQKQGQGQW  3
```

>Methylocella silvestris strain TVC NODE_58_length_1453_cov_3.56561, whole genome shotgun sequence
Sequence ID: NZ_PDZR01000058.1 Length: 1453
Range 1: 379 to 1413

```
Score:647 bits(1670), Expect:0.0,
Method:Compositional matrix adjust.,
Identities:322/345(93%), Positives:336/345(97%), Gaps:0/345(0%)

Query  1     PISPIETVPVKLKPGMDGPKVKQWPLTEEKIKALVEICTEMEKEGKISKIGPENPYNTPV  60
             PISPIETVPVKLKPGMDGPKVKQWPLTEEKIKALVEICTEMEKEGKI+KIGPENPYNTP+
Sbjct  379   PISPIETVPVKLKPGMDGPKVKQWPLTEEKIKALVEICTEMEKEGKITKIGPENPYNTPI  558

Query  61    FAIKKKDSTKWRKLVDFRELNKRTQDFWEVQLGIPHPAGLKKKKSVTVLDVGDAYFSVPL  120
             FAIKKK+S KWRKLVDFRELNKRTQ+FWEVQLGIPHPAGLKKKKSVTVLDVGDAYFSVPL
Sbjct  559   FAIKKKNSNKWRKLVDFRELNKRTQEFWEVQLGIPHPAGLKKKKSVTVLDVGDAYFSVPL  738

Query  121   DEDFRKYTAFTIPSINNETPGIRYQYNVLPQGWKGSPAIFQSSMTKILEPFRKQNPDIVI  180
             D DFRKYTAFTIPS NNETPGIRYQYNVLPQGWKGSPAIFQSSMTKILEPFRKQNPD+VI
Sbjct  739   DPDFRKYTAFTIPSRNNETPGIRYQYNVLPQGWKGSPAIFQSSMTKILEPFRKQNPDMVI  918

Query  181   YQYMDDLYVGSDLEIGQHRTKIEELRQHLLRWGLTTPDKKHQKEPPFLWMGYELHPDKWT  240
             YQYMDDLYVGSDLE+ QHRTK+EELR HLL+ G +TPD+KHQKEPPFLWMGYELHPDKWT
Sbjct  919   YQYMDDLYVGSDLELEQHRTKVEELRAHLLK*GFSTPDQKHQKEPPFLWMGYELHPDKWT  1098

Query  241   VQPIVLPEKDSWTVNDIQKLVGKLNWASQIYPGIKVRQLCKLLRGTKALTEVIPLTeeae  300
             VQPIVLPEKDSWTVNDIQKLVGKLNWASQIY GIKV+QLCKLLRGTKALTEV+PL+EEAE
Sbjct  1099  VQPIVLPEKDSWTVNDIQKLVGKLNWASQIYAGIKVKQLCKLLRGTKALTEVVPLSEEAE  1278

Query  301   lelaeNREILKEPVHGVYYDPSKDLIAEIQKQGQGQWTYQIYQEP  345
             LELAENREILKEPVHGVYYDPSKDL+AEIQKQ GQGQWTYQIYQEP
Sbjct  1279  LELAENREILKEPVHGVYYDPSKDLVAEIQKQEQGQWTYQIYQEP  1413



>Escherichia coli strain 8374wH11 NODE_36_length_1397_cov_3.54567, whole genome shotgun sequence
Sequence ID: NZ_WSJI01000036.1 Length: 1397
Range 1: 385 to 1395

Score:644 bits(1662), Expect:0.0,
Method:Compositional matrix adjust.,
Identities:320/337(95%), Positives:329/337(97%), Gaps:0/337(0%)

Query  1     PISPIETVPVKLKPGMDGPKVKQWPLTEEKIKALVEICTEMEKEGKISKIGPENPYNTPV  60
             PISPI+TVPVKLKPGMDGPKVKQWPLTEEKIKALVEICTEMEKEGKISKIGPENPYNTPV
Sbjct  385   PISPIDTVPVKLKPGMDGPKVKQWPLTEEKIKALVEICTEMEKEGKISKIGPENPYNTPV  564

Query  61    FAIKKKDSTKWRKLVDFRELNKRTQDFWEVQLGIPHPAGLKKKKSVTVLDVGDAYFSVPL  120
             FAIKKKDSTKWRKLVDFRELNKRTQDFWEVQLGIPHPAGLKKKKSVTVLDVGDAYFSVPL
Sbjct  565   FAIKKKDSTKWRKLVDFRELNKRTQDFWEVQLGIPHPAGLKKKKSVTVLDVGDAYFSVPL  744

Query  121   DEDFRKYTAFTIPSINNETPGIRYQYNVLPQGWKGSPAIFQSSMTKILEPFRKQNPDIVI  180
             D++FRKYTAFTIPS NNETPGIRYQYNVLPQGWKGSPAIFQ SMTKILEPFRKQNPDIVI
Sbjct  745   DKEFRKYTAFTIPSTNNETPGIRYQYNVLPQGWKGSPAIFQCSMTKILEPFRKQNPDIVI  924

Query  181   YQYMDDLYVGSDLEIGQHRTKIEELRQHLLRWGLTTPDKKHQKEPPFLWMGYELHPDKWT  240
             YQY+DDLYVGSDLEI QHRTK+EELRQHLL WG TTPDKKHQKEPPFLWMGYELHPDKWT
Sbjct  925   YQYVDDLYVGSDLEIEQHRTKVEELRQHLLTWGFTTPDKKHQKEPPFLWMGYELHPDKWT  1104

Query  241   VQPIVLPEKDSWTVNDIQKLVGKLNWASQIYPGIKVRQLCKLLRGTKALTEVIPLTeeae  300
             VQPIVLPEKDSWTVNDIQKLVGKLNWASQIYPGIKVRQLCKLLRG KALTEV+ LT+EAE
Sbjct  1105  VQPIVLPEKDSWTVNDIQKLVGKLNWASQIYPGIKVRQLCKLLRGAKALTEVVTLTKEAE  1284

Query  301   lelaeNREILKEPVHGVYYDPSKDLIAEIQKQGQGQW  337
             LELAENREI+KEPVHGVYYDP+KDLIAEIQKQGQ QW
Sbjct  1285  LELAENREIIKEPVHGVYYDPTKDLIAEIQKQGQDQW  1395



>Staphylococcus epidermidis strain PR246B0 NODE_70_length_1113_cov_1.18763_ID_139, whole genome shotgun sequence
Sequence ID: NZ_PCFD01000070.1 Length: 1113
Range 1: 3 to 1103

Score:644 bits(1662), Expect:0.0,
Method:Compositional matrix adjust.,
Identities:329/367(90%), Positives:349/367(95%), Gaps:0/367(0%)

Query  72    RKLVDFRELNKRTQDFWEVQLGIPHPAGLKKKKSVTVLDVGDAYFSVPLDEDFRKYTAFT  131
             +K+VDFRELNKRTQDFWEVQLGIPHPAGLKKKKSVTVLDVGDA+FSVPLDE FRKYTAFT
Sbjct  1103  KKVVDFRELNKRTQDFWEVQLGIPHPAGLKKKKSVTVLDVGDAFFSVPLDESFRKYTAFT  924

Query  132   IPSINNETPGIRYQYNVLPQGWKGSPAIFQSSMTKILEPFRKQNPDIVIYQYMDDLYVGS  191
             IPSINN TPGIRYQYNVLPQGWKGSPAIFQSSMTKILEPFR +NP++VIYQY+DDLYV S
Sbjct  923   IPSINNATPGIRYQYNVLPQGWKGSPAIFQSSMTKILEPFRTKNPEMVIYQYVDDLYVES  744

Query  192   DLEIGQHRTKIEELRQHLLRWGLTTPDKKHQKEPPFLWMGYELHPDKWTVQPIVLPEKDS  251
             DLEI QHR K+EELR+HLL+WG TTPD+KHQKEPPFLWMGYELHPDKWTVQPI LPEKD
Sbjct  743   DLEIMQHRAKVEELREHLLKWGFTTPDEKHQKEPPFLWMGYELHPDKWTVQPIQLPEKDI  564

Query  252   WTVNDIQKLVGKLNWASQIYPGIKVRQLCKLLRGTKALTEVIPLTeeaelelaeNREILK  311
             WTVNDIQKLVGKLNWASQIYP IKV+QLCKLLRGTKALT+++ LTEEAELELAENREILK
Sbjct  563   WTVNDIQKLVGKLNWASQIYPEIKVKQLCKLLRGTKALTDIVTLTEEAELELAENREILK  384

Query  312   EPVHGVYYDPSKDLIAEIQKQGQGQWTYQIYQEPFKNLKTGKYARMRGAHTNDVKQLTEA  371
             EPVHGVYYDPSKDLIAEIQKQG  QWTYQIYQEPFKNLKTGKYA+MR AHTNDV+QLTEA
Sbjct  383   EPVHGVYYDPSKDLIAEIQKQGNDQWTYQIYQEPFKNLKTGKYAKMRTAHTNDVRQLTEA  204

Query  372   VQKITTESIVIWGKTPKFKLPIQKetwetwwteYWQATWIPEWEFVNTPPLVKLWYQLEK  431
             VQKI  ESIVIWGK PKF+LPIQKETWETWWT+YWQATWIP+WEFVNTPPLVKLWYQLEK
Sbjct  203   VQKIARESIVIWGKIPKFRLPIQKETWETWWTDYWQATWIPDWEFVNTPPLVKLWYQLEK  24

Query  432   EPIVGAE  438
             EPI GAE
Sbjct  23    EPIAGAE  3



>Helicobacter acinonychis isolate 212_10, whole genome shotgun sequence
Sequence ID: NZ_FZLW01000041.1 Length: 1526
Range 1: 103 to 1104

Score:642 bits(1656), Expect:0.0,
Method:Compositional matrix adjust.,
Identities:318/334(95%), Positives:328/334(98%), Gaps:0/334(0%)

Query  1     PISPIETVPVKLKPGMDGPKVKQWPLTEEKIKALVEICTEMEKEGKISKIGPENPYNTPV  60
             PISPIETVPVKLKPGMDGPKVKQWPLTEEKIKAL+EICTEMEKEGKISKIGPENPYNTPV
Sbjct  1104  PISPIETVPVKLKPGMDGPKVKQWPLTEEKIKALIEICTEMEKEGKISKIGPENPYNTPV  925

Query  61    FAIKKKDSTKWRKLVDFRELNKRTQDFWEVQLGIPHPAGLKKKKSVTVLDVGDAYFSVPL  120
             FAIKKKDSTKWRKLVDFRELNK+TQDFWEVQLGIPHP+GLKKKKSVTVLDVGDAYFSVPL
Sbjct  924   FAIKKKDSTKWRKLVDFRELNKKTQDFWEVQLGIPHPSGLKKKKSVTVLDVGDAYFSVPL  745

Query  121   DEDFRKYTAFTIPSINNETPGIRYQYNVLPQGWKGSPAIFQSSMTKILEPFRKQNPDIVI  180
             D+DFRKYTAFTIPS+NNETPGIRYQYNVLPQGWKGSPAIFQ SMTKILEPFRKQNP+IVI
```

```
Sbjct  744   DKDFRKYTAFTIPSMNNETPGIRYQYNVLPQGWKGSPAIFQHSMTKILEPFRKQNPNIVI  565

Query  181   YQYMDDLYVGSDLEIGQHRTKIEELRQHLLRWGLTTPDKKHQKEPPFLWMGYELHPDKWT  240
             YQYMDDLYVGSDLEI QHRTKIEELRQHLL WG +TPDKKHQKEPPFLWMGYELHPDKWT
Sbjct  564   YQYMDDLYVGSDLEIEQHRTKIEELRQHLLGWGFSTPDKKHQKEPPFLWMGYELHPDKWT  385

Query  241   VQPIVLPEKDSWTVNDIQKLVGKLNWASQIYPGIKVRQLCKLLRGTKALTEVIPLTeeae  300
             VQPI+LPEKDSWTVNDIQKLVGKLNWASQIYPGIKVRQLCKLLRGTKALTEV+PLT EAE
Sbjct  384   VQPIMLPEKDSWTVNDIQKLVGKLNWASQIYPGIKVRQLCKLLRGTKALTEVVPLTAEAE  205

Query  301   lelaeNREILKEPVHGVYYDPSKDLIAEIQKQGQ  334
             LELAENREILKEPVHGVYYDPSKDLIAE+QKQ Q
Sbjct  204   LELAENREILKEPVHGVYYDPSKDLIAELQKQEQ  103



>Klebsiella aerogenes strain AUH-KAM5 NODE_25_length_1427_cov_1.66077, whole genome shotgun sequence
Sequence ID: NZ_PYHJ01000025.1 Length: 1427
Range 1: 381 to 1397

Score:635 bits(1637), Expect:0.0,
Method:Compositional matrix adjust.,
Identities:317/339(94%), Positives:329/339(97%), Gaps:0/339(0%)

Query  1     PISPIETVPVKLKPGMDGPKVKQWPLTEEKIKALVEICTEMEKEGKISKIGPENPYNTPV  60
             PISPIETVPVKLKPGMDGPKVKQWPLTEEKIKALVEICTEMEKEGKISKIGPENPYNTPV
Sbjct  381   PISPIETVPVKLKPGMDGPKVKQWPLTEEKIKALVEICTEMEKEGKISKIGPENPYNTPV  560

Query  61    FAIKKKDSTKWRKLVDFRELNKRTQDFWEVQLGIPHPAGLKKKKSVTVLDVGDAYFSVPL  120
             FAIKKKDSTKWRKLVDFRELNKRTQDFWEVQLGIPHPAGLKKKKSVTVLDVGDAYFSVPL
Sbjct  561   FAIKKKDSTKWRKLVDFRELNKRTQDFWEVQLGIPHPAGLKKKKSVTVLDVGDAYFSVPL  740

Query  121   DEDFRKYTAFTIPSINNETPGIRYQYNVLPQGWKGSPAIFQSSMTKILEPFRKQNPDIVI  180
             D++FRKYTAFTIPS NNETPGIRYQYNVLPQGWKGSPAIFQSSMTKILEPFRKQNPDIVI
Sbjct  741   DKEFRKYTAFTIPSRNNETPGIRYQYNVLPQGWKGSPAIFQSSMTKILEPFRKQNPDIVI  920

Query  181   YQYMDDLYVGSDLEIGQHRTKIEELRQHLLRWGLTTPDKKHQKEPPFLWMGYELHPDKWT  240
             YQY+DDLYVGSDLE+ QHRTK+EELRQHLL WG +TPD+KHQKEPPFLWMGYELHPDKWT
Sbjct  921   YQYIDDLYVGSDLELEQHRTKVEELRQHLLTWGFSTPDQKHQKEPPFLWMGYELHPDKWT  1100

Query  241   VQPIVLPEKDSWTVNDIQKLVGKLNWASQIYPGIKVRQLCKLLRGTKALTEVIPLTeeae  300
             VQPIVLPEKDSWTVNDIQKLVGKLNWASQIY GIKV+QLCKLL+GTKALTEV+PL+EEAE
Sbjct  1101  VQPIVLPEKDSWTVNDIQKLVGKLNWASQIYAGIKVKQLCKLLKGTKALTEVVPLSEEAE  1280

Query  301   lelaeNREILKEPVHGVYYDPSKDLIAEIQKQGQGQWTY  339
             LELAENREILKEPVHGVYYDPSKDL+AEIQKQ  G   Y
Sbjct  1281  LELAENREILKEPVHGVYYDPSKDLVAEIQKQEHGPVPY  1397



>Enterobacter hormaechei strain AUH-ENM6 NODE_27_length_1394_cov_0.980268, whole genome shotgun sequence
Sequence ID: NZ_PYHB01000027.1 Length: 1394
Range 1: 394 to 1392

Score:629 bits(1623), Expect:0.0,
Method:Compositional matrix adjust.,
Identities:314/333(94%), Positives:325/333(97%), Gaps:0/333(0%)

Query  1     PISPIETVPVKLKPGMDGPKVKQWPLTEEKIKALVEICTEMEKEGKISKIGPENPYNTPV  60
             PISPIETVPVKLKPGMDGPKVKQWPLTEEKIKALVEICTEMEKEGKISKIGPENPYNTP+
Sbjct  394   PISPIETVPVKLKPGMDGPKVKQWPLTEEKIKALVEICTEMEKEGKISKIGPENPYNTPI  573

Query  61    FAIKKKDSTKWRKLVDFRELNKRTQDFWEVQLGIPHPAGLKKKKSVTVLDVGDAYFSVPL  120
             FAIKKK+S KWRKLVDFRELNKRTQ+FWEVQLGIPHPAGLKKKKSVTVLDVGDAYFSVPL
Sbjct  574   FAIKKKNSDKWRKLVDFRELNKRTQEFWEVQLGIPHPAGLKKKKSVTVLDVGDAYFSVPL  753

Query  121   DEDFRKYTAFTIPSINNETPGIRYQYNVLPQGWKGSPAIFQSSMTKILEPFRKQNPDIVI  180
             D DFRKYTAFTIPS NNETPGIRYQYNVLPQGWKGSPAIFQSSMTKILEPFRKQNPDIVI
Sbjct  754   DPDFRKYTAFTIPSRNNETPGIRYQYNVLPQGWKGSPAIFQSSMTKILEPFRKQNPDIVI  933

Query  181   YQYMDDLYVGSDLEIGQHRTKIEELRQHLLRWGLTTPDKKHQKEPPFLWMGYELHPDKWT  240
             YQYMDDLYVGSDLE+ QHRTK+EELR HLL+WG +TPD+KHQKEPPFLWMGYELHPDKWT
Sbjct  934   YQYMDDLYVGSDLELEQHRTKVEELRAHLLKWGFSTPDQKHQKEPPFLWMGYELHPDKWT  1113

Query  241   VQPIVLPEKDSWTVNDIQKLVGKLNWASQIYPGIKVRQLCKLLRGTKALTEVIPLTeeae  300
             VQPIVLPEKDSWTVNDIQKLVGKLNWASQIY  IKV+QLCKLLRGTKALTEV+PL+EEAE
Sbjct  1114  VQPIVLPEKDSWTVNDIQKLVGKLNWASQIYAEIKVKQLCKLLRGTKALTEVVPLSEEAE  1293

Query  301   lelaeNREILKEPVHGVYYDPSKDLIAEIQKQG  333
             LELAENREILKEPVHGVYYDPSKDLIAEIQKQG
Sbjct  1294  LELAENREILKEPVHGVYYDPSKDLIAEIQKQG  1392



>Acinetobacter baumannii strain ACM-35 NODE_65_length_1409_cov_2.30031, whole genome shotgun sequence
Sequence ID: NZ_QEHN01000065.1 Length: 1409
Range 1: 405 to 1409

Score:626 bits(1615), Expect:0.0,
Method:Compositional matrix adjust.,
Identities:314/335(94%), Positives:325/335(97%), Gaps:0/335(0%)

Query  1     PISPIETVPVKLKPGMDGPKVKQWPLTEEKIKALVEICTEMEKEGKISKIGPENPYNTPV  60
             PISPIETVPVKLKPGMDGPKVKQWPLTEEKIKALVEICTEMEKEGKISKIGPENPYNTP+
Sbjct  405   PISPIETVPVKLKPGMDGPKVKQWPLTEEKIKALVEICTEMEKEGKISKIGPENPYNTPI  584

Query  61    FAIKKKDSTKWRKLVDFRELNKRTQDFWEVQLGIPHPAGLKKKKSVTVLDVGDAYFSVPL  120
             FAIKKK+S KWRKLVDFRELNKRTQ+FWEVQLGIPHPAGLKKKKSVTVLDVGDAYFSVPL
Sbjct  585   FAIKKKNSDKWRKLVDFRELNKRTQEFWEVQLGIPHPAGLKKKKSVTVLDVGDAYFSVPL  764

Query  121   DEDFRKYTAFTIPSINNETPGIRYQYNVLPQGWKGSPAIFQSSMTKILEPFRKQNPDIVI  180
             D DFRKYTAFTIPS NNETPGIRYQYNVLPQGWKGSPAIFQSSMTKILEPFRKQNPD+VI
Sbjct  765   DPDFRKYTAFTIPSRNNETPGIRYQYNVLPQGWKGSPAIFQSSMTKILEPFRKQNPDMVI  944

Query  181   YQYMDDLYVGSDLEIGQHRTKIEELRQHLLRWGLTTPDKKHQKEPPFLWMGYELHPDKWT  240
             YQYMDDLYVGSDLEI QHRTK+EELR HLL+WG +TPD+KHQKEPPFLWMGYELHPDKWT
Sbjct  945   YQYMDDLYVGSDLEIEQHRTKVEELRAHLLKWGFSTPDQKHQKEPPFLWMGYELHPDKWT  1124

Query  241   VQPIVLPEKDSWTVNDIQKLVGKLNWASQIYPGIKVRQLCKLLRGTKALTEVIPLTeeae  300
             VQPIVLPEKDSWTVNDIQKLVGKLNWASQIY GIKV+QLCKLLRG KALTEV+ LTEEAE
Sbjct  1125  VQPIVLPEKDSWTVNDIQKLVGKLNWASQIYAGIKVKQLCKLLRGAKALTEVVTLTEEAE  1304

Query  301   lelaeNREILKEPVHGVYYDPSKDLIAEIQKQGQG  335
             LELAENREILKEPVHGVYYDPSKDL+AEIQKQG G
Sbjct  1305  LELAENREILKEPVHGVYYDPSKDLVAEIQKQGLG  1409



>Staphylococcus aureus strain PN235B0 NODE_123_length_1117_cov_0.929293_ID_245, whole genome shotgun sequence
Sequence ID: NZ_PDUY01000159.1 Length: 1083
```

```
Range 1: 30 to 1082

Score:614 bits(1584), Expect:0.0,
Method:Compositional matrix adjust.,
Identities:311/351(89%), Positives:331/351(94%), Gaps:0/351(0%)

Query  91    QLGIPHPAGLKKKKSVTVLDVGDAYFSVPLDEDFRKYTAFTIPSINNETPGIRYQYNVLP  150
             Q+GIPHPAG+KK KSVTVLDVGDAYFSVPL EDFRKYTAFTIPS+NNE PGIRYQYNVLP
Sbjct  30    QIGIPHPAGIKKNKSVTVLDVGDAYFSVPLHEDFRKYTAFTIPSVNNEAPGIRYQYNVLP  209

Query  151   QGWKGSPAIFQSSMTKILEPFRKQNPDIVIYQYMDDLYVGSDLEIGQHRTKIEELRQHLL  210
             QGWKGSPAIFQSSMTKILEPFR +NP++VIYQY+DDLY+ SDLEI QHR K+EELR+HLL
Sbjct  210   QGWKGSPAIFQSSMTKILEPFRTKNPEVVIYQYVDDLYIESDLEIMQHRAKVEELREHLL  389

Query  211   RWGLTTPDKKHQKEPPFLWMGYELHPDKWTVQPIVLPEKDSWTVNDIQKLVGKLNWASQI  270
             +WG TTPD+KHQKEPPFLWMGYELHPDKWTVQPI LPEKD WTVNDIQKLVGKLNWASQI
Sbjct  390   KWGFTTPDEKHQKEPPFLWMGYELHPDKWTVQPIQLPEKDIWTVNDIQKLVGKLNWASQI  569

Query  271   YPGIKVRQLCKLLRGTKALTEVIPLTeeaelelaeNREILKEPVHGVYYDPSKDLIAEIQ  330
             YP IKV+QLCKLLRGTKALT+++PLTEEAELELAENREILKEPVHG YYDPSKDLIAEIQ
Sbjct  570   YPEIKVKQLCKLLRGTKALTDIVPLTEEAELELAENREILKEPVHGAYYDPSKDLIAEIQ  749

Query  331   KQGQGQWTYQIYQEPFKNLKTGKYARMRGAHTNDVKQLTEAVQKITTESIVIWGKTPKFK  390
             KQG  QWTYQIYQEPFKNLKTGKYA+MR AHTNDVKQLTEAVQKI  E IVIWGK PKF+
Sbjct  750   KQGNDQWTYQIYQEPFKNLKTGKYAKMRTAHTNDVKQLTEAVQKIARECIVIWGKIPKFR  929

Query  391   LPIQKetwetwwteYWQATWIPEWEFVNTPPLVKLWYQLEKEPIVGAETFY  441
             LPIQKETWETWWT+YWQATWIP+WEFVNTPPLVKLWYQLEKEPI GAETFY
Sbjct  930   LPIQKETWETWWTDYWQATWIPDWEFVNTPPLVKLWYQLEKEPIAGAETFY  1082



>Klebsiella pneumoniae strain P7701 NODE_50_length_1305_cov_481.863, whole genome shotgun sequence
Sequence ID: NZ_JACAWM010000050.1 Length: 1305
Range 1: 21 to 908

Score:601 bits(1550), Expect:0.0,
Method:Compositional matrix adjust.,
Identities:290/296(98%), Positives:293/296(98%), Gaps:0/296(0%)

Query  1    PISPIETVPVKLKPGMDGPKVKQWPLTEEKIKALVEICTEMEKEGKISKIGPENPYNTPV  60
            PISPIETVPVKLKPGMDGPKVKQWPLTEEKIKALVEICTEMEKEGKISKIGPENPYNTPV
Sbjct  908  PISPIETVPVKLKPGMDGPKVKQWPLTEEKIKALVEICTEMEKEGKISKIGPENPYNTPV  729

Query  61   FAIKKKDSTKWRKLVDFRELNKRTQDFWEVQLGIPHPAGLKKKKSVTVLDVGDAYFSVPL  120
            FAIKKKDSTKWRKLVDFRELNKRTQDFWEVQLGIPHPAGLK+KKSVTVLDVGDAYFSVPL
Sbjct  728  FAIKKKDSTKWRKLVDFRELNKRTQDFWEVQLGIPHPAGLKQKKSVTVLDVGDAYFSVPL  549

Query  121  DEDFRKYTAFTIPSINNETPGIRYQYNVLPQGWKGSPAIFQSSMTKILEPFRKQNPDIVI  180
            D+DFRKYTAFTIPSINNETPGIRYQYNVLPQGWKGSPAIFQ SMTKILEPFRKQNPDIVI
Sbjct  548  DKDFRKYTAFTIPSINNETPGIRYQYNVLPQGWKGSPAIFQCSMTKILEPFRKQNPDIVI  369

Query  181  YQYMDDLYVGSDLEIGQHRTKIEELRQHLLRWGLTTPDKKHQKEPPFLWMGYELHPDKWT  240
            YQYMDDLYVGSDLEIGQHRTKIEELRQHLLRWG TTPDKKHQKEPPFLWMGYELHPDKWT
Sbjct  368  YQYMDDLYVGSDLEIGQHRTKIEELRQHLLRWGFTTPDKKHQKEPPFLWMGYELHPDKWT  189

Query  241  VQPIVLPEKDSWTVNDIQKLVGKLNWASQIYPGIKVRQLCKLLRGTKALTEVIPLT  296
            VQPIVLPEKDSWTVNDIQKLVGKLNWASQIY GIKVRQLCKLLRGTKALTEV+PLT
Sbjct  188  VQPIVLPEKDSWTVNDIQKLVGKLNWASQIYAGIKVRQLCKLLRGTKALTEVVPLT  21



>Salirhabdus euzebyi strain Marseille-Q1438, whole genome shotgun sequence
Sequence ID: NZ_CADDWK010000035.1 Length: 1279
Range 1: 9 to 896

Score:600 bits(1547), Expect:0.0,
Method:Compositional matrix adjust.,
Identities:288/296(97%), Positives:295/296(99%), Gaps:0/296(0%)

Query  1    PISPIETVPVKLKPGMDGPKVKQWPLTEEKIKALVEICTEMEKEGKISKIGPENPYNTPV  60
            PISPIETVPVKLKPGMDGPKVKQWPLTEEKIKALVEICTEMEKEGKIS++GPENPYNTP+
Sbjct  896  PISPIETVPVKLKPGMDGPKVKQWPLTEEKIKALVEICTEMEKEGKISRVGPENPYNTPI  717

Query  61   FAIKKKDSTKWRKLVDFRELNKRTQDFWEVQLGIPHPAGLKKKKSVTVLDVGDAYFSVPL  120
            FAIKKKDSTKWRKLVDFRELNK+TQDFWEVQLGIPHPAGLKKKKSVTVLDVGDAYFSVPL
Sbjct  716  FAIKKKDSTKWRKLVDFRELNKKTQDFWEVQLGIPHPAGLKKKKSVTVLDVGDAYFSVPL  537

Query  121  DEDFRKYTAFTIPSINNETPGIRYQYNVLPQGWKGSPAIFQSSMTKILEPFRKQNPDIVI  180
            D+DFRKYTAFTIPS+NNETPGIRYQYNVLPQGWKGSPAIFQSSMTKILEPFRKQNPDIVI
Sbjct  536  DKDFRKYTAFTIPSVNNETPGIRYQYNVLPQGWKGSPAIFQSSMTKILEPFRKQNPDIVI  357

Query  181  YQYMDDLYVGSDLEIGQHRTKIEELRQHLLRWGLTTPDKKHQKEPPFLWMGYELHPDKWT  240
            YQYMDDLYVGSDLEIGQHRTKIEELRQHLLRWG TTPDKKHQKEPPFLWMGYELHPDKWT
Sbjct  356  YQYMDDLYVGSDLEIGQHRTKIEELRQHLLRWGFTTPDKKHQKEPPFLWMGYELHPDKWT  177

Query  241  VQPIVLPEKDSWTVNDIQKLVGKLNWASQIYPGIKVRQLCKLLRGTKALTEVIPLT  296
            VQPIVLPEKDSWTVNDIQKLVGKLNWASQIYPGIKV+QLCKLLRGTKALTEVIPLT
Sbjct  176  VQPIVLPEKDSWTVNDIQKLVGKLNWASQIYPGIKVKQLCKLLRGTKALTEVIPLT  9



>Methylobacterium radiotolerans strain MAMP 4754 MP-M-radiotolerans_contig_704, whole genome shotgun sequence
Sequence ID: NZ_NKQS01000691.1 Length: 1636
Range 1: 618 to 1634

Score:588 bits(1517), Expect:0.0,
Method:Compositional matrix adjust.,
Identities:305/339(90%), Positives:321/339(94%), Gaps:0/339(0%)

Query  222   QKEPPFLWMGYELHPDKWTVQPIVLPEKDSWTVNDIQKLVGKLNWASQIYPGIKVRQLCK  281
             QKEPPFLWMGYELHPDKWTVQPIVLPEK+SWTVNDIQKLVGKLNWASQIYPGIKVRQ+CK
Sbjct  1634  QKEPPFLWMGYELHPDKWTVQPIVLPEKESWTVNDIQKLVGKLNWASQIYPGIKVRQMCK  1455

Query  282   LLRGTKALTEVIPLTeeaelelaeNREILKEPVHGVYYDPSKDLIAEIQKQGQGQWTYQI  341
             LLRGTKALT+++PLT EAELELAENREILKEPVHG YYDPSKDLIAEIQKQG  QWTYQI
Sbjct  1454  LLRGTKALTDIVPLTTEAELELAENREILKEPVHGAYYDPSKDLIAEIQKQGHDQWTYQI  1275

Query  342   YQEPFKNLKTGKYARMRGAHTNDVKQLTEAVQKITTESIVIWGKTPKFKLPIQKetwetw  401
             YQEPFKNLKTGKYA M+ A+ NDVKQLTEAVQKI  ESIVIWGKTPKF+LPIQKETWETW
Sbjct  1274  YQEPFKNLKTGKYA*MKSANINDVKQLTEAVQKIALESIVIWGKTPKFRLPIQKETWETW  1095

Query  402   wteYWQATWIPEWEFVNTPPLVKLWYQLEKEPIVGAETFYVDGAANRETKLGKAGYVTNR  461
             WT+YWQATWIPEWEFVNTPPLVKLWYQLEKEPI GAETFYVDGAANRETK GKAGYVT+R
Sbjct  1094  WTDYWQATWIPEWEFVNTPPLVKLWYQLEKEPIAGAETFYVDGAANRETKRGKAGYVTDR  915

Query  462   GRQKVVTLTDTTNQKTELQAIYLALQDSGLEVNIVTDSQYALGIIQAQPDQSESELVNQI  521
             GRQK+V L +TTNQK ELQAI LALQDSG EVNIVTDSQYALGIIQAQPD+SESELVNQI
Sbjct  914   GRQKIVPLNETTNQKAELQAIQLALQDSGSEVNIVTDSQYALGIIQAQPDKSESELVNQI  735

Query  522   IEQLIKKEKVYLAWVPAHKGIGGNEQVDKLVSAGIRKVL  560
```

```
           IEQLI KE++YL+WVPAHKGIGGNEQVDKLVS GIRKVL
Sbjct  734   IEQLINKERIYLSWVPAHKGIGGNEQVDKLVSTGIRKVL  618
```

>Mycobacterium tuberculosis variant bovis strain CSURQ1132, whole genome shotgun sequence
Sequence ID: NZ_CAJEUK010000155.1 Length: 1524
Range 1: 123 to 1010

Score:578 bits(1490), Expect:0.0,
Method:Compositional matrix adjust.,
Identities:273/296(92%), Positives:289/296(97%), Gaps:0/296(0%)

```
Query  1     PISPIETVPVKLKPGMDGPKVKQWPLTEEKIKALVEICTEMEKEGKISKIGPENPYNTPV  60
             PISPIETVPVKLKPGMDGPKVKQWPLTEEKIKAL EICTEMEKEGKISKIGPENPYNTPV
Sbjct  1010  PISPIETVPVKLKPGMDGPKVKQWPLTEEKIKALTEICTEMEKEGKISKIGPENPYNTPV  831

Query  61    FAIKKKDSTKWRKLVDFRELNKRTQDFWEVQLGIPHPAGLKKKKSVTVLDVGDAYFSVPL  120
             FAI+KKDSTKWRKLVDFRELNKRTQDFWE+QLGIPHPAGLKKKKSVTVLDVGDAYFSVPL
Sbjct  830   FAIRKKDSTKWRKLVDFRELNKRTQDFWEIQLGIPHPAGLKKKKSVTVLDVGDAYFSVPL  651

Query  121   DEDFRKYTAFTIPSINNETPGIRYQYNVLPQGWKGSPAIFQSSMTKILEPFRKQNPDIVI  180
             D+DFRKYTAFTIPSINNETPGIRYQYNVLPQGWKGSPAIFQ+SMTKILEPFR +NP++VI
Sbjct  650   DKDFRKYTAFTIPSINNETPGIRYQYNVLPQGWKGSPAIFQASMTKILEPFRTKNPELVI  471

Query  181   YQYMDDLYVGSDLEIGQHRTKIEELRQHLLRWGLTTPDKKHQKEPPFLWMGYELHPDKWT  240
             YQYMDDLY+GSDLEIGQHRTKIEELR HLLRWG TTPDKKHQKEPPFLWMGYELHPDKWT
Sbjct  470   YQYMDDLYIGSDLEIGQHRTKIEELRAHLLRWGFTTPDKKHQKEPPFLWMGYELHPDKWT  291

Query  241   VQPIVLPEKDSWTVNDIQKLVGKLNWASQIYPGIKVRQLCKLLRGTKALTEVIPLT  296
             VQP+ LP+K++WTVNDIQKLVGKLNWASQIY GIKV+QLCKLLRG KALT+++PLT
Sbjct  290   VQPVQLPDKENWTVNDIQKLVGKLNWASQIYAGIKVKQLCKLLRGAKALTDIVPLT  123
```

Range 2: 123 to 1010

Score:578 bits(1490), Expect:0.0,
Method:Compositional matrix adjust.,
Identities:273/296(92%), Positives:289/296(97%), Gaps:0/296(0%)

```
Query  1     PISPIETVPVKLKPGMDGPKVKQWPLTEEKIKALVEICTEMEKEGKISKIGPENPYNTPV  60
             PISPIETVPVKLKPGMDGPKVKQWPLTEEKIKAL EICTEMEKEGKISKIGPENPYNTPV
Sbjct  1010  PISPIETVPVKLKPGMDGPKVKQWPLTEEKIKALTEICTEMEKEGKISKIGPENPYNTPV  831

Query  61    FAIKKKDSTKWRKLVDFRELNKRTQDFWEVQLGIPHPAGLKKKKSVTVLDVGDAYFSVPL  120
             FAI+KKDSTKWRKLVDFRELNKRTQDFWE+QLGIPHPAGLKKKKSVTVLDVGDAYFSVPL
Sbjct  830   FAIRKKDSTKWRKLVDFRELNKRTQDFWEIQLGIPHPAGLKKKKSVTVLDVGDAYFSVPL  651

Query  121   DEDFRKYTAFTIPSINNETPGIRYQYNVLPQGWKGSPAIFQSSMTKILEPFRKQNPDIVI  180
             D+DFRKYTAFTIPSINNETPGIRYQYNVLPQGWKGSPAIFQ+SMTKILEPFR +NP++VI
Sbjct  650   DKDFRKYTAFTIPSINNETPGIRYQYNVLPQGWKGSPAIFQASMTKILEPFRTKNPELVI  471

Query  181   YQYMDDLYVGSDLEIGQHRTKIEELRQHLLRWGLTTPDKKHQKEPPFLWMGYELHPDKWT  240
             YQYMDDLY+GSDLEIGQHRTKIEELR HLLRWG TTPDKKHQKEPPFLWMGYELHPDKWT
Sbjct  470   YQYMDDLYIGSDLEIGQHRTKIEELRAHLLRWGFTTPDKKHQKEPPFLWMGYELHPDKWT  291

Query  241   VQPIVLPEKDSWTVNDIQKLVGKLNWASQIYPGIKVRQLCKLLRGTKALTEVIPLT  296
             VQP+ LP+K++WTVNDIQKLVGKLNWASQIY GIKV+QLCKLLRG KALT+++PLT
Sbjct  290   VQPVQLPDKENWTVNDIQKLVGKLNWASQIYAGIKVKQLCKLLRGAKALTDIVPLT  123
```

>Mycobacterium tuberculosis variant bovis strain CSURQ1134, whole genome shotgun sequence
Sequence ID: NZ_CAJEUT010000109.1 Length: 1697
Range 1: 85 to 972

Score:578 bits(1490), Expect:0.0,
Method:Compositional matrix adjust.,
Identities:273/296(92%), Positives:289/296(97%), Gaps:0/296(0%)

```
Query  1    PISPIETVPVKLKPGMDGPKVKQWPLTEEKIKALVEICTEMEKEGKISKIGPENPYNTPV  60
            PISPIETVPVKLKPGMDGPKVKQWPLTEEKIKAL EICTEMEKEGKISKIGPENPYNTPV
Sbjct  972  PISPIETVPVKLKPGMDGPKVKQWPLTEEKIKALTEICTEMEKEGKISKIGPENPYNTPV  793

Query  61   FAIKKKDSTKWRKLVDFRELNKRTQDFWEVQLGIPHPAGLKKKKSVTVLDVGDAYFSVPL  120
            FAI+KKDSTKWRKLVDFRELNKRTQDFWE+QLGIPHPAGLKKKKSVTVLDVGDAYFSVPL
Sbjct  792  FAIRKKDSTKWRKLVDFRELNKRTQDFWEIQLGIPHPAGLKKKKSVTVLDVGDAYFSVPL  613

Query  121  DEDFRKYTAFTIPSINNETPGIRYQYNVLPQGWKGSPAIFQSSMTKILEPFRKQNPDIVI  180
            D+DFRKYTAFTIPSINNETPGIRYQYNVLPQGWKGSPAIFQ+SMTKILEPFR +NP++VI
Sbjct  612  DKDFRKYTAFTIPSINNETPGIRYQYNVLPQGWKGSPAIFQASMTKILEPFRTKNPELVI  433

Query  181  YQYMDDLYVGSDLEIGQHRTKIEELRQHLLRWGLTTPDKKHQKEPPFLWMGYELHPDKWT  240
            YQYMDDLY+GSDLEIGQHRTKIEELR HLLRWG TTPDKKHQKEPPFLWMGYELHPDKWT
Sbjct  432  YQYMDDLYIGSDLEIGQHRTKIEELRAHLLRWGFTTPDKKHQKEPPFLWMGYELHPDKWT  253

Query  241  VQPIVLPEKDSWTVNDIQKLVGKLNWASQIYPGIKVRQLCKLLRGTKALTEVIPLT  296
            VQP+ LP+K++WTVNDIQKLVGKLNWASQIY GIKV+QLCKLLRG KALT+++PLT
Sbjct  252  VQPVQLPDKENWTVNDIQKLVGKLNWASQIYAGIKVKQLCKLLRGAKALTDIVPLT  85
```

Range 2: 85 to 972

Score:578 bits(1490), Expect:0.0,
Method:Compositional matrix adjust.,
Identities:273/296(92%), Positives:289/296(97%), Gaps:0/296(0%)

```
Query  1    PISPIETVPVKLKPGMDGPKVKQWPLTEEKIKALVEICTEMEKEGKISKIGPENPYNTPV  60
            PISPIETVPVKLKPGMDGPKVKQWPLTEEKIKAL EICTEMEKEGKISKIGPENPYNTPV
Sbjct  972  PISPIETVPVKLKPGMDGPKVKQWPLTEEKIKALTEICTEMEKEGKISKIGPENPYNTPV  793

Query  61   FAIKKKDSTKWRKLVDFRELNKRTQDFWEVQLGIPHPAGLKKKKSVTVLDVGDAYFSVPL  120
            FAI+KKDSTKWRKLVDFRELNKRTQDFWE+QLGIPHPAGLKKKKSVTVLDVGDAYFSVPL
Sbjct  792  FAIRKKDSTKWRKLVDFRELNKRTQDFWEIQLGIPHPAGLKKKKSVTVLDVGDAYFSVPL  613

Query  121  DEDFRKYTAFTIPSINNETPGIRYQYNVLPQGWKGSPAIFQSSMTKILEPFRKQNPDIVI  180
            D+DFRKYTAFTIPSINNETPGIRYQYNVLPQGWKGSPAIFQ+SMTKILEPFR +NP++VI
Sbjct  612  DKDFRKYTAFTIPSINNETPGIRYQYNVLPQGWKGSPAIFQASMTKILEPFRTKNPELVI  433

Query  181  YQYMDDLYVGSDLEIGQHRTKIEELRQHLLRWGLTTPDKKHQKEPPFLWMGYELHPDKWT  240
            YQYMDDLY+GSDLEIGQHRTKIEELR HLLRWG TTPDKKHQKEPPFLWMGYELHPDKWT
Sbjct  432  YQYMDDLYIGSDLEIGQHRTKIEELRAHLLRWGFTTPDKKHQKEPPFLWMGYELHPDKWT  253

Query  241  VQPIVLPEKDSWTVNDIQKLVGKLNWASQIYPGIKVRQLCKLLRGTKALTEVIPLT  296
            VQP+ LP+K++WTVNDIQKLVGKLNWASQIY GIKV+QLCKLLRG KALT+++PLT
Sbjct  252  VQPVQLPDKENWTVNDIQKLVGKLNWASQIYAGIKVKQLCKLLRGAKALTDIVPLT  85
```

```
>Escherichia coli strain 8374wA11 NODE_37_length_940_cov_2.75892, whole genome shotgun sequence
Sequence ID: NZ_WSPR01000036.1 Length: 940
Range 1: 2 to 883

Score:569 bits(1467), Expect:0.0,
Method:Compositional matrix adjust.,
Identities:270/294(92%), Positives:286/294(97%), Gaps:0/294(0%)

Query  1    PISPIETVPVKLKPGMDGPKVKQWPLTEEKIKALVEICTEMEKEGKISKIGPENPYNTPV  60
            PISPIETVPVKLKPGMDGP+VKQWPLTEEKIKAL  IC EMEKEGKI+KIGPENPYNTPV
Sbjct  883  PISPIETVPVKLKPGMDGPRVKQWPLTEEKIKALQAICEEMEKEGKITKIGPENPYNTPV  704

Query  61   FAIKKKDSTKWRKLVDFRELNKRTQDFWEVQLGIPHPAGLKKKKSVTVLDVGDAYFSVPL  120
            FAIKKKDSTKWRKLVDFRELNKRTQDFWEVQLGIPHPAGLKKKKSVTVLDVGDAYFSVPL
Sbjct  703  FAIKKKDSTKWRKLVDFRELNKRTQDFWEVQLGIPHPAGLKKKKSVTVLDVGDAYFSVPL  524

Query  121  DEDFRKYTAFTIPSINNETPGIRYQYNVLPQGWKGSPAIFQSSMTKILEPFRKQNPDIVI  180
            DE+FRKYTAFTIPSINN TPGIRYQYNVLPQGWKGSPAIFQS+MTKIL+PFR +NPDIVI
Sbjct  523  DENFRKYTAFTIPSINNATPGIRYQYNVLPQGWKGSPAIFQSTMTKILQPFRTRNPDIVI  344

Query  181  YQYMDDLYVGSDLEIGQHRTKIEELRQHLLRWGLTTPDKKHQKEPPFLWMGYELHPDKWT  240
            YQYMDDLYVGSDLEIGQHR KIEELR+HLL+WGLTTPDKKHQKEPPFLWMGYELHPDKWT
Sbjct  343  YQYMDDLYVGSDLEIGQHRAKIEELREHLLKWGLTTPDKKHQKEPPFLWMGYELHPDKWT  164

Query  241  VQPIVLPEKDSWTVNDIQKLVGKLNWASQIYPGIKVRQLCKLLRGTKALTEVIP  294
            VQPI LP+K+SWTVNDIQKLVGKLNWASQIYPGI+V++LC+LLRG KALT++IP
Sbjct  163  VQPIQLPDKESWTVNDIQKLVGKLNWASQIYPGIRVKRLCRLLRGAKALTDIIP  2



>Escherichia coli strain TzEc080 NODE_37_length_940_cov_2.75892, whole genome shotgun sequence
Sequence ID: NZ_WSFZ01000036.1 Length: 940
Range 1: 2 to 883

Score:569 bits(1467), Expect:0.0,
Method:Compositional matrix adjust.,
Identities:270/294(92%), Positives:286/294(97%), Gaps:0/294(0%)

Query  1    PISPIETVPVKLKPGMDGPKVKQWPLTEEKIKALVEICTEMEKEGKISKIGPENPYNTPV  60
            PISPIETVPVKLKPGMDGP+VKQWPLTEEKIKAL  IC EMEKEGKI+KIGPENPYNTPV
Sbjct  883  PISPIETVPVKLKPGMDGPRVKQWPLTEEKIKALQAICEEMEKEGKITKIGPENPYNTPV  704

Query  61   FAIKKKDSTKWRKLVDFRELNKRTQDFWEVQLGIPHPAGLKKKKSVTVLDVGDAYFSVPL  120
            FAIKKKDSTKWRKLVDFRELNKRTQDFWEVQLGIPHPAGLKKKKSVTVLDVGDAYFSVPL
Sbjct  703  FAIKKKDSTKWRKLVDFRELNKRTQDFWEVQLGIPHPAGLKKKKSVTVLDVGDAYFSVPL  524

Query  121  DEDFRKYTAFTIPSINNETPGIRYQYNVLPQGWKGSPAIFQSSMTKILEPFRKQNPDIVI  180
            DE+FRKYTAFTIPSINN TPGIRYQYNVLPQGWKGSPAIFQS+MTKIL+PFR +NPDIVI
Sbjct  523  DENFRKYTAFTIPSINNATPGIRYQYNVLPQGWKGSPAIFQSTMTKILQPFRTRNPDIVI  344

Query  181  YQYMDDLYVGSDLEIGQHRTKIEELRQHLLRWGLTTPDKKHQKEPPFLWMGYELHPDKWT  240
            YQYMDDLYVGSDLEIGQHR KIEELR+HLL+WGLTTPDKKHQKEPPFLWMGYELHPDKWT
Sbjct  343  YQYMDDLYVGSDLEIGQHRAKIEELREHLLKWGLTTPDKKHQKEPPFLWMGYELHPDKWT  164

Query  241  VQPIVLPEKDSWTVNDIQKLVGKLNWASQIYPGIKVRQLCKLLRGTKALTEVIP  294
            VQPI LP+K+SWTVNDIQKLVGKLNWASQIYPGI+V++LC+LLRG KALT++IP
Sbjct  163  VQPIQLPDKESWTVNDIQKLVGKLNWASQIYPGIRVKRLCRLLRGAKALTDIIP  2



>Mycobacterium tuberculosis strain 32846_S30 184, whole genome shotgun sequence
Sequence ID: NZ_QTBW01000041.1 Length: 932
Range 1: 4 to 930

Score:552 bits(1422), Expect:0.0,
Method:Compositional matrix adjust.,
Identities:283/309(92%), Positives:297/309(96%), Gaps:0/309(0%)

Query  126  KYTAFTIPSINNETPGIRYQYNVLPQGWKGSPAIFQSSMTKILEPFRKQNPDIVIYQYMD  185
            KYTAFTIPSINNETPGIRYQYNVLPQGWKGSPAIFQ SMTKILEPFR +NP+IVIYQYMD
Sbjct  930  KYTAFTIPSINNETPGIRYQYNVLPQGWKGSPAIFQYSMTKILEPFRAKNPEIVIYQYMD  751

Query  186  DLYVGSDLEIGQHRTKIEELRQHLLRWGLTTPDKKHQKEPPFLWMGYELHPDKWTVQPIV  245
            DLYVGSDLEIGQHR KIEELR HLL+WG TTPDKKHQKEPPFLWMGYELHPDKWTVQPI
Sbjct  750  DLYVGSDLEIGQHRVKIEELRGHLLKWGFTTPDKKHQKEPPFLWMGYELHPDKWTVQPIQ  571

Query  246  LPEKDSWTVNDIQKLVGKLNWASQIYPGIKVRQLCKLLRGTKALTEVIPLTeeaelelae  305
            LPEKDSWTVNDIQKLVGKLNWASQIYPGIKV+QLC+LLRG KALT+++PLTEEAELELAE
Sbjct  570  LPEKDSWTVNDIQKLVGKLNWASQIYPGIKVKQLCRLLRGAKALTDIVPLTEEAELELAE  391

Query  306  NREILKEPVHGVYYDPSKDLIAEIQKQGQGQWTYQIYQEPFKNLKTGKYARMRGAHTNDV  365
            NREILKEPVHGVYYDP+KDLIAEIQKQ  GQWTYQIYQEPFKNLKTGKYA+MR AHTNDV
Sbjct  390  NREILKEPVHGVYYDPAKDLIAEIQKQDHGQWTYQIYQEPFKNLKTGKYAKMRTAHTNDV  211

Query  366  KQLTEAVQKITTESIVIWGKTPKFKLPIQKetwetwwteYWQATWIPEWEFVNTPPLVKL  425
            KQLTEAVQKI  ESIV+WGKTPKF+LPIQKETWETWWTEYWQATWIPEWEFVNTPPLVKL
Sbjct  210  KQLTEAVQKIAMESIVVWGKTPKFRLPIQKETWETWWTEYWQATWIPEWEFVNTPPLVKL  31

Query  426  WYQLEKEPI  434
            WYQLEK+P+
Sbjct  30   WYQLEKDPL  4



>Acinetobacter baumannii strain ACM-36 NODE_62_length_953_cov_1.8184, whole genome shotgun sequence
Sequence ID: NZ_QAGO01000062.1 Length: 953
Range 1: 69 to 953

Score:550 bits(1416), Expect:0.0,
Method:Compositional matrix adjust.,
Identities:272/295(92%), Positives:285/295(96%), Gaps:0/295(0%)

Query  42   EKEGKISKIGPENPYNTPVFAIKKKDSTKWRKLVDFRELNKRTQDFWEVQLGIPHPAGLK  101
            EKEGKISKIG E PYNTP+FAIKKK+S KWRKLVDFRELNKRTQ+FWEVQLGIPHPAGLK
Sbjct  953  EKEGKISKIGSETPYNTPIFAIKKKNSDKWRKLVDFRELNKRTQEFWEVQLGIPHPAGLK  774

Query  102  KKKSVTVLDVGDAYFSVPLDEDFRKYTAFTIPSINNETPGIRYQYNVLPQGWKGSPAIFQ  161
            KKKSVTVLDVGDAYFSVPLD DFRKYTAFTIPS NNETPGIRYQYNVLPQGWKGSPAIFQ
Sbjct  773  KKKSVTVLDVGDAYFSVPLDPDFRKYTAFTIPSRNNETPGIRYQYNVLPQGWKGSPAIFQ  594

Query  162  SSMTKILEPFRKQNPDIVIYQYMDDLYVGSDLEIGQHRTKIEELRQHLLRWGLTTPDKKH  221
            SSMTKILEPFRKQNPD+VIYQYMDDLYVGSDLE+ QHRTK+EELR HLL+WG +TPD+KH
Sbjct  593  SSMTKILEPFRKQNPDMVIYQYMDDLYVGSDLELEQHRTKVEELRAHLLKWGFSTPDQKH  414

Query  222  QKEPPFLWMGYELHPDKWTVQPIVLPEKDSWTVNDIQKLVGKLNWASQIYPGIKVRQLCK  281
            QKEPPFLWMGYELHPDKWTVQPIVLPEKDSWTVNDIQKLVGKLNWASQIY GIKV+QLCK
Sbjct  413  QKEPPFLWMGYELHPDKWTVQPIVLPEKDSWTVNDIQKLVGKLNWASQIYAGIKVKQLCK  234

Query  282  LLRGTKALTEVIPLTeeaelelaeNREILKEPVHGVYYDPSKDLIAEIQKQGQGQ  336
```

```
             LLRGTKALTEV+PL+EEAELELAENREILKEPVHGVYYDPSKDL+AEIQKQG GQ
Sbjct  233  LLRGTKALTEVVPLSEEAELELAENREILKEPVHGVYYDPSKDLVAEIQKQGLGQ  69




>Providencia rettgeri strain AFE652_S1 38_contig_28, whole genome shotgun sequence
Sequence ID: NZ_SHDK01000028.1 Length: 1127
Range 1: 1 to 810

Score:533 bits(1373), Expect:0.0,
Method:Compositional matrix adjust.,
Identities:255/270(94%), Positives:263/270(97%), Gaps:0/270(0%)

Query  1    PISPIETVPVKLKPGMDGPKVKQWPLTEEKIKALVEICTEMEKEGKISKIGPENPYNTPV  60
            PISPIETVPVKLKPGMDGP+VKQWPLTEEKIKAL EIC EMEKEGKI +IGPENPYNTPV
Sbjct  810  PISPIETVPVKLKPGMDGPRVKQWPLTEEKIKALTEICEEMEKEGKIERIGPENPYNTPV  631

Query  61   FAIKKKDSTKWRKLVDFRELNKRTQDFWEVQLGIPHPAGLKKKKSVTVLDVGDAYFSVPL  120
            FAIKKKDSTKWRKLVDFRELNKRTQDFWEVQLGIPHPAGLKKKKSVTVLDVGDAYFSVPL
Sbjct  630  FAIKKKDSTKWRKLVDFRELNKRTQDFWEVQLGIPHPAGLKKKKSVTVLDVGDAYFSVPL  451

Query  121  DEDFRKYTAFTIPSINNETPGIRYQYNVLPQGWKGSPAIFQSSMTKILEPFRKQNPDIVI  180
            DEDFRKYTAFTIPSINNETPGIRYQYNVLPQGWKGSPAIFQSSM +ILEPFR +NP+IVI
Sbjct  450  DEDFRKYTAFTIPSINNETPGIRYQYNVLPQGWKGSPAIFQSSMIRILEPFRAKNPEIVI  271

Query  181  YQYMDDLYVGSDLEIGQHRTKIEELRQHLLRWGLTTPDKKHQKEPPFLWMGYELHPDKWT  240
            YQYMDDLYVGSDLEIGQHR KIE+LR+HLL+WGLTTPDKKHQKEPPFLWMGYELHPDKWT
Sbjct  270  YQYMDDLYVGSDLEIGQHRAKIEKLREHLLKWGLTTPDKKHQKEPPFLWMGYELHPDKWT  91

Query  241  VQPIVLPEKDSWTVNDIQKLVGKLNWASQI  270
            VQPI LPEKDSWTVNDIQKLVGKLNWASQI
Sbjct  90   VQPIQLPEKDSWTVNDIQKLVGKLNWASQI  1




>Mycobacterium tuberculosis variant bovis strain CSURQ1134, whole genome shotgun sequence
Sequence ID: NZ_CAJEUT010000134.1 Length: 910
Range 1: 54 to 890

Score:518 bits(1334), Expect:5e-179,
Method:Compositional matrix adjust.,
Identities:251/279(90%), Positives:260/279(93%), Gaps:0/279(0%)

Query  1    PISPIETVPVKLKPGMDGPKVKQWPLTEEKIKALVEICTEMEKEGKISKIGPENPYNTPV  60
            P  P ET+ VKLK  M GP+ KQWPLT   I +L EICTEMEKEGKISKIGPENPYNTPV
Sbjct  890  PSRPTETLRVKLKR*MVGPQAKQWPLTA*TINSLTEICTEMEKEGKISKIGPENPYNTPV  711

Query  61   FAIKKKDSTKWRKLVDFRELNKRTQDFWEVQLGIPHPAGLKKKKSVTVLDVGDAYFSVPL  120
            FAIKKK+STKWRKLVDFRELNK+TQDFWEVQLGIPHP+GLKKKKSVTVLDVGDAYFSVPL
Sbjct  710  FAIKKKNSTKWRKLVDFRELNKKTQDFWEVQLGIPHPSGLKKKKSVTVLDVGDAYFSVPL  531

Query  121  DEDFRKYTAFTIPSINNETPGIRYQYNVLPQGWKGSPAIFQSSMTKILEPFRKQNPDIVI  180
            DEDFRKYTAFTIPS+NN TPGIRYQYNVLPQGWKGSPAIFQSSMTKILEPFRKQNPDIVI
Sbjct  530  DEDFRKYTAFTIPSVNNATPGIRYQYNVLPQGWKGSPAIFQSSMTKILEPFRKQNPDIVI  351

Query  181  YQYMDDLYVGSDLEIGQHRTKIEELRQHLLRWGLTTPDKKHQKEPPFLWMGYELHPDKWT  240
            YQY+DDLYVGSDLEI QHRTK+EELRQHL RWG  TPDKKHQKEPPFLWMGYELHPDKWT
Sbjct  350  YQYVDDLYVGSDLEIEQHRTKVEELRQHLWRWGFDTPDKKHQKEPPFLWMGYELHPDKWT  171

Query  241  VQPIVLPEKDSWTVNDIQKLVGKLNWASQIYPGIKVRQL  279
            VQPIVLPEKDSWTVNDIQKLVGKLNWASQIYPGIKV QL
Sbjct  170  VQPIVLPEKDSWTVNDIQKLVGKLNWASQIYPGIKVNQL  54




Range 2: 54 to 890

Score:518 bits(1334), Expect:5e-179,
Method:Compositional matrix adjust.,
Identities:251/279(90%), Positives:260/279(93%), Gaps:0/279(0%)

Query  1    PISPIETVPVKLKPGMDGPKVKQWPLTEEKIKALVEICTEMEKEGKISKIGPENPYNTPV  60
            P  P ET+ VKLK  M GP+ KQWPLT   I +L EICTEMEKEGKISKIGPENPYNTPV
Sbjct  890  PSRPTETLRVKLKR*MVGPQAKQWPLTA*TINSLTEICTEMEKEGKISKIGPENPYNTPV  711

Query  61   FAIKKKDSTKWRKLVDFRELNKRTQDFWEVQLGIPHPAGLKKKKSVTVLDVGDAYFSVPL  120
            FAIKKK+STKWRKLVDFRELNK+TQDFWEVQLGIPHP+GLKKKKSVTVLDVGDAYFSVPL
Sbjct  710  FAIKKKNSTKWRKLVDFRELNKKTQDFWEVQLGIPHPSGLKKKKSVTVLDVGDAYFSVPL  531

Query  121  DEDFRKYTAFTIPSINNETPGIRYQYNVLPQGWKGSPAIFQSSMTKILEPFRKQNPDIVI  180
            DEDFRKYTAFTIPS+NN TPGIRYQYNVLPQGWKGSPAIFQSSMTKILEPFRKQNPDIVI
Sbjct  530  DEDFRKYTAFTIPSVNNATPGIRYQYNVLPQGWKGSPAIFQSSMTKILEPFRKQNPDIVI  351

Query  181  YQYMDDLYVGSDLEIGQHRTKIEELRQHLLRWGLTTPDKKHQKEPPFLWMGYELHPDKWT  240
            YQY+DDLYVGSDLEI QHRTK+EELRQHL RWG  TPDKKHQKEPPFLWMGYELHPDKWT
Sbjct  350  YQYVDDLYVGSDLEIEQHRTKVEELRQHLWRWGFDTPDKKHQKEPPFLWMGYELHPDKWT  171

Query  241  VQPIVLPEKDSWTVNDIQKLVGKLNWASQIYPGIKVRQL  279
            VQPIVLPEKDSWTVNDIQKLVGKLNWASQIYPGIKV QL
Sbjct  170  VQPIVLPEKDSWTVNDIQKLVGKLNWASQIYPGIKVNQL  54




>Candidatus Tokpelaia sp. strain RhiAcro1 isolate RhiAcro1_RAo1 NODE_51_length_826_cov_3.34846_ID_101, whole genome shotgun sequence
Sequence ID: NZ_QLZD01000026.1 Length: 826
Range 1: 2 to 826

Score:511 bits(1316), Expect:8e-177,
Method:Compositional matrix adjust.,
Identities:255/275(93%), Positives:267/275(97%), Gaps:0/275(0%)

Query  62   AIKKKDSTKWRKLVDFRELNKRTQDFWEVQLGIPHPAGLKKKKSVTVLDVGDAYFSVPLD  121
            AIKKKDSTKWRKLVDFRELNKRTQDFWEVQLGIPHPAGLKKKKSVTVLDVGDAYFSVPLD
Sbjct  826  AIKKKDSTKWRKLVDFRELNKRTQDFWEVQLGIPHPAGLKKKKSVTVLDVGDAYFSVPLD  647

Query  122  EDFRKYTAFTIPSINNETPGIRYQYNVLPQGWKGSPAIFQSSMTKILEPFRKQNPDIVIY  181
            ++FRKYT FTIPSINNETPGIRYQYNVLPQGWKGSPAIFQ SMTKILEPFRKQNP++VIY
Sbjct  646  KEFRKYTTFTIPSINNETPGIRYQYNVLPQGWKGSPAIFQCSMTKILEPFRKQNPEMVIY  467

Query  182  QYMDDLYVGSDLEIGQHRTKIEELRQHLLRWGLTTPDKKHQKEPPFLWMGYELHPDKWTV  241
            QYMDDLYVGSDLE+GQHRTKIEELR+HLL WG TTPDKKHQKEPPFLWMGYELHPDKWTV
Sbjct  466  QYMDDLYVGSDLELGQHRTKIEELRKHLLAWGFTTPDKKHQKEPPFLWMGYELHPDKWTV  287

Query  242  QPIVLPEKDSWTVNDIQKLVGKLNWASQIYPGIKVRQLCKLLRGTKALTEVIPLTeeael  301
            QPI+LPEKDSWTVNDIQKLVGKLNWASQIY GIKV+QLCKLLRG K+LTEVI LTEEAEL
Sbjct  286  QPIMLPEKDSWTVNDIQKLVGKLNWASQIYAGIKVKQLCKLLRGAKSLTEVITLTEEAEL  107

Query  302  elaeNREILKEPVHGVYYDPSKDLIAEIQKQGQGQ  336
            ELAENREILKEPVHG YYDP+KDL+AE+QKQGQGQ
Sbjct  106  ELAENREILKEPVHGAYYDPTKDLVAELQKQGQGQ  2
```

```
>Mycobacterium tuberculosis variant bovis strain CSURQ1139, whole genome shotgun sequence
Sequence ID: NZ_CAJEUO010000157.1 Length: 848
Range 1: 1 to 777

Score:510 bits(1313), Expect:4e-176,
Method:Compositional matrix adjust.,
Identities:241/259(93%), Positives:254/259(98%), Gaps:0/259(0%)

Query  1    PISPIETVPVKLKPGMDGPKVKQWPLTEEKIKALVEICTEMEKEGKISKIGPENPYNTPV  60
            PISPIETVPVKLKPGMDGPKVKQWPLTEEKIKAL EICTEMEKEGKISKIGPENPYNTPV
Sbjct  777  PISPIETVPVKLKPGMDGPKVKQWPLTEEKIKALTEICTEMEKEGKISKIGPENPYNTPV  598

Query  61   FAIKKKDSTKWRKLVDFRELNKRTQDFWEVQLGIPHPAGLKKKKSVTVLDVGDAYFSVPL  120
            FAI+KKDSTKWRKLVDFRELNKRTQDFWE+QLGIPHPAGLKKKKSVTVLDVGDAYFSVPL
Sbjct  597  FAIRKKDSTKWRKLVDFRELNKRTQDFWEIQLGIPHPAGLKKKKSVTVLDVGDAYFSVPL  418

Query  121  DEDFRKYTAFTIPSINNETPGIRYQYNVLPQGWKGSPAIFQSSMTKILEPFRKQNPDIVI  180
            D+DFRKYTAFTIPSINNETPGIRYQYNVLPQGWKGSPAIFQ+SMTKILEPFR +NP++VI
Sbjct  417  DKDFRKYTAFTIPSINNETPGIRYQYNVLPQGWKGSPAIFQASMTKILEPFRTKNPELVI  238

Query  181  YQYMDDLYVGSDLEIGQHRTKIEELRQHLLRWGLTTPDKKHQKEPPFLWMGYELHPDKWT  240
            YQYMDDLY+GSDLEIGQHRTKIEELR HLLRWG TTPDKKHQKEPPFLWMGYELHPDKWT
Sbjct  237  YQYMDDLYIGSDLEIGQHRTKIEELRAHLLRWGFTTPDKKHQKEPPFLWMGYELHPDKWT  58

Query  241  VQPIVLPEKDSWTVNDIQK  259
            VQP+ LP+K++WTVNDI+K
Sbjct  57   VQPVQLPDKENWTVNDIRK  1
```

```
Range 2: 1 to 777

Score:510 bits(1313), Expect:4e-176,
Method:Compositional matrix adjust.,
Identities:241/259(93%), Positives:254/259(98%), Gaps:0/259(0%)

Query  1    PISPIETVPVKLKPGMDGPKVKQWPLTEEKIKALVEICTEMEKEGKISKIGPENPYNTPV  60
            PISPIETVPVKLKPGMDGPKVKQWPLTEEKIKAL EICTEMEKEGKISKIGPENPYNTPV
Sbjct  777  PISPIETVPVKLKPGMDGPKVKQWPLTEEKIKALTEICTEMEKEGKISKIGPENPYNTPV  598

Query  61   FAIKKKDSTKWRKLVDFRELNKRTQDFWEVQLGIPHPAGLKKKKSVTVLDVGDAYFSVPL  120
            FAI+KKDSTKWRKLVDFRELNKRTQDFWE+QLGIPHPAGLKKKKSVTVLDVGDAYFSVPL
Sbjct  597  FAIRKKDSTKWRKLVDFRELNKRTQDFWEIQLGIPHPAGLKKKKSVTVLDVGDAYFSVPL  418

Query  121  DEDFRKYTAFTIPSINNETPGIRYQYNVLPQGWKGSPAIFQSSMTKILEPFRKQNPDIVI  180
            D+DFRKYTAFTIPSINNETPGIRYQYNVLPQGWKGSPAIFQ+SMTKILEPFR +NP++VI
Sbjct  417  DKDFRKYTAFTIPSINNETPGIRYQYNVLPQGWKGSPAIFQASMTKILEPFRTKNPELVI  238

Query  181  YQYMDDLYVGSDLEIGQHRTKIEELRQHLLRWGLTTPDKKHQKEPPFLWMGYELHPDKWT  240
            YQYMDDLY+GSDLEIGQHRTKIEELR HLLRWG TTPDKKHQKEPPFLWMGYELHPDKWT
Sbjct  237  YQYMDDLYIGSDLEIGQHRTKIEELRAHLLRWGFTTPDKKHQKEPPFLWMGYELHPDKWT  58

Query  241  VQPIVLPEKDSWTVNDIQK  259
            VQP+ LP+K++WTVNDI+K
Sbjct  57   VQPVQLPDKENWTVNDIRK  1
```

```
>Levilactobacillus brevis strain CRL 2013 scaffold12, whole genome shotgun sequence
Sequence ID: NZ_MZMW01000016.1 Length: 1126
Range 1: 359 to 1126

Score:513 bits(1322), Expect:4e-176,
Method:Compositional matrix adjust.,
Identities:248/256(97%), Positives:249/256(97%), Gaps:0/256(0%)

Query  1     PISPIETVPVKLKPGMDGPKVKQWPLTEEKIKALVEICTEMEKEGKISKIGPENPYNTPV  60
             PISPIETVPVKLKPGMDGPKVKQWPLTEEKIKAL EICTEMEKEGKISKIGPENPYNTP
Sbjct  359   PISPIETVPVKLKPGMDGPKVKQWPLTEEKIKALXEICTEMEKEGKISKIGPENPYNTPX  538

Query  61    FAIKKKDSTKWRKLVDFRELNKRTQDFWEVQLGIPHPAGLKKKKSVTVLDVGDAYFSVPL  120
             FAIKKKDSTKWRKLVDFRELNKRTQDFWEVQLGIPHPAGLK+ KSVTVLDVGDAYFSVPL
Sbjct  539   FAIKKKDSTKWRKLVDFRELNKRTQDFWEVQLGIPHPAGLKQNKSVTVLDVGDAYFSVPL  718

Query  121   DEDFRKYTAFTIPSINNETPGIRYQYNVLPQGWKGSPAIFQSSMTKILEPFRKQNPDIVI  180
             DE FRKYTAFTIPSINNETPGIRYQYNVLPQGWKGSPAIFQSSMTKILEPFRKQNPDIVI
Sbjct  719   DEAFRKYTAFTIPSINNETPGIRYQYNVLPQGWKGSPAIFQSSMTKILEPFRKQNPDIVI  898

Query  181   YQYMDDLYVGSDLEIGQHRTKIEELRQHLLRWGLTTPDKKHQKEPPFLWMGYELHPDKWT  240
             YQYMDDLYVGSDLEIGQHR KIEELRQHLLRWG TTPDKKHQKE PFLWMGYELHPDKWT
Sbjct  899   YQYMDDLYVGSDLEIGQHRAKIEELRQHLLRWGFTTPDKKHQKEHPFLWMGYELHPDKWT  1078

Query  241   VQPIVLPEKDSWTVND  256
             VQPIVLPEKDSWTVND
Sbjct  1079  VQPIVLPEKDSWTVND  1126
```

```
>Pseudomonas aeruginosa strain VET-42 NODE_85_length_1204_cov_6.50932_ID_169, whole genome shotgun sequence
Sequence ID: NZ_RACX01000100.1 Length: 1166
Range 1: 325 to 1104

Score:510 bits(1313), Expect:1e-174,
Method:Compositional matrix adjust.,
Identities:243/260(93%), Positives:252/260(96%), Gaps:0/260(0%)

Query  1     PISPIETVPVKLKPGMDGPKVKQWPLTEEKIKALVEICTEMEKEGKISKIGPENPYNTPV  60
             PISPIETVPVKLKPGMDGPKVKQWPLTEEKIKAL EICTEMEKEGKISKIGPENPYNTP+
Sbjct  325   PISPIETVPVKLKPGMDGPKVKQWPLTEEKIKALTEICTEMEKEGKISKIGPENPYNTPI  504

Query  61    FAIKKKDSTKWRKLVDFRELNKRTQDFWEVQLGIPHPAGLKKKKSVTVLDVGDAYFSVPL  120
             FAIKKKDSTKWRKLVDFRELNKRTQDFWEVQLGIPHPAGLKKKKSVT+LDVGDAYFSVPL
Sbjct  505   FAIKKKDSTKWRKLVDFRELNKRTQDFWEVQLGIPHPAGLKKKKSVTILDVGDAYFSVPL  684

Query  121   DEDFRKYTAFTIPSINNETPGIRYQYNVLPQGWKGSPAIFQSSMTKILEPFRKQNPDIVI  180
             D DFRKYTAFTIPSINNETPGIRYQYNVLPQGWKGSPAIFQ SMTKILEPFRKQNP++VI
Sbjct  685   DPDFRKYTAFTIPSINNETPGIRYQYNVLPQGWKGSPAIFQCSMTKILEPFRKQNPELVI  864

Query  181   YQYMDDLYVGSDLEIGQHRTKIEELRQHLLRWGLTTPDKKHQKEPPFLWMGYELHPDKWT  240
             YQYMDDLY+GSDLEIGQHR KIEELR+HLL+WG TTPDKKHQKEPPFLWMGYELHPDKWT
Sbjct  865   YQYMDDLYIGSDLEIGQHRAKIEELREHLLKWGFTTPDKKHQKEPPFLWMGYELHPDKWT  1044

Query  241   VQPIVLPEKDSWTVNDIQKL  260
             VQPI LP+K+SWTVNDI  L
Sbjct  1045  VQPIELPDKESWTVNDICLL  1104
```

```
>Pseudomonas aeruginosa strain VET-121 NODE_73_length_1192_cov_8.02063_ID_145, whole genome shotgun sequence
Sequence ID: NZ_QZXB01000185.1 Length: 1158
Range 1: 57 to 824

Score:508 bits(1307), Expect:1e-173,
Method:Compositional matrix adjust.,
Identities:241/256(94%), Positives:250/256(97%), Gaps:0/256(0%)

Query  1    PISPIETVPVKLKPGMDGPKVKQWPLTEEKIKALVEICTEMEKEGKISKIGPENPYNTPV  60
            PISPIETVPVKLKPGMDGPKVKQWPLTEEKIKAL EICTEMEKEGKISKIGPENPYNTP+
Sbjct  824  PISPIETVPVKLKPGMDGPKVKQWPLTEEKIKALTEICTEMEKEGKISKIGPENPYNTPI  645

Query  61   FAIKKKDSTKWRKLVDFRELNKRTQDFWEVQLGIPHPAGLKKKKSVTVLDVGDAYFSVPL  120
            FAIKKKDSTKWRKLVDFRELNKRTQDFWEVQLGIPHPAGLKKKKSVT+LDVGDAYFSVPL
Sbjct  644  FAIKKKDSTKWRKLVDFRELNKRTQDFWEVQLGIPHPAGLKKKKSVTILDVGDAYFSVPL  465

Query  121  DEDFRKYTAFTIPSINNETPGIRYQYNVLPQGWKGSPAIFQSSMTKILEPFRKQNPDIVI  180
            D DFRKYTAFTIPSINNETPGIRYQYNVLPQGWKGSPAIFQ SMTKILEPFRKQNP++VI
Sbjct  464  DPDFRKYTAFTIPSINNETPGIRYQYNVLPQGWKGSPAIFQCSMTKILEPFRKQNPELVI  285

Query  181  YQYMDDLYVGSDLEIGQHRTKIEELRQHLLRWGLTTPDKKHQKEPPFLWMGYELHPDKWT  240
            YQYMDDLY+GSDLEIGQHR KIEELR+HLL+WG TTPDKKHQKEPPFLWMGYELHPDKWT
Sbjct  284  YQYMDDLYIGSDLEIGQHRAKIEELREHLLKWGFTTPDKKHQKEPPFLWMGYELHPDKWT  105

Query  241  VQPIVLPEKDSWTVND  256
            VQPI LP+K+SWTVND
Sbjct  104  VQPIELPDKESWTVND  57



>Acinetobacter baumannii strain ACM-27 NODE_68_length_895_cov_1.29167, whole genome shotgun sequence
Sequence ID: NZ_PYDM01000068.1 Length: 895
Range 1: 15 to 890

Score:504 bits(1297), Expect:2e-173,
Method:Compositional matrix adjust.,
Identities:241/292(83%), Positives:267/292(91%), Gaps:0/292(0%)

Query  5    IETVPVKLKPGMDGPKVKQWPLTEEKIKALVEICTEMEKEGKISKIGPENPYNTPVFAIK  64
            IET  VKL  GM GPK+KQW      I AL EI TE++++GKISKIGPENP NTP+FAIK
Sbjct  890  IETNKVKL*SGMYGPKIKQWS*ITHIIVALTEIFTEVKQDGKISKIGPENPCNTPIFAIK  711

Query  65   KKDSTKWRKLVDFRELNKRTQDFWEVQLGIPHPAGLKKKKSVTVLDVGDAYFSVPLDEDF  124
            KK+ST WRKL+DFRELNKRTQDFWEVQLGIPHPAGL+KKKSVTVLDVGDAYFS+PLDEDF
Sbjct  710  KKNSTGWRKLMDFRELNKRTQDFWEVQLGIPHPAGLEKKKSVTVLDVGDAYFSIPLDEDF  531

Query  125  RKYTAFTIPSINNETPGIRYQYNVLPQGWKGSPAIFQSSMTKILEPFRKQNPDIVIYQYM  184
            RKYTAFTIPS NNETPG+RYQYNVLPQGWKGSPAIFQSSMT+ILEPFR +NP+IVIYQ +
Sbjct  530  RKYTAFTIPSTNNETPGVRYQYNVLPQGWKGSPAIFQSSMTRILEPFRAKNPEIVIYQSV  351

Query  185  DDLYVGSDLEIGQHRTKIEELRQHLLRWGLTTPDKKHQKEPPFLWMGYELHPDKWTVQPI  244
            DDLYVGSDLEIGQHR KI+ELR++L +WG  TPDKKHQ+EPPFLWMGY+LHPDKWTVQPI
Sbjct  350  DDLYVGSDLEIGQHRAKIKELREYLWKWGFYTPDKKHQEEPPFLWMGYKLHPDKWTVQPI  171

Query  245  VLPEKDSWTVNDIQKLVGKLNWASQIYPGIKVRQLCKLLRGTKALTEVIPLT  296
             LP K+SWTVNDIQKLVGKLNWASQIYPGIKV+QLCKLLRGTKALT+++PLT
Sbjct  170  QLPNKESWTVNDIQKLVGKLNWASQIYPGIKVKQLCKLLRGTKALTDIVPLT  15



>Klebsiella pneumoniae strain EuSCAPE_DE086, whole genome shotgun sequence
Sequence ID: NZ_UKLW02000059.1 Length: 995
Range 1: 2 to 994

Score:495 bits(1275), Expect:1e-169,
Method:Compositional matrix adjust.,
Identities:267/331(81%), Positives:289/331(87%), Gaps:0/331(0%)

Query  163  SMTKILEPFRKQNPDIVIYQYMDDLYVGSDLEIGQHRTKIEELRQHLLRWGLTTPDKKHQ  222
            SMTKILEPF+ QNP+IVIYQY+DDLYV SDL+I QHR KIEELR+HLLR   TTPDKKHQ
Sbjct  2    SMTKILEPFKAQNPEIVIYQYIDDLYVRSDLKIRQHRAKIEELREHLLR*RFTTPDKKHQ  181

Query  223  KEPPFLWMGYELHPDKWTVQPIVLPEKDSWTVNDIQKLVGKLNWASQIYPGIKVRQLCKL  282
            KEPPFL + YELHPDK TVQPI LPEK+S TVNDIQKLV KLN ASQIY  IKV+QLCKL
Sbjct  182  KEPPFL*IKYELHPDK*TVQPIKLPEKES*TVNDIQKLVRKLN*ASQIYSKIKVKQLCKL  361

Query  283  LRGTKALTEVIPLTeeaelelaeNREILKEPVHGVYYDPSKDLIAEIQKQGQGQWTYQIY  342
            LR  KALT+++PLTEEAELELAENREILKEPVH  YYDPSKDLIAE+QKQG  QWTYQIY
Sbjct  362  LREAKALTDIVPLTEEAELELAENREILKEPVHEAYYDPSKDLIAELQKQGHDQWTYQIY  541

Query  343  QEPFKNLKTGKYARMRGAHTNDVKQLTEAVQKITTESIVIWGKTPKFKLPIQKetwetww  402
            Q+PFKNLKTGKYA+MR AHTNDVKQLTEAVQKI  E IVI  KTPKFKLPIQKET E  W
Sbjct  542  QKPFKNLKTGKYAKMRTAHTNDVKQLTEAVQKIAIECIVI*RKTPKFKLPIQKET*EL*W  721

Query  403  teYWQATWIPEWEFVNTPPLVKLWYQLEKEPIVGAETFYVDGAANRETKLGKAGYVTNRG  462
            T+YWQAT IPE EFVNTPPLVKL YQLEKEPIV AETFYVD AAN+ETK+ KA YVT++
Sbjct  722  TDYWQAT*IPE*EFVNTPPLVKL*YQLEKEPIVRAETFYVDRAANKETKIRKARYVTDKR  901

Query  463  RQKVVTLTDTTNQKTELQAIYLALQDSGLEV  493
            RQK+V+LT+TTNQKTELQ I LALQDSG EV
Sbjct  902  RQKIVSLTETTNQKTELQTIQLALQDSGSEV  994



>Providencia rettgeri strain APW139_S1 24_contig_64, whole genome shotgun sequence
Sequence ID: NZ_SHDG01000064.1 Length: 902
Range 1: 135 to 902

Score:490 bits(1262), Expect:4e-168,
Method:Compositional matrix adjust.,
Identities:230/256(90%), Positives:245/256(95%), Gaps:0/256(0%)

Query  1    PISPIETVPVKLKPGMDGPKVKQWPLTEEKIKALVEICTEMEKEGKISKIGPENPYNTPV  60
            PISPI+TVPVKLKPGMDGPKVKQWPLTEEKIKAL  IC EMEKEGKI+KIGPENPYNTP+
Sbjct  135  PISPIKTVPVKLKPGMDGPKVKQWPLTEEKIKALTAICEEMEKEGKITKIGPENPYNTPI  314

Query  61   FAIKKKDSTKWRKLVDFRELNKRTQDFWEVQLGIPHPAGLKKKKSVTVLDVGDAYFSVPL  120
            FAIKKK STKWRK+VDFRELNKRTQDFWEVQLGIPHPAGLK+ KS+TVLDVGDA+FSVPL
Sbjct  315  FAIKKKGSTKWRKVVDFRELNKRTQDFWEVQLGIPHPAGLKRNKSMTVLDVGDAFFSVPL  494

Query  121  DEDFRKYTAFTIPSINNETPGIRYQYNVLPQGWKGSPAIFQSSMTKILEPFRKQNPDIVI  180
            D+DFRKYTAFTIPS+NN TPGIRYQYNVLPQGWKGSPAIFQ SMTKILEPFR QNPDIVI
Sbjct  495  DKDFRKYTAFTIPSVNNATPGIRYQYNVLPQGWKGSPAIFQCSMTKILEPFRTQNPDIVI  674

Query  181  YQYMDDLYVGSDLEIGQHRTKIEELRQHLLRWGLTTPDKKHQKEPPFLWMGYELHPDKWT  240
            YQY+DDLYVGSDLEIGQHR K+EELR+HLL+WG TTPDKKHQKEPPFLWMGYELHPDKWT
Sbjct  675  YQYVDDLYVGSDLEIGQHRAKVEELRKHLLKWGFTTPDKKHQKEPPFLWMGYELHPDKWT  854

Query  241  VQPIVLPEKDSWTVND  256
```

```
            VQPI LP+K+SWTVND
Sbjct  855  VQPIQLPDKESWTVND  902
```

>Levilactobacillus brevis strain CRL 2013 scaffold13, whole genome shotgun sequence
Sequence ID: NZ_MZMW01000017.1 Length: 1030
Range 1: 298 to 1029

Score:488 bits(1255), Expect:2e-166,
Method:Compositional matrix adjust.,
Identities:234/244(96%), Positives:238/244(97%), Gaps:0/244(0%)

```
Query  1     PISPIETVPVKLKPGMDGPKVKQWPLTEEKIKALVEICTEMEKEGKISKIGPENPYNTPV  60
             PISPIETVPVKLKPGMDGPKVKQWPLTEEKI AL EIC EMEKEGKISKIGPENPYNTPV
Sbjct  298   PISPIETVPVKLKPGMDGPKVKQWPLTEEKIXALTEICXEMEKEGKISKIGPENPYNTPV  477

Query  61    FAIKKKDSTKWRKLVDFRELNKRTQDFWEVQLGIPHPAGLKKKKSVTVLDVGDAYFSVPL  120
             FAIKKKDSTKWRKLVDFRELNKRTQDFWEVQLGIPHPAGLK+KKSVTVLDVGDAYFSVPL
Sbjct  478   FAIKKKDSTKWRKLVDFRELNKRTQDFWEVQLGIPHPAGLKRKKSVTVLDVGDAYFSVPL  657

Query  121   DEDFRKYTAFTIPSINNETPGIRYQYNVLPQGWKGSPAIFQSSMTKILEPFRKQNPDIVI  180
             DE+FRKYTAFTIPS NNETPGIRYQYNVLPQGWKGSPAIFQSSMTKILEPFRKQNPDI I
Sbjct  658   DENFRKYTAFTIPSTNNETPGIRYQYNVLPQGWKGSPAIFQSSMTKILEPFRKQNPDIDI  837

Query  181   YQYMDDLYVGSDLEIGQHRTKIEELRQHLLRWGLTTPDKKHQKEPPFLWMGYELHPDKWT  240
             YQYMDDLYVGSDLEIGQHR KIEELR+HLLRWGLTTPDKKHQK+PPFLWMGYELHPDKWT
Sbjct  838   YQYMDDLYVGSDLEIGQHRAKIEELREHLLRWGLTTPDKKHQKKPPFLWMGYELHPDKWT  1017

Query  241   VQPI  244
             VQPI
Sbjct  1018  VQPI  1029
```

>Providencia rettgeri strain AFE96_S1 41_contig_60, whole genome shotgun sequence
Sequence ID: NZ_SHDP01000060.1 Length: 1101
Range 1: 354 to 1100

Score:488 bits(1257), Expect:2e-166,
Method:Compositional matrix adjust.,
Identities:234/249(94%), Positives:240/249(96%), Gaps:0/249(0%)

```
Query  1     PISPIETVPVKLKPGMDGPKVKQWPLTEEKIKALVEICTEMEKEGKISKIGPENPYNTPV  60
             PISPIETVPVKLKPGMDGPKVKQWPLTEEKIKAL  IC EMEKEGKI+KIGPENPYNTP+
Sbjct  354   PISPIETVPVKLKPGMDGPKVKQWPLTEEKIKALTAICKEMEKEGKITKIGPENPYNTPI  533

Query  61    FAIKKKDSTKWRKLVDFRELNKRTQDFWEVQLGIPHPAGLKKKKSVTVLDVGDAYFSVPL  120
             FAIKKKDSTKWRKLVDFRELNKRTQDFWEVQLGIPHPAGLKKKKSVTVLDVGDAYFSVPL
Sbjct  534   FAIKKKDSTKWRKLVDFRELNKRTQDFWEVQLGIPHPAGLKKKKSVTVLDVGDAYFSVPL  713

Query  121   DEDFRKYTAFTIPSINNETPGIRYQYNVLPQGWKGSPAIFQSSMTKILEPFRKQNPDIVI  180
             DE FRKYTAFTIPSINNETPGIRYQYNVLPQGWKGSPAIFQSSMTKILEPFR QNP+IVI
Sbjct  714   DEGFRKYTAFTIPSINNETPGIRYQYNVLPQGWKGSPAIFQSSMTKILEPFRAQNPEIVI  893

Query  181   YQYMDDLYVGSDLEIGQHRTKIEELRQHLLRWGLTTPDKKHQKEPPFLWMGYELHPDKWT  240
             YQY+DDLYVGSDLEIGQHR KIEELR+HLL+WG TTPDKKHQKEPPFLWMGYELHPDKWT
Sbjct  894   YQYVDDLYVGSDLEIGQHRAKIEELREHLLKWGFTTPDKKHQKEPPFLWMGYELHPDKWT  1073

Query  241   VQPIVLPEK  249
             VQPI LP K
Sbjct  1074  VQPIQLPTK  1100
```

>Mycobacterium tuberculosis strain 32845_S29 549, whole genome shotgun sequence
Sequence ID: NZ_QTBV01000198.1 Length: 825
Range 1: 2 to 823

Score:481 bits(1237), Expect:7e-165,
Method:Compositional matrix adjust.,
Identities:249/274(91%), Positives:262/274(95%), Gaps:0/274(0%)

```
Query  163  SMTKILEPFRKQNPDIVIYQYMDDLYVGSDLEIGQHRTKIEELRQHLLRWGLTTPDKKHQ  222
            SMTKILEPFR +NP+IVIYQYMDDLYVGSDLEIGQHR KIEELR HLL+WG TTPDKKHQ
Sbjct  823  SMTKILEPFRTENPEIVIYQYMDDLYVGSDLEIGQHRVKIEELRGHLLKWGFTTPDKKHQ  644

Query  223  KEPPFLWMGYELHPDKWTVQPIVLPEKDSWTVNDIQKLVGKLNWASQIYPGIKVRQLCKL  282
            KEPPFLWMGYELHPDKWTVQPI LPEKDSWTVNDIQKLVGKLNWASQIYPGIKV+QLC+L
Sbjct  643  KEPPFLWMGYELHPDKWTVQPIQLPEKDSWTVNDIQKLVGKLNWASQIYPGIKVKQLCRL  464

Query  283  LRGTKALTEVIPLTeeaelelaeNREILKEPVHGVYYDPSKDLIAEIQKQGQGQWTYQIY  342
            LRG KALT+++PLTEEAELELAENREILKEPVHGVYYDP+KDLIAEIQKQ  GQWTYQIY
Sbjct  463  LRGAKALTDIVPLTEEAELELAENREILKEPVHGVYYDPAKDLIAEIQKQDHGQWTYQIY  284

Query  343  QEPFKNLKTGKYARMRGAHTNDVKQLTEAVQKITTESIVIWGKTPKFKLPIQKetwetww  402
            QEPFKNLKTGKYA+MR AHTNDVKQLTEAVQKI  ESIV+WGKTPKF+LPIQKETWETWW
Sbjct  283  QEPFKNLKTGKYAKMRTAHTNDVKQLTEAVQKIAMESIVVWGKTPKFRLPIQKETWETWW  104

Query  403  teYWQATWIPEWEFVNTPPLVKLWYQLEKEPIVG  436
            TEYWQATWIPEWEFVNTPPLVKLWYQLEK+PI G
Sbjct  103  TEYWQATWIPEWEFVNTPPLVKLWYQLEKDPIAG  2
```

>Enterococcus durans strain E115_10 NODE_63_length_955_cov_1.30556, whole genome shotgun sequence
Sequence ID: NZ_JAANZI010000063.1 Length: 955
Range 1: 3 to 734

Score:476 bits(1225), Expect:2e-162,
Method:Compositional matrix adjust.,
Identities:227/244(93%), Positives:235/244(96%), Gaps:0/244(0%)

```
Query  1    PISPIETVPVKLKPGMDGPKVKQWPLTEEKIKALVEICTEMEKEGKISKIGPENPYNTPV  60
            PISPIETVPVKLKPGMDGPKVKQWPLTEEKIKAL EIC EMEKEGKI+KIGPENPYNTPV
Sbjct  734  PISPIETVPVKLKPGMDGPKVKQWPLTEEKIKALTEICEEMEKEGKITKIGPENPYNTPV  555

Query  61   FAIKKKDSTKWRKLVDFRELNKRTQDFWEVQLGIPHPAGLKKKKSVTVLDVGDAYFSVPL  120
            FAIKKKDSTKWRKLVDFRELNKRTQDFWEVQLGIPHPAGLK+ KSVTVLDVGDAYFSVPL
Sbjct  554  FAIKKKDSTKWRKLVDFRELNKRTQDFWEVQLGIPHPAGLKQNKSVTVLDVGDAYFSVPL  375

Query  121  DEDFRKYTAFTIPSINNETPGIRYQYNVLPQGWKGSPAIFQSSMTKILEPFRKQNPDIVI  180
            DE FRKYTAFTIPSINNETPGIRYQYNVLPQGWKGSPAIFQ SMTKILEPFR +NP++VI
Sbjct  374  DEGFRKYTAFTIPSINNETPGIRYQYNVLPQGWKGSPAIFQCSMTKILEPFRAKNPELVI  195

Query  181  YQYMDDLYVGSDLEIGQHRTKIEELRQHLLRWGLTTPDKKHQKEPPFLWMGYELHPDKWT  240
            YQY+DDLYVGSDLEIGQHR KIEELR+HLL+WG TTPDKKHQKE PFLWMGYELHPDKWT
Sbjct  194  YQYVDDLYVGSDLEIGQHRAKIEELREHLLKWGFTTPDKKHQKEHPFLWMGYELHPDKWT  15

Query  241  VQPI  244
```

```
          VQPI
Sbjct  14   VQPI  3
```

```
>Enterobacter hormaechei strain AUH-ENM4 NODE_27_length_809_cov_0.64956, whole genome shotgun sequence
Sequence ID: NZ_PYGZ01000027.1 Length: 809
Range 1: 3 to 734

Score:471 bits(1211), Expect:5e-161,
Method:Compositional matrix adjust.,
Identities:224/244(92%), Positives:236/244(96%), Gaps:0/244(0%)

Query  1    PISPIETVPVKLKPGMDGPKVKQWPLTEEKIKALVEICTEMEKEGKISKIGPENPYNTPV  60
            PISPIETVPVKLKPGMDGPKVKQWPLTEEKIKALVEICTEMEKEGKISKIGPENPYNTP+
Sbjct  734  PISPIETVPVKLKPGMDGPKVKQWPLTEEKIKALVEICTEMEKEGKISKIGPENPYNTPI  555

Query  61   FAIKKKDSTKWRKLVDFRELNKRTQDFWEVQLGIPHPAGLKKKKSVTVLDVGDAYFSVPL  120
            FAIKKK+S KWRKLV FRELNKRTQ+FWE+QLGIPHPAGLKKKKSVTVLDVGDAYFSVPL
Sbjct  554  FAIKKKNSDKWRKLVGFRELNKRTQEFWEIQLGIPHPAGLKKKKSVTVLDVGDAYFSVPL  375

Query  121  DEDFRKYTAFTIPSINNETPGIRYQYNVLPQGWKGSPAIFQSSMTKILEPFRKQNPDIVI  180
            D++FRKYTAFTIPS NN+TPGIRYQYNVLPQGWKGSPAIFQSSMTKILEPFRKQNPD+VI
Sbjct  374  DKEFRKYTAFTIPSTNNKTPGIRYQYNVLPQGWKGSPAIFQSSMTKILEPFRKQNPDMVI  195

Query  181  YQYMDDLYVGSDLEIGQHRTKIEELRQHLLRWGLTTPDKKHQKEPPFLWMGYELHPDKWT  240
             QYMDDLYVGSDLEI Q+R K+EELR HLL+WG TTPD+KHQKEPPFLWMGYELHPDKWT
Sbjct  194  CQYMDDLYVGSDLEIEQNRIKVEELRAHLLKWGFTTPDQKHQKEPPFLWMGYELHPDKWT  15

Query  241  VQPI  244
            VQPI
Sbjct  14   VQPI  3
```

```
>Klebsiella oxytoca strain WCHKO085006 NODE_152_length_682_cov_23.205, whole genome shotgun sequence
Sequence ID: NZ_NWES01000143.1 Length: 682
Range 1: 4 to 678

Score:462 bits(1188), Expect:4e-158,
Method:Compositional matrix adjust.,
Identities:221/225(98%), Positives:223/225(99%), Gaps:0/225(0%)

Query  24   WPLTEEKIKALVEICTEMEKEGKISKIGPENPYNTPVFAIKKKDSTKWRKLVDFRELNKR  83
            WPLTEEKIKALVEICTEMEKEGKISKIGPENPYNTPVFAIKKKDSTKWRKLVDFRELNKR
Sbjct  4    WPLTEEKIKALVEICTEMEKEGKISKIGPENPYNTPVFAIKKKDSTKWRKLVDFRELNKR  183

Query  84   TQDFWEVQLGIPHPAGLKKKKSVTVLDVGDAYFSVPLDEDFRKYTAFTIPSINNETPGIR  143
            TQDFWEVQLGIPHPAGLK+KKSVTVLDVGDAYFSVPLD+DFRKYTAFTIPSINNETPGIR
Sbjct  184  TQDFWEVQLGIPHPAGLKQKKSVTVLDVGDAYFSVPLDKDFRKYTAFTIPSINNETPGIR  363

Query  144  YQYNVLPQGWKGSPAIFQSSMTKILEPFRKQNPDIVIYQYMDDLYVGSDLEIGQHRTKIE  203
            YQYNVLPQGWKGSPAIFQ SMTKILEPFRKQNPDIVIYQYMDDLYVGSDLEIGQHRTKIE
Sbjct  364  YQYNVLPQGWKGSPAIFQCSMTKILEPFRKQNPDIVIYQYMDDLYVGSDLEIGQHRTKIE  543

Query  204  ELRQHLLRWGLTTPDKKHQKEPPFLWMGYELHPDKWTVQPIVLPE  248
            ELRQHLLRWG TTPDKKHQKEPPFLWMGYELHPDKWTVQPIVLPE
Sbjct  544  ELRQHLLRWGFTTPDKKHQKEPPFLWMGYELHPDKWTVQPIVLPE  678
```

```
>Escherichia coli strain 8374wB4 NODE_74_length_738_cov_3.41735, whole genome shotgun sequence
Sequence ID: NZ_WSPX01000074.1 Length: 738
Range 1: 6 to 737

Score:458 bits(1178), Expect:3e-156,
Method:Compositional matrix adjust.,
Identities:214/244(88%), Positives:235/244(96%), Gaps:0/244(0%)

Query  1    PISPIETVPVKLKPGMDGPKVKQWPLTEEKIKALVEICTEMEKEGKISKIGPENPYNTPV  60
            PISPIETVPVKLKPGMDGPK+KQWPLT+EKI+AL EICTE+E++GKISKIGPENPYNTP+
Sbjct  6    PISPIETVPVKLKPGMDGPKIKQWPLTKEKIEALTEICTELEQDGKISKIGPENPYNTPI  185

Query  61   FAIKKKDSTKWRKLVDFRELNKRTQDFWEVQLGIPHPAGLKKKKSVTVLDVGDAYFSVPL  120
            FAIKKK+ST WRKL+DFRELNKRTQDFWEVQLGIPHPAGL+KKKSVTVLDVGDAYFS+PL
Sbjct  186  FAIKKKNSTGWRKLMDFRELNKRTQDFWEVQLGIPHPAGLEKKKSVTVLDVGDAYFSIPL  365

Query  121  DEDFRKYTAFTIPSINNETPGIRYQYNVLPQGWKGSPAIFQSSMTKILEPFRKQNPDIVI  180
            DEDFRKYTAFTIPS NNETPG+RYQYNVLPQGWKGSPAIFQSSMT+ILEPFR +NP+IVI
Sbjct  366  DEDFRKYTAFTIPSTNNETPGVRYQYNVLPQGWKGSPAIFQSSMTRILEPFRAKNPEIVI  545

Query  181  YQYMDDLYVGSDLEIGQHRTKIEELRQHLLRWGLTTPDKKHQKEPPFLWMGYELHPDKWT  240
            YQY+DDLYVGSDLEIGQHR KI+ELR++L +WG  TPDKKHQ+ PPFLWMGYELHPDKWT
Sbjct  546  YQYVDDLYVGSDLEIGQHRAKIKELREYLWKWGFYTPDKKHQEGPPFLWMGYELHPDKWT  725

Query  241  VQPI  244
            VQPI
Sbjct  726  VQPI  737
```

```
>Acinetobacter baumannii strain ACM-17 NODE_68_length_757_cov_0.560317, whole genome shotgun sequence
Sequence ID: NZ_PYDD01000068.1 Length: 757
Range 1: 8 to 757

Score:455 bits(1171), Expect:3e-155,
Method:Compositional matrix adjust.,
Identities:226/250(90%), Positives:237/250(94%), Gaps:0/250(0%)

Query  106  VTVLDVGDAYFSVPLDEDFRKYTAFTIPSINNETPGIRYQYNVLPQGWKGSPAIFQSSMT  165
            VTV+DVGDAYFSVPLD DFRKYTAFTIPS NNETPGIRYQYNVLPQGWKGSPAIFQSSMT
Sbjct  757  VTVMDVGDAYFSVPLDPDFRKYTAFTIPSKNNETPGIRYQYNVLPQGWKGSPAIFQSSMT  578

Query  166  KILEPFRKQNPDIVIYQYMDDLYVGSDLEIGQHRTKIEELRQHLLRWGLTTPDKKHQKEP  225
            KILEPFRKQNPDIVI QYMDDLYVGSDLEI QHRTK+EELR +LL+WG +TPDKKHQKEP
Sbjct  577  KILEPFRKQNPDIVICQYMDDLYVGSDLEIEQHRTKVEELRAYLLKWGFSTPDKKHQKEP  398

Query  226  PFLWMGYELHPDKWTVQPIVLPEKDSWTVNDIQKLVGKLNWASQIYPGIKVRQLCKLLRG  285
            PFLWMGYELHPDKWTVQPIVLPEKDSWTVNDIQKLVGKLNWASQIY GIKV+QLCKLLRG
Sbjct  397  PFLWMGYELHPDKWTVQPIVLPEKDSWTVNDIQKLVGKLNWASQIYAGIKVKQLCKLLRG  218

Query  286  TKALTEVIPLTeeaelelaeNREILKEPVHGVYYDPSKDLIAEIQKQGQGQWTYQIYQEP  345
            TKALTEV+PL+EEAELELAENREILKEPVHGVYYDPSKDL+AEIQKQGQGQWTYQIYQEP
Sbjct  217  TKALTEVVPLSEEAELELAENREILKEPVHGVYYDPSKDLVAEIQKQGQGQWTYQIYQEP  38

Query  346  FKNLKTGKYA  355
                ++  YA
Sbjct  37   AHETRSSSYA  8
```

>Acinetobacter baumannii strain ACM-22 NODE_78_length_726_cov_1.48748, whole genome shotgun sequence
Sequence ID: NZ_PYDI01000078.1 Length: 726
Range 1: 1 to 726

Score:449 bits(1156), Expect:5e-153,
Method:Compositional matrix adjust.,
Identities:223/242(92%), Positives:232/242(95%), Gaps:0/242(0%)

```
Query  79   ELNKRTQDFWEVQLGIPHPAGLKKKKSVTVLDVGDAYFSVPLDEDFRKYTAFTIPSINNE  138
            ELNKRTQDFWEVQLGIPHPAGLKKKKS+TVLDVGDAYFSVPLDE FRKYTAFTIPS NNE
Sbjct  1    ELNKRTQDFWEVQLGIPHPAGLKKKKSMTVLDVGDAYFSVPLDEGFRKYTAFTIPSRNNE  180

Query  139  TPGIRYQYNVLPQGWKGSPAIFQSSMTKILEPFRKQNPDIVIYQYMDDLYVGSDLEIGQH  198
            TPGIRYQYNVLPQGWKGSPAIFQ SMTKILEPFR +NP+I+I QYMDDLYVGSDLEIGQH
Sbjct  181  TPGIRYQYNVLPQGWKGSPAIFQCSMTKILEPFRAKNPEIIICQYMDDLYVGSDLEIGQH  360

Query  199  RTKIEELRQHLLRWGLTTPDKKHQKEPPFLWMGYELHPDKWTVQPIVLPEKDSWTVNDIQ  258
            R KIEELR HLL+WG TTPDKKHQKEPPFLWMGYELHPDKWTVQPI LPEKDSWTVNDIQ
Sbjct  361  RAKIEELRDHLLKWGFTTPDKKHQKEPPFLWMGYELHPDKWTVQPIQLPEKDSWTVNDIQ  540

Query  259  KLVGKLNWASQIYPGIKVRQLCKLLRGTKALTEVIPLTeeaelelaeNREILKEPVHGVY  318
            KLVGKLNWASQIYPGIKV+QLCKLLRG KALT+++PLTEEAELELAENREILKEPVHGVY
Sbjct  541  KLVGKLNWASQIYPGIKVKQLCKLLRGAKALTDIVPLTEEAELELAENREILKEPVHGVY  720

Query  319  YD  320
            YD
Sbjct  721  YD  726
```

>Acinetobacter baumannii strain ACM-13 NODE_69_length_701_cov_0.510453, whole genome shotgun sequence
Sequence ID: NZ_PYCZ01000069.1 Length: 701
Range 1: 3 to 701

Score:435 bits(1118), Expect:2e-147,
Method:Compositional matrix adjust.,
Identities:208/233(89%), Positives:219/233(93%), Gaps:0/233(0%)

```
Query  43   KEGKISKIGPENPYNTPVFAIKKKDSTKWRKLVDFRELNKRTQDFWEVQLGIPHPAGLKK  102
            KE K SK GPENPYNTP+FAIKKK+S KWRKLVDFRELNKRTQ+FWEVQLGIPHPAGLKK
Sbjct  3    KEWKTSKSGPENPYNTPIFAIKKKNSDKWRKLVDFRELNKRTQEFWEVQLGIPHPAGLKK  182

Query  103  KKSVTVLDVGDAYFSVPLDEDFRKYTAFTIPSINNETPGIRYQYNVLPQGWKGSPAIFQS  162
            K SVTVL VGDAYFSVP D DFRKYTAF+IPS NN+TPGIRYQYNVLPQGWKGSPAIFQS
Sbjct  183  KISVTVLCVGDAYFSVP*DPDFRKYTAFSIPSRNNKTPGIRYQYNVLPQGWKGSPAIFQS  362

Query  163  SMTKILEPFRKQNPDIVIYQYMDDLYVGSDLEIGQHRTKIEELRQHLLRWGLTTPDKKHQ  222
            SMTKILEPFRKQNPD+VI QYMDDLYVGSDLEI QHR K+EELR HLL+WG +TPD+KHQ
Sbjct  363  SMTKILEPFRKQNPDMVICQYMDDLYVGSDLEIEQHRIKVEELRAHLLKWGFSTPDQKHQ  542

Query  223  KEPPFLWMGYELHPDKWTVQPIVLPEKDSWTVNDIQKLVGKLNWASQIYPGIK  275
            KEPPFLWMGYELHPDKWTVQPIVLPEKDSWTVNDIQKLVGKLNWASQIYPG+K
Sbjct  543  KEPPFLWMGYELHPDKWTVQPIVLPEKDSWTVNDIQKLVGKLNWASQIYPGVK  701
```

>Escherichia coli strain VRES0669, whole genome shotgun sequence
Sequence ID: NZ_UINV01000086.1 Length: 5544
Range 1: 3408 to 4259

Score:474 bits(1221), Expect:4e-146,
Method:Compositional matrix adjust.,
Identities:251/284(88%), Positives:267/284(94%), Gaps:0/284(0%)

```
Query  277   RQLCKLLRGTKALTEVIPLTeeaelelaeNREILKEPVHGVYYDPSKDLIAEIQKQGQGQ  336
             RQLC+LLRGTKALT+++PLTEEAELELAENREILKEPVHGVYYDPSKDLIAEIQKQG  Q
Sbjct  4259  RQLCRLLRGTKALTDIVPLTEEAELELAENREILKEPVHGVYYDPSKDLIAEIQKQGNDQ  4080

Query  337   WTYQIYQEPFKNLKTGKYARMRGAHTNDVKQLTEAVQKITTESIVIWGKTPKFKLPIQKe  396
             WTYQIYQEPFKNLKTGKYARMR AHTNDVKQLTEAVQKI  ESIVIWGK PKF+LPIQKE
Sbjct  4079  WTYQIYQEPFKNLKTGKYARMRTAHTNDVKQLTEAVQKIVMESIVIWGKPPKFRLPIQKE  3900

Query  397   twetwwteYWQATWIPEWEFVNTPPLVKLWYQLEKEPIVGAETFYVDGAANRETKLGKAG  456
             TWETWW++YWQATWIPEWEFVNTPPLVKLWYQLEKEPI G ETFYVDGAANRETK GKAG
Sbjct  3899  TWETWWSDYWQATWIPEWEFVNTPPLVKLWYQLEKEPIEGVETFYVDGAANRETKAGKAG  3720

Query  457   YVTNRGRQKVVTLTDTTNQKTELQAIYLALQDSGLEVNIVTDSQYALGIIQAQPDQSESE  516
             YVT+RGRQKV++ T+TTNQK ELQAI LALQDSG EVNIVTDSQYALGII A PD+SESE
Sbjct  3719  YVTDRGRQKVISFTETTNQKAELQAIQLALQDSGSEVNIVTDSQYALGIIHALPDKSESE  3540

Query  517   LVNQIIEQLIKKEKVYLAWVPAHKGIGGNEQVDKLVSAGIRKVL  560
             LVN+IIE LI+KE+VYL+WVPAHKGIGGNEQVDKLVS GIRKVL
Sbjct  3539  LVNKIIELLIQKERVYLSWVPAHKGIGGNEQVDKLVSTGIRKVL  3408
```

Range 2: 4301 to 4876

Score:379 bits(973), Expect:1e-111,
Method:Compositional matrix adjust.,
Identities:181/192(94%), Positives:188/192(97%), Gaps:0/192(0%)

```
Query  1     PISPIETVPVKLKPGMDGPKVKQWPLTEEKIKALVEICTEMEKEGKISKIGPENPYNTPV  60
             PISPIETVPVKLKPGMDGPKVKQWPLTEEKIKAL EIC EMEKEGKI+KIGP+NPYNTPV
Sbjct  4876  PISPIETVPVKLKPGMDGPKVKQWPLTEEKIKALTEICEEMEKEGKITKIGPDNPYNTPV  4697

Query  61    FAIKKKDSTKWRKLVDFRELNKRTQDFWEVQLGIPHPAGLKKKKSVTVLDVGDAYFSVPL  120
             FAIKKKDSTKWRKL+DFRELNKRTQDFWEVQLGIPHPAGLKKKKSVTVLDVGDAYFSVPL
Sbjct  4696  FAIKKKDSTKWRKLIDFRELNKRTQDFWEVQLGIPHPAGLKKKKSVTVLDVGDAYFSVPL  4517

Query  121   DEDFRKYTAFTIPSINNETPGIRYQYNVLPQGWKGSPAIFQSSMTKILEPFRKQNPDIVI  180
             DE+FRKYTAFTIPS NNETPGIRYQYNVLPQGWKGSPAIFQSSMT+ILEPFR QNP+IVI
Sbjct  4516  DENFRKYTAFTIPSTNNETPGIRYQYNVLPQGWKGSPAIFQSSMTRILEPFRAQNPEIVI  4337

Query  181   YQYMDDLYVGSD  192
             YQY+DDLYVGSD
Sbjct  4336  YQYVDDLYVGSD  4301
```

>Acinetobacter baumannii strain ACM-41 NODE_63_length_718_cov_0.852792, whole genome shotgun sequence
Sequence ID: NZ_QAGT01000063.1 Length: 718
Range 1: 2 to 682

Score:427 bits(1097), Expect:4e-144,
Method:Compositional matrix adjust.,
Identities:204/227(90%), Positives:213/227(93%), Gaps:0/227(0%)

```
Query  1    PISPIETVPVKLKPGMDGPKVKQWPLTEEKIKALVEICTEMEKEGKISKIGPENPYNTPV  60
            PISPIETVPVKLKPGMDGPKVKQWPL+EEKIKAL EIC EMEKEGKISKIGPENPYNTPV
Sbjct  682  PISPIETVPVKLKPGMDGPKVKQWPLSEEKIKALTEICEEMEKEGKISKIGPENPYNTPV  503

Query  61   FAIKKKDSTKWRKLVDFRELNKRTQDFWEVQLGIPHPAGLKKKKSVTVLDVGDAYFSVPL  120
            FAIK++DSTKWRKLVDFRELNKRTQDFWEVQLGIPHPAGLKKKKS+TVLDVGDAYFSVPL
Sbjct  502  FAIKRRDSTKWRKLVDFRELNKRTQDFWEVQLGIPHPAGLKKKKSMTVLDVGDAYFSVPL  323

Query  121  DEDFRKYTAFTIPSINNETPGIRYQYNVLPQGWKGSPAIFQSSMTKILEPFRKQNPDIVI  180
            DE FRKYTAFTIPS NNETPGIR QYNVLPQGWKGSPAIF+ SMTK LEP R +NP+I+I
Sbjct  322  DEGFRKYTAFTIPSRNNETPGIRKQYNVLPQGWKGSPAIFKYSMTKTLEPIRAKNPEIII  143

Query  181  YQYMDDLYVGSDLEIGQHRTKIEELRQHLLRWGLTTPDKKHQKEPPF  227
             QYMDDLYVGSDLEIGQHR KIEELR HLL+WG TTPDKKHQKEPP
Sbjct  142  CQYMDDLYVGSDLEIGQHRAKIEELRDHLLKWGFTTPDKKHQKEPPL  2



>Escherichia coli strain ECM-2 NODE_127_length_636_cov_2.0668, whole genome shotgun sequence
Sequence ID: NZ_PXIA01000127.1 Length: 636
Range 1: 3 to 635

Score:417 bits(1072), Expect:8e-141,
Method:Compositional matrix adjust.,
Identities:201/211(95%), Positives:206/211(97%), Gaps:0/211(0%)

Query  4    PIETVPVKLKPGMDGPKVKQWPLTEEKIKALVEICTEMEKEGKISKIGPENPYNTPVFAI  63
            PIETVPVKLKPGMDGPKVKQWPLTEEKIKALVEICTEMEKEGKISKIGPENPYNTP+FAI
Sbjct  3    PIETVPVKLKPGMDGPKVKQWPLTEEKIKALVEICTEMEKEGKISKIGPENPYNTPIFAI  182

Query  64   KKKDSTKWRKLVDFRELNKRTQDFWEVQLGIPHPAGLKKKKSVTVLDVGDAYFSVPLDED  123
            KKK+S KWRKLVDFRELNKRTQ+FWEVQLGIPHPAGLKKKKSVTVLDVGDAYFSVPLD D
Sbjct  183  KKKNSDKWRKLVDFRELNKRTQEFWEVQLGIPHPAGLKKKKSVTVLDVGDAYFSVPLDPD  362

Query  124  FRKYTAFTIPSINNETPGIRYQYNVLPQGWKGSPAIFQSSMTKILEPFRKQNPDIVIYQY  183
            FRKYTAFTIPS NNETPGIRYQYNVLPQGWKGSPAIFQSSMTKILEPFRKQNPDIVIYQY
Sbjct  363  FRKYTAFTIPSRNNETPGIRYQYNVLPQGWKGSPAIFQSSMTKILEPFRKQNPDIVIYQY  542

Query  184  MDDLYVGSDLEIGQHRTKIEELRQHLLRWGL  214
            MDDLYVGSDLEI QHRTK+EELRQHLL+WG
Sbjct  543  MDDLYVGSDLEIEQHRTKVEELRQHLLKWGF  635



>Acinetobacter baumannii strain ACM-14 NODE_75_length_665_cov_0.98513, whole genome shotgun sequence
Sequence ID: NZ_PYDA01000075.1 Length: 665
Range 1: 1 to 657

Score:412 bits(1059), Expect:1e-138,
Method:Compositional matrix adjust.,
Identities:207/219(95%), Positives:213/219(97%), Gaps:0/219(0%)

Query  114  AYFSVPLDEDFRKYTAFTIPSINNETPGIRYQYNVLPQGWKGSPAIFQSSMTKILEPFRK  173
            AYFSVPLD DFRKYTAFTIPS NNETPGIRYQYNVLPQGWKGSPAIFQSSMTKILEPFRK
Sbjct  1    AYFSVPLDPDFRKYTAFTIPSRNNETPGIRYQYNVLPQGWKGSPAIFQSSMTKILEPFRK  180

Query  174  QNPDIVIYQYMDDLYVGSDLEIGQHRTKIEELRQHLLRWGLTTPDKKHQKEPPFLWMGYE  233
            QNPDIVIYQY+DDLYVGSDLEI QHRTK+EELRQHLL WG TTPDKKHQKEPPFLWMGYE
Sbjct  181  QNPDIVIYQYVDDLYVGSDLEIEQHRTKVEELRQHLLTWGFTTPDKKHQKEPPFLWMGYE  360

Query  234  LHPDKWTVQPIVLPEKDSWTVNDIQKLVGKLNWASQIYPGIKVRQLCKLLRGTKALTEVI  293
            LHPDKWTVQPIVLPEKDSWTVNDIQKLVGKLNWASQIY GIKV+QLCKLLRGTKALTEV+
Sbjct  361  LHPDKWTVQPIVLPEKDSWTVNDIQKLVGKLNWASQIYAGIKVKQLCKLLRGTKALTEVV  540

Query  294  PLTeeaelelaeNREILKEPVHGVYYDPSKDLIAEIQKQ  332
            PL+EEAELELAENREILKEPVHGVYYDPSKDL+AEIQKQ
Sbjct  541  PLSEEAELELAENREILKEPVHGVYYDPSKDLVAEIQKQ  657



>Acinetobacter baumannii strain ACM-11 NODE_65_length_706_cov_0.799655, whole genome shotgun sequence
Sequence ID: NZ_PYCW01000065.1 Length: 706
Range 1: 2 to 706

Score:410 bits(1054), Expect:1e-137,
Method:Compositional matrix adjust.,
Identities:206/235(88%), Positives:221/235(94%), Gaps:0/235(0%)

Query  90   VQLGIPHPAGLKKKKSVTVLDVGDAYFSVPLDEDFRKYTAFTIPSINNETPGIRYQYNVL  149
            V+LG      LK K S+++LDVGDAYFSVPLDE+FRKYTAFTIPS+NNETPGIRYQYNVL
Sbjct  2    VKLGKLQQPRLKHK*SLSILDVGDAYFSVPLDENFRKYTAFTIPSVNNETPGIRYQYNVL  181

Query  150  PQGWKGSPAIFQSSMTKILEPFRKQNPDIVIYQYMDDLYVGSDLEIGQHRTKIEELRQHL  209
            PQGWKGSPAIFQ+SMTKILEPFR +NPDIVIYQYMDDLYVGSDLEIGQHR KIEELR+HL
Sbjct  182  PQGWKGSPAIFQASMTKILEPFRLKNPDIVIYQYMDDLYVGSDLEIGQHRAKIEELREHL  361

Query  210  LRWGLTTPDKKHQKEPPFLWMGYELHPDKWTVQPIVLPEKDSWTVNDIQKLVGKLNWASQ  269
            L+WGLTTPDKKHQKEPPFLWMGYELHPDKWTVQPI LPEKDSWTVNDIQKLVGKL WASQ
Sbjct  362  LKWGLTTPDKKHQKEPPFLWMGYELHPDKWTVQPIQLPEKDSWTVNDIQKLVGKLIWASQ  541

Query  270  IYPGIKVRQLCKLLRGTKALTEVIPLTeeaelelaeNREILKEPVHGVYYDPSKD  324
            IY GIKV+QLCKLLRG KALT+VIPLTEEAELELAENREIL+EPVHGVYYDP++D
Sbjct  542  IYAGIKVKQLCKLLRGAKALTDVIPLTEEAELELAENREILREPVHGVYYDPTRD  706



>Providencia rettgeri strain APW139_S1 24_contig_65, whole genome shotgun sequence
Sequence ID: NZ_SHDG01000065.1 Length: 941
Range 1: 303 to 941

Score:410 bits(1053), Expect:2e-136,
Method:Compositional matrix adjust.,
Identities:194/213(91%), Positives:205/213(96%), Gaps:0/213(0%)

Query  1    PISPIETVPVKLKPGMDGPKVKQWPLTEEKIKALVEICTEMEKEGKISKIGPENPYNTPV  60
            PIS IETVPVKLKPGMDGP+VKQWPLTEEKIKAL EIC EMEKEGKI+KIGPENPYNTPV
Sbjct  303  PISSIETVPVKLKPGMDGPRVKQWPLTEEKIKALTEICEEMEKEGKITKIGPENPYNTPV  482

Query  61   FAIKKKDSTKWRKLVDFRELNKRTQDFWEVQLGIPHPAGLKKKKSVTVLDVGDAYFSVPL  120
            FAIKKKDSTKWRK+VDFRELNKRTQDFWEVQLGIPHPAG+KK KSVTVLDVGDA+FSVPL
Sbjct  483  FAIKKKDSTKWRKVVDFRELNKRTQDFWEVQLGIPHPAGIKKNKSVTVLDVGDAFFSVPL  662

Query  121  DEDFRKYTAFTIPSINNETPGIRYQYNVLPQGWKGSPAIFQSSMTKILEPFRKQNPDIVI  180
            DE FRKYTAFTIPSINNE PGIRYQYNVLPQGWKGSPAIFQSSMT+ILEPFR +NP+IVI
Sbjct  663  DESFRKYTAFTIPSINNEAPGIRYQYNVLPQGWKGSPAIFQSSMTRILEPFRAKNPEIVI  842

Query  181  YQYMDDLYVGSDLEIGQHRTKIEELRQHLLRWG  213
            YQY+DDLYVGSDLEIGQHR KIEELR+HLL+WG
Sbjct  843  YQYVDDLYVGSDLEIGQHRAKIEELREHLLKWG  941
```

```
>Providencia rettgeri strain AFE97_S1 21_contig_71, whole genome shotgun sequence
Sequence ID: NZ_SHDO01000071.1 Length: 1157
Range 1: 364 to 804

Score:279 bits(714), Expect:5e-136,
Method:Compositional matrix adjust.,
Identities:133/147(90%), Positives:139/147(94%), Gaps:0/147(0%)

Query  1    PISPIETVPVKLKPGMDGPKVKQWPLTEEKIKALVEICTEMEKEGKISKIGPENPYNTPV  60
            PISPIETVPVKLKPGMDGPKVKQWPLTEEKIKAL  IC EMEKEGKI+KIGPENPYNTP+
Sbjct  364  PISPIETVPVKLKPGMDGPKVKQWPLTEEKIKALTAICEEMEKEGKITKIGPENPYNTPI  543

Query  61   FAIKKKDSTKWRKLVDFRELNKRTQDFWEVQLGIPHPAGLKKKKSVTVLDVGDAYFSVPL  120
            FAIKKKDSTKWRKLVDFRELNKRTQDFWEVQLGIPHPAGLK+ KS+TVLDVGDAYFSVPL
Sbjct  544  FAIKKKDSTKWRKLVDFRELNKRTQDFWEVQLGIPHPAGLKRNKSMTVLDVGDAYFSVPL  723

Query  121  DEDFRKYTAFTIPSINNETPGIRYQYN  147
            DE FRKYTAFTIPSINNETPGI +  +
Sbjct  724  DEGFRKYTAFTIPSINNETPGISFHRD  804



Range 2: 792 to 1157

Score:240 bits(612), Expect:5e-136,
Method:Compositional matrix adjust.,
Identities:113/122(93%), Positives:116/122(95%), Gaps:0/122(0%)

Query  149   LPQGWKGSPAIFQSSMTKILEPFRKQNPDIVIYQYMDDLYVGSDLEIGQHRTKIEELRQH  208
             LPQGWKGSPAIFQSSMTKILEPFR QNP++VIYQYMDDLYVGSDLEIGQHR KIEELR H
Sbjct  792   LPQGWKGSPAIFQSSMTKILEPFRTQNPEMVIYQYMDDLYVGSDLEIGQHRAKIEELRAH  971

Query  209   LLRWGLTTPDKKHQKEPPFLWMGYELHPDKWTVQPIVLPEKDSWTVNDIQKLVGKLNWAS  268
             LL+WG TTPDKKHQKEPPFLWMGYELHPDKWTVQPI LPEKDSWTVNDIQKL GKLNWAS
Sbjct  972   LLKWGFTTPDKKHQKEPPFLWMGYELHPDKWTVQPIKLPEKDSWTVNDIQKLGGKLNWAS  1151

Query  269   QI  270
             QI
Sbjct  1152  QI  1157



>Mycobacterium tuberculosis strain 32847_S31 312, whole genome shotgun sequence
Sequence ID: NZ_QTBX01000084.1 Length: 704
Range 1: 3 to 704

Score:404 bits(1039), Expect:2e-135,
Method:Compositional matrix adjust.,
Identities:213/234(91%), Positives:224/234(95%), Gaps:0/234(0%)

Query  202  IEELRQHLLRWGLTTPDKKHQKEPPFLWMGYELHPDKWTVQPIVLPEKDSWTVNDIQKLV  261
            IEELR HLL+WG TTPDKKHQKEPPFLWMGYELHPDKWTVQPI LPEKDSWTVNDIQKLV
Sbjct  704  IEELRGHLLKWGFTTPDKKHQKEPPFLWMGYELHPDKWTVQPIQLPEKDSWTVNDIQKLV  525

Query  262  GKLNWASQIYPGIKVRQLCKLLRGTKALTEVIPLTeeaelelaeNREILKEPVHGVYYDP  321
            GKLNWASQIYPGIKV+QLC+LLRG KALT+++PLTEEAELELAENREILKEPVHGVYYDP
Sbjct  524  GKLNWASQIYPGIKVKQLCRLLRGAKALTDIVPLTEEAELELAENREILKEPVHGVYYDP  345

Query  322  SKDLIAEIQKQGQGQWTYQIYQEPFKNLKTGKYARMRGAHTNDVKQLTEAVQKITTESIV  381
            +KDLIAEIQKQ  GQWTYQIYQEPFKNLKTGKYA+MR AHTNDVKQLTEAVQKI  ESIV
Sbjct  344  AKDLIAEIQKQDHGQWTYQIYQEPFKNLKTGKYAKMRTAHTNDVKQLTEAVQKIAMESIV  165

Query  382  IWGKTPKFKLPIQKetwetwwteYWQATWIPEWEFVNTPPLVKLWYQLEKEPIV  435
            +WGKTPKF+LPIQKETWETWWTEYWQATWIPEWEFVNTPPLVKLWYQLEK+PI
Sbjct  164  VWGKTPKFRLPIQKETWETWWTEYWQATWIPEWEFVNTPPLVKLWYQLEKDPIA  3



>Acinetobacter baumannii strain ACM-29 NODE_109_length_888_cov_0.827858, whole genome shotgun sequence
Sequence ID: NZ_PYDO01000109.1 Length: 888
Range 1: 2 to 625

Score:402 bits(1033), Expect:1e-133,
Method:Compositional matrix adjust.,
Identities:193/208(93%), Positives:202/208(97%), Gaps:0/208(0%)

Query  1    PISPIETVPVKLKPGMDGPKVKQWPLTEEKIKALVEICTEMEKEGKISKIGPENPYNTPV  60
            PISPIETVPVKLKPGMDGPKVKQWPLTEEKIKALVEICTEMEKEGKI+KIGPENPYNTP+
Sbjct  625  PISPIETVPVKLKPGMDGPKVKQWPLTEEKIKALVEICTEMEKEGKITKIGPENPYNTPI  446

Query  61   FAIKKKDSTKWRKLVDFRELNKRTQDFWEVQLGIPHPAGLKKKKSVTVLDVGDAYFSVPL  120
            FAIKKK+S KWRKLVDFRELNKRTQ+FWEVQLGIPHPAGLKKKKSVTVLDVGDAYFSVPL
Sbjct  445  FAIKKKNSDKWRKLVDFRELNKRTQEFWEVQLGIPHPAGLKKKKSVTVLDVGDAYFSVPL  266

Query  121  DEDFRKYTAFTIPSINNETPGIRYQYNVLPQGWKGSPAIFQSSMTKILEPFRKQNPDIVI  180
            D DFRKYTAFT PS NNETPGIRYQYNVLPQGWKGSPAIFQSSMTKILEPFRKQNPD+VI
Sbjct  265  DPDFRKYTAFTTPSRNNETPGIRYQYNVLPQGWKGSPAIFQSSMTKILEPFRKQNPDMVI  86

Query  181  YQYMDDLYVGSDLEIGQHRTKIEELRQH  208
            YQY+DDLYVGSDLE+ QHRTK+EELR +
Sbjct  85   YQYIDDLYVGSDLELEQHRTKVEELRAY  2



>Acinetobacter baumannii strain ACM-25 NODE_71_length_740_cov_1.2447, whole genome shotgun sequence
Sequence ID: NZ_PXYP01000071.1 Length: 655
Range 1: 1 to 654

Score:397 bits(1021), Expect:6e-133,
Method:Compositional matrix adjust.,
Identities:198/218(91%), Positives:206/218(94%), Gaps:0/218(0%)

Query  126  KYTAFTIPSINNETPGIRYQYNVLPQGWKGSPAIFQSSMTKILEPFRKQNPDIVIYQYMD  185
            KYTAFTIPS NNETPGIRYQYNVLPQGWKGSPAIFQ SMTKILEPFR +NP+I+I QYMD
Sbjct  654  KYTAFTIPSRNNETPGIRYQYNVLPQGWKGSPAIFQCSMTKILEPFRAKNPEIIICQYMD  475

Query  186  DLYVGSDLEIGQHRTKIEELRQHLLRWGLTTPDKKHQKEPPFLWMGYELHPDKWTVQPIV  245
            DLYVGSDLEIGQHR KIEELR HLL+WG TTPDKKHQKEPPFLWMGYELHPDKWTVQPI
Sbjct  474  DLYVGSDLEIGQHRAKIEELRDHLLKWGFTTPDKKHQKEPPFLWMGYELHPDKWTVQPIQ  295

Query  246  LPEKDSWTVNDIQKLVGKLNWASQIYPGIKVRQLCKLLRGTKALTEVIPLTeeaelelae  305
            LPEKDSWTVNDIQKLVGKLNWASQIYPGIKV+QLCKLLRG KALT+++PLTEEAELELAE
Sbjct  294  LPEKDSWTVNDIQKLVGKLNWASQIYPGIKVKQLCKLLRGAKALTDIVPLTEEAELELAE  115

Query  306  NREILKEPVHGVYYDPSKDLIAEIQKQGQGQWTYQIYQ  343
            NREILKEPVHG YYDPSKDLIAEIQKQG  QWTYQIYQ
Sbjct  114  NREILKEPVHGAYYDPSKDLIAEIQKQGNDQWTYQIYQ  1
```

```
>Levilactobacillus brevis strain CRL 2013 scaffold15, whole genome shotgun sequence
Sequence ID: NZ_MZMW01000019.1 Length: 566
Range 1: 3 to 566

Score:391 bits(1005), Expect:5e-131,
Method:Compositional matrix adjust.,
Identities:186/188(99%), Positives:187/188(99%), Gaps:0/188(0%)

Query  54   NPYNTPVFAIKKKDSTKWRKLVDFRELNKRTQDFWEVQLGIPHPAGLKKKKSVTVLDVGD  113
            NPYNTPVFAIKKKDSTKWRKLVDFRELNKRTQDFWEVQLGIPHPAGLKKKKSVTVLDVGD
Sbjct  566  NPYNTPVFAIKKKDSTKWRKLVDFRELNKRTQDFWEVQLGIPHPAGLKKKKSVTVLDVGD  387

Query  114  AYFSVPLDEDFRKYTAFTIPSINNETPGIRYQYNVLPQGWKGSPAIFQSSMTKILEPFRK  173
            AYFSVPLD+DFRKYTAFTIPSINNETPGIRYQYNVLPQGWKGSPAIFQSSMTKILEPFRK
Sbjct  386  AYFSVPLDKDFRKYTAFTIPSINNETPGIRYQYNVLPQGWKGSPAIFQSSMTKILEPFRK  207

Query  174  QNPDIVIYQYMDDLYVGSDLEIGQHRTKIEELRQHLLRWGLTTPDKKHQKEPPFLWMGYE  233
            QNPDIVIYQYMDDLYVGSDLEIGQHRTKIEELRQHLLRWG TTPDKKHQKEPPFLWMGYE
Sbjct  206  QNPDIVIYQYMDDLYVGSDLEIGQHRTKIEELRQHLLRWGFTTPDKKHQKEPPFLWMGYE  27

Query  234  LHPDKWTV  241
            LHPDKWTV
Sbjct  26   LHPDKWTV  3



>Pseudomonas aeruginosa strain ENV-297 NODE_95_length_638_cov_6.84848_ID_189, whole genome shotgun sequence
Sequence ID: NZ_RADD01000130.1 Length: 638
Range 1: 2 to 589

Score:390 bits(1001), Expect:5e-130,
Method:Compositional matrix adjust.,
Identities:183/196(93%), Positives:191/196(97%), Gaps:0/196(0%)

Query  61   FAIKKKDSTKWRKLVDFRELNKRTQDFWEVQLGIPHPAGLKKKKSVTVLDVGDAYFSVPL  120
            FAIKKKDSTKWRKLVDFRELNKRTQDFWEVQLGIPHPAGLKKKKSVT+LDVGDAYFSVPL
Sbjct  2    FAIKKKDSTKWRKLVDFRELNKRTQDFWEVQLGIPHPAGLKKKKSVTILDVGDAYFSVPL  181

Query  121  DEDFRKYTAFTIPSINNETPGIRYQYNVLPQGWKGSPAIFQSSMTKILEPFRKQNPDIVI  180
            D DFRKYTAFTIPSINNETPGIRYQYNVLPQGWKGSPAIFQ SMTKILEPFRKQNP++VI
Sbjct  182  DPDFRKYTAFTIPSINNETPGIRYQYNVLPQGWKGSPAIFQCSMTKILEPFRKQNPELVI  361

Query  181  YQYMDDLYVGSDLEIGQHRTKIEELRQHLLRWGLTTPDKKHQKEPPFLWMGYELHPDKWT  240
            YQYMDDLY+GSDLEIGQHR KIEELR+HLL+WG TTPDKKHQKEPPFLWMGYELHPDKWT
Sbjct  362  YQYMDDLYIGSDLEIGQHRAKIEELREHLLKWGFTTPDKKHQKEPPFLWMGYELHPDKWT  541

Query  241  VQPIVLPEKDSWTVND  256
            VQPI LP+K+SWTVND
Sbjct  542  VQPIELPDKESWTVND  589



>Providencia rettgeri strain AFE422_S1 40_contig_34, whole genome shotgun sequence
Sequence ID: NZ_SHDL01000034.1 Length: 1058
Range 1: 303 to 743

Score:278 bits(712), Expect:6e-129,
Method:Compositional matrix adjust.,
Identities:133/147(90%), Positives:139/147(94%), Gaps:0/147(0%)

Query  1    PISPIETVPVKLKPGMDGPKVKQWPLTEEKIKALVEICTEMEKEGKISKIGPENPYNTPV  60
            PISPIETVPVKLKPGMDGPKVKQWPLTEEKIKAL  IC EMEKEGKI+KIGPENPYNTP+
Sbjct  303  PISPIETVPVKLKPGMDGPKVKQWPLTEEKIKALTAICEEMEKEGKITKIGPENPYNTPI  482

Query  61   FAIKKKDSTKWRKLVDFRELNKRTQDFWEVQLGIPHPAGLKKKKSVTVLDVGDAYFSVPL  120
            FAIKKKDSTKWRKLVDFRELNKRTQDFWEVQLGIPHPAGLK+ KS+TVLDVGDAYFSVPL
Sbjct  483  FAIKKKDSTKWRKLVDFRELNKRTQDFWEVQLGIPHPAGLKRNKSMTVLDVGDAYFSVPL  662

Query  121  DEDFRKYTAFTIPSINNETPGIRYQYN  147
            DE FRKYTAFTIPSINNETPGI +  +
Sbjct  663  DEGFRKYTAFTIPSINNETPGISFHRD  743



Range 2: 731 to 1057

Score:217 bits(553), Expect:6e-129,
Method:Compositional matrix adjust.,
Identities:101/109(93%), Positives:104/109(95%), Gaps:0/109(0%)

Query  149   LPQGWKGSPAIFQSSMTKILEPFRKQNPDIVIYQYMDDLYVGSDLEIGQHRTKIEELRQH  208
             LPQGWKGSPAIFQSSMTKILEPFR QNP++VIYQYMDDLYVGSDLEIGQHR KIEELR H
Sbjct  731   LPQGWKGSPAIFQSSMTKILEPFRTQNPEMVIYQYMDDLYVGSDLEIGQHRAKIEELRAH  910

Query  209   LLRWGLTTPDKKHQKEPPFLWMGYELHPDKWTVQPIVLPEKDSWTVNDI  257
             LL+WG TTPDKKHQKEPPFLWMGYELHPDKWTVQPI LPEKDSWTVNDI
Sbjct  911   LLKWGFTTPDKKHQKEPPFLWMGYELHPDKWTVQPIKLPEKDSWTVNDI  1057



>Providencia rettgeri strain AMW697_S1 18_contig_57, whole genome shotgun sequence
Sequence ID: NZ_SHDI01000057.1 Length: 585
Range 1: 2 to 583

Score:382 bits(980), Expect:4e-127,
Method:Compositional matrix adjust.,
Identities:181/194(93%), Positives:186/194(95%), Gaps:0/194(0%)

Query  23   QWPLTEEKIKALVEICTEMEKEGKISKIGPENPYNTPVFAIKKKDSTKWRKLVDFRELNK  82
            QWPLTEEKIKAL EIC EMEKEGKI+KIGPENPYNTP+FAIKKKDSTKWRKLVDFRELNK
Sbjct  2    QWPLTEEKIKALTEICEEMEKEGKITKIGPENPYNTPIFAIKKKDSTKWRKLVDFRELNK  181

Query  83   RTQDFWEVQLGIPHPAGLKKKKSVTVLDVGDAYFSVPLDEDFRKYTAFTIPSINNETPGI  142
            RTQDFWEVQLGIPHPAGLKK KSVTVLDVGDAYFSVPLDE FRKYTAFTIPSINNETPGI
Sbjct  182  RTQDFWEVQLGIPHPAGLKKNKSVTVLDVGDAYFSVPLDEGFRKYTAFTIPSINNETPGI  361

Query  143  RYQYNVLPQGWKGSPAIFQSSMTKILEPFRKQNPDIVIYQYMDDLYVGSDLEIGQHRTKI  202
            RYQYNVLPQGWKGSPAIFQSSMTKILEPFR QNP++VIYQYMDDLYVGSDLEIGQHR KI
Sbjct  362  RYQYNVLPQGWKGSPAIFQSSMTKILEPFRAQNPELVIYQYMDDLYVGSDLEIGQHRAKI  541

Query  203  EELRQHLLRWGLTT  216
            EELR HLL+WG TT
Sbjct  542  EELRNHLLKWGFTT  583
```

```
>Levilactobacillus brevis strain CRL 2013 scaffold16, whole genome shotgun sequence
Sequence ID: NZ_MZMW01000020.1 Length: 547
Range 1: 2 to 547

Score:372 bits(956), Expect:1e-123,
Method:Compositional matrix adjust.,
Identities:177/182(97%), Positives:179/182(98%), Gaps:0/182(0%)

Query  63    IKKDSTKWRKLVDFRELNKRTQDFWEVQLGIPHPAGLKKKKSVTVLDVGDAYFSVPLDE  122
             IKKDSTKWRKLVDFRELNKRTQDFWEVQLGIPHPAGLKKKK+VTVLDVGDAYFSVPLD+
Sbjct  547   IKKDSTKWRKLVDFRELNKRTQDFWEVQLGIPHPAGLKKKKAVTVLDVGDAYFSVPLDK  368

Query  123   DFRKYTAFTIPSINNETPGIRYQYNVLPQGWKGSPAIFQSSMTKILEPFRKQNPDIVIYQ  182
             DFRKYTAFTIPSINNETPGIRYQYNVLPQGWKGSPAIFQSSMTKILEPFRKQNPDIVIYQ
Sbjct  367   DFRKYTAFTIPSINNETPGIRYQYNVLPQGWKGSPAIFQSSMTKILEPFRKQNPDIVIYQ  188

Query  183   YMDDLYVGSDLEIGQHRTKIEELRQHLLRWGLTTPDKKHQKEPPFLWMGYELHPDKWTVQ  242
             YMDDLYVGSDLEIGQHR KIEELRQHLL WG TTPDKKHQKEPPFLWMGYELHPDKWTVQ
Sbjct  187   YMDDLYVGSDLEIGQHRAKIEELRQHLLEWGFTTPDKKHQKEPPFLWMGYELHPDKWTVQ  8

Query  243   PI  244
             PI
Sbjct  7     PI  2



>Providencia rettgeri strain AFE97_S1 21_contig_72, whole genome shotgun sequence
Sequence ID: NZ_SHDO01000072.1 Length: 854
Range 1: 291 to 854

Score:369 bits(947), Expect:9e-121,
Method:Compositional matrix adjust.,
Identities:176/188(94%), Positives:184/188(97%), Gaps:0/188(0%)

Query  3     SPIETVPVKLKPGMDGPKVKQWPLTEEKIKALVEICTEMEKEGKISKIGPENPYNTPVFA  62
             SPI+TVPVKLKPGMDGPKVKQWPLTEEKIKAL EIC EMEKEGKI+KIGPENPYNTP+FA
Sbjct  291   SPIKTVPVKLKPGMDGPKVKQWPLTEEKIKALTEICDEMEKEGKITKIGPENPYNTPIFA  470

Query  63    IKKKDSTKWRKLVDFRELNKRTQDFWEVQLGIPHPAGLKKKKSVTVLDVGDAYFSVPLDE  122
             IK+KDSTKWRKLVDFRELNKRTQDFWEVQLGIPHPAGLKKKKS+TVLDVGDAYFSVPLDE
Sbjct  471   IKRKDSTKWRKLVDFRELNKRTQDFWEVQLGIPHPAGLKKKKSMTVLDVGDAYFSVPLDE  650

Query  123   DFRKYTAFTIPSINNETPGIRYQYNVLPQGWKGSPAIFQSSMTKILEPFRKQNPDIVIYQ  182
              FRKYTAFTIPSINNETPGIRYQYNVLPQGWKGSPAIFQSSMTKILEPFR +NP+IVIYQ
Sbjct  651   SFRKYTAFTIPSINNETPGIRYQYNVLPQGWKGSPAIFQSSMTKILEPFRAKNPEIVIYQ  830

Query  183   YMDDLYVG  190
             Y+DDLYVG
Sbjct  831   YVDDLYVG  854



>Cuspidothrix issatschenkoi CHARLIE-1 contig_175, whole genome shotgun sequence
Sequence ID: NZ_PGEM01000173.1 Length: 2237
Range 1: 286 to 1899

Score:385 bits(988), Expect:1e-120,
Method:Compositional matrix adjust.,
Identities:233/543(43%), Positives:321/543(59%), Gaps:7/543(1%)

Query  10    VKLKPGMDGPKVKQWPLTEEKIKALVEICTEMEKEGKISKIGPENPYNTPVFAIKKKDST  69
             ++LK G  GPK+ QWPLT+EK++   EI   +  EGKIS+    NPYN+P+F IKK+ S
Sbjct  286   IELKEGTMGPKIPQWPLTKEKLEGAKEIVQRLLSEGKISEASDNNPYNSPIFVIKKR-SG  462

Query  70    KWRKLVDFRELNKRTQDFWEVQLGIPHPAGLKKKKSVTVLDVGDAYFSVPLDEDFRKYTA  129
             KWR L D RELNK  Q   E+  G+PHP GL K K +TVLD+GDAYF++PLD +FR YTA
Sbjct  463   KWRLLQDLRELNKTVQVGTEISRGLPHPGGLIKCKHMTVLDIGDAYFTIPLDPEFRPYTA  642

Query  130   FTIPSINNETPGIRYQYNVLPQGWKGSPAIFQSSMTKILEPFRKQNPDIVIYQYMDDLYV  189
             FTIPSIN++ P  RY +N LPQG+  SP I+Q ++ +IL+PFR++ P++ +YQYMDDL+V
Sbjct  643   FTIPSINHQEPDKRYVWNCLPQGFVLSPYIYQKTLQEILQPFRERYPEVQLYQYMDDLFV  822

Query  190   GSDLEIGQHRTKIEELRQHLLRWGLTTPDKKHQKEPPFLWMGYELHPDKWTVQPIVLPEK  249
             GS+    QH+  I ELR  LL  G  TPD K Q+ PP+ W+GY+L P+ W VQ + L
Sbjct  823   GSNGSKKQHKELIIELRAILLEKGFETPDDKLQEVPPYSWLGYQLCPENWKVQKMQLDMV  1002

Query  250   DSWTVNDIQKLVGKLNWASQIYPGIKVRQLCKLLRGTKALTEVIPLTeeaelelaeNREI  309
              + T+ND+QKL+G + W S   PG+ V+ +    +G   L + +  TEEA+ EL EN E
Sbjct  1003  KNPTLNDVQKLMGNITWMSSGVPGLTVKHIAATTKGCLELNQKVIWTEEAQKELEENNEK  1182

Query  310   LKEPVHGVYYDPSKDLIAEIQKQGQGQWTYQIYQEPFKNLKTGKYARMRGAHTNDVKQLT  369
             +K      YY+P ++++ E++     + TY I Q     L  GK         + VK L
Sbjct  1183  IKNAQGLQYYNPEEEMLCEVEITKNYEATYVIKQSQ-GILWAGKKIMKANKGWSTVKNLM  1359

Query  370   EAVQKITTESIVIWGKTPKFKLPIQKetwetwwteYWQATWIPEWEFVNTPPLVKLWYQL  429
               +Q + TESI   GK P FK+P  KE       + W  +W+PE  + +         +L
Sbjct  1360  LLLQHVATESITRVGKCPTFKVPFTKEQVMWEMQKGWYYSWLPEIVYTHQVVHDDWRMKL  1539

Query  430   EKEPIVGAETFYVDGAANRETKLGKAGYVTNRGRQKVVTLTDTTNQKTELQAIYLALQDS  489
              +EP  G  T Y DG   ++   G A YVT+ GR K   L   T+Q  E  AI +AL+D+
Sbjct  1540  VEEPTSGI-TIYTDGG--KQNGEGIAAYVTSNGRTKQKRLGPVTHQVAERMAIQMALEDT  1710

Query  490   -GLEVNIVTDSQYAL-GIIQAQPDQSESELVNQIIEQLIKKEKVYLAWVPAHKGIGGNEQ  547
                +VNIVTDS Y    I +    +        II+ + +KE VY AWVP HKGI GN+
Sbjct  1711  RDKQVNIVTDSYYCWKNITEGLGLEGPQSPWWPIIQNIREKEIVYFAWVPGHKGICGNQL  1890

Query  548   VDK  550
              D+
Sbjct  1891  ADE  1899



>Enterococcus faecalis strain 44.1 NODE_36_length_611_cov_1.15083, whole genome shotgun sequence
Sequence ID: NZ_NSJR01000036.1 Length: 611
Range 1: 2 to 610

Score:364 bits(934), Expect:5e-120,
Method:Compositional matrix adjust.,
Identities:182/203(90%), Positives:194/203(95%), Gaps:0/203(0%)

Query  127   YTAFTIPSINNETPGIRYQYNVLPQGWKGSPAIFQSSMTKILEPFRKQNPDIVIYQYMDD  186
             YTAFTI S  NET GI+Y YN+LPQGWKGSPAIFQSSMTKILEPFRKQNPDIVIYQY+DD
Sbjct  610   YTAFTILSRKNETSGIKY*YNILPQGWKGSPAIFQSSMTKILEPFRKQNPDIVIYQYVDD  431

Query  187   LYVGSDLEIGQHRTKIEELRQHLLRWGLTTPDKKHQKEPPFLWMGYELHPDKWTVQPIVL  246
             LYVGSDLE+ QHRTK+EELR HLL+WG +TPD+KHQKEPPFLWMGYELHPDKWTVQPIVL
Sbjct  430   LYVGSDLELEQHRTKVEELRAHLLKWGFSTPDQKHQKEPPFLWMGYELHPDKWTVQPIVL  251

Query  247   PEKDSWTVNDIQKLVGKLNWASQIYPGIKVRQLCKLLRGTKALTEVIPLTeeaelelaeN  306
             PEKDSWTVNDIQKLVGKLNWASQIY GIKV+QLCKLLRGTKALTEV+PL+EEAELELAEN
```

```
Sbjct  250  PEKDSWTVNDIQKLVGKLNWASQIYAGIKVKQLCKLLRGTKALTEVVPLSEEAELELAEN  71

Query  307  REILKEPVHGVYYDPSKDLIAEI  329
            REILKEPVHGVYYDPSKDL+AEI
Sbjct  70   REILKEPVHGVYYDPSKDLVAEI  2
```

```
>Enterobacter hormaechei strain 2011WA-SCV PROKKA_contig000046, whole genome shotgun sequence
Sequence ID: NZ_SJEC01000046.1 Length: 544
Range 1: 4 to 543

Score:349 bits(896), Expect:1e-114,
Method:Compositional matrix adjust.,
Identities:166/180(92%), Positives:172/180(95%), Gaps:0/180(0%)

Query  66   KDSTKWRKLVDFRELNKRTQDFWEVQLGIPHPAGLKKKKSVTVLDVGDAYFSVPLDEDFR  125
            KDST+W KL DFRELNKR QDFWEVQLGIPHPAGLKKKKSVTVLDVGDAYFSVPLD+DFR
Sbjct  4    KDSTQWGKLGDFRELNKRIQDFWEVQLGIPHPAGLKKKKSVTVLDVGDAYFSVPLDKDFR  183

Query  126  KYTAFTIPSINNETPGIRYQYNVLPQGWKGSPAIFQSSMTKILEPFRKQNPDIVIYQYMD  185
            KYTAFTIPS NNETPG+RYQYNVLPQGWKGSPAIFQSSM KILEPFRKQNP+I I QYMD
Sbjct  184  KYTAFTIPSRNNETPGMRYQYNVLPQGWKGSPAIFQSSMIKILEPFRKQNPNIEINQYMD  363

Query  186  DLYVGSDLEIGQHRTKIEELRQHLLRWGLTTPDKKHQKEPPFLWMGYELHPDKWTVQPIV  245
            DLYVGSDLEIGQHRTK+EELRQHLL+WGLTTPDKKHQKEPPFLWMGYELHPDKWTVQPI
Sbjct  364  DLYVGSDLEIGQHRTKVEELRQHLLKWGLTTPDKKHQKEPPFLWMGYELHPDKWTVQPIT  543
```

```
>Levilactobacillus brevis strain CRL 2013 scaffold14, whole genome shotgun sequence
Sequence ID: NZ_MZMW01000018.1 Length: 588
Range 1: 62 to 586

Score:348 bits(893), Expect:6e-114,
Method:Compositional matrix adjust.,
Identities:165/175(94%), Positives:167/175(95%), Gaps:0/175(0%)

Query  82   KRTQDFWEVQLGIPHPAGLKKKKSVTVLDVGDAYFSVPLDEDFRKYTAFTIPSINNETPG  141
            KRTQDFWEVQLGIPHPAGLKKKKSVTVLDVGDAYFSVPLD DFRKYTAFTIPS+NNETPG
Sbjct  586  KRTQDFWEVQLGIPHPAGLKKKKSVTVLDVGDAYFSVPLDXDFRKYTAFTIPSVNNETPG  407

Query  142  IRYQYNVLPQGWKGSPAIFQSSMTKILEPFRKQNPDIVIYQYMDDLYVGSDLEIGQHRTK  201
            +RYQYNVLPQGWKGSPAIFQSSMTKILEPFRK  PDIVIYQYMDDLYVGSDLEI QHR K
Sbjct  406  VRYQYNVLPQGWKGSPAIFQSSMTKILEPFRKXXPDIVIYQYMDDLYVGSDLEIEQHRAK  227

Query  202  IEELRQHLLRWGLTTPDKKHQKEPPFLWMGYELHPDKWTVQPIVLPEKDSWTVND  256
            IEELR HLL WG TTPDKKHQKEPPFLWMGYELHPDKWTVQPIVLPEKDSWTVND
Sbjct  226  IEELRXHLLXWGFTTPDKKHQKEPPFLWMGYELHPDKWTVQPIVLPEKDSWTVND  62
```

```
Range 2: 2 to 70

Score:49.7 bits(117), Expect:0.034,
Method:Compositional matrix adjust.,
Identities:22/23(96%), Positives:22/23(95%), Gaps:0/23(0%)

Query  234  LHPDKWTVQPIVLPEKDSWTVND  256
            LHPDKWTVQPIVLPEKDSWTV D
Sbjct  2    LHPDKWTVQPIVLPEKDSWTVID  70
```

```
>Acinetobacter baumannii strain ACM-38 NODE_62_length_542_cov_0.942169, whole genome shotgun sequence
Sequence ID: NZ_QAGQ01000062.1 Length: 542
Range 1: 2 to 541

Score:347 bits(890), Expect:7e-114,
Method:Compositional matrix adjust.,
Identities:164/180(91%), Positives:172/180(95%), Gaps:0/180(0%)

Query  98   AGLKKKKSVTVLDVGDAYFSVPLDEDFRKYTAFTIPSINNETPGIRYQYNVLPQGWKGSP  157
            A LKKK S+TVLDVGDAYFSVPLD DFRKYTAFTIPS NNETPGIRYQYNVLPQGWKGSP
Sbjct  541  AELKKKNSLTVLDVGDAYFSVPLDPDFRKYTAFTIPSRNNETPGIRYQYNVLPQGWKGSP  362

Query  158  AIFQSSMTKILEPFRKQNPDIVIYQYMDDLYVGSDLEIGQHRTKIEELRQHLLRWGLTTP  217
            AIFQSSMTKILEPFRKQNPD+VIYQYMDDLYVGSDLE+ QHRTK+EELR HLL+WG +TP
Sbjct  361  AIFQSSMTKILEPFRKQNPDMVIYQYMDDLYVGSDLELEQHRTKVEELRAHLLKWGFSTP  182

Query  218  DKKHQKEPPFLWMGYELHPDKWTVQPIVLPEKDSWTVNDIQKLVGKLNWASQIYPGIKVR  277
            D+KHQKEPPFLWMGYELHPDKWTVQPIVLPEKDSWTVNDIQKLVGKLNWASQIY GIKV+
Sbjct  181  DQKHQKEPPFLWMGYELHPDKWTVQPIVLPEKDSWTVNDIQKLVGKLNWASQIYAGIKVK  2
```

```
>Cellulophaga baltica strain 14 NODE_54_length_581_cov_63.513216, whole genome shotgun sequence
Sequence ID: NZ_VIQP01000054.1 Length: 581
Range 1: 1 to 579

Score:342 bits(877), Expect:1e-111,
Method:Compositional matrix adjust.,
Identities:173/193(90%), Positives:184/193(95%), Gaps:0/193(0%)

Query  151  QGWKGSPAIFQSSMTKILEPFRKQNPDIVIYQYMDDLYVGSDLEIGQHRTKIEELRQHLL  210
            QGWKGSPAIFQ+SMTKIL+PFR++NP+IVIYQYMDDLYVGSDLEIGQHR KIEELR HLL
Sbjct  1    QGWKGSPAIFQASMTKILDPFRRKNPEIVIYQYMDDLYVGSDLEIGQHRAKIEELRAHLL  180

Query  211  RWGLTTPDKKHQKEPPFLWMGYELHPDKWTVQPIVLPEKDSWTVNDIQKLVGKLNWASQI  270
             WG TTPDKKHQKEPPFLWMGYELHPDKWTVQPI LP+KDSWTVNDIQKLVGKLNWASQI
Sbjct  181  NWGFTTPDKKHQKEPPFLWMGYELHPDKWTVQPIQLPDKDSWTVNDIQKLVGKLNWASQI  360

Query  271  YPGIKVRQLCKLLRGTKALTEVIPLTeeaelelaeNREILKEPVHGVYYDPSKDLIAEIQ  330
            YPGIKV +LCKLLRG KALT+++ LTEEAELELAENREILKEPVHGVYYDPSKDL+AEIQ
Sbjct  361  YPGIKVTRLCKLLRGAKALTDIVTLTEEAELELAENREILKEPVHGVYYDPSKDLVAEIQ  540

Query  331  KQGQGQWTYQIYQ  343
            KQGQ QWTYQIYQ
Sbjct  541  KQGQDQWTYQIYQ  579
```

```
>Shigella sonnei strain 07401ss_1, whole genome shotgun sequence
Sequence ID: NZ_CWST01000362.1 Length: 509
Range 1: 3 to 509

Score:338 bits(867), Expect:2e-110,
Method:Compositional matrix adjust.,
```

```
Identities:158/169(93%), Positives:165/169(97%), Gaps:0/169(0%)

Query  80   LNKRTQDFWEVQLGIPHPAGLKKKKSVTVLDVGDAYFSVPLDEDFRKYTAFTIPSINNET  139
            LNKRTQDFWEVQLGIPHPAGLKKKKSVTVLDVGDAYFSVPLD+DFRKYTAFTIPS+NNET
Sbjct  509  LNKRTQDFWEVQLGIPHPAGLKKKKSVTVLDVGDAYFSVPLDKDFRKYTAFTIPSVNNET  330

Query  140  PGIRYQYNVLPQGWKGSPAIFQSSMTKILEPFRKQNPDIVIYQYMDDLYVGSDLEIGQHR  199
            PGIRYQYNVLPQGWKGSPAIFQ+SMTKILEPFR +NP+IVIYQYMDDLYVGSDLEIGQHR
Sbjct  329  PGIRYQYNVLPQGWKGSPAIFQASMTKILEPFRIKNPEIVIYQYMDDLYVGSDLEIGQHR  150

Query  200  TKIEELRQHLLRWGLTTPDKKHQKEPPFLWMGYELHPDKWTVQPIVLPE  248
             KIEELR+HLL+WG TTPDKKHQKEPPFLWMGYELHPDKWTVQPI LPE
Sbjct  149  AKIEELREHLLKWGFTTPDKKHQKEPPFLWMGYELHPDKWTVQPIALPE  3



>Mycobacterium tuberculosis variant bovis strain CSURQ1131, whole genome shotgun sequence
Sequence ID: NZ_CAJEUJ010000145.1 Length: 888
Range 1: 114 to 701

Score:341 bits(875), Expect:1e-109,
Method:Compositional matrix adjust.,
Identities:166/205(81%), Positives:177/205(86%), Gaps:9/205(4%)

Query  41   MEKEGKISKIGPENPYNTPVFAIKKKDSTKWRKLVDFRELNKRTQDFWEVQLGIPHPAGL  100
            +E EG +S IGPENPYNTPVFAIKKK+STKWRKLVDFRELNK+TQDFWEVQLGIPHP+GL
Sbjct  114  VEHEGSLSNIGPENPYNTPVFAIKKKNSTKWRKLVDFRELNKKTQDFWEVQLGIPHPSGL  293

Query  101  KKKKSVTVLDVGDAYFSVPLDEDFRKYTAFTIPSINNETPGIRYQYNVLPQGWKGSPAIF  160
            KKKKSVTVLDVGDAYFSVPLDEDFRKYTAFTIPS+NN TPGIRYQYNVLPQGWKGSPAIF
Sbjct  294  KKKKSVTVLDVGDAYFSVPLDEDFRKYTAFTIPSVNNTTPGIRYQYNVLPQGWKGSPAIF  473

Query  161  QSSMTKILEPFRKQNPDIVIYQYMDDLYVGSDLEIGQHRTKIEELRQHLLRWGLTTPDKK  220
            QSSMTKILEPFRKQNPDIVIYQY+DDLYVGSDLEI QHR K+EELRQHL RWGLT   K
Sbjct  474  QSSMTKILEPFRKQNPDIVIYQYVDDLYVGSDLEIEQHRAKVEELRQHLWRWGLTHQTKN  653

Query  221  HQKEPPFLWMGYELHPDKWTVQPIV  245
             +K          LH   W +  I+
Sbjct  654  IRK---------NLHSFGWVMNSIL  701



Range 2: 629 to 769

Score:88.2 bits(217), Expect:1e-14,
Method:Compositional matrix adjust.,
Identities:39/47(83%), Positives:43/47(91%), Gaps:0/47(0%)

Query  213  GLTTPDKKHQKEPPFLWMGYELHPDKWTVQPIVLPEKDSWTVNDIQK  259
            G+ TPDKKHQKEPPFLWMGYELHPDKWTVQPIVLPEKDSW V+ + +
Sbjct  629  GIDTPDKKHQKEPPFLWMGYELHPDKWTVQPIVLPEKDSWNVHRLMQ  769



Range 3: 114 to 701

Score:341 bits(875), Expect:1e-109,
Method:Compositional matrix adjust.,
Identities:166/205(81%), Positives:177/205(86%), Gaps:9/205(4%)

Query  41   MEKEGKISKIGPENPYNTPVFAIKKKDSTKWRKLVDFRELNKRTQDFWEVQLGIPHPAGL  100
            +E EG +S IGPENPYNTPVFAIKKK+STKWRKLVDFRELNK+TQDFWEVQLGIPHP+GL
Sbjct  114  VEHEGSLSNIGPENPYNTPVFAIKKKNSTKWRKLVDFRELNKKTQDFWEVQLGIPHPSGL  293

Query  101  KKKKSVTVLDVGDAYFSVPLDEDFRKYTAFTIPSINNETPGIRYQYNVLPQGWKGSPAIF  160
            KKKKSVTVLDVGDAYFSVPLDEDFRKYTAFTIPS+NN TPGIRYQYNVLPQGWKGSPAIF
Sbjct  294  KKKKSVTVLDVGDAYFSVPLDEDFRKYTAFTIPSVNNTTPGIRYQYNVLPQGWKGSPAIF  473

Query  161  QSSMTKILEPFRKQNPDIVIYQYMDDLYVGSDLEIGQHRTKIEELRQHLLRWGLTTPDKK  220
            QSSMTKILEPFRKQNPDIVIYQY+DDLYVGSDLEI QHR K+EELRQHL RWGLT   K
Sbjct  474  QSSMTKILEPFRKQNPDIVIYQYVDDLYVGSDLEIEQHRAKVEELRQHLWRWGLTHQTKN  653

Query  221  HQKEPPFLWMGYELHPDKWTVQPIV  245
             +K          LH   W +  I+
Sbjct  654  IRK---------NLHSFGWVMNSIL  701



Range 4: 629 to 769

Score:88.2 bits(217), Expect:1e-14,
Method:Compositional matrix adjust.,
Identities:39/47(83%), Positives:43/47(91%), Gaps:0/47(0%)

Query  213  GLTTPDKKHQKEPPFLWMGYELHPDKWTVQPIVLPEKDSWTVNDIQK  259
            G+ TPDKKHQKEPPFLWMGYELHPDKWTVQPIVLPEKDSW V+ + +
Sbjct  629  GIDTPDKKHQKEPPFLWMGYELHPDKWTVQPIVLPEKDSWNVHRLMQ  769



>Klebsiella aerogenes strain AUH-KAM3 NODE_59_length_558_cov_0.865429, whole genome shotgun sequence
Sequence ID: NZ_PYHH01000059.1 Length: 558
Range 1: 1 to 558

Score:336 bits(861), Expect:3e-109,
Method:Compositional matrix adjust.,
Identities:168/186(90%), Positives:179/186(96%), Gaps:0/186(0%)

Query  135  INNETPGIRYQYNVLPQGWKGSPAIFQSSMTKILEPFRKQNPDIVIYQYMDDLYVGSDLE  194
            +NNETPGIRYQYNVLPQGWKGSPAIFQ+SMTKILEPFR +NPDIVIYQYMDDLYVGSDLE
Sbjct  558  VNNETPGIRYQYNVLPQGWKGSPAIFQASMTKILEPFRLKNPDIVIYQYMDDLYVGSDLE  379

Query  195  IGQHRTKIEELRQHLLRWGLTTPDKKHQKEPPFLWMGYELHPDKWTVQPIVLPEKDSWTV  254
            IGQHRTK+EELR HLL+ G +TPD+KHQKEPPFLW+GYELHPDKWTVQPI LPEKDSWTV
Sbjct  378  IGQHRTKVEELRAHLLK*GFSTPDQKHQKEPPFLWIGYELHPDKWTVQPIQLPEKDSWTV  199

Query  255  NDIQKLVGKLNWASQIYPGIKVRQLCKLLRGTKALTEVIPLTeeaelelaeNREILKEPV  314
            NDIQKLVGKLNWASQIY GIKV+QLCKLLRG KALT+VIPLTEEAELELAENREIL+EPV
Sbjct  198  NDIQKLVGKLNWASQIYAGIKVKQLCKLLRGAKALTDVIPLTEEAELELAENREILREPV  19

Query  315  HGVYYD  320
            HGVYYD
Sbjct  18   HGVYYD  1



>Enterococcus faecium strain INF91_9 NODE_617_length_515_cov_1.76289, whole genome shotgun sequence
Sequence ID: NZ_JAANZG010000617.1 Length: 515
```

```
Range 1: 2 to 514

Score:334 bits(857), Expect:6e-109,
Method:Compositional matrix adjust.,
Identities:158/171(92%), Positives:162/171(94%), Gaps:0/171(0%)

Query  87   FWEVQLGIPHPAGLKKKKSVTVLDVGDAYFSVPLDEDFRKYTAFTIPSINNETPGIRYQY  146
            FWEVQLGIPHP GLKKKKSVTVLDVGDAYFSVPLDE FRKYTAFTIPS NNETPGIRYQY
Sbjct  2    FWEVQLGIPHPGGLKKKKSVTVLDVGDAYFSVPLDEGFRKYTAFTIPSTNNETPGIRYQY  181

Query  147  NVLPQGWKGSPAIFQSSMTKILEPFRKQNPDIVIYQYMDDLYVGSDLEIGQHRTKIEELR  206
            NVLPQGWKGSPAIFQSSMTKILEPFR +NPDIVIYQY+DDL VGSDLEIGQHR KIEELR
Sbjct  182  NVLPQGWKGSPAIFQSSMTKILEPFRAKNPDIVIYQYVDDLLVGSDLEIGQHRAKIEELR  361

Query  207  QHLLRWGLTTPDKKHQKEPPFLWMGYELHPDKWTVQPIVLPEKDSWTVNDI  257
            +HLLRWG  TPDKKHQKEPPFLWMGYELHPDKWT+QPI LPEKDSWTVNDI
Sbjct  362  EHLLRWGFYTPDKKHQKEPPFLWMGYELHPDKWTIQPIQLPEKDSWTVNDI  514



>Neisseria gonorrhoeae strain GC-206 NODE_117_length_504_cov_1.30067_ID_20406, whole genome shotgun sequence
Sequence ID: NZ_SAWN01000117.1 Length: 504
Range 1: 2 to 502

Score:329 bits(843), Expect:7e-107,
Method:Compositional matrix adjust.,
Identities:153/167(92%), Positives:160/167(95%), Gaps:0/167(0%)

Query  114  AYFSVPLDEDFRKYTAFTIPSINNETPGIRYQYNVLPQGWKGSPAIFQSSMTKILEPFRK  173
            AYFSVPLDE+FRKYTAFTIPSINNETPGIRYQYNVLPQGWKGSP+IFQSSM KILEPFR
Sbjct  502  AYFSVPLDENFRKYTAFTIPSINNETPGIRYQYNVLPQGWKGSPSIFQSSMIKILEPFRA  323

Query  174  QNPDIVIYQYMDDLYVGSDLEIGQHRTKIEELRQHLLRWGLTTPDKKHQKEPPFLWMGYE  233
            +NP++VIYQY+DDLYVGSDLEIGQHR KIEELR HLL+WG TTPDKKHQKEPPFLWMGYE
Sbjct  322  ENPELVIYQYIDDLYVGSDLEIGQHRAKIEELRDHLLKWGFTTPDKKHQKEPPFLWMGYE  143

Query  234  LHPDKWTVQPIVLPEKDSWTVNDIQKLVGKLNWASQIYPGIKVRQLC  280
            LHPDKWTVQPI LP KDSWTVNDIQKLVGKLNWASQIYPGIKVRQLC
Sbjct  142  LHPDKWTVQPIQLPTKDSWTVNDIQKLVGKLNWASQIYPGIKVRQLC  2



>Providencia rettgeri strain AFE96_S1 41_contig_58, whole genome shotgun sequence
Sequence ID: NZ_SHDP01000058.1 Length: 546
Range 1: 20 to 544

Score:328 bits(841), Expect:2e-106,
Method:Compositional matrix adjust.,
Identities:157/175(90%), Positives:165/175(94%), Gaps:0/175(0%)

Query  40   EMEKEGKISKIGPENPYNTPVFAIKKKDSTKWRKLVDFRELNKRTQDFWEVQLGIPHPAG  99
            EMEKE KI+K G ENPYNTPVFAIKKKDSTKWRKLVD RELNKRT+D WEVQLGIPHPAG
Sbjct  20   EMEKEEKITKNGTENPYNTPVFAIKKKDSTKWRKLVDVRELNKRTKDMWEVQLGIPHPAG  199

Query  100  LKKKKSVTVLDVGDAYFSVPLDEDFRKYTAFTIPSINNETPGIRYQYNVLPQGWKGSPAI  159
            LKK KSVT+LDVGDAYFSVPLDE+FRKYTAFTIPSINN TPGIRYQYNVLPQGWKGSPAI
Sbjct  200  LKKNKSVTILDVGDAYFSVPLDENFRKYTAFTIPSINNATPGIRYQYNVLPQGWKGSPAI  379

Query  160  FQSSMTKILEPFRKQNPDIVIYQYMDDLYVGSDLEIGQHRTKIEELRQHLLRWGL  214
            FQSSMTKILEPF  +NPDIVIYQY+DDLYVGSDLEIGQHR KIEELR+HLL+WGL
Sbjct  380  FQSSMTKILEPFIAKNPDIVIYQYVDDLYVGSDLEIGQHRRKIEELREHLLKWGL  544



>Providencia rettgeri strain AFE422_S1 40_contig_33, whole genome shotgun sequence
Sequence ID: NZ_SHDL01000033.1 Length: 501
Range 1: 1 to 501

Score:327 bits(838), Expect:4e-106,
Method:Compositional matrix adjust.,
Identities:153/167(92%), Positives:162/167(97%), Gaps:0/167(0%)

Query  59   PVFAIKKKDSTKWRKLVDFRELNKRTQDFWEVQLGIPHPAGLKKKKSVTVLDVGDAYFSV  118
            P+FAIK+KDSTKWRKLVDFRELNKRTQDFWEVQLGIPHPAGLKKKKS+TVLDVGDAYFSV
Sbjct  501  PIFAIKRKDSTKWRKLVDFRELNKRTQDFWEVQLGIPHPAGLKKKKSMTVLDVGDAYFSV  322

Query  119  PLDEDFRKYTAFTIPSINNETPGIRYQYNVLPQGWKGSPAIFQSSMTKILEPFRKQNPDI  178
            PLDE FRKYTAFTIPSINNETPGIRYQYNVLPQGWKGSPAIFQSSMT+ILEPFR +NP+I
Sbjct  321  PLDEGFRKYTAFTIPSINNETPGIRYQYNVLPQGWKGSPAIFQSSMTRILEPFRAKNPEI  142

Query  179  VIYQYMDDLYVGSDLEIGQHRTKIEELRQHLLRWGLTTPDKKHQKEP  225
            VIYQY+DDLYVGSDLEIGQHR K+EELR+HLL WG TTPDKKHQKEP
Sbjct  141  VIYQYVDDLYVGSDLEIGQHRAKVEELRKHLLMWGFTTPDKKHQKEP  1



>Providencia rettgeri strain AFE220_S1 26_contig_66, whole genome shotgun sequence
Sequence ID: NZ_SHDM01000066.1 Length: 502
Range 1: 5 to 502

Score:323 bits(829), Expect:9e-105,
Method:Compositional matrix adjust.,
Identities:152/166(92%), Positives:160/166(96%), Gaps:0/166(0%)

Query  90   VQLGIPHPAGLKKKKSVTVLDVGDAYFSVPLDEDFRKYTAFTIPSINNETPGIRYQYNVL  149
            VQLGIPHPAGLKKKKSVTVLDVGDAYFSVPLDE+FRKYTAFTIPSINNETPGIRYQYNVL
Sbjct  502  VQLGIPHPAGLKKKKSVTVLDVGDAYFSVPLDENFRKYTAFTIPSINNETPGIRYQYNVL  323

Query  150  PQGWKGSPAIFQSSMTKILEPFRKQNPDIVIYQYMDDLYVGSDLEIGQHRTKIEELRQHL  209
            PQGWKGSPAIFQ SMTKILEPFR +NP+IVIYQYMDDLYVGSDLEIGQHR KIEELR+HL
Sbjct  322  PQGWKGSPAIFQCSMTKILEPFRAKNPEIVIYQYMDDLYVGSDLEIGQHRAKIEELREHL  143

Query  210  LRWGLTTPDKKHQKEPPFLWMGYELHPDKWTVQPIVLPEKDSWTVN  255
            L+WGLTTPDKKHQKEPPFLWMGYELHPDKWTVQPI LPEK+ + +N
Sbjct  142  LKWGLTTPDKKHQKEPPFLWMGYELHPDKWTVQPIQLPEKELYLIN  5



>Escherichia coli strain 8374wG10 NODE_51_length_497_cov_1.16802, whole genome shotgun sequence
Sequence ID: NZ_WSJQ01000051.1 Length: 497
Range 1: 3 to 497

Score:321 bits(823), Expect:7e-104,
Method:Compositional matrix adjust.,
Identities:152/165(92%), Positives:160/165(96%), Gaps:0/165(0%)

Query  20   KVKQWPLTEEKIKALVEICTEMEKEGKISKIGPENPYNTPVFAIKKKDSTKWRKLVDFRE  79
            ++KQWPLTEEKIKAL  IC EMEKEGKI+KIGPENPYNTPVFAIKKKDSTKWRKLVDFRE
Sbjct  497  RIKQWPLTEEKIKALQAICEEMEKEGKITKIGPENPYNTPVFAIKKKDSTKWRKLVDFRE  318
```

```
Query  80   LNKRTQDFWEVQLGIPHPAGLKKKKSVTVLDVGDAYFSVPLDEDFRKYTAFTIPSINNET  139
            LNKRTQDFWEVQLGIPHPAGLKKKKSVTVLDVGDAYFSVPLDE+FRKYTAFTIPSINN T
Sbjct  317  LNKRTQDFWEVQLGIPHPAGLKKKKSVTVLDVGDAYFSVPLDENFRKYTAFTIPSINNAT  138

Query  140  PGIRYQYNVLPQGWKGSPAIFQSSMTKILEPFRKQNPDIVIYQYM  184
            PGIRYQYNVLPQGWKGSPAIFQS+MTKIL+PFR +NPD+VIYQYM
Sbjct  137  PGIRYQYNVLPQGWKGSPAIFQSTMTKILQPFRTRNPDLVIYQYM  3



>Klebsiella oxytoca strain WCHKO085010 NODE_171_length_472_cov_19.6304, whole genome shotgun sequence
Sequence ID: NZ_NWER01000158.1 Length: 472
Range 1: 4 to 471

Score:320 bits(820), Expect:1e-103,
Method:Compositional matrix adjust.,
Identities:151/156(97%), Positives:154/156(98%), Gaps:0/156(0%)

Query  94   IPHPAGLKKKKSVTVLDVGDAYFSVPLDEDFRKYTAFTIPSINNETPGIRYQYNVLPQGW  153
            IPHPAGLK+KKSVTVLDVGDAYFSVPLD+DFRKYTAFTIPSINNETPGIRYQYNVLPQGW
Sbjct  4    IPHPAGLKQKKSVTVLDVGDAYFSVPLDKDFRKYTAFTIPSINNETPGIRYQYNVLPQGW  183

Query  154  KGSPAIFQSSMTKILEPFRKQNPDIVIYQYMDDLYVGSDLEIGQHRTKIEELRQHLLRWG  213
            KGSPAIFQ SMTKILEPFRKQNPDIVIYQYMDDLYVGSDLEIGQHRTKIEELRQHLLRWG
Sbjct  184  KGSPAIFQCSMTKILEPFRKQNPDIVIYQYMDDLYVGSDLEIGQHRTKIEELRQHLLRWG  363

Query  214  LTTPDKKHQKEPPFLWMGYELHPDKWTVQPIVLPEK  249
             TTPDKKHQKEPPFLWMGYELHPDKWTVQPIVLPE+
Sbjct  364  FTTPDKKHQKEPPFLWMGYELHPDKWTVQPIVLPER  471



>Escherichia coli strain 8374wH11 NODE_48_length_500_cov_2.00268, whole genome shotgun sequence
Sequence ID: NZ_WSJI01000048.1 Length: 500
Range 1: 1 to 495

Score:319 bits(817), Expect:5e-103,
Method:Compositional matrix adjust.,
Identities:151/165(92%), Positives:160/165(96%), Gaps:0/165(0%)

Query  26   LTEEKIKALVEICTEMEKEGKISKIGPENPYNTPVFAIKKKDSTKWRKLVDFRELNKRTQ  85
            L+EEKIKAL  IC EMEKEGKISKIGPENPYNTPVFAIK+KDSTKWRKLVDFRELNKRTQ
Sbjct  495  LSEEKIKALTGICEEMEKEGKISKIGPENPYNTPVFAIKRKDSTKWRKLVDFRELNKRTQ  316

Query  86   DFWEVQLGIPHPAGLKKKKSVTVLDVGDAYFSVPLDEDFRKYTAFTIPSINNETPGIRYQ  145
            DFWEVQLGIPHPAGLKKKKS+TVLDVGDAYFSVPLDE+FRKYTAFTIPSINN TPGIRYQ
Sbjct  315  DFWEVQLGIPHPAGLKKKKSMTVLDVGDAYFSVPLDENFRKYTAFTIPSINNATPGIRYQ  136

Query  146  YNVLPQGWKGSPAIFQSSMTKILEPFRKQNPDIVIYQYMDDLYVG  190
            YNVLPQGWKGSPAIFQS+MTKIL+PFR +NPDIVIYQY++DLYVG
Sbjct  135  YNVLPQGWKGSPAIFQSTMTKILQPFRTRNPDIVIYQYVEDLYVG  1



>Mycobacterium tuberculosis strain 32846_S30 476, whole genome shotgun sequence
Sequence ID: NZ_QTBW01000113.1 Length: 861
Range 1: 1 to 483

Score:322 bits(824), Expect:3e-102,
Method:Compositional matrix adjust.,
Identities:155/161(96%), Positives:156/161(96%), Gaps:0/161(0%)

Query  1    PISPIETVPVKLKPGMDGPKVKQWPLTEEKIKALVEICTEMEKEGKISKIGPENPYNTPV  60
            PISPIETVPVKLKPGMDGPKVKQWPLTEEKIKAL  IC EMEKEGKI+KIGPENPYNTPV
Sbjct  483  PISPIETVPVKLKPGMDGPKVKQWPLTEEKIKALTAICEEMEKEGKITKIGPENPYNTPV  304

Query  61   FAIKKKDSTKWRKLVDFRELNKRTQDFWEVQLGIPHPAGLKKKKSVTVLDVGDAYFSVPL  120
            FAIKKKDSTKWRKLVDFRELNKRTQDFWEVQLGIPHPAGLKK KSVTVLDVGDAYFSVPL
Sbjct  303  FAIKKKDSTKWRKLVDFRELNKRTQDFWEVQLGIPHPAGLKKNKSVTVLDVGDAYFSVPL  124

Query  121  DEDFRKYTAFTIPSINNETPGIRYQYNVLPQGWKGSPAIFQ  161
            DE FRKYTAFTIPSINNETPGIRYQYNVLPQGWKGSPAIFQ
Sbjct  123  DESFRKYTAFTIPSINNETPGIRYQYNVLPQGWKGSPAIFQ  1



>Escherichia coli strain 8374wH11 NODE_51_length_464_cov_1.98813, whole genome shotgun sequence
Sequence ID: NZ_WSJI01000051.1 Length: 464
Range 1: 1 to 462

Score:306 bits(784), Expect:3e-98,
Method:Compositional matrix adjust.,
Identities:143/154(93%), Positives:150/154(97%), Gaps:0/154(0%)

Query  103  KKSVTVLDVGDAYFSVPLDEDFRKYTAFTIPSINNETPGIRYQYNVLPQGWKGSPAIFQS  162
            KK+VTVLDVGDAYFSVPLDE FRKYTAFTIPS+NNETPGIRYQYNVLPQGWKGSPAIFQ+
Sbjct  462  KKAVTVLDVGDAYFSVPLDESFRKYTAFTIPSVNNETPGIRYQYNVLPQGWKGSPAIFQA  283

Query  163  SMTKILEPFRKQNPDIVIYQYMDDLYVGSDLEIGQHRTKIEELRQHLLRWGLTTPDKKHQ  222
            SMTKILEPFR +NPDIVIYQYM+DLYVGSDLEIGQHR KIEELR+HLL+WGLTTPDKKHQ
Sbjct  282  SMTKILEPFRLKNPDIVIYQYMEDLYVGSDLEIGQHRAKIEELREHLLKWGLTTPDKKHQ  103

Query  223  KEPPFLWMGYELHPDKWTVQPIVLPEKDSWTVND  256
            KEPPFLWMGYELHPDKWTVQPI LPEKDSWTVND
Sbjct  102  KEPPFLWMGYELHPDKWTVQPIQLPEKDSWTVND  1



>Escherichia coli strain P010 NODE_181_length_556_cov_0.773893_ID_361, whole genome shotgun sequence
Sequence ID: NZ_VKOQ01000154.1 Length: 556
Range 1: 2 to 556

Score:298 bits(762), Expect:2e-94,
Method:Compositional matrix adjust.,
Identities:163/185(88%), Positives:173/185(93%), Gaps:0/185(0%)

Query  241  VQPIVLPEKDSWTVNDIQKLVGKLNWASQIYPGIKVRQLCKLLRGTKALTEVIPLTeeae  300
            VQPI LPEKD WTVNDIQKLVGKLNWASQIYP IKV+QLCKLLRGTKALT+++PLTEEAE
Sbjct  556  VQPIQLPEKDIWTVNDIQKLVGKLNWASQIYPEIKVKQLCKLLRGTKALTDIVPLTEEAE  377

Query  301  lelaeNREILKEPVHGVYYDPSKDLIAEIQKQGQGQWTYQIYQEPFKNLKTGKYARMRGA  360
            L LAE REILKEPVHG YYDPS+DLIAEIQKQG  +WTYQIYQEPFKNLKTGKYA+MR A
Sbjct  376  LALAEIREILKEPVHGAYYDPSEDLIAEIQKQGNDKWTYQIYQEPFKNLKTGKYAKMRTA  197

Query  361  HTNDVKQLTEAVQKITTESIVIWGKTPKFKLPIQKetwetwwteYWQATWIPEWEFVNTP  420
            HTNDVKQLTEAVQKI  ESIVIWGK PKF+LPIQKETWETWWT+YWQATWIP+WEFVNTP
Sbjct  196  HTNDVKQLTEAVQKIARESIVIWGKIPKFRLPIQKETWETWWTDYWQATWIPDWEFVNTP  17
```

```
Query  421  PLVKL  425
            PLVKL
Sbjct  16   PLVKL  2
```

```
>Acinetobacter baumannii strain ACM-35 NODE_89_length_466_cov_2.51917, whole genome shotgun sequence
Sequence ID: NZ_QEHN01000089.1 Length: 466
Range 1: 29 to 466

Score:295 bits(754), Expect:1e-93,
Method:Compositional matrix adjust.,
Identities:143/146(98%), Positives:145/146(99%), Gaps:0/146(0%)

Query  1    PISPIETVPVKLKPGMDGPKVKQWPLTEEKIKALVEICTEMEKEGKISKIGPENPYNTPV  60
            PISPIETVPVKLKPGMDGPKVKQWPLTEEKIKALVEICTEMEKEGKISKIGPENPYNTPV
Sbjct  29   PISPIETVPVKLKPGMDGPKVKQWPLTEEKIKALVEICTEMEKEGKISKIGPENPYNTPV  208

Query  61   FAIKKKDSTKWRKLVDFRELNKRTQDFWEVQLGIPHPAGLKKKKSVTVLDVGDAYFSVPL  120
            FAIKKKDSTKWRKLVDFRELNKRTQDFWEVQLGIPHPAGLKKKKSVTVLDVGDAYFSVPL
Sbjct  209  FAIKKKDSTKWRKLVDFRELNKRTQDFWEVQLGIPHPAGLKKKKSVTVLDVGDAYFSVPL  388

Query  121  DEDFRKYTAFTIPSINNETPGIRYQY  146
            D++FRKYTAFTIPS NNETPGIRYQY
Sbjct  389  DKEFRKYTAFTIPSTNNETPGIRYQY  466
```

```
>Klebsiella pneumoniae strain EuSCAPE_IL021, whole genome shotgun sequence
Sequence ID: NZ_UKJX02000112.1 Length: 426
Range 1: 1 to 426

Score:289 bits(740), Expect:8e-92,
Method:Compositional matrix adjust.,
Identities:139/142(98%), Positives:139/142(97%), Gaps:0/142(0%)

Query  64   KKKDSTKWRKLVDFRELNKRTQDFWEVQLGIPHPAGLKKKKSVTVLDVGDAYFSVPLDED  123
            KKKDSTKWRKLVDFRELNKRTQDFWEVQLGIPHPAGLKKKKSVTVLDVGDAYFSVPLDED
Sbjct  1    KKKDSTKWRKLVDFRELNKRTQDFWEVQLGIPHPAGLKKKKSVTVLDVGDAYFSVPLDED  180

Query  124  FRKYTAFTIPSINNETPGIRYQYNVLPQGWKGSPAIFQSSMTKILEPFRKQNPDIVIYQY  183
            FRKYTAFTIPSINNETPGIRYQYNVLPQGWKGSPAIFQSSMTKILEPFR QNP IVIYQY
Sbjct  181  FRKYTAFTIPSINNETPGIRYQYNVLPQGWKGSPAIFQSSMTKILEPFRAQNPQIVIYQY  360

Query  184  MDDLYVGSDLEIGQHRTKIEEL  205
            MDDLYVGSDLEIGQHR KIEEL
Sbjct  361  MDDLYVGSDLEIGQHRAKIEEL  426
```

```
>Escherichia coli strain 8374wA11 NODE_51_length_444_cov_2.55205, whole genome shotgun sequence
Sequence ID: NZ_WSPR01000049.1 Length: 444
Range 1: 1 to 444

Score:288 bits(738), Expect:2e-91,
Method:Compositional matrix adjust.,
Identities:135/148(91%), Positives:139/148(93%), Gaps:0/148(0%)

Query  112  GDAYFSVPLDEDFRKYTAFTIPSINNETPGIRYQYNVLPQGWKGSPAIFQSSMTKILEPF  171
            GDAYFSVPLDE FRKYTAFTIPS NNETPGIRYQYNVLPQGWKGSPAIFQ SMTKILEPF
Sbjct  444  GDAYFSVPLDEGFRKYTAFTIPSRNNETPGIRYQYNVLPQGWKGSPAIFQCSMTKILEPF  265

Query  172  RKQNPDIVIYQYMDDLYVGSDLEIGQHRTKIEELRQHLLRWGLTTPDKKHQKEPPFLWMG  231
            R +NP+I+I QYMDDLYVGSDLEIGQHR KIEELR HLL+WG TTPDKKHQKEPPFLWMG
Sbjct  264  RAKNPEIIICQYMDDLYVGSDLEIGQHRAKIEELRDHLLKWGFTTPDKKHQKEPPFLWMG  85

Query  232  YELHPDKWTVQPIVLPEKDSWTVNDIQK  259
            YELHPDKWTVQPI LPEKDSWTVNDIQK
Sbjct  84   YELHPDKWTVQPIQLPEKDSWTVNDIQK  1
```

```
>Escherichia coli strain TzEc080 NODE_51_length_444_cov_2.55205, whole genome shotgun sequence
Sequence ID: NZ_WSFZ01000049.1 Length: 444
Range 1: 1 to 444

Score:288 bits(738), Expect:2e-91,
Method:Compositional matrix adjust.,
Identities:135/148(91%), Positives:139/148(93%), Gaps:0/148(0%)

Query  112  GDAYFSVPLDEDFRKYTAFTIPSINNETPGIRYQYNVLPQGWKGSPAIFQSSMTKILEPF  171
            GDAYFSVPLDE FRKYTAFTIPS NNETPGIRYQYNVLPQGWKGSPAIFQ SMTKILEPF
Sbjct  444  GDAYFSVPLDEGFRKYTAFTIPSRNNETPGIRYQYNVLPQGWKGSPAIFQCSMTKILEPF  265

Query  172  RKQNPDIVIYQYMDDLYVGSDLEIGQHRTKIEELRQHLLRWGLTTPDKKHQKEPPFLWMG  231
            R +NP+I+I QYMDDLYVGSDLEIGQHR KIEELR HLL+WG TTPDKKHQKEPPFLWMG
Sbjct  264  RAKNPEIIICQYMDDLYVGSDLEIGQHRAKIEELRDHLLKWGFTTPDKKHQKEPPFLWMG  85

Query  232  YELHPDKWTVQPIVLPEKDSWTVNDIQK  259
            YELHPDKWTVQPI LPEKDSWTVNDIQK
Sbjct  84   YELHPDKWTVQPIQLPEKDSWTVNDIQK  1
```

```
>Klebsiella pneumoniae strain B2 isolate B2, whole genome shotgun sequence
Sequence ID: NZ_CABWNT010000058.1 Length: 7306
Range 1: 4178 to 5398

Score:315 bits(807), Expect:2e-89,
Method:Compositional matrix adjust.,
Identities:185/412(45%), Positives:256/412(62%), Gaps:5/412(1%)

Query  2     ISPIETVPVKLKPGMDGPKVKQWPLTEEKIKALVEICTEMEKEGKISKIGPENPYNTPVF  61
             I+P E   +KLK G  GPKV QWPLT+EK+    EI  ++  EGKIS+   +NPYN+P+F
Sbjct  4178  ITPRE---IKLKTGTVGPKVPQWPLTKEKLLGAKEIVKKLLDEGKISEASDDNPYNSPIF  4348

Query  62    AIKKKDSTKWRKLVDFRELNKRTQDFWEVQLGIPHPAGLKKKKSVTVLDVGDAYFSVPLD  121
              IKKK S KWR L D RELNK  Q   E+  G+PHP GL K   +TVLD+GDAYF++PLD
Sbjct  4349  VIKKK-SGKWRLLQDLRELNKVVQVGTEISRGLPHPGGLIKCNHMTVLDIGDAYFTIPLD  4525

Query  122   EDFRKYTAFTIPSINNETPGIRYQYNVLPQGWKGSPAIFQSSMTKILEPFRKQNPDIVIY  181
               FR+YTAFT+PSIN++ P  RY +N LPQG+  SP I+Q ++  IL+ FR+++PD+ +Y
Sbjct  4526  PKFRQYTAFTVPSINHQEPDKRYVWNCLPQGFVLSPYIYQKTLQDILQAFRERHPDVQLY  4705

Query  182   QYMDDLYVGSDLEIGQHRTKIEELRQHLLRWGLTTPDKKHQKEPPFLWMGYELHPDKWTV  241
             QYMDDL++GS+    QH+  +EELR  LL  G  TP+ K Q+E P+ W+GY+L P  W V
Sbjct  4706  QYMDDLFIGSNESKRQHKELVEELRAILLEKGFETPEDKLQEEAPYNWLGYQLSPGNWKV  4885

Query  242   QPIVLPEKDSWTVNDIQKLVGKLNWASQIYPGIKVRQLCKLLRGTKALTEVIPLTeeael  301
```

```
             Q + L      T+ND+QKL+G + W S   PG+ V+Q+    +G   L + +  TEEA+
Sbjct  4886  QKMQLELVKEPTLNDVQKLMGNITWMSSGVPGLTVKQIAATTKGCLDLNQKVVWTEEAQK  5065

Query  302   elaeNREILKEPVHGVYYDPSKDLIAEIQKQGQGQWTYQIYQEPFKNLKTGKYARMRGAH  361
             EL EN + ++E     YY+P +++I EI+     + TY I Q     L  GK
Sbjct  5066  ELEENNKKIQEAQGLQYYNPEEEVICEIEITKNYEATYIIKQSQ-GILWAGKKIMRANKG  5242

Query  362   TNDVKQLTEAVQKITTESIVIWGKTPKFKLPIQKetwetwwteYWQATWIPE  413
              +  K L   +Q + TESIV  G  PKFK+P  KE  +    + W  +W+P+
Sbjct  5243  WSAAKNLMLLLQHVATESIVRIGTCPKFKVPFTKEQVKWEMEKGWYYSWLPD  5398
```

Range 2: 5654 to 5953

Score:60.5 bits(145), Expect:1e-04,
Method:Compositional matrix adjust.,
Identities:42/100(42%), Positives:54/100(54%), Gaps:2/100(2%)

```
Query  453   GKAGYVTNRGRQKVVTLTDTTNQKTELQAIYLALQDSGLE-VNIVTDSQYAL-GIIQAQP  510
             G A YVT+ G+ K   L   T+Q  E  AI +AL+D+    VNIVTDS Y    I +
Sbjct  5654  GVAAYVTSNGKTKQKRLGPVTHQTAERIAIQMALEDTEETLVNIVTDSYYCWKNITEGLG  5833

Query  511   DQSESELVNQIIEQLIKKEKVYLAWVPAHKGIGGNEQVDK  550
              +        II+ +  KE VY AWVP HKGI GN+  D+
Sbjct  5834  LEGPDSPWWPIIQNIRAKEMVYFAWVPGHKGIYGNQLADE  5953
```

>Burkholderia pseudomallei strain 2604STDY5518971 ERS240105SCcontig000116, whole genome shotgun sequence
Sequence ID: NZ_UVFZ02000116.1 Length: 427
Range 1: 1 to 426

Score:283 bits(724), Expect:2e-89,
Method:Compositional matrix adjust.,
Identities:135/142(95%), Positives:139/142(97%), Gaps:0/142(0%)

```
Query  26    LTEEKIKALVEICTEMEKEGKISKIGPENPYNTPVFAIKKKDSTKWRKLVDFRELNKRTQ  85
             LTEEKIKAL EIC +MEKEGKISKIGPENPYNTP+FAIKKKDSTKWRKLVDFRELNKRTQ
Sbjct  426   LTEEKIKALTEICNDMEKEGKISKIGPENPYNTPIFAIKKKDSTKWRKLVDFRELNKRTQ  247

Query  86    DFWEVQLGIPHPAGLKKKKSVTVLDVGDAYFSVPLDEDFRKYTAFTIPSINNETPGIRYQ  145
             DFWEVQLGIPHPAGLKKKKSVTVLDVGDAYFSVPLDE+FRKYTAFTIPS NNETPG+RYQ
Sbjct  246   DFWEVQLGIPHPAGLKKKKSVTVLDVGDAYFSVPLDENFRKYTAFTIPSRNNETPGVRYQ  67

Query  146   YNVLPQGWKGSPAIFQSSMTKI  167
             YNVLPQGWKGSPAIFQSSMTKI
Sbjct  66    YNVLPQGWKGSPAIFQSSMTKI  1
```

>Escherichia coli strain 8374wB11 NODE_47_length_476_cov_2.59885, whole genome shotgun sequence
Sequence ID: NZ_WSJO01000047.1 Length: 476
Range 1: 1 to 474

Score:281 bits(720), Expect:2e-88,
Method:Compositional matrix adjust.,
Identities:141/158(89%), Positives:153/158(96%), Gaps:0/158(0%)

```
Query  167   ILEPFRKQNPDIVIYQYMDDLYVGSDLEIGQHRTKIEELRQHLLRWGLTTPDKKHQKEPP  226
             IL+PFR +NPDIVIYQY+DDLYVGSDLEIGQHR KIEELR+HLL+WGLTTPDKKHQKEPP
Sbjct  474   ILQPFRTRNPDIVIYQYVDDLYVGSDLEIGQHRAKIEELREHLLKWGLTTPDKKHQKEPP  295

Query  227   FLWMGYELHPDKWTVQPIVLPEKDSWTVNDIQKLVGKLNWASQIYPGIKVRQLCKLLRGT  286
             FLWMGYELHPDKWTVQPI LP+K+SWTVNDIQKLVGKLNWASQIYPGI+V+ LCKLLRG
Sbjct  294   FLWMGYELHPDKWTVQPIQLPDKESWTVNDIQKLVGKLNWASQIYPGIRVKHLCKLLRGA  115

Query  287   KALTEVIPLTeeaelelaeNREILKEPVHGVYYDPSKD  324
             KALT++IPLTEEAELELAENREILKEPVHGVYYDPS+D
Sbjct  114   KALTDIIPLTEEAELELAENREILKEPVHGVYYDPSRD  1
```

>Escherichia coli strain 8374wB11 NODE_56_length_419_cov_1.43493, whole genome shotgun sequence
Sequence ID: NZ_WSJO01000056.1 Length: 419
Range 1: 2 to 418

Score:278 bits(712), Expect:1e-87,
Method:Compositional matrix adjust.,
Identities:131/139(94%), Positives:136/139(97%), Gaps:0/139(0%)

```
Query  65    KKDSTKWRKLVDFRELNKRTQDFWEVQLGIPHPAGLKKKKSVTVLDVGDAYFSVPLDEDF  124
             K+DSTKWRKLVDFRELNKRTQDFWEVQLGIPHPAGLK+KKSVTVLDVGDAYFSVPLDE F
Sbjct  418   KRDSTKWRKLVDFRELNKRTQDFWEVQLGIPHPAGLKQKKSVTVLDVGDAYFSVPLDESF  239

Query  125   RKYTAFTIPSINNETPGIRYQYNVLPQGWKGSPAIFQSSMTKILEPFRKQNPDIVIYQYM  184
             RKYTAFTIPS+NNETPGIRYQYNVLPQGWKGSPAIFQ+SMTKILEPFR +NPDIVIYQYM
Sbjct  238   RKYTAFTIPSVNNETPGIRYQYNVLPQGWKGSPAIFQASMTKILEPFRLKNPDIVIYQYM  59

Query  185   DDLYVGSDLEIGQHRTKIE  203
             DDLYVGSDLEIGQHR KIE
Sbjct  58    DDLYVGSDLEIGQHRAKIE  2
```

>Burkholderia pseudomallei strain 2604STDY5518971 ERS240105SCcontig000113, whole genome shotgun sequence
Sequence ID: NZ_UVFZ02000113.1 Length: 457
Range 1: 2 to 457

Score:278 bits(712), Expect:2e-87,
Method:Compositional matrix adjust.,
Identities:141/152(93%), Positives:146/152(96%), Gaps:0/152(0%)

```
Query  188   YVGSDLEIGQHRTKIEELRQHLLRWGLTTPDKKHQKEPPFLWMGYELHPDKWTVQPIVLP  247
             YVGSDLEI QHRTKIEELR HLL+WG TTPDKKHQKEPPFLWMGYELHPDKWTVQPI LP
Sbjct  2     YVGSDLEIEQHRTKIEELRHHLLKWGFTTPDKKHQKEPPFLWMGYELHPDKWTVQPIKLP  181

Query  248   EKDSWTVNDIQKLVGKLNWASQIYPGIKVRQLCKLLRGTKALTEVIPLTeeaelelaeNR  307
             EKDSWTVNDIQKLVGKLNWASQIYPGIKV+QLCKL+RGTKALTEV+PLT+EAELELAENR
Sbjct  182   EKDSWTVNDIQKLVGKLNWASQIYPGIKVKQLCKLIRGTKALTEVVPLTKEAELELAENR  361

Query  308   EILKEPVHGVYYDPSKDLIAEIQKQGQGQWTY  339
             EILKEPVHG YYDPSKDLIAEIQKQG GQWTY
Sbjct  362   EILKEPVHGTYYDPSKDLIAEIQKQGNGQWTY  457
```

>Cronobacter sakazakii strain cro4215W contig98, whole genome shotgun sequence

```
Sequence ID: NZ_NRKF01000098.1 Length: 429
Range 1: 3 to 401

Score:272 bits(695), Expect:4e-85,
Method:Compositional matrix adjust.,
Identities:133/133(100%), Positives:133/133(100%), Gaps:0/133(0%)

Query  1    PISPIETVPVKLKPGMDGPKVKQWPLTEEKIKALVEICTEMEKEGKISKIGPENPYNTPV  60
            PISPIETVPVKLKPGMDGPKVKQWPLTEEKIKALVEICTEMEKEGKISKIGPENPYNTPV
Sbjct  401  PISPIETVPVKLKPGMDGPKVKQWPLTEEKIKALVEICTEMEKEGKISKIGPENPYNTPV  222

Query  61   FAIKKKDSTKWRKLVDFRELNKRTQDFWEVQLGIPHPAGLKKKKSVTVLDVGDAYFSVPL  120
            FAIKKKDSTKWRKLVDFRELNKRTQDFWEVQLGIPHPAGLKKKKSVTVLDVGDAYFSVPL
Sbjct  221  FAIKKKDSTKWRKLVDFRELNKRTQDFWEVQLGIPHPAGLKKKKSVTVLDVGDAYFSVPL  42

Query  121  DEDFRKYTAFTIP  133
            DEDFRKYTAFTIP
Sbjct  41   DEDFRKYTAFTIP  3
```

```
>Levilactobacillus brevis strain CRL 2013 scaffold20, whole genome shotgun sequence
Sequence ID: NZ_MZMW01000021.1 Length: 398
Range 1: 2 to 397

Score:270 bits(689), Expect:3e-84,
Method:Compositional matrix adjust.,
Identities:128/132(97%), Positives:129/132(97%), Gaps:0/132(0%)

Query  85   QDFWEVQLGIPHPAGLKKKKSVTVLDVGDAYFSVPLDEDFRKYTAFTIPSINNETPGIRY  144
            QDFWEVQLGIPHPAGLKKKKSVTVLDVGDAYFSVPLD+DFRKYTAFTIPSINNETPGIRY
Sbjct  2    QDFWEVQLGIPHPAGLKKKKSVTVLDVGDAYFSVPLDKDFRKYTAFTIPSINNETPGIRY  181

Query  145  QYNVLPQGWKGSPAIFQSSMTKILEPFRKQNPDIVIYQYMDDLYVGSDLEIGQHRTKIEE  204
            QYNVLPQGWKGSPAIFQSSMTKILEPFRKQNPDIVIYQYMDDLYVGSDLEIGQHR KIEE
Sbjct  182  QYNVLPQGWKGSPAIFQSSMTKILEPFRKQNPDIVIYQYMDDLYVGSDLEIGQHRAKIEE  361

Query  205  LRQHLLRWGLTT  216
            LRQHLL WG TT
Sbjct  362  LRQHLLTWGFTT  397
```

```
>Cronobacter sakazakii strain cro4114A1 contig119, whole genome shotgun sequence
Sequence ID: NZ_NRKJ01000119.1 Length: 429
Range 1: 3 to 401

Score:270 bits(690), Expect:3e-84,
Method:Compositional matrix adjust.,
Identities:132/133(99%), Positives:133/133(100%), Gaps:0/133(0%)

Query  1    PISPIETVPVKLKPGMDGPKVKQWPLTEEKIKALVEICTEMEKEGKISKIGPENPYNTPV  60
            PISPIETVPVKLKPGMDGPKVKQWPLTEEKIKALVEICTEMEKEGKISKIGPENPYNTPV
Sbjct  401  PISPIETVPVKLKPGMDGPKVKQWPLTEEKIKALVEICTEMEKEGKISKIGPENPYNTPV  222

Query  61   FAIKKKDSTKWRKLVDFRELNKRTQDFWEVQLGIPHPAGLKKKKSVTVLDVGDAYFSVPL  120
            FAIKKKDSTKWRKLVDFRELNKRTQDFWEVQLGIPHPAGLKKKKSVTVLDVGDAYFSVPL
Sbjct  221  FAIKKKDSTKWRKLVDFRELNKRTQDFWEVQLGIPHPAGLKKKKSVTVLDVGDAYFSVPL  42

Query  121  DEDFRKYTAFTIP  133
            DE+FRKYTAFTIP
Sbjct  41   DEEFRKYTAFTIP  3
```

```
>Cronobacter sakazakii strain cro4114B1 contig94, whole genome shotgun sequence
Sequence ID: NZ_NRKI01000094.1 Length: 429
Range 1: 29 to 427

Score:269 bits(687), Expect:8e-84,
Method:Compositional matrix adjust.,
Identities:130/133(98%), Positives:133/133(100%), Gaps:0/133(0%)

Query  1    PISPIETVPVKLKPGMDGPKVKQWPLTEEKIKALVEICTEMEKEGKISKIGPENPYNTPV  60
            PISPIETVPVKLKPGMDGPKVKQWPLTEEKIKALVEICTEMEKEGKISKIGPENPYNTPV
Sbjct  29   PISPIETVPVKLKPGMDGPKVKQWPLTEEKIKALVEICTEMEKEGKISKIGPENPYNTPV  208

Query  61   FAIKKKDSTKWRKLVDFRELNKRTQDFWEVQLGIPHPAGLKKKKSVTVLDVGDAYFSVPL  120
            FAIKKKDSTKWRKLVDFRELNKRTQDFWEVQLGIPHPAGLK+KKSVTVLDVGDAYFS+PL
Sbjct  209  FAIKKKDSTKWRKLVDFRELNKRTQDFWEVQLGIPHPAGLKRKKSVTVLDVGDAYFSIPL  388

Query  121  DEDFRKYTAFTIP  133
            DE+FRKYTAFTIP
Sbjct  389  DEEFRKYTAFTIP  427
```

```
>Escherichia coli strain 8374wB11 NODE_55_length_422_cov_0.938983, whole genome shotgun sequence
Sequence ID: NZ_WSJO01000055.1 Length: 422
Range 1: 2 to 421

Score:268 bits(686), Expect:1e-83,
Method:Compositional matrix adjust.,
Identities:122/140(87%), Positives:134/140(95%), Gaps:0/140(0%)

Query  105  SVTVLDVGDAYFSVPLDEDFRKYTAFTIPSINNETPGIRYQYNVLPQGWKGSPAIFQSSM  164
            SVTV+DVGDAYFS+PLDEDFRKYTAFTIPS NNETPG+RYQYNVLPQGWKGSPAIFQSSM
Sbjct  421  SVTVMDVGDAYFSIPLDEDFRKYTAFTIPSTNNETPGVRYQYNVLPQGWKGSPAIFQSSM  242

Query  165  TKILEPFRKQNPDIVIYQYMDDLYVGSDLEIGQHRTKIEELRQHLLRWGLTTPDKKHQKE  224
            T+ILEPFR +NP+IVIYQY+DDLYVGSDLEIGQHR KI+ELR++L +WG  TPDKKHQ+E
Sbjct  241  TRILEPFRAKNPEIVIYQYVDDLYVGSDLEIGQHRAKIKELREYLWKWGFYTPDKKHQEE  62

Query  225  PPFLWMGYELHPDKWTVQPI  244
            PPFLWMGYELHPDKWTVQPI
Sbjct  61   PPFLWMGYELHPDKWTVQPI  2
```

```
>Escherichia coli strain 8374wB4 NODE_86_length_485_cov_1.49162, whole genome shotgun sequence
Sequence ID: NZ_WSPX01000086.1 Length: 485
Range 1: 78 to 485

Score:267 bits(683), Expect:5e-83,
Method:Compositional matrix adjust.,
Identities:130/136(96%), Positives:133/136(97%), Gaps:0/136(0%)

Query  1    PISPIETVPVKLKPGMDGPKVKQWPLTEEKIKALVEICTEMEKEGKISKIGPENPYNTPV  60
            PISPIETVPVKLKPGMDGP+VKQWPLTEEKIKAL  IC EMEKEGKI+KIGPENPYNTPV
```

```
Sbjct  78   PISPIETVPVKLKPGMDGPRVKQWPLTEEKIKALQAICEEMEKEGKITKIGPENPYNTPV  257

Query  61   FAIKKKDSTKWRKLVDFRELNKRTQDFWEVQLGIPHPAGLKKKKSVTVLDVGDAYFSVPL  120
            FAIKKKDSTKWRKLVDFRELNKRTQDFWEVQLGIPHPAGLKKKKSVTVLDVGDAYFSVPL
Sbjct  258  FAIKKKDSTKWRKLVDFRELNKRTQDFWEVQLGIPHPAGLKKKKSVTVLDVGDAYFSVPL  437

Query  121  DEDFRKYTAFTIPSIN  136
            DE+FRKYTAFTIPSIN
Sbjct  438  DENFRKYTAFTIPSIN  485
```

>Klebsiella pneumoniae strain EuSCAPE_IL018, whole genome shotgun sequence
Sequence ID: NZ_UKJU02000100.1 Length: 392
Range 1: 3 to 392

Score:264 bits(674), Expect:4e-82,
Method:Compositional matrix adjust.,
Identities:123/130(95%), Positives:126/130(96%), Gaps:0/130(0%)

```
Query  115  YFSVPLDEDFRKYTAFTIPSINNETPGIRYQYNVLPQGWKGSPAIFQSSMTKILEPFRKQ  174
            YFSVPL EDFRKYTAFTIPSINNETPGIRYQYNVLPQGWKGSPAIFQSSMTKILEPFR Q
Sbjct  392  YFSVPLYEDFRKYTAFTIPSINNETPGIRYQYNVLPQGWKGSPAIFQSSMTKILEPFRTQ  213

Query  175  NPDIVIYQYMDDLYVGSDLEIGQHRTKIEELRQHLLRWGLTTPDKKHQKEPPFLWMGYEL  234
            NP+IVIYQYMDDLYVGSDLEIGQHR KIEELR+HLL+WG TTPDKKHQKEPPFLWMGYEL
Sbjct  212  NPEIVIYQYMDDLYVGSDLEIGQHRAKIEELREHLLKWGFTTPDKKHQKEPPFLWMGYEL  33

Query  235  HPDKWTVQPI  244
            HPDKWTVQPI
Sbjct  32   HPDKWTVQPI  3
```

>Cronobacter malonaticus strain cro3263A1 contig63, whole genome shotgun sequence
Sequence ID: NZ_NRMM01000063.1 Length: 429
Range 1: 29 to 427

Score:264 bits(674), Expect:6e-82,
Method:Compositional matrix adjust.,
Identities:127/133(95%), Positives:132/133(99%), Gaps:0/133(0%)

```
Query  1    PISPIETVPVKLKPGMDGPKVKQWPLTEEKIKALVEICTEMEKEGKISKIGPENPYNTPV  60
            PISPI+TVPVKLKPGMDGPKVKQWPLTEEKIKAL+EICTEMEKEGKISKIGPENPYNTPV
Sbjct  29   PISPIDTVPVKLKPGMDGPKVKQWPLTEEKIKALIEICTEMEKEGKISKIGPENPYNTPV  208

Query  61   FAIKKKDSTKWRKLVDFRELNKRTQDFWEVQLGIPHPAGLKKKKSVTVLDVGDAYFSVPL  120
            FAIKKKDSTKWRKLVDFRELNKRTQDF EVQLGIPHPAGLK+KKSVTVLDVGDAYFS+PL
Sbjct  209  FAIKKKDSTKWRKLVDFRELNKRTQDFSEVQLGIPHPAGLKRKKSVTVLDVGDAYFSIPL  388

Query  121  DEDFRKYTAFTIP  133
            DE+FRKYTAFTIP
Sbjct  389  DEEFRKYTAFTIP  427
```

>Cronobacter dublinensis strain cro981C2 contig293, whole genome shotgun sequence
Sequence ID: NZ_NROB01000293.1 Length: 429
Range 1: 29 to 427

Score:263 bits(673), Expect:1e-81,
Method:Compositional matrix adjust.,
Identities:128/133(96%), Positives:130/133(97%), Gaps:0/133(0%)

```
Query  1    PISPIETVPVKLKPGMDGPKVKQWPLTEEKIKALVEICTEMEKEGKISKIGPENPYNTPV  60
            PISPI+TVPVKLKPGMDGPKVKQWPLTEEKIKAL EIC EMEKEGKISKIGPENPYNTPV
Sbjct  29   PISPIDTVPVKLKPGMDGPKVKQWPLTEEKIKALTEICKEMEKEGKISKIGPENPYNTPV  208

Query  61   FAIKKKDSTKWRKLVDFRELNKRTQDFWEVQLGIPHPAGLKKKKSVTVLDVGDAYFSVPL  120
            FAIKKKDS KWRKLVDFRELNKRTQDFWEVQLGIPHPAGLKKKKSVTVLDVGDAYFSVPL
Sbjct  209  FAIKKKDSNKWRKLVDFRELNKRTQDFWEVQLGIPHPAGLKKKKSVTVLDVGDAYFSVPL  388

Query  121  DEDFRKYTAFTIP  133
            DE+FRKYTAFTIP
Sbjct  389  DENFRKYTAFTIP  427
```

>Cronobacter sakazakii strain cro3040W contig78, whole genome shotgun sequence
Sequence ID: NZ_NRJO01000078.1 Length: 429
Range 1: 29 to 427

Score:263 bits(671), Expect:2e-81,
Method:Compositional matrix adjust.,
Identities:128/133(96%), Positives:129/133(96%), Gaps:0/133(0%)

```
Query  1    PISPIETVPVKLKPGMDGPKVKQWPLTEEKIKALVEICTEMEKEGKISKIGPENPYNTPV  60
            PISPI+TVPV LKPGMDGPKVKQWPLTEEKIKAL EIC EMEKEGKISKIGPENPYNTPV
Sbjct  29   PISPIDTVPVTLKPGMDGPKVKQWPLTEEKIKALTEICREMEKEGKISKIGPENPYNTPV  208

Query  61   FAIKKKDSTKWRKLVDFRELNKRTQDFWEVQLGIPHPAGLKKKKSVTVLDVGDAYFSVPL  120
            FAIKKKDSTKWRKLVDFRELNKRTQDFWEVQLGIPHPAGLKKKKSVTVLDVGDAYFSVPL
Sbjct  209  FAIKKKDSTKWRKLVDFRELNKRTQDFWEVQLGIPHPAGLKKKKSVTVLDVGDAYFSVPL  388

Query  121  DEDFRKYTAFTIP  133
            DE FRKYTAFTIP
Sbjct  389  DESFRKYTAFTIP  427
```

>Cronobacter sakazakii strain cro4215W contig97, whole genome shotgun sequence
Sequence ID: NZ_NRKF01000097.1 Length: 429
Range 1: 29 to 427

Score:262 bits(669), Expect:4e-81,
Method:Compositional matrix adjust.,
Identities:126/133(95%), Positives:129/133(96%), Gaps:0/133(0%)

```
Query  1    PISPIETVPVKLKPGMDGPKVKQWPLTEEKIKALVEICTEMEKEGKISKIGPENPYNTPV  60
            PISPI+T+PV LKPGMDGPKVKQWPLTEEKIKAL EIC EMEKEGKISKIGPENPYNTPV
Sbjct  29   PISPIDTIPVTLKPGMDGPKVKQWPLTEEKIKALTEICKEMEKEGKISKIGPENPYNTPV  208

Query  61   FAIKKKDSTKWRKLVDFRELNKRTQDFWEVQLGIPHPAGLKKKKSVTVLDVGDAYFSVPL  120
            FAIKKKDSTKWRKLVDFRELNKRTQDFWEVQLGIPHPAGLKKKKSVTVLD+GDAYFSVPL
Sbjct  209  FAIKKKDSTKWRKLVDFRELNKRTQDFWEVQLGIPHPAGLKKKKSVTVLDIGDAYFSVPL  388

Query  121  DEDFRKYTAFTIP  133
            DE FRKYTAFTIP
Sbjct  389  DESFRKYTAFTIP  427
```

```
>Cronobacter sakazakii strain cro4114B1 contig95, whole genome shotgun sequence
Sequence ID: NZ_NRKI01000095.1 Length: 429
Range 1: 29 to 427

Score:262 bits(669), Expect:4e-81,
Method:Compositional matrix adjust.,
Identities:126/133(95%), Positives:129/133(96%), Gaps:0/133(0%)

Query  1    PISPIETVPVKLKPGMDGPKVKQWPLTEEKIKALVEICTEMEKEGKISKIGPENPYNTPV  60
            PISPI+T+PV LKPGMDGPKVKQWPLTEEKIKAL EIC EMEKEGKISKIGPENPYNTPV
Sbjct  29   PISPIDTIPVTLKPGMDGPKVKQWPLTEEKIKALTEICKEMEKEGKISKIGPENPYNTPV  208

Query  61   FAIKKKDSTKWRKLVDFRELNKRTQDFWEVQLGIPHPAGLKKKKSVTVLDVGDAYFSVPL  120
            FAIKKKDSTKWRKLVDFRELNKRTQDFWEVQLGIPHPAGLKKKKSVTVLD+GDAYFSVPL
Sbjct  209  FAIKKKDSTKWRKLVDFRELNKRTQDFWEVQLGIPHPAGLKKKKSVTVLDIGDAYFSVPL  388

Query  121  DEDFRKYTAFTIP  133
            DE FRKYTAFTIP
Sbjct  389  DESFRKYTAFTIP  427



>Cronobacter sakazakii strain cro4114A1 contig120, whole genome shotgun sequence
Sequence ID: NZ_NRKJ01000120.1 Length: 429
Range 1: 29 to 427

Score:262 bits(669), Expect:4e-81,
Method:Compositional matrix adjust.,
Identities:126/133(95%), Positives:129/133(96%), Gaps:0/133(0%)

Query  1    PISPIETVPVKLKPGMDGPKVKQWPLTEEKIKALVEICTEMEKEGKISKIGPENPYNTPV  60
            PISPI+T+PV LKPGMDGPKVKQWPLTEEKIKAL EIC EMEKEGKISKIGPENPYNTPV
Sbjct  29   PISPIDTIPVTLKPGMDGPKVKQWPLTEEKIKALTEICKEMEKEGKISKIGPENPYNTPV  208

Query  61   FAIKKKDSTKWRKLVDFRELNKRTQDFWEVQLGIPHPAGLKKKKSVTVLDVGDAYFSVPL  120
            FAIKKKDSTKWRKLVDFRELNKRTQDFWEVQLGIPHPAGLKKKKSVTVLD+GDAYFSVPL
Sbjct  209  FAIKKKDSTKWRKLVDFRELNKRTQDFWEVQLGIPHPAGLKKKKSVTVLDIGDAYFSVPL  388

Query  121  DEDFRKYTAFTIP  133
            DE FRKYTAFTIP
Sbjct  389  DESFRKYTAFTIP  427



>Cronobacter sakazakii strain cro4191B1-2 contig80, whole genome shotgun sequence
Sequence ID: NZ_NRKG01000080.1 Length: 429
Range 1: 29 to 427

Score:262 bits(669), Expect:4e-81,
Method:Compositional matrix adjust.,
Identities:127/133(95%), Positives:129/133(96%), Gaps:0/133(0%)

Query  1    PISPIETVPVKLKPGMDGPKVKQWPLTEEKIKALVEICTEMEKEGKISKIGPENPYNTPV  60
            PISPI+TVPV LKPGMDGPKVKQWPLTEEKIKAL EIC EMEKEGKI+KIGPENPYNTPV
Sbjct  29   PISPIDTVPVTLKPGMDGPKVKQWPLTEEKIKALTEICREMEKEGKITKIGPENPYNTPV  208

Query  61   FAIKKKDSTKWRKLVDFRELNKRTQDFWEVQLGIPHPAGLKKKKSVTVLDVGDAYFSVPL  120
            FAIKKKDSTKWRKLVDFRELNKRTQDFWEVQLGIPHPAGLKKKKSVTVLDVGDAYFSVPL
Sbjct  209  FAIKKKDSTKWRKLVDFRELNKRTQDFWEVQLGIPHPAGLKKKKSVTVLDVGDAYFSVPL  388

Query  121  DEDFRKYTAFTIP  133
            DE FRKYTAFTIP
Sbjct  389  DESFRKYTAFTIP  427



>Acinetobacter baumannii strain ACM-28 NODE_88_length_395_cov_2.15672, whole genome shotgun sequence
Sequence ID: NZ_PYDN01000087.1 Length: 395
Range 1: 3 to 395

Score:260 bits(665), Expect:1e-80,
Method:Compositional matrix adjust.,
Identities:122/131(93%), Positives:127/131(96%), Gaps:0/131(0%)

Query  135  INNETPGIRYQYNVLPQGWKGSPAIFQSSMTKILEPFRKQNPDIVIYQYMDDLYVGSDLE  194
            +NNETPGIRYQYNVLPQGWKGSPAIFQ+SMTKILEP R +NPDIVIYQYMDDLYVGSDLE
Sbjct  3    VNNETPGIRYQYNVLPQGWKGSPAIFQASMTKILEPLRLKNPDIVIYQYMDDLYVGSDLE  182

Query  195  IGQHRTKIEELRQHLLRWGLTTPDKKHQKEPPFLWMGYELHPDKWTVQPIVLPEKDSWTV  254
            IGQHR KIEELR+HLL+WGLTTPDKKHQKEPPFLWMGYELHPDKWTVQPI LPEKDSWTV
Sbjct  183  IGQHRAKIEELREHLLKWGLTTPDKKHQKEPPFLWMGYELHPDKWTVQPIQLPEKDSWTV  362

Query  255  NDIQKLVGKLN  265
            NDIQKLVGKLN
Sbjct  363  NDIQKLVGKLN  395



>Mycobacterium tuberculosis strain 32847_S31 462, whole genome shotgun sequence
Sequence ID: NZ_QTBX01000137.1 Length: 893
Range 1: 491 to 892

Score:266 bits(680), Expect:2e-80,
Method:Compositional matrix adjust.,
Identities:128/134(96%), Positives:130/134(97%), Gaps:0/134(0%)

Query  1    PISPIETVPVKLKPGMDGPKVKQWPLTEEKIKALVEICTEMEKEGKISKIGPENPYNTPV  60
            PISPIETVPVKLKPGMDGPKVKQWPLTEEKIKAL  IC EMEKEGKI+KIGPENPYNTPV
Sbjct  491  PISPIETVPVKLKPGMDGPKVKQWPLTEEKIKALTAICEEMEKEGKITKIGPENPYNTPV  670

Query  61   FAIKKKDSTKWRKLVDFRELNKRTQDFWEVQLGIPHPAGLKKKKSVTVLDVGDAYFSVPL  120
            FAIKKKDSTKWRKLVDFRELNKRTQDFWE+QLGIPHPAGLKKKKSVTVLDVGDAYFSVPL
Sbjct  671  FAIKKKDSTKWRKLVDFRELNKRTQDFWEIQLGIPHPAGLKKKKSVTVLDVGDAYFSVPL  850

Query  121  DEDFRKYTAFTIPS  134
            DE FRKYTAFTIPS
Sbjct  851  DESFRKYTAFTIPS  892



>Acinetobacter baumannii strain ACM-14 NODE_74_length_787_cov_0.637879, whole genome shotgun sequence
Sequence ID: NZ_PYDA01000074.1 Length: 787
Range 1: 3 to 404

Score:263 bits(673), Expect:6e-80,
```

```
Method:Compositional matrix adjust.,
Identities:128/134(96%), Positives:132/134(98%), Gaps:0/134(0%)

Query  1    PISPIETVPVKLKPGMDGPKVKQWPLTEEKIKALVEICTEMEKEGKISKIGPENPYNTPV  60
            PISPIETVPVKLKPGMDGPKVKQWPLTEEKIKALVEICTEMEKEGKISKIGPENPYNTP+
Sbjct  404  PISPIETVPVKLKPGMDGPKVKQWPLTEEKIKALVEICTEMEKEGKISKIGPENPYNTPI  225

Query  61   FAIKKKDSTKWRKLVDFRELNKRTQDFWEVQLGIPHPAGLKKKKSVTVLDVGDAYFSVPL  120
            FAIKKK+S KWRKLVDFRELNKRTQDFWEVQLGIPHPAGLKKKKSVTVLDVGDAYFSVPL
Sbjct  224  FAIKKKNSNKWRKLVDFRELNKRTQDFWEVQLGIPHPAGLKKKKSVTVLDVGDAYFSVPL  45

Query  121  DEDFRKYTAFTIPS  134
            D++FRKYTA TIPS
Sbjct  44   DKEFRKYTASTIPS  3
```

```
>Klebsiella pneumoniae strain EuSCAPE_MT012, whole genome shotgun sequence
Sequence ID: NZ_UKIX02000064.1 Length: 452
Range 1: 3 to 452

Score:259 bits(661), Expect:9e-80,
Method:Compositional matrix adjust.,
Identities:136/150(91%), Positives:147/150(98%), Gaps:0/150(0%)

Query  391  LPIQKetwetwwteYWQATWIPEWEFVNTPPLVKLWYQLEKEPIVGAETFYVDGAANRET  450
            LPIQKETWETWWT+YWQATWIPEWEFVNTPPLVKLWYQLEKEPI+GAETFYVDGAANR+T
Sbjct  452  LPIQKETWETWWTDYWQATWIPEWEFVNTPPLVKLWYQLEKEPIIGAETFYVDGAANRDT  273

Query  451  KLGKAGYVTNRGRQKVVTLTDTTNQKTELQAIYLALQDSGLEVNIVTDSQYALGIIQAQP  510
            K+GKAGYVT++GRQKVV+LT+TTNQKTEL AI LALQDSG EVNIVTDSQYALGIIQAQP
Sbjct  272  KMGKAGYVTDKGRQKVVSLTETTNQKTELHAIQLALQDSGPEVNIVTDSQYALGIIQAQP  93

Query  511  DQSESELVNQIIEQLIKKEKVYLAWVPAHK  540
            D+SESELVNQIIEQLIKKE+VYL+WVPAHK
Sbjct  92   DKSESELVNQIIEQLIKKERVYLSWVPAHK  3
```

```
>Cronobacter malonaticus strain cro4164A3 contig124, whole genome shotgun sequence
Sequence ID: NZ_NRNF01000124.1 Length: 429
Range 1: 29 to 427

Score:258 bits(660), Expect:9e-80,
Method:Compositional matrix adjust.,
Identities:125/133(94%), Positives:129/133(96%), Gaps:0/133(0%)

Query  1    PISPIETVPVKLKPGMDGPKVKQWPLTEEKIKALVEICTEMEKEGKISKIGPENPYNTPV  60
            PISPI+TVPVKLKPGMDGPKVKQWPLT+EKI+AL  IC EMEKEGKI+KIGPENPYNTPV
Sbjct  29   PISPIDTVPVKLKPGMDGPKVKQWPLTKEKIEALTAICDEMEKEGKITKIGPENPYNTPV  208

Query  61   FAIKKKDSTKWRKLVDFRELNKRTQDFWEVQLGIPHPAGLKKKKSVTVLDVGDAYFSVPL  120
            FAIKKKDSTKWRKLVDFRELNKRTQDFWEVQLGIPHPAGLKKKKSVTVLDVGDAYFSVPL
Sbjct  209  FAIKKKDSTKWRKLVDFRELNKRTQDFWEVQLGIPHPAGLKKKKSVTVLDVGDAYFSVPL  388

Query  121  DEDFRKYTAFTIP  133
            DE FRKYTAFTIP
Sbjct  389  DESFRKYTAFTIP  427
```

```
>Burkholderia pseudomallei strain 2604STDY5518971 ERS240105SCcontig000097, whole genome shotgun sequence
Sequence ID: NZ_UVFZ02000097.1 Length: 1345
Range 1: 205 to 567

Score:219 bits(557), Expect:1e-79,
Method:Compositional matrix adjust.,
Identities:107/121(88%), Positives:118/121(97%), Gaps:0/121(0%)

Query  440  FYVDGAANRETKLGKAGYVTNRGRQKVVTLTDTTNQKTELQAIYLALQDSGLEVNIVTDS  499
            +YVDGAA+RETKLGKAGYVT+RGRQKVV+LT+TTNQKTEL AI+LALQDSG EVNIVTDS
Sbjct  205  YYVDGAASRETKLGKAGYVTDRGRQKVVSLTETTNQKTELHAIHLALQDSGSEVNIVTDS  384

Query  500  QYALGIIQAQPDQSESELVNQIIEQLIKKEKVYLAWVPAHKGIGGNEQVDKLVSAGIRKV  559
            QYALGIIQAQPD SESE+VNQIIE+LIKKE+VYL+WVPAHKGIGGNEQVDKLVS+GIRKV
Sbjct  385  QYALGIIQAQPDMSESEVVNQIIEELIKKERVYLSWVPAHKGIGGNEQVDKLVSSGIRKV  564

Query  560  L  560
            L
Sbjct  565  L  567
```

```
Range 2: 2 to 208

Score:112 bits(281), Expect:1e-79,
Method:Compositional matrix adjust.,
Identities:61/69(88%), Positives:67/69(97%), Gaps:0/69(0%)

Query  351  TGKYARMRGAHTNDVKQLTEAVQKITTESIVIWGKTPKFKLPIQKetwetwwteYWQATW  410
            TGKYA++RGAHTNDVK LTEAVQKI TESIVIWGKTP+F+LPIQ+ETW+TWWTEYWQATW
Sbjct  2    TGKYAKIRGAHTNDVKLLTEAVQKIATESIVIWGKTPRFRLPIQRETWDTWWTEYWQATW  181

Query  411  IPEWEFVNT  419
            IPEWEFVNT
Sbjct  182  IPEWEFVNT  208
```

```
>Cronobacter sakazakii strain cro3040W contig77, whole genome shotgun sequence
Sequence ID: NZ_NRJO01000077.1 Length: 429
Range 1: 3 to 401

Score:258 bits(658), Expect:2e-79,
Method:Compositional matrix adjust.,
Identities:125/133(94%), Positives:129/133(96%), Gaps:0/133(0%)

Query  1    PISPIETVPVKLKPGMDGPKVKQWPLTEEKIKALVEICTEMEKEGKISKIGPENPYNTPV  60
            PISPIETVPVKLKPGMDGPKVKQWPLT+EKI+AL  IC EMEKEGKI+KIGPENPYNTP+
Sbjct  401  PISPIETVPVKLKPGMDGPKVKQWPLTKEKIEALTAICDEMEKEGKITKIGPENPYNTPI  222

Query  61   FAIKKKDSTKWRKLVDFRELNKRTQDFWEVQLGIPHPAGLKKKKSVTVLDVGDAYFSVPL  120
            FAIKKKDSTKWRKLVDFRELNKRTQDFWEVQLGIPHPAGLKKKKSVTVLDVGDAYFSVPL
Sbjct  221  FAIKKKDSTKWRKLVDFRELNKRTQDFWEVQLGIPHPAGLKKKKSVTVLDVGDAYFSVPL  42

Query  121  DEDFRKYTAFTIP  133
             EDFRKYTAFTIP
Sbjct  41   YEDFRKYTAFTIP  3
```

>Escherichia coli strain 8374wB11 NODE_49_length_463_cov_1.77083, whole genome shotgun sequence
Sequence ID: NZ_WSJO01000049.1 Length: 463
Range 1: 58 to 459

Score:254 bits(649), Expect:6e-78,
Method:Compositional matrix adjust.,
Identities:120/134(90%), Positives:131/134(97%), Gaps:0/134(0%)

```
Query  1    PISPIETVPVKLKPGMDGPKVKQWPLTEEKIKALVEICTEMEKEGKISKIGPENPYNTPV  60
            PISPIETVPVKLKPGMDGPK+KQWPLT+EKI+AL EICTE+E++GKISKIGPENPYNTP+
Sbjct  58   PISPIETVPVKLKPGMDGPKIKQWPLTKEKIEALTEICTELEQDGKISKIGPENPYNTPI  237

Query  61   FAIKKKDSTKWRKLVDFRELNKRTQDFWEVQLGIPHPAGLKKKKSVTVLDVGDAYFSVPL  120
            FAIKKK+ST WRKL+DFRELNKRTQDFWEVQLGIPHPAGL+KKKSVTVLDVGDAYFS+PL
Sbjct  238  FAIKKKNSTGWRKLMDFRELNKRTQDFWEVQLGIPHPAGLEKKKSVTVLDVGDAYFSIPL  417

Query  121  DEDFRKYTAFTIPS  134
            DEDFRK TAFTIPS
Sbjct  418  DEDFRKDTAFTIPS  459
```

>Enterococcus faecalis strain 44.1 NODE_33_length_806_cov_1.20619, whole genome shotgun sequence
Sequence ID: NZ_NSJR01000033.1 Length: 806
Range 1: 414 to 806

Score:258 bits(659), Expect:7e-78,
Method:Compositional matrix adjust.,
Identities:125/131(95%), Positives:127/131(96%), Gaps:0/131(0%)

```
Query  1    PISPIETVPVKLKPGMDGPKVKQWPLTEEKIKALVEICTEMEKEGKISKIGPENPYNTPV  60
            PISPIETVPVKLKPGMDGPKVKQWPLTEEKIKALVEICTEMEKEGKISKIGPENPYNTP+
Sbjct  414  PISPIETVPVKLKPGMDGPKVKQWPLTEEKIKALVEICTEMEKEGKISKIGPENPYNTPI  593

Query  61   FAIKKKDSTKWRKLVDFRELNKRTQDFWEVQLGIPHPAGLKKKKSVTVLDVGDAYFSVPL  120
            FAIKKK+S KWRKLVDFR LNKRTQ FWEVQLGIPHPAGLKKKKSVTVLDVGDAYFSVPL
Sbjct  594  FAIKKKNSNKWRKLVDFRGLNKRTQGFWEVQLGIPHPAGLKKKKSVTVLDVGDAYFSVPL  773

Query  121  DEDFRKYTAFT  131
            D DFRKYTAFT
Sbjct  774  DPDFRKYTAFT  806
```

>Acinetobacter baumannii strain ACM-29 NODE_131_length_447_cov_1.05937, whole genome shotgun sequence
Sequence ID: NZ_PYDO01000131.1 Length: 447
Range 1: 29 to 442

Score:251 bits(642), Expect:5e-77,
Method:Compositional matrix adjust.,
Identities:128/138(93%), Positives:135/138(97%), Gaps:0/138(0%)

```
Query  208  HLLRWGLTTPDKKHQKEPPFLWMGYELHPDKWTVQPIVLPEKDSWTVNDIQKLVGKLNWA  267
            HLL+WG +TPD+KHQKEPPFLWMGYELHPDKWTVQPIVLPEKDSWTVNDIQKLVGKLNWA
Sbjct  442  HLLKWGFSTPDQKHQKEPPFLWMGYELHPDKWTVQPIVLPEKDSWTVNDIQKLVGKLNWA  263

Query  268  SQIYPGIKVRQLCKLLRGTKALTEVIPLTeeaelelaeNREILKEPVHGVYYDPSKDLIA  327
            SQIY GIKV+QLCKLLRGTKALTEV+PL+EEAELELAENREILKEPVHGVYYDPSKDL+A
Sbjct  262  SQIYAGIKVKQLCKLLRGTKALTEVVPLSEEAELELAENREILKEPVHGVYYDPSKDLVA  83

Query  328  EIQKQGQGQWTYQIYQEP  345
            EIQKQG GQWTYQIYQEP
Sbjct  82   EIQKQGLGQWTYQIYQEP  29
```

>Escherichia coli strain 8374wH11 NODE_62_length_437_cov_1.95484, whole genome shotgun sequence
Sequence ID: NZ_WSJI01000059.1 Length: 437
Range 1: 2 to 436

Score:250 bits(639), Expect:1e-76,
Method:Compositional matrix adjust.,
Identities:128/145(88%), Positives:139/145(95%), Gaps:0/145(0%)

```
Query  189  VGSDLEIGQHRTKIEELRQHLLRWGLTTPDKKHQKEPPFLWMGYELHPDKWTVQPIVLPE  248
            VGSDLEIGQHR KI+ELR++L +WG  TPDKKHQ+EPPFLWMGYELHPDKWTVQPI LP
Sbjct  436  VGSDLEIGQHRAKIKELREYLWKWGFYTPDKKHQEEPPFLWMGYELHPDKWTVQPIQLPN  257

Query  249  KDSWTVNDIQKLVGKLNWASQIYPGIKVRQLCKLLRGTKALTEVIPLTeeaelelaeNRE  308
            K+SWTVNDIQKLVGKLNWASQIYPGIKV+QLCKLLRGTKALT+++PLTEEAELELAENRE
Sbjct  256  KESWTVNDIQKLVGKLNWASQIYPGIKVKQLCKLLRGTKALTDIVPLTEEAELELAENRE  77

Query  309  ILKEPVHGVYYDPSKDLIAEIQKQG  333
            ILKEPVHGVYYDPSKDLIAE+QKQG
Sbjct  76   ILKEPVHGVYYDPSKDLIAEVQKQG  2
```

>Escherichia coli strain 8374wB4 NODE_91_length_444_cov_2.17035, whole genome shotgun sequence
Sequence ID: NZ_WSPX01000090.1 Length: 444
Range 1: 78 to 443

Score:245 bits(625), Expect:2e-74,
Method:Compositional matrix adjust.,
Identities:113/122(93%), Positives:118/122(96%), Gaps:0/122(0%)

```
Query  112  GDAYFSVPLDEDFRKYTAFTIPSINNETPGIRYQYNVLPQGWKGSPAIFQSSMTKILEPF  171
            GDAYFSVPLDE FRKYTAFTIPS+NNETPGIRYQYNVLPQGWKGSPAIFQ+SMTKILEPF
Sbjct  78   GDAYFSVPLDESFRKYTAFTIPSVNNETPGIRYQYNVLPQGWKGSPAIFQASMTKILEPF  257

Query  172  RKQNPDIVIYQYMDDLYVGSDLEIGQHRTKIEELRQHLLRWGLTTPDKKHQKEPPFLWMG  231
            R +NPDIVIYQYMDDLYVGSDLEIGQHR KIEELR+HLL+WGLTTPDKKHQKEPPFLWMG
Sbjct  258  RLKNPDIVIYQYMDDLYVGSDLEIGQHRAKIEELREHLLKWGLTTPDKKHQKEPPFLWMG  437

Query  232  YE  233
             E
Sbjct  438  CE  443
```

Range 2: 2 to 103

Score:58.2 bits(139), Expect:2e-05,
Method:Composition-based stats.,
Identities:28/34(82%), Positives:30/34(88%), Gaps:1/34(2%)

```
Query  87   FWEVQLGIPHPAGLKKKKSVTVLDVGDA-YFSVP  119
            FWEVQLGIPHPAGLK+KKSVTVLDVG   +FS P
Sbjct  2    FWEVQLGIPHPAGLKQKKSVTVLDVGGCIFFSSP  103
```

>Klebsiella aerogenes strain AUH-KAM4 NODE_62_length_381_cov_0.925197, whole genome shotgun sequence
Sequence ID: NZ_PYHI01000061.1 Length: 381
Range 1: 2 to 379

Score:243 bits(621), Expect:4e-74,
Method:Compositional matrix adjust.,
Identities:115/126(91%), Positives:119/126(94%), Gaps:0/126(0%)

```
Query  91   QLGIPHPAGLKKKKSVTVLDVGDAYFSVPLDEDFRKYTAFTIPSINNETPGIRYQYNVLP  150
            QLGIPHPAGLKKKKSVTVLDVGDAYFSVPLDE+FRKYTAFTIPSINN TPGIRYQYNVLP
Sbjct  379  QLGIPHPAGLKKKKSVTVLDVGDAYFSVPLDENFRKYTAFTIPSINNATPGIRYQYNVLP  200

Query  151  QGWKGSPAIFQSSMTKILEPFRKQNPDIVIYQYMDDLYVGSDLEIGQHRTKIEELRQHLL  210
            QGWKGSPAIFQSSMTKILEPFRKQNPDIVIYQY DDLYVGS LE+ QHR K+EELR HLL
Sbjct  199  QGWKGSPAIFQSSMTKILEPFRKQNPDIVIYQYRDDLYVGSGLELEQHRIKVEELRAHLL  20

Query  211  RWGLTT  216
            +WG TT
Sbjct  19   KWGFTT  2
```

>Escherichia coli strain 8374wA11 NODE_52_length_437_cov_2.24516, whole genome shotgun sequence
Sequence ID: NZ_WSPR01000050.1 Length: 437
Range 1: 1 to 435

Score:244 bits(623), Expect:4e-74,
Method:Compositional matrix adjust.,
Identities:125/145(86%), Positives:137/145(94%), Gaps:0/145(0%)

```
Query  193  LEIGQHRTKIEELRQHLLRWGLTTPDKKHQKEPPFLWMGYELHPDKWTVQPIVLPEKDSW  252
            LEIGQHR KI+ELR++L +WG  TPDKKHQ+EPPFLWMGYELHPDKWTVQPI LP K+SW
Sbjct  1    LEIGQHRAKIKELREYLWKWGFYTPDKKHQEEPPFLWMGYELHPDKWTVQPIQLPNKESW  180

Query  253  TVNDIQKLVGKLNWASQIYPGIKVRQLCKLLRGTKALTEVIPLTeeaelelaeNREILKE  312
            TVNDIQKLVGKLNWASQIYPGIKV+QLCKLLRGTKALT+++PLTEEAELELAENREILKE
Sbjct  181  TVNDIQKLVGKLNWASQIYPGIKVKQLCKLLRGTKALTDIVPLTEEAELELAENREILKE  360

Query  313  PVHGVYYDPSKDLIAEIQKQGQGQW  337
            PVHGVYYDPSK+LIAE+QKQG  QW
Sbjct  361  PVHGVYYDPSKELIAEVQKQGLDQW  435
```

>Escherichia coli strain TzEc080 NODE_52_length_437_cov_2.24516, whole genome shotgun sequence
Sequence ID: NZ_WSFZ01000050.1 Length: 437
Range 1: 1 to 435

Score:244 bits(623), Expect:4e-74,
Method:Compositional matrix adjust.,
Identities:125/145(86%), Positives:137/145(94%), Gaps:0/145(0%)

```
Query  193  LEIGQHRTKIEELRQHLLRWGLTTPDKKHQKEPPFLWMGYELHPDKWTVQPIVLPEKDSW  252
            LEIGQHR KI+ELR++L +WG  TPDKKHQ+EPPFLWMGYELHPDKWTVQPI LP K+SW
Sbjct  1    LEIGQHRAKIKELREYLWKWGFYTPDKKHQEEPPFLWMGYELHPDKWTVQPIQLPNKESW  180

Query  253  TVNDIQKLVGKLNWASQIYPGIKVRQLCKLLRGTKALTEVIPLTeeaelelaeNREILKE  312
            TVNDIQKLVGKLNWASQIYPGIKV+QLCKLLRGTKALT+++PLTEEAELELAENREILKE
Sbjct  181  TVNDIQKLVGKLNWASQIYPGIKVKQLCKLLRGTKALTDIVPLTEEAELELAENREILKE  360

Query  313  PVHGVYYDPSKDLIAEIQKQGQGQW  337
            PVHGVYYDPSK+LIAE+QKQG  QW
Sbjct  361  PVHGVYYDPSKELIAEVQKQGLDQW  435
```

>Acinetobacter baumannii strain ACM-24 NODE_72_length_503_cov_0.531915, whole genome shotgun sequence
Sequence ID: NZ_PYDK01000072.1 Length: 503
Range 1: 1 to 369

Score:242 bits(618), Expect:3e-73,
Method:Compositional matrix adjust.,
Identities:117/123(95%), Positives:121/123(98%), Gaps:0/123(0%)

```
Query  1    PISPIETVPVKLKPGMDGPKVKQWPLTEEKIKALVEICTEMEKEGKISKIGPENPYNTPV  60
            PISPIETVPV LKPGMDGPKVKQWPLTEEKIKAL +ICTEME+EGKISKIGPENPYNTPV
Sbjct  369  PISPIETVPVTLKPGMDGPKVKQWPLTEEKIKALTDICTEMEQEGKISKIGPENPYNTPV  190

Query  61   FAIKKKDSTKWRKLVDFRELNKRTQDFWEVQLGIPHPAGLKKKKSVTVLDVGDAYFSVPL  120
            FAIKKKDSTKWRKLVDFRELNKRTQDFWEVQLGIPHPAGLK+KKSVTVLDVGDAYFSVPL
Sbjct  189  FAIKKKDSTKWRKLVDFRELNKRTQDFWEVQLGIPHPAGLKQKKSVTVLDVGDAYFSVPL  10

Query  121  DED  123
            DE+
Sbjct  9    DEN  1
```

>Klebsiella pneumoniae strain EuSCAPE_MT020, whole genome shotgun sequence
Sequence ID: NZ_UKJB02000092.1 Length: 368
Range 1: 1 to 366

Score:240 bits(613), Expect:4e-73,
Method:Compositional matrix adjust.,
Identities:117/122(96%), Positives:119/122(97%), Gaps:0/122(0%)

```
Query  4    PIETVPVKLKPGMDGPKVKQWPLTEEKIKALVEICTEMEKEGKISKIGPENPYNTPVFAI  63
            PIETVPVKLKPGMDGPKVKQWPLTEEKIKAL  IC EMEKEGKI+KIGPENPYNTPVFAI
Sbjct  366  PIETVPVKLKPGMDGPKVKQWPLTEEKIKALTAICEEMEKEGKITKIGPENPYNTPVFAI  187

Query  64   KKKDSTKWRKLVDFRELNKRTQDFWEVQLGIPHPAGLKKKKSVTVLDVGDAYFSVPLDED  123
            KKKDSTKWRKLVDFRELNKRTQDFWEVQLGIPHPAGLKKKKSVTVLDVGDAYFSVPLDED
Sbjct  186  KKKDSTKWRKLVDFRELNKRTQDFWEVQLGIPHPAGLKKKKSVTVLDVGDAYFSVPLDED  7

Query  124  FR  125
            F+
Sbjct  6    FK  1
```

>Providencia rettgeri strain AFE85_S2 19_contig_46, whole genome shotgun sequence
Sequence ID: NZ_SHDQ01000046.1 Length: 846

Range 1: 3 to 374

Score:246 bits(629), Expect:4e-73,
Method:Compositional matrix adjust.,
Identities:120/124(97%), Positives:121/124(97%), Gaps:0/124(0%)

```
Query  1    PISPIETVPVKLKPGMDGPKVKQWPLTEEKIKALVEICTEMEKEGKISKIGPENPYNTPV  60
            PISPIETVPVKLKPGMDGP+VKQWPLTEEKIKAL EIC EMEKEGKI KIGPENPYNTPV
Sbjct  374  PISPIETVPVKLKPGMDGPRVKQWPLTEEKIKALTEICEEMEKEGKIEKIGPENPYNTPV  195

Query  61   FAIKKKDSTKWRKLVDFRELNKRTQDFWEVQLGIPHPAGLKKKKSVTVLDVGDAYFSVPL  120
            FAIKKKDSTKWRKLVDFRELNKRTQDFWEVQLGIPHPAGLKKKKSVTVLDVGDAYFSVPL
Sbjct  194  FAIKKKDSTKWRKLVDFRELNKRTQDFWEVQLGIPHPAGLKKKKSVTVLDVGDAYFSVPL  15

Query  121  DEDF  124
            DEDF
Sbjct  14   DEDF  3
```

Range 2: 680 to 844

Score:77.4 bits(189), Expect:5e-11,
Method:Compositional matrix adjust.,
Identities:39/55(71%), Positives:42/55(76%), Gaps:0/55(0%)

```
Query  13   KPGMDGPKVKQWPLTEEKIKALVEICTEMEKEGKISKIGPENPYNTPVFAIKKKD  67
            KP     K +  PLTEEKIKA+ EIC EMEKEGKI+KIGPEN YNTP FAIKK D
Sbjct  680  KPAFA*KKKRH*PLTEEKIKAITEICKEMEKEGKITKIGPENQYNTPGFAIKKND  844
```

>Escherichia coli strain 8374wH11 NODE_67_length_417_cov_0.882759, whole genome shotgun sequence
Sequence ID: NZ_WSJI01000064.1 Length: 417
Range 1: 3 to 413

Score:241 bits(614), Expect:6e-73,
Method:Compositional matrix adjust.,
Identities:121/137(88%), Positives:133/137(97%), Gaps:0/137(0%)

```
Query  189  VGSDLEIGQHRTKIEELRQHLLRWGLTTPDKKHQKEPPFLWMGYELHPDKWTVQPIVLPE  248
            VGSDLEIGQHR KIEE+R+HLL+WGLTTPD+KHQKEPPFLWMGYELHPDKWTVQPI LP+
Sbjct  413  VGSDLEIGQHRAKIEEIREHLLKWGLTTPDRKHQKEPPFLWMGYELHPDKWTVQPIQLPD  234

Query  249  KDSWTVNDIQKLVGKLNWASQIYPGIKVRQLCKLLRGTKALTEVIPLTeeaelelaeNRE  308
            K+SWTVNDIQKLVGKLNWASQIYPGI+V+ LCKLLRG KALT+++PLTEEAELELAENRE
Sbjct  233  KESWTVNDIQKLVGKLNWASQIYPGIRVKHLCKLLRGAKALTDIMPLTEEAELELAENRE  54

Query  309  ILKEPVHGVYYDPSKDL  325
            ILKEPVHGVYYDPSK+L
Sbjct  53   ILKEPVHGVYYDPSKEL  3
```

>Acinetobacter baumannii strain ACM-37 NODE_88_length_418_cov_0.570447, whole genome shotgun sequence
Sequence ID: NZ_QAGP01000088.1 Length: 418
Range 1: 1 to 417

Score:238 bits(606), Expect:9e-72,
Method:Compositional matrix adjust.,
Identities:116/139(83%), Positives:124/139(89%), Gaps:0/139(0%)

```
Query  5    IETVPVKLKPGMDGPKVKQWPLTEEKIKALVEICTEMEKEGKISKIGPENPYNTPVFAIK  64
            IET  ++LK G DGP++KQW LTE KIK+L  IC EMEK GKI+KIGPENPYNTPVFAIK
Sbjct  417  IETETIRLKLGRDGPRIKQWALTEGKIKSLQAICEEMEKVGKITKIGPENPYNTPVFAIK  238

Query  65   KKDSTKWRKLVDFRELNKRTQDFWEVQLGIPHPAGLKKKKSVTVLDVGDAYFSVPLDEDF  124
            KK STKWRKLVDFRELNKRTQDFWEVQLGIPHPAGLKKKKSVTVLDV DAYFSV LDE+F
Sbjct  237  KKASTKWRKLVDFRELNKRTQDFWEVQLGIPHPAGLKKKKSVTVLDVVDAYFSVLLDENF  58

Query  125  RKYTAFTIPSINNETPGIR  143
             KYTAFTIPSINN TPG+R
Sbjct  57   MKYTAFTIPSINNATPGVR  1
```

>Mycobacterium tuberculosis strain 32845_S29 230, whole genome shotgun sequence
Sequence ID: NZ_QTBV01000049.1 Length: 601
Range 1: 1 to 366

Score:239 bits(611), Expect:1e-71,
Method:Compositional matrix adjust.,
Identities:116/122(95%), Positives:118/122(96%), Gaps:0/122(0%)

```
Query  1    PISPIETVPVKLKPGMDGPKVKQWPLTEEKIKALVEICTEMEKEGKISKIGPENPYNTPV  60
            PISPIETVPVKLKPGMDGPKVKQWPLTEEKIKAL  IC EMEKEGKI+KIGPENPYNTPV
Sbjct  366  PISPIETVPVKLKPGMDGPKVKQWPLTEEKIKALTAICEEMEKEGKITKIGPENPYNTPV  187

Query  61   FAIKKKDSTKWRKLVDFRELNKRTQDFWEVQLGIPHPAGLKKKKSVTVLDVGDAYFSVPL  120
            FAIKKKDSTKWRKLVDFRELNKRTQDFWE+QLGIPHPAGLKK KSVTVLDVGDAYFSVPL
Sbjct  186  FAIKKKDSTKWRKLVDFRELNKRTQDFWEIQLGIPHPAGLKKNKSVTVLDVGDAYFSVPL  7

Query  121  DE  122
            DE
Sbjct  6    DE  1
```

>Escherichia coli strain 8374wA11 NODE_70_length_381_cov_1.25984, whole genome shotgun sequence
Sequence ID: NZ_WSPR01000067.1 Length: 381
Range 1: 1 to 378

Score:236 bits(602), Expect:2e-71,
Method:Compositional matrix adjust.,
Identities:114/126(90%), Positives:118/126(93%), Gaps:0/126(0%)

```
Query  14   PGMDGPKVKQWPLTEEKIKALVEICTEMEKEGKISKIGPENPYNTPVFAIKKKDSTKWRK  73
            PGMDGP VKQWP +EE IKAL E C +MEKEGKISKIGPENPYNTPVFAIK+KDSTKWRK
Sbjct  1    PGMDGP*VKQWPWSEETIKALTESCEDMEKEGKISKIGPENPYNTPVFAIKRKDSTKWRK  180

Query  74   LVDFRELNKRTQDFWEVQLGIPHPAGLKKKKSVTVLDVGDAYFSVPLDEDFRKYTAFTIP  133
            LVDFRELNKRTQDFWEVQLGIPHPAGLKKKKS+TVLDVGDAYFSVPLDE FRKYTAFTIP
Sbjct  181  LVDFRELNKRTQDFWEVQLGIPHPAGLKKKKSMTVLDVGDAYFSVPLDEGFRKYTAFTIP  360

Query  134  SINNET  139
            S NNET
Sbjct  361  SRNNET  378
```

```
>Escherichia coli strain TzEc080 NODE_70_length_381_cov_1.25984, whole genome shotgun sequence
Sequence ID: NZ_WSFZ01000067.1 Length: 381
Range 1: 1 to 378

Score:236 bits(602), Expect:2e-71,
Method:Compositional matrix adjust.,
Identities:114/126(90%), Positives:118/126(93%), Gaps:0/126(0%)

Query  14   PGMDGPKVKQWPLTEEKIKALVEICTEMEKEGKISKIGPENPYNTPVFAIKKKDSTKWRK  73
            PGMDGP VKQWP +EE IKAL E C +MEKEGKISKIGPENPYNTPVFAIK+KDSTKWRK
Sbjct  1    PGMDGP*VKQWPWSEETIKALTESCEDMEKEGKISKIGPENPYNTPVFAIKRKDSTKWRK  180

Query  74   LVDFRELNKRTQDFWEVQLGIPHPAGLKKKKSVTVLDVGDAYFSVPLDEDFRKYTAFTIP  133
            LVDFRELNKRTQDFWEVQLGIPHPAGLKKKKS+TVLDVGDAYFSVPLDE FRKYTAFTIP
Sbjct  181  LVDFRELNKRTQDFWEVQLGIPHPAGLKKKKSMTVLDVGDAYFSVPLDEGFRKYTAFTIP  360

Query  134  SINNET  139
            S NNET
Sbjct  361  SRNNET  378



>Mycobacterium tuberculosis strain 32845_S29 147, whole genome shotgun sequence
Sequence ID: NZ_QTBV01000025.1 Length: 357
Range 1: 3 to 356

Score:234 bits(598), Expect:7e-71,
Method:Compositional matrix adjust.,
Identities:109/118(92%), Positives:112/118(94%), Gaps:0/118(0%)

Query  115  YFSVPLDEDFRKYTAFTIPSINNETPGIRYQYNVLPQGWKGSPAIFQSSMTKILEPFRKQ  174
            YFSVPLDE FRKYTAFTIPSINNETPGIRYQYNVLPQGWKGSPAIFQ SMTKILEPFR +
Sbjct  3    YFSVPLDESFRKYTAFTIPSINNETPGIRYQYNVLPQGWKGSPAIFQYSMTKILEPFRAK  182

Query  175  NPDIVIYQYMDDLYVGSDLEIGQHRTKIEELRQHLLRWGLTTPDKKHQKEPPFLWMGY  232
            NP+IVIYQYMDDLYVGSDLEIGQHR KIEELR HLL+WG TTPDKKHQKEPPFLWMGY
Sbjct  183  NPEIVIYQYMDDLYVGSDLEIGQHRVKIEELRGHLLKWGFTTPDKKHQKEPPFLWMGY  356



>Mycobacterium tuberculosis strain 32847_S31 147, whole genome shotgun sequence
Sequence ID: NZ_QTBX01000037.1 Length: 357
Range 1: 2 to 355

Score:234 bits(598), Expect:7e-71,
Method:Compositional matrix adjust.,
Identities:109/118(92%), Positives:112/118(94%), Gaps:0/118(0%)

Query  115  YFSVPLDEDFRKYTAFTIPSINNETPGIRYQYNVLPQGWKGSPAIFQSSMTKILEPFRKQ  174
            YFSVPLDE FRKYTAFTIPSINNETPGIRYQYNVLPQGWKGSPAIFQ SMTKILEPFR +
Sbjct  355  YFSVPLDESFRKYTAFTIPSINNETPGIRYQYNVLPQGWKGSPAIFQYSMTKILEPFRAK  176

Query  175  NPDIVIYQYMDDLYVGSDLEIGQHRTKIEELRQHLLRWGLTTPDKKHQKEPPFLWMGY  232
            NP+IVIYQYMDDLYVGSDLEIGQHR KIEELR HLL+WG TTPDKKHQKEPPFLWMGY
Sbjct  175  NPEIVIYQYMDDLYVGSDLEIGQHRVKIEELRGHLLKWGFTTPDKKHQKEPPFLWMGY  2



>Klebsiella aerogenes strain AUH-KAM2 NODE_49_length_404_cov_0.407942, whole genome shotgun sequence
Sequence ID: NZ_PYHG01000048.1 Length: 404
Range 1: 3 to 404

Score:231 bits(589), Expect:3e-69,
Method:Compositional matrix adjust.,
Identities:118/134(88%), Positives:126/134(94%), Gaps:0/134(0%)

Query  192  DLEIGQHRTKIEELRQHLLRWGLTTPDKKHQKEPPFLWMGYELHPDKWTVQPIVLPEKDS  251
            DLEIGQHR KIEELR HLL+WG TTPDKKHQKEPPFLWMGYELHPDKWTV+PI LPEKDS
Sbjct  404  DLEIGQHRAKIEELRDHLLKWGFTTPDKKHQKEPPFLWMGYELHPDKWTVKPIQLPEKDS  225

Query  252  WTVNDIQKLVGKLNWASQIYPGIKVRQLCKLLRGTKALTEVIPLTeeaelelaeNREILK  311
            WTVN IQKLVGKLNWASQIYPGIK++QLCKLLRG KALT+++ LTEEAELELAENREILK
Sbjct  224  WTVNGIQKLVGKLNWASQIYPGIKIKQLCKLLRGAKALTDIVQLTEEAELELAENREILK  45

Query  312  EPVHGVYYDPSKDL  325
            EP HGVYYDPSK+L
Sbjct  44   EPAHGVYYDPSKEL  3



>Methylobacterium radiotolerans strain MAMP 4754 MP-M-radiotolerans_contig_1065, whole genome shotgun sequence
Sequence ID: NZ_NKQS01001052.1 Length: 539
Range 1: 166 to 519

Score:231 bits(589), Expect:1e-68,
Method:Compositional matrix adjust.,
Identities:107/118(91%), Positives:113/118(95%), Gaps:0/118(0%)

Query  179  VIYQYMDDLYVGSDLEIGQHRTKIEELRQHLLRWGLTTPDKKHQKEPPFLWMGYELHPDK  238
             +YQYMDDLYVGSDLEIGQHR KIEELR+HLLRWGLTTPDKKHQKEPPFLWMGYELHPDK
Sbjct  519  CVYQYMDDLYVGSDLEIGQHRAKIEELRKHLLRWGLTTPDKKHQKEPPFLWMGYELHPDK  340

Query  239  WTVQPIVLPEKDSWTVNDIQKLVGKLNWASQIYPGIKVRQLCKLLRGTKALTEVIPLT  296
            WTVQPIVLPEK+SWTVNDIQKLVGKLNWASQIY GIKV Q+CKLLR TKALT++IPLT
Sbjct  339  WTVQPIVLPEKESWTVNDIQKLVGKLNWASQIYTGIKVTQMCKLLR*TKALTDIIPLT  166



>Mycobacterium tuberculosis variant bovis strain CSURQ1132, whole genome shotgun sequence
Sequence ID: NZ_CAJEUK010000179.1 Length: 862
Range 1: 377 to 739

Score:229 bits(584), Expect:2e-66,
Method:Compositional matrix adjust.,
Identities:113/121(93%), Positives:118/121(97%), Gaps:0/121(0%)

Query  1    PISPIETVPVKLKPGMDGPKVKQWPLTEEKIKALVEICTEMEKEGKISKIGPENPYNTPV  60
            PISPIETVPVKLKPGMDGPKVKQWPLTEEKIKAL+EICTEMEKEGKISKIGPENPYNTPV
Sbjct  377  PISPIETVPVKLKPGMDGPKVKQWPLTEEKIKALIEICTEMEKEGKISKIGPENPYNTPV  556

Query  61   FAIKKKDSTKWRKLVDFRELNKRTQDFWEVQLGIPHPAGLKKKKSVTVLDVGDAYFSVPL  120
            FAIKKK+STKWRKLVDFRELNK+TQDFWEVQLGIPHP+GLKKKKSVTVLDVGDA  SV +
Sbjct  557  FAIKKKNSTKWRKLVDFRELNKKTQDFWEVQLGIPHPSGLKKKKSVTVLDVGDA*SSVHV  736

Query  121  D  121
            D
Sbjct  737  D  739
```

Range 2: 377 to 739

Score:229 bits(584), Expect:2e-66,
Method:Compositional matrix adjust.,
Identities:113/121(93%), Positives:118/121(97%), Gaps:0/121(0%)

```
Query  1    PISPIETVPVKLKPGMDGPKVKQWPLTEEKIKALVEICTEMEKEGKISKIGPENPYNTPV  60
            PISPIETVPVKLKPGMDGPKVKQWPLTEEKIKAL+EICTEMEKEGKISKIGPENPYNTPV
Sbjct  377  PISPIETVPVKLKPGMDGPKVKQWPLTEEKIKALIEICTEMEKEGKISKIGPENPYNTPV  556

Query  61   FAIKKKDSTKWRKLVDFRELNKRTQDFWEVQLGIPHPAGLKKKKSVTVLDVGDAYFSVPL  120
            FAIKKK+STKWRKLVDFRELNK+TQDFWEVQLGIPHP+GLKKKKSVTVLDVGDA  SV +
Sbjct  557  FAIKKKNSTKWRKLVDFRELNKKTQDFWEVQLGIPHPSGLKKKKSVTVLDVGDA*SSVHV  736

Query  121  D  121
            D
Sbjct  737  D  739
```

>Escherichia coli strain ECM-2 NODE_147_length_382_cov_1.52941, whole genome shotgun sequence
Sequence ID: NZ_PXIA01000147.1 Length: 382
Range 1: 1 to 375

Score:222 bits(565), Expect:8e-66,
Method:Compositional matrix adjust.,
Identities:115/125(92%), Positives:121/125(96%), Gaps:0/125(0%)

```
Query  213  GLTTPDKKHQKEPPFLWMGYELHPDKWTVQPIVLPEKDSWTVNDIQKLVGKLNWASQIYP  272
            GL+TPD+KHQKEPPFLWMGYEL PDKWTVQPIVLPEKDSWTVNDIQKLVGKLNWASQIY
Sbjct  375  GLSTPDQKHQKEPPFLWMGYELRPDKWTVQPIVLPEKDSWTVNDIQKLVGKLNWASQIYA  196

Query  273  GIKVRQLCKLLRGTKALTEVIPLTeeaelelaeNREILKEPVHGVYYDPSKDLIAEIQKQ  332
            GIKV+QLCKLLRG KALTEV+PL+EEAELELAENREILKEPVHGVYYDPSKDL+AEIQKQ
Sbjct  195  GIKVKQLCKLLRGAKALTEVVPLSEEAELELAENREILKEPVHGVYYDPSKDLVAEIQKQ  16

Query  333  GQGQW  337
            G GQW
Sbjct  15   GLGQW  1
```

>Escherichia coli strain 8374wA11 NODE_44_length_523_cov_1.7197, whole genome shotgun sequence
Sequence ID: NZ_WSPR01000043.1 Length: 523
Range 1: 196 to 522

Score:205 bits(521), Expect:1e-58,
Method:Compositional matrix adjust.,
Identities:97/109(89%), Positives:107/109(98%), Gaps:0/109(0%)

```
Query  1    PISPIETVPVKLKPGMDGPKVKQWPLTEEKIKALVEICTEMEKEGKISKIGPENPYNTPV  60
            PISPIETVPVKLKPGMDGPK+KQWPLT+EKI+AL EICTE+E++GKISKIGPENPYNTP+
Sbjct  196  PISPIETVPVKLKPGMDGPKIKQWPLTKEKIEALTEICTELEQDGKISKIGPENPYNTPI  375

Query  61   FAIKKKDSTKWRKLVDFRELNKRTQDFWEVQLGIPHPAGLKKKKSVTVL  109
            FAIKKK+ST WRKL+DFRELNKRTQDFWEVQLGIPHPAGL+KKKSVTVL
Sbjct  376  FAIKKKNSTGWRKLMDFRELNKRTQDFWEVQLGIPHPAGLEKKKSVTVL  522
```

>Escherichia coli strain TzEc080 NODE_44_length_523_cov_1.7197, whole genome shotgun sequence
Sequence ID: NZ_WSFZ01000043.1 Length: 523
Range 1: 196 to 522

Score:205 bits(521), Expect:1e-58,
Method:Compositional matrix adjust.,
Identities:97/109(89%), Positives:107/109(98%), Gaps:0/109(0%)

```
Query  1    PISPIETVPVKLKPGMDGPKVKQWPLTEEKIKALVEICTEMEKEGKISKIGPENPYNTPV  60
            PISPIETVPVKLKPGMDGPK+KQWPLT+EKI+AL EICTE+E++GKISKIGPENPYNTP+
Sbjct  196  PISPIETVPVKLKPGMDGPKIKQWPLTKEKIEALTEICTELEQDGKISKIGPENPYNTPI  375

Query  61   FAIKKKDSTKWRKLVDFRELNKRTQDFWEVQLGIPHPAGLKKKKSVTVL  109
            FAIKKK+ST WRKL+DFRELNKRTQDFWEVQLGIPHPAGL+KKKSVTVL
Sbjct  376  FAIKKKNSTGWRKLMDFRELNKRTQDFWEVQLGIPHPAGLEKKKSVTVL  522
```

>Acinetobacter baumannii strain ACM-32 NODE_81_length_654_cov_1.01708, whole genome shotgun sequence
Sequence ID: NZ_PYDR01000081.1 Length: 654
Range 1: 318 to 653

Score:204 bits(520), Expect:7e-58,
Method:Compositional matrix adjust.,
Identities:101/112(90%), Positives:107/112(95%), Gaps:0/112(0%)

```
Query  1    PISPIETVPVKLKPGMDGPKVKQWPLTEEKIKALVEICTEMEKEGKISKIGPENPYNTPV  60
            PISPIETVPVKLKPGMDGP+VKQWPLTEEKIKAL  IC EMEKEGKI+KIGPENPYNTPV
Sbjct  318  PISPIETVPVKLKPGMDGPRVKQWPLTEEKIKALQAICEEMEKEGKITKIGPENPYNTPV  497

Query  61   FAIKKKDSTKWRKLVDFRELNKRTQDFWEVQLGIPHPAGLKKKKSVTVLDVG  112
            FAIKKK+STK RKL+DFR+LNK +QDFWEVQLGIPHPAGLKKKKSVTVLDVG
Sbjct  498  FAIKKKNSTK*RKLIDFRKLNKISQDFWEVQLGIPHPAGLKKKKSVTVLDVG  653
```

>Klebsiella pneumoniae strain EuSCAPE_LT004, whole genome shotgun sequence
Sequence ID: NZ_UKJE02000070.1 Length: 335
Range 1: 1 to 333

Score:200 bits(508), Expect:1e-57,
Method:Compositional matrix adjust.,
Identities:96/111(86%), Positives:103/111(92%), Gaps:0/111(0%)

```
Query  74   LVDFRELNKRTQDFWEVQLGIPHPAGLKKKKSVTVLDVGDAYFSVPLDEDFRKYTAFTIP  133
            LVDF+ELNKRTQDFWEVQLGIPHPA LKKKKSVTVLDV DAYFSVPLD+ F+KYTAFTIP
Sbjct  333  LVDFKELNKRTQDFWEVQLGIPHPAELKKKKSVTVLDVKDAYFSVPLDKGFKKYTAFTIP  154

Query  134  SINNETPGIRYQYNVLPQGWKGSPAIFQSSMTKILEPFRKQNPDIVIYQYM  184
            SINNETP IRYQYNVLPQ  K SPAIFQSSMT+ILEPFR +NP+IVIYQY+
Sbjct  153  SINNETPEIRYQYNVLPQR*KRSPAIFQSSMTRILEPFRAKNPEIVIYQYI  1
```

>Klebsiella pneumoniae strain EuSCAPE_MT006, whole genome shotgun sequence
Sequence ID: NZ_UKIU02000071.1 Length: 455
Range 1: 3 to 305

Score:201 bits(510), Expect:2e-57,
Method:Compositional matrix adjust.,
Identities:97/101(96%), Positives:98/101(97%), Gaps:0/101(0%)

```
Query  1    PISPIETVPVKLKPGMDGPKVKQWPLTEEKIKALVEICTEMEKEGKISKIGPENPYNTPV  60
            PISPIETVPVKLKPGMDGPKVKQWPLTEEKIKAL  IC EMEKEGKI+KIGPENPYNTPV
Sbjct  305  PISPIETVPVKLKPGMDGPKVKQWPLTEEKIKALTAICEEMEKEGKITKIGPENPYNTPV  126

Query  61   FAIKKKDSTKWRKLVDFRELNKRTQDFWEVQLGIPHPAGLK  101
            FAIKKKDSTKWRKLVDFRELNKRTQDFWEVQLGIPHPAGLK
Sbjct  125  FAIKKKDSTKWRKLVDFRELNKRTQDFWEVQLGIPHPAGLK  3
```

>Klebsiella pneumoniae strain EuSCAPE_IL005, whole genome shotgun sequence
Sequence ID: NZ_UKJL02000063.1 Length: 298
Range 1: 3 to 296

Score:198 bits(503), Expect:4e-57,
Method:Compositional matrix adjust.,
Identities:95/98(97%), Positives:96/98(97%), Gaps:0/98(0%)

```
Query  37   ICTEMEKEGKISKIGPENPYNTPVFAIKKKDSTKWRKLVDFRELNKRTQDFWEVQLGIPH  96
            IC EMEKEGKI+KIGPENPYNTPVFAIKKKDSTKWRKLVDFRELNKRTQDFWEVQLGIPH
Sbjct  3    ICEEMEKEGKITKIGPENPYNTPVFAIKKKDSTKWRKLVDFRELNKRTQDFWEVQLGIPH  182

Query  97   PAGLKKKKSVTVLDVGDAYFSVPLDEDFRKYTAFTIPS  134
            PAGLKKKKSVTVLDVGDAYFSVPLDE FRKYTAFTIPS
Sbjct  183  PAGLKKKKSVTVLDVGDAYFSVPLDEGFRKYTAFTIPS  296
```

>Acinetobacter baumannii strain ACM-38 NODE_58_length_652_cov_1.55048, whole genome shotgun sequence
Sequence ID: NZ_QAGQ01000058.1 Length: 652
Range 1: 1 to 303

Score:202 bits(513), Expect:6e-57,
Method:Compositional matrix adjust.,
Identities:97/101(96%), Positives:100/101(99%), Gaps:0/101(0%)

```
Query  1    PISPIETVPVKLKPGMDGPKVKQWPLTEEKIKALVEICTEMEKEGKISKIGPENPYNTPV  60
            PISPIETVPVKLKPGMDGPKVKQWPLTEEKIKALVEICTEMEKEGKISKIGPENPYNTP+
Sbjct  303  PISPIETVPVKLKPGMDGPKVKQWPLTEEKIKALVEICTEMEKEGKISKIGPENPYNTPI  124

Query  61   FAIKKKDSTKWRKLVDFRELNKRTQDFWEVQLGIPHPAGLK  101
            FAIKKK+S KWRKLVDFRELNKRTQ+FWEVQLGIPHPAGLK
Sbjct  123  FAIKKKNSDKWRKLVDFRELNKRTQEFWEVQLGIPHPAGLK  1
```

>Klebsiella pneumoniae strain EuSCAPE_LT004, whole genome shotgun sequence
Sequence ID: NZ_UKJE02000066.1 Length: 367
Range 1: 1 to 366

Score:197 bits(501), Expect:2e-56,
Method:Compositional matrix adjust.,
Identities:102/122(84%), Positives:113/122(92%), Gaps:0/122(0%)

```
Query  260  LVGKLNWASQIYPGIKVRQLCKLLRGTKALTEVIPLTeeaelelaeNREILKEPVHGVYY  319
            LVGKLNWASQIYPGIKVRQLC+LLRG KALT+V+PLTEEAEL+LAEN+EILKEPVH VYY
Sbjct  1    LVGKLNWASQIYPGIKVRQLCRLLRGAKALTDVVPLTEEAELKLAENKEILKEPVHKVYY  180

Query  320  DPSKDLIAEIQKQGQGQWTYQIYQEPFKNLKTGKYARMRGAHTNDVKQLTEAVQKITTES  379
            DPSK+LIAEIQKQ   Q TYQ++QEPFKNLKTGKYA+M+ AHTNDVKQLTEAVQKI  E+
Sbjct  181  DPSKELIAEIQKQKNDQ*TYQVFQEPFKNLKTGKYAKMKSAHTNDVKQLTEAVQKIALEA  360

Query  380  IV  381
            IV
Sbjct  361  IV  366
```

>Klebsiella pneumoniae strain EuSCAPE_MT015, whole genome shotgun sequence
Sequence ID: NZ_UKIW02000099.1 Length: 435
Range 1: 3 to 299

Score:198 bits(503), Expect:2e-56,
Method:Compositional matrix adjust.,
Identities:95/99(96%), Positives:96/99(96%), Gaps:0/99(0%)

```
Query  1    PISPIETVPVKLKPGMDGPKVKQWPLTEEKIKALVEICTEMEKEGKISKIGPENPYNTPV  60
            PISPIETVPVKLKPGMDGPKVKQWPLTEEKIKAL  IC EMEKEGKI+KIGPENPYNTPV
Sbjct  299  PISPIETVPVKLKPGMDGPKVKQWPLTEEKIKALTAICEEMEKEGKITKIGPENPYNTPV  120

Query  61   FAIKKKDSTKWRKLVDFRELNKRTQDFWEVQLGIPHPAG  99
            FAIKKKDSTKWRKLVDFRELNKRTQDFWEVQLGIPHPAG
Sbjct  119  FAIKKKDSTKWRKLVDFRELNKRTQDFWEVQLGIPHPAG  3
```

>Klebsiella pneumoniae strain EuSCAPE_IL005, whole genome shotgun sequence
Sequence ID: NZ_UKJL02000064.1 Length: 283
Range 1: 5 to 283

Score:186 bits(471), Expect:2e-52,
Method:Compositional matrix adjust.,
Identities:86/93(92%), Positives:88/93(94%), Gaps:0/93(0%)

```
Query  124  FRKYTAFTIPSINNETPGIRYQYNVLPQGWKGSPAIFQSSMTKILEPFRKQNPDIVIYQY  183
            F KYTAFTIPS NNETPGIRYQYNVLPQGWKGSPAIFQSSMTKILEPFR QNPDIVIYQY
Sbjct  5    FMKYTAFTIPSTNNETPGIRYQYNVLPQGWKGSPAIFQSSMTKILEPFRAQNPDIVIYQY  184

Query  184  MDDLYVGSDLEIGQHRTKIEELRQHLLRWGLTT  216
            MDDLYVGSDLEIGQHR KIEELR+HLL+WG TT
Sbjct  185  MDDLYVGSDLEIGQHRAKIEELREHLLKWGFTT  283
```

>Klebsiella oxytoca strain WCHK085010 NODE_190_length_275_cov_14.1515, whole genome shotgun sequence
Sequence ID: NZ_NWER01000176.1 Length: 275
Range 1: 1 to 273

Score:183 bits(465), Expect:1e-51,
Method:Compositional matrix adjust.,

```
Identities:89/91(98%), Positives:90/91(98%), Gaps:0/91(0%)

Query  23   QWPLTEEKIKALVEICTEMEKEGKISKIGPENPYNTPVFAIKKKDSTKWRKLVDFRELNK  82
             WPLTEEKIKALVEICTEMEKEGKISKIGPENPYNTPVFAIKKKDSTKWRKLVDFRELNK
Sbjct  1    NWPLTEEKIKALVEICTEMEKEGKISKIGPENPYNTPVFAIKKKDSTKWRKLVDFRELNK  180

Query  83   RTQDFWEVQLGIPHPAGLKKKKSVTVLDVGD  113
            RTQDFWEVQLGIPHPAGLK+KKSVTVLDVGD
Sbjct  181  RTQDFWEVQLGIPHPAGLKQKKSVTVLDVGD  273
```

```
>Klebsiella pneumoniae strain EuSCAPE_DE082, whole genome shotgun sequence
Sequence ID: NZ_UKLV02000054.1 Length: 295
Range 1: 3 to 293

Score:177 bits(448), Expect:4e-49,
Method:Compositional matrix adjust.,
Identities:86/97(89%), Positives:89/97(91%), Gaps:0/97(0%)

Query  33   ALVEICTEMEKEGKISKIGPENPYNTPVFAIKKKDSTKWRKLVDFRELNKRTQDFWEVQL  92
            AL  IC EME EGKI+KI PENPYNTPVFAIKKKDSTKWRKLVDFRELNKRTQDF EVQL
Sbjct  3    ALTAICEEMENEGKITKIRPENPYNTPVFAIKKKDSTKWRKLVDFRELNKRTQDF*EVQL  182

Query  93   GIPHPAGLKKKKSVTVLDVGDAYFSVPLDEDFRKYTA  129
             IPHPAGLKKKKSVTVLDV DAYFSVPLD++FRKYTA
Sbjct  183  RIPHPAGLKKKKSVTVLDVEDAYFSVPLDKNFRKYTA  293
```

```
>Klebsiella oxytoca strain WCHKO085010 NODE_191_length_262_cov_12.3243, whole genome shotgun sequence
Sequence ID: NZ_NWER01000177.1 Length: 262
Range 1: 4 to 258

Score:174 bits(440), Expect:6e-48,
Method:Compositional matrix adjust.,
Identities:83/85(98%), Positives:83/85(97%), Gaps:0/85(0%)

Query  94   IPHPAGLKKKKSVTVLDVGDAYFSVPLDEDFRKYTAFTIPSINNETPGIRYQYNVLPQGW  153
            IPHPAGLKKKKSVTVLDVGDAYFSVPL EDFRKYTAFTIPSINNE PGIRYQYNVLPQGW
Sbjct  4    IPHPAGLKKKKSVTVLDVGDAYFSVPLYEDFRKYTAFTIPSINNEAPGIRYQYNVLPQGW  183

Query  154  KGSPAIFQSSMTKILEPFRKQNPDI  178
            KGSPAIFQSSMTKILEPFRKQNPDI
Sbjct  184  KGSPAIFQSSMTKILEPFRKQNPDI  258
```

```
>Escherichia coli strain 8374wB11 NODE_46_length_516_cov_2.15167, whole genome shotgun sequence
Sequence ID: NZ_WSJO01000046.1 Length: 516
Range 1: 181 to 441

Score:168 bits(425), Expect:1e-44,
Method:Compositional matrix adjust.,
Identities:81/87(93%), Positives:84/87(96%), Gaps:0/87(0%)

Query  1    PISPIETVPVKLKPGMDGPKVKQWPLTEEKIKALVEICTEMEKEGKISKIGPENPYNTPV  60
            PISPIETVPVKLKPGMDGP+VKQWPLTEEKIKAL  IC EMEKEGKI+KIGPENPYNTPV
Sbjct  441  PISPIETVPVKLKPGMDGPRVKQWPLTEEKIKALQAICEEMEKEGKITKIGPENPYNTPV  262

Query  61   FAIKKKDSTKWRKLVDFRELNKRTQDF  87
            FAIKKK+STKWRKLVDFRELNKRTQDF
Sbjct  261  FAIKKKNSTKWRKLVDFRELNKRTQDF  181
```

```
Range 2: 2 to 226

Score:129 bits(325), Expect:2e-30,
Method:Compositional matrix adjust.,
Identities:61/75(81%), Positives:65/75(86%), Gaps:0/75(0%)

Query  73   KLVDFRELNKRTQDFWEVQLGIPHPAGLKKKKSVTVLDVGDAYFSVPLDEDFRKYTAFTI  132
            K+  F+    +   FWEVQLGIPHPAGLKKKKSVTVLDVGDAYFSVPLDE+FRKYTAFTI
Sbjct  226  KISRFQGTQ*KNSRFWEVQLGIPHPAGLKKKKSVTVLDVGDAYFSVPLDENFRKYTAFTI  47

Query  133  PSINNETPGIRYQYN  147
            PSINN TPGIRYQYN
Sbjct  46   PSINNATPGIRYQYN  2
```

```
>Acinetobacter baumannii strain ACM-23 NODE_77_length_505_cov_0.960317, whole genome shotgun sequence
Sequence ID: NZ_PYDJ01000077.1 Length: 505
Range 1: 2 to 235

Score:158 bits(399), Expect:5e-41,
Method:Compositional matrix adjust.,
Identities:77/78(99%), Positives:78/78(100%), Gaps:0/78(0%)

Query  1    PISPIETVPVKLKPGMDGPKVKQWPLTEEKIKALVEICTEMEKEGKISKIGPENPYNTPV  60
            PISPIETVPVKLKPGMDGPKVKQWPLTEEKIKALVEICTEMEK+GKISKIGPENPYNTPV
Sbjct  235  PISPIETVPVKLKPGMDGPKVKQWPLTEEKIKALVEICTEMEKKGKISKIGPENPYNTPV  56

Query  61   FAIKKKDSTKWRKLVDFR  78
            FAIKKKDSTKWRKLVDFR
Sbjct  55   FAIKKKDSTKWRKLVDFR  2
```

```
>Mycobacterium tuberculosis strain 32847_S31 146, whole genome shotgun sequence
Sequence ID: NZ_QTBX01000036.1 Length: 235
Range 1: 2 to 235

Score:153 bits(386), Expect:3e-40,
Method:Compositional matrix adjust.,
Identities:70/78(90%), Positives:73/78(93%), Gaps:0/78(0%)

Query  155  GSPAIFQSSMTKILEPFRKQNPDIVIYQYMDDLYVGSDLEIGQHRTKIEELRQHLLRWGL  214
            GSPAIFQ SMTKILEPFR +NP+IVIYQYMDDLYVGSDLEIGQHR KIEELR HLL+WG
Sbjct  235  GSPAIFQHSMTKILEPFRTENPEIVIYQYMDDLYVGSDLEIGQHRVKIEELRGHLLKWGF  56

Query  215  TTPDKKHQKEPPFLWMGY  232
            TTPDKKHQKEPPFLWMGY
Sbjct  55   TTPDKKHQKEPPFLWMGY  2
```

```
>Chlamydia abortus strain 09_01, whole genome shotgun sequence
Sequence ID: NZ_FQLJ01000007.1 Length: 847
Range 1: 52 to 777

Score:156 bits(394), Expect:7e-39,
Method:Compositional matrix adjust.,
Identities:95/247(38%), Positives:139/247(56%), Gaps:6/247(2%)

Query  21   VKQWPLTEEKIKALVEICTEMEKEGKISKIGPENPYNTPVFAIKKKDSTKWRKLVDFREL  80
            V QW L+ EK++A++++  E  K G I      +P+NTP+F IKKK S KWR L D R +
Sbjct  777  VPQWHLSSEKLEAVIQLVEEQLKLGHIEP--STSPWNTPIFVIKKK-SGKWRLLHDLRAI  607

Query  81   NKRTQDFWEVQLGIPHPAGLKKKKSVTVLDVGDAYFSVPLDEDFRKYTAFTIPSINNETP  140
            N++   F  VQ G+P  + L +  ++ ++D+ D +FS+PL    R   AFTIPSIN+  P
Sbjct  606  NEQMNLFGPVQRGLPVLSALPRGWNLIIIDIKDCFFSIPLCPRDRPRFAFTIPSINHMEP  427

Query  141  GIRYQYNVLPQGWKGSPAIFQSSMTKILEPFRKQNPDIVIYQYMDD-LYVGSDLEIGQHR  199
              RYQ+ VLPQG   SP + Q  + + L P R+Q P +++  YMDD L    DL + Q
Sbjct  426  DKRYQWKVLPQGMSNSPTMCQLYVQEALLPVREQFPSLILLLYMDDILLCHKDLTMLQKA  247

Query  200  TKIEELRQHLLRWGLTTPDKKHQKEPPFLWMGYELHPDKWTVQPIVLPEKDSWTVNDIQK  259
                  L + L +WGL    +K Q      ++G  + PDK   Q + +      T+ND QK
Sbjct  246  YPF--LLKTLSQWGLQIATEKVQISDTGQFLGSVVSPDKIVPQKVEIRRDHLHTLNDFQK  73

Query  260  LVGKLNW  266
            L+G +NW
Sbjct  72   LLGDINW  52



>Chlamydia trachomatis strain R33512extract, whole genome shotgun sequence
Sequence ID: NZ_CVNW01000006.1 Length: 1732
Range 1: 596 to 1384

Score:159 bits(403), Expect:1e-37,
Method:Compositional matrix adjust.,
Identities:100/268(37%), Positives:147/268(54%), Gaps:8/268(2%)

Query  21    VKQWPLTEEKIKALVEICTEMEKEGKISKIGPENPYNTPVFAIKKKDSTKWRKLVDFREL  80
             V QW L+ EK++A++++  E  K G I      +P+NTP+F IKKK S KWR L D R +
Sbjct  596   VPQWHLSSEKLEAVIQLVEEQLKLGHIEP--STSPWNTPIFVIKKK-SGKWRLLHDLRAI  766

Query  81    NKRTQDFWEVQLGIPHPAGLKKKKSVTVLDVGDAYFSVPLDEDFRKYTAFTIPSINNETP  140
             N++   F  VQ G+P  + L +  ++ ++D+ D +FS+PL    R   AFTIPSIN+  P
Sbjct  767   NEQMNLFGPVQRGLPVLSALPRGWNLIIIDIKDCFFSIPLCPRDRPRFAFTIPSINHMEP  946

Query  141   GIRYQYNVLPQGWKGSPAIFQSSMTKILEPFRKQNPDIVIYQYMDD-LYVGSDLEIGQHR  199
               RYQ+ VLPQG   SP + Q  + + L P R+Q P +++  YMDD L    DL + Q
Sbjct  947   DKRYQWKVLPQGMSNSPTMCQLYVQEALLPVREQFPSLILLLYMDDILLCHKDLTMLQKA  1126

Query  200   TKIEELRQHLLRWGLTTPDKKHQKEPPFLWMGYELHPDKWTVQPIVLPEKDSWTVNDIQK  259
                   L + L +WGL    +K Q      ++G  + PDK   Q + +      T+ND QK
Sbjct  1127  YPF--LLKTLSQWGLQIATEKVQISDTGQFLGSVVSPDKIVPQKVEIRRDHLHTLNDFQK  1300

Query  260   LVGKLNWASQIY--PGIKVRQLCKLLRG  285
             L+G +NW       P  ++R L  +L G
Sbjct  1301  LLGDINWLRPFLKIPSAELRPLFGILEG  1384



>Chlamydia trachomatis strain R4175extract, whole genome shotgun sequence
Sequence ID: NZ_CVNX01000006.1 Length: 2328
Range 1: 1202 to 2023

Score:160 bits(405), Expect:4e-37,
Method:Compositional matrix adjust.,
Identities:102/279(37%), Positives:151/279(54%), Gaps:8/279(2%)

Query  21    VKQWPLTEEKIKALVEICTEMEKEGKISKIGPENPYNTPVFAIKKKDSTKWRKLVDFREL  80
             V QW L+ EK++A++++  E  K G I      +P+NTP+F IKKK S KWR L D R +
Sbjct  2023  VPQWHLSSEKLEAVIQLVEEQLKLGHIEP--STSPWNTPIFVIKKK-SGKWRLLHDLRAI  1853

Query  81    NKRTQDFWEVQLGIPHPAGLKKKKSVTVLDVGDAYFSVPLDEDFRKYTAFTIPSINNETP  140
             N++   F  VQ G+P  + L +  ++ ++D+ D +FS+PL    R   AFTIPSIN+  P
Sbjct  1852  NEQMNLFGPVQRGLPVLSALPRGWNLIIIDIKDCFFSIPLCPRDRPRFAFTIPSINHMEP  1673

Query  141   GIRYQYNVLPQGWKGSPAIFQSSMTKILEPFRKQNPDIVIYQYMDD-LYVGSDLEIGQHR  199
               RYQ+ VLPQG   SP + Q  + + L P R+Q P +++  YMDD L    DL + Q
Sbjct  1672  DKRYQWKVLPQGMSNSPTMCQLYVQEALLPVREQFPSLILLLYMDDILLCHKDLTMLQKA  1493

Query  200   TKIEELRQHLLRWGLTTPDKKHQKEPPFLWMGYELHPDKWTVQPIVLPEKDSWTVNDIQK  259
                   L + L +WGL    +K Q      ++G  + PDK   Q + +      T+ND QK
Sbjct  1492  YPF--LLKTLSQWGLQIATEKVQISDTGQFLGSVVSPDKIVPQKVEIRRDHLHTLNDFQK  1319

Query  260   LVGKLNWASQIY--PGIKVRQLCKLLRGTKALTEVIPLT  296
             L+G +NW       P  ++R L  +L G   ++    LT
Sbjct  1318  LLGDINWLRPFLKIPSAELRPLFGILEGDPHISSPRTLT  1202



>Chlamydia abortus strain 09_03, whole genome shotgun sequence
Sequence ID: NZ_FQLL01000003.1 Length: 4440
Range 1: 80 to 901

Score:161 bits(407), Expect:4e-37,
Method:Compositional matrix adjust.,
Identities:102/279(37%), Positives:151/279(54%), Gaps:8/279(2%)

Query  21   VKQWPLTEEKIKALVEICTEMEKEGKISKIGPENPYNTPVFAIKKKDSTKWRKLVDFREL  80
            V QW L+ EK++A++++  E  K G I      +P+NTP+F IKKK S KWR L D R +
Sbjct  80   VPQWHLSSEKLEAVIQLVEEQLKLGHIEP--STSPWNTPIFVIKKK-SGKWRLLHDLRAI  250

Query  81   NKRTQDFWEVQLGIPHPAGLKKKKSVTVLDVGDAYFSVPLDEDFRKYTAFTIPSINNETP  140
            N++   F  VQ G+P  + L +  ++ ++D+ D +FS+PL    R   AFTIPSIN+  P
Sbjct  251  NEQMNLFGPVQRGLPVLSALPRGWNLIIIDIKDCFFSIPLCPRDRPRFAFTIPSINHMEP  430

Query  141  GIRYQYNVLPQGWKGSPAIFQSSMTKILEPFRKQNPDIVIYQYMDD-LYVGSDLEIGQHR  199
              RYQ+ VLPQG   SP + Q  + + L P R+Q P +++  YMDD L    DL + Q
Sbjct  431  DKRYQWKVLPQGMSNSPTMCQLYVQEALLPVREQFPSLILLLYMDDILLCHKDLTMLQKA  610

Query  200  TKIEELRQHLLRWGLTTPDKKHQKEPPFLWMGYELHPDKWTVQPIVLPEKDSWTVNDIQK  259
                  L + L +WGL    +K Q      ++G  + PDK   Q + +      T+ND QK
Sbjct  611  YPF--LLKTLSQWGLQIATEKVQISDTGQFLGSVVSPDKIVPQKVEIRRDHLHTLNDFQK  784

Query  260  LVGKLNWASQIY--PGIKVRQLCKLLRGTKALTEVIPLT  296
            L+G +NW       P  ++R L  +L G   ++    LT
Sbjct  785  LLGDINWLRPFLKIPSAELRPLFSILEGDPHISSPRTLT  901
```

>Chlamydia abortus strain 15a, whole genome shotgun sequence
Sequence ID: NZ_FQLM01000004.1 Length: 3222
Range 1: 1236 to 2057

Score:160 bits(406), Expect:4e-37,
Method:Compositional matrix adjust.,
Identities:102/279(37%), Positives:151/279(54%), Gaps:8/279(2%)

```
Query  21    VKQWPLTEEKIKALVEICTEMEKEGKISKIGPENPYNTPVFAIKKKDSTKWRKLVDFREL  80
             V QW L+ EK++A++++  E  K G I       +P+NTP+F IKKK S KWR L D R +
Sbjct  2057  VPQWHLSSEKLEAVIQLVEEQLKLGHIEP--STSPWNTPIFVIKKK-SGKWRLLHDLRAI  1887

Query  81    NKRTQDFWEVQLGIPHPAGLKKKKSVTVLDVGDAYFSVPLDEDFRKYTAFTIPSINNETP  140
             N++   F  VQ G+P  + L +  ++ ++D+ D +FS+PL    R   AFTIPSIN+  P
Sbjct  1886  NEQMNLFGPVQRGLPVLSALPRGWNLIIIDIKDCFFSIPLCPRDRPRFAFTIPSINHMEP  1707

Query  141   GIRYQYNVLPQGWKGSPAIFQSSMTKILEPFRKQNPDIVIYQYMDD-LYVGSDLEIGQHR  199
               RYQ+ VLPQG   SP + Q  + + L P R+Q P +++  YMDD L    DL + Q
Sbjct  1706  DKRYQWKVLPQGMSNSPTMCQLYVQEALLPVREQFPSLILLLYMDDILLCHKDLTMLQKA  1527

Query  200   TKIEELRQHLLRWGLTTPDKKHQKEPPFLWMGYELHPDKWTVQPIVLPEKDSWTVNDIQK  259
                  L + L +WGL    +K Q       ++G  + PDK   Q + +       T+ND QK
Sbjct  1526  YPF--LLKTLSQWGLQIATEKVQISDTGQFLGSVVSPDKIVPQKVEIRRDHLHTLNDFQK  1353

Query  260   LVGKLNWASQIY--PGIKVRQLCKLLRGTKALTEVIPLT  296
             L+G +NW        P  ++R L  +L G   ++    LT
Sbjct  1352  LLGDINWLRPFLKIPSAELRPLFSILEGDPHISSPRTLT  1236
```

>Chlamydia abortus strain 11_01, whole genome shotgun sequence
Sequence ID: NZ_FQLK01000002.1 Length: 5240
Range 1: 2674 to 3495

Score:160 bits(406), Expect:5e-37,
Method:Compositional matrix adjust.,
Identities:102/279(37%), Positives:151/279(54%), Gaps:8/279(2%)

```
Query  21    VKQWPLTEEKIKALVEICTEMEKEGKISKIGPENPYNTPVFAIKKKDSTKWRKLVDFREL  80
             V QW L+ EK++A++++  E  K G I       +P+NTP+F IKKK S KWR L D R +
Sbjct  3495  VPQWHLSSEKLEAVIQLVEEQLKLGHIEP--STSPWNTPIFVIKKK-SGKWRLLHDLRAI  3325

Query  81    NKRTQDFWEVQLGIPHPAGLKKKKSVTVLDVGDAYFSVPLDEDFRKYTAFTIPSINNETP  140
             N++   F  VQ G+P  + L +  ++ ++D+ D +FS+PL    R   AFTIPSIN+  P
Sbjct  3324  NEQMNLFGPVQRGLPVLSALPRGWNLIIIDIKDCFFSIPLCPRDRPRFAFTIPSINHMEP  3145

Query  141   GIRYQYNVLPQGWKGSPAIFQSSMTKILEPFRKQNPDIVIYQYMDD-LYVGSDLEIGQHR  199
               RYQ+ VLPQG   SP + Q  + + L P R+Q P +++  YMDD L    DL + Q
Sbjct  3144  DKRYQWKVLPQGMSNSPTMCQLYVQEALLPVREQFPSLILLLYMDDILLCHKDLTMLQKA  2965

Query  200   TKIEELRQHLLRWGLTTPDKKHQKEPPFLWMGYELHPDKWTVQPIVLPEKDSWTVNDIQK  259
                  L + L +WGL    +K Q       ++G  + PDK   Q + +       T+ND QK
Sbjct  2964  YPF--LLKTLSQWGLQIATEKVQISDTGQFLGSVVSPDKIVPQKVEIRRDHLHTLNDFQK  2791

Query  260   LVGKLNWASQIY--PGIKVRQLCKLLRGTKALTEVIPLT  296
             L+G +NW        P  ++R L  +L G   ++    LT
Sbjct  2790  LLGDINWLRPFLKIPSAELRPLFSILEGDPHISSPRTLT  2674
```

>Chlamydia abortus strain 09_02, whole genome shotgun sequence
Sequence ID: NZ_FQLN01000004.1 Length: 4507
Range 1: 2412 to 3233

Score:160 bits(405), Expect:8e-37,
Method:Compositional matrix adjust.,
Identities:102/279(37%), Positives:151/279(54%), Gaps:8/279(2%)

```
Query  21    VKQWPLTEEKIKALVEICTEMEKEGKISKIGPENPYNTPVFAIKKKDSTKWRKLVDFREL  80
             V QW L+ EK++A++++  E  K G I       +P+NTP+F IKKK S KWR L D R +
Sbjct  2412  VPQWHLSSEKLEAVIQLVEEQLKLGHIEP--STSPWNTPIFVIKKK-SGKWRLLHDLRAI  2582

Query  81    NKRTQDFWEVQLGIPHPAGLKKKKSVTVLDVGDAYFSVPLDEDFRKYTAFTIPSINNETP  140
             N++   F  VQ G+P  + L +  ++ ++D+ D +FS+PL    R   AFTIPSIN+  P
Sbjct  2583  NEQMNLFGPVQRGLPVLSALPRGWNLIIIDIKDCFFSIPLCPRDRPRFAFTIPSINHMEP  2762

Query  141   GIRYQYNVLPQGWKGSPAIFQSSMTKILEPFRKQNPDIVIYQYMDD-LYVGSDLEIGQHR  199
               RYQ+ VLPQG   SP + Q  + + L P R+Q P +++  YMDD L    DL + Q
Sbjct  2763  DKRYQWKVLPQGMSNSPTMCQLYVQEALLPVREQFPSLILLLYMDDILLCHKDLTMLQKA  2942

Query  200   TKIEELRQHLLRWGLTTPDKKHQKEPPFLWMGYELHPDKWTVQPIVLPEKDSWTVNDIQK  259
                  L + L +WGL    +K Q       ++G  + PDK   Q + +       T+ND QK
Sbjct  2943  YPF--LLKTLSQWGLQIATEKVQISDTGQFLGSVVSPDKIVPQKVEIRRDHLHTLNDFQK  3116

Query  260   LVGKLNWASQIY--PGIKVRQLCKLLRGTKALTEVIPLT  296
             L+G +NW        P  ++R L  +L G   ++    LT
Sbjct  3117  LLGDINWLRPFLKIPSAELRPLFGILEGDPHISSPRTLT  3233
```

>Chlamydia abortus strain 09_05, whole genome shotgun sequence
Sequence ID: NZ_FQLG01000005.1 Length: 4831
Range 1: 3076 to 3801

Score:157 bits(396), Expect:1e-35,
Method:Compositional matrix adjust.,
Identities:95/247(38%), Positives:139/247(56%), Gaps:6/247(2%)

```
Query  21    VKQWPLTEEKIKALVEICTEMEKEGKISKIGPENPYNTPVFAIKKKDSTKWRKLVDFREL  80
             V QW L+ EK++A++++  E  K G I       +P+NTP+F IKKK S KWR L D R +
Sbjct  3801  VPQWHLSSEKLEAVIQLVEEQLKLGHIEP--STSPWNTPIFVIKKK-SGKWRLLHDLRAI  3631

Query  81    NKRTQDFWEVQLGIPHPAGLKKKKSVTVLDVGDAYFSVPLDEDFRKYTAFTIPSINNETP  140
             N++   F  VQ G+P  + L +  ++ ++D+ D +FS+PL    R   AFTIPSIN+  P
Sbjct  3630  NEQMNLFGPVQRGLPVLSALPRGWNLIIIDIKDCFFSIPLCPRDRPRFAFTIPSINHMEP  3451

Query  141   GIRYQYNVLPQGWKGSPAIFQSSMTKILEPFRKQNPDIVIYQYMDD-LYVGSDLEIGQHR  199
               RYQ+ VLPQG   SP + Q  + + L P R+Q P +++  YMDD L    DL + Q
Sbjct  3450  DKRYQWKVLPQGMSNSPTMCQLYVQEALLPVREQFPSLILLLYMDDILLCHKDLTMLQKA  3271

Query  200   TKIEELRQHLLRWGLTTPDKKHQKEPPFLWMGYELHPDKWTVQPIVLPEKDSWTVNDIQK  259
                  L + L +WGL    +K Q       ++G  + PDK   Q + +       T+ND QK
Sbjct  3270  YPF--LLKTLSQWGLQIATEKVQISDTGQFLGSVVSPDKIVPQKVEIRRDHLHTLNDFQK  3097

Query  260   LVGKLNW  266
             L+G +NW
Sbjct  3096  LLGDINW  3076
```

>Providencia rettgeri strain AFE220_S1 26_contig_67, whole genome shotgun sequence
Sequence ID: NZ_SHDM01000067.1 Length: 512
Range 1: 1 to 219

Score:142 bits(358), Expect:4e-35,
Method:Compositional matrix adjust.,
Identities:68/73(93%), Positives:71/73(97%), Gaps:0/73(0%)

```
Query  1    PISPIETVPVKLKPGMDGPKVKQWPLTEEKIKALVEICTEMEKEGKISKIGPENPYNTPV  60
            PISPIETVPVKLKPGMDGPKVKQWPLTEEKIKAL EIC EME+EGKI+KIGPENPYNTP+
Sbjct  219  PISPIETVPVKLKPGMDGPKVKQWPLTEEKIKALTEICDEMEREGKITKIGPENPYNTPI  40

Query  61   FAIKKKDSTKWRK  73
            FAIKKKDSTKWRK
Sbjct  39   FAIKKKDSTKWRK  1
```

>Pseudomonas aromaticivorans strain MAP12 34, whole genome shotgun sequence
Sequence ID: NZ_JAHRGL010000035.1 Length: 7031
Range 1: 3 to 806

Score:143 bits(360), Expect:5e-31,
Method:Compositional matrix adjust.,
Identities:91/272(33%), Positives:147/272(54%), Gaps:6/272(2%)

```
Query  21   VKQWPLTEEKIKALVEICTEMEKEGKISKIGPENPYNTPVFAIKKKDSTKWRKLVDFREL  80
            V QWP+  E++ +L ++         + +     P+N+ VF IKKK S KWR L D R +
Sbjct  3    VGQWPMNSEELVSLEKLVH*QLDA*HVEETT--FPWNSTVFVIKKK-SGKWRMLTDLRAI  173

Query  81   NKRTQDFWEVQLGIPHPAGLKKKKSVTVLDVGDAYFSVPLDEDFRKYTAFTIPSINNETP  140
            NK  Q    +Q G+P P+ + K   + V+D+ D +F++PL E+ R+  AFT+P++NN  P
Sbjct  174  NKVIQPMGSLQPGMPLPSLIPKGWPIIVIDLKDCFFTIPLQENDREKFAFTVPTLNNSQP  353

Query  141  GIRYQYNVLPQGWKGSPAIFQSSMTKILEPFRKQNPDIVIYQYMDDLYVGSDLEIGQHRT  200
              RYQ+ VLPQG   SP + Q  + + LE  RK+ P  ++Y YMDD+ + SD
Sbjct  354  VRRYQWRVLPQGMLNSPTLCQHFVQQPLEIIRKKLPQSLVYHYMDDILL-SDSNKETLER  530

Query  201  KIEELRQHLLRWGLTTPDKKHQKEPPFLWMGYELHPDKWTVQPIVLPEKDSWTVNDIQKL  260
              E +++ L RWGL    +K Q+     ++GY++   +   Q + +      T+N +QKL
Sbjct  531  MFEMVKEVLPRWGLQIAPEKIQRGDSINYLGYKIDLQRIRPQKVQIRRDRYQTLNSLQKL  710

Query  261  VGKLNWASQI--YPGIKVRQLCKLLRGTKALT  290
            +G+++    I    G  ++ L   L+G K L
Sbjct  711  LGEISQLQTIIGVEGHDLKHLKMALKGDKDLN  806
```

>Neisseria gonorrhoeae strain GC-206 NODE_177_length_223_cov_0.803571_ID_20526, whole genome shotgun sequence
Sequence ID: NZ_SAWN01000176.1 Length: 223
Range 1: 26 to 223

Score:125 bits(314), Expect:4e-30,
Method:Compositional matrix adjust.,
Identities:61/66(92%), Positives:63/66(95%), Gaps:0/66(0%)

```
Query  1    PISPIETVPVKLKPGMDGPKVKQWPLTEEKIKALVEICTEMEKEGKISKIGPENPYNTPV  60
            PISPIETVPVKLKPGMDGPKVKQWPLTEEKIKAL  IC EMEKEGKI+KIGPENPYNTPV
Sbjct  26   PISPIETVPVKLKPGMDGPKVKQWPLTEEKIKALTAICEEMEKEGKITKIGPENPYNTPV  205

Query  61   FAIKKK  66
            FAI+KK
Sbjct  206  FAIRKK  223
```

>Enterococcus faecium strain E21_6 NODE_88_length_518_cov_1.29923, whole genome shotgun sequence
Sequence ID: NZ_JAANZJ010000088.1 Length: 518
Range 1: 305 to 514

Score:128 bits(322), Expect:4e-30,
Method:Compositional matrix adjust.,
Identities:61/70(87%), Positives:67/70(95%), Gaps:0/70(0%)

```
Query  1    PISPIETVPVKLKPGMDGPKVKQWPLTEEKIKALVEICTEMEKEGKISKIGPENPYNTPV  60
            PIS IETVPVKLKPGMDGP+VKQWPLTEEKIKAL EIC EME+EGKI+KIGPENPYNTP+
Sbjct  305  PISTIETVPVKLKPGMDGPRVKQWPLTEEKIKALTEICAEMEREGKITKIGPENPYNTPI  484

Query  61   FAIKKKDSTK  70
            FAIKKK+ST+
Sbjct  485  FAIKKKNSTE  514
```

>Methylobacterium radiotolerans strain MAMP 4754 MP-M-radiotolerans_contig_206, whole genome shotgun sequence
Sequence ID: NZ_NKQS01000204.1 Length: 483
Range 1: 3 to 191

Score:122 bits(307), Expect:4e-28,
Method:Compositional matrix adjust.,
Identities:59/63(94%), Positives:61/63(96%), Gaps:0/63(0%)

```
Query  1    PISPIETVPVKLKPGMDGPKVKQWPLTEEKIKALVEICTEMEKEGKISKIGPENPYNTPV  60
            PISPI+TVPVKLKPGMDGPKVKQWPLTEEKIKAL EIC +MEKEGKISKIGPENPYNTPV
Sbjct  191  PISPIDTVPVKLKPGMDGPKVKQWPLTEEKIKALTEICEDMEKEGKISKIGPENPYNTPV  12

Query  61   FAI  63
            FAI
Sbjct  11   FAI  3
```

>Acinetobacter baumannii strain ACM-36 NODE_67_length_567_cov_1.13409, whole genome shotgun sequence
Sequence ID: NZ_QAGO01000067.1 Length: 567
Range 1: 387 to 566

Score:122 bits(305), Expect:2e-27,
Method:Compositional matrix adjust.,
Identities:58/60(97%), Positives:60/60(100%), Gaps:0/60(0%)

```
Query  1    PISPIETVPVKLKPGMDGPKVKQWPLTEEKIKALVEICTEMEKEGKISKIGPENPYNTPV  60
            PISPIETVPVKLKPGMDGPKVKQWPLTEEKIKALVEICTEMEKEGKI+KIGPENPYNTP+
Sbjct  387  PISPIETVPVKLKPGMDGPKVKQWPLTEEKIKALVEICTEMEKEGKITKIGPENPYNTPI  566
```

>Escherichia coli strain 8374wG10 NODE_65_length_433_cov_1.76797, whole genome shotgun sequence

```
Sequence ID: NZ_WSJQ01000065.1 Length: 433
Range 1: 232 to 432

Score:112 bits(280), Expect:2e-24,
Method:Compositional matrix adjust.,
Identities:55/67(82%), Positives:59/67(88%), Gaps:0/67(0%)

Query  1    PISPIETVPVKLKPGMDGPKVKQWPLTEEKIKALVEICTEMEKEGKISKIGPENPYNTPV  60
            PISPIETVPVKLKPGMDGPK+KQWPLT+EKI+AL E C E EKEGKISKIGPENP N PV
Sbjct  232  PISPIETVPVKLKPGMDGPKIKQWPLTKEKIEALTESCEEREKEGKISKIGPENPSNPPV  411

Query  61   FAIKKKD  67
            F  K+KD
Sbjct  412  FDRKRKD  432




>Pseudomonas aeruginosa strain VET-42 NODE_86_length_1154_cov_8.30919_ID_171, whole genome shotgun sequence
Sequence ID: NZ_RACX01000118.1 Length: 1154
Range 1: 939 to 1103

Score:104 bits(260), Expect:1e-19,
Method:Compositional matrix adjust.,
Identities:50/55(91%), Positives:55/55(100%), Gaps:0/55(0%)

Query  506   IQAQPDQSESELVNQIIEQLIKKEKVYLAWVPAHKGIGGNEQVDKLVSAGIRKVL  560
             +QAQPD+SESE+VNQIIEQLIKKEKVYL+WVPAHKGIGGNEQVDKLVS+GIRKVL
Sbjct  1103  VQAQPDKSESEVVNQIIEQLIKKEKVYLSWVPAHKGIGGNEQVDKLVSSGIRKVL  939




>Burkholderia pseudomallei strain A-175-05-2-02, whole genome shotgun sequence
Sequence ID: NZ_UVFW01000138.1 Length: 455
Range 1: 1 to 138

Score:89.7 bits(221), Expect:2e-16,
Method:Compositional matrix adjust.,
Identities:43/46(93%), Positives:44/46(95%), Gaps:0/46(0%)

Query  1    PISPIETVPVKLKPGMDGPKVKQWPLTEEKIKALVEICTEMEKEGK  46
            PISPIETVPVKLKPGMDGPKVKQWPLTEEKIKAL EIC +MEKEGK
Sbjct  138  PISPIETVPVKLKPGMDGPKVKQWPLTEEKIKALTEICNDMEKEGK  1




>Tumebacillus flagellatus strain GST4 Contig077, whole genome shotgun sequence
Sequence ID: NZ_JMIR01000077.1 Length: 2091
Range 1: 1046 to 1819

Score:87.8 bits(216), Expect:2e-13,
Method:Compositional matrix adjust.,
Identities:71/267(27%), Positives:128/267(47%), Gaps:21/267(7%)

Query  20    KVKQWPLTEEKIKALVEICTEMEKEGKISKIGPENPYNTPVFAIKKKDSTKWRKLVDFRE  79
              +KQ+P++++    +      +  +G +  +  ++P+NTP+  +KK  +  +R + D RE
Sbjct  1819  SIKQYPMSQKARLGIKPHIQRLLDQGIL--VPCQSPWNTPLLPVKKPGTNDYRPVQDLRE  1646

Query  80    LNKRTQDFWEVQLGIPHP----AGLKKKKS-VTVLDVGDAYFSVPLDEDFRKYTAFTIPS  134
             +NKR +D   +   +P+P    +GL       TVLD+ DA+F + L    +   AF
Sbjct  1645  VNKRVED---IHPTVPNPYNLLSGLPPSHQWYTVLDLKDAFFCLRLHPTSQPLFAF---E  1484

Query  135   INNETPGIRYQ--YNVLPQGWKGSPAIFQSSMTKILEPFRKQNPDIVIYQYMDDLYVGSD  192
               +   GI  Q  +  LPQG+K SP +F  ++ + L  FR Q+PD+++ QY+DDL + +
Sbjct  1483  WRDPEMGISGQLTWTRLPQGFKNSPTLFDEALHRDLADFRIQHPDLILLQYVDDLLLAAT  1304

Query  193   LEIGQHRTKIEELRQHLLRWGLTTPDKKHQ-KEPPFLWMGYEL-HPDKWTVQP---IVLP  247
              E+   +     L Q L   G     KK Q  +    ++GY L    +W  +     V+
Sbjct  1303  SELDCQQ-GTRALLQTLGNLGYRASAKKAQICQKQVKYLGYLLKEGQRWLTEARKETVMG  1127

Query  248   EKDSWTVNDIQKLVGKLNWASQIYPGI  274
             +    T   +++ +G   +     PG
Sbjct  1126  QPTPKTPRQLRRFLGTAGFCRLFIPGF  1046




>Klebsiella pneumoniae strain BL2026 NODE_25_length_1934_cov_0.482568, whole genome shotgun sequence
Sequence ID: NZ_VICY01000025.1 Length: 1934
Range 1: 1208 to 1894

Score:81.6 bits(200), Expect:2e-11,
Method:Compositional matrix adjust.,
Identities:70/240(29%), Positives:121/240(50%), Gaps:22/240(9%)

Query  6     ETVPVKLKPGMDGPKVKQWPLTEEKIKALVEICTEMEKEGKISKIGPENPYNTPVFAIKK  65
             E V + LK       + KQ+PL  E  + L ++     K+ ++  +  E+ +NTP+  +KK
Sbjct  1208  EPVKIFLKESTKLVRQKQYPLKLEARRGLEKMIENFLKKYRLL-VECESEFNTPILPVKK  1384

Query  66    KDSTKWRKLVDFRELNKRTQDFWEVQLGIPHP--AGLKKK-KSVTVLDVGDAYFSVPLDE  122
              +S ++R + DF   NK     +    G P      LK+  K  TVL++ DA+F + LD+
Sbjct  1385  PESEEYRLVQDFNS-NKFNSSRYSPLSGKPVSLLTALKENHKWFTVLNL*DAFFCILLDK  1561

Query  123   DFRKYTAFTIPSINNETPGIRYQYN--VLPQGWKGSPAIFQSSMTKILEPFRKQNPDIVI  180
             + + Y AF    + N T G + Q    VLPQG+K SP IF + + K LE  +   P+ ++
Sbjct  1562  NSQAYFAF---ELQNPTTGQKTQLTWAVLPQGFKNSPMIFGNQLAKELETCKIIYPERLV  1732

Query  181   YQYMDDLYVGSDLEIGQHRTKIEELRQHLLRWGLTTPDKKHQKEPP------FLWMGYEL  234
              QY+DD+ + ++      R +   L  +LL + L   D +  K+         +++G+E+
Sbjct  1733  LQYVDDILIATE-----TREEYFNLTTNLLNF-LGQSDNRFSKKKAQIGKETVMYLGFEI  1894




>Streptococcus pneumoniae strain NN2021-1 12_contig_107, whole genome shotgun sequence
Sequence ID: NZ_JAGYHD010000101.1 Length: 1657
Range 1: 54 to 641

Score:79.3 bits(194), Expect:6e-11,
Method:Compositional matrix adjust.,
Identities:64/208(31%), Positives:102/208(49%), Gaps:20/208(9%)

Query  6    ETVPVKLKPGMDGPKVKQWPLTEEKIKALVEICTEMEKEGKISKIGPENPYNTPVFAIKK  65
            E V + L+     P  KQ+PL  E  + L  +  +    G +  +   +P NTP+ ++KK
Sbjct  54   ELVIIVLQDPTRFPNQKQYPLKREAREGLQPLINKFLACGLL--VPTSSPCNTPILSVKK  227

Query  66   KDSTKWRKLVDFRELNKRTQDFWEVQLGIPHP-----AGLKKKKSVTVLDVGDAYFSVPL  120
            KD T WR + D R +N+           +P+P       L   K  TVLD+ DA+F +PL
Sbjct  228  KDRT-WRMVQDLRIINEAVVPLHPT---VPNPYVILGEILPSAKWFTVLDLKDAFFCIPL  395

Query  121  DEDFRKYTAFTIPSINNETPGIRYQ---YNVLPQGWKGSPAIFQSSMTKILEPFRKQNPD  177
             ++ +   AF       E PG ++Q   + VLPQG++ SP +F  ++++ L       P+
```

```
Sbjct  396  AKESQYLFAFEW-----EAPGEKHQQMTWTVLPQGFRDSPHLFGQALSRDLL-DLDLGPN  557

Query  178  IVIYQYMDDLYVGSDLEIGQHRTKIEEL  205
              I QY+DDL + S  E    +  I+ L
Sbjct  558  GKILQYVDDLLICSPDEKSAQQHAIQVL  641
```

>Klebsiella pneumoniae strain BL1427 NODE_43_length_1114_cov_0.430598, whole genome shotgun sequence
Sequence ID: NZ_VICX01000043.1 Length: 1114
Range 1: 49 to 609

Score:76.6 bits(187), Expect:2e-10,
Method:Compositional matrix adjust.,
Identities:59/192(31%), Positives:100/192(52%), Gaps:10/192(5%)

```
Query  6    ETVPVKLKPGMDGPKVKQWPLTEEKIKALVEICTEMEKEGKISKIGPENPYNTPVFAIKK  65
            E V + LK      +  Q+P   E  + L ++     K+ ++  +  E+ +NTP+  +KK
Sbjct  609  EPVKIFLKESTKLVRQNQYPSKLEARRGLEKMIENFLKKYRLL-VECESEFNTPILPVKK  433

Query  66   KDSTKWRKLVDFRELNKRTQDFWEVQLGIPHP--AGLKKK-KSVTVLDVGDAYFSVPLDE  122
             +S ++R + DF   NK     +    G P      LK+  K  TVL++ DA+F + LD+
Sbjct  432  PESEEYRLVQDFNS-NKFNSSRYSPLSGKPVSLLTALKENHKWFTVLNL*DAFFCILLDK  256

Query  123  DFRKYTAFTIPSINNETPGIRYQYN--VLPQGWKGSPAIFQSSMTKILEPFRKQNPDIVI  180
            + + Y AF    + N T G + Q    VLPQG+K SP IF + + K LE  +   P+ ++
Sbjct  255  NSQAYFAF---ELQNPTTGQKTQLTWAVLPQGFKNSPMIFGNQLAKELETCKIIYPERLV  85

Query  181  YQYMDDLYVGSD  192
             QY+DD+ + ++
Sbjct  84   LQYVDDILIATE  49
```

>Escherichia coli B7A gcontig_1112495760658, whole genome shotgun sequence
Sequence ID: NZ_AAJT02000281.1 Length: 1060
Range 1: 213 to 923

Score:76.3 bits(186), Expect:3e-10,
Method:Compositional matrix adjust.,
Identities:67/256(26%), Positives:120/256(46%), Gaps:34/256(13%)

```
Query  36   EICTEMEKEGKISKIGP-ENPYNTPVFAIKKKDSTKWRKLVDFRELNKRTQDFWEVQLGI  94
            EI T++++   +  I P ++PY++ +  ++KK + KWR  VDFR++N ++     ++
Sbjct  213  EINTKVDELLTMGCIEPSQSPYSSLIKMVRKKQTGKWRLCVDFRQINAKS-----IKDAY  377

Query  95   PHPA------GLKKKKSVTVLDVGDAYFSVPLDEDFRKYTAFTIPSINNETPGIRYQYNV  148
            P P        L++ K ++ LD+ D Y+ +PL+E+ R  TAFT+P       G+ +Q  V
Sbjct  378  PMPRIDYILNQLREAKFISSLDLKDGYWQIPLEENSRPMTAFTVPG-----KGL-FQXKV  539

Query  149  LPQGWKGSPAIFQSSMTKILEPFRKQNPDIVIYQYMDDLYVGSDLEIGQHRTKIEELRQH  208
            +P G   + A FQ ++ +++ P    N     + Y DD+ V S   I      ++E+
Sbjct  540  MPFGLHSASATFQRALDQVIGPEMIPNA----FAYQDDIVVVSK-TISXXMXHLQEVFSR  704

Query  209  LLRWGLTTPDKKHQ---KEPPFLWM-----GYELHPDKWTVQPIVLPEKDSWTVNDIQKL  260
            L    L    +K Q   KE  FL       G    PDK      +   K      ++++
Sbjct  705  LKEANLRINAEKCQFFRKELKFLGHLVTENGIATDPDKVAA---IAQLKPPTCTKELRQY  875

Query  261  VGKLNWASQIYPGIKV  276
            +G  +W  +  P   +
Sbjct  876  LGVASWYRRFVPNFAM  923
```

>Thalassovita mangrovi strain GS-10 Scaffold30_1, whole genome shotgun sequence
Sequence ID: NZ_WWEN01000028.1 Length: 722
Range 1: 569 to 706

Score:73.9 bits(180), Expect:4e-10,
Method:Compositional matrix adjust.,
Identities:33/46(72%), Positives:41/46(89%), Gaps:0/46(0%)

```
Query  515  SELVNQIIEQLIKKEKVYLAWVPAHKGIGGNEQVDKLVSAGIRKVL  560
            ++  +++IEQ+ KK + Y+AWVPAHKGIGGNEQVDKLVSAGIRKVL
Sbjct  706  AKAASKVIEQITKKAEAYMAWVPAHKGIGGNEQVDKLVSAGIRKVL  569
```

>Rickettsia endosymbiont of Ceutorhynchus assimilis isolate Rickettsia_endosymbiont_Ceutorhynchus_assimilis chromosome 1
Sequence ID: NZ_OU906081.1 Length: 4063491
Range 1: 14359 to 15096

Score:77.4 bits(189), Expect:6e-10,
Method:Compositional matrix adjust.,
Identities:70/264(27%), Positives:121/264(45%), Gaps:29/264(10%)

```
Query  22     KQWPLTEEKIKALVEICTEMEKEGKISKIGPENPYNTPVFAIKKKDSTKWRKLVDFRELN  81
              + +P++ EK K L     +M + G I      + Y +P   + + D  + R  VD+R +N
Sbjct  15096  RCYPMSPEKRKILYGQLNQMLQAGVIEPT--TSAYASPPVLVTRPDK-EPRFCVDYRAVN  14926

Query  82     KRTQD----FWEVQLGIPHPAGLKKKKSVTVLDVGDAYFSVPLDEDFRKYTAFTIPSINN  137
                T D       +Q  +     L + K  T+LD+   Y+ +P+DE  +KYTAF+ P
Sbjct  14925  DITLDESTTLPRIQDCV---KPLAQAKIFTLLDLKSGYWQIPMDEQSKKYTAFSTPD---  14764

Query  138    ETPGIRYQYNVLPQGWKGSPAIFQSSMTKILEPFRKQNPDIVIYQYMDDLYVGSDL---E  194
                 G  YQ+ V+P G K +P+ FQ  MT+ +     Q+    ++ Y+DD+ + S+
Sbjct  14763  ---GASYQFTVMPFGLKNAPSTFQKLMTQEVLVGHLQH---FVHVYLDDIIIYSETMEEH  14602

Query  195    IGQHRTKIEELRQHLLRWGLTTPDKKHQKEPPFLWMGYELHPDKWTVQPIVLPEKDSWT-  253
              +   R   E LR H L+           Q+     ++GY +  D    Q   L + ++++
Sbjct  14601  VNHVRKVFERLRIHKLQVSAAKCQIATQE---LDYLGYHIKGDITVPQTKHLKQIETFSR  14431

Query  254    -VN--DIQKLVGKLNWASQIYPGI  274
               VN   +Q  +G  NW  +  P +
Sbjct  14430  PVNKKQLQSFIGTCNWLREHVPHM  14359
```

Range 2: 43090 to 43827

Score:77.0 bits(188), Expect:7e-10,
Method:Compositional matrix adjust.,
Identities:70/264(27%), Positives:121/264(45%), Gaps:29/264(10%)

```
Query  22     KQWPLTEEKIKALVEICTEMEKEGKISKIGPENPYNTPVFAIKKKDSTKWRKLVDFRELN  81
              + +P++ EK K L     +M + G I      + Y +P   + + D  + R  VD+R +N
Sbjct  43090  RCYPMSPEKRKILYGQLNQMLQAGVIEPT--TSAYASPPVLVTRPDK-EPRFCVDYRAVN  43260

Query  82     KRTQD----FWEVQLGIPHPAGLKKKKSVTVLDVGDAYFSVPLDEDFRKYTAFTIPSINN  137
```

```
                T D       +Q  +     L + K  T+LD+   Y+ +P+DE  +KYTAF+ P
Sbjct  43261  DITLDESTTLPRIQDCV---KPLAQAKIFTLLDLKSGYWQIPMDEQSKKYTAFSTPD---  43422

Query  138    ETPGIRYQYNVLPQGWKGSPAIFQSSMTKILEPFRKQNPDIVIYQYMDDLYVGSDL---E  194
                G  YQ+ V+P G K +P+ FQ  MT+ +     Q+    ++ Y+DD+ + S+
Sbjct  43423  ---GASYQFTVMPFGLKNAPSTFQKLMTQEVLVGHLQH---FVHVYLDDIIIYSETMEEH  43584

Query  195    IGQHRTKIEELRQHLLRWGLTTPDKKHQKEPPFLWMGYELHPDKWTVQPIVLPEKDSWT-  253
              +   R   E LR H L+          Q+     ++GY +   D    Q   L + ++++
Sbjct  43585  VNHVRKVFERLRIHKLQVSAAKCQIATQE---LDYLGYHIKGDITVPQNKHLKQIETFSR  43755

Query  254    -VN--DIQKLVGKLNWASQIYPGI  274
               VN   +Q  +G  NW  +  P +
Sbjct  43756  PVNKKQLQSFIGTCNWLREHVPHM  43827



Range 3: 168432 to 168887

Score:53.1 bits(126), Expect:0.022,
Method:Compositional matrix adjust.,
Identities:47/172(27%), Positives:83/172(48%), Gaps:26/172(15%)

Query  40      EMEKEGKISKIGPENPYNTPVFAIKKKDSTKWRKLVDFRELNKRTQDFWEVQLGIPHP--  97
               +M +EG I +    +P++ PV  + KKD +  R  VD+R+LN  T+     +   P P
Sbjct  168432  KMLQEGIIEE--SNSPWSAPVVLVTKKDGS-LRFCVDYRKLNDVTK-----KDSFPLPRI  168587

Query  98      ----AGLKKKKSVTVLDVGDAYFSVPLDEDFRKYTAFTIPSINNETPGIRYQYNVLPQGW  153
                     L   K  + LD+   Y+ V + +D ++ TAF+  S      G+ YQ+NV+P G
Sbjct  168588  DDTLTTLSGSKWFSTLDLKSGYWQVAIHDDEKEKTAFSTGS------GL-YQFNVMPFGL  168746

Query  154     KGSPAIFQSSMTKILEPFRKQNPDIVIYQYMDDLYVGSDLEIGQHRTKIEEL  205
                 +PA F+  M  +L     +   +    Y+DD+ +  +    +H   ++E+
Sbjct  168747  CNAPATFERLMHLVLRGLTWRTCLV----YLDDVMIMGN-SFQEHLNNLKEV  168887




>Klebsiella pneumoniae strain EuSCAPE_NO017, whole genome shotgun sequence
Sequence ID: NZ_UKIO02000069.1 Length: 458
Range 1: 354 to 455

Score:70.9 bits(172), Expect:9e-10,
Method:Compositional matrix adjust.,
Identities:34/34(100%), Positives:34/34(100%), Gaps:0/34(0%)

Query  1    PISPIETVPVKLKPGMDGPKVKQWPLTEEKIKAL  34
            PISPIETVPVKLKPGMDGPKVKQWPLTEEKIKAL
Sbjct  354  PISPIETVPVKLKPGMDGPKVKQWPLTEEKIKAL  455




>Pseudomonas aeruginosa strain ENV-297 NODE_79_length_1100_cov_9.24438_ID_157, whole genome shotgun sequence
Sequence ID: NZ_RADD01000036.1 Length: 1100
Range 1: 945 to 1055

Score:74.7 bits(182), Expect:1e-09,
Method:Compositional matrix adjust.,
Identities:35/37(95%), Positives:37/37(100%), Gaps:0/37(0%)

Query  524   QLIKKEKVYLAWVPAHKGIGGNEQVDKLVSAGIRKVL  560
             QLIKKEKVYL+WVPAHKGIGGNEQVDKLVS+GIRKVL
Sbjct  1055  QLIKKEKVYLSWVPAHKGIGGNEQVDKLVSSGIRKVL  945




>Pseudomonas aeruginosa strain P60_London_6_VIM_2_11_13, whole genome shotgun sequence
Sequence ID: NZ_CVXD01000084.1 Length: 395
Range 1: 3 to 101

Score:68.6 bits(166), Expect:3e-09,
Method:Compositional matrix adjust.,
Identities:33/33(100%), Positives:33/33(100%), Gaps:0/33(0%)

Query  1    PISPIETVPVKLKPGMDGPKVKQWPLTEEKIKA  33
            PISPIETVPVKLKPGMDGPKVKQWPLTEEKIKA
Sbjct  101  PISPIETVPVKLKPGMDGPKVKQWPLTEEKIKA  3




>Listeria monocytogenes strain 1827/2,1 NODE_48_length_4618_cov_6.857588, whole genome shotgun sequence
Sequence ID: NZ_JAENKW010000028.1 Length: 4618
Range 1: 2144 to 2731

Score:73.9 bits(180), Expect:6e-09,
Method:Compositional matrix adjust.,
Identities:62/208(30%), Positives:101/208(48%), Gaps:20/208(9%)

Query  6     ETVPVKLKPGMDGPKVKQWPLTEEKIKALVEICTEMEKEGKISKIGPENPYNTPVFAIKK  65
             E V + L+     P  KQ+PL  E  + L  +  +    G +  +   +P NTP+ ++KK
Sbjct  2144  EPVIIVLRDPTRFPNRKQYPLKREAREGLQPLINKFLACGLL--VPTSSPCNTPILSVKK  2317

Query  66    KDSTKWRKLVDFRELNKRTQDFWEVQLGIPHPAGL-----KKKKSVTVLDVGDAYFSVPL  120
             KD T W  + D R +N+            +P+P  +         K  TVLD+ DA+F +PL
Sbjct  2318  KDGT-W*MVQDLRIINEAVVPLHPT---VPNPYVILGEIPPSAKWFTVLDLKDAFFCIPL  2485

Query  121   DEDFRKYTAFTIPSINNETPGIRYQ---YNVLPQGWKGSPAIFQSSMTKILEPFRKQNPD  177
              ++ +   AF       E PG ++Q   + VLPQG++ SP +F  ++++ L       P+
Sbjct  2486  AKESQYLFAF-----EWEAPGEKHQQMTWTVLPQGFRDSPHLFGQALSQDLL-DLDLGPN  2647

Query  178   IVIYQYMDDLYVGSDLEIGQHRTKIEEL  205
               I QY+DDL + S  E    +  I+ L
Sbjct  2648  GKILQYVDDLLICSPDEKSAQQHAIQVL  2731




>Klebsiella pneumoniae strain BL2026 NODE_43_length_659_cov_0.466165, whole genome shotgun sequence
Sequence ID: NZ_VICY01000043.1 Length: 659
Range 1: 198 to 650

Score:66.2 bits(160), Expect:1e-07,
Method:Compositional matrix adjust.,
Identities:51/160(32%), Positives:84/160(52%), Gaps:16/160(10%)

Query  8    VPVKLKPGMDGPKVKQWPLTEEKIKALVEICTEMEKEGKISKIGPENPYNTPVFAIKKKD  67
            V VK+K G    ++KQ+PL +E  + +  +  + G + +   E  +NTP+   ++K D
Sbjct  198  VEVKVKEGRRPVRIKQYPLKKEDREGIRPVIEKFLQLGLLKE*VSE--FNTPILPVRKPD  371

Query  68   STKWRKLVDFRELNKRTQDFWEVQLGIPHPAGLKKKKS-----VTVLDVGDAYFSVPLDE  122
```

```
            + +R + D R +NK T+D + V   + +P  L    +      TVLD+ DA+F +PL E
Sbjct  372  GS-YRVVQDLRAVNKITEDLYPV---VANPYTLLTVLTPELIWFTVLDLKDAFFCLPLHE  539

Query  123  DFRKYTAFTIPSINNETPGIRYQ--YNVLPQGWKGSPAIF  160
             +K  AF      N   G + Q  +  LPQG+K SP +F
Sbjct  540  ASQKLFAF---EWENPKSGRKTQLTWTALPQGFKNSPTLF  650



>Thalassovita mangrovi strain GS-10 Scaffold19_1, whole genome shotgun sequence
Sequence ID: NZ_WWEN01000019.1 Length: 2900
Range 1: 2800 to 2898

Score:68.2 bits(165), Expect:3e-07,
Method:Compositional matrix adjust.,
Identities:32/33(97%), Positives:33/33(100%), Gaps:0/33(0%)

Query  1     PISPIETVPVKLKPGMDGPKVKQWPLTEEKIKA  33
             PISPIETVPVKLKPGMDGP+VKQWPLTEEKIKA
Sbjct  2800  PISPIETVPVKLKPGMDGPQVKQWPLTEEKIKA  2898



>Sphingobacterium bovisgrunnientis strain KCTC 52685 scaffold_128, whole genome shotgun sequence
Sequence ID: NZ_WUQY01000124.1 Length: 503
Range 1: 11 to 448

Score:63.5 bits(153), Expect:4e-07,
Method:Compositional matrix adjust.,
Identities:43/164(26%), Positives:81/164(49%), Gaps:22/164(13%)

Query  54   NPYNTPVFAIKKKDSTK---WRKLVDFRELNKRT-QDFWEVQLGIPHPAGLKKKKSVTVL  109
            +PYN+P F ++  +  K    R ++D+RELN +T  + +++ +        ++  K  +
Sbjct  448  SPYNSPAFMVRNHNEIKRGKARMVIDYRELNSKTIPNSYKLPIKDLLIKQIQGSKIYSKF  269

Query  110  DVGDAYFSVPLDEDFRKYTAFTIPSINNETPGIRYQYNVLPQGWKGSPAIFQSSMTKILE  169
            D    Y+ + LDE  +  T+FT P+     P  ++++NV+  G   +PAIFQ  M KI+
Sbjct  268  DCKSGYWQIKLDEISKPLTSFTTPN-----PLGQFEWNVMSFGLMNAPAIFQQRMDKIIR  104

Query  170  PFRKQNPDIVIYQYMDDLYVGSDLEIGQHRTKIEELRQHLLRWG  213
            P +            Y+DD+ +         H   ++E  +HL ++
Sbjct  103  PIQT-----FCIAYVDDILI--------HSKNLKEHIEHLKKFN  11



>Acinetobacter baumannii strain ACM-15 NODE_85_length_432_cov_0.596721, whole genome shotgun sequence
Sequence ID: NZ_PYDB01000084.1 Length: 432
Range 1: 309 to 431

Score:62.8 bits(151), Expect:5e-07,
Method:Compositional matrix adjust.,
Identities:33/41(80%), Positives:35/41(85%), Gaps:0/41(0%)

Query  1    PISPIETVPVKLKPGMDGPKVKQWPLTEEKIKALVEICTEM  41
            PISPIETVPVKLK GMDGPKVKQW LTE+K+KA VE   EM
Sbjct  309  PISPIETVPVKLKQGMDGPKVKQWLLTEDKLKA*VET*KEM  431



>Proteus mirabilis strain PM2-4 PM2-4_scaffold37, whole genome shotgun sequence
Sequence ID: NZ_JAMKNT010000037.1 Length: 5277
Range 1: 2288 to 2986

Score:66.6 bits(161), Expect:1e-06,
Method:Compositional matrix adjust.,
Identities:61/248(25%), Positives:104/248(41%), Gaps:31/248(12%)

Query  44    EGKISKIGPENPYNTPVFAIKKKD-----STKWRKLVDFRELNKRTQDFWEVQLGIPHPA  98
             +G I    P   +N+P+  ++KKD     S   R  +D R +N         +  +P  A
Sbjct  2986  DGVIEIAPPNTAFNSPLLTVRKKDANGNYSDDVRVCIDPRHINAILIKDRTDRFQLPLIA  2807

Query  99    GLKKKKSV----TVLDVGDAYFSVPLDEDFRKYTAFTIPSINNETPGIRYQYNVLPQGWK  154
              L +K S     TVLD+  A+  +PL E+ R  TAFT         G +YQ+   P G
Sbjct  2806  ELHQKMSAATLFTVLDLKQAFHRLPLAEEHRALTAFTF-------NGQQYQFCRAPFGLT  2648

Query  155   GSPAIFQSSMTKILE--PFRKQNPDIVIYQYMDDLYVGSDLEIGQHRTKIEELRQHLLRW  212
               P  FQ  +T +L   P+        +  ++DDL +G+  +I +H   +  + + L
Sbjct  2647  PIPNHFQRVLTTLLHDLPY--------VTCFIDDLTIGTGPDIEEHIKCVTTVIERLTSA  2492

Query  213   G-LTTPDKKHQKEPPFLWMGYELHPDKWTVQPIVLPEKDSWTV----NDIQKLVGKLNWA  267
               +   DK H  +     +G+ +  +  ++    +     +W V     DIQ+ +G  N+
Sbjct  2491  KFILNVDKCHFMQTSVNILGFTISHNSLSLDSRKVANALAWPVPKTGKDIQRFLGLANYF  2312

Query  268   SQIYPGIK  275
                 P
Sbjct  2311  RSHLPNFS  2288



>Klebsiella pneumoniae strain c10 contig85, whole genome shotgun sequence
Sequence ID: NZ_JAIBKQ010000085.1 Length: 6264
Range 1: 3409 to 4095

Score:66.2 bits(160), Expect:2e-06,
Method:Compositional matrix adjust.,
Identities:66/245(27%), Positives:113/245(46%), Gaps:23/245(9%)

Query  37    ICTEMEKEGKISKIGPENPYNTPVFAIKKKDSTKWRKLVDFRELNKRT-QDFWEVQLGIP  95
             I  EM KEG I   G  +P+ +PV  +KKD +  R  VD+R+LN  T +D + +
Sbjct  4095  IVQEMAKEGVIEPSG--SPWASPVVLVKKKDGST-RFCVDYRQLNNVTKKDSYPLPRIDD  3925

Query  96    HPAGLKKKKSVTVLDVGDAYFSVPLDEDFRKYTAFTIPSINNETPGIRYQYNVLPQGWKG  155
                L   +  + LD+    Y+ V L  + ++ TAFT+ +      G+ +Q+ V+P G
Sbjct  3924  TLDTLAGSRIFSTLDLKSGYWQVELAPEDKEKTAFTVGT------GL-WQFTVMPFGLCN  3766

Query  156   SPAIFQSSMTKILEPFRKQNPDIVIYQYMDD-LYVGSDLEIGQHRTKIEELRQHLLRWGL  214
             +PA F+  M  +L+    +   +    Y+DD + VG   E   H   +EE+ Q L   GL
Sbjct  3765  APATFERLMDTVLKGLSWKTCLV----YLDDIIVVGKTFE--DHLRNLEEVFQRLRNSGL  3604

Query  215   TTPDKK-HQKEPPFLWMGYELHPDKWTVQPIVLPEKDSWTV----NDIQKLVGKLNWASQ  269
                KK H  +    ++G+ +  +   V    +     W V    ++++  +G   +
Sbjct  3603  RLSSKKCHLFQREVQYLGHVVSSEGVAVDKKKVESVKEWPVPTDKHELRSFLGLCTYYRR  3424

Query  270   IYPGI  274
               PG
Sbjct  3423  FVPGF  3409
```

```
>Streptococcus anginosus strain San49 NODE_781_length_269_cov_0.734375, whole genome shotgun sequence
Sequence ID: NZ_JAXIIG010000228.1 Length: 269
Range 1: 6 to 245

Score:57.4 bits(137), Expect:1e-05,
Method:Composition-based stats.,
Identities:29/81(36%), Positives:49/81(60%), Gaps:1/81(1%)

Query  54   NPYNTPVFAIKKKDSTKWRKLVDFRELNKRTQDFWEVQLGIPHPAGLKKKKSVTVLDVGD  113
            +P+NTP+F IKK+ S KWR L D + +N + Q    +Q G+P  + +  +  + ++++ D
Sbjct  245  SPWNTPIFVIKKQ-SGKWRILHDLKAINSQMQIMGPIQRGLPLLSSIPTEWLIIIINIKD  69

Query  114  AYFSVPLDEDFRKYTAFTIPS  134
             +FS+ L    R   AFT+PS
Sbjct  68   CFFSISLAPQDRVRFAFTLPS  6



>Klebsiella pneumoniae strain BL198 NODE_37_length_611_cov_0.421488, whole genome shotgun sequence
Sequence ID: NZ_VICV01000037.1 Length: 611
Range 1: 65 to 448

Score:60.1 bits(144), Expect:1e-05,
Method:Compositional matrix adjust.,
Identities:40/137(29%), Positives:74/137(54%), Gaps:17/137(12%)

Query  106  VTVLDVGDAYFSVPLDEDFRKYTAFTI--PSINNETPGIRYQYNVLPQGWKGSPAIFQSS  163
             TVL++ DA+F + LD++ + Y AF +  P+   +T   +  + VLPQG+K SP IF +
Sbjct  65   FTVLNL*DAFFCILLDKNSQAYFAFELQNPTTRQKT---QLTWAVLPQGFKNSPMIFGNQ  235

Query  164  MTKILEPFRKQNPDIVIYQYMDDLYVGSDLEIGQHRTKIEELRQHLLRWGLTTPDKKHQK  223
            + K LE  +   P+ ++ QY+DD+ + ++       R +   L  +LL + L   D +  K
Sbjct  236  LAKELETCKIIYPERLVLQYVDDILIATE-----TREEYFNLTTNLLNF-LGQSDNRFSK  397

Query  224  EPP------FLWMGYEL  234
            +         +++G+E+
Sbjct  398  KKAQIGKETVMYLGFEI  448



>MAG: Thalassolituus oleivorans isolate SRR17458625_maxbin2_1_1705322953 ERZ21854034.981, whole genome shotgun sequence
Sequence ID: NZ_CAWUIR010000113.1 Length: 10375
Range 1: 4275 to 5099

Score:62.8 bits(151), Expect:2e-05,
Method:Compositional matrix adjust.,
Identities:80/299(27%), Positives:124/299(41%), Gaps:45/299(15%)

Query  2     ISPIETVPVKLKPGMDGPK-------VKQWPLTEEKIKALVEICTEMEKEGKISKIGPEN  54
             +S ++ + V+LKPG + P         +Q     +KIK L++I    E    I
Sbjct  5099  MSKLDPIEVELKPGHEPPNSYGRTLGYEQRMFLNDKIKDLIKIGILKESANPI-------  4941

Query  55    PYNTPVFAIKKKDSTKWRKLVDFRELNKRT-QDFWEVQLGIPHPAGLKKKKSVTVLDVGD  113
              Y +P+F + KK   KWR +VD R LN  T +   E+ L       +K  K     DV
Sbjct  4940  -YGSPIFIVPKKGPKKWRMVVDMRALNTITIKTAMEMPLLDEQIDWIKGAKVFGTFDVLS  4764

Query  114   AYFSVPLDEDFRKYTAFTIPSINNETPGIRYQYNVLPQGWKGSPAIFQSSMTK-ILEPFR  172
              +  +P  E+ +KY  FT+  I  E     Y     P GW  +PA+F   M + IL+P
Sbjct  4763  GFDFLPTAENSQKY--FTL--ITKEGA---YTMCGAPMGWVNTPALFMDRMVRNILKP--  4611

Query  173   KQNPDIVIY--------QYMDDLYVGSDLEIGQHRTKIEELRQHLLRWGLTTPDKKHQKE  224
                  I IY        Q++DD  + +D      +   + L Q + R    + DK H
Sbjct  4610  -----IGIYGIATNGAIQWLDDTCLYADSFDRYAKLLDKFLAQMIKRAVRLSIDKCHLYG  4446

Query  225   PPFLWMGYELHPDKWTVQP----IVLPEKDSWTVNDIQKLVGKLNWASQIYPGIKVRQL  279
                 W G  ++   W  Q      +L  +    V ++ +++   NW S   P  K+ QL
Sbjct  4445  EEVEWCGRRINAKGWNFQSKFYDKILTVQKPQRVWEMAQVIYLCNWLSPAIP--KLSQL  4275



>Polyangium sp. y55x31 NODE_26_length_169233_cov_17.164843, whole genome shotgun sequence
Sequence ID: NZ_JARZHG010000026.1 Length: 169233
Range 1: 66608 to 66976

Score:62.4 bits(150), Expect:3e-05,
Method:Compositional matrix adjust.,
Identities:41/124(33%), Positives:66/124(53%), Gaps:8/124(6%)

Query  441    YVDGAANRETKLGKAGYVTNRGRQKVVT---LTDTTNQKTELQAIYLAL---QDSGLEVN  494
              Y DGA           G V   G +++     L D TN   EL AI  A+   +D+G  +
Sbjct  66976  YADGACTGNPGPAGLGVVVVDGDRRIEISEYLGDGTNNIAELTAILRAVGAVEDTGKPLL  66797

Query  495    IVTDSQYALGIIQAQ-PDQSESELVNQIIEQLIKKEKVYLAWVPAHKGIGGNEQVDKLVS  553
              + TDSQYA+G++Q     ++ ++LV Q ++ ++K+ K  L +VP H G+  NE+ D+L
Sbjct  66796  VRTDSQYAIGVLQKGWKAKANTDLVAQ-VKDVLKRNKTRLEYVPGHSGVALNERADELAR  66620

Query  554    AGIR  557
                +R
Sbjct  66619  EAVR  66608



>Polyangium spumosum strain DSM 14734 Scaffold16, whole genome shotgun sequence
Sequence ID: NZ_WJIE01000016.1 Length: 180314
Range 1: 106389 to 106757

Score:61.2 bits(147), Expect:6e-05,
Method:Compositional matrix adjust.,
Identities:41/124(33%), Positives:66/124(53%), Gaps:8/124(6%)

Query  441     YVDGAANRETKLGKAGYVTNRGRQKVVT---LTDTTNQKTELQAIYLAL---QDSGLEVN  494
               Y DGA           G V   G +++     L D TN   EL AI  A+   Q++G  +
Sbjct  106389  YADGACTGNPGPAGLGVVVVDGDRRIEVSEYLGDGTNNIAELTAILRAVEAVQETGKPLL  106568

Query  495     IVTDSQYALGIIQAQ-PDQSESELVNQIIEQLIKKEKVYLAWVPAHKGIGGNEQVDKLVS  553
               + TDSQYA+G++Q     ++ ++LV Q+ + ++K+ K  L +VP H G+  NE+ D+L
Sbjct  106569  VRTDSQYAIGVLQKGWKAKANTDLVAQV-KDVLKRNKTRLEYVPGHSGVALNERADELAR  106745

Query  554     AGIR  557
                 +R
Sbjct  106746  EAVR  106757



>Streptomyces albidochromogenes strain DSM 41800 NODE_804_length_521_cov_1.81218, whole genome shotgun sequence
Sequence ID: NZ_VBVX01000736.1 Length: 521
Range 1: 4 to 498
```

```
Score:57.4 bits(137), Expect:7e-05,
Method:Compositional matrix adjust.,
Identities:52/186(28%), Positives:82/186(44%), Gaps:27/186(14%)

Query  10   VKLKPGMDGPKVKQWPLTEEKIKALVEICTEMEKEGKISKIGPENPYNTPVFAIKKKDST  69
            + L PG        + +++ + + L +   E+ ++G I K    +PY  PV  +KKKD
Sbjct  498  IHLIPGAKPTARPPYRISQFETEELKKQIEELLQQGFIQK--SNSPYAAPVLFVKKKDKA  325

Query  70   KWRKLVDFRELNKRTQDFWEVQLGIPHP------AGLKKKKSVTVLDVGDAYFSVPLDED  123
              R  VDFR LN  T     ++   P P        +   K  + LD+   YF + +D
Sbjct  324  -LRLCVDFRLLNINT-----IKNAFPIPLIEDLFMKIGTAKVFSKLDLMSGYFQIRMDPR  163

Query  124  FRKYTAFTIPSINNETPGIRYQYNVLPQGWKGSPAIFQSSMTKILEPFRKQNPDIVIYQY  183
              + T F        TP   Y + V+P G    PA FQS M  IL       P++++  Y
Sbjct  162  DEEKTGFV-------TPFGHYHWRVMPFGVVNGPATFQSFMNDIL----GDTPNVLV--Y  22

Query  184  MDDLYV  189
            +DD+ +
Sbjct  21   LDDILI  4




>Klebsiella pneumoniae strain BL3427 NODE_36_length_966_cov_0.293206, whole genome shotgun sequence
Sequence ID: NZ_VIDH01000036.1 Length: 966
Range 1: 228 to 599

Score:58.2 bits(139), Expect:2e-04,
Method:Compositional matrix adjust.,
Identities:40/133(30%), Positives:70/133(52%), Gaps:17/133(12%)

Query  110  DVGDAYFSVPLDEDFRKYTAFTIPSINNETPGIRYQYN--VLPQGWKGSPAIFQSSMTKI  167
             V DA+F + LD++ + Y AF    + N T G + Q +  VLPQG+K SP IF + + K
Sbjct  228  SV*DAFFCILLDKNSQAYFAF---ELQNPTTGQKTQLSWAVLPQGFKNSPMIFGNQLAKE  398

Query  168  LEPFRKQNPDIVIYQYMDDLYVGSDLEIGQHRTKIEELRQHLLRWGLTTPDKKHQKEPP-  226
            LE  +   P+ ++ QY+DD+ + ++      R +   L  +LL + L   D +  K+
Sbjct  399  LETCKIIYPERLVLQYVDDILIATE-----TREEYFNLTTNLLNF-LGQSDNRFSKKKAQ  560

Query  227  -----FLWMGYEL  234
                  +++G+E+
Sbjct  561  IGKETVMYLGFEI  599




>MAG: Thiolapillus sp. isolate Ifr-SOX-238 NODE_5703_length_2294_cov_6.676024, whole genome shotgun sequence
Sequence ID: NZ_JANEJA010000497.1 Length: 2294
Range 1: 1495 to 2115

Score:59.3 bits(142), Expect:2e-04,
Method:Compositional matrix adjust.,
Identities:66/233(28%), Positives:108/233(46%), Gaps:35/233(15%)

Query  20    KVKQWPLTEEKIKALVEICTEMEKEGKISKIGPE-NPYNTPVFAIKKKDSTKWRKLVDFR  78
             ++K +P+   K +   EI  E++    +  I P  + Y++P+  +KKKD T  R  +DFR
Sbjct  1495  RLKSYPIPFAKRE---EIRKEVKSMLDLGVIEPSTSSYSSPIVLVKKKDGTN-RFCIDFR  1662

Query  79    ELNKRTQDFWEVQLGIPHP------AGLKKKKSVTVLDVGDAYFSVPLDEDFRKYTAFTI  132
              LNK T     V    P P      A L K +  T +D+   Y+ +P+ ++ R  TAF
Sbjct  1663  ALNKVT-----VFDAEPIPDTEELFARLTKARFFTKIDLAKGYWQIPVKQEDRHKTAF--  1821

Query  133   PSINNETPGIRYQYNVLPQGWKGSPAIFQSSMTKI-LEPFRKQNPDIVIYQYMDDLYVGS  191
                   +TP   YQ+  +P G   +PA F   M K+ LE     N       + DD+ + +
Sbjct  1822  -----QTPLGLYQWTKMPFGLVSAPASFARMMRKLKLEENSALN-------FFDDILIAT  1965

Query  192   DLEIGQHRTKIEELRQHLLRWGLTT-PDKKHQKEPPFLWMGYELHPDKWTVQP  243
             +     H   ++ + Q L ++GLT  P K          ++G+ +   K T+QP
Sbjct  1966  E-RWDDHLESVKGVLQKLQKFGLTARPTKIFAGFQTLEFLGHTV--GKGTMQP  2115




>Polyangium sp. 15x6 NODE_112_length_36528_cov_15.618939, whole genome shotgun sequence
Sequence ID: NZ_JASBQE010000112.1 Length: 36528
Range 1: 5134 to 5502

Score:58.5 bits(140), Expect:4e-04,
Method:Compositional matrix adjust.,
Identities:40/124(32%), Positives:66/124(53%), Gaps:8/124(6%)

Query  441   YVDGAANRETKLGKAGYVTNRGRQKVVT---LTDTTNQKTELQAIYLAL---QDSGLEVN  494
             Y DGA           G V   G +++     L + TN   EL AI  A+   +D+G  +
Sbjct  5502  YADGACTGNPGPAGLGVVVVDGDRRIEISEYLGNGTNNIAELTAILRAVGAVEDTGKPLL  5323

Query  495   IVTDSQYALGIIQAQ-PDQSESELVNQIIEQLIKKEKVYLAWVPAHKGIGGNEQVDKLVS  553
             + TDSQYA+G++Q     ++ +ELV + ++ ++K+ K  L +VP H G+  NE+ D+L
Sbjct  5322  VRTDSQYAIGVLQKGWKAKANTELVAE-VKDVLKRNKTRLEYVPGHSGVALNERADELAR  5146

Query  554   AGIR  557
               +R
Sbjct  5145  EAVR  5134




>Penaeicola halotolerans strain LMIT005 NODE_33_length_336_cov_0.594595, whole genome shotgun sequence
Sequence ID: NZ_JAEAGO010000029.1 Length: 336
Range 1: 3 to 176

Score:53.1 bits(126), Expect:6e-04,
Method:Compositional matrix adjust.,
Identities:28/61(46%), Positives:39/61(63%), Gaps:3/61(4%)

Query  22   KQWPLTEEKIKALVEICTEMEKEGKISKIGPENPYNTPVFAIKKKDSTKWRKLVDFRELN  81
            KQWPL EEK++AL ++  E    G I +    +P+N+PVF +KKK   KWR + D R +N
Sbjct  176  KQWPLAEEKLQALEQLVQEQLDAGHIEE--STSPWNSPVFVVKKKFG-KWRMVTDLRAIN  6

Query  82   K  82
            K
Sbjct  5    K  3




>Piscirickettsia salmonis strain FIUCHILE-89L01, whole genome shotgun sequence
Sequence ID: NZ_LELB01000266.1 Length: 2765
Range 1: 476 to 865

Score:57.8 bits(138), Expect:6e-04,
Method:Compositional matrix adjust.,
Identities:41/141(29%), Positives:67/141(47%), Gaps:12/141(8%)

Query  53   ENPYNTPVFAIKKKDSTKWRKLVDFRELNKRT-QDFWEVQLGIPHPAGLKKKKSVTVLDV  111
```

```
            E+ Y +PV  +KK+D +  R  VDFR LN+ T +D + + L   H   L   K  + LD+
Sbjct  865  ESEYASPVLLVKKRDGSD-RLCVDFRALNRITVKDRYPLPLIDDHIDRLGSSKYFSCLDM  689

Query  112  GDAYFSVPLDEDFRKYTAFTIPSINNETPGIRYQYNVLPQGWKGSPAIFQSSMTKILEPF  171
               +  +P+ ED    T F        TP   ++Y  +P G   SP ++Q  +   L
Sbjct  688  ASGFHQIPIHEDSIHKTGFV-------TPEGHFEYLKMPFGLCNSPTVYQRIINGTLRRL  530

Query  172  RKQNPDIVIYQYMDDLYVGSD  192
             +    +V   Y+DD+ + S+
Sbjct  529  IETGNVLV---YVDDVLLVSE  476
```

>Escherichia coli strain 8551F 237, whole genome shotgun sequence
Sequence ID: NZ_JAFCNN010000170.1 Length: 689
Range 1: 93 to 665

Score:55.5 bits(132), Expect:7e-04,
Method:Compositional matrix adjust.,
Identities:60/216(28%), Positives:96/216(44%), Gaps:32/216(14%)

```
Query  29   EKIKALVEICTEMEKEGKISKIGPENPYNTPVFAIKKKDSTKWRKLVDFRELNKRTQDFW  88
            E++K  +E   E+ ++G I K    +P   PV  +KKD    R  VDFR LN  T
Sbjct  665  EELKKQIE---ELLQQGFIQK--SNSPXAAPVLFVKKKDKA-LRLCVDFRLLNINT----  516

Query  89   EVQLGIPHP------AGLKKKKSVTVLDVGDAYFSVPLDEDFRKYTAFTIPSINNETPGI  142
             ++   P P        +   K  + LD+   YF + +D    + T F        TP
Sbjct  515  -IKNAFPIPLIEDLFMKIGTAKVFSKLDLMSGYFQIRMDPRDEEKTGFV-------TPFG  360

Query  143  RYQYNVLPQGWKGSPAIFQSSMTKILEPFRKQNPDIVIYQYMDDLYVGSDLEIGQHRTKI  202
             Y + V+P G    PA FQS M  IL       P++++  Y+DD+ + S     +H   +
Sbjct  359  HYHWRVMPFGVVNGPATFQSFMNDIL----GDTPNVLV--YLDDILIYSH-NRKEHERDL  201

Query  203  EELRQHLLRWGLTTPDKKHQKEPPFL-WMGYELHPD  237
            + + + L    L   +KK +     L ++G+ + PD
Sbjct  200  KRVLRVLRENKLIAKEKKCEFFKDTLDFLGHTITPD  93
```

>Candidatus Erwinia dacicola strain IL contig_608, whole genome shotgun sequence
Sequence ID: NZ_MAYS01000566.1 Length: 1576
Range 1: 725 to 1012

Score:57.0 bits(136), Expect:7e-04,
Method:Compositional matrix adjust.,
Identities:34/107(32%), Positives:58/107(54%), Gaps:17/107(15%)

```
Query  70    KWRKLVDFRELNKRTQDFWEVQLGIPHPA------GLKKKKSVTVLDVGDAYFSVPLDED  123
             K RK  D+R+LN R+     V    P P        L++ + ++ LD+ + Y+ +PL++
Sbjct  725   KRRKCTDYRQLNARS-----VPDAYPLPRIQHILDKLRQARYISSLDLKNGYWQIPLEQG  889

Query  124   FRKYTAFTIPSINNETPGIRYQYNVLPQGWKGSPAIFQSSMTKILEP  170
              R YTAFT+P       G+ +Q+ V+P G   +PA FQ  + +++ P
Sbjct  890   SRPYTAFTVPG-----RGL-FQWRVMPFGLHSAPATFQRVLDQVIGP  1012
```

>Polyangium sp. 6x1 NODE_22_length_166486_cov_22.101912, whole genome shotgun sequence
Sequence ID: NZ_JARZHH010000022.1 Length: 166486
Range 1: 110104 to 110472

Score:57.8 bits(138), Expect:7e-04,
Method:Compositional matrix adjust.,
Identities:41/124(33%), Positives:62/124(50%), Gaps:8/124(6%)

```
Query  441     YVDGAANRETKLGKAGYVTNRGRQKVVT---LTDTTNQKTELQAIYLALQ---DSGLEVN  494
               Y DGA          G +   G +++     L   TN   EL AI  A++    +G  +
Sbjct  110104  YADGACTGNPGPAGLGVLVEDGDRRIEISEYLGQGTNNIAELTAILRAVEAVEGTGKSLL  110283

Query  495     IVTDSQYALGIIQAQ-PDQSESELVNQIIEQLIKKEKVYLAWVPAHKGIGGNEQVDKLVS  553
               + TDSQYA+G++Q     ++ +ELV  + + L KK K  L +VP H G+ GNE+ D L
Sbjct  110284  VRTDSQYAIGVLQKGWKAKANTELVAAVKDAL-KKSKARLEYVPGHSGVRGNERADALAR  110460

Query  554     AGIR  557
                 +R
Sbjct  110461  EAVR  110472
```

>Pseudomonas aeruginosa isolate PSA-13, whole genome shotgun sequence
Sequence ID: NZ_FNME01000065.1 Length: 505
Range 1: 74 to 448

Score:53.5 bits(127), Expect:0.001,
Method:Composition-based stats.,
Identities:42/137(31%), Positives:68/137(49%), Gaps:13/137(9%)

```
Query  54   NPYNTPVFAIKKKDSTKWRKLVDFRELNKRT-QDFWEVQLGIPHPAGLKKKKSVTVLDVG  112
            +P     F +KKKD T  R  VD+R LN  T ++ + + L       L+  +  T LD+
Sbjct  448  SPAGAGFFFVKKKDGT-LRPCVDYRGLNDITVKNRYPLLLMSSAFEILQGARFFTKLDLR  272

Query  113  DAYFSVPLDEDFRKYTAFTIPSINNETPGIRYQYNVLPQGWKGSPAIFQSSMTKILEPFR  172
            +AY  V + E     TAF        TP   ++Y VLP G   +PA+FQ+ +  +L
Sbjct  271  NAYHLVRIREGDEWKTAFN-------TPLGHFEYRVLPFGLANAPAVFQALVNDVL----  125

Query  173  KQNPDIVIYQYMDDLYV  189
            +   +I ++ Y+DD+ +
Sbjct  124  RDMLNIFVFVYLDDILI  74
```

>Enterobacter cloacae complex sp. GF14B NODE_151_length_4351_cov_0.721117, whole genome shotgun sequence
Sequence ID: NZ_QFRV01000155.1 Length: 4351
Range 1: 3283 to 3981

Score:56.2 bits(134), Expect:0.002,
Method:Compositional matrix adjust.,
Identities:71/263(27%), Positives:109/263(41%), Gaps:52/263(19%)

```
Query  6     ETVPVKLKPGMDGPKVKQWPLTEEKIKALVEICTEMEKEGKISKIGPENPYNTPVFAIKK  65
             E + ++LK G    + +   + +E+++AL E   ++ K G I  +     + +PV    K
Sbjct  3283  EEMHIELKEGAKPVRQRLRRMGQEQMEALREEVDKLLKAGFIVPV-ETTEWVSPVVVSPK  3459

Query  66    KDSTKWRKLVDFRELNKRTQDFWEVQLGIPHP-----------AGLKKKKSVTVLDVGDA  114
             KD  KWR  +DF+ LN  T+         P+P           AG ++    +V D
Sbjct  3460  KDG-KWRICIDFKPLNAATKKN-------PYPLPFIDQILDSVAGYER---YSVCDGFSG  3606

Query  115   YFSVPLDEDFRKYTAFTIPSINNETPGIRYQYNVLPQGWKGSPAIFQSSMTKILEPFRKQ  174
             YF + +    R  T F        TP   Y Y VLP G    P+ +Q     +L PF
```

```
Sbjct  3607  YFQLKIALKDRTKTTFI-------TPWGCYCYTVLPFGLSNGPSFYQERANWVLAPFIGS  3765

Query  175   NPDIVIYQYMDDLYVGSDLEIGQHRTKIEELRQHLLRW----GLTTPDKKHQKEPPFLWM  230
                  +  +MDD  V S     +H    E+L+  L R+    G   P K H  +P    +
Sbjct  3766  ----CVKAFMDDFCVYSSR--AEHH---EKLQMVLARYDECGGQLNPKKCHLAQPRVKLL  3918

Query  231   GY-------ELHPDKWTVQPIVL  246
             G+       E  PDK  V+ I+L
Sbjct  3919  GHMVSENGIEADPDK--VKSIIL  3981
```

>Methylobacterium radiotolerans strain MAMP 4754 MP-M-radiotolerans_contig_818, whole genome shotgun sequence
Sequence ID: NZ_NKQS01000805.1 Length: 253
Range 1: 12 to 155

Score:50.1 bits(118), Expect:0.003,
Method:Compositional matrix adjust.,
Identities:27/48(56%), Positives:33/48(68%), Gaps:0/48(0%)

```
Query  97   PAGLKKKKSVTVLDVGDAYFSVPLDEDFRKYTAFTIPSINNETPGIRY  144
             AGLKKKK+ TVLDVGD   S  + E+FR+ T FT P+ +NET  I Y
Sbjct  12   AAGLKKKKTETVLDVGDT*MSDTVTENFRQDTEFTRPNRHNETTWIIY  155
```

>Polyangium jinanense strain SDU14 Scaffold9, whole genome shotgun sequence
Sequence ID: NZ_JAGTJJ010000009.1 Length: 231060
Range 1: 11612 to 11965

Score:55.1 bits(131), Expect:0.004,
Method:Compositional matrix adjust.,
Identities:38/119(32%), Positives:64/119(53%), Gaps:8/119(6%)

```
Query  441    YVDGAANRETKLGKAGYVTNRGRQKVVT---LTDTTNQKTELQAIYLAL---QDSGLEVN  494
              Y DGA           G V   G +++     L + TN   EL AI  A+   +D+G  +
Sbjct  11612  YADGACTGNPGPAGLGVVVVDGDRRIEISEYLGNGTNNIAELTAILRAVGAVEDTGKPLL  11791

Query  495    IVTDSQYALGIIQAQ-PDQSESELVNQIIEQLIKKEKVYLAWVPAHKGIGGNEQVDKLV  552
              + TDSQYA+G++Q     ++ ++LV + ++ ++K+ K  L +VP H G+  NE+ D+L
Sbjct  11792  VRTDSQYAIGVLQKGWKAKANTDLVAE-VKDVLKRNKTRLEYVPGHSGVALNERADELA  11965
```

>Polyangium jinanense strain SDU13 Scaffold15, whole genome shotgun sequence
Sequence ID: NZ_JAGTJK010000015.1 Length: 231263
Range 1: 11765 to 12118

Score:55.1 bits(131), Expect:0.004,
Method:Compositional matrix adjust.,
Identities:38/119(32%), Positives:64/119(53%), Gaps:8/119(6%)

```
Query  441    YVDGAANRETKLGKAGYVTNRGRQKVVT---LTDTTNQKTELQAIYLAL---QDSGLEVN  494
              Y DGA           G V   G +++     L + TN   EL AI  A+   +D+G  +
Sbjct  11765  YADGACTGNPGPAGLGVVVVDGDRRIEISEYLGNGTNNIAELTAILRAVGAVEDTGKPLL  11944

Query  495    IVTDSQYALGIIQAQ-PDQSESELVNQIIEQLIKKEKVYLAWVPAHKGIGGNEQVDKLV  552
              + TDSQYA+G++Q     ++ ++LV + ++ ++K+ K  L +VP H G+  NE+ D+L
Sbjct  11945  VRTDSQYAIGVLQKGWKAKANTDLVAE-VKDVLKRNKTRLEYVPGHSGVALNERADELA  12118
```

>Proteus terrae subsp. cibarius strain pt-OS9-1 NODE_353_length_267_cov_0.768421, whole genome shotgun sequence
Sequence ID: NZ_JBBEZU010000351.1 Length: 267
Range 1: 13 to 267

Score:49.3 bits(116), Expect:0.006,
Method:Composition-based stats.,
Identities:31/94(33%), Positives:49/94(52%), Gaps:12/94(12%)

```
Query  78   RELNKRTQDFWE---VQLGIPHPAGLKKKKSVTVLDVGDAYFSVPLDEDFRKYTAFTIPS  134
            RE+NK      E   +QL + +    +   K ++++D+ +AYF +P+ +  +KYT F+
Sbjct  267  REINKHMISETEKPPLQLDVLN--SFQGAKFISIVDLNNAYFQIPISDSSKKYTGFSF--  100

Query  135  INNETPGIRYQYNVLPQGWKGSPAIFQSSMTKIL  168
                  G  Y YNVLPQG K S   F  +M  I+
Sbjct  99   -----NGKSYVYNVLPQGLKTSVGSFSRAMDFII  13
```

>Streptococcus mutans strain PKUSS-9 C209, whole genome shotgun sequence
Sequence ID: NZ_NBFI01000031.1 Length: 307
Range 1: 12 to 230

Score:49.7 bits(117), Expect:0.006,
Method:Composition-based stats.,
Identities:29/75(39%), Positives:43/75(57%), Gaps:3/75(4%)

```
Query  149  LPQGWKGSPAIFQSSMTKILEPFRKQNPDIVIYQYMDDLYVGSDLEIG-QHRTKIEELRQ  207
            LPQG+K SP +F  ++ + L  FR Q+PD+++ QY+DDL + +  E+  Q  T+   L Q
Sbjct  12   LPQGFKNSPTLFDEALHRDLADFRIQHPDLILLQYVDDLLLAATSELDCQQGTR--ALLQ  185

Query  208  HLLRWGLTTPDKKHQ  222
             L   G     KK Q
Sbjct  186  TLGNLGYRASAKKAQ  230
```

>Bradyrhizobium sp. 33ap4 NODE_290_length_534_cov_0.849015, whole genome shotgun sequence
Sequence ID: NZ_JAUMTJ010000287.1 Length: 534
Range 1: 107 to 481

Score:51.2 bits(121), Expect:0.008,
Method:Composition-based stats.,
Identities:42/143(29%), Positives:72/143(50%), Gaps:25/143(17%)

```
Query  54   NPYNTPVFAIKKKDSTKWRKLVDFRELNKRTQDFWEVQLGI-PHPA------GLKKKKSV  106
            +P+++PV  +KKD T  R  VD+R+LN  T+       G+ P P        L++ + +
Sbjct  107  SPWSSPVVLVKKKDGT-LRFCVDYRKLNSVTKK------GVCPLPRIDDTLDRLRRSRYL  265

Query  107  TVLDVGDAYFSVPLDEDFRKYTAFTIPSINNETPGIRYQYNVLPQGWKGSPAIFQSSMTK  166
            + +D+   ++ + +DE  R+ TAF        TP   Y++ V+P G    PA FQ  +
Sbjct  266  SSMDLKSGHWQIEVDERDREKTAFI-------TPYGLYEFKVMPSGLCTPPATFQRVIDT  424

Query  167  ILEPFRKQNPDIVIYQYMDDLYV  189
            +L   + Q   +    Y+DD+ V
Sbjct  425  VLAGPKWQTCLV----YLDDVVV  481
```

```
>MAG: Breoghania sp. isolate MO_035.B22 MO_035.c_2803, whole genome shotgun sequence
Sequence ID: NZ_JASJAV010000047.1 Length: 6353
Range 1: 3126 to 3500

Score:53.9 bits(128), Expect:0.010,
Method:Compositional matrix adjust.,
Identities:40/145(28%), Positives:66/145(45%), Gaps:31/145(21%)

Query  56    YNTPVFAIKKKDSTKWRKLVDFRELNKRTQDFWEVQLGIPH---------PAGLKKKKSV  106
             + + VF + KK+  K R +++ R LN+         L  PH          A ++
Sbjct  3500  FYSNVFLVPKKNG-KLRPVINLRCLNR--------VLHCPHFQMETVGSISAAIQPGDWA  3348

Query  107   TVLDVGDAYFSVPLDEDFRKYTAFTIPSINNETPGIRYQYNVLPQGWKGSPAIFQSSMTK  166
             T LD+ DAYF +P+   FRKY  F +        G  +Q+  LP G   +P +     T+
Sbjct  3347  TSLDLTDAYFHIPIAPWFRKYLRFVV-------AGQVFQFRALPFGLAPAPLV----CTR  3201

Query  167   ILEPF--RKQNPDIVIYQYMDDLYV  189
             +L P      +  I+ ++Y+DDL +
Sbjct  3200  LLAPLAIHLHSQGILFHRYLDDLLI  3126



>Klebsiella pneumoniae strain BL198 NODE_55_length_484_cov_0.434174, whole genome shotgun sequence
Sequence ID: NZ_VICV01000055.1 Length: 484
Range 1: 12 to 482

Score:50.4 bits(119), Expect:0.011,
Method:Compositional matrix adjust.,
Identities:40/160(25%), Positives:73/160(45%), Gaps:9/160(5%)

Query  114  AYFSVPLDEDFRKYTAFTIPSINNETPGIRYQ--YNVLPQGWKGSPAIFQSSMTKILEPF  171
            A+F +PL E  +K  AF     +N   G + Q  + VLPQG+K SP IF + + K LE +
Sbjct  482  AFFCLPLHEASQKLFAF---EWDNPKSGRKTQLTWTVLPQGFKNSPTIFGNQLAKDLESW  312

Query  172  RKQNPDIVIYQYMDDLYVGSDLEIGQHRTKIEELRQHLLRWGLTTPDKKHQKEPPFLWMG  231
               + +  + QY+DD+ + +  E       +  L    L+    +  K    +    ++G
Sbjct  311  ETPSGEGKLLQYVDDILIATKTEEDCMAWTVSLLNFLGLQGYRVSKKKAQVMQRKVNYLG  132

Query  232  YELHPDKWTV----QPIVLPEKDSWTVNDIQKLVGKLNWA  267
            YE+   + T+    +  +   +   TV +++  +G   W
Sbjct  131  YEISAGQRTLGQARKEAICQTQRPQTVKELRTFLGMTGWC  12



>MAG: Breoghania sp. isolate MO_035.B22 MO_035.c_6474, whole genome shotgun sequence
Sequence ID: NZ_JASJAV010000052.1 Length: 5920
Range 1: 2426 to 3088

Score:53.9 bits(128), Expect:0.011,
Method:Compositional matrix adjust.,
Identities:63/247(26%), Positives:104/247(42%), Gaps:50/247(20%)

Query  15    GMDGPKVKQWPLTEEKIKA-----LVEICTEMEKEG--KISKIGPEN-PYNTPVFAIKKK  66
             G   P  ++W   E           + EI T + K+   K+S+  PE+  + + +F + KK
Sbjct  3088  GTRPPLTRRWSAHESATSGNRAVLIQEIQTLLRKQAVEKVSR--PESLGFYSHIFLVPKK  2915

Query  67    DSTKWRKLVDFRELNKRTQDFWEVQLGIPH---------PAGLKKKKSVTVLDVGDAYFS  117
             +  + R +++ R LN+         L  PH          A ++      T +D+ DAYF
Sbjct  2914  NG-QLRPVINLRPLNR--------VLLCPHFQMETVGTIAAAIQPGDWATSVDLTDAYFH  2762

Query  118   VPLDEDFRKYTAFTIPSINNETPGIRYQYNVLPQGWKGSPAIFQSSMTKILEPF--RKQN  175
             +P+   FRKY  F +        G  +Q+  LP G   +P +     T++L P
Sbjct  2761  IPIAPWFRKYLRFVV-------AGEVFQFRALPFGLAPAPLV----CTRLLAPLAIHLHT  2615

Query  176   PDIVIYQYMDDLYV---GSDLEIGQHRTKIEELRQHLLRWGLTTPDKKHQKEPP--FLWM  230
               ++ ++Y+DDL +    SDL       T I      L R GL      K + EP   F+++
Sbjct  2614  QGVLFHRYLDDLLIRAPSSDLCRRWTDTTISL----LYRMGLGVNLDKSELEPSRDFVYV  2447

Query  231   GYELHPD  237
             G     D
Sbjct  2446  GVRFRTD  2426



>Wolbachia endosymbiont of Corcyra cephalonica isolate SYLI2103 plasmid unnamed3, complete sequence
Sequence ID: NZ_CP087956.1 Length: 5332
Range 1: 1719 to 2189

Score:53.5 bits(127), Expect:0.012,
Method:Compositional matrix adjust.,
Identities:49/169(29%), Positives:84/169(49%), Gaps:14/169(8%)

Query  54    NPYNTPVFAIKKKDSTKWRKLVDFRELNKRTQ-DFWEVQLGIPHPAGLKKKKSVTVLDVG  112
             +P+++PV  +KKD T  R  VD+R LN  T+ D + +            L   K  + +D+
Sbjct  1719  SPWSSPVVLVKKKDGTT-RFCVDYRMLNNVTKKDSYPLPRIDDTLDMLSGMKWFSTIDLK  1895

Query  113   DAYFSVPLDEDFRKYTAFTIPSINNETPGIRYQYNVLPQGWKGSPAIFQSSMTKILEPFR  172
               Y+ V +  + ++ TAF+   I N   G+ +Q+NV+P G   +PA F+  M +++
Sbjct  1896  SGYWQVEISPEDKEKTAFST-GIGN---GL-WQFNVMPFGLCNAPATFERLMERVMVGLL  2060

Query  173   KQNPDIVIYQYMDDLYV-GSDLEIGQHRTKIEELRQHLLRWGLTTPDKK  220
                  V   Y+DD+ + G D E  +H   +E++ Q +    L    KK
Sbjct  2061  GD----VCLVYLDDVIIMGRDFE--EHLKNLEKVFQRIRGANLKLSTKK  2189



>Klebsiella pneumoniae strain A13 NODE_160_length_467_cov_0.764103, whole genome shotgun sequence
Sequence ID: NZ_JARWSQ010000155.1 Length: 333
Range 1: 5 to 316

Score:49.3 bits(116), Expect:0.012,
Method:Composition-based stats.,
Identities:34/112(30%), Positives:56/112(50%), Gaps:9/112(8%)

Query  54   NPYNTPVFAIKKKDSTKWRKLVDFRELNKRT-QDFWEVQLGIPHPAGLKKKKSVTVLDVG  112
            +PY++PV  ++KKD T WR  +D++ LN  T +D + + +        LK  +  T LD+
Sbjct  316  SPYSSPVLLVRKKDGT-WRMCIDYQALNSMTVKDKYPIPVVDELLDELKGARVFTKLDLR  140

Query  113  DAYFSVPLDEDFRKYTAFTIPSINNETPGIRYQYNVLPQGWKGSPAIFQSSM  164
              Y  + + +D    TAF        T    Y++ V+  G   +P+ FQS M
Sbjct  139  SGYHQIRVCDDGIPKTAF-------RTHDGHYEFLVMSFGLTNAPSTFQSLM  5



>Clostridium paraputrificum strain 28-X28 Scaffold60_1, whole genome shotgun sequence
Sequence ID: NZ_RAYJ01000063.1 Length: 680
```

```
Range 1: 1 to 363

Score:51.6 bits(122), Expect:0.013,
Method:Compositional matrix adjust.,
Identities:41/135(30%), Positives:61/135(45%), Gaps:15/135(11%)

Query  59   PVFAIKKKDSTKWRKLVDFRELNKRTQ-DFWEVQLGIPHPAGLKKKKSVTVLDVGDAYFS  117
            P   + K D T +R   DFR++N  T+ D + +         +   K V+  D+   Y+
Sbjct  1    PCILVPKPDGT-YRLCTDFRKVNSVTKTDSYPIPRIDDCIDKIGSAKFVSKFDLLKGYWQ  177

Query  118  VPLDEDFRKYTAFTIPSINNETPGIRYQYNVLPQGWKGSPAIFQSSMTKILEPFRKQNPD  177
            VPL E  R+ +AF        TP   YQY V+P G K +PA FQ  +  +L
Sbjct  178  VPLTERAREISAFA-------TPDGLYQYKVMPFGMKNAPATFQRMIHSLLNHLEGCE--  330

Query  178  IVIYQYMDDLYVGSD  192
                 Y+DD+ + SD
Sbjct  331  ----AYIDDVIIYSD  363
```

```
>Campylobacter sp. JMF_08 NE1 NODE_13_length_3482_cov_0.037836, whole genome shotgun sequence
Sequence ID: NZ_JAOVUC010000013.1 Length: 3482
Range 1: 873 to 1355

Score:53.5 bits(127), Expect:0.014,
Method:Compositional matrix adjust.,
Identities:53/181(29%), Positives:86/181(47%), Gaps:25/181(13%)

Query  6     ETVPVKLKPGMDGPKVKQWPLTEEKIKALVEICTEMEKEGKISKIGP-ENPYNTPVFAIK  64
             +T P+K  P    P VKQ  L +E +K L+E             I P ++ Y++P   +
Sbjct  1355  DTQPIKQHPYRMNP-VKQQCLRDE-VKYLLEN----------DFIEPSQSNYSSPCILVP  1212

Query  65    KKDSTKWRKLVDFRELNKRTQ-DFWEVQLGIPHPAGLKKKKSVTVLDVGDAYFSVPLDED  123
             K + T +R   D+R++N  T+ D + +         + K K VT  D+   ++ VPL +
Sbjct  1211  KSNGT-YRMCTDYRKINAVTKTDSFPIPRIDDCIDNVGKAKYVTKFDLLKGFWQVPLTDR  1035

Query  124   FRKYTAFTIPSINNETPGIRYQYNVLPQGWKGSPAIFQSSMTKI---LEPFRKQNPDIVI  180
              ++ +AF        TP   +QY V+P G K SPA FQ  +  I   L+  +    D++I
Sbjct  1034  AKEISAFV-------TPDGLFQYKVMPFGMKNSPATFQRLINSILAGLDNCKAYIDDVII  876

Query  181   Y  181
             Y
Sbjct  875   Y  873
```

```
>Campylobacter sp. JMF_10 EL2 NODE_13_length_1688_cov_0.027690, whole genome shotgun sequence
Sequence ID: NZ_JAOVTV010000013.1 Length: 1688
Range 1: 349 to 831

Score:52.8 bits(125), Expect:0.015,
Method:Compositional matrix adjust.,
Identities:53/181(29%), Positives:86/181(47%), Gaps:25/181(13%)

Query  6    ETVPVKLKPGMDGPKVKQWPLTEEKIKALVEICTEMEKEGKISKIGP-ENPYNTPVFAIK  64
            +T P+K  P    P VKQ  L +E +K L+E             I P ++ Y++P   +
Sbjct  831  DTQPIKQHPYRMNP-VKQQCLKDE-VKYLLEN----------DFIEPSQSNYSSPCILVP  688

Query  65   KKDSTKWRKLVDFRELNKRTQ-DFWEVQLGIPHPAGLKKKKSVTVLDVGDAYFSVPLDED  123
            K + T +R   D+R++N  T+ D + +         + K K VT  D+   ++ VPL +
Sbjct  687  KSNGT-YRMCTDYRKINAVTKTDSFPIPRIDDCIDNVGKAKYVTKFDLLKGFWQVPLTDR  511

Query  124  FRKYTAFTIPSINNETPGIRYQYNVLPQGWKGSPAIFQSSMTKI---LEPFRKQNPDIVI  180
             ++ +AF        TP   +QY V+P G K SPA FQ  +  I   L+  +    D++I
Sbjct  510  AKEISAFV-------TPDGLFQYKVMPFGMKNSPATFQRLINSILAGLDNCKAYIDDVII  352

Query  181  Y  181
            Y
Sbjct  351  Y  349
```

```
>Campylobacter sp. CS_ED1 strain Cs ED1 NODE_9_length_5400_cov_0.175250, whole genome shotgun sequence
Sequence ID: NZ_JAOVUL010000009.1 Length: 5400
Range 1: 3715 to 4197

Score:53.1 bits(126), Expect:0.016,
Method:Compositional matrix adjust.,
Identities:53/181(29%), Positives:86/181(47%), Gaps:25/181(13%)

Query  6     ETVPVKLKPGMDGPKVKQWPLTEEKIKALVEICTEMEKEGKISKIGP-ENPYNTPVFAIK  64
             +T P+K  P    P VKQ  L +E +K L+E             I P ++ Y++P   +
Sbjct  3715  DTQPIKQHPYRMNP-VKQQCLKDE-VKYLLEN----------DFIEPSQSNYSSPCILVP  3858

Query  65    KKDSTKWRKLVDFRELNKRTQ-DFWEVQLGIPHPAGLKKKKSVTVLDVGDAYFSVPLDED  123
             K + T +R   D+R++N  T+ D + +         + K K VT  D+   ++ VPL +
Sbjct  3859  KSNGT-YRMCTDYRKINAVTKTDSFPIPRIDDCIDNVGKAKYVTKFDLLKGFWQVPLTDR  4035

Query  124   FRKYTAFTIPSINNETPGIRYQYNVLPQGWKGSPAIFQSSMTKI---LEPFRKQNPDIVI  180
              ++ +AF        TP   +QY V+P G K SPA FQ  +  I   L+  +    D++I
Sbjct  4036  AKEISAFV-------TPDGLFQYKVMPFGMKNSPATFQRLINSILAGLDNCKAYIDDVII  4194

Query  181   Y  181
             Y
Sbjct  4195  Y  4197
```

```
>Campylobacter sp. CN_NE4 strain Cn NE4 NODE_12_length_5576_cov_0.042351, whole genome shotgun sequence
Sequence ID: NZ_JAOVUF010000012.1 Length: 5576
Range 1: 3483 to 3965

Score:53.1 bits(126), Expect:0.017,
Method:Compositional matrix adjust.,
Identities:53/181(29%), Positives:86/181(47%), Gaps:25/181(13%)

Query  6     ETVPVKLKPGMDGPKVKQWPLTEEKIKALVEICTEMEKEGKISKIGP-ENPYNTPVFAIK  64
             +T P+K  P    P VKQ  L +E +K L+E             I P ++ Y++P   +
Sbjct  3483  DTQPIKQHPYRMNP-VKQQCLKDE-VKYLLEN----------DFIEPSQSNYSSPCILVP  3626

Query  65    KKDSTKWRKLVDFRELNKRTQ-DFWEVQLGIPHPAGLKKKKSVTVLDVGDAYFSVPLDED  123
             K + T +R   D+R++N  T+ D + +         + K K VT  D+   ++ VPL +
Sbjct  3627  KSNGT-YRMCTDYRKINAVTKTDSFPIPRIDDCIDNVGKAKYVTKFDLLKGFWQVPLTDR  3803

Query  124   FRKYTAFTIPSINNETPGIRYQYNVLPQGWKGSPAIFQSSMTKI---LEPFRKQNPDIVI  180
              ++ +AF        TP   +QY V+P G K SPA FQ  +  I   L+  +    D++I
Sbjct  3804  AKEISAFV-------TPDGLFQYKVMPFGMKNSPATFQRLINSILAGLDNCKAYIDDVII  3962

Query  181   Y  181
             Y
```

```
Sbjct  3963  Y  3965
```

>Klebsiella pneumoniae strain BL1427 NODE_183_length_321_cov_0.118557, whole genome shotgun sequence
Sequence ID: NZ_VICX01000183.1 Length: 321
Range 1: 11 to 316

Score:48.1 bits(113), Expect:0.026,
Method:Compositional matrix adjust.,
Identities:27/104(26%), Positives:57/104(54%), Gaps:2/104(1%)

```
Query  164  MTKILEPFRKQNPDIVIYQYMDDLYVGSDLEIGQHRTKIEELRQHLLRWGLTTPDKKHQK  223
            +T+ + P R  + + +IY YMDD+ + +  +    R  + ++   L R+ L    +K Q
Sbjct  316  VTQAIAPVRHLHKNSLIYHYMDDILIAAPNDSLGDRV-LNDITNALSRFKLVIAQEKIQT  140

Query  224  EPPFLWMGYELHPDKWTVQPIVLPEKDSWTVNDIQKLVGKLNWA  267
              P+ ++G+++       Q + +P+   +++ND+QKL+G +NW
Sbjct  139  VSPWQYLGWQIMERTIRPQKLKIPQT-VFSLNDLQKLLGAINWV  11
```

>Pseudomonas aromaticivorans strain MAP12 47, whole genome shotgun sequence
Sequence ID: NZ_JAHRGL010000047.1 Length: 2177
Range 1: 5 to 367

Score:52.4 bits(124), Expect:0.028,
Method:Compositional matrix adjust.,
Identities:35/126(28%), Positives:66/126(52%), Gaps:10/126(7%)

```
Query  10   VKLKPGMDGPKVKQWPLTEEKIKALVEICTEMEKEGKISKIGPENPYNTPVFAIKKKDST  69
            V LK  +    V+Q+P+ +E  + +      + ++G +  +  ++P+NTP+  ++K  +
Sbjct  367  VTLKAAILPASVRQYPMPKEAREGIRPHIKRLLEQGIL--VPCKSPWNTPLLPVRKPGTN  194

Query  70   KWRKLVDFRELNKRTQDFWEVQLGIPHP----AGLKKKKS-VTVLDVGDAYFSVPLDEDF  124
             +R + D RE+NKR +D       +P+P    +GL    +  TVLD+ DA+F + L
Sbjct  193  DYRPVQDLREVNKRIEDMHPT---VPNPYNLLSGLPPNYTWYTVLDLKDAFFCLRLHPTN  23

Query  125  RKYTAF  130
            +   AF
Sbjct  22   QPIFAF  5
```

>Lachnoclostridium phocaeense strain Marseille-P3177T chromosome contig00001
Sequence ID: NZ_LT635479.1 Length: 3500754
Range 1: 1458059 to 1458487

Score:52.4 bits(124), Expect:0.030,
Method:Compositional matrix adjust.,
Identities:41/143(29%), Positives:72/143(50%), Gaps:8/143(5%)

```
Query  422      LVKLWYQLEKEPIVGAETFYVDGAAN-RETKLGKAGYVTNRGRQKVV-----TLTDTTNQ  475
                +++L+ +L KE + G  T Y DG          G  G V      KV+         +TN
Sbjct  1458487  ILELFPELAKEKLPGKYTVYTDGGCGFNPGGPGGIGAVILDVEGKVIQEISKGFHASTNN  1458308

Query  476      KTELQAIYLALQD--SGLEVNIVTDSQYALGIIQAQPDQSESELVNQIIEQLIKKEKVYL  533
                + EL A   AL+   +G EV + +DSQY +  ++    ++++  + + +++ ++ +KV
Sbjct  1458307  RMELLAAIEALKTIPAGAEVKVKSDSQYLVQTMEEGWSRNKNHDLWKQLDEALRGKKVSF  1458128

Query  534      AWVPAHKGIGGNEQVDKLVSAGI  556
                 WV  H G   NE+ D+L +AGI
Sbjct  1458127  KWVRGHNGNQWNERCDELATAGI  1458059
```

>Kribbella italica strain DSM 28967 Ga0415174_01, whole genome shotgun sequence
Sequence ID: NZ_JACHMY010000001.1 Length: 8885108
Range 1: 163423 to 163851

Score:52.4 bits(124), Expect:0.032,
Method:Compositional matrix adjust.,
Identities:41/145(28%), Positives:62/145(42%), Gaps:20/145(13%)

```
Query  433     PIVGAETF---YVDGAANRETKLGKAGYVTNRGRQKVVTLTDTTNQKTELQAIYLALQDS  489
               P VGA      Y DGA +  T  G   +V   G +      DTTNQ+ ELQA Y A Q
Sbjct  163423  PEVGASDLIEVYTDGACSGGT--GGWAWVVADGPEGSGATNDTTNQRMELQAAYEAAQSL  163596

Query  490     GLEVNIVTDSQYALGIIQ---------------AQPDQSESELVNQIIEQLIKKEKVYLA  534
                  + IV+DS Y +   +                 A+   +  ++   +IE ++ +  +
Sbjct  163597  FGPLTIVSDSSYVVNCFKQGWWKGWHARNWVNSARKPVANKDIWEPLIELVLHRGDIDFR  163776

Query  535     WVPAHKGIGGNEQVDKLVSAGIRKV  559
               W+  H G   NE+ D+L  A  + V
Sbjct  163777  WIKGHSGHPLNERADRLAVAARQGV  163851
```

>Campylobacter sp. JMF_03 NE3 NODE_16_length_1924_cov_0.035616, whole genome shotgun sequence
Sequence ID: NZ_JAOVTS010000016.1 Length: 1924
Range 1: 1428 to 1820

Score:52.0 bits(123), Expect:0.035,
Method:Compositional matrix adjust.,
Identities:40/139(29%), Positives:69/139(49%), Gaps:13/139(9%)

```
Query  53    ENPYNTPVFAIKKKDSTKWRKLVDFRELNKRTQ-DFWEVQLGIPHPAGLKKKKSVTVLDV  111
             ++ Y++P   + K + T +R   D+R++N  T+ D + +         + K K VT  D+
Sbjct  1820  QSNYSSPCILVPKSNGT-YRMCTDYRKINAVTKTDSFPIPRIDDCIDNVGKAKYVTKFDL  1644

Query  112   GDAYFSVPLDEDFRKYTAFTIPSINNETPGIRYQYNVLPQGWKGSPAIFQSSMTKI---L  168
               ++ VPL +  ++ +AF         TP   +QY V+P G K SPA FQ  +  I   L
Sbjct  1643  LKGFWQVPLTDRAKEISAFV-------TPDGLFQYKVMPFGMKNSPATFQRLINSILAGL  1485

Query  169   EPFRKQNPDIVIY-QYMDD  186
             +  +    D++IY +  DD
Sbjct  1484  DNCKAYIDDVIIYTETFDD  1428
```

>Campylobacter jejuni strain CJJD2010009 CJJD2010009_contig_657, whole genome shotgun sequence
Sequence ID: NZ_JAJUND010000202.1 Length: 251
Range 1: 5 to 133

Score:47.0 bits(110), Expect:0.035,
Method:Compositional matrix adjust.,
Identities:23/44(52%), Positives:30/44(68%), Gaps:1/44(2%)

```
Query  55   PYNTPVFAIKKKDSTKWRKLVDFRELNKRTQDFWEVQLGIPHPA  98
            P+N+PVF IKKK   KWR L D +E+NK  +    +QLG+P PA
Sbjct  133  PWNSPVFVIKKK-YEKWRILTDLQEVNKYIEPMGALQLGLPSPA  5
```

>Saccharospirillum mangrovi strain HK-33 Scaffold2, whole genome shotgun sequence
Sequence ID: NZ_QAPE01000002.1 Length: 517067
Range 1: 16487 to 16972

Score:52.4 bits(124), Expect:0.035,
Method:Compositional matrix adjust.,
Identities:43/162(27%), Positives:71/162(43%), Gaps:21/162(12%)

```
Query  420    PPLVKLWYQLEKEP---IVGAETFYVDGAANRETKLGKAGYVTNRGRQKVVTL---TDTT  473
              PP  ++W     +    +  A T Y DGA       G  G +   G  +        +TT
Sbjct  16972  PPSSQIWPNRSDDQGIFMSDAVTLYTDGACRGNPGPGGWGVLMRSGAHEKTLHGGEAETT  16793

Query  474    NQKTELQAIYLALQD--SGLEVNIVTDSQYAL-GIIQ------------AQPDQSESELV  518
              N + EL A  + L+      +V++ TDSQY   GI +             A     +++ +
Sbjct  16792  NNRMELMAAIVGLESLKRRCQVDLYTDSQYVRKGITEWIHGWKRNGWKTASKQPVKNQDL  16613

Query  519    NQIIEQLIKKEKVYLAWVPAHKGIGGNEQVDKLVSAGIRKVL  560
               Q ++Q I K +V   WV  H G+ GNE+ D L + G+ +++
Sbjct  16612  WQRLDQQIGKHQVEWHWVKGHAGVDGNERADALANQGVDEII  16487
```

>Saccharospirillum mangrovi strain HK-33 chromosome, complete genome
Sequence ID: NZ_CP031415.1 Length: 3686911
Range 1: 1731954 to 1732439

Score:52.4 bits(124), Expect:0.035,
Method:Compositional matrix adjust.,
Identities:43/162(27%), Positives:71/162(43%), Gaps:21/162(12%)

```
Query  420      PPLVKLWYQLEKEP---IVGAETFYVDGAANRETKLGKAGYVTNRGRQKVVTL---TDTT  473
                PP  ++W     +    +  A T Y DGA       G  G +   G  +        +TT
Sbjct  1731954  PPSSQIWPNRSDDQGIFMSDAVTLYTDGACRGNPGPGGWGVLMRSGAHEKTLHGGEAETT  1732133

Query  474      NQKTELQAIYLALQD--SGLEVNIVTDSQYAL-GIIQ------------AQPDQSESELV  518
                N + EL A  + L+      +V++ TDSQY   GI +             A     +++ +
Sbjct  1732134  NNRMELMAAIVGLESLKRRCQVDLYTDSQYVRKGITEWIHGWKRNGWKTASKQPVKNQDL  1732313

Query  519      NQIIEQLIKKEKVYLAWVPAHKGIGGNEQVDKLVSAGIRKVL  560
                 Q ++Q I K +V   WV  H G+ GNE+ D L + G+ +++
Sbjct  1732314  WQRLDQQIGKHQVEWHWVKGHAGVDGNERADALANQGVDEII  1732439
```

>Pseudomonas morbosilactucae strain MAFF 302046 302046_contig_42, whole genome shotgun sequence
Sequence ID: NZ_JALQCX010000042.1 Length: 73784
Range 1: 43168 to 43593

Score:52.4 bits(124), Expect:0.035,
Method:Compositional matrix adjust.,
Identities:44/142(31%), Positives:61/142(42%), Gaps:25/142(17%)

```
Query  439    TFYVDGAANRE----TKLGKAGYVTNRGRQK------VVTLTDTTNQKTELQAIYLALQD  488
              T YVDGA         K G   Y+TN           V      TN + EL A  +AL+
Sbjct  43593  TIYVDGACPNNGKPNAKAGWGAYLTNPHGDTLELAGPVPANQPQTNGRAELLAPLMALRR  43414

Query  489    S--GLEVNIVTDSQY-------------ALGIIQAQPDQSESELVNQIIEQLIKKEKVYL  533
                 G  +NIV+DSQY             A G  +A   + E   + ++I+Q ++K ++
Sbjct  43413  CKPGAAINIVSDSQYVVKGAMEWLQGWKARGWRKADKKEVEHRDLWELIDQEMQKREIRF  43234

Query  534    AWVPAHKGIGGNEQVDKLVSAG  555
               WV  H G  GNE+ D L   G
Sbjct  43233  QWVRGHSGDTGNERADTLACRG  43168
```

Query #11: sp|P04585|p15_RNAseH_HV1H2 Gag-Pol polyprotein OS=Human immunodeficiency virus type 1 group M subtype B (isolate HXB2) OX=11706 GN=gag-pol PE=1 SV=4 Query ID: lcl|Query_2387719 Length: 120

Sequences producing significant alignments:

| Description | Scientific Name | Common Name | Taxid | Max Score | Total Score | Query cover | E Value | Per. Ident | Acc. Len | Accession |
|---|---|---|---|---|---|---|---|---|---|---|
| Klebsiella pneumoniae strain ARGID_31501, whole genome shotgun... | Klebsiella p... | NA | 573 | 230 | 230 | 100% | 1e-67 | 95.00 | 2655 | NZ_CAKLPD0100... |
| Burkholderia pseudomallei strain 2604STDY5518971... | Burkholderia... | NA | 28450 | 220 | 220 | 100% | 5e-67 | 89.17 | 1345 | NZ_UVFZ020000... |
| Actinoplanes campanulatus strain JCM 3059 sequence058, whole... | Actinoplanes... | NA | 113559 | 231 | 231 | 100% | 5e-67 | 95.83 | 4754 | NZ_BMPW010000... |
| Enterococcus alcedinis strain CCM 8433 sequence23, whole genom... | Enterococcus... | NA | 1274384 | 231 | 231 | 100% | 5e-67 | 95.83 | 6075 | NZ_BMDT010000... |
| Deinococcus aquiradiocola strain JCM 14371 sequence30, whole... | Deinococcus ... | NA | 393059 | 231 | 231 | 100% | 5e-67 | 95.83 | 6128 | NZ_BMOE010000... |
| Methylobacterium radiotolerans strain MAMP 4754... | Methylobacte... | NA | 31998 | 214 | 214 | 100% | 8e-64 | 87.50 | 1636 | NZ_NKQS010006... |
| Escherichia coli strain VRES0669, whole genome shotgun sequence | Escherichia ... | NA | 562 | 207 | 207 | 100% | 9e-59 | 85.00 | 5544 | NZ_UINV010000... |
| Thalassovita mangrovi strain GS-10 Scaffold20_1, whole genome... | Thalassovita... | NA | 2692236 | 199 | 199 | 85% | 2e-58 | 95.15 | 1488 | NZ_WWEN010000... |
| Klebsiella pneumoniae strain EuSCAPE_MT012, whole genome shotg... | Klebsiella p... | NA | 573 | 182 | 182 | 83% | 4e-56 | 88.00 | 452 | NZ_UKIX020000... |
| Pseudomonas aeruginosa strain VET-42... | Pseudomonas ... | NA | 287 | 107 | 107 | 45% | 4e-24 | 90.91 | 1154 | NZ_RACX010001... |
| Klebsiella pneumoniae strain EuSCAPE_DE086, whole genome shotg... | Klebsiella p... | NA | 573 | 79.7 | 79.7 | 44% | 3e-14 | 73.58 | 995 | NZ_UKLW020000... |
| Pseudomonas aeruginosa strain ENV-297... | Pseudomonas ... | NA | 287 | 76.3 | 76.3 | 30% | 6e-13 | 94.59 | 1100 | NZ_RADD010000... |
| Thalassovita mangrovi strain GS-10 Scaffold30_1, whole genome... | Thalassovita... | NA | 2692236 | 75.1 | 75.1 | 38% | 7e-13 | 71.74 | 722 | NZ_WWEN010000... |
| Cuspidothrix issatschenkoi CHARLIE-1 contig_175, whole genome... | Cuspidothrix... | NA | 2052836 | 69.3 | 69.3 | 91% | 3e-10 | 41.07 | 2237 | NZ_PGEM010001... |
| Klebsiella pneumoniae strain B2 isolate B2, whole genome shotg... | Klebsiella p... | NA | 573 | 64.3 | 64.3 | 81% | 2e-08 | 42.31 | 7306 | NZ_CABWNT0100... |
| Polyangium sp. y55x31 NODE_26_length_169233_cov_17.164843, who... | Polyangium s... | NA | 3042688 | 61.2 | 61.2 | 97% | 2e-07 | 33.06 | 169233 | NZ_JARZHG0100... |
| Polyangium sp. 15x6 NODE_112_length_36528_cov_15.618939, whole... | Polyangium s... | NA | 3042687 | 58.2 | 58.2 | 97% | 3e-06 | 32.26 | 36528 | NZ_JASBQE0100... |
| Polyangium sp. 6x1 NODE_22_length_166486_cov_22.101912, whole... | Polyangium s... | NA | 3042689 | 56.6 | 56.6 | 97% | 9e-06 | 33.06 | 166486 | NZ_JARZHH0100... |
| Polyangium jinanense strain SDU14 Scaffold9, whole genome... | Polyangium j... | NA | 2829994 | 54.7 | 54.7 | 93% | 4e-05 | 31.93 | 231060 | NZ_JAGTJJ0100... |
| Polyangium jinanense strain SDU13 Scaffold15, whole genome... | Polyangium j... | NA | 2829994 | 54.7 | 54.7 | 93% | 4e-05 | 31.93 | 231263 | NZ_JAGTJK0100... |
| Roseiconus lacunae strain JC742 soapdenovo_K_105_82, whole... | Roseiconus l... | NA | 2605694 | 50.8 | 50.8 | 100% | 0.001 | 28.28 | 35968 | NZ_JAJMQV0100... |
| Kribbella italica strain DSM 28967 Ga0415174_01, whole genome... | Kribbella it... | NA | 1540520 | 50.1 | 50.1 | 99% | 0.002 | 27.61 | 8885108 | NZ_JACHMY0100... |
| Stutzerimonas stutzeri strain 24a75 44Pst24A75gv17C029, whole... | Stutzerimona... | NA | 316 | 49.7 | 49.7 | 72% | 0.003 | 33.98 | 55843 | NZ_JAMOHU0100... |
| Stutzerimonas stutzeri strain 24a75... | Stutzerimona... | NA | 316 | 49.7 | 49.7 | 72% | 0.003 | 33.98 | 75459 | NZ_POUT010000... |
| Pseudomonas sp. BAY1663 contig000002, whole genome shotgun... | Pseudomonas ... | NA | 1439940 | 49.7 | 49.7 | 72% | 0.003 | 33.98 | 124554 | NZ_AZSV010000... |
| Nitrosomonas sp. isolate RBC_AOB3 BC1.188.fa-20, whole genome... | Nitrosomonas... | NA | 42353 | 49.3 | 49.3 | 100% | 0.003 | 30.43 | 55000 | NZ_JALDQX0100... |
| Deinococcus aerius strain TR0125 DAERI_Contig_05, whole genome... | Deinococcus ... | NA | 200253 | 49.3 | 49.3 | 93% | 0.003 | 33.08 | 248739 | NZ_BFAG010000... |
| Nitrosomonas sp. Nm51, whole genome shotgun sequence | Nitrosomonas... | NA | 133720 | 48.9 | 48.9 | 100% | 0.004 | 28.99 | 20055 | NZ_FOGH010000... |
| Stutzerimonas stutzeri strain GD04160 15, whole genome shotgun... | Stutzerimona... | NA | 316 | 48.9 | 48.9 | 96% | 0.004 | 32.59 | 112086 | NZ_JAODYR0100... |
| Stutzerimonas frequens strain HI00D01 HI00D01_c19, whole genom... | Stutzerimona... | NA | 2968969 | 48.9 | 48.9 | 96% | 0.004 | 32.59 | 149482 | NZ_LWEF010000... |
| Stutzerimonas frequens strain GOM2 scaffold_3, whole genome... | Stutzerimona... | NA | 2968969 | 48.9 | 48.9 | 96% | 0.004 | 32.59 | 232832 | NZ_WOUM010000... |
| Stutzerimonas frequens strain DNSP21 27DNSP21gv5C003, whole... | Stutzerimona... | NA | 2968969 | 48.9 | 48.9 | 96% | 0.004 | 32.59 | 433620 | NZ_JAMOHM0100... |
| Stutzerimonas frequens strain S18 3, whole genome shotgun... | Stutzerimona... | NA | 2968969 | 48.9 | 48.9 | 96% | 0.004 | 32.59 | 681260 | NZ_JASSZI0100... |
| Stutzerimonas frequens strain DNSP21... | Stutzerimona... | NA | 2968969 | 48.9 | 48.9 | 96% | 0.004 | 32.59 | 722045 | NZ_POUJ010000... |
| Stutzerimonas frequens strain KOL14.W.20.12... | Stutzerimona... | NA | 2968969 | 48.9 | 48.9 | 96% | 0.004 | 32.59 | 879884 | NZ_WIFY010000... |
| Stutzerimonas frequens strain KOL14.W.90.4... | Stutzerimona... | NA | 2968969 | 48.9 | 48.9 | 96% | 0.004 | 32.59 | 1280911 | NZ_WIGH010000... |
| Stutzerimonas frequens strain KOL14.W.90.9... | Stutzerimona... | NA | 2968969 | 48.9 | 48.9 | 96% | 0.004 | 32.59 | 1283483 | NZ_WIFU010000... |
| Stutzerimonas frequens strain KOL14.W.45.16... | Stutzerimona... | NA | 2968969 | 48.9 | 48.9 | 96% | 0.004 | 32.59 | 1368612 | NZ_WIGI010000... |
| Stutzerimonas frequens strain CAM01 k141_91720, whole genome... | Stutzerimona... | NA | 2968969 | 48.9 | 48.9 | 96% | 0.004 | 32.59 | 1835512 | NZ_JAQBSX0100... |
| Stutzerimonas frequens strain FDAARGOS_877 chromosome, complet... | Stutzerimona... | NA | 2968969 | 48.9 | 48.9 | 96% | 0.004 | 32.59 | 4394686 | NZ_CP065720.1 |
| Stutzerimonas frequens strain TPA3 chromosome, complete genome | Stutzerimona... | NA | 2968969 | 48.9 | 48.9 | 96% | 0.004 | 32.59 | 4415884 | NZ_CP071899.1 |

| | | | | | | | | | | |
|---|---|---|---|---|---|---|---|---|---|---|
| Stutzerimonas frequens strain THAF7b chromosome, complete genome | Stutzerimona... | NA | 2968969 | 48.9 | 48.9 | 96% | 0.004 | 32.59 | 4522538 | NZ_CP045416.1 |
| Stutzerimonas frequens strain L27 chromosome, complete genome | Stutzerimona... | NA | 2968969 | 48.9 | 48.9 | 96% | 0.004 | 32.59 | 4544032 | NZ_CP136501.1 |
| Stutzerimonas frequens strain KOL14.W.5.3... | Stutzerimona... | NA | 2968969 | 48.9 | 48.9 | 96% | 0.004 | 32.59 | 4554507 | NZ_WIGG010000... |
| Stutzerimonas frequens strain TF18 chromosome, complete genome | Stutzerimona... | NA | 2968969 | 48.9 | 48.9 | 96% | 0.004 | 32.59 | 4918844 | NZ_CP113257.1 |
| Stutzerimonas stutzeri strain PS_027... | Stutzerimona... | NA | 316 | 48.9 | 48.9 | 72% | 0.004 | 33.98 | 26947 | NZ_RHRQ010000... |
| Stutzerimonas stutzeri strain GD03740 30, whole genome shotgun... | Stutzerimona... | NA | 316 | 48.9 | 48.9 | 72% | 0.004 | 33.98 | 38162 | NZ_JAOCHW0100... |
| Stutzerimonas stutzeri strain INSAq93 Aq93_contig_23, whole... | Stutzerimona... | NA | 316 | 48.9 | 48.9 | 72% | 0.004 | 33.98 | 38538 | NZ_JAKCMP0100... |
| Stutzerimonas stutzeri XLDN-R contig000128, whole genome shotg... | Stutzerimona... | NA | 1200345 | 48.9 | 48.9 | 72% | 0.004 | 33.98 | 40595 | NZ_AKYE010001... |
| Stutzerimonas stutzeri strain 1223_PMEN 439_42450_436684, whol... | Stutzerimona... | NA | 316 | 48.9 | 48.9 | 72% | 0.004 | 33.98 | 42450 | NZ_JVYC010001... |
| Stutzerimonas stutzeri strain GD03749 27, whole genome shotgun... | Stutzerimona... | NA | 316 | 48.9 | 48.9 | 72% | 0.004 | 33.98 | 62832 | NZ_JAOCHN0100... |
| Stutzerimonas stutzeri strain PSB00024 PSB00024_c0024, whole... | Stutzerimona... | NA | 316 | 48.9 | 48.9 | 72% | 0.004 | 33.98 | 63007 | NZ_JADTWB0100... |
| Stutzerimonas stutzeri strain A655/78 11PstA665_78gv1C022, who... | Stutzerimona... | NA | 316 | 48.9 | 48.9 | 72% | 0.004 | 33.98 | 63676 | NZ_JAMOJY0100... |
| Stutzerimonas stutzeri strain INSAq87 Aq87_contig_65, whole... | Stutzerimona... | NA | 316 | 48.9 | 48.9 | 72% | 0.004 | 33.98 | 64028 | NZ_JAKCMO0100... |
| Stutzerimonas stutzeri strain SD93936 09PstSD93936gv1c014, who... | Stutzerimona... | NA | 316 | 48.9 | 48.9 | 72% | 0.004 | 33.98 | 64350 | NZ_JAMOJZ0100... |
| Stutzerimonas stutzeri strain GD04104 21, whole genome shotgun... | Stutzerimona... | NA | 316 | 48.9 | 48.9 | 72% | 0.004 | 33.98 | 65673 | NZ_JAOEAT0100... |
| Stutzerimonas stutzeri strain LH-42 Contig004, whole genome... | Stutzerimona... | NA | 316 | 48.9 | 48.9 | 72% | 0.004 | 33.98 | 68185 | NZ_NCKS020000... |
| Stutzerimonas stutzeri strain A160/74 10PstA160_74gv1c016, who... | Stutzerimona... | NA | 316 | 48.9 | 48.9 | 72% | 0.004 | 33.98 | 82616 | NZ_JAMOHN0100... |
| Stutzerimonas stutzeri strain NT-I, whole genome shotgun sequence | Stutzerimona... | NA | 316 | 48.9 | 48.9 | 72% | 0.004 | 33.98 | 97403 | NZ_BCAJ010000... |
| Stutzerimonas stutzeri strain GD04106 14, whole genome shotgun... | Stutzerimona... | NA | 316 | 48.9 | 48.9 | 72% | 0.004 | 33.98 | 109530 | NZ_JAOEAR0100... |
| Stutzerimonas stutzeri strain GD04164 8, whole genome shotgun... | Stutzerimona... | NA | 316 | 48.9 | 48.9 | 72% | 0.004 | 33.98 | 111383 | NZ_JAODYN0100... |
| Stutzerimonas stutzeri strain PS_197... | Stutzerimona... | NA | 316 | 48.9 | 48.9 | 72% | 0.004 | 33.98 | 111597 | NZ_RHRB010000... |
| Stutzerimonas stutzeri strain PS_234... | Stutzerimona... | NA | 316 | 48.9 | 48.9 | 72% | 0.004 | 33.98 | 111675 | NZ_RHQZ010000... |
| Stutzerimonas stutzeri strain GD04161 18, whole genome shotgun... | Stutzerimona... | NA | 316 | 48.9 | 48.9 | 72% | 0.004 | 33.98 | 112063 | NZ_JAODYQ0100... |
| Stutzerimonas stutzeri strain GD03774 16, whole genome shotgun... | Stutzerimona... | NA | 316 | 48.9 | 48.9 | 72% | 0.004 | 33.98 | 112070 | NZ_JAOCGO0100... |
| Stutzerimonas stutzeri strain CM14... | Stutzerimona... | NA | 316 | 48.9 | 48.9 | 72% | 0.004 | 33.98 | 115186 | NZ_SRXR010000... |
| Stutzerimonas stutzeri strain CM... | Stutzerimona... | NA | 316 | 48.9 | 48.9 | 72% | 0.004 | 33.98 | 115190 | NZ_QLAB010000... |
| Stutzerimonas stutzeri strain 40D2... | Stutzerimona... | NA | 316 | 48.9 | 48.9 | 72% | 0.004 | 33.98 | 115488 | NZ_MWUI010000... |
| Stutzerimonas stutzeri strain PS_138... | Stutzerimona... | NA | 316 | 48.9 | 48.9 | 72% | 0.004 | 33.98 | 125664 | NZ_RHRE010000... |
| Stutzerimonas stutzeri strain NGHE31... | Stutzerimona... | NA | 316 | 48.9 | 48.9 | 72% | 0.004 | 33.98 | 130955 | NZ_JANAJI0100... |
| Stutzerimonas stutzeri strain PS_001... | Stutzerimona... | NA | 316 | 48.9 | 48.9 | 72% | 0.004 | 33.98 | 130975 | NZ_RXOW010000... |
| Stutzerimonas stutzeri strain PS_211... | Stutzerimona... | NA | 316 | 48.9 | 48.9 | 72% | 0.004 | 33.98 | 130975 | NZ_RHRA010000... |
| Stutzerimonas stutzeri strain PS_072... | Stutzerimona... | NA | 316 | 48.9 | 48.9 | 72% | 0.004 | 33.98 | 135183 | NZ_RHRO010000... |
| Stutzerimonas stutzeri strain PS_075... | Stutzerimona... | NA | 316 | 48.9 | 48.9 | 72% | 0.004 | 33.98 | 135224 | NZ_RHRN010000... |
| Stutzerimonas stutzeri strain GD04143 6, whole genome shotgun... | Stutzerimona... | NA | 316 | 48.9 | 48.9 | 72% | 0.004 | 33.98 | 166685 | NZ_JAODZI0100... |
| Stutzerimonas stutzeri strain GD04147 6, whole genome shotgun... | Stutzerimona... | NA | 316 | 48.9 | 48.9 | 72% | 0.004 | 33.98 | 166685 | NZ_JAODZE0100... |
| Stutzerimonas stutzeri strain PS_257... | Stutzerimona... | NA | 316 | 48.9 | 48.9 | 72% | 0.004 | 33.98 | 183596 | NZ_RHQY010000... |
| Stutzerimonas stutzeri strain GD04144 10, whole genome shotgun... | Stutzerimona... | NA | 316 | 48.9 | 48.9 | 72% | 0.004 | 33.98 | 184200 | NZ_JAODZH0100... |
| Stutzerimonas stutzeri strain GD03747 7, whole genome shotgun... | Stutzerimona... | NA | 316 | 48.9 | 48.9 | 72% | 0.004 | 33.98 | 199808 | NZ_JAOCHP0100... |
| Stutzerimonas stutzeri strain 109A1... | Stutzerimona... | NA | 316 | 48.9 | 48.9 | 72% | 0.004 | 33.98 | 231087 | NZ_JAAMRD0100... |
| Stutzerimonas stutzeri strain PAH8... | Stutzerimona... | NA | 316 | 48.9 | 48.9 | 72% | 0.004 | 33.98 | 231702 | NZ_JAGFOX0100... |
| Stutzerimonas stutzeri strain SDU10 Scaffold8, whole genome... | Stutzerimona... | NA | 316 | 48.9 | 48.9 | 72% | 0.004 | 33.98 | 233818 | NZ_JAJHZH0100... |
| Stutzerimonas stutzeri strain AR9-4 contig_1, whole genome... | Stutzerimona... | NA | 316 | 48.9 | 48.9 | 72% | 0.004 | 33.98 | 233982 | NZ_MDGV010000... |
| Stutzerimonas stutzeri strain DIA-8 S.stutzeri8_contig_8, whol... | Stutzerimona... | NA | 316 | 48.9 | 48.9 | 72% | 0.004 | 33.98 | 234537 | NZ_JAZGZT0100... |
| Stutzerimonas stutzeri isolate PSEUDT2.31D-1, whole genome... | Stutzerimona... | NA | 316 | 48.9 | 48.9 | 72% | 0.004 | 33.98 | 237644 | NZ_CAJFAG0100... |
| Stutzerimonas stutzeri strain GD04155 4, whole genome shotgun... | Stutzerimona... | NA | 316 | 48.9 | 48.9 | 72% | 0.004 | 33.98 | 238078 | NZ_JAODYW0100... |
| Stutzerimonas stutzeri strain GD03864 1, whole genome shotgun... | Stutzerimona... | NA | 316 | 48.9 | 48.9 | 72% | 0.004 | 33.98 | 249048 | NZ_JAOCDG0100... |
| Stutzerimonas stutzeri strain PS_366... | Stutzerimona... | NA | 316 | 48.9 | 48.9 | 72% | 0.004 | 33.98 | 258367 | NZ_RHQX010000... |
| Stutzerimonas stutzeri strain 4MP-B contig_5, whole genome... | Stutzerimona... | NA | 316 | 48.9 | 48.9 | 72% | 0.004 | 33.98 | 276576 | NZ_RBEA010000... |
| Stutzerimonas stutzeri strain PS_134... | Stutzerimona... | NA | 316 | 48.9 | 48.9 | 72% | 0.004 | 33.98 | 278675 | NZ_RHRF010000... |
| Stutzerimonas stutzeri strain PS_133... | Stutzerimona... | NA | 316 | 48.9 | 48.9 | 72% | 0.004 | 33.98 | 278690 | NZ_RHRG010000... |
| Stutzerimonas stutzeri strain PS_130... | Stutzerimona... | NA | 316 | 48.9 | 48.9 | 72% | 0.004 | 33.98 | 278690 | NZ_RHRI010000... |
| Stutzerimonas stutzeri strain PS_131... | Stutzerimona... | NA | 316 | 48.9 | 48.9 | 72% | 0.004 | 33.98 | 278962 | NZ_RHRH010000... |
| Stutzerimonas stutzeri strain PS_050... | Stutzerimona... | NA | 316 | 48.9 | 48.9 | 72% | 0.004 | 33.98 | 280932 | NZ_RIAN010000... |
| Stutzerimonas stutzeri strain G434 17141_contig_1, whole genom... | Stutzerimona... | NA | 316 | 48.9 | 48.9 | 72% | 0.004 | 33.98 | 328625 | NZ_JASGXY0100... |
| Stutzerimonas stutzeri strain GD04107 5, whole genome shotgun... | Stutzerimona... | NA | 316 | 48.9 | 48.9 | 72% | 0.004 | 33.98 | 336744 | NZ_JAOEAQ0100... |
| Stutzerimonas stutzeri strain PS_125... | Stutzerimona... | NA | 316 | 48.9 | 48.9 | 72% | 0.004 | 33.98 | 336950 | NZ_RHRK010000... |
| Stutzerimonas stutzeri strain PS_087... | Stutzerimona... | NA | 316 | 48.9 | 48.9 | 72% | 0.004 | 33.98 | 352814 | NZ_RHRM010000... |
| Stutzerimonas stutzeri strain PS_066... | Stutzerimona... | NA | 316 | 48.9 | 48.9 | 72% | 0.004 | 33.98 | 359430 | NZ_RHRP010000... |
| Stutzerimonas stutzeri strain GD04118 4, whole genome shotgun... | Stutzerimona... | NA | 316 | 48.9 | 48.9 | 72% | 0.004 | 33.98 | 364296 | NZ_JAOEAF0100... |
| Stutzerimonas stutzeri strain G432 17142_contig_21, whole geno... | Stutzerimona... | NA | 316 | 48.9 | 48.9 | 72% | 0.004 | 33.98 | 367318 | NZ_JASGXX0100... |
| Stutzerimonas stutzeri strain GD03946 3, whole genome shotgun... | Stutzerimona... | NA | 316 | 48.9 | 48.9 | 72% | 0.004 | 33.98 | 401344 | NZ_JAOCAF0100... |
| Stutzerimonas stutzeri strain GD03845 1, whole genome shotgun... | Stutzerimona... | NA | 316 | 48.9 | 48.9 | 72% | 0.004 | 33.98 | 423848 | NZ_JAOCDX0100... |
| Stutzerimonas stutzeri strain NT0124 211_430101_56647634_15539... | Stutzerimona... | NA | 316 | 48.9 | 48.9 | 72% | 0.004 | 33.98 | 430101 | NZ_JXTL010000... |
| Stutzerimonas stutzeri strain G431 17143_contig_2, whole genom... | Stutzerimona... | NA | 316 | 48.9 | 48.9 | 72% | 0.004 | 33.98 | 463823 | NZ_JASGXW0100... |
| Stutzerimonas stutzeri strain 69C1... | Stutzerimona... | NA | 316 | 48.9 | 48.9 | 72% | 0.004 | 33.98 | 470705 | NZ_JAAMQV0100... |
| Stutzerimonas stutzeri strain MP4687... | Stutzerimona... | NA | 316 | 48.9 | 48.9 | 72% | 0.004 | 33.98 | 579614 | NZ_WWNS010000... |
| Stutzerimonas stutzeri strain PS_377... | Stutzerimona... | NA | 316 | 48.9 | 48.9 | 72% | 0.004 | 33.98 | 617223 | NZ_RHQV010000... |
| Stutzerimonas stutzeri strain MM2021_5... | Stutzerimona... | NA | 316 | 48.9 | 48.9 | 72% | 0.004 | 33.98 | 633422 | NZ_JAFMNW0100... |
| Stutzerimonas stutzeri strain GD04140 1, whole genome shotgun... | Stutzerimona... | NA | 316 | 48.9 | 48.9 | 72% | 0.004 | 33.98 | 687846 | NZ_JAODZL0100... |
| Stutzerimonas stutzeri strain NBRC 105234 sequence01, whole... | Stutzerimona... | NA | 316 | 48.9 | 48.9 | 72% | 0.004 | 33.98 | 710836 | NZ_BSTP010000... |
| Stutzerimonas stutzeri strain ODKF13 contig_3, whole genome... | Stutzerimona... | NA | 316 | 48.9 | 48.9 | 72% | 0.004 | 33.98 | 719382 | NZ_LSVE010000... |
| Stutzerimonas stutzeri strain KJ3... | Stutzerimona... | NA | 316 | 48.9 | 48.9 | 72% | 0.004 | 33.98 | 767050 | NZ_JBBELA0100... |
| Stutzerimonas stutzeri strain KOL14.W.90.5... | Stutzerimona... | NA | 316 | 48.9 | 48.9 | 72% | 0.004 | 33.98 | 1379714 | NZ_WIFW010000... |
| Stutzerimonas stutzeri strain KOL14.W25.495.90... | Stutzerimona... | NA | 316 | 48.9 | 48.9 | 72% | 0.004 | 33.98 | 2608624 | NZ_WIGJ010000... |
| Stutzerimonas stutzeri strain XX1 chromosome, complete genome | Stutzerimona... | NA | 316 | 48.9 | 48.9 | 72% | 0.004 | 33.98 | 4424182 | NZ_CP091174.1 |
| Stutzerimonas stutzeri strain GD04095 DMICFCLD_1, whole genome... | Stutzerimona... | NA | 316 | 48.9 | 48.9 | 72% | 0.004 | 33.98 | 4437022 | NZ_JAOEBC0100... |
| Stutzerimonas stutzeri strain NCTC10450 chromosome 1, complete... | Stutzerimona... | NA | 316 | 48.9 | 48.9 | 72% | 0.004 | 33.98 | 4438731 | NZ_LR134319.1 |
| Stutzerimonas stutzeri strain SB-1 chromosome, complete genome | Stutzerimona... | NA | 316 | 48.9 | 48.9 | 72% | 0.004 | 33.98 | 4473665 | NZ_CP088004.1 |
| Stutzerimonas stutzeri strain YWX-1 chromosome, complete genome | Stutzerimona... | NA | 316 | 48.9 | 48.9 | 72% | 0.004 | 33.98 | 4488441 | NZ_CP078087.1 |
| Stutzerimonas stutzeri strain ZDHY95 chromosome, complete genome | Stutzerimona... | NA | 316 | 48.9 | 48.9 | 72% | 0.004 | 33.98 | 4500524 | NZ_CP063358.1 |
| Stutzerimonas stutzeri strain SGAir0442 chromosome, complete... | Stutzerimona... | NA | 316 | 48.9 | 48.9 | 72% | 0.004 | 33.98 | 4524655 | NZ_CP025149.2 |
| Stutzerimonas stutzeri strain F2a chromosome, complete genome | Stutzerimona... | NA | 316 | 48.9 | 48.9 | 72% | 0.004 | 33.98 | 4545686 | NZ_AP024722.1 |
| Stutzerimonas stutzeri A1501, complete sequence | Stutzerimona... | NA | 379731 | 48.9 | 48.9 | 72% | 0.004 | 33.98 | 4567418 | NC_009434.1 |
| Stutzerimonas stutzeri strain NRCB010 chromosome, complete genome | Stutzerimona... | NA | 316 | 48.9 | 48.9 | 72% | 0.004 | 33.98 | 4570338 | NZ_CP098731.1 |
| Stutzerimonas stutzeri strain MJL19 chromosome, complete genome | Stutzerimona... | NA | 316 | 48.9 | 48.9 | 72% | 0.004 | 33.98 | 4572525 | NZ_CP140298.1 |
| Stutzerimonas stutzeri strain 2020WEIHUA_G chromosome, complet... | Stutzerimona... | NA | 316 | 48.9 | 48.9 | 72% | 0.004 | 33.98 | 4591234 | NZ_CP062162.1 |
| Stutzerimonas stutzeri strain SLG510A3-8 chromosome, complete... | Stutzerimona... | NA | 316 | 48.9 | 48.9 | 72% | 0.004 | 33.98 | 4650155 | NZ_CP011854.1 |
| Stutzerimonas stutzeri strain J15 chromosome, complete genome | Stutzerimona... | NA | 316 | 48.9 | 48.9 | 72% | 0.004 | 33.98 | 4683655 | NZ_CP101739.1 |
| Stutzerimonas stutzeri DSM 4166, complete sequence | Stutzerimona... | NA | 996285 | 48.9 | 48.9 | 72% | 0.004 | 33.98 | 4689946 | NC_017532.1 |
| Stutzerimonas stutzeri strain H46 chromosome, complete genome | Stutzerimona... | NA | 316 | 48.9 | 48.9 | 72% | 0.004 | 33.98 | 4720414 | NZ_CP139417.1 |
| Stutzerimonas stutzeri strain GD04120 chromosome, complete genome | Stutzerimona... | NA | 316 | 48.9 | 48.9 | 72% | 0.004 | 33.98 | 4775048 | NZ_CP104580.1 |
| Stutzerimonas stutzeri strain GD04113 PNKOAAFP_1, whole genome... | Stutzerimona... | NA | 316 | 48.9 | 48.9 | 72% | 0.004 | 33.98 | 4814082 | NZ_JAOEAK0100... |
| Stutzerimonas frequens strain XN05-1 chromosome, complete genome | Stutzerimona... | NA | 2968969 | 48.9 | 48.9 | 96% | 0.005 | 32.59 | 4411446 | NZ_CP142122.1 |
| Stutzerimonas frequens strain R2A2 chromosome, complete genome | Stutzerimona... | NA | 2968969 | 48.9 | 48.9 | 96% | 0.005 | 32.59 | 4559447 | NZ_CP029772.1 |
| Stutzerimonas stutzeri strain C2 contig309, whole genome shotg... | Stutzerimona... | NA | 316 | 48.9 | 48.9 | 71% | 0.005 | 34.31 | 16666 | NZ_LDWB010000... |
| Stutzerimonas stutzeri strain 267_PSTU... | Stutzerimona... | NA | 316 | 48.9 | 48.9 | 71% | 0.005 | 34.31 | 88813 | NZ_JVNQ010000... |
| Stutzerimonas stutzeri isolate BB1462, whole genome shotgun... | Stutzerimona... | NA | 316 | 48.9 | 48.9 | 71% | 0.005 | 34.31 | 108117 | NZ_CAHPQS0100... |
| Stutzerimonas stutzeri isolate BB1470, whole genome shotgun... | Stutzerimona... | NA | 316 | 48.9 | 48.9 | 71% | 0.005 | 34.31 | 108117 | NZ_CAHPRE0100... |
| Stutzerimonas stutzeri strain PS_376... | Stutzerimona... | NA | 316 | 48.9 | 48.9 | 71% | 0.005 | 34.31 | 135172 | NZ_RHQW010000... |
| Stutzerimonas stutzeri strain GD03947 6, whole genome shotgun... | Stutzerimona... | NA | 316 | 48.9 | 48.9 | 71% | 0.005 | 34.31 | 284342 | NZ_JAOCAE0100... |
| Stutzerimonas stutzeri strain AK6 Contig_22_len_290503, whole... | Stutzerimona... | NA | 316 | 48.9 | 48.9 | 71% | 0.005 | 34.31 | 290503 | NZ_PZYR010000... |
| Stutzerimonas sp. FeSN7 contig_1_pilon_pilon_pilon, whole geno... | Stutzerimona... | NA | 3035479 | 48.9 | 48.9 | 71% | 0.005 | 34.31 | 4789579 | NZ_JARPMN0100... |
| Stutzerimonas stutzeri strain GD03838 14, whole genome shotgun... | Stutzerimona... | NA | 316 | 48.9 | 48.9 | 72% | 0.005 | 33.98 | 117613 | NZ_JAOCEE0100... |
| Stutzerimonas stutzeri strain PS_151... | Stutzerimona... | NA | 316 | 48.9 | 48.9 | 72% | 0.005 | 33.98 | 167014 | NZ_RHRD010000... |
| Stutzerimonas stutzeri strain PS_128... | Stutzerimona... | NA | 316 | 48.9 | 48.9 | 72% | 0.005 | 33.98 | 364407 | NZ_RHRJ010000... |
| Stutzerimonas stutzeri strain S116 chromosome, complete genome | Stutzerimona... | NA | 316 | 48.9 | 48.9 | 72% | 0.005 | 33.98 | 4440952 | NZ_CP078509.1 |
| Stutzerimonas stutzeri strain KOL14.W.495.96... | Stutzerimona... | NA | 316 | 48.5 | 48.5 | 96% | 0.005 | 31.85 | 212774 | NZ_WIFO010000... |
| Stutzerimonas stutzeri KOS6 scaffold2, whole genome shotgun... | Stutzerimona... | NA | 1218352 | 48.5 | 48.5 | 96% | 0.005 | 31.85 | 2709949 | NZ_KK020676.1 |
| Stutzerimonas sp. R40042 chromosome, complete genome | Stutzerimona... | NA | 2998559 | 48.5 | 48.5 | 72% | 0.005 | 34.95 | 4666542 | NZ_CP113258.1 |
| Pseudomonas morbosilactucae strain MAFF 302046 302046_contig_4... | Pseudomonas ... | NA | 2938197 | 48.5 | 48.5 | 95% | 0.005 | 30.71 | 73784 | NZ_JALQCX0100... |
| Deinococcus aluminii strain NBRC 112889 112889_is_8, whole... | Deinococcus ... | NA | 1656885 | 48.5 | 48.5 | 93% | 0.005 | 32.33 | 134314 | NZ_BAABRV0100... |
| Deinococcus metallilatus strain MA1002... | Deinococcus ... | NA | 1211322 | 48.5 | 48.5 | 93% | 0.006 | 32.33 | 246168 | NZ_SDEC010000... |
| Deinococcus metallilatus strain MA1002-m5... | Deinococcus ... | NA | 1211322 | 48.5 | 48.5 | 93% | 0.006 | 32.33 | 246233 | NZ_VBRC010000... |
| Deinococcus metallilatus strain NBRC 110141 sequence15, whole... | Deinococcus ... | NA | 1211322 | 48.5 | 48.5 | 93% | 0.006 | 32.33 | 279199 | NZ_BSUI010000... |
| Deinococcus metallilatus strain DSM 105434 Ga0416734_07, whole... | Deinococcus ... | NA | 1211322 | 48.5 | 48.5 | 93% | 0.006 | 32.33 | 279933 | NZ_JACHFV0100... |
| Deinococcus metallilatus strain MA1002 chromosome | Deinococcus ... | NA | 1211322 | 48.5 | 48.5 | 93% | 0.006 | 32.33 | 3062136 | NZ_CP038512.1 |
| Pseudomonas lopnurensis strain AL-54 46, whole genome shotgun... | Pseudomonas ... | NA | 1477517 | 48.5 | 48.5 | 72% | 0.006 | 33.98 | 22361 | NZ_JADDIX0100... |
| Pseudomonas lopnurensis isolate CTOTU48819... | Pseudomonas ... | NA | 1477517 | 48.5 | 48.5 | 72% | 0.006 | 33.98 | 48550 | NZ_DAMBEU0100... |
| Pseudomonas lopnurensis isolate CTOTU48911... | Pseudomonas ... | NA | 1477517 | 48.5 | 48.5 | 72% | 0.006 | 33.98 | 57616 | NZ_DAMBHS0100... |
| Stutzerimonas stutzeri strain CBC1 contig_18, whole genome... | Stutzerimona... | NA | 316 | 48.5 | 48.5 | 72% | 0.006 | 33.98 | 94832 | NZ_JAXBVD0100... |
| Stutzerimonas stutzeri strain PS_110... | Stutzerimona... | NA | 316 | 48.5 | 48.5 | 72% | 0.006 | 33.98 | 115677 | NZ_RHRL010000... |
| Stutzerimonas stutzeri strain FDAARGOS_875 chromosome, complet... | Stutzerimona... | NA | 316 | 48.5 | 48.5 | 72% | 0.006 | 33.98 | 4535380 | NZ_CP065750.1 |
| Stutzerimonas stutzeri, complete sequence | Stutzerimona... | NA | 316 | 48.5 | 48.5 | 72% | 0.006 | 33.98 | 4547930 | NC_015740.1 |
| Stutzerimonas nitrititolerans isolate CTOTU48809... | Stutzerimona... | NA | 2482751 | 48.5 | 48.5 | 70% | 0.006 | 34.00 | 140689 | NZ_DAMBEK0100... |

| | | | | | | | | | | |
|---|---|---|---|---|---|---|---|---|---|---|
| Saccharospirillum mangrovi strain HK-33 Scaffold2, whole genom... | Saccharospir... | NA | 2161747 | 48.1 | 48.1 | 100% | 0.007 | 26.81 | 517067 | NZ_QAPE010000... |
| Saccharospirillum mangrovi strain HK-33 chromosome, complete... | Saccharospir... | NA | 2161747 | 48.1 | 48.1 | 100% | 0.007 | 26.81 | 3686911 | NZ_CP031415.1 |
| Nitrosomonas sp. isolate HKST-UBA152 k127_817706, whole genome... | Nitrosomonas... | NA | 42353 | 47.8 | 47.8 | 100% | 0.010 | 29.71 | 19510 | NZ_JAGRFN0100... |
| Deinococcus hopiensis KR-140, whole genome shotgun sequence | Deinococcus ... | NA | 695939 | 47.8 | 47.8 | 93% | 0.011 | 31.58 | 3128235 | NZ_FWWU010000... |
| Hydrogenophilus thiooxidans strain SS56 contig-100_16, whole... | Hydrogenophi... | NA | 2820326 | 47.8 | 47.8 | 96% | 0.012 | 30.60 | 46787 | NZ_JAHLSY0100... |
| Deinococcus pimensis DSM 21231 DeipiDRAFT_Scaffold38.32, whole... | Deinococcus ... | NA | 926554 | 47.4 | 47.4 | 99% | 0.017 | 28.26 | 52679 | NZ_KI912645.1 |
| Methylomonas sp. SURF-1 SURF1.00003, whole genome shotgun... | Methylomonas... | NA | 2952224 | 47.0 | 47.0 | 71% | 0.018 | 30.69 | 328747 | NZ_JANIBM0100... |
| Brucella anthropi isolate UBA809 UBA809_contig_1384, whole... | Brucella ant... | NA | 529 | 47.0 | 47.0 | 96% | 0.019 | 27.01 | 11382 | NZ_DBIF010000... |
| uncultured Halovibrio sp. isolate... | uncultured H... | NA | 985049 | 46.6 | 46.6 | 100% | 0.024 | 30.94 | 14964 | NZ_CAJXMA0100... |
| Halomonas sp. 22501_18_FS NODE_5_length_234959_cov_13.5081,... | Halomonas sp... | NA | 2665505 | 46.6 | 46.6 | 100% | 0.024 | 30.94 | 234959 | NZ_WMFD010000... |
| Vreelandella utahensis strain 22507_15_FS... | Vreelandella... | NA | 86177 | 46.6 | 46.6 | 100% | 0.024 | 30.94 | 234959 | NZ_WMEX010000... |
| Chishuiella changwenlii strain DSM 27989, whole genome shotgun... | Chishuiella ... | NA | 1434701 | 46.6 | 46.6 | 69% | 0.024 | 32.63 | 305184 | NZ_FRBH010000... |
| Chishuiella changwenlii strain CGMCC 1.12707 sequence01, whole... | Chishuiella ... | NA | 1434701 | 46.6 | 46.6 | 69% | 0.024 | 32.63 | 305393 | NZ_BMFL010000... |
| Chishuiella changwenlii isolate CTOTU47302... | Chishuiella ... | NA | 1434701 | 46.6 | 46.6 | 69% | 0.024 | 32.63 | 371829 | NZ_DAINVI0100... |
| Halospina sp. K52047b contig02, whole genome shotgun sequence | Halospina sp... | NA | 2614160 | 46.6 | 46.6 | 100% | 0.024 | 30.94 | 478842 | NZ_VYJH010000... |
| Halovibrio salipaludis strain YL5-2 contig2, whole genome... | Halovibrio s... | NA | 2032626 | 46.6 | 46.6 | 100% | 0.024 | 30.94 | 639556 | NZ_NSKD010000... |
| Lachnoclostridium phocaeense strain Marseille-P3177T chromosom... | Lachnoclostr... | NA | 1871021 | 46.6 | 46.6 | 70% | 0.025 | 31.03 | 3500754 | NZ_LT635479.1 |
| Deinococcus aetherius strain JCM 11751 sequence18, whole genom... | Deinococcus ... | NA | 200252 | 46.6 | 46.6 | 93% | 0.025 | 30.60 | 486419 | NZ_BAABDW0100... |
| Deinococcus aetherius strain ST0316 chromosome, complete genome | Deinococcus ... | NA | 200252 | 46.6 | 46.6 | 93% | 0.025 | 30.60 | 3135460 | NZ_AP026560.1 |
| Nitrosospira sp. Nsp2 Ga0181081_101, whole genome shotgun... | Nitrosospira... | NA | 136548 | 46.2 | 46.2 | 100% | 0.034 | 28.26 | 921569 | NZ_QAOU010000... |
| Nitrosospira sp. Is2 chromosome, complete genome | Nitrosospira... | NA | 3080532 | 46.2 | 46.2 | 100% | 0.034 | 28.26 | 3574273 | NZ_CP137134.1 |
| Brucella intermedia isolate CTOTU48914... | Brucella int... | NA | 94625 | 46.2 | 46.2 | 73% | 0.037 | 27.88 | 124729 | NZ_DAMBHV0100... |
| Chitinophaga sp. Cy-1792 NODE_2_length_2808195_cov_51.575576/0... | Chitinophaga... | NA | 2608339 | 46.2 | 46.2 | 95% | 0.042 | 29.77 | 2808195 | NZ_VWWO010000... |
| Pseudomonas sp. isolate RS261_metabat.bin.8 RS261_k141_142368,... | Pseudomonas sp. | NA | 306 | 45.8 | 45.8 | 70% | 0.046 | 33.66 | 165529 | NZ_JANJWQ0100... |
| Pseudomonas sp. FIP_A4 k141_127, whole genome shotgun sequence | Pseudomonas ... | NA | 3070684 | 45.8 | 45.8 | 70% | 0.046 | 33.66 | 170970 | NZ_JAYDXP0100... |

Alignments:

>Klebsiella pneumoniae strain ARGID_31501, whole genome shotgun sequence
Sequence ID: NZ_CAKLPD010000006.1 Length: 2655
Range 1: 1544 to 1903

Score:230 bits(587), Expect:1e-67,
Method:Compositional matrix adjust.,
Identities:114/120(95%), Positives:117/120(97%), Gaps:0/120(0%)

```
Query  1     YVDGAANRETKLGKAGYVTNRGRQKVVTLTDTTNQKTELQAIYLALQDSGLEVNIVTDSQ  60
             YVDGAANRETK+GKAGYVTNRGRQKVV LTDTTNQKTELQAIYLALQDS LEVNIVTDSQ
Sbjct  1544  YVDGAANRETKIGKAGYVTNRGRQKVVPLTDTTNQKTELQAIYLALQDSELEVNIVTDSQ  1723

Query  61    YALGIIQAQPDQSESELVNQIIEQLIKKEKVYLAWVPAHKGIGGNEQVDKLVSAGIRKVL  120
             YALGIIQAQPD+SESELVNQIIEQLIKKEK+YLAWVPAHKGIGGNEQVDKLVS GIRKVL
Sbjct  1724  YALGIIQAQPDKSESELVNQIIEQLIKKEKIYLAWVPAHKGIGGNEQVDKLVSTGIRKVL  1903
```

>Burkholderia pseudomallei strain 2604STDY5518971 ERS240105SCcontig000097, whole genome shotgun sequence
Sequence ID: NZ_UVFZ02000097.1 Length: 1345
Range 1: 208 to 567

Score:220 bits(560), Expect:5e-67,
Method:Compositional matrix adjust.,
Identities:107/120(89%), Positives:117/120(97%), Gaps:0/120(0%)

```
Query  1    YVDGAANRETKLGKAGYVTNRGRQKVVTLTDTTNQKTELQAIYLALQDSGLEVNIVTDSQ  60
            YVDGAA+RETKLGKAGYVT+RGRQKVV+LT+TTNQKTEL AI+LALQDSG EVNIVTDSQ
Sbjct  208  YVDGAASRETKLGKAGYVTDRGRQKVVSLTETTNQKTELHAIHLALQDSGSEVNIVTDSQ  387

Query  61   YALGIIQAQPDQSESELVNQIIEQLIKKEKVYLAWVPAHKGIGGNEQVDKLVSAGIRKVL  120
            YALGIIQAQPD SESE+VNQIIE+LIKKE+VYL+WVPAHKGIGGNEQVDKLVS+GIRKVL
Sbjct  388  YALGIIQAQPDMSESEVVNQIIEELIKKERVYLSWVPAHKGIGGNEQVDKLVSSGIRKVL  567
```

>Actinoplanes campanulatus strain JCM 3059 sequence058, whole genome shotgun sequence
Sequence ID: NZ_BMPW01000058.1 Length: 4754
Range 1: 3177 to 3536

Score:231 bits(589), Expect:5e-67,
Method:Compositional matrix adjust.,
Identities:115/120(96%), Positives:119/120(99%), Gaps:0/120(0%)

```
Query  1     YVDGAANRETKLGKAGYVTNRGRQKVVTLTDTTNQKTELQAIYLALQDSGLEVNIVTDSQ  60
             YVDGAANRETKLGKAGYVT+RGRQKVV LTDTTNQKTELQAI+LALQDSGLEVNIVTDSQ
Sbjct  3177  YVDGAANRETKLGKAGYVTDRGRQKVVPLTDTTNQKTELQAIHLALQDSGLEVNIVTDSQ  3356

Query  61    YALGIIQAQPDQSESELVNQIIEQLIKKEKVYLAWVPAHKGIGGNEQVDKLVSAGIRKVL  120
             YALGIIQAQPD+SESELV+QIIEQLIKKEKVYLAWVPAHKGIGGNEQVDKLVSAGIRKVL
Sbjct  3357  YALGIIQAQPDKSESELVSQIIEQLIKKEKVYLAWVPAHKGIGGNEQVDKLVSAGIRKVL  3536
```

>Enterococcus alcedinis strain CCM 8433 sequence23, whole genome shotgun sequence
Sequence ID: NZ_BMDT01000023.1 Length: 6075
Range 1: 3135 to 3494

Score:231 bits(589), Expect:5e-67,
Method:Compositional matrix adjust.,
Identities:115/120(96%), Positives:119/120(99%), Gaps:0/120(0%)

```
Query  1     YVDGAANRETKLGKAGYVTNRGRQKVVTLTDTTNQKTELQAIYLALQDSGLEVNIVTDSQ  60
             YVDGAANRETKLGKAGYVT+RGRQKVV LTDTTNQKTELQAI+LALQDSGLEVNIVTDSQ
Sbjct  3135  YVDGAANRETKLGKAGYVTDRGRQKVVPLTDTTNQKTELQAIHLALQDSGLEVNIVTDSQ  3314

Query  61    YALGIIQAQPDQSESELVNQIIEQLIKKEKVYLAWVPAHKGIGGNEQVDKLVSAGIRKVL  120
             YALGIIQAQPD+SESELV+QIIEQLIKKEKVYLAWVPAHKGIGGNEQVDKLVSAGIRKVL
Sbjct  3315  YALGIIQAQPDKSESELVSQIIEQLIKKEKVYLAWVPAHKGIGGNEQVDKLVSAGIRKVL  3494
```

>Deinococcus aquiradiocola strain JCM 14371 sequence30, whole genome shotgun sequence
Sequence ID: NZ_BMOE01000030.1 Length: 6128
Range 1: 3188 to 3547

Score:231 bits(589), Expect:5e-67,
Method:Compositional matrix adjust.,
Identities:115/120(96%), Positives:119/120(99%), Gaps:0/120(0%)

```
Query  1     YVDGAANRETKLGKAGYVTNRGRQKVVTLTDTTNQKTELQAIYLALQDSGLEVNIVTDSQ  60
             YVDGAANRETKLGKAGYVT+RGRQKVV LTDTTNQKTELQAI+LALQDSGLEVNIVTDSQ
Sbjct  3188  YVDGAANRETKLGKAGYVTDRGRQKVVPLTDTTNQKTELQAIHLALQDSGLEVNIVTDSQ  3367

Query  61    YALGIIQAQPDQSESELVNQIIEQLIKKEKVYLAWVPAHKGIGGNEQVDKLVSAGIRKVL  120
             YALGIIQAQPD+SESELV+QIIEQLIKKEKVYLAWVPAHKGIGGNEQVDKLVSAGIRKVL
Sbjct  3368  YALGIIQAQPDKSESELVSQIIEQLIKKEKVYLAWVPAHKGIGGNEQVDKLVSAGIRKVL  3547
```

>Methylobacterium radiotolerans strain MAMP 4754 MP-M-radiotolerans_contig_704, whole genome shotgun sequence
Sequence ID: NZ_NKQS01000691.1 Length: 1636
Range 1: 618 to 977

Score:214 bits(545), Expect:8e-64,
Method:Compositional matrix adjust.,
Identities:105/120(88%), Positives:112/120(93%), Gaps:0/120(0%)

```
Query  1    YVDGAANRETKLGKAGYVTNRGRQKVVTLTDTTNQKTELQAIYLALQDSGLEVNIVTDSQ  60
            YVDGAANRETK GKAGYVT+RGRQK+V L +TTNQK ELQAI LALQDSG EVNIVTDSQ
Sbjct  977  YVDGAANRETKRGKAGYVTDRGRQKIVPLNETTNQKAELQAIQLALQDSGSEVNIVTDSQ  798

Query  61   YALGIIQAQPDQSESELVNQIIEQLIKKEKVYLAWVPAHKGIGGNEQVDKLVSAGIRKVL  120
            YALGIIQAQPD+SESELVNQIIEQLI KE++YL+WVPAHKGIGGNEQVDKLVS GIRKVL
Sbjct  797  YALGIIQAQPDKSESELVNQIIEQLINKERIYLSWVPAHKGIGGNEQVDKLVSTGIRKVL  618
```

>Escherichia coli strain VRES0669, whole genome shotgun sequence
Sequence ID: NZ_UINV01000086.1 Length: 5544
Range 1: 3408 to 3767

Score:207 bits(528), Expect:9e-59,
Method:Compositional matrix adjust.,
Identities:102/120(85%), Positives:111/120(92%), Gaps:0/120(0%)

```
Query  1     YVDGAANRETKLGKAGYVTNRGRQKVVTLTDTTNQKTELQAIYLALQDSGLEVNIVTDSQ  60
             YVDGAANRETK GKAGYVT+RGRQKV++ T+TTNQK ELQAI LALQDSG EVNIVTDSQ
Sbjct  3767  YVDGAANRETKAGKAGYVTDRGRQKVISFTETTNQKAELQAIQLALQDSGSEVNIVTDSQ  3588

Query  61    YALGIIQAQPDQSESELVNQIIEQLIKKEKVYLAWVPAHKGIGGNEQVDKLVSAGIRKVL  120
             YALGII A PD+SESELVN+IIE LI+KE+VYL+WVPAHKGIGGNEQVDKLVS GIRKVL
Sbjct  3587  YALGIIHALPDKSESELVNKIIELLIQKERVYLSWVPAHKGIGGNEQVDKLVSTGIRKVL  3408
```

>Thalassovita mangrovi strain GS-10 Scaffold20_1, whole genome shotgun sequence
Sequence ID: NZ_WWEN01000020.1 Length: 1488
Range 1: 2 to 310

Score:199 bits(505), Expect:2e-58,
Method:Compositional matrix adjust.,
Identities:98/103(95%), Positives:102/103(99%), Gaps:0/103(0%)

```
Query  1    YVDGAANRETKLGKAGYVTNRGRQKVVTLTDTTNQKTELQAIYLALQDSGLEVNIVTDSQ  60
            YVDGAANRETKLGKAGYVT+RGRQKVV LTDTTNQKTELQAI+LALQDSGLEVNIVTDSQ
Sbjct  310  YVDGAANRETKLGKAGYVTDRGRQKVVPLTDTTNQKTELQAIHLALQDSGLEVNIVTDSQ  131

Query  61   YALGIIQAQPDQSESELVNQIIEQLIKKEKVYLAWVPAHKGIG  103
            YALGIIQAQPD+SESELV+QIIEQLIKKEKVYLAWVPAHKGIG
Sbjct  130  YALGIIQAQPDKSESELVSQIIEQLIKKEKVYLAWVPAHKGIG  2
```

>Klebsiella pneumoniae strain EuSCAPE_MT012, whole genome shotgun sequence
Sequence ID: NZ_UKIX02000064.1 Length: 452
Range 1: 3 to 302

Score:182 bits(463), Expect:4e-56,
Method:Compositional matrix adjust.,
Identities:88/100(88%), Positives:97/100(97%), Gaps:0/100(0%)

```
Query  1    YVDGAANRETKLGKAGYVTNRGRQKVVTLTDTTNQKTELQAIYLALQDSGLEVNIVTDSQ  60
            YVDGAANR+TK+GKAGYVT++GRQKVV+LT+TTNQKTEL AI LALQDSG EVNIVTDSQ
Sbjct  302  YVDGAANRDTKMGKAGYVTDKGRQKVVSLTETTNQKTELHAIQLALQDSGPEVNIVTDSQ  123

Query  61   YALGIIQAQPDQSESELVNQIIEQLIKKEKVYLAWVPAHK  100
            YALGIIQAQPD+SESELVNQIIEQLIKKE+VYL+WVPAHK
Sbjct  122  YALGIIQAQPDKSESELVNQIIEQLIKKERVYLSWVPAHK  3
```

>Pseudomonas aeruginosa strain VET-42 NODE_86_length_1154_cov_8.30919_ID_171, whole genome shotgun sequence
Sequence ID: NZ_RACX01000118.1 Length: 1154
Range 1: 939 to 1103

Score:107 bits(266), Expect:4e-24,
Method:Compositional matrix adjust.,
Identities:50/55(91%), Positives:55/55(100%), Gaps:0/55(0%)

```
Query  66    IQAQPDQSESELVNQIIEQLIKKEKVYLAWVPAHKGIGGNEQVDKLVSAGIRKVL  120
             +QAQPD+SESE+VNQIIEQLIKKEKVYL+WVPAHKGIGGNEQVDKLVS+GIRKVL
Sbjct  1103  VQAQPDKSESEVVNQIIEQLIKKEKVYLSWVPAHKGIGGNEQVDKLVSSGIRKVL  939
```

>Klebsiella pneumoniae strain EuSCAPE_DE086, whole genome shotgun sequence
Sequence ID: NZ_UKLW02000059.1 Length: 995
Range 1: 836 to 994

Score:79.7 bits(195), Expect:3e-14,
Method:Compositional matrix adjust.,
Identities:39/53(74%), Positives:46/53(86%), Gaps:0/53(0%)

```
Query  1    YVDGAANRETKLGKAGYVTNRGRQKVVTLTDTTNQKTELQAIYLALQDSGLEV  53
            YVD AAN+ETK+ KA YVT++ RQK+V+LT+TTNQKTELQ I LALQDSG EV
Sbjct  836  YVDRAANKETKIRKARYVTDKRRQKIVSLTETTNQKTELQTIQLALQDSGSEV  994
```

>Pseudomonas aeruginosa strain ENV-297 NODE_79_length_1100_cov_9.24438_ID_157, whole genome shotgun sequence
Sequence ID: NZ_RADD01000036.1 Length: 1100
Range 1: 945 to 1055

Score:76.3 bits(186), Expect:6e-13,
Method:Compositional matrix adjust.,
Identities:35/37(95%), Positives:37/37(100%), Gaps:0/37(0%)

```
Query  84    QLIKKEKVYLAWVPAHKGIGGNEQVDKLVSAGIRKVL  120
             QLIKKEKVYL+WVPAHKGIGGNEQVDKLVS+GIRKVL
Sbjct  1055  QLIKKEKVYLSWVPAHKGIGGNEQVDKLVSSGIRKVL  945
```

>Thalassovita mangrovi strain GS-10 Scaffold30_1, whole genome shotgun sequence
Sequence ID: NZ_WWEN01000028.1 Length: 722
Range 1: 569 to 706

```
Score:75.1 bits(183), Expect:7e-13,
Method:Compositional matrix adjust.,
Identities:33/46(72%), Positives:41/46(89%), Gaps:0/46(0%)

Query  75   SELVNQIIEQLIKKEKVYLAWVPAHKGIGGNEQVDKLVSAGIRKVL  120
            ++  +++IEQ+ KK + Y+AWVPAHKGIGGNEQVDKLVSAGIRKVL
Sbjct  706  AKAASKVIEQITKKAEAYMAWVPAHKGIGGNEQVDKLVSAGIRKVL  569
```

```
>Cuspidothrix issatschenkoi CHARLIE-1 contig_175, whole genome shotgun sequence
Sequence ID: NZ_PGEM01000173.1 Length: 2237
Range 1: 1570 to 1899

Score:69.3 bits(168), Expect:3e-10,
Method:Compositional matrix adjust.,
Identities:46/112(41%), Positives:61/112(54%), Gaps:4/112(3%)

Query  1     YVDGAANRETKLGKAGYVTNRGRQKVVTLTDTTNQKTELQAIYLALQDS-GLEVNIVTDS  59
             Y DG   ++   G A YVT+ GR K   L   T+Q  E  AI +AL+D+   +VNIVTDS
Sbjct  1570  YTDGG--KQNGEGIAAYVTSNGRTKQKRLGPVTHQVAERMAIQMALEDTRDKQVNIVTDS  1743

Query  60    QYAL-GIIQAQPDQSESELVNQIIEQLIKKEKVYLAWVPAHKGIGGNEQVDK  110
              Y    I +    +        II+ + +KE VY AWVP HKGI GN+  D+
Sbjct  1744  YYCWKNITEGLGLEGPQSPWWPIIQNIREKEIVYFAWVPGHKGICGNQLADE  1899
```

```
>Klebsiella pneumoniae strain B2 isolate B2, whole genome shotgun sequence
Sequence ID: NZ_CABWNT010000058.1 Length: 7306
Range 1: 5654 to 5953

Score:64.3 bits(155), Expect:2e-08,
Method:Compositional matrix adjust.,
Identities:44/104(42%), Positives:55/104(52%), Gaps:10/104(9%)

Query  13    GKAGYVTNRGRQKVVTLTDTTNQKTELQAIYLALQDSGLE-VNIVTDSQYAL-----GII  66
             G A YVT+ G+ K   L   T+Q  E  AI +AL+D+    VNIVTDS Y       G+
Sbjct  5654  GVAAYVTSNGKTKQKRLGPVTHQTAERIAIQMALEDTEETLVNIVTDSYYCWKNITEGLG  5833

Query  67    QAQPDQSESELVNQIIEQLIKKEKVYLAWVPAHKGIGGNEQVDK  110
                PD         II+ +  KE VY AWVP HKGI GN+  D+
Sbjct  5834  LEGPDSPWWP----IIQNIRAKEMVYFAWVPGHKGIYGNQLADE  5953
```

```
>Polyangium sp. y55x31 NODE_26_length_169233_cov_17.164843, whole genome shotgun sequence
Sequence ID: NZ_JARZHG010000026.1 Length: 169233
Range 1: 66608 to 66976

Score:61.2 bits(147), Expect:2e-07,
Method:Composition-based stats.,
Identities:41/124(33%), Positives:66/124(53%), Gaps:8/124(6%)

Query  1      YVDGAANRETKLGKAGYVTNRGRQKVVT---LTDTTNQKTELQAIYLAL---QDSGLEVN  54
              Y DGA          G V   G +++     L D TN   EL AI  A+   +D+G  +
Sbjct  66976  YADGACTGNPGPAGLGVVVVDGDRRIEISEYLGDGTNNIAELTAILRAVGAVEDTGKPLL  66797

Query  55     IVTDSQYALGIIQ-AQPDQSESELVNQIIEQLIKKEKVYLAWVPAHKGIGGNEQVDKLVS  113
              + TDSQYA+G++Q     ++ ++LV Q ++ ++K+ K  L +VP H G+  NE+ D+L
Sbjct  66796  VRTDSQYAIGVLQKGWKAKANTDLVAQ-VKDVLKRNKTRLEYVPGHSGVALNERADELAR  66620

Query  114    AGIR  117
                +R
Sbjct  66619  EAVR  66608
```

```
>Polyangium sp. 15x6 NODE_112_length_36528_cov_15.618939, whole genome shotgun sequence
Sequence ID: NZ_JASBQE010000112.1 Length: 36528
Range 1: 5134 to 5502

Score:58.2 bits(139), Expect:3e-06,
Method:Compositional matrix adjust.,
Identities:40/124(32%), Positives:66/124(53%), Gaps:8/124(6%)

Query  1     YVDGAANRETKLGKAGYVTNRGRQKVVT---LTDTTNQKTELQAIYLAL---QDSGLEVN  54
             Y DGA          G V   G +++     L + TN   EL AI  A+   +D+G  +
Sbjct  5502  YADGACTGNPGPAGLGVVVVDGDRRIEISEYLGNGTNNIAELTAILRAVGAVEDTGKPLL  5323

Query  55    IVTDSQYALGIIQ-AQPDQSESELVNQIIEQLIKKEKVYLAWVPAHKGIGGNEQVDKLVS  113
             + TDSQYA+G++Q     ++ +ELV + ++ ++K+ K  L +VP H G+  NE+ D+L
Sbjct  5322  VRTDSQYAIGVLQKGWKAKANTELVAE-VKDVLKRNKTRLEYVPGHSGVALNERADELAR  5146

Query  114   AGIR  117
               +R
Sbjct  5145  EAVR  5134
```

```
>Polyangium sp. 6x1 NODE_22_length_166486_cov_22.101912, whole genome shotgun sequence
Sequence ID: NZ_JARZHH010000022.1 Length: 166486
Range 1: 110104 to 110472

Score:56.6 bits(135), Expect:9e-06,
Method:Compositional matrix adjust.,
Identities:41/124(33%), Positives:62/124(50%), Gaps:8/124(6%)

Query  1       YVDGAANRETKLGKAGYVTNRGRQKVVT---LTDTTNQKTELQAIYLALQ---DSGLEVN  54
               Y DGA          G +   G +++     L   TN   EL AI  A++    +G  +
Sbjct  110104  YADGACTGNPGPAGLGVLVEDGDRRIEISEYLGQGTNNIAELTAILRAVEAVEGTGKSLL  110283

Query  55      IVTDSQYALGIIQ-AQPDQSESELVNQIIEQLIKKEKVYLAWVPAHKGIGGNEQVDKLVS  113
               + TDSQYA+G++Q     ++ +ELV  + + L KK K  L +VP H G+ GNE+ D L
Sbjct  110284  VRTDSQYAIGVLQKGWKAKANTELVAAVKDAL-KKSKARLEYVPGHSGVRGNERADALAR  110460

Query  114     AGIR  117
                 +R
Sbjct  110461  EAVR  110472
```

```
>Polyangium jinanense strain SDU14 Scaffold9, whole genome shotgun sequence
Sequence ID: NZ_JAGTJJ010000009.1 Length: 231060
Range 1: 11612 to 11965

Score:54.7 bits(130), Expect:4e-05,
```

```
Method:Compositional matrix adjust.,
Identities:38/119(32%), Positives:64/119(53%), Gaps:8/119(6%)

Query  1      YVDGAANRETKLGKAGYVTNRGRQKVVT---LTDTTNQKTELQAIYLAL---QDSGLEVN  54
              Y DGA          G V   G +++     L + TN   EL AI  A+   +D+G  +
Sbjct  11612  YADGACTGNPGPAGLGVVVVDGDRRIEISEYLGNGTNNIAELTAILRAVGAVEDTGKPLL  11791

Query  55     IVTDSQYALGIIQ-AQPDQSESELVNQIIEQLIKKEKVYLAWVPAHKGIGGNEQVDKLV  112
              + TDSQYA+G++Q     ++ ++LV + ++ ++K+ K  L +VP H G+  NE+ D+L
Sbjct  11792  VRTDSQYAIGVLQKGWKAKANTDLVAE-VKDVLKRNKTRLEYVPGHSGVALNERADELA  11965
```

>Polyangium jinanense strain SDU13 Scaffold15, whole genome shotgun sequence
Sequence ID: NZ_JAGTJK010000015.1 Length: 231263
Range 1: 11765 to 12118

```
Score:54.7 bits(130), Expect:4e-05,
Method:Compositional matrix adjust.,
Identities:38/119(32%), Positives:64/119(53%), Gaps:8/119(6%)

Query  1      YVDGAANRETKLGKAGYVTNRGRQKVVT---LTDTTNQKTELQAIYLAL---QDSGLEVN  54
              Y DGA          G V   G +++     L + TN   EL AI  A+   +D+G  +
Sbjct  11765  YADGACTGNPGPAGLGVVVVDGDRRIEISEYLGNGTNNIAELTAILRAVGAVEDTGKPLL  11944

Query  55     IVTDSQYALGIIQ-AQPDQSESELVNQIIEQLIKKEKVYLAWVPAHKGIGGNEQVDKLV  112
              + TDSQYA+G++Q     ++ ++LV + ++ ++K+ K  L +VP H G+  NE+ D+L
Sbjct  11945  VRTDSQYAIGVLQKGWKAKANTDLVAE-VKDVLKRNKTRLEYVPGHSGVALNERADELA  12118
```

>Roseiconus lacunae strain JC742 soapdenovo_K_105_82, whole genome shotgun sequence
Sequence ID: NZ_JAJMQV010000082.1 Length: 35968
Range 1: 13245 to 13676

```
Score:50.8 bits(120), Expect:0.001,
Method:Composition-based stats.,
Identities:41/145(28%), Positives:69/145(47%), Gaps:26/145(17%)

Query  1      YVDGAANRETKLGKAGYVT-------NRGRQKVVTLTDTTNQKTELQAIYLALQDSGLE-  52
              Y DG+++ ++ +G   Y+         +   Q+  +    +N + EL+A+   L+  G E
Sbjct  13245  YTDGSSDPDSGVGGWAYLIINEDSPLDLSDQQAGSHPRASNNRMELEAVLQGLRAIGEEP  13424

Query  53     --VNIVTDSQYA-------LGI-IQAQPDQ-------SESELVNQIIEQLIKKEKVYLAW  95
                + I TDS+Y        L I + ++PD        S   L  ++IE LI + +VY  W
Sbjct  13425  ACITIKTDSEYVERGLNEYLPIWLSSRPDSRFKDPIPSHPRLWRRVIE-LINRHEVYCIW  13601

Query  96     VPAHKGIGGNEQVDKLVSAGIRKVL  120
              +P H G   NE VD+L  + +R  L
Sbjct  13602  IPGHSGHTENEIVDRLAGSAVRDHL  13676
```

>Kribbella italica strain DSM 28967 Ga0415174_01, whole genome shotgun sequence
Sequence ID: NZ_JACHMY010000001.1 Length: 8885108
Range 1: 163456 to 163851

```
Score:50.1 bits(118), Expect:0.002,
Method:Composition-based stats.,
Identities:37/134(28%), Positives:58/134(43%), Gaps:17/134(12%)

Query  1       YVDGAANRETKLGKAGYVTNRGRQKVVTLTDTTNQKTELQAIYLALQDSGLEVNIVTDSQ  60
               Y DGA +  T  G   +V   G +       DTTNQ+ ELQA Y A Q     + IV+DS
Sbjct  163456  YTDGACSGGT--GGWAWVVADGPEGSGATNDTTNQRMELQAAYEAAQSLFGPLTIVSDSS  163629

Query  61      YALGIIQ---------------AQPDQSESELVNQIIEQLIKKEKVYLAWVPAHKGIGGN  105
               Y +   +                A+   +  ++   +IE ++ +  +   W+  H G   N
Sbjct  163630  YVVNCFKQGWWKGWHARNWVNSARKPVANKDIWEPLIELVLHRGDIDFRWIKGHSGHPLN  163809

Query  106     EQVDKLVSAGIRKV  119
               E+ D+L  A  + V
Sbjct  163810  ERADRLAVAARQGV  163851
```

>Stutzerimonas stutzeri strain 24a75 44Pst24A75gv17C029, whole genome shotgun sequence
Sequence ID: NZ_JAMOHU010000029.1 Length: 55843
Range 1: 1033 to 1338

```
Score:49.7 bits(117), Expect:0.003,
Method:Compositional matrix adjust.,
Identities:35/103(34%), Positives:50/103(48%), Gaps:17/103(16%)

Query  30    TDTTNQKTELQAIYLALQD--SGLEVNIVTDSQYAL-GIIQAQPDQSE------------  74
              DTTN + EL A   AL +     +V +VTDSQY + GI    P+  +
Sbjct  1033  ADTTNNRMELMAAIRALAELKRPCKVRLVTDSQYVMQGINDWMPNWKKRGWKTASKQPVK  1212

Query  75    -SELVNQIIEQLIKKEKVYLAWVPAHKGIGGNEQVDKLVSAGI  116
              ++L  Q+ EQ + + +V   WV  H G  GNEQ D L + G+
Sbjct  1213  NADLWQQLDEQ-VNRHQVSWEWVRGHNGHPGNEQADLLANRGV  1338
```

>Stutzerimonas stutzeri strain 24a75 NODE_16_length_75459_cov_15.8451, whole genome shotgun sequence
Sequence ID: NZ_POUT01000016.1 Length: 75459
Range 1: 62540 to 62845

```
Score:49.7 bits(117), Expect:0.003,
Method:Compositional matrix adjust.,
Identities:35/103(34%), Positives:50/103(48%), Gaps:17/103(16%)

Query  30     TDTTNQKTELQAIYLALQD--SGLEVNIVTDSQYAL-GIIQAQPDQSE------------  74
               DTTN + EL A   AL +     +V +VTDSQY + GI    P+  +
Sbjct  62845  ADTTNNRMELMAAIRALAELKRPCKVRLVTDSQYVMQGINDWMPNWKKRGWKTASKQPVK  62666

Query  75     -SELVNQIIEQLIKKEKVYLAWVPAHKGIGGNEQVDKLVSAGI  116
               ++L  Q+ EQ + + +V   WV  H G  GNEQ D L + G+
Sbjct  62665  NADLWQQLDEQ-VNRHQVSWEWVRGHNGHPGNEQADLLANRGV  62540
```

>Pseudomonas sp. BAY1663 contig000002, whole genome shotgun sequence
Sequence ID: NZ_AZSV01000002.1 Length: 124554
Range 1: 62477 to 62782

```
Score:49.7 bits(117), Expect:0.003,
Method:Compositional matrix adjust.,
Identities:35/103(34%), Positives:50/103(48%), Gaps:17/103(16%)
```

```
Query  30     TDTTNQKTELQAIYLALQD--SGLEVNIVTDSQYAL-GIIQAQPDQSE------------  74
               DTTN + EL A   AL +     +V +VTDSQY + GI    P+  +
Sbjct  62477  ADTTNNRMELMAAIRALAELKRPCKVRLVTDSQYVMQGINDWMPNWKKRGWKTASKQPVK  62656

Query  75     -SELVNQIIEQLIKKEKVYLAWVPAHKGIGGNEQVDKLVSAGI  116
               ++L  Q+ EQ + + +V   WV  H G  GNEQ D L + G+
Sbjct  62657  NADLWQQLDEQ-VNRHQVSWEWVRGHNGHPGNEQADLLANRGV  62782
```

```
>MAG: Nitrosomonas sp. isolate RBC_AOB3 BC1.188.fa-20, whole genome shotgun sequence
Sequence ID: NZ_JALDQX010000020.1 Length: 55000
Range 1: 35481 to 35894

Score:49.3 bits(116), Expect:0.003,
Method:Compositional matrix adjust.,
Identities:42/138(30%), Positives:60/138(43%), Gaps:18/138(13%)

Query  1      YVDGAANRETKLGKAGYVTNRGRQKVVTL---TDTTNQKTELQAIYLALQD--SGLEVNI  55
              Y DGA      +G  G V   G  +         TTN + EL A   AL+       V +
Sbjct  35481  YTDGACKGNPGVGGWGAVLKYGVHEQEIFGGEKHTTNNRMELLAAIRALESLKRACRVRL  35660

Query  56     VTDSQYAL-GIIQ------------AQPDQSESELVNQIIEQLIKKEKVYLAWVPAHKGI  102
               TDSQY   GI Q             A     ++E + + ++QL +K  +   WV  H G
Sbjct  35661  YTDSQYVHNGIGQWMQAWKARNWRTANKKPVKNEDLWRTLDQLSQKHDIEWLWVRGHAGH  35840

Query  103    GGNEQVDKLVSAGIRKVL  120
               GNE+ D L + G++K+L
Sbjct  35841  EGNERADMLANRGVKKIL  35894
```

```
>Deinococcus aerius strain TR0125 DAERI_Contig_05, whole genome shotgun sequence
Sequence ID: NZ_BFAG01000005.1 Length: 248739
Range 1: 101310 to 101702

Score:49.3 bits(116), Expect:0.003,
Method:Compositional matrix adjust.,
Identities:44/133(33%), Positives:68/133(51%), Gaps:23/133(17%)

Query  1       YVDGAANRETKLGKAGYVT---NRGRQKVVT--LTDTTNQKTELQAIYLALQ--DSGLEV  53
               Y DGA +  T  G  G+ T    RG++ V++    DTTN + EL+ +   L+      +V
Sbjct  101310  YSDGACD--TAAGHGGWATILNYRGKELVLSGHERDTTNNRMELRGLLEGLKVLKRPCQV  101483

Query  54      NIVTDSQYAL------GIIQAQPDQSES----ELVNQ-IIEQLIKKEKVY---LAWVPAH  99
                +VTDSQY          I++ Q +  ++     + NQ + E+LI + K++     WV  H
Sbjct  101484  RVVTDSQYLRKAFTDGWILKWQRNGWKTAGGEPVKNQDLWEELIAQAKIHALTFIWVRGH  101663

Query  100     KGIGGNEQVDKLV  112
                G G NE+VDKL
Sbjct  101664  NGHGENERVDKLA  101702
```

```
>Nitrosomonas sp. Nm51, whole genome shotgun sequence
Sequence ID: NZ_FOGH01000046.1 Length: 20055
Range 1: 4147 to 4560

Score:48.9 bits(115), Expect:0.004,
Method:Compositional matrix adjust.,
Identities:40/138(29%), Positives:63/138(45%), Gaps:18/138(13%)

Query  1     YVDGAANRETKLGKAGYVTNRGRQKVVTLTD---TTNQKTELQAIYLALQ--DSGLEVNI  55
             Y DGA      +G  G V   G  +         TTN + EL A   AL+      +V +
Sbjct  4560  YTDGACKGNPGVGGWGAVLKYGAHEQEIFGGEKYTTNNRMELLAAIRALELLKRSCKVRL  4381

Query  56    VTDSQYAL-GIIQAQP------------DQSESELVNQIIEQLIKKEKVYLAWVPAHKGI  102
              TDSQY   GI Q  P              Q ++E + +++++L ++  +   WV  H G
Sbjct  4380  YTDSQYVHNGISQWIPAWKARDWKTADKKQVKNEDLWRMLDRLSQQHDIEWLWVRGHAGD  4201

Query  103   GGNEQVDKLVSAGIRKVL  120
              GNE+ D L + G++K++
Sbjct  4200  EGNERADMLANRGVKKII  4147
```

```
>Stutzerimonas stutzeri strain GD04160 15, whole genome shotgun sequence
Sequence ID: NZ_JAODYR010000015.1 Length: 112086
Range 1: 38343 to 38744

Score:48.9 bits(115), Expect:0.004,
Method:Compositional matrix adjust.,
Identities:44/135(33%), Positives:61/135(45%), Gaps:20/135(14%)

Query  1      YVDGAANRETKLGKAG-YVTNRG--RQKVVTLTDTTNQKTELQAIYLALQD--SGLEVNI  55
              Y DGA       G  G  +  +G  R+      DTTN + EL A   AL +     +V +
Sbjct  38744  YTDGACKGNPGPGGWGALLVYKGVKRELWGGEPDTTNNRMELMAAIRALAELKRPCKVRL  38565

Query  56     VTDSQYAL-GIIQAQPDQSE-------------SELVNQIIEQLIKKEKVYLAWVPAHKG  101
              VTDSQY + GI    P+  +              ++L  Q+ EQ+ + E V   WV  H G
Sbjct  38564  VTDSQYVMQGINDWMPNWKKRGWKTASKQPVKNADLWQQLDEQVSRHE-VSWQWVRGHTG  38388

Query  102    IGGNEQVDKLVSAGI  116
               GNEQ D L + G+
Sbjct  38387  HPGNEQADLLANRGV  38343
```

```
>Stutzerimonas frequens strain HI00D01 HI00D01_c19, whole genome shotgun sequence
Sequence ID: NZ_LWEF01000079.1 Length: 149482
Range 1: 120550 to 120951

Score:48.9 bits(115), Expect:0.004,
Method:Compositional matrix adjust.,
Identities:44/135(33%), Positives:61/135(45%), Gaps:20/135(14%)

Query  1       YVDGAANRETKLGKAG-YVTNRG--RQKVVTLTDTTNQKTELQAIYLALQD--SGLEVNI  55
               Y DGA       G  G  +  +G  R+      DTTN + EL A   AL +     +V +
Sbjct  120951  YTDGACKGNPGPGGWGALLVYKGVKRELWGGEPDTTNNRMELMAAIRALAELKRPCKVRL  120772

Query  56      VTDSQYAL-GIIQAQPDQSE-------------SELVNQIIEQLIKKEKVYLAWVPAHKG  101
               VTDSQY + GI    P+  +              ++L  Q+ EQ+ + E V   WV  H G
Sbjct  120771  VTDSQYVMQGINDWMPNWKKRGWKTASKQPVKNADLWQQLDEQVSRHE-VSWQWVRGHTG  120595

Query  102     IGGNEQVDKLVSAGI  116
                GNEQ D L + G+
Sbjct  120594  HPGNEQADLLANRGV  120550
```

```
>Stutzerimonas frequens strain GOM2 scaffold_3, whole genome shotgun sequence
Sequence ID: NZ_WOUM01000018.1 Length: 232832
Range 1: 155361 to 155762

Score:48.9 bits(115), Expect:0.004,
Method:Compositional matrix adjust.,
Identities:44/135(33%), Positives:61/135(45%), Gaps:20/135(14%)

Query  1       YVDGAANRETKLGKAG-YVTNRG--RQKVVTLTDTTNQKTELQAIYLALQD--SGLEVNI  55
               Y DGA       G  G  +  +G  R+      DTTN + EL A   AL +     +V +
Sbjct  155762  YTDGACKGNPGPGGWGALLVYKGVKRELWGGEPDTTNNRMELMAAIRALAELKRPCKVRL  155583

Query  56      VTDSQYAL-GIIQAQPDQSE-------------SELVNQIIEQLIKKEKVYLAWVPAHKG  101
               VTDSQY + GI    P+  +              ++L  Q+ EQ+ + E V   WV  H G
Sbjct  155582  VTDSQYVMQGINDWMPNWKKRGWKTASKQPVKNADLWQQLDEQVSRHE-VSWQWVRGHTG  155406

Query  102     IGGNEQVDKLVSAGI  116
                 GNEQ D L + G+
Sbjct  155405  HPGNEQADLLANRGV  155361
```

```
>Stutzerimonas frequens strain DNSP21 27DNSP21gv5C003, whole genome shotgun sequence
Sequence ID: NZ_JAMOHM010000003.1 Length: 433620
Range 1: 361739 to 362140

Score:48.9 bits(115), Expect:0.004,
Method:Compositional matrix adjust.,
Identities:44/135(33%), Positives:61/135(45%), Gaps:20/135(14%)

Query  1       YVDGAANRETKLGKAG-YVTNRG--RQKVVTLTDTTNQKTELQAIYLALQD--SGLEVNI  55
               Y DGA       G  G  +  +G  R+      DTTN + EL A   AL +     +V +
Sbjct  362140  YTDGACKGNPGPGGWGALLVYKGVKRELWGGEPDTTNNRMELMAAIRALAELKRPCKVRL  361961

Query  56      VTDSQYAL-GIIQAQPDQSE-------------SELVNQIIEQLIKKEKVYLAWVPAHKG  101
               VTDSQY + GI    P+  +              ++L  Q+ EQ+ + E V   WV  H G
Sbjct  361960  VTDSQYVMQGINDWMPNWKKRGWKTASKQPVKNADLWQQLDEQVSRHE-VSWQWVRGHTG  361784

Query  102     IGGNEQVDKLVSAGI  116
                 GNEQ D L + G+
Sbjct  361783  HPGNEQADLLANRGV  361739
```

```
>Stutzerimonas frequens strain S18 3, whole genome shotgun sequence
Sequence ID: NZ_JASSZI010000003.1 Length: 681260
Range 1: 71528 to 71929

Score:48.9 bits(115), Expect:0.004,
Method:Compositional matrix adjust.,
Identities:44/135(33%), Positives:61/135(45%), Gaps:20/135(14%)

Query  1      YVDGAANRETKLGKAG-YVTNRG--RQKVVTLTDTTNQKTELQAIYLALQD--SGLEVNI  55
              Y DGA       G  G  +  +G  R+      DTTN + EL A   AL +     +V +
Sbjct  71528  YTDGACKGNPGPGGWGALLVYKGVKRELWGGEPDTTNNRMELMAAIRALAELKRPCKVRL  71707

Query  56     VTDSQYAL-GIIQAQPDQSE-------------SELVNQIIEQLIKKEKVYLAWVPAHKG  101
              VTDSQY + GI    P+  +              ++L  Q+ EQ+ + E V   WV  H G
Sbjct  71708  VTDSQYVMQGINDWMPNWKKRGWKTASKQPVKNADLWQQLDEQVSRHE-VSWQWVRGHTG  71884

Query  102    IGGNEQVDKLVSAGI  116
                GNEQ D L + G+
Sbjct  71885  HPGNEQADLLANRGV  71929
```

```
>Stutzerimonas frequens strain DNSP21 NODE_4_length_722045_cov_18.4055, whole genome shotgun sequence
Sequence ID: NZ_POUJ01000004.1 Length: 722045
Range 1: 71788 to 72189

Score:48.9 bits(115), Expect:0.004,
Method:Compositional matrix adjust.,
Identities:44/135(33%), Positives:61/135(45%), Gaps:20/135(14%)

Query  1      YVDGAANRETKLGKAG-YVTNRG--RQKVVTLTDTTNQKTELQAIYLALQD--SGLEVNI  55
              Y DGA       G  G  +  +G  R+      DTTN + EL A   AL +     +V +
Sbjct  71788  YTDGACKGNPGPGGWGALLVYKGVKRELWGGEPDTTNNRMELMAAIRALAELKRPCKVRL  71967

Query  56     VTDSQYAL-GIIQAQPDQSE-------------SELVNQIIEQLIKKEKVYLAWVPAHKG  101
              VTDSQY + GI    P+  +              ++L  Q+ EQ+ + E V   WV  H G
Sbjct  71968  VTDSQYVMQGINDWMPNWKKRGWKTASKQPVKNADLWQQLDEQVSRHE-VSWQWVRGHTG  72144

Query  102    IGGNEQVDKLVSAGI  116
                GNEQ D L + G+
Sbjct  72145  HPGNEQADLLANRGV  72189
```

```
>Stutzerimonas frequens strain KOL14.W.20.12 Scaffold_2_length_878494_pilon, whole genome shotgun sequence
Sequence ID: NZ_WIFY01000002.1 Length: 879884
Range 1: 173732 to 174133

Score:48.9 bits(115), Expect:0.004,
Method:Compositional matrix adjust.,
Identities:44/135(33%), Positives:61/135(45%), Gaps:20/135(14%)

Query  1       YVDGAANRETKLGKAG-YVTNRG--RQKVVTLTDTTNQKTELQAIYLALQD--SGLEVNI  55
               Y DGA       G  G  +  +G  R+      DTTN + EL A   AL +     +V +
Sbjct  173732  YTDGACKGNPGPGGWGALLVYKGVKRELWGGEPDTTNNRMELMAAIRALAELKRPCKVRL  173911

Query  56      VTDSQYAL-GIIQAQPDQSE-------------SELVNQIIEQLIKKEKVYLAWVPAHKG  101
               VTDSQY + GI    P+  +              ++L  Q+ EQ+ + E V   WV  H G
Sbjct  173912  VTDSQYVMQGINDWMPNWKKRGWKTASKQPVKNADLWQQLDEQVSRHE-VSWQWVRGHTG  174088

Query  102     IGGNEQVDKLVSAGI  116
                 GNEQ D L + G+
Sbjct  174089  HPGNEQADLLANRGV  174133
```

```
>Stutzerimonas frequens strain KOL14.W.90.4 Scaffold_2_length_1279239_pilon, whole genome shotgun sequence
Sequence ID: NZ_WIGH01000002.1 Length: 1280911
Range 1: 568007 to 568408

Score:48.9 bits(115), Expect:0.004,
Method:Compositional matrix adjust.,
```

```
Identities:44/135(33%), Positives:61/135(45%), Gaps:20/135(14%)

Query  1       YVDGAANRETKLGKAG-YVTNRG--RQKVVTLTDTTNQKTELQAIYLALQD--SGLEVNI  55
               Y DGA       G  G  +  +G  R+      DTTN + EL A   AL +     +V +
Sbjct  568007  YTDGACKGNPGPGGWGALLVYKGVKRELWGGEPDTTNNRMELMAAIRALAELKRPCKVRL  568186

Query  56      VTDSQYAL-GIIQAQPDQSE-------------SELVNQIIEQLIKKEKVYLAWVPAHKG  101
               VTDSQY + GI    P+  +              ++L  Q+ EQ+ + E V   WV  H G
Sbjct  568187  VTDSQYVMQGINDWMPNWKKRGWKTASKQPVKNADLWQQLDEQVSRHE-VSWQWVRGHTG  568363

Query  102     IGGNEQVDKLVSAGI  116
                 GNEQ D L + G+
Sbjct  568364  HPGNEQADLLANRGV  568408
```

>Stutzerimonas frequens strain KOL14.W.90.9 Scaffold_3_length_1279107_pilon, whole genome shotgun sequence
Sequence ID: NZ_WIFU01000002.1 Length: 1283483
Range 1: 567985 to 568386

Score:48.9 bits(115), Expect:0.004,
Method:Compositional matrix adjust.,
Identities:44/135(33%), Positives:61/135(45%), Gaps:20/135(14%)

```
Query  1       YVDGAANRETKLGKAG-YVTNRG--RQKVVTLTDTTNQKTELQAIYLALQD--SGLEVNI  55
               Y DGA       G  G  +  +G  R+      DTTN + EL A   AL +     +V +
Sbjct  567985  YTDGACKGNPGPGGWGALLVYKGVKRELWGGEPDTTNNRMELMAAIRALAELKRPCKVRL  568164

Query  56      VTDSQYAL-GIIQAQPDQSE-------------SELVNQIIEQLIKKEKVYLAWVPAHKG  101
               VTDSQY + GI    P+  +              ++L  Q+ EQ+ + E V   WV  H G
Sbjct  568165  VTDSQYVMQGINDWMPNWKKRGWKTASKQPVKNADLWQQLDEQVSRHE-VSWQWVRGHTG  568341

Query  102     IGGNEQVDKLVSAGI  116
                 GNEQ D L + G+
Sbjct  568342  HPGNEQADLLANRGV  568386
```

>Stutzerimonas frequens strain KOL14.W.45.16 Scaffold_5_length_1368366_pilon, whole genome shotgun sequence
Sequence ID: NZ_WIGI01000005.1 Length: 1368612
Range 1: 691303 to 691704

Score:48.9 bits(115), Expect:0.004,
Method:Compositional matrix adjust.,
Identities:44/135(33%), Positives:61/135(45%), Gaps:20/135(14%)

```
Query  1       YVDGAANRETKLGKAG-YVTNRG--RQKVVTLTDTTNQKTELQAIYLALQD--SGLEVNI  55
               Y DGA       G  G  +  +G  R+      DTTN + EL A   AL +     +V +
Sbjct  691303  YTDGACKGNPGPGGWGALLVYKGVKRELWGGEPDTTNNRMELMAAIRALAELKRPCKVRL  691482

Query  56      VTDSQYAL-GIIQAQPDQSE-------------SELVNQIIEQLIKKEKVYLAWVPAHKG  101
               VTDSQY + GI    P+  +              ++L  Q+ EQ+ + E V   WV  H G
Sbjct  691483  VTDSQYVMQGINDWMPNWKKRGWKTASKQPVKNADLWQQLDEQVSRHE-VSWQWVRGHTG  691659

Query  102     IGGNEQVDKLVSAGI  116
                 GNEQ D L + G+
Sbjct  691660  HPGNEQADLLANRGV  691704
```

>Stutzerimonas frequens strain CAM01 k141_91720, whole genome shotgun sequence
Sequence ID: NZ_JAQBSX010000067.1 Length: 1835512
Range 1: 518882 to 519283

Score:48.9 bits(115), Expect:0.004,
Method:Compositional matrix adjust.,
Identities:44/135(33%), Positives:61/135(45%), Gaps:20/135(14%)

```
Query  1       YVDGAANRETKLGKAG-YVTNRG--RQKVVTLTDTTNQKTELQAIYLALQD--SGLEVNI  55
               Y DGA       G  G  +  +G  R+      DTTN + EL A   AL +     +V +
Sbjct  519283  YTDGACKGNPGPGGWGALLVYKGVKRELWGGEPDTTNNRMELMAAIRALAELKRPCKVRL  519104

Query  56      VTDSQYAL-GIIQAQPDQSE-------------SELVNQIIEQLIKKEKVYLAWVPAHKG  101
               VTDSQY + GI    P+  +              ++L  Q+ EQ+ + E V   WV  H G
Sbjct  519103  VTDSQYVMQGINDWMPNWKKRGWKTASKQPVKNADLWQQLDEQVSRHE-VSWQWVRGHTG  518927

Query  102     IGGNEQVDKLVSAGI  116
                 GNEQ D L + G+
Sbjct  518926  HPGNEQADLLANRGV  518882
```

>Stutzerimonas frequens strain FDAARGOS_877 chromosome, complete genome
Sequence ID: NZ_CP065720.1 Length: 4394686
Range 1: 3314036 to 3314437

Score:48.9 bits(115), Expect:0.004,
Method:Compositional matrix adjust.,
Identities:44/135(33%), Positives:61/135(45%), Gaps:20/135(14%)

```
Query  1        YVDGAANRETKLGKAG-YVTNRG--RQKVVTLTDTTNQKTELQAIYLALQD--SGLEVNI  55
                Y DGA       G  G  +  +G  R+      DTTN + EL A   AL +     +V +
Sbjct  3314437  YTDGACKGNPGPGGWGALLVYKGVKRELWGGEPDTTNNRMELMAAIRALAELKRPCKVRL  3314258

Query  56       VTDSQYAL-GIIQAQPDQSE-------------SELVNQIIEQLIKKEKVYLAWVPAHKG  101
                VTDSQY + GI    P+  +              ++L  Q+ EQ+ + E V   WV  H G
Sbjct  3314257  VTDSQYVMQGINDWMPNWKKRGWKTASKQPVKNADLWQQLDEQVSRHE-VSWQWVRGHTG  3314081

Query  102      IGGNEQVDKLVSAGI  116
                  GNEQ D L + G+
Sbjct  3314080  HPGNEQADLLANRGV  3314036
```

>Stutzerimonas frequens strain TPA3 chromosome, complete genome
Sequence ID: NZ_CP071899.1 Length: 4415884
Range 1: 2305661 to 2306062

Score:48.9 bits(115), Expect:0.004,
Method:Compositional matrix adjust.,
Identities:44/135(33%), Positives:61/135(45%), Gaps:20/135(14%)

```
Query  1        YVDGAANRETKLGKAG-YVTNRG--RQKVVTLTDTTNQKTELQAIYLALQD--SGLEVNI  55
                Y DGA       G  G  +  +G  R+      DTTN + EL A   AL +     +V +
Sbjct  2306062  YTDGACKGNPGPGGWGALLVYKGVKRELWGGEPDTTNNRMELMAAIRALAELKRPCKVRL  2305883

Query  56       VTDSQYAL-GIIQAQPDQSE-------------SELVNQIIEQLIKKEKVYLAWVPAHKG  101
                VTDSQY + GI    P+  +              ++L  Q+ EQ+ + E V   WV  H G
Sbjct  2305882  VTDSQYVMQGINDWMPNWKKRGWKTASKQPVKNADLWQQLDEQVSRHE-VSWQWVRGHTG  2305706
```

```
Query  102      IGGNEQVDKLVSAGI  116
                  GNEQ D L + G+
Sbjct  2305705  HPGNEQADLLANRGV  2305661
```

>Stutzerimonas frequens strain THAF7b chromosome, complete genome
Sequence ID: NZ_CP045416.1 Length: 4522538
Range 1: 2419478 to 2419879

Score:48.9 bits(115), Expect:0.004,
Method:Compositional matrix adjust.,
Identities:44/135(33%), Positives:61/135(45%), Gaps:20/135(14%)

```
Query  1        YVDGAANRETKLGKAG-YVTNRG--RQKVVTLTDTTNQKTELQAIYLALQD--SGLEVNI  55
                Y DGA       G  G  +  +G  R+      DTTN + EL A   AL +     +V +
Sbjct  2419879  YTDGACKGNPGPGGWGALLVYKGVKRELWGGEPDTTNNRMELMAAIRALAELKRPCKVRL  2419700

Query  56       VTDSQYAL-GIIQAQPDQSE------------SELVNQIIEQLIKKEKVYLAWVPAHKG  101
                VTDSQY + GI    P+  +             ++L  Q+ EQ+ + E V   WV  H G
Sbjct  2419699  VTDSQYVMQGINDWMPNWKKRGWKTASKQPVKNADLWQQLDEQVSRHE-VSWQWVRGHTG  2419523

Query  102      IGGNEQVDKLVSAGI  116
                  GNEQ D L + G+
Sbjct  2419522  HPGNEQADLLANRGV  2419478
```

>Stutzerimonas frequens strain L27 chromosome, complete genome
Sequence ID: NZ_CP136501.1 Length: 4544032
Range 1: 3530417 to 3530818

Score:48.9 bits(115), Expect:0.004,
Method:Compositional matrix adjust.,
Identities:44/135(33%), Positives:61/135(45%), Gaps:20/135(14%)

```
Query  1        YVDGAANRETKLGKAG-YVTNRG--RQKVVTLTDTTNQKTELQAIYLALQD--SGLEVNI  55
                Y DGA       G  G  +  +G  R+      DTTN + EL A   AL +     +V +
Sbjct  3530818  YTDGACKGNPGPGGWGALLVYKGVKRELWGGEPDTTNNRMELMAAIRALAELKRPCKVRL  3530639

Query  56       VTDSQYAL-GIIQAQPDQSE------------SELVNQIIEQLIKKEKVYLAWVPAHKG  101
                VTDSQY + GI    P+  +             ++L  Q+ EQ+ + E V   WV  H G
Sbjct  3530638  VTDSQYVMQGINDWMPNWKKRGWKTASKQPVKNADLWQQLDEQVSRHE-VSWQWVRGHTG  3530462

Query  102      IGGNEQVDKLVSAGI  116
                  GNEQ D L + G+
Sbjct  3530461  HPGNEQADLLANRGV  3530417
```

>Stutzerimonas frequens strain KOL14.W.5.3 Scaffold_1_length_4537025_pilon, whole genome shotgun sequence
Sequence ID: NZ_WIGG01000001.1 Length: 4554507
Range 1: 709425 to 709826

Score:48.9 bits(115), Expect:0.004,
Method:Compositional matrix adjust.,
Identities:44/135(33%), Positives:61/135(45%), Gaps:20/135(14%)

```
Query  1       YVDGAANRETKLGKAG-YVTNRG--RQKVVTLTDTTNQKTELQAIYLALQD--SGLEVNI  55
               Y DGA       G  G  +  +G  R+      DTTN + EL A   AL +     +V +
Sbjct  709826  YTDGACKGNPGPGGWGALLVYKGVKRELWGGEPDTTNNRMELMAAIRALAELKRPCKVRL  709647

Query  56      VTDSQYAL-GIIQAQPDQSE------------SELVNQIIEQLIKKEKVYLAWVPAHKG  101
               VTDSQY + GI    P+  +             ++L  Q+ EQ+ + E V   WV  H G
Sbjct  709646  VTDSQYVMQGINDWMPNWKKRGWKTASKQPVKNADLWQQLDEQVSRHE-VSWQWVRGHTG  709470

Query  102     IGGNEQVDKLVSAGI  116
                 GNEQ D L + G+
Sbjct  709469  HPGNEQADLLANRGV  709425
```

>Stutzerimonas frequens strain TF18 chromosome, complete genome
Sequence ID: NZ_CP113257.1 Length: 4918844
Range 1: 2659503 to 2659904

Score:48.9 bits(115), Expect:0.004,
Method:Compositional matrix adjust.,
Identities:44/135(33%), Positives:61/135(45%), Gaps:20/135(14%)

```
Query  1        YVDGAANRETKLGKAG-YVTNRG--RQKVVTLTDTTNQKTELQAIYLALQD--SGLEVNI  55
                Y DGA       G  G  +  +G  R+      DTTN + EL A   AL +     +V +
Sbjct  2659904  YTDGACKGNPGPGGWGALLVYKGVKRELWGGEPDTTNNRMELMAAIRALAELKRPCKVRL  2659725

Query  56       VTDSQYAL-GIIQAQPDQSE------------SELVNQIIEQLIKKEKVYLAWVPAHKG  101
                VTDSQY + GI    P+  +             ++L  Q+ EQ+ + E V   WV  H G
Sbjct  2659724  VTDSQYVMQGINDWMPNWKKRGWKTASKQPVKNADLWQQLDEQVSRHE-VSWQWVRGHTG  2659548

Query  102      IGGNEQVDKLVSAGI  116
                  GNEQ D L + G+
Sbjct  2659547  HPGNEQADLLANRGV  2659503
```

>Stutzerimonas stutzeri strain PS_027 NODE_44_length_26947_cov_4.29568, whole genome shotgun sequence
Sequence ID: NZ_RHRQ01000044.1 Length: 26947
Range 1: 3697 to 4002

Score:48.9 bits(115), Expect:0.004,
Method:Compositional matrix adjust.,
Identities:35/103(34%), Positives:50/103(48%), Gaps:17/103(16%)

```
Query  30    TDTTNQKTELQAIYLALQD--SGLEVNIVTDSQYAL-GIIQAQPDQSE------------  74
              DTTN + EL A   AL +     +V +VTDSQY + GI    P+  +
Sbjct  3697  PDTTNNRMELMAAIRALAELKRPCKVRLVTDSQYVMQGINDWMPNWKKRGWKTASKQPVK  3876

Query  75    -SELVNQIIEQLIKKEKVYLAWVPAHKGIGGNEQVDKLVSAGI  116
              ++L  Q+ EQ + + +V   WV  H G  GNEQ D L + G+
Sbjct  3877  NADLWQQLDEQ-VNRHEVSWQWVRGHTGHPGNEQADLLANRGV  4002
```

>Stutzerimonas stutzeri strain GD03740 30, whole genome shotgun sequence
Sequence ID: NZ_JAOCHW010000030.1 Length: 38162
Range 1: 13681 to 13986

Score:48.9 bits(115), Expect:0.004,

```
Method:Compositional matrix adjust.,
Identities:35/103(34%), Positives:50/103(48%), Gaps:17/103(16%)

Query  30     TDTTNQKTELQAIYLALQD--SGLEVNIVTDSQYAL-GIIQAQPDQSE------------  74
               DTTN + EL A   AL +     +V +VTDSQY + GI    P+  +
Sbjct  13986  PDTTNNRMELMAAIRALAELKRPCKVRLVTDSQYVMQGINDWMPNWKKRGWKTASKQPVK  13807

Query  75     -SELVNQIIEQLIKKEKVYLAWVPAHKGIGGNEQVDKLVSAGI  116
               ++L  Q+ EQ + + +V   WV  H G  GNEQ D L + G+
Sbjct  13806  NADLWQQLDEQ-VNRHEVSWQWVRGHTGHPGNEQADLLANRGV  13681
```

>Stutzerimonas stutzeri strain INSAq93 Aq93_contig_23, whole genome shotgun sequence
Sequence ID: NZ_JAKCMP010000023.1 Length: 38538
Range 1: 13814 to 14119

```
Score:48.9 bits(115), Expect:0.004,
Method:Compositional matrix adjust.,
Identities:35/103(34%), Positives:50/103(48%), Gaps:17/103(16%)

Query  30     TDTTNQKTELQAIYLALQD--SGLEVNIVTDSQYAL-GIIQAQPDQSE------------  74
               DTTN + EL A   AL +     +V +VTDSQY + GI    P+  +
Sbjct  14119  PDTTNNRMELMAAIRALAELKRPCKVRLVTDSQYVMQGINDWMPNWKKRGWKTASKQPVK  13940

Query  75     -SELVNQIIEQLIKKEKVYLAWVPAHKGIGGNEQVDKLVSAGI  116
               ++L  Q+ EQ + + +V   WV  H G  GNEQ D L + G+
Sbjct  13939  NADLWQQLDEQ-VNRHEVSWQWVRGHTGHPGNEQADLLANRGV  13814
```

>Stutzerimonas stutzeri XLDN-R contig000128, whole genome shotgun sequence
Sequence ID: NZ_AKYE01000128.1 Length: 40595
Range 1: 38507 to 38812

```
Score:48.9 bits(115), Expect:0.004,
Method:Compositional matrix adjust.,
Identities:35/103(34%), Positives:50/103(48%), Gaps:17/103(16%)

Query  30     TDTTNQKTELQAIYLALQD--SGLEVNIVTDSQYAL-GIIQAQPDQSE------------  74
               DTTN + EL A   AL +     +V +VTDSQY + GI    P+  +
Sbjct  38812  PDTTNNRMELMAAIRALAELKRPCKVRLVTDSQYVMQGINDWMPNWKKRGWKTASKQPVK  38633

Query  75     -SELVNQIIEQLIKKEKVYLAWVPAHKGIGGNEQVDKLVSAGI  116
               ++L  Q+ EQ + + +V   WV  H G  GNEQ D L + G+
Sbjct  38632  NADLWQQLDEQ-VNRHEVSWQWVRGHTGHPGNEQADLLANRGV  38507
```

>Stutzerimonas stutzeri strain 1223_PMEN 439_42450_436684, whole genome shotgun sequence
Sequence ID: NZ_JVYC01000128.1 Length: 42450
Range 1: 3728 to 4033

```
Score:48.9 bits(115), Expect:0.004,
Method:Compositional matrix adjust.,
Identities:35/103(34%), Positives:50/103(48%), Gaps:17/103(16%)

Query  30    TDTTNQKTELQAIYLALQD--SGLEVNIVTDSQYAL-GIIQAQPDQSE------------  74
              DTTN + EL A   AL +     +V +VTDSQY + GI    P+  +
Sbjct  3728  PDTTNNRMELMAAIRALAELKRPCKVRLVTDSQYVMQGINDWMPNWKKRGWKTASKQPVK  3907

Query  75    -SELVNQIIEQLIKKEKVYLAWVPAHKGIGGNEQVDKLVSAGI  116
              ++L  Q+ EQ + + +V   WV  H G  GNEQ D L + G+
Sbjct  3908  NADLWQQLDEQ-VNRHEVSWQWVRGHTGHPGNEQADLLANRGV  4033
```

>Stutzerimonas stutzeri strain GD03749 27, whole genome shotgun sequence
Sequence ID: NZ_JAOCHN010000027.1 Length: 62832
Range 1: 24138 to 24443

```
Score:48.9 bits(115), Expect:0.004,
Method:Compositional matrix adjust.,
Identities:35/103(34%), Positives:50/103(48%), Gaps:17/103(16%)

Query  30     TDTTNQKTELQAIYLALQD--SGLEVNIVTDSQYAL-GIIQAQPDQSE------------  74
               DTTN + EL A   AL +     +V +VTDSQY + GI    P+  +
Sbjct  24138  PDTTNNRMELMAAIRALAELKRPCKVRLVTDSQYVMQGINDWMPNWKKRGWKTASKQPVK  24317

Query  75     -SELVNQIIEQLIKKEKVYLAWVPAHKGIGGNEQVDKLVSAGI  116
               ++L  Q+ EQ + + +V   WV  H G  GNEQ D L + G+
Sbjct  24318  NADLWQQLDEQ-VNRHEVSWQWVRGHTGHPGNEQADLLANRGV  24443
```

>Stutzerimonas stutzeri strain PSB00024 PSB00024_c0024, whole genome shotgun sequence
Sequence ID: NZ_JADTWB010000024.1 Length: 63007
Range 1: 38486 to 38791

```
Score:48.9 bits(115), Expect:0.004,
Method:Compositional matrix adjust.,
Identities:35/103(34%), Positives:50/103(48%), Gaps:17/103(16%)

Query  30     TDTTNQKTELQAIYLALQD--SGLEVNIVTDSQYAL-GIIQAQPDQSE------------  74
               DTTN + EL A   AL +     +V +VTDSQY + GI    P+  +
Sbjct  38791  PDTTNNRMELMAAIRALAELKRPCKVRLVTDSQYVMQGINDWMPNWKKRGWKTASKQPVK  38612

Query  75     -SELVNQIIEQLIKKEKVYLAWVPAHKGIGGNEQVDKLVSAGI  116
               ++L  Q+ EQ + + +V   WV  H G  GNEQ D L + G+
Sbjct  38611  NADLWQQLDEQ-VNRHEVSWQWVRGHTGHPGNEQADLLANRGV  38486
```

>Stutzerimonas stutzeri strain A655/78 11PstA665_78gv1C022, whole genome shotgun sequence
Sequence ID: NZ_JAMOJY010000022.1 Length: 63676
Range 1: 13351 to 13656

```
Score:48.9 bits(115), Expect:0.004,
Method:Compositional matrix adjust.,
Identities:35/103(34%), Positives:50/103(48%), Gaps:17/103(16%)

Query  30     TDTTNQKTELQAIYLALQD--SGLEVNIVTDSQYAL-GIIQAQPDQSE------------  74
               DTTN + EL A   AL +     +V +VTDSQY + GI    P+  +
Sbjct  13656  PDTTNNRMELMAAIRALAELKRPCKVRLVTDSQYVMQGINDWMPNWKKRGWKTASKQPVK  13477

Query  75     -SELVNQIIEQLIKKEKVYLAWVPAHKGIGGNEQVDKLVSAGI  116
               ++L  Q+ EQ + + +V   WV  H G  GNEQ D L + G+
Sbjct  13476  NADLWQQLDEQ-VNRHEVSWQWVRGHTGHPGNEQADLLANRGV  13351
```

```
>Stutzerimonas stutzeri strain INSAq87 Aq87_contig_65, whole genome shotgun sequence
Sequence ID: NZ_JAKCMO010000065.1 Length: 64028
Range 1: 57114 to 57419

Score:48.9 bits(115), Expect:0.004,
Method:Compositional matrix adjust.,
Identities:35/103(34%), Positives:50/103(48%), Gaps:17/103(16%)

Query  30     TDTTNQKTELQAIYLALQD--SGLEVNIVTDSQYAL-GIIQAQPDQSE------------  74
               DTTN + EL A   AL +     +V +VTDSQY + GI    P+  +
Sbjct  57114  PDTTNNRMELMAAIRALAELKRPCKVRLVTDSQYVMQGINDWMPNWKKRGWKTASKQPVK  57293

Query  75     -SELVNQIIEQLIKKEKVYLAWVPAHKGIGGNEQVDKLVSAGI  116
               ++L  Q+ EQ + + +V   WV  H G  GNEQ D L + G+
Sbjct  57294  NADLWQQLDEQ-VNRHEVSWQWVRGHTGHPGNEQADLLANRGV  57419



>Stutzerimonas stutzeri strain SD93936 09PstSD93936gv1c014, whole genome shotgun sequence
Sequence ID: NZ_JAMOJZ010000014.1 Length: 64350
Range 1: 40993 to 41298

Score:48.9 bits(115), Expect:0.004,
Method:Compositional matrix adjust.,
Identities:35/103(34%), Positives:50/103(48%), Gaps:17/103(16%)

Query  30     TDTTNQKTELQAIYLALQD--SGLEVNIVTDSQYAL-GIIQAQPDQSE------------  74
               DTTN + EL A   AL +     +V +VTDSQY + GI    P+  +
Sbjct  40993  PDTTNNRMELMAAIRALAELKRPCKVRLVTDSQYVMQGINDWMPNWKKRGWKTASKQPVK  41172

Query  75     -SELVNQIIEQLIKKEKVYLAWVPAHKGIGGNEQVDKLVSAGI  116
               ++L  Q+ EQ + + +V   WV  H G  GNEQ D L + G+
Sbjct  41173  NADLWQQLDEQ-VNRHEVSWQWVRGHTGHPGNEQADLLANRGV  41298



>Stutzerimonas stutzeri strain GD04104 21, whole genome shotgun sequence
Sequence ID: NZ_JAOEAT010000021.1 Length: 65673
Range 1: 41192 to 41497

Score:48.9 bits(115), Expect:0.004,
Method:Compositional matrix adjust.,
Identities:35/103(34%), Positives:50/103(48%), Gaps:17/103(16%)

Query  30     TDTTNQKTELQAIYLALQD--SGLEVNIVTDSQYAL-GIIQAQPDQSE------------  74
               DTTN + EL A   AL +     +V +VTDSQY + GI    P+  +
Sbjct  41497  PDTTNNRMELMAAIRALAELKRPCKVRLVTDSQYVMQGINDWMPNWKKRGWKTASKQPVK  41318

Query  75     -SELVNQIIEQLIKKEKVYLAWVPAHKGIGGNEQVDKLVSAGI  116
               ++L  Q+ EQ + + +V   WV  H G  GNEQ D L + G+
Sbjct  41317  NADLWQQLDEQ-VNRHEVSWQWVRGHTGHPGNEQADLLANRGV  41192



>Stutzerimonas stutzeri strain LH-42 Contig004, whole genome shotgun sequence
Sequence ID: NZ_NCKS02000004.1 Length: 68185
Range 1: 64207 to 64512

Score:48.9 bits(115), Expect:0.004,
Method:Compositional matrix adjust.,
Identities:35/103(34%), Positives:50/103(48%), Gaps:17/103(16%)

Query  30     TDTTNQKTELQAIYLALQD--SGLEVNIVTDSQYAL-GIIQAQPDQSE------------  74
               DTTN + EL A   AL +     +V +VTDSQY + GI    P+  +
Sbjct  64512  PDTTNNRMELMAAIRALAELKRPCKVRLVTDSQYVMQGINDWMPNWKKRGWKTASKQPVK  64333

Query  75     -SELVNQIIEQLIKKEKVYLAWVPAHKGIGGNEQVDKLVSAGI  116
               ++L  Q+ EQ + + +V   WV  H G  GNEQ D L + G+
Sbjct  64332  NADLWQQLDEQ-VNRHEVSWQWVRGHTGHPGNEQADLLANRGV  64207



>Stutzerimonas stutzeri strain A160/74 10PstA160_74gv1c016, whole genome shotgun sequence
Sequence ID: NZ_JAMOHN010000016.1 Length: 82616
Range 1: 8902 to 9207

Score:48.9 bits(115), Expect:0.004,
Method:Compositional matrix adjust.,
Identities:35/103(34%), Positives:50/103(48%), Gaps:17/103(16%)

Query  30    TDTTNQKTELQAIYLALQD--SGLEVNIVTDSQYAL-GIIQAQPDQSE------------  74
              DTTN + EL A   AL +     +V +VTDSQY + GI    P+  +
Sbjct  9207  PDTTNNRMELMAAIRALAELKRPCKVRLVTDSQYVMQGINDWMPNWKKRGWKTASKQPVK  9028

Query  75    -SELVNQIIEQLIKKEKVYLAWVPAHKGIGGNEQVDKLVSAGI  116
              ++L  Q+ EQ + + +V   WV  H G  GNEQ D L + G+
Sbjct  9027  NADLWQQLDEQ-VNRHEVSWQWVRGHTGHPGNEQADLLANRGV  8902



>Stutzerimonas stutzeri strain NT-I, whole genome shotgun sequence
Sequence ID: NZ_BCAJ01000037.1 Length: 97403
Range 1: 22481 to 22786

Score:48.9 bits(115), Expect:0.004,
Method:Compositional matrix adjust.,
Identities:35/103(34%), Positives:50/103(48%), Gaps:17/103(16%)

Query  30     TDTTNQKTELQAIYLALQD--SGLEVNIVTDSQYAL-GIIQAQPDQSE------------  74
               DTTN + EL A   AL +     +V +VTDSQY + GI    P+  +
Sbjct  22786  PDTTNNRMELMAAIRALAELKRPCKVRLVTDSQYVMQGINDWMPNWKKRGWKTASKQPVK  22607

Query  75     -SELVNQIIEQLIKKEKVYLAWVPAHKGIGGNEQVDKLVSAGI  116
               ++L  Q+ EQ + + +V   WV  H G  GNEQ D L + G+
Sbjct  22606  NADLWQQLDEQ-VNRHEVSWQWVRGHTGHPGNEQADLLANRGV  22481



>Stutzerimonas stutzeri strain GD04106 14, whole genome shotgun sequence
Sequence ID: NZ_JAOEAR010000014.1 Length: 109530
Range 1: 41192 to 41497

Score:48.9 bits(115), Expect:0.004,
Method:Compositional matrix adjust.,
```

```
Identities:35/103(34%), Positives:50/103(48%), Gaps:17/103(16%)

Query  30     TDTTNQKTELQAIYLALQD--SGLEVNIVTDSQYAL-GIIQAQPDQSE------------  74
               DTTN + EL A   AL +     +V +VTDSQY + GI    P+  +
Sbjct  41497  PDTTNNRMELMAAIRALAELKRPCKVRLVTDSQYVMQGINDWMPNWKKRGWKTASKQPVK  41318

Query  75     -SELVNQIIEQLIKKEKVYLAWVPAHKGIGGNEQVDKLVSAGI  116
               ++L  Q+ EQ + + +V   WV  H G  GNEQ D L + G+
Sbjct  41317  NADLWQQLDEQ-VNRHEVSWQWVRGHTGHPGNEQADLLANRGV  41192
```

```
>Stutzerimonas stutzeri strain GD04164 8, whole genome shotgun sequence
Sequence ID: NZ_JAODYN010000008.1 Length: 111383
Range 1: 73385 to 73690

Score:48.9 bits(115), Expect:0.004,
Method:Compositional matrix adjust.,
Identities:35/103(34%), Positives:50/103(48%), Gaps:17/103(16%)

Query  30     TDTTNQKTELQAIYLALQD--SGLEVNIVTDSQYAL-GIIQAQPDQSE------------  74
               DTTN + EL A   AL +     +V +VTDSQY + GI    P+  +
Sbjct  73385  PDTTNNRMELMAAIRALAELKRPCKVRLVTDSQYVMQGINDWMPNWKKRGWKTASKQPVK  73564

Query  75     -SELVNQIIEQLIKKEKVYLAWVPAHKGIGGNEQVDKLVSAGI  116
               ++L  Q+ EQ + + +V   WV  H G  GNEQ D L + G+
Sbjct  73565  NADLWQQLDEQ-VNRHEVSWQWVRGHTGHPGNEQADLLANRGV  73690
```

```
>Stutzerimonas stutzeri strain PS_197 NODE_16_length_111597_cov_35.5926, whole genome shotgun sequence
Sequence ID: NZ_RHRB01000016.1 Length: 111597
Range 1: 73518 to 73823

Score:48.9 bits(115), Expect:0.004,
Method:Compositional matrix adjust.,
Identities:35/103(34%), Positives:50/103(48%), Gaps:17/103(16%)

Query  30     TDTTNQKTELQAIYLALQD--SGLEVNIVTDSQYAL-GIIQAQPDQSE------------  74
               DTTN + EL A   AL +     +V +VTDSQY + GI    P+  +
Sbjct  73518  PDTTNNRMELMAAIRALAELKRPCKVRLVTDSQYVMQGINDWMPNWKKRGWKTASKQPVK  73697

Query  75     -SELVNQIIEQLIKKEKVYLAWVPAHKGIGGNEQVDKLVSAGI  116
               ++L  Q+ EQ + + +V   WV  H G  GNEQ D L + G+
Sbjct  73698  NADLWQQLDEQ-VNRHEVSWQWVRGHTGHPGNEQADLLANRGV  73823
```

```
>Stutzerimonas stutzeri strain PS_234 NODE_10_length_111675_cov_16.9529, whole genome shotgun sequence
Sequence ID: NZ_RHQZ01000010.1 Length: 111675
Range 1: 37822 to 38127

Score:48.9 bits(115), Expect:0.004,
Method:Compositional matrix adjust.,
Identities:35/103(34%), Positives:50/103(48%), Gaps:17/103(16%)

Query  30     TDTTNQKTELQAIYLALQD--SGLEVNIVTDSQYAL-GIIQAQPDQSE------------  74
               DTTN + EL A   AL +     +V +VTDSQY + GI    P+  +
Sbjct  38127  PDTTNNRMELMAAIRALAELKRPCKVRLVTDSQYVMQGINDWMPNWKKRGWKTASKQPVK  37948

Query  75     -SELVNQIIEQLIKKEKVYLAWVPAHKGIGGNEQVDKLVSAGI  116
               ++L  Q+ EQ + + +V   WV  H G  GNEQ D L + G+
Sbjct  37947  NADLWQQLDEQ-VNRHEVSWQWVRGHTGHPGNEQADLLANRGV  37822
```

```
>Stutzerimonas stutzeri strain GD04161 18, whole genome shotgun sequence
Sequence ID: NZ_JAODYQ010000018.1 Length: 112063
Range 1: 38338 to 38643

Score:48.9 bits(115), Expect:0.004,
Method:Compositional matrix adjust.,
Identities:35/103(34%), Positives:50/103(48%), Gaps:17/103(16%)

Query  30     TDTTNQKTELQAIYLALQD--SGLEVNIVTDSQYAL-GIIQAQPDQSE------------  74
               DTTN + EL A   AL +     +V +VTDSQY + GI    P+  +
Sbjct  38643  PDTTNNRMELMAAIRALAELKRPCKVRLVTDSQYVMQGINDWMPNWKKRGWKTASKQPVK  38464

Query  75     -SELVNQIIEQLIKKEKVYLAWVPAHKGIGGNEQVDKLVSAGI  116
               ++L  Q+ EQ + + +V   WV  H G  GNEQ D L + G+
Sbjct  38463  NADLWQQLDEQ-VNRHEVSWQWVRGHTGHPGNEQADLLANRGV  38338
```

```
>Stutzerimonas stutzeri strain GD03774 16, whole genome shotgun sequence
Sequence ID: NZ_JAOCGO010000016.1 Length: 112070
Range 1: 73392 to 73697

Score:48.9 bits(115), Expect:0.004,
Method:Compositional matrix adjust.,
Identities:35/103(34%), Positives:50/103(48%), Gaps:17/103(16%)

Query  30     TDTTNQKTELQAIYLALQD--SGLEVNIVTDSQYAL-GIIQAQPDQSE------------  74
               DTTN + EL A   AL +     +V +VTDSQY + GI    P+  +
Sbjct  73392  PDTTNNRMELMAAIRALAELKRPCKVRLVTDSQYVMQGINDWMPNWKKRGWKTASKQPVK  73571

Query  75     -SELVNQIIEQLIKKEKVYLAWVPAHKGIGGNEQVDKLVSAGI  116
               ++L  Q+ EQ + + +V   WV  H G  GNEQ D L + G+
Sbjct  73572  NADLWQQLDEQ-VNRHEVSWQWVRGHTGHPGNEQADLLANRGV  73697
```

```
>Stutzerimonas stutzeri strain CM14 NODE_16_length_115186_cov_71.345649, whole genome shotgun sequence
Sequence ID: NZ_SRXR01000016.1 Length: 115186
Range 1: 73512 to 73817

Score:48.9 bits(115), Expect:0.004,
Method:Compositional matrix adjust.,
Identities:35/103(34%), Positives:50/103(48%), Gaps:17/103(16%)

Query  30     TDTTNQKTELQAIYLALQD--SGLEVNIVTDSQYAL-GIIQAQPDQSE------------  74
               DTTN + EL A   AL +     +V +VTDSQY + GI    P+  +
Sbjct  73512  PDTTNNRMELMAAIRALAELKRPCKVRLVTDSQYVMQGINDWMPNWKKRGWKTASKQPVK  73691

Query  75     -SELVNQIIEQLIKKEKVYLAWVPAHKGIGGNEQVDKLVSAGI  116
               ++L  Q+ EQ + + +V   WV  H G  GNEQ D L + G+
Sbjct  73692  NADLWQQLDEQ-VNRHEVSWQWVRGHTGHPGNEQADLLANRGV  73817
```

```
>Stutzerimonas stutzeri strain CM NODE_17_length_115190_cov_58.8358, whole genome shotgun sequence
Sequence ID: NZ_QLAB01000017.1 Length: 115190
Range 1: 73516 to 73821

Score:48.9 bits(115), Expect:0.004,
Method:Compositional matrix adjust.,
Identities:35/103(34%), Positives:50/103(48%), Gaps:17/103(16%)

Query  30     TDTTNQKTELQAIYLALQD--SGLEVNIVTDSQYAL-GIIQAQPDQSE------------  74
               DTTN + EL A   AL +     +V +VTDSQY + GI    P+  +
Sbjct  73516  PDTTNNRMELMAAIRALAELKRPCKVRLVTDSQYVMQGINDWMPNWKKRGWKTASKQPVK  73695

Query  75     -SELVNQIIEQLIKKEKVYLAWVPAHKGIGGNEQVDKLVSAGI  116
               ++L  Q+ EQ + + +V   WV  H G  GNEQ D L + G+
Sbjct  73696  NADLWQQLDEQ-VNRHEVSWQWVRGHTGHPGNEQADLLANRGV  73821
```

```
>Stutzerimonas stutzeri strain 40D2 NODE_16_length_115488_cov_67.7601_ID_31, whole genome shotgun sequence
Sequence ID: NZ_MWUI01000016.1 Length: 115488
Range 1: 41676 to 41981

Score:48.9 bits(115), Expect:0.004,
Method:Compositional matrix adjust.,
Identities:35/103(34%), Positives:50/103(48%), Gaps:17/103(16%)

Query  30     TDTTNQKTELQAIYLALQD--SGLEVNIVTDSQYAL-GIIQAQPDQSE------------  74
               DTTN + EL A   AL +     +V +VTDSQY + GI    P+  +
Sbjct  41981  PDTTNNRMELMAAIRALAELKRPCKVRLVTDSQYVMQGINDWMPNWKKRGWKTASKQPVK  41802

Query  75     -SELVNQIIEQLIKKEKVYLAWVPAHKGIGGNEQVDKLVSAGI  116
               ++L  Q+ EQ + + +V   WV  H G  GNEQ D L + G+
Sbjct  41801  NADLWQQLDEQ-VNRHEVSWQWVRGHTGHPGNEQADLLANRGV  41676
```

```
>Stutzerimonas stutzeri strain PS_138 NODE_12_length_125664_cov_16.9743, whole genome shotgun sequence
Sequence ID: NZ_RHRE01000012.1 Length: 125664
Range 1: 68096 to 68401

Score:48.9 bits(115), Expect:0.004,
Method:Compositional matrix adjust.,
Identities:35/103(34%), Positives:50/103(48%), Gaps:17/103(16%)

Query  30     TDTTNQKTELQAIYLALQD--SGLEVNIVTDSQYAL-GIIQAQPDQSE------------  74
               DTTN + EL A   AL +     +V +VTDSQY + GI    P+  +
Sbjct  68096  PDTTNNRMELMAAIRALAELKRPCKVRLVTDSQYVMQGINDWMPNWKKRGWKTASKQPVK  68275

Query  75     -SELVNQIIEQLIKKEKVYLAWVPAHKGIGGNEQVDKLVSAGI  116
               ++L  Q+ EQ + + +V   WV  H G  GNEQ D L + G+
Sbjct  68276  NADLWQQLDEQ-VNRHEVSWQWVRGHTGHPGNEQADLLANRGV  68401
```

```
>Stutzerimonas stutzeri strain NGHE31 NODE_11_length_130955_cov_43.307594, whole genome shotgun sequence
Sequence ID: NZ_JANAJI010000011.1 Length: 130955
Range 1: 57239 to 57544

Score:48.9 bits(115), Expect:0.004,
Method:Compositional matrix adjust.,
Identities:35/103(34%), Positives:50/103(48%), Gaps:17/103(16%)

Query  30     TDTTNQKTELQAIYLALQD--SGLEVNIVTDSQYAL-GIIQAQPDQSE------------  74
               DTTN + EL A   AL +     +V +VTDSQY + GI    P+  +
Sbjct  57544  PDTTNNRMELMAAIRALAELKRPCKVRLVTDSQYVMQGINDWMPNWKKRGWKTASKQPVK  57365

Query  75     -SELVNQIIEQLIKKEKVYLAWVPAHKGIGGNEQVDKLVSAGI  116
               ++L  Q+ EQ + + +V   WV  H G  GNEQ D L + G+
Sbjct  57364  NADLWQQLDEQ-VNRHEVSWQWVRGHTGHPGNEQADLLANRGV  57239
```

```
>Stutzerimonas stutzeri strain PS_001 NODE_7_length_130975_cov_100.528633, whole genome shotgun sequence
Sequence ID: NZ_RXOW01000007.1 Length: 130975
Range 1: 57263 to 57568

Score:48.9 bits(115), Expect:0.004,
Method:Compositional matrix adjust.,
Identities:35/103(34%), Positives:50/103(48%), Gaps:17/103(16%)

Query  30     TDTTNQKTELQAIYLALQD--SGLEVNIVTDSQYAL-GIIQAQPDQSE------------  74
               DTTN + EL A   AL +     +V +VTDSQY + GI    P+  +
Sbjct  57568  PDTTNNRMELMAAIRALAELKRPCKVRLVTDSQYVMQGINDWMPNWKKRGWKTASKQPVK  57389

Query  75     -SELVNQIIEQLIKKEKVYLAWVPAHKGIGGNEQVDKLVSAGI  116
               ++L  Q+ EQ + + +V   WV  H G  GNEQ D L + G+
Sbjct  57388  NADLWQQLDEQ-VNRHEVSWQWVRGHTGHPGNEQADLLANRGV  57263
```

```
>Stutzerimonas stutzeri strain PS_211 NODE_13_length_130975_cov_25.6613, whole genome shotgun sequence
Sequence ID: NZ_RHRA01000013.1 Length: 130975
Range 1: 73408 to 73713

Score:48.9 bits(115), Expect:0.004,
Method:Compositional matrix adjust.,
Identities:35/103(34%), Positives:50/103(48%), Gaps:17/103(16%)

Query  30     TDTTNQKTELQAIYLALQD--SGLEVNIVTDSQYAL-GIIQAQPDQSE------------  74
               DTTN + EL A   AL +     +V +VTDSQY + GI    P+  +
Sbjct  73408  PDTTNNRMELMAAIRALAELKRPCKVRLVTDSQYVMQGINDWMPNWKKRGWKTASKQPVK  73587

Query  75     -SELVNQIIEQLIKKEKVYLAWVPAHKGIGGNEQVDKLVSAGI  116
               ++L  Q+ EQ + + +V   WV  H G  GNEQ D L + G+
Sbjct  73588  NADLWQQLDEQ-VNRHEVSWQWVRGHTGHPGNEQADLLANRGV  73713
```

```
>Stutzerimonas stutzeri strain PS_072 NODE_10_length_135183_cov_17.9828, whole genome shotgun sequence
Sequence ID: NZ_RHRO01000010.1 Length: 135183
Range 1: 73540 to 73845

Score:48.9 bits(115), Expect:0.004,
Method:Compositional matrix adjust.,
Identities:35/103(34%), Positives:50/103(48%), Gaps:17/103(16%)
```

```
Query  30     TDTTNQKTELQAIYLALQD--SGLEVNIVTDSQYAL-GIIQAQPDQSE------------  74
               DTTN + EL A   AL +     +V +VTDSQY + GI     P+  +
Sbjct  73540  PDTTNNRMELMAAIRALAELKRPCKVRLVTDSQYVMQGINDWMPNWKKRGWKTASKQPVK  73719

Query  75     -SELVNQIIEQLIKKEKVYLAWVPAHKGIGGNEQVDKLVSAGI  116
               ++L  Q+ EQ + + +V   WV  H G  GNEQ D L + G+
Sbjct  73720  NADLWQQLDEQ-VNRHEVSWQWVRGHTGHPGNEQADLLANRGV  73845



>Stutzerimonas stutzeri strain PS_075 NODE_9_length_135224_cov_14.5207, whole genome shotgun sequence
Sequence ID: NZ_RHRN01000009.1 Length: 135224
Range 1: 61396 to 61701

Score:48.9 bits(115), Expect:0.004,
Method:Compositional matrix adjust.,
Identities:35/103(34%), Positives:50/103(48%), Gaps:17/103(16%)

Query  30     TDTTNQKTELQAIYLALQD--SGLEVNIVTDSQYAL-GIIQAQPDQSE------------  74
               DTTN + EL A   AL +     +V +VTDSQY + GI     P+  +
Sbjct  61701  PDTTNNRMELMAAIRALAELKRPCKVRLVTDSQYVMQGINDWMPNWKKRGWKTASKQPVK  61522

Query  75     -SELVNQIIEQLIKKEKVYLAWVPAHKGIGGNEQVDKLVSAGI  116
               ++L  Q+ EQ + + +V   WV  H G  GNEQ D L + G+
Sbjct  61521  NADLWQQLDEQ-VNRHEVSWQWVRGHTGHPGNEQADLLANRGV  61396



>Stutzerimonas stutzeri strain GD04143 6, whole genome shotgun sequence
Sequence ID: NZ_JAODZI010000006.1 Length: 166685
Range 1: 93085 to 93390

Score:48.9 bits(115), Expect:0.004,
Method:Compositional matrix adjust.,
Identities:35/103(34%), Positives:50/103(48%), Gaps:17/103(16%)

Query  30     TDTTNQKTELQAIYLALQD--SGLEVNIVTDSQYAL-GIIQAQPDQSE------------  74
               DTTN + EL A   AL +     +V +VTDSQY + GI     P+  +
Sbjct  93390  PDTTNNRMELMAAIRALAELKRPCKVRLVTDSQYVMQGINDWMPNWKKRGWKTASKQPVK  93211

Query  75     -SELVNQIIEQLIKKEKVYLAWVPAHKGIGGNEQVDKLVSAGI  116
               ++L  Q+ EQ + + +V   WV  H G  GNEQ D L + G+
Sbjct  93210  NADLWQQLDEQ-VNRHEVSWQWVRGHTGHPGNEQADLLANRGV  93085



>Stutzerimonas stutzeri strain GD04147 6, whole genome shotgun sequence
Sequence ID: NZ_JAODZE010000006.1 Length: 166685
Range 1: 93085 to 93390

Score:48.9 bits(115), Expect:0.004,
Method:Compositional matrix adjust.,
Identities:35/103(34%), Positives:50/103(48%), Gaps:17/103(16%)

Query  30     TDTTNQKTELQAIYLALQD--SGLEVNIVTDSQYAL-GIIQAQPDQSE------------  74
               DTTN + EL A   AL +     +V +VTDSQY + GI     P+  +
Sbjct  93390  PDTTNNRMELMAAIRALAELKRPCKVRLVTDSQYVMQGINDWMPNWKKRGWKTASKQPVK  93211

Query  75     -SELVNQIIEQLIKKEKVYLAWVPAHKGIGGNEQVDKLVSAGI  116
               ++L  Q+ EQ + + +V   WV  H G  GNEQ D L + G+
Sbjct  93210  NADLWQQLDEQ-VNRHEVSWQWVRGHTGHPGNEQADLLANRGV  93085



>Stutzerimonas stutzeri strain PS_257 NODE_3_length_183596_cov_42.2268, whole genome shotgun sequence
Sequence ID: NZ_RHQY01000003.1 Length: 183596
Range 1: 73489 to 73794

Score:48.9 bits(115), Expect:0.004,
Method:Compositional matrix adjust.,
Identities:35/103(34%), Positives:50/103(48%), Gaps:17/103(16%)

Query  30     TDTTNQKTELQAIYLALQD--SGLEVNIVTDSQYAL-GIIQAQPDQSE------------  74
               DTTN + EL A   AL +     +V +VTDSQY + GI     P+  +
Sbjct  73489  PDTTNNRMELMAAIRALAELKRPCKVRLVTDSQYVMQGINDWMPNWKKRGWKTASKQPVK  73668

Query  75     -SELVNQIIEQLIKKEKVYLAWVPAHKGIGGNEQVDKLVSAGI  116
               ++L  Q+ EQ + + +V   WV  H G  GNEQ D L + G+
Sbjct  73669  NADLWQQLDEQ-VNRHEVSWQWVRGHTGHPGNEQADLLANRGV  73794



>Stutzerimonas stutzeri strain GD04144 10, whole genome shotgun sequence
Sequence ID: NZ_JAODZH010000010.1 Length: 184200
Range 1: 110508 to 110813

Score:48.9 bits(115), Expect:0.004,
Method:Compositional matrix adjust.,
Identities:35/103(34%), Positives:50/103(48%), Gaps:17/103(16%)

Query  30      TDTTNQKTELQAIYLALQD--SGLEVNIVTDSQYAL-GIIQAQPDQSE------------  74
                DTTN + EL A   AL +     +V +VTDSQY + GI     P+  +
Sbjct  110813  PDTTNNRMELMAAIRALAELKRPCKVRLVTDSQYVMQGINDWMPNWKKRGWKTASKQPVK  110634

Query  75      -SELVNQIIEQLIKKEKVYLAWVPAHKGIGGNEQVDKLVSAGI  116
                ++L  Q+ EQ + + +V   WV  H G  GNEQ D L + G+
Sbjct  110633  NADLWQQLDEQ-VNRHEVSWQWVRGHTGHPGNEQADLLANRGV  110508



>Stutzerimonas stutzeri strain GD03747 7, whole genome shotgun sequence
Sequence ID: NZ_JAOCHP010000007.1 Length: 199808
Range 1: 175368 to 175673

Score:48.9 bits(115), Expect:0.004,
Method:Compositional matrix adjust.,
Identities:35/103(34%), Positives:50/103(48%), Gaps:17/103(16%)

Query  30      TDTTNQKTELQAIYLALQD--SGLEVNIVTDSQYAL-GIIQAQPDQSE------------  74
                DTTN + EL A   AL +     +V +VTDSQY + GI     P+  +
Sbjct  175673  PDTTNNRMELMAAIRALAELKRPCKVRLVTDSQYVMQGINDWMPNWKKRGWKTASKQPVK  175494

Query  75      -SELVNQIIEQLIKKEKVYLAWVPAHKGIGGNEQVDKLVSAGI  116
                ++L  Q+ EQ + + +V   WV  H G  GNEQ D L + G+
Sbjct  175493  NADLWQQLDEQ-VNRHEVSWQWVRGHTGHPGNEQADLLANRGV  175368
```

```
>Stutzerimonas stutzeri strain 109A1 NODE_6_length_231087_cov_45.093358, whole genome shotgun sequence
Sequence ID: NZ_JAAMRD010000006.1 Length: 231087
Range 1: 157253 to 157558

Score:48.9 bits(115), Expect:0.004,
Method:Compositional matrix adjust.,
Identities:35/103(34%), Positives:50/103(48%), Gaps:17/103(16%)

Query  30      TDTTNQKTELQAIYLALQD--SGLEVNIVTDSQYAL-GIIQAQPDQSE------------  74
                DTTN + EL A   AL +     +V +VTDSQY + GI    P+  +
Sbjct  157558  PDTTNNRMELMAAIRALAELKRPCKVRLVTDSQYVMQGINDWMPNWKKRGWKTASKQPVK  157379

Query  75      -SELVNQIIEQLIKKEKVYLAWVPAHKGIGGNEQVDKLVSAGI  116
                ++L  Q+ EQ + + +V   WV  H G  GNEQ D L + G+
Sbjct  157378  NADLWQQLDEQ-VNRHEVSWQWVRGHTGHPGNEQADLLANRGV  157253
```

```
>Stutzerimonas stutzeri strain PAH8 NODE_11_length_231702_cov_23.415533, whole genome shotgun sequence
Sequence ID: NZ_JAGFOX010000011.1 Length: 231702
Range 1: 72678 to 72983

Score:48.9 bits(115), Expect:0.004,
Method:Compositional matrix adjust.,
Identities:35/103(34%), Positives:50/103(48%), Gaps:17/103(16%)

Query  30     TDTTNQKTELQAIYLALQD--SGLEVNIVTDSQYAL-GIIQAQPDQSE------------  74
               DTTN + EL A   AL +     +V +VTDSQY + GI    P+  +
Sbjct  72678  PDTTNNRMELMAAIRALAELKRPCKVRLVTDSQYVMQGINDWMPNWKKRGWKTASKQPVK  72857

Query  75     -SELVNQIIEQLIKKEKVYLAWVPAHKGIGGNEQVDKLVSAGI  116
               ++L  Q+ EQ + + +V   WV  H G  GNEQ D L + G+
Sbjct  72858  NADLWQQLDEQ-VNRHEVSWQWVRGHTGHPGNEQADLLANRGV  72983
```

```
>Stutzerimonas stutzeri strain SDU10 Scaffold8, whole genome shotgun sequence
Sequence ID: NZ_JAJHZH010000008.1 Length: 233818
Range 1: 73400 to 73705

Score:48.9 bits(115), Expect:0.004,
Method:Compositional matrix adjust.,
Identities:35/103(34%), Positives:50/103(48%), Gaps:17/103(16%)

Query  30     TDTTNQKTELQAIYLALQD--SGLEVNIVTDSQYAL-GIIQAQPDQSE------------  74
               DTTN + EL A   AL +     +V +VTDSQY + GI    P+  +
Sbjct  73400  PDTTNNRMELMAAIRALAELKRPCKVRLVTDSQYVMQGINDWMPNWKKRGWKTASKQPVK  73579

Query  75     -SELVNQIIEQLIKKEKVYLAWVPAHKGIGGNEQVDKLVSAGI  116
               ++L  Q+ EQ + + +V   WV  H G  GNEQ D L + G+
Sbjct  73580  NADLWQQLDEQ-VNRHEVSWQWVRGHTGHPGNEQADLLANRGV  73705
```

```
>Stutzerimonas stutzeri strain AR9-4 contig_1, whole genome shotgun sequence
Sequence ID: NZ_MDGV01000001.1 Length: 233982
Range 1: 160174 to 160479

Score:48.9 bits(115), Expect:0.004,
Method:Compositional matrix adjust.,
Identities:35/103(34%), Positives:50/103(48%), Gaps:17/103(16%)

Query  30      TDTTNQKTELQAIYLALQD--SGLEVNIVTDSQYAL-GIIQAQPDQSE------------  74
                DTTN + EL A   AL +     +V +VTDSQY + GI    P+  +
Sbjct  160479  PDTTNNRMELMAAIRALAELKRPCKVRLVTDSQYVMQGINDWMPNWKKRGWKTASKQPVK  160300

Query  75      -SELVNQIIEQLIKKEKVYLAWVPAHKGIGGNEQVDKLVSAGI  116
                ++L  Q+ EQ + + +V   WV  H G  GNEQ D L + G+
Sbjct  160299  NADLWQQLDEQ-VNRHEVSWQWVRGHTGHPGNEQADLLANRGV  160174
```

```
>Stutzerimonas stutzeri strain DIA-8 S.stutzeri8_contig_8, whole genome shotgun sequence
Sequence ID: NZ_JAZGZT010000008.1 Length: 234537
Range 1: 73370 to 73675

Score:48.9 bits(115), Expect:0.004,
Method:Compositional matrix adjust.,
Identities:35/103(34%), Positives:50/103(48%), Gaps:17/103(16%)

Query  30     TDTTNQKTELQAIYLALQD--SGLEVNIVTDSQYAL-GIIQAQPDQSE------------  74
               DTTN + EL A   AL +     +V +VTDSQY + GI    P+  +
Sbjct  73370  PDTTNNRMELMAAIRALAELKRPCKVRLVTDSQYVMQGINDWMPNWKKRGWKTASKQPVK  73549

Query  75     -SELVNQIIEQLIKKEKVYLAWVPAHKGIGGNEQVDKLVSAGI  116
               ++L  Q+ EQ + + +V   WV  H G  GNEQ D L + G+
Sbjct  73550  NADLWQQLDEQ-VNRHEVSWQWVRGHTGHPGNEQADLLANRGV  73675
```

```
>Stutzerimonas stutzeri isolate PSEUDT2.31D-1, whole genome shotgun sequence
Sequence ID: NZ_CAJFAG010000045.1 Length: 237644
Range 1: 166235 to 166540

Score:48.9 bits(115), Expect:0.004,
Method:Compositional matrix adjust.,
Identities:35/103(34%), Positives:50/103(48%), Gaps:17/103(16%)

Query  30      TDTTNQKTELQAIYLALQD--SGLEVNIVTDSQYAL-GIIQAQPDQSE------------  74
                DTTN + EL A   AL +     +V +VTDSQY + GI    P+  +
Sbjct  166540  PDTTNNRMELMAAIRALAELKRPCKVRLVTDSQYVMQGINDWMPNWKKRGWKTASKQPVK  166361

Query  75      -SELVNQIIEQLIKKEKVYLAWVPAHKGIGGNEQVDKLVSAGI  116
                ++L  Q+ EQ + + +V   WV  H G  GNEQ D L + G+
Sbjct  166360  NADLWQQLDEQ-VNRHEVSWQWVRGHTGHPGNEQADLLANRGV  166235
```

```
>Stutzerimonas stutzeri strain GD04155 4, whole genome shotgun sequence
Sequence ID: NZ_JAODYW010000004.1 Length: 238078
Range 1: 73422 to 73727

Score:48.9 bits(115), Expect:0.004,
Method:Compositional matrix adjust.,
Identities:35/103(34%), Positives:50/103(48%), Gaps:17/103(16%)
```

```
Query  30     TDTTNQKTELQAIYLALQD--SGLEVNIVTDSQYAL-GIIQAQPDQSE------------  74
               DTTN + EL A   AL +     +V +VTDSQY + GI    P+  +
Sbjct  73422  PDTTNNRMELMAAIRALAELKRPCKVRLVTDSQYVMQGINDWMPNWKKRGWKTASKQPVK  73601

Query  75     -SELVNQIIEQLIKKEKVYLAWVPAHKGIGGNEQVDKLVSAGI  116
               ++L  Q+ EQ + + +V   WV  H G  GNEQ D L + G+
Sbjct  73602  NADLWQQLDEQ-VNRHEVSWQWVRGHTGHPGNEQADLLANRGV  73727
```

>Stutzerimonas stutzeri strain GD03864 1, whole genome shotgun sequence
Sequence ID: NZ_JAOCDG010000001.1 Length: 249048
Range 1: 175321 to 175626

Score:48.9 bits(115), Expect:0.004,
Method:Compositional matrix adjust.,
Identities:35/103(34%), Positives:50/103(48%), Gaps:17/103(16%)

```
Query  30      TDTTNQKTELQAIYLALQD--SGLEVNIVTDSQYAL-GIIQAQPDQSE------------  74
                DTTN + EL A   AL +     +V +VTDSQY + GI    P+  +
Sbjct  175626  PDTTNNRMELMAAIRALAELKRPCKVRLVTDSQYVMQGINDWMPNWKKRGWKTASKQPVK  175447

Query  75      -SELVNQIIEQLIKKEKVYLAWVPAHKGIGGNEQVDKLVSAGI  116
                ++L  Q+ EQ + + +V   WV  H G  GNEQ D L + G+
Sbjct  175446  NADLWQQLDEQ-VNRHEVSWQWVRGHTGHPGNEQADLLANRGV  175321
```

>Stutzerimonas stutzeri strain PS_366 NODE_4_length_258367_cov_46.287870, whole genome shotgun sequence
Sequence ID: NZ_RHQX01000004.1 Length: 258367
Range 1: 184574 to 184879

Score:48.9 bits(115), Expect:0.004,
Method:Compositional matrix adjust.,
Identities:35/103(34%), Positives:50/103(48%), Gaps:17/103(16%)

```
Query  30      TDTTNQKTELQAIYLALQD--SGLEVNIVTDSQYAL-GIIQAQPDQSE------------  74
                DTTN + EL A   AL +     +V +VTDSQY + GI    P+  +
Sbjct  184879  PDTTNNRMELMAAIRALAELKRPCKVRLVTDSQYVMQGINDWMPNWKKRGWKTASKQPVK  184700

Query  75      -SELVNQIIEQLIKKEKVYLAWVPAHKGIGGNEQVDKLVSAGI  116
                ++L  Q+ EQ + + +V   WV  H G  GNEQ D L + G+
Sbjct  184699  NADLWQQLDEQ-VNRHEVSWQWVRGHTGHPGNEQADLLANRGV  184574
```

>Stutzerimonas stutzeri strain 4MP-B contig_5, whole genome shotgun sequence
Sequence ID: NZ_RBEA01000005.1 Length: 276576
Range 1: 202395 to 202700

Score:48.9 bits(115), Expect:0.004,
Method:Compositional matrix adjust.,
Identities:35/103(34%), Positives:50/103(48%), Gaps:17/103(16%)

```
Query  30      TDTTNQKTELQAIYLALQD--SGLEVNIVTDSQYAL-GIIQAQPDQSE------------  74
                DTTN + EL A   AL +     +V +VTDSQY + GI    P+  +
Sbjct  202700  PDTTNNRMELMAAIRALAELKRPCKVRLVTDSQYVMQGINDWMPNWKKRGWKTASKQPVK  202521

Query  75      -SELVNQIIEQLIKKEKVYLAWVPAHKGIGGNEQVDKLVSAGI  116
                ++L  Q+ EQ + + +V   WV  H G  GNEQ D L + G+
Sbjct  202520  NADLWQQLDEQ-VNRHEVSWQWVRGHTGHPGNEQADLLANRGV  202395
```

>Stutzerimonas stutzeri strain PS_134 NODE_7_length_278675_cov_20.2503, whole genome shotgun sequence
Sequence ID: NZ_RHRF01000007.1 Length: 278675
Range 1: 73502 to 73807

Score:48.9 bits(115), Expect:0.004,
Method:Compositional matrix adjust.,
Identities:35/103(34%), Positives:50/103(48%), Gaps:17/103(16%)

```
Query  30     TDTTNQKTELQAIYLALQD--SGLEVNIVTDSQYAL-GIIQAQPDQSE------------  74
               DTTN + EL A   AL +     +V +VTDSQY + GI    P+  +
Sbjct  73502  PDTTNNRMELMAAIRALAELKRPCKVRLVTDSQYVMQGINDWMPNWKKRGWKTASKQPVK  73681

Query  75     -SELVNQIIEQLIKKEKVYLAWVPAHKGIGGNEQVDKLVSAGI  116
               ++L  Q+ EQ + + +V   WV  H G  GNEQ D L + G+
Sbjct  73682  NADLWQQLDEQ-VNRHEVSWQWVRGHTGHPGNEQADLLANRGV  73807
```

>Stutzerimonas stutzeri strain PS_133 NODE_6_length_278690_cov_22.099, whole genome shotgun sequence
Sequence ID: NZ_RHRG01000006.1 Length: 278690
Range 1: 73502 to 73807

Score:48.9 bits(115), Expect:0.004,
Method:Compositional matrix adjust.,
Identities:35/103(34%), Positives:50/103(48%), Gaps:17/103(16%)

```
Query  30     TDTTNQKTELQAIYLALQD--SGLEVNIVTDSQYAL-GIIQAQPDQSE------------  74
               DTTN + EL A   AL +     +V +VTDSQY + GI    P+  +
Sbjct  73502  PDTTNNRMELMAAIRALAELKRPCKVRLVTDSQYVMQGINDWMPNWKKRGWKTASKQPVK  73681

Query  75     -SELVNQIIEQLIKKEKVYLAWVPAHKGIGGNEQVDKLVSAGI  116
               ++L  Q+ EQ + + +V   WV  H G  GNEQ D L + G+
Sbjct  73682  NADLWQQLDEQ-VNRHEVSWQWVRGHTGHPGNEQADLLANRGV  73807
```

>Stutzerimonas stutzeri strain PS_130 NODE_7_length_278690_cov_19.9998, whole genome shotgun sequence
Sequence ID: NZ_RHRI01000007.1 Length: 278690
Range 1: 73502 to 73807

Score:48.9 bits(115), Expect:0.004,
Method:Compositional matrix adjust.,
Identities:35/103(34%), Positives:50/103(48%), Gaps:17/103(16%)

```
Query  30     TDTTNQKTELQAIYLALQD--SGLEVNIVTDSQYAL-GIIQAQPDQSE------------  74
               DTTN + EL A   AL +     +V +VTDSQY + GI    P+  +
Sbjct  73502  PDTTNNRMELMAAIRALAELKRPCKVRLVTDSQYVMQGINDWMPNWKKRGWKTASKQPVK  73681

Query  75     -SELVNQIIEQLIKKEKVYLAWVPAHKGIGGNEQVDKLVSAGI  116
               ++L  Q+ EQ + + +V   WV  H G  GNEQ D L + G+
Sbjct  73682  NADLWQQLDEQ-VNRHEVSWQWVRGHTGHPGNEQADLLANRGV  73807
```

```
>Stutzerimonas stutzeri strain PS_131 NODE_5_length_278962_cov_20.0832, whole genome shotgun sequence
Sequence ID: NZ_RHRH01000005.1 Length: 278962
Range 1: 73776 to 74081

Score:48.9 bits(115), Expect:0.004,
Method:Compositional matrix adjust.,
Identities:35/103(34%), Positives:50/103(48%), Gaps:17/103(16%)

Query  30     TDTTNQKTELQAIYLALQD--SGLEVNIVTDSQYAL-GIIQAQPDQSE------------  74
               DTTN + EL A   AL +     +V +VTDSQY + GI    P+  +
Sbjct  73776  PDTTNNRMELMAAIRALAELKRPCKVRLVTDSQYVMQGINDWMPNWKKRGWKTASKQPVK  73955

Query  75     -SELVNQIIEQLIKKEKVYLAWVPAHKGIGGNEQVDKLVSAGI  116
               ++L  Q+ EQ + + +V   WV  H G  GNEQ D L + G+
Sbjct  73956  NADLWQQLDEQ-VNRHEVSWQWVRGHTGHPGNEQADLLANRGV  74081
```

```
>Stutzerimonas stutzeri strain PS_050 NODE_6_length_280932_cov_15.3034, whole genome shotgun sequence
Sequence ID: NZ_RIAN01000006.1 Length: 280932
Range 1: 73755 to 74060

Score:48.9 bits(115), Expect:0.004,
Method:Compositional matrix adjust.,
Identities:35/103(34%), Positives:50/103(48%), Gaps:17/103(16%)

Query  30     TDTTNQKTELQAIYLALQD--SGLEVNIVTDSQYAL-GIIQAQPDQSE------------  74
               DTTN + EL A   AL +     +V +VTDSQY + GI    P+  +
Sbjct  73755  PDTTNNRMELMAAIRALAELKRPCKVRLVTDSQYVMQGINDWMPNWKKRGWKTASKQPVK  73934

Query  75     -SELVNQIIEQLIKKEKVYLAWVPAHKGIGGNEQVDKLVSAGI  116
               ++L  Q+ EQ + + +V   WV  H G  GNEQ D L + G+
Sbjct  73935  NADLWQQLDEQ-VNRHEVSWQWVRGHTGHPGNEQADLLANRGV  74060
```

```
>Stutzerimonas stutzeri strain G434 17141_contig_1, whole genome shotgun sequence
Sequence ID: NZ_JASGXY010000001.1 Length: 328625
Range 1: 73411 to 73716

Score:48.9 bits(115), Expect:0.004,
Method:Compositional matrix adjust.,
Identities:35/103(34%), Positives:50/103(48%), Gaps:17/103(16%)

Query  30     TDTTNQKTELQAIYLALQD--SGLEVNIVTDSQYAL-GIIQAQPDQSE------------  74
               DTTN + EL A   AL +     +V +VTDSQY + GI    P+  +
Sbjct  73411  PDTTNNRMELMAAIRALAELKRPCKVRLVTDSQYVMQGINDWMPNWKKRGWKTASKQPVK  73590

Query  75     -SELVNQIIEQLIKKEKVYLAWVPAHKGIGGNEQVDKLVSAGI  116
               ++L  Q+ EQ + + +V   WV  H G  GNEQ D L + G+
Sbjct  73591  NADLWQQLDEQ-VNRHEVSWQWVRGHTGHPGNEQADLLANRGV  73716
```

```
>Stutzerimonas stutzeri strain GD04107 5, whole genome shotgun sequence
Sequence ID: NZ_JAOEAQ010000005.1 Length: 336744
Range 1: 73387 to 73692

Score:48.9 bits(115), Expect:0.004,
Method:Compositional matrix adjust.,
Identities:35/103(34%), Positives:50/103(48%), Gaps:17/103(16%)

Query  30     TDTTNQKTELQAIYLALQD--SGLEVNIVTDSQYAL-GIIQAQPDQSE------------  74
               DTTN + EL A   AL +     +V +VTDSQY + GI    P+  +
Sbjct  73387  PDTTNNRMELMAAIRALAELKRPCKVRLVTDSQYVMQGINDWMPNWKKRGWKTASKQPVK  73566

Query  75     -SELVNQIIEQLIKKEKVYLAWVPAHKGIGGNEQVDKLVSAGI  116
               ++L  Q+ EQ + + +V   WV  H G  GNEQ D L + G+
Sbjct  73567  NADLWQQLDEQ-VNRHEVSWQWVRGHTGHPGNEQADLLANRGV  73692
```

```
>Stutzerimonas stutzeri strain PS_125 NODE_5_length_336950_cov_18.2593, whole genome shotgun sequence
Sequence ID: NZ_RHRK01000005.1 Length: 336950
Range 1: 263130 to 263435

Score:48.9 bits(115), Expect:0.004,
Method:Compositional matrix adjust.,
Identities:35/103(34%), Positives:50/103(48%), Gaps:17/103(16%)

Query  30      TDTTNQKTELQAIYLALQD--SGLEVNIVTDSQYAL-GIIQAQPDQSE------------  74
                DTTN + EL A   AL +     +V +VTDSQY + GI    P+  +
Sbjct  263435  PDTTNNRMELMAAIRALAELKRPCKVRLVTDSQYVMQGINDWMPNWKKRGWKTASKQPVK  263256

Query  75      -SELVNQIIEQLIKKEKVYLAWVPAHKGIGGNEQVDKLVSAGI  116
                ++L  Q+ EQ + + +V   WV  H G  GNEQ D L + G+
Sbjct  263255  NADLWQQLDEQ-VNRHEVSWQWVRGHTGHPGNEQADLLANRGV  263130
```

```
>Stutzerimonas stutzeri strain PS_087 NODE_4_length_352814_cov_14.3544, whole genome shotgun sequence
Sequence ID: NZ_RHRM01000004.1 Length: 352814
Range 1: 73540 to 73845

Score:48.9 bits(115), Expect:0.004,
Method:Compositional matrix adjust.,
Identities:35/103(34%), Positives:50/103(48%), Gaps:17/103(16%)

Query  30     TDTTNQKTELQAIYLALQD--SGLEVNIVTDSQYAL-GIIQAQPDQSE------------  74
               DTTN + EL A   AL +     +V +VTDSQY + GI    P+  +
Sbjct  73540  PDTTNNRMELMAAIRALAELKRPCKVRLVTDSQYVMQGINDWMPNWKKRGWKTASKQPVK  73719

Query  75     -SELVNQIIEQLIKKEKVYLAWVPAHKGIGGNEQVDKLVSAGI  116
               ++L  Q+ EQ + + +V   WV  H G  GNEQ D L + G+
Sbjct  73720  NADLWQQLDEQ-VNRHEVSWQWVRGHTGHPGNEQADLLANRGV  73845
```

```
>Stutzerimonas stutzeri strain PS_066 NODE_2_length_359430_cov_15.0875, whole genome shotgun sequence
Sequence ID: NZ_RHRP01000002.1 Length: 359430
Range 1: 291030 to 291335

Score:48.9 bits(115), Expect:0.004,
Method:Compositional matrix adjust.,
Identities:35/103(34%), Positives:50/103(48%), Gaps:17/103(16%)

Query  30      TDTTNQKTELQAIYLALQD--SGLEVNIVTDSQYAL-GIIQAQPDQSE------------  74
```

```
                DTTN + EL A   AL +     +V +VTDSQY + GI    P+  +
Sbjct  291335  PDTTNNRMELMAAIRALAELKRPCKVRLVTDSQYVMQGINDWMPNWKKRGWKTASKQPVK  291156

Query  75      -SELVNQIIEQLIKKEKVYLAWVPAHKGIGGNEQVDKLVSAGI  116
                ++L  Q+ EQ + + +V   WV  H G  GNEQ D L + G+
Sbjct  291155  NADLWQQLDEQ-VNRHEVSWQWVRGHTGHPGNEQADLLANRGV  291030
```

```
>Stutzerimonas stutzeri strain GD04118 4, whole genome shotgun sequence
Sequence ID: NZ_JAOEAF010000004.1 Length: 364296
Range 1: 71319 to 71624

Score:48.9 bits(115), Expect:0.004,
Method:Compositional matrix adjust.,
Identities:35/103(34%), Positives:50/103(48%), Gaps:17/103(16%)

Query  30     TDTTNQKTELQAIYLALQD--SGLEVNIVTDSQYAL-GIIQAQPDQSE------------  74
               DTTN + EL A   AL +     +V +VTDSQY + GI    P+  +
Sbjct  71319  PDTTNNRMELMAAIRALAELKRPCKVRLVTDSQYVMQGINDWMPNWKKRGWKTASKQPVK  71498

Query  75     -SELVNQIIEQLIKKEKVYLAWVPAHKGIGGNEQVDKLVSAGI  116
               ++L  Q+ EQ + + +V   WV  H G  GNEQ D L + G+
Sbjct  71499  NADLWQQLDEQ-VNRHEVSWQWVRGHTGHPGNEQADLLANRGV  71624
```

```
>Stutzerimonas stutzeri strain G432 17142_contig_21, whole genome shotgun sequence
Sequence ID: NZ_JASGXX010000021.1 Length: 367318
Range 1: 73586 to 73891

Score:48.9 bits(115), Expect:0.004,
Method:Compositional matrix adjust.,
Identities:35/103(34%), Positives:50/103(48%), Gaps:17/103(16%)

Query  30     TDTTNQKTELQAIYLALQD--SGLEVNIVTDSQYAL-GIIQAQPDQSE------------  74
               DTTN + EL A   AL +     +V +VTDSQY + GI    P+  +
Sbjct  73586  PDTTNNRMELMAAIRALAELKRPCKVRLVTDSQYVMQGINDWMPNWKKRGWKTASKQPVK  73765

Query  75     -SELVNQIIEQLIKKEKVYLAWVPAHKGIGGNEQVDKLVSAGI  116
               ++L  Q+ EQ + + +V   WV  H G  GNEQ D L + G+
Sbjct  73766  NADLWQQLDEQ-VNRHEVSWQWVRGHTGHPGNEQADLLANRGV  73891
```

```
>Stutzerimonas stutzeri strain GD03946 3, whole genome shotgun sequence
Sequence ID: NZ_JAOCAF010000003.1 Length: 401344
Range 1: 67992 to 68297

Score:48.9 bits(115), Expect:0.004,
Method:Compositional matrix adjust.,
Identities:35/103(34%), Positives:50/103(48%), Gaps:17/103(16%)

Query  30     TDTTNQKTELQAIYLALQD--SGLEVNIVTDSQYAL-GIIQAQPDQSE------------  74
               DTTN + EL A   AL +     +V +VTDSQY + GI    P+  +
Sbjct  67992  PDTTNNRMELMAAIRALAELKRPCKVRLVTDSQYVMQGINDWMPNWKKRGWKTASKQPVK  68171

Query  75     -SELVNQIIEQLIKKEKVYLAWVPAHKGIGGNEQVDKLVSAGI  116
               ++L  Q+ EQ + + +V   WV  H G  GNEQ D L + G+
Sbjct  68172  NADLWQQLDEQ-VNRHEVSWQWVRGHTGHPGNEQADLLANRGV  68297
```

```
>Stutzerimonas stutzeri strain GD03845 1, whole genome shotgun sequence
Sequence ID: NZ_JAOCDX010000001.1 Length: 423848
Range 1: 73296 to 73601

Score:48.9 bits(115), Expect:0.004,
Method:Compositional matrix adjust.,
Identities:35/103(34%), Positives:50/103(48%), Gaps:17/103(16%)

Query  30     TDTTNQKTELQAIYLALQD--SGLEVNIVTDSQYAL-GIIQAQPDQSE------------  74
               DTTN + EL A   AL +     +V +VTDSQY + GI    P+  +
Sbjct  73296  PDTTNNRMELMAAIRALAELKRPCKVRLVTDSQYVMQGINDWMPNWKKRGWKTASKQPVK  73475

Query  75     -SELVNQIIEQLIKKEKVYLAWVPAHKGIGGNEQVDKLVSAGI  116
               ++L  Q+ EQ + + +V   WV  H G  GNEQ D L + G+
Sbjct  73476  NADLWQQLDEQ-VNRHEVSWQWVRGHTGHPGNEQADLLANRGV  73601
```

```
>Stutzerimonas stutzeri strain NT0124 211_430101_56647634_15539_, whole genome shotgun sequence
Sequence ID: NZ_JXTL01000024.1 Length: 430101
Range 1: 350811 to 351116

Score:48.9 bits(115), Expect:0.004,
Method:Compositional matrix adjust.,
Identities:35/103(34%), Positives:50/103(48%), Gaps:17/103(16%)

Query  30      TDTTNQKTELQAIYLALQD--SGLEVNIVTDSQYAL-GIIQAQPDQSE------------  74
                DTTN + EL A   AL +     +V +VTDSQY + GI    P+  +
Sbjct  351116  PDTTNNRMELMAAIRALAELKRPCKVRLVTDSQYVMQGINDWMPNWKKRGWKTASKQPVK  350937

Query  75      -SELVNQIIEQLIKKEKVYLAWVPAHKGIGGNEQVDKLVSAGI  116
                ++L  Q+ EQ + + +V   WV  H G  GNEQ D L + G+
Sbjct  350936  NADLWQQLDEQ-VNRHEVSWQWVRGHTGHPGNEQADLLANRGV  350811
```

```
>Stutzerimonas stutzeri strain G431 17143_contig_2, whole genome shotgun sequence
Sequence ID: NZ_JASGXW010000002.1 Length: 463823
Range 1: 389933 to 390238

Score:48.9 bits(115), Expect:0.004,
Method:Compositional matrix adjust.,
Identities:35/103(34%), Positives:50/103(48%), Gaps:17/103(16%)

Query  30      TDTTNQKTELQAIYLALQD--SGLEVNIVTDSQYAL-GIIQAQPDQSE------------  74
                DTTN + EL A   AL +     +V +VTDSQY + GI    P+  +
Sbjct  390238  PDTTNNRMELMAAIRALAELKRPCKVRLVTDSQYVMQGINDWMPNWKKRGWKTASKQPVK  390059

Query  75      -SELVNQIIEQLIKKEKVYLAWVPAHKGIGGNEQVDKLVSAGI  116
                ++L  Q+ EQ + + +V   WV  H G  GNEQ D L + G+
Sbjct  390058  NADLWQQLDEQ-VNRHEVSWQWVRGHTGHPGNEQADLLANRGV  389933
```

```
>Stutzerimonas stutzeri strain 69C1 NODE_5_length_470705_cov_45.205883, whole genome shotgun sequence
Sequence ID: NZ_JAAMQV010000005.1 Length: 470705
Range 1: 396857 to 397162

Score:48.9 bits(115), Expect:0.004,
Method:Compositional matrix adjust.,
Identities:35/103(34%), Positives:50/103(48%), Gaps:17/103(16%)

Query  30      TDTTNQKTELQAIYLALQD--SGLEVNIVTDSQYAL-GIIQAQPDQSE------------  74
                DTTN + EL A   AL +     +V +VTDSQY + GI    P+  +
Sbjct  397162  PDTTNNRMELMAAIRALAELKRPCKVRLVTDSQYVMQGINDWMPNWKKRGWKTASKQPVK  396983

Query  75      -SELVNQIIEQLIKKEKVYLAWVPAHKGIGGNEQVDKLVSAGI  116
                ++L  Q+ EQ + + +V   WV  H G  GNEQ D L + G+
Sbjct  396982  NADLWQQLDEQ-VNRHEVSWQWVRGHTGHPGNEQADLLANRGV  396857



>Stutzerimonas stutzeri strain MP4687 NODE_3_length_579614_cov_54.2791, whole genome shotgun sequence
Sequence ID: NZ_WWNS01000003.1 Length: 579614
Range 1: 73510 to 73815

Score:48.9 bits(115), Expect:0.004,
Method:Compositional matrix adjust.,
Identities:35/103(34%), Positives:50/103(48%), Gaps:17/103(16%)

Query  30     TDTTNQKTELQAIYLALQD--SGLEVNIVTDSQYAL-GIIQAQPDQSE------------  74
               DTTN + EL A   AL +     +V +VTDSQY + GI    P+  +
Sbjct  73510  PDTTNNRMELMAAIRALAELKRPCKVRLVTDSQYVMQGINDWMPNWKKRGWKTASKQPVK  73689

Query  75     -SELVNQIIEQLIKKEKVYLAWVPAHKGIGGNEQVDKLVSAGI  116
               ++L  Q+ EQ + + +V   WV  H G  GNEQ D L + G+
Sbjct  73690  NADLWQQLDEQ-VNRHEVSWQWVRGHTGHPGNEQADLLANRGV  73815



>Stutzerimonas stutzeri strain PS_377 NODE_2_length_617223_cov_44.615388, whole genome shotgun sequence
Sequence ID: NZ_RHQV01000002.1 Length: 617223
Range 1: 73381 to 73686

Score:48.9 bits(115), Expect:0.004,
Method:Compositional matrix adjust.,
Identities:35/103(34%), Positives:50/103(48%), Gaps:17/103(16%)

Query  30     TDTTNQKTELQAIYLALQD--SGLEVNIVTDSQYAL-GIIQAQPDQSE------------  74
               DTTN + EL A   AL +     +V +VTDSQY + GI    P+  +
Sbjct  73381  PDTTNNRMELMAAIRALAELKRPCKVRLVTDSQYVMQGINDWMPNWKKRGWKTASKQPVK  73560

Query  75     -SELVNQIIEQLIKKEKVYLAWVPAHKGIGGNEQVDKLVSAGI  116
               ++L  Q+ EQ + + +V   WV  H G  GNEQ D L + G+
Sbjct  73561  NADLWQQLDEQ-VNRHEVSWQWVRGHTGHPGNEQADLLANRGV  73686



>Stutzerimonas stutzeri strain MM2021_5 NODE_3_length_633422_cov_23.013727, whole genome shotgun sequence
Sequence ID: NZ_JAFMNW010000003.1 Length: 633422
Range 1: 559572 to 559877

Score:48.9 bits(115), Expect:0.004,
Method:Compositional matrix adjust.,
Identities:35/103(34%), Positives:50/103(48%), Gaps:17/103(16%)

Query  30      TDTTNQKTELQAIYLALQD--SGLEVNIVTDSQYAL-GIIQAQPDQSE------------  74
                DTTN + EL A   AL +     +V +VTDSQY + GI    P+  +
Sbjct  559877  PDTTNNRMELMAAIRALAELKRPCKVRLVTDSQYVMQGINDWMPNWKKRGWKTASKQPVK  559698

Query  75      -SELVNQIIEQLIKKEKVYLAWVPAHKGIGGNEQVDKLVSAGI  116
                ++L  Q+ EQ + + +V   WV  H G  GNEQ D L + G+
Sbjct  559697  NADLWQQLDEQ-VNRHEVSWQWVRGHTGHPGNEQADLLANRGV  559572



>Stutzerimonas stutzeri strain GD04140 1, whole genome shotgun sequence
Sequence ID: NZ_JAODZL010000001.1 Length: 687846
Range 1: 67992 to 68297

Score:48.9 bits(115), Expect:0.004,
Method:Compositional matrix adjust.,
Identities:35/103(34%), Positives:50/103(48%), Gaps:17/103(16%)

Query  30     TDTTNQKTELQAIYLALQD--SGLEVNIVTDSQYAL-GIIQAQPDQSE------------  74
               DTTN + EL A   AL +     +V +VTDSQY + GI    P+  +
Sbjct  67992  PDTTNNRMELMAAIRALAELKRPCKVRLVTDSQYVMQGINDWMPNWKKRGWKTASKQPVK  68171

Query  75     -SELVNQIIEQLIKKEKVYLAWVPAHKGIGGNEQVDKLVSAGI  116
               ++L  Q+ EQ + + +V   WV  H G  GNEQ D L + G+
Sbjct  68172  NADLWQQLDEQ-VNRHEVSWQWVRGHTGHPGNEQADLLANRGV  68297



>Stutzerimonas stutzeri strain NBRC 105234 sequence01, whole genome shotgun sequence
Sequence ID: NZ_BSTP01000001.1 Length: 710836
Range 1: 638230 to 638535

Score:48.9 bits(115), Expect:0.004,
Method:Compositional matrix adjust.,
Identities:35/103(34%), Positives:50/103(48%), Gaps:17/103(16%)

Query  30      TDTTNQKTELQAIYLALQD--SGLEVNIVTDSQYAL-GIIQAQPDQSE------------  74
                DTTN + EL A   AL +     +V +VTDSQY + GI    P+  +
Sbjct  638535  PDTTNNRMELMAAIRALAELKRPCKVRLVTDSQYVMQGINDWMPNWKKRGWKTASKQPVK  638356

Query  75      -SELVNQIIEQLIKKEKVYLAWVPAHKGIGGNEQVDKLVSAGI  116
                ++L  Q+ EQ + + +V   WV  H G  GNEQ D L + G+
Sbjct  638355  NADLWQQLDEQ-VNRHEVSWQWVRGHTGHPGNEQADLLANRGV  638230



>Stutzerimonas stutzeri strain ODKF13 contig_3, whole genome shotgun sequence
Sequence ID: NZ_LSVE01000023.1 Length: 719382
Range 1: 645569 to 645874

Score:48.9 bits(115), Expect:0.004,
Method:Compositional matrix adjust.,
Identities:35/103(34%), Positives:50/103(48%), Gaps:17/103(16%)

Query  30      TDTTNQKTELQAIYLALQD--SGLEVNIVTDSQYAL-GIIQAQPDQSE------------  74
                DTTN + EL A   AL +     +V +VTDSQY + GI    P+  +
```

```
Sbjct  645874  PDTTNNRMELMAAIRALAELKRPCKVRLVTDSQYVMQGINDWMPNWKKRGWKTASKQPVK  645695

Query  75      -SELVNQIIEQLIKKEKVYLAWVPAHKGIGGNEQVDKLVSAGI  116
                ++L  Q+ EQ + + +V   WV  H G  GNEQ D L + G+
Sbjct  645694  NADLWQQLDEQ-VNRHEVSWQWVRGHTGHPGNEQADLLANRGV  645569




>Stutzerimonas stutzeri strain KJ3 NODE_1_length_767050_cov_139.532478, whole genome shotgun sequence
Sequence ID: NZ_JBBELA010000001.1 Length: 767050
Range 1: 693279 to 693584

Score:48.9 bits(115), Expect:0.004,
Method:Compositional matrix adjust.,
Identities:35/103(34%), Positives:50/103(48%), Gaps:17/103(16%)

Query  30      TDTTNQKTELQAIYLALQD--SGLEVNIVTDSQYAL-GIIQAQPDQSE------------  74
                DTTN + EL A   AL +     +V +VTDSQY + GI    P+  +
Sbjct  693584  PDTTNNRMELMAAIRALAELKRPCKVRLVTDSQYVMQGINDWMPNWKKRGWKTASKQPVK  693405

Query  75      -SELVNQIIEQLIKKEKVYLAWVPAHKGIGGNEQVDKLVSAGI  116
                ++L  Q+ EQ + + +V   WV  H G  GNEQ D L + G+
Sbjct  693404  NADLWQQLDEQ-VNRHEVSWQWVRGHTGHPGNEQADLLANRGV  693279




>Stutzerimonas stutzeri strain KOL14.W.90.5 Scaffold_2_length_1379524_pilon, whole genome shotgun sequence
Sequence ID: NZ_WIFW01000002.1 Length: 1379714
Range 1: 73715 to 74020

Score:48.9 bits(115), Expect:0.004,
Method:Compositional matrix adjust.,
Identities:35/103(34%), Positives:50/103(48%), Gaps:17/103(16%)

Query  30     TDTTNQKTELQAIYLALQD--SGLEVNIVTDSQYAL-GIIQAQPDQSE------------  74
               DTTN + EL A   AL +     +V +VTDSQY + GI    P+  +
Sbjct  73715  PDTTNNRMELMAAIRALAELKRPCKVRLVTDSQYVMQGINDWMPNWKKRGWKTASKQPVK  73894

Query  75     -SELVNQIIEQLIKKEKVYLAWVPAHKGIGGNEQVDKLVSAGI  116
               ++L  Q+ EQ + + +V   WV  H G  GNEQ D L + G+
Sbjct  73895  NADLWQQLDEQ-VNRHEVSWQWVRGHTGHPGNEQADLLANRGV  74020




>Stutzerimonas stutzeri strain KOL14.W25.495.90 Scaffold_4_length_2608715_pilon, whole genome shotgun sequence
Sequence ID: NZ_WIGJ01000004.1 Length: 2608624
Range 1: 1830939 to 1831244

Score:48.9 bits(115), Expect:0.004,
Method:Compositional matrix adjust.,
Identities:35/103(34%), Positives:50/103(48%), Gaps:17/103(16%)

Query  30       TDTTNQKTELQAIYLALQD--SGLEVNIVTDSQYAL-GIIQAQPDQSE------------  74
                 DTTN + EL A   AL +     +V +VTDSQY + GI    P+  +
Sbjct  1831244  PDTTNNRMELMAAIRALAELKRPCKVRLVTDSQYVMQGINDWMPNWKKRGWKTASKQPVK  1831065

Query  75       -SELVNQIIEQLIKKEKVYLAWVPAHKGIGGNEQVDKLVSAGI  116
                 ++L  Q+ EQ + + +V   WV  H G  GNEQ D L + G+
Sbjct  1831064  NADLWQQLDEQ-VNRHEVSWQWVRGHTGHPGNEQADLLANRGV  1830939




>Stutzerimonas stutzeri strain XX1 chromosome, complete genome
Sequence ID: NZ_CP091174.1 Length: 4424182
Range 1: 2784177 to 2784482

Score:48.9 bits(115), Expect:0.004,
Method:Compositional matrix adjust.,
Identities:35/103(34%), Positives:50/103(48%), Gaps:17/103(16%)

Query  30       TDTTNQKTELQAIYLALQD--SGLEVNIVTDSQYAL-GIIQAQPDQSE------------  74
                 DTTN + EL A   AL +     +V +VTDSQY + GI    P+  +
Sbjct  2784482  PDTTNNRMELMAAIRALAELKRPCKVRLVTDSQYVMQGINDWMPNWKKRGWKTASKQPVK  2784303

Query  75       -SELVNQIIEQLIKKEKVYLAWVPAHKGIGGNEQVDKLVSAGI  116
                 ++L  Q+ EQ + + +V   WV  H G  GNEQ D L + G+
Sbjct  2784302  NADLWQQLDEQ-VNRHEVSWQWVRGHTGHPGNEQADLLANRGV  2784177




>Stutzerimonas stutzeri strain GD04095 DMICFCLD_1, whole genome shotgun sequence
Sequence ID: NZ_JAOEBC010000001.1 Length: 4437022
Range 1: 2486179 to 2486484

Score:48.9 bits(115), Expect:0.004,
Method:Compositional matrix adjust.,
Identities:35/103(34%), Positives:50/103(48%), Gaps:17/103(16%)

Query  30       TDTTNQKTELQAIYLALQD--SGLEVNIVTDSQYAL-GIIQAQPDQSE------------  74
                 DTTN + EL A   AL +     +V +VTDSQY + GI    P+  +
Sbjct  2486484  PDTTNNRMELMAAIRALAELKRPCKVRLVTDSQYVMQGINDWMPNWKKRGWKTASKQPVK  2486305

Query  75       -SELVNQIIEQLIKKEKVYLAWVPAHKGIGGNEQVDKLVSAGI  116
                 ++L  Q+ EQ + + +V   WV  H G  GNEQ D L + G+
Sbjct  2486304  NADLWQQLDEQ-VNRHEVSWQWVRGHTGHPGNEQADLLANRGV  2486179




>Stutzerimonas stutzeri strain NCTC10450 chromosome 1, complete sequence
Sequence ID: NZ_LR134319.1 Length: 4438731
Range 1: 2460817 to 2461122

Score:48.9 bits(115), Expect:0.004,
Method:Compositional matrix adjust.,
Identities:35/103(34%), Positives:50/103(48%), Gaps:17/103(16%)

Query  30       TDTTNQKTELQAIYLALQD--SGLEVNIVTDSQYAL-GIIQAQPDQSE------------  74
                 DTTN + EL A   AL +     +V +VTDSQY + GI    P+  +
Sbjct  2461122  PDTTNNRMELMAAIRALAELKRPCKVRLVTDSQYVMQGINDWMPNWKKRGWKTASKQPVK  2460943

Query  75       -SELVNQIIEQLIKKEKVYLAWVPAHKGIGGNEQVDKLVSAGI  116
                 ++L  Q+ EQ + + +V   WV  H G  GNEQ D L + G+
Sbjct  2460942  NADLWQQLDEQ-VNRHEVSWQWVRGHTGHPGNEQADLLANRGV  2460817




>Stutzerimonas stutzeri strain SB-1 chromosome, complete genome
```

```
Sequence ID: NZ_CP088004.1 Length: 4473665
Range 1: 2532932 to 2533237

Score:48.9 bits(115), Expect:0.004,
Method:Compositional matrix adjust.,
Identities:35/103(34%), Positives:50/103(48%), Gaps:17/103(16%)

Query  30       TDTTNQKTELQAIYLALQD--SGLEVNIVTDSQYAL-GIIQAQPDQSE------------  74
                 DTTN + EL A   AL +     +V +VTDSQY + GI    P+  +
Sbjct  2533237  PDTTNNRMELMAAIRALAELKRPCKVRLVTDSQYVMQGINDWMPNWKKRGWKTASKQPVK  2533058

Query  75       -SELVNQIIEQLIKKEKVYLAWVPAHKGIGGNEQVDKLVSAGI  116
                 ++L  Q+ EQ + + +V   WV  H G  GNEQ D L + G+
Sbjct  2533057  NADLWQQLDEQ-VNRHEVSWQWVRGHTGHPGNEQADLLANRGV  2532932
```

```
>Stutzerimonas stutzeri strain YWX-1 chromosome, complete genome
Sequence ID: NZ_CP078087.1 Length: 4488441
Range 1: 2436930 to 2437235

Score:48.9 bits(115), Expect:0.004,
Method:Compositional matrix adjust.,
Identities:35/103(34%), Positives:50/103(48%), Gaps:17/103(16%)

Query  30       TDTTNQKTELQAIYLALQD--SGLEVNIVTDSQYAL-GIIQAQPDQSE------------  74
                 DTTN + EL A   AL +     +V +VTDSQY + GI    P+  +
Sbjct  2437235  PDTTNNRMELMAAIRALAELKRPCKVRLVTDSQYVMQGINDWMPNWKKRGWKTASKQPVK  2437056

Query  75       -SELVNQIIEQLIKKEKVYLAWVPAHKGIGGNEQVDKLVSAGI  116
                 ++L  Q+ EQ + + +V   WV  H G  GNEQ D L + G+
Sbjct  2437055  NADLWQQLDEQ-VNRHEVSWQWVRGHTGHPGNEQADLLANRGV  2436930
```

```
>Stutzerimonas stutzeri strain ZDHY95 chromosome, complete genome
Sequence ID: NZ_CP063358.1 Length: 4500524
Range 1: 1987285 to 1987590

Score:48.9 bits(115), Expect:0.004,
Method:Compositional matrix adjust.,
Identities:35/103(34%), Positives:50/103(48%), Gaps:17/103(16%)

Query  30       TDTTNQKTELQAIYLALQD--SGLEVNIVTDSQYAL-GIIQAQPDQSE------------  74
                 DTTN + EL A   AL +     +V +VTDSQY + GI    P+  +
Sbjct  1987285  PDTTNNRMELMAAIRALAELKRPCKVRLVTDSQYVMQGINDWMPNWKKRGWKTASKQPVK  1987464

Query  75       -SELVNQIIEQLIKKEKVYLAWVPAHKGIGGNEQVDKLVSAGI  116
                 ++L  Q+ EQ + + +V   WV  H G  GNEQ D L + G+
Sbjct  1987465  NADLWQQLDEQ-VNRHEVSWQWVRGHTGHPGNEQADLLANRGV  1987590
```

```
>Stutzerimonas stutzeri strain SGAir0442 chromosome, complete genome
Sequence ID: NZ_CP025149.2 Length: 4524655
Range 1: 1803318 to 1803623

Score:48.9 bits(115), Expect:0.004,
Method:Compositional matrix adjust.,
Identities:35/103(34%), Positives:50/103(48%), Gaps:17/103(16%)

Query  30       TDTTNQKTELQAIYLALQD--SGLEVNIVTDSQYAL-GIIQAQPDQSE------------  74
                 DTTN + EL A   AL +     +V +VTDSQY + GI    P+  +
Sbjct  1803318  PDTTNNRMELMAAIRALAELKRPCKVRLVTDSQYVMQGINDWMPNWKKRGWKTASKQPVK  1803497

Query  75       -SELVNQIIEQLIKKEKVYLAWVPAHKGIGGNEQVDKLVSAGI  116
                 ++L  Q+ EQ + + +V   WV  H G  GNEQ D L + G+
Sbjct  1803498  NADLWQQLDEQ-VNRHEVSWQWVRGHTGHPGNEQADLLANRGV  1803623
```

```
>Stutzerimonas stutzeri strain F2a chromosome, complete genome
Sequence ID: NZ_AP024722.1 Length: 4545686
Range 1: 2125415 to 2125720

Score:48.9 bits(115), Expect:0.004,
Method:Compositional matrix adjust.,
Identities:35/103(34%), Positives:50/103(48%), Gaps:17/103(16%)

Query  30       TDTTNQKTELQAIYLALQD--SGLEVNIVTDSQYAL-GIIQAQPDQSE------------  74
                 DTTN + EL A   AL +     +V +VTDSQY + GI    P+  +
Sbjct  2125415  PDTTNNRMELMAAIRALAELKRPCKVRLVTDSQYVMQGINDWMPNWKKRGWKTASKQPVK  2125594

Query  75       -SELVNQIIEQLIKKEKVYLAWVPAHKGIGGNEQVDKLVSAGI  116
                 ++L  Q+ EQ + + +V   WV  H G  GNEQ D L + G+
Sbjct  2125595  NADLWQQLDEQ-VNRHEVSWQWVRGHTGHPGNEQADLLANRGV  2125720
```

```
>Stutzerimonas stutzeri A1501, complete sequence
Sequence ID: NC_009434.1 Length: 4567418
Range 1: 2455637 to 2455942

Score:48.9 bits(115), Expect:0.004,
Method:Compositional matrix adjust.,
Identities:35/103(34%), Positives:50/103(48%), Gaps:17/103(16%)

Query  30       TDTTNQKTELQAIYLALQD--SGLEVNIVTDSQYAL-GIIQAQPDQSE------------  74
                 DTTN + EL A   AL +     +V +VTDSQY + GI    P+  +
Sbjct  2455942  PDTTNNRMELMAAIRALAELKRPCKVRLVTDSQYVMQGINDWMPNWKKRGWKTASKQPVK  2455763

Query  75       -SELVNQIIEQLIKKEKVYLAWVPAHKGIGGNEQVDKLVSAGI  116
                 ++L  Q+ EQ + + +V   WV  H G  GNEQ D L + G+
Sbjct  2455762  NADLWQQLDEQ-VNRHEVSWQWVRGHTGHPGNEQADLLANRGV  2455637
```

```
>Stutzerimonas stutzeri strain NRCB010 chromosome, complete genome
Sequence ID: NZ_CP098731.1 Length: 4570338
Range 1: 2457746 to 2458051

Score:48.9 bits(115), Expect:0.004,
Method:Compositional matrix adjust.,
Identities:35/103(34%), Positives:50/103(48%), Gaps:17/103(16%)

Query  30       TDTTNQKTELQAIYLALQD--SGLEVNIVTDSQYAL-GIIQAQPDQSE------------  74
                 DTTN + EL A   AL +     +V +VTDSQY + GI    P+  +
Sbjct  2458051  PDTTNNRMELMAAIRALAELKRPCKVRLVTDSQYVMQGINDWMPNWKKRGWKTASKQPVK  2457872
```

```
Query  75       -SELVNQIIEQLIKKEKVYLAWVPAHKGIGGNEQVDKLVSAGI  116
                 ++L  Q+ EQ + + +V   WV  H G  GNEQ D L + G+
Sbjct  2457871  NADLWQQLDEQ-VNRHEVSWQWVRGHTGHPGNEQADLLANRGV  2457746
```

>Stutzerimonas stutzeri strain MJL19 chromosome, complete genome
Sequence ID: NZ_CP140298.1 Length: 4572525
Range 1: 2406043 to 2406348

Score:48.9 bits(115), Expect:0.004,
Method:Compositional matrix adjust.,
Identities:35/103(34%), Positives:50/103(48%), Gaps:17/103(16%)

```
Query  30       TDTTNQKTELQAIYLALQD--SGLEVNIVTDSQYAL-GIIQAQPDQSE------------  74
                 DTTN + EL A   AL +     +V +VTDSQY + GI    P+  +
Sbjct  2406348  PDTTNNRMELMAAIRALAELKRPCKVRLVTDSQYVMQGINDWMPNWKKRGWKTASKQPVK  2406169

Query  75       -SELVNQIIEQLIKKEKVYLAWVPAHKGIGGNEQVDKLVSAGI  116
                 ++L  Q+ EQ + + +V   WV  H G  GNEQ D L + G+
Sbjct  2406168  NADLWQQLDEQ-VNRHEVSWQWVRGHTGHPGNEQADLLANRGV  2406043
```

>Stutzerimonas stutzeri strain 2020WEIHUA_G chromosome, complete genome
Sequence ID: NZ_CP062162.1 Length: 4591234
Range 1: 2102282 to 2102587

Score:48.9 bits(115), Expect:0.004,
Method:Compositional matrix adjust.,
Identities:35/103(34%), Positives:50/103(48%), Gaps:17/103(16%)

```
Query  30       TDTTNQKTELQAIYLALQD--SGLEVNIVTDSQYAL-GIIQAQPDQSE------------  74
                 DTTN + EL A   AL +     +V +VTDSQY + GI    P+  +
Sbjct  2102282  PDTTNNRMELMAAIRALAELKRPCKVRLVTDSQYVMQGINDWMPNWKKRGWKTASKQPVK  2102461

Query  75       -SELVNQIIEQLIKKEKVYLAWVPAHKGIGGNEQVDKLVSAGI  116
                 ++L  Q+ EQ + + +V   WV  H G  GNEQ D L + G+
Sbjct  2102462  NADLWQQLDEQ-VNRHEVSWQWVRGHTGHPGNEQADLLANRGV  2102587
```

>Stutzerimonas stutzeri strain SLG510A3-8 chromosome, complete genome
Sequence ID: NZ_CP011854.1 Length: 4650155
Range 1: 2220903 to 2221208

Score:48.9 bits(115), Expect:0.004,
Method:Compositional matrix adjust.,
Identities:35/103(34%), Positives:50/103(48%), Gaps:17/103(16%)

```
Query  30       TDTTNQKTELQAIYLALQD--SGLEVNIVTDSQYAL-GIIQAQPDQSE------------  74
                 DTTN + EL A   AL +     +V +VTDSQY + GI    P+  +
Sbjct  2220903  PDTTNNRMELMAAIRALAELKRPCKVRLVTDSQYVMQGINDWMPNWKKRGWKTASKQPVK  2221082

Query  75       -SELVNQIIEQLIKKEKVYLAWVPAHKGIGGNEQVDKLVSAGI  116
                 ++L  Q+ EQ + + +V   WV  H G  GNEQ D L + G+
Sbjct  2221083  NADLWQQLDEQ-VNRHEVSWQWVRGHTGHPGNEQADLLANRGV  2221208
```

>Stutzerimonas stutzeri strain J15 chromosome, complete genome
Sequence ID: NZ_CP101739.1 Length: 4683655
Range 1: 1802706 to 1803011

Score:48.9 bits(115), Expect:0.004,
Method:Compositional matrix adjust.,
Identities:35/103(34%), Positives:50/103(48%), Gaps:17/103(16%)

```
Query  30       TDTTNQKTELQAIYLALQD--SGLEVNIVTDSQYAL-GIIQAQPDQSE------------  74
                 DTTN + EL A   AL +     +V +VTDSQY + GI    P+  +
Sbjct  1802706  PDTTNNRMELMAAIRALAELKRPCKVRLVTDSQYVMQGINDWMPNWKKRGWKTASKQPVK  1802885

Query  75       -SELVNQIIEQLIKKEKVYLAWVPAHKGIGGNEQVDKLVSAGI  116
                 ++L  Q+ EQ + + +V   WV  H G  GNEQ D L + G+
Sbjct  1802886  NADLWQQLDEQ-VNRHEVSWQWVRGHTGHPGNEQADLLANRGV  1803011
```

>Stutzerimonas stutzeri DSM 4166, complete sequence
Sequence ID: NC_017532.1 Length: 4689946
Range 1: 2447283 to 2447588

Score:48.9 bits(115), Expect:0.004,
Method:Compositional matrix adjust.,
Identities:35/103(34%), Positives:50/103(48%), Gaps:17/103(16%)

```
Query  30       TDTTNQKTELQAIYLALQD--SGLEVNIVTDSQYAL-GIIQAQPDQSE------------  74
                 DTTN + EL A   AL +     +V +VTDSQY + GI    P+  +
Sbjct  2447588  PDTTNNRMELMAAIRALAELKRPCKVRLVTDSQYVMQGINDWMPNWKKRGWKTASKQPVK  2447409

Query  75       -SELVNQIIEQLIKKEKVYLAWVPAHKGIGGNEQVDKLVSAGI  116
                 ++L  Q+ EQ + + +V   WV  H G  GNEQ D L + G+
Sbjct  2447408  NADLWQQLDEQ-VNRHEVSWQWVRGHTGHPGNEQADLLANRGV  2447283
```

>Stutzerimonas stutzeri strain H46 chromosome, complete genome
Sequence ID: NZ_CP139417.1 Length: 4720414
Range 1: 2530679 to 2530984

Score:48.9 bits(115), Expect:0.004,
Method:Compositional matrix adjust.,
Identities:35/103(34%), Positives:50/103(48%), Gaps:17/103(16%)

```
Query  30       TDTTNQKTELQAIYLALQD--SGLEVNIVTDSQYAL-GIIQAQPDQSE------------  74
                 DTTN + EL A   AL +     +V +VTDSQY + GI    P+  +
Sbjct  2530984  PDTTNNRMELMAAIRALAELKRPCKVRLVTDSQYVMQGINDWMPNWKKRGWKTASKQPVK  2530805

Query  75       -SELVNQIIEQLIKKEKVYLAWVPAHKGIGGNEQVDKLVSAGI  116
                 ++L  Q+ EQ + + +V   WV  H G  GNEQ D L + G+
Sbjct  2530804  NADLWQQLDEQ-VNRHEVSWQWVRGHTGHPGNEQADLLANRGV  2530679
```

>Stutzerimonas stutzeri strain GD04120 chromosome, complete genome
Sequence ID: NZ_CP104580.1 Length: 4775048

```
Range 1: 2866218 to 2866523

Score:48.9 bits(115), Expect:0.004,
Method:Compositional matrix adjust.,
Identities:35/103(34%), Positives:50/103(48%), Gaps:17/103(16%)

Query  30       TDTTNQKTELQAIYLALQD--SGLEVNIVTDSQYAL-GIIQAQPDQSE------------  74
                 DTTN + EL A   AL +     +V +VTDSQY + GI    P+  +
Sbjct  2866523  PDTTNNRMELMAAIRALAELKRPCKVRLVTDSQYVMQGINDWMPNWKKRGWKTASKQPVK  2866344

Query  75       -SELVNQIIEQLIKKEKVYLAWVPAHKGIGGNEQVDKLVSAGI  116
                 ++L  Q+ EQ + + +V   WV  H G  GNEQ D L + G+
Sbjct  2866343  NADLWQQLDEQ-VNRHEVSWQWVRGHTGHPGNEQADLLANRGV  2866218
```

```
>Stutzerimonas stutzeri strain GD04113 PNKOAAFP_1, whole genome shotgun sequence
Sequence ID: NZ_JAOEAK010000001.1 Length: 4814082
Range 1: 2063339 to 2063644

Score:48.9 bits(115), Expect:0.004,
Method:Compositional matrix adjust.,
Identities:35/103(34%), Positives:50/103(48%), Gaps:17/103(16%)

Query  30       TDTTNQKTELQAIYLALQD--SGLEVNIVTDSQYAL-GIIQAQPDQSE------------  74
                 DTTN + EL A   AL +     +V +VTDSQY + GI    P+  +
Sbjct  2063339  PDTTNNRMELMAAIRALAELKRPCKVRLVTDSQYVMQGINDWMPNWKKRGWKTASKQPVK  2063518

Query  75       -SELVNQIIEQLIKKEKVYLAWVPAHKGIGGNEQVDKLVSAGI  116
                 ++L  Q+ EQ + + +V   WV  H G  GNEQ D L + G+
Sbjct  2063519  NADLWQQLDEQ-VNRHEVSWQWVRGHTGHPGNEQADLLANRGV  2063644
```

```
>Stutzerimonas frequens strain XN05-1 chromosome, complete genome
Sequence ID: NZ_CP142122.1 Length: 4411446
Range 1: 2379444 to 2379845

Score:48.9 bits(115), Expect:0.005,
Method:Compositional matrix adjust.,
Identities:44/135(33%), Positives:61/135(45%), Gaps:20/135(14%)

Query  1        YVDGAANRETKLGKAG-YVTNRG--RQKVVTLTDTTNQKTELQAIYLALQD--SGLEVNI  55
                Y DGA       G  G  +  +G  R+      DTTN + EL A   AL +     +V +
Sbjct  2379845  YTDGACKGNPGPGGWGALLVYKGVKRELWGGEPDTTNNRMELMAAIRALAELKRPCKVRL  2379666

Query  56       VTDSQYAL-GIIQAQPDQSE-------------SELVNQIIEQLIKKEKVYLAWVPAHKG  101
                VTDSQY + GI    P+  +             ++L  Q+ EQ+ + E V   WV  H G
Sbjct  2379665  VTDSQYVMQGINDWMPNWKKRGWKTASKQPVKNADLWQQLDEQVSRHE-VSWQWVRGHTG  2379489

Query  102      IGGNEQVDKLVSAGI  116
                  GNEQ D L + G+
Sbjct  2379488  DPGNEQADLLANRGV  2379444
```

```
>Stutzerimonas frequens strain R2A2 chromosome, complete genome
Sequence ID: NZ_CP029772.1 Length: 4559447
Range 1: 1751505 to 1751906

Score:48.9 bits(115), Expect:0.005,
Method:Compositional matrix adjust.,
Identities:44/135(33%), Positives:61/135(45%), Gaps:20/135(14%)

Query  1        YVDGAANRETKLGKAG-YVTNRG--RQKVVTLTDTTNQKTELQAIYLALQD--SGLEVNI  55
                Y DGA       G  G  +  +G  R+      DTTN + EL A   AL +     +V +
Sbjct  1751505  YTDGACKGNPGPGGWGALLVYKGVKRELWGGEPDTTNNRMELMAAIRALAELKRPCKVRL  1751684

Query  56       VTDSQYAL-GIIQAQPDQSE-------------SELVNQIIEQLIKKEKVYLAWVPAHKG  101
                VTDSQY + GI    P+  +             ++L  Q+ EQ+ + E V   WV  H G
Sbjct  1751685  VTDSQYVMQGINDWMPNWKKRGWKTASKQPVKNADLWQQLDEQVSRHE-VSWQWVRGHTG  1751861

Query  102      IGGNEQVDKLVSAGI  116
                  GNEQ D L + G+
Sbjct  1751862  DPGNEQADLLANRGV  1751906
```

```
>Stutzerimonas stutzeri strain C2 contig309, whole genome shotgun sequence
Sequence ID: NZ_LDWB01000066.1 Length: 16666
Range 1: 13616 to 13918

Score:48.9 bits(115), Expect:0.005,
Method:Compositional matrix adjust.,
Identities:35/102(34%), Positives:50/102(49%), Gaps:17/102(16%)

Query  31     DTTNQKTELQAIYLALQD--SGLEVNIVTDSQYAL-GIIQAQPDQSE------------  74
              DTTN + EL A   AL +     +V +VTDSQY + GI    P+  +
Sbjct  13918  DTTNNRMELMAAIRALAELKRPCKVRLVTDSQYVMQGINDWMPNWKKRGWKTASKQPVKN  13739

Query  75     SELVNQIIEQLIKKEKVYLAWVPAHKGIGGNEQVDKLVSAGI  116
              ++L  Q+ EQ + + +V   WV  H G  GNEQ D L + G+
Sbjct  13738  ADLWQQLDEQ-VNRHEVSWQWVRGHTGHPGNEQADLLANRGV  13616
```

```
>Stutzerimonas stutzeri strain 267_PSTU 1208_88756_1251158_409+,...,270+, whole genome shotgun sequence
Sequence ID: NZ_JVNQ01000084.1 Length: 88813
Range 1: 11642 to 11944

Score:48.9 bits(115), Expect:0.005,
Method:Compositional matrix adjust.,
Identities:35/102(34%), Positives:50/102(49%), Gaps:17/102(16%)

Query  31     DTTNQKTELQAIYLALQD--SGLEVNIVTDSQYAL-GIIQAQPDQSE------------  74
              DTTN + EL A   AL +     +V +VTDSQY + GI    P+  +
Sbjct  11944  DTTNNRMELMAAIRALAELKRPCKVRLVTDSQYVMQGINDWMPNWKKRGWKTASKQPVKN  11765

Query  75     SELVNQIIEQLIKKEKVYLAWVPAHKGIGGNEQVDKLVSAGI  116
              ++L  Q+ EQ + + +V   WV  H G  GNEQ D L + G+
Sbjct  11764  ADLWQQLDEQ-VNRHEVSWQWVRGHTGHPGNEQADLLANRGV  11642
```

```
>Stutzerimonas stutzeri isolate BB1462, whole genome shotgun sequence
Sequence ID: NZ_CAHPQS010000015.1 Length: 108117
Range 1: 41312 to 41614
```

```
Score:48.9 bits(115), Expect:0.005,
Method:Compositional matrix adjust.,
Identities:35/102(34%), Positives:50/102(49%), Gaps:17/102(16%)

Query  31     DTTNQKTELQAIYLALQD--SGLEVNIVTDSQYAL-GIIQAQPDQSE-------------  74
              DTTN + EL A   AL +     +V +VTDSQY + GI    P+  +
Sbjct  41614  DTTNNRMELMAAIRALAELKRPCKVRLVTDSQYVMQGINDWMPNWKKRGWKTASKQPVKN  41435

Query  75     SELVNQIIEQLIKKEKVYLAWVPAHKGIGGNEQVDKLVSAGI  116
              ++L  Q+ EQ + + +V   WV  H G  GNEQ D L + G+
Sbjct  41434  ADLWQQLDEQ-VNRHEVSWQWVRGHTGHPGNEQADLLANRGV  41312
```

>Stutzerimonas stutzeri isolate BB1470, whole genome shotgun sequence
Sequence ID: NZ_CAHPRE010000013.1 Length: 108117
Range 1: 66504 to 66806

```
Score:48.9 bits(115), Expect:0.005,
Method:Compositional matrix adjust.,
Identities:35/102(34%), Positives:50/102(49%), Gaps:17/102(16%)

Query  31     DTTNQKTELQAIYLALQD--SGLEVNIVTDSQYAL-GIIQAQPDQSE-------------  74
              DTTN + EL A   AL +     +V +VTDSQY + GI    P+  +
Sbjct  66504  DTTNNRMELMAAIRALAELKRPCKVRLVTDSQYVMQGINDWMPNWKKRGWKTASKQPVKN  66683

Query  75     SELVNQIIEQLIKKEKVYLAWVPAHKGIGGNEQVDKLVSAGI  116
              ++L  Q+ EQ + + +V   WV  H G  GNEQ D L + G+
Sbjct  66684  ADLWQQLDEQ-VNRHEVSWQWVRGHTGHPGNEQADLLANRGV  66806
```

>Stutzerimonas stutzeri strain PS_376 NODE_10_length_135172_cov_44.059425, whole genome shotgun sequence
Sequence ID: NZ_RHQW01000010.1 Length: 135172
Range 1: 61360 to 61662

```
Score:48.9 bits(115), Expect:0.005,
Method:Compositional matrix adjust.,
Identities:35/102(34%), Positives:50/102(49%), Gaps:17/102(16%)

Query  31     DTTNQKTELQAIYLALQD--SGLEVNIVTDSQYAL-GIIQAQPDQSE-------------  74
              DTTN + EL A   AL +     +V +VTDSQY + GI    P+  +
Sbjct  61662  DTTNNRMELMAAIRALAELKRPCKVRLVTDSQYVMQGINDWMPNWKKRGWKTASKQPVKN  61483

Query  75     SELVNQIIEQLIKKEKVYLAWVPAHKGIGGNEQVDKLVSAGI  116
              ++L  Q+ EQ + + +V   WV  H G  GNEQ D L + G+
Sbjct  61482  ADLWQQLDEQ-VNRHEVSWQWVRGHTGHPGNEQADLLANRGV  61360
```

>Stutzerimonas stutzeri strain GD03947 6, whole genome shotgun sequence
Sequence ID: NZ_JAOCAE010000006.1 Length: 284342
Range 1: 22133 to 22435

```
Score:48.9 bits(115), Expect:0.005,
Method:Compositional matrix adjust.,
Identities:35/102(34%), Positives:50/102(49%), Gaps:17/102(16%)

Query  31     DTTNQKTELQAIYLALQD--SGLEVNIVTDSQYAL-GIIQAQPDQSE-------------  74
              DTTN + EL A   AL +     +V +VTDSQY + GI    P+  +
Sbjct  22133  DTTNNRMELMAAIRALAELKRPCKVRLVTDSQYVMQGINDWMPNWKKRGWKTASKQPVKN  22312

Query  75     SELVNQIIEQLIKKEKVYLAWVPAHKGIGGNEQVDKLVSAGI  116
              ++L  Q+ EQ + + +V   WV  H G  GNEQ D L + G+
Sbjct  22313  ADLWQQLDEQ-VNRHEVSWQWVRGHTGHPGNEQADLLANRGV  22435
```

>Stutzerimonas stutzeri strain AK6 Contig_22_len_290503, whole genome shotgun sequence
Sequence ID: NZ_PZYR01000022.1 Length: 290503
Range 1: 61395 to 61697

```
Score:48.9 bits(115), Expect:0.005,
Method:Compositional matrix adjust.,
Identities:35/102(34%), Positives:50/102(49%), Gaps:17/102(16%)

Query  31     DTTNQKTELQAIYLALQD--SGLEVNIVTDSQYAL-GIIQAQPDQSE-------------  74
              DTTN + EL A   AL +     +V +VTDSQY + GI    P+  +
Sbjct  61697  DTTNNRMELMAAIRALAELKRPCKVRLVTDSQYVMQGINDWMPNWKKRGWKTASKQPVKN  61518

Query  75     SELVNQIIEQLIKKEKVYLAWVPAHKGIGGNEQVDKLVSAGI  116
              ++L  Q+ EQ + + +V   WV  H G  GNEQ D L + G+
Sbjct  61517  ADLWQQLDEQ-VNRHEVSWQWVRGHTGHPGNEQADLLANRGV  61395
```

>Stutzerimonas sp. FeSN7 contig_1_pilon_pilon_pilon, whole genome shotgun sequence
Sequence ID: NZ_JARPMN010000001.1 Length: 4789579
Range 1: 1863530 to 1863832

```
Score:48.9 bits(115), Expect:0.005,
Method:Compositional matrix adjust.,
Identities:35/102(34%), Positives:50/102(49%), Gaps:17/102(16%)

Query  31       DTTNQKTELQAIYLALQD--SGLEVNIVTDSQYAL-GIIQAQPDQSE-------------  74
                DTTN + EL A   AL +     +V +VTDSQY + GI    P+  +
Sbjct  1863530  DTTNNRMELMAAIRALAELKRPCKVRLVTDSQYVMQGINDWMPNWKKRGWKTASKQPVKN  1863709

Query  75       SELVNQIIEQLIKKEKVYLAWVPAHKGIGGNEQVDKLVSAGI  116
                ++L  Q+ EQ + + +V   WV  H G  GNEQ D L + G+
Sbjct  1863710  ADLWQQLDEQ-VNRHEVSWQWVRGHTGHPGNEQADLLANRGV  1863832
```

>Stutzerimonas stutzeri strain GD03838 14, whole genome shotgun sequence
Sequence ID: NZ_JAOCEE010000014.1 Length: 117613
Range 1: 24134 to 24439

```
Score:48.9 bits(115), Expect:0.005,
Method:Compositional matrix adjust.,
Identities:35/103(34%), Positives:50/103(48%), Gaps:17/103(16%)

Query  30     TDTTNQKTELQAIYLALQD--SGLEVNIVTDSQYAL-GIIQAQPDQSE------------  74
               DTTN + EL A   AL +     +V +VTDSQY + GI    P+  +
Sbjct  24134  PDTTNNRMELMAAIRALAELKRPCKVRLVTDSQYVMQGINDWMPNWKKRGWKTASKQPVK  24313

Query  75     -SELVNQIIEQLIKKEKVYLAWVPAHKGIGGNEQVDKLVSAGI  116
               ++L  Q+ EQ + + +V   WV  H G  GNEQ D L + G+
```

```
Sbjct  24314  NADLWQQLDEQ-VNRHEVSWQWVRGHTGHPGNEQADLLANRGV  24439



>Stutzerimonas stutzeri strain PS_151 NODE_10_length_167014_cov_15.9041, whole genome shotgun sequence
Sequence ID: NZ_RHRD01000010.1 Length: 167014
Range 1: 93196 to 93501

Score:48.9 bits(115), Expect:0.005,
Method:Compositional matrix adjust.,
Identities:35/103(34%), Positives:50/103(48%), Gaps:17/103(16%)

Query  30     TDTTNQKTELQAIYLALQD--SGLEVNIVTDSQYAL-GIIQAQPDQSE------------  74
               DTTN + EL A   AL +      +V +VTDSQY + GI     P+  +
Sbjct  93501  PDTTNNRMELMAAIRALAELKRPCKVRLVTDSQYVMQGINDWMPNWKKRGWKTASKQPVK  93322

Query  75     -SELVNQIIEQLIKKEKVYLAWVPAHKGIGGNEQVDKLVSAGI  116
               ++L  Q+ EQ + + +V   WV  H G  GNEQ D L + G+
Sbjct  93321  NADLWQQLDEQ-VNRHEVSWQWVRGHTGHPGNEQADLLANRGV  93196



>Stutzerimonas stutzeri strain PS_128 NODE_4_length_364407_cov_26.9692, whole genome shotgun sequence
Sequence ID: NZ_RHRJ01000004.1 Length: 364407
Range 1: 290568 to 290873

Score:48.9 bits(115), Expect:0.005,
Method:Compositional matrix adjust.,
Identities:35/103(34%), Positives:50/103(48%), Gaps:17/103(16%)

Query  30      TDTTNQKTELQAIYLALQD--SGLEVNIVTDSQYAL-GIIQAQPDQSE------------  74
                DTTN + EL A   AL +      +V +VTDSQY + GI     P+  +
Sbjct  290873  PDTTNNRMELMAAIRALAELKRPCKVRLVTDSQYVMQGINDWMPNWKKRGWKTASKQPVK  290694

Query  75      -SELVNQIIEQLIKKEKVYLAWVPAHKGIGGNEQVDKLVSAGI  116
                ++L  Q+ EQ + + +V   WV  H G  GNEQ D L + G+
Sbjct  290693  NADLWQQLDEQ-VNRHEVSWQWVRGHTGHPGNEQADLLANRGV  290568



>Stutzerimonas stutzeri strain S116 chromosome, complete genome
Sequence ID: NZ_CP078509.1 Length: 4440952
Range 1: 1621738 to 1622043

Score:48.9 bits(115), Expect:0.005,
Method:Compositional matrix adjust.,
Identities:35/103(34%), Positives:50/103(48%), Gaps:17/103(16%)

Query  30       TDTTNQKTELQAIYLALQD--SGLEVNIVTDSQYAL-GIIQAQPDQSE------------  74
                 DTTN + EL A   AL +      +V +VTDSQY + GI     P+  +
Sbjct  1622043  PDTTNNRMELMAAIRALAELKRPCKVRLVTDSQYVMQGINDWMPNWKKRGWKTASKQPVK  1621864

Query  75       -SELVNQIIEQLIKKEKVYLAWVPAHKGIGGNEQVDKLVSAGI  116
                 ++L  Q+ EQ + + +V   WV  H G  GNEQ D L + G+
Sbjct  1621863  NADLWQQLDEQ-VNRHEVSWQWVRGHTGHPGNEQADLLANRGV  1621738



>Stutzerimonas stutzeri strain KOL14.W.495.96 Scaffold_7_length_212774_pilon, whole genome shotgun sequence
Sequence ID: NZ_WIFO01000006.1 Length: 212774
Range 1: 68749 to 69150

Score:48.5 bits(114), Expect:0.005,
Method:Compositional matrix adjust.,
Identities:43/135(32%), Positives:61/135(45%), Gaps:20/135(14%)

Query  1      YVDGAANRETKLGKAG-YVTNRG--RQKVVTLTDTTNQKTELQAIYLALQD--SGLEVNI  55
              Y DGA       G  G  +  +G  R+      DTTN + EL A   AL +      +V +
Sbjct  68749  YTDGACKGNPGPGGWGALLVYKGVKRELWGGEPDTTNNRMELMAAIRALAELKRPCKVRL  68928

Query  56     VTDSQYAL-GIIQAQPDQSE-------------SELVNQIIEQLIKKEKVYLAWVPAHKG  101
              VTDSQY + GI     P+  +              ++L  Q+ EQ + + +V   WV  H G
Sbjct  68929  VTDSQYVMQGINDWMPNWKKRGWKTASKQPVKNADLWQQLDEQ-VGRHQVSWEWVRGHTG  69105

Query  102    IGGNEQVDKLVSAGI  116
                GNEQ D L + G+
Sbjct  69106  HPGNEQADLLANRGV  69150



>Stutzerimonas stutzeri KOS6 scaffold2, whole genome shotgun sequence
Sequence ID: NZ_KK020676.1 Length: 2709949
Range 1: 1879963 to 1880364

Score:48.5 bits(114), Expect:0.005,
Method:Compositional matrix adjust.,
Identities:43/135(32%), Positives:61/135(45%), Gaps:20/135(14%)

Query  1        YVDGAANRETKLGKAG-YVTNRG--RQKVVTLTDTTNQKTELQAIYLALQD--SGLEVNI  55
                Y DGA       G  G  +  +G  R+      DTTN + EL A   AL +      +V +
Sbjct  1879963  YTDGACKGNPGPGGWGALLVYKGVKRELWGGEPDTTNNRMELMAAIRALAELKRPCKVRL  1880142

Query  56       VTDSQYAL-GIIQAQPDQSE-------------SELVNQIIEQLIKKEKVYLAWVPAHKG  101
                VTDSQY + GI     P+  +              ++L  Q+ EQ + + +V   WV  H G
Sbjct  1880143  VTDSQYVMQGINDWMPNWKKRGWKTASKQPVKNADLWQQLDEQ-VGRHQVSWEWVRGHTG  1880319

Query  102      IGGNEQVDKLVSAGI  116
                  GNEQ D L + G+
Sbjct  1880320  HPGNEQADLLANRGV  1880364



>Stutzerimonas sp. R40042 chromosome, complete genome
Sequence ID: NZ_CP113258.1 Length: 4666542
Range 1: 2584241 to 2584546

Score:48.5 bits(114), Expect:0.005,
Method:Compositional matrix adjust.,
Identities:36/103(35%), Positives:50/103(48%), Gaps:17/103(16%)

Query  30       TDTTNQKTELQAIYLALQD--SGLEVNIVTDSQYAL-GIIQAQPDQSE------------  74
                 DTTN + EL A   AL +      +V +VTDSQY + GI     P+  +
Sbjct  2584546  PDTTNNRMELMAAIRALAELKRPCKVRLVTDSQYVMQGINDWMPNWKKRGWKTASKQPVK  2584367

Query  75       -SELVNQIIEQLIKKEKVYLAWVPAHKGIGGNEQVDKLVSAGI  116
                 ++L  Q+ EQ+ + E V   WV  H G  GNEQ D L + G+
Sbjct  2584366  NADLWQQLDEQVSRHE-VSWQWVRGHTGDPGNEQADLLANRGV  2584241
```

>Pseudomonas morbosilactucae strain MAFF 302046 302046_contig_42, whole genome shotgun sequence
Sequence ID: NZ_JALQCX010000042.1 Length: 73784
Range 1: 43168 to 43587

Score:48.5 bits(114), Expect:0.005,
Method:Composition-based stats.,
Identities:43/140(31%), Positives:60/140(42%), Gaps:25/140(17%)

```
Query  1      YVDGA----ANRETKLGKAGYVTNRGRQK------VVTLTDTTNQKTELQAIYLALQ--D  48
              YVDGA         K G   Y+TN           V      TN + EL A  +AL+
Sbjct  43587  YVDGACPNNGKPNAKAGWGAYLTNPHGDTLELAGPVPANQPQTNGRAELLAPLMALRRCK  43408

Query  49     SGLEVNIVTDSQY-------------ALGIIQAQPDQSESELVNQIIEQLIKKEKVYLAW  95
               G  +NIV+DSQY             A G  +A   + E   + ++I+Q ++K ++   W
Sbjct  43407  PGAAINIVSDSQYVVKGAMEWLQGWKARGWRKADKKEVEHRDLWELIDQEMQKREIRFQW  43228

Query  96     VPAHKGIGGNEQVDKLVSAG  115
              V  H G  GNE+ D L   G
Sbjct  43227  VRGHSGDTGNERADTLACRG  43168
```

>Deinococcus aluminii strain NBRC 112889 112889_is_8, whole genome shotgun sequence
Sequence ID: NZ_BAABRV010000008.1 Length: 134314
Range 1: 92267 to 92659

Score:48.5 bits(114), Expect:0.005,
Method:Compositional matrix adjust.,
Identities:43/133(32%), Positives:68/133(51%), Gaps:23/133(17%)

```
Query  1      YVDGAANRETKLGKAGYVT---NRGRQKVVTLT--DTTNQKTELQAIYLALQ--DSGLEV  53
              Y DGA   +T+ G  G+ T    RG++ V++    DTTN + EL+ +   L+      +V
Sbjct  92267  YSDGAC--DTQAGHGGWATILRYRGKELVLSGNERDTTNNRMELRGLLEGLKVLKRPCQV  92440

Query  54     NIVTDSQYAL------GIIQAQPDQSES----ELVNQ-IIEQLIKKEKVY---LAWVPAH  99
               +VTDSQY         I++ Q +  ++     + NQ + E+LI + + +     WV  H
Sbjct  92441  RVVTDSQYLRKAFTDGWILKWQRNGWKTAGGDPVKNQDLWEELIAQARTHALTFVWVRGH  92620

Query  100    KGIGGNEQVDKLV  112
               G G NE+VDKL
Sbjct  92621  NGHGENERVDKLA  92659
```

>Deinococcus metallilatus strain MA1002 497.DSPMA1002.2contig00006, whole genome shotgun sequence
Sequence ID: NZ_SDEC01000029.1 Length: 246168
Range 1: 132436 to 132828

Score:48.5 bits(114), Expect:0.006,
Method:Compositional matrix adjust.,
Identities:43/133(32%), Positives:68/133(51%), Gaps:23/133(17%)

```
Query  1       YVDGAANRETKLGKAGYVT---NRGRQKVVTLT--DTTNQKTELQAIYLALQ--DSGLEV  53
               Y DGA   +T+ G  G+ T    RG++ V++    DTTN + EL+ +   L+      +V
Sbjct  132436  YSDGAC--DTQAGHGGWATILRYRGKELVLSGNERDTTNNRMELRGLLEGLKVLKRPCQV  132609

Query  54      NIVTDSQYAL------GIIQAQPDQSES----ELVNQ-IIEQLIKKEKVY---LAWVPAH  99
                +VTDSQY         I++ Q +  ++     + NQ + E+LI + + +     WV  H
Sbjct  132610  RVVTDSQYLRKAFTDGWILKWQRNGWKTAGGDPVKNQDLWEELIAQARTHALTFVWVRGH  132789

Query  100     KGIGGNEQVDKLV  112
                G G NE+VDKL
Sbjct  132790  NGHGENERVDKLA  132828
```

>Deinococcus metallilatus strain MA1002-m5 497.DSPMA1002.3_contig00006, whole genome shotgun sequence
Sequence ID: NZ_VBRC01000006.1 Length: 246233
Range 1: 113378 to 113770

Score:48.5 bits(114), Expect:0.006,
Method:Compositional matrix adjust.,
Identities:43/133(32%), Positives:68/133(51%), Gaps:23/133(17%)

```
Query  1       YVDGAANRETKLGKAGYVT---NRGRQKVVTLT--DTTNQKTELQAIYLALQ--DSGLEV  53
               Y DGA   +T+ G  G+ T    RG++ V++    DTTN + EL+ +   L+      +V
Sbjct  113770  YSDGAC--DTQAGHGGWATILRYRGKELVLSGNERDTTNNRMELRGLLEGLKVLKRPCQV  113597

Query  54      NIVTDSQYAL------GIIQAQPDQSES----ELVNQ-IIEQLIKKEKVY---LAWVPAH  99
                +VTDSQY         I++ Q +  ++     + NQ + E+LI + + +     WV  H
Sbjct  113596  RVVTDSQYLRKAFTDGWILKWQRNGWKTAGGDPVKNQDLWEELIAQARTHALTFVWVRGH  113417

Query  100     KGIGGNEQVDKLV  112
                G G NE+VDKL
Sbjct  113416  NGHGENERVDKLA  113378
```

>Deinococcus metallilatus strain NBRC 110141 sequence15, whole genome shotgun sequence
Sequence ID: NZ_BSUI01000015.1 Length: 279199
Range 1: 146381 to 146773

Score:48.5 bits(114), Expect:0.006,
Method:Compositional matrix adjust.,
Identities:43/133(32%), Positives:68/133(51%), Gaps:23/133(17%)

```
Query  1       YVDGAANRETKLGKAGYVT---NRGRQKVVTLT--DTTNQKTELQAIYLALQ--DSGLEV  53
               Y DGA   +T+ G  G+ T    RG++ V++    DTTN + EL+ +   L+      +V
Sbjct  146773  YSDGAC--DTQAGHGGWATILRYRGKELVLSGNERDTTNNRMELRGLLEGLKVLKRPCQV  146600

Query  54      NIVTDSQYAL------GIIQAQPDQSES----ELVNQ-IIEQLIKKEKVY---LAWVPAH  99
                +VTDSQY         I++ Q +  ++     + NQ + E+LI + + +     WV  H
Sbjct  146599  RVVTDSQYLRKAFTDGWILKWQRNGWKTAGGDPVKNQDLWEELIAQARTHALTFVWVRGH  146420

Query  100     KGIGGNEQVDKLV  112
                G G NE+VDKL
Sbjct  146419  NGHGENERVDKLA  146381
```

>Deinococcus metallilatus strain DSM 105434 Ga0416734_07, whole genome shotgun sequence
Sequence ID: NZ_JACHFV010000007.1 Length: 279933
Range 1: 146684 to 147076

Score:48.5 bits(114), Expect:0.006,
Method:Compositional matrix adjust.,

Identities:43/133(32%), Positives:68/133(51%), Gaps:23/133(17%)

```
Query  1       YVDGAANRETKLGKAGYVT---NRGRQKVVTLT--DTTNQKTELQAIYLALQ--DSGLEV  53
               Y DGA   +T+ G  G+ T    RG++ V++    DTTN + EL+ +   L+       +V
Sbjct  147076  YSDGAC--DTQAGHGGWATILRYRGKELVLSGNERDTTNNRMELRGLLEGLKVLKRPCQV  146903

Query  54      NIVTDSQYAL------GIIQAQPDQSES----ELVNQ-IIEQLIKKEKVY---LAWVPAH  99
                +VTDSQY          I++ Q +  ++     + NQ + E+LI + + +      WV  H
Sbjct  146902  RVVTDSQYLRKAFTDGWILKWQRNGWKTAGGDPVKNQDLWEELIAQARTHALTFVWVRGH  146723

Query  100     KGIGGNEQVDKLV  112
                G G NE+VDKL
Sbjct  146722  NGHGENERVDKLA  146684
```

>Deinococcus metallilatus strain MA1002 chromosome
Sequence ID: NZ_CP038512.1 Length: 3062136
Range 1: 547044 to 547436

Score:48.5 bits(114), Expect:0.006,
Method:Compositional matrix adjust.,
Identities:43/133(32%), Positives:68/133(51%), Gaps:23/133(17%)

```
Query  1       YVDGAANRETKLGKAGYVT---NRGRQKVVTLT--DTTNQKTELQAIYLALQ--DSGLEV  53
               Y DGA   +T+ G  G+ T    RG++ V++    DTTN + EL+ +   L+       +V
Sbjct  547044  YSDGAC--DTQAGHGGWATILRYRGKELVLSGNERDTTNNRMELRGLLEGLKVLKRPCQV  547217

Query  54      NIVTDSQYAL------GIIQAQPDQSES----ELVNQ-IIEQLIKKEKVY---LAWVPAH  99
                +VTDSQY          I++ Q +  ++     + NQ + E+LI + + +      WV  H
Sbjct  547218  RVVTDSQYLRKAFTDGWILKWQRNGWKTAGGDPVKNQDLWEELIAQARTHALTFVWVRGH  547397

Query  100     KGIGGNEQVDKLV  112
                G G NE+VDKL
Sbjct  547398  NGHGENERVDKLA  547436
```

>Pseudomonas lopnurensis strain AL-54 46, whole genome shotgun sequence
Sequence ID: NZ_JADDIX010000046.1 Length: 22361
Range 1: 9406 to 9711

Score:48.5 bits(114), Expect:0.006,
Method:Compositional matrix adjust.,
Identities:35/103(34%), Positives:50/103(48%), Gaps:17/103(16%)

```
Query  30    TDTTNQKTELQAIYLALQD--SGLEVNIVTDSQYAL-GIIQAQPDQSE------------  74
              DTTN + EL A   AL +     +V ++TDSQY + GI    P+  +
Sbjct  9711  ADTTNNRMELMAAIRALAELKRPCKVRLITDSQYVMQGINDWMPNWKKRGWKTASKQPVK  9532

Query  75    -SELVNQIIEQLIKKEKVYLAWVPAHKGIGGNEQVDKLVSAGI  116
              ++L  Q+ EQ+ + E V   WV  H G  GNEQ D L + G+
Sbjct  9531  NADLWRQLDEQVGRHE-VSWEWVRGHTGHPGNEQADLLANRGV  9406
```

>MAG: Pseudomonas lopnurensis isolate CTOTU48819 NODE_78_length_48550_cov_72.279493, whole genome shotgun sequence
Sequence ID: NZ_DAMBEU010000041.1 Length: 48550
Range 1: 22239 to 22544

Score:48.5 bits(114), Expect:0.006,
Method:Compositional matrix adjust.,
Identities:35/103(34%), Positives:50/103(48%), Gaps:17/103(16%)

```
Query  30     TDTTNQKTELQAIYLALQD--SGLEVNIVTDSQYAL-GIIQAQPDQSE------------  74
               DTTN + EL A   AL +     +V ++TDSQY + GI    P+  +
Sbjct  22239  ADTTNNRMELMAAIRALAELKRPCKVRLITDSQYVMQGINDWMPNWKKRGWKTASKQPVK  22418

Query  75     -SELVNQIIEQLIKKEKVYLAWVPAHKGIGGNEQVDKLVSAGI  116
               ++L  Q+ EQ+ + E V   WV  H G  GNEQ D L + G+
Sbjct  22419  NADLWRQLDEQVGRHE-VSWEWVRGHTGHPGNEQADLLANRGV  22544
```

>MAG: Pseudomonas lopnurensis isolate CTOTU48911 NODE_32_length_57616_cov_49.862372, whole genome shotgun sequence
Sequence ID: NZ_DAMBHS010000032.1 Length: 57616
Range 1: 26064 to 26369

Score:48.5 bits(114), Expect:0.006,
Method:Compositional matrix adjust.,
Identities:35/103(34%), Positives:50/103(48%), Gaps:17/103(16%)

```
Query  30     TDTTNQKTELQAIYLALQD--SGLEVNIVTDSQYAL-GIIQAQPDQSE------------  74
               DTTN + EL A   AL +     +V ++TDSQY + GI    P+  +
Sbjct  26369  ADTTNNRMELMAAIRALAELKRPCKVRLITDSQYVMQGINDWMPNWKKRGWKTASKQPVK  26190

Query  75     -SELVNQIIEQLIKKEKVYLAWVPAHKGIGGNEQVDKLVSAGI  116
               ++L  Q+ EQ+ + E V   WV  H G  GNEQ D L + G+
Sbjct  26189  NADLWRQLDEQVGRHE-VSWEWVRGHTGHPGNEQADLLANRGV  26064
```

>Stutzerimonas stutzeri strain CBC1 contig_18, whole genome shotgun sequence
Sequence ID: NZ_JAXBVD010000018.1 Length: 94832
Range 1: 73595 to 73900

Score:48.5 bits(114), Expect:0.006,
Method:Compositional matrix adjust.,
Identities:35/103(34%), Positives:50/103(48%), Gaps:17/103(16%)

```
Query  30     TDTTNQKTELQAIYLALQD--SGLEVNIVTDSQYAL-GIIQAQPDQSE------------  74
               DTTN + EL A   AL +     +V +VTDSQY + GI    P+  +
Sbjct  73595  PDTTNNRMELMAAIRALAELKRPCKVRLVTDSQYVMQGINDWMPNWKKRGWKTASKQPVK  73774

Query  75     -SELVNQIIEQLIKKEKVYLAWVPAHKGIGGNEQVDKLVSAGI  116
               ++L  Q+ EQ + + +V   WV  H G  GNEQ D L + G+
Sbjct  73775  NADLWQQLDEQ-VDRHEVSWQWVRGHTGHPGNEQADLLANRGV  73900
```

>Stutzerimonas stutzeri strain PS_110 NODE_12_length_115677_cov_22.2726, whole genome shotgun sequence
Sequence ID: NZ_RHRL01000012.1 Length: 115677
Range 1: 73495 to 73800

Score:48.5 bits(114), Expect:0.006,
Method:Compositional matrix adjust.,
Identities:35/103(34%), Positives:50/103(48%), Gaps:17/103(16%)

```
Query  30     TDTTNQKTELQAIYLALQD--SGLEVNIVTDSQYAL-GIIQAQPDQSE------------  74
               DTTN + EL A   AL +     +V +VTDSQY + GI    P+  +
Sbjct  73495  PDTTNNRMELMAAIRALAELKRPCKVQLVTDSQYVMQGINDWMPNWKKRGWKTASKQPVK  73674

Query  75     -SELVNQIIEQLIKKEKVYLAWVPAHKGIGGNEQVDKLVSAGI  116
               ++L  Q+ EQ + + +V   WV  H G  GNEQ D L + G+
Sbjct  73675  NADLWQQLDEQ-VNRHEVSWQWVRGHTGHPGNEQADLLANRGV  73800
```

>Stutzerimonas stutzeri strain FDAARGOS_875 chromosome, complete genome
Sequence ID: NZ_CP065750.1 Length: 4535380
Range 1: 218646 to 218951

Score:48.5 bits(114), Expect:0.006,
Method:Compositional matrix adjust.,
Identities:35/103(34%), Positives:50/103(48%), Gaps:17/103(16%)

```
Query  30      TDTTNQKTELQAIYLALQD--SGLEVNIVTDSQYAL-GIIQAQPDQSE------------  74
                DTTN + EL A   AL +     +V +VTDSQY + GI    P+  +
Sbjct  218951  PDTTNNRMELMAAIRALAELKRPCKVQLVTDSQYVMQGINDWMPNWKKRGWKTASKQPVK  218772

Query  75      -SELVNQIIEQLIKKEKVYLAWVPAHKGIGGNEQVDKLVSAGI  116
                ++L  Q+ EQ + + +V   WV  H G  GNEQ D L + G+
Sbjct  218771  NADLWQQLDEQ-VNRHEVSWQWVRGHTGHPGNEQADLLANRGV  218646
```

>Stutzerimonas stutzeri, complete sequence
Sequence ID: NC_015740.1 Length: 4547930
Range 1: 2286049 to 2286354

Score:48.5 bits(114), Expect:0.006,
Method:Compositional matrix adjust.,
Identities:35/103(34%), Positives:50/103(48%), Gaps:17/103(16%)

```
Query  30       TDTTNQKTELQAIYLALQD--SGLEVNIVTDSQYAL-GIIQAQPDQSE------------  74
                 DTTN + EL A   AL +     +V +VTDSQY + GI    P+  +
Sbjct  2286354  PDTTNNRMELMAAIRALAELKRPCKVQLVTDSQYVMQGINDWMPNWKKRGWKTASKQPVK  2286175

Query  75       -SELVNQIIEQLIKKEKVYLAWVPAHKGIGGNEQVDKLVSAGI  116
                 ++L  Q+ EQ + + +V   WV  H G  GNEQ D L + G+
Sbjct  2286174  NADLWQQLDEQ-VNRHEVSWQWVRGHTGHPGNEQADLLANRGV  2286049
```

>MAG: Stutzerimonas nitrititolerans isolate CTOTU48809 NODE_12_length_140689_cov_50.709387, whole genome shotgun sequence
Sequence ID: NZ_DAMBEK010000011.1 Length: 140689
Range 1: 137 to 436

Score:48.5 bits(114), Expect:0.006,
Method:Compositional matrix adjust.,
Identities:34/100(34%), Positives:49/100(49%), Gaps:15/100(15%)

```
Query  33   TNQKTELQAIYLALQDSGLEVNIV--TDSQYALGIIQA-----------QPDQSESELVN  79
            TN + EL A+  AL+       I+  TDS+Y    +Q             + D+   E ++
Sbjct  137  TNSRAELMAVAEALKQCKQPAPIILHTDSEYIAKALQGWLASWKAKGWKKSDRKPPEHLD  316

Query  80   --QIIEQLIKKEKVYLAWVPAHKGIGGNEQVDKLVSAGIR  117
              Q I+QL++ + V   WV AH GI GNE+ D L S G R
Sbjct  317  LWQRIDQLLQGKNVTTHWVKAHSGIAGNERADTLASLGAR  436
```

>Saccharospirillum mangrovi strain HK-33 Scaffold2, whole genome shotgun sequence
Sequence ID: NZ_QAPE01000002.1 Length: 517067
Range 1: 16487 to 16900

Score:48.1 bits(113), Expect:0.007,
Method:Compositional matrix adjust.,
Identities:37/138(27%), Positives:60/138(43%), Gaps:18/138(13%)

```
Query  1      YVDGAANRETKLGKAGYVTNRGRQKVVTL---TDTTNQKTELQAIYLALQD--SGLEVNI  55
              Y DGA       G  G +   G  +        +TTN + EL A  + L+      +V++
Sbjct  16900  YTDGACRGNPGPGGWGVLMRSGAHEKTLHGGEAETTNNRMELMAAIVGLESLKRRCQVDL  16721

Query  56     VTDSQYAL-------------GIIQAQPDQSESELVNQIIEQLIKKEKVYLAWVPAHKGI  102
               TDSQY                G   A     +++ + Q ++Q  I  K +V   WV  H G+
Sbjct  16720  YTDSQYVRKGITEWIHGWKRNGWKTASKQPVKNQDLWQRLDQQIGKHQVEWHWVKGHAGV  16541

Query  103    GGNEQVDKLVSAGIRKVL  120
               GNE+ D L + G+ +++
Sbjct  16540  DGNERADALANQGVDEII  16487
```

>Saccharospirillum mangrovi strain HK-33 chromosome, complete genome
Sequence ID: NZ_CP031415.1 Length: 3686911
Range 1: 1732026 to 1732439

Score:48.1 bits(113), Expect:0.007,
Method:Compositional matrix adjust.,
Identities:37/138(27%), Positives:60/138(43%), Gaps:18/138(13%)

```
Query  1        YVDGAANRETKLGKAGYVTNRGRQKVVTL---TDTTNQKTELQAIYLALQD--SGLEVNI  55
                Y DGA       G  G +   G  +        +TTN + EL A  + L+      +V++
Sbjct  1732026  YTDGACRGNPGPGGWGVLMRSGAHEKTLHGGEAETTNNRMELMAAIVGLESLKRRCQVDL  1732205

Query  56       VTDSQYAL-------------GIIQAQPDQSESELVNQIIEQLIKKEKVYLAWVPAHKGI  102
                 TDSQY                G   A     +++ + Q ++Q  I  K +V   WV  H G+
Sbjct  1732206  YTDSQYVRKGITEWIHGWKRNGWKTASKQPVKNQDLWQRLDQQIGKHQVEWHWVKGHAGV  1732385

Query  103      GGNEQVDKLVSAGIRKVL  120
                 GNE+ D L + G+ +++
Sbjct  1732386  DGNERADALANQGVDEII  1732439
```

>MAG: Nitrosomonas sp. isolate HKST-UBA152 k127_817706, whole genome shotgun sequence
Sequence ID: NZ_JAGRFN010000156.1 Length: 19510
Range 1: 12912 to 13325

Score:47.8 bits(112), Expect:0.010,
Method:Compositional matrix adjust.,
Identities:41/138(30%), Positives:60/138(43%), Gaps:18/138(13%)

```
Query  1      YVDGAANRETKLGKAGYVTNRGRQKVVTL---TDTTNQKTELQAIYLALQD--SGLEVNI  55
              Y DGA      +G  G V   G  +         TTN + EL A   AL+       V +
```

```
Sbjct  13325  YTDGACKGNPGVGGWGAVLKYGVHEQEIFGGEKHTTNNRMELLAAIRALESLKRACRVRL  13146

Query  56     VTDSQYAL-GIIQ------------AQPDQSESELVNQIIEQLIKKEKVYLAWVPAHKGI  102
               TDSQY   GI Q            A     ++E + + +++L +K  +   WV  H G
Sbjct  13145  YTDSQYVHNGIGQWMQAWKARNWRTANKKPVKNEDLWRTLDRLSQKHDIEWLWVRGHAGH  12966

Query  103    GGNEQVDKLVSAGIRKVL  120
               GNE+ D L + G++K+L
Sbjct  12965  EGNERADMLANRGVKKIL  12912




>Deinococcus hopiensis KR-140, whole genome shotgun sequence
Sequence ID: NZ_FWWU01000009.1 Length: 3128235
Range 1: 514444 to 514836

Score:47.8 bits(112), Expect:0.011,
Method:Compositional matrix adjust.,
Identities:42/133(32%), Positives:69/133(51%), Gaps:23/133(17%)

Query  1       YVDGAANRETKLGKAGYVT---NRGRQKVVTLTD--TTNQKTELQAIYLALQ--DSGLEV  53
               Y DGA   +T+ G  G+ T    RG++ V++  +  TTN + EL+ +   L+      ++
Sbjct  514836  YSDGAC--DTQAGHGGWATILNYRGKELVLSGNEEGTTNNRMELRGLLEGLKVLKRPCQI  514663

Query  54      NIVTDSQYAL------GIIQAQPDQSES----ELVNQ-IIEQLIKKEKVY---LAWVPAH  99
                +VTDSQY         I++ Q +  ++     + NQ + E+LI + K++     WV  H
Sbjct  514662  RVVTDSQYLRKAFTDGWILKWQRNGWKTAGGEPVKNQDLWEELIGQAKIHALTFVWVRGH  514483

Query  100     KGIGGNEQVDKLV  112
                G G NE+VDKL
Sbjct  514482  NGHGENERVDKLA  514444




>Hydrogenophilus thiooxidans strain SS56 contig-100_16, whole genome shotgun sequence
Sequence ID: NZ_JAHLSY010000017.1 Length: 46787
Range 1: 10691 to 11092

Score:47.8 bits(112), Expect:0.012,
Method:Composition-based stats.,
Identities:41/134(31%), Positives:62/134(46%), Gaps:18/134(13%)

Query  1      YVDGAANRETKLGKAGYVTN-RGRQKVVTLTD--TTNQKTELQAIYLALQ--DSGLEVNI  55
              Y DGA +     G  G V   RGR++ ++ ++  TTN + ELQA+  AL+     + V I
Sbjct  10691  YTDGACSGNPGPGGWGAVLRWRGRERQLSGSEAHTTNNRMELQAVIAALEALKRPVPVTI  10870

Query  56     VTDSQYAL-GIIQAQP--------DQSESELVN----QIIEQLIKKEKVYLAWVPAHKGI  102
               TDSQY   G+ +  P           S   + N    + ++ L  +  +   WV  H G+
Sbjct  10871  YTDSQYVQKGVTEWLPRWKARQWRTASGHAVANRDLWERLDDLATQLPITWCWVKGHDGV  11050

Query  103    GGNEQVDKLVSAGI  116
               GNE  D+L    I
Sbjct  11051  AGNELADRLARQAI  11092




>Deinococcus pimensis DSM 21231 DeipiDRAFT_Scaffold38.32, whole genome shotgun sequence
Sequence ID: NZ_KI912645.1 Length: 52679
Range 1: 42732 to 43145

Score:47.4 bits(111), Expect:0.017,
Method:Composition-based stats.,
Identities:39/138(28%), Positives:65/138(47%), Gaps:19/138(13%)

Query  1      YVDGAANRETKLGKAGYVTN-RGRQKVVT--LTDTTNQKTELQAIYLALQ--DSGLEVNI  55
              Y DGA + +  LG    + N RGR+ V++    DTTN + EL+A+   L+       V +
Sbjct  43145  YSDGACDTQAGLGGWATILNCRGRELVLSGGERDTTNNRMELRALLEGLRALKRPCNVRV  42966

Query  56     VTDSQYAL--------------GIIQAQPDQSESELVNQIIEQLIKKEKVYLAWVPAHKG  101
              VTDSQY                 G   +  D  +++ + + + +  ++  +   WV  H G
Sbjct  42965  VTDSQYLRKAFTDSWILKWQRNGWKTSGNDSVKNQDLWEALVEEARRHSLTFVWVKGHAG  42786

Query  102    IGGNEQVDKLVSAGIRKV  119
               G NE+VD+L     +K+
Sbjct  42785  HGENERVDQLAVTERKKL  42732




>Methylomonas sp. SURF-1 SURF1.00003, whole genome shotgun sequence
Sequence ID: NZ_JANIBM010000003.1 Length: 328747
Range 1: 296644 to 296946

Score:47.0 bits(110), Expect:0.018,
Method:Compositional matrix adjust.,
Identities:31/101(31%), Positives:50/101(49%), Gaps:15/101(14%)

Query  31      DTTNQKTELQAIYLALQ--DSGLEVNIVTDSQYAL-GIIQAQPDQSESEL---VNQIIEQ  84
               +TTN + EL A   AL+  +    V + TDS+Y L GI +  P+  +       NQ ++
Sbjct  296644  ETTNNRMELMAAIQALETLNKPCNVRLYTDSKYVLQGITEWMPNWKKRAWKTAANQPVKN  296823

Query  85      L---------IKKEKVYLAWVPAHKGIGGNEQVDKLVSAGI  116
               +         I+  ++   WV  H G+ GNE+ D+L + GI
Sbjct  296824  VELWRRLDAAIQLHRIDWVWVKGHAGVAGNERADQLANQGI  296946




>MAG: Brucella anthropi isolate UBA809 UBA809_contig_1384, whole genome shotgun sequence
Sequence ID: NZ_DBIF01000027.1 Length: 11382
Range 1: 164 to 574

Score:47.0 bits(110), Expect:0.019,
Method:Compositional matrix adjust.,
Identities:37/137(27%), Positives:63/137(45%), Gaps:21/137(15%)

Query  1    YVDGAANRETKLGKAGYVTNRGRQKVVT----LTDTTNQKTELQAIYLALQDS---GLEV  53
            + DG        G  G+   R   +V +    + DTTN   EL  + ++++ +   G  V
Sbjct  574  FSDGGCEPNPGDGGWGFAVYRDGVEVHSDCGGMLDTTNNVMELTGMLMSIRWAVVQGEPV  395

Query  54   NIVTDSQYAL------------GIIQAQPDQ-SESELVNQIIEQLIKKEKVYLAWVPAH  99
            +I+ DSQY +              G  + + + + S+L  QI   L  K+ + + WV  H
Sbjct  394  HILCDSQYVVNGCNDWRHGWKKKGWKRGETKELANSDLWQQIDAVLDGKDHITIGWVKGH  215

Query  100  KGIGGNEQVDKLVSAGI  116
             GI GNE+ D+L + G+
Sbjct  214  VGITGNERADELATMGM  164
```

```
>MAG: uncultured Halovibrio sp. isolate SRR6201609_bin.19_MetaBAT_v2.12.1_MAG, whole genome shotgun sequence
Sequence ID: NZ_CAJXMA010000070.1 Length: 14964
Range 1: 14038 to 14451

Score:46.6 bits(109), Expect:0.024,
Method:Compositional matrix adjust.,
Identities:43/139(31%), Positives:60/139(43%), Gaps:20/139(14%)

Query  1      YVDGAANRETKLGKAGYVTNRGRQKVVTL---TDTTNQKTELQAIY--LALQDSGLEVNI  55
              Y DGA       G  G V   G  +         TTN + EL+A    LA      +V+I
Sbjct  14451  YTDGACKGNPGPGGWGVVLRYGDHEKHLWGGEPGTTNNRMELRAAIEGLAALRRQCDVDI  14272

Query  56     VTDSQYAL-GIIQ-------------AQPDQSESELVNQIIEQLIKKEKVYLAWVPAHKG  101
               TDSQY   GI Q             A+     +EL  Q +++L++  ++   WV  H G
Sbjct  14271  YTDSQYVRNGINQWMANWKRNGWKTSARKPVKNAELW-QRLDELVQSHRIQWHWVRGHSG  14095

Query  102    IGGNEQVDKLVSAGIRKVL  120
                GNEQ D L + G  +V+
Sbjct  14094  NDGNEQADALANRGAAEVM  14038



>Halomonas sp. 22501_18_FS NODE_5_length_234959_cov_13.5081, whole genome shotgun sequence
Sequence ID: NZ_WMFD01000005.1 Length: 234959
Range 1: 205025 to 205438

Score:46.6 bits(109), Expect:0.024,
Method:Compositional matrix adjust.,
Identities:43/139(31%), Positives:60/139(43%), Gaps:20/139(14%)

Query  1       YVDGAANRETKLGKAGYVTNRGRQKVVTL---TDTTNQKTELQAIY--LALQDSGLEVNI  55
               Y DGA       G  G V   G  +         TTN + EL+A    LA      +V+I
Sbjct  205025  YTDGACKGNPGPGGWGVVLRYGDHEKHLWGGEPGTTNNRMELRAAIEGLAALRRQCDVDI  205204

Query  56      VTDSQYAL-GIIQ-------------AQPDQSESELVNQIIEQLIKKEKVYLAWVPAHKG  101
                TDSQY   GI Q             A+     +EL  Q +++L++  ++   WV  H G
Sbjct  205205  YTDSQYVRNGINQWMANWKRNGWKTSARKPVKNAELW-QRLDELVQSHRIQWHWVRGHSG  205381

Query  102     IGGNEQVDKLVSAGIRKVL  120
                 GNEQ D L + G  +V+
Sbjct  205382  NDGNEQADALANRGAAEVM  205438



>Vreelandella utahensis strain 22507_15_FS NODE_6_length_234959_cov_92.6278, whole genome shotgun sequence
Sequence ID: NZ_WMEX01000006.1 Length: 234959
Range 1: 205025 to 205438

Score:46.6 bits(109), Expect:0.024,
Method:Compositional matrix adjust.,
Identities:43/139(31%), Positives:60/139(43%), Gaps:20/139(14%)

Query  1       YVDGAANRETKLGKAGYVTNRGRQKVVTL---TDTTNQKTELQAIY--LALQDSGLEVNI  55
               Y DGA       G  G V   G  +         TTN + EL+A    LA      +V+I
Sbjct  205025  YTDGACKGNPGPGGWGVVLRYGDHEKHLWGGEPGTTNNRMELRAAIEGLAALRRQCDVDI  205204

Query  56      VTDSQYAL-GIIQ-------------AQPDQSESELVNQIIEQLIKKEKVYLAWVPAHKG  101
                TDSQY   GI Q             A+     +EL  Q +++L++  ++   WV  H G
Sbjct  205205  YTDSQYVRNGINQWMANWKRNGWKTSARKPVKNAELW-QRLDELVQSHRIQWHWVRGHSG  205381

Query  102     IGGNEQVDKLVSAGIRKVL  120
                 GNEQ D L + G  +V+
Sbjct  205382  NDGNEQADALANRGAAEVM  205438



>Chishuiella changwenlii strain DSM 27989, whole genome shotgun sequence
Sequence ID: NZ_FRBH01000007.1 Length: 305184
Range 1: 88186 to 88470

Score:46.6 bits(109), Expect:0.024,
Method:Compositional matrix adjust.,
Identities:31/95(33%), Positives:46/95(48%), Gaps:12/95(12%)

Query  32     TTNQKTELQAIYLALQD---SGLEVNIVTDSQYA---------LGIIQAQPDQSESELVN  79
              TTN + EL A+ +AL+    +  E+ IVTDS+Y           G +Q      ++E +
Sbjct  88186  TTNNRMELMAVIVALETLKFTNTEITIVTDSKYVVDAVEKRWVFGWLQKGFKGKKNEDLW  88365

Query  80     QIIEQLIKKEKVYLAWVPAHKGIGGNEQVDKLVSA  114
              +   Q+ K  KV   W+  H G   NE+ DKL  A
Sbjct  88366  RRFLQVYKIHKVKFQWIKGHAGHHWNEEADKLAVA  88470



>Chishuiella changwenlii strain CGMCC 1.12707 sequence01, whole genome shotgun sequence
Sequence ID: NZ_BMFL01000001.1 Length: 305393
Range 1: 216756 to 217040

Score:46.6 bits(109), Expect:0.024,
Method:Compositional matrix adjust.,
Identities:31/95(33%), Positives:46/95(48%), Gaps:12/95(12%)

Query  32      TTNQKTELQAIYLALQD---SGLEVNIVTDSQYA---------LGIIQAQPDQSESELVN  79
               TTN + EL A+ +AL+    +  E+ IVTDS+Y           G +Q      ++E +
Sbjct  217040  TTNNRMELMAVIVALETLKFTNTEITIVTDSKYVVDAVEKRWVFGWLQKGFKGKKNEDLW  216861

Query  80      QIIEQLIKKEKVYLAWVPAHKGIGGNEQVDKLVSA  114
               +   Q+ K  KV   W+  H G   NE+ DKL  A
Sbjct  216860  RRFLQVYKIHKVKFQWIKGHAGHHWNEEADKLAVA  216756



>MAG: Chishuiella changwenlii isolate CTOTU47302 NODE_1_length_371829_cov_5.156235, whole genome shotgun sequence
Sequence ID: NZ_DAINVI010000001.1 Length: 371829
Range 1: 163189 to 163473

Score:46.6 bits(109), Expect:0.024,
Method:Compositional matrix adjust.,
Identities:31/95(33%), Positives:46/95(48%), Gaps:12/95(12%)

Query  32      TTNQKTELQAIYLALQD---SGLEVNIVTDSQYA---------LGIIQAQPDQSESELVN  79
               TTN + EL A+ +AL+    +  E+ IVTDS+Y           G +Q      ++E +
Sbjct  163189  TTNNRMELMAVIVALETLKFTNTEITIVTDSKYVVDAVEKRWVFGWLQKGFKGKKNEDLW  163368

Query  80      QIIEQLIKKEKVYLAWVPAHKGIGGNEQVDKLVSA  114
               +   Q+ K  KV   W+  H G   NE+ DKL  A
Sbjct  163369  RRFLQVYKIHKVKFQWIKGHAGHHWNEEADKLAVA  163473
```

```
>Halospina sp. K52047b contig02, whole genome shotgun sequence
Sequence ID: NZ_VYJH01000002.1 Length: 478842
Range 1: 422564 to 422977

Score:46.6 bits(109), Expect:0.024,
Method:Compositional matrix adjust.,
Identities:43/139(31%), Positives:60/139(43%), Gaps:20/139(14%)

Query  1       YVDGAANRETKLGKAGYVTNRGRQKVVTL---TDTTNQKTELQAIY--LALQDSGLEVNI  55
               Y DGA       G  G V   G  +          TTN + EL+A    LA      +V+I
Sbjct  422977  YTDGACKGNPGPGGWGVVLRYGDHEKHLWGGEPGTTNNRMELRAAIEGLAALRRQCDVDI  422798

Query  56      VTDSQYAL-GIIQ-------------AQPDQSESELVNQIIEQLIKKEKVYLAWVPAHKG  101
                TDSQY   GI Q              A+     +EL  Q +++L++  ++   WV  H G
Sbjct  422797  YTDSQYVRNGINQWMANWKRNGWKTSARKPVKNAELW-QRLDELVQSHRIQWHWVRGHSG  422621

Query  102     IGGNEQVDKLVSAGIRKVL  120
                 GNEQ D L + G  +V+
Sbjct  422620  NDGNEQADALANRGAAEVM  422564



>Halovibrio salipaludis strain YL5-2 contig2, whole genome shotgun sequence
Sequence ID: NZ_NSKD01000002.1 Length: 639556
Range 1: 173229 to 173642

Score:46.6 bits(109), Expect:0.024,
Method:Compositional matrix adjust.,
Identities:43/139(31%), Positives:60/139(43%), Gaps:20/139(14%)

Query  1       YVDGAANRETKLGKAGYVTNRGRQKVVTL---TDTTNQKTELQAIY--LALQDSGLEVNI  55
               Y DGA       G  G V   G  +          TTN + EL+A    LA      +V+I
Sbjct  173229  YTDGACKGNPGPGGWGVVLRYGDHEKHLWGGEPGTTNNRMELRAAIEGLAALRRQCDVDI  173408

Query  56      VTDSQYAL-GIIQ-------------AQPDQSESELVNQIIEQLIKKEKVYLAWVPAHKG  101
                TDSQY   GI Q              A+     +EL  Q +++L++  ++   WV  H G
Sbjct  173409  YTDSQYVRNGINQWMANWKRNGWKTSARKPVKNAELW-QRLDELVQSHRIQWHWVRGHSG  173585

Query  102     IGGNEQVDKLVSAGIRKVL  120
                 GNEQ D L + G  +V+
Sbjct  173586  NDGNEQADALANRGAAEVM  173642



>Lachnoclostridium phocaeense strain Marseille-P3177T chromosome contig00001
Sequence ID: NZ_LT635479.1 Length: 3500754
Range 1: 1458059 to 1458319

Score:46.6 bits(109), Expect:0.025,
Method:Composition-based stats.,
Identities:27/87(31%), Positives:51/87(58%), Gaps:2/87(2%)

Query  32       TTNQKTELQAIYLALQD--SGLEVNIVTDSQYALGIIQAQPDQSESELVNQIIEQLIKKE  89
                +TN + EL A   AL+   +G EV + +DSQY +  ++    ++++  + + +++ ++ +
Sbjct  1458319  STNNRMELLAAIEALKTIPAGAEVKVKSDSQYLVQTMEEGWSRNKNHDLWKQLDEALRGK  1458140

Query  90       KVYLAWVPAHKGIGGNEQVDKLVSAGI  116
                KV   WV  H G   NE+ D+L +AGI
Sbjct  1458139  KVSFKWVRGHNGNQWNERCDELATAGI  1458059



>Deinococcus aetherius strain JCM 11751 sequence18, whole genome shotgun sequence
Sequence ID: NZ_BAABDW010000018.1 Length: 486419
Range 1: 328488 to 328880

Score:46.6 bits(109), Expect:0.025,
Method:Compositional matrix adjust.,
Identities:41/134(31%), Positives:65/134(48%), Gaps:25/134(18%)

Query  1       YVDGAANRETKLGKAGYVT---NRGRQKVVTLTD--TTNQKTELQAIYLALQ--DSGLEV  53
               Y DGA +  T  G  G+ T    RG++ V++  +  TTN + EL+ +   L+      +V
Sbjct  328488  YSDGACD--TAAGHGGWATILKYRGQEFVLSGNEEGTTNNRMELRGLLEGLRALKRPCQV  328661

Query  54      NIVTDSQYAL------GIIQAQPDQSES---------ELVNQIIEQLIKKEKVYLAWVPA  98
                ++TDSQY          I++ Q +  ++          EL  ++IEQ  KK  +   WV
Sbjct  328662  RVITDSQYLRKAFTDGWILKWQRNGWKTAGGEPVKNRELWEELIEQ-AKKHALTFVWVRG  328838

Query  99      HKGIGGNEQVDKLV  112
               H G   NE+VD+L
Sbjct  328839  HDGHAENERVDRLA  328880



>Deinococcus aetherius strain ST0316 chromosome, complete genome
Sequence ID: NZ_AP026560.1 Length: 3135460
Range 1: 2367705 to 2368097

Score:46.6 bits(109), Expect:0.025,
Method:Compositional matrix adjust.,
Identities:41/134(31%), Positives:65/134(48%), Gaps:25/134(18%)

Query  1        YVDGAANRETKLGKAGYVT---NRGRQKVVTLTD--TTNQKTELQAIYLALQ--DSGLEV  53
                Y DGA +  T  G  G+ T    RG++ V++  +  TTN + EL+ +   L+      +V
Sbjct  2368097  YSDGACD--TAAGHGGWATILKYRGQEFVLSGNEEGTTNNRMELRGLLEGLRALKRPCQV  2367924

Query  54       NIVTDSQYAL------GIIQAQPDQSES---------ELVNQIIEQLIKKEKVYLAWVPA  98
                 ++TDSQY          I++ Q +  ++          EL  ++IEQ  KK  +   WV
Sbjct  2367923  RVITDSQYLRKAFTDGWILKWQRNGWKTAGGEPVKNRELWEELIEQ-AKKHALTFVWVRG  2367747

Query  99       HKGIGGNEQVDKLV  112
                H G   NE+VD+L
Sbjct  2367746  HDGHAENERVDRLA  2367705



>Nitrosospira sp. Nsp2 Ga0181081_101, whole genome shotgun sequence
Sequence ID: NZ_QAOU01000001.1 Length: 921569
Range 1: 834128 to 834541

Score:46.2 bits(108), Expect:0.034,
Method:Compositional matrix adjust.,
Identities:39/138(28%), Positives:63/138(45%), Gaps:18/138(13%)

Query  1       YVDGAANRETKLGKAGYV---TNRGRQKVVTLTDTTNQKTELQAIYLALQ--DSGLEVNI  55
               + DGA      +G  G +     R R+       TTN + EL A+  AL+      EV +
Sbjct  834128  FTDGACKGNPGVGGWGALLRYNGRTRELCGGEKLTTNNRMELLAVIRALEALTKPCEVRV  834307
```

```
Query  56      VTDSQYA-LGIIQ------------AQPDQSESELVNQIIEQLIKKEKVYLAWVPAHKGI  102
                TDS Y   GI +            A     +++ + + +++L ++ K+   WV  H G
Sbjct  834308  HTDSTYVQKGISEWIHSWKKRDWRTADKKPVKNDDLWKELDRLTQRHKIEWLWVRGHSGH  834487

Query  103     GGNEQVDKLVSAGIRKVL  120
                GNEQ D+L + G++ VL
Sbjct  834488  DGNEQADRLANQGVQSVL  834541
```

>Nitrosospira sp. Is2 chromosome, complete genome
Sequence ID: NZ_CP137134.1 Length: 3574273
Range 1: 197306 to 197719

Score:46.2 bits(108), Expect:0.034,
Method:Compositional matrix adjust.,
Identities:39/138(28%), Positives:63/138(45%), Gaps:18/138(13%)

```
Query  1       YVDGAANRETKLGKAGYV---TNRGRQKVVTLTDTTNQKTELQAIYLALQ--DSGLEVNI  55
               + DGA      +G  G +     R R+       TTN + EL A+  AL+      EV +
Sbjct  197719  FTDGACKGNPGVGGWGALLRYNGRTRELCGGEKLTTNNRMELLAVIRALEALTKPCEVRV  197540

Query  56      VTDSQYA-LGIIQ------------AQPDQSESELVNQIIEQLIKKEKVYLAWVPAHKGI  102
                TDS Y   GI +            A     +++ + + +++L ++ K+   WV  H G
Sbjct  197539  HTDSTYVQKGISEWIHSWKKRDWRTADKKPVKNDDLWKELDRLTQRHKIEWLWVRGHSGH  197360

Query  103     GGNEQVDKLVSAGIRKVL  120
                GNEQ D+L + G++ VL
Sbjct  197359  DGNEQADRLANQGVQSVL  197306
```

>MAG: Brucella intermedia isolate CTOTU48914 NODE_45_length_124729_cov_3.484375, whole genome shotgun sequence
Sequence ID: NZ_DAMBHV010000011.1 Length: 124729
Range 1: 107320 to 107631

Score:46.2 bits(108), Expect:0.037,
Method:Composition-based stats.,
Identities:29/104(28%), Positives:53/104(50%), Gaps:16/104(15%)

```
Query  29      LTDTTNQKTELQAIYLALQDSGLE---VNIVTDSQYAL-------------GIIQAQPDQ  72
               + D+TN   EL  + +A++ + ++   V+I+ DSQY +                G  + +
Sbjct  107320  MLDSTNNIMELTGMLMAIRWAAVQGEPVHILCDSQYVVNGCNDWRHSWKKAGWKRGKKAL  107499

Query  73      SESELVNQIIEQLIKKEKVYLAWVPAHKGIGGNEQVDKLVSAGI  116
               + ++L  QI    L  K+ + + WV  H GI GNE+ D+L + G+
Sbjct  107500  ANADLWQQIDAVLDGKDHITIGWVKGHAGITGNERADELATTGM  107631
```

>Chitinophaga sp. Cy-1792 NODE_2_length_2808195_cov_51.575576/0, whole genome shotgun sequence
Sequence ID: NZ_VWWO01000002.1 Length: 2808195
Range 1: 2205257 to 2205646

Score:46.2 bits(108), Expect:0.042,
Method:Composition-based stats.,
Identities:39/131(30%), Positives:59/131(45%), Gaps:17/131(12%)

```
Query  1        YVDGAANRETKLGKAGYVT---NRGRQKVVTLTDTTNQKTELQAIYL---ALQDSGLEVN  54
                Y DG++      G  G +    N  ++       TTN + EL A+ +   AL+  GLEVN
Sbjct  2205646  YTDGSSRGNPGPGGYGVILIWGNIRKEMSQGYRKTTNNRMELLAVIVGLEALKRDGLEVN  2205467

Query  55       IVTDSQYALGIIQAQ----------PDQSESELVNQIIEQLIKKEKVYLAWVPAHKGIGG  104
                I TDS+Y +  I+               D+   +L  + I  L +K KV + WV  H
Sbjct  2205466  IFTDSKYVVDSIEKGWIWGWVKTGFKDKKNKDLWMRFIP-LFRKHKVKMNWVKGHATNPF  2205290

Query  105      NEQVDKLVSAG  115
                N + D+L +A
Sbjct  2205289  NNRCDELATAA  2205257
```

>MAG: Pseudomonas sp. isolate RS261_metabat.bin.8 RS261_k141_142368, whole genome shotgun sequence
Sequence ID: NZ_JANJWQ010000069.1 Length: 165529
Range 1: 68514 to 68813

Score:45.8 bits(107), Expect:0.046,
Method:Compositional matrix adjust.,
Identities:34/101(34%), Positives:49/101(48%), Gaps:17/101(16%)

```
Query  32     TTNQKTELQAIYLALQD--SGLEVNIVTDSQYAL-GIIQAQPDQSE-------------S  75
              TTN + EL A   AL +     +V +VTDSQY + GI    P+  +             +
Sbjct  68813  TTNNRMELMAAIRALAELKRPCQVRLVTDSQYVMQGINDWLPNWKKRGWKTASKQPVKNA  68634

Query  76     ELVNQIIEQLIKKEKVYLAWVPAHKGIGGNEQVDKLVSAGI  116
              +L  Q+ EQ + + +V   WV  H G  GNEQ D L + G+
Sbjct  68633  DLWQQLDEQ-VNRHQVSWKWVRGHTGHPGNEQADLLANRGV  68514
```

>Pseudomonas sp. FIP_A4 k141_127, whole genome shotgun sequence
Sequence ID: NZ_JAYDXP010000130.1 Length: 170970
Range 1: 103434 to 103733

Score:45.8 bits(107), Expect:0.046,
Method:Compositional matrix adjust.,
Identities:34/101(34%), Positives:49/101(48%), Gaps:17/101(16%)

```
Query  32      TTNQKTELQAIYLALQD--SGLEVNIVTDSQYAL-GIIQAQPDQSE-------------S  75
               TTN + EL A   AL +     +V +VTDSQY + GI    P+  +             +
Sbjct  103733  TTNNRMELMAAIRALAELKRPCQVRLVTDSQYVMQGINDWLPNWKKRGWKTASKQPVKNA  103554

Query  76      ELVNQIIEQLIKKEKVYLAWVPAHKGIGGNEQVDKLVSAGI  116
               +L  Q+ EQ + + +V   WV  H G  GNEQ D L + G+
Sbjct  103553  DLWQQLDEQ-VNRHQVSWKWVRGHTGHPGNEQADLLANRGV  103434
```

Query #12: sp|P04585|Integrase_HV1H2 Gag-Pol polyprotein OS=Human immunodeficiency virus type 1 group M subtype B (isolate HXB2) OX=11706 GN=gag-pol PE=1 SV=4 Query ID: lcl|Query_2387720 Length: 288

Sequences producing significant alignments:

| Description | Scientific Name | Common Name | Taxid | Max Score | Total Score | Query cover | E Value | Per. Ident | Acc. Len | Accession |
|---|---|---|---|---|---|---|---|---|---|---|
| Actinoplanes campanulatus strain JCM 3059 sequence058, whole... | Actinoplanes... | NA | 113559 | 591 | 591 | 100% | 0.0 | 97.22 | 4754 | NZ_BMPW010000... |
| Escherichia coli strain VRES0669, whole genome shotgun sequence | Escherichia ... | NA | 562 | 575 | 575 | 100% | 0.0 | 93.75 | 5544 | NZ_UINV010000... |

| | | | | | | | | | | |
|---|---|---|---|---|---|---|---|---|---|---|
| Pseudomonas aeruginosa strain ENV-297... | Pseudomonas ... | NA | 287 | 560 | 560 | 100% | 0.0 | 90.62 | 1100 | NZ_RADD010000... |
| Pseudomonas aeruginosa strain VET-42... | Pseudomonas ... | NA | 287 | 559 | 559 | 100% | 0.0 | 90.28 | 1154 | NZ_RACX010001... |
| Burkholderia pseudomallei strain 2604STDY5518971... | Burkholderia... | NA | 28450 | 518 | 518 | 89% | 0.0 | 93.05 | 1345 | NZ_UVFZ020000... |
| Klebsiella pneumoniae strain P7701... | Klebsiella p... | NA | 573 | 482 | 482 | 82% | 2e-170 | 97.05 | 711 | NZ_JACAWM0100... |
| Klebsiella pneumoniae strain ARGID_31501, whole genome shotgun... | Klebsiella p... | NA | 573 | 504 | 504 | 86% | 7e-170 | 94.40 | 2655 | NZ_CAKLPD0100... |
| Enterococcus alcedinis strain CCM 8433 sequence23, whole genom... | Enterococcus... | NA | 1274384 | 455 | 455 | 77% | 8e-143 | 96.86 | 6075 | NZ_BMDT010000... |
| Deinococcus aquiradiocola strain JCM 14371 sequence30, whole... | Deinococcus ... | NA | 393059 | 455 | 455 | 77% | 8e-143 | 96.86 | 6128 | NZ_BMOE010000... |
| Thalassovita mangrovi strain GS-10 Scaffold30_1, whole genome... | Thalassovita... | NA | 2692236 | 392 | 392 | 65% | 6e-135 | 96.83 | 722 | NZ_WWEN010000... |
| Methylobacterium radiotolerans strain MAMP 4754... | Methylobacte... | NA | 31998 | 350 | 350 | 65% | 8e-114 | 85.42 | 1636 | NZ_NKQS010006... |
| Nocardioides luteus strain JCM 3358 sequence39, whole genome... | Nocardioides... | NA | 1844 | 370 | 370 | 63% | 1e-112 | 95.08 | 5754 | NZ_BMRK010000... |
| Streptomyces griseoflavus strain JCM 4479 sequence37, whole... | Streptomyces... | NA | 35619 | 370 | 370 | 63% | 1e-112 | 95.08 | 5775 | NZ_BMUC010000... |
| Saccharothrix coeruleofusca strain JCM 3313 sequence45, whole... | Saccharothri... | NA | 33919 | 370 | 370 | 63% | 1e-112 | 95.08 | 5775 | NZ_BMRG010000... |
| Actinoplanes ianthinogenes strain JCM 3249 sequence037, whole... | Actinoplanes... | NA | 122358 | 370 | 370 | 63% | 1e-112 | 95.08 | 5815 | NZ_BMQZ010000... |
| Sinomonas sp. KCTC 49339 sequence28, whole genome shotgun... | Sinomonas sp... | NA | 2769422 | 370 | 370 | 63% | 1e-112 | 95.08 | 5849 | NZ_BNCM010000... |
| Saccharothrix mutabilis subsp. mutabilis strain JCM 3380... | Saccharothri... | NA | 66855 | 370 | 370 | 63% | 1e-112 | 95.08 | 5899 | NZ_BAAABU0100... |
| Klebsiella pneumoniae strain EuSCAPE_IL016, whole genome shotg... | Klebsiella p... | NA | 573 | 295 | 295 | 60% | 3e-97 | 83.91 | 523 | NZ_UKKA020000... |
| Klebsiella pneumoniae strain EuSCAPE_LT003, whole genome shotg... | Klebsiella p... | NA | 573 | 261 | 261 | 47% | 1e-84 | 91.91 | 422 | NZ_UKJH020000... |
| Klebsiella pneumoniae strain EuSCAPE_IL004, whole genome shotg... | Klebsiella p... | NA | 573 | 254 | 254 | 44% | 7e-82 | 92.91 | 382 | NZ_UKJP020001... |
| Klebsiella pneumoniae strain EuSCAPE_MT010, whole genome shotg... | Klebsiella p... | NA | 573 | 233 | 233 | 42% | 2e-73 | 92.62 | 367 | NZ_UKIT020000... |
| Neisseria gonorrhoeae strain GC-206... | Neisseria go... | NA | 485 | 226 | 226 | 38% | 3e-71 | 92.79 | 344 | NZ_SAWN010001... |
| Klebsiella pneumoniae strain EuSCAPE_IL007, whole genome shotg... | Klebsiella p... | NA | 573 | 220 | 220 | 39% | 1e-68 | 92.92 | 343 | NZ_UKJT020000... |
| Thalassovita mangrovi strain GS-10 Scaffold21_1, whole genome... | Thalassovita... | NA | 2692236 | 227 | 227 | 39% | 1e-66 | 97.35 | 1440 | NZ_WWEN010000... |
| Klebsiella pneumoniae strain EuSCAPE_LT003, whole genome shotg... | Klebsiella p... | NA | 573 | 197 | 197 | 34% | 7e-60 | 92.86 | 297 | NZ_UKJH020000... |
| Deinococcus aquiradiocola strain JCM 14371 sequence36, whole... | Deinococcus ... | NA | 393059 | 203 | 203 | 35% | 1e-59 | 98.02 | 869 | NZ_BMOE010000... |
| Klebsiella pneumoniae strain EuSCAPE_IL005, whole genome shotg... | Klebsiella p... | NA | 573 | 196 | 196 | 35% | 4e-58 | 92.23 | 636 | NZ_UKJL020000... |
| Salmonella enterica subsp. enterica serovar Anatum strain... | Salmonella e... | NA | 58712 | 164 | 164 | 30% | 4e-47 | 89.66 | 264 | NZ_CALPAM0100... |
| Klebsiella pneumoniae strain EuSCAPE_IL004, whole genome shotg... | Klebsiella p... | NA | 573 | 162 | 162 | 28% | 2e-46 | 95.06 | 244 | NZ_UKJP020001... |
| Klebsiella pneumoniae strain B2 isolate B2, whole genome shotg... | Klebsiella p... | NA | 573 | 175 | 175 | 93% | 1e-44 | 36.06 | 7306 | NZ_CABWNT0100... |
| Klebsiella pneumoniae strain EuSCAPE_LT003, whole genome shotg... | Klebsiella p... | NA | 573 | 153 | 153 | 25% | 1e-42 | 93.24 | 305 | NZ_UKJH020000... |
| Methylobacterium radiotolerans strain MAMP 4754... | Methylobacte... | NA | 31998 | 150 | 150 | 28% | 9e-42 | 88.89 | 289 | NZ_NKQS010005... |
| Methylobacterium radiotolerans strain MAMP 4754... | Methylobacte... | NA | 31998 | 135 | 135 | 29% | 1e-35 | 80.95 | 279 | NZ_NKQS010012... |
| Klebsiella pneumoniae strain EuSCAPE_MT012, whole genome shotg... | Klebsiella p... | NA | 573 | 104 | 104 | 18% | 3e-23 | 96.23 | 385 | NZ_UKIX020000... |
| Acinetobacter baumannii strain P4 contig_P4_92, whole genome... | Acinetobacte... | NA | 470 | 103 | 103 | 18% | 2e-20 | 92.31 | 1226 | NZ_JBBXLU0100... |
| Acinetobacter baumannii strain E12 contig_E12_75, whole genome... | Acinetobacte... | NA | 470 | 82.8 | 82.8 | 14% | 1e-14 | 90.48 | 514 | NZ_JBBXKS0100... |
| Chlamydia abortus strain 09_05, whole genome shotgun sequence | Chlamydia ab... | NA | 83555 | 80.5 | 80.5 | 92% | 3e-12 | 26.06 | 4831 | NZ_FQLG010000... |
| Uncultured Roseobacter sp. isolate 53.fasta NODE_38, whole... | uncultured R... | NA | 114847 | 73.9 | 73.9 | 12% | 5e-12 | 100.00 | 315 | NZ_CANMKI0100... |
| Chlamydia abortus strain 09_03, whole genome shotgun sequence | Chlamydia ab... | NA | 83555 | 79.3 | 152 | 92% | 9e-12 | 25.70 | 4440 | NZ_FQLL010000... |
| Chlamydia abortus strain 11_01, whole genome shotgun sequence | Chlamydia ab... | NA | 83555 | 79.0 | 79.0 | 92% | 1e-11 | 25.35 | 5240 | NZ_FQLK010000... |
| Uncultured Dokdonia sp. isolate 746.fasta NODE_102, whole geno... | uncultured D... | NA | 575653 | 73.9 | 73.9 | 12% | 1e-11 | 100.00 | 440 | NZ_CANLOB0100... |
| Chlamydia trachomatis strain R33512extract, whole genome shotg... | Chlamydia tr... | NA | 813 | 78.2 | 78.2 | 87% | 1e-11 | 26.30 | 1310 | NZ_CVNW010000... |
| Cellulophaga baltica isolate 925.fasta NODE_42, whole genome... | Cellulophaga... | NA | 76594 | 72.0 | 72.0 | 11% | 2e-11 | 100.00 | 316 | NZ_CANLPS0100... |
| Acinetobacter baumannii strain E1 contig_E1_51, whole genome... | Acinetobacte... | NA | 470 | 69.3 | 69.3 | 12% | 6e-10 | 88.57 | 412 | NZ_JBBXKX0100... |
| Salmonella enterica subsp. enterica serovar Typhi strain 7-sc-... | Salmonella e... | NA | 90370 | 67.8 | 67.8 | 11% | 7e-10 | 100.00 | 289 | NZ_JAEPKN0100... |
| Corynebacterium glutamicum strain 39 Scaffolds_32_9166, whole... | Corynebacter... | NA | 1718 | 72.4 | 72.4 | 12% | 2e-09 | 94.59 | 8862 | NZ_JACEQW0100... |
| Corynebacterium glutamicum strain 37 Scaffolds_33_8928, whole... | Corynebacter... | NA | 1718 | 72.4 | 72.4 | 12% | 2e-09 | 94.59 | 8928 | NZ_JACENO0100... |
| Pseudomonas aromaticivorans strain MAP12 34, whole genome... | Pseudomonas ... | NA | 2849492 | 72.4 | 72.4 | 84% | 2e-09 | 25.59 | 7031 | NZ_JAHRGL0100... |
| Chlamydia abortus strain 09_01, whole genome shotgun sequence | Chlamydia ab... | NA | 83555 | 68.2 | 68.2 | 57% | 3e-08 | 28.09 | 1528 | NZ_FQLJ010000... |
| Enterococcus faecalis strain 15224... | Enterococcus... | NA | 1351 | 63.5 | 63.5 | 45% | 3e-07 | 29.66 | 632 | NZ_QNGP010000... |
| Uncultured Roseobacter sp. isolate 930.fasta NODE_31, whole... | uncultured R... | NA | 114847 | 57.0 | 57.0 | 9% | 5e-06 | 100.00 | 257 | NZ_CANLQB0100... |
| Brachybacterium paraconglomeratum isolate 859.fasta NODE_71,... | Brachybacter... | NA | 173362 | 57.0 | 57.0 | 9% | 5e-06 | 100.00 | 258 | NZ_CANLNX0100... |
| Enterococcus faecalis strain AKSZ-144... | Enterococcus... | NA | 1351 | 56.2 | 56.2 | 9% | 6e-06 | 96.15 | 234 | NZ_JAOXSP0100... |
| Roseobacter litoralis isolate 289-G.fasta NODE_32, whole genom... | Roseobacter ... | NA | 42443 | 56.2 | 56.2 | 9% | 8e-06 | 96.15 | 249 | NZ_CANMVH0100... |
| Uncultured Pseudoteredinibacter sp. isolate 208.fasta NODE_42,... | uncultured P... | NA | 1641701 | 56.2 | 56.2 | 9% | 8e-06 | 96.15 | 253 | NZ_CANMSS0100... |
| Bacillus subtilis strain RS3017... | Bacillus sub... | NA | 1423 | 56.2 | 56.2 | 8% | 8e-06 | 100.00 | 232 | NZ_JASDBS0100... |
| Uncultured Tateyamaria sp. isolate 203.fasta NODE_54, whole... | uncultured T... | NA | 455651 | 56.2 | 56.2 | 9% | 8e-06 | 96.15 | 263 | NZ_CANLVZ0100... |
| Uncultured Roseobacter sp. isolate 908.fasta NODE_28, whole... | uncultured R... | NA | 114847 | 56.2 | 56.2 | 8% | 9e-06 | 100.00 | 244 | NZ_CANMNK0100... |
| Uncultured Roseibium sp. isolate 190.fasta NODE_26, whole geno... | uncultured R... | NA | 1936171 | 56.2 | 56.2 | 8% | 9e-06 | 100.00 | 248 | NZ_CANNEN0100... |
| Uncultured Roseobacter sp. isolate 407-3Z.fasta NODE_45, whole... | uncultured R... | NA | 114847 | 56.2 | 56.2 | 8% | 1e-05 | 100.00 | 252 | NZ_CANMKD0100... |
| Epibacterium ulvae isolate 20.fasta NODE_62, whole genome... | Epibacterium... | NA | 1156985 | 56.2 | 56.2 | 8% | 1e-05 | 100.00 | 257 | NZ_CANLDO0100... |
| Roseibium album isolate 321-3Z.fasta NODE_38, whole genome... | Roseibium album | NA | 311410 | 56.2 | 56.2 | 8% | 1e-05 | 100.00 | 255 | NZ_CANMGD0100... |
| Uncultured Ruegeria sp. isolate 409.fasta NODE_44, whole genom... | uncultured R... | NA | 259304 | 56.2 | 56.2 | 8% | 1e-05 | 100.00 | 256 | NZ_CANLNV0100... |
| Uncultured Winogradskyella sp. isolate 408-3Z.fasta NODE_32,... | uncultured W... | NA | 395353 | 56.2 | 56.2 | 8% | 1e-05 | 100.00 | 256 | NZ_CANMJY0100... |
| Pseudomonas sp. QS1027... | Pseudomonas ... | NA | 2056244 | 55.8 | 55.8 | 8% | 1e-05 | 100.00 | 240 | NZ_PHSU010000... |
| Pseudomonas nitroreducens strain FY43... | Pseudomonas ... | NA | 46680 | 56.2 | 56.2 | 9% | 2e-05 | 96.15 | 306 | NZ_JAJISQ0100... |
| Proteus mirabilis strain PM2-4 PM2-4_scaffold43, whole genome... | Proteus mira... | NA | 584 | 60.5 | 60.5 | 52% | 2e-05 | 28.31 | 3726 | NZ_JAMKNT0100... |
| Uncultured Tateyamaria sp. isolate 159-G.fasta NODE_34, whole... | uncultured T... | NA | 455651 | 55.5 | 55.5 | 9% | 2e-05 | 96.15 | 252 | NZ_CANNDY0100... |
| Uncultured Tenacibaculum sp. isolate 838.fasta NODE_42, whole... | uncultured T... | NA | 174713 | 55.5 | 55.5 | 9% | 2e-05 | 96.15 | 262 | NZ_CANMMT0100... |
| Escherichia coli strain w183 NODE_188_length_269_cov_26.989,... | Escherichia ... | NA | 562 | 54.3 | 54.3 | 8% | 4e-05 | 100.00 | 269 | NZ_JAJNDO0100... |
| Klebsiella pneumoniae isolate ena-yuan-GCF_010597935.1... | Klebsiella p... | NA | 573 | 52.8 | 52.8 | 7% | 1e-04 | 100.00 | 234 | NZ_CAWPJY0100... |
| Klebsiella pneumoniae strain AS012315 AS012315_632, whole geno... | Klebsiella p... | NA | 573 | 52.8 | 52.8 | 7% | 1e-04 | 100.00 | 234 | NZ_VKXG010006... |
| Escherichia coli strain w189 NODE_129_length_265_cov_25.6333,... | Escherichia ... | NA | 562 | 52.8 | 52.8 | 7% | 2e-04 | 100.00 | 265 | NZ_JAJNDQ0100... |
| Escherichia coli strain w190 NODE_198_length_265_cov_25.5778,... | Escherichia ... | NA | 562 | 52.8 | 52.8 | 7% | 2e-04 | 100.00 | 265 | NZ_JAJNDR0100... |
| Escherichia coli strain w176 NODE_127_length_264_cov_65.95,... | Escherichia ... | NA | 562 | 52.8 | 52.8 | 7% | 2e-04 | 100.00 | 264 | NZ_JAJNDN0100... |
| Escherichia coli strain w202 NODE_136_length_268_cov_26.95,... | Escherichia ... | NA | 562 | 52.8 | 52.8 | 7% | 2e-04 | 100.00 | 268 | NZ_JAJNDS0100... |
| Escherichia coli strain w184 NODE_200_length_268_cov_29.5537,... | Escherichia ... | NA | 562 | 52.8 | 52.8 | 7% | 2e-04 | 100.00 | 268 | NZ_JAJNDP0100... |
| Escherichia coli strain w208 NODE_126_length_276_cov_44.5, who... | Escherichia ... | NA | 562 | 52.8 | 52.8 | 7% | 2e-04 | 100.00 | 276 | NZ_JAJNDT0100... |
| Enterococcus faecalis strain 17838... | Enterococcus... | NA | 1351 | 52.0 | 52.0 | 31% | 4e-04 | 33.70 | 288 | NZ_QNGS010001... |
| Devosia insulae DS-56 contig234, whole genome shotgun sequence | Devosia insu... | NA | 1116389 | 56.2 | 56.2 | 37% | 5e-04 | 30.53 | 24799 | NZ_LAJE020001... |
| Eikenella corrodens strain 491_NMEN, whole genome shotgun... | Eikenella co... | NA | 539 | 52.8 | 52.8 | 42% | 0.002 | 28.46 | 659 | NZ_JVFA010000... |
| Escherichia coli strain RF108-1... | Escherichia ... | NA | 562 | 52.4 | 52.4 | 18% | 0.002 | 41.51 | 640 | NZ_JAAQGD0100... |
| Eikenella corrodens strain EI_05 NODE_86_length_856_cov_136.29... | Eikenella co... | NA | 539 | 52.8 | 52.8 | 42% | 0.002 | 28.46 | 856 | NZ_VYYO010000... |
| Rhodovulum sulfidophilum strain SE1... | Rhodovulum s... | NA | 35806 | 51.2 | 51.2 | 30% | 0.002 | 30.00 | 467 | NZ_JAAEAJ0100... |
| Devosia sp. Root105 contig_7, whole genome shotgun sequence | Devosia sp. ... | NA | 1736423 | 53.5 | 53.5 | 37% | 0.003 | 30.53 | 620452 | NZ_LMCR010000... |
| Devosia sp. Root413D1 contig_4, whole genome shotgun sequence | Devosia sp. ... | NA | 1736531 | 53.5 | 53.5 | 37% | 0.003 | 30.53 | 575790 | NZ_LMEA010000... |
| Escherichia coli strain 90 NODE_134_length_1268_cov_0.816121,... | Escherichia ... | NA | 562 | 52.8 | 52.8 | 23% | 0.004 | 33.77 | 1268 | NZ_JACAMA0100... |
| Albimonas sp. CAU 1670 contig2, whole genome shotgun sequence | Albimonas sp... | NA | 3032599 | 53.1 | 53.1 | 34% | 0.004 | 36.27 | 847010 | NZ_JARHUC0100... |
| Agreia sp. PsM10 NODE_68_length_253_cov_0.818182, whole genome... | Agreia sp. P... | NA | 3030533 | 48.5 | 48.5 | 19% | 0.004 | 43.86 | 253 | NZ_JARUXC0100... |
| Enterococcus faecalis strain 15407... | Enterococcus... | NA | 1351 | 48.9 | 48.9 | 30% | 0.005 | 32.58 | 285 | NZ_QNGQ010002... |
| Eikenella corrodens strain EI_05... | Eikenella co... | NA | 539 | 52.4 | 52.4 | 42% | 0.008 | 28.46 | 165032 | NZ_VYYO010000... |
| Eikenella corrodens strain EI_07... | Eikenella co... | NA | 539 | 52.4 | 52.4 | 42% | 0.008 | 26.92 | 123520 | NZ_VYYM010000... |
| Eikenella corrodens ATCC 23834 Scfld2, whole genome shotgun... | Eikenella co... | NA | 546274 | 52.0 | 52.0 | 42% | 0.009 | 28.46 | 131618 | NZ_EQ973318.1 |
| Eikenella corrodens strain FDAARGOS_1474 chromosome, complete... | Eikenella co... | NA | 539 | 52.0 | 52.0 | 42% | 0.009 | 28.46 | 2192571 | NZ_CP082861.1 |
| Eikenella corrodens strain NCTC10596 chromosome 1, complete... | Eikenella co... | NA | 539 | 52.0 | 52.0 | 42% | 0.009 | 28.46 | 2192577 | NZ_LT906482.1 |
| Eikenella corrodens strain KCOM 1378 B230_1_2, whole genome... | Eikenella co... | NA | 539 | 52.0 | 52.0 | 42% | 0.009 | 28.46 | 572658 | NZ_NHRS010000... |
| Eikenella corrodens strain NML01-0040 Eikcor_contig000008, who... | Eikenella co... | NA | 539 | 52.0 | 52.0 | 42% | 0.009 | 28.46 | 65735 | NZ_LXSE010000... |
| Eikenella corrodens strain EI_08... | Eikenella co... | NA | 539 | 52.0 | 52.0 | 42% | 0.009 | 28.46 | 16570 | NZ_VYYL010000... |
| Escherichia coli strain E-4H4 NODE_99_length_2231_cov_0.479087... | Escherichia ... | NA | 562 | 50.1 | 50.1 | 24% | 0.010 | 37.84 | 590 | NZ_JALSFQ0100... |
| Klebsiella pneumoniae strain CL1616... | Klebsiella p... | NA | 573 | 51.2 | 51.2 | 26% | 0.010 | 35.44 | 1091 | NZ_RPET010001... |
| Klebsiella pneumoniae strain CL1549... | Klebsiella p... | NA | 573 | 51.2 | 51.2 | 26% | 0.010 | 35.44 | 1225 | NZ_RPES010001... |
| Klebsiella pneumoniae strain CL2074... | Klebsiella p... | NA | 573 | 51.6 | 51.6 | 26% | 0.010 | 35.44 | 2903 | NZ_RPFA010000... |
| Staphylococcus aureus strain 17PQ_1_195... | Staphylococc... | NA | 1280 | 50.8 | 50.8 | 27% | 0.016 | 34.83 | 1338 | NZ_JAOBRT0100... |
| Eikenella corrodens strain KCOM 1378 B230_1_3, whole genome... | Eikenella co... | NA | 539 | 51.2 | 51.2 | 42% | 0.017 | 28.46 | 129678 | NZ_NHRS010000... |
| Acinetobacter bereziniae strain A005.1... | Acinetobacte... | NA | 106648 | 50.4 | 50.4 | 27% | 0.017 | 33.71 | 855 | NZ_JACDXR0100... |
| Haemophilus sp. SZY H51 NODE_22_length_1889_cov_0.990066, whol... | Haemophilus ... | NA | 3041427 | 50.8 | 50.8 | 34% | 0.019 | 27.93 | 1889 | NZ_JASGBG0100... |
| Rickettsia endosymbiont of Ceutorhynchus assimilis isolate... | Rickettsia e... | NA | 2884482 | 51.2 | 51.2 | 35% | 0.020 | 31.82 | 4063491 | NZ_OU906081.1 |
| Escherichia coli strain E-4H5 NODE_187_length_488_cov_0.063712... | Escherichia ... | NA | 562 | 48.1 | 48.1 | 28% | 0.027 | 35.71 | 488 | NZ_JALYCP0100... |
| Mycobacterium innocens strain MK13 isolate MK13, whole genome... | Mycobacteriu... | NA | 2341083 | 50.1 | 50.1 | 40% | 0.027 | 34.38 | 1438 | NZ_UPHQ010003... |
| Eikenella corrodens strain NML01-0040 Eikcor_contig000002, who... | Eikenella co... | NA | 539 | 50.8 | 50.8 | 42% | 0.028 | 27.69 | 226986 | NZ_LXSE010000... |
| Eikenella corrodens strain NML140207 Eikcor_contig000050, whol... | Eikenella co... | NA | 539 | 50.4 | 50.4 | 43% | 0.031 | 27.41 | 5581 | NZ_LXSJ010000... |
| Eikenella corrodens strain NML120819 Eikcor_contig000019, whol... | Eikenella co... | NA | 539 | 50.4 | 50.4 | 42% | 0.032 | 27.69 | 95946 | NZ_LXSH010000... |
| Acinetobacter bereziniae strain A005.2... | Acinetobacte... | NA | 106648 | 49.3 | 49.3 | 22% | 0.035 | 37.88 | 954 | NZ_JABWPZ0100... |
| Klebsiella pneumoniae strain KP231... | Klebsiella p... | NA | 573 | 45.8 | 45.8 | 17% | 0.036 | 46.94 | 233 | NZ_JAUUHQ0100... |
| Pseudomonas reactans strain IPO3751... | Pseudomonas ... | NA | 117680 | 48.5 | 48.5 | 19% | 0.038 | 38.60 | 594 | NZ_JACAQP0100... |
| Corynebacterium sp. c3Ub_189 c3Ub189_contig_53, whole genome... | Corynebacter... | NA | 3032331 | 49.7 | 49.7 | 38% | 0.039 | 31.25 | 1136 | NZ_JASBOF0100... |
| Aeromonas jandaei strain 82 NODE_15_length_4445_cov_0.106043,... | Aeromonas ja... | NA | 650 | 50.1 | 50.1 | 30% | 0.039 | 28.43 | 4445 | NZ_RPDF010000... |
| Aeromonas jandaei isolate ena-yuan-GCF_009749285.1... | Aeromonas ja... | NA | 650 | 50.1 | 50.1 | 30% | 0.039 | 28.43 | 4445 | NZ_CAWPET0100... |
| Corynebacterium tuberculostearicum strain CTNIH9... | Corynebacter... | NA | 38304 | 49.7 | 49.7 | 38% | 0.040 | 31.50 | 1392 | NZ_JAVBHZ0100... |
| Corynebacterium sp. ACRPF... | Corynebacter... | NA | 2918197 | 49.7 | 49.7 | 38% | 0.040 | 31.50 | 1392 | NZ_JAKRNB0100... |
| Corynebacterium marquesiae strain MSK087... | Corynebacter... | NA | 2913503 | 48.9 | 48.9 | 38% | 0.044 | 31.50 | 882 | NZ_JASPIN0100... |
| Pseudarthrobacter sp. W1I19 Ga0505363_01, whole genome shotgun... | Pseudarthrob... | NA | 3042288 | 50.1 | 50.1 | 44% | 0.045 | 26.67 | 4812144 | NZ_JAUSZR0100... |
| Corynebacterium marquesiae strain MSK093... | Corynebacter... | NA | 2913503 | 48.5 | 48.5 | 38% | 0.046 | 31.50 | 702 | NZ_JASPII0100... |
| Corynebacterium marquesiae strain MSK097... | Corynebacter... | NA | 2913503 | 48.5 | 48.5 | 38% | 0.046 | 31.50 | 702 | NZ_JASPIF0100... |
| Escherichia coli strain E-4H5 NODE_89_length_1734_cov_0.210330... | Escherichia ... | NA | 562 | 49.7 | 49.7 | 27% | 0.047 | 33.71 | 1734 | NZ_JALYCP0100... |

Alignments:

```
>Actinoplanes campanulatus strain JCM 3059 sequence058, whole genome shotgun sequence
Sequence ID: NZ_BMPW01000058.1 Length: 4754
Range 1: 3537 to 4400

Score:591 bits(1523), Expect:0.0,
Method:Compositional matrix adjust.,
Identities:280/288(97%), Positives:286/288(99%), Gaps:0/288(0%)

Query  1     FLDGIDKAQDEHEKYHSNWRAMASDFNLPPVVAKEIVASCDKCQLKGEAMHGQVDCSPGI  60
             FLDGIDKAQ+EHEKYHSNWRAMASDFNLPPVVAKEIVASCDKCQLKGEAMHGQVDCSPGI
Sbjct  3537  FLDGIDKAQEEHEKYHSNWRAMASDFNLPPVVAKEIVASCDKCQLKGEAMHGQVDCSPGI  3716

Query  61    WQLDCTHLEGKVILVAVHVASGYIEAEVIPAETGQETAYFLLKLAGRWPVKTIHTDNGSN  120
             WQLDCTHLEGKVILVAVHVASGYIEAEVIPAETGQETAYFLLKLAGRWPVKT+HTDNGSN
Sbjct  3717  WQLDCTHLEGKVILVAVHVASGYIEAEVIPAETGQETAYFLLKLAGRWPVKTVHTDNGSN  3896

Query  121   FTGATVRAACWWAGIKQEFGIPYNPQSQGVVESMNKELKKIIGQVRDQAEHLKTAVQMAV  180
             FT  TV+AACWWAGIKQEFGIPYNPQSQGV+ESMNKELKKIIGQVRDQAEHLKTAVQMAV
Sbjct  3897  FTSTTVKAACWWAGIKQEFGIPYNPQSQGVIESMNKELKKIIGQVRDQAEHLKTAVQMAV  4076

Query  181   FIHNFKRKGGIGGYSAGERIVDIIATDIQTKELQKQITKIQNFRVYYRDSRNPLWKGPAK  240
             FIHNFKRKGGIGGYSAGERIVDIIATDIQTKELQKQITKIQNFRVYYRDSR+P+WKGPAK
Sbjct  4077  FIHNFKRKGGIGGYSAGERIVDIIATDIQTKELQKQITKIQNFRVYYRDSRDPVWKGPAK  4256

Query  241   LLWKGEGAVVIQDNSDIKVVPRRKAKIIRDYGKQMAGDDCVASRQDED  288
             LLWKGEGAVVIQDNSDIKVVPRRKAKIIRDYGKQMAGDDCVASRQDED
Sbjct  4257  LLWKGEGAVVIQDNSDIKVVPRRKAKIIRDYGKQMAGDDCVASRQDED  4400
```

```
>Escherichia coli strain VRES0669, whole genome shotgun sequence
Sequence ID: NZ_UINV01000086.1 Length: 5544
Range 1: 2544 to 3407

Score:575 bits(1481), Expect:0.0,
Method:Compositional matrix adjust.,
Identities:270/288(94%), Positives:282/288(97%), Gaps:0/288(0%)

Query  1     FLDGIDKAQDEHEKYHSNWRAMASDFNLPPVVAKEIVASCDKCQLKGEAMHGQVDCSPGI  60
             FLDGIDKAQ+EHEKYHSNWRAMAS+FNLPP+VAKEIVASCDKCQLKGEAMHGQVDCSPGI
Sbjct  3407  FLDGIDKAQEEHEKYHSNWRAMASEFNLPPIVAKEIVASCDKCQLKGEAMHGQVDCSPGI  3228

Query  61    WQLDCTHLEGKVILVAVHVASGYIEAEVIPAETGQETAYFLLKLAGRWPVKTIHTDNGSN  120
             WQLDCTHLEGKVILVAVHVASGY+EAEVIPAETGQETAYF+LKLAGRWPVKTIHTDNGSN
Sbjct  3227  WQLDCTHLEGKVILVAVHVASGYMEAEVIPAETGQETAYFILKLAGRWPVKTIHTDNGSN  3048

Query  121   FTGATVRAACWWAGIKQEFGIPYNPQSQGVVESMNKELKKIIGQVRDQAEHLKTAVQMAV  180
             FT   V+AACWWAGI+QEFGIPYNPQSQGVVESMNKELKKII Q+RDQAEHLKTAVQMAV
Sbjct  3047  FTSTVVKAACWWAGIQQEFGIPYNPQSQGVVESMNKELKKIIEQLRDQAEHLKTAVQMAV  2868

Query  181   FIHNFKRKGGIGGYSAGERIVDIIATDIQTKELQKQITKIQNFRVYYRDSRNPLWKGPAK  240
             F+HNFKRKGGIGGYSAGERI+DIIATDIQTKELQKQITKIQNFRVYYRDSR+P+WKGPAK
Sbjct  2867  FVHNFKRKGGIGGYSAGERIIDIIATDIQTKELQKQITKIQNFRVYYRDSRDPIWKGPAK  2688

Query  241   LLWKGEGAVVIQDNSDIKVVPRRKAKIIRDYGKQMAGDDCVASRQDED  288
             LLWKGEGAVVIQDNSDIKVVPRRKAKIIRDYGKQMAG DCVA RQDED
Sbjct  2687  LLWKGEGAVVIQDNSDIKVVPRRKAKIIRDYGKQMAGTDCVAGRQDED  2544
```

```
>Pseudomonas aeruginosa strain ENV-297 NODE_79_length_1100_cov_9.24438_ID_157, whole genome shotgun sequence
Sequence ID: NZ_RADD01000036.1 Length: 1100
Range 1: 81 to 944

Score:560 bits(1443), Expect:0.0,
Method:Compositional matrix adjust.,
Identities:261/288(91%), Positives:278/288(96%), Gaps:0/288(0%)

Query  1    FLDGIDKAQDEHEKYHSNWRAMASDFNLPPVVAKEIVASCDKCQLKGEAMHGQVDCSPGI  60
            FLDGIDKAQ+EHE+YH+NWRAMASDFNLPP+VAKEIVASCDKCQ KGEAMHGQVDCSPGI
Sbjct  944  FLDGIDKAQEEHERYHNNWRAMASDFNLPPIVAKEIVASCDKCQQKGEAMHGQVDCSPGI  765

Query  61   WQLDCTHLEGKVILVAVHVASGYIEAEVIPAETGQETAYFLLKLAGRWPVKTIHTDNGSN  120
            WQ+DCTHLEGK+I+VAVHVASGYIEAEVIPAETGQETAYF+LKLA RWPVK IHTDNGSN
Sbjct  764  WQIDCTHLEGKIIIVAVHVASGYIEAEVIPAETGQETAYFILKLAARWPVKVIHTDNGSN  585

Query  121  FTGATVRAACWWAGIKQEFGIPYNPQSQGVVESMNKELKKIIGQVRDQAEHLKTAVQMAV  180
            FT A V+AACWWA + QEFGIPYNPQSQGVVESMNKELKKIIGQ+RDQAEHLKTAVQMAV
Sbjct  584  FTSAAVKAACWWANVTQEFGIPYNPQSQGVVESMNKELKKIIGQIRDQAEHLKTAVQMAV  405

Query  181  FIHNFKRKGGIGGYSAGERIVDIIATDIQTKELQKQITKIQNFRVYYRDSRNPLWKGPAK  240
            FIHNFKRKGGIG YSAGERI+DIIA+DIQTKELQKQI KIQNFRVYYRDSR+P+WKGPAK
Sbjct  404  FIHNFKRKGGIGRYSAGERIIDIIASDIQTKELQKQIIKIQNFRVYYRDSRDPIWKGPAK  225

Query  241  LLWKGEGAVVIQDNSDIKVVPRRKAKIIRDYGKQMAGDDCVASRQDED  288
            LLWKGEGAVVIQDNS+IKV+PRRKAKIIRDYGKQMAGDDCVA RQDED
Sbjct  224  LLWKGEGAVVIQDNSEIKVIPRRKAKIIRDYGKQMAGDDCVAGRQDED  81
```

```
>Pseudomonas aeruginosa strain VET-42 NODE_86_length_1154_cov_8.30919_ID_171, whole genome shotgun sequence
Sequence ID: NZ_RACX01000118.1 Length: 1154
Range 1: 75 to 938

Score:559 bits(1440), Expect:0.0,
Method:Compositional matrix adjust.,
Identities:260/288(90%), Positives:278/288(96%), Gaps:0/288(0%)

Query  1    FLDGIDKAQDEHEKYHSNWRAMASDFNLPPVVAKEIVASCDKCQLKGEAMHGQVDCSPGI  60
            FLDGIDKAQ+EHE+YH+NWRAMASDFNLPP+VAKEIVASCDKCQ KGEAMHGQVDCSPGI
Sbjct  938  FLDGIDKAQEEHERYHNNWRAMASDFNLPPIVAKEIVASCDKCQQKGEAMHGQVDCSPGI  759

Query  61   WQLDCTHLEGKVILVAVHVASGYIEAEVIPAETGQETAYFLLKLAGRWPVKTIHTDNGSN  120
            WQ+DCTHLEGK+I+VAVHVASGYIEAEVIPAETGQETAYF+LKLA RWPVK IHTDNGSN
Sbjct  758  WQIDCTHLEGKIIIVAVHVASGYIEAEVIPAETGQETAYFILKLAARWPVKVIHTDNGSN  579

Query  121  FTGATVRAACWWAGIKQEFGIPYNPQSQGVVESMNKELKKIIGQVRDQAEHLKTAVQMAV  180
            FT A V+AACWWA + QEFGIPYNPQSQGVVESMNKELKKIIGQ+RDQAEHLKTAVQMAV
Sbjct  578  FTSAAVKAACWWANVTQEFGIPYNPQSQGVVESMNKELKKIIGQIRDQAEHLKTAVQMAV  399

Query  181  FIHNFKRKGGIGGYSAGERIVDIIATDIQTKELQKQITKIQNFRVYYRDSRNPLWKGPAK  240
            FIHNFKRKGGIG YSAGERI+DIIA+DIQTKELQKQI KIQNFRVYYRDSR+P+WKGPAK
Sbjct  398  FIHNFKRKGGIGRYSAGERIIDIIASDIQTKELQKQIIKIQNFRVYYRDSRDPIWKGPAK  219

Query  241  LLWKGEGAVVIQDNSDIKVVPRRKAKIIRDYGKQMAGDDCVASRQDED  288
            LLWKGEGAVVIQDNS+IKV+PRRKAKIIRDYGKQMAGDDCVA RQDE+
Sbjct  218  LLWKGEGAVVIQDNSEIKVIPRRKAKIIRDYGKQMAGDDCVAGRQDEN  75
```

```
>Burkholderia pseudomallei strain 2604STDY5518971 ERS240105SCcontig000097, whole genome shotgun sequence
Sequence ID: NZ_UVFZ02000097.1 Length: 1345
Range 1: 568 to 1344

Score:518 bits(1335), Expect:0.0,
Method:Compositional matrix adjust.,
Identities:241/259(93%), Positives:253/259(97%), Gaps:0/259(0%)

Query  1     FLDGIDKAQDEHEKYHSNWRAMASDFNLPPVVAKEIVASCDKCQLKGEAMHGQVDCSPGI  60
             FLDGIDKAQ++HE+YHSNWR MASDFNLPP+VAKEIVA+CDKCQLKGEAMHGQVDCSPGI
Sbjct  568   FLDGIDKAQEDHERYHSNWRTMASDFNLPPIVAKEIVANCDKCQLKGEAMHGQVDCSPGI  747

Query  61    WQLDCTHLEGKVILVAVHVASGYIEAEVIPAETGQETAYFLLKLAGRWPVKTIHTDNGSN  120
             WQLDCTHLEGKVILVAVHVASGYIEAEVIPAETGQETAYFLLKLAGRWPVK IHTDNGSN
Sbjct  748   WQLDCTHLEGKVILVAVHVASGYIEAEVIPAETGQETAYFLLKLAGRWPVKVIHTDNGSN  927

Query  121   FTGATVRAACWWAGIKQEFGIPYNPQSQGVVESMNKELKKIIGQVRDQAEHLKTAVQMAV  180
             FT   V+AACWWA ++QEFGIPYNPQSQGVVESMNKELKKIIGQVR+QAEHLKTAVQMAV
Sbjct  928   FTSTAVKAACWWANVQQEFGIPYNPQSQGVVESMNKELKKIIGQVREQAEHLKTAVQMAV  1107

Query  181   FIHNFKRKGGIGGYSAGERIVDIIATDIQTKELQKQITKIQNFRVYYRDSRNPLWKGPAK  240
             FIHNFKRKGGIGGYSAGERI+DIIATDIQTKELQKQITKIQNFRVYYRDSR+P+WKGPAK
Sbjct  1108  FIHNFKRKGGIGGYSAGERIIDIIATDIQTKELQKQITKIQNFRVYYRDSRDPIWKGPAK  1287

Query  241   LLWKGEGAVVIQDNSDIKV  259
             LLWKGEGAVVIQDNSDIKV
Sbjct  1288  LLWKGEGAVVIQDNSDIKV  1344
```

```
>Klebsiella pneumoniae strain P7701 NODE_60_length_711_cov_294.76, whole genome shotgun sequence
Sequence ID: NZ_JACAWM010000058.1 Length: 711
Range 1: 1 to 711

Score:482 bits(1241), Expect:2e-170,
Method:Compositional matrix adjust.,
Identities:230/237(97%), Positives:235/237(99%), Gaps:0/237(0%)

Query  50   MHGQVDCSPGIWQLDCTHLEGKVILVAVHVASGYIEAEVIPAETGQETAYFLLKLAGRWP  109
            MHGQVDCSPGIWQLDCTHLEGKVILVAVHVASGYIEAEVIPAETGQETAYFLLKLAGRWP
Sbjct  1    MHGQVDCSPGIWQLDCTHLEGKVILVAVHVASGYIEAEVIPAETGQETAYFLLKLAGRWP  180

Query  110  VKTIHTDNGSNFTGATVRAACWWAGIKQEFGIPYNPQSQGVVESMNKELKKIIGQVRDQA  169
            VKT+HTDNGSNFT  TV+AACWWAGIKQEFGIPYNPQSQGV+ESMNKELKKIIGQVRDQA
Sbjct  181  VKTVHTDNGSNFTSTTVKAACWWAGIKQEFGIPYNPQSQGVIESMNKELKKIIGQVRDQA  360

Query  170  EHLKTAVQMAVFIHNFKRKGGIGGYSAGERIVDIIATDIQTKELQKQITKIQNFRVYYRD  229
            EHLKTAVQMAVFIHNFKRKGGIGGYSAGERIVDIIATDIQTKELQKQITKIQNFRVYYRD
Sbjct  361  EHLKTAVQMAVFIHNFKRKGGIGGYSAGERIVDIIATDIQTKELQKQITKIQNFRVYYRD  540

Query  230  SRNPLWKGPAKLLWKGEGAVVIQDNSDIKVVPRRKAKIIRDYGKQMAGDDCVASRQD  286
            SR+P+WKGPAKLLWKGEGAVVIQDNSDIKVVPRRKAKIIRDYGKQMAGDDCVASRQD
Sbjct  541  SRDPVWKGPAKLLWKGEGAVVIQDNSDIKVVPRRKAKIIRDYGKQMAGDDCVASRQD  711
```

```
>Klebsiella pneumoniae strain ARGID_31501, whole genome shotgun sequence
Sequence ID: NZ_CAKLPD010000006.1 Length: 2655
Range 1: 1904 to 2653

Score:504 bits(1299), Expect:7e-170,
Method:Compositional matrix adjust.,
Identities:236/250(94%), Positives:246/250(98%), Gaps:0/250(0%)

Query  1     FLDGIDKAQDEHEKYHSNWRAMASDFNLPPVVAKEIVASCDKCQLKGEAMHGQVDCSPGI  60
             FLDGIDKAQ++H+KYHSNWRAMASDFNLPP+VAKEIVASCDKCQLKGEAMHGQVDCSPGI
Sbjct  1904  FLDGIDKAQEDHDKYHSNWRAMASDFNLPPIVAKEIVASCDKCQLKGEAMHGQVDCSPGI  2083

Query  61    WQLDCTHLEGKVILVAVHVASGYIEAEVIPAETGQETAYFLLKLAGRWPVKTIHTDNGSN  120
             WQLDCTHLEGK+ILVAVHVASGYIEAEVIPAETGQETAYF+LKLAGRWPV TIHTDNG N
Sbjct  2084  WQLDCTHLEGKIILVAVHVASGYIEAEVIPAETGQETAYFILKLAGRWPVTTIHTDNGPN  2263

Query  121   FTGATVRAACWWAGIKQEFGIPYNPQSQGVVESMNKELKKIIGQVRDQAEHLKTAVQMAV  180
             F  +TV+AACWWAGIKQEFGIPYNPQSQGVVESMNKELKKIIGQVRDQAEHL+TAVQMAV
Sbjct  2264  FISSTVKAACWWAGIKQEFGIPYNPQSQGVVESMNKELKKIIGQVRDQAEHLRTAVQMAV  2443

Query  181   FIHNFKRKGGIGGYSAGERIVDIIATDIQTKELQKQITKIQNFRVYYRDSRNPLWKGPAK  240
             FIHNFKRKGGIGGYSAGERIVDIIATDIQTKELQKQITKIQNFRVYYRDSR+PLWKGPAK
Sbjct  2444  FIHNFKRKGGIGGYSAGERIVDIIATDIQTKELQKQITKIQNFRVYYRDSRDPLWKGPAK  2623

Query  241   LLWKGEGAVV  250
             LLWKGEGAVV
Sbjct  2624  LLWKGEGAVV  2653
```

```
>Enterococcus alcedinis strain CCM 8433 sequence23, whole genome shotgun sequence
Sequence ID: NZ_BMDT01000023.1 Length: 6075
Range 1: 3495 to 4163

Score:455 bits(1171), Expect:8e-143,
Method:Compositional matrix adjust.,
Identities:216/223(97%), Positives:220/223(98%), Gaps:0/223(0%)

Query  1     FLDGIDKAQDEHEKYHSNWRAMASDFNLPPVVAKEIVASCDKCQLKGEAMHGQVDCSPGI  60
             FLDGIDKAQ+EHEKYHSNWRAMASDFNLPPVVAKEIVASCDKCQLKGEAMHGQVDCSPGI
Sbjct  3495  FLDGIDKAQEEHEKYHSNWRAMASDFNLPPVVAKEIVASCDKCQLKGEAMHGQVDCSPGI  3674

Query  61    WQLDCTHLEGKVILVAVHVASGYIEAEVIPAETGQETAYFLLKLAGRWPVKTIHTDNGSN  120
             WQLDCTHLEGKVILVAVHVASGYIEAEVIPAETGQETAYFLLKLAGRWPVKT+HTDNGSN
Sbjct  3675  WQLDCTHLEGKVILVAVHVASGYIEAEVIPAETGQETAYFLLKLAGRWPVKTVHTDNGSN  3854

Query  121   FTGATVRAACWWAGIKQEFGIPYNPQSQGVVESMNKELKKIIGQVRDQAEHLKTAVQMAV  180
             FT  TV+AACWWAGIKQEFGIPYNPQSQGV+ESMNKELKKIIGQVRDQAEHLKTAVQMAV
Sbjct  3855  FTSTTVKAACWWAGIKQEFGIPYNPQSQGVIESMNKELKKIIGQVRDQAEHLKTAVQMAV  4034

Query  181   FIHNFKRKGGIGGYSAGERIVDIIATDIQTKELQKQITKIQNF  223
             FIHNFKRKGGIGGYSAGERIVDIIATDIQTKELQKQITKIQ F
Sbjct  4035  FIHNFKRKGGIGGYSAGERIVDIIATDIQTKELQKQITKIQIF  4163
```

```
>Deinococcus aquiradiocola strain JCM 14371 sequence30, whole genome shotgun sequence
Sequence ID: NZ_BMOE01000030.1 Length: 6128
Range 1: 3548 to 4216

Score:455 bits(1171), Expect:8e-143,
```

```
Method:Compositional matrix adjust.,
Identities:216/223(97%), Positives:220/223(98%), Gaps:0/223(0%)

Query  1     FLDGIDKAQDEHEKYHSNWRAMASDFNLPPVVAKEIVASCDKCQLKGEAMHGQVDCSPGI  60
             FLDGIDKAQ+EHEKYHSNWRAMASDFNLPPVVAKEIVASCDKCQLKGEAMHGQVDCSPGI
Sbjct  3548  FLDGIDKAQEEHEKYHSNWRAMASDFNLPPVVAKEIVASCDKCQLKGEAMHGQVDCSPGI  3727

Query  61    WQLDCTHLEGKVILVAVHVASGYIEAEVIPAETGQETAYFLLKLAGRWPVKTIHTDNGSN  120
             WQLDCTHLEGKVILVAVHVASGYIEAEVIPAETGQETAYFLLKLAGRWPVKT+HTDNGSN
Sbjct  3728  WQLDCTHLEGKVILVAVHVASGYIEAEVIPAETGQETAYFLLKLAGRWPVKTVHTDNGSN  3907

Query  121   FTGATVRAACWWAGIKQEFGIPYNPQSQGVVESMNKELKKIIGQVRDQAEHLKTAVQMAV  180
             FT  TV+AACWWAGIKQEFGIPYNPQSQGV+ESMNKELKKIIGQVRDQAEHLKTAVQMAV
Sbjct  3908  FTSTTVKAACWWAGIKQEFGIPYNPQSQGVIESMNKELKKIIGQVRDQAEHLKTAVQMAV  4087

Query  181   FIHNFKRKGGIGGYSAGERIVDIIATDIQTKELQKQITKIQNF  223
             FIHNFKRKGGIGGYSAGERIVDIIATDIQTKELQKQITKIQ F
Sbjct  4088  FIHNFKRKGGIGGYSAGERIVDIIATDIQTKELQKQITKIQIF  4216
```

>Thalassovita mangrovi strain GS-10 Scaffold30_1, whole genome shotgun sequence
Sequence ID: NZ_WWEN01000028.1 Length: 722
Range 1: 2 to 568

```
Score:392 bits(1008), Expect:6e-135,
Method:Compositional matrix adjust.,
Identities:183/189(97%), Positives:187/189(98%), Gaps:0/189(0%)

Query  1    FLDGIDKAQDEHEKYHSNWRAMASDFNLPPVVAKEIVASCDKCQLKGEAMHGQVDCSPGI  60
            FLDGIDKAQ+EHEKYHSNWRAMASDFNLPPVVAKEIVASCDKCQLKGEAMHGQVDCSPGI
Sbjct  568  FLDGIDKAQEEHEKYHSNWRAMASDFNLPPVVAKEIVASCDKCQLKGEAMHGQVDCSPGI  389

Query  61   WQLDCTHLEGKVILVAVHVASGYIEAEVIPAETGQETAYFLLKLAGRWPVKTIHTDNGSN  120
            WQLDCTHLEGKVILVAVHVASGYIEAEVIPAETGQETAYFLLKLAGRWPVKT+HTDNGSN
Sbjct  388  WQLDCTHLEGKVILVAVHVASGYIEAEVIPAETGQETAYFLLKLAGRWPVKTVHTDNGSN  209

Query  121  FTGATVRAACWWAGIKQEFGIPYNPQSQGVVESMNKELKKIIGQVRDQAEHLKTAVQMAV  180
            FT  TV+AACWWAGIKQEFGIPYNPQSQGV+ESMNKELKKIIGQVRDQAEHLKTAVQMAV
Sbjct  208  FTSTTVKAACWWAGIKQEFGIPYNPQSQGVIESMNKELKKIIGQVRDQAEHLKTAVQMAV  29

Query  181  FIHNFKRKG  189
            FIHNFKRKG
Sbjct  28   FIHNFKRKG  2
```

>Methylobacterium radiotolerans strain MAMP 4754 MP-M-radiotolerans_contig_704, whole genome shotgun sequence
Sequence ID: NZ_NKQS01000691.1 Length: 1636
Range 1: 42 to 617

```
Score:350 bits(898), Expect:8e-114,
Method:Compositional matrix adjust.,
Identities:164/192(85%), Positives:178/192(92%), Gaps:2/192(1%)

Query  1    FLDGIDKAQDEHEKYHSNWRAMASDFNLPPVVAKEIVASCDKCQLKGEAMHGQVDCSPGI  60
            FLDGIDKAQ+EHEKYHSNWRAMAS+FNLPP+VAKEIVASCDKCQLKGEAMHGQVDCSPGI
Sbjct  617  FLDGIDKAQEEHEKYHSNWRAMASEFNLPPIVAKEIVASCDKCQLKGEAMHGQVDCSPGI  438

Query  61   WQLDCTHLEGKVILVAVHVASGYIEAEVIPAETGQETAYFLLKLAGRWPVKTIHTDNGSN  120
            WQLDCTHLEGK+ILVAVHVASGYIEAEVIPAETGQETAYF+LKLAGRWPVK IHTDNGSN
Sbjct  437  WQLDCTHLEGKIILVAVHVASGYIEAEVIPAETGQETAYFILKLAGRWPVKVIHTDNGSN  258

Query  121  FTGATVRAACWWAGIKQEFGIPYNPQSQGVVESMNKELKKIIGQVRDQAEHLKTAVQMAV  180
            FT A V+AACWWAGI+QEFGI YNPQ +GVV+SM+ EL KIIGQ RD+ EH+KTAV MAV
Sbjct  257  FTSAAVKAACWWAGIQQEFGIHYNPQRKGVVDSMHTELTKIIGQERDKDEHIKTAVLMAV  78

Query  181  FIHN--FKRKGG  190
            +IHN   KR+ G
Sbjct  77   YIHNCSSKREAG  42
```

>Nocardioides luteus strain JCM 3358 sequence39, whole genome shotgun sequence
Sequence ID: NZ_BMRK01000039.1 Length: 5754
Range 1: 1433 to 1981

```
Score:370 bits(950), Expect:1e-112,
Method:Compositional matrix adjust.,
Identities:174/183(95%), Positives:181/183(98%), Gaps:0/183(0%)

Query  53    QVDCSPGIWQLDCTHLEGKVILVAVHVASGYIEAEVIPAETGQETAYFLLKLAGRWPVKT  112
             ++DCSPGIWQLDCTHLEGKVILVAVHVASGYIEAEVIPAETGQETAYFLLKLAGRWPVKT
Sbjct  1433  RLDCSPGIWQLDCTHLEGKVILVAVHVASGYIEAEVIPAETGQETAYFLLKLAGRWPVKT  1612

Query  113   IHTDNGSNFTGATVRAACWWAGIKQEFGIPYNPQSQGVVESMNKELKKIIGQVRDQAEHL  172
             +HTDNGSNFT  TV+AACWWAGIKQEFGIPYNPQSQGV+ESMNKELKKIIGQVRDQAEHL
Sbjct  1613  VHTDNGSNFTSTTVKAACWWAGIKQEFGIPYNPQSQGVIESMNKELKKIIGQVRDQAEHL  1792

Query  173   KTAVQMAVFIHNFKRKGGIGGYSAGERIVDIIATDIQTKELQKQITKIQNFRVYYRDSRN  232
             KTAVQMAVFIHNFKRKGGIGGYSAGERIVDIIATDIQTKELQKQITKIQNFRVYYRDSR+
Sbjct  1793  KTAVQMAVFIHNFKRKGGIGGYSAGERIVDIIATDIQTKELQKQITKIQNFRVYYRDSRD  1972

Query  233   PLW  235
             P+W
Sbjct  1973  PVW  1981
```

>Streptomyces griseoflavus strain JCM 4479 sequence37, whole genome shotgun sequence
Sequence ID: NZ_BMUC01000037.1 Length: 5775
Range 1: 3767 to 4315

```
Score:370 bits(950), Expect:1e-112,
Method:Compositional matrix adjust.,
Identities:174/183(95%), Positives:181/183(98%), Gaps:0/183(0%)

Query  53    QVDCSPGIWQLDCTHLEGKVILVAVHVASGYIEAEVIPAETGQETAYFLLKLAGRWPVKT  112
             ++DCSPGIWQLDCTHLEGKVILVAVHVASGYIEAEVIPAETGQETAYFLLKLAGRWPVKT
Sbjct  4315  RLDCSPGIWQLDCTHLEGKVILVAVHVASGYIEAEVIPAETGQETAYFLLKLAGRWPVKT  4136

Query  113   IHTDNGSNFTGATVRAACWWAGIKQEFGIPYNPQSQGVVESMNKELKKIIGQVRDQAEHL  172
             +HTDNGSNFT  TV+AACWWAGIKQEFGIPYNPQSQGV+ESMNKELKKIIGQVRDQAEHL
Sbjct  4135  VHTDNGSNFTSTTVKAACWWAGIKQEFGIPYNPQSQGVIESMNKELKKIIGQVRDQAEHL  3956

Query  173   KTAVQMAVFIHNFKRKGGIGGYSAGERIVDIIATDIQTKELQKQITKIQNFRVYYRDSRN  232
             KTAVQMAVFIHNFKRKGGIGGYSAGERIVDIIATDIQTKELQKQITKIQNFRVYYRDSR+
Sbjct  3955  KTAVQMAVFIHNFKRKGGIGGYSAGERIVDIIATDIQTKELQKQITKIQNFRVYYRDSRD  3776

Query  233   PLW  235
```

```
           P+W
Sbjct  3775  PVW  3767
```

>Saccharothrix coeruleofusca strain JCM 3313 sequence45, whole genome shotgun sequence
Sequence ID: NZ_BMRG01000045.1 Length: 5775
Range 1: 3767 to 4315

Score:370 bits(950), Expect:1e-112,
Method:Compositional matrix adjust.,
Identities:174/183(95%), Positives:181/183(98%), Gaps:0/183(0%)

```
Query  53    QVDCSPGIWQLDCTHLEGKVILVAVHVASGYIEAEVIPAETGQETAYFLLKLAGRWPVKT  112
             ++DCSPGIWQLDCTHLEGKVILVAVHVASGYIEAEVIPAETGQETAYFLLKLAGRWPVKT
Sbjct  4315  RLDCSPGIWQLDCTHLEGKVILVAVHVASGYIEAEVIPAETGQETAYFLLKLAGRWPVKT  4136

Query  113   IHTDNGSNFTGATVRAACWWAGIKQEFGIPYNPQSQGVVESMNKELKKIIGQVRDQAEHL  172
             +HTDNGSNFT  TV+AACWWAGIKQEFGIPYNPQSQGV+ESMNKELKKIIGQVRDQAEHL
Sbjct  4135  VHTDNGSNFTSTTVKAACWWAGIKQEFGIPYNPQSQGVIESMNKELKKIIGQVRDQAEHL  3956

Query  173   KTAVQMAVFIHNFKRKGGIGGYSAGERIVDIIATDIQTKELQKQITKIQNFRVYYRDSRN  232
             KTAVQMAVFIHNFKRKGGIGGYSAGERIVDIIATDIQTKELQKQITKIQNFRVYYRDSR+
Sbjct  3955  KTAVQMAVFIHNFKRKGGIGGYSAGERIVDIIATDIQTKELQKQITKIQNFRVYYRDSRD  3776

Query  233   PLW  235
             P+W
Sbjct  3775  PVW  3767
```

>Actinoplanes ianthinogenes strain JCM 3249 sequence037, whole genome shotgun sequence
Sequence ID: NZ_BMQZ01000037.1 Length: 5815
Range 1: 3787 to 4335

Score:370 bits(950), Expect:1e-112,
Method:Compositional matrix adjust.,
Identities:174/183(95%), Positives:181/183(98%), Gaps:0/183(0%)

```
Query  53    QVDCSPGIWQLDCTHLEGKVILVAVHVASGYIEAEVIPAETGQETAYFLLKLAGRWPVKT  112
             ++DCSPGIWQLDCTHLEGKVILVAVHVASGYIEAEVIPAETGQETAYFLLKLAGRWPVKT
Sbjct  4335  RLDCSPGIWQLDCTHLEGKVILVAVHVASGYIEAEVIPAETGQETAYFLLKLAGRWPVKT  4156

Query  113   IHTDNGSNFTGATVRAACWWAGIKQEFGIPYNPQSQGVVESMNKELKKIIGQVRDQAEHL  172
             +HTDNGSNFT  TV+AACWWAGIKQEFGIPYNPQSQGV+ESMNKELKKIIGQVRDQAEHL
Sbjct  4155  VHTDNGSNFTSTTVKAACWWAGIKQEFGIPYNPQSQGVIESMNKELKKIIGQVRDQAEHL  3976

Query  173   KTAVQMAVFIHNFKRKGGIGGYSAGERIVDIIATDIQTKELQKQITKIQNFRVYYRDSRN  232
             KTAVQMAVFIHNFKRKGGIGGYSAGERIVDIIATDIQTKELQKQITKIQNFRVYYRDSR+
Sbjct  3975  KTAVQMAVFIHNFKRKGGIGGYSAGERIVDIIATDIQTKELQKQITKIQNFRVYYRDSRD  3796

Query  233   PLW  235
             P+W
Sbjct  3795  PVW  3787
```

>Sinomonas sp. KCTC 49339 sequence28, whole genome shotgun sequence
Sequence ID: NZ_BNCM01000028.1 Length: 5849
Range 1: 3758 to 4306

Score:370 bits(950), Expect:1e-112,
Method:Compositional matrix adjust.,
Identities:174/183(95%), Positives:181/183(98%), Gaps:0/183(0%)

```
Query  53    QVDCSPGIWQLDCTHLEGKVILVAVHVASGYIEAEVIPAETGQETAYFLLKLAGRWPVKT  112
             ++DCSPGIWQLDCTHLEGKVILVAVHVASGYIEAEVIPAETGQETAYFLLKLAGRWPVKT
Sbjct  4306  RLDCSPGIWQLDCTHLEGKVILVAVHVASGYIEAEVIPAETGQETAYFLLKLAGRWPVKT  4127

Query  113   IHTDNGSNFTGATVRAACWWAGIKQEFGIPYNPQSQGVVESMNKELKKIIGQVRDQAEHL  172
             +HTDNGSNFT  TV+AACWWAGIKQEFGIPYNPQSQGV+ESMNKELKKIIGQVRDQAEHL
Sbjct  4126  VHTDNGSNFTSTTVKAACWWAGIKQEFGIPYNPQSQGVIESMNKELKKIIGQVRDQAEHL  3947

Query  173   KTAVQMAVFIHNFKRKGGIGGYSAGERIVDIIATDIQTKELQKQITKIQNFRVYYRDSRN  232
             KTAVQMAVFIHNFKRKGGIGGYSAGERIVDIIATDIQTKELQKQITKIQNFRVYYRDSR+
Sbjct  3946  KTAVQMAVFIHNFKRKGGIGGYSAGERIVDIIATDIQTKELQKQITKIQNFRVYYRDSRD  3767

Query  233   PLW  235
             P+W
Sbjct  3766  PVW  3758
```

>Saccharothrix mutabilis subsp. mutabilis strain JCM 3380 sequence46, whole genome shotgun sequence
Sequence ID: NZ_BAAABU010000046.1 Length: 5899
Range 1: 1481 to 2029

Score:370 bits(950), Expect:1e-112,
Method:Compositional matrix adjust.,
Identities:174/183(95%), Positives:181/183(98%), Gaps:0/183(0%)

```
Query  53    QVDCSPGIWQLDCTHLEGKVILVAVHVASGYIEAEVIPAETGQETAYFLLKLAGRWPVKT  112
             ++DCSPGIWQLDCTHLEGKVILVAVHVASGYIEAEVIPAETGQETAYFLLKLAGRWPVKT
Sbjct  1481  RLDCSPGIWQLDCTHLEGKVILVAVHVASGYIEAEVIPAETGQETAYFLLKLAGRWPVKT  1660

Query  113   IHTDNGSNFTGATVRAACWWAGIKQEFGIPYNPQSQGVVESMNKELKKIIGQVRDQAEHL  172
             +HTDNGSNFT  TV+AACWWAGIKQEFGIPYNPQSQGV+ESMNKELKKIIGQVRDQAEHL
Sbjct  1661  VHTDNGSNFTSTTVKAACWWAGIKQEFGIPYNPQSQGVIESMNKELKKIIGQVRDQAEHL  1840

Query  173   KTAVQMAVFIHNFKRKGGIGGYSAGERIVDIIATDIQTKELQKQITKIQNFRVYYRDSRN  232
             KTAVQMAVFIHNFKRKGGIGGYSAGERIVDIIATDIQTKELQKQITKIQNFRVYYRDSR+
Sbjct  1841  KTAVQMAVFIHNFKRKGGIGGYSAGERIVDIIATDIQTKELQKQITKIQNFRVYYRDSRD  2020

Query  233   PLW  235
             P+W
Sbjct  2021  PVW  2029
```

>Klebsiella pneumoniae strain EuSCAPE_IL016, whole genome shotgun sequence
Sequence ID: NZ_UKKA02000066.1 Length: 523
Range 1: 2 to 523

Score:295 bits(754), Expect:3e-97,
Method:Compositional matrix adjust.,
Identities:146/174(84%), Positives:159/174(91%), Gaps:0/174(0%)

```
Query  9    QDEHEKYHSNWRAMASDFNLPPVVAKEIVASCDKCQLKGEAMHGQVDCSPGIWQLDCTHL  68
            Q+EHEKYHSN RAMAS+FNLPP+VAKEIVAS DKCQLK EA+ GQVDCSP IWQLDCTHL
```

```
Sbjct  523  QEEHEKYHSN*RAMASEFNLPPIVAKEIVASYDKCQLKKEAIQGQVDCSPRIWQLDCTHL  344

Query  69   EGKVILVAVHVASGYIEAEVIPAETGQETAYFLLKLAGRWPVKTIHTDNGSNFTGATVRA  128
            E K+ILVAVHVASGY+EAEVIPA+TGQETAY++LKLAGRWPVK IHTDN SNFT A V+A
Sbjct  343  ERKIILVAVHVASGYLEAEVIPAKTGQETAYYILKLAGRWPVKVIHTDNSSNFTSAAVKA  164

Query  129  ACWWAGIKQEFGIPYNPQSQGVVESMNKELKKIIGQVRDQAEHLKTAVQMAVFI  182
            AC  AGI+QEF IPYNPQSQ VVESMNKELKKII QV++QAEHLKTAVQMAVFI
Sbjct  163  AC**AGIQQEFRIPYNPQSQRVVESMNKELKKIIRQVKEQAEHLKTAVQMAVFI  2
```

```
>Klebsiella pneumoniae strain EuSCAPE_LT003, whole genome shotgun sequence
Sequence ID: NZ_UKJH02000060.1 Length: 422
Range 1: 15 to 422

Score:261 bits(668), Expect:1e-84,
Method:Compositional matrix adjust.,
Identities:125/136(92%), Positives:133/136(97%), Gaps:0/136(0%)

Query  153  SMNKELKKIIGQVRDQAEHLKTAVQMAVFIHNFKRKGGIGGYSAGERIVDIIATDIQTKE  212
            SMNKELKKIIGQVRDQAE LKTAVQMAVFIHNFKRKGGIGGYSAGERI+DIIA+DIQTKE
Sbjct  422  SMNKELKKIIGQVRDQAEQLKTAVQMAVFIHNFKRKGGIGGYSAGERIIDIIASDIQTKE  243

Query  213  LQKQITKIQNFRVYYRDSRNPLWKGPAKLLWKGEGAVVIQDNSDIKVVPRRKAKIIRDYG  272
            LQKQITK+QNFRVYYRDSR+P+WKGPAKLLWKGEGAVV+QDNSDIKVVPRRKAKII+DYG
Sbjct  242  LQKQITKLQNFRVYYRDSRDPIWKGPAKLLWKGEGAVVLQDNSDIKVVPRRKAKIIKDYG  63

Query  273  KQMAGDDCVASRQDED  288
            KQMAG DCVA RQ+ED
Sbjct  62   KQMAGADCVAGRQNED  15
```

```
>Klebsiella pneumoniae strain EuSCAPE_IL004, whole genome shotgun sequence
Sequence ID: NZ_UKJP02000105.1 Length: 382
Range 1: 2 to 382

Score:254 bits(648), Expect:7e-82,
Method:Compositional matrix adjust.,
Identities:118/127(93%), Positives:123/127(96%), Gaps:0/127(0%)

Query  28   LPPVVAKEIVASCDKCQLKGEAMHGQVDCSPGIWQLDCTHLEGKVILVAVHVASGYIEAE  87
            LPPVVAKEIVASCDKCQLKG AMHGQVDCSPGIWQLDCTHLEGK+ILVAVHVASGYIEAE
Sbjct  382  LPPVVAKEIVASCDKCQLKGGAMHGQVDCSPGIWQLDCTHLEGKIILVAVHVASGYIEAE  203

Query  88   VIPAETGQETAYFLLKLAGRWPVKTIHTDNGSNFTGATVRAACWWAGIKQEFGIPYNPQS  147
            VIPAETGQETAY++LKLAGRWPVK IHTDNGSNFT A V+AACWWAGI+QEFGIPYNPQS
Sbjct  202  VIPAETGQETAYYILKLAGRWPVKVIHTDNGSNFTSAAVKAACWWAGIQQEFGIPYNPQS  23

Query  148  QGVVESM  154
            QGVVESM
Sbjct  22   QGVVESM  2
```

```
>Klebsiella pneumoniae strain EuSCAPE_MT010, whole genome shotgun sequence
Sequence ID: NZ_UKIT02000046.1 Length: 367
Range 1: 1 to 366

Score:233 bits(593), Expect:2e-73,
Method:Compositional matrix adjust.,
Identities:113/122(93%), Positives:119/122(97%), Gaps:0/122(0%)

Query  155  NKELKKIIGQVRDQAEHLKTAVQMAVFIHNFKRKGGIGGYSAGERIVDIIATDIQTKELQ  214
            NKELKKII QV+DQAEHLKTAVQMAVFIHNFKRKGGI GYSAGERI+DIIATD+QTKELQ
Sbjct  1    NKELKKIIRQVKDQAEHLKTAVQMAVFIHNFKRKGGIRGYSAGERIIDIIATDLQTKELQ  180

Query  215  KQITKIQNFRVYYRDSRNPLWKGPAKLLWKGEGAVVIQDNSDIKVVPRRKAKIIRDYGKQ  274
            KQI KIQNFRVYYRDSR+P+WKGPAKLLWKGEGAVVIQDNSDIKVVPRRKAKII+DYGKQ
Sbjct  181  KQIIKIQNFRVYYRDSRDPIWKGPAKLLWKGEGAVVIQDNSDIKVVPRRKAKIIKDYGKQ  360

Query  275  MA  276
            MA
Sbjct  361  MA  366
```

```
>Neisseria gonorrhoeae strain GC-206 NODE_134_length_344_cov_1.3045_ID_20440, whole genome shotgun sequence
Sequence ID: NZ_SAWN01000134.1 Length: 344
Range 1: 1 to 333

Score:226 bits(577), Expect:3e-71,
Method:Compositional matrix adjust.,
Identities:103/111(93%), Positives:111/111(100%), Gaps:0/111(0%)

Query  1    FLDGIDKAQDEHEKYHSNWRAMASDFNLPPVVAKEIVASCDKCQLKGEAMHGQVDCSPGI  60
            FLDGIDKAQ+EHE+YHSNWRAMAS+FNLPP+VAKEIVASCDKCQLKGEA+HGQVDCSPGI
Sbjct  333  FLDGIDKAQEEHERYHSNWRAMASEFNLPPIVAKEIVASCDKCQLKGEAIHGQVDCSPGI  154

Query  61   WQLDCTHLEGKVILVAVHVASGYIEAEVIPAETGQETAYFLLKLAGRWPVK  111
            WQLDCTHLEGK+ILVAVHVASGY+EAEVIPAETGQETAYF+LKLAGRWPVK
Sbjct  153  WQLDCTHLEGKIILVAVHVASGYMEAEVIPAETGQETAYFILKLAGRWPVK  1
```

```
>Klebsiella pneumoniae strain EuSCAPE_IL007, whole genome shotgun sequence
Sequence ID: NZ_UKJT02000028.1 Length: 343
Range 1: 3 to 341

Score:220 bits(560), Expect:1e-68,
Method:Compositional matrix adjust.,
Identities:105/113(93%), Positives:110/113(97%), Gaps:0/113(0%)

Query  169  AEHLKTAVQMAVFIHNFKRKGGIGGYSAGERIVDIIATDIQTKELQKQITKIQNFRVYYR  228
            AEHL+TAVQMAVFIHNFKRKGGIGGYSAGERI+DIIATDIQTKELQKQI KIQNFRVYYR
Sbjct  3    AEHLRTAVQMAVFIHNFKRKGGIGGYSAGERIIDIIATDIQTKELQKQIIKIQNFRVYYR  182

Query  229  DSRNPLWKGPAKLLWKGEGAVVIQDNSDIKVVPRRKAKIIRDYGKQMAGDDCV  281
            DSR+P+WKGPAKLLWKGEGAVVIQDNSDIKVVPRRK KII+DYGKQMAG DCV
Sbjct  183  DSRDPIWKGPAKLLWKGEGAVVIQDNSDIKVVPRRKVKIIKDYGKQMAGADCV  341
```

```
>Thalassovita mangrovi strain GS-10 Scaffold21_1, whole genome shotgun sequence
Sequence ID: NZ_WWEN01000021.1 Length: 1440
Range 1: 1101 to 1439
```

```
Score:227 bits(578), Expect:1e-66,
Method:Compositional matrix adjust.,
Identities:110/113(97%), Positives:112/113(99%), Gaps:0/113(0%)

Query  176   VQMAVFIHNFKRKGGIGGYSAGERIVDIIATDIQTKELQKQITKIQNFRVYYRDSRNPLW  235
             VQMAV IHNFKRKGGIGGYSAGERIVDIIATDIQTKELQKQITKIQNFRVYYRDSR+P+W
Sbjct  1439  VQMAVLIHNFKRKGGIGGYSAGERIVDIIATDIQTKELQKQITKIQNFRVYYRDSRDPVW  1260

Query  236   KGPAKLLWKGEGAVVIQDNSDIKVVPRRKAKIIRDYGKQMAGDDCVASRQDED  288
             KGPAKLLWKGEGAVVIQDNSDIKVVPRRKAKIIRDYGKQMAGDDCVASRQDED
Sbjct  1259  KGPAKLLWKGEGAVVIQDNSDIKVVPRRKAKIIRDYGKQMAGDDCVASRQDED  1101




>Klebsiella pneumoniae strain EuSCAPE_LT003, whole genome shotgun sequence
Sequence ID: NZ_UKJH02000070.1 Length: 297
Range 1: 2 to 295

Score:197 bits(501), Expect:7e-60,
Method:Compositional matrix adjust.,
Identities:91/98(93%), Positives:94/98(95%), Gaps:0/98(0%)

Query  53   QVDCSPGIWQLDCTHLEGKVILVAVHVASGYIEAEVIPAETGQETAYFLLKLAGRWPVKT  112
            QVDCSPGIWQLDCTHLEGKVILVAVHVASGYIEAEVIPAETGQETAYF+LKLAGRWPVK
Sbjct  295  QVDCSPGIWQLDCTHLEGKVILVAVHVASGYIEAEVIPAETGQETAYFILKLAGRWPVKV  116

Query  113  IHTDNGSNFTGATVRAACWWAGIKQEFGIPYNPQSQGV  150
            IHTDNGSNFT   V+AACWWAGI+QEFGIPYNPQSQGV
Sbjct  115  IHTDNGSNFTSNAVKAACWWAGIQQEFGIPYNPQSQGV  2




>Deinococcus aquiradiocola strain JCM 14371 sequence36, whole genome shotgun sequence
Sequence ID: NZ_BMOE01000036.1 Length: 869
Range 1: 2 to 304

Score:203 bits(516), Expect:1e-59,
Method:Compositional matrix adjust.,
Identities:99/101(98%), Positives:101/101(100%), Gaps:0/101(0%)

Query  188  KGGIGGYSAGERIVDIIATDIQTKELQKQITKIQNFRVYYRDSRNPLWKGPAKLLWKGEG  247
            KGGIGGYSAGERIVDIIATDIQTKELQKQITKIQNFRVYYRDSR+P+WKGPAKLLWKGEG
Sbjct  2    KGGIGGYSAGERIVDIIATDIQTKELQKQITKIQNFRVYYRDSRDPVWKGPAKLLWKGEG  181

Query  248  AVVIQDNSDIKVVPRRKAKIIRDYGKQMAGDDCVASRQDED  288
            AVVIQDNSDIKVVPRRKAKIIRDYGKQMAGDDCVASRQDED
Sbjct  182  AVVIQDNSDIKVVPRRKAKIIRDYGKQMAGDDCVASRQDED  304




>Klebsiella pneumoniae strain EuSCAPE_IL005, whole genome shotgun sequence
Sequence ID: NZ_UKJL02000050.1 Length: 636
Range 1: 326 to 634

Score:196 bits(499), Expect:4e-58,
Method:Compositional matrix adjust.,
Identities:95/103(92%), Positives:99/103(96%), Gaps:0/103(0%)

Query  186  KRKGGIGGYSAGERIVDIIATDIQTKELQKQITKIQNFRVYYRDSRNPLWKGPAKLLWKG  245
            KRKGGIGGYSAGERI+DIIATDIQTKELQKQI KIQNFRVYYRDSR+P+WKGPAKLLWKG
Sbjct  634  KRKGGIGGYSAGERIIDIIATDIQTKELQKQIIKIQNFRVYYRDSRDPIWKGPAKLLWKG  455

Query  246  EGAVVIQDNSDIKVVPRRKAKIIRDYGKQMAGDDCVASRQDED  288
            EGAVV+QDNSDIKVVPRRK KIIRDYGKQMAG DCVA RQDED
Sbjct  454  EGAVVLQDNSDIKVVPRRKVKIIRDYGKQMAGADCVAGRQDED  326




>Salmonella enterica subsp. enterica serovar Anatum strain 2014LSAL01667 2014LSAL01667_120, whole genome shotgun sequence
Sequence ID: NZ_CALPAM010000120.1 Length: 264
Range 1: 2 to 262

Score:164 bits(415), Expect:4e-47,
Method:Compositional matrix adjust.,
Identities:78/87(90%), Positives:85/87(97%), Gaps:0/87(0%)

Query  199  RIVDIIATDIQTKELQKQITKIQNFRVYYRDSRNPLWKGPAKLLWKGEGAVVIQDNSDIK  258
            RI+DIIA+DIQTKELQKQITKIQ FRVYYRDSR+P+WKGPAKLLWKGEGAVVI DNSDIK
Sbjct  262  RIIDIIASDIQTKELQKQITKIQKFRVYYRDSRDPIWKGPAKLLWKGEGAVVIPDNSDIK  83

Query  259  VVPRRKAKIIRDYGKQMAGDDCVASRQ  285
            VVPRRKAKI+RDYG+QMAG+DCVASRQ
Sbjct  82   VVPRRKAKILRDYGQQMAGNDCVASRQ  2




>Klebsiella pneumoniae strain EuSCAPE_IL004, whole genome shotgun sequence
Sequence ID: NZ_UKJP02000116.1 Length: 244
Range 1: 2 to 244

Score:162 bits(410), Expect:2e-46,
Method:Compositional matrix adjust.,
Identities:77/81(95%), Positives:81/81(100%), Gaps:0/81(0%)

Query  157  ELKKIIGQVRDQAEHLKTAVQMAVFIHNFKRKGGIGGYSAGERIVDIIATDIQTKELQKQ  216
            ELKKIIGQVRDQAEHLKTAVQMAVFIHNFKR+GGIGGYSAGERI+DIIATDIQTKELQKQ
Sbjct  2    ELKKIIGQVRDQAEHLKTAVQMAVFIHNFKRRGGIGGYSAGERIIDIIATDIQTKELQKQ  181

Query  217  ITKIQNFRVYYRDSRNPLWKG  237
            ITKIQNFRVYYRDSR+P+WKG
Sbjct  182  ITKIQNFRVYYRDSRDPIWKG  244




>Klebsiella pneumoniae strain B2 isolate B2, whole genome shotgun sequence
Sequence ID: NZ_CABWNT010000058.1 Length: 7306
Range 1: 6380 to 7180

Score:175 bits(443), Expect:1e-44,
Method:Compositional matrix adjust.,
Identities:97/269(36%), Positives:150/269(55%), Gaps:3/269(1%)

Query  1     FLDGIDKAQDEHEKYHSNWRAMASDFNLPPVVAKEIVASCDKCQLKGEAMHGQVDCSPGI  60
             +++ I +AQDEHE +H++ + +A  + LP  VAK+I   C  C  +G    G V  SP
Sbjct  6380  WVENIQEAQDEHENWHTSPKILAKRYGLPLTVAKQITQECPHCTKQGSGPAGCVMRSPNH  6559

Query  61    WQLDCTHLEGKVILVAVHVASGYIEAEVIPAETGQETAYFLLKLAGRWPVKTIHTDNGSN  120
             WQ DCTHLE +VI+  V   SGYI A ++  E    T+  +L+    +  K++HTDNG+N
```

```
Sbjct  6560  WQADCTHLENRVIMTFVESNSGYIHATLLSKENALCTSLAILEWVRLFSPKSLHTDNGTN  6739

Query  121   FTGATVRAACWWAGIKQEFGIPYNPQSQGVVESMNKELKKIIGQVRDQAEHLKTAVQMAV  180
             F   +V     +  +    GIPY+P+SQG+VE  N+ LK+ I   R+  + L+ A+Q+A+
Sbjct  6740  FVAESVANLLKFLKVTHTTGIPYHPESQGIVERANRTLKEKIKSHRENTQTLEAALQLAL  6919

Query  181   FIHNFKRKGGIGGYSAGERIVDIIATDIQTKELQKQITKIQNFRVYYRDSRNPLWKGPAK  240
                N K +  +GG +  E  +   A  I  + L +Q    + F  +Y+      WKGP +
Sbjct  6920  ITCN-KGRESMGGQTPWEVFITNQAQTIHEELLLQQAQSSKKF-CFYKIPGEHNWKGPTR  7093

Query  241   LLWKGEGAVVIQDNSD-IKVVPRRKAKII  268
             +LWKG+GAVV+ D    I  VP  + K++
Sbjct  7094  VLWKGDGAVVVNDEKKGIIAVPLTRTKLL  7180




>Klebsiella pneumoniae strain EuSCAPE_LT003, whole genome shotgun sequence
Sequence ID: NZ_UKJH02000069.1 Length: 305
Range 1: 84 to 305

Score:153 bits(386), Expect:1e-42,
Method:Compositional matrix adjust.,
Identities:69/74(93%), Positives:73/74(98%), Gaps:0/74(0%)

Query  1    FLDGIDKAQDEHEKYHSNWRAMASDFNLPPVVAKEIVASCDKCQLKGEAMHGQVDCSPGI  60
            FLD IDKAQ+EHEKYH+NWRA+ASDFNLPP+VAKEIVASCDKCQLKGEAMHGQVDCSPGI
Sbjct  84   FLDRIDKAQEEHEKYHNNWRAIASDFNLPPIVAKEIVASCDKCQLKGEAMHGQVDCSPGI  263

Query  61   WQLDCTHLEGKVIL  74
            WQLDCTHLEGKVIL
Sbjct  264  WQLDCTHLEGKVIL  305




>Methylobacterium radiotolerans strain MAMP 4754 MP-M-radiotolerans_contig_571, whole genome shotgun sequence
Sequence ID: NZ_NKQS01000559.1 Length: 289
Range 1: 2 to 244

Score:150 bits(380), Expect:9e-42,
Method:Compositional matrix adjust.,
Identities:72/81(89%), Positives:76/81(93%), Gaps:0/81(0%)

Query  208  IQTKELQKQITKIQNFRVYYRDSRNPLWKGPAKLLWKGEGAVVIQDNSDIKVVPRRKAKI  267
            IQTK +QKQI KIQNFRVYYRDSR+P+WKGPAKLLWKGEGAVVIQDNSDIKVVPRRK KI
Sbjct  2    IQTKAVQKQIIKIQNFRVYYRDSRDPIWKGPAKLLWKGEGAVVIQDNSDIKVVPRRKVKI  181

Query  268  IRDYGKQMAGDDCVASRQDED  288
            I+DYGKQMAG DCVA RQDED
Sbjct  182  IKDYGKQMAGADCVAGRQDED  244




>Methylobacterium radiotolerans strain MAMP 4754 MP-M-radiotolerans_contig_1227, whole genome shotgun sequence
Sequence ID: NZ_NKQS01001213.1 Length: 279
Range 1: 3 to 254

Score:135 bits(339), Expect:1e-35,
Method:Compositional matrix adjust.,
Identities:68/84(81%), Positives:74/84(88%), Gaps:0/84(0%)

Query  168  QAEHLKTAVQMAVFIHNFKRKGGIGGYSAGERIVDIIATDIQTKELQKQITKIQNFRVYY  227
            QAEHLKTAV MAVFIHNFKRKGGIGGYSAGERI+DIIATDIQTKELQKQ  KIQN++V Y
Sbjct  3    QAEHLKTAVLMAVFIHNFKRKGGIGGYSAGERIIDIIATDIQTKELQKQNIKIQNYQVDY  182

Query  228  RDSRNPLWKGPAKLLWKGEGAVVI  251
            RDS +   KG AKL+WKG+GAV I
Sbjct  183  RDSIDHSRKGSAKLIWKGKGAVGI  254




>Klebsiella pneumoniae strain EuSCAPE_MT012, whole genome shotgun sequence
Sequence ID: NZ_UKIX02000067.1 Length: 385
Range 1: 226 to 384

Score:104 bits(259), Expect:3e-23,
Method:Compositional matrix adjust.,
Identities:51/53(96%), Positives:51/53(96%), Gaps:0/53(0%)

Query  236  KGPAKLLWKGEGAVVIQDNSDIKVVPRRKAKIIRDYGKQMAGDDCVASRQDED  288
            KGPAKLLWKGEGAVVIQDNSDIKVVPRRKAKIIRDYGKQMAG DCVA RQDED
Sbjct  384  KGPAKLLWKGEGAVVIQDNSDIKVVPRRKAKIIRDYGKQMAGADCVAGRQDED  226




>Acinetobacter baumannii strain P4 contig_P4_92, whole genome shotgun sequence
Sequence ID: NZ_JBBXLU010000071.1 Length: 1226
Range 1: 333 to 488

Score:103 bits(257), Expect:2e-20,
Method:Compositional matrix adjust.,
Identities:48/52(92%), Positives:52/52(100%), Gaps:0/52(0%)

Query  185  FKRKGGIGGYSAGERIVDIIATDIQTKELQKQITKIQNFRVYYRDSRNPLWK  236
            FKRKGGIGGYSAGERIVDIIATDIQTKELQKQITKIQNFRVYYRDSR+P+++
Sbjct  333  FKRKGGIGGYSAGERIVDIIATDIQTKELQKQITKIQNFRVYYRDSRDPVYR  488




>Acinetobacter baumannii strain E12 contig_E12_75, whole genome shotgun sequence
Sequence ID: NZ_JBBXKS010000057.1 Length: 514
Range 1: 2 to 127

Score:82.8 bits(203), Expect:1e-14,
Method:Compositional matrix adjust.,
Identities:38/42(90%), Positives:42/42(100%), Gaps:0/42(0%)

Query  195  SAGERIVDIIATDIQTKELQKQITKIQNFRVYYRDSRNPLWK  236
            SAGERIVDIIATDIQTKELQKQITKIQNFRVYYRDSR+P+++
Sbjct  2    SAGERIVDIIATDIQTKELQKQITKIQNFRVYYRDSRDPVYR  127




>Chlamydia abortus strain 09_05, whole genome shotgun sequence
Sequence ID: NZ_FQLG01000005.1 Length: 4831
Range 1: 1318 to 2118

Score:80.5 bits(197), Expect:3e-12,
```

```
Method:Compositional matrix adjust.,
Identities:74/284(26%), Positives:122/284(42%), Gaps:36/284(12%)

Query  5     IDKAQDEHEKYHSNWRAMASDFNLPPVVAKEIVASCDKCQLKGEAMHGQVDCSP------  58
             ++ A++ H  +H     + S F+L    A++IV  C  C       H  V  +P
Sbjct  2118  VEAARNFHNNFHVTAETLRSRFSLTRKEARDIVTQCQSCCEFLPVPH--VGINPRGIRPL  1945

Query  59    GIWQLDCTHLE--GKV--ILVAVHVASGYIEAEVIPAETGQETAYFLLKLAGRW-PVKTI  113
              +WQ+D TH+   GK+  + V++   SG + A  +  E         L+    W   + +
Sbjct  1944  QVWQMDVTHVSSFGKLQYLHVSIDTCSGIMFASPLTGEKASHVIQHCLEAWSAWGKPRLL  1765

Query  114   HTDNGSNFTGATVRAACWWAGIKQEFGIPYNPQSQGVVESMNKELKK-IIGQVRDQAEHL  172
              TDNG  +T    +  C    +    G+PYNPQ QG+VE  ++ LK  +I Q R   E
Sbjct  1764  KTDNGPAYTSQKFQQFCRQMDVTHLTGLPYNPQGQGIVERAHRTLKAYLIKQKRGTFEET  1585

Query  173   -----KTAVQMAVFIHNFKRKGGIGGYSAGERIVDIIATDIQTKELQKQITKIQNFRVYY  227
                   + +V MA+F  NF       G++A ER            E  +      N  V +
Sbjct  1584  VPRAPRVSVSMALFTLNFLNIDA-HGHTAAER---------HCSEPDR-----PNEMVKW  1450

Query  228   RDSRNPLWKGPAKLLWKGEGAVVI--QDNSDIKVVPRRKAKIIR  269
             ++  +  W GP  +L +  GA+ +  Q+  +   VP R  + I+
Sbjct  1449  KNVLDNKWYGPDPILIRSRGAICVFPQNEDNPFWVPERLTRKIQ  1318
```

```
>Uncultured Roseobacter sp. isolate 53.fasta NODE_38, whole genome shotgun sequence
Sequence ID: NZ_CANMKI010000038.1 Length: 315
Range 1: 210 to 314

Score:73.9 bits(180), Expect:5e-12,
Method:Compositional matrix adjust.,
Identities:35/35(100%), Positives:35/35(100%), Gaps:0/35(0%)

Query  177  QMAVFIHNFKRKGGIGGYSAGERIVDIIATDIQTK  211
            QMAVFIHNFKRKGGIGGYSAGERIVDIIATDIQTK
Sbjct  210  QMAVFIHNFKRKGGIGGYSAGERIVDIIATDIQTK  314
```

```
>Chlamydia abortus strain 09_03, whole genome shotgun sequence
Sequence ID: NZ_FQLL01000003.1 Length: 4440
Range 1: 2393 to 3193

Score:79.3 bits(194), Expect:9e-12,
Method:Compositional matrix adjust.,
Identities:73/284(26%), Positives:122/284(42%), Gaps:36/284(12%)

Query  5     IDKAQDEHEKYHSNWRAMASDFNLPPVVAKEIVASCDKCQLKGEAMHGQVDCSP------  58
             ++ A++ H  +H     + S F+L    A++IV  C  C       H  V  +P
Sbjct  2393  VEAARNFHNNFHVTAETLRSRFSLTRKEARDIVTQCQSCCEFLPVPH--VGINPRGIRPL  2566

Query  59    GIWQLDCTHLE--GKV--ILVAVHVASGYIEAEVIPAETGQETAYFLLKLAGRW-PVKTI  113
              +WQ+D TH+   GK+  + V++   SG + A  +  E         L+    W   + +
Sbjct  2567  QVWQMDVTHVSSFGKLQYLHVSIDTCSGIMFASPLTGEKASHVIQHCLEAWSAWGKPRLL  2746

Query  114   HTDNGSNFTGATVRAACWWAGIKQEFGIPYNPQSQGVVESMNKELKK-IIGQVRDQAEHL  172
              TDNG  +T    +  C    +    G+PYNPQ QG+VE  ++ LK  +I Q R   E
Sbjct  2747  KTDNGPAYTSQKFQQFCRQMDVTHLTGLPYNPQGQGIVERAHRTLKAYLIKQKRGTFEET  2926

Query  173   -----KTAVQMAVFIHNFKRKGGIGGYSAGERIVDIIATDIQTKELQKQITKIQNFRVYY  227
                   + +V +A+F  NF       G++A ER            E  +      N  V +
Sbjct  2927  VPRAPRVSVSLALFTLNFLNIDA-HGHTAAER---------HCSEPDR-----PNEMVKW  3061

Query  228   RDSRNPLWKGPAKLLWKGEGAVVI--QDNSDIKVVPRRKAKIIR  269
             ++  +  W GP  +L +  GA+ +  Q+  +   VP R  + I+
Sbjct  3062  KNVLDNKWYGPDPILIRSRGAICVFPQNEDNPFWVPERLTRKIQ  3193
```

```
Range 2: 1757 to 2413

Score:72.8 bits(177), Expect:1e-09,
Method:Compositional matrix adjust.,
Identities:61/231(26%), Positives:98/231(42%), Gaps:24/231(10%)

Query  5     IDKAQDEHEKYHSNWRAMASDFNLPPVVAKEIVASCDKCQLKGEAMHGQVDCSP------  58
             ++ A++ H  +H     + S F+L    A++IV  C  C       H  V  +P
Sbjct  1757  VEAARNFHNNFHVTAETLRSRFSLTRKEARDIVTQCQSCCEFLPVPH--VGINPRGIRPL  1930

Query  59    GIWQLDCTHLE--GKV--ILVAVHVASGYIEAEVIPAETGQETAYFLLKLAGRW-PVKTI  113
              +WQ+D TH+   GK+  + V++   SG + A  +  E         L+    W   K +
Sbjct  1931  QVWQMDVTHVSSFGKLQYLHVSIDTCSGIMFASPLTGEKASHVIQHCLEAWSAWGKPKLL  2110

Query  114   HTDNGSNFTGATVRAACWWAGIKQEFGIPYNPQSQGVVESMNKELKKIIGQVRDQAEHLK  173
              TDNG  +T    +  C    +    G+PYNPQ QG+VE  ++ LK  +         LK
Sbjct  2111  KTDNGPAYTSQKFQQFCRQMDVTHLTGLPYNPQGQGIVERAHRTLKSYL---------LK  2263

Query  174   TAVQMAVFIHNFKRKGGIGGYSA-GERIVDIIATDIQTKELQKQITKIQNF  223
                    V+I + +   G+ G  A G  + D  AT +    L   +   +NF
Sbjct  2264  QKRGXPVYITHVRAHSGLPGPMALGNDLAD-KATKVVAAALSSPVEAARNF  2413
```

```
>Chlamydia abortus strain 11_01, whole genome shotgun sequence
Sequence ID: NZ_FQLK01000002.1 Length: 5240
Range 1: 1018 to 1818

Score:79.0 bits(193), Expect:1e-11,
Method:Compositional matrix adjust.,
Identities:72/284(25%), Positives:122/284(42%), Gaps:36/284(12%)

Query  5     IDKAQDEHEKYHSNWRAMASDFNLPPVVAKEIVASCDKCQLKGEAMHGQVDCSP------  58
             ++ A++ H  +H     + S F+L    A++IV  C  C       H  V  +P
Sbjct  1818  VEAARNFHNNFHVTAETLRSRFSLTRKEARDIVTQCQSCCEFLPVPH--VGINPRGIRPL  1645

Query  59    GIWQLDCTHLE--GKV--ILVAVHVASGYIEAEVIPAETGQETAYFLLKLAGRW-PVKTI  113
              +WQ+D TH+   GK+  + V++   SG + A  +  E         L+    W   + +
Sbjct  1644  QVWQMDVTHVSSFGKLQYLHVSIDTCSGIMFASPLTGEKASHVIQHCLEAWSAWGKPRLL  1465

Query  114   HTDNGSNFTGATVRAACWWAGIKQEFGIPYNPQSQGVVESMNKELKK-IIGQVRDQAEHL  172
              TDNG  +T    +  C    +    G+PYNPQ QG+VE  ++ LK  +I Q R   E
Sbjct  1464  KTDNGPAYTSQKFQQFCRQMDVTHLTGLPYNPQGQGIVERAHRTLKAYLIKQKRGTFEET  1285

Query  173   -----KTAVQMAVFIHNFKRKGGIGGYSAGERIVDIIATDIQTKELQKQITKIQNFRVYY  227
                   + +V +A+F  NF       G++A ER            E  +      N  V +
Sbjct  1284  VPRAPRVSVSLALFTLNFLNIDA-HGHTAAER---------HCSEPDR-----PNEMVKW  1150

Query  228   RDSRNPLWKGPAKLLWKGEGAVVI--QDNSDIKVVPRRKAKIIR  269
             ++  +  W GP  +L +  GA+ +  Q+  +   +P R  + I+
Sbjct  1149  KNVLDNKWYGPDPILIRSRGAICVFPQNEDNPFWIPERLTRKIQ  1018
```

```
>Uncultured Dokdonia sp. isolate 746.fasta NODE_102, whole genome shotgun sequence
Sequence ID: NZ_CANLOB010000102.1 Length: 440
Range 1: 2 to 106

Score:73.9 bits(180), Expect:1e-11,
Method:Compositional matrix adjust.,
Identities:35/35(100%), Positives:35/35(100%), Gaps:0/35(0%)

Query  177  QMAVFIHNFKRKGGIGGYSAGERIVDIIATDIQTK  211
            QMAVFIHNFKRKGGIGGYSAGERIVDIIATDIQTK
Sbjct  106  QMAVFIHNFKRKGGIGGYSAGERIVDIIATDIQTK  2
```

```
>Chlamydia trachomatis strain R33512extract, whole genome shotgun sequence
Sequence ID: NZ_CVNW01000009.1 Length: 1310
Range 1: 13 to 771

Score:78.2 bits(191), Expect:1e-11,
Method:Compositional matrix adjust.,
Identities:71/270(26%), Positives:115/270(42%), Gaps:35/270(12%)

Query  5    IDKAQDEHEKYHSNWRAMASDFNLPPVVAKEIVASCDKCQLKGEAMHGQVDCSP------  58
            ++ A++ H  +H     + S F+L    A++IV  C  C       H  V  +P
Sbjct  771  VEAARNFHNNFHVTAETLRSRFSLTRKEARDIVTQCQSCCEFLPVPH--VGINPRGIRPL  598

Query  59   GIWQLDCTHLE--GKV--ILVAVHVASGYIEAEVIPAETGQETAYFLLKLAGRW-PVKTI  113
             +WQ+D TH+   GK+  + V++   SG + A  +  E         L+    W   + +
Sbjct  597  QVWQMDVTHVSSFGKLQYLHVSIDTCSGIMFASPLTGEKASHVIQHCLEAWSAWGKPRLL  418

Query  114  HTDNGSNFTGATVRAACWWAGIKQEFGIPYNPQSQGVVESMNKELKK-IIGQVRDQAEHL  172
             TDNG  +T    +  C    +    G+PYNPQ QG+VE  ++ LK  +I Q R   E
Sbjct  417  KTDNGPAYTSQKFQQFCRQMDVTHLTGLPYNPQGQGIVERAHRTLKAYLIKQKRGTFEET  238

Query  173  -----KTAVQMAVFIHNFKRKGGIGGYSAGERIVDIIATDIQTKELQKQITKIQNFRVYY  227
                 + +V +A+F  NF       G++A ER            E  +      N  V +
Sbjct  237  VPRAPRVSVSLALFTLNFLNIDA-HGHTAAER---------HCSEPDR-----PNEMVKW  103

Query  228  RDSRNPLWKGPAKLLWKGEGAV-VIQDNSD  256
            ++  +  W GP  +L +  GA+ V   N D
Sbjct  102  KNVLDNKWYGPDPILIRSRGAICVFPQNED  13
```

```
>Cellulophaga baltica isolate 925.fasta NODE_42, whole genome shotgun sequence
Sequence ID: NZ_CANLPS010000042.1 Length: 316
Range 1: 35 to 136

Score:72.0 bits(175), Expect:2e-11,
Method:Compositional matrix adjust.,
Identities:34/34(100%), Positives:34/34(100%), Gaps:0/34(0%)

Query  177  QMAVFIHNFKRKGGIGGYSAGERIVDIIATDIQT  210
            QMAVFIHNFKRKGGIGGYSAGERIVDIIATDIQT
Sbjct  136  QMAVFIHNFKRKGGIGGYSAGERIVDIIATDIQT  35
```

```
>Acinetobacter baumannii strain E1 contig_E1_51, whole genome shotgun sequence
Sequence ID: NZ_JBBXKX010000035.1 Length: 412
Range 1: 2 to 106

Score:69.3 bits(168), Expect:6e-10,
Method:Compositional matrix adjust.,
Identities:31/35(89%), Positives:35/35(100%), Gaps:0/35(0%)

Query  202  DIIATDIQTKELQKQITKIQNFRVYYRDSRNPLWK  236
            DIIATDIQTKELQKQITKIQNFRVYYRDSR+P+++
Sbjct  2    DIIATDIQTKELQKQITKIQNFRVYYRDSRDPVYR  106
```

```
>Salmonella enterica subsp. enterica serovar Typhi strain 7-sc-cl NODE_243_length_289_cov_0.314815, whole genome shotgun sequence
Sequence ID: NZ_JAEPKN010000213.1 Length: 289
Range 1: 1 to 96

Score:67.8 bits(164), Expect:7e-10,
Method:Compositional matrix adjust.,
Identities:32/32(100%), Positives:32/32(100%), Gaps:0/32(0%)

Query  177  QMAVFIHNFKRKGGIGGYSAGERIVDIIATDI  208
            QMAVFIHNFKRKGGIGGYSAGERIVDIIATDI
Sbjct  96   QMAVFIHNFKRKGGIGGYSAGERIVDIIATDI  1
```

```
>Corynebacterium glutamicum strain 39 Scaffolds_32_9166, whole genome shotgun sequence
Sequence ID: NZ_JACEQW010000032.1 Length: 8862
Range 1: 3976 to 4086

Score:72.4 bits(176), Expect:2e-09,
Method:Compositional matrix adjust.,
Identities:35/37(95%), Positives:36/37(97%), Gaps:0/37(0%)

Query  185   FKRKGGIGGYSAGERIVDIIATDIQTKELQKQITKIQ  221
             FKRKGGIGGYSAGERIVDIIATDIQTKELQKQITK +
Sbjct  4086  FKRKGGIGGYSAGERIVDIIATDIQTKELQKQITKFK  3976
```

```
>Corynebacterium glutamicum strain 37 Scaffolds_33_8928, whole genome shotgun sequence
Sequence ID: NZ_JACENO010000033.1 Length: 8928
Range 1: 3950 to 4060

Score:72.4 bits(176), Expect:2e-09,
Method:Compositional matrix adjust.,
Identities:35/37(95%), Positives:36/37(97%), Gaps:0/37(0%)

Query  185   FKRKGGIGGYSAGERIVDIIATDIQTKELQKQITKIQ  221
             FKRKGGIGGYSAGERIVDIIATDIQTKELQKQITK +
Sbjct  4060  FKRKGGIGGYSAGERIVDIIATDIQTKELQKQITKFK  3950
```

>Pseudomonas aromaticivorans strain MAP12 34, whole genome shotgun sequence
Sequence ID: NZ_JAHRGL010000035.1 Length: 7031
Range 1: 1653 to 2372

Score:72.4 bits(176), Expect:2e-09,
Method:Compositional matrix adjust.,
Identities:65/254(26%), Positives:104/254(40%), Gaps:25/254(9%)

```
Query  1     FLDGIDKAQDEHEKYHSNWRAMASDFNLPPVVAKEIVASCDKCQLKGEAMHGQVDCSPG-  59
              +  + +A + H+K+H N + +  DF++    AKEIV +C  C    +       C+P
Sbjct  1653  LIGSVLEASEFHKKHHVNSKGLKKDFSITWQQAKEIVRNCPTCSFYNQTP-LPAGCNPKG  1829

Query  60    -----IWQLDCTHLEGKVILVAVH----VASGYIEAEVIPAETGQETAYFLLKLAGRWPV  110
                  +WQ+D  H      L  VH      SG+  A  + +E        LL++     +
Sbjct  1830  IRRNEVWQMDVFHFAEFGNLKYVHHTIDTFSGFQWATALNSEKADSVITHLLEVMAVMGI  2009

Query  111   KT-IHTDNGSNFTGATVRAACWWAGIKQEFGIPYNPQSQGVVESMNKELKKIIGQVRDQA  169
                 I TDN   +    +     +  IK   GIP+NP  Q VVE  N+ LK+++ +   +
Sbjct  2010  PAQIKTDNAPAYVSTKLEQFFKYYNIKHVTGIPHNPTGQAVVERSNRTLKEMLNKQAWKT  2189

Query  170   EHLKTAVQMAVFIHNFKRKGGIGGYSAGERIVDIIATDIQTKELQKQITKIQNFRVYYRD  229
             +  K  +  A+   NF      G  +A         T  +T EL        N  VY++D
Sbjct  2190  KPPKHRLHNALLTLNFLNANEKGQTAAKRH-----WTREKTAEL--------NQPVYFKD  2330

Query  230   SRNPLWKGPAKLLW  243
                 +WK    L W
Sbjct  2331  VLTSVWKSGHVL*W  2372
```

>Chlamydia abortus strain 09_01, whole genome shotgun sequence
Sequence ID: NZ_FQLJ01000005.1 Length: 1528
Range 1: 750 to 1277

Score:68.2 bits(165), Expect:3e-08,
Method:Compositional matrix adjust.,
Identities:50/178(28%), Positives:80/178(44%), Gaps:14/178(7%)

```
Query  5     IDKAQDEHEKYHSNWRAMASDFNLPPVVAKEIVASCDKCQLKGEAMHGQVDCSP------  58
             ++ A++ H  +H     + S F+L    A++IV  C  C       H  V  +P
Sbjct  750   VEAARNFHNNFHVTAETLRSRFSLTRKEARDIVTQCQSCCEFLPVPH--VGINPRGIRPL  923

Query  59    GIWQLDCTHLE--GKV--ILVAVHVASGYIEAEVIPAETGQETAYFLLKLAGRW-PVKTI  113
              +WQ+D TH+   GK+  + V++   SG + A  +  E         L+    W   + +
Sbjct  924   QVWQMDVTHVSSFGKLQYLHVSIDTCSGIMFASPLTGEKASHVIQHCLEAWSAWGKPRLL  1103

Query  114   HTDNGSNFTGATVRAACWWAGIKQEFGIPYNPQSQGVVESMNKELKK-IIGQVRDQAE  170
              TDNG  +T    +  C    +    G+PYNPQ QG+VE  ++ LK  +I Q R   E
Sbjct  1104  KTDNGPAYTSQKFQQFCRQMDVTHLTGLPYNPQGQGIVERAHRTLKAYLIKQKRGTFE  1277
```

>Enterococcus faecalis strain 15224 NODE_126_length_632_cov_0.918812, whole genome shotgun sequence
Sequence ID: NZ_QNGP01000071.1 Length: 632
Range 1: 177 to 611

Score:63.5 bits(153), Expect:3e-07,
Method:Compositional matrix adjust.,
Identities:43/145(30%), Positives:68/145(46%), Gaps:15/145(10%)

```
Query  33   AKEIVASCDKCQLKGEAMHGQVDCSPGIWQLDCTHLEG-------------KVILVAVHV  79
            ++E V +CD CQ  G   HG      GI +++   + G               ILVAV
Sbjct  177  SREFVRNCDSCQRAGNLPHGYAMPQQGILEIELFDVWGIDFMGPFPPSYSNTYILVAVDY  356

Query  80   ASGYIEAEVIPAETGQETAYFLLK-LAGRWPV-KTIHTDNGSNFTGATVRAACWWAGIKQ  137
             S ++EA   P    +    FL K +  R+ V + + +D G++F    + +A    GI
Sbjct  357  VSKWVEAIATPTNDTKTVLKFLQKHIFSRFGVPRALISDGGTHFCNKQLYSAMVRYGISH  536

Query  138  EFGIPYNPQSQGVVESMNKELKKII  162
            +   PY+PQ+ G  E  N+ELK+I+
Sbjct  537  KVATPYHPQTNGQAEVSNRELKRIL  611
```

>Uncultured Roseobacter sp. isolate 930.fasta NODE_31, whole genome shotgun sequence
Sequence ID: NZ_CANLQB010000031.1 Length: 257
Range 1: 180 to 257

Score:57.0 bits(136), Expect:5e-06,
Method:Compositional matrix adjust.,
Identities:26/26(100%), Positives:26/26(100%), Gaps:0/26(0%)

```
Query  177  QMAVFIHNFKRKGGIGGYSAGERIVD  202
            QMAVFIHNFKRKGGIGGYSAGERIVD
Sbjct  180  QMAVFIHNFKRKGGIGGYSAGERIVD  257
```

>Brachybacterium paraconglomeratum isolate 859.fasta NODE_71, whole genome shotgun sequence
Sequence ID: NZ_CANLNX010000071.1 Length: 258
Range 1: 181 to 258

Score:57.0 bits(136), Expect:5e-06,
Method:Compositional matrix adjust.,
Identities:26/26(100%), Positives:26/26(100%), Gaps:0/26(0%)

```
Query  177  QMAVFIHNFKRKGGIGGYSAGERIVD  202
            QMAVFIHNFKRKGGIGGYSAGERIVD
Sbjct  181  QMAVFIHNFKRKGGIGGYSAGERIVD  258
```

>Enterococcus faecalis strain AKSZ-144 NODE_42_length_234_cov_1622.847134, whole genome shotgun sequence
Sequence ID: NZ_JAOXSP010000040.1 Length: 234
Range 1: 153 to 230

Score:56.2 bits(134), Expect:6e-06,
Method:Composition-based stats.,
Identities:25/26(96%), Positives:26/26(100%), Gaps:0/26(0%)

```
Query  177  QMAVFIHNFKRKGGIGGYSAGERIVD  202
            QMAVFIHNFKRKGGIGGYSAGERIV+
Sbjct  153  QMAVFIHNFKRKGGIGGYSAGERIVE  230
```

>Roseobacter litoralis isolate 289-G.fasta NODE_32, whole genome shotgun sequence

Sequence ID: NZ_CANMVH010000032.1 Length: 249
Range 1: 2 to 79

Score:56.2 bits(134), Expect:8e-06,
Method:Composition-based stats.,
Identities:25/26(96%), Positives:26/26(100%), Gaps:0/26(0%)

```
Query  177  QMAVFIHNFKRKGGIGGYSAGERIVD  202
            QMAVFIHNFKRKGGIGGYSAGERIV+
Sbjct  79   QMAVFIHNFKRKGGIGGYSAGERIVE  2
```

>Uncultured Pseudoteredinibacter sp. isolate 208.fasta NODE_42, whole genome shotgun sequence
Sequence ID: NZ_CANMSS010000042.1 Length: 253
Range 1: 174 to 251

Score:56.2 bits(134), Expect:8e-06,
Method:Composition-based stats.,
Identities:25/26(96%), Positives:26/26(100%), Gaps:0/26(0%)

```
Query  177  QMAVFIHNFKRKGGIGGYSAGERIVD  202
            QMAVFIHNFKRKGGIGGYSAGERIV+
Sbjct  174  QMAVFIHNFKRKGGIGGYSAGERIVE  251
```

>Bacillus subtilis strain RS3017 NODE_116_length_232_cov_0.748387, whole genome shotgun sequence
Sequence ID: NZ_JASDBS010000113.1 Length: 232
Range 1: 156 to 230

Score:56.2 bits(134), Expect:8e-06,
Method:Composition-based stats.,
Identities:25/25(100%), Positives:25/25(100%), Gaps:0/25(0%)

```
Query  177  QMAVFIHNFKRKGGIGGYSAGERIV  201
            QMAVFIHNFKRKGGIGGYSAGERIV
Sbjct  156  QMAVFIHNFKRKGGIGGYSAGERIV  230
```

>Uncultured Tateyamaria sp. isolate 203.fasta NODE_54, whole genome shotgun sequence
Sequence ID: NZ_CANLVZ010000054.1 Length: 263
Range 1: 185 to 262

Score:56.2 bits(134), Expect:8e-06,
Method:Composition-based stats.,
Identities:25/26(96%), Positives:26/26(100%), Gaps:0/26(0%)

```
Query  177  QMAVFIHNFKRKGGIGGYSAGERIVD  202
            QMAVFIHNFKRKGGIGGYSAGERIV+
Sbjct  185  QMAVFIHNFKRKGGIGGYSAGERIVE  262
```

>Uncultured Roseobacter sp. isolate 908.fasta NODE_28, whole genome shotgun sequence
Sequence ID: NZ_CANMNK010000028.1 Length: 244
Range 1: 168 to 242

Score:56.2 bits(134), Expect:9e-06,
Method:Composition-based stats.,
Identities:25/25(100%), Positives:25/25(100%), Gaps:0/25(0%)

```
Query  177  QMAVFIHNFKRKGGIGGYSAGERIV  201
            QMAVFIHNFKRKGGIGGYSAGERIV
Sbjct  168  QMAVFIHNFKRKGGIGGYSAGERIV  242
```

>Uncultured Roseibium sp. isolate 190.fasta NODE_26, whole genome shotgun sequence
Sequence ID: NZ_CANNEN010000026.1 Length: 248
Range 1: 172 to 246

Score:56.2 bits(134), Expect:9e-06,
Method:Composition-based stats.,
Identities:25/25(100%), Positives:25/25(100%), Gaps:0/25(0%)

```
Query  177  QMAVFIHNFKRKGGIGGYSAGERIV  201
            QMAVFIHNFKRKGGIGGYSAGERIV
Sbjct  172  QMAVFIHNFKRKGGIGGYSAGERIV  246
```

>Uncultured Roseobacter sp. isolate 407-3Z.fasta NODE_45, whole genome shotgun sequence
Sequence ID: NZ_CANMKD010000045.1 Length: 252
Range 1: 3 to 77

Score:56.2 bits(134), Expect:1e-05,
Method:Composition-based stats.,
Identities:25/25(100%), Positives:25/25(100%), Gaps:0/25(0%)

```
Query  177  QMAVFIHNFKRKGGIGGYSAGERIV  201
            QMAVFIHNFKRKGGIGGYSAGERIV
Sbjct  77   QMAVFIHNFKRKGGIGGYSAGERIV  3
```

>Epibacterium ulvae isolate 20.fasta NODE_62, whole genome shotgun sequence
Sequence ID: NZ_CANLDO010000062.1 Length: 257
Range 1: 181 to 255

Score:56.2 bits(134), Expect:1e-05,
Method:Composition-based stats.,
Identities:25/25(100%), Positives:25/25(100%), Gaps:0/25(0%)

```
Query  177  QMAVFIHNFKRKGGIGGYSAGERIV  201
            QMAVFIHNFKRKGGIGGYSAGERIV
Sbjct  181  QMAVFIHNFKRKGGIGGYSAGERIV  255
```

>Roseibium album isolate 321-3Z.fasta NODE_38, whole genome shotgun sequence
Sequence ID: NZ_CANMGD010000038.1 Length: 255
Range 1: 3 to 77

Score:56.2 bits(134), Expect:1e-05,

```
Method:Composition-based stats.,
Identities:25/25(100%), Positives:25/25(100%), Gaps:0/25(0%)

Query  177  QMAVFIHNFKRKGGIGGYSAGERIV  201
            QMAVFIHNFKRKGGIGGYSAGERIV
Sbjct  77   QMAVFIHNFKRKGGIGGYSAGERIV  3
```

```
>Uncultured Ruegeria sp. isolate 409.fasta NODE_44, whole genome shotgun sequence
Sequence ID: NZ_CANLNV010000044.1 Length: 256
Range 1: 3 to 77

Score:56.2 bits(134), Expect:1e-05,
Method:Composition-based stats.,
Identities:25/25(100%), Positives:25/25(100%), Gaps:0/25(0%)

Query  177  QMAVFIHNFKRKGGIGGYSAGERIV  201
            QMAVFIHNFKRKGGIGGYSAGERIV
Sbjct  77   QMAVFIHNFKRKGGIGGYSAGERIV  3
```

```
>Uncultured Winogradskyella sp. isolate 408-3Z.fasta NODE_32, whole genome shotgun sequence
Sequence ID: NZ_CANMJY010000032.1 Length: 256
Range 1: 180 to 254

Score:56.2 bits(134), Expect:1e-05,
Method:Composition-based stats.,
Identities:25/25(100%), Positives:25/25(100%), Gaps:0/25(0%)

Query  177  QMAVFIHNFKRKGGIGGYSAGERIV  201
            QMAVFIHNFKRKGGIGGYSAGERIV
Sbjct  180  QMAVFIHNFKRKGGIGGYSAGERIV  254
```

```
>Pseudomonas sp. QS1027 NG-9811_QS116_lib127709_4633_4663_4674_contig_78, whole genome shotgun sequence
Sequence ID: NZ_PHSU01000078.1 Length: 240
Range 1: 9 to 83

Score:55.8 bits(133), Expect:1e-05,
Method:Composition-based stats.,
Identities:25/25(100%), Positives:25/25(100%), Gaps:0/25(0%)

Query  177  QMAVFIHNFKRKGGIGGYSAGERIV  201
            QMAVFIHNFKRKGGIGGYSAGERIV
Sbjct  83   QMAVFIHNFKRKGGIGGYSAGERIV  9
```

```
>Pseudomonas nitroreducens strain FY43 NODE_35_length_306_cov_242.462882, whole genome shotgun sequence
Sequence ID: NZ_JAJISQ010000035.1 Length: 306
Range 1: 161 to 238

Score:56.2 bits(134), Expect:2e-05,
Method:Compositional matrix adjust.,
Identities:25/26(96%), Positives:26/26(100%), Gaps:0/26(0%)

Query  177  QMAVFIHNFKRKGGIGGYSAGERIVD  202
            QMAVFIHNFKRKGGIGGYSAGERIV+
Sbjct  161  QMAVFIHNFKRKGGIGGYSAGERIVE  238
```

```
>Proteus mirabilis strain PM2-4 PM2-4_scaffold43, whole genome shotgun sequence
Sequence ID: NZ_JAMKNT010000043.1 Length: 3726
Range 1: 2922 to 3395

Score:60.5 bits(145), Expect:2e-05,
Method:Compositional matrix adjust.,
Identities:47/166(28%), Positives:71/166(42%), Gaps:22/166(13%)

Query  19    WRAMASDFNLPPVVAKEIVASCDKCQLKG------EAMHGQVDCSP-GIWQLDCTHLEG-  70
             W  M  D        +E    C +CQ+ G      E     V   P     LD  H++
Sbjct  3395  WENMTHDI-------QEYTRKCSRCQMHGSKSLREELYPIPVSTKPFDRIALDVKHVQAS  3237

Query  71    ----KVILVAVHVASGYIEAEVIPAETGQETAYFLL-KLAGRWPVKTI-HTDNGSNFTGA  124
                 + I+  +   + Y+EA  +  +T  E A FL  ++  R    TI  +DNG  F
Sbjct  3236  RAGYRYIIAGIDYLTKYVEARPMRFQTSSEIALFLYDEIICRHGCPTIIVSDNGKPFLSD  3057

Query  125   TVRAACWWAGIKQEFGIPYNPQSQGVVESMNKELKKIIGQVRDQAE  170
              +R  C    I  +   PYNPQS G++E  N+ L +I+ Q R+ +E
Sbjct  3056  LIRQVCRNYSIIHKTTTPYNPQSNGLIERFNRTLGQIL-QRREDSE  2922
```

```
>Uncultured Tateyamaria sp. isolate 159-G.fasta NODE_34, whole genome shotgun sequence
Sequence ID: NZ_CANNDY010000034.1 Length: 252
Range 1: 168 to 245

Score:55.5 bits(132), Expect:2e-05,
Method:Compositional matrix adjust.,
Identities:25/26(96%), Positives:26/26(100%), Gaps:0/26(0%)

Query  177  QMAVFIHNFKRKGGIGGYSAGERIVD  202
            QMAVFIHNFKRKGGIGGYSAGERIV+
Sbjct  168  QMAVFIHNFKRKGGIGGYSAGERIVE  245
```

```
>Uncultured Tenacibaculum sp. isolate 838.fasta NODE_42, whole genome shotgun sequence
Sequence ID: NZ_CANMMT010000042.1 Length: 262
Range 1: 180 to 257

Score:55.5 bits(132), Expect:2e-05,
Method:Compositional matrix adjust.,
Identities:25/26(96%), Positives:26/26(100%), Gaps:0/26(0%)

Query  177  QMAVFIHNFKRKGGIGGYSAGERIVD  202
            QMAVFIHNFKRKGGIGGYSAGERIV+
Sbjct  180  QMAVFIHNFKRKGGIGGYSAGERIVE  257
```

```
>Escherichia coli strain w183 NODE_188_length_269_cov_26.989, whole genome shotgun sequence
```

```
Sequence ID: NZ_JAJNDO010000188.1 Length: 269
Range 1: 198 to 269

Score:54.3 bits(129), Expect:4e-05,
Method:Composition-based stats.,
Identities:24/24(100%), Positives:24/24(100%), Gaps:0/24(0%)

Query  177  QMAVFIHNFKRKGGIGGYSAGERI  200
            QMAVFIHNFKRKGGIGGYSAGERI
Sbjct  198  QMAVFIHNFKRKGGIGGYSAGERI  269



>Klebsiella pneumoniae isolate ena-yuan-GCF_010597935.1 NZ_VKXG01000632.1, whole genome shotgun sequence
Sequence ID: NZ_CAWPJY010000593.1 Length: 234
Range 1: 166 to 234

Score:52.8 bits(125), Expect:1e-04,
Method:Composition-based stats.,
Identities:23/23(100%), Positives:23/23(100%), Gaps:0/23(0%)

Query  177  QMAVFIHNFKRKGGIGGYSAGER  199
            QMAVFIHNFKRKGGIGGYSAGER
Sbjct  166  QMAVFIHNFKRKGGIGGYSAGER  234



>Klebsiella pneumoniae strain AS012315 AS012315_632, whole genome shotgun sequence
Sequence ID: NZ_VKXG01000632.1 Length: 234
Range 1: 166 to 234

Score:52.8 bits(125), Expect:1e-04,
Method:Composition-based stats.,
Identities:23/23(100%), Positives:23/23(100%), Gaps:0/23(0%)

Query  177  QMAVFIHNFKRKGGIGGYSAGER  199
            QMAVFIHNFKRKGGIGGYSAGER
Sbjct  166  QMAVFIHNFKRKGGIGGYSAGER  234



>Escherichia coli strain w189 NODE_129_length_265_cov_25.6333, whole genome shotgun sequence
Sequence ID: NZ_JAJNDQ010000122.1 Length: 265
Range 1: 195 to 263

Score:52.8 bits(125), Expect:2e-04,
Method:Composition-based stats.,
Identities:23/23(100%), Positives:23/23(100%), Gaps:0/23(0%)

Query  177  QMAVFIHNFKRKGGIGGYSAGER  199
            QMAVFIHNFKRKGGIGGYSAGER
Sbjct  195  QMAVFIHNFKRKGGIGGYSAGER  263



>Escherichia coli strain w190 NODE_198_length_265_cov_25.5778, whole genome shotgun sequence
Sequence ID: NZ_JAJNDR010000198.1 Length: 265
Range 1: 3 to 71

Score:52.8 bits(125), Expect:2e-04,
Method:Composition-based stats.,
Identities:23/23(100%), Positives:23/23(100%), Gaps:0/23(0%)

Query  177  QMAVFIHNFKRKGGIGGYSAGER  199
            QMAVFIHNFKRKGGIGGYSAGER
Sbjct  71   QMAVFIHNFKRKGGIGGYSAGER  3



>Escherichia coli strain w176 NODE_127_length_264_cov_65.95, whole genome shotgun sequence
Sequence ID: NZ_JAJNDN010000127.1 Length: 264
Range 1: 3 to 71

Score:52.8 bits(125), Expect:2e-04,
Method:Composition-based stats.,
Identities:23/23(100%), Positives:23/23(100%), Gaps:0/23(0%)

Query  177  QMAVFIHNFKRKGGIGGYSAGER  199
            QMAVFIHNFKRKGGIGGYSAGER
Sbjct  71   QMAVFIHNFKRKGGIGGYSAGER  3



>Escherichia coli strain w202 NODE_136_length_268_cov_26.95, whole genome shotgun sequence
Sequence ID: NZ_JAJNDS010000125.1 Length: 268
Range 1: 198 to 266

Score:52.8 bits(125), Expect:2e-04,
Method:Composition-based stats.,
Identities:23/23(100%), Positives:23/23(100%), Gaps:0/23(0%)

Query  177  QMAVFIHNFKRKGGIGGYSAGER  199
            QMAVFIHNFKRKGGIGGYSAGER
Sbjct  198  QMAVFIHNFKRKGGIGGYSAGER  266



>Escherichia coli strain w184 NODE_200_length_268_cov_29.5537, whole genome shotgun sequence
Sequence ID: NZ_JAJNDP010000200.1 Length: 268
Range 1: 3 to 71

Score:52.8 bits(125), Expect:2e-04,
Method:Composition-based stats.,
Identities:23/23(100%), Positives:23/23(100%), Gaps:0/23(0%)

Query  177  QMAVFIHNFKRKGGIGGYSAGER  199
            QMAVFIHNFKRKGGIGGYSAGER
Sbjct  71   QMAVFIHNFKRKGGIGGYSAGER  3



>Escherichia coli strain w208 NODE_126_length_276_cov_44.5, whole genome shotgun sequence
Sequence ID: NZ_JAJNDT010000124.1 Length: 276
Range 1: 3 to 71

Score:52.8 bits(125), Expect:2e-04,
```

```
Method:Composition-based stats.,
Identities:23/23(100%), Positives:23/23(100%), Gaps:0/23(0%)

Query  177  QMAVFIHNFKRKGGIGGYSAGER  199
            QMAVFIHNFKRKGGIGGYSAGER
Sbjct  71   QMAVFIHNFKRKGGIGGYSAGER  3




>Enterococcus faecalis strain 17838 NODE_651_length_288_cov_1.37267, whole genome shotgun sequence
Sequence ID: NZ_QNGS01000142.1 Length: 288
Range 1: 6 to 281

Score:52.0 bits(123), Expect:4e-04,
Method:Compositional matrix adjust.,
Identities:31/92(34%), Positives:50/92(54%), Gaps:2/92(2%)

Query  73   ILVAVHVASGYIEAEVIPAETGQETAYFLLK-LAGRWPV-KTIHTDNGSNFTGATVRAAC  130
            ILVAV   S ++EA   P    +    FL K +  R+ V + + +D G++F    + +A
Sbjct  6    ILVAVDYVSKWVEAIATPNNDTKTVLKFLQKHIFSRFGVPRVLISDGGTHFCNKQLYSAM  185

Query  131  WWAGIKQEFGIPYNPQSQGVVESMNKELKKII  162
               GI+ +   PY+PQ+ G  E  N+ELK+I+
Sbjct  186  VRYGIRHKVATPYHPQTNGQAEVSNRELKRIL  281




>Devosia insulae DS-56 contig234, whole genome shotgun sequence
Sequence ID: NZ_LAJE02000151.1 Length: 24799
Range 1: 24261 to 24653

Score:56.2 bits(134), Expect:5e-04,
Method:Compositional matrix adjust.,
Identities:40/131(31%), Positives:59/131(45%), Gaps:24/131(18%)

Query  52     GQVDCSP-GIWQLDCTHL---EG--KVILVAVHVASGYIEAEVIPAETGQETAYFLLKL-  104
              G+ + +P G   +D  HL   EG    + VA+  A+ ++  E+I        A  L++L
Sbjct  24653  GRFEPAPVGFIHIDLKHLTRLEGHPSYVFVAIDRATRFVHIEIIERRDASTIAACLVRLL  24474

Query  105    -AGRWPVKTIHTDNGSNFT---GAT-------------VRAACWWAGIKQEFGIPYNPQS  147
               A  +PV TI TDNG+ FT   GA                    C W GI       P++PQ+
Sbjct  24473  TAFPYPVHTILTDNGAEFTDRFGAARWRNPSRPTGKHAFDLICRWYGIDHRLTRPFHPQT  24294

Query  148    QGVVESMNKEL  158
               G+ E  N+ L
Sbjct  24293  NGMAERFNRRL  24261




>Eikenella corrodens strain 491_NMEN, whole genome shotgun sequence
Sequence ID: NZ_JVFA01000031.1 Length: 659
Range 1: 60 to 449

Score:52.8 bits(125), Expect:0.002,
Method:Compositional matrix adjust.,
Identities:37/130(28%), Positives:60/130(46%), Gaps:9/130(6%)

Query  73   ILVAVHVASGYIEAEVIPAETGQETAYFLLKLAGR---WPVKTIHTDNGSNFTGATVRA-  128
            + VA+   S  + A ++P +T    A FL +       + ++ +++DNGS + G    A
Sbjct  449  LFVAIDDFSRELYAAILPDKTADSAAKFLAEQVIEPCPYLIECVYSDNGSEYKGGANHAF  270

Query  129  --ACWWAGIKQEFGIPYNPQSQGVVESMNKELKKI--IGQVRDQAEHL-KTAVQMAVFIH  183
               CW  GI Q+F  P  PQ+ G  E + + L ++    Q  D  EH  K   +   F +
Sbjct  269  GVTCWENGINQKFTRPARPQTNGKAERVIRTLMEMWHEKQSFDSPEHRQKELCRFVNFYN  90

Query  184  NFKRKGGIGG  193
              K  G +GG
Sbjct  89   TVKPHGSLGG  60




>Escherichia coli strain RF108-1 NODE_139_length_640_cov_0.833037, whole genome shotgun sequence
Sequence ID: NZ_JAAQGD010000139.1 Length: 640
Range 1: 128 to 286

Score:52.4 bits(124), Expect:0.002,
Method:Compositional matrix adjust.,
Identities:22/53(42%), Positives:32/53(60%), Gaps:0/53(0%)

Query  110  VKTIHTDNGSNFTGATVRAACWWAGIKQEFGIPYNPQSQGVVESMNKELKKII  162
            VK + +DNG  +T    +  C   GIK+EF  PYNPQ  GV E MN+ +++ +
Sbjct  286  VKCLRSDNGGEYTSKAFQDFCDMKGIKREFTAPYNPQQNGVAERMNRTIQEKV  128




>Eikenella corrodens strain EI_05 NODE_86_length_856_cov_136.295, whole genome shotgun sequence
Sequence ID: NZ_VYYO01000085.1 Length: 856
Range 1: 408 to 797

Score:52.8 bits(125), Expect:0.002,
Method:Compositional matrix adjust.,
Identities:37/130(28%), Positives:61/130(46%), Gaps:9/130(6%)

Query  73   ILVAVHVASGYIEAEVIPAETGQETAYFLLKLAGR---WPVKTIHTDNGSNFTGATVRA-  128
            + VA+   S  + A ++P +T    A FL +       + ++ +++DNGS + G    A
Sbjct  408  LFVAIDDFSRELYAAILPDKTADSAAKFLAEQVIEPCPYLIECVYSDNGSEYKGGANHAF  587

Query  129  --ACWWAGIKQEFGIPYNPQSQGVVESMNKELKKI--IGQVRDQAEHL-KTAVQMAVFIH  183
               CW  GI Q+F  P  PQ+ G  E + + L ++    Q  D +EH  K   +   F +
Sbjct  588  GVTCWENGINQKFTRPARPQTNGKAERVIRTLMEMWHEKQSFDSSEHRQKELCRFVNFYN  767

Query  184  NFKRKGGIGG  193
              K  G +GG
Sbjct  768  TVKPHGSLGG  797




>Rhodovulum sulfidophilum strain SE1 Rhodovulum_DR_SE_assembly_contig_147, whole genome shotgun sequence
Sequence ID: NZ_JAAEAJ010000147.1 Length: 467
Range 1: 198 to 467

Score:51.2 bits(121), Expect:0.002,
Method:Compositional matrix adjust.,
Identities:27/90(30%), Positives:43/90(47%), Gaps:2/90(2%)

Query  75   VAVHVASGYIEAEVIPAETGQETAYFLLKLAGRWPVKT--IHTDNGSNFTGATVRAACWW  132
            VA+  A+ ++   + PA+T      FL  L    P+K   + TDNG  FT       C
Sbjct  467  VAIDRATRWVFVRIYPAQTAANARRFLRDLQRAAPMKITRVLTDNGKAFTDRLFDLLCTD  288
```

```
Query  133  AGIKQEFGIPYNPQSQGVVESMNKELKKII  162
             GI+     P  PQ+ G+VE  N  +++++
Sbjct  287  LGIEHRLAPPMRPQTNGMVERFNGRIEEVL  198
```

>Devosia sp. Root105 contig_7, whole genome shotgun sequence
Sequence ID: NZ_LMCR01000019.1 Length: 620452
Range 1: 183 to 575

Score:53.5 bits(127), Expect:0.003,
Method:Compositional matrix adjust.,
Identities:40/131(31%), Positives:59/131(45%), Gaps:24/131(18%)

```
Query  52   GQVDCSP-GIWQLDCTHL---EG--KVILVAVHVASGYIEAEVIPAETGQETAYFLLKL-  104
            G+ + +P G   +D  HL   EG    + VA+  A+ ++  E+I        A  L++L
Sbjct  575  GRFEPAPVGFIHIDLKHLTRLEGHPSYVFVAIDRATRFVHIEIIERRDASTIAACLVRLL  396

Query  105  -AGRWPVKTIHTDNGSNFT---GAT------------VRAACWWAGIKQEFGIPYNPQS  147
             A  +PV TI TDNG+ FT   GA                   C W GI      P++PQ+
Sbjct  395  TAFPYPVHTILTDNGAEFTDRFGAARWRNPSKPTGKHAFDLVCRWYGIDHRLTRPFHPQT  216

Query  148  QGVVESMNKEL  158
             G+ E  N+ L
Sbjct  215  NGMAERFNRRL  183
```

>Devosia sp. Root413D1 contig_4, whole genome shotgun sequence
Sequence ID: NZ_LMEA01000009.1 Length: 575790
Range 1: 574957 to 575349

Score:53.5 bits(127), Expect:0.003,
Method:Compositional matrix adjust.,
Identities:40/131(31%), Positives:59/131(45%), Gaps:24/131(18%)

```
Query  52      GQVDCSP-GIWQLDCTHL---EG--KVILVAVHVASGYIEAEVIPAETGQETAYFLLKL-  104
               G+ + +P G   +D  HL   EG    + VA+  A+ ++  E+I        A  L++L
Sbjct  575349  GRFEPAPVGFIHIDLKHLTRLEGHPSYVFVAIDRATRFVHIEIIERRDASTIAACLVRLL  575170

Query  105     -AGRWPVKTIHTDNGSNFT---GAT------------VRAACWWAGIKQEFGIPYNPQS  147
                A  +PV TI TDNG+ FT   GA                   C W GI      P++PQ+
Sbjct  575169  TAFPYPVHTILTDNGAEFTDRFGAARWRNPSKPTGKHAFDLVCRWYGIDHRLTRPFHPQT  574990

Query  148     QGVVESMNKEL  158
                G+ E  N+ L
Sbjct  574989  NGMAERFNRRL  574957
```

>Escherichia coli strain 90 NODE_134_length_1268_cov_0.816121, whole genome shotgun sequence
Sequence ID: NZ_JACAMA010000134.1 Length: 1268
Range 1: 537 to 767

Score:52.8 bits(125), Expect:0.004,
Method:Compositional matrix adjust.,
Identities:26/77(34%), Positives:40/77(51%), Gaps:8/77(10%)

```
Query  110  VKTIHTDNGSNFTGATVRAACWWAGIKQEFGIPYNPQSQGVVESMNKELKKIIGQVRDQ-  168
            VK + +DNG  +T    +  C   GIK+EF  PYNPQ  G+ E MN+ +++ +  +
Sbjct  767  VKCLRSDNGGEYTSKAFQDFCDMKGIKREFTAPYNPQQNGIAERMNRTIQEKVRSMLSNA  588

Query  169  -------AEHLKTAVQM  178
                   AE L TA+ +
Sbjct  587  SLPNGFWAEALATAIHL  537
```

>Albimonas sp. CAU 1670 contig2, whole genome shotgun sequence
Sequence ID: NZ_JARHUC010000002.1 Length: 847010
Range 1: 590130 to 590423

Score:53.1 bits(126), Expect:0.004,
Method:Compositional matrix adjust.,
Identities:37/102(36%), Positives:46/102(45%), Gaps:6/102(5%)

```
Query  53      QVDCSPGIWQLDCTHLEGKV-ILVAVHVASGYIEAEVIPAETGQETAYFLLKLAGRWP-V  110
               Q+D SP     D     G+  I +AV V S      V P           + K    W
Sbjct  590423  QIDASPA----DIMLTSGRASIYMAVDVYSRRAIVLVTPTPRASAVGLLVRKCIKTWGWP  590256

Query  111     KTIHTDNGSNFTGATVRAACWWAGIKQEFGIPYNPQSQGVVE  152
               + IHTDNGS+FT    R     AGI  E   PYNP+S+G+VE
Sbjct  590255  ERIHTDNGSDFTAHQTRQLFDSAGIHVELSPPYNPKSKGIVE  590130
```

>Agreia sp. PsM10 NODE_68_length_253_cov_0.818182, whole genome shotgun sequence
Sequence ID: NZ_JARUXC010000068.1 Length: 253
Range 1: 78 to 245

Score:48.5 bits(114), Expect:0.004,
Method:Composition-based stats.,
Identities:25/57(44%), Positives:33/57(57%), Gaps:1/57(1%)

```
Query  109  PVKTIHTDNGSNFTGATVRAACWWAGIKQEFGIPYNPQSQGVVESMNKELKKIIGQV  165
            P K I TDNGSNF    +R  C    I+     PY P+  GVVE+ NK +KKI+ ++
Sbjct  245  PSKII-TDNGSNFNNKMMRELCEEFKIEHHNSSPYRPKMNGVVEAANKNIKKIVQKM  78
```

>Enterococcus faecalis strain 15407 NODE_710_length_285_cov_0.436709, whole genome shotgun sequence
Sequence ID: NZ_QNGQ01000207.1 Length: 285
Range 1: 17 to 283

Score:48.9 bits(115), Expect:0.005,
Method:Compositional matrix adjust.,
Identities:29/89(33%), Positives:46/89(51%), Gaps:2/89(2%)

```
Query  76   AVHVASGYIEAEVIPAETGQETAYFLLK-LAGRWPV-KTIHTDNGSNFTGATVRAACWWA  133
            AV   S ++EA   P    +    FL K +  R+ V + + +D GS+F    + A
Sbjct  283  AVDYVSKWVEAIATPTNDNKVVMNFLRKNIFSRFGVPRALISDGGSHFCNRPLEALLLRY  104

Query  134  GIKQEFGIPYNPQSQGVVESMNKELKKII  162
            G+K +   PY+PQ+ G  E  N+ELK+I+
Sbjct  103  GVKHKVATPYHPQTSGQTEISNRELKRIL  17
```

>Eikenella corrodens strain EI_05 NODE_1_length_165032_cov_49.9348, whole genome shotgun sequence
Sequence ID: NZ_VYYO01000001.1 Length: 165032
Range 1: 50396 to 50785

Score:52.4 bits(124), Expect:0.008,
Method:Compositional matrix adjust.,
Identities:37/130(28%), Positives:60/130(46%), Gaps:9/130(6%)

```
Query  73     ILVAVHVASGYIEAEVIPAETGQETAYFLLKLAGR---WPVKTIHTDNGSNFTGATVRA-  128
              + VA+   S  + A ++P +T    A FL +       + ++ +++DNGS + G    A
Sbjct  50396  LFVAIDDFSRELYAAILPDKTADSAAKFLAEQVIEPCPYLIECVYSDNGSEYKGGANHAF  50575

Query  129    --ACWWAGIKQEFGIPYNPQSQGVVESMNKELKKI--IGQVRDQAEHL-KTAVQMAVFIH  183
                CW  GI Q+F  P  PQ+ G  E + + L ++    Q  D  EH  K   +   F +
Sbjct  50576  GVTCWENGINQKFTRPARPQTNGKAERVIRTLMEMWHEKQSFDSPEHRQKELCRFVNFYN  50755

Query  184    NFKRKGGIGG  193
                K  G +GG
Sbjct  50756  TVKPHGSLGG  50785
```

>Eikenella corrodens strain EI_07 NODE_3_length_123520_cov_80.810171, whole genome shotgun sequence
Sequence ID: NZ_VYYM01000003.1 Length: 123520
Range 1: 8861 to 9250

Score:52.4 bits(124), Expect:0.008,
Method:Compositional matrix adjust.,
Identities:35/130(27%), Positives:61/130(46%), Gaps:9/130(6%)

```
Query  73    ILVAVHVASGYIEAEVIPAETGQETAYFLLKLAGR---WPVKTIHTDNGSNFTGATVRA-  128
             + VA+   S  + A ++P +T      FL +       + ++ +++DNGS + G    A
Sbjct  8861  LFVAIDDFSRELYAAILPDKTADSATKFLAEQVIEPCPYLIECVYSDNGSEYKGGANHAF  9040

Query  129   --ACWWAGIKQEFGIPYNPQSQGVVESMNKELKKI--IGQVRDQAEHLKTAV-QMAVFIH  183
               CW  GI Q+F  P  PQ+ G  E + + L ++    Q+ D  EH +  + +   F +
Sbjct  9041  GVTCWENGINQKFTRPARPQTNGKTERVIRTLMEMWHEKQLFDSPEHRRKELCRFVNFYN  9220

Query  184   NFKRKGGIGG  193
               K  G +GG
Sbjct  9221  TVKPHGSLGG  9250
```

>Eikenella corrodens ATCC 23834 Scfld2, whole genome shotgun sequence
Sequence ID: NZ_EQ973318.1 Length: 131618
Range 1: 59282 to 59671

Score:52.0 bits(123), Expect:0.009,
Method:Compositional matrix adjust.,
Identities:37/130(28%), Positives:60/130(46%), Gaps:9/130(6%)

```
Query  73     ILVAVHVASGYIEAEVIPAETGQETAYFLLKLAGR---WPVKTIHTDNGSNFTGATVRA-  128
              + VA+   S  + A ++P +T    A FL +       + ++ +++DNGS + G    A
Sbjct  59282  LFVAIDDFSRELYAAILPDKTADSAAKFLAEQVIEPCPYLIECVYSDNGSEYKGGANHAF  59461

Query  129    --ACWWAGIKQEFGIPYNPQSQGVVESMNKELKKI--IGQVRDQAEHL-KTAVQMAVFIH  183
                CW  GI Q+F  P  PQ+ G  E + + L ++    Q  D  EH  K   +   F +
Sbjct  59462  GVTCWENGINQKFTRPARPQTNGKAERVIRTLMEMWHEKQSFDSPEHRQKELCRFVNFYN  59641

Query  184    NFKRKGGIGG  193
                K  G +GG
Sbjct  59642  TVKPHGSLGG  59671
```

>Eikenella corrodens strain FDAARGOS_1474 chromosome, complete genome
Sequence ID: NZ_CP082861.1 Length: 2192571
Range 1: 344094 to 344483

Score:52.0 bits(123), Expect:0.009,
Method:Compositional matrix adjust.,
Identities:37/130(28%), Positives:60/130(46%), Gaps:9/130(6%)

```
Query  73      ILVAVHVASGYIEAEVIPAETGQETAYFLLKLAGR---WPVKTIHTDNGSNFTGATVRA-  128
               + VA+   S  + A ++P +T    A FL +       + ++ +++DNGS + G    A
Sbjct  344483  LFVAIDDFSRELYAAILPDKTADSAAKFLAEQVIEPCPYLIECVYSDNGSEYKGGANHAF  344304

Query  129     --ACWWAGIKQEFGIPYNPQSQGVVESMNKELKKI--IGQVRDQAEHL-KTAVQMAVFIH  183
                 CW  GI Q+F  P  PQ+ G  E + + L ++    Q  D  EH  K   +   F +
Sbjct  344303  GVTCWENGINQKFTRPARPQTNGKAERVIRTLMEMWHEKQSFDSPEHRQKELCRFVNFYN  344124

Query  184     NFKRKGGIGG  193
                 K  G +GG
Sbjct  344123  TVKPHGSLGG  344094
```

>Eikenella corrodens strain NCTC10596 chromosome 1, complete sequence
Sequence ID: NZ_LT906482.1 Length: 2192577
Range 1: 716530 to 716919

Score:52.0 bits(123), Expect:0.009,
Method:Compositional matrix adjust.,
Identities:37/130(28%), Positives:60/130(46%), Gaps:9/130(6%)

```
Query  73      ILVAVHVASGYIEAEVIPAETGQETAYFLLKLAGR---WPVKTIHTDNGSNFTGATVRA-  128
               + VA+   S  + A ++P +T    A FL +       + ++ +++DNGS + G    A
Sbjct  716530  LFVAIDDFSRELYAAILPDKTADSAAKFLAEQVIEPCPYLIECVYSDNGSEYKGGANHAF  716709

Query  129     --ACWWAGIKQEFGIPYNPQSQGVVESMNKELKKI--IGQVRDQAEHL-KTAVQMAVFIH  183
                 CW  GI Q+F  P  PQ+ G  E + + L ++    Q  D  EH  K   +   F +
Sbjct  716710  GVTCWENGINQKFTRPARPQTNGKAERVIRTLMEMWHEKQSFDSPEHRQKELCRFVNFYN  716889

Query  184     NFKRKGGIGG  193
                 K  G +GG
Sbjct  716890  TVKPHGSLGG  716919
```

>Eikenella corrodens strain KCOM 1378 B230_1_2, whole genome shotgun sequence
Sequence ID: NZ_NHRS01000002.1 Length: 572658
Range 1: 511602 to 511991

Score:52.0 bits(123), Expect:0.009,
Method:Compositional matrix adjust.,
Identities:37/130(28%), Positives:60/130(46%), Gaps:9/130(6%)

```
Query  73      ILVAVHVASGYIEAEVIPAETGQETAYFLLKLAGR---WPVKTIHTDNGSNFTGATVRA-  128
               + VA+   S  + A ++P +T    A FL +       + ++ +++DNGS + G    A
Sbjct  511602  LFVAIDDFSRELYAAILPDKTADSAAKFLAEQVIEPCPYLIECVYSDNGSEYKGGANHAF  511781

Query  129     --ACWWAGIKQEFGIPYNPQSQGVVESMNKELKKI--IGQVRDQAEHL-KTAVQMAVFIH  183
                 CW  GI Q+F  P  PQ+ G  E + + L ++    Q  D  EH  K   +   F +
Sbjct  511782  GVTCWENGINQKFTRPARPQTNGKAERVIRTLMEMWHEKQSFDSPEHRQKELCRFVNFYN  511961

Query  184     NFKRKGGIGG  193
                 K  G +GG
Sbjct  511962  TVKPHGSLGG  511991
```

>Eikenella corrodens strain NML01-0040 Eikcor_contig000008, whole genome shotgun sequence
Sequence ID: NZ_LXSE01000008.1 Length: 65735
Range 1: 45607 to 45996

Score:52.0 bits(123), Expect:0.009,
Method:Compositional matrix adjust.,
Identities:37/130(28%), Positives:60/130(46%), Gaps:9/130(6%)

```
Query  73     ILVAVHVASGYIEAEVIPAETGQETAYFLLKLAGR---WPVKTIHTDNGSNFTGATVRA-  128
              + VA+   S  + A ++P +T    A FL +       + ++ +++DNGS + G    A
Sbjct  45607  LFVAIDDFSRELYAAILPDKTADSAAKFLAEQVIEPCPYLIECVYSDNGSEYKGGANHAF  45786

Query  129    --ACWWAGIKQEFGIPYNPQSQGVVESMNKELKKI--IGQVRDQAEHL-KTAVQMAVFIH  183
                CW  GI Q+F  P  PQ+ G  E + + L ++    Q  D  EH  K   +   F +
Sbjct  45787  GVTCWENGINQKFTRPARPQTNGKAERVIRTLMEMWHEKQSFDSPEHRQKELCRFVNFYN  45966

Query  184    NFKRKGGIGG  193
                K  G +GG
Sbjct  45967  TVKPHGSLGG  45996
```

>Eikenella corrodens strain EI_08 NODE_51_length_16570_cov_73.383, whole genome shotgun sequence
Sequence ID: NZ_VYYL01000051.1 Length: 16570
Range 1: 11273 to 11662

Score:52.0 bits(123), Expect:0.009,
Method:Compositional matrix adjust.,
Identities:37/130(28%), Positives:60/130(46%), Gaps:9/130(6%)

```
Query  73     ILVAVHVASGYIEAEVIPAETGQETAYFLLKLAGR---WPVKTIHTDNGSNFTGATVRA-  128
              + VA+   S  + A ++P +T    A FL +       + ++ +++DNGS + G    A
Sbjct  11662  LFVAIDDFSRELYAAILPDKTADSAAKFLAEQVIEPCPYLIECVYSDNGSEYKGGANHAF  11483

Query  129    --ACWWAGIKQEFGIPYNPQSQGVVESMNKELKKI--IGQVRDQAEHL-KTAVQMAVFIH  183
                CW  GI Q+F  P  PQ+ G  E + + L ++    Q  D  EH  K   +   F +
Sbjct  11482  GVTCWENGINQKFTRPARPQTNGKAERVIRTLMEMWHEKQSFDSPEHRQKELCRFVNFYN  11303

Query  184    NFKRKGGIGG  193
                K  G +GG
Sbjct  11302  TVKPHGSLGG  11273
```

>Escherichia coli strain E-4H4 NODE_99_length_2231_cov_0.479087, whole genome shotgun sequence
Sequence ID: NZ_JALSFQ010000093.1 Length: 590
Range 1: 25 to 246

Score:50.1 bits(118), Expect:0.010,
Method:Compositional matrix adjust.,
Identities:28/74(38%), Positives:35/74(47%), Gaps:4/74(5%)

```
Query  110  VKTIHTDNGSNFTGATVRAACWWAGIKQEFGIPYNPQSQGVVESMNKEL----KKIIGQV  165
            VK I +DNGS F    V       GIK EF  PY PQ  GVVE  N+ L    + ++G+
Sbjct  246  VKKIRSDNGSEFKNLQVEEYLEEEGIKHEFSAPYTPQQNGVVERKNRTLIDMARTMLGEF  67

Query  166  RDQAEHLKTAVQMA  179
            +        AV  A
Sbjct  66   KTPERFWSEAVNTA  25
```

>Klebsiella pneumoniae strain CL1616 NODE_145_length_1091_cov_1.30913, whole genome shotgun sequence
Sequence ID: NZ_RPET01000142.1 Length: 1091
Range 1: 670 to 906

Score:51.2 bits(121), Expect:0.010,
Method:Compositional matrix adjust.,
Identities:28/79(35%), Positives:40/79(50%), Gaps:4/79(5%)

```
Query  110  VKTIHTDNGSNFTGATVRAACWWAGIKQEFGIPYNPQSQGVVESMNKELKKIIGQVRDQA  169
            VK + +DNG  +T   +   C   GI+Q+F  PY PQ  GV E MN+ L +    + + A
Sbjct  670  VKVLRSDNGGEYTSGAMAKLCADRGIEQKFTPPYTPQLNGVAERMNRTLVECARCMLEHA  849

Query  170  ----EHLKTAVQMAVFIHN  184
                E+   AV  A F+ N
Sbjct  850  GLSKEYWGEAVMTATFLRN  906
```

>Klebsiella pneumoniae strain CL1549 NODE_149_length_1225_cov_0.87796, whole genome shotgun sequence
Sequence ID: NZ_RPES01000124.1 Length: 1225
Range 1: 171 to 407

Score:51.2 bits(121), Expect:0.010,
Method:Compositional matrix adjust.,
Identities:28/79(35%), Positives:40/79(50%), Gaps:4/79(5%)

```
Query  110  VKTIHTDNGSNFTGATVRAACWWAGIKQEFGIPYNPQSQGVVESMNKELKKIIGQVRDQA  169
            VK + +DNG  +T   +   C   GI+Q+F  PY PQ  GV E MN+ L +    + + A
Sbjct  407  VKVLRSDNGGEYTSGAMAKLCADRGIEQKFTPPYTPQLNGVAERMNRTLVECARCMLEHA  228

Query  170  ----EHLKTAVQMAVFIHN  184
                E+   AV  A F+ N
Sbjct  227  GLSKEYWGEAVMTATFLRN  171
```

>Klebsiella pneumoniae strain CL2074 NODE_44_length_2903_cov_2.16679, whole genome shotgun sequence
Sequence ID: NZ_RPFA01000043.1 Length: 2903
Range 1: 2109 to 2345

Score:51.6 bits(122), Expect:0.010,

```
Method:Compositional matrix adjust.,
Identities:28/79(35%), Positives:40/79(50%), Gaps:4/79(5%)

Query  110   VKTIHTDNGSNFTGATVRAACWWAGIKQEFGIPYNPQSQGVVESMNKELKKIIGQVRDQA  169
             VK + +DNG  +T   +   C   GI+Q+F  PY PQ  GV E MN+ L +    + + A
Sbjct  2109  VKVLRSDNGGEYTSGAMAKLCADRGIEQKFTPPYTPQLNGVAERMNRTLVECARCMLEHA  2288

Query  170   ----EHLKTAVQMAVFIHN  184
                 E+   AV  A F+ N
Sbjct  2289  GLSKEYWGEAVMTATFLRN  2345



>Staphylococcus aureus strain 17PQ_1_195 17PQ-1-195NODE_43_length_1998_cov_0.982301, whole genome shotgun sequence
Sequence ID: NZ_JAOBRT010000041.1 Length: 1338
Range 1: 511 to 777

Score:50.8 bits(120), Expect:0.016,
Method:Compositional matrix adjust.,
Identities:31/89(35%), Positives:41/89(46%), Gaps:10/89(11%)

Query  110  VKTIHTDNGSNFTGATVRAACWWAGIKQEFGIPYNPQSQGVVESMNKEL----KKIIGQV  165
            VK I +DNGS F    V       GIK EF  PY PQ  GVVE  N+ L    + ++G+
Sbjct  511  VKKIRSDNGSEFKNLQVEEFLEEEGIKHEFSAPYTPQQNGVVERKNRTLIDMARTMLGEF  690

Query  166  RDQAEHLKTAVQMA------VFIHNFKRK  188
            +        AV  A      V++H   +K
Sbjct  691  KTPERFWSEAVNTACHAINRVYLHRLLKK  777



>Eikenella corrodens strain KCOM 1378 B230_1_3, whole genome shotgun sequence
Sequence ID: NZ_NHRS01000003.1 Length: 129678
Range 1: 85685 to 86074

Score:51.2 bits(121), Expect:0.017,
Method:Compositional matrix adjust.,
Identities:37/130(28%), Positives:60/130(46%), Gaps:9/130(6%)

Query  73     ILVAVHVASGYIEAEVIPAETGQETAYFLLKLAGR---WPVKTIHTDNGSNFTGATVRA-  128
              + VA+   S  + A ++P +T    A FL +       + ++ +++DNGS + G    A
Sbjct  85685  LFVAIDDFSRELYAAILPDKTADSAAKFLAEQVIEPCPYLIECVYSDNGSEYKGGANHAF  85864

Query  129    --ACWWAGIKQEFGIPYNPQSQGVVESMNKELKKI--IGQVRDQAEHL-KTAVQMAVFIH  183
                 CW  GI Q+F  P  PQ+ G  E + + L ++    Q  D  EH  K   +   F +
Sbjct  85865  GVTCWENGINQKFTRPACPQTNGKAERVIRTLMEMWHEKQSFDSPEHRQKELCRFVNFYN  86044

Query  184    NFKRKGGIGG  193
                K  G +GG
Sbjct  86045  TVKPHGSLGG  86074



>Acinetobacter bereziniae strain A005.1 NODE_166_length_855_cov_1.037275, whole genome shotgun sequence
Sequence ID: NZ_JACDXR010000165.1 Length: 855
Range 1: 431 to 697

Score:50.4 bits(119), Expect:0.017,
Method:Compositional matrix adjust.,
Identities:30/89(34%), Positives:41/89(46%), Gaps:10/89(11%)

Query  110  VKTIHTDNGSNFTGATVRAACWWAGIKQEFGIPYNPQSQGVVESMNKEL----KKIIGQV  165
            VK I +DNGS F    V       GIK EF  PY PQ  GVVE  N+ L    + ++G+
Sbjct  431  VKKIRSDNGSEFKNLQVEEYLEEEGIKHEFSAPYTPQQNGVVERKNRTLIDMARTMLGEF  610

Query  166  RDQAEHLKTAVQMA------VFIHNFKRK  188
            +        AV  A      +++H   +K
Sbjct  611  KTLERFWSEAVNTACHAINRLYLHRLLKK  697



>Haemophilus sp. SZY H51 NODE_22_length_1889_cov_0.990066, whole genome shotgun sequence
Sequence ID: NZ_JASGBG010000022.1 Length: 1889
Range 1: 1146 to 1478

Score:50.8 bits(120), Expect:0.019,
Method:Compositional matrix adjust.,
Identities:31/111(28%), Positives:52/111(46%), Gaps:13/111(11%)

Query  91    AETGQETAYFLLKLAGRWPV---KTIHTDNGSNFTGATVRAACWWAGIKQEFGIPYNPQS  147
              ET       +L+L   +P+   + I +DNGS F  +     C   G++ +F  PY P
Sbjct  1146  TETFSFARNLILRLQNEFPLNSMRAIRSDNGSEFKNSDFEKFCNSLGLEHQFSSPYTPPQ  1325

Query  148   QGVVESMNKELKKIIGQVRDQ--------AEHLKTAVQMA--VFIHNFKRK  188
              GVVE  N+ L ++   + D+        AE + TA  ++  +F+  F +K
Sbjct  1326  NGVVERKNRTLVEMARTMLDEHRTPRRYWAEAMNTACYISNRIFLRAFLKK  1478



>Rickettsia endosymbiont of Ceutorhynchus assimilis isolate Rickettsia_endosymbiont_Ceutorhynchus_assimilis chromosome 1
Sequence ID: NZ_OU906081.1 Length: 4063491
Range 1: 170223 to 170552

Score:51.2 bits(121), Expect:0.020,
Method:Compositional matrix adjust.,
Identities:35/110(32%), Positives:55/110(50%), Gaps:7/110(6%)

Query  70      GKVILVAVHVASGYIEAEVIPAETGQETAYFLL-KLAGRWPVKT-IHTDNGSNFTGATVR  127
                K ILV ++  S + E   IP +     A  L+ ++  R  V   +H+D G NF  A  +
Sbjct  170223  NKYILVIMNYFSKWPEVFAIPNQESITVAQVLVDEVFSRHGVPLELHSDQGRNFESAIFK  170402

Query  128     AACWWAGIKQEFGIPYNPQSQGVVESMNKELKKIIGQV-----RDQAEHL  172
                 C   GI++    P +PQS G+VE  NK +++ + +V     RD  +HL
Sbjct  170403  KVCEIMGIRKTRTTPLHPQSDGMVERFNKTIEEHLTKVVAKNQRDWDKHL  170552



>Escherichia coli strain E-4H5 NODE_187_length_488_cov_0.063712, whole genome shotgun sequence
Sequence ID: NZ_JALYCP010000174.1 Length: 488
Range 1: 79 to 321

Score:48.1 bits(113), Expect:0.027,
Method:Compositional matrix adjust.,
Identities:30/84(36%), Positives:40/84(47%), Gaps:6/84(7%)

Query  95   QETAYFLLKLAG---RWPVKTIHTDNGSNFTGATVRAACWWAGIKQEFGIPYNPQSQGVV  151
            QET    L+ A    R  +K I +DNG+ F  + +       GIK EF  PY PQ  GVV
Sbjct  321  QETLKGFLRRAQNEFRLRIKKIRSDNGT*FKNSQIEGFLEDEGIKHEFSSPYTPQQNGVV  142
```

```
Query  152  ESMNKELKKIIGQVRDQAEHLKTA  175
            E  N+ L   +   R   +  KT+
Sbjct  141  ERKNRTL---LDTARTMLDEYKTS  79
```

>Mycobacterium innocens strain MK13 isolate MK13, whole genome shotgun sequence
Sequence ID: NZ_UPHQ01000351.1 Length: 1438
Range 1: 736 to 1107

Score:50.1 bits(118), Expect:0.027,
Method:Compositional matrix adjust.,
Identities:44/128(34%), Positives:59/128(46%), Gaps:15/128(11%)

```
Query  61    WQLDCTHL---EGKVILVAV-HVASGYIEAEVIPAETGQETAYFLLKLA-----GRWPVK  111
             W  D T +   EGK+ L +V  +AS  I     I       E AY  L +A     GR  +
Sbjct  736   WYGDGTEVDTDEGKLQLDSVLDMASRRIIGFAIGEHHDAELAYGALSMAVAVRGGRQQIA  915

Query  112   --TIHTDNGSNFTGATVRAACWWAGIKQEFGIPYNPQSQGVVESMNKELKKIIGQVRDQA  169
               T+HTD GS +T    RAAC   GI+Q  G P +     V+ES +  L+  +  +
Sbjct  916   GVTLHTDQGSEYTAGAFRAACRRCGIRQSMGRPGSALDNAVIESWHSTLEFELRML----  1083

Query  170   EHLKTAVQ  177
             EH  T VQ
Sbjct  1084  EHFDTKVQ  1107
```

>Eikenella corrodens strain NML01-0040 Eikcor_contig000002, whole genome shotgun sequence
Sequence ID: NZ_LXSE01000002.1 Length: 226986
Range 1: 61932 to 62321

Score:50.8 bits(120), Expect:0.028,
Method:Compositional matrix adjust.,
Identities:36/130(28%), Positives:60/130(46%), Gaps:9/130(6%)

```
Query  73     ILVAVHVASGYIEAEVIPAETGQETAYFLLKLAGR---WPVKTIHTDNGSNFTGATVRA-  128
              + VA+   S  + A ++P +T    A FL +       + ++ +++DNGS + G    A
Sbjct  61932  LFVAIDDFSRELYAAILPDKTADSAAKFLAEQVIEPCPYLIECVYSDNGSEYKGGANHAF  62111

Query  129    --ACWWAGIKQEFGIPYNPQSQGVVESMNKELKKI-IGQVRDQAEHL-KTAVQMAVFIH  183
                 CW  GI Q+F  P  PQ+ G  E + + + ++    Q  D  EH  K   +   F +
Sbjct  62112  GVTCWENGINQKFTRPARPQTNGKAERVIRTIMEMWHEKQSFDSPEHRQKELCRFVNFYN  62291

Query  184    NFKRKGGIGG  193
                K  G +GG
Sbjct  62292  TVKPHGSLGG  62321
```

>Eikenella corrodens strain NML140207 Eikcor_contig000050, whole genome shotgun sequence
Sequence ID: NZ_LXSJ01000050.1 Length: 5581
Range 1: 408 to 812

Score:50.4 bits(119), Expect:0.031,
Method:Compositional matrix adjust.,
Identities:37/135(27%), Positives:62/135(45%), Gaps:9/135(6%)

```
Query  73   ILVAVHVASGYIEAEVIPAETGQETAYFLLKLAGR---WPVKTIHTDNGSNFTGATVRA-  128
            + VA+   S  + A ++P +T    A FL +       + ++ +++DNGS + G    A
Sbjct  408  LFVAIDDFSRELYAAILPDKTADSAAKFLAEQVIEPCPYLIECVYSDNGSEYKGGANHAF  587

Query  129  --ACWWAGIKQEFGIPYNPQSQGVVESMNKELKKI--IGQVRDQAEHL-KTAVQMAVFIH  183
               CW  GI Q+F  P  PQ+ G  E + + L ++    Q  D +EH  K   +   F +
Sbjct  588  GVTCWENGINQKFTRPARPQTNGKAERVIRTLMEMWHEKQSFDSSEHRQKELCRFVNFYN  767

Query  184  NFKRKGGIGGYSAGE  198
              K  G + G +  E
Sbjct  768  TVKPHGSLDGNTPFE  812
```

>Eikenella corrodens strain NML120819 Eikcor_contig000019, whole genome shotgun sequence
Sequence ID: NZ_LXSH01000019.1 Length: 95946
Range 1: 93863 to 94252

Score:50.4 bits(119), Expect:0.032,
Method:Compositional matrix adjust.,
Identities:36/130(28%), Positives:59/130(45%), Gaps:9/130(6%)

```
Query  73     ILVAVHVASGYIEAEVIPAETGQETAYFLLKLAGR---WPVKTIHTDNGSNFTGATVRA-  128
              + VA+   S  + A ++P +T      FL +       + ++ +++DNGS + G    A
Sbjct  94252  LFVAIDDFSRELYAAILPDKTADSATKFLAEQVIEPCPYLIECVYSDNGSEYKGGANHAF  94073

Query  129    --ACWWAGIKQEFGIPYNPQSQGVVESMNKELKKI--IGQVRDQAEHL-KTAVQMAVFIH  183
                 CW  GI Q+F  P  PQ+ G  E + + L ++    Q  D  EH  K   +   F +
Sbjct  94072  GVTCWENGINQKFTRPARPQTNGKAERVIRTLMEMWHEKQSFDNPEHRQKELCRFVNFYN  93893

Query  184    NFKRKGGIGG  193
                K  G +GG
Sbjct  93892  TVKPHGSLGG  93863
```

>Acinetobacter bereziniae strain A005.2 NODE_137_length_954_cov_0.819124, whole genome shotgun sequence
Sequence ID: NZ_JABWPZ010000137.1 Length: 954
Range 1: 178 to 366

Score:49.3 bits(116), Expect:0.035,
Method:Compositional matrix adjust.,
Identities:25/66(38%), Positives:33/66(50%), Gaps:3/66(4%)

```
Query  110  VKTIHTDNGSNFTGATVRAACWWAGIKQEFGIPYNPQSQGVVESMNKELKKIIGQVRDQA  169
            +K I +DNG+ F  + +       GIK EF  PY PQ  GVVE  N+ L   +   R
Sbjct  178  IKRIRSDNGTEFKNSQIEGFLEEEGIKHEFSSPYTPQQNGVVERKNRTL---MDMARTML  348

Query  170  EHLKTA  175
            E  KT+
Sbjct  349  EEYKTS  366
```

>Klebsiella pneumoniae strain KP231 NODE_2205_length_233_cov_2.807692, whole genome shotgun sequence
Sequence ID: NZ_JAUUHQ010001574.1 Length: 233
Range 1: 22 to 168

Score:45.8 bits(107), Expect:0.036,

```
Method:Composition-based stats.,
Identities:23/49(47%), Positives:26/49(53%), Gaps:0/49(0%)

Query  110  VKTIHTDNGSNFTGATVRAACWWAGIKQEFGIPYNPQSQGVVESMNKEL  158
            VK I +DNGS F    V       GIK +F  PY PQ  GVVE  N+ L
Sbjct  22   VKKIRSDNGSEFKNLQVEEYLEEEGIKHKFSAPYTPQQNGVVERKNRTL  168
```

>Pseudomonas reactans strain IPO3751 IPO_3751_S34_L003_contig_216, whole genome shotgun sequence
Sequence ID: NZ_JACAQP010000157.1 Length: 594
Range 1: 397 to 567

```
Score:48.5 bits(114), Expect:0.038,
Method:Compositional matrix adjust.,
Identities:22/57(39%), Positives:31/57(54%), Gaps:0/57(0%)

Query  102  LKLAGRWPVKTIHTDNGSNFTGATVRAACWWAGIKQEFGIPYNPQSQGVVESMNKEL  158
            ++L  + PV+ I +DNG  F   T+ +      GI+  F  PY PQ  GVVE  N+ L
Sbjct  567  IELKLKRPVRRIRSDNGLEFKNITLDSFLKNKGIEHNFSAPYTPQQNGVVERRNRTL  397
```

>Corynebacterium sp. c3Ub_189 c3Ub189_contig_53, whole genome shotgun sequence
Sequence ID: NZ_JASBOF010000053.1 Length: 1136
Range 1: 279 to 659

```
Score:49.7 bits(117), Expect:0.039,
Method:Compositional matrix adjust.,
Identities:40/128(31%), Positives:57/128(44%), Gaps:18/128(14%)

Query  53   QVDCSPGIWQLDCTHL-----EGKVILVAVHVASGYIEA-EVIPAETGQETAYFLLKLAG  106
            Q D     WQLD +H      +   IL  +   S Y+ A + +  ETG       +
Sbjct  279  QADQPNETWQLDYSHWYLNNNQRVAILTIIDDHSRYVIACKAVEHETGTNVIDTFISAGN  458

Query  107  R--WPVKTIHTDNGSNFTGATVRAA---------CWWAGIKQEFGIPYNPQSQGVVESMN  155
            +  +P  T+ TDNGS FT    R A             GI Q+ G PY+PQ+QG VE  +
Sbjct  459  KHGFPKSTL-TDNGSVFTTKLYRKADTRNGFEQLLLQMGIHQKNGTPYHPQTQGKVERFH  635

Query  156  KELKKIIG  163
            + L+K +
Sbjct  636  QTLQKALN  659
```

>Aeromonas jandaei strain 82 NODE_15_length_4445_cov_0.106043, whole genome shotgun sequence
Sequence ID: NZ_RPDF01000015.1 Length: 4445
Range 1: 78 to 383

```
Score:50.1 bits(118), Expect:0.039,
Method:Compositional matrix adjust.,
Identities:29/102(28%), Positives:50/102(49%), Gaps:13/102(12%)

Query  100  FLLKLAGRWP---VKTIHTDNGSNFTGATVRAACWWAGIKQEFGIPYNPQSQGVVESMNK  156
             +L+L   +P   ++ I +DNGS F  +     C   G++ +F  PY P   GVVE  N+
Sbjct  78   LILRLQNEFPKNSMRAIRSDNGSEFKNSDFEKFCNSLGLEHQFSSPYTPPQNGVVERKNR  257

Query  157  ELKKIIGQVRDQ--------AEHLKTAVQMA--VFIHNFKRK  188
             L ++   + D+        AE + TA  ++  +F+  F +K
Sbjct  258  TLVEMARTMLDEHRTPRRYWAEAMSTACYISNRIFLRAFLKK  383
```

>Aeromonas jandaei isolate ena-yuan-GCF_009749285.1 NZ_RPDF01000015.1, whole genome shotgun sequence
Sequence ID: NZ_CAWPET010000058.1 Length: 4445
Range 1: 78 to 383

```
Score:50.1 bits(118), Expect:0.039,
Method:Compositional matrix adjust.,
Identities:29/102(28%), Positives:50/102(49%), Gaps:13/102(12%)

Query  100  FLLKLAGRWP---VKTIHTDNGSNFTGATVRAACWWAGIKQEFGIPYNPQSQGVVESMNK  156
             +L+L   +P   ++ I +DNGS F  +     C   G++ +F  PY P   GVVE  N+
Sbjct  78   LILRLQNEFPKNSMRAIRSDNGSEFKNSDFEKFCNSLGLEHQFSSPYTPPQNGVVERKNR  257

Query  157  ELKKIIGQVRDQ--------AEHLKTAVQMA--VFIHNFKRK  188
             L ++   + D+        AE + TA  ++  +F+  F +K
Sbjct  258  TLVEMARTMLDEHRTPRRYWAEAMSTACYISNRIFLRAFLKK  383
```

>Corynebacterium tuberculostearicum strain CTNIH9 NODE_36_length_1392_cov_16.726987_pilon, whole genome shotgun sequence
Sequence ID: NZ_JAVBHZ010000036.1 Length: 1392
Range 1: 405 to 782

```
Score:49.7 bits(117), Expect:0.040,
Method:Compositional matrix adjust.,
Identities:40/127(31%), Positives:56/127(44%), Gaps:18/127(14%)

Query  53   QVDCSPGIWQLDCTHL-----EGKVILVAVHVASGY-IEAEVIPAETGQETAYFLLKLAG  106
            Q D     WQLD +H      +   IL  +   S Y I  + +  ETG       +
Sbjct  782  QADQPNETWQLDYSHWYLNNNQRVAILTIIDDHSRYVISCKAVEHETGNNVIDTFISAGN  603

Query  107  R--WPVKTIHTDNGSNFTGATVRAA---------CWWAGIKQEFGIPYNPQSQGVVESMN  155
            +  +P  T+ TDNGS FT    R A             GI Q+ G PY+PQ+QG VE  +
Sbjct  602  KHGFPKSTL-TDNGSVFTTKLYRKADTRNGFEQLLLQMGIHQKNGTPYHPQTQGKVERFH  426

Query  156  KELKKII  162
            + L+K +
Sbjct  425  QTLQKAL  405
```

>Corynebacterium sp. ACRPF NODE_37_length_1392_cov_16.843430_pilon, whole genome shotgun sequence
Sequence ID: NZ_JAKRNB010000037.1 Length: 1392
Range 1: 405 to 782

```
Score:49.7 bits(117), Expect:0.040,
Method:Compositional matrix adjust.,
Identities:40/127(31%), Positives:56/127(44%), Gaps:18/127(14%)

Query  53   QVDCSPGIWQLDCTHL-----EGKVILVAVHVASGY-IEAEVIPAETGQETAYFLLKLAG  106
            Q D     WQLD +H      +   IL  +   S Y I  + +  ETG       +
Sbjct  782  QADQPNETWQLDYSHWYLNNNQRVAILTIIDDHSRYVISCKAVEHETGNNVIDTFISAGN  603

Query  107  R--WPVKTIHTDNGSNFTGATVRAA---------CWWAGIKQEFGIPYNPQSQGVVESMN  155
            +  +P  T+ TDNGS FT    R A             GI Q+ G PY+PQ+QG VE  +
Sbjct  602  KHGFPKSTL-TDNGSVFTTKLYRKADTRNGFEQLLLQMGIHQKNGTPYHPQTQGKVERFH  426
```

```
Query  156  KELKKII  162
            + L+K +
Sbjct  425  QTLQKAL  405
```

>Corynebacterium marquesiae strain MSK087 NODE_41_length_882_cov_36.411180, whole genome shotgun sequence
Sequence ID: NZ_JASPIN010000041.1 Length: 882
Range 1: 233 to 610

Score:48.9 bits(115), Expect:0.044,
Method:Compositional matrix adjust.,
Identities:40/127(31%), Positives:57/127(44%), Gaps:18/127(14%)

```
Query  53   QVDCSPGIWQLDCTHL-----EGKVILVAVHVASGYIEA-EVIPAETGQETAYFLLKLAG  106
            Q D     WQLD +H      +   IL  +   S Y+ A + +  ETG       +
Sbjct  610  QADQPNETWQLDYSHWYLNNNQRVAILTIIDDHSRYVIACKAVEHETGTNVIDTFISAGN  431

Query  107  R--WPVKTIHTDNGSNFTGATVRAA---------CWWAGIKQEFGIPYNPQSQGVVESMN  155
            +  +P  T+ TDNGS FT    R A              GI Q+ G PY+PQ+QG VE  +
Sbjct  430  KHGFPKSTL-TDNGSVFTTKLYRKADTRNGFEQLLLQMGIHQKNGTPYHPQTQGKVERFH  254

Query  156  KELKKII  162
            + L+K +
Sbjct  253  QTLQKAL  233
```

>Pseudarthrobacter sp. W1I19 Ga0505363_01, whole genome shotgun sequence
Sequence ID: NZ_JAUSZR010000001.1 Length: 4812144
Range 1: 3313123 to 3313527

Score:50.1 bits(118), Expect:0.045,
Method:Compositional matrix adjust.,
Identities:36/135(27%), Positives:62/135(45%), Gaps:8/135(5%)

```
Query  76       AVHVASGYIEAEVIPAETGQETAYFLLK----LAGRW-PVKTIHTDNGSNFTGATVRAAC  130
                AV   S ++ +E++P ET +   + F+L      A R   ++ + TDNG+ +   T  A
Sbjct  3313527  AVDDHSRFVYSEILPDETKETASAFMLNAIAAFAARGVKIQRVLTDNGACYRSRTFAAVL  3313348

Query  131      WWAGIKQEFGIPYNPQSQGVVESMNKELKK---IIGQVRDQAEHLKTAVQMAVFIHNFKR  187
                  AGIK +   PY PQ+ G VE  N+ L++         R +AE          + +  +
Sbjct  3313347  AEAGIKHKRTRPYRPQTNGKVERFNRILQEEWAYAKTYRSEAERQACYPDFIKYYNQRRP  3313168

Query  188      KGGIGGYSAGERIVD  202
                   + G S   R+++
Sbjct  3313167  HTALKGASPASRVIN  3313123
```

>Corynebacterium marquesiae strain MSK093 NODE_39_length_702_cov_24.545600, whole genome shotgun sequence
Sequence ID: NZ_JASPII010000039.1 Length: 702
Range 1: 53 to 430

Score:48.5 bits(114), Expect:0.046,
Method:Compositional matrix adjust.,
Identities:40/127(31%), Positives:57/127(44%), Gaps:18/127(14%)

```
Query  53   QVDCSPGIWQLDCTHL-----EGKVILVAVHVASGYIEA-EVIPAETGQETAYFLLKLAG  106
            Q D     WQLD +H      +   IL  +   S Y+ A + +  ETG       +
Sbjct  430  QADQPNETWQLDYSHWYLNNNQRVAILTIIDDHSRYVIACKAVEHETGTNVIDTFISAGN  251

Query  107  R--WPVKTIHTDNGSNFTGATVRAA---------CWWAGIKQEFGIPYNPQSQGVVESMN  155
            +  +P  T+ TDNGS FT    R A              GI Q+ G PY+PQ+QG VE  +
Sbjct  250  KHGFPKSTL-TDNGSVFTTKLYRKADTRNGFEQLLLQMGIHQKNGTPYHPQTQGKVERFH  74

Query  156  KELKKII  162
            + L+K +
Sbjct  73   QTLQKAL  53
```

>Corynebacterium marquesiae strain MSK097 NODE_40_length_702_cov_29.779200, whole genome shotgun sequence
Sequence ID: NZ_JASPIF010000040.1 Length: 702
Range 1: 53 to 430

Score:48.5 bits(114), Expect:0.046,
Method:Compositional matrix adjust.,
Identities:40/127(31%), Positives:57/127(44%), Gaps:18/127(14%)

```
Query  53   QVDCSPGIWQLDCTHL-----EGKVILVAVHVASGYIEA-EVIPAETGQETAYFLLKLAG  106
            Q D     WQLD +H      +   IL  +   S Y+ A + +  ETG       +
Sbjct  430  QADQPNETWQLDYSHWYLNNNQRVAILTIIDDHSRYVIACKAVEHETGTNVIDTFISAGN  251

Query  107  R--WPVKTIHTDNGSNFTGATVRAA---------CWWAGIKQEFGIPYNPQSQGVVESMN  155
            +  +P  T+ TDNGS FT    R A              GI Q+ G PY+PQ+QG VE  +
Sbjct  250  KHGFPKSTL-TDNGSVFTTKLYRKADTRNGFEQLLLQMGIHQKNGTPYHPQTQGKVERFH  74

Query  156  KELKKII  162
            + L+K +
Sbjct  73   QTLQKAL  53
```

>Escherichia coli strain E-4H5 NODE_89_length_1734_cov_0.210330, whole genome shotgun sequence
Sequence ID: NZ_JALYCP010000088.1 Length: 1734
Range 1: 888 to 1154

Score:49.7 bits(117), Expect:0.047,
Method:Compositional matrix adjust.,
Identities:30/89(34%), Positives:41/89(46%), Gaps:10/89(11%)

```
Query  110   VKTIHTDNGSNFTGATVRAACWWAGIKQEFGIPYNPQSQGVVESMNKEL----KKIIGQV  165
             VK I ++NGS F    V       GIK EF  PY PQ  GVVE  N+ L    + ++G+
Sbjct  1154  VKKIRSNNGSEFKNLQVEEFLEEEGIKHEFSAPYTPQQNGVVERKNRTLIDMARTMLGEF  975

Query  166   RDQAEHLKTAVQMA------VFIHNFKRK  188
             +         AV  A      V++H   +K
Sbjct  974   KTPERFWSEAVNTACHAINRVYLHRLLKK  888
```

Query #13: sp|P04585|Protease_HV1H2 Gag-Pol polyprotein OS=Human immunodeficiency virus type 1 group M subtype B (isolate HXB2) OX=11706 GN=gag-pol PE=1 SV=4 Query ID: lcl|Query_2387717 Length: 99

Sequences producing significant alignments:

| Description | Scientific Name | Common Name | Taxid | Max Score | Total Score | Query cover | E Value | Per. Ident | Acc. Len | Accession |
|---|---|---|---|---|---|---|---|---|---|---|
| Klebsiella pneumoniae strain P7701... | Klebsiella p... | NA | 573 | 199 | 199 | 100% | 2e-59 | 97.98 | 1305 | NZ_JACAWM0100... |
| Acinetobacter baumannii strain ACM-36... | Acinetobacte... | NA | 470 | 189 | 189 | 100% | 2e-58 | 89.90 | 567 | NZ_QAGO010000... |
| Salirhabdus euzebyi strain Marseille-Q1438, whole genome shotg... | Salirhabdus ... | NA | 394506 | 195 | 195 | 100% | 7e-58 | 94.95 | 1279 | NZ_CADDWK0100... |
| Escherichia coli strain 8374wH11... | Escherichia ... | NA | 562 | 195 | 195 | 100% | 1e-57 | 94.95 | 1397 | NZ_WSJI010000... |
| Acinetobacter baumannii strain ACM-15... | Acinetobacte... | NA | 470 | 183 | 183 | 100% | 8e-57 | 89.90 | 432 | NZ_PYDB010000... |
| Enterococcus faecalis strain 44.1... | Enterococcus... | NA | 1351 | 187 | 187 | 100% | 9e-57 | 88.89 | 806 | NZ_NSJR010000... |
| Mycobacterium tuberculosis strain 32847_S31 462, whole genome... | Mycobacteriu... | NA | 1773 | 188 | 188 | 100% | 9e-57 | 88.89 | 893 | NZ_QTBX010001... |
| Mycobacterium tuberculosis strain 32846_S30 476, whole genome... | Mycobacteriu... | NA | 1773 | 187 | 187 | 100% | 1e-56 | 88.89 | 861 | NZ_QTBW010001... |
| Acinetobacter baumannii strain ACM-35... | Acinetobacte... | NA | 470 | 192 | 192 | 100% | 1e-56 | 91.92 | 1409 | NZ_QEHN010000... |
| Acinetobacter baumannii strain ACM-14... | Acinetobacte... | NA | 470 | 186 | 186 | 100% | 1e-56 | 89.90 | 787 | NZ_PYDA010000... |
| Actinoplanes campanulatus strain JCM 3059 sequence058, whole... | Actinoplanes... | NA | 113559 | 199 | 199 | 100% | 3e-56 | 97.98 | 4754 | NZ_BMPW010000... |
| Enterococcus alcedinis strain CCM 8433 sequence23, whole genom... | Enterococcus... | NA | 1274384 | 199 | 199 | 100% | 3e-56 | 97.98 | 6075 | NZ_BMDT010000... |
| Deinococcus aquiradiocola strain JCM 14371 sequence30, whole... | Deinococcus ... | NA | 393059 | 199 | 199 | 100% | 3e-56 | 97.98 | 6128 | NZ_BMOE010000... |
| Enterobacter hormaechei strain AUH-ENM6... | Enterobacter... | NA | 158836 | 191 | 191 | 100% | 5e-56 | 90.91 | 1394 | NZ_PYHB010000... |
| Thalassovita mangrovi strain GS-10 Scaffold19_1, whole genome... | Thalassovita... | NA | 2692236 | 198 | 198 | 100% | 6e-56 | 97.98 | 2900 | NZ_WWEN010000... |
| Escherichia coli strain 8374wG10... | Escherichia ... | NA | 562 | 190 | 190 | 100% | 7e-56 | 92.93 | 1373 | NZ_WSJQ010000... |
| Providencia rettgeri strain AFE220_S1 26_contig_67, whole geno... | Providencia ... | NA | 587 | 181 | 181 | 97% | 1e-55 | 88.66 | 512 | NZ_SHDM010000... |
| Klebsiella pneumoniae strain EuSCAPE_NO017, whole genome shotg... | Klebsiella p... | NA | 573 | 181 | 181 | 100% | 1e-55 | 85.86 | 458 | NZ_UKIO020000... |
| Klebsiella aerogenes strain AUH-KAM5... | Klebsiella a... | NA | 548 | 190 | 190 | 100% | 1e-55 | 92.93 | 1427 | NZ_PYHJ010000... |
| Helicobacter acinonychis isolate 212_10, whole genome shotgun... | Helicobacter... | NA | 212 | 190 | 241 | 100% | 2e-55 | 89.90 | 1526 | NZ_FZLW010000... |
| Providencia rettgeri strain AFE652_S1 38_contig_28, whole geno... | Providencia ... | NA | 587 | 187 | 187 | 100% | 2e-55 | 88.89 | 1127 | NZ_SHDK010000... |
| Escherichia coli strain 8374wB4 NODE_60_length_1385_cov_3.2655... | Escherichia ... | NA | 562 | 189 | 189 | 100% | 2e-55 | 92.93 | 1385 | NZ_WSPX010000... |
| Providencia rettgeri strain AFE422_S1 40_contig_34, whole geno... | Providencia ... | NA | 587 | 186 | 186 | 100% | 4e-55 | 88.89 | 1058 | NZ_SHDL010000... |
| Methylocella silvestris strain TVC... | Methylocella... | NA | 199596 | 189 | 189 | 100% | 6e-55 | 89.90 | 1453 | NZ_PDZR010000... |
| Providencia rettgeri strain AFE97_S1 21_contig_71, whole genom... | Providencia ... | NA | 587 | 186 | 186 | 100% | 1e-54 | 88.89 | 1157 | NZ_SHDO010000... |
| Listeria monocytogenes strain YA00082404... | Listeria mon... | NA | 1639 | 189 | 189 | 100% | 3e-54 | 89.90 | 1816 | NZ_QEXE010000... |
| Pseudomonas aeruginosa strain P60_London_6_VIM_2_11_13, whole... | Pseudomonas ... | NA | 287 | 175 | 175 | 97% | 1e-53 | 83.51 | 395 | NZ_CVXD010000... |
| Enterococcus faecium strain E21_6... | Enterococcus... | NA | 1352 | 176 | 176 | 100% | 1e-53 | 84.16 | 518 | NZ_JAANZJ0100... |
| Methylobacterium radiotolerans strain MAMP 4754... | Methylobacte... | NA | 31998 | 175 | 175 | 97% | 2e-53 | 84.54 | 483 | NZ_NKQS010002... |
| Mycobacterium tuberculosis variant bovis strain CSURQ1132, who... | Mycobacteriu... | NA | 1765 | 184 | 369 | 100% | 4e-53 | 90.91 | 1524 | NZ_CAJEUK0100... |
| Levilactobacillus brevis strain CRL 2013 scaffold12, whole... | Levilactobac... | NA | 1580 | 181 | 181 | 100% | 6e-53 | 86.87 | 1126 | NZ_MZMW010000... |
| Klebsiella pneumoniae strain NUBRI-K 73, whole genome shotgun... | Klebsiella p... | NA | 573 | 185 | 185 | 100% | 6e-53 | 85.86 | 1717 | NZ_SOYT010000... |
| Enterococcus faecium strain A209R1B0C1... | Enterococcus... | NA | 1352 | 186 | 186 | 100% | 7e-53 | 88.89 | 1819 | NZ_PGCR010000... |
| Acinetobacter baumannii strain ACM-32... | Acinetobacte... | NA | 470 | 176 | 176 | 100% | 8e-53 | 82.83 | 654 | NZ_PYDR010000... |
| Mycobacterium tuberculosis variant bovis strain CSURQ1134, who... | Mycobacteriu... | NA | 1765 | 184 | 369 | 100% | 1e-52 | 90.91 | 1697 | NZ_CAJEUT0100... |
| Providencia rettgeri strain APW139_S1 24_contig_65, whole geno... | Providencia ... | NA | 587 | 177 | 177 | 98% | 4e-52 | 85.71 | 941 | NZ_SHDG010000... |
| Levilactobacillus brevis strain CRL 2013 scaffold13, whole... | Levilactobac... | NA | 1580 | 177 | 177 | 100% | 1e-51 | 86.87 | 1030 | NZ_MZMW010000... |
| Escherichia coli strain VRES0669, whole genome shotgun sequence | Escherichia ... | NA | 562 | 186 | 186 | 100% | 1e-51 | 88.89 | 5544 | NZ_UINV010000... |
| Enterococcus faecium strain A206R2B0... | Enterococcus... | NA | 1352 | 181 | 181 | 100% | 2e-51 | 86.87 | 1761 | NZ_PGCU010000... |
| Pseudomonas aeruginosa strain VET-121... | Pseudomonas ... | NA | 287 | 174 | 174 | 98% | 3e-50 | 85.71 | 1158 | NZ_QZXB010001... |
| Acinetobacter baumannii strain ACM-23... | Acinetobacte... | NA | 470 | 167 | 167 | 90% | 4e-50 | 88.89 | 505 | NZ_PYDJ010000... |
| Acinetobacter baumannii strain ACM-38... | Acinetobacte... | NA | 470 | 167 | 167 | 100% | 2e-49 | 81.82 | 652 | NZ_QAGQ010000... |
| Pseudomonas aeruginosa strain VET-42... | Pseudomonas ... | NA | 287 | 170 | 170 | 98% | 1e-48 | 83.67 | 1166 | NZ_RACX010001... |
| Mycobacterium tuberculosis variant bovis strain CSURQ1132, who... | Mycobacteriu... | NA | 1765 | 166 | 333 | 98% | 2e-48 | 84.69 | 862 | NZ_CAJEUK0100... |
| Escherichia coli strain YA00028237 contig_187, whole genome... | Escherichia ... | NA | 562 | 173 | 173 | 100% | 5e-48 | 75.76 | 1828 | NZ_NGBP020001... |
| Acinetobacter baumannii strain ACM-29... | Acinetobacte... | NA | 470 | 163 | 163 | 87% | 6e-47 | 88.51 | 888 | NZ_PYDO010001... |
| Escherichia coli strain 8374wB11... | Escherichia ... | NA | 562 | 166 | 166 | 95% | 8e-47 | 82.11 | 1310 | NZ_WSJO010000... |
| Providencia rettgeri strain AFE85_S2 19_contig_46, whole genom... | Providencia ... | NA | 587 | 160 | 160 | 86% | 7e-46 | 88.37 | 846 | NZ_SHDQ010000... |
| Providencia rettgeri strain AFE97_S1 21_contig_72, whole genom... | Providencia ... | NA | 587 | 160 | 160 | 87% | 7e-46 | 87.36 | 854 | NZ_SHDO010000... |
| Providencia rettgeri strain AFE96_S1 41_contig_60, whole genom... | Providencia ... | NA | 587 | 162 | 162 | 86% | 1e-45 | 88.37 | 1101 | NZ_SHDP010000... |
| Klebsiella pneumoniae strain EuSCAPE_IL004, whole genome shotg... | Klebsiella p... | NA | 573 | 151 | 151 | 80% | 1e-44 | 90.00 | 356 | NZ_UKJP020001... |
| Acinetobacter baumannii strain ACM-5... | Acinetobacte... | NA | 470 | 157 | 157 | 85% | 2e-43 | 87.06 | 1274 | NZ_PYCX010000... |
| Mycobacterium tuberculosis strain 32845_S29 230, whole genome... | Mycobacteriu... | NA | 1773 | 150 | 150 | 78% | 6e-43 | 92.31 | 601 | NZ_QTBV010000... |
| Cellulophaga baltica strain 14 NODE_63_length_422_cov_10.20678... | Cellulophaga... | NA | 76594 | 144 | 144 | 82% | 2e-41 | 85.37 | 422 | NZ_VIQP010000... |
| Enterococcus faecium strain A201R2B0... | Enterococcus... | NA | 1352 | 151 | 151 | 85% | 3e-40 | 84.71 | 1589 | NZ_PGCT010000... |
| Klebsiella pneumoniae strain EuSCAPE_IL005, whole genome shotg... | Klebsiella p... | NA | 573 | 135 | 135 | 73% | 3e-38 | 86.30 | 323 | NZ_UKJL020000... |
| Klebsiella pneumoniae strain EuSCAPE_LT007, whole genome shotg... | Klebsiella p... | NA | 573 | 134 | 134 | 72% | 2e-37 | 87.50 | 392 | NZ_UKJI020000... |
| Enterococcus durans strain E115_10... | Enterococcus... | NA | 53345 | 134 | 134 | 73% | 3e-35 | 80.82 | 955 | NZ_JAANZI0100... |
| Klebsiella pneumoniae strain ARGID_31501, whole genome shotgun... | Klebsiella p... | NA | 573 | 139 | 139 | 74% | 4e-35 | 90.54 | 2655 | NZ_CAKLPD0100... |
| Escherichia coli strain 8374wG10 NODE_65_length_433_cov_1.7679... | Escherichia ... | NA | 562 | 125 | 125 | 77% | 7e-34 | 72.73 | 433 | NZ_WSJQ010000... |
| Escherichia coli strain 8374wA11 NODE_44_length_523_cov_1.7197... | Escherichia ... | NA | 562 | 104 | 104 | 64% | 3e-25 | 71.88 | 523 | NZ_WSPR010000... |
| Escherichia coli strain TzEc080 NODE_44_length_523_cov_1.7197,... | Escherichia ... | NA | 562 | 104 | 104 | 64% | 3e-25 | 71.88 | 523 | NZ_WSFZ010000... |
| Klebsiella pneumoniae strain EuSCAPE_MT006, whole genome shotg... | Klebsiella p... | NA | 573 | 95.9 | 95.9 | 50% | 4e-22 | 90.00 | 455 | NZ_UKIU020000... |
| Klebsiella pneumoniae strain EuSCAPE_MT015, whole genome shotg... | Klebsiella p... | NA | 573 | 88.2 | 88.2 | 45% | 4e-19 | 93.33 | 435 | NZ_UKIW020000... |
| Providencia rettgeri strain APW139_S1 24_contig_64, whole geno... | Providencia ... | NA | 587 | 85.9 | 85.9 | 44% | 5e-17 | 93.18 | 902 | NZ_SHDG010000... |
| Acinetobacter baumannii strain ACM-24... | Acinetobacte... | NA | 470 | 82.4 | 82.4 | 44% | 1e-16 | 88.64 | 503 | NZ_PYDK010000... |
| Klebsiella pneumoniae strain EuSCAPE_IL017, whole genome shotg... | Klebsiella p... | NA | 573 | 76.6 | 76.6 | 41% | 4e-15 | 82.93 | 325 | NZ_UKKG020000... |
| Burkholderia pseudomallei strain A-175-05-2-02, whole genome... | Burkholderia... | NA | 28450 | 72.0 | 72.0 | 38% | 1e-12 | 89.47 | 455 | NZ_UVFW010001... |
| Mycobacterium tuberculosis strain 32845_S29 214, whole genome... | Mycobacteriu... | NA | 1773 | 66.6 | 66.6 | 34% | 1e-11 | 85.29 | 223 | NZ_QTBV010000... |
| Enterobacter hormaechei strain AUH-ENM4... | Enterobacter... | NA | 158836 | 53.1 | 53.1 | 25% | 5e-05 | 96.00 | 809 | NZ_PYGZ010000... |
| Escherichia coli strain 8374wB11 NODE_46_length_516_cov_2.1516... | Escherichia ... | NA | 562 | 51.6 | 51.6 | 25% | 1e-04 | 88.00 | 516 | NZ_WSJO010000... |
| Mycobacterium tuberculosis variant bovis strain CSURQ1139, who... | Mycobacteriu... | NA | 1765 | 51.2 | 102 | 23% | 3e-04 | 100.00 | 848 | NZ_CAJEUO0100... |
| Pseudomonas aromaticivorans strain MAP12 34, whole genome... | Pseudomonas ... | NA | 2849492 | 48.9 | 48.9 | 93% | 0.002 | 32.32 | 7031 | NZ_JAHRGL0100... |

Alignments:

```
>Klebsiella pneumoniae strain P7701 NODE_50_length_1305_cov_481.863, whole genome shotgun sequence
Sequence ID: NZ_JACAWM010000050.1 Length: 1305
Range 1: 909 to 1205

Score:199 bits(506), Expect:2e-59,
Method:Compositional matrix adjust.,
Identities:97/99(98%), Positives:99/99(100%), Gaps:0/99(0%)

Query  1     PQVTLWQRPLVTIKIGGQLKEALLDTGADDTVLEEMSLPGRWKPKMIGGIGGFIKVRQYD  60
             PQ+TLWQRPLVTIKIGGQLKEALLDTGADDTVLEEM+LPGRWKPKMIGGIGGFIKVRQYD
Sbjct  1205  PQITLWQRPLVTIKIGGQLKEALLDTGADDTVLEEMNLPGRWKPKMIGGIGGFIKVRQYD  1026

Query  61    QILIEICGHKAIGTVLVGPTPVNIIGRNLLTQIGCTLNF  99
             QILIEICGHKAIGTVLVGPTPVNIIGRNLLTQIGCTLNF
Sbjct  1025  QILIEICGHKAIGTVLVGPTPVNIIGRNLLTQIGCTLNF  909
```

```
>Acinetobacter baumannii strain ACM-36 NODE_67_length_567_cov_1.13409, whole genome shotgun sequence
Sequence ID: NZ_QAGO01000067.1 Length: 567
Range 1: 90 to 386

Score:189 bits(479), Expect:2e-58,
Method:Compositional matrix adjust.,
Identities:89/99(90%), Positives:97/99(97%), Gaps:0/99(0%)

Query  1    PQVTLWQRPLVTIKIGGQLKEALLDTGADDTVLEEMSLPGRWKPKMIGGIGGFIKVRQYD  60
            PQ+TLWQRP+VTIK+GGQLKEALLDTGADDTVLE+M LPGRWKPKMIGGIGGFIKVRQYD
Sbjct  90   PQITLWQRPVVTIKVGGQLKEALLDTGADDTVLEDMQLPGRWKPKMIGGIGGFIKVRQYD  269

Query  61   QILIEICGHKAIGTVLVGPTPVNIIGRNLLTQIGCTLNF  99
            ++ IEICGHKAIGTVL+GPTPVNIIGRNLLTQ+GCTLNF
Sbjct  270  EVPIEICGHKAIGTVLIGPTPVNIIGRNLLTQLGCTLNF  386
```

```
>Salirhabdus euzebyi strain Marseille-Q1438, whole genome shotgun sequence
Sequence ID: NZ_CADDWK010000035.1 Length: 1279
Range 1: 897 to 1193

Score:195 bits(495), Expect:7e-58,
Method:Compositional matrix adjust.,
Identities:94/99(95%), Positives:97/99(97%), Gaps:0/99(0%)
```

```
Query  1     PQVTLWQRPLVTIKIGGQLKEALLDTGADDTVLEEMSLPGRWKPKMIGGIGGFIKVRQYD  60
             PQ+TLWQRPLV IKIGGQLKEALLDTGADDTVLEEM+LPGRWKPKMIGGIGGFIKVRQYD
Sbjct  1193  PQITLWQRPLVAIKIGGQLKEALLDTGADDTVLEEMNLPGRWKPKMIGGIGGFIKVRQYD  1014

Query  61    QILIEICGHKAIGTVLVGPTPVNIIGRNLLTQIGCTLNF  99
             QI IEICGHKAIGTVL+GPTPVNIIGRNLLTQIGCTLNF
Sbjct  1013  QIPIEICGHKAIGTVLIGPTPVNIIGRNLLTQIGCTLNF  897
```

>Escherichia coli strain 8374wH11 NODE_36_length_1397_cov_3.54567, whole genome shotgun sequence
Sequence ID: NZ_WSJI01000036.1 Length: 1397
Range 1: 88 to 384

Score:195 bits(496), Expect:1e-57,
Method:Compositional matrix adjust.,
Identities:94/99(95%), Positives:98/99(98%), Gaps:0/99(0%)

```
Query  1    PQVTLWQRPLVTIKIGGQLKEALLDTGADDTVLEEMSLPGRWKPKMIGGIGGFIKVRQYD  60
            PQ+TLWQRPLVTIKIGGQLKEALLDTGADDTVLEEM+LPGRWKPKMIGGIGGFIKVRQYD
Sbjct  88   PQITLWQRPLVTIKIGGQLKEALLDTGADDTVLEEMNLPGRWKPKMIGGIGGFIKVRQYD  267

Query  61   QILIEICGHKAIGTVLVGPTPVNIIGRNLLTQIGCTLNF  99
            QI IEICGHKAIGTVLVGPTPVNIIGRN+LTQ+GCTLNF
Sbjct  268  QIPIEICGHKAIGTVLVGPTPVNIIGRNMLTQLGCTLNF  384
```

>Acinetobacter baumannii strain ACM-15 NODE_85_length_432_cov_0.596721, whole genome shotgun sequence
Sequence ID: NZ_PYDB01000084.1 Length: 432
Range 1: 12 to 308

Score:183 bits(465), Expect:8e-57,
Method:Compositional matrix adjust.,
Identities:89/99(90%), Positives:96/99(96%), Gaps:0/99(0%)

```
Query  1    PQVTLWQRPLVTIKIGGQLKEALLDTGADDTVLEEMSLPGRWKPKMIGGIGGFIKVRQYD  60
            PQ+TLWQRPLV+IK+GGQ+KEALLDTGADDTVLEE+SLPG+WKPKMIGGIGGFIKVRQY
Sbjct  12   PQITLWQRPLVSIKVGGQIKEALLDTGADDTVLEEISLPGKWKPKMIGGIGGFIKVRQYG  191

Query  61   QILIEICGHKAIGTVLVGPTPVNIIGRNLLTQIGCTLNF  99
            QI IEICG KAIGTVLVGPTPVNIIGRNLLTQ+GCTLNF
Sbjct  192  QIPIEICGKKAIGTVLVGPTPVNIIGRNLLTQLGCTLNF  308
```

>Enterococcus faecalis strain 44.1 NODE_33_length_806_cov_1.20619, whole genome shotgun sequence
Sequence ID: NZ_NSJR01000033.1 Length: 806
Range 1: 117 to 413

Score:187 bits(476), Expect:9e-57,
Method:Compositional matrix adjust.,
Identities:88/99(89%), Positives:96/99(96%), Gaps:0/99(0%)

```
Query  1    PQVTLWQRPLVTIKIGGQLKEALLDTGADDTVLEEMSLPGRWKPKMIGGIGGFIKVRQYD  60
            PQ+TLWQRP+VTIK+GGQLKEALLDTGADDTVLEEM+LPGRWKPKMIGGIGGFIKVRQYD
Sbjct  117  PQITLWQRPVVTIKVGGQLKEALLDTGADDTVLEEMNLPGRWKPKMIGGIGGFIKVRQYD  296

Query  61   QILIEICGHKAIGTVLVGPTPVNIIGRNLLTQIGCTLNF  99
            ++ IEICGHK IGTVL+G TPVNIIGRNLLTQ+GCTLNF
Sbjct  297  EVPIEICGHKVIGTVLIGSTPVNIIGRNLLTQLGCTLNF  413
```

>Mycobacterium tuberculosis strain 32847_S31 462, whole genome shotgun sequence
Sequence ID: NZ_QTBX01000137.1 Length: 893
Range 1: 194 to 490

Score:188 bits(478), Expect:9e-57,
Method:Compositional matrix adjust.,
Identities:88/99(89%), Positives:98/99(98%), Gaps:0/99(0%)

```
Query  1    PQVTLWQRPLVTIKIGGQLKEALLDTGADDTVLEEMSLPGRWKPKMIGGIGGFIKVRQYD  60
            PQ+TLWQRPLV+IK+GGQ+EALLDTGADDTVLE+++LPG+WKPKMIGGIGGFIKVRQYD
Sbjct  194  PQITLWQRPLVSIKVGGQIREALLDTGADDTVLEDINLPGKWKPKMIGGIGGFIKVRQYD  373

Query  61   QILIEICGHKAIGTVLVGPTPVNIIGRNLLTQIGCTLNF  99
            QILIEICG KAIGTVLVGPTPVNIIGRN+LTQIGCTLNF
Sbjct  374  QILIEICGKKAIGTVLVGPTPVNIIGRNMLTQIGCTLNF  490
```

>Mycobacterium tuberculosis strain 32846_S30 476, whole genome shotgun sequence
Sequence ID: NZ_QTBW01000113.1 Length: 861
Range 1: 484 to 780

Score:187 bits(476), Expect:1e-56,
Method:Compositional matrix adjust.,
Identities:88/99(89%), Positives:98/99(98%), Gaps:0/99(0%)

```
Query  1    PQVTLWQRPLVTIKIGGQLKEALLDTGADDTVLEEMSLPGRWKPKMIGGIGGFIKVRQYD  60
            PQ+TLWQRPLV+IK+GGQ++EALLDTGADDTVLE+++LPG+WKPKMIGGIGGFIKVRQYD
Sbjct  780  PQITLWQRPLVSIKVGGQIREALLDTGADDTVLEDINLPGKWKPKMIGGIGGFIKVRQYD  601

Query  61   QILIEICGHKAIGTVLVGPTPVNIIGRNLLTQIGCTLNF  99
            QILIEICG KAIGTVLVGPTPVNIIGRN+LTQIGCTLNF
Sbjct  600  QILIEICGKKAIGTVLVGPTPVNIIGRNMLTQIGCTLNF  484
```

>Acinetobacter baumannii strain ACM-35 NODE_65_length_1409_cov_2.30031, whole genome shotgun sequence
Sequence ID: NZ_QEHN01000065.1 Length: 1409
Range 1: 108 to 404

Score:192 bits(489), Expect:1e-56,
Method:Compositional matrix adjust.,
Identities:91/99(92%), Positives:97/99(97%), Gaps:0/99(0%)

```
Query  1    PQVTLWQRPLVTIKIGGQLKEALLDTGADDTVLEEMSLPGRWKPKMIGGIGGFIKVRQYD  60
            PQ+TLWQRP+VTIK+GGQLKEALLDTGADDTVLEEM+LPGRWKPKMIGGIGGFIKVRQYD
Sbjct  108  PQITLWQRPVVTIKVGGQLKEALLDTGADDTVLEEMNLPGRWKPKMIGGIGGFIKVRQYD  287

Query  61   QILIEICGHKAIGTVLVGPTPVNIIGRNLLTQIGCTLNF  99
            QI IEICGHK IGTVL+GPTPVNIIGRNLLTQ+GCTLNF
Sbjct  288  QIPIEICGHKVIGTVLIGPTPVNIIGRNLLTQLGCTLNF  404
```

```
>Acinetobacter baumannii strain ACM-14 NODE_74_length_787_cov_0.637879, whole genome shotgun sequence
Sequence ID: NZ_PYDA01000074.1 Length: 787
Range 1: 405 to 701

Score:186 bits(473), Expect:1e-56,
Method:Compositional matrix adjust.,
Identities:89/99(90%), Positives:95/99(95%), Gaps:0/99(0%)

Query  1    PQVTLWQRPLVTIKIGGQLKEALLDTGADDTVLEEMSLPGRWKPKMIGGIGGFIKVRQYD  60
            PQ+TLWQRPLVTIKIGGQLKEALLDTGADDTVLE+M LP RWKPKMIGGI GFIKVRQYD
Sbjct  701  PQITLWQRPLVTIKIGGQLKEALLDTGADDTVLEDMQLPERWKPKMIGGIRGFIKVRQYD  522

Query  61   QILIEICGHKAIGTVLVGPTPVNIIGRNLLTQIGCTLNF  99
            ++ IEICGHKAIGTVL+GPTPVNIIGRNLLTQ+GCTLNF
Sbjct  521  EVPIEICGHKAIGTVLIGPTPVNIIGRNLLTQLGCTLNF  405



>Actinoplanes campanulatus strain JCM 3059 sequence058, whole genome shotgun sequence
Sequence ID: NZ_BMPW01000058.1 Length: 4754
Range 1: 1560 to 1856

Score:199 bits(507), Expect:3e-56,
Method:Compositional matrix adjust.,
Identities:97/99(98%), Positives:99/99(100%), Gaps:0/99(0%)

Query  1     PQVTLWQRPLVTIKIGGQLKEALLDTGADDTVLEEMSLPGRWKPKMIGGIGGFIKVRQYD  60
             PQ+TLWQRPLVTIKIGGQLKEALLDTGADDTVLEEM+LPGRWKPKMIGGIGGFIKVRQYD
Sbjct  1560  PQITLWQRPLVTIKIGGQLKEALLDTGADDTVLEEMNLPGRWKPKMIGGIGGFIKVRQYD  1739

Query  61    QILIEICGHKAIGTVLVGPTPVNIIGRNLLTQIGCTLNF  99
             QILIEICGHKAIGTVLVGPTPVNIIGRNLLTQIGCTLNF
Sbjct  1740  QILIEICGHKAIGTVLVGPTPVNIIGRNLLTQIGCTLNF  1856



>Enterococcus alcedinis strain CCM 8433 sequence23, whole genome shotgun sequence
Sequence ID: NZ_BMDT01000023.1 Length: 6075
Range 1: 1518 to 1814

Score:199 bits(507), Expect:3e-56,
Method:Compositional matrix adjust.,
Identities:97/99(98%), Positives:99/99(100%), Gaps:0/99(0%)

Query  1     PQVTLWQRPLVTIKIGGQLKEALLDTGADDTVLEEMSLPGRWKPKMIGGIGGFIKVRQYD  60
             PQ+TLWQRPLVTIKIGGQLKEALLDTGADDTVLEEM+LPGRWKPKMIGGIGGFIKVRQYD
Sbjct  1518  PQITLWQRPLVTIKIGGQLKEALLDTGADDTVLEEMNLPGRWKPKMIGGIGGFIKVRQYD  1697

Query  61    QILIEICGHKAIGTVLVGPTPVNIIGRNLLTQIGCTLNF  99
             QILIEICGHKAIGTVLVGPTPVNIIGRNLLTQIGCTLNF
Sbjct  1698  QILIEICGHKAIGTVLVGPTPVNIIGRNLLTQIGCTLNF  1814



>Deinococcus aquiradiocola strain JCM 14371 sequence30, whole genome shotgun sequence
Sequence ID: NZ_BMOE01000030.1 Length: 6128
Range 1: 1571 to 1867

Score:199 bits(507), Expect:3e-56,
Method:Compositional matrix adjust.,
Identities:97/99(98%), Positives:99/99(100%), Gaps:0/99(0%)

Query  1     PQVTLWQRPLVTIKIGGQLKEALLDTGADDTVLEEMSLPGRWKPKMIGGIGGFIKVRQYD  60
             PQ+TLWQRPLVTIKIGGQLKEALLDTGADDTVLEEM+LPGRWKPKMIGGIGGFIKVRQYD
Sbjct  1571  PQITLWQRPLVTIKIGGQLKEALLDTGADDTVLEEMNLPGRWKPKMIGGIGGFIKVRQYD  1750

Query  61    QILIEICGHKAIGTVLVGPTPVNIIGRNLLTQIGCTLNF  99
             QILIEICGHKAIGTVLVGPTPVNIIGRNLLTQIGCTLNF
Sbjct  1751  QILIEICGHKAIGTVLVGPTPVNIIGRNLLTQIGCTLNF  1867



>Enterobacter hormaechei strain AUH-ENM6 NODE_27_length_1394_cov_0.980268, whole genome shotgun sequence
Sequence ID: NZ_PYHB01000027.1 Length: 1394
Range 1: 97 to 393

Score:191 bits(485), Expect:5e-56,
Method:Compositional matrix adjust.,
Identities:90/99(91%), Positives:97/99(97%), Gaps:0/99(0%)

Query  1    PQVTLWQRPLVTIKIGGQLKEALLDTGADDTVLEEMSLPGRWKPKMIGGIGGFIKVRQYD  60
            PQ+TLWQRP+VTIK+GGQLKEALLDTGADDTVLE+M LPGRWKPKMIGGIGGFIKVRQYD
Sbjct  97   PQITLWQRPVVTIKVGGQLKEALLDTGADDTVLEDMHLPGRWKPKMIGGIGGFIKVRQYD  276

Query  61   QILIEICGHKAIGTVLVGPTPVNIIGRNLLTQIGCTLNF  99
            ++ IEICGHKAIGTVLVGPTPVNIIGRNLLTQ+GCTLNF
Sbjct  277  EVPIEICGHKAIGTVLVGPTPVNIIGRNLLTQLGCTLNF  393



>Thalassovita mangrovi strain GS-10 Scaffold19_1, whole genome shotgun sequence
Sequence ID: NZ_WWEN01000019.1 Length: 2900
Range 1: 2503 to 2799

Score:198 bits(503), Expect:6e-56,
Method:Compositional matrix adjust.,
Identities:97/99(98%), Positives:99/99(100%), Gaps:0/99(0%)

Query  1     PQVTLWQRPLVTIKIGGQLKEALLDTGADDTVLEEMSLPGRWKPKMIGGIGGFIKVRQYD  60
             PQ+TLWQRPLVTIKIGGQLKEALLDTGADDTVLEEM+LPGRWKPKMIGGIGGFIKVRQYD
Sbjct  2503  PQITLWQRPLVTIKIGGQLKEALLDTGADDTVLEEMNLPGRWKPKMIGGIGGFIKVRQYD  2682

Query  61    QILIEICGHKAIGTVLVGPTPVNIIGRNLLTQIGCTLNF  99
             QILIEICGHKAIGTVLVGPTPVNIIGRNLLTQIGCTLNF
Sbjct  2683  QILIEICGHKAIGTVLVGPTPVNIIGRNLLTQIGCTLNF  2799



>Escherichia coli strain 8374wG10 NODE_34_length_1373_cov_2.57865, whole genome shotgun sequence
Sequence ID: NZ_WSJQ01000034.1 Length: 1373
Range 1: 1014 to 1310

Score:190 bits(483), Expect:7e-56,
Method:Compositional matrix adjust.,
Identities:92/99(93%), Positives:96/99(96%), Gaps:0/99(0%)
```

```
Query  1     PQVTLWQRPLVTIKIGGQLKEALLDTGADDTVLEEMSLPGRWKPKMIGGIGGFIKVRQYD  60
             PQ+TLWQRPLVTIKIGGQLKEALLDTGADDTVLEEM+LPGRWKPKMIGGIGGFIKVRQYD
Sbjct  1310  PQITLWQRPLVTIKIGGQLKEALLDTGADDTVLEEMNLPGRWKPKMIGGIGGFIKVRQYD  1131

Query  61    QILIEICGHKAIGTVLVGPTPVNIIGRNLLTQIGCTLNF  99
             QI IEICGHK IGTVL+G TPVNIIGRNLLTQ+GCTLNF
Sbjct  1130  QIPIEICGHKVIGTVLIGSTPVNIIGRNLLTQLGCTLNF  1014
```

>Providencia rettgeri strain AFE220_S1 26_contig_67, whole genome shotgun sequence
Sequence ID: NZ_SHDM01000067.1 Length: 512
Range 1: 220 to 510

Score:181 bits(460), Expect:1e-55,
Method:Compositional matrix adjust.,
Identities:86/97(89%), Positives:96/97(98%), Gaps:0/97(0%)

```
Query  3    VTLWQRPLVTIKIGGQLKEALLDTGADDTVLEEMSLPGRWKPKMIGGIGGFIKVRQYDQI  62
            +TLWQRPLV+IK+GGQ+KEALLDTGADDTVLE+++LPG+WKPKMIGGIGGFIKVRQYDQI
Sbjct  510  ITLWQRPLVSIKVGGQIKEALLDTGADDTVLEDINLPGKWKPKMIGGIGGFIKVRQYDQI  331

Query  63   LIEICGHKAIGTVLVGPTPVNIIGRNLLTQIGCTLNF  99
            LIEICG KAIGTVLVGPTPVNIIGRN+LTQ+GCTLNF
Sbjct  330  LIEICGKKAIGTVLVGPTPVNIIGRNMLTQLGCTLNF  220
```

>Klebsiella pneumoniae strain EuSCAPE_NO017, whole genome shotgun sequence
Sequence ID: NZ_UKIO02000069.1 Length: 458
Range 1: 57 to 353

Score:181 bits(458), Expect:1e-55,
Method:Compositional matrix adjust.,
Identities:85/99(86%), Positives:97/99(97%), Gaps:0/99(0%)

```
Query  1    PQVTLWQRPLVTIKIGGQLKEALLDTGADDTVLEEMSLPGRWKPKMIGGIGGFIKVRQYD  60
            PQ+TLWQRPLV+IK+GGQ++EALLDTGADDTVLE+++LPG+WKPKMIGGIGGFIKVRQY+
Sbjct  57   PQITLWQRPLVSIKVGGQIREALLDTGADDTVLEDINLPGKWKPKMIGGIGGFIKVRQYE  236

Query  61   QILIEICGHKAIGTVLVGPTPVNIIGRNLLTQIGCTLNF  99
             ILIEICG KAIGTVLVGPTPVNIIGRN+LTQ+GCTLNF
Sbjct  237  DILIEICGKKAIGTVLVGPTPVNIIGRNMLTQLGCTLNF  353
```

>Klebsiella aerogenes strain AUH-KAM5 NODE_25_length_1427_cov_1.66077, whole genome shotgun sequence
Sequence ID: NZ_PYHJ01000025.1 Length: 1427
Range 1: 84 to 380

Score:190 bits(483), Expect:1e-55,
Method:Compositional matrix adjust.,
Identities:92/99(93%), Positives:96/99(96%), Gaps:0/99(0%)

```
Query  1    PQVTLWQRPLVTIKIGGQLKEALLDTGADDTVLEEMSLPGRWKPKMIGGIGGFIKVRQYD  60
            PQ+TLWQRPLVTIKIGGQLKEALLDTGADDTVLEEM+LPGRWKPKMIGGIGGFIKVRQYD
Sbjct  84   PQITLWQRPLVTIKIGGQLKEALLDTGADDTVLEEMNLPGRWKPKMIGGIGGFIKVRQYD  263

Query  61   QILIEICGHKAIGTVLVGPTPVNIIGRNLLTQIGCTLNF  99
            QI IEICGHK IGTVL+G TPVNIIGRNLLTQ+GCTLNF
Sbjct  264  QIPIEICGHKVIGTVLIGSTPVNIIGRNLLTQLGCTLNF  380
```

>Helicobacter acinonychis isolate 212_10, whole genome shotgun sequence
Sequence ID: NZ_FZLW01000041.1 Length: 1526
Range 1: 1105 to 1401

Score:190 bits(483), Expect:2e-55,
Method:Compositional matrix adjust.,
Identities:89/99(90%), Positives:97/99(97%), Gaps:0/99(0%)

```
Query  1     PQVTLWQRPLVTIKIGGQLKEALLDTGADDTVLEEMSLPGRWKPKMIGGIGGFIKVRQYD  60
             PQ+TLWQRP+VTIKIGGQLKEALLDTGADDTVLEEMSLPG+WKPKMIGGIGGFIKVRQY+
Sbjct  1401  PQITLWQRPIVTIKIGGQLKEALLDTGADDTVLEEMSLPGKWKPKMIGGIGGFIKVRQYE  1222

Query  61    QILIEICGHKAIGTVLVGPTPVNIIGRNLLTQIGCTLNF  99
             ++ IEICGHKA GTVL+GPTPVNIIGRNLLTQ+GCTLNF
Sbjct  1221  EVPIEICGHKATGTVLIGPTPVNIIGRNLLTQLGCTLNF  1105
```

Range 2: 1 to 75

Score:51.2 bits(121), Expect:3e-04,
Method:Compositional matrix adjust.,
Identities:23/25(92%), Positives:25/25(100%), Gaps:0/25(0%)

```
Query  1   PQVTLWQRPLVTIKIGGQLKEALLD  25
           PQ+TLWQRP+VTIKIGGQLKEALLD
Sbjct  75  PQITLWQRPIVTIKIGGQLKEALLD  1
```

>Providencia rettgeri strain AFE652_S1 38_contig_28, whole genome shotgun sequence
Sequence ID: NZ_SHDK01000028.1 Length: 1127
Range 1: 811 to 1107

Score:187 bits(475), Expect:2e-55,
Method:Compositional matrix adjust.,
Identities:88/99(89%), Positives:97/99(97%), Gaps:0/99(0%)

```
Query  1     PQVTLWQRPLVTIKIGGQLKEALLDTGADDTVLEEMSLPGRWKPKMIGGIGGFIKVRQYD  60
             PQ+TLWQRPLV+IK+GGQ+KEALLDTGADDTVLEE++LPG+WKPKMIGGIGGFIKVRQYD
Sbjct  1107  PQITLWQRPLVSIKVGGQIKEALLDTGADDTVLEEINLPGKWKPKMIGGIGGFIKVRQYD  928

Query  61    QILIEICGHKAIGTVLVGPTPVNIIGRNLLTQIGCTLNF  99
              ILIEICG KAIGTVLVGPTPVNIIGRN+LTQ+GCTLNF
Sbjct  927   HILIEICGKKAIGTVLVGPTPVNIIGRNMLTQLGCTLNF  811
```

>Escherichia coli strain 8374wB4 NODE_60_length_1385_cov_3.2655, whole genome shotgun sequence
Sequence ID: NZ_WSPX01000060.1 Length: 1385
Range 1: 66 to 362

Score:189 bits(480), Expect:2e-55,

```
Method:Compositional matrix adjust.,
Identities:92/99(93%), Positives:97/99(97%), Gaps:0/99(0%)

Query  1    PQVTLWQRPLVTIKIGGQLKEALLDTGADDTVLEEMSLPGRWKPKMIGGIGGFIKVRQYD  60
            PQ+TLWQRPLVTIKIGGQLKEALLDTGADDTVLEE++LPG+WKPKMIGGIGGFIKVRQYD
Sbjct  66   PQITLWQRPLVTIKIGGQLKEALLDTGADDTVLEEINLPGKWKPKMIGGIGGFIKVRQYD  245

Query  61   QILIEICGHKAIGTVLVGPTPVNIIGRNLLTQIGCTLNF  99
            QI IEICG KAIGTVLVGPTPVNIIGRNLLTQ+GCTLNF
Sbjct  246  QIPIEICGKKAIGTVLVGPTPVNIIGRNLLTQLGCTLNF  362
```

```
>Providencia rettgeri strain AFE422_S1 40_contig_34, whole genome shotgun sequence
Sequence ID: NZ_SHDL01000034.1 Length: 1058
Range 1: 6 to 302

Score:186 bits(471), Expect:4e-55,
Method:Compositional matrix adjust.,
Identities:88/99(89%), Positives:97/99(97%), Gaps:0/99(0%)

Query  1    PQVTLWQRPLVTIKIGGQLKEALLDTGADDTVLEEMSLPGRWKPKMIGGIGGFIKVRQYD  60
            PQ+TLWQRPLV+IK+GGQ+KEALLDTGADDTVLEE++LPG+WKPKMIGGIGGFIKVRQYD
Sbjct  6    PQITLWQRPLVSIKVGGQVKEALLDTGADDTVLEEINLPGKWKPKMIGGIGGFIKVRQYD  185

Query  61   QILIEICGHKAIGTVLVGPTPVNIIGRNLLTQIGCTLNF  99
            QI IEICG KAIGTVLVGPTPVNIIGRN+LTQ+GCTLNF
Sbjct  186  QIPIEICGKKAIGTVLVGPTPVNIIGRNMLTQLGCTLNF  302
```

```
>Methylocella silvestris strain TVC NODE_58_length_1453_cov_3.56561, whole genome shotgun sequence
Sequence ID: NZ_PDZR01000058.1 Length: 1453
Range 1: 82 to 378

Score:189 bits(479), Expect:6e-55,
Method:Compositional matrix adjust.,
Identities:89/99(90%), Positives:96/99(96%), Gaps:0/99(0%)

Query  1    PQVTLWQRPLVTIKIGGQLKEALLDTGADDTVLEEMSLPGRWKPKMIGGIGGFIKVRQYD  60
            PQ+TLWQRP+V I+IGGQLKEALLDTGADDTVLE+M LPGRWKPKMIGGIGGFIKVRQYD
Sbjct  82   PQITLWQRPVVEIRIGGQLKEALLDTGADDTVLEDMQLPGRWKPKMIGGIGGFIKVRQYD  261

Query  61   QILIEICGHKAIGTVLVGPTPVNIIGRNLLTQIGCTLNF  99
            ++ IEICGHKAIGTVLVGPTPVNIIGRNLLTQ+GCTLNF
Sbjct  262  EVPIEICGHKAIGTVLVGPTPVNIIGRNLLTQLGCTLNF  378
```

```
>Providencia rettgeri strain AFE97_S1 21_contig_71, whole genome shotgun sequence
Sequence ID: NZ_SHDO01000071.1 Length: 1157
Range 1: 67 to 363

Score:186 bits(471), Expect:1e-54,
Method:Compositional matrix adjust.,
Identities:88/99(89%), Positives:97/99(97%), Gaps:0/99(0%)

Query  1    PQVTLWQRPLVTIKIGGQLKEALLDTGADDTVLEEMSLPGRWKPKMIGGIGGFIKVRQYD  60
            PQ+TLWQRPLV+IK+GGQ+KEALLDTGADDTVLEE++LPG+WKPKMIGGIGGFIKVRQYD
Sbjct  67   PQITLWQRPLVSIKVGGQVKEALLDTGADDTVLEEINLPGKWKPKMIGGIGGFIKVRQYD  246

Query  61   QILIEICGHKAIGTVLVGPTPVNIIGRNLLTQIGCTLNF  99
            QI IEICG KAIGTVLVGPTPVNIIGRN+LTQ+GCTLNF
Sbjct  247  QIPIEICGKKAIGTVLVGPTPVNIIGRNMLTQLGCTLNF  363
```

```
>Listeria monocytogenes strain YA00082404 NODE_73_length_1816_cov_5.664890, whole genome shotgun sequence
Sequence ID: NZ_QEXE01000073.1 Length: 1816
Range 1: 202 to 498

Score:189 bits(481), Expect:3e-54,
Method:Compositional matrix adjust.,
Identities:89/99(90%), Positives:97/99(97%), Gaps:0/99(0%)

Query  1    PQVTLWQRPLVTIKIGGQLKEALLDTGADDTVLEEMSLPGRWKPKMIGGIGGFIKVRQYD  60
            PQ+TLWQRP+VTIK+GGQ+KEALLDTGADDTVLE+M LPGRWKPKMIGGIGGFIKVRQYD
Sbjct  202  PQITLWQRPMVTIKVGGQVKEALLDTGADDTVLEDMELPGRWKPKMIGGIGGFIKVRQYD  381

Query  61   QILIEICGHKAIGTVLVGPTPVNIIGRNLLTQIGCTLNF  99
            QI+IEICG KAIGTVLVGPTPVNIIGRN+LTQ+GCTLNF
Sbjct  382  QIVIEICGKKAIGTVLVGPTPVNIIGRNMLTQLGCTLNF  498
```

```
>Pseudomonas aeruginosa strain P60_London_6_VIM_2_11_13, whole genome shotgun sequence
Sequence ID: NZ_CVXD01000084.1 Length: 395
Range 1: 102 to 392

Score:175 bits(443), Expect:1e-53,
Method:Compositional matrix adjust.,
Identities:81/97(84%), Positives:94/97(96%), Gaps:0/97(0%)

Query  3    VTLWQRPLVTIKIGGQLKEALLDTGADDTVLEEMSLPGRWKPKMIGGIGGFIKVRQYDQI  62
            +TLWQRP+V +K+GGQL+EALLDTGADDTVLE+++LPG+WKPKMIGGIGGFIKV+QYDQI
Sbjct  392  ITLWQRPVVKVKVGGQLREALLDTGADDTVLEDINLPGKWKPKMIGGIGGFIKVKQYDQI  213

Query  63   LIEICGHKAIGTVLVGPTPVNIIGRNLLTQIGCTLNF  99
             I+ICGH+A GTVLVGPTPVNIIGRN+LTQIGCTLNF
Sbjct  212  TIDICGHRATGTVLVGPTPVNIIGRNMLTQIGCTLNF  102
```

```
>Enterococcus faecium strain E21_6 NODE_88_length_518_cov_1.29923, whole genome shotgun sequence
Sequence ID: NZ_JAANZJ010000088.1 Length: 518
Range 1: 2 to 304

Score:176 bits(446), Expect:1e-53,
Method:Compositional matrix adjust.,
Identities:85/101(84%), Positives:96/101(95%), Gaps:2/101(1%)

Query  1    PQVTLWQRPLVTIKIGGQLKEALLDTGADDTVLEEM--SLPGRWKPKMIGGIGGFIKVRQ  58
            PQ+TLWQRPLV+IKIGGQ+KEALLDTGADDTVLEE+   LPG+WKPKMIGGIGGFIKV+Q
Sbjct  2    PQITLWQRPLVSIKIGGQIKEALLDTGADDTVLEELGLELPGKWKPKMIGGIGGFIKVKQ  181

Query  59   YDQILIEICGHKAIGTVLVGPTPVNIIGRNLLTQIGCTLNF  99
            Y++I IEICG +AIGTVLVGPTPVNIIGRN+LTQ+GCTLNF
Sbjct  182  YEEIPIEICGKRAIGTVLVGPTPVNIIGRNMLTQLGCTLNF  304
```

>Methylobacterium radiotolerans strain MAMP 4754 MP-M-radiotolerans_contig_206, whole genome shotgun sequence
Sequence ID: NZ_NKQS01000204.1 Length: 483
Range 1: 192 to 482

Score:175 bits(443), Expect:2e-53,
Method:Compositional matrix adjust.,
Identities:82/97(85%), Positives:94/97(96%), Gaps:0/97(0%)

```
Query  3    VTLWQRPLVTIKIGGQLKEALLDTGADDTVLEEMSLPGRWKPKMIGGIGGFIKVRQYDQI  62
            +TLWQRPLV+IK+ GQ+KEALLDTGADDTVLE+M+LPG+WKP+MIGGIGGFIKVRQY+QI
Sbjct  482  ITLWQRPLVSIKVEGQIKEALLDTGADDTVLEDMNLPGKWKPRMIGGIGGFIKVRQYEQI  303

Query  63   LIEICGHKAIGTVLVGPTPVNIIGRNLLTQIGCTLNF  99
             IEICG KAIGTVLVGPTP+NIIGRN+LTQ+GCTLNF
Sbjct  302  NIEICGKKAIGTVLVGPTPINIIGRNMLTQLGCTLNF  192
```

>Mycobacterium tuberculosis variant bovis strain CSURQ1132, whole genome shotgun sequence
Sequence ID: NZ_CAJEUK010000155.1 Length: 1524
Range 1: 1011 to 1307

Score:184 bits(468), Expect:4e-53,
Method:Compositional matrix adjust.,
Identities:90/99(91%), Positives:95/99(95%), Gaps:0/99(0%)

```
Query  1     PQVTLWQRPLVTIKIGGQLKEALLDTGADDTVLEEMSLPGRWKPKMIGGIGGFIKVRQYD  60
             PQ+TLWQRPLVT++I GQL EALLDTGADDTVLEE++LPGRWKPKMIGGIGGFIKVRQYD
Sbjct  1307  PQITLWQRPLVTVRIEGQLIEALLDTGADDTVLEEINLPGRWKPKMIGGIGGFIKVRQYD  1128

Query  61    QILIEICGHKAIGTVLVGPTPVNIIGRNLLTQIGCTLNF  99
             QI IEICG KAIGTVLVGPTPVNIIGRNLLTQIGCTLNF
Sbjct  1127  QISIEICGKKAIGTVLVGPTPVNIIGRNLLTQIGCTLNF  1011
```

Range 2: 1011 to 1307

Score:184 bits(468), Expect:4e-53,
Method:Compositional matrix adjust.,
Identities:90/99(91%), Positives:95/99(95%), Gaps:0/99(0%)

```
Query  1     PQVTLWQRPLVTIKIGGQLKEALLDTGADDTVLEEMSLPGRWKPKMIGGIGGFIKVRQYD  60
             PQ+TLWQRPLVT++I GQL EALLDTGADDTVLEE++LPGRWKPKMIGGIGGFIKVRQYD
Sbjct  1307  PQITLWQRPLVTVRIEGQLIEALLDTGADDTVLEEINLPGRWKPKMIGGIGGFIKVRQYD  1128

Query  61    QILIEICGHKAIGTVLVGPTPVNIIGRNLLTQIGCTLNF  99
             QI IEICG KAIGTVLVGPTPVNIIGRNLLTQIGCTLNF
Sbjct  1127  QISIEICGKKAIGTVLVGPTPVNIIGRNLLTQIGCTLNF  1011
```

>Levilactobacillus brevis strain CRL 2013 scaffold12, whole genome shotgun sequence
Sequence ID: NZ_MZMW01000016.1 Length: 1126
Range 1: 62 to 358

Score:181 bits(459), Expect:6e-53,
Method:Compositional matrix adjust.,
Identities:86/99(87%), Positives:94/99(94%), Gaps:0/99(0%)

```
Query  1    PQVTLWQRPLVTIKIGGQLKEALLDTGADDTVLEEMSLPGRWKPKMIGGIGGFIKVRQYD  60
            PQ+TLWQRPLVT K+GGQL EALLDTGADDTVLE+++L G+WKPKMIGGIGGFIKV+QYD
Sbjct  62   PQITLWQRPLVTXKVGGQLXEALLDTGADDTVLEDINLXGKWKPKMIGGIGGFIKVKQYD  241

Query  61   QILIEICGHKAIGTVLVGPTPVNIIGRNLLTQIGCTLNF  99
             ILIEICGHKAIGTVLVGPTP NIIGRNLLTQ+GCTLNF
Sbjct  242  NILIEICGHKAIGTVLVGPTPXNIIGRNLLTQLGCTLNF  358
```

>Klebsiella pneumoniae strain NUBRI-K 73, whole genome shotgun sequence
Sequence ID: NZ_SOYT01000073.1 Length: 1717
Range 1: 155 to 451

Score:185 bits(470), Expect:6e-53,
Method:Compositional matrix adjust.,
Identities:85/99(86%), Positives:98/99(98%), Gaps:0/99(0%)

```
Query  1    PQVTLWQRPLVTIKIGGQLKEALLDTGADDTVLEEMSLPGRWKPKMIGGIGGFIKVRQYD  60
            PQ+TLWQRPLV+IK+GGQ++EALLDTGADDTVLE+++LPG+WKPKMIGGIGGFIKVRQYD
Sbjct  155  PQITLWQRPLVSIKVGGQIREALLDTGADDTVLEDINLPGKWKPKMIGGIGGFIKVRQYD  334

Query  61   QILIEICGHKAIGTVLVGPTPVNIIGRNLLTQIGCTLNF  99
            +ILIEICG KAIG+VLVGPTPVNIIGRN+LTQ+GCTLNF
Sbjct  335  EILIEICGKKAIGSVLVGPTPVNIIGRNMLTQLGCTLNF  451
```

>Enterococcus faecium strain A209R1B0C1 NODE_70_length_1819_cov_3.01832_ID_139, whole genome shotgun sequence
Sequence ID: NZ_PGCR01000088.1 Length: 1819
Range 1: 189 to 485

Score:186 bits(471), Expect:7e-53,
Method:Compositional matrix adjust.,
Identities:88/99(89%), Positives:97/99(97%), Gaps:0/99(0%)

```
Query  1    PQVTLWQRPLVTIKIGGQLKEALLDTGADDTVLEEMSLPGRWKPKMIGGIGGFIKVRQYD  60
            PQ+TLWQRPLVTI++GGQ+KEALLDTGADDTVLE+++LPG+WKPKMIGGIGGFIKVRQYD
Sbjct  189  PQITLWQRPLVTIRVGGQIKEALLDTGADDTVLEDINLPGKWKPKMIGGIGGFIKVRQYD  368

Query  61   QILIEICGHKAIGTVLVGPTPVNIIGRNLLTQIGCTLNF  99
            QI IEICG KAIGTVLVGPTPVNIIGRNLLTQ+GCTLNF
Sbjct  369  QIPIEICGKKAIGTVLVGPTPVNIIGRNLLTQLGCTLNF  485
```

>Acinetobacter baumannii strain ACM-32 NODE_81_length_654_cov_1.01708, whole genome shotgun sequence
Sequence ID: NZ_PYDR01000081.1 Length: 654
Range 1: 21 to 317

Score:176 bits(445), Expect:8e-53,
Method:Compositional matrix adjust.,
Identities:82/99(83%), Positives:95/99(95%), Gaps:0/99(0%)

```
Query  1    PQVTLWQRPLVTIKIGGQLKEALLDTGADDTVLEEMSLPGRWKPKMIGGIGGFIKVRQYD  60
            PQ+T WQRPLV IK+GGQ+KEALLDTGADDTVLE+++LPG+WKP+MIGGIGGFIKVRQY+
Sbjct  21   PQITPWQRPLVPIKVGGQIKEALLDTGADDTVLEDINLPGKWKPRMIGGIGGFIKVRQYE  200

Query  61   QILIEICGHKAIGTVLVGPTPVNIIGRNLLTQIGCTLNF  99
            QI IEICG KAIGT+LVGPTP+NIIGRN+LTQ+GCTLNF
Sbjct  201  QIPIEICGKKAIGTILVGPTPINIIGRNMLTQLGCTLNF  317
```

>Mycobacterium tuberculosis variant bovis strain CSURQ1134, whole genome shotgun sequence
Sequence ID: NZ_CAJEUT010000109.1 Length: 1697
Range 1: 973 to 1269

Score:184 bits(468), Expect:1e-52,
Method:Compositional matrix adjust.,
Identities:90/99(91%), Positives:95/99(95%), Gaps:0/99(0%)

```
Query  1     PQVTLWQRPLVTIKIGGQLKEALLDTGADDTVLEEMSLPGRWKPKMIGGIGGFIKVRQYD  60
             PQ+TLWQRPLVT++I GQL EALLDTGADDTVLEE++LPGRWKPKMIGGIGGFIKVRQYD
Sbjct  1269  PQITLWQRPLVTVRIEGQLIEALLDTGADDTVLEEINLPGRWKPKMIGGIGGFIKVRQYD  1090

Query  61    QILIEICGHKAIGTVLVGPTPVNIIGRNLLTQIGCTLNF  99
             QI IEICG KAIGTVLVGPTPVNIIGRNLLTQIGCTLNF
Sbjct  1089  QISIEICGKKAIGTVLVGPTPVNIIGRNLLTQIGCTLNF  973
```

Range 2: 973 to 1269

Score:184 bits(468), Expect:1e-52,
Method:Compositional matrix adjust.,
Identities:90/99(91%), Positives:95/99(95%), Gaps:0/99(0%)

```
Query  1     PQVTLWQRPLVTIKIGGQLKEALLDTGADDTVLEEMSLPGRWKPKMIGGIGGFIKVRQYD  60
             PQ+TLWQRPLVT++I GQL EALLDTGADDTVLEE++LPGRWKPKMIGGIGGFIKVRQYD
Sbjct  1269  PQITLWQRPLVTVRIEGQLIEALLDTGADDTVLEEINLPGRWKPKMIGGIGGFIKVRQYD  1090

Query  61    QILIEICGHKAIGTVLVGPTPVNIIGRNLLTQIGCTLNF  99
             QI IEICG KAIGTVLVGPTPVNIIGRNLLTQIGCTLNF
Sbjct  1089  QISIEICGKKAIGTVLVGPTPVNIIGRNLLTQIGCTLNF  973
```

>Providencia rettgeri strain APW139_S1 24_contig_65, whole genome shotgun sequence
Sequence ID: NZ_SHDG01000065.1 Length: 941
Range 1: 9 to 302

Score:177 bits(448), Expect:4e-52,
Method:Compositional matrix adjust.,
Identities:84/98(86%), Positives:93/98(94%), Gaps:0/98(0%)

```
Query  2    QVTLWQRPLVTIKIGGQLKEALLDTGADDTVLEEMSLPGRWKPKMIGGIGGFIKVRQYDQ  61
            ++TLW+RPL TIK+G Q+KEALLDTGADDTVLEE++LPG+WKPKMIGGIGGFIKVRQYD
Sbjct  9    RITLWRRPLDTIKVGSQIKEALLDTGADDTVLEEINLPGKWKPKMIGGIGGFIKVRQYDH  188

Query  62   ILIEICGHKAIGTVLVGPTPVNIIGRNLLTQIGCTLNF  99
            I IEICG KAIGTVLVGPTPVNIIGRNLLTQ+GCTLNF
Sbjct  189  IPIEICGKKAIGTVLVGPTPVNIIGRNLLTQLGCTLNF  302
```

>Levilactobacillus brevis strain CRL 2013 scaffold13, whole genome shotgun sequence
Sequence ID: NZ_MZMW01000017.1 Length: 1030
Range 1: 1 to 297

Score:177 bits(448), Expect:1e-51,
Method:Compositional matrix adjust.,
Identities:86/99(87%), Positives:91/99(91%), Gaps:0/99(0%)

```
Query  1    PQVTLWQRPLVTIKIGGQLKEALLDTGADDTVLEEMSLPGRWKPKMIGGIGGFIKVRQYD  60
            PQ+TLWQRPLVTIK+G QLKEALLDTGADDTVLE ++LPG WKPKMIGGIGGFIKV QYD
Sbjct  1    PQITLWQRPLVTIKVGXQLKEALLDTGADDTVLEXINLPGXWKPKMIGGIGGFIKVXQYD  180

Query  61   QILIEICGHKAIGTVLVGPTPVNIIGRNLLTQIGCTLNF  99
             I IEICGH A GTVLVGPTPVNIIGRNLLTQ+GCTLNF
Sbjct  181  XIPIEICGHXAXGTVLVGPTPVNIIGRNLLTQLGCTLNF  297
```

>Escherichia coli strain VRES0669, whole genome shotgun sequence
Sequence ID: NZ_UINV01000086.1 Length: 5544
Range 1: 4877 to 5173

Score:186 bits(472), Expect:1e-51,
Method:Compositional matrix adjust.,
Identities:88/99(89%), Positives:97/99(97%), Gaps:0/99(0%)

```
Query  1     PQVTLWQRPLVTIKIGGQLKEALLDTGADDTVLEEMSLPGRWKPKMIGGIGGFIKVRQYD  60
             PQ+TLWQRPLV+IK+GGQ+KEALLDTGADDTVLEE++LPG+WKPKMIGGIGGFIKVRQYD
Sbjct  5173  PQITLWQRPLVSIKVGGQIKEALLDTGADDTVLEEINLPGKWKPKMIGGIGGFIKVRQYD  4994

Query  61    QILIEICGHKAIGTVLVGPTPVNIIGRNLLTQIGCTLNF  99
             QI IEICG KAIGTVLVGPTPVNIIGRN+LTQ+GCTLNF
Sbjct  4993  QIPIEICGKKAIGTVLVGPTPVNIIGRNMLTQLGCTLNF  4877
```

>Enterococcus faecium strain A206R2B0 NODE_116_length_1761_cov_4.01102_ID_231, whole genome shotgun sequence
Sequence ID: NZ_PGCU01000052.1 Length: 1761
Range 1: 173 to 469

Score:181 bits(460), Expect:2e-51,
Method:Compositional matrix adjust.,
Identities:86/99(87%), Positives:96/99(96%), Gaps:0/99(0%)

```
Query  1    PQVTLWQRPLVTIKIGGQLKEALLDTGADDTVLEEMSLPGRWKPKMIGGIGGFIKVRQYD  60
            PQ+TLWQRPLV+IKI GQ+KEALLDTGADDTVLEE++LPG+WKPKMIGGIGGFIKVRQY+
Sbjct  173  PQITLWQRPLVSIKIEGQVKEALLDTGADDTVLEEINLPGKWKPKMIGGIGGFIKVRQYE  352

Query  61   QILIEICGHKAIGTVLVGPTPVNIIGRNLLTQIGCTLNF  99
            +ILIEICG KA GTVLVGPTPVNIIGRN+LTQ+GCTLNF
Sbjct  353  EILIEICGKKARGTVLVGPTPVNIIGRNMLTQLGCTLNF  469
```

>Pseudomonas aeruginosa strain VET-121 NODE_73_length_1192_cov_8.02063_ID_145, whole genome shotgun sequence

```
Sequence ID: NZ_QZXB01000185.1 Length: 1158
Range 1: 825 to 1118

Score:174 bits(441), Expect:3e-50,
Method:Compositional matrix adjust.,
Identities:84/98(86%), Positives:92/98(93%), Gaps:0/98(0%)

Query  2     QVTLWQRPLVTIKIGGQLKEALLDTGADDTVLEEMSLPGRWKPKMIGGIGGFIKVRQYDQ  61
             Q+TLWQRPLVT++I  QL EALLDTGADDTVLE++ LPG+WKPKMIGGIGGFIKVRQYDQ
Sbjct  1118  QITLWQRPLVTVRIEEQLIEALLDTGADDTVLEDIDLPGKWKPKMIGGIGGFIKVRQYDQ  939

Query  62    ILIEICGHKAIGTVLVGPTPVNIIGRNLLTQIGCTLNF  99
             I IEICG KAIGTVLVGPTPVNIIGRN+LT+IGCTLNF
Sbjct  938   IHIEICGKKAIGTVLVGPTPVNIIGRNILTKIGCTLNF  825
```

```
>Acinetobacter baumannii strain ACM-23 NODE_77_length_505_cov_0.960317, whole genome shotgun sequence
Sequence ID: NZ_PYDJ01000077.1 Length: 505
Range 1: 236 to 505

Score:167 bits(423), Expect:4e-50,
Method:Compositional matrix adjust.,
Identities:80/90(89%), Positives:87/90(96%), Gaps:0/90(0%)

Query  10   LVTIKIGGQLKEALLDTGADDTVLEEMSLPGRWKPKMIGGIGGFIKVRQYDQILIEICGH  69
            +V I+IGGQLKEALLDTGADDTVLE+M LPGRWKPKMIGGIGGFIKVRQYD++ IEICGH
Sbjct  505  VVEIRIGGQLKEALLDTGADDTVLEDMQLPGRWKPKMIGGIGGFIKVRQYDEVPIEICGH  326

Query  70   KAIGTVLVGPTPVNIIGRNLLTQIGCTLNF  99
            KAIGTVL+GPTPVNIIGRNLLTQ+GCTLNF
Sbjct  325  KAIGTVLIGPTPVNIIGRNLLTQLGCTLNF  236
```

```
>Acinetobacter baumannii strain ACM-38 NODE_58_length_652_cov_1.55048, whole genome shotgun sequence
Sequence ID: NZ_QAGQ01000058.1 Length: 652
Range 1: 304 to 600

Score:167 bits(423), Expect:2e-49,
Method:Compositional matrix adjust.,
Identities:81/99(82%), Positives:90/99(90%), Gaps:0/99(0%)

Query  1    PQVTLWQRPLVTIKIGGQLKEALLDTGADDTVLEEMSLPGRWKPKMIGGIGGFIKVRQYD  60
            PQ+T+W  P+VTI++G QLK ALLDTG  DTVLE+M LPGRWKPKMIGGI GFIKVRQYD
Sbjct  600  PQITIWLLPVVTIQVGRQLKVALLDTGEYDTVLEDMQLPGRWKPKMIGGIRGFIKVRQYD  421

Query  61   QILIEICGHKAIGTVLVGPTPVNIIGRNLLTQIGCTLNF  99
            ++ IEICGHKAIGTVLVGPTPVNIIGRNLLTQ+GCTLNF
Sbjct  420  EVPIEICGHKAIGTVLVGPTPVNIIGRNLLTQLGCTLNF  304
```

```
>Pseudomonas aeruginosa strain VET-42 NODE_85_length_1204_cov_6.50932_ID_169, whole genome shotgun sequence
Sequence ID: NZ_RACX01000100.1 Length: 1166
Range 1: 31 to 324

Score:170 bits(431), Expect:1e-48,
Method:Compositional matrix adjust.,
Identities:82/98(84%), Positives:90/98(91%), Gaps:0/98(0%)

Query  2    QVTLWQRPLVTIKIGGQLKEALLDTGADDTVLEEMSLPGRWKPKMIGGIGGFIKVRQYDQ  61
            +  LWQRPLVT++I  QL EALLDTGADDTVLE++ LPG+WKPKMIGGIGGFIKVRQYDQ
Sbjct  31   ETALWQRPLVTVRIEEQLIEALLDTGADDTVLEDIDLPGKWKPKMIGGIGGFIKVRQYDQ  210

Query  62   ILIEICGHKAIGTVLVGPTPVNIIGRNLLTQIGCTLNF  99
            I IEICG KAIGTVLVGPTPVNIIGRN+LT+IGCTLNF
Sbjct  211  IHIEICGKKAIGTVLVGPTPVNIIGRNILTKIGCTLNF  324
```

```
>Mycobacterium tuberculosis variant bovis strain CSURQ1132, whole genome shotgun sequence
Sequence ID: NZ_CAJEUK010000179.1 Length: 862
Range 1: 83 to 376

Score:166 bits(421), Expect:2e-48,
Method:Compositional matrix adjust.,
Identities:83/98(85%), Positives:90/98(91%), Gaps:0/98(0%)

Query  2    QVTLWQRPLVTIKIGGQLKEALLDTGADDTVLEEMSLPGRWKPKMIGGIGGFIKVRQYDQ  61
            Q T+ +  +V IK GGQ+KEALLDTGADDTVLEE+ LPGRWKPKMIGGIGGFIKVRQYDQ
Sbjct  83   QDTVEEGEIV*IKRGGQVKEALLDTGADDTVLEEIDLPGRWKPKMIGGIGGFIKVRQYDQ  262

Query  62   ILIEICGHKAIGTVLVGPTPVNIIGRNLLTQIGCTLNF  99
            + +EICGHKAIGTVLVGPTPVNIIGRNLLTQIGCTLNF
Sbjct  263  VQVEICGHKAIGTVLVGPTPVNIIGRNLLTQIGCTLNF  376
```

```
Range 2: 83 to 376

Score:166 bits(421), Expect:2e-48,
Method:Compositional matrix adjust.,
Identities:83/98(85%), Positives:90/98(91%), Gaps:0/98(0%)

Query  2    QVTLWQRPLVTIKIGGQLKEALLDTGADDTVLEEMSLPGRWKPKMIGGIGGFIKVRQYDQ  61
            Q T+ +  +V IK GGQ+KEALLDTGADDTVLEE+ LPGRWKPKMIGGIGGFIKVRQYDQ
Sbjct  83   QDTVEEGEIV*IKRGGQVKEALLDTGADDTVLEEIDLPGRWKPKMIGGIGGFIKVRQYDQ  262

Query  62   ILIEICGHKAIGTVLVGPTPVNIIGRNLLTQIGCTLNF  99
            + +EICGHKAIGTVLVGPTPVNIIGRNLLTQIGCTLNF
Sbjct  263  VQVEICGHKAIGTVLVGPTPVNIIGRNLLTQIGCTLNF  376
```

```
>Escherichia coli strain YA00028237 contig_187, whole genome shotgun sequence
Sequence ID: NZ_NGBP02000187.1 Length: 1828
Range 1: 1326 to 1622

Score:173 bits(438), Expect:5e-48,
Method:Compositional matrix adjust.,
Identities:75/99(76%), Positives:94/99(94%), Gaps:0/99(0%)

Query  1     PQVTLWQRPLVTIKIGGQLKEALLDTGADDTVLEEMSLPGRWKPKMIGGIGGFIKVRQYD  60
             PQ+TLWQRP VT+K+GGQ++EAL+DTGADDTVLE+++LPG+W PK+IGGIGGF+KV+QYD
Sbjct  1622  PQITLWQRPFVTVKVGGQIREALIDTGADDTVLEDINLPGKWTPKIIGGIGGFVKVKQYD  1443

Query  61    QILIEICGHKAIGTVLVGPTPVNIIGRNLLTQIGCTLNF  99
```

```
             I+IE+CG KAIG+VLVGPTP NIIGRN+L+Q+GCTLNF
Sbjct  1442  HIVIEMCGKKAIGSVLVGPTPANIIGRNMLSQLGCTLNF  1326
```

```
>Acinetobacter baumannii strain ACM-29 NODE_109_length_888_cov_0.827858, whole genome shotgun sequence
Sequence ID: NZ_PYDO01000109.1 Length: 888
Range 1: 626 to 886

Score:163 bits(413), Expect:6e-47,
Method:Compositional matrix adjust.,
Identities:77/87(89%), Positives:84/87(96%), Gaps:0/87(0%)

Query  13   IKIGGQLKEALLDTGADDTVLEEMSLPGRWKPKMIGGIGGFIKVRQYDQILIEICGHKAI  72
            IK+GGQLKEALLDTGA+DTVLE+M LPGRWKPKMIGGI GFIKVRQYD++ IEICGHKAI
Sbjct  886  IKVGGQLKEALLDTGAEDTVLEDMQLPGRWKPKMIGGIRGFIKVRQYDEVPIEICGHKAI  707

Query  73   GTVLVGPTPVNIIGRNLLTQIGCTLNF  99
            GTVL+GPTPVNIIGRNLLTQ+GCTLNF
Sbjct  706  GTVLIGPTPVNIIGRNLLTQLGCTLNF  626
```

```
>Escherichia coli strain 8374wB11 NODE_34_length_1310_cov_3.21048, whole genome shotgun sequence
Sequence ID: NZ_WSJO01000034.1 Length: 1310
Range 1: 1024 to 1308

Score:166 bits(421), Expect:8e-47,
Method:Compositional matrix adjust.,
Identities:78/95(82%), Positives:86/95(90%), Gaps:0/95(0%)

Query  5     LWQRPLVTIKIGGQLKEALLDTGADDTVLEEMSLPGRWKPKMIGGIGGFIKVRQYDQILI  64
             LW RP V ++IGGQ +EALLD GADD VLEE++LPG+WKPKMIGGIGGFIKVRQYDQI I
Sbjct  1308  LWHRPFVPVRIGGQPREALLDNGADDPVLEEINLPGKWKPKMIGGIGGFIKVRQYDQIPI  1129

Query  65    EICGHKAIGTVLVGPTPVNIIGRNLLTQIGCTLNF  99
             EICGHK IGTVL+G TPVNIIGRNLLTQ+GCTLNF
Sbjct  1128  EICGHKVIGTVLIGSTPVNIIGRNLLTQLGCTLNF  1024
```

```
>Providencia rettgeri strain AFE85_S2 19_contig_46, whole genome shotgun sequence
Sequence ID: NZ_SHDQ01000046.1 Length: 846
Range 1: 375 to 632

Score:160 bits(405), Expect:7e-46,
Method:Compositional matrix adjust.,
Identities:76/86(88%), Positives:84/86(97%), Gaps:0/86(0%)

Query  14   KIGGQLKEALLDTGADDTVLEEMSLPGRWKPKMIGGIGGFIKVRQYDQILIEICGHKAIG  73
            ++GGQ+KEALLDTGADDTVLEE++LPG+WKPKMIGGIGGFIKVRQYD ILIEICG KAIG
Sbjct  632  EVGGQIKEALLDTGADDTVLEEINLPGKWKPKMIGGIGGFIKVRQYDHILIEICGKKAIG  453

Query  74   TVLVGPTPVNIIGRNLLTQIGCTLNF  99
            TVLVGPTPVNIIGRN+LTQ+GCTLNF
Sbjct  452  TVLVGPTPVNIIGRNMLTQLGCTLNF  375
```

```
>Providencia rettgeri strain AFE97_S1 21_contig_72, whole genome shotgun sequence
Sequence ID: NZ_SHDO01000072.1 Length: 854
Range 1: 3 to 263

Score:160 bits(405), Expect:7e-46,
Method:Compositional matrix adjust.,
Identities:76/87(87%), Positives:85/87(97%), Gaps:0/87(0%)

Query  13   IKIGGQLKEALLDTGADDTVLEEMSLPGRWKPKMIGGIGGFIKVRQYDQILIEICGHKAI  72
            IKIGGQ+KEALLDTGADDTVLEE++LPG+WKPKMIGGIGGFIKVRQY++I IEICG KAI
Sbjct  3    IKIGGQIKEALLDTGADDTVLEEINLPGKWKPKMIGGIGGFIKVRQYEEIPIEICGKKAI  182

Query  73   GTVLVGPTPVNIIGRNLLTQIGCTLNF  99
            G+VLVGPTPVNIIGRN+LTQ+GCTLNF
Sbjct  183  GSVLVGPTPVNIIGRNMLTQLGCTLNF  263
```

```
>Providencia rettgeri strain AFE96_S1 41_contig_60, whole genome shotgun sequence
Sequence ID: NZ_SHDP01000060.1 Length: 1101
Range 1: 96 to 353

Score:162 bits(409), Expect:1e-45,
Method:Compositional matrix adjust.,
Identities:76/86(88%), Positives:85/86(98%), Gaps:0/86(0%)

Query  14   KIGGQLKEALLDTGADDTVLEEMSLPGRWKPKMIGGIGGFIKVRQYDQILIEICGHKAIG  73
            K+GGQ+KEALLDTGADDTVLE+++LPG+WKP+MIGGIGGFIKVRQYDQILIEICG KAIG
Sbjct  96   KVGGQVKEALLDTGADDTVLEDINLPGKWKPRMIGGIGGFIKVRQYDQILIEICGKKAIG  275

Query  74   TVLVGPTPVNIIGRNLLTQIGCTLNF  99
            TVLVGPTPVNIIGRN+LTQ+GCTLNF
Sbjct  276  TVLVGPTPVNIIGRNMLTQLGCTLNF  353
```

```
>Klebsiella pneumoniae strain EuSCAPE_IL004, whole genome shotgun sequence
Sequence ID: NZ_UKJP02000108.1 Length: 356
Range 1: 1 to 240

Score:151 bits(382), Expect:1e-44,
Method:Compositional matrix adjust.,
Identities:72/80(90%), Positives:78/80(97%), Gaps:0/80(0%)

Query  1    PQVTLWQRPLVTIKIGGQLKEALLDTGADDTVLEEMSLPGRWKPKMIGGIGGFIKVRQYD  60
            PQ+TLWQRPLV+IK+GGQ KEALLDTGADDTVLEE++LPG+WKPKMIGGIGGFIKVRQYD
Sbjct  240  PQITLWQRPLVSIKVGGQTKEALLDTGADDTVLEEINLPGKWKPKMIGGIGGFIKVRQYD  61

Query  61   QILIEICGHKAIGTVLVGPT  80
            QILIEICG KAIGTVLVGPT
Sbjct  60   QILIEICGKKAIGTVLVGPT  1
```

```
>Acinetobacter baumannii strain ACM-5 NODE_55_length_1274_cov_1.90584, whole genome shotgun sequence
Sequence ID: NZ_PYCX01000055.1 Length: 1274
Range 1: 1019 to 1273
```

```
Score:157 bits(397), Expect:2e-43,
Method:Compositional matrix adjust.,
Identities:74/85(87%), Positives:81/85(95%), Gaps:0/85(0%)

Query  15    IGGQLKEALLDTGADDTVLEEMSLPGRWKPKMIGGIGGFIKVRQYDQILIEICGHKAIGT  74
             + GQLKEALLDTGA+DTVLE+M LPGRWKPKMIGGI GFIKVRQYD++ IEICGHKAIGT
Sbjct  1273  VEGQLKEALLDTGAEDTVLEDMQLPGRWKPKMIGGIRGFIKVRQYDEVPIEICGHKAIGT  1094

Query  75    VLVGPTPVNIIGRNLLTQIGCTLNF  99
             VL+GPTPVNIIGRNLLTQ+GCTLNF
Sbjct  1093  VLIGPTPVNIIGRNLLTQLGCTLNF  1019




>Mycobacterium tuberculosis strain 32845_S29 230, whole genome shotgun sequence
Sequence ID: NZ_QTBV01000049.1 Length: 601
Range 1: 367 to 600

Score:150 bits(379), Expect:6e-43,
Method:Compositional matrix adjust.,
Identities:72/78(92%), Positives:77/78(98%), Gaps:0/78(0%)

Query  22   ALLDTGADDTVLEEMSLPGRWKPKMIGGIGGFIKVRQYDQILIEICGHKAIGTVLVGPTP  81
            ALLDTGADDTVLE+++LPG+WKPKMIGGIGGFIKVRQYDQILIEICG KAIGTVLVGPTP
Sbjct  600  ALLDTGADDTVLEDINLPGKWKPKMIGGIGGFIKVRQYDQILIEICGKKAIGTVLVGPTP  421

Query  82   VNIIGRNLLTQIGCTLNF  99
            VNIIGRN+LTQIGCTLNF
Sbjct  420  VNIIGRNMLTQIGCTLNF  367




>Cellulophaga baltica strain 14 NODE_63_length_422_cov_10.206780, whole genome shotgun sequence
Sequence ID: NZ_VIQP01000062.1 Length: 422
Range 1: 1 to 246

Score:144 bits(363), Expect:2e-41,
Method:Compositional matrix adjust.,
Identities:70/82(85%), Positives:78/82(95%), Gaps:0/82(0%)

Query  1    PQVTLWQRPLVTIKIGGQLKEALLDTGADDTVLEEMSLPGRWKPKMIGGIGGFIKVRQYD  60
            PQ+TLWQRPLVT++I GQL EALLDTGADDTVLE+++LPG+WKPKMIGGIGGFIKVRQYD
Sbjct  246  PQITLWQRPLVTVRIEGQLIEALLDTGADDTVLEDINLPGKWKPKMIGGIGGFIKVRQYD  67

Query  61   QILIEICGHKAIGTVLVGPTPV  82
            QIL+EI G KAIGTVLVGPTPV
Sbjct  66   QILMEIHGKKAIGTVLVGPTPV  1




>Enterococcus faecium strain A201R2B0 NODE_60_length_1589_cov_4.20178_ID_119, whole genome shotgun sequence
Sequence ID: NZ_PGCT01000032.1 Length: 1589
Range 1: 1331 to 1585

Score:151 bits(381), Expect:3e-40,
Method:Compositional matrix adjust.,
Identities:72/85(85%), Positives:81/85(95%), Gaps:0/85(0%)

Query  15    IGGQLKEALLDTGADDTVLEEMSLPGRWKPKMIGGIGGFIKVRQYDQILIEICGHKAIGT  74
             I GQ+KEALL+TG DDTVLEE++LPG+WKPKMIGGIGGFIKVRQY++ILIEICG KA GT
Sbjct  1585  IEGQVKEALLETGEDDTVLEEINLPGKWKPKMIGGIGGFIKVRQYEEILIEICGKKARGT  1406

Query  75    VLVGPTPVNIIGRNLLTQIGCTLNF  99
             VLVGPTPVNIIGRN+LTQ+GCTLNF
Sbjct  1405  VLVGPTPVNIIGRNMLTQLGCTLNF  1331




>Klebsiella pneumoniae strain EuSCAPE_IL005, whole genome shotgun sequence
Sequence ID: NZ_UKJL02000059.1 Length: 323
Range 1: 103 to 321

Score:135 bits(340), Expect:3e-38,
Method:Compositional matrix adjust.,
Identities:63/73(86%), Positives:70/73(95%), Gaps:0/73(0%)

Query  1    PQVTLWQRPLVTIKIGGQLKEALLDTGADDTVLEEMSLPGRWKPKMIGGIGGFIKVRQYD  60
            PQ+TLWQRPLV+IK+GGQ KEALLDTGADDTVLEE++LPG+WKPKMIGGIGGFIKVRQYD
Sbjct  103  PQITLWQRPLVSIKVGGQTKEALLDTGADDTVLEEINLPGKWKPKMIGGIGGFIKVRQYD  282

Query  61   QILIEICGHKAIG  73
            +I IEICG KAIG
Sbjct  283  EITIEICGKKAIG  321




>Klebsiella pneumoniae strain EuSCAPE_LT007, whole genome shotgun sequence
Sequence ID: NZ_UKJI02000072.1 Length: 392
Range 1: 1 to 216

Score:134 bits(336), Expect:2e-37,
Method:Compositional matrix adjust.,
Identities:63/72(88%), Positives:70/72(97%), Gaps:0/72(0%)

Query  1    PQVTLWQRPLVTIKIGGQLKEALLDTGADDTVLEEMSLPGRWKPKMIGGIGGFIKVRQYD  60
            PQ+TLWQRP+VTIK+GGQ+KEALLDTGADDTVLEE++LPG+WKPKMIGGIGGFIKVRQYD
Sbjct  216  PQITLWQRPVVTIKVGGQVKEALLDTGADDTVLEEINLPGKWKPKMIGGIGGFIKVRQYD  37

Query  61   QILIEICGHKAI  72
            QI IEICG KAI
Sbjct  36   QIPIEICGKKAI  1




>Enterococcus durans strain E115_10 NODE_63_length_955_cov_1.30556, whole genome shotgun sequence
Sequence ID: NZ_JAANZI010000063.1 Length: 955
Range 1: 735 to 953

Score:134 bits(336), Expect:3e-35,
Method:Compositional matrix adjust.,
Identities:59/73(81%), Positives:70/73(95%), Gaps:0/73(0%)

Query  27   GADDTVLEEMSLPGRWKPKMIGGIGGFIKVRQYDQILIEICGHKAIGTVLVGPTPVNIIG  86
            GADDT+LE+++LPG+WKPKM GGIGGFIKV+QYDQ+L+EICG K IGTVLVGPTP+NIIG
Sbjct  953  GADDTILEDINLPGKWKPKMAGGIGGFIKVKQYDQVLVEICGQKVIGTVLVGPTPINIIG  774

Query  87   RNLLTQIGCTLNF  99
            RNL+TQ+GCTLNF
```

```
Sbjct  773  RNLMTQLGCTLNF  735
```

>Klebsiella pneumoniae strain ARGID_31501, whole genome shotgun sequence
Sequence ID: NZ_CAKLPD010000006.1 Length: 2655
Range 1: 2 to 223

Score:139 bits(350), Expect:4e-35,
Method:Compositional matrix adjust.,
Identities:67/74(91%), Positives:70/74(94%), Gaps:0/74(0%)

```
Query  26   TGADDTVLEEMSLPGRWKPKMIGGIGGFIKVRQYDQILIEICGHKAIGTVLVGPTPVNII  85
            TGAD TVLE+M L G+WKPKMIGGIGGFIKVRQYDQI IEICGHKAIGTVLVGPTPVNII
Sbjct  2    TGADGTVLEDMQLSGKWKPKMIGGIGGFIKVRQYDQIPIEICGHKAIGTVLVGPTPVNII  181

Query  86   GRNLLTQIGCTLNF  99
            GRNLLTQ+GCTLNF
Sbjct  182  GRNLLTQLGCTLNF  223
```

>Escherichia coli strain 8374wG10 NODE_65_length_433_cov_1.76797, whole genome shotgun sequence
Sequence ID: NZ_WSJQ01000065.1 Length: 433
Range 1: 1 to 231

Score:125 bits(314), Expect:7e-34,
Method:Compositional matrix adjust.,
Identities:56/77(73%), Positives:68/77(88%), Gaps:0/77(0%)

```
Query  23   LLDTGADDTVLEEMSLPGRWKPKMIGGIGGFIKVRQYDQILIEICGHKAIGTVLVGPTPV  82
            +LDTGADDTV E ++LPG+W PK+IGG+GG ++V+QYD ILIE  G KAIG+VLVGPTP
Sbjct  1    ILDTGADDTVFENINLPGKWTPKLIGGVGGLVRVKQYDNILIEFEGRKAIGSVLVGPTPS  180

Query  83   NIIGRNLLTQIGCTLNF  99
            NIIGRN++TQIGCTLNF
Sbjct  181  NIIGRNMMTQIGCTLNF  231
```

>Escherichia coli strain 8374wA11 NODE_44_length_523_cov_1.7197, whole genome shotgun sequence
Sequence ID: NZ_WSPR01000043.1 Length: 523
Range 1: 4 to 195

Score:104 bits(259), Expect:3e-25,
Method:Compositional matrix adjust.,
Identities:46/64(72%), Positives:57/64(89%), Gaps:0/64(0%)

```
Query  36   MSLPGRWKPKMIGGIGGFIKVRQYDQILIEICGHKAIGTVLVGPTPVNIIGRNLLTQIGC  95
            ++LPG+W PK+IGG+GG ++V+QYD ILIE  G KAIG+VLVGPTP NIIGRN++TQIGC
Sbjct  4    INLPGKWTPKLIGGVGGLVRVKQYDNILIEFEGRKAIGSVLVGPTPSNIIGRNMMTQIGC  183

Query  96   TLNF  99
            TLNF
Sbjct  184  TLNF  195
```

>Escherichia coli strain TzEc080 NODE_44_length_523_cov_1.7197, whole genome shotgun sequence
Sequence ID: NZ_WSFZ01000043.1 Length: 523
Range 1: 4 to 195

Score:104 bits(259), Expect:3e-25,
Method:Compositional matrix adjust.,
Identities:46/64(72%), Positives:57/64(89%), Gaps:0/64(0%)

```
Query  36   MSLPGRWKPKMIGGIGGFIKVRQYDQILIEICGHKAIGTVLVGPTPVNIIGRNLLTQIGC  95
            ++LPG+W PK+IGG+GG ++V+QYD ILIE  G KAIG+VLVGPTP NIIGRN++TQIGC
Sbjct  4    INLPGKWTPKLIGGVGGLVRVKQYDNILIEFEGRKAIGSVLVGPTPSNIIGRNMMTQIGC  183

Query  96   TLNF  99
            TLNF
Sbjct  184  TLNF  195
```

>Klebsiella pneumoniae strain EuSCAPE_MT006, whole genome shotgun sequence
Sequence ID: NZ_UKIU02000071.1 Length: 455
Range 1: 306 to 455

Score:95.9 bits(237), Expect:4e-22,
Method:Compositional matrix adjust.,
Identities:45/50(90%), Positives:48/50(96%), Gaps:0/50(0%)

```
Query  50   IGGFIKVRQYDQILIEICGHKAIGTVLVGPTPVNIIGRNLLTQIGCTLNF  99
            IGGFIKVRQYDQI IEICG KAIGTVL+GPTPVNIIGRN+LTQ+GCTLNF
Sbjct  455  IGGFIKVRQYDQIPIEICGKKAIGTVLIGPTPVNIIGRNMLTQLGCTLNF  306
```

>Klebsiella pneumoniae strain EuSCAPE_MT015, whole genome shotgun sequence
Sequence ID: NZ_UKIW02000099.1 Length: 435
Range 1: 300 to 434

Score:88.2 bits(217), Expect:4e-19,
Method:Compositional matrix adjust.,
Identities:42/45(93%), Positives:43/45(95%), Gaps:0/45(0%)

```
Query  55   KVRQYDQILIEICGHKAIGTVLVGPTPVNIIGRNLLTQIGCTLNF  99
            KVRQYDQI IEICG KAIGTVLVGPTPVNIIGRNLLTQ+GCTLNF
Sbjct  434  KVRQYDQITIEICGKKAIGTVLVGPTPVNIIGRNLLTQLGCTLNF  300
```

>Providencia rettgeri strain APW139_S1 24_contig_64, whole genome shotgun sequence
Sequence ID: NZ_SHDG01000064.1 Length: 902
Range 1: 3 to 134

Score:85.9 bits(211), Expect:5e-17,
Method:Compositional matrix adjust.,
Identities:41/44(93%), Positives:42/44(95%), Gaps:0/44(0%)

```
Query  56   VRQYDQILIEICGHKAIGTVLVGPTPVNIIGRNLLTQIGCTLNF  99
            VRQYDQI IEICG KAIGTVLVGPTPVNIIGRNLLTQ+GCTLNF
Sbjct  3    VRQYDQIPIEICGKKAIGTVLVGPTPVNIIGRNLLTQLGCTLNF  134
```

>Acinetobacter baumannii strain ACM-24 NODE_72_length_503_cov_0.531915, whole genome shotgun sequence
Sequence ID: NZ_PYDK01000072.1 Length: 503
Range 1: 370 to 501

Score:82.4 bits(202), Expect:1e-16,
Method:Compositional matrix adjust.,
Identities:39/44(89%), Positives:41/44(93%), Gaps:0/44(0%)

```
Query  56   VRQYDQILIEICGHKAIGTVLVGPTPVNIIGRNLLTQIGCTLNF  99
            VRQY+QI IEICG K IGTVLVGPTPVNIIGRN+LTQIGCTLNF
Sbjct  501  VRQYNQIPIEICGKKTIGTVLVGPTPVNIIGRNILTQIGCTLNF  370
```

>Klebsiella pneumoniae strain EuSCAPE_IL017, whole genome shotgun sequence
Sequence ID: NZ_UKKG02000044.1 Length: 325
Range 1: 202 to 324

Score:76.6 bits(187), Expect:4e-15,
Method:Compositional matrix adjust.,
Identities:34/41(83%), Positives:40/41(97%), Gaps:0/41(0%)

```
Query  1    PQVTLWQRPLVTIKIGGQLKEALLDTGADDTVLEEMSLPGR  41
            PQ+TLWQRPLV+IK+GGQ+KEALLDTGADDTVLE+M LPG+
Sbjct  202  PQITLWQRPLVSIKVGGQIKEALLDTGADDTVLEDMHLPGK  324
```

>Burkholderia pseudomallei strain A-175-05-2-02, whole genome shotgun sequence
Sequence ID: NZ_UVFW01000138.1 Length: 455
Range 1: 139 to 252

Score:72.0 bits(175), Expect:1e-12,
Method:Compositional matrix adjust.,
Identities:34/38(89%), Positives:36/38(94%), Gaps:0/38(0%)

```
Query  62   ILIEICGHKAIGTVLVGPTPVNIIGRNLLTQIGCTLNF  99
            + IEICG KAIGTVLVGPTPVNIIGRN+LTQIGCTLNF
Sbjct  252  LCIEICGKKAIGTVLVGPTPVNIIGRNMLTQIGCTLNF  139
```

>Mycobacterium tuberculosis strain 32845_S29 214, whole genome shotgun sequence
Sequence ID: NZ_QTBV01000045.1 Length: 223
Range 1: 121 to 222

Score:66.6 bits(161), Expect:1e-11,
Method:Compositional matrix adjust.,
Identities:29/34(85%), Positives:34/34(100%), Gaps:0/34(0%)

```
Query  1    PQVTLWQRPLVTIKIGGQLKEALLDTGADDTVLE  34
            PQ+TLWQRPLV+IK+GGQ++EALLDTGADDTVLE
Sbjct  121  PQITLWQRPLVSIKVGGQIREALLDTGADDTVLE  222
```

>Enterobacter hormaechei strain AUH-ENM4 NODE_27_length_809_cov_0.64956, whole genome shotgun sequence
Sequence ID: NZ_PYGZ01000027.1 Length: 809
Range 1: 735 to 809

Score:53.1 bits(126), Expect:5e-05,
Method:Compositional matrix adjust.,
Identities:24/25(96%), Positives:25/25(100%), Gaps:0/25(0%)

```
Query  75   VLVGPTPVNIIGRNLLTQIGCTLNF  99
            VLVGPTPVNIIGRNLLTQ+GCTLNF
Sbjct  809  VLVGPTPVNIIGRNLLTQLGCTLNF  735
```

>Escherichia coli strain 8374wB11 NODE_46_length_516_cov_2.15167, whole genome shotgun sequence
Sequence ID: NZ_WSJO01000046.1 Length: 516
Range 1: 442 to 516

Score:51.6 bits(122), Expect:1e-04,
Method:Compositional matrix adjust.,
Identities:22/25(88%), Positives:25/25(100%), Gaps:0/25(0%)

```
Query  75   VLVGPTPVNIIGRNLLTQIGCTLNF  99
            VLVGPTP+NIIGRN+LTQ+GCTLNF
Sbjct  516  VLVGPTPINIIGRNMLTQLGCTLNF  442
```

>Mycobacterium tuberculosis variant bovis strain CSURQ1139, whole genome shotgun sequence
Sequence ID: NZ_CAJEUO010000157.1 Length: 848
Range 1: 778 to 846

Score:51.2 bits(121), Expect:3e-04,
Method:Compositional matrix adjust.,
Identities:23/23(100%), Positives:23/23(100%), Gaps:0/23(0%)

```
Query  77   VGPTPVNIIGRNLLTQIGCTLNF  99
            VGPTPVNIIGRNLLTQIGCTLNF
Sbjct  846  VGPTPVNIIGRNLLTQIGCTLNF  778
```

Range 2: 778 to 846

Score:51.2 bits(121), Expect:3e-04,
Method:Compositional matrix adjust.,
Identities:23/23(100%), Positives:23/23(100%), Gaps:0/23(0%)

```
Query  77   VGPTPVNIIGRNLLTQIGCTLNF  99
            VGPTPVNIIGRNLLTQIGCTLNF
Sbjct  846  VGPTPVNIIGRNLLTQIGCTLNF  778
```

>Pseudomonas aromaticivorans strain MAP12 34, whole genome shotgun sequence
Sequence ID: NZ_JAHRGL010000035.1 Length: 7031
Range 1: 6356 to 6652

```
Score:48.9 bits(115), Expect:0.002,
Method:Compositional matrix adjust.,
Identities:32/99(32%), Positives:50/99(50%), Gaps:6/99(6%)

Query  7     QRPLVTIKIGGQLKEALLDTGADDTVLEEMSLPGRWKPKMIG----GIGGFIKVRQYDQI  62
             +RP + +KI  ++   L+DTGAD T++ + S P +W  +       GIG   +VRQ
Sbjct  6356  KRPQLKVKINNKVISGLVDTGADVTIITQKSWPQKWPLREANVQFLGIGTLSRVRQSVNS  6535

Query  63    LIEI--CGHKAIGTVLVGPTPVNIIGRNLLTQIGCTLNF  99
             ++ I   G K I    V    +N+ GR+LL Q    +N
Sbjct  6536  VVCIGPEGQKGILKPYVADIAINLWGRDLLQQWNTQINI  6652
```

Query #14: sp|P04585|p6_pol_HV1H2 Gag-Pol polyprotein OS=Human immunodeficiency virus type 1 group M subtype B (isolate HXB2) OX=11706 GN=gag-pol PE=1 SV=4 Query ID: lcl|Query_2387716 Length: 48

Sequences producing significant alignments:

| Description | Scientific Name | Common Name | Taxid | Max Score | Total Score | Query cover | E Value | Per. Ident | Acc. Len | Accession |
|---|---|---|---|---|---|---|---|---|---|---|
| Thalassovita mangrovi strain GS-10 Scaffold19_1, whole genome... | Thalassovita... | NA | 2692236 | 92.8 | 92.8 | 97% | 9e-20 | 95.74 | 2900 | NZ_WWEN010000... |
| Deinococcus aquiradiocola strain JCM 14371 sequence30, whole... | Deinococcus ... | NA | 393059 | 92.4 | 92.4 | 97% | 1e-19 | 95.74 | 6128 | NZ_BMOE010000... |
| Enterococcus alcedinis strain CCM 8433 sequence23, whole genom... | Enterococcus... | NA | 1274384 | 92.4 | 92.4 | 97% | 1e-19 | 95.74 | 6075 | NZ_BMDT010000... |
| Actinoplanes campanulatus strain JCM 3059 sequence058, whole... | Actinoplanes... | NA | 113559 | 92.4 | 92.4 | 97% | 1e-19 | 95.74 | 4754 | NZ_BMPW010000... |
| Klebsiella pneumoniae strain P7701... | Klebsiella p... | NA | 573 | 67.8 | 67.8 | 68% | 5e-11 | 93.94 | 1305 | NZ_JACAWM0100... |
| Helicobacter acinonychis isolate 212_10, whole genome shotgun... | Helicobacter... | NA | 212 | 66.2 | 66.2 | 83% | 2e-10 | 75.00 | 1526 | NZ_FZLW010000... |
| Acinetobacter baumannii strain ACM-35... | Acinetobacte... | NA | 470 | 61.6 | 61.6 | 72% | 8e-09 | 82.86 | 1409 | NZ_QEHN010000... |
| Acinetobacter baumannii strain ACM-36... | Acinetobacte... | NA | 470 | 55.8 | 55.8 | 60% | 4e-07 | 86.21 | 567 | NZ_QAGO010000... |
| Enterobacter hormaechei strain AUH-ENM6... | Enterobacter... | NA | 158836 | 55.8 | 55.8 | 66% | 1e-06 | 81.25 | 1394 | NZ_PYHB010000... |
| Klebsiella pneumoniae strain EuSCAPE_MT015, whole genome shotg... | Klebsiella p... | NA | 573 | 52.0 | 52.0 | 93% | 2e-06 | 61.54 | 281 | NZ_UKIW020001... |
| Klebsiella pneumoniae strain EuSCAPE_IL017, whole genome shotg... | Klebsiella p... | NA | 573 | 52.0 | 52.0 | 93% | 3e-06 | 62.22 | 325 | NZ_UKKG020000... |
| Klebsiella pneumoniae strain NUBRI-K 73, whole genome shotgun... | Klebsiella p... | NA | 573 | 54.3 | 54.3 | 93% | 4e-06 | 63.46 | 1717 | NZ_SOYT010000... |
| Escherichia coli strain YA00028237 contig_187, whole genome... | Escherichia ... | NA | 562 | 54.3 | 54.3 | 93% | 4e-06 | 63.46 | 1828 | NZ_NGBP020001... |
| Klebsiella pneumoniae strain EuSCAPE_LT007, whole genome shotg... | Klebsiella p... | NA | 573 | 52.4 | 52.4 | 97% | 4e-06 | 65.96 | 392 | NZ_UKJI020000... |
| Burkholderia pseudomallei strain A-175-05-2-02, whole genome... | Burkholderia... | NA | 28450 | 50.8 | 50.8 | 91% | 2e-05 | 70.45 | 455 | NZ_UVFW010001... |
| Burkholderia pseudomallei strain 2604STDY5519032... | Burkholderia... | NA | 28450 | 51.2 | 51.2 | 93% | 3e-05 | 62.22 | 656 | NZ_UVID020000... |
| Acinetobacter baumannii strain ACM-14... | Acinetobacte... | NA | 470 | 50.8 | 50.8 | 58% | 5e-05 | 78.57 | 787 | NZ_PYDA010000... |
| Escherichia coli strain 8374wH11... | Escherichia ... | NA | 562 | 50.8 | 50.8 | 58% | 6e-05 | 78.57 | 1397 | NZ_WSJI010000... |
| Enterococcus faecium strain A209R1B0C1... | Enterococcus... | NA | 1352 | 48.5 | 48.5 | 93% | 4e-04 | 62.22 | 1819 | NZ_PGCR010000... |
| Listeria monocytogenes strain YA00082404... | Listeria mon... | NA | 1639 | 47.8 | 47.8 | 93% | 6e-04 | 57.69 | 1816 | NZ_QEXE010000... |
| Methylocella silvestris strain TVC... | Methylocella... | NA | 199596 | 47.8 | 47.8 | 56% | 7e-04 | 81.48 | 1453 | NZ_PDZR010000... |
| Enterococcus faecalis strain 44.1... | Enterococcus... | NA | 1351 | 47.0 | 47.0 | 52% | 0.001 | 84.00 | 806 | NZ_NSJR010000... |
| Cellulophaga baltica strain 14 NODE_63_length_422_cov_10.20678... | Cellulophaga... | NA | 76594 | 45.8 | 45.8 | 97% | 0.001 | 60.42 | 422 | NZ_VIQP010000... |
| Mycobacterium tuberculosis variant bovis strain CSURQ1132, who... | Mycobacteriu... | NA | 1765 | 46.6 | 93.2 | 97% | 0.002 | 60.42 | 1524 | NZ_CAJEUK0100... |
| Mycobacterium tuberculosis variant bovis strain CSURQ1134, who... | Mycobacteriu... | NA | 1765 | 46.6 | 93.2 | 97% | 0.002 | 60.42 | 1697 | NZ_CAJEUT0100... |
| Salirhabdus euzebyi strain Marseille-Q1438, whole genome shotg... | Salirhabdus ... | NA | 394506 | 45.8 | 45.8 | 58% | 0.004 | 78.57 | 1279 | NZ_CADDWK0100... |
| Mycobacterium tuberculosis strain 32847_S31 462, whole genome... | Mycobacteriu... | NA | 1773 | 43.9 | 43.9 | 93% | 0.015 | 48.98 | 893 | NZ_QTBX010001... |
| Pseudomonas sp. MPR-R2A3 NODE_254_length_244_cov_0.832335, who... | Pseudomonas ... | NA | 2070623 | 42.0 | 42.0 | 45% | 0.017 | 95.45 | 244 | NZ_POFW010002... |
| Klebsiella pneumoniae strain EuSCAPE_IL004, whole genome shotg... | Klebsiella p... | NA | 573 | 42.0 | 42.0 | 77% | 0.034 | 62.16 | 356 | NZ_UKJP020001... |

Alignments:

>Thalassovita mangrovi strain GS-10 Scaffold19_1, whole genome shotgun sequence
Sequence ID: NZ_WWEN01000019.1 Length: 2900
Range 1: 2362 to 2502

```
Score:92.8 bits(229), Expect:9e-20,
Method:Compositional matrix adjust.,
Identities:45/47(96%), Positives:46/47(97%), Gaps:0/47(0%)

Query  2     QGKAREFSSEQTRANSPTRRELQVWGRDNNSPSEAGADRQGTVSFNF  48
             QGKAREFSSEQTRANSPTRRELQVWGRDNNS SEAGADRQGTVSF+F
Sbjct  2362  QGKAREFSSEQTRANSPTRRELQVWGRDNNSLSEAGADRQGTVSFSF  2502
```

>Deinococcus aquiradiocola strain JCM 14371 sequence30, whole genome shotgun sequence
Sequence ID: NZ_BMOE01000030.1 Length: 6128
Range 1: 1430 to 1570

```
Score:92.4 bits(228), Expect:1e-19,
Method:Composition-based stats.,
Identities:45/47(96%), Positives:46/47(97%), Gaps:0/47(0%)

Query  2     QGKAREFSSEQTRANSPTRRELQVWGRDNNSPSEAGADRQGTVSFNF  48
             QGKAREFSSEQTRANSPTRRELQVWGRDNNS SEAGADRQGTVSF+F
Sbjct  1430  QGKAREFSSEQTRANSPTRRELQVWGRDNNSLSEAGADRQGTVSFSF  1570
```

>Enterococcus alcedinis strain CCM 8433 sequence23, whole genome shotgun sequence
Sequence ID: NZ_BMDT01000023.1 Length: 6075
Range 1: 1377 to 1517

```
Score:92.4 bits(228), Expect:1e-19,
Method:Composition-based stats.,
Identities:45/47(96%), Positives:46/47(97%), Gaps:0/47(0%)

Query  2     QGKAREFSSEQTRANSPTRRELQVWGRDNNSPSEAGADRQGTVSFNF  48
             QGKAREFSSEQTRANSPTRRELQVWGRDNNS SEAGADRQGTVSF+F
Sbjct  1377  QGKAREFSSEQTRANSPTRRELQVWGRDNNSLSEAGADRQGTVSFSF  1517
```

>Actinoplanes campanulatus strain JCM 3059 sequence058, whole genome shotgun sequence
Sequence ID: NZ_BMPW01000058.1 Length: 4754
Range 1: 1419 to 1559

```
Score:92.4 bits(228), Expect:1e-19,
Method:Composition-based stats.,
Identities:45/47(96%), Positives:46/47(97%), Gaps:0/47(0%)

Query  2     QGKAREFSSEQTRANSPTRRELQVWGRDNNSPSEAGADRQGTVSFNF  48
             QGKAREFSSEQTRANSPTRRELQVWGRDNNS SEAGADRQGTVSF+F
Sbjct  1419  QGKAREFSSEQTRANSPTRRELQVWGRDNNSLSEAGADRQGTVSFSF  1559
```

>Klebsiella pneumoniae strain P7701 NODE_50_length_1305_cov_481.863, whole genome shotgun sequence
Sequence ID: NZ_JACAWM010000050.1 Length: 1305
Range 1: 1206 to 1304

```
Score:67.8 bits(164), Expect:5e-11,
Method:Composition-based stats.,
Identities:31/33(94%), Positives:32/33(96%), Gaps:0/33(0%)

Query  16    NSPTRRELQVWGRDNNSPSEAGADRQGTVSFNF  48
```

```
             NSPTRRELQVWGRDNNS SEAGADRQGTVSF+F
Sbjct  1304  NSPTRRELQVWGRDNNSLSEAGADRQGTVSFSF  1206
```

>Helicobacter acinonychis isolate 212_10, whole genome shotgun sequence
Sequence ID: NZ_FZLW01000041.1 Length: 1526
Range 1: 1402 to 1521

Score:66.2 bits(160), Expect:2e-10,
Method:Composition-based stats.,
Identities:30/40(75%), Positives:35/40(87%), Gaps:0/40(0%)

```
Query  9     SSEQTRANSPTRRELQVWGRDNNSPSEAGADRQGTVSFNF  48
             +S  +RANSPT RELQVWG D+NSPSEAGADRQG +SF+F
Sbjct  1521  NSPTSRANSPTSRELQVWGGDSNSPSEAGADRQGDISFSF  1402
```

>Acinetobacter baumannii strain ACM-35 NODE_65_length_1409_cov_2.30031, whole genome shotgun sequence
Sequence ID: NZ_QEHN01000065.1 Length: 1409
Range 1: 3 to 107

Score:61.6 bits(148), Expect:8e-09,
Method:Composition-based stats.,
Identities:29/35(83%), Positives:31/35(88%), Gaps:0/35(0%)

```
Query  14   RANSPTRRELQVWGRDNNSPSEAGADRQGTVSFNF  48
            RANSPTRRE Q WG D+NS SEAGADRQGTVSF+F
Sbjct  3    RANSPTRREPQAWGGDSNSLSEAGADRQGTVSFSF  107
```

>Acinetobacter baumannii strain ACM-36 NODE_67_length_567_cov_1.13409, whole genome shotgun sequence
Sequence ID: NZ_QAGO01000067.1 Length: 567
Range 1: 3 to 89

Score:55.8 bits(133), Expect:4e-07,
Method:Composition-based stats.,
Identities:25/29(86%), Positives:26/29(89%), Gaps:0/29(0%)

```
Query  20  RRELQVWGRDNNSPSEAGADRQGTVSFNF  48
           RRELQ WG DNNS SEAGADRQGTVSF+F
Sbjct  3   RRELQAWGGDNNSLSEAGADRQGTVSFSF  89
```

>Enterobacter hormaechei strain AUH-ENM6 NODE_27_length_1394_cov_0.980268, whole genome shotgun sequence
Sequence ID: NZ_PYHB01000027.1 Length: 1394
Range 1: 1 to 96

Score:55.8 bits(133), Expect:1e-06,
Method:Composition-based stats.,
Identities:26/32(81%), Positives:27/32(84%), Gaps:0/32(0%)

```
Query  17  SPTRRELQVWGRDNNSPSEAGADRQGTVSFNF  48
           SPTRRE Q WG D NS SEAGADRQGTVSF+F
Sbjct  1   SPTRREPQAWGGDRNSLSEAGADRQGTVSFSF  96
```

>Klebsiella pneumoniae strain EuSCAPE_MT015, whole genome shotgun sequence
Sequence ID: NZ_UKIW02000113.1 Length: 281
Range 1: 95 to 244

Score:52.0 bits(123), Expect:2e-06,
Method:Compositional matrix adjust.,
Identities:32/52(62%), Positives:36/52(69%), Gaps:9/52(17%)

```
Query  2    QGKAREFSSEQTRAN-------SPTRRELQVWGRDNNSPSEAGADRQGTVSF  46
            QG+AREF  EQTRAN       SPT RELQV  R NN  SEAG +RQGT+F
Sbjct  95   QGEAREFPPEQTRANSPTSRANSPTSRELQV--RGNNPGSEAGGERQGTLNF  244
```

>Klebsiella pneumoniae strain EuSCAPE_IL017, whole genome shotgun sequence
Sequence ID: NZ_UKKG02000044.1 Length: 325
Range 1: 73 to 201

Score:52.0 bits(123), Expect:3e-06,
Method:Compositional matrix adjust.,
Identities:28/45(62%), Positives:34/45(75%), Gaps:2/45(4%)

```
Query  2    QGKAREFSSEQTRANSPTRRELQVWGRDNNSPSEAGADRQGTVSF  46
            QG+AREF  EQ RA SPT RELQV  R +N  SEAG +RQG+++F
Sbjct  73   QGEAREFPPEQARAISPTSRELQV--RGDNPRSEAGVERQGSLNF  201
```

>Klebsiella pneumoniae strain NUBRI-K 73, whole genome shotgun sequence
Sequence ID: NZ_SOYT01000073.1 Length: 1717
Range 1: 5 to 154

Score:54.3 bits(129), Expect:4e-06,
Method:Composition-based stats.,
Identities:33/52(63%), Positives:37/52(71%), Gaps:9/52(17%)

```
Query  2    QGKAREFSSEQ-------TRANSPTRRELQVWGRDNNSPSEAGADRQGTVSF  46
            QGKAREF  EQ       +RANSPT RELQV  R +NS SEAGA RQGT+F
Sbjct  5    QGKAREFPPEQARTNSPTSRANSPTSRELQV--RGDNSRSEAGAKRQGTLNF  154
```

>Escherichia coli strain YA00028237 contig_187, whole genome shotgun sequence
Sequence ID: NZ_NGBP02000187.1 Length: 1828
Range 1: 1623 to 1772

Score:54.3 bits(129), Expect:4e-06,
Method:Composition-based stats.,
Identities:33/52(63%), Positives:37/52(71%), Gaps:9/52(17%)

```
Query  2     QGKAREFSSEQTRANSPTR-------RELQVWGRDNNSPSEAGADRQGTVSF  46
             QG+AREF SEQTRANSPT        RELQV  R +NS SEAG +RQGT +F
Sbjct  1772  QGEAREFPSEQTRANSPTSRANSPTGRELQV--RGDNSHSEAGEERQGTFNF  1623
```

>Klebsiella pneumoniae strain EuSCAPE_LT007, whole genome shotgun sequence
Sequence ID: NZ_UKJI02000072.1 Length: 392
Range 1: 217 to 345

Score:52.4 bits(124), Expect:4e-06,
Method:Compositional matrix adjust.,
Identities:31/47(66%), Positives:34/47(72%), Gaps:4/47(8%)

```
Query  2    QGKAREFSSEQTRANSPTRRELQVWGRDNNSPSEAGADRQGTVSFNF  48
            QG+AREF SEQ RA SPT RELQV  R +N  SEAG + QG  SFNF
Sbjct  345  QGEAREFPSEQARAISPTSRELQV--RGDNPCSEAGVEGQG--SFNF  217
```

>Burkholderia pseudomallei strain A-175-05-2-02, whole genome shotgun sequence
Sequence ID: NZ_UVFW01000138.1 Length: 455
Range 1: 273 to 401

Score:50.8 bits(120), Expect:2e-05,
Method:Compositional matrix adjust.,
Identities:31/44(70%), Positives:33/44(75%), Gaps:1/44(2%)

```
Query  2    QGKAREFSSEQTRANSPTRRELQVWGRDNNSPSEAGADRQGTVS  45
            Q KAREFSSEQ RANSPT REL   GRD N  +EAGA+ QGT S
Sbjct  401  QRKAREFSSEQNRANSPTSRELGDGGRD-NLLTEAGAEGQGTSS  273
```

>Burkholderia pseudomallei strain 2604STDY5519032 ERS240165SCcontig000074, whole genome shotgun sequence
Sequence ID: NZ_UVID02000074.1 Length: 656
Range 1: 26 to 154

Score:51.2 bits(121), Expect:3e-05,
Method:Compositional matrix adjust.,
Identities:28/45(62%), Positives:33/45(73%), Gaps:2/45(4%)

```
Query  2    QGKAREFSSEQTRANSPTRRELQVWGRDNNSPSEAGADRQGTVSF  46
            QG+AREF  EQTRANSPT RELQV  R  N  +EAG + +GT +F
Sbjct  154  QGEAREFHPEQTRANSPTSRELQV--RGGNPCAEAGVEGEGTFNF  26
```

>Acinetobacter baumannii strain ACM-14 NODE_74_length_787_cov_0.637879, whole genome shotgun sequence
Sequence ID: NZ_PYDA01000074.1 Length: 787
Range 1: 702 to 785

Score:50.8 bits(120), Expect:5e-05,
Method:Composition-based stats.,
Identities:22/28(79%), Positives:24/28(85%), Gaps:0/28(0%)

```
Query  21   RELQVWGRDNNSPSEAGADRQGTVSFNF  48
            RELQVWG DNNSPSEAG +RQG VSF+
Sbjct  785  RELQVWGGDNNSPSEAGGERQGDVSFSL  702
```

>Escherichia coli strain 8374wH11 NODE_36_length_1397_cov_3.54567, whole genome shotgun sequence
Sequence ID: NZ_WSJI01000036.1 Length: 1397
Range 1: 4 to 87

Score:50.8 bits(120), Expect:6e-05,
Method:Composition-based stats.,
Identities:22/28(79%), Positives:24/28(85%), Gaps:0/28(0%)

```
Query  21  RELQVWGRDNNSPSEAGADRQGTVSFNF  48
           RELQVWG DNNSPSEAG +RQG VSF+
Sbjct  4   RELQVWGGDNNSPSEAGGERQGDVSFSL  87
```

>Enterococcus faecium strain A209R1B0C1 NODE_70_length_1819_cov_3.01832_ID_139, whole genome shotgun sequence
Sequence ID: NZ_PGCR01000088.1 Length: 1819
Range 1: 66 to 188

Score:48.5 bits(114), Expect:4e-04,
Method:Composition-based stats.,
Identities:28/45(62%), Positives:33/45(73%), Gaps:4/45(8%)

```
Query  2    QGKAREFSSEQTRANSPTRRELQVWGRDNNSPSEAGADRQGTVSF  46
            QG+AREF SEQTRANSPT  ELQV  R +N  SE G   +GT++F
Sbjct  66   QGEAREFPSEQTRANSPTSGELQV--RGDNPCSETGV--EGTLNF  188
```

>Listeria monocytogenes strain YA00082404 NODE_73_length_1816_cov_5.664890, whole genome shotgun sequence
Sequence ID: NZ_QEXE01000073.1 Length: 1816
Range 1: 52 to 201

Score:47.8 bits(112), Expect:6e-04,
Method:Composition-based stats.,
Identities:30/52(58%), Positives:35/52(67%), Gaps:9/52(17%)

```
Query  2    QGKAREFSSEQ-------TRANSPTRRELQVWGRDNNSPSEAGADRQGTVSF  46
            QG+AREF SEQ       +RANSPT RELQV  R +N  SE GA+ QGT +F
Sbjct  52   QGEAREFPSEQARTNSPTSRANSPTGRELQV--RGDNPRSETGAEGQGTFNF  201
```

>Methylocella silvestris strain TVC NODE_58_length_1453_cov_3.56561, whole genome shotgun sequence
Sequence ID: NZ_PDZR01000058.1 Length: 1453
Range 1: 1 to 81

Score:47.8 bits(112), Expect:7e-04,
Method:Composition-based stats.,
Identities:22/27(81%), Positives:23/27(85%), Gaps:0/27(0%)

```
Query  22  ELQVWGRDNNSPSEAGADRQGTVSFNF  48
           E Q WG DNNS SEAGADRQGTVSF+F
Sbjct  1   EPQAWGGDNNSLSEAGADRQGTVSFSF  81
```

>Enterococcus faecalis strain 44.1 NODE_33_length_806_cov_1.20619, whole genome shotgun sequence
Sequence ID: NZ_NSJR01000033.1 Length: 806

Range 1: 42 to 116

Score:47.0 bits(110), Expect:0.001,
Method:Composition-based stats.,
Identities:21/25(84%), Positives:22/25(88%), Gaps:0/25(0%)

```
Query  24   QVWGRDNNSPSEAGADRQGTVSFNF  48
            Q WG DNNS SEAGADRQGTVSF+F
Sbjct  42   QAWGGDNNSLSEAGADRQGTVSFSF  116
```

>Cellulophaga baltica strain 14 NODE_63_length_422_cov_10.206780, whole genome shotgun sequence
Sequence ID: NZ_VIQP01000062.1 Length: 422
Range 1: 247 to 387

Score:45.8 bits(107), Expect:0.001,
Method:Compositional matrix adjust.,
Identities:29/48(60%), Positives:33/48(68%), Gaps:2/48(4%)

```
Query  2    QGKAREFSSEQTRANSPTRRELQVWGRDNNSPSEAGADRQGTV-SFNF  48
            Q +AR+FSSEQTR N PT RE    GRDN   S AGA+ QG + SFNF
Sbjct  387  QREARKFSSEQTRTNCPTSREHGDGGRDNLL-SAAGAEGQGAISSFNF  247
```

>Mycobacterium tuberculosis variant bovis strain CSURQ1132, whole genome shotgun sequence
Sequence ID: NZ_CAJEUK010000155.1 Length: 1524
Range 1: 1308 to 1448

Score:46.6 bits(109), Expect:0.002,
Method:Composition-based stats.,
Identities:29/48(60%), Positives:32/48(66%), Gaps:2/48(4%)

```
Query  2     QGKAREFSSEQTRANSPTRRELQVWGRDNNSPSEAGADRQGTV-SFNF  48
             QG+AR+FSS QT  NSPT REL   GRD   P EAG   QG + SFNF
Sbjct  1448  QGEARKFSSGQTGTNSPTSRELWNGGRDTILP-EAGTGGQGPISSFNF  1308
```

Range 2: 1308 to 1448

Score:46.6 bits(109), Expect:0.002,
Method:Composition-based stats.,
Identities:29/48(60%), Positives:32/48(66%), Gaps:2/48(4%)

```
Query  2     QGKAREFSSEQTRANSPTRRELQVWGRDNNSPSEAGADRQGTV-SFNF  48
             QG+AR+FSS QT  NSPT REL   GRD   P EAG   QG + SFNF
Sbjct  1448  QGEARKFSSGQTGTNSPTSRELWNGGRDTILP-EAGTGGQGPISSFNF  1308
```

>Mycobacterium tuberculosis variant bovis strain CSURQ1134, whole genome shotgun sequence
Sequence ID: NZ_CAJEUT010000109.1 Length: 1697
Range 1: 1270 to 1410

Score:46.6 bits(109), Expect:0.002,
Method:Composition-based stats.,
Identities:29/48(60%), Positives:32/48(66%), Gaps:2/48(4%)

```
Query  2     QGKAREFSSEQTRANSPTRRELQVWGRDNNSPSEAGADRQGTV-SFNF  48
             QG+AR+FSS QT  NSPT REL   GRD   P EAG   QG + SFNF
Sbjct  1410  QGEARKFSSGQTGTNSPTSRELWNGGRDTILP-EAGTGGQGPISSFNF  1270
```

Range 2: 1270 to 1410

Score:46.6 bits(109), Expect:0.002,
Method:Composition-based stats.,
Identities:29/48(60%), Positives:32/48(66%), Gaps:2/48(4%)

```
Query  2     QGKAREFSSEQTRANSPTRRELQVWGRDNNSPSEAGADRQGTV-SFNF  48
             QG+AR+FSS QT  NSPT REL   GRD   P EAG   QG + SFNF
Sbjct  1410  QGEARKFSSGQTGTNSPTSRELWNGGRDTILP-EAGTGGQGPISSFNF  1270
```

>Salirhabdus euzebyi strain Marseille-Q1438, whole genome shotgun sequence
Sequence ID: NZ_CADDWK010000035.1 Length: 1279
Range 1: 1194 to 1277

Score:45.8 bits(107), Expect:0.004,
Method:Composition-based stats.,
Identities:22/28(79%), Positives:24/28(85%), Gaps:0/28(0%)

```
Query  21    RELQVWGRDNNSPSEAGADRQGTVSFNF  48
             R+ QV GRDNNS SEAGADRQG VSF+F
Sbjct  1277  RKPQVGGRDNNSLSEAGADRQGHVSFSF  1194
```

>Mycobacterium tuberculosis strain 32847_S31 462, whole genome shotgun sequence
Sequence ID: NZ_QTBX01000137.1 Length: 893
Range 1: 47 to 193

Score:43.9 bits(102), Expect:0.015,
Method:Composition-based stats.,
Identities:24/49(49%), Positives:32/49(65%), Gaps:4/49(8%)

```
Query  2    QGKAREFSSEQTRANSPTRRELQVWGRDNNSPS----EAGADRQGTVSF  46
            QG+AREF  EQ RANSPT R+L       +SP+    +AGA+RQG ++
Sbjct  47   QGEAREFYPEQARANSPTSRKLPPEQARADSPTSRELQAGAERQGALNL  193
```

>Pseudomonas sp. MPR-R2A3 NODE_254_length_244_cov_0.832335, whole genome shotgun sequence
Sequence ID: NZ_POFW01000253.1 Length: 244
Range 1: 58 to 123

Score:42.0 bits(97), Expect:0.017,
Method:Compositional matrix adjust.,
Identities:21/22(95%), Positives:21/22(95%), Gaps:0/22(0%)

```
Query  2    QGKAREFSSEQTRANSPTRREL  23
            QGKAREFSSEQTRANSPTRR L
```

```
Sbjct  58   QGKAREFSSEQTRANSPTRRWL  123
```

>Klebsiella pneumoniae strain EuSCAPE_IL004, whole genome shotgun sequence
Sequence ID: NZ_UKJP02000108.1 Length: 356
Range 1: 241 to 345

Score:42.0 bits(97), Expect:0.034,
Method:Compositional matrix adjust.,
Identities:23/37(62%), Positives:28/37(75%), Gaps:2/37(5%)

```
Query  10   SEQTRANSPTRRELQVWGRDNNSPSEAGADRQGTVSF  46
            S  +RANSPT RELQV  R +N  SEAGA+ QGT+F
Sbjct  345  SPTSRANSPTSRELQV--RGDNPCSEAGAEGQGTLNF  241
```

Query #15: sp|P04601|NEF_HV1H2 Protein Nef OS=Human immunodeficiency virus type 1 group M subtype B (isolate HXB2) OX=11706 GN=nef PE=1 SV=5 Query ID: lcl|Query_2387711
Length: 206

Sequences producing significant alignments:

| Description | Scientific Name | Common Name | Taxid | Max Score | Total Score | Query cover | E Value | Per. Ident | Acc. Len | Accession |
|---|---|---|---|---|---|---|---|---|---|---|
| Photobacterium rosenbergii strain DP6N14-7 DP6N14-7_ctg0054,... | Photobacteri... | NA | 294936 | 324 | 324 | 100% | 5e-108 | 73.30 | 960 | NZ_JAHVIB0100... |
| Bacillus velezensis strain TN14-11 TN14-11_ctg0036, whole geno... | Bacillus vel... | NA | 492670 | 325 | 325 | 100% | 9e-108 | 73.79 | 1101 | NZ_JAHVIU0100... |
| Acinetobacter baumannii strain P2 contig_P2_66, whole genome... | Acinetobacte... | NA | 470 | 323 | 323 | 84% | 3e-98 | 89.08 | 4185 | NZ_JBBXLW0100... |
| Qipengyuania pelagi strain JCM 17468 JCM17468_9, whole genome... | Qipengyuania... | NA | 994320 | 304 | 304 | 81% | 4e-96 | 86.23 | 1945 | NZ_WTYD010000... |
| Klebsiella pneumoniae strain EuSCAPE_NO002, whole genome shotg... | Klebsiella p... | NA | 573 | 269 | 269 | 92% | 1e-87 | 65.97 | 676 | NZ_UKMC020000... |
| Klebsiella pneumoniae strain EuSCAPE_IL007, whole genome shotg... | Klebsiella p... | NA | 573 | 260 | 260 | 83% | 4e-84 | 69.54 | 697 | NZ_UKJT020000... |
| Acinetobacter baumannii strain XH798 XH798_contig_187, whole... | Acinetobacte... | NA | 470 | 253 | 253 | 61% | 7e-82 | 92.13 | 588 | NZ_LYGA010000... |
| Klebsiella pneumoniae strain EuSCAPE_MT009, whole genome shotg... | Klebsiella p... | NA | 573 | 251 | 251 | 87% | 2e-80 | 66.30 | 670 | NZ_UKIV020000... |
| Klebsiella pneumoniae strain EuSCAPE_DE088, whole genome shotg... | Klebsiella p... | NA | 573 | 241 | 241 | 70% | 2e-77 | 76.03 | 535 | NZ_UKLX020000... |
| Sphingomonas sp. UV9 NODE_33_length_614_cov_0.261122, whole... | Sphingomonas... | NA | 1851410 | 239 | 239 | 65% | 4e-76 | 85.82 | 614 | NZ_SCIN010000... |
| Klebsiella pneumoniae strain EuSCAPE_IL022, whole genome shotg... | Klebsiella p... | NA | 573 | 224 | 224 | 82% | 1e-70 | 62.84 | 594 | NZ_UKKD020000... |
| Klebsiella pneumoniae strain EuSCAPE_IL021, whole genome shotg... | Klebsiella p... | NA | 573 | 218 | 218 | 66% | 1e-68 | 72.99 | 415 | NZ_UKJX020001... |
| Acinetobacter baumannii strain XH797 XH797_contig_168, whole... | Acinetobacte... | NA | 470 | 215 | 215 | 51% | 1e-67 | 92.52 | 418 | NZ_LYGB010000... |
| Klebsiella pneumoniae strain EuSCAPE_IL001, whole genome shotg... | Klebsiella p... | NA | 573 | 213 | 213 | 70% | 4e-66 | 66.88 | 519 | NZ_UKJN020000... |
| Klebsiella pneumoniae strain EuSCAPE_MT003, whole genome shotg... | Klebsiella p... | NA | 573 | 216 | 216 | 72% | 1e-65 | 66.88 | 964 | NZ_UKIK020000... |
| Klebsiella pneumoniae strain EuSCAPE_LT010, whole genome shotg... | Klebsiella p... | NA | 573 | 202 | 202 | 59% | 2e-62 | 74.59 | 494 | NZ_UKJK020000... |
| Klebsiella pneumoniae strain EuSCAPE_LT004, whole genome shotg... | Klebsiella p... | NA | 573 | 195 | 195 | 53% | 4e-60 | 83.04 | 348 | NZ_UKJE020000... |
| Klebsiella pneumoniae strain EuSCAPE_DE083, whole genome shotg... | Klebsiella p... | NA | 573 | 194 | 194 | 54% | 9e-60 | 76.99 | 349 | NZ_UKLY020000... |
| Klebsiella pneumoniae strain EuSCAPE_MT011, whole genome shotg... | Klebsiella p... | NA | 573 | 188 | 188 | 52% | 8e-57 | 75.23 | 431 | NZ_UKIY020000... |
| Acinetobacter baumannii strain XH744 XH744_contig_84, whole... | Acinetobacte... | NA | 470 | 184 | 184 | 46% | 3e-55 | 88.42 | 397 | NZ_LYHI010000... |
| Klebsiella pneumoniae strain EuSCAPE_IL020, whole genome shotg... | Klebsiella p... | NA | 573 | 179 | 179 | 64% | 2e-53 | 66.90 | 422 | NZ_UKKB020000... |
| Klebsiella pneumoniae strain EuSCAPE_IL005, whole genome shotg... | Klebsiella p... | NA | 573 | 173 | 173 | 48% | 1e-51 | 77.00 | 307 | NZ_UKJL020000... |
| Pseudomonas aeruginosa strain ENV-297... | Pseudomonas ... | NA | 287 | 174 | 174 | 52% | 2e-51 | 71.30 | 378 | NZ_RADD010002... |
| Klebsiella pneumoniae strain EuSCAPE_MT019, whole genome shotg... | Klebsiella p... | NA | 573 | 172 | 172 | 53% | 3e-51 | 76.36 | 332 | NZ_UKJD020000... |
| Klebsiella pneumoniae strain EuSCAPE_MT015, whole genome shotg... | Klebsiella p... | NA | 573 | 171 | 171 | 60% | 2e-50 | 67.97 | 372 | NZ_UKIW020001... |
| Acinetobacter baumannii strain XH821 XH821_contig_233, whole... | Acinetobacte... | NA | 470 | 172 | 172 | 47% | 6e-50 | 84.54 | 572 | NZ_LYFC010000... |
| Klebsiella pneumoniae strain EuSCAPE_MT014, whole genome shotg... | Klebsiella p... | NA | 573 | 166 | 166 | 45% | 2e-48 | 75.53 | 388 | NZ_UKJF020000... |
| Escherichia coli strain EN5308 contig00099, whole genome shotg... | Escherichia ... | NA | 562 | 168 | 168 | 58% | 1e-43 | 67.21 | 2827 | NZ_JARBDE0100... |
| Klebsiella pneumoniae strain EuSCAPE_MT009, whole genome shotg... | Klebsiella p... | NA | 573 | 149 | 149 | 40% | 3e-42 | 78.57 | 262 | NZ_UKIV020000... |
| Klebsiella pneumoniae strain EuSCAPE_IL005, whole genome shotg... | Klebsiella p... | NA | 573 | 150 | 150 | 50% | 5e-42 | 72.12 | 442 | NZ_UKJL020000... |
| Acinetobacter baumannii strain XH821 XH821_contig_145, whole... | Acinetobacte... | NA | 470 | 142 | 142 | 35% | 8e-39 | 87.84 | 427 | NZ_LYFC010000... |
| Acinetobacter baumannii strain P25 contig_P25_82, whole genome... | Acinetobacte... | NA | 470 | 150 | 150 | 43% | 2e-37 | 81.11 | 2628 | NZ_JBBXLB0100... |
| Acinetobacter baumannii strain E13 contig_E13_79, whole genome... | Acinetobacte... | NA | 470 | 150 | 150 | 43% | 2e-37 | 81.11 | 2763 | NZ_JBBXKR0100... |
| Klebsiella pneumoniae strain EuSCAPE_LT003, whole genome shotg... | Klebsiella p... | NA | 573 | 126 | 126 | 41% | 6e-33 | 69.77 | 350 | NZ_UKJH020000... |
| Klebsiella pneumoniae strain EuSCAPE_MT002, whole genome shotg... | Klebsiella p... | NA | 573 | 121 | 121 | 47% | 1e-30 | 69.23 | 418 | NZ_UKIM020000... |
| Acinetobacter baumannii strain XH744 XH744_contig_122, whole... | Acinetobacte... | NA | 470 | 121 | 121 | 36% | 2e-30 | 78.95 | 489 | NZ_LYHI010000... |
| Escherichia coli strain NBE0009 contig00151, whole genome... | Escherichia ... | NA | 562 | 124 | 124 | 44% | 3e-30 | 68.75 | 802 | NZ_JARBDL0100... |
| Escherichia coli strain NBE0003 contig00096, whole genome... | Escherichia ... | NA | 562 | 127 | 127 | 46% | 2e-29 | 64.58 | 2837 | NZ_JARBDG0100... |
| Escherichia coli strain EN5169 contig00082, whole genome shotg... | Escherichia ... | NA | 562 | 127 | 127 | 46% | 3e-29 | 64.58 | 2837 | NZ_JAPMNJ0100... |
| Escherichia coli strain EN5317 contig00057, whole genome shotg... | Escherichia ... | NA | 562 | 124 | 124 | 46% | 4e-28 | 65.66 | 2867 | NZ_JARBDO0100... |
| Klebsiella pneumoniae strain EuSCAPE_IL015, whole genome shotg... | Klebsiella p... | NA | 573 | 109 | 109 | 32% | 2e-26 | 71.21 | 301 | NZ_UKJV020000... |
| Escherichia coli strain EN5090 contig00081, whole genome shotg... | Escherichia ... | NA | 562 | 116 | 116 | 44% | 1e-25 | 63.04 | 2798 | NZ_JAODTD0100... |
| Escherichia coli strain NBE0006 contig00145, whole genome... | Escherichia ... | NA | 562 | 110 | 110 | 39% | 2e-25 | 65.85 | 738 | NZ_JARBDI0100... |
| Escherichia coli strain EN5076 contig00066, whole genome shotg... | Escherichia ... | NA | 562 | 115 | 115 | 43% | 2e-25 | 62.64 | 2732 | NZ_JAODTC0100... |
| Escherichia coli strain EN5374 contig00094, whole genome shotg... | Escherichia ... | NA | 562 | 115 | 115 | 42% | 4e-25 | 64.04 | 2720 | NZ_JAPTGI0100... |
| Klebsiella pneumoniae strain EuSCAPE_LT003, whole genome shotg... | Klebsiella p... | NA | 573 | 104 | 104 | 43% | 2e-24 | 55.56 | 289 | NZ_UKJH020000... |
| Escherichia coli strain EN5379 contig00092, whole genome shotg... | Escherichia ... | NA | 562 | 112 | 112 | 39% | 7e-24 | 64.71 | 2592 | NZ_JAPTGJ0100... |
| Escherichia coli strain EN5383 contig00096, whole genome shotg... | Escherichia ... | NA | 562 | 111 | 111 | 41% | 1e-23 | 63.22 | 2210 | NZ_JAPTZN0100... |
| Escherichia coli strain NBE0009 contig00153, whole genome... | Escherichia ... | NA | 562 | 103 | 103 | 37% | 2e-22 | 64.56 | 753 | NZ_JARBDL0100... |
| Escherichia coli strain NBE0010 contig00080, whole genome... | Escherichia ... | NA | 562 | 106 | 180 | 42% | 5e-22 | 60.23 | 2743 | NZ_JARBDM0100... |
| Escherichia coli strain EN5383 contig00089, whole genome shotg... | Escherichia ... | NA | 562 | 103 | 103 | 40% | 6e-21 | 61.90 | 1970 | NZ_JAPTZN0100... |
| Escherichia coli strain NBE0009 contig00116, whole genome... | Escherichia ... | NA | 562 | 102 | 102 | 42% | 8e-21 | 55.88 | 1713 | NZ_JARBDL0100... |
| Klebsiella pneumoniae strain EuSCAPE_NO017, whole genome shotg... | Klebsiella p... | NA | 573 | 96.7 | 96.7 | 36% | 3e-20 | 62.96 | 607 | NZ_UKIO020000... |
| Escherichia coli strain EN5192 contig00135, whole genome shotg... | Escherichia ... | NA | 562 | 100 | 100 | 37% | 3e-20 | 62.82 | 1341 | NZ_JAPMNL0100... |
| Acinetobacter baumannii strain XH798 XH798_contig_106, whole... | Acinetobacte... | NA | 470 | 92.8 | 92.8 | 29% | 4e-19 | 75.00 | 507 | NZ_LYGA010000... |
| Escherichia coli strain EN5286 contig00120, whole genome shotg... | Escherichia ... | NA | 562 | 94.7 | 94.7 | 42% | 6e-18 | 54.90 | 3033 | NZ_JAPMNM0100... |
| Escherichia coli strain EN5177 contig00087, whole genome shotg... | Escherichia ... | NA | 562 | 94.7 | 94.7 | 42% | 6e-18 | 54.90 | 3403 | NZ_JAPMNK0100... |
| Escherichia coli strain EN5308 contig00153, whole genome shotg... | Escherichia ... | NA | 562 | 90.9 | 90.9 | 40% | 1e-17 | 57.65 | 786 | NZ_JARBDE0100... |
| Escherichia coli strain EN5308 contig00096, whole genome shotg... | Escherichia ... | NA | 562 | 93.2 | 93.2 | 42% | 2e-17 | 54.46 | 3024 | NZ_JARBDE0100... |
| Escherichia coli strain EN5143 contig00071, whole genome shotg... | Escherichia ... | NA | 562 | 91.7 | 91.7 | 42% | 7e-17 | 54.46 | 2774 | NZ_JAPMNI0100... |
| Klebsiella pneumoniae strain EuSCAPE_IL015, whole genome shotg... | Klebsiella p... | NA | 573 | 85.1 | 85.1 | 32% | 3e-16 | 61.76 | 532 | NZ_UKJV020000... |
| Escherichia coli strain EN5095 contig00078, whole genome shotg... | Escherichia ... | NA | 562 | 88.6 | 88.6 | 36% | 8e-16 | 57.14 | 1972 | NZ_JAPYLY0100... |
| Escherichia coli strain NBE0006 contig00147, whole genome... | Escherichia ... | NA | 562 | 85.1 | 85.1 | 33% | 1e-15 | 58.57 | 694 | NZ_JARBDI0100... |
| Escherichia coli strain NBE0002 contig00097, whole genome... | Escherichia ... | NA | 562 | 87.4 | 87.4 | 41% | 2e-15 | 53.00 | 2717 | NZ_JARBDF0100... |
| Escherichia coli strain NBE0007 contig00093, whole genome... | Escherichia ... | NA | 562 | 85.5 | 85.5 | 39% | 9e-15 | 53.12 | 3013 | NZ_JARBDJ0100... |
| Escherichia coli strain EN5132 contig00077, whole genome shotg... | Escherichia ... | NA | 562 | 85.5 | 85.5 | 39% | 1e-14 | 57.14 | 2498 | NZ_JAPYLZ0100... |
| Acinetobacter baumannii strain XH797 XH797_contig_152, whole... | Acinetobacte... | NA | 470 | 78.6 | 78.6 | 25% | 2e-14 | 71.70 | 304 | NZ_LYGB010000... |
| Bacillus cereus strain SJ-S17... | Bacillus cereus | NA | 1396 | 77.0 | 77.0 | 17% | 3e-14 | 97.30 | 219 | NZ_NMYP010000... |
| Klebsiella pneumoniae strain EuSCAPE_LT007, whole genome shotg... | Klebsiella p... | NA | 573 | 75.9 | 75.9 | 23% | 1e-13 | 67.35 | 253 | NZ_UKJI020000... |
| Escherichia coli strain NBE0004 contig00110, whole genome... | Escherichia ... | NA | 562 | 80.1 | 80.1 | 43% | 6e-13 | 53.54 | 1390 | NZ_JARBDH0100... |
| Acinetobacter baumannii strain XH822 XH822_contig_193, whole... | Acinetobacte... | NA | 470 | 77.8 | 77.8 | 20% | 7e-13 | 88.37 | 753 | NZ_LYFB010000... |
| Acinetobacter baumannii strain XH851 XH851_contig_81, whole... | Acinetobacte... | NA | 470 | 77.8 | 77.8 | 20% | 1e-12 | 88.37 | 882 | NZ_LYJM010000... |
| Acinetobacter baumannii strain XH840 XH840_contig_135, whole... | Acinetobacte... | NA | 470 | 77.8 | 77.8 | 20% | 3e-12 | 88.37 | 1251 | NZ_LYEX010000... |
| Streptomyces sp. AA4 supercont1.1, whole genome shotgun sequence | Streptomyces... | NA | 591158 | 76.6 | 76.6 | 19% | 1e-11 | 82.61 | 8475969 | NZ_GG657746.1 |
| Nocardioides luteus strain JCM 3358 sequence39, whole genome... | Nocardioides... | NA | 1844 | 76.3 | 76.3 | 19% | 2e-11 | 90.00 | 5754 | NZ_BMRK010000... |
| Streptomyces griseoflavus strain JCM 4479 sequence37, whole... | Streptomyces... | NA | 35619 | 76.3 | 76.3 | 19% | 2e-11 | 90.00 | 5775 | NZ_BMUC010000... |
| Saccharothrix coeruleofusca strain JCM 3313 sequence45, whole... | Saccharothri... | NA | 33919 | 76.3 | 76.3 | 19% | 2e-11 | 90.00 | 5775 | NZ_BMRG010000... |
| Actinoplanes ianthinogenes strain JCM 3249 sequence037, whole... | Actinoplanes... | NA | 122358 | 76.3 | 76.3 | 19% | 2e-11 | 90.00 | 5815 | NZ_BMQZ010000... |
| Sinomonas sp. KCTC 49339 sequence28, whole genome shotgun... | Sinomonas sp... | NA | 2769422 | 76.3 | 76.3 | 19% | 2e-11 | 90.00 | 5849 | NZ_BNCM010000... |
| Saccharothrix mutabilis subsp. mutabilis strain JCM 3380... | Saccharothri... | NA | 66855 | 76.3 | 76.3 | 19% | 2e-11 | 90.00 | 5899 | NZ_BAAABU0100... |
| Enterococcus alcedinis strain CCM 8433 sequence23, whole genom... | Enterococcus... | NA | 1274384 | 75.5 | 75.5 | 23% | 3e-11 | 74.07 | 6075 | NZ_BMDT010000... |
| Deinococcus aquiradiocola strain JCM 14371 sequence30, whole... | Deinococcus ... | NA | 393059 | 75.5 | 75.5 | 23% | 3e-11 | 74.07 | 6128 | NZ_BMOE010000... |
| Escherichia coli strain EN5192 contig00108, whole genome shotg... | Escherichia ... | NA | 562 | 73.9 | 73.9 | 37% | 8e-11 | 51.65 | 3032 | NZ_JAPMNL0100... |
| Escherichia coli strain EN5143 contig00069, whole genome shotg... | Escherichia ... | NA | 562 | 68.9 | 68.9 | 39% | 4e-09 | 56.04 | 2688 | NZ_JAPMNI0100... |
| Acinetobacter baumannii strain XH799 XH799_contig_168, whole... | Acinetobacte... | NA | 470 | 63.2 | 63.2 | 16% | 1e-08 | 88.57 | 292 | NZ_LYFZ010000... |
| Escherichia coli strain EN5177 contig00148, whole genome shotg... | Escherichia ... | NA | 562 | 66.2 | 66.2 | 35% | 2e-08 | 51.81 | 850 | NZ_JAPMNK0100... |
| Acinetobacter baumannii strain E16 contig_E16_244, whole genom... | Acinetobacte... | NA | 470 | 63.2 | 63.2 | 24% | 1e-07 | 62.75 | 651 | NZ_JBBXKO0100... |
| Acinetobacter baumannii strain E13 contig_E13_197, whole genom... | Acinetobacte... | NA | 470 | 63.2 | 63.2 | 24% | 2e-07 | 62.75 | 703 | NZ_JBBXKR0100... |
| Acinetobacter baumannii strain E11 contig_E11_176, whole genom... | Acinetobacte... | NA | 470 | 63.2 | 63.2 | 24% | 3e-07 | 62.75 | 888 | NZ_JBBXKT0100... |
| Acinetobacter baumannii strain P11 contig_P11_393, whole genom... | Acinetobacte... | NA | 470 | 63.2 | 63.2 | 24% | 4e-07 | 62.75 | 1140 | NZ_JBBXLN0100... |
| Acinetobacter baumannii strain P28 contig_P28_100, whole genom... | Acinetobacte... | NA | 470 | 63.2 | 63.2 | 24% | 5e-07 | 62.75 | 1822 | NZ_JBBXKY0100... |
| Acinetobacter baumannii strain P18 contig_P18_196, whole genom... | Acinetobacte... | NA | 470 | 63.2 | 63.2 | 24% | 5e-07 | 62.75 | 2259 | NZ_JBBXLH0100... |
| Acinetobacter baumannii strain P1 contig_P1_68, whole genome... | Acinetobacte... | NA | 470 | 63.2 | 63.2 | 24% | 5e-07 | 62.75 | 2273 | NZ_JBBXLX0100... |
| Acinetobacter baumannii strain E2 contig_E2_954, whole genome... | Acinetobacte... | NA | 470 | 63.2 | 63.2 | 24% | 5e-07 | 62.75 | 2573 | NZ_JBBXKW0100... |
| Acinetobacter baumannii strain P27 contig_P27_23, whole genome... | Acinetobacte... | NA | 470 | 63.2 | 63.2 | 24% | 5e-07 | 62.75 | 2573 | NZ_JBBXKZ0100... |
| Acinetobacter baumannii strain P16 contig_P16_190, whole genom... | Acinetobacte... | NA | 470 | 63.2 | 63.2 | 24% | 5e-07 | 62.75 | 2599 | NZ_JBBXLI0100... |
| Acinetobacter baumannii strain P5 contig_P5_54, whole genome... | Acinetobacte... | NA | 470 | 63.2 | 63.2 | 24% | 5e-07 | 62.75 | 2679 | NZ_JBBXLT0100... |

| | | | | | | | | | | |
|---|---|---|---|---|---|---|---|---|---|---|
| Acinetobacter baumannii strain P7 contig_P7_199, whole genome... | Acinetobacte... | NA | 470 | 63.2 | 63.2 | 24% | 5e-07 | 62.75 | 2736 | NZ_JBBXLR0100... |
| Acinetobacter baumannii strain P19 contig_P19_138, whole genom... | Acinetobacte... | NA | 470 | 63.2 | 63.2 | 24% | 5e-07 | 62.75 | 2736 | NZ_JBBXLG0100... |
| Acinetobacter baumannii strain P15 contig_P15_245, whole genom... | Acinetobacte... | NA | 470 | 63.2 | 63.2 | 24% | 5e-07 | 62.75 | 2736 | NZ_JBBXLJ0100... |
| Acinetobacter baumannii strain P21 contig_P21_632, whole genom... | Acinetobacte... | NA | 470 | 63.2 | 63.2 | 24% | 5e-07 | 62.75 | 2804 | NZ_JBBXLE0100... |
| Acinetobacter baumannii strain E17 contig_E17_94, whole genome... | Acinetobacte... | NA | 470 | 63.2 | 63.2 | 24% | 5e-07 | 62.75 | 2965 | NZ_JBBXKN0100... |
| Acinetobacter baumannii strain P23 contig_P23_230, whole genom... | Acinetobacte... | NA | 470 | 63.2 | 63.2 | 24% | 5e-07 | 62.75 | 3261 | NZ_JBBXLC0100... |
| Klebsiella pneumoniae strain CL2087... | Klebsiella p... | NA | 573 | 62.0 | 62.0 | 15% | 1e-06 | 90.32 | 2194 | NZ_RPFC010000... |
| Klebsiella pneumoniae strain EuSCAPE_LT003, whole genome shotg... | Klebsiella p... | NA | 573 | 53.5 | 53.5 | 15% | 6e-05 | 71.88 | 317 | NZ_UKJH020000... |
| Klebsiella pneumoniae strain EuSCAPE_MT017, whole genome shotg... | Klebsiella p... | NA | 573 | 54.7 | 54.7 | 19% | 6e-05 | 60.87 | 513 | NZ_UKJA020000... |

Alignments:

>Photobacterium rosenbergii strain DP6N14-7 DP6N14-7_ctg0054, whole genome shotgun sequence
Sequence ID: NZ_JAHVIB010000049.1 Length: 960
Range 1: 222 to 839

Score:324 bits(830), Expect:5e-108,
Method:Compositional matrix adjust.,
Identities:162/206(79%), Positives:182/206(88%), Gaps:0/206(0%)

```
Query  1    MGGKWSKSSVIGWPTVRERMRRAEPAADRVGAASRDLEKHGaitssntaatnaacaWLEA  60
            MGGKWSKSS+IGWP +RERMRR EPAAD VGA S+DLEK GA+TS+NTA TNA CAWLEA
Sbjct  222  MGGKWSKSSIIGWPKIRERMRRTEPAADGVGAVSQDLEKRGAVTSNNTAETNADCAWLEA  401

Query  61   QEEEEVGFPVTPQVPLRPMTYKAAVDLSHFLKEKGGLEGLIHSQRRQDILDLWIYHTQGY  120
            QEEEEVGFPV PQVPLRPMT+K A DLS FLKEKGGLEGLI+S++RQ+ILDLW+YHTQGY
Sbjct  402  QEEEEVGFPVRPQVPLRPMTFKGAFDLSFFLKEKGGLEGLIYSKKRQEILDLWVYHTQGY  581

Query  121  FPDWQNYTPGPGVRYPLTFGWCYKLVPVEPDKIEEANKGENTSLLHPVSLHGMDDPEREV  180
            FPDWQNYTPGPGVRYPLTFGWC+KLVPV+ + +EEANKGE+  LLHPV  HGMDD  +EV
Sbjct  582  FPDWQNYTPGPGVRYPLTFGWCFKLVPVDQEAVEEANKGEDNCLLHPVCQHGMDDEHKEV  761

Query  181  LEWRFDSRLAFHHVARELHPEYFKNC  206
            L W+FDS LA  H ARELHPE++KNC
Sbjct  762  LMWKFDSSLARTHRARELHPEFYKNC  839
```

>Bacillus velezensis strain TN14-11 TN14-11_ctg0036, whole genome shotgun sequence
Sequence ID: NZ_JAHVIU010000019.1 Length: 1101
Range 1: 363 to 980

Score:325 bits(833), Expect:9e-108,
Method:Compositional matrix adjust.,
Identities:163/206(79%), Positives:182/206(88%), Gaps:0/206(0%)

```
Query  1    MGGKWSKSSVIGWPTVRERMRRAEPAADRVGAASRDLEKHGaitssntaatnaacaWLEA  60
            MGGKWSKSS+IGWP +RERMRR EPAAD VGA S+DLEK GA+TS+NTA TNA CAWLEA
Sbjct  363  MGGKWSKSSIIGWPKIRERMRRTEPAADGVGAVSQDLEKRGAVTSNNTAETNADCAWLEA  542

Query  61   QEEEEVGFPVTPQVPLRPMTYKAAVDLSHFLKEKGGLEGLIHSQRRQDILDLWIYHTQGY  120
            QEEEEVGFPV PQVPLRPMT+K A DLS FLKEKGGLEGLI+S++RQ+ILDLW+YHTQGY
Sbjct  543  QEEEEVGFPVRPQVPLRPMTFKGAFDLSFFLKEKGGLEGLIYSKKRQEILDLWVYHTQGY  722

Query  121  FPDWQNYTPGPGVRYPLTFGWCYKLVPVEPDKIEEANKGENTSLLHPVSLHGMDDPEREV  180
            FPDWQNYTPGPGVRYPLTFGWC+KLVPVE + +EEANKGE+  LLHPV  HGMDD  +EV
Sbjct  723  FPDWQNYTPGPGVRYPLTFGWCFKLVPVEQEAVEEANKGEDNCLLHPVCQHGMDDEHKEV  902

Query  181  LEWRFDSRLAFHHVARELHPEYFKNC  206
            L W+FDS LA  H ARELHPE++KNC
Sbjct  903  LMWKFDSSLARTHRARELHPEFYKNC  980
```

>Acinetobacter baumannii strain P2 contig_P2_66, whole genome shotgun sequence
Sequence ID: NZ_JBBXLW010000045.1 Length: 4185
Range 1: 2473 to 2994

Score:323 bits(828), Expect:3e-98,
Method:Compositional matrix adjust.,
Identities:169/174(97%), Positives:170/174(97%), Gaps:0/174(0%)

```
Query  33    ASRDLEKHGaitssntaatnaacaWLEAQEEEEVGFPVTPQVPLRPMTYKAAVDLSHFLK  92
              SRDLEKHGAITSSNTAATNA CAWLEAQEEEEVGFPVTPQVPLRPMTYKAAVDLSHFLK
Sbjct  2994  CSRDLEKHGAITSSNTAATNADCAWLEAQEEEEVGFPVTPQVPLRPMTYKAAVDLSHFLK  2815

Query  93    EKGGLEGLIHSQRRQDILDLWIYHTQGYFPDWQNYTPGPGVRYPLTFGWCYKLVPVEPDK  152
             EK GLEGLIHSQRRQDILDLWIYHTQGYFPD QNYTPGPGVRYPLTFGWCYKLVPVEPDK
Sbjct  2814  EKRGLEGLIHSQRRQDILDLWIYHTQGYFPD*QNYTPGPGVRYPLTFGWCYKLVPVEPDK  2635

Query  153   IEEANKGENTSLLHPVSLHGMDDPEREVLEWRFDSRLAFHHVARELHPEYFKNC  206
             +EEANKGENTSLLHPVSLHGMDDPEREVLEWRFDSRLAFHHVARELHPEYFKNC
Sbjct  2634  VEEANKGENTSLLHPVSLHGMDDPEREVLEWRFDSRLAFHHVARELHPEYFKNC  2473
```

>Qipengyuania pelagi strain JCM 17468 JCM17468_9, whole genome shotgun sequence
Sequence ID: NZ_WTYD01000009.1 Length: 1945
Range 1: 1416 to 1916

Score:304 bits(779), Expect:4e-96,
Method:Compositional matrix adjust.,
Identities:158/167(95%), Positives:164/167(98%), Gaps:0/167(0%)

```
Query  35    RDLEKHGaitssntaatnaacaWLEAQEEEEVGFPVTPQVPLRPMTYKAAVDLSHFLKEK  94
             RDLEKHGAITSSNTAATNA CAWLEAQEEEEVGFPVTPQVPLRPMTYKAAVDLSHFLKEK
Sbjct  1416  RDLEKHGAITSSNTAATNADCAWLEAQEEEEVGFPVTPQVPLRPMTYKAAVDLSHFLKEK  1595

Query  95    GGLEGLIHSQRRQDILDLWIYHTQGYFPDWQNYTPGPGVRYPLTFGWCYKLVPVEPDKIE  154
             GGLEGLIHSQRRQDILDLWIYHTQGYFPDWQNYTPGPG+RYPLTFGWCYKLVPVE +K+E
Sbjct  1596  GGLEGLIHSQRRQDILDLWIYHTQGYFPDWQNYTPGPGIRYPLTFGWCYKLVPVEQEKVE  1775

Query  155   EANKGENTSLLHPVSLHGMDDPEREVLEWRFDSRLAFHHVARELHPE  201
             EAN+GENT LLHPVSLHGMDDPEREVLEWRFDSRLAFHH+ARELHP+
Sbjct  1776  EANEGENTRLLHPVSLHGMDDPEREVLEWRFDSRLAFHHMARELHPD  1916
```

>Klebsiella pneumoniae strain EuSCAPE_NO002, whole genome shotgun sequence
Sequence ID: NZ_UKMC02000074.1 Length: 676
Range 1: 1 to 573

Score:269 bits(687), Expect:1e-87,
Method:Compositional matrix adjust.,
Identities:139/191(73%), Positives:168/191(87%), Gaps:1/191(0%)

```
Query  17   RERMRRAEPAADRVGAASRDLEKHGaitssntaatnaacaWLEAQEE-EEVGFPVTPQVP  75
            RER+RR +PAA++V AAS+DL+K+ A+TSSNTAATNA CAWLEAQEE +EV FPV PQVP
Sbjct  1    RERIRRTKPAAEKVRAASQDLDKYRALTSSNTAATNADCAWLEAQEEADEVSFPVRPQVP  180
```

```
Query  76   LRPMTYKAAVDLSHFLKEKGGLEGLIHSQRRQDILDLWIYHTQGYFPDWQNYTPGPGVRY  135
            LRPMTYKAAVDLS FLKEKGGL+ LI+S+++QDILDL +YHTQG+FPDWQNYTP P VRY
Sbjct  181  LRPMTYKAAVDLSFFLKEKGGLDELIYSKKKQDILDL*VYHTQGFFPDWQNYTPRPKVRY  360

Query  136  PLTFGWCYKLVPVEPDKIEEANKGENTSLLHPVSLHGMDDPEREVLEWRFDSRLAFHHVA  195
            PLTF WCYKLVPV+P +++EANK ENT LLHP+S H ++D  REVL+W+FDS LA  H+A
Sbjct  361  PLTFRWCYKLVPVDPKEVKEANKEENTCLLHPISQHKIEDAHREVLQWQFDSHLARRHMA  540

Query  196  RELHPEYFKNC  206
            RELHP+Y+K+C
Sbjct  541  RELHPKYYKDC  573
```

>Klebsiella pneumoniae strain EuSCAPE_IL007, whole genome shotgun sequence
Sequence ID: NZ_UKJT02000022.1 Length: 697
Range 1: 104 to 625

Score:260 bits(664), Expect:4e-84,
Method:Compositional matrix adjust.,
Identities:133/174(76%), Positives:156/174(89%), Gaps:1/174(0%)

```
Query  34   SRDLEKHGaitssntaatnaacaWLEAQEE-EEVGFPVTPQVPLRPMTYKAAVDLSHFLK  92
            S+DL +HGAIT+SNTAATNA  AW+EAQEE EEVGFPV PQVPLRPMTYK A DL  FLK
Sbjct  625  SQDLARHGAITTSNTAATNADVAWVEAQEEGEEVGFPVRPQVPLRPMTYKGAFDLGFFLK  446

Query  93   EKGGLEGLIHSQRRQDILDLWIYHTQGYFPDWQNYTPGPGVRYPLTFGWCYKLVPVEPDK  152
            EKGGL+GLI S++RQ+ILDLW+Y+TQGYFPDWQNYTPGPGVRYPLTFGWCYKLVPV+P +
Sbjct  445  EKGGLDGLIWSKKRQEILDLWVYNTQGYFPDWQNYTPGPGVRYPLTFGWCYKLVPVDPRE  266

Query  153  IEEANKGENTSLLHPVSLHGMDDPEREVLEWRFDSRLAFHHVARELHPEYFKNC  206
            +EEAN+GEN  LLHP+SLHGM+D  +EVL+W FDS+LA  H+ARELHP+Y+K+C
Sbjct  265  VEEANEGENNCLLHPMSLHGMEDEHKEVLKWVFDSQLARRHIARELHPDYYKDC  104
```

>Acinetobacter baumannii strain XH798 XH798_contig_187, whole genome shotgun sequence
Sequence ID: NZ_LYGA01000021.1 Length: 588
Range 1: 206 to 586

Score:253 bits(646), Expect:7e-82,
Method:Compositional matrix adjust.,
Identities:117/127(92%), Positives:125/127(98%), Gaps:0/127(0%)

```
Query  75   PLRPMTYKAAVDLSHFLKEKGGLEGLIHSQRRQDILDLWIYHTQGYFPDWQNYTPGPGVR  134
            PLRPMTY++AVDLSHFLKEKGGLEGLIHSQRRQDILDLWIYHTQGYFPDWQNYTPGPG+R
Sbjct  586  PLRPMTYQSAVDLSHFLKEKGGLEGLIHSQRRQDILDLWIYHTQGYFPDWQNYTPGPGIR  407

Query  135  YPLTFGWCYKLVPVEPDKIEEANKGENTSLLHPVSLHGMDDPEREVLEWRFDSRLAFHHV  194
            YPLTFGWCYKLVPVE +K+EEAN+GENT LLHPVSLHGMDDPEREVLEWRFDSRLAFHH+
Sbjct  406  YPLTFGWCYKLVPVEQEKVEEANEGENTRLLHPVSLHGMDDPEREVLEWRFDSRLAFHHM  227

Query  195  ARELHPE  201
            ARELHP+
Sbjct  226  ARELHPD  206
```

>Klebsiella pneumoniae strain EuSCAPE_MT009, whole genome shotgun sequence
Sequence ID: NZ_UKIV02000085.1 Length: 670
Range 1: 2 to 544

Score:251 bits(640), Expect:2e-80,
Method:Compositional matrix adjust.,
Identities:129/181(71%), Positives:155/181(85%), Gaps:1/181(0%)

```
Query  26   AADRVGAASRDLEKHGaitssntaatnaacaWLEAQEE-EEVGFPVTPQVPLRPMTYKAA  84
            AA+RV AASRDLE+H A+T+SNTA  N  CAWL AQ+E EEVGFPV PQVP+RPMTYK A
Sbjct  2    AAERVRAASRDLERHEALTTSNTATNNPNCAWLAAQKEKEEVGFPVRPQVPVRPMTYKRA  181

Query  85   VDLSHFLKEKGGLEGLIHSQRRQDILDLWIYHTQGYFPDWQNYTPGPGVRYPLTFGWCYK  144
             DLS FLKEKGGL+GLI+SQ+RQDILDLW+Y+TQG+FPDWQNYTPGP VRYPLTF  CYK
Sbjct  182  FDLSFFLKEKGGLDGLIYSQQRQDILDLWVYNTQGFFPDWQNYTPGPEVRYPLTFK*CYK  361

Query  145  LVPVEPDKIEEANKGENTSLLHPVSLHGMDDPEREVLEWRFDSRLAFHHVARELHPEYFK  204
            LVPV+P ++E+ NKG+N SLLHP+S H ++D  REVL+ +FD+ LA  H+ARELHPEY+K
Sbjct  362  LVPVDPSEVEKDNKGKNNSLLHPISQHRIEDTNREVLK*KFDTNLARRHLARELHPEYYK  541

Query  205  N  205
            +
Sbjct  542  D  544
```

>Klebsiella pneumoniae strain EuSCAPE_DE088, whole genome shotgun sequence
Sequence ID: NZ_UKLX02000048.1 Length: 535
Range 1: 91 to 528

Score:241 bits(615), Expect:2e-77,
Method:Compositional matrix adjust.,
Identities:111/146(76%), Positives:131/146(89%), Gaps:0/146(0%)

```
Query  61   QEEEEVGFPVTPQVPLRPMTYKAAVDLSHFLKEKGGLEGLIHSQRRQDILDLWIYHTQGY  120
            +E+EEVGFPV PQVPLRPMTYK A DLS FLKEKGGL+ LI+S++RQDILDLW+YHTQGY
Sbjct  528  EEDEEVGFPVRPQVPLRPMTYKGAFDLSFFLKEKGGLDRLIYSKKRQDILDLWVYHTQGY  349

Query  121  FPDWQNYTPGPGVRYPLTFGWCYKLVPVEPDKIEEANKGENTSLLHPVSLHGMDDPEREV  180
            FPDWQNYTPGPGVRYPLTF W +KLVPV+P +IEEAN+G+N  LLHP++ HGM+D +REV
Sbjct  348  FPDWQNYTPGPGVRYPLTFRWPFKLVPVDPREIEEANEGDNACLLHPINQHGMEDADREV  169

Query  181  LEWRFDSRLAFHHVARELHPEYFKNC  206
            L+W+FDS LA  HVARELHPEY+K+C
Sbjct  168  LKWQFDSHLARRHVARELHPEYYKDC  91
```

>Sphingomonas sp. UV9 NODE_33_length_614_cov_0.261122, whole genome shotgun sequence
Sequence ID: NZ_SCIN01000017.1 Length: 614
Range 1: 2 to 403

Score:239 bits(609), Expect:4e-76,
Method:Compositional matrix adjust.,
Identities:129/134(96%), Positives:130/134(97%), Gaps:0/134(0%)

```
Query  33   ASRDLEKHGaitssntaatnaacaWLEAQEEEEVGFPVTPQVPLRPMTYKAAVDLSHFLK  92
             SRDLEKHGAITSSNTAATNA CAWLEAQEEEEVGFPVTPQVPLRPMTYKAAVDLSHFLK
Sbjct  403  CSRDLEKHGAITSSNTAATNADCAWLEAQEEEEVGFPVTPQVPLRPMTYKAAVDLSHFLK  224
```

```
Query  93   EKGGLEGLIHSQRRQDILDLWIYHTQGYFPDWQNYTPGPGVRYPLTFGWCYKLVPVEPDK  152
            EK GLEGLIHSQRRQDILDLWIYHTQGYFPD QNYTPGPGVRYPLTFGWCYKLVPVEPDK
Sbjct  223  EKRGLEGLIHSQRRQDILDLWIYHTQGYFPD*QNYTPGPGVRYPLTFGWCYKLVPVEPDK  44

Query  153  IEEANKGENTSLLH  166
            +EEANKGENTSLLH
Sbjct  43   VEEANKGENTSLLH  2
```

>Klebsiella pneumoniae strain EuSCAPE_IL022, whole genome shotgun sequence
Sequence ID: NZ_UKKD02000021.1 Length: 594
Range 1: 3 to 551

Score:224 bits(572), Expect:1e-70,
Method:Compositional matrix adjust.,
Identities:127/183(69%), Positives:150/183(81%), Gaps:13/183(7%)

```
Query  12   GWPTVRERMRRAEPAADRVGAA------------SRDLEKHGaitssntaatnaacaWLE  59
            GW  + E++R+ +PAA+  GAA            SRDL+K+GA+TSSNT ATNA CAWLE
Sbjct  551  GWSAMMEKIRKTDPAAEGAGAAEGVGAAAGVGAASRDLDKYGALTSSNTGATNADCAWLE  372

Query  60   AQEE-EEVGFPVTPQVPLRPMTYKAAVDLSHFLKEKGGLEGLIHSQRRQDILDLWIYHTQ  118
            AQEE EEVGFPV PQVPLRPMTYK A D+  FLKEKGGL+GLIHS++RQDILDLW+Y+TQ
Sbjct  371  AQEESEEVGFPVRPQVPLRPMTYKGAFDIGFFLKEKGGLDGLIHSKKRQDILDLWVYNTQ  192

Query  119  GYFPDWQNYTPGPGVRYPLTFGWCYKLVPVEPDKIEEANKGENTSLLHPVSLHGMDDPER  178
            GYFPDWQNYTPGPGVRYPLTFGW +KLVPV+P ++E AN+GEN  LLHP+S HGM+D +R
Sbjct  191  GYFPDWQNYTPGPGVRYPLTFGWPFKLVPVDPREVEGANEGENNCLLHPISQHGMEDDDR  12

Query  179  EVL  181
            EVL
Sbjct  11   EVL  3
```

>Klebsiella pneumoniae strain EuSCAPE_IL021, whole genome shotgun sequence
Sequence ID: NZ_UKJX02000113.1 Length: 415
Range 1: 2 to 412

Score:218 bits(554), Expect:1e-68,
Method:Compositional matrix adjust.,
Identities:100/137(73%), Positives:120/137(87%), Gaps:0/137(0%)

```
Query  61   QEEEEVGFPVTPQVPLRPMTYKAAVDLSHFLKEKGGLEGLIHSQRRQDILDLWIYHTQGY  120
            + E EVGFPV PQVPLRPMTYK AVDL  FLKEKGGL+GLI+S++R++ILDLW+Y+TQGY
Sbjct  2    ETEGEVGFPVRPQVPLRPMTYKGAVDLGFFLKEKGGLDGLIYSEKRKEILDLWVYNTQGY  181

Query  121  FPDWQNYTPGPGVRYPLTFGWCYKLVPVEPDKIEEANKGENTSLLHPVSLHGMDDPEREV  180
            FPDWQNYTPGPG+RYPLTFGW +KLVPV+P ++EEAN+GEN  LLHP+  HGM+D  REV
Sbjct  182  FPDWQNYTPGPGIRYPLTFGWPFKLVPVDPREVEEANEGENNCLLHPMHQHGMEDEHREV  361

Query  181  LEWRFDSRLAFHHVARE  197
            L W+FDS+LA  H+ARE
Sbjct  362  LMWKFDSQLARRHIARE  412
```

>Acinetobacter baumannii strain XH797 XH797_contig_168, whole genome shotgun sequence
Sequence ID: NZ_LYGB01000013.1 Length: 418
Range 1: 3 to 323

Score:215 bits(548), Expect:1e-67,
Method:Compositional matrix adjust.,
Identities:99/107(93%), Positives:105/107(98%), Gaps:0/107(0%)

```
Query  95   GGLEGLIHSQRRQDILDLWIYHTQGYFPDWQNYTPGPGVRYPLTFGWCYKLVPVEPDKIE  154
            GGLEGLIHSQRRQDILDLWIYHTQGYFPDWQNYTPGPG+RYPLTFGWCYKLVPVE +K+E
Sbjct  3    GGLEGLIHSQRRQDILDLWIYHTQGYFPDWQNYTPGPGIRYPLTFGWCYKLVPVEQEKVE  182

Query  155  EANKGENTSLLHPVSLHGMDDPEREVLEWRFDSRLAFHHVARELHPE  201
            EAN+GENT LLHPVSLHGMDDPEREVLEWRFDSRLAFHH+ARELHP+
Sbjct  183  EANEGENTRLLHPVSLHGMDDPEREVLEWRFDSRLAFHHMARELHPD  323
```

>Klebsiella pneumoniae strain EuSCAPE_IL001, whole genome shotgun sequence
Sequence ID: NZ_UKJN02000091.1 Length: 519
Range 1: 50 to 517

Score:213 bits(541), Expect:4e-66,
Method:Compositional matrix adjust.,
Identities:118/160(74%), Positives:132/160(82%), Gaps:19/160(11%)

```
Query  1    MGGKWSKSSVIGWPTVRERMRRAEPAADR--------------VGAASRDLEKHGaitss  46
            MGGKWSK    GWP VR+RMR+A+PAA+R              VGAASRDLEKHGA+T+S
Sbjct  50   MGGKWSK----GWPAVRDRMRQADPAAERLGARRRQTEPAAVGVGAASRDLEKHGALTTS  217

Query  47   ntaatnaacaWLEAQEEEE-VGFPVTPQVPLRPMTYKAAVDLSHFLKEKGGLEGLIHSQR  105
            NT A NAACAWL+AQEE+E VGFPV PQVPLRPMTYK A DL  FLKEKGGL GLI+S++
Sbjct  218  NTPANNAACAWLQAQEEDEDVGFPVRPQVPLRPMTYKGAFDLGFFLKEKGGLHGLIYSKK  397

Query  106  RQDILDLWIYHTQGYFPDWQNYTPGPGVRYPLTFGWCYKL  145
            RQDILDLW+Y+TQG+FPDWQNYTPGPGVRYPLTFGWCYKL
Sbjct  398  RQDILDLWVYNTQGFFPDWQNYTPGPGVRYPLTFGWCYKL  517
```

>Klebsiella pneumoniae strain EuSCAPE_MT003, whole genome shotgun sequence
Sequence ID: NZ_UKIK02000085.1 Length: 964
Range 1: 2 to 475

Score:216 bits(551), Expect:1e-65,
Method:Compositional matrix adjust.,
Identities:118/160(74%), Positives:135/160(84%), Gaps:12/160(7%)

```
Query  1    MGGKWSKSSVIGWPTVRERMRRA---------EPAADRVGAASRDLEKHGaitssntaat  51
            MGGK SKS  IGW  +RERMR+          E AA+ VGAASRDL+KHGA+TSSNTA+
Sbjct  475  MGGKMSKS--IGWSAIRERMRKTPVGERARENESAAEGVGAASRDLDKHGALTSSNTASN  302

Query  52   naacaWLEAQEE-EEVGFPVTPQVPLRPMTYKAAVDLSHFLKEKGGLEGLIHSQRRQDIL  110
            NA CAWLEAQEE ++VGFPV PQVPLRPMTYK AVDL  FLKEKGGL+GL+HSQ+RQ+IL
Sbjct  301  NADCAWLEAQEEADDVGFPVCPQVPLRPMTYKGAVDLGFFLKEKGGLDGLVHSQKRQEIL  122

Query  111  DLWIYHTQGYFPDWQNYTPGPGVRYPLTFGWCYKLVPVEP  150
            DLW+YHTQG+FPDWQNYTPGPG+RYPLTFGW +KLVPV+P
Sbjct  121  DLWVYHTQGFFPDWQNYTPGPGIRYPLTFGWPFKLVPVDP  2
```

```
>Klebsiella pneumoniae strain EuSCAPE_LT010, whole genome shotgun sequence
Sequence ID: NZ_UKJK02000057.1 Length: 494
Range 1: 2 to 367

Score:202 bits(515), Expect:2e-62,
Method:Compositional matrix adjust.,
Identities:91/122(75%), Positives:109/122(89%), Gaps:0/122(0%)

Query  84   AVDLSHFLKEKGGLEGLIHSQRRQDILDLWIYHTQGYFPDWQNYTPGPGVRYPLTFGWCY  143
            A DL  FLKEKGGL+GLI+S++RQ+ILDLW+YHTQG+FPDWQNYTPGPGVRYPLTFGWCY
Sbjct  2    AFDLGFFLKEKGGLDGLIYSKKRQEILDLWVYHTQGFFPDWQNYTPGPGVRYPLTFGWCY  181

Query  144  KLVPVEPDKIEEANKGENTSLLHPVSLHGMDDPEREVLEWRFDSRLAFHHVARELHPEYF  203
            KLVPV+P ++EEAN+GEN  LLHP+S HGM+D   EVL+W+FDS LA  H+ARELHPEY+
Sbjct  182  KLVPVDPQEVEEANQGENNCLLHPMSQHGMEDDCGEVLKWKFDSSLARRHMARELHPEYY  361

Query  204  KN  205
            K+
Sbjct  362  KD  367




>Klebsiella pneumoniae strain EuSCAPE_LT004, whole genome shotgun sequence
Sequence ID: NZ_UKJE02000067.1 Length: 348
Range 1: 1 to 336

Score:195 bits(496), Expect:4e-60,
Method:Compositional matrix adjust.,
Identities:93/112(83%), Positives:102/112(91%), Gaps:1/112(0%)

Query  57   WLEAQE-EEEVGFPVTPQVPLRPMTYKAAVDLSHFLKEKGGLEGLIHSQRRQDILDLWIY  115
            WLEAQE EEEVGFPV PQVPLRPMTYK AVDL  FLKEKGGLEGLI+S++RQ+ILDL +Y
Sbjct  336  WLEAQEQEEEVGFPVRPQVPLRPMTYKGAVDLRFFLKEKGGLEGLIYSKKRQEILDL*VY  157

Query  116  HTQGYFPDWQNYTPGPGVRYPLTFGWCYKLVPVEPDKIEEANKGENTSLLHP  167
            HTQGYFPDW NYTPGPGVRYPLTFGWCYKLVPV+P ++EEAN+GEN  LLHP
Sbjct  156  HTQGYFPDWHNYTPGPGVRYPLTFGWCYKLVPVDPQELEEANEGENNCLLHP  1




>Klebsiella pneumoniae strain EuSCAPE_DE083, whole genome shotgun sequence
Sequence ID: NZ_UKLY02000035.1 Length: 349
Range 1: 10 to 348

Score:194 bits(493), Expect:9e-60,
Method:Compositional matrix adjust.,
Identities:87/113(77%), Positives:103/113(91%), Gaps:0/113(0%)

Query  61   QEEEEVGFPVTPQVPLRPMTYKAAVDLSHFLKEKGGLEGLIHSQRRQDILDLWIYHTQGY  120
            +EEE+VGFPV PQVPLRPMTYKAA DLS FLKEKGGL+GLI+S++RQDILDLW+Y+TQG+
Sbjct  10   KEEEDVGFPVRPQVPLRPMTYKAAFDLSFFLKEKGGLDGLIYSKKRQDILDLWVYNTQGF  189

Query  121  FPDWQNYTPGPGVRYPLTFGWCYKLVPVEPDKIEEANKGENTSLLHPVSLHGM  173
            FPDWQNYTPGPG R PLTFGWCYKLVPV+P ++EEA +GE+  LLHP++ HGM
Sbjct  190  FPDWQNYTPGPGTRLPLTFGWCYKLVPVDPREVEEATEGEDNCLLHPMNQHGM  348




>Klebsiella pneumoniae strain EuSCAPE_MT011, whole genome shotgun sequence
Sequence ID: NZ_UKIY02000084.1 Length: 431
Range 1: 103 to 429

Score:188 bits(477), Expect:8e-57,
Method:Compositional matrix adjust.,
Identities:82/109(75%), Positives:100/109(91%), Gaps:0/109(0%)

Query  98   EGLIHSQRRQDILDLWIYHTQGYFPDWQNYTPGPGVRYPLTFGWCYKLVPVEPDKIEEAN  157
            EGL++S++RQDILDLW+Y+TQGYFPDWQNYTPGPGVRYPLT GWC+KLVPV+P ++EEAN
Sbjct  429  EGLVYSKQRQDILDLWVYNTQGYFPDWQNYTPGPGVRYPLTLGWCFKLVPVDPREVEEAN  250

Query  158  KGENTSLLHPVSLHGMDDPEREVLEWRFDSRLAFHHVARELHPEYFKNC  206
            +GEN+ LLHPVS HGM+D +REVL W+FDS LA  H+ARELHPEY+K+C
Sbjct  249  QGENSCLLHPVSQHGMEDEDREVLVWKFDSNLARRHMARELHPEYYKDC  103




>Acinetobacter baumannii strain XH744 XH744_contig_84, whole genome shotgun sequence
Sequence ID: NZ_LYHI01000082.1 Length: 397
Range 1: 112 to 396

Score:184 bits(466), Expect:3e-55,
Method:Compositional matrix adjust.,
Identities:84/95(88%), Positives:91/95(95%), Gaps:0/95(0%)

Query  107  QDILDLWIYHTQGYFPDWQNYTPGPGVRYPLTFGWCYKLVPVEPDKIEEANKGENTSLLH  166
            Q +LDLWI HTQGYFPDWQNYTPGPG+RYPLTFGWCYKLVPVE +K+EEAN+GENT LLH
Sbjct  396  QGMLDLWI*HTQGYFPDWQNYTPGPGIRYPLTFGWCYKLVPVEQEKVEEANEGENTRLLH  217

Query  167  PVSLHGMDDPEREVLEWRFDSRLAFHHVARELHPE  201
            PVSLHGMDDPEREVLEWRFDSRLAFHH+ARELHP+
Sbjct  216  PVSLHGMDDPEREVLEWRFDSRLAFHHMARELHPD  112




>Klebsiella pneumoniae strain EuSCAPE_IL020, whole genome shotgun sequence
Sequence ID: NZ_UKKB02000091.1 Length: 422
Range 1: 1 to 420

Score:179 bits(455), Expect:2e-53,
Method:Compositional matrix adjust.,
Identities:107/142(75%), Positives:120/142(84%), Gaps:11/142(7%)

Query  4    KWSKSSVIGWPTVRERMRRA--------EPAADRVGAASRDLEKHGaitssntaatnaac  55
            KWSKS+   W  +RER+RRA        EPAA  VGAAS+DL+K+GA+TSSNT ATNA C
Sbjct  1    KWSKSN--SWAAIRERIRRAVPAVPAGAEPAAVGVGAASQDLDKYGALTSSNTPATNATC  174

Query  56   aWLEAQEEEE-VGFPVTPQVPLRPMTYKAAVDLSHFLKEKGGLEGLIHSQRRQDILDLWI  114
            AW+EAQEEEE VGFPV PQVPLRPMTYK AVDL  FLKEKGGL+G+IHS++RQDILDLW+
Sbjct  175  AWVEAQEEEEEVGFPVRPQVPLRPMTYKGAVDLGFFLKEKGGLDGMIHSKKRQDILDLWV  354

Query  115  YHTQGYFPDWQNYTPGPGVRYP  136
            YHTQGYFPDWQNYTPGPGVRYP
Sbjct  355  YHTQGYFPDWQNYTPGPGVRYP  420
```

```
>Klebsiella pneumoniae strain EuSCAPE_IL005, whole genome shotgun sequence
Sequence ID: NZ_UKJL02000062.1 Length: 307
Range 1: 8 to 307

Score:173 bits(439), Expect:1e-51,
Method:Compositional matrix adjust.,
Identities:77/100(77%), Positives:93/100(93%), Gaps:0/100(0%)

Query  91   LKEKGGLEGLIHSQRRQDILDLWIYHTQGYFPDWQNYTPGPGVRYPLTFGWCYKLVPVEP  150
            LKEKGGL+GLI+S++RQDILDLW+Y+TQG+FPDWQNYTPGPGVRYPLTFGWC+KLVPV+P
Sbjct  307  LKEKGGLDGLIYSKQRQDILDLWVYNTQGFFPDWQNYTPGPGVRYPLTFGWCFKLVPVDP  128

Query  151  DKIEEANKGENTSLLHPVSLHGMDDPEREVLEWRFDSRLA  190
             ++EEANKGEN  LLHP+S HGM+D   EVL+W+FDS+LA
Sbjct  127  REVEEANKGENNCLLHPMSQHGMEDEHGEVLKWKFDSQLA  8



>Pseudomonas aeruginosa strain ENV-297 NODE_110_length_378_cov_2.35548_ID_219, whole genome shotgun sequence
Sequence ID: NZ_RADD01000248.1 Length: 378
Range 1: 31 to 351

Score:174 bits(440), Expect:2e-51,
Method:Compositional matrix adjust.,
Identities:77/108(71%), Positives:92/108(85%), Gaps:1/108(0%)

Query  99   GLIHSQRRQDILDLWIYHTQGYFPDWQNYTPGPGVRYPLTFGWCYKLVPVEPDKIEEANK  158
            GLI+S++RQ+ILDLW+YHTQGYFPDWQNYTPGPG R+PLTFGWCYKLVPV+P ++EEA
Sbjct  351  GLIYSKKRQEILDLWVYHTQGYFPDWQNYTPGPGTRFPLTFGWCYKLVPVDPREVEEATT  172

Query  159  GENTSLLHPVSLHGMDDPEREVLEWRFDSRLAFHHVARELHPEYFKNC  206
            GEN  LLHP+ LHG +D E EVL W+FDS LA  H+ARE HPE++ NC
Sbjct  171  GENNCLLHPICLHGAEDEEGEVLMWKFDSSLARRHIAREKHPEFY-NC  31



>Klebsiella pneumoniae strain EuSCAPE_MT019, whole genome shotgun sequence
Sequence ID: NZ_UKJD02000054.1 Length: 332
Range 1: 1 to 330

Score:172 bits(437), Expect:3e-51,
Method:Compositional matrix adjust.,
Identities:84/110(76%), Positives:93/110(84%), Gaps:0/110(0%)

Query  90   FLKEKGGLEGLIHSQRRQDILDLWIYHTQGYFPDWQNYTPGPGVRYPLTFGWCYKLVPVE  149
            FLKEKGGLEGL  SQRRQDILDL +Y+TQGYFPDWQNYTPGPGVRYPLTFGW +KLVPV+
Sbjct  1    FLKEKGGLEGLP*SQRRQDILDL*VYNTQGYFPDWQNYTPGPGVRYPLTFGWPFKLVPVD  180

Query  150  PDKIEEANKGENTSLLHPVSLHGMDDPEREVLEWRFDSRLAFHHVARELH  199
            P ++EEAN GEN  LLHPVS H MDD +REVL  +FDS L   H+ARELH
Sbjct  181  PKEVEEANNGENNCLLHPVSQHRMDDADREVLI*KFDSHLVHRHMARELH  330



>Klebsiella pneumoniae strain EuSCAPE_MT015, whole genome shotgun sequence
Sequence ID: NZ_UKIW02000106.1 Length: 372
Range 1: 1 to 372

Score:171 bits(433), Expect:2e-50,
Method:Compositional matrix adjust.,
Identities:99/128(77%), Positives:112/128(87%), Gaps:7/128(5%)

Query  4    KWSKSSVIGWPTVRERMRRA--EPAADRVGAASRDLEKHGaitssntaatnaacaWLEAQ  61
            KWSKS    WP VR+RMRR   EPAA+ VGAAS+DL+KHGA+TSSNTAATN  CAWLEAQ
Sbjct  372  KWSKS----WPAVRDRMRRTRTEPAAEGVGAASQDLDKHGALTSSNTAATNNDCAWLEAQ  205

Query  62   EE-EEVGFPVTPQVPLRPMTYKAAVDLSHFLKEKGGLEGLIHSQRRQDILDLWIYHTQGY  120
            EE E+VGFPV PQVPLRPMTYKAA DL  FLKEKGGL+GLI+S++RQ+ILDLW+YHTQG+
Sbjct  204  EESEDVGFPVRPQVPLRPMTYKAAFDLGWFLKEKGGLDGLIYSKKRQEILDLWVYHTQGF  25

Query  121  FPDWQNYT  128
            FPDWQNYT
Sbjct  24   FPDWQNYT  1



>Acinetobacter baumannii strain XH821 XH821_contig_233, whole genome shotgun sequence
Sequence ID: NZ_LYFC01000027.1 Length: 572
Range 1: 280 to 570

Score:172 bits(436), Expect:6e-50,
Method:Compositional matrix adjust.,
Identities:96/97(99%), Positives:96/97(98%), Gaps:0/97(0%)

Query  35   RDLEKHGaitssntaatnaacaWLEAQEEEEVGFPVTPQVPLRPMTYKAAVDLSHFLKEK  94
            RDLEKHGAITSSNTAATNA CAWLEAQEEEEVGFPVTPQVPLRPMTYKAAVDLSHFLKEK
Sbjct  280  RDLEKHGAITSSNTAATNADCAWLEAQEEEEVGFPVTPQVPLRPMTYKAAVDLSHFLKEK  459

Query  95   GGLEGLIHSQRRQDILDLWIYHTQGYFPDWQNYTPGP  131
            GGLEGLIHSQRRQDILDLWIYHTQGYFPDWQNYTPGP
Sbjct  460  GGLEGLIHSQRRQDILDLWIYHTQGYFPDWQNYTPGP  570



>Klebsiella pneumoniae strain EuSCAPE_MT014, whole genome shotgun sequence
Sequence ID: NZ_UKJF02000036.1 Length: 388
Range 1: 2 to 283

Score:166 bits(420), Expect:2e-48,
Method:Compositional matrix adjust.,
Identities:71/94(76%), Positives:84/94(89%), Gaps:0/94(0%)

Query  113  WIYHTQGYFPDWQNYTPGPGVRYPLTFGWCYKLVPVEPDKIEEANKGENTSLLHPVSLHG  172
            W+YHTQG+FPDWQNYTPGPGVRYPLTFGWC+KLVPV+P  +E+AN+GEN  LLHP+S HG
Sbjct  2    WVYHTQGFFPDWQNYTPGPGVRYPLTFGWCFKLVPVDPKDVEKANEGENNCLLHPMSQHG  181

Query  173  MDDPEREVLEWRFDSRLAFHHVARELHPEYFKNC  206
            MDDPE+EVL W+FDS LA  H+ARE HPEY+K+C
Sbjct  182  MDDPEKEVLMWKFDSSLARRHIAREQHPEYYKDC  283



>Escherichia coli strain EN5308 contig00099, whole genome shotgun sequence
Sequence ID: NZ_JARBDE010000099.1 Length: 2827
Range 1: 3 to 365

Score:168 bits(426), Expect:1e-43,
Method:Compositional matrix adjust.,
```

```
Identities:94/122(77%), Positives:109/122(89%), Gaps:2/122(1%)

Query  1    MGGKWSKSSVIGWPTVRERMRRAEPAADRVGAASRDLEKHGaitssntaatnaacaWLEA  60
            MGGKWS+S ++GWP VR+RMRR EPAA+ VGAAS+DL+K+GA+TSSNTA TNA CAWLEA
Sbjct  365  MGGKWSRSKIVGWPAVRDRMRRTEPAAEGVGAASQDLDKYGALTSSNTADTNADCAWLEA  186

Query  61   QEE-EEVGFPVTPQVPLRPMTYKAAVDLSHFLKEKGGLEGLIHSQRRQDILDLWIYHTQG  119
            QEE  EVGFPV PQV LRPMT+K A DLS FLKEKGGL+GLI+S+RRQ+ILDLW+YHTQG
Sbjct  185  QEEANEVGFPVRPQV-LRPMTFKGAFDLSFFLKEKGGLDGLIYSKRRQEILDLWVYHTQG  9

Query  120  YF  121
            YF
Sbjct  8    YF  3
```

```
>Klebsiella pneumoniae strain EuSCAPE_MT009, whole genome shotgun sequence
Sequence ID: NZ_UKIV02000098.1 Length: 262
Range 1: 10 to 261

Score:149 bits(375), Expect:3e-42,
Method:Compositional matrix adjust.,
Identities:66/84(79%), Positives:77/84(91%), Gaps:0/84(0%)

Query  122  PDWQNYTPGPGVRYPLTFGWCYKLVPVEPDKIEEANKGENTSLLHPVSLHGMDDPEREVL  181
            PDWQNYTPGPGVRYPLTFGWCYKLVPV+P +IEEAN+GEN SLLHP+S HGM+DP REVL
Sbjct  261  PDWQNYTPGPGVRYPLTFGWCYKLVPVDPSEIEEANEGENNSLLHPMSQHGMEDPNREVL  82

Query  182  EWRFDSRLAFHHVARELHPEYFKN  205
            +W+FD+ LA  H+ARELHPEY+K+
Sbjct  81   KWKFDTSLARRHLARELHPEYYKD  10
```

```
>Klebsiella pneumoniae strain EuSCAPE_IL005, whole genome shotgun sequence
Sequence ID: NZ_UKJL02000056.1 Length: 442
Range 1: 2 to 313

Score:150 bits(379), Expect:5e-42,
Method:Compositional matrix adjust.,
Identities:88/104(85%), Positives:96/104(92%), Gaps:1/104(0%)

Query  1    MGGKWSKSSVIGWPTVRERMRRAEPAADRVGAASRDLEKHGaitssntaatnaacaWLEA  60
            MGGKWSKSS++GWP VRER+RR EPAA+ VGAAS+DL+KHGA+TSSNTAATNA CAWLEA
Sbjct  313  MGGKWSKSSIVGWPAVRERIRRTEPAAEGVGAASQDLDKHGALTSSNTAATNADCAWLEA  134

Query  61   QEE-EEVGFPVTPQVPLRPMTYKAAVDLSHFLKEKGGLEGLIHS  103
            QEE EEVGFPV PQVPLRPMTYKAA DLS FLKEKGGLEGLI+S
Sbjct  133  QEESEEVGFPVRPQVPLRPMTYKAAFDLSFFLKEKGGLEGLIYS  2
```

```
>Acinetobacter baumannii strain XH821 XH821_contig_145, whole genome shotgun sequence
Sequence ID: NZ_LYFC01000009.1 Length: 427
Range 1: 1 to 222

Score:142 bits(358), Expect:8e-39,
Method:Compositional matrix adjust.,
Identities:65/74(88%), Positives:71/74(95%), Gaps:0/74(0%)

Query  128  TPGPGVRYPLTFGWCYKLVPVEPDKIEEANKGENTSLLHPVSLHGMDDPEREVLEWRFDS  187
            TPGP +RYPLTFGWCYKLVPVE +K+EEAN+GENT LLHPVSLHGMDDPEREVLEWRFDS
Sbjct  1    TPGPEIRYPLTFGWCYKLVPVEQEKVEEANEGENTRLLHPVSLHGMDDPEREVLEWRFDS  180

Query  188  RLAFHHVARELHPE  201
            RLAFHH+ARELHP+
Sbjct  181  RLAFHHMARELHPD  222
```

```
>Acinetobacter baumannii strain P25 contig_P25_82, whole genome shotgun sequence
Sequence ID: NZ_JBBXLB010000056.1 Length: 2628
Range 1: 2357 to 2626

Score:150 bits(380), Expect:2e-37,
Method:Compositional matrix adjust.,
Identities:87/90(97%), Positives:87/90(96%), Gaps:0/90(0%)

Query  33    ASRDLEKHGaitssntaatnaacaWLEAQEEEEVGFPVTPQVPLRPMTYKAAVDLSHFLK  92
              SRDLEKHGAITSSNTAATNA CAWLEAQEEEEVGFPVTPQVPLRPMTYKAAVDLSHFLK
Sbjct  2357  CSRDLEKHGAITSSNTAATNADCAWLEAQEEEEVGFPVTPQVPLRPMTYKAAVDLSHFLK  2536

Query  93    EKGGLEGLIHSQRRQDILDLWIYHTQGYFP  122
             EK GLEGLIHSQRRQDILDLWIYHTQGYFP
Sbjct  2537  EKRGLEGLIHSQRRQDILDLWIYHTQGYFP  2626
```

```
>Acinetobacter baumannii strain E13 contig_E13_79, whole genome shotgun sequence
Sequence ID: NZ_JBBXKR010000053.1 Length: 2763
Range 1: 2492 to 2761

Score:150 bits(380), Expect:2e-37,
Method:Compositional matrix adjust.,
Identities:87/90(97%), Positives:87/90(96%), Gaps:0/90(0%)

Query  33    ASRDLEKHGaitssntaatnaacaWLEAQEEEEVGFPVTPQVPLRPMTYKAAVDLSHFLK  92
              SRDLEKHGAITSSNTAATNA CAWLEAQEEEEVGFPVTPQVPLRPMTYKAAVDLSHFLK
Sbjct  2492  CSRDLEKHGAITSSNTAATNADCAWLEAQEEEEVGFPVTPQVPLRPMTYKAAVDLSHFLK  2671

Query  93    EKGGLEGLIHSQRRQDILDLWIYHTQGYFP  122
             EK GLEGLIHSQRRQDILDLWIYHTQGYFP
Sbjct  2672  EKRGLEGLIHSQRRQDILDLWIYHTQGYFP  2761
```

```
>Klebsiella pneumoniae strain EuSCAPE_LT003, whole genome shotgun sequence
Sequence ID: NZ_UKJH02000066.1 Length: 350
Range 1: 1 to 246

Score:126 bits(316), Expect:6e-33,
Method:Compositional matrix adjust.,
Identities:60/86(70%), Positives:72/86(83%), Gaps:4/86(4%)

Query  121  FPDWQNYTPGPGVRYPLTFGWCYKLVPVEPDKIEEANKGENTSLLHPVSLHGMDDPEREV  180
            FPDWQ YTPGPGVRYPLTFGWC+KLVPV+P ++EEAN+GEN  LLH  S HGM+D  +EV
Sbjct  1    FPDWQCYTPGPGVRYPLTFGWCFKLVPVDPSEVEEANEGEN--LLH--SQHGMEDEHKEV  168
```

```
Query  181  LEWRFDSRLAFHHVARELHPEYFKNC  206
            L W+FDS LA  H+ARE HPE++K+C
Sbjct  169  LVWKFDSMLARRHLARERHPEFYKDC  246
```

>Klebsiella pneumoniae strain EuSCAPE_MT002, whole genome shotgun sequence
Sequence ID: NZ_UKIM02000084.1 Length: 418
Range 1: 3 to 314

Score:121 bits(303), Expect:1e-30,
Method:Compositional matrix adjust.,
Identities:81/104(78%), Positives:88/104(84%), Gaps:7/104(6%)

```
Query  1    MGGKWSKSSVIGWPTVRERMRRAEPAADR-------VGAASRDLEKHGaitssntaatna  53
            MG KWSKSSV+GWP VRERMRRAEPAA+R       VGAASRDLE+HGA+T+SNT+  N
Sbjct  314  MGSKWSKSSVVGWPAVRERMRRAEPAAERAEPAAEGVGAASRDLERHGALTTSNTSHNNE  135

Query  54   acaWLEAQEEEEVGFPVTPQVPLRPMTYKAAVDLSHFLKEKGGL  97
            ACAWLEA E+EEVGFPV PQVPLRPMTYKAA DLS FLKEKGGL
Sbjct  134  ACAWLEAHEDEEVGFPVRPQVPLRPMTYKAAFDLSFFLKEKGGL  3
```

>Acinetobacter baumannii strain XH744 XH744_contig_122, whole genome shotgun sequence
Sequence ID: NZ_LYHI01000009.1 Length: 489
Range 1: 3 to 230

Score:121 bits(303), Expect:2e-30,
Method:Compositional matrix adjust.,
Identities:74/76(97%), Positives:74/76(97%), Gaps:0/76(0%)

```
Query  35   RDLEKHGaitssntaatnaacaWLEAQEEEEVGFPVTPQVPLRPMTYKAAVDLSHFLKEK  94
            RDLEKHGAITSSNTAATNA CAWLEA EEEEVGFPVTPQVPLRPMTYKAAVDLSHFLKEK
Sbjct  230  RDLEKHGAITSSNTAATNADCAWLEAPEEEEVGFPVTPQVPLRPMTYKAAVDLSHFLKEK  51

Query  95   GGLEGLIHSQRRQDIL  110
            GGLEGLIHSQRRQDIL
Sbjct  50   GGLEGLIHSQRRQDIL  3
```

>Escherichia coli strain NBE0009 contig00151, whole genome shotgun sequence
Sequence ID: NZ_JARBDL010000151.1 Length: 802
Range 1: 1 to 288

Score:124 bits(310), Expect:3e-30,
Method:Compositional matrix adjust.,
Identities:79/96(82%), Positives:84/96(87%), Gaps:4/96(4%)

```
Query  1    MGGKWSKSSVIGWPTVRERMRR---AEPAADRVGAASRDLEKHGaitssntaatnaacaW  57
            MGGKWSKSS IGWP VR+RMRR   AEPAA+ VGAAS+DL+K+GAITSSNTA TNA CAW
Sbjct  288  MGGKWSKSSKIGWPAVRDRMRRTQPAEPAAEGVGAASQDLDKYGAITSSNTADTNADCAW  109

Query  58   LEAQEE-EEVGFPVTPQVPLRPMTYKAAVDLSHFLK  92
            LEAQEE EEVGFPV PQVPLRPMTYK AVDLS FLK
Sbjct  108  LEAQEEAEEVGFPVRPQVPLRPMTYKGAVDLSFFLK  1
```

>Escherichia coli strain NBE0003 contig00096, whole genome shotgun sequence
Sequence ID: NZ_JARBDG010000096.1 Length: 2837
Range 1: 1 to 288

Score:127 bits(320), Expect:2e-29,
Method:Compositional matrix adjust.,
Identities:74/96(77%), Positives:85/96(88%), Gaps:1/96(1%)

```
Query  1    MGGKWSKSSVIGWPTVRERMRRAEPAADRVGAASRDLEKHGaitssntaatnaacaWLEA  60
            MGGKWS+S ++GWP VR+RMRR EPAA+ VGAAS+DL+K+GA+TSSNTA TNA CAWL+A
Sbjct  288  MGGKWSRSKIVGWPAVRDRMRRTEPAAEGVGAASQDLDKYGALTSSNTADTNADCAWLKA  109

Query  61   QEE-EEVGFPVTPQVPLRPMTYKAAVDLSHFLKEKG  95
            QEE  EVGFPV PQVPLRPMTYK+A DLS FLKEKG
Sbjct  108  QEEANEVGFPVRPQVPLRPMTYKSAFDLSFFLKEKG  1
```

>Escherichia coli strain EN5169 contig00082, whole genome shotgun sequence
Sequence ID: NZ_JAPMNJ010000082.1 Length: 2837
Range 1: 1 to 288

Score:127 bits(319), Expect:3e-29,
Method:Compositional matrix adjust.,
Identities:74/96(77%), Positives:84/96(87%), Gaps:1/96(1%)

```
Query  1    MGGKWSKSSVIGWPTVRERMRRAEPAADRVGAASRDLEKHGaitssntaatnaacaWLEA  60
            MGGKWS+S ++GWP VR+RMRR EPAA+ VGAAS+DL+K+GA+TSSNTA TNA CAWLEA
Sbjct  288  MGGKWSRSKIVGWPAVRDRMRRTEPAAEGVGAASQDLDKYGALTSSNTADTNADCAWLEA  109

Query  61   QEE-EEVGFPVTPQVPLRPMTYKAAVDLSHFLKEKG  95
            QEE  EVGFPV PQVPLRPMT+K A DLS FLKEKG
Sbjct  108  QEEANEVGFPVRPQVPLRPMTFKGAFDLSFFLKEKG  1
```

>Escherichia coli strain EN5317 contig00057, whole genome shotgun sequence
Sequence ID: NZ_JARBDO010000057.1 Length: 2867
Range 1: 1 to 297

Score:124 bits(310), Expect:4e-28,
Method:Compositional matrix adjust.,
Identities:76/99(77%), Positives:83/99(83%), Gaps:4/99(4%)

```
Query  1    MGGKWSKSSVIGWPTVRERMRR---AEPAADRVGAASRDLEKHGaitssntaatnaacaW  57
            MGGKWSKSS IGWP VR+RMRR   AEPAA+ VGAAS+DL+K+GAITSSNT   NA CAW
Sbjct  297  MGGKWSKSSKIGWPAVRDRMRRTQPAEPAAEGVGAASQDLDKYGAITSSNTDTNNADCAW  118

Query  58   LEAQEEEE-VGFPVTPQVPLRPMTYKAAVDLSHFLKEKG  95
            L AQEE+E VGFPV PQVPLRPMT+K A DLS FLKEKG
Sbjct  117  LRAQEEDEDVGFPVKPQVPLRPMTFKGAFDLSFFLKEKG  1
```

>Klebsiella pneumoniae strain EuSCAPE_IL015, whole genome shotgun sequence
Sequence ID: NZ_UKJV02000049.1 Length: 301
Range 1: 104 to 301

```
Score:109 bits(272), Expect:2e-26,
Method:Compositional matrix adjust.,
Identities:47/66(71%), Positives:59/66(89%), Gaps:0/66(0%)

Query  141  WCYKLVPVEPDKIEEANKGENTSLLHPVSLHGMDDPEREVLEWRFDSRLAFHHVARELHP  200
            WC+KLVPV+P ++EEAN+GEN  LLHP+S HGM+DPEREVL+W+FDS LA  H+ARELHP
Sbjct  301  WCFKLVPVDPGEVEEANQGENNCLLHPISQHGMEDPEREVLKWQFDSHLARRHMARELHP  122

Query  201  EYFKNC  206
            EY+K+C
Sbjct  121  EYYKDC  104
```

```
>Escherichia coli strain EN5090 contig00081, whole genome shotgun sequence
Sequence ID: NZ_JAODTD010000081.1 Length: 2798
Range 1: 2 to 277

Score:116 bits(291), Expect:1e-25,
Method:Compositional matrix adjust.,
Identities:69/92(75%), Positives:78/92(84%), Gaps:1/92(1%)

Query  1    MGGKWSKSSVIGWPTVRERMRRAEPAADRVGAASRDLEKHGaitssntaatnaacaWLEA  60
            M  KWSKSS++GWP VRER+RR EPAA+ VGAAS+DL+K+GA+TSSNT   NAACAWL A
Sbjct  277  MESKWSKSSIVGWPAVRERIRRTEPAAEGVGAASQDLDKYGALTSSNTDTNNAACAWLRA  98

Query  61   Q-EEEEVGFPVTPQVPLRPMTYKAAVDLSHFL  91
            Q EEEEVGFPV PQVPLRPMT+K A DLS FL
Sbjct  97   QEEEEEVGFPVRPQVPLRPMTFKGAFDLSFFL  2
```

```
>Escherichia coli strain NBE0006 contig00145, whole genome shotgun sequence
Sequence ID: NZ_JARBDI010000145.1 Length: 738
Range 1: 491 to 736

Score:110 bits(276), Expect:2e-25,
Method:Compositional matrix adjust.,
Identities:64/82(78%), Positives:72/82(87%), Gaps:1/82(1%)

Query  1    MGGKWSKSSVIGWPTVRERMRRAEPAADRVGAASRDLEKHGaitssntaatnaacaWLEA  60
            MGGKWSKSS++GWP VRER+RR +PAA+ VGAAS+DL+KHGA+TSSNT   NA CAWL A
Sbjct  491  MGGKWSKSSIVGWPAVRERIRRTQPAAEGVGAASQDLDKHGALTSSNTDTNNADCAWLRA  670

Query  61   QEEE-EVGFPVTPQVPLRPMTY  81
            QEEE EVGFPV PQVPLRPMTY
Sbjct  671  QEEEGEVGFPVKPQVPLRPMTY  736
```

```
>Escherichia coli strain EN5076 contig00066, whole genome shotgun sequence
Sequence ID: NZ_JAODTC010000066.1 Length: 2732
Range 1: 2 to 274

Score:115 bits(289), Expect:2e-25,
Method:Compositional matrix adjust.,
Identities:68/91(75%), Positives:77/91(84%), Gaps:1/91(1%)

Query  1    MGGKWSKSSVIGWPTVRERMRRAEPAADRVGAASRDLEKHGaitssntaatnaacaWLEA  60
            MG KWSKSS++GWP VRER+RR EPAA+ VGAAS+DL+K+GA+TSSNT   NAACAWL A
Sbjct  274  MGSKWSKSSIVGWPAVRERIRRTEPAAEGVGAASQDLDKYGALTSSNTDTNNAACAWLRA  95

Query  61   Q-EEEEVGFPVTPQVPLRPMTYKAAVDLSHF  90
            Q EEEEVGFPV PQVPL PMT+K A DLS F
Sbjct  94   QEEEEEVGFPVRPQVPLTPMTFKGAFDLSFF  2
```

```
>Escherichia coli strain EN5374 contig00094, whole genome shotgun sequence
Sequence ID: NZ_JAPTGI010000094.1 Length: 2720
Range 1: 2453 to 2719

Score:115 bits(288), Expect:4e-25,
Method:Compositional matrix adjust.,
Identities:68/89(76%), Positives:77/89(86%), Gaps:1/89(1%)

Query  1     MGGKWSKSSVIGWPTVRERMRRAEPAADRVGAASRDLEKHGaitssntaatnaacaWLEA  60
             MG KWSKSS++GWP VRER+RR EPAA+ VGAAS+DL+K+GA+TSSNT   NAACAWL A
Sbjct  2453  MGSKWSKSSIVGWPAVRERIRRTEPAAEGVGAASQDLDKYGALTSSNTDTNNAACAWLRA  2632

Query  61    Q-EEEEVGFPVTPQVPLRPMTYKAAVDLS  88
             Q EEEEVGFPV PQVPLRPMT+K A DLS
Sbjct  2633  QEEEEEVGFPVRPQVPLRPMTFKGAFDLS  2719
```

```
>Klebsiella pneumoniae strain EuSCAPE_LT003, whole genome shotgun sequence
Sequence ID: NZ_UKJH02000071.1 Length: 289
Range 1: 2 to 271

Score:104 bits(259), Expect:2e-24,
Method:Compositional matrix adjust.,
Identities:59/90(66%), Positives:73/90(81%), Gaps:1/90(1%)

Query  26   AADRVGAASRDLEKHGaitssntaatnaacaWLEAQEEE-EVGFPVTPQVPLRPMTYKAA  84
             A+RVGA ++D  +HGA  SSNT  TN A A LE Q+EE +VGFPV PQVPLRPMTYK A
Sbjct  271  GAERVGAVTQDSNRHGASASSNTPTTNQAYARLEEQDEEGDVGFPVRPQVPLRPMTYKGA  92

Query  85   VDLSHFLKEKGGLEGLIHSQRRQDILDLWI  114
             DLS FLKEKGGL+GL++S++R++ILDLW+
Sbjct  91   FDLSFFLKEKGGLDGLVYSEKRKEILDLWV  2
```

```
>Escherichia coli strain EN5379 contig00092, whole genome shotgun sequence
Sequence ID: NZ_JAPTGJ010000092.1 Length: 2592
Range 1: 1 to 255

Score:112 bits(279), Expect:7e-24,
Method:Compositional matrix adjust.,
Identities:66/85(78%), Positives:74/85(87%), Gaps:3/85(3%)

Query  1    MGGKWSKSSVIGWPTVRERMRRAEPAADRVGAASRDLEKHGaitssntaatnaacaWLEA  60
            MGGKWSKSS++GWP VRER+RRAEPAA+ VGAAS+DL+K+GAITSSNT   NA CAWL A
Sbjct  255  MGGKWSKSSIVGWPAVRERIRRAEPAAEGVGAASQDLDKYGAITSSNTDTNNADCAWLRA  76

Query  61   QEEEE---VGFPVTPQVPLRPMTYK  82
```

```
            QEE+E   VGFPV PQVPLRPMTYK
Sbjct  75   QEEDEDGDVGFPVKPQVPLRPMTYK  1
```

>Escherichia coli strain EN5383 contig00096, whole genome shotgun sequence
Sequence ID: NZ_JAPTZN010000096.1 Length: 2210
Range 1: 1949 to 2209

Score:111 bits(277), Expect:1e-23,
Method:Compositional matrix adjust.,
Identities:66/87(76%), Positives:75/87(86%), Gaps:1/87(1%)

```
Query  1     MGGKWSKSSVIGWPTVRERMRRAEPAADRVGAASRDLEKHGaitssntaatnaacaWLEA  60
             MG KWSKSS++GWP VRER+RR EPAA+ VGAAS+DL+K+GA+TSSNT   NAACAWL A
Sbjct  1949  MGSKWSKSSLVGWPAVRERIRRTEPAAEGVGAASQDLDKYGALTSSNTDTNNAACAWLRA  2128

Query  61    Q-EEEEVGFPVTPQVPLRPMTYKAAVD  86
             Q EEEEVGFPV PQVPLRPMT+K A D
Sbjct  2129  QEEEEEVGFPVRPQVPLRPMTFKGAFD  2209
```

>Escherichia coli strain NBE0009 contig00153, whole genome shotgun sequence
Sequence ID: NZ_JARBDL010000153.1 Length: 753
Range 1: 515 to 751

Score:103 bits(257), Expect:2e-22,
Method:Compositional matrix adjust.,
Identities:62/79(78%), Positives:70/79(88%), Gaps:1/79(1%)

```
Query  1    MGGKWSKSSVIGWPTVRERMRRAEPAADRVGAASRDLEKHGaitssntaatnaacaWLEA  60
            MGGKWSKSS++GWP VRER+RRAEPAA+ VGAAS+DL+K+GAITSSNT   NA CAWL A
Sbjct  515  MGGKWSKSSIVGWPAVRERIRRAEPAAEGVGAASQDLDKYGAITSSNTDTNNADCAWLRA  694

Query  61   QEE-EEVGFPVTPQVPLRP  78
            QEE E+VGFPV PQVPLRP
Sbjct  695  QEEDEDVGFPVKPQVPLRP  751
```

>Escherichia coli strain NBE0010 contig00080, whole genome shotgun sequence
Sequence ID: NZ_JARBDM010000080.1 Length: 2743
Range 1: 2154 to 2417

Score:106 bits(264), Expect:5e-22,
Method:Compositional matrix adjust.,
Identities:64/88(73%), Positives:73/88(82%), Gaps:1/88(1%)

```
Query  1     MGGKWSKSSVIGWPTVRERMRRAEPAADRVGAASRDLEKHGaitssntaatnaacaWLEA  60
             MG KWSKSS++GWP VRER+RR EPAA+ VGAAS+DL+K+GA+TSSNT   NAACAWL A
Sbjct  2154  MGSKWSKSSIVGWPAVRERIRRTEPAAEGVGAASQDLDKYGALTSSNTDTNNAACAWLRA  2333

Query  61    Q-EEEEVGFPVTPQVPLRPMTYKAAVDL  87
             Q EEEEVGFPV PQVPLRPMT    + L
Sbjct  2334  QEEEEEVGFPVRPQVPLRPMTLLKVIGL  2417
```

Range 2: 2384 to 2683

Score:73.9 bits(180), Expect:1e-10,
Method:Compositional matrix adjust.,
Identities:54/100(54%), Positives:69/100(69%), Gaps:13/100(13%)

```
Query  1     MGGKWSKSSVIGWPTV-----------RERMRRAEPAADRVGAASRDLEKHGaitssnta  49
             MGGKWSK S++  P +            +++R+A+PAA+RV AAS++L K+ AITSSNT
Sbjct  2683  MGGKWSKQSIVE*PAIKKKIKRAKPAAEQKIRQAKPAAERVKAASQNLNKYRAITSSNTD  2504

Query  50    atnaacaWLEAQEEEE--VGFPVTPQVPLRPMTYKAAVDL  87
              +NA CA L AQEEEE  V FPV PQVPLRPMT+K  + L
Sbjct  2503  TSNADCA*LRAQEEEEEDVSFPVKPQVPLRPMTFKRVIGL  2384
```

>Escherichia coli strain EN5383 contig00089, whole genome shotgun sequence
Sequence ID: NZ_JAPTZN010000089.1 Length: 1970
Range 1: 1725 to 1970

Score:103 bits(257), Expect:6e-21,
Method:Compositional matrix adjust.,
Identities:55/84(65%), Positives:66/84(78%), Gaps:3/84(3%)

```
Query  1     MGGKWSKSSVIGWPTVRERMRRAEPAADRVGAASRDLEKHGaitssntaatnaacaWLEA  60
             MGGKWSKSS++GWPT+RER+RR EPAA+ VGAAS+DL++ G    +     N AC WL A
Sbjct  1725  MGGKWSKSSIVGWPTIRERIRRTEPAAEGVGAASQDLDRRG--AITINNLNNEACTWLRA  1898

Query  61    QEEEE-VGFPVTPQVPLRPMTYKA  83
             QEE+E VGFPV PQVPLRPMT+K
Sbjct  1899  QEEDEGVGFPVRPQVPLRPMTFKG  1970
```

>Escherichia coli strain NBE0009 contig00116, whole genome shotgun sequence
Sequence ID: NZ_JARBDL010000116.1 Length: 1713
Range 1: 1 to 300

Score:102 bits(255), Expect:8e-21,
Method:Compositional matrix adjust.,
Identities:60/102(59%), Positives:71/102(69%), Gaps:16/102(15%)

```
Query  1    MGGKWSKSSVIGWPTVRERMRRAEPAADR-----------VGAASRDLEKHGaitssnta  49
            MGGKWSK S++GWP +RER+RRAEPAA+            VGAAS+DL+KH     +++
Sbjct  300  MGGKWSKRSIVGWPAIRERIRRAEPAAEHRTRQGEPAAEGVGAASQDLDKH--EAITSSN  127

Query  50   atnaacaWLEAQEEEE---VGFPVTPQVPLRPMTYKAAVDLS  88
              NA C WL AQEE+E   VGFPV PQVPLRPMTYK A DLS
Sbjct  126  TNNADCTWLRAQEEDEDGDVGFPVKPQVPLRPMTYKGAFDLS  1
```

>Klebsiella pneumoniae strain EuSCAPE_NO017, whole genome shotgun sequence
Sequence ID: NZ_UKIO02000061.1 Length: 607
Range 1: 365 to 607

Score:96.7 bits(239), Expect:3e-20,
Method:Compositional matrix adjust.,

```
Identities:64/81(79%), Positives:71/81(87%), Gaps:5/81(6%)

Query  1    MGGKWSKSSVIGWPTVRERMRR----AEPAADRVGAASRDLEKHGaitssntaatnaaca  56
            MG KWSKSS++GWP VRER+RR    AEPAA+ VGAAS+DL+KHGA+TSSNTAATNA CA
Sbjct  365  MGNKWSKSSIVGWPAVRERIRRTRPSAEPAAEGVGAASQDLDKHGALTSSNTAATNADCA  544

Query  57   WLEAQEE-EEVGFPVTPQVPL  76
            WLEAQEE EEVGFPV PQVPL
Sbjct  545  WLEAQEETEEVGFPVRPQVPL  607
```

```
>Escherichia coli strain EN5192 contig00135, whole genome shotgun sequence
Sequence ID: NZ_JAPMNL010000135.1 Length: 1341
Range 1: 2 to 235

Score:100 bits(249), Expect:3e-20,
Method:Compositional matrix adjust.,
Identities:59/78(76%), Positives:68/78(87%), Gaps:1/78(1%)

Query  1    MGGKWSKSSVIGWPTVRERMRRAEPAADRVGAASRDLEKHGaitssntaatnaacaWLEA  60
            MGGKWSKSS++GWP VRER+RR +PAA+ VGAAS+DL+KHGA+TSSNT   NA CAWL A
Sbjct  235  MGGKWSKSSIVGWPAVRERIRRTQPAAEGVGAASQDLDKHGALTSSNTDTNNADCAWLRA  56

Query  61   QEEE-EVGFPVTPQVPLR  77
            QEEE EVGFPV PQVP+R
Sbjct  55   QEEEGEVGFPVKPQVPVR  2
```

```
>Acinetobacter baumannii strain XH798 XH798_contig_106, whole genome shotgun sequence
Sequence ID: NZ_LYGA01000003.1 Length: 507
Range 1: 328 to 507

Score:92.8 bits(229), Expect:4e-19,
Method:Compositional matrix adjust.,
Identities:59/60(98%), Positives:59/60(98%), Gaps:0/60(0%)

Query  35   RDLEKHGaitssntaatnaacaWLEAQEEEEVGFPVTPQVPLRPMTYKAAVDLSHFLKEK  94
            RDLEKHGAITSSNTAATNA CAWLEAQEEEEVGFPVTPQVPLRPMTYKAAVDLSHFLKEK
Sbjct  328  RDLEKHGAITSSNTAATNADCAWLEAQEEEEVGFPVTPQVPLRPMTYKAAVDLSHFLKEK  507
```

```
>Escherichia coli strain EN5286 contig00120, whole genome shotgun sequence
Sequence ID: NZ_JAPMNM010000120.1 Length: 3033
Range 1: 2734 to 3033

Score:94.7 bits(234), Expect:6e-18,
Method:Compositional matrix adjust.,
Identities:67/102(66%), Positives:72/102(70%), Gaps:16/102(15%)

Query  1     MGGKWSKSSVIGWPTVRERMRRA------------EPAADRVGAASRDLEKHGaitssnt  48
             MG K SK S  GWP VRERMRR             E AA+ VGAAS+DL +HGA+TSSNT
Sbjct  2734  MGSKLSKCS--GWPAVRERMRRTRLTEEGTAEGATEGAAEGVGAASQDLARHGALTSSNT  2907

Query  49    aatnaacaWLEAQEEEE--VGFPVTPQVPLRPMTYKAAVDLS  88
                NAACAWL AQEEEE  VGFPV PQVPLRPMTYK A+DLS
Sbjct  2908  VTNNAACAWLAAQEEEEEDVGFPVRPQVPLRPMTYKGALDLS  3033
```

```
>Escherichia coli strain EN5177 contig00087, whole genome shotgun sequence
Sequence ID: NZ_JAPMNK010000087.1 Length: 3403
Range 1: 3104 to 3403

Score:94.7 bits(234), Expect:6e-18,
Method:Compositional matrix adjust.,
Identities:67/102(66%), Positives:72/102(70%), Gaps:16/102(15%)

Query  1     MGGKWSKSSVIGWPTVRERMRRA------------EPAADRVGAASRDLEKHGaitssnt  48
             MG K SK S  GWP VRERMRR             E AA+ VGAAS+DL +HGA+TSSNT
Sbjct  3104  MGSKLSKCS--GWPAVRERMRRTRLTEEGTAEGATEGAAEGVGAASQDLARHGALTSSNT  3277

Query  49    aatnaacaWLEAQEEEE--VGFPVTPQVPLRPMTYKAAVDLS  88
                NAACAWL AQEEEE  VGFPV PQVPLRPMTYK A+DLS
Sbjct  3278  VTNNAACAWLAAQEEEEEDVGFPVRPQVPLRPMTYKGALDLS  3403
```

```
>Escherichia coli strain EN5308 contig00153, whole genome shotgun sequence
Sequence ID: NZ_JARBDE010000153.1 Length: 786
Range 1: 3 to 254

Score:90.9 bits(224), Expect:1e-17,
Method:Compositional matrix adjust.,
Identities:61/85(72%), Positives:69/85(81%), Gaps:2/85(2%)

Query  1    MGGKWSKSSVIGWPTVRERMRRAEPAADRVGAASRDLEKHGaitssntaatnaacaWLEA  60
            M GKWSK  +  WP +RE++RR EPAA+RVGAAS+DL K+ A+TSSNT ATNA CA L A
Sbjct  254  MRGKWSKG-IARWPAIREKIRRTEPAAERVGAASQDLNKYRALTSSNTEATNADCARLRA  78

Query  61   QEEEE-VGFPVTPQVPLRPMTYKAA  84
            QEEEE VGFPV PQVPLRPMTYK A
Sbjct  77   QEEEEEVGFPVRPQVPLRPMTYKGA  3
```

```
>Escherichia coli strain EN5308 contig00096, whole genome shotgun sequence
Sequence ID: NZ_JARBDE010000096.1 Length: 3024
Range 1: 1 to 297

Score:93.2 bits(230), Expect:2e-17,
Method:Compositional matrix adjust.,
Identities:66/101(65%), Positives:71/101(70%), Gaps:16/101(15%)

Query  1    MGGKWSKSSVIGWPTVRERMRRA------------EPAADRVGAASRDLEKHGaitssnt  48
            MG K SK S  GWP VRERMRR             E AA+ VGAAS+DL +HGA+TSSNT
Sbjct  297  MGSKLSKCS--GWPAVRERMRRTRLTAEGTAEGATEGAAEGVGAASQDLARHGALTSSNT  124

Query  49   aatnaacaWLEAQEEEE--VGFPVTPQVPLRPMTYKAAVDL  87
               NAACAWL AQEEEE  VGFPV PQVPLRPMTYK A+DL
Sbjct  123  VTNNAACAWLAAQEEEEEDVGFPVRPQVPLRPMTYKGALDL  1
```

```
>Escherichia coli strain EN5143 contig00071, whole genome shotgun sequence
```

```
Sequence ID: NZ_JAPMNI010000071.1 Length: 2774
Range 1: 2478 to 2774

Score:91.7 bits(226), Expect:7e-17,
Method:Compositional matrix adjust.,
Identities:66/101(65%), Positives:70/101(69%), Gaps:16/101(15%)

Query  1     MGGKWSKSSVIGWPTVRERMRRA------------EPAADRVGAASRDLEKHGaitssnt  48
             MG K SK S  GWP VRERMRR             E AA  VGAAS+DL +HGA+TSSNT
Sbjct  2478  MGSKLSKCS--GWPAVRERMRRTRLTEEGTAEGATEGAAVGVGAASQDLARHGALTSSNT  2651

Query  49    aatnaacaWLEAQEEEE--VGFPVTPQVPLRPMTYKAAVDL  87
                NAACAWL AQEEEE  VGFPV PQVPLRPMTYK A+DL
Sbjct  2652  VTNNAACAWLAAQEEEEEDVGFPVRPQVPLRPMTYKGALDL  2774
```

```
>Klebsiella pneumoniae strain EuSCAPE_IL015, whole genome shotgun sequence
Sequence ID: NZ_UKJV02000040.1 Length: 532
Range 1: 329 to 532

Score:85.1 bits(209), Expect:3e-16,
Method:Compositional matrix adjust.,
Identities:51/68(75%), Positives:60/68(88%), Gaps:1/68(1%)

Query  1    MGGKWSKSSVIGWPTVRERMRRAEPAADRVGAASRDLEKHGaitssntaatnaacaWLEA  60
            MGGKWSKSS++GWP +RER+RR EPAA+ VGAASRDL+K+GA+T+SNTA  NA  AWLEA
Sbjct  329  MGGKWSKSSIVGWPAIRERIRRTEPAAEGVGAASRDLDKYGALTNSNTAQNNADVAWLEA  508

Query  61   QEE-EEVG  67
            QEE EEVG
Sbjct  509  QEEAEEVG  532
```

```
>Escherichia coli strain EN5095 contig00078, whole genome shotgun sequence
Sequence ID: NZ_JAPYLY010000078.1 Length: 1972
Range 1: 1 to 231

Score:88.6 bits(218), Expect:8e-16,
Method:Compositional matrix adjust.,
Identities:56/77(73%), Positives:67/77(87%), Gaps:1/77(1%)

Query  1    MGGKWSKSSVIGWPTVRERMRRAEPAADRVGAASRDLEKHGaitssntaatnaacaWLEA  60
            MGGKWSKSS++ WP VRER+RR  PAA+ VGAAS+DL+K+GA+T+SNTAATNA CAWLE+
Sbjct  231  MGGKWSKSSIVRWPAVRERLRRTAPAAEGVGAASQDLDKYGALTNSNTAATNADCAWLES  52

Query  61   QEEEE-VGFPVTPQVPL  76
            QEE + VGFPV PQV L
Sbjct  51   QEEADGVGFPVKPQVLL  1
```

```
>Escherichia coli strain NBE0006 contig00147, whole genome shotgun sequence
Sequence ID: NZ_JARBDI010000147.1 Length: 694
Range 1: 2 to 211

Score:85.1 bits(209), Expect:1e-15,
Method:Compositional matrix adjust.,
Identities:53/70(76%), Positives:62/70(88%), Gaps:1/70(1%)

Query  1    MGGKWSKSSVIGWPTVRERMRRAEPAADRVGAASRDLEKHGaitssntaatnaacaWLEA  60
            MGGKWS+S ++GWP VR+RMRR EPAA+ VGAAS+DL+K+GA+TSSNTA TNA CAWLEA
Sbjct  211  MGGKWSRSKIVGWPAVRDRMRRTEPAAEGVGAASQDLDKYGALTSSNTADTNADCAWLEA  32

Query  61   QEE-EEVGFP  69
            QEE  EVGFP
Sbjct  31   QEEANEVGFP  2
```

```
>Escherichia coli strain NBE0002 contig00097, whole genome shotgun sequence
Sequence ID: NZ_JARBDF010000097.1 Length: 2717
Range 1: 2 to 295

Score:87.4 bits(215), Expect:2e-15,
Method:Compositional matrix adjust.,
Identities:64/100(64%), Positives:68/100(68%), Gaps:16/100(16%)

Query  1    MGGKWSKSSVIGWPTVRERMRRA------------EPAADRVGAASRDLEKHGaitssnt  48
            MG K SK S  GWP VRERMRR             E AA  VGAAS+DL +HGA+TSSNT
Sbjct  295  MGSKLSKCS--GWPAVRERMRRTRLTEEGTAEGATEGAAVGVGAASQDLARHGALTSSNT  122

Query  49   aatnaacaWLEAQEEEE--VGFPVTPQVPLRPMTYKAAVD  86
               NAACAWL AQEEEE  VGFPV PQVPLRPMT+K A D
Sbjct  121  VTNNAACAWLAAQEEEEEDVGFPVRPQVPLRPMTFKGAFD  2
```

```
>Escherichia coli strain NBE0007 contig00093, whole genome shotgun sequence
Sequence ID: NZ_JARBDJ010000093.1 Length: 3013
Range 1: 3 to 284

Score:85.5 bits(210), Expect:9e-15,
Method:Compositional matrix adjust.,
Identities:62/96(65%), Positives:67/96(69%), Gaps:16/96(16%)

Query  1    MGGKWSKSSVIGWPTVRERMRRA------------EPAADRVGAASRDLEKHGaitssnt  48
            MG K SK S  GWP VRERMRR             E AA+ VGAAS+DL +HGA+TSSNT
Sbjct  284  MGSKLSKCS--GWPAVRERMRRTRLTEEGTAEGATEGAAEGVGAASQDLARHGALTSSNT  111

Query  49   aatnaacaWLEAQEEEE--VGFPVTPQVPLRPMTYK  82
               NAACAWL AQEEEE  VGFPV PQVPLRPMT+K
Sbjct  110  VTNNAACAWLAAQEEEEEDVGFPVRPQVPLRPMTFK  3
```

```
>Escherichia coli strain EN5132 contig00077, whole genome shotgun sequence
Sequence ID: NZ_JAPYLZ010000077.1 Length: 2498
Range 1: 2236 to 2496

Score:85.5 bits(210), Expect:1e-14,
Method:Compositional matrix adjust.,
Identities:63/91(69%), Positives:70/91(76%), Gaps:13/91(14%)

Query  1     MGGKWSKSSVIGWPTVRERMRRAEPAADRVGAA-------SRDLEKHGaitssntaatna  53
             MG KWSKS    WP VR+R+RR +PAA+RVGAA       S+DL+KHGA+TSSNT   NA
Sbjct  2236  MGNKWSKS----WPAVRDRIRRTQPAAERVGAAAEGVGAVSQDLDKHGALTSSNTDTNNA  2403
```

```
Query  54    acaWLEAQEEEE--VGFPVTPQVPLRPMTYK  82
             ACAWL AQEEEE  VGFPV PQVPLRPMTYK
Sbjct  2404  ACAWLRAQEEEEEEVGFPVRPQVPLRPMTYK  2496
```

>Acinetobacter baumannii strain XH797 XH797_contig_152, whole genome shotgun sequence
Sequence ID: NZ_LYGB01000010.1 Length: 304
Range 1: 3 to 161

Score:78.6 bits(192), Expect:2e-14,
Method:Compositional matrix adjust.,
Identities:52/53(98%), Positives:52/53(98%), Gaps:0/53(0%)

```
Query  35   RDLEKHGaitssntaatnaacaWLEAQEEEEVGFPVTPQVPLRPMTYKAAVDL  87
            RDLEKHGAITSSNTAATNA CAWLEAQEEEEVGFPVTPQVPLRPMTYKAAVDL
Sbjct  161  RDLEKHGAITSSNTAATNADCAWLEAQEEEEVGFPVTPQVPLRPMTYKAAVDL  3
```

>Bacillus cereus strain SJ-S17 s_1_17_trimmed_nodup_SPAdes_89_len:290, whole genome shotgun sequence
Sequence ID: NZ_NMYP01000066.1 Length: 219
Range 1: 6 to 116

Score:77.0 bits(188), Expect:3e-14,
Method:Compositional matrix adjust.,
Identities:36/37(97%), Positives:37/37(100%), Gaps:0/37(0%)

```
Query  74   VPLRPMTYKAAVDLSHFLKEKGGLEGLIHSQRRQDIL  110
            VPLRPMTYKAAVDLSHFLKEKGGLEGLIHSQRRQD+L
Sbjct  116  VPLRPMTYKAAVDLSHFLKEKGGLEGLIHSQRRQDLL  6
```

>Klebsiella pneumoniae strain EuSCAPE_LT007, whole genome shotgun sequence
Sequence ID: NZ_UKJI02000075.1 Length: 253
Range 1: 3 to 149

Score:75.9 bits(185), Expect:1e-13,
Method:Compositional matrix adjust.,
Identities:33/49(67%), Positives:40/49(81%), Gaps:0/49(0%)

```
Query  158  KGENTSLLHPVSLHGMDDPEREVLEWRFDSRLAFHHVARELHPEYFKNC  206
             GEN  LLHP+S HG +D EREVL W+FDS+LA  H+ARELHPEY+K+C
Sbjct  3    NGENNCLLHPMSQHGAEDEEREVLRWKFDSQLARRHMARELHPEYYKDC  149
```

>Escherichia coli strain NBE0004 contig00110, whole genome shotgun sequence
Sequence ID: NZ_JARBDH010000110.1 Length: 1390
Range 1: 2 to 286

Score:80.1 bits(196), Expect:6e-13,
Method:Compositional matrix adjust.,
Identities:64/99(65%), Positives:71/99(71%), Gaps:13/99(13%)

```
Query  1    MGGKWSKSSVIGWPTVRERMRRAEPAADRVGAA-------SRDLEKHGaitssntaatna  53
            M  KWSK+    WP VR+R+RR  P A+RV AA       S+DL+KHGA+TSSNT   NA
Sbjct  286  MRNKWSKT----WPAVRDRIRRTRPTAERVEAAAEGVGAASQDLDKHGALTSSNTDTNNA  119

Query  54   acaWLEAQEEEE--VGFPVTPQVPLRPMTYKAAVDLSHF  90
            ACAWL AQEEEE  VGFPV PQVPLRPMTYK A DLS F
Sbjct  118  ACAWLRAQEEEEEEVGFPVRPQVPLRPMTYKGAFDLSFF  2
```

>Acinetobacter baumannii strain XH822 XH822_contig_193, whole genome shotgun sequence
Sequence ID: NZ_LYFB01000021.1 Length: 753
Range 1: 529 to 657

Score:77.8 bits(190), Expect:7e-13,
Method:Compositional matrix adjust.,
Identities:38/43(88%), Positives:39/43(90%), Gaps:0/43(0%)

```
Query  68   FPVTPQVPLRPMTYKAAVDLSHFLKEKGGLEGLIHSQRRQDIL  110
            F  T  VPLRPMTYKAAVDLSHFLKEKGGLEGLIHSQRRQD+L
Sbjct  529  F*STLDVPLRPMTYKAAVDLSHFLKEKGGLEGLIHSQRRQDLL  657
```

>Acinetobacter baumannii strain XH851 XH851_contig_81, whole genome shotgun sequence
Sequence ID: NZ_LYJM01000091.1 Length: 882
Range 1: 652 to 780

Score:77.8 bits(190), Expect:1e-12,
Method:Compositional matrix adjust.,
Identities:38/43(88%), Positives:39/43(90%), Gaps:0/43(0%)

```
Query  68   FPVTPQVPLRPMTYKAAVDLSHFLKEKGGLEGLIHSQRRQDIL  110
            F  T  VPLRPMTYKAAVDLSHFLKEKGGLEGLIHSQRRQD+L
Sbjct  652  F*STLDVPLRPMTYKAAVDLSHFLKEKGGLEGLIHSQRRQDLL  780
```

>Acinetobacter baumannii strain XH840 XH840_contig_135, whole genome shotgun sequence
Sequence ID: NZ_LYEX01000008.1 Length: 1251
Range 1: 925 to 1053

Score:77.8 bits(190), Expect:3e-12,
Method:Compositional matrix adjust.,
Identities:38/43(88%), Positives:39/43(90%), Gaps:0/43(0%)

```
Query  68    FPVTPQVPLRPMTYKAAVDLSHFLKEKGGLEGLIHSQRRQDIL  110
             F  T  VPLRPMTYKAAVDLSHFLKEKGGLEGLIHSQRRQD+L
Sbjct  925   F*STLDVPLRPMTYKAAVDLSHFLKEKGGLEGLIHSQRRQDLL  1053
```

>Streptomyces sp. AA4 supercont1.1, whole genome shotgun sequence
Sequence ID: NZ_GG657746.1 Length: 8475969
Range 1: 2393665 to 2393802

Score:76.6 bits(187), Expect:1e-11,
Method:Compositional matrix adjust.,
Identities:38/46(83%), Positives:40/46(86%), Gaps:5/46(10%)

```
Query  74       VPLRPMTYKAAVDLSHFLKEKGGLEGLIHSQRRQDIL-----DLWI  114
                VPLRPMTYKAAVDLSHFLKEKGGLEGLIHSQRRQD+L      LW+
Sbjct  2393665  VPLRPMTYKAAVDLSHFLKEKGGLEGLIHSQRRQDLLFACTGSLWL  2393802
```

>Nocardioides luteus strain JCM 3358 sequence39, whole genome shotgun sequence
Sequence ID: NZ_BMRK01000039.1 Length: 5754
Range 1: 5574 to 5693

Score:76.3 bits(186), Expect:2e-11,
Method:Compositional matrix adjust.,
Identities:36/40(90%), Positives:38/40(95%), Gaps:0/40(0%)

```
Query  71    TPQVPLRPMTYKAAVDLSHFLKEKGGLEGLIHSQRRQDIL  110
             +  VPLRPMTYKAAVDLSHFLKEKGGLEGLIHSQRRQD+L
Sbjct  5574  SSSVPLRPMTYKAAVDLSHFLKEKGGLEGLIHSQRRQDLL  5693
```

>Streptomyces griseoflavus strain JCM 4479 sequence37, whole genome shotgun sequence
Sequence ID: NZ_BMUC01000037.1 Length: 5775
Range 1: 55 to 174

Score:76.3 bits(186), Expect:2e-11,
Method:Compositional matrix adjust.,
Identities:36/40(90%), Positives:38/40(95%), Gaps:0/40(0%)

```
Query  71   TPQVPLRPMTYKAAVDLSHFLKEKGGLEGLIHSQRRQDIL  110
            +  VPLRPMTYKAAVDLSHFLKEKGGLEGLIHSQRRQD+L
Sbjct  174  SSSVPLRPMTYKAAVDLSHFLKEKGGLEGLIHSQRRQDLL  55
```

>Saccharothrix coeruleofusca strain JCM 3313 sequence45, whole genome shotgun sequence
Sequence ID: NZ_BMRG01000045.1 Length: 5775
Range 1: 55 to 174

Score:76.3 bits(186), Expect:2e-11,
Method:Compositional matrix adjust.,
Identities:36/40(90%), Positives:38/40(95%), Gaps:0/40(0%)

```
Query  71   TPQVPLRPMTYKAAVDLSHFLKEKGGLEGLIHSQRRQDIL  110
            +  VPLRPMTYKAAVDLSHFLKEKGGLEGLIHSQRRQD+L
Sbjct  174  SSSVPLRPMTYKAAVDLSHFLKEKGGLEGLIHSQRRQDLL  55
```

>Actinoplanes ianthinogenes strain JCM 3249 sequence037, whole genome shotgun sequence
Sequence ID: NZ_BMQZ01000037.1 Length: 5815
Range 1: 75 to 194

Score:76.3 bits(186), Expect:2e-11,
Method:Compositional matrix adjust.,
Identities:36/40(90%), Positives:38/40(95%), Gaps:0/40(0%)

```
Query  71   TPQVPLRPMTYKAAVDLSHFLKEKGGLEGLIHSQRRQDIL  110
            +  VPLRPMTYKAAVDLSHFLKEKGGLEGLIHSQRRQD+L
Sbjct  194  SSSVPLRPMTYKAAVDLSHFLKEKGGLEGLIHSQRRQDLL  75
```

>Sinomonas sp. KCTC 49339 sequence28, whole genome shotgun sequence
Sequence ID: NZ_BNCM01000028.1 Length: 5849
Range 1: 46 to 165

Score:76.3 bits(186), Expect:2e-11,
Method:Compositional matrix adjust.,
Identities:36/40(90%), Positives:38/40(95%), Gaps:0/40(0%)

```
Query  71   TPQVPLRPMTYKAAVDLSHFLKEKGGLEGLIHSQRRQDIL  110
            +  VPLRPMTYKAAVDLSHFLKEKGGLEGLIHSQRRQD+L
Sbjct  165  SSSVPLRPMTYKAAVDLSHFLKEKGGLEGLIHSQRRQDLL  46
```

>Saccharothrix mutabilis subsp. mutabilis strain JCM 3380 sequence46, whole genome shotgun sequence
Sequence ID: NZ_BAAABU010000046.1 Length: 5899
Range 1: 5671 to 5790

Score:76.3 bits(186), Expect:2e-11,
Method:Compositional matrix adjust.,
Identities:36/40(90%), Positives:38/40(95%), Gaps:0/40(0%)

```
Query  71    TPQVPLRPMTYKAAVDLSHFLKEKGGLEGLIHSQRRQDIL  110
             +  VPLRPMTYKAAVDLSHFLKEKGGLEGLIHSQRRQD+L
Sbjct  5671  SSSVPLRPMTYKAAVDLSHFLKEKGGLEGLIHSQRRQDLL  5790
```

>Enterococcus alcedinis strain CCM 8433 sequence23, whole genome shotgun sequence
Sequence ID: NZ_BMDT01000023.1 Length: 6075
Range 1: 5907 to 6068

Score:75.5 bits(184), Expect:3e-11,
Method:Compositional matrix adjust.,
Identities:40/54(74%), Positives:40/54(74%), Gaps:5/54(9%)

```
Query  60    AQEEEEVGFPVTPQ-----VPLRPMTYKAAVDLSHFLKEKGGLEGLIHSQRRQD  108
             A     VG P  P      VPLRPMTYKAAVDLSHFLKEKGGLEGLIHSQRRQD
Sbjct  5907  AFALRRVGSPFGPPPRLVLVPLRPMTYKAAVDLSHFLKEKGGLEGLIHSQRRQD  6068
```

>Deinococcus aquiradiocola strain JCM 14371 sequence30, whole genome shotgun sequence
Sequence ID: NZ_BMOE01000030.1 Length: 6128
Range 1: 5960 to 6121

Score:75.5 bits(184), Expect:3e-11,
Method:Compositional matrix adjust.,
Identities:40/54(74%), Positives:40/54(74%), Gaps:5/54(9%)

```
Query  60    AQEEEEVGFPVTPQ-----VPLRPMTYKAAVDLSHFLKEKGGLEGLIHSQRRQD  108
             A     VG P  P      VPLRPMTYKAAVDLSHFLKEKGGLEGLIHSQRRQD
Sbjct  5960  AFALRRVGSPFGPPPRLVLVPLRPMTYKAAVDLSHFLKEKGGLEGLIHSQRRQD  6121
```

```
>Escherichia coli strain EN5192 contig00108, whole genome shotgun sequence
Sequence ID: NZ_JAPMNL010000108.1 Length: 3032
Range 1: 2765 to 3031

Score:73.9 bits(180), Expect:8e-11,
Method:Compositional matrix adjust.,
Identities:58/91(64%), Positives:61/91(67%), Gaps:16/91(17%)

Query  1     MGGKWSKSSVIGWPTVRERMRRA------------EPAADRVGAASRDLEKHGaitssnt  48
             MG K SK S  GWP VRERMRR             E AA  VGAAS+DL +HGA+TSSNT
Sbjct  2765  MGSKLSKCS--GWPAVRERMRRTRLTEEGTAEGATEGAAVGVGAASQDLARHGALTSSNT  2938

Query  49    aatnaacaWLEAQEEEE--VGFPVTPQVPLR  77
                NAACAWL AQEEEE  VGFPV PQVPLR
Sbjct  2939  VTNNAACAWLAAQEEEEEDVGFPVRPQVPLR  3031



>Escherichia coli strain EN5143 contig00069, whole genome shotgun sequence
Sequence ID: NZ_JAPMNI010000069.1 Length: 2688
Range 1: 2426 to 2686

Score:68.9 bits(167), Expect:4e-09,
Method:Compositional matrix adjust.,
Identities:62/91(68%), Positives:68/91(74%), Gaps:13/91(14%)

Query  1     MGGKWSKSSVIGWPTVRERMRRAEPAADRVGAA-------SRDLEKHGaitssntaatna  53
             MG KWSKS    WP VR+R+RR  PAA+RV AA       S+DL+KHGA+TSSNT   NA
Sbjct  2426  MGNKWSKS----WPAVRDRIRRTRPAAERVEAAAEGVGAASQDLDKHGALTSSNTDTNNA  2593

Query  54    acaWLEAQEEEE--VGFPVTPQVPLRPMTYK  82
             ACAWL AQEEEE  VGFPV PQVPLRPMTYK
Sbjct  2594  ACAWLRAQEEEEEEVGFPVRPQVPLRPMTYK  2686



>Acinetobacter baumannii strain XH799 XH799_contig_168, whole genome shotgun sequence
Sequence ID: NZ_LYFZ01000011.1 Length: 292
Range 1: 1 to 105

Score:63.2 bits(152), Expect:1e-08,
Method:Compositional matrix adjust.,
Identities:31/35(89%), Positives:31/35(88%), Gaps:0/35(0%)

Query  68   FPVTPQVPLRPMTYKAAVDLSHFLKEKGGLEGLIH  102
            F  T  VPLRPMTYKAAVDLSHFLKEKGGLEGLIH
Sbjct  105  F*STLDVPLRPMTYKAAVDLSHFLKEKGGLEGLIH  1



>Escherichia coli strain EN5177 contig00148, whole genome shotgun sequence
Sequence ID: NZ_JAPMNK010000148.1 Length: 850
Range 1: 2 to 250

Score:66.2 bits(160), Expect:2e-08,
Method:Compositional matrix adjust.,
Identities:54/83(65%), Positives:62/83(74%), Gaps:9/83(10%)

Query  12   GWPTVRERMRRAEPAADRVGAA-------SRDLEKHGaitssntaatnaacaWLEAQ--E  62
             WP V++R+RR  PAA++V AA       S+DL+KH A+TSSNT   NAACAW  AQ  E
Sbjct  250  SWPAVKDRIRRTRPAAEKVEAAAEGVGAASQDLDKHRALTSSNTDTNNAACAWPRAQKEE  71

Query  63   EEEVGFPVTPQVPLRPMTYKAAV  85
            EEEVGFPVTPQVPLRPMTYK AV
Sbjct  70   EEEVGFPVTPQVPLRPMTYKGAV  2



>Acinetobacter baumannii strain E16 contig_E16_244, whole genome shotgun sequence
Sequence ID: NZ_JBBXKO010000176.1 Length: 651
Range 1: 500 to 649

Score:63.2 bits(152), Expect:1e-07,
Method:Compositional matrix adjust.,
Identities:32/51(63%), Positives:37/51(72%), Gaps:1/51(1%)

Query  72   PQVPLRPMTYKAAVDLSHFLKEKGGLEGLIHSQRRQDILDLWIYHTQGYFP  122
            P +PL    +K  V + + L +   LEGLIHSQRRQDILDLWIYHTQGYFP
Sbjct  500  PVIPLNYFHFKEIVFVKYVL-QT**LEGLIHSQRRQDILDLWIYHTQGYFP  649



>Acinetobacter baumannii strain E13 contig_E13_197, whole genome shotgun sequence
Sequence ID: NZ_JBBXKR010000127.1 Length: 703
Range 1: 3 to 152

Score:63.2 bits(152), Expect:2e-07,
Method:Compositional matrix adjust.,
Identities:32/51(63%), Positives:37/51(72%), Gaps:1/51(1%)

Query  72   PQVPLRPMTYKAAVDLSHFLKEKGGLEGLIHSQRRQDILDLWIYHTQGYFP  122
            P +PL    +K  V + + L +   LEGLIHSQRRQDILDLWIYHTQGYFP
Sbjct  152  PVIPLNYFHFKEIVFVKYVL-QT**LEGLIHSQRRQDILDLWIYHTQGYFP  3



>Acinetobacter baumannii strain E11 contig_E11_176, whole genome shotgun sequence
Sequence ID: NZ_JBBXKT010000132.1 Length: 888
Range 1: 737 to 886

Score:63.2 bits(152), Expect:3e-07,
Method:Compositional matrix adjust.,
Identities:32/51(63%), Positives:37/51(72%), Gaps:1/51(1%)

Query  72   PQVPLRPMTYKAAVDLSHFLKEKGGLEGLIHSQRRQDILDLWIYHTQGYFP  122
            P +PL    +K  V + + L +   LEGLIHSQRRQDILDLWIYHTQGYFP
Sbjct  737  PVIPLNYFHFKEIVFVKYVL-QT**LEGLIHSQRRQDILDLWIYHTQGYFP  886



>Acinetobacter baumannii strain P11 contig_P11_393, whole genome shotgun sequence
Sequence ID: NZ_JBBXLN010000030.1 Length: 1140
Range 1: 3 to 152
```

```
Score:63.2 bits(152), Expect:4e-07,
Method:Compositional matrix adjust.,
Identities:32/51(63%), Positives:37/51(72%), Gaps:1/51(1%)

Query  72   PQVPLRPMTYKAAVDLSHFLKEKGGLEGLIHSQRRQDILDLWIYHTQGYFP  122
            P +PL    +K  V + + L +   LEGLIHSQRRQDILDLWIYHTQGYFP
Sbjct  152  PVIPLNYFHFKEIVFVKYVL-QT**LEGLIHSQRRQDILDLWIYHTQGYFP  3
```

>Acinetobacter baumannii strain P28 contig_P28_100, whole genome shotgun sequence
Sequence ID: NZ_JBBXKY010000067.1 Length: 1822
Range 1: 3 to 152

```
Score:63.2 bits(152), Expect:5e-07,
Method:Compositional matrix adjust.,
Identities:32/51(63%), Positives:37/51(72%), Gaps:1/51(1%)

Query  72   PQVPLRPMTYKAAVDLSHFLKEKGGLEGLIHSQRRQDILDLWIYHTQGYFP  122
            P +PL    +K  V + + L +   LEGLIHSQRRQDILDLWIYHTQGYFP
Sbjct  152  PVIPLNYFHFKEIVFVKYVL-QT**LEGLIHSQRRQDILDLWIYHTQGYFP  3
```

>Acinetobacter baumannii strain P18 contig_P18_196, whole genome shotgun sequence
Sequence ID: NZ_JBBXLH010000065.1 Length: 2259
Range 1: 2108 to 2257

```
Score:63.2 bits(152), Expect:5e-07,
Method:Compositional matrix adjust.,
Identities:32/51(63%), Positives:37/51(72%), Gaps:1/51(1%)

Query  72    PQVPLRPMTYKAAVDLSHFLKEKGGLEGLIHSQRRQDILDLWIYHTQGYFP  122
             P +PL    +K  V + + L +   LEGLIHSQRRQDILDLWIYHTQGYFP
Sbjct  2108  PVIPLNYFHFKEIVFVKYVL-QT**LEGLIHSQRRQDILDLWIYHTQGYFP  2257
```

>Acinetobacter baumannii strain P1 contig_P1_68, whole genome shotgun sequence
Sequence ID: NZ_JBBXLX010000052.1 Length: 2273
Range 1: 3 to 152

```
Score:63.2 bits(152), Expect:5e-07,
Method:Compositional matrix adjust.,
Identities:32/51(63%), Positives:37/51(72%), Gaps:1/51(1%)

Query  72   PQVPLRPMTYKAAVDLSHFLKEKGGLEGLIHSQRRQDILDLWIYHTQGYFP  122
            P +PL    +K  V + + L +   LEGLIHSQRRQDILDLWIYHTQGYFP
Sbjct  152  PVIPLNYFHFKEIVFVKYVL-QT**LEGLIHSQRRQDILDLWIYHTQGYFP  3
```

>Acinetobacter baumannii strain E2 contig_E2_954, whole genome shotgun sequence
Sequence ID: NZ_JBBXKW010000145.1 Length: 2573
Range 1: 2422 to 2571

```
Score:63.2 bits(152), Expect:5e-07,
Method:Compositional matrix adjust.,
Identities:32/51(63%), Positives:37/51(72%), Gaps:1/51(1%)

Query  72    PQVPLRPMTYKAAVDLSHFLKEKGGLEGLIHSQRRQDILDLWIYHTQGYFP  122
             P +PL    +K  V + + L +   LEGLIHSQRRQDILDLWIYHTQGYFP
Sbjct  2422  PVIPLNYFHFKEIVFVKYVL-QT**LEGLIHSQRRQDILDLWIYHTQGYFP  2571
```

>Acinetobacter baumannii strain P27 contig_P27_23, whole genome shotgun sequence
Sequence ID: NZ_JBBXKZ010000016.1 Length: 2573
Range 1: 2422 to 2571

```
Score:63.2 bits(152), Expect:5e-07,
Method:Compositional matrix adjust.,
Identities:32/51(63%), Positives:37/51(72%), Gaps:1/51(1%)

Query  72    PQVPLRPMTYKAAVDLSHFLKEKGGLEGLIHSQRRQDILDLWIYHTQGYFP  122
             P +PL    +K  V + + L +   LEGLIHSQRRQDILDLWIYHTQGYFP
Sbjct  2422  PVIPLNYFHFKEIVFVKYVL-QT**LEGLIHSQRRQDILDLWIYHTQGYFP  2571
```

>Acinetobacter baumannii strain P16 contig_P16_190, whole genome shotgun sequence
Sequence ID: NZ_JBBXLI010000090.1 Length: 2599
Range 1: 3 to 152

```
Score:63.2 bits(152), Expect:5e-07,
Method:Compositional matrix adjust.,
Identities:32/51(63%), Positives:37/51(72%), Gaps:1/51(1%)

Query  72   PQVPLRPMTYKAAVDLSHFLKEKGGLEGLIHSQRRQDILDLWIYHTQGYFP  122
            P +PL    +K  V + + L +   LEGLIHSQRRQDILDLWIYHTQGYFP
Sbjct  152  PVIPLNYFHFKEIVFVKYVL-QT**LEGLIHSQRRQDILDLWIYHTQGYFP  3
```

>Acinetobacter baumannii strain P5 contig_P5_54, whole genome shotgun sequence
Sequence ID: NZ_JBBXLT010000030.1 Length: 2679
Range 1: 2528 to 2677

```
Score:63.2 bits(152), Expect:5e-07,
Method:Compositional matrix adjust.,
Identities:32/51(63%), Positives:37/51(72%), Gaps:1/51(1%)

Query  72    PQVPLRPMTYKAAVDLSHFLKEKGGLEGLIHSQRRQDILDLWIYHTQGYFP  122
             P +PL    +K  V + + L +   LEGLIHSQRRQDILDLWIYHTQGYFP
Sbjct  2528  PVIPLNYFHFKEIVFVKYVL-QT**LEGLIHSQRRQDILDLWIYHTQGYFP  2677
```

>Acinetobacter baumannii strain P7 contig_P7_199, whole genome shotgun sequence
Sequence ID: NZ_JBBXLR010000069.1 Length: 2736
Range 1: 3 to 152

```
Score:63.2 bits(152), Expect:5e-07,
Method:Compositional matrix adjust.,
Identities:32/51(63%), Positives:37/51(72%), Gaps:1/51(1%)
```

```
Query  72   PQVPLRPMTYKAAVDLSHFLKEKGGLEGLIHSQRRQDILDLWIYHTQGYFP  122
            P +PL    +K  V + + L +   LEGLIHSQRRQDILDLWIYHTQGYFP
Sbjct  152  PVIPLNYFHFKEIVFVKYVL-QT**LEGLIHSQRRQDILDLWIYHTQGYFP  3
```

>Acinetobacter baumannii strain P19 contig_P19_138, whole genome shotgun sequence
Sequence ID: NZ_JBBXLG010000094.1 Length: 2736
Range 1: 3 to 152

Score:63.2 bits(152), Expect:5e-07,
Method:Compositional matrix adjust.,
Identities:32/51(63%), Positives:37/51(72%), Gaps:1/51(1%)

```
Query  72   PQVPLRPMTYKAAVDLSHFLKEKGGLEGLIHSQRRQDILDLWIYHTQGYFP  122
            P +PL    +K  V + + L +   LEGLIHSQRRQDILDLWIYHTQGYFP
Sbjct  152  PVIPLNYFHFKEIVFVKYVL-QT**LEGLIHSQRRQDILDLWIYHTQGYFP  3
```

>Acinetobacter baumannii strain P15 contig_P15_245, whole genome shotgun sequence
Sequence ID: NZ_JBBXLJ010000096.1 Length: 2736
Range 1: 2585 to 2734

Score:63.2 bits(152), Expect:5e-07,
Method:Compositional matrix adjust.,
Identities:32/51(63%), Positives:37/51(72%), Gaps:1/51(1%)

```
Query  72    PQVPLRPMTYKAAVDLSHFLKEKGGLEGLIHSQRRQDILDLWIYHTQGYFP  122
             P +PL    +K  V + + L +   LEGLIHSQRRQDILDLWIYHTQGYFP
Sbjct  2585  PVIPLNYFHFKEIVFVKYVL-QT**LEGLIHSQRRQDILDLWIYHTQGYFP  2734
```

>Acinetobacter baumannii strain P21 contig_P21_632, whole genome shotgun sequence
Sequence ID: NZ_JBBXLE010000174.1 Length: 2804
Range 1: 2653 to 2802

Score:63.2 bits(152), Expect:5e-07,
Method:Compositional matrix adjust.,
Identities:32/51(63%), Positives:37/51(72%), Gaps:1/51(1%)

```
Query  72    PQVPLRPMTYKAAVDLSHFLKEKGGLEGLIHSQRRQDILDLWIYHTQGYFP  122
             P +PL    +K  V + + L +   LEGLIHSQRRQDILDLWIYHTQGYFP
Sbjct  2653  PVIPLNYFHFKEIVFVKYVL-QT**LEGLIHSQRRQDILDLWIYHTQGYFP  2802
```

>Acinetobacter baumannii strain E17 contig_E17_94, whole genome shotgun sequence
Sequence ID: NZ_JBBXKN010000051.1 Length: 2965
Range 1: 3 to 152

Score:63.2 bits(152), Expect:5e-07,
Method:Compositional matrix adjust.,
Identities:32/51(63%), Positives:37/51(72%), Gaps:1/51(1%)

```
Query  72   PQVPLRPMTYKAAVDLSHFLKEKGGLEGLIHSQRRQDILDLWIYHTQGYFP  122
            P +PL    +K  V + + L +   LEGLIHSQRRQDILDLWIYHTQGYFP
Sbjct  152  PVIPLNYFHFKEIVFVKYVL-QT**LEGLIHSQRRQDILDLWIYHTQGYFP  3
```

>Acinetobacter baumannii strain P23 contig_P23_230, whole genome shotgun sequence
Sequence ID: NZ_JBBXLC010000084.1 Length: 3261
Range 1: 3110 to 3259

Score:63.2 bits(152), Expect:5e-07,
Method:Compositional matrix adjust.,
Identities:32/51(63%), Positives:37/51(72%), Gaps:1/51(1%)

```
Query  72    PQVPLRPMTYKAAVDLSHFLKEKGGLEGLIHSQRRQDILDLWIYHTQGYFP  122
             P +PL    +K  V + + L +   LEGLIHSQRRQDILDLWIYHTQGYFP
Sbjct  3110  PVIPLNYFHFKEIVFVKYVL-QT**LEGLIHSQRRQDILDLWIYHTQGYFP  3259
```

>Klebsiella pneumoniae strain CL2087 NODE_71_length_2194_cov_4.05709, whole genome shotgun sequence
Sequence ID: NZ_RPFC01000071.1 Length: 2194
Range 1: 15 to 107

Score:62.0 bits(149), Expect:1e-06,
Method:Compositional matrix adjust.,
Identities:28/31(90%), Positives:30/31(96%), Gaps:0/31(0%)

```
Query  79   MTYKAAVDLSHFLKEKGGLEGLIHSQRRQDI  109
            MTYKAAVDLSHFLKEKGGLEGLIHSQR Q++
Sbjct  107  MTYKAAVDLSHFLKEKGGLEGLIHSQRCQEL  15
```

>Klebsiella pneumoniae strain EuSCAPE_LT003, whole genome shotgun sequence
Sequence ID: NZ_UKJH02000068.1 Length: 317
Range 1: 23 to 118

Score:53.5 bits(127), Expect:6e-05,
Method:Compositional matrix adjust.,
Identities:23/32(72%), Positives:28/32(87%), Gaps:0/32(0%)

```
Query  1    MGGKWSKSSVIGWPTVRERMRRAEPAADRVGA  32
            MGGKWSKSS++GWP +RER+RR  PAA+RV A
Sbjct  118  MGGKWSKSSLVGWPAIRERIRRTNPAAERVRA  23
```

>Klebsiella pneumoniae strain EuSCAPE_MT017, whole genome shotgun sequence
Sequence ID: NZ_UKJA02000086.1 Length: 513
Range 1: 1 to 138

Score:54.7 bits(130), Expect:6e-05,
Method:Compositional matrix adjust.,
Identities:28/46(61%), Positives:33/46(71%), Gaps:5/46(10%)

```
Query  1    MGGKWSKSSVIGWPTVRERMRRAE-----PAADRVGAASRDLEKHG  41
            MGGKWSK S + WP VRER+R+ E     P A+ VGAAS+DL KHG
Sbjct  138  MGGKWSKKSKVEWPAVRERIRKTEPVAEGPVAEGVGAASQDLAKHG  1
```

Query #16: sp|P04618|REV_HV1H2 Protein Rev OS=Human immunodeficiency virus type 1 group M subtype B (isolate HXB2) OX=11706 GN=rev PE=1 SV=1 Query ID: lcl|Query_2387709
Length: 116

Sequences producing significant alignments:

| Description | Scientific Name | Common Name | Taxid | Max Score | Total Score | Query cover | E Value | Per. Ident | Acc. Len | Accession |
|---|---|---|---|---|---|---|---|---|---|---|
| Actinoplanes campanulatus strain JCM 3059 sequence095, whole... | Actinoplanes... | NA | 113559 | 180 | 180 | 100% | 2e-54 | 78.45 | 582 | NZ_BMPW010000... |
| Deinococcus aquiradiocola strain JCM 14371 sequence41, whole... | Deinococcus ... | NA | 393059 | 180 | 180 | 100% | 5e-54 | 78.45 | 690 | NZ_BMOE010000... |
| Enterococcus alcedinis strain CCM 8433 sequence26, whole genom... | Enterococcus... | NA | 1274384 | 180 | 180 | 100% | 2e-53 | 78.45 | 853 | NZ_BMDT010000... |
| Corynebacterium glutamicum strain 38 Scaffolds_31_4109, whole... | Corynebacter... | NA | 1718 | 180 | 180 | 100% | 3e-49 | 78.45 | 3118 | NZ_JACENN0100... |
| Lactiplantibacillus plantarum strain MH-301 Scaffolds_50_4236,... | Lactiplantib... | NA | 1590 | 180 | 180 | 100% | 3e-49 | 78.45 | 4236 | NZ_JAOYUG0100... |
| Corynebacterium glutamicum strain 51 Scaffolds_32_4236, whole... | Corynebacter... | NA | 1718 | 180 | 180 | 100% | 3e-49 | 78.45 | 4236 | NZ_JACENK0100... |
| Aggregatibacter actinomycetemcomitans serotype d str. SA269... | Aggregatibac... | NA | 1434259 | 126 | 126 | 84% | 1e-33 | 69.39 | 504 | NZ_AZTX010004... |
| Klebsiella pneumoniae strain EuSCAPE_MT015, whole genome shotg... | Klebsiella p... | NA | 573 | 92.8 | 92.8 | 84% | 4e-21 | 54.08 | 328 | NZ_UKIW020001... |
| Klebsiella pneumoniae strain EuSCAPE_IL022, whole genome shotg... | Klebsiella p... | NA | 573 | 80.1 | 80.1 | 56% | 9e-16 | 64.62 | 411 | NZ_UKKD020000... |
| Klebsiella pneumoniae strain EuSCAPE_IL006, whole genome shotg... | Klebsiella p... | NA | 573 | 77.0 | 77.0 | 84% | 4e-14 | 48.57 | 560 | NZ_UKJR020001... |
| Escherichia coli strain EN5090 contig00081, whole genome shotg... | Escherichia ... | NA | 562 | 80.1 | 80.1 | 79% | 4e-14 | 50.00 | 2798 | NZ_JAODTD0100... |
| Escherichia coli strain EN5383 contig00096, whole genome shotg... | Escherichia ... | NA | 562 | 79.7 | 79.7 | 79% | 6e-14 | 50.00 | 2210 | NZ_JAPTZN0100... |
| Escherichia coli strain EN5374 contig00094, whole genome shotg... | Escherichia ... | NA | 562 | 79.0 | 79.0 | 86% | 1e-13 | 47.00 | 2720 | NZ_JAPTGI0100... |
| Escherichia coli strain EN5076 contig00066, whole genome shotg... | Escherichia ... | NA | 562 | 79.0 | 79.0 | 86% | 1e-13 | 47.00 | 2732 | NZ_JAODTC0100... |
| Escherichia coli strain NBE0010 contig00080, whole genome... | Escherichia ... | NA | 562 | 79.0 | 79.0 | 86% | 1e-13 | 47.00 | 2743 | NZ_JARBDM0100... |
| Escherichia coli strain EN5143 contig00069, whole genome shotg... | Escherichia ... | NA | 562 | 78.6 | 124 | 79% | 2e-13 | 63.89 | 2688 | NZ_JAPMNI0100... |
| Escherichia coli strain EN5132 contig00077, whole genome shotg... | Escherichia ... | NA | 562 | 78.2 | 124 | 99% | 2e-13 | 50.00 | 2498 | NZ_JAPYLZ0100... |
| Klebsiella pneumoniae strain EuSCAPE_MT003, whole genome shotg... | Klebsiella p... | NA | 573 | 77.0 | 77.0 | 75% | 3e-13 | 50.00 | 964 | NZ_UKIK020000... |
| Escherichia coli strain EN5379 contig00092, whole genome shotg... | Escherichia ... | NA | 562 | 77.0 | 77.0 | 79% | 4e-13 | 51.52 | 2592 | NZ_JAPTGJ0100... |
| Escherichia coli strain NBE0009 contig00116, whole genome... | Escherichia ... | NA | 562 | 77.0 | 77.0 | 84% | 5e-13 | 50.48 | 1713 | NZ_JARBDL0100... |
| Escherichia coli strain NBE0007 contig00093, whole genome... | Escherichia ... | NA | 562 | 75.5 | 75.5 | 79% | 1e-12 | 51.52 | 3013 | NZ_JARBDJ0100... |
| Escherichia coli strain NBE0006 contig00145, whole genome... | Escherichia ... | NA | 562 | 73.6 | 73.6 | 79% | 2e-12 | 49.49 | 738 | NZ_JARBDI0100... |
| Escherichia coli strain EN5177 contig00148, whole genome shotg... | Escherichia ... | NA | 562 | 73.2 | 73.2 | 78% | 5e-12 | 48.98 | 850 | NZ_JAPMNK0100... |
| Escherichia coli strain NBE0006 contig00147, whole genome... | Escherichia ... | NA | 562 | 71.6 | 71.6 | 79% | 9e-12 | 48.48 | 694 | NZ_JARBDI0100... |
| Escherichia coli strain NBE0004 contig00110, whole genome... | Escherichia ... | NA | 562 | 73.2 | 73.2 | 78% | 1e-11 | 48.98 | 1390 | NZ_JARBDH0100... |
| Escherichia coli strain EN5095 contig00078, whole genome shotg... | Escherichia ... | NA | 562 | 73.2 | 73.2 | 79% | 1e-11 | 49.49 | 1972 | NZ_JAPYLY0100... |
| Escherichia coli strain EN5286 contig00120, whole genome shotg... | Escherichia ... | NA | 562 | 73.2 | 73.2 | 84% | 1e-11 | 47.62 | 3033 | NZ_JAPMNM0100... |
| Escherichia coli strain NBE0009 contig00153, whole genome... | Escherichia ... | NA | 562 | 71.6 | 71.6 | 84% | 1e-11 | 47.62 | 753 | NZ_JARBDL0100... |
| Escherichia coli strain EN5192 contig00108, whole genome shotg... | Escherichia ... | NA | 562 | 72.8 | 72.8 | 84% | 1e-11 | 47.62 | 3032 | NZ_JAPMNL0100... |
| Escherichia coli strain EN5192 contig00135, whole genome shotg... | Escherichia ... | NA | 562 | 72.4 | 72.4 | 79% | 2e-11 | 48.48 | 1341 | NZ_JAPMNL0100... |
| Escherichia coli strain EN5308 contig00099, whole genome shotg... | Escherichia ... | NA | 562 | 71.6 | 117 | 100% | 4e-11 | 48.48 | 2827 | NZ_JARBDE0100... |
| Escherichia coli strain EN5169 contig00082, whole genome shotg... | Escherichia ... | NA | 562 | 71.6 | 117 | 100% | 4e-11 | 48.48 | 2837 | NZ_JAPMNJ0100... |
| Escherichia coli strain NBE0003 contig00096, whole genome... | Escherichia ... | NA | 562 | 71.6 | 117 | 100% | 4e-11 | 48.48 | 2837 | NZ_JARBDG0100... |
| Escherichia coli strain NBE0002 contig00097, whole genome... | Escherichia ... | NA | 562 | 71.6 | 117 | 100% | 4e-11 | 46.67 | 2717 | NZ_JARBDF0100... |
| Escherichia coli strain EN5177 contig00087, whole genome shotg... | Escherichia ... | NA | 562 | 69.3 | 69.3 | 84% | 2e-10 | 45.71 | 3403 | NZ_JAPMNK0100... |
| Escherichia coli strain EN5143 contig00071, whole genome shotg... | Escherichia ... | NA | 562 | 69.3 | 69.3 | 84% | 3e-10 | 45.71 | 2774 | NZ_JAPMNI0100... |
| Escherichia coli strain EN5308 contig00096, whole genome shotg... | Escherichia ... | NA | 562 | 69.3 | 69.3 | 84% | 3e-10 | 45.71 | 3024 | NZ_JARBDE0100... |
| Klebsiella pneumoniae strain EuSCAPE_IL009, whole genome shotg... | Klebsiella p... | NA | 573 | 63.9 | 63.9 | 51% | 5e-10 | 58.33 | 274 | NZ_UKJO020000... |
| Klebsiella pneumoniae strain EuSCAPE_DE091, whole genome shotg... | Klebsiella p... | NA | 573 | 65.9 | 65.9 | 53% | 9e-10 | 60.87 | 618 | NZ_UKMA020000... |
| Klebsiella pneumoniae strain EuSCAPE_IL001, whole genome shotg... | Klebsiella p... | NA | 573 | 65.1 | 65.1 | 79% | 2e-09 | 44.44 | 599 | NZ_UKJN020000... |
| Escherichia coli strain NBE0009 contig00151, whole genome... | Escherichia ... | NA | 562 | 65.5 | 65.5 | 84% | 2e-09 | 43.81 | 802 | NZ_JARBDL0100... |
| Escherichia coli strain EN5317 contig00057, whole genome shotg... | Escherichia ... | NA | 562 | 66.6 | 66.6 | 84% | 2e-09 | 44.76 | 2867 | NZ_JARBDO0100... |
| Klebsiella pneumoniae strain EuSCAPE_IL005, whole genome shotg... | Klebsiella p... | NA | 573 | 63.5 | 63.5 | 73% | 3e-09 | 46.74 | 455 | NZ_UKJL020000... |
| Klebsiella pneumoniae strain EuSCAPE_MT017, whole genome shotg... | Klebsiella p... | NA | 573 | 62.8 | 62.8 | 50% | 8e-09 | 57.58 | 513 | NZ_UKJA020000... |
| Klebsiella pneumoniae strain EuSCAPE_IL018, whole genome shotg... | Klebsiella p... | NA | 573 | 61.2 | 61.2 | 78% | 2e-08 | 43.88 | 388 | NZ_UKJU020001... |
| Escherichia coli strain EN5308 contig00153, whole genome shotg... | Escherichia ... | NA | 562 | 63.2 | 63.2 | 79% | 2e-08 | 42.42 | 786 | NZ_JARBDE0100... |
| Escherichia coli strain NBE0010 contig00098, whole genome... | Escherichia ... | NA | 562 | 63.5 | 63.5 | 64% | 2e-08 | 50.67 | 1912 | NZ_JARBDM0100... |
| Klebsiella pneumoniae strain EuSCAPE_MT009, whole genome shotg... | Klebsiella p... | NA | 573 | 57.4 | 57.4 | 34% | 5e-07 | 67.50 | 398 | NZ_UKIV020000... |
| Klebsiella pneumoniae strain OK8 jcf7180000003506, whole genom... | Klebsiella p... | NA | 573 | 55.5 | 55.5 | 56% | 6e-06 | 51.35 | 577 | NZ_MZXR010000... |
| Klebsiella pneumoniae strain EuSCAPE_IL015, whole genome shotg... | Klebsiella p... | NA | 573 | 55.5 | 55.5 | 45% | 6e-06 | 55.00 | 532 | NZ_UKJV020000... |
| Photobacterium rosenbergii strain DP6N14-7 DP6N14-7_ctg0065,... | Photobacteri... | NA | 294936 | 52.8 | 52.8 | 49% | 7e-05 | 52.63 | 564 | NZ_JAHVIB0100... |
| Klebsiella pneumoniae strain EuSCAPE_NO017, whole genome shotg... | Klebsiella p... | NA | 573 | 52.8 | 52.8 | 56% | 8e-05 | 55.56 | 607 | NZ_UKIO020000... |
| Escherichia coli strain NBE0006 contig00106, whole genome... | Escherichia ... | NA | 562 | 53.5 | 53.5 | 62% | 8e-05 | 43.84 | 2090 | NZ_JARBDI0100... |
| Bacillus velezensis strain TN14-11 TN14-11_ctg0032, whole geno... | Bacillus vel... | NA | 492670 | 52.8 | 52.8 | 49% | 2e-04 | 52.63 | 1125 | NZ_JAHVIU0100... |
| Klebsiella pneumoniae strain EuSCAPE_LT010, whole genome shotg... | Klebsiella p... | NA | 573 | 46.2 | 46.2 | 22% | 0.007 | 76.92 | 384 | NZ_UKJK020000... |
| Klebsiella pneumoniae strain EuSCAPE_MT011, whole genome shotg... | Klebsiella p... | NA | 573 | 45.4 | 45.4 | 23% | 0.013 | 74.07 | 368 | NZ_UKIY020000... |
| Klebsiella pneumoniae strain EuSCAPE_IL004, whole genome shotg... | Klebsiella p... | NA | 573 | 44.7 | 44.7 | 22% | 0.037 | 73.08 | 411 | NZ_UKJP020001... |

Alignments:

>Actinoplanes campanulatus strain JCM 3059 sequence095, whole genome shotgun sequence
Sequence ID: NZ_BMPW01000095.1 Length: 582
Range 1: 35 to 382

Score:180 bits(456), Expect:2e-54,
Method:Compositional matrix adjust.,
Identities:108/116(93%), Positives:112/116(96%), Gaps:0/116(0%)

```
Query  1    MAGRSGDSDEELIRTVRLIKLLYQSNPPPNPEGTrqarrnrrrrwrerqrqIHSISERIL  60
            MAGRSGDSDE+L++ VRLIK LYQSNPPPNPEGTRQARRNRRRRWRERQRQIHSISERIL
Sbjct  35   MAGRSGDSDEDLLKAVRLIKFLYQSNPPPNPEGTRQARRNRRRRWRERQRQIHSISERIL  214

Query  61   GTYLGRSAEPVPLQLPPLERLTLDCNEDCGTSGTQGVGSPQILVESPTVLESGTKE  116
             TYLGRSAEPVPLQLPPLERLTLDCNEDCGTSGTQGVGSPQILVESPT+LESG KE
Sbjct  215  STYLGRSAEPVPLQLPPLERLTLDCNEDCGTSGTQGVGSPQILVESPTILESGAKE  382
```

>Deinococcus aquiradiocola strain JCM 14371 sequence41, whole genome shotgun sequence
Sequence ID: NZ_BMOE01000041.1 Length: 690
Range 1: 31 to 378

Score:180 bits(456), Expect:5e-54,
Method:Compositional matrix adjust.,
Identities:108/116(93%), Positives:112/116(96%), Gaps:0/116(0%)

```
Query  1    MAGRSGDSDEELIRTVRLIKLLYQSNPPPNPEGTrqarrnrrrrwrerqrqIHSISERIL  60
            MAGRSGDSDE+L++ VRLIK LYQSNPPPNPEGTRQARRNRRRRWRERQRQIHSISERIL
Sbjct  31   MAGRSGDSDEDLLKAVRLIKFLYQSNPPPNPEGTRQARRNRRRRWRERQRQIHSISERIL  210

Query  61   GTYLGRSAEPVPLQLPPLERLTLDCNEDCGTSGTQGVGSPQILVESPTVLESGTKE  116
             TYLGRSAEPVPLQLPPLERLTLDCNEDCGTSGTQGVGSPQILVESPT+LESG KE
Sbjct  211  STYLGRSAEPVPLQLPPLERLTLDCNEDCGTSGTQGVGSPQILVESPTILESGAKE  378
```

>Enterococcus alcedinis strain CCM 8433 sequence26, whole genome shotgun sequence
Sequence ID: NZ_BMDT01000026.1 Length: 853
Range 1: 53 to 400

Score:180 bits(456), Expect:2e-53,
Method:Compositional matrix adjust.,
Identities:108/116(93%), Positives:112/116(96%), Gaps:0/116(0%)

```
Query  1    MAGRSGDSDEELIRTVRLIKLLYQSNPPPNPEGTrqarrnrrrrwrerqrqIHSISERIL  60
            MAGRSGDSDE+L++ VRLIK LYQSNPPPNPEGTRQARRNRRRRWRERQRQIHSISERIL
Sbjct  53   MAGRSGDSDEDLLKAVRLIKFLYQSNPPPNPEGTRQARRNRRRRWRERQRQIHSISERIL  232

Query  61   GTYLGRSAEPVPLQLPPLERLTLDCNEDCGTSGTQGVGSPQILVESPTVLESGTKE  116
             TYLGRSAEPVPLQLPPLERLTLDCNEDCGTSGTQGVGSPQILVESPT+LESG KE
```

```
Sbjct  233  STYLGRSAEPVPLQLPPLERLTLDCNEDCGTSGTQGVGSPQILVESPTILESGAKE  400
```

>Corynebacterium glutamicum strain 38 Scaffolds_31_4109, whole genome shotgun sequence
Sequence ID: NZ_JACENN010000031.1 Length: 3118
Range 1: 2305 to 2652

Score:180 bits(456), Expect:3e-49,
Method:Compositional matrix adjust.,
Identities:108/116(93%), Positives:112/116(96%), Gaps:0/116(0%)

```
Query  1     MAGRSGDSDEELIRTVRLIKLLYQSNPPPNPEGTrqarrnrrrrwrerqrqIHSISERIL  60
             MAGRSGDSDE+L++ VRLIK LYQSNPPPNPEGTRQARRNRRRRWRERQRQIHSISERIL
Sbjct  2305  MAGRSGDSDEDLLKAVRLIKFLYQSNPPPNPEGTRQARRNRRRRWRERQRQIHSISERIL  2484

Query  61    GTYLGRSAEPVPLQLPPLERLTLDCNEDCGTSGTQGVGSPQILVESPTVLESGTKE  116
              TYLGRSAEPVPLQLPPLERLTLDCNEDCGTSGTQGVGSPQILVESPT+LESG KE
Sbjct  2485  STYLGRSAEPVPLQLPPLERLTLDCNEDCGTSGTQGVGSPQILVESPTILESGAKE  2652
```

>Lactiplantibacillus plantarum strain MH-301 Scaffolds_50_4236, whole genome shotgun sequence
Sequence ID: NZ_JAOYUG010000050.1 Length: 4236
Range 1: 120 to 467

Score:180 bits(456), Expect:3e-49,
Method:Compositional matrix adjust.,
Identities:108/116(93%), Positives:112/116(96%), Gaps:0/116(0%)

```
Query  1    MAGRSGDSDEELIRTVRLIKLLYQSNPPPNPEGTrqarrnrrrrwrerqrqIHSISERIL  60
            MAGRSGDSDE+L++ VRLIK LYQSNPPPNPEGTRQARRNRRRRWRERQRQIHSISERIL
Sbjct  120  MAGRSGDSDEDLLKAVRLIKFLYQSNPPPNPEGTRQARRNRRRRWRERQRQIHSISERIL  299

Query  61   GTYLGRSAEPVPLQLPPLERLTLDCNEDCGTSGTQGVGSPQILVESPTVLESGTKE  116
             TYLGRSAEPVPLQLPPLERLTLDCNEDCGTSGTQGVGSPQILVESPT+LESG KE
Sbjct  300  STYLGRSAEPVPLQLPPLERLTLDCNEDCGTSGTQGVGSPQILVESPTILESGAKE  467
```

>Corynebacterium glutamicum strain 51 Scaffolds_32_4236, whole genome shotgun sequence
Sequence ID: NZ_JACENK010000032.1 Length: 4236
Range 1: 2887 to 3234

Score:180 bits(456), Expect:3e-49,
Method:Compositional matrix adjust.,
Identities:108/116(93%), Positives:112/116(96%), Gaps:0/116(0%)

```
Query  1     MAGRSGDSDEELIRTVRLIKLLYQSNPPPNPEGTrqarrnrrrrwrerqrqIHSISERIL  60
             MAGRSGDSDE+L++ VRLIK LYQSNPPPNPEGTRQARRNRRRRWRERQRQIHSISERIL
Sbjct  2887  MAGRSGDSDEDLLKAVRLIKFLYQSNPPPNPEGTRQARRNRRRRWRERQRQIHSISERIL  3066

Query  61    GTYLGRSAEPVPLQLPPLERLTLDCNEDCGTSGTQGVGSPQILVESPTVLESGTKE  116
              TYLGRSAEPVPLQLPPLERLTLDCNEDCGTSGTQGVGSPQILVESPT+LESG KE
Sbjct  3067  STYLGRSAEPVPLQLPPLERLTLDCNEDCGTSGTQGVGSPQILVESPTILESGAKE  3234
```

>Aggregatibacter actinomycetemcomitans serotype d str. SA269 contig0497, whole genome shotgun sequence
Sequence ID: NZ_AZTX01000497.1 Length: 504
Range 1: 99 to 392

Score:126 bits(316), Expect:1e-33,
Method:Compositional matrix adjust.,
Identities:85/98(87%), Positives:89/98(90%), Gaps:0/98(0%)

```
Query  19   IKLLYQSNPPPNPEGTrqarrnrrrrwrerqrqIHSISERILGTYLGRSAEPVPLQLPPL  78
            I  +  S+PPP  EG+RQARRNRRRRWRERQRQIHSIS+RIL TYLGRSAEPVPLQLPPL
Sbjct  99   ILTIIVSDPPPTSEGSRQARRNRRRRWRERQRQIHSISDRILSTYLGRSAEPVPLQLPPL  278

Query  79   ERLTLDCNEDCGTSGTQGVGSPQILVESPTVLESGTKE  116
            ERLTLDCNEDCGT GTQ VGSPQILVESPTVLESGTKE
Sbjct  279  ERLTLDCNEDCGTPGTQRVGSPQILVESPTVLESGTKE  392
```

>Klebsiella pneumoniae strain EuSCAPE_MT015, whole genome shotgun sequence
Sequence ID: NZ_UKIW02000110.1 Length: 328
Range 1: 12 to 305

Score:92.8 bits(229), Expect:4e-21,
Method:Compositional matrix adjust.,
Identities:68/98(69%), Positives:77/98(78%), Gaps:0/98(0%)

```
Query  19   IKLLYQSNPPPNPEGTrqarrnrrrrwrerqrqIHSISERILGTYLGRSAEPVPLQLPPL  78
            I  L  S+P P PEGTRQAR+NRRRRWR RQRQIHS+SERIL T LGR AEPVP QLPP+
Sbjct  12   ILTLVVSDPYPKPEGTRQARKNRRRRWRARQRQIHSLSERILSTCLGRPAEPVPFQLPPI  191

Query  79   ERLTLDCNEDCGTSGTQGVGSPQILVESPTVLESGTKE  116
            ERL +DC+E  GTSGTQGVG P +  +S  VL SGTK+
Sbjct  192  ERLHIDCSEGSGTSGTQGVGHP*VSGKSWAVLGSGTKK  305
```

>Klebsiella pneumoniae strain EuSCAPE_IL022, whole genome shotgun sequence
Sequence ID: NZ_UKKD02000023.1 Length: 411
Range 1: 185 to 379

Score:80.1 bits(196), Expect:9e-16,
Method:Compositional matrix adjust.,
Identities:42/65(65%), Positives:51/65(78%), Gaps:0/65(0%)

```
Query  52   IHSISERILGTYLGRSAEPVPLQLPPLERLTLDCNEDCGTSGTQGVGSPQILVESPTVLE  111
            I+SISERIL T LGRS EPVP QLPP+ERLT+DC+E  GTSGT+GVG+  I  +S T+L
Sbjct  379  INSISERILSTCLGRSEEPVPFQLPPIERLTIDCDESGGTSGTEGVGNS*ISGKSYTILG  200

Query  112  SGTKE  116
            S  K+
Sbjct  199  SRIKK  185
```

>Klebsiella pneumoniae strain EuSCAPE_IL006, whole genome shotgun sequence
Sequence ID: NZ_UKJR02000114.1 Length: 560
Range 1: 99 to 413

Score:77.0 bits(188), Expect:4e-14,

Method:Compositional matrix adjust.,
Identities:67/105(64%), Positives:74/105(70%), Gaps:7/105(6%)

```
Query  19   IKLLYQSNPPPNPEGTrqarrnrrrrwrerqrqIHSISERILGTYLGRSAEPVPLQLPPL  78
            I  L  ++P P PEGTRQARRNRRRRWR RQRQIHSISERIL   LGRS EPVP QLPP+
Sbjct  413  ILTLVVADPYPKPEGTRQARRNRRRRWRARQRQIHSISERILSDCLGRSTEPVPFQLPPI  234

Query  79   ERLTLDCNEDCGTSGTQ-------GVGSPQILVESPTVLESGTKE  116
            ERL +  +E  GTSGTQ       GVGSP I  +   VL SGTK+
Sbjct  233  ERLNIGDSESSGTSGTQLPQGSTEGVGSP*ISGKYCAVLGSGTKK  99
```

>Escherichia coli strain EN5090 contig00081, whole genome shotgun sequence
Sequence ID: NZ_JAODTD010000081.1 Length: 2798
Range 1: 424 to 699

Score:80.1 bits(196), Expect:4e-14,
Method:Compositional matrix adjust.,
Identities:60/92(65%), Positives:73/92(79%), Gaps:0/92(0%)

```
Query  25   SNPPPNPEGTrqarrnrrrrwrerqrqIHSISERILGTYLGRSAEPVPLQLPPLERLTLD  84
            S+P P P+GTRQA++NRRRRWR RQRQIHSISERIL + LGR AEPVPLQLPP+E L +
Sbjct  699  SDPYPKPKGTRQAQKNRRRRWRARQRQIHSISERILSSCLGRPAEPVPLQLPPIEELHIG  520

Query  85   CNEDCGTSGTQGVGSPQILVESPTVLESGTKE  116
             ++  GTSGTQG+GSP I  +S  +L SG K+
Sbjct  519  DSKSGGTSGTQGMGSP*ISRKSCAILGSGAKK  424
```

>Escherichia coli strain EN5383 contig00096, whole genome shotgun sequence
Sequence ID: NZ_JAPTZN010000096.1 Length: 2210
Range 1: 1527 to 1802

Score:79.7 bits(195), Expect:6e-14,
Method:Compositional matrix adjust.,
Identities:61/92(66%), Positives:74/92(80%), Gaps:0/92(0%)

```
Query  25    SNPPPNPEGTrqarrnrrrrwrerqrqIHSISERILGTYLGRSAEPVPLQLPPLERLTLD  84
             S+P P P+GTRQAR+NRRRRWR RQRQIHSISERIL + LGR AEPVPLQLPP+E+L +
Sbjct  1527  SDPYPEPKGTRQARKNRRRRWRARQRQIHSISERILSSCLGRPAEPVPLQLPPIEKLHIG  1706

Query  85    CNEDCGTSGTQGVGSPQILVESPTVLESGTKE  116
              ++  GTSGTQG+GSP I  +S  +L SG K+
Sbjct  1707  DSKSGGTSGTQGMGSP*ISRKSCAILGSGAKK  1802
```

>Escherichia coli strain EN5374 contig00094, whole genome shotgun sequence
Sequence ID: NZ_JAPTGI010000094.1 Length: 2720
Range 1: 2010 to 2306

Score:79.0 bits(193), Expect:1e-13,
Method:Compositional matrix adjust.,
Identities:62/100(62%), Positives:77/100(77%), Gaps:1/100(1%)

```
Query  17    RLIKLLYQSNPPPNPEGTrqarrnrrrrwrerqrqIHSISERILGTYLGRSAEPVPLQLP  76
             RL+  +  S+P P P+GTRQAR+NRRRRWR RQRQIHSISERIL + LGR AEPVPLQLP
Sbjct  2010  RLLTFVV-SDPYPEPKGTRQARKNRRRRWRARQRQIHSISERILSSCLGRPAEPVPLQLP  2186

Query  77    PLERLTLDCNEDCGTSGTQGVGSPQILVESPTVLESGTKE  116
             P+E+L +  ++  GTSG QG+GSP I  +S  +L SG K+
Sbjct  2187  PIEKLHIGDSKSGGTSGAQGMGSP*ISRKSCAILGSGAKK  2306
```

>Escherichia coli strain EN5076 contig00066, whole genome shotgun sequence
Sequence ID: NZ_JAODTC010000066.1 Length: 2732
Range 1: 421 to 717

Score:79.0 bits(193), Expect:1e-13,
Method:Compositional matrix adjust.,
Identities:62/100(62%), Positives:77/100(77%), Gaps:1/100(1%)

```
Query  17   RLIKLLYQSNPPPNPEGTrqarrnrrrrwrerqrqIHSISERILGTYLGRSAEPVPLQLP  76
            RL+  +  S+P P P+GTRQAR+NRRRRWR RQRQIHSISERIL + LGR AEPVPLQLP
Sbjct  717  RLLTFVV-SDPYPEPKGTRQARKNRRRRWRARQRQIHSISERILSSCLGRPAEPVPLQLP  541

Query  77   PLERLTLDCNEDCGTSGTQGVGSPQILVESPTVLESGTKE  116
            P+E+L +  ++  GTSG QG+GSP I  +S  +L SG K+
Sbjct  540  PIEKLHIGDSKSGGTSGAQGMGSP*ISRKSCAILGSGAKK  421
```

>Escherichia coli strain NBE0010 contig00080, whole genome shotgun sequence
Sequence ID: NZ_JARBDM010000080.1 Length: 2743
Range 1: 1711 to 2007

Score:79.0 bits(193), Expect:1e-13,
Method:Compositional matrix adjust.,
Identities:62/100(62%), Positives:77/100(77%), Gaps:1/100(1%)

```
Query  17    RLIKLLYQSNPPPNPEGTrqarrnrrrrwrerqrqIHSISERILGTYLGRSAEPVPLQLP  76
             RL+  +  S+P P P+GTRQAR+NRRRRWR RQRQIHSISERIL + LGR AEPVPLQLP
Sbjct  1711  RLLTFVV-SDPYPEPKGTRQARKNRRRRWRARQRQIHSISERILSSCLGRPAEPVPLQLP  1887

Query  77    PLERLTLDCNEDCGTSGTQGVGSPQILVESPTVLESGTKE  116
             P+E+L +  ++  GTSG QG+GSP I  +S  +L SG K+
Sbjct  1888  PIEKLHIGDSKSGGTSGAQGMGSP*ISRKSCAILGSGAKK  2007
```

>Escherichia coli strain EN5143 contig00069, whole genome shotgun sequence
Sequence ID: NZ_JAPMNI010000069.1 Length: 2688
Range 1: 2064 to 2279

Score:78.6 bits(192), Expect:2e-13,
Method:Compositional matrix adjust.,
Identities:46/72(64%), Positives:52/72(72%), Gaps:7/72(9%)

```
Query  52    IHSISERILGTYLGRSAEPVPLQLPPLERLTLDCNEDCGTSGTQ-------GVGSPQILV  104
             IHSISERIL + LGRSAEPVPLQLPP+ERL +  +E  GTSGTQ       GVGSP I
Sbjct  2064  IHSISERILSSCLGRSAEPVPLQLPPIERLHISDSESAGTSGTQQSQGTAEGVGSP*ISG  2243

Query  105   ESPTVLESGTKE  116
             +S  VL SGTK+
Sbjct  2244  KSCAVLGSGTKK  2279
```

```
Range 2: 103 to 183

Score:45.8 bits(107), Expect:0.045,
Method:Compositional matrix adjust.,
Identities:19/27(70%), Positives:24/27(88%), Gaps:0/27(0%)

Query  1    MAGRSGDSDEELIRTVRLIKLLYQSNP  27
            MAGRSGDSDE L++ VR+IK+LYQS+
Sbjct  103  MAGRSGDSDEALLKAVRIIKILYQSSK  183
```

```
>Escherichia coli strain EN5132 contig00077, whole genome shotgun sequence
Sequence ID: NZ_JAPYLZ010000077.1 Length: 2498
Range 1: 1775 to 2086

Score:78.2 bits(191), Expect:2e-13,
Method:Compositional matrix adjust.,
Identities:66/104(63%), Positives:74/104(71%), Gaps:7/104(6%)

Query  19    IKLLYQSNPPPNPEGTrqarrnrrrrwrerqrqIHSISERILGTYLGRSAEPVPLQLPPL  78
             I     S+P P P GTRQAR+NRRRRWR RQ+QIHSISERIL + LGRSAEPVPLQLPP+
Sbjct  1775  ILTFVVSDPYPEPRGTRQARKNRRRRWRARQKQIHSISERILSSCLGRSAEPVPLQLPPI  1954

Query  79    ERLTLDCNEDCGTSGTQ-------GVGSPQILVESPTVLESGTK  115
             ERL +  +E  GTSGTQ       GVGSP I  +   VL SGTK
Sbjct  1955  ERLHIGDSESAGTSGTQQSQGTAEGVGSP*ISGKPCAVLGSGTK  2086
```

```
Range 2: 103 to 183

Score:45.8 bits(107), Expect:0.044,
Method:Compositional matrix adjust.,
Identities:19/27(70%), Positives:24/27(88%), Gaps:0/27(0%)

Query  1    MAGRSGDSDEELIRTVRLIKLLYQSNP  27
            MAGRSGDSDE L++ VR+IK+LYQS+
Sbjct  103  MAGRSGDSDEALLKAVRIIKILYQSSK  183
```

```
>Klebsiella pneumoniae strain EuSCAPE_MT003, whole genome shotgun sequence
Sequence ID: NZ_UKIK02000085.1 Length: 964
Range 1: 634 to 897

Score:77.0 bits(188), Expect:3e-13,
Method:Compositional matrix adjust.,
Identities:59/88(67%), Positives:69/88(78%), Gaps:0/88(0%)

Query  25   SNPPPNPEGTrqarrnrrrrwrerqrqIHSISERILGTYLGRSAEPVPLQLPPLERLTLD  84
            ++P P P GTRQAR+NRRRRWR RQRQIHSIS+RIL T LGR  EPVP QLPP+ERL +D
Sbjct  897  ADPYPEPRGTRQARKNRRRRWRARQRQIHSISKRILFTCLGRLTEPVPFQLPPIERLHID  718

Query  85   CNEDCGTSGTQGVGSPQILVESPTVLES  112
            C+E  G SGTQGVG+P +  +  TVL S
Sbjct  717  CSESGGNSGTQGVGNP*VSGKPYTVLGS  634
```

```
>Escherichia coli strain EN5379 contig00092, whole genome shotgun sequence
Sequence ID: NZ_JAPTGJ010000092.1 Length: 2592
Range 1: 402 to 698

Score:77.0 bits(188), Expect:4e-13,
Method:Compositional matrix adjust.,
Identities:65/99(66%), Positives:74/99(74%), Gaps:7/99(7%)

Query  25   SNPPPNPEGTrqarrnrrrrwrerqrqIHSISERILGTYLGRSAEPVPLQLPPLERLTLD  84
            S+P P P+GTRQA++NRRRRWR RQRQIHSISERIL T LGRS EPVPLQLPP+ERL +
Sbjct  698  SDPYPKPKGTRQAQKNRRRRWRARQRQIHSISERILITCLGRSTEPVPLQLPPIERLHIT  519

Query  85   CNEDCGTSGTQ-------GVGSPQILVESPTVLESGTKE  116
             +E  GTSGTQ       GVGSP I  +   VL SGTK+
Sbjct  518  DSESGGTSGTQQSPGTTEGVGSP*ISGKPCAVLGSGTKK  402
```

```
>Escherichia coli strain NBE0009 contig00116, whole genome shotgun sequence
Sequence ID: NZ_JARBDL010000116.1 Length: 1713
Range 1: 447 to 761

Score:77.0 bits(188), Expect:5e-13,
Method:Compositional matrix adjust.,
Identities:67/105(64%), Positives:74/105(70%), Gaps:7/105(6%)

Query  19   IKLLYQSNPPPNPEGTrqarrnrrrrwrerqrqIHSISERILGTYLGRSAEPVPLQLPPL  78
            I     S+P P P GTRQA+RNRRRRWR RQ+QIHSISERIL T LGRSAEPVPLQLPPL
Sbjct  761  ILTFVVSDPYPEPRGTRQAQRNRRRRWRARQKQIHSISERILSTCLGRSAEPVPLQLPPL  582

Query  79   ERLTLDCNEDCGTSGTQ-------GVGSPQILVESPTVLESGTKE  116
            ERL +  +E  GT GTQ       GVGSP I  +   VL SGTK+
Sbjct  581  ERLHITDSESGGTFGTQQSPGTTEGVGSP*ISGKPCAVLGSGTKK  447
```

```
>Escherichia coli strain NBE0007 contig00093, whole genome shotgun sequence
Sequence ID: NZ_JARBDJ010000093.1 Length: 3013
Range 1: 431 to 727

Score:75.5 bits(184), Expect:1e-12,
Method:Compositional matrix adjust.,
Identities:66/99(67%), Positives:73/99(73%), Gaps:7/99(7%)

Query  25   SNPPPNPEGTrqarrnrrrrwrerqrqIHSISERILGTYLGRSAEPVPLQLPPLERLTLD  84
            S+P P P GTRQAR+NRRRRWR RQRQIHS+SERIL T LGRSAE VPLQLPP+ERL +
Sbjct  727  SDPCPEPRGTRQARKNRRRRWRARQRQIHSLSERILSTCLGRSAELVPLQLPPIERLHIS  548

Query  85   CNEDCGTSGTQ-------GVGSPQILVESPTVLESGTKE  116
             +E  GTSGTQ       GVGSP I  +S  VL SG KE
Sbjct  547  DSESGGTSGTQQPQGTTEGVGSP*ISGKSCAVLGSGIKE  431
```

>Escherichia coli strain NBE0006 contig00145, whole genome shotgun sequence
Sequence ID: NZ_JARBDI010000145.1 Length: 738
Range 1: 63 to 359

Score:73.6 bits(179), Expect:2e-12,
Method:Compositional matrix adjust.,
Identities:63/99(64%), Positives:71/99(71%), Gaps:7/99(7%)

```
Query  19   IKLLYQSNPPPNPEGTrqarrnrrrrwrerqrqIHSISERILGTYLGRSAEPVPLQLPPL  78
            I     S+P P P GTRQAR+NRRRRWR RQ+QIHSISERIL + LGRSAEPVPLQLPP+
Sbjct  63   ILTFVVSDPYPEPRGTRQARKNRRRRWRARQKQIHSISERILSSCLGRSAEPVPLQLPPI  242

Query  79   ERLTLDCNEDCGTSGTQ-------GVGSPQILVESPTVL  110
            ERL +  +E  GTSGTQ       GVGSP I  +S  VL
Sbjct  243  ERLHISDSESGGTSGTQQSQGTAEGVGSP*ISGKSCAVL  359
```

>Escherichia coli strain EN5177 contig00148, whole genome shotgun sequence
Sequence ID: NZ_JAPMNK010000148.1 Length: 850
Range 1: 421 to 714

Score:73.2 bits(178), Expect:5e-12,
Method:Compositional matrix adjust.,
Identities:61/98(62%), Positives:70/98(71%), Gaps:7/98(7%)

```
Query  25   SNPPPNPEGTrqarrnrrrrwrerqrqIHSISERILGTYLGRSAEPVPLQLPPLERLTLD  84
            S+P P P+GTRQAR+NRRRR R RQ+QIHSISERIL + LGRSAEPVPLQLPP+ERL +
Sbjct  714  SDPYPEPKGTRQARKNRRRR*RARQKQIHSISERILSSCLGRSAEPVPLQLPPIERLHIS  535

Query  85   CNEDCGTSGTQ-------GVGSPQILVESPTVLESGTK  115
             +E  GTS TQ        V SP I  +S  VL S TK
Sbjct  534  GSESSGTSRTQQSQRTAEKVRSP*ISRKSCAVLRSRTK  421
```

>Escherichia coli strain NBE0006 contig00147, whole genome shotgun sequence
Sequence ID: NZ_JARBDI010000147.1 Length: 694
Range 1: 358 to 654

Score:71.6 bits(174), Expect:9e-12,
Method:Compositional matrix adjust.,
Identities:63/99(64%), Positives:71/99(71%), Gaps:7/99(7%)

```
Query  25   SNPPPNPEGTrqarrnrrrrwrerqrqIHSISERILGTYLGRSAEPVPLQLPPLERLTLD  84
            S+P P P+GTRQAR+NRRRRWR RQRQI SISERIL T LGR  EPVPLQLPP+ERL +
Sbjct  654  SDPYPKPKGTRQARKNRRRRWRARQRQIDSISERILSTCLGRPVEPVPLQLPPIERLNIG  475

Query  85   CNEDCGTSGTQ-------GVGSPQILVESPTVLESGTKE  116
             +E  GTSGTQ       GVGSP +      VL SGTK+
Sbjct  474  DSESGGTSGTQQSQGTTEGVGSP*VSGRYCAVLGSGTKK  358
```

>Escherichia coli strain NBE0004 contig00110, whole genome shotgun sequence
Sequence ID: NZ_JARBDH010000110.1 Length: 1390
Range 1: 436 to 729

Score:73.2 bits(178), Expect:1e-11,
Method:Compositional matrix adjust.,
Identities:61/98(62%), Positives:70/98(71%), Gaps:7/98(7%)

```
Query  25   SNPPPNPEGTrqarrnrrrrwrerqrqIHSISERILGTYLGRSAEPVPLQLPPLERLTLD  84
            S+P P P+GTRQAR+NRRRR R RQ+QIHSISERIL + LGRSAEPVPLQLPP+ERL +
Sbjct  729  SDPYPEPKGTRQARKNRRRR*RARQKQIHSISERILSSCLGRSAEPVPLQLPPIERLHIS  550

Query  85   CNEDCGTSGTQ-------GVGSPQILVESPTVLESGTK  115
             +E  GTS TQ        V SP I  +S  VL S TK
Sbjct  549  GSESSGTSRTQQSQRTAEKVRSP*ISRKSCAVLRSRTK  436
```

>Escherichia coli strain EN5095 contig00078, whole genome shotgun sequence
Sequence ID: NZ_JAPYLY010000078.1 Length: 1972
Range 1: 378 to 674

Score:73.2 bits(178), Expect:1e-11,
Method:Compositional matrix adjust.,
Identities:64/99(65%), Positives:71/99(71%), Gaps:7/99(7%)

```
Query  25   SNPPPNPEGTrqarrnrrrrwrerqrqIHSISERILGTYLGRSAEPVPLQLPPLERLTLD  84
            S+P P P GTRQAR+NRRRRWR RQRQI SIS RIL T LGRS EPVPLQLPP+ERLT+
Sbjct  674  SDPYPEPRGTRQARKNRRRRWRARQRQIDSISTRILSTCLGRSEEPVPLQLPPIERLTIG  495

Query  85   CNEDCGTSGTQ-------GVGSPQILVESPTVLESGTKE  116
             NE  G SGTQ       GVGSP +  +   VL SGTK+
Sbjct  494  DNESGGNSGTQQSQGTTEGVGSP*VSGKYCAVLGSGTKK  378
```

>Escherichia coli strain EN5286 contig00120, whole genome shotgun sequence
Sequence ID: NZ_JAPMNM010000120.1 Length: 3033
Range 1: 2273 to 2587

Score:73.2 bits(178), Expect:1e-11,
Method:Compositional matrix adjust.,
Identities:64/105(61%), Positives:72/105(68%), Gaps:7/105(6%)

```
Query  19    IKLLYQSNPPPNPEGTrqarrnrrrrwrerqrqIHSISERILGTYLGRSAEPVPLQLPPL  78
             I     S+P P P GTRQAR+NRRRRWR RQ+Q+HSISERIL T LGR AEPVP  LPP+
Sbjct  2273  ILTFVISDPCPEPRGTRQARKNRRRRWRARQKQLHSISERILSTCLGRPAEPVPFLLPPI  2452

Query  79    ERLTLDCNEDCGTSGTQ-------GVGSPQILVESPTVLESGTKE  116
             ERL +  +E  GTSGTQ       GVGSP I  +S  VL SG KE
Sbjct  2453  ERLNISDSESGGTSGTQQPQGTTEGVGSP*ISGKSCAVLGSGIKE  2587
```

>Escherichia coli strain NBE0009 contig00153, whole genome shotgun sequence
Sequence ID: NZ_JARBDL010000153.1 Length: 753
Range 1: 54 to 368

Score:71.6 bits(174), Expect:1e-11,
Method:Compositional matrix adjust.,
Identities:65/105(62%), Positives:72/105(68%), Gaps:7/105(6%)

```
Query  19   IKLLYQSNPPPNPEGTrqarrnrrrrwrerqrqIHSISERILGTYLGRSAEPVPLQLPPL  78
```

```
            I     S+P P P GTRQAR+NRRRRWR RQRQIHSISERIL T LGRS EPVP QLPP+
Sbjct  54   ILTFVVSDPYPEPRGTRQARKNRRRRWRARQRQIHSISERILITCLGRSTEPVPFQLPPI  233

Query  79   ERLTLDCNEDCGTSGTQ-------GVGSPQILVESPTVLESGTKE  116
            ERL +  +E  GTSGTQ       GV SP I  +   VL SGTK+
Sbjct  234  ERLHIGDSESGGTSGTQQSQGTTEGVESP*ISEKPCAVLGSGTKK  368
```

```
>Escherichia coli strain EN5192 contig00108, whole genome shotgun sequence
Sequence ID: NZ_JAPMNL010000108.1 Length: 3032
Range 1: 2304 to 2618

Score:72.8 bits(177), Expect:1e-11,
Method:Compositional matrix adjust.,
Identities:64/105(61%), Positives:71/105(67%), Gaps:7/105(6%)

Query  19    IKLLYQSNPPPNPEGTrqarrnrrrrwrerqrqIHSISERILGTYLGRSAEPVPLQLPPL  78
             I     S+P P P GTRQAR+NRRRRWR RQ+Q+HSISERIL T LGR AEPVP  LPP+
Sbjct  2304  ILTFVISDPCPEPRGTRQARKNRRRRWRARQKQLHSISERILSTCLGRPAEPVPFLLPPI  2483

Query  79    ERLTLDCNEDCGTSGTQ-------GVGSPQILVESPTVLESGTKE  116
             ERL +  NE  GTSGTQ       GVGSP I  +   VL SG KE
Sbjct  2484  ERLNISDNESGGTSGTQQPQGTAEGVGSP*ISGKPCAVLGSGIKE  2618
```

```
>Escherichia coli strain EN5192 contig00135, whole genome shotgun sequence
Sequence ID: NZ_JAPMNL010000135.1 Length: 1341
Range 1: 367 to 663

Score:72.4 bits(176), Expect:2e-11,
Method:Compositional matrix adjust.,
Identities:62/99(63%), Positives:71/99(71%), Gaps:7/99(7%)

Query  19   IKLLYQSNPPPNPEGTrqarrnrrrrwrerqrqIHSISERILGTYLGRSAEPVPLQLPPL  78
            I     S+P P P GTRQAR+NRRRRWR RQ+QIHS+SERIL + LGRSAEPVPLQLPP+
Sbjct  663  ILTFVVSDPYPEPRGTRQARKNRRRRWRARQKQIHSLSERILSSCLGRSAEPVPLQLPPI  484

Query  79   ERLTLDCNEDCGTSGTQ-------GVGSPQILVESPTVL  110
            ERL +  +E  GTSGTQ       GVGSP I  +S  VL
Sbjct  483  ERLHISDSESGGTSGTQQSQGTAEGVGSP*ISGKSCAVL  367
```

```
>Escherichia coli strain EN5308 contig00099, whole genome shotgun sequence
Sequence ID: NZ_JARBDE010000099.1 Length: 2827
Range 1: 512 to 808

Score:71.6 bits(174), Expect:4e-11,
Method:Compositional matrix adjust.,
Identities:63/99(64%), Positives:71/99(71%), Gaps:7/99(7%)

Query  25   SNPPPNPEGTrqarrnrrrrwrerqrqIHSISERILGTYLGRSAEPVPLQLPPLERLTLD  84
            S+P P P+GTRQAR+NRRRRWR RQRQI SISERIL T LGR  EPVPLQLPP+ERL +
Sbjct  808  SDPYPKPKGTRQARKNRRRRWRARQRQIDSISERILSTCLGRPVEPVPLQLPPIERLNIG  629

Query  85   CNEDCGTSGTQ-------GVGSPQILVESPTVLESGTKE  116
             +E  GTSGTQ       GVGSP +      VL SGTK+
Sbjct  628  DSESGGTSGTQQSQGTTEGVGSP*VSGRYCAVLGSGTKK  512
```

```
Range 2: 2699 to 2779

Score:45.8 bits(107), Expect:0.045,
Method:Compositional matrix adjust.,
Identities:19/27(70%), Positives:24/27(88%), Gaps:0/27(0%)

Query  1     MAGRSGDSDEELIRTVRLIKLLYQSNP  27
             MAGRSGDSDE L++ VR+IK+LYQS+
Sbjct  2779  MAGRSGDSDEALLKAVRIIKILYQSSK  2699
```

```
>Escherichia coli strain EN5169 contig00082, whole genome shotgun sequence
Sequence ID: NZ_JAPMNJ010000082.1 Length: 2837
Range 1: 435 to 731

Score:71.6 bits(174), Expect:4e-11,
Method:Compositional matrix adjust.,
Identities:63/99(64%), Positives:71/99(71%), Gaps:7/99(7%)

Query  25   SNPPPNPEGTrqarrnrrrrwrerqrqIHSISERILGTYLGRSAEPVPLQLPPLERLTLD  84
            S+P P P+GTRQAR+NRRRRWR RQRQI SISERIL T LGR  EPVPLQLPP+ERL +
Sbjct  731  SDPYPKPKGTRQARKNRRRRWRARQRQIDSISERILSTCLGRPVEPVPLQLPPIERLNIG  552

Query  85   CNEDCGTSGTQ-------GVGSPQILVESPTVLESGTKE  116
             +E  GTSGTQ       GVGSP +      VL SGTK+
Sbjct  551  DSESGGTSGTQQSQGTTEGVGSP*VSGRYCAVLGSGTKK  435
```

```
Range 2: 2655 to 2735

Score:45.8 bits(107), Expect:0.049,
Method:Compositional matrix adjust.,
Identities:19/27(70%), Positives:24/27(88%), Gaps:0/27(0%)

Query  1     MAGRSGDSDEELIRTVRLIKLLYQSNP  27
             MAGRSGDSDE L+R VR+I++LYQS+
Sbjct  2735  MAGRSGDSDEALLRAVRIIRILYQSSK  2655
```

```
>Escherichia coli strain NBE0003 contig00096, whole genome shotgun sequence
Sequence ID: NZ_JARBDG010000096.1 Length: 2837
Range 1: 435 to 731

Score:71.6 bits(174), Expect:4e-11,
Method:Compositional matrix adjust.,
Identities:63/99(64%), Positives:71/99(71%), Gaps:7/99(7%)

Query  25   SNPPPNPEGTrqarrnrrrrwrerqrqIHSISERILGTYLGRSAEPVPLQLPPLERLTLD  84
            S+P P P+GTRQAR+NRRRRWR RQRQI SISERIL T LGR  EPVPLQLPP+ERL +
Sbjct  731  SDPYPKPKGTRQARKNRRRRWRARQRQIDSISERILSTCLGRPVEPVPLQLPPIERLNIG  552
```

```
Query  85   CNEDCGTSGTQ-------GVGSPQILVESPTVLESGTKE  116
             +E  GTSGTQ       GVGSP +      VL SGTK+
Sbjct  551  DSESGGTSGTQQSQGTTEGVGSP*VSGRYCAVLGSGTKK  435
```

Range 2: 2655 to 2735

Score:45.8 bits(107), Expect:0.049,
Method:Compositional matrix adjust.,
Identities:19/27(70%), Positives:24/27(88%), Gaps:0/27(0%)

```
Query  1     MAGRSGDSDEELIRTVRLIKLLYQSNP  27
             MAGRSGDSDE L+R VR+I++LYQS+
Sbjct  2735  MAGRSGDSDEALLRAVRIIRILYQSSK  2655
```

>Escherichia coli strain NBE0002 contig00097, whole genome shotgun sequence
Sequence ID: NZ_JARBDF010000097.1 Length: 2717
Range 1: 442 to 756

Score:71.6 bits(174), Expect:4e-11,
Method:Compositional matrix adjust.,
Identities:63/105(60%), Positives:71/105(67%), Gaps:7/105(6%)

```
Query  19   IKLLYQSNPPPNPEGTrqarrnrrrrwrerqrqIHSISERILGTYLGRSAEPVPLQLPPL  78
            I     S+P P P GTRQAR+NRRRRWR RQ+Q+HSISERIL T LGR AEPVP  LPP+
Sbjct  756  ILTFVISDPCPEPRGTRQARKNRRRRWRARQKQLHSISERILSTCLGRPAEPVPFLLPPI  577

Query  79   ERLTLDCNEDCGTSGTQ-------GVGSPQILVESPTVLESGTKE  116
            ERL +  +E  G SGTQ       GVGSP I  +S  VL SG KE
Sbjct  576  ERLNISDSESGGASGTQQPQGTTEGVGSP*ISGKSCAVLGSGIKE  442
```

Range 2: 2535 to 2615

Score:45.8 bits(107), Expect:0.045,
Method:Compositional matrix adjust.,
Identities:19/27(70%), Positives:24/27(88%), Gaps:0/27(0%)

```
Query  1     MAGRSGDSDEELIRTVRLIKLLYQSNP  27
             MAGRSGDSDE L++ VR+IK+LYQS+
Sbjct  2615  MAGRSGDSDEALLKAVRIIKILYQSSK  2535
```

>Escherichia coli strain EN5177 contig00087, whole genome shotgun sequence
Sequence ID: NZ_JAPMNK010000087.1 Length: 3403
Range 1: 2643 to 2957

Score:69.3 bits(168), Expect:2e-10,
Method:Compositional matrix adjust.,
Identities:62/105(59%), Positives:71/105(67%), Gaps:7/105(6%)

```
Query  19    IKLLYQSNPPPNPEGTrqarrnrrrrwrerqrqIHSISERILGTYLGRSAEPVPLQLPPL  78
             I     S+P P P GTRQAR+NRRRRWR RQ+Q+HSISERIL T LGR AEPVP  LPP+
Sbjct  2643  ILTFVISDPCPEPRGTRQARKNRRRRWRARQKQLHSISERILSTCLGRPAEPVPFLLPPI  2822

Query  79    ERLTLDCNEDCGTSGTQ-------GVGSPQILVESPTVLESGTKE  116
             ERL +  +E  GTSGTQ       GVGSP I  +   VL SG K+
Sbjct  2823  ERLNISDSESGGTSGTQQPQGTTEGVGSP*ISGKPCAVLGSGIKK  2957
```

>Escherichia coli strain EN5143 contig00071, whole genome shotgun sequence
Sequence ID: NZ_JAPMNI010000071.1 Length: 2774
Range 1: 2017 to 2331

Score:69.3 bits(168), Expect:3e-10,
Method:Compositional matrix adjust.,
Identities:62/105(59%), Positives:70/105(66%), Gaps:7/105(6%)

```
Query  19    IKLLYQSNPPPNPEGTrqarrnrrrrwrerqrqIHSISERILGTYLGRSAEPVPLQLPPL  78
             I     S+P P P GTRQAR+NRRRRWR RQ+Q+HSISERIL T LGR AEPVP  LPP+
Sbjct  2017  ILTFVISDPCPEPRGTRQARKNRRRRWRARQKQLHSISERILSTCLGRPAEPVPFLLPPI  2196

Query  79    ERLTLDCNEDCGTSGTQ-------GVGSPQILVESPTVLESGTKE  116
             ERL +  +E  G SGTQ       GVGSP I  +   VL SG KE
Sbjct  2197  ERLNISDSESGGASGTQQPQGTTEGVGSP*ISGKPCAVLGSGIKE  2331
```

>Escherichia coli strain EN5308 contig00096, whole genome shotgun sequence
Sequence ID: NZ_JARBDE010000096.1 Length: 3024
Range 1: 444 to 758

Score:69.3 bits(168), Expect:3e-10,
Method:Compositional matrix adjust.,
Identities:62/105(59%), Positives:70/105(66%), Gaps:7/105(6%)

```
Query  19   IKLLYQSNPPPNPEGTrqarrnrrrrwrerqrqIHSISERILGTYLGRSAEPVPLQLPPL  78
            I     S+P P P GTRQAR+NRRRRWR RQ+Q+HSISERIL T LGR AEPVP  LPP+
Sbjct  758  ILTFVISDPCPEPRGTRQARKNRRRRWRARQKQLHSISERILSTCLGRPAEPVPFLLPPI  579

Query  79   ERLTLDCNEDCGTSGTQ-------GVGSPQILVESPTVLESGTKE  116
            ERL +  +E  G SGTQ       GVGSP I  +   VL SG KE
Sbjct  578  ERLNISDSESGGASGTQQPQGTTEGVGSP*ISGKPCAVLGSGIKE  444
```

>Klebsiella pneumoniae strain EuSCAPE_IL009, whole genome shotgun sequence
Sequence ID: NZ_UKJO02000079.1 Length: 274
Range 1: 3 to 182

Score:63.9 bits(154), Expect:5e-10,
Method:Compositional matrix adjust.,
Identities:51/60(85%), Positives:55/60(91%), Gaps:0/60(0%)

```
Query  25   SNPPPNPEGTrqarrnrrrrwrerqrqIHSISERILGTYLGRSAEPVPLQLPPLERLTLD  84
            S+P PNPEGTRQARRNRRRRWR RQRQIHSISERIL + LGRSAEPVPLQLPP+ERL +D
Sbjct  182  SDPYPNPEGTRQARRNRRRRWRARQRQIHSISERILSSCLGRSAEPVPLQLPPIERLHID  3
```

```
>Klebsiella pneumoniae strain EuSCAPE_DE091, whole genome shotgun sequence
Sequence ID: NZ_UKMA02000049.1 Length: 618
Range 1: 356 to 562

Score:65.9 bits(159), Expect:9e-10,
Method:Compositional matrix adjust.,
Identities:42/69(61%), Positives:47/69(68%), Gaps:7/69(10%)

Query  55   ISERILGTYLGRSAEPVPLQLPPLERLTLDCNEDCGTSG-------TQGVGSPQILVESP  107
            ISERIL T L RS EPVP QLPP+ERLTLD +E  GTSG       T+GVGS  I  +S
Sbjct  562  ISERILDTCLERSEEPVPFQLPPIERLTLDNSESGGTSGTLQPQGTTEGVGSS*ISGKSC  383

Query  108  TVLESGTKE  116
             VL SGTK+
Sbjct  382  AVLGSGTKK  356



>Klebsiella pneumoniae strain EuSCAPE_IL001, whole genome shotgun sequence
Sequence ID: NZ_UKJN02000084.1 Length: 599
Range 1: 193 to 489

Score:65.1 bits(157), Expect:2e-09,
Method:Compositional matrix adjust.,
Identities:60/99(61%), Positives:70/99(70%), Gaps:7/99(7%)

Query  25   SNPPPNPEGTrqarrnrrrrwrerqrqIHSISERILGTYLGRSAEPVPLQLPPLERLTLD  84
            S+P P PEG+RQARRNRRRRWR RQRQI+ ISERILG+ LGR  EPVP  LPP+ERL +
Sbjct  193  SDPYPKPEGSRQARRNRRRRWRARQRQINQISERILGSILGRPEEPVPFLLPPIERLHIG  372

Query  85   CNEDCGTSGTQ-------GVGSPQILVESPTVLESGTKE  116
             +E  GTSGTQ       GVG+P I  +   VL SG K+
Sbjct  373  DSESSGTSGTQQPQGTTEGVGNP*ISGKPCAVLGSGIKK  489



>Escherichia coli strain NBE0009 contig00151, whole genome shotgun sequence
Sequence ID: NZ_JARBDL010000151.1 Length: 802
Range 1: 435 to 749

Score:65.5 bits(158), Expect:2e-09,
Method:Compositional matrix adjust.,
Identities:60/105(57%), Positives:70/105(66%), Gaps:7/105(6%)

Query  19   IKLLYQSNPPPNPEGTrqarrnrrrrwrerqrqIHSISERILGTYLGRSAEPVPLQLPPL  78
            I     S+P P+P GTRQA+RNRRRRWR RQ QI +ISERIL   LGR AEPVPLQLPP+
Sbjct  749  ILTFVVSDPYPDPRGTRQAQRNRRRRWRARQSQIRAISERILSDCLGRPAEPVPLQLPPI  570

Query  79   ERLTLDCNEDCGTSGTQ-------GVGSPQILVESPTVLESGTKE  116
            ERLT+  +E  GTSGTQ       G GS +       VL SGT++
Sbjct  569  ERLTIGDSESGGTSGTQLSSGTTEGEGSTEASGGHCAVLGSGTEK  435



>Escherichia coli strain EN5317 contig00057, whole genome shotgun sequence
Sequence ID: NZ_JARBDO010000057.1 Length: 2867
Range 1: 444 to 758

Score:66.6 bits(161), Expect:2e-09,
Method:Compositional matrix adjust.,
Identities:61/105(58%), Positives:70/105(66%), Gaps:7/105(6%)

Query  19   IKLLYQSNPPPNPEGTrqarrnrrrrwrerqrqIHSISERILGTYLGRSAEPVPLQLPPL  78
            I     S+P P+P GTRQA+RNRRRRWR RQ QI +ISERIL   LGR AEPVPLQLPPL
Sbjct  758  ILTFVVSDPYPDPRGTRQAQRNRRRRWRARQSQIRAISERILSDCLGRPAEPVPLQLPPL  579

Query  79   ERLTLDCNEDCGTSGTQ-------GVGSPQILVESPTVLESGTKE  116
            ERLT+  +E  GTSGTQ       G GS +       VL SGT++
Sbjct  578  ERLTIGDSESGGTSGTQLSSGTTEGEGSTEASGGHCAVLGSGTEK  444



>Klebsiella pneumoniae strain EuSCAPE_IL005, whole genome shotgun sequence
Sequence ID: NZ_UKJL02000055.1 Length: 455
Range 1: 179 to 454

Score:63.5 bits(153), Expect:3e-09,
Method:Compositional matrix adjust.,
Identities:57/92(62%), Positives:67/92(72%), Gaps:7/92(7%)

Query  26   NPPPNPEGTrqarrnrrrrwrerqrqIHSISERILGTYLGRSAEPVPLQLPPLERLTLDC  85
            +P P PEGTRQAR+NRRRRWR RQ+QI+SISERIL T LGR AEPVP  LPP+ERL +
Sbjct  179  DPYPKPEGTRQARKNRRRRWRARQKQINSISERILSTCLGRPAEPVPFLLPPIERLHIGD  358

Query  86   NEDCGTSGTQ-------GVGSPQILVESPTVL  110
            +E  GTSGTQ       GVG+P +  +S  VL
Sbjct  359  SESSGTSGTQQPQGTPEGVGNP*VSGKSCAVL  454



>Klebsiella pneumoniae strain EuSCAPE_MT017, whole genome shotgun sequence
Sequence ID: NZ_UKJA02000086.1 Length: 513
Range 1: 303 to 500

Score:62.8 bits(151), Expect:8e-09,
Method:Compositional matrix adjust.,
Identities:38/66(58%), Positives:45/66(68%), Gaps:7/66(10%)

Query  52   IHSISERILGTYLGRSAEPVPLQLPPLERLTLDCNEDCGTSGTQ-------GVGSPQILV  104
            I S+S RILG  LGRS EPVPLQLPP+ERL +  +E  GTSGTQ       GVG+P +
Sbjct  500  IDSLSARILGACLGRSEEPVPLQLPPIERLNIGDSESGGTSGTQQPQGTTEGVGNP*VPG  321

Query  105  ESPTVL  110
            +S TVL
Sbjct  320  KSYTVL  303



>Klebsiella pneumoniae strain EuSCAPE_IL018, whole genome shotgun sequence
Sequence ID: NZ_UKJU02000101.1 Length: 388
Range 1: 67 to 360

Score:61.2 bits(147), Expect:2e-08,
Method:Compositional matrix adjust.,
Identities:58/98(59%), Positives:69/98(70%), Gaps:7/98(7%)

Query  26   NPPPNPEGTrqarrnrrrrwrerqrqIHSISERILGTYLGRSAEPVPLQLPPLERLTLDC  85
            +P P PEGTRQAR+NRRRRWR RQRQI +ISERIL T LGRSAEPVP  LPP+ERL +
```

```
Sbjct  360   DPYPKPEGTRQARKNRRRRWRARQRQIRAISERILSTCLGRSAEPVPFLLPPIERLHIGD  181

Query  86    NEDCGTSGTQ-------GVGSPQILVESPTVLESGTKE  116
             +E  GTSGTQ       GVG+  +  +S  VL +  K+
Sbjct  180   SESSGTSGTQQSQGTPEGVGNS*VSGKSCAVLGARAKK  67
```

>Escherichia coli strain EN5308 contig00153, whole genome shotgun sequence
Sequence ID: NZ_JARBDE010000153.1 Length: 786
Range 1: 401 to 697

Score:63.2 bits(152), Expect:2e-08,
Method:Compositional matrix adjust.,
Identities:55/99(56%), Positives:69/99(69%), Gaps:7/99(7%)

```
Query  25    SNPPPNPEGTrqarrnrrrrwrerqrqIHSISERILGTYLGRSAEPVPLQLPPLERLTLD  84
             S+P P P  TRQA++NRRR+WR RQRQIHSISERIL + LGR AEPVPLQLPP++RL +
Sbjct  697   SDPYPEPRRTRQAQKNRRRKWRARQRQIHSISERILSSCLGRPAEPVPLQLPPIKRLHIS  518

Query  85    CNEDCGTSG-------TQGVGSPQILVESPTVLESGTKE  116
              +E+ GTS        T+ V S  +  +S  VL S TK+
Sbjct  517   DSENGGTSKTPQSQRTTEEVRSS*VSKKSCAVLRSRTKK  401
```

>Escherichia coli strain NBE0010 contig00098, whole genome shotgun sequence
Sequence ID: NZ_JARBDM010000098.1 Length: 1912
Range 1: 826 to 1050

Score:63.5 bits(153), Expect:2e-08,
Method:Compositional matrix adjust.,
Identities:52/75(69%), Positives:58/75(77%), Gaps:0/75(0%)

```
Query  19     IKLLYQSNPPPNPEGTrqarrnrrrrwrerqrqIHSISERILGTYLGRSAEPVPLQLPPL  78
              I     S+P P P GTRQA+RNRRRRWR RQ+QIHSISERIL T LGRSAEPVPLQLPP+
Sbjct  1050  ILTFVVSDPYPEPRGTRQAQRNRRRRWRARQKQIHSISERILSTCLGRSAEPVPLQLPPI  871

Query  79     ERLTLDCNEDCGTSG  93
              ERL +  +E  GT G
Sbjct  870    ERLHIADSESGGTFG  826
```

>Klebsiella pneumoniae strain EuSCAPE_MT009, whole genome shotgun sequence
Sequence ID: NZ_UKIV02000092.1 Length: 398
Range 1: 3 to 122

Score:57.4 bits(137), Expect:5e-07,
Method:Compositional matrix adjust.,
Identities:27/40(68%), Positives:33/40(82%), Gaps:0/40(0%)

```
Query  56    SERILGTYLGRSAEPVPLQLPPLERLTLDCNEDCGTSGTQ  95
             S++IL T LGRSAEPVPLQLPP++RL +DC+E   TS TQ
Sbjct  3     SKQILSTCLGRSAEPVPLQLPPIKRLHIDCSESSRTSKTQ  122
```

>Klebsiella pneumoniae strain OK8 jcf7180000003506, whole genome shotgun sequence
Sequence ID: NZ_MZXR01000036.1 Length: 577
Range 1: 195 to 410

Score:55.5 bits(132), Expect:6e-06,
Method:Compositional matrix adjust.,
Identities:38/74(51%), Positives:45/74(60%), Gaps:11/74(14%)

```
Query  52    IHSISERILGTYLGRSAEPVPLQLPPLERLTL---------DCNEDCGTSGTQGVGSPQI  102
             IH++SERIL + LGR AEPVPLQLPPLE L +            +  GT  T GVGSP I
Sbjct  195   IHTLSERILSSCLGRPAEPVPLQLPPLEGLHISGSESGGTSGTQQSQGT--TDGVGSP*I  368

Query  103   LVESPTVLESGTKE  116
               +S  VL SG K+
Sbjct  369   SGKSCAVLGSGAKK  410
```

>Klebsiella pneumoniae strain EuSCAPE_IL015, whole genome shotgun sequence
Sequence ID: NZ_UKJV02000040.1 Length: 532
Range 1: 3 to 182

Score:55.5 bits(132), Expect:6e-06,
Method:Compositional matrix adjust.,
Identities:33/60(55%), Positives:39/60(65%), Gaps:7/60(11%)

```
Query  64    LGRSAEPVPLQLPPLERLTLDCNEDCGTSGTQ-------GVGSPQILVESPTVLESGTKE  116
             LGRSAEPV  QLPP+ERL +  +E  GTSGTQ       GVG P +  +S  VL SGTK+
Sbjct  3     LGRSAEPVSFQLPPIERLNIGDSESGGTSGTQQSQGTTEGVGRP*VSGKSCAVLGSGTKK  182
```

>Photobacterium rosenbergii strain DP6N14-7 DP6N14-7_ctg0065, whole genome shotgun sequence
Sequence ID: NZ_JAHVIB010000057.1 Length: 564
Range 1: 277 to 447

Score:52.8 bits(125), Expect:7e-05,
Method:Compositional matrix adjust.,
Identities:45/57(79%), Positives:50/57(87%), Gaps:0/57(0%)

```
Query  25    SNPPPNPEGTrqarrnrrrrwrerqrqIHSISERILGTYLGRSAEPVPLQLPPLERL  81
             ++P P P GTRQAR+NRRRRWR RQRQI +ISERIL + LGRSAEPVPLQLPPLERL
Sbjct  277   TDPHPEPSGTRQARKNRRRRWRARQRQISAISERILSSCLGRSAEPVPLQLPPLERL  447
```

>Klebsiella pneumoniae strain EuSCAPE_NO017, whole genome shotgun sequence
Sequence ID: NZ_UKIO02000061.1 Length: 607
Range 1: 3 to 218

Score:52.8 bits(125), Expect:8e-05,
Method:Compositional matrix adjust.,
Identities:40/72(56%), Positives:48/72(66%), Gaps:7/72(9%)

```
Query  52    IHSISERILGTYLGRSAEPVPLQLPPLERLTLDCNEDCGTSGT-------QGVGSPQILV  104
             I+SISERIL T LGR AEPVP QLPP+ERL +  +E  GTSGT       +GVG+P I
Sbjct  3     INSISERILSTCLGRPAEPVPFQLPPIERLHIGDSESSGTSGTQSSQGTPEGVGNP*ISR  182

Query  105   ESPTVLESGTKE  116
```

```
            +S TVL    K+
Sbjct  183  KSCTVLGDRVKK  218
```

>Escherichia coli strain NBE0006 contig00106, whole genome shotgun sequence
Sequence ID: NZ_JARBDI010000106.1 Length: 2090
Range 1: 2 to 220

Score:53.5 bits(127), Expect:8e-05,
Method:Compositional matrix adjust.,
Identities:46/73(63%), Positives:53/73(72%), Gaps:0/73(0%)

```
Query  19   IKLLYQSNPPPNPEGTrqarrnrrrrwrerqrqIHSISERILGTYLGRSAEPVPLQLPPL  78
            I     S+P P P GTRQAR+NRRRRWR RQ+Q+HSISERIL T LGR AEPVP  LPP+
Sbjct  220  ILTFVISDPCPEPRGTRQARKNRRRRWRARQKQLHSISERILSTCLGRPAEPVPFLLPPI  41

Query  79   ERLTLDCNEDCGT  91
            ERL +  +E  G
Sbjct  40   ERLNISDSESGGA  2
```

>Bacillus velezensis strain TN14-11 TN14-11_ctg0032, whole genome shotgun sequence
Sequence ID: NZ_JAHVIU010000017.1 Length: 1125
Range 1: 277 to 447

Score:52.8 bits(125), Expect:2e-04,
Method:Compositional matrix adjust.,
Identities:45/57(79%), Positives:50/57(87%), Gaps:0/57(0%)

```
Query  25   SNPPPNPEGTrqarrnrrrrwrerqrqIHSISERILGTYLGRSAEPVPLQLPPLERL  81
            ++P P P GTRQAR+NRRRRWR RQRQI +ISERIL + LGRSAEPVPLQLPPLERL
Sbjct  277  TDPHPEPSGTRQARKNRRRRWRARQRQISAISERILSSCLGRSAEPVPLQLPPLERL  447
```

>Klebsiella pneumoniae strain EuSCAPE_LT010, whole genome shotgun sequence
Sequence ID: NZ_UKJK02000062.1 Length: 384
Range 1: 304 to 381

Score:46.2 bits(108), Expect:0.007,
Method:Compositional matrix adjust.,
Identities:20/26(77%), Positives:24/26(92%), Gaps:0/26(0%)

```
Query  1    MAGRSGDSDEELIRTVRLIKLLYQSN  26
            MAGRSGDSDE L+R VR+IK+LYQS+
Sbjct  381  MAGRSGDSDEALLRAVRIIKILYQSS  304
```

>Klebsiella pneumoniae strain EuSCAPE_MT011, whole genome shotgun sequence
Sequence ID: NZ_UKIY02000086.1 Length: 368
Range 1: 91 to 171

Score:45.4 bits(106), Expect:0.013,
Method:Compositional matrix adjust.,
Identities:20/27(74%), Positives:23/27(85%), Gaps:0/27(0%)

```
Query  1    MAGRSGDSDEELIRTVRLIKLLYQSNP  27
            MAGRSGDSDE L+R VR IK+LYQS+
Sbjct  171  MAGRSGDSDEALLRAVRTIKILYQSSK  91
```

>Klebsiella pneumoniae strain EuSCAPE_IL004, whole genome shotgun sequence
Sequence ID: NZ_UKJP02000103.1 Length: 411
Range 1: 4 to 81

Score:44.7 bits(104), Expect:0.037,
Method:Compositional matrix adjust.,
Identities:19/26(73%), Positives:24/26(92%), Gaps:0/26(0%)

```
Query  1  MAGRSGDSDEELIRTVRLIKLLYQSN  26
          MAGRSGDSDE L++ VR+IK+LYQS+
Sbjct  4  MAGRSGDSDEALLQAVRIIKILYQSS  81
```

Query #17: sp|P04608|TAT_HV1H2 Protein Tat OS=Human immunodeficiency virus type 1 group M subtype B (isolate HXB2) OX=11706 GN=tat PE=1 SV=2 Query ID: lcl|Query_2387707
Length: 86

Sequences producing significant alignments:

| Description | Scientific Name | Common Name | Taxid | Max Score | Total Score | Query cover | E Value | Per. Ident | Acc. Len | Accession |
|---|---|---|---|---|---|---|---|---|---|---|
| Klebsiella pneumoniae strain EuSCAPE_IL018, whole genome shotg... | Klebsiella p... | NA | 573 | 97.4 | 97.4 | 83% | 5e-23 | 63.89 | 409 | NZ_UKJU020000... |
| Klebsiella pneumoniae strain EuSCAPE_MT011, whole genome shotg... | Klebsiella p... | NA | 573 | 96.3 | 96.3 | 83% | 7e-23 | 63.89 | 368 | NZ_UKIY020000... |
| Escherichia coli strain VRES0669, whole genome shotgun sequence | Escherichia ... | NA | 562 | 94.0 | 94.0 | 93% | 2e-19 | 56.25 | 5544 | NZ_UINV010000... |
| Klebsiella pneumoniae strain EuSCAPE_IL013, whole genome shotg... | Klebsiella p... | NA | 573 | 79.7 | 79.7 | 53% | 2e-16 | 76.09 | 329 | NZ_UKJW020000... |
| Klebsiella pneumoniae strain EuSCAPE_IL020, whole genome shotg... | Klebsiella p... | NA | 573 | 73.6 | 73.6 | 51% | 6e-14 | 72.73 | 355 | NZ_UKKB020000... |
| Escherichia coli strain EN5374 contig00094, whole genome shotg... | Escherichia ... | NA | 562 | 78.2 | 78.2 | 68% | 6e-14 | 62.71 | 2720 | NZ_JAPTGI0100... |
| Escherichia coli strain EN5076 contig00066, whole genome shotg... | Escherichia ... | NA | 562 | 78.2 | 78.2 | 68% | 6e-14 | 62.71 | 2732 | NZ_JAODTC0100... |
| Escherichia coli strain NBE0009 contig00112, whole genome... | Escherichia ... | NA | 562 | 75.5 | 75.5 | 68% | 6e-13 | 61.02 | 2003 | NZ_JARBDL0100... |
| Escherichia coli strain EN5192 contig00108, whole genome shotg... | Escherichia ... | NA | 562 | 70.1 | 70.1 | 69% | 4e-11 | 56.67 | 3032 | NZ_JAPMNL0100... |
| Escherichia coli strain EN5090 contig00081, whole genome shotg... | Escherichia ... | NA | 562 | 70.1 | 70.1 | 68% | 4e-11 | 57.63 | 2798 | NZ_JAODTD0100... |
| Escherichia coli strain NBE0004 contig00090, whole genome... | Escherichia ... | NA | 562 | 69.7 | 69.7 | 68% | 7e-11 | 55.93 | 2818 | NZ_JARBDH0100... |
| Escherichia coli strain EN5143 contig00069, whole genome shotg... | Escherichia ... | NA | 562 | 68.9 | 68.9 | 68% | 1e-10 | 55.93 | 2688 | NZ_JAPMNI0100... |
| Escherichia coli strain EN5132 contig00077, whole genome shotg... | Escherichia ... | NA | 562 | 68.9 | 68.9 | 68% | 1e-10 | 55.93 | 2498 | NZ_JAPYLZ0100... |
| Escherichia coli strain NBE0002 contig00097, whole genome... | Escherichia ... | NA | 562 | 68.9 | 68.9 | 68% | 1e-10 | 55.93 | 2717 | NZ_JARBDF0100... |
| Escherichia coli strain NBE0009 contig00111, whole genome... | Escherichia ... | NA | 562 | 68.6 | 68.6 | 68% | 1e-10 | 57.63 | 2027 | NZ_JARBDL0100... |
| Escherichia coli strain EN5317 contig00057, whole genome shotg... | Escherichia ... | NA | 562 | 68.6 | 68.6 | 68% | 1e-10 | 57.63 | 2867 | NZ_JARBDO0100... |
| Escherichia coli strain EN5169 contig00082, whole genome shotg... | Escherichia ... | NA | 562 | 68.6 | 68.6 | 68% | 2e-10 | 55.93 | 2837 | NZ_JAPMNJ0100... |
| Escherichia coli strain NBE0003 contig00096, whole genome... | Escherichia ... | NA | 562 | 68.6 | 68.6 | 68% | 2e-10 | 55.93 | 2837 | NZ_JARBDG0100... |
| Escherichia coli strain EN5383 contig00089, whole genome shotg... | Escherichia ... | NA | 562 | 68.6 | 68.6 | 68% | 2e-10 | 55.93 | 1970 | NZ_JAPTZN0100... |
| Escherichia coli strain NBE0002 contig00132, whole genome... | Escherichia ... | NA | 562 | 68.2 | 68.2 | 68% | 2e-10 | 55.93 | 1381 | NZ_JARBDF0100... |
| Escherichia coli strain EN5308 contig00096, whole genome shotg... | Escherichia ... | NA | 562 | 68.2 | 68.2 | 68% | 2e-10 | 55.93 | 3024 | NZ_JARBDE0100... |
| Escherichia coli strain NBE0007 contig00093, whole genome... | Escherichia ... | NA | 562 | 67.8 | 67.8 | 68% | 3e-10 | 55.93 | 3013 | NZ_JARBDJ0100... |
| Escherichia coli strain EN5286 contig00120, whole genome shotg... | Escherichia ... | NA | 562 | 67.8 | 67.8 | 68% | 3e-10 | 55.93 | 3033 | NZ_JAPMNM0100... |
| Escherichia coli strain EN5177 contig00087, whole genome shotg... | Escherichia ... | NA | 562 | 67.8 | 67.8 | 68% | 3e-10 | 55.93 | 3403 | NZ_JAPMNK0100... |
| Klebsiella pneumoniae strain EuSCAPE_NO002, whole genome shotg... | Klebsiella p... | NA | 573 | 62.4 | 62.4 | 41% | 2e-08 | 77.78 | 907 | NZ_UKMC020000... |
| Burkholderia pseudomallei strain 2604STDY5518971... | Burkholderia... | NA | 28450 | 53.1 | 53.1 | 30% | 2e-05 | 88.46 | 603 | NZ_UVFZ020001... |
| Actinoplanes campanulatus strain JCM 3059 sequence095, whole... | Actinoplanes... | NA | 113559 | 49.7 | 49.7 | 48% | 3e-04 | 66.67 | 582 | NZ_BMPW010000... |
| Enterococcus alcedinis strain CCM 8433 sequence26, whole genom... | Enterococcus... | NA | 1274384 | 50.4 | 50.4 | 48% | 3e-04 | 66.67 | 853 | NZ_BMDT010000... |
| Deinococcus aquiradiocola strain JCM 14371 sequence41, whole... | Deinococcus ... | NA | 393059 | 49.7 | 49.7 | 48% | 5e-04 | 66.67 | 690 | NZ_BMOE010000... |
| Corynebacterium glutamicum strain 38 Scaffolds_31_4109, whole... | Corynebacter... | NA | 1718 | 49.3 | 49.3 | 48% | 0.001 | 66.67 | 3118 | NZ_JACENN0100... |
| Lactiplantibacillus plantarum strain MH-301 Scaffolds_50_4236,... | Lactiplantib... | NA | 1590 | 49.3 | 49.3 | 48% | 0.001 | 66.67 | 4236 | NZ_JAOYUG0100... |
| Corynebacterium glutamicum strain 51 Scaffolds_32_4236, whole... | Corynebacter... | NA | 1718 | 49.3 | 49.3 | 48% | 0.001 | 66.67 | 4236 | NZ_JACENK0100... |

Alignments:

>Klebsiella pneumoniae strain EuSCAPE_IL018, whole genome shotgun sequence
Sequence ID: NZ_UKJU02000098.1 Length: 409
Range 1: 10 to 225

Score:97.4 bits(241), Expect:5e-23,
Method:Compositional matrix adjust.,
Identities:54/72(75%), Positives:58/72(80%), Gaps:0/72(0%)

```
Query  1    MEPVDPRLEPWKHPGSQPKTACTNCYCKKCCFHCQVCFITKALGISYGrkkrrqrrrAHQ  60
            MEPVDP LEPW HPGSQPKTAC NCYCKKC +HC VCF  K LGISYGRKKRRQRR A
Sbjct  225  MEPVDPNLEPWNHPGSQPKTACNNCYCKKCSYHCLVCFQKKGLGISYGRKKRRQRRSAPP  46

Query  61   NSQTHQASLSKQ  72
            +S+ HQ  +SKQ
Sbjct  45   SSEDHQNLVSKQ  10
```

>Klebsiella pneumoniae strain EuSCAPE_MT011, whole genome shotgun sequence
Sequence ID: NZ_UKIY02000086.1 Length: 368
Range 1: 95 to 310

Score:96.3 bits(238), Expect:7e-23,
Method:Compositional matrix adjust.,
Identities:54/72(75%), Positives:58/72(80%), Gaps:0/72(0%)

```
Query  1    MEPVDPRLEPWKHPGSQPKTACTNCYCKKCCFHCQVCFITKALGISYGrkkrrqrrrAHQ  60
            MEPVDP LEPWKHPGSQPKTAC  CYCK C +HC VCF TK LGISYGRKKRRQRR A
Sbjct  310  MEPVDPNLEPWKHPGSQPKTACNKCYCKHCSYHCLVCFQTKGLGISYGRKKRRQRRSAPP  131

Query  61   NSQTHQASLSKQ  72
            +S+ HQ  +SKQ
Sbjct  130  SSEDHQDPISKQ  95
```

>Escherichia coli strain VRES0669, whole genome shotgun sequence
Sequence ID: NZ_UINV01000086.1 Length: 5544
Range 1: 1568 to 1807

Score:94.0 bits(232), Expect:2e-19,
Method:Compositional matrix adjust.,
Identities:53/80(66%), Positives:60/80(75%), Gaps:0/80(0%)

```
Query  1     MEPVDPRLEPWKHPGSQPKTACTNCYCKKCCFHCQVCFITKALGISYGrkkrrqrrrAHQ  60
             M+PVDP+LEPW HPGSQPKT C  CYCK+C +HC VCF TK LGI YGRKKRRQRR A
Sbjct  1807  MDPVDPKLEPWNHPGSQPKTPCNKCYCKQCSYHCLVCFQTKGLGIYYGRKKRRQRRSAPP  1628

Query  61    NSQTHQASLSKQPTSQPRGD  80
             +S+ HQ  LSKQ     RG+
Sbjct  1627  SSEDHQDPLSKQ*VPYGRGN  1568
```

>Klebsiella pneumoniae strain EuSCAPE_IL013, whole genome shotgun sequence
Sequence ID: NZ_UKJW02000092.1 Length: 329
Range 1: 3 to 140

Score:79.7 bits(195), Expect:2e-16,
Method:Compositional matrix adjust.,
Identities:35/46(76%), Positives:37/46(80%), Gaps:0/46(0%)

```
Query  1    MEPVDPRLEPWKHPGSQPKTACTNCYCKKCCFHCQVCFITKALGIS  46
            MEP+DP LEPW HPGSQPKTAC  CYCK C +HC VCF TK LGIS
Sbjct  140  MEPIDPNLEPWNHPGSQPKTACNKCYCKACSYHCLVCFQTKGLGIS  3
```

>Klebsiella pneumoniae strain EuSCAPE_IL020, whole genome shotgun sequence
Sequence ID: NZ_UKKB02000095.1 Length: 355
Range 1: 224 to 355

Score:73.6 bits(179), Expect:6e-14,
Method:Compositional matrix adjust.,
Identities:32/44(73%), Positives:34/44(77%), Gaps:0/44(0%)

```
Query  1    MEPVDPRLEPWKHPGSQPKTACTNCYCKKCCFHCQVCFITKALG  44
            MEPVDP LEPW HPGSQP T C  CYCK C +HC VCF+TK LG
Sbjct  224  MEPVDPNLEPWNHPGSQPATPCNKCYCKICSYHCPVCFLTKGLG  355
```

>Escherichia coli strain EN5374 contig00094, whole genome shotgun sequence
Sequence ID: NZ_JAPTGI010000094.1 Length: 2720
Range 1: 3 to 179

Score:78.2 bits(191), Expect:6e-14,
Method:Compositional matrix adjust.,
Identities:45/59(76%), Positives:51/59(86%), Gaps:0/59(0%)

```
Query  14   PGSQPKTACTNCYCKKCCFHCQVCFITKALGISYGrkkrrqrrrAHQNSQTHQASLSKQ  72
            PGSQPKTACTNCYCK+C +HCQVCF+TK LGISYGRKKRRQRR A  +S+ HQ  +SKQ
Sbjct  3    PGSQPKTACTNCYCKRCSYHCQVCFLTKGLGISYGRKKRRQRRSAPPSSEDHQNLISKQ  179
```

>Escherichia coli strain EN5076 contig00066, whole genome shotgun sequence
Sequence ID: NZ_JAODTC010000066.1 Length: 2732
Range 1: 2554 to 2730

Score:78.2 bits(191), Expect:6e-14,
Method:Compositional matrix adjust.,
Identities:45/59(76%), Positives:51/59(86%), Gaps:0/59(0%)

```
Query  14    PGSQPKTACTNCYCKKCCFHCQVCFITKALGISYGrkkrrqrrrAHQNSQTHQASLSKQ  72
             PGSQPKTACTNCYCK+C +HCQVCF+TK LGISYGRKKRRQRR A  +S+ HQ  +SKQ
Sbjct  2730  PGSQPKTACTNCYCKRCSYHCQVCFLTKGLGISYGRKKRRQRRSAPPSSEDHQNLISKQ  2554
```

>Escherichia coli strain NBE0009 contig00112, whole genome shotgun sequence
Sequence ID: NZ_JARBDL010000112.1 Length: 2003
Range 1: 1825 to 2001

Score:75.5 bits(184), Expect:6e-13,
Method:Compositional matrix adjust.,
Identities:44/59(75%), Positives:50/59(84%), Gaps:0/59(0%)

```
Query  14    PGSQPKTACTNCYCKKCCFHCQVCFITKALGISYGrkkrrqrrrAHQNSQTHQASLSKQ  72
             PGSQPKTACTNCYCK+C +HC VCF+TK LGISYGRKKRRQRR A  +S+ HQ  +SKQ
Sbjct  2001  PGSQPKTACTNCYCKRCSYHCPVCFLTKGLGISYGRKKRRQRRSAPPSSEDHQNLISKQ  1825
```

>Escherichia coli strain EN5192 contig00108, whole genome shotgun sequence
Sequence ID: NZ_JAPMNL010000108.1 Length: 3032
Range 1: 1 to 180

Score:70.1 bits(170), Expect:4e-11,
Method:Compositional matrix adjust.,
Identities:42/60(70%), Positives:47/60(78%), Gaps:0/60(0%)

```
Query  13   HPGSQPKTACTNCYCKKCCFHCQVCFITKALGISYGrkkrrqrrrAHQNSQTHQASLSKQ  72
            HPGSQPKTAC  CYCK+C +HC VCF  K LGISYGRKKRRQRR A  +S+ HQ  +SKQ
Sbjct  1    HPGSQPKTACNQCYCKRCSYHCLVCFQKKGLGISYGRKKRRQRRSAPPSSEDHQNHISKQ  180
```

>Escherichia coli strain EN5090 contig00081, whole genome shotgun sequence
Sequence ID: NZ_JAODTD010000081.1 Length: 2798
Range 1: 2620 to 2796

Score:70.1 bits(170), Expect:4e-11,
Method:Compositional matrix adjust.,
Identities:42/59(71%), Positives:47/59(79%), Gaps:0/59(0%)

```
Query  14    PGSQPKTACTNCYCKKCCFHCQVCFITKALGISYGrkkrrqrrrAHQNSQTHQASLSKQ  72
             PGSQP+TAC NCYCKKC +HC VCF  K LGISYGRKKRRQRR A  +S+ HQ  +SKQ
Sbjct  2796  PGSQPRTACNNCYCKKCSYHCLVCFQKKGLGISYGRKKRRQRRSAPPSSEDHQNPISKQ  2620
```

>Escherichia coli strain NBE0004 contig00090, whole genome shotgun sequence
Sequence ID: NZ_JARBDH010000090.1 Length: 2818
Range 1: 2640 to 2816

Score:69.7 bits(169), Expect:7e-11,
Method:Compositional matrix adjust.,
Identities:41/59(69%), Positives:47/59(79%), Gaps:0/59(0%)

```
Query  14    PGSQPKTACTNCYCKKCCFHCQVCFITKALGISYGrkkrrqrrrAHQNSQTHQASLSKQ  72
             PGSQP+TAC NCYCK+C +HC VCF  K LGISYGRKKRRQRR A  +S+ HQ  +SKQ
Sbjct  2816  PGSQPRTACNNCYCKRCSYHCLVCFQKKGLGISYGRKKRRQRRSAPPSSEDHQNHISKQ  2640
```

>Escherichia coli strain EN5143 contig00069, whole genome shotgun sequence
Sequence ID: NZ_JAPMNI010000069.1 Length: 2688
Range 1: 3 to 179

Score:68.9 bits(167), Expect:1e-10,
Method:Compositional matrix adjust.,
Identities:41/59(69%), Positives:46/59(77%), Gaps:0/59(0%)

```
Query  14   PGSQPKTACTNCYCKKCCFHCQVCFITKALGISYGrkkrrqrrrAHQNSQTHQASLSKQ  72
            PGSQP+TAC NCYCK C +HC VCF  K LGISYGRKKRRQRR   Q+S+ HQ  +SKQ
Sbjct  3    PGSQPRTACNNCYCKGCSYHCLVCFQKKGLGISYGRKKRRQRRSTPQSSEDHQNLVSKQ  179
```

>Escherichia coli strain EN5132 contig00077, whole genome shotgun sequence
Sequence ID: NZ_JAPYLZ010000077.1 Length: 2498
Range 1: 3 to 179

Score:68.9 bits(167), Expect:1e-10,
Method:Compositional matrix adjust.,
Identities:41/59(69%), Positives:46/59(77%), Gaps:0/59(0%)

```
Query  14   PGSQPKTACTNCYCKKCCFHCQVCFITKALGISYGrkkrrqrrrAHQNSQTHQASLSKQ  72
            PGSQPKTAC  CYCK+C +HC VCF  K LGISYGRKKRRQRR   Q+S+ HQ  +SKQ
Sbjct  3    PGSQPKTACNQCYCKRCSYHCLVCFQKKGLGISYGRKKRRQRRSTPQSSEDHQNLVSKQ  179
```

>Escherichia coli strain NBE0002 contig00097, whole genome shotgun sequence
Sequence ID: NZ_JARBDF010000097.1 Length: 2717
Range 1: 2539 to 2715

Score:68.9 bits(167), Expect:1e-10,
Method:Compositional matrix adjust.,
Identities:41/59(69%), Positives:46/59(77%), Gaps:0/59(0%)

```
Query  14    PGSQPKTACTNCYCKKCCFHCQVCFITKALGISYGrkkrrqrrrAHQNSQTHQASLSKQ  72
             PGSQPKTAC  CYCK+C +HC VCF  K LGISYGRKKRRQRR   Q+S+ HQ  +SKQ
Sbjct  2715  PGSQPKTACNQCYCKRCSYHCLVCFQKKGLGISYGRKKRRQRRSTPQSSEDHQNLVSKQ  2539
```

>Escherichia coli strain NBE0009 contig00111, whole genome shotgun sequence
Sequence ID: NZ_JARBDL010000111.1 Length: 2027
Range 1: 3 to 179

Score:68.6 bits(166), Expect:1e-10,
Method:Compositional matrix adjust.,
Identities:41/59(69%), Positives:45/59(76%), Gaps:0/59(0%)

```
Query  14   PGSQPKTACTNCYCKKCCFHCQVCFITKALGISYGrkkrrqrrrAHQNSQTHQASLSKQ  72
            PGSQP TACTNCYCK+C FHC  CF  K LGISYGRKKR QRR A  +S+ HQ  +SKQ
Sbjct  3    PGSQPATACTNCYCKRCSFHCIACFQKKGLGISYGRKKRGQRRSAPPSSEDHQNPISKQ  179
```

>Escherichia coli strain EN5317 contig00057, whole genome shotgun sequence
Sequence ID: NZ_JARBDO010000057.1 Length: 2867
Range 1: 2689 to 2865

Score:68.6 bits(166), Expect:1e-10,
Method:Compositional matrix adjust.,
Identities:41/59(69%), Positives:45/59(76%), Gaps:0/59(0%)

```
Query  14    PGSQPKTACTNCYCKKCCFHCQVCFITKALGISYGrkkrqrrrAHQNSQTHQASLSKQ  72
             PGSQP TACTNCYCK+C FHC  CF  K LGISYGRKKR QRR A  +S+ HQ  +SKQ
Sbjct  2865  PGSQPATACTNCYCKRCSFHCIACFQKKGLGISYGRKKRGQRRSAPPSSEDHQNPISKQ  2689
```

>Escherichia coli strain EN5169 contig00082, whole genome shotgun sequence
Sequence ID: NZ_JAPMNJ010000082.1 Length: 2837
Range 1: 2659 to 2835

Score:68.6 bits(166), Expect:2e-10,
Method:Compositional matrix adjust.,
Identities:41/59(69%), Positives:47/59(79%), Gaps:0/59(0%)

```
Query  14    PGSQPKTACTNCYCKKCCFHCQVCFITKALGISYGrkkrqrrrAHQNSQTHQASLSKQ  72
             PGSQP+TAC NCYCK+  +HC VCF  K LGISYGRKKRRQRR A Q+S+ HQ  +SKQ
Sbjct  2835  PGSQPETACNNCYCKRYSYHCLVCFQRKGLGISYGRKKRRQRRSAPQSSEDHQNFVSKQ  2659
```

>Escherichia coli strain NBE0003 contig00096, whole genome shotgun sequence
Sequence ID: NZ_JARBDG010000096.1 Length: 2837
Range 1: 2659 to 2835

Score:68.6 bits(166), Expect:2e-10,
Method:Compositional matrix adjust.,
Identities:41/59(69%), Positives:47/59(79%), Gaps:0/59(0%)

```
Query  14    PGSQPKTACTNCYCKKCCFHCQVCFITKALGISYGrkkrqrrrAHQNSQTHQASLSKQ  72
             PGSQP+TAC NCYCK+  +HC VCF  K LGISYGRKKRRQRR A Q+S+ HQ  +SKQ
Sbjct  2835  PGSQPETACNNCYCKRYSYHCLVCFQRKGLGISYGRKKRRQRRSAPQSSEDHQNFVSKQ  2659
```

>Escherichia coli strain EN5383 contig00089, whole genome shotgun sequence
Sequence ID: NZ_JAPTZN010000089.1 Length: 1970
Range 1: 3 to 179

Score:68.6 bits(166), Expect:2e-10,
Method:Compositional matrix adjust.,
Identities:41/59(69%), Positives:46/59(77%), Gaps:0/59(0%)

```
Query  14   PGSQPKTACTNCYCKKCCFHCQVCFITKALGISYGrkkrqrrrAHQNSQTHQASLSKQ  72
            PGSQP+TAC NCYCKKC +HC VCF  K LGISYGRKKRRQRR    +S+ HQ  +SKQ
Sbjct  3    PGSQPRTACNNCYCKKCSYHCLVCFQKKGLGISYGRKKRRQRRSTPPSSEDHQNLISKQ  179
```

>Escherichia coli strain NBE0002 contig00132, whole genome shotgun sequence
Sequence ID: NZ_JARBDF010000132.1 Length: 1381
Range 1: 1203 to 1379

Score:68.2 bits(165), Expect:2e-10,
Method:Compositional matrix adjust.,
Identities:41/59(69%), Positives:47/59(79%), Gaps:0/59(0%)

```
Query  14    PGSQPKTACTNCYCKKCCFHCQVCFITKALGISYGrkkrqrrrAHQNSQTHQASLSKQ  72
             PGSQP+TAC NC+CKKC +HC VCF  K LGISYGRKKRRQRR A  +S+ HQ  +SKQ
Sbjct  1379  PGSQPETACNNCFCKKCSYHCLVCFQKKGLGISYGRKKRRQRRSAPPSSEDHQNLISKQ  1203
```

>Escherichia coli strain EN5308 contig00096, whole genome shotgun sequence
Sequence ID: NZ_JARBDE010000096.1 Length: 3024
Range 1: 2846 to 3022

Score:68.2 bits(165), Expect:2e-10,
Method:Compositional matrix adjust.,
Identities:41/59(69%), Positives:47/59(79%), Gaps:0/59(0%)

```
Query  14    PGSQPKTACTNCYCKKCCFHCQVCFITKALGISYGrkkrqrrrAHQNSQTHQASLSKQ  72
             PGSQP+TAC NC+CKKC +HC VCF  K LGISYGRKKRRQRR A  +S+ HQ  +SKQ
Sbjct  3022  PGSQPETACNNCFCKKCSYHCLVCFQKKGLGISYGRKKRRQRRSAPPSSEDHQNHISKQ  2846
```

>Escherichia coli strain NBE0007 contig00093, whole genome shotgun sequence
Sequence ID: NZ_JARBDJ010000093.1 Length: 3013
Range 1: 2835 to 3011

Score:67.8 bits(164), Expect:3e-10,
Method:Compositional matrix adjust.,
Identities:41/59(69%), Positives:46/59(77%), Gaps:0/59(0%)

```
Query  14    PGSQPKTACTNCYCKKCCFHCQVCFITKALGISYGrkkrqrrrAHQNSQTHQASLSKQ  72
             PGSQP+TAC NCYCK C +HC VCF  K LGISYGRKKRRQRR A  +S+ HQ  +SKQ
Sbjct  3011  PGSQPRTACNNCYCKGCSYHCLVCFQKKGLGISYGRKKRRQRRSAPPSSEDHQNHISKQ  2835
```

>Escherichia coli strain EN5286 contig00120, whole genome shotgun sequence
Sequence ID: NZ_JAPMNM010000120.1 Length: 3033
Range 1: 3 to 179

Score:67.8 bits(164), Expect:3e-10,
Method:Compositional matrix adjust.,
Identities:41/59(69%), Positives:46/59(77%), Gaps:0/59(0%)

```
Query  14   PGSQPKTACTNCYCKKCCFHCQVCFITKALGISYGrkkrqrrrAHQNSQTHQASLSKQ  72
            PGSQP+TAC NCYCK C +HC VCF  K LGISYGRKKRRQRR A  +S+ HQ  +SKQ
Sbjct  3    PGSQPRTACNNCYCKGCSYHCLVCFQKKGLGISYGRKKRRQRRSAPPSSEDHQNHISKQ  179
```

>Escherichia coli strain EN5177 contig00087, whole genome shotgun sequence
Sequence ID: NZ_JAPMNK010000087.1 Length: 3403
Range 1: 3 to 179

Score:67.8 bits(164), Expect:3e-10,
Method:Compositional matrix adjust.,
Identities:41/59(69%), Positives:46/59(77%), Gaps:0/59(0%)

```
Query  14   PGSQPKTACTNCYCKKCCFHCQVCFITKALGISYGrkkrqrrrAHQNSQTHQASLSKQ  72
            PGSQPKTAC  CYCK+C +HC VCF  K LGISYGRKKRRQRR A  +S+ HQ  +SKQ
Sbjct  3    PGSQPKTACNQCYCKRCSYHCLVCFQKKGLGISYGRKKRRQRRSAPPSSEDHQNHISKQ  179
```

```
>Klebsiella pneumoniae strain EuSCAPE_NO002, whole genome shotgun sequence
Sequence ID: NZ_UKMC02000071.1 Length: 907
Range 1: 3 to 110

Score:62.4 bits(150), Expect:2e-08,
Method:Compositional matrix adjust.,
Identities:28/36(78%), Positives:31/36(86%), Gaps:0/36(0%)

Query  1    MEPVDPRLEPWKHPGSQPKTACTNCYCKKCCFHCQV  36
            +EPVDP LEP  HPGSQPKTAC +CYCKKC +HCQV
Sbjct  110  IEPVDPNLEP*NHPGSQPKTACNSCYCKKCSYHCQV  3
```

```
>Burkholderia pseudomallei strain 2604STDY5518971 ERS240105SCcontig000108, whole genome shotgun sequence
Sequence ID: NZ_UVFZ02000108.1 Length: 603
Range 1: 526 to 603

Score:53.1 bits(126), Expect:2e-05,
Method:Compositional matrix adjust.,
Identities:23/26(88%), Positives:24/26(92%), Gaps:0/26(0%)

Query  7    RLEPWKHPGSQPKTACTNCYCKKCCF  32
             LEPWKHPGSQP TACTNCYCKKCC+
Sbjct  526  NLEPWKHPGSQPTTACTNCYCKKCCW  603
```

```
>Actinoplanes campanulatus strain JCM 3059 sequence095, whole genome shotgun sequence
Sequence ID: NZ_BMPW01000095.1 Length: 582
Range 1: 22 to 147

Score:49.7 bits(117), Expect:3e-04,
Method:Compositional matrix adjust.,
Identities:37/42(88%), Positives:37/42(88%), Gaps:0/42(0%)

Query  43   LGISYGrkkrrqrrrAHQNSQTHQASLSKQPTSQPRGDPTGP  84
            LGISYGRKKRRQRRR  Q SQTHQ SLSKQPTSQ RGDPTGP
Sbjct  22   LGISYGRKKRRQRRRPPQGSQTHQVSLSKQPTSQSRGDPTGP  147
```

```
>Enterococcus alcedinis strain CCM 8433 sequence26, whole genome shotgun sequence
Sequence ID: NZ_BMDT01000026.1 Length: 853
Range 1: 40 to 165

Score:50.4 bits(119), Expect:3e-04,
Method:Compositional matrix adjust.,
Identities:37/42(88%), Positives:37/42(88%), Gaps:0/42(0%)

Query  43   LGISYGrkkrrqrrrAHQNSQTHQASLSKQPTSQPRGDPTGP  84
            LGISYGRKKRRQRRR  Q SQTHQ SLSKQPTSQ RGDPTGP
Sbjct  40   LGISYGRKKRRQRRRPPQGSQTHQVSLSKQPTSQSRGDPTGP  165
```

```
>Deinococcus aquiradiocola strain JCM 14371 sequence41, whole genome shotgun sequence
Sequence ID: NZ_BMOE01000041.1 Length: 690
Range 1: 18 to 143

Score:49.7 bits(117), Expect:5e-04,
Method:Compositional matrix adjust.,
Identities:37/42(88%), Positives:37/42(88%), Gaps:0/42(0%)

Query  43   LGISYGrkkrrqrrrAHQNSQTHQASLSKQPTSQPRGDPTGP  84
            LGISYGRKKRRQRRR  Q SQTHQ SLSKQPTSQ RGDPTGP
Sbjct  18   LGISYGRKKRRQRRRPPQGSQTHQVSLSKQPTSQSRGDPTGP  143
```

```
>Corynebacterium glutamicum strain 38 Scaffolds_31_4109, whole genome shotgun sequence
Sequence ID: NZ_JACENN010000031.1 Length: 3118
Range 1: 2292 to 2417

Score:49.3 bits(116), Expect:0.001,
Method:Compositional matrix adjust.,
Identities:37/42(88%), Positives:37/42(88%), Gaps:0/42(0%)

Query  43    LGISYGrkkrrqrrrAHQNSQTHQASLSKQPTSQPRGDPTGP  84
             LGISYGRKKRRQRRR  Q SQTHQ SLSKQPTSQ RGDPTGP
Sbjct  2292  LGISYGRKKRRQRRRPPQGSQTHQVSLSKQPTSQSRGDPTGP  2417
```

```
>Lactiplantibacillus plantarum strain MH-301 Scaffolds_50_4236, whole genome shotgun sequence
Sequence ID: NZ_JAOYUG010000050.1 Length: 4236
Range 1: 107 to 232

Score:49.3 bits(116), Expect:0.001,
Method:Compositional matrix adjust.,
Identities:37/42(88%), Positives:37/42(88%), Gaps:0/42(0%)

Query  43   LGISYGrkkrrqrrrAHQNSQTHQASLSKQPTSQPRGDPTGP  84
            LGISYGRKKRRQRRR  Q SQTHQ SLSKQPTSQ RGDPTGP
Sbjct  107  LGISYGRKKRRQRRRPPQGSQTHQVSLSKQPTSQSRGDPTGP  232
```

```
>Corynebacterium glutamicum strain 51 Scaffolds_32_4236, whole genome shotgun sequence
Sequence ID: NZ_JACENK010000032.1 Length: 4236
Range 1: 2874 to 2999

Score:49.3 bits(116), Expect:0.001,
Method:Compositional matrix adjust.,
Identities:37/42(88%), Positives:37/42(88%), Gaps:0/42(0%)

Query  43    LGISYGrkkrrqrrrAHQNSQTHQASLSKQPTSQPRGDPTGP  84
             LGISYGRKKRRQRRR  Q SQTHQ SLSKQPTSQ RGDPTGP
Sbjct  2874  LGISYGRKKRRQRRRPPQGSQTHQVSLSKQPTSQSRGDPTGP  2999
```

Query #18: sp|P05919|VPU_HV1H2 Protein Vpu OS=Human immunodeficiency virus type 1 group M subtype B (isolate HXB2) OX=11706 GN=vpu PE=1 SV=2 Query ID: lcl|Query_2387710
Length: 82

Sequences producing significant alignments:

| Description | Scientific Name | Common Name | Taxid | Max Score | Total Score | Query cover | E Value | Per. Ident | Acc. Len | Accession |
|---|---|---|---|---|---|---|---|---|---|---|
| Burkholderia pseudomallei strain 2604STDY5519032... | Burkholderia... | NA | 28450 | 62.4 | 62.4 | 65% | 1e-09 | 64.81 | 356 | NZ_UVID020000... |
| Escherichia coli strain EN5169 contig00082, whole genome shotg... | Escherichia ... | NA | 562 | 63.5 | 63.5 | 65% | 7e-09 | 66.67 | 2837 | NZ_JAPMNJ0100... |
| Escherichia coli strain NBE0003 contig00096, whole genome... | Escherichia ... | NA | 562 | 63.5 | 63.5 | 65% | 7e-09 | 66.67 | 2837 | NZ_JARBDG0100... |
| Escherichia coli strain VRES0669, whole genome shotgun sequence | Escherichia ... | NA | 562 | 63.2 | 63.2 | 54% | 1e-08 | 73.33 | 5544 | NZ_UINV010000... |
| Escherichia coli strain EN5379 contig00092, whole genome shotg... | Escherichia ... | NA | 562 | 62.4 | 62.4 | 65% | 2e-08 | 62.96 | 2592 | NZ_JAPTGJ0100... |
| Klebsiella pneumoniae strain EuSCAPE_IL004, whole genome shotg... | Klebsiella p... | NA | 573 | 58.9 | 58.9 | 65% | 4e-08 | 64.81 | 411 | NZ_UKJP020001... |
| Klebsiella pneumoniae strain EuSCAPE_LT010, whole genome shotg... | Klebsiella p... | NA | 573 | 58.5 | 58.5 | 65% | 4e-08 | 59.26 | 384 | NZ_UKJK020000... |
| Escherichia coli strain EN5090 contig00081, whole genome shotg... | Escherichia ... | NA | 562 | 61.2 | 61.2 | 65% | 5e-08 | 62.96 | 2798 | NZ_JAODTD0100... |
| Escherichia coli strain EN5308 contig00099, whole genome shotg... | Escherichia ... | NA | 562 | 61.2 | 61.2 | 65% | 5e-08 | 61.11 | 2827 | NZ_JARBDE0100... |
| Escherichia coli strain NBE0004 contig00090, whole genome... | Escherichia ... | NA | 562 | 60.5 | 60.5 | 65% | 8e-08 | 61.11 | 2818 | NZ_JARBDH0100... |
| Escherichia coli strain NBE0007 contig00093, whole genome... | Escherichia ... | NA | 562 | 60.5 | 60.5 | 65% | 8e-08 | 61.11 | 3013 | NZ_JARBDJ0100... |
| Escherichia coli strain EN5286 contig00120, whole genome shotg... | Escherichia ... | NA | 562 | 60.5 | 60.5 | 65% | 8e-08 | 61.11 | 3033 | NZ_JAPMNM0100... |
| Escherichia coli strain EN5177 contig00087, whole genome shotg... | Escherichia ... | NA | 562 | 60.5 | 60.5 | 65% | 8e-08 | 61.11 | 3403 | NZ_JAPMNK0100... |
| Escherichia coli strain EN5374 contig00094, whole genome shotg... | Escherichia ... | NA | 562 | 60.5 | 60.5 | 65% | 9e-08 | 61.11 | 2720 | NZ_JAPTGI0100... |
| Escherichia coli strain EN5132 contig00077, whole genome shotg... | Escherichia ... | NA | 562 | 59.3 | 59.3 | 65% | 2e-07 | 59.26 | 2498 | NZ_JAPYLZ0100... |
| Escherichia coli strain EN5143 contig00071, whole genome shotg... | Escherichia ... | NA | 562 | 58.9 | 58.9 | 48% | 3e-07 | 72.09 | 2774 | NZ_JAPMNI0100... |
| Escherichia coli strain EN5383 contig00089, whole genome shotg... | Escherichia ... | NA | 562 | 58.9 | 58.9 | 65% | 3e-07 | 61.11 | 1970 | NZ_JAPTZN0100... |
| Escherichia coli strain EN5308 contig00096, whole genome shotg... | Escherichia ... | NA | 562 | 58.5 | 58.5 | 65% | 4e-07 | 59.26 | 3024 | NZ_JARBDE0100... |
| Pseudomonas aeruginosa strain VET-42... | Pseudomonas ... | NA | 287 | 57.8 | 57.8 | 65% | 7e-07 | 53.70 | 963 | NZ_RACX010002... |
| Escherichia coli strain EN5192 contig00149, whole genome shotg... | Escherichia ... | NA | 562 | 57.4 | 57.4 | 65% | 1e-06 | 59.26 | 1397 | NZ_JAPMNL0100... |
| Escherichia coli strain EN5076 contig00066, whole genome shotg... | Escherichia ... | NA | 562 | 57.4 | 57.4 | 65% | 1e-06 | 59.26 | 2732 | NZ_JAODTC0100... |
| Escherichia coli strain NBE0009 contig00112, whole genome... | Escherichia ... | NA | 562 | 57.0 | 57.0 | 65% | 2e-06 | 61.11 | 2003 | NZ_JARBDL0100... |
| Escherichia coli strain EN5143 contig00069, whole genome shotg... | Escherichia ... | NA | 562 | 56.6 | 56.6 | 65% | 2e-06 | 59.26 | 2688 | NZ_JAPMNI0100... |
| Escherichia coli strain NBE0002 contig00097, whole genome... | Escherichia ... | NA | 562 | 56.6 | 56.6 | 65% | 2e-06 | 59.26 | 2717 | NZ_JARBDF0100... |
| Escherichia coli strain NBE0006 contig00124, whole genome... | Escherichia ... | NA | 562 | 56.2 | 56.2 | 65% | 2e-06 | 59.26 | 1306 | NZ_JARBDI0100... |
| Escherichia coli strain EN5177 contig00105, whole genome shotg... | Escherichia ... | NA | 562 | 55.8 | 55.8 | 65% | 4e-06 | 57.41 | 1783 | NZ_JAPMNK0100... |
| Escherichia coli strain EN5192 contig00108, whole genome shotg... | Escherichia ... | NA | 562 | 55.5 | 55.5 | 48% | 5e-06 | 69.77 | 3032 | NZ_JAPMNL0100... |
| Escherichia coli strain NBE0009 contig00111, whole genome... | Escherichia ... | NA | 562 | 53.5 | 53.5 | 65% | 2e-05 | 55.56 | 2027 | NZ_JARBDL0100... |
| Escherichia coli strain EN5317 contig00057, whole genome shotg... | Escherichia ... | NA | 562 | 52.0 | 52.0 | 65% | 9e-05 | 53.70 | 2867 | NZ_JARBDO0100... |
| Escherichia coli strain NBE0002 contig00132, whole genome... | Escherichia ... | NA | 562 | 50.4 | 50.4 | 65% | 3e-04 | 51.85 | 1381 | NZ_JARBDF0100... |
| Escherichia coli strain NBE0010 contig00080, whole genome... | Escherichia ... | NA | 562 | 44.3 | 44.3 | 37% | 0.050 | 70.59 | 2743 | NZ_JARBDM0100... |

Alignments:

>Burkholderia pseudomallei strain 2604STDY5519032 ERS240165SCcontig000088, whole genome shotgun sequence
Sequence ID: NZ_UVID02000088.1 Length: 356
Range 1: 64 to 222

Score:62.4 bits(150), Expect:1e-09,
Method:Compositional matrix adjust.,
Identities:35/54(65%), Positives:41/54(75%), Gaps:1/54(1%)

```
Query  29   EYRKILRQRKIDRLIDRLIERAEDSGNESEGEISALVEMGVEMGHHAPWDVDDL  82
            EYRK++RQRKID LI+R+ ERAEDSGNESEG+   L  M V+MGH    DV DL
Sbjct  222  EYRKLVRQRKIDWLIERIRERAEDSGNESEGDTEELATM-VDMGHLRLLDVHDL  64
```

>Escherichia coli strain EN5169 contig00082, whole genome shotgun sequence
Sequence ID: NZ_JAPMNJ010000082.1 Length: 2837
Range 1: 2401 to 2559

Score:63.5 bits(153), Expect:7e-09,
Method:Compositional matrix adjust.,
Identities:36/54(67%), Positives:41/54(75%), Gaps:1/54(1%)

```
Query  29    EYRKILRQRKIDRLIDRLIERAEDSGNESEGEISALVEMGVEMGHHAPWDVDDL  82
             EYRK+LRQRKID LI R+ ERAEDSGNESEG+   L  M V+MGH    DV+DL
Sbjct  2559  EYRKLLRQRKIDWLIKRIRERAEDSGNESEGDTEELSTM-VDMGHLRLLDVNDL  2401
```

>Escherichia coli strain NBE0003 contig00096, whole genome shotgun sequence
Sequence ID: NZ_JARBDG010000096.1 Length: 2837
Range 1: 2401 to 2559

Score:63.5 bits(153), Expect:7e-09,
Method:Compositional matrix adjust.,
Identities:36/54(67%), Positives:41/54(75%), Gaps:1/54(1%)

```
Query  29    EYRKILRQRKIDRLIDRLIERAEDSGNESEGEISALVEMGVEMGHHAPWDVDDL  82
             EYRK+LRQRKID LI R+ ERAEDSGNESEG+   L  M V+MGH    DV+DL
Sbjct  2559  EYRKLLRQRKIDWLIKRIRERAEDSGNESEGDTEELSTM-VDMGHLRLLDVNDL  2401
```

>Escherichia coli strain VRES0669, whole genome shotgun sequence
Sequence ID: NZ_UINV01000086.1 Length: 5544
Range 1: 1339 to 1470

Score:63.2 bits(152), Expect:1e-08,
Method:Compositional matrix adjust.,
Identities:33/45(73%), Positives:38/45(84%), Gaps:1/45(2%)

```
Query  29    EYRKILRQRKIDRLIDRLIERAEDSGNESEGEISALVEMGVEMGH  73
             EYRKI+RQRKIDRLI+R+ ERAEDSGNESEG+   L  M V+MGH
Sbjct  1470  EYRKIVRQRKIDRLIERIRERAEDSGNESEGDTEELATM-VDMGH  1339
```

>Escherichia coli strain EN5379 contig00092, whole genome shotgun sequence
Sequence ID: NZ_JAPTGJ010000092.1 Length: 2592
Range 1: 2355 to 2513

Score:62.4 bits(150), Expect:2e-08,
Method:Compositional matrix adjust.,
Identities:34/54(63%), Positives:42/54(77%), Gaps:1/54(1%)

```
Query  29    EYRKILRQRKIDRLIDRLIERAEDSGNESEGEISALVEMGVEMGHHAPWDVDDL  82
             EYRK++ QRKIDRLI+R+ ERAEDSGNESEG+   L  M V+MGH    DV++L
Sbjct  2513  EYRKLVSQRKIDRLIERIRERAEDSGNESEGDTEELSTM-VDMGHLRLLDVNEL  2355
```

>Klebsiella pneumoniae strain EuSCAPE_IL004, whole genome shotgun sequence
Sequence ID: NZ_UKJP02000103.1 Length: 411
Range 1: 192 to 350

Score:58.9 bits(141), Expect:4e-08,
Method:Compositional matrix adjust.,
Identities:35/54(65%), Positives:40/54(74%), Gaps:1/54(1%)

```
Query  29   EYRKILRQRKIDRLIDRLIERAEDSGNESEGEISALVEMGVEMGHHAPWDVDDL  82
            EYRKI+RQRKID LI R+ ERAEDSGNESEG+   L  M V+M H    DV+DL
Sbjct  192  EYRKIVRQRKIDWLIKRIRERAEDSGNESEGDNEDLATM-VDMEHLRLLDVNDL  350
```

>Klebsiella pneumoniae strain EuSCAPE_LT010, whole genome shotgun sequence
Sequence ID: NZ_UKJK02000062.1 Length: 384
Range 1: 35 to 193

Score:58.5 bits(140), Expect:4e-08,
Method:Compositional matrix adjust.,
Identities:32/54(59%), Positives:41/54(75%), Gaps:1/54(1%)

```
Query  29   EYRKILRQRKIDRLIDRLIERAEDSGNESEGEISALVEMGVEMGHHAPWDVDDL  82
            EYRK+LRQ++ID+LI R+ ERAEDSGNES+G+   L  M V+MGH    D +DL
Sbjct  193  EYRKLLRQKRIDQLIKRIRERAEDSGNESDGDTEELSTM-VDMGHLRLLDGNDL  35
```

>Escherichia coli strain EN5090 contig00081, whole genome shotgun sequence
Sequence ID: NZ_JAODTD010000081.1 Length: 2798
Range 1: 2362 to 2520

Score:61.2 bits(147), Expect:5e-08,
Method:Compositional matrix adjust.,
Identities:34/54(63%), Positives:42/54(77%), Gaps:1/54(1%)

```
Query  29    EYRKILRQRKIDRLIDRLIERAEDSGNESEGEISALVEMGVEMGHHAPWDVDDL  82
             EYRK+LRQ+KID LI+R+ ERAEDSGNESEG+   L  M V+MGH    DV++L
Sbjct  2520  EYRKLLRQKKIDWLIERIRERAEDSGNESEGDTEELSTM-VDMGHLRLLDVNEL  2362
```

>Escherichia coli strain EN5308 contig00099, whole genome shotgun sequence
Sequence ID: NZ_JARBDE010000099.1 Length: 2827
Range 1: 2445 to 2603

Score:61.2 bits(147), Expect:5e-08,
Method:Compositional matrix adjust.,
Identities:33/54(61%), Positives:41/54(75%), Gaps:1/54(1%)

```
Query  29    EYRKILRQRKIDRLIDRLIERAEDSGNESEGEISALVEMGVEMGHHAPWDVDDL  82
             EYRK+++QRKIDR+I R+ ERAEDSGNESEG+I  L  M V+MG     D +DL
Sbjct  2603  EYRKLVKQRKIDRIIKRIRERAEDSGNESEGDIEELSTM-VDMGQLRLLDANDL  2445
```

>Escherichia coli strain NBE0004 contig00090, whole genome shotgun sequence
Sequence ID: NZ_JARBDH010000090.1 Length: 2818
Range 1: 2388 to 2546

Score:60.5 bits(145), Expect:8e-08,
Method:Compositional matrix adjust.,
Identities:33/54(61%), Positives:42/54(77%), Gaps:1/54(1%)

```
Query  29    EYRKILRQRKIDRLIDRLIERAEDSGNESEGEISALVEMGVEMGHHAPWDVDDL  82
             EYRK+ +QRKIDRLI+R+ ERAEDSGNESEG+   L  + V+MG+    DV+DL
Sbjct  2546  EYRKLAKQRKIDRLIERIRERAEDSGNESEGDTEELSTL-VDMGNLRLLDVNDL  2388
```

>Escherichia coli strain NBE0007 contig00093, whole genome shotgun sequence
Sequence ID: NZ_JARBDJ010000093.1 Length: 3013
Range 1: 2583 to 2741

Score:60.5 bits(145), Expect:8e-08,
Method:Compositional matrix adjust.,
Identities:33/54(61%), Positives:42/54(77%), Gaps:1/54(1%)

```
Query  29    EYRKILRQRKIDRLIDRLIERAEDSGNESEGEISALVEMGVEMGHHAPWDVDDL  82
             EYRK+ +QRKIDRLI+R+ ERAEDSGNESEG+   L  + V+MG+    DV+DL
Sbjct  2741  EYRKLAKQRKIDRLIERIRERAEDSGNESEGDTEELSTL-VDMGNLRLLDVNDL  2583
```

>Escherichia coli strain EN5286 contig00120, whole genome shotgun sequence
Sequence ID: NZ_JAPMNM010000120.1 Length: 3033
Range 1: 273 to 431

Score:60.5 bits(145), Expect:8e-08,
Method:Compositional matrix adjust.,
Identities:33/54(61%), Positives:42/54(77%), Gaps:1/54(1%)

```
Query  29   EYRKILRQRKIDRLIDRLIERAEDSGNESEGEISALVEMGVEMGHHAPWDVDDL  82
            EYRK+ +QRKIDRLI+R+ ERAEDSGNESEG+   L  + V+MG+    DV+DL
Sbjct  273  EYRKLAKQRKIDRLIERIRERAEDSGNESEGDTEELSTL-VDMGNLRLLDVNDL  431
```

>Escherichia coli strain EN5177 contig00087, whole genome shotgun sequence
Sequence ID: NZ_JAPMNK010000087.1 Length: 3403
Range 1: 273 to 431

Score:60.5 bits(145), Expect:8e-08,
Method:Compositional matrix adjust.,
Identities:33/54(61%), Positives:42/54(77%), Gaps:1/54(1%)

```
Query  29   EYRKILRQRKIDRLIDRLIERAEDSGNESEGEISALVEMGVEMGHHAPWDVDDL  82
            EYRK+ +QRKIDRLI+R+ ERAEDSGNESEG+   L  + V+MG+    DV+DL
Sbjct  273  EYRKLAKQRKIDRLIERIRERAEDSGNESEGDTEELSTL-VDMGNLRLLDVNDL  431
```

>Escherichia coli strain EN5374 contig00094, whole genome shotgun sequence
Sequence ID: NZ_JAPTGI010000094.1 Length: 2720
Range 1: 273 to 431

Score:60.5 bits(145), Expect:9e-08,
Method:Compositional matrix adjust.,
Identities:33/54(61%), Positives:42/54(77%), Gaps:1/54(1%)

```
Query  29   EYRKILRQRKIDRLIDRLIERAEDSGNESEGEISALVEMGVEMGHHAPWDVDDL  82
            EYRK++ QRKIDRLI+R+ ERAEDSGNESEG++  L  M V+MG     DV++L
Sbjct  273  EYRKLVSQRKIDRLIERIRERAEDSGNESEGDVEELSTM-VDMGRLRLLDVNEL  431
```

```
>Escherichia coli strain EN5132 contig00077, whole genome shotgun sequence
Sequence ID: NZ_JAPYLZ010000077.1 Length: 2498
Range 1: 279 to 437

Score:59.3 bits(142), Expect:2e-07,
Method:Compositional matrix adjust.,
Identities:32/54(59%), Positives:41/54(75%), Gaps:1/54(1%)

Query  29   EYRKILRQRKIDRLIDRLIERAEDSGNESEGEISALVEMGVEMGHHAPWDVDDL  82
            EYRK+++QRKID+LI R+ ER EDSGNESEG+I  L  M V+MG+    D +DL
Sbjct  279  EYRKLVKQRKIDQLIKRIRERTEDSGNESEGDIEELSTM-VDMGNLRLLDANDL  437
```

```
>Escherichia coli strain EN5143 contig00071, whole genome shotgun sequence
Sequence ID: NZ_JAPMNI010000071.1 Length: 2774
Range 1: 30 to 158

Score:58.9 bits(141), Expect:3e-07,
Method:Compositional matrix adjust.,
Identities:31/43(72%), Positives:37/43(86%), Gaps:3/43(6%)

Query  29   EYRKILRQRKIDRLIDRLIERAEDSGNESEG---EISALVEMG  68
            EYRK+ +QRKIDRLI+R+ ERAEDSGNESEG   E+S LV+MG
Sbjct  30   EYRKLAKQRKIDRLIERIRERAEDSGNESEGDTEELSTLVDMG  158
```

```
>Escherichia coli strain EN5383 contig00089, whole genome shotgun sequence
Sequence ID: NZ_JAPTZN010000089.1 Length: 1970
Range 1: 279 to 437

Score:58.9 bits(141), Expect:3e-07,
Method:Compositional matrix adjust.,
Identities:33/54(61%), Positives:41/54(75%), Gaps:1/54(1%)

Query  29   EYRKILRQRKIDRLIDRLIERAEDSGNESEGEISALVEMGVEMGHHAPWDVDDL  82
            EYRK+++QRKID LI R+ ERAEDSGNESEG+   L  M V+MG+    DV+DL
Sbjct  279  EYRKLVKQRKIDWLIKRIRERAEDSGNESEGDTEELSTM-VDMGNLRLLDVNDL  437
```

```
>Escherichia coli strain EN5308 contig00096, whole genome shotgun sequence
Sequence ID: NZ_JARBDE010000096.1 Length: 3024
Range 1: 2594 to 2752

Score:58.5 bits(140), Expect:4e-07,
Method:Compositional matrix adjust.,
Identities:32/54(59%), Positives:40/54(74%), Gaps:1/54(1%)

Query  29    EYRKILRQRKIDRLIDRLIERAEDSGNESEGEISALVEMGVEMGHHAPWDVDDL  82
             EYRK+ +QRKIDRLI R+ ER EDSGNESEG+   L  + V+MG+    DV+DL
Sbjct  2752  EYRKLAKQRKIDRLIKRIRERTEDSGNESEGDTEELSTL-VDMGNLRLLDVNDL  2594
```

```
>Pseudomonas aeruginosa strain VET-42 NODE_89_length_1008_cov_1.38346_ID_177, whole genome shotgun sequence
Sequence ID: NZ_RACX01000220.1 Length: 963
Range 1: 128 to 286

Score:57.8 bits(138), Expect:7e-07,
Method:Compositional matrix adjust.,
Identities:29/54(54%), Positives:42/54(77%), Gaps:1/54(1%)

Query  29   EYRKILRQRKIDRLIDRLIERAEDSGNESEGEISALVEMGVEMGHHAPWDVDDL  82
            +Y++I +QRKID+L+DR+ ERAEDSGNESEG+   L ++ +EMG+   W  D+L
Sbjct  128  QYKEIRKQRKIDKLLDRIRERAEDSGNESEGDTDELAKL-MEMGNFDAWVGDNL  286
```

```
>Escherichia coli strain EN5192 contig00149, whole genome shotgun sequence
Sequence ID: NZ_JAPMNL010000149.1 Length: 1397
Range 1: 144 to 302

Score:57.4 bits(137), Expect:1e-06,
Method:Compositional matrix adjust.,
Identities:32/54(59%), Positives:40/54(74%), Gaps:1/54(1%)

Query  29   EYRKILRQRKIDRLIDRLIERAEDSGNESEGEISALVEMGVEMGHHAPWDVDDL  82
            EYRK+++QRKID LI R+ ER EDSGNESEG+I  L  M V+MG+    D +DL
Sbjct  144  EYRKLVKQRKIDWLIKRIRERTEDSGNESEGDIEELSTM-VDMGNLRLLDANDL  302
```

```
>Escherichia coli strain EN5076 contig00066, whole genome shotgun sequence
Sequence ID: NZ_JAODTC010000066.1 Length: 2732
Range 1: 2296 to 2454

Score:57.4 bits(137), Expect:1e-06,
Method:Compositional matrix adjust.,
Identities:32/54(59%), Positives:40/54(74%), Gaps:1/54(1%)

Query  29    EYRKILRQRKIDRLIDRLIERAEDSGNESEGEISALVEMGVEMGHHAPWDVDDL  82
             EYR+++ QRKID LI R+ ERAEDSGNESEG+   L  M V+MGH    DV++L
Sbjct  2454  EYRRLVNQRKIDWLIKRIRERAEDSGNESEGDTEELSTM-VDMGHLRLLDVNEL  2296
```

```
>Escherichia coli strain NBE0009 contig00112, whole genome shotgun sequence
Sequence ID: NZ_JARBDL010000112.1 Length: 2003
Range 1: 1567 to 1725

Score:57.0 bits(136), Expect:2e-06,
Method:Compositional matrix adjust.,
Identities:33/54(61%), Positives:40/54(74%), Gaps:1/54(1%)

Query  29    EYRKILRQRKIDRLIDRLIERAEDSGNESEGEISALVEMGVEMGHHAPWDVDDL  82
             EYRK++ QRKID LI R+ ERAEDSGNESEG+I  L  M V+MG     DV++L
Sbjct  1725  EYRKLVSQRKIDWLIKRIRERAEDSGNESEGDIEELSTM-VDMGRLRLLDVNEL  1567
```

>Escherichia coli strain EN5143 contig00069, whole genome shotgun sequence
Sequence ID: NZ_JAPMNI010000069.1 Length: 2688
Range 1: 279 to 437

Score:56.6 bits(135), Expect:2e-06,
Method:Compositional matrix adjust.,
Identities:32/54(59%), Positives:39/54(72%), Gaps:1/54(1%)

```
Query  29   EYRKILRQRKIDRLIDRLIERAEDSGNESEGEISALVEMGVEMGHHAPWDVDDL  82
            EYRK+++QRKID LI R+ ER EDSGNESEG+I  L  M V+MG     D +DL
Sbjct  279  EYRKLVKQRKIDWLIKRIRERTEDSGNESEGDIEELSTM-VDMGRIRLLDANDL  437
```

>Escherichia coli strain NBE0002 contig00097, whole genome shotgun sequence
Sequence ID: NZ_JARBDF010000097.1 Length: 2717
Range 1: 2281 to 2439

Score:56.6 bits(135), Expect:2e-06,
Method:Compositional matrix adjust.,
Identities:32/54(59%), Positives:39/54(72%), Gaps:1/54(1%)

```
Query  29    EYRKILRQRKIDRLIDRLIERAEDSGNESEGEISALVEMGVEMGHHAPWDVDDL  82
             EYRK+++QRKID LI R+ ER EDSGNESEG+I  L  M V+MG     D +DL
Sbjct  2439  EYRKLVKQRKIDWLIKRIRERTEDSGNESEGDIEELSTM-VDMGRIRLLDANDL  2281
```

>Escherichia coli strain NBE0006 contig00124, whole genome shotgun sequence
Sequence ID: NZ_JARBDI010000124.1 Length: 1306
Range 1: 1073 to 1231

Score:56.2 bits(134), Expect:2e-06,
Method:Compositional matrix adjust.,
Identities:32/54(59%), Positives:39/54(72%), Gaps:1/54(1%)

```
Query  29    EYRKILRQRKIDRLIDRLIERAEDSGNESEGEISALVEMGVEMGHHAPWDVDDL  82
             EYRK+++QRKID LI R+ ER EDSGNESEG+I  L  M V+MG     D +DL
Sbjct  1231  EYRKLVKQRKIDWLIKRIRERTEDSGNESEGDIEELSTM-VDMGRIRLLDANDL  1073
```

>Escherichia coli strain EN5177 contig00105, whole genome shotgun sequence
Sequence ID: NZ_JAPMNK010000105.1 Length: 1783
Range 1: 97 to 255

Score:55.8 bits(133), Expect:4e-06,
Method:Compositional matrix adjust.,
Identities:31/54(57%), Positives:39/54(72%), Gaps:1/54(1%)

```
Query  29   EYRKILRQRKIDRLIDRLIERAEDSGNESEGEISALVEMGVEMGHHAPWDVDDL  82
            EYRK+++QRKIDRLI R+ ER EDSGNESEG+I  L  M V++      D +DL
Sbjct  97   EYRKLVKQRKIDRLIKRIKERTEDSGNESEGDIEELSTM-VDIRRIRLLDANDL  255
```

>Escherichia coli strain EN5192 contig00108, whole genome shotgun sequence
Sequence ID: NZ_JAPMNL010000108.1 Length: 3032
Range 1: 274 to 402

Score:55.5 bits(132), Expect:5e-06,
Method:Compositional matrix adjust.,
Identities:30/43(70%), Positives:36/43(83%), Gaps:3/43(6%)

```
Query  29   EYRKILRQRKIDRLIDRLIERAEDSGNESEG---EISALVEMG  68
            EYRK+ +QRKIDRLI+R+ ERAEDSGNESE    E+S LV+MG
Sbjct  274  EYRKLAKQRKIDRLIERIRERAEDSGNESERDTEELSTLVDMG  402
```

>Escherichia coli strain NBE0009 contig00111, whole genome shotgun sequence
Sequence ID: NZ_JARBDL010000111.1 Length: 2027
Range 1: 279 to 437

Score:53.5 bits(127), Expect:2e-05,
Method:Compositional matrix adjust.,
Identities:30/54(56%), Positives:40/54(74%), Gaps:1/54(1%)

```
Query  29   EYRKILRQRKIDRLIDRLIERAEDSGNESEGEISALVEMGVEMGHHAPWDVDDL  82
            EY+K+ +Q++ID LI+R+ ERAEDSGNESEG+   L  M V+MG     DV+DL
Sbjct  279  EYKKLKKQKRIDWLIERIRERAEDSGNESEGDTEELSTM-VDMGRLRLLDVNDL  437
```

>Escherichia coli strain EN5317 contig00057, whole genome shotgun sequence
Sequence ID: NZ_JARBDO010000057.1 Length: 2867
Range 1: 2431 to 2589

Score:52.0 bits(123), Expect:9e-05,
Method:Compositional matrix adjust.,
Identities:29/54(54%), Positives:40/54(74%), Gaps:1/54(1%)

```
Query  29    EYRKILRQRKIDRLIDRLIERAEDSGNESEGEISALVEMGVEMGHHAPWDVDDL  82
             EY+K+ +Q++ID LI+R+ ERAEDSGNESEG+   L  M V+MG     DV++L
Sbjct  2589  EYKKLKKQKRIDWLIERIRERAEDSGNESEGDTEELSTM-VDMGRLRLLDVNEL  2431
```

>Escherichia coli strain NBE0002 contig00132, whole genome shotgun sequence
Sequence ID: NZ_JARBDF010000132.1 Length: 1381
Range 1: 945 to 1103

Score:50.4 bits(119), Expect:3e-04,
Method:Compositional matrix adjust.,
Identities:28/54(52%), Positives:39/54(72%), Gaps:1/54(1%)

```
Query  29    EYRKILRQRKIDRLIDRLIERAEDSGNESEGEISALVEMGVEMGHHAPWDVDDL  82
             EY+K+ +QRKIDR+I R+ ERAEDSGN+SE +I  L  M V++      DV++L
Sbjct  1103  EYKKLAKQRKIDRIIKRIKERAEDSGNKSEEDIEELSTM-VDIRRLRLLDVNEL  945
```

>Escherichia coli strain NBE0010 contig00080, whole genome shotgun sequence
Sequence ID: NZ_JARBDM010000080.1 Length: 2743
Range 1: 1 to 102

Score:44.3 bits(103), Expect:0.050,
Method:Compositional matrix adjust.,
Identities:24/34(71%), Positives:27/34(79%), Gaps:3/34(8%)

```
Query  38   KIDRLIDRLIERAEDSGNESEG---EISALVEMG  68
            KIDRLI R+ ERAEDSGNESEG   E+S +V MG
Sbjct  1    KIDRLIKRIRERAEDSGNESEGDIEELSTMVNMG  102
```

Query #19: sp|P69723|VIF_HV1H2 Virion infectivity factor OS=Human immunodeficiency virus type 1 group M subtype B (isolate HXB2) OX=11706 GN=vif PE=1 SV=1 Query ID: lcl|Query_2387721 Length: 192

Sequences producing significant alignments:

| Description | Scientific Name | Common Name | Taxid | Max Score | Total Score | Query cover | E Value | Per. Ident | Acc. Len | Accession |
|---|---|---|---|---|---|---|---|---|---|---|
| Aeromicrobium phoceense strain Marseille-Q0843... | Aeromicrobiu... | NA | 2754045 | 333 | 400 | 98% | 7e-112 | 83.68 | 944 | NZ_JACEOG0100... |
| Pseudomonas sp. Marseille-Q0931, whole genome shotgun sequence | Pseudomonas ... | NA | 2697507 | 333 | 333 | 98% | 7e-112 | 83.68 | 944 | NZ_CALNWT0100... |
| Streptomyces mexicanus strain Marseille-Q0842, whole genome... | Streptomyces... | NA | 178566 | 332 | 412 | 99% | 1e-111 | 84.13 | 946 | NZ_LR732595.1 |
| Streptomyces mexicanus strain Marseille-Q0842... | Streptomyces... | NA | 178566 | 313 | 407 | 98% | 3e-104 | 83.33 | 944 | NZ_JACMHY0100... |
| Escherichia coli strain VRES0669, whole genome shotgun sequence | Escherichia ... | NA | 562 | 311 | 311 | 100% | 1e-93 | 74.48 | 5544 | NZ_UINV010000... |
| Klebsiella pneumoniae strain EuSCAPE_NO002, whole genome shotg... | Klebsiella p... | NA | 573 | 275 | 275 | 100% | 3e-89 | 70.83 | 907 | NZ_UKMC020000... |
| Anaerococcus sp. Marseille-P9784, whole genome shotgun sequence | Anaerococcus... | NA | 2614127 | 243 | 414 | 98% | 2e-76 | 86.92 | 1004 | NZ_CABVOZ0100... |
| Klebsiella pneumoniae strain EuSCAPE_IL005, whole genome shotg... | Klebsiella p... | NA | 573 | 226 | 226 | 65% | 4e-71 | 83.33 | 636 | NZ_UKJL020000... |
| Klebsiella pneumoniae strain EuSCAPE_LT003, whole genome shotg... | Klebsiella p... | NA | 573 | 214 | 214 | 70% | 2e-67 | 77.21 | 410 | NZ_UKJH020000... |
| Klebsiella pneumoniae strain EuSCAPE_LT010, whole genome shotg... | Klebsiella p... | NA | 573 | 207 | 207 | 59% | 8e-65 | 82.46 | 391 | NZ_UKJK020000... |
| Klebsiella pneumoniae strain EuSCAPE_IL009, whole genome shotg... | Klebsiella p... | NA | 573 | 204 | 204 | 68% | 8e-64 | 74.05 | 394 | NZ_UKJO020000... |
| Klebsiella pneumoniae strain EuSCAPE_IL007, whole genome shotg... | Klebsiella p... | NA | 573 | 194 | 194 | 58% | 9e-60 | 80.53 | 359 | NZ_UKJT020000... |
| Burkholderia pseudomallei strain 2604STDY5518971... | Burkholderia... | NA | 28450 | 195 | 195 | 67% | 5e-59 | 70.54 | 603 | NZ_UVFZ020001... |
| Klebsiella pneumoniae strain EuSCAPE_IL001, whole genome shotg... | Klebsiella p... | NA | 573 | 184 | 184 | 53% | 5e-56 | 81.55 | 311 | NZ_UKJN020001... |
| Klebsiella pneumoniae strain EuSCAPE_NO001, whole genome shotg... | Klebsiella p... | NA | 573 | 177 | 177 | 59% | 2e-53 | 72.81 | 351 | NZ_UKMF020000... |
| Klebsiella pneumoniae strain EuSCAPE_MT012, whole genome shotg... | Klebsiella p... | NA | 573 | 157 | 157 | 46% | 4e-45 | 78.89 | 385 | NZ_UKIX020000... |
| Klebsiella pneumoniae strain EuSCAPE_LT006, whole genome shotg... | Klebsiella p... | NA | 573 | 151 | 151 | 44% | 2e-43 | 81.18 | 258 | NZ_UKJC020000... |
| Klebsiella pneumoniae strain EuSCAPE_DE083, whole genome shotg... | Klebsiella p... | NA | 573 | 151 | 151 | 43% | 3e-43 | 83.13 | 263 | NZ_UKLY020000... |
| Klebsiella pneumoniae strain EuSCAPE_MT014, whole genome shotg... | Klebsiella p... | NA | 573 | 145 | 145 | 42% | 6e-41 | 80.49 | 296 | NZ_UKJF020000... |
| Klebsiella pneumoniae strain EuSCAPE_IL005, whole genome shotg... | Klebsiella p... | NA | 573 | 129 | 129 | 43% | 2e-34 | 71.43 | 307 | NZ_UKJL020000... |
| Klebsiella pneumoniae strain EuSCAPE_MT009, whole genome shotg... | Klebsiella p... | NA | 573 | 93.2 | 93.2 | 33% | 3e-20 | 70.31 | 318 | NZ_UKIV020000... |
| Pseudomonas aeruginosa strain ENV-297... | Pseudomonas ... | NA | 287 | 81.3 | 81.3 | 22% | 1e-13 | 81.82 | 1100 | NZ_RADD010000... |
| Methylobacterium radiotolerans strain MAMP 4754... | Methylobacte... | NA | 31998 | 59.7 | 59.7 | 16% | 2e-07 | 81.25 | 289 | NZ_NKQS010005... |
| Klebsiella pneumoniae strain EuSCAPE_IL004, whole genome shotg... | Klebsiella p... | NA | 573 | 58.5 | 58.5 | 23% | 4e-07 | 60.87 | 244 | NZ_UKJP020001... |
| Pseudomonas aeruginosa strain VET-42... | Pseudomonas ... | NA | 287 | 59.7 | 59.7 | 15% | 4e-06 | 93.10 | 1154 | NZ_RACX010001... |
| Streptomyces mexicanus strain Marseille-Q0842, whole genome... | Streptomyces... | NA | 178566 | 54.3 | 54.3 | 27% | 4e-04 | 50.94 | 7159421 | NZ_LR732544.1 |
| Klebsiella pneumoniae strain EuSCAPE_MT014, whole genome shotg... | Klebsiella p... | NA | 573 | 50.4 | 50.4 | 14% | 5e-04 | 78.57 | 265 | NZ_UKJF020000... |
| Klebsiella pneumoniae strain EuSCAPE_DE084, whole genome shotg... | Klebsiella p... | NA | 573 | 48.5 | 48.5 | 23% | 0.002 | 54.35 | 250 | NZ_UKLZ020000... |
| Klebsiella pneumoniae strain EuSCAPE_IL009, whole genome shotg... | Klebsiella p... | NA | 573 | 48.5 | 48.5 | 14% | 0.002 | 75.00 | 261 | NZ_UKJO020000... |
| Klebsiella pneumoniae strain EuSCAPE_LT010, whole genome shotg... | Klebsiella p... | NA | 573 | 48.5 | 48.5 | 14% | 0.003 | 75.00 | 325 | NZ_UKJK020000... |

Alignments:

>Aeromicrobium phoceense strain Marseille-Q0843 NODE_5_length_944_cov_66.0355, whole genome shotgun sequence
Sequence ID: NZ_JACEOG010000003.1 Length: 944
Range 1: 359 to 928

Score:333 bits(854), Expect:7e-112,
Method:Compositional matrix adjust.,
Identities:169/190(89%), Positives:180/190(94%), Gaps:0/190(0%)

```
Query  3    NRWQVMIVWQVDRMRIRTWKSLVKHHMYVSGKARGWFYRHHYESPHPRISSEVHIPLGDA  62
            +RWQVMIVWQVDRMRI TWK LVKHHMY+S KA+ WFYRHHYES +P+ISSEVHIPLGDA
Sbjct  928  SRWQVMIVWQVDRMRINTWKRLVKHHMYISRKAKDWFYRHHYESTNPKISSEVHIPLGDA  749

Query  63   RLVITTYWGLHTGERDWHLGQGVSIEWRKKRYSTQVDPELADQLIHLYYFDCFSDSAIRK  122
            +LVITTYWGLHTGERDWHLGQGVSIEWRKKRYSTQVDP+LADQLIHL+YFDCFS+SAIR
Sbjct  748  KLVITTYWGLHTGERDWHLGQGVSIEWRKKRYSTQVDPDLADQLIHLHYFDCFSESAIRN  569

Query  123  ALLGHIVSPRCEYQAGHNKVGSLQYLALAAlitpkkikpplpSVTKLTEDRWNKPQKTKG  182
             +LG IVSPRCEYQAGHNKVGSLQYLALAALI PK+IKPPLPSV KLTEDRWNKPQKTKG
Sbjct  568  TILGRIVSPRCEYQAGHNKVGSLQYLALAALIKPKQIKPPLPSVRKLTEDRWNKPQKTKG  389

Query  183  HRGSHTMNGH  192
            HRGSHTMNGH
Sbjct  388  HRGSHTMNGH  359
```

Range 2: 1 to 111

Score:66.6 bits(161), Expect:1e-08,
Method:Compositional matrix adjust.,
Identities:30/37(81%), Positives:33/37(89%), Gaps:0/37(0%)

```
Query  3    NRWQVMIVWQVDRMRIRTWKSLVKHHMYVSGKARGWF  39
            +RWQVMIVWQVDRMRI TWK LVKHHMY+S KA+ WF
Sbjct  111  SRWQVMIVWQVDRMRINTWKRLVKHHMYISRKAKDWF  1
```

>Pseudomonas sp. Marseille-Q0931, whole genome shotgun sequence
Sequence ID: NZ_CALNWT010000029.1 Length: 944
Range 1: 252 to 821

Score:333 bits(854), Expect:7e-112,
Method:Compositional matrix adjust.,
Identities:169/190(89%), Positives:180/190(94%), Gaps:0/190(0%)

```
Query  3    NRWQVMIVWQVDRMRIRTWKSLVKHHMYVSGKARGWFYRHHYESPHPRISSEVHIPLGDA  62
            +RWQVMIVWQVDRMRI TWK LVKHHMY+S KA+ WFYRHHYES +P+ISSEVHIPLGDA
Sbjct  821  SRWQVMIVWQVDRMRINTWKRLVKHHMYISRKAKDWFYRHHYESTNPKISSEVHIPLGDA  642

Query  63   RLVITTYWGLHTGERDWHLGQGVSIEWRKKRYSTQVDPELADQLIHLYYFDCFSDSAIRK  122
            +LVITTYWGLHTGERDWHLGQGVSIEWRKKRYSTQVDP+LADQLIHL+YFDCFS+SAIR
Sbjct  641  KLVITTYWGLHTGERDWHLGQGVSIEWRKKRYSTQVDPDLADQLIHLHYFDCFSESAIRN  462

Query  123  ALLGHIVSPRCEYQAGHNKVGSLQYLALAAlitpkkikpplpSVTKLTEDRWNKPQKTKG  182
             +LG IVSPRCEYQAGHNKVGSLQYLALAALI PK+IKPPLPSV KLTEDRWNKPQKTKG
Sbjct  461  TILGRIVSPRCEYQAGHNKVGSLQYLALAALIKPKQIKPPLPSVRKLTEDRWNKPQKTKG  282

Query  183  HRGSHTMNGH  192
            HRGSHTMNGH
Sbjct  281  HRGSHTMNGH  252
```

>Streptomyces mexicanus strain Marseille-Q0842, whole genome shotgun sequence
Sequence ID: NZ_LR732595.1 Length: 946
Range 1: 2 to 568

Score:332 bits(852), Expect:1e-111,

```
Method:Compositional matrix adjust.,
Identities:169/189(89%), Positives:179/189(94%), Gaps:0/189(0%)

Query  4    RWQVMIVWQVDRMRIRTWKSLVKHHMYVSGKARGWFYRHHYESPHPRISSEVHIPLGDAR  63
            RWQVMIVWQVDRMRI TWK LVKHHMY+S KA+ WFYRHHYES +P+ISSEVHIPLGDA+
Sbjct  2    RWQVMIVWQVDRMRINTWKRLVKHHMYISRKAKDWFYRHHYESTNPKISSEVHIPLGDAK  181

Query  64   LVITTYWGLHTGERDWHLGQGVSIEWRKKRYSTQVDPELADQLIHLYYFDCFSDSAIRKA  123
            LVITTYWGLHTGERDWHLGQGVSIEWRKKRYSTQVDP+LADQLIHL+YFDCFS+SAIR
Sbjct  182  LVITTYWGLHTGERDWHLGQGVSIEWRKKRYSTQVDPDLADQLIHLHYFDCFSESAIRNT  361

Query  124  LLGHIVSPRCEYQAGHNKVGSLQYLALAAlitpkkikpplpSVTKLTEDRWNKPQKTKGH  183
            +LG IVSPRCEYQAGHNKVGSLQYLALAALI PK+IKPPLPSV KLTEDRWNKPQKTKGH
Sbjct  362  ILGRIVSPRCEYQAGHNKVGSLQYLALAALIKPKQIKPPLPSVRKLTEDRWNKPQKTKGH  541

Query  184  RGSHTMNGH  192
            RGSHTMNGH
Sbjct  542  RGSHTMNGH  568




Range 2: 815 to 946

Score:80.1 bits(196), Expect:2e-13,
Method:Compositional matrix adjust.,
Identities:36/44(82%), Positives:40/44(90%), Gaps:0/44(0%)

Query  2    ENRWQVMIVWQVDRMRIRTWKSLVKHHMYVSGKARGWFYRHHYE  45
            ++RWQVMIVWQVDRMRI TWK LVKHHMY+S KA+ WFYRHHYE
Sbjct  815  KHRWQVMIVWQVDRMRINTWKRLVKHHMYISRKAKDWFYRHHYE  946




>Streptomyces mexicanus strain Marseille-Q0842 NODE_30_length_944_cov_25.2289, whole genome shotgun sequence
Sequence ID: NZ_JACMHY010000029.1 Length: 944
Range 1: 405 to 944

Score:313 bits(803), Expect:3e-104,
Method:Compositional matrix adjust.,
Identities:160/180(89%), Positives:170/180(94%), Gaps:0/180(0%)

Query  13   VDRMRIRTWKSLVKHHMYVSGKARGWFYRHHYESPHPRISSEVHIPLGDARLVITTYWGL  72
            VDRMRI TWK LVKHHMY+S KA+ WFYRHHYES +P+ISSEVHIPLGDA+LVITTYWGL
Sbjct  944  VDRMRINTWKRLVKHHMYISRKAKDWFYRHHYESTNPKISSEVHIPLGDAKLVITTYWGL  765

Query  73   HTGERDWHLGQGVSIEWRKKRYSTQVDPELADQLIHLYYFDCFSDSAIRKALLGHIVSPR  132
            HTGERDWHLGQGVSIEWRKKRYSTQVDP+LADQLIHL+YFDCFS+SAIR  +LG IVSPR
Sbjct  764  HTGERDWHLGQGVSIEWRKKRYSTQVDPDLADQLIHLHYFDCFSESAIRNTILGRIVSPR  585

Query  133  CEYQAGHNKVGSLQYLALAAlitpkkikpplpSVTKLTEDRWNKPQKTKGHRGSHTMNGH  192
            CEYQAGHNKVGSLQYLALAALI PK+IKPPLPSV KLTEDRWNKPQKTKGHRGSHTMNGH
Sbjct  584  CEYQAGHNKVGSLQYLALAALIKPKQIKPPLPSVRKLTEDRWNKPQKTKGHRGSHTMNGH  405




Range 2: 2 to 157

Score:93.2 bits(230), Expect:3e-18,
Method:Compositional matrix adjust.,
Identities:42/52(81%), Positives:47/52(90%), Gaps:0/52(0%)

Query  3    NRWQVMIVWQVDRMRIRTWKSLVKHHMYVSGKARGWFYRHHYESPHPRISSE  54
            +RWQVMIVWQVDRMRI TWK LVKHHMY+S KA+ WFYRHHYES +P+ISSE
Sbjct  157  SRWQVMIVWQVDRMRINTWKRLVKHHMYISRKAKDWFYRHHYESTNPKISSE  2




>Escherichia coli strain VRES0669, whole genome shotgun sequence
Sequence ID: NZ_UINV01000086.1 Length: 5544
Range 1: 2021 to 2596

Score:311 bits(797), Expect:1e-93,
Method:Compositional matrix adjust.,
Identities:153/192(80%), Positives:174/192(90%), Gaps:0/192(0%)

Query  1     MENRWQVMIVWQVDRMRIRTWKSLVKHHMYVSGKARGWFYRHHYESPHPRISSEVHIPLG  60
             MENRWQ +IVWQVDRM+IRTW SLVKHHMY+S +A GWFY+HHYES HP++SSEVHIPLG
Sbjct  2596  MENRWQALIVWQVDRMKIRTWNSLVKHHMYISERASGWFYKHHYESRHPKVSSEVHIPLG  2417

Query  61    DARLVITTYWGLHTGERDWHLGQGVSIEWRKKRYSTQVDPELADQLIHLYYFDCFSDSAI  120
             +A+LVITTYWGL TGERDWHLG GVSIEWR +RYSTQVDP++ADQLIH++YFDCF+DSAI
Sbjct  2416  EAKLVITTYWGLQTGERDWHLGHGVSIEWRLRRYSTQVDPDMADQLIHIHYFDCFADSAI  2237

Query  121   RKALLGHIVSPRCEYQAGHNKVGSLQYLALAAlitpkkikpplpSVTKLTEDRWNKPQKT  180
             RKA+LGHIV PRC+Y AGHNKVGSLQYLAL ALI PKK KPPLPS+ KL EDRWN+PQKT
Sbjct  2236  RKAILGHIVIPRCDYPAGHNKVGSLQYLALTALIKPKKRKPPLPSIRKLVEDRWNRPQKT  2057

Query  181   KGHRGSHTMNGH  192
             +G RG+ T NGH
Sbjct  2056  RGRRGNRTTNGH  2021




>Klebsiella pneumoniae strain EuSCAPE_NO002, whole genome shotgun sequence
Sequence ID: NZ_UKMC02000071.1 Length: 907
Range 1: 324 to 899

Score:275 bits(704), Expect:3e-89,
Method:Compositional matrix adjust.,
Identities:146/192(76%), Positives:165/192(85%), Gaps:0/192(0%)

Query  1    MENRWQVMIVWQVDRMRIRTWKSLVKHHMYVSGKARGWFYRHHYESPHPRISSEVHIPLG  60
            +ENRWQV+IVWQVDRM+IRT  SLVKHHMYVS +A   FYRHHYES +P+ISSEVHIP+
Sbjct  899  IENRWQVLIVWQVDRMKIRT*NSLVKHHMYVSKRASR*FYRHHYESRNPKISSEVHIPVE  720

Query  61   DARLVITTYWGLHTGERDWHLGQGVSIEWRKKRYSTQVDPELADQLIHLYYFDCFSDSAI  120
            +ARLVI TY GL TGER+ HLG  VSIEWR +RYSTQVDP LADQLIH++YFDCF+DSAI
Sbjct  719  EARLVIITY*GLQTGERE*HLGHRVSIEWRLRRYSTQVDPGLADQLIHMHYFDCFADSAI  540

Query  121  RKALLGHIVSPRCEYQAGHNKVGSLQYLALAAlitpkkikpplpSVTKLTEDRWNKPQKT  180
            RKA+L HIV+PRC+Y AGH++V SLQYLAL ALI PK+IKPPLPS  KL EDR NKPQKT
Sbjct  539  RKAILRHIVTPRCDYPAGHSQVRSLQYLALTALIKPKRIKPPLPSAKKLVEDR*NKPQKT  360

Query  181  KGHRGSHTMNGH  192
            KG R SHTMNGH
Sbjct  359  KGRRRSHTMNGH  324
```

```
>Anaerococcus sp. Marseille-P9784, whole genome shotgun sequence
Sequence ID: NZ_CABVOZ010000006.1 Length: 1004
Range 1: 615 to 1004

Score:243 bits(621), Expect:2e-76,
Method:Compositional matrix adjust.,
Identities:113/130(87%), Positives:122/130(93%), Gaps:0/130(0%)

Query  4     RWQVMIVWQVDRMRIRTWKSLVKHHMYVSGKARGWFYRHHYESPHPRISSEVHIPLGDAR  63
             RWQVMIVWQVDRMRI TWK LVKHHMY+S KA+ WFYRHHYES +P+ISSEVHIPLGDA+
Sbjct  615   RWQVMIVWQVDRMRINTWKRLVKHHMYISRKAKDWFYRHHYESTNPKISSEVHIPLGDAK  794

Query  64    LVITTYWGLHTGERDWHLGQGVSIEWRKKRYSTQVDPELADQLIHLYYFDCFSDSAIRKA  123
             LVITTYWGLHTGERDWHLGQGVSIEWRKKRYSTQVDP+LADQLIHL+YFDCFS+SAIR
Sbjct  795   LVITTYWGLHTGERDWHLGQGVSIEWRKKRYSTQVDPDLADQLIHLHYFDCFSESAIRNT  974

Query  124   LLGHIVSPRC  133
             +LG IVSPRC
Sbjct  975   ILGRIVSPRC  1004



Range 2: 2 to 304

Score:171 bits(432), Expect:7e-48,
Method:Compositional matrix adjust.,
Identities:91/101(90%), Positives:96/101(95%), Gaps:0/101(0%)

Query  92   KRYSTQVDPELADQLIHLYYFDCFSDSAIRKALLGHIVSPRCEYQAGHNKVGSLQYLALA  151
            KRYSTQVDP+LADQLIHL+YFDCFS+SAIR  +LG IVSPRCEYQAGHNKVGSLQYLALA
Sbjct  2    KRYSTQVDPDLADQLIHLHYFDCFSESAIRNTILGRIVSPRCEYQAGHNKVGSLQYLALA  181

Query  152  AlitpkkikpplpSVTKLTEDRWNKPQKTKGHRGSHTMNGH  192
            ALI PK+IKPPLPSV KLTEDRWNKPQKTKGHRGSHTMNGH
Sbjct  182  ALIKPKQIKPPLPSVRKLTEDRWNKPQKTKGHRGSHTMNGH  304
```

```
>Klebsiella pneumoniae strain EuSCAPE_IL005, whole genome shotgun sequence
Sequence ID: NZ_UKJL02000050.1 Length: 636
Range 1: 1 to 378

Score:226 bits(576), Expect:4e-71,
Method:Compositional matrix adjust.,
Identities:105/126(83%), Positives:115/126(91%), Gaps:0/126(0%)

Query  1    MENRWQVMIVWQVDRMRIRTWKSLVKHHMYVSGKARGWFYRHHYESPHPRISSEVHIPLG  60
            MENRWQV+IVWQVDRM+IRTW SLVKHHMYVS KA  W YRHHYES HPRISSEVHIPLG
Sbjct  378  MENRWQVLIVWQVDRMKIRTWNSLVKHHMYVSKKANRWVYRHHYESRHPRISSEVHIPLG  199

Query  61   DARLVITTYWGLHTGERDWHLGQGVSIEWRKKRYSTQVDPELADQLIHLYYFDCFSDSAI  120
            DARL+I TYWGLHTGER+WHLG GVSIEWR + YSTQVDP LADQLIH++YFDCF+DSAI
Sbjct  198  DARLIIKTYWGLHTGEREWHLGHGVSIEWRLREYSTQVDPGLADQLIHMHYFDCFADSAI  19

Query  121  RKALLG  126
            +KA+LG
Sbjct  18   KKAILG  1
```

```
>Klebsiella pneumoniae strain EuSCAPE_LT003, whole genome shotgun sequence
Sequence ID: NZ_UKJH02000063.1 Length: 410
Range 1: 3 to 410

Score:214 bits(544), Expect:2e-67,
Method:Compositional matrix adjust.,
Identities:114/136(84%), Positives:124/136(91%), Gaps:0/136(0%)

Query  35   ARGWFYRHHYESPHPRISSEVHIPLGDARLVITTYWGLHTGERDWHLGQGVSIEWRKKRY  94
            A G FYRHHYES HP+ISSEVHIPLGDARLVITTYWGL+TGER+WHLG GVSIEWR +RY
Sbjct  3    ANG*FYRHHYESRHPKISSEVHIPLGDARLVITTYWGLNTGEREWHLGHGVSIEWRLRRY  182

Query  95   STQVDPELADQLIHLYYFDCFSDSAIRKALLGHIVSPRCEYQAGHNKVGSLQYLALAAli  154
            STQVDP LADQLIH++YFDCF+DSAIRKA+LGHIV  RCEYQAGHNKVGSLQYLAL ALI
Sbjct  183  STQVDPGLADQLIHMHYFDCFTDSAIRKAILGHIVFHRCEYQAGHNKVGSLQYLALTALI  362

Query  155  tpkkikpplpSVTKLT  170
             P+K+KPPLPSV KL
Sbjct  363  KPRKLKPPLPSVRKLV  410
```

```
>Klebsiella pneumoniae strain EuSCAPE_LT010, whole genome shotgun sequence
Sequence ID: NZ_UKJK02000060.1 Length: 391
Range 1: 1 to 342

Score:207 bits(527), Expect:8e-65,
Method:Compositional matrix adjust.,
Identities:94/114(82%), Positives:107/114(93%), Gaps:0/114(0%)

Query  39   FYRHHYESPHPRISSEVHIPLGDARLVITTYWGLHTGERDWHLGQGVSIEWRKKRYSTQV  98
            FYRHHY+S HP++SSEVHIPLG+ARLVITTYWGL TGERDWHLG GVSIEWR ++YSTQV
Sbjct  1    FYRHHYQSRHPKVSSEVHIPLGEARLVITTYWGLQTGERDWHLGHGVSIEWRLRKYSTQV  180

Query  99   DPELADQLIHLYYFDCFSDSAIRKALLGHIVSPRCEYQAGHNKVGSLQYLALAA  152
            DP+LADQLIH++YFDCF+DSAIR+A+LGHIV PRC+YQAGH KVGSLQYLAL A
Sbjct  181  DPDLADQLIHMHYFDCFADSAIRQAILGHIVFPRCDYQAGHRKVGSLQYLALTA  342
```

```
>Klebsiella pneumoniae strain EuSCAPE_IL009, whole genome shotgun sequence
Sequence ID: NZ_UKJO02000078.1 Length: 394
Range 1: 2 to 394

Score:204 bits(520), Expect:8e-64,
Method:Compositional matrix adjust.,
Identities:107/131(82%), Positives:119/131(90%), Gaps:0/131(0%)

Query  44   YESPHPRISSEVHIPLGDARLVITTYWGLHTGERDWHLGQGVSIEWRKKRYSTQVDPELA  103
            +ES HP++SSE+HIPLG+ARLV+TTYWGL TGER+WHLG GVSIEWR KRYSTQVDP LA
Sbjct  394  FESRHPKVSSEIHIPLGEARLVVTTYWGLQTGEREWHLGHGVSIEWRWKRYSTQVDPGLA  215

Query  104  DQLIHLYYFDCFSDSAIRKALLGHIVSPRCEYQAGHNKVGSLQYLALAAlitpkkikppl  163
            DQLIH +YFDCF+DSAIR+A+LGHIV PRC+YQAGHNKVGSLQYLAL ALI PKK KPPL
Sbjct  214  DQLIHTHYFDCFADSAIRQAILGHIVYPRCDYQAGHNKVGSLQYLALTALIKPKKRKPPL  35

Query  164  pSVTKLTEDRW  174
            PSV KL EDRW
```

```
Sbjct  34   PSVRKLVEDRW  2
```

>Klebsiella pneumoniae strain EuSCAPE_IL007, whole genome shotgun sequence
Sequence ID: NZ_UKJT02000027.1 Length: 359
Range 1: 18 to 356

Score:194 bits(492), Expect:9e-60,
Method:Compositional matrix adjust.,
Identities:91/113(81%), Positives:101/113(89%), Gaps:0/113(0%)

```
Query  40   YRHHYESPHPRISSEVHIPLGDARLVITTYWGLHTGERDWHLGQGVSIEWRKKRYSTQVD  99
            Y HHYES HP+ISSEVHIPLG+ARLVITTYWGL  GERDWHLG GVSI+WR + YSTQVD
Sbjct  356  YSHHYESRHPKISSEVHIPLGEARLVITTYWGLQPGERDWHLGHGVSIDWRLRGYSTQVD  177

Query  100  PELADQLIHLYYFDCFSDSAIRKALLGHIVSPRCEYQAGHNKVGSLQYLALAA  152
            P LADQLIH++YFDCF++SAIR A+LGH V PRCEYQAGHN+VGSLQYLAL A
Sbjct  176  PGLADQLIHIHYFDCFAESAIRNAILGHRVIPRCEYQAGHNRVGSLQYLALTA  18
```

>Burkholderia pseudomallei strain 2604STDY5518971 ERS240105SCcontig000108, whole genome shotgun sequence
Sequence ID: NZ_UVFZ02000108.1 Length: 603
Range 1: 2 to 388

Score:195 bits(495), Expect:5e-59,
Method:Compositional matrix adjust.,
Identities:101/129(78%), Positives:114/129(88%), Gaps:0/129(0%)

```
Query  64   LVITTYWGLHTGERDWHLGQGVSIEWRKKRYSTQVDPELADQLIHLYYFDCFSDSAIRKA  123
            L+I TYWGL TGE+DWHLG GVSIEW ++ YSTQ+DP+LADQLIHL YFDCFSDSAIRKA
Sbjct  2    LIIRTYWGLQTGEKDWHLGHGVSIEWSQRNYSTQIDPDLADQLIHLQYFDCFSDSAIRKA  181

Query  124  LLGHIVSPRCEYQAGHNKVGSLQYLALAAlitpkkikpplpSVTKLTEDRWNKPQKTKGH  183
            +LG +VS RCEY +GHNKVGSLQYLAL AL TPKKI+PPLPSV KLTED+WNKPQK +GH
Sbjct  182  ILGQVVSHRCEYPSGHNKVGSLQYLALKALATPKKIRPPLPSVKKLTEDKWNKPQKIRGH  361

Query  184  RGSHTMNGH  192
            R + TMNGH
Sbjct  362  RENPTMNGH  388
```

>Klebsiella pneumoniae strain EuSCAPE_IL001, whole genome shotgun sequence
Sequence ID: NZ_UKJN02000104.1 Length: 311
Range 1: 2 to 310

Score:184 bits(466), Expect:5e-56,
Method:Compositional matrix adjust.,
Identities:84/103(82%), Positives:95/103(92%), Gaps:0/103(0%)

```
Query  42   HHYESPHPRISSEVHIPLGDARLVITTYWGLHTGERDWHLGQGVSIEWRKKRYSTQVDPE  101
            HH+ES HP++SSEVHIPLGDARLVITTYWGL  GERDW+LG GVSIEWR ++YSTQVDP
Sbjct  2    HHFESRHPKVSSEVHIPLGDARLVITTYWGLQPGERDWYLGHGVSIEWRLRKYSTQVDPG  181

Query  102  LADQLIHLYYFDCFSDSAIRKALLGHIVSPRCEYQAGHNKVGS  144
            LADQLIH++YFDCF+DSAIRKA+LGHIV PRC+Y AGHNKVGS
Sbjct  182  LADQLIHMHYFDCFADSAIRKAILGHIVIPRCDYPAGHNKVGS  310
```

>Klebsiella pneumoniae strain EuSCAPE_NO001, whole genome shotgun sequence
Sequence ID: NZ_UKMF02000043.1 Length: 351
Range 1: 2 to 343

Score:177 bits(450), Expect:2e-53,
Method:Compositional matrix adjust.,
Identities:92/114(81%), Positives:104/114(91%), Gaps:0/114(0%)

```
Query  79   WHLGQGVSIEWRKKRYSTQVDPELADQLIHLYYFDCFSDSAIRKALLGHIVSPRCEYQAG  138
            WHLG GVSIEWR +RYSTQVDP LADQLIH++YFDCF+DSAIRKALLGHIV+PRC+Y AG
Sbjct  2    WHLGHGVSIEWRLRRYSTQVDPGLADQLIHMHYFDCFADSAIRKALLGHIVTPRCDYPAG  181

Query  139  HNKVGSLQYLALAAlitpkkikpplpSVTKLTEDRWNKPQKTKGHRGSHTMNGH  192
            H++VGSLQYLAL ALI PK+I+PPLPS  KL EDRWNKPQKT+G RG+HTMNGH
Sbjct  182  HSQVGSLQYLALTALIKPKRIRPPLPSTRKLVEDRWNKPQKTRGRRGNHTMNGH  343
```

>Klebsiella pneumoniae strain EuSCAPE_MT012, whole genome shotgun sequence
Sequence ID: NZ_UKIX02000067.1 Length: 385
Range 1: 9 to 278

Score:157 bits(397), Expect:4e-45,
Method:Compositional matrix adjust.,
Identities:71/90(79%), Positives:80/90(88%), Gaps:0/90(0%)

```
Query  1    MENRWQVMIVWQVDRMRIRTWKSLVKHHMYVSGKARGWFYRHHYESPHPRISSEVHIPLG  60
            MENRWQ +IVWQVDRMRIRTW SLVKHHMY+S +A  W YRHHYES HPR+S+EVHIPLG
Sbjct  278  MENRWQALIVWQVDRMRIRTWNSLVKHHMYISKRAEKWVYRHHYESRHPRVSAEVHIPLG  99

Query  61   DARLVITTYWGLHTGERDWHLGQGVSIEWR  90
            +ARLV+ +YWGL TGER+WHLG GVSIEWR
Sbjct  98   EARLVVISYWGLQTGEREWHLGHGVSIEWR  9
```

>Klebsiella pneumoniae strain EuSCAPE_LT006, whole genome shotgun sequence
Sequence ID: NZ_UKJC02000059.1 Length: 258
Range 1: 3 to 257

Score:151 bits(382), Expect:2e-43,
Method:Compositional matrix adjust.,
Identities:69/85(81%), Positives:78/85(91%), Gaps:0/85(0%)

```
Query  16   MRIRTWKSLVKHHMYVSGKARGWFYRHHYESPHPRISSEVHIPLGDARLVITTYWGLHTG  75
            MRIRTW SLVKHHMYVS +A+GWFYRHH+ES HP++SSEVHIPLG+A+LVI TYWGL TG
Sbjct  257  MRIRTWNSLVKHHMYVSKRAKGWFYRHHFESRHPKVSSEVHIPLGEAKLVIITYWGLQTG  78

Query  76   ERDWHLGQGVSIEWRKKRYSTQVDP  100
            ER+WHLG GVSIEWR +RYSTQVDP
Sbjct  77   EREWHLGHGVSIEWRLRRYSTQVDP  3
```

>Klebsiella pneumoniae strain EuSCAPE_DE083, whole genome shotgun sequence

Sequence ID: NZ_UKLY02000037.1 Length: 263
Range 1: 2 to 250

Score:151 bits(381), Expect:3e-43,
Method:Compositional matrix adjust.,
Identities:69/83(83%), Positives:76/83(91%), Gaps:0/83(0%)

```
Query  70   WGLHTGERDWHLGQGVSIEWRKKRYSTQVDPELADQLIHLYYFDCFSDSAIRKALLGHIV  129
            WGL TGER+WHLG GVSIEWR +RYSTQVDP LADQLIH++YFDCF+DSAIR A+LGH V
Sbjct  2    WGLQTGEREWHLGNGVSIEWRLRRYSTQVDPGLADQLIHMHYFDCFADSAIRTAILGHRV  181

Query  130  SPRCEYQAGHNKVGSLQYLALAA  152
            SPRC+YQAGHNKVGSLQYLAL A
Sbjct  182  SPRCDYQAGHNKVGSLQYLALTA  250
```

>Klebsiella pneumoniae strain EuSCAPE_MT014, whole genome shotgun sequence
Sequence ID: NZ_UKJF02000040.1 Length: 296
Range 1: 3 to 248

Score:145 bits(366), Expect:6e-41,
Method:Compositional matrix adjust.,
Identities:66/82(80%), Positives:75/82(91%), Gaps:0/82(0%)

```
Query  71   GLHTGERDWHLGQGVSIEWRKKRYSTQVDPELADQLIHLYYFDCFSDSAIRKALLGHIVS  130
            GL TGER+WHLG GVSIEWR  RY TQVDP+LADQLIH++YFDCF+DSAIRKA+LGHIV+
Sbjct  3    GLQTGEREWHLGHGVSIEWRLGRYKTQVDPDLADQLIHIHYFDCFADSAIRKAILGHIVT  182

Query  131  PRCEYQAGHNKVGSLQYLALAA  152
            PRC+YQAGH+KVGS QYLAL A
Sbjct  183  PRCDYQAGHSKVGSPQYLALTA  248
```

>Klebsiella pneumoniae strain EuSCAPE_IL005, whole genome shotgun sequence
Sequence ID: NZ_UKJL02000061.1 Length: 307
Range 1: 56 to 307

Score:129 bits(323), Expect:2e-34,
Method:Compositional matrix adjust.,
Identities:69/84(82%), Positives:76/84(90%), Gaps:0/84(0%)

```
Query  109  LYYFDCFSDSAIRKALLGHIVSPRCEYQAGHNKVGSLQYLALAAlitpkkikpplpSVTK  168
            ++YFDCF+DSAIRKA+LGHIV PRC+YQAGHNKVGSLQYLAL AL+ PKK KPPLPSV K
Sbjct  307  MHYFDCFADSAIRKAILGHIVIPRCDYQAGHNKVGSLQYLALTALLKPKKRKPPLPSVRK  128

Query  169  LTEDRWNKPQKTKGHRGSHTMNGH  192
            L EDRWNKPQKT+ HR SHTMNGH
Sbjct  127  LVEDRWNKPQKTRDHRESHTMNGH  56
```

>Klebsiella pneumoniae strain EuSCAPE_MT009, whole genome shotgun sequence
Sequence ID: NZ_UKIV02000096.1 Length: 318
Range 1: 122 to 313

Score:93.2 bits(230), Expect:3e-20,
Method:Compositional matrix adjust.,
Identities:55/64(86%), Positives:58/64(90%), Gaps:0/64(0%)

```
Query  129  VSPRCEYQAGHNKVGSLQYLALAAlitpkkikpplpSVTKLTEDRWNKPQKTKGHRGSHT  188
            VSPRCEYQ GHNKVGSLQYLAL ALI PKKI+PPLPSV KL EDRWNKPQKT+G RG+HT
Sbjct  313  VSPRCEYQTGHNKVGSLQYLALTALIKPKKIRPPLPSVWKLVEDRWNKPQKTRGRRGNHT  134

Query  189  MNGH  192
            MNGH
Sbjct  133  MNGH  122
```

>Pseudomonas aeruginosa strain ENV-297 NODE_79_length_1100_cov_9.24438_ID_157, whole genome shotgun sequence
Sequence ID: NZ_RADD01000036.1 Length: 1100
Range 1: 2 to 133

Score:81.3 bits(199), Expect:1e-13,
Method:Compositional matrix adjust.,
Identities:36/44(82%), Positives:40/44(90%), Gaps:0/44(0%)

```
Query  1    MENRWQVMIVWQVDRMRIRTWKSLVKHHMYVSGKARGWFYRHHY  44
            MENRWQVMIVWQVDRMRI  WKSLVK+HM++S KA+ WFYRHHY
Sbjct  133  MENRWQVMIVWQVDRMRINKWKSLVKYHMHISKKAKNWFYRHHY  2
```

>Methylobacterium radiotolerans strain MAMP 4754 MP-M-radiotolerans_contig_571, whole genome shotgun sequence
Sequence ID: NZ_NKQS01000559.1 Length: 289
Range 1: 192 to 287

Score:59.7 bits(143), Expect:2e-07,
Method:Compositional matrix adjust.,
Identities:26/32(81%), Positives:31/32(96%), Gaps:0/32(0%)

```
Query  1    MENRWQVMIVWQVDRMRIRTWKSLVKHHMYVS  32
            MENRWQV+IVWQVDRM+IRTW SLVK+HM++S
Sbjct  192  MENRWQVLIVWQVDRMKIRTWNSLVKYHMHIS  287
```

>Klebsiella pneumoniae strain EuSCAPE_IL004, whole genome shotgun sequence
Sequence ID: NZ_UKJP02000115.1 Length: 244
Range 1: 107 to 244

Score:58.5 bits(140), Expect:4e-07,
Method:Compositional matrix adjust.,
Identities:38/46(83%), Positives:40/46(86%), Gaps:0/46(0%)

```
Query  147  YLALAAlitpkkikpplpSVTKLTEDRWNKPQKTKGHRGSHTMNGH  192
            YLAL ALI PKK KPPLPSV KL EDRWNKPQKT+G RG+HTMNGH
Sbjct  244  YLALTALIKPKKRKPPLPSVQKLVEDRWNKPQKTRGRRGNHTMNGH  107
```

>Pseudomonas aeruginosa strain VET-42 NODE_86_length_1154_cov_8.30919_ID_171, whole genome shotgun sequence
Sequence ID: NZ_RACX01000118.1 Length: 1154
Range 1: 41 to 127

Score:59.7 bits(143), Expect:4e-06,
Method:Compositional matrix adjust.,
Identities:27/29(93%), Positives:28/29(96%), Gaps:0/29(0%)

```
Query  1    MENRWQVMIVWQVDRMRIRTWKSLVKHHM  29
            MENRWQVMIVWQVDRMRI TWKSLVK+HM
Sbjct  127  MENRWQVMIVWQVDRMRINTWKSLVKYHM  41
```

>Streptomyces mexicanus strain Marseille-Q0842, whole genome shotgun sequence
Sequence ID: NZ_LR732544.1 Length: 7159421
Range 1: 453356 to 453514

Score:54.3 bits(129), Expect:4e-04,
Method:Compositional matrix adjust.,
Identities:32/53(60%), Positives:41/53(77%), Gaps:0/53(0%)

```
Query  137     AGHNKVGSLQYLALAAlitpkkikpplpSVTKLTEDRWNKPQKTKGHRGSHTM  189
               AGH K GS++ LA+A  I  K+IK  L SV K+TE+RWNKPQKTKG+RGS+ +
Sbjct  453514  AGHKKGGSIK*LAIAE*IKTKQIKTTLQSVRKMTEERWNKPQKTKGYRGSNKL  453356
```

>Klebsiella pneumoniae strain EuSCAPE_MT014, whole genome shotgun sequence
Sequence ID: NZ_UKJF02000041.1 Length: 265
Range 1: 7 to 90

Score:50.4 bits(119), Expect:5e-04,
Method:Compositional matrix adjust.,
Identities:22/28(79%), Positives:24/28(85%), Gaps:0/28(0%)

```
Query  165  SVTKLTEDRWNKPQKTKGHRGSHTMNGH  192
            SV KL EDRWNKPQKT+  RG+HTMNGH
Sbjct  7    SVRKLVEDRWNKPQKTRDRRGNHTMNGH  90
```

>Klebsiella pneumoniae strain EuSCAPE_DE084, whole genome shotgun sequence
Sequence ID: NZ_UKLZ02000063.1 Length: 250
Range 1: 3 to 140

Score:48.5 bits(114), Expect:0.002,
Method:Compositional matrix adjust.,
Identities:36/46(78%), Positives:39/46(84%), Gaps:0/46(0%)

```
Query  147  YLALAAlitpkkikpplpSVTKLTEDRWNKPQKTKGHRGSHTMNGH  192
            YLAL ALI PKKIKPPLPSV KL EDR NKPQKT+G R +HTM+GH
Sbjct  3    YLALTALIKPKKIKPPLPSVRKLVEDR*NKPQKTRGRRKNHTMSGH  140
```

>Klebsiella pneumoniae strain EuSCAPE_IL009, whole genome shotgun sequence
Sequence ID: NZ_UKJO02000080.1 Length: 261
Range 1: 156 to 239

Score:48.5 bits(114), Expect:0.002,
Method:Compositional matrix adjust.,
Identities:21/28(75%), Positives:23/28(82%), Gaps:0/28(0%)

```
Query  165  SVTKLTEDRWNKPQKTKGHRGSHTMNGH  192
            SV KL EDRWNKPQKT+  RG+H MNGH
Sbjct  239  SVRKLVEDRWNKPQKTRDRRGNHIMNGH  156
```

>Klebsiella pneumoniae strain EuSCAPE_LT010, whole genome shotgun sequence
Sequence ID: NZ_UKJK02000067.1 Length: 325
Range 1: 21 to 104

Score:48.5 bits(114), Expect:0.003,
Method:Compositional matrix adjust.,
Identities:21/28(75%), Positives:24/28(85%), Gaps:0/28(0%)

```
Query  165  SVTKLTEDRWNKPQKTKGHRGSHTMNGH  192
            SV KL EDRWNKP+KT+G R +HTMNGH
Sbjct  21   SVKKLVEDRWNKPRKTRGRRENHTMNGH  104
```

Query #20: sp|P69726|VPR_HV1H2 Protein Vpr OS=Human immunodeficiency virus type 1 group M subtype B (isolate HXB2) OX=11706 GN=vpr PE=1 SV=2 Query ID: lcl|Query_2387708
Length: 96

Sequences producing significant alignments:

| Description | Scientific Name | Common Name | Taxid | Max Score | Total Score | Query cover | E Value | Per. Ident | Acc. Len | Accession |
|---|---|---|---|---|---|---|---|---|---|---|
| Aeromicrobium phoceense strain Marseille-Q0843... | Aeromicrobiu... | NA | 2754045 | 193 | 193 | 100% | 2e-58 | 96.88 | 944 | NZ_JACEOG0100... |
| Streptomyces mexicanus strain Marseille-Q0842... | Streptomyces... | NA | 178566 | 193 | 193 | 100% | 2e-58 | 96.88 | 944 | NZ_JACMHY0100... |
| Pseudomonas sp. Marseille-Q0931, whole genome shotgun sequence | Pseudomonas ... | NA | 2697507 | 193 | 264 | 100% | 2e-58 | 96.88 | 944 | NZ_CALNWT0100... |
| Streptomyces mexicanus strain Marseille-Q0842, whole genome... | Streptomyces... | NA | 178566 | 193 | 193 | 100% | 2e-58 | 96.88 | 946 | NZ_LR732595.1 |
| Anaerococcus sp. Marseille-P9784, whole genome shotgun sequence | Anaerococcus... | NA | 2614127 | 193 | 193 | 100% | 3e-58 | 96.88 | 1004 | NZ_CABVOZ0100... |
| Klebsiella pneumoniae strain EuSCAPE_LT010, whole genome shotg... | Klebsiella p... | NA | 573 | 162 | 162 | 95% | 4e-49 | 91.30 | 325 | NZ_UKJK020000... |
| Escherichia coli strain VRES0669, whole genome shotgun sequence | Escherichia ... | NA | 562 | 170 | 170 | 100% | 3e-46 | 83.33 | 5544 | NZ_UINV010000... |
| Klebsiella pneumoniae strain EuSCAPE_NO002, whole genome shotg... | Klebsiella p... | NA | 573 | 147 | 147 | 100% | 9e-41 | 79.17 | 907 | NZ_UKMC020000... |
| Klebsiella pneumoniae strain EuSCAPE_IL020, whole genome shotg... | Klebsiella p... | NA | 573 | 140 | 140 | 83% | 3e-40 | 81.25 | 355 | NZ_UKKB020000... |
| Klebsiella pneumoniae strain EuSCAPE_IL009, whole genome shotg... | Klebsiella p... | NA | 573 | 136 | 136 | 73% | 5e-39 | 88.73 | 261 | NZ_UKJO020000... |
| Klebsiella pneumoniae strain EuSCAPE_MT014, whole genome shotg... | Klebsiella p... | NA | 573 | 134 | 134 | 80% | 3e-38 | 89.61 | 265 | NZ_UKJF020000... |
| Klebsiella pneumoniae strain EuSCAPE_IL013, whole genome shotg... | Klebsiella p... | NA | 573 | 127 | 127 | 70% | 3e-35 | 89.71 | 329 | NZ_UKJW020000... |
| Klebsiella pneumoniae strain EuSCAPE_IL018, whole genome shotg... | Klebsiella p... | NA | 573 | 121 | 121 | 68% | 2e-32 | 87.88 | 409 | NZ_UKJU020000... |
| Klebsiella pneumoniae strain EuSCAPE_MT009, whole genome shotg... | Klebsiella p... | NA | 573 | 99.4 | 99.4 | 61% | 4e-24 | 89.83 | 318 | NZ_UKIV020000... |
| Klebsiella pneumoniae strain EuSCAPE_IL004, whole genome shotg... | Klebsiella p... | NA | 573 | 93.2 | 93.2 | 56% | 7e-22 | 88.89 | 244 | NZ_UKJP020001... |
| Klebsiella pneumoniae strain EuSCAPE_DE084, whole genome shotg... | Klebsiella p... | NA | 573 | 90.1 | 90.1 | 58% | 1e-20 | 85.71 | 250 | NZ_UKLZ020000... |
| Klebsiella pneumoniae strain EuSCAPE_IL005, whole genome shotg... | Klebsiella p... | NA | 573 | 60.1 | 60.1 | 38% | 1e-08 | 91.89 | 307 | NZ_UKJL020000... |
| Klebsiella pneumoniae strain EuSCAPE_NO001, whole genome shotg... | Klebsiella p... | NA | 573 | 45.8 | 45.8 | 22% | 0.005 | 90.91 | 351 | NZ_UKMF020000... |
| Klebsiella pneumoniae strain EuSCAPE_MT011, whole genome shotg... | Klebsiella p... | NA | 573 | 45.8 | 45.8 | 26% | 0.006 | 84.00 | 368 | NZ_UKIY020000... |

Alignments:

>Aeromicrobium phoceense strain Marseille-Q0843 NODE_5_length_944_cov_66.0355, whole genome shotgun sequence
Sequence ID: NZ_JACEOG010000003.1 Length: 944
Range 1: 129 to 416

Score:193 bits(490), Expect:2e-58,
Method:Compositional matrix adjust.,
Identities:93/96(97%), Positives:95/96(98%), Gaps:0/96(0%)

```
Query  1    MEQAPEDQGPQREPHNEWTLELLEELKNEAVRHFPRIWLHGLGQHIYETYGDTWAGVEAI  60
            MEQAPEDQGPQREP+NEWTLELLEELK+EAVRHFPRIWLH LGQHIYETYGDTWAGVEAI
Sbjct  416  MEQAPEDQGPQREPYNEWTLELLEELKSEAVRHFPRIWLHNLGQHIYETYGDTWAGVEAI  237

Query  61   IRILQQLLFIHFRIGCRHSRIGVTRQRRARNGASRS  96
            IRILQQLLFIHFRIGCRHSRIGVTRQRRARNGASRS
Sbjct  236  IRILQQLLFIHFRIGCRHSRIGVTRQRRARNGASRS  129
```

```
>Streptomyces mexicanus strain Marseille-Q0842 NODE_30_length_944_cov_25.2289, whole genome shotgun sequence
Sequence ID: NZ_JACMHY010000029.1 Length: 944
Range 1: 175 to 462

Score:193 bits(490), Expect:2e-58,
Method:Compositional matrix adjust.,
Identities:93/96(97%), Positives:95/96(98%), Gaps:0/96(0%)

Query  1    MEQAPEDQGPQREPHNEWTLELLEELKNEAVRHFPRIWLHGLGQHIYETYGDTWAGVEAI  60
            MEQAPEDQGPQREP+NEWTLELLEELK+EAVRHFPRIWLH LGQHIYETYGDTWAGVEAI
Sbjct  462  MEQAPEDQGPQREPYNEWTLELLEELKSEAVRHFPRIWLHNLGQHIYETYGDTWAGVEAI  283

Query  61   IRILQQLLFIHFRIGCRHSRIGVTRQRRARNGASRS  96
            IRILQQLLFIHFRIGCRHSRIGVTRQRRARNGASRS
Sbjct  282  IRILQQLLFIHFRIGCRHSRIGVTRQRRARNGASRS  175
```

```
>Pseudomonas sp. Marseille-Q0931, whole genome shotgun sequence
Sequence ID: NZ_CALNWT010000029.1 Length: 944
Range 1: 22 to 309

Score:193 bits(490), Expect:2e-58,
Method:Compositional matrix adjust.,
Identities:93/96(97%), Positives:95/96(98%), Gaps:0/96(0%)

Query  1    MEQAPEDQGPQREPHNEWTLELLEELKNEAVRHFPRIWLHGLGQHIYETYGDTWAGVEAI  60
            MEQAPEDQGPQREP+NEWTLELLEELK+EAVRHFPRIWLH LGQHIYETYGDTWAGVEAI
Sbjct  309  MEQAPEDQGPQREPYNEWTLELLEELKSEAVRHFPRIWLHNLGQHIYETYGDTWAGVEAI  130

Query  61   IRILQQLLFIHFRIGCRHSRIGVTRQRRARNGASRS  96
            IRILQQLLFIHFRIGCRHSRIGVTRQRRARNGASRS
Sbjct  129  IRILQQLLFIHFRIGCRHSRIGVTRQRRARNGASRS  22
```

```
Range 2: 839 to 943

Score:71.2 bits(173), Expect:1e-11,
Method:Compositional matrix adjust.,
Identities:35/35(100%), Positives:35/35(100%), Gaps:0/35(0%)

Query  62   RILQQLLFIHFRIGCRHSRIGVTRQRRARNGASRS  96
            RILQQLLFIHFRIGCRHSRIGVTRQRRARNGASRS
Sbjct  943  RILQQLLFIHFRIGCRHSRIGVTRQRRARNGASRS  839
```

```
>Streptomyces mexicanus strain Marseille-Q0842, whole genome shotgun sequence
Sequence ID: NZ_LR732595.1 Length: 946
Range 1: 511 to 798

Score:193 bits(490), Expect:2e-58,
Method:Compositional matrix adjust.,
Identities:93/96(97%), Positives:95/96(98%), Gaps:0/96(0%)

Query  1    MEQAPEDQGPQREPHNEWTLELLEELKNEAVRHFPRIWLHGLGQHIYETYGDTWAGVEAI  60
            MEQAPEDQGPQREP+NEWTLELLEELK+EAVRHFPRIWLH LGQHIYETYGDTWAGVEAI
Sbjct  511  MEQAPEDQGPQREPYNEWTLELLEELKSEAVRHFPRIWLHNLGQHIYETYGDTWAGVEAI  690

Query  61   IRILQQLLFIHFRIGCRHSRIGVTRQRRARNGASRS  96
            IRILQQLLFIHFRIGCRHSRIGVTRQRRARNGASRS
Sbjct  691  IRILQQLLFIHFRIGCRHSRIGVTRQRRARNGASRS  798
```

```
>Anaerococcus sp. Marseille-P9784, whole genome shotgun sequence
Sequence ID: NZ_CABVOZ010000006.1 Length: 1004
Range 1: 247 to 534

Score:193 bits(490), Expect:3e-58,
Method:Compositional matrix adjust.,
Identities:93/96(97%), Positives:95/96(98%), Gaps:0/96(0%)

Query  1    MEQAPEDQGPQREPHNEWTLELLEELKNEAVRHFPRIWLHGLGQHIYETYGDTWAGVEAI  60
            MEQAPEDQGPQREP+NEWTLELLEELK+EAVRHFPRIWLH LGQHIYETYGDTWAGVEAI
Sbjct  247  MEQAPEDQGPQREPYNEWTLELLEELKSEAVRHFPRIWLHNLGQHIYETYGDTWAGVEAI  426

Query  61   IRILQQLLFIHFRIGCRHSRIGVTRQRRARNGASRS  96
            IRILQQLLFIHFRIGCRHSRIGVTRQRRARNGASRS
Sbjct  427  IRILQQLLFIHFRIGCRHSRIGVTRQRRARNGASRS  534
```

```
>Klebsiella pneumoniae strain EuSCAPE_LT010, whole genome shotgun sequence
Sequence ID: NZ_UKJK02000067.1 Length: 325
Range 1: 47 to 322

Score:162 bits(411), Expect:4e-49,
Method:Compositional matrix adjust.,
Identities:84/92(91%), Positives:88/92(95%), Gaps:0/92(0%)

Query  1    MEQAPEDQGPQREPHNEWTLELLEELKNEAVRHFPRIWLHGLGQHIYETYGDTWAGVEAI  60
            MEQ PEDQGPQREP+NEWTLELLEELK EAVRHFPR WLHGLGQHIYETYGDTWAGVEA+
Sbjct  47   MEQTPEDQGPQREPYNEWTLELLEELKQEAVRHFPRPWLHGLGQHIYETYGDTWAGVEAL  226

Query  61   IRILQQLLFIHFRIGCRHSRIGVTRQRRARNG  92
            IRILQQLLFIHFRIGC+HSRIG+ RQRRARNG
Sbjct  227  IRILQQLLFIHFRIGCQHSRIGILRQRRARNG  322
```

```
>Escherichia coli strain VRES0669, whole genome shotgun sequence
Sequence ID: NZ_UINV01000086.1 Length: 5544
Range 1: 1791 to 2078

Score:170 bits(431), Expect:3e-46,
```

```
Method:Compositional matrix adjust.,
Identities:80/96(83%), Positives:89/96(92%), Gaps:0/96(0%)

Query  1     MEQAPEDQGPQREPHNEWTLELLEELKNEAVRHFPRIWLHGLGQHIYETYGDTWAGVEAI  60
             MEQAPEDQGPQREP+NEWTLE+L+ELK EAVRHFPR WLH LGQHI++TYGDTW GVEA+
Sbjct  2078  MEQAPEDQGPQREPYNEWTLEILKELKQEAVRHFPRPWLHLLGQHIHDTYGDTWTGVEAM  1899

Query  61    IRILQQLLFIHFRIGCRHSRIGVTRQRRARNGASRS  96
             IR LQQLLFIHFRIGC+HSRIG+  QRRARNG+SRS
Sbjct  1898  IRTLQQLLFIHFRIGCQHSRIGILPQRRARNGSSRS  1791
```

```
>Klebsiella pneumoniae strain EuSCAPE_NO002, whole genome shotgun sequence
Sequence ID: NZ_UKMC02000071.1 Length: 907
Range 1: 94 to 381

Score:147 bits(372), Expect:9e-41,
Method:Compositional matrix adjust.,
Identities:76/96(79%), Positives:86/96(89%), Gaps:0/96(0%)

Query  1    MEQAPEDQGPQREPHNEWTLELLEELKNEAVRHFPRIWLHGLGQHIYETYGDTWAGVEAI  60
            +EQAPEDQ PQ+EP+NEWTLELLEELK EAVRHFPR  LH LGQ++YETY DTW GVEAI
Sbjct  381  VEQAPEDQRPQKEPYNEWTLELLEELKQEAVRHFPRP*LHSLGQYVYETYRDTWTGVEAI  202

Query  61   IRILQQLLFIHFRIGCRHSRIGVTRQRRARNGASRS  96
            IRILQQLLF+HFRI C+HSRIG+ RQRRA++ ASRS
Sbjct  201  IRILQQLLFVHFRISCQHSRIGILRQRRAKDRASRS  94
```

```
>Klebsiella pneumoniae strain EuSCAPE_IL020, whole genome shotgun sequence
Sequence ID: NZ_UKKB02000095.1 Length: 355
Range 1: 1 to 240

Score:140 bits(353), Expect:3e-40,
Method:Compositional matrix adjust.,
Identities:65/80(81%), Positives:72/80(90%), Gaps:0/80(0%)

Query  17   EWTLELLEELKNEAVRHFPRIWLHGLGQHIYETYGDTWAGVEAIIRILQQLLFIHFRIGC  76
            EW LE+LEELK EAVRHFPR+WLH LGQ+IY TYGDTW GVEA+IR LQQLLFIHFRIGC
Sbjct  1    EWALEILEELKQEAVRHFPRVWLHNLGQYIYATYGDTWTGVEALIRTLQQLLFIHFRIGC  180

Query  77   RHSRIGVTRQRRARNGASRS  96
            +HSRIG+ +QRRARNGA RS
Sbjct  181  QHSRIGILQQRRARNGARRS  240
```

```
>Klebsiella pneumoniae strain EuSCAPE_IL009, whole genome shotgun sequence
Sequence ID: NZ_UKJO02000080.1 Length: 261
Range 1: 1 to 213

Score:136 bits(343), Expect:5e-39,
Method:Compositional matrix adjust.,
Identities:63/71(89%), Positives:68/71(95%), Gaps:0/71(0%)

Query  1    MEQAPEDQGPQREPHNEWTLELLEELKNEAVRHFPRIWLHGLGQHIYETYGDTWAGVEAI  60
            MEQAPEDQGPQREP+NEW LE+LEELK EAVRHFPR+WLH LGQ+IYETYGDTWAGVEA+
Sbjct  213  MEQAPEDQGPQREPYNEWALEVLEELKQEAVRHFPRLWLHSLGQYIYETYGDTWAGVEAL  34

Query  61   IRILQQLLFIH  71
            IRILQQLLFIH
Sbjct  33   IRILQQLLFIH  1
```

```
>Klebsiella pneumoniae strain EuSCAPE_MT014, whole genome shotgun sequence
Sequence ID: NZ_UKJF02000041.1 Length: 265
Range 1: 33 to 263

Score:134 bits(338), Expect:3e-38,
Method:Compositional matrix adjust.,
Identities:69/77(90%), Positives:73/77(94%), Gaps:0/77(0%)

Query  1    MEQAPEDQGPQREPHNEWTLELLEELKNEAVRHFPRIWLHGLGQHIYETYGDTWAGVEAI  60
            MEQAPEDQGPQREP+NEWTLELLEELK EAVRHFPR WLH LGQ+IYETYGDTW GVEAI
Sbjct  33   MEQAPEDQGPQREPYNEWTLELLEELKQEAVRHFPRPWLHSLGQYIYETYGDTWTGVEAI  212

Query  61   IRILQQLLFIHFRIGCR  77
            IRILQQLLF+HFRIGC+
Sbjct  213  IRILQQLLFVHFRIGCQ  263
```

```
>Klebsiella pneumoniae strain EuSCAPE_IL013, whole genome shotgun sequence
Sequence ID: NZ_UKJW02000092.1 Length: 329
Range 1: 124 to 327

Score:127 bits(320), Expect:3e-35,
Method:Compositional matrix adjust.,
Identities:61/68(90%), Positives:64/68(94%), Gaps:0/68(0%)

Query  29   EAVRHFPRIWLHGLGQHIYETYGDTWAGVEAIIRILQQLLFIHFRIGCRHSRIGVTRQRR  88
            EAVRHFPR WLH LGQHIYETYGDTW GVEAIIRILQQLLFIHFRIGC+HSRIG+ RQRR
Sbjct  327  EAVRHFPRPWLHSLGQHIYETYGDTWTGVEAIIRILQQLLFIHFRIGCQHSRIGILRQRR  148

Query  89   ARNGASRS  96
            ARNGA+RS
Sbjct  147  ARNGANRS  124
```

```
>Klebsiella pneumoniae strain EuSCAPE_IL018, whole genome shotgun sequence
Sequence ID: NZ_UKJU02000098.1 Length: 409
Range 1: 212 to 409

Score:121 bits(303), Expect:2e-32,
Method:Compositional matrix adjust.,
Identities:58/66(88%), Positives:60/66(90%), Gaps:0/66(0%)

Query  30   AVRHFPRIWLHGLGQHIYETYGDTWAGVEAIIRILQQLLFIHFRIGCRHSRIGVTRQRRA  89
            AVRHFPR WLH LGQ+IY TYGDTW GVEAIIRILQQLLFIHFRIGCRHSRIG+ RQRR
Sbjct  409  AVRHFPREWLHNLGQYIYNTYGDTWRGVEAIIRILQQLLFIHFRIGCRHSRIGIMRQRRT  230

Query  90   RNGASR  95
            RNGASR
Sbjct  229  RNGASR  212
```

>Klebsiella pneumoniae strain EuSCAPE_MT009, whole genome shotgun sequence
Sequence ID: NZ_UKIV02000096.1 Length: 318
Range 1: 3 to 179

Score:99.4 bits(246), Expect:4e-24,
Method:Compositional matrix adjust.,
Identities:53/59(90%), Positives:54/59(91%), Gaps:0/59(0%)

```
Query  1    MEQAPEDQGPQREPHNEWTLELLEELKNEAVRHFPRIWLHGLGQHIYETYGDTWAGVEA  59
            MEQAPEDQGPQREPHNEWTLELLEELK EAVRHFPR WLH L QHIY+TYGDTW GVEA
Sbjct  179  MEQAPEDQGPQREPHNEWTLELLEELKQEAVRHFPRPWLHNLRQHIYDTYGDTWTGVEA  3
```

>Klebsiella pneumoniae strain EuSCAPE_IL004, whole genome shotgun sequence
Sequence ID: NZ_UKJP02000115.1 Length: 244
Range 1: 3 to 164

Score:93.2 bits(230), Expect:7e-22,
Method:Compositional matrix adjust.,
Identities:48/54(89%), Positives:51/54(94%), Gaps:0/54(0%)

```
Query  1    MEQAPEDQGPQREPHNEWTLELLEELKNEAVRHFPRIWLHGLGQHIYETYGDTW  54
            MEQAPEDQGPQREP+NEW LELLEELK EAVRHFPR+WLH LGQHIY+TYGDTW
Sbjct  164  MEQAPEDQGPQREPYNEWALELLEELKQEAVRHFPRMWLHSLGQHIYDTYGDTW  3
```

>Klebsiella pneumoniae strain EuSCAPE_DE084, whole genome shotgun sequence
Sequence ID: NZ_UKLZ02000063.1 Length: 250
Range 1: 83 to 250

Score:90.1 bits(222), Expect:1e-20,
Method:Compositional matrix adjust.,
Identities:48/56(86%), Positives:53/56(94%), Gaps:0/56(0%)

```
Query  1    MEQAPEDQGPQREPHNEWTLELLEELKNEAVRHFPRIWLHGLGQHIYETYGDTWAG  56
            +EQAPEDQGPQ+EP+NEWTLELLEELK EAVRHFPR+WLHGLGQ+IYETY DT AG
Sbjct  83   IEQAPEDQGPQKEPYNEWTLELLEELKQEAVRHFPRVWLHGLGQYIYETYRDT*AG  250
```

>Klebsiella pneumoniae strain EuSCAPE_IL005, whole genome shotgun sequence
Sequence ID: NZ_UKJL02000061.1 Length: 307
Range 1: 3 to 113

Score:60.1 bits(144), Expect:1e-08,
Method:Compositional matrix adjust.,
Identities:34/37(92%), Positives:36/37(97%), Gaps:0/37(0%)

```
Query  1    MEQAPEDQGPQREPHNEWTLELLEELKNEAVRHFPRI  37
            MEQAPEDQGPQREP+NEWTLELLEELK EAVRHFPR+
Sbjct  113  MEQAPEDQGPQREPYNEWTLELLEELKQEAVRHFPRV  3
```

>Klebsiella pneumoniae strain EuSCAPE_NO001, whole genome shotgun sequence
Sequence ID: NZ_UKMF02000043.1 Length: 351
Range 1: 286 to 351

Score:45.8 bits(107), Expect:0.005,
Method:Compositional matrix adjust.,
Identities:20/22(91%), Positives:21/22(95%), Gaps:0/22(0%)

```
Query  1    MEQAPEDQGPQREPHNEWTLEL  22
            MEQAPEDQGPQREP+NEW LEL
Sbjct  286  MEQAPEDQGPQREPYNEWALEL  351
```

>Klebsiella pneumoniae strain EuSCAPE_MT011, whole genome shotgun sequence
Sequence ID: NZ_UKIY02000086.1 Length: 368
Range 1: 294 to 368

Score:45.8 bits(107), Expect:0.006,
Method:Compositional matrix adjust.,
Identities:21/25(84%), Positives:24/25(96%), Gaps:0/25(0%)

```
Query  72   FRIGCRHSRIGVTRQRRARNGASRS  96
            FRIGC+HSRIG+ +QRRARNGASRS
Sbjct  368  FRIGCQHSRIGILQQRRARNGASRS  294
```
