## Supplementary File 2 for "HIV-1 protein coding sequences are present in relevant bacteria"

Suppementary File 2: Multiple Sequence Alignments of HIV-1 proteins and top TBLASTN hits to bacterial genomes

### HIV-1 gp120

```
                  NC_001802.1|GP120 KLWVTVYYGVPVWKEATTTLFCASDAKAYDTEVHNVWATHACVPTDPNPQEVVLVNVTEN 60
NZ_JARBDL010000112.1|Escherichia_coli -LWVTVYYGVPVWTEAKTTLFCASDAKAYEKEVHNVWATHACVPTDPNPQEMVLENVTEN 59
NZ_JARBDL010000111.1|Escherichia_coli -LWVTVYYGVPVWKEAKTTLFCASDAKGYETEVHNVWATHACVPTDPSPQEMRLDNVTEN 59
NZ_JAPMNK010000087.1|Escherichia_coli -LWVTVYYGVPVWRDANATLFCASDAKVYKTEVHNVWATHACVPTDPNPQELVLENVTEN 59
NZ_JARBDH010000090.1|Escherichia_coli -LWVTVYYGVPVWRDANATLFCASDAKVYKTEVHNVWATHACVPTDPNPQELVLENVTEN 59
NZ_JAODTC010000066.1|Escherichia_coli -LWVTVYYGVPVWKEAKTTLFCASDAKAYKTEVHNVWATHACVPTDPNPQEMVLENVTEN 59
NZ_JARBDM010000080.1|Escherichia_coli -LWVTVYYGVPVWRDAKTTLFCASDAKAYEKEVHNVWATHACVPTDPNPQEMVLENVTEN 59
NZ_JAPTGI010000094.1|Escherichia_coli -LWVTVYYGVPVWRDAKTTLFCASDAKAYEKEVHNVWATHACVPTDPNPQEMVLENVTEN 59
NZ_JAPMNK010000105.1|Escherichia_coli ---VTVYYEVPVWKEAKATLFCASDAKAHEREVHNVWATHACVPTDPNPQEIVLENVTEN 57
NZ_JAODTD010000081.1|Escherichia_coli -LWVTVYYGVPVWKEAKTTLFCASDAKAYKTEVHNVWATHACVPTDPNPQEMVLENVTEN 59

                  NC_001802.1|GP120 FNMWKNDMVEQMHEDIISLWDQSLKPCVKLTPLCVSLKCTD------LKNDTNTNSSSGR 114
NZ_JARBDL010000112.1|Escherichia_coli FNMWKNDMVNQMHEDVISLWDQSLKPCVKLTPLCVTLECNS------STYNQSSVR---- 109
NZ_JARBDL010000111.1|Escherichia_coli FNMWENNMVDQMHEDIISLWDQSLKPCIKLTPLCVTLECEK------A--NRTVTNILNE 111
NZ_JAPMNK010000087.1|Escherichia_coli FNMWENDMVEQMHQDVISIWDQGLKPCVKLTPLCVTLNCTTYEPKKVTADNNNTCAPSQN 119
NZ_JARBDH010000090.1|Escherichia_coli FNMWENDMVEQMHQDVISIWDQGLKPCVKLTPLCVTLNCTTYEPKKVTADNNNTCAPSQN 119
NZ_JAODTC010000066.1|Escherichia_coli FNMWKNDMVNQMHEDVISLWDQSLKPCVKLTPLCVTLKCNA------TTHNGTSDAADND 113
NZ_JARBDM010000080.1|Escherichia_coli FNMWKNDMVNQMHEDVISLWDQSLKPCVKLTPLCVTLKCNA------TTHNGTSDAADND 113
NZ_JAPTGI010000094.1|Escherichia_coli FNMWKNDMVNQMHEDVISLWDQSLKPCVKLTPLCVTLKCNA------TTHNGTSDAADND 113
NZ_JAPMNK010000105.1|Escherichia_coli FNMWKSDMVDQMHEDVISL-DQSLKPCVKLTPLCVTLECD----------NETVSVNHTD 106
NZ_JAODTD010000081.1|Escherichia_coli FNMWKNDMVNQMHEDVISLWDQSLKPCVKLTPLCVTLKCTA------TTHNGTRDAADND 113

                  NC_001802.1|GP120 MI--------MEKGEIKNCSFNISTSIRGKVQKEYAFFYKLDIIPIDN----------DT 156
NZ_JARBDL010000112.1|Escherichia_coli ------------VHEVKNCSFNATTELRDKKHKVQALFHRLDVVPLDNRSSNENST---- 153
NZ_JARBDL010000111.1|Escherichia_coli TY--------DSKNEIQNCSFNATTEIEDKKKKEYALFYRLDIVSLDN-ENNS--SGNDS 160
NZ_JAPMNK010000087.1|Escherichia_coli TTVKNLTIDPEAVKEISNCSFNTTTELRDKTRKVHALFYKIDMVELGE-NSTA----PNF 174
NZ_JARBDH010000090.1|Escherichia_coli TTVKNLTIDPEAVKEISNCSFNTTTELRDKTRKVHALFYKIDMVELGE-NSTA----PNF 174
NZ_JAODTC010000066.1|Escherichia_coli TS--------SGGEEMKNCSFNITTELRDRKQKVYALFYKLDVVPLDN-TNGS----NSS 160
NZ_JARBDM010000080.1|Escherichia_coli TS--------SGGEEMKNCSFNITTELRDRKQKVYALFYKLDVVPLDN-TNGS----NSS 160
NZ_JAPTGI010000094.1|Escherichia_coli TS--------SGGEEMKNCSFNITTELRDRKQKVYALFYKLDVVPLDN-TNGS----NSS 160
NZ_JAPMNK010000105.1|Escherichia_coli E-----------SKELKRCFFNASTEVEDKKQKVSALFYSLDIRPLNK-SSSRNSSINSN 154
NZ_JAODTD010000081.1|Escherichia_coli TS--------SGGEEMKNCSFNITTELRDRKQKVYALFYKLDVVPLDN-TNGS----NSS 160

                  NC_001802.1|GP120 TSYKLTSCNTSVITQACPKVSFEPIPIHYCAPAGFAILKCNNKTFNGTGPCTNVSTVQCT 216
NZ_JARBDL010000112.1|Escherichia_coli GYYRLINCNTSVITQACPKVTFDPIPIHYCTPAGYAILQCNDKTFNGTGPCNNVSTVQCT 213
NZ_JARBDL010000111.1|Escherichia_coli RPYRLINCNTSAITQACPKVSFDPIPIHYCAPAGYAILKCNNKTFNGTGPCNNVSTVQCT 220
NZ_JAPMNK010000087.1|Escherichia_coli TLYRLINCNTSAITQACPKVSFEPIPIHYCAPAGYALLKCNNNTFNGTGPCRNISTVQCT 234
NZ_JARBDH010000090.1|Escherichia_coli TLYRLINCNTSAITQACPKVSFEPIPIHYCAPAGYALLKCNNNTFNGTGPCRNISTVQCT 234
NZ_JAODTC010000066.1|Escherichia_coli RSYRLINCNTSAITQACPKVSFDPIPIHYCAPAGYAILRCNDKTFNGTGPCHNVSTVQCT 220
NZ_JARBDM010000080.1|Escherichia_coli RSYRLINCNTSAITQACPKVSFDPIPIHYCAPAGYAILRCNDKTFNGTGPCHNVSTVQCT 220
NZ_JAPTGI010000094.1|Escherichia_coli RSYRLINCNTSAITQACPKVSFDPIPIHYCAPAGYAILRCNDKTFNGTGPCHNVSTVQCT 220
NZ_JAPMNK010000105.1|Escherichia_coli ETYVLISCNTSVIKQACPKVTFDPIPIHYCAPAGYAILKCNNETFNRTGPCNNVSTVQCT 214
NZ_JAODTD010000081.1|Escherichia_coli RPYRLINCNTSAITQACPKVSFDPIPIHYCAPAGYAILRCNDKTFNGTGPCHNVSTVQCT 220

                  NC_001802.1|GP120 HGIRPVVSTQLLLNGSLAEEEVVIRSVNFTDNAKTIIVQLNTSVEINCTRPNNNTRKRIR 276
NZ_JARBDL010000112.1|Escherichia_coli HGIKPVVSTQLLLNGSLAEEEIVIRSENLTNNAKTIIVHLNKSVDIVCTRPNNNTRKSIR 273
NZ_JARBDL010000111.1|Escherichia_coli HGIKPVVSTQLLVNGSLAEEEIIIRSENLTDNAKTIIVQLNKSVPITCIRPNNNTRKSIR 280
NZ_JAPMNK010000087.1|Escherichia_coli HGIRPVVSTQLLLNGSLAEGNVTIRSENLTDNAKTIIAHLNQSVEILCTRPNNNTRQSVR 294
NZ_JARBDH010000090.1|Escherichia_coli HGIRPVVSTQLLLNGSLAEGNVTIRSENLTDNAKTIIAHLNQSVEILCTRPNNNTRQSVR 294
NZ_JAODTC010000066.1|Escherichia_coli HGIKPVVSTQLLLNGSLAEEEIIIRSQNLSDNAKTIIVHLNQSVEIVCTRPHNNTRKSIR 280
NZ_JARBDM010000080.1|Escherichia_coli HGIKPVVSTQLLLNGSLAEEEIIIRSQNLSDNAKTIIVHLNQSVEIVCTRPHNNTRKSIR 280
NZ_JAPTGI010000094.1|Escherichia_coli HGIKPVVSTQLLLNGSLAEEEIIIRSQNLSDNAKTIIVHLNQSVEIVCTRPHNNTRKSIR 280
NZ_JAPMNK010000105.1|Escherichia_coli HRIKPVVSTQLLLNSSLAKGEIIIRSENLTDNAKTIIVHLNESVEINCTRPNNNTRKSIR 274
NZ_JAODTD010000081.1|Escherichia_coli HGIKPVVSTQLLLNGSLAEEEIIIRSQNLSDNAKTIIVHLNQSVEIVCTRPHNNTRKSIR 280

                  NC_001802.1|GP120 IQRGPGRAFVTIGK-IGNMRQAHCNISRAKWNNTLKQIASKLREQFGNNKTIIFKQSSGG 335
NZ_JARBDL010000112.1|Escherichia_coli I--GPGQTFYATGDIIGDIRKAHCNISLNRWNDTILKVREKLAEHFPN-RTIEFQPSSGG 330
NZ_JARBDL010000111.1|Escherichia_coli I--GPGQTVYATGDSIGDIRQAHCNVSRKDWEGTLYNVNKKLKEHFPNITTIKFKPSSGG 338
NZ_JAPMNK010000087.1|Escherichia_coli I--GPGQTFFATGDIIGDIRRAHCNVSAKHWHTALRNVSIELAKLFPNNKTINFTSPSGG 352
NZ_JARBDH010000090.1|Escherichia_coli I--GPGQTFFATGDIIGDIRRAHCNVSAKHWHTALRNVSIELAKLFPNNKTINFTSPSGG 352
NZ_JAODTC010000066.1|Escherichia_coli I--GPGQTFYATGAIIGNIRQAYCSINISKWSETLHNVSKKLAERFPN-KTINFAASSGG 337
NZ_JARBDM010000080.1|Escherichia_coli I--GPGQTFYATGAIIGNIRQAYCSINISKWSETLHNVSKKLAERFPN-KTINFAASSGG 337
NZ_JAPTGI010000094.1|Escherichia_coli I--GPGQTFYATGAIIGNIRQAYCSINISKWSETLHNVSKKLAERFPN-KTINFAASSGG 337
NZ_JAPMNK010000105.1|Escherichia_coli I--GPGQTFYATGEIVRNIRQAHCNIS-EKENKTLQKVGIKLAEHFPN-KTINFTSHSGG 330
NZ_JAODTD010000081.1|Escherichia_coli I--GPGQTFYATGAIIGNIRQAYCSINISKWSETLHNVSKKLAERFPN-KTINFAASSGG 337
```

>>>

### HIV-1 gp120 (continuation)

```
                     NC_001802.1|GP120 DPEIVTHSFNCGGEFFYCNSTQLFNSTW--FNSTWST----------EGS---------- 373
NZ_JARBDL010000112.1|Escherichia_coli DLEITTHSFNCGGEFFYCNTSNLFNFTYYYDNGTYHY----------NGTYY---NS--- 374
NZ_JARBDL010000111.1|Escherichia_coli DLEITTHSFNCRGEFFYCNTSDLFNGTY---NSTY------------KGTYN---NTG-- 378
NZ_JAPMNK010000087.1|Escherichia_coli DLEITTHSFNCRGEFFYCNTSQLFNSLYMPNNDTYIWYNENHTYVWHNGTQYVINGSTQY 412
NZ_JARBDH010000090.1|Escherichia_coli DLEITTHSFNCRGEFFYCNTSQLFNSLYMPNNDTYIWYNENHTYVWHNGTQYVINGSTQY 412
NZ_JAODTC010000066.1|Escherichia_coli DLEVTTHSFNCRGEFFYCNTSELFNSTYIYLNRTYMF----------NGT---------- 377
NZ_JARBDM010000080.1|Escherichia_coli DLEVTTHSFNCRGEFFYCNTSELFNSTYIYLNRTYMF----------NGT---------- 377
NZ_JAPTGI010000094.1|Escherichia_coli DLEVTTHSFNCRGEFFYCNTSELFNSTYIYLNRTYMF----------NGT---------- 377
NZ_JAPMNK010000105.1|Escherichia_coli DLEIMTHSFNCRGEFFYCNTSSLFNSTY---NSTYIR----------N------------ 365
NZ_JAODTD010000081.1|Escherichia_coli DLEVTTHSFNCRGEFFYCNTSELFNSTYIYPDRTYMF----------NGT---------- 377

                     NC_001802.1|GP120 ------------NNTEGSDTITLPCRIKQIINMWQKVGKAMYAPPISGQIRCSSNITGLL 421
NZ_JARBDL010000112.1|Escherichia_coli ------TQY----NS-T-ANITIPCRIKQIINMWQEVGRAMYAPPIAGNITCRSNITGLL 422
NZ_JARBDL010000111.1|Escherichia_coli ------GNS----SA-ADPNITLQCRIKQIINMWQQVGKAMYAPPIAGNITCISNITGLL 427
NZ_JAPMNK010000087.1|Escherichia_coli VTLKNSAQNVTINNI-TDFNITLPCRIKQIINMWQEVGKAMYAPPIAGNITCLSNITGLL 471
NZ_JARBDH010000090.1|Escherichia_coli VTLKNSAQNVTINNI-TDFNITLPCRIKQIINMWQEVGKAMYAPPIAGNITCLSNITGLL 471
NZ_JAODTC010000066.1|Escherichia_coli ------------NGT-STNNFTIPCRIKQIINMWQEVGRAMYAPPIEGNITCKSNITGLL 424
NZ_JARBDM010000080.1|Escherichia_coli ------------NGT-STNNFTIPCRIKQIINMWQEVGRAMYAPPIEGNITCKSNITGLL 424
NZ_JAPTGI010000094.1|Escherichia_coli ------------NGT-STNNFTIPCRIKQIINMWQEVGRAMYAPPIEGNITCKSNITGLL 424
NZ_JAPMNK010000105.1|Escherichia_coli -------------NS-S-SIITIQCRIKQIINMWQEVGRAMYAPPIAGNITCQSNITGLL 410
NZ_JAODTD010000081.1|Escherichia_coli ------------NGT-STNNFTIPCRIKQIINMWQKVGRAMYAPPIEGNITCKSNITGLL 424

                     NC_001802.1|GP120 LTRDGGNSN---NESEIFRPGGGDMRDNWRSELYKYKVVKIEPLGVAPTKAKRRVVQREK 478
NZ_JARBDL010000112.1|Escherichia_coli LVRDGGPSS---NETETFRPGGGDMRDNWRSELYKYKVVEIKPLGIAPTGAKRRVVGREK 479
NZ_JARBDL010000111.1|Escherichia_coli LVRDGGT-NRTDNDTETFRPGGGNMKDNWRSELYKYKVVEIQPLGVAPTGAKRRVVEREK 486
NZ_JAPMNK010000087.1|Escherichia_coli LTRDGGKGDEIRNDTEIFRPGGGNMKDNWRSELYKYKVVEIKPLGVAPT----------- 520
NZ_JARBDH010000090.1|Escherichia_coli LTRDGGKGDEIRNDTEIFRPGGGNMKDNWRSELYKYKVVEIKPLGVAPT----------- 520
NZ_JAODTC010000066.1|Escherichia_coli LERDGGE---ENRTTEIFRP---------------------------------------- 441
NZ_JARBDM010000080.1|Escherichia_coli LERDGGE---ENRTTEIFRP---------------------------------------- 441
NZ_JAPTGI010000094.1|Escherichia_coli LERDGGE---ENRTTEIFRP---------------------------------------- 441
NZ_JAPMNK010000105.1|Escherichia_coli LVRDRGHRNETSNSTETFRPKGGDMKNN-RSELYKYKVVEIKPLGIAPTAAKRRVV---- 465
NZ_JAODTD010000081.1|Escherichia_coli LERDGGE--EENRTTEIFRP---------------------------------------- 442

                     NC_001802.1|GP120 R                                                            479
NZ_JARBDL010000112.1|Escherichia_coli R                                                            480
NZ_JARBDL010000111.1|Escherichia_coli R                                                            487
NZ_JAPMNK010000087.1|Escherichia_coli -                                                            520
NZ_JARBDH010000090.1|Escherichia_coli -                                                            520
NZ_JAODTC010000066.1|Escherichia_coli -                                                            441
NZ_JARBDM010000080.1|Escherichia_coli -                                                            441
NZ_JAPTGI010000094.1|Escherichia_coli -                                                            441
NZ_JAPMNK010000105.1|Escherichia_coli -                                                            465
NZ_JAODTD010000081.1|Escherichia_coli -                                                            442
```

### HIV-1 gp41

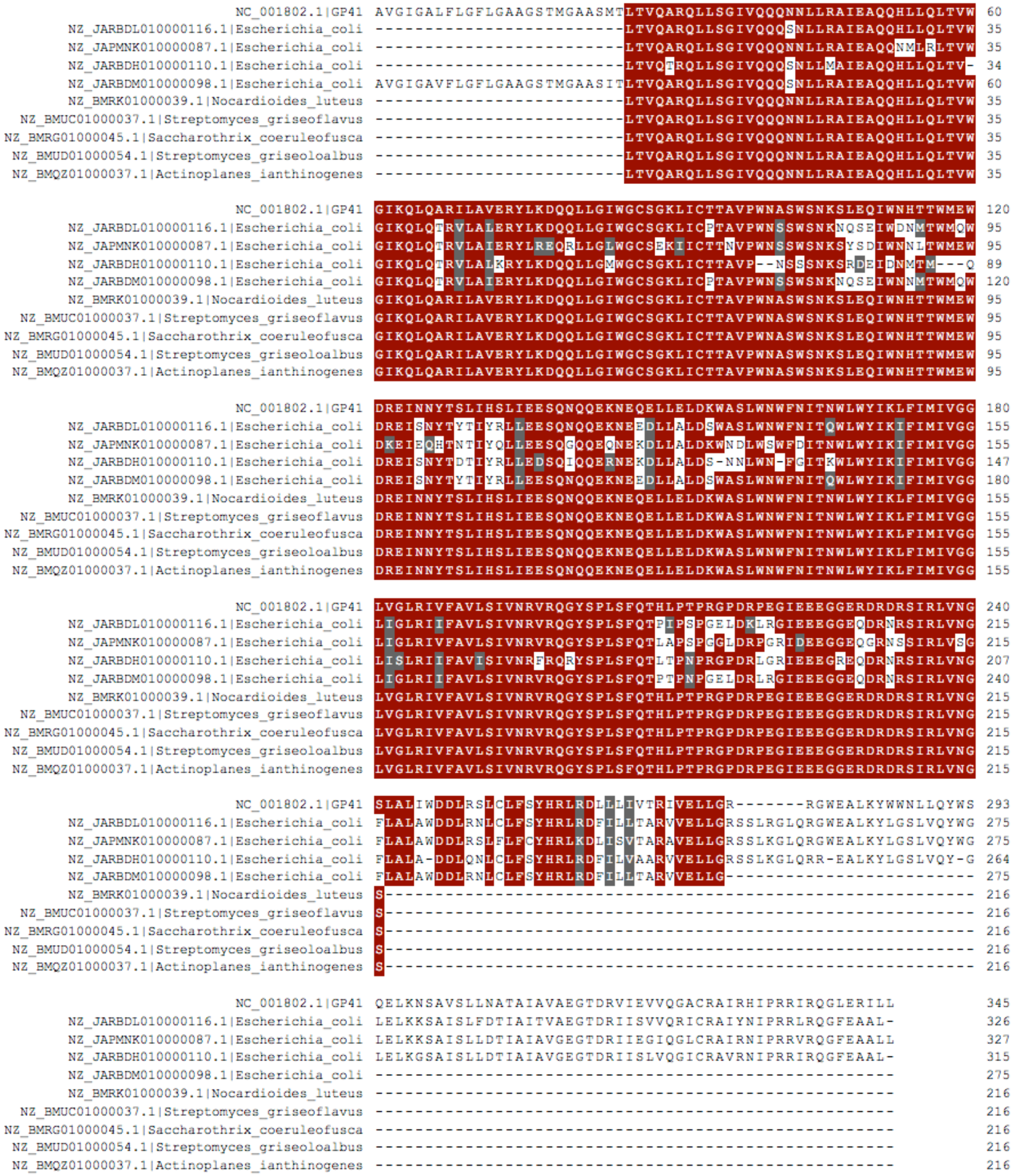
NC_001802.1|GP41 AVGIGALFLGFLGAAGSTMGAASMTLTVQARQLLSGIVQQQNNLLRAIEAQQHLLQLTVW 60
NZ_JARBDL010000116.1|Escherichia_coli -------------------------LTVQARQLLSGIVQQQSNLLRAIEAQQHLLQLTVW 35
NZ_JAPMNK010000087.1|Escherichia_coli -------------------------LTVQARQLLSGIVQQQNNLLRAIEAQQNMLRLTVW 35
NZ_JARBDH010000110.1|Escherichia_coli -------------------------LTVQTRQLLSGIVQQQSNLLMAIEAQQHLLQLTV- 34
NZ_JARBDM010000098.1|Escherichia_coli AVGIGAVFLGFLGAAGSTMGAASITLTVQARQLLSGIVQQQSNLLRAIEAQQHLLQLTVW 60
NZ_BMRK01000039.1|Nocardioides_luteus -------------------------LTVQARQLLSGIVQQQNNLLRAIEAQQHLLQLTVW 35
NZ_BMUC01000037.1|Streptomyces_griseoflavus -------------------------LTVQARQLLSGIVQQQNNLLRAIEAQQHLLQLTVW 35
NZ_BMRG01000045.1|Saccharothrix_coeruleofusca -------------------------LTVQARQLLSGIVQQQNNLLRAIEAQQHLLQLTVW 35
NZ_BMUD01000054.1|Streptomyces_griseoloalbus -------------------------LTVQARQLLSGIVQQQNNLLRAIEAQQHLLQLTVW 35
NZ_BMQZ01000037.1|Actinoplanes_ianthinogenes -------------------------LTVQARQLLSGIVQQQNNLLRAIEAQQHLLQLTVW 35
NC_001802.1|GP41 GIKQLQARILAVERYLKDQQLLGIWGCSGKLICTTAVPWNASWSNKSLEQIWNHTTWMEW 120
NZ_JARBDL010000116.1|Escherichia_coli GIKQLQTRVLALERYLKDQQLLGIWGCSGKLICPTAVPWNSSWSNKNQSEIWDNMTWMQW 95
NZ_JAPMNK010000087.1|Escherichia_coli GIKQLQTRVLAIERYLREQRLLGLWGCSEKIICTTNVPWNSSWSNKSYSDIWNNLTWMEW 95
NZ_JARBDH010000110.1|Escherichia_coli GIKQLQTRVLALKRYLKDQQLLGMWGCSGKLICTTAVP--NSSSNKSRDEIDNMTM---Q 89
NZ_JARBDM010000098.1|Escherichia_coli GIKQLQTRVLAIERYLKDQQLLGIWGCSGKLICPTAVPWNSSWSNKNQSEIWNNMTWMQW 120
NZ_BMRK01000039.1|Nocardioides_luteus GIKQLQARILAVERYLKDQQLLGIWGCSGKLICTTAVPWNASWSNKSLEQIWNHTTWMEW 95
NZ_BMUC01000037.1|Streptomyces_griseoflavus GIKQLQARILAVERYLKDQQLLGIWGCSGKLICTTAVPWNASWSNKSLEQIWNHTTWMEW 95
NZ_BMRG01000045.1|Saccharothrix_coeruleofusca GIKQLQARILAVERYLKDQQLLGIWGCSGKLICTTAVPWNASWSNKSLEQIWNHTTWMEW 95
NZ_BMUD01000054.1|Streptomyces_griseoloalbus GIKQLQARILAVERYLKDQQLLGIWGCSGKLICTTAVPWNASWSNKSLEQIWNHTTWMEW 95
NZ_BMQZ01000037.1|Actinoplanes_ianthinogenes GIKQLQARILAVERYLKDQQLLGIWGCSGKLICTTAVPWNASWSNKSLEQIWNHTTWMEW 95
NC_001802.1|GP41 DREINNYTSLIHSLIEESQNQQEKNEQELLELDKWASLWNWFNITNWLWYIKLFIMIVGG 180
NZ_JARBDL010000116.1|Escherichia_coli DREISNYTYTIYRLLEESQNQQEKNEEDLLALDSWASLWNWFNITQWLWYIKIFIMIVGG 155
NZ_JAPMNK010000087.1|Escherichia_coli DKEIEQHTNTIYQLLEESQGQQEQNEKDLLALDKWNDLWSWFDITNWLWYIKIFIMIVGG 155
NZ_JARBDH010000110.1|Escherichia_coli DREISNYTDTIYRLLEDSQIQQERNEKDLLALDS-NNLWN-FGITKWLWYIKIFIMIVGG 147
NZ_JARBDM010000098.1|Escherichia_coli DREISNYTYTIYRLLEESQNQQEKNEEDLLALDSWASLWNWFNITQWLWYIKIFIMIVGG 180
NZ_BMRK01000039.1|Nocardioides_luteus DREINNYTSLIHSLIEESQNQQEKNEQELLELDKWASLWNWFNITNWLWYIKLFIMIVGG 155
NZ_BMUC01000037.1|Streptomyces_griseoflavus DREINNYTSLIHSLIEESQNQQEKNEQELLELDKWASLWNWFNITNWLWYIKLFIMIVGG 155
NZ_BMRG01000045.1|Saccharothrix_coeruleofusca DREINNYTSLIHSLIEESQNQQEKNEQELLELDKWASLWNWFNITNWLWYIKLFIMIVGG 155
NZ_BMUD01000054.1|Streptomyces_griseoloalbus DREINNYTSLIHSLIEESQNQQEKNEQELLELDKWASLWNWFNITNWLWYIKLFIMIVGG 155
NZ_BMQZ01000037.1|Actinoplanes_ianthinogenes DREINNYTSLIHSLIEESQNQQEKNEQELLELDKWASLWNWFNITNWLWYIKLFIMIVGG 155
NC_001802.1|GP41 LVGLRIVFAVLSIVNRVRQGYSPLSFQTHLPTPRGPDRPEGIEEEGGERDRDRSIRLVNG 240
NZ_JARBDL010000116.1|Escherichia_coli LIGLRIIFAVLSIVNRVRQGYSPLSFQTPIPSPGELDKLRGIEEEGGEQDRNRSIRLVNG 215
NZ_JAPMNK010000087.1|Escherichia_coli LIGLRIVFAVLSIVNRVRQGYSPLSFQTLAPSPGGLDRPGRIDEEGGEQGRNSSIRLVSG 215
NZ_JARBDH010000110.1|Escherichia_coli LISLRIIFAVISIVNRFRQRYSPLSFQTLTPNPRGPDRLGRIEEEGREQDRNRSIRLVNG 207
NZ_JARBDM010000098.1|Escherichia_coli LIGLRIIFAVLSIVNRVRQGYSPLSFQTPTPNPGELDRLRGIEEEGGEQDRNRSIRLVNG 240
NZ_BMRK01000039.1|Nocardioides_luteus LVGLRIVFAVLSIVNRVRQGYSPLSFQTHLPTPRGPDRPEGIEEEGGERDRDRSIRLVNG 215
NZ_BMUC01000037.1|Streptomyces_griseoflavus LVGLRIVFAVLSIVNRVRQGYSPLSFQTHLPTPRGPDRPEGIEEEGGERDRDRSIRLVNG 215
NZ_BMRG01000045.1|Saccharothrix_coeruleofusca LVGLRIVFAVLSIVNRVRQGYSPLSFQTHLPTPRGPDRPEGIEEEGGERDRDRSIRLVNG 215
NZ_BMUD01000054.1|Streptomyces_griseoloalbus LVGLRIVFAVLSIVNRVRQGYSPLSFQTHLPTPRGPDRPEGIEEEGGERDRDRSIRLVNG 215
NZ_BMQZ01000037.1|Actinoplanes_ianthinogenes LVGLRIVFAVLSIVNRVRQGYSPLSFQTHLPTPRGPDRPEGIEEEGGERDRDRSIRLVNG 215
NC_001802.1|GP41 SLALIWDDLRSLCLFSYHRLRDLLLIVTRIVELLGR-------RGWEALKYWWNLLQYWS 293
NZ_JARBDL010000116.1|Escherichia_coli FLALAWDDLRNLCLFSYHRLRDFILLTARVVELLGRSSLRGLQRGWEALKYLGSLVQYWG 275
NZ_JAPMNK010000087.1|Escherichia_coli FLALAWDDLRSLFLFCYHRLKDLISVTARAVELLGRSSLKGLQRGWEALKYLGSLVQYWG 275
NZ_JARBDH010000110.1|Escherichia_coli FLALA-DDLQNLCLFSYHRLRDFILVAARVVELLGRSSLKGLQRR-EALKYLGSLVQY-G 264
NZ_JARBDM010000098.1|Escherichia_coli FLALAWDDLRNLCLFSYHRLRDFILLTARVVELLG------------------------- 275
NZ_BMRK01000039.1|Nocardioides_luteus S----------------------------------------------------------- 216
NZ_BMUC01000037.1|Streptomyces_griseoflavus S----------------------------------------------------------- 216
NZ_BMRG01000045.1|Saccharothrix_coeruleofusca S----------------------------------------------------------- 216
NZ_BMUD01000054.1|Streptomyces_griseoloalbus S----------------------------------------------------------- 216
NZ_BMQZ01000037.1|Actinoplanes_ianthinogenes S----------------------------------------------------------- 216
NC_001802.1|GP41 QELKNSAVSLLNATAIAVAEGTDRVIEVVQGACRAIRHIPRRIRQGLERILL 345
NZ_JARBDL010000116.1|Escherichia_coli LELKKSAISLFDTIAITVAEGTDRIISVVQRICRAIYNIPRRLRQGFEAAL- 326
NZ_JAPMNK010000087.1|Escherichia_coli LELKKSAISLLDTIAIAVGEGTDRIIEGIQGLCRAIRNIPRRVRQGFEAALL 327
NZ_JARBDH010000110.1|Escherichia_coli LELKGSAISLLDTIAIAVGEGTDRIISLVQGICRAVRNIPRRIRQGFEAAL- 315
NZ_JARBDM010000098.1|Escherichia_coli ---------------------------------------------------- 275
NZ_BMRK01000039.1|Nocardioides_luteus ---------------------------------------------------- 216
NZ_BMUC01000037.1|Streptomyces_griseoflavus ---------------------------------------------------- 216
NZ_BMRG01000045.1|Saccharothrix_coeruleofusca ---------------------------------------------------- 216
NZ_BMUD01000054.1|Streptomyces_griseoloalbus ---------------------------------------------------- 216
NZ_BMQZ01000037.1|Actinoplanes_ianthinogenes ---------------------------------------------------- 216

### HIV-1 Gag p6

```
                      NC_001802.1|GAG_p6 LQSRPEPTAPPEESFRSGVETTTPPQKQEPIDKELYPLTSLRSLFGNDPSSQ 52
NZ_BMOE01000030.1|Deinococcus_aquiradiocola LQSRPEPTAPPEESFRFGEETTTPSQKQEPIDKELYPLASLRSLFGSDPSSQ 52
   NZ_BMDT01000023.1|Enterococcus_alcedinis LQSRPEPTAPPEESFRFGEETTTPSQKQEPIDKELYPLASLRSLFGSDPSSQ 52
NZ_BMPW01000058.1|Actinoplanes_campanulatus LQSRPEPTAPPEESFRFGEETTTPSQKQEPIDKELYPLASLRSLFGSDPSSQ 52
     NZ_WWEN01000019.1|Thalassovita_mangrovi LQSRPEPTAPPEESFRFGEETTTPSQKQEPIDKELYPLASLRSLFGSDPSSQ 52
  NZ_JACAWM010000050.1|Klebsiella_pneumoniae -------TAPPEESFRFGEETTTPSQKQEPIDKELYPLASLRSLFGSDPSSQ 45
    NZ_QEHN01000065.1|Acinetobacter_baumannii ----PEPTAPPEESLRLGGETATPSQKQEPTDKELYPLASLRSLFGND---- 44
    NZ_QAGO01000067.1|Acinetobacter_baumannii ----------PEESFRLGGETTTPSQKQEPTDKELYPLASLRSLFGND---- 38
   NZ_CADDWK010000035.1|Salirhabdus_euzebyi -----------EESLRLGEETTTPSQKQEQIDKDMYPLASLRSLFGNDPSSQ 41
    NZ_PYHB01000027.1|Enterobacter_hormaechei --------APPEESLRLGGETATPSQKQEPTDKELYPLASLRSLFGND---- 40
```

### HIV-1 Gag p24

```
NC_001802.1|GAG_p24                          PIVQNIQGQMVHQAISPRTLNAWVKVVEEKAFSPEVIPMFSALSEGATPQDLNTMLNTVG 60
NZ_WWEN01000019.1|Thalassovita_mangrovi      PIVQNLQGQMVHQAISPRTLNAWVKVVEEKAFSPEVIPMFSALSEGATPQDLNTMLNTVG 60
NZ_BMPW01000058.1|Actinoplanes_campanulatus  PIVQNLQGQMVHQAISPRTLNAWVKVVEEKAFSPEVIPMFSALSEGATPQDLNTMLNTVG 60
NZ_BMDT01000023.1|Enterococcus_alcedinis     PIVQNLQGQMVHQAISPRTLNAWVKVVEEKAFSPEVIPMFSALSEGATPQDLNTMLNTVG 60
NZ_BMOE01000030.1|Deinococcus_aquiradiocola  PIVQNLQGQMVHQAISPRTLNAWVKVVEEKAFSPEVIPMFSALSEGATPQDLNTMLNTVG 60
NZ_UKKG02000043.1|Klebsiella_pneumoniae      ------------------------------------------------------------ 0
NZ_UKKB02000090.1|Klebsiella_pneumoniae      ------------------------------------------------------------ 0
NZ_UKIK02000113.1|Klebsiella_pneumoniae      ------------------------------------------------------------ 0
NZ_UKJZ02000061.1|Klebsiella_pneumoniae      ------------------------------------------------------------ 0
NZ_UVFZ02000120.1|Burkholderia_pseudomallei  ------------------------------------------------------------ 0

NC_001802.1|GAG_p24                          GHQAAMQMLKETINEEAAEWDRVHPVHAGPIAPGQMREPRGSDIAGTTSTLQEQIGWMTN 120
NZ_WWEN01000019.1|Thalassovita_mangrovi      GHQAAMQMLKETINEEAAEWDRLHPVHAGPIAPGQMREPRGSDIAGTTSTLQEQIGWMTH 120
NZ_BMPW01000058.1|Actinoplanes_campanulatus  GHQAAMQMLKETINEEAAEWDRLHPVHAGPIAPGQMREPRGSDIAGTTSTLQEQIGWMTH 120
NZ_BMDT01000023.1|Enterococcus_alcedinis     GHQAAMQMLKETINEEAAEWDRLHPVHAGPIAPGQMREPRGSDIAGTTSTLQEQIGWMTH 120
NZ_BMOE01000030.1|Deinococcus_aquiradiocola  GHQAAMQMLKETINEEAAEWDRLHPVHAGPIAPGQMREPRGSDIAGTTSTLQEQIGWMTH 120
NZ_UKKG02000043.1|Klebsiella_pneumoniae      ----------------------------------------GSDIAGTTSTLQEQITWMTS 20
NZ_UKKB02000090.1|Klebsiella_pneumoniae      ----------------------------------------GSDIAGTTSTLQEQVAWITG 20
NZ_UKIK02000113.1|Klebsiella_pneumoniae      ----------------------------------------GSDIAGTTSSLQEQIGWMTS 20
NZ_UKJZ02000061.1|Klebsiella_pneumoniae      ----------------------------------------GSDIAGTTSTLQEQIAWMTS 20
NZ_UVFZ02000120.1|Burkholderia_pseudomallei  ------------------------------------------------------------ 0

NC_001802.1|GAG_p24                          NPPIPVGEIYKRWIILGLNKIVRMYSPTSILDIRQGPKEPFRDYVDRFYKTLRAEQASQE 180
NZ_WWEN01000019.1|Thalassovita_mangrovi      NPPIPVGEIYKRWIILGLNKIVRMYSPTSILDIRQGPKEPFRDYVDRFYKTLRAEQASQE 180
NZ_BMPW01000058.1|Actinoplanes_campanulatus  NPPIPVGEIYKRWIILGLNKIVRMYSPTSILDIRQGPKEPFRDYVDRFYKTLRAEQASQE 180
NZ_BMDT01000023.1|Enterococcus_alcedinis     NPPIPVGEIYKRWIILGLNKIVRMYSPTSILDIRQGPKEPFRDYVDRFYKTLRAEQASQE 180
NZ_BMOE01000030.1|Deinococcus_aquiradiocola  NPPIPVGEIYKRWIILGLNKIVRMYSPTSILDIRQGPKEPFRDYVDRFYKTLRAEQASQE 180
NZ_UKKG02000043.1|Klebsiella_pneumoniae      NPPVPVGDIYKRWIILGLNKIVRMYSPVSILDIRQGPKEPFRDYVDRFFKTLRAEQATQD 80
NZ_UKKB02000090.1|Klebsiella_pneumoniae      NPAIPVGEIYKRWIILGLNKIVRMYSPVSILDIKQGPKEPFRDYVDRFFKCLRAEQATQE 80
NZ_UKIK02000113.1|Klebsiella_pneumoniae      NPPIPVGDIYKRWIILGLNKIVRMYSPVSILDIKQGPKEPFRDYVDRFFKTLRAEQATQD 80
NZ_UKJZ02000061.1|Klebsiella_pneumoniae      NPPIPVGDIYKRWIILGLNKIVRMYSPVSILDIRQGPKEPFRDYVDRFFKTLRAEQATQE 80
NZ_UVFZ02000120.1|Burkholderia_pseudomallei  --------IYKRWIILGLNKIVRMYSPVSILDIKQGPKEPFRDYVDRFYRTLRAEQATQE 52

NC_001802.1|GAG_p24                          VKNWMTETLLVQNANPDCKTILKALGPAATLEEMMTACQGVGGPGHKARVL 231
NZ_WWEN01000019.1|Thalassovita_mangrovi      VKNWMTETLLVQNANPDCKTILKALGPGATLEEMMTACQGVGGPGHKARVL 231
NZ_BMPW01000058.1|Actinoplanes_campanulatus  VKNWMTETLLVQNANPDCKTILKALGPGATLEEMMTACQGVGGPGHKARVL 231
NZ_BMDT01000023.1|Enterococcus_alcedinis     VKNWMTETLLVQNANPDCKTILKALGPGATLEEMMTACQGVGGPGHKARVL 231
NZ_BMOE01000030.1|Deinococcus_aquiradiocola  VKNWMTETLLVQNANPDCKTILKALGPGATLEEMMTACQGVGGPGHKARVL 231
NZ_UKKG02000043.1|Klebsiella_pneumoniae      VKNWMTDTLLVQNANPDCKTILRALGPGATLEEMMTACQGVGGPGHKARVL 131
NZ_UKKB02000090.1|Klebsiella_pneumoniae      VKNWMTDTLLVQNANPDCKAILRALGPGANLEEMMTACQGVGGPSHKARVL 131
NZ_UKIK02000113.1|Klebsiella_pneumoniae      VKNWMTDTLLVQNANPDCKTILRAL-------------------------- 105
NZ_UKJZ02000061.1|Klebsiella_pneumoniae      VKNWMTETLLVQNANPDCKTILR---------------------------- 103
NZ_UVFZ02000120.1|Burkholderia_pseudomallei  VKNWMTETLLVQNANPDCKSILKALGTGATLEEMMTACQGVGGPSHKARVL 103
```

### HIV-1 Gag p7

```
          NC_001802.1|GAG_NCPp7 MQRGNFRNQRKIVKCFNCGKEGHTARNCRAPRKKGCWKCGKEGHQMKDCTERQAN 55
 NZ_UKJZ02000062.1|Klebsiella_pneumoniae MQKGNFKGPKRIIKCFNCGKEGHIARNCRAPRKKGCWKCGKEGHQMKDCTERQAN 55
 NZ_WWEN01000019.1|Thalassovita_mangrovi IQKGNFRNQRKTVKCFNCGKEGHIAKNCRAPRKKGCWKCGKEGHQMKDCTERQAN 55
NZ_BMPW01000058.1|Actinoplanes_campanulatus IQKGNFRNQRKTVKCFNCGKEGHIAKNCRAPRKKGCWKCGKEGHQMKDCTERQAN 55
 NZ_BMDT01000023.1|Enterococcus_alcedinis IQKGNFRNQRKTVKCFNCGKEGHIAKNCRAPRKKGCWKCGKEGHQMKDCTERQAN 55
NZ_BMOE01000030.1|Deinococcus_aquiradiocola IQKGNFRNQRKTVKCFNCGKEGHIAKNCRAPRKKGCWKCGKEGHQMKDCTERQAN 55
NZ_UVID02000074.1|Burkholderia_pseudomallei MQKNNFKGPRRTVKCFNCGKEGHIAKNCRAPRKKGCWKCGKEGHQMKDCTERQAN 55
 NZ_UKIO02000074.1|Klebsiella_pneumoniae MQKSNFKGPRRAVKCFNCGKEGHIARNCRAPRKKGCWKCGREGHQMKD------- 48
 NZ_UKJH02000061.1|Klebsiella_pneumoniae MQKSNFKN-RKIVKCFNCGKEGHIARNCRAPRKKGCWKCGREGHQM--------- 45
 NZ_SAWN01000136.1|Neisseria_gonorrhoeae MQRGNFKGPKRIIKCFNCGKEGHIAKNCRAPRKKGCWKCGKEGHQ---------- 45
```

### HIV-1 Gag p17, Matrix Protein

```
                  NC_001802.1|GAG_MPp17 GARASVLSGGELDRWEKIRLRPGGKKKYKLKHIVWASRELERFAVNPGLLETSEGCRQIL 60
     NZ_WWEN01000019.1|Thalassovita_mangrovi GARASVLSGGELDKWEKIRLRPGGKKQYKLKHIVWASRELERFAVNPGLLETSEGCRQIL 60
 NZ_BMPW01000058.1|Actinoplanes_campanulatus GARASVLSGGELDKWEKIRLRPGGKKQYKLKHIVWASRELERFAVNPGLLETSEGCRQIL 60
    NZ_BMDT01000023.1|Enterococcus_alcedinis GARASVLSGGELDKWEKIRLRPGGKKQYKLKHIVWASRELERFAVNPGLLETSEGCRQIL 60
 NZ_BMOE01000030.1|Deinococcus_aquiradiocola GARASVLSGGELDKWEKIRLRPGGKKQYKLKHIVWASRELERFAVNPGLLETSEGCRQIL 60
   NZ_JAHVIU010000020.1|Bacillus_velezensis GARASILRGGKLDKWEKIRLRPGGKKHYMLKHLVWASRELERFAVNPGLLETAEGCKQII 60
NZ_JAHVIB010000060.1|Photobacterium_rosenbergii GARASILRGGKLDKWEKIRLRPGGKKHYMLKHLVWASRELERFAVNPGLLETAEGCKQII 60
  NZ_UVID02000086.1|Burkholderia_pseudomallei GARASILRGGKLDTWEKIRLRPGGKKHYMLKHLVWASRELERFALNPGLLETSEGCKQII 60
NZ_JAHVIB010000058.1|Photobacterium_rosenbergii GARASILRGGKLGKWEKIKLRPVGKKHYMLKHLVWASRELERFAVNPGLLETAEGCRQII 60
   NZ_JAHVKH010000037.1|Vibrio_hangzhouensis GARASILRGGKLGKWEKIKLRPVGKKHYMLKHLVWASRELERFAVNPGLLETAEGCRQII 60

                  NC_001802.1|GAG_MPp17 GQLQPSLQTGSEELRSLYNTVATLYCVHQRIEIKDTKEALDKIEEEQNKSKKKAQQAAAD 120
     NZ_WWEN01000019.1|Thalassovita_mangrovi GQLQPSLQTGSEELRSLYNTIAVLYCVHQRIDVKDTKEALDKIEEEQNKSKKKAQQAAAD 120
 NZ_BMPW01000058.1|Actinoplanes_campanulatus GQLQPSLQTGSEELRSLYNTIAVLYCVHQRIDVKDTKEALDKIEEEQNKSKKKAQQAAAD 120
    NZ_BMDT01000023.1|Enterococcus_alcedinis GQLQPSLQTGSEELRSLYNTIAVLYCVHQRIDVKDTKEALDKIEEEQNKSKKKAQQAAAD 120
 NZ_BMOE01000030.1|Deinococcus_aquiradiocola GQLQPSLQTGSEELRSLYNTIAVLYCVHQRIDVKDTKEALDKIEEEQNKSKKKAQQAAAD 120
   NZ_JAHVIU010000020.1|Bacillus_velezensis NQLQPALQTGTEELRSLYNTVAVLYCVHKGIEVRDTKEALDKI----------------- 103
NZ_JAHVIB010000060.1|Photobacterium_rosenbergii NQLQPALQTGTEELRSLYNTVAVLYCVHKGIEVRDTKEALDKI----------------- 103
  NZ_UVID02000086.1|Burkholderia_pseudomallei QQLQPALQTGTEELRSLFNTVATLYCVHKDIKVADTKEALDKI----------------- 103
NZ_JAHVIB010000058.1|Photobacterium_rosenbergii NQLQPALQTGTEELRSLYNTVAVLYCVHKEIEVRDTKEALDKI----------------- 103
   NZ_JAHVKH010000037.1|Vibrio_hangzhouensis NQLQPALQTGTEELRSLYNTVAVLYCVHKEIEVRDTKEALDKI----------------- 103

                  NC_001802.1|GAG_MPp17 TGHSNQVSQNY 131
     NZ_WWEN01000019.1|Thalassovita_mangrovi TGNNSQVSQNY 131
 NZ_BMPW01000058.1|Actinoplanes_campanulatus TGNNSQVSQNY 131
    NZ_BMDT01000023.1|Enterococcus_alcedinis TGNNSQVSQNY 131
 NZ_BMOE01000030.1|Deinococcus_aquiradiocola TGNNSQVSQNY 131
   NZ_JAHVIU010000020.1|Bacillus_velezensis ----------- 103
NZ_JAHVIB010000060.1|Photobacterium_rosenbergii ----------- 103
  NZ_UVID02000086.1|Burkholderia_pseudomallei ----------- 103
NZ_JAHVIB010000058.1|Photobacterium_rosenbergii ----------- 103
   NZ_JAHVKH010000037.1|Vibrio_hangzhouensis ----------- 103
```

### HIV-1 Reverse Transcriptase, p51

```
                    NC_001802.1|POL_RTp51 PISPIETVPVKLKPGMDGPKVKQWPLTEEKIKALVEICTEMEKEGKISKIGPENPYNTPV 60
  NZ_BMDT01000023.1|Enterococcus_alcedinis PISPIETVPVKLKPGMDGPKVKQWPLTEEKIKALVEICTEMEKEGKISKIGPENPYNTPV 60
NZ_BMOE01000030.1|Deinococcus_aquiradiocola PISPIETVPVKLKPGMDGPKVKQWPLTEEKIKALVEICTEMEKEGKISKIGPENPYNTPV 60
NZ_BMPW01000058.1|Actinoplanes_campanulatus PISPIETVPVKLKPGMDGPKVKQWPLTEEKIKALVEICTEMEKEGKISKIGPENPYNTPV 60
 NZ_CAKLPD010000006.1|Klebsiella_pneumoniae PISPIETVPVKLKPGMDGPKVKQWPLTEEKIKALVEICTEMEKEGKISKIGPENPYNTPI 60
    NZ_WWEN01000020.1|Thalassovita_mangrovi ------------------------------------------------KIGPENPYNTPV 12
     NZ_PGCT01000032.1|Enterococcus_faecium PISPIGTVPVKLKPGMDGPKVKQWPLTEEKIKALTAICEEMEKEGKISKIGPENPYNTPV 60
     NZ_PGCR01000088.1|Enterococcus_faecium PISPIETVPVKLKPGMDGPKVKQWPLTEEKIKALTEICNEMEKEGKITKIGPENPYNTPI 60
  NZ_QEXE01000073.1|Listeria_monocytogenes PISPIETIPVKLKPGMDGPRVKQWPLTEEKIKALTEICEDMEKEGKITRIGPENPYNTPI 60
       NZ_NGBP02000187.1|Escherichia_coli PISPIETVPVKLKPGMDGPRVKQWPLTEEKIKALTEICDEMKKEGKITKIGPENPYNTPI 60

                    NC_001802.1|POL_RTp51 FAIKKKDSTKWRKLVDFRELNKRTQDFWEVQLGIPHPAGLKKKKSVTVLDVGDAYFSVPL 120
  NZ_BMDT01000023.1|Enterococcus_alcedinis FAIKKKDSTKWRKLVDFRELNKRTQDFWEVQLGIPHPAGLKQKKSVTVLDVGDAYFSVPL 120
NZ_BMOE01000030.1|Deinococcus_aquiradiocola FAIKKKDSTKWRKLVDFRELNKRTQDFWEVQLGIPHPAGLKQKKSVTVLDVGDAYFSVPL 120
NZ_BMPW01000058.1|Actinoplanes_campanulatus FAIKKKDSTKWRKLVDFRELNKRTQDFWEVQLGIPHPAGLKQKKSVTVLDVGDAYFSVPL 120
 NZ_CAKLPD010000006.1|Klebsiella_pneumoniae FAIKKKDSTKWRKLVDFRELNKRTQDFWEVQLGIPHPAGLKKKKSVTVLDVGDAYFSVPL 120
    NZ_WWEN01000020.1|Thalassovita_mangrovi FAIKKKDSTKWRKLVDFRELNKRTQDFWEVQLGIPHPAGLKQKKSVTVLDVGDAYFSVPL 72
     NZ_PGCT01000032.1|Enterococcus_faecium FAIKKKDSTKWRKLVDFRELNKRTQDFWEVQLGIPHPAGLKKKKSVTVLDVGDAYFSVPL 120
     NZ_PGCR01000088.1|Enterococcus_faecium FAIKKKDSTKWRKLVDFRELNKRTQDFWEVQLGIPHPAGLKKKKSVTVLDVGDAYFSVPL 120
  NZ_QEXE01000073.1|Listeria_monocytogenes FAIKRKDGTKWRKLVDFRELNKRTQDFWEVQLGIPHPAGLKKKKSMTVLDVGDAYFSVPL 120
       NZ_NGBP02000187.1|Escherichia_coli FAIKKKNSTKWRKLVDFRELNKRTQDFWEVQLGIPHPAGLKKNKSVTVLDVGDAYFSVPL 120

                    NC_001802.1|POL_RTp51 DEDFRKYTAFTIPSINNETPGIRYQYNVLPQGWKGSPAIFQSSMTKILEPFRKQNPDIVI 180
  NZ_BMDT01000023.1|Enterococcus_alcedinis DKDFRKYTAFTIPSINNETPGIRYQYNVLPQGWKGSPAIFQCSMTKILEPFRKQNPDIVI 180
NZ_BMOE01000030.1|Deinococcus_aquiradiocola DKDFRKYTAFTIPSINNETPGIRYQYNVLPQGWKGSPAIFQCSMTKILEPFRKQNPDIVI 180
NZ_BMPW01000058.1|Actinoplanes_campanulatus DKDFRKYTAFTIPSINNETPGIRYQYNVLPQGWKGSPAIFQCSMTKILEPFRKQNPDIVI 180
 NZ_CAKLPD010000006.1|Klebsiella_pneumoniae DQDFRKYTAFTIPSINNETPGVRYQYNVLPQGWKGSPAIFQSSMTKILEPFRKQNPDIVI 180
    NZ_WWEN01000020.1|Thalassovita_mangrovi DKDFRKYTAFTIPSINNETPGIRYQYNVLPQGWKGSPAIFQCSMTKILEPFRKQNPDIVI 132
     NZ_PGCT01000032.1|Enterococcus_faecium DEGFRRYTAFTIPSINNETPGIRYQYNVLPQGWKGSPAIFQSSMTKILEPFRTRNPDIII 180
     NZ_PGCR01000088.1|Enterococcus_faecium DKEFRKYTAFTIPSINNETPGIRYQYNVLPQGWKGSPSIFQSSMIKILEPFRAKNPEMVI 180
  NZ_QEXE01000073.1|Listeria_monocytogenes DESFRKYTAFTIPSINNSTPGIRYQYNVLPQGWKGSPAIFQSSMTKILQPFRDENPDIVI 180
       NZ_NGBP02000187.1|Escherichia_coli DENFRKYTAFTIPSVNNETPGIRYQYNVLPQGWKGSPAIFQSSMTKILEPFRAQNPEIVI 180

                    NC_001802.1|POL_RTp51 YQYMDDLYVGSDLEIGQHRTKIEELRQHLLRWGLTTPDKKHQKEPPFLWMGYELHPDKWT 240
  NZ_BMDT01000023.1|Enterococcus_alcedinis YQYMDDLYVGSDLEIGQHRTKIEELRQHLLRWGFTTPDKKHQKEPPFLWMGYELHPDKWT 240
NZ_BMOE01000030.1|Deinococcus_aquiradiocola YQYMDDLYVGSDLEIGQHRTKIEELRQHLLRWGFTTPDKKHQKEPPFLWMGYELHPDKWT 240
NZ_BMPW01000058.1|Actinoplanes_campanulatus YQYMDDLYVGSDLEIGQHRTKIEELRQHLLRWGFTTPDKKHQKEPPFLWMGYELHPDKWT 240
 NZ_CAKLPD010000006.1|Klebsiella_pneumoniae YQYMDDLYVGSDLEIGQHRIKIEELRQHLLKWGFTTPDKKHQKEPPFLWMGYELHPDKWT 240
    NZ_WWEN01000020.1|Thalassovita_mangrovi YQYMDDLYVGSDLEIGQHRTKIEELRQHLLRWGFTTPDKKHQKEPPFLWMGYELHPDKWT 192
     NZ_PGCT01000032.1|Enterococcus_faecium YQYMDDLYVGSDLEIGQHRAKIEELREHLLRWGLTTPDKKHQKEPPFLWMGYELHPDKWT 240
     NZ_PGCR01000088.1|Enterococcus_faecium YQYMDDLYVGSDLEIGQHRAKIEELREHLLRWGFTTPDKKHQKEPPFLWMGYELHPDKWT 240
  NZ_QEXE01000073.1|Listeria_monocytogenes YQYVDDLLVGSDLEIGQHRAKIEQLREHLLRWGLTTPDKKHQKEPPFLWMGYELHPDKWT 240
       NZ_NGBP02000187.1|Escherichia_coli YQYVDDLYVGSDLEIGQHRAKVEKLRDHLLKWGFTTPDKKHQKEPPFLWMGYELHPDKWT 240

                    NC_001802.1|POL_RTp51 VQPIVLPEKDSWTVNDIQKLVGKLNWASQIYPGIKVRQLCKLLRGTKALTEVIPLTEEAE 300
  NZ_BMDT01000023.1|Enterococcus_alcedinis VQPIVLPEKDSWTVNDIQKLVGKLNWASQIYAGIKVRQLCKLLRGTKALTEVVPLTEEAE 300
NZ_BMOE01000030.1|Deinococcus_aquiradiocola VQPIVLPEKDSWTVNDIQKLVGKLNWASQIYAGIKVRQLCKLLRGTKALTEVVPLTEEAE 300
NZ_BMPW01000058.1|Actinoplanes_campanulatus VQPIVLPEKDSWTVNDIQKLVGKLNWASQIYAGIKVRQLCKLLRGTKALTEVVPLTEEAE 300
 NZ_CAKLPD010000006.1|Klebsiella_pneumoniae VQPIKLPEKDSWTVNDIQKLVGKLNWASQIYAGIKVRQLCKLLRGAKALTEVVPLTKEAE 300
    NZ_WWEN01000020.1|Thalassovita_mangrovi VQPIVLPEKDSWTVNDIQKLVGKLNWASQIYAGIKVRQLCKLLRGTKALTEVVPLTEEAE 252
     NZ_PGCT01000032.1|Enterococcus_faecium VQPIVLPEKDSWTVNDIQKLVGKLNWASQIYQGIKVKQLCKLLRGAKALTDIVPLTEEAE 300
     NZ_PGCR01000088.1|Enterococcus_faecium VQPIQLPDKDSWTVNDIQKLVGKLNWASQIYPGIKVRQLCKLLRGAKALTDIVPLTEEAE 300
  NZ_QEXE01000073.1|Listeria_monocytogenes VQPIQLPEKDSWTVNDIQKLVGKLNWASQIYPGICTKQLCRLLRGTKALTDIVPLTEEAE 300
       NZ_NGBP02000187.1|Escherichia_coli VQQIQLPEKESWTVNDIQKLVGKLNWASQIYAGIKVKQLCRLLRGAKALTDIVPLTEEAE 300

                    NC_001802.1|POL_RTp51 LELAENREILKEPVHGVYYDPSKDLIAEIQKQGQGQWTYQIYQEPFKNLKTGKYARMRGA 360
  NZ_BMDT01000023.1|Enterococcus_alcedinis LELAENREILKEPVHGVYYDPSKDLIAEIQKQGQGQWTYQIYQEPFKNLKTGKYARMKGA 360
NZ_BMOE01000030.1|Deinococcus_aquiradiocola LELAENREILKEPVHGVYYDPSKDLIAEIQKQGQGQWTYQIYQEPFKNLKTGKYARMKGA 360
NZ_BMPW01000058.1|Actinoplanes_campanulatus LELAENREILKEPVHGVYYDPSKDLIAEIQKQGQGQWTYQIYQEPFKNLKTGKYARMKGA 360
 NZ_CAKLPD010000006.1|Klebsiella_pneumoniae LELAENREILKEPVHGVYYDPSKELIAEIQKQGEGQWTYQIYQEQFKNLKTGKYARMRGT 360
    NZ_WWEN01000020.1|Thalassovita_mangrovi LELAENREILKEPVHGVYYDPSKDLIAEIQKQGQGQWTYQIYQEPFKNLKTGKYARMKGA 312
     NZ_PGCT01000032.1|Enterococcus_faecium LELAENREILKEPVHGVYYDPSKDLIAEIQKQGHDQWTYQIYQEPFKNLKTGKYAKMRTA 360
     NZ_PGCR01000088.1|Enterococcus_faecium LELAENREILKEPVHGVYYDPSKDLIAEIQKQGHDQWTYQIYQDPFKNLKTGKYAKMRTA 360
  NZ_QEXE01000073.1|Listeria_monocytogenes LELAENREILKEPVHGVYYDPSKDLIAEIQKQGDDQWTYQIYQEPFKNLKTGKYAKMRTA 360
       NZ_NGBP02000187.1|Escherichia_coli LELAENREILKEPVHGVYYDPSKDLVAEIQKQGNDQWTYQIYQEPFKNLKTGKYAKRRTA 360
```

>>>

### HIV-1 Reverse Transcriptase, Pol p51 (continuation)

```
NC_001802.1|POL_RTp51                     HTNDVKQLTEAVQKITTESIVIWGKTPKFKLPIQKETWETWWTEYWQATWIPEWEFVNTP 420
NZ_BMDT01000023.1|Enterococcus_alcedinis  HTNDVKQLTEAVQKIATESIVIWGKTPKFKLPIQKETWEAWWTEYWQATWIPEWEFVNTP 420
NZ_BMOE01000030.1|Deinococcus_aquiradiocola HTNDVKQLTEAVQKIATESIVIWGKTPKFKLPIQKETWEAWWTEYWQATWIPEWEFVNTP 420
NZ_BMPW01000058.1|Actinoplanes_campanulatus HTNDVKQLTEAVQKIATESIVIWGKTPKFKLPIQKETWEAWWTEYWQATWIPEWEFVNTP 420
NZ_CAKLPD010000006.1|Klebsiella_pneumoniae HTNDVKQLTEAVQKIATESIVIWGKTPKFRLPIHKETWETWWTDYWQATWIPEWEFVNTP 420
NZ_WWEN01000020.1|Thalassovita_mangrovi   HTNDVKQLTEAVQKIATESIVIWGKTPKFKLPIQKETWEAWWTEYWQATWIPEWEFVNTP 372
NZ_PGCT01000032.1|Enterococcus_faecium    HTNDIKQLTEAVQKIAMESIVIWGKTPKFRLPIQKGTWETWWTDYWQATWIPDWEFVNTP 420
NZ_PGCR01000088.1|Enterococcus_faecium    HTNDVKQLAEAVQKIAIESIVIWGKTPKFRLPIQKETWETWWTDYWQATWIPDWEFVNTP 420
NZ_QEXE01000073.1|Listeria_monocytogenes  HTNDVKQLTEAVQKIALESIVIWGKTPKFRLPIQKETWEIWWTDYWQATWIPEWEFVNTP 420
NZ_NGBP02000187.1|Escherichia_coli        HTNDVQQLTEVVQKISRESIVIWGKTPKFRLPIQKETWETWWADYWQATWIPDWEFVNTP 420

NC_001802.1|POL_RTp51                     PLVKLWYQLEKEPIVGAETFYVDGAANRETKLGKAGYVTNRGRQKVVTLTDTTNQKTELQ 480
NZ_BMDT01000023.1|Enterococcus_alcedinis  PLVKLWYQLEKEPIIGAETFYVDGAANRETKLGKAGYVTDRGRQKVVPLTDTTNQKTELQ 480
NZ_BMOE01000030.1|Deinococcus_aquiradiocola PLVKLWYQLEKEPIIGAETFYVDGAANRETKLGKAGYVTDRGRQKVVPLTDTTNQKTELQ 480
NZ_BMPW01000058.1|Actinoplanes_campanulatus PLVKLWYQLEKEPIIGAETFYVDGAANRETKLGKAGYVTDRGRQKVVPLTDTTNQKTELQ 480
NZ_CAKLPD010000006.1|Klebsiella_pneumoniae PLVKLWYQLEKEPIEGAETFYVDGAANRETKIGKAGYVTNRGRQKVVPLTDTTNQKTELQ 480
NZ_WWEN01000020.1|Thalassovita_mangrovi   PLVKLWYQLEKEPIIGAETFYVDGAANRETKLGKAGYVTDRGRQKVVPLTDTTNQKTELQ 432
NZ_PGCT01000032.1|Enterococcus_faecium    PLVKLWYQLEKEPIAGAETFYVD------------------------------------- 443
NZ_PGCR01000088.1|Enterococcus_faecium    PLVKLWYQLVKEPIIGAETFYVDG------------------------------------ 444
NZ_QEXE01000073.1|Listeria_monocytogenes  PLVKLWYQLEKEPIAEAET----------------------------------------- 439
NZ_NGBP02000187.1|Escherichia_coli        PLVKLWYQLEKDPIAGAETFY--------------------------------------- 441

NC_001802.1|POL_RTp51                     AIYLALQDSGLEVNIVTDSQYALGIIQAQPDQSESELVNQIIEQLIKKEKVYLAWVPAHK 540
NZ_BMDT01000023.1|Enterococcus_alcedinis  AIHLALQDSGLEVNIVTDSQYALGIIQAQPDKSESELVSQIIEQLIKKEKVYLAWVPAHK 540
NZ_BMOE01000030.1|Deinococcus_aquiradiocola AIHLALQDSGLEVNIVTDSQYALGIIQAQPDKSESELVSQIIEQLIKKEKVYLAWVPAHK 540
NZ_BMPW01000058.1|Actinoplanes_campanulatus AIHLALQDSGLEVNIVTDSQYALGIIQAQPDKSESELVSQIIEQLIKKEKVYLAWVPAHK 540
NZ_CAKLPD010000006.1|Klebsiella_pneumoniae AIYLALQDSELEVNIVTDSQYALGIIQAQPDKSESELVNQIIEQLIKKEKIYLAWVPAHK 540
NZ_WWEN01000020.1|Thalassovita_mangrovi   AIHLALQDSGLEVNIVTDSQYALGIIQAQPDKSESELVSQIIEQLIKKEKVYLAWVPAHK 492
NZ_PGCT01000032.1|Enterococcus_faecium    ------------------------------------------------------------ 443
NZ_PGCR01000088.1|Enterococcus_faecium    ------------------------------------------------------------ 444
NZ_QEXE01000073.1|Listeria_monocytogenes  ------------------------------------------------------------ 439
NZ_NGBP02000187.1|Escherichia_coli        ------------------------------------------------------------ 441

NC_001802.1|POL_RTp51                     GIGGNEQVDKLVSAGIRKVL 560
NZ_BMDT01000023.1|Enterococcus_alcedinis  GIGGNEQVDKLVSAGIRKVL 560
NZ_BMOE01000030.1|Deinococcus_aquiradiocola GIGGNEQVDKLVSAGIRKVL 560
NZ_BMPW01000058.1|Actinoplanes_campanulatus GIGGNEQVDKLVSAGIRKVL 560
NZ_CAKLPD010000006.1|Klebsiella_pneumoniae GIGGNEQVDKLVSTGIRKVL 560
NZ_WWEN01000020.1|Thalassovita_mangrovi   GIG----------------- 495
NZ_PGCT01000032.1|Enterococcus_faecium    -------------------- 443
NZ_PGCR01000088.1|Enterococcus_faecium    -------------------- 444
NZ_QEXE01000073.1|Listeria_monocytogenes  -------------------- 439
NZ_NGBP02000187.1|Escherichia_coli        -------------------- 441
```

### HIV-1 RNAse H, p15

```
                NC_001802.1|POL_RNASEp15 YVDGAANRETKLGKAGYVTNRGRQKVVTLTDTTNQKTELQAIYLALQDSGLEVNIVTDSQ  60
  NZ_CAKLPD010000006.1|Klebsiella_pneumoniae YVDGAANRETKIGKAGYVTNRGRQKVVPLTDTTNQKTELQAIYLALQDSELEVNIVTDSQ  60
   NZ_UVFZ02000097.1|Burkholderia_pseudomallei YVDGAASRETKLGKAGYVTDRGRQKVVSLTETTNQKTELHAIHLALQDSGSEVNIVTDSQ  60
   NZ_BMPW01000058.1|Actinoplanes_campanulatus YVDGAANRETKLGKAGYVTDRGRQKVVPLTDTTNQKTELQAIHLALQDSGLEVNIVTDSQ  60
      NZ_BMDT01000023.1|Enterococcus_alcedinis YVDGAANRETKLGKAGYVTDRGRQKVVPLTDTTNQKTELQAIHLALQDSGLEVNIVTDSQ  60
   NZ_BMOE01000030.1|Deinococcus_aquiradiocola YVDGAANRETKLGKAGYVTDRGRQKVVPLTDTTNQKTELQAIHLALQDSGLEVNIVTDSQ  60
NZ_NKQS01000691.1|Methylobacterium_radiotolerans YVDGAANRETKRGKAGYVTDRGRQKIVPLNETTNQKAELQAIQLALQDSGSEVNIVTDSQ  60
          NZ_UINV01000086.1|Escherichia_coli YVDGAANRETKAGKAGYVTDRGRQKVISFTETTNQKAELQAIQLALQDSGSEVNIVTDSQ  60
      NZ_WWEN01000020.1|Thalassovita_mangrovi YVDGAANRETKLGKAGYVTDRGRQKVVPLTDTTNQKTELQAIHLALQDSGLEVNIVTDSQ  60
      NZ_UKIX02000064.1|Klebsiella_pneumoniae YVDGAANRDTKMGKAGYVTDKGRQKVVSLTETTNQKTELHAIQLALQDSGPEVNIVTDSQ  60

                NC_001802.1|POL_RNASEp15 YALGIIQAQPDQSESELVNQIIEQLIKKEKVYLAWVPAHKGIGGNEQVDKLVSAGIRKVL  120
  NZ_CAKLPD010000006.1|Klebsiella_pneumoniae YALGIIQAQPDKSESELVNQIIEQLIKKEKIYLAWVPAHKGIGGNEQVDKLVSTGIRKVL  120
   NZ_UVFZ02000097.1|Burkholderia_pseudomallei YALGIIQAQPDMSESEVVNQIIEELIKKERVYLSWVPAHKGIGGNEQVDKLVSSGIRKVL  120
   NZ_BMPW01000058.1|Actinoplanes_campanulatus YALGIIQAQPDKSESELVSQIIEQLIKKEKVYLAWVPAHKGIGGNEQVDKLVSAGIRKVL  120
      NZ_BMDT01000023.1|Enterococcus_alcedinis YALGIIQAQPDKSESELVSQIIEQLIKKEKVYLAWVPAHKGIGGNEQVDKLVSAGIRKVL  120
   NZ_BMOE01000030.1|Deinococcus_aquiradiocola YALGIIQAQPDKSESELVSQIIEQLIKKEKVYLAWVPAHKGIGGNEQVDKLVSAGIRKVL  120
NZ_NKQS01000691.1|Methylobacterium_radiotolerans YALGIIQAQPDKSESELVNQIIEQLINKERIYLSWVPAHKGIGGNEQVDKLVSTGIRKVL  120
          NZ_UINV01000086.1|Escherichia_coli YALGIIHALPDKSESELVNKIIELLIQKERVYLSWVPAHKGIGGNEQVDKLVSTGIRKVL  120
      NZ_WWEN01000020.1|Thalassovita_mangrovi YALGIIQAQPDKSESELVSQIIEQLIKKEKVYLAWVPAHKGIG-----------------  103
      NZ_UKIX02000064.1|Klebsiella_pneumoniae YALGIIQAQPDKSESELVNQIIEQLIKKERVYLSWVPAHK--------------------  100
```

### HIV-1 Integrase. p31

```
                  NC_001802.1|POL_INTEGp31  FLDGIDKAQDEHEKYHSNWRAMASDFNLPPVVAKEIVASCDKCQLKGEAMHGQVDCSPGI  60
NZ_BMPW01000058.1|Actinoplanes_campanulatus  FLDGIDKAQEEHEKYHSNWRAMASDFNLPPVVAKEIVASCDKCQLKGEAMHGQVDCSPGI  60
       NZ_UINV01000086.1|Escherichia_coli  FLDGIDKAQEEHEKYHSNWRAMASEFNLPPIVAKEIVASCDKCQLKGEAMHGQVDCSPGI  60
  NZ_RADD01000036.1|Pseudomonas_aeruginosa  FLDGIDKAQEEHERYHNNWRAMASDFNLPPIVAKEIVASCDKCQQKGEAMHGQVDCSPGI  60
  NZ_RACX01000118.1|Pseudomonas_aeruginosa  FLDGIDKAQEEHERYHNNWRAMASDFNLPPIVAKEIVASCDKCQQKGEAMHGQVDCSPGI  60
NZ_UVFZ02000097.1|Burkholderia_pseudomallei  FLDGIDKAQEDHERYHSNWRTMASDFNLPPIVAKEIVANCDKCQLKGEAMHGQVDCSPGI  60
 NZ_JACAWM010000058.1|Klebsiella_pneumoniae  -------------------------------------------------MHGQVDCSPGI  11
 NZ_CAKLPD010000006.1|Klebsiella_pneumoniae  FLDGIDKAQEDHDKYHSNWRAMASDFNLPPIVAKEIVASCDKCQLKGEAMHGQVDCSPGI  60
  NZ_BMDT01000023.1|Enterococcus_alcedinis  FLDGIDKAQEEHEKYHSNWRAMASDFNLPPVVAKEIVASCDKCQLKGEAMHGQVDCSPGI  60
NZ_BMOE01000030.1|Deinococcus_aquiradiocola  FLDGIDKAQEEHEKYHSNWRAMASDFNLPPVVAKEIVASCDKCQLKGEAMHGQVDCSPGI  60

                  NC_001802.1|POL_INTEGp31  WQLDCTHLEGKVILVAVHVASGYIEAEVIPAETGQETAYFLLKLAGRWPVKTIHTDNGSN  120
NZ_BMPW01000058.1|Actinoplanes_campanulatus  WQLDCTHLEGKVILVAVHVASGYIEAEVIPAETGQETAYFLLKLAGRWPVKTVHTDNGSN  120
       NZ_UINV01000086.1|Escherichia_coli  WQLDCTHLEGKVILVAVHVASGYMEAEVIPAETGQETAYFILKLAGRWPVKTIHTDNGSN  120
  NZ_RADD01000036.1|Pseudomonas_aeruginosa  WQIDCTHLEGKIIIVAVHVASGYIEAEVIPAETGQETAYFILKLAARWPVKVIHTDNGSN  120
  NZ_RACX01000118.1|Pseudomonas_aeruginosa  WQIDCTHLEGKIIIVAVHVASGYIEAEVIPAETGQETAYFILKLAARWPVKVIHTDNGSN  120
NZ_UVFZ02000097.1|Burkholderia_pseudomallei  WQLDCTHLEGKVILVAVHVASGYIEAEVIPAETGQETAYFLLKLAGRWPVKVIHTDNGSN  120
 NZ_JACAWM010000058.1|Klebsiella_pneumoniae  WQLDCTHLEGKVILVAVHVASGYIEAEVIPAETGQETAYFLLKLAGRWPVKTVHTDNGSN  71
 NZ_CAKLPD010000006.1|Klebsiella_pneumoniae  WQLDCTHLEGKIILVAVHVASGYIEAEVIPAETGQETAYFILKLAGRWPVTTIHTDNGPN  120
  NZ_BMDT01000023.1|Enterococcus_alcedinis  WQLDCTHLEGKVILVAVHVASGYIEAEVIPAETGQETAYFLLKLAGRWPVKTVHTDNGSN  120
NZ_BMOE01000030.1|Deinococcus_aquiradiocola  WQLDCTHLEGKVILVAVHVASGYIEAEVIPAETGQETAYFLLKLAGRWPVKTVHTDNGSN  120

                  NC_001802.1|POL_INTEGp31  FTGATVRAACWWAGIKQEFGIPYNPQSQGVVESMNKELKKIIGQVRDQAEHLKTAVQMAV  180
NZ_BMPW01000058.1|Actinoplanes_campanulatus  FTSTTVKAACWWAGIKQEFGIPYNPQSQGVIESMNKELKKIIGQVRDQAEHLKTAVQMAV  180
       NZ_UINV01000086.1|Escherichia_coli  FTSTVVKAACWWAGIQQEFGIPYNPQSQGVVESMNKELKKIIEQLRDQAEHLKTAVQMAV  180
  NZ_RADD01000036.1|Pseudomonas_aeruginosa  FTSAAVKAACWWANVTQEFGIPYNPQSQGVVESMNKELKKIIGQIRDQAEHLKTAVQMAV  180
  NZ_RACX01000118.1|Pseudomonas_aeruginosa  FTSAAVKAACWWANVTQEFGIPYNPQSQGVVESMNKELKKIIGQIRDQAEHLKTAVQMAV  180
NZ_UVFZ02000097.1|Burkholderia_pseudomallei  FTSTAVKAACWWANVQQEFGIPYNPQSQGVVESMNKELKKIIGQVREQAEHLKTAVQMAV  180
 NZ_JACAWM010000058.1|Klebsiella_pneumoniae  FTSTTVKAACWWAGIKQEFGIPYNPQSQGVIESMNKELKKIIGQVRDQAEHLKTAVQMAV  131
 NZ_CAKLPD010000006.1|Klebsiella_pneumoniae  FISSTVKAACWWAGIKQEFGIPYNPQSQGVVESMNKELKKIIGQVRDQAEHLRTAVQMAV  180
  NZ_BMDT01000023.1|Enterococcus_alcedinis  FTSTTVKAACWWAGIKQEFGIPYNPQSQGVIESMNKELKKIIGQVRDQAEHLKTAVQMAV  180
NZ_BMOE01000030.1|Deinococcus_aquiradiocola  FTSTTVKAACWWAGIKQEFGIPYNPQSQGVIESMNKELKKIIGQVRDQAEHLKTAVQMAV  180

                  NC_001802.1|POL_INTEGp31  FIHNFKRKGGIGGYSAGERIVDIIATDIQTKELQKQITKIQNFRVYYRDSRNPLWKGPAK  240
NZ_BMPW01000058.1|Actinoplanes_campanulatus  FIHNFKRKGGIGGYSAGERIVDIIATDIQTKELQKQITKIQNFRVYYRDSRDPVWKGPAK  240
       NZ_UINV01000086.1|Escherichia_coli  FVHNFKRKGGIGGYSAGERIIDIIATDIQTKELQKQITKIQNFRVYYRDSRDPIWKGPAK  240
  NZ_RADD01000036.1|Pseudomonas_aeruginosa  FIHNFKRKGGIGRYSAGERIIDIIASDIQTKELQKQIIKIQNFRVYYRDSRDPIWKGPAK  240
  NZ_RACX01000118.1|Pseudomonas_aeruginosa  FIHNFKRKGGIGRYSAGERIIDIIASDIQTKELQKQIIKIQNFRVYYRDSRDPIWKGPAK  240
NZ_UVFZ02000097.1|Burkholderia_pseudomallei  FIHNFKRKGGIGGYSAGERIIDIIATDIQTKELQKQITKIQNFRVYYRDSRDPIWKGPAK  240
 NZ_JACAWM010000058.1|Klebsiella_pneumoniae  FIHNFKRKGGIGGYSAGERIVDIIATDIQTKELQKQITKIQNFRVYYRDSRDPVWKGPAK  191
 NZ_CAKLPD010000006.1|Klebsiella_pneumoniae  FIHNFKRKGGIGGYSAGERIVDIIATDIQTKELQKQITKIQNFRVYYRDSRDPLWKGPAK  240
  NZ_BMDT01000023.1|Enterococcus_alcedinis  FIHNFKRKGGIGGYSAGERIVDIIATDIQTKELQKQITKIQIF-----------------  223
NZ_BMOE01000030.1|Deinococcus_aquiradiocola  FIHNFKRKGGIGGYSAGERIVDIIATDIQTKELQKQITKIQIF-----------------  223

                  NC_001802.1|POL_INTEGp31  LLWKGEGAVVIQDNSDIKVVPRRKAKIIRDYGKQMAGDDCVASRQDED  288
NZ_BMPW01000058.1|Actinoplanes_campanulatus  LLWKGEGAVVIQDNSDIKVVPRRKAKIIRDYGKQMAGDDCVASRQDED  288
       NZ_UINV01000086.1|Escherichia_coli  LLWKGEGAVVIQDNSDIKVVPRRKAKIIRDYGKQMAGTDCVAGRQDED  288
  NZ_RADD01000036.1|Pseudomonas_aeruginosa  LLWKGEGAVVIQDNSEIKVIPRRKAKIIRDYGKQMAGDDCVAGRQDED  288
  NZ_RACX01000118.1|Pseudomonas_aeruginosa  LLWKGEGAVVIQDNSEIKVIPRRKAKIIRDYGKQMAGDDCVAGRQDEN  288
NZ_UVFZ02000097.1|Burkholderia_pseudomallei  LLWKGEGAVVIQDNSDIKV-----------------------------  259
 NZ_JACAWM010000058.1|Klebsiella_pneumoniae  LLWKGEGAVVIQDNSDIKVVPRRKAKIIRDYGKQMAGDDCVASRQD--  237
 NZ_CAKLPD010000006.1|Klebsiella_pneumoniae  LLWKGEGAVV--------------------------------------  250
  NZ_BMDT01000023.1|Enterococcus_alcedinis  ------------------------------------------------  223
NZ_BMOE01000030.1|Deinococcus_aquiradiocola  ------------------------------------------------  223
```

### HIV-1 Protease

```
                 NC_001802.1|POL_Protease PQVTLWQRPLVTIKIGGQLKEALLDTGADDTVLEEMSLPGRWKPKMIGGIGGFIKVRQYD  60
  NZ_JACAWM010000050.1|Klebsiella_pneumoniae PQITLWQRPLVTIKIGGQLKEALLDTGADDTVLEEMNLPGRWKPKMIGGIGGFIKVRQYD  60
   NZ_QAGO01000067.1|Acinetobacter_baumannii PQITLWQRPVVTIKVGGQLKEALLDTGADDTVLEDMQLPGRWKPKMIGGIGGFIKVRQYD  60
    NZ_CADDWK010000035.1|Salirhabdus_euzebyi PQITLWQRPLVAIKIGGQLKEALLDTGADDTVLEEMNLPGRWKPKMIGGIGGFIKVRQYD  60
         NZ_WSJI01000036.1|Escherichia_coli PQITLWQRPLVTIKIGGQLKEALLDTGADDTVLEEMNLPGRWKPKMIGGIGGFIKVRQYD  60
   NZ_PYDB01000084.1|Acinetobacter_baumannii PQITLWQRPLVSIKVGGQIKEALLDTGADDTVLEEISLPGKWKPKMIGGIGGFIKVRQYG  60
     NZ_NSJR01000033.1|Enterococcus_faecalis PQITLWQRPVVTIKVGGQLKEALLDTGADDTVLEEMNLPGRWKPKMIGGIGGFIKVRQYD  60
NZ_QTBX01000137.1|Mycobacterium_tuberculosis PQITLWQRPLVSIKVGGQIREALLDTGADDTVLEDINLPGKWKPKMIGGIGGFIKVRQYD  60
NZ_QTBW01000113.1|Mycobacterium_tuberculosis PQITLWQRPLVSIKVGGQIREALLDTGADDTVLEDINLPGKWKPKMIGGIGGFIKVRQYD  60
   NZ_QEHN01000065.1|Acinetobacter_baumannii PQITLWQRPVVTIKVGGQLKEALLDTGADDTVLEEMNLPGRWKPKMIGGIGGFIKVRQYD  60

                 NC_001802.1|POL_Protease QILIEICGHKAIGTVLVGPTPVNIIGRNLLTQIGCTLNF                      99
  NZ_JACAWM010000050.1|Klebsiella_pneumoniae QILIEICGHKAIGTVLVGPTPVNIIGRNLLTQIGCTLNF                      99
   NZ_QAGO01000067.1|Acinetobacter_baumannii EVPIEICGHKAIGTVLIGPTPVNIIGRNLLTQLGCTLNF                      99
    NZ_CADDWK010000035.1|Salirhabdus_euzebyi QIPIEICGHKAIGTVLIGPTPVNIIGRNLLTQIGCTLNF                      99
         NZ_WSJI01000036.1|Escherichia_coli QIPIEICGHKAIGTVLVGPTPVNIIGRNMLTQLGCTLNF                      99
   NZ_PYDB01000084.1|Acinetobacter_baumannii QIPIEICGKKAIGTVLVGPTPVNIIGRNLLTQLGCTLNF                      99
     NZ_NSJR01000033.1|Enterococcus_faecalis EVPIEICGHKVIGTVLIGSTPVNIIGRNLLTQLGCTLNF                      99
NZ_QTBX01000137.1|Mycobacterium_tuberculosis QILIEICGKKAIGTVLVGPTPVNIIGRNMLTQIGCTLNF                      99
NZ_QTBW01000113.1|Mycobacterium_tuberculosis QILIEICGKKAIGTVLVGPTPVNIIGRNMLTQIGCTLNF                      99
   NZ_QEHN01000065.1|Acinetobacter_baumannii QIPIEICGHKVIGTVLIGPTPVNIIGRNLLTQLGCTLNF                      99
```

### HIV-1 p6pol (POL)

```
             NC_001802.1|POL_p6 LQGKAREFSSEQTRANSPTRRELQVWGRDNNSPSEAGADRQGTVSFNF 48
   NZ_WWEN01000019.1|Thalassovita_mangrovi -QGKAREFSSEQTRANSPTRRELQVWGRDNNSLSEAGADRQGTVSFSF 47
NZ_BMOE01000030.1|Deinococcus_aquiradiocola -QGKAREFSSEQTRANSPTRRELQVWGRDNNSLSEAGADRQGTVSFSF 47
  NZ_BMDT01000023.1|Enterococcus_alcedinis -QGKAREFSSEQTRANSPTRRELQVWGRDNNSLSEAGADRQGTVSFSF 47
NZ_BMPW01000058.1|Actinoplanes_campanulatus -QGKAREFSSEQTRANSPTRRELQVWGRDNNSLSEAGADRQGTVSFSF 47
 NZ_JACAWM010000050.1|Klebsiella_pneumoniae ----------------NSPTRRELQVWGRDNNSLSEAGADRQGTVSFSF 33
   NZ_FZLW01000041.1|Helicobacter_acinonychis ---------NSPTSRANSPTSRELQVWGGDSNSPSEAGADRQGDISFSF 40
     NZ_QEHN01000065.1|Acinetobacter_baumannii --------------RANSPTRREPQAWGGDSNSLSEAGADRQGTVSFSF 35
     NZ_QAGO01000067.1|Acinetobacter_baumannii --------------------RRELQAWGGDNNSLSEAGADRQGTVSFSF 29
     NZ_PYHB01000027.1|Enterobacter_hormaechei -----------------SPTRREPQAWGGDRNSLSEAGADRQGTVSFSF 32
```

### HIV-1 Nef

```
                                NC_001802.1|NEF  MGGKWSKSSVIGWPTVRERMRRAEPAADRVGAASRDLEKHGAITSSNTAATNAACAWLEA  60
NZ_JAHVIB010000049.1|Photobacterium_rosenbergii  MGGKWSKSSIIGWPKIRERMRRTEPAADGVGAVSQDLEKRGAVTSNNTAETNADCAWLEA  60
     NZ_JAHVIU010000019.1|Bacillus_velezensis  MGGKWSKSSIIGWPKIRERMRRTEPAADGVGAVSQDLEKRGAVTSNNTAETNADCAWLEA  60
   NZ_JBBXLW010000045.1|Acinetobacter_baumannii  --------------------------------CSRDLEKHGAITSSNTAATNADCAWLEA  28
         NZ_WTYD01000009.1|Qipengyuania_pelagi  ----------------------------------RDLEKHGAITSSNTAATNADCAWLEA  26
      NZ_UKMC02000074.1|Klebsiella_pneumoniae  ----------------RERIRRTKPAAEKVRAASQDLDKYRALTSSNTAATNADCAWLEA  44
      NZ_UKJT02000022.1|Klebsiella_pneumoniae  ---------------------------------SQDLARHGAITTSNTAATNADVAWVEA  27
    NZ_LYGA01000021.1|Acinetobacter_baumannii  ------------------------------------------------------------  0
      NZ_UKIV02000085.1|Klebsiella_pneumoniae  -------------------------AAERVRAASRDLERHEALTTSNTATNNPNCAWLAA  35
      NZ_UKLX02000048.1|Klebsiella_pneumoniae  ------------------------------------------------------------  0

                                NC_001802.1|NEF  QEE-EEVGFPVTPQVPLRPMTYKAAVDLSHFLKEKGGLEGLIHSQRRQDILDLWIYHTQG  119
NZ_JAHVIB010000049.1|Photobacterium_rosenbergii  QEE-EEVGFPVRPQVPLRPMTFKGAFDLSFFLKEKGGLEGLIYSKKRQEILDLWVYHTQG  119
     NZ_JAHVIU010000019.1|Bacillus_velezensis  QEE-EEVGFPVRPQVPLRPMTFKGAFDLSFFLKEKGGLEGLIYSKKRQEILDLWVYHTQG  119
   NZ_JBBXLW010000045.1|Acinetobacter_baumannii  QEE-EEVGFPVTPQVPLRPMTYKAAVDLSHFLKEKRGLEGLIHSQRRQDILDLWIYHTQG  87
         NZ_WTYD01000009.1|Qipengyuania_pelagi  QEE-EEVGFPVTPQVPLRPMTYKAAVDLSHFLKEKGGLEGLIHSQRRQDILDLWIYHTQG  85
      NZ_UKMC02000074.1|Klebsiella_pneumoniae  QEEADEVSFPVRPQVPLRPMTYKAAVDLSFFLKEKGGLDELIYSKKKQDILDL-VYHTQG  103
      NZ_UKJT02000022.1|Klebsiella_pneumoniae  QEEGEEVGFPVRPQVPLRPMTYKGAFDLGFFLKEKGGLDGLIWSKKRQEILDLWVYNTQG  87
    NZ_LYGA01000021.1|Acinetobacter_baumannii  ---------------PLRPMTYQSAVDLSHFLKEKGGLEGLIHSQRRQDILDLWIYHTQG  45
      NZ_UKIV02000085.1|Klebsiella_pneumoniae  QKEKEEVGFPVRPQVPVRPMTYKRAFDLSFFLKEKGGLDGLIYSQQRQDILDLWVYNTQG  95
      NZ_UKLX02000048.1|Klebsiella_pneumoniae  EED-EEVGFPVRPQVPLRPMTYKGAFDLSFFLKEKGGLDRLIYSKKRQDILDLWVYHTQG  59

                                NC_001802.1|NEF  YFPD-QNYTPGPGVRYPLTFGWCYKLVPVEPDKIEEANKGENTSLLHPVSLHGMDDPERE  178
NZ_JAHVIB010000049.1|Photobacterium_rosenbergii  YFPDWQNYTPGPGVRYPLTFGWCFKLVPVDQEAVEEANKGEDNCLLHPVCQHGMDDEHKE  179
     NZ_JAHVIU010000019.1|Bacillus_velezensis  YFPDWQNYTPGPGVRYPLTFGWCFKLVPVEQEAVEEANKGEDNCLLHPVCQHGMDDEHKE  179
   NZ_JBBXLW010000045.1|Acinetobacter_baumannii  YFPD-QNYTPGPGVRYPLTFGWCYKLVPVEPDKVEEANKGENTSLLHPVSLHGMDDPERE  146
         NZ_WTYD01000009.1|Qipengyuania_pelagi  YFPDWQNYTPGPGIRYPLTFGWCYKLVPVEQEKVEEANEGENTRLLHPVSLHGMDDPERE  145
      NZ_UKMC02000074.1|Klebsiella_pneumoniae  FFPDWQNYTPRPKVRYPLTFRWCYKLVPVDPKEVKEANKEENTCLLHPISQHKIEDAHRE  163
      NZ_UKJT02000022.1|Klebsiella_pneumoniae  YFPDWQNYTPGPGVRYPLTFGWCYKLVPVDPREVEEANEGENNCLLHPMSLHGMEDEHKE  147
    NZ_LYGA01000021.1|Acinetobacter_baumannii  YFPDWQNYTPGPGIRYPLTFGWCYKLVPVEQEKVEEANEGENTRLLHPVSLHGMDDPERE  105
      NZ_UKIV02000085.1|Klebsiella_pneumoniae  FFPDWQNYTPGPEVRYPLTFK-CYKLVPVDPSEVEKDNKGKNNSLLHPISQHRIEDTNRE  154
      NZ_UKLX02000048.1|Klebsiella_pneumoniae  YFPDWQNYTPGPGVRYPLTFRWPFKLVPVDPREIEEANEGDNACLLHPINQHGMEDADRE  119

                                NC_001802.1|NEF  VLEWRFDSRLAFHHVARELHPEYFKNC  205
NZ_JAHVIB010000049.1|Photobacterium_rosenbergii  VLMWKFDSSLARTHRARELHPEFYKNC  206
     NZ_JAHVIU010000019.1|Bacillus_velezensis  VLMWKFDSSLARTHRARELHPEFYKNC  206
   NZ_JBBXLW010000045.1|Acinetobacter_baumannii  VLEWRFDSRLAFHHVARELHPEYFKNC  173
         NZ_WTYD01000009.1|Qipengyuania_pelagi  VLEWRFDSRLAFHHMARELHPD-----  167
      NZ_UKMC02000074.1|Klebsiella_pneumoniae  VLQWQFDSHLARRHMARELHPKYYKDC  190
      NZ_UKJT02000022.1|Klebsiella_pneumoniae  VLKWVFDSQLARRHIARELHPDYYKDC  174
    NZ_LYGA01000021.1|Acinetobacter_baumannii  VLEWRFDSRLAFHHMARELHPD-----  127
      NZ_UKIV02000085.1|Klebsiella_pneumoniae  VLK-KFDTNLARRHLARELHPEYYKD-  179
      NZ_UKLX02000048.1|Klebsiella_pneumoniae  VLKWQFDSHLARRHVARELHPEYYKDC  146
```

### HIV-1 Rev

```
                           NC_001802.1|REV MAGRSGDSDEELIRTVRLIKLLYQSNPPPNPEGTRQARRNRRRRWRERQRQIHSISERIL 60
   NZ_BMPW01000095.1|Actinoplanes_campanulatus MAGRSGDSDEDLLKAVRLIKFLYQSNPPPNPEGTRQARRNRRRRWRERQRQIHSISERIL 60
   NZ_BMOE01000041.1|Deinococcus_aquiradiocola MAGRSGDSDEDLLKAVRLIKFLYQSNPPPNPEGTRQARRNRRRRWRERQRQIHSISERIL 60
      NZ_BMDT01000026.1|Enterococcus_alcedinis MAGRSGDSDEDLLKAVRLIKFLYQSNPPPNPEGTRQARRNRRRRWRERQRQIHSISERIL 60
  NZ_JACENN010000031.1|Corynebacterium_glutamicum MAGRSGDSDEDLLKAVRLIKFLYQSNPPPNPEGTRQARRNRRRRWRERQRQIHSISERIL 60
NZ_JAOYUG010000050.1|Lactiplantibacillus_plantarum MAGRSGDSDEDLLKAVRLIKFLYQSNPPPNPEGTRQARRNRRRRWRERQRQIHSISERIL 60
  NZ_JACENK010000032.1|Corynebacterium_glutamicum MAGRSGDSDEDLLKAVRLIKFLYQSNPPPNPEGTRQARRNRRRRWRERQRQIHSISERIL 60
NZ_AZTX01000497.1|Aggregatibacter_actinomycetemcomitans ------------------ILTIIVSDPPPTSEGSRQARRNRRRRWRERQRQIHSISDRIL 42
       NZ_UKIW02000110.1|Klebsiella_pneumoniae ------------------ILTLVVSDPYPKPEGTRQARKNRRRRWRARQRQIHSLSERIL 42
       NZ_UKKD02000023.1|Klebsiella_pneumoniae ---------------------------------------------------INSISERIL 9

                           NC_001802.1|REV GTYLGRSAEPVPLQLPPLERLTLDCNEDCGTSGTQGVGSPQILVESPTVLESGTKE 116
   NZ_BMPW01000095.1|Actinoplanes_campanulatus STYLGRSAEPVPLQLPPLERLTLDCNEDCGTSGTQGVGSPQILVESPTILESGAKE 116
   NZ_BMOE01000041.1|Deinococcus_aquiradiocola STYLGRSAEPVPLQLPPLERLTLDCNEDCGTSGTQGVGSPQILVESPTILESGAKE 116
      NZ_BMDT01000026.1|Enterococcus_alcedinis STYLGRSAEPVPLQLPPLERLTLDCNEDCGTSGTQGVGSPQILVESPTILESGAKE 116
  NZ_JACENN010000031.1|Corynebacterium_glutamicum STYLGRSAEPVPLQLPPLERLTLDCNEDCGTSGTQGVGSPQILVESPTILESGAKE 116
NZ_JAOYUG010000050.1|Lactiplantibacillus_plantarum STYLGRSAEPVPLQLPPLERLTLDCNEDCGTSGTQGVGSPQILVESPTILESGAKE 116
  NZ_JACENK010000032.1|Corynebacterium_glutamicum STYLGRSAEPVPLQLPPLERLTLDCNEDCGTSGTQGVGSPQILVESPTILESGAKE 116
NZ_AZTX01000497.1|Aggregatibacter_actinomycetemcomitans STYLGRSAEPVPLQLPPLERLTLDCNEDCGTPGTQRVGSPQILVESPTVLESGTKE 98
       NZ_UKIW02000110.1|Klebsiella_pneumoniae STCLGRPAEPVPFQLPPIERLHIDCSEGSGTSGTQGVGHP-VSGKSWAVLGSGTKK 97
       NZ_UKKD02000023.1|Klebsiella_pneumoniae STCLGRSEEPVPFQLPPIERLTIDCDESGGTSGTEGVGNS-ISGKSYTILGSRIKK 64
```

### HIV-1 Tat

```
                      NC_001802.1|TAT MEPVDPRLEPWKHPGSQPKTACTNCYCKKCCFHCQVCFITKALGISYGRKKRRQRRRAHQ 60
NZ_UKJU02000098.1|Klebsiella_pneumoniae MEPVDPNLEPWNHPGSQPKTACNNCYCKKCSYHCLVCFQKKGLGISYGRKKRRQRRSAPP 60
NZ_UKIY02000086.1|Klebsiella_pneumoniae MEPVDPNLEPWKHPGSQPKTACNKCYCKHCSYHCLVCFQTKGLGISYGRKKRRQRRSAPP 60
      NZ_UINV01000086.1|Escherichia_coli MDPVDPKLEPWNHPGSQPKTPCNKCYCKQCSYHCLVCFQTKGLGIYYGRKKRRQRRSAPP 60
NZ_UKJW02000092.1|Klebsiella_pneumoniae MEPIDPNLEPWNHPGSQPKTACNKCYCKACSYHCLVCFQTKGLGIS-------------- 46
NZ_UKKB02000095.1|Klebsiella_pneumoniae MEPVDPNLEPWNHPGSQPATPCNKCYCKICSYHCPVCFLTKGLG---------------- 44
   NZ_JAPTGI010000094.1|Escherichia_coli -------------PGSQPKTACTNCYCKRCSYHCQVCFLTKGLGISYGRKKRRQRRSAPP 47
   NZ_JAODTC010000066.1|Escherichia_coli -------------PGSQPKTACTNCYCKRCSYHCQVCFLTKGLGISYGRKKRRQRRSAPP 47
   NZ_JARBDL010000112.1|Escherichia_coli -------------PGSQPKTACTNCYCKRCSYHCPVCFLTKGLGISYGRKKRRQRRSAPP 47
   NZ_JAPMNL010000108.1|Escherichia_coli ------------HPGSQPKTACNQCYCKRCSYHCLVCFQKKGLGISYGRKKRRQRRSAPP 48

                      NC_001802.1|TAT NSQTHQASLSKQ 72
NZ_UKJU02000098.1|Klebsiella_pneumoniae SSEDHQNLVSKQ 72
NZ_UKIY02000086.1|Klebsiella_pneumoniae SSEDHQDPISKQ 72
      NZ_UINV01000086.1|Escherichia_coli SSEDHQDPLSKQ 72
NZ_UKJW02000092.1|Klebsiella_pneumoniae ------------ 46
NZ_UKKB02000095.1|Klebsiella_pneumoniae ------------ 44
   NZ_JAPTGI010000094.1|Escherichia_coli SSEDHQNLISKQ 59
   NZ_JAODTC010000066.1|Escherichia_coli SSEDHQNLISKQ 59
   NZ_JARBDL010000112.1|Escherichia_coli SSEDHQNLISKQ 59
   NZ_JAPMNL010000108.1|Escherichia_coli SSEDHQNHISKQ 60
```

### HIV-1 Vpu

```
                             NC_001802.1|VPU  EYRKILRQRKIDRLIDRLIERAEDSGNESEGEISALVEMGVEMGHHAPWDVDDL  54
NZ_UVID02000088.1|Burkholderia_pseudomallei  EYRKLVRQRKIDWLIERIRERAEDSGNESEGDTEELATM-VDMGHLRLLDVHDL  53
       NZ_JAPMNJ010000082.1|Escherichia_coli  EYRKLLRQRKIDWLIKRIRERAEDSGNESEGDTEELSTM-VDMGHLRLLDVNDL  53
       NZ_JARBDG010000096.1|Escherichia_coli  EYRKLLRQRKIDWLIKRIRERAEDSGNESEGDTEELSTM-VDMGHLRLLDVNDL  53
          NZ_UINV01000086.1|Escherichia_coli  EYRKIVRQRKIDRLIERIRERAEDSGNESEGDTEELATM-VDMGH---------  44
       NZ_JAPTGJ010000092.1|Escherichia_coli  EYRKLVSQRKIDRLIERIRERAEDSGNESEGDTEELSTM-VDMGHLRLLDVNEL  53
    NZ_UKJP02000103.1|Klebsiella_pneumoniae  EYRKIVRQRKIDWLIKRIRERAEDSGNESEGDNEDLATM-VDMEHLRLLDVNDL  53
    NZ_UKJK02000062.1|Klebsiella_pneumoniae  EYRKLLRQKRIDQLIKRIRERAEDSGNESDGDTEELSTM-VDMGHLRLLDGNDL  53
       NZ_JAODTD010000081.1|Escherichia_coli  EYRKLLRQKKIDWLIERIRERAEDSGNESEGDTEELSTM-VDMGHLRLLDVNEL  53
       NZ_JARBDE010000099.1|Escherichia_coli  EYRKLVKQRKIDRIIKRIRERAEDSGNESEGDIEELSTM-VDMGQLRLLDANDL  53
```

### HIV-1 Vif

```
                       NC_001802.1|VIF MENRWQVMIVWQVDRMRIRTWKSLVKHHMYVSGKARGWFYRHHYESPHPRISSEVHIPLG  60
 NZ_JACEOG010000003.1|Aeromicrobium_phoceense --SRWQVMIVWQVDRMRINTWKRLVKHHMYISRKAKDWFYRHHYESTNPKISSEVHIPLG  58
   NZ_CALNWT010000029.1|Pseudomonas_sp --SRWQVMIVWQVDRMRINTWKRLVKHHMYISRKAKDWFYRHHYESTNPKISSEVHIPLG  58
  NZ_LR732595.1|Streptomyces_mexicanus ---RWQVMIVWQVDRMRINTWKRLVKHHMYISRKAKDWFYRHHYESTNPKISSEVHIPLG  57
NZ_JACMHY010000029.1|Streptomyces_mexicanus ------------VDRMRINTWKRLVKHHMYISRKAKDWFYRHHYESTNPKISSEVHIPLG  48
      NZ_UINV01000086.1|Escherichia_coli MENRWQALIVWQVDRMKIRTWNSLVKHHMYISERASGWFYKHHYESRHPKVSSEVHIPLG  60
  NZ_UKMC02000071.1|Klebsiella_pneumoniae IENRWQVLIVWQVDRMKIRT-NSLVKHHMYVSKRASR-FYRHHYESRNPKISSEVHIPVE  58
  NZ_CABVOZ010000006.1|Anaerococcus_sp ---RWQVMIVWQVDRMRINTWKRLVKHHMYISRKAKDWFYRHHYESTNPKISSEVHIPLG  57
  NZ_UKJL02000050.1|Klebsiella_pneumoniae MENRWQVLIVWQVDRMKIRTWNSLVKHHMYVSKKANRWVYRHHYESRHPRISSEVHIPLG  60
  NZ_UKJH02000063.1|Klebsiella_pneumoniae -----------------------------------ANGFYRHHYESRHPKISSEVHIPLG  25

                       NC_001802.1|VIF DARLVITTYWGLHTGERDWHLGQGVSIEWRKKRYSTQVDPELADQLIHLYYFDCFSDSAI 120
 NZ_JACEOG010000003.1|Aeromicrobium_phoceense DAKLVITTYWGLHTGERDWHLGQGVSIEWRKKRYSTQVDPDLADQLIHLHYFDCFSESAI 118
   NZ_CALNWT010000029.1|Pseudomonas_sp DAKLVITTYWGLHTGERDWHLGQGVSIEWRKKRYSTQVDPDLADQLIHLHYFDCFSESAI 118
  NZ_LR732595.1|Streptomyces_mexicanus DAKLVITTYWGLHTGERDWHLGQGVSIEWRKKRYSTQVDPDLADQLIHLHYFDCFSESAI 117
NZ_JACMHY010000029.1|Streptomyces_mexicanus DAKLVITTYWGLHTGERDWHLGQGVSIEWRKKRYSTQVDPDLADQLIHLHYFDCFSESAI 108
      NZ_UINV01000086.1|Escherichia_coli EAKLVITTYWGLQTGERDWHLGHGVSIEWRLRRYSTQVDPDMADQLIHIHYFDCFADSAI 120
  NZ_UKMC02000071.1|Klebsiella_pneumoniae EARLVIITY-GLQTGERE-HLGHRVSIEWRLRRYSTQVDPGLADQLIHMHYFDCFADSAI 116
  NZ_CABVOZ010000006.1|Anaerococcus_sp DAKLVITTYWGLHTGERDWHLGQGVSIEWRKKRYSTQVDPDLADQLIHLHYFDCFSESAI 117
  NZ_UKJL02000050.1|Klebsiella_pneumoniae DARLIIKTYWGLHTGEREWHLGHGVSIEWRLREYSTQVDPGLADQLIHMHYFDCFADSAI 120
  NZ_UKJH02000063.1|Klebsiella_pneumoniae DARLVITTYWGLNTGEREWHLGHGVSIEWRLRRYSTQVDPGLADQLIHMHYFDCFTDSAI  85

                       NC_001802.1|VIF RKALLGHIVSPRCEYQAGHNKVGSLQYLALAALITPKKIKPPLPSVTKLTEDRWNKPQKT 180
 NZ_JACEOG010000003.1|Aeromicrobium_phoceense RNTILGRIVSPRCEYQAGHNKVGSLQYLALAALIKPKQIKPPLPSVRKLTEDRWNKPQKT 178
   NZ_CALNWT010000029.1|Pseudomonas_sp RNTILGRIVSPRCEYQAGHNKVGSLQYLALAALIKPKQIKPPLPSVRKLTEDRWNKPQKT 178
  NZ_LR732595.1|Streptomyces_mexicanus RNTILGRIVSPRCEYQAGHNKVGSLQYLALAALIKPKQIKPPLPSVRKLTEDRWNKPQKT 177
NZ_JACMHY010000029.1|Streptomyces_mexicanus RNTILGRIVSPRCEYQAGHNKVGSLQYLALAALIKPKQIKPPLPSVRKLTEDRWNKPQKT 168
      NZ_UINV01000086.1|Escherichia_coli RKAILGHIVIPRCDYPAGHNKVGSLQYLALTALIKPKKRKPPLPSIRKLVEDRWNRPQKT 180
  NZ_UKMC02000071.1|Klebsiella_pneumoniae RKAILRHIVTPRCDYPAGHSQVRSLQYLALTALIKPKRIKPPLPSAKKLVEDR-NKPQKT 175
  NZ_CABVOZ010000006.1|Anaerococcus_sp RNTILGRIVSPRC----------------------------------------------- 130
  NZ_UKJL02000050.1|Klebsiella_pneumoniae KKAILG------------------------------------------------------ 126
  NZ_UKJH02000063.1|Klebsiella_pneumoniae RKAILGHIVFHRCEYQAGHNKVGSLQYLALTALIKPRKLKPPLPSVRKLV---------- 135

                       NC_001802.1|VIF KGHRGSHTMNGH 192
 NZ_JACEOG010000003.1|Aeromicrobium_phoceense KGHRGSHTMNGH 190
   NZ_CALNWT010000029.1|Pseudomonas_sp KGHRGSHTMNGH 190
  NZ_LR732595.1|Streptomyces_mexicanus KGHRGSHTMNGH 189
NZ_JACMHY010000029.1|Streptomyces_mexicanus KGHRGSHTMNGH 180
      NZ_UINV01000086.1|Escherichia_coli RGRRGNRTTNGH 192
  NZ_UKMC02000071.1|Klebsiella_pneumoniae KGRRRSHTMNGH 187
  NZ_CABVOZ010000006.1|Anaerococcus_sp ------------ 130
  NZ_UKJL02000050.1|Klebsiella_pneumoniae ------------ 126
  NZ_UKJH02000063.1|Klebsiella_pneumoniae ------------ 135
```

### HIV-1 Vpr

```
NC_001802.1|VPR                              MEQAPEDQGPQREPHNEWTLELLEELKNEAVRHFPRIWLHGLGQHIYETYGDTWAGVEAI 60
NZ_JACEOG010000003.1|Aeromicrobium_phoceense MEQAPEDQGPQREPYNEWTLELLEELKSEAVRHFPRIWLHNLGQHIYETYGDTWAGVEAI 60
NZ_JACMHY010000029.1|Streptomyces_mexicanus  MEQAPEDQGPQREPYNEWTLELLEELKSEAVRHFPRIWLHNLGQHIYETYGDTWAGVEAI 60
NZ_CALNWT010000029.1|Pseudomonas_sp          MEQAPEDQGPQREPYNEWTLELLEELKSEAVRHFPRIWLHNLGQHIYETYGDTWAGVEAI 60
NZ_LR732595.1|Streptomyces_mexicanus         MEQAPEDQGPQREPYNEWTLELLEELKSEAVRHFPRIWLHNLGQHIYETYGDTWAGVEAI 60
NZ_CABVOZ010000006.1|Anaerococcus_sp         MEQAPEDQGPQREPYNEWTLELLEELKSEAVRHFPRIWLHNLGQHIYETYGDTWAGVEAI 60
NZ_UKJK02000067.1|Klebsiella_pneumoniae      MEQTPEDQGPQREPYNEWTLELLEELKQEAVRHFPRPWLHGLGQHIYETYGDTWAGVEAL 60
NZ_UINV01000086.1|Escherichia_coli           MEQAPEDQGPQREPYNEWTLEILKELKQEAVRHFPRPWLHLLGQHIHDTYGDTWTGVEAM 60
NZ_UKMC02000071.1|Klebsiella_pneumoniae      VEQAPEDQRPQKEPYNEWTLELLEELKQEAVRHFPRP-LHSLGQYVYETYRDTWTGVEAI 59
NZ_UKKB02000095.1|Klebsiella_pneumoniae      ----------------EWALEILEELKQEAVRHFPRVWLHNLGQYIYATYGDTWTGVEAL 44

NC_001802.1|VPR                              IRILQQLLFIHFRIGCRHSRIGVTRQRRARNGASRS                         96
NZ_JACEOG010000003.1|Aeromicrobium_phoceense IRILQQLLFIHFRIGCRHSRIGVTRQRRARNGASRS                         96
NZ_JACMHY010000029.1|Streptomyces_mexicanus  IRILQQLLFIHFRIGCRHSRIGVTRQRRARNGASRS                         96
NZ_CALNWT010000029.1|Pseudomonas_sp          IRILQQLLFIHFRIGCRHSRIGVTRQRRARNGASRS                         96
NZ_LR732595.1|Streptomyces_mexicanus         IRILQQLLFIHFRIGCRHSRIGVTRQRRARNGASRS                         96
NZ_CABVOZ010000006.1|Anaerococcus_sp         IRILQQLLFIHFRIGCRHSRIGVTRQRRARNGASRS                         96
NZ_UKJK02000067.1|Klebsiella_pneumoniae      IRILQQLLFIHFRIGCQHSRIGILRQRRARNG----                         92
NZ_UINV01000086.1|Escherichia_coli           IRTLQQLLFIHFRIGCQHSRIGILPQRRARNGSSRS                         96
NZ_UKMC02000071.1|Klebsiella_pneumoniae      IRILQQLLFVHFRISCQHSRIGILRQRRAKDRASRS                         95
NZ_UKKB02000095.1|Klebsiella_pneumoniae      IRTLQQLLFIHFRIGCQHSRIGILQQRRARNGARRS                         80
```
