## Supplementary Table S1 for "HIV-1 protein coding sequences are present in relevant bacteria"

**Table S1.** HIV-1 proteome coverage statistics in *E. coli* and *K. pneumoniae.*

| Bacteria | *E. coli* | | | | *K. pneumoniae* | | | |
|---|---|---|---|---|---|---|---|---|
| **Viral protein** | **RBS [1]** | **E-value** | **Q-coverage** | **Identity** | **RBS [1]** | **E-value** | **Q-coverage** | **Identity** |
| **gp120** | 70.37% | 0 | 99% | 70.88% | 47.15% | $1\times10^{-160}$ | 70% | 68.24% |
| **gp41** | 79.49% | $4\times10^{-147}$ | 92% | 67.79% | 43.91% | $2\times10^{-75}$ | 52% | 64.55% |
| **p24** | 18.22% | $3\times10^{-15}$ | 18%% | 93.02% | 54.15% | $3\times10^{-85}$ | 56% | 91.6% |
| **p17** | – | – | – | – | – | – | – | – |
| **p7** | 39.5% | $2\times10^{-4}$ | 67% | 48.65% | 85.83% | $1\times10^{-26}$ | 100% | 85.45% |
| **p6gag** | 58.11% | $9\times10^{-9}$ | 76% | 75% | 74.53% | $7\times10^{-15}$ | 86% | 88.89% |
| **RT p51** | 72.35% | 0 | 78% | 86.17% | 96.26% | 0 | 100% | 92.5% |
| **Protease** | 96.53% | $1\times10^{-57}$ | 100% | 94.95% | 98.51% | $2\times10^{-59}$ | 100% | 97.98% |
| **Integrase** | 95.51% | 0 | 100% | 93.75% | 83.72% | $7\times10^{-107}$ | 86% | 94.4% |
| **p15** | 86.25% | $9\times10^{-59}$ | 100% | 85% | 95.83% | $1\times10^{-67}$ | 100% | 95% |
| **p6pol** | 54.41% | $4\times10^{-6}$ | 93% | 63.46% | 67.94% | $5\times10^{-11}$ | 68% | 94.94% |
| **Nef** | 46.27% | $5\times10^{-55}$ | 42% | 60.23% | 69.15% | $1\times10^{-87}$ | 92% | 65.97% |
| **Rev** | 63.91% | $2\times10^{-13}$ | 99% | 50% | 47.83% | $4\times10^{-21}$ | 84% | 54.08% |
| **Tat** | 72.31% | $2\times10^{-19}$ | 93% | 56.25% | 74.92% | $5\times10^{-23}$ | 83% | 63.89% |
| **Vpu** | 58.26% | $7\times10^{-9}$ | 65% | 66.67% | 54.04% | $4\times10^{-8}$ | 65% | 64.81% |
| **Vif** | 83.6% | $1\times10^{-93}$ | 100% | 74.48% | 73.92% | $3\times10^{-89}$ | 100% | 70.83% |
| **Vpr** | 88% | $3\times10^{-46}$ | 100% | 83.33% | 84% | $3\times10^{-49}$ | 95% | 91.3% |
| **VPC [2]** | **63.71%** | | | | **67.74%** | | | |

Table shows TBLASTN statistics of best hits between HIV-1 proteins and E. coli and K. pneumoniae. [1] Relative Bit-Score, RBS, computed dividing the bit-score of the relevant hit by the maximum bit-score of the query obtained from a self-hit. [2]Viral Proteome Coverage, VPC, is the sum of RBS values divided by the total number of proteins. RBS and VPC are in percentage (details in Methods). – No hit.
