## Supplementary Table S2 for "HIV-1 protein coding sequences are present in relevant bacteria"

**Table S2.** IAV proteome coverage in *L. monocytogenes*.

| Viral protein | Bit-Score | mBS [1] | RBS [2] | E-value | Q-coverage | Identity |
|---|---|---|---|---|---|---|
| Polymerase PB2 | 1476 | 1529 | 96.53% | 0 | 97% | 96.22% |
| Polymerase PB1 | 1577 | 1590 | 99.18% | 0 | 99% | 100% |
| PB1-F2 protein | 158 | 184 | 85.86% | $4\times10^{-42}$ | 100% | 87.78% |
| Polymerase PA | 1212 | 1454 | 83.36% | 0 | 84% | 96.38 |
| PA-X protein | 399 | 400 | 99.75% | $9\times10^{-137}$ | 100% | 99.48% |
| Hemagglutinin | 1155 | 1170 | 98.72% | 0 | 100% | 98.21% |
| Nucleocapsid protein | 1032 | 1032 | 100% | 0 | 100 | 99.8 |
| Matrix protein 2 | 161 | 162 | 99.38 | $2\times10^{-45}$ | 97% | 80% |
| Matrix protein 1 | 524 | 524 | 100% | 0 | 100% | 100% |
| Nuclear export protein | 200 | 200 | 100% | $2\times10^{-61}$ | 93% | 89.38 |
| Nonstructural protein 1 | 444 | 444 | 100% | $2\times10^{-155}$ | 100% | 99.54 % |
| Neuraminidase | 886 | 892 | 99.33% | 0 | 100% | 92.04 |
| **VPC [3]** | 96.84% | | | | | |

Table shows TBLASTN statistics of best hits between IAV proteins and *L. monocytogenes*. [1] Maximum Bit-Score, mBS, obtained from a self-hit alignment. [2] Relative Bit-Score, RBS, computed dividing the bit-score of the relevant hit by the mBS. [3] Viral Proteome Coverage, VPC, is the sum of RBS values divided by the total number of proteins. RBS and VPC are in percentage (details in Methods).
