## Supplementary Table S3 for "HIV-1 protein coding sequences are present in relevant bacteria"

**Table S3.** HBV proteome coverage in *Streptomyces sp. 2BBP J2*.

| Viral protein | Bit-Score | mBS [1] | RBS [2] | E-value | Q-coverage | Identity |
|---|---|---|---|---|---|---|
| MEP | 461 | 461 | 100% | $4\times10^{-151}$ | 100% | 84.7% |
| LEP | 677 | 677 | 100% | 0 | 99% | 88.69% |
| Pol | 1568 | 1568 | 100% | 0 | 100% | 92.43% |
| C0 uORF | 48.9 | 48.9 | 100% | $7\times10^{-5}$ | 100% | 100% |
| Capsid protein | 305 | 305 | 100% | $2\times10^{-93}$ | 81% | 100% |
| X | 275 | 285 | 96.49% | $1\times10^{-82}$ | 97% | 90% |
| SEP | 357 | 357 | 100% | $3\times10^{-112}$ | 100% | 81.86% |
| Precapsid prot | 362 | 366 | 98.91% | $3\times10^{-147}$ | 83% | 100% |
| **VPC [3]** | 99.43% | | | | | |

Table shows TBLASTN statistics of best hits between HBV proteins and *Streptomyces sp. 2BBP J2*. [1] Maximum Bit-Score, mBS, obtained from a self-hit alignment. [2] Relative Bit-Score, RBS, computed dividing the bit-score of the relevant hit by the mBS. [3] Viral Proteome Coverage, VPC, is the sum of RBS values divided by the total number of proteins. RBS and VPC are in percentage (details in Methods).
