## Supplementary Table S4 for "HIV-1 protein coding sequences are present in relevant bacteria"

**Table S4.** IAV proteome coverage in *E. coli.*

| Viral protein | Bit-Score | mBS [1] | RBS [2] | E-value | Q-coverage | Identity |
|---|---|---|---|---|---|---|
| Polymerase PB2 | 668 | 1529 | 43.69% | 0 | 43% | 98.17% |
| Polymerase PB1 | 622 | 1590 | 39.12% | 0 | 40% | 98.36% |
| PB1-F2 protein | – | 184 | – | – | – | – |
| Polymerase PA | 1079 | 1454 | 74.27% | 0 | 64% | 99.18% |
| PA-X protein | 209 | 400 | 52.25% | $9{\times}10^{-74}$ | 76% | 99.75 |
| Hemagglutinin | 553 | 1170 | 47.27% | 0 | 48% | 95.93% |
| Nucleocapsid protein | – | 1032 | – | – | – | – |
| Matrix protein 2 | 143 | 162 | 88.27% | $1{\times}10^{-38}$ | 96% | 70.21% |
| Matrix protein 1 | 499. | 524 | 95.23% | $2{\times}10^{-176}$ | 100% | 93.65% |
| Nuclear export protein | 191 | 200 | 95.5% | $1{\times}10^{-57}$ | 93% | 85.84% |
| Nonstructural protein 1 | 398 | 444 | 89.64% | $2{\times}10^{-137}$ | 100% | 88.48% |
| Neuraminidase | 284 | 892 | 31.81% | $1{\times}10^{-90}$ | 45% | 62.86% |
| **VPC [3]** | 54.75% | | | | | |

Table shows TBLASTN statistics of best hits between IAV proteins and *E. coli*. [1] Maximum Bit-Score, mBS, obtained from a self-hit alignment. [2] Relative Bit-Score, RBS, computed dividing the bit-score of the relevant hit by the mBS. [3] Viral Proteome Coverage, VPC, is the sum of RBS values divided by the total number of proteins. RBS and VPC are in percentage (details in Methods). – No hit.

**Table S5.** HBV proteome coverage in *K. pneumonia.*

| Viral protein | Bit-Score | mBS [1] | RBS [2] | E-value | Q-coverage | Identity |
|---|---|---|---|---|---|---|
| MEP | 335 | 461 | 72.66% | $1\times10^{-100}$ | 100% | 74.78 |
| LEP | 500 | 677 | 73.90% | 0 | 83% | 87.42 % |
| Pol | 1372 | 1568 | 87.5% | 0 | 100% | 80.43% |
| C0 uORF | - | 48.9 | - | - | - | - |
| Capsid protein | 295 | 305 | 96.72% | $4\times10^{-88}$ | 81% | 94.63% |
| X | - | 285 | - | - | - | - |
| SEP | 334 | 357 | 93.48% | $1\times10^{-99}$ | 100% | 74.78% |
| Precapsid prot | 357 | 366 | 97.54% | $7\times10^{-110}$ | 83% | 95.51% |
| **VPC [3]** | **65.22%** | | | | | |

Table shows TBLASTN statistics of best hits between HBV proteins and *K. pneumoniae*. [1] Maximum Bit-Score, mBS, obtained from a self-hit alignment. [2] Relative Bit-Score, RBS, computed dividing the bit-score of the relevant hit by the mBS. [3] Viral Proteome Coverage, VPC, is the sum of RBS values divided by the total number of proteins. RBS and VPC are in percentage (details in Methods).
